# C1qa□ muscularis macrophages maintain enteric synaptic homeostasis to regulate gastrointestinal motility

**DOI:** 10.64898/2026.06.03.729640

**Authors:** Mario D’Ambrosio, Juan Sebastian Ortiz Colmenares, Khaled Warasnhe, Yuanhang Liu, Mohamed H. Eldesouki, Dokic Vladimir, Natalie R Wertish, Xin-yi Chai, Sara Traserra, Trace A. Christensen, Elisabetta Bigagli, Cristina Luceri, Keith A. Sharkey, Madhusudan Grover, Marcel Jiménez, Arthur Beyder, Gianrico Farrugia, Gianluca Cipriani

## Abstract

The enteric nervous system (ENS) is a complex peripheral neural network that coordinates gastrointestinal motility through highly organized synaptic communication. Although tissue-resident muscularis macrophages (MMs) closely associate with enteric neurons, whether they regulate enteric synaptic organization remains unknown. In the central nervous system (CNS), microglia sculpt neural circuits through complement-dependent synaptic remodeling, raising the possibility that analogous neuroimmune mechanisms operate in the gut. Here, we identify a previously unrecognized role for C1qa□ MMs in regulating enteric synaptic homeostasis and gastrointestinal motility. Using macrophage-specific constitutive and inducible C1qa deletion models, single-cell RNA sequencing, enteric synaptosome proteomics, physiology, and advanced imaging, we demonstrate that loss of MMs-derived C1qa increases enteric synaptic density without altering neuronal numbers. C1qa deficiency induced broad transcriptional changes in enteric neurons and macrophages, including altered synapse-associated, lysosomal, and endocytic programs. Proteomic analysis revealed that enteric synapses share a conserved molecular architecture with brain synapses while exhibiting distinct gastrointestinal-specific complement-associated synaptic networks enriched for structural and receptor-localization pathways.

Functionally, macrophage-specific C1qa deletion altered excitatory and inhibitory enteric neurotransmission, enhanced cholinergic signaling, reduced nitrergic responses, and accelerated gastrointestinal transit, while smooth muscle responsiveness remained preserved. C1qa□ MMs displayed transcriptional and functional features consistent with a phagocytic synapse-remodeling phenotype, including enrichment of complement, lysosomal, and engulfment pathways. Loss of C1qa impaired macrophage phagocytic activity both *in vitro* and *in vivo* and was associated with synapse accumulation and altered macrophage morphology. Importantly, inducible deletion of C1qa in adulthood recapitulated the synaptic and motility phenotypes, demonstrating that C1qa□ MMs continuously regulate enteric synaptic organization beyond development.

Together, these findings identify a complement-dependent neuroimmune mechanism that regulates enteric circuit organization and gut motility, establishing MMs as active modulators of adult ENS synaptic homeostasis.

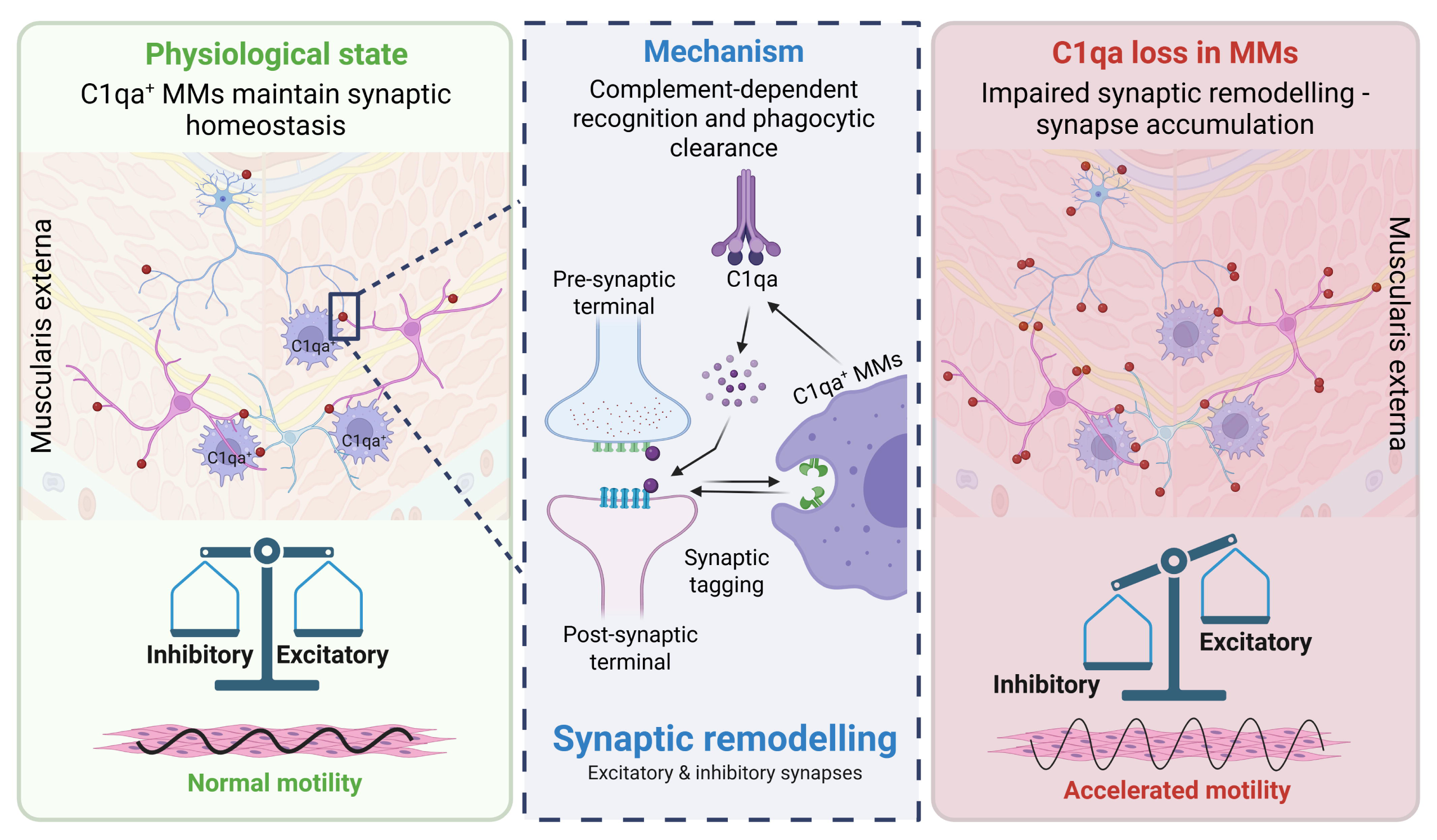

## INTRODUCTION

The gastrointestinal (GI) tract is innervated by the enteric nervous system (ENS), a complex, semi-autonomous neural network often referred to as the “second brain” due to its ability to independently control digestive functions including motility, secretion, and local blood flow (**1–4**). This intricate control relies on a highly organized synaptic architecture that enables precise communication between diverse neuronal populations and other cell types in the gut wall (**2,5**). The ENS operates within a dynamic microenvironment shared with specialized immune cells, called muscularis macrophages (MMs), establishing pivotal neuro-immune interactions that are critical for maintaining gut homeostasis and preventing disease (**6–11**). However, the precise mechanisms by which MMs dynamically regulate the structural and functional plasticity of ENS circuitry remain largely unknown. In the central nervous system (CNS), the role of local immune cells, microglia, in shaping neuronal circuits through synaptic pruning is well-established (**12,13**). This process involves components of the classical complement pathway, where the primary component C1q tags weak synapses, leading to their opsonization by C3 fragments and subsequent phagocytic engulfment by microglia (**13–16**). This tightly regulated mechanism is vital for synaptic refinement during development, and its dysregulation contributes to neurodevelopmental disorders and neurodegenerative diseases, highlighting complement signaling as a key regulator of synaptic homeostasis (**13,16,17**). Given the parallels between the ENS and CNS, particularly in their complex neuronal network organization and intimate association with tissue-resident immune cells, we hypothesized that a similar complement-mediated synaptic pruning mechanism might operate within the gut, orchestrated by resident MMs.

Independent studies, including our own, have identified a subset of MMs characterized by the expression of C1qa, the initiating component of the classical complement cascade, and demonstrated their association with enteric neurons (**10,11**). These studies showed that macrophage-specific C1qa deletion accelerates GI transit and alters neuronal gene expression, suggesting a regulatory role for C1qa^+^ MMs in ENS function. However, the precise molecular and cellular mechanisms by which C1qa^+^ MMs regulate ENS circuitry, and whether this involves synaptic pruning, remain unknown.

To address this critical gap, we investigated the role of C1qa^+^ MMs in maintaining enteric synaptic homeostasis. We first developed and validated a method to isolate gut synaptosomes, enabling the characterization of the complex proteomic landscape of C1q-dependent enteric synapses and their comparison with brain synapses. We then used genetic mouse models, single□cell RNA sequencing, *ex vivo* and *in vivo* functional assays, and advanced imaging to dissect the effects of C1qa deficiency on ENS synaptic architecture and GI motility. Our findings support a model in which C1qa^+^ MMs regulate ENS synaptic density through a phagocytic synaptic remodeling mechanism that persists in adulthood. These findings establish a novel neuro-immune axis in the gut, providing a mechanistic framework for enteric circuit regulation in health and disease, with potential implications for the development of therapeutic strategies aimed at targeting complement-dependent pathways.

## METHODS

### Transgenic mouse models

All procedures were carried out in accordance with the NIH Guide for the Care and Use of Laboratory Animals and approved by the Mayo Clinic Institutional Animal Care and Use Committee. We generated a constitutive C1qa knockout in macrophages (***C1qa^CKO^***) by crossing C1qa flox mice (***C1qa^FLX^***; B6(SJL) - C1qatm1c(EUCOMM)Wtsi/TennJ - Jackson# 031261) with Lyz2^Cre^ (B6.129P2-Lyz2tm1(cre)Ifo/J - Jackson# 004781) mice, and an inducible C1qa knockout from macrophages (***iC1qa^CKO^***) by crossing C1qa flox mice (B6(SJL) - C1qatm1c(EUCOMM)Wtsi/TennJ - Jackson# 031261) with Lyz2^CreERT2^ (Lyz2tm1(cre/ERT2)Grtn/J - Jackson# 031674) mice. We also generated a constitutive mouse model (Lyz2^eGFP^) in which all macrophages express the Enhanced Green Fluorescent Protein (EGFP) gene inserted into the *Gt(ROSA)26Sor* locus by crossing Lyz2^Cre^ (B6.129P2-Lyz2tm1(cre)Ifo/J - Jackson# 004781) mice with ROSA26-EGFP^flox^ (B6;129-Gt(ROSA)26Sortm2Sho/J - Jackson# 004077). Wild-type (WT - C57BL/6J - Jackson# 000664) mice were used for synaptosome isolation. Mice were genotyped using the automated genotyping system Transnetyx (Cordova, TN, USA). Male mice, 3-4 months of age, were used for the experiments. Male mice were used to minimize biological variability and maintain consistency across experimental groups. Water and food were available *ad libitum*, the lighting regime was a standard 12 h light and 12 h dark, temperature was maintained constant at 21 ± 2°C. Mice were humanely euthanized by carbon dioxide exposure followed by cervical dislocation in accordance with established institutional protocols.

### Tamoxifen-induced *Cre* activity

Temporal control of cell-specific recombination (***iC1qa^CKO^***) was obtained by administering tamoxifen (10mg/ml in corn oil; Sigma-Aldrich, St. Louis, MO, USA) to 3-4-month-old Lyz2^CreERT2^*C1qa mice via daily intraperitoneal doses (75mg/kg) for three consecutive days. All experiments were performed after a 2□week tamoxifen washout. Age-matched corn-oil injected Lyz2^CreERT2^*C1qa mice were used as controls (Vehicle).

### Whole gut transit assay

Whole Gut transit was performed as described previously (**18**). Briefly, carmine red solution (6% w/v; Sigma Aldrich) prepared in methylcellulose (0.5% w/v: Sigma Aldrich) was administered by gastric gavage. Mice were singly housed to follow pellets output until the appearance of the first red pellet.

### Single-cell sequencing

Single-cell sequencing was performed on the muscularis externa obtained from the colon, of age-matched C1qa^FL^, C1qa^CKO^ and iC1qa^CKO^ mice. Cells were assessed for viability using the Vi-Cell XR Cell Viability Analyzer (Beckman-Coulter, Indianapolis, IN, USA), and their count was determined. Following this, barcoded Gel Beads were thawed, and a cDNA master mix was prepared according to Chromium Single Cell 3’ v3 library kit instructions. Cells were mixed with the master mix and Gel Beads, then processed using the Chromium Controller to capture individual cells in uniquely labeled Gel Beads-In-Emulsion (GEMs). After collecting the GEMs, reverse transcription, dissolution, and cDNA clean-up were performed, resulting in a pool of barcoded cDNA. This cDNA was then used to construct libraries, incorporating Illumina sequencing primers and unique i7 Sample index, and assessed for quality using Qubit High Sensitivity assays (Thermo Fisher Scientific, Waltham, MA, USA), Agilent Bioanalyzer High Sensitivity chips (Agilent, Santa Clara, CA, USA), and Kapa DNA Quantification reagents (Kapa Biosystems, Wilmington, MA, USA). Sequencing was conducted at a rate of 60,000 fragment reads per cell according to Illumina’s standard protocol, using the Illumina cBot and HiSeq 3000/4000 PE Cluster Kit. The flow cells were sequenced as 100 X 2 paired end reads on an Illumina HiSeq 4000 using HiSeq 3000/4000 sequencing kit and HCS v3.3.52 collection software. Base-calling was performed using Illumina’s RTA version 2.7.3. Raw BCL files were demultiplexed into FASTQ sequencing files using the Cell Ranger version 3.1 suite from 10xGenomics.

### Single cell RNA-seq preprocessing and quality control

We used 10X Genomics Cell Ranger Single Cell Software Suite (v7.2.0) to perform alignment to mm10 reference genome, filtering, barcode counting, and UMI counting. For subsequent analysis, we followed the standard integrated analysis workflow in the Seurat package (v4) (**19**). Cell doublet identification was performed using scDblFinder (**20**). Predicted doublets were removed from downstream analysis. The remaining cells that met any of the criteria below were further removed from downstream analysis for each sample: 1). Cells expressing fewer than 50 genes; 2) Cells with a total UMI lower than 200; 3). Cells with percentage of mitochondrial reads > 40%; 4). Cells with a percentage of hemoglobin concentration higher than 20%. Genes expressed in fewer than 10 cells were removed from downstream analysis. Each sample was normalized using SCTransform and scaled for each gene across all cells (**21**). Sample C1qa^FL^ and C1qa^CKO^ were first integrated and clustered on the low-dimensional space with resolution 0.3 to identify 23 unsupervised clusters. Sample iC1qa^CKO^ was then mapped to the integrated object using Seurat’s FindTransferAnchors.

### Cell-type annotation of macrophage and enteric neuron populations

Canonical marker genes for macrophage and enteric neuron/glia population (**Supplementary Table 1**) were used to calculate cell type activity score using add module score function from Seurat, and sctype (**22**). Cluster 3 and 22 showed high cell type activity scores for Enteric Neuron (EN). Cluster 22 was not included for downstream analysis due to the limited number of cells. Cluster 5 showed high cell type activity scores for macrophage population. The two cell types were further subclustered to identify four subclusters in each population (EN: 1-4, MM: 1-4).

### Differential expression and pathway analysis

Differential expression analysis was performed using Wilcoxon Rank Sum test as implemented in Presto. Significant differentially expressed genes were identified with Bonferroni adjusted p-value < 0.01 and fold change >1.5 in either direction or percentage of expressing cells > 10% in at least one of the groups. Pathway analysis was performed using GSEA preranked with log2 fold change as the ranking metric (**23**). Biological process from Gene Ontology, and KEGG terms were used as pathway databases for GSEA analysis. Significant altered pathways were identified with FDR < 0.25 (https://mforgeweb.mayo.edu/bsi/archive/PI/Cipriani_Gianluca_m101167/tertiary/s310567.c1qa_scrnaseq/mrnaseq/122223_subcluster/Seurat_report.html).

### Immunohistochemistry

Small intestines and colons were collected, and the muscularis externa separated from the mucosa. Tissues were then fixed with 4% paraformaldehyde (PFA, Sigma Aldrich) in 0.1 M phosphate buffer overnight at 4°C. After fixation, following three washes to remove residual PFA, tissues were incubated with a blocking solution containing 5% Bovine Serum Albumin (BSA, Sigma Aldrich) in 1% Phosphate-buffered saline (PBS, Sigma Aldrich) and 1% TritonX-100 (Sigma Aldrich) overnight at room temperature (RT). After blocking, tissues were labeled in the presence of the primary antibodies overnight at RT (**Table 1**). After washing, the tissues were incubated with the appropriate secondary antibodies (**Table 2**) overnight at RT, in the dark, and then mounted using SlowFade™ Gold Antifade Mountant (Thermo Fisher). For all immunohistochemical studies, controls omitting the primary antibodies and controls in double labeling experiments with a mis-matched secondary antibody were performed. For quantitative analysis, labeled tissues were examined with a laser scanning confocal microscope (Nikon AX-R) using a 20X (NA 0.75 WI) Nikon Apo LWD objective (Nikon, Melville, NY, USA) using identical fluorescence acquisition settings (laser power, gain, and exposure) across all groups. Stacks of confocal images across the full thickness of the muscularis propria were collected from 6 to 8 mice for each antibody and condition, based on the experiment. For labeling quantification, 5 non-adjacent fields of view selected randomly from each mouse. All the confocal image stacks were flattened into projections using NIS elements AR (Nikon) for quantification of the labeling, ensuring that signal from the entire z-depth was included in the analysis. PSD-95 immunoreactivity was quantified as the number of positive perisomatic regions surrounding neuronal cell bodies whereas Synaptophysin (SYP) and Vesicular Acetylcholine Transporter (VAChT) were quantified by mean fluorescence intensity. Labeling quantification has been performed using FIJI image analysis software (https://imagej.net/software/fiji/). For representative micrographs, images were acquired using a 20X (NA 0.75 WI) Nikon Apo LWD objective (Nikon) at a field of view (FOV) from 5 to 7 μm with a pixel size from 0.173 to 0.093 μm/pixel to better illustrate morphological details.

**Table 1:**
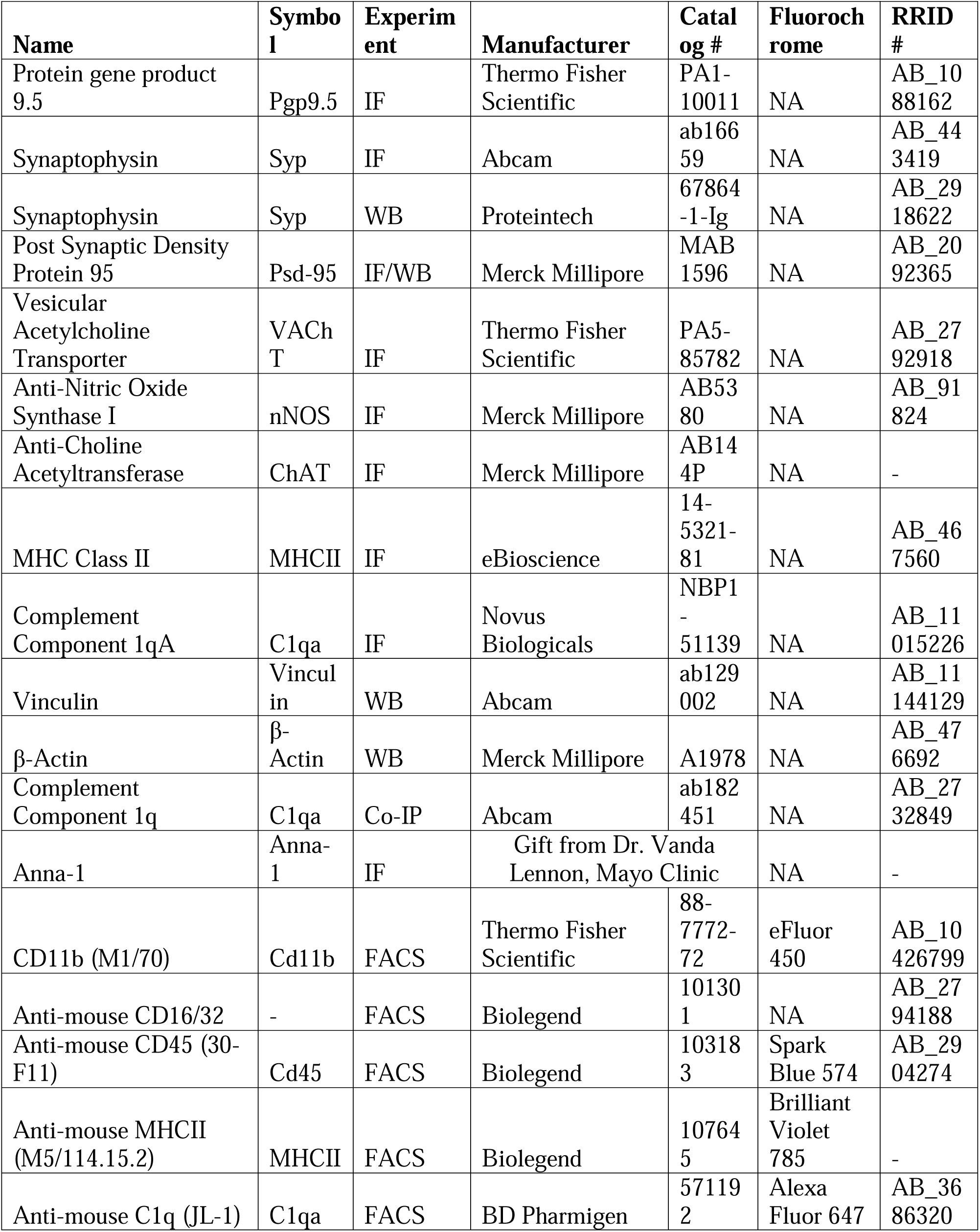
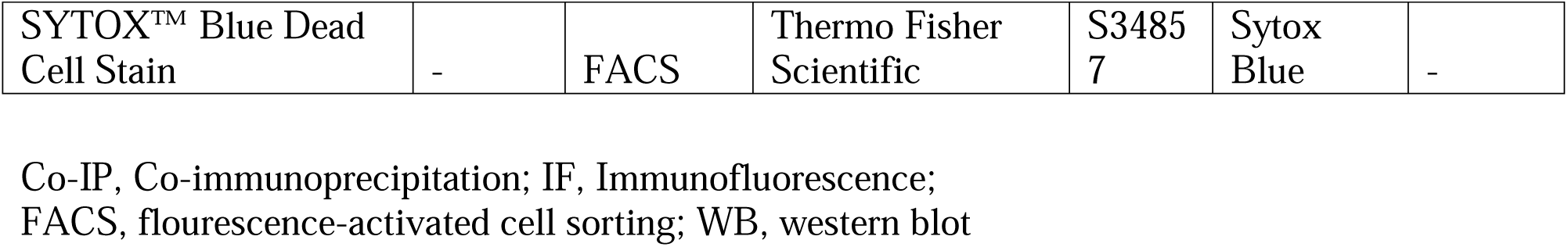
Primary antibody.

**Table 2:**
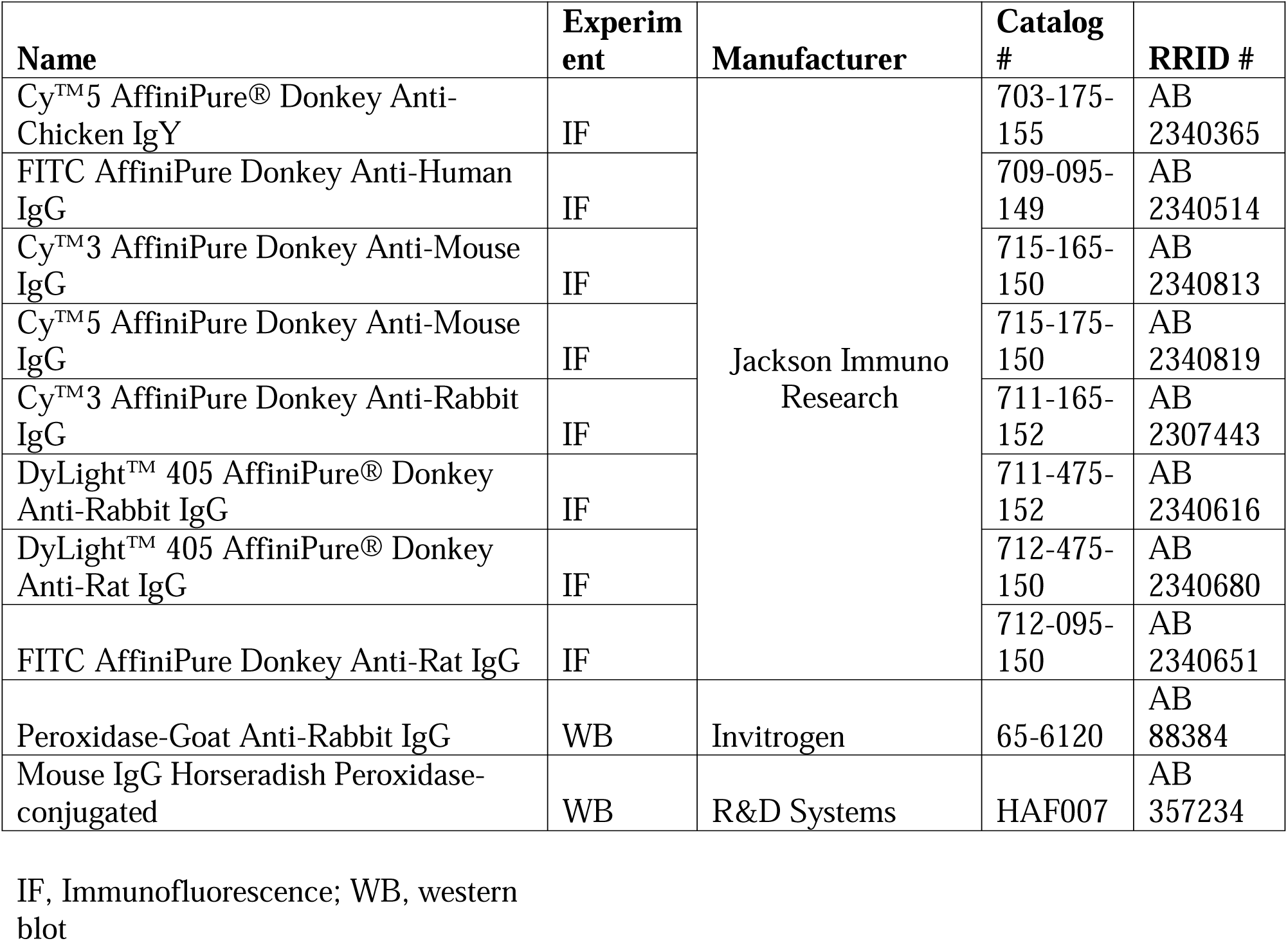
Secondary antibody.

### NADPH diaphorase (NADPH-d) histochemistry

NADPH□d histochemistry was performed as previously described (**24**). Briefly, small intestines and colons were fixed as described above. Wholemount preparations were labeled with neuronal nitric oxide synthase (nNOS, **Table 1**) prior to NADPH-d staining. Following staining, tissues were washed in PBS and incubated with 1% Triton X-100 in PBS for 10 min followed by a 10 min PBS wash. Wholemounts were then incubated in NADPH-d solution (0.1M Tris buffer, pH 7.6, 0.5% Triton X-100, 0.25 mg/mL nitroblue tetrazolium (Thermo Fisher Scientific), 1 mg/mL β-NADPH (Sigma-Aldrich) for 20-30 min at 37°C, with the color reaction monitored throughout. Following this, wholemount preparations were washed in PBS (3x 10 min) and mounted with SlowFade™ Gold Antifade Mountant (Thermo Fisher). For quantification, five randomly selected, non-adjacent fields of view were analyzed per tissue. Images were acquired sequentially from 6 different mice for each condition, first for nNOS and then for NADPH-d staining. Imaging was performed using Zeiss Axio Imager.Z2 motorized upright microscope (Zeiss, White Plains, NY, USA) equipped with a Plan-Apochromat 63x/1.4 Oil objective at 20X magnification for quantitation and 40X magnification for representative images. NADPH-d/nNOS ratio quantification was conducted on FIJI image analysis software (https://imagej.net/software/fiji/).

### Western blot

For western blot analysis, protein extraction was performed on muscularis separated from the mucosa of small intestine and colon. The total protein content was evaluated by using the Bio-Rad DC protein assay kit (Bio-Rad, Hercules, CA, USA), using a BSA standard solution (ranging from 0.2 to 2 mg/mL) for the calibration curve. Twenty micrograms of protein were separated on 4–20% SDS-PAGE (Thermo Fisher Scientific) and transferred into PVDF membranes (60 min at 398 mA) using standard procedures (**25**). Blots were incubated overnight at 4°C with primary antibodies (**Table 1**) diluted in PBS containing 5% BSA or 5% non-fat dry milk and 0.05% Tween 20. The antigen-antibody complexes were visualized using appropriate secondary antibodies (**Table 2**), diluted in PBS containing 5% BSA or 5% non-fat dry milk and 0.05% Tween 20. Blot development was performed using the enhanced chemiluminescence detection system SuperSignal™ Western Blot Enhancer (Thermo Fisher Scientific). Exposure and developing times were standardized for all blots. Densitometric analysis was performed using the public Quantity One 1-D Analysis Software (Bio-Rad). Each gel was loaded with all the experimental groups to standardize image acquisition and densitometric analysis. Data are presented as mean ± SEM and are reported as arbitrary units (AU), consisting of the ratio between the level of the target protein expression and that of β-Actin.

### Synaptosome isolation

Synaptosome isolation was performed on muscularis externa from small intestines and colons, taken from WT male mice (3-4 months old) according to Scott-Hewitt *et al.* (**26**), with the following modification: the synaptic fraction was isolated using the Syn-PER™ Synaptic Protein Extraction Reagent (Thermo Fisher) according to manufacturer’s instructions. Protein content was evaluated by using the Bio-Rad DC protein assay kit (Bio-Rad, Hercules), using a BSA standard solution (ranging from 0.2 to 2 mg/mL) for the calibration curve. Synaptic fraction purity was validated by western blot analysis, by comparison with the relative cytosolic fraction. β-Actin was used as a loading control.

### C1q Co-Immunoprecipitation (Co-IP)

Synaptosome lysates were used immediately for C1q co-IP with a highly specific monoclonal anti-C1q (**Table 1**) antibody, at a concentration of 2μg/mg of protein. Co-IP was performed using the Dynabeads® Protein G Immunoprecipitation kit (Thermo Fisher) according to the manufacturer’s instructions. Protein content was evaluated by using the Bio-Rad DC protein assay kit (Bio-Rad, Hercules), using a BSA standard solution (ranging from 0.2 to 2 mg/mL) for the calibration curve.

### Sample processing for mass spectrometry

The eluted proteins were lyophilized and processed using an S-trap micro cartridge (https://www.Protifi.com) following the vendor’s recommended protocol. Briefly, the proteins were solubilized in 5% SDS / 50mM triethylammonium bicarbonate (TEAB) pH 8.5 and reduced using 5mM TCEP at 90°C for 10 minutes, cooled to RT, then alkylated in the dark with 10mM iodoacetamide at RT for 20 minutes. The solution was acidified with phosphoric acid to a final concentration of 1% and 90% MeOH/100mM TEAB buffer was added to create a protein suspension. The suspension solution was then transferred to the S-trap mini cartridge, centrifuged at 4000 x g for 30 seconds followed by 3 repeated additions and spins with 90% MeOH/100mM TEAB buffer. Protein digestion was performed by adding 125ul of trypsin/lys-C mix (Promega, Madison, WI, USA) in 50mM TEAB pH 8.5 with 1:10 enzyme to protein ratio to the cartridge and incubating at 37°C overnight. Peptides were extracted from the cartridge with 3 elution steps into a clean tube using 50mM TEAB pH 8.5, 0.2% formic acid, then 50% acetonitrile / 0.1% trifluoroacetic acid (TFA) and lyophilized.

### NanoLC-tandem mass spectrometry data acquisition

The protein relative quantitation data was acquired by nanoLC-tandem mass spectrometry using a Thermo Scientific Orbitrap Exploris 480 mass spectrometer coupled to a Thermo Vanquish Neo HPLC system (Thermo Fisher) with 0.1% formic acid in 98% water / 2% acetonitrile for the A solvent and 0.1% formic acid in 80% acetonitrile / 10% isopropanol / 10% water for the B solvent. The peptides were solubilized in 0.2% formic acid and pumped onto a Halo C18 2.7µm EXP stem trap (Optimize Technologies, Oregon City, OR, USA) with A solvent at a flow rate of 8µl / min. The trap was placed in line with a Bruker PepSep C18 1.5µm, 40cm x 100µm column (Bruker, Billerica, MA, USA) and the peptides separated with a gradient of 3%B to 35%B over 120minutes at a flow rate of 350nl / minute. The mass spectrometer data was acquired in data dependent acquisition mode with a 3 second cycle time. The MS1 survey scan range was 350-1600 m/z at resolution of 120,000 (200m/z) with the automatic gain control (AGC) set to 2x106 ions (200%) and ion inject time set to 50ms. Precursor ions with positive charge states from 2-5, were sequentially selected with an isolation window of 1.2 m/z and fragmented with high energy collisional dissociation (HCD) with a normal collision energy (NCE) setting of 30% and scanned at resolution 15,000 with the AGC setting at 80% (8x104 ions) and the ion inject time set to 60ms. The dynamic exclusion feature was used to prevent any ion within a 10ppm window of the selected ion, from being chosen again for MS/MS fragmentation for at least 30 seconds.

### Protein identification and analysis

The mass spectrometry raw data files were searched with MSFragger against the Swissprot mouse database (ver. 2024_06) with parameters set for full trypsin specificity with oxidized Met and N-term protein acetylation allowed as variable modifications and carbamidomethyl cysteine as a fixed modification. Mass tolerances were set at 20ppm for MS1 and 20ppm for MS2 and protein identifications with a minimum of 1 peptide were filtered at 1% false discovery rate (FDR) at the peptide and protein level. Protein intensities were used to determine relative quantitation using an in-house R-script. Samples were normalized by median subtraction and equal variance t-test comparisons determined with protein level log2 transformed protein intensity values. The p-values were adjusted for multiple hypothesis testing using Benjamini-Hochberg procedure.

### Data analysis of synaptic proteomes

Synaptic proteome analysis was performed according to the workflow described previously (**26**), and the resulting GI synaptosome dataset was compared to the reference brain synaptic proteome reported (**27**). All analyses were conducted in R v4.4.1. The significance of overlapping between GI synaptosomes and Brain synaptosomes is calculated using hypergeometric tests. Synapse function for GI synaptosome was conducted using SynGO with default parameters (**28**). STRING interaction networks were constructed in Cytoscape (v3.10.2) using the STRINGapp plugin. Protein complexes were annotated using the CORUM database to identify conserved synaptic protein assemblies shared with the brain dataset. Enrichment analysis between defined groups of synaptosomes (brain_specific, GI_specific, shared) and excitatory and inhibitory modules identified from the reference brain synaptosome were calculated using Fisher’s Exact Test. Heatmaps were visualized using ComplexHeatmap (**29**).

### Muscle bath

After removal of mucosal layer, colon tissue was cut into 5 mm long ring segments for measuring circular muscle responses. Colonic rings were immersed into 30 mL organ filled with conventional Krebs solution (aerated with 95% O_2_ / 5% CO_2_ and heated to 37°C) and mounted between two hooks with the upper one connected to the force transducer (Radnoti 4-Unit Tissue Organ Bath System Model 159920, Radnoti, Dublin, Ireland). Platinum electrodes were used for electrical stimulation. Contractile activity was registered using a PowerLab data acquisition system (ADInstruments, Dunedin, New Zealand) coupled with LabChart software. All colonic rings were equilibrated for 60 min. During equilibration period, the Krebs solution was changed every 15-20 min, and tissues were gradually stretched to the optimal resting tension of 1g. Electrical field stimulation (EFS) was applied with trains of electrical pulses (pulse duration=0.4 ms; train frequency of the pulse=50Hz and duration=300 ms) every 100 s, without renewal of Krebs solution. Voltage-response curves were created by measuring the amplitude of the contraction at voltages of 5, 10, 15, 20, 25, and 30 V. After the generation of basal voltage-response curve, the circular muscle rings were washed three times. Maximal response was obtained with the muscarinic receptor agonist Carbachol (CCh) 10^-5^M (Thermo Fisher, Waltham, MA, USA). Following the basal response, the tissue was incubated with Atropine (10^-6^M) and another voltage-response to EFS was performed. The muscarinic response was obtained by performing the subtraction (i.e. basal response-atropine response for each voltage). The nitrergic contribution was evaluated performing additional voltage-response curves in the presence of MRS2500 (10^-6^M) and Atropine (10^-6^M) for the basal response and repeated in the presence of L-NAME (10^-4^M). To calculate the nitrergic response, the subtraction between basal and L-NAME responses was performed for each voltage (**30**).

### Flow cytometry and fluorescence-activated cell sorting (FACS)

Mucosal-free muscularis from small intestines and colons were dissociated as previously described (**10**). Cell sorting for CD11b^+^ cells was performed using a BD FACSMelody™ Cell Sorter with BD FACSChorus™ Software (BD Bioscience, Franklin Lakes, NJ, USA), located at the Mayo Clinic Flow Cytometry Core Facility (Rochester, MN). Aliquots of cells were either unstained or stained with individual fluorescently labeled antibodies (**Table 1**) to establish instrument voltages, compensation, and appropriate gates. Briefly, forward and side scatter parameters were established using the unstained cell populations, and the baseline voltage levels for each fluorochrome were set between the 1st and 2nd logs on the fluorescence scale. Each positive control tube was initially run without storing the data to ensure that the positive signals were on scale. For separation of C1qa^+^ and C1qa^-^, cells were incubated with anti-mouse CD16/32 antibody for 10 minutes to block nonspecific Fc receptor binding. Cells were then stained on ice for 30 minutes with a myeloid cell panel consisting of fluorophore-conjugated surface marker antibodies (**Table 1**). Following staining, cells were washed with FACS buffer (1% BSA in Hanks’ Balanced Salt Solution (HBSS)) and centrifuged at 300 x g for 5 minutes at 4°C. The supernatant was discarded, and cells were resuspended in 100µl FACS buffer then incubated with SYTOX^TM^ Blue dead cell stain (Thermo Fisher) for 5 minutes on ice. Cells were subsequently washed with 3 mL of FACS buffer and centrifuged again at 300 x g for 5 minutes at 4°C. Samples were resuspended in 300 µL of FACS buffer and data were acquired on the Cytek Aurora spectral flow cytometer (Cytek® Bioscience, Bethesda, MD, USA). Reference controls included an unstained sample and single-stain controls for each fluorophore using UltraComp eBeads™ (Thermo Fisher). Data were spectrally unmixed using SpectroFlo v3.3.0 software (Cytek® Bioscience). The resulting unmixed FCS files were analyzed in FlowJo v10.10.0 using manual gating. Briefly, single and live cells were first identified, followed by gating on CD45^+^ cells. Within this population, CD11b^+^MHCII^+^ cells were selected, and C1q positive and negative subsets were then defined.

For *in vitro* phagocytic assay, CD11b^+^ cells were sorted directly into culture media DMEM/F-12 supplemented with 10% Fetal Bovine Serum (FBS), 10mM L-Glutamine, 100 IU/mL Penicillin, 100 μg/mL Streptomycin and 20 ng/mL mouse Colony-Stimulating Factor 1 (CSF1) for following *in vitro* evaluations.

### Phagocytic activity

#### In vitro phagocytosis assay

FACS-sorted CD11b□ cells from the muscularis of small intestine and colon as described above were seeded in 24-well plates (50,000 cells/well) in DMEM/F-12 supplemented with 10% FBS, 10 mM L-glutamine, 100 IU/mL penicillin, 100 μg/mL streptomycin, and 20 ng/mL mouse CSF-1 and maintained for 7 days. Cells were then incubated with pHrodo™ Red BioParticles™ (Thermo Fisher) for 6 h at 37 °C in a humidified 5% CO_2_ incubator. Cells were subsequently lysed according to the manufacturer’s instructions, and fluorescence intensity was measured using a Synergy Mx Multi-Mode Microplate Reader (BioTek, Winooski, VT, USA). Fluorescence values were normalized to total protein concentration determined by the Bio-Rad DC protein assay kit (Bio-Rad, Segrate, Milan, Italy), using a BSA standard solution (ranging from 0.2 to 2 mg/mL) for the calibration curve. For representative micrographs, images were taken using a laser scanning confocal microscope (Nikon AX-R) using a 20X (NA 0.75 WI) Nikon Apo LWD objective (Nikon).

#### In vivo phagocytosis assay

pHrodo™ Red BioParticles™ (Thermo Fisher) were administered intraperitoneally to age-matched C1qa^FL^ and C1qa^CKO^ mice and allowed to circulate for 6 hours. The muscularis of the small intestine and colon were carefully separated from the mucosa and imaged immediately *ex vivo*. Labeled tissues were examined with a laser-scanning confocal microscope (Nikon AX-R) using a 20X (NA 0.75 WI) Nikon Apo LWD objective (Nikon). Stacks of confocal images across the full thickness of the muscularis propria were collected from 6 mice. For quantification, 5 non-adjacent different fields of view were randomly selected from each tissue and the fluorescence produced by the internalized pHrodo™ Red BioParticles™ quantified using FIJI image analysis software (https://imagej.net/software/fiji/). All the confocal image stacks were flattened into projections using NIS elements AR (Nikon) for quantification of the labeling, ensuring that signal from the entire z-depth was included in the analysis. To verify that internalized particles were localized within MMs, parallel experiments were performed in Lyz2^eGFP^ mice, in which eGFP labels myeloid cells.

### Macrophage morphology determination

MMs’ morphology was evaluated according to Young *et al.* (**31**). Briefly, the standard fluorescence immunohistochemistry protocol using a pan-macrophages marker MHCII (**Table 1**) was performed and labeled tissues were examined with a laser-scanning confocal microscope (Nikon AX-R) using a 20X (NA 0.75 WI) Nikon Apo LWD objective (Nikon). Stacks of confocal images across the full thickness of the muscularis propria were collected from 6 different mice. For quantification, 5 non-adjacent different fields of view were randomly selected from each tissue, and the skeleton analysis was performed as reported in the paper using FIJI image analysis software (https://imagej.net/software/fiji/).

### Transmission electron microscopy (TEM)

Small intestines and colons were dissected and placed immediately into fixative (2% paraformaldehyde + 2% glutaraldehyde in 0.15 M cacodylate buffer containing 2 mM calcium chloride, pH 7.2) for minimum of 12 hour. Following fixation, tissue was processed using a protocol developed to enhance staining (**32**). Briefly, fixed tissue was rinsed in 0.1 M cacodylate buffer and placed into 2% osmium tetroxide + 1.5% potassium ferrocyanide in 0.1 M cacodylate, washed with nH_2_O, incubated at 50°C in 1% thiocarbohydrazide, incubated again in 2% osmium tetroxide in nH_2_O, rinsed in nH_2_O and placed in 2% uranyl acetate overnight. The next day sample is rinsed again in nH_2_O, incubated with Walton’s lead aspartate, dehydrated through an ethanol series, and embedded in Embed 812 resin. Following a 24-hour polymerization at 60°C, 0.1 mm ultrathin sections were prepared and post-stained with lead citrate. Micrographs were acquired using a JEOL 1400+ transmission electron microscope (JEOL, Inc., Peabody, MA) operating at 80 kV equipped with a NanoSprint12 CMOS camera (AMT, Woburn, MA).

### Realtime PCR

Total RNA was extracted from the muscularis of the small intestine and colon using the RNeasy plus universal kit (Qiagen, Hilden, Germany) according to the manufacturer’s specifications. RNA concentration was determined by using a NanoDrop ND-1000 spectrophotometer (Thermo Fisher Scientific). For first-strand cDNA synthesis, 500 ng of total RNA from each sample was reverse-transcribed using the SuperScript VILO complementary DNA Synthesis Kit (Invitrogen, Waltham, MA, USA). Real-time PCR was performed using the LightCycler® 480 SYBR Green I Master Mix (Roche Molecular Systems, San Francisco, CA, USA). Primers were designed based on the mouse GenBank sequences (**Table 3**). The relative expression of mRNA was normalized by glyceraldehyde-3-phosphate dehydrogenase (Gapdh, Qiagen, catalog # PPM02946E-200) and calculated by the 2^−ΔΔCt^ method.

**Table 3:**
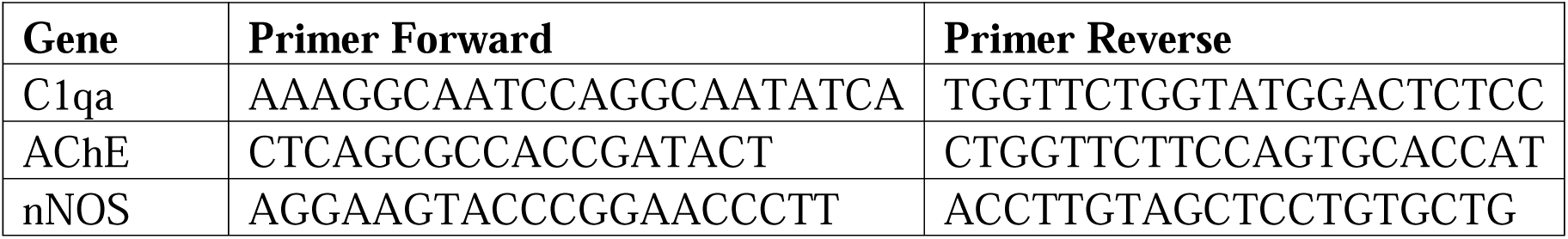
Primer sequences.

### Statistical analysis

Data points represent biological replicates. Results are presented as means ± SEM. Statistical analyses were conducted using the GraphPad Prism 8.0.2 software (GraphPad, San Diego, CA, USA). Specific statistical analyses used are described in figure legends. Statistical significance was assigned at p < 0.05.

## RESULTS

### C1qa^+^ MMs modulate ENS synaptic homeostasis

To determine the impact of C1qa^+^ MMs on enteric synapses, we compared pre- and post-synaptic protein distribution between C1qa^CKO^ and C1qa^FL^ mice (**Figure 1A-E**). Immunostaining for the presynaptic marker SYP revealed a significant increase in synaptic density in C1qa^CKO^ mice compared to controls (**Figure 1B,D**). Consistent with this finding, the postsynaptic marker PSD-95 was also significantly increased, as reflected by a higher number of PSD-95-positive perisomatic neuronal cell bodies in C1qa^CKO^ tissues (**Figure 1C,E**), indicating a coordinated upregulation of pre- and postsynaptic components. These observations were further confirmed by quantitative western blot analysis, which showed increased SYP and PSD-95 protein levels in both the small intestine and colon of C1qa^CKO^ mice (**Supplementary Figure 1A,B**).

**Figure 1.**
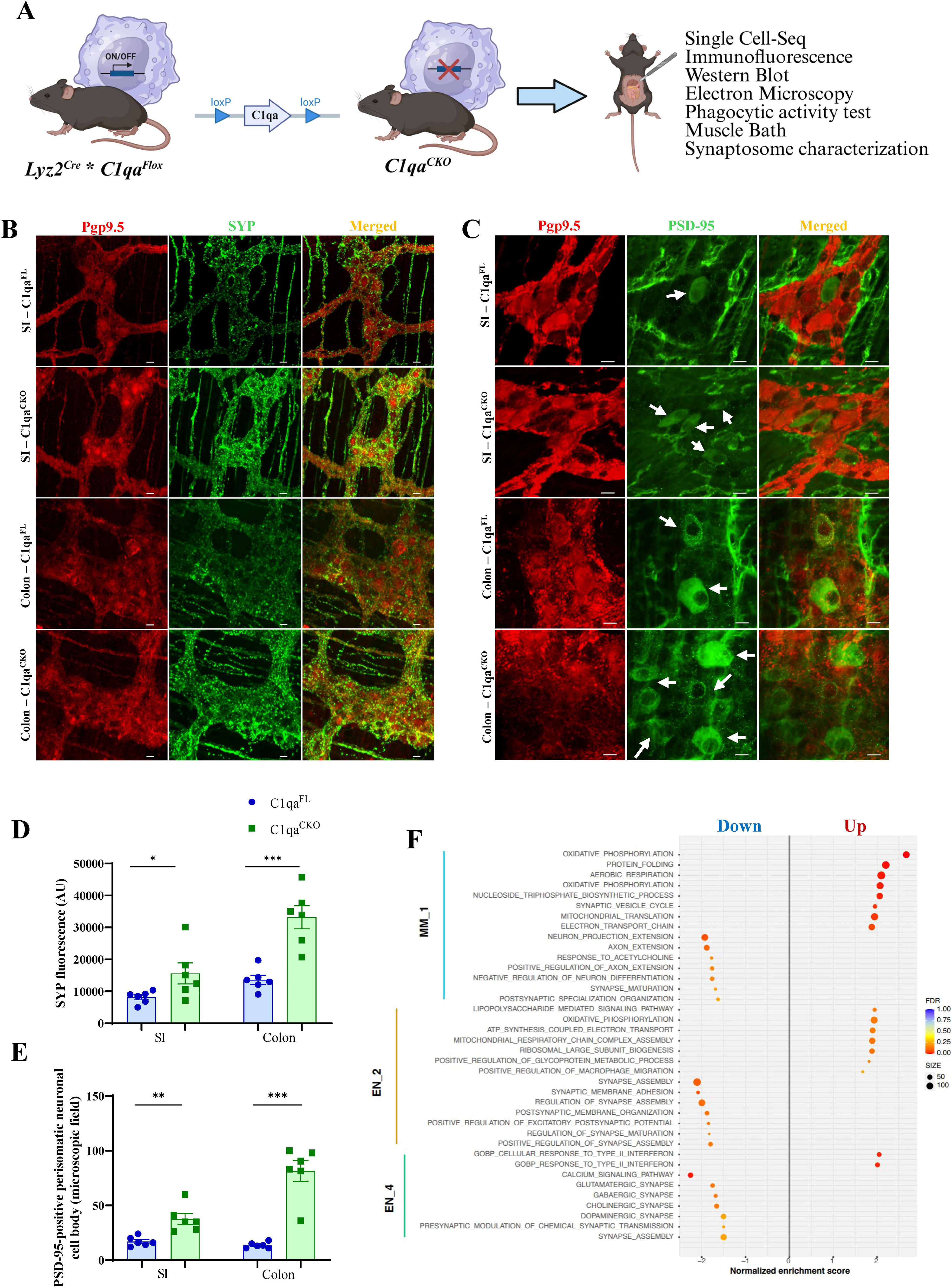
C1qa^+^ muscularis macrophages modulate ENS synaptic homeostasis. **A**. Schematic of C1qa conditional deletion strategy and experimental workflow. **B**. Representative immunofluorescence micrographs showing staining for the pan-neuronal marker Pgp9.5 (red) and the pre-synaptic marker synaptophysin (SYP, green) in the small intestinal (SI) and colonic myenteric plexus of C1qa^FL^ and C1qa^CKO^ mice. Scale bar: 50 μm. **C**. Representative immunofluorescence micrographs showing staining for the pan-neuronal marker Pgp9.5 (red) and the post-synaptic marker PSD-95 (green) in the small intestinal (SI) and colon myenteric plexus of C1qa^FL^ and C1qa^CKO^ mice. Scale bar: 20 μm. White arrows indicate positive PSD-95 perisomatic neuronal cell bodies. **D**. Quantification of mean fluorescence intensity of SYP in C1qa^FL^ (blue dots/bars) and C1qa^CKO^ (green dots/bars) mice. Data are presented as mean ± SEM. * p<0.05 and *** p<0.001 by one-way ANOVA and Tukey’s multiple comparisons test. **E.** Quantification of the post-synaptic marker PSD-95 (expressed as PSD-95 positive perisomatic neuronal cell bodies) in C1qa^FL^ (blue dots/bars) and C1qa^CKO^ (green dots/bars) mice. Data are presented as mean ± SEM. ** p<0.01 and *** p<0.001 by one-way ANOVA and Tukey’s multiple comparisons test. **F.** scRNA-seq pathway enrichment analysis comparing C1qa^CKO^ and C1qa^FL^ mice. Selected MMs (MM_1) and enteric neurons (EN_2 and _4) subclusters show enrichment of synapse-related GO terms and upregulation of synaptic genes. Visualization summarizes key upregulated synaptic genes and enriched pathways (heatmap/bubble plot).

In line with these findings, whole-mount immunofluorescence demonstrated an extensive punctate distribution across the small intestine and colon, with overlapping SYP and PSD-95 signals, supporting the presence of organized pre- and post-synaptic compartments indicative of synaptic junctions (**Supplementary Figure 1C**). This synaptic architecture was further confirmed by electron microscopy, which revealed presynaptic terminals, filled with synaptic vesicles, converging onto post-synaptic terminals (**Supplementary Figure 1D**). Synaptic markers were detected in both nitrergic (nNOS□) and non-nitrergic enteric neurons (nNOS^-^), consistent with the heterogeneous neuronal composition of the ENS and its dual inhibitory and excitatory control of gut motility (**Supplementary Figure 1E**) (**2,33–35**).

To characterize cellular effects of macrophage□specific C1qa deletion, we performed single-cell RNA sequencing on dissociated cells from the colonic muscularis propria of C1qa^FL^ and C1qa^CKO^ mice. Unsupervised clustering identified enteric neuron and MM populations, validated by canonical marker expression (**Supplementary Table 1; Supplementary Figure 2A-C**). Additionally, unbiased clustering uncovered heterogeneity within both populations, revealing four distinct subtypes for each cell type (**Supplementary Figure 2D,E**), defined by the enrichment of distinct marker genes (**Supplementary Figure 2G,H; Supplementary Table 2**) Notably, neither the number of clusters nor the proportion of each subtype differed between C1qa^FL^ and C1qa^CKO^ mice, with stable cellular proportions between genotypes (**Supplementary Figure 2F**).

Differential gene expression analysis revealed transcriptional differences between C1qa^CKO^ and C1qa^FL^ mice (**Supplementary Figure 3A,B; Supplementary Table 3**). Specifically, C1qa^FL^ MMs were enriched for complement and phagocytic genes (*C1qa/b/c, C3*) and genes involved in opsonization and debris recognition (*Apoe, Lrp1*). Moreover, genes involved in phagosome formation and lysosomal processing (*Cd68, Lamp1, Ctss*) were enriched in C1qa^FL^ MMs, suggesting altered phagocytic capacity with C1qa loss (**Supplementary Figure 3A**). Similarly, enteric neurons from C1qa^FL^ mice were enriched for genes linked to synaptic function and neuronal signaling, including key regulators of neurotransmission and synaptic organization (*Snap25, Stx1a, Grin2a,* and *Kcnj* family members). By contrast, enteric neurons from C1qa^CKO^ mice showed increased expression of activity-dependent genes associated with neuronal excitability and remodeling (*Egr1, Fos*; **Supplementary Figure 3B**). Pathway enrichment analysis was performed to define biological processes associated with these transcriptional changes. Among the different MMs populations, we found that MM_1, characterized by a complement-enriched program (*C1qa, C1qb, C1qc*), coupled with a phagocytic/lysosomal signature (*Cd68, Ctsd, Fcgrt*) and a neuro-interaction/vesicle trafficking module, was the most affected MMs population, where C1qa deletion altered pathways associated with oxidative phosphorylation, synaptic vesicle cycle, neuron projection extension, axon extension, synapse maturation and postsynaptic specialization organization (**Figure 1F; Supplementary Table 4**). Among enteric neuron populations, EN_2 and EN_4, characterized by neuronal signaling components (*Gphn, Ntng1, Prkca*), extracellular matrix-associated program (*Efemp1, Col14a1*) alongside complement-related genes (*Cfh*) and neuronal regulators (*Negr1, Cacna1d, Epha7*), were the subclasses of enteric neurons most affected, showing altered pathways in synapse assembly, synaptic membrane adhesion, postsynaptic membrane organization, regulation of synapse maturation and a broad neurotransmitter synapse pathway (e.g. cholinergic and glutamatergic) (**Figure 1F; Supplementary Table 4**). These findings collectively demonstrate that C1qa deficiency in MMs induces molecular changes in subsets of MMs and enteric neurons, resulting in alteration of synapse-related pathways.

### Unbiased proteomic analysis uncovers complement-specific protein interactions in the ENS

The ENS exhibits extensive neurochemical and functional heterogeneity and is composed of diverse neuronal subtypes that govern GI function (**1,5**). In the brain, C1q-dependent synaptic interactions are well recognized as modulators of synapse organization and maintenance (**12,13**); however, whether similar complement-mediated mechanisms contribute to synaptic diversity in the ENS remains largely unexplored. To address this gap, we adapted an established brain synaptosome isolation method to enrich synapses from the GI tract (**26**). Using this approach, we generated a comprehensive proteomic profile of gut C1q-dependent synaptic proteins and compared it with brain C1q proteomes to identify complement-regulated synapse classes in the GI (**Supplementary Figure 4A**). Western blot analysis confirmed the successful enrichment of synaptic components in the isolated synaptosome fraction. The presynaptic marker SYP (33 kDa) showed strong enrichment in the synaptosome fraction, with an absent signal in the cytosolic fraction. Conversely, vinculin (114 and 124 kDa), a cytosolic marker, was absent in the synaptosome fraction, confirming effective separation of synaptic structures from non-synaptic cellular material (**Supplementary Figure 4B**). Mass spectrometry-based proteomic analysis yielded a high number of quantifiable proteins across replicates (**Supplementary Figure 4C**), demonstrating substantial proteomic depth. The reproducibility was high, with most protein groups exhibiting a low coefficient of variation (**Supplementary Figure 4D**).

The proteomic analysis identified a set of C1q-dependent synaptic proteins within the GI tract (2,727 proteins). Comparative analysis with a reference brain synaptic proteome (**26**) revealed, together with a core of shared proteins, a remarkable enrichment of GI-specific proteins. Specifically, 700 proteins were identified as common to both GI and brain synaptosomes, demonstrating a highly significant similarity (p < 2.22E-16, Fold of enrichment = 3.99). This shared core machinery was supplemented by 2,027 proteins unique to GI synaptosomes and 677 unique to brain synaptosomes (**Figure 2A**). This extensive shared proteome highlights that C1q-dependent ENS synapses share a fundamental molecular architecture with CNS synapses, while also possessing distinct components that are likely to confer their specialized functions within the gut.

**Figure 2:**
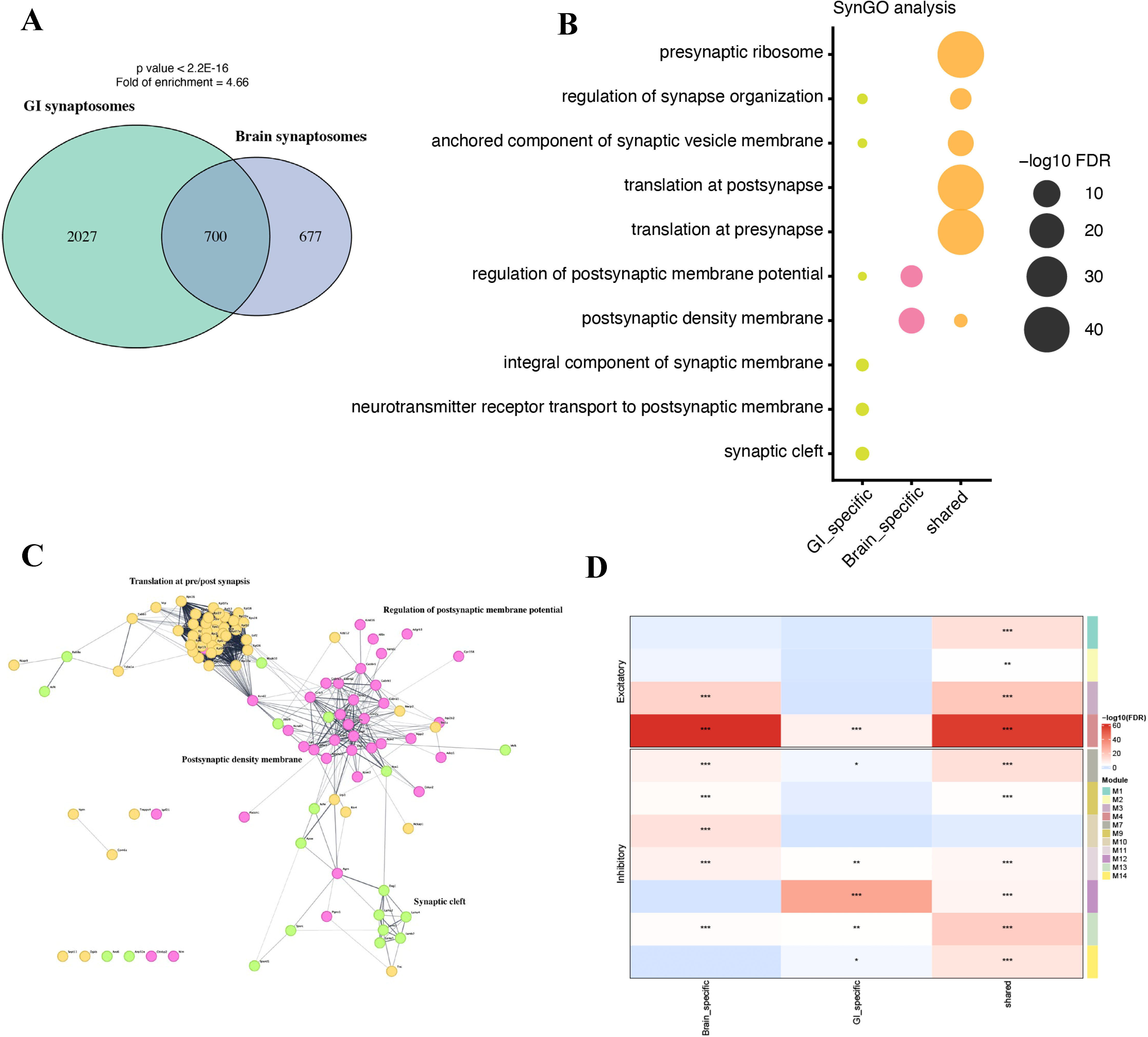
C1qa^+^ muscularis macrophages regulate the synaptic proteome with significant overlap to brain synapses. **A**. Venn diagram illustrating the overlap of identified proteins between C1q-dependent GI synaptosomes and brain synaptosomes. The left circle (green) represents proteins identified in GI synaptosomes, the right circle (blue) represents proteins identified in brain synaptosomes, and the overlapping region shows the number of proteins common to both. p value < 2.22E-16, fold enrichment = 3.99. **B**. SynGO enrichment analysis of proteins identified in GI, brain and shared between them. The dot plot displays significantly enriched Gene Ontology (GO) terms related to synaptic components and processes. The size of each dot represents the number of genes in that term, and the color intensity reflects the significance (-log10 adjusted p-value). **C**. Network analysis of shared C1q-associated synaptosome proteomes, performed using STRING, revealed distinct functional modules. Nodes represent proteins, and edges indicate protein-protein interactions. Node colors denote functionally related clusters, while network connectivity reflects the degree of interaction between proteins. Distinct clusters highlight major synaptic modules, including translation at pre- and postsynaptic compartments, post-synaptic density membrane organization, regulation of post-synaptic membrane potential, and synaptic cleft components. **D**. Heatmap illustrating the distribution of excitatory and inhibitory synaptic modules within brain-specific, GI-specific, and shared C1q-associated synaptosome proteomes. Rows represent individual modules (from module 01 to 14, color-coded) or functional group (excitatory on top and inhibitory on bottom), columns represent different groups (brain specific, GI specific and shared). Color intensity indicates Pearson’s correlation coefficient, with red denoting positive correlation and blue denoting negative correlation. Asterisks indicate statistical significance (*p < 0.05, **p < 0.01, ***p < 0.001). This analysis provides insights into the organization and functional relationships of different synaptic components.

SynGO ontology analysis, based on these three sets, revealed that the C1q-associated synaptic proteome in the GI tract is enriched for pathways supporting synaptic organization and maintenance (**Figure 2B; Supplementary Table 5**). In detail, a core set of highly significant pathways, including translation at pre- and post-synapse, regulation of synapse organization, and anchored components of the synaptic vesicle membrane, was shared across the GI tract and brain (**orange dots**), indicating a broad association of C1q with the maintenance of synaptic machinery. Notably, GI-specific proteins (**green dots**) were predominantly associated with structural organization and receptor localization (neurotransmitter receptor localization and regulation of synapse organization), suggesting a specialized role in maintaining synaptic stability and spatial organization within the ENS. Importantly, the identification of these GI-specific synaptic features provides, for the first time, a framework to understand how neuroimmune regulation in the ENS diverges from canonical brain mechanisms, with potential implications for GI motility disorders. Functionally, these data support a model in which C1qa□ MMs contribute to the structural integrity of enteric synapses by interacting with postsynaptic and synaptic cleft components enriched in the gut, thereby influencing synaptic assembly and maintenance. Consistent with these enrichment patterns, protein-protein interaction network analysis (**Figure 2C and Supplementary Figure 4F,G**) showed that the C1q-associated synaptic proteome revealed a highly organized architecture. While the global network showed a dense core of interconnected proteins associated with translation at pre- and post-synaptic compartments and postsynaptic density organization, a more defined GI-specific subnetwork emerged, highlighting features unique to enteric synapses. In detail, this GI-enriched module was centered on synaptic cleft and postsynaptic membrane components, including extracellular matrix proteins (*Lamc1, Lama2, Lama4*), integrins (*Itgb1, Itgb5, Itgb3*), and signaling-associated molecules (*Egfr, Ncam2*), forming a tightly interconnected cluster. Notably, the presence of proteins involved in neurotransmission (*Nos1, Ache*), within this network suggests that these structural clusters are functionally linked to the regulation of inhibitory and excitatory signaling in the ENS. Together, these data indicate that, beyond a conserved synaptic core, C1q-associated proteins in the GI tract preferentially assemble into networks that support synaptic structure and stability, highlighting a tissue-specific mechanism for regulating enteric synaptic architecture.

We next asked whether C1q□dependent regulation preferentially affects excitatory or inhibitory circuitry. To address this, based on the work of Van Oostrum *et al.*, we extracted the protein-protein network based on excitatory and inhibitory synaptic modules and integrated our GI synaptic proteome alongside the reference brain synaptic proteome within it (**27**). As shown in **Figure 2D** and **Supplementary Figure 4E**, both excitatory and inhibitory modules showed a C1q□dependent regulation of synaptic proteins shared between brain and GI tract. Notably, two tissue□specific modules emerged: module 10 (brain□selective) and module 12 (GI□selective). Module 10 comprises genes involved in synaptic adhesion and organization (*Nlgn2, Nrxn3* and *Caskin1*), vesicle release machinery (*Vamp1, Sv2a, Rab3b, Rab3gap1* and *Cadps*) and synaptic signaling regulators (*Necab2* and *Iqsec3*), while module 12 is a network enriched for mitochondrial & metabolic machinery (*Nduf, Uqcr, Cox, Atp5, Idh3a/g, Ogdh, Acot9* and *Mecr*), transport & trafficking (*Slc25, Slc7a5, Slc12a2, Snx1* and *Snx6*), membrane regulators (*Magi1, Clasp1* and *Fnbp1*) and protein folding (*Timm, Hspd1* and *Hspe1*). The different compositions of modules 10 and 12 further highlight functional differences between the brain and GI tract, as module 10 (brain) is directly involved in synaptic transmission, whereas module 12 (GI) is related to mitochondrial and metabolic support, consistent with a more structural and supportive role.

### C1qa^+^ MMs regulate enteric neurotransmission

Proteomic analysis revealed that complement-mediated modulation of synaptic processes affects both inhibitory and excitatory proteins. To examine the functional implications, we used *ex vivo* colonic preparations (**Figure 3A**) and dissected C1qa-dependent effects at the circuit level using electrical field stimulation (EFS) in an organ bath setup. By isolating the cholinergic component (atropine-sensitive), we found that EFS-induced contractions were increased in colonic tissues isolated from C1qa^CKO^ compared to C1qa^FL^ mice across stimulation voltages (**Figure 3B**). To confirm that the observed functional changes were due to neuronal alterations rather than intrinsic smooth muscle dysfunction, we tested responses to the muscarinic receptor agonist carbachol. There was no difference between C1qa^CKO^ and C1qa^FL^ mice (**Figure 3C**), indicating that the increased EFS-dependent response in C1qa^CKO^ mice was primarily neuronal.

**Figure 3.**
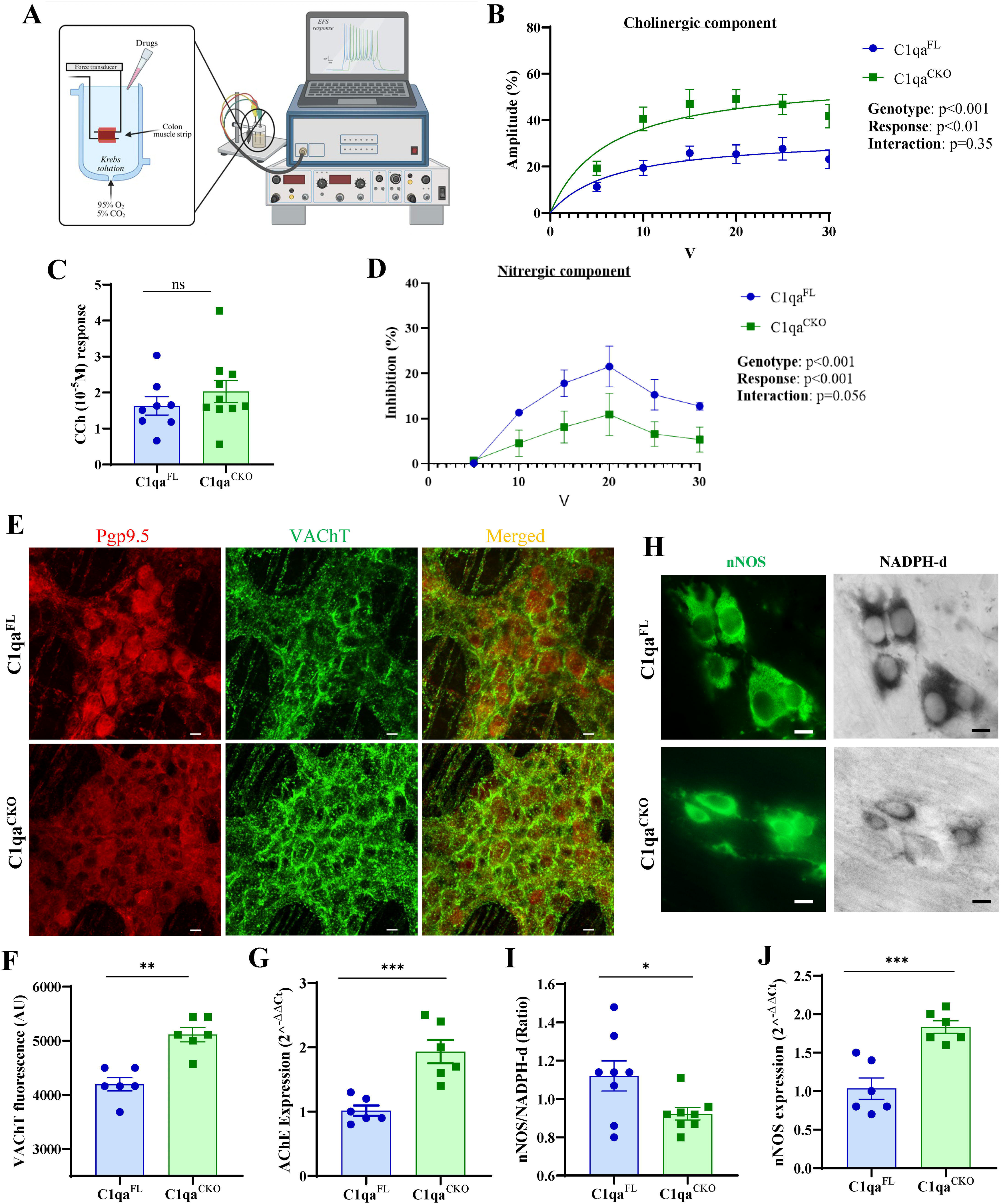
C1qa^+^ muscularis macrophages modulate enteric neurotransmission. **A**. Schematic of *ex vivo* colonic muscle strip preparation and recording setup. **B-D**. Evoked contractile responses of colonic muscle strips from C1qa^FL^ (blue) and C1qa^CKO^ (green) mice. **B.** Cholinergic component evoked amplitude (percent of maximum response obtained in the presence of CCh), determined as atropine-sensitive fraction (difference between basal and atropine-treated EFS responses) across stimulation voltages. **C.** Representative single□pulse contractile responses (CCh responses) and quantification. ns = not significant. **D.** Nitrergic component evoked amplitude determined as the L-NAME-sensitive fraction of the EFS response, calculated from recordings obtained in the presence of atropine and MRS2500 (both 10□□ M) before and after addition of L-NAME (10□□ M). Data are plotted as mean ± SEM, n=8. p=0.35, p=0.056, p<0.01 and p<0.001 by two-way ANOVA (genotype x stimulation intensity) and Sidak’s multiple comparison test. **E.** Representative immunostaining for the pan-neuronal marker Pgp9.5 (red) and the Vesicular Acetylcholine Transporter (VAChT, green) in the colonic myenteric plexus of C1qa^FL^ and C1qa^CKO^ mice. Scale bar: 50 μm. **F.** Quantification of mean fluorescence intensity of VAChT in the colon of C1qa^FL^ (blue dots/bars) and C1qa^CKO^ (green dots/bars) mice. **G.** AChE mRNA expression in colonic tissues from C1qa^FL^ (blue dots/bars) and C1qa^CKO^ (green dots/bars). **H.** Representative immunohistochemistry for the nitrergic neuronal marker nNOS (green) and the histochemical marker of nitric oxide synthase activity (NADPH-d, gray scale) in the colonic myenteric plexus of C1qa^FL^ and C1qa^CKO^ mice. Scale bar: 10 μm. **I.** Ratio quantification between nNOS+/NADPH-d+ neurons in the colon myenteric plexus of C1qa^FL^ (blue dots/bars) and C1qa^CKO^ (green dots/bars) mice. **J.** nNOS mRNA expression in colonic tissues from C1qa^FL^ (blue dots/bars) and C1qa^CKO^ (green dots/bars). Data are expressed as mean ± SEM. * p<0.05, ** p<0.01 and *** p<0.001 by unpaired t-test.

We then examined the nitrergic component by repeating the EFS protocol after L-NAME treatment, in the presence of MRS2500 and atropine. C1qa^CKO^ mice showed a reduced inhibitory response compared with C1qa^FL^ mice (**Figure 3D**). These findings indicate that C1qa deficiency alters both cholinergic and nitrergic neurotransmission, producing an imbalance that favors hypermotility, which may drive the accelerated GI transit previously observed (**10,11**). To determine whether the functional differences corresponded to changes in neuron numbers or solely to synaptic alterations, we quantified enteric neuron subtypes by immunohistochemistry. Consistent with previous reports (**11**), total enteric neuron number did not differ between genotypes (**Supplementary Figure 5A,B**). We next quantified enteric neuron subtypes to assess potential shifts between inhibitory (nitrergic) and excitatory (cholinergic) populations. We found no change in the number of nNOS^+^ and ChAT^+^ neurons in C1qa^CKO^ mice compared with C1qa^FL^ mice (**Supplementary Figure 5A,B**), suggesting that the functional phenotype observed in C1qa^CKO^ mice may reflect synaptic modifications rather than changes in neuronal number.

Despite unchanged neuronal numbers, immunostaining for the vesicular acetylcholine transporter (VAChT) was increased (**Figure 3E,F**), as was the neurotransmitter-related gene acetylcholinesterase (AChE) (**Figure 3G**). The proportion of nNOS□/NADPH-d^+^ neurons, a marker for nitrergic neuron identity and functional capacity, was reduced in C1qa^CKO^ mice compared with C1qa^FL^ mice (**Figure 3H,I**). Interestingly, nNOS gene expression was increased in C1qa^CKO^ compared to C1qa^FL^ mice (**Figure 3J**). Together, these data indicate that loss of C1qa MMs is associated with altered cholinergic/nitrergic synaptic neurotransmission driving the enhanced GI motility previously reported (**10,11**).

### C1qa^+^ MMs regulate synaptic pruning via phagocytosis

C1qa^+^ MMs represent a subpopulation of tissue-resident MMs (**Figure 4A**, **Supplementary Figure 6A,B**). To phenotypically characterize these cells, we separated C1qa^+^ and C1qa^−^ MMs from the single-cell dataset and performed differential expression and gene set enrichment analyses (**Figure 4A,B; Supplementary Table 6**). C1qa^+^ MMs showed upregulation of lysosomal, endocytic, complement activation, and synapse□remodeling pathways, consistent with a phagocytic, degradative phenotype. By contrast, C1qa^−^ MMs were enriched for ribosomal and cytoplasmic translation programs, indicating a transcriptional state biased toward protein synthesis rather than cellular clearance.

**Figure 4:**
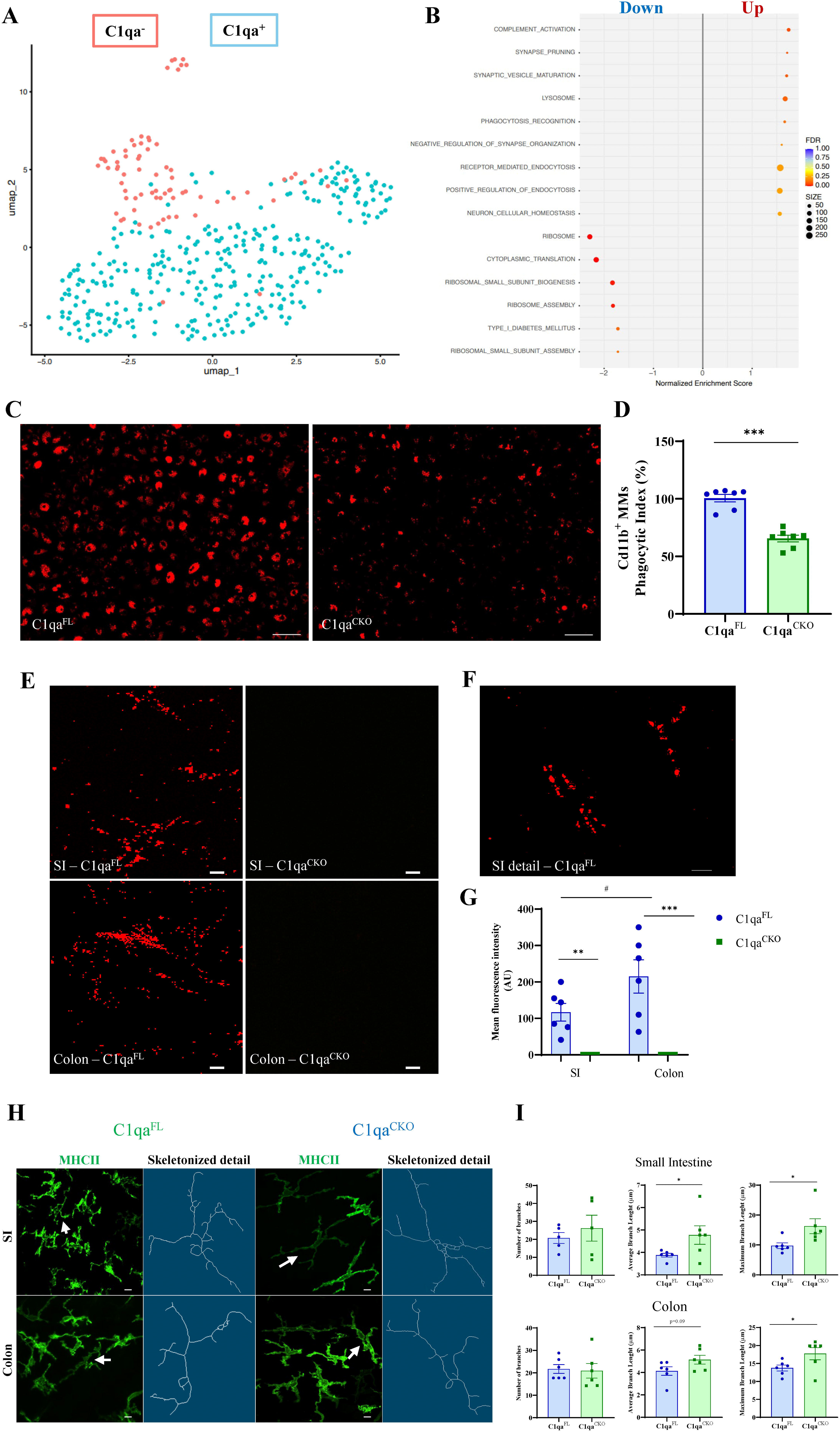
C1qa^+^ muscularis macrophages regulate synapses through phagocytosis. **A**. UMAP of MMs divided by C1qa expression (C1qa^+^ pink dots and C1qa^−^ cyan dots). **B.** Pathway enrichment plot showing phagocytosis□ and synapse□related pathways upregulated in C1qa^+^ MMs (right - red = up, left - blue = down). **C-D.** *In vitro* phagocytosis assay. **C.** Representative images of fluorescent red bioparticles internalized by FACS-sorted CD11b^+^ MMs from C1qa^FL^ and C1qa^CKO^ mice. Scale bar: 50 μm. **D.** Quantification of phagocytic index (fluorescence intensity) of internalized bioparticles following cell lysis and expressed as a percentage relative to CD11b□ MMs from C1qa^FL^ mice. Data are expressed as mean ± SEM. ***p < 0.001 by unpaired t-test. **E-G.** *In vivo* phagocytosis assay. **E.** Representative high□magnification images of pHrodo bioparticles accumulation in the small intestine (SI) and colon from C1qa^FL^ and C1qa^CKO^ mice. Scale bar: 50 μm. **F.** Representative high□magnification detail in the SI of C1qa^FL^ showing accumulation of red bioparticles in MMs. Scale bar: 20 μm. **G.** Quantification of the phagocytic index quantified as mean fluorescence intensity (arbitrary units). Data are expressed as mean ± SEM. **p < 0.01, ***p < 0.001 vs C1qa^FL^ and ^#^ p<0.05 in C1qa^FL^ SI vs Colon, by one-way ANOVA with Tukey’s multiple comparison test/ **H–I.** MMs’ morphology and branches’ analysis. **H.** Representative MHCII immunofluorescence and corresponding skeletonized images (SI and colon) showing altered MMs process complexity in C1qa^CKO^ vs C1qa^FL^ (arrows). Scale bar: 50 μm. **I.** Quantification of shape parameters (number of branches, average branch length and maximum branch length). Data are expressed as mean ± SEM. *p < 0.05 by unpaired t-test.

We next tested whether the synaptic changes observed in C1qa^CKO^ mice depend on deficits in phagocytic activity. We isolated CD11b^+^ cells from the gut muscularis externa of both C1qa^CKO^ and C1qa^FL^ mice (**Supplementary Figure 6C,D**) to evaluate their phagocytic activity following incubation with pHrodo bioparticles. MMs isolated from C1qa^CKO^ mice exhibited a reduced phagocytic index compared to MMs isolated from C1qa^FL^ mice (**Figure 4 C,D**). This finding was replicated *in vivo*, where we observed accumulation of bioparticles in C1qa^FL^ mice within the muscularis externa taken up by MMs (**Figure 4E-G and Supplementary Figure 6E**), while C1qa^CKO^ mice showed no accumulation (**Figure 4E-G**) 6 hours post injection. This functional impairment aligns with the molecular reprogramming revealed by the single-cell RNA sequencing data and underscores the critical role of C1qa in maintaining effective macrophage clearance functions. To provide direct evidence of MMs phagocytic activity on synapses, we performed immunohistochemistry and 3D reconstruction of labeled tissues at high magnification. Images revealed SYP within MMs’ cytoplasm (**Supplementary Figure 6F**). These observations, particularly from 3D reconstructions, support the possibility that MMs internalize synaptic material.

Macrophage morphology often reflects their activation state and phagocytic capacity (**36**). We observed significant changes in the morphology of MMs from C1qa^CKO^ mice (**Figure 4H,I**) compared with C1qa^FL^ mice. Specifically, although the overall number of branches did not change, MMs from C1qa^CKO^ mice showed increased average and maximum branch length compared with those from C1qa^FL^ mice. These morphological changes further support the notion that C1qa deficiency fundamentally alters MM functional state, contributing to their reduced ability to clear synaptic components.

### C1qa^+^ MMs regulate synaptic density and GI motility in adulthood

To distinguish whether regulation of enteric synapses occurs throughout the lifespan we next assessed whether C1qa^+^ MMs can regulate enteric synapses in adulthood. To accomplish this, we utilized an inducible mouse model in which tamoxifen-induced C1qa deletion in MMs was achieved in adult mice (**Figure 5A**), allowing the assessment of C1qa function independently of constitutive deletion. qPCR analysis confirmed a significant reduction of C1qa mRNA expression in small intestines and colon tissues of tamoxifen-treated iC1qa^CKO^ mice (**Figure 5B**), validating the inducible genetic deletion. Furthermore, no significant differences were observed in small intestine or colon length or in body weight (**Supplementary Figure 7A-B**) between tamoxifen- and vehicle-treated mice, indicating that the observed changes are specific to C1qa deletion rather than developmental abnormalities or systemic distress.

**Figure 5.**
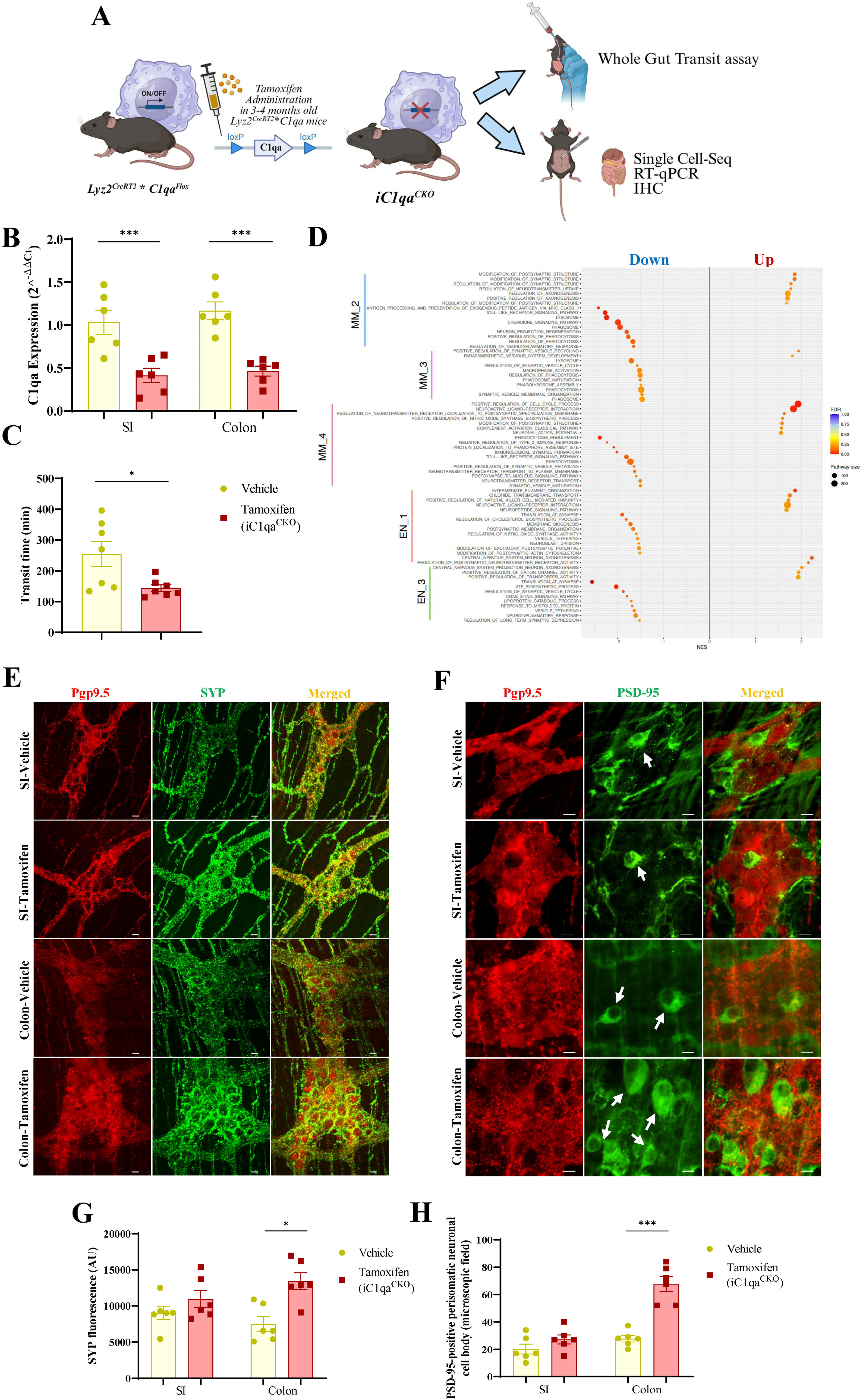
C1qa^+^ Muscularis macrophages regulate synapses in adulthood. **A.** Experimental timeline and schematic of inducible, macrophage□specific C1qa deletion (Lyz2^CreERT2^ * C1qa^fl/fl^ → iC1qa^CKO^). Tamoxifen (75 mg/kg) or vehicle (corn oil) was administered, followed by washout and functional/tissue analyses (whole□gut transit, single□cell RNA□seq, qPCR, IHC). **B.** qPCR validation of C1qa deletion (2−ddCt) in the small intestine (SI) and colon. Data are expressed as mean ± SEM. ***p < 0.001 by one-way ANOVA with Tukey’s multiple comparison test. **C.** Whole□gut transit time (minutes) assay performed in vehicle and tamoxifen (iC1qa^CKO^)-treated mice. Data are expressed as mean ± SEM. *p < 0.05 by unpaired t-test. **D.** GSEA pathway enrichment comparing iC1qa^CKO^ to C1qa^FL^ mice**. E.** Representative immunofluorescence micrographs showing staining for the pan-neuronal marker Pgp9.5 (red) and the pre-synaptic marker SYP (green) in the small intestine (SI) and colonic myenteric plexus of vehicle and tamoxifen (iC1qa^CKO^) treated mice. Scale bar: 50 μm. **F.** Representative immunofluorescence micrographs showing staining for the pan-neuronal marker Pgp9.5 (red) and the post-synaptic marker PSD-95 (green) in the small intestine (SI) and colonic myenteric plexus of vehicle and tamoxifen (iC1qa^CKO^) treated mice. Scale bar: 20 μm. White arrows indicate positive PSD-95 perisomatic neuronal cell bodies. **G.** Quantification of mean fluorescence intensity of SYP in vehicle (yellow dots/bars) and tamoxifen (iC1qa^CKO^ red dots/bars) treated mice. Data are presented as mean ± SEM. * p<0.05 by one-way ANOVA with Tukey’s multiple comparisons test. **H.** Quantification of the post-synaptic marker PSD-95 (expressed as PSD-95 positive perisomatic neuronal cell bodies) in vehicle (yellow dots/bars) and tamoxifen (iC1qa^CKO^ red dots/bars) treated mice. Data are presented as mean ± SEM. *** p<0.001 by one-way ANOVA and Tukey’s multiple comparisons test.

Consistent with the constitutive C1qa^CKO^ model, iC1qa^CKO^ mice exhibited a significantly accelerated whole gut transit time compared to controls (**Figure 5C**). These findings indicate that C1qa^+^ MMs regulate GI motility independently of developmental effects.

To identify the underlying mechanisms regulating these changes, we performed single-cell RNA-seq on colonic muscularis from iC1qa^CKO^ and C1qa^FL^ mice. Following the workflow used for the constitutive model, we identified MMs and enteric neurons by marker expression (**Supplementary Table 1**) and further divided each into four distinct subgroups (**Supplementary Table 2**), which were not different between the mouse models (**Supplementary Figure 7C-H**). Pathway analysis across MM and enteric neuron clusters revealed dysregulation of synapse-pruning, endocytic/lysosomal, and synaptic□maintenance pathways in iC1qa^CKO^ tissue (**Figure 5D**). C1qa^+^ MMs-associated programs (lysosome, receptor□mediated endocytosis, complement activation, synapse remodeling) were reduced in the inducible knockout model, while translation/ribosomal signatures were relatively increased, recapitulating the transcriptional shift observed in the constitutive mouse model. Parallel changes in enteric neuron clusters (EN_1, EN_3) indicate altered synaptic maintenance signatures in neurons as well, consistent with impaired complement□dependent synapse remodeling driving the observed hypercontractility. Together, these data indicate that adult C1qa^+^ MMs actively maintain ENS synaptic organization, and that loss of C1qa in adulthood is sufficient to perturb synapse□related programs and alter gut motility.

As with the constitutive mouse model, we first assessed whether the synaptic changes were reflected in altered neuronal numbers. Immunohistochemistry showed no change in total enteric neuron number or in the ChAT^+^ or nNOS^+^ neuronal subpopulations (**Supplementary Figure 8A,B**), consistent with the constitutive C1qa^CKO^ model. We then investigated the molecular impact of C1qa deletion on synaptic components by immunohistochemical analysis. Tamoxifen-induced iC1qa^CKO^ mice showed a significant increase in the pre-synaptic marker SYP and in the post-synaptic marker PSD-95 in the colon, compared to vehicle-treated mice, while the expression of both SYP and PSD-95 was only slightly increased in the small intestine (**Figure 5E-H**). In contrast, cholinergic and nitrergic neurons (VAChT and NADPH-d/nNOS) that were altered in the constitutive C1qa^CKO^ model were unchanged in the iC1qa^CKO^ mice (**Supplementary Figure 8C-F**).

To further define how C1qa in MMs affects different stages of life, we compared C1qa^CKO^ and iC1qa^CKO^ mice to identify stage-specific transcriptional programs (**Supplementary Figure 9A-C; Supplementary Tables 3,4**). Gene set enrichment analysis revealed distinct pathway shifts between groups: C1qa^CKO^ MMs were enriched for lysosome, proteasome, translation at synapse, regulation of phagocytosis/autophagosome assembly, and neuropeptide signaling pathways. In contrast, iC1qa^CKO^ MMs showed enrichment in synaptic transmission, neurotransmitter receptor interaction, antigen processing and presentation, toll□like receptor signaling, and regulation of membrane invagination, NOS biosynthetic process and cholinergic synaptic transmission. This profile indicates that, after adult (inducible) C1qa deletion, MMs not only engage degradative/proteostatic programs but also actively regulate synaptic function, participating in synaptic protein translation/turnover, neuropeptide signaling, nitric□oxide–related pathways, and neurotransmitter handling, consistent with a role in real time modulation of neuron□to□neuron communication. Finally, consistent with the ability of C1qa^+^ MMs to regulate synapses in adulthood, we found changes to MM morphology, consistent with the phenotype observed in the C1qa^CKO^ mouse model (**Supplementary Figure 10A-B**). This suggests that C1qa loss drives a conserved morphological and transcriptional shift that impairs MM□mediated synaptic maintenance.

## DISCUSSION

In this study, we identified a previously unrecognized role of C1qa^+^ MMs in regulating enteric synaptic organization and GI function. Integrating proteomic, transcriptomic, imaging, and functional studies, we show that C1qa-dependent mechanisms modulate synaptic dynamics and neuronally driven contractility. Complementing these anatomical and functional changes, C1qa^+^ MMs exhibit phagocytic activity *in vitro* and *in vivo*, supporting a model in which C1qa^+^ MMs contribute to ongoing ENS synaptic turnover.

Our proteomic analysis revealed that C1q-associated synapses in the GI tract substantially overlap with those in the brain while also displaying distinct features, indicating both conservation and tissue-specific specialization of synaptic organization. C1q-dependent regulation affects both excitatory and inhibitory synapses, indicating that complement-mediated mechanisms are not restricted to a specific neurotransmitter class but broadly influence synaptic composition. However, network and module analyses identify C1q□sensitive synapse categories and highlight GI-modules enriched for bioenergetic, trafficking, and local□translation machinery. These organ□specific signatures suggest the different ways MMs might selectively target synapse subtypes and support the idea that peripheral synapses are specialized, immune□accessible structures, potentially amenable to pharmacological modulation. This resource complements emerging studies of enteric neuron diversity (**37–40**) and advances classification of GI-specific ENS synapse classes.

Our results suggest synaptic dysregulation, rather than enteric neuron loss, may drive GI motility dysfunction. After macrophage-specific C1qa depletion, we observed clear increases in the number of synapses without changes in total neuronal count, suggesting that synaptic remodeling may be sufficient to alter circuit output. This opens an important area of research: 1) to determine whether disorders already linked to ENS alterations (**41**) also exhibit synaptic deficits, and 2) to test whether interventions that restore synaptic balance can rescue motility. Together, these future studies could establish synapses as functionally relevant targets for therapeutic intervention in GI motility disorders.

Multiple lines of evidence point to phagocytic pruning as a major mechanism contributing to the functions of MMs (**42,43**). C1qa^+^ MMs are transcriptionally enriched for lysosomal, endocytic, and complement related programs, and loss of C1qa shifts MMs toward a less phagocytic, more translation biased state. Functionally, impaired clearance by C1qa-deficient MMs coincides with synapse accumulation, and synaptic proteins co associate with C1q in synaptosome preparations. Together, these findings support a model in which complement dependent engulfment contributes substantially to synapse turnover in the gut.

These molecular and proteomic changes alter enteric circuit behavior. *Ex vivo* organ bath recordings show that macrophage-specific C1qa deficiency modifies neurogenic contractile responses, by altering both cholinergic and nitrergic signaling, while smooth muscle responsiveness is preserved. Thus, the phenotype arises from neural-circuit changes rather than intrinsic muscle dysfunction, consistent with a model in which disrupted synaptic organization leads to altered neurotransmission and, ultimately, accelerated GI transit.

Another critical question is whether MMs regulate synapses across the lifespan. In the brain, microglia shape circuits during development and continue to modulate synapses in adulthood (**44–47**). Gut studies have largely focused on developmental neuroimmune interactions, with limited data on adult MM-ENS communication (**48**). Using an inducible, macrophage-targeted C1qa deletion, we provide the first evidence that C1qa^+^ MMs actively regulate ENS synaptic organization in adult animals. Acute loss of C1qa in MMs is sufficient to alter synaptic composition and produce GI dysmotility, indicating ongoing, lifelong regulation of enteric synapses by MMs. This is particularly relevant given the continuous mechanical and microbial challenges faced by the gut, which may necessitate dynamic remodeling of enteric circuits (**48–50**).

In conclusion, our data identify C1qa^+^ MMs as active, adult regulators of enteric synapse homeostasis and establish a mechanistic framework in which complement-dependent neuro-immune interactions contribute to the maintenance of ENS circuits and gut motility in disease and aging. While these findings suggest that such pathways may represent upstream mechanisms through which enteric circuitry could be modulated, the essential physiological roles of complement signaling warrant caution, and future studies will be required to determine whether selective or context-dependent therapeutic modulation is feasible. Together, this work positions neuroimmune regulation of synaptic organization as a potential avenue for modulating GI motility in health and disease.

## Funding

DK129297 (GC); DK127992-1 (GC); DK115255 (GC); ANMS Young Investigator grant (GC); AGA grant#36 (GC); P30DK084567 (GC); Robert and Arlene Kogod Center on Aging Fundamental Mechanisms of Aging Award (GC and AB); ANMS Discovery Grants Program (MD).

## Author contributions

**MD**: Conceptualization of the work, data analysis, writing of the original draft, experimental supervision, review of the original draft and editing. **JSOC**: Performance of experiments, review of the original draft and editing. **KW**: Performance of experiments, visualization, review of the original draft and editing. **YL**: Data curation and analysis, visualization, review of the original draft and editing. **MHE**: Performance of experiments, review of the original draft and editing. **VD**: Performance of experiments, data curation, review of the original draft and editing. **NW**: Performance of analysis, review of the original draft and editing. **XC**: Performance of experiments, review of the original draft and editing. **ST**: Performance of analysis, review of the original draft and editing. **TAC**: Performance of experiments, review of the original draft and editing. **EB**: Intellectual inputs, review of the original draft and editing. **CL**: Intellectual inputs, review of the original draft and editing. **KAS**: Data curation, intellectual input, review of the original draft and editing. **MG**: Intellectual input, review of the original draft and editing. **MJ**: Data curation, intellectual input, review of the original draft and editing. **AB**: Intellectual input, review of the original draft and editing. **GF**: Intellectual input, review of the original draft and editing. **GC**: Conceptualization of the work, provision of resources, writing of the original draft, study supervision, review of the original draft and editing.

## Acknowledgements

The authors thank the Mayo Clinic Flow Cytometry Core, Genome Analysis Core and Proteomics Core for their technical support, data acquisition, and expert guidance throughout this project.

## Data Sharing

All data generated or analyzed in this study are included in the article and its supplementary materials; additional information can be obtained from the corresponding authors upon request.

## Conflicts of interest

The authors report no commercial or financial relationships that could be perceived as a potential conflict of interest.

## SUPPLEMENTARY FIGURE LEGENDS

**Supplementary Figure 1.**
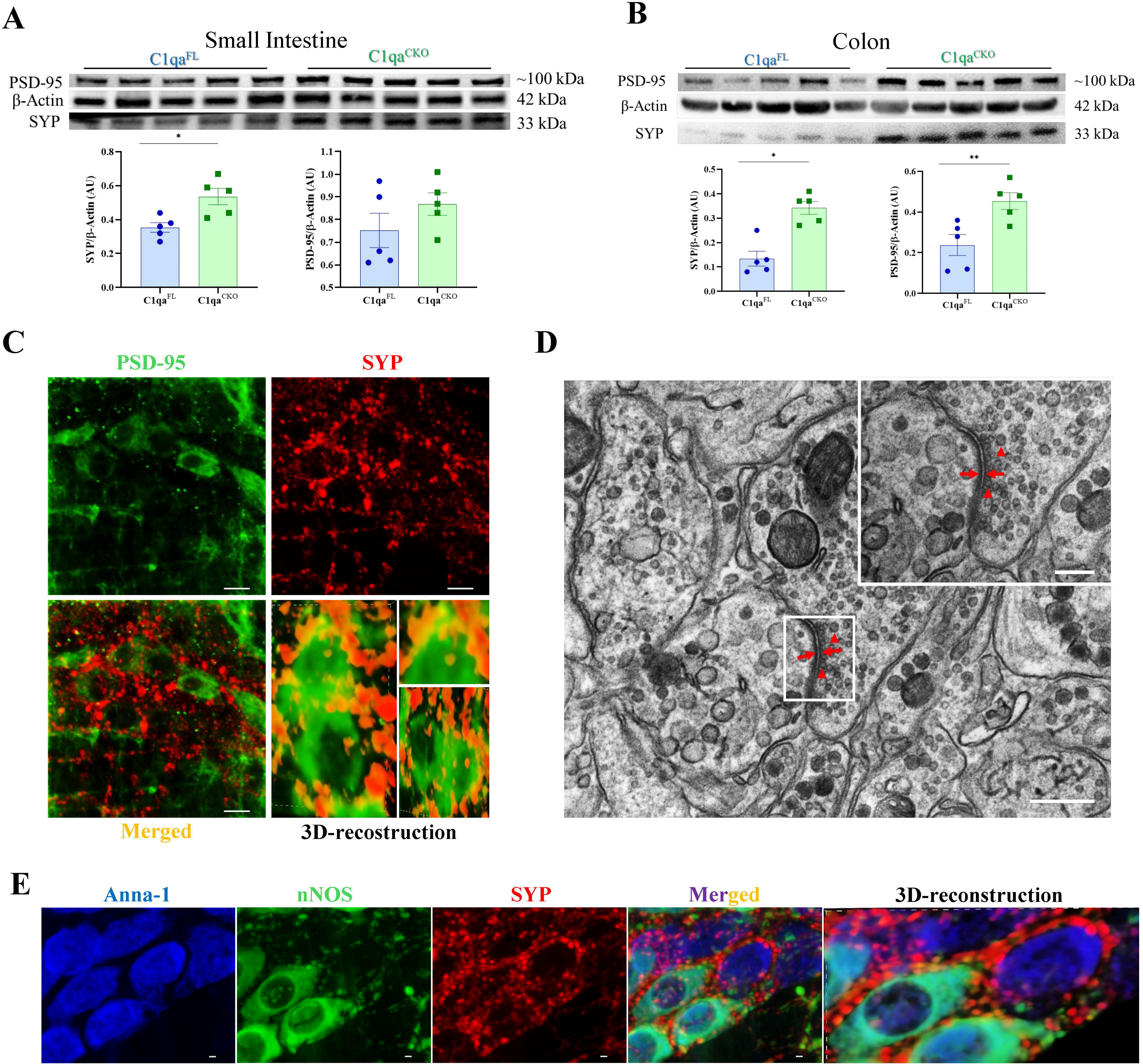
Characterization of ENS synapses. **A-B**. Western blot analysis quantifying the expression levels of the presynaptic marker SYP (**A**) and the postsynaptic marker PSD-95 (**B**) in the small intestine (SI) and colon of C1qa^CKO^ and C1qa^FL^ mice. Densiometric analysis is expressed as mean ± SEM. * p<0.05 and ** p<0.01 by unpaired t-test. **C**. Representative immunofluorescence images of colonic muscularis whole mounts from C1qa^FL^ mice showing distribution of the presynaptic marker SYP (red) and the postsynaptic marked PSD-95 (green). Merged image and 3D-reconstruction (yellow) indicate colocalization of SYP and PSD-95 in the synaptic puncta. Scale bars: 50 µm. **D**. Representative TEM image of the colonic myenteric plexus, illustrating the ultrastructural features of chemical synapses. Highlighted are pre-synaptic terminals containing synaptic vesicles (red arrows heads), the synaptic cleft, and post-synaptic densities (thin red arrows). Scale bars: 400 nm in the overview image and 100 nm in the magnified inset. **E**. Representative immunofluorescence micrographs showing the pan-neuronal marker Anna-1 (blue), nNOS neurons (green) and the presynaptic marker SYP (red). Merged image and 3D-recostruction highlighted (white arrows) the equal co-localization of SYP on both nNOS^+^ and nNOS^-^ enteric neurons. Scale bars: 10 µm..

**Supplementary Figure 2.**
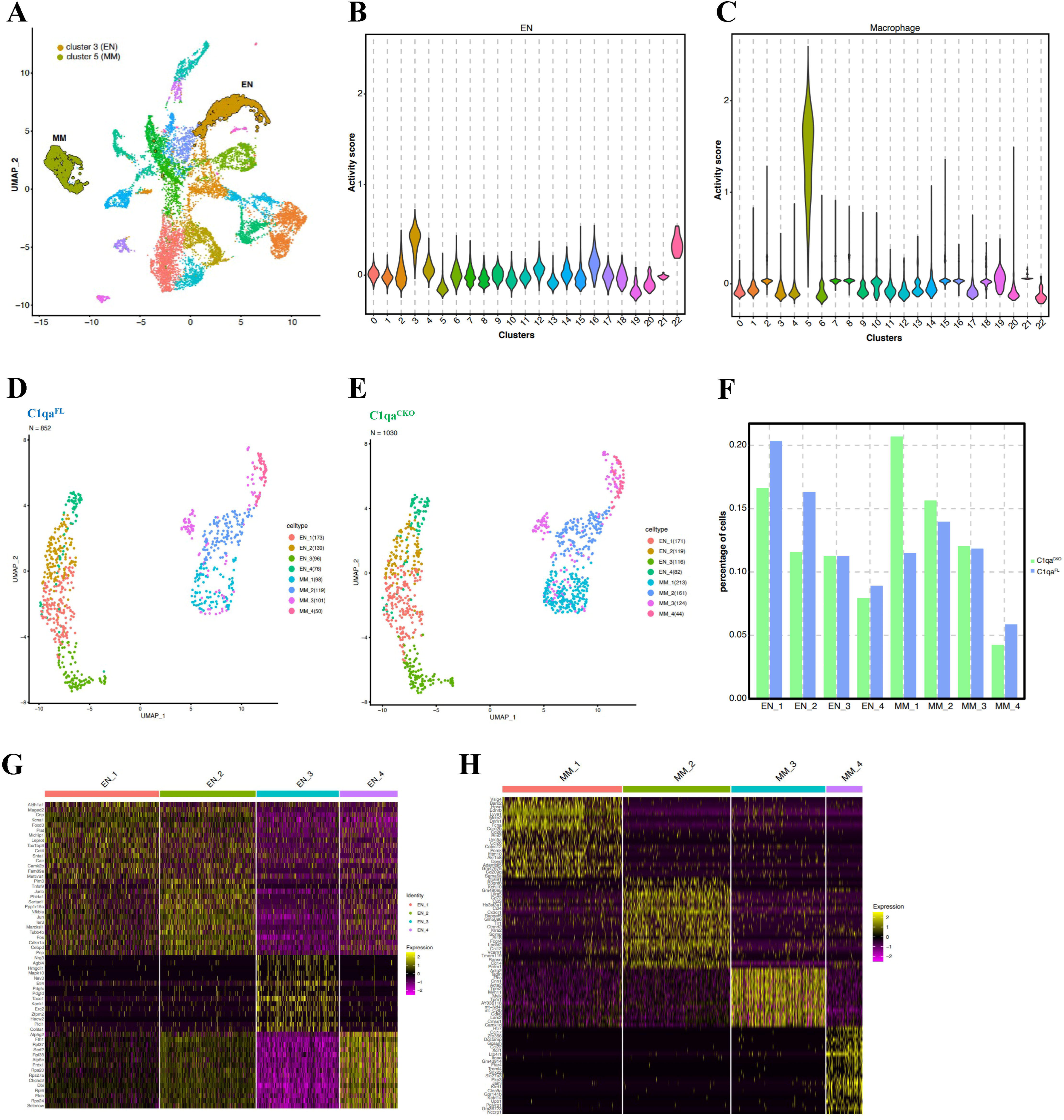
Identification of discrete populations of enteric neurons and muscularis macrophages (MMs) by single-cell RNA sequencing. **A**. UMAP plot displaying the unsupervised clustering of single-cell RNA sequencing (scRNA-seq) data from the colon muscularis externa. Each dot represents an individual cell, colored according to its assigned cluster. Cell populations of interest, including MMs and enteric neurons (EN), are highlighted (e.g., cluster 5 as MM, cluster 3 as EN). **B**. Violin plot illustrates the distribution of a calculated “activity score” for EN marker genes across all identified cell clusters. High activity scores in specific clusters (e.g., cluster 3) confirm their identity as enteric neurons. **C**. Violin plot illustrates the distribution of a calculated “activity score” for macrophage marker genes across all identified cell clusters. High activity scores in specific clusters (e.g., cluster 5) confirm their identity as MMs. **D**. UMAP plot highlights the subclusters identified in MMs and enteric neurons in C1qa^FL^ mice. The legend provides examples of specific sub-clusters (e.g., EN_1, MM_1) within these populations. **E**. UMAP plots showing the sub-clustering of the identified MMs and ENs populations in C1qa^CKO^ mice. Each sub-cluster (colored differently, e.g., 0-7) represents a transcriptionally distinct subtype within these main cell lineages, revealing their inherent heterogeneity. **F**. Bar graph displaying the percentage of cells within each identified cluster for both C1qa^CKO^ (green bars) and C1qa^FL^ (cyan bars) samples. This panel demonstrates that the overall proportions of identified cell clusters remain largely consistent between genotypes, confirming that C1qa deletion does not lead to major shifts in cell type abundance. **G.** Heatmap showing the expression of representative marker genes across enteric neuron (EN) subclusters. The heatmap highlights the distinct transcriptional signatures that define each EN population and illustrates the molecular heterogeneity of enteric neurons. **H**. Heatmap showing the expression of representative marker genes across muscularis macrophages (MM) subclusters. The heatmap highlights the distinct transcriptional signatures that define each MM population and illustrates the molecular heterogeneity of MMs.

**Supplementary Figure 3.**
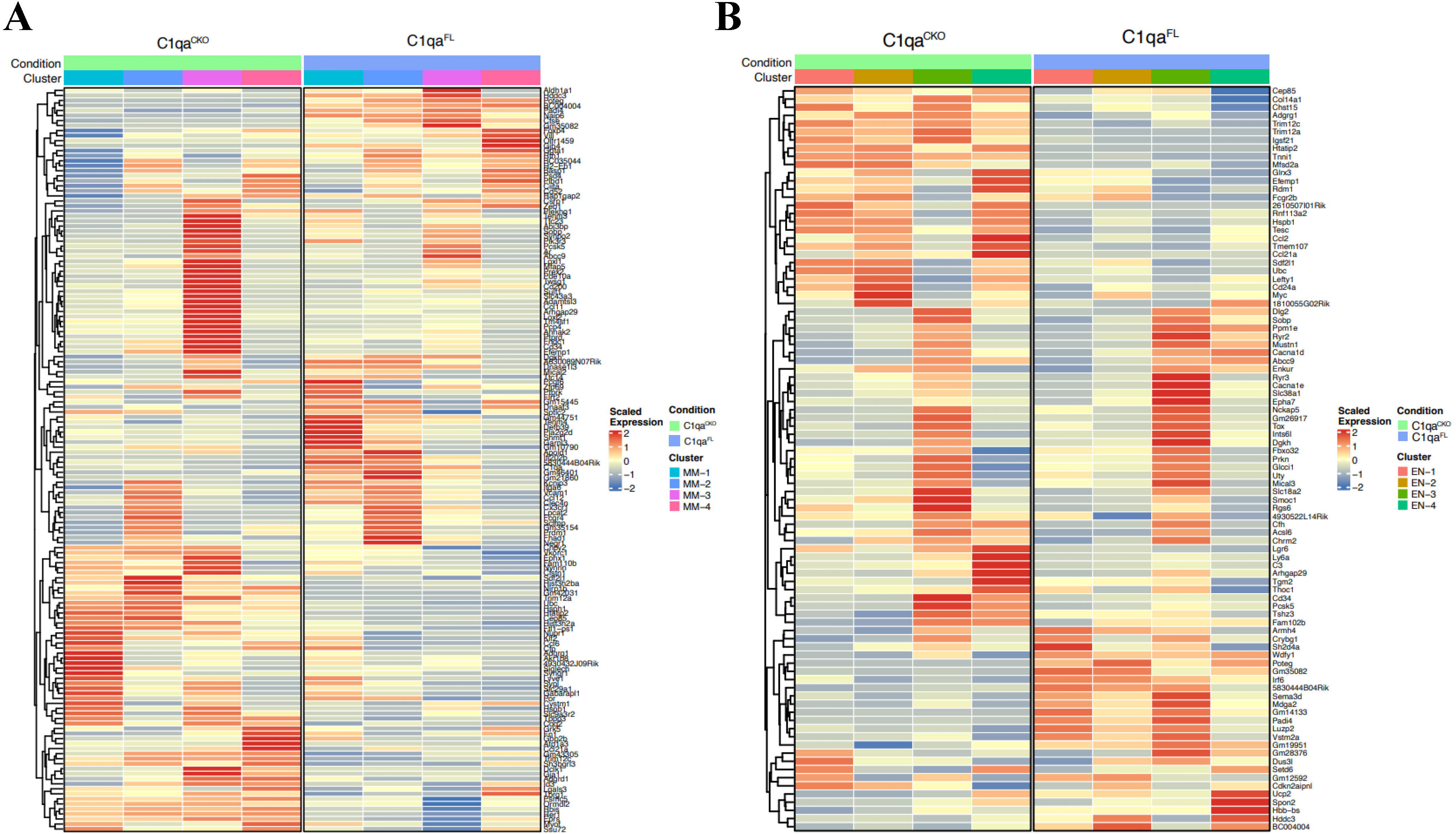
C1qa deficiency drives molecular reprogramming in muscularis macrophages (MMs) and enteric neurons. **A**. Heatmap specifically showing differentially expressed genes (DEGs) between C1qa^CKO^ and C1qa^FL^ MMs. This panel may focus on key genes or a subset of the DEGs from panel A, providing a more granular view of the molecular changes within the MMs population. **B**. Heatmap specifically showing differentially expressed genes (DEGs) between C1qa^CKO^ and C1qa^FL^ enteric neurons. This panel highlights key genes or a subset of the DEGs from panel B, providing a more granular view of the molecular changes within the enteric neuron population.

**Supplementary Figure 4.**
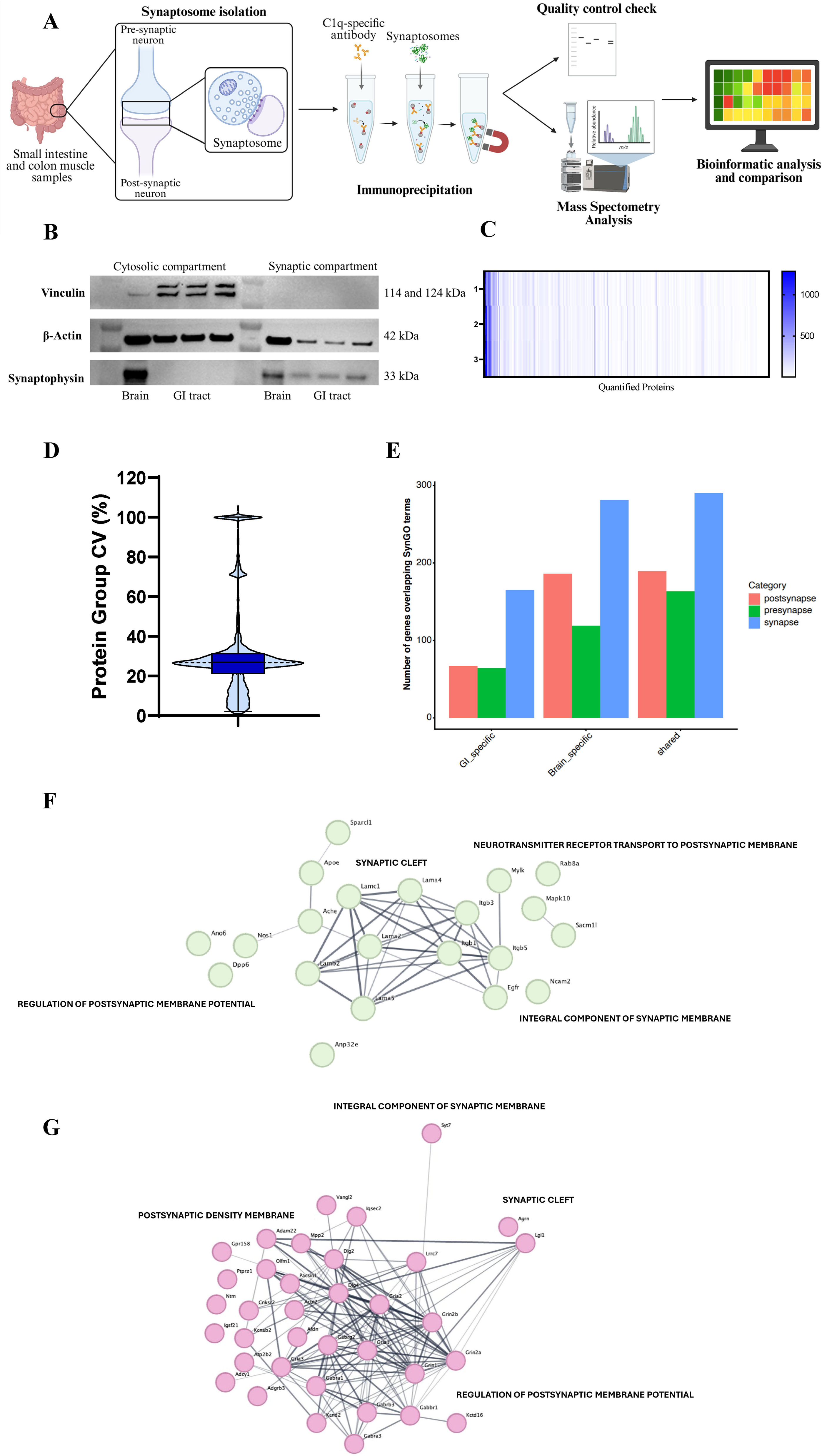
Characterization of the workflow used to isolate synaptosomes from intestinal muscularis externa. **A**. Schematic representation of the synaptosome isolation and proteomic analysis workflow from mouse gastrointestinal (GI) tissue. The process begins with dissection of GI tissue, followed by homogenization and differential centrifugation to isolate synaptosomes. An immunoprecipitation step (using a highly selective C1q antibody) is then performed to further purify the synaptosomes. Quality control checks (western blot analysis) ensure purity before mass spectrometry for proteomic analysis. Bioinformatic analysis then identifies and quantifies synaptic proteins. **B**. Western blots validating fractionation and synaptosome enrichment. Representative gel images loaded with proteins obtained from 3 different samples for GI and from 1 brain as control, show Vinculin (cytosolic marker; 114/124 kDa), β-Actin (loading control; 42 kDa), and Synaptophysin (synaptic marker; 33 kDa) across cytosolic and synaptic fractions, demonstrating enrichment of synaptic protein in the synaptosome compartment. **C**. Quality control heatmap of quantified proteins from mass spectrometry of enriched synaptosome preparations. Intensity scale (right) indicates protein abundance across samples and demonstrates reproducible synaptic protein recovery. **D**. Coefficient of variation (CV) across quantified protein groups (%). Violin/box plot (with median and IQR) summarizes reproducibility across biological replicates (n = 3). Most protein groups show low-to-moderate CV, indicating consistent preparation quality. **E.** Summary bar plot showing classification of quantified synaptic proteins into GI□specific, brain□specific, and shared synapse categories (postsynaptic, presynaptic, synapse). Numbers indicate gene count overlapping SynGO terms per category**. F-G.** Network analysis of GI-specific (**F**) and brain-specific (**G**) C1q-associated synaptic proteome, performed using STRING, revealed distinct functional modules. Nodes represent proteins, and edges indicate protein-protein interactions. Network connectivity reflects the degree of interaction between proteins, while spatial organization highlights functionally related synaptic modules.

**Supplementary Figure 5.**
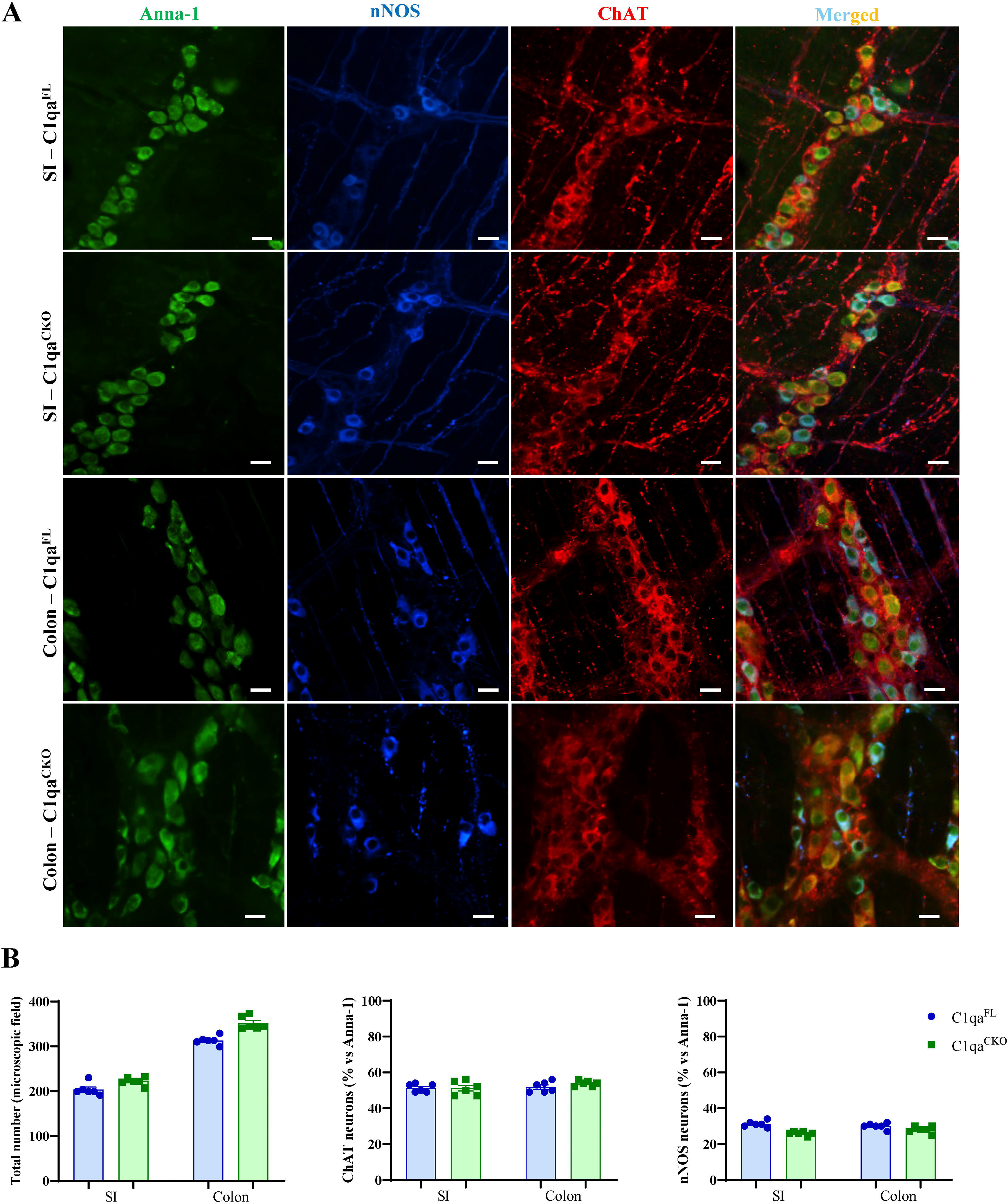
Subpopulations of enteric neurons are not different in C1qa^CKO^ mice. **A.** Representative immunostaining for the pan-neuronal marker Anna-1 (green), the nitrergic neurons marker (nNOS, blue) and the cholinergic neurons marker (ChAT, red) images used for quantification of enteric neurons in the small intestine (SI) and colon of C1qa^FL^ and C1qa^CKO^ mice. Merged image highlights that no difference in enteric neuronal subtypes were observed between the 2 genetic models. Scale bars: 20 µm. **B.** Quantification of enteric neuronal populations in the small intestine (SI) and colon of C1qa^FL^ (blue dots/bars) and C1qa^CKO^ (green dots/bars) mice. Data are presented as mean ± SEM.

**Supplementary Figure 6.**
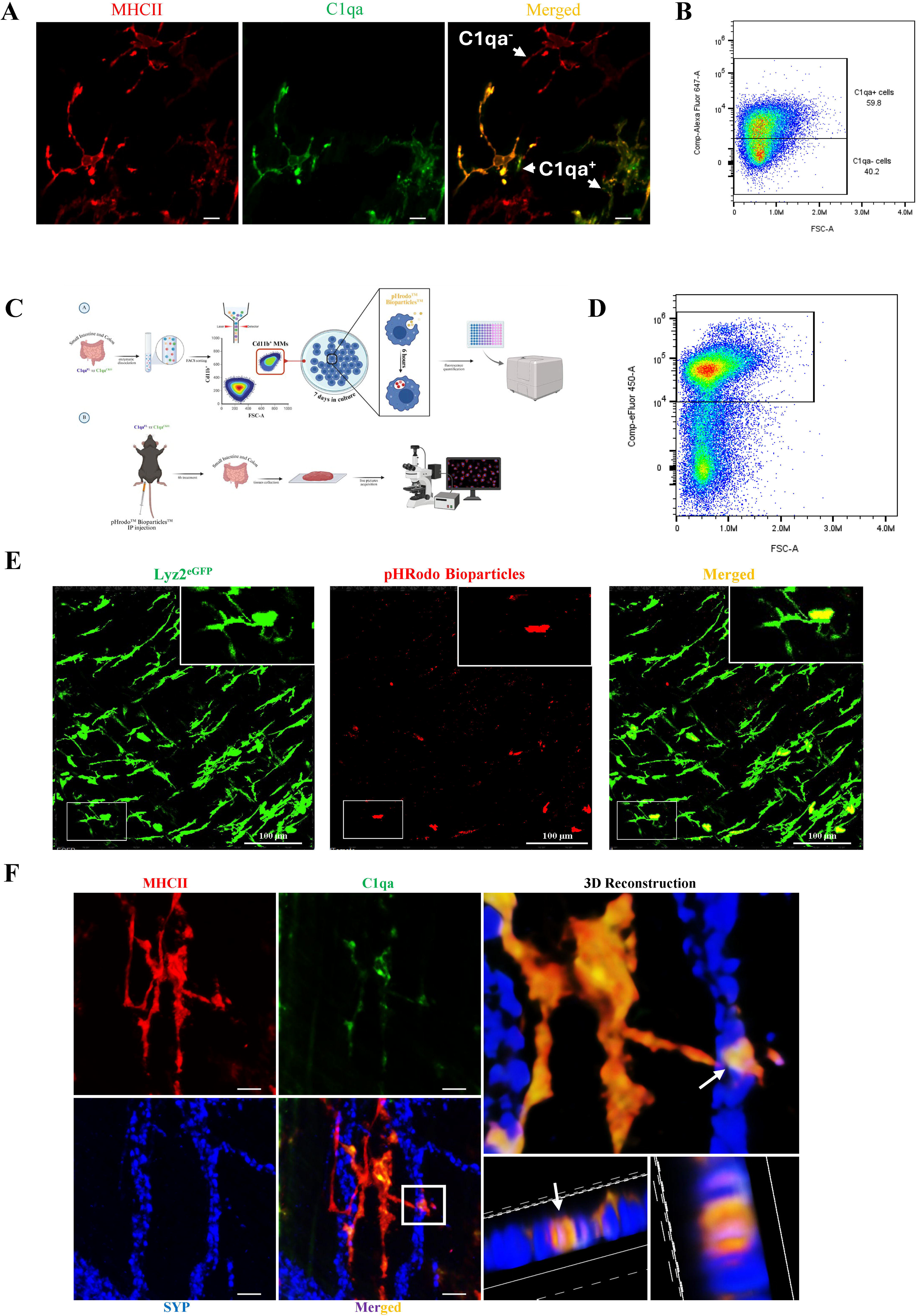
Characterization of C1qa^+^ muscularis macrophages (MMs) and their interactions with enteric neurons. **A.** Representative confocal micrographs showing immunostaining for the pan-macrophage marker MHCII (red) and C1qa (green). Merged channels shown as C1qa^+^ MMs represent a sub-population of tissue resident MMs. Scale bars: 50 µm. **B.** Flow cytometry plots used to gate and quantify C1qa^+^ versus C1qa^−^ MMs populations from intestinal tissue. Gates shown were applied consistently across samples; percentages indicated on plots. **C-D.** Experimental schematic summarizing isolation of CD11b^+^ for *in vitro* phagocytic assay evaluation (top), and histology/imaging pipeline for tissue validation (bottom). **E.** Validation of the effective pHRodo Bioparticle internalization in MMs performed by using a Lyz2^eGFP^ mouse model, in which myeloid cells are genetically tagged in green, following the same pipeline used for experiments in the genetic mouse model. Scale bars: 100 µm. **F.** High-resolution confocal validation of close apposition between C1qa^+^ MMs (yellow staining obtained by using a pan-macrophage marker MHCII - red and a C1qa marker - green) and enteric synaptic structures (presynaptic marker SYP - blue). Scale bars: 20 µm.

**Supplementary Figure 7.**
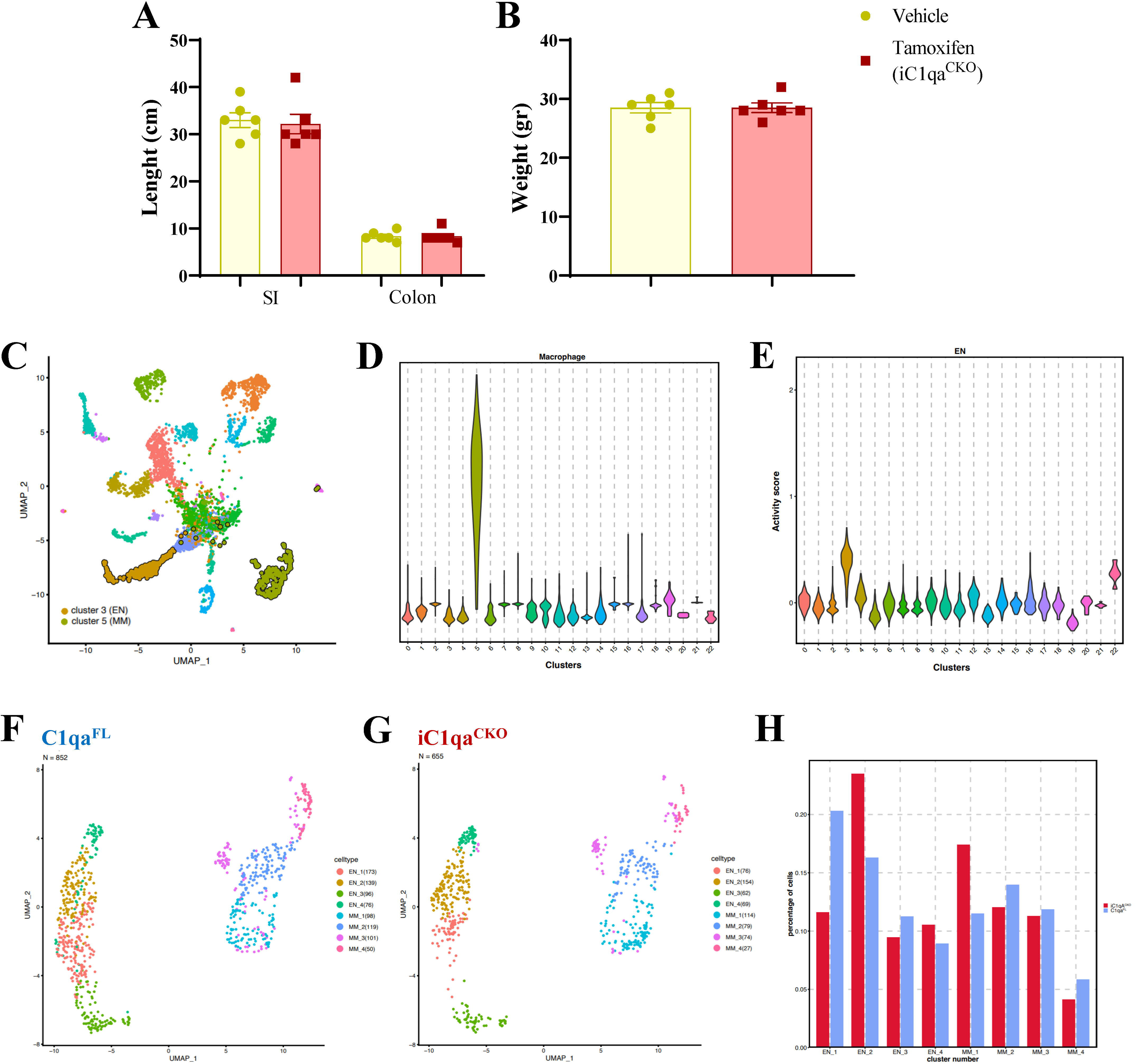
Characterization of iC1qa^CKO^ mouse model. **A.** Small intestine and colon lengths measured at harvest in vehicle (yellow dots/bars) and tamoxifen (iC1qa^CKO^ red dots/bars) treated mice. **B.** Body weights at time of tissue collection in vehicle (yellow dots/bars) and tamoxifen (iC1qa^CKO^ red dots/bars) treated mice. **C.** UMAP projection showing identified cell clusters from the scRNA-seq dataset; each dot represents a single cell colored by cluster identity. **D.** Violin plot illustrates the distribution and density of expression for a macrophage marker across the clusters. **E.** Violin plot of an enteric neuron activity score across the same clusters. **F**. UMAP showing the different subgroups of enteric neurons and MMs identified in the C1qa^FL^ mice. **G**. UMAP showing the different subgroups of ENs and muscularis macrophages (MMs) identified in the iC1qa^CKO^ mice. **H**. Bar chart showing relative abundance of selected clusters in C1qa^FL^ (blue) versus iC1qA^CKO^ (red) samples; values represent fraction of total cells per sample.

**Supplementary Figure 8.**
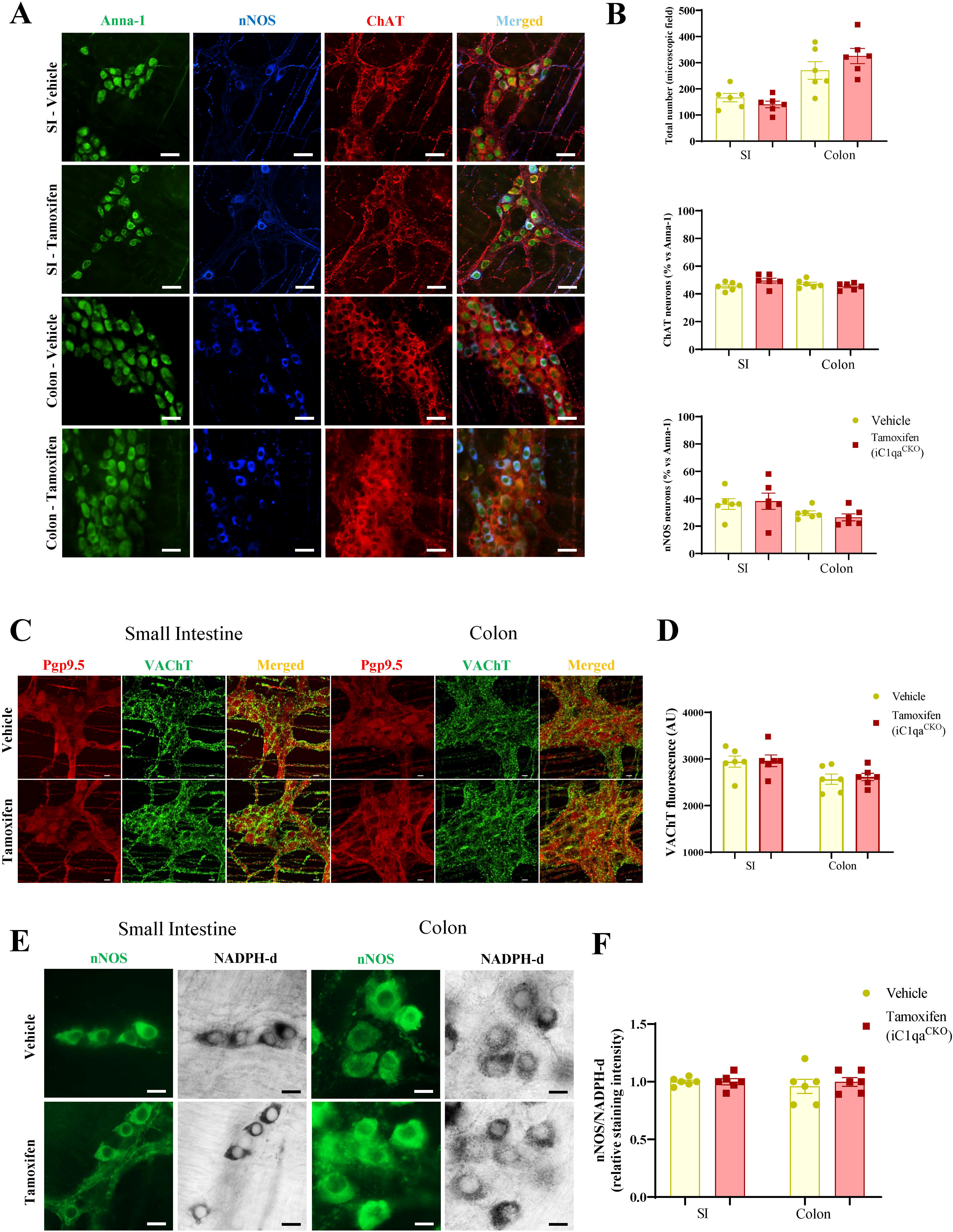
Enteric neurons are not changed in the iC1qa^CKO^ mouse model. **A.** Representative immunostaining for the pan-neuronal marker Anna-1 (green), the nitrergic neuronal marker nNOS (blue) and the cholinergic neuronal marker ChAT (red). Images were used for the quantification of enteric neurons in the small intestine (SI) and colon of vehicle and tamoxifen (iC1qa^CKO^) treated mice. Merged image highlights no differences in enteric neuronal subtypes numbers between the 2 genetic models. Scale bars: 50 μm. **B.** Quantification of enteric neuronal populations in vehicle (yellow dots/bars) and tamoxifen (iC1qa^CKO^ red dots/bars) mice in the small intestine (SI) and colon. Data are presented as mean ± SEM. **C**. Representative immunostaining for the pan-neuronal marker Pgp9.5 (red) and the Vesicular Acetylcholine Transporter (VAChT, green) in the small intestine (SI) and colon myenteric plexus of vehicle and tamoxifen (iC1qa^CKO^) treated mice. Scale bars: 50 μm. **D**. Quantification of mean fluorescence intensity of VAChT in the small intestine (SI) and colon ENS of vehicle (yellow dots/bars) and tamoxifen (iC1qa^CKO^ red dots/bars) mice. **E.** Representative immunohistochemistry for the nitrergic neuronal marker nNOS (green) and the histochemical marker of nitric oxide synthase activity (NADPH-d, gray scale) in the small intestine (SI) and colon myenteric plexus of vehicle and tamoxifen (iC1qa^CKO^) treated mice. Scale bars: 10 μm. **F**. Ratio quantification between nNOS^+^/NADPH-d^+^ neurons in the small intestine (SI) and colon myenteric plexus of vehicle (yellow dots/bars) and tamoxifen (iC1qa^CKO^ red dots/bars) treated mice. Data are expressed as mean ± SEM.

**Supplementary Figure 9.**
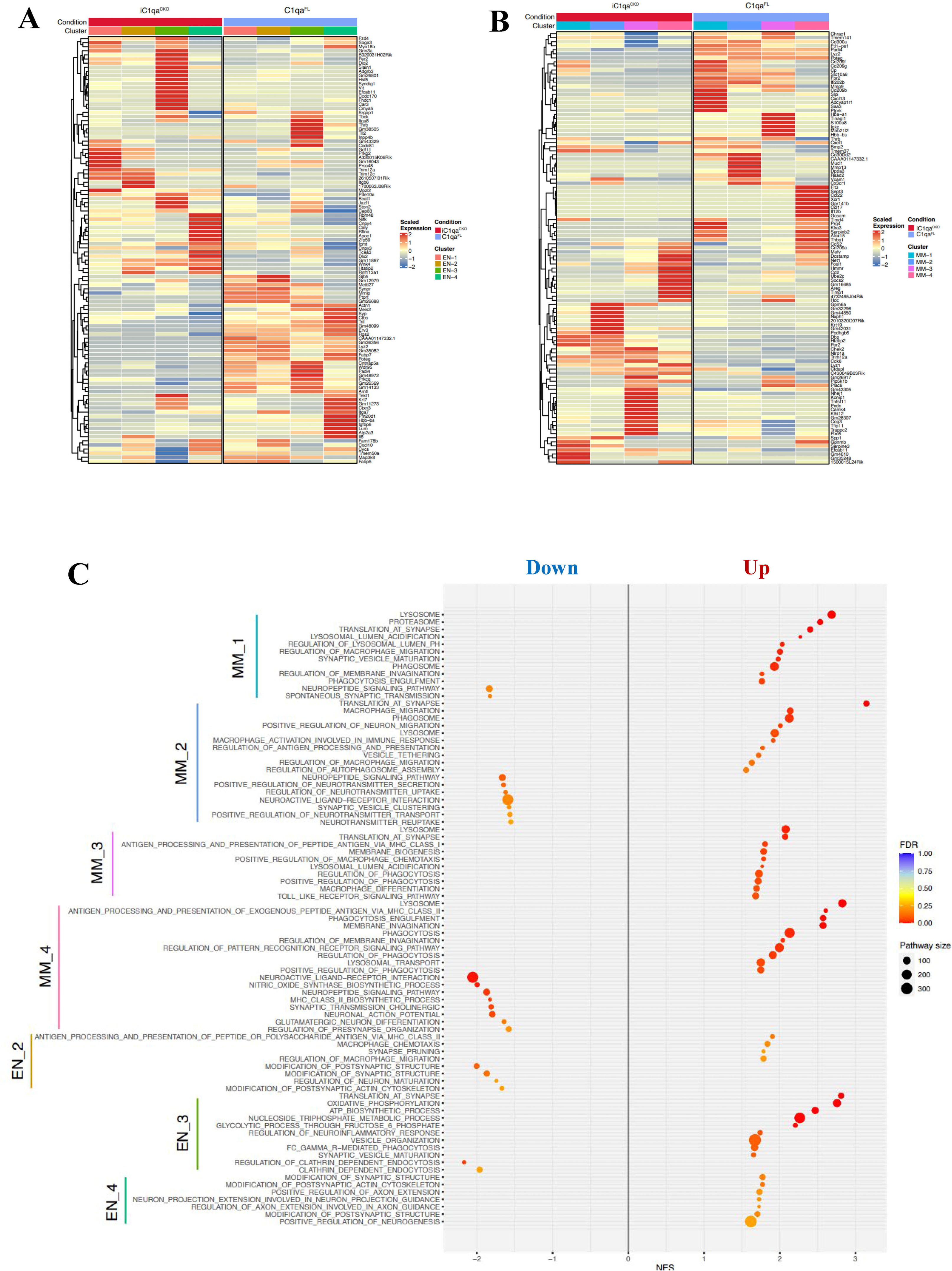
Transcriptional differences between C1qa^CKO^ and iC1qa^CKO^ A-B. Heatmaps of differentially expressed genes between iC1qa^CKO^ (left columns - red) and C1qa^FL^ (right columns - blue) conditions. Rows represent genes (labeled at right) and columns represent individual cells or pseudo□bulk samples grouped by annotated subclusters (color bar at top: EN_1-4 = enteric neuron subclusters; MM_1-4 = muscularis macrophage subclusters). **A**. Heatmap highlighting genes enriched in EN clusters. **B.** Heatmap highlighting genes enriched in MMs clusters. Heatmaps display representative differentially expressed genes per cluster. **C.** GSEA dot plot comparing C1qa^CKO^ versus inducible iC1qa^CKO^ models across selected subclusters.

**Supplementary Figure 10.**
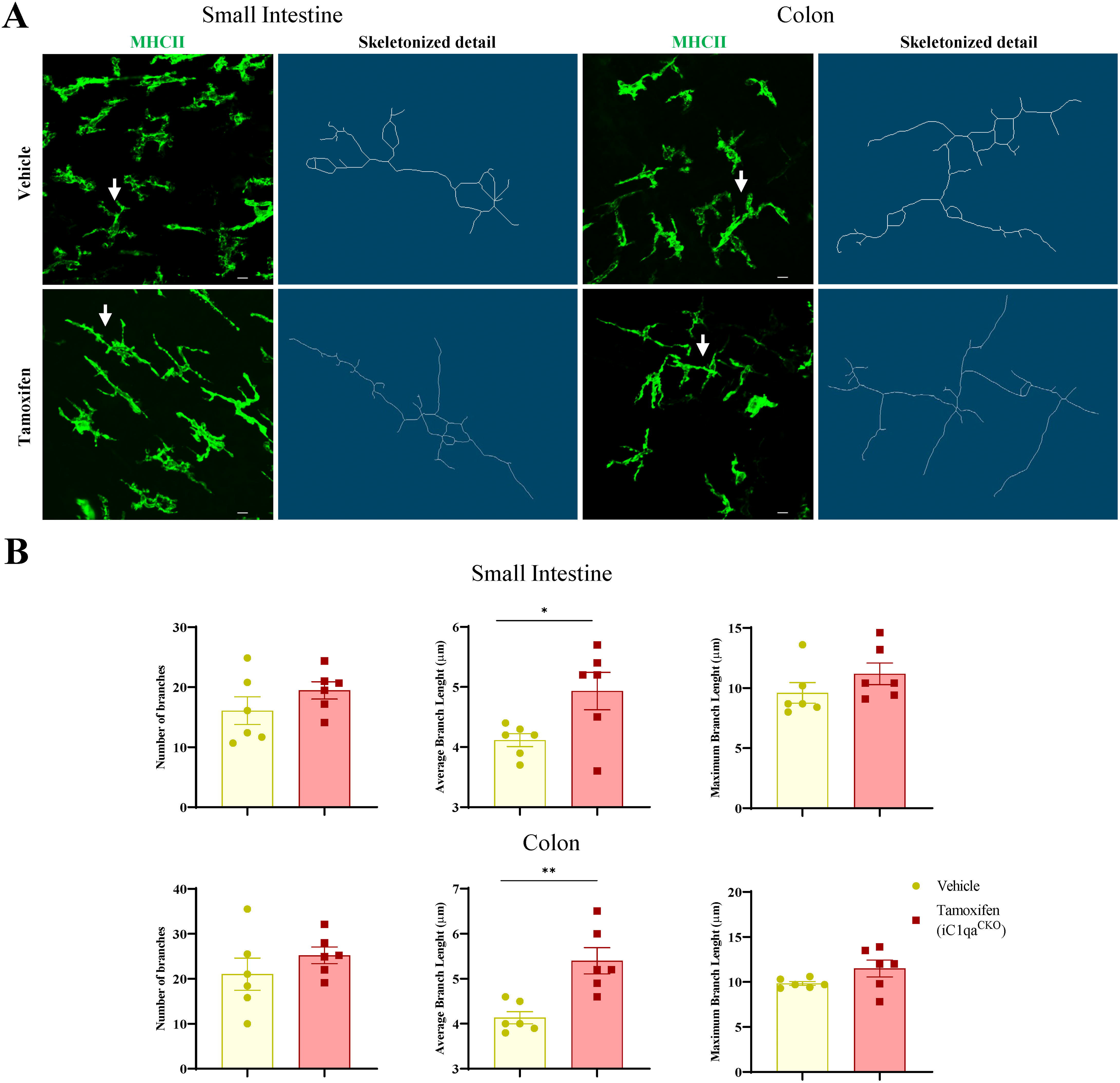
Inducible C1qa deletion alters muscularis macrophage (MMs) morphology. A-B. Muscularis macrophage morphology and branches analysis. **A.** Representative MHCII immunofluorescence and corresponding skeletonized images (small intestine and colon) showing altered MMs process complexity in vehicle and tamoxifen (iC1qa^CKO^) treated mice. Scale bars: 50 μm. **B.** Quantification of shape parameters (number of branches, average branch length and maximum branches length). Data are expressed as mean ± SEM. *p < 0.05 and ** p<0.01 by unpaired t-test.

## SUPPLEMENTARY TABLE LEGENDS

**Supplementary Table 1:** Canonical marker genes for macrophage and enteric neuron/glia population

**Supplementary Table 2:** Enriched marker genes defining EN and MM subclusters

**Supplementary Table 3:** Differentially expressed genes (DEGs) across EN and MM subclusters following C1qa deletion

**Supplementary Table 4:** Significantly enriched GSEA pathways identified in the C1qa^CKO^ vs C1qa^FL^, C1qa^CKO^ vs iC1qa^CKO^, and C1qa^FL^ vs iC1qa^CKO^ differential expression comparisons

**Supplementary Table 5:** Proteins associated with selected SynGO terms

**Supplementary Table 6:** Significantly enriched GSEA pathways in C1qa□ versus C1qa□ populations in C1qa^FL^ mice

## Notes

### Competing Interest Statement

The authors have declared no competing interest.

