## Supplementary Table 1 for "C1qa□ muscularis macrophages maintain enteric synaptic homeostasis to regulate gastrointestinal motility"

| celltype | gene |
|---|---|
| EN/Glia | Tubb3 |
| EN/Glia | Elavl4 |
| EN/Glia | Ret |
| EN/Glia | Phox2b |
| EN/Glia | Chrnb4 |
| EN/Glia | Eml5 |
| EN/Glia | Smpd3 |
| EN/Glia | Tagln3 |
| EN/Glia | Snap25 |
| EN/Glia | Gpr22 |
| EN/Glia | Gdap1l1 |
| EN/Glia | Stmn3 |
| EN/Glia | Chrna3 |
| EN/Glia | Scg3 |
| EN/Glia | Syt4 |
| EN/Glia | Ncan |
| EN/Glia | Crmp1 |
| EN/Glia | Adcyap1r1 |
| EN/Glia | Elavl3 |
| EN/Glia | Dlg2 |
| EN/Glia | Cacna2d |
| EN/Glia | Erbb3 |
| EN/Glia | Sox10 |
| EN/Glia | Fabp7 |
| EN/Glia | Plp1 |
| EN/Glia | Gas7 |
| EN/Glia | Nid1 |
| EN/Glia | Qk |
| EN/Glia | Sparc |
| EN/Glia | Mest |
| EN/Glia | Nfia |
| EN/Glia | Wwtr1 |
| EN/Glia | Gpm6b |
| EN/Glia | Rasa3 |
| EN/Glia | Flrt1 |
| EN/Glia | Itpripl1 |
| EN/Glia | Itga4 |
| EN/Glia | Lama4 |
| EN/Glia | Postn |
| EN/Glia | Ptprz1 |
| EN/Glia | Pdpn |
| EN/Glia | Col18a1 |
| EN/Glia | Nrcam |
| Macrophage | H2-Ab1 |
| Macrophage | H2-Eb1 |
| Macrophage | Itgam |

| | |
|---|---|
| Macrophage | Adgre1 |
| Macrophage | Mrc1 |
| Macrophage | Retnla |
| Macrophage | Cd68 |
| Macrophage | Csf1r |
| Macrophage | Cx3cr1 |
| Macrophage | Ccr2 |
| Macrophage | Cd163 |
