## Supplementary Table 2 for "C1qa□ muscularis macrophages maintain enteric synaptic homeostasis to regulate gastrointestinal motility"

| KO_p_val | KO_avg_log2FC | KO_pct.1 | KO_pct.2 | KO_p_val_adj | WT_p_val | WT_avg_log2FC | WT_pct.1 | WT_pct.2 | WT_p_val_adj | max_pval | minimump_p_val | gene | cluster |
|---|---|---|---|---|---|---|---|---|---|---|---|---|---|
| | | | | | | EN | | | | | | | |
| 3.39708E-23 | 0.857267775 | 1 | 0.785 | 8.02119E-19 | 2.35639E-20 | 0.759991126 | 1 | 0.796 | 5.56391E-16 | 2.35639E-20 | 6.79416E-23 | Kcna1 | EN_1 |
| 6.14962E-11 | 0.935073478 | 0.635 | 0.319 | 1.45205E-06 | 4.38195E-18 | 1.204896952 | 0.756 | 0.384 | 1.03467E-13 | 6.14962E-11 | 8.7639E-18 | Aldh1a1 | EN_1 |
| 7.70136E-18 | 0.827470251 | 0.927 | 0.51 | 1.81845E-13 | 2.55908E-16 | 0.79379416 | 0.956 | 0.551 | 6.04249E-12 | 2.55908E-16 | 1.54027E-17 | Cnp | EN_1 |
| 2.01497E-17 | 0.645399571 | 1 | 0.807 | 4.75774E-13 | 5.92533E-14 | 0.594291485 | 0.994 | 0.831 | 1.39909E-09 | 5.92533E-14 | 4.02993E-17 | H2-D1 | EN_1 |
| 6.05488E-17 | 0.628075121 | 1 | 0.826 | 1.42968E-12 | 3.55207E-16 | 0.633288727 | 0.994 | 0.831 | 8.38715E-12 | 3.55207E-16 | 1.21098E-16 | Plp1 | EN_1 |
| 8.55971E-12 | 0.655049054 | 0.91 | 0.578 | 2.02112E-07 | 6.58421E-17 | 0.943502681 | 0.922 | 0.586 | 1.55466E-12 | 8.55971E-12 | 1.31684E-16 | Foxd3 | EN_1 |
| 8.46229E-16 | 0.844564009 | 0.781 | 0.343 | 1.99812E-11 | 1.10744E-11 | 0.678871434 | 0.789 | 0.43 | 2.61488E-07 | 1.10744E-11 | 1.69246E-15 | Tax1bp3 | EN_1 |
| 9.16003E-13 | 0.740158462 | 0.854 | 0.469 | 2.16287E-08 | 1.58881E-15 | 0.849946625 | 0.85 | 0.457 | 3.75151E-11 | 9.16003E-13 | 3.17763E-15 | Plat | EN_1 |
| 1.86016E-15 | 0.821008168 | 0.949 | 0.567 | 4.39221E-11 | 1.75507E-11 | 0.616794771 | 0.894 | 0.524 | 4.14408E-07 | 1.75507E-11 | 3.72032E-15 | Calr | EN_1 |
| 1.21842E-13 | 0.818828442 | 0.854 | 0.504 | 2.87692E-09 | 9.0175E-15 | 0.749075544 | 0.844 | 0.462 | 2.12921E-10 | 1.21842E-13 | 1.8035E-14 | Mid1ip1 | EN_1 |
| 7.00911E-14 | 0.872124028 | 0.719 | 0.319 | 1.65499E-09 | 2.18528E-07 | 0.601731143 | 0.65 | 0.401 | 0.005159878 | 2.18528E-07 | 1.40182E-13 | Cct4 | EN_1 |
| 1.94742E-13 | 0.680842922 | 0.938 | 0.564 | 4.59824E-09 | 5.98337E-08 | 0.606992258 | 0.772 | 0.487 | 0.001412793 | 5.98337E-08 | 3.89483E-13 | Ccnd1 | EN_1 |
| 7.7832E-12 | 0.624252789 | 0.983 | 0.722 | 1.83777E-07 | 8.76773E-13 | 0.66583682 | 0.961 | 0.707 | 2.07024E-08 | 7.7832E-12 | 1.75355E-12 | Aspa | EN_1 |
| 1.39519E-12 | 0.632820677 | 0.86 | 0.45 | 3.29431E-08 | 2.68449E-11 | 0.621990451 | 0.833 | 0.476 | 6.33862E-07 | 2.68449E-11 | 2.79037E-12 | Ajap1 | EN_1 |
| 1.56247E-10 | 0.670251983 | 0.848 | 0.485 | 3.6893E-06 | 1.40882E-12 | 0.687002033 | 0.822 | 0.46 | 3.32651E-08 | 1.56247E-10 | 2.81765E-12 | Mettl7a1 | EN_1 |
| 1.56954E-12 | 1.009962922 | 0.624 | 0.27 | 3.706E-08 | 7.48006E-10 | 0.670738669 | 0.683 | 0.339 | 1.76619E-05 | 7.48006E-10 | 3.13909E-12 | Maged2 | EN_1 |
| 1.95633E-12 | 0.707503853 | 0.888 | 0.49 | 4.61928E-08 | 3.38348E-11 | 0.620211615 | 0.878 | 0.516 | 7.98907E-07 | 3.38348E-11 | 3.91266E-12 | Tspan15 | EN_1 |
| 5.46452E-11 | 0.697595666 | 0.5 | 0.196 | 1.29028E-06 | 1.2174E-08 | 0.72538895 | 0.622 | 0.333 | 0.000287452 | 1.2174E-08 | 1.0929E-10 | Camk2b | EN_1 |
| 1.11886E-10 | 0.634891855 | 0.719 | 0.322 | 2.64186E-06 | 7.45312E-09 | 0.603067091 | 0.644 | 0.344 | 0.000175983 | 7.45312E-09 | 2.23773E-10 | Hsd17b12 | EN_1 |
| 1.44078E-10 | 0.606087769 | 0.657 | 0.316 | 3.40196E-06 | 2.02994E-09 | 0.641318608 | 0.528 | 0.237 | 4.79308E-05 | 2.02994E-09 | 2.88155E-10 | Pdp1 | EN_1 |
| 1.48516E-10 | 0.691317453 | 0.494 | 0.202 | 3.50676E-06 | 1.27915E-07 | 0.714288162 | 0.506 | 0.253 | 0.003020324 | 1.27915E-07 | 2.97032E-10 | Fam89a | EN_1 |
| 1.63375E-10 | 0.777379571 | 0.64 | 0.308 | 3.85761E-06 | 1.57071E-09 | 0.687676263 | 0.628 | 0.323 | 3.70876E-05 | 1.57071E-09 | 3.2675E-10 | Snta1 | EN_1 |
| 2.74385E-09 | 0.929187406 | 0.517 | 0.237 | 6.47878E-05 | 8.75698E-09 | 0.626837075 | 0.572 | 0.288 | 0.00020677 | 8.75698E-09 | 5.4877E-09 | Leprot | EN_1 |
| 3.74673E-18 | 1.046913189 | 0.971 | 0.77 | 8.84677E-14 | 4.43215E-33 | 1.485101702 | 0.982 | 0.796 | 1.04652E-28 | 3.74673E-18 | 8.86431E-33 | Jun | EN_2 |
| 5.42568E-24 | 1.43744415 | 0.963 | 0.716 | 1.28111E-19 | 6.44367E-31 | 1.477680701 | 0.97 | 0.706 | 1.52148E-26 | 5.42568E-24 | 1.28873E-30 | Junb | EN_2 |
| 1.42512E-20 | 1.134810819 | 0.985 | 0.714 | 3.36499E-16 | 2.37504E-28 | 1.326363971 | 0.994 | 0.794 | 5.60795E-24 | 1.42512E-20 | 4.75008E-28 | Fos | EN_2 |
| 2.93812E-23 | 0.789002719 | 1 | 0.856 | 6.93748E-19 | 1.95476E-25 | 0.793133487 | 1 | 0.897 | 4.61557E-21 | 2.93812E-23 | 3.90951E-25 | Eif1 | EN_2 |
| 8.00789E-18 | 1.155032038 | 0.868 | 0.526 | 1.89082E-13 | 3.02284E-25 | 1.366657996 | 0.896 | 0.487 | 7.13752E-21 | 8.00789E-18 | 6.04567E-25 | Ier3 | EN_2 |
| 3.97787E-18 | 0.72980251 | 1 | 0.861 | 9.39255E-14 | 4.83615E-25 | 0.83201892 | 1 | 0.884 | 1.14191E-20 | 3.97787E-18 | 9.67231E-25 | Rplp1 | EN_2 |
| 1.48199E-20 | 0.814416562 | 0.985 | 0.797 | 3.49928E-16 | 1.28469E-24 | 0.882301434 | 0.988 | 0.784 | 3.0334E-20 | 1.48199E-20 | 2.56937E-24 | Rpl17 | EN_2 |
| 1.56574E-24 | 0.951455651 | 1 | 0.778 | 3.69702E-20 | 7.02876E-18 | 0.728653254 | 0.982 | 0.791 | 1.65963E-13 | 7.02876E-18 | 3.13148E-24 | Selenof | EN_2 |
| 1.58712E-24 | 0.995882545 | 1 | 0.919 | 3.74751E-20 | 2.01982E-20 | 0.838745059 | 1 | 0.915 | 4.7692E-16 | 2.01982E-20 | 3.17424E-24 | Vim | EN_2 |
| 6.52851E-18 | 0.830246262 | 0.985 | 0.782 | 1.54151E-13 | 1.65897E-24 | 0.827086709 | 0.994 | 0.835 | 3.91716E-20 | 6.52851E-18 | 3.31794E-24 | Rps23 | EN_2 |
| 8.97525E-19 | 0.774691152 | 1 | 0.924 | 2.11924E-14 | 1.87093E-24 | 0.77493834 | 1 | 0.936 | 4.41764E-20 | 8.97525E-19 | 3.74186E-24 | Tpt1 | EN_2 |
| 2.71376E-19 | 0.80197223 | 0.985 | 0.844 | 6.40772E-15 | 2.80901E-24 | 0.800898605 | 1 | 0.848 | 6.63263E-20 | 2.71376E-19 | 5.61802E-24 | Rps27 | EN_2 |
| 5.89095E-24 | 0.764863825 | 1 | 0.88 | 1.39097E-19 | 2.03787E-19 | 0.625405454 | 1 | 0.881 | 4.81181E-15 | 2.03787E-19 | 1.17819E-23 | Ubb | EN_2 |
| 8.4124E-18 | 0.812361144 | 0.993 | 0.751 | 1.98634E-13 | 9.92786E-23 | 0.865238386 | 0.988 | 0.768 | 2.34417E-18 | 8.4124E-18 | 1.98557E-22 | Rpl3 | EN_2 |
| 2.3411E-18 | 0.837164614 | 0.993 | 0.807 | 5.52781E-14 | 1.30821E-22 | 0.85063905 | 0.982 | 0.786 | 3.08895E-18 | 2.3411E-18 | 2.61642E-22 | Rpl21 | EN_2 |
| 3.22063E-17 | 0.980194912 | 0.985 | 0.787 | 7.60455E-13 | 1.34829E-22 | 1.08756057 | 0.988 | 0.796 | 3.18359E-18 | 3.22063E-17 | 2.69658E-22 | Mt1 | EN_2 |
| 4.40854E-14 | 0.672724737 | 0.963 | 0.79 | 1.04094E-09 | 1.95469E-22 | 0.85210101 | 0.988 | 0.776 | 4.61541E-18 | 4.40854E-14 | 3.90937E-22 | Rps19 | EN_2 |
| 1.13614E-17 | 0.736941144 | 0.993 | 0.826 | 2.68265E-13 | 2.7669E-22 | 0.707229318 | 1 | 0.866 | 6.5332E-18 | 1.13614E-17 | 5.53379E-22 | Rps10 | EN_2 |
| 1.30724E-14 | 0.675554322 | 0.978 | 0.812 | 3.08667E-10 | 3.67359E-22 | 0.812259737 | 0.994 | 0.802 | 8.67407E-18 | 1.30724E-14 | 7.34717E-22 | Rps12 | EN_2 |
| 5.1753E-22 | 0.971409144 | 0.978 | 0.716 | 1.22199E-17 | 6.43078E-18 | 0.679608735 | 0.976 | 0.784 | 1.51844E-13 | 6.43078E-18 | 1.03506E-21 | Rpl7 | EN_2 |
| 1.42563E-21 | 0.904324826 | 1 | 0.826 | 3.3662E-17 | 2.61742E-21 | 0.741412303 | 0.994 | 0.822 | 6.18026E-17 | 2.61742E-21 | 2.85126E-21 | Ftl1 | EN_2 |
| 1.20052E-15 | 1.076100202 | 0.912 | 0.621 | 2.83468E-11 | 3.37586E-21 | 1.111976149 | 0.951 | 0.683 | 7.97107E-17 | 1.20052E-15 | 6.75171E-21 | Egr1 | EN_2 |
| 2.8206E-16 | 0.708444951 | 0.993 | 0.812 | 6.66E-12 | 5.48678E-21 | 0.727539825 | 1 | 0.822 | 1.29554E-16 | 2.8206E-16 | 1.09736E-20 | Rpl27a | EN_2 |
| 9.71828E-21 | 0.896173824 | 1 | 0.8 | 2.29468E-16 | 2.53568E-18 | 0.776252723 | 1 | 0.82 | 5.98725E-14 | 2.53568E-18 | 1.94366E-20 | Arpc1b | EN_2 |
| 1.10792E-20 | 0.750060004 | 1 | 0.878 | 2.61602E-16 | 3.18004E-18 | 0.59338963 | 1 | 0.897 | 7.50871E-14 | 3.18004E-18 | 2.21584E-20 | Rpl13 | EN_2 |
| 1.62306E-20 | 0.805819407 | 0.993 | 0.802 | 3.83238E-16 | 2.90276E-20 | 0.608871979 | 0.994 | 0.771 | 6.854E-16 | 2.90276E-20 | 3.24613E-20 | Rps4x | EN_2 |
| 1.29131E-18 | 1.148379361 | 0.985 | 0.692 | 3.04904E-14 | 2.67025E-20 | 1.213993396 | 0.951 | 0.686 | 6.30499E-16 | 1.29131E-18 | 5.3405E-20 | Cebpd | EN_2 |
| 2.7098E-20 | 1.022065884 | 0.978 | 0.731 | 6.39838E-16 | 1.64257E-14 | 0.720340117 | 0.951 | 0.716 | 3.87843E-10 | 1.64257E-14 | 5.4196E-20 | Tuba1a | EN_2 |
| 3.09794E-20 | 1.225765889 | 0.949 | 0.597 | 7.31487E-16 | 9.69721E-10 | 0.687200085 | 0.805 | 0.562 | 2.2897E-05 | 9.69721E-10 | 6.19589E-20 | Stmn1 | EN_2 |
| 8.41958E-18 | 0.854707009 | 0.978 | 0.731 | 1.98803E-13 | 6.084E-20 | 0.807450415 | 1 | 0.771 | 1.43655E-15 | 8.41958E-18 | 1.2168E-19 | Rpl10 | EN_2 |
| 8.37902E-17 | 0.671953111 | 0.985 | 0.844 | 1.97845E-12 | 7.71357E-20 | 0.611656532 | 1 | 0.83 | 1.82133E-15 | 8.37902E-17 | 1.54271E-19 | Rpl19 | EN_2 |
| 2.98273E-16 | 0.699011819 | 1 | 0.949 | 7.04283E-12 | 1.01054E-19 | 0.687489968 | 1 | 0.964 | 2.3861E-15 | 2.98273E-16 | 2.02109E-19 | Tmsb4x | EN_2 |
| 1.86776E-19 | 0.842028537 | 0.978 | 0.785 | 4.41016E-15 | 5.46803E-16 | 0.643578071 | 0.988 | 0.768 | 1.29111E-11 | 5.46803E-16 | 3.73553E-19 | Rpl18 | EN_2 |
| 2.63035E-13 | 1.177492904 | 0.904 | 0.621 | 6.21079E-09 | 3.03635E-19 | 1.104541391 | 0.939 | 0.655 | 7.16943E-15 | 2.63035E-13 | 6.0727E-19 | Socs3 | EN_2 |
| 5.428E-19 | 0.760087761 | 1 | 0.824 | 1.28166E-14 | 2.99641E-18 | 0.717714209 | 0.994 | 0.871 | 7.07513E-14 | 2.99641E-18 | 1.0856E-18 | H3f3b | EN_2 |
| 3.68245E-17 | 0.630735881 | 1 | 0.934 | 8.695E-13 | 5.44182E-19 | 0.595264814 | 1 | 0.91 | 1.28492E-14 | 3.68245E-17 | 1.08836E-18 | Rps29 | EN_2 |
| 2.26958E-16 | 1.257536869 | 0.875 | 0.55 | 5.35894E-12 | 5.48693E-19 | 1.35280017 | 0.848 | 0.531 | 1.29558E-14 | 2.26958E-16 | 1.09739E-18 | Nfkbia | EN_2 |
| 6.69725E-19 | 0.874598593 | 0.993 | 0.758 | 1.58136E-14 | 5.86128E-18 | 0.777375866 | 0.982 | 0.82 | 1.38397E-13 | 5.86128E-18 | 1.33945E-18 | Rps13 | EN_2 |
| 4.65945E-16 | 0.848322733 | 0.978 | 0.822 | 1.10019E-11 | 8.66094E-19 | 0.803970267 | 0.97 | 0.812 | 2.04502E-14 | 4.65945E-16 | 1.73219E-18 | Hspa8 | EN_2 |
| 2.45794E-17 | 0.838393376 | 1 | 0.841 | 5.80368E-13 | 9.5885E-19 | 0.955103434 | 0.982 | 0.827 | 2.26404E-14 | 2.45794E-17 | 1.9177E-18 | Jund | EN_2 |
| 1.40614E-16 | 0.783900392 | 0.993 | 0.77 | 3.32018E-12 | 9.93032E-19 | 0.762335966 | 0.994 | 0.789 | 2.34475E-14 | 1.40614E-16 | 1.98606E-18 | Rps15a | EN_2 |
| 4.04892E-17 | 0.766966119 | 0.985 | 0.814 | 9.56031E-13 | 1.31359E-18 | 0.710800167 | 1 | 0.825 | 3.10164E-14 | 4.04892E-17 | 2.62717E-18 | Rps9 | EN_2 |
| 8.52577E-17 | 0.809291914 | 0.978 | 0.724 | 2.0131E-12 | 1.55841E-18 | 0.747443958 | 0.988 | 0.771 | 3.67971E-14 | 8.52577E-17 | 3.11681E-18 | Rps18 | EN_2 |
| 9.69467E-13 | 1.293177701 | 0.588 | 0.269 | 2.2891E-08 | 1.605E-18 | 1.910463598 | 0.537 | 0.191 | 3.78973E-14 | 9.69467E-13 | 3.21001E-18 | Pim3 | EN_2 |
| 1.71373E-17 | 0.78492349 | 0.978 | 0.782 | 4.04645E-13 | 1.66821E-18 | 0.791604785 | 0.988 | 0.809 | 3.93898E-14 | 1.71373E-17 | 3.33643E-18 | Rpl8 | EN_2 |
| 1.72482E-15 | 0.723344913 | 1 | 0.797 | 4.07265E-11 | 1.69631E-18 | 0.659019059 | 0.988 | 0.807 | 4.00533E-14 | 1.72482E-15 | 3.39263E-18 | Rpl36 | EN_2 |
| 1.1552E-12 | 0.846718716 | 0.897 | 0.589 | 2.72766E-08 | 3.6188E-18 | 1.015306362 | 0.896 | 0.557 | 8.5447E-14 | 1.1552E-12 | 7.23759E-18 | Atf3 | EN_2 |
| 1.92306E-17 | 0.622002351 | 1 | 0.863 | 4.54073E-13 | 5.83967E-18 | 0.603097271 | 1 | 0.869 | 1.37886E-13 | 1.92306E-17 | 1.16793E-17 | Rps21 | EN_2 |
| 5.91072E-13 | 0.703126531 | 0.971 | 0.689 | 1.39564E-08 | 5.87865E-18 | 0.800234897 | 0.963 | 0.701 | 1.38807E-13 | 5.91072E-13 | 1.17573E-17 | Rps6 | EN_2 |
| 3.06532E-16 | 0.83395388 | 0.956 | 0.726 | 7.23785E-12 | 7.33097E-18 | 0.676003825 | 0.982 | 0.812 | 1.73099E-13 | 3.06532E-16 | 1.46619E-17 | Rps7 | EN_2 |
| 1.48003E-17 | 1.222608951 | 0.919 | 0.499 | 3.49464E-13 | 7.48581E-13 | 0.884917584 | 0.848 | 0.528 | 1.76755E-08 | 7.48581E-13 | 2.96005E-17 | Fabp5 | EN_2 |
| 1.98625E-17 | 1.54885307 | 0.713 | 0.367 | 4.68994E-13 | 1.22692E-11 | 1.155290163 | 0.61 | 0.33 | 2.89701E-07 | 1.22692E-11 | 3.97251E-17 | Ppp1r15a | EN_2 |
| 8.6896E-12 | 0.952240566 | 0.794 | 0.494 | 2.05179E-07 | 2.02289E-17 | 1.074213484 | 0.884 | 0.546 | 4.77644E-13 | 8.6896E-12 | 4.04577E-17 | Zfp36 | EN_2 |
| 2.69474E-16 | 0.603396984 | 1 | 0.875 | 6.36283E-12 | 2.35389E-17 | 0.643323867 | 1 | 0.879 | 5.558E-13 | 2.69474E-16 | 4.70777E-17 | Ptma | EN_2 |
| 1.8658E-13 | 0.701152888 | 0.949 | 0.655 | 4.40553E-09 | 3.23025E-17 | 0.679835319 | 0.97 | 0.686 | 7.62726E-13 | 1.8658E-13 | 6.46049E-17 | Btf3 | EN_2 |
| 5.83941E-17 | 1.364179957 | 0.853 | 0.523 | 1.3788E-12 | 5.4996E-13 | 1.115526438 | 0.762 | 0.474 | 1.29856E-08 | 5.4996E-13 | 1.16788E-16 | Tubb4b | EN_2 |

| | | | | | | | | | | | | | |
|---|---|---|---|---|---|---|---|---|---|---|---|---|---|
| 7.56749E-17 | 0.728557961 | 1 | 0.785 | 1.78684E-12 | 3.56319E-13 | 0.597670006 | 0.988 | 0.755 | 8.41341E-09 | 3.56319E-13 | 1.5135E-16 | Rpl15 | EN_2 |
| 8.0113E-16 | 0.756431364 | 0.985 | 0.773 | 1.89163E-11 | 8.46068E-17 | 0.661570709 | 0.982 | 0.802 | 1.99774E-12 | 8.0113E-16 | 1.69214E-16 | Rps5 | EN_2 |
| 1.87967E-14 | 0.709448567 | 0.978 | 0.748 | 4.43827E-10 | 1.01949E-16 | 0.684385314 | 0.97 | 0.735 | 2.40722E-12 | 1.87967E-14 | 2.03898E-16 | Rps3 | EN_2 |
| 1.30531E-16 | 1.093894946 | 0.853 | 0.491 | 3.08209E-12 | 5.04268E-14 | 1.010575051 | 0.78 | 0.503 | 1.19068E-09 | 5.04268E-14 | 2.61061E-16 | Eif3f | EN_2 |
| 1.54022E-16 | 0.796916517 | 0.993 | 0.765 | 3.63677E-12 | 2.9037E-15 | 0.654418016 | 0.994 | 0.758 | 6.85622E-11 | 2.9037E-15 | 3.08044E-16 | Rplp0 | EN_2 |
| 5.35942E-14 | 0.927827725 | 0.868 | 0.528 | 1.26547E-09 | 3.27706E-16 | 0.855177464 | 0.896 | 0.557 | 7.7378E-12 | 5.35942E-14 | 6.55412E-16 | Ier2 | EN_2 |
| 1.57212E-12 | 0.751658939 | 0.971 | 0.765 | 3.7121E-08 | 3.43711E-16 | 0.81579996 | 0.951 | 0.673 | 8.11571E-12 | 1.57212E-12 | 6.87423E-16 | Anxa2 | EN_2 |
| 1.20449E-12 | 1.133254734 | 0.831 | 0.531 | 2.84403E-08 | 4.80583E-16 | 1.21573977 | 0.787 | 0.438 | 1.13475E-11 | 1.20449E-12 | 9.61165E-16 | Pnp | EN_2 |
| 3.02619E-15 | 0.645357023 | 0.993 | 0.851 | 7.14545E-11 | 5.15507E-16 | 0.672084925 | 0.988 | 0.851 | 1.21721E-11 | 3.02619E-15 | 1.03101E-15 | Rps8 | EN_2 |
| 2.76321E-11 | 0.616867138 | 0.971 | 0.692 | 6.52448E-07 | 6.51672E-16 | 0.672279983 | 0.963 | 0.704 | 1.53873E-11 | 2.76321E-11 | 1.30334E-15 | Rpl36a | EN_2 |
| 9.70658E-16 | 1.147270556 | 0.772 | 0.435 | 2.29192E-11 | 1.31215E-07 | 0.873427298 | 0.634 | 0.405 | 0.003098244 | 1.31215E-07 | 1.94132E-15 | Slc25a5 | EN_2 |
| 3.47745E-14 | 0.632745203 | 0.993 | 0.802 | 8.21095E-10 | 1.03582E-15 | 0.586825089 | 0.994 | 0.807 | 2.44578E-11 | 3.47745E-14 | 2.07164E-15 | Rpl18a | EN_2 |
| 1.14341E-15 | 0.821981998 | 0.978 | 0.797 | 2.69982E-11 | 1.0648E-14 | 0.753995071 | 0.982 | 0.802 | 2.5142E-10 | 1.0648E-14 | 2.28682E-15 | Rsrp1 | EN_2 |
| 1.46024E-15 | 0.719749926 | 0.993 | 0.812 | 3.44791E-11 | 9.99699E-14 | 0.640157935 | 0.97 | 0.773 | 2.36049E-09 | 9.99699E-14 | 2.92048E-15 | Laptm4a | EN_2 |
| 3.03723E-10 | 0.707545831 | 0.853 | 0.614 | 7.1715E-06 | 2.91658E-15 | 0.783995854 | 0.89 | 0.603 | 6.88664E-11 | 3.03723E-10 | 5.83317E-15 | Gas5 | EN_2 |
| 5.89204E-15 | 1.223744948 | 0.816 | 0.504 | 1.39123E-10 | 3.56805E-15 | 1.233654669 | 0.799 | 0.482 | 8.42488E-11 | 5.89204E-15 | 7.1361E-15 | Cdkn1a | EN_2 |
| 4.10992E-15 | 1.382647972 | 0.699 | 0.355 | 9.70434E-11 | 2.21791E-08 | 1.112800428 | 0.579 | 0.369 | 0.000523692 | 2.21791E-08 | 8.21984E-15 | Marcksl1 | EN_2 |
| 1.47326E-09 | 0.669279716 | 0.743 | 0.472 | 3.47867E-05 | 5.83536E-15 | 1.061742794 | 0.768 | 0.454 | 1.37784E-10 | 1.47326E-09 | 1.16707E-14 | Nudc | EN_2 |
| 7.96365E-15 | 0.819813597 | 0.993 | 0.721 | 1.88038E-10 | 1.22973E-11 | 0.593522532 | 0.957 | 0.747 | 2.90363E-07 | 1.22973E-11 | 1.59273E-14 | Ctsl | EN_2 |
| 5.70415E-12 | 0.73147381 | 0.912 | 0.606 | 1.34686E-07 | 9.81732E-15 | 0.796523645 | 0.902 | 0.577 | 2.31807E-10 | 5.70415E-12 | 1.96346E-14 | Hmgn1 | EN_2 |
| 1.57065E-14 | 1.342793096 | 0.632 | 0.291 | 3.70861E-10 | 1.25134E-13 | 1.401269994 | 0.567 | 0.263 | 2.95466E-09 | 1.25134E-13 | 3.14129E-14 | Sertad1 | EN_2 |
| 2.03755E-14 | 0.877051336 | 0.949 | 0.709 | 4.81107E-10 | 7.22311E-12 | 0.695662487 | 0.921 | 0.709 | 1.70552E-07 | 7.22311E-12 | 4.0751E-14 | Dynll1 | EN_2 |
| 3.08561E-13 | 0.88090001 | 0.941 | 0.667 | 7.28574E-09 | 2.47517E-14 | 1.041756446 | 0.884 | 0.572 | 5.84436E-10 | 3.08561E-13 | 4.95033E-14 | Fcgr2b | EN_2 |
| 3.7866E-11 | 0.778076266 | 0.787 | 0.511 | 8.94092E-07 | 4.99064E-14 | 0.808313694 | 0.835 | 0.552 | 1.17839E-09 | 3.7866E-11 | 9.98129E-14 | Mt2 | EN_2 |
| 1.07427E-08 | 0.793302226 | 0.728 | 0.472 | 0.000253656 | 5.22064E-14 | 1.147539766 | 0.793 | 0.49 | 1.2327E-09 | 1.07427E-08 | 1.04413E-13 | Btg2 | EN_2 |
| 1.55009E-12 | 0.797107566 | 0.853 | 0.55 | 3.66007E-08 | 6.70263E-14 | 0.852478315 | 0.841 | 0.503 | 1.58263E-09 | 1.55009E-12 | 1.34053E-13 | Mrfap1 | EN_2 |
| 7.5183E-14 | 0.813828105 | 0.904 | 0.577 | 1.77522E-09 | 7.20184E-11 | 0.697118171 | 0.878 | 0.629 | 1.7005E-06 | 7.20184E-11 | 1.50366E-13 | Cntf | EN_2 |
| 8.35265E-13 | 0.732052743 | 0.993 | 0.702 | 1.97223E-08 | 1.05484E-13 | 0.730830603 | 0.97 | 0.693 | 2.49068E-09 | 8.35265E-13 | 2.10967E-13 | Nrn1 | EN_2 |
| 1.15437E-13 | 1.014392602 | 0.772 | 0.445 | 2.7257E-09 | 1.47958E-07 | 0.590656852 | 0.805 | 0.575 | 0.00349359 | 1.47958E-07 | 2.30874E-13 | S100a4 | EN_2 |
| 2.63253E-11 | 0.62568079 | 0.956 | 0.726 | 6.21593E-07 | 1.31475E-13 | 0.600495008 | 0.957 | 0.758 | 3.10439E-09 | 2.63253E-11 | 2.6295E-13 | Rps25 | EN_2 |
| 1.40018E-09 | 0.584998379 | 0.941 | 0.677 | 3.30611E-05 | 1.34376E-13 | 0.673417407 | 0.976 | 0.714 | 3.17289E-09 | 1.40018E-09 | 2.68752E-13 | Prdx6 | EN_2 |
| 2.29215E-10 | 1.050575154 | 0.772 | 0.521 | 5.41223E-06 | 1.64936E-13 | 1.029598517 | 0.811 | 0.526 | 3.89448E-09 | 2.29215E-10 | 3.29873E-13 | Dusp1 | EN_2 |
| 8.07862E-13 | 1.051359258 | 0.581 | 0.257 | 1.90752E-08 | 1.71457E-13 | 1.043415611 | 0.616 | 0.309 | 4.04844E-09 | 8.07862E-13 | 3.42913E-13 | Cdkn2d | EN_2 |
| 1.50849E-11 | 0.603954867 | 0.926 | 0.699 | 3.56185E-07 | 2.77277E-13 | 0.694833392 | 0.951 | 0.706 | 6.54706E-09 | 1.50849E-11 | 5.54554E-13 | Rpl7a | EN_2 |
| 1.92324E-11 | 1.011285629 | 0.75 | 0.484 | 4.54115E-07 | 2.93104E-13 | 0.875409147 | 0.713 | 0.448 | 6.92078E-09 | 1.92324E-11 | 5.86209E-13 | Ube2s | EN_2 |
| 2.95686E-13 | 0.814017682 | 0.912 | 0.636 | 6.98173E-09 | 9.76898E-12 | 0.666599312 | 0.878 | 0.595 | 2.30665E-07 | 9.76898E-12 | 5.91372E-13 | Npm1 | EN_2 |
| 2.00141E-10 | 0.750943848 | 0.728 | 0.443 | 4.72573E-06 | 3.00532E-13 | 0.786075356 | 0.762 | 0.454 | 7.09616E-09 | 2.00141E-10 | 6.01064E-13 | Gm10076 | EN_2 |
| 3.7135E-09 | 0.921053404 | 0.713 | 0.477 | 8.76832E-05 | 3.03241E-13 | 0.957034794 | 0.756 | 0.436 | 7.16013E-09 | 3.7135E-09 | 6.06482E-13 | Hmgb2 | EN_2 |
| 6.90089E-11 | 0.857946205 | 0.728 | 0.435 | 1.62944E-06 | 4.19143E-13 | 1.086725051 | 0.695 | 0.397 | 9.8968E-09 | 6.90089E-11 | 8.38286E-13 | Klf2 | EN_2 |
| 9.73613E-13 | 1.002777871 | 0.816 | 0.523 | 2.2989E-08 | 3.21212E-10 | 0.787904242 | 0.762 | 0.523 | 7.58446E-06 | 3.21212E-10 | 1.94723E-12 | Gatm | EN_2 |
| 1.53473E-12 | 0.766183427 | 0.949 | 0.699 | 3.62381E-08 | 9.60528E-11 | 0.608640369 | 0.945 | 0.701 | 2.268E-06 | 9.60528E-11 | 3.06947E-12 | Cfl1 | EN_2 |
| 2.12773E-12 | 0.645633553 | 0.941 | 0.689 | 5.02399E-08 | 1.2059E-11 | 0.585279575 | 0.957 | 0.68 | 2.84737E-07 | 1.2059E-11 | 4.25545E-12 | Sumo2 | EN_2 |
| 2.89492E-07 | 0.907102532 | 0.574 | 0.367 | 0.006835482 | 2.71879E-12 | 0.958293695 | 0.604 | 0.299 | 6.41962E-08 | 2.89492E-07 | 5.43759E-12 | Il11ra1 | EN_2 |
| 3.92588E-08 | 0.843303205 | 0.706 | 0.487 | 0.000926978 | 2.79166E-12 | 0.963813501 | 0.695 | 0.423 | 6.59167E-08 | 3.92588E-08 | 5.58333E-12 | Pgp | EN_2 |
| 3.42545E-12 | 0.75891772 | 0.86 | 0.616 | 8.08818E-08 | 2.85968E-08 | 0.594956418 | 0.817 | 0.582 | 0.000675229 | 2.85968E-08 | 6.85091E-12 | Tle5 | EN_2 |
| 1.36867E-09 | 0.715646738 | 0.765 | 0.516 | 3.23171E-05 | 5.15081E-12 | 0.802915977 | 0.817 | 0.521 | 1.21621E-07 | 1.36867E-09 | 1.03016E-11 | Syf2 | EN_2 |
| 8.74897E-09 | 0.631457211 | 0.882 | 0.599 | 0.000206581 | 8.45306E-12 | 1.038837432 | 0.829 | 0.593 | 1.99594E-07 | 8.74897E-09 | 1.69061E-11 | G0s2 | EN_2 |
| 9.17742E-12 | 0.676631014 | 0.934 | 0.746 | 2.16697E-07 | 6.75531E-10 | 0.614201835 | 0.963 | 0.74 | 1.59506E-05 | 6.75531E-10 | 1.83548E-11 | Gnai2 | EN_2 |
| 1.25162E-11 | 1.439709278 | 0.551 | 0.257 | 2.95533E-07 | 1.8061E-09 | 1.35778212 | 0.518 | 0.294 | 4.26456E-05 | 1.8061E-09 | 2.50325E-11 | Phlda1 | EN_2 |
| 1.42386E-11 | 1.128322483 | 0.61 | 0.308 | 3.36202E-07 | 5.77418E-08 | 1.147683265 | 0.482 | 0.265 | 0.0013634 | 5.77418E-08 | 2.84773E-11 | Gadd45b | EN_2 |
| 1.43506E-11 | 0.822345406 | 0.897 | 0.567 | 3.38847E-07 | 4.95125E-11 | 0.755456297 | 0.848 | 0.577 | 1.16909E-06 | 4.95125E-11 | 2.87012E-11 | Hnrnpa0 | EN_2 |
| 1.48583E-11 | 0.884804226 | 0.816 | 0.57 | 3.50833E-07 | 3.04318E-08 | 0.739753313 | 0.768 | 0.572 | 0.000718555 | 3.04318E-08 | 2.97165E-11 | Eif4a1 | EN_2 |
| 4.68571E-11 | 0.636761493 | 0.875 | 0.572 | 1.10639E-06 | 1.87904E-11 | 0.609901671 | 0.817 | 0.554 | 4.43679E-07 | 4.68571E-11 | 3.75808E-11 | Clta | EN_2 |
| 2.00688E-07 | 0.589181676 | 0.735 | 0.457 | 0.004738636 | 2.96994E-11 | 0.720584184 | 0.707 | 0.42 | 7.01263E-07 | 2.00688E-07 | 5.93989E-11 | Cdkn2c | EN_2 |
| 1.71263E-10 | 0.700718862 | 0.912 | 0.663 | 4.04387E-06 | 3.64375E-11 | 0.697947451 | 0.909 | 0.649 | 8.60362E-07 | 1.71263E-10 | 7.2875E-11 | Tuba1b | EN_2 |
| 3.77454E-11 | 0.664807398 | 0.956 | 0.665 | 8.91244E-07 | 5.67287E-08 | 0.633192059 | 0.884 | 0.673 | 0.001339478 | 5.67287E-08 | 7.54908E-11 | Tmem176a | EN_2 |
| 7.33468E-08 | 0.644883302 | 0.824 | 0.54 | 0.001731864 | 5.88579E-11 | 0.759555413 | 0.817 | 0.5 | 1.38975E-06 | 7.33468E-08 | 1.17716E-10 | Fosb | EN_2 |
| 7.57934E-11 | 0.778952705 | 0.978 | 0.795 | 1.78963E-06 | 4.44284E-10 | 0.610634842 | 0.89 | 0.698 | 1.04904E-05 | 4.44284E-10 | 1.51587E-10 | Ubc | EN_2 |
| 2.26265E-07 | 1.826308336 | 0.294 | 0.117 | 0.005342581 | 1.80119E-10 | 1.128032639 | 0.323 | 0.098 | 4.25297E-06 | 2.26265E-07 | 3.60238E-10 | Tnfsf9 | EN_2 |
| 2.25118E-10 | 0.806640175 | 0.721 | 0.43 | 5.3155E-06 | 4.48411E-10 | 1.326204471 | 0.598 | 0.343 | 1.05879E-05 | 4.48411E-10 | 4.50237E-10 | Lgals3 | EN_2 |
| 4.72424E-08 | 0.793887078 | 0.735 | 0.509 | 0.001115488 | 2.27525E-10 | 0.880959724 | 0.732 | 0.454 | 5.37232E-06 | 4.72424E-08 | 4.5505E-10 | Ppp1r10 | EN_2 |
| 7.30456E-09 | 0.707263832 | 0.647 | 0.367 | 0.000172475 | 7.23579E-10 | 0.909036105 | 0.634 | 0.387 | 1.70851E-05 | 7.30456E-09 | 1.44716E-09 | Pfdn2 | EN_2 |
| 7.61356E-10 | 0.766486288 | 0.64 | 0.364 | 1.79771E-05 | 3.62367E-07 | 0.660779935 | 0.537 | 0.335 | 0.008556215 | 3.62367E-07 | 1.52271E-09 | Tmem208 | EN_2 |
| 3.18325E-09 | 0.82308268 | 0.676 | 0.413 | 7.51628E-05 | 8.37528E-10 | 0.738842732 | 0.671 | 0.412 | 1.97757E-05 | 3.18325E-09 | 1.67506E-09 | Anp32b | EN_2 |
| 1.08314E-09 | 0.79959162 | 0.684 | 0.423 | 2.55751E-05 | 1.07309E-08 | 0.640089936 | 0.652 | 0.392 | 0.000253377 | 1.07309E-08 | 2.16628E-09 | Lysmd2 | EN_2 |
| 1.42491E-09 | 0.606004868 | 0.846 | 0.545 | 3.3645E-05 | 3.95423E-09 | 0.596513509 | 0.817 | 0.564 | 9.33674E-05 | 3.95423E-09 | 2.84983E-09 | Polr2f | EN_2 |
| 1.37609E-08 | 0.656360607 | 0.904 | 0.704 | 0.000324921 | 4.22412E-09 | 0.606410796 | 0.921 | 0.755 | 9.974E-05 | 1.37609E-08 | 8.44824E-09 | Lmna | EN_2 |
| 2.11614E-08 | 0.628575189 | 0.956 | 0.79 | 0.000499664 | 6.22673E-09 | 0.612584665 | 0.976 | 0.807 | 0.000147026 | 2.11614E-08 | 1.24535E-08 | Sdc4 | EN_2 |
| 7.93438E-09 | 0.631812651 | 0.816 | 0.518 | 0.000187347 | 1.01148E-08 | 0.673810146 | 0.811 | 0.528 | 0.00023883 | 1.01148E-08 | 1.58688E-08 | Id3 | EN_2 |
| 1.0119E-08 | 0.673311319 | 0.809 | 0.565 | 0.000238931 | 2.80005E-08 | 0.644319328 | 0.805 | 0.567 | 0.000661148 | 2.80005E-08 | 2.02381E-08 | Sdf4 | EN_2 |
| 1.2536E-08 | 0.75847838 | 0.647 | 0.389 | 0.000296 | 1.35673E-07 | 0.911497825 | 0.537 | 0.32 | 0.003203511 | 1.35673E-07 | 2.5072E-08 | Tm2d2 | EN_2 |
| 1.58923E-07 | 0.782236679 | 0.75 | 0.504 | 0.003752491 | 2.2832E-08 | 0.932239217 | 0.701 | 0.464 | 0.000539108 | 1.58923E-07 | 4.56639E-08 | Tnfaip6 | EN_2 |
| 2.67578E-07 | 0.795207321 | 0.662 | 0.438 | 0.006318043 | 4.31331E-08 | 0.611436373 | 0.665 | 0.392 | 0.001018458 | 2.67578E-07 | 8.62661E-08 | Rcn1 | EN_2 |
| 9.56812E-08 | 0.651443844 | 0.574 | 0.347 | 0.002259223 | 5.86907E-08 | 1.136243906 | 0.5 | 0.294 | 0.001385805 | 9.56812E-08 | 1.17381E-07 | Gtf2b | EN_2 |
| 1.51116E-45 | 1.3576092 | 0.993 | 0.916 | 3.56815E-41 | 1.64848E-28 | 0.954510657 | 0.992 | 0.922 | 3.8924E-24 | 1.64848E-28 | 3.02232E-45 | Zbtb20 | EN_3 |
| 5.06991E-42 | 2.162713459 | 0.957 | 0.798 | 1.19711E-37 | 2.12999E-35 | 1.816805552 | 0.975 | 0.825 | 5.02933E-31 | 2.12999E-35 | 1.01398E-41 | Rora | EN_3 |
| 1.66782E-39 | 1.965112592 | 0.943 | 0.743 | 3.93806E-35 | 6.85095E-25 | 1.402967619 | 0.949 | 0.765 | 1.61765E-20 | 6.85095E-25 | 3.33564E-39 | Pard3b | EN_3 |
| 3.78513E-38 | 3.669795477 | 0.721 | 0.215 | 8.93744E-34 | 4.6611E-23 | 2.708390501 | 0.661 | 0.251 | 1.10058E-18 | 4.6611E-23 | 7.57025E-38 | Gm26917 | EN_3 |
| 1.95274E-36 | 1.553861989 | 0.971 | 0.874 | 4.61081E-32 | 3.71025E-25 | 1.264809526 | 0.966 | 0.876 | 8.76065E-21 | 3.71025E-25 | 3.90548E-36 | Plce1 | EN_3 |
| 2.46462E-36 | 1.662298523 | 0.993 | 0.884 | 5.81946E-32 | 2.66372E-21 | 1.356172676 | 0.975 | 0.917 | 6.28957E-17 | 2.66372E-21 | 4.92924E-36 | Prkca | EN_3 |
| 2.65673E-36 | 1.764364669 | 0.979 | 0.872 | 6.27307E-32 | 1.23956E-20 | 1.320874232 | 0.975 | 0.876 | 2.92685E-16 | 1.23956E-20 | 5.31346E-36 | Stard13 | EN_3 |

| | | | | | | | | | | | | | |
|---|---|---|---|---|---|---|---|---|---|---|---|---|---|
| 5.1929E-36 | 2.035402108 | 0.914 | 0.756 | 1.22615E-31 | 3.30993E-21 | 1.614064241 | 0.915 | 0.73 | 7.81541E-17 | 3.30993E-21 | 1.03858E-35 | Mir100hg | EN_3 |
| 1.50527E-35 | 2.61265832 | 0.893 | 0.612 | 3.55425E-31 | 5.10547E-18 | 2.238939485 | 0.856 | 0.657 | 1.2055E-13 | 5.10547E-18 | 3.01054E-35 | Mir99ahg | EN_3 |
| 2.17311E-35 | 1.519574675 | 1 | 0.886 | 5.13114E-31 | 2.09542E-15 | 1.192027713 | 0.958 | 0.889 | 4.9477E-11 | 2.09542E-15 | 4.34622E-35 | Celf2 | EN_3 |
| 2.72801E-35 | 1.532779987 | 0.986 | 0.896 | 6.44138E-31 | 2.47001E-23 | 1.196265177 | 0.992 | 0.894 | 5.8322E-19 | 2.47001E-23 | 5.45602E-35 | Nkain2 | EN_3 |
| 7.84289E-35 | 2.048878689 | 0.936 | 0.743 | 1.85186E-30 | 5.56891E-27 | 1.798642711 | 0.949 | 0.774 | 1.31493E-22 | 5.56891E-27 | 1.56858E-34 | Maml2 | EN_3 |
| 2.95723E-34 | 1.801108091 | 0.95 | 0.805 | 6.98261E-30 | 6.33573E-29 | 1.500200784 | 0.983 | 0.841 | 1.49599E-24 | 6.33573E-29 | 5.91446E-34 | Zswim6 | EN_3 |
| 1.54575E-32 | 1.942413156 | 0.886 | 0.647 | 3.64981E-28 | 2.46313E-20 | 1.690817256 | 0.839 | 0.638 | 5.81594E-16 | 2.46313E-20 | 3.09149E-32 | Rbms3 | EN_3 |
| 1.64508E-32 | 1.745665339 | 0.95 | 0.785 | 3.88435E-28 | 1.68411E-14 | 1.196993216 | 0.932 | 0.8 | 3.97651E-10 | 1.68411E-14 | 3.29015E-32 | Sox6 | EN_3 |
| 1.98808E-32 | 1.910073483 | 0.914 | 0.778 | 4.69426E-28 | 9.26806E-21 | 1.366908406 | 0.941 | 0.82 | 2.18838E-16 | 9.26806E-21 | 3.97617E-32 | Agap1 | EN_3 |
| 5.38283E-32 | 1.636530013 | 0.943 | 0.795 | 1.27099E-27 | 1.70986E-23 | 1.476612526 | 0.949 | 0.836 | 4.03732E-19 | 1.70986E-23 | 1.07657E-31 | Dmd | EN_3 |
| 5.48567E-32 | 1.85931036 | 0.907 | 0.652 | 1.29528E-27 | 3.7215E-21 | 1.530950512 | 0.873 | 0.668 | 8.78721E-17 | 3.7215E-21 | 1.09713E-31 | Prkce | EN_3 |
| 2.24587E-31 | 1.580599086 | 0.943 | 0.793 | 5.30294E-27 | 2.10621E-19 | 1.241438547 | 0.949 | 0.836 | 4.97319E-15 | 2.10621E-19 | 4.49173E-31 | Utrn | EN_3 |
| 2.08196E-30 | 1.85272443 | 0.9 | 0.721 | 4.91592E-26 | 9.86358E-18 | 1.508628669 | 0.881 | 0.684 | 2.32899E-13 | 9.86358E-18 | 4.16392E-30 | Zeb1 | EN_3 |
| 2.25832E-30 | 1.579887323 | 0.943 | 0.815 | 5.33235E-26 | 1.57119E-20 | 1.222744021 | 0.915 | 0.809 | 3.70989E-16 | 1.57119E-20 | 4.51664E-30 | Sorbs1 | EN_3 |
| 2.26747E-30 | 2.15181048 | 0.821 | 0.543 | 5.35395E-26 | 6.5877E-21 | 1.824835179 | 0.831 | 0.588 | 1.55549E-16 | 6.5877E-21 | 4.53494E-30 | Zcchc7 | EN_3 |
| 2.57344E-30 | 3.826918477 | 0.5 | 0.081 | 6.07641E-26 | 1.39289E-20 | 3.920421247 | 0.39 | 0.071 | 3.2889E-16 | 1.39289E-20 | 5.14688E-30 | Taco1 | EN_3 |
| 5.43767E-30 | 2.020890213 | 0.9 | 0.642 | 1.28394E-25 | 3.08031E-15 | 2.080294954 | 0.763 | 0.539 | 7.27324E-11 | 3.08031E-15 | 1.08753E-29 | Sgip1 | EN_3 |
| 8.21126E-30 | 1.502386643 | 0.964 | 0.886 | 1.93884E-25 | 4.98266E-20 | 1.293189118 | 0.983 | 0.871 | 1.17651E-15 | 4.98266E-20 | 1.64225E-29 | Slc35f1 | EN_3 |
| 3.90403E-26 | 2.062001732 | 0.85 | 0.647 | 9.21819E-22 | 1.95581E-29 | 1.904045554 | 0.941 | 0.714 | 4.61806E-25 | 3.90403E-26 | 3.91162E-29 | Prkg1 | EN_3 |
| 1.96252E-29 | 1.315732256 | 0.95 | 0.906 | 4.6339E-25 | 3.12272E-14 | 1.029803727 | 0.932 | 0.908 | 7.37337E-10 | 3.12272E-14 | 3.92504E-29 | Sgcd | EN_3 |
| 3.65003E-29 | 1.865665488 | 0.893 | 0.726 | 8.61846E-25 | 2.13268E-18 | 1.460872119 | 0.89 | 0.728 | 5.03568E-14 | 2.13268E-18 | 7.30007E-29 | Ldlrad4 | EN_3 |
| 8.90258E-29 | 1.50727911 | 0.929 | 0.763 | 2.10208E-24 | 1.56808E-14 | 1.086383206 | 0.89 | 0.8 | 3.70254E-10 | 1.56808E-14 | 1.78052E-28 | Plekha5 | EN_3 |
| 4.08317E-28 | 1.557270779 | 0.943 | 0.867 | 9.64118E-24 | 7.04086E-18 | 1.450410378 | 0.932 | 0.857 | 1.66249E-13 | 7.04086E-18 | 8.16634E-28 | Sorbs2 | EN_3 |
| 1.63523E-27 | 1.806630605 | 0.921 | 0.714 | 3.86111E-23 | 4.28057E-21 | 1.479263686 | 0.907 | 0.714 | 1.01073E-16 | 4.28057E-21 | 3.27046E-27 | Il1rapl1 | EN_3 |
| 6.23653E-27 | 2.049179698 | 0.836 | 0.615 | 1.47257E-22 | 3.09473E-19 | 1.79554688 | 0.814 | 0.611 | 7.30728E-15 | 3.09473E-19 | 1.24731E-26 | Lncpint | EN_3 |
| 1.34475E-26 | 1.596863562 | 0.879 | 0.701 | 3.17521E-22 | 1.52642E-16 | 1.450360594 | 0.831 | 0.694 | 3.60418E-12 | 1.52642E-16 | 2.68949E-26 | Aopep | EN_3 |
| 1.70475E-26 | 1.912184035 | 0.857 | 0.664 | 4.02526E-22 | 6.52907E-18 | 1.484923423 | 0.864 | 0.689 | 1.54164E-13 | 6.52907E-18 | 3.4095E-26 | Dip2c | EN_3 |
| 2.03358E-26 | 1.183790967 | 0.993 | 0.953 | 4.80168E-22 | 1.30397E-15 | 0.80697617 | 0.992 | 0.947 | 3.07894E-11 | 1.30397E-15 | 4.06715E-26 | Ank3 | EN_3 |
| 2.23539E-26 | 1.861318884 | 0.921 | 0.753 | 5.2782E-22 | 2.48007E-15 | 1.674753332 | 0.915 | 0.783 | 5.85594E-11 | 2.48007E-15 | 4.47078E-26 | Kcnq5 | EN_3 |
| 3.80207E-26 | 2.135821863 | 0.821 | 0.551 | 8.97745E-22 | 3.10111E-17 | 1.758031451 | 0.78 | 0.597 | 7.32233E-13 | 3.10111E-17 | 7.60414E-26 | 2610307P | EN_3 |
| 7.86635E-26 | 2.025609058 | 0.85 | 0.694 | 1.8574E-21 | 1.2156E-20 | 1.482890625 | 0.907 | 0.747 | 2.87028E-16 | 1.2156E-20 | 1.57327E-25 | Sox5 | EN_3 |
| 8.69368E-26 | 1.692472972 | 0.857 | 0.714 | 2.05275E-21 | 8.10781E-15 | 1.325521311 | 0.839 | 0.737 | 1.91442E-10 | 8.10781E-15 | 1.73874E-25 | Msi2 | EN_3 |
| 2.32581E-25 | 1.60774087 | 0.9 | 0.778 | 5.4917E-21 | 4.70051E-15 | 1.254614906 | 0.881 | 0.786 | 1.10988E-10 | 4.70051E-15 | 4.65161E-25 | Piezo2 | EN_3 |
| 3.09613E-25 | 2.019682279 | 0.864 | 0.664 | 7.31057E-21 | 7.3843E-16 | 2.005937951 | 0.788 | 0.599 | 1.74358E-11 | 7.3843E-16 | 6.19225E-25 | Sorcs1 | EN_3 |
| 5.28424E-25 | 1.564249555 | 0.879 | 0.699 | 1.24772E-20 | 1.27188E-13 | 1.225130872 | 0.831 | 0.747 | 3.00316E-09 | 1.27188E-13 | 1.05685E-24 | Gnaq | EN_3 |
| 1.95447E-24 | 1.028352191 | 0.957 | 0.877 | 4.6149E-20 | 7.85713E-16 | 0.735543904 | 0.992 | 0.908 | 1.85522E-11 | 7.85713E-16 | 3.90894E-24 | Gphn | EN_3 |
| 3.11757E-24 | 1.383169547 | 0.886 | 0.807 | 7.3612E-20 | 4.71326E-13 | 1.024452857 | 0.873 | 0.802 | 1.1129E-08 | 4.71326E-13 | 6.23513E-24 | Nfia | EN_3 |
| 4.53854E-24 | 1.229604038 | 0.986 | 0.904 | 1.07164E-19 | 6.01098E-12 | 0.843684486 | 0.983 | 0.901 | 1.41931E-07 | 6.01098E-12 | 9.07707E-24 | Cdh19 | EN_3 |
| 4.88557E-24 | 2.273530821 | 0.743 | 0.444 | 1.15358E-19 | 1.21308E-13 | 1.854501606 | 0.703 | 0.486 | 2.86434E-09 | 1.21308E-13 | 9.77113E-24 | Fbxl17 | EN_3 |
| 9.38009E-24 | 1.738096393 | 0.821 | 0.644 | 2.21483E-19 | 6.83033E-19 | 1.417220889 | 0.864 | 0.687 | 1.61278E-14 | 6.83033E-19 | 1.87602E-23 | Rere | EN_3 |
| 1.09087E-23 | 1.600668729 | 0.879 | 0.733 | 2.57575E-19 | 4.39728E-08 | 0.950687692 | 0.881 | 0.751 | 0.001038285 | 4.39728E-08 | 2.18173E-23 | Tmtc2 | EN_3 |
| 2.0928E-19 | 2.129848985 | 0.857 | 0.741 | 4.94152E-15 | 1.25615E-23 | 2.197288056 | 0.881 | 0.7 | 2.96602E-19 | 2.0928E-19 | 2.5123E-23 | Plcb1 | EN_3 |
| 4.40882E-23 | 1.944002323 | 0.786 | 0.546 | 1.04101E-18 | 4.8301E-17 | 1.729646119 | 0.78 | 0.532 | 1.14048E-12 | 4.8301E-17 | 8.81765E-23 | Exoc4 | EN_3 |
| 6.76306E-23 | 1.816795076 | 0.786 | 0.558 | 1.59689E-18 | 1.22962E-16 | 1.630078307 | 0.788 | 0.541 | 2.90338E-12 | 1.22962E-16 | 1.35261E-22 | Pard3 | EN_3 |
| 1.02107E-22 | 1.704365514 | 0.85 | 0.704 | 2.41094E-18 | 1.66165E-15 | 1.350864192 | 0.89 | 0.728 | 3.92349E-11 | 1.66165E-15 | 2.04213E-22 | Sntb1 | EN_3 |
| 1.60401E-22 | 1.918258844 | 0.807 | 0.551 | 3.78739E-18 | 3.15028E-13 | 1.371958702 | 0.797 | 0.645 | 7.43845E-09 | 3.15028E-13 | 3.20802E-22 | Ptprm | EN_3 |
| 1.73866E-22 | 1.538925082 | 0.9 | 0.81 | 4.10533E-18 | 2.17917E-19 | 1.636325172 | 0.881 | 0.77 | 5.14545E-15 | 2.17917E-19 | 3.47732E-22 | Dtna | EN_3 |
| 2.9499E-22 | 1.898442051 | 0.821 | 0.63 | 6.96531E-18 | 3.70293E-16 | 2.300542858 | 0.763 | 0.53 | 8.74336E-12 | 3.70293E-16 | 5.89981E-22 | Adgrl3 | EN_3 |
| 3.79477E-22 | 1.872422152 | 0.793 | 0.625 | 8.96021E-18 | 1.56965E-17 | 1.606387881 | 0.831 | 0.634 | 3.70627E-13 | 1.56965E-17 | 7.58954E-22 | Zfhx4 | EN_3 |
| 1.05524E-21 | 1.559056316 | 0.829 | 0.679 | 2.49164E-17 | 1.37668E-15 | 1.316498169 | 0.856 | 0.694 | 3.25061E-11 | 1.37668E-15 | 2.11049E-21 | Fndc3b | EN_3 |
| 1.5432E-21 | 1.713588234 | 0.836 | 0.627 | 3.6438E-17 | 7.53254E-12 | 1.134900119 | 0.822 | 0.68 | 1.77858E-07 | 7.53254E-12 | 3.0864E-21 | Dpyd | EN_3 |
| 1.76801E-21 | 1.568814181 | 0.829 | 0.733 | 4.17462E-17 | 1.12817E-17 | 1.324883566 | 0.89 | 0.76 | 2.66383E-13 | 1.12817E-17 | 3.53601E-21 | Igf1r | EN_3 |
| 1.91206E-21 | 1.627763459 | 0.936 | 0.83 | 4.51477E-17 | 2.60114E-11 | 1.217667488 | 0.932 | 0.827 | 6.14181E-07 | 2.60114E-11 | 3.82413E-21 | Frmd4a | EN_3 |
| 4.8837E-21 | 1.688185036 | 0.9 | 0.837 | 1.15314E-16 | 1.93884E-21 | 1.696787576 | 0.924 | 0.795 | 4.57799E-17 | 4.8837E-21 | 3.87768E-21 | Grik2 | EN_3 |
| 2.0129E-21 | 1.822953865 | 0.786 | 0.546 | 4.75286E-17 | 1.49022E-14 | 1.440733854 | 0.805 | 0.62 | 3.5187E-10 | 1.49022E-14 | 4.0258E-21 | Akt3 | EN_3 |
| 3.35818E-21 | 2.235945111 | 0.671 | 0.336 | 7.92933E-17 | 1.37249E-08 | 1.50598943 | 0.602 | 0.385 | 0.000324072 | 1.37249E-08 | 6.71636E-21 | Cdkal1 | EN_3 |
| 4.04461E-21 | 1.283095995 | 0.971 | 0.83 | 9.55014E-17 | 1.52938E-08 | 0.853719947 | 0.924 | 0.825 | 0.000361117 | 1.52938E-08 | 8.08922E-21 | Zfp536 | EN_3 |
| 5.15484E-21 | 1.811350312 | 0.779 | 0.474 | 1.21716E-16 | 1.32515E-13 | 1.521178143 | 0.737 | 0.551 | 3.12894E-09 | 1.32515E-13 | 1.03097E-20 | Arid1b | EN_3 |
| 6.84483E-21 | 2.110591992 | 0.779 | 0.583 | 1.6162E-16 | 4.17705E-10 | 1.677550155 | 0.729 | 0.571 | 9.86286E-06 | 4.17705E-10 | 1.36897E-20 | Maml3 | EN_3 |
| 7.92579E-21 | 0.962359475 | 0.971 | 0.844 | 1.87144E-16 | 8.40879E-12 | 0.589330211 | 1 | 0.878 | 1.98548E-07 | 8.40879E-12 | 1.58516E-20 | Lars2 | EN_3 |
| 8.40972E-21 | 1.807579387 | 0.921 | 0.788 | 1.9857E-16 | 5.53565E-15 | 1.627908699 | 0.975 | 0.829 | 1.30708E-10 | 5.53565E-15 | 1.68194E-20 | Csmd1 | EN_3 |
| 1.28649E-20 | 0.909994418 | 0.957 | 0.874 | 3.03765E-16 | 3.37459E-16 | 0.778758807 | 0.983 | 0.929 | 7.96809E-12 | 3.37459E-16 | 2.57297E-20 | Dst | EN_3 |
| 3.63593E-20 | 1.328234969 | 0.836 | 0.751 | 8.58515E-16 | 5.23174E-09 | 0.720920048 | 0.873 | 0.779 | 0.000123532 | 5.23174E-09 | 7.27185E-20 | Clasp2 | EN_3 |
| 4.97446E-20 | 2.02409907 | 0.7 | 0.435 | 1.17457E-15 | 5.874E-14 | 1.458566118 | 0.754 | 0.5 | 1.38697E-09 | 5.874E-14 | 9.94892E-20 | 4930402H | EN_3 |
| 6.1339E-20 | 1.87783836 | 0.736 | 0.454 | 1.44834E-15 | 4.77255E-10 | 1.479336401 | 0.669 | 0.465 | 1.1269E-05 | 4.77255E-10 | 1.22678E-19 | Clasp1 | EN_3 |
| 6.18313E-20 | 1.831223109 | 0.764 | 0.531 | 1.45996E-15 | 7.14948E-15 | 1.658604028 | 0.797 | 0.576 | 1.68813E-10 | 7.14948E-15 | 1.23663E-19 | Bcas3 | EN_3 |
| 7.86506E-20 | 1.913314379 | 0.771 | 0.523 | 1.8571E-15 | 1.84041E-09 | 1.404411353 | 0.72 | 0.548 | 4.34557E-05 | 1.84041E-09 | 1.57301E-19 | Gab2 | EN_3 |
| 1.79648E-19 | 1.824607007 | 0.8 | 0.6 | 4.24184E-15 | 3.36042E-10 | 1.332847046 | 0.754 | 0.613 | 7.93462E-06 | 3.36042E-10 | 3.59295E-19 | Fign | EN_3 |
| 1.86287E-19 | 1.586789135 | 0.85 | 0.756 | 4.39862E-15 | 7.07026E-15 | 1.327457799 | 0.89 | 0.781 | 1.66943E-10 | 7.07026E-15 | 3.72575E-19 | Adam11 | EN_3 |
| 1.95429E-19 | 0.756596257 | 1 | 0.919 | 4.61447E-15 | 3.9421E-16 | 0.647963259 | 1 | 0.922 | 9.30809E-12 | 3.9421E-16 | 3.90858E-19 | Camk1d | EN_3 |
| 1.97901E-19 | 1.417317888 | 0.814 | 0.664 | 4.67284E-15 | 2.1002E-10 | 1.118200705 | 0.814 | 0.765 | 4.959E-06 | 2.1002E-10 | 3.95802E-19 | Nf1 | EN_3 |
| 2.16633E-19 | 2.066117463 | 0.7 | 0.472 | 5.11515E-15 | 8.43826E-18 | 1.826994624 | 0.763 | 0.514 | 1.99244E-13 | 8.43826E-18 | 4.33267E-19 | Dennd1a | EN_3 |
| 2.17071E-19 | 1.653116381 | 0.8 | 0.588 | 5.12549E-15 | 2.65827E-10 | 1.291415298 | 0.771 | 0.618 | 6.2767E-06 | 2.65827E-10 | 4.34142E-19 | Arhgap32 | EN_3 |
| 2.262E-19 | 1.986679962 | 0.771 | 0.523 | 5.34103E-15 | 5.58412E-09 | 1.357813337 | 0.686 | 0.569 | 0.000131852 | 5.58412E-09 | 4.524E-19 | Ehbp1 | EN_3 |
| 2.95454E-19 | 1.733226751 | 0.764 | 0.556 | 6.97626E-15 | 2.11502E-11 | 1.338533057 | 0.771 | 0.548 | 4.99399E-07 | 2.11502E-11 | 5.90908E-19 | Ncoa2 | EN_3 |
| 3.03767E-19 | 2.106813026 | 0.757 | 0.506 | 7.17255E-15 | 4.82682E-15 | 2.231392331 | 0.703 | 0.484 | 1.13971E-10 | 4.82682E-15 | 6.07534E-19 | Pde7b | EN_3 |
| 3.20447E-19 | 1.688587188 | 0.786 | 0.617 | 7.56638E-15 | 1.77932E-12 | 1.311688038 | 0.797 | 0.638 | 4.20132E-08 | 1.77932E-12 | 6.40893E-19 | Bach2 | EN_3 |
| 3.41054E-19 | 2.030383986 | 0.793 | 0.59 | 8.05296E-15 | 3.06944E-18 | 1.97269729 | 0.814 | 0.594 | 7.24756E-14 | 3.06944E-18 | 6.82108E-19 | Klhl29 | EN_3 |
| 3.68679E-19 | 1.430511965 | 0.857 | 0.654 | 8.70524E-15 | 7.19894E-14 | 1.246803542 | 0.822 | 0.673 | 1.69981E-09 | 7.19894E-14 | 7.37357E-19 | Zfp608 | EN_3 |
| 1.12689E-18 | 2.351733301 | 0.757 | 0.546 | 2.66082E-14 | 2.17386E-11 | 1.852114317 | 0.695 | 0.576 | 5.13292E-07 | 2.17386E-11 | 2.25378E-18 | Pbx1 | EN_3 |
| 1.3593E-18 | 1.763666554 | 0.779 | 0.573 | 3.20958E-14 | 1.45849E-08 | 1.33348234 | 0.746 | 0.638 | 0.000344378 | 1.45849E-08 | 2.7186E-18 | Ptprj | EN_3 |

| | | | | | | | | | | | | | |
|---|---|---|---|---|---|---|---|---|---|---|---|---|---|
| 1.70836E-18 | 1.434684795 | 0.857 | 0.706 | 4.03378E-14 | 2.87067E-13 | 1.371939018 | 0.814 | 0.65 | 6.77824E-09 | 2.87067E-13 | 3.41672E-18 | Col5a2 | EN_3 |
| 2.48991E-18 | 1.458888417 | 0.879 | 0.763 | 5.87919E-14 | 2.65076E-13 | 1.148326935 | 0.941 | 0.781 | 6.25897E-09 | 2.65076E-13 | 4.97983E-18 | Gpcpd1 | EN_3 |
| 6.42228E-18 | 2.086735857 | 0.707 | 0.491 | 1.51643E-13 | 8.3064E-14 | 1.822049513 | 0.703 | 0.498 | 1.96131E-09 | 8.3064E-14 | 1.28446E-17 | Dock10 | EN_3 |
| 6.50411E-18 | 0.864231234 | 0.921 | 0.874 | 1.53575E-13 | 8.72534E-13 | 0.784454191 | 0.924 | 0.894 | 2.06023E-08 | 8.72534E-13 | 1.30082E-17 | Mbnl2 | EN_3 |
| 8.38157E-18 | 1.159225887 | 0.907 | 0.847 | 1.97906E-13 | 8.65213E-13 | 1.095043605 | 0.924 | 0.853 | 2.04294E-08 | 8.65213E-13 | 1.67631E-17 | Gulp1 | EN_3 |
| 1.02444E-17 | 1.428541853 | 0.871 | 0.743 | 2.41891E-13 | 5.53216E-12 | 1.234521081 | 0.89 | 0.8 | 1.30625E-07 | 5.53216E-12 | 2.04888E-17 | Lrrc4c | EN_3 |
| 1.28876E-17 | 1.337118759 | 0.829 | 0.709 | 3.04301E-13 | 4.05871E-07 | 0.721813092 | 0.847 | 0.779 | 0.009583431 | 4.05871E-07 | 2.57751E-17 | Pdzd2 | EN_3 |
| 1.39769E-17 | 1.300945275 | 0.829 | 0.672 | 3.30023E-13 | 4.66473E-11 | 1.17950043 | 0.805 | 0.698 | 1.10144E-06 | 4.66473E-11 | 2.79538E-17 | Stag1 | EN_3 |
| 1.43668E-17 | 0.896163841 | 0.971 | 0.946 | 3.39229E-13 | 1.29166E-10 | 0.667567229 | 0.958 | 0.933 | 3.04987E-06 | 1.29166E-10 | 2.87336E-17 | Ncam1 | EN_3 |
| 1.86266E-17 | 1.427508344 | 0.857 | 0.684 | 4.39812E-13 | 7.99853E-11 | 1.580280427 | 0.814 | 0.643 | 1.88861E-06 | 7.99853E-11 | 3.72532E-17 | Adamts12 | EN_3 |
| 3.00607E-17 | 3.215000573 | 0.35 | 0.072 | 7.09793E-13 | 2.03575E-12 | 2.942659274 | 0.331 | 0.092 | 4.80681E-08 | 2.03575E-12 | 6.01214E-17 | Foxp2 | EN_3 |
| 1.05068E-16 | 1.96040608 | 0.714 | 0.496 | 2.48087E-12 | 3.12457E-17 | 1.954466654 | 0.754 | 0.484 | 7.37774E-13 | 1.05068E-16 | 6.24915E-17 | Ablim2 | EN_3 |
| 3.69361E-17 | 1.608620046 | 0.757 | 0.543 | 8.72135E-13 | 3.66617E-13 | 1.33778773 | 0.814 | 0.601 | 8.65655E-09 | 3.66617E-13 | 7.38722E-17 | Nedd4l | EN_3 |
| 4.55605E-17 | 1.525714852 | 0.8 | 0.59 | 1.07577E-12 | 5.79224E-09 | 0.973331025 | 0.763 | 0.671 | 0.000136766 | 5.79224E-09 | 9.1121E-17 | Lama4 | EN_3 |
| 4.79252E-17 | 1.713755335 | 0.779 | 0.607 | 1.13161E-12 | 2.37611E-12 | 1.307652615 | 0.78 | 0.682 | 5.61047E-08 | 2.37611E-12 | 9.58505E-17 | Ldb2 | EN_3 |
| 5.93478E-17 | 1.406810432 | 0.814 | 0.654 | 1.40132E-12 | 1.55982E-10 | 1.047626507 | 0.78 | 0.661 | 3.68305E-06 | 1.55982E-10 | 1.18696E-16 | Macf1 | EN_3 |
| 9.60294E-17 | 2.034685087 | 0.671 | 0.412 | 2.26745E-12 | 2.28391E-13 | 1.825803474 | 0.669 | 0.406 | 5.39277E-09 | 2.28391E-13 | 1.92059E-16 | Vps13b | EN_3 |
| 1.25381E-16 | 1.671169749 | 0.736 | 0.531 | 2.96051E-12 | 9.33211E-09 | 1.355412002 | 0.703 | 0.569 | 0.00022035 | 9.33211E-09 | 2.50763E-16 | Dock5 | EN_3 |
| 1.34812E-16 | 1.522511507 | 0.779 | 0.598 | 3.18319E-12 | 1.32717E-09 | 1.57886253 | 0.737 | 0.59 | 3.13372E-05 | 1.32717E-09 | 2.69625E-16 | Lpp | EN_3 |
| 1.59298E-16 | 1.06922358 | 0.964 | 0.904 | 3.76134E-12 | 3.13911E-11 | 0.854678719 | 0.966 | 0.917 | 7.41207E-07 | 3.13911E-11 | 3.18595E-16 | Zeb2 | EN_3 |
| 1.27508E-12 | 1.379421487 | 0.821 | 0.741 | 3.01071E-08 | 1.9251E-16 | 1.35354556 | 0.924 | 0.744 | 4.54556E-12 | 1.27508E-12 | 3.85021E-16 | P3h2 | EN_3 |
| 2.3957E-16 | 1.547066833 | 0.743 | 0.56 | 5.65673E-12 | 7.76511E-13 | 1.399595805 | 0.737 | 0.578 | 1.8335E-08 | 7.76511E-13 | 4.79141E-16 | Kmt2c | EN_3 |
| 2.45489E-16 | 1.604613563 | 0.736 | 0.521 | 5.79648E-12 | 2.47948E-10 | 1.24950332 | 0.737 | 0.606 | 5.85456E-06 | 2.47948E-10 | 4.90977E-16 | Fam172a | EN_3 |
| 2.4609E-16 | 3.62194472 | 0.35 | 0.079 | 5.81068E-12 | 4.57923E-11 | 3.316366563 | 0.28 | 0.071 | 1.08125E-06 | 4.57923E-11 | 4.9218E-16 | Plcl1 | EN_3 |
| 2.88051E-16 | 1.515710518 | 0.786 | 0.605 | 6.80145E-12 | 8.15438E-12 | 1.381935071 | 0.788 | 0.622 | 1.92541E-07 | 8.15438E-12 | 5.76101E-16 | Zfp462 | EN_3 |
| 4.8609E-16 | 1.799414693 | 0.757 | 0.615 | 1.14776E-11 | 8.54762E-13 | 1.46094957 | 0.805 | 0.643 | 2.01826E-08 | 8.54762E-13 | 9.72181E-16 | Fmnl2 | EN_3 |
| 6.56507E-16 | 2.04191272 | 0.614 | 0.363 | 1.55014E-11 | 1.42793E-10 | 1.795205315 | 0.593 | 0.355 | 3.37163E-06 | 1.42793E-10 | 1.31301E-15 | Nr3c2 | EN_3 |
| 7.85284E-16 | 1.772845335 | 0.686 | 0.474 | 1.85421E-11 | 6.62828E-14 | 1.44140588 | 0.746 | 0.495 | 1.56507E-09 | 6.62828E-14 | 1.57057E-15 | Mbd5 | EN_3 |
| 8.8664E-16 | 1.618428776 | 0.764 | 0.56 | 2.09353E-11 | 1.38616E-09 | 1.3302916 | 0.754 | 0.608 | 3.27301E-05 | 1.38616E-09 | 1.77328E-15 | Srgap2 | EN_3 |
| 8.8958E-16 | 1.266746558 | 0.964 | 0.812 | 2.10048E-11 | 2.56752E-08 | 0.810071164 | 0.941 | 0.855 | 0.000606242 | 2.56752E-08 | 1.77916E-15 | Des | EN_3 |
| 1.06357E-15 | 1.738394545 | 0.686 | 0.474 | 2.51129E-11 | 4.03687E-13 | 1.634337146 | 0.729 | 0.516 | 9.53186E-09 | 4.03687E-13 | 2.12713E-15 | Fto | EN_3 |
| 3.43817E-14 | 1.714802236 | 0.807 | 0.765 | 8.1182E-10 | 1.09404E-15 | 1.635448345 | 0.847 | 0.698 | 2.58324E-11 | 3.43817E-14 | 2.18808E-15 | Ppp2r2b | EN_3 |
| 1.23256E-15 | 1.843261023 | 0.643 | 0.38 | 2.91032E-11 | 2.55001E-07 | 1.387361994 | 0.576 | 0.422 | 0.006021085 | 2.55001E-07 | 2.46512E-15 | Lrba | EN_3 |
| 1.37078E-15 | 1.115687136 | 0.914 | 0.758 | 3.23669E-11 | 8.75415E-08 | 0.759015542 | 0.898 | 0.825 | 0.00206703 | 8.75415E-08 | 2.74156E-15 | Dlgap1 | EN_3 |
| 1.50936E-15 | 1.992260576 | 0.65 | 0.363 | 3.56391E-11 | 1.16974E-10 | 1.763313632 | 0.653 | 0.424 | 2.762E-06 | 1.16974E-10 | 3.01873E-15 | Prkn | EN_3 |
| 2.12551E-15 | 1.016182055 | 0.9 | 0.842 | 5.01874E-11 | 6.21677E-11 | 0.852513443 | 0.898 | 0.823 | 1.4679E-06 | 6.21677E-11 | 4.25101E-15 | Foxp1 | EN_3 |
| 2.22238E-15 | 1.682460089 | 0.707 | 0.499 | 5.24748E-11 | 7.38524E-14 | 1.759707155 | 0.737 | 0.537 | 1.7438E-09 | 7.38524E-14 | 4.44476E-15 | Ldlrad3 | EN_3 |
| 2.36604E-15 | 2.440031214 | 0.536 | 0.257 | 5.5867E-11 | 1.2963E-09 | 1.900405159 | 0.525 | 0.288 | 3.06083E-05 | 1.2963E-09 | 4.73208E-15 | Aff3 | EN_3 |
| 2.56819E-15 | 2.351707333 | 0.579 | 0.309 | 6.06401E-11 | 6.42506E-08 | 1.836020205 | 0.559 | 0.396 | 0.001517085 | 6.42506E-08 | 5.13638E-15 | Tox | EN_3 |
| 2.60809E-15 | 1.542044273 | 0.714 | 0.496 | 6.15821E-11 | 2.40929E-10 | 1.300895977 | 0.703 | 0.514 | 5.68881E-06 | 2.40929E-10 | 5.21617E-15 | Psme4 | EN_3 |
| 3.56267E-15 | 1.477243866 | 0.736 | 0.536 | 8.41218E-11 | 1.62661E-09 | 1.30779773 | 0.712 | 0.528 | 3.84076E-05 | 1.62661E-09 | 7.12534E-15 | Setd5 | EN_3 |
| 3.96265E-15 | 1.469339241 | 0.807 | 0.644 | 9.3566E-11 | 2.48758E-08 | 1.182603403 | 0.771 | 0.703 | 0.000587368 | 2.48758E-08 | 7.92529E-15 | Rai14 | EN_3 |
| 5.18706E-15 | 2.692124355 | 0.564 | 0.281 | 1.22477E-10 | 5.24969E-10 | 2.179895413 | 0.517 | 0.27 | 1.23956E-05 | 5.24969E-10 | 1.03741E-14 | Dock4 | EN_3 |
| 5.65048E-15 | 0.955373157 | 0.929 | 0.906 | 1.33419E-10 | 2.35852E-07 | 0.659102885 | 0.907 | 0.878 | 0.005568947 | 2.35852E-07 | 1.1301E-14 | Adam23 | EN_3 |
| 4.14782E-07 | 0.917084297 | 0.693 | 0.637 | 0.009793843 | 5.97835E-15 | 1.327595891 | 0.805 | 0.647 | 1.41161E-10 | 4.14782E-07 | 1.19567E-14 | Chd9 | EN_3 |
| 7.18596E-15 | 1.529230477 | 0.843 | 0.751 | 1.69675E-10 | 9.34054E-12 | 1.014898235 | 0.898 | 0.823 | 2.20549E-07 | 9.34054E-12 | 1.43719E-14 | Rbms1 | EN_3 |
| 8.24663E-15 | 1.953084004 | 0.743 | 0.612 | 1.94719E-10 | 2.34349E-11 | 1.510218826 | 0.737 | 0.576 | 5.53344E-07 | 2.34349E-11 | 1.64933E-14 | Ext1 | EN_3 |
| 1.22807E-14 | 1.085993583 | 0.95 | 0.886 | 2.89973E-10 | 9.02953E-13 | 1.377297969 | 0.975 | 0.871 | 2.13205E-08 | 9.02953E-13 | 2.45615E-14 | Acta2 | EN_3 |
| 1.24472E-12 | 7.041351953 | 0.171 | 0.015 | 2.93903E-08 | 1.86057E-14 | 6.852120868 | 0.186 | 0.014 | 4.39317E-10 | 1.24472E-12 | 3.72114E-14 | Nrg3 | EN_3 |
| 1.94969E-14 | 2.039861685 | 0.579 | 0.314 | 4.6036E-10 | 3.29439E-10 | 1.734943624 | 0.568 | 0.343 | 7.77871E-06 | 3.29439E-10 | 3.89937E-14 | Stk3 | EN_3 |
| 2.20502E-14 | 1.80691202 | 0.693 | 0.516 | 5.20649E-10 | 5.47921E-14 | 1.520413487 | 0.746 | 0.546 | 1.29375E-09 | 5.47921E-14 | 4.41004E-14 | Dock1 | EN_3 |
| 2.21545E-14 | 1.224497038 | 0.836 | 0.748 | 5.23111E-10 | 2.07967E-11 | 0.971572459 | 0.864 | 0.818 | 4.91052E-07 | 2.07967E-11 | 4.43089E-14 | Dock7 | EN_3 |
| 3.78981E-14 | 1.205670984 | 0.843 | 0.723 | 8.9485E-10 | 2.84188E-11 | 1.137599538 | 0.873 | 0.719 | 6.71025E-07 | 2.84188E-11 | 7.57962E-14 | St6gal1 | EN_3 |
| 4.06139E-14 | 2.556504043 | 0.4 | 0.131 | 9.58976E-10 | 2.0627E-11 | 2.813631631 | 0.39 | 0.141 | 4.87044E-07 | 2.0627E-11 | 8.12278E-14 | A330023F[illegible] | EN_3 |
| 4.28375E-14 | 1.04315955 | 0.943 | 0.889 | 1.01148E-09 | 5.82928E-09 | 0.785706264 | 0.941 | 0.901 | 0.000137641 | 5.82928E-09 | 8.5675E-14 | Ptprz1 | EN_3 |
| 4.4379E-14 | 1.844149058 | 0.671 | 0.479 | 1.04788E-09 | 2.44486E-09 | 1.844866064 | 0.568 | 0.341 | 5.7728E-05 | 2.44486E-09 | 8.8758E-14 | Elmo1 | EN_3 |
| 1.68607E-12 | 1.528791373 | 0.693 | 0.528 | 3.98114E-08 | 4.46881E-14 | 1.706755272 | 0.763 | 0.592 | 1.05517E-09 | 1.68607E-12 | 8.93761E-14 | Diaph2 | EN_3 |
| 1.14213E-13 | 1.434821757 | 0.9 | 0.79 | 2.69679E-09 | 4.9983E-14 | 1.726598723 | 0.924 | 0.767 | 1.1802E-09 | 1.14213E-13 | 9.9966E-14 | Myh11 | EN_3 |
| 5.35796E-14 | 1.743545993 | 0.643 | 0.41 | 1.26512E-09 | 3.01682E-08 | 1.427927816 | 0.636 | 0.468 | 0.000712332 | 3.01682E-08 | 1.07159E-13 | Ogt | EN_3 |
| 6.51296E-14 | 1.933829742 | 0.707 | 0.509 | 1.53784E-09 | 2.07236E-12 | 1.485996922 | 0.763 | 0.567 | 4.89325E-08 | 2.07236E-12 | 1.30259E-13 | Lrmda | EN_3 |
| 6.81544E-14 | 1.432446449 | 0.743 | 0.595 | 1.60926E-09 | 6.30757E-09 | 1.194472547 | 0.788 | 0.643 | 0.000148934 | 6.30757E-09 | 1.36309E-13 | Ssbp2 | EN_3 |
| 1.10029E-10 | 1.592477106 | 0.679 | 0.523 | 2.59801E-06 | 7.69938E-14 | 1.667222059 | 0.788 | 0.592 | 1.81798E-09 | 1.10029E-10 | 1.53988E-13 | Tenm3 | EN_3 |
| 8.25819E-14 | 1.478743184 | 0.743 | 0.6 | 1.94992E-09 | 1.15931E-12 | 1.503477883 | 0.78 | 0.647 | 2.73736E-08 | 1.15931E-12 | 1.65164E-13 | Phf21a | EN_3 |
| 8.26309E-14 | 1.313269874 | 0.764 | 0.61 | 1.95108E-09 | 8.20613E-08 | 0.971249486 | 0.754 | 0.638 | 0.001937631 | 8.20613E-08 | 1.65262E-13 | Kansl1 | EN_3 |
| 8.98402E-14 | 2.26347861 | 0.686 | 0.484 | 2.12131E-09 | 4.47752E-11 | 1.974128694 | 0.678 | 0.495 | 1.05723E-06 | 4.47752E-11 | 1.7968E-13 | Nav2 | EN_3 |
| 9.86429E-14 | 1.101954844 | 0.857 | 0.78 | 2.32916E-09 | 6.40414E-08 | 0.740178609 | 0.873 | 0.767 | 0.001512145 | 6.40414E-08 | 1.97286E-13 | Akap13 | EN_3 |
| 1.18917E-13 | 1.55292287 | 0.8 | 0.674 | 2.80787E-09 | 2.24599E-11 | 1.563837655 | 0.771 | 0.634 | 5.30324E-07 | 2.24599E-11 | 2.37834E-13 | Tanc2 | EN_3 |
| 9.88316E-12 | 2.038500968 | 0.586 | 0.365 | 2.33361E-07 | 1.20585E-13 | 1.843956093 | 0.636 | 0.369 | 2.84725E-09 | 9.88316E-12 | 2.4117E-13 | Neat1 | EN_3 |
| 7.15598E-10 | 2.255861057 | 0.379 | 0.158 | 1.68967E-05 | 1.68934E-13 | 2.833269551 | 0.424 | 0.143 | 3.98888E-09 | 7.15598E-10 | 3.37869E-13 | Greb1 | EN_3 |
| 1.13155E-10 | 5.094520624 | 0.157 | 0.017 | 2.67181E-06 | 1.71493E-13 | 5.338469895 | 0.144 | 0.005 | 4.0493E-09 | 1.13155E-10 | 3.42986E-13 | Mapk10 | EN_3 |
| 1.7213E-13 | 1.441547164 | 0.743 | 0.59 | 4.06433E-09 | 1.53931E-08 | 1.169368804 | 0.72 | 0.59 | 0.000363463 | 1.53931E-08 | 3.44259E-13 | Baz2b | EN_3 |
| 1.77382E-13 | 0.966120129 | 0.936 | 0.84 | 4.18834E-09 | 3.49451E-13 | 1.20487452 | 0.975 | 0.862 | 8.25123E-09 | 3.49451E-13 | 3.54764E-13 | Actg2 | EN_3 |
| 1.89229E-13 | 1.751738556 | 0.736 | 0.583 | 4.46808E-09 | 2.88969E-07 | 1.246252339 | 0.695 | 0.583 | 0.006823144 | 2.88969E-07 | 3.78459E-13 | Fam78b | EN_3 |
| 2.00005E-13 | 1.92927503 | 0.721 | 0.565 | 4.72251E-09 | 1.65547E-08 | 1.536964461 | 0.72 | 0.569 | 0.000390891 | 1.65547E-08 | 4.0001E-13 | Prickle2 | EN_3 |
| 3.7918E-11 | 1.553694885 | 0.729 | 0.644 | 8.9532E-07 | 2.25533E-13 | 1.242682015 | 0.805 | 0.581 | 5.32528E-09 | 3.7918E-11 | 4.51066E-13 | Samd4 | EN_3 |
| 2.26412E-13 | 1.958367004 | 0.586 | 0.333 | 5.34604E-09 | 3.62824E-11 | 1.546432991 | 0.593 | 0.339 | 8.56699E-07 | 3.62824E-11 | 4.52824E-13 | Vwa8 | EN_3 |
| 6.02916E-13 | 1.067087588 | 0.843 | 0.778 | 1.4236E-08 | 2.30248E-13 | 1.113495448 | 0.873 | 0.804 | 5.43661E-09 | 6.02916E-13 | 4.60496E-13 | Tcf12 | EN_3 |
| 1.61159E-11 | 0.752275863 | 0.914 | 0.906 | 3.80528E-07 | 2.44112E-13 | 0.697588014 | 0.958 | 0.915 | 5.76398E-09 | 1.61159E-11 | 4.88224E-13 | Son | EN_3 |
| 5.55941E-13 | 1.525374512 | 0.786 | 0.63 | 1.31269E-08 | 3.33768E-13 | 1.535712811 | 0.822 | 0.636 | 7.88093E-09 | 5.55941E-13 | 6.67536E-13 | Frmd5 | EN_3 |
| 6.24753E-09 | 1.910011914 | 0.486 | 0.299 | 0.000147517 | 3.45785E-13 | 2.106193927 | 0.585 | 0.297 | 8.16467E-09 | 6.24753E-09 | 6.91569E-13 | Pxdn | EN_3 |
| 2.27971E-12 | 0.775468477 | 0.993 | 0.948 | 5.38286E-08 | 4.45345E-13 | 0.740208867 | 0.983 | 0.952 | 1.05155E-08 | 2.27971E-12 | 8.9069E-13 | Chl1 | EN_3 |

| | | | | | | | | | | | | | |
|---|---|---|---|---|---|---|---|---|---|---|---|---|---|
| 5.98478E-13 | 2.230152763 | 0.486 | 0.212 | 1.41313E-08 | 1.43057E-12 | 2.411496694 | 0.475 | 0.203 | 3.37787E-08 | 1.43057E-12 | 1.19696E-12 | Gm4258 | EN_3 |
| 6.59019E-13 | 1.693705627 | 0.636 | 0.415 | 1.55608E-08 | 8.90441E-13 | 1.724750504 | 0.678 | 0.412 | 2.10251E-08 | 8.90441E-13 | 1.31804E-12 | Ambra1 | EN_3 |
| 9.25404E-13 | 1.116496217 | 0.85 | 0.793 | 2.18506E-08 | 4.09587E-11 | 0.926147782 | 0.873 | 0.839 | 9.67116E-07 | 4.09587E-11 | 1.85081E-12 | Itga1 | EN_3 |
| 3.19684E-08 | 1.032120807 | 0.75 | 0.662 | 0.000754837 | 9.77789E-13 | 1.222696226 | 0.797 | 0.682 | 2.30876E-08 | 3.19684E-08 | 1.95558E-12 | Numb | EN_3 |
| 3.52963E-09 | 1.140015737 | 0.721 | 0.602 | 8.33416E-05 | 1.02142E-12 | 1.309186331 | 0.78 | 0.594 | 2.41177E-08 | 3.52963E-09 | 2.04284E-12 | Susd6 | EN_3 |
| 1.03757E-12 | 2.124220293 | 0.564 | 0.314 | 2.44992E-08 | 2.57788E-07 | 1.499912351 | 0.568 | 0.373 | 0.006086888 | 2.57788E-07 | 2.07515E-12 | Pknox2 | EN_3 |
| 1.09118E-12 | 2.06540757 | 0.636 | 0.407 | 2.5765E-08 | 4.35277E-09 | 2.031414245 | 0.559 | 0.348 | 0.000102778 | 4.35277E-09 | 2.18236E-12 | Arhgap42 | EN_3 |
| 1.24454E-12 | 2.558504069 | 0.457 | 0.2 | 2.93861E-08 | 1.16785E-07 | 1.879603584 | 0.398 | 0.205 | 0.002757523 | 1.16785E-07 | 2.48908E-12 | Kcnq1ot1 | EN_3 |
| 1.25332E-12 | 1.712229795 | 0.643 | 0.435 | 2.95933E-08 | 5.29502E-09 | 1.354796977 | 0.669 | 0.512 | 0.000125026 | 5.29502E-09 | 2.50663E-12 | Pan3 | EN_3 |
| 1.63704E-12 | 2.638169722 | 0.45 | 0.207 | 3.86537E-08 | 4.42382E-09 | 2.277412448 | 0.432 | 0.214 | 0.000104455 | 4.42382E-09 | 3.27408E-12 | Prdm5 | EN_3 |
| 1.89098E-12 | 1.433383365 | 0.736 | 0.637 | 4.46498E-08 | 1.5491E-07 | 1.176202404 | 0.678 | 0.576 | 0.003657733 | 1.5491E-07 | 3.78196E-12 | Strn3 | EN_3 |
| 7.25004E-12 | 1.411809186 | 0.7 | 0.56 | 1.71188E-07 | 2.11537E-12 | 1.375287443 | 0.737 | 0.562 | 4.99481E-08 | 7.25004E-12 | 4.23074E-12 | Ddx17 | EN_3 |
| 2.41982E-12 | 1.500641288 | 0.729 | 0.573 | 5.71369E-08 | 2.25932E-07 | 1.20239586 | 0.763 | 0.618 | 0.005334711 | 2.25932E-07 | 4.83965E-12 | Cep112 | EN_3 |
| 2.97515E-12 | 2.837330344 | 0.586 | 0.375 | 7.02492E-08 | 3.92852E-07 | 1.993696857 | 0.551 | 0.389 | 0.009276017 | 3.92852E-07 | 5.95029E-12 | Tshz2 | EN_3 |
| 3.60354E-12 | 1.953120745 | 0.6 | 0.39 | 8.50869E-08 | 4.5991E-10 | 1.649831487 | 0.593 | 0.366 | 1.08594E-05 | 4.5991E-10 | 7.20709E-12 | Smyd3 | EN_3 |
| 1.36732E-08 | 1.223835211 | 0.671 | 0.593 | 0.000322851 | 5.58375E-12 | 1.205206798 | 0.763 | 0.571 | 1.31843E-07 | 1.36732E-08 | 1.11675E-11 | Mpdz | EN_3 |
| 6.24614E-12 | 3.74369601 | 0.3 | 0.084 | 1.47484E-07 | 4.86797E-08 | 3.549599663 | 0.212 | 0.058 | 0.001149426 | 4.86797E-08 | 1.24923E-11 | Erc2 | EN_3 |
| 1.00534E-08 | 1.914877939 | 0.521 | 0.338 | 0.000237381 | 6.48644E-12 | 2.257965043 | 0.576 | 0.323 | 1.53158E-07 | 1.00534E-08 | 1.29729E-11 | Pde4d | EN_3 |
| 7.36174E-12 | 1.353011446 | 0.693 | 0.531 | 1.73825E-07 | 4.24487E-11 | 1.309651613 | 0.754 | 0.594 | 1.0023E-06 | 4.24487E-11 | 1.47235E-11 | Mycbp2 | EN_3 |
| 5.36554E-10 | 3.60087493 | 0.193 | 0.035 | 1.26691E-05 | 7.56561E-12 | 4.522570255 | 0.161 | 0.014 | 1.78639E-07 | 5.36554E-10 | 1.51312E-11 | Pdgfc | EN_3 |
| 7.91037E-12 | 1.625030635 | 0.65 | 0.437 | 1.8678E-07 | 8.06478E-09 | 1.504572089 | 0.653 | 0.486 | 0.000190426 | 8.06478E-09 | 1.58207E-11 | Gm10863 | EN_3 |
| 1.56746E-11 | 2.055790432 | 0.464 | 0.212 | 3.70108E-07 | 8.27316E-12 | 2.113338499 | 0.534 | 0.265 | 1.95346E-07 | 1.56746E-11 | 1.65463E-11 | Snx29 | EN_3 |
| 9.68079E-12 | 1.739081871 | 0.671 | 0.521 | 2.28583E-07 | 7.50119E-11 | 1.37837924 | 0.737 | 0.558 | 1.77118E-06 | 7.50119E-11 | 1.93616E-11 | Tnik | EN_3 |
| 1.06175E-11 | 2.310668949 | 0.321 | 0.094 | 2.50701E-07 | 4.59278E-08 | 2.639041136 | 0.271 | 0.092 | 0.001084446 | 4.59278E-08 | 2.1235E-11 | Carmn | EN_3 |
| 1.10926E-11 | 1.348808601 | 0.736 | 0.642 | 2.61919E-07 | 8.64769E-10 | 1.241837389 | 0.763 | 0.647 | 2.04189E-05 | 8.64769E-10 | 2.21853E-11 | Cep85l | EN_3 |
| 1.15154E-11 | 6.096923456 | 0.121 | 0.002 | 2.71901E-07 | 6.32564E-11 | 4.480820793 | 0.136 | 0.009 | 1.49361E-06 | 6.32564E-11 | 2.30307E-11 | Hmgcll1 | EN_3 |
| 1.16926E-11 | 1.313298407 | 0.793 | 0.696 | 2.76085E-07 | 1.32418E-11 | 1.195566949 | 0.847 | 0.707 | 3.12666E-07 | 1.32418E-11 | 2.33851E-11 | Akap6 | EN_3 |
| 6.13476E-09 | 2.452680324 | 0.429 | 0.23 | 0.000144854 | 1.27793E-11 | 2.725345642 | 0.441 | 0.171 | 3.01746E-07 | 6.13476E-09 | 2.55587E-11 | Dlg2 | EN_3 |
| 1.31082E-11 | 1.397465592 | 0.743 | 0.605 | 3.09511E-07 | 4.78214E-10 | 1.348784095 | 0.788 | 0.629 | 1.12916E-05 | 4.78214E-10 | 2.62164E-11 | Plekhg1 | EN_3 |
| 1.53181E-08 | 2.109809259 | 0.479 | 0.299 | 0.000361691 | 1.37084E-11 | 1.957790245 | 0.559 | 0.302 | 3.23684E-07 | 1.53181E-08 | 2.74169E-11 | Large1 | EN_3 |
| 6.20584E-10 | 3.47223353 | 0.157 | 0.02 | 1.46532E-05 | 1.50332E-11 | 3.253508105 | 0.254 | 0.053 | 3.54963E-07 | 6.20584E-10 | 3.00663E-11 | Gucy1a2 | EN_3 |
| 1.79377E-11 | 1.887066716 | 0.643 | 0.479 | 4.23546E-07 | 6.61917E-09 | 1.567030302 | 0.661 | 0.498 | 0.000156292 | 6.61917E-09 | 3.58755E-11 | Galnt17 | EN_3 |
| 1.89498E-11 | 5.672786519 | 0.179 | 0.022 | 4.47442E-07 | 1.84418E-08 | 4.930850513 | 0.153 | 0.025 | 0.000435448 | 1.84418E-08 | 3.78995E-11 | Agbl4 | EN_3 |
| 3.7457E-11 | 1.06301105 | 0.8 | 0.753 | 8.84434E-07 | 1.96345E-11 | 0.951692367 | 0.864 | 0.753 | 4.6361E-07 | 3.7457E-11 | 3.9269E-11 | Luc7l2 | EN_3 |
| 2.8804E-08 | 1.935934812 | 0.429 | 0.24 | 0.00068012 | 1.98601E-11 | 2.039939441 | 0.517 | 0.249 | 4.68937E-07 | 2.8804E-08 | 3.97202E-11 | Garem1 | EN_3 |
| 3.0071E-07 | 1.315330207 | 0.579 | 0.437 | 0.007100368 | 2.0418E-11 | 1.659769436 | 0.669 | 0.461 | 4.82111E-07 | 3.0071E-07 | 4.08361E-11 | Slmap | EN_3 |
| 2.09314E-11 | 1.416592014 | 0.729 | 0.57 | 4.94232E-07 | 1.07505E-07 | 1.113169937 | 0.78 | 0.647 | 0.002538416 | 1.07505E-07 | 4.18628E-11 | Exoc6b | EN_3 |
| 2.52158E-11 | 1.976447515 | 0.657 | 0.489 | 5.95396E-07 | 8.25404E-10 | 1.550938025 | 0.686 | 0.505 | 1.94894E-05 | 8.25404E-10 | 5.04317E-11 | Ank2 | EN_3 |
| 5.36895E-09 | 1.894601084 | 0.5 | 0.301 | 0.000126772 | 2.71013E-11 | 1.668573696 | 0.585 | 0.313 | 6.39915E-07 | 5.36895E-09 | 5.42026E-11 | Cog5 | EN_3 |
| 8.67624E-10 | 1.551870372 | 0.607 | 0.42 | 2.04863E-05 | 2.71855E-11 | 1.565558407 | 0.695 | 0.44 | 6.41905E-07 | 8.67624E-10 | 5.43711E-11 | Itsn2 | EN_3 |
| 3.34048E-11 | 3.334256592 | 0.179 | 0.022 | 7.88754E-07 | 9.13978E-08 | 2.31851784 | 0.195 | 0.046 | 0.002158085 | 9.13978E-08 | 6.68096E-11 | Robo2 | EN_3 |
| 3.34793E-11 | 2.181790684 | 0.493 | 0.267 | 7.90514E-07 | 7.43614E-08 | 2.413701769 | 0.356 | 0.154 | 0.001755822 | 7.43614E-08 | 6.69587E-11 | Sulf1 | EN_3 |
| 3.36499E-11 | 4.345707392 | 0.143 | 0.01 | 7.94542E-07 | 3.46518E-08 | 2.613067021 | 0.127 | 0.016 | 0.000818198 | 3.46518E-08 | 6.72998E-11 | Hecw2 | EN_3 |
| 4.58173E-08 | 2.24539332 | 0.4 | 0.202 | 0.001081838 | 3.39494E-11 | 2.474719416 | 0.449 | 0.2 | 8.01613E-07 | 4.58173E-08 | 6.78988E-11 | Cacna2d1 | EN_3 |
| 3.67106E-11 | 1.126286063 | 0.779 | 0.681 | 8.6681E-07 | 8.75744E-08 | 0.863265061 | 0.797 | 0.707 | 0.002067806 | 8.75744E-08 | 7.34212E-11 | Nipbl | EN_3 |
| 4.91056E-11 | 1.196350461 | 0.757 | 0.672 | 1.15948E-06 | 1.46845E-08 | 0.950981834 | 0.797 | 0.724 | 0.000346731 | 1.46845E-08 | 9.82111E-11 | Tnrc6c | EN_3 |
| 4.9745E-11 | 1.7408746 | 0.707 | 0.617 | 1.17458E-06 | 2.31514E-09 | 1.370415697 | 0.737 | 0.585 | 5.46652E-05 | 2.31514E-09 | 9.94901E-11 | Agmo | EN_3 |
| 1.06109E-09 | 1.625136967 | 0.586 | 0.383 | 2.50545E-05 | 6.10962E-11 | 1.67942185 | 0.593 | 0.355 | 1.4426E-06 | 1.06109E-09 | 1.22192E-10 | Herc1 | EN_3 |
| 6.71495E-11 | 2.527556877 | 0.421 | 0.193 | 1.58553E-06 | 1.94713E-09 | 2.695793418 | 0.415 | 0.194 | 4.59757E-05 | 1.94713E-09 | 1.34299E-10 | Ptprg | EN_3 |
| 7.68943E-11 | 1.532051685 | 0.714 | 0.612 | 1.81563E-06 | 1.4757E-07 | 1.140410155 | 0.814 | 0.659 | 0.00348442 | 1.4757E-07 | 1.53789E-10 | Afap1l2 | EN_3 |
| 7.72394E-11 | 3.487007187 | 0.221 | 0.044 | 1.82378E-06 | 1.09904E-10 | 3.83867252 | 0.203 | 0.035 | 2.59505E-06 | 1.09904E-10 | 1.54479E-10 | Kank1 | EN_3 |
| 7.75731E-11 | 1.376232814 | 0.764 | 0.716 | 1.83166E-06 | 3.76775E-10 | 1.290547472 | 0.814 | 0.698 | 8.8964E-06 | 3.76775E-10 | 1.55146E-10 | Plxdc2 | EN_3 |
| 7.79482E-11 | 1.761547225 | 0.679 | 0.523 | 1.84051E-06 | 2.319E-07 | 1.357041989 | 0.72 | 0.56 | 0.005475618 | 2.319E-07 | 1.55896E-10 | Xkr4 | EN_3 |
| 7.92089E-11 | 2.743144661 | 0.414 | 0.188 | 1.87028E-06 | 7.99701E-08 | 2.531619222 | 0.373 | 0.173 | 0.001888255 | 7.99701E-08 | 1.58418E-10 | Plcb4 | EN_3 |
| 8.67378E-11 | 1.280961286 | 0.743 | 0.642 | 2.04805E-06 | 7.95871E-11 | 1.078783339 | 0.814 | 0.65 | 1.87921E-06 | 8.67378E-11 | 1.59174E-10 | Tnrc6b | EN_3 |
| 8.34525E-11 | 3.505146721 | 0.314 | 0.101 | 1.97048E-06 | 1.81067E-08 | 3.275616893 | 0.271 | 0.088 | 0.000427534 | 1.81067E-08 | 1.66905E-10 | Col8a1 | EN_3 |
| 9.04575E-11 | 4.280178523 | 0.179 | 0.025 | 2.13588E-06 | 2.20814E-08 | 3.710055857 | 0.169 | 0.032 | 0.000521386 | 2.20814E-08 | 1.80915E-10 | Pdgfd | EN_3 |
| 1.01002E-10 | 2.088324978 | 0.5 | 0.277 | 2.38486E-06 | 7.73265E-08 | 1.986095148 | 0.483 | 0.286 | 0.001825834 | 7.73265E-08 | 2.02004E-10 | Lncppara | EN_3 |
| 1.0169E-10 | 2.264307264 | 0.45 | 0.21 | 2.4011E-06 | 4.58896E-09 | 1.870747912 | 0.441 | 0.203 | 0.000108354 | 4.58896E-09 | 2.0338E-10 | Atxn7l1 | EN_3 |
| 1.04668E-10 | 1.019736792 | 0.914 | 0.842 | 2.47142E-06 | 6.29112E-09 | 1.055850057 | 0.975 | 0.887 | 0.000148546 | 6.29112E-09 | 2.09336E-10 | Tpm1 | EN_3 |
| 1.26175E-10 | 1.545743838 | 0.629 | 0.435 | 2.97925E-06 | 1.43605E-09 | 1.371217872 | 0.686 | 0.502 | 3.3908E-05 | 1.43605E-09 | 2.5235E-10 | Ksr1 | EN_3 |
| 1.27451E-10 | 2.445234524 | 0.521 | 0.314 | 3.00937E-06 | 1.20591E-08 | 1.77902696 | 0.559 | 0.371 | 0.00028474 | 1.20591E-08 | 2.54902E-10 | Efna5 | EN_3 |
| 1.42279E-10 | 2.257588873 | 0.579 | 0.4 | 3.35948E-06 | 1.35898E-10 | 2.12385243 | 0.568 | 0.348 | 3.20883E-06 | 1.42279E-10 | 2.71797E-10 | Airn | EN_3 |
| 4.45546E-09 | 1.530940641 | 0.593 | 0.43 | 0.000105202 | 1.78144E-10 | 1.365618128 | 0.644 | 0.426 | 4.20633E-06 | 4.45546E-09 | 3.56288E-10 | Rbm6 | EN_3 |
| 3.33351E-10 | 1.216612391 | 0.707 | 0.551 | 7.87107E-06 | 1.87439E-10 | 1.46405629 | 0.703 | 0.535 | 4.42581E-06 | 3.33351E-10 | 3.74878E-10 | Mkln1 | EN_3 |
| 2.02579E-10 | 1.093069908 | 0.95 | 0.847 | 4.78331E-06 | 1.58655E-08 | 0.915099681 | 0.966 | 0.908 | 0.000374615 | 1.58655E-08 | 4.05159E-10 | Tpm2 | EN_3 |
| 2.27676E-10 | 1.863173524 | 0.564 | 0.346 | 5.37589E-06 | 1.2158E-07 | 1.343127591 | 0.602 | 0.44 | 0.002870746 | 1.2158E-07 | 4.55352E-10 | Cdk14 | EN_3 |
| 2.45309E-10 | 1.388939817 | 0.729 | 0.64 | 5.79223E-06 | 3.95162E-10 | 1.171953682 | 0.788 | 0.659 | 9.33058E-06 | 3.95162E-10 | 4.90617E-10 | Foxo1 | EN_3 |
| 2.97106E-10 | 1.462128326 | 0.636 | 0.506 | 7.01526E-06 | 4.16255E-07 | 1.008948782 | 0.653 | 0.518 | 0.009828615 | 4.16255E-07 | 5.94212E-10 | Zfand3 | EN_3 |
| 9.01298E-10 | 1.877651097 | 0.686 | 0.553 | 2.12815E-05 | 3.20981E-10 | 2.043728346 | 0.627 | 0.433 | 7.579E-06 | 9.01298E-10 | 6.41962E-10 | Ltbp1 | EN_3 |
| 4.84779E-10 | 1.708776267 | 0.55 | 0.368 | 1.14466E-05 | 1.66645E-09 | 1.342905512 | 0.602 | 0.385 | 3.93482E-05 | 1.66645E-09 | 9.69558E-10 | Ppp6r3 | EN_3 |
| 5.86127E-10 | 1.726080402 | 0.586 | 0.393 | 1.38396E-05 | 2.66516E-08 | 1.402950149 | 0.576 | 0.364 | 0.000629299 | 2.66516E-08 | 1.17225E-09 | Arl15 | EN_3 |
| 6.68763E-10 | 2.870397638 | 0.343 | 0.133 | 1.57908E-05 | 1.08724E-07 | 2.642175672 | 0.339 | 0.143 | 0.002567193 | 1.08724E-07 | 1.33753E-09 | Gm4876 | EN_3 |
| 6.89394E-10 | 1.529484908 | 0.614 | 0.449 | 1.6278E-05 | 9.21457E-08 | 1.483923733 | 0.593 | 0.415 | 0.002175743 | 9.21457E-08 | 1.37879E-09 | Ptbp2 | EN_3 |
| 7.72064E-10 | 0.873929459 | 0.836 | 0.807 | 1.823E-05 | 3.51678E-07 | 0.732714308 | 0.847 | 0.79 | 0.008303832 | 3.51678E-07 | 1.54413E-09 | Nfib | EN_3 |
| 8.503E-10 | 2.241717601 | 0.493 | 0.294 | 2.00773E-05 | 1.45435E-09 | 2.347468133 | 0.483 | 0.256 | 3.43402E-05 | 1.45435E-09 | 1.7006E-09 | Setbp1 | EN_3 |
| 8.75497E-10 | 1.145352555 | 0.743 | 0.672 | 2.06722E-05 | 1.6505E-07 | 0.892078611 | 0.788 | 0.705 | 0.003897154 | 1.6505E-07 | 1.75099E-09 | Frmd6 | EN_3 |
| 1.045E-09 | 1.292352414 | 0.736 | 0.612 | 2.46744E-05 | 1.68863E-08 | 0.89806186 | 0.78 | 0.641 | 0.000398718 | 1.68863E-08 | 2.08999E-09 | Tead1 | EN_3 |
| 1.0998E-09 | 1.406533715 | 0.693 | 0.56 | 2.59684E-05 | 3.47133E-09 | 1.383948506 | 0.72 | 0.569 | 8.19651E-05 | 3.47133E-09 | 2.1996E-09 | Crppa | EN_3 |
| 1.24501E-09 | 1.599171503 | 0.614 | 0.462 | 2.93973E-05 | 1.88506E-08 | 1.172258603 | 0.678 | 0.498 | 0.0004451 | 1.88506E-08 | 2.49003E-09 | Dyrk1a | EN_3 |
| 5.80207E-08 | 1.527661915 | 0.564 | 0.407 | 0.001369985 | 1.26311E-09 | 1.329563178 | 0.61 | 0.385 | 2.98246E-05 | 5.80207E-08 | 2.52622E-09 | Braf | EN_3 |

| | | | | | | | | | | | | | |
|---|---|---|---|---|---|---|---|---|---|---|---|---|---|
| 1.5386E-08 | 1.563302661 | 0.579 | 0.4 | 0.000363293 | 1.30584E-09 | 1.571123784 | 0.602 | 0.406 | 3.08336E-05 | 1.5386E-08 | 2.61169E-09 | Ubr3 | EN_3 |
| 1.38508E-09 | 1.959528628 | 0.486 | 0.264 | 3.27046E-05 | 3.43837E-07 | 1.466857657 | 0.525 | 0.334 | 0.008118683 | 3.43837E-07 | 2.77017E-09 | Fbxl20 | EN_3 |
| 2.71917E-08 | 1.403515318 | 0.65 | 0.511 | 0.00064205 | 1.47519E-09 | 1.484307983 | 0.678 | 0.486 | 3.48322E-05 | 2.71917E-08 | 2.95038E-09 | Gsap | EN_3 |
| 2.92522E-07 | 1.305188548 | 0.636 | 0.519 | 0.006907018 | 1.72628E-09 | 1.360383069 | 0.678 | 0.495 | 4.07609E-05 | 2.92522E-07 | 3.45255E-09 | Rarb | EN_3 |
| 1.8725E-09 | 2.188819094 | 0.45 | 0.242 | 4.42135E-05 | 2.97675E-08 | 1.697581004 | 0.568 | 0.353 | 0.000702869 | 2.97675E-08 | 3.745E-09 | Thsd4 | EN_3 |
| 7.36277E-09 | 1.62732892 | 0.579 | 0.43 | 0.00017385 | 1.98457E-09 | 1.513853356 | 0.576 | 0.385 | 4.68597E-05 | 7.36277E-09 | 3.96915E-09 | Faf1 | EN_3 |
| 2.11523E-09 | 1.648330933 | 0.621 | 0.514 | 4.99449E-05 | 3.44251E-07 | 1.48622218 | 0.619 | 0.488 | 0.008128466 | 3.44251E-07 | 4.23047E-09 | Atxn1 | EN_3 |
| 1.83034E-08 | 1.429024084 | 0.593 | 0.427 | 0.000432181 | 2.28419E-09 | 1.344858482 | 0.686 | 0.525 | 5.39343E-05 | 1.83034E-08 | 4.56838E-09 | Daam1 | EN_3 |
| 2.4297E-09 | 3.383008746 | 0.207 | 0.047 | 5.73702E-05 | 6.04453E-09 | 3.796400459 | 0.212 | 0.051 | 0.000142723 | 6.04453E-09 | 4.85941E-09 | Zfpm2 | EN_3 |
| 1.04073E-08 | 1.53910789 | 0.593 | 0.44 | 0.000245736 | 2.74616E-09 | 1.468312753 | 0.61 | 0.463 | 6.48424E-05 | 1.04073E-08 | 5.49233E-09 | Rbfox2 | EN_3 |
| 2.771E-09 | 1.886596861 | 0.5 | 0.319 | 6.54289E-05 | 3.31647E-09 | 1.552405891 | 0.559 | 0.341 | 7.83086E-05 | 3.31647E-09 | 5.54201E-09 | Uty | EN_3 |
| 1.05559E-08 | 4.750000266 | 0.393 | 0.198 | 0.000249247 | 2.97199E-09 | 3.946884228 | 0.449 | 0.242 | 7.01746E-05 | 1.05559E-08 | 5.94398E-09 | Etl4 | EN_3 |
| 3.77779E-08 | 1.505733169 | 0.55 | 0.388 | 0.000892011 | 3.19862E-09 | 1.507402086 | 0.619 | 0.44 | 7.55258E-05 | 3.77779E-08 | 6.39724E-09 | Trio | EN_3 |
| 1.09102E-08 | 1.58781651 | 0.529 | 0.331 | 0.000257612 | 3.50286E-09 | 1.622051149 | 0.559 | 0.371 | 8.27095E-05 | 1.09102E-08 | 7.00572E-09 | Nbea | EN_3 |
| 4.5403E-09 | 1.590139513 | 0.664 | 0.541 | 0.000107206 | 8.71728E-08 | 1.287650723 | 0.729 | 0.535 | 0.002058325 | 8.71728E-08 | 9.0806E-09 | Slc22a23 | EN_3 |
| 5.36161E-09 | 2.050350947 | 0.4 | 0.185 | 0.000126598 | 2.95904E-07 | 1.785338312 | 0.415 | 0.21 | 0.006986882 | 2.95904E-07 | 1.07232E-08 | Tmem135 | EN_3 |
| 5.42365E-09 | 1.05175131 | 0.714 | 0.607 | 0.000128063 | 3.93943E-07 | 0.994912486 | 0.737 | 0.654 | 0.009301784 | 3.93943E-07 | 1.08473E-08 | Syne1 | EN_3 |
| 5.93251E-09 | 1.054224326 | 0.714 | 0.612 | 0.000140078 | 2.59815E-07 | 1.085873577 | 0.72 | 0.601 | 0.006134748 | 2.59815E-07 | 1.1865E-08 | Nsd1 | EN_3 |
| 7.88251E-09 | 1.253299457 | 0.836 | 0.881 | 0.000186122 | 7.83798E-08 | 1.364858563 | 0.831 | 0.809 | 0.001850705 | 7.83798E-08 | 1.5765E-08 | Hand2os1 | EN_3 |
| 1.00647E-08 | 1.792251196 | 0.586 | 0.44 | 0.000237649 | 1.34923E-07 | 1.55854179 | 0.619 | 0.468 | 0.003185813 | 1.34923E-07 | 2.01295E-08 | Grik3 | EN_3 |
| 1.02946E-08 | 1.428924508 | 0.671 | 0.585 | 0.000243077 | 1.32488E-07 | 1.173600444 | 0.737 | 0.615 | 0.003128301 | 1.32488E-07 | 2.05893E-08 | Atp8a1 | EN_3 |
| 1.09104E-08 | 0.999589712 | 0.664 | 0.558 | 0.000257616 | 5.51569E-08 | 1.10935492 | 0.678 | 0.541 | 0.001302364 | 5.51569E-08 | 2.18207E-08 | Tnrc6a | EN_3 |
| 1.16541E-08 | 5.159519393 | 0.121 | 0.012 | 0.000275176 | 3.64688E-07 | 4.159969666 | 0.127 | 0.021 | 0.008611009 | 3.64688E-07 | 2.33081E-08 | Nav3 | EN_3 |
| 1.32782E-08 | 1.935551336 | 0.457 | 0.254 | 0.000313525 | 7.5567E-08 | 2.293091121 | 0.475 | 0.3 | 0.001784287 | 7.5567E-08 | 2.65564E-08 | Pcdh7 | EN_3 |
| 1.35955E-08 | 1.579212076 | 0.493 | 0.296 | 0.000321018 | 4.871E-08 | 1.588357884 | 0.576 | 0.378 | 0.001150141 | 4.871E-08 | 2.71911E-08 | Atad2b | EN_3 |
| 1.92498E-07 | 1.623875569 | 0.486 | 0.306 | 0.004545251 | 1.45208E-08 | 1.677759268 | 0.602 | 0.394 | 0.000342865 | 1.92498E-07 | 2.90416E-08 | Btbd9 | EN_3 |
| 2.64919E-08 | 3.068652104 | 0.221 | 0.062 | 0.000625527 | 1.96314E-08 | 2.444999942 | 0.322 | 0.12 | 0.000463537 | 2.64919E-08 | 3.92628E-08 | B3galt1 | EN_3 |
| 2.66574E-08 | 1.300614639 | 0.671 | 0.511 | 0.000629435 | 1.14007E-07 | 1.324580018 | 0.686 | 0.546 | 0.002691931 | 1.14007E-07 | 5.33149E-08 | Megf10 | EN_3 |
| 5.56119E-08 | 1.970253348 | 0.5 | 0.341 | 0.001313108 | 3.22069E-08 | 1.719067863 | 0.568 | 0.403 | 0.000760469 | 5.56119E-08 | 6.44138E-08 | Rabgap1l | EN_3 |
| 4.84922E-08 | 1.474844904 | 0.571 | 0.459 | 0.001144998 | 7.97895E-08 | 1.664891723 | 0.61 | 0.445 | 0.00188399 | 7.97895E-08 | 9.69844E-08 | Fars2 | EN_3 |
| 4.92436E-08 | 1.465701416 | 0.614 | 0.496 | 0.001162741 | 1.55326E-07 | 1.424213637 | 0.61 | 0.445 | 0.003667559 | 1.55326E-07 | 9.84873E-08 | Phlpp1 | EN_3 |
| 1.28912E-07 | 1.826748781 | 0.514 | 0.343 | 0.003043874 | 5.20706E-08 | 2.323598457 | 0.466 | 0.274 | 0.00122949 | 1.28912E-07 | 1.04141E-07 | Adam12 | EN_3 |
| 7.10934E-08 | 1.530510338 | 0.529 | 0.36 | 0.001678659 | 5.31638E-08 | 1.29983405 | 0.61 | 0.426 | 0.001255304 | 7.10934E-08 | 1.06328E-07 | Rictor | EN_3 |
| 1.84996E-07 | 1.22048833 | 0.664 | 0.61 | 0.004368136 | 9.04113E-08 | 1.078305127 | 0.771 | 0.615 | 0.002134792 | 1.84996E-07 | 1.80823E-07 | Sgms1 | EN_3 |
| 1.08643E-07 | 2.600602975 | 0.279 | 0.106 | 0.002565272 | 9.60554E-08 | 2.295430272 | 0.271 | 0.092 | 0.00226806 | 1.08643E-07 | 1.92111E-07 | Plekha6 | EN_3 |
| 2.49202E-07 | 1.3511388 | 0.529 | 0.385 | 0.005884156 | 1.34494E-07 | 1.365516921 | 0.61 | 0.417 | 0.00317567 | 2.49202E-07 | 2.68988E-07 | Atrn | EN_3 |
| 1.39546E-07 | 1.834136924 | 0.471 | 0.289 | 0.003294952 | 2.83353E-07 | 1.927372524 | 0.466 | 0.272 | 0.006690532 | 2.83353E-07 | 2.79091E-07 | Pde8b | EN_3 |
| 2.86756E-07 | 0.929491142 | 0.764 | 0.654 | 0.006770875 | 1.5196E-07 | 1.057372891 | 0.771 | 0.675 | 0.003588073 | 2.86756E-07 | 3.03919E-07 | Jmjd1c | EN_3 |
| 3.90887E-07 | 3.184558554 | 0.193 | 0.057 | 0.00922962 | 1.55142E-07 | 2.166954538 | 0.254 | 0.083 | 0.003663207 | 3.90887E-07 | 3.10283E-07 | Rnf150 | EN_3 |
| 2.85278E-07 | 2.653161234 | 0.271 | 0.106 | 0.006735975 | 1.57591E-07 | 2.099797299 | 0.254 | 0.083 | 0.003721032 | 2.85278E-07 | 3.15181E-07 | Ppm1l | EN_3 |
| 1.64064E-07 | 0.909377594 | 0.807 | 0.785 | 0.003873874 | 2.27987E-07 | 0.852001617 | 0.831 | 0.772 | 0.005383237 | 2.27987E-07 | 3.28128E-07 | Adgrg6 | EN_3 |
| 3.21221E-07 | 1.404561078 | 0.707 | 0.669 | 0.007584675 | 3.59212E-07 | 1.206005823 | 0.746 | 0.684 | 0.008481714 | 3.59212E-07 | 6.42442E-07 | Palld | EN_3 |
| 4.73573E-23 | 1.548558926 | 0.956 | 0.892 | 1.1182E-18 | 3.55648E-15 | 1.458387736 | 0.889 | 0.913 | 8.39756E-11 | 3.55648E-15 | 9.47146E-23 | Rps24 | EN_4 |
| 1.67655E-22 | 1.682201362 | 0.956 | 0.881 | 3.95867E-18 | 1.33778E-15 | 1.789627349 | 0.878 | 0.911 | 3.15877E-11 | 1.33778E-15 | 3.3531E-22 | Rpl38 | EN_4 |
| 1.69923E-21 | 1.560504597 | 0.923 | 0.868 | 4.01221E-17 | 1.68348E-17 | 1.674571422 | 0.9 | 0.892 | 3.97503E-13 | 1.68348E-17 | 3.39845E-21 | Rps27a | EN_4 |
| 1.11105E-20 | 1.582053839 | 0.923 | 0.879 | 2.6234E-16 | 3.23965E-15 | 1.793639955 | 0.889 | 0.9 | 7.64946E-11 | 3.23965E-15 | 2.22209E-20 | Rps20 | EN_4 |
| 2.75699E-20 | 1.818775698 | 0.945 | 0.903 | 6.5098E-16 | 3.68188E-15 | 1.774804268 | 0.9 | 0.911 | 8.69366E-11 | 3.68188E-15 | 5.51398E-20 | Rpl37 | EN_4 |
| 3.04406E-20 | 2.104508704 | 1 | 0.978 | 7.18764E-16 | 2.99028E-12 | 1.817345017 | 0.989 | 0.978 | 7.06066E-08 | 2.99028E-12 | 6.08813E-20 | Fth1 | EN_4 |
| 4.57995E-19 | 1.276822852 | 0.967 | 0.941 | 1.08142E-14 | 2.50943E-17 | 1.316533727 | 0.967 | 0.955 | 5.92527E-13 | 2.50943E-17 | 9.1599E-19 | Rpl41 | EN_4 |
| 7.4396E-19 | 1.357461127 | 0.901 | 0.85 | 1.75664E-14 | 3.59169E-09 | 1.031551294 | 0.889 | 0.877 | 8.48069E-05 | 3.59169E-09 | 1.48792E-18 | Rpl11 | EN_4 |
| 9.59126E-19 | 1.434244918 | 0.923 | 0.85 | 2.26469E-14 | 6.0065E-12 | 1.36805341 | 0.867 | 0.861 | 1.41825E-07 | 6.0065E-12 | 1.91825E-18 | Rpl9 | EN_4 |
| 1.06231E-12 | 1.623354126 | 0.868 | 0.841 | 2.50834E-08 | 2.53724E-17 | 1.962811548 | 0.9 | 0.857 | 5.99092E-13 | 1.06231E-12 | 5.07447E-17 | Serf2 | EN_4 |
| 1.12857E-16 | 1.320666159 | 0.901 | 0.874 | 2.66477E-12 | 1.96205E-08 | 1.11555419 | 0.822 | 0.877 | 0.00046328 | 1.96205E-08 | 2.25714E-16 | Rps28 | EN_4 |
| 1.24634E-16 | 1.496295913 | 0.868 | 0.859 | 2.94287E-12 | 8.49089E-13 | 1.337864265 | 0.856 | 0.894 | 2.00487E-08 | 8.49089E-13 | 2.49269E-16 | Rpl30 | EN_4 |
| 4.13125E-16 | 1.275001383 | 0.89 | 0.857 | 9.7547E-12 | 6.21289E-09 | 1.100993917 | 0.844 | 0.913 | 0.000146699 | 6.21289E-09 | 8.2625E-16 | Rpl39 | EN_4 |
| 9.06543E-16 | 1.322487605 | 0.912 | 0.857 | 2.14053E-11 | 1.05317E-08 | 1.239603148 | 0.822 | 0.89 | 0.000248674 | 1.05317E-08 | 1.81309E-15 | Rps3a1 | EN_4 |
| 1.1449E-15 | 1.156803231 | 0.967 | 0.883 | 2.70334E-11 | 7.22702E-09 | 0.933271816 | 0.878 | 0.913 | 0.000170644 | 7.22702E-09 | 2.2898E-15 | Rps21 | EN_4 |
| 2.79616E-15 | 1.581937067 | 0.868 | 0.855 | 6.60228E-11 | 6.35906E-11 | 1.527720417 | 0.867 | 0.872 | 1.5015E-06 | 6.35906E-11 | 5.59231E-15 | Rpl6 | EN_4 |
| 3.09579E-15 | 1.162457871 | 0.934 | 0.921 | 7.30978E-11 | 6.84222E-08 | 0.950065679 | 0.867 | 0.931 | 0.001615585 | 6.84222E-08 | 6.19158E-15 | Rpl37a | EN_4 |
| 4.808E-15 | 1.213031897 | 0.901 | 0.846 | 1.13527E-10 | 5.66258E-09 | 1.165358666 | 0.8 | 0.861 | 0.000133705 | 5.66258E-09 | 9.616E-15 | Rpsa | EN_4 |
| 1.67941E-14 | 1.433325865 | 0.967 | 0.877 | 3.96542E-10 | 7.1454E-08 | 1.730374236 | 0.967 | 0.9 | 0.001687171 | 7.1454E-08 | 3.35882E-14 | Dbi | EN_4 |
| 2.22464E-14 | 1.025657679 | 0.978 | 0.945 | 5.25282E-10 | 7.59911E-11 | 0.916956912 | 0.944 | 0.935 | 1.7943E-06 | 7.59911E-11 | 4.44928E-14 | Rps29 | EN_4 |
| 2.79693E-14 | 1.388513499 | 0.912 | 0.868 | 6.6041E-10 | 1.63396E-12 | 1.327398215 | 0.9 | 0.913 | 3.85811E-08 | 1.63396E-12 | 5.59385E-14 | Fau | EN_4 |
| 9.0624E-13 | 1.239222126 | 0.879 | 0.852 | 2.13981E-08 | 3.01902E-14 | 1.4799716 | 0.856 | 0.881 | 7.1285E-10 | 9.0624E-13 | 6.03803E-14 | Cox8a | EN_4 |
| 6.30472E-14 | 1.619634652 | 0.857 | 0.824 | 1.48867E-09 | 3.71864E-11 | 1.781405967 | 0.811 | 0.833 | 8.78046E-07 | 3.71864E-11 | 1.26094E-13 | Prdx1 | EN_4 |
| 7.22575E-14 | 1.191627833 | 0.923 | 0.879 | 1.70614E-09 | 1.90002E-11 | 1.195882627 | 0.878 | 0.913 | 4.48633E-07 | 1.90002E-11 | 1.44515E-13 | Rpl35a | EN_4 |
| 9.70145E-14 | 1.068611441 | 0.912 | 0.866 | 2.29071E-09 | 2.55965E-08 | 0.839134167 | 0.889 | 0.905 | 0.000604385 | 2.55965E-08 | 1.94029E-13 | Rps16 | EN_4 |
| 1.85843E-13 | 1.406665235 | 0.846 | 0.839 | 4.38811E-09 | 8.06005E-10 | 1.534072632 | 0.811 | 0.857 | 1.90314E-05 | 8.06005E-10 | 3.71685E-13 | Rps2 | EN_4 |
| 3.3938E-13 | 0.956604246 | 0.934 | 0.971 | 8.01344E-09 | 8.7515E-13 | 0.972688715 | 0.867 | 0.981 | 2.0664E-08 | 8.7515E-13 | 6.7876E-13 | Myl6 | EN_4 |
| 4.24203E-13 | 1.656292965 | 0.813 | 0.775 | 1.00163E-08 | 8.89992E-11 | 1.776353508 | 0.756 | 0.788 | 2.10145E-06 | 8.89992E-11 | 8.48406E-13 | Atp5e | EN_4 |
| 2.77411E-11 | 1.440299553 | 0.835 | 0.811 | 6.55023E-07 | 4.45579E-13 | 1.535016406 | 0.856 | 0.855 | 1.0521E-08 | 2.77411E-11 | 8.91158E-13 | Selenow | EN_4 |
| 1.37513E-07 | 1.206232359 | 0.802 | 0.8 | 0.003246947 | 4.72116E-13 | 1.820157921 | 0.811 | 0.794 | 1.11476E-08 | 1.37513E-07 | 9.44231E-13 | Elob | EN_4 |
| 1.10622E-12 | 1.224469245 | 0.912 | 0.85 | 2.61201E-08 | 3.6838E-09 | 1.288906687 | 0.822 | 0.866 | 8.69818E-05 | 3.6838E-09 | 2.21245E-12 | Cox7c | EN_4 |
| 4.65399E-12 | 1.043469587 | 0.923 | 0.899 | 1.0989E-07 | 1.49479E-12 | 1.02155056 | 0.956 | 0.907 | 3.5295E-08 | 4.65399E-12 | 2.98959E-12 | Ppia | EN_4 |
| 3.3542E-12 | 1.509851484 | 0.879 | 0.828 | 7.91993E-08 | 4.20809E-11 | 1.664017635 | 0.789 | 0.855 | 9.93613E-07 | 4.20809E-11 | 6.70839E-12 | Chchd2 | EN_4 |
| 4.81593E-09 | 1.240775755 | 0.868 | 0.835 | 0.000113714 | 7.00306E-12 | 1.312624887 | 0.9 | 0.877 | 1.65356E-07 | 4.81593E-09 | 1.40061E-11 | S100b | EN_4 |
| 1.36775E-11 | 0.916306421 | 0.934 | 0.903 | 3.22954E-07 | 2.89293E-09 | 0.81414914 | 0.911 | 0.931 | 6.83078E-05 | 2.89293E-09 | 2.73551E-11 | Rpl13 | EN_4 |
| 1.40163E-11 | 1.399733179 | 0.89 | 0.833 | 3.30953E-07 | 2.64022E-11 | 1.38160314 | 0.856 | 0.879 | 6.23409E-07 | 2.64022E-11 | 2.80326E-11 | Selenom | EN_4 |
| 1.56757E-11 | 1.049755706 | 0.857 | 0.806 | 3.70136E-07 | 1.05717E-07 | 0.988972618 | 0.833 | 0.874 | 0.002496191 | 1.05717E-07 | 3.13515E-11 | Rpl34 | EN_4 |
| 3.36475E-11 | 1.133220425 | 0.857 | 0.824 | 7.94485E-07 | 1.76678E-11 | 1.149030417 | 0.856 | 0.872 | 4.17171E-07 | 3.36475E-11 | 3.53355E-11 | Rplp2 | EN_4 |
| 3.53596E-11 | 0.938542987 | 0.912 | 0.885 | 8.3491E-07 | 3.96946E-10 | 1.009834821 | 0.844 | 0.903 | 9.37269E-06 | 3.96946E-10 | 7.07191E-11 | Rpl23 | EN_4 |

| | | | | | | | | | | | | | |
|---|---|---|---|---|---|---|---|---|---|---|---|---|---|
| 4.01484E-11 | 1.097476105 | 0.978 | 0.896 | 9.47985E-07 | 4.6146E-08 | 0.936610093 | 0.956 | 0.909 | 0.001089599 | 4.6146E-08 | 8.02968E-11 | Ubb | EN_4 |
| 7.46335E-11 | 1.378957211 | 0.791 | 0.775 | 1.76225E-06 | 1.44128E-08 | 1.520411066 | 0.744 | 0.799 | 0.000340316 | 1.44128E-08 | 1.49267E-10 | Ndufa4 | EN_4 |
| 1.32111E-09 | 1.998007751 | 0.692 | 0.65 | 3.1194E-05 | 1.09223E-10 | 1.925123274 | 0.711 | 0.719 | 2.57897E-06 | 1.32111E-09 | 2.18445E-10 | Atp5g2 | EN_4 |
| 6.44492E-10 | 1.17182943 | 0.835 | 0.852 | 1.52177E-05 | 5.39241E-09 | 1.296407248 | 0.822 | 0.84 | 0.000127326 | 5.39241E-09 | 1.28898E-09 | Rps4x | EN_4 |
| 6.93654E-10 | 0.987821846 | 0.89 | 0.877 | 1.63786E-05 | 2.53436E-09 | 1.057115214 | 0.822 | 0.892 | 5.98413E-05 | 2.53436E-09 | 1.38731E-09 | Rpl19 | EN_4 |
| 8.83491E-10 | 0.960286478 | 0.912 | 0.883 | 2.0861E-05 | 2.62105E-08 | 0.983945479 | 0.833 | 0.887 | 0.000618882 | 2.62105E-08 | 1.76698E-09 | Ifitm3 | EN_4 |
| 4.89228E-09 | 0.926349959 | 0.89 | 0.863 | 0.000115517 | 2.55845E-08 | 0.791456434 | 0.9 | 0.907 | 0.0006041 | 2.55845E-08 | 9.78457E-09 | Rps10 | EN_4 |
| 7.21394E-09 | 1.024174493 | 0.813 | 0.833 | 0.000170336 | 2.83364E-08 | 1.42406502 | 0.778 | 0.857 | 0.00066908 | 2.83364E-08 | 1.44279E-08 | Cox6c | EN_4 |
| | | | | | | MM | | | | | | | |
| 4.36593E-46 | 3.740905585 | 0.835 | 0.117 | 1.03088E-41 | 1.51985E-86 | 3.453990671 | 0.986 | 0.183 | 3.58866E-82 | 4.36593E-46 | 3.03969E-86 | Fcna | MM_1 |
| 2.66202E-49 | 4.015164829 | 0.826 | 0.083 | 6.28556E-45 | 3.1306E-85 | 3.459543987 | 0.977 | 0.191 | 7.39198E-81 | 2.66202E-49 | 6.26121E-85 | Ednrb | MM_1 |
| 3.97099E-55 | 4.013921325 | 0.954 | 0.159 | 9.37631E-51 | 2.50995E-80 | 2.999341502 | 1 | 0.316 | 5.92649E-76 | 3.97099E-55 | 5.0199E-80 | Lyve1 | MM_1 |
| 3.48426E-41 | 2.275408385 | 0.982 | 0.514 | 8.22704E-37 | 6.00195E-76 | 2.406947334 | 1 | 0.616 | 1.41718E-71 | 3.48426E-41 | 1.20039E-75 | F13a1 | MM_1 |
| 7.21621E-47 | 2.776643879 | 0.945 | 0.193 | 1.70389E-42 | 7.40107E-73 | 2.333290984 | 0.991 | 0.33 | 1.74754E-68 | 7.21621E-47 | 1.48021E-72 | Folr2 | MM_1 |
| 2.87794E-43 | 2.284290565 | 0.908 | 0.169 | 6.79539E-39 | 1.19381E-70 | 2.694429001 | 0.932 | 0.21 | 2.81882E-66 | 2.87794E-43 | 2.38762E-70 | Plekhg5 | MM_1 |
| 1.08642E-34 | 2.012101993 | 1 | 0.641 | 2.56526E-30 | 4.38549E-69 | 2.026524178 | 1 | 0.703 | 1.0355E-64 | 1.08642E-34 | 8.77097E-69 | Cd36 | MM_1 |
| 1.07354E-38 | 2.731948453 | 0.899 | 0.217 | 2.53485E-34 | 3.00745E-68 | 2.610620892 | 0.959 | 0.243 | 7.10118E-64 | 1.07354E-38 | 6.01489E-68 | Ccl24 | MM_1 |
| 1.33E-36 | 1.748651685 | 1 | 0.783 | 3.1404E-32 | 3.6591E-65 | 1.660852889 | 1 | 0.828 | 8.63987E-61 | 1.33E-36 | 7.3182E-65 | Selenop | MM_1 |
| 3.08171E-30 | 1.643822718 | 0.936 | 0.414 | 7.27653E-26 | 3.08994E-64 | 2.08246562 | 0.986 | 0.499 | 7.29595E-60 | 3.08171E-30 | 6.17987E-64 | Ninj1 | MM_1 |
| 1.15984E-23 | 1.488547273 | 0.954 | 0.562 | 2.73862E-19 | 1.10688E-60 | 1.756392609 | 1 | 0.635 | 2.61357E-56 | 1.15984E-23 | 2.21377E-60 | Fcgrt | MM_1 |
| 4.18151E-21 | 1.156608679 | 0.972 | 0.61 | 9.87337E-17 | 2.06844E-60 | 1.751282746 | 1 | 0.744 | 4.88399E-56 | 4.18151E-21 | 4.13687E-60 | Ctsd | MM_1 |
| 8.3586E-27 | 2.138847324 | 0.716 | 0.166 | 1.97363E-22 | 1.00314E-59 | 2.693435001 | 0.856 | 0.193 | 2.36861E-55 | 8.3586E-27 | 2.00627E-59 | Tfrc | MM_1 |
| 5.0324E-30 | 2.388224307 | 0.826 | 0.248 | 1.18825E-25 | 5.71825E-59 | 2.027419838 | 0.937 | 0.292 | 1.35019E-54 | 5.0324E-30 | 1.14365E-58 | Clec10a | MM_1 |
| 7.21071E-27 | 1.558584727 | 0.972 | 0.6 | 1.70259E-22 | 1.10385E-58 | 1.868308514 | 0.991 | 0.676 | 2.6064E-54 | 7.21071E-27 | 2.20769E-58 | Mtss1 | MM_1 |
| 1.46114E-27 | 2.194315322 | 0.743 | 0.166 | 3.45005E-23 | 1.87179E-58 | 2.052727966 | 0.892 | 0.199 | 4.41966E-54 | 1.46114E-27 | 3.74357E-58 | Pdgfc | MM_1 |
| 7.47259E-34 | 1.884363327 | 0.982 | 0.479 | 1.76443E-29 | 3.9685E-58 | 1.663510944 | 1 | 0.621 | 9.37043E-54 | 7.47259E-34 | 7.937E-58 | Gas6 | MM_1 |
| 6.38749E-32 | 2.331770364 | 0.963 | 0.51 | 1.50822E-27 | 3.28511E-57 | 2.299919118 | 0.995 | 0.627 | 7.75681E-53 | 6.38749E-32 | 6.57023E-57 | Wfdc17 | MM_1 |
| 3.1741E-20 | 1.236364552 | 0.917 | 0.448 | 7.49469E-16 | 4.88831E-57 | 1.678956629 | 0.995 | 0.575 | 1.15423E-52 | 3.1741E-20 | 9.77661E-57 | Serpinb6a | MM_1 |
| 2.27099E-19 | 1.313518237 | 0.927 | 0.59 | 5.36226E-15 | 3.05105E-56 | 1.637869635 | 0.995 | 0.673 | 7.20414E-52 | 2.27099E-19 | 6.1021E-56 | Pepd | MM_1 |
| 2.60094E-30 | 1.722094318 | 0.945 | 0.307 | 6.14133E-26 | 3.87213E-56 | 1.745562876 | 0.995 | 0.447 | 9.14286E-52 | 2.60094E-30 | 7.74425E-56 | Cbr2 | MM_1 |
| 2.4661E-32 | 2.497365151 | 0.817 | 0.186 | 5.82295E-28 | 5.92841E-56 | 2.3433665 | 0.856 | 0.174 | 1.39982E-51 | 2.4661E-32 | 1.18568E-55 | C4b | MM_1 |
| 1.16148E-26 | 1.651100211 | 0.972 | 0.641 | 2.74248E-22 | 7.27752E-56 | 1.618740739 | 0.991 | 0.708 | 1.71837E-51 | 1.16148E-26 | 1.4555E-55 | Snx2 | MM_1 |
| 2.78711E-23 | 1.529773221 | 0.642 | 0.131 | 6.58094E-19 | 7.3235E-56 | 2.020885759 | 0.847 | 0.177 | 1.72922E-51 | 2.78711E-23 | 1.4647E-55 | Mgst1 | MM_1 |
| 1.03937E-23 | 1.522495792 | 0.881 | 0.338 | 2.45416E-19 | 1.05561E-55 | 1.998851233 | 0.968 | 0.428 | 2.49249E-51 | 1.03937E-23 | 2.11121E-55 | Ltc4s | MM_1 |
| 1.5547E-28 | 3.736694749 | 0.45 | 0.021 | 3.67097E-24 | 5.7918E-55 | 3.056772099 | 0.662 | 0.054 | 1.36756E-50 | 1.5547E-28 | 1.15836E-54 | Coro2b | MM_1 |
| 4.34493E-28 | 2.056914004 | 0.872 | 0.348 | 1.02592E-23 | 1.75881E-54 | 1.750325997 | 0.977 | 0.392 | 4.1529E-50 | 4.34493E-28 | 3.51762E-54 | Sulf2 | MM_1 |
| 2.43735E-23 | 4.082276222 | 0.431 | 0.041 | 5.75507E-19 | 2.18437E-54 | 2.944773149 | 0.707 | 0.093 | 5.15773E-50 | 2.43735E-23 | 4.36873E-54 | Hpse | MM_1 |
| 6.0377E-28 | 4.231339736 | 0.486 | 0.038 | 1.42562E-23 | 3.37861E-54 | 3.495510673 | 0.622 | 0.038 | 7.97756E-50 | 6.0377E-28 | 6.75721E-54 | Barx2 | MM_1 |
| 5.64454E-27 | 1.319926089 | 1 | 0.779 | 1.33279E-22 | 4.14154E-53 | 1.381278432 | 1 | 0.798 | 9.779E-49 | 5.64454E-27 | 8.28308E-53 | Grn | MM_1 |
| 5.34192E-18 | 3.839661427 | 0.339 | 0.031 | 1.26134E-13 | 3.03374E-52 | 3.361447026 | 0.644 | 0.057 | 7.16326E-48 | 5.34192E-18 | 6.06747E-52 | Dnm1 | MM_1 |
| 1.39587E-22 | 1.056862368 | 1 | 0.776 | 3.29592E-18 | 5.49348E-52 | 1.107807146 | 1 | 0.804 | 1.29712E-47 | 1.39587E-22 | 1.0987E-51 | Lamp1 | MM_1 |
| 1.32521E-26 | 1.127176004 | 1 | 0.869 | 3.12909E-22 | 7.79472E-52 | 1.188297363 | 1 | 0.891 | 1.84049E-47 | 1.32521E-26 | 1.55894E-51 | Ctsb | MM_1 |
| 1.06137E-24 | 1.544004249 | 0.991 | 0.662 | 2.50611E-20 | 3.64394E-51 | 1.843850304 | 0.995 | 0.782 | 8.60407E-47 | 1.06137E-24 | 7.28788E-51 | Ccl6 | MM_1 |
| 2.77209E-23 | 1.604023915 | 0.807 | 0.259 | 6.54547E-19 | 4.55017E-51 | 1.861739599 | 0.928 | 0.292 | 1.07439E-46 | 2.77209E-23 | 9.10035E-51 | Fgfr1 | MM_1 |
| 2.3168E-22 | 1.960386872 | 0.78 | 0.286 | 5.47043E-18 | 9.05969E-51 | 1.801490348 | 0.977 | 0.433 | 2.13917E-46 | 2.3168E-22 | 1.81194E-50 | Ecm1 | MM_1 |
| 1.16053E-24 | 2.1238118 | 0.789 | 0.276 | 2.74025E-20 | 1.25206E-50 | 2.050559919 | 0.919 | 0.332 | 2.95637E-46 | 1.16053E-24 | 2.50413E-50 | Fxyd2 | MM_1 |
| 4.60645E-21 | 1.241590435 | 0.963 | 0.528 | 1.08767E-16 | 1.59882E-50 | 1.715158615 | 0.982 | 0.613 | 3.77512E-46 | 4.60645E-21 | 3.19763E-50 | Arhgef3 | MM_1 |
| 1.42369E-19 | 1.588808814 | 0.817 | 0.328 | 3.36161E-15 | 7.52034E-50 | 1.792969921 | 0.919 | 0.294 | 1.7757E-45 | 1.42369E-19 | 1.50407E-49 | Rnasel | MM_1 |
| 1.01246E-23 | 1.454291325 | 0.927 | 0.476 | 2.39062E-19 | 1.78453E-49 | 1.545754018 | 0.973 | 0.526 | 4.21364E-45 | 1.01246E-23 | 3.56907E-49 | Stard8 | MM_1 |
| 4.37809E-30 | 1.993734905 | 0.835 | 0.221 | 1.03375E-25 | 1.89732E-49 | 1.578771997 | 0.901 | 0.275 | 4.47994E-45 | 4.37809E-30 | 3.79463E-49 | Reps2 | MM_1 |
| 4.91359E-24 | 1.455547204 | 0.963 | 0.466 | 1.1602E-19 | 3.16145E-49 | 1.517693276 | 0.995 | 0.591 | 7.46481E-45 | 4.91359E-24 | 6.32289E-49 | Pmp22 | MM_1 |
| 6.61268E-23 | 1.3042757 | 0.982 | 0.707 | 1.56139E-18 | 7.08774E-49 | 1.488378001 | 0.995 | 0.725 | 1.67356E-44 | 6.61268E-23 | 1.41755E-48 | Mctp1 | MM_1 |
| 2.13106E-24 | 2.664683458 | 0.56 | 0.09 | 5.03185E-20 | 7.41846E-49 | 1.922764187 | 0.806 | 0.183 | 1.75165E-44 | 2.13106E-24 | 1.48369E-48 | Selenbp1 | MM_1 |
| 3.83765E-22 | 1.450822394 | 0.89 | 0.448 | 9.06146E-18 | 8.63778E-49 | 1.440959079 | 0.991 | 0.537 | 2.03955E-44 | 3.83765E-22 | 1.72756E-48 | Tns1 | MM_1 |
| 1.32961E-22 | 1.228523534 | 1 | 0.652 | 3.13949E-18 | 1.25135E-48 | 1.229154583 | 1 | 0.747 | 2.9547E-44 | 1.32961E-22 | 2.50271E-48 | Timp2 | MM_1 |
| 9.81727E-21 | 1.717239165 | 0.798 | 0.307 | 2.31805E-16 | 2.20757E-47 | 1.769819727 | 0.919 | 0.376 | 5.21252E-43 | 9.81727E-21 | 4.41515E-47 | Tcn2 | MM_1 |
| 9.62992E-22 | 2.38154395 | 0.459 | 0.055 | 2.27382E-17 | 5.50028E-47 | 1.72865918 | 0.703 | 0.104 | 1.29873E-42 | 9.62992E-22 | 1.10006E-46 | Ms4a8a | MM_1 |
| 2.23835E-24 | 1.673091057 | 0.917 | 0.566 | 5.28519E-20 | 9.80896E-47 | 1.725902287 | 0.955 | 0.553 | 2.31609E-42 | 2.23835E-24 | 1.96179E-46 | Cfh | MM_1 |
| 6.15254E-22 | 1.354993678 | 0.936 | 0.472 | 1.45274E-17 | 9.86403E-47 | 1.506548305 | 0.973 | 0.51 | 2.32909E-42 | 6.15254E-22 | 1.97281E-46 | Mpp1 | MM_1 |
| 5.41265E-19 | 1.240325934 | 0.963 | 0.472 | 1.27804E-14 | 2.89057E-46 | 1.474028957 | 1 | 0.572 | 6.82523E-42 | 5.41265E-19 | 5.78115E-46 | Ccl9 | MM_1 |
| 3.28316E-26 | 1.38965184 | 0.991 | 0.648 | 7.7522E-22 | 9.86614E-46 | 1.314532868 | 1 | 0.706 | 2.32959E-41 | 3.28316E-26 | 1.97323E-45 | Maf | MM_1 |
| 3.76125E-18 | 1.119894101 | 0.991 | 0.686 | 8.88107E-14 | 2.87524E-45 | 1.295065733 | 0.995 | 0.73 | 6.78903E-41 | 3.76125E-18 | 5.75049E-45 | Tgfbi | MM_1 |
| 6.9643E-26 | 1.954800236 | 0.596 | 0.093 | 1.64441E-21 | 4.97407E-45 | 1.87510713 | 0.73 | 0.134 | 1.17448E-40 | 6.9643E-26 | 9.94814E-45 | Egfr | MM_1 |
| 1.5513E-23 | 2.213696969 | 0.55 | 0.083 | 3.66292E-19 | 5.72948E-45 | 3.042379403 | 0.635 | 0.093 | 1.35284E-40 | 1.5513E-23 | 1.1459E-44 | Adamtsl5 | MM_1 |
| 2.78552E-13 | 0.996347703 | 0.927 | 0.555 | 6.57717E-09 | 2.09671E-44 | 1.380827089 | 0.977 | 0.591 | 4.95074E-40 | 2.78552E-13 | 4.19341E-44 | Snx9 | MM_1 |
| 1.41423E-18 | 1.158796811 | 0.963 | 0.59 | 3.33927E-14 | 1.00967E-43 | 1.571428942 | 0.986 | 0.619 | 2.38403E-39 | 1.41423E-18 | 2.01933E-43 | Sgpl1 | MM_1 |
| 2.28826E-15 | 2.155874494 | 0.303 | 0.028 | 5.40303E-11 | 3.59855E-43 | 3.530397526 | 0.554 | 0.049 | 8.4969E-39 | 2.28826E-15 | 7.1971E-43 | Ifitm10 | MM_1 |
| 7.21471E-18 | 1.056831433 | 0.844 | 0.379 | 1.70354E-13 | 4.44361E-43 | 1.220710021 | 0.95 | 0.441 | 1.04922E-38 | 7.21471E-18 | 8.88721E-43 | Idh2 | MM_1 |
| 8.91572E-18 | 2.332047223 | 0.413 | 0.062 | 2.10518E-13 | 6.34251E-43 | 1.908994242 | 0.68 | 0.114 | 1.49759E-38 | 8.91572E-18 | 1.2685E-42 | Gypc | MM_1 |
| 2.68836E-17 | 0.669094213 | 0.835 | 0.324 | 6.34776E-13 | 8.87418E-43 | 1.763404135 | 0.856 | 0.292 | 2.09537E-38 | 2.68836E-17 | 1.77484E-42 | Tbc1d4 | MM_1 |
| 2.93001E-21 | 1.30140606 | 0.972 | 0.586 | 6.91834E-17 | 1.50034E-42 | 1.289571914 | 0.995 | 0.678 | 3.54261E-38 | 2.93001E-21 | 3.00069E-42 | Pltp | MM_1 |
| 4.26056E-19 | 1.748207359 | 0.651 | 0.186 | 1.006E-14 | 4.6284E-42 | 1.910817452 | 0.77 | 0.199 | 1.09286E-37 | 4.26056E-19 | 9.2568E-42 | Fam43a | MM_1 |
| 2.56884E-15 | 1.465760601 | 0.569 | 0.19 | 6.06556E-11 | 1.05214E-41 | 2.041671598 | 0.784 | 0.223 | 2.48431E-37 | 2.56884E-15 | 2.10428E-41 | Rcn3 | MM_1 |
| 8.43362E-23 | 1.289887456 | 0.991 | 0.559 | 1.99135E-18 | 1.14776E-41 | 1.31653205 | 0.991 | 0.665 | 2.71009E-37 | 8.43362E-23 | 2.29552E-41 | Psd3 | MM_1 |
| 9.3057E-17 | 1.573943203 | 0.651 | 0.214 | 2.19726E-12 | 1.35943E-41 | 2.081618552 | 0.766 | 0.207 | 3.20988E-37 | 9.3057E-17 | 2.71885E-41 | Mfsd6 | MM_1 |
| 1.65886E-19 | 3.884385099 | 0.303 | 0.01 | 3.91691E-15 | 1.74468E-41 | 2.699396399 | 0.5 | 0.027 | 4.11953E-37 | 1.65886E-19 | 3.48936E-41 | Mctp2 | MM_1 |
| 6.23201E-14 | 0.966958079 | 0.881 | 0.476 | 1.4715E-09 | 2.38199E-41 | 1.172073716 | 0.937 | 0.466 | 5.62436E-37 | 6.23201E-14 | 4.76398E-41 | Cd164 | MM_1 |
| 2.50842E-17 | 1.011959412 | 0.963 | 0.562 | 5.92288E-13 | 2.62149E-41 | 1.353679109 | 0.973 | 0.567 | 6.18985E-37 | 2.50842E-17 | 5.24297E-41 | Ctnnd1 | MM_1 |
| 6.61323E-31 | 1.489850536 | 0.982 | 0.766 | 1.56152E-26 | 2.88476E-41 | 1.128554716 | 1 | 0.801 | 6.81149E-37 | 6.61323E-31 | 5.76951E-41 | Mrc1 | MM_1 |
| 5.7184E-22 | 1.374303185 | 0.798 | 0.259 | 1.35023E-17 | 9.34086E-41 | 1.399229253 | 0.919 | 0.422 | 2.20556E-36 | 5.7184E-22 | 1.86817E-40 | Vat1 | MM_1 |
| 1.29984E-12 | 1.488552339 | 0.55 | 0.19 | 3.06918E-08 | 1.42147E-40 | 1.768582982 | 0.815 | 0.24 | 3.35637E-36 | 1.29984E-12 | 2.84293E-40 | S1pr1 | MM_1 |

| | | | | | | | | | | | | | |
|---|---|---|---|---|---|---|---|---|---|---|---|---|---|
| 3.20749E-27 | 1.74143761 | 0.991 | 0.548 | 7.57353E-23 | 1.8599E-40 | 1.282141178 | 1 | 0.586 | 4.39159E-36 | 3.20749E-27 | 3.7198E-40 | Rgl1 | MM_1 |
| 5.05527E-20 | 1.889707508 | 0.679 | 0.203 | 1.19365E-15 | 2.3857E-40 | 1.705473944 | 0.833 | 0.259 | 5.63312E-36 | 5.05527E-20 | 4.77141E-40 | Npl | MM_1 |
| 1.07488E-26 | 1.683477682 | 0.835 | 0.221 | 2.53801E-22 | 2.39426E-40 | 1.443878429 | 0.865 | 0.292 | 5.65334E-36 | 1.07488E-26 | 4.78853E-40 | Etv1 | MM_1 |
| 1.25981E-24 | 1.511735788 | 0.991 | 0.552 | 2.97467E-20 | 1.17568E-39 | 1.224955785 | 0.991 | 0.61 | 2.77601E-35 | 1.25981E-24 | 2.35136E-39 | Itsn1 | MM_1 |
| 3.85143E-22 | 1.524757985 | 0.78 | 0.224 | 9.094E-18 | 2.44956E-39 | 1.32650395 | 0.878 | 0.302 | 5.78391E-35 | 3.85143E-22 | 4.89913E-39 | Chp2 | MM_1 |
| 3.20058E-16 | 2.716885946 | 0.394 | 0.062 | 7.5572E-12 | 3.04073E-39 | 2.674418853 | 0.541 | 0.063 | 7.17977E-35 | 3.20058E-16 | 6.08146E-39 | Gprc5c | MM_1 |
| 1.86764E-12 | 1.070906232 | 0.78 | 0.403 | 4.40986E-08 | 3.37221E-39 | 1.502885909 | 0.932 | 0.45 | 7.96247E-35 | 1.86764E-12 | 6.74443E-39 | Slc43a2 | MM_1 |
| 7.78369E-23 | 1.255030794 | 0.982 | 0.683 | 1.83789E-18 | 4.21122E-39 | 1.099571042 | 1 | 0.692 | 9.94352E-35 | 7.78369E-23 | 8.42243E-39 | Wwp1 | MM_1 |
| 2.62801E-18 | 1.967130941 | 0.514 | 0.107 | 6.20525E-14 | 1.59447E-38 | 1.87452778 | 0.721 | 0.174 | 3.76487E-34 | 2.62801E-18 | 3.18895E-38 | Epb41l1 | MM_1 |
| 1.95437E-21 | 1.939310354 | 0.734 | 0.262 | 4.61465E-17 | 1.79227E-38 | 1.453611715 | 0.878 | 0.351 | 4.23192E-34 | 1.95437E-21 | 3.58455E-38 | Cln8 | MM_1 |
| 6.46301E-13 | 1.286300405 | 0.688 | 0.31 | 1.52605E-08 | 5.55534E-38 | 1.630201904 | 0.869 | 0.354 | 1.31173E-33 | 6.46301E-13 | 1.11107E-37 | Idh1 | MM_1 |
| 1.09728E-25 | 2.02672603 | 0.899 | 0.376 | 2.59089E-21 | 6.96175E-38 | 1.571454875 | 0.914 | 0.45 | 1.64381E-33 | 1.09728E-25 | 1.39235E-37 | Pros1 | MM_1 |
| 1.09491E-22 | 1.646394261 | 0.853 | 0.341 | 2.58531E-18 | 7.51338E-38 | 1.304804007 | 0.91 | 0.428 | 1.77406E-33 | 1.09491E-22 | 1.50268E-37 | Rab11fip5 | MM_1 |
| 2.03884E-15 | 0.944985913 | 0.991 | 0.631 | 4.81412E-11 | 8.82323E-38 | 1.03147728 | 1 | 0.698 | 2.08334E-33 | 2.03884E-15 | 1.76465E-37 | Ctsa | MM_1 |
| 2.569E-26 | 2.649288006 | 0.45 | 0.028 | 6.06593E-22 | 1.01505E-37 | 2.997758302 | 0.532 | 0.06 | 2.39674E-33 | 2.569E-26 | 2.0301E-37 | Cd209g | MM_1 |
| 2.00717E-24 | 1.497092925 | 0.945 | 0.462 | 4.73932E-20 | 1.04666E-37 | 1.219099489 | 0.986 | 0.518 | 2.47138E-33 | 2.00717E-24 | 2.09333E-37 | Igf1 | MM_1 |
| 3.73456E-10 | 0.843442831 | 0.826 | 0.483 | 8.81805E-06 | 1.57183E-37 | 1.493270899 | 0.946 | 0.471 | 3.71142E-33 | 3.73456E-10 | 3.14367E-37 | Lima1 | MM_1 |
| 1.46945E-09 | 1.289967357 | 0.642 | 0.324 | 3.46967E-05 | 1.62816E-37 | 1.567509617 | 0.856 | 0.351 | 3.84441E-33 | 1.46945E-09 | 3.25632E-37 | B3gnt2 | MM_1 |
| 1.05717E-16 | 1.265599262 | 0.908 | 0.5 | 2.49618E-12 | 2.27719E-37 | 1.3075761 | 0.977 | 0.567 | 5.37691E-33 | 1.05717E-16 | 4.55439E-37 | Rin2 | MM_1 |
| 4.83237E-17 | 1.311730171 | 0.872 | 0.483 | 1.14102E-12 | 3.54257E-37 | 1.465338797 | 0.919 | 0.485 | 8.36472E-33 | 4.83237E-17 | 7.08514E-37 | Mfsd1 | MM_1 |
| 1.92923E-17 | 1.099243063 | 0.963 | 0.666 | 4.5553E-13 | 4.69469E-37 | 1.111570657 | 0.986 | 0.687 | 1.10851E-32 | 1.92923E-17 | 9.38938E-37 | Aplp2 | MM_1 |
| 1.09034E-10 | 0.871369294 | 0.45 | 0.138 | 2.57451E-06 | 5.91514E-37 | 1.937865745 | 0.671 | 0.153 | 1.39668E-32 | 1.09034E-10 | 1.18303E-36 | Mindy1 | MM_1 |
| 4.63455E-17 | 1.096123161 | 0.835 | 0.448 | 1.09431E-12 | 6.1038E-37 | 1.108660591 | 0.964 | 0.646 | 1.44123E-32 | 4.63455E-17 | 1.22076E-36 | Aldh2 | MM_1 |
| 5.00696E-13 | 0.814100225 | 0.908 | 0.552 | 1.18224E-08 | 6.8422E-37 | 1.282662071 | 0.968 | 0.575 | 1.61558E-32 | 5.00696E-13 | 1.36844E-36 | Zdhhc14 | MM_1 |
| 2.93281E-22 | 2.156798953 | 0.716 | 0.241 | 6.92495E-18 | 1.22409E-36 | 1.875664297 | 0.829 | 0.322 | 2.89032E-32 | 2.93281E-22 | 2.44818E-36 | Rasgrp3 | MM_1 |
| 2.18592E-14 | 0.891521199 | 0.917 | 0.528 | 5.16139E-10 | 1.30609E-36 | 1.166447809 | 0.959 | 0.537 | 3.08393E-32 | 2.18592E-14 | 2.61217E-36 | Lrp6 | MM_1 |
| 2.3073E-12 | 1.187459979 | 0.752 | 0.341 | 5.44799E-08 | 1.60523E-36 | 1.683650203 | 0.874 | 0.365 | 3.79028E-32 | 2.3073E-12 | 3.21047E-36 | Fam219a | MM_1 |
| 2.96655E-17 | 1.269250743 | 0.752 | 0.303 | 7.00462E-13 | 1.68749E-36 | 1.555547062 | 0.838 | 0.305 | 3.98451E-32 | 2.96655E-17 | 3.37499E-36 | Pikfyve | MM_1 |
| 2.79361E-10 | 1.186709246 | 0.798 | 0.497 | 6.59628E-06 | 2.61595E-36 | 1.151351733 | 0.955 | 0.594 | 6.17677E-32 | 2.79361E-10 | 5.23189E-36 | Ap2m1 | MM_1 |
| 9.44729E-18 | 1.788254413 | 0.523 | 0.114 | 2.23069E-13 | 3.07699E-36 | 2.775870979 | 0.586 | 0.112 | 7.26539E-32 | 9.44729E-18 | 6.15398E-36 | Timd4 | MM_1 |
| 3.29082E-21 | 1.323063535 | 0.945 | 0.448 | 7.77029E-17 | 6.00964E-36 | 1.031657684 | 0.955 | 0.518 | 1.419E-31 | 3.29082E-21 | 1.20193E-35 | Tpp1 | MM_1 |
| 9.19979E-10 | 0.704151687 | 0.523 | 0.2 | 2.17225E-05 | 7.22013E-36 | 1.749533654 | 0.784 | 0.286 | 1.70482E-31 | 9.19979E-10 | 1.44403E-35 | Prxl2b | MM_1 |
| 1.56829E-11 | 2.078211679 | 0.413 | 0.124 | 3.70304E-07 | 8.22E-36 | 2.687342568 | 0.568 | 0.095 | 1.94091E-31 | 1.56829E-11 | 1.644E-35 | Fam118a | MM_1 |
| 2.14672E-25 | 1.402466079 | 0.945 | 0.369 | 5.06882E-21 | 8.57363E-36 | 1.271554147 | 0.937 | 0.471 | 2.02441E-31 | 2.14672E-25 | 1.71473E-35 | Serpinb8 | MM_1 |
| 1.41795E-24 | 2.475057535 | 0.514 | 0.066 | 3.34806E-20 | 1.45742E-35 | 2.504914492 | 0.59 | 0.104 | 3.44126E-31 | 1.41795E-24 | 2.91484E-35 | Cd209f | MM_1 |
| 1.36361E-11 | 0.780685879 | 0.55 | 0.19 | 3.21977E-07 | 1.5046E-35 | 1.459700951 | 0.811 | 0.264 | 3.55265E-31 | 1.36361E-11 | 3.00919E-35 | Emilin2 | MM_1 |
| 1.15265E-17 | 1.386207456 | 0.881 | 0.466 | 2.72164E-13 | 2.48287E-35 | 1.366321326 | 0.946 | 0.441 | 5.86255E-31 | 1.15265E-17 | 4.96574E-35 | Nrp2 | MM_1 |
| 5.33989E-22 | 1.514544532 | 0.954 | 0.507 | 1.26086E-17 | 2.55109E-35 | 1.160272708 | 0.977 | 0.564 | 6.02364E-31 | 5.33989E-22 | 5.10219E-35 | Stab1 | MM_1 |
| 5.58054E-18 | 1.516296943 | 0.78 | 0.297 | 1.31768E-13 | 3.93124E-35 | 1.172202449 | 0.883 | 0.392 | 9.28244E-31 | 5.58054E-18 | 7.86248E-35 | Cmah | MM_1 |
| 1.23266E-18 | 1.708927793 | 0.807 | 0.345 | 2.91056E-14 | 4.2986E-35 | 1.369213914 | 0.865 | 0.417 | 1.01498E-30 | 1.23266E-18 | 8.59719E-35 | Klhl9 | MM_1 |
| 5.67884E-12 | 1.267764912 | 0.303 | 0.052 | 1.34089E-07 | 4.32793E-35 | 2.891367093 | 0.536 | 0.079 | 1.02191E-30 | 5.67884E-12 | 8.65587E-35 | Sema6a | MM_1 |
| 4.07197E-16 | 0.918700661 | 0.945 | 0.538 | 9.61473E-12 | 6.86949E-35 | 1.280433769 | 0.959 | 0.556 | 1.62202E-30 | 4.07197E-16 | 1.3739E-34 | Snx8 | MM_1 |
| 1.31693E-19 | 1.332255492 | 0.872 | 0.393 | 3.10954E-15 | 7.08957E-35 | 1.127708663 | 0.982 | 0.537 | 1.67399E-30 | 1.31693E-19 | 1.41791E-34 | Rnase4 | MM_1 |
| 1.03405E-13 | 0.662657204 | 0.817 | 0.403 | 2.4416E-09 | 7.24116E-35 | 1.262464077 | 0.887 | 0.392 | 1.70978E-30 | 1.03405E-13 | 1.44823E-34 | Usp24 | MM_1 |
| 8.50685E-16 | 1.1436723 | 0.945 | 0.669 | 2.00864E-11 | 9.79915E-35 | 1.341587077 | 0.973 | 0.695 | 2.31377E-30 | 8.50685E-16 | 1.95983E-34 | Osbpl9 | MM_1 |
| 1.00679E-21 | 1.444695651 | 0.917 | 0.441 | 2.37724E-17 | 1.24093E-34 | 1.184033426 | 0.973 | 0.531 | 2.93008E-30 | 1.00679E-21 | 2.48185E-34 | Cfp | MM_1 |
| 6.20185E-16 | 0.838828554 | 0.982 | 0.569 | 1.46438E-11 | 2.13946E-34 | 1.190069382 | 0.995 | 0.657 | 5.0517E-30 | 6.20185E-16 | 4.27892E-34 | Ctsl | MM_1 |
| 1.82787E-14 | 2.343372051 | 0.532 | 0.162 | 4.31597E-10 | 3.08225E-34 | 2.023597134 | 0.811 | 0.327 | 7.2778E-30 | 1.82787E-14 | 6.16449E-34 | Retnla | MM_1 |
| 1.92531E-14 | 1.005801502 | 0.927 | 0.648 | 4.54604E-10 | 3.51627E-34 | 1.131920682 | 0.977 | 0.668 | 8.30261E-30 | 1.92531E-14 | 7.03254E-34 | Eps15 | MM_1 |
| 5.55022E-16 | 1.396591889 | 0.945 | 0.545 | 1.31052E-11 | 3.57494E-34 | 1.577687707 | 0.968 | 0.627 | 8.44114E-30 | 5.55022E-16 | 7.14987E-34 | Rhob | MM_1 |
| 6.6718E-16 | 1.025953173 | 0.982 | 0.717 | 1.57535E-11 | 4.85651E-34 | 1.213253743 | 0.995 | 0.779 | 1.14672E-29 | 6.6718E-16 | 9.71301E-34 | Gas7 | MM_1 |
| 8.40839E-11 | 1.472882863 | 0.45 | 0.141 | 1.98539E-06 | 5.22829E-34 | 2.040590889 | 0.644 | 0.15 | 1.2345E-29 | 8.40839E-11 | 1.04566E-33 | Cdc14b | MM_1 |
| 3.04009E-23 | 2.027968682 | 0.716 | 0.197 | 7.17827E-19 | 7.94069E-34 | 1.558347915 | 0.775 | 0.245 | 1.87496E-29 | 3.04009E-23 | 1.58814E-33 | Pde2a | MM_1 |
| 3.73698E-16 | 0.950635761 | 0.927 | 0.424 | 8.82376E-12 | 9.50974E-34 | 1.389780238 | 0.95 | 0.474 | 2.24544E-29 | 3.73698E-16 | 1.90195E-33 | Mertk | MM_1 |
| 6.85203E-13 | 1.215812602 | 0.725 | 0.359 | 1.6179E-08 | 1.23642E-33 | 1.54783926 | 0.815 | 0.316 | 2.91944E-29 | 6.85203E-13 | 2.47284E-33 | Agfg1 | MM_1 |
| 6.65301E-12 | 0.832267629 | 0.945 | 0.648 | 1.57091E-07 | 1.4021E-33 | 1.032072415 | 0.986 | 0.662 | 3.31064E-29 | 6.65301E-12 | 2.8042E-33 | Tut7 | MM_1 |
| 6.49184E-09 | 0.868635327 | 0.578 | 0.29 | 0.000153285 | 2.0603E-33 | 1.501115794 | 0.802 | 0.281 | 4.86479E-29 | 6.49184E-09 | 4.12061E-33 | Jup | MM_1 |
| 2.10571E-17 | 2.245484339 | 0.495 | 0.103 | 4.97201E-13 | 2.96406E-33 | 1.909781083 | 0.64 | 0.134 | 6.99875E-29 | 2.10571E-17 | 5.92813E-33 | Slc40a1 | MM_1 |
| 1.52863E-09 | 0.729111257 | 0.927 | 0.617 | 3.60941E-05 | 3.95231E-33 | 0.952802795 | 0.991 | 0.676 | 9.3322E-29 | 1.52863E-09 | 7.90463E-33 | St13 | MM_1 |
| 9.13637E-16 | 2.033973189 | 0.514 | 0.131 | 2.15728E-11 | 5.32211E-33 | 1.468637328 | 0.698 | 0.191 | 1.25666E-28 | 9.13637E-16 | 1.06442E-32 | Tmod1 | MM_1 |
| 2.79735E-17 | 1.859121504 | 0.624 | 0.183 | 6.6051E-13 | 5.80054E-33 | 1.685016621 | 0.802 | 0.316 | 1.36962E-28 | 2.79735E-17 | 1.16011E-32 | Alox5 | MM_1 |
| 1.97526E-13 | 0.821501252 | 0.936 | 0.603 | 4.66398E-09 | 6.77957E-33 | 1.063038635 | 0.973 | 0.638 | 1.60079E-28 | 1.97526E-13 | 1.35591E-32 | Fkbp1a | MM_1 |
| 1.1806E-11 | 0.874707706 | 0.853 | 0.552 | 2.78762E-07 | 8.5229E-33 | 1.332550078 | 0.901 | 0.469 | 2.01243E-28 | 1.1806E-11 | 1.70458E-32 | Arap1 | MM_1 |
| 6.06212E-24 | 1.526491206 | 0.963 | 0.555 | 1.43139E-19 | 1.17948E-32 | 1.216390758 | 0.986 | 0.635 | 2.78499E-28 | 6.06212E-24 | 2.35896E-32 | Glul | MM_1 |
| 2.55835E-15 | 1.224511808 | 0.752 | 0.303 | 6.04078E-11 | 1.61265E-32 | 1.348105822 | 0.869 | 0.381 | 3.80779E-28 | 2.55835E-15 | 3.2253E-32 | Mfsd11 | MM_1 |
| 4.42864E-12 | 0.866615381 | 0.89 | 0.541 | 1.04569E-07 | 2.42475E-32 | 0.95543954 | 0.959 | 0.599 | 5.72531E-28 | 4.42864E-12 | 4.84949E-32 | Snx6 | MM_1 |
| 1.04722E-14 | 1.042249653 | 0.835 | 0.41 | 2.4727E-10 | 2.64961E-32 | 1.350624197 | 0.923 | 0.466 | 6.25626E-28 | 1.04722E-14 | 5.29922E-32 | Tmem106a | MM_1 |
| 5.22015E-12 | 1.768399965 | 0.477 | 0.148 | 1.23258E-07 | 3.00995E-32 | 1.299960381 | 0.685 | 0.177 | 7.1071E-28 | 5.22015E-12 | 6.01991E-32 | Tslp | MM_1 |
| 3.67766E-20 | 1.076283507 | 1 | 0.745 | 8.6837E-16 | 3.70483E-32 | 1.008402812 | 1 | 0.796 | 8.74786E-28 | 3.67766E-20 | 7.40967E-32 | Lgmn | MM_1 |
| 1.09247E-11 | 0.97609486 | 0.917 | 0.597 | 2.57955E-07 | 6.12792E-32 | 1.134755893 | 0.977 | 0.629 | 1.44693E-27 | 1.09247E-11 | 1.22558E-31 | Eps8 | MM_1 |
| 1.56256E-25 | 2.556399113 | 0.743 | 0.2 | 3.68951E-21 | 9.86364E-32 | 1.967627908 | 0.721 | 0.207 | 2.329E-27 | 1.56256E-25 | 1.97273E-31 | Ccl8 | MM_1 |
| 4.38375E-24 | 1.884686636 | 0.945 | 0.562 | 1.03509E-19 | 1.12329E-31 | 1.236559005 | 0.995 | 0.73 | 2.65232E-27 | 4.38375E-24 | 2.24659E-31 | Klf2 | MM_1 |
| 1.17548E-10 | 0.808943635 | 0.899 | 0.614 | 2.77553E-06 | 1.15351E-31 | 1.012567093 | 0.991 | 0.654 | 2.72367E-27 | 1.17548E-10 | 2.30702E-31 | Scpep1 | MM_1 |
| 7.33625E-16 | 2.148963836 | 0.367 | 0.055 | 1.73223E-11 | 1.87655E-31 | 2.119240855 | 0.559 | 0.114 | 4.43092E-27 | 7.33625E-16 | 3.75311E-31 | Cacna1a | MM_1 |
| 1.78765E-16 | 0.910375231 | 0.954 | 0.538 | 4.221E-12 | 1.9077E-31 | 1.152596488 | 0.964 | 0.575 | 4.50446E-27 | 1.78765E-16 | 3.8154E-31 | Dse | MM_1 |
| 2.01429E-14 | 1.825329855 | 0.413 | 0.083 | 4.75615E-10 | 2.22079E-31 | 2.570670765 | 0.514 | 0.084 | 5.24373E-27 | 2.01429E-14 | 4.44158E-31 | Mcoln2 | MM_1 |
| 1.8073E-08 | 1.085795929 | 0.431 | 0.162 | 0.00042674 | 2.25701E-31 | 1.33161423 | 0.784 | 0.243 | 5.32926E-27 | 1.8073E-08 | 4.51402E-31 | Nupr1 | MM_1 |
| 6.91968E-17 | 1.534084126 | 0.78 | 0.355 | 1.63387E-12 | 2.87738E-31 | 1.669094184 | 0.829 | 0.444 | 6.79406E-27 | 6.91968E-17 | 5.75476E-31 | Cd163 | MM_1 |
| 5.52834E-12 | 0.911746017 | 0.789 | 0.445 | 1.30535E-07 | 2.95092E-31 | 1.299192889 | 0.887 | 0.45 | 6.96772E-27 | 5.52834E-12 | 5.90185E-31 | Cpeb4 | MM_1 |
| 1.99464E-16 | 1.29748553 | 0.67 | 0.214 | 4.70975E-12 | 3.41583E-31 | 1.81012151 | 0.689 | 0.207 | 8.06546E-27 | 1.99464E-16 | 6.83166E-31 | Ralgds | MM_1 |
| 1.87664E-21 | 1.009041636 | 1 | 0.831 | 4.43113E-17 | 6.19244E-31 | 0.950761171 | 1 | 0.858 | 1.46216E-26 | 1.87664E-21 | 1.23849E-30 | Rbpj | MM_1 |

| | | | | | | | | | | | | | |
|---|---|---|---|---|---|---|---|---|---|---|---|---|---|
| 2.59534E-19 | 1.069744499 | 0.991 | 0.7 | 6.12811E-15 | 7.64398E-31 | 0.897840981 | 1 | 0.755 | 1.8049E-26 | 2.59534E-19 | 1.5288E-30 | Dab2 | MM_1 |
| 8.95478E-10 | 0.693044627 | 0.982 | 0.745 | 2.1144E-05 | 1.03424E-30 | 0.91936551 | 1 | 0.703 | 2.44204E-26 | 8.95478E-10 | 2.06847E-30 | Akr1a1 | MM_1 |
| 8.40488E-17 | 1.392130921 | 0.743 | 0.317 | 1.98456E-12 | 1.75376E-30 | 0.992559051 | 0.856 | 0.376 | 4.14097E-26 | 8.40488E-17 | 3.50751E-30 | Tpcn1 | MM_1 |
| 4.74838E-15 | 1.754642548 | 0.716 | 0.321 | 1.12119E-10 | 2.06496E-30 | 1.806729862 | 0.815 | 0.36 | 4.87579E-26 | 4.74838E-15 | 4.12993E-30 | Emp1 | MM_1 |
| 3.39056E-12 | 1.056061246 | 0.706 | 0.29 | 8.0058E-08 | 2.43773E-30 | 1.607803271 | 0.752 | 0.264 | 5.75597E-26 | 3.39056E-12 | 4.87546E-30 | Cmklr1 | MM_1 |
| 8.92614E-17 | 1.031299401 | 0.908 | 0.5 | 2.10764E-12 | 2.5578E-30 | 1.064690054 | 0.95 | 0.559 | 6.03948E-26 | 8.92614E-17 | 5.1156E-30 | Kif1b | MM_1 |
| 3.33154E-07 | 0.903803501 | 0.697 | 0.403 | 0.007866421 | 2.60438E-30 | 1.418675517 | 0.847 | 0.401 | 6.14946E-26 | 3.33154E-07 | 5.20875E-30 | Col4a3bp | MM_1 |
| 1.35231E-11 | 0.811801999 | 0.789 | 0.383 | 3.19307E-07 | 2.8163E-30 | 1.213838063 | 0.901 | 0.444 | 6.64986E-26 | 1.35231E-11 | 5.63261E-30 | Evi5 | MM_1 |
| 1.61668E-10 | 0.703099079 | 0.321 | 0.066 | 3.8173E-06 | 3.18678E-30 | 1.965453498 | 0.495 | 0.079 | 7.52462E-26 | 1.61668E-10 | 6.37356E-30 | Enpep | MM_1 |
| 7.06314E-10 | 1.019015778 | 0.789 | 0.462 | 1.66775E-05 | 3.37768E-30 | 1.297273218 | 0.874 | 0.455 | 7.97537E-26 | 7.06314E-10 | 6.75535E-30 | Slc23a2 | MM_1 |
| 1.35475E-08 | 1.833411551 | 0.358 | 0.114 | 0.000319884 | 3.46479E-30 | 1.47536737 | 0.563 | 0.123 | 8.18106E-26 | 1.35475E-08 | 6.92958E-30 | Bst1 | MM_1 |
| 2.22873E-09 | 0.701428551 | 0.927 | 0.7 | 5.26249E-05 | 3.58837E-30 | 1.028794346 | 0.973 | 0.728 | 8.47287E-26 | 2.22873E-09 | 7.17675E-30 | Cd68 | MM_1 |
| 5.06935E-14 | 0.82843069 | 0.817 | 0.366 | 1.19697E-09 | 4.51767E-30 | 1.20074721 | 0.887 | 0.42 | 1.06671E-25 | 5.06935E-14 | 9.03533E-30 | Arhgap10 | MM_1 |
| 4.52342E-14 | 0.808199233 | 0.991 | 0.679 | 1.06807E-09 | 7.65444E-30 | 1.064649363 | 0.977 | 0.67 | 1.80737E-25 | 4.52342E-14 | 1.53089E-29 | BC005537 | MM_1 |
| 2.64726E-07 | 0.636609767 | 0.587 | 0.31 | 0.00625071 | 8.47267E-30 | 1.443163564 | 0.802 | 0.362 | 2.00057E-25 | 2.64726E-07 | 1.69453E-29 | Stom | MM_1 |
| 1.51974E-16 | 0.903510571 | 1 | 0.707 | 3.58841E-12 | 8.62077E-30 | 0.993129062 | 0.991 | 0.714 | 2.03554E-25 | 1.51974E-16 | 1.72415E-29 | Frmd4b | MM_1 |
| 1.7999E-09 | 1.201381928 | 0.486 | 0.186 | 4.24993E-05 | 8.73015E-30 | 1.551729458 | 0.689 | 0.204 | 2.06136E-25 | 1.7999E-09 | 1.74603E-29 | P2ry13 | MM_1 |
| 1.86676E-12 | 1.079932306 | 0.826 | 0.483 | 4.40779E-08 | 9.69579E-30 | 1.193172332 | 0.91 | 0.507 | 2.28937E-25 | 1.86676E-12 | 1.93916E-29 | Adam15 | MM_1 |
| 1.04453E-07 | 0.668851686 | 0.771 | 0.466 | 0.002466346 | 1.06043E-29 | 1.103818685 | 0.91 | 0.482 | 2.50389E-25 | 1.04453E-07 | 2.12086E-29 | Rybp | MM_1 |
| 9.38173E-21 | 2.474402636 | 0.459 | 0.059 | 2.21521E-16 | 1.08987E-29 | 1.804390673 | 0.509 | 0.09 | 2.57339E-25 | 9.38173E-21 | 2.17973E-29 | Rgs7bp | MM_1 |
| 6.92463E-25 | 1.049433913 | 1 | 0.886 | 1.63504E-20 | 1.54615E-29 | 0.784186201 | 1 | 0.877 | 3.65078E-25 | 6.92463E-25 | 3.09231E-29 | Serinc3 | MM_1 |
| 2.61496E-11 | 0.758924274 | 0.817 | 0.441 | 6.17443E-07 | 1.94003E-29 | 1.24637417 | 0.878 | 0.411 | 4.5808E-25 | 2.61496E-11 | 3.88006E-29 | Anxa6 | MM_1 |
| 5.90123E-17 | 1.04868951 | 0.881 | 0.466 | 1.3934E-12 | 2.21874E-29 | 1.377789268 | 0.856 | 0.422 | 5.23888E-25 | 5.90123E-17 | 4.43748E-29 | Clcn5 | MM_1 |
| 1.89475E-18 | 1.064346889 | 1 | 0.659 | 4.47389E-14 | 2.64229E-29 | 0.974974913 | 1 | 0.76 | 6.23897E-25 | 1.89475E-18 | 5.28458E-29 | Pf4 | MM_1 |
| 5.2588E-18 | 1.113948155 | 0.963 | 0.714 | 1.24171E-13 | 2.87974E-29 | 0.993852767 | 0.991 | 0.738 | 6.79964E-25 | 5.2588E-18 | 5.75948E-29 | Rbms1 | MM_1 |
| 2.36203E-12 | 0.768724178 | 0.835 | 0.507 | 5.57723E-08 | 3.09856E-29 | 1.031658763 | 0.959 | 0.594 | 7.31632E-25 | 2.36203E-12 | 6.19712E-29 | Lipa | MM_1 |
| 9.75996E-10 | 0.669290023 | 0.927 | 0.621 | 2.30452E-05 | 3.69127E-29 | 1.04573763 | 0.991 | 0.67 | 8.71583E-25 | 9.75996E-10 | 7.38254E-29 | Atp6v1a | MM_1 |
| 7.88027E-15 | 1.060017266 | 0.89 | 0.5 | 1.86069E-10 | 3.69447E-29 | 1.043351807 | 0.955 | 0.64 | 8.72339E-25 | 7.88027E-15 | 7.38895E-29 | Blvrb | MM_1 |
| 6.30365E-22 | 1.407302802 | 0.972 | 0.562 | 1.48842E-17 | 4.01691E-29 | 1.154057091 | 0.995 | 0.662 | 9.48474E-25 | 6.30365E-22 | 8.03383E-29 | Mafb | MM_1 |
| 6.1647E-13 | 1.733687895 | 0.505 | 0.152 | 1.45561E-08 | 4.81659E-29 | 1.430475864 | 0.622 | 0.174 | 1.13729E-24 | 6.1647E-13 | 9.63318E-29 | Maoa | MM_1 |
| 2.93442E-22 | 1.425895982 | 0.945 | 0.497 | 6.92875E-18 | 5.50796E-29 | 1.039116301 | 0.964 | 0.529 | 1.30054E-24 | 2.93442E-22 | 1.10159E-28 | Igfbp4 | MM_1 |
| 9.08611E-13 | 1.270510765 | 0.642 | 0.262 | 2.14541E-08 | 5.90195E-29 | 1.541141189 | 0.752 | 0.281 | 1.39357E-24 | 9.08611E-13 | 1.18039E-28 | Mtmr10 | MM_1 |
| 4.46974E-13 | 0.738885587 | 0.963 | 0.703 | 1.0554E-08 | 6.58492E-29 | 0.854137803 | 0.991 | 0.695 | 1.55483E-24 | 4.46974E-13 | 1.31698E-28 | Myo5a | MM_1 |
| 1.03841E-10 | 1.555066932 | 0.477 | 0.166 | 2.45189E-06 | 9.01392E-29 | 1.505952583 | 0.595 | 0.15 | 2.12837E-24 | 1.03841E-10 | 1.80278E-28 | Prkar2b | MM_1 |
| 1.45092E-08 | 0.894578182 | 0.706 | 0.403 | 0.000342592 | 9.37907E-29 | 1.104863895 | 0.901 | 0.414 | 2.21459E-24 | 1.45092E-08 | 1.87581E-28 | Add3 | MM_1 |
| 8.33887E-12 | 0.878102614 | 0.706 | 0.31 | 1.96897E-07 | 1.13913E-28 | 0.874390856 | 0.865 | 0.33 | 2.68971E-24 | 8.33887E-12 | 2.27826E-28 | Gda | MM_1 |
| 2.56144E-13 | 0.939978458 | 0.697 | 0.303 | 6.04807E-09 | 1.37025E-28 | 1.014928644 | 0.829 | 0.33 | 3.23544E-24 | 2.56144E-13 | 2.74051E-28 | Abcc1 | MM_1 |
| 4.6274E-09 | 0.871330832 | 0.587 | 0.252 | 0.000109262 | 1.51448E-28 | 1.370396741 | 0.739 | 0.267 | 3.57598E-24 | 4.6274E-09 | 3.02895E-28 | Dusp7 | MM_1 |
| 2.91365E-13 | 2.379182696 | 0.385 | 0.083 | 6.87971E-09 | 1.81476E-28 | 1.174273911 | 0.68 | 0.215 | 4.285E-24 | 2.91365E-13 | 3.62951E-28 | Dpp7 | MM_1 |
| 1.90339E-10 | 0.978523917 | 0.697 | 0.348 | 4.49428E-06 | 2.16779E-28 | 1.271339395 | 0.833 | 0.411 | 5.11858E-24 | 1.90339E-10 | 4.33557E-28 | Renbp | MM_1 |
| 3.04107E-13 | 1.64523084 | 0.807 | 0.462 | 7.18057E-09 | 2.19132E-28 | 1.523228244 | 0.941 | 0.567 | 5.17415E-24 | 3.04107E-13 | 4.38264E-28 | Klf4 | MM_1 |
| 1.44991E-17 | 0.975042808 | 0.982 | 0.707 | 3.42353E-13 | 2.50958E-28 | 0.818613471 | 0.991 | 0.774 | 5.92562E-24 | 1.44991E-17 | 5.01916E-28 | Hexa | MM_1 |
| 2.04286E-14 | 1.010458964 | 0.89 | 0.531 | 4.82361E-10 | 2.55683E-28 | 0.889811638 | 0.941 | 0.515 | 6.03718E-24 | 2.04286E-14 | 5.11366E-28 | Snx29 | MM_1 |
| 6.95635E-09 | 0.91180754 | 0.294 | 0.066 | 0.000164253 | 2.73847E-28 | 1.781938938 | 0.509 | 0.093 | 6.46608E-24 | 6.95635E-09 | 5.47694E-28 | Ankrd33b | MM_1 |
| 7.13251E-11 | 1.446804097 | 0.642 | 0.297 | 1.68413E-06 | 3.08393E-28 | 1.168068868 | 0.838 | 0.395 | 7.28177E-24 | 7.13251E-11 | 6.16785E-28 | Gabarapl1 | MM_1 |
| 2.40229E-15 | 1.325871307 | 0.908 | 0.541 | 5.67228E-11 | 3.89758E-28 | 0.945881854 | 0.964 | 0.537 | 9.20296E-24 | 2.40229E-15 | 7.79515E-28 | Fchsd2 | MM_1 |
| 8.22029E-14 | 0.676292122 | 0.734 | 0.293 | 1.94098E-09 | 4.03219E-28 | 0.655853994 | 0.892 | 0.39 | 9.52081E-24 | 8.22029E-14 | 8.06438E-28 | Gstm1 | MM_1 |
| 1.9032E-11 | 1.334360466 | 0.431 | 0.124 | 4.49385E-07 | 5.39117E-28 | 1.793667143 | 0.599 | 0.15 | 1.27296E-23 | 1.9032E-11 | 1.07823E-27 | Thrb | MM_1 |
| 1.09579E-16 | 0.940461106 | 0.972 | 0.59 | 2.58739E-12 | 5.4428E-28 | 1.008412785 | 0.995 | 0.632 | 1.28515E-23 | 1.09579E-16 | 1.08856E-27 | Abca1 | MM_1 |
| 1.62935E-15 | 0.970590306 | 0.78 | 0.3 | 3.84721E-11 | 5.53129E-28 | 1.249117439 | 0.833 | 0.354 | 1.30605E-23 | 1.62935E-15 | 1.10626E-27 | Samd4 | MM_1 |
| 3.61594E-07 | 1.089095571 | 0.147 | 0.017 | 0.008537954 | 6.91358E-28 | 3.33928097 | 0.36 | 0.025 | 1.63243E-23 | 3.61594E-07 | 1.38272E-27 | Akr1b8 | MM_1 |
| 9.4862E-13 | 1.296895544 | 0.734 | 0.366 | 2.23988E-08 | 7.55904E-28 | 1.155047365 | 0.865 | 0.414 | 1.78484E-23 | 9.4862E-13 | 1.51181E-27 | Dnase2a | MM_1 |
| 7.00834E-14 | 0.891946822 | 0.936 | 0.621 | 1.65481E-09 | 7.81588E-28 | 0.873259908 | 0.991 | 0.687 | 1.84549E-23 | 7.00834E-14 | 1.56318E-27 | Iqgap2 | MM_1 |
| 1.45636E-20 | 1.983766113 | 0.642 | 0.183 | 3.43876E-16 | 8.30211E-28 | 1.237790732 | 0.712 | 0.267 | 1.9603E-23 | 1.45636E-20 | 1.66042E-27 | Agtrap | MM_1 |
| 4.7768E-13 | 0.790041017 | 0.835 | 0.469 | 1.1279E-08 | 8.43256E-28 | 1.116146811 | 0.923 | 0.523 | 1.9911E-23 | 4.7768E-13 | 1.68651E-27 | Sema4a | MM_1 |
| 4.9378E-09 | 0.761771893 | 0.826 | 0.534 | 0.000116591 | 1.12394E-27 | 0.955601282 | 0.941 | 0.567 | 2.65384E-23 | 4.9378E-09 | 2.24788E-27 | Rab11a | MM_1 |
| 1.82341E-10 | 0.94339859 | 0.642 | 0.303 | 4.30544E-06 | 1.53686E-27 | 0.974555928 | 0.847 | 0.42 | 3.62884E-23 | 1.82341E-10 | 3.07372E-27 | Npc1 | MM_1 |
| 4.9639E-13 | 0.933541996 | 0.862 | 0.441 | 1.17208E-08 | 1.8797E-27 | 0.754415226 | 0.959 | 0.531 | 4.43835E-23 | 4.9639E-13 | 3.7594E-27 | Hacd4 | MM_1 |
| 3.62235E-19 | 1.044917268 | 1 | 0.814 | 8.55308E-15 | 2.15248E-27 | 0.817283779 | 1 | 0.82 | 5.08243E-23 | 3.62235E-19 | 4.30495E-27 | Cltc | MM_1 |
| 9.15074E-21 | 1.534935474 | 0.927 | 0.441 | 2.16067E-16 | 2.32015E-27 | 1.097094568 | 0.964 | 0.55 | 5.47835E-23 | 9.15074E-21 | 4.64031E-27 | Hmox1 | MM_1 |
| 3.43188E-14 | 1.105706696 | 0.862 | 0.452 | 8.10336E-10 | 2.92681E-27 | 1.059617784 | 0.901 | 0.477 | 6.91078E-23 | 3.43188E-14 | 5.85362E-27 | Rell1 | MM_1 |
| 5.51878E-15 | 1.030896399 | 0.881 | 0.39 | 1.30309E-10 | 3.57615E-27 | 1.031427676 | 0.932 | 0.46 | 8.444E-23 | 5.51878E-15 | 7.15229E-27 | Mir99ahg | MM_1 |
| 1.67157E-13 | 1.491035808 | 0.578 | 0.2 | 3.9469E-09 | 4.01218E-27 | 1.869356882 | 0.644 | 0.204 | 9.47355E-23 | 1.67157E-13 | 8.02436E-27 | Rasal2 | MM_1 |
| 2.05925E-12 | 1.014559285 | 0.706 | 0.31 | 4.86231E-08 | 5.21534E-27 | 0.874718376 | 0.86 | 0.387 | 1.23145E-22 | 2.05925E-12 | 1.04307E-26 | Xdh | MM_1 |
| 1.67919E-09 | 1.256348799 | 0.266 | 0.048 | 3.9649E-05 | 5.3496E-27 | 2.334405725 | 0.455 | 0.071 | 1.26315E-22 | 1.67919E-09 | 1.06992E-26 | Cela1 | MM_1 |
| 3.00909E-09 | 0.622455401 | 0.89 | 0.534 | 7.10507E-05 | 5.48969E-27 | 0.926511624 | 0.946 | 0.531 | 1.29623E-22 | 3.00909E-09 | 1.09794E-26 | Dusp3 | MM_1 |
| 1.87286E-13 | 2.165792623 | 0.44 | 0.107 | 4.4222E-09 | 7.50642E-27 | 1.922641982 | 0.604 | 0.169 | 1.77242E-22 | 1.87286E-13 | 1.50128E-26 | Cnrip1 | MM_1 |
| 3.12457E-18 | 1.651352918 | 0.725 | 0.231 | 7.37774E-14 | 1.39369E-26 | 1.370971817 | 0.806 | 0.332 | 3.29077E-22 | 3.12457E-18 | 2.78737E-26 | Mt2 | MM_1 |
| 1.50671E-08 | 0.593121127 | 0.844 | 0.548 | 0.000355765 | 1.3971E-26 | 0.863477661 | 0.941 | 0.608 | 3.29883E-22 | 1.50671E-08 | 2.7942E-26 | Creg1 | MM_1 |
| 7.25508E-11 | 0.934029255 | 0.844 | 0.472 | 1.71307E-06 | 1.46152E-26 | 0.948426156 | 0.968 | 0.553 | 3.45093E-22 | 7.25508E-11 | 2.92303E-26 | Cndp2 | MM_1 |
| 1.20085E-08 | 0.732317022 | 0.321 | 0.09 | 0.000283546 | 1.58728E-26 | 1.820173526 | 0.581 | 0.15 | 3.74788E-22 | 1.20085E-08 | 3.17456E-26 | Mamdc2 | MM_1 |
| 8.40123E-15 | 0.988402787 | 0.927 | 0.497 | 1.9837E-10 | 1.73012E-26 | 0.921571124 | 0.955 | 0.55 | 4.08517E-22 | 8.40123E-15 | 3.46025E-26 | Cd93 | MM_1 |
| 1.31254E-13 | 0.825203485 | 0.963 | 0.714 | 3.09916E-09 | 1.75065E-26 | 0.791668395 | 0.986 | 0.719 | 4.13362E-22 | 1.31254E-13 | 3.50129E-26 | Nav1 | MM_1 |
| 1.62323E-16 | 1.133767628 | 0.908 | 0.483 | 3.83277E-12 | 1.77876E-26 | 1.026585702 | 0.946 | 0.561 | 4.2E-22 | 1.62323E-16 | 3.55751E-26 | Man1c1 | MM_1 |
| 4.56863E-12 | 0.951884015 | 0.789 | 0.41 | 1.07874E-07 | 2.35626E-26 | 1.032433568 | 0.932 | 0.51 | 5.5636E-22 | 4.56863E-12 | 4.71252E-26 | Ms4a6d | MM_1 |
| 1.59146E-11 | 1.175491297 | 0.771 | 0.424 | 3.75775E-07 | 3.35688E-26 | 1.062898587 | 0.91 | 0.49 | 7.92626E-22 | 1.59146E-11 | 6.71376E-26 | Ap3s1 | MM_1 |
| 3.13602E-08 | 0.659037947 | 0.725 | 0.397 | 0.000740477 | 5.42802E-26 | 1.110331037 | 0.896 | 0.501 | 1.28166E-21 | 3.13602E-08 | 1.0856E-25 | Card19 | MM_1 |
| 5.59317E-15 | 0.772565946 | 0.954 | 0.648 | 1.32066E-10 | 7.71601E-26 | 0.958376158 | 0.941 | 0.621 | 1.8219E-21 | 5.59317E-15 | 1.5432E-25 | Wdfy3 | MM_1 |
| 3.74023E-12 | 0.810268418 | 0.954 | 0.6 | 8.83143E-08 | 7.7893E-26 | 0.884677412 | 0.973 | 0.643 | 1.83921E-21 | 3.74023E-12 | 1.55786E-25 | Rab31 | MM_1 |
| 3.6857E-09 | 0.831778783 | 0.431 | 0.145 | 8.70268E-05 | 8.43558E-26 | 1.663361514 | 0.626 | 0.196 | 1.99181E-21 | 3.6857E-09 | 1.68712E-25 | Aldh7a1 | MM_1 |
| 2.16079E-12 | 1.362492657 | 0.651 | 0.269 | 5.10206E-08 | 1.04683E-25 | 1.343874437 | 0.73 | 0.281 | 2.47178E-21 | 2.16079E-12 | 2.09366E-25 | Snx27 | MM_1 |

| | | | | | | | | | | | | | |
|---|---|---|---|---|---|---|---|---|---|---|---|---|---|
| 4.1256E-10 | 1.197289432 | 0.55 | 0.228 | 9.74137E-06 | 1.22669E-25 | 1.400197688 | 0.667 | 0.221 | 2.89646E-21 | 4.1256E-10 | 2.45338E-25 | Ptk2 | MM_1 |
| 9.73632E-15 | 1.183304326 | 0.881 | 0.5 | 2.29894E-10 | 1.23921E-25 | 0.97858793 | 0.896 | 0.515 | 2.92603E-21 | 9.73632E-15 | 2.47843E-25 | Dnajc13 | MM_1 |
| 1.77131E-17 | 1.060248782 | 0.807 | 0.324 | 4.18241E-13 | 1.35226E-25 | 0.997548707 | 0.82 | 0.392 | 3.19297E-21 | 1.77131E-17 | 2.70453E-25 | Zcchc24 | MM_1 |
| 4.09999E-09 | 0.900586245 | 0.376 | 0.11 | 9.68091E-05 | 1.44748E-25 | 1.80153846 | 0.608 | 0.183 | 3.41778E-21 | 4.09999E-09 | 2.89495E-25 | Cped1 | MM_1 |
| 3.13549E-13 | 0.884210332 | 0.972 | 0.645 | 7.40351E-09 | 1.66068E-25 | 0.934203771 | 0.964 | 0.635 | 3.9212E-21 | 3.13549E-13 | 3.32137E-25 | Ddx3x | MM_1 |
| 6.20283E-14 | 0.905605448 | 0.991 | 0.731 | 1.46461E-09 | 1.72429E-25 | 0.805786977 | 0.986 | 0.722 | 4.0714E-21 | 6.20283E-14 | 3.44858E-25 | Slc9a9 | MM_1 |
| 1.87488E-20 | 1.662785369 | 0.789 | 0.255 | 4.42696E-16 | 2.77307E-25 | 1.155189446 | 0.716 | 0.243 | 6.54778E-21 | 1.87488E-20 | 5.54615E-25 | Cp | MM_1 |
| 6.04123E-08 | 0.786387868 | 0.798 | 0.51 | 0.001426455 | 3.96992E-25 | 0.99462101 | 0.932 | 0.537 | 9.37378E-21 | 6.04123E-08 | 7.93984E-25 | Atp6v1b2 | MM_1 |
| 1.16087E-16 | 1.207782535 | 0.78 | 0.3 | 2.74104E-12 | 5.21978E-25 | 0.995803953 | 0.829 | 0.349 | 1.2325E-20 | 1.16087E-16 | 1.04396E-24 | P2ry12 | MM_1 |
| 8.59453E-13 | 1.610401276 | 0.459 | 0.124 | 2.02934E-08 | 5.49142E-25 | 1.032789235 | 0.712 | 0.262 | 1.29663E-20 | 8.59453E-13 | 1.09828E-24 | Gdf15 | MM_1 |
| 8.68877E-25 | 2.724963146 | 0.422 | 0.024 | 2.05159E-20 | 1.17044E-19 | 2.394287593 | 0.297 | 0.033 | 2.76364E-15 | 1.17044E-19 | 1.73775E-24 | Slc7a13 | MM_1 |
| 2.20946E-17 | 1.088645241 | 0.899 | 0.438 | 5.21699E-13 | 8.74743E-25 | 1.108389008 | 0.932 | 0.529 | 2.06544E-20 | 2.20946E-17 | 1.74949E-24 | Lifr | MM_1 |
| 2.12515E-08 | 1.010667204 | 0.606 | 0.303 | 0.000501791 | 1.00907E-24 | 1.253921998 | 0.793 | 0.357 | 2.38262E-20 | 2.12515E-08 | 2.01814E-24 | Tmem165 | MM_1 |
| 1.00604E-23 | 1.510019759 | 0.761 | 0.214 | 2.37546E-19 | 1.161E-24 | 1.016708567 | 0.676 | 0.223 | 2.74136E-20 | 1.00604E-23 | 2.32201E-24 | Ddx60 | MM_1 |
| 7.95668E-15 | 1.295029906 | 0.844 | 0.448 | 1.87873E-10 | 1.36452E-24 | 1.066492831 | 0.91 | 0.474 | 3.22191E-20 | 7.95668E-15 | 2.72905E-24 | Gpr34 | MM_1 |
| 1.33851E-12 | 0.756190793 | 0.789 | 0.41 | 3.16048E-08 | 1.46584E-24 | 0.749122085 | 0.851 | 0.447 | 3.46115E-20 | 1.33851E-12 | 2.93169E-24 | Grina | MM_1 |
| 1.01045E-09 | 0.928664226 | 0.495 | 0.172 | 2.38588E-05 | 1.54066E-24 | 1.618408822 | 0.599 | 0.188 | 3.6378E-20 | 1.01045E-09 | 3.08132E-24 | Tgfbrap1 | MM_1 |
| 1.97973E-13 | 0.898997843 | 0.844 | 0.476 | 4.67455E-09 | 1.64995E-24 | 1.011961948 | 0.914 | 0.499 | 3.89585E-20 | 1.97973E-13 | 3.29989E-24 | Herc1 | MM_1 |
| 2.68291E-17 | 1.085217351 | 0.982 | 0.638 | 6.33488E-13 | 1.68586E-24 | 0.907763379 | 0.986 | 0.629 | 3.98065E-20 | 2.68291E-17 | 3.37172E-24 | Nrp1 | MM_1 |
| 6.6129E-10 | 1.309563046 | 0.661 | 0.324 | 1.56144E-05 | 1.86871E-24 | 0.869729252 | 0.838 | 0.398 | 4.41239E-20 | 6.6129E-10 | 3.73741E-24 | Sypl | MM_1 |
| 2.95564E-10 | 0.598429528 | 0.927 | 0.628 | 6.97887E-06 | 1.93996E-24 | 0.84846476 | 0.973 | 0.665 | 4.58063E-20 | 2.95564E-10 | 3.87992E-24 | Atp1a1 | MM_1 |
| 6.9668E-11 | 1.693523094 | 0.459 | 0.152 | 1.645E-06 | 2.89745E-24 | 1.538440653 | 0.568 | 0.153 | 6.84145E-20 | 6.9668E-11 | 5.79489E-24 | Slc12a7 | MM_1 |
| 2.56385E-12 | 0.70580489 | 0.982 | 0.703 | 6.05377E-08 | 3.51922E-24 | 0.724410855 | 0.995 | 0.774 | 8.30959E-20 | 2.56385E-12 | 7.03845E-24 | Snx3 | MM_1 |
| 6.28666E-11 | 0.652161237 | 0.862 | 0.572 | 1.48441E-06 | 3.55499E-24 | 0.993865115 | 0.887 | 0.504 | 8.39405E-20 | 6.28666E-11 | 7.10998E-24 | Tns3 | MM_1 |
| 1.22317E-08 | 0.822861347 | 0.587 | 0.259 | 0.000288815 | 3.61853E-24 | 0.751861878 | 0.815 | 0.368 | 8.54408E-20 | 1.22317E-08 | 7.23706E-24 | Slc29a1 | MM_1 |
| 3.03305E-08 | 0.699672154 | 0.972 | 0.772 | 0.000716163 | 4.91087E-24 | 0.901136858 | 1 | 0.82 | 1.15955E-19 | 3.03305E-08 | 9.82173E-24 | Lgals1 | MM_1 |
| 8.58111E-19 | 1.314894506 | 0.798 | 0.279 | 2.02617E-14 | 5.35037E-24 | 0.951958817 | 0.775 | 0.297 | 1.26333E-19 | 8.58111E-19 | 1.07007E-23 | Cd38 | MM_1 |
| 5.43766E-13 | 1.048490949 | 0.771 | 0.359 | 1.28394E-08 | 5.47766E-24 | 0.769868506 | 0.887 | 0.414 | 1.29339E-19 | 5.43766E-13 | 1.09553E-23 | Atxn1 | MM_1 |
| 9.06387E-13 | 1.042553515 | 0.807 | 0.476 | 2.14016E-08 | 9.24184E-24 | 0.895930276 | 0.914 | 0.531 | 2.18218E-19 | 9.06387E-13 | 1.84837E-23 | 2610507B1 | MM_1 |
| 5.56337E-12 | 0.945171728 | 0.651 | 0.276 | 1.31362E-07 | 1.31798E-23 | 1.20425179 | 0.721 | 0.294 | 3.11202E-19 | 5.56337E-12 | 2.63596E-23 | Gm10134 | MM_1 |
| 4.35836E-09 | 1.445255591 | 0.468 | 0.193 | 0.00010291 | 1.47328E-23 | 1.564816706 | 0.694 | 0.264 | 3.47871E-19 | 4.35836E-09 | 2.94656E-23 | Trmt1 | MM_1 |
| 3.96918E-13 | 1.152657933 | 0.734 | 0.334 | 9.37203E-09 | 1.6197E-23 | 0.934714534 | 0.797 | 0.351 | 3.82443E-19 | 3.96918E-13 | 3.23939E-23 | Tlr4 | MM_1 |
| 1.03358E-09 | 1.042834047 | 0.688 | 0.345 | 2.4405E-05 | 1.70944E-23 | 1.229013169 | 0.802 | 0.384 | 4.03632E-19 | 1.03358E-09 | 3.41887E-23 | Dennd4c | MM_1 |
| 2.20072E-16 | 2.525992989 | 0.358 | 0.045 | 5.19633E-12 | 1.73453E-23 | 2.207970667 | 0.432 | 0.082 | 4.09557E-19 | 2.20072E-16 | 3.46905E-23 | Tppp | MM_1 |
| 1.34846E-11 | 0.83717932 | 0.936 | 0.572 | 3.18399E-07 | 1.77907E-23 | 0.83323142 | 0.964 | 0.673 | 4.20075E-19 | 1.34846E-11 | 3.55815E-23 | Dhrs3 | MM_1 |
| 9.90082E-18 | 1.446224118 | 0.761 | 0.269 | 2.33778E-13 | 3.33215E-23 | 0.963601352 | 0.856 | 0.406 | 7.86787E-19 | 9.90082E-18 | 6.6643E-23 | Sh3bp5 | MM_1 |
| 2.44347E-15 | 0.826971862 | 0.872 | 0.403 | 5.76953E-11 | 3.38632E-23 | 0.950551789 | 0.91 | 0.512 | 7.99577E-19 | 2.44347E-15 | 6.77263E-23 | Tceal9 | MM_1 |
| 1.07291E-15 | 0.856187544 | 0.78 | 0.338 | 2.53335E-11 | 3.45283E-23 | 0.94866932 | 0.829 | 0.387 | 8.15282E-19 | 1.07291E-15 | 6.90566E-23 | Pstpip2 | MM_1 |
| 7.96105E-14 | 0.967542121 | 0.844 | 0.428 | 1.87976E-09 | 3.51998E-23 | 0.800671034 | 0.919 | 0.49 | 8.31139E-19 | 7.96105E-14 | 7.03997E-23 | Arhgef12 | MM_1 |
| 5.54725E-12 | 1.138337724 | 0.688 | 0.283 | 1.30982E-07 | 3.77875E-23 | 1.06986927 | 0.757 | 0.343 | 8.92238E-19 | 5.54725E-12 | 7.5575E-23 | Twf1 | MM_1 |
| 5.50833E-17 | 0.838187365 | 1 | 0.776 | 1.30063E-12 | 4.15245E-23 | 0.661540671 | 1 | 0.807 | 9.80476E-19 | 5.50833E-17 | 8.3049E-23 | Lamp2 | MM_1 |
| 2.5097E-14 | 0.917837503 | 0.936 | 0.583 | 5.92591E-10 | 5.66672E-23 | 0.837076171 | 0.973 | 0.61 | 1.33803E-18 | 2.5097E-14 | 1.13334E-22 | Sbf2 | MM_1 |
| 2.0749E-09 | 1.541386182 | 0.422 | 0.138 | 4.89925E-05 | 6.5073E-23 | 1.126577771 | 0.59 | 0.193 | 1.5365E-18 | 2.0749E-09 | 1.30146E-22 | Rusc2 | MM_1 |
| 2.69527E-14 | 0.983586878 | 0.917 | 0.483 | 6.36407E-10 | 6.98758E-23 | 0.861824591 | 0.919 | 0.518 | 1.64991E-18 | 2.69527E-14 | 1.39752E-22 | Ophn1 | MM_1 |
| 5.43718E-10 | 1.156053718 | 0.385 | 0.107 | 1.28383E-05 | 7.6899E-23 | 1.311736353 | 0.545 | 0.15 | 1.81574E-18 | 5.43718E-10 | 1.53798E-22 | Trpv4 | MM_1 |
| 3.04037E-14 | 1.620062456 | 0.477 | 0.114 | 7.17893E-10 | 7.72164E-23 | 1.637395534 | 0.536 | 0.15 | 1.82323E-18 | 3.04037E-14 | 1.54433E-22 | Myo7a | MM_1 |
| 9.94258E-11 | 0.774122031 | 0.734 | 0.393 | 2.34764E-06 | 7.90409E-23 | 1.131122581 | 0.82 | 0.428 | 1.86631E-18 | 9.94258E-11 | 1.58082E-22 | Osbpl11 | MM_1 |
| 1.65543E-12 | 0.761103688 | 0.972 | 0.659 | 3.90881E-08 | 9.28347E-23 | 0.765743496 | 0.991 | 0.7 | 2.19201E-18 | 1.65543E-12 | 1.85669E-22 | Rab1a | MM_1 |
| 2.18304E-08 | 1.083137143 | 0.651 | 0.331 | 0.00051546 | 9.30806E-23 | 1.007180357 | 0.793 | 0.36 | 2.19782E-18 | 2.18304E-08 | 1.86161E-22 | Il16 | MM_1 |
| 1.06369E-11 | 0.763150831 | 1 | 0.728 | 2.5116E-07 | 1.01609E-22 | 0.922699973 | 0.995 | 0.752 | 2.39919E-18 | 1.06369E-11 | 2.03218E-22 | Zfp36l2 | MM_1 |
| 1.93978E-13 | 1.380982421 | 0.862 | 0.459 | 4.5802E-09 | 1.07261E-22 | 1.298273491 | 0.856 | 0.49 | 2.53266E-18 | 1.93978E-13 | 2.14523E-22 | P2rx7 | MM_1 |
| 2.68202E-10 | 0.690755515 | 0.927 | 0.621 | 6.33279E-06 | 1.29953E-22 | 0.856517329 | 0.937 | 0.627 | 3.06844E-18 | 2.68202E-10 | 2.59905E-22 | Mat2a | MM_1 |
| 1.61968E-16 | 1.108655757 | 0.798 | 0.328 | 3.82438E-12 | 1.45104E-22 | 1.249486131 | 0.77 | 0.36 | 3.4262E-18 | 1.61968E-16 | 2.90209E-22 | Rhoq | MM_1 |
| 1.35127E-13 | 1.371549829 | 0.505 | 0.138 | 3.19063E-09 | 1.73248E-22 | 1.36473861 | 0.626 | 0.204 | 4.09073E-18 | 1.35127E-13 | 3.46496E-22 | 5033421B0 | MM_1 |
| 1.77554E-16 | 0.854183883 | 1 | 0.707 | 4.1924E-12 | 2.04748E-22 | 0.774678711 | 0.986 | 0.741 | 4.83451E-18 | 1.77554E-16 | 4.09496E-22 | Tcf4 | MM_1 |
| 2.30756E-22 | 1.913541526 | 0.505 | 0.069 | 5.44862E-18 | 1.54998E-21 | 1.493583998 | 0.428 | 0.084 | 3.65981E-17 | 1.54998E-21 | 4.61513E-22 | Afap1l1 | MM_1 |
| 7.31902E-10 | 0.796930682 | 0.936 | 0.559 | 1.72817E-05 | 2.57361E-22 | 0.824812313 | 0.973 | 0.613 | 6.07682E-18 | 7.31902E-10 | 5.14723E-22 | Sash1 | MM_1 |
| 1.28567E-10 | 0.819777302 | 0.817 | 0.5 | 3.03572E-06 | 2.83212E-22 | 0.643773654 | 0.91 | 0.55 | 6.68721E-18 | 1.28567E-10 | 5.66425E-22 | Dmxl1 | MM_1 |
| 5.26074E-11 | 0.879318453 | 0.523 | 0.172 | 1.24217E-06 | 3.11348E-22 | 1.851202635 | 0.486 | 0.123 | 7.35154E-18 | 5.26074E-11 | 6.22695E-22 | Tfec | MM_1 |
| 4.98873E-13 | 0.76051831 | 0.954 | 0.659 | 1.17794E-08 | 3.55673E-22 | 0.763960789 | 0.959 | 0.692 | 8.39814E-18 | 4.98873E-13 | 7.11345E-22 | Rreb1 | MM_1 |
| 5.26716E-09 | 0.873736196 | 0.587 | 0.259 | 0.000124368 | 3.82919E-22 | 1.052819276 | 0.698 | 0.313 | 9.04148E-18 | 5.26716E-09 | 7.65837E-22 | Uap1l1 | MM_1 |
| 1.9708E-13 | 1.32887543 | 0.385 | 0.072 | 4.65346E-09 | 3.95303E-22 | 0.720775397 | 0.532 | 0.134 | 9.33389E-18 | 1.9708E-13 | 7.90605E-22 | Gimap6 | MM_1 |
| 1.27755E-10 | 0.773021511 | 0.945 | 0.679 | 3.01656E-06 | 4.46264E-22 | 0.781469266 | 0.995 | 0.747 | 1.05372E-17 | 1.27755E-10 | 8.92528E-22 | Tmcc1 | MM_1 |
| 6.15889E-11 | 0.959532146 | 0.624 | 0.255 | 1.45424E-06 | 4.533E-22 | 1.346889096 | 0.676 | 0.253 | 1.07033E-17 | 6.15889E-11 | 9.06599E-22 | Slc38a6 | MM_1 |
| 2.90044E-11 | 1.80395381 | 0.376 | 0.093 | 6.84852E-07 | 7.93412E-22 | 1.812402069 | 0.473 | 0.117 | 1.8734E-17 | 2.90044E-11 | 1.58682E-21 | Optn | MM_1 |
| 2.74557E-10 | 0.888354881 | 0.927 | 0.686 | 6.48284E-06 | 9.22688E-22 | 0.998708779 | 0.964 | 0.703 | 2.17865E-17 | 2.74557E-10 | 1.84538E-21 | Fnip1 | MM_1 |
| 1.23477E-07 | 0.699617233 | 0.734 | 0.41 | 0.002915549 | 1.12602E-21 | 0.765281954 | 0.887 | 0.49 | 2.65876E-17 | 1.23477E-07 | 2.25204E-21 | Washc2 | MM_1 |
| 4.33391E-12 | 0.955682325 | 0.56 | 0.179 | 1.02332E-07 | 1.37965E-21 | 0.756621691 | 0.694 | 0.243 | 3.25762E-17 | 4.33391E-12 | 2.75929E-21 | Lpar1 | MM_1 |
| 5.39223E-08 | 0.937504721 | 0.468 | 0.186 | 0.001273213 | 1.46358E-21 | 1.278832994 | 0.581 | 0.204 | 3.4558E-17 | 5.39223E-08 | 2.92715E-21 | Sord | MM_1 |
| 1.0269E-07 | 0.680829012 | 0.862 | 0.545 | 0.002424708 | 1.59589E-21 | 0.931263218 | 0.901 | 0.542 | 3.76822E-17 | 1.0269E-07 | 3.19179E-21 | Pdcd6ip | MM_1 |
| 5.19157E-11 | 1.113749609 | 0.394 | 0.1 | 1.22583E-06 | 2.04423E-21 | 1.897744931 | 0.514 | 0.153 | 4.82683E-17 | 5.19157E-11 | 4.08846E-21 | Epn2 | MM_1 |
| 1.00989E-07 | 1.001476193 | 0.422 | 0.169 | 0.002384545 | 2.09863E-21 | 0.903678718 | 0.613 | 0.204 | 4.95528E-17 | 1.00989E-07 | 4.19726E-21 | Hrh1 | MM_1 |
| 4.00091E-10 | 0.946471076 | 0.615 | 0.266 | 9.44694E-06 | 2.19203E-21 | 0.885569488 | 0.698 | 0.281 | 5.17582E-17 | 4.00091E-10 | 4.38406E-21 | Ccdc152 | MM_1 |
| 4.51602E-10 | 0.684491194 | 0.927 | 0.676 | 1.06632E-05 | 2.34119E-21 | 0.829397745 | 0.991 | 0.684 | 5.52801E-17 | 4.51602E-10 | 4.68237E-21 | Wdr26 | MM_1 |
| 5.25495E-16 | 1.503298148 | 0.596 | 0.183 | 1.2408E-11 | 2.71769E-21 | 1.299696657 | 0.631 | 0.232 | 6.41702E-17 | 5.25495E-16 | 5.43539E-21 | Nhsl2 | MM_1 |
| 4.11252E-08 | 1.164709017 | 0.44 | 0.169 | 0.000971048 | 3.02541E-21 | 1.676979601 | 0.554 | 0.18 | 7.1436E-17 | 4.11252E-08 | 6.05082E-21 | Cemip2 | MM_1 |
| 2.56235E-10 | 0.833737375 | 0.945 | 0.628 | 6.05022E-06 | 3.03491E-21 | 0.779343294 | 0.991 | 0.695 | 7.16603E-17 | 2.56235E-10 | 6.06982E-21 | Ptafr | MM_1 |
| 6.88571E-12 | 1.366222872 | 0.67 | 0.297 | 1.62585E-07 | 3.06145E-21 | 0.950622546 | 0.811 | 0.387 | 7.2287E-17 | 6.88571E-12 | 6.12291E-21 | Ly96 | MM_1 |
| 2.20002E-09 | 1.081832149 | 0.633 | 0.321 | 5.19469E-05 | 3.46617E-21 | 1.056650463 | 0.797 | 0.384 | 8.18432E-17 | 2.20002E-09 | 6.93234E-21 | Tmem87b | MM_1 |
| 3.40945E-08 | 1.288815073 | 0.55 | 0.283 | 0.000805038 | 3.87534E-21 | 1.00969217 | 0.752 | 0.351 | 9.15046E-17 | 3.40945E-08 | 7.75069E-21 | Gnpda1 | MM_1 |
| 1.63047E-15 | 1.024741301 | 0.945 | 0.579 | 3.84987E-11 | 3.90409E-21 | 0.839804029 | 0.95 | 0.651 | 9.21834E-17 | 1.63047E-15 | 7.80819E-21 | Ap2a2 | MM_1 |

| | | | | | | | | | | | | | |
|---|---|---|---|---|---|---|---|---|---|---|---|---|---|
| 2.24021E-11 | 0.790507377 | 0.881 | 0.5 | 5.28957E-07 | 4.24364E-21 | 0.968521699 | 0.851 | 0.496 | 1.00201E-16 | 2.24021E-11 | 8.48729E-21 | Ano6 | MM_1 |
| 2.76951E-08 | 0.771657145 | 0.505 | 0.197 | 0.000653936 | 5.07925E-21 | 0.951156237 | 0.707 | 0.278 | 1.19931E-16 | 2.76951E-08 | 1.01585E-20 | Dpysl3 | MM_1 |
| 2.77899E-11 | 1.238069601 | 0.807 | 0.469 | 6.56175E-07 | 5.90816E-21 | 0.968933917 | 0.874 | 0.488 | 1.39503E-16 | 2.77899E-11 | 1.18163E-20 | Abl2 | MM_1 |
| 4.18379E-12 | 1.33161174 | 0.532 | 0.176 | 9.87875E-08 | 6.47999E-21 | 1.156604653 | 0.59 | 0.196 | 1.53006E-16 | 4.18379E-12 | 1.296E-20 | Lpin1 | MM_1 |
| 1.2908E-08 | 0.979833036 | 0.633 | 0.328 | 0.000304785 | 6.59514E-21 | 0.930958031 | 0.815 | 0.411 | 1.55724E-16 | 1.2908E-08 | 1.31903E-20 | Hsd17b4 | MM_1 |
| 2.69249E-13 | 0.79283626 | 0.991 | 0.738 | 6.35751E-09 | 6.637E-21 | 0.709703471 | 1 | 0.777 | 1.56713E-16 | 2.69249E-13 | 1.3274E-20 | Csf1r | MM_1 |
| 3.6285E-08 | 1.130723841 | 0.367 | 0.114 | 0.000856761 | 7.23363E-21 | 1.223777354 | 0.464 | 0.112 | 1.70801E-16 | 3.6285E-08 | 1.44673E-20 | Sgsh | MM_1 |
| 2.49328E-20 | 4.721715728 | 0.376 | 0.034 | 5.88713E-16 | 7.43897E-21 | 2.036050214 | 0.369 | 0.063 | 1.75649E-16 | 2.49328E-20 | 1.48779E-20 | Vsig4 | MM_1 |
| 1.23188E-07 | 0.722045552 | 0.844 | 0.569 | 0.002908722 | 7.54733E-21 | 0.859951077 | 0.901 | 0.534 | 1.78208E-16 | 1.23188E-07 | 1.50947E-20 | 5031439G | MM_1 |
| 4.00326E-10 | 2.643099182 | 0.229 | 0.031 | 9.4525E-06 | 7.57166E-21 | 2.761423553 | 0.365 | 0.063 | 1.78782E-16 | 4.00326E-10 | 1.51433E-20 | Cd209d | MM_1 |
| 3.52138E-14 | 0.938371238 | 0.844 | 0.428 | 8.31468E-10 | 8.06394E-21 | 1.124238366 | 0.824 | 0.452 | 1.90406E-16 | 3.52138E-14 | 1.61279E-20 | Tbc1d14 | MM_1 |
| 1.30342E-12 | 0.773746666 | 0.963 | 0.741 | 3.07765E-08 | 8.9745E-21 | 0.734181169 | 0.995 | 0.747 | 2.11906E-16 | 1.30342E-12 | 1.7949E-20 | Ehd4 | MM_1 |
| 6.61992E-12 | 0.725003898 | 0.872 | 0.534 | 1.5631E-07 | 9.14523E-21 | 0.685511404 | 0.932 | 0.616 | 2.15937E-16 | 6.61992E-12 | 1.82905E-20 | Scamp2 | MM_1 |
| 3.87775E-09 | 0.837450591 | 0.945 | 0.738 | 9.15615E-05 | 9.46089E-21 | 0.925122625 | 0.977 | 0.768 | 2.23391E-16 | 3.87775E-09 | 1.89218E-20 | Neat1 | MM_1 |
| 1.55734E-14 | 0.84333935 | 0.991 | 0.772 | 3.67718E-10 | 9.79341E-21 | 0.649981426 | 1 | 0.785 | 2.31242E-16 | 1.55734E-14 | 1.95868E-20 | Qk | MM_1 |
| 2.69482E-10 | 0.738232024 | 0.862 | 0.472 | 6.36301E-06 | 1.091E-20 | 1.055811457 | 0.878 | 0.512 | 2.57607E-16 | 2.69482E-10 | 2.182E-20 | Fnip2 | MM_1 |
| 5.75432E-09 | 0.644050059 | 0.945 | 0.634 | 0.000135871 | 1.20247E-20 | 0.796786439 | 0.973 | 0.689 | 2.83927E-16 | 5.75432E-09 | 2.40494E-20 | Gns | MM_1 |
| 2.12605E-12 | 0.970001457 | 0.853 | 0.455 | 5.02003E-08 | 1.22524E-20 | 0.903254988 | 0.901 | 0.52 | 2.89305E-16 | 2.12605E-12 | 2.45049E-20 | Ap1b1 | MM_1 |
| 1.11927E-09 | 0.772980733 | 0.761 | 0.434 | 2.64283E-05 | 1.52392E-20 | 0.757605727 | 0.842 | 0.466 | 3.59828E-16 | 1.11927E-09 | 3.04784E-20 | Vps13c | MM_1 |
| 4.02306E-17 | 1.196537023 | 0.679 | 0.214 | 9.49925E-13 | 1.54134E-20 | 1.185086624 | 0.721 | 0.305 | 3.6394E-16 | 4.02306E-17 | 3.08267E-20 | Rhobtb1 | MM_1 |
| 1.45828E-13 | 1.147010478 | 0.862 | 0.472 | 3.44329E-09 | 1.96722E-20 | 0.784984442 | 0.865 | 0.474 | 4.64501E-16 | 1.45828E-13 | 3.93445E-20 | Raph1 | MM_1 |
| 1.02983E-07 | 3.10699246 | 0.11 | 0.003 | 0.002431632 | 2.07508E-20 | 3.780532321 | 0.225 | 0.003 | 4.89968E-16 | 1.02983E-07 | 4.15016E-20 | Ccl26 | MM_1 |
| 2.17397E-15 | 1.297323101 | 0.679 | 0.231 | 5.13318E-11 | 2.26096E-20 | 1.35544472 | 0.604 | 0.21 | 5.33857E-16 | 2.17397E-15 | 4.52191E-20 | Gm21188 | MM_1 |
| 4.92245E-15 | 0.808185686 | 0.982 | 0.683 | 1.16229E-10 | 2.32071E-20 | 0.668689874 | 0.982 | 0.695 | 5.47967E-16 | 4.92245E-15 | 4.64143E-20 | Tgfbr2 | MM_1 |
| 5.67984E-11 | 0.786240925 | 0.844 | 0.507 | 1.34112E-06 | 2.39453E-20 | 0.728699311 | 0.919 | 0.542 | 5.65397E-16 | 5.67984E-11 | 4.78906E-20 | Ehd1 | MM_1 |
| 3.05948E-11 | 1.116416151 | 0.752 | 0.359 | 7.22404E-07 | 3.39999E-20 | 1.048088907 | 0.743 | 0.324 | 8.02805E-16 | 3.05948E-11 | 6.79997E-20 | Dusp6 | MM_1 |
| 9.64504E-14 | 1.385226045 | 0.569 | 0.186 | 2.27739E-09 | 3.40367E-20 | 1.069425302 | 0.617 | 0.234 | 8.03674E-16 | 9.64504E-14 | 6.80733E-20 | Wdr36 | MM_1 |
| 2.07818E-09 | 1.112082791 | 0.33 | 0.083 | 4.907E-05 | 3.57683E-20 | 1.69500711 | 0.432 | 0.101 | 8.4456E-16 | 2.07818E-09 | 7.15365E-20 | Ikbke | MM_1 |
| 4.75216E-11 | 0.771641454 | 0.853 | 0.466 | 1.12208E-06 | 4.01411E-20 | 0.867172934 | 0.878 | 0.523 | 9.47811E-16 | 4.75216E-11 | 8.02821E-20 | Snx30 | MM_1 |
| 6.00387E-11 | 1.060084924 | 0.55 | 0.19 | 1.41763E-06 | 4.66462E-20 | 1.653747464 | 0.505 | 0.15 | 1.10141E-15 | 6.00387E-11 | 9.32924E-20 | Mmp9 | MM_1 |
| 9.01098E-12 | 0.852193435 | 0.89 | 0.552 | 2.12767E-07 | 5.89194E-20 | 0.832230483 | 0.923 | 0.578 | 1.3912E-15 | 9.01098E-12 | 1.17839E-19 | Atp6ap1 | MM_1 |
| 6.15731E-09 | 0.71712639 | 0.853 | 0.507 | 0.000145387 | 5.90576E-20 | 0.704259947 | 0.959 | 0.556 | 1.39447E-15 | 6.15731E-09 | 1.18115E-19 | Hpgds | MM_1 |
| 2.17141E-08 | 0.799761026 | 0.771 | 0.472 | 0.000512714 | 6.27654E-20 | 0.854266806 | 0.865 | 0.548 | 1.48202E-15 | 2.17141E-08 | 1.25531E-19 | Dtnbp1 | MM_1 |
| 2.1133E-10 | 0.726731012 | 0.982 | 0.607 | 4.98993E-06 | 7.28244E-20 | 0.731691106 | 0.986 | 0.73 | 1.71953E-15 | 2.1133E-10 | 1.45649E-19 | Cd63 | MM_1 |
| 1.48386E-08 | 2.835194907 | 0.156 | 0.014 | 0.00035037 | 7.32667E-20 | 2.572858096 | 0.297 | 0.033 | 1.72997E-15 | 1.48386E-08 | 1.46533E-19 | Tox2 | MM_1 |
| 1.80467E-09 | 1.84616902 | 0.22 | 0.031 | 4.26119E-05 | 7.42351E-20 | 3.209938595 | 0.311 | 0.041 | 1.75284E-15 | 1.80467E-09 | 1.4847E-19 | Dpyd | MM_1 |
| 5.32452E-10 | 1.28835561 | 0.55 | 0.224 | 1.25723E-05 | 9.5678E-20 | 0.895803706 | 0.716 | 0.302 | 2.25915E-15 | 5.32452E-10 | 1.91356E-19 | Plod1 | MM_1 |
| 1.3528E-07 | 0.940219276 | 0.633 | 0.334 | 0.003194242 | 1.02984E-19 | 0.830876945 | 0.788 | 0.398 | 2.43166E-15 | 1.3528E-07 | 2.05969E-19 | Anxa11 | MM_1 |
| 1.14224E-12 | 0.78830797 | 0.927 | 0.621 | 2.69706E-08 | 1.04655E-19 | 0.717844407 | 0.919 | 0.624 | 2.4711E-15 | 1.14224E-12 | 2.09309E-19 | Dip2b | MM_1 |
| 1.02753E-08 | 0.835592431 | 0.606 | 0.262 | 0.000242621 | 1.41557E-19 | 1.261115798 | 0.577 | 0.193 | 3.34245E-15 | 1.02753E-08 | 2.83114E-19 | Gas2l3 | MM_1 |
| 2.30954E-11 | 1.293637558 | 0.624 | 0.266 | 5.45328E-07 | 1.53058E-19 | 1.142531041 | 0.685 | 0.297 | 3.614E-15 | 2.30954E-11 | 3.06115E-19 | Fgd6 | MM_1 |
| 3.52059E-09 | 0.605572667 | 0.862 | 0.559 | 8.31282E-05 | 1.54187E-19 | 0.8833905 | 0.892 | 0.575 | 3.64067E-15 | 3.52059E-09 | 3.08375E-19 | Bin1 | MM_1 |
| 2.41224E-08 | 1.004209424 | 0.596 | 0.297 | 0.000569579 | 1.82848E-19 | 1.082843508 | 0.676 | 0.297 | 4.3174E-15 | 2.41224E-08 | 3.65695E-19 | Glb1 | MM_1 |
| 1.74406E-10 | 0.661439001 | 0.853 | 0.5 | 4.11808E-06 | 1.90847E-19 | 0.774525866 | 0.932 | 0.529 | 4.50628E-15 | 1.74406E-10 | 3.81694E-19 | Dclre1c | MM_1 |
| 8.64956E-11 | 1.164792889 | 0.734 | 0.379 | 2.04234E-06 | 2.70846E-19 | 1.265648174 | 0.779 | 0.401 | 6.39522E-15 | 8.64956E-11 | 5.41692E-19 | Pdpk1 | MM_1 |
| 2.12089E-11 | 0.835758072 | 0.817 | 0.407 | 5.00784E-07 | 2.77185E-19 | 0.708866015 | 0.878 | 0.444 | 6.54489E-15 | 2.12089E-11 | 5.5437E-19 | Msr1 | MM_1 |
| 4.87462E-11 | 0.927738218 | 0.697 | 0.324 | 1.151E-06 | 2.83323E-19 | 0.811485359 | 0.752 | 0.349 | 6.68982E-15 | 4.87462E-11 | 5.66645E-19 | Tspan14 | MM_1 |
| 1.14127E-08 | 0.751363195 | 0.679 | 0.355 | 0.000269477 | 3.53989E-19 | 0.797782158 | 0.824 | 0.444 | 8.35839E-15 | 1.14127E-08 | 7.07978E-19 | Rpn1 | MM_1 |
| 1.146E-12 | 0.929696019 | 0.752 | 0.348 | 2.70593E-08 | 3.80798E-19 | 0.702516117 | 0.797 | 0.376 | 8.99141E-15 | 1.146E-12 | 7.61596E-19 | Prune2 | MM_1 |
| 5.4717E-14 | 1.200175606 | 0.862 | 0.493 | 1.29198E-09 | 4.3884E-19 | 0.64591002 | 0.941 | 0.589 | 1.03619E-14 | 5.4717E-14 | 8.77679E-19 | Pirb | MM_1 |
| 3.58962E-14 | 1.112381714 | 0.835 | 0.434 | 8.47582E-10 | 4.78584E-19 | 0.895653449 | 0.842 | 0.466 | 1.13003E-14 | 3.58962E-14 | 9.57167E-19 | Vwa5a | MM_1 |
| 1.12469E-07 | 1.14303308 | 0.44 | 0.179 | 0.002655622 | 5.22213E-19 | 1.394884525 | 0.568 | 0.213 | 1.23305E-14 | 1.12469E-07 | 1.04443E-18 | Rfk | MM_1 |
| 1.0861E-08 | 0.747204119 | 0.798 | 0.452 | 0.000256451 | 5.71461E-19 | 0.631764498 | 0.838 | 0.471 | 1.34933E-14 | 1.0861E-08 | 1.14292E-18 | Ctdsp2 | MM_1 |
| 1.63637E-08 | 0.728805489 | 0.798 | 0.49 | 0.000386379 | 5.78636E-19 | 0.812394556 | 0.878 | 0.542 | 1.36627E-14 | 1.63637E-08 | 1.15727E-18 | Hip1 | MM_1 |
| 8.0751E-16 | 3.379307834 | 0.284 | 0.021 | 1.90669E-11 | 6.08283E-19 | 1.916319492 | 0.338 | 0.054 | 1.43628E-14 | 8.0751E-16 | 1.21657E-18 | Cd28 | MM_1 |
| 6.40229E-10 | 0.777921691 | 0.734 | 0.362 | 1.51171E-05 | 7.14662E-19 | 0.794239153 | 0.865 | 0.469 | 1.68746E-14 | 6.40229E-10 | 1.42932E-18 | Slc11a1 | MM_1 |
| 1.89198E-07 | 0.62537315 | 0.67 | 0.383 | 0.004467344 | 8.6756E-19 | 1.055689099 | 0.82 | 0.447 | 2.04848E-14 | 1.89198E-07 | 1.73512E-18 | Atp6v1d | MM_1 |
| 1.74589E-10 | 0.659284644 | 0.642 | 0.269 | 4.12239E-06 | 8.84166E-19 | 1.022266457 | 0.694 | 0.313 | 2.08769E-14 | 1.74589E-10 | 1.76833E-18 | Rgs18 | MM_1 |
| 2.70351E-11 | 0.866280836 | 0.853 | 0.497 | 6.38352E-07 | 1.12252E-18 | 0.635172652 | 0.892 | 0.616 | 2.6505E-14 | 2.70351E-11 | 2.24504E-18 | Tm9sf2 | MM_1 |
| 8.39198E-08 | 1.271852529 | 0.358 | 0.117 | 0.001981515 | 1.20392E-18 | 1.152147358 | 0.554 | 0.18 | 2.84269E-14 | 8.39198E-08 | 2.40784E-18 | Sorbs3 | MM_1 |
| 1.30342E-07 | 0.805499604 | 0.679 | 0.393 | 0.003077644 | 1.30562E-18 | 0.657139759 | 0.793 | 0.384 | 3.08282E-14 | 1.30342E-07 | 2.61123E-18 | Pik3cg | MM_1 |
| 7.92336E-16 | 1.499778977 | 0.661 | 0.217 | 1.87086E-11 | 1.53617E-18 | 0.888240333 | 0.622 | 0.229 | 3.6272E-14 | 7.92336E-16 | 3.07233E-18 | Gab1 | MM_1 |
| 2.11944E-12 | 0.988258873 | 0.89 | 0.607 | 5.00442E-08 | 1.66218E-18 | 0.631392324 | 0.914 | 0.589 | 3.92475E-14 | 2.11944E-12 | 3.32437E-18 | G3bp2 | MM_1 |
| 8.13071E-11 | 0.603893391 | 0.385 | 0.1 | 1.91982E-06 | 1.79697E-18 | 1.602129977 | 0.486 | 0.15 | 4.24301E-14 | 8.13071E-11 | 3.59395E-18 | Kifc3 | MM_1 |
| 1.32947E-09 | 0.74975239 | 0.725 | 0.417 | 3.13915E-05 | 1.80937E-18 | 0.74711832 | 0.829 | 0.469 | 4.27227E-14 | 1.32947E-09 | 3.61873E-18 | Ap1s2 | MM_1 |
| 6.52767E-08 | 0.737805752 | 0.679 | 0.386 | 0.001541314 | 2.19158E-18 | 0.608233974 | 0.851 | 0.485 | 5.17475E-14 | 6.52767E-08 | 4.38315E-18 | Cd37 | MM_1 |
| 8.79076E-10 | 0.813520159 | 0.945 | 0.683 | 2.07567E-05 | 2.20886E-18 | 0.756514739 | 0.959 | 0.684 | 5.21556E-14 | 8.79076E-10 | 4.41772E-18 | Srgap2 | MM_1 |
| 8.59929E-11 | 0.740355487 | 0.835 | 0.479 | 2.03046E-06 | 2.50109E-18 | 0.692929781 | 0.887 | 0.488 | 5.90557E-14 | 8.59929E-11 | 5.00218E-18 | Rab3il1 | MM_1 |
| 3.54604E-07 | 0.893870405 | 0.67 | 0.372 | 0.008372902 | 2.53732E-18 | 0.822574723 | 0.779 | 0.433 | 5.99112E-14 | 3.54604E-07 | 5.07464E-18 | Atp6v1h | MM_1 |
| 6.61626E-08 | 0.998321209 | 0.624 | 0.31 | 0.001562231 | 2.67218E-18 | 1.242865951 | 0.734 | 0.395 | 6.30954E-14 | 6.61626E-08 | 5.34435E-18 | Mospd2 | MM_1 |
| 3.15102E-18 | 1.280575555 | 0.798 | 0.286 | 7.4402E-14 | 3.957E-18 | 0.702930163 | 0.788 | 0.368 | 9.34326E-14 | 3.957E-18 | 6.30205E-18 | Fer | MM_1 |
| 3.57614E-14 | 1.277037155 | 0.541 | 0.162 | 8.44399E-10 | 3.22861E-18 | 1.537516791 | 0.586 | 0.221 | 7.6234E-14 | 3.57614E-14 | 6.45722E-18 | Otud7b | MM_1 |
| 1.53703E-10 | 1.413689515 | 0.606 | 0.266 | 3.62923E-06 | 4.52292E-18 | 0.872143226 | 0.707 | 0.338 | 1.06795E-13 | 1.53703E-10 | 9.04584E-18 | Agpat3 | MM_1 |
| 6.8011E-11 | 1.049404731 | 0.651 | 0.286 | 1.60587E-06 | 4.77497E-18 | 0.914464358 | 0.698 | 0.302 | 1.12747E-13 | 6.8011E-11 | 9.54994E-18 | 2610203C | MM_1 |
| 2.15044E-11 | 0.871351434 | 0.55 | 0.197 | 5.07761E-07 | 8.12423E-18 | 0.781437944 | 0.613 | 0.229 | 1.91829E-13 | 2.15044E-11 | 1.62485E-17 | Prkca | MM_1 |
| 4.82106E-10 | 0.832530368 | 0.706 | 0.334 | 1.13835E-05 | 9.08111E-18 | 0.860708194 | 0.698 | 0.308 | 2.14423E-13 | 4.82106E-10 | 1.81622E-17 | Prkn | MM_1 |
| 2.50086E-11 | 0.844557881 | 0.743 | 0.359 | 5.90503E-07 | 1.02643E-17 | 0.798025606 | 0.829 | 0.436 | 2.4236E-13 | 2.50086E-11 | 2.05286E-17 | Anxa4 | MM_1 |
| 8.65455E-11 | 0.973646523 | 0.853 | 0.51 | 2.04351E-06 | 1.23769E-17 | 0.784601065 | 0.883 | 0.553 | 2.92243E-13 | 8.65455E-11 | 2.47538E-17 | Plekhm2 | MM_1 |
| 3.69744E-08 | 0.711941389 | 0.872 | 0.579 | 0.000873039 | 1.30197E-17 | 0.788493826 | 0.896 | 0.569 | 3.07422E-13 | 3.69744E-08 | 2.60395E-17 | Gapvd1 | MM_1 |
| 1.06733E-10 | 0.604761253 | 0.917 | 0.576 | 2.52018E-06 | 1.43037E-17 | 0.727330359 | 0.955 | 0.651 | 3.37738E-13 | 1.06733E-10 | 2.86073E-17 | Mitf | MM_1 |
| 1.75913E-13 | 0.861851924 | 0.899 | 0.493 | 4.15365E-09 | 1.53804E-17 | 0.718088918 | 0.923 | 0.553 | 3.63163E-13 | 1.75913E-13 | 3.07609E-17 | Abca9 | MM_1 |

| | | | | | | | | | | | | | |
|---|---|---|---|---|---|---|---|---|---|---|---|---|---|
| 1.93774E-11 | 0.677595426 | 0.872 | 0.545 | 4.57538E-07 | 1.70941E-17 | 0.63966139 | 0.892 | 0.575 | 4.03626E-13 | 1.93774E-11 | 3.41882E-17 | Serinc1 | MM_1 |
| 1.9998E-08 | 0.767439592 | 0.651 | 0.297 | 0.000472194 | 1.81439E-17 | 0.728116627 | 0.707 | 0.313 | 4.28414E-13 | 1.9998E-08 | 3.62878E-17 | Acp2 | MM_1 |
| 1.55854E-07 | 0.784237891 | 0.688 | 0.397 | 0.003680014 | 1.93295E-17 | 0.795814639 | 0.82 | 0.496 | 4.56407E-13 | 1.55854E-07 | 3.86589E-17 | Degs1 | MM_1 |
| 2.8887E-07 | 0.670660656 | 0.817 | 0.528 | 0.006820808 | 2.55144E-17 | 0.905553951 | 0.842 | 0.54 | 6.02446E-13 | 2.8887E-07 | 5.10288E-17 | Rragc | MM_1 |
| 1.05881E-14 | 1.264894474 | 0.771 | 0.341 | 2.50007E-10 | 3.00389E-17 | 0.606351383 | 0.752 | 0.346 | 7.09279E-13 | 1.05881E-14 | 6.00779E-17 | Pde4d | MM_1 |
| 9.4937E-14 | 1.468146941 | 0.56 | 0.2 | 2.24165E-09 | 3.04183E-17 | 1.035754261 | 0.554 | 0.213 | 7.18238E-13 | 9.4937E-14 | 6.08367E-17 | Pgm2l1 | MM_1 |
| 8.91574E-08 | 0.647616662 | 0.697 | 0.372 | 0.002105183 | 3.15291E-17 | 0.747730294 | 0.766 | 0.392 | 7.44464E-13 | 8.91574E-08 | 6.30581E-17 | Lats2 | MM_1 |
| 3.30552E-14 | 2.21386734 | 0.606 | 0.231 | 7.80499E-10 | 3.18147E-17 | 1.74591069 | 0.613 | 0.272 | 7.51209E-13 | 3.30552E-14 | 6.36294E-17 | Hgsnat | MM_1 |
| 3.6219E-07 | 0.818374275 | 0.44 | 0.183 | 0.008552037 | 3.30619E-17 | 0.772178071 | 0.586 | 0.213 | 7.80658E-13 | 3.6219E-07 | 6.61238E-17 | Ampd3 | MM_1 |
| 6.25638E-12 | 1.192031121 | 0.661 | 0.293 | 1.47726E-07 | 3.37163E-17 | 0.800086157 | 0.748 | 0.343 | 7.96109E-13 | 6.25638E-12 | 6.74326E-17 | Gpr160 | MM_1 |
| 1.13923E-09 | 1.007471915 | 0.569 | 0.231 | 2.68996E-05 | 4.00539E-17 | 1.025199948 | 0.626 | 0.256 | 9.45753E-13 | 1.13923E-09 | 8.01079E-17 | Tbc1d2b | MM_1 |
| 2.98167E-10 | 3.236660688 | 0.211 | 0.024 | 7.04033E-06 | 4.34814E-17 | 1.47427785 | 0.293 | 0.044 | 1.02668E-12 | 2.98167E-10 | 8.69628E-17 | Btnl2 | MM_1 |
| 1.08164E-08 | 0.585700645 | 0.817 | 0.5 | 0.000255397 | 4.50773E-17 | 0.673470873 | 0.842 | 0.471 | 1.06436E-12 | 1.08164E-08 | 9.01545E-17 | Lrrc58 | MM_1 |
| 4.54719E-17 | 2.870085139 | 0.459 | 0.1 | 1.07368E-12 | 2.27498E-14 | 2.042446489 | 0.419 | 0.158 | 5.37168E-10 | 2.27498E-14 | 9.09438E-17 | Colec12 | MM_1 |
| 3.00441E-12 | 2.122278814 | 0.404 | 0.103 | 7.09401E-08 | 4.67267E-17 | 1.351216246 | 0.455 | 0.131 | 1.10331E-12 | 3.00441E-12 | 9.34534E-17 | Cxcl12 | MM_1 |
| 6.17459E-15 | 1.160120852 | 0.872 | 0.428 | 1.45795E-10 | 5.04828E-17 | 0.742894636 | 0.905 | 0.493 | 1.192E-12 | 6.17459E-15 | 1.00966E-16 | Cd33 | MM_1 |
| 7.03289E-08 | 0.621162347 | 0.734 | 0.414 | 0.001660605 | 5.08779E-17 | 0.761993366 | 0.851 | 0.469 | 1.20133E-12 | 7.03289E-08 | 1.01756E-16 | Fgd4 | MM_1 |
| 7.73217E-12 | 0.838939735 | 0.826 | 0.483 | 1.82572E-07 | 6.35567E-17 | 0.666241994 | 0.838 | 0.447 | 1.5007E-12 | 7.73217E-12 | 1.27113E-16 | Prkacb | MM_1 |
| 5.68208E-08 | 1.168167489 | 0.495 | 0.217 | 0.001341653 | 6.97233E-17 | 1.1335439 | 0.572 | 0.229 | 1.64631E-12 | 5.68208E-08 | 1.39447E-16 | Ticam2 | MM_1 |
| 1.2563E-11 | 0.865243701 | 0.908 | 0.576 | 2.96638E-07 | 7.27532E-17 | 0.81665654 | 0.919 | 0.599 | 1.71785E-12 | 1.2563E-11 | 1.45506E-16 | Il6ra | MM_1 |
| 1.9148E-08 | 0.998801469 | 0.532 | 0.214 | 0.000452124 | 7.76631E-17 | 0.980116162 | 0.707 | 0.322 | 1.83378E-12 | 1.9148E-08 | 1.55326E-16 | Tnfsf13 | MM_1 |
| 9.81385E-11 | 1.232571744 | 0.697 | 0.355 | 2.31725E-06 | 9.31289E-17 | 0.809969336 | 0.779 | 0.392 | 2.19896E-12 | 9.81385E-11 | 1.86258E-16 | Parvb | MM_1 |
| 4.81659E-09 | 0.907222912 | 0.743 | 0.393 | 0.000113729 | 1.02392E-16 | 0.780477537 | 0.752 | 0.376 | 2.41768E-12 | 4.81659E-09 | 2.04784E-16 | Slc16a7 | MM_1 |
| 5.42269E-09 | 0.807821656 | 0.798 | 0.51 | 0.00012804 | 1.05881E-16 | 0.65704413 | 0.838 | 0.485 | 2.50006E-12 | 5.42269E-09 | 2.11762E-16 | Tmem30a | MM_1 |
| 2.2495E-08 | 0.759896396 | 0.743 | 0.434 | 0.000531151 | 1.06571E-16 | 0.713947592 | 0.829 | 0.477 | 2.51636E-12 | 2.2495E-08 | 2.13143E-16 | Pcna | MM_1 |
| 7.2569E-11 | 1.219292724 | 0.422 | 0.117 | 1.7135E-06 | 1.09402E-16 | 1.141679195 | 0.505 | 0.174 | 2.58321E-12 | 7.2569E-11 | 2.18805E-16 | Gcnt1 | MM_1 |
| 2.03622E-07 | 0.773689823 | 0.367 | 0.131 | 0.004807924 | 1.11746E-16 | 1.434470684 | 0.5 | 0.18 | 2.63855E-12 | 2.03622E-07 | 2.23493E-16 | Kank2 | MM_1 |
| 2.24787E-08 | 0.725629774 | 0.56 | 0.245 | 0.000530767 | 1.38949E-16 | 0.848058687 | 0.617 | 0.256 | 3.28087E-12 | 2.24787E-08 | 2.77899E-16 | Gab3 | MM_1 |
| 1.64409E-10 | 0.958028028 | 0.807 | 0.5 | 3.88202E-06 | 1.42892E-16 | 0.688346231 | 0.91 | 0.575 | 3.37397E-12 | 1.64409E-10 | 2.85784E-16 | Pura | MM_1 |
| 1.45533E-08 | 0.978831172 | 0.624 | 0.279 | 0.000343633 | 1.53337E-16 | 0.921863039 | 0.712 | 0.338 | 3.6206E-12 | 1.45533E-08 | 3.06675E-16 | Tom1 | MM_1 |
| 1.21882E-10 | 0.643945428 | 0.991 | 0.755 | 2.87788E-06 | 1.60316E-16 | 0.659490681 | 0.995 | 0.747 | 3.78539E-12 | 1.21882E-10 | 3.20632E-16 | Trf | MM_1 |
| 2.62009E-07 | 0.808280944 | 0.514 | 0.231 | 0.006186564 | 1.78702E-16 | 0.752224736 | 0.64 | 0.253 | 4.21951E-12 | 2.62009E-07 | 3.57404E-16 | Eef2k | MM_1 |
| 6.3993E-10 | 0.728677539 | 0.844 | 0.507 | 1.511E-05 | 1.86997E-16 | 0.781084038 | 0.923 | 0.572 | 4.41538E-12 | 6.3993E-10 | 3.73994E-16 | Ptpn12 | MM_1 |
| 1.0493E-07 | 2.367011459 | 0.165 | 0.021 | 0.002477613 | 1.94325E-16 | 3.018292909 | 0.248 | 0.027 | 4.58841E-12 | 1.0493E-07 | 3.8865E-16 | Gm47675 | MM_1 |
| 1.16552E-07 | 0.85891779 | 0.661 | 0.39 | 0.002752029 | 3.13734E-16 | 0.689786094 | 0.811 | 0.422 | 7.40789E-12 | 1.16552E-07 | 6.27469E-16 | Fkbp15 | MM_1 |
| 1.54904E-10 | 1.902438633 | 0.339 | 0.079 | 3.65759E-06 | 3.75855E-16 | 1.998669313 | 0.378 | 0.098 | 8.87468E-12 | 1.54904E-10 | 7.51709E-16 | Hbegf | MM_1 |
| 3.83968E-16 | 1.726075561 | 0.642 | 0.234 | 9.06624E-12 | 1.11046E-14 | 0.777378369 | 0.658 | 0.322 | 2.62203E-10 | 1.11046E-14 | 7.67935E-16 | Scamp1 | MM_1 |
| 6.12892E-10 | 0.699257381 | 0.706 | 0.372 | 1.44716E-05 | 4.33439E-16 | 0.836897503 | 0.784 | 0.417 | 1.02344E-11 | 6.12892E-10 | 8.66879E-16 | Arhgap12 | MM_1 |
| 1.16821E-07 | 1.091285603 | 0.532 | 0.255 | 0.002758384 | 4.45413E-16 | 0.964381099 | 0.586 | 0.256 | 1.05171E-11 | 1.16821E-07 | 8.90827E-16 | Map3k11 | MM_1 |
| 1.07326E-11 | 0.790696195 | 0.945 | 0.672 | 2.53419E-07 | 4.45684E-16 | 0.630441786 | 0.946 | 0.665 | 1.05235E-11 | 1.07326E-11 | 8.91368E-16 | Fcho2 | MM_1 |
| 2.32223E-10 | 2.697670162 | 0.248 | 0.038 | 5.48325E-06 | 5.03055E-16 | 0.922162476 | 0.284 | 0.044 | 1.18781E-11 | 2.32223E-10 | 1.00611E-15 | Kcnt2 | MM_1 |
| 9.61655E-11 | 2.758807863 | 0.22 | 0.024 | 2.27066E-06 | 5.15372E-16 | 1.071440358 | 0.333 | 0.068 | 1.2169E-11 | 9.61655E-11 | 1.03074E-15 | A4galt | MM_1 |
| 1.49005E-14 | 1.472389857 | 0.633 | 0.228 | 3.51831E-10 | 5.66855E-16 | 0.857986496 | 0.599 | 0.234 | 1.33846E-11 | 1.49005E-14 | 1.13371E-15 | Gbp7 | MM_1 |
| 1.02238E-07 | 1.392683018 | 0.165 | 0.021 | 0.002414049 | 6.13246E-16 | 1.919070004 | 0.252 | 0.03 | 1.448E-11 | 1.02238E-07 | 1.22649E-15 | Wfdc18 | MM_1 |
| 4.56711E-08 | 1.019420614 | 0.56 | 0.272 | 0.001078387 | 6.55881E-16 | 1.033295396 | 0.658 | 0.289 | 1.54867E-11 | 4.56711E-08 | 1.31176E-15 | Mob3c | MM_1 |
| 2.95104E-07 | 0.868980785 | 0.55 | 0.293 | 0.006967991 | 7.0437E-16 | 0.815701758 | 0.703 | 0.335 | 1.66316E-11 | 2.95104E-07 | 1.40874E-15 | Pdxk | MM_1 |
| 2.73426E-12 | 0.963733254 | 0.817 | 0.466 | 6.45613E-08 | 7.26703E-16 | 0.849160213 | 0.878 | 0.534 | 1.71589E-11 | 2.73426E-12 | 1.45341E-15 | Tsc22d3 | MM_1 |
| 4.23806E-09 | 0.719230771 | 0.963 | 0.728 | 0.000100069 | 7.69788E-16 | 0.653192568 | 0.968 | 0.782 | 1.81762E-11 | 4.23806E-09 | 1.53958E-15 | Txnip | MM_1 |
| 3.64397E-10 | 0.761441722 | 0.67 | 0.334 | 8.60413E-06 | 7.79754E-16 | 0.811231746 | 0.649 | 0.3 | 1.84116E-11 | 3.64397E-10 | 1.55951E-15 | Vcpip1 | MM_1 |
| 2.57562E-10 | 0.734546935 | 0.55 | 0.197 | 6.08156E-06 | 7.82997E-16 | 0.908202625 | 0.662 | 0.283 | 1.84881E-11 | 2.57562E-10 | 1.56599E-15 | Bmp2 | MM_1 |
| 3.76885E-10 | 1.246018078 | 0.385 | 0.11 | 8.89902E-06 | 8.15316E-16 | 1.156484333 | 0.432 | 0.131 | 1.92512E-11 | 3.76885E-10 | 1.63063E-15 | Kif3a | MM_1 |
| 4.31288E-09 | 0.642798085 | 0.587 | 0.259 | 0.000101836 | 9.44991E-16 | 0.941841135 | 0.649 | 0.313 | 2.23131E-11 | 4.31288E-09 | 1.88998E-15 | Arfgef2 | MM_1 |
| 3.77197E-10 | 0.755601724 | 0.624 | 0.283 | 8.90637E-06 | 1.14336E-15 | 0.769371753 | 0.725 | 0.362 | 2.69969E-11 | 3.77197E-10 | 2.28671E-15 | Insr | MM_1 |
| 8.60086E-12 | 0.992445661 | 0.817 | 0.448 | 2.03083E-07 | 1.28214E-15 | 0.779797284 | 0.856 | 0.512 | 3.02738E-11 | 8.60086E-12 | 2.56427E-15 | Man2a1 | MM_1 |
| 1.21728E-09 | 1.35901577 | 0.413 | 0.134 | 2.87424E-05 | 1.28657E-15 | 0.586699022 | 0.505 | 0.18 | 3.03785E-11 | 1.21728E-09 | 2.57314E-15 | Gas2l1 | MM_1 |
| 3.3145E-07 | 1.806532317 | 0.303 | 0.1 | 0.007826208 | 1.34653E-15 | 1.069458041 | 0.437 | 0.136 | 3.17943E-11 | 3.3145E-07 | 2.69306E-15 | Tuft1 | MM_1 |
| 4.2648E-11 | 1.022945277 | 0.706 | 0.317 | 1.007E-06 | 1.57124E-15 | 0.767802622 | 0.739 | 0.403 | 3.71002E-11 | 4.2648E-11 | 3.14249E-15 | Neu1 | MM_1 |
| 3.43106E-09 | 0.94406329 | 0.734 | 0.4 | 8.10141E-05 | 1.62185E-15 | 0.953393514 | 0.788 | 0.441 | 3.82952E-11 | 3.43106E-09 | 3.24371E-15 | Mylip | MM_1 |
| 3.5123E-08 | 0.649470887 | 0.936 | 0.538 | 0.000829323 | 1.78195E-15 | 0.628564591 | 0.982 | 0.621 | 4.20754E-11 | 3.5123E-08 | 3.5639E-15 | C3ar1 | MM_1 |
| 1.34962E-10 | 0.7715798 | 0.761 | 0.4 | 3.18672E-06 | 1.78457E-15 | 0.72007947 | 0.788 | 0.458 | 4.21372E-11 | 1.34962E-10 | 3.56913E-15 | Micu1 | MM_1 |
| 1.24668E-09 | 0.727985795 | 0.651 | 0.307 | 2.94367E-05 | 1.80528E-15 | 0.88320046 | 0.644 | 0.297 | 4.26264E-11 | 1.24668E-09 | 3.61057E-15 | Cd200r1 | MM_1 |
| 1.69249E-10 | 0.673065262 | 0.936 | 0.593 | 3.99631E-06 | 1.8084E-15 | 0.764528167 | 0.919 | 0.651 | 4.26999E-11 | 1.69249E-10 | 3.61679E-15 | Pitpnc1 | MM_1 |
| 4.41905E-08 | 0.689968744 | 0.697 | 0.366 | 0.001043427 | 2.04489E-15 | 0.787200713 | 0.73 | 0.401 | 4.8284E-11 | 4.41905E-08 | 4.08978E-15 | Cers2 | MM_1 |
| 3.91496E-10 | 0.810559434 | 0.752 | 0.4 | 9.24401E-06 | 2.28565E-15 | 0.600633487 | 0.766 | 0.395 | 5.39688E-11 | 3.91496E-10 | 4.5713E-15 | Sh3pxd2a | MM_1 |
| 1.18238E-07 | 0.689250941 | 0.706 | 0.414 | 0.002791835 | 2.67452E-15 | 1.00191262 | 0.73 | 0.436 | 6.31509E-11 | 1.18238E-07 | 5.34905E-15 | Stx4a | MM_1 |
| 9.73676E-14 | 2.853968306 | 0.33 | 0.052 | 2.29904E-09 | 3.23363E-15 | 1.854606946 | 0.306 | 0.06 | 7.63525E-11 | 9.73676E-14 | 6.46726E-15 | Pomk | MM_1 |
| 3.6898E-07 | 0.717077952 | 0.716 | 0.421 | 0.008712347 | 4.28099E-15 | 0.62377212 | 0.82 | 0.482 | 1.01083E-10 | 3.6898E-07 | 8.56198E-15 | Rap2b | MM_1 |
| 1.30697E-08 | 1.012651216 | 0.45 | 0.169 | 0.000308602 | 5.22448E-15 | 1.010329683 | 0.541 | 0.204 | 1.23361E-10 | 1.30697E-08 | 1.0449E-14 | Scamp5 | MM_1 |
| 1.68943E-11 | 0.814405659 | 0.817 | 0.445 | 3.98907E-07 | 5.58643E-15 | 0.72656185 | 0.838 | 0.49 | 1.31907E-10 | 1.68943E-11 | 1.11729E-14 | Klf7 | MM_1 |
| 3.81033E-07 | 0.801585971 | 0.606 | 0.307 | 0.008996962 | 5.79531E-15 | 0.661515415 | 0.748 | 0.411 | 1.36839E-10 | 3.81033E-07 | 1.15906E-14 | Sdhc | MM_1 |
| 6.62401E-11 | 1.207343392 | 0.633 | 0.276 | 1.56406E-06 | 7.55334E-15 | 0.658499241 | 0.676 | 0.322 | 1.7835E-10 | 6.62401E-11 | 1.51067E-14 | Notch1 | MM_1 |
| 7.0797E-09 | 0.7026703 | 0.798 | 0.414 | 0.000167166 | 8.85845E-15 | 0.704093205 | 0.815 | 0.441 | 2.09166E-10 | 7.0797E-09 | 1.77169E-14 | Sesn1 | MM_1 |
| 5.79518E-09 | 0.920564033 | 0.67 | 0.341 | 0.000136836 | 1.03809E-14 | 0.876350876 | 0.779 | 0.428 | 2.45114E-10 | 5.79518E-09 | 2.07618E-14 | Nck1 | MM_1 |
| 7.66507E-14 | 1.388547065 | 0.716 | 0.293 | 1.80988E-09 | 1.14691E-14 | 0.847151673 | 0.761 | 0.39 | 2.70809E-10 | 7.66507E-14 | 2.29383E-14 | Ccl7 | MM_1 |
| 1.08967E-10 | 0.860817314 | 0.954 | 0.641 | 2.57292E-06 | 1.18742E-14 | 0.685554117 | 0.973 | 0.668 | 2.80374E-10 | 1.08967E-10 | 2.37484E-14 | Ifi207 | MM_1 |
| 3.09615E-07 | 0.673967642 | 0.826 | 0.524 | 0.007310628 | 1.36343E-14 | 0.601299593 | 0.914 | 0.567 | 3.21932E-10 | 3.09615E-07 | 2.72685E-14 | Tiparp | MM_1 |
| 1.92834E-07 | 1.073648384 | 0.532 | 0.266 | 0.004553199 | 1.68403E-14 | 0.944816877 | 0.586 | 0.259 | 3.97632E-10 | 1.92834E-07 | 3.36805E-14 | Tdrd7 | MM_1 |
| 6.24301E-08 | 0.846866062 | 0.633 | 0.341 | 0.0014741 | 2.21546E-14 | 0.628409974 | 0.707 | 0.368 | 5.23114E-10 | 6.24301E-08 | 4.43091E-14 | Kif13b | MM_1 |
| 2.33337E-07 | 0.742139548 | 0.422 | 0.179 | 0.005509557 | 2.26891E-14 | 1.030259197 | 0.532 | 0.218 | 5.35735E-10 | 2.33337E-07 | 4.53782E-14 | Vezt | MM_1 |
| 1.76389E-10 | 1.448960467 | 0.477 | 0.166 | 4.1649E-06 | 2.36495E-14 | 1.168676611 | 0.509 | 0.207 | 5.58411E-10 | 1.76389E-10 | 4.7299E-14 | Slc38a7 | MM_1 |
| 6.81313E-08 | 0.666508461 | 0.431 | 0.166 | 0.001608717 | 2.76708E-14 | 0.828403097 | 0.532 | 0.207 | 6.53363E-10 | 6.81313E-08 | 5.53416E-14 | Vps33a | MM_1 |

| | | | | | | | | | | | | | |
|---|---|---|---|---|---|---|---|---|---|---|---|---|---|
| 5.75566E-09 | 0.825745743 | 0.413 | 0.134 | 0.000135903 | 3.63761E-14 | 0.84797195 | 0.514 | 0.193 | 8.58914E-10 | 5.75566E-09 | 7.27523E-14 | Tmem104 | MM_1 |
| 3.75663E-08 | 0.786845122 | 0.642 | 0.328 | 0.000887015 | 3.6476E-14 | 0.835617935 | 0.685 | 0.365 | 8.61271E-10 | 3.75663E-08 | 7.2952E-14 | Stxbp5 | MM_1 |
| 8.48373E-13 | 3.187107782 | 0.22 | 0.014 | 2.00318E-08 | 4.9279E-14 | 2.769884925 | 0.216 | 0.027 | 1.16358E-09 | 8.48373E-13 | 9.8558E-14 | Unc5a | MM_1 |
| 3.30473E-08 | 1.173146893 | 0.587 | 0.29 | 0.000780312 | 5.65973E-14 | 0.765579956 | 0.658 | 0.324 | 1.33638E-09 | 3.30473E-08 | 1.13195E-13 | Samd8 | MM_1 |
| 2.89374E-08 | 0.901046723 | 0.651 | 0.372 | 0.00068327 | 5.99395E-14 | 0.590777962 | 0.788 | 0.45 | 1.41529E-09 | 2.89374E-08 | 1.19879E-13 | Psmd1 | MM_1 |
| 9.95422E-08 | 0.819914557 | 0.67 | 0.407 | 0.00235039 | 9.02632E-14 | 0.585780159 | 0.811 | 0.493 | 2.13129E-09 | 9.95422E-08 | 1.80526E-13 | 9930111J2 | MM_1 |
| 1.30236E-11 | 0.968718673 | 0.991 | 0.724 | 3.07514E-07 | 1.06459E-13 | 0.648396283 | 1 | 0.826 | 2.51372E-09 | 1.30236E-11 | 2.12919E-13 | Mt1 | MM_1 |
| 1.74435E-07 | 1.053759869 | 0.514 | 0.241 | 0.004118765 | 1.30946E-13 | 0.764944447 | 0.653 | 0.324 | 3.0919E-09 | 1.74435E-07 | 2.61892E-13 | Acot13 | MM_1 |
| 5.43419E-08 | 0.741265672 | 0.881 | 0.652 | 0.00128312 | 1.38379E-13 | 0.593826797 | 0.95 | 0.725 | 3.26741E-09 | 5.43419E-08 | 2.76759E-13 | Wsb1 | MM_1 |
| 2.70845E-07 | 2.01371028 | 0.211 | 0.045 | 0.006395204 | 1.70337E-13 | 1.900928424 | 0.284 | 0.063 | 4.02199E-09 | 2.70845E-07 | 3.40673E-13 | Adcy3 | MM_1 |
| 2.80962E-09 | 1.010614786 | 0.771 | 0.431 | 6.63409E-05 | 1.93151E-13 | 0.961350264 | 0.775 | 0.463 | 4.56067E-09 | 2.80962E-09 | 3.86301E-13 | Kansl1l | MM_1 |
| 2.77568E-10 | 1.156224065 | 0.532 | 0.207 | 6.55392E-06 | 2.20625E-13 | 0.704946603 | 0.563 | 0.234 | 5.2094E-09 | 2.77568E-10 | 4.4125E-13 | Tmem106l | MM_1 |
| 2.00615E-08 | 2.280691172 | 0.174 | 0.021 | 0.000473692 | 5.52774E-13 | 2.039660526 | 0.203 | 0.025 | 1.30521E-08 | 2.00615E-08 | 1.10555E-12 | Cd209b | MM_1 |
| 1.92701E-11 | 1.246261936 | 0.321 | 0.062 | 4.55006E-07 | 5.78606E-13 | 0.856711448 | 0.36 | 0.106 | 1.3662E-08 | 1.92701E-11 | 1.15721E-12 | Uaca | MM_1 |
| 1.07728E-12 | 1.291411178 | 0.477 | 0.134 | 2.54368E-08 | 1.24506E-11 | 0.746245988 | 0.459 | 0.188 | 2.93984E-07 | 1.24506E-11 | 2.15457E-12 | Abcd2 | MM_1 |
| 8.29108E-11 | 0.911077492 | 0.716 | 0.366 | 1.95769E-06 | 1.23491E-12 | 0.713215396 | 0.667 | 0.343 | 2.91588E-08 | 8.29108E-11 | 2.46983E-12 | D1Ertd622 | MM_1 |
| 1.76532E-08 | 0.649870767 | 0.853 | 0.583 | 0.000416827 | 1.3198E-12 | 0.603412069 | 0.919 | 0.605 | 3.11632E-08 | 1.76532E-08 | 2.6396E-12 | Rabep1 | MM_1 |
| 6.14801E-12 | 2.139462503 | 0.376 | 0.09 | 1.45167E-07 | 1.60801E-12 | 0.790952934 | 0.446 | 0.177 | 3.79683E-08 | 6.14801E-12 | 3.21602E-12 | Bmi1 | MM_1 |
| 8.10771E-09 | 0.615005325 | 0.642 | 0.3 | 0.000191439 | 1.68731E-12 | 0.748695977 | 0.68 | 0.343 | 3.98407E-08 | 8.10771E-09 | 3.37462E-12 | Camkk2 | MM_1 |
| 1.07357E-07 | 1.179887491 | 0.486 | 0.21 | 0.002534921 | 1.78459E-12 | 0.635341532 | 0.554 | 0.245 | 4.21378E-08 | 1.07357E-07 | 3.56918E-12 | Tk2 | MM_1 |
| 2.18591E-07 | 0.781737283 | 0.56 | 0.272 | 0.00516136 | 4.30037E-12 | 0.964507677 | 0.586 | 0.281 | 1.0154E-07 | 2.18591E-07 | 8.60075E-12 | Slc41a2 | MM_1 |
| 3.93316E-08 | 1.653988928 | 0.33 | 0.1 | 0.000928698 | 5.13659E-12 | 1.217110631 | 0.378 | 0.125 | 1.21285E-07 | 3.93316E-08 | 1.02732E-11 | Siglec1 | MM_1 |
| 3.41227E-08 | 0.602943785 | 0.771 | 0.441 | 0.000805704 | 5.44903E-12 | 0.819412384 | 0.739 | 0.403 | 1.28662E-07 | 3.41227E-08 | 1.08981E-11 | Zfp703 | MM_1 |
| 4.47288E-08 | 0.946947857 | 0.523 | 0.228 | 0.001056135 | 9.08002E-12 | 0.616648807 | 0.55 | 0.256 | 2.14398E-07 | 4.47288E-08 | 1.816E-11 | Cdip1 | MM_1 |
| 7.83799E-10 | 1.227272354 | 0.312 | 0.069 | 1.85071E-05 | 9.97465E-12 | 1.732050165 | 0.333 | 0.109 | 2.35521E-07 | 7.83799E-10 | 1.99493E-11 | Poglut3 | MM_1 |
| 5.92221E-08 | 0.832809712 | 0.761 | 0.445 | 0.001398352 | 1.00586E-11 | 0.622044055 | 0.811 | 0.518 | 2.37504E-07 | 5.92221E-08 | 2.01172E-11 | Arl8a | MM_1 |
| 2.09616E-08 | 0.919264945 | 0.642 | 0.307 | 0.000494944 | 2.01579E-11 | 0.600351065 | 0.721 | 0.392 | 4.75969E-07 | 2.09616E-08 | 4.03159E-11 | Acadl | MM_1 |
| 4.53167E-09 | 0.961798509 | 0.569 | 0.245 | 0.000107002 | 2.92825E-11 | 0.608991528 | 0.55 | 0.248 | 6.91418E-07 | 4.53167E-09 | 5.8565E-11 | Pkp4 | MM_1 |
| 1.16389E-08 | 0.727972865 | 0.596 | 0.269 | 0.000274817 | 8.41382E-11 | 0.760408799 | 0.59 | 0.308 | 1.98667E-06 | 1.16389E-08 | 1.68276E-10 | Dusp22 | MM_1 |
| 4.01453E-10 | 1.032727275 | 0.67 | 0.321 | 9.47912E-06 | 8.97138E-11 | 0.781058924 | 0.653 | 0.351 | 2.11832E-06 | 4.01453E-10 | 1.79428E-10 | Spred2 | MM_1 |
| 6.12382E-09 | 2.39936658 | 0.193 | 0.024 | 0.000144596 | 9.37272E-11 | 1.382582223 | 0.234 | 0.052 | 2.21309E-06 | 6.12382E-09 | 1.87454E-10 | Atp1b1 | MM_1 |
| 1.87845E-07 | 0.738290188 | 0.523 | 0.245 | 0.004435403 | 9.76277E-11 | 0.84163387 | 0.55 | 0.275 | 2.30518E-06 | 1.87845E-07 | 1.95255E-10 | Zcchc2 | MM_1 |
| 1.0903E-10 | 1.614997676 | 0.248 | 0.034 | 2.57442E-06 | 7.60509E-09 | 1.25655041 | 0.243 | 0.071 | 0.000179571 | 7.60509E-09 | 2.18061E-10 | Flrt2 | MM_1 |
| 1.59534E-10 | 1.266350259 | 0.486 | 0.162 | 3.76693E-06 | 1.11407E-10 | 0.931216897 | 0.437 | 0.177 | 2.63055E-06 | 1.59534E-10 | 2.22815E-10 | Slc28a2 | MM_1 |
| 1.70833E-10 | 1.940819693 | 0.422 | 0.131 | 4.0337E-06 | 9.04042E-10 | 0.985057627 | 0.41 | 0.174 | 2.13462E-05 | 9.04042E-10 | 3.41665E-10 | Prps2 | MM_1 |
| 1.98006E-08 | 2.654541817 | 0.183 | 0.024 | 0.000467532 | 2.23613E-10 | 1.07449149 | 0.216 | 0.044 | 5.27994E-06 | 1.98006E-08 | 4.47225E-10 | Trpm1 | MM_1 |
| 3.94907E-07 | 1.216544335 | 0.312 | 0.1 | 0.009324555 | 2.89906E-10 | 0.695105392 | 0.324 | 0.109 | 6.84527E-06 | 3.94907E-07 | 5.79812E-10 | Tmem8 | MM_1 |
| 3.82726E-09 | 1.15660812 | 0.229 | 0.038 | 9.03692E-05 | 4.53131E-10 | 1.006479771 | 0.266 | 0.074 | 1.06993E-05 | 3.82726E-09 | 9.06262E-10 | Cobll1 | MM_1 |
| 2.364E-07 | 1.24432194 | 0.33 | 0.107 | 0.005581876 | 4.66704E-10 | 1.233554139 | 0.311 | 0.098 | 1.10198E-05 | 2.364E-07 | 9.33407E-10 | A930007I1 | MM_1 |
| 1.01549E-08 | 0.863711374 | 0.679 | 0.369 | 0.000239777 | 7.09219E-10 | 0.610403738 | 0.631 | 0.362 | 1.67461E-05 | 1.01549E-08 | 1.41844E-09 | Hexim1 | MM_1 |
| 3.93954E-07 | 1.102059794 | 0.257 | 0.069 | 0.00930204 | 7.09567E-10 | 1.027060564 | 0.311 | 0.104 | 1.67543E-05 | 3.93954E-07 | 1.41913E-09 | Dennd2a | MM_1 |
| 3.08454E-07 | 0.795955614 | 0.826 | 0.503 | 0.007283224 | 8.25444E-10 | 0.599478061 | 0.761 | 0.482 | 1.94904E-05 | 3.08454E-07 | 1.65089E-09 | Tsc22d2 | MM_1 |
| 3.56229E-07 | 1.0745451 | 0.596 | 0.341 | 0.00841129 | 1.93852E-09 | 0.593895422 | 0.649 | 0.384 | 4.57723E-05 | 3.56229E-07 | 3.87704E-09 | Rffl | MM_1 |
| 2.31038E-09 | 1.585140964 | 0.257 | 0.048 | 5.45528E-05 | 3.46833E-08 | 0.978116536 | 0.284 | 0.104 | 0.000818943 | 3.46833E-08 | 4.62077E-09 | Trim2 | MM_1 |
| 2.62531E-09 | 1.148188633 | 0.541 | 0.221 | 6.19887E-05 | 6.45944E-09 | 0.866752644 | 0.473 | 0.243 | 0.00015252 | 6.45944E-09 | 5.25061E-09 | Ank2 | MM_1 |
| 3.65938E-08 | 1.424731502 | 0.321 | 0.097 | 0.000864054 | 3.06234E-09 | 1.96018097 | 0.18 | 0.035 | 7.2308E-05 | 3.65938E-08 | 6.12468E-09 | Gm44751 | MM_1 |
| 3.17663E-08 | 0.7006314 | 0.771 | 0.448 | 0.000750065 | 3.51982E-09 | 0.592411597 | 0.797 | 0.515 | 8.31101E-05 | 3.17663E-08 | 7.03965E-09 | Ankrd12 | MM_1 |
| 7.56746E-08 | 1.28990242 | 0.624 | 0.352 | 0.001786828 | 5.73578E-09 | 0.65301848 | 0.631 | 0.365 | 0.000135433 | 7.56746E-08 | 1.14716E-08 | Il4ra | MM_1 |
| 2.06247E-08 | 0.805449538 | 0.725 | 0.393 | 0.000486989 | 6.03988E-09 | 0.606649451 | 0.662 | 0.387 | 0.000142614 | 2.06247E-08 | 1.20798E-08 | Sfmbt1 | MM_1 |
| 1.15383E-08 | 1.345164187 | 0.431 | 0.152 | 0.000272443 | 8.28033E-09 | 0.715316734 | 0.392 | 0.174 | 0.000195515 | 1.15383E-08 | 1.65606E-08 | Lrp5 | MM_1 |
| 3.3438E-08 | 1.163809891 | 0.486 | 0.207 | 0.000789539 | 8.75977E-09 | 0.738093734 | 0.536 | 0.278 | 0.000206836 | 3.3438E-08 | 1.75195E-08 | Cyp20a1 | MM_1 |
| 2.5036E-07 | 0.905291887 | 0.532 | 0.259 | 0.005911492 | 8.79828E-09 | 0.594179287 | 0.577 | 0.316 | 0.000207745 | 2.5036E-07 | 1.75966E-08 | Brox | MM_1 |
| 4.08873E-07 | 0.756850615 | 0.413 | 0.159 | 0.009654321 | 1.35127E-08 | 0.67498729 | 0.473 | 0.243 | 0.000319062 | 4.08873E-07 | 2.70254E-08 | Acadvl | MM_1 |
| 1.29679E-07 | 1.068838825 | 0.514 | 0.228 | 0.003061973 | 1.65415E-08 | 0.852103858 | 0.491 | 0.264 | 0.000390577 | 1.29679E-07 | 3.30829E-08 | Slc25a28 | MM_1 |
| 2.37969E-08 | 1.532190628 | 0.523 | 0.228 | 0.000561893 | 1.30308E-07 | 0.645521389 | 0.482 | 0.259 | 0.003076821 | 1.30308E-07 | 4.75939E-08 | Gm17268 | MM_1 |
| 2.43956E-07 | 2.587114795 | 0.211 | 0.045 | 0.005760284 | 5.33031E-08 | 1.192115303 | 0.207 | 0.06 | 0.001258593 | 2.43956E-07 | 1.06606E-07 | Gpt2 | MM_1 |
| 4.1854E-07 | 0.953669834 | 0.486 | 0.214 | 0.009882557 | 8.01309E-08 | 0.652721279 | 0.473 | 0.256 | 0.00189205 | 4.1854E-07 | 1.60262E-07 | Fhit | MM_1 |
| 2.12388E-07 | 1.352709226 | 0.339 | 0.117 | 0.005014904 | 8.13014E-08 | 0.857493733 | 0.342 | 0.153 | 0.001919688 | 2.12388E-07 | 1.62603E-07 | 2210408I2 | MM_1 |
| 9.62741E-37 | 2.400887685 | 0.92 | 0.292 | 2.27322E-32 | 1.28907E-73 | 3.65667068 | 0.895 | 0.144 | 3.04375E-69 | 9.62741E-37 | 2.57814E-73 | Cx3cr1 | MM_2 |
| 5.19516E-35 | 2.65559771 | 0.768 | 0.15 | 1.22668E-30 | 4.14217E-67 | 3.05427372 | 0.808 | 0.101 | 9.78049E-63 | 5.19516E-35 | 8.28433E-67 | Lilra5 | MM_2 |
| 1.21923E-25 | 1.665561144 | 0.832 | 0.299 | 2.87886E-21 | 2.99102E-56 | 3.345798964 | 0.75 | 0.125 | 7.06239E-52 | 1.21923E-25 | 5.98203E-56 | Lpcat2 | MM_2 |
| 2.39281E-22 | 1.210859049 | 0.992 | 0.814 | 5.6499E-18 | 3.56173E-55 | 2.367058166 | 0.994 | 0.655 | 8.40996E-51 | 2.39281E-22 | 7.12346E-55 | H2-Eb1 | MM_2 |
| 3.14085E-22 | 1.013942 | 0.952 | 0.387 | 7.41617E-18 | 4.3612E-55 | 2.447592777 | 0.884 | 0.23 | 1.02977E-50 | 3.14085E-22 | 8.72241E-55 | Basp1 | MM_2 |
| 6.18741E-24 | 1.45338242 | 0.968 | 0.511 | 1.46097E-19 | 8.18382E-54 | 2.319691964 | 0.977 | 0.523 | 1.93236E-49 | 6.18741E-24 | 1.63676E-53 | Cxcl16 | MM_2 |
| 2.83536E-22 | 1.135788046 | 0.992 | 0.828 | 6.69486E-18 | 3.45371E-53 | 2.172388613 | 0.988 | 0.691 | 8.15489E-49 | 2.83536E-22 | 6.90741E-53 | H2-Aa | MM_2 |
| 4.16535E-25 | 1.85952552 | 0.864 | 0.401 | 9.83523E-21 | 4.24268E-52 | 2.090546889 | 0.948 | 0.381 | 1.00178E-47 | 4.16535E-25 | 8.48537E-52 | Cd300c2 | MM_2 |
| 4.43821E-22 | 1.026038669 | 1 | 0.901 | 1.04795E-17 | 4.52781E-50 | 1.964980537 | 1 | 0.808 | 1.06911E-45 | 4.43821E-22 | 9.05563E-50 | Cd74 | MM_2 |
| 1.70108E-25 | 1.766857588 | 0.704 | 0.179 | 4.01659E-21 | 1.33791E-49 | 3.586956562 | 0.634 | 0.077 | 3.15906E-45 | 1.70108E-25 | 2.67581E-49 | Fcgr4 | MM_2 |
| 3.48347E-19 | 1.062635924 | 0.984 | 0.81 | 8.22516E-15 | 1.78182E-49 | 2.080104453 | 1 | 0.743 | 4.20723E-45 | 3.48347E-19 | 3.56364E-49 | H2-Ab1 | MM_2 |
| 5.87429E-27 | 1.603936614 | 0.968 | 0.453 | 1.38704E-22 | 1.43463E-45 | 2.284929629 | 0.936 | 0.432 | 3.38744E-41 | 5.87429E-27 | 2.86925E-45 | Bcl2a1b | MM_2 |
| 1.53951E-23 | 1.330225002 | 0.848 | 0.296 | 3.6351E-19 | 6.96718E-45 | 2.021121524 | 0.802 | 0.206 | 1.64509E-40 | 1.53951E-23 | 1.39344E-44 | Axl | MM_2 |
| 3.93946E-14 | 0.837203398 | 0.984 | 0.646 | 9.30184E-10 | 4.1202E-43 | 1.452060075 | 0.971 | 0.64 | 9.72861E-39 | 3.93946E-14 | 8.24039E-43 | Aif1 | MM_2 |
| 1.77228E-25 | 2.630662227 | 0.64 | 0.142 | 4.1847E-21 | 7.96646E-42 | 2.358682451 | 0.709 | 0.18 | 1.88104E-37 | 1.77228E-25 | 1.59329E-41 | Cd72 | MM_2 |
| 2.98898E-23 | 2.043507347 | 0.968 | 0.544 | 7.05757E-19 | 1.97558E-41 | 2.666135146 | 0.977 | 0.645 | 4.66474E-37 | 2.98898E-23 | 3.95116E-41 | Cd14 | MM_2 |
| 4.68872E-23 | 1.408870889 | 0.992 | 0.529 | 1.1071E-18 | 2.43842E-39 | 1.454481664 | 0.994 | 0.64 | 5.7576E-35 | 4.68872E-23 | 4.87685E-39 | Ms4a7 | MM_2 |
| 6.34704E-17 | 0.986548338 | 0.976 | 0.609 | 1.49866E-12 | 4.21799E-39 | 1.247749455 | 1 | 0.763 | 9.95953E-35 | 6.34704E-17 | 8.43599E-39 | Tmem176l | MM_2 |
| 1.91015E-11 | 1.452599476 | 0.672 | 0.314 | 4.51024E-07 | 4.31321E-39 | 3.225670614 | 0.744 | 0.24 | 1.01844E-34 | 1.91015E-11 | 8.62643E-39 | Ccl12 | MM_2 |
| 3.90577E-24 | 1.835394168 | 0.856 | 0.347 | 9.2223E-20 | 4.28842E-38 | 2.335507732 | 0.82 | 0.343 | 1.01258E-33 | 3.90577E-24 | 8.57684E-38 | Gpr65 | MM_2 |
| 1.1407E-20 | 1.558808339 | 0.808 | 0.339 | 2.69342E-16 | 4.5653E-37 | 2.095292424 | 0.802 | 0.293 | 1.07796E-32 | 1.1407E-20 | 9.1306E-37 | Slamf9 | MM_2 |
| 1.90046E-22 | 1.464731232 | 0.56 | 0.102 | 4.48737E-18 | 2.78701E-36 | 2.285564731 | 0.512 | 0.058 | 6.5807E-32 | 1.90046E-22 | 5.57403E-36 | Itga6 | MM_2 |
| 1.83722E-14 | 0.761182444 | 0.936 | 0.518 | 4.33805E-10 | 3.33292E-36 | 1.383854181 | 0.971 | 0.614 | 7.86968E-32 | 1.83722E-14 | 6.66583E-36 | Lst1 | MM_2 |

| | | | | | | | | | | | | | |
|---|---|---|---|---|---|---|---|---|---|---|---|---|---|
| 4.88234E-23 | 1.384597781 | 0.88 | 0.365 | 1.15282E-18 | 3.98039E-36 | 1.613850887 | 0.93 | 0.458 | 9.3985E-32 | 4.88234E-23 | 7.96078E-36 | Creb5 | MM_2 |
| 3.8274E-17 | 1.66604444 | 0.592 | 0.179 | 9.03725E-13 | 1.24888E-35 | 2.397805092 | 0.581 | 0.108 | 2.94886E-31 | 3.8274E-17 | 2.49776E-35 | Serpinf1 | MM_2 |
| 2.56865E-20 | 1.9683858 | 0.52 | 0.091 | 6.06511E-16 | 3.71479E-35 | 1.81526413 | 0.616 | 0.122 | 8.77135E-31 | 2.56865E-20 | 7.42957E-35 | Col14a1 | MM_2 |
| 1.7242E-21 | 1.490904968 | 0.776 | 0.303 | 4.07118E-17 | 2.00636E-34 | 1.52194494 | 0.907 | 0.436 | 4.73743E-30 | 1.7242E-21 | 4.01273E-34 | Zeb2os | MM_2 |
| 7.25059E-16 | 0.839448869 | 1 | 0.726 | 1.71201E-11 | 1.32555E-32 | 1.210503524 | 0.988 | 0.782 | 3.12989E-28 | 7.25059E-16 | 2.6511E-32 | Ly86 | MM_2 |
| 2.2533E-13 | 1.727876541 | 0.496 | 0.15 | 5.32049E-09 | 2.30789E-32 | 3.808894473 | 0.442 | 0.053 | 5.44939E-28 | 2.2533E-13 | 4.61578E-32 | Scimp | MM_2 |
| 7.85514E-16 | 1.217332941 | 0.752 | 0.321 | 1.85476E-11 | 2.45473E-32 | 1.747274041 | 0.837 | 0.386 | 5.79612E-28 | 7.85514E-16 | 4.90947E-32 | Fcgr1 | MM_2 |
| 2.58727E-16 | 1.084949884 | 0.96 | 0.547 | 6.10907E-12 | 8.30265E-32 | 1.220594875 | 0.971 | 0.695 | 1.96042E-27 | 2.58727E-16 | 1.66053E-31 | Tmem176a | MM_2 |
| 6.8287E-23 | 2.200131679 | 0.56 | 0.099 | 1.61239E-18 | 1.65106E-31 | 2.375009178 | 0.512 | 0.084 | 3.89848E-27 | 6.8287E-23 | 3.30212E-31 | Ctnnd2 | MM_2 |
| 6.02013E-22 | 1.607253334 | 0.872 | 0.361 | 1.42147E-17 | 2.07536E-31 | 1.747847321 | 0.843 | 0.4 | 4.90034E-27 | 6.02013E-22 | 4.15072E-31 | St3gal6 | MM_2 |
| 1.35387E-12 | 1.443064866 | 0.624 | 0.241 | 3.19677E-08 | 2.3318E-31 | 1.763443291 | 0.866 | 0.408 | 5.50584E-27 | 1.35387E-12 | 4.66359E-31 | G530011O | MM_2 |
| 9.13399E-20 | 1.383065228 | 0.832 | 0.332 | 2.15672E-15 | 4.47529E-31 | 1.670418223 | 0.814 | 0.372 | 1.05671E-26 | 9.13399E-20 | 8.95059E-31 | Itgb5 | MM_2 |
| 1.23397E-16 | 1.28304602 | 0.832 | 0.427 | 2.91365E-12 | 5.71156E-31 | 1.521566471 | 0.901 | 0.516 | 1.34861E-26 | 1.23397E-16 | 1.14231E-30 | Lacc1 | MM_2 |
| 3.90603E-15 | 0.928307464 | 0.984 | 0.792 | 9.22291E-11 | 7.25069E-31 | 1.329440197 | 0.977 | 0.882 | 1.71203E-26 | 3.90603E-15 | 1.45014E-30 | Apoe | MM_2 |
| 5.75732E-13 | 1.02710895 | 0.888 | 0.526 | 1.35942E-08 | 1.19186E-30 | 1.783389979 | 0.89 | 0.516 | 2.81423E-26 | 5.75732E-13 | 2.38373E-30 | Skil | MM_2 |
| 6.1008E-12 | 1.175764079 | 0.664 | 0.292 | 1.44052E-07 | 1.86636E-30 | 1.509693967 | 0.756 | 0.242 | 4.40686E-26 | 6.1008E-12 | 3.73273E-30 | Ccnd2 | MM_2 |
| 5.73611E-15 | 0.634493401 | 1 | 0.836 | 1.35441E-10 | 3.18906E-30 | 0.754127481 | 1 | 0.88 | 7.53001E-26 | 5.73611E-15 | 6.37812E-30 | Fcer1g | MM_2 |
| 3.49215E-16 | 1.011650665 | 0.872 | 0.412 | 8.24567E-12 | 3.56634E-30 | 1.585151463 | 0.866 | 0.42 | 8.42084E-26 | 3.49215E-16 | 7.13268E-30 | Fcrls | MM_2 |
| 1.80572E-13 | 1.137016146 | 0.768 | 0.434 | 4.26367E-09 | 6.22476E-30 | 1.58509212 | 0.872 | 0.422 | 1.46979E-25 | 1.80572E-13 | 1.24495E-29 | Plxna4 | MM_2 |
| 6.23934E-25 | 1.516070524 | 0.968 | 0.507 | 1.47323E-20 | 2.82614E-29 | 1.371562985 | 0.953 | 0.624 | 6.67309E-25 | 6.23934E-25 | 5.65228E-29 | Apobec1 | MM_2 |
| 1.5026E-17 | 0.920023572 | 1 | 0.85 | 3.54793E-13 | 3.23694E-29 | 1.059107102 | 1 | 0.911 | 7.64305E-25 | 1.5026E-17 | 6.47387E-29 | H3f3b | MM_2 |
| 3.57116E-17 | 1.011368565 | 0.664 | 0.226 | 8.43222E-13 | 4.81247E-29 | 1.95043532 | 0.616 | 0.18 | 1.13632E-24 | 3.57116E-17 | 9.62493E-29 | Pdgfb | MM_2 |
| 1.51359E-23 | 1.606615226 | 0.792 | 0.288 | 3.57389E-19 | 5.89312E-29 | 1.815367748 | 0.738 | 0.312 | 1.39148E-24 | 1.51359E-23 | 1.17862E-28 | Rcbtb2 | MM_2 |
| 4.5429E-17 | 2.279299693 | 0.552 | 0.15 | 1.07267E-12 | 4.12458E-28 | 2.049325809 | 0.564 | 0.144 | 9.73897E-24 | 4.5429E-17 | 8.24917E-28 | Gm5086 | MM_2 |
| 5.12016E-12 | 0.715595447 | 0.52 | 0.161 | 1.20897E-07 | 4.38825E-28 | 1.847694922 | 0.512 | 0.096 | 1.03615E-23 | 5.12016E-12 | 8.77649E-28 | Cadm1 | MM_2 |
| 2.34932E-18 | 1.366496516 | 0.824 | 0.431 | 5.54722E-14 | 7.51445E-28 | 2.141315462 | 0.756 | 0.372 | 1.77431E-23 | 2.34932E-18 | 1.50289E-27 | Tgfbr1 | MM_2 |
| 2.80346E-20 | 1.071008134 | 1 | 0.661 | 6.61953E-16 | 9.02685E-28 | 0.844402119 | 1 | 0.839 | 2.13142E-23 | 2.80346E-20 | 1.80537E-27 | C1qb | MM_2 |
| 5.64933E-22 | 1.398034688 | 0.872 | 0.423 | 1.33392E-17 | 2.44437E-27 | 1.562656063 | 0.831 | 0.424 | 5.77166E-23 | 5.64933E-22 | 4.88875E-27 | Slc15a3 | MM_2 |
| 7.17498E-15 | 0.974779781 | 0.904 | 0.485 | 1.69416E-10 | 3.0272E-27 | 1.255843967 | 0.924 | 0.53 | 7.14783E-23 | 7.17498E-15 | 6.0544E-27 | Clec4a2 | MM_2 |
| 1.24223E-19 | 0.875680513 | 1 | 0.734 | 2.93316E-15 | 4.49175E-27 | 1.075830465 | 0.988 | 0.635 | 1.06059E-22 | 1.24223E-19 | 8.9835E-27 | C1qa | MM_2 |
| 1.96098E-13 | 1.222719473 | 0.704 | 0.328 | 4.63027E-09 | 7.49666E-27 | 2.006182255 | 0.669 | 0.259 | 1.77011E-22 | 1.96098E-13 | 1.49933E-26 | Inpp4b | MM_2 |
| 2.19703E-13 | 1.426911993 | 0.728 | 0.391 | 5.18764E-09 | 1.35642E-26 | 1.533427886 | 0.773 | 0.326 | 3.20277E-22 | 2.19703E-13 | 2.71283E-26 | Stap1 | MM_2 |
| 3.8926E-19 | 2.420057982 | 0.792 | 0.391 | 9.1912E-15 | 2.72833E-26 | 2.553563945 | 0.808 | 0.439 | 6.44214E-22 | 3.8926E-19 | 5.45666E-26 | Ccl4 | MM_2 |
| 6.81786E-15 | 1.520809094 | 0.616 | 0.23 | 1.60983E-10 | 3.72277E-26 | 1.821396058 | 0.686 | 0.276 | 8.7902E-22 | 6.81786E-15 | 7.44553E-26 | Lpar6 | MM_2 |
| 3.58977E-20 | 1.589553467 | 0.68 | 0.208 | 8.47617E-16 | 5.50665E-26 | 1.849174614 | 0.599 | 0.175 | 1.30023E-21 | 3.58977E-20 | 1.10133E-25 | Gm35154 | MM_2 |
| 8.46234E-26 | 1.741501248 | 0.792 | 0.237 | 1.99813E-21 | 7.36661E-18 | 0.587816097 | 0.733 | 0.362 | 1.7394E-13 | 7.36661E-18 | 1.69247E-25 | Adap2os | MM_2 |
| 2.55467E-19 | 1.266631056 | 0.584 | 0.139 | 6.03209E-15 | 1.06882E-25 | 1.220281148 | 0.483 | 0.091 | 2.5237E-21 | 2.55467E-19 | 2.13764E-25 | Gngt2 | MM_2 |
| 4.25386E-12 | 0.78867784 | 0.984 | 0.642 | 1.00442E-07 | 1.74725E-25 | 0.961715686 | 0.965 | 0.607 | 4.12562E-21 | 4.25386E-12 | 3.49451E-25 | Mpeg1 | MM_2 |
| 4.1709E-20 | 1.166431353 | 0.896 | 0.405 | 9.84832E-16 | 1.88813E-25 | 1.426273537 | 0.837 | 0.47 | 4.45826E-21 | 4.1709E-20 | 3.77627E-25 | Tmcc3 | MM_2 |
| 1.27659E-17 | 1.073606327 | 0.928 | 0.54 | 3.01429E-13 | 1.97507E-25 | 1.091427997 | 0.924 | 0.554 | 4.66354E-21 | 1.27659E-17 | 3.95014E-25 | Otulinl | MM_2 |
| 1.71914E-09 | 0.848178926 | 0.576 | 0.259 | 4.05923E-05 | 3.40379E-25 | 1.380174112 | 0.622 | 0.182 | 8.03703E-21 | 1.71914E-09 | 6.80758E-25 | Ciita | MM_2 |
| 1.04757E-19 | 1.716353687 | 0.832 | 0.423 | 2.47352E-15 | 1.19626E-24 | 1.332947086 | 0.797 | 0.444 | 2.8246E-20 | 1.04757E-19 | 2.39251E-24 | Clec12a | MM_2 |
| 3.00272E-14 | 0.699556433 | 1 | 0.854 | 7.09003E-10 | 1.3158E-24 | 0.807059566 | 1 | 0.894 | 3.10687E-20 | 3.00272E-14 | 2.6316E-24 | Ctss | MM_2 |
| 2.53453E-12 | 0.733584214 | 0.984 | 0.73 | 5.98453E-08 | 2.4864E-24 | 0.833692646 | 0.994 | 0.82 | 5.87088E-20 | 2.53453E-12 | 4.9728E-24 | Ctsh | MM_2 |
| 9.04298E-12 | 0.759457851 | 0.896 | 0.533 | 2.13523E-07 | 2.9344E-24 | 1.22471003 | 0.907 | 0.525 | 6.92871E-20 | 9.04298E-12 | 5.8688E-24 | Gatm | MM_2 |
| 7.63796E-08 | 1.212701567 | 0.4 | 0.161 | 0.001803476 | 5.96694E-24 | 2.555874535 | 0.43 | 0.082 | 1.40891E-19 | 7.63796E-08 | 1.19339E-23 | Prdm1 | MM_2 |
| 2.86421E-18 | 2.093157382 | 0.456 | 0.08 | 6.76296E-14 | 9.1354E-24 | 2.796739103 | 0.413 | 0.074 | 2.15705E-19 | 2.86421E-18 | 1.82708E-23 | Klra2 | MM_2 |
| 2.65072E-14 | 0.92365194 | 0.888 | 0.511 | 6.25887E-10 | 1.52403E-23 | 1.070410609 | 0.895 | 0.542 | 3.59855E-19 | 2.65072E-14 | 3.04807E-23 | Arrb2 | MM_2 |
| 2.56489E-09 | 0.802307264 | 0.696 | 0.365 | 6.05622E-05 | 1.96E-23 | 1.175677771 | 0.791 | 0.381 | 4.62794E-19 | 2.56489E-09 | 3.91999E-23 | Fgl2 | MM_2 |
| 2.92341E-07 | 0.649204584 | 0.776 | 0.474 | 0.006902749 | 3.25283E-23 | 1.285204852 | 0.785 | 0.386 | 7.68058E-19 | 2.92341E-07 | 6.50566E-23 | Ccnd1 | MM_2 |
| 6.48718E-17 | 1.23811971 | 0.912 | 0.485 | 1.53175E-12 | 3.62E-23 | 1.150756824 | 0.913 | 0.566 | 8.54755E-19 | 6.48718E-17 | 7.24E-23 | Adap2 | MM_2 |
| 5.15201E-14 | 2.082684468 | 0.48 | 0.146 | 1.21649E-09 | 4.41825E-23 | 1.907594295 | 0.494 | 0.127 | 1.04324E-18 | 5.15201E-14 | 8.83651E-23 | Tmem273 | MM_2 |
| 1.65686E-09 | 0.767858169 | 0.976 | 0.759 | 3.91218E-05 | 6.31193E-23 | 1.192021328 | 0.977 | 0.791 | 1.49037E-18 | 1.65686E-09 | 1.26239E-22 | Cd83 | MM_2 |
| 2.23414E-14 | 0.891223463 | 0.984 | 0.686 | 5.27524E-10 | 6.60019E-23 | 1.007260158 | 0.977 | 0.698 | 1.55844E-18 | 2.23414E-14 | 1.32004E-22 | Fyb | MM_2 |
| 1.90287E-11 | 1.72658394 | 0.632 | 0.31 | 4.49305E-07 | 7.75558E-23 | 1.75020591 | 0.721 | 0.326 | 1.83125E-18 | 1.90287E-11 | 1.55112E-22 | Ccrl2 | MM_2 |
| 3.20836E-07 | 0.732013733 | 0.464 | 0.204 | 0.007575583 | 1.45832E-22 | 1.77870114 | 0.512 | 0.127 | 3.44338E-18 | 3.20836E-07 | 2.91664E-22 | Sema4d | MM_2 |
| 7.05608E-15 | 0.993094241 | 0.856 | 0.438 | 1.66608E-10 | 1.97737E-22 | 1.468555946 | 0.756 | 0.381 | 4.66898E-18 | 7.05608E-15 | 3.95475E-22 | Dock4 | MM_2 |
| 1.87305E-14 | 1.277355058 | 0.592 | 0.215 | 4.42264E-10 | 2.64987E-22 | 1.60004743 | 0.622 | 0.245 | 6.25688E-18 | 1.87305E-14 | 5.29974E-22 | Milr1 | MM_2 |
| 1.67003E-13 | 0.919822782 | 0.72 | 0.354 | 3.94329E-09 | 3.60836E-22 | 1.605441209 | 0.715 | 0.333 | 8.52005E-18 | 1.67003E-13 | 7.21671E-22 | Arl4c | MM_2 |
| 3.1718E-07 | 0.782052402 | 0.6 | 0.339 | 0.007489261 | 4.33076E-22 | 1.497049595 | 0.674 | 0.326 | 1.02258E-17 | 3.1718E-07 | 8.66152E-22 | Ntpcr | MM_2 |
| 1.01779E-18 | 1.31399372 | 0.848 | 0.376 | 2.4032E-14 | 4.38085E-22 | 1.177423782 | 0.872 | 0.518 | 1.03441E-17 | 1.01779E-18 | 8.76169E-22 | Lair1 | MM_2 |
| 6.11675E-08 | 0.661408015 | 0.576 | 0.27 | 0.001444286 | 1.90143E-21 | 2.294633443 | 0.378 | 0.067 | 4.48967E-17 | 6.11675E-08 | 3.80287E-21 | Rtn1 | MM_2 |
| 6.93553E-11 | 0.736726819 | 0.816 | 0.423 | 1.63762E-06 | 2.1392E-21 | 1.244622019 | 0.843 | 0.482 | 5.05107E-17 | 6.93553E-11 | 4.27839E-21 | Ccr5 | MM_2 |
| 2.11183E-12 | 1.632873463 | 0.632 | 0.288 | 4.98644E-08 | 2.3254E-21 | 2.031120945 | 0.616 | 0.247 | 5.49074E-17 | 2.11183E-12 | 4.6508E-21 | E230029C0 | MM_2 |
| 2.75854E-21 | 1.438301224 | 0.896 | 0.442 | 6.51347E-17 | 8.88084E-20 | 1.115299724 | 0.849 | 0.511 | 2.09694E-15 | 8.88084E-20 | 5.51708E-21 | Cdk6 | MM_2 |
| 6.60144E-10 | 0.91131986 | 0.536 | 0.226 | 1.55873E-05 | 3.22666E-21 | 1.450884386 | 0.506 | 0.144 | 7.61879E-17 | 6.60144E-10 | 6.45332E-21 | Tle1 | MM_2 |
| 5.68267E-21 | 2.462997676 | 0.72 | 0.263 | 1.34179E-16 | 8.0572E-17 | 2.218731448 | 0.692 | 0.369 | 1.90247E-12 | 8.0572E-17 | 1.13653E-20 | Ccl3 | MM_2 |
| 5.4794E-16 | 1.290880806 | 0.832 | 0.427 | 1.2938E-11 | 6.6491E-21 | 1.022705834 | 0.831 | 0.492 | 1.56999E-16 | 5.4794E-16 | 1.32982E-20 | Retreg1 | MM_2 |
| 1.23125E-13 | 2.443218311 | 0.36 | 0.069 | 2.90722E-09 | 7.65761E-21 | 1.530850658 | 0.401 | 0.082 | 1.80812E-16 | 1.23125E-13 | 1.53152E-20 | Hs3st3a1 | MM_2 |
| 1.78156E-13 | 1.915342706 | 0.304 | 0.04 | 4.20663E-09 | 1.12555E-20 | 1.540895842 | 0.291 | 0.031 | 2.65765E-16 | 1.78156E-13 | 2.2511E-20 | Arap3 | MM_2 |
| 5.93751E-15 | 0.897585559 | 0.992 | 0.686 | 1.40197E-10 | 1.16534E-20 | 0.929510699 | 0.994 | 0.758 | 2.75161E-16 | 5.93751E-15 | 2.33069E-20 | Cybb | MM_2 |
| 4.0813E-12 | 2.911161396 | 0.344 | 0.077 | 9.63677E-08 | 1.19263E-20 | 3.13310148 | 0.326 | 0.046 | 2.81603E-16 | 4.0813E-12 | 2.38526E-20 | Gm48065 | MM_2 |
| 1.58407E-07 | 0.749383491 | 0.624 | 0.365 | 0.003740306 | 1.34866E-20 | 1.246167981 | 0.773 | 0.42 | 3.18445E-16 | 1.58407E-07 | 2.69732E-20 | Hmgcl | MM_2 |
| 2.81125E-11 | 0.67216801 | 0.992 | 0.821 | 6.63793E-07 | 1.54408E-20 | 0.754220225 | 0.994 | 0.911 | 3.64589E-16 | 2.81125E-11 | 3.08817E-20 | Ubb | MM_2 |
| 2.30566E-20 | 1.205159905 | 0.96 | 0.504 | 5.44411E-16 | 1.55172E-10 | 0.585057191 | 0.971 | 0.669 | 3.66392E-06 | 1.55172E-10 | 4.61131E-20 | C3ar1 | MM_2 |
| 7.3599E-10 | 1.72278969 | 0.304 | 0.073 | 1.73782E-05 | 2.31652E-20 | 1.693834735 | 0.349 | 0.06 | 5.46977E-16 | 7.3599E-10 | 4.63304E-20 | Zmynd15 | MM_2 |
| 2.99964E-11 | 1.197159691 | 0.552 | 0.234 | 7.08276E-07 | 3.45791E-20 | 2.215970685 | 0.552 | 0.216 | 8.16481E-16 | 2.99964E-11 | 6.91581E-20 | AU020206 | MM_2 |
| 1.75186E-07 | 0.854460937 | 0.744 | 0.493 | 0.004136488 | 4.21414E-20 | 1.66979209 | 0.849 | 0.537 | 9.95042E-16 | 1.75186E-07 | 8.42827E-20 | Tnf | MM_2 |
| 4.97388E-14 | 1.122865896 | 0.728 | 0.336 | 1.17443E-09 | 8.02119E-20 | 1.082701939 | 0.767 | 0.432 | 1.89396E-15 | 4.97388E-14 | 1.60424E-19 | Bmyc | MM_2 |
| 1.40486E-07 | 1.385636387 | 0.216 | 0.044 | 0.003317144 | 1.99713E-19 | 2.730173694 | 0.227 | 0.012 | 4.71563E-15 | 1.40486E-07 | 3.99427E-19 | Rapsn | MM_2 |
| 3.30357E-13 | 0.864862127 | 0.96 | 0.522 | 7.8004E-09 | 2.44977E-19 | 1.043891329 | 0.971 | 0.612 | 5.78441E-15 | 3.30357E-13 | 4.89955E-19 | Tanc2 | MM_2 |

| | | | | | | | | | | | | | |
|---|---|---|---|---|---|---|---|---|---|---|---|---|---|
| 4.30431E-09 | 0.937428024 | 0.48 | 0.186 | 0.000101633 | 3.96515E-19 | 1.13043654 | 0.517 | 0.163 | 9.36251E-15 | 4.30431E-09 | 7.9303E-19 | Gm20513 | MM_2 |
| 4.73466E-14 | 1.210943051 | 0.832 | 0.449 | 1.11795E-09 | 5.18103E-19 | 1.167760285 | 0.814 | 0.506 | 1.22334E-14 | 4.73466E-14 | 1.03621E-18 | Pou2f2 | MM_2 |
| 1.24177E-14 | 0.696049841 | 0.992 | 0.719 | 2.93207E-10 | 8.68025E-19 | 0.650633291 | 1 | 0.832 | 2.04958E-14 | 1.24177E-14 | 1.73605E-18 | Cd81 | MM_2 |
| 4.35026E-12 | 0.780404606 | 0.856 | 0.551 | 1.02718E-07 | 9.05462E-19 | 1.027099996 | 0.86 | 0.535 | 2.13798E-14 | 4.35026E-12 | 1.81092E-18 | Cttnbp2nl | MM_2 |
| 3.29465E-08 | 2.015644413 | 0.216 | 0.04 | 0.000777934 | 1.16446E-18 | 3.788046445 | 0.221 | 0.012 | 2.74952E-14 | 3.29465E-08 | 2.32892E-18 | St18 | MM_2 |
| 5.81097E-18 | 1.469890993 | 0.736 | 0.299 | 1.37209E-13 | 2.06741E-12 | 0.853290215 | 0.61 | 0.302 | 4.88156E-08 | 2.06741E-12 | 1.16219E-17 | Glcci1 | MM_2 |
| 7.48922E-18 | 1.842054623 | 0.528 | 0.139 | 1.76835E-13 | 4.03202E-10 | 0.906363549 | 0.488 | 0.254 | 9.52041E-06 | 4.03202E-10 | 1.49784E-17 | Lysmd4 | MM_2 |
| 1.22339E-11 | 0.667239648 | 0.984 | 0.781 | 2.88866E-07 | 1.14036E-17 | 0.758189448 | 0.994 | 0.794 | 2.69261E-13 | 1.22339E-11 | 2.28072E-17 | Mef2c | MM_2 |
| 1.59373E-17 | 1.105191391 | 0.888 | 0.416 | 3.76312E-13 | 6.86354E-11 | 0.630591943 | 0.831 | 0.592 | 1.62062E-06 | 6.86354E-11 | 3.18746E-17 | Cd300ld | MM_2 |
| 1.80658E-16 | 1.00716163 | 0.968 | 0.697 | 4.26571E-12 | 2.60503E-17 | 0.734115877 | 0.977 | 0.751 | 6.151E-13 | 1.80658E-16 | 5.21006E-17 | Tpd52 | MM_2 |
| 2.65434E-17 | 1.235995586 | 0.752 | 0.303 | 6.26742E-13 | 5.7629E-12 | 0.815786954 | 0.587 | 0.297 | 1.36074E-07 | 5.7629E-12 | 5.30867E-17 | Maml3 | MM_2 |
| 1.41827E-14 | 0.993853088 | 0.832 | 0.445 | 3.34883E-10 | 4.66319E-17 | 0.976465767 | 0.744 | 0.408 | 1.10107E-12 | 1.41827E-14 | 9.32638E-17 | Dst | MM_2 |
| 2.33853E-09 | 2.071936424 | 0.432 | 0.168 | 5.52173E-05 | 4.6743E-17 | 1.413715339 | 0.564 | 0.228 | 1.1037E-12 | 2.33853E-09 | 9.34861E-17 | Acsl1 | MM_2 |
| 4.7472E-17 | 2.24020084 | 0.48 | 0.109 | 1.12091E-12 | 9.83779E-15 | 1.935457443 | 0.378 | 0.115 | 2.3229E-10 | 9.83779E-15 | 9.4944E-17 | Tlr1 | MM_2 |
| 1.14409E-08 | 0.994156819 | 0.744 | 0.471 | 0.000270143 | 5.03203E-17 | 1.217402992 | 0.808 | 0.487 | 1.18816E-12 | 1.14409E-08 | 1.00641E-16 | Bcl6 | MM_2 |
| 6.519E-16 | 1.043147636 | 0.976 | 0.558 | 1.53927E-11 | 5.32679E-17 | 0.793057886 | 1 | 0.722 | 1.25776E-12 | 6.519E-16 | 1.06536E-16 | C5ar1 | MM_2 |
| 1.00612E-16 | 1.887193852 | 0.544 | 0.161 | 2.37566E-12 | 3.00703E-12 | 1.392856368 | 0.488 | 0.235 | 7.10021E-08 | 3.00703E-12 | 2.01225E-16 | Mvb12b | MM_2 |
| 1.21034E-16 | 1.010429089 | 0.936 | 0.577 | 2.85785E-12 | 2.49067E-14 | 0.659072893 | 0.977 | 0.7 | 5.88096E-10 | 2.49067E-14 | 2.42067E-16 | Lilrb4a | MM_2 |
| 2.41655E-11 | 0.601822739 | 1 | 0.898 | 5.70597E-07 | 1.28087E-16 | 0.619317904 | 1 | 0.94 | 3.02438E-12 | 2.41655E-11 | 2.56173E-16 | Lyz2 | MM_2 |
| 9.03338E-08 | 1.337569485 | 0.4 | 0.15 | 0.002132962 | 1.60967E-16 | 2.921609664 | 0.349 | 0.082 | 3.80074E-12 | 9.03338E-08 | 3.21933E-16 | Vcam1 | MM_2 |
| 3.78702E-10 | 1.039442699 | 0.776 | 0.482 | 8.94192E-06 | 2.37414E-16 | 1.012253235 | 0.901 | 0.571 | 5.60582E-12 | 3.78702E-10 | 4.74828E-16 | Neurl3 | MM_2 |
| 3.43711E-08 | 0.667531343 | 0.448 | 0.172 | 0.000811571 | 4.12648E-16 | 1.238699363 | 0.471 | 0.149 | 9.74344E-12 | 3.43711E-08 | 8.25295E-16 | Ramp1 | MM_2 |
| 5.15492E-13 | 0.800348931 | 0.816 | 0.427 | 1.21718E-08 | 4.54698E-16 | 1.548725913 | 0.686 | 0.379 | 1.07363E-11 | 5.15492E-13 | 9.09396E-16 | Itgav | MM_2 |
| 3.45398E-07 | 0.69142838 | 0.576 | 0.318 | 0.008155531 | 5.06442E-16 | 1.311016294 | 0.674 | 0.369 | 1.19581E-11 | 3.45398E-07 | 1.01288E-15 | Pdcd2l | MM_2 |
| 8.23798E-11 | 0.933417114 | 0.48 | 0.168 | 1.94515E-06 | 5.11887E-16 | 1.487575349 | 0.5 | 0.211 | 1.20867E-11 | 8.23798E-11 | 1.02377E-15 | Dcakd | MM_2 |
| 3.59728E-07 | 0.806510627 | 0.584 | 0.328 | 0.008493891 | 5.52924E-16 | 1.297933656 | 0.552 | 0.237 | 1.30556E-11 | 3.59728E-07 | 1.10585E-15 | Disc1 | MM_2 |
| 2.81275E-12 | 0.804801341 | 0.968 | 0.591 | 6.64148E-08 | 9.1177E-16 | 0.788713795 | 1 | 0.755 | 2.15287E-11 | 2.81275E-12 | 1.82354E-15 | Adgre1 | MM_2 |
| 6.07628E-09 | 1.137869472 | 0.544 | 0.263 | 0.000143473 | 1.34392E-15 | 2.163997695 | 0.558 | 0.254 | 3.17326E-11 | 6.07628E-09 | 2.68784E-15 | Pmepa1 | MM_2 |
| 2.01277E-09 | 0.702873446 | 0.944 | 0.624 | 4.75255E-05 | 1.73116E-15 | 0.709378396 | 0.983 | 0.739 | 4.08762E-11 | 2.01277E-09 | 3.46232E-15 | Ifngr1 | MM_2 |
| 2.26721E-15 | 1.442500566 | 0.84 | 0.456 | 5.35333E-11 | 2.04545E-13 | 1.182950868 | 0.89 | 0.609 | 4.82971E-09 | 2.04545E-13 | 4.53441E-15 | Gpr183 | MM_2 |
| 7.05715E-12 | 0.876353738 | 0.792 | 0.515 | 1.66633E-07 | 2.468E-15 | 0.892228729 | 0.849 | 0.58 | 5.82743E-11 | 7.05715E-12 | 4.93599E-15 | Ifngr2 | MM_2 |
| 8.21681E-12 | 0.87414808 | 0.856 | 0.442 | 1.94015E-07 | 2.93788E-15 | 0.851979389 | 0.878 | 0.552 | 6.93692E-11 | 8.21681E-12 | 5.87576E-15 | Slco2b1 | MM_2 |
| 6.27572E-08 | 1.0996868 | 0.736 | 0.522 | 0.001481824 | 2.9457E-15 | 1.346838705 | 0.785 | 0.52 | 6.95538E-11 | 6.27572E-08 | 5.8914E-15 | Gadd45b | MM_2 |
| 1.10187E-10 | 2.307805005 | 0.32 | 0.077 | 2.60173E-06 | 3.07195E-15 | 1.470960846 | 0.355 | 0.091 | 7.25348E-11 | 1.10187E-10 | 6.14389E-15 | Rapgef5 | MM_2 |
| 9.81372E-11 | 1.403023152 | 0.6 | 0.281 | 2.31721E-06 | 4.31532E-15 | 1.418197111 | 0.692 | 0.398 | 1.01893E-10 | 9.81372E-11 | 8.63065E-15 | Rab20 | MM_2 |
| 6.96923E-11 | 0.667720857 | 1 | 0.668 | 1.64558E-06 | 4.59648E-15 | 0.612701769 | 1 | 0.832 | 1.08532E-10 | 6.96923E-11 | 9.19296E-15 | C1qc | MM_2 |
| 4.34263E-09 | 0.919023729 | 0.8 | 0.478 | 0.000102538 | 4.98454E-15 | 0.881709099 | 0.797 | 0.465 | 1.17695E-10 | 4.34263E-09 | 9.96908E-15 | Rgs2 | MM_2 |
| 5.32549E-15 | 1.367090784 | 0.44 | 0.095 | 1.25745E-10 | 3.44959E-09 | 0.853116181 | 0.401 | 0.173 | 8.14518E-05 | 3.44959E-09 | 1.0651E-14 | Pparg | MM_2 |
| 1.18594E-09 | 1.115738569 | 0.608 | 0.31 | 2.80024E-05 | 6.3706E-15 | 1.121097455 | 0.669 | 0.386 | 1.50423E-10 | 1.18594E-09 | 1.27412E-14 | H2-M3 | MM_2 |
| 4.83521E-10 | 1.902202187 | 0.48 | 0.193 | 1.14169E-05 | 6.47365E-15 | 1.60196031 | 0.488 | 0.189 | 1.52856E-10 | 4.83521E-10 | 1.29473E-14 | Rgs1 | MM_2 |
| 2.45388E-14 | 1.055902696 | 0.808 | 0.365 | 5.7941E-10 | 7.48914E-15 | 0.776450545 | 0.866 | 0.518 | 1.76834E-10 | 2.45388E-14 | 1.49783E-14 | Trem2 | MM_2 |
| 4.83415E-12 | 0.82470291 | 0.904 | 0.511 | 1.14144E-07 | 7.78893E-15 | 0.885612008 | 0.878 | 0.559 | 1.83912E-10 | 4.83415E-12 | 1.55779E-14 | P2ry6 | MM_2 |
| 8.9035E-08 | 0.621765158 | 0.512 | 0.23 | 0.002102295 | 9.68188E-15 | 1.002002962 | 0.413 | 0.127 | 2.28608E-10 | 8.9035E-08 | 1.93638E-14 | Grk3 | MM_2 |
| 5.40112E-13 | 0.83643946 | 0.952 | 0.617 | 1.27531E-08 | 1.04006E-14 | 0.845533177 | 0.936 | 0.681 | 2.45579E-10 | 5.40112E-13 | 2.08012E-14 | Gab2 | MM_2 |
| 7.45443E-12 | 0.740287791 | 0.904 | 0.609 | 1.76014E-07 | 1.1525E-14 | 0.787755631 | 0.942 | 0.724 | 2.72129E-10 | 7.45443E-12 | 2.30501E-14 | Ifnar2 | MM_2 |
| 1.59813E-14 | 1.231917267 | 0.856 | 0.493 | 3.77352E-10 | 3.01331E-12 | 0.84802154 | 0.901 | 0.691 | 7.11502E-08 | 3.01331E-12 | 3.19627E-14 | Metrnl | MM_2 |
| 1.68934E-07 | 0.758415658 | 0.744 | 0.467 | 0.003988879 | 1.6642E-14 | 0.905129293 | 0.831 | 0.544 | 3.92951E-10 | 1.68934E-07 | 3.3284E-14 | Lrrc25 | MM_2 |
| 1.35687E-11 | 1.021901421 | 0.696 | 0.31 | 3.20384E-07 | 1.91097E-14 | 1.097420706 | 0.68 | 0.357 | 4.51218E-10 | 1.35687E-11 | 3.82194E-14 | Rnf180 | MM_2 |
| 6.94984E-12 | 1.630019212 | 0.48 | 0.179 | 1.641E-07 | 2.10392E-14 | 1.35676878 | 0.552 | 0.261 | 4.96777E-10 | 6.94984E-12 | 4.20783E-14 | Susd3 | MM_2 |
| 8.75213E-08 | 0.787581128 | 0.456 | 0.19 | 0.002066553 | 2.28649E-14 | 1.702183512 | 0.465 | 0.182 | 5.39886E-10 | 8.75213E-08 | 4.57298E-14 | Icosl | MM_2 |
| 1.32212E-07 | 1.106476452 | 0.184 | 0.029 | 0.003121782 | 2.87727E-14 | 2.821014351 | 0.209 | 0.024 | 6.79381E-10 | 1.32212E-07 | 5.75454E-14 | Tmem119 | MM_2 |
| 2.71479E-08 | 1.158630721 | 0.752 | 0.478 | 0.000641015 | 3.02292E-14 | 1.104127555 | 0.837 | 0.544 | 7.13772E-10 | 2.71479E-08 | 6.04584E-14 | Rasgef1b | MM_2 |
| 1.11033E-08 | 0.701792593 | 0.872 | 0.569 | 0.000262172 | 5.09044E-14 | 0.824925749 | 0.878 | 0.614 | 1.20195E-09 | 1.11033E-08 | 1.01809E-13 | Il10ra | MM_2 |
| 2.3609E-10 | 1.53925294 | 0.472 | 0.182 | 5.57456E-06 | 5.60543E-14 | 1.325025824 | 0.413 | 0.144 | 1.32355E-09 | 2.3609E-10 | 1.12109E-13 | Tifab | MM_2 |
| 2.08444E-13 | 0.939473656 | 0.912 | 0.504 | 4.92179E-09 | 6.64137E-14 | 0.945669239 | 0.872 | 0.568 | 1.56816E-09 | 2.08444E-13 | 1.32827E-13 | Hpgd | MM_2 |
| 1.53688E-10 | 0.983892321 | 0.408 | 0.113 | 3.62889E-06 | 6.64384E-14 | 1.754237855 | 0.331 | 0.084 | 1.56874E-09 | 1.53688E-10 | 1.32877E-13 | Bcl2a1a | MM_2 |
| 1.04665E-13 | 0.872889997 | 0.864 | 0.467 | 2.47136E-09 | 9.94109E-12 | 0.669687649 | 0.808 | 0.508 | 2.34729E-07 | 9.94109E-12 | 2.09331E-13 | Arhgap22 | MM_2 |
| 1.11873E-10 | 0.769671183 | 0.864 | 0.595 | 2.64155E-06 | 1.3767E-13 | 0.642672958 | 0.942 | 0.691 | 3.25066E-09 | 1.11873E-10 | 2.7534E-13 | Plekho1 | MM_2 |
| 1.82746E-13 | 0.800635308 | 0.896 | 0.449 | 4.31499E-09 | 1.40832E-10 | 0.648256786 | 0.919 | 0.633 | 3.32532E-06 | 1.40832E-10 | 3.65491E-13 | Aoah | MM_2 |
| 2.27001E-10 | 1.4729604 | 0.456 | 0.168 | 5.35995E-06 | 1.87179E-13 | 1.412819406 | 0.384 | 0.127 | 4.41967E-09 | 2.27001E-10 | 3.74358E-13 | Ildr1 | MM_2 |
| 7.54885E-12 | 1.184135544 | 0.672 | 0.339 | 1.78243E-07 | 2.08756E-13 | 0.614886197 | 0.762 | 0.46 | 4.92914E-09 | 7.54885E-12 | 4.17512E-13 | Rhoc | MM_2 |
| 2.98251E-08 | 0.730877529 | 0.592 | 0.296 | 0.00070423 | 2.40367E-13 | 1.159464499 | 0.471 | 0.182 | 5.67555E-09 | 2.98251E-08 | 4.80734E-13 | Ggta1 | MM_2 |
| 9.33876E-11 | 0.863185481 | 0.768 | 0.423 | 2.20507E-06 | 5.72946E-13 | 0.683262179 | 0.802 | 0.564 | 1.35284E-08 | 9.33876E-11 | 1.14589E-12 | Nmt1 | MM_2 |
| 3.57216E-08 | 1.518220709 | 0.44 | 0.19 | 0.000843459 | 1.03072E-12 | 1.725940783 | 0.442 | 0.187 | 2.43374E-08 | 3.57216E-08 | 2.06145E-12 | Siglece | MM_2 |
| 2.58191E-08 | 0.720744594 | 0.888 | 0.624 | 0.000609641 | 1.14572E-12 | 0.619602704 | 0.913 | 0.652 | 2.70528E-08 | 2.58191E-08 | 2.29144E-12 | Mir142hg | MM_2 |
| 9.82538E-08 | 0.677960226 | 0.712 | 0.398 | 0.002319968 | 1.46286E-12 | 1.159086428 | 0.68 | 0.398 | 3.45411E-08 | 9.82538E-08 | 2.92573E-12 | Bcl2l1 | MM_2 |
| 3.33193E-07 | 0.74860689 | 0.8 | 0.504 | 0.007867357 | 1.81064E-12 | 0.708432168 | 0.89 | 0.619 | 4.27529E-08 | 3.33193E-07 | 3.62129E-12 | Clec4a3 | MM_2 |
| 4.66707E-08 | 1.086855658 | 0.408 | 0.157 | 0.001101989 | 2.06389E-12 | 1.303745211 | 0.453 | 0.199 | 4.87325E-08 | 4.66707E-08 | 4.12778E-12 | 4933406I1 | MM_2 |
| 3.48009E-08 | 2.062073754 | 0.288 | 0.08 | 0.000821719 | 4.30028E-12 | 2.434286277 | 0.262 | 0.062 | 1.01538E-07 | 3.48009E-08 | 8.60057E-12 | Negr1 | MM_2 |
| 6.83035E-12 | 3.199940585 | 0.248 | 0.029 | 1.61278E-07 | 2.57483E-08 | 1.450794022 | 0.291 | 0.113 | 0.000607969 | 2.57483E-08 | 1.36607E-11 | B3gnt8 | MM_2 |
| 1.24605E-10 | 0.807752024 | 0.904 | 0.58 | 2.94217E-06 | 6.91484E-12 | 0.76113277 | 0.93 | 0.659 | 1.63273E-07 | 1.24605E-10 | 1.38297E-11 | Pik3ap1 | MM_2 |
| 1.93463E-08 | 0.649168572 | 0.72 | 0.42 | 0.000456805 | 1.0489E-11 | 0.804220422 | 0.727 | 0.458 | 2.47667E-07 | 1.93463E-08 | 2.09781E-11 | Pde3b | MM_2 |
| 6.03259E-10 | 1.471846965 | 0.264 | 0.047 | 1.42442E-05 | 1.09479E-11 | 2.065172847 | 0.198 | 0.031 | 2.58501E-07 | 6.03259E-10 | 2.18957E-11 | Abi3 | MM_2 |
| 1.35561E-11 | 1.728269485 | 0.448 | 0.15 | 3.20086E-07 | 2.32941E-08 | 1.39920006 | 0.326 | 0.132 | 0.000550021 | 2.32941E-08 | 2.71122E-11 | Adam22 | MM_2 |
| 1.49774E-11 | 1.255880478 | 0.592 | 0.259 | 3.53646E-07 | 7.8802E-11 | 1.103863591 | 0.599 | 0.35 | 1.86067E-06 | 7.8802E-11 | 2.99548E-11 | Mgat4a | MM_2 |
| 3.77961E-07 | 0.932356662 | 0.856 | 0.62 | 0.008924412 | 1.92553E-11 | 0.814133004 | 0.965 | 0.839 | 4.54656E-07 | 3.77961E-07 | 3.85106E-11 | Hspa1a | MM_2 |
| 2.21591E-11 | 1.206744785 | 0.92 | 0.748 | 5.23221E-07 | 1.81253E-07 | 0.671485619 | 0.942 | 0.842 | 0.004279748 | 1.81253E-07 | 4.43182E-11 | Fos | MM_2 |
| 7.73376E-09 | 1.377738606 | 0.576 | 0.299 | 0.00018261 | 2.28062E-11 | 1.250156112 | 0.547 | 0.276 | 5.38499E-07 | 7.73376E-09 | 4.56123E-11 | Gm46224 | MM_2 |
| 7.93292E-09 | 0.705395785 | 0.552 | 0.241 | 0.000187312 | 4.20222E-11 | 0.836995672 | 0.57 | 0.321 | 9.92229E-07 | 7.93292E-09 | 8.40444E-11 | Il21r | MM_2 |
| 2.57988E-07 | 1.397684584 | 0.624 | 0.38 | 0.006091616 | 5.70557E-11 | 1.713261141 | 0.744 | 0.518 | 1.3472E-06 | 2.57988E-07 | 1.14111E-10 | Cxcl2 | MM_2 |
| 2.20358E-08 | 0.942744644 | 0.72 | 0.431 | 0.000520309 | 6.25731E-11 | 0.829254268 | 0.785 | 0.571 | 1.47748E-06 | 2.20358E-08 | 1.25146E-10 | Aopep | MM_2 |

| | | | | | | | | | | | | | |
|---|---|---|---|---|---|---|---|---|---|---|---|---|---|
| 3.62747E-08 | 1.126019772 | 0.376 | 0.135 | 0.000856518 | 6.6624E-11 | 1.222852314 | 0.407 | 0.173 | 1.57313E-06 | 3.62747E-08 | 1.33248E-10 | Gm44710 | MM_2 |
| 8.23579E-11 | 0.863759982 | 0.872 | 0.536 | 1.94464E-06 | 1.52426E-08 | 0.585785863 | 0.924 | 0.698 | 0.000359909 | 1.52426E-08 | 1.64716E-10 | Lilr4b | MM_2 |
| 1.23885E-10 | 0.784607215 | 0.752 | 0.372 | 2.92517E-06 | 2.18058E-08 | 0.693356926 | 0.692 | 0.448 | 0.000514878 | 2.18058E-08 | 2.4777E-10 | Ets2 | MM_2 |
| 1.46199E-10 | 0.968829623 | 0.68 | 0.347 | 3.45205E-06 | 3.26433E-09 | 0.858554432 | 0.68 | 0.477 | 7.70774E-05 | 3.26433E-09 | 2.92398E-10 | Mknk1 | MM_2 |
| 7.07715E-08 | 0.887015027 | 0.584 | 0.332 | 0.001671055 | 2.14377E-10 | 0.760529013 | 0.68 | 0.41 | 5.06188E-06 | 7.07715E-08 | 4.28754E-10 | Igsf6 | MM_2 |
| 1.38754E-08 | 2.057514287 | 0.408 | 0.157 | 0.000327625 | 3.27645E-10 | 1.560140661 | 0.512 | 0.261 | 7.73637E-06 | 1.38754E-08 | 6.55291E-10 | Cxcl10 | MM_2 |
| 1.44048E-07 | 0.960366427 | 0.512 | 0.237 | 0.003401271 | 4.39248E-10 | 0.696003466 | 0.488 | 0.221 | 1.03715E-05 | 1.44048E-07 | 8.78495E-10 | Clec4b1 | MM_2 |
| 6.56426E-09 | 0.824275891 | 0.536 | 0.245 | 0.000154995 | 5.01245E-10 | 0.797194792 | 0.517 | 0.278 | 1.18354E-05 | 6.56426E-09 | 1.00249E-09 | 2510009E | MM_2 |
| 1.65406E-07 | 0.791829463 | 0.584 | 0.31 | 0.003905565 | 5.85986E-10 | 1.000387892 | 0.61 | 0.396 | 1.38363E-05 | 1.65406E-07 | 1.17197E-09 | Slc29a3 | MM_2 |
| 7.20746E-08 | 1.101922985 | 0.4 | 0.15 | 0.001701826 | 5.87861E-10 | 1.135457122 | 0.547 | 0.293 | 1.38806E-05 | 7.20746E-08 | 1.17572E-09 | Gm15726 | MM_2 |
| 2.58946E-08 | 0.601563783 | 0.88 | 0.62 | 0.000611423 | 6.12774E-10 | 0.64861056 | 0.872 | 0.638 | 1.44688E-05 | 2.58946E-08 | 1.22555E-09 | Epb41l2 | MM_2 |
| 9.26931E-10 | 1.631921124 | 0.304 | 0.073 | 2.18867E-05 | 9.70967E-08 | 2.482710767 | 0.198 | 0.06 | 0.002292647 | 9.70967E-08 | 1.85386E-09 | Gm33370 | MM_2 |
| 5.03892E-08 | 0.612796269 | 0.808 | 0.453 | 0.001189789 | 9.78342E-10 | 0.692245297 | 0.849 | 0.59 | 2.31006E-05 | 5.03892E-08 | 1.95668E-09 | Ms4a6b | MM_2 |
| 1.18655E-09 | 4.290139578 | 0.144 | 0.004 | 2.80168E-05 | 3.4151E-09 | 1.862432227 | 0.134 | 0.017 | 8.06372E-05 | 3.4151E-09 | 2.3731E-09 | Zfp691 | MM_2 |
| 1.34888E-09 | 0.831383835 | 0.592 | 0.259 | 3.18496E-05 | 1.52736E-09 | 1.154719007 | 0.512 | 0.309 | 3.60641E-05 | 1.52736E-09 | 2.69775E-09 | AW11201 | MM_2 |
| 2.28975E-08 | 0.593515894 | 0.624 | 0.314 | 0.000540656 | 1.35149E-09 | 0.937439417 | 0.663 | 0.42 | 3.19114E-05 | 2.28975E-08 | 2.70298E-09 | Ptgs1 | MM_2 |
| 1.63113E-09 | 1.098258346 | 0.408 | 0.135 | 3.85144E-05 | 1.51667E-08 | 1.153449636 | 0.43 | 0.218 | 0.000358115 | 1.51667E-08 | 3.26227E-09 | Fmnl3 | MM_2 |
| 2.98107E-08 | 0.715308828 | 0.848 | 0.555 | 0.000703891 | 2.26343E-09 | 0.671817926 | 0.878 | 0.652 | 5.34442E-05 | 2.98107E-08 | 4.52686E-09 | Slc3a2 | MM_2 |
| 4.53628E-09 | 1.217110192 | 0.352 | 0.102 | 0.000107111 | 1.50104E-07 | 0.989705012 | 0.372 | 0.177 | 0.003544253 | 1.50104E-07 | 9.07257E-09 | Slc25a33 | MM_2 |
| 4.82935E-09 | 0.643309006 | 0.88 | 0.562 | 0.000114031 | 1.53867E-08 | 0.719533726 | 0.808 | 0.576 | 0.000363311 | 1.53867E-08 | 9.65871E-09 | Lrmda | MM_2 |
| 5.18542E-09 | 1.247117693 | 0.432 | 0.172 | 0.000122438 | 7.07247E-08 | 0.914310416 | 0.477 | 0.271 | 0.001669951 | 7.07247E-08 | 1.03708E-08 | Arl11 | MM_2 |
| 7.90596E-09 | 1.105949781 | 0.304 | 0.077 | 0.000186675 | 2.47692E-07 | 0.686359543 | 0.262 | 0.096 | 0.005848503 | 2.47692E-07 | 1.58119E-08 | Stard13 | MM_2 |
| 2.83248E-07 | 1.025604775 | 0.48 | 0.237 | 0.006688062 | 9.35869E-09 | 1.740402154 | 0.407 | 0.197 | 0.000220977 | 2.83248E-07 | 1.87174E-08 | Clec4n | MM_2 |
| 1.5028E-07 | 0.939797972 | 0.552 | 0.288 | 0.003548421 | 9.96781E-09 | 0.845721021 | 0.599 | 0.357 | 0.00023536 | 1.5028E-07 | 1.99356E-08 | Tagap | MM_2 |
| 1.25096E-08 | 0.784563867 | 0.808 | 0.485 | 0.000295376 | 1.86263E-07 | 0.711688659 | 0.756 | 0.571 | 0.004398044 | 1.86263E-07 | 2.50191E-08 | Lpxn | MM_2 |
| 2.49314E-08 | 0.836917326 | 0.688 | 0.401 | 0.00058868 | 3.56232E-08 | 0.83677006 | 0.663 | 0.475 | 0.000841134 | 3.56232E-08 | 4.98628E-08 | Tifa | MM_2 |
| 3.4202E-07 | 1.998306642 | 0.272 | 0.084 | 0.008075778 | 3.56905E-08 | 1.74235115 | 0.227 | 0.07 | 0.000842723 | 3.4202E-07 | 7.13809E-08 | 2010013B | MM_2 |
| 6.39097E-08 | 1.891606203 | 0.248 | 0.062 | 0.001509036 | 3.8655E-08 | 1.339941507 | 0.273 | 0.098 | 0.000912722 | 6.39097E-08 | 7.73101E-08 | A930006K | MM_2 |
| 5.78242E-08 | 3.188235563 | 0.16 | 0.018 | 0.001365344 | 4.07842E-08 | 2.721653661 | 0.116 | 0.014 | 0.000962996 | 5.78242E-08 | 8.15683E-08 | Kcnj10 | MM_2 |
| 1.04662E-07 | 1.096419636 | 0.288 | 0.08 | 0.002471275 | 3.23736E-07 | 1.660470042 | 0.308 | 0.139 | 0.007644055 | 3.23736E-07 | 2.09324E-07 | 4930430E | MM_2 |
| 1.30056E-07 | 0.672026004 | 0.424 | 0.172 | 0.003070872 | 1.56935E-07 | 0.66197475 | 0.372 | 0.17 | 0.003705561 | 1.56935E-07 | 2.60111E-07 | Zbtb16 | MM_2 |
| 2.28018E-20 | 2.589491533 | 0.858 | 0.92 | 5.38396E-16 | 4.93054E-38 | 3.217047437 | 0.946 | 0.95 | 1.1642E-33 | 2.28018E-20 | 9.86107E-38 | Tagln | MM_3 |
| 3.32681E-28 | 2.253713055 | 0.947 | 0.958 | 7.85526E-24 | 9.39151E-35 | 2.899761237 | 0.918 | 0.959 | 2.21752E-30 | 3.32681E-28 | 1.8783E-34 | Tpm2 | MM_3 |
| 3.54148E-23 | 2.373800861 | 0.885 | 0.962 | 8.36213E-19 | 8.34402E-33 | 2.93908422 | 0.918 | 0.971 | 1.97019E-28 | 3.54148E-23 | 1.6688E-32 | Acta2 | MM_3 |
| 2.52839E-29 | 1.616826516 | 1 | 1 | 5.97004E-25 | 4.67057E-32 | 1.502486113 | 0.993 | 1 | 1.10281E-27 | 2.52839E-29 | 9.34113E-32 | mt-Cytb | MM_3 |
| 1.08276E-11 | 2.120120574 | 0.805 | 0.958 | 2.55661E-07 | 2.06621E-31 | 2.745487585 | 0.898 | 0.952 | 4.87874E-27 | 1.08276E-11 | 4.13242E-31 | Tpm1 | MM_3 |
| 9.17321E-21 | 1.447355 | 1 | 1 | 2.16598E-16 | 2.92761E-29 | 1.468550533 | 0.993 | 1 | 6.91268E-25 | 9.17321E-21 | 5.85523E-29 | mt-Nd2 | MM_3 |
| 7.37072E-25 | 2.453371061 | 0.903 | 0.958 | 1.74037E-20 | 6.6347E-29 | 3.084566439 | 0.884 | 0.941 | 1.56659E-24 | 7.37072E-25 | 1.32694E-28 | Des | MM_3 |
| 8.62177E-23 | 1.330448585 | 1 | 1 | 2.03577E-18 | 4.06964E-28 | 1.315699917 | 1 | 1 | 9.60923E-24 | 8.62177E-23 | 8.13928E-28 | mt-Atp6 | MM_3 |
| 8.87051E-26 | 1.436462258 | 1 | 1 | 2.09451E-21 | 5.65075E-28 | 1.332323645 | 1 | 1 | 1.33426E-23 | 8.87051E-26 | 1.13015E-27 | mt-Co3 | MM_3 |
| 2.09859E-16 | 1.63940982 | 0.929 | 0.969 | 4.95518E-12 | 1.1631E-27 | 2.159719313 | 0.912 | 0.968 | 2.74632E-23 | 2.09859E-16 | 2.32621E-27 | AY036118 | MM_3 |
| 1.22945E-26 | 1.285486819 | 1 | 1 | 2.90297E-22 | 1.30265E-27 | 1.18393677 | 1 | 1 | 3.07582E-23 | 1.22945E-26 | 2.6053E-27 | mt-Co2 | MM_3 |
| 2.20093E-10 | 2.229407161 | 0.752 | 0.871 | 5.19684E-06 | 1.30405E-27 | 2.851022201 | 0.857 | 0.896 | 3.07911E-23 | 2.20093E-10 | 2.60809E-27 | Mylk | MM_3 |
| 1.86539E-17 | 2.729804532 | 0.841 | 0.944 | 4.40455E-13 | 2.31398E-26 | 3.001439412 | 0.891 | 0.941 | 5.46378E-22 | 1.86539E-17 | 4.62797E-26 | Actg2 | MM_3 |
| 4.59887E-10 | 0.961525497 | 0.973 | 0.993 | 1.08588E-05 | 2.69699E-23 | 1.652892631 | 0.98 | 0.998 | 6.36814E-19 | 4.59887E-10 | 5.39398E-23 | Camk1d | MM_3 |
| 1.10022E-18 | 1.25448553 | 0.991 | 1 | 2.59784E-14 | 3.18281E-23 | 1.16720135 | 1 | 1 | 7.51525E-19 | 1.10022E-18 | 6.36562E-23 | mt-Nd1 | MM_3 |
| 2.43204E-22 | 1.351447963 | 1 | 1 | 5.74254E-18 | 5.51949E-22 | 1.279893939 | 0.986 | 1 | 1.30326E-17 | 5.51949E-22 | 4.86409E-22 | mt-Nd4 | MM_3 |
| 2.6011E-17 | 1.522748657 | 0.938 | 0.986 | 6.14172E-13 | 8.25649E-20 | 1.872510011 | 0.925 | 0.989 | 1.94952E-15 | 2.6011E-17 | 1.6513E-19 | Cdk8 | MM_3 |
| 2.31486E-09 | 1.315402917 | 0.938 | 1 | 5.46584E-05 | 1.51894E-19 | 1.869557119 | 0.952 | 0.995 | 3.58652E-15 | 2.31486E-09 | 3.03788E-19 | Cmss1 | MM_3 |
| 6.24467E-13 | 2.252981252 | 0.788 | 0.888 | 1.47449E-08 | 2.12841E-19 | 2.798400538 | 0.816 | 0.925 | 5.02559E-15 | 6.24467E-13 | 4.25681E-19 | Myh11 | MM_3 |
| 4.21798E-14 | 1.130408759 | 1 | 1 | 9.95949E-10 | 9.08201E-19 | 1.448135137 | 0.993 | 1 | 2.14445E-14 | 4.21798E-14 | 1.8164E-18 | Gm42418 | MM_3 |
| 3.80245E-14 | 0.894031747 | 1 | 1 | 8.97836E-10 | 1.70008E-14 | 0.778584136 | 1 | 1 | 4.01423E-10 | 3.80245E-14 | 3.40016E-14 | mt-Co1 | MM_3 |
| 1.30866E-11 | 1.469375209 | 0.876 | 0.99 | 3.09E-07 | 1.66385E-13 | 1.759878756 | 0.871 | 0.982 | 3.92869E-09 | 1.30866E-11 | 3.3277E-13 | Lars2 | MM_3 |
| 1.35821E-12 | 2.423099612 | 0.743 | 0.815 | 3.20701E-08 | 3.04865E-11 | 2.709459034 | 0.728 | 0.837 | 7.19848E-07 | 3.04865E-11 | 2.71643E-12 | Cnn1 | MM_3 |
| 1.90207E-10 | 1.338740285 | 0.929 | 1 | 4.49116E-06 | 3.49996E-10 | 1.083123137 | 0.884 | 0.993 | 8.26411E-06 | 3.49996E-10 | 3.80413E-10 | mt-Nd3 | MM_3 |
| 1.19518E-07 | 1.150042536 | 0.841 | 0.986 | 0.00282206 | 4.99023E-09 | 1.393728329 | 0.816 | 0.986 | 0.000117829 | 1.19518E-07 | 9.98046E-09 | mt-Nd5 | MM_3 |
| 1.4219E-07 | 1.63311658 | 0.752 | 0.948 | 0.003357393 | 1.92184E-07 | 1.494752961 | 0.741 | 0.952 | 0.004537858 | 1.92184E-07 | 2.8438E-07 | mt-Nd4l | MM_3 |
| 1.24478E-58 | 5.119108444 | 0.865 | 0.032 | 2.93917E-54 | 3.82059E-74 | 4.21222113 | 0.854 | 0.031 | 9.02117E-70 | 1.24478E-58 | 7.64117E-74 | Napsa | MM_4 |
| 1.34527E-52 | 6.762636867 | 0.673 | 0.009 | 3.17644E-48 | 8.27156E-66 | 4.059905057 | 0.771 | 0.03 | 1.95308E-61 | 1.34527E-52 | 1.65431E-65 | Ltb4r1 | MM_4 |
| 2.22617E-44 | 5.292458414 | 0.615 | 0.014 | 5.25643E-40 | 3.66718E-65 | 5.196775173 | 0.771 | 0.033 | 8.65895E-61 | 2.22617E-44 | 7.33436E-65 | Cd226 | MM_4 |
| 1.07685E-40 | 3.447457386 | 0.827 | 0.084 | 2.54266E-36 | 2.08712E-56 | 4.313241353 | 0.917 | 0.087 | 4.92811E-52 | 1.07685E-40 | 4.17425E-56 | Rab11fip1 | MM_4 |
| 6.90826E-29 | 3.554607143 | 0.558 | 0.043 | 1.63118E-24 | 9.77888E-56 | 3.216836506 | 0.792 | 0.046 | 2.30899E-51 | 6.90826E-29 | 1.95578E-55 | Gm15987 | MM_4 |
| 1.42977E-33 | 4.904136905 | 0.462 | 0.009 | 3.37597E-29 | 4.48363E-55 | 7.426523484 | 0.458 | 0.002 | 1.05867E-50 | 1.42977E-33 | 8.96726E-55 | Klrd1 | MM_4 |
| 7.33921E-55 | 4.99001265 | 0.808 | 0.029 | 1.73293E-50 | 2.25573E-52 | 4.445795018 | 0.75 | 0.05 | 5.32622E-48 | 2.25573E-52 | 1.46784E-54 | Itgb7 | MM_4 |
| 1.13784E-34 | 4.003737427 | 0.538 | 0.02 | 2.68668E-30 | 6.29928E-51 | 3.053909621 | 0.667 | 0.033 | 1.48739E-46 | 1.13784E-34 | 1.25986E-50 | Dna2 | MM_4 |
| 1.64419E-50 | 3.934771046 | 0.904 | 0.069 | 3.88227E-46 | 2.33753E-42 | 3.798511451 | 0.792 | 0.092 | 5.51937E-38 | 2.33753E-42 | 3.28839E-50 | Bhlhe40 | MM_4 |
| 1.67105E-21 | 2.840774115 | 0.596 | 0.086 | 3.94569E-17 | 6.41444E-50 | 3.587497914 | 0.708 | 0.043 | 1.51458E-45 | 1.67105E-21 | 1.28289E-49 | Kcnn4 | MM_4 |
| 2.24303E-32 | 4.598131729 | 0.462 | 0.012 | 5.29624E-28 | 1.27224E-49 | 5.974811934 | 0.438 | 0.004 | 3.00401E-45 | 2.24303E-32 | 2.54447E-49 | Gm36723 | MM_4 |
| 1.20328E-34 | 2.993042844 | 0.827 | 0.104 | 2.84118E-30 | 3.29176E-49 | 3.495664841 | 0.833 | 0.078 | 7.7725E-45 | 1.20328E-34 | 6.58351E-49 | Coro2a | MM_4 |
| 1.21685E-25 | 4.567452795 | 0.346 | 0.006 | 2.87322E-21 | 5.37885E-49 | 4.560474386 | 0.5 | 0.011 | 1.27006E-44 | 1.21685E-25 | 1.07577E-48 | Klri1 | MM_4 |
| 6.44544E-49 | 4.918841654 | 0.673 | 0.014 | 1.5219E-44 | 1.34872E-33 | 4.898375725 | 0.438 | 0.022 | 3.18459E-29 | 1.34872E-33 | 1.28909E-48 | Klrb1b | MM_4 |
| 3.11819E-35 | 4.039459258 | 0.596 | 0.032 | 7.36267E-31 | 1.50382E-48 | 7.507172697 | 0.458 | 0.007 | 3.55081E-44 | 3.11819E-35 | 3.00763E-48 | Jaml | MM_4 |
| 1.03894E-33 | 3.584450579 | 0.615 | 0.04 | 2.45315E-29 | 5.95392E-48 | 4.129151113 | 0.604 | 0.028 | 1.40584E-43 | 1.03894E-33 | 1.19078E-47 | Gpr35 | MM_4 |
| 5.98209E-35 | 2.840897897 | 0.731 | 0.066 | 1.41249E-30 | 4.69803E-46 | 3.478819547 | 0.688 | 0.046 | 1.1093E-41 | 5.98209E-35 | 9.39605E-46 | Adam8 | MM_4 |
| 2.66788E-10 | 4.508608881 | 0.135 | 0.003 | 6.29939E-06 | 8.60049E-45 | 6.743332453 | 0.396 | 0.004 | 2.03075E-40 | 2.66788E-10 | 1.7201E-44 | Upb1 | MM_4 |
| 2.98481E-29 | 3.227219355 | 0.635 | 0.058 | 7.04773E-25 | 2.69283E-44 | 3.889039716 | 0.625 | 0.037 | 6.35832E-40 | 2.98481E-29 | 5.38567E-44 | H2-DMb2 | MM_4 |
| 2.2777E-43 | 4.377077749 | 0.769 | 0.052 | 5.3781E-39 | 2.67716E-42 | 3.707094679 | 0.688 | 0.059 | 6.32132E-38 | 2.67716E-42 | 4.55539E-43 | Cd24a | MM_4 |
| 1.13699E-39 | 3.972574292 | 0.942 | 0.153 | 2.68465E-35 | 1.55767E-42 | 3.570558437 | 0.917 | 0.135 | 3.67797E-38 | 1.13699E-39 | 3.11534E-42 | Cytip | MM_4 |
| 3.54814E-33 | 4.263174921 | 0.519 | 0.02 | 8.37787E-29 | 2.57731E-42 | 4.582895187 | 0.521 | 0.022 | 6.08554E-38 | 3.54814E-33 | 5.15461E-42 | Ear2 | MM_4 |
| 4.99815E-41 | 4.7473753 | 0.615 | 0.02 | 1.18016E-36 | 1.93116E-37 | 2.229275734 | 0.5 | 0.024 | 4.55985E-33 | 1.93116E-37 | 9.99629E-41 | P2ry10 | MM_4 |
| 5.57658E-28 | 3.2335823 | 0.558 | 0.043 | 1.31674E-23 | 2.64527E-40 | 3.736785825 | 0.646 | 0.054 | 6.246E-36 | 5.57658E-28 | 5.29053E-40 | Ryr1 | MM_4 |

| | | | | | | | | | | | | | |
|---|---|---|---|---|---|---|---|---|---|---|---|---|---|
| 5.02615E-40 | 5.054506116 | 0.5 | 0.003 | 1.18678E-35 | 5.66049E-24 | 5.191970473 | 0.208 | 0.002 | 1.33656E-19 | 5.66049E-24 | 1.00523E-39 | Sept3 | MM_4 |
| 1.14268E-39 | 4.088432731 | 0.538 | 0.009 | 2.69809E-35 | 2.08112E-29 | 3.311431682 | 0.333 | 0.011 | 4.91395E-25 | 2.08112E-29 | 2.28535E-39 | Flt3 | MM_4 |
| 4.14638E-39 | 2.921294494 | 0.692 | 0.04 | 9.79043E-35 | 1.9506E-34 | 3.798592177 | 0.5 | 0.031 | 4.60575E-30 | 1.9506E-34 | 8.29276E-39 | Dpp4 | MM_4 |
| 6.43161E-39 | 8.542593314 | 0.5 | 0.006 | 1.51863E-34 | 8.89036E-32 | 8.228138912 | 0.271 | 0.002 | 2.09919E-27 | 8.89036E-32 | 1.28632E-38 | Htr7 | MM_4 |
| 4.5241E-18 | 2.670617878 | 0.269 | 0.009 | 1.06823E-13 | 9.20331E-39 | 3.689670577 | 0.458 | 0.017 | 2.17309E-34 | 4.5241E-18 | 1.84066E-38 | Wnt11 | MM_4 |
| 2.73279E-38 | 3.323152104 | 0.635 | 0.029 | 6.45266E-34 | 6.21857E-38 | 3.784283455 | 0.458 | 0.018 | 1.46833E-33 | 6.21857E-38 | 5.46557E-38 | Cd244a | MM_4 |
| 5.43922E-34 | 3.490722544 | 0.654 | 0.046 | 1.28431E-29 | 1.25496E-37 | 4.064002902 | 0.542 | 0.033 | 2.9632E-33 | 5.43922E-34 | 2.50991E-37 | Itgax | MM_4 |
| 2.2908E-23 | 1.788404565 | 0.481 | 0.04 | 5.40905E-19 | 1.19114E-36 | 3.963345326 | 0.5 | 0.028 | 2.81251E-32 | 2.2908E-23 | 2.38227E-36 | Nup210 | MM_4 |
| 6.37023E-12 | 4.101569242 | 0.25 | 0.023 | 1.50414E-07 | 6.12522E-36 | 3.296239897 | 0.5 | 0.028 | 1.44629E-31 | 6.37023E-12 | 1.22504E-35 | Skint3 | MM_4 |
| 7.86178E-19 | 2.567050567 | 0.365 | 0.026 | 1.85632E-14 | 7.73087E-36 | 4.566518287 | 0.417 | 0.015 | 1.82541E-31 | 7.86178E-19 | 1.54617E-35 | Lmo1 | MM_4 |
| 2.3214E-34 | 4.228713436 | 0.538 | 0.02 | 5.48128E-30 | 1.0805E-35 | 4.862158194 | 0.479 | 0.026 | 2.55128E-31 | 2.3214E-34 | 2.161E-35 | Uck2 | MM_4 |
| 1.84136E-35 | 6.871793169 | 0.462 | 0.006 | 4.34781E-31 | 3.84989E-24 | 7.553204464 | 0.208 | 0.002 | 9.09035E-20 | 3.84989E-24 | 3.68271E-35 | Xcr1 | MM_4 |
| 6.2093E-29 | 3.405035318 | 0.558 | 0.043 | 1.46614E-24 | 3.78501E-35 | 5.005419204 | 0.5 | 0.031 | 8.93717E-31 | 6.2093E-29 | 7.57003E-35 | Itgal | MM_4 |
| 5.05269E-15 | 7.986681808 | 0.173 | 0 | 1.19304E-10 | 4.07209E-35 | 5.162501008 | 0.354 | 0.007 | 9.61503E-31 | 5.05269E-15 | 8.14419E-35 | Dcstamp | MM_4 |
| 4.49736E-35 | 4.517083631 | 0.558 | 0.023 | 1.06192E-30 | 1.37585E-34 | 5.461952816 | 0.375 | 0.011 | 3.24865E-30 | 1.37585E-34 | 8.99471E-35 | H2-Oa | MM_4 |
| 6.4845E-35 | 3.261586779 | 0.615 | 0.035 | 1.53112E-30 | 5.2486E-28 | 2.727056543 | 0.396 | 0.022 | 1.2393E-23 | 5.2486E-28 | 1.2969E-34 | Ap1s3 | MM_4 |
| 1.38439E-33 | 5.702516237 | 0.423 | 0.003 | 3.26881E-29 | 1.63317E-34 | 4.599995861 | 0.333 | 0.006 | 3.85624E-30 | 1.38439E-33 | 3.26634E-34 | Pkp3 | MM_4 |
| 1.29799E-23 | 1.755081022 | 0.942 | 0.207 | 3.0648E-19 | 2.87533E-34 | 2.753884276 | 0.958 | 0.187 | 6.78923E-30 | 1.29799E-23 | 5.75066E-34 | Ccr2 | MM_4 |
| 1.37921E-33 | 4.941315091 | 0.462 | 0.009 | 3.25658E-29 | 1.50949E-27 | 7.384593478 | 0.271 | 0.006 | 3.56421E-23 | 1.50949E-27 | 2.75841E-33 | Clec9a | MM_4 |
| 1.08614E-24 | 3.49441144 | 0.404 | 0.017 | 2.5646E-20 | 3.93944E-33 | 3.508213458 | 0.396 | 0.015 | 9.30181E-29 | 1.08614E-24 | 7.87889E-33 | Klrk1 | MM_4 |
| 4.20736E-33 | 2.569907654 | 0.942 | 0.19 | 9.93441E-29 | 1.28674E-27 | 3.121958193 | 0.875 | 0.246 | 3.03825E-23 | 1.28674E-27 | 8.41471E-33 | Agpat4 | MM_4 |
| 4.85459E-33 | 2.827393746 | 0.654 | 0.055 | 1.14627E-28 | 5.59039E-24 | 2.938176911 | 0.625 | 0.102 | 1.32E-19 | 5.59039E-24 | 9.70919E-33 | Bri3bp | MM_4 |
| 5.51029E-33 | 5.685433945 | 0.558 | 0.032 | 1.30109E-28 | 4.8243E-12 | 5.32471688 | 0.25 | 0.031 | 1.13911E-07 | 4.8243E-12 | 1.10206E-32 | Ifi205 | MM_4 |
| 8.28637E-33 | 3.155524069 | 0.673 | 0.061 | 1.95658E-28 | 2.40537E-23 | 3.63773638 | 0.604 | 0.102 | 5.67955E-19 | 2.40537E-23 | 1.65727E-32 | Nectin1 | MM_4 |
| 2.6772E-32 | 8.06820743 | 0.385 | 0 | 6.32142E-28 | 1.60613E-07 | 3.497093371 | 0.104 | 0.007 | 0.003792387 | 1.60613E-07 | 5.35441E-32 | Zfp366 | MM_4 |
| 2.3355E-21 | 3.593580143 | 0.346 | 0.014 | 5.51457E-17 | 2.854E-32 | 5.359558442 | 0.312 | 0.006 | 6.73885E-28 | 2.3355E-21 | 5.70799E-32 | Mcemp1 | MM_4 |
| 2.50148E-30 | 5.340724513 | 0.385 | 0.003 | 5.90651E-26 | 2.97285E-29 | 7.132467347 | 0.229 | 0 | 7.01948E-25 | 2.97285E-29 | 5.00297E-30 | Gpr141b | MM_4 |
| 6.44993E-30 | 3.253457827 | 1 | 0.55 | 1.52296E-25 | 2.17764E-27 | 3.360323166 | 0.979 | 0.571 | 5.14184E-23 | 2.17764E-27 | 1.28999E-29 | Gm2a | MM_4 |
| 6.85759E-30 | 2.567347644 | 0.846 | 0.133 | 1.61921E-25 | 2.55784E-09 | 0.946625853 | 0.854 | 0.383 | 6.03956E-05 | 2.55784E-09 | 1.37152E-29 | St3gal5 | MM_4 |
| 7.58481E-30 | 3.084001558 | 0.846 | 0.156 | 1.79093E-25 | 2.11032E-29 | 3.339217514 | 0.833 | 0.174 | 4.98288E-25 | 2.11032E-29 | 1.51696E-29 | Tnip3 | MM_4 |
| 1.16466E-29 | 2.236150796 | 0.942 | 0.202 | 2.74999E-25 | 1.20963E-26 | 2.425822388 | 0.833 | 0.194 | 2.85617E-22 | 1.20963E-26 | 2.32932E-29 | Itpr1 | MM_4 |
| 1.26063E-29 | 3.68397727 | 0.577 | 0.043 | 2.97659E-25 | 1.42057E-19 | 4.443620267 | 0.521 | 0.091 | 3.35426E-15 | 1.42057E-19 | 2.52125E-29 | Tnni2 | MM_4 |
| 1.27349E-22 | 3.765507388 | 0.346 | 0.012 | 3.00696E-18 | 1.54219E-29 | 3.07960018 | 0.458 | 0.031 | 3.64142E-25 | 1.27349E-22 | 3.08438E-29 | Mefv | MM_4 |
| 1.72306E-29 | 3.018281549 | 0.981 | 0.366 | 4.06849E-25 | 1.45147E-20 | 2.107117781 | 0.979 | 0.56 | 3.42722E-16 | 1.45147E-20 | 3.44612E-29 | Cd52 | MM_4 |
| 1.27566E-16 | 3.451059692 | 0.308 | 0.02 | 3.01209E-12 | 2.78076E-29 | 2.928607388 | 0.479 | 0.037 | 6.56593E-25 | 1.27566E-16 | 5.56152E-29 | Slc2a6 | MM_4 |
| 3.4547E-29 | 4.306686928 | 0.442 | 0.014 | 8.15724E-25 | 6.57023E-17 | 3.377663518 | 0.25 | 0.017 | 1.55136E-12 | 6.57023E-17 | 6.9094E-29 | Btla | MM_4 |
| 3.02675E-16 | 1.837742277 | 0.577 | 0.107 | 7.14677E-12 | 4.10489E-29 | 2.891857456 | 0.646 | 0.085 | 9.69247E-25 | 3.02675E-16 | 8.20978E-29 | Map4k1 | MM_4 |
| 4.27531E-29 | 7.516772823 | 0.346 | 0 | 1.00949E-24 | 3.73308E-24 | 7.696186878 | 0.188 | 0 | 8.81454E-20 | 3.73308E-24 | 8.55062E-29 | Gcsam | MM_4 |
| 5.14785E-29 | 3.067594099 | 0.538 | 0.035 | 1.21551E-24 | 1.34407E-20 | 2.638307966 | 0.417 | 0.044 | 3.17361E-16 | 1.34407E-20 | 1.02957E-28 | Grap2 | MM_4 |
| 1.18481E-11 | 1.73135379 | 0.442 | 0.095 | 2.79756E-07 | 6.32802E-29 | 3.397102573 | 0.604 | 0.076 | 1.49417E-24 | 1.18481E-11 | 1.2656E-28 | Fgr | MM_4 |
| 3.60688E-27 | 2.981628756 | 0.577 | 0.052 | 8.51655E-23 | 7.35653E-29 | 3.984467623 | 0.521 | 0.048 | 1.73702E-24 | 3.60688E-27 | 1.47131E-28 | Hepacam2 | MM_4 |
| 1.39118E-28 | 4.481504303 | 0.404 | 0.009 | 3.28487E-24 | 1.6206E-18 | 2.105649561 | 0.271 | 0.017 | 3.82656E-14 | 1.6206E-18 | 2.78237E-28 | Gpr171 | MM_4 |
| 3.61291E-28 | 3.908424211 | 0.769 | 0.147 | 8.53081E-24 | 3.35571E-22 | 2.954532196 | 0.688 | 0.146 | 7.9235E-18 | 3.35571E-22 | 7.22582E-28 | Nr4a2 | MM_4 |
| 7.6095E-10 | 1.497611294 | 0.327 | 0.055 | 1.79675E-05 | 4.01033E-28 | 3.435622957 | 0.604 | 0.076 | 9.46918E-24 | 7.6095E-10 | 8.02065E-28 | Unc119b | MM_4 |
| 4.25619E-28 | 3.028090665 | 0.635 | 0.066 | 1.00497E-23 | 4.7709E-15 | 2.468117186 | 0.354 | 0.046 | 1.1265E-10 | 4.7709E-15 | 8.51238E-28 | Tspan33 | MM_4 |
| 4.05622E-24 | 3.240405065 | 0.673 | 0.11 | 9.57756E-20 | 4.28882E-28 | 2.844055125 | 0.688 | 0.107 | 1.01268E-23 | 4.05622E-24 | 8.57764E-28 | Sept6 | MM_4 |
| 2.46068E-25 | 2.655471607 | 0.942 | 0.3 | 5.81015E-21 | 4.53595E-28 | 2.619165724 | 0.938 | 0.27 | 1.07103E-23 | 2.46068E-25 | 9.07191E-28 | Gpr132 | MM_4 |
| 1.32113E-17 | 2.335193614 | 0.808 | 0.256 | 3.11946E-13 | 7.90683E-28 | 2.641198386 | 0.875 | 0.213 | 1.86696E-23 | 1.32113E-17 | 1.58137E-27 | Rin3 | MM_4 |
| 8.2489E-28 | 3.076337697 | 0.846 | 0.187 | 1.94773E-23 | 4.48774E-24 | 2.823247534 | 0.792 | 0.211 | 1.05965E-19 | 4.48774E-24 | 1.64978E-27 | Unc119 | MM_4 |
| 1.58428E-15 | 4.816515688 | 0.25 | 0.012 | 3.74081E-11 | 1.07652E-27 | 3.367804894 | 0.292 | 0.007 | 2.54187E-23 | 1.58428E-15 | 2.15303E-27 | Ramp3 | MM_4 |
| 1.66586E-27 | 6.256904579 | 0.327 | 0 | 3.93343E-23 | 4.43952E-19 | 6.088091801 | 0.146 | 0 | 1.04826E-14 | 4.43952E-19 | 3.33172E-27 | Gm43914 | MM_4 |
| 1.95218E-27 | 7.329461436 | 0.385 | 0.009 | 4.60949E-23 | 1.79994E-14 | 2.471531173 | 0.146 | 0.004 | 4.25001E-10 | 1.79994E-14 | 3.90437E-27 | Ccl22 | MM_4 |
| 3.78916E-27 | 2.83027062 | 0.981 | 0.45 | 8.94697E-23 | 6.45085E-27 | 2.574215524 | 1 | 0.606 | 1.52317E-22 | 6.45085E-27 | 7.57833E-27 | Lsp1 | MM_4 |
| 3.87543E-27 | 3.34763792 | 1 | 0.602 | 9.15067E-23 | 1.04146E-25 | 3.362106799 | 0.979 | 0.641 | 2.45911E-21 | 1.04146E-25 | 7.75086E-27 | Tmsb10 | MM_4 |
| 6.6578E-23 | 1.724807767 | 0.654 | 0.101 | 1.57204E-18 | 4.7042E-27 | 3.337459907 | 0.583 | 0.074 | 1.11076E-22 | 6.6578E-23 | 9.4084E-27 | Srgap3 | MM_4 |
| 8.00461E-26 | 4.146068587 | 0.654 | 0.095 | 1.89005E-21 | 7.91628E-27 | 3.570581712 | 0.688 | 0.122 | 1.86919E-22 | 8.00461E-26 | 1.58326E-26 | Avpi1 | MM_4 |
| 3.12532E-26 | 4.7997677 | 0.404 | 0.014 | 7.37951E-22 | 8.99801E-27 | 2.622802851 | 0.312 | 0.011 | 2.12461E-22 | 3.12532E-26 | 1.7996E-26 | Strip2 | MM_4 |
| 9.2135E-27 | 3.666200868 | 0.769 | 0.153 | 2.17549E-22 | 2.86411E-20 | 2.940819697 | 0.688 | 0.174 | 6.76273E-16 | 2.86411E-20 | 1.8427E-26 | Traf1 | MM_4 |
| 9.75631E-27 | 3.76745034 | 0.558 | 0.049 | 2.30366E-22 | 3.48706E-14 | 2.245398985 | 0.375 | 0.057 | 8.23366E-10 | 3.48706E-14 | 1.95126E-26 | Cyfip2 | MM_4 |
| 1.0632E-26 | 3.458451264 | 0.692 | 0.11 | 2.51042E-22 | 3.44839E-15 | 2.412126867 | 0.646 | 0.192 | 8.14234E-11 | 3.44839E-15 | 2.12639E-26 | Plp2 | MM_4 |
| 6.23648E-09 | 3.923419789 | 0.135 | 0.006 | 0.000147256 | 1.29349E-26 | 6.953683247 | 0.229 | 0.002 | 3.05419E-22 | 6.23648E-09 | 2.58699E-26 | Kctd14 | MM_4 |
| 3.958E-14 | 2.969418989 | 0.288 | 0.023 | 9.34563E-10 | 1.7183E-26 | 2.997557848 | 0.333 | 0.015 | 4.05725E-22 | 3.958E-14 | 3.4366E-26 | Gm9530 | MM_4 |
| 3.93159E-12 | 3.197572148 | 0.288 | 0.032 | 9.28326E-08 | 2.98973E-26 | 2.15099449 | 0.333 | 0.015 | 7.05935E-22 | 3.93159E-12 | 5.97946E-26 | Clec2g | MM_4 |
| 1.36695E-25 | 2.677908077 | 0.846 | 0.19 | 3.22764E-21 | 6.54347E-26 | 2.729851259 | 0.875 | 0.229 | 1.54504E-21 | 1.36695E-25 | 1.30869E-25 | Tiam1 | MM_4 |
| 3.01797E-10 | 1.320485863 | 0.558 | 0.161 | 7.12604E-06 | 7.96836E-26 | 2.451878166 | 0.708 | 0.116 | 1.88149E-21 | 3.01797E-10 | 1.59367E-25 | B4galnt1 | MM_4 |
| 1.16035E-25 | 5.259134392 | 0.365 | 0.009 | 2.73981E-21 | 1.33198E-08 | 3.293679645 | 0.125 | 0.009 | 0.000314508 | 1.33198E-08 | 2.32069E-25 | Ly75 | MM_4 |
| 1.95686E-25 | 2.385524272 | 0.981 | 0.427 | 4.62054E-21 | 1.70292E-23 | 2.361565337 | 1 | 0.625 | 4.02095E-19 | 1.70292E-23 | 3.91372E-25 | Anxa2 | MM_4 |
| 1.9563E-21 | 2.131939878 | 0.846 | 0.239 | 4.61921E-17 | 2.1848E-25 | 2.599961962 | 0.854 | 0.237 | 5.15875E-21 | 1.9563E-21 | 4.3696E-25 | Olfm1 | MM_4 |
| 3.23632E-25 | 2.55720017 | 1 | 0.847 | 7.64161E-21 | 6.56343E-21 | 2.0193288 | 0.979 | 0.941 | 1.54976E-16 | 6.56343E-21 | 6.47265E-25 | Crip1 | MM_4 |
| 2.62013E-16 | 3.032673112 | 0.308 | 0.02 | 6.18665E-12 | 4.11727E-25 | 2.825355232 | 0.312 | 0.013 | 9.72169E-21 | 2.62013E-16 | 8.23454E-25 | Siglecg | MM_4 |
| 3.86319E-19 | 2.295969782 | 0.481 | 0.055 | 9.12177E-15 | 9.01052E-25 | 2.748188046 | 0.604 | 0.089 | 2.12756E-20 | 3.86319E-19 | 1.8021E-24 | Gramd4 | MM_4 |
| 2.26508E-17 | 2.674295625 | 0.481 | 0.066 | 5.3483E-13 | 9.06342E-25 | 3.189496288 | 0.521 | 0.063 | 2.14005E-20 | 2.26508E-17 | 1.81268E-24 | Lsr | MM_4 |
| 2.1175E-21 | 2.191514458 | 0.981 | 0.392 | 4.99984E-17 | 1.28299E-24 | 2.635194004 | 0.958 | 0.37 | 3.0294E-20 | 2.1175E-21 | 2.56598E-24 | Alcam | MM_4 |
| 2.62567E-24 | 2.598367243 | 0.75 | 0.144 | 6.19974E-20 | 1.49431E-24 | 2.476905822 | 0.708 | 0.137 | 3.52836E-20 | 2.62567E-24 | 2.98862E-24 | Crybg1 | MM_4 |
| 1.81895E-24 | 6.476857876 | 0.346 | 0.009 | 4.2949E-20 | 9.35919E-12 | 5.725767648 | 0.146 | 0.007 | 2.20989E-07 | 9.35919E-12 | 3.6379E-24 | Itgae | MM_4 |
| 2.18212E-24 | 2.782323097 | 0.942 | 0.297 | 5.15243E-20 | 3.00508E-18 | 2.427045776 | 0.812 | 0.292 | 7.09558E-14 | 3.00508E-18 | 4.36424E-24 | Bcl2a1d | MM_4 |
| 1.40705E-16 | 2.353036522 | 0.385 | 0.04 | 3.32232E-12 | 3.73264E-24 | 2.824962049 | 0.5 | 0.057 | 8.81351E-20 | 1.40705E-16 | 7.46528E-24 | Inf2 | MM_4 |
| 2.44139E-13 | 1.40858266 | 0.827 | 0.314 | 5.76462E-09 | 4.15222E-24 | 2.004891748 | 0.958 | 0.336 | 9.80423E-20 | 2.44139E-13 | 8.30445E-24 | Ezr | MM_4 |
| 4.50345E-24 | 2.815476758 | 0.885 | 0.277 | 1.06335E-19 | 6.69953E-24 | 2.941925887 | 0.833 | 0.231 | 1.58189E-19 | 6.69953E-24 | 9.0069E-24 | Fyn | MM_4 |
| 2.08485E-16 | 5.428929129 | 0.212 | 0.003 | 4.92274E-12 | 5.66049E-24 | 4.692380898 | 0.208 | 0.002 | 1.33656E-19 | 2.08485E-16 | 1.1321E-23 | Dhrs9 | MM_4 |
| 3.25919E-10 | 2.963741675 | 0.192 | 0.014 | 7.6956E-06 | 8.02823E-24 | 4.506721655 | 0.271 | 0.009 | 1.89562E-19 | 3.25919E-10 | 1.60565E-23 | Gm30978 | MM_4 |

| | | | | | | | | | | | | | |
|---|---|---|---|---|---|---|---|---|---|---|---|---|---|
| 1.01549E-23 | 2.829437493 | 0.923 | 0.493 | 2.39778E-19 | 9.37885E-23 | 2.679392367 | 0.958 | 0.409 | 2.21453E-18 | 9.37885E-23 | 2.03098E-23 | Plbd1 | MM_4 |
| 1.27819E-23 | 3.207818056 | 0.904 | 0.343 | 3.01806E-19 | 7.60488E-18 | 2.696238332 | 0.896 | 0.366 | 1.79566E-13 | 7.60488E-18 | 2.55638E-23 | Naaa | MM_4 |
| 1.64389E-22 | 4.841523457 | 0.327 | 0.009 | 3.88155E-18 | 1.29618E-23 | 2.609462117 | 0.354 | 0.022 | 3.06055E-19 | 1.64389E-22 | 2.59237E-23 | Haao | MM_4 |
| 1.4636E-23 | 2.009265543 | 0.865 | 0.21 | 3.45584E-19 | 5.84034E-21 | 2.609987429 | 0.792 | 0.213 | 1.37902E-16 | 5.84034E-21 | 2.92719E-23 | Sowahc | MM_4 |
| 1.47289E-23 | 3.607558654 | 0.404 | 0.02 | 3.47779E-19 | 1.6207E-18 | 2.344028718 | 0.354 | 0.035 | 3.82681E-14 | 1.6207E-18 | 2.94578E-23 | Gpr68 | MM_4 |
| 1.83779E-23 | 2.799519751 | 0.962 | 0.539 | 4.33938E-19 | 3.02002E-22 | 2.499023771 | 0.958 | 0.749 | 7.13086E-18 | 3.02002E-22 | 3.67558E-23 | S100a11 | MM_4 |
| 2.42549E-23 | 3.209232781 | 0.673 | 0.118 | 5.72708E-19 | 1.29561E-22 | 2.874737302 | 0.75 | 0.172 | 3.0592E-18 | 1.29561E-22 | 4.85099E-23 | Rnase6 | MM_4 |
| 4.08889E-23 | 3.749135964 | 0.442 | 0.032 | 9.65468E-19 | 2.49867E-17 | 2.005984614 | 0.417 | 0.059 | 5.89986E-13 | 2.49867E-17 | 8.17777E-23 | Dock5 | MM_4 |
| 5.6168E-23 | 2.39327596 | 0.788 | 0.176 | 1.32624E-18 | 4.76911E-22 | 2.885940793 | 0.75 | 0.196 | 1.12608E-17 | 4.76911E-22 | 1.12336E-22 | 4930523C | MM_4 |
| 1.84409E-16 | 2.75448365 | 0.442 | 0.061 | 4.35427E-12 | 6.24865E-23 | 2.551634353 | 0.562 | 0.083 | 1.47543E-18 | 1.84409E-16 | 1.24973E-22 | Glipr2 | MM_4 |
| 8.93776E-23 | 5.947926232 | 0.269 | 0 | 2.11038E-18 | 1.42969E-11 | 5.749629319 | 0.104 | 0.002 | 3.37577E-07 | 1.42969E-11 | 1.78755E-22 | Treml4 | MM_4 |
| 8.93776E-23 | 6.098124842 | 0.269 | 0 | 2.11038E-18 | 2.08359E-19 | 5.834398495 | 0.188 | 0.004 | 4.91976E-15 | 2.08359E-19 | 1.78755E-22 | Ffar4 | MM_4 |
| 1.14668E-22 | 2.691850856 | 0.75 | 0.156 | 2.70754E-18 | 2.53231E-18 | 2.535703773 | 0.688 | 0.172 | 5.9793E-14 | 2.53231E-18 | 2.29336E-22 | Tmtc2 | MM_4 |
| 1.16701E-22 | 5.495342149 | 0.288 | 0.003 | 2.75554E-18 | 8.81786E-10 | 3.877824004 | 0.104 | 0.004 | 2.08207E-05 | 8.81786E-10 | 2.33401E-22 | Tnfrsf18 | MM_4 |
| 1.23309E-22 | 4.180466959 | 0.346 | 0.012 | 2.91157E-18 | 4.1522E-22 | 2.467957346 | 0.292 | 0.015 | 9.80418E-18 | 4.1522E-22 | 2.46618E-22 | Gm9844 | MM_4 |
| 1.2925E-22 | 3.297927575 | 0.423 | 0.029 | 3.05185E-18 | 2.82227E-17 | 5.242116116 | 0.25 | 0.017 | 6.66395E-13 | 2.82227E-17 | 2.585E-22 | Ccser1 | MM_4 |
| 1.22573E-11 | 2.476317827 | 0.673 | 0.285 | 2.89419E-07 | 1.72291E-22 | 2.692331133 | 0.917 | 0.327 | 4.06813E-18 | 1.22573E-11 | 3.44581E-22 | S100a4 | MM_4 |
| 1.94337E-22 | 2.182277783 | 1 | 0.715 | 4.58868E-18 | 5.27705E-15 | 2.079948489 | 0.938 | 0.725 | 1.24602E-10 | 5.27705E-15 | 3.88674E-22 | Btg1 | MM_4 |
| 2.24619E-22 | 1.757235705 | 1 | 0.888 | 5.30369E-18 | 2.64018E-20 | 1.538377446 | 1 | 0.911 | 6.23398E-16 | 2.64018E-20 | 4.49237E-22 | Actg1 | MM_4 |
| 3.73509E-22 | 2.386559131 | 0.692 | 0.127 | 8.8193E-18 | 1.02869E-16 | 2.956605663 | 0.562 | 0.128 | 2.42895E-12 | 1.02869E-16 | 7.47019E-22 | Dusp5 | MM_4 |
| 1.78529E-21 | 3.36972244 | 0.5 | 0.055 | 4.21542E-17 | 1.32065E-21 | 2.686687444 | 0.708 | 0.172 | 3.11831E-17 | 1.78529E-21 | 2.64129E-21 | Hopx | MM_4 |
| 2.8182E-21 | 1.94713082 | 0.962 | 0.435 | 6.65433E-17 | 4.92047E-19 | 2.24191471 | 0.938 | 0.536 | 1.16182E-14 | 4.92047E-19 | 5.6364E-21 | Zyx | MM_4 |
| 6.34637E-19 | 1.941793133 | 0.731 | 0.167 | 1.49851E-14 | 2.85227E-21 | 2.122882294 | 0.812 | 0.233 | 6.73479E-17 | 6.34637E-19 | 5.70455E-21 | Me2 | MM_4 |
| 3.26681E-21 | 8.49237277 | 0.25 | 0 | 7.71359E-17 | 1.73477E-16 | 4.015549589 | 0.146 | 0.002 | 4.09613E-12 | 1.73477E-16 | 6.53362E-21 | Ccl17 | MM_4 |
| 3.29296E-21 | 2.028557316 | 0.981 | 0.438 | 7.77534E-17 | 3.0064E-14 | 1.543712994 | 0.979 | 0.651 | 7.09872E-10 | 3.0064E-14 | 6.58592E-21 | Lgals3 | MM_4 |
| 4.13211E-21 | 2.048972404 | 0.923 | 0.329 | 9.75675E-17 | 8.48806E-19 | 1.989471751 | 0.875 | 0.372 | 2.0042E-14 | 8.48806E-19 | 8.26423E-21 | Myo1g | MM_4 |
| 1.28527E-20 | 2.499111041 | 0.596 | 0.095 | 3.03478E-16 | 7.74021E-11 | 2.186051504 | 0.479 | 0.133 | 1.82762E-06 | 7.74021E-11 | 2.57054E-20 | H2-Ob | MM_4 |
| 4.18045E-10 | 1.766290465 | 0.673 | 0.219 | 9.87089E-06 | 1.63711E-20 | 2.536037685 | 0.729 | 0.183 | 3.86555E-16 | 4.18045E-10 | 3.27423E-20 | Chn2 | MM_4 |
| 2.81286E-20 | 3.901895342 | 0.346 | 0.017 | 6.64172E-16 | 5.15375E-16 | 3.053944742 | 0.271 | 0.022 | 1.2169E-11 | 5.15375E-16 | 5.62571E-20 | Gm10552 | MM_4 |
| 3.03225E-20 | 2.485803622 | 0.558 | 0.081 | 7.15974E-16 | 3.09326E-12 | 3.273891618 | 0.333 | 0.057 | 7.30382E-08 | 3.09326E-12 | 6.0645E-20 | Gfod1 | MM_4 |
| 1.88386E-08 | 1.233805999 | 0.788 | 0.343 | 0.000444817 | 3.52441E-20 | 2.25286323 | 0.896 | 0.325 | 8.32184E-16 | 1.88386E-08 | 7.04882E-20 | Pfkp | MM_4 |
| 3.87404E-20 | 3.02337558 | 0.519 | 0.063 | 9.14737E-16 | 8.07366E-11 | 3.036071318 | 0.271 | 0.041 | 1.90635E-06 | 8.07366E-11 | 7.74807E-20 | Card11 | MM_4 |
| 2.8791E-16 | 1.705251108 | 0.788 | 0.21 | 6.79833E-12 | 4.17452E-20 | 2.054477131 | 0.771 | 0.192 | 9.85688E-16 | 2.8791E-16 | 8.34904E-20 | Sema4d | MM_4 |
| 5.44422E-15 | 1.866551216 | 0.923 | 0.513 | 1.28549E-10 | 4.63944E-20 | 2.021287066 | 1 | 0.529 | 1.09546E-15 | 5.44422E-15 | 9.27887E-20 | Arhgap26 | MM_4 |
| 5.44388E-20 | 3.454365269 | 0.5 | 0.063 | 1.28541E-15 | 1.10562E-14 | 2.806344097 | 0.458 | 0.089 | 2.6106E-10 | 1.10562E-14 | 1.08878E-19 | Nod2 | MM_4 |
| 7.48829E-20 | 1.496250673 | 1 | 0.919 | 1.76813E-15 | 1.03519E-16 | 1.054182398 | 1 | 0.941 | 2.44429E-12 | 1.03519E-16 | 1.49766E-19 | Rpsa | MM_4 |
| 8.74473E-20 | 4.608641263 | 0.308 | 0.012 | 2.06481E-15 | 6.5852E-14 | 4.578368376 | 0.208 | 0.015 | 1.5549E-09 | 6.5852E-14 | 1.74895E-19 | 4931406G | MM_4 |
| 2.28677E-16 | 1.720877548 | 0.596 | 0.112 | 5.39952E-12 | 1.2403E-19 | 2.411422197 | 0.729 | 0.183 | 2.92859E-15 | 2.28677E-16 | 2.4806E-19 | Pira2 | MM_4 |
| 1.28734E-19 | 1.698292851 | 0.827 | 0.219 | 3.03966E-15 | 2.68126E-15 | 2.119282632 | 0.75 | 0.238 | 6.33099E-11 | 2.68126E-15 | 2.57467E-19 | 1700025G | MM_4 |
| 1.84054E-19 | 2.734895817 | 0.635 | 0.124 | 4.34588E-15 | 1.04461E-08 | 2.205389523 | 0.375 | 0.1 | 0.000246653 | 1.04461E-08 | 3.68108E-19 | Lmnb1 | MM_4 |
| 1.84704E-19 | 3.038305513 | 0.885 | 0.36 | 4.36124E-15 | 4.35394E-17 | 2.896326545 | 0.812 | 0.348 | 1.02805E-12 | 4.35394E-17 | 3.69409E-19 | Jak2 | MM_4 |
| 2.25516E-19 | 2.241696876 | 0.731 | 0.156 | 5.32488E-15 | 2.19985E-15 | 2.364148156 | 0.562 | 0.129 | 5.19429E-11 | 2.19985E-15 | 4.51031E-19 | Runx3 | MM_4 |
| 5.49501E-18 | 5.581888874 | 0.231 | 0.003 | 1.29748E-13 | 2.31002E-19 | 4.899700991 | 0.188 | 0.004 | 5.45441E-15 | 5.49501E-18 | 4.62003E-19 | 6430531B | MM_4 |
| 1.48428E-15 | 1.760603236 | 0.981 | 0.582 | 3.50469E-11 | 2.72508E-19 | 2.105392382 | 0.979 | 0.689 | 6.43446E-15 | 1.48428E-15 | 5.45016E-19 | S100a6 | MM_4 |
| 3.66736E-19 | 1.706624631 | 0.981 | 0.562 | 8.65937E-15 | 5.40892E-17 | 1.803273066 | 0.938 | 0.617 | 1.27716E-12 | 5.40892E-17 | 7.33472E-19 | H2afy | MM_4 |
| 1.4713E-14 | 1.577352829 | 1 | 0.689 | 3.47403E-10 | 6.11055E-19 | 2.066901912 | 1 | 0.75 | 1.44282E-14 | 1.4713E-14 | 1.22211E-18 | Cd44 | MM_4 |
| 6.98092E-19 | 1.6615758 | 0.962 | 0.666 | 1.64833E-14 | 1.06352E-11 | 0.905617899 | 0.958 | 0.882 | 2.51118E-07 | 1.06352E-11 | 1.39618E-18 | Sh3bgrl3 | MM_4 |
| 1.15578E-14 | 1.195249756 | 1 | 0.634 | 2.72903E-10 | 9.83191E-19 | 1.497443399 | 1 | 0.686 | 2.32151E-14 | 1.15578E-14 | 1.96638E-18 | Fxyd5 | MM_4 |
| 3.35667E-12 | 2.766096423 | 0.25 | 0.02 | 7.92577E-08 | 1.05945E-18 | 2.452923395 | 0.292 | 0.02 | 2.50157E-14 | 3.35667E-12 | 2.1189E-18 | Gm15228 | MM_4 |
| 3.96006E-18 | 2.608993208 | 0.288 | 0.012 | 9.35048E-14 | 1.14252E-18 | 3.360217338 | 0.25 | 0.013 | 2.69771E-14 | 3.96006E-18 | 2.28504E-18 | Gm13373 | MM_4 |
| 1.34215E-18 | 1.442522478 | 1 | 0.89 | 3.16908E-14 | 1.52221E-14 | 1.097615859 | 1 | 0.919 | 3.59424E-10 | 1.52221E-14 | 2.68429E-18 | Rplp0 | MM_4 |
| 8.18336E-18 | 3.811109979 | 0.538 | 0.098 | 1.93226E-13 | 2.68966E-18 | 3.14365199 | 0.521 | 0.092 | 6.35083E-14 | 8.18336E-18 | 5.37933E-18 | Lrrk2 | MM_4 |
| 6.17229E-15 | 2.020018435 | 0.385 | 0.046 | 1.4574E-10 | 2.93744E-18 | 2.983506083 | 0.5 | 0.085 | 6.93589E-14 | 6.17229E-15 | 5.87488E-18 | Acot7 | MM_4 |
| 4.99944E-12 | 2.700808126 | 0.308 | 0.037 | 1.18047E-07 | 3.13492E-18 | 1.851451844 | 0.417 | 0.052 | 7.40218E-14 | 4.99944E-12 | 6.26984E-18 | Ccdc88c | MM_4 |
| 3.64878E-18 | 3.570714135 | 0.558 | 0.104 | 8.6155E-14 | 1.07455E-07 | 2.312292592 | 0.458 | 0.165 | 0.002537217 | 1.07455E-07 | 7.29756E-18 | Slc25a20 | MM_4 |
| 3.89732E-18 | 2.010136043 | 0.962 | 0.536 | 9.20234E-14 | 2.19917E-14 | 1.537006134 | 1 | 0.774 | 5.19267E-10 | 2.19917E-14 | 7.79463E-18 | S100a10 | MM_4 |
| 4.1751E-18 | 5.906543995 | 0.212 | 0 | 9.85825E-14 | 3.53593E-12 | 4.98182342 | 0.125 | 0.004 | 8.34904E-08 | 3.53593E-12 | 8.3502E-18 | Snx22 | MM_4 |
| 5.70889E-18 | 2.470303177 | 0.904 | 0.49 | 1.34798E-13 | 6.09023E-09 | 1.426311153 | 0.812 | 0.53 | 0.000143803 | 6.09023E-09 | 1.14178E-17 | Dapp1 | MM_4 |
| 6.84321E-18 | 3.018089493 | 0.596 | 0.121 | 1.61582E-13 | 2.30176E-08 | 2.251743851 | 0.5 | 0.174 | 0.000543491 | 2.30176E-08 | 1.36864E-17 | Havcr2 | MM_4 |
| 1.55317E-17 | 2.432928785 | 0.846 | 0.3 | 3.66733E-13 | 7.3192E-18 | 2.251562114 | 0.812 | 0.264 | 1.72821E-13 | 1.55317E-17 | 1.46384E-17 | Pmaip1 | MM_4 |
| 9.95736E-12 | 2.332745083 | 0.231 | 0.017 | 2.35113E-07 | 9.41735E-18 | 3.11763376 | 0.229 | 0.011 | 2.22362E-13 | 9.95736E-12 | 1.88347E-17 | 5830428M | MM_4 |
| 1.18035E-17 | 2.183619812 | 0.885 | 0.354 | 2.78703E-13 | 1.10082E-13 | 1.811669192 | 0.917 | 0.573 | 2.59926E-09 | 1.10082E-13 | 2.36069E-17 | Anxa1 | MM_4 |
| 3.86079E-16 | 1.531364623 | 0.981 | 0.726 | 9.1161E-12 | 1.19471E-17 | 1.974008108 | 1 | 0.75 | 2.82095E-13 | 3.86079E-16 | 2.38942E-17 | Etv6 | MM_4 |
| 9.27732E-15 | 4.309582227 | 0.192 | 0.003 | 2.19056E-10 | 1.38006E-17 | 4.824804171 | 0.188 | 0.006 | 3.2586E-13 | 9.27732E-15 | 2.76012E-17 | Mycl | MM_4 |
| 1.27645E-10 | 2.914779548 | 0.212 | 0.017 | 3.01395E-06 | 1.38279E-17 | 2.649420282 | 0.25 | 0.015 | 3.26505E-13 | 1.27645E-10 | 2.76558E-17 | Slc36a3os | MM_4 |
| 1.52506E-17 | 1.886129877 | 1 | 0.752 | 3.60097E-13 | 5.50928E-13 | 1.34632924 | 1 | 0.878 | 1.30085E-08 | 5.50928E-13 | 3.05012E-17 | Vim | MM_4 |
| 1.57855E-17 | 2.452593439 | 0.827 | 0.282 | 3.72727E-13 | 9.1043E-12 | 2.363631311 | 0.688 | 0.262 | 2.14971E-07 | 9.1043E-12 | 3.15709E-17 | Phlpp1 | MM_4 |
| 1.64289E-17 | 3.048605534 | 0.615 | 0.144 | 3.87919E-13 | 2.78591E-12 | 2.836508637 | 0.521 | 0.153 | 6.57809E-08 | 2.78591E-12 | 3.28578E-17 | Zbtb46 | MM_4 |
| 2.81744E-09 | 1.04103533 | 0.981 | 0.579 | 6.65254E-05 | 1.69103E-17 | 1.888937606 | 1 | 0.625 | 3.99287E-13 | 2.81744E-09 | 3.38207E-17 | Itga4 | MM_4 |
| 5.79386E-15 | 2.764668583 | 0.635 | 0.17 | 1.36805E-10 | 1.76176E-17 | 1.980998875 | 0.708 | 0.19 | 4.15986E-13 | 5.79386E-15 | 3.52351E-17 | Qpct | MM_4 |
| 1.8665E-17 | 1.120689157 | 1 | 0.922 | 4.40718E-13 | 5.76433E-12 | 1.018586792 | 1 | 0.895 | 1.36107E-07 | 5.76433E-12 | 3.733E-17 | Psap | MM_4 |
| 1.9075E-17 | 2.111642297 | 0.731 | 0.173 | 4.50399E-13 | 1.59629E-10 | 1.538476463 | 0.562 | 0.166 | 3.76916E-06 | 1.59629E-10 | 3.815E-17 | Tspan13 | MM_4 |
| 9.12559E-12 | 3.303462981 | 0.269 | 0.029 | 2.15473E-07 | 2.06851E-17 | 2.310625577 | 0.562 | 0.111 | 4.88418E-13 | 9.12559E-12 | 4.13703E-17 | Gm14548 | MM_4 |
| 7.92686E-10 | 2.994870462 | 0.269 | 0.037 | 1.87169E-05 | 2.22361E-17 | 5.025713503 | 0.375 | 0.048 | 5.25038E-13 | 7.92686E-10 | 4.44722E-17 | Il6 | MM_4 |
| 1.237E-07 | 1.775245713 | 0.269 | 0.052 | 0.0029208 | 2.68095E-17 | 2.618205817 | 0.5 | 0.091 | 6.33027E-13 | 1.237E-07 | 5.36191E-17 | Kcnq1 | MM_4 |
| 2.73057E-17 | 2.926809463 | 0.308 | 0.017 | 6.44743E-13 | 4.63647E-13 | 6.091407517 | 0.146 | 0.006 | 1.09476E-08 | 4.63647E-13 | 5.46115E-17 | Pglyrp1 | MM_4 |
| 2.81941E-17 | 2.721106502 | 0.519 | 0.084 | 6.6572E-13 | 1.6318E-14 | 2.383727218 | 0.521 | 0.118 | 3.85301E-10 | 1.6318E-14 | 5.63883E-17 | Krt80 | MM_4 |
| 5.38008E-17 | 2.45237122 | 0.769 | 0.3 | 1.27034E-12 | 3.19579E-17 | 1.6115485 | 0.917 | 0.36 | 7.54589E-13 | 5.38008E-17 | 6.39157E-17 | Cbfa2t3 | MM_4 |
| 5.05269E-15 | 5.76816777 | 0.173 | 0 | 1.19304E-10 | 5.22174E-17 | 3.721678561 | 0.208 | 0.009 | 1.23296E-12 | 5.05269E-15 | 1.04435E-16 | Slc27a3 | MM_4 |
| 5.45076E-17 | 1.897136207 | 0.75 | 0.21 | 1.28703E-12 | 7.08292E-12 | 1.31363651 | 0.667 | 0.226 | 1.67242E-07 | 7.08292E-12 | 1.09015E-16 | Tes | MM_4 |
| 2.05709E-15 | 2.430177003 | 0.769 | 0.274 | 4.8572E-11 | 7.76695E-17 | 2.715509345 | 0.708 | 0.205 | 1.83393E-12 | 2.05709E-15 | 1.55339E-16 | Il1b | MM_4 |

| | | | | | | | | | | | | | |
|---|---|---|---|---|---|---|---|---|---|---|---|---|---|
| 9.44232E-17 | 2.537419033 | 0.75 | 0.254 | 2.22952E-12 | 3.3891E-08 | 2.189237567 | 0.667 | 0.327 | 0.000800234 | 3.3891E-08 | 1.88846E-16 | Naga | MM_4 |
| 1.06319E-16 | 1.585104102 | 0.981 | 0.729 | 2.51041E-12 | 1.70739E-13 | 1.387854077 | 0.979 | 0.784 | 4.03149E-09 | 1.70739E-13 | 2.12639E-16 | Atox1 | MM_4 |
| 1.31595E-16 | 2.53119702 | 0.558 | 0.104 | 3.10722E-12 | 3.88335E-13 | 2.125264496 | 0.521 | 0.128 | 9.16936E-09 | 3.88335E-13 | 2.6319E-16 | Mthfd2 | MM_4 |
| 1.46174E-16 | 5.62961848 | 0.192 | 0 | 3.45147E-12 | 1.50704E-16 | 5.771517829 | 0.125 | 0 | 3.55842E-12 | 1.50704E-16 | 2.92349E-16 | Nccrp1 | MM_4 |
| 1.76083E-07 | 3.185678116 | 0.115 | 0.006 | 0.004157679 | 1.50704E-16 | 5.611469057 | 0.125 | 0 | 3.55842E-12 | 1.76083E-07 | 3.01408E-16 | Mbnl3 | MM_4 |
| 1.5304E-16 | 2.061602529 | 0.788 | 0.231 | 3.61359E-12 | 5.65964E-11 | 1.876891942 | 0.625 | 0.222 | 1.33636E-06 | 5.65964E-11 | 3.06081E-16 | Ldlr | MM_4 |
| 1.81462E-16 | 1.596901091 | 0.923 | 0.432 | 4.28469E-12 | 1.75964E-12 | 1.494045982 | 0.875 | 0.553 | 4.15486E-08 | 1.75964E-12 | 3.62925E-16 | Selplg | MM_4 |
| 7.05631E-12 | 1.512736398 | 0.885 | 0.53 | 1.66614E-07 | 2.57648E-16 | 1.599270778 | 0.958 | 0.573 | 6.08359E-12 | 7.05631E-12 | 5.15296E-16 | Ccdc88a | MM_4 |
| 2.90132E-16 | 4.242898994 | 0.288 | 0.017 | 6.8506E-12 | 7.45517E-09 | 4.43128098 | 0.167 | 0.018 | 0.000176031 | 7.45517E-09 | 5.80265E-16 | 5730522E[illegible] | MM_4 |
| 4.15584E-16 | 1.669207848 | 0.962 | 0.732 | 9.81278E-12 | 1.37453E-10 | 1.296091257 | 0.979 | 0.821 | 3.24554E-06 | 1.37453E-10 | 8.31169E-16 | Sub1 | MM_4 |
| 5.54961E-15 | 2.261065109 | 0.769 | 0.268 | 1.31037E-10 | 4.6466E-16 | 2.134051808 | 0.75 | 0.235 | 1.09716E-11 | 5.54961E-15 | 9.2932E-16 | Gpd2 | MM_4 |
| 4.65622E-16 | 3.133805044 | 0.423 | 0.055 | 1.09943E-11 | 3.42661E-12 | 2.273080276 | 0.354 | 0.065 | 8.09091E-08 | 3.42661E-12 | 9.31243E-16 | Net1 | MM_4 |
| 4.79844E-16 | 2.047160543 | 0.885 | 0.38 | 1.13301E-11 | 2.06465E-09 | 1.568229335 | 0.833 | 0.49 | 4.87504E-05 | 2.06465E-09 | 9.59688E-16 | BC028528 | MM_4 |
| 7.56501E-15 | 1.422784516 | 1 | 0.689 | 1.78625E-10 | 4.81994E-16 | 1.699700725 | 0.958 | 0.693 | 1.13808E-11 | 7.56501E-15 | 9.63988E-16 | Coro1a | MM_4 |
| 5.10248E-16 | 3.615155905 | 0.327 | 0.026 | 1.2048E-11 | 1.23631E-11 | 4.574674004 | 0.208 | 0.02 | 2.91917E-07 | 1.23631E-11 | 1.0205E-15 | Gpr141 | MM_4 |
| 5.70393E-16 | 1.633968808 | 0.981 | 0.545 | 1.34681E-11 | 5.50567E-16 | 1.889705326 | 0.979 | 0.566 | 1.3E-11 | 5.70393E-16 | 1.10113E-15 | Lrrfip1 | MM_4 |
| 1.44372E-13 | 2.802404546 | 0.212 | 0.009 | 3.40891E-09 | 5.9603E-16 | 4.6541646 | 0.229 | 0.015 | 1.40735E-11 | 1.44372E-13 | 1.19206E-15 | Hr | MM_4 |
| 7.17975E-16 | 1.492963971 | 0.981 | 0.594 | 1.69528E-11 | 2.38666E-11 | 1.413656381 | 0.896 | 0.597 | 5.63538E-07 | 2.38666E-11 | 1.43595E-15 | Pim1 | MM_4 |
| 5.24202E-08 | 1.895896024 | 0.231 | 0.035 | 0.001237746 | 7.41338E-16 | 2.30747613 | 0.375 | 0.05 | 1.75045E-11 | 5.24202E-08 | 1.48268E-15 | Dgkg | MM_4 |
| 7.92398E-16 | 2.126537668 | 0.654 | 0.176 | 1.87101E-11 | 1.35674E-08 | 1.764708525 | 0.667 | 0.305 | 0.000320353 | 1.35674E-08 | 1.5848E-15 | Pkib | MM_4 |
| 1.97891E-11 | 2.24058543 | 0.288 | 0.035 | 4.67261E-07 | 8.4606E-16 | 3.402409308 | 0.312 | 0.033 | 1.99772E-11 | 1.97891E-11 | 1.69212E-15 | Il7r | MM_4 |
| 8.57434E-16 | 2.67396438 | 0.462 | 0.069 | 2.02457E-11 | 1.98183E-08 | 1.610074795 | 0.292 | 0.061 | 0.00046795 | 1.98183E-08 | 1.71487E-15 | Fndc7 | MM_4 |
| 1.19441E-15 | 3.704286119 | 0.558 | 0.124 | 2.82025E-11 | 5.9205E-09 | 2.22887888 | 0.438 | 0.126 | 0.000139795 | 5.9205E-09 | 2.38883E-15 | Zc3h12c | MM_4 |
| 5.68396E-15 | 1.551574686 | 0.865 | 0.308 | 1.3421E-10 | 1.52348E-15 | 1.82546993 | 0.875 | 0.325 | 3.59723E-11 | 5.68396E-15 | 3.04695E-15 | Atp8b4 | MM_4 |
| 3.28208E-13 | 1.380299303 | 0.923 | 0.712 | 7.74965E-09 | 1.68652E-15 | 1.646828746 | 0.958 | 0.632 | 3.98221E-11 | 3.28208E-13 | 3.37304E-15 | H2-DMa | MM_4 |
| 3.04038E-11 | 0.864971353 | 1 | 0.81 | 7.17895E-07 | 1.74271E-15 | 1.072398744 | 1 | 0.841 | 4.11488E-11 | 3.04038E-11 | 3.48541E-15 | Actr3 | MM_4 |
| 2.01825E-15 | 1.453729951 | 1 | 0.784 | 4.76549E-11 | 4.24478E-13 | 1.331562518 | 1 | 0.823 | 1.00228E-08 | 4.24478E-13 | 4.0365E-15 | Srgn | MM_4 |
| 3.6316E-13 | 0.889187166 | 1 | 0.888 | 8.57494E-09 | 2.02357E-15 | 0.871553929 | 1 | 0.904 | 4.77804E-11 | 3.6316E-13 | 4.04713E-15 | Rack1 | MM_4 |
| 8.6052E-10 | 1.159996053 | 0.635 | 0.202 | 2.03186E-05 | 2.16425E-15 | 1.622081725 | 0.708 | 0.201 | 5.11023E-11 | 8.6052E-10 | 4.3285E-15 | Ramp1 | MM_4 |
| 1.94798E-07 | 0.961179796 | 0.904 | 0.522 | 0.004599568 | 2.32225E-15 | 1.399855965 | 0.979 | 0.547 | 5.4833E-11 | 1.94798E-07 | 4.6445E-15 | Fem1c | MM_4 |
| 2.67185E-15 | 1.840575392 | 0.615 | 0.133 | 6.30876E-11 | 1.4376E-10 | 1.553278243 | 0.542 | 0.161 | 3.39447E-06 | 1.4376E-10 | 5.34369E-15 | Amz1 | MM_4 |
| 3.17512E-15 | 3.063428953 | 0.288 | 0.02 | 7.49709E-11 | 7.84914E-10 | 2.860401775 | 0.208 | 0.026 | 1.85334E-05 | 7.84914E-10 | 6.35024E-15 | Clcf1 | MM_4 |
| 1.00094E-14 | 3.550184188 | 0.212 | 0.006 | 2.36341E-10 | 3.4166E-15 | 3.615933536 | 0.167 | 0.006 | 8.06728E-11 | 1.00094E-14 | 6.83321E-15 | Cd300c | MM_4 |
| 4.12946E-15 | 2.653428498 | 0.442 | 0.066 | 9.75047E-11 | 4.28683E-08 | 1.953184021 | 0.396 | 0.113 | 0.001012206 | 4.28683E-08 | 8.25891E-15 | Asap2 | MM_4 |
| 4.19814E-15 | 1.542958154 | 0.942 | 0.519 | 9.91264E-11 | 2.32308E-07 | 1.160757192 | 0.854 | 0.623 | 0.005485264 | 2.32308E-07 | 8.39627E-15 | Nr4a1 | MM_4 |
| 3.06014E-10 | 4.363680794 | 0.135 | 0.003 | 7.22561E-06 | 4.21899E-15 | 3.358945111 | 0.167 | 0.006 | 9.96188E-11 | 3.06014E-10 | 8.43798E-15 | Klri2 | MM_4 |
| 1.9954E-14 | 1.468816497 | 0.942 | 0.637 | 4.71155E-10 | 4.75631E-15 | 1.703558991 | 0.938 | 0.593 | 1.12306E-10 | 1.9954E-14 | 9.51263E-15 | H2-DMb1 | MM_4 |
| 5.00841E-15 | 1.928984623 | 0.923 | 0.778 | 1.18259E-10 | 2.85808E-09 | 1.354090643 | 0.958 | 0.843 | 6.7485E-05 | 2.85808E-09 | 1.00168E-14 | H2afz | MM_4 |
| 6.95413E-15 | 3.130619232 | 0.615 | 0.167 | 1.64201E-10 | 2.48158E-09 | 2.602326222 | 0.458 | 0.144 | 5.85952E-05 | 2.48158E-09 | 1.39083E-14 | Dusp2 | MM_4 |
| 6.95736E-15 | 4.669185579 | 0.212 | 0.006 | 1.64277E-10 | 1.24847E-10 | 4.241829312 | 0.125 | 0.006 | 2.94788E-06 | 1.24847E-10 | 1.39147E-14 | Tlr11 | MM_4 |
| 7.43587E-15 | 3.492196372 | 0.365 | 0.043 | 1.75576E-10 | 1.10835E-11 | 2.784973007 | 0.25 | 0.03 | 2.61704E-07 | 1.10835E-11 | 1.48717E-14 | Chsy3 | MM_4 |
| 8.11314E-15 | 1.088194565 | 1 | 0.896 | 1.91567E-10 | 1.19645E-10 | 0.733075637 | 1 | 0.928 | 2.82505E-06 | 1.19645E-10 | 1.62263E-14 | Rps5 | MM_4 |
| 8.1659E-15 | 1.758098767 | 0.827 | 0.311 | 1.92813E-10 | 9.12231E-09 | 1.636503813 | 0.688 | 0.311 | 0.000215396 | 9.12231E-09 | 1.63318E-14 | Vrk1 | MM_4 |
| 8.21028E-15 | 4.406642172 | 0.212 | 0.006 | 1.93861E-10 | 1.46478E-08 | 4.450035548 | 0.104 | 0.006 | 0.000345863 | 1.46478E-08 | 1.64206E-14 | Mmp25 | MM_4 |
| 8.09929E-10 | 1.608836211 | 0.692 | 0.256 | 1.9124E-05 | 8.22395E-15 | 1.293744843 | 0.771 | 0.237 | 1.94184E-10 | 8.09929E-10 | 1.64479E-14 | Xylt1 | MM_4 |
| 8.80279E-09 | 4.521096938 | 0.115 | 0.003 | 0.000207851 | 8.76039E-15 | 3.649467913 | 0.188 | 0.009 | 2.0685E-10 | 8.80279E-09 | 1.75208E-14 | Ccdc170 | MM_4 |
| 6.57464E-08 | 3.522375245 | 0.135 | 0.009 | 0.001552403 | 8.81691E-15 | 3.193422634 | 0.229 | 0.017 | 2.08185E-10 | 6.57464E-08 | 1.76338E-14 | Sigirr | MM_4 |
| 9.98849E-15 | 0.867266833 | 1 | 0.928 | 2.35848E-10 | 3.33322E-12 | 0.665857588 | 1 | 0.963 | 7.87039E-08 | 3.33322E-12 | 1.9977E-14 | Rpl41 | MM_4 |
| 7.70211E-09 | 0.88375142 | 1 | 0.81 | 0.000181862 | 1.04573E-14 | 1.308035094 | 1 | 0.852 | 2.46918E-10 | 7.70211E-09 | 2.09146E-14 | Ptprc | MM_4 |
| 3.63242E-09 | 1.214033579 | 0.365 | 0.078 | 8.57686E-05 | 1.06535E-14 | 2.579671453 | 0.458 | 0.089 | 2.51551E-10 | 3.63242E-09 | 2.1307E-14 | Aldh1a2 | MM_4 |
| 2.57413E-13 | 4.137635114 | 0.192 | 0.006 | 6.07804E-09 | 1.09289E-14 | 4.208975612 | 0.208 | 0.013 | 2.58053E-10 | 2.57413E-13 | 2.18578E-14 | Tbc1d10c | MM_4 |
| 1.46416E-10 | 2.106319199 | 0.538 | 0.161 | 3.45717E-06 | 1.18205E-14 | 2.31676248 | 0.625 | 0.166 | 2.79106E-10 | 1.46416E-10 | 2.3641E-14 | Nfil3 | MM_4 |
| 1.21446E-14 | 2.034156599 | 0.577 | 0.133 | 2.86758E-10 | 2.14299E-09 | 1.227486312 | 0.625 | 0.22 | 5.06003E-05 | 2.14299E-09 | 2.42892E-14 | Apobr | MM_4 |
| 2.20848E-09 | 0.936848096 | 0.827 | 0.415 | 5.21465E-05 | 1.24728E-14 | 1.384807709 | 0.938 | 0.494 | 2.94507E-10 | 2.20848E-09 | 2.49456E-14 | H13 | MM_4 |
| 1.29551E-14 | 1.90909641 | 0.865 | 0.429 | 3.05895E-10 | 1.18883E-12 | 1.591152049 | 0.854 | 0.473 | 2.80706E-08 | 1.18883E-12 | 2.59102E-14 | Gng10 | MM_4 |
| 1.37649E-14 | 2.988699244 | 0.269 | 0.017 | 3.25016E-10 | 2.50988E-10 | 3.287247091 | 0.188 | 0.018 | 5.92634E-06 | 2.50988E-10 | 2.75297E-14 | Aldoc | MM_4 |
| 1.52966E-14 | 2.340653489 | 0.596 | 0.144 | 3.61184E-10 | 1.46536E-07 | 1.587248966 | 0.583 | 0.264 | 0.003460016 | 1.46536E-07 | 3.05933E-14 | Cdkn2d | MM_4 |
| 1.5334E-14 | 1.017636522 | 0.423 | 0.058 | 3.62066E-10 | 1.0956E-08 | 1.281662538 | 0.271 | 0.052 | 0.000258693 | 1.0956E-08 | 3.0668E-14 | Bcl11a | MM_4 |
| 2.52908E-10 | 1.67089556 | 0.769 | 0.334 | 5.97167E-06 | 1.55473E-14 | 1.694564411 | 0.896 | 0.368 | 3.67102E-10 | 2.52908E-10 | 3.10945E-14 | Lrrc8c | MM_4 |
| 1.5927E-14 | 1.625066939 | 0.865 | 0.375 | 3.76069E-10 | 2.17077E-10 | 1.327076655 | 0.833 | 0.416 | 5.12563E-06 | 2.17077E-10 | 3.18541E-14 | Cd9 | MM_4 |
| 1.65959E-14 | 2.175907614 | 0.673 | 0.21 | 3.91863E-10 | 8.54929E-09 | 2.118334678 | 0.542 | 0.19 | 0.000201866 | 8.54929E-09 | 3.31919E-14 | Pik3cb | MM_4 |
| 1.76162E-14 | 1.162695685 | 1 | 0.908 | 4.15953E-10 | 6.90496E-11 | 0.878595704 | 1 | 0.915 | 1.6304E-06 | 6.90496E-11 | 3.52323E-14 | Rpl17 | MM_4 |
| 4.20951E-07 | 1.062374445 | 0.25 | 0.046 | 0.009939491 | 1.80341E-14 | 2.649998043 | 0.417 | 0.072 | 4.25821E-10 | 4.20951E-07 | 3.60682E-14 | Gm17749 | MM_4 |
| 3.89469E-08 | 1.007294901 | 0.808 | 0.438 | 0.000919615 | 1.87899E-14 | 1.570412042 | 0.958 | 0.429 | 4.43668E-10 | 3.89469E-08 | 3.75798E-14 | Stk10 | MM_4 |
| 1.18114E-11 | 1.490147235 | 0.865 | 0.438 | 2.78892E-07 | 1.92357E-14 | 1.861838369 | 0.896 | 0.495 | 4.54192E-10 | 1.18114E-11 | 3.84713E-14 | St3gal1 | MM_4 |
| 5.31693E-14 | 2.937949936 | 0.712 | 0.251 | 1.25543E-09 | 2.04888E-14 | 3.38236979 | 0.667 | 0.214 | 4.83782E-10 | 5.31693E-14 | 4.09776E-14 | Malt1 | MM_4 |
| 1.0524E-07 | 2.18519003 | 0.288 | 0.061 | 0.00248492 | 2.33734E-14 | 3.379762919 | 0.375 | 0.057 | 5.51894E-10 | 1.0524E-07 | 4.67469E-14 | Gm35853 | MM_4 |
| 2.48046E-14 | 2.697500098 | 0.615 | 0.161 | 5.85686E-10 | 4.98826E-08 | 1.402583973 | 0.583 | 0.222 | 0.001177829 | 4.98826E-08 | 4.96092E-14 | Sgk1 | MM_4 |
| 2.93868E-14 | 1.341782354 | 1 | 0.686 | 6.9388E-10 | 7.19117E-10 | 1.217842551 | 0.958 | 0.715 | 1.69798E-05 | 7.19117E-10 | 5.87735E-14 | Rel | MM_4 |
| 3.01999E-14 | 1.758580042 | 0.846 | 0.34 | 7.13081E-10 | 4.99184E-14 | 1.7402237 | 0.917 | 0.42 | 1.17867E-09 | 4.99184E-14 | 6.03998E-14 | Mbp | MM_4 |
| 2.89414E-13 | 1.182222662 | 0.923 | 0.504 | 6.83365E-09 | 3.066E-14 | 1.410143819 | 0.958 | 0.649 | 7.23944E-10 | 2.89414E-13 | 6.132E-14 | Ldha | MM_4 |
| 3.38502E-14 | 1.231136373 | 1 | 0.882 | 7.9927E-10 | 3.34813E-13 | 0.895233542 | 0.979 | 0.904 | 7.90562E-09 | 3.34813E-13 | 6.77003E-14 | Rps7 | MM_4 |
| 3.49963E-14 | 1.285695645 | 0.981 | 0.55 | 8.26333E-10 | 2.72969E-07 | 0.958915291 | 0.875 | 0.625 | 0.006445349 | 2.72969E-07 | 6.99926E-14 | Irf5 | MM_4 |
| 1.96408E-12 | 0.821653059 | 1 | 0.988 | 4.63759E-08 | 3.53734E-14 | 0.792285576 | 1 | 1 | 8.35237E-10 | 1.96408E-12 | 7.07468E-14 | Actb | MM_4 |
| 4.13489E-14 | 1.694845939 | 0.75 | 0.213 | 9.7633E-10 | 3.24768E-08 | 1.512354737 | 0.646 | 0.307 | 0.000766843 | 3.24768E-08 | 8.26978E-14 | Lactb | MM_4 |
| 4.65015E-14 | 2.393500155 | 0.865 | 0.507 | 1.09799E-09 | 3.16399E-12 | 2.185598166 | 0.917 | 0.525 | 7.47081E-08 | 3.16399E-12 | 9.3003E-14 | Wdfy4 | MM_4 |
| 1.52144E-13 | 3.328517746 | 0.404 | 0.066 | 3.59243E-09 | 4.92486E-14 | 2.371695538 | 0.396 | 0.065 | 1.16286E-09 | 1.52144E-13 | 9.84973E-14 | Mir155hg | MM_4 |
| 7.05869E-12 | 1.269362092 | 1 | 0.686 | 1.6667E-07 | 5.1628E-14 | 1.64310698 | 1 | 0.734 | 1.21904E-09 | 7.05869E-12 | 1.03256E-13 | Jarid2 | MM_4 |
| 7.9588E-09 | 4.979638121 | 0.115 | 0.003 | 0.000187923 | 5.36435E-14 | 3.874228888 | 0.125 | 0.002 | 1.26663E-09 | 7.9588E-09 | 1.07287E-13 | Klrb1a | MM_4 |
| 1.28843E-07 | 1.16346439 | 0.654 | 0.288 | 0.00304223 | 6.45047E-14 | 2.015711062 | 0.812 | 0.362 | 1.52309E-09 | 1.28843E-07 | 1.29009E-13 | Adss | MM_4 |
| 6.69199E-14 | 3.275012253 | 0.423 | 0.072 | 1.58011E-09 | 4.36481E-10 | 2.337013229 | 0.375 | 0.089 | 1.03062E-05 | 4.36481E-10 | 1.3384E-13 | Grasp | MM_4 |

| | | | | | | | | | | | | | |
|---|---|---|---|---|---|---|---|---|---|---|---|---|---|
| 1.56958E-07 | 3.181195601 | 0.115 | 0.006 | 0.003706091 | 7.38918E-14 | 4.23469091 | 0.167 | 0.007 | 1.74473E-09 | 1.56958E-07 | 1.47784E-13 | Vdr | MM_4 |
| 1.60011E-11 | 2.401242008 | 0.5 | 0.135 | 3.77818E-07 | 1.13582E-13 | 2.095551313 | 0.604 | 0.17 | 2.6819E-09 | 1.60011E-11 | 2.27165E-13 | Ppm1m | MM_4 |
| 1.18572E-13 | 3.035365122 | 0.385 | 0.055 | 2.79971E-09 | 1.22185E-11 | 2.044747558 | 0.417 | 0.089 | 2.88503E-07 | 1.22185E-11 | 2.37143E-13 | Mical1 | MM_4 |
| 2.06936E-07 | 2.313877994 | 0.442 | 0.15 | 0.004886173 | 1.34133E-13 | 2.392347868 | 0.771 | 0.309 | 3.16714E-09 | 2.06936E-07 | 2.68265E-13 | Ifitm6 | MM_4 |
| 8.89263E-13 | 2.639233177 | 0.231 | 0.014 | 2.09973E-08 | 1.4238E-13 | 2.689788092 | 0.188 | 0.011 | 3.36188E-09 | 8.89263E-13 | 2.8476E-13 | Adgrg5 | MM_4 |
| 1.436E-13 | 1.606851373 | 0.75 | 0.239 | 3.39067E-09 | 2.16451E-12 | 1.54153586 | 0.729 | 0.268 | 5.11084E-08 | 2.16451E-12 | 2.87199E-13 | Sms | MM_4 |
| 1.61783E-10 | 1.026683018 | 0.769 | 0.288 | 3.82003E-06 | 1.43948E-13 | 1.586411946 | 0.854 | 0.377 | 3.3989E-09 | 1.61783E-10 | 2.87896E-13 | Sik1 | MM_4 |
| 1.55681E-13 | 1.973788822 | 0.808 | 0.343 | 3.67595E-09 | 2.59529E-11 | 1.356730302 | 0.896 | 0.458 | 6.128E-07 | 2.59529E-11 | 3.11363E-13 | Lmo4 | MM_4 |
| 1.72637E-13 | 5.289401592 | 0.154 | 0 | 4.07631E-09 | 4.00708E-12 | 3.599203768 | 0.125 | 0.004 | 9.46152E-08 | 4.00708E-12 | 3.45274E-13 | Sv2b | MM_4 |
| 1.90363E-13 | 0.993385872 | 1 | 0.899 | 4.49484E-09 | 1.66053E-10 | 0.689482658 | 0.979 | 0.928 | 3.92084E-06 | 1.66053E-10 | 3.80725E-13 | Rps19 | MM_4 |
| 8.27712E-12 | 0.801614093 | 1 | 0.87 | 1.95439E-07 | 2.03821E-13 | 0.799824405 | 1 | 0.882 | 4.81261E-09 | 8.27712E-12 | 4.07641E-13 | Arpc2 | MM_4 |
| 2.6393E-13 | 0.97007415 | 1 | 0.919 | 6.23191E-09 | 2.59084E-12 | 0.792892458 | 1 | 0.928 | 6.11748E-08 | 2.59084E-12 | 5.2786E-13 | Rps26 | MM_4 |
| 2.93052E-13 | 4.863638226 | 0.192 | 0.006 | 6.91955E-09 | 9.44608E-09 | 2.476170794 | 0.208 | 0.03 | 0.000223041 | 9.44608E-09 | 5.86105E-13 | Gm10974 | MM_4 |
| 2.99922E-13 | 1.391950555 | 0.904 | 0.467 | 7.08177E-09 | 5.24041E-10 | 1.004227853 | 0.958 | 0.58 | 1.23736E-05 | 5.24041E-10 | 5.99845E-13 | Riok3 | MM_4 |
| 8.86405E-13 | 1.388024865 | 0.808 | 0.369 | 2.09298E-08 | 3.20243E-13 | 1.577609646 | 0.854 | 0.394 | 7.56158E-09 | 8.86405E-13 | 6.40487E-13 | Cnn2 | MM_4 |
| 3.91344E-13 | 1.799682372 | 0.962 | 0.674 | 9.24041E-09 | 3.31299E-13 | 1.82648601 | 0.979 | 0.678 | 7.82263E-09 | 3.91344E-13 | 6.62598E-13 | Dennd4a | MM_4 |
| 1.44059E-11 | 0.97699939 | 0.981 | 0.824 | 3.40151E-07 | 3.5625E-13 | 0.870701016 | 1 | 0.896 | 8.41178E-09 | 1.44059E-11 | 7.125E-13 | Calm1 | MM_4 |
| 1.95616E-07 | 0.585962249 | 1 | 0.844 | 0.004618877 | 4.30517E-13 | 0.86705109 | 1 | 0.872 | 1.01654E-08 | 1.95616E-07 | 8.61035E-13 | Arpc1b | MM_4 |
| 5.28475E-13 | 0.85232139 | 1 | 0.914 | 1.24783E-08 | 5.92474E-11 | 0.689742304 | 0.979 | 0.941 | 1.39895E-06 | 5.92474E-11 | 1.05695E-12 | Rpl19 | MM_4 |
| 1.80439E-12 | 1.631235805 | 0.808 | 0.291 | 4.26052E-08 | 6.52723E-13 | 1.931080608 | 0.792 | 0.388 | 1.54121E-08 | 1.80439E-12 | 1.30545E-12 | Batf3 | MM_4 |
| 6.95847E-13 | 1.756688152 | 0.577 | 0.135 | 1.64303E-08 | 5.76802E-11 | 1.487455555 | 0.5 | 0.133 | 1.36194E-06 | 5.76802E-11 | 1.39169E-12 | Cyp51 | MM_4 |
| 1.80207E-11 | 1.516401957 | 0.827 | 0.424 | 4.25504E-07 | 7.14222E-13 | 1.391860276 | 0.917 | 0.519 | 1.68642E-08 | 1.80207E-11 | 1.42844E-12 | Syngr2 | MM_4 |
| 7.74791E-13 | 1.089617932 | 0.904 | 0.375 | 1.82944E-08 | 3.6283E-11 | 1.371880263 | 0.812 | 0.405 | 8.56715E-07 | 3.6283E-11 | 1.54958E-12 | Sp140 | MM_4 |
| 7.85735E-13 | 1.114518473 | 0.865 | 0.378 | 1.85528E-08 | 4.38291E-11 | 1.173528352 | 0.938 | 0.543 | 1.03489E-06 | 4.38291E-11 | 1.57147E-12 | Pgam1 | MM_4 |
| 1.14701E-12 | 2.8413273 | 0.327 | 0.04 | 2.70833E-08 | 1.71753E-07 | 2.065720714 | 0.312 | 0.078 | 0.00405543 | 1.71753E-07 | 2.29403E-12 | Tmem238 | MM_4 |
| 1.28191E-12 | 1.195039331 | 1 | 0.7 | 3.02685E-08 | 5.82696E-09 | 1.071143013 | 0.979 | 0.815 | 0.000137586 | 5.82696E-09 | 2.56382E-12 | Spag9 | MM_4 |
| 1.46015E-12 | 1.319251585 | 0.712 | 0.228 | 3.44772E-08 | 1.01831E-08 | 1.272073959 | 0.667 | 0.261 | 0.000240442 | 1.01831E-08 | 2.92031E-12 | Mob3a | MM_4 |
| 3.06014E-10 | 4.197579236 | 0.135 | 0.003 | 7.22561E-06 | 1.70561E-12 | 2.621439333 | 0.229 | 0.022 | 4.02728E-08 | 3.06014E-10 | 3.41121E-12 | Tmem154 | MM_4 |
| 1.75169E-12 | 1.142571944 | 1 | 0.89 | 4.13609E-08 | 3.69355E-10 | 0.803456751 | 0.979 | 0.896 | 8.7212E-06 | 3.69355E-10 | 3.50338E-12 | Rpl18 | MM_4 |
| 1.7832E-12 | 2.511641679 | 0.404 | 0.069 | 4.21048E-08 | 2.5406E-09 | 1.844550469 | 0.312 | 0.061 | 5.99886E-05 | 2.5406E-09 | 3.56639E-12 | D930030I0 | MM_4 |
| 1.7872E-12 | 1.329236412 | 0.923 | 0.522 | 4.21993E-08 | 2.40149E-09 | 1.179999006 | 0.854 | 0.591 | 5.6704E-05 | 2.40149E-09 | 3.57439E-12 | Slk | MM_4 |
| 1.18799E-09 | 0.946752815 | 0.923 | 0.478 | 2.80509E-05 | 2.17715E-12 | 1.288151112 | 0.979 | 0.603 | 5.14069E-08 | 1.18799E-09 | 4.3543E-12 | Csf2ra | MM_4 |
| 2.18047E-11 | 3.007608256 | 0.212 | 0.014 | 5.14852E-07 | 2.30837E-12 | 2.659471181 | 0.229 | 0.022 | 5.45053E-08 | 2.18047E-11 | 4.61674E-12 | Gm45238 | MM_4 |
| 2.4221E-12 | 1.472559226 | 0.865 | 0.473 | 5.71907E-08 | 6.56188E-11 | 1.337351564 | 0.875 | 0.512 | 1.54939E-06 | 6.56188E-11 | 4.84421E-12 | Snx20 | MM_4 |
| 2.73058E-12 | 0.912012678 | 1 | 0.911 | 6.44745E-08 | 1.57912E-10 | 0.687925966 | 1 | 0.952 | 3.72863E-06 | 1.57912E-10 | 5.46117E-12 | Rps16 | MM_4 |
| 1.50236E-08 | 1.750785535 | 0.615 | 0.259 | 0.000354736 | 2.75265E-12 | 2.005587643 | 0.708 | 0.288 | 6.49956E-08 | 1.50236E-08 | 5.50531E-12 | Plaur | MM_4 |
| 7.70904E-12 | 1.296495968 | 1 | 0.7 | 1.82026E-07 | 2.83847E-12 | 1.226814581 | 1 | 0.793 | 6.7022E-08 | 7.70904E-12 | 5.67695E-12 | Uvrag | MM_4 |
| 3.98602E-09 | 1.207996947 | 0.808 | 0.38 | 9.4118E-05 | 3.28439E-12 | 1.535670351 | 0.833 | 0.399 | 7.7551E-08 | 3.98602E-09 | 6.56878E-12 | Sept9 | MM_4 |
| 3.50165E-12 | 0.98596709 | 1 | 0.914 | 8.2681E-08 | 1.56209E-11 | 0.783946739 | 0.979 | 0.945 | 3.68841E-07 | 1.56209E-11 | 7.0033E-12 | Rpl28 | MM_4 |
| 3.53198E-12 | 1.515144166 | 0.731 | 0.248 | 8.33972E-08 | 1.61547E-08 | 1.432289112 | 0.646 | 0.298 | 0.000381445 | 1.61547E-08 | 7.06397E-12 | Bloc1s2 | MM_4 |
| 3.63644E-12 | 0.903293122 | 1 | 0.934 | 8.58636E-08 | 6.77446E-11 | 0.685833838 | 0.979 | 0.946 | 1.59959E-06 | 6.77446E-11 | 7.27288E-12 | Rps27a | MM_4 |
| 8.81297E-09 | 2.241574249 | 0.25 | 0.037 | 0.000208092 | 3.68052E-12 | 3.148016484 | 0.333 | 0.057 | 8.69045E-08 | 8.81297E-09 | 7.36104E-12 | Adora2b | MM_4 |
| 3.82635E-12 | 1.27022641 | 0.885 | 0.45 | 9.03478E-08 | 1.16096E-07 | 0.859116661 | 0.938 | 0.645 | 0.002741265 | 1.16096E-07 | 7.6527E-12 | Rnh1 | MM_4 |
| 3.9455E-12 | 1.086656609 | 1 | 0.735 | 9.31611E-08 | 4.50852E-11 | 0.908074566 | 0.979 | 0.823 | 1.06455E-06 | 4.50852E-11 | 7.891E-12 | Alox5ap | MM_4 |
| 3.97993E-12 | 1.160440766 | 0.981 | 0.695 | 9.3974E-08 | 6.07655E-09 | 1.041221354 | 0.979 | 0.736 | 0.000143479 | 6.07655E-09 | 7.95985E-12 | Asap1 | MM_4 |
| 7.34682E-12 | 1.178406397 | 0.923 | 0.64 | 1.73473E-07 | 4.19413E-12 | 1.407460037 | 0.917 | 0.619 | 9.90317E-08 | 7.34682E-12 | 8.38825E-12 | Rab43 | MM_4 |
| 4.40611E-12 | 1.116184582 | 0.942 | 0.585 | 1.04037E-07 | 4.2804E-09 | 0.989201404 | 0.938 | 0.684 | 0.000101069 | 4.2804E-09 | 8.81223E-12 | Tpm4 | MM_4 |
| 4.42159E-12 | 0.91909656 | 0.981 | 0.818 | 1.04403E-07 | 8.09514E-09 | 0.671883149 | 1 | 0.835 | 0.000191143 | 8.09514E-09 | 8.84318E-12 | Rpl4 | MM_4 |
| 4.77664E-12 | 1.000539004 | 1 | 0.853 | 1.12786E-07 | 7.86349E-09 | 0.677830869 | 0.958 | 0.895 | 0.000185673 | 7.86349E-09 | 9.55329E-12 | Rpl14 | MM_4 |
| 4.91171E-12 | 1.388340014 | 0.788 | 0.297 | 1.15975E-07 | 6.4264E-10 | 1.112981548 | 0.812 | 0.423 | 1.5174E-05 | 6.4264E-10 | 9.82342E-12 | Elovl5 | MM_4 |
| 6.4625E-09 | 0.822992861 | 1 | 0.986 | 0.000152593 | 5.0741E-12 | 0.828252251 | 1 | 0.983 | 1.1981E-07 | 6.4625E-09 | 1.01482E-11 | Tmsb4x | MM_4 |
| 1.76966E-08 | 0.883869149 | 0.962 | 0.911 | 0.000417852 | 5.30188E-12 | 0.757773303 | 1 | 0.932 | 1.25188E-07 | 1.76966E-08 | 1.06038E-11 | Rpl32 | MM_4 |
| 5.45861E-12 | 1.346475276 | 0.75 | 0.28 | 1.28889E-07 | 4.41735E-10 | 1.518369122 | 0.729 | 0.29 | 1.04302E-05 | 4.41735E-10 | 1.09172E-11 | Dot1l | MM_4 |
| 2.24533E-11 | 1.339416277 | 0.75 | 0.288 | 5.30167E-07 | 5.86289E-12 | 1.753045715 | 0.75 | 0.307 | 1.38435E-07 | 2.24533E-11 | 1.17258E-11 | Prkar2a | MM_4 |
| 1.71203E-07 | 1.133343549 | 0.827 | 0.438 | 0.004042453 | 5.88313E-12 | 1.810822001 | 0.958 | 0.547 | 1.38912E-07 | 1.71203E-07 | 1.17663E-11 | Sept11 | MM_4 |
| 6.71532E-12 | 1.146075143 | 1 | 0.905 | 1.58562E-07 | 5.17248E-08 | 0.768166502 | 0.979 | 0.932 | 0.001221326 | 5.17248E-08 | 1.34306E-11 | Rps11 | MM_4 |
| 3.98302E-08 | 1.061874134 | 0.712 | 0.305 | 0.00094047 | 7.15268E-12 | 1.302035224 | 0.75 | 0.272 | 1.68889E-07 | 3.98302E-08 | 1.43054E-11 | Ciita | MM_4 |
| 7.61147E-12 | 1.776264051 | 0.462 | 0.098 | 1.79722E-07 | 3.51732E-08 | 2.023930505 | 0.396 | 0.111 | 0.000830511 | 3.51732E-08 | 1.52229E-11 | Cyb561a3 | MM_4 |
| 1.79425E-10 | 1.368000337 | 0.692 | 0.274 | 4.23659E-06 | 7.9531E-12 | 2.912169976 | 0.604 | 0.214 | 1.87788E-07 | 1.79425E-10 | 1.59062E-11 | Nr4a3 | MM_4 |
| 1.66711E-08 | 0.92304476 | 0.923 | 0.68 | 0.000393637 | 8.14443E-12 | 0.897924941 | 0.958 | 0.767 | 1.92306E-07 | 1.66711E-08 | 1.62889E-11 | Myl12b | MM_4 |
| 1.62992E-07 | 1.211369464 | 0.462 | 0.141 | 0.003848568 | 8.40478E-12 | 1.984025125 | 0.5 | 0.122 | 1.98454E-07 | 1.62992E-07 | 1.68096E-11 | Slc2a1 | MM_4 |
| 8.80217E-12 | 2.407612087 | 0.346 | 0.052 | 2.07837E-07 | 1.0818E-08 | 1.631102711 | 0.292 | 0.057 | 0.000255435 | 1.0818E-08 | 1.76043E-11 | Ccdc102a | MM_4 |
| 6.40227E-08 | 2.068806417 | 0.135 | 0.009 | 0.001511704 | 9.45611E-12 | 2.216068211 | 0.229 | 0.024 | 2.23278E-07 | 6.40227E-08 | 1.89122E-11 | Adora2a | MM_4 |
| 1.0534E-11 | 1.049065735 | 0.981 | 0.718 | 2.4873E-07 | 2.14636E-10 | 0.911046099 | 0.979 | 0.813 | 5.06798E-06 | 2.14636E-10 | 2.10681E-11 | Ywhaz | MM_4 |
| 4.60577E-09 | 1.009400378 | 0.865 | 0.493 | 0.000108751 | 1.20778E-11 | 1.274093657 | 0.854 | 0.436 | 2.85181E-07 | 4.60577E-09 | 2.41556E-11 | Vasp | MM_4 |
| 1.21268E-11 | 1.041145777 | 1 | 0.836 | 2.86337E-07 | 3.76765E-11 | 0.793387848 | 0.979 | 0.898 | 8.89619E-07 | 3.76765E-11 | 2.42535E-11 | Gpx1 | MM_4 |
| 1.27484E-11 | 1.032576701 | 1 | 0.879 | 3.01015E-07 | 5.19199E-10 | 0.721387588 | 1 | 0.928 | 1.22593E-05 | 5.19199E-10 | 2.54967E-11 | Rps15a | MM_4 |
| 3.28426E-09 | 2.72302775 | 0.154 | 0.009 | 7.75479E-05 | 1.28126E-11 | 3.843161078 | 0.146 | 0.007 | 3.02532E-07 | 3.28426E-09 | 2.56252E-11 | Gm37982 | MM_4 |
| 1.35601E-11 | 1.277151289 | 0.75 | 0.254 | 3.20181E-07 | 1.51004E-10 | 1.432108489 | 0.583 | 0.177 | 3.5655E-06 | 1.51004E-10 | 2.71202E-11 | Grk3 | MM_4 |
| 1.36402E-11 | 3.983996198 | 0.212 | 0.014 | 3.22072E-07 | 1.48743E-10 | 4.040671016 | 0.146 | 0.009 | 3.51213E-06 | 1.48743E-10 | 2.72804E-11 | F2rl2 | MM_4 |
| 1.76977E-11 | 0.843825788 | 1 | 0.957 | 4.17878E-07 | 2.15866E-10 | 0.654564179 | 1 | 0.961 | 5.09703E-06 | 2.15866E-10 | 3.53954E-11 | Rpl13 | MM_4 |
| 2.38655E-09 | 1.92025558 | 0.385 | 0.089 | 5.63512E-05 | 1.78169E-11 | 0.928768465 | 0.396 | 0.078 | 4.20693E-07 | 2.38655E-09 | 3.56338E-11 | Plcb2 | MM_4 |
| 1.98332E-11 | 1.17601571 | 0.731 | 0.259 | 4.68302E-07 | 2.15282E-10 | 1.471190259 | 0.875 | 0.436 | 5.08324E-06 | 2.15282E-10 | 3.96664E-11 | Ybx3 | MM_4 |
| 2.00288E-11 | 1.288182909 | 0.481 | 0.112 | 4.7292E-07 | 2.05998E-11 | 2.654433103 | 0.417 | 0.096 | 4.86402E-07 | 2.05998E-11 | 4.00576E-11 | Chst11 | MM_4 |
| 1.92801E-10 | 2.391801258 | 0.269 | 0.035 | 4.55241E-06 | 2.10941E-11 | 2.841814622 | 0.354 | 0.065 | 4.98073E-07 | 1.92801E-10 | 4.21881E-11 | Prr7 | MM_4 |
| 2.17904E-10 | 1.347682832 | 0.827 | 0.458 | 5.14516E-06 | 2.23818E-11 | 1.424088021 | 0.938 | 0.519 | 5.28479E-07 | 2.17904E-10 | 4.47636E-11 | Cycs | MM_4 |
| 2.80865E-08 | 1.053474852 | 0.923 | 0.617 | 0.000663179 | 2.35328E-11 | 1.205181932 | 0.979 | 0.645 | 5.55657E-07 | 2.80865E-08 | 4.70657E-11 | Baz1a | MM_4 |
| 2.58373E-11 | 0.947604137 | 1 | 0.89 | 6.10071E-07 | 7.29956E-11 | 0.735318314 | 1 | 0.922 | 1.72357E-06 | 7.29956E-11 | 5.16747E-11 | Rpl39 | MM_4 |
| 3.04367E-07 | 0.786396658 | 0.327 | 0.078 | 0.007186716 | 2.6894E-11 | 1.856237532 | 0.542 | 0.157 | 6.35022E-07 | 3.04367E-07 | 5.37881E-11 | Lrrc42 | MM_4 |
| 2.36483E-10 | 1.240717453 | 0.942 | 0.66 | 5.58383E-06 | 2.76564E-11 | 1.402624736 | 0.979 | 0.693 | 6.53022E-07 | 2.36483E-10 | 5.53127E-11 | Fbxo11 | MM_4 |
| 3.01302E-11 | 0.974445494 | 1 | 0.85 | 7.11435E-07 | 4.03117E-08 | 0.627893206 | 0.979 | 0.876 | 0.000951839 | 4.03117E-08 | 6.02604E-11 | Rps2 | MM_4 |

| | | | | | | | | | | | | | |
|---|---|---|---|---|---|---|---|---|---|---|---|---|---|
| 3.18442E-11 | 1.118292256 | 0.942 | 0.597 | 7.51905E-07 | 3.00862E-10 | 1.388705293 | 0.938 | 0.678 | 7.10394E-06 | 3.00862E-10 | 6.36884E-11 | Taldo1 | MM_4 |
| 2.39333E-07 | 0.717281189 | 0.885 | 0.493 | 0.005651121 | 3.24071E-11 | 1.12835263 | 0.875 | 0.516 | 7.65196E-07 | 2.39333E-07 | 6.48142E-11 | Furin | MM_4 |
| 9.58679E-11 | 0.754715082 | 1 | 0.896 | 2.26363E-06 | 3.57774E-11 | 0.685440366 | 0.979 | 0.935 | 8.44776E-07 | 9.58679E-11 | 7.15548E-11 | Rpl6 | MM_4 |
| 9.64558E-11 | 1.163941996 | 0.865 | 0.421 | 2.27751E-06 | 4.24599E-11 | 1.398955077 | 0.854 | 0.449 | 1.00256E-06 | 9.64558E-11 | 8.49198E-11 | Plcg2 | MM_4 |
| 5.20588E-10 | 0.712833597 | 1 | 0.896 | 1.22921E-05 | 4.26336E-11 | 0.652104455 | 1 | 0.911 | 1.00667E-06 | 5.20588E-10 | 8.52673E-11 | H3f3a | MM_4 |
| 3.42435E-09 | 0.994629683 | 0.885 | 0.476 | 8.08557E-05 | 4.29874E-11 | 1.388349109 | 0.938 | 0.658 | 1.01502E-06 | 3.42435E-09 | 8.59747E-11 | Itgb2 | MM_4 |
| 5.10786E-11 | 1.464602813 | 0.865 | 0.432 | 1.20607E-06 | 8.35823E-09 | 1.368686909 | 0.875 | 0.503 | 0.000197355 | 8.35823E-09 | 1.02157E-10 | Arih2 | MM_4 |
| 5.89217E-11 | 1.166428181 | 0.827 | 0.363 | 1.39126E-06 | 2.06993E-09 | 0.907249002 | 0.812 | 0.386 | 4.88753E-05 | 2.06993E-09 | 1.17843E-10 | Nuak2 | MM_4 |
| 5.98762E-11 | 2.851141123 | 0.25 | 0.026 | 1.4138E-06 | 7.69426E-10 | 2.241281108 | 0.271 | 0.044 | 1.81677E-05 | 7.69426E-10 | 1.19752E-10 | Ptgir | MM_4 |
| 6.63293E-11 | 1.191492881 | 0.904 | 0.573 | 1.56617E-06 | 2.95307E-07 | 0.775704349 | 0.958 | 0.726 | 0.006972783 | 2.95307E-07 | 1.32659E-10 | Anp32b | MM_4 |
| 6.70224E-11 | 1.282180475 | 0.923 | 0.542 | 1.58253E-06 | 1.09742E-08 | 1.214085381 | 0.854 | 0.538 | 0.000259122 | 1.09742E-08 | 1.34045E-10 | Pde4b | MM_4 |
| 7.20979E-11 | 0.804614996 | 0.962 | 0.669 | 1.70238E-06 | 5.49084E-09 | 0.790452155 | 0.979 | 0.773 | 0.00012965 | 5.49084E-09 | 1.44196E-10 | Arf5 | MM_4 |
| 8.65329E-11 | 2.066568268 | 0.404 | 0.081 | 2.04321E-06 | 2.65628E-10 | 2.846443159 | 0.375 | 0.083 | 6.27201E-06 | 2.65628E-10 | 1.73066E-10 | Egr3 | MM_4 |
| 9.06559E-11 | 1.328297963 | 0.962 | 0.816 | 2.14057E-06 | 4.24549E-09 | 0.877905619 | 0.958 | 0.863 | 0.000100244 | 4.24549E-09 | 1.81312E-10 | Eef1b2 | MM_4 |
| 5.5696E-10 | 1.159844477 | 0.75 | 0.297 | 1.31509E-05 | 1.0736E-10 | 1.072405575 | 0.812 | 0.34 | 2.53499E-06 | 5.5696E-10 | 2.1472E-10 | Esyt1 | MM_4 |
| 1.16047E-10 | 1.236068782 | 0.692 | 0.242 | 2.74011E-06 | 4.23282E-07 | 0.781920751 | 0.625 | 0.27 | 0.009994536 | 4.23282E-07 | 2.32094E-10 | Glipr1 | MM_4 |
| 1.67106E-07 | 1.431942731 | 0.404 | 0.118 | 0.003945715 | 1.23128E-10 | 1.870844755 | 0.438 | 0.107 | 2.90731E-06 | 1.67106E-07 | 2.46257E-10 | Prr5l | MM_4 |
| 1.25654E-10 | 1.871123878 | 0.808 | 0.398 | 2.96694E-06 | 2.42154E-07 | 1.593559757 | 0.792 | 0.464 | 0.005717735 | 2.42154E-07 | 2.51308E-10 | Lrrk1 | MM_4 |
| 1.37605E-10 | 0.814731436 | 0.962 | 0.879 | 3.24913E-06 | 2.14962E-07 | 0.592292818 | 1 | 0.9 | 0.005075681 | 2.14962E-07 | 2.7521E-10 | Rpl3 | MM_4 |
| 1.52571E-10 | 0.66509311 | 1 | 0.957 | 3.60251E-06 | 3.16188E-10 | 0.657718067 | 1 | 0.943 | 7.46584E-06 | 3.16188E-10 | 3.05142E-10 | Rps9 | MM_4 |
| 7.77415E-09 | 0.613118058 | 0.981 | 0.885 | 0.000183563 | 1.59169E-10 | 0.659892119 | 0.979 | 0.9 | 3.7583E-06 | 7.77415E-09 | 3.18339E-10 | Pfn1 | MM_4 |
| 3.01409E-09 | 1.499896993 | 0.327 | 0.061 | 7.11687E-05 | 1.69629E-10 | 2.184662934 | 0.312 | 0.054 | 4.00528E-06 | 3.01409E-09 | 3.39258E-10 | Traf3ip3 | MM_4 |
| 1.80113E-10 | 0.996638335 | 0.981 | 0.62 | 4.25283E-06 | 8.79016E-09 | 1.051299638 | 0.958 | 0.701 | 0.000207553 | 8.79016E-09 | 3.60226E-10 | Diaph1 | MM_4 |
| 1.92238E-10 | 1.009476228 | 0.885 | 0.386 | 4.53912E-06 | 1.46814E-07 | 1.130159568 | 0.771 | 0.41 | 0.003466581 | 1.46814E-07 | 3.84476E-10 | Slc38a1 | MM_4 |
| 1.96023E-10 | 5.635668193 | 0.115 | 0 | 4.6285E-06 | 1.6966E-08 | 3.02933777 | 0.125 | 0.009 | 0.000400601 | 1.6966E-08 | 3.92046E-10 | Spint1 | MM_4 |
| 2.38547E-10 | 0.825796474 | 1 | 0.893 | 5.63258E-06 | 1.08992E-08 | 0.617604147 | 1 | 0.948 | 0.000257352 | 1.08992E-08 | 4.77095E-10 | Rps10 | MM_4 |
| 2.9055E-10 | 1.671131039 | 0.577 | 0.17 | 6.86048E-06 | 1.1273E-07 | 1.562955734 | 0.562 | 0.22 | 0.002661781 | 1.1273E-07 | 5.81101E-10 | Atp6v0a2 | MM_4 |
| 2.96991E-10 | 3.069720186 | 0.231 | 0.023 | 7.01256E-06 | 1.62886E-07 | 2.40379035 | 0.146 | 0.017 | 0.003846075 | 1.62886E-07 | 5.93982E-10 | Gm47922 | MM_4 |
| 3.06014E-10 | 4.239599467 | 0.135 | 0.003 | 7.22561E-06 | 7.37865E-10 | 4.641288704 | 0.104 | 0.004 | 1.74225E-05 | 7.37865E-10 | 6.12028E-10 | Kcne3 | MM_4 |
| 2.85596E-08 | 1.149494202 | 0.865 | 0.539 | 0.00067435 | 3.09598E-10 | 0.916938526 | 0.958 | 0.645 | 7.31022E-06 | 2.85596E-08 | 6.19195E-10 | Sfr1 | MM_4 |
| 1.52069E-07 | 1.593100601 | 0.673 | 0.32 | 0.003590664 | 3.48543E-10 | 1.826766775 | 0.771 | 0.355 | 8.22981E-06 | 1.52069E-07 | 6.97087E-10 | Grk5 | MM_4 |
| 1.97014E-07 | 1.836120697 | 0.731 | 0.369 | 0.004651902 | 3.50879E-10 | 2.326368856 | 0.729 | 0.436 | 8.28496E-06 | 1.97014E-07 | 7.01759E-10 | Samsn1 | MM_4 |
| 4.15229E-07 | 0.769374726 | 0.75 | 0.357 | 0.009804394 | 3.5167E-10 | 1.301583481 | 0.812 | 0.381 | 8.30364E-06 | 4.15229E-07 | 7.03341E-10 | Nfkbie | MM_4 |
| 3.61713E-10 | 0.936557352 | 0.942 | 0.769 | 8.54077E-06 | 3.71663E-09 | 0.685754291 | 1 | 0.848 | 8.77571E-05 | 3.71663E-09 | 7.23426E-10 | Btf3 | MM_4 |
| 5.33856E-08 | 1.034662621 | 0.827 | 0.401 | 0.001260541 | 4.08267E-10 | 1.277971446 | 0.854 | 0.438 | 9.63999E-06 | 5.33856E-08 | 8.16533E-10 | Rilpl2 | MM_4 |
| 4.18003E-10 | 1.575447276 | 0.5 | 0.138 | 9.86989E-06 | 1.6216E-08 | 0.950216031 | 0.5 | 0.148 | 0.000382892 | 1.6216E-08 | 8.36006E-10 | Slco3a1 | MM_4 |
| 1.05069E-09 | 0.975880954 | 0.846 | 0.372 | 2.48088E-05 | 4.75926E-10 | 1.542546354 | 0.771 | 0.39 | 1.12376E-05 | 1.05069E-09 | 9.51852E-10 | Pik3r5 | MM_4 |
| 4.98362E-10 | 1.866142677 | 0.692 | 0.311 | 1.17673E-05 | 1.74267E-09 | 2.016540436 | 0.625 | 0.251 | 4.11479E-05 | 1.74267E-09 | 9.96725E-10 | Ahr | MM_4 |
| 5.02896E-10 | 1.011930846 | 0.962 | 0.85 | 1.18744E-05 | 3.34727E-07 | 0.945506877 | 0.958 | 0.806 | 0.007903566 | 3.34727E-07 | 1.00579E-09 | H2-Ab1 | MM_4 |
| 5.04805E-10 | 1.398453874 | 0.731 | 0.303 | 1.19194E-05 | 8.06368E-09 | 1.346396842 | 0.667 | 0.275 | 0.0001904 | 8.06368E-09 | 1.00961E-09 | Lyst | MM_4 |
| 5.92994E-10 | 0.784120711 | 0.962 | 0.507 | 1.40018E-05 | 7.39445E-09 | 1.183481457 | 0.833 | 0.562 | 0.000174598 | 7.39445E-09 | 1.18599E-09 | Lrrfip2 | MM_4 |
| 5.96669E-10 | 1.040864757 | 0.923 | 0.66 | 1.40886E-05 | 1.4836E-09 | 0.938714859 | 0.938 | 0.717 | 3.50309E-05 | 1.4836E-09 | 1.19334E-09 | Ndufa6 | MM_4 |
| 1.15278E-08 | 1.633831351 | 0.327 | 0.066 | 0.000272195 | 6.18311E-10 | 2.348686413 | 0.333 | 0.067 | 1.45995E-05 | 1.15278E-08 | 1.23662E-09 | Zfp971 | MM_4 |
| 3.97793E-07 | 1.643245497 | 0.346 | 0.092 | 0.009392686 | 6.41491E-10 | 2.046154876 | 0.542 | 0.172 | 1.51469E-05 | 3.97793E-07 | 1.28298E-09 | Mmp19 | MM_4 |
| 1.66255E-07 | 2.408230867 | 0.115 | 0.006 | 0.003925611 | 7.83087E-10 | 4.339442881 | 0.104 | 0.004 | 1.84903E-05 | 1.66255E-07 | 1.56617E-09 | Rab11fip4 | MM_4 |
| 8.38164E-10 | 0.858220211 | 0.962 | 0.876 | 1.97907E-05 | 5.62486E-09 | 0.776055824 | 0.958 | 0.908 | 0.000132814 | 5.62486E-09 | 1.67633E-09 | Rps18 | MM_4 |
| 8.39303E-10 | 0.786889252 | 0.981 | 0.899 | 1.98176E-05 | 1.51235E-07 | 0.618586467 | 0.979 | 0.93 | 0.003570958 | 1.51235E-07 | 1.67861E-09 | Rps4x | MM_4 |
| 3.93534E-08 | 0.668573236 | 0.962 | 0.611 | 0.000929212 | 8.44828E-10 | 0.835551112 | 0.938 | 0.734 | 1.99481E-05 | 3.93534E-08 | 1.68966E-09 | Smdt1 | MM_4 |
| 1.02981E-09 | 1.471088195 | 0.673 | 0.242 | 2.43159E-05 | 3.64025E-09 | 0.974323626 | 0.667 | 0.246 | 8.59537E-05 | 3.64025E-09 | 2.05962E-09 | Slc1a5 | MM_4 |
| 4.46412E-09 | 1.324138307 | 0.231 | 0.029 | 0.000105407 | 1.04503E-09 | 2.548316341 | 0.188 | 0.02 | 2.46752E-05 | 4.46412E-09 | 2.09005E-09 | Ass1 | MM_4 |
| 1.07758E-09 | 1.173682032 | 0.923 | 0.467 | 2.54438E-05 | 3.90866E-08 | 0.678511758 | 0.938 | 0.656 | 0.000922912 | 3.90866E-08 | 2.15516E-09 | Adgre5 | MM_4 |
| 5.38458E-09 | 1.369055472 | 0.538 | 0.176 | 0.000127141 | 1.16452E-09 | 1.222199837 | 0.667 | 0.253 | 2.74967E-05 | 5.38458E-09 | 2.32904E-09 | Bak1 | MM_4 |
| 1.41929E-09 | 1.026616265 | 0.673 | 0.248 | 3.35123E-05 | 9.13604E-08 | 1.021109097 | 0.75 | 0.346 | 0.002157201 | 9.13604E-08 | 2.83858E-09 | Fam129b | MM_4 |
| 1.45436E-09 | 0.838291064 | 1 | 0.89 | 3.43405E-05 | 2.00492E-09 | 0.684492922 | 1 | 0.906 | 4.73402E-05 | 2.00492E-09 | 2.90873E-09 | Rps13 | MM_4 |
| 8.27632E-09 | 0.812796705 | 0.962 | 0.706 | 0.000195421 | 1.47571E-09 | 0.851677002 | 0.979 | 0.782 | 3.48445E-05 | 8.27632E-09 | 2.95142E-09 | Eif5a | MM_4 |
| 1.18805E-08 | 1.317612447 | 0.173 | 0.014 | 0.000280522 | 1.50262E-09 | 1.859957578 | 0.146 | 0.011 | 3.54799E-05 | 1.18805E-08 | 3.00524E-09 | Lamb3 | MM_4 |
| 1.50686E-09 | 1.087310813 | 0.885 | 0.53 | 3.55801E-05 | 3.22117E-08 | 0.973504552 | 0.896 | 0.617 | 0.000760584 | 3.22117E-08 | 3.01373E-09 | Tkt | MM_4 |
| 1.72027E-09 | 1.148699539 | 0.769 | 0.317 | 4.06189E-05 | 2.07133E-09 | 1.343901595 | 0.771 | 0.36 | 4.89083E-05 | 2.07133E-09 | 3.44053E-09 | Prkd3 | MM_4 |
| 2.58456E-09 | 2.626902069 | 0.269 | 0.04 | 6.10265E-05 | 1.74608E-09 | 1.232953154 | 0.271 | 0.046 | 4.12283E-05 | 2.58456E-09 | 3.49215E-09 | Lrrc1 | MM_4 |
| 1.79831E-09 | 1.149183648 | 0.635 | 0.21 | 4.24616E-05 | 3.76326E-09 | 1.501245925 | 0.646 | 0.264 | 8.88581E-05 | 3.76326E-09 | 3.59661E-09 | Pqlc1 | MM_4 |
| 1.81262E-09 | 1.588342599 | 0.538 | 0.187 | 4.27996E-05 | 2.21773E-09 | 1.778961504 | 0.625 | 0.246 | 5.23651E-05 | 2.21773E-09 | 3.62524E-09 | Gfpt1 | MM_4 |
| 1.94714E-08 | 1.020294473 | 0.615 | 0.231 | 0.000459759 | 2.11738E-09 | 1.614092093 | 0.708 | 0.303 | 4.99957E-05 | 1.94714E-08 | 4.23477E-09 | Phlda1 | MM_4 |
| 2.63129E-09 | 3.969025094 | 0.154 | 0.009 | 6.21301E-05 | 1.3029E-08 | 3.968663046 | 0.125 | 0.009 | 0.00030764 | 1.3029E-08 | 5.26259E-09 | Olfr433 | MM_4 |
| 2.75466E-08 | 0.953718561 | 0.731 | 0.317 | 0.000650431 | 2.86667E-09 | 1.195596202 | 0.75 | 0.36 | 6.76877E-05 | 2.75466E-08 | 5.73333E-09 | Nudt21 | MM_4 |
| 2.96106E-09 | 0.885275126 | 0.25 | 0.035 | 6.99165E-05 | 1.22423E-08 | 2.31451592 | 0.271 | 0.054 | 0.000289064 | 1.22423E-08 | 5.92212E-09 | Gcnt2 | MM_4 |
| 2.86839E-08 | 2.542124921 | 0.231 | 0.035 | 0.000677285 | 3.72247E-09 | 2.47072995 | 0.292 | 0.057 | 8.7895E-05 | 2.86839E-08 | 7.44494E-09 | Cd300lg | MM_4 |
| 3.79366E-09 | 1.480753632 | 0.885 | 0.594 | 8.95759E-05 | 4.1522E-08 | 0.949414339 | 0.938 | 0.601 | 0.000980417 | 4.1522E-08 | 7.58732E-09 | Map4k4 | MM_4 |
| 5.67109E-09 | 0.720581994 | 1 | 0.89 | 0.000133906 | 1.17027E-08 | 0.593362554 | 0.979 | 0.924 | 0.000276325 | 1.17027E-08 | 1.13422E-08 | Rpl24 | MM_4 |
| 6.0752E-09 | 1.072270166 | 0.692 | 0.251 | 0.000143448 | 2.81208E-07 | 0.967857127 | 0.688 | 0.325 | 0.006639891 | 2.81208E-07 | 1.21504E-08 | Kmt5a | MM_4 |
| 6.24938E-09 | 1.285648964 | 0.981 | 0.778 | 0.00014756 | 3.94234E-07 | 0.99703359 | 0.958 | 0.837 | 0.009308653 | 3.94234E-07 | 1.24988E-08 | Fnbp1 | MM_4 |
| 2.92124E-08 | 0.854117781 | 0.981 | 0.859 | 0.000689763 | 6.27743E-09 | 0.664219679 | 0.979 | 0.902 | 0.000148223 | 2.92124E-08 | 1.25549E-08 | Rpl15 | MM_4 |
| 1.18255E-07 | 1.56091061 | 0.635 | 0.303 | 0.00279223 | 6.3289E-09 | 1.457734777 | 0.688 | 0.292 | 0.000149438 | 1.18255E-07 | 1.26578E-08 | Id2 | MM_4 |
| 6.81722E-09 | 1.027961736 | 0.981 | 0.784 | 0.000160968 | 4.27997E-08 | 1.096723029 | 0.958 | 0.847 | 0.001010586 | 4.27997E-08 | 1.36344E-08 | Rab8b | MM_4 |
| 7.1403E-09 | 1.897365373 | 0.385 | 0.095 | 0.000168597 | 2.71625E-07 | 1.548028518 | 0.396 | 0.118 | 0.006413613 | 2.71625E-07 | 1.42806E-08 | Pak4 | MM_4 |
| 1.00617E-08 | 1.261900446 | 0.615 | 0.213 | 0.000237576 | 9.05188E-09 | 1.22364544 | 0.646 | 0.246 | 0.000213733 | 1.00617E-08 | 1.81038E-08 | Tmem51 | MM_4 |
| 9.13875E-09 | 1.210762016 | 0.731 | 0.311 | 0.000215784 | 1.12165E-08 | 0.984986899 | 0.792 | 0.36 | 0.000264843 | 1.12165E-08 | 1.82775E-08 | Map2k3 | MM_4 |
| 9.23618E-09 | 0.667408015 | 0.962 | 0.712 | 0.000218085 | 9.43769E-09 | 1.279002215 | 0.979 | 0.688 | 0.000222843 | 9.43769E-09 | 1.84724E-08 | Dock10 | MM_4 |
| 2.4473E-07 | 1.443525921 | 0.577 | 0.262 | 0.005778554 | 1.24578E-08 | 1.851388828 | 0.604 | 0.253 | 0.000294153 | 2.4473E-07 | 2.49156E-08 | Emb | MM_4 |
| 1.26245E-08 | 1.150889449 | 0.75 | 0.329 | 0.00029809 | 2.97049E-08 | 0.802543334 | 0.854 | 0.425 | 0.000701392 | 2.97049E-08 | 2.5249E-08 | Cs | MM_4 |
| 1.33165E-08 | 1.012472013 | 0.692 | 0.256 | 0.00031443 | 1.71078E-07 | 1.052168814 | 0.771 | 0.381 | 0.004039494 | 1.71078E-07 | 2.66331E-08 | Dipk1a | MM_4 |
| 2.55406E-08 | 1.603297008 | 0.692 | 0.343 | 0.000603065 | 1.33542E-08 | 1.321538792 | 0.833 | 0.468 | 0.00031532 | 2.55406E-08 | 2.67084E-08 | Tuba1a | MM_4 |

| | | | | | | | | | | | | | |
|---|---|---|---|---|---|---|---|---|---|---|---|---|---|
| 7.53925E-08 | 0.809508243 | 0.962 | 0.692 | 0.001780168 | 1.82797E-08 | 0.826181726 | 0.979 | 0.782 | 0.000431619 | 7.53925E-08 | 3.65593E-08 | Cd47 | MM_4 |
| 2.69757E-08 | 1.398104485 | 0.769 | 0.424 | 0.00063695 | 7.52783E-08 | 1.281602622 | 0.833 | 0.471 | 0.00177747 | 7.52783E-08 | 5.39514E-08 | Fgl2 | MM_4 |
| 2.94304E-08 | 0.966085417 | 0.788 | 0.432 | 0.00069491 | 3.81339E-07 | 1.093845977 | 0.75 | 0.451 | 0.009004172 | 3.81339E-07 | 5.88608E-08 | Got1 | MM_4 |
| 2.88936E-07 | 0.816022557 | 0.962 | 0.718 | 0.006822364 | 3.25146E-08 | 0.828384096 | 0.979 | 0.688 | 0.000767735 | 2.88936E-07 | 6.50292E-08 | Mpeg1 | MM_4 |
| 3.57959E-08 | 1.859317717 | 0.404 | 0.112 | 0.000845212 | 1.5787E-07 | 1.812771576 | 0.333 | 0.089 | 0.003727618 | 1.5787E-07 | 7.15917E-08 | Gm10634 | MM_4 |
| 2.2768E-07 | 0.931551106 | 0.558 | 0.205 | 0.005375976 | 4.38968E-08 | 1.0893734 | 0.625 | 0.24 | 0.001036491 | 2.2768E-07 | 8.77936E-08 | Utp18 | MM_4 |
| 2.56991E-07 | 1.315443525 | 0.538 | 0.19 | 0.006068079 | 5.4722E-08 | 1.128963148 | 0.604 | 0.238 | 0.001292096 | 2.56991E-07 | 1.09444E-07 | Snhg15 | MM_4 |
| 9.38772E-08 | 1.320856008 | 0.558 | 0.199 | 0.002216629 | 6.49589E-08 | 1.358432451 | 0.625 | 0.259 | 0.00153381 | 9.38772E-08 | 1.29918E-07 | Glrx | MM_4 |
| 7.63315E-08 | 1.038802966 | 0.808 | 0.432 | 0.00180234 | 1.25898E-07 | 1.051442066 | 0.812 | 0.494 | 0.002972703 | 1.25898E-07 | 1.52663E-07 | Gpsm3 | MM_4 |
| 1.71101E-07 | 3.148567998 | 0.115 | 0.006 | 0.004040034 | 1.11748E-07 | 3.648481512 | 0.146 | 0.017 | 0.002638588 | 1.71101E-07 | 2.23496E-07 | Cyp2s1 | MM_4 |
| 1.941E-07 | 1.926454112 | 0.404 | 0.121 | 0.004583091 | 3.89877E-07 | 2.767616104 | 0.354 | 0.109 | 0.009205774 | 3.89877E-07 | 3.882E-07 | Morrbid | MM_4 |
| 3.63602E-07 | 1.328838869 | 0.538 | 0.205 | 0.008585382 | 2.96527E-07 | 1.953123984 | 0.521 | 0.213 | 0.007001592 | 3.63602E-07 | 5.93054E-07 | Ezh2 | MM_4 |
