## Supplementary Table 3 for "C1qa□ muscularis macrophages maintain enteric synaptic homeostasis to regulate gastrointestinal motility"

| gene_symbol | ensembl_gene_id | gene_biotype | mgi_description | p_val | avg_log2FC | pct.1 | pct.2 | p_val_adj | Cluster |
|---|---|---|---|---|---|---|---|---|---|
| C1qaCKOvsC1qaFL | | | | | | | | | |
| Trim12a | ENSMUSG00000066258 | protein_coding | tripartite motif-containing 12A | 6.04E-14 | 4.282906206 | 0.309 | 0.017 | 1.33746E-09 | EN_1 |
| Rgs6 | ENSMUSG00000021219 | protein_coding | regulator of G-protein signaling 6 | 2.243E-09 | 4.010095878 | 0.281 | 0.05 | 4.96779E-05 | EN_1 |
| Tnni1 | ENSMUSG00000026418 | protein_coding | troponin I, skeletal, slow 1 | 2.119E-10 | 3.475551284 | 0.253 | 0.022 | 4.6922E-06 | EN_1 |
| Trim12c | ENSMUSG00000057143 | protein_coding | tripartite motif-containing 12C | 8.405E-09 | 2.196691911 | 0.303 | 0.067 | 0.000186139 | EN_1 |
| Hspb1 | ENSMUSG00000004951 | protein_coding | heat shock protein 1 | 9.016E-24 | 2.017350109 | 0.921 | 0.667 | 1.99667E-19 | EN_1 |
| Chst15 | ENSMUSG00000030930 | protein_coding | carbohydrate sulfotransferase 15 | 6.051E-08 | 1.890588783 | 0.371 | 0.133 | 0.001339935 | EN_1 |
| Igsf21 | ENSMUSG00000040972 | protein_coding | immunoglobulin superfamily, member 21 | 8.58E-17 | 1.861609716 | 0.685 | 0.283 | 1.89996E-12 | EN_1 |
| Ubc | ENSMUSG00000008348 | protein_coding | ubiquitin C | 7.163E-39 | 1.739233752 | 0.994 | 0.922 | 1.58614E-34 | EN_1 |
| Sdf2l1 | ENSMUSG00000022769 | protein_coding | stromal cell-derived factor 2-like 1 | 1.491E-09 | 1.524266569 | 0.517 | 0.233 | 3.30265E-05 | EN_1 |
| Kcnj12 | ENSMUSG00000042529 | protein_coding | potassium inwardly-rectifying channel, subfamily J, member 12 | 1.076E-07 | 1.410398604 | 0.416 | 0.161 | 0.002382356 | EN_1 |
| Gm26802 | ENSMUSG00000097266 | lncRNA | predicted gene, 26802 | 3.36E-07 | 1.386488115 | 0.393 | 0.156 | 0.00744052 | EN_1 |
| Serpinh1 | ENSMUSG00000070436 | protein_coding | serine (or cysteine) peptidase inhibitor, clade H, member 1 | 5.95E-19 | 1.117318388 | 0.933 | 0.756 | 1.31755E-14 | EN_1 |
| Hspa5 | ENSMUSG00000026864 | protein_coding | heat shock protein 5 | 9.655E-17 | 1.103393487 | 0.972 | 0.861 | 2.13811E-12 | EN_1 |
| Msmo1 | ENSMUSG00000031604 | protein_coding | methylsterol monoxygenase 1 | 2.398E-07 | 1.042837004 | 0.652 | 0.394 | 0.005310108 | EN_1 |
| Hist1h2bc | ENSMUSG00000018102 | protein_coding | histone cluster 1, H2bc | 1.485E-09 | 1.036157419 | 0.736 | 0.456 | 3.28916E-05 | EN_1 |
| Sgip1 | ENSMUSG00000028524 | protein_coding | SH3-domain GRB2-like (endophilin) interacting protein 1 | 2.376E-08 | 0.888029192 | 0.865 | 0.706 | 0.000526139 | EN_1 |
| Cryab | ENSMUSG00000032060 | protein_coding | crystallin, alpha B | 2.362E-14 | 0.784100349 | 0.994 | 1 | 5.23149E-10 | EN_1 |
| Pdia6 | ENSMUSG00000020571 | protein_coding | protein disulfide isomerase associated 6 | 2.26E-07 | 0.706600357 | 0.876 | 0.733 | 0.005005654 | EN_1 |
| Hsp90aa1 | ENSMUSG00000021270 | protein_coding | heat shock protein 90, alpha (cytosolic), class A member 1 | 4.5E-10 | 0.694628155 | 0.944 | 0.894 | 9.96523E-06 | EN_1 |
| Rpl26 | ENSMUSG00000060938 | protein_coding | ribosomal protein L26 | 5.182E-12 | 0.689862761 | 0.978 | 0.967 | 1.1475E-07 | EN_1 |
| Atp5md | ENSMUSG00000071528 | protein_coding | ATP synthase membrane subunit DAPIT | 1.635E-08 | 0.685971064 | 0.91 | 0.767 | 0.000362037 | EN_1 |
| Vim | ENSMUSG00000026728 | protein_coding | vimentin | 1.384E-18 | 0.679688488 | 1 | 1 | 3.0643E-14 | EN_1 |
| Hsp90ab1 | ENSMUSG00000023944 | protein_coding | heat shock protein 90 alpha (cytosolic), class B member 1 | 4.216E-17 | 0.624447615 | 1 | 0.989 | 9.33683E-13 | EN_1 |
| Gnb2 | ENSMUSG00000029713 | protein_coding | guanine nucleotide binding protein (G protein), beta 2 | 3.245E-15 | -0.770113474 | 0.882 | 0.972 | 7.18602E-11 | EN_1 |
| Wdfy1 | ENSMUSG00000073643 | protein_coding | WD repeat and FYVE domain containing 1 | 3.713E-13 | -1.505417455 | 0.236 | 0.622 | 8.22241E-09 | EN_1 |
| Luzp2 | ENSMUSG00000063297 | protein_coding | leucine zipper protein 2 | 2.454E-28 | -1.870344215 | 0.646 | 0.939 | 5.43535E-24 | EN_1 |
| Ucp2 | ENSMUSG00000033685 | protein_coding | uncoupling protein 2 (mitochondrial, proton carrier) | 8.664E-10 | -2.665704375 | 0.062 | 0.311 | 1.91864E-05 | EN_1 |
| Crybg1 | ENSMUSG00000019866 | protein_coding | crystallin beta-gamma domain containing 1 | 2.666E-07 | -4.89077093 | 0 | 0.139 | 0.005904786 | EN_1 |
| Gm14133 | ENSMUSG00000087029 | lncRNA | predicted gene 14133 | 1.579E-22 | -6.060695932 | 0.006 | 0.439 | 3.49575E-18 | EN_1 |
| Poteg | ENSMUSG00000063932 | protein_coding | POTE ankyrin domain family, member G | 4.46E-33 | -7.530774795 | 0 | 0.589 | 9.87727E-29 | EN_1 |
| Trim12a | ENSMUSG00000066258 | protein_coding | tripartite motif-containing 12A | 1.221E-10 | 5.793651119 | 0.228 | 0 | 2.70457E-06 | EN_2 |
| Trim12c | ENSMUSG00000057143 | protein_coding | tripartite motif-containing 12C | 1.09E-07 | 3.81440968 | 0.191 | 0.012 | 0.002414621 | EN_2 |
| Igsf21 | ENSMUSG00000040972 | protein_coding | immunoglobulin superfamily, member 21 | 1.36E-11 | 1.92216586 | 0.529 | 0.165 | 3.01164E-07 | EN_2 |
| Lefty1 | ENSMUSG00000038793 | protein_coding | left right determination factor 1 | 3.697E-07 | 1.644484678 | 0.397 | 0.146 | 0.008188087 | EN_2 |
| Ubc | ENSMUSG00000008348 | protein_coding | ubiquitin C | 9.519E-24 | 1.564068667 | 0.985 | 0.909 | 2.10805E-19 | EN_2 |
| Hspb1 | ENSMUSG00000004951 | protein_coding | heat shock protein 1 | 9.48E-14 | 1.178530211 | 0.941 | 0.768 | 2.09936E-09 | EN_2 |
| Serpinh1 | ENSMUSG00000070436 | protein_coding | serine (or cysteine) peptidase inhibitor, clade H, member 1 | 3.226E-08 | 0.853463667 | 0.882 | 0.707 | 0.000714301 | EN_2 |
| Hspa5 | ENSMUSG00000026864 | protein_coding | heat shock protein 5 | 1.086E-09 | 0.846100063 | 0.971 | 0.866 | 2.40401E-05 | EN_2 |
| Pdia3 | ENSMUSG00000027248 | protein_coding | protein disulfide isomerase associated 3 | 5.396E-09 | 0.745753022 | 0.963 | 0.841 | 0.000119494 | EN_2 |
| Cryab | ENSMUSG00000032060 | protein_coding | crystallin, alpha B | 5.176E-11 | 0.713870064 | 1 | 0.988 | 1.14618E-06 | EN_2 |
| Tomm6 | ENSMUSG00000033475 | protein_coding | translocase of outer mitochondrial membrane 6 | 1.603E-08 | 0.703048571 | 0.919 | 0.811 | 0.000355026 | EN_2 |
| Vim | ENSMUSG00000026728 | protein_coding | vimentin | 1.404E-13 | 0.700562599 | 1 | 1 | 3.11015E-09 | EN_2 |
| Hsp90b1 | ENSMUSG00000020048 | protein_coding | heat shock protein 90, beta (Grp94), member 1 | 1.562E-07 | 0.596241844 | 0.963 | 0.945 | 0.003458807 | EN_2 |
| Stard13 | ENSMUSG00000016128 | protein_coding | StAR-related lipid transfer (START) domain containing 13 | 2.024E-07 | -0.590347112 | 0.971 | 0.97 | 0.004483144 | EN_2 |
| Gm42418 | ENSMUSG00000098178 | lncRNA | predicted gene, 42418 | 1.903E-08 | -0.605362068 | 1 | 1 | 0.00042137 | EN_2 |
| Gphn | ENSMUSG00000047454 | protein_coding | gephyrin | 4.246E-08 | -0.632321009 | 0.956 | 0.982 | 0.000940258 | EN_2 |
| Kmt2e | ENSMUSG00000029004 | protein_coding | lysine (K)-specific methyltransferase 2E | 1.393E-07 | -0.675463053 | 0.735 | 0.878 | 0.003083764 | EN_2 |
| Prkca | ENSMUSG00000050965 | protein_coding | protein kinase C, alpha | 4.189E-10 | -0.70893927 | 0.971 | 0.994 | 9.27577E-06 | EN_2 |
| Rbms1 | ENSMUSG00000026970 | protein_coding | RNA binding motif, single stranded interacting protein 1 | 1.164E-07 | -0.812635759 | 0.801 | 0.921 | 0.002578759 | EN_2 |
| Lars2 | ENSMUSG00000035202 | protein_coding | leucyl-tRNA synthetase, mitochondrial | 5.08E-08 | -0.854238972 | 0.919 | 0.921 | 0.00112492 | EN_2 |
| Lrrc8b | ENSMUSG00000070639 | protein_coding | leucine rich repeat containing 8 family, member B | 4.619E-09 | -0.903778708 | 0.676 | 0.89 | 0.000102282 | EN_2 |
| Ntng1 | ENSMUSG00000059857 | protein_coding | netrin G1 | 1.284E-08 | -1.006995504 | 0.684 | 0.86 | 0.000284438 | EN_2 |
| Il31ra | ENSMUSG00000050377 | protein_coding | interleukin 31 receptor A | 4.191E-10 | -1.192882813 | 0.5 | 0.78 | 9.28035E-06 | EN_2 |
| BC004004 | ENSMUSG00000052712 | protein_coding | cDNA sequence BC004004 | 4.542E-13 | -1.450133167 | 0.39 | 0.756 | 1.00577E-08 | EN_2 |
| Luzp2 | ENSMUSG00000063297 | protein_coding | leucine zipper protein 2 | 1.886E-15 | -1.610197281 | 0.419 | 0.817 | 4.17554E-11 | EN_2 |
| Wdfy1 | ENSMUSG00000073643 | protein_coding | WD repeat and FYVE domain containing 1 | 1.361E-09 | -1.742734877 | 0.14 | 0.463 | 3.01498E-05 | EN_2 |
| Gm14133 | ENSMUSG00000087029 | lncRNA | predicted gene 14133 | 3.985E-07 | -3.344620681 | 0.022 | 0.22 | 0.008824021 | EN_2 |
| Ucp2 | ENSMUSG00000033685 | protein_coding | uncoupling protein 2 (mitochondrial, proton carrier) | 6.854E-10 | -3.354401702 | 0.029 | 0.305 | 1.51772E-05 | EN_2 |
| Poteg | ENSMUSG00000063932 | protein_coding | POTE ankyrin domain family, member G | 4.007E-31 | -8.051838932 | 0 | 0.671 | 8.873E-27 | EN_2 |
| Trim12a | ENSMUSG00000066258 | protein_coding | tripartite motif-containing 12A | 1.872E-10 | 5.797754152 | 0.293 | 0 | 4.14575E-06 | EN_3 |
| Igsf21 | ENSMUSG00000040972 | protein_coding | immunoglobulin superfamily, member 21 | 1.885E-08 | 2.240625859 | 0.45 | 0.136 | 0.000417443 | EN_3 |
| Luzp2 | ENSMUSG00000063297 | protein_coding | leucine zipper protein 2 | 6.242E-15 | -2.230258826 | 0.3 | 0.737 | 1.38231E-10 | EN_3 |
| Gm14133 | ENSMUSG00000087029 | lncRNA | predicted gene 14133 | 8.293E-14 | -5.785798779 | 0.007 | 0.356 | 1.83657E-09 | EN_3 |
| Poteg | ENSMUSG00000063932 | protein_coding | POTE ankyrin domain family, member G | 5.66E-10 | -6.129283017 | 0 | 0.246 | 1.25334E-05 | EN_3 |
| Ly6a | ENSMUSG00000075602 | protein_coding | lymphocyte antigen 6 complex, locus A | 1.118E-08 | 3.042952145 | 0.363 | 0.022 | 0.000247628 | EN_4 |
| Cfh | ENSMUSG00000026365 | protein_coding | complement component factor h | 4.479E-07 | 2.305986551 | 0.385 | 0.067 | 0.009917931 | EN_4 |
| Efemp1 | ENSMUSG00000020467 | protein_coding | epidermal growth factor-containing fibulin-like extracellular matrix protein 1 | 3.417E-11 | 1.921322701 | 0.78 | 0.233 | 7.56668E-07 | EN_4 |
| Col14a1 | ENSMUSG00000022371 | protein_coding | collagen, type XIV, alpha 1 | 3.544E-07 | 1.4604965 | 0.626 | 0.233 | 0.0078491 | EN_4 |
| Ubc | ENSMUSG00000008348 | protein_coding | ubiquitin C | 4.86E-10 | 1.291083728 | 1 | 0.933 | 1.07629E-05 | EN_4 |
| Myom1 | ENSMUSG00000024049 | protein_coding | myomesin 1 | 1.887E-09 | -0.600904045 | 0.077 | 0.522 | 4.17893E-05 | EN_4 |
| Macrod2 | ENSMUSG00000068205 | protein_coding | MACRO domain containing 2 | 2.41E-07 | -0.941940962 | 0.242 | 0.678 | 0.005336629 | EN_4 |
| BC004004 | ENSMUSG00000052712 | protein_coding | cDNA sequence BC004004 | 4.504E-08 | -1.092562825 | 0.209 | 0.722 | 0.000997355 | EN_4 |
| Inpp4b | ENSMUSG00000037940 | protein_coding | inositol polyphosphate-4-phosphatase, type II | 6.549E-08 | -1.185866545 | 0.022 | 0.333 | 0.001450166 | EN_4 |
| Hdac9 | ENSMUSG00000004698 | protein_coding | histone deacetylase 9 | 1.542E-07 | -1.400605394 | 0.077 | 0.444 | 0.003414955 | EN_4 |
| Negr1 | ENSMUSG00000040037 | protein_coding | neuronal growth regulator 1 | 1.131E-10 | -1.402964667 | 0.066 | 0.544 | 2.50437E-06 | EN_4 |
| Cacna1d | ENSMUSG00000015968 | protein_coding | calcium channel, voltage-dependent, L type, alpha 1D subunit | 3.05E-09 | -1.479342065 | 0.22 | 0.678 | 6.75472E-05 | EN_4 |
| Prkn | ENSMUSG00000023826 | protein_coding | parkin RBR E3 ubiquitin protein ligase | 2.453E-08 | -1.505326385 | 0.187 | 0.611 | 0.000543259 | EN_4 |
| Epha7 | ENSMUSG00000028289 | protein_coding | Eph receptor A7 | 2.17E-13 | -1.637429921 | 0.088 | 0.633 | 4.8046E-09 | EN_4 |
| Dgkh | ENSMUSG00000034731 | protein_coding | diacylglycerol kinase, eta | 6.116E-10 | -1.793549123 | 0.154 | 0.633 | 1.35447E-05 | EN_4 |
| Tox | ENSMUSG00000041272 | protein_coding | thymocyte selection-associated high mobility group box | 1.185E-11 | -1.994567893 | 0.055 | 0.533 | 2.62513E-07 | EN_4 |
| Uty | ENSMUSG00000068457 | protein_coding | ubiquitously transcribed tetratricopeptide repeat gene, Y chromosome | 2.354E-09 | -2.752907138 | 0.077 | 0.478 | 5.21343E-05 | EN_4 |
| Ppm1e | ENSMUSG00000046442 | protein_coding | protein phosphatase 1E (PP2C domain containing) | 1.254E-07 | -2.979415668 | 0.022 | 0.322 | 0.002777938 | EN_4 |
| Poteg | ENSMUSG00000063932 | protein_coding | POTE ankyrin domain family, member G | 1.478E-07 | -6.555100355 | 0 | 0.267 | 0.003272546 | EN_4 |
| Trim12a | ENSMUSG00000066258 | protein_coding | tripartite motif-containing 12A | 2.347E-07 | 4.806658473 | 0.212 | 0 | 0.005197145 | MM_1 |
| Htatip2 | ENSMUSG00000039745 | protein_coding | HIV-1 Tat interactive protein 2 | 3.033E-07 | 3.19616088 | 0.23 | 0.009 | 0.006717525 | MM_1 |
| Siglech | ENSMUSG00000051504 | protein_coding | sialic acid binding Ig-like lectin H | 2.903E-10 | 2.620131504 | 0.5 | 0.165 | 6.42791E-06 | MM_1 |
| Hspb1 | ENSMUSG00000004951 | protein_coding | heat shock protein 1 | 2.659E-08 | 2.414749225 | 0.613 | 0.303 | 0.000588803 | MM_1 |
| Sh3bgrl3 | ENSMUSG00000028843 | protein_coding | SH3 domain binding glutamic acid-rich protein-like 3 | 5.091E-34 | 2.079438918 | 0.959 | 0.771 | 1.12735E-29 | MM_1 |
| Cystm1 | ENSMUSG00000046727 | protein_coding | cysteine-rich transmembrane module containing 1 | 1.461E-08 | 1.973768458 | 0.446 | 0.138 | 0.000323492 | MM_1 |
| Ftl1-ps1 | ENSMUSG00000062382 | protein_coding | ferritin light polypeptide 1, pseudogene 1 | 2.469E-19 | 1.649333508 | 0.887 | 0.514 | 5.46852E-15 | MM_1 |
| Lgals3 | ENSMUSG00000050335 | protein_coding | lectin, galactose binding, soluble 3 | 2.875E-09 | 1.628884186 | 0.797 | 0.569 | 6.36564E-05 | MM_1 |
| Ccl6 | ENSMUSG00000018927 | protein_coding | chemokine (C-C motif) ligand 6 | 1.061E-22 | 1.590497206 | 0.995 | 0.991 | 2.34891E-18 | MM_1 |
| Tppp3 | ENSMUSG00000014846 | protein_coding | tubulin polymerization-promoting protein family member 3 | 9.863E-15 | 1.489660566 | 0.775 | 0.312 | 2.18426E-10 | MM_1 |
| Pdia4 | ENSMUSG00000025823 | protein_coding | protein disulfide isomerase associated 4 | 4.641E-08 | 1.367682666 | 0.613 | 0.321 | 0.001027651 | MM_1 |
| Hspa1b | ENSMUSG00000090877 | protein_coding | heat shock protein 1B | 5.024E-12 | 1.357074478 | 0.928 | 0.826 | 1.11254E-07 | MM_1 |
| Pdia6 | ENSMUSG00000020571 | protein_coding | protein disulfide isomerase associated 6 | 3.028E-10 | 1.324623739 | 0.77 | 0.477 | 6.70603E-06 | MM_1 |
| St3gal5 | ENSMUSG00000056091 | protein_coding | ST3 beta-galactoside alpha-2,3-sialyltransferase 5 | 7.407E-09 | 1.316160656 | 0.64 | 0.275 | 0.000164018 | MM_1 |
| Fcna | ENSMUSG00000026938 | protein_coding | ficolin A | 1.489E-18 | 1.314839641 | 0.986 | 0.826 | 3.29831E-14 | MM_1 |
| Hspe1 | ENSMUSG00000073676 | protein_coding | heat shock protein 1 (chaperonin 10) | 1.055E-11 | 1.306936304 | 0.928 | 0.679 | 2.33573E-07 | MM_1 |
| Tmem37 | ENSMUSG00000050777 | protein_coding | transmembrane protein 37 | 5.4E-08 | 1.239358569 | 0.626 | 0.339 | 0.001195909 | MM_1 |
| S100a10 | ENSMUSG00000041959 | protein_coding | S100 calcium binding protein A10 (calpactin) | 1.442E-12 | 1.206214544 | 0.946 | 0.752 | 3.19289E-08 | MM_1 |
| Anxa2 | ENSMUSG00000032231 | protein_coding | annexin A2 | 6.16E-10 | 1.197516159 | 0.838 | 0.56 | 1.36407E-05 | MM_1 |
| Crip1 | ENSMUSG00000006360 | protein_coding | cysteine-rich protein 1 (intestinal) | 5.425E-17 | 1.182790116 | 0.991 | 0.945 | 1.20145E-12 | MM_1 |
| Hspa1a | ENSMUSG00000091971 | protein_coding | heat shock protein 1A | 5.331E-10 | 1.173949601 | 0.955 | 0.862 | 1.18053E-05 | MM_1 |
| Hsp90aa1 | ENSMUSG00000021270 | protein_coding | heat shock protein 90, alpha (cytosolic), class A member 1 | 2.332E-07 | 1.105041044 | 0.991 | 0.899 | 0.005164755 | MM_1 |
| Prxl2b | ENSMUSG00000029059 | protein_coding | peroxiredoxin like 2B | 5.212E-08 | 1.079563122 | 0.739 | 0.468 | 0.001154307 | MM_1 |
| C1qb | ENSMUSG00000036905 | protein_coding | complement component 1, q subcomponent, beta polypeptide | 1.913E-22 | 1.063238088 | 1 | 0.991 | 4.23612E-18 | MM_1 |
| Card19 | ENSMUSG00000037960 | protein_coding | caspase recruitment domain family, member 19 | 1.765E-09 | 1.05038974 | 0.838 | 0.633 | 3.90874E-05 | MM_1 |
| Wfdc17 | ENSMUSG00000069792 | protein_coding | WAP four-disulfide core domain 17 | 4.764E-10 | 1.048058515 | 0.991 | 0.963 | 1.0549E-05 | MM_1 |
| Aldh2 | ENSMUSG00000029455 | protein_coding | aldehyde dehydrogenase 2, mitochondrial | 1.602E-15 | 1.032984501 | 0.95 | 0.761 | 3.54758E-11 | MM_1 |
| Pdia3 | ENSMUSG00000027248 | protein_coding | protein disulfide isomerase associated 3 | 8.225E-15 | 1.028333932 | 0.968 | 0.835 | 1.8215E-10 | MM_1 |
| Dbi | ENSMUSG00000026385 | protein_coding | diazepam binding inhibitor | 6.401E-10 | 1.011243164 | 0.874 | 0.651 | 1.41752E-05 | MM_1 |
| Cox5b | ENSMUSG00000061518 | protein_coding | cytochrome c oxidase subunit 5B | 3.69E-10 | 0.961695626 | 0.905 | 0.697 | 8.17135E-06 | MM_1 |
| Cbr2 | ENSMUSG00000025150 | protein_coding | carbonyl reductase 2 | 1.891E-18 | 0.958787067 | 0.995 | 0.945 | 4.18863E-14 | MM_1 |
| Lyve1 | ENSMUSG00000030787 | protein_coding | lymphatic vessel endothelial hyaluronan receptor 1 | 5.234E-16 | 0.953062169 | 1 | 0.954 | 1.159E-11 | MM_1 |
| Ctsd | ENSMUSG00000007891 | protein_coding | cathepsin D | 5.292E-18 | 0.949260756 | 1 | 0.972 | 1.17183E-13 | MM_1 |
| Ndufs7 | ENSMUSG00000020153 | protein_coding | NADH:ubiquinone oxidoreductase core subunit S7 | 3.14E-09 | 0.948142651 | 0.784 | 0.505 | 6.95388E-05 | MM_1 |
| Ubc | ENSMUSG00000008348 | protein_coding | ubiquitin C | 5.957E-14 | 0.946671535 | 1 | 0.917 | 1.31913E-09 | MM_1 |
| Plin2 | ENSMUSG00000028494 | protein_coding | perilipin 2 | 2.185E-09 | 0.945423833 | 0.95 | 0.817 | 4.8379E-05 | MM_1 |

| | | | | | | | | | |
|---|---|---|---|---|---|---|---|---|---|
| Ltc4s | ENSMUSG00000020377 | protein_coding | leukotriene C4 synthase | 3.462E-10 | 0.925078966 | 0.964 | 0.881 | 7.66594E-06 | MM_1 |
| Nme2 | ENSMUSG00000020857 | protein_coding | NME/NM23 nucleoside diphosphate kinase 2 | 5.359E-11 | 0.918552846 | 0.941 | 0.789 | 1.18671E-06 | MM_1 |
| Hsp90ab1 | ENSMUSG00000023944 | protein_coding | heat shock protein 90 alpha (cytosolic), class B member 1 | 1.06E-21 | 0.914890278 | 1 | 1 | 2.34712E-17 | MM_1 |
| Cyb5a | ENSMUSG00000024646 | protein_coding | cytochrome b5 type A (microsomal) | 3.914E-11 | 0.904698066 | 0.905 | 0.688 | 8.6684E-07 | MM_1 |
| Vat1 | ENSMUSG00000034993 | protein_coding | vesicle amine transport 1 | 3.003E-08 | 0.877195199 | 0.856 | 0.679 | 0.000664916 | MM_1 |
| Klf2 | ENSMUSG00000055148 | protein_coding | Kruppel-like factor 2 (lung) | 9.542E-10 | 0.870691197 | 0.991 | 0.89 | 2.11302E-05 | MM_1 |
| Fcgrt | ENSMUSG00000003420 | protein_coding | Fc receptor, IgG, alpha chain transporter | 1.745E-16 | 0.8684586 | 0.995 | 0.927 | 3.86419E-12 | MM_1 |
| Cstb | ENSMUSG00000005054 | protein_coding | cystatin B | 1.468E-10 | 0.855675293 | 0.982 | 0.817 | 3.25028E-06 | MM_1 |
| Emp3 | ENSMUSG00000040212 | protein_coding | epithelial membrane protein 3 | 4.98E-09 | 0.851284452 | 0.982 | 0.908 | 0.000110279 | MM_1 |
| Serpinb6a | ENSMUSG00000060147 | protein_coding | serine (or cysteine) peptidase inhibitor, clade B, member 6a | 2.1E-12 | 0.843587467 | 0.986 | 0.872 | 4.6498E-08 | MM_1 |
| Ednrb | ENSMUSG00000022122 | protein_coding | endothelin receptor type B | 1.082E-11 | 0.837284783 | 0.973 | 0.817 | 2.39556E-07 | MM_1 |
| Prdx1 | ENSMUSG00000028691 | protein_coding | peroxiredoxin 1 | 4.668E-13 | 0.836451157 | 0.995 | 0.982 | 1.03369E-08 | MM_1 |
| Cd68 | ENSMUSG00000018774 | protein_coding | CD68 antigen | 8.641E-12 | 0.833862889 | 0.973 | 0.927 | 1.91346E-07 | MM_1 |
| Folr2 | ENSMUSG00000032725 | protein_coding | folate receptor 2 (fetal) | 1.336E-14 | 0.829758156 | 0.991 | 0.945 | 2.95967E-10 | MM_1 |
| C1qc | ENSMUSG00000036896 | protein_coding | complement component 1, q subcomponent, C chain | 1.345E-28 | 0.821629396 | 1 | 1 | 2.97766E-24 | MM_1 |
| Hsp90b1 | ENSMUSG00000020048 | protein_coding | heat shock protein 90, beta (Grp94), member 1 | 7.384E-11 | 0.782825676 | 0.982 | 0.899 | 1.63526E-06 | MM_1 |
| Ninj1 | ENSMUSG00000037966 | protein_coding | ninjurin 1 | 6.137E-13 | 0.782518412 | 0.982 | 0.936 | 1.35898E-08 | MM_1 |
| Tagln2 | ENSMUSG00000026547 | protein_coding | transgelin 2 | 1.297E-12 | 0.78112338 | 0.991 | 0.954 | 2.87226E-08 | MM_1 |
| Ost4 | ENSMUSG00000038803 | protein_coding | oligosaccharyltransferase complex subunit 4 (non-catalytic) | 6.461E-08 | 0.776395812 | 0.892 | 0.688 | 0.001430766 | MM_1 |
| Ap2m1 | ENSMUSG00000022841 | protein_coding | adaptor-related protein complex 2, mu 1 subunit | 1.212E-09 | 0.75907936 | 0.914 | 0.706 | 2.68495E-05 | MM_1 |
| Gapdh | ENSMUSG00000057666 | protein_coding | glyceraldehyde-3-phosphate dehydrogenase | 3.262E-10 | 0.758500472 | 0.955 | 0.817 | 7.22353E-06 | MM_1 |
| Ccl9 | ENSMUSG00000019122 | protein_coding | chemokine (C-C motif) ligand 9 | 1.105E-12 | 0.757074915 | 1 | 0.945 | 2.44747E-08 | MM_1 |
| Lgals1 | ENSMUSG00000068220 | protein_coding | lectin, galactose binding, soluble 1 | 3.319E-09 | 0.742115375 | 0.995 | 0.963 | 7.34932E-05 | MM_1 |
| Alox5ap | ENSMUSG00000060063 | protein_coding | arachidonate 5-lipoxygenase activating protein | 3.574E-10 | 0.741987706 | 0.982 | 0.927 | 7.91519E-06 | MM_1 |
| Gpx1 | ENSMUSG00000063856 | protein_coding | glutathione peroxidase 1 | 1.513E-13 | 0.739303205 | 0.995 | 0.972 | 3.35074E-09 | MM_1 |
| Txn1 | ENSMUSG00000028367 | protein_coding | thioredoxin 1 | 1.867E-07 | 0.702051809 | 0.883 | 0.679 | 0.004133896 | MM_1 |
| Fth1 | ENSMUSG00000024661 | protein_coding | ferritin heavy polypeptide 1 | 1.176E-17 | 0.673991985 | 1 | 1 | 2.60337E-13 | MM_1 |
| Ecm1 | ENSMUSG00000028108 | protein_coding | extracellular matrix protein 1 | 1.87E-08 | 0.664083959 | 0.95 | 0.743 | 0.000414153 | MM_1 |
| Aldoa | ENSMUSG00000030695 | protein_coding | aldolase A, fructose-bisphosphate | 1.116E-07 | 0.651840364 | 0.923 | 0.734 | 0.002471138 | MM_1 |
| Pepd | ENSMUSG00000063931 | protein_coding | peptidase D | 5.321E-11 | 0.637442935 | 0.995 | 0.908 | 1.17838E-06 | MM_1 |
| Ppia | ENSMUSG00000071866 | protein_coding | peptidylprolyl isomerase A | 4.039E-14 | 0.617990172 | 1 | 1 | 8.94532E-10 | MM_1 |
| Tomm6 | ENSMUSG00000033475 | protein_coding | translocase of outer mitochondrial membrane 6 | 2.172E-07 | 0.605462009 | 0.91 | 0.743 | 0.004809348 | MM_1 |
| Blvrb | ENSMUSG00000040466 | protein_coding | biliverdin reductase B (flavin reductase (NADPH)) | 5.969E-08 | 0.604617904 | 0.95 | 0.89 | 0.00132175 | MM_1 |
| St13 | ENSMUSG00000022403 | protein_coding | suppression of tumorigenicity 13 | 1.165E-08 | 0.596698809 | 0.968 | 0.826 | 0.000258017 | MM_1 |
| Ptafr | ENSMUSG00000056529 | protein_coding | platelet-activating factor receptor | 8.006E-10 | 0.587202898 | 0.991 | 0.945 | 1.77298E-05 | MM_1 |
| Ndufa4 | ENSMUSG00000029632 | protein_coding | Ndufa4, mitochondrial complex associated | 3.866E-10 | 0.586620356 | 0.959 | 0.899 | 8.56118E-06 | MM_1 |
| Marcks | ENSMUSG00000069662 | protein_coding | myristoylated alanine rich protein kinase C substrate | 1.353E-10 | -0.633914119 | 0.973 | 0.982 | 2.99652E-06 | MM_1 |
| Slc8a1 | ENSMUSG00000054640 | protein_coding | solute carrier family 8 (sodium/calcium exchanger), member 1 | 3.833E-09 | -0.790972944 | 0.914 | 0.954 | 8.48719E-05 | MM_1 |
| Tsc22d1 | ENSMUSG00000022010 | protein_coding | TSC22 domain family, member 1 | 3.813E-07 | -0.866451098 | 0.315 | 0.606 | 0.008444244 | MM_1 |
| Gnb2 | ENSMUSG00000029713 | protein_coding | guanine nucleotide binding protein (G protein), beta 2 | 1.093E-16 | -0.892606977 | 0.928 | 0.972 | 2.42138E-12 | MM_1 |
| BC004004 | ENSMUSG00000052712 | protein_coding | cDNA sequence BC004004 | 1.116E-11 | -1.514169588 | 0.311 | 0.651 | 2.4709E-07 | MM_1 |
| H2-Eb1 | ENSMUSG00000060586 | protein_coding | histocompatibility 2, class II antigen E beta | 7.288E-08 | -1.5247102 | 0.518 | 0.725 | 0.001613955 | MM_1 |
| BC035044 | ENSMUSG00000090164 | protein_coding | cDNA sequence BC035044 | 6.764E-08 | -1.971783758 | 0.108 | 0.349 | 0.001497931 | MM_1 |
| Pla2g2d | ENSMUSG00000041202 | protein_coding | phospholipase A2, group IID | 8.504E-09 | -2.036499877 | 0.126 | 0.394 | 0.000188323 | MM_1 |
| Zeb1 | ENSMUSG00000024238 | protein_coding | zinc finger E-box binding homeobox 1 | 2.561E-09 | -2.053712278 | 0.09 | 0.358 | 5.67229E-05 | MM_1 |
| Basp1 | ENSMUSG00000045763 | protein_coding | brain abundant, membrane attached signal protein 1 | 5.177E-08 | -2.348159636 | 0.099 | 0.339 | 0.001146472 | MM_1 |
| Lpcat2 | ENSMUSG00000033192 | protein_coding | lysophosphatidylcholine acyltransferase 2 | 6.075E-10 | -2.456865896 | 0.081 | 0.349 | 1.34528E-05 | MM_1 |
| Cx3cr1 | ENSMUSG00000052336 | protein_coding | chemokine (C-X3-C motif) receptor 1 | 1.949E-07 | -2.566799923 | 0.086 | 0.303 | 0.004316848 | MM_1 |
| Ifi202b | ENSMUSG00000026535 | protein_coding | interferon activated gene 202B | 1.983E-23 | -3.157476075 | 0.086 | 0.578 | 4.39073E-19 | MM_1 |
| Gm21860 | ENSMUSG00000095366 | lncRNA | predicted gene, 21860 | 3.212E-07 | -3.348159636 | 0.014 | 0.156 | 0.007113795 | MM_1 |
| C1qa | ENSMUSG00000036887 | protein_coding | complement component 1, q subcomponent, alpha polypeptide | 9.37E-50 | -3.466041569 | 0.932 | 1 | 2.07495E-45 | MM_1 |
| Rtn1 | ENSMUSG00000021087 | protein_coding | reticulon 1 | 4.569E-11 | -4.222628754 | 0.023 | 0.248 | 1.01173E-06 | MM_1 |
| Naip6 | ENSMUSG00000078942 | protein_coding | NLR family, apoptosis inhibitory protein 6 | 4.686E-12 | -5.670087731 | 0 | 0.202 | 1.03768E-07 | MM_1 |
| Poteg | ENSMUSG00000063932 | protein_coding | POTE ankyrin domain family, member G | 3.942E-14 | -6.383783546 | 0 | 0.239 | 8.72909E-10 | MM_1 |
| Trim12a | ENSMUSG00000066258 | protein_coding | tripartite motif-containing 12A | 7.14E-11 | 4.879369533 | 0.302 | 0.008 | 1.58122E-06 | MM_2 |
| Htatip2 | ENSMUSG00000039745 | protein_coding | HIV-1 Tat interactive protein 2 | 3.217E-08 | 4.861447625 | 0.215 | 0 | 0.000712503 | MM_2 |
| Gm42031 | ENSMUSG00000110386 | lncRNA | predicted gene, 42031 | 2.675E-09 | 4.267439985 | 0.262 | 0.008 | 5.92434E-05 | MM_2 |
| Gm43305 | | | | 1.308E-10 | 2.665050412 | 0.395 | 0.064 | 2.89716E-06 | MM_2 |
| Nlrp1b | ENSMUSG00000070390 | protein_coding | NLR family, pyrin domain containing 1B | 2.807E-11 | 2.424876964 | 0.471 | 0.112 | 6.21617E-07 | MM_2 |
| Sh3bgrl3 | ENSMUSG00000028843 | protein_coding | SH3 domain binding glutamic acid-rich protein-like 3 | 6.195E-37 | 2.016943928 | 1 | 0.856 | 1.37178E-32 | MM_2 |
| Klf2 | ENSMUSG00000055148 | protein_coding | Kruppel-like factor 2 (lung) | 1.937E-09 | 1.981907517 | 0.779 | 0.576 | 4.29043E-05 | MM_2 |
| Sdf2l1 | ENSMUSG00000022769 | protein_coding | stromal cell-derived factor 2-like 1 | 1.684E-11 | 1.783645473 | 0.762 | 0.512 | 3.72997E-07 | MM_2 |
| Cd52 | ENSMUSG00000000682 | protein_coding | CD52 antigen | 1.078E-22 | 1.714847112 | 0.977 | 0.68 | 2.38643E-18 | MM_2 |
| Ubc | ENSMUSG00000008348 | protein_coding | ubiquitin C | 1.009E-17 | 1.486080271 | 0.988 | 0.904 | 2.23439E-13 | MM_2 |
| Ftl1-ps1 | ENSMUSG00000062382 | protein_coding | ferritin light polypeptide 1, pseudogene 1 | 5.418E-15 | 1.367338555 | 0.843 | 0.536 | 1.19978E-10 | MM_2 |
| Hspa1b | ENSMUSG00000090877 | protein_coding | heat shock protein 1B | 9.194E-13 | 1.296823012 | 0.924 | 0.76 | 2.03592E-08 | MM_2 |
| Hspa1a | ENSMUSG00000091971 | protein_coding | heat shock protein 1A | 1.932E-12 | 1.229385762 | 0.965 | 0.816 | 4.27765E-08 | MM_2 |
| Pdia3 | ENSMUSG00000027248 | protein_coding | protein disulfide isomerase associated 3 | 3.341E-17 | 1.225245008 | 0.959 | 0.824 | 7.39812E-13 | MM_2 |
| Aldh2 | ENSMUSG00000029455 | protein_coding | aldehyde dehydrogenase 2, mitochondrial | 4.771E-10 | 1.216542584 | 0.779 | 0.536 | 1.05649E-05 | MM_2 |
| G530011O06Rik | | | | 3.045E-10 | 1.1614086 | 0.837 | 0.608 | 6.74239E-06 | MM_2 |
| Pdia6 | ENSMUSG00000020571 | protein_coding | protein disulfide isomerase associated 6 | 8.827E-09 | 1.038479513 | 0.837 | 0.68 | 0.000195475 | MM_2 |
| C1qb | ENSMUSG00000036905 | protein_coding | complement component 1, q subcomponent, beta polypeptide | 9.978E-28 | 0.976317396 | 1 | 1 | 2.20963E-23 | MM_2 |
| Slamf7 | ENSMUSG00000038179 | protein_coding | SLAM family member 7 | 4.179E-08 | 0.841639144 | 0.82 | 0.504 | 0.000925441 | MM_2 |
| Hsp90ab1 | ENSMUSG00000023944 | protein_coding | heat shock protein 90 alpha (cytosolic), class B member 1 | 4.854E-13 | 0.812909627 | 1 | 0.992 | 1.07494E-08 | MM_2 |
| C1qc | ENSMUSG00000036896 | protein_coding | complement component 1, q subcomponent, C chain | 7.654E-24 | 0.800578092 | 1 | 1 | 1.69491E-19 | MM_2 |
| Jund | ENSMUSG00000071076 | protein_coding | jun D proto-oncogene | 5.112E-08 | 0.777133405 | 0.988 | 0.944 | 0.0011321 | MM_2 |
| Sec61g | ENSMUSG00000078974 | protein_coding | SEC61, gamma subunit | 9.765E-08 | 0.711344064 | 0.948 | 0.88 | 0.002162445 | MM_2 |
| Hspa8 | ENSMUSG00000015656 | protein_coding | heat shock protein 8 | 3.053E-07 | 0.679236146 | 0.994 | 0.992 | 0.006761754 | MM_2 |
| Selenof | ENSMUSG00000037072 | protein_coding | selenoprotein F | 8.19E-09 | 0.668417263 | 0.936 | 0.808 | 0.000181366 | MM_2 |
| Sec61b | ENSMUSG00000053317 | protein_coding | Sec61 beta subunit | 3.319E-07 | 0.657960987 | 0.924 | 0.76 | 0.007349498 | MM_2 |
| Ppib | ENSMUSG00000032383 | protein_coding | peptidylprolyl isomerase B | 3.818E-09 | 0.636932032 | 0.959 | 0.88 | 8.45478E-05 | MM_2 |
| Ubb | ENSMUSG00000019505 | protein_coding | ubiquitin B | 2.228E-10 | 0.619090944 | 0.994 | 1 | 4.93386E-06 | MM_2 |
| Uba52 | ENSMUSG00000090137 | protein_coding | ubiquitin A-52 residue ribosomal protein fusion product 1 | 1.604E-08 | 0.595579489 | 0.965 | 0.888 | 0.000355163 | MM_2 |
| AY036118 | ENSMUSG00000105361 | lncRNA | cDNA sequence AY036118 | 3.581E-11 | -0.603683634 | 0.977 | 1 | 7.93113E-07 | MM_2 |
| Dab2 | ENSMUSG00000022150 | protein_coding | disabled 2, mitogen-responsive phosphoprotein | 4.206E-08 | -0.613719882 | 0.988 | 0.984 | 0.000931526 | MM_2 |
| Gphn | ENSMUSG00000047454 | protein_coding | gephyrin | 5.847E-15 | -0.815100728 | 1 | 1 | 1.29477E-10 | MM_2 |
| Lars2 | ENSMUSG00000035202 | protein_coding | leucyl-tRNA synthetase, mitochondrial | 1.205E-16 | -0.849219482 | 0.983 | 1 | 2.66801E-12 | MM_2 |
| Gnb2 | ENSMUSG00000029713 | protein_coding | guanine nucleotide binding protein (G protein), beta 2 | 3.391E-17 | -0.878520457 | 0.907 | 1 | 7.50992E-13 | MM_2 |
| BC005537 | ENSMUSG00000019132 | protein_coding | cDNA sequence BC005537 | 2.53E-07 | -0.898881962 | 0.703 | 0.88 | 0.005603699 | MM_2 |
| Gm42418 | ENSMUSG00000098178 | lncRNA | predicted gene, 42418 | 1.518E-30 | -1.012266517 | 1 | 1 | 3.36173E-26 | MM_2 |
| Il31ra | ENSMUSG00000050377 | protein_coding | interleukin 31 receptor A | 8.777E-10 | -1.037909298 | 0.483 | 0.784 | 1.94365E-05 | MM_2 |
| Gigyf1 | ENSMUSG00000029714 | protein_coding | GRB10 interacting GYF protein 1 | 3.575E-07 | -1.279908224 | 0.262 | 0.528 | 0.007917108 | MM_2 |
| BC004004 | ENSMUSG00000052712 | protein_coding | cDNA sequence BC004004 | 4.719E-18 | -1.52390585 | 0.424 | 0.848 | 1.04495E-13 | MM_2 |
| Ctse | ENSMUSG00000004552 | protein_coding | cathepsin E | 1.79E-08 | -2.154377342 | 0.076 | 0.328 | 0.00039633 | MM_2 |
| Gm21860 | ENSMUSG00000095366 | lncRNA | predicted gene, 21860 | 6.183E-08 | -2.931786189 | 0.058 | 0.28 | 0.001369234 | MM_2 |
| C1qa | ENSMUSG00000036887 | protein_coding | complement component 1, q subcomponent, alpha polypeptide | 2.229E-47 | -3.15980129 | 0.983 | 1 | 4.93654E-43 | MM_2 |
| Ifi202b | ENSMUSG00000026535 | protein_coding | interferon activated gene 202B | 4.852E-24 | -3.575957687 | 0.076 | 0.616 | 1.07446E-19 | MM_2 |
| Naip6 | ENSMUSG00000078942 | protein_coding | NLR family, apoptosis inhibitory protein 6 | 9.685E-10 | -5.318461465 | 0 | 0.2 | 2.14472E-05 | MM_2 |
| Poteg | ENSMUSG00000063932 | protein_coding | POTE ankyrin domain family, member G | 3.877E-27 | -7.52656966 | 0 | 0.536 | 8.5849E-23 | MM_2 |
| Arhgap29 | ENSMUSG00000039831 | protein_coding | Rho GTPase activating protein 29 | 5.568E-12 | 3.924287366 | 0.381 | 0.018 | 1.23299E-07 | MM_3 |
| Id3 | ENSMUSG00000007872 | protein_coding | inhibitor of DNA binding 3 | 2.079E-12 | 3.413064538 | 0.408 | 0.027 | 4.60462E-08 | MM_3 |
| Adgrd1 | ENSMUSG00000044017 | protein_coding | adhesion G protein-coupled receptor D1 | 5.497E-09 | 3.320946336 | 0.299 | 0.018 | 0.000121736 | MM_3 |
| Sulf1 | ENSMUSG00000016918 | protein_coding | sulfatase 1 | 1.005E-09 | 2.983076697 | 0.367 | 0.044 | 2.2256E-05 | MM_3 |
| Adamtsl3 | ENSMUSG00000070469 | protein_coding | ADAMTS-like 3 | 9.137E-13 | 2.707969459 | 0.524 | 0.097 | 2.02343E-08 | MM_3 |
| Tm4sf1 | ENSMUSG00000027800 | protein_coding | transmembrane 4 superfamily member 1 | 1.21E-10 | 2.640406175 | 0.367 | 0.027 | 2.67982E-06 | MM_3 |
| Prex2 | ENSMUSG00000048960 | protein_coding | phosphatidylinositol-3,4,5-trisphosphate-dependent Rac exchange factor 2 | 2.046E-08 | 2.364667713 | 0.32 | 0.035 | 0.000453164 | MM_3 |
| Efemp1 | ENSMUSG00000020467 | protein_coding | epidermal growth factor-containing fibulin-like extracellular matrix protein 1 | 3.744E-16 | 2.28347163 | 0.66 | 0.133 | 8.29028E-12 | MM_3 |
| Dclk1 | ENSMUSG00000027797 | protein_coding | doublecortin-like kinase 1 | 9.815E-08 | 2.24853784 | 0.34 | 0.062 | 0.00217361 | MM_3 |
| Fndc1 | ENSMUSG00000071984 | protein_coding | fibronectin type III domain containing 1 | 8.389E-13 | 2.164827134 | 0.565 | 0.124 | 1.85785E-08 | MM_3 |
| Pcsk5 | ENSMUSG00000024713 | protein_coding | proprotein convertase subtilisin/kexin type 5 | 5.692E-09 | 2.144068574 | 0.34 | 0.035 | 0.000126051 | MM_3 |
| Slc29a1 | ENSMUSG00000023942 | protein_coding | solute carrier family 29 (nucleoside transporters), member 1 | 8.985E-09 | 2.13507979 | 0.429 | 0.088 | 0.00019897 | MM_3 |
| Sh3bgrl3 | ENSMUSG00000028843 | protein_coding | SH3 domain binding glutamic acid-rich protein-like 3 | 2.594E-16 | 2.127466606 | 0.85 | 0.31 | 5.74445E-12 | MM_3 |
| Cd200 | ENSMUSG00000022661 | protein_coding | CD200 antigen | 6.812E-08 | 2.05791193 | 0.32 | 0.044 | 0.001508429 | MM_3 |
| Ephx1 | ENSMUSG00000038776 | protein_coding | epoxide hydrolase 1, microsomal | 6.073E-11 | 2.035544117 | 0.463 | 0.071 | 1.34496E-06 | MM_3 |
| Sypl | ENSMUSG00000020570 | protein_coding | synaptophysin-like protein | 2.937E-08 | 1.854303802 | 0.51 | 0.168 | 0.000650393 | MM_3 |
| Mfap5 | ENSMUSG00000030116 | protein_coding | microfibrillar associated protein 5 | 2.539E-09 | 1.837737334 | 0.374 | 0.044 | 5.62269E-05 | MM_3 |
| Rer1 | ENSMUSG00000029048 | protein_coding | retention in endoplasmic reticulum sorting receptor 1 | 2.932E-12 | 1.817808058 | 0.558 | 0.115 | 6.4922E-08 | MM_3 |
| Gabarapl1 | ENSMUSG00000030161 | protein_coding | gamma-aminobutyric acid (GABA) A receptor-associated protein-like 1 | 1.014E-07 | 1.68138886 | 0.49 | 0.168 | 0.002246175 | MM_3 |
| Ptprk | ENSMUSG00000019889 | protein_coding | protein tyrosine phosphatase, receptor type, K | 1.055E-07 | 1.5103237 | 0.395 | 0.097 | 0.002336368 | MM_3 |
| Cd34 | ENSMUSG00000016494 | protein_coding | CD34 antigen | 5.955E-09 | 1.485577037 | 0.469 | 0.115 | 0.000131871 | MM_3 |
| Myof | ENSMUSG00000048612 | protein_coding | myoferlin | 1.884E-07 | 1.455875916 | 0.571 | 0.23 | 0.004172942 | MM_3 |

| | | | | | | | | | |
|---|---|---|---|---|---|---|---|---|---|
| Vat1 | ENSMUSG00000034993 | protein_coding | vesicle amine transport 1 | 1.128E-08 | 1.448325642 | 0.469 | 0.106 | 0.000249713 | MM_3 |
| Ly6a | ENSMUSG00000075602 | protein_coding | lymphocyte antigen 6 complex, locus A | 1.605E-11 | 1.42786154 | 0.497 | 0.08 | 3.55342E-07 | MM_3 |
| Hspa1a | ENSMUSG00000091971 | protein_coding | heat shock protein 1A | 3.29E-08 | 1.407657969 | 0.939 | 0.779 | 0.000728675 | MM_3 |
| Hspa1b | ENSMUSG00000090877 | protein_coding | heat shock protein 1B | 1.232E-08 | 1.404502524 | 0.918 | 0.779 | 0.000272889 | MM_3 |
| S100a10 | ENSMUSG00000041959 | protein_coding | S100 calcium binding protein A10 (calpactin) | 7.334E-12 | 1.378533827 | 0.952 | 0.832 | 1.62414E-07 | MM_3 |
| Pros1 | ENSMUSG00000022912 | protein_coding | protein S (alpha) | 2.474E-08 | 1.363348778 | 0.571 | 0.195 | 0.0005479 | MM_3 |
| AC149090.1 | | | | 4.863E-09 | 1.334638577 | 0.884 | 0.504 | 0.000107701 | MM_3 |
| Atf3 | ENSMUSG00000026628 | protein_coding | activating transcription factor 3 | 1.259E-07 | 1.293538873 | 0.871 | 0.655 | 0.002788729 | MM_3 |
| Gm47283 | ENSMUSG00000096768 | lncRNA | predicted gene, 47283 | 2.983E-09 | 1.28347163 | 0.605 | 0.186 | 6.6056E-05 | MM_3 |
| Fkbp2 | ENSMUSG00000056629 | protein_coding | FK506 binding protein 2 | 2.958E-08 | 1.243943266 | 0.476 | 0.115 | 0.000655104 | MM_3 |
| C3 | ENSMUSG00000024164 | protein_coding | complement component 3 | 2.605E-09 | 1.141622636 | 0.755 | 0.363 | 5.76944E-05 | MM_3 |
| Kras | ENSMUSG00000030265 | protein_coding | Kirsten rat sarcoma viral oncogene homolog | 4.326E-07 | 1.113838643 | 0.585 | 0.221 | 0.00958027 | MM_3 |
| Ubc | ENSMUSG00000008348 | protein_coding | ubiquitin C | 2.626E-10 | 1.112659308 | 0.973 | 0.903 | 5.81542E-06 | MM_3 |
| Crip1 | ENSMUSG00000006360 | protein_coding | cysteine-rich protein 1 (intestinal) | 3.89E-09 | 1.106080441 | 0.993 | 0.956 | 8.61495E-05 | MM_3 |
| Fbln2 | ENSMUSG00000064080 | protein_coding | fibulin 2 | 1.648E-07 | 1.089059626 | 0.633 | 0.292 | 0.003649971 | MM_3 |
| Pdlim1 | ENSMUSG00000055044 | protein_coding | PDZ and LIM domain 1 (elfin) | 3.611E-09 | 1.015807899 | 0.531 | 0.124 | 7.99694E-05 | MM_3 |
| Sema6a | ENSMUSG00000019647 | protein_coding | sema domain, transmembrane domain (TM), and cytoplasmic domain, (semaphorin) 6A | 2.464E-08 | 0.978058622 | 0.388 | 0.071 | 0.000545756 | MM_3 |
| Gpx3 | ENSMUSG00000018339 | protein_coding | glutathione peroxidase 3 | 2.412E-07 | 0.749239932 | 0.694 | 0.336 | 0.005341664 | MM_3 |
| Akap6 | ENSMUSG00000061603 | protein_coding | A kinase (PRKA) anchor protein 6 | 4.43E-08 | -1.21599465 | 0.061 | 0.345 | 0.000980971 | MM_3 |
| Gnb2 | ENSMUSG00000029713 | protein_coding | guanine nucleotide binding protein (G protein), beta 2 | 4.521E-10 | -1.281128253 | 0.837 | 0.938 | 1.00126E-05 | MM_3 |
| Tenm3 | ENSMUSG00000031561 | protein_coding | teneurin transmembrane protein 3 | 3.504E-07 | -1.508776399 | 0.048 | 0.283 | 0.007758782 | MM_3 |
| Abcc9 | ENSMUSG00000030249 | protein_coding | ATP-binding cassette, sub-family C (CFTR/MRP), member 9 | 3.321E-07 | -1.71652837 | 0.068 | 0.319 | 0.007354214 | MM_3 |
| Negr1 | ENSMUSG00000040037 | protein_coding | neuronal growth regulator 1 | 1.868E-13 | -2.453493964 | 0.075 | 0.487 | 4.13577E-09 | MM_3 |
| BC004004 | ENSMUSG00000052712 | protein_coding | cDNA sequence BC004004 | 2.535E-17 | -2.497888083 | 0.15 | 0.673 | 5.61294E-13 | MM_3 |
| C1qa | ENSMUSG00000036887 | protein_coding | complement component 1, q subcomponent, alpha polypeptide | 4.508E-21 | -3.638227651 | 0.32 | 0.77 | 9.98393E-17 | MM_3 |
| Poteg | ENSMUSG00000063932 | protein_coding | POTE ankyrin domain family, member G | 1.213E-07 | -6.423887502 | 0 | 0.177 | 0.002685159 | MM_3 |
| Sh3bgrl3 | ENSMUSG00000028843 | protein_coding | SH3 domain binding glutamic acid-rich protein-like 3 | 7.906E-11 | 1.228935267 | 0.958 | 0.962 | 1.75083E-06 | MM_4 |
| BC004004 | ENSMUSG00000052712 | protein_coding | cDNA sequence BC004004 | 3.578E-11 | -2.783995907 | 0.229 | 0.846 | 7.92448E-07 | MM_4 |
| Poteg | ENSMUSG00000063932 | protein_coding | POTE ankyrin domain family, member G | 1.913E-11 | -5.639410285 | 0 | 0.654 | 4.23536E-07 | MM_4 |
| | | | **C1qaCKOvsiC1qaCKO** | | | | | | |
| Gulp1 | ENSMUSG00000056870 | protein_coding | GULP, engulfment adaptor PTB domain containing 1 | 9.072E-09 | 0.728169459 | 1 | 0.961 | 0.000214212 | EN_1 |
| Per2 | ENSMUSG00000055866 | protein_coding | period circadian clock 2 | 3.312E-13 | -3.382807493 | 0.058 | 0.421 | 7.82061E-09 | EN_1 |
| Dbi | ENSMUSG00000026385 | protein_coding | diazepam binding inhibitor | 1.421E-35 | 3.004548224 | 1 | 0.908 | 3.35447E-31 | EN_1 |
| Gpr37l1 | ENSMUSG00000026424 | protein_coding | G protein-coupled receptor 37-like 1 | 1.682E-10 | 0.728228345 | 1 | 0.934 | 3.97211E-06 | EN_1 |
| Gm2000 | ENSMUSG00000078193 | protein_coding | predicted gene 2000 | 2.913E-09 | 2.309087181 | 0.503 | 0.079 | 6.87851E-05 | EN_1 |
| Prdx6 | ENSMUSG00000026701 | protein_coding | peroxiredoxin 6 | 7.948E-11 | 1.075321784 | 0.947 | 0.75 | 1.87664E-06 | EN_1 |
| G0s2 | ENSMUSG00000009633 | protein_coding | G0/G1 switch gene 2 | 1.411E-08 | -0.951579842 | 0.895 | 0.947 | 0.000333194 | EN_1 |
| Camk1d | ENSMUSG00000039145 | protein_coding | calcium/calmodulin-dependent protein kinase ID | 2.255E-21 | -0.978521839 | 1 | 1 | 5.32557E-17 | EN_1 |
| Nebl | ENSMUSG00000053702 | protein_coding | nebulette | 4.415E-08 | 0.906352601 | 0.836 | 0.461 | 0.001042557 | EN_1 |
| Hspa5 | ENSMUSG00000026864 | protein_coding | heat shock protein 5 | 4.556E-11 | 1.091067237 | 0.977 | 0.829 | 1.07585E-06 | EN_1 |
| Gsn | ENSMUSG00000026879 | protein_coding | gelsolin | 6.301E-11 | 0.785737858 | 1 | 1 | 1.48787E-06 | EN_1 |
| Rpl35 | ENSMUSG00000062997 | protein_coding | ribosomal protein L35 | 8.318E-08 | 0.767754514 | 0.965 | 0.868 | 0.001963962 | EN_1 |
| Cd59a | ENSMUSG00000032679 | protein_coding | CD59a antigen | 6.021E-16 | 1.04002246 | 1 | 0.895 | 1.42176E-11 | EN_1 |
| Atp5e | ENSMUSG00000016252 | protein_coding | ATP synthase, H+ transporting, mitochondrial F1 complex, epsilon subunit | 4.129E-09 | 0.864123993 | 0.959 | 0.75 | 9.74838E-05 | EN_1 |
| Fdps | ENSMUSG00000059743 | protein_coding | farnesyl diphosphate synthetase | 1.803E-09 | 1.892844225 | 0.626 | 0.171 | 4.25762E-05 | EN_1 |
| Slc39a1 | ENSMUSG00000052310 | protein_coding | solute carrier family 39 (zinc transporter), member 1 | 1.206E-07 | 1.525188029 | 0.614 | 0.237 | 0.00284788 | EN_1 |
| S100a16 | ENSMUSG00000074457 | protein_coding | S100 calcium binding protein A16 | 7.071E-08 | 0.906468462 | 0.953 | 0.737 | 0.001669718 | EN_1 |
| Rhoc | ENSMUSG00000002233 | protein_coding | ras homolog family member C | 4.256E-16 | 1.592193219 | 0.959 | 0.513 | 1.00493E-11 | EN_1 |
| Gm11808 | ENSMUSG00000068240 | protein_coding | predicted gene 11808 | 7.364E-09 | 2.100679179 | 0.509 | 0.092 | 0.000173875 | EN_1 |
| Bach2 | ENSMUSG00000040270 | protein_coding | BTB and CNC homology, basic leucine zipper transcription factor 2 | 2.896E-07 | 0.972068771 | 0.871 | 0.513 | 0.006838051 | EN_1 |
| Foxd3 | ENSMUSG00000067261 | protein_coding | forkhead box D3 | 4.528E-11 | 1.223725634 | 0.906 | 0.579 | 1.06907E-06 | EN_1 |
| Sgip1 | ENSMUSG00000028524 | protein_coding | SH3-domain GRB2-like (endophilin) interacting protein 1 | 6.091E-08 | 1.197433725 | 0.865 | 0.566 | 0.001438271 | EN_1 |
| Prdx1 | ENSMUSG00000028691 | protein_coding | peroxiredoxin 1 | 5.124E-09 | 0.75537278 | 0.977 | 0.803 | 0.00012099 | EN_1 |
| Insig1 | ENSMUSG00000045294 | protein_coding | insulin induced gene 1 | 1.531E-12 | 2.541720715 | 0.667 | 0.145 | 3.61531E-08 | EN_1 |
| Atp5k | ENSMUSG00000050856 | protein_coding | ATP synthase, H+ transporting, mitochondrial F1F0 complex, subunit E | 1.549E-09 | 1.178295549 | 0.889 | 0.579 | 3.65788E-05 | EN_1 |
| Ubc | ENSMUSG00000008348 | protein_coding | ubiquitin C | 7.226E-16 | 1.203817767 | 0.994 | 0.921 | 1.7061E-11 | EN_1 |
| Actb | ENSMUSG00000029580 | protein_coding | actin, beta | 5.19E-26 | 1.170300153 | 1 | 0.987 | 1.22549E-21 | EN_1 |
| Cdk8 | ENSMUSG00000029635 | protein_coding | cyclin-dependent kinase 8 | 1.235E-34 | -2.632505253 | 0.994 | 1 | 2.9156E-30 | EN_1 |
| Hsph1 | ENSMUSG00000029657 | protein_coding | heat shock 105kDa/110kDa protein 1 | 1.849E-07 | -1.575008428 | 0.45 | 0.697 | 0.004365602 | EN_1 |
| Gadd45a | ENSMUSG00000036390 | protein_coding | growth arrest and DNA-damage-inducible 45 alpha | 1.502E-09 | 2.333202552 | 0.591 | 0.171 | 3.54695E-05 | EN_1 |
| Actg2 | ENSMUSG00000059430 | protein_coding | actin, gamma 2, smooth muscle, enteric | 8.238E-11 | 1.225284881 | 0.971 | 0.645 | 1.94505E-06 | EN_1 |
| Gapdh | ENSMUSG00000057666 | protein_coding | glyceraldehyde-3-phosphate dehydrogenase | 2.374E-09 | 0.982620178 | 0.936 | 0.684 | 5.6058E-05 | EN_1 |
| Kcna1 | ENSMUSG00000047976 | protein_coding | potassium voltage-gated channel, shaker-related subfamily, member 1 | 1.348E-16 | 1.001830691 | 1 | 0.921 | 3.18385E-12 | EN_1 |
| Ldhb | ENSMUSG00000030246 | protein_coding | lactate dehydrogenase B | 4.026E-07 | 0.986302823 | 0.842 | 0.553 | 0.00950642 | EN_1 |
| Ndufa3 | ENSMUSG00000035674 | protein_coding | NADH:ubiquinone oxidoreductase subunit A3 | 8.558E-09 | 1.249207804 | 0.801 | 0.395 | 0.000202062 | EN_1 |
| Aldoa | ENSMUSG00000030695 | protein_coding | aldolase A, fructose-bisphosphate | 6.971E-11 | 1.153845313 | 0.918 | 0.684 | 1.64607E-06 | EN_1 |
| Cd81 | ENSMUSG00000037706 | protein_coding | CD81 antigen | 3.2E-14 | 0.843812175 | 1 | 0.947 | 7.55547E-10 | EN_1 |
| Msmo1 | ENSMUSG00000031604 | protein_coding | methylsterol monoxygenase 1 | 1.624E-09 | 1.853444105 | 0.649 | 0.184 | 3.83391E-05 | EN_1 |
| Uba52 | ENSMUSG00000090137 | protein_coding | ubiquitin A-52 residue ribosomal protein fusion product 1 | 2.511E-07 | 1.104649395 | 0.76 | 0.395 | 0.005930124 | EN_1 |
| Tecr | ENSMUSG00000031708 | protein_coding | trans-2,3-enoyl-CoA reductase | 3.162E-07 | 0.812595841 | 0.912 | 0.724 | 0.007465119 | EN_1 |
| Lpcat2 | ENSMUSG00000033192 | protein_coding | lysophosphatidylcholine acyltransferase 2 | 1.597E-10 | 2.117097792 | 0.573 | 0.105 | 3.77065E-06 | EN_1 |
| Rora | ENSMUSG00000032238 | protein_coding | RAR-related orphan receptor alpha | 1.638E-11 | -0.880174883 | 1 | 0.987 | 3.8666E-07 | EN_1 |
| Anxa2 | ENSMUSG00000032231 | protein_coding | annexin A2 | 7.083E-10 | 0.865803787 | 0.971 | 0.829 | 1.67237E-05 | EN_1 |
| Lars2 | ENSMUSG00000035202 | protein_coding | leucyl-tRNA synthetase, mitochondrial | 4.645E-33 | -2.211265511 | 0.994 | 1 | 1.09689E-28 | EN_1 |
| Ccn2 | ENSMUSG00000019997 | protein_coding | cellular communication network factor 2 | 3.766E-10 | 2.033791395 | 0.836 | 0.566 | 8.89173E-06 | EN_1 |
| Fabp7 | ENSMUSG00000019874 | protein_coding | fatty acid binding protein 7, brain | 1.311E-14 | 3.97948042 | 0.596 | 0.053 | 3.09656E-10 | EN_1 |
| Arid5b | ENSMUSG00000019947 | protein_coding | AT rich interactive domain 5B (MRF1-like) | 4.348E-10 | 1.193854082 | 0.895 | 0.5 | 1.0267E-05 | EN_1 |
| Cirbp | ENSMUSG00000045193 | protein_coding | cold inducible RNA binding protein | 8.319E-10 | 1.405962793 | 0.778 | 0.316 | 1.96423E-05 | EN_1 |
| Gadd45b | ENSMUSG00000015312 | protein_coding | growth arrest and DNA-damage-inducible 45 beta | 2.732E-07 | 1.978357931 | 0.468 | 0.105 | 0.006450972 | EN_1 |
| Tle5 | ENSMUSG00000054452 | protein_coding | TLE family member 5, transcriptional modulator | 9.549E-08 | 0.980622595 | 0.877 | 0.592 | 0.002254641 | EN_1 |
| Hsp90b1 | ENSMUSG00000020048 | protein_coding | heat shock protein 90, beta (Grp94), member 1 | 2.184E-08 | 0.706052653 | 0.994 | 0.921 | 0.000515637 | EN_1 |
| Dcn | ENSMUSG00000019929 | protein_coding | decorin | 8.031E-11 | 2.023527511 | 0.632 | 0.132 | 1.89622E-06 | EN_1 |
| Lyz2 | ENSMUSG00000069516 | protein_coding | lysozyme 2 | 5.207E-10 | 6.620648077 | 0.386 | 0 | 1.2294E-05 | EN_1 |
| Rpl41 | ENSMUSG00000093674 | protein_coding | ribosomal protein L41 | 7.765E-17 | 0.886573771 | 1 | 0.987 | 1.83351E-12 | EN_1 |
| Rps26 | ENSMUSG00000025362 | protein_coding | ribosomal protein S26 | 3.997E-07 | 0.587576584 | 0.994 | 0.895 | 0.009437972 | EN_1 |
| Bloc1s1 | ENSMUSG00000090247 | protein_coding | biogenesis of lysosomal organelles complex-1, subunit 1 | 1.386E-07 | 2.160306121 | 0.485 | 0.118 | 0.003273259 | EN_1 |
| Rtn4 | ENSMUSG00000020458 | protein_coding | reticulon 4 | 2.877E-07 | 0.77071554 | 0.965 | 0.803 | 0.00679325 | EN_1 |
| Sqstm1 | ENSMUSG00000015837 | protein_coding | sequestosome 1 | 1.787E-08 | 1.062922471 | 0.865 | 0.461 | 0.000422009 | EN_1 |
| Irf1 | ENSMUSG00000018899 | protein_coding | interferon regulatory factor 1 | 3.315E-07 | 1.318881703 | 0.591 | 0.237 | 0.007827037 | EN_1 |
| Pdlim4 | ENSMUSG00000020388 | protein_coding | PDZ and LIM domain 4 | 2.792E-09 | 0.867276074 | 0.982 | 0.882 | 6.59149E-05 | EN_1 |
| Nr1d1 | ENSMUSG00000020889 | protein_coding | nuclear receptor subfamily 1, group D, member 1 | 9.687E-17 | 2.563356717 | 0.795 | 0.184 | 2.28729E-12 | EN_1 |
| Rpl38 | ENSMUSG00000057322 | protein_coding | ribosomal protein L38 | 7.338E-16 | 1.054217148 | 1 | 0.895 | 1.73263E-11 | EN_1 |
| Jpt1 | ENSMUSG00000020737 | protein_coding | Jupiter microtubule associated homolog 1 | 1.555E-07 | 0.986219702 | 0.889 | 0.658 | 0.003670772 | EN_1 |
| Socs3 | ENSMUSG00000053113 | protein_coding | suppressor of cytokine signaling 3 | 2.213E-07 | 1.186004936 | 0.877 | 0.579 | 0.005225587 | EN_1 |
| Klhl29 | ENSMUSG00000020627 | protein_coding | kelch-like 29 | 6.397E-12 | -1.289323905 | 0.789 | 0.934 | 1.51054E-07 | EN_1 |
| Pdia6 | ENSMUSG00000020571 | protein_coding | protein disulfide isomerase associated 6 | 6.079E-11 | 1.334207217 | 0.901 | 0.526 | 1.4354E-06 | EN_1 |
| Asap2 | ENSMUSG00000052632 | protein_coding | ArfGAP with SH3 domain, ankyrin repeat and PH domain 2 | 2.253E-08 | 0.813345665 | 0.982 | 0.868 | 0.000532076 | EN_1 |
| Nfkbia | ENSMUSG00000021025 | protein_coding | nuclear factor of kappa light polypeptide gene enhancer in B cells inhibitor, alpha | 8.372E-10 | 1.510061953 | 0.865 | 0.513 | 1.97679E-05 | EN_1 |
| Rps29 | ENSMUSG00000034892 | protein_coding | ribosomal protein S29 | 1.54E-10 | 0.690467717 | 0.982 | 0.974 | 3.6361E-06 | EN_1 |
| Frmd6 | ENSMUSG00000048285 | protein_coding | FERM domain containing 6 | 8.321E-09 | 1.150217849 | 0.871 | 0.487 | 0.000196479 | EN_1 |
| Gphn | ENSMUSG00000047454 | protein_coding | gephyrin | 6.553E-28 | -1.498931751 | 1 | 1 | 1.5473E-23 | EN_1 |
| Sptlc2 | ENSMUSG00000021036 | protein_coding | serine palmitoyltransferase, long chain base subunit 2 | 1.875E-07 | 1.513392898 | 0.608 | 0.211 | 0.004428211 | EN_1 |
| Gm19951 | ENSMUSG00000113136 | lncRNA | predicted gene, 19951 | 2.505E-09 | -1.939004436 | 0.316 | 0.605 | 5.91577E-05 | EN_1 |
| Ckb | ENSMUSG00000001270 | protein_coding | creatine kinase, brain | 8.896E-27 | 1.9950557 | 1 | 0.855 | 2.10061E-22 | EN_1 |
| Atp5mpl | ENSMUSG00000021290 | protein_coding | ATP synthase membrane subunit 6.8PL | 4.541E-09 | 1.021129732 | 0.924 | 0.592 | 0.000107221 | EN_1 |
| Crip1 | ENSMUSG00000006360 | protein_coding | cysteine-rich protein 1 (intestinal) | 3.513E-08 | 0.617965765 | 1 | 1 | 0.000829467 | EN_1 |
| Hist1h4d | ENSMUSG00000061482 | protein_coding | histone cluster 1, H4d | 6.482E-12 | -1.78727723 | 0.608 | 0.789 | 1.53045E-07 | EN_1 |
| Gm48099 | ENSMUSG00000114304 | lncRNA | predicted gene, 48099 | 5.435E-08 | 2.381685953 | 0.45 | 0.079 | 0.001283259 | EN_1 |
| Plac9b | ENSMUSG00000072674 | protein_coding | placenta specific 9b | 1.605E-10 | 2.131228614 | 0.637 | 0.184 | 3.78947E-06 | EN_1 |
| Arhgef3 | ENSMUSG00000021895 | protein_coding | Rho guanine nucleotide exchange factor (GEF) 3 | 1.277E-09 | 1.823300119 | 0.684 | 0.224 | 3.01631E-05 | EN_1 |
| Pnp | ENSMUSG00000115338 | protein_coding | purine-nucleoside phosphorylase | 1.367E-08 | 1.523307349 | 0.801 | 0.474 | 0.00032289 | EN_1 |
| Dad1 | ENSMUSG00000022174 | protein_coding | defender against cell death 1 | 2.328E-07 | 0.683834686 | 0.942 | 0.803 | 0.005497239 | EN_1 |
| Pabpn1 | ENSMUSG00000022194 | protein_coding | poly(A) binding protein, nuclear 1 | 2.751E-07 | 1.087982018 | 0.813 | 0.513 | 0.0064952 | EN_1 |
| Cmtm5 | ENSMUSG00000040759 | protein_coding | CKLF-like MARVEL transmembrane domain containing 5 | 1.936E-07 | 0.884911285 | 0.942 | 0.697 | 0.004570146 | EN_1 |
| Spry2 | ENSMUSG00000022114 | protein_coding | sprouty RTK signaling antagonist 2 | 2.43E-10 | 1.732402941 | 0.749 | 0.25 | 5.73802E-06 | EN_1 |
| Tuba1b | ENSMUSG00000023004 | protein_coding | tubulin, alpha 1B | 5.003E-10 | 1.071207904 | 0.93 | 0.737 | 1.18138E-05 | EN_1 |
| Tuba1a | ENSMUSG00000072235 | protein_coding | tubulin, alpha 1A | 4.367E-12 | 1.021347534 | 0.988 | 0.855 | 1.03121E-07 | EN_1 |
| Tmbim6 | ENSMUSG00000023010 | protein_coding | transmembrane BAX inhibitor motif containing 6 | 1.606E-07 | 0.749988198 | 0.959 | 0.789 | 0.00379311 | EN_1 |
| Cebpd | ENSMUSG00000071637 | protein_coding | CCAAT/enhancer binding protein (C/EBP), delta | 1.274E-07 | 1.038057274 | 0.936 | 0.763 | 0.003007254 | EN_1 |
| Sdf2l1 | ENSMUSG00000022769 | protein_coding | stromal cell-derived factor 2-like 1 | 5.598E-09 | 2.251460454 | 0.526 | 0.118 | 0.000132187 | EN_1 |
| Comt | ENSMUSG00000000326 | protein_coding | catechol-O-methyltransferase | 1.564E-08 | 0.900403881 | 0.971 | 0.711 | 0.000369212 | EN_1 |

| | | | | | | | | | |
|---|---|---|---|---|---|---|---|---|---|
| Gm15564 | ENSMUSG00000086324 | lncRNA | predicted gene 15564 | 2.46E-18 | -2.499868599 | 0.433 | 0.842 | 5.80973E-14 | EN_1 |
| Cmss1 | ENSMUSG00000022748 | protein_coding | cms small ribosomal subunit 1 | 6.15E-32 | -2.190086654 | 1 | 1 | 1.45214E-27 | EN_1 |
| Fndc1 | ENSMUSG00000071984 | protein_coding | fibronectin type III domain containing 1 | 3.532E-10 | -1.415750902 | 0.772 | 0.934 | 8.34063E-06 | EN_1 |
| Ppp1r10 | ENSMUSG00000039220 | protein_coding | protein phosphatase 1, regulatory subunit 10 | 1.186E-07 | 1.245607884 | 0.76 | 0.434 | 0.002799269 | EN_1 |
| Gm42418 | ENSMUSG00000098178 | lncRNA | predicted gene, 42418 | 1.045E-34 | -1.958916796 | 1 | 1 | 2.46724E-30 | EN_1 |
| Dtna | ENSMUSG00000024302 | protein_coding | dystrobrevin alpha | 2.117E-08 | 0.932033679 | 0.971 | 0.75 | 0.000499823 | EN_1 |
| Ndfip1 | ENSMUSG00000024425 | protein_coding | Nedd4 family interacting protein 1 | 2.407E-10 | 0.648979331 | 0.994 | 0.961 | 5.68269E-06 | EN_1 |
| Gstp1 | ENSMUSG00000060803 | protein_coding | glutathione S-transferase, pi 1 | 2.556E-12 | 2.251558942 | 0.661 | 0.145 | 6.03473E-08 | EN_1 |
| Banf1 | ENSMUSG00000024844 | protein_coding | barrier to autointegration factor 1 | 1.621E-07 | 1.702645014 | 0.509 | 0.132 | 0.003827135 | EN_1 |
| Cfl1 | ENSMUSG00000056201 | protein_coding | cofilin 1, non-muscle | 3.562E-11 | 0.909216964 | 0.994 | 0.803 | 8.41044E-07 | EN_1 |
| Acta2 | ENSMUSG00000035783 | protein_coding | actin, alpha 2, smooth muscle, aorta | 1.362E-10 | 0.957203312 | 0.959 | 0.711 | 3.21691E-06 | EN_1 |
| Pdlim1 | ENSMUSG00000055044 | protein_coding | PDZ and LIM domain 1 (elfin) | 7.186E-08 | 0.86795896 | 0.965 | 0.711 | 0.001696827 | EN_1 |
| Scd2 | ENSMUSG00000025203 | protein_coding | stearoyl-Coenzyme A desaturase 2 | 2.25E-14 | 1.139738804 | 1 | 0.882 | 5.31217E-10 | EN_1 |
| Rpl10 | ENSMUSG00000008682 | protein_coding | ribosomal protein L10 | 1.953E-08 | 0.690827892 | 0.982 | 0.921 | 0.000461073 | EN_1 |
| Sat1 | ENSMUSG00000025283 | protein_coding | spermidine/spermine N1-acetyl transferase 1 | 5.824E-12 | 0.958779442 | 0.988 | 0.908 | 1.3751E-07 | EN_1 |
| Gm47283 | ENSMUSG00000096768 | lncRNA | predicted gene, 47283 | 1.531E-11 | 2.479643599 | 0.608 | 0.105 | 3.61535E-07 | EN_1 |
| mt-Co3 | ENSMUSG00000064358 | protein_coding | mitochondrially encoded cytochrome c oxidase III | 1.682E-13 | -0.641267408 | 1 | 1 | 3.9715E-09 | EN_1 |
| mt-Nd3 | ENSMUSG00000064360 | protein_coding | mitochondrially encoded NADH dehydrogenase 3 | 2.752E-12 | -0.822184203 | 0.994 | 1 | 6.49721E-08 | EN_1 |
| AC149090.1 | ENSMUSG00000095041 | protein_coding | | 2.437E-07 | 0.974577933 | 0.813 | 0.395 | 0.005754298 | EN_1 |
| Coq10b | ENSMUSG00000025981 | protein_coding | coenzyme Q10B | 1.994E-08 | -1.303506046 | 0.345 | 0.662 | 0.000470735 | EN_2 |
| Des | ENSMUSG00000026208 | protein_coding | desmin | 1.563E-08 | 1.058411203 | 0.832 | 0.63 | 0.000368978 | EN_2 |
| Itm2c | ENSMUSG00000026223 | protein_coding | integral membrane protein 2C | 7.193E-08 | 0.802733953 | 0.882 | 0.714 | 0.001698335 | EN_2 |
| Per2 | ENSMUSG00000055866 | protein_coding | period circadian clock 2 | 3.108E-08 | -3.816412431 | 0.025 | 0.273 | 0.000733922 | EN_2 |
| Cdh19 | ENSMUSG00000047216 | protein_coding | cadherin 19, type 2 | 1.231E-09 | -0.610204988 | 0.983 | 1 | 2.90647E-05 | EN_2 |
| Dbi | ENSMUSG00000026385 | protein_coding | diazepam binding inhibitor | 2.778E-44 | 2.74747494 | 1 | 0.948 | 6.55855E-40 | EN_2 |
| Gpr37l1 | ENSMUSG00000026424 | protein_coding | G protein-coupled receptor 37-like 1 | 7.721E-10 | 0.622943493 | 0.975 | 0.948 | 1.82302E-05 | EN_2 |
| Csrp1 | ENSMUSG00000026421 | protein_coding | cysteine and glycine-rich protein 1 | 1.977E-08 | 0.604386648 | 0.992 | 0.948 | 0.000466716 | EN_2 |
| Gm2000 | ENSMUSG00000078193 | protein_coding | predicted gene 2000 | 2.472E-10 | 1.919491577 | 0.487 | 0.149 | 5.83726E-06 | EN_2 |
| Prdx6 | ENSMUSG00000026701 | protein_coding | peroxiredoxin 6 | 7.066E-08 | 0.662918625 | 0.933 | 0.857 | 0.001668366 | EN_2 |
| Traf5 | ENSMUSG00000026637 | protein_coding | TNF receptor-associated factor 5 | 5.011E-08 | 1.675643082 | 0.462 | 0.162 | 0.001183242 | EN_2 |
| G0s2 | ENSMUSG00000009633 | protein_coding | G0/G1 switch gene 2 | 1.788E-15 | -1.475636376 | 0.891 | 0.981 | 4.22179E-11 | EN_2 |
| Camk1d | ENSMUSG00000039145 | protein_coding | calcium/calmodulin-dependent protein kinase ID | 2.446E-31 | -1.377080289 | 0.95 | 1 | 5.7754E-27 | EN_2 |
| Gsn | ENSMUSG00000026879 | protein_coding | gelsolin | 5.429E-09 | 0.668587766 | 0.983 | 0.987 | 0.000128185 | EN_2 |
| Rpl35 | ENSMUSG00000062997 | protein_coding | ribosomal protein L35 | 6.719E-15 | 0.885626712 | 0.975 | 0.922 | 1.58641E-10 | EN_2 |
| Cd59a | ENSMUSG00000032679 | protein_coding | CD59a antigen | 1.931E-12 | 0.80427096 | 0.975 | 0.961 | 4.55935E-08 | EN_2 |
| Gatm | ENSMUSG00000027199 | protein_coding | glycine amidinotransferase (L-arginine:glycine amidinotransferase) | 3.544E-08 | 1.048297085 | 0.824 | 0.656 | 0.000836819 | EN_2 |
| Fkbp1a | ENSMUSG00000032966 | protein_coding | FK506 binding protein 1a | 3.614E-12 | 1.207030993 | 0.866 | 0.597 | 8.53297E-08 | EN_2 |
| Myl9 | ENSMUSG00000067818 | protein_coding | myosin, light polypeptide 9, regulatory | 5.646E-09 | 1.021722482 | 0.798 | 0.539 | 0.000133324 | EN_2 |
| Atp5e | ENSMUSG00000016252 | protein_coding | ATP synthase, H+ transporting, mitochondrial F1 complex, epsilon subunit | 4.275E-11 | 0.719992208 | 0.95 | 0.922 | 1.00948E-06 | EN_2 |
| Anxa5 | ENSMUSG00000027712 | protein_coding | annexin A5 | 8.111E-15 | 0.689692968 | 1 | 0.994 | 1.91522E-10 | EN_2 |
| Fdps | ENSMUSG00000059743 | protein_coding | farnesyl diphosphate synthetase | 2.076E-12 | 2.102408513 | 0.58 | 0.188 | 4.90299E-08 | EN_2 |
| S100a16 | ENSMUSG00000074457 | protein_coding | S100 calcium binding protein A16 | 2.457E-17 | 1.198167711 | 0.958 | 0.721 | 5.80212E-13 | EN_2 |
| Rhoc | ENSMUSG00000002233 | protein_coding | ras homolog family member C | 3.265E-18 | 1.394599765 | 0.941 | 0.649 | 7.71046E-14 | EN_2 |
| Gstm5 | ENSMUSG00000004032 | protein_coding | glutathione S-transferase, mu 5 | 3.76E-07 | 0.935499526 | 0.891 | 0.61 | 0.00887907 | EN_2 |
| Gm11808 | ENSMUSG00000068240 | protein_coding | predicted gene 11808 | 1.537E-11 | 2.285938552 | 0.529 | 0.156 | 3.62859E-07 | EN_2 |
| Ccl21a | ENSMUSG00000094686 | protein_coding | chemokine (C-C motif) ligand 21A (serine) | 9.937E-10 | 3.80717955 | 0.269 | 0.019 | 2.34638E-05 | EN_2 |
| Foxd3 | ENSMUSG00000067261 | protein_coding | forkhead box D3 | 3.895E-08 | 0.988674827 | 0.857 | 0.701 | 0.00091977 | EN_2 |
| Prdx1 | ENSMUSG00000028691 | protein_coding | peroxiredoxin 1 | 2.427E-11 | 0.776412609 | 0.958 | 0.916 | 5.7311E-07 | EN_2 |
| Vwa1 | ENSMUSG00000042116 | protein_coding | von Willebrand factor A domain containing 1 | 6.655E-08 | 1.063945482 | 0.849 | 0.688 | 0.001571459 | EN_2 |
| Insig1 | ENSMUSG00000045294 | protein_coding | insulin induced gene 1 | 8.967E-15 | 2.375775595 | 0.622 | 0.188 | 2.11737E-10 | EN_2 |
| Hadhb | ENSMUSG00000059447 | protein_coding | hydroxyacyl-Coenzyme A dehydrogenase/3-ketoacyl-Coenzyme A thiolase/enoyl-Coenzyme A hydratase (trifunctional protein), beta subunit | 4.115E-07 | 1.107680134 | 0.647 | 0.357 | 0.009715258 | EN_2 |
| Rpl6 | ENSMUSG00000029614 | protein_coding | ribosomal protein L6 | 5.906E-10 | 0.599538085 | 0.992 | 0.961 | 1.39443E-05 | EN_2 |
| AI480526 | ENSMUSG00000090086 | lncRNA | expressed sequence AI480526 | 3.56E-07 | 3.956399107 | 0.176 | 0.006 | 0.008404701 | EN_2 |
| Actb | ENSMUSG00000029580 | protein_coding | actin, beta | 1.97E-24 | 0.902293595 | 1 | 1 | 4.65158E-20 | EN_2 |
| Cdk8 | ENSMUSG00000029635 | protein_coding | cyclin-dependent kinase 8 | 7.781E-35 | -2.727562415 | 0.866 | 0.994 | 1.83735E-30 | EN_2 |
| Hsph1 | ENSMUSG00000029657 | protein_coding | heat shock 105kDa/110kDa protein 1 | 2.719E-12 | -2.325641021 | 0.328 | 0.675 | 6.41928E-08 | EN_2 |
| Gapdh | ENSMUSG00000057666 | protein_coding | glyceraldehyde-3-phosphate dehydrogenase | 6.869E-08 | 0.805569592 | 0.924 | 0.87 | 0.00162196 | EN_2 |
| Kcna1 | ENSMUSG00000047976 | protein_coding | potassium voltage-gated channel, shaker-related subfamily, member 1 | 4.857E-10 | 0.742628321 | 0.95 | 0.935 | 1.14691E-05 | EN_2 |
| Zfp580 | ENSMUSG00000055633 | protein_coding | zinc finger protein 580 | 1.117E-07 | 3.004856731 | 0.235 | 0.026 | 0.002638336 | EN_2 |
| Sirt2 | ENSMUSG00000015149 | protein_coding | sirtuin 2 | 1.601E-09 | 0.90935988 | 0.882 | 0.643 | 3.78017E-05 | EN_2 |
| Mapk3 | ENSMUSG00000063065 | protein_coding | mitogen-activated protein kinase 3 | 4.808E-10 | 1.108952183 | 0.84 | 0.565 | 1.13529E-05 | EN_2 |
| Aldoa | ENSMUSG00000030695 | protein_coding | aldolase A, fructose-bisphosphate | 4.21E-08 | 0.911829973 | 0.874 | 0.74 | 0.000994115 | EN_2 |
| Ifitm3 | ENSMUSG00000025492 | protein_coding | interferon induced transmembrane protein 3 | 5.368E-12 | 0.697873675 | 1 | 0.987 | 1.26747E-07 | EN_2 |
| Cd151 | ENSMUSG00000025510 | protein_coding | CD151 antigen | 2.172E-07 | 1.014961111 | 0.723 | 0.474 | 0.005127819 | EN_2 |
| Ctsd | ENSMUSG00000007891 | protein_coding | cathepsin D | 1.586E-08 | 0.995823828 | 0.782 | 0.571 | 0.00037441 | EN_2 |
| Cd81 | ENSMUSG00000037706 | protein_coding | CD81 antigen | 1.801E-18 | 0.824802889 | 1 | 0.994 | 4.25168E-14 | EN_2 |
| Plat | ENSMUSG00000031538 | protein_coding | plasminogen activator, tissue | 1.378E-07 | 1.106841896 | 0.681 | 0.429 | 0.003253096 | EN_2 |
| Msmo1 | ENSMUSG00000031604 | protein_coding | methylsterol monoxygenase 1 | 9.787E-10 | 1.755779009 | 0.571 | 0.234 | 2.31097E-05 | EN_2 |
| Uba52 | ENSMUSG00000090137 | protein_coding | ubiquitin A-52 residue ribosomal protein fusion product 1 | 2.81E-15 | 2.017000042 | 0.689 | 0.279 | 6.63382E-11 | EN_2 |
| Tecr | ENSMUSG00000031708 | protein_coding | trans-2,3-enoyl-CoA reductase | 5.888E-11 | 0.960119313 | 0.933 | 0.773 | 1.39036E-06 | EN_2 |
| Rora | ENSMUSG00000032238 | protein_coding | RAR-related orphan receptor alpha | 3.916E-14 | -1.217200855 | 0.866 | 0.961 | 9.24696E-10 | EN_2 |
| Col12a1 | ENSMUSG00000032332 | protein_coding | collagen, type XII, alpha 1 | 8.192E-08 | -0.724525596 | 0.891 | 0.961 | 0.001934189 | EN_2 |
| Gnai2 | ENSMUSG00000032562 | protein_coding | guanine nucleotide binding protein (G protein), alpha inhibiting 2 | 5.958E-08 | 0.601045061 | 0.924 | 0.935 | 0.001406736 | EN_2 |
| Lars2 | ENSMUSG00000035202 | protein_coding | leucyl-tRNA synthetase, mitochondrial | 3.066E-34 | -2.487256239 | 0.866 | 0.961 | 7.23951E-30 | EN_2 |
| Ccn2 | ENSMUSG00000019997 | protein_coding | cellular communication network factor 2 | 1.48E-09 | 1.724996134 | 0.773 | 0.513 | 3.49363E-05 | EN_2 |
| Fabp7 | ENSMUSG00000019874 | protein_coding | fatty acid binding protein 7, brain | 1.219E-24 | 4.501261518 | 0.639 | 0.065 | 2.87779E-20 | EN_2 |
| Cirbp | ENSMUSG00000045193 | protein_coding | cold inducible RNA binding protein | 6.696E-14 | 1.897641611 | 0.756 | 0.364 | 1.58096E-09 | EN_2 |
| Cry1 | ENSMUSG00000020038 | protein_coding | cryptochrome 1 (photolyase-like) | 1.688E-07 | -2.012051851 | 0.059 | 0.312 | 0.003986279 | EN_2 |
| Hsp90b1 | ENSMUSG00000020048 | protein_coding | heat shock protein 90, beta (Grp94), member 1 | 6.022E-08 | 0.636734369 | 0.958 | 0.929 | 0.001421873 | EN_2 |
| Dcn | ENSMUSG00000019929 | protein_coding | decorin | 1.401E-11 | 1.804569485 | 0.462 | 0.097 | 3.30813E-07 | EN_2 |
| Lyz2 | ENSMUSG00000069516 | protein_coding | lysozyme 2 | 6.942E-16 | 8.716531059 | 0.361 | 0 | 1.63915E-11 | EN_2 |
| Rpl41 | ENSMUSG00000093674 | protein_coding | ribosomal protein L41 | 4.447E-19 | 0.863839057 | 1 | 1 | 1.04996E-14 | EN_2 |
| Bloc1s1 | ENSMUSG00000090247 | protein_coding | biogenesis of lysosomal organelles complex-1, subunit 1 | 1.389E-11 | 2.369151095 | 0.479 | 0.117 | 3.2806E-07 | EN_2 |
| Pdlim4 | ENSMUSG00000020388 | protein_coding | PDZ and LIM domain 4 | 5.385E-09 | 0.904657176 | 0.975 | 0.916 | 0.000127153 | EN_2 |
| Ywhae | ENSMUSG00000020849 | protein_coding | tyrosine 3-monooxygenase/tryptophan 5-monooxygenase activation protein, epsilon polypeptide | 5.646E-08 | 0.622531536 | 0.975 | 0.883 | 0.001333153 | EN_2 |
| Nr1d1 | ENSMUSG00000020889 | protein_coding | nuclear receptor subfamily 1, group D, member 1 | 1.384E-11 | 1.588668673 | 0.689 | 0.26 | 3.26792E-07 | EN_2 |
| Cnp | ENSMUSG00000006782 | protein_coding | 2',3'-cyclic nucleotide 3' phosphodiesterase | 3.512E-07 | 1.089445293 | 0.84 | 0.688 | 0.008293499 | EN_2 |
| Abca8a | ENSMUSG00000041828 | protein_coding | ATP-binding cassette, sub-family A (ABC1), member 8a | 4.326E-08 | -0.748182115 | 0.874 | 0.955 | 0.001021394 | EN_2 |
| Rpl38 | ENSMUSG00000057322 | protein_coding | ribosomal protein L38 | 1.655E-20 | 1.003399179 | 0.983 | 0.955 | 3.90807E-16 | EN_2 |
| Jpt1 | ENSMUSG00000020737 | protein_coding | Jupiter microtubule associated homolog 1 | 2.807E-09 | 0.940908048 | 0.874 | 0.675 | 6.62888E-05 | EN_2 |
| Ubald2 | ENSMUSG00000050628 | protein_coding | UBA-like domain containing 2 | 4.86E-08 | 1.676851727 | 0.496 | 0.195 | 0.001147654 | EN_2 |
| Klhl29 | ENSMUSG00000020627 | protein_coding | kelch-like 29 | 1.939E-08 | -1.27189154 | 0.546 | 0.753 | 0.000457869 | EN_2 |
| Cys1 | ENSMUSG00000062563 | protein_coding | cystin 1 | 2.508E-10 | -1.769468586 | 0.235 | 0.591 | 5.92097E-06 | EN_2 |
| Rps29 | ENSMUSG00000034892 | protein_coding | ribosomal protein S29 | 1.866E-16 | 0.720491931 | 1 | 1 | 4.40667E-12 | EN_2 |
| Gphn | ENSMUSG00000047454 | protein_coding | gephyrin | 5.265E-28 | -1.604079988 | 0.908 | 0.987 | 1.24306E-23 | EN_2 |
| Slirp | ENSMUSG00000021040 | protein_coding | SRA stem-loop interacting RNA binding protein | 1.441E-07 | 1.148239146 | 0.639 | 0.351 | 0.003403563 | EN_2 |
| Gm19951 | ENSMUSG00000113136 | lncRNA | predicted gene, 19951 | 3.769E-11 | -2.40256928 | 0.134 | 0.5 | 8.89835E-07 | EN_2 |
| Ckb | ENSMUSG00000001270 | protein_coding | creatine kinase, brain | 1.366E-35 | 2.068458928 | 0.992 | 0.935 | 3.22594E-31 | EN_2 |
| Atp5mpl | ENSMUSG00000021290 | protein_coding | ATP synthase membrane subunit 6.8PL | 8.032E-11 | 0.901454609 | 0.916 | 0.695 | 1.89644E-06 | EN_2 |
| Idi1 | ENSMUSG00000058258 | protein_coding | isopentenyl-diphosphate delta isomerase | 3.967E-08 | 1.401265457 | 0.546 | 0.221 | 0.000936719 | EN_2 |
| Hist1h4d | ENSMUSG00000061482 | protein_coding | histone cluster 1, H4d | 2.099E-21 | -2.164984484 | 0.462 | 0.844 | 4.95655E-17 | EN_2 |
| Cox7c | ENSMUSG00000017778 | protein_coding | cytochrome c oxidase subunit 7C | 2.354E-11 | 0.635652919 | 0.983 | 0.968 | 5.55743E-07 | EN_2 |
| Hmgcs1 | ENSMUSG00000093930 | protein_coding | 3-hydroxy-3-methylglutaryl-Coenzyme A synthase 1 | 4.686E-09 | 1.723868855 | 0.563 | 0.24 | 0.000110646 | EN_2 |
| Fam107a | ENSMUSG00000021750 | protein_coding | family with sequence similarity 107, member A | 3.86E-11 | -1.617337083 | 0.328 | 0.695 | 9.11384E-07 | EN_2 |
| Gm48099 | ENSMUSG00000114304 | lncRNA | predicted gene, 48099 | 9.011E-11 | 5.047352765 | 0.261 | 0.006 | 2.12771E-06 | EN_2 |
| Plac9b | ENSMUSG00000072674 | protein_coding | placenta specific 9b | 3.384E-16 | 2.945586978 | 0.555 | 0.11 | 7.99136E-12 | EN_2 |
| Dad1 | ENSMUSG00000022174 | protein_coding | defender against cell death 1 | 1.019E-12 | 0.80898127 | 0.95 | 0.838 | 2.40543E-08 | EN_2 |
| Rgcc | ENSMUSG00000022018 | protein_coding | regulator of cell cycle | 3.048E-10 | -1.023000623 | 0.782 | 0.909 | 7.19737E-06 | EN_2 |
| Rpl37 | ENSMUSG00000041841 | protein_coding | ribosomal protein L37 | 7.471E-12 | 0.673145561 | 0.992 | 0.994 | 1.7641E-07 | EN_2 |
| Cox6c | ENSMUSG00000014313 | protein_coding | cytochrome c oxidase subunit 6C | 3.435E-11 | 0.639319881 | 0.983 | 0.955 | 8.10983E-07 | EN_2 |
| Azin1 | ENSMUSG00000037458 | protein_coding | antizyme inhibitor 1 | 3.531E-08 | -0.733804134 | 0.832 | 0.929 | 0.000833671 | EN_2 |
| Pdgfb | ENSMUSG00000000489 | protein_coding | platelet derived growth factor, B polypeptide | 4.783E-08 | -0.866652674 | 0.655 | 0.844 | 0.001129288 | EN_2 |
| Tuba1b | ENSMUSG00000023004 | protein_coding | tubulin, alpha 1B | 3.33E-12 | 1.076518099 | 0.908 | 0.799 | 7.8623E-08 | EN_2 |
| Tuba1a | ENSMUSG00000072235 | protein_coding | tubulin, alpha 1A | 1.953E-18 | 1.148456047 | 0.983 | 0.955 | 4.61028E-14 | EN_2 |
| Tmbim6 | ENSMUSG00000023010 | protein_coding | transmembrane BAX inhibitor motif containing 6 | 2.873E-07 | 0.727595619 | 0.916 | 0.76 | 0.006784184 | EN_2 |
| Igfbp6 | ENSMUSG00000023046 | protein_coding | insulin-like growth factor binding protein 6 | 2.659E-07 | 2.452865443 | 0.244 | 0.032 | 0.006277377 | EN_2 |
| Gm15564 | ENSMUSG00000086324 | lncRNA | predicted gene 15564 | 1.316E-17 | -2.601430408 | 0.176 | 0.675 | 3.10647E-13 | EN_2 |
| Cmss1 | ENSMUSG00000022748 | protein_coding | cms small ribosomal subunit 1 | 3.147E-31 | -2.353512568 | 0.924 | 1 | 7.42997E-27 | EN_2 |
| Elob | ENSMUSG00000055839 | protein_coding | elongin B | 1.168E-10 | 0.650869947 | 1 | 0.935 | 2.75722E-06 | EN_2 |
| Cuta | ENSMUSG00000024194 | protein_coding | cutA divalent cation tolerance homolog | 2.945E-07 | 0.999470288 | 0.756 | 0.513 | 0.006953537 | EN_2 |
| Rps28 | ENSMUSG00000067288 | protein_coding | ribosomal protein S28 | 1.754E-10 | 0.698481144 | 0.992 | 0.981 | 4.14099E-06 | EN_2 |

| | | | | | | | | | |
|---|---|---|---|---|---|---|---|---|---|
| Ndufa7 | ENSMUSG00000041881 | protein_coding | NADH:ubiquinone oxidoreductase subunit A7 | 2.142E-09 | 0.756123017 | 0.941 | 0.838 | 5.05686E-05 | EN_2 |
| Gm42418 | ENSMUSG00000098178 | lncRNA | predicted gene, 42418 | 1.411E-37 | -2.063192349 | 1 | 1 | 3.33165E-33 | EN_2 |
| Tomm6 | ENSMUSG00000033475 | protein_coding | translocase of outer mitochondrial membrane 6 | 1.104E-08 | 0.767092972 | 0.908 | 0.792 | 0.000260565 | EN_2 |
| Rpl36 | ENSMUSG00000057863 | protein_coding | ribosomal protein L36 | 3.189E-09 | 0.617460676 | 1 | 0.987 | 7.52935E-05 | EN_2 |
| Gstp1 | ENSMUSG00000060803 | protein_coding | glutathione S-transferase, pi 1 | 1.16E-10 | 1.820749943 | 0.639 | 0.292 | 2.73811E-06 | EN_2 |
| Cntf | ENSMUSG00000079415 | protein_coding | ciliary neurotrophic factor | 1.16E-09 | 1.078653601 | 0.924 | 0.721 | 2.7399E-05 | EN_2 |
| Acta2 | ENSMUSG00000035783 | protein_coding | actin, alpha 2, smooth muscle, aorta | 9.08E-13 | 1.220990139 | 0.891 | 0.63 | 2.144E-08 | EN_2 |
| Plce1 | ENSMUSG00000024998 | protein_coding | phospholipase C, epsilon 1 | 1.747E-09 | -0.615104792 | 0.966 | 1 | 4.12481E-05 | EN_2 |
| Scd2 | ENSMUSG00000025203 | protein_coding | stearoyl-Coenzyme A desaturase 2 | 4.224E-07 | 0.751668055 | 0.908 | 0.909 | 0.009973447 | EN_2 |
| Atp5md | ENSMUSG00000071528 | protein_coding | ATP synthase membrane subunit DAPIT | 3.03E-07 | 0.794326325 | 0.866 | 0.675 | 0.007155476 | EN_2 |
| Cox7b | ENSMUSG00000031231 | protein_coding | cytochrome c oxidase subunit 7B | 3.626E-07 | 0.766549208 | 0.874 | 0.779 | 0.00856242 | EN_2 |
| Pgk1 | ENSMUSG00000062070 | protein_coding | phosphoglycerate kinase 1 | 9.935E-08 | 0.972592047 | 0.748 | 0.487 | 0.002345814 | EN_2 |
| Sat1 | ENSMUSG00000025283 | protein_coding | spermidine/spermine N1-acetyl transferase 1 | 5.437E-16 | 0.859358529 | 1 | 0.974 | 1.28386E-11 | EN_2 |
| Gm47283 | ENSMUSG00000096768 | lncRNA | predicted gene, 47283 | 2.523E-07 | 1.952870474 | 0.412 | 0.143 | 0.005956179 | EN_2 |
| mt-Nd3 | ENSMUSG00000064360 | protein_coding | mitochondrially encoded NADH dehydrogenase 3 | 6.65E-11 | -0.710960829 | 0.992 | 0.994 | 1.57027E-06 | EN_2 |
| Dbi | ENSMUSG00000026385 | protein_coding | diazepam binding inhibitor | 2.461E-08 | 3.666434264 | 0.595 | 0.21 | 0.000581099 | EN_3 |
| Frmd4a | ENSMUSG00000026657 | protein_coding | FERM domain containing 4A | 2.751E-08 | 1.32505409 | 0.931 | 0.79 | 0.000649543 | EN_3 |
| Camk1d | ENSMUSG00000039145 | protein_coding | calcium/calmodulin-dependent protein kinase ID | 9.54E-11 | -0.844786742 | 1 | 1 | 2.2525E-06 | EN_3 |
| S100a6 | ENSMUSG00000001025 | protein_coding | S100 calcium binding protein A6 (calcyclin) | 1.296E-08 | 1.881231033 | 0.784 | 0.339 | 0.000305924 | EN_3 |
| Pappa | ENSMUSG00000028370 | protein_coding | pregnancy-associated plasma protein A | 1.045E-08 | 2.188715544 | 0.741 | 0.323 | 0.000246735 | EN_3 |
| Sgip1 | ENSMUSG00000028524 | protein_coding | SH3-domain GRB2-like (endophilin) interacting protein 1 | 2.138E-07 | 1.053271119 | 0.897 | 0.532 | 0.00504849 | EN_3 |
| Actb | ENSMUSG00000029580 | protein_coding | actin, beta | 2.205E-11 | 1.307093621 | 0.974 | 0.79 | 5.20735E-07 | EN_3 |
| Cdk8 | ENSMUSG00000029635 | protein_coding | cyclin-dependent kinase 8 | 9.626E-28 | -2.797974263 | 0.966 | 1 | 2.27298E-23 | EN_3 |
| Tmsb10 | ENSMUSG00000079523 | protein_coding | thymosin, beta 10 | 5.466E-08 | 2.844804199 | 0.526 | 0.097 | 0.001290581 | EN_3 |
| Ifitm2 | ENSMUSG00000060591 | protein_coding | interferon induced transmembrane protein 2 | 1.44E-07 | 3.006095772 | 0.517 | 0.113 | 0.003400366 | EN_3 |
| Lars2 | ENSMUSG00000035202 | protein_coding | leucyl-tRNA synthetase, mitochondrial | 3.111E-24 | -2.073127637 | 0.966 | 1 | 7.34577E-20 | EN_3 |
| Ank3 | ENSMUSG00000069601 | protein_coding | ankyrin 3, epithelial | 1.926E-09 | 0.870373234 | 0.991 | 0.984 | 4.54837E-05 | EN_3 |
| Rpl41 | ENSMUSG00000093674 | protein_coding | ribosomal protein L41 | 3.623E-08 | 1.781298017 | 0.784 | 0.419 | 0.000855554 | EN_3 |
| Ppia | ENSMUSG00000071866 | protein_coding | peptidylprolyl isomerase A | 1.874E-07 | 1.963552248 | 0.655 | 0.226 | 0.004424779 | EN_3 |
| Rpl19 | ENSMUSG00000017404 | protein_coding | ribosomal protein L19 | 3.724E-08 | 2.281040797 | 0.595 | 0.145 | 0.000879293 | EN_3 |
| Rpl38 | ENSMUSG00000057322 | protein_coding | ribosomal protein L38 | 3.626E-07 | 2.173404768 | 0.586 | 0.194 | 0.008562122 | EN_3 |
| Rps29 | ENSMUSG00000034892 | protein_coding | ribosomal protein S29 | 3.734E-08 | 1.649252341 | 0.828 | 0.468 | 0.000881571 | EN_3 |
| Frmd6 | ENSMUSG00000048285 | protein_coding | FERM domain containing 6 | 5.166E-08 | 1.357194833 | 0.707 | 0.242 | 0.001219806 | EN_3 |
| Gphn | ENSMUSG00000047454 | protein_coding | gephyrin | 7.357E-21 | -1.635315967 | 0.957 | 1 | 1.73717E-16 | EN_3 |
| Gm19951 | ENSMUSG00000113136 | lncRNA | predicted gene, 19951 | 1.842E-10 | -2.634228098 | 0.19 | 0.613 | 4.34961E-06 | EN_3 |
| Ckb | ENSMUSG00000001270 | protein_coding | creatine kinase, brain | 1.29E-07 | 1.677481706 | 0.707 | 0.306 | 0.003046704 | EN_3 |
| Gm15564 | ENSMUSG00000086324 | lncRNA | predicted gene 15564 | 1.133E-15 | -2.908477508 | 0.284 | 0.79 | 2.67424E-11 | EN_3 |
| Cmss1 | ENSMUSG00000022748 | protein_coding | cms small ribosomal subunit 1 | 2.542E-27 | -2.467419789 | 1 | 1 | 6.00316E-23 | EN_3 |
| Gm26917 | ENSMUSG00000097971 | lncRNA | predicted gene, 26917 | 3.954E-08 | -1.548542695 | 0.69 | 0.871 | 0.000933542 | EN_3 |
| Gm42418 | ENSMUSG00000098178 | lncRNA | predicted gene, 42418 | 1.91E-26 | -2.306936303 | 1 | 1 | 4.50879E-22 | EN_3 |
| AY036118 | ENSMUSG00000105361 | lncRNA | cDNA sequence AY036118 | 5.001E-08 | -0.859352873 | 0.991 | 1 | 0.001180749 | EN_3 |
| Fth1 | ENSMUSG00000024661 | protein_coding | ferritin heavy polypeptide 1 | 6.177E-08 | 1.443632369 | 0.931 | 0.597 | 0.001458576 | EN_3 |
| Gm47283 | ENSMUSG00000096768 | lncRNA | predicted gene, 47283 | 7.391E-08 | 3.070189091 | 0.5 | 0.097 | 0.001745157 | EN_3 |
| Dbi | ENSMUSG00000026385 | protein_coding | diazepam binding inhibitor | 2.549E-15 | 2.28377273 | 0.963 | 0.884 | 6.01768E-11 | EN_4 |
| G0s2 | ENSMUSG00000009633 | protein_coding | G0/G1 switch gene 2 | 2.446E-08 | -1.766706823 | 0.512 | 0.797 | 0.000577552 | EN_4 |
| Actb | ENSMUSG00000029580 | protein_coding | actin, beta | 6.867E-10 | 0.904654483 | 0.988 | 0.942 | 1.62137E-05 | EN_4 |
| Actg2 | ENSMUSG00000059430 | protein_coding | actin, gamma 2, smooth muscle, enteric | 1.904E-07 | 2.737285913 | 0.598 | 0.232 | 0.004495601 | EN_4 |
| Rpl32 | ENSMUSG00000057841 | protein_coding | ribosomal protein L32 | 9.011E-08 | -0.947799509 | 0.854 | 0.971 | 0.002127665 | EN_4 |
| Tagln | ENSMUSG00000032085 | protein_coding | transgelin | 2.834E-08 | 1.801281432 | 0.793 | 0.493 | 0.000669203 | EN_4 |
| Ckb | ENSMUSG00000001270 | protein_coding | creatine kinase, brain | 1.486E-11 | 1.777672343 | 0.89 | 0.594 | 3.50908E-07 | EN_4 |
| Mylk | ENSMUSG00000022836 | protein_coding | myosin, light polypeptide kinase | 2.708E-07 | 2.787615683 | 0.512 | 0.116 | 0.006394416 | EN_4 |
| Cox5b | ENSMUSG00000061518 | protein_coding | cytochrome c oxidase subunit 5B | 4.738E-14 | 1.113332654 | 0.93 | 0.702 | 1.11882E-09 | MM_1 |
| Slc40a1 | ENSMUSG00000025993 | protein_coding | solute carrier family 40 (iron-regulated transporter), member 1 | 4.551E-10 | 1.469624028 | 0.634 | 0.246 | 1.07452E-05 | MM_1 |
| Hspe1 | ENSMUSG00000073676 | protein_coding | heat shock protein 1 (chaperonin 10) | 1.284E-12 | 1.143518079 | 0.953 | 0.754 | 3.03099E-08 | MM_1 |
| Ndufb3 | ENSMUSG00000026032 | protein_coding | NADH:ubiquinone oxidoreductase subunit B3 | 5.28E-09 | 0.780790965 | 0.887 | 0.64 | 0.00012466 | MM_1 |
| Slc11a1 | ENSMUSG00000026177 | protein_coding | solute carrier family 11 (proton-coupled divalent metal ion transporters), member 1 | 5.237E-08 | 0.786885643 | 0.869 | 0.64 | 0.001236558 | MM_1 |
| Des | ENSMUSG00000026208 | protein_coding | desmin | 1.425E-09 | 0.595121488 | 0.958 | 0.693 | 3.36383E-05 | MM_1 |
| Per2 | ENSMUSG00000055866 | protein_coding | period circadian clock 2 | 6.322E-19 | -4.221717829 | 0.023 | 0.395 | 1.49272E-14 | MM_1 |
| Cops9 | ENSMUSG00000073616 | protein_coding | COP9 signalosome subunit 9 | 1.366E-10 | 0.933708416 | 0.859 | 0.561 | 3.22647E-06 | MM_1 |
| Fam174a | ENSMUSG00000051185 | protein_coding | family with sequence similarity 174, member A | 1.817E-08 | 0.979980072 | 0.798 | 0.439 | 0.000428934 | MM_1 |
| Dbi | ENSMUSG00000026385 | protein_coding | diazepam binding inhibitor | 8.377E-12 | 0.932184268 | 0.948 | 0.667 | 1.978E-07 | MM_1 |
| Adipor1 | ENSMUSG00000026457 | protein_coding | adiponectin receptor 1 | 1.456E-07 | 0.961127764 | 0.845 | 0.561 | 0.003437194 | MM_1 |
| Gm2000 | ENSMUSG00000078193 | protein_coding | predicted gene 2000 | 8.903E-21 | 2.134286107 | 0.789 | 0.281 | 2.10216E-16 | MM_1 |
| Creg1 | ENSMUSG00000040713 | protein_coding | cellular repressor of E1A-stimulated genes 1 | 4.854E-10 | 0.769636477 | 0.939 | 0.746 | 1.14624E-05 | MM_1 |
| Sdhc | ENSMUSG00000058076 | protein_coding | succinate dehydrogenase complex, subunit C, integral membrane protein | 2.112E-09 | 1.097964135 | 0.751 | 0.386 | 4.98667E-05 | MM_1 |
| Ndufs2 | ENSMUSG00000013593 | protein_coding | NADH:ubiquinone oxidoreductase core subunit S2 | 3.673E-07 | 0.807650323 | 0.704 | 0.36 | 0.008673117 | MM_1 |
| Pex19 | ENSMUSG00000003464 | protein_coding | peroxisomal biogenesis factor 19 | 4.801E-08 | 2.309197961 | 0.338 | 0.061 | 0.001133707 | MM_1 |
| Nenf | ENSMUSG00000037499 | protein_coding | neuron derived neurotrophic factor | 8.548E-09 | 0.907418778 | 0.831 | 0.482 | 0.000201836 | MM_1 |
| Phyh | ENSMUSG00000026664 | protein_coding | phytanoyl-CoA hydroxylase | 2.936E-07 | 1.394574144 | 0.502 | 0.202 | 0.006931388 | MM_1 |
| Camk1d | ENSMUSG00000039145 | protein_coding | calcium/calmodulin-dependent protein kinase ID | 1.307E-15 | -0.945509238 | 0.995 | 1 | 3.0867E-11 | MM_1 |
| Dpp7 | ENSMUSG00000026958 | protein_coding | dipeptidylpeptidase 7 | 1.022E-13 | 1.649344707 | 0.685 | 0.211 | 2.41275E-09 | MM_1 |
| Tmem141 | ENSMUSG00000026939 | protein_coding | transmembrane protein 141 | 2.684E-08 | 1.305319698 | 0.62 | 0.272 | 0.000633689 | MM_1 |
| Fcna | ENSMUSG00000026938 | protein_coding | ficolin A | 3.244E-19 | 1.396465701 | 0.986 | 0.807 | 7.66019E-15 | MM_1 |
| Cdk9 | ENSMUSG00000009555 | protein_coding | cyclin-dependent kinase 9 (CDC2-related kinase) | 1.734E-07 | 1.181290831 | 0.681 | 0.333 | 0.004095435 | MM_1 |
| Stxbp1 | ENSMUSG00000026797 | protein_coding | syntaxin binding protein 1 | 5.197E-08 | 1.048477532 | 0.488 | 0.158 | 0.001227061 | MM_1 |
| Gsn | ENSMUSG00000026879 | protein_coding | gelsolin | 1.039E-11 | 1.013491807 | 0.878 | 0.491 | 2.45323E-07 | MM_1 |
| Rpl35 | ENSMUSG00000062997 | protein_coding | ribosomal protein L35 | 5.555E-13 | 0.712956591 | 0.977 | 0.868 | 1.31155E-08 | MM_1 |
| Tanc1 | ENSMUSG00000035168 | protein_coding | tetratricopeptide repeat, ankyrin repeat and coiled-coil containing 1 | 1.273E-07 | 2.24794665 | 0.338 | 0.07 | 0.00300518 | MM_1 |
| Atp5g3 | ENSMUSG00000018770 | protein_coding | ATP synthase, H+ transporting, mitochondrial F0 complex, subunit C3 (subunit 9) | 2.201E-14 | 1.02416928 | 0.934 | 0.605 | 5.19688E-10 | MM_1 |
| Ndufs3 | ENSMUSG00000005510 | protein_coding | NADH:ubiquinone oxidoreductase core subunit S3 | 3.301E-07 | 1.011967009 | 0.638 | 0.307 | 0.007794506 | MM_1 |
| Psmc3 | ENSMUSG00000002102 | protein_coding | proteasome (prosome, macropain) 26S subunit, ATPase 3 | 5.405E-09 | 0.755117599 | 0.779 | 0.404 | 0.000127632 | MM_1 |
| Lmo2 | ENSMUSG00000032698 | protein_coding | LIM domain only 2 | 3.366E-10 | 0.918034236 | 0.897 | 0.649 | 7.94676E-06 | MM_1 |
| Pdia3 | ENSMUSG00000027248 | protein_coding | protein disulfide isomerase associated 3 | 1.35E-14 | 0.944712755 | 0.981 | 0.816 | 3.18688E-10 | MM_1 |
| B2m | ENSMUSG00000060802 | protein_coding | beta-2 microglobulin | 3.319E-21 | 0.805823456 | 0.995 | 0.982 | 7.83789E-17 | MM_1 |
| Blvra | ENSMUSG00000001999 | protein_coding | biliverdin reductase A | 5.443E-10 | 1.10533018 | 0.761 | 0.351 | 1.2853E-05 | MM_1 |
| Snrpb | ENSMUSG00000027404 | protein_coding | small nuclear ribonucleoprotein B | 2.271E-08 | 0.61193026 | 0.944 | 0.719 | 0.000536266 | MM_1 |
| Idh3b | ENSMUSG00000027406 | protein_coding | isocitrate dehydrogenase 3 (NAD+) beta | 1.71E-07 | 1.147024989 | 0.582 | 0.246 | 0.004036944 | MM_1 |
| Dstn | ENSMUSG00000015932 | protein_coding | destrin | 9.845E-10 | 0.621903307 | 0.981 | 0.842 | 2.3247E-05 | MM_1 |
| Rin2 | ENSMUSG00000001768 | protein_coding | Ras and Rab interactor 2 | 2.745E-13 | 0.936911946 | 0.977 | 0.746 | 6.48224E-09 | MM_1 |
| Cst3 | ENSMUSG00000027447 | protein_coding | cystatin C | 1.334E-18 | 0.985576042 | 1 | 0.956 | 3.1501E-14 | MM_1 |
| Abhd12 | ENSMUSG00000032046 | protein_coding | abhydrolase domain containing 12 | 3.011E-07 | 0.601324802 | 0.911 | 0.711 | 0.007110744 | MM_1 |
| Fkbp1a | ENSMUSG00000032966 | protein_coding | FK506 binding protein 1a | 1.472E-10 | 0.679361319 | 0.972 | 0.781 | 3.47479E-06 | MM_1 |
| Ergic3 | ENSMUSG00000005881 | protein_coding | ERGIC and golgi 3 | 1.56E-10 | 1.265987218 | 0.761 | 0.333 | 3.68245E-06 | MM_1 |
| Romo1 | ENSMUSG00000067847 | protein_coding | reactive oxygen species modulator 1 | 1.611E-12 | 0.897722798 | 0.92 | 0.632 | 3.80289E-08 | MM_1 |
| Scand1 | ENSMUSG00000046229 | protein_coding | SCAN domain-containing 1 | 2.626E-07 | 0.79754259 | 0.854 | 0.675 | 0.006199823 | MM_1 |
| Myl9 | ENSMUSG00000067818 | protein_coding | myosin, light polypeptide 9, regulatory | 1.335E-11 | 0.842909257 | 0.784 | 0.377 | 3.15135E-07 | MM_1 |
| Rpn2 | ENSMUSG00000027642 | protein_coding | ribophorin II | 7.45E-08 | 0.740016547 | 0.84 | 0.535 | 0.001759125 | MM_1 |
| Gm826 | ENSMUSG00000074623 | protein_coding | predicted gene 826 | 2.918E-07 | -2.404180072 | 0.08 | 0.289 | 0.006889972 | MM_1 |
| Ctsa | ENSMUSG00000017760 | protein_coding | cathepsin A | 2.037E-19 | 0.953773099 | 1 | 0.825 | 4.80877E-15 | MM_1 |
| Pltp | ENSMUSG00000017754 | protein_coding | phospholipid transfer protein | 5.232E-15 | 0.865007833 | 0.995 | 0.877 | 1.23532E-10 | MM_1 |
| Pcif1 | ENSMUSG00000039849 | protein_coding | PDX1 C-terminal inhibiting factor 1 | 2.969E-08 | 1.026043921 | 0.587 | 0.237 | 0.00070115 | MM_1 |
| Sulf2 | ENSMUSG00000006800 | protein_coding | sulfatase 2 | 4.191E-08 | 0.653658485 | 0.977 | 0.763 | 0.000989543 | MM_1 |
| 5031425F14Rik | ENSMUSG00000085129 | lncRNA | RIKEN cDNA 5031425F14 gene | 5.052E-09 | 1.594715064 | 0.526 | 0.158 | 0.000119278 | MM_1 |
| Ube2v1 | ENSMUSG00000078923 | protein_coding | ubiquitin-conjugating enzyme E2 variant 1 | 1.83E-08 | 0.883049526 | 0.718 | 0.333 | 0.00043219 | MM_1 |
| Tmem189 | ENSMUSG00000090213 | protein_coding | transmembrane protein 189 | 1.129E-08 | 0.950197634 | 0.826 | 0.491 | 0.000266677 | MM_1 |
| Gnas | ENSMUSG00000027523 | protein_coding | GNAS (guanine nucleotide binding protein, alpha stimulating) complex locus | 6.107E-15 | 0.943762801 | 0.972 | 0.833 | 1.44207E-10 | MM_1 |
| Ctsz | ENSMUSG00000016256 | protein_coding | cathepsin Z | 4.296E-18 | 0.982524632 | 0.986 | 0.895 | 1.0144E-13 | MM_1 |
| Atp5e | ENSMUSG00000016252 | protein_coding | ATP synthase, H+ transporting, mitochondrial F1 complex, epsilon subunit | 2.6E-14 | 0.941608129 | 0.944 | 0.798 | 6.13854E-10 | MM_1 |
| Gyg | ENSMUSG00000019528 | protein_coding | glycogenin | 8.997E-09 | 1.64549429 | 0.559 | 0.228 | 0.000212436 | MM_1 |
| Sec62 | ENSMUSG00000027706 | protein_coding | SEC62 homolog (S. cerevisiae) | 1.812E-08 | 0.600838847 | 0.953 | 0.746 | 0.000427861 | MM_1 |
| Dnajc19 | ENSMUSG00000027679 | protein_coding | DnaJ heat shock protein family (Hsp40) member C19 | 2.978E-07 | 0.920899492 | 0.732 | 0.421 | 0.00703177 | MM_1 |
| Anxa5 | ENSMUSG00000027712 | protein_coding | annexin A5 | 1.024E-10 | 0.678644537 | 0.995 | 0.912 | 2.41772E-06 | MM_1 |
| Ndufc1 | ENSMUSG00000037152 | protein_coding | NADH:ubiquinone oxidoreductase subunit C1 | 1.142E-14 | 1.352749161 | 0.854 | 0.491 | 2.69735E-10 | MM_1 |
| P2ry13 | ENSMUSG00000036362 | protein_coding | purinergic receptor P2Y, G-protein coupled 13 | 2.683E-11 | 1.413079856 | 0.69 | 0.281 | 6.33413E-07 | MM_1 |
| Mfsd1 | ENSMUSG00000027775 | protein_coding | major facilitator superfamily domain containing 1 | 5.764E-10 | 0.861991836 | 0.92 | 0.675 | 1.36111E-05 | MM_1 |
| Glmp | ENSMUSG00000001418 | protein_coding | glycosylated lysosomal membrane protein | 3.068E-08 | 0.959656761 | 0.746 | 0.456 | 0.000724302 | MM_1 |
| Sema4a | ENSMUSG00000028064 | protein_coding | sema domain, immunoglobulin domain (Ig), transmembrane domain (TM) and short cytoplasmic domain, (semaphorin) 4A | 2.347E-09 | 0.791601323 | 0.925 | 0.632 | 5.54193E-05 | MM_1 |
| Rit1 | ENSMUSG00000028057 | protein_coding | Ras-like without CAAX 1 | 1.041E-07 | 1.218914278 | 0.526 | 0.193 | 0.00245867 | MM_1 |
| Dpm3 | ENSMUSG00000042737 | protein_coding | dolichyl-phosphate mannosyltransferase polypeptide 3 | 1.853E-09 | 1.111349317 | 0.793 | 0.491 | 4.37644E-05 | MM_1 |
| Snapin | ENSMUSG00000001018 | protein_coding | SNAP-associated protein | 7.981E-12 | 1.656325176 | 0.62 | 0.193 | 1.8845E-07 | MM_1 |
| S100a1 | ENSMUSG00000044080 | protein_coding | S100 calcium binding protein A1 | 1.672E-07 | 1.155177674 | 0.671 | 0.404 | 0.003948853 | MM_1 |

| | | | | | | | | | |
|---|---|---|---|---|---|---|---|---|---|
| S100a11 | ENSMUSG00000027907 | protein_coding | S100 calcium binding protein A11 | 7.593E-08 | 0.720304097 | 0.869 | 0.57 | 0.001792752 | MM_1 |
| Psmb4 | ENSMUSG00000005779 | protein_coding | proteasome (prosome, macropain) subunit, beta type 4 | 4.455E-10 | 0.980597815 | 0.897 | 0.649 | 1.05181E-05 | MM_1 |
| Ctss | ENSMUSG00000038642 | protein_coding | cathepsin S | 3.32E-14 | 0.63092017 | 1 | 0.939 | 7.83869E-10 | MM_1 |
| Ecm1 | ENSMUSG00000028108 | protein_coding | extracellular matrix protein 1 | 2.697E-07 | 0.681946136 | 0.981 | 0.825 | 0.006367095 | MM_1 |
| Txnip | ENSMUSG00000038393 | protein_coding | thioredoxin interacting protein | 3.995E-08 | 0.640907495 | 0.967 | 0.842 | 0.000943193 | MM_1 |
| Gstm1 | ENSMUSG00000058135 | protein_coding | glutathione S-transferase, mu 1 | 6.886E-12 | 1.015506248 | 0.887 | 0.491 | 1.62581E-07 | MM_1 |
| Sars | ENSMUSG00000068739 | protein_coding | seryl-aminoacyl-tRNA synthetase | 4.069E-07 | 1.010755371 | 0.615 | 0.325 | 0.009607037 | MM_1 |
| S1pr1 | ENSMUSG00000045092 | protein_coding | sphingosine-1-phosphate receptor 1 | 1.677E-07 | 0.853462547 | 0.817 | 0.465 | 0.00396026 | MM_1 |
| 1810037I17Rik | ENSMUSG00000054091 | protein_coding | RIKEN cDNA 1810037I17 gene | 4.881E-10 | 0.89965404 | 0.897 | 0.667 | 1.15246E-05 | MM_1 |
| Dnajb14 | ENSMUSG00000074212 | protein_coding | DnaJ heat shock protein family (Hsp40) member B14 | 1.021E-07 | 0.907388061 | 0.761 | 0.465 | 0.002410859 | MM_1 |
| Selenof | ENSMUSG00000037072 | protein_coding | selenoprotein F | 1.626E-10 | 0.691690937 | 0.967 | 0.807 | 3.83931E-06 | MM_1 |
| Gng5 | ENSMUSG00000068523 | protein_coding | guanine nucleotide binding protein (G protein), gamma 5 | 1.501E-12 | 0.682786716 | 0.981 | 0.842 | 3.54458E-08 | MM_1 |
| Wls | ENSMUSG00000028173 | protein_coding | wntless WNT ligand secretion mediator | 2.249E-07 | 0.846338504 | 0.859 | 0.614 | 0.00531019 | MM_1 |
| Gm11808 | ENSMUSG00000068240 | protein_coding | predicted gene 11808 | 4.818E-25 | 2.904512779 | 0.775 | 0.149 | 1.13774E-20 | MM_1 |
| Wwp1 | ENSMUSG00000041058 | protein_coding | WW domain containing E3 ubiquitin protein ligase 1 | 2.172E-12 | 0.593055958 | 1 | 0.974 | 5.12812E-08 | MM_1 |
| Ndufb6 | ENSMUSG00000071014 | protein_coding | NADH:ubiquinone oxidoreductase subunit B6 | 2.493E-12 | 1.383417551 | 0.737 | 0.325 | 5.88657E-08 | MM_1 |
| Bag1 | ENSMUSG00000028416 | protein_coding | BCL2-associated athanogene 1 | 2.07E-08 | 0.769101161 | 0.854 | 0.465 | 0.000488815 | MM_1 |
| Ccl21a | ENSMUSG00000094686 | protein_coding | chemokine (C-C motif) ligand 21A (serine) | 1.693E-14 | 3.710164191 | 0.441 | 0.026 | 3.99786E-10 | MM_1 |
| Tpm2 | ENSMUSG00000028464 | protein_coding | tropomyosin 2, beta | 1.947E-09 | 0.745932678 | 0.953 | 0.693 | 4.59689E-05 | MM_1 |
| Txn1 | ENSMUSG00000028367 | protein_coding | thioredoxin 1 | 2.428E-10 | 0.806277067 | 0.944 | 0.693 | 5.7329E-06 | MM_1 |
| Atp6v1g1 | ENSMUSG00000039105 | protein_coding | ATPase, H+ transporting, lysosomal V1 subunit G1 | 1.907E-10 | 0.769125055 | 0.967 | 0.781 | 4.50276E-06 | MM_1 |
| Plin2 | ENSMUSG00000028494 | protein_coding | perilipin 2 | 5.155E-20 | 1.446844936 | 0.977 | 0.658 | 1.21709E-15 | MM_1 |
| Hacd4 | ENSMUSG00000028497 | protein_coding | 3-hydroxyacyl-CoA dehydratase 4 | 2.782E-07 | 0.64824087 | 0.958 | 0.754 | 0.006569792 | MM_1 |
| Akr1a1 | ENSMUSG00000028692 | protein_coding | aldo-keto reductase family 1, member A1 (aldehyde reductase) | 1.657E-21 | 1.076501227 | 1 | 0.816 | 3.9121E-17 | MM_1 |
| Prdx1 | ENSMUSG00000028691 | protein_coding | peroxiredoxin 1 | 1.6E-19 | 1.079586511 | 0.995 | 0.877 | 3.77815E-15 | MM_1 |
| Zmpste24 | ENSMUSG00000043207 | protein_coding | zinc metallopeptidase, STE24 | 6.825E-08 | 1.458921491 | 0.549 | 0.202 | 0.001611582 | MM_1 |
| Ndufs5 | ENSMUSG00000028648 | protein_coding | NADH:ubiquinone oxidoreductase core subunit S5 | 1.221E-09 | 0.910178838 | 0.864 | 0.57 | 2.8829E-05 | MM_1 |
| Fhl3 | ENSMUSG00000032643 | protein_coding | four and a half LIM domains 3 | 8.964E-08 | 1.356893461 | 0.413 | 0.114 | 0.002116691 | MM_1 |
| Psmb2 | ENSMUSG00000028837 | protein_coding | proteasome (prosome, macropain) subunit, beta type 2 | 2.051E-07 | 0.769804728 | 0.85 | 0.544 | 0.004842133 | MM_1 |
| Ptp4a2 | ENSMUSG00000028788 | protein_coding | protein tyrosine phosphatase 4a2 | 1.703E-12 | 0.618033357 | 0.995 | 0.921 | 4.02201E-08 | MM_1 |
| Atpif1 | ENSMUSG00000054428 | protein_coding | ATPase inhibitory factor 1 | 9.384E-11 | 0.779523703 | 0.981 | 0.781 | 2.21575E-06 | MM_1 |
| Ptafr | ENSMUSG00000056529 | protein_coding | platelet-activating factor receptor | 4.516E-11 | 0.743624487 | 0.991 | 0.93 | 1.06621E-06 | MM_1 |
| Themis2 | ENSMUSG00000037731 | protein_coding | thymocyte selection associated family member 2 | 5.514E-08 | 1.210350195 | 0.671 | 0.342 | 0.001302079 | MM_1 |
| Man1c1 | ENSMUSG00000037306 | protein_coding | mannosidase, alpha, class 1C, member 1 | 1.671E-18 | 1.458478659 | 0.948 | 0.64 | 3.94623E-14 | MM_1 |
| C1qb | ENSMUSG00000036905 | protein_coding | complement component 1, q subcomponent, beta polypeptide | 1.954E-12 | 0.654052359 | 1 | 0.974 | 4.61433E-08 | MM_1 |
| Micos10 | ENSMUSG00000050608 | protein_coding | mitochondrial contact site and cristae organizing system subunit 10 | 1.822E-08 | 0.585186462 | 0.925 | 0.754 | 0.000430315 | MM_1 |
| Dhrs3 | ENSMUSG00000066026 | protein_coding | dehydrogenase/reductase (SDR family) member 3 | 5.908E-19 | 1.122128903 | 0.962 | 0.798 | 1.39502E-14 | MM_1 |
| Agtrap | ENSMUSG00000029007 | protein_coding | angiotensin II, type I receptor-associated protein | 1.634E-08 | 1.115343699 | 0.718 | 0.368 | 0.000385922 | MM_1 |
| Eno1 | ENSMUSG00000063524 | protein_coding | enolase 1, alpha non-neuron | 9.661E-17 | 1.610031632 | 0.826 | 0.421 | 2.28113E-12 | MM_1 |
| Per3 | ENSMUSG00000028957 | protein_coding | period circadian clock 3 | 7.161E-08 | -2.754304096 | 0.099 | 0.316 | 0.001690915 | MM_1 |
| Prxl2b | ENSMUSG00000029059 | protein_coding | peroxiredoxin like 2B | 5.655E-08 | 0.884287122 | 0.784 | 0.491 | 0.001335246 | MM_1 |
| Cd36 | ENSMUSG00000002944 | protein_coding | CD36 molecule | 2.509E-23 | 0.982812417 | 1 | 0.965 | 5.92426E-19 | MM_1 |
| Psmc2 | ENSMUSG00000028932 | protein_coding | proteasome (prosome, macropain) 26S subunit, ATPase 2 | 9.136E-09 | 1.058686712 | 0.7 | 0.377 | 0.000215725 | MM_1 |
| Tomm7 | ENSMUSG00000028998 | protein_coding | translocase of outer mitochondrial membrane 7 | 6.422E-08 | 0.643284731 | 0.977 | 0.719 | 0.00151639 | MM_1 |
| Ost4 | ENSMUSG00000038803 | protein_coding | oligosaccharyltransferase complex subunit 4 (non-catalytic) | 5.001E-14 | 1.027656851 | 0.93 | 0.649 | 1.18075E-09 | MM_1 |
| Man2b2 | ENSMUSG00000029119 | protein_coding | mannosidase 2, alpha B2 | 3.015E-08 | 1.39872556 | 0.446 | 0.123 | 0.000711927 | MM_1 |
| Wdr1 | ENSMUSG00000005103 | protein_coding | WD repeat domain 1 | 1.327E-07 | 0.695735263 | 0.906 | 0.64 | 0.0031343 | MM_1 |
| Med28 | ENSMUSG00000015804 | protein_coding | mediator complex subunit 28 | 1.822E-09 | 1.109943038 | 0.784 | 0.439 | 4.30201E-05 | MM_1 |
| Igfbp7 | ENSMUSG00000036256 | protein_coding | insulin-like growth factor binding protein 7 | 2.099E-10 | 2.794772534 | 0.385 | 0.053 | 4.95526E-06 | MM_1 |
| Pf4 | ENSMUSG00000029373 | protein_coding | platelet factor 4 | 2.577E-16 | 0.904857023 | 1 | 0.912 | 6.08411E-12 | MM_1 |
| Scarb2 | ENSMUSG00000029426 | protein_coding | scavenger receptor class B, member 2 | 1.199E-08 | 0.997096466 | 0.864 | 0.57 | 0.000283048 | MM_1 |
| Hnrnpd | ENSMUSG00000000568 | protein_coding | heterogeneous nuclear ribonucleoprotein D | 1.232E-07 | 0.657762592 | 0.883 | 0.596 | 0.002908895 | MM_1 |
| Hpse | ENSMUSG00000035273 | protein_coding | heparanase | 6.203E-09 | 1.069872686 | 0.709 | 0.368 | 0.000146456 | MM_1 |
| Atp5k | ENSMUSG00000050856 | protein_coding | ATP synthase, H+ transporting, mitochondrial F1F0 complex, subunit E | 1.523E-15 | 1.16038467 | 0.93 | 0.623 | 3.59614E-11 | MM_1 |
| Cox6a1 | ENSMUSG00000041697 | protein_coding | cytochrome c oxidase subunit 6A1 | 1.186E-18 | 1.200802197 | 0.953 | 0.702 | 2.80067E-14 | MM_1 |
| Pebp1 | ENSMUSG00000032959 | protein_coding | phosphatidylethanolamine binding protein 1 | 6.212E-11 | 1.088061463 | 0.854 | 0.535 | 1.46684E-06 | MM_1 |
| Tpcn1 | ENSMUSG00000032741 | protein_coding | two pore channel 1 | 2.532E-07 | 0.701509776 | 0.854 | 0.544 | 0.005977438 | MM_1 |
| Rpl6 | ENSMUSG00000029614 | protein_coding | ribosomal protein L6 | 8.021E-11 | 0.588278254 | 1 | 0.904 | 1.89401E-06 | MM_1 |
| Aldh2 | ENSMUSG00000029455 | protein_coding | aldehyde dehydrogenase 2, mitochondrial | 2.515E-12 | 0.808411359 | 0.962 | 0.825 | 5.93825E-08 | MM_1 |
| Hvcn1 | ENSMUSG00000064267 | protein_coding | hydrogen voltage-gated channel 1 | 2.751E-12 | 1.99440887 | 0.516 | 0.105 | 6.4953E-08 | MM_1 |
| Ccl26 | ENSMUSG00000070464 | protein_coding | chemokine (C-C motif) ligand 26 | 3.383E-08 | 4.152255249 | 0.23 | 0 | 0.000798848 | MM_1 |
| Hspb1 | ENSMUSG00000004951 | protein_coding | heat shock protein 1 | 1.098E-10 | 1.397205022 | 0.775 | 0.368 | 2.59152E-06 | MM_1 |
| Znhit1 | ENSMUSG00000059518 | protein_coding | zinc finger, HIT domain containing 1 | 4.097E-10 | 1.388129574 | 0.62 | 0.228 | 9.67421E-06 | MM_1 |
| Tsc22d4 | ENSMUSG00000029723 | protein_coding | TSC22 domain family, member 4 | 3.048E-08 | 0.778603383 | 0.892 | 0.684 | 0.000719592 | MM_1 |
| Actb | ENSMUSG00000029580 | protein_coding | actin, beta | 1.239E-23 | 0.732814843 | 1 | 1 | 2.92597E-19 | MM_1 |
| Atp5j2 | ENSMUSG00000038690 | protein_coding | ATP synthase, H+ transporting, mitochondrial F0 complex, subunit F2 | 4.889E-14 | 0.998600611 | 0.948 | 0.728 | 1.15447E-09 | MM_1 |
| Cdk8 | ENSMUSG00000029635 | protein_coding | cyclin-dependent kinase 8 | 3.873E-48 | -3.147047071 | 0.991 | 0.991 | 9.14493E-44 | MM_1 |
| Alox5ap | ENSMUSG00000060063 | protein_coding | arachidonate 5-lipoxygenase activating protein | 7.218E-12 | 0.81942282 | 0.981 | 0.851 | 1.70438E-07 | MM_1 |
| Ndufa4 | ENSMUSG00000029632 | protein_coding | Ndufa4, mitochondrial complex associated | 4.148E-11 | 0.665168629 | 0.977 | 0.816 | 9.7935E-07 | MM_1 |
| Ndufa5 | ENSMUSG00000023089 | protein_coding | NADH:ubiquinone oxidoreductase subunit A5 | 1.632E-16 | 1.418591167 | 0.864 | 0.447 | 3.85356E-12 | MM_1 |
| Atp6v1f | ENSMUSG00000004285 | protein_coding | ATPase, H+ transporting, lysosomal V1 subunit F | 2.876E-09 | 0.653036717 | 0.981 | 0.789 | 6.7906E-05 | MM_1 |
| Mkrn1 | ENSMUSG00000029922 | protein_coding | makorin, ring finger protein, 1 | 6.929E-08 | 0.828367432 | 0.709 | 0.325 | 0.001636003 | MM_1 |
| Ndufb2 | ENSMUSG00000002416 | protein_coding | NADH:ubiquinone oxidoreductase subunit B2 | 2.541E-08 | 1.076084114 | 0.746 | 0.482 | 0.000600085 | MM_1 |
| Mrps33 | ENSMUSG00000029918 | protein_coding | mitochondrial ribosomal protein S33 | 2.416E-07 | 0.90988834 | 0.77 | 0.43 | 0.005705168 | MM_1 |
| Rarres2 | ENSMUSG00000009281 | protein_coding | retinoic acid receptor responder (tazarotene induced) 2 | 9.891E-12 | 2.083315272 | 0.455 | 0.07 | 2.33551E-07 | MM_1 |
| Hpgds | ENSMUSG00000029919 | protein_coding | hematopoietic prostaglandin D synthase | 8.116E-09 | 0.639238762 | 0.958 | 0.737 | 0.000191638 | MM_1 |
| St3gal5 | ENSMUSG00000056091 | protein_coding | ST3 beta-galactoside alpha-2,3-sialyltransferase 5 | 1.835E-15 | 2.180814183 | 0.723 | 0.289 | 4.33226E-11 | MM_1 |
| Vamp8 | ENSMUSG00000050732 | protein_coding | vesicle-associated membrane protein 8 | 1.849E-13 | 0.821590974 | 0.972 | 0.737 | 4.36518E-09 | MM_1 |
| Htra2 | ENSMUSG00000068329 | protein_coding | HtrA serine peptidase 2 | 1.93E-08 | 1.53864392 | 0.54 | 0.202 | 0.000455646 | MM_1 |
| Actg2 | ENSMUSG00000059430 | protein_coding | actin, gamma 2, smooth muscle, enteric | 2.556E-13 | 0.968283348 | 0.934 | 0.614 | 6.03612E-09 | MM_1 |
| Nagk | ENSMUSG00000034744 | protein_coding | N-acetylglucosamine kinase | 1.551E-07 | 0.848121176 | 0.746 | 0.412 | 0.003661578 | MM_1 |
| Rab11fip5 | ENSMUSG00000051343 | protein_coding | RAB11 family interacting protein 5 (class I) | 4.551E-09 | 0.756457303 | 0.911 | 0.658 | 0.000107449 | MM_1 |
| Camk1 | ENSMUSG00000030272 | protein_coding | calcium/calmodulin-dependent protein kinase I | 2.44E-09 | 0.673354935 | 0.944 | 0.754 | 5.76048E-05 | MM_1 |
| Arpc4 | ENSMUSG00000079426 | protein_coding | actin related protein 2/3 complex, subunit 4 | 2.28E-11 | 0.813886089 | 0.958 | 0.702 | 5.38265E-07 | MM_1 |
| Lpcat3 | ENSMUSG00000004270 | protein_coding | lysophosphatidylcholine acyltransferase 3 | 2.404E-07 | 1.168679927 | 0.465 | 0.149 | 0.005676469 | MM_1 |
| Grcc10 | ENSMUSG00000072772 | protein_coding | gene rich cluster, C10 gene | 1.078E-09 | 0.881022406 | 0.892 | 0.579 | 2.54481E-05 | MM_1 |
| Gapdh | ENSMUSG00000057666 | protein_coding | glyceraldehyde-3-phosphate dehydrogenase | 5.159E-19 | 1.232887695 | 0.967 | 0.746 | 1.21817E-14 | MM_1 |
| Ltbr | ENSMUSG00000030339 | protein_coding | lymphotoxin B receptor | 2.493E-07 | 1.079617337 | 0.526 | 0.193 | 0.005886102 | MM_1 |
| Gabarapl1 | ENSMUSG00000030161 | protein_coding | gamma-aminobutyric acid (GABA) A receptor-associated protein-like 1 | 3.761E-17 | 1.702623328 | 0.836 | 0.351 | 8.881E-13 | MM_1 |
| Klra3 | ENSMUSG00000067591 | protein_coding | killer cell lectin-like receptor, subfamily A, member 3 | 7.543E-09 | 4.871049499 | 0.249 | 0 | 0.000178098 | MM_1 |
| Mgp | ENSMUSG00000030218 | protein_coding | matrix Gla protein | 4.389E-08 | 1.776538573 | 0.376 | 0.079 | 0.001036254 | MM_1 |
| Mgst1 | ENSMUSG00000008540 | protein_coding | microsomal glutathione S-transferase 1 | 5.088E-13 | 1.358846163 | 0.85 | 0.474 | 1.20149E-08 | MM_1 |
| Ndufa3 | ENSMUSG00000035674 | protein_coding | NADH:ubiquinone oxidoreductase subunit A3 | 9.945E-21 | 1.494665226 | 0.925 | 0.474 | 2.34822E-16 | MM_1 |
| Chmp2a | ENSMUSG00000033916 | protein_coding | charged multivesicular body protein 2A | 1.426E-10 | 0.972072193 | 0.878 | 0.623 | 3.36807E-06 | MM_1 |
| Selenow | ENSMUSG00000041571 | protein_coding | selenoprotein W | 2.249E-12 | 0.933104576 | 0.944 | 0.684 | 5.30951E-08 | MM_1 |
| Tmem160 | ENSMUSG00000019158 | protein_coding | transmembrane protein 160 | 2.595E-07 | 1.071703499 | 0.681 | 0.36 | 0.006126636 | MM_1 |
| Ap2s1 | ENSMUSG00000008036 | protein_coding | adaptor-related protein complex 2, sigma 1 subunit | 1.16E-09 | 0.989191062 | 0.84 | 0.509 | 2.73785E-05 | MM_1 |
| Dmpk | ENSMUSG00000030409 | protein_coding | dystrophia myotonica-protein kinase | 1.002E-10 | 1.928986275 | 0.521 | 0.14 | 2.36619E-06 | MM_1 |
| Snrpd2 | ENSMUSG00000040824 | protein_coding | small nuclear ribonucleoprotein D2 | 7.37E-10 | 0.942953654 | 0.845 | 0.579 | 1.74017E-05 | MM_1 |
| Clptm1 | ENSMUSG00000002981 | protein_coding | cleft lip and palate associated transmembrane protein 1 | 3.986E-10 | 1.191181057 | 0.685 | 0.281 | 9.41062E-06 | MM_1 |
| Gsk3a | ENSMUSG00000057177 | protein_coding | glycogen synthase kinase 3 alpha | 1.051E-07 | 1.038170059 | 0.577 | 0.211 | 0.002481072 | MM_1 |
| Mrps12 | ENSMUSG00000045948 | protein_coding | mitochondrial ribosomal protein S12 | 3.965E-07 | 1.123646398 | 0.531 | 0.202 | 0.009363276 | MM_1 |
| Sirt2 | ENSMUSG00000015149 | protein_coding | sirtuin 2 | 5.128E-09 | 0.796133576 | 0.85 | 0.561 | 0.000121071 | MM_1 |
| Rinl | ENSMUSG00000051735 | protein_coding | Ras and Rab interactor-like | 4.507E-08 | 1.933051988 | 0.376 | 0.079 | 0.001064098 | MM_1 |
| Hnrnpl | ENSMUSG00000015165 | protein_coding | heterogeneous nuclear ribonucleoprotein L | 1.321E-08 | 0.722128505 | 0.85 | 0.465 | 0.000312023 | MM_1 |
| Yif1b | ENSMUSG00000030588 | protein_coding | Yip1 interacting factor homolog B (S. cerevisiae) | 2.407E-07 | 1.185319665 | 0.531 | 0.211 | 0.00568247 | MM_1 |
| Capns1 | ENSMUSG00000001794 | protein_coding | calpain, small subunit 1 | 1.052E-10 | 0.86333964 | 0.911 | 0.64 | 2.48382E-06 | MM_1 |
| Tbcb | ENSMUSG00000006095 | protein_coding | tubulin folding cofactor B | 1.5E-07 | 0.869182208 | 0.775 | 0.482 | 0.003542894 | MM_1 |
| Psenen | ENSMUSG00000036835 | protein_coding | presenilin enhancer gamma secretase subunit | 1.769E-09 | 0.765503102 | 0.93 | 0.658 | 4.17626E-05 | MM_1 |
| Cox6b1 | ENSMUSG00000036751 | protein_coding | cytochrome c oxidase, subunit 6B1 | 6.699E-17 | 0.929470545 | 0.977 | 0.807 | 1.58175E-12 | MM_1 |
| Cd33 | ENSMUSG00000004609 | protein_coding | CD33 antigen | 1.943E-09 | 0.849511278 | 0.911 | 0.658 | 4.58696E-05 | MM_1 |
| Rcn3 | ENSMUSG00000019539 | protein_coding | reticulocalbin 3, EF-hand calcium binding domain | 1.638E-11 | 1.171252828 | 0.784 | 0.404 | 3.8672E-07 | MM_1 |
| Fcgrt | ENSMUSG00000003420 | protein_coding | Fc receptor, IgG, alpha chain transporter | 2.913E-18 | 0.928170355 | 1 | 0.877 | 6.87893E-14 | MM_1 |
| Cd37 | ENSMUSG00000030798 | protein_coding | CD37 antigen | 6.299E-14 | 1.20542014 | 0.85 | 0.404 | 1.48726E-09 | MM_1 |
| Ftl1 | ENSMUSG00000050708 | protein_coding | ferritin light polypeptide 1 | 1.003E-18 | 0.830582591 | 1 | 0.956 | 2.36897E-14 | MM_1 |
| Dbp | ENSMUSG00000059824 | protein_coding | D site albumin promoter binding protein | 7.914E-17 | -2.334008336 | 0.263 | 0.649 | 1.86877E-12 | MM_1 |
| Tmem86a | ENSMUSG00000010307 | protein_coding | transmembrane protein 86A | 1.585E-08 | 1.403260429 | 0.577 | 0.246 | 0.000374249 | MM_1 |
| Idh2 | ENSMUSG00000030541 | protein_coding | isocitrate dehydrogenase 2 (NADP+), mitochondrial | 1.261E-09 | 0.85746901 | 0.953 | 0.623 | 2.97633E-05 | MM_1 |
| Cpeb1 | ENSMUSG00000025586 | protein_coding | cytoplasmic polyadenylation element binding protein 1 | 1.524E-07 | 2.42523436 | 0.343 | 0.079 | 0.003597415 | MM_1 |
| Ndufc2 | ENSMUSG00000030647 | protein_coding | NADH:ubiquinone oxidoreductase subunit C2 | 3.86E-11 | 1.013988815 | 0.883 | 0.579 | 9.11388E-07 | MM_1 |
| Folr2 | ENSMUSG00000032725 | protein_coding | folate receptor 2 (fetal) | 5.128E-17 | 0.914349873 | 0.991 | 0.86 | 1.21089E-12 | MM_1 |

| | | | | | | | | | |
|---|---|---|---|---|---|---|---|---|---|
| Taf10 | ENSMUSG00000043866 | protein_coding | TATA-box binding protein associated factor 10 | 3.841E-08 | 0.785809845 | 0.803 | 0.456 | 0.000906958 | MM_1 |
| Tpp1 | ENSMUSG00000030894 | protein_coding | tripeptidyl peptidase I | 4.219E-09 | 0.681473174 | 0.953 | 0.746 | 9.96212E-05 | MM_1 |
| Arntl | ENSMUSG00000055116 | protein_coding | aryl hydrocarbon receptor nuclear translocator-like | 1.031E-21 | 3.807333988 | 0.624 | 0.053 | 2.43373E-17 | MM_1 |
| Chp2 | ENSMUSG00000030865 | protein_coding | calcineurin-like EF hand protein 2 | 1.152E-11 | 1.115098921 | 0.878 | 0.491 | 2.71947E-07 | MM_1 |
| Cln3 | ENSMUSG00000030720 | protein_coding | ceroid lipofuscinosis, neuronal 3, juvenile (Batten, Spielmeyer-Vogt disease) | 7.383E-13 | 1.308956526 | 0.784 | 0.36 | 1.74319E-08 | MM_1 |
| Bola2 | ENSMUSG00000047721 | protein_coding | bolA-like 2 (E. coli) | 4.788E-10 | 1.342191505 | 0.718 | 0.368 | 1.13043E-05 | MM_1 |
| Mapk3 | ENSMUSG00000063065 | protein_coding | mitogen-activated protein kinase 3 | 6.797E-11 | 1.306646506 | 0.77 | 0.36 | 1.60488E-06 | MM_1 |
| Aldoa | ENSMUSG00000030695 | protein_coding | aldolase A, fructose-bisphosphate | 1.58E-23 | 1.517790471 | 0.977 | 0.561 | 3.73165E-19 | MM_1 |
| Fuom | ENSMUSG00000025466 | protein_coding | fucose mutarotase | 1.76E-09 | 2.465119672 | 0.39 | 0.07 | 4.155E-05 | MM_1 |
| Ifitm2 | ENSMUSG00000060591 | protein_coding | interferon induced transmembrane protein 2 | 6.957E-18 | 1.002473279 | 0.991 | 0.868 | 1.64265E-13 | MM_1 |
| Ifitm3 | ENSMUSG00000025492 | protein_coding | interferon induced transmembrane protein 3 | 1.24E-16 | 1.034495577 | 0.991 | 0.877 | 2.92886E-12 | MM_1 |
| Ifitm6 | ENSMUSG00000059108 | protein_coding | interferon induced transmembrane protein 6 | 1.443E-09 | 1.72347865 | 0.549 | 0.175 | 3.40702E-05 | MM_1 |
| Ctsd | ENSMUSG00000007891 | protein_coding | cathepsin D | 2.404E-28 | 1.270574433 | 1 | 0.965 | 5.67534E-24 | MM_1 |
| Mrpl23 | ENSMUSG00000037772 | protein_coding | mitochondrial ribosomal protein L23 | 1.151E-07 | 0.789465736 | 0.84 | 0.623 | 0.002716672 | MM_1 |
| Cd81 | ENSMUSG00000037706 | protein_coding | CD81 antigen | 1.601E-19 | 0.829179199 | 0.995 | 0.886 | 3.78097E-15 | MM_1 |
| Tssc4 | ENSMUSG00000045752 | protein_coding | tumor-suppressing subchromosomal transferable fragment 4 | 8.001E-08 | 0.911935698 | 0.469 | 0.149 | 0.001889151 | MM_1 |
| Pet100 | ENSMUSG00000087687 | protein_coding | PET100 homolog | 3.233E-11 | 1.217461036 | 0.798 | 0.447 | 7.63424E-07 | MM_1 |
| Stxbp2 | ENSMUSG00000004626 | protein_coding | syntaxin binding protein 2 | 1.757E-12 | 1.980325638 | 0.549 | 0.123 | 4.14848E-08 | MM_1 |
| Cd209b | ENSMUSG00000065987 | protein_coding | CD209b antigen | 3.026E-07 | 5.904742296 | 0.202 | 0 | 0.007145915 | MM_1 |
| Cd209f | ENSMUSG00000051906 | protein_coding | CD209f antigen | 2.164E-08 | 1.963632819 | 0.596 | 0.316 | 0.000511026 | MM_1 |
| Cln8 | ENSMUSG00000026317 | protein_coding | ceroid-lipofuscinosis, neuronal 8 | 2.45E-08 | 0.80226871 | 0.883 | 0.588 | 0.000578508 | MM_1 |
| Slc25a4 | ENSMUSG00000031633 | protein_coding | solute carrier family 25 (mitochondrial carrier, adenine nucleotide translocator), member 4 | 8.059E-08 | 0.82201896 | 0.812 | 0.518 | 0.00190296 | MM_1 |
| Atp6v1b2 | ENSMUSG00000006273 | protein_coding | ATPase, H+ transporting, lysosomal V1 subunit B2 | 1.567E-07 | 0.675943074 | 0.93 | 0.675 | 0.003700325 | MM_1 |
| Ndufa13 | ENSMUSG00000036199 | protein_coding | NADH:ubiquinone oxidoreductase subunit A13 | 2.218E-10 | 0.874756558 | 0.911 | 0.702 | 5.23721E-06 | MM_1 |
| Cope | ENSMUSG00000055681 | protein_coding | coatomer protein complex, subunit epsilon | 1.44E-07 | 0.998889254 | 0.714 | 0.377 | 0.003399194 | MM_1 |
| Uba52 | ENSMUSG00000090137 | protein_coding | ubiquitin A-52 residue ribosomal protein fusion product 1 | 5.563E-22 | 1.900842152 | 0.836 | 0.474 | 1.3135E-17 | MM_1 |
| Fkbp8 | ENSMUSG00000019428 | protein_coding | FK506 binding protein 8 | 8.915E-09 | 0.840841021 | 0.779 | 0.456 | 0.000210511 | MM_1 |
| Gdf15 | ENSMUSG00000038508 | protein_coding | growth differentiation factor 15 | 2.822E-08 | 1.129889393 | 0.718 | 0.333 | 0.000666325 | MM_1 |
| Ifi30 | ENSMUSG00000031838 | lncRNA | interferon gamma inducible protein 30 | 2.583E-09 | 0.740903698 | 0.953 | 0.825 | 6.09781E-05 | MM_1 |
| Babam1 | ENSMUSG00000031820 | protein_coding | BRISC and BRCA1 A complex member 1 | 1.174E-08 | 1.630266921 | 0.488 | 0.149 | 0.000277152 | MM_1 |
| Sin3b | ENSMUSG00000031622 | protein_coding | transcriptional regulator, SIN3B (yeast) | 3.175E-09 | 0.941052093 | 0.845 | 0.491 | 7.49763E-05 | MM_1 |
| Ndufb7 | ENSMUSG00000033938 | protein_coding | NADH:ubiquinone oxidoreductase subunit B7 | 2.288E-09 | 0.907061679 | 0.883 | 0.561 | 5.4033E-05 | MM_1 |
| Trmt1 | ENSMUSG00000001909 | protein_coding | tRNA methyltransferase 1 | 2.766E-10 | 1.409669958 | 0.695 | 0.298 | 6.53105E-06 | MM_1 |
| Lyl1 | ENSMUSG00000034041 | protein_coding | lymphoblastomic leukemia 1 | 2.759E-07 | 0.953584025 | 0.709 | 0.377 | 0.006515612 | MM_1 |
| Dnase2a | ENSMUSG00000003812 | protein_coding | deoxyribonuclease II alpha | 3.754E-08 | 0.665363188 | 0.859 | 0.561 | 0.000886339 | MM_1 |
| Wdr83os | ENSMUSG00000059355 | protein_coding | WD repeat domain 83 opposite strand | 4.939E-10 | 1.219969856 | 0.779 | 0.421 | 1.1662E-05 | MM_1 |
| Pla2g15 | ENSMUSG00000031903 | protein_coding | phospholipase A2, group XV | 4.993E-08 | 1.126307384 | 0.718 | 0.386 | 0.001178996 | MM_1 |
| Cox4i1 | ENSMUSG00000031818 | protein_coding | cytochrome c oxidase subunit 4I1 | 1.353E-12 | 0.696359597 | 0.995 | 0.904 | 3.1937E-08 | MM_1 |
| Cyba | ENSMUSG00000006519 | protein_coding | cytochrome b-245, alpha polypeptide | 1.628E-07 | 0.604206783 | 0.972 | 0.895 | 0.003844409 | MM_1 |
| Ubl5 | ENSMUSG00000084786 | protein_coding | ubiquitin-like 5 | 2.718E-11 | 0.770614006 | 0.939 | 0.737 | 6.41812E-07 | MM_1 |
| Cnn1 | ENSMUSG00000001349 | protein_coding | calponin 1 | 8.313E-08 | 0.850831854 | 0.808 | 0.509 | 0.001962852 | MM_1 |
| Atp5l | ENSMUSG00000038717 | protein_coding | ATP synthase, H+ transporting, mitochondrial F0 complex, subunit G | 9.277E-16 | 0.904056774 | 0.972 | 0.816 | 2.19053E-11 | MM_1 |
| Fxyd2 | ENSMUSG00000059412 | protein_coding | FXYD domain-containing ion transport regulator 2 | 6.925E-11 | 1.139181315 | 0.92 | 0.649 | 1.63509E-06 | MM_1 |
| Tagln | ENSMUSG00000032085 | protein_coding | transgelin | 4.065E-09 | 0.611782666 | 0.934 | 0.754 | 9.59839E-05 | MM_1 |
| Rexo2 | ENSMUSG00000032026 | protein_coding | RNA exonuclease 2 | 2.035E-07 | 0.808722267 | 0.831 | 0.561 | 0.004805284 | MM_1 |
| Rpl10-ps3 | ENSMUSG00000058443 | protein_coding | ribosomal protein L10, pseudogene 3 | 2.372E-11 | 2.131806269 | 0.535 | 0.158 | 5.60079E-07 | MM_1 |
| Timm8b | ENSMUSG00000039016 | protein_coding | translocase of inner mitochondrial membrane 8B | 5.696E-10 | 1.730360866 | 0.573 | 0.202 | 1.34498E-05 | MM_1 |
| Tspan3 | ENSMUSG00000032324 | protein_coding | tetraspanin 3 | 4.141E-07 | 0.945289702 | 0.728 | 0.404 | 0.00977859 | MM_1 |
| Commd4 | ENSMUSG00000032299 | protein_coding | COMM domain containing 4 | 2.218E-07 | 1.061873205 | 0.62 | 0.272 | 0.005235996 | MM_1 |
| Cox5a | ENSMUSG00000000088 | protein_coding | cytochrome c oxidase subunit 5A | 2.809E-07 | 0.678958852 | 0.939 | 0.772 | 0.006633679 | MM_1 |
| Scamp2 | ENSMUSG00000040188 | protein_coding | secretory carrier membrane protein 2 | 2.817E-13 | 0.978229808 | 0.93 | 0.702 | 6.65188E-09 | MM_1 |
| Hexa | ENSMUSG00000025232 | protein_coding | hexosaminidase A | 3.525E-12 | 0.627309668 | 0.991 | 0.912 | 8.32311E-08 | MM_1 |
| Anp32a | ENSMUSG00000032249 | protein_coding | acidic (leucine-rich) nuclear phosphoprotein 32 family, member A | 2.579E-08 | 0.647759595 | 0.953 | 0.789 | 0.000608885 | MM_1 |
| Rps27l | ENSMUSG00000036781 | protein_coding | ribosomal protein S27-like | 3.585E-08 | 0.703234734 | 0.925 | 0.711 | 0.000846372 | MM_1 |
| Arpp19 | ENSMUSG00000007656 | protein_coding | cAMP-regulated phosphoprotein 19 | 8.489E-08 | 0.805008232 | 0.826 | 0.561 | 0.002004377 | MM_1 |
| Cox7a2 | ENSMUSG00000032330 | protein_coding | cytochrome c oxidase subunit 7A2 | 2.179E-10 | 0.802052314 | 0.939 | 0.719 | 5.14499E-06 | MM_1 |
| Trf | ENSMUSG00000032554 | protein_coding | transferrin | 1.108E-12 | 0.658330141 | 0.995 | 0.939 | 2.61734E-08 | MM_1 |
| Inka1 | ENSMUSG00000042106 | protein_coding | inka box actin regulator 1 | 4.187E-09 | 1.962128832 | 0.437 | 0.105 | 9.8858E-05 | MM_1 |
| Rhoa | ENSMUSG00000007815 | protein_coding | ras homolog family member A | 1.805E-15 | 0.691943571 | 1 | 0.956 | 4.2629E-11 | MM_1 |
| Gpx1 | ENSMUSG00000063856 | protein_coding | glutathione peroxidase 1 | 3.411E-26 | 1.234234137 | 0.995 | 0.842 | 8.05297E-22 | MM_1 |
| Uqcrc1 | ENSMUSG00000025651 | protein_coding | ubiquinol-cytochrome c reductase core protein 1 | 5.923E-08 | 0.823726459 | 0.728 | 0.368 | 0.001398452 | MM_1 |
| Shisa5 | ENSMUSG00000025647 | protein_coding | shisa family member 5 | 1.167E-07 | 0.74925621 | 0.869 | 0.526 | 0.002755618 | MM_1 |
| Tma7 | ENSMUSG00000091537 | protein_coding | translational machinery associated 7 | 2.619E-07 | 0.595398685 | 0.887 | 0.632 | 0.006184464 | MM_1 |
| Cmtm6 | ENSMUSG00000032434 | protein_coding | CKLF-like MARVEL transmembrane domain containing 6 | 1.357E-10 | 1.051718249 | 0.831 | 0.404 | 3.2033E-06 | MM_1 |
| Cmtm7 | ENSMUSG00000032436 | protein_coding | CKLF-like MARVEL transmembrane domain containing 7 | 4.774E-09 | 1.102963479 | 0.742 | 0.439 | 0.000112727 | MM_1 |
| Rps27rt | ENSMUSG00000050621 | protein_coding | ribosomal protein S27, retrogene | 1.44E-10 | 1.235531787 | 0.667 | 0.237 | 3.40022E-06 | MM_1 |
| Lars2 | ENSMUSG00000035202 | protein_coding | leucyl-tRNA synthetase, mitochondrial | 8.034E-41 | -1.95624604 | 0.972 | 0.991 | 1.8969E-36 | MM_1 |
| Mthfd1l | ENSMUSG00000040675 | protein_coding | methylenetetrahydrofolate dehydrogenase (NADP+ dependent) 1-like | 7.526E-22 | -3.630984311 | 0.08 | 0.535 | 1.77699E-17 | MM_1 |
| Hivep2 | ENSMUSG00000015501 | protein_coding | human immunodeficiency virus type I enhancer binding protein 2 | 4.518E-08 | -0.851633485 | 0.915 | 0.921 | 0.001066769 | MM_1 |
| Tspyl1 | ENSMUSG00000047514 | protein_coding | testis-specific protein, Y-encoded-like 1 | 3.208E-10 | 1.432781144 | 0.648 | 0.219 | 7.57584E-06 | MM_1 |
| Snx3 | ENSMUSG00000019804 | protein_coding | sorting nexin 3 | 3.759E-15 | 0.756112271 | 0.995 | 0.912 | 8.87538E-11 | MM_1 |
| Pkib | ENSMUSG00000019876 | protein_coding | protein kinase inhibitor beta, cAMP dependent, testis specific | 1.417E-09 | 1.627482041 | 0.474 | 0.132 | 3.34672E-05 | MM_1 |
| Snrpd3 | ENSMUSG00000020180 | protein_coding | small nuclear ribonucleoprotein D3 | 4.903E-13 | 1.270467528 | 0.826 | 0.412 | 1.15781E-08 | MM_1 |
| Mif | ENSMUSG00000033307 | protein_coding | macrophage migration inhibitory factor (glycosylation-inhibiting factor) | 2.043E-10 | 1.040597098 | 0.859 | 0.509 | 4.82354E-06 | MM_1 |
| Sumo3 | ENSMUSG00000020265 | protein_coding | small ubiquitin-like modifier 3 | 1.19E-09 | 0.879603021 | 0.793 | 0.439 | 2.80987E-05 | MM_1 |
| Agpat3 | ENSMUSG00000001211 | protein_coding | 1-acylglycerol-3-phosphate O-acyltransferase 3 | 9.55E-10 | 1.293993605 | 0.7 | 0.36 | 2.25484E-05 | MM_1 |
| Cstb | ENSMUSG00000005054 | protein_coding | cystatin B | 1.443E-09 | 0.739443075 | 0.986 | 0.825 | 3.4064E-05 | MM_1 |
| Gpx4 | ENSMUSG00000075706 | protein_coding | glutathione peroxidase 4 | 1.37E-16 | 1.125016562 | 0.962 | 0.728 | 3.23598E-12 | MM_1 |
| Sbno2 | ENSMUSG00000035673 | protein_coding | strawberry notch 2 | 2.776E-09 | -1.472496337 | 0.69 | 0.772 | 6.5555E-05 | MM_1 |
| Atp5d | ENSMUSG00000003072 | protein_coding | ATP synthase, H+ transporting, mitochondrial F1 complex, delta subunit | 1.257E-07 | 0.724912993 | 0.854 | 0.649 | 0.002968321 | MM_1 |
| Cirbp | ENSMUSG00000045193 | protein_coding | cold inducible RNA binding protein | 3.464E-07 | 1.069662648 | 0.521 | 0.202 | 0.008179649 | MM_1 |
| Ndufs7 | ENSMUSG00000020153 | protein_coding | NADH:ubiquinone oxidoreductase core subunit S7 | 1.303E-17 | 1.29948879 | 0.915 | 0.535 | 3.07575E-13 | MM_1 |
| Adamtsl5 | ENSMUSG00000043822 | protein_coding | ADAMTS-like 5 | 4.202E-07 | 0.743806179 | 0.634 | 0.298 | 0.00992249 | MM_1 |
| Uqcr11 | ENSMUSG00000020163 | protein_coding | ubiquinol-cytochrome c reductase, complex III subunit XI | 9.153E-10 | 0.649385576 | 0.967 | 0.719 | 2.16114E-05 | MM_1 |
| Lsm7 | ENSMUSG00000035215 | protein_coding | LSM7 homolog, U6 small nuclear RNA and mRNA degradation associated | 3.67E-07 | 1.257281466 | 0.54 | 0.219 | 0.008665584 | MM_1 |
| Nfic | ENSMUSG00000055053 | protein_coding | nuclear factor I/C | 4.558E-08 | 0.839137738 | 0.859 | 0.553 | 0.001076238 | MM_1 |
| Tle5 | ENSMUSG00000054452 | protein_coding | TLE family member 5, transcriptional modulator | 1.444E-10 | 0.894293737 | 0.873 | 0.579 | 3.40891E-06 | MM_1 |
| Igf1 | ENSMUSG00000020053 | protein_coding | insulin-like growth factor 1 | 1.131E-09 | 0.716308428 | 0.986 | 0.842 | 2.67141E-05 | MM_1 |
| Uhrf1bp1l | ENSMUSG00000019951 | protein_coding | UHRF1 (ICBP90) binding protein 1-like | 3.706E-12 | 1.215031478 | 0.784 | 0.421 | 8.75135E-08 | MM_1 |
| Slc25a3 | ENSMUSG00000061904 | protein_coding | solute carrier family 25 (mitochondrial carrier, phosphate carrier), member 3 | 1.283E-09 | 0.592266215 | 0.991 | 0.895 | 3.03019E-05 | MM_1 |
| Ndufa12 | ENSMUSG00000020022 | protein_coding | NADH:ubiquinone oxidoreductase subunit A12 | 1.248E-11 | 1.380515677 | 0.761 | 0.395 | 2.94697E-07 | MM_1 |
| Dcn | ENSMUSG00000019929 | protein_coding | decorin | 1.509E-17 | 1.757845797 | 0.728 | 0.211 | 3.56416E-13 | MM_1 |
| Lyz2 | ENSMUSG00000069516 | protein_coding | lysozyme 2 | 2.298E-52 | 12.97393227 | 1 | 0 | 5.42611E-48 | MM_1 |
| Lyz1 | ENSMUSG00000069515 | protein_coding | lysozyme 1 | 2.294E-10 | -2.519645522 | 0.3 | 0.57 | 5.41749E-06 | MM_1 |
| Rxylt1 | ENSMUSG00000034620 | protein_coding | ribitol xylosyltransferase 1 | 2.246E-08 | 1.502373309 | 0.531 | 0.193 | 0.000530353 | MM_1 |
| Ctdsp2 | ENSMUSG00000078429 | protein_coding | CTD (carboxy-terminal domain, RNA polymerase II, polypeptide A) small phosphatase 2 | 4.116E-09 | 0.849641304 | 0.831 | 0.5 | 9.71785E-05 | MM_1 |
| Dctn2 | ENSMUSG00000025410 | protein_coding | dynactin 2 | 2.865E-09 | 1.222225928 | 0.676 | 0.316 | 6.76585E-05 | MM_1 |
| Atp5b | ENSMUSG00000025393 | protein_coding | ATP synthase, H+ transporting mitochondrial F1 complex, beta subunit | 6.481E-13 | 0.865764844 | 0.962 | 0.816 | 1.53024E-08 | MM_1 |
| Myl6 | ENSMUSG00000090841 | protein_coding | myosin, light polypeptide 6, alkali, smooth muscle and non-muscle | 1.803E-16 | 0.659985908 | 1 | 0.956 | 4.25716E-12 | MM_1 |
| Rpl41 | ENSMUSG00000093674 | protein_coding | ribosomal protein L41 | 5.877E-21 | 0.75279536 | 0.995 | 0.912 | 1.38775E-16 | MM_1 |
| Rps26 | ENSMUSG00000025362 | protein_coding | ribosomal protein S26 | 1.385E-14 | 0.734519542 | 0.986 | 0.904 | 3.26921E-10 | MM_1 |
| Bloc1s1 | ENSMUSG00000090247 | protein_coding | biogenesis of lysosomal organelles complex-1, subunit 1 | 1.355E-07 | 1.539266551 | 0.469 | 0.158 | 0.003198343 | MM_1 |
| Smtn | ENSMUSG00000020439 | protein_coding | smoothelin | 1.416E-09 | 1.386927377 | 0.469 | 0.105 | 3.34255E-05 | MM_1 |
| Tcn2 | ENSMUSG00000020432 | protein_coding | transcobalamin 2 | 1.78E-10 | 0.791945579 | 0.92 | 0.658 | 4.20198E-06 | MM_1 |
| Ap1b1 | ENSMUSG00000009090 | protein_coding | adaptor protein complex AP-1, beta 1 subunit | 1.413E-22 | 1.903844891 | 0.901 | 0.404 | 3.33557E-18 | MM_1 |
| Ppia | ENSMUSG00000071866 | protein_coding | peptidylprolyl isomerase A | 2.536E-23 | 0.970998641 | 1 | 0.904 | 5.98699E-19 | MM_1 |
| Sec61g | ENSMUSG00000078974 | protein_coding | SEC61, gamma subunit | 2.171E-10 | 0.871537726 | 0.934 | 0.711 | 5.12693E-06 | MM_1 |
| Mdh1 | ENSMUSG00000020321 | protein_coding | malate dehydrogenase 1, NAD (soluble) | 2.883E-11 | 1.248225334 | 0.751 | 0.351 | 6.80633E-07 | MM_1 |
| Commd1 | ENSMUSG00000051355 | protein_coding | COMM domain containing 1 | 3.955E-07 | 0.931841007 | 0.746 | 0.465 | 0.009338867 | MM_1 |
| Ltc4s | ENSMUSG00000020377 | protein_coding | leukotriene C4 synthase | 5.555E-18 | 1.296643263 | 0.967 | 0.763 | 1.31175E-13 | MM_1 |
| 0610009B22Rik | ENSMUSG00000007777 | protein_coding | RIKEN cDNA 0610009B22 gene | 9.89E-10 | 1.985398284 | 0.446 | 0.096 | 2.33532E-05 | MM_1 |
| Uqcrq | ENSMUSG00000044894 | protein_coding | ubiquinol-cytochrome c reductase, complex III subunit VII | 2.968E-13 | 1.049103633 | 0.915 | 0.711 | 7.00894E-09 | MM_1 |
| Hint1 | ENSMUSG00000020267 | protein_coding | histidine triad nucleotide binding protein 1 | 2.849E-09 | 0.67398981 | 0.958 | 0.798 | 6.72739E-05 | MM_1 |
| Anxa6 | ENSMUSG00000018340 | protein_coding | annexin A6 | 6.918E-12 | 1.169833738 | 0.883 | 0.553 | 1.6335E-07 | MM_1 |
| Rnf187 | ENSMUSG00000020496 | protein_coding | ring finger protein 187 | 1.581E-07 | 0.993747854 | 0.653 | 0.325 | 0.003734213 | MM_1 |
| Ubb | ENSMUSG00000019505 | protein_coding | ubiquitin B | 8.903E-08 | 0.611919672 | 0.981 | 0.895 | 0.00210226 | MM_1 |
| Cd68 | ENSMUSG00000018774 | protein_coding | CD68 antigen | 8.14E-19 | 1.120198136 | 0.972 | 0.816 | 1.92203E-14 | MM_1 |
| 0610010K14Rik | ENSMUSG00000020831 | protein_coding | RIKEN cDNA 0610010K14 gene | 4.084E-07 | 0.938081896 | 0.681 | 0.36 | 0.009643997 | MM_1 |
| Rnasek | ENSMUSG00000093989 | protein_coding | ribonuclease, RNase K | 1.421E-14 | 1.071668933 | 0.911 | 0.605 | 3.35529E-10 | MM_1 |
| Slc43a2 | ENSMUSG00000038178 | protein_coding | solute carrier family 43, member 2 | 4.515E-11 | 0.941269547 | 0.934 | 0.649 | 1.06607E-06 | MM_1 |
| 2610507B11Rik | ENSMUSG00000010277 | protein_coding | RIKEN cDNA 2610507B11 gene | 1.557E-07 | 0.605818914 | 0.92 | 0.702 | 0.003675407 | MM_1 |

| | | | | | | | | | |
|---|---|---|---|---|---|---|---|---|---|
| Lgals9 | ENSMUSG00000001123 | protein_coding | lectin, galactose binding, soluble 9 | 4.825E-10 | 1.080691671 | 0.808 | 0.439 | 1.13932E-05 | MM_1 |
| Evi2a | ENSMUSG00000078771 | protein_coding | ecotropic viral integration site 2a | 4.142E-10 | 0.961452481 | 0.915 | 0.596 | 9.77981E-06 | MM_1 |
| Ccl6 | ENSMUSG00000018927 | protein_coding | chemokine (C-C motif) ligand 6 | 1.111E-17 | 1.297835286 | 0.995 | 0.904 | 2.62317E-13 | MM_1 |
| Wfdc17 | ENSMUSG00000069792 | protein_coding | WAP four-disulfide core domain 17 | 6.955E-09 | 0.89718516 | 0.995 | 0.895 | 0.000164223 | MM_1 |
| Nme2 | ENSMUSG00000020857 | protein_coding | NME/NM23 nucleoside diphosphate kinase 2 | 2.138E-08 | 0.761326991 | 0.967 | 0.763 | 0.000504913 | MM_1 |
| Nme1 | ENSMUSG00000037601 | protein_coding | NME/NM23 nucleoside diphosphate kinase 1 | 3.85E-10 | 0.787832868 | 0.85 | 0.544 | 9.09087E-06 | MM_1 |
| Phb | ENSMUSG00000038845 | protein_coding | prohibitin | 8.491E-13 | 1.780053748 | 0.554 | 0.114 | 2.00481E-08 | MM_1 |
| Atp5g1 | ENSMUSG00000006057 | protein_coding | ATP synthase, H+ transporting, mitochondrial F0 complex, subunit C1 (subunit 9) | 1.155E-09 | 1.001267796 | 0.845 | 0.588 | 2.72777E-05 | MM_1 |
| Psmb3 | ENSMUSG00000069744 | protein_coding | proteasome (prosome, macropain) subunit, beta type 3 | 4.055E-10 | 0.999793143 | 0.826 | 0.491 | 9.57356E-06 | MM_1 |
| Rab5c | ENSMUSG00000019173 | protein_coding | RAB5C, member RAS oncogene family | 9.123E-13 | 0.796643007 | 0.962 | 0.754 | 2.15418E-08 | MM_1 |
| Naglu | ENSMUSG00000001751 | protein_coding | alpha-N-acetylglucosaminidase (Sanfilippo disease IIIB) | 1.05E-07 | 1.679959403 | 0.432 | 0.132 | 0.002478954 | MM_1 |
| Vat1 | ENSMUSG00000034993 | protein_coding | vesicle amine transport 1 | 3.231E-11 | 1.009110637 | 0.925 | 0.596 | 7.62841E-07 | MM_1 |
| Grn | ENSMUSG00000034708 | protein_coding | granulin | 1.32E-15 | 0.743088237 | 1 | 0.956 | 3.11726E-11 | MM_1 |
| Arf2 | ENSMUSG00000062421 | protein_coding | ADP-ribosylation factor 2 | 6.963E-08 | 0.944845483 | 0.812 | 0.482 | 0.001644055 | MM_1 |
| Rpl38 | ENSMUSG00000057322 | protein_coding | ribosomal protein L38 | 3.432E-31 | 1.089424672 | 0.995 | 0.939 | 8.10308E-27 | MM_1 |
| Gprc5c | ENSMUSG00000051043 | protein_coding | G protein-coupled receptor, family C, group 5, member C | 1.419E-08 | 1.347397512 | 0.545 | 0.202 | 0.000335164 | MM_1 |
| Cd300a | ENSMUSG00000034652 | protein_coding | CD300A molecule | 3.535E-14 | 1.368680257 | 0.864 | 0.439 | 8.3457E-10 | MM_1 |
| Atp5h | ENSMUSG00000034566 | protein_coding | ATP synthase, H+ transporting, mitochondrial F0 complex, subunit D | 5.475E-16 | 0.884706476 | 0.962 | 0.798 | 1.29274E-11 | MM_1 |
| Jpt1 | ENSMUSG00000020737 | protein_coding | Jupiter microtubule associated homolog 1 | 5.806E-08 | 0.707017487 | 0.925 | 0.684 | 0.001370889 | MM_1 |
| Mif4gd | ENSMUSG00000020743 | protein_coding | MIF4G domain containing | 1.127E-08 | 0.950979747 | 0.474 | 0.132 | 0.000266055 | MM_1 |
| Ubald2 | ENSMUSG00000050628 | protein_coding | UBA-like domain containing 2 | 1.826E-11 | 1.259517647 | 0.737 | 0.298 | 4.31256E-07 | MM_1 |
| Socs3 | ENSMUSG00000053113 | protein_coding | suppressor of cytokine signaling 3 | 1.051E-07 | -1.646347621 | 0.667 | 0.711 | 0.002481144 | MM_1 |
| Gaa | ENSMUSG00000025579 | protein_coding | glucosidase, alpha, acid | 1.618E-07 | 1.194244442 | 0.559 | 0.228 | 0.0038204 | MM_1 |
| Arhgdia | ENSMUSG00000025132 | protein_coding | Rho GDP dissociation inhibitor (GDI) alpha | 1.34E-08 | 0.697905092 | 0.977 | 0.816 | 0.0003163 | MM_1 |
| Cbr2 | ENSMUSG00000025150 | protein_coding | carbonyl reductase 2 | 5.894E-25 | 1.166115489 | 0.995 | 0.904 | 1.39181E-20 | MM_1 |
| Laptm4a | ENSMUSG00000020585 | protein_coding | lysosomal-associated protein transmembrane 4A | 1.438E-09 | 0.615238755 | 0.986 | 0.833 | 3.39625E-05 | MM_1 |
| Ywhaq | ENSMUSG00000076432 | protein_coding | tyrosine 3-monooxygenase/tryptophan 5-monooxygenase activation protein theta | 1.689E-07 | 0.707011486 | 0.948 | 0.719 | 0.00398777 | MM_1 |
| Hbp1 | ENSMUSG00000002996 | protein_coding | high mobility group box transcription factor 1 | 2.415E-07 | 1.050340363 | 0.657 | 0.351 | 0.005703085 | MM_1 |
| Psma6 | ENSMUSG00000021024 | protein_coding | proteasome (prosome, macropain) subunit, alpha type 6 | 1.876E-07 | 0.800983024 | 0.789 | 0.456 | 0.004428559 | MM_1 |
| Rps29 | ENSMUSG00000034892 | protein_coding | ribosomal protein S29 | 3.947E-20 | 0.774596675 | 1 | 0.956 | 9.32051E-16 | MM_1 |
| Churc1 | ENSMUSG00000090258 | protein_coding | churchill domain containing 1 | 1.412E-07 | 1.046864746 | 0.648 | 0.298 | 0.003333096 | MM_1 |
| Gphn | ENSMUSG00000047454 | protein_coding | gephyrin | 3.218E-29 | -1.402769217 | 0.991 | 0.982 | 7.59892E-25 | MM_1 |
| Susd6 | ENSMUSG00000021133 | protein_coding | sushi domain containing 6 | 2.447E-13 | -1.503487899 | 0.803 | 0.895 | 5.77719E-09 | MM_1 |
| Srsf5 | ENSMUSG00000021134 | protein_coding | serine/arginine-rich splicing factor 5 | 4.007E-11 | 0.820049887 | 0.939 | 0.693 | 9.46172E-07 | MM_1 |
| Tmed10 | ENSMUSG00000021248 | protein_coding | transmembrane p24 trafficking protein 10 | 1.42E-07 | 0.599829195 | 0.944 | 0.789 | 0.003353452 | MM_1 |
| Slirp | ENSMUSG00000021040 | protein_coding | SRA stem-loop interacting RNA binding protein | 4.669E-10 | 1.161594622 | 0.718 | 0.377 | 1.10233E-05 | MM_1 |
| Gm19951 | ENSMUSG00000113136 | lncRNA | predicted gene, 19951 | 1.126E-13 | -1.900787151 | 0.521 | 0.746 | 2.65891E-09 | MM_1 |
| Ndufb1-ps | ENSMUSG00000113902 | protein_coding | NADH:ubiquinone oxidoreductase subunit B1 | 2.508E-19 | 1.257997712 | 0.953 | 0.728 | 5.92177E-15 | MM_1 |
| Ckb | ENSMUSG00000001270 | protein_coding | creatine kinase, brain | 1.104E-07 | 0.735519962 | 0.911 | 0.667 | 0.002605966 | MM_1 |
| Atp5mpl | ENSMUSG00000021290 | protein_coding | ATP synthase membrane subunit 6.8PL | 1.988E-21 | 1.387461653 | 0.934 | 0.667 | 4.69456E-17 | MM_1 |
| Pld4 | ENSMUSG00000052160 | protein_coding | phospholipase D family, member 4 | 8.172E-14 | 0.977722654 | 0.958 | 0.737 | 1.92958E-09 | MM_1 |
| Crip1 | ENSMUSG00000006360 | protein_coding | cysteine-rich protein 1 (intestinal) | 3.923E-10 | 0.768797243 | 1 | 0.939 | 9.26264E-06 | MM_1 |
| Ighm | ENSMUSG00000076617 | IG_C_gene | immunoglobulin heavy constant mu | 1.061E-15 | 1.433675721 | 0.897 | 0.544 | 2.50418E-11 | MM_1 |
| Psma2 | ENSMUSG00000015671 | protein_coding | proteasome (prosome, macropain) subunit, alpha type 2 | 4.437E-08 | 0.784491014 | 0.864 | 0.632 | 0.001047585 | MM_1 |
| Cmah | ENSMUSG00000016756 | protein_coding | cytidine monophospho-N-acetylneuraminic acid hydroxylase | 1.607E-15 | 1.282084392 | 0.892 | 0.544 | 3.79327E-11 | MM_1 |
| Ninj1 | ENSMUSG00000037966 | protein_coding | ninjurin 1 | 8.758E-18 | 1.03715357 | 0.986 | 0.789 | 2.068E-13 | MM_1 |
| Higd2a | ENSMUSG00000025868 | protein_coding | HIG1 domain family, member 2A | 2.027E-07 | 0.801349477 | 0.775 | 0.43 | 0.004786469 | MM_1 |
| Prelid1 | ENSMUSG00000021486 | protein_coding | PRELI domain containing 1 | 1.058E-07 | 0.920890507 | 0.737 | 0.404 | 0.002498151 | MM_1 |
| Uqcrb | ENSMUSG00000021520 | protein_coding | ubiquinol-cytochrome c reductase binding protein | 5.235E-15 | 1.070141028 | 0.948 | 0.728 | 1.23598E-10 | MM_1 |
| Sdha | ENSMUSG00000021577 | protein_coding | succinate dehydrogenase complex, subunit A, flavoprotein (Fp) | 1.766E-07 | 0.854046832 | 0.826 | 0.518 | 0.004169264 | MM_1 |
| Ftl1-ps1 | ENSMUSG00000062382 | protein_coding | ferritin light polypeptide 1, pseudogene 1 | 2.181E-32 | 2.703524965 | 0.93 | 0.333 | 5.14965E-28 | MM_1 |
| Cox7c | ENSMUSG00000017778 | protein_coding | cytochrome c oxidase subunit 7C | 4.002E-23 | 1.093669625 | 0.995 | 0.825 | 9.4495E-19 | MM_1 |
| Tbca | ENSMUSG00000042043 | protein_coding | tubulin cofactor A | 1.711E-07 | 0.698302745 | 0.906 | 0.667 | 0.004039667 | MM_1 |
| Naip6 | ENSMUSG00000078942 | protein_coding | NLR family, apoptosis inhibitory protein 6 | 6.381E-08 | -4.417686938 | 0 | 0.132 | 0.001506628 | MM_1 |
| Mrps36 | ENSMUSG00000061474 | protein_coding | mitochondrial ribosomal protein S36 | 3.458E-08 | 1.249336158 | 0.624 | 0.289 | 0.000816472 | MM_1 |
| Map3k1 | ENSMUSG00000021754 | protein_coding | mitogen-activated protein kinase kinase kinase 1 | 3.081E-07 | 0.683643385 | 0.883 | 0.649 | 0.007274248 | MM_1 |
| Il31ra | ENSMUSG00000050377 | protein_coding | interleukin 31 receptor A | 7.963E-12 | -1.286840785 | 0.864 | 0.868 | 1.88022E-07 | MM_1 |
| Ndufs4 | ENSMUSG00000021764 | protein_coding | NADH:ubiquinone oxidoreductase core subunit S4 | 4.975E-09 | 1.038705762 | 0.784 | 0.395 | 0.00011748 | MM_1 |
| Psmd6 | ENSMUSG00000021737 | protein_coding | proteasome (prosome, macropain) 26S subunit, non-ATPase, 6 | 7.231E-08 | 1.013944901 | 0.69 | 0.36 | 0.001707281 | MM_1 |
| Nr1d2 | ENSMUSG00000021775 | protein_coding | nuclear receptor subfamily 1, group D, member 2 | 1.946E-08 | -1.36459755 | 0.559 | 0.702 | 0.000459551 | MM_1 |
| Gm48099 | ENSMUSG00000114304 | lncRNA | predicted gene, 48099 | 1.499E-20 | 2.71370564 | 0.737 | 0.167 | 3.54033E-16 | MM_1 |
| Spcs1 | ENSMUSG00000021917 | protein_coding | signal peptidase complex subunit 1 homolog (S. cerevisiae) | 2.862E-09 | 0.943405199 | 0.84 | 0.491 | 6.75718E-05 | MM_1 |
| Smim4 | ENSMUSG00000058351 | protein_coding | small integral membrane protein 4 | 5.817E-08 | 1.126077709 | 0.62 | 0.263 | 0.001373525 | MM_1 |
| Stab1 | ENSMUSG00000042286 | protein_coding | stabilin 1 | 7.159E-09 | 0.616155459 | 0.977 | 0.921 | 0.000169048 | MM_1 |
| Nisch | ENSMUSG00000021910 | protein_coding | nischarin | 1.121E-07 | 0.725833613 | 0.93 | 0.667 | 0.002645845 | MM_1 |
| Dad1 | ENSMUSG00000022174 | protein_coding | defender against cell death 1 | 3.196E-12 | 0.909995461 | 0.93 | 0.588 | 7.54665E-08 | MM_1 |
| Mrpl52 | ENSMUSG00000010406 | protein_coding | mitochondrial ribosomal protein L52 | 1.181E-11 | 0.996489541 | 0.864 | 0.57 | 2.78812E-07 | MM_1 |
| Slc7a8 | ENSMUSG00000022180 | protein_coding | solute carrier family 7 (cationic amino acid transporter, y+ system), member 8 | 1.083E-07 | 0.654903015 | 0.812 | 0.474 | 0.002556165 | MM_1 |
| Psme1 | ENSMUSG00000022216 | protein_coding | proteasome (prosome, macropain) activator subunit 1 (PA28 alpha) | 7.382E-12 | 1.045724496 | 0.878 | 0.588 | 1.74301E-07 | MM_1 |
| Mrpl57 | ENSMUSG00000021967 | protein_coding | mitochondrial ribosomal protein L57 | 1.375E-07 | 1.294896532 | 0.587 | 0.272 | 0.003246133 | MM_1 |
| Dnajc15 | ENSMUSG00000022013 | protein_coding | DnaJ heat shock protein family (Hsp40) member C15 | 6.77E-09 | 1.116419788 | 0.77 | 0.421 | 0.000159851 | MM_1 |
| Gpr183 | ENSMUSG00000051212 | protein_coding | G protein-coupled receptor 183 | 1.276E-09 | 0.881368199 | 0.873 | 0.465 | 3.01332E-05 | MM_1 |
| Selenop | ENSMUSG00000064373 | protein_coding | selenoprotein P | 7.772E-16 | 0.660330415 | 1 | 0.974 | 1.83507E-11 | MM_1 |
| Ccdc152 | ENSMUSG00000091119 | protein_coding | coiled-coil domain containing 152 | 4.393E-11 | 1.599794618 | 0.695 | 0.316 | 1.03737E-06 | MM_1 |
| Cox6c | ENSMUSG00000014313 | protein_coding | cytochrome c oxidase subunit 6C | 6.83E-23 | 1.137386019 | 0.981 | 0.763 | 1.61271E-18 | MM_1 |
| Ly6e | ENSMUSG00000022587 | protein_coding | lymphocyte antigen 6 complex, locus E | 4.354E-12 | 1.311595428 | 0.911 | 0.632 | 1.028E-07 | MM_1 |
| Vps28 | ENSMUSG00000115987 | protein_coding | vacuolar protein sorting 28 | 1.173E-09 | 1.168510585 | 0.737 | 0.368 | 2.76999E-05 | MM_1 |
| Rac2 | ENSMUSG00000033220 | protein_coding | Rac family small GTPase 2 | 5.661E-09 | 1.015374944 | 0.793 | 0.465 | 0.000133666 | MM_1 |
| Lgals1 | ENSMUSG00000068220 | protein_coding | lectin, galactose binding, soluble 1 | 5.416E-17 | 1.09951116 | 1 | 0.868 | 1.27891E-12 | MM_1 |
| Rbx1 | ENSMUSG00000022400 | protein_coding | ring-box 1 | 2.328E-08 | 0.673891596 | 0.944 | 0.789 | 0.000549745 | MM_1 |
| Tef | ENSMUSG00000022389 | protein_coding | thyrotroph embryonic factor | 3.449E-10 | -2.117624324 | 0.239 | 0.535 | 8.14349E-06 | MM_1 |
| Snu13 | ENSMUSG00000063480 | protein_coding | SNU13 homolog, small nuclear ribonucleoprotein (U4/U6.U5) | 5.946E-09 | 0.8055592 | 0.869 | 0.596 | 0.000140398 | MM_1 |
| Ndufa6 | ENSMUSG00000022450 | protein_coding | NADH:ubiquinone oxidoreductase subunit A6 | 1.453E-07 | 0.731915434 | 0.864 | 0.623 | 0.003431295 | MM_1 |
| Cerk | ENSMUSG00000035891 | protein_coding | ceramide kinase | 3.096E-07 | 0.730316716 | 0.897 | 0.632 | 0.007309842 | MM_1 |
| Slc48a1 | ENSMUSG00000081534 | protein_coding | solute carrier family 48 (heme transporter), member 1 | 1.686E-07 | 1.40131382 | 0.474 | 0.158 | 0.003981048 | MM_1 |
| Tuba1b | ENSMUSG00000023004 | protein_coding | tubulin, alpha 1B | 4.897E-20 | 1.217574249 | 0.977 | 0.807 | 1.15625E-15 | MM_1 |
| Cela1 | ENSMUSG00000023031 | protein_coding | chymotrypsin-like elastase family, member 1 | 8.513E-09 | 1.758812565 | 0.441 | 0.114 | 0.000201021 | MM_1 |
| Igfbp6 | ENSMUSG00000023046 | protein_coding | insulin-like growth factor binding protein 6 | 2.999E-09 | 3.076219977 | 0.324 | 0.035 | 7.08165E-05 | MM_1 |
| Prr13 | ENSMUSG00000023048 | protein_coding | proline rich 13 | 5.999E-09 | 0.949273743 | 0.817 | 0.456 | 0.000141651 | MM_1 |
| Atp5g2 | ENSMUSG00000062683 | protein_coding | ATP synthase, H+ transporting, mitochondrial F0 complex, subunit C2 (subunit 9) | 4.516E-16 | 0.836120419 | 0.977 | 0.807 | 1.06637E-11 | MM_1 |
| Hmox2 | ENSMUSG00000004070 | protein_coding | heme oxygenase 2 | 6.008E-10 | 0.645860433 | 0.925 | 0.693 | 1.41855E-05 | MM_1 |
| Ranbp1 | ENSMUSG00000005732 | protein_coding | RAN binding protein 1 | 4.321E-09 | 0.91727127 | 0.84 | 0.526 | 0.000102028 | MM_1 |
| Ap2m1 | ENSMUSG00000022841 | protein_coding | adaptor-related protein complex 2, mu 1 subunit | 5.2E-13 | 0.914817587 | 0.958 | 0.737 | 1.22776E-08 | MM_1 |
| Tfrc | ENSMUSG00000022797 | protein_coding | transferrin receptor | 5.016E-08 | 0.839360193 | 0.854 | 0.57 | 0.001184366 | MM_1 |
| Mylk | ENSMUSG00000022836 | protein_coding | myosin, light polypeptide kinase | 1.3E-07 | 0.658872543 | 0.906 | 0.561 | 0.003068393 | MM_1 |
| Gm15564 | ENSMUSG00000086324 | lncRNA | predicted gene 15564 | 9.613E-30 | -2.274801109 | 0.737 | 0.93 | 2.26985E-25 | MM_1 |
| Ndufb4 | ENSMUSG00000022820 | protein_coding | NADH:ubiquinone oxidoreductase subunit B4 | 6.313E-11 | 1.028844155 | 0.84 | 0.535 | 1.49063E-06 | MM_1 |
| Cd200r1 | ENSMUSG00000022667 | protein_coding | CD200 receptor 1 | 1.521E-07 | 1.10192329 | 0.648 | 0.36 | 0.003591331 | MM_1 |
| Cmss1 | ENSMUSG00000022748 | protein_coding | cms small ribosomal subunit 1 | 2.231E-40 | -2.330101496 | 0.991 | 0.982 | 5.26899E-36 | MM_1 |
| Mir99ahg | ENSMUSG00000090386 | lncRNA | Mir99a and Mirlet7c-1 host gene (non-protein coding) | 9.865E-08 | 0.747703305 | 0.93 | 0.763 | 0.00232936 | MM_1 |
| Atp5j | ENSMUSG00000022890 | protein_coding | ATP synthase, H+ transporting, mitochondrial F0 complex, subunit F | 3.416E-14 | 0.966865241 | 0.939 | 0.728 | 8.06478E-10 | MM_1 |
| Cryzl1 | ENSMUSG00000058240 | protein_coding | crystallin, zeta (quinone reductase)-like 1 | 1.293E-08 | 0.642861478 | 0.554 | 0.193 | 0.000305208 | MM_1 |
| Atp5o | ENSMUSG00000022956 | protein_coding | ATP synthase, H+ transporting, mitochondrial F1 complex, O subunit | 2.953E-12 | 1.173369404 | 0.859 | 0.509 | 6.97188E-08 | MM_1 |
| Rnaset2b | ENSMUSG00000094724 | protein_coding | ribonuclease T2B | 6.72E-12 | 1.740468739 | 0.662 | 0.281 | 1.58666E-07 | MM_1 |
| Rnaset2a | ENSMUSG00000095687 | protein_coding | ribonuclease T2A | 1.216E-16 | 1.500848001 | 0.883 | 0.439 | 2.8703E-12 | MM_1 |
| Elob | ENSMUSG00000055839 | protein_coding | elongin B | 1.63E-14 | 1.025504562 | 0.944 | 0.702 | 3.84953E-10 | MM_1 |
| Atp6v0c | ENSMUSG00000024121 | protein_coding | ATPase, H+ transporting, lysosomal V0 subunit C | 9.205E-15 | 0.789704253 | 0.995 | 0.912 | 2.17338E-10 | MM_1 |
| Ndufb10 | ENSMUSG00000040048 | protein_coding | NADH:ubiquinone oxidoreductase subunit B10 | 4.624E-08 | 0.766320981 | 0.84 | 0.561 | 0.001091873 | MM_1 |
| Haghl | ENSMUSG00000061046 | protein_coding | hydroxyacylglutathione hydrolase-like | 1.013E-07 | 1.084615964 | 0.305 | 0.044 | 0.002391406 | MM_1 |
| Fam173a | ENSMUSG00000057411 | protein_coding | family with sequence similarity 173, member A | 5.044E-09 | 1.264301103 | 0.606 | 0.254 | 0.000119098 | MM_1 |
| Stub1 | ENSMUSG00000039615 | protein_coding | STIP1 homology and U-Box containing protein 1 | 1.978E-13 | 1.370196131 | 0.789 | 0.298 | 4.67067E-09 | MM_1 |
| Fam234a | ENSMUSG00000024187 | protein_coding | family with sequence similarity 234, member A | 3.826E-07 | 0.750022626 | 0.789 | 0.43 | 0.009033779 | MM_1 |
| Cuta | ENSMUSG00000024194 | protein_coding | cutA divalent cation tolerance homolog | 1.714E-08 | 0.906626737 | 0.822 | 0.5 | 0.00040482 | MM_1 |
| Uqcc2 | ENSMUSG00000024208 | protein_coding | ubiquinol-cytochrome c reductase complex assembly factor 2 | 2.463E-08 | 0.998971719 | 0.742 | 0.395 | 0.000581499 | MM_1 |
| BC004004 | ENSMUSG00000052712 | protein_coding | cDNA sequence BC004004 | 1.499E-10 | 1.758872946 | 0.521 | 0.132 | 3.53904E-06 | MM_1 |
| Mtch1 | ENSMUSG00000024012 | protein_coding | mitochondrial carrier 1 | 2.515E-11 | 1.34032478 | 0.723 | 0.281 | 5.93936E-07 | MM_1 |
| Glo1 | ENSMUSG00000024026 | protein_coding | glyoxalase 1 | 3.87E-07 | 1.025365063 | 0.601 | 0.281 | 0.009138561 | MM_1 |
| Ndufv3 | ENSMUSG00000024038 | protein_coding | NADH:ubiquinone oxidoreductase core subunit V3 | 1.828E-07 | 0.855356478 | 0.789 | 0.491 | 0.004315908 | MM_1 |
| Rps28 | ENSMUSG00000067288 | protein_coding | ribosomal protein S28 | 5.305E-24 | 0.799626072 | 1 | 0.947 | 1.2527E-19 | MM_1 |
| Ndufa7 | ENSMUSG00000041881 | protein_coding | NADH:ubiquinone oxidoreductase subunit A7 | 3.321E-09 | 0.866546274 | 0.897 | 0.64 | 7.84162E-05 | MM_1 |
| Rnf5 | ENSMUSG00000015478 | protein_coding | ring finger protein 5 | 1.208E-11 | 1.931296098 | 0.502 | 0.105 | 2.85176E-07 | MM_1 |

| | | | | | | | | | |
|---|---|---|---|---|---|---|---|---|---|
| Hspa1b | ENSMUSG00000090877 | protein_coding | heat shock protein 1B | 1.172E-14 | 1.542510355 | 0.958 | 0.851 | 2.76729E-10 | MM_1 |
| Hspa1a | ENSMUSG00000091971 | protein_coding | heat shock protein 1A | 4.636E-09 | 1.092500174 | 0.986 | 0.877 | 0.000109459 | MM_1 |
| Lst1 | ENSMUSG00000073412 | protein_coding | leukocyte specific transcript 1 | 8.845E-08 | 0.874229229 | 0.836 | 0.544 | 0.002088476 | MM_1 |
| H2-D1 | ENSMUSG00000073411 | protein_coding | histocompatibility 2, D region locus 1 | 3.289E-13 | 0.636944483 | 0.995 | 0.947 | 7.76554E-09 | MM_1 |
| Gm42418 | ENSMUSG00000098178 | lncRNA | predicted gene, 42418 | 6.063E-47 | -2.000346381 | 1 | 1 | 1.43171E-42 | MM_1 |
| AY036118 | ENSMUSG00000105361 | lncRNA | cDNA sequence AY036118 | 5.032E-23 | -1.414364272 | 0.948 | 0.965 | 1.18825E-18 | MM_1 |
| Slc29a1 | ENSMUSG00000023942 | protein_coding | solute carrier family 29 (nucleoside transporters), member 1 | 5.299E-11 | 1.201154456 | 0.812 | 0.439 | 1.25108E-06 | MM_1 |
| Yipf3 | ENSMUSG00000071074 | protein_coding | Yip1 domain family, member 3 | 1.439E-09 | 1.470243743 | 0.498 | 0.14 | 3.39828E-05 | MM_1 |
| Tomm6 | ENSMUSG00000033475 | protein_coding | translocase of outer mitochondrial membrane 6 | 4.605E-18 | 1.155279948 | 0.934 | 0.675 | 1.08735E-13 | MM_1 |
| Ubxn6 | ENSMUSG00000019578 | protein_coding | UBX domain protein 6 | 3.048E-07 | 1.381214931 | 0.488 | 0.175 | 0.007195939 | MM_1 |
| Ndufa11 | ENSMUSG00000002379 | protein_coding | NADH:ubiquinone oxidoreductase subunit A11 | 5.499E-14 | 1.438777024 | 0.803 | 0.412 | 1.29834E-09 | MM_1 |
| Rfx2 | ENSMUSG00000024206 | protein_coding | regulatory factor X, 2 (influences HLA class II expression) | 7.804E-09 | -2.394456633 | 0.103 | 0.351 | 0.00018426 | MM_1 |
| Ndufv2 | ENSMUSG00000024099 | protein_coding | NADH:ubiquinone oxidoreductase core subunit V2 | 9.837E-08 | 0.885553277 | 0.765 | 0.421 | 0.002322819 | MM_1 |
| Rasgrp3 | ENSMUSG00000071042 | protein_coding | RAS, guanyl releasing protein 3 | 2.114E-08 | 1.101542367 | 0.831 | 0.544 | 0.000499123 | MM_1 |
| Galm | ENSMUSG00000035473 | protein_coding | galactose mutarotase | 1.148E-07 | 1.392941906 | 0.394 | 0.096 | 0.002711787 | MM_1 |
| Mcfd2 | ENSMUSG00000024150 | protein_coding | multiple coagulation factor deficiency 2 | 1.208E-07 | 0.934105749 | 0.671 | 0.307 | 0.002852828 | MM_1 |
| Tslp | ENSMUSG00000024379 | protein_coding | thymic stromal lymphopoietin | 9.474E-09 | 1.088982223 | 0.676 | 0.298 | 0.000223707 | MM_1 |
| Cystm1 | ENSMUSG00000046727 | protein_coding | cysteine-rich transmembrane module containing 1 | 7.729E-11 | 1.929529198 | 0.582 | 0.202 | 1.82496E-06 | MM_1 |
| Ndufa2 | ENSMUSG00000014294 | protein_coding | NADH:ubiquinone oxidoreductase subunit A2 | 1.256E-08 | 0.706646178 | 0.944 | 0.719 | 0.000296525 | MM_1 |
| Commd10 | ENSMUSG00000042705 | protein_coding | COMM domain containing 10 | 9.175E-10 | 1.221609026 | 0.714 | 0.333 | 2.16639E-05 | MM_1 |
| Sema6a | ENSMUSG00000019647 | protein_coding | sema domain, transmembrane domain (TM), and cytoplasmic domain, (semaphorin) 6A | 1.714E-08 | 1.702896025 | 0.531 | 0.193 | 0.000404752 | MM_1 |
| Hsd17b4 | ENSMUSG00000024507 | protein_coding | hydroxysteroid (17-beta) dehydrogenase 4 | 9.88E-08 | 0.849510042 | 0.808 | 0.465 | 0.002332786 | MM_1 |
| Aldh7a1 | ENSMUSG00000053644 | protein_coding | aldehyde dehydrogenase family 7, member A1 | 1.873E-09 | 1.025976953 | 0.629 | 0.219 | 4.42281E-05 | MM_1 |
| Csf1r | ENSMUSG00000024621 | protein_coding | colony stimulating factor 1 receptor | 4.839E-16 | 0.687504253 | 1 | 0.939 | 1.14262E-11 | MM_1 |
| Atp5a1 | ENSMUSG00000025428 | protein_coding | ATP synthase, H+ transporting, mitochondrial F1 complex, alpha subunit 1 | 2.711E-10 | 0.709177931 | 0.948 | 0.868 | 6.40088E-06 | MM_1 |
| Cyb5a | ENSMUSG00000024646 | protein_coding | cytochrome b5 type A (microsomal) | 1.175E-13 | 1.014175162 | 0.939 | 0.64 | 2.77377E-09 | MM_1 |
| Ndufs8 | ENSMUSG00000059734 | protein_coding | NADH:ubiquinone oxidoreductase core subunit S8 | 9.237E-11 | 1.29705072 | 0.723 | 0.316 | 2.18109E-06 | MM_1 |
| Ndufv1 | ENSMUSG00000037916 | protein_coding | NADH:ubiquinone oxidoreductase core subunit V1 | 2.583E-08 | 1.420030974 | 0.484 | 0.149 | 0.000609984 | MM_1 |
| Gstp1 | ENSMUSG00000060803 | protein_coding | glutathione S-transferase, pi 1 | 1.604E-10 | 2.34876135 | 0.404 | 0.053 | 3.7882E-06 | MM_1 |
| Banf1 | ENSMUSG00000024844 | protein_coding | barrier to autointegration factor 1 | 2.342E-08 | 1.032358128 | 0.69 | 0.342 | 0.000553088 | MM_1 |
| Drap1 | ENSMUSG00000024914 | protein_coding | Dr1 associated protein 1 (negative cofactor 2 alpha) | 1.069E-07 | 0.70563409 | 0.883 | 0.579 | 0.002525028 | MM_1 |
| Cfl1 | ENSMUSG00000056201 | protein_coding | cofilin 1, non-muscle | 1.545E-16 | 0.740315012 | 1 | 0.921 | 3.64752E-12 | MM_1 |
| Fam89b | ENSMUSG00000024939 | protein_coding | family with sequence similarity 89, member B | 3.523E-09 | 1.137113624 | 0.714 | 0.351 | 8.31832E-05 | MM_1 |
| Fkbp2 | ENSMUSG00000056629 | protein_coding | FK506 binding protein 2 | 1.1E-07 | 0.775137544 | 0.845 | 0.526 | 0.002598025 | MM_1 |
| Otub1 | ENSMUSG00000024767 | protein_coding | OTU domain, ubiquitin aldehyde binding 1 | 1.598E-07 | 1.245288195 | 0.554 | 0.228 | 0.003772884 | MM_1 |
| Cox8a | ENSMUSG00000035885 | protein_coding | cytochrome c oxidase subunit 8A | 7.38E-18 | 0.946483061 | 0.981 | 0.798 | 1.74266E-13 | MM_1 |
| Tmem179b | ENSMUSG00000118346 | protein_coding | transmembrane protein 179B | 5.032E-08 | 1.22503772 | 0.549 | 0.202 | 0.001188224 | MM_1 |
| Fth1 | ENSMUSG00000024661 | protein_coding | ferritin heavy polypeptide 1 | 1.159E-25 | 0.924671969 | 1 | 0.982 | 2.73775E-21 | MM_1 |
| Tmem258 | ENSMUSG00000036372 | protein_coding | transmembrane protein 258 | 1.945E-12 | 1.079827536 | 0.892 | 0.526 | 4.5921E-08 | MM_1 |
| Ostf1 | ENSMUSG00000024725 | protein_coding | osteoclast stimulating factor 1 | 3.856E-09 | 0.705311067 | 0.977 | 0.746 | 9.10434E-05 | MM_1 |
| Gda | ENSMUSG00000058624 | protein_coding | guanine deaminase | 1.544E-07 | -1.058490922 | 0.869 | 0.833 | 0.003645255 | MM_1 |
| Rpl9-ps6 | ENSMUSG00000062456 | protein_coding | ribosomal protein L9, pseudogene 6 | 4.454E-10 | 2.446019428 | 0.399 | 0.061 | 1.0517E-05 | MM_1 |
| Acta2 | ENSMUSG00000035783 | protein_coding | actin, alpha 2, smooth muscle, aorta | 7.036E-18 | 1.114321731 | 0.972 | 0.684 | 1.66134E-13 | MM_1 |
| Lipa | ENSMUSG00000024781 | protein_coding | lysosomal acid lipase A | 7.35E-16 | 1.074040252 | 0.967 | 0.667 | 1.7354E-11 | MM_1 |
| Ndufb8 | ENSMUSG00000025204 | protein_coding | NADH:ubiquinone oxidoreductase subunit B8 | 2.45E-09 | 0.882621066 | 0.892 | 0.596 | 5.78524E-05 | MM_1 |
| Lzts2 | ENSMUSG00000035342 | protein_coding | leucine zipper, putative tumor suppressor 2 | 5.899E-08 | 0.846489345 | 0.507 | 0.167 | 0.001392971 | MM_1 |
| Atp5md | ENSMUSG00000071528 | protein_coding | ATP synthase membrane subunit DAPIT | 5.413E-18 | 1.207703648 | 0.925 | 0.667 | 1.27816E-13 | MM_1 |
| Gpr34 | ENSMUSG00000040229 | protein_coding | G protein-coupled receptor 34 | 1.848E-18 | 1.549589177 | 0.906 | 0.579 | 4.36321E-14 | MM_1 |
| Ndufb11 | ENSMUSG00000031059 | protein_coding | NADH:ubiquinone oxidoreductase subunit B11 | 5.528E-08 | 0.778047899 | 0.854 | 0.57 | 0.001305277 | MM_1 |
| Cfp | ENSMUSG00000001128 | protein_coding | complement factor properdin | 5.431E-18 | 1.057885717 | 0.972 | 0.833 | 1.28226E-13 | MM_1 |
| Slc25a5 | ENSMUSG00000016319 | protein_coding | solute carrier family 25 (mitochondrial carrier, adenine nucleotide translocator), member 5 | 3.792E-16 | 0.991264158 | 0.972 | 0.772 | 8.95411E-12 | MM_1 |
| Ndufa1 | ENSMUSG00000016427 | protein_coding | NADH:ubiquinone oxidoreductase subunit A1 | 6.226E-13 | 1.224976976 | 0.84 | 0.5 | 1.47002E-08 | MM_1 |
| Rtl8a | ENSMUSG00000067925 | protein_coding | retrotransposon Gag like 8A | 8.165E-08 | 1.613527635 | 0.408 | 0.114 | 0.001927812 | MM_1 |
| Renbp | ENSMUSG00000031387 | protein_coding | renin binding protein | 2.709E-09 | 0.842399564 | 0.84 | 0.5 | 6.39648E-05 | MM_1 |
| Cox7b | ENSMUSG00000031231 | protein_coding | cytochrome c oxidase subunit 7B | 7.722E-15 | 1.161246069 | 0.925 | 0.667 | 1.82322E-10 | MM_1 |
| Bex3 | ENSMUSG00000046432 | protein_coding | brain expressed X-linked 3 | 6.627E-10 | 1.429918764 | 0.643 | 0.272 | 1.56465E-05 | MM_1 |
| Sat1 | ENSMUSG00000025283 | protein_coding | spermidine/spermine N1-acetyl transferase 1 | 1.941E-11 | 0.7889031 | 0.991 | 0.877 | 4.58398E-07 | MM_1 |
| Reps2 | ENSMUSG00000040855 | protein_coding | RALBP1 associated Eps domain containing protein 2 | 3.205E-13 | 1.006616283 | 0.897 | 0.518 | 7.56733E-09 | MM_1 |
| G530011O06Rik | ENSMUSG00000072844 | lncRNA | RIKEN cDNA G530011O06 gene | 5.4E-09 | 1.81381948 | 0.624 | 0.289 | 0.000127498 | MM_1 |
| mt-Nd3 | ENSMUSG00000064360 | protein_coding | mitochondrially encoded NADH dehydrogenase 3 | 5.015E-12 | -0.782702635 | 1 | 0.982 | 1.18421E-07 | MM_1 |
| AC149090.1 | ENSMUSG00000095041 | protein_coding | | 4.576E-18 | 1.337942711 | 0.944 | 0.693 | 1.08049E-13 | MM_1 |
| Per2 | ENSMUSG00000055866 | protein_coding | period circadian clock 2 | 1.906E-08 | -3.636124587 | 0.037 | 0.278 | 0.000450004 | MM_2 |
| Gm2000 | ENSMUSG00000078193 | protein_coding | predicted gene 2000 | 4.253E-21 | 3.07433004 | 0.82 | 0.165 | 1.00431E-16 | MM_2 |
| Camk1d | ENSMUSG00000039145 | protein_coding | calcium/calmodulin-dependent protein kinase ID | 1.633E-22 | -1.02603555 | 1 | 1 | 3.85612E-18 | MM_2 |
| Rpl35 | ENSMUSG00000062997 | protein_coding | ribosomal protein L35 | 1.605E-13 | 0.749697101 | 0.994 | 0.975 | 3.78934E-09 | MM_2 |
| B2m | ENSMUSG00000060802 | protein_coding | beta-2 microglobulin | 2.365E-17 | 0.770175014 | 1 | 0.987 | 5.58422E-13 | MM_2 |
| Cst3 | ENSMUSG00000027447 | protein_coding | cystatin C | 3.241E-11 | 0.722376324 | 1 | 0.987 | 7.65177E-07 | MM_2 |
| Myl9 | ENSMUSG00000067818 | protein_coding | myosin, light polypeptide 9, regulatory | 1.307E-09 | 1.313840708 | 0.807 | 0.342 | 3.08686E-05 | MM_2 |
| Selenof | ENSMUSG00000037072 | protein_coding | selenoprotein F | 9.969E-09 | 0.773278411 | 0.944 | 0.797 | 0.000235395 | MM_2 |
| Gm11808 | ENSMUSG00000068240 | protein_coding | predicted gene 11808 | 1.985E-25 | 2.96472635 | 0.919 | 0.215 | 4.6862E-21 | MM_2 |
| Ptafr | ENSMUSG00000056529 | protein_coding | platelet-activating factor receptor | 3.86E-07 | 1.076202186 | 0.882 | 0.595 | 0.009113995 | MM_2 |
| Ost4 | ENSMUSG00000038803 | protein_coding | oligosaccharyltransferase complex subunit 4 (non-catalytic) | 4.192E-08 | 0.815015652 | 0.938 | 0.696 | 0.000989912 | MM_2 |
| Cdk8 | ENSMUSG00000029635 | protein_coding | cyclin-dependent kinase 8 | 4.612E-33 | -2.728195582 | 0.988 | 0.987 | 1.08898E-28 | MM_2 |
| Actg2 | ENSMUSG00000059430 | protein_coding | actin, gamma 2, smooth muscle, enteric | 1.367E-10 | 1.14890292 | 0.932 | 0.582 | 3.22729E-06 | MM_2 |
| Gapdh | ENSMUSG00000057666 | protein_coding | glyceraldehyde-3-phosphate dehydrogenase | 1.459E-11 | 0.921329502 | 0.975 | 0.848 | 3.44449E-07 | MM_2 |
| Ftl1 | ENSMUSG00000050708 | protein_coding | ferritin light polypeptide 1 | 2.994E-16 | 0.7565342 | 1 | 0.987 | 7.0704E-12 | MM_2 |
| Dbp | ENSMUSG00000059824 | protein_coding | D site albumin promoter binding protein | 9.213E-09 | -2.007306345 | 0.273 | 0.57 | 0.000217544 | MM_2 |
| Ifitm2 | ENSMUSG00000060591 | protein_coding | interferon induced transmembrane protein 2 | 3.585E-09 | 0.785884096 | 0.994 | 0.899 | 8.46486E-05 | MM_2 |
| Uba52 | ENSMUSG00000090137 | protein_coding | ubiquitin A-52 residue ribosomal protein fusion product 1 | 2.094E-23 | 1.890776434 | 0.969 | 0.557 | 4.94526E-19 | MM_2 |
| Cnn1 | ENSMUSG00000001349 | protein_coding | calponin 1 | 1.123E-07 | 0.983757048 | 0.87 | 0.506 | 0.002651097 | MM_2 |
| Gpx1 | ENSMUSG00000063856 | protein_coding | glutathione peroxidase 1 | 1.465E-12 | 1.021113694 | 0.981 | 0.899 | 3.45945E-08 | MM_2 |
| Lars2 | ENSMUSG00000035202 | protein_coding | leucyl-tRNA synthetase, mitochondrial | 2.693E-30 | -2.200585919 | 0.988 | 0.987 | 6.35901E-26 | MM_2 |
| Cirbp | ENSMUSG00000045193 | protein_coding | cold inducible RNA binding protein | 2.788E-07 | 1.52728442 | 0.547 | 0.152 | 0.006582842 | MM_2 |
| Tle5 | ENSMUSG00000054452 | protein_coding | TLE family member 5, transcriptional modulator | 2.835E-07 | 0.904801358 | 0.876 | 0.57 | 0.006694855 | MM_2 |
| Lyz2 | ENSMUSG00000069516 | protein_coding | lysozyme 2 | 1.398E-37 | 12.83527697 | 1 | 0 | 3.30098E-33 | MM_2 |
| Lyz1 | ENSMUSG00000069515 | protein_coding | lysozyme 1 | 4.418E-09 | -4.337004659 | 0.143 | 0.443 | 0.000104315 | MM_2 |
| Rpl41 | ENSMUSG00000093674 | protein_coding | ribosomal protein L41 | 2.064E-17 | 0.73535075 | 1 | 0.987 | 4.87383E-13 | MM_2 |
| Bloc1s1 | ENSMUSG00000090247 | protein_coding | biogenesis of lysosomal organelles complex-1, subunit 1 | 1.434E-08 | 1.976864261 | 0.503 | 0.101 | 0.000338622 | MM_2 |
| Smtn | ENSMUSG00000020439 | protein_coding | smoothelin | 6.98E-08 | 1.652503747 | 0.484 | 0.101 | 0.001648091 | MM_2 |
| Ubb | ENSMUSG00000019505 | protein_coding | ubiquitin B | 3.501E-08 | 0.625963209 | 0.994 | 0.975 | 0.00082667 | MM_2 |
| Cd68 | ENSMUSG00000018774 | protein_coding | CD68 antigen | 1.526E-07 | 0.699243676 | 0.975 | 0.911 | 0.003602152 | MM_2 |
| Lgals9 | ENSMUSG00000001123 | protein_coding | lectin, galactose binding, soluble 9 | 9.203E-09 | 1.494779769 | 0.658 | 0.253 | 0.000217297 | MM_2 |
| Evi2a | ENSMUSG00000078771 | protein_coding | ecotropic viral integration site 2a | 3.816E-10 | 1.048570319 | 0.894 | 0.532 | 9.00982E-06 | MM_2 |
| Rpl38 | ENSMUSG00000057322 | protein_coding | ribosomal protein L38 | 6.19E-25 | 1.015729419 | 1 | 0.937 | 1.46163E-20 | MM_2 |
| Rps29 | ENSMUSG00000034892 | protein_coding | ribosomal protein S29 | 5.442E-21 | 0.766785972 | 1 | 1 | 1.28496E-16 | MM_2 |
| Gphn | ENSMUSG00000047454 | protein_coding | gephyrin | 3.812E-24 | -1.436260814 | 1 | 0.987 | 9.00139E-20 | MM_2 |
| Gm19951 | ENSMUSG00000113136 | lncRNA | predicted gene, 19951 | 5.2E-14 | -2.13020207 | 0.354 | 0.709 | 1.22789E-09 | MM_2 |
| Ndufb1-ps | ENSMUSG00000113902 | protein_coding | NADH:ubiquinone oxidoreductase subunit B1 | 4.263E-08 | 0.749692648 | 0.957 | 0.722 | 0.00100663 | MM_2 |
| Ckb | ENSMUSG00000001270 | protein_coding | creatine kinase, brain | 1.118E-07 | 1.035022077 | 0.938 | 0.734 | 0.002640741 | MM_2 |
| Pld4 | ENSMUSG00000052160 | protein_coding | phospholipase D family, member 4 | 1.026E-07 | 0.782169369 | 0.932 | 0.709 | 0.002422611 | MM_2 |
| Hist1h4d | ENSMUSG00000061482 | protein_coding | histone cluster 1, H4d | 9.122E-08 | -2.394574834 | 0.224 | 0.506 | 0.002153853 | MM_2 |
| Ftl1-ps1 | ENSMUSG00000062382 | protein_coding | ferritin light polypeptide 1, pseudogene 1 | 4.281E-20 | 2.459509697 | 0.876 | 0.228 | 1.01078E-15 | MM_2 |
| Phf11b | ENSMUSG00000091649 | protein_coding | PHD finger protein 11B | 4.223E-07 | 1.473755627 | 0.565 | 0.203 | 0.009970913 | MM_2 |
| Tmbim6 | ENSMUSG00000023010 | protein_coding | transmembrane BAX inhibitor motif containing 6 | 2.149E-08 | 0.727119236 | 0.957 | 0.861 | 0.00050731 | MM_2 |
| Gm15564 | ENSMUSG00000086324 | lncRNA | predicted gene 15564 | 8.201E-19 | -2.582823953 | 0.534 | 0.873 | 1.93637E-14 | MM_2 |
| Cmss1 | ENSMUSG00000022748 | protein_coding | cms small ribosomal subunit 1 | 3.147E-33 | -2.163099663 | 1 | 0.987 | 7.43089E-29 | MM_2 |
| Rnaset2a | ENSMUSG00000095687 | protein_coding | ribonuclease T2A | 5.747E-12 | 1.324313213 | 0.876 | 0.443 | 1.35699E-07 | MM_2 |
| Atp6v0c | ENSMUSG00000024121 | protein_coding | ATPase, H+ transporting, lysosomal V0 subunit C | 2.858E-07 | 0.648656674 | 0.981 | 0.924 | 0.006747746 | MM_2 |
| Rps28 | ENSMUSG00000067288 | protein_coding | ribosomal protein S28 | 4.154E-16 | 0.698025028 | 1 | 0.975 | 9.80879E-12 | MM_2 |
| H2-Ab1 | ENSMUSG00000073421 | protein_coding | histocompatibility 2, class II antigen A, beta 1 | 5.515E-13 | 0.965565995 | 1 | 0.949 | 1.30213E-08 | MM_2 |
| H2-Aa | ENSMUSG00000036594 | protein_coding | histocompatibility 2, class II antigen A, alpha | 9.282E-13 | 0.868867007 | 0.988 | 0.962 | 2.19159E-08 | MM_2 |
| H2-Eb1 | ENSMUSG00000060586 | protein_coding | histocompatibility 2, class II antigen E beta | 1.293E-07 | 0.639067079 | 0.994 | 0.937 | 0.003051869 | MM_2 |
| Hspa1b | ENSMUSG00000090877 | protein_coding | heat shock protein 1B | 9.165E-09 | 1.291751943 | 0.938 | 0.873 | 0.0002164 | MM_2 |
| Hspa1a | ENSMUSG00000091971 | protein_coding | heat shock protein 1A | 8.163E-09 | 1.221408421 | 0.963 | 0.835 | 0.000192746 | MM_2 |
| Lst1 | ENSMUSG00000073412 | protein_coding | leukocyte specific transcript 1 | 1.511E-10 | 0.926887316 | 0.975 | 0.81 | 3.56687E-06 | MM_2 |
| H2-D1 | ENSMUSG00000073411 | protein_coding | histocompatibility 2, D region locus 1 | 2.029E-17 | 0.948637531 | 1 | 0.962 | 4.79124E-13 | MM_2 |
| H2-T23 | ENSMUSG00000067212 | protein_coding | histocompatibility 2, T region locus 23 | 7.472E-08 | 1.095497375 | 0.882 | 0.532 | 0.001764369 | MM_2 |
| Gm42418 | ENSMUSG00000098178 | lncRNA | predicted gene, 42418 | 6.371E-36 | -2.401195676 | 1 | 1 | 1.50423E-31 | MM_2 |
| AY036118 | ENSMUSG00000105361 | lncRNA | cDNA sequence AY036118 | 1.229E-27 | -1.671761545 | 0.988 | 0.987 | 2.90304E-23 | MM_2 |
| Gm49980 | ENSMUSG00000117465 | lncRNA | predicted gene, 49980 | 1.674E-07 | 4.279744878 | 0.286 | 0 | 0.003952432 | MM_2 |
| Cd74 | ENSMUSG00000024610 | protein_coding | CD74 antigen (invariant polypeptide of major histocompatibility complex, class II antigen-associated) | 2.015E-19 | 1.21112008 | 1 | 1 | 4.75688E-15 | MM_2 |

| | | | | | | | | | |
|---|---|---|---|---|---|---|---|---|---|
| Csf1r | ENSMUSG00000024621 | protein_coding | colony stimulating factor 1 receptor | 3.902E-09 | 0.659324433 | 1 | 0.949 | 9.21346E-05 | MM_2 |
| Fth1 | ENSMUSG00000024661 | protein_coding | ferritin heavy polypeptide 1 | 7.439E-11 | 0.624731038 | 1 | 0.987 | 1.7564E-06 | MM_2 |
| Acta2 | ENSMUSG00000035783 | protein_coding | actin, alpha 2, smooth muscle, aorta | 2.906E-11 | 0.854709283 | 0.975 | 0.658 | 6.86125E-07 | MM_2 |
| Atp5md | ENSMUSG00000071528 | protein_coding | ATP synthase membrane subunit DAPIT | 2.48E-07 | 0.875486461 | 0.901 | 0.684 | 0.005856546 | MM_2 |
| Rpl39 | ENSMUSG00000079641 | protein_coding | ribosomal protein L39 | 7.975E-15 | 0.687186115 | 0.994 | 0.949 | 1.88308E-10 | MM_2 |
| G530011O06Rik | ENSMUSG00000072844 | lncRNA | RIKEN cDNA G530011O06 gene | 6.477E-17 | 2.554243072 | 0.857 | 0.316 | 1.52937E-12 | MM_2 |
| Gm15726 | ENSMUSG00000087263 | lncRNA | predicted gene 15726 | 8.391E-08 | 1.528289311 | 0.528 | 0.139 | 0.001981191 | MM_2 |
| mt-Nd3 | ENSMUSG00000064360 | protein_coding | mitochondrially encoded NADH dehydrogenase 3 | 8.643E-10 | -0.753766869 | 0.981 | 0.987 | 2.04071E-05 | MM_2 |
| Actb | ENSMUSG00000029580 | protein_coding | actin, beta | 4.56E-11 | 1.175054462 | 1 | 0.892 | 1.07675E-06 | MM_3 |
| Cdk8 | ENSMUSG00000029635 | protein_coding | cyclin-dependent kinase 8 | 6.776E-28 | -2.54339974 | 0.919 | 1 | 1.59989E-23 | MM_3 |
| Mt1 | ENSMUSG00000031765 | protein_coding | metallothionein 1 | 1.955E-07 | 2.270259835 | 0.573 | 0.189 | 0.004616981 | MM_3 |
| Lars2 | ENSMUSG00000035202 | protein_coding | leucyl-tRNA synthetase, mitochondrial | 2.989E-18 | -1.833085103 | 0.879 | 0.986 | 7.05794E-14 | MM_3 |
| Lyz2 | ENSMUSG00000069516 | protein_coding | lysozyme 2 | 4.651E-24 | 11.78132661 | 0.806 | 0 | 1.09826E-19 | MM_3 |
| Rpl38 | ENSMUSG00000057322 | protein_coding | ribosomal protein L38 | 1.148E-07 | 1.537954691 | 0.871 | 0.486 | 0.00271091 | MM_3 |
| Gphn | ENSMUSG00000047454 | protein_coding | gephyrin | 4.761E-14 | -1.52460754 | 0.887 | 0.946 | 1.12421E-09 | MM_3 |
| Crip1 | ENSMUSG00000006360 | protein_coding | cysteine-rich protein 1 (intestinal) | 1.106E-11 | 1.885262782 | 0.79 | 0.324 | 2.61037E-07 | MM_3 |
| Gm15564 | ENSMUSG00000086324 | lncRNA | predicted gene 15564 | 2.156E-12 | -2.330229164 | 0.298 | 0.73 | 5.09148E-08 | MM_3 |
| Cmss1 | ENSMUSG00000022748 | protein_coding | cms small ribosomal subunit 1 | 2.322E-19 | -1.849497092 | 0.96 | 0.973 | 5.48298E-15 | MM_3 |
| Gm42418 | ENSMUSG00000098178 | lncRNA | predicted gene, 42418 | 4.149E-23 | -1.733125589 | 0.992 | 1 | 9.79637E-19 | MM_3 |
| mt-Nd3 | ENSMUSG00000064360 | protein_coding | mitochondrially encoded NADH dehydrogenase 3 | 3.867E-07 | -1.138223363 | 0.863 | 0.932 | 0.009129881 | MM_3 |
| Gm11808 | ENSMUSG00000068240 | protein_coding | predicted gene 11808 | 3.176E-09 | 3.075955115 | 0.909 | 0.37 | 7.50004E-05 | MM_4 |
| Cdk8 | ENSMUSG00000029635 | protein_coding | cyclin-dependent kinase 8 | 2.632E-10 | -2.834189869 | 0.977 | 1 | 6.21482E-06 | MM_4 |
| Lyz2 | ENSMUSG00000069516 | protein_coding | lysozyme 2 | 4.791E-13 | 10.49818644 | 1 | 0 | 1.13119E-08 | MM_4 |
| Cmss1 | ENSMUSG00000022748 | protein_coding | cms small ribosomal subunit 1 | 3.76E-08 | -1.990493955 | 1 | 1 | 0.00088784 | MM_4 |
| Gm42418 | ENSMUSG00000098178 | lncRNA | predicted gene, 42418 | 2.187E-08 | -2.075695142 | 1 | 1 | 0.000516391 | MM_4 |
| **iC1qaCKOvsC1qaFL** | | | | | | | | | |
| Gulp1 | ENSMUSG00000056870 | protein_coding | GULP, engulfment adaptor PTB domain containing 1 | 9.192E-10 | -0.924385889 | 0.961 | 0.988 | 2.17048E-05 | EN_1 |
| Per2 | ENSMUSG00000055866 | protein_coding | period circadian clock 2 | 3.119E-14 | 3.518121027 | 0.421 | 0.046 | 7.36482E-10 | EN_1 |
| Dbi | ENSMUSG00000026385 | protein_coding | diazepam binding inhibitor | 5.238E-35 | -2.82938143 | 0.908 | 1 | 1.23671E-30 | EN_1 |
| Prdx6 | ENSMUSG00000026701 | protein_coding | peroxiredoxin 6 | 7.638E-10 | -0.953270056 | 0.75 | 0.925 | 1.80353E-05 | EN_1 |
| G0s2 | ENSMUSG00000009633 | protein_coding | G0/G1 switch gene 2 | 2.489E-13 | 1.429285968 | 0.947 | 0.821 | 5.87622E-09 | EN_1 |
| Frmd4a | ENSMUSG00000026657 | protein_coding | FERM domain containing 4A | 5.69E-08 | -0.774407819 | 0.961 | 0.994 | 0.001343451 | EN_1 |
| Camk1d | ENSMUSG00000039145 | protein_coding | calcium/calmodulin-dependent protein kinase ID | 1.823E-22 | 0.971154809 | 1 | 1 | 4.30369E-18 | EN_1 |
| Gsn | ENSMUSG00000026879 | protein_coding | gelsolin | 2.909E-11 | -0.768788614 | 1 | 1 | 6.86798E-07 | EN_1 |
| Zeb2 | ENSMUSG00000026872 | protein_coding | zinc finger E-box binding homeobox 2 | 3.466E-09 | -0.70624615 | 0.987 | 0.994 | 8.18336E-05 | EN_1 |
| Cd59a | ENSMUSG00000032679 | protein_coding | CD59a antigen | 3.251E-16 | -1.03928976 | 0.895 | 0.994 | 7.67666E-12 | EN_1 |
| Gm14133 | ENSMUSG00000087029 | lncRNA | predicted gene 14133 | 5.728E-12 | -5.847082553 | 0 | 0.451 | 1.35246E-07 | EN_1 |
| Col20a1 | ENSMUSG00000016356 | protein_coding | collagen, type XX, alpha 1 | 9.561E-08 | -1.582051238 | 0.329 | 0.699 | 0.002257483 | EN_1 |
| Rhoc | ENSMUSG00000002233 | protein_coding | ras homolog family member C | 3.213E-13 | -1.425317374 | 0.513 | 0.942 | 7.58621E-09 | EN_1 |
| Kcna2 | ENSMUSG00000040724 | protein_coding | potassium voltage-gated channel, shaker-related subfamily, member 2 | 2.771E-17 | -1.089632893 | 0.921 | 1 | 6.5431E-13 | EN_1 |
| Ntng1 | ENSMUSG00000059857 | protein_coding | netrin G1 | 6.061E-10 | -1.179528483 | 0.737 | 0.948 | 1.43106E-05 | EN_1 |
| Gm11808 | ENSMUSG00000068240 | protein_coding | predicted gene 11808 | 2.733E-07 | -1.889313669 | 0.092 | 0.451 | 0.006454216 | EN_1 |
| Dnaja1 | ENSMUSG00000028410 | protein_coding | DnaJ heat shock protein family (Hsp40) member A1 | 1.143E-07 | 1.077327503 | 0.868 | 0.775 | 0.002699458 | EN_1 |
| Tpm2 | ENSMUSG00000028464 | protein_coding | tropomyosin 2, beta | 7.234E-09 | -0.843948618 | 0.776 | 0.977 | 0.000170808 | EN_1 |
| Col27a1 | ENSMUSG00000045672 | protein_coding | collagen, type XXVII, alpha 1 | 2.823E-07 | -1.069356996 | 0.461 | 0.803 | 0.006664654 | EN_1 |
| Foxd3 | ENSMUSG00000067261 | protein_coding | forkhead box D3 | 1.856E-13 | -1.406044883 | 0.579 | 0.925 | 4.38171E-09 | EN_1 |
| Atp6v0b | ENSMUSG00000033379 | protein_coding | ATPase, H+ transporting, lysosomal V0 subunit B | 1.363E-07 | 1.068377578 | 0.829 | 0.751 | 0.003219003 | EN_1 |
| Lrrc8b | ENSMUSG00000070639 | protein_coding | leucine rich repeat containing 8 family, member B | 1.674E-10 | -1.051980037 | 0.75 | 0.954 | 3.9517E-06 | EN_1 |
| Atp5k | ENSMUSG00000050856 | protein_coding | ATP synthase, H+ transporting, mitochondrial F1F0 complex, subunit E | 3.258E-09 | -1.028018488 | 0.579 | 0.902 | 7.69351E-05 | EN_1 |
| Hspb1 | ENSMUSG00000004951 | protein_coding | heat shock protein 1 | 2.842E-10 | 1.799376445 | 0.855 | 0.688 | 6.71026E-06 | EN_1 |
| Gnb2 | ENSMUSG00000029713 | protein_coding | guanine nucleotide binding protein (G protein), beta 2 | 5.492E-19 | -1.421432956 | 0.711 | 0.971 | 1.2968E-14 | EN_1 |
| Actb | ENSMUSG00000029580 | protein_coding | actin, beta | 4.401E-21 | -1.014967216 | 0.987 | 1 | 1.03926E-16 | EN_1 |
| Cdk8 | ENSMUSG00000029635 | protein_coding | cyclin-dependent kinase 8 | 8.824E-34 | 2.310439756 | 1 | 1 | 2.08348E-29 | EN_1 |
| Hsph1 | ENSMUSG00000029657 | protein_coding | heat shock 105kDa/110kDa protein 1 | 1.064E-08 | 1.9650965 | 0.697 | 0.41 | 0.000251279 | EN_1 |
| Stard13 | ENSMUSG00000016128 | protein_coding | StAR-related lipid transfer (START) domain containing 13 | 6.218E-11 | -0.682751551 | 1 | 1 | 1.4683E-06 | EN_1 |
| Actg2 | ENSMUSG00000059430 | protein_coding | actin, gamma 2, smooth muscle, enteric | 2.772E-11 | -1.318437711 | 0.645 | 0.931 | 6.54595E-07 | EN_1 |
| Foxp1 | ENSMUSG00000030067 | protein_coding | forkhead box P1 | 4.228E-08 | -0.641064162 | 0.895 | 0.983 | 0.0009983 | EN_1 |
| Kcna1 | ENSMUSG00000047976 | protein_coding | potassium voltage-gated channel, shaker-related subfamily, member 1 | 5.74E-16 | -1.000063402 | 0.921 | 1 | 1.35543E-11 | EN_1 |
| Sox5 | ENSMUSG00000041540 | protein_coding | SRY (sex determining region Y)-box 5 | 4.233E-08 | -0.951261194 | 0.724 | 0.948 | 0.000999551 | EN_1 |
| Capns1 | ENSMUSG00000001794 | protein_coding | calpain, small subunit 1 | 2.901E-07 | -0.845574254 | 0.724 | 0.902 | 0.00684958 | EN_1 |
| Luzp2 | ENSMUSG00000063297 | protein_coding | leucine zipper protein 2 | 3.793E-19 | -2.020952796 | 0.434 | 0.936 | 8.95514E-15 | EN_1 |
| Igf1r | ENSMUSG00000005533 | protein_coding | insulin-like growth factor I receptor | 5.132E-08 | -0.74794303 | 0.816 | 0.948 | 0.001211854 | EN_1 |
| Trim12a | ENSMUSG00000066258 | protein_coding | tripartite motif-containing 12A | 2.189E-09 | 4.100478119 | 0.25 | 0.017 | 5.16864E-05 | EN_1 |
| Sox6 | ENSMUSG00000051910 | protein_coding | SRY (sex determining region Y)-box 6 | 2.085E-09 | -0.815221545 | 0.921 | 0.988 | 4.92209E-05 | EN_1 |
| Cd81 | ENSMUSG00000037706 | protein_coding | CD81 antigen | 1.517E-10 | -0.698663706 | 0.947 | 1 | 3.58095E-06 | EN_1 |
| Poteg | ENSMUSG00000063932 | protein_coding | POTE ankyrin domain family, member G | 2.11E-17 | -6.377701726 | 0 | 0.607 | 4.983E-13 | EN_1 |
| Lpcat2 | ENSMUSG00000033192 | protein_coding | lysophosphatidylcholine acyltransferase 2 | 3.001E-08 | -2.043413683 | 0.105 | 0.491 | 0.000708519 | EN_1 |
| Col5a3 | ENSMUSG00000004098 | protein_coding | collagen, type V, alpha 3 | 7.644E-12 | -1.317003194 | 0.618 | 0.931 | 1.80501E-07 | EN_1 |
| Col12a1 | ENSMUSG00000032332 | protein_coding | collagen, type XII, alpha 1 | 1.555E-07 | 0.806113454 | 0.987 | 0.954 | 0.003671605 | EN_1 |
| Lars2 | ENSMUSG00000035202 | protein_coding | leucyl-tRNA synthetase, mitochondrial | 2.087E-31 | 1.763187571 | 1 | 1 | 4.92703E-27 | EN_1 |
| Ccn2 | ENSMUSG00000019997 | protein_coding | cellular communication network factor 2 | 3.024E-10 | -1.943842082 | 0.566 | 0.815 | 7.14099E-06 | EN_1 |
| Marcks | ENSMUSG00000069662 | protein_coding | myristoylated alanine rich protein kinase C substrate | 3.241E-14 | -0.788439646 | 0.974 | 0.994 | 7.65213E-10 | EN_1 |
| Fabp7 | ENSMUSG00000019874 | protein_coding | fatty acid binding protein 7, brain | 3.218E-18 | -4.780915867 | 0.053 | 0.676 | 7.59734E-14 | EN_1 |
| Arid5b | ENSMUSG00000019947 | protein_coding | AT rich interactive domain 5B (MRF1-like) | 2.812E-07 | -0.918873817 | 0.5 | 0.855 | 0.006638757 | EN_1 |
| Cirbp | ENSMUSG00000045193 | protein_coding | cold inducible RNA binding protein | 3.495E-11 | -1.618180389 | 0.316 | 0.78 | 8.25193E-07 | EN_1 |
| Nr1d1 | ENSMUSG00000020889 | protein_coding | nuclear receptor subfamily 1, group D, member 1 | 5.412E-15 | -2.484811925 | 0.184 | 0.74 | 1.27783E-10 | EN_1 |
| Cnp | ENSMUSG00000006782 | protein_coding | 2',3'-cyclic nucleotide 3' phosphodiesterase | 8.357E-09 | -0.977508854 | 0.789 | 0.96 | 0.000197325 | EN_1 |
| Rpl38 | ENSMUSG00000057322 | protein_coding | ribosomal protein L38 | 1.342E-11 | -0.873063378 | 0.895 | 0.988 | 3.168E-07 | EN_1 |
| Klhl29 | ENSMUSG00000020627 | protein_coding | kelch-like 29 | 1.468E-07 | 0.909402133 | 0.934 | 0.78 | 0.00346703 | EN_1 |
| Asap2 | ENSMUSG00000052632 | protein_coding | ArfGAP with SH3 domain, ankyrin repeat and PH domain 2 | 1.337E-13 | -1.11276858 | 0.868 | 0.988 | 3.15712E-09 | EN_1 |
| Arhgap5 | ENSMUSG00000035133 | protein_coding | Rho GTPase activating protein 5 | 2.957E-08 | -0.800245142 | 0.921 | 0.988 | 0.0006982 | EN_1 |
| Frmd6 | ENSMUSG00000048285 | protein_coding | FERM domain containing 6 | 4.196E-09 | -1.247554292 | 0.487 | 0.832 | 9.90704E-05 | EN_1 |
| Gphn | ENSMUSG00000047454 | protein_coding | gephyrin | 1.314E-21 | 1.035500597 | 1 | 1 | 3.1015E-17 | EN_1 |
| Ckb | ENSMUSG00000001270 | protein_coding | creatine kinase, brain | 1.715E-22 | -1.667812479 | 0.855 | 1 | 4.05064E-18 | EN_1 |
| Hist1h4d | ENSMUSG00000061482 | protein_coding | histone cluster 1, H4d | 5.183E-12 | 1.761657546 | 0.789 | 0.538 | 1.22383E-07 | EN_1 |
| Hist1h1e | ENSMUSG00000051627 | protein_coding | histone cluster 1, H1e | 1.544E-08 | 1.49107292 | 0.697 | 0.376 | 0.000364483 | EN_1 |
| Iqgap2 | ENSMUSG00000021676 | protein_coding | IQ motif containing GTPase activating protein 2 | 6.607E-08 | 0.612174227 | 1 | 0.983 | 0.001560128 | EN_1 |
| Hexb | ENSMUSG00000021665 | protein_coding | hexosaminidase B | 1.2E-12 | -0.944556885 | 0.868 | 0.983 | 2.83256E-08 | EN_1 |
| Itga2 | ENSMUSG00000015533 | protein_coding | integrin alpha 2 | 1.101E-08 | -1.051347445 | 0.671 | 0.913 | 0.000260083 | EN_1 |
| Fam107a | ENSMUSG00000021750 | protein_coding | family with sequence similarity 107, member A | 3.477E-08 | 1.603320013 | 0.645 | 0.393 | 0.000820891 | EN_1 |
| Gm48099 | ENSMUSG00000114304 | lncRNA | predicted gene, 48099 | 2.801E-12 | -2.91765366 | 0.079 | 0.584 | 6.6128E-08 | EN_1 |
| Arhgef3 | ENSMUSG00000021895 | protein_coding | Rho guanine nucleotide exchange factor (GEF) 3 | 1.944E-07 | -1.938969982 | 0.224 | 0.595 | 0.004589288 | EN_1 |
| Spry2 | ENSMUSG00000022114 | protein_coding | sprouty RTK signaling antagonist 2 | 3.671E-07 | -1.388168869 | 0.25 | 0.665 | 0.008667485 | EN_1 |
| Azin1 | ENSMUSG00000037458 | protein_coding | antizyme inhibitor 1 | 1.238E-07 | 0.728847788 | 0.934 | 0.867 | 0.002923398 | EN_1 |
| Sntb1 | ENSMUSG00000060429 | protein_coding | syntrophin, basic 1 | 1.777E-07 | -1.052034817 | 0.803 | 0.908 | 0.004196829 | EN_1 |
| Gm15564 | ENSMUSG00000086324 | lncRNA | predicted gene 15564 | 4.792E-16 | 2.240460994 | 0.842 | 0.451 | 1.13156E-11 | EN_1 |
| Zbtb20 | ENSMUSG00000022708 | protein_coding | zinc finger and BTB domain containing 20 | 1.074E-09 | -0.594799759 | 1 | 1 | 2.5364E-05 | EN_1 |
| Cmss1 | ENSMUSG00000022748 | protein_coding | cms small ribosomal subunit 1 | 3.406E-28 | 1.718602667 | 1 | 1 | 8.04276E-24 | EN_1 |
| Fndc1 | ENSMUSG00000071984 | protein_coding | fibronectin type III domain containing 1 | 2.377E-07 | 1.075603195 | 0.934 | 0.746 | 0.005613118 | EN_1 |
| BC004004 | ENSMUSG00000052712 | protein_coding | cDNA sequence BC004004 | 1.591E-07 | -1.529417891 | 0.276 | 0.665 | 0.003757308 | EN_1 |
| Gm42418 | ENSMUSG00000098178 | lncRNA | predicted gene, 42418 | 1.945E-33 | 1.527626609 | 1 | 1 | 4.59157E-29 | EN_1 |
| Zfp608 | ENSMUSG00000052713 | protein_coding | zinc finger protein 608 | 5.104E-09 | -1.20189291 | 0.566 | 0.867 | 0.000120519 | EN_1 |
| Gstp1 | ENSMUSG00000060803 | protein_coding | glutathione S-transferase, pi 1 | 4.371E-12 | -2.179307466 | 0.145 | 0.659 | 1.03205E-07 | EN_1 |
| Cntf | ENSMUSG00000079415 | protein_coding | ciliary neurotrophic factor | 2.94E-08 | -1.186691455 | 0.605 | 0.89 | 0.000694189 | EN_1 |
| Acta2 | ENSMUSG00000035783 | protein_coding | actin, alpha 2, smooth muscle, aorta | 4.241E-08 | -0.926530823 | 0.711 | 0.936 | 0.001001381 | EN_1 |
| Scd2 | ENSMUSG00000025203 | protein_coding | stearoyl-Coenzyme A desaturase 2 | 5.905E-13 | -1.171943237 | 0.882 | 0.988 | 1.39424E-08 | EN_1 |
| L1cam | ENSMUSG00000031391 | protein_coding | L1 cell adhesion molecule | 1.305E-08 | -0.740274281 | 0.842 | 0.965 | 0.000308186 | EN_1 |
| Sat1 | ENSMUSG00000025283 | protein_coding | spermidine/spermine N1-acetyl transferase 1 | 1.832E-09 | -0.828712946 | 0.908 | 1 | 4.32528E-05 | EN_1 |
| Hspe1 | ENSMUSG00000073676 | protein_coding | heat shock protein 1 (chaperonin 10) | 1.2E-07 | 0.783231693 | 0.916 | 0.777 | 0.002833187 | EN_2 |
| Des | ENSMUSG00000026208 | protein_coding | desmin | 1.576E-11 | -1.249246769 | 0.63 | 0.842 | 3.72032E-07 | EN_2 |
| Itm2c | ENSMUSG00000026223 | protein_coding | integral membrane protein 2C | 8.443E-09 | -0.862066453 | 0.714 | 0.935 | 0.00019936 | EN_2 |
| Per2 | ENSMUSG00000055866 | protein_coding | period circadian clock 2 | 3.479E-07 | 2.365757877 | 0.273 | 0.05 | 0.008214184 | EN_2 |
| Dbi | ENSMUSG00000026385 | protein_coding | diazepam binding inhibitor | 4.843E-47 | -2.592267435 | 0.948 | 1 | 1.14358E-42 | EN_2 |
| Prdx6 | ENSMUSG00000026701 | protein_coding | peroxiredoxin 6 | 1.417E-10 | -0.704940509 | 0.857 | 0.978 | 3.34687E-06 | EN_2 |
| G0s2 | ENSMUSG00000009633 | protein_coding | G0/G1 switch gene 2 | 1.157E-14 | 1.279809985 | 0.981 | 0.842 | 2.73166E-10 | EN_2 |
| Camk1d | ENSMUSG00000039145 | protein_coding | calcium/calmodulin-dependent protein kinase ID | 1.588E-27 | 1.026074469 | 1 | 0.971 | 3.74873E-23 | EN_2 |
| Gsn | ENSMUSG00000026879 | protein_coding | gelsolin | 6.175E-08 | -0.619936209 | 0.987 | 0.993 | 0.001458032 | EN_2 |
| Rpl35 | ENSMUSG00000062997 | protein_coding | ribosomal protein L35 | 5.745E-13 | -0.795732405 | 0.922 | 0.986 | 1.35649E-08 | EN_2 |
| Cd59a | ENSMUSG00000032679 | protein_coding | CD59a antigen | 7.893E-17 | -0.80318062 | 0.961 | 1 | 1.8638E-12 | EN_2 |
| Gatm | ENSMUSG00000027199 | protein_coding | glycine amidinotransferase (L-arginine:glycine amidinotransferase) | 2.252E-07 | -0.968933253 | 0.656 | 0.799 | 0.005316363 | EN_2 |
| Gm14133 | ENSMUSG00000087029 | lncRNA | predicted gene 14133 | 2.694E-09 | -5.47509692 | 0 | 0.209 | 6.36188E-05 | EN_2 |

| | | | | | | | | | |
|---|---|---|---|---|---|---|---|---|---|
| Myl9 | ENSMUSG00000067818 | protein_coding | myosin, light polypeptide 9, regulatory | 4.966E-12 | -1.204694568 | 0.539 | 0.892 | 1.17253E-07 | EN_2 |
| Atp9a | ENSMUSG00000027546 | protein_coding | ATPase, class II, type 9A | 6.07E-09 | -1.640222956 | 0.169 | 0.496 | 0.000143319 | EN_2 |
| Col20a1 | ENSMUSG00000016356 | protein_coding | collagen, type XX, alpha 1 | 1.588E-08 | -1.981513751 | 0.221 | 0.511 | 0.000374875 | EN_2 |
| Fdps | ENSMUSG00000059743 | protein_coding | farnesyl diphosphate synthetase | 6.044E-08 | -1.690453091 | 0.188 | 0.489 | 0.001427209 | EN_2 |
| S100a16 | ENSMUSG00000074457 | protein_coding | S100 calcium binding protein A16 | 3.916E-08 | -0.838450217 | 0.721 | 0.878 | 0.000924647 | EN_2 |
| Rhoc | ENSMUSG00000002233 | protein_coding | ras homolog family member C | 5.533E-14 | -1.121078093 | 0.649 | 0.928 | 1.30637E-09 | EN_2 |
| Kcna2 | ENSMUSG00000040724 | protein_coding | potassium voltage-gated channel, shaker-related subfamily, member 2 | 4.842E-16 | -1.059067855 | 0.909 | 0.978 | 1.14321E-11 | EN_2 |
| Ntng1 | ENSMUSG00000059857 | protein_coding | netrin G1 | 1.306E-11 | -1.260812611 | 0.63 | 0.871 | 3.08399E-07 | EN_2 |
| Gm11808 | ENSMUSG00000068240 | protein_coding | predicted gene 11808 | 5.15E-10 | -1.993315551 | 0.156 | 0.496 | 1.216E-05 | EN_2 |
| Tpm2 | ENSMUSG00000028464 | protein_coding | tropomyosin 2, beta | 2.003E-10 | -0.913552696 | 0.74 | 0.942 | 4.72949E-06 | EN_2 |
| Gnb1 | ENSMUSG00000029064 | protein_coding | guanine nucleotide binding protein (G protein), beta 1 | 2.508E-07 | -0.764716637 | 0.669 | 0.82 | 0.005921049 | EN_2 |
| Insig1 | ENSMUSG00000045294 | protein_coding | insulin induced gene 1 | 1.682E-08 | -1.625811946 | 0.188 | 0.489 | 0.000397264 | EN_2 |
| Sparcl1 | ENSMUSG00000029309 | protein_coding | SPARC-like 1 | 1.342E-07 | -1.114001152 | 0.669 | 0.827 | 0.003168664 | EN_2 |
| Lrrc8b | ENSMUSG00000070639 | protein_coding | leucine rich repeat containing 8 family, member B | 1.958E-07 | -0.753334725 | 0.682 | 0.885 | 0.004624184 | EN_2 |
| Atp5k | ENSMUSG00000050856 | protein_coding | ATP synthase, H+ transporting, mitochondrial F1F0 complex, subunit E | 9.932E-08 | -0.738776087 | 0.708 | 0.885 | 0.002345108 | EN_2 |
| Ubc | ENSMUSG00000008348 | protein_coding | ubiquitin C | 3.008E-09 | 0.845341544 | 0.961 | 0.892 | 7.10132E-05 | EN_2 |
| Hspb1 | ENSMUSG00000004951 | protein_coding | heat shock protein 1 | 3.545E-08 | 1.40468319 | 0.857 | 0.777 | 0.000836997 | EN_2 |
| Gigyf1 | ENSMUSG00000029714 | protein_coding | GRB10 interacting GYF protein 1 | 4.796E-11 | -2.268880837 | 0.104 | 0.446 | 1.13236E-06 | EN_2 |
| Gnb2 | ENSMUSG00000029713 | protein_coding | guanine nucleotide binding protein (G protein), beta 2 | 8.825E-18 | -1.083829192 | 0.838 | 0.95 | 2.0838E-13 | EN_2 |
| Actb | ENSMUSG00000029580 | protein_coding | actin, beta | 1.608E-20 | -0.798140458 | 1 | 1 | 3.79733E-16 | EN_2 |
| Cdk8 | ENSMUSG00000029635 | protein_coding | cyclin-dependent kinase 8 | 2.283E-33 | 2.307553547 | 0.994 | 0.928 | 5.38947E-29 | EN_2 |
| Hsph1 | ENSMUSG00000029657 | protein_coding | heat shock 105kDa/110kDa protein 1 | 3.404E-13 | 2.294310493 | 0.675 | 0.317 | 8.0378E-09 | EN_2 |
| Actg2 | ENSMUSG00000059430 | protein_coding | actin, gamma 2, smooth muscle, enteric | 8.189E-21 | -1.547203474 | 0.604 | 0.935 | 1.93361E-16 | EN_2 |
| Mgll | ENSMUSG00000033174 | protein_coding | monoglyceride lipase | 5.265E-12 | -1.95694009 | 0.188 | 0.554 | 1.24311E-07 | EN_2 |
| Kcna1 | ENSMUSG00000047976 | protein_coding | potassium voltage-gated channel, shaker-related subfamily, member 1 | 7.612E-15 | -0.955045525 | 0.935 | 0.978 | 1.79728E-10 | EN_2 |
| Luzp2 | ENSMUSG00000063297 | protein_coding | leucine zipper protein 2 | 1.612E-18 | -1.958280472 | 0.39 | 0.806 | 3.80535E-14 | EN_2 |
| Igf1r | ENSMUSG00000005533 | protein_coding | insulin-like growth factor I receptor | 8.614E-09 | -1.02659366 | 0.571 | 0.82 | 0.000203393 | EN_2 |
| Serpinh1 | ENSMUSG00000070436 | protein_coding | serine (or cysteine) peptidase inhibitor, clade H, member 1 | 1.226E-07 | 0.83680626 | 0.838 | 0.683 | 0.002895959 | EN_2 |
| Ucp2 | ENSMUSG00000033685 | protein_coding | uncoupling protein 2 (mitochondrial, proton carrier) | 3.607E-12 | -3.869965648 | 0.013 | 0.309 | 8.5174E-08 | EN_2 |
| Hbb-bs | ENSMUSG00000052305 | protein_coding | hemoglobin, beta adult s chain | 4.053E-07 | -3.067156191 | 0.026 | 0.216 | 0.009571024 | EN_2 |
| Trim12a | ENSMUSG00000066258 | protein_coding | tripartite motif-containing 12A | 7.89E-08 | 5.379378885 | 0.188 | 0 | 0.001863104 | EN_2 |
| Trim12c | ENSMUSG00000057143 | protein_coding | tripartite motif-containing 12C | 2.432E-07 | 5.445450487 | 0.175 | 0 | 0.00574307 | EN_2 |
| Sox6 | ENSMUSG00000051910 | protein_coding | SRY (sex determining region Y)-box 6 | 2.336E-08 | -0.900308086 | 0.747 | 0.878 | 0.00055157 | EN_2 |
| Mapk3 | ENSMUSG00000063065 | protein_coding | mitogen-activated protein kinase 3 | 6.374E-09 | -1.0230681 | 0.565 | 0.82 | 0.0001505 | EN_2 |
| Aldoa | ENSMUSG00000030695 | protein_coding | aldolase A, fructose-bisphosphate | 3.74E-07 | -0.765955619 | 0.74 | 0.849 | 0.008830251 | EN_2 |
| Poteg | ENSMUSG00000063932 | protein_coding | POTE ankyrin domain family, member G | 1.715E-33 | -8.479966378 | 0 | 0.676 | 4.04975E-29 | EN_2 |
| Rfxank | ENSMUSG00000036120 | protein_coding | regulatory factor X-associated ankyrin-containing protein | 3.054E-07 | -2.707466953 | 0.026 | 0.223 | 0.007211847 | EN_2 |
| Uba52 | ENSMUSG00000090137 | protein_coding | ubiquitin A-52 residue ribosomal protein fusion product 1 | 1.5E-09 | -1.53380809 | 0.279 | 0.619 | 3.54116E-05 | EN_2 |
| Gm26887 | ENSMUSG00000097120 | lncRNA | predicted gene, 26887 | 1.348E-09 | 1.754114663 | 0.552 | 0.23 | 3.18279E-05 | EN_2 |
| Col5a3 | ENSMUSG00000004098 | protein_coding | collagen, type V, alpha 3 | 2.659E-11 | -1.461394734 | 0.448 | 0.748 | 6.2783E-07 | EN_2 |
| Fez1 | ENSMUSG00000032118 | protein_coding | fasciculation and elongation protein zeta 1 (zygin I) | 2.099E-08 | -0.771121482 | 0.818 | 0.906 | 0.000495702 | EN_2 |
| Mcam | ENSMUSG00000032135 | protein_coding | melanoma cell adhesion molecule | 8.557E-10 | -1.352772339 | 0.383 | 0.712 | 2.02037E-05 | EN_2 |
| Tagln | ENSMUSG00000032085 | protein_coding | transgelin | 3.443E-10 | -0.854811377 | 0.74 | 0.942 | 8.12996E-06 | EN_2 |
| Cryab | ENSMUSG00000032060 | protein_coding | crystallin, alpha B | 1.494E-08 | 0.686063174 | 1 | 0.993 | 0.000352845 | EN_2 |
| Rora | ENSMUSG00000032238 | protein_coding | RAR-related orphan receptor alpha | 1.125E-09 | 0.82690176 | 0.961 | 0.871 | 2.65662E-05 | EN_2 |
| Col12a1 | ENSMUSG00000032332 | protein_coding | collagen, type XII, alpha 1 | 5.238E-09 | 0.80328764 | 0.961 | 0.842 | 0.000123675 | EN_2 |
| Me1 | ENSMUSG00000032418 | protein_coding | malic enzyme 1, NADP(+)-dependent, cytosolic | 2.697E-07 | -2.298648105 | 0.032 | 0.237 | 0.006367782 | EN_2 |
| Lars2 | ENSMUSG00000035202 | protein_coding | leucyl-tRNA synthetase, mitochondrial | 6.3E-27 | 1.601703291 | 0.961 | 0.892 | 1.48745E-22 | EN_2 |
| Ccn2 | ENSMUSG00000019997 | protein_coding | cellular communication network factor 2 | 3.89E-10 | -1.668749467 | 0.513 | 0.77 | 9.18536E-06 | EN_2 |
| Marcks | ENSMUSG00000069662 | protein_coding | myristoylated alanine rich protein kinase C substrate | 4.637E-15 | -0.664384584 | 0.994 | 1 | 1.09499E-10 | EN_2 |
| Fabp7 | ENSMUSG00000019874 | protein_coding | fatty acid binding protein 7, brain | 1.088E-35 | -4.730751002 | 0.065 | 0.784 | 2.56894E-31 | EN_2 |
| Cirbp | ENSMUSG00000045193 | protein_coding | cold inducible RNA binding protein | 4.656E-14 | -1.706987405 | 0.364 | 0.741 | 1.09939E-09 | EN_2 |
| Nudt4 | ENSMUSG00000020029 | protein_coding | nudix (nucleoside diphosphate linked moiety X)-type motif 4 | 1.469E-07 | 0.933281053 | 0.825 | 0.568 | 0.003467742 | EN_2 |
| Dcn | ENSMUSG00000019929 | protein_coding | decorin | 3.79E-09 | -1.768401813 | 0.097 | 0.403 | 8.94839E-05 | EN_2 |
| Lyz2 | ENSMUSG00000069516 | protein_coding | lysozyme 2 | 9.358E-12 | -7.276006889 | 0 | 0.266 | 2.20952E-07 | EN_2 |
| Bloc1s1 | ENSMUSG00000090247 | protein_coding | biogenesis of lysosomal organelles complex-1, subunit 1 | 3.553E-09 | -2.079383181 | 0.117 | 0.417 | 8.3889E-05 | EN_2 |
| Pdlim4 | ENSMUSG00000020388 | protein_coding | PDZ and LIM domain 4 | 2.899E-07 | -0.742539721 | 0.916 | 0.957 | 0.006845149 | EN_2 |
| Rpl26 | ENSMUSG00000060938 | protein_coding | ribosomal protein L26 | 3.52E-09 | 0.598649123 | 0.994 | 0.957 | 8.31251E-05 | EN_2 |
| Thra | ENSMUSG00000058756 | protein_coding | thyroid hormone receptor alpha | 2.362E-10 | -0.918907734 | 0.558 | 0.835 | 5.5781E-06 | EN_2 |
| Nr1d1 | ENSMUSG00000020889 | protein_coding | nuclear receptor subfamily 1, group D, member 1 | 9.904E-19 | -2.081599313 | 0.26 | 0.755 | 2.33845E-14 | EN_2 |
| Cnp | ENSMUSG00000006782 | protein_coding | 2',3'-cyclic nucleotide 3' phosphodiesterase | 2.368E-11 | -1.208941471 | 0.688 | 0.827 | 5.5913E-07 | EN_2 |
| Ramp2 | ENSMUSG00000001240 | protein_coding | receptor (calcitonin) activity modifying protein 2 | 3.551E-08 | 1.131263177 | 0.695 | 0.468 | 0.000838346 | EN_2 |
| Rnd2 | ENSMUSG00000001313 | protein_coding | Rho family GTPase 2 | 4.026E-09 | -2.455259583 | 0.097 | 0.381 | 9.50617E-05 | EN_2 |
| Rpl38 | ENSMUSG00000057322 | protein_coding | ribosomal protein L38 | 1.275E-18 | -0.890537494 | 0.955 | 0.993 | 3.01093E-14 | EN_2 |
| Jpt1 | ENSMUSG00000020737 | protein_coding | Jupiter microtubule associated homolog 1 | 6.053E-08 | -0.845824654 | 0.675 | 0.871 | 0.001429173 | EN_2 |
| Asap2 | ENSMUSG00000052632 | protein_coding | ArfGAP with SH3 domain, ankyrin repeat and PH domain 2 | 1.625E-10 | -0.9242598 | 0.812 | 0.906 | 3.83762E-06 | EN_2 |
| Cys1 | ENSMUSG00000062563 | protein_coding | cystin 1 | 1.198E-07 | 1.346327369 | 0.591 | 0.324 | 0.002828602 | EN_2 |
| Frmd6 | ENSMUSG00000048285 | protein_coding | FERM domain containing 6 | 8.967E-08 | -1.038155664 | 0.455 | 0.727 | 0.002117353 | EN_2 |
| Gphn | ENSMUSG00000047454 | protein_coding | gephyrin | 1.086E-18 | 1.013354202 | 0.987 | 0.942 | 2.56315E-14 | EN_2 |
| Ifi27l2a | ENSMUSG00000079017 | protein_coding | interferon, alpha-inducible protein 27 like 2A | 3.097E-08 | -1.446423476 | 0.571 | 0.82 | 0.000731255 | EN_2 |
| Ckb | ENSMUSG00000001270 | protein_coding | creatine kinase, brain | 1.204E-27 | -1.500500374 | 0.935 | 0.986 | 2.84261E-23 | EN_2 |
| Elmo1 | ENSMUSG00000041112 | protein_coding | engulfment and cell motility 1 | 3.88E-09 | 1.667291779 | 0.545 | 0.223 | 9.16065E-05 | EN_2 |
| Hist1h4d | ENSMUSG00000061482 | protein_coding | histone cluster 1, H4d | 9.612E-20 | 1.848890053 | 0.844 | 0.511 | 2.26947E-15 | EN_2 |
| Hist1h1e | ENSMUSG00000051627 | protein_coding | histone cluster 1, H1e | 1.967E-07 | 1.226787569 | 0.604 | 0.338 | 0.004645284 | EN_2 |
| Hexb | ENSMUSG00000021665 | protein_coding | hexosaminidase B | 1.577E-11 | -1.021709838 | 0.727 | 0.871 | 3.72289E-07 | EN_2 |
| Fam107a | ENSMUSG00000021750 | protein_coding | family with sequence similarity 107, member A | 2.521E-14 | 1.802123978 | 0.695 | 0.273 | 5.95285E-10 | EN_2 |
| Gm48099 | ENSMUSG00000114304 | lncRNA | predicted gene, 48099 | 7.471E-17 | -5.944871517 | 0.006 | 0.388 | 1.76395E-12 | EN_2 |
| Plac9b | ENSMUSG00000072674 | protein_coding | placenta specific 9b | 1.429E-14 | -2.797937686 | 0.11 | 0.518 | 3.37488E-10 | EN_2 |
| Dpysl2 | ENSMUSG00000022048 | protein_coding | dihydropyrimidinase-like 2 | 3.222E-10 | -0.69317631 | 0.857 | 0.957 | 7.60697E-06 | EN_2 |
| Rgcc | ENSMUSG00000022018 | protein_coding | regulator of cell cycle | 3.23E-10 | 0.940976934 | 0.909 | 0.791 | 7.62637E-06 | EN_2 |
| Azin1 | ENSMUSG00000037458 | protein_coding | antizyme inhibitor 1 | 1.75E-07 | 0.70045456 | 0.929 | 0.842 | 0.004132941 | EN_2 |
| Deptor | ENSMUSG00000022419 | protein_coding | DEP domain containing MTOR-interacting protein | 2.132E-09 | -1.128993659 | 0.656 | 0.835 | 5.0347E-05 | EN_2 |
| Tuba1b | ENSMUSG00000023004 | protein_coding | tubulin, alpha 1B | 2.8E-10 | -0.796032761 | 0.799 | 0.914 | 6.61194E-06 | EN_2 |
| Tuba1a | ENSMUSG00000072235 | protein_coding | tubulin, alpha 1A | 2.21E-09 | -0.695686245 | 0.955 | 0.942 | 5.21759E-05 | EN_2 |
| Mylk | ENSMUSG00000022836 | protein_coding | myosin, light polypeptide kinase | 9.726E-10 | -1.197536978 | 0.455 | 0.77 | 2.2965E-05 | EN_2 |
| Gm15564 | ENSMUSG00000086324 | lncRNA | predicted gene 15564 | 5.842E-14 | 1.86977543 | 0.675 | 0.317 | 1.37949E-09 | EN_2 |
| Cmss1 | ENSMUSG00000022748 | protein_coding | cms small ribosomal subunit 1 | 8.571E-26 | 1.796581109 | 1 | 0.986 | 2.02368E-21 | EN_2 |
| BC004004 | ENSMUSG00000052712 | protein_coding | cDNA sequence BC004004 | 1.773E-21 | -2.298284003 | 0.221 | 0.755 | 4.1867E-17 | EN_2 |
| H2-Q4 | ENSMUSG00000035929 | protein_coding | histocompatibility 2, Q region locus 4 | 8.136E-09 | -1.131136761 | 0.519 | 0.755 | 0.000192102 | EN_2 |
| Gm42418 | ENSMUSG00000098178 | lncRNA | predicted gene, 42418 | 4.193E-31 | 1.513594238 | 1 | 1 | 9.89954E-27 | EN_2 |
| B4galt6 | ENSMUSG00000056124 | protein_coding | UDP-Gal:betaGlcNAc beta 1,4-galactosyltransferase, polypeptide 6 | 1.83E-08 | 1.412915064 | 0.63 | 0.396 | 0.000432016 | EN_2 |
| Gfra3 | ENSMUSG00000024366 | protein_coding | glial cell line derived neurotrophic factor family receptor alpha 3 | 1.643E-09 | -2.350346436 | 0.11 | 0.41 | 3.87847E-05 | EN_2 |
| Kcnn2 | ENSMUSG00000054477 | protein_coding | potassium intermediate/small conductance calcium-activated channel, subfamily N, member 2 | 3.097E-08 | -1.774597472 | 0.169 | 0.475 | 0.000731328 | EN_2 |
| Gstp1 | ENSMUSG00000060803 | protein_coding | glutathione S-transferase, pi 1 | 2.532E-10 | -1.872920617 | 0.292 | 0.633 | 5.97902E-06 | EN_2 |
| Cntf | ENSMUSG00000079415 | protein_coding | ciliary neurotrophic factor | 5.875E-10 | -1.209685123 | 0.721 | 0.892 | 1.3873E-05 | EN_2 |
| Acta2 | ENSMUSG00000035783 | protein_coding | actin, alpha 2, smooth muscle, aorta | 3.04E-13 | -1.125268754 | 0.63 | 0.899 | 7.17919E-09 | EN_2 |
| Scd2 | ENSMUSG00000025203 | protein_coding | stearoyl-Coenzyme A desaturase 2 | 1.647E-08 | -0.743438346 | 0.909 | 0.942 | 0.000389003 | EN_2 |
| Bgn | ENSMUSG00000031375 | protein_coding | biglycan | 4.648E-08 | -0.742698463 | 0.805 | 0.935 | 0.001097422 | EN_2 |
| Sat1 | ENSMUSG00000025283 | protein_coding | spermidine/spermine N1-acetyl transferase 1 | 2.875E-12 | -0.720784506 | 0.974 | 0.986 | 6.78755E-08 | EN_2 |
| mt-Atp8 | ENSMUSG00000064356 | protein_coding | mitochondrially encoded ATP synthase 8 | 2.877E-07 | 0.862598442 | 0.799 | 0.691 | 0.006792932 | EN_2 |
| mt-Nd6 | ENSMUSG00000064368 | protein_coding | mitochondrially encoded NADH dehydrogenase 6 | 1.018E-08 | 1.404401879 | 0.604 | 0.309 | 0.000240457 | EN_2 |
| Dbi | ENSMUSG00000026385 | protein_coding | diazepam binding inhibitor | 1.974E-08 | -3.454274678 | 0.21 | 0.594 | 0.000466081 | EN_3 |
| Frmd4a | ENSMUSG00000026657 | protein_coding | FERM domain containing 4A | 3.646E-07 | -1.238497055 | 0.79 | 0.927 | 0.008609222 | EN_3 |
| Camk1d | ENSMUSG00000039145 | protein_coding | calcium/calmodulin-dependent protein kinase ID | 3.388E-11 | 0.870844868 | 1 | 1 | 8.00025E-07 | EN_3 |
| Gm14133 | ENSMUSG00000087029 | lncRNA | predicted gene 14133 | 1.075E-07 | -6.508584396 | 0 | 0.365 | 0.002539386 | EN_3 |
| Tmem50a | ENSMUSG00000028822 | protein_coding | transmembrane protein 50A | 1.211E-07 | -4.825139209 | 0.016 | 0.385 | 0.002858402 | EN_3 |
| Actb | ENSMUSG00000029580 | protein_coding | actin, beta | 1.164E-10 | -1.37427565 | 0.79 | 0.958 | 2.74941E-06 | EN_3 |
| Cdk8 | ENSMUSG00000029635 | protein_coding | cyclin-dependent kinase 8 | 3.891E-25 | 2.564974652 | 1 | 0.958 | 9.18668E-21 | EN_3 |
| Luzp2 | ENSMUSG00000063297 | protein_coding | leucine zipper protein 2 | 6.026E-09 | -2.355875152 | 0.258 | 0.698 | 0.000142278 | EN_3 |
| Lars2 | ENSMUSG00000035202 | protein_coding | leucyl-tRNA synthetase, mitochondrial | 8.672E-23 | 1.898919476 | 1 | 1 | 2.04768E-18 | EN_3 |
| Ccn2 | ENSMUSG00000019997 | protein_coding | cellular communication network factor 2 | 5.674E-08 | -2.749037289 | 0.242 | 0.635 | 0.001339644 | EN_3 |
| Ank3 | ENSMUSG00000069601 | protein_coding | ankyrin 3, epithelial | 1.167E-07 | -0.749683923 | 0.984 | 0.99 | 0.002754876 | EN_3 |
| Rpl41 | ENSMUSG00000093674 | protein_coding | ribosomal protein L41 | 2.089E-09 | -1.81049939 | 0.419 | 0.802 | 4.93338E-05 | EN_3 |
| Rpl19 | ENSMUSG00000017404 | protein_coding | ribosomal protein L19 | 4.158E-07 | -2.126677831 | 0.145 | 0.562 | 0.009817799 | EN_3 |
| Rpl38 | ENSMUSG00000057322 | protein_coding | ribosomal protein L38 | 2.122E-08 | -2.194339569 | 0.194 | 0.635 | 0.000500955 | EN_3 |
| H3f3b | ENSMUSG00000016559 | protein_coding | H3 histone, family 3B | 4.808E-09 | -2.453862949 | 0.177 | 0.646 | 0.000113532 | EN_3 |
| Frmd6 | ENSMUSG00000048285 | protein_coding | FERM domain containing 6 | 6.384E-08 | -1.37929377 | 0.242 | 0.74 | 0.001507456 | EN_3 |
| Gphn | ENSMUSG00000047454 | protein_coding | gephyrin | 1.448E-18 | 1.517727849 | 1 | 0.99 | 3.41822E-14 | EN_3 |
| Ckb | ENSMUSG00000001270 | protein_coding | creatine kinase, brain | 8.375E-08 | -1.885590484 | 0.306 | 0.719 | 0.001977618 | EN_3 |
| Gm48099 | ENSMUSG00000114304 | lncRNA | predicted gene, 48099 | 1.494E-07 | -3.565163084 | 0.016 | 0.396 | 0.003528746 | EN_3 |
| Gm15564 | ENSMUSG00000086324 | lncRNA | predicted gene 15564 | 5.58E-11 | 1.85317609 | 0.79 | 0.354 | 1.31744E-06 | EN_3 |

| | | | | | | | | | |
|---|---|---|---|---|---|---|---|---|---|
| Cmss1 | ENSMUSG00000022748 | protein_coding | cms small ribosomal subunit 1 | 1.14E-24 | 2.206559168 | 1 | 0.99 | 2.69107E-20 | EN_3 |
| Gm26917 | ENSMUSG00000097971 | lncRNA | predicted gene, 26917 | 2.82E-09 | 1.837783033 | 0.871 | 0.625 | 6.65819E-05 | EN_3 |
| Gm42418 | ENSMUSG00000098178 | lncRNA | predicted gene, 42418 | 5.875E-23 | 1.899794528 | 1 | 1 | 1.3871E-18 | EN_3 |
| Rpl39 | ENSMUSG00000079641 | protein_coding | ribosomal protein L39 | 2.509E-07 | -1.482133511 | 0.21 | 0.635 | 0.005925199 | EN_3 |
| Tmsb4x | ENSMUSG00000049775 | protein_coding | thymosin, beta 4, X chromosome | 2.332E-08 | -1.528479482 | 0.484 | 0.875 | 0.000550673 | EN_3 |
| Dbi | ENSMUSG00000026385 | protein_coding | diazepam binding inhibitor | 2.806E-12 | -2.599798378 | 0.884 | 0.961 | 6.62493E-08 | EN_4 |
| G0s2 | ENSMUSG00000009633 | protein_coding | G0/G1 switch gene 2 | 3.195E-08 | 1.823628755 | 0.797 | 0.421 | 0.000754367 | EN_4 |
| Tpm2 | ENSMUSG00000028464 | protein_coding | tropomyosin 2, beta | 1.163E-07 | -2.096277585 | 0.232 | 0.645 | 0.002744998 | EN_4 |
| Fabp7 | ENSMUSG00000019874 | protein_coding | fatty acid binding protein 7, brain | 7.78E-08 | -3.937768924 | 0.058 | 0.447 | 0.001837002 | EN_4 |
| Ckb | ENSMUSG00000001270 | protein_coding | creatine kinase, brain | 2.784E-07 | -1.174695425 | 0.594 | 0.842 | 0.006573437 | EN_4 |
| Rgcc | ENSMUSG00000022018 | protein_coding | regulator of cell cycle | 1.623E-07 | 2.134439722 | 0.71 | 0.329 | 0.003832129 | EN_4 |
| Rpl37a | ENSMUSG00000046330 | protein_coding | ribosomal protein L37a | 1.016E-14 | -0.774879999 | 0.965 | 1 | 2.39796E-10 | MM_1 |
| Des | ENSMUSG00000026208 | protein_coding | desmin | 2.294E-08 | -0.815439037 | 0.693 | 0.939 | 0.000541615 | MM_1 |
| Per2 | ENSMUSG00000055866 | protein_coding | period circadian clock 2 | 2.139E-09 | 3.338648583 | 0.395 | 0.051 | 5.05054E-05 | MM_1 |
| Ifi202b | ENSMUSG00000026535 | protein_coding | interferon activated gene 202B | 1.324E-16 | -3.281104133 | 0.079 | 0.612 | 3.12621E-12 | MM_1 |
| Camk1d | ENSMUSG00000039145 | protein_coding | calcium/calmodulin-dependent protein kinase ID | 5.395E-08 | 0.677008736 | 1 | 0.98 | 0.001273982 | MM_1 |
| Cdk9 | ENSMUSG00000009555 | protein_coding | cyclin-dependent kinase 9 (CDC2-related kinase) | 2.564E-07 | -1.256376791 | 0.333 | 0.694 | 0.006052978 | MM_1 |
| Rpl35 | ENSMUSG00000062997 | protein_coding | ribosomal protein L35 | 3.308E-08 | -0.730936973 | 0.868 | 0.949 | 0.00078117 | MM_1 |
| Lmo2 | ENSMUSG00000032698 | protein_coding | LIM domain only 2 | 3.784E-10 | -1.154953876 | 0.649 | 0.898 | 8.93521E-06 | MM_1 |
| B2m | ENSMUSG00000060802 | protein_coding | beta-2 microglobulin | 2.348E-15 | -0.775129692 | 0.982 | 1 | 5.54484E-11 | MM_1 |
| Cst3 | ENSMUSG00000027447 | protein_coding | cystatin C | 6.929E-14 | -0.969815319 | 0.956 | 0.99 | 1.63607E-09 | MM_1 |
| Abhd12 | ENSMUSG00000032046 | protein_coding | abhydrolase domain containing 12 | 4.028E-07 | -0.818189154 | 0.711 | 0.918 | 0.009511557 | MM_1 |
| Myl9 | ENSMUSG00000067818 | protein_coding | myosin, light polypeptide 9, regulatory | 4.861E-09 | -1.039946532 | 0.377 | 0.776 | 0.000114774 | MM_1 |
| Serinc3 | ENSMUSG00000017707 | protein_coding | serine incorporator 3 | 2.451E-18 | -0.781663267 | 0.974 | 1 | 5.78732E-14 | MM_1 |
| Cp | ENSMUSG00000003617 | protein_coding | ceruloplasmin | 3.262E-08 | -1.250314409 | 0.421 | 0.806 | 0.000770298 | MM_1 |
| Gm11808 | ENSMUSG00000068240 | protein_coding | predicted gene 11808 | 7.407E-14 | -2.734387498 | 0.149 | 0.622 | 1.74897E-09 | MM_1 |
| Wwp1 | ENSMUSG00000041058 | protein_coding | WW domain containing E3 ubiquitin protein ligase 1 | 8.244E-09 | -0.697750322 | 0.974 | 0.99 | 0.000194658 | MM_1 |
| Tpm2 | ENSMUSG00000028464 | protein_coding | tropomyosin 2, beta | 3.904E-07 | -0.960972882 | 0.693 | 0.898 | 0.009218417 | MM_1 |
| Akr1a1 | ENSMUSG00000028692 | protein_coding | aldo-keto reductase family 1, member A1 (aldehyde reductase) | 1.644E-07 | -0.84756347 | 0.816 | 0.98 | 0.003882505 | MM_1 |
| Ptp4a2 | ENSMUSG00000028788 | protein_coding | protein tyrosine phosphatase 4a2 | 3.95E-08 | -0.664184463 | 0.921 | 0.99 | 0.00093273 | MM_1 |
| Cd52 | ENSMUSG00000000682 | protein_coding | CD52 antigen | 4.153E-10 | 3.42203867 | 0.447 | 0.071 | 9.80654E-06 | MM_1 |
| Sh3bgrl3 | ENSMUSG00000028843 | protein_coding | SH3 domain binding glutamic acid-rich protein-like 3 | 1.162E-15 | 1.658492665 | 0.895 | 0.786 | 2.74461E-11 | MM_1 |
| Man1c1 | ENSMUSG00000037306 | protein_coding | mannosidase, alpha, class 1C, member 1 | 3.148E-15 | -1.616893011 | 0.64 | 0.918 | 7.4336E-11 | MM_1 |
| C1qa | ENSMUSG00000036887 | protein_coding | complement component 1, q subcomponent, alpha polypeptide | 1.898E-31 | -2.500862228 | 0.842 | 1 | 4.48136E-27 | MM_1 |
| Dhrs3 | ENSMUSG00000066026 | protein_coding | dehydrogenase/reductase (SDR family) member 3 | 1.57E-07 | -0.895179344 | 0.798 | 0.929 | 0.003706137 | MM_1 |
| 5830444B04Rik | ENSMUSG00000084803 | lncRNA | RIKEN cDNA 5830444B04 gene | 2.698E-07 | -3.820814124 | 0.009 | 0.235 | 0.006370879 | MM_1 |
| Cd36 | ENSMUSG00000002944 | protein_coding | CD36 molecule | 8.57E-08 | -0.750437542 | 0.965 | 1 | 0.002023663 | MM_1 |
| Pf4 | ENSMUSG00000029373 | protein_coding | platelet factor 4 | 5.665E-08 | -0.73112586 | 0.912 | 1 | 0.001337668 | MM_1 |
| Gigyf1 | ENSMUSG00000029714 | protein_coding | GRB10 interacting GYF protein 1 | 4.279E-10 | -1.265304608 | 0.246 | 0.663 | 1.01035E-05 | MM_1 |
| Gnb2 | ENSMUSG00000029713 | protein_coding | guanine nucleotide binding protein (G protein), beta 2 | 3.39E-15 | -1.060228231 | 0.877 | 0.99 | 8.00446E-11 | MM_1 |
| Pcolce | ENSMUSG00000029718 | protein_coding | procollagen C-endopeptidase enhancer protein | 1.119E-07 | -2.804433126 | 0.035 | 0.306 | 0.002642707 | MM_1 |
| Lrch4 | ENSMUSG00000093445 | protein_coding | leucine-rich repeats and calponin homology (CH) domain containing 4 | 8.962E-09 | -1.786127412 | 0.219 | 0.602 | 0.000211605 | MM_1 |
| Actb | ENSMUSG00000029580 | protein_coding | actin, beta | 3.65E-13 | -0.664009753 | 1 | 1 | 8.61902E-09 | MM_1 |
| Cdk8 | ENSMUSG00000029635 | protein_coding | cyclin-dependent kinase 8 | 3.222E-34 | 3.004958432 | 0.991 | 0.98 | 7.60759E-30 | MM_1 |
| Actg2 | ENSMUSG00000059430 | protein_coding | actin, gamma 2, smooth muscle, enteric | 1.317E-11 | -1.203141206 | 0.614 | 0.939 | 3.10953E-07 | MM_1 |
| Cd33 | ENSMUSG00000004609 | protein_coding | CD33 antigen | 8.857E-10 | -1.278579723 | 0.658 | 0.867 | 2.09143E-05 | MM_1 |
| Dbp | ENSMUSG00000059824 | protein_coding | D site albumin promoter binding protein | 4.959E-11 | 2.169083829 | 0.649 | 0.276 | 1.171E-06 | MM_1 |
| Gm44751 | ENSMUSG00000109244 | lncRNA | predicted gene 44751 | 4.207E-07 | -2.262209288 | 0.053 | 0.327 | 0.009932943 | MM_1 |
| Tenm4 | ENSMUSG00000048078 | protein_coding | teneurin transmembrane protein 4 | 4.546E-09 | -2.403496854 | 0.123 | 0.48 | 0.000107349 | MM_1 |
| Arntl | ENSMUSG00000055116 | protein_coding | aryl hydrocarbon receptor nuclear translocator-like | 5.329E-13 | -3.746688725 | 0.053 | 0.48 | 1.25838E-08 | MM_1 |
| Mapk3 | ENSMUSG00000063065 | protein_coding | mitogen-activated protein kinase 3 | 8.723E-08 | -1.365725513 | 0.36 | 0.735 | 0.002059638 | MM_1 |
| Poteg | ENSMUSG00000063932 | protein_coding | POTE ankyrin domain family, member G | 8.171E-13 | -5.469164223 | 0 | 0.378 | 1.92944E-08 | MM_1 |
| Hpgd | ENSMUSG00000031613 | protein_coding | hydroxyprostaglandin dehydrogenase 15 (NAD) | 5.925E-09 | -1.444048876 | 0.632 | 0.867 | 0.00013989 | MM_1 |
| Uba52 | ENSMUSG00000090137 | protein_coding | ubiquitin A-52 residue ribosomal protein fusion product 1 | 1.397E-09 | -1.42144143 | 0.474 | 0.755 | 3.29953E-05 | MM_1 |
| Mast3 | ENSMUSG00000031833 | protein_coding | microtubule associated serine/threonine kinase 3 | 2.694E-09 | -1.819835567 | 0.158 | 0.571 | 6.36133E-05 | MM_1 |
| Maf | ENSMUSG00000055435 | protein_coding | avian musculoaponeurotic fibrosarcoma oncogene homolog | 2.843E-13 | -0.890943946 | 0.974 | 1 | 6.71308E-09 | MM_1 |
| Scamp2 | ENSMUSG00000040188 | protein_coding | secretory carrier membrane protein 2 | 1.928E-07 | -0.917344707 | 0.702 | 0.878 | 0.004552119 | MM_1 |
| Trf | ENSMUSG00000032554 | protein_coding | transferrin | 1.769E-08 | -0.588232743 | 0.939 | 1 | 0.000417626 | MM_1 |
| Cmtm6 | ENSMUSG00000032434 | protein_coding | CKLF-like MARVEL transmembrane domain containing 6 | 3.582E-08 | -1.223061256 | 0.404 | 0.816 | 0.000845797 | MM_1 |
| Cx3cr1 | ENSMUSG00000052336 | protein_coding | chemokine (C-X3-C motif) receptor 1 | 1.838E-09 | -4.04893892 | 0.026 | 0.337 | 4.33886E-05 | MM_1 |
| Lars2 | ENSMUSG00000035202 | protein_coding | leucyl-tRNA synthetase, mitochondrial | 3.603E-23 | 1.503264494 | 0.991 | 0.969 | 8.50819E-19 | MM_1 |
| Mthfd1l | ENSMUSG00000040675 | protein_coding | methylenetetrahydrofolate dehydrogenase (NADP+ dependent) 1-like | 5.088E-12 | 2.842603592 | 0.535 | 0.092 | 1.20142E-07 | MM_1 |
| Marcks | ENSMUSG00000069662 | protein_coding | myristoylated alanine rich protein kinase C substrate | 5.893E-13 | -0.978535882 | 0.939 | 1 | 1.39143E-08 | MM_1 |
| Sumo3 | ENSMUSG00000020265 | protein_coding | small ubiquitin-like modifier 3 | 4.968E-08 | -1.154813208 | 0.439 | 0.745 | 0.001173108 | MM_1 |
| Igf1 | ENSMUSG00000020053 | protein_coding | insulin-like growth factor 1 | 8.307E-08 | -0.876684979 | 0.842 | 0.949 | 0.001961406 | MM_1 |
| Lyz2 | ENSMUSG00000069516 | protein_coding | lysozyme 2 | 2.053E-42 | -12.99784748 | 0 | 1 | 4.84846E-38 | MM_1 |
| Ctdsp2 | ENSMUSG00000078429 | protein_coding | CTD (carboxy-terminal domain, RNA polymerase II, polypeptide A) small phosphatase 2 | 5.063E-08 | -1.181104886 | 0.5 | 0.806 | 0.0011954 | MM_1 |
| Myl6 | ENSMUSG00000090841 | protein_coding | myosin, light polypeptide 6, alkali, smooth muscle and non-muscle | 2.535E-10 | -0.635632318 | 0.956 | 0.99 | 5.98517E-06 | MM_1 |
| Rpl41 | ENSMUSG00000093674 | protein_coding | ribosomal protein L41 | 1.101E-08 | -0.592433926 | 0.912 | 0.99 | 0.000260004 | MM_1 |
| Ap1b1 | ENSMUSG00000009090 | protein_coding | adaptor protein complex AP-1, beta 1 subunit | 1.096E-17 | -2.128677677 | 0.404 | 0.867 | 2.589E-13 | MM_1 |
| Plscr3 | ENSMUSG00000019461 | protein_coding | phospholipid scramblase 3 | 4.141E-07 | -1.836399981 | 0.167 | 0.5 | 0.009777463 | MM_1 |
| Evi2a | ENSMUSG00000078771 | protein_coding | ecotropic viral integration site 2a | 6.312E-08 | -1.161281053 | 0.596 | 0.888 | 0.001490377 | MM_1 |
| Rpl38 | ENSMUSG00000057322 | protein_coding | ribosomal protein L38 | 9.163E-20 | -1.017949091 | 0.939 | 0.98 | 2.16349E-15 | MM_1 |
| Rps29 | ENSMUSG00000034892 | protein_coding | ribosomal protein S29 | 8.277E-11 | -0.665617551 | 0.956 | 1 | 1.9543E-06 | MM_1 |
| Gphn | ENSMUSG00000047454 | protein_coding | gephyrin | 3.395E-12 | 0.960824722 | 0.982 | 0.969 | 8.01581E-08 | MM_1 |
| Actn1 | ENSMUSG00000015143 | protein_coding | actinin, alpha 1 | 2.034E-07 | -1.997270031 | 0.14 | 0.469 | 0.004803115 | MM_1 |
| Srsf5 | ENSMUSG00000021134 | protein_coding | serine/arginine-rich splicing factor 5 | 1.275E-09 | -0.980975337 | 0.693 | 0.918 | 3.00959E-05 | MM_1 |
| Gm19951 | ENSMUSG00000113136 | lncRNA | predicted gene, 19951 | 8.543E-10 | 1.838802913 | 0.746 | 0.469 | 2.01716E-05 | MM_1 |
| Ighm | ENSMUSG00000076617 | IG_C_gene | immunoglobulin heavy constant mu | 7.326E-14 | -1.437535989 | 0.544 | 0.939 | 1.72976E-09 | MM_1 |
| Cox7c | ENSMUSG00000017778 | protein_coding | cytochrome c oxidase subunit 7C | 2.092E-07 | -0.684498344 | 0.825 | 0.99 | 0.004938509 | MM_1 |
| Nr1d2 | ENSMUSG00000021775 | protein_coding | nuclear receptor subfamily 1, group D, member 2 | 8.261E-08 | 1.569643547 | 0.702 | 0.459 | 0.001950545 | MM_1 |
| Gm48099 | ENSMUSG00000114304 | lncRNA | predicted gene, 48099 | 3.104E-24 | -3.613677318 | 0.167 | 0.847 | 7.3303E-20 | MM_1 |
| Stab1 | ENSMUSG00000042286 | protein_coding | stabilin 1 | 2.903E-08 | -0.909628023 | 0.921 | 0.969 | 0.000685531 | MM_1 |
| Cryl1 | ENSMUSG00000021947 | protein_coding | crystallin, lambda 1 | 3.424E-07 | -1.83223661 | 0.228 | 0.551 | 0.008085157 | MM_1 |
| Lcp1 | ENSMUSG00000021998 | protein_coding | lymphocyte cytosolic protein 1 | 1.215E-11 | -0.743870396 | 0.947 | 1 | 2.86791E-07 | MM_1 |
| Dab2 | ENSMUSG00000022150 | protein_coding | disabled 2, mitogen-responsive phosphoprotein | 9.67E-09 | -0.660190879 | 1 | 0.99 | 0.000228319 | MM_1 |
| Fyb | ENSMUSG00000022148 | protein_coding | FYN binding protein | 1.227E-07 | -0.851367676 | 0.833 | 0.939 | 0.002897821 | MM_1 |
| Ly6e | ENSMUSG00000022587 | protein_coding | lymphocyte antigen 6 complex, locus E | 6.09E-12 | -1.45835379 | 0.632 | 0.918 | 1.43796E-07 | MM_1 |
| Tef | ENSMUSG00000022389 | protein_coding | thyrotroph embryonic factor | 7.104E-08 | 1.963441675 | 0.535 | 0.194 | 0.001677425 | MM_1 |
| Dazap2 | ENSMUSG00000000346 | protein_coding | DAZ associated protein 2 | 7.426E-08 | -0.704946461 | 0.816 | 0.98 | 0.001753337 | MM_1 |
| Snx29 | ENSMUSG00000071669 | protein_coding | sorting nexin 29 | 1.722E-07 | -0.81492578 | 0.763 | 0.898 | 0.004065767 | MM_1 |
| Mylk | ENSMUSG00000022836 | protein_coding | myosin, light polypeptide kinase | 5.901E-08 | -1.038283801 | 0.561 | 0.867 | 0.001393333 | MM_1 |
| Gm15564 | ENSMUSG00000086324 | lncRNA | predicted gene 15564 | 7.396E-14 | 1.455791232 | 0.93 | 0.776 | 1.74646E-09 | MM_1 |
| Cmss1 | ENSMUSG00000022748 | protein_coding | cms small ribosomal subunit 1 | 3.77E-26 | 1.996304066 | 0.982 | 1 | 8.90224E-22 | MM_1 |
| St3gal6 | ENSMUSG00000022747 | protein_coding | ST3 beta-galactoside alpha-2,3-sialyltransferase 6 | 5.27E-08 | -1.368821028 | 0.316 | 0.694 | 0.001244442 | MM_1 |
| Rnaset2b | ENSMUSG00000094724 | protein_coding | ribonuclease T2B | 3.657E-07 | -1.703212038 | 0.281 | 0.602 | 0.008634567 | MM_1 |
| BC004004 | ENSMUSG00000052712 | protein_coding | cDNA sequence BC004004 | 3.346E-22 | -3.175571454 | 0.132 | 0.786 | 7.89976E-18 | MM_1 |
| Rps28 | ENSMUSG00000067288 | protein_coding | ribosomal protein S28 | 1.289E-13 | -0.755554111 | 0.947 | 0.99 | 3.04368E-09 | MM_1 |
| Aif1 | ENSMUSG00000024397 | protein_coding | allograft inflammatory factor 1 | 1.22E-09 | -1.181668811 | 0.64 | 0.898 | 2.8818E-05 | MM_1 |
| Gm42418 | ENSMUSG00000098178 | lncRNA | predicted gene, 42418 | 5.387E-29 | 1.490010976 | 1 | 1 | 1.27187E-24 | MM_1 |
| AY036118 | ENSMUSG00000105361 | lncRNA | cDNA sequence AY036118 | 5.508E-09 | 0.640809013 | 0.965 | 0.939 | 0.00013006 | MM_1 |
| Slc8a1 | ENSMUSG00000054640 | protein_coding | solute carrier family 8 (sodium/calcium exchanger), member 1 | 1.432E-11 | -1.007926932 | 0.895 | 0.98 | 3.38064E-07 | MM_1 |
| Csf1r | ENSMUSG00000024621 | protein_coding | colony stimulating factor 1 receptor | 2.029E-14 | -0.928058676 | 0.939 | 0.99 | 4.7912E-10 | MM_1 |
| Unc93b1 | ENSMUSG00000036908 | protein_coding | unc-93 homolog B1, TLR signaling regulator | 1.796E-08 | -0.669953579 | 0.886 | 0.99 | 0.000424117 | MM_1 |
| Gda | ENSMUSG00000058624 | protein_coding | guanine deaminase | 4.183E-08 | 1.293067668 | 0.833 | 0.694 | 0.000987795 | MM_1 |
| Acta2 | ENSMUSG00000035783 | protein_coding | actin, alpha 2, smooth muscle, aorta | 4.376E-14 | -1.423482766 | 0.684 | 0.959 | 1.03315E-09 | MM_1 |
| Gpr34 | ENSMUSG00000040229 | protein_coding | G protein-coupled receptor 34 | 1.528E-10 | -1.663911461 | 0.579 | 0.847 | 3.60706E-06 | MM_1 |
| Reps2 | ENSMUSG00000040855 | protein_coding | RALBP1 associated Eps domain containing protein 2 | 1.108E-08 | -0.967519315 | 0.518 | 0.847 | 0.000261626 | MM_1 |
| Des | ENSMUSG00000026208 | protein_coding | desmin | 9.567E-08 | -0.839511241 | 0.722 | 0.966 | 0.002259072 | MM_2 |
| Per2 | ENSMUSG00000055866 | protein_coding | period circadian clock 2 | 6.314E-09 | 4.57499828 | 0.278 | 0.008 | 0.000149081 | MM_2 |
| Csrp1 | ENSMUSG00000026421 | protein_coding | cysteine and glycine-rich protein 1 | 6.686E-10 | -1.307336567 | 0.405 | 0.849 | 1.57862E-05 | MM_2 |
| Gm2000 | ENSMUSG00000078193 | protein_coding | predicted gene 2000 | 3.295E-11 | -2.327396843 | 0.165 | 0.647 | 7.77949E-07 | MM_2 |
| Ifi202b | ENSMUSG00000026535 | protein_coding | interferon activated gene 202B | 6.463E-16 | -4.367127588 | 0.038 | 0.63 | 1.52607E-11 | MM_2 |
| Camk1d | ENSMUSG00000039145 | protein_coding | calcium/calmodulin-dependent protein kinase ID | 6.32E-10 | 0.602082759 | 1 | 1 | 1.49238E-05 | MM_2 |
| Rpl35 | ENSMUSG00000062997 | protein_coding | ribosomal protein L35 | 1.914E-09 | -0.679035244 | 0.975 | 0.992 | 4.51916E-05 | MM_2 |
| B2m | ENSMUSG00000060802 | protein_coding | beta-2 microglobulin | 2.951E-13 | -0.654186261 | 0.987 | 1 | 6.96867E-09 | MM_2 |
| Myl9 | ENSMUSG00000067818 | protein_coding | myosin, light polypeptide 9, regulatory | 6.426E-08 | -1.305540193 | 0.342 | 0.79 | 0.00151738 | MM_2 |
| Gm11808 | ENSMUSG00000068240 | protein_coding | predicted gene 11808 | 2.456E-15 | -2.52396131 | 0.215 | 0.79 | 5.79988E-11 | MM_2 |
| Dnaja1 | ENSMUSG00000028410 | protein_coding | DnaJ heat shock protein family (Hsp40) member A1 | 1.157E-09 | 1.361203803 | 0.949 | 0.874 | 2.73116E-05 | MM_2 |
| Ptp4a2 | ENSMUSG00000028788 | protein_coding | protein tyrosine phosphatase 4a2 | 9.341E-08 | -0.697172921 | 0.886 | 1 | 0.002205613 | MM_2 |
| Cd52 | ENSMUSG00000000682 | protein_coding | CD52 antigen | 2.892E-11 | 1.352669957 | 0.924 | 0.689 | 6.82803E-07 | MM_2 |

| | | | | | | | | | |
|---|---|---|---|---|---|---|---|---|---|
| Sh3bgrl3 | ENSMUSG00000028843 | protein_coding | SH3 domain binding glutamic acid-rich protein-like 3 | 1.882E-19 | 1.676870437 | 0.962 | 0.891 | 4.4448E-15 | MM_2 |
| C1qa | ENSMUSG00000036887 | protein_coding | complement component 1, q subcomponent, alpha polypeptide | 1.31E-17 | -1.264764564 | 0.924 | 1 | 3.09387E-13 | MM_2 |
| Gigyf1 | ENSMUSG00000029714 | protein_coding | GRB10 interacting GYF protein 1 | 9.907E-08 | -1.713916503 | 0.177 | 0.588 | 0.00233918 | MM_2 |
| Gnb2 | ENSMUSG00000029713 | protein_coding | guanine nucleotide binding protein (G protein), beta 2 | 3.403E-15 | -1.046551255 | 0.835 | 1 | 8.03591E-11 | MM_2 |
| Cdk8 | ENSMUSG00000029635 | protein_coding | cyclin-dependent kinase 8 | 1.32E-29 | 2.560047976 | 0.987 | 0.992 | 3.11703E-25 | MM_2 |
| Actg2 | ENSMUSG00000059430 | protein_coding | actin, gamma 2, smooth muscle, enteric | 5.884E-09 | -1.099368549 | 0.582 | 0.941 | 0.00013893 | MM_2 |
| Dbp | ENSMUSG00000059824 | protein_coding | D site albumin promoter binding protein | 9.547E-09 | 2.25620355 | 0.57 | 0.244 | 0.000225419 | MM_2 |
| Poteg | ENSMUSG00000063932 | protein_coding | POTE ankyrin domain family, member G | 1.852E-15 | -6.333443462 | 0 | 0.563 | 4.37186E-11 | MM_2 |
| Uba52 | ENSMUSG00000090137 | protein_coding | ubiquitin A-52 residue ribosomal protein fusion product 1 | 2.342E-11 | -1.328974148 | 0.557 | 0.899 | 5.53027E-07 | MM_2 |
| Gm42031 | ENSMUSG00000110386 | lncRNA | predicted gene, 42031 | 5.437E-12 | 5.062728419 | 0.367 | 0.008 | 1.28373E-07 | MM_2 |
| Cx3cr1 | ENSMUSG00000052336 | protein_coding | chemokine (C-X3-C motif) receptor 1 | 5.92E-08 | -1.094077748 | 0.595 | 0.924 | 0.001397779 | MM_2 |
| Lars2 | ENSMUSG00000035202 | protein_coding | leucyl-tRNA synthetase, mitochondrial | 1.134E-20 | 1.401322758 | 0.987 | 1 | 2.67727E-16 | MM_2 |
| Psap | ENSMUSG00000004207 | protein_coding | prosaposin | 1.637E-08 | -0.588570546 | 0.975 | 1 | 0.000386478 | MM_2 |
| Lyz2 | ENSMUSG00000069516 | protein_coding | lysozyme 2 | 9.001E-35 | -12.6371382 | 0 | 1 | 2.12538E-30 | MM_2 |
| Lgals9 | ENSMUSG00000001123 | protein_coding | lectin, galactose binding, soluble 9 | 2.517E-08 | -1.515383127 | 0.253 | 0.664 | 0.000594409 | MM_2 |
| Rpl38 | ENSMUSG00000057322 | protein_coding | ribosomal protein L38 | 7.832E-15 | -0.770682569 | 0.937 | 1 | 1.84931E-10 | MM_2 |
| Cd300ld | ENSMUSG00000034641 | protein_coding | CD300 molecule like family member d | 7.485E-08 | -0.960407964 | 0.633 | 0.882 | 0.001767383 | MM_2 |
| Laptm4a | ENSMUSG00000020585 | protein_coding | lysosomal-associated protein transmembrane 4A | 3.321E-07 | -0.754214402 | 0.797 | 0.95 | 0.007840757 | MM_2 |
| Gphn | ENSMUSG00000047454 | protein_coding | gephyrin | 9.289E-08 | 0.653753834 | 0.987 | 0.992 | 0.00219323 | MM_2 |
| Gm19951 | ENSMUSG00000113136 | lncRNA | predicted gene, 19951 | 7.985E-09 | 1.549590817 | 0.709 | 0.42 | 0.000188533 | MM_2 |
| Ckb | ENSMUSG00000001270 | protein_coding | creatine kinase, brain | 9.012E-10 | -1.264590932 | 0.734 | 0.933 | 2.12786E-05 | MM_2 |
| Hist1h4d | ENSMUSG00000061482 | protein_coding | histone cluster 1, H4d | 1.208E-07 | 1.987942363 | 0.506 | 0.193 | 0.002852185 | MM_2 |
| Gm48099 | ENSMUSG00000114304 | lncRNA | predicted gene, 48099 | 7.897E-10 | -2.111679583 | 0.152 | 0.622 | 1.86474E-05 | MM_2 |
| Gm15564 | ENSMUSG00000086324 | lncRNA | predicted gene 15564 | 1.741E-11 | 1.812072375 | 0.873 | 0.689 | 4.10996E-07 | MM_2 |
| Cmss1 | ENSMUSG00000022748 | protein_coding | cms small ribosomal subunit 1 | 5.618E-29 | 1.762915431 | 0.987 | 1 | 1.32654E-24 | MM_2 |
| BC004004 | ENSMUSG00000052712 | protein_coding | cDNA sequence BC004004 | 4.034E-18 | -2.426217244 | 0.215 | 0.866 | 9.52419E-14 | MM_2 |
| H2-Aa | ENSMUSG00000036594 | protein_coding | histocompatibility 2, class II antigen A, alpha | 1.984E-07 | -0.59444459 | 0.962 | 0.992 | 0.004684396 | MM_2 |
| Gm42418 | ENSMUSG00000098178 | lncRNA | predicted gene, 42418 | 5.096E-28 | 1.396867996 | 1 | 1 | 1.20316E-23 | MM_2 |
| AY036118 | ENSMUSG00000105361 | lncRNA | cDNA sequence AY036118 | 6.626E-18 | 1.064103655 | 0.987 | 0.983 | 1.56456E-13 | MM_2 |
| Hsp90ab1 | ENSMUSG00000023944 | protein_coding | heat shock protein 90 alpha (cytosolic), class B member 1 | 2.857E-10 | 0.811201005 | 1 | 0.992 | 6.74683E-06 | MM_2 |
| Cd74 | ENSMUSG00000024610 | protein_coding | CD74 antigen (invariant polypeptide of major histocompatibility complex, class II antigen-associated) | 4.037E-15 | -0.930500412 | 1 | 1 | 9.53161E-11 | MM_2 |
| Csf1r | ENSMUSG00000024621 | protein_coding | colony stimulating factor 1 receptor | 2.915E-11 | -0.867255354 | 0.949 | 0.992 | 6.8821E-07 | MM_2 |
| Unc93b1 | ENSMUSG00000036908 | protein_coding | unc-93 homolog B1, TLR signaling regulator | 3.814E-07 | -0.58502398 | 0.949 | 1 | 0.009006063 | MM_2 |
| Acta2 | ENSMUSG00000035783 | protein_coding | actin, alpha 2, smooth muscle, aorta | 1.076E-09 | -0.87470086 | 0.658 | 0.966 | 2.54102E-05 | MM_2 |
| Camk1d | ENSMUSG00000039145 | protein_coding | calcium/calmodulin-dependent protein kinase ID | 3.539E-11 | 0.978146081 | 0.986 | 0.97 | 8.35742E-07 | MM_3 |
| C1qa | ENSMUSG00000036887 | protein_coding | complement component 1, q subcomponent, alpha polypeptide | 5.875E-17 | -3.624378134 | 0.162 | 0.792 | 1.3873E-12 | MM_3 |
| Cdk8 | ENSMUSG00000029635 | protein_coding | cyclin-dependent kinase 8 | 2.749E-27 | 2.862860818 | 1 | 0.941 | 6.49024E-23 | MM_3 |
| Lars2 | ENSMUSG00000035202 | protein_coding | leucyl-tRNA synthetase, mitochondrial | 1.207E-16 | 1.755201873 | 0.986 | 0.871 | 2.85007E-12 | MM_3 |
| Lyz2 | ENSMUSG00000069516 | protein_coding | lysozyme 2 | 1.795E-22 | -11.89928589 | 0 | 0.792 | 4.23889E-18 | MM_3 |
| Gphn | ENSMUSG00000047454 | protein_coding | gephyrin | 2.938E-15 | 1.731753263 | 0.946 | 0.842 | 6.9361E-11 | MM_3 |
| Gm15564 | ENSMUSG00000086324 | lncRNA | predicted gene 15564 | 2.044E-10 | 1.991661388 | 0.73 | 0.337 | 4.82677E-06 | MM_3 |
| Cmss1 | ENSMUSG00000022748 | protein_coding | cms small ribosomal subunit 1 | 3.915E-19 | 2.173649673 | 0.973 | 0.931 | 9.24405E-15 | MM_3 |
| Gm42418 | ENSMUSG00000098178 | lncRNA | predicted gene, 42418 | 7.209E-21 | 1.605759153 | 1 | 1 | 1.7021E-16 | MM_3 |
| Cd74 | ENSMUSG00000024610 | protein_coding | CD74 antigen (invariant polypeptide of major histocompatibility complex, class II antigen-associated) | 1.467E-07 | -1.596021037 | 0.635 | 0.871 | 0.003463512 | MM_3 |
| Gm11808 | ENSMUSG00000068240 | protein_coding | predicted gene 11808 | 2.754E-09 | -3.07438697 | 0.37 | 0.88 | 6.50172E-05 | MM_4 |
| Gnb2 | ENSMUSG00000029713 | protein_coding | guanine nucleotide binding protein (G protein), beta 2 | 1.782E-08 | -1.372290064 | 0.889 | 0.96 | 0.000420815 | MM_4 |
| Cdk8 | ENSMUSG00000029635 | protein_coding | cyclin-dependent kinase 8 | 1.789E-10 | 2.689172684 | 1 | 0.98 | 4.22426E-06 | MM_4 |
| Poteg | ENSMUSG00000063932 | protein_coding | POTE ankyrin domain family, member G | 4.661E-10 | -4.774915212 | 0 | 0.82 | 1.10055E-05 | MM_4 |
| Lyz2 | ENSMUSG00000069516 | protein_coding | lysozyme 2 | 1.83E-13 | -9.264960678 | 0 | 1 | 4.32119E-09 | MM_4 |
| Cmss1 | ENSMUSG00000022748 | protein_coding | cms small ribosomal subunit 1 | 3.443E-07 | 1.606880783 | 1 | 1 | 0.008130244 | MM_4 |
| BC004004 | ENSMUSG00000052712 | protein_coding | cDNA sequence BC004004 | 1.006E-09 | -3.002654696 | 0.259 | 0.92 | 2.37427E-05 | MM_4 |
