## Supplementary Table 4 for "C1qa□ muscularis macrophages maintain enteric synaptic homeostasis to regulate gastrointestinal motility"

| NAME | GS<br> follow link to MSigDB | SIZE | ES | NES | NOM p-val | FDR q-val | FWER p-val | RANK AT MAX | LEADING EDGE | cluster |
|---|---|---|---|---|---|---|---|---|---|---|
| C1qaCKOvsC1qaFL | | | | | | | | | | |
| GOBP_REGULATION_OF_ | GOBP_REGULATION_OF_BIOLOGICAL_PROC | 33 | 0.656166 | 2.021311 | 0 | 0.05333728 | 0.058 | 431 | tags=18%, list=4%, sig | EN_1 |
| GOBP_MODULATION_BY_ | GOBP_MODULATION_BY_SYMBIONT_OF_EN | 31 | 0.638483 | 1.90094 | 0 | 0.24495755 | 0.434 | 431 | tags=16%, list=4%, sig | EN_1 |
| GOBP_RESPONSE_TO_CH | GOBP_RESPONSE_TO_CHEMOKINE | 23 | 0.673789 | 1.891698 | 0 | 0.19154216 | 0.485 | 425 | tags=26%, list=4%, sig | EN_1 |
| GOBP_NEUTROPHIL_CHE | GOBP_NEUTROPHIL_CHEMOTAXIS | 29 | 0.628571 | 1.878582 | 0.00135318 | 0.1729305 | 0.561 | 1264 | tags=34%, list=13%, si | EN_1 |
| GOBP_CHAPERONE_COFA | GOBP_CHAPERONE_COFACTOR_DEPENDEN | 28 | 0.640999 | 1.877758 | 0.001416431 | 0.14000615 | 0.565 | 1700 | tags=50%, list=18%, si | EN_1 |
| GOBP_CELL_KILLING | GOBP_CELL_KILLING | 75 | 0.536621 | 1.876182 | 0 | 0.12128147 | 0.577 | 1439 | tags=35%, list=15%, si | EN_1 |
| GOBP_SUPPRESSION_OF_ | GOBP_SUPPRESSION_OF_VIRAL_RELEASE_B | 16 | 0.729872 | 1.862977 | 0 | 0.12791929 | 0.658 | 11 | tags=13%, list=0%, sig | EN_1 |
| GOBP_NEGATIVE_REGUL | GOBP_NEGATIVE_REGULATION_OF_VIRAL_ | 21 | 0.673884 | 1.850042 | 0 | 0.13427623 | 0.72 | 52 | tags=14%, list=1%, sig | EN_1 |
| GOBP_ANTIMICROBIAL_H | GOBP_ANTIMICROBIAL_HUMORAL_RESPON | 26 | 0.647128 | 1.846359 | 0 | 0.1265225 | 0.74 | 1313 | tags=42%, list=14%, si | EN_1 |
| GOBP_CHAPERONE_MED | GOBP_CHAPERONE_MEDIATED_PROTEIN_F | 51 | 0.560703 | 1.838004 | 0 | 0.13024849 | 0.787 | 1700 | tags=35%, list=18%, si | EN_1 |
| GOBP_HUMORAL_IMMUI | GOBP_HUMORAL_IMMUNE_RESPONSE | 47 | 0.563482 | 1.832237 | 0 | 0.1282987 | 0.811 | 1328 | tags=34%, list=14%, si | EN_1 |
| GOBP_ADENYLATE_CYCLA | GOBP_ADENYLATE_CYCLASE_ACTIVATING_ | 38 | 0.574305 | 1.812828 | 0.001329787 | 0.15723775 | 0.884 | 1564 | tags=26%, list=16%, si | EN_1 |
| GOBP_DE_NOVO_PROTEI | GOBP_DE_NOVO_PROTEIN_FOLDING | 35 | 0.591371 | 1.802504 | 0.001308901 | 0.16873987 | 0.913 | 1896 | tags=46%, list=20%, si | EN_1 |
| GOBP_NEUTROPHIL_MIG | GOBP_NEUTROPHIL_MIGRATION | 38 | 0.573191 | 1.782773 | 0.001303781 | 0.20637633 | 0.955 | 985 | tags=26%, list=10%, si | EN_1 |
| GOBP_CHONDROCYTE_D | GOBP_CHONDROCYTE_DEVELOPMENT | 17 | 0.688055 | 1.778238 | 0.001459854 | 0.20417714 | 0.962 | 821 | tags=41%, list=9%, sig | EN_1 |
| GOBP_ANTIMICROBIAL_H | GOBP_ANTIMICROBIAL_HUMORAL_IMMUN | 19 | 0.652627 | 1.776813 | 0.001451379 | 0.19406271 | 0.963 | 1313 | tags=42%, list=14%, si | EN_1 |
| LEGIONELLOSIS | LEGIONELLOSIS | 33 | 0.582184 | 1.771632 | 0.00405954 | 0.19522305 | 0.97 | 2493 | tags=55%, list=26%, si | EN_1 |
| GOBP_REGULATION_OF_ | GOBP_REGULATION_OF_T_HELPER_CELL_D | 17 | 0.66766 | 1.767686 | 0.002898551 | 0.19476463 | 0.974 | 821 | tags=29%, list=9%, sig | EN_1 |
| GOBP_NEGATIVE_REGUL | GOBP_NEGATIVE_REGULATION_OF_IMMUN | 52 | 0.540275 | 1.763314 | 0.00130039 | 0.195613 | 0.978 | 1713 | tags=35%, list=18%, si | EN_1 |
| GOBP_TRIGLYCERIDE_BIO | GOBP_TRIGLYCERIDE_BIOSYNTHETIC_PROCE | 31 | 0.593943 | 1.759881 | 0.001338688 | 0.19393928 | 0.984 | 230 | tags=16%, list=2%, sig | EN_1 |
| GOBP_POSITIVE_REGULA | GOBP_POSITIVE_REGULATION_OF_COLLAGE | 21 | 0.647346 | 1.756962 | 0.004149378 | 0.19128744 | 0.986 | 585 | tags=29%, list=6%, sig | EN_1 |
| GOBP_PROTEIN_MATURA | GOBP_PROTEIN_MATURATION_BY_IRON_SU | 15 | 0.690686 | 1.745861 | 0.005952381 | 0.2087238 | 0.993 | 1559 | tags=60%, list=16%, si | EN_1 |
| IL-17_SIGNALING_PATHW | IL-17_SIGNALING_PATHWAY | 40 | 0.552442 | 1.737879 | 0.002680965 | 0.22160003 | 0.996 | 1693 | tags=38%, list=18%, si | EN_1 |
| GOBP_NEGATIVE_REGUL | GOBP_NEGATIVE_REGULATION_OF_LYMPH | 31 | 0.588885 | 1.734582 | 0 | 0.2206561 | 0.998 | 833 | tags=32%, list=9%, sig | EN_1 |
| ANTIGEN_PROCESSING_A | ANTIGEN_PROCESSING_AND_PRESENTATIO | 38 | 0.558255 | 1.732529 | 0.002617801 | 0.21647292 | 0.998 | 1110 | tags=37%, list=12%, si | EN_1 |
| GOBP_SYMBIONT_ENTRY | GOBP_SYMBIONT_ENTRY_INTO_HOST | 53 | 0.523119 | 1.731208 | 0 | 0.21196881 | 0.998 | 52 | tags=9%, list=1%, sign | EN_1 |
| GOBP_KILLING_OF_CELLS | GOBP_KILLING_OF_CELLS_OF_ANOTHER_OI | 23 | 0.619598 | 1.730507 | 0.005494506 | 0.20579086 | 0.998 | 1313 | tags=57%, list=14%, si | EN_1 |
| GOBP_DISRUPTION_OF_A | GOBP_DISRUPTION_OF_ANATOMICAL_STRU | 25 | 0.607399 | 1.729069 | 0.006756757 | 0.20215544 | 0.998 | 1313 | tags=52%, list=14%, si | EN_1 |
| GOBP_POSITIVE_REGULA | GOBP_POSITIVE_REGULATION_OF_MYELOID | 33 | 0.57224 | 1.72897 | 0.006877579 | 0.19550444 | 0.998 | 1437 | tags=36%, list=15%, si | EN_1 |
| GOBP_POSITIVE_REGULA | GOBP_POSITIVE_REGULATION_OF_CANONI | 98 | 0.481187 | 1.728659 | 0 | 0.18969359 | 0.998 | 1334 | tags=22%, list=14%, si | EN_1 |
| GOBP_GRANULOCYTE_CH | GOBP_GRANULOCYTE_CHEMOTAXIS | 42 | 0.536495 | 1.724564 | 0 | 0.19368245 | 0.998 | 985 | tags=26%, list=10%, si | EN_1 |
| GOBP_NEGATIVE_REGUL | GOBP_NEGATIVE_REGULATION_OF_LEUKO | 27 | 0.591152 | 1.72328 | 0.001340483 | 0.1902535 | 0.998 | 2025 | tags=48%, list=21%, si | EN_1 |
| GOBP_POSITIVE_REGULA | GOBP_POSITIVE_REGULATION_OF_LEUKOCY | 19 | 0.653309 | 1.720347 | 0.002949853 | 0.19102582 | 0.998 | 441 | tags=26%, list=5%, sig | EN_1 |
| GOBP_STEROL_BIOSYNTH | GOBP_STEROL_BIOSYNTHETIC_PROCESS | 40 | 0.541203 | 1.719146 | 0.003957784 | 0.1882808 | 0.999 | 3017 | tags=65%, list=31%, si | EN_1 |
| GOBP_REGULATION_OF_ | GOBP_REGULATION_OF_LEUKOCYTE_DIFFE | 176 | 0.450217 | 1.71595 | 0 | 0.18948427 | 0.999 | 1437 | tags=26%, list=15%, si | EN_1 |
| GOBP_MONOCYTE_DIFFE | GOBP_MONOCYTE_DIFFERENTIATION | 17 | 0.657004 | 1.714101 | 0.008645534 | 0.18826763 | 0.999 | 1122 | tags=41%, list=12%, si | EN_1 |
| CORTISOL_SYNTHESIS_AN | CORTISOL_SYNTHESIS_AND_SECRETION | 35 | 0.569465 | 1.711382 | 0.00130719 | 0.1888109 | 0.999 | 2207 | tags=46%, list=23%, si | EN_1 |
| GOBP_REGULATION_OF_ | GOBP_REGULATION_OF_B_CELL_DIFFERENT | 18 | 0.648611 | 1.708055 | 0.005722461 | 0.19142543 | 0.999 | 1619 | tags=28%, list=17%, si | EN_1 |
| GOBP_REGULATION_OF_ | GOBP_REGULATION_OF_CELL_KILLING | 42 | 0.546846 | 1.707016 | 0.002583979 | 0.18956526 | 0.999 | 1439 | tags=31%, list=15%, si | EN_1 |
| GLYCINE,_SERINE_AND_T | GLYCINE,_SERINE_AND_THREONINE_METAE | 16 | 0.652623 | 1.698277 | 0.005722461 | 0.20503889 | 0.999 | 1330 | tags=44%, list=14%, si | EN_1 |
| ESTROGEN_SIGNALING_P | ESTROGEN_SIGNALING_PATHWAY | 74 | 0.491111 | 1.697314 | 0.001231527 | 0.20246185 | 1 | 1564 | tags=31%, list=16%, si | EN_1 |
| GOBP_MONOCARBOXYLI | GOBP_MONOCARBOXYLIC_ACID_BIOSYNTH | 106 | 0.472286 | 1.697079 | 0 | 0.19807965 | 1 | 2669 | tags=45%, list=28%, si | EN_1 |
| GOBP_NEGATIVE_REGUL | GOBP_NEGATIVE_REGULATION_OF_ADAPTI | 17 | 0.648827 | 1.69646 | 0.011396011 | 0.19488584 | 1 | 2478 | tags=53%, list=26%, si | EN_1 |
| GOBP_NUCLEOSIDE_BISP | GOBP_NUCLEOSIDE_BISPHOSPHATE_BIOSYN | 36 | 0.553577 | 1.695028 | 0.009150327 | 0.1933476 | 1 | 2674 | tags=50%, list=28%, si | EN_1 |
| GOBP_LIPOPOLYSACCHAI | GOBP_LIPOPOLYSACCHARIDE_MEDIATED_SI | 30 | 0.569578 | 1.694674 | 0.011034483 | 0.18997198 | 1 | 1334 | tags=27%, list=14%, si | EN_1 |
| GOBP_NEGATIVE_REGUL | GOBP_NEGATIVE_REGULATION_OF_LYMPH | 21 | 0.612 | 1.691655 | 0.005649718 | 0.19232939 | 1 | 2025 | tags=43%, list=21%, si | EN_1 |
| GOBP_PROTEIN_REFOLDI | GOBP_PROTEIN_REFOLDING | 22 | 0.605453 | 1.690079 | 0.005449591 | 0.19216011 | 1 | 2740 | tags=64%, list=29%, si | EN_1 |
| GOBP_REGULATION_OF_ | GOBP_REGULATION_OF_B_CELL_APOPTOTI | 15 | 0.676297 | 1.688605 | 0.007429421 | 0.19095859 | 1 | 52 | tags=20%, list=1%, sig | EN_1 |
| GOBP_RESPONSE_TO_INT | GOBP_RESPONSE_TO_INTERFERON_BETA | 27 | 0.579797 | 1.686948 | 0.012693935 | 0.19076411 | 1 | 1882 | tags=48%, list=20%, si | EN_1 |
| GOBP_SMALL_MOLECULE | GOBP_SMALL_MOLECULE_BIOSYNTHETIC_P | 291 | 0.421362 | 1.685492 | 0 | 0.19006248 | 1 | 2671 | tags=42%, list=28%, si | EN_1 |
| GOBP_T_CELL_ACTIVATIO | GOBP_T_CELL_ACTIVATION_INVOLVED_IN_ | 47 | 0.520328 | 1.682009 | 0.009079118 | 0.19414654 | 1 | 972 | tags=28%, list=10%, si | EN_1 |
| GOBP_CANONICAL_NF_K | GOBP_CANONICAL_NF_KAPPAB_SIGNAL_TR | 163 | 0.442657 | 1.677141 | 0 | 0.20088813 | 1 | 1837 | tags=27%, list=19%, si | EN_1 |
| GOBP_MALE_MEIOTIC_N | GOBP_MALE_MEIOTIC_NUCLEAR_DIVISION | 15 | 0.671044 | 1.674871 | 0.011412269 | 0.20215303 | 1 | 970 | tags=40%, list=10%, si | EN_1 |
| GOBP_RNA_METHYLATIO | GOBP_RNA_METHYLATION | 46 | 0.523966 | 1.672727 | 0.003797468 | 0.2027409 | 1 | 2655 | tags=59%, list=28%, si | EN_1 |
| FERROPTOSIS | FERROPTOSIS | 30 | 0.570335 | 1.672385 | 0.004126547 | 0.19970769 | 1 | 1699 | tags=30%, list=18%, si | EN_1 |
| GOBP_POSITIVE_REGULA | GOBP_POSITIVE_REGULATION_OF_MYELOID | 20 | 0.601764 | 1.662695 | 0.009762901 | 0.21854946 | 1 | 2378 | tags=45%, list=25%, si | EN_1 |
| GOBP_POSITIVE_REGULA | GOBP_POSITIVE_REGULATION_OF_HEMOPO | 104 | 0.457569 | 1.661058 | 0.001175088 | 0.21900083 | 1 | 1238 | tags=23%, list=13%, si | EN_1 |
| GOBP_THIOESTER_BIOSYI | GOBP_THIOESTER_BIOSYNTHETIC_PROCESS | 25 | 0.581199 | 1.658967 | 0.005369128 | 0.21983135 | 1 | 2674 | tags=52%, list=28%, si | EN_1 |
| GOBP_REGULATION_OF_ | GOBP_REGULATION_OF_G_PROTEIN_COUP | 68 | 0.474286 | 1.658553 | 0.001242236 | 0.21723051 | 1 | 1297 | tags=24%, list=14%, si | EN_1 |
| GOBP_RESPONSE_TO_PR | GOBP_RESPONSE_TO_PROTOZOAN | 15 | 0.64863 | 1.658275 | 0.015492958 | 0.2142241 | 1 | 2073 | tags=67%, list=22%, si | EN_1 |
| GOBP_BIOLOGICAL_PROC | GOBP_BIOLOGICAL_PROCESS_INVOLVED_IN | 83 | 0.473245 | 1.655449 | 0.001183432 | 0.21700978 | 1 | 1845 | tags=24%, list=19%, si | EN_1 |
| GOBP_FATTY_ACID_BIOS | GOBP_FATTY_ACID_BIOSYNTHETIC_PROCES | 82 | 0.472856 | 1.653288 | 0.001204819 | 0.21819292 | 1 | 2453 | tags=43%, list=26%, si | EN_1 |
| GOBP_DEFENSE_RESPON | GOBP_DEFENSE_RESPONSE_TO_GRAM_NE | 26 | 0.568106 | 1.650139 | 0.013947002 | 0.22231817 | 1 | 2378 | tags=46%, list=25%, si | EN_1 |
| GOBP_PYROPTOSIS | GOBP_PYROPTOSIS | 15 | 0.638032 | 1.643538 | 0.015363129 | 0.23474723 | 1 | 1407 | tags=47%, list=15%, si | EN_1 |
| GOBP_REGULATION_OF_ | GOBP_REGULATION_OF_FATTY_ACID_OXID | 27 | 0.557378 | 1.640692 | 0.005256242 | 0.2387201 | 1 | 2593 | tags=59%, list=27%, si | EN_1 |
| GOBP_NEGATIVE_REGUL | GOBP_NEGATIVE_REGULATION_OF_HEMOP | 60 | 0.486383 | 1.638933 | 0.003764116 | 0.23961233 | 1 | 833 | tags=25%, list=9%, sig | EN_1 |
| GOBP_POSITIVE_REGULA | GOBP_POSITIVE_REGULATION_OF_FATTY_A | 25 | 0.575106 | 1.636517 | 0.012261581 | 0.24219997 | 1 | 2593 | tags=48%, list=27%, si | EN_1 |
| GOBP_CELLULAR_RESPON | GOBP_CELLULAR_RESPONSE_TO_BIOTIC_ST | 123 | 0.44227 | 1.636414 | 0.001146789 | 0.23881441 | 1 | 1388 | tags=23%, list=14%, si | EN_1 |
| GOBP_VERY_LONG_CHAI | GOBP_VERY_LONG_CHAIN_FATTY_ACID_M | 23 | 0.577242 | 1.635761 | 0.010899182 | 0.23697124 | 1 | 2567 | tags=57%, list=27%, si | EN_1 |
| GOBP_POSITIVE_REGULA | GOBP_POSITIVE_REGULATION_OF_RESPON | 37 | 0.535416 | 1.635747 | 0.003875969 | 0.23359904 | 1 | 470 | tags=22%, list=5%, sig | EN_1 |
| GOBP_RESPONSE_TO_TY | GOBP_RESPONSE_TO_TYPE_II_INTERFERON | 71 | 0.478225 | 1.63482 | 0.007434944 | 0.2325548 | 1 | 1929 | tags=35%, list=20%, si | EN_1 |
| GOBP_RESPONSE_TO_HE | GOBP_RESPONSE_TO_HEAT | 57 | 0.488602 | 1.634543 | 0.006337136 | 0.22992617 | 1 | 3622 | tags=58%, list=38%, si | EN_1 |
| GOBP_NUCLEOSIDE_MON | GOBP_NUCLEOSIDE_MONOPHOSPHATE_ME | 38 | 0.531087 | 1.633677 | 0.006451613 | 0.22860521 | 1 | 3064 | tags=55%, list=32%, si | EN_1 |
| GOBP_INTERLEUKIN_2_P | GOBP_INTERLEUKIN_2_PRODUCTION | 31 | 0.550938 | 1.633546 | 0.004021448 | 0.22584002 | 1 | 2319 | tags=48%, list=24%, si | EN_1 |
| GOBP_REGULATION_OF_ | GOBP_REGULATION_OF_LYMPHOCYTE_DIFF | 114 | 0.443399 | 1.630957 | 0 | 0.2291958 | 1 | 1231 | tags=22%, list=13%, si | EN_1 |
| CHEMOKINE_SIGNALING_ | CHEMOKINE_SIGNALING_PATHWAY | 93 | 0.455132 | 1.630155 | 0.004807693 | 0.22815762 | 1 | 1622 | tags=26%, list=17%, si | EN_1 |
| GOBP_REGULATION_OF_ | GOBP_REGULATION_OF_LYMPHOCYTE_APO | 39 | 0.521287 | 1.629111 | 0.006443299 | 0.2273374 | 1 | 1744 | tags=36%, list=18%, si | EN_1 |
| GOBP_RESPONSE_TO_TO | GOBP_RESPONSE_TO_TOPOLOGICALLY_INC | 107 | 0.448122 | 1.627591 | 0.001173709 | 0.22762226 | 1 | 2908 | tags=48%, list=30%, si | EN_1 |
| GOBP_POSITIVE_REGULA | GOBP_POSITIVE_REGULATION_OF_NF_KAPI | 74 | 0.467004 | 1.623614 | 0.00477327 | 0.23426783 | 1 | 1587 | tags=26%, list=17%, si | EN_1 |
| GOBP_GRANULOCYTE_M | GOBP_GRANULOCYTE_MIGRATION | 52 | 0.497828 | 1.622178 | 0.007662835 | 0.23500006 | 1 | 985 | tags=23%, list=10%, si | EN_1 |

| | | | | | | | | | | |
|---|---|---|---|---|---|---|---|---|---|---|
| GOBP_REGULATION_OF_ | GOBP_REGULATION_OF_MYELOID_LEUKOC | 70 | 0.47329 | 1.621967 | 0.006016847 | 0.23273516 | 1 | 1684 | tags=33%, list=18%, si | EN_1 |
| GOBP_REGULATION_OF_ | GOBP_REGULATION_OF_ADAPTIVE_IMMUN | 92 | 0.451844 | 1.620371 | 0.00243309 | 0.23381913 | 1 | 2568 | tags=40%, list=27%, si | EN_1 |
| CHEMICAL_CARCINOGEN | CHEMICAL_CARCINOGENESIS | 16 | 0.627183 | 1.618049 | 0.008559201 | 0.23670745 | 1 | 2455 | tags=56%, list=26%, si | EN_1 |
| GOBP_RIBOSE_PHOSPHA | GOBP_RIBOSE_PHOSPHATE_BIOSYNTHETIC_ | 146 | 0.428181 | 1.613994 | 0.001146789 | 0.24345183 | 1 | 2593 | tags=38%, list=27%, si | EN_1 |
| OXIDATIVE_PHOSPHORYL | OXIDATIVE_PHOSPHORYLATION | 99 | 0.6117 | 2.666369 | 0 | 0 | 0 | 2562 | tags=77%, list=30%, si | MM_1 |
| PARKINSON_DISEASE | PARKINSON_DISEASE | 99 | 0.544024 | 2.360052 | 0 | 0 | 0 | 2562 | tags=73%, list=30%, si | MM_1 |
| GOBP_PROTEIN_FOLDING | GOBP_PROTEIN_FOLDING | 128 | 0.485299 | 2.199288 | 0 | 0.00430829 | 0.014 | 1523 | tags=39%, list=18%, si | MM_1 |
| GOBP_PROTEIN_MATURA | GOBP_PROTEIN_MATURATION_BY_IRON_SU | 15 | 0.759709 | 2.155382 | 0 | 0.007371451 | 0.032 | 1366 | tags=53%, list=16%, si | MM_1 |
| GOBP_RESPONSE_TO_HE | GOBP_RESPONSE_TO_HEAT | 56 | 0.564211 | 2.14588 | 0 | 0.006448926 | 0.035 | 1196 | tags=36%, list=14%, si | MM_1 |
| GOBP_ESTABLISHMENT_( | GOBP_ESTABLISHMENT_OF_PROTEIN_LOCA | 21 | 0.692427 | 2.121259 | 0 | 0.008309053 | 0.053 | 1169 | tags=67%, list=13%, si | MM_1 |
| GOBP_PROTON_MOTIVE_ | GOBP_PROTON_MOTIVE_FORCE_DRIVEN_A | 52 | 0.549513 | 2.1059 | 0 | 0.008946934 | 0.066 | 2515 | tags=67%, list=29%, si | MM_1 |
| GOBP_AEROBIC_RESPIRA | GOBP_AEROBIC_RESPIRATION | 145 | 0.459113 | 2.0968 | 0 | 0.009085899 | 0.077 | 2139 | tags=50%, list=25%, si | MM_1 |
| GOBP_OXIDATIVE_PHOSF | GOBP_OXIDATIVE_PHOSPHORYLATION | 106 | 0.481051 | 2.069118 | 0 | 0.012646558 | 0.119 | 2139 | tags=53%, list=25%, si | MM_1 |
| GOBP_NUCLEOSIDE_TRIP | GOBP_NUCLEOSIDE_TRIPHOSPHATE_BIOSYN | 86 | 0.489349 | 2.061838 | 0 | 0.012760043 | 0.132 | 2607 | tags=62%, list=30%, si | MM_1 |
| GOBP_CELLULAR_RESPIR | GOBP_CELLULAR_RESPIRATION | 177 | 0.432111 | 2.020794 | 0 | 0.022579469 | 0.239 | 2139 | tags=48%, list=25%, si | MM_1 |
| GOBP_NUCLEOSIDE_TRIP | GOBP_NUCLEOSIDE_TRIPHOSPHATE_METAE | 179 | 0.430923 | 2.019704 | 0 | 0.020852841 | 0.24 | 1851 | tags=39%, list=21%, si | MM_1 |
| GOBP_ATP_BIOSYNTHETI | GOBP_ATP_BIOSYNTHETIC_PROCESS | 74 | 0.499799 | 2.011386 | 0 | 0.021237766 | 0.264 | 2354 | tags=59%, list=27%, si | MM_1 |
| PROTEASOME | PROTEASOME | 42 | 0.552707 | 2.00628 | 0 | 0.021038555 | 0.277 | 1884 | tags=52%, list=22%, si | MM_1 |
| GOBP_ATP_METABOLIC_ | GOBP_ATP_METABOLIC_PROCESS | 147 | 0.442128 | 1.997441 | 0 | 0.022005763 | 0.306 | 2088 | tags=44%, list=24%, si | MM_1 |
| GOBP_PROTEIN_REFOLDI | GOBP_PROTEIN_REFOLDING | 18 | 0.67188 | 1.9737 | 0 | 0.02746964 | 0.392 | 950 | tags=56%, list=11%, si | MM_1 |
| SYNAPTIC_VESICLE_CYCL | SYNAPTIC_VESICLE_CYCLE | 33 | 0.565123 | 1.952635 | 0 | 0.033044104 | 0.47 | 2140 | tags=55%, list=25%, si | MM_1 |
| GOBP_MITOCHONDRIAL_ | GOBP_MITOCHONDRIAL_TRANSLATION | 113 | 0.442895 | 1.943932 | 0 | 0.03416131 | 0.501 | 1849 | tags=47%, list=21%, si | MM_1 |
| GOBP_SUPPRESSION_OF_ | GOBP_SUPPRESSION_OF_VIRAL_RELEASE_B | 19 | 0.649308 | 1.929679 | 0.002475248 | 0.037926324 | 0.561 | 268 | tags=21%, list=3%, sig | MM_1 |
| GOBP_RESPONSE_TO_TE | GOBP_RESPONSE_TO_TEMPERATURE_STIM | 90 | 0.458755 | 1.924064 | 0 | 0.038982164 | 0.585 | 1444 | tags=34%, list=17%, si | MM_1 |
| GOBP_NADH_DEHYDROG | GOBP_NADH_DEHYDROGENASE_COMPLEX_ | 52 | 0.510317 | 1.916536 | 0 | 0.040700153 | 0.618 | 2424 | tags=62%, list=28%, si | MM_1 |
| GOBP_CHAPERONE_MED | GOBP_CHAPERONE_MEDIATED_PROTEIN_F | 51 | 0.510017 | 1.91125 | 0 | 0.040900473 | 0.636 | 950 | tags=31%, list=11%, si | MM_1 |
| GOBP_CELLULAR_RESPON | GOBP_CELLULAR_RESPONSE_TO_HEAT | 37 | 0.545486 | 1.902954 | 0.002808989 | 0.043725695 | 0.675 | 1196 | tags=32%, list=14%, si | MM_1 |
| HUNTINGTON_DISEASE | HUNTINGTON_DISEASE | 138 | 0.415739 | 1.887181 | 0 | 0.050105743 | 0.737 | 2562 | tags=64%, list=30%, si | MM_1 |
| GOBP_ELECTRON_TRANS | GOBP_ELECTRON_TRANSPORT_CHAIN | 88 | 0.451179 | 1.879558 | 0 | 0.05273961 | 0.767 | 2139 | tags=48%, list=25%, si | MM_1 |
| GOBP_ANTIMICROBIAL_H | GOBP_ANTIMICROBIAL_HUMORAL_IMMUN | 31 | 0.557609 | 1.874823 | 0 | 0.053755656 | 0.785 | 642 | tags=32%, list=7%, sig | MM_1 |
| GOBP_MITOCHONDRIAL_ | GOBP_MITOCHONDRIAL_RESPIRATORY_CHA | 81 | 0.458043 | 1.868368 | 0 | 0.055454858 | 0.805 | 2424 | tags=56%, list=28%, si | MM_1 |
| THERMOGENESIS | THERMOGENESIS | 160 | 0.397743 | 1.848958 | 0 | 0.0654638 | 0.867 | 2354 | tags=52%, list=27%, si | MM_1 |
| GOBP_ATP_SYNTHESIS_C | GOBP_ATP_SYNTHESIS_COUPLED_ELECTROI | 66 | 0.465184 | 1.840977 | 0 | 0.068874955 | 0.892 | 2139 | tags=53%, list=25%, si | MM_1 |
| GOBP_MITOCHONDRIAL_ | GOBP_MITOCHONDRIAL_GENE_EXPRESSION | 136 | 0.404423 | 1.827641 | 0 | 0.0768482 | 0.926 | 1849 | tags=45%, list=21%, si | MM_1 |
| GOBP_ENERGY_DERIVATI | GOBP_ENERGY_DERIVATION_BY_OXIDATIOI | 231 | 0.372773 | 1.824244 | 0 | 0.07707594 | 0.934 | 2139 | tags=42%, list=25%, si | MM_1 |
| GOBP_AEROBIC_ELECTRO | GOBP_AEROBIC_ELECTRON_TRANSPORT_CH | 56 | 0.46512 | 1.822783 | 0 | 0.07576019 | 0.937 | 2139 | tags=55%, list=25%, si | MM_1 |
| GOBP_CARBOHYDRATE_D | GOBP_CARBOHYDRATE_DERIVATIVE_CATAB | 144 | 0.402071 | 1.810302 | 0 | 0.08342316 | 0.954 | 1097 | tags=26%, list=13%, si | MM_1 |
| RIBOSOME | RIBOSOME | 126 | 0.398184 | 1.788741 | 0 | 0.099868946 | 0.975 | 1877 | tags=27%, list=22%, si | MM_1 |
| GOBP_GTP_METABOLIC_ | GOBP_GTP_METABOLIC_PROCESS | 19 | 0.595779 | 1.778483 | 0.007177034 | 0.10729631 | 0.98 | 1946 | tags=53%, list=22%, si | MM_1 |
| PROTEIN_PROCESSING_IN | PROTEIN_PROCESSING_IN_ENDOPLASMIC_F | 142 | 0.392304 | 1.765774 | 0 | 0.117272854 | 0.989 | 1336 | tags=25%, list=15%, si | MM_1 |
| GOBP_PROTEIN_MATURA | GOBP_PROTEIN_MATURATION | 308 | 0.350695 | 1.763936 | 0 | 0.116016306 | 0.989 | 1415 | tags=28%, list=16%, si | MM_1 |
| GOBP_RESPONSE_TO_FO | GOBP_RESPONSE_TO_FOOD | 15 | 0.62447 | 1.743189 | 0.012406948 | 0.13658114 | 0.996 | 1834 | tags=73%, list=21%, si | MM_1 |
| GOBP_MITOCHONDRIAL_ | GOBP_MITOCHONDRIAL_ELECTRON_TRANS | 27 | 0.518292 | 1.707323 | 0.005263158 | 0.1831214 | 1 | 2088 | tags=59%, list=24%, si | MM_1 |
| PPAR_SIGNALING_PATHV | PPAR_SIGNALING_PATHWAY | 30 | 0.497595 | 1.707195 | 0.007772021 | 0.17868574 | 1 | 1632 | tags=50%, list=19%, si | MM_1 |
| GOBP_GENERATION_OF_ | GOBP_GENERATION_OF_PRECURSOR_META | 305 | 0.341888 | 1.705181 | 0 | 0.17763023 | 1 | 2088 | tags=38%, list=24%, si | MM_1 |
| GOBP_AUTOPHAGOSOMI | GOBP_AUTOPHAGOSOME_MATURATION | 55 | 0.430655 | 1.669019 | 0.005405406 | 0.23185882 | 1 | 2416 | tags=49%, list=28%, si | MM_1 |
| GOBP_ANTEROGRADE_A) | GOBP_ANTEROGRADE_AXONAL_TRANSPOR | 33 | 0.484746 | 1.6643 | 0.010230179 | 0.23520125 | 1 | 168 | tags=12%, list=2%, sig | MM_1 |
| GOBP_NUCLEOSIDE_PHO | GOBP_NUCLEOSIDE_PHOSPHATE_CATABOLI | 99 | 0.385403 | 1.658229 | 0.006802721 | 0.24112931 | 1 | 1768 | tags=34%, list=20%, si | MM_1 |
| GOBP_RIBOSE_PHOSPHA | GOBP_RIBOSE_PHOSPHATE_BIOSYNTHETIC_ | 139 | 0.36775 | 1.658199 | 0 | 0.23581122 | 1 | 2354 | tags=45%, list=27%, si | MM_1 |
| METABOLISM_OF_XENOE | METABOLISM_OF_XENOBIOTICS_BY_CYTOC | 15 | 0.605165 | 1.657271 | 0.01891253 | 0.23234086 | 1 | 1189 | tags=47%, list=14%, si | MM_1 |
| GOBP_GLYCOPROTEIN_C/ | GOBP_GLYCOPROTEIN_CATABOLIC_PROCES | 22 | 0.530607 | 1.657236 | 0.01173709 | 0.22743599 | 1 | 774 | tags=32%, list=9%, sig | MM_1 |
| ALZHEIMER_DISEASE | ALZHEIMER_DISEASE | 129 | 0.368134 | 1.649656 | 0 | 0.23657265 | 1 | 2092 | tags=50%, list=24%, si | MM_1 |
| PROTEIN_EXPORT | PROTEIN_EXPORT | 22 | 0.534928 | 1.644993 | 0.016990291 | 0.24075815 | 1 | 2863 | tags=64%, list=33%, si | MM_1 |
| GOBP_REGULATION_OF_ | GOBP_REGULATION_OF_SPROUTING_ANGI( | 20 | 0.549076 | 1.64197 | 0.02184466 | 0.2413497 | 1 | 1697 | tags=50%, list=20%, si | MM_1 |
| GOBP_PROTEIN_TARGETI | GOBP_PROTEIN_TARGETING_TO_MITOCHO | 62 | 0.412111 | 1.641016 | 0.005509642 | 0.23811588 | 1 | 1419 | tags=39%, list=16%, si | MM_1 |
| GOBP_REGULATION_OF_ | GOBP_REGULATION_OF_LIPOPOLYSACCHAR | 22 | 0.5342 | 1.635486 | 0.012048192 | 0.24342449 | 1 | 268 | tags=18%, list=3%, sig | MM_1 |
| GOBP_POSITIVE_REGULA | GOBP_POSITIVE_REGULATION_OF_ERAD_P/ | 15 | 0.579519 | 1.635112 | 0.036057692 | 0.23947738 | 1 | 1174 | tags=27%, list=14%, si | MM_1 |
| AGE-RAGE_SIGNALING_P | AGE-RAGE_SIGNALING_PATHWAY_IN_DIABI | 63 | -0.60111 | -2.15101 | 0 | 0.019849846 | 0.019 | 1132 | tags=35%, list=13%, si | MM_1 |
| ECM-RECEPTOR_INTERAC | ECM-RECEPTOR_INTERACTION | 17 | -0.7811 | -2.10231 | 0 | 0.026178258 | 0.049 | 387 | tags=35%, list=4%, sig | MM_1 |
| FOCAL_ADHESION | FOCAL_ADHESION | 110 | -0.5379 | -2.10152 | 0 | 0.018489292 | 0.052 | 1730 | tags=37%, list=20%, si | MM_1 |
| GOBP_ANTIGEN_PROCES | GOBP_ANTIGEN_PROCESSING_AND_PRESEN | 16 | -0.78464 | -2.10124 | 0 | 0.013866969 | 0.052 | 544 | tags=44%, list=6%, sig | MM_1 |
| HYPERTROPHIC_CARDIOM | HYPERTROPHIC_CARDIOMYOPATHY_(HCM) | 30 | -0.67617 | -2.08 | 0 | 0.01737006 | 0.082 | 638 | tags=30%, list=7%, sig | MM_1 |
| GOBP_ANTIGEN_PROCES | GOBP_ANTIGEN_PROCESSING_AND_PRESEN | 24 | -0.69007 | -2.02667 | 0 | 0.03662361 | 0.189 | 1264 | tags=42%, list=15%, si | MM_1 |
| HEDGEHOG_SIGNALING_ | HEDGEHOG_SIGNALING_PATHWAY | 27 | -0.67334 | -2.02193 | 0 | 0.03348901 | 0.199 | 1296 | tags=44%, list=15%, si | MM_1 |
| GOBP_ANTIGEN_PROCES | GOBP_ANTIGEN_PROCESSING_AND_PRESEN | 28 | -0.65433 | -2.00146 | 0 | 0.042080063 | 0.278 | 1264 | tags=36%, list=15%, si | MM_1 |
| GOBP_ANTIGEN_PROCES | GOBP_ANTIGEN_PROCESSING_AND_PRESEN | 19 | -0.72212 | -1.9866 | 0 | 0.047386833 | 0.331 | 544 | tags=37%, list=6%, sig | MM_1 |
| DILATED_CARDIOMYOPAT | DILATED_CARDIOMYOPATHY_(DCM) | 31 | -0.63567 | -1.96692 | 0 | 0.055800114 | 0.409 | 638 | tags=26%, list=7%, sig | MM_1 |
| GOBP_REGULATION_OF_ | GOBP_REGULATION_OF_ENDOTHELIAL_CEL | 27 | -0.63717 | -1.96329 | 0 | 0.053779654 | 0.427 | 1019 | tags=33%, list=12%, si | MM_1 |
| CELL_ADHESION_MOLECU | CELL_ADHESION_MOLECULES_(CAMS) | 40 | -0.60325 | -1.92878 | 0 | 0.077777326 | 0.577 | 696 | tags=28%, list=8%, sig | MM_1 |
| GOBP_NEURON_PROJECT | GOBP_NEURON_PROJECTION_EXTENSION | 95 | -0.50315 | -1.92869 | 0 | 0.07179446 | 0.577 | 2650 | tags=48%, list=31%, si | MM_1 |
| ARRHYTHMOGENIC_RIGH | ARRHYTHMOGENIC_RIGHT_VENTRICULAR_( | 22 | -0.66687 | -1.91705 | 0 | 0.0775591 | 0.63 | 487 | tags=27%, list=6%, sig | MM_1 |
| GOBP_REGULATION_OF_ | GOBP_REGULATION_OF_EPITHELIAL_CELL_I | 67 | -0.51764 | -1.89316 | 0 | 0.10094123 | 0.759 | 1365 | tags=28%, list=16%, si | MM_1 |
| GOBP_AXON_EXTENSION | GOBP_AXON_EXTENSION | 66 | -0.51619 | -1.88504 | 0 | 0.10546733 | 0.8 | 2728 | tags=53%, list=31%, si | MM_1 |
| GOBP_NEGATIVE_REGUL | GOBP_NEGATIVE_REGULATION_OF_EPITHEI | 22 | -0.63787 | -1.87819 | 0.003502627 | 0.1064042 | 0.819 | 1787 | tags=55%, list=21%, si | MM_1 |
| GOBP_DICARBOXYLIC_AC | GOBP_DICARBOXYLIC_ACID_TRANSPORT | 41 | -0.563 | -1.8682 | 0 | 0.11356614 | 0.851 | 427 | tags=22%, list=5%, sig | MM_1 |
| GOBP_TRANSFORMING_( | GOBP_TRANSFORMING_GROWTH_FACTOR_ | 15 | -0.71407 | -1.85253 | 0.003683241 | 0.12929493 | 0.905 | 1019 | tags=47%, list=12%, si | MM_1 |
| GOBP_REGULATION_OF_ | GOBP_REGULATION_OF_EXTENT_OF_CELL_ | 58 | -0.51473 | -1.85163 | 0 | 0.12424463 | 0.908 | 2728 | tags=50%, list=31%, si | MM_1 |
| TRANSCRIPTIONAL_MISRI | TRANSCRIPTIONAL_MISREGULATION_IN_CA | 107 | -0.46772 | -1.84919 | 0 | 0.12254668 | 0.915 | 1962 | tags=38%, list=23%, si | MM_1 |
| GOBP_FORMATION_OF_F | GOBP_FORMATION_OF_PRIMARY_GERM_L/ | 50 | -0.54349 | -1.83416 | 0.003039514 | 0.13968651 | 0.952 | 1577 | tags=38%, list=18%, si | MM_1 |
| GOBP_DETECTION_OF_BI | GOBP_DETECTION_OF_BIOTIC_STIMULUS | 23 | -0.62368 | -1.82097 | 0 | 0.15454485 | 0.966 | 503 | tags=22%, list=6%, sig | MM_1 |
| GOBP_ENDODERMAL_CE | GOBP_ENDODERMAL_CELL_DIFFERENTIATI( | 15 | -0.68472 | -1.81787 | 0.005244755 | 0.15334393 | 0.966 | 969 | tags=27%, list=11%, si | MM_1 |
| GOBP_REGULATION_OF_ | GOBP_REGULATION_OF_SODIUM_ION_TRA | 21 | -0.62961 | -1.80599 | 0.006745363 | 0.1669305 | 0.982 | 865 | tags=33%, list=10%, si | MM_1 |

| | | | | | | | | | | |
|---|---|---|---|---|---|---|---|---|---|---|
| GOBP_ARTERY_MORPHO | GOBP_ARTERY_MORPHOGENESIS | 33 | -0.55811 | -1.80537 | 0.001650165 | 0.1619174 | 0.982 | 1510 | tags=39%, list=17%, si | MM_1 |
| GOBP_MESODERM_DEVE | GOBP_MESODERM_DEVELOPMENT | 46 | -0.53857 | -1.80103 | 0 | 0.1635741 | 0.982 | 1738 | tags=48%, list=20%, si | MM_1 |
| GOBP_ANTERIOR_POSTEI | GOBP_ANTERIOR_POSTERIOR_PATTERN_SP | 77 | -0.47419 | -1.78925 | 0 | 0.17970334 | 0.992 | 2062 | tags=42%, list=24%, si | MM_1 |
| GOBP_C4_DICARBOXYLAT | GOBP_C4_DICARBOXYLATE_TRANSPORT | 15 | -0.68167 | -1.7863 | 0.006932409 | 0.17956287 | 0.993 | 427 | tags=40%, list=5%, sig | MM_1 |
| GOBP_REGULATION_OF_ | GOBP_REGULATION_OF_EPITHELIAL_TO_MI | 41 | -0.54337 | -1.78535 | 0.003125 | 0.17538866 | 0.993 | 1787 | tags=46%, list=21%, si | MM_1 |
| GOBP_ENDOTHELIAL_CEL | GOBP_ENDOTHELIAL_CELL_DIFFERENTIATIO | 59 | -0.5072 | -1.78331 | 0.003081664 | 0.173673 | 0.994 | 1793 | tags=41%, list=21%, si | MM_1 |
| SIGNALING_PATHWAYS_I | SIGNALING_PATHWAYS_REGULATING_PLUR | 64 | -0.49989 | -1.78297 | 0.001569859 | 0.16886368 | 0.994 | 2076 | tags=39%, list=24%, si | MM_1 |
| GOBP_RESPONSE_TO_AC | GOBP_RESPONSE_TO_ACETYLCHOLINE | 18 | -0.6461 | -1.77513 | 0.006688963 | 0.17860094 | 0.996 | 85 | tags=22%, list=1%, sig | MM_1 |
| GRAFT-VERSUS-HOST_DIS | GRAFT-VERSUS-HOST_DISEASE | 19 | -0.62333 | -1.77251 | 0.006546645 | 0.1785383 | 0.996 | 154 | tags=21%, list=2%, sig | MM_1 |
| MALARIA | MALARIA | 18 | -0.64549 | -1.76904 | 0.012750455 | 0.18023363 | 0.997 | 927 | tags=33%, list=11%, si | MM_1 |
| GOBP_DETECTION_OF_EX | GOBP_DETECTION_OF_EXTERNAL_BIOTIC_S | 18 | -0.6452 | -1.76506 | 0.010067114 | 0.18298076 | 0.997 | 503 | tags=22%, list=6%, sig | MM_1 |
| GOBP_POSITIVE_REGULA | GOBP_POSITIVE_REGULATION_OF_SMOOTH | 40 | -0.53026 | -1.76438 | 0 | 0.17936678 | 0.998 | 1510 | tags=43%, list=17%, si | MM_1 |
| SMALL_CELL_LUNG_CAN( | SMALL_CELL_LUNG_CANCER | 58 | -0.50048 | -1.76353 | 0 | 0.17632572 | 0.999 | 1720 | tags=36%, list=20%, si | MM_1 |
| GOBP_POSITIVE_REGULA | GOBP_POSITIVE_REGULATION_OF_AXON_E | 32 | -0.55882 | -1.76255 | 0.006535948 | 0.17383999 | 0.999 | 2476 | tags=53%, list=29%, si | MM_1 |
| GOBP_NEGATIVE_REGUL | GOBP_NEGATIVE_REGULATION_OF_NEURO | 35 | -0.55024 | -1.76026 | 0 | 0.17363048 | 0.999 | 346 | tags=20%, list=4%, sig | MM_1 |
| STAPHYLOCOCCUS_AURE | STAPHYLOCOCCUS_AUREUS_INFECTION | 24 | -0.58745 | -1.75117 | 0.00805153 | 0.1861771 | 1 | 1015 | tags=38%, list=12%, si | MM_1 |
| TYPE_I_DIABETES_MELLIT | TYPE_I_DIABETES_MELLITUS | 20 | -0.60729 | -1.7488 | 0.015625 | 0.18629786 | 1 | 154 | tags=20%, list=2%, sig | MM_1 |
| GOBP_ASTROCYTE_ACTIV | GOBP_ASTROCYTE_ACTIVATION | 16 | -0.67128 | -1.74859 | 0.003424658 | 0.18232945 | 1 | 153 | tags=19%, list=2%, sig | MM_1 |
| GOBP_REGULATION_OF_ | GOBP_REGULATION_OF_SODIUM_ION_TRA | 27 | -0.59364 | -1.74773 | 0.007987221 | 0.17958917 | 1 | 865 | tags=30%, list=10%, si | MM_1 |
| GOBP_REGULATION_OF_ | GOBP_REGULATION_OF_TISSUE_REMODELI | 40 | -0.53015 | -1.7394 | 0.001694915 | 0.19096169 | 1 | 254 | tags=15%, list=3%, sig | MM_1 |
| GOBP_DEVELOPMENTAL_ | GOBP_DEVELOPMENTAL_GROWTH_INVOLV | 128 | -0.43389 | -1.7353 | 0 | 0.19558492 | 1 | 2650 | tags=45%, list=31%, si | MM_1 |
| GOBP_EMBRYONIC_PATT | GOBP_EMBRYONIC_PATTERN_SPECIFICATIO | 23 | -0.60379 | -1.73422 | 0.005067567 | 0.19351342 | 1 | 1590 | tags=43%, list=18%, si | MM_1 |
| GOBP_ENTRAINMENT_OI | GOBP_ENTRAINMENT_OF_CIRCADIAN_CLO( | 16 | -0.66091 | -1.73367 | 0.010169491 | 0.19046378 | 1 | 2087 | tags=50%, list=24%, si | MM_1 |
| GOBP_REGULATION_OF_ | GOBP_REGULATION_OF_ASTROCYTE_DIFFEI | 18 | -0.62883 | -1.73312 | 0.012589928 | 0.18783768 | 1 | 604 | tags=28%, list=7%, sig | MM_1 |
| GOBP_FATTY_ACID_DERI | GOBP_FATTY_ACID_DERIVATIVE_METABOLI | 23 | -0.59287 | -1.73307 | 0.010273973 | 0.18412241 | 1 | 1186 | tags=39%, list=14%, si | MM_1 |
| GOBP_SMOOTH_MUSCLE | GOBP_SMOOTH_MUSCLE_CONTRACTION | 37 | -0.52962 | -1.72974 | 0.00660066 | 0.18684702 | 1 | 1581 | tags=46%, list=18%, si | MM_1 |
| VASCULAR_SMOOTH_MU | VASCULAR_SMOOTH_MUSCLE_CONTRACTI( | 59 | -0.48812 | -1.7293 | 0.001512859 | 0.18395643 | 1 | 1520 | tags=37%, list=18%, si | MM_1 |
| GOBP_POSITIVE_REGULA | GOBP_POSITIVE_REGULATION_OF_AXONO( | 50 | -0.49893 | -1.72882 | 0.004651163 | 0.1813716 | 1 | 2567 | tags=46%, list=30%, si | MM_1 |
| GOBP_CELL_MORPHOGEI | GOBP_CELL_MORPHOGENESIS | 471 | -0.37054 | -1.71707 | 0 | 0.19966878 | 1 | 1867 | tags=29%, list=22%, si | MM_1 |
| CORTISOL_SYNTHESIS_AI | CORTISOL_SYNTHESIS_AND_SECRETION | 28 | -0.56997 | -1.71658 | 0.006525286 | 0.19714203 | 1 | 346 | tags=21%, list=4%, sig | MM_1 |
| GOBP_SODIUM_ION_TRA | GOBP_SODIUM_ION_TRANSMEMBRANE_TR | 41 | -0.52469 | -1.71538 | 0.003194888 | 0.19613692 | 1 | 1269 | tags=29%, list=15%, si | MM_1 |
| GOBP_DEVELOPMENTAL_ | GOBP_DEVELOPMENTAL_CELL_GROWTH | 137 | -0.42229 | -1.71389 | 0 | 0.19579273 | 1 | 2728 | tags=42%, list=31%, si | MM_1 |
| GOBP_SMOOTH_MUSCLE | GOBP_SMOOTH_MUSCLE_CELL_MIGRATION | 68 | -0.47466 | -1.71318 | 0.001488095 | 0.19371587 | 1 | 1510 | tags=38%, list=17%, si | MM_1 |
| GOBP_NATURAL_KILLER_ | GOBP_NATURAL_KILLER_CELL_DIFFERENTIA | 22 | -0.59193 | -1.71196 | 0.013536379 | 0.19271828 | 1 | 700 | tags=27%, list=8%, sig | MM_1 |
| GOBP_ASTROCYTE_DEVEI | GOBP_ASTROCYTE_DEVELOPMENT | 21 | -0.5999 | -1.71034 | 0.010380623 | 0.19272648 | 1 | 153 | tags=14%, list=2%, sig | MM_1 |
| GOBP_PROGRAMMED_CI | GOBP_PROGRAMMED_CELL_DEATH_INVOLV | 15 | -0.65071 | -1.70771 | 0.006908463 | 0.19467103 | 1 | 1070 | tags=40%, list=12%, si | MM_1 |
| GOBP_CELL_MATRIX_ADI | GOBP_CELL_MATRIX_ADHESION | 111 | -0.43098 | -1.70243 | 0 | 0.20159802 | 1 | 747 | tags=18%, list=9%, sig | MM_1 |
| GOBP_MRNA_MODIFICAT | GOBP_MRNA_MODIFICATION | 17 | -0.62588 | -1.69956 | 0.015544041 | 0.20366828 | 1 | 1434 | tags=41%, list=17%, si | MM_1 |
| GOBP_POSITIVE_REGULA | GOBP_POSITIVE_REGULATION_OF_NEURO( | 154 | -0.40795 | -1.6965 | 0 | 0.20621733 | 1 | 1183 | tags=20%, list=14%, si | MM_1 |
| GOBP_MESODERM_MOR | GOBP_MESODERM_MORPHOGENESIS | 29 | -0.54365 | -1.69528 | 0.011235955 | 0.20560084 | 1 | 2062 | tags=52%, list=24%, si | MM_1 |
| GOBP_RELAXATION_OF_I | GOBP_RELAXATION_OF_MUSCLE | 16 | -0.63791 | -1.69507 | 0.008 | 0.20286585 | 1 | 1582 | tags=56%, list=18%, si | MM_1 |
| GOBP_RESPONSE_TO_PU | GOBP_RESPONSE_TO_PURINE_CONTAINING | 49 | -0.48627 | -1.69094 | 0.0048 | 0.20820782 | 1 | 1511 | tags=29%, list=17%, si | MM_1 |
| GOBP_SYNAPSE_MATUR/ | GOBP_SYNAPSE_MATURATION | 17 | -0.6206 | -1.68564 | 0.011945393 | 0.21533805 | 1 | 275 | tags=18%, list=3%, sig | MM_1 |
| GOBP_CELL_FATE_SPECIF | GOBP_CELL_FATE_SPECIFICATION | 32 | -0.53729 | -1.68451 | 0.008605852 | 0.2144446 | 1 | 2266 | tags=44%, list=26%, si | MM_1 |
| GOBP_REGULATION_OF_ | GOBP_REGULATION_OF_AXONOGENESIS | 79 | -0.45094 | -1.68413 | 0.003076923 | 0.21218736 | 1 | 2367 | tags=39%, list=27%, si | MM_1 |
| GOBP_CELL_MORPHOGEI | GOBP_CELL_MORPHOGENESIS_INVOLVED_I | 271 | -0.38292 | -1.6834 | 0 | 0.21059562 | 1 | 2650 | tags=43%, list=31%, si | MM_1 |
| GOBP_POSITIVE_REGULA | GOBP_POSITIVE_REGULATION_OF_JNK_CAS | 57 | -0.47865 | -1.68151 | 0.003053435 | 0.21143138 | 1 | 418 | tags=11%, list=5%, sig | MM_1 |
| GOBP_CELL_PROJECTION | GOBP_CELL_PROJECTION_MORPHOGENESIS | 306 | -0.37807 | -1.67995 | 0 | 0.21136996 | 1 | 2730 | tags=44%, list=31%, si | MM_1 |
| GOBP_CELL_SUBSTRATE_ | GOBP_CELL_SUBSTRATE_ADHESION | 165 | -0.40166 | -1.67957 | 0.001385042 | 0.2092065 | 1 | 1182 | tags=22%, list=14%, si | MM_1 |
| GOBP_ASTROCYTE_DIFFE | GOBP_ASTROCYTE_DIFFERENTIATION | 48 | -0.49055 | -1.67867 | 0.00974026 | 0.20776942 | 1 | 907 | tags=19%, list=10%, si | MM_1 |
| AMOEBIASIS | AMOEBIASIS | 45 | -0.49665 | -1.67228 | 0.00621118 | 0.21751158 | 1 | 491 | tags=20%, list=6%, sig | MM_1 |
| VIRAL_MYOCARDITIS | VIRAL_MYOCARDITIS | 37 | -0.51843 | -1.67077 | 0.014492754 | 0.21735883 | 1 | 238 | tags=16%, list=3%, sig | MM_1 |
| GOBP_INTERLEUKIN_1_M | GOBP_INTERLEUKIN_1_MEDIATED_SIGNALI | 21 | -0.59874 | -1.66955 | 0.008532424 | 0.2170796 | 1 | 552 | tags=19%, list=6%, sig | MM_1 |
| GOBP_T_CELL_DIFFERENT | GOBP_T_CELL_DIFFERENTIATION_IN_THYM | 61 | -0.46847 | -1.66427 | 0.004594181 | 0.22441849 | 1 | 997 | tags=20%, list=11%, si | MM_1 |
| GOBP_POSITIVE_REGULA | GOBP_POSITIVE_REGULATION_OF_NERVOU | 165 | -0.40054 | -1.66279 | 0.002688172 | 0.2244731 | 1 | 1183 | tags=21%, list=14%, si | MM_1 |
| GOBP_POSITIVE_REGULA | GOBP_POSITIVE_REGULATION_OF_CELL_CY | 16 | -0.63222 | -1.66103 | 0.019097222 | 0.22552596 | 1 | 865 | tags=38%, list=10%, si | MM_1 |
| PROTEOGLYCANS_IN_CAI | PROTEOGLYCANS_IN_CANCER | 125 | -0.41421 | -1.65921 | 0.001362398 | 0.2261868 | 1 | 1038 | tags=20%, list=12%, si | MM_1 |
| GOBP_NEGATIVE_REGUL | GOBP_NEGATIVE_REGULATION_OF_SMOOT | 25 | -0.54705 | -1.65752 | 0.015410959 | 0.22687611 | 1 | 1302 | tags=48%, list=15%, si | MM_1 |
| GOBP_REGULATION_OF_ | GOBP_REGULATION_OF_DENDRITIC_SPINE_ | 30 | -0.52184 | -1.65164 | 0.013490725 | 0.23577948 | 1 | 1866 | tags=33%, list=22%, si | MM_1 |
| GOBP_RESPONSE_TO_OR | GOBP_RESPONSE_TO_ORGANOPHOSPHORU | 45 | -0.49055 | -1.65136 | 0.014173228 | 0.233595 | 1 | 767 | tags=20%, list=9%, sig | MM_1 |
| GOBP_REGULATION_OF_ | GOBP_REGULATION_OF_DENDRITE_EXTENS | 16 | -0.61174 | -1.65057 | 0.010909091 | 0.23233779 | 1 | 1679 | tags=44%, list=19%, si | MM_1 |
| GOBP_STRIATED_MUSCLI | GOBP_STRIATED_MUSCLE_CELL_DEVELOPM | 28 | -0.54445 | -1.6475 | 0.016393442 | 0.23555173 | 1 | 1520 | tags=39%, list=18%, si | MM_1 |
| INFLAMMATORY_BOWEL | INFLAMMATORY_BOWEL_DISEASE_(IBD) | 29 | -0.53318 | -1.64617 | 0.018242123 | 0.23585677 | 1 | 1201 | tags=41%, list=14%, si | MM_1 |
| GOBP_POSITIVE_REGULA | GOBP_POSITIVE_REGULATION_OF_PHOSPH | 86 | -0.43711 | -1.64551 | 0.002894356 | 0.2347131 | 1 | 1942 | tags=37%, list=22%, si | MM_1 |
| GOBP_PATTERN_SPECIFI( | GOBP_PATTERN_SPECIFICATION_PROCESS | 144 | -0.40656 | -1.64445 | 0.002801121 | 0.23445256 | 1 | 2062 | tags=35%, list=24%, si | MM_1 |
| GOBP_VASCULAR_ASSOC | GOBP_VASCULAR_ASSOCIATED_SMOOTH_M | 27 | -0.54373 | -1.64315 | 0.016949153 | 0.23443691 | 1 | 1705 | tags=44%, list=20%, si | MM_1 |
| GOBP_MUSCLE_CONTRA( | GOBP_MUSCLE_CONTRACTION | 104 | -0.42123 | -1.64265 | 0.00443787 | 0.23279445 | 1 | 1582 | tags=34%, list=18%, si | MM_1 |
| ENDOMETRIAL_CANCER | ENDOMETRIAL_CANCER | 46 | -0.47879 | -1.64195 | 0.012383901 | 0.23151663 | 1 | 1720 | tags=33%, list=20%, si | MM_1 |
| REGULATION_OF_ACTIN_ | REGULATION_OF_ACTIN_CYTOSKELETON | 129 | -0.41009 | -1.64057 | 0.001447178 | 0.23172367 | 1 | 1720 | tags=27%, list=20%, si | MM_1 |
| GOBP_CARDIAC_MUSCLE | GOBP_CARDIAC_MUSCLE_CONTRACTION | 49 | -0.47288 | -1.64054 | 0.013953488 | 0.22932863 | 1 | 1582 | tags=39%, list=18%, si | MM_1 |
| PROTEIN_DIGESTION_AN | PROTEIN_DIGESTION_AND_ABSORPTION | 16 | -0.61386 | -1.63905 | 0.024193548 | 0.22987437 | 1 | 487 | tags=31%, list=6%, sig | MM_1 |
| GOBP_POSITIVE_REGULA | GOBP_POSITIVE_REGULATION_OF_EPITHELI | 22 | -0.5676 | -1.63784 | 0.014010508 | 0.22980805 | 1 | 1754 | tags=45%, list=20%, si | MM_1 |
| GOBP_ACUTE_INFLAMM/ | GOBP_ACUTE_INFLAMMATORY_RESPONSE | 42 | -0.49496 | -1.62736 | 0.015649453 | 0.24970378 | 1 | 142 | tags=14%, list=2%, sig | MM_1 |
| GOBP_DETECTION_OF_ST | GOBP_DETECTION_OF_STIMULUS | 62 | -0.46129 | -1.62733 | 0.003139718 | 0.24720229 | 1 | 1046 | tags=24%, list=12%, si | MM_1 |
| GOBP_POSTSYNAPTIC_SP | GOBP_POSTSYNAPTIC_SPECIALIZATION_OR( | 23 | -0.56495 | -1.62608 | 0.021207178 | 0.24744515 | 1 | 2020 | tags=48%, list=23%, si | MM_1 |
| GOBP_ACTOMYOSIN_STR | GOBP_ACTOMYOSIN_STRUCTURE_ORGANIZ | 109 | -0.4148 | -1.62597 | 0.002932551 | 0.24522254 | 1 | 1520 | tags=30%, list=18%, si | MM_1 |
| GOBP_POSITIVE_REGULA | GOBP_POSITIVE_REGULATION_OF_ORGANI( | 24 | -0.55753 | -1.62532 | 0.02173913 | 0.24416311 | 1 | 1246 | tags=33%, list=14%, si | MM_1 |
| GOBP_NEGATIVE_REGUL | GOBP_NEGATIVE_REGULATION_OF_ORGAN | 25 | -0.54399 | -1.62457 | 0.008547009 | 0.24354626 | 1 | 2271 | tags=52%, list=26%, si | MM_1 |
| GOBP_PYROPTOSIS | GOBP_PYROPTOSIS | 27 | -0.54081 | -1.62456 | 0.025641026 | 0.24120447 | 1 | 571 | tags=22%, list=7%, sig | MM_1 |
| GOBP_POSTSYNAPTIC_DE | GOBP_POSTSYNAPTIC_DENSITY_ORGANIZAT | 18 | -0.58734 | -1.62353 | 0.023569023 | 0.24104618 | 1 | 2020 | tags=56%, list=23%, si | MM_1 |
| GOBP_ARTERY_DEVELOPI | GOBP_ARTERY_DEVELOPMENT | 54 | -0.47022 | -1.62247 | 0.010802469 | 0.24104846 | 1 | 1754 | tags=35%, list=20%, si | MM_1 |
| GOBP_REGULATION_OF_ | GOBP_REGULATION_OF_POTASSIUM_ION_ | 31 | -0.5223 | -1.62219 | 0.013400335 | 0.23941222 | 1 | 778 | tags=29%, list=9%, sig | MM_1 |

| | | | | | | | | | | |
|---|---|---|---|---|---|---|---|---|---|---|
| GOBP_ENDOTHELIUM_DE | GOBP_ENDOTHELIUM_DEVELOPMENT | 72 | -0.44468 | -1.62198 | 0.008810572 | 0.2376516 | 1 | 1793 | tags=36%, list=21%, si | MM_1 |
| GOBP_ENDODERM_FORM | GOBP_ENDODERM_FORMATION | 19 | -0.58711 | -1.62084 | 0.025510205 | 0.23769458 | 1 | 1577 | tags=32%, list=18%, si | MM_1 |
| TYPE_II_DIABETES_MELLI | TYPE_II_DIABETES_MELLITUS | 25 | -0.55073 | -1.62023 | 0.02631579 | 0.23678729 | 1 | 1720 | tags=36%, list=20%, si | MM_1 |
| GOBP_TRIGLYCERIDE_BIO | GOBP_TRIGLYCERIDE_BIOSYNTHETIC_PROCE | 29 | -0.52402 | -1.62005 | 0.013536379 | 0.23506741 | 1 | 2331 | tags=55%, list=27%, si | MM_1 |
| TOXOPLASMOSIS | TOXOPLASMOSIS | 72 | -0.43799 | -1.61948 | 0.004587156 | 0.23422763 | 1 | 1235 | tags=24%, list=14%, si | MM_1 |
| GOBP_GASTRULATION | GOBP_GASTRULATION | 95 | -0.4196 | -1.61947 | 0.004484305 | 0.2322199 | 1 | 1962 | tags=32%, list=23%, si | MM_1 |
| GOBP_CELL_JUNCTION_A | GOBP_CELL_JUNCTION_ASSEMBLY | 188 | -0.38263 | -1.61767 | 0 | 0.23377527 | 1 | 1679 | tags=27%, list=19%, si | MM_1 |
| GOBP_CARDIOCYTE_DIFF | GOBP_CARDIOCYTE_DIFFERENTIATION | 91 | -0.42143 | -1.61576 | 0.010086455 | 0.23565258 | 1 | 1581 | tags=26%, list=18%, si | MM_1 |
| GOBP_MEMBRANE_PROT | GOBP_MEMBRANE_PROTEIN_ECTODOMAIN | 31 | -0.52353 | -1.6146 | 0.019704433 | 0.23597114 | 1 | 806 | tags=19%, list=9%, sig | MM_1 |
| GOBP_REGULATION_OF_ | GOBP_REGULATION_OF_T_HELPER_1_TYPE_ | 15 | -0.62176 | -1.61383 | 0.029010238 | 0.23556186 | 1 | 1409 | tags=60%, list=16%, si | MM_1 |
| HERPES_SIMPLEX_VIRUS_ | HERPES_SIMPLEX_VIRUS_1_INFECTION | 187 | -0.38064 | -1.60784 | 0 | 0.24539864 | 1 | 1809 | tags=31%, list=21%, si | MM_1 |
| GOBP_CELL_DIFFERENTIA | GOBP_CELL_DIFFERENTIATION_INVOLVED_I | 17 | -0.59252 | -1.60532 | 0.03760684 | 0.2481706 | 1 | 1302 | tags=41%, list=15%, si | MM_1 |
| GOBP_ACIDIC_AMINO_AC | GOBP_ACIDIC_AMINO_ACID_TRANSPORT | 29 | -0.52031 | -1.60385 | 0.031719532 | 0.2491342 | 1 | 2261 | tags=38%, list=26%, si | MM_1 |
| GOBP_ANTIMICROBIAL_H | GOBP_ANTIMICROBIAL_HUMORAL_RESPON | 24 | 0.628788 | 1.995796 | 0 | 0.22358476 | 0.457 | 1288 | tags=42%, list=15%, si | EN_2 |
| GOBP_NEGATIVE_REGUL | GOBP_NEGATIVE_REGULATION_OF_VIRAL_I | 18 | 0.667654 | 1.975182 | 0 | 0.18769376 | 0.531 | 359 | tags=17%, list=4%, sig | EN_2 |
| OXIDATIVE_PHOSPHORYL | OXIDATIVE_PHOSPHORYLATION | 99 | 0.447466 | 1.962044 | 0 | 0.16713284 | 0.581 | 2970 | tags=60%, list=34%, si | EN_2 |
| GOBP_LIPOPOLYSACCHAR | GOBP_LIPOPOLYSACCHARIDE_MEDIATED_SI | 27 | 0.585027 | 1.944645 | 0 | 0.16006272 | 0.64 | 186 | tags=11%, list=2%, sig | EN_2 |
| GOBP_OXIDATIVE_PHOSP | GOBP_OXIDATIVE_PHOSPHORYLATION | 105 | 0.437873 | 1.933865 | 0 | 0.14981705 | 0.677 | 2708 | tags=51%, list=31%, si | EN_2 |
| GOBP_ANTIMICROBIAL_H | GOBP_ANTIMICROBIAL_HUMORAL_IMMUN | 18 | 0.64493 | 1.926945 | 0.004807693 | 0.13676547 | 0.699 | 1288 | tags=44%, list=15%, si | EN_2 |
| GOBP_ATP_SYNTHESIS_C | GOBP_ATP_SYNTHESIS_COUPLED_ELECTRO | 68 | 0.465026 | 1.900023 | 0 | 0.15593947 | 0.799 | 2708 | tags=51%, list=31%, si | EN_2 |
| PARKINSON_DISEASE | PARKINSON_DISEASE | 104 | 0.428164 | 1.896725 | 0 | 0.14277504 | 0.807 | 2708 | tags=56%, list=31%, si | EN_2 |
| GOBP_MITOCHONDRIAL_ | GOBP_MITOCHONDRIAL_RESPIRATORY_CHA | 82 | 0.445254 | 1.890217 | 0 | 0.13823135 | 0.834 | 2569 | tags=49%, list=29%, si | EN_2 |
| GOBP_RIBOSOMAL_LARG | GOBP_RIBOSOMAL_LARGE_SUBUNIT_BIOGE | 54 | 0.48131 | 1.885331 | 0 | 0.1314573 | 0.851 | 1637 | tags=33%, list=19%, si | EN_2 |
| GOBP_REGULATION_OF_ | GOBP_REGULATION_OF_NEUTROPHIL_MIGI | 16 | 0.635566 | 1.855002 | 0.002450981 | 0.16446304 | 0.928 | 246 | tags=19%, list=3%, sig | EN_2 |
| FERROPTOSIS | FERROPTOSIS | 28 | 0.545245 | 1.838131 | 0.002617801 | 0.17745897 | 0.946 | 2112 | tags=43%, list=24%, si | EN_2 |
| GOBP_MODULATION_BY_ | GOBP_MODULATION_BY_SYMBIONT_OF_EN | 26 | 0.565087 | 1.836827 | 0.005524862 | 0.16682714 | 0.949 | 359 | tags=12%, list=4%, sig | EN_2 |
| GOBP_KILLING_OF_CELLS | GOBP_KILLING_OF_CELLS_OF_ANOTHER_OI | 23 | 0.550132 | 1.822953 | 0.005012531 | 0.177751 | 0.967 | 1031 | tags=35%, list=12%, si | EN_2 |
| GOBP_POSITIVE_REGULA | GOBP_POSITIVE_REGULATION_OF_GLYCOPI | 16 | 0.61046 | 1.817894 | 0.01173709 | 0.17384052 | 0.972 | 64 | tags=13%, list=1%, sig | EN_2 |
| PROTEASOME | PROTEASOME | 41 | 0.490725 | 1.803909 | 0 | 0.18432865 | 0.982 | 3120 | tags=59%, list=35%, si | EN_2 |
| GOBP_MATURATION_OF_ | GOBP_MATURATION_OF_5_8S_RRNA_FROM | 17 | 0.601748 | 1.796965 | 0.007407407 | 0.18454315 | 0.987 | 3009 | tags=94%, list=34%, si | EN_2 |
| GOBP_REGULATION_OF_ | GOBP_REGULATION_OF_GRANULOCYTE_CH | 21 | 0.580803 | 1.790742 | 0.00486618 | 0.18440858 | 0.989 | 246 | tags=19%, list=3%, sig | EN_2 |
| GOBP_MITOCHONDRIAL_ | GOBP_MITOCHONDRIAL_TRANSLATION | 104 | 0.405716 | 1.777062 | 0 | 0.19575003 | 0.994 | 2613 | tags=55%, list=30%, si | EN_2 |
| GOBP_REGULATION_OF_ | GOBP_REGULATION_OF_CELL_ADHESION_M | 18 | 0.615597 | 1.775993 | 0.010282776 | 0.1881039 | 0.994 | 557 | tags=22%, list=6%, sig | EN_2 |
| GOBP_DISRUPTION_OF_A | GOBP_DISRUPTION_OF_ANATOMICAL_STRU | 25 | 0.538163 | 1.76454 | 0.005102041 | 0.19803308 | 0.999 | 1031 | tags=32%, list=12%, si | EN_2 |
| GOBP_RIBOSE_PHOSPHA | GOBP_RIBOSE_PHOSPHATE_BIOSYNTHETIC_ | 139 | 0.382355 | 1.75449 | 0.003236246 | 0.2046862 | 0.999 | 1599 | tags=29%, list=18%, si | EN_2 |
| GOBP_PROTEIN_NEDDYL | GOBP_PROTEIN_NEDDYLATION | 17 | 0.599936 | 1.748266 | 0.007537689 | 0.20680986 | 1 | 2685 | tags=59%, list=30%, si | EN_2 |
| DRUG_METABOLISM | DRUG_METABOLISM | 36 | 0.49125 | 1.735699 | 0.002427185 | 0.21921122 | 1 | 845 | tags=31%, list=10%, si | EN_2 |
| GOBP_MITOCHONDRIAL_ | GOBP_MITOCHONDRIAL_GENE_EXPRESSION | 124 | 0.382947 | 1.731855 | 0 | 0.21713012 | 1 | 2567 | tags=49%, list=29%, si | EN_2 |
| GOBP_REGULATION_OF_ | GOBP_REGULATION_OF_BIOLOGICAL_PROC | 29 | 0.512977 | 1.728152 | 0.01591512 | 0.21567054 | 1 | 2 | tags=7%, list=0%, signi | EN_2 |
| GOBP_RESPIRATORY_CHA | GOBP_RESPIRATORY_CHAIN_COMPLEX_IV_ | 18 | 0.584794 | 1.727956 | 0.007712083 | 0.20823753 | 1 | 2029 | tags=56%, list=23%, si | EN_2 |
| GOBP_ELECTRON_TRANS | GOBP_ELECTRON_TRANSPORT_CHAIN | 90 | 0.405522 | 1.704201 | 0.00625 | 0.2418085 | 1 | 2708 | tags=49%, list=31%, si | EN_2 |
| GOBP_TAIL_ANCHORED_ | GOBP_TAIL_ANCHORED_MEMBRANE_PROT | 15 | 0.608122 | 1.701108 | 0.019184653 | 0.23947755 | 1 | 1413 | tags=47%, list=16%, si | EN_2 |
| GOBP_RESPONSE_TO_CH | GOBP_RESPONSE_TO_CHEMOKINE | 22 | 0.536608 | 1.7003 | 0.022160664 | 0.23308271 | 1 | 557 | tags=27%, list=6%, sig | EN_2 |
| GOBP_SYNAPSE_ASSEMB | GOBP_SYNAPSE_ASSEMBLY | 131 | -0.50558 | -2.10504 | 0 | 0.095668964 | 0.08 | 2990 | tags=62%, list=34%, si | EN_2 |
| GOBP_SYNAPTIC_MEMBR | GOBP_SYNAPTIC_MEMBRANE_ADHESION | 22 | -0.70019 | -2.0827 | 0 | 0.07230292 | 0.118 | 549 | tags=36%, list=6%, sig | EN_2 |
| GOBP_GLANDULAR_EPITI | GOBP_GLANDULAR_EPITHELIAL_CELL_DIFFE | 34 | -0.61011 | -2.01027 | 0 | 0.13659716 | 0.309 | 2892 | tags=74%, list=33%, si | EN_2 |
| GOBP_REGULATION_OF_ | GOBP_REGULATION_OF_SYNAPSE_ASSEMBI | 98 | -0.50641 | -1.99501 | 0 | 0.12688667 | 0.362 | 2990 | tags=64%, list=34%, si | EN_2 |
| GOBP_CELL_JUNCTION_A | GOBP_CELL_JUNCTION_ASSEMBLY | 263 | -0.44269 | -1.98907 | 0 | 0.110602014 | 0.381 | 2547 | tags=46%, list=29%, si | EN_2 |
| GOBP_HIPPO_SIGNALING | GOBP_HIPPO_SIGNALING | 29 | -0.63565 | -1.97291 | 0 | 0.118541524 | 0.462 | 2339 | tags=62%, list=26%, si | EN_2 |
| GOBP_REGULATION_OF_ | GOBP_REGULATION_OF_CARDIAC_MUSCLE_ | 25 | -0.6371 | -1.96105 | 0 | 0.1202718 | 0.522 | 2959 | tags=84%, list=33%, si | EN_2 |
| MELANOGENESIS | MELANOGENESIS | 49 | -0.56374 | -1.94733 | 0 | 0.12343237 | 0.576 | 2344 | tags=51%, list=26%, si | EN_2 |
| GOBP_COLUMNAR_CUBO | GOBP_COLUMNAR_CUBOIDAL_EPITHELIAL_ | 27 | -0.61837 | -1.91984 | 0.001623377 | 0.15757832 | 0.709 | 2486 | tags=67%, list=28%, si | EN_2 |
| GOBP_REGULATION_OF_ | GOBP_REGULATION_OF_CELL_JUNCTION_A | 161 | -0.44573 | -1.91956 | 0 | 0.14239174 | 0.709 | 2547 | tags=46%, list=29%, si | EN_2 |
| GOBP_COLUMNAR_CUBO | GOBP_COLUMNAR_CUBOIDAL_EPITHELIAL_ | 64 | -0.51197 | -1.91235 | 0 | 0.14069307 | 0.747 | 2945 | tags=66%, list=33%, si | EN_2 |
| GOBP_NEGATIVE_REGUL | GOBP_NEGATIVE_REGULATION_OF_HORMO | 34 | -0.58718 | -1.90499 | 0.001639344 | 0.14147663 | 0.783 | 745 | tags=29%, list=8%, sig | EN_2 |
| GOBP_SECRETION_BY_TIS | GOBP_SECRETION_BY_TISSUE | 19 | -0.66853 | -1.90485 | 0.001706485 | 0.13077058 | 0.783 | 1405 | tags=47%, list=16%, si | EN_2 |
| GOBP_REGULATION_OF_ | GOBP_REGULATION_OF_CIRCADIAN_RHYTH | 68 | -0.50762 | -1.90371 | 0 | 0.12273164 | 0.789 | 2945 | tags=65%, list=33%, si | EN_2 |
| GOBP_NEURON_PROJECT | GOBP_NEURON_PROJECTION_GUIDANCE | 112 | -0.46727 | -1.89643 | 0 | 0.12434542 | 0.814 | 2990 | tags=53%, list=34%, si | EN_2 |
| ECM-RECEPTOR_INTERAC | ECM-RECEPTOR_INTERACTION | 36 | -0.56601 | -1.88775 | 0.001620746 | 0.12916176 | 0.846 | 2825 | tags=67%, list=32%, si | EN_2 |
| GOBP_POSTSYNAPTIC_M | GOBP_POSTSYNAPTIC_MEMBRANE_ORGAN | 35 | -0.57771 | -1.87886 | 0 | 0.13340977 | 0.869 | 1582 | tags=40%, list=18%, si | EN_2 |
| GOBP_NEGATIVE_REGUL | GOBP_NEGATIVE_REGULATION_OF_INSULIN | 27 | -0.60559 | -1.86905 | 0.001584786 | 0.1413639 | 0.893 | 731 | tags=30%, list=8%, sig | EN_2 |
| GOBP_GLANDULAR_EPITI | GOBP_GLANDULAR_EPITHELIAL_CELL_DEVE | 17 | -0.67665 | -1.86269 | 0.003407155 | 0.14313507 | 0.905 | 2420 | tags=71%, list=27%, si | EN_2 |
| CELL_ADHESION_MOLECU | CELL_ADHESION_MOLECULES_(CAMS) | 51 | -0.53021 | -1.86066 | 0.003149606 | 0.13899358 | 0.912 | 471 | tags=25%, list=5%, sig | EN_2 |
| GOBP_FACULTATIVE_HET | GOBP_FACULTATIVE_HETEROCHROMATIN_I | 37 | -0.55674 | -1.8494 | 0 | 0.1498622 | 0.94 | 2100 | tags=54%, list=24%, si | EN_2 |
| GOBP_POSITIVE_REGULA | GOBP_POSITIVE_REGULATION_OF_EXCITAT | 18 | -0.65161 | -1.84394 | 0.001683502 | 0.15221624 | 0.953 | 617 | tags=28%, list=7%, sig | EN_2 |
| GOBP_CELL_CELL_ADHES | GOBP_CELL_CELL_ADHESION_VIA_PLASMA_ | 75 | -0.48962 | -1.84383 | 0 | 0.1457959 | 0.953 | 2589 | tags=53%, list=29%, si | EN_2 |
| GOBP_CARDIAC_MUSCLE | GOBP_CARDIAC_MUSCLE_CELL_PROLIFERAT | 33 | -0.57086 | -1.83723 | 0 | 0.15056951 | 0.964 | 2959 | tags=76%, list=33%, si | EN_2 |
| GOBP_RHYTHMIC_PROCE | GOBP_RHYTHMIC_PROCESS | 166 | -0.4319 | -1.83667 | 0 | 0.14568281 | 0.964 | 2945 | tags=57%, list=33%, si | EN_2 |
| GOBP_CENTRAL_NERVOU | GOBP_CENTRAL_NERVOUS_SYSTEM_NEURO | 47 | -0.52619 | -1.83274 | 0 | 0.14663795 | 0.968 | 1812 | tags=51%, list=20%, si | EN_2 |
| GOBP_REGULATION_OF_ | GOBP_REGULATION_OF_SYNAPSE_MATURA | 15 | -0.67828 | -1.82848 | 0.003395586 | 0.14853859 | 0.974 | 323 | tags=27%, list=4%, sig | EN_2 |
| GOBP_REGULATION_OF_ | GOBP_REGULATION_OF_FAT_CELL_DIFFERE | 77 | -0.48609 | -1.82594 | 0.004615385 | 0.1468489 | 0.977 | 2297 | tags=53%, list=26%, si | EN_2 |
| GOBP_RESPONSE_TO_TR | GOBP_RESPONSE_TO_TRANSFORMING_GRO | 171 | -0.43017 | -1.82591 | 0 | 0.14190315 | 0.977 | 2661 | tags=49%, list=30%, si | EN_2 |
| GOBP_POSTSYNAPSE_AS | GOBP_POSTSYNAPSE_ASSEMBLY | 48 | -0.51763 | -1.82025 | 0.003072197 | 0.14449298 | 0.983 | 2420 | tags=52%, list=27%, si | EN_2 |
| GOBP_RESPONSE_TO_EST | GOBP_RESPONSE_TO_ESTRADIOL | 25 | -0.58907 | -1.81091 | 0.003333333 | 0.15385167 | 0.985 | 1641 | tags=52%, list=19%, si | EN_2 |
| GOBP_CELL_JUNCTION_O | GOBP_CELL_JUNCTION_ORGANIZATION | 462 | -0.3831 | -1.8086 | 0 | 0.15206364 | 0.987 | 2587 | tags=44%, list=29%, si | EN_2 |
| GOBP_CELLULAR_RESPON | GOBP_CELLULAR_RESPONSE_TO_CARBOHYI | 88 | -0.46193 | -1.80561 | 0 | 0.1514922 | 0.989 | 1746 | tags=34%, list=20%, si | EN_2 |
| GOBP_REGULATION_OF_ | GOBP_REGULATION_OF_PEPTIDE_TRANSPO | 118 | -0.43643 | -1.80513 | 0 | 0.1479067 | 0.99 | 1731 | tags=31%, list=20%, si | EN_2 |
| GOBP_POSITIVE_REGULA | GOBP_POSITIVE_REGULATION_OF_CELL_JU | 65 | -0.49123 | -1.80258 | 0 | 0.14767714 | 0.99 | 1860 | tags=38%, list=21%, si | EN_2 |
| GOBP_INSULIN_SECRETIO | GOBP_INSULIN_SECRETION_INVOLVED_IN_ | 53 | -0.51021 | -1.79898 | 0.003081664 | 0.14827783 | 0.991 | 1746 | tags=34%, list=20%, si | EN_2 |
| GOBP_POSITIVE_REGULA | GOBP_POSITIVE_REGULATION_OF_SYNAPSE | 38 | -0.54299 | -1.79766 | 0 | 0.14639276 | 0.992 | 1577 | tags=42%, list=18%, si | EN_2 |
| GOBP_DNA_METHYLATIO | GOBP_DNA_METHYLATION_DEPENDENT_HI | 26 | -0.59627 | -1.79114 | 0.001584786 | 0.15196882 | 0.994 | 1734 | tags=54%, list=20%, si | EN_2 |
| GOBP_CARBOHYDRATE_H | GOBP_CARBOHYDRATE_HOMEOSTASIS | 149 | -0.42434 | -1.78973 | 0 | 0.14984956 | 0.994 | 1746 | tags=32%, list=20%, si | EN_2 |

| | | | | | | | | | | |
|---|---|---|---|---|---|---|---|---|---|---|
| GOBP_REGULATION_OF_ | GOBP_REGULATION_OF_HORMONE_SECRET | 137 | -0.42834 | -1.78427 | 0 | 0.15500593 | 0.995 | 1731 | tags=31%, list=20%, si | EN_2 |
| GOBP_CELL_PROJECTION | GOBP_CELL_PROJECTION_MORPHOGENESIS | 411 | -0.38264 | -1.77863 | 0 | 0.15907131 | 0.996 | 2076 | tags=35%, list=23%, si | EN_2 |
| GOBP_BASEMENT_MEME | GOBP_BASEMENT_MEMBRANE_ORGANIZAT | 31 | -0.55766 | -1.77038 | 0.006472492 | 0.1676656 | 0.998 | 2114 | tags=55%, list=24%, si | EN_2 |
| GOBP_CELL_MORPHOGEI | GOBP_CELL_MORPHOGENESIS_INVOLVED_I | 361 | -0.38036 | -1.7634 | 0 | 0.17600892 | 0.999 | 2668 | tags=45%, list=30%, si | EN_2 |
| GOBP_EXCITATORY_SYNA | GOBP_EXCITATORY_SYNAPSE_ASSEMBLY | 20 | -0.61829 | -1.76338 | 0.003401361 | 0.17200872 | 0.999 | 2049 | tags=60%, list=23%, si | EN_2 |
| GOBP_DENDRITE_MORPH | GOBP_DENDRITE_MORPHOGENESIS | 111 | -0.43743 | -1.76234 | 0.001481482 | 0.16990426 | 0.999 | 2820 | tags=48%, list=32%, si | EN_2 |
| GOBP_POSTSYNAPTIC_SP | GOBP_POSTSYNAPTIC_SPECIALIZATION_ASS | 20 | -0.60979 | -1.75685 | 0.004769475 | 0.17491713 | 0.999 | 2049 | tags=55%, list=23%, si | EN_2 |
| GOBP_REGULATION_OF_ | GOBP_REGULATION_OF_NERVOUS_SYSTEM | 308 | -0.38311 | -1.75684 | 0 | 0.1711955 | 0.999 | 2807 | tags=46%, list=32%, si | EN_2 |
| GOBP_NEGATIVE_REGUL | GOBP_NEGATIVE_REGULATION_OF_PEPTID | 30 | -0.55924 | -1.75527 | 0.008431703 | 0.17023475 | 1 | 731 | tags=27%, list=8%, sig | EN_2 |
| GOBP_CIRCADIAN_RHYTH | GOBP_CIRCADIAN_RHYTHM | 123 | -0.43239 | -1.75512 | 0 | 0.16701527 | 1 | 2945 | tags=59%, list=33%, si | EN_2 |
| ADHERENS_JUNCTION | ADHERENS_JUNCTION | 54 | -0.49267 | -1.75435 | 0 | 0.16481084 | 1 | 2598 | tags=43%, list=29%, si | EN_2 |
| GOBP_REGULATION_OF_ | GOBP_REGULATION_OF_HIPPO_SIGNALING | 15 | -0.6372 | -1.75189 | 0.006837607 | 0.16536434 | 1 | 2275 | tags=60%, list=26%, si | EN_2 |
| GOBP_REGULATION_OF_ | GOBP_REGULATION_OF_STEM_CELL_PROLI | 50 | -0.48978 | -1.74423 | 0.00304878 | 0.17366998 | 1 | 3018 | tags=62%, list=34%, si | EN_2 |
| GOBP_REGULATION_OF_ | GOBP_REGULATION_OF_PROTEIN_LOCALIZA | 21 | -0.59407 | -1.7432 | 0.005163511 | 0.17206335 | 1 | 2063 | tags=57%, list=23%, si | EN_2 |
| GOBP_OUTFLOW_TRACT_ | GOBP_OUTFLOW_TRACT_MORPHOGENESIS | 46 | -0.49866 | -1.73956 | 0.006369427 | 0.17453377 | 1 | 2959 | tags=61%, list=33%, si | EN_2 |
| GOBP_EPITHELIAL_CELL_ | GOBP_EPITHELIAL_CELL_MORPHOGENESIS | 23 | -0.57097 | -1.73831 | 0.013029316 | 0.17327876 | 1 | 3195 | tags=78%, list=36%, si | EN_2 |
| GOBP_TRANSFORMING_C | GOBP_TRANSFORMING_GROWTH_FACTOR_ | 138 | -0.41693 | -1.73563 | 0 | 0.17445615 | 1 | 2661 | tags=50%, list=30%, si | EN_2 |
| GOBP_POSITIVE_REGULA | GOBP_POSITIVE_REGULATION_OF_HORMOI | 87 | -0.44113 | -1.73477 | 0 | 0.17289096 | 1 | 1662 | tags=33%, list=19%, si | EN_2 |
| GOBP_REGULATION_OF_ | GOBP_REGULATION_OF_BEHAVIOR | 36 | -0.53475 | -1.73402 | 0.008169935 | 0.17120145 | 1 | 2200 | tags=53%, list=25%, si | EN_2 |
| GOBP_STRIATED_MUSCLI | GOBP_STRIATED_MUSCLE_CELL_PROLIFERA | 45 | -0.49539 | -1.73286 | 0.001686341 | 0.16999535 | 1 | 2959 | tags=69%, list=33%, si | EN_2 |
| GOBP_SYNAPSE_ORGANI | GOBP_SYNAPSE_ORGANIZATION | 323 | -0.37889 | -1.7323 | 0 | 0.16799551 | 1 | 2630 | tags=46%, list=30%, si | EN_2 |
| GOBP_LUNG_CELL_DIFFE | GOBP_LUNG_CELL_DIFFERENTIATION | 23 | -0.58948 | -1.73208 | 0.006410257 | 0.165578 | 1 | 2618 | tags=70%, list=30%, si | EN_2 |
| GOBP_CENTRAL_NERVOU | GOBP_CENTRAL_NERVOUS_SYSTEM_NEURC | 87 | -0.44544 | -1.7305 | 0.002980626 | 0.16527349 | 1 | 1965 | tags=41%, list=22%, si | EN_2 |
| GOBP_MORPHOGENESIS_ | GOBP_MORPHOGENESIS_OF_AN_EPITHELIU | 293 | -0.38029 | -1.7304 | 0 | 0.16273975 | 1 | 2589 | tags=43%, list=29%, si | EN_2 |
| GOBP_POSITIVE_REGULA | GOBP_POSITIVE_REGULATION_OF_NEUROB | 21 | -0.59396 | -1.72918 | 0.004958678 | 0.16186927 | 1 | 1563 | tags=38%, list=18%, si | EN_2 |
| GOBP_REGULATION_OF_ | GOBP_REGULATION_OF_NEURON_PROJECT | 28 | -0.54688 | -1.72421 | 0.008431703 | 0.16744164 | 1 | 2599 | tags=54%, list=29%, si | EN_2 |
| GOBP_PRESYNAPSE_ASSE | GOBP_PRESYNAPSE_ASSEMBLY | 28 | -0.54214 | -1.72024 | 0.013355592 | 0.17119896 | 1 | 2049 | tags=46%, list=23%, si | EN_2 |
| GOBP_MODULATION_OF | GOBP_MODULATION_OF_EXCITATORY_POS | 29 | -0.55031 | -1.7198 | 0.009852217 | 0.1691687 | 1 | 617 | tags=28%, list=7%, sig | EN_2 |
| GOBP_REGULATION_OF_ | GOBP_REGULATION_OF_SYNAPSE_STRUCTU | 202 | -0.3957 | -1.71798 | 0 | 0.16947244 | 1 | 2630 | tags=44%, list=30%, si | EN_2 |
| GOBP_EMBRYONIC_EYE_ | GOBP_EMBRYONIC_EYE_MORPHOGENESIS | 18 | -0.6289 | -1.71755 | 0.006944445 | 0.16769087 | 1 | 1614 | tags=44%, list=18%, si | EN_2 |
| GOBP_REGULATION_OF_ | GOBP_REGULATION_OF_INSULIN_SECRETIO | 105 | -0.43517 | -1.71732 | 0 | 0.16562036 | 1 | 1731 | tags=30%, list=20%, si | EN_2 |
| GOBP_MAMMARY_GLAN | GOBP_MAMMARY_GLAND_MORPHOGENES | 33 | -0.52426 | -1.71506 | 0.006389776 | 0.16687879 | 1 | 2838 | tags=61%, list=32%, si | EN_2 |
| GOBP_ERYTHROCYTE_DE | GOBP_ERYTHROCYTE_DEVELOPMENT | 31 | -0.53719 | -1.71332 | 0.011456628 | 0.16706157 | 1 | 1478 | tags=42%, list=17%, si | EN_2 |
| GOBP_BODY_FLUID_SECF | GOBP_BODY_FLUID_SECRETION | 39 | -0.51827 | -1.70889 | 0.003120125 | 0.17149194 | 1 | 1416 | tags=38%, list=16%, si | EN_2 |
| GOBP_PROTEIN_PALMITO | GOBP_PROTEIN_PALMITOYLATION | 17 | -0.61705 | -1.70666 | 0.0224 | 0.1725365 | 1 | 1582 | tags=41%, list=18%, si | EN_2 |
| FC_EPSILON_RI_SIGNALIN | FC_EPSILON_RI_SIGNALING_PATHWAY | 32 | -0.5336 | -1.70507 | 0.006329114 | 0.17300878 | 1 | 2551 | tags=53%, list=29%, si | EN_2 |
| CIRCADIAN_RHYTHM | CIRCADIAN_RHYTHM | 22 | -0.58099 | -1.70467 | 0.010471204 | 0.17119555 | 1 | 1867 | tags=50%, list=21%, si | EN_2 |
| GOBP_REGULATION_OF_ | GOBP_REGULATION_OF_NERVOUS_SYSTEM | 62 | -0.466 | -1.70289 | 0.003100775 | 0.1715853 | 1 | 1139 | tags=29%, list=13%, si | EN_2 |
| SIGNALING_PATHWAYS_I | SIGNALING_PATHWAYS_REGULATING_PLUR | 72 | -0.45615 | -1.70263 | 0 | 0.16970599 | 1 | 2598 | tags=51%, list=29%, si | EN_2 |
| GOBP_REGULATION_OF_ | GOBP_REGULATION_OF_SIGNALING_RECEP | 46 | -0.49222 | -1.70235 | 0.006420546 | 0.1680475 | 1 | 677 | tags=24%, list=8%, sig | EN_2 |
| FOCAL_ADHESION | FOCAL_ADHESION | 124 | -0.41236 | -1.69823 | 0 | 0.17237036 | 1 | 2623 | tags=45%, list=30%, si | EN_2 |
| GOBP_AXON_DEVELOPM | GOBP_AXON_DEVELOPMENT | 303 | -0.37273 | -1.69584 | 0 | 0.17403403 | 1 | 2668 | tags=45%, list=30%, si | EN_2 |
| GOBP_LUNG_EPITHELIUN | GOBP_LUNG_EPITHELIUM_DEVELOPMENT | 33 | -0.52905 | -1.69556 | 0.008250825 | 0.17234087 | 1 | 2618 | tags=61%, list=30%, si | EN_2 |
| GOBP_HEAD_DEVELOPM | GOBP_HEAD_DEVELOPMENT | 413 | -0.36072 | -1.69344 | 0 | 0.1734435 | 1 | 2619 | tags=44%, list=30%, si | EN_2 |
| GOBP_WNT_SIGNALING_ | GOBP_WNT_SIGNALING_PATHWAY_PLANA | 21 | -0.58087 | -1.69259 | 0.010186757 | 0.17285483 | 1 | 1982 | tags=52%, list=22%, si | EN_2 |
| GOBP_MESONEPHROS_D | GOBP_MESONEPHROS_DEVELOPMENT | 43 | -0.49944 | -1.6892 | 0.006430868 | 0.17566724 | 1 | 1700 | tags=37%, list=19%, si | EN_2 |
| GOBP_RHYTHMIC_BEHAV | GOBP_RHYTHMIC_BEHAVIOR | 24 | -0.55588 | -1.68783 | 0.015570934 | 0.17558326 | 1 | 2705 | tags=71%, list=31%, si | EN_2 |
| GOBP_REGULATION_OF_ | GOBP_REGULATION_OF_SYNAPTIC_TRANSN | 15 | -0.6331 | -1.68634 | 0.010434783 | 0.17597543 | 1 | 1770 | tags=47%, list=20%, si | EN_2 |
| GOBP_REGULATION_OF_ | GOBP_REGULATION_OF_BROWN_FAT_CELL | 16 | -0.6134 | -1.68618 | 0.015517241 | 0.17424715 | 1 | 1637 | tags=63%, list=19%, si | EN_2 |
| GOBP_TOOTH_MINERALI | GOBP_TOOTH_MINERALIZATION | 17 | -0.60579 | -1.68395 | 0.014084507 | 0.17596976 | 1 | 596 | tags=29%, list=7%, sig | EN_2 |
| GOBP_REGULATION_OF_ | GOBP_REGULATION_OF_CHROMATIN_ORG/ | 28 | -0.53735 | -1.6832 | 0.004830918 | 0.17517932 | 1 | 1734 | tags=46%, list=20%, si | EN_2 |
| GOBP_EPITHELIAL_CELL_ | GOBP_EPITHELIAL_CELL_DEVELOPMENT | 119 | -0.41664 | -1.68136 | 0.002828854 | 0.17613852 | 1 | 3136 | tags=56%, list=35%, si | EN_2 |
| GOBP_EPITHELIAL_TUBE_ | GOBP_EPITHELIAL_TUBE_MORPHOGENESIS | 207 | -0.38454 | -1.68045 | 0 | 0.17567013 | 1 | 2980 | tags=52%, list=34%, si | EN_2 |
| GOBP_REGULATION_OF_ | GOBP_REGULATION_OF_POSTSYNAPTIC_DE | 16 | -0.60991 | -1.6786 | 0.024752475 | 0.17678925 | 1 | 955 | tags=38%, list=11%, si | EN_2 |
| GOBP_REGULATION_OF_ | GOBP_REGULATION_OF_CELLULAR_RESPON | 97 | -0.42598 | -1.67776 | 0.002915452 | 0.17627731 | 1 | 2661 | tags=49%, list=30%, si | EN_2 |
| GOBP_FOREBRAIN_NEUR | GOBP_FOREBRAIN_NEURON_DIFFERENTIAT | 21 | -0.58466 | -1.67718 | 0.015358361 | 0.17536871 | 1 | 1805 | tags=52%, list=20%, si | EN_2 |
| GOBP_RENAL_SYSTEM_D | GOBP_RENAL_SYSTEM_DEVELOPMENT | 170 | -0.39312 | -1.67663 | 0 | 0.17434914 | 1 | 2648 | tags=46%, list=30%, si | EN_2 |
| GOBP_POSITIVE_REGULA | GOBP_POSITIVE_REGULATION_OF_PEPTIDE | 73 | -0.45316 | -1.67609 | 0.003007519 | 0.17339534 | 1 | 1662 | tags=34%, list=19%, si | EN_2 |
| GOBP_POSITIVE_REGULA | GOBP_POSITIVE_REGULATION_OF_NERVOU | 208 | -0.38167 | -1.67298 | 0 | 0.17667973 | 1 | 1679 | tags=30%, list=19%, si | EN_2 |
| GOBP_MIDBRAIN_DEVEL | GOBP_MIDBRAIN_DEVELOPMENT | 23 | -0.55595 | -1.67151 | 0.014729951 | 0.1773885 | 1 | 2014 | tags=57%, list=23%, si | EN_2 |
| GOBP_POSITIVE_REGULA | GOBP_POSITIVE_REGULATION_OF_FAT_CEL | 34 | -0.51 | -1.66738 | 0.011146497 | 0.18184525 | 1 | 2102 | tags=53%, list=24%, si | EN_2 |
| GNRH_SIGNALING_PATH | GNRH_SIGNALING_PATHWAY | 54 | -0.46999 | -1.66691 | 0.006410257 | 0.18076265 | 1 | 2344 | tags=41%, list=26%, si | EN_2 |
| GOBP_REGULATION_OF_ | GOBP_REGULATION_OF_DENDRITE_MORPH | 57 | -0.45996 | -1.66515 | 0.004559271 | 0.18152174 | 1 | 3172 | tags=56%, list=36%, si | EN_2 |
| GOBP_PIGMENTATION | GOBP_PIGMENTATION | 67 | -0.4399 | -1.66383 | 0.003134796 | 0.18152824 | 1 | 1659 | tags=37%, list=19%, si | EN_2 |
| GOBP_CORPUS_CALLOSU | GOBP_CORPUS_CALLOSUM_DEVELOPMENT | 17 | -0.59827 | -1.66186 | 0.013223141 | 0.1827536 | 1 | 1217 | tags=41%, list=14%, si | EN_2 |
| GOBP_GLAND_MORPHOC | GOBP_GLAND_MORPHOGENESIS | 79 | -0.43069 | -1.66136 | 0.005865103 | 0.18171626 | 1 | 2860 | tags=52%, list=32%, si | EN_2 |
| GOBP_REGULATION_OF_ | GOBP_REGULATION_OF_PRESYNAPSE_ORG/ | 22 | -0.56198 | -1.65962 | 0.012965964 | 0.18268706 | 1 | 472 | tags=23%, list=5%, sig | EN_2 |
| GOBP_LENS_DEVELOPME | GOBP_LENS_DEVELOPMENT_IN_CAMERA_T | 52 | -0.46662 | -1.65939 | 0.003039514 | 0.18134092 | 1 | 2026 | tags=42%, list=23%, si | EN_2 |
| GOBP_KIDNEY_EPITHELIU | GOBP_KIDNEY_EPITHELIUM_DEVELOPMENT | 63 | -0.45258 | -1.65872 | 0.004651163 | 0.18061952 | 1 | 2860 | tags=56%, list=32%, si | EN_2 |
| GOBP_DNA_INTEGRITY_C | GOBP_DNA_INTEGRITY_CHECKPOINT_SIGN/ | 78 | -0.43213 | -1.65804 | 0.002967359 | 0.1799438 | 1 | 2210 | tags=40%, list=25%, si | EN_2 |
| MTOR_SIGNALING_PATH | MTOR_SIGNALING_PATHWAY | 99 | -0.41465 | -1.65532 | 0.001468429 | 0.1827541 | 1 | 2551 | tags=44%, list=29%, si | EN_2 |
| GOBP_REGULATION_OF_ | GOBP_REGULATION_OF_AXON_REGENERAT | 24 | -0.53827 | -1.65515 | 0.019354839 | 0.18133263 | 1 | 2599 | tags=54%, list=29%, si | EN_2 |
| GOBP_SMAD_PROTEIN_S | GOBP_SMAD_PROTEIN_SIGNAL_TRANSDUC | 49 | -0.47272 | -1.65463 | 0.007751938 | 0.18039425 | 1 | 2838 | tags=53%, list=32%, si | EN_2 |
| GOBP_CARDIAC_SEPTUM | GOBP_CARDIAC_SEPTUM_MORPHOGENESIS | 44 | -0.48223 | -1.65438 | 0.00624025 | 0.17920013 | 1 | 2860 | tags=64%, list=32%, si | EN_2 |
| GOBP_EMBRYONIC_ORG/ | GOBP_EMBRYONIC_ORGAN_MORPHOGENE | 139 | -0.39975 | -1.65367 | 0.0013947 | 0.17870143 | 1 | 2813 | tags=47%, list=32%, si | EN_2 |
| GOBP_ENDOCRINE_SYST | GOBP_ENDOCRINE_SYSTEM_DEVELOPMENT | 50 | -0.47988 | -1.65297 | 0.01236476 | 0.17813477 | 1 | 2598 | tags=52%, list=29%, si | EN_2 |
| GOBP_NEGATIVE_REGUL | GOBP_NEGATIVE_REGULATION_OF_PROTEI | 41 | -0.50175 | -1.65284 | 0.011272142 | 0.17674677 | 1 | 745 | tags=27%, list=8%, sig | EN_2 |
| GOBP_EPITHELIAL_CELL_ | GOBP_EPITHELIAL_CELL_DIFFERENTIATION | 294 | -0.36142 | -1.64959 | 0 | 0.18044342 | 1 | 2648 | tags=44%, list=30%, si | EN_2 |
| GOBP_RECEPTOR_CLUST | GOBP_RECEPTOR_CLUSTERING | 50 | -0.47187 | -1.64956 | 0.007788162 | 0.17897166 | 1 | 1010 | tags=26%, list=11%, si | EN_2 |
| GOBP_INSULIN_SECRETIO | GOBP_INSULIN_SECRETION | 122 | -0.39938 | -1.64675 | 0.001459854 | 0.18175052 | 1 | 1746 | tags=30%, list=20%, si | EN_2 |
| GOBP_TISSUE_MORPHOC | GOBP_TISSUE_MORPHOGENESIS | 337 | -0.35443 | -1.63859 | 0 | 0.19322282 | 1 | 2598 | tags=42%, list=29%, si | EN_2 |
| GOBP_CARDIAC_SEPTUM | GOBP_CARDIAC_SEPTUM_DEVELOPMENT | 73 | -0.4237 | -1.63684 | 0.004658385 | 0.19476826 | 1 | 2860 | tags=55%, list=32%, si | EN_2 |

| | | | | | | | | | | |
|---|---|---|---|---|---|---|---|---|---|---|
| VEGF_SIGNALING_PATHW | VEGF_SIGNALING_PATHWAY | 33 | -0.50529 | -1.6362 | 0.019292604 | 0.19452122 | 1 | 2699 | tags=64%, list=31%, si | EN_2 |
| SPHINGOLIPID_SIGNALIN | SPHINGOLIPID_SIGNALING_PATHWAY | 84 | -0.42025 | -1.63546 | 0.005706134 | 0.19385636 | 1 | 2632 | tags=43%, list=30%, si | EN_2 |
| GOBP_REGULATION_OF_ | GOBP_REGULATION_OF_CELLULAR_RESPON | 175 | -0.38094 | -1.63488 | 0 | 0.19300891 | 1 | 2661 | tags=43%, list=30%, si | EN_2 |
| GOBP_POSTSYNAPSE_OR | GOBP_POSTSYNAPSE_ORGANIZATION | 171 | -0.38278 | -1.63425 | 0 | 0.19259465 | 1 | 2605 | tags=43%, list=29%, si | EN_2 |
| GOBP_NEURON_MIGRAT | GOBP_NEURON_MIGRATION | 106 | -0.40265 | -1.63415 | 0.001472754 | 0.1912564 | 1 | 1980 | tags=42%, list=22%, si | EN_2 |
| ABC_TRANSPORTERS | ABC_TRANSPORTERS | 19 | -0.57292 | -1.63364 | 0.0237691 | 0.1906539 | 1 | 1487 | tags=42%, list=17%, si | EN_2 |
| GOBP_NEGATIVE_REGUL | GOBP_NEGATIVE_REGULATION_OF_MAPK_ | 94 | -0.41448 | -1.63331 | 0.002816901 | 0.18968844 | 1 | 1905 | tags=31%, list=22%, si | EN_2 |
| GOBP_NEGATIVE_REGUL | GOBP_NEGATIVE_REGULATION_OF_BMP_S | 30 | -0.51706 | -1.63312 | 0.009950249 | 0.18849309 | 1 | 2860 | tags=60%, list=32%, si | EN_2 |
| HEPATITIS_B | HEPATITIS_B | 104 | -0.40976 | -1.63302 | 0.002980626 | 0.18714802 | 1 | 2128 | tags=38%, list=24%, si | EN_2 |
| GOBP_EMBRYONIC_CAM | GOBP_EMBRYONIC_CAMERA_TYPE_EYE_M | 16 | -0.60475 | -1.63274 | 0.02586207 | 0.18618041 | 1 | 1614 | tags=44%, list=18%, si | EN_2 |
| GOBP_BROWN_FAT_CELL | GOBP_BROWN_FAT_CELL_DIFFERENTIATION | 31 | -0.51824 | -1.63169 | 0.01875 | 0.186407 | 1 | 1863 | tags=48%, list=21%, si | EN_2 |
| GOBP_NEURON_PROJECT | GOBP_NEURON_PROJECTION_REGENERATI | 44 | -0.47514 | -1.62998 | 0.010852713 | 0.18756388 | 1 | 2599 | tags=45%, list=29%, si | EN_2 |
| GOBP_NEUROTRANSMITT | GOBP_NEUROTRANSMITTER_RECEPTOR_LO | 21 | -0.56174 | -1.627 | 0.018242123 | 0.19066143 | 1 | 2284 | tags=52%, list=26%, si | EN_2 |
| GOBP_RESPONSE_TO_CO | GOBP_RESPONSE_TO_COLD | 25 | -0.53299 | -1.62641 | 0.011475409 | 0.19007485 | 1 | 122 | tags=12%, list=1%, sig | EN_2 |
| GOBP_FAT_CELL_DIFFERE | GOBP_FAT_CELL_DIFFERENTIATION | 145 | -0.38939 | -1.6259 | 0.00405954 | 0.18953156 | 1 | 2674 | tags=48%, list=30%, si | EN_2 |
| GOBP_BRANCHING_INVO | GOBP_BRANCHING_INVOLVED_IN_MAMMA | 15 | -0.61044 | -1.62587 | 0.022184301 | 0.18816462 | 1 | 903 | tags=40%, list=10%, si | EN_2 |
| GOBP_PRESYNAPSE_ORG | GOBP_PRESYNAPSE_ORGANIZATION | 34 | -0.50471 | -1.62559 | 0.02247191 | 0.1872133 | 1 | 2764 | tags=56%, list=31%, si | EN_2 |
| GOBP_METANEPHROS_D | GOBP_METANEPHROS_DEVELOPMENT | 30 | -0.51096 | -1.62512 | 0.026578072 | 0.18652874 | 1 | 1563 | tags=40%, list=18%, si | EN_2 |
| GOBP_PIGMENT_CELL_D | GOBP_PIGMENT_CELL_DIFFERENTIATION | 27 | -0.52743 | -1.62398 | 0.013513514 | 0.18667656 | 1 | 1545 | tags=44%, list=17%, si | EN_2 |
| GOBP_RESPONSE_TO_GR | GOBP_RESPONSE_TO_GROWTH_FACTOR | 390 | -0.34934 | -1.62379 | 0 | 0.18563442 | 1 | 2661 | tags=42%, list=30%, si | EN_2 |
| GOBP_REGULATION_OF_ | GOBP_REGULATION_OF_CELL_MORPHOGEN | 172 | -0.37898 | -1.6231 | 0.001349528 | 0.18544054 | 1 | 2411 | tags=40%, list=27%, si | EN_2 |
| GOBP_CELL_PROLIFERATI | GOBP_CELL_PROLIFERATION_IN_FOREBRAIN | 19 | -0.57572 | -1.62202 | 0.019769358 | 0.18580216 | 1 | 2797 | tags=74%, list=32%, si | EN_2 |
| GOBP_NEGATIVE_REGUL | GOBP_NEGATIVE_REGULATION_OF_CELLUL | 55 | -0.44865 | -1.62196 | 0.006060606 | 0.18459108 | 1 | 2860 | tags=47%, list=32%, si | EN_2 |
| GOBP_PROTEIN_LOCALIZ | GOBP_PROTEIN_LOCALIZATION_TO_MICRO | 30 | -0.51019 | -1.61944 | 0.021630615 | 0.18717583 | 1 | 2304 | tags=47%, list=26%, si | EN_2 |
| GOBP_NEGATIVE_REGUL | GOBP_NEGATIVE_REGULATION_OF_TRANSL | 17 | -0.58954 | -1.61801 | 0.023728814 | 0.18796022 | 1 | 2190 | tags=53%, list=25%, si | EN_2 |
| GOBP_NATURAL_KILLER_ | GOBP_NATURAL_KILLER_CELL_MEDIATED_I | 21 | -0.54977 | -1.61723 | 0.016722407 | 0.18784933 | 1 | 1723 | tags=48%, list=19%, si | EN_2 |
| GOBP_FOREBRAIN_GENE | GOBP_FOREBRAIN_GENERATION_OF_NEUR | 28 | -0.51839 | -1.61689 | 0.02733119 | 0.18708818 | 1 | 1805 | tags=50%, list=20%, si | EN_2 |
| GOBP_RESPONSE_TO_EL | GOBP_RESPONSE_TO_ELECTRICAL_STIMULU | 15 | -0.59542 | -1.61558 | 0.024263432 | 0.18774936 | 1 | 1770 | tags=47%, list=20%, si | EN_2 |
| GOBP_ESTABLISHMENT_ | GOBP_ESTABLISHMENT_OF_MITOCHONDRI | 24 | -0.54017 | -1.6153 | 0.014681893 | 0.18689215 | 1 | 1812 | tags=42%, list=20%, si | EN_2 |
| GOBP_NEGATIVE_REGUL | GOBP_NEGATIVE_REGULATION_OF_SECRET | 80 | -0.42211 | -1.61465 | 0.008797654 | 0.18659534 | 1 | 904 | tags=23%, list=10%, si | EN_2 |
| GOBP_RESPIRATORY_GAS | GOBP_RESPIRATORY_GASEOUS_EXCHANGE | 36 | -0.48322 | -1.61388 | 0.010971786 | 0.18635572 | 1 | 2114 | tags=44%, list=24%, si | EN_2 |
| HIPPO_SIGNALING_PATH | HIPPO_SIGNALING_PATHWAY | 93 | -0.41048 | -1.61267 | 0.002915452 | 0.18692835 | 1 | 2625 | tags=43%, list=30%, si | EN_2 |
| GOBP_CRANIAL_NERVE_ | GOBP_CRANIAL_NERVE_DEVELOPMENT | 28 | -0.51489 | -1.61225 | 0.020249221 | 0.18636295 | 1 | 1934 | tags=54%, list=22%, si | EN_2 |
| GOBP_REGULATION_OF_ | GOBP_REGULATION_OF_HETEROCHROMATI | 18 | -0.58627 | -1.61207 | 0.017636685 | 0.18547496 | 1 | 2100 | tags=61%, list=24%, si | EN_2 |
| GOBP_AXON_REGENERAT | GOBP_AXON_REGENERATION | 36 | -0.49345 | -1.60995 | 0.018092105 | 0.18762134 | 1 | 2599 | tags=47%, list=29%, si | EN_2 |
| GOBP_DENDRITE_DEVEL | GOBP_DENDRITE_DEVELOPMENT | 192 | -0.37134 | -1.60891 | 0.00136612 | 0.18799034 | 1 | 2668 | tags=43%, list=30%, si | EN_2 |
| GOBP_PEPTIDE_TRANSP | GOBP_PEPTIDE_TRANSPORT | 143 | -0.38392 | -1.60567 | 0 | 0.19181249 | 1 | 1746 | tags=30%, list=20%, si | EN_2 |
| GOBP_POSITIVE_REGULA | GOBP_POSITIVE_REGULATION_OF_CELL_M | 44 | -0.46882 | -1.60473 | 0.009615385 | 0.19182214 | 1 | 2795 | tags=50%, list=32%, si | EN_2 |
| GOBP_PERIPHERAL_NERV | GOBP_PERIPHERAL_NERVOUS_SYSTEM_AX | 23 | -0.53471 | -1.60441 | 0.027350428 | 0.19105588 | 1 | 2835 | tags=65%, list=32%, si | EN_2 |
| GOBP_POSITIVE_REGULA | GOBP_POSITIVE_REGULATION_OF_DEFENSE | 16 | -0.59654 | -1.60318 | 0.017006803 | 0.19172016 | 1 | 2131 | tags=56%, list=24%, si | EN_2 |
| GOBP_DIGESTIVE_SYSTEM | GOBP_DIGESTIVE_SYSTEM_DEVELOPMENT | 61 | -0.44385 | -1.60188 | 0.009036144 | 0.19246046 | 1 | 2088 | tags=41%, list=24%, si | EN_2 |
| GOBP_CELL_SUBSTRATE_ | GOBP_CELL_SUBSTRATE_JUNCTION_ORGAN | 82 | -0.41981 | -1.59825 | 0.004405286 | 0.1968836 | 1 | 3300 | tags=57%, list=37%, si | EN_2 |
| GOBP_PROTEIN_LOCALIZ | GOBP_PROTEIN_LOCALIZATION_TO_CHROM | 29 | -0.52059 | -1.59756 | 0.01923077 | 0.1968051 | 1 | 3256 | tags=72%, list=37%, si | EN_2 |
| PI3K-AKT_SIGNALING_PA | PI3K-AKT_SIGNALING_PATHWAY | 176 | -0.36871 | -1.59735 | 0.001373626 | 0.19596396 | 1 | 2632 | tags=43%, list=30%, si | EN_2 |
| GOBP_T_HELPER_17_TYP | GOBP_T_HELPER_17_TYPE_IMMUNE_RESP | 19 | -0.55855 | -1.59727 | 0.02451839 | 0.19489941 | 1 | 3336 | tags=79%, list=38%, si | EN_2 |
| GOBP_DEVELOPMENTAL_ | GOBP_DEVELOPMENTAL_GROWTH_INVOLV | 163 | -0.38027 | -1.59724 | 0.001367989 | 0.19377343 | 1 | 2885 | tags=50%, list=33%, si | EN_2 |
| GOBP_TRANSFORMING_ | GOBP_TRANSFORMING_GROWTH_FACTOR_ | 19 | -0.56284 | -1.5965 | 0.03482587 | 0.19378482 | 1 | 931 | tags=32%, list=11%, si | EN_2 |
| GOBP_POSITIVE_REGULA | GOBP_POSITIVE_REGULATION_OF_CELLULA | 21 | -0.55065 | -1.59443 | 0.021775544 | 0.19591738 | 1 | 2303 | tags=52%, list=26%, si | EN_2 |
| GOBP_SKIN_DEVELOPME | GOBP_SKIN_DEVELOPMENT | 138 | -0.38354 | -1.59299 | 0.0013947 | 0.1970813 | 1 | 2200 | tags=39%, list=25%, si | EN_2 |
| GOBP_REGULATION_OF_ | GOBP_REGULATION_OF_PROTEIN_SECRETI | 159 | -0.37417 | -1.59202 | 0 | 0.19744201 | 1 | 1731 | tags=27%, list=20%, si | EN_2 |
| GOBP_CELL_CYCLE_PHAS | GOBP_CELL_CYCLE_PHASE_TRANSITION | 317 | -0.34725 | -1.59113 | 0 | 0.19774213 | 1 | 2139 | tags=34%, list=24%, si | EN_2 |
| GOBP_INTRACELLULAR_ | GOBP_INTRACELLULAR_GLUCOSE_HOMEOS | 94 | -0.40466 | -1.59015 | 0.005952381 | 0.19830222 | 1 | 1746 | tags=30%, list=20%, si | EN_2 |
| GOBP_REGULATION_OF_ | GOBP_REGULATION_OF_NEUROBLAST_PRO | 31 | -0.4972 | -1.58967 | 0.0112 | 0.19789633 | 1 | 2860 | tags=55%, list=32%, si | EN_2 |
| GOBP_REGULATION_OF_ | GOBP_REGULATION_OF_NEUROGENESIS | 253 | -0.35583 | -1.58814 | 0 | 0.19902954 | 1 | 2795 | tags=44%, list=32%, si | EN_2 |
| GOBP_EMBRYONIC_CRAN | GOBP_EMBRYONIC_CRANIAL_SKELETON_M | 25 | -0.5282 | -1.58792 | 0.026936026 | 0.19831172 | 1 | 2959 | tags=64%, list=33%, si | EN_2 |
| GOBP_EMBRYONIC_SKEL | GOBP_EMBRYONIC_SKELETAL_SYSTEM_MO | 46 | -0.45327 | -1.58718 | 0.010869565 | 0.19835345 | 1 | 2793 | tags=54%, list=32%, si | EN_2 |
| GOBP_RESPONSE_TO_AX | GOBP_RESPONSE_TO_AXON_INJURY | 47 | -0.45549 | -1.58633 | 0.01910828 | 0.19847405 | 1 | 2609 | tags=45%, list=29%, si | EN_2 |
| GOBP_REGULATION_OF_ | GOBP_REGULATION_OF_CELL_SHAPE | 91 | -0.40421 | -1.58564 | 0.007440476 | 0.19848776 | 1 | 2198 | tags=40%, list=25%, si | EN_2 |
| GOBP_REGULATION_OF_ | GOBP_REGULATION_OF_CALCIUM_ION_IM | 20 | -0.54735 | -1.58521 | 0.023809524 | 0.19807376 | 1 | 1563 | tags=50%, list=18%, si | EN_2 |
| GOBP_REPRODUCTIVE_B | GOBP_REPRODUCTIVE_BEHAVIOR | 24 | -0.5321 | -1.57722 | 0.022653721 | 0.21040514 | 1 | 2611 | tags=58%, list=30%, si | EN_2 |
| LYSINE_DEGRADATION | LYSINE_DEGRADATION | 42 | -0.4636 | -1.57591 | 0.026101142 | 0.2115747 | 1 | 1176 | tags=31%, list=13%, si | EN_2 |
| GOBP_PHOSPHATIDYLIN | GOBP_PHOSPHATIDYLINOSITOL_3_KINASE_ | 164 | -0.36996 | -1.57474 | 0.004115226 | 0.21220198 | 1 | 1934 | tags=37%, list=22%, si | EN_2 |
| GOBP_BRANCHING_MOR | GOBP_BRANCHING_MORPHOGENESIS_OF_ | 84 | -0.40567 | -1.57416 | 0.004545454 | 0.21191534 | 1 | 2860 | tags=58%, list=32%, si | EN_2 |
| GOBP_REGULATION_OF_ | GOBP_REGULATION_OF_POSTSYNAPSE_OR | 110 | -0.38915 | -1.57139 | 0.00736377 | 0.21525623 | 1 | 2587 | tags=44%, list=29%, si | EN_2 |
| GOBP_REGULATION_OF_ | GOBP_REGULATION_OF_SMAD_PROTEIN_S | 34 | -0.47649 | -1.57134 | 0.023622047 | 0.21422137 | 1 | 2838 | tags=56%, list=32%, si | EN_2 |
| GOBP_POSITIVE_REGULA | GOBP_POSITIVE_REGULATION_OF_COLD_IN | 48 | -0.44299 | -1.57074 | 0.013803681 | 0.21403678 | 1 | 1377 | tags=27%, list=16%, si | EN_2 |
| GOBP_REGULATION_OF_ | GOBP_REGULATION_OF_RESPONSE_TO_NU | 16 | -0.56939 | -1.57009 | 0.032367975 | 0.21397619 | 1 | 660 | tags=31%, list=7%, sig | EN_2 |
| UBIQUITIN_MEDIATED_P | UBIQUITIN_MEDIATED_PROTEOLYSIS | 113 | -0.38829 | -1.57004 | 0.004297994 | 0.21291023 | 1 | 2263 | tags=35%, list=26%, si | EN_2 |
| GOBP_REGULATION_OF_ | GOBP_REGULATION_OF_ORGAN_GROWTH | 66 | -0.42224 | -1.56995 | 0.005934718 | 0.211957 | 1 | 1770 | tags=35%, list=20%, si | EN_2 |
| GOBP_REGULATION_OF_ | GOBP_REGULATION_OF_CELL_CYCLE_PHAS | 255 | -0.35177 | -1.56853 | 0 | 0.21321853 | 1 | 2170 | tags=35%, list=25%, si | EN_2 |
| GOBP_POSITIVE_REGULA | GOBP_POSITIVE_REGULATION_OF_STEM_C | 26 | -0.51133 | -1.56796 | 0.027350428 | 0.2131442 | 1 | 3259 | tags=69%, list=37%, si | EN_2 |
| GOBP_KERATINOCYTE_DI | GOBP_KERATINOCYTE_DIFFERENTIATION | 46 | -0.45369 | -1.56685 | 0.014610389 | 0.21367618 | 1 | 2502 | tags=52%, list=28%, si | EN_2 |
| GOBP_NEGATIVE_REGUL | GOBP_NEGATIVE_REGULATION_OF_TRANSL | 84 | -0.40129 | -1.5667 | 0.009146341 | 0.21276826 | 1 | 2290 | tags=38%, list=26%, si | EN_2 |
| CIRCADIAN_ENTRAINMEN | CIRCADIAN_ENTRAINMENT | 45 | -0.45757 | -1.56645 | 0.0203125 | 0.21206832 | 1 | 2262 | tags=38%, list=26%, si | EN_2 |
| GOBP_POSITIVE_REGULA | GOBP_POSITIVE_REGULATION_OF_CELLULA | 334 | -0.3387 | -1.56595 | 0 | 0.21179423 | 1 | 1779 | tags=29%, list=20%, si | EN_2 |
| GOBP_EMBRYONIC_CAM | GOBP_EMBRYONIC_CAMERA_TYPE_EYE_DE | 21 | -0.52825 | -1.56576 | 0.029565217 | 0.21108793 | 1 | 2980 | tags=67%, list=34%, si | EN_2 |
| GOBP_FOREBRAIN_DEVE | GOBP_FOREBRAIN_DEVELOPMENT | 214 | -0.35511 | -1.56477 | 0 | 0.2117825 | 1 | 2668 | tags=48%, list=30%, si | EN_2 |
| GOBP_RESPONSE_TO_INS | GOBP_RESPONSE_TO_INSULIN | 172 | -0.3636 | -1.56392 | 0.001371742 | 0.21198638 | 1 | 2720 | tags=43%, list=31%, si | EN_2 |
| GOBP_T_CELL_CYTOKINE | GOBP_T_CELL_CYTOKINE_PRODUCTION | 16 | -0.56742 | -1.56287 | 0.031088082 | 0.21282688 | 1 | 2386 | tags=56%, list=27%, si | EN_2 |
| GOBP_HORMONE_TRANS | GOBP_HORMONE_TRANSPORT | 166 | -0.37139 | -1.56197 | 0.002766252 | 0.21324901 | 1 | 1746 | tags=29%, list=20%, si | EN_2 |
| HUMAN_PAPILLOMAVIRU | HUMAN_PAPILLOMAVIRUS_INFECTION | 198 | -0.35799 | -1.56115 | 0.00270636 | 0.21361232 | 1 | 2632 | tags=42%, list=30%, si | EN_2 |
| GOBP_AMEBOIDAL_TYPE | GOBP_AMEBOIDAL_TYPE_CELL_MIGRATION | 268 | -0.34766 | -1.55898 | 0 | 0.2162877 | 1 | 2026 | tags=34%, list=23%, si | EN_2 |

| | | | | | | | | | | |
|---|---|---|---|---|---|---|---|---|---|---|
| GOBP_HOMOPHILIC_CELI | GOBP_HOMOPHILIC_CELL_ADHESION_VIA_I | 31 | -0.48235 | -1.55848 | 0.020933978 | 0.2161202 | 1 | 2500 | tags=55%, list=28%, si | EN_2 |
| GOBP_NEGATIVE_REGUL | GOBP_NEGATIVE_REGULATION_OF_MITOTI | 131 | -0.3731 | -1.55847 | 0.008784773 | 0.21507148 | 1 | 1653 | tags=31%, list=19%, si | EN_2 |
| GOBP_REGULATION_OF_ | GOBP_REGULATION_OF_SECRETION | 302 | -0.34118 | -1.55773 | 0 | 0.21517566 | 1 | 1696 | tags=26%, list=19%, si | EN_2 |
| GOBP_REGULATION_OF_ | GOBP_REGULATION_OF_EPITHELIAL_CELL_I | 77 | -0.41145 | -1.55543 | 0.007530121 | 0.21802129 | 1 | 2583 | tags=47%, list=29%, si | EN_2 |
| INSULIN_RESISTANCE | INSULIN_RESISTANCE | 59 | -0.42676 | -1.55458 | 0.017713366 | 0.21819218 | 1 | 2815 | tags=49%, list=32%, si | EN_2 |
| GOBP_TUBE_FORMATION | GOBP_TUBE_FORMATION | 110 | -0.38869 | -1.5528 | 0.00862069 | 0.22006038 | 1 | 2871 | tags=47%, list=32%, si | EN_2 |
| GOBP_PRIMARY_NEURAL | GOBP_PRIMARY_NEURAL_TUBE_FORMATIO | 85 | -0.40022 | -1.55249 | 0.009090909 | 0.21962386 | 1 | 3186 | tags=54%, list=36%, si | EN_2 |
| GOBP_AXO_DENDRITIC_T | GOBP_AXO_DENDRITIC_TRANSPORT | 68 | -0.41301 | -1.55198 | 0.013574661 | 0.21944065 | 1 | 2599 | tags=46%, list=29%, si | EN_2 |
| GOBP_POSITIVE_REGULA | GOBP_POSITIVE_REGULATION_OF_CELL_DIF | 494 | -0.32399 | -1.55149 | 0 | 0.21924767 | 1 | 2668 | tags=41%, list=30%, si | EN_2 |
| GOBP_MIRNA_METABOL | GOBP_MIRNA_METABOLIC_PROCESS | 71 | -0.41617 | -1.55057 | 0.015432099 | 0.21988796 | 1 | 2955 | tags=51%, list=33%, si | EN_2 |
| GOBP_LEUKOCYTE_DEGR | GOBP_LEUKOCYTE_DEGRANULATION | 23 | -0.51828 | -1.54938 | 0.042301185 | 0.2209634 | 1 | 1545 | tags=43%, list=17%, si | EN_2 |
| GOBP_ANIMAL_ORGAN_ | GOBP_ANIMAL_ORGAN_MORPHOGENESIS | 495 | -0.32471 | -1.5488 | 0 | 0.22082514 | 1 | 2656 | tags=43%, list=30%, si | EN_2 |
| GOBP_DEVELOPMENTAL_ | GOBP_DEVELOPMENTAL_PIGMENTATION | 33 | -0.4808 | -1.54817 | 0.024271844 | 0.22071452 | 1 | 1545 | tags=36%, list=17%, si | EN_2 |
| GOBP_ESTABLISHMENT_( | GOBP_ESTABLISHMENT_OF_TISSUE_POLARI | 21 | -0.52406 | -1.54812 | 0.035117056 | 0.21976066 | 1 | 1742 | tags=38%, list=20%, si | EN_2 |
| GOBP_NEURAL_TUBE_FO | GOBP_NEURAL_TUBE_FORMATION | 94 | -0.39308 | -1.54798 | 0.001451379 | 0.21898764 | 1 | 2447 | tags=40%, list=28%, si | EN_2 |
| COMPLEMENT_AND_COA | COMPLEMENT_AND_COAGULATION_CASCA | 15 | -0.59185 | -1.54698 | 0.043405674 | 0.21966639 | 1 | 266 | tags=27%, list=3%, sig | EN_2 |
| GOBP_SALIVARY_GLAND_ | GOBP_SALIVARY_GLAND_DEVELOPMENT | 18 | -0.53889 | -1.54577 | 0.029616725 | 0.22062337 | 1 | 2623 | tags=61%, list=30%, si | EN_2 |
| GOBP_MITOTIC_CELL_CY | GOBP_MITOTIC_CELL_CYCLE_PHASE_TRANS | 261 | -0.34283 | -1.54467 | 0.001269036 | 0.22135757 | 1 | 2266 | tags=36%, list=26%, si | EN_2 |
| GOBP_NUCLEAR_TRANSC | GOBP_NUCLEAR_TRANSCRIBED_MRNA_POL | 23 | -0.5283 | -1.5439 | 0.041597337 | 0.22151364 | 1 | 2189 | tags=52%, list=25%, si | EN_2 |
| GOBP_MAINTENANCE_OF | GOBP_MAINTENANCE_OF_CELL_NUMBER | 142 | -0.36606 | -1.54174 | 0.001404494 | 0.2240629 | 1 | 2649 | tags=39%, list=30%, si | EN_2 |
| GOBP_RESPONSE_TO_CA | GOBP_RESPONSE_TO_CARBOHYDRATE | 114 | -0.38061 | -1.54143 | 0.008583691 | 0.22366194 | 1 | 1746 | tags=30%, list=20%, si | EN_2 |
| AXON_GUIDANCE | AXON_GUIDANCE | 103 | -0.38887 | -1.541 | 0.005839416 | 0.22334562 | 1 | 2710 | tags=46%, list=31%, si | EN_2 |
| GOBP_MULTICELLULAR_( | GOBP_MULTICELLULAR_ORGANISMAL_REPF | 388 | -0.33012 | -1.5406 | 0 | 0.2230475 | 1 | 2343 | tags=36%, list=26%, si | EN_2 |
| GOBP_CHEMICAL_SYNAP | GOBP_CHEMICAL_SYNAPTIC_TRANSMISSIOI | 45 | -0.44316 | -1.53888 | 0.028239202 | 0.22502273 | 1 | 854 | tags=22%, list=10%, si | EN_2 |
| GOBP_NEGATIVE_REGUL | GOBP_NEGATIVE_REGULATION_OF_MAP_K | 26 | -0.50065 | -1.53827 | 0.02538071 | 0.22507882 | 1 | 1797 | tags=31%, list=20%, si | EN_2 |
| GOBP_COLLAGEN_FIBRIL | GOBP_COLLAGEN_FIBRIL_ORGANIZATION | 43 | -0.45274 | -1.53778 | 0.017628206 | 0.22491008 | 1 | 1379 | tags=30%, list=16%, si | EN_2 |
| GOBP_PROTEIN_ACYLATI | GOBP_PROTEIN_ACYLATION | 69 | -0.4106 | -1.53753 | 0.012084592 | 0.2243675 | 1 | 1677 | tags=30%, list=19%, si | EN_2 |
| GOBP_CENTRAL_NERVOU | GOBP_CENTRAL_NERVOUS_SYSTEM_PROJE( | 15 | -0.57671 | -1.53612 | 0.0411449 | 0.22573516 | 1 | 1715 | tags=47%, list=19%, si | EN_2 |
| GOBP_STEM_CELL_DIFFE | GOBP_STEM_CELL_DIFFERENTIATION | 151 | -0.36523 | -1.5359 | 0.002728513 | 0.22518358 | 1 | 2656 | tags=42%, list=30%, si | EN_2 |
| GOBP_REGULATION_OF_ | GOBP_REGULATION_OF_SYNAPTIC_TRANSN | 32 | -0.48257 | -1.5355 | 0.043902438 | 0.22479174 | 1 | 759 | tags=25%, list=9%, sig | EN_2 |
| GOBP_MAMMARY_GLAN | GOBP_MAMMARY_GLAND_DEVELOPMENT | 74 | -0.4092 | -1.53511 | 0.018045112 | 0.22445329 | 1 | 2838 | tags=54%, list=32%, si | EN_2 |
| GOBP_MULTICELLULAR_( | GOBP_MULTICELLULAR_ORGANISMAL_LEVE | 430 | -0.32371 | -1.53455 | 0 | 0.22441784 | 1 | 2207 | tags=35%, list=25%, si | EN_2 |
| GOBP_TEMPERATURE_H( | GOBP_TEMPERATURE_HOMEOSTASIS | 88 | -0.39609 | -1.53447 | 0.012326657 | 0.22357713 | 1 | 2392 | tags=40%, list=27%, si | EN_2 |
| GOBP_EAR_DEVELOPMEI | GOBP_EAR_DEVELOPMENT | 98 | -0.38408 | -1.53439 | 0.007320644 | 0.222744 | 1 | 2551 | tags=45%, list=29%, si | EN_2 |
| GOBP_SENSORY_ORGAN_ | GOBP_SENSORY_ORGAN_MORPHOGENESIS | 125 | -0.37361 | -1.53422 | 0.008333334 | 0.2220988 | 1 | 2647 | tags=46%, list=30%, si | EN_2 |
| GOBP_MITOCHONDRION_ | GOBP_MITOCHONDRION_DISTRIBUTION | 20 | -0.54819 | -1.53379 | 0.03448276 | 0.22193636 | 1 | 2472 | tags=60%, list=28%, si | EN_2 |
| GOBP_REGULATION_OF_ | GOBP_REGULATION_OF_TRANSMEMBRANE | 147 | -0.36774 | -1.53322 | 0.005509642 | 0.22196542 | 1 | 2890 | tags=49%, list=33%, si | EN_2 |
| GOBP_EPIDERMIS_DEVEL | GOBP_EPIDERMIS_DEVELOPMENT | 153 | -0.36353 | -1.53264 | 0.002832861 | 0.22197467 | 1 | 2648 | tags=46%, list=30%, si | EN_2 |
| GOBP_NERVE_DEVELOPN | GOBP_NERVE_DEVELOPMENT | 46 | -0.44781 | -1.53156 | 0.01369863 | 0.22271316 | 1 | 2056 | tags=48%, list=23%, si | EN_2 |
| GOBP_SIGNAL_RELEASE | GOBP_SIGNAL_RELEASE | 244 | -0.34384 | -1.53046 | 0.001293661 | 0.22368829 | 1 | 1891 | tags=30%, list=21%, si | EN_2 |
| GOBP_NEGATIVE_REGUL | GOBP_NEGATIVE_REGULATION_OF_ORGAN | 26 | -0.50411 | -1.52902 | 0.035117056 | 0.22510691 | 1 | 1770 | tags=46%, list=20%, si | EN_2 |
| GOBP_REGULATION_OF_ | GOBP_REGULATION_OF_SYNAPTIC_PLASTIC | 124 | -0.37044 | -1.52882 | 0.008510638 | 0.22452676 | 1 | 2579 | tags=42%, list=29%, si | EN_2 |
| GOBP_BRANCHING_INVO | GOBP_BRANCHING_INVOLVED_IN_BLOOD_ | 20 | -0.53003 | -1.52843 | 0.05892256 | 0.22424741 | 1 | 2656 | tags=80%, list=30%, si | EN_2 |
| GOBP_FOREBRAIN_NEUR | GOBP_FOREBRAIN_NEURON_DEVELOPMEN | 17 | -0.54998 | -1.52795 | 0.04730832 | 0.22424679 | 1 | 2512 | tags=65%, list=28%, si | EN_2 |
| GOBP_DEVELOPMENTAL_ | GOBP_DEVELOPMENTAL_CELL_GROWTH | 168 | -0.35511 | -1.52739 | 0 | 0.22422785 | 1 | 2628 | tags=42%, list=30%, si | EN_2 |
| GOBP_SPINAL_CORD_DE | GOBP_SPINAL_CORD_DEVELOPMENT | 36 | -0.45913 | -1.52733 | 0.032258064 | 0.22342318 | 1 | 2148 | tags=44%, list=24%, si | EN_2 |
| GOBP_CENTRAL_NERVOU | GOBP_CENTRAL_NERVOUS_SYSTEM_NEUR( | 24 | -0.51094 | -1.5268 | 0.03242321 | 0.22345231 | 1 | 1775 | tags=50%, list=20%, si | EN_2 |
| GOBP_POSITIVE_REGULA | GOBP_POSITIVE_REGULATION_OF_SMAD_P | 22 | -0.52268 | -1.52548 | 0.036392406 | 0.22471866 | 1 | 2838 | tags=59%, list=32%, si | EN_2 |
| GOBP_ERYTHROCYTE_MA | GOBP_ERYTHROCYTE_MATURATION | 15 | -0.58101 | -1.52543 | 0.042345278 | 0.22390355 | 1 | 1454 | tags=40%, list=16%, si | EN_2 |
| GOBP_MITOTIC_G2_DNA | GOBP_MITOTIC_G2_DNA_DAMAGE_CHECKI | 24 | -0.50842 | -1.52513 | 0.034305315 | 0.22349045 | 1 | 1998 | tags=46%, list=23%, si | EN_2 |
| GOBP_POSITIVE_REGULA | GOBP_POSITIVE_REGULATION_OF_INSULIN_ | 62 | -0.41774 | -1.52387 | 0.024767801 | 0.22457658 | 1 | 1615 | tags=31%, list=18%, si | EN_2 |
| GOBP_DIENCEPHALON_D | GOBP_DIENCEPHALON_DEVELOPMENT | 27 | -0.4854 | -1.52222 | 0.033492822 | 0.22647728 | 1 | 2807 | tags=56%, list=32%, si | EN_2 |
| GOBP_MONOATOMIC_AI | GOBP_MONOATOMIC_ANION_TRANSMEME | 24 | -0.50968 | -1.52148 | 0.03691275 | 0.22678794 | 1 | 2708 | tags=58%, list=31%, si | EN_2 |
| GOBP_EYE_MORPHOGEN | GOBP_EYE_MORPHOGENESIS | 82 | -0.39259 | -1.52124 | 0.007800312 | 0.22639251 | 1 | 2878 | tags=51%, list=33%, si | EN_2 |
| GOBP_ENDOCRINE_PANC | GOBP_ENDOCRINE_PANCREAS_DEVELOPME | 15 | -0.57382 | -1.5185 | 0.05821918 | 0.23013176 | 1 | 2420 | tags=53%, list=27%, si | EN_2 |
| GOBP_GLAND_DEVELOPN | GOBP_GLAND_DEVELOPMENT | 230 | -0.34175 | -1.51826 | 0.001285347 | 0.22965966 | 1 | 2860 | tags=47%, list=32%, si | EN_2 |
| GOBP_EXPORT_FROM_CE | GOBP_EXPORT_FROM_CELL | 460 | -0.32167 | -1.51805 | 0 | 0.2290347 | 1 | 1631 | tags=25%, list=18%, si | EN_2 |
| INSULIN_SIGNALING_PAT | INSULIN_SIGNALING_PATHWAY | 91 | -0.38865 | -1.51764 | 0.010542168 | 0.22877713 | 1 | 2551 | tags=40%, list=29%, si | EN_2 |
| GOBP_POSITIVE_REGULA | GOBP_POSITIVE_REGULATION_OF_CELL_DE | 286 | -0.33613 | -1.51689 | 0.00131579 | 0.22929937 | 1 | 2871 | tags=46%, list=32%, si | EN_2 |
| RAP1_SIGNALING_PATHV | RAP1_SIGNALING_PATHWAY | 102 | -0.38141 | -1.5164 | 0.00761035 | 0.22933227 | 1 | 1790 | tags=34%, list=20%, si | EN_2 |
| GOBP_THYMUS_DEVELOF | GOBP_THYMUS_DEVELOPMENT | 32 | -0.47011 | -1.5164 | 0.028938906 | 0.2284764 | 1 | 1700 | tags=44%, list=19%, si | EN_2 |
| GOBP_POSITIVE_REGULA | GOBP_POSITIVE_REGULATION_OF_RECEPTO | 17 | -0.55834 | -1.51581 | 0.046434496 | 0.22865847 | 1 | 1917 | tags=41%, list=22%, si | EN_2 |
| GOBP_CIRCADIAN_REGUI | GOBP_CIRCADIAN_REGULATION_OF_GENE_ | 54 | -0.42292 | -1.51535 | 0.015527951 | 0.22849528 | 1 | 2945 | tags=54%, list=33%, si | EN_2 |
| GOBP_FOCAL_ADHESION_ | GOBP_FOCAL_ADHESION_ASSEMBLY | 67 | -0.40925 | -1.51373 | 0.013533834 | 0.23033647 | 1 | 3548 | tags=60%, list=40%, si | EN_2 |
| GOBP_EXOCRINE_SYSTEN | GOBP_EXOCRINE_SYSTEM_DEVELOPMENT | 24 | -0.51052 | -1.51339 | 0.033707865 | 0.2300316 | 1 | 2623 | tags=54%, list=30%, si | EN_2 |
| B_CELL_RECEPTOR_SIGN | B_CELL_RECEPTOR_SIGNALING_PATHWAY | 41 | -0.45013 | -1.51334 | 0.038961038 | 0.22926535 | 1 | 2699 | tags=51%, list=31%, si | EN_2 |
| GOBP_REGULATION_OF_ | GOBP_REGULATION_OF_MITOTIC_CELL_CY( | 203 | -0.34724 | -1.51318 | 0.002717391 | 0.22872713 | 1 | 2266 | tags=37%, list=26%, si | EN_2 |
| GOBP_GAMETE_GENERAT | GOBP_GAMETE_GENERATION | 316 | -0.32786 | -1.51211 | 0 | 0.22967175 | 1 | 2318 | tags=35%, list=26%, si | EN_2 |
| GOBP_REGULATION_OF_ | GOBP_REGULATION_OF_POSTSYNAPTIC_ME | 45 | -0.44091 | -1.51183 | 0.041335452 | 0.22930047 | 1 | 854 | tags=22%, list=10%, si | EN_2 |
| GOBP_MESODERM_DEVE | GOBP_MESODERM_DEVELOPMENT | 55 | -0.42169 | -1.51167 | 0.024390243 | 0.22868153 | 1 | 2598 | tags=44%, list=29%, si | EN_2 |
| GOBP_POSITIVE_REGULA | GOBP_POSITIVE_REGULATION_OF_VASOCO | 16 | -0.55294 | -1.51156 | 0.032871973 | 0.22803845 | 1 | 1422 | tags=38%, list=16%, si | EN_2 |
| GOBP_AMIDE_TRANSPOF | GOBP_AMIDE_TRANSPORT | 158 | -0.3619 | -1.50995 | 0.008486563 | 0.22978322 | 1 | 1746 | tags=28%, list=20%, si | EN_2 |
| GOBP_NON_CANONICAL_ | GOBP_NON_CANONICAL_WNT_SIGNALING_ | 31 | -0.46741 | -1.50978 | 0.049441785 | 0.22927287 | 1 | 1982 | tags=42%, list=22%, si | EN_2 |
| GOBP_CHROMOSOME_O | GOBP_CHROMOSOME_ORGANIZATION_INV | 19 | -0.52909 | -1.50621 | 0.050653595 | 0.23441042 | 1 | 1811 | tags=37%, list=20%, si | EN_2 |
| ERBB_SIGNALING_PATHW | ERBB_SIGNALING_PATHWAY | 62 | -0.41401 | -1.50595 | 0.01674277 | 0.23404053 | 1 | 2551 | tags=47%, list=29%, si | EN_2 |
| GOBP_POSITIVE_REGULA | GOBP_POSITIVE_REGULATION_OF_DENDRIT | 52 | -0.42237 | -1.5042 | 0.029320987 | 0.23613608 | 1 | 3172 | tags=52%, list=36%, si | EN_2 |
| GOBP_MITOTIC_G2_M_T | GOBP_MITOTIC_G2_M_TRANSITION_CHECK | 35 | -0.45929 | -1.50278 | 0.046623792 | 0.23778051 | 1 | 2059 | tags=43%, list=23%, si | EN_2 |
| ALDOSTERONE-REGULATI | ALDOSTERONE-REGULATED_SODIUM_REAB | 19 | -0.52564 | -1.50131 | 0.049833886 | 0.23948954 | 1 | 2651 | tags=53%, list=30%, si | EN_2 |
| GOBP_RESPONSE_TO_OR | GOBP_RESPONSE_TO_ORGANOPHOSPHORU | 41 | -0.43579 | -1.4996 | 0.03125 | 0.24179223 | 1 | 1538 | tags=34%, list=17%, si | EN_2 |
| CHOLINERGIC_SYNAPSE | CHOLINERGIC_SYNAPSE | 59 | -0.41271 | -1.4988 | 0.031397175 | 0.242287 | 1 | 2262 | tags=44%, list=26%, si | EN_2 |
| GOBP_CELL_DIFFERENTIA | GOBP_CELL_DIFFERENTIATION_INVOLVED_I | 24 | -0.50374 | -1.49767 | 0.048622366 | 0.24332066 | 1 | 2959 | tags=71%, list=33%, si | EN_2 |
| GOBP_MELANOCYTE_DIF | GOBP_MELANOCYTE_DIFFERENTIATION | 19 | -0.53237 | -1.49695 | 0.057660624 | 0.24378751 | 1 | 2088 | tags=58%, list=24%, si | EN_2 |

| | | | | | | | | | | |
|---|---|---|---|---|---|---|---|---|---|---|
| GOBP_CELL_CELL_ADHES | GOBP_CELL_CELL_ADHESION | 374 | -0.32088 | -1.4956 | 0.001201923 | 0.24525394 | 1 | 1810 | tags=30%, list=20%, si | EN_2 |
| GOBP_EAR_MORPHOGEN | GOBP_EAR_MORPHOGENESIS | 49 | -0.4221 | -1.49559 | 0.02733119 | 0.24441533 | 1 | 2551 | tags=49%, list=29%, si | EN_2 |
| GOBP_LENS_FIBER_CELL_ | GOBP_LENS_FIBER_CELL_DIFFERENTIATION | 22 | -0.50548 | -1.49558 | 0.0375817 | 0.2435866 | 1 | 1374 | tags=36%, list=16%, si | EN_2 |
| CUSHING_SYNDROME | CUSHING_SYNDROME | 75 | -0.39096 | -1.4939 | 0.023291925 | 0.2456337 | 1 | 2454 | tags=43%, list=28%, si | EN_2 |
| GOBP_POSITIVE_REGULA | GOBP_POSITIVE_REGULATION_OF_MYOBLA | 31 | -0.47321 | -1.49386 | 0.03785489 | 0.24484167 | 1 | 3402 | tags=61%, list=38%, si | EN_2 |
| GOBP_MOLTING_CYCLE | GOBP_MOLTING_CYCLE | 67 | -0.39596 | -1.49228 | 0.031770047 | 0.24670711 | 1 | 2114 | tags=39%, list=24%, si | EN_2 |
| GOBP_CELL_SURFACE_RE | GOBP_CELL_SURFACE_RECEPTOR_PROTEIN_ | 338 | -0.32231 | -1.49226 | 0 | 0.24587761 | 1 | 2632 | tags=39%, list=30%, si | EN_2 |
| GOBP_REGULATION_OF_ | GOBP_REGULATION_OF_HORMONE_LEVELS | 226 | -0.33438 | -1.49185 | 0.001290323 | 0.24572116 | 1 | 1746 | tags=28%, list=20%, si | EN_2 |
| GOBP_ESTABLISHMENT_( | GOBP_ESTABLISHMENT_OF_EPITHELIAL_CEI | 24 | -0.49149 | -1.49144 | 0.055462185 | 0.2456707 | 1 | 694 | tags=29%, list=8%, sig | EN_2 |
| GOBP_HEART_GROWTH | GOBP_HEART_GROWTH | 66 | -0.40145 | -1.49085 | 0.032012194 | 0.24589483 | 1 | 2138 | tags=39%, list=24%, si | EN_2 |
| GOBP_REGULATION_OF_ | GOBP_REGULATION_OF_MUSCLE_CELL_DIF | 97 | -0.37713 | -1.49008 | 0.013867488 | 0.24652019 | 1 | 1746 | tags=33%, list=20%, si | EN_2 |
| GOBP_RESPONSE_TO_PE | GOBP_RESPONSE_TO_PEPTIDE_HORMONE | 244 | -0.33708 | -1.48968 | 0.003916449 | 0.24634105 | 1 | 2720 | tags=42%, list=31%, si | EN_2 |
| GOBP_NEURON_PROJECT | GOBP_NEURON_PROJECTION_EXTENSION | 115 | -0.37068 | -1.48914 | 0.011644833 | 0.24646224 | 1 | 2885 | tags=47%, list=33%, si | EN_2 |
| GOBP_DOPAMINE_META | GOBP_DOPAMINE_METABOLIC_PROCESS | 19 | -0.52647 | -1.48905 | 0.046901174 | 0.24579345 | 1 | 778 | tags=32%, list=9%, sig | EN_2 |
| GOBP_PROTEIN_LOCALIZ | GOBP_PROTEIN_LOCALIZATION_TO_EXTRAC | 220 | -0.33889 | -1.48891 | 0.002617801 | 0.24519037 | 1 | 1746 | tags=27%, list=20%, si | EN_2 |
| GOBP_LENS_MORPHOGE | GOBP_LENS_MORPHOGENESIS_IN_CAMERA | 17 | -0.54604 | -1.48818 | 0.052117262 | 0.24576272 | 1 | 2647 | tags=59%, list=30%, si | EN_2 |
| GOBP_SCHWANN_CELL_I | GOBP_SCHWANN_CELL_DIFFERENTIATION | 43 | -0.43855 | -1.48806 | 0.040387724 | 0.24511105 | 1 | 2835 | tags=56%, list=32%, si | EN_2 |
| GOBP_REGULATION_OF_ | GOBP_REGULATION_OF_BODY_FLUID_LEVE | 118 | -0.36834 | -1.48748 | 0.011764706 | 0.24534558 | 1 | 2088 | tags=35%, list=24%, si | EN_2 |
| GOBP_NEGATIVE_REGUL | GOBP_NEGATIVE_REGULATION_OF_TRANSI | 90 | -0.38199 | -1.48671 | 0.016200295 | 0.24576318 | 1 | 2661 | tags=47%, list=30%, si | EN_2 |
| GOBP_NEGATIVE_REGUL | GOBP_NEGATIVE_REGULATION_OF_SIGNAL | 15 | -0.56009 | -1.48669 | 0.051813472 | 0.24498357 | 1 | 654 | tags=33%, list=7%, sig | EN_2 |
| GOBP_STEM_CELL_PROLI | GOBP_STEM_CELL_PROLIFERATION | 66 | -0.40271 | -1.48657 | 0.023622047 | 0.2443766 | 1 | 3018 | tags=52%, list=34%, si | EN_2 |
| GOBP_POSITIVE_REGULA | GOBP_POSITIVE_REGULATION_OF_PROTEIN | 51 | -0.41745 | -1.48657 | 0.015360983 | 0.24358061 | 1 | 1630 | tags=31%, list=18%, si | EN_2 |
| GOBP_SECRETION | GOBP_SECRETION | 468 | -0.31357 | -1.48651 | 0 | 0.24286719 | 1 | 1891 | tags=28%, list=21%, si | EN_2 |
| GOBP_NEGATIVE_REGUL | GOBP_NEGATIVE_REGULATION_OF_TRANSF | 67 | -0.39997 | -1.48633 | 0.03343949 | 0.24240538 | 1 | 2661 | tags=49%, list=30%, si | EN_2 |
| GOBP_RESPONSE_TO_GR | GOBP_RESPONSE_TO_GROWTH_HORMONE | 17 | -0.52551 | -1.48492 | 0.0539629 | 0.24402502 | 1 | 1723 | tags=41%, list=19%, si | EN_2 |
| GOBP_HEPATICOBILIARY_ | GOBP_HEPATICOBILIARY_SYSTEM_DEVELOP | 81 | -0.38924 | -1.48431 | 0.012066365 | 0.24421024 | 1 | 2860 | tags=54%, list=32%, si | EN_2 |
| GOBP_MORPHOGENESIS_ | GOBP_MORPHOGENESIS_OF_EMBRYONIC_E | 109 | -0.36564 | -1.48382 | 0.019089574 | 0.24429129 | 1 | 2447 | tags=41%, list=28%, si | EN_2 |
| GOBP_VESICLE_TRANSPO | GOBP_VESICLE_TRANSPORT_ALONG_MICRO | 33 | -0.45288 | -1.48138 | 0.042207792 | 0.2474642 | 1 | 2599 | tags=52%, list=29%, si | EN_2 |
| GOBP_RESPIRATORY_SYS | GOBP_RESPIRATORY_SYSTEM_DEVELOPMEI | 144 | -0.34869 | -1.48066 | 0.012820513 | 0.24786356 | 1 | 2942 | tags=49%, list=33%, si | EN_2 |
| GOBP_HETEROPHILIC_CE | GOBP_HETEROPHILIC_CELL_CELL_ADHESION | 23 | -0.49561 | -1.48061 | 0.03583618 | 0.24717405 | 1 | 2434 | tags=74%, list=28%, si | EN_2 |
| GOBP_EXCITATORY_POST | GOBP_EXCITATORY_POSTSYNAPTIC_POTENT | 41 | -0.43956 | -1.47989 | 0.03497615 | 0.24763806 | 1 | 854 | tags=22%, list=10%, si | EN_2 |
| INSULIN_SECRETION | INSULIN_SECRETION | 33 | -0.45549 | -1.47901 | 0.055462185 | 0.24829157 | 1 | 1416 | tags=27%, list=16%, si | EN_2 |
| GOBP_SERTOLI_CELL_DIF | GOBP_SERTOLI_CELL_DIFFERENTIATION | 15 | -0.55352 | -1.47846 | 0.07321131 | 0.24847503 | 1 | 1718 | tags=47%, list=19%, si | EN_2 |
| GOBP_REGULATION_OF_ | GOBP_REGULATION_OF_NEURONAL_SYNAF | 37 | -0.4532 | -1.47798 | 0.048154093 | 0.24852675 | 1 | 958 | tags=27%, list=11%, si | EN_2 |
| GOBP_RESPONSE_TO_NE | GOBP_RESPONSE_TO_NERVE_GROWTH_FA( | 24 | -0.48674 | -1.47773 | 0.05050505 | 0.24818806 | 1 | 1628 | tags=42%, list=18%, si | EN_2 |
| GOBP_NEGATIVE_REGUL | GOBP_NEGATIVE_REGULATION_OF_STEM_( | 23 | -0.49336 | -1.47717 | 0.050161812 | 0.24838877 | 1 | 2505 | tags=52%, list=28%, si | EN_2 |
| GOBP_ADAPTIVE_THERM | GOBP_ADAPTIVE_THERMOGENESIS | 84 | -0.37328 | -1.47646 | 0.024709303 | 0.2489269 | 1 | 2392 | tags=37%, list=27%, si | EN_2 |
| GOBP_CELLULAR_RESPOI | GOBP_CELLULAR_RESPONSE_TO_NITROGEN | 334 | -0.3199 | -1.47625 | 0.001253133 | 0.24852926 | 1 | 2603 | tags=39%, list=29%, si | EN_2 |
| GOBP_NEUROMUSCULAR | GOBP_NEUROMUSCULAR_JUNCTION_DEVE | 39 | -0.44608 | -1.47563 | 0.041733548 | 0.24886648 | 1 | 1917 | tags=46%, list=22%, si | EN_2 |
| GOBP_SOMATIC_STEM_C | GOBP_SOMATIC_STEM_CELL_POPULATION_ | 47 | -0.41969 | -1.47552 | 0.03599374 | 0.24831733 | 1 | 2718 | tags=45%, list=31%, si | EN_2 |
| GOBP_POSITIVE_REGULA | GOBP_POSITIVE_REGULATION_OF_NEUROG | 182 | -0.33956 | -1.47486 | 0.005494506 | 0.24867508 | 1 | 2795 | tags=43%, list=32%, si | EN_2 |
| GOBP_SENSORY_SYSTEM | GOBP_SENSORY_SYSTEM_DEVELOPMENT | 212 | -0.33474 | -1.4747 | 0.00270636 | 0.24817511 | 1 | 2647 | tags=44%, list=30%, si | EN_2 |
| GOBP_CELLULAR_RESPOI | GOBP_CELLULAR_RESPONSE_TO_PEPTIDE_H | 190 | -0.34078 | -1.474 | 0.00130039 | 0.24867079 | 1 | 2648 | tags=40%, list=30%, si | EN_2 |
| GOBP_TELENCEPHALON_ | GOBP_TELENCEPHALON_DEVELOPMENT | 147 | -0.34802 | -1.47329 | 0.004213483 | 0.24901731 | 1 | 2668 | tags=45%, list=30%, si | EN_2 |
| GOBP_REGULATION_OF_ | GOBP_REGULATION_OF_NEURON_PROJECT | 328 | -0.32123 | -1.47328 | 0.002484472 | 0.24830386 | 1 | 2605 | tags=40%, list=29%, si | EN_2 |
| GOBP_REGULATION_OF_ | GOBP_REGULATION_OF_AXONOGENESIS | 100 | -0.37365 | -1.47326 | 0.01459854 | 0.2475844 | 1 | 2795 | tags=49%, list=32%, si | EN_2 |
| GOBP_MAMMARY_GLAN | GOBP_MAMMARY_GLAND_DUCT_MORPHO | 22 | -0.50458 | -1.47188 | 0.064080946 | 0.24915047 | 1 | 2838 | tags=59%, list=32%, si | EN_2 |
| GOBP_BODY_MORPHOGI | GOBP_BODY_MORPHOGENESIS | 46 | -0.4303 | -1.47064 | 0.031152649 | 0.24992794 | 1 | 2977 | tags=48%, list=34%, si | EN_2 |
| GOBP_POSITIVE_REGULA | GOBP_POSITIVE_REGULATION_OF_INSULIN | 25 | -0.48432 | -1.47036 | 0.041935485 | 0.24965055 | 1 | 1159 | tags=28%, list=13%, si | EN_2 |
| LONG-TERM_DEPRESSION | LONG-TERM_DEPRESSION | 34 | -0.45459 | -1.46979 | 0.057971016 | 0.24994318 | 1 | 2551 | tags=47%, list=29%, si | EN_2 |
| GOBP_REGULATION_OF_ | GOBP_REGULATION_OF_DEFENSE_RESPONS | 22 | -0.50788 | -1.4697 | 0.046075087 | 0.24937725 | 1 | 2619 | tags=55%, list=30%, si | EN_2 |
| GOBP_TISSUE_MIGRATIO | GOBP_TISSUE_MIGRATION | 186 | -0.33883 | -1.46939 | 0.005563282 | 0.2491577 | 1 | 2720 | tags=45%, list=31%, si | EN_2 |
| GOBP_STEM_CELL_DIVISI | GOBP_STEM_CELL_DIVISION | 25 | -0.47855 | -1.46629 | 0.040518638 | 0.24998277 | 1 | 2373 | tags=44%, list=27%, si | EN_2 |
| GOBP_EPIDERMIS_MORP | GOBP_EPIDERMIS_MORPHOGENESIS | 23 | -0.49374 | -1.46592 | 0.05059022 | 0.2498988 | 1 | 1563 | tags=35%, list=18%, si | EN_2 |
| GOBP_EMBRYONIC_ORG/ | GOBP_EMBRYONIC_ORGAN_DEVELOPMENT | 237 | -0.32923 | -1.46575 | 0.001293661 | 0.2495316 | 1 | 2656 | tags=41%, list=30%, si | EN_2 |
| GOBP_NEURON_RECOGN | GOBP_NEURON_RECOGNITION | 21 | -0.50966 | -1.46528 | 0.05263158 | 0.24965523 | 1 | 2857 | tags=52%, list=32%, si | EN_2 |
| GOBP_REGULATION_OF_ | GOBP_REGULATION_OF_HEART_GROWTH | 50 | -0.41249 | -1.46501 | 0.04043546 | 0.24948443 | 1 | 2959 | tags=54%, list=33%, si | EN_2 |
| GOBP_BIOLOGICAL_PROC | GOBP_BIOLOGICAL_PROCESS_INVOLVED_IN | 28 | -0.4627 | -1.4644 | 0.058419243 | 0.24988107 | 1 | 2190 | tags=43%, list=25%, si | EN_2 |
| GOBP_VIRAL_RNA_GENO | GOBP_VIRAL_RNA_GENOME_REPLICATION | 22 | -0.50374 | -1.46281 | 0.06632653 | 0.24986206 | 1 | 2190 | tags=45%, list=25%, si | EN_2 |
| GOBP_MIRNA_TRANSCRI | GOBP_MIRNA_TRANSCRIPTION | 50 | -0.41446 | -1.45917 | 0.03834356 | 0.24972105 | 1 | 2955 | tags=52%, list=33%, si | EN_2 |
| GOBP_CHAPERONE_MED | GOBP_CHAPERONE_MEDIATED_PROTEIN_F( | 52 | 0.633156 | 2.460414 | 0 | 0 | 0 | 819 | tags=33%, list=9%, sig | MM_2 |
| GOBP_PROTEIN_REFOLDI | GOBP_PROTEIN_REFOLDING | 20 | 0.77359 | 2.3369 | 0 | 0.001744138 | 0.004 | 606 | tags=50%, list=7%, sig | MM_2 |
| GOBP_ANTIVIRAL_INNAT | GOBP_ANTIVIRAL_INNATE_IMMUNE_RESPO | 32 | 0.655779 | 2.306262 | 0 | 0.003862584 | 0.013 | 883 | tags=41%, list=10%, si | MM_2 |
| GOBP_DE_NOVO_PROTEI | GOBP_DE_NOVO_PROTEIN_FOLDING | 34 | 0.633534 | 2.293525 | 0 | 0.003122408 | 0.014 | 737 | tags=32%, list=8%, sig | MM_2 |
| GOBP_RESPONSE_TO_IN1 | GOBP_RESPONSE_TO_INTERFERON_ALPHA | 20 | 0.759658 | 2.287174 | 0 | 0.002865391 | 0.016 | 792 | tags=45%, list=9%, sig | MM_2 |
| GOBP_PROTEIN_FOLDING | GOBP_PROTEIN_FOLDING | 127 | 0.493849 | 2.277178 | 0 | 0.002538928 | 0.017 | 1480 | tags=35%, list=17%, si | MM_2 |
| GOBP_NEGATIVE_REGUL | GOBP_NEGATIVE_REGULATION_OF_VIRAL_ | 26 | 0.660875 | 2.246924 | 0 | 0.003443754 | 0.026 | 1409 | tags=54%, list=16%, si | MM_2 |
| GOBP_ANTIMICROBIAL_H | GOBP_ANTIMICROBIAL_HUMORAL_IMMUN | 32 | 0.627862 | 2.244529 | 0 | 0.003013285 | 0.026 | 1396 | tags=47%, list=16%, si | MM_2 |
| RIBOSOME | RIBOSOME | 125 | 0.495584 | 2.231257 | 0 | 0.003472065 | 0.033 | 3262 | tags=72%, list=37%, si | MM_2 |
| GOBP_CHAPERONE_COF/ | GOBP_CHAPERONE_COFACTOR_DEPENDEN | 29 | 0.66749 | 2.229142 | 0 | 0.003124859 | 0.033 | 737 | tags=34%, list=8%, sig | MM_2 |
| TYPE_I_DIABETES_MELLIT | TYPE_I_DIABETES_MELLITUS | 20 | 0.701897 | 2.221739 | 0 | 0.003081312 | 0.036 | 1873 | tags=75%, list=21%, si | MM_2 |
| GOBP_ANTIMICROBIAL_H | GOBP_ANTIMICROBIAL_HUMORAL_RESPON | 40 | 0.60067 | 2.179699 | 0 | 0.005820497 | 0.073 | 1396 | tags=48%, list=16%, si | MM_2 |
| ANTIGEN_PROCESSING_A | ANTIGEN_PROCESSING_AND_PRESENTATIO | 45 | 0.581644 | 2.168375 | 0 | 0.006125909 | 0.083 | 1805 | tags=67%, list=21%, si | MM_2 |
| GOBP_REGULATION_OF_ | GOBP_REGULATION_OF_VIRAL_LIFE_CYCLE | 92 | 0.496895 | 2.162444 | 0 | 0.00594842 | 0.087 | 1409 | tags=34%, list=16%, si | MM_2 |
| ALLOGRAFT_REJECTION | ALLOGRAFT_REJECTION | 18 | 0.707494 | 2.140156 | 0 | 0.007278068 | 0.114 | 1727 | tags=72%, list=20%, si | MM_2 |
| GRAFT-VERSUS-HOST_DIS | GRAFT-VERSUS-HOST_DISEASE | 18 | 0.703424 | 2.113979 | 0 | 0.009609495 | 0.159 | 1873 | tags=78%, list=21%, si | MM_2 |
| GOBP_SUPPRESSION_OF_ | GOBP_SUPPRESSION_OF_VIRAL_RELEASE_B | 17 | 0.714192 | 2.100774 | 0 | 0.010786854 | 0.188 | 220 | tags=18%, list=3%, sig | MM_2 |
| GOBP_REGULATION_OF_ | GOBP_REGULATION_OF_LIPOPOLYSACCHAR | 22 | 0.680564 | 2.100764 | 0 | 0.010187583 | 0.188 | 1041 | tags=41%, list=12%, si | MM_2 |
| OXIDATIVE_PHOSPHORYL | OXIDATIVE_PHOSPHORYLATION | 99 | 0.474371 | 2.096108 | 0 | 0.010124686 | 0.197 | 2448 | tags=61%, list=28%, si | MM_2 |
| GOBP_DEFENSE_RESPON | GOBP_DEFENSE_RESPONSE_TO_VIRUS | 191 | 0.416619 | 2.079145 | 0 | 0.011943352 | 0.238 | 803 | tags=18%, list=9%, sig | MM_2 |
| GOBP_RESPONSE_TO_HE | GOBP_RESPONSE_TO_HEAT | 59 | 0.513776 | 2.077749 | 0 | 0.0114572 | 0.239 | 819 | tags=25%, list=9%, sig | MM_2 |
| GOBP_NEGATIVE_REGUL | GOBP_NEGATIVE_REGULATION_OF_VIRAL_I | 69 | 0.502103 | 2.056995 | 0 | 0.013627192 | 0.295 | 1462 | tags=38%, list=17%, si | MM_2 |

| | | | | | | | | | | |
|---|---|---|---|---|---|---|---|---|---|---|
| AUTOIMMUNE_THYROID_ | AUTOIMMUNE_THYROID_DISEASE | 17 | 0.698755 | 2.044551 | 0 | 0.015599017 | 0.343 | 1727 | tags=71%, list=20%, si | MM_2 |
| GOBP_REGULATION_OF_ | GOBP_REGULATION_OF_OXIDATIVE_PHOSP | 18 | 0.675554 | 2.028236 | 0 | 0.018350992 | 0.405 | 453 | tags=28%, list=5%, sig | MM_2 |
| PARKINSON_DISEASE | PARKINSON_DISEASE | 99 | 0.467024 | 2.026913 | 0 | 0.017800653 | 0.407 | 2448 | tags=56%, list=28%, si | MM_2 |
| GOBP_TRANSLATION_AT_ | GOBP_TRANSLATION_AT_SYNAPSE | 48 | 0.535883 | 2.022965 | 0 | 0.017700505 | 0.419 | 3417 | tags=83%, list=39%, si | MM_2 |
| GOBP_REGULATION_OF_ | GOBP_REGULATION_OF_BIOLOGICAL_PROC | 37 | 0.549523 | 2.002138 | 0 | 0.021977656 | 0.505 | 1829 | tags=49%, list=21%, si | MM_2 |
| GOBP_MODULATION_BY_ | GOBP_MODULATION_BY_SYMBIONT_OF_EN | 36 | 0.530457 | 1.939883 | 0 | 0.039489828 | 0.713 | 1829 | tags=47%, list=21%, si | MM_2 |
| GOBP_NEGATIVE_REGUL | GOBP_NEGATIVE_REGULATION_OF_VIRAL_ | 36 | 0.534826 | 1.930208 | 0.00304878 | 0.041633822 | 0.746 | 1350 | tags=39%, list=15%, si | MM_2 |
| GOBP_REGULATION_OF_ | GOBP_REGULATION_OF_VIRAL_GENOME_R | 57 | 0.484105 | 1.924845 | 0.003436426 | 0.042216185 | 0.764 | 1368 | tags=37%, list=16%, si | MM_2 |
| GOBP_OXIDATIVE_PHOSP | GOBP_OXIDATIVE_PHOSPHORYLATION | 107 | 0.427999 | 1.913955 | 0 | 0.045567486 | 0.799 | 2690 | tags=58%, list=31%, si | MM_2 |
| CYTOKINE-CYTOKINE_REC | CYTOKINE-CYTOKINE_RECEPTOR_INTERACTI | 87 | 0.428767 | 1.867484 | 0 | 0.06635569 | 0.91 | 1327 | tags=33%, list=15%, si | MM_2 |
| GOBP_RESPONSE_TO_VIR | GOBP_RESPONSE_TO_VIRUS | 227 | 0.37674 | 1.866838 | 0 | 0.06475516 | 0.912 | 803 | tags=15%, list=9%, sig | MM_2 |
| GOBP_REGULATION_OF_ | GOBP_REGULATION_OF_VIRAL_PROCESS | 114 | 0.417108 | 1.864146 | 0 | 0.06424947 | 0.916 | 1462 | tags=29%, list=17%, si | MM_2 |
| GOBP_RESPONSE_TO_FO | GOBP_RESPONSE_TO_FOOD | 15 | 0.620428 | 1.835191 | 0 | 0.081234574 | 0.962 | 916 | tags=33%, list=10%, si | MM_2 |
| GOBP_LIPOPOLYSACCHAR | GOBP_LIPOPOLYSACCHARIDE_MEDIATED_SI | 46 | 0.493353 | 1.834556 | 0.00304878 | 0.07942115 | 0.964 | 669 | tags=20%, list=8%, sig | MM_2 |
| GOBP_HUMORAL_IMMU | GOBP_HUMORAL_IMMUNE_RESPONSE | 74 | 0.428706 | 1.812442 | 0.003703704 | 0.091849856 | 0.985 | 1010 | tags=34%, list=12%, si | MM_2 |
| GOBP_MYELOID_DENDRI | GOBP_MYELOID_DENDRITIC_CELL_ACTIVAT | 24 | 0.541493 | 1.801877 | 0.01058201 | 0.09779382 | 0.991 | 1199 | tags=38%, list=14%, si | MM_2 |
| GOBP_MITOCHONDRIAL_ | GOBP_MITOCHONDRIAL_TRANSLATION | 112 | 0.398596 | 1.796377 | 0 | 0.09923519 | 0.993 | 2317 | tags=54%, list=26%, si | MM_2 |
| GOBP_NEGATIVE_REGUL | GOBP_NEGATIVE_REGULATION_OF_CYTOKI | 18 | 0.593686 | 1.794455 | 0.014204546 | 0.09879498 | 0.994 | 411 | tags=28%, list=5%, sig | MM_2 |
| GOBP_DNA_REPLICATION | GOBP_DNA_REPLICATION_INITIATION | 20 | 0.575504 | 1.780337 | 0.008426966 | 0.10714395 | 0.997 | 985 | tags=35%, list=11%, si | MM_2 |
| INTESTINAL_IMMUNE_NE | INTESTINAL_IMMUNE_NETWORK_FOR_IGA_ | 20 | 0.580685 | 1.779624 | 0.005899705 | 0.10523374 | 0.997 | 1809 | tags=55%, list=21%, si | MM_2 |
| GOBP_POSITIVE_REGULA | GOBP_POSITIVE_REGULATION_OF_CALCIUM | 36 | 0.485281 | 1.773507 | 0.006134969 | 0.107791506 | 0.998 | 884 | tags=31%, list=10%, si | MM_2 |
| GOBP_NADH_DEHYDROG | GOBP_NADH_DEHYDROGENASE_COMPLEX_ | 52 | 0.451881 | 1.772569 | 0 | 0.10621295 | 0.998 | 2545 | tags=52%, list=29%, si | MM_2 |
| GOBP_INTRINSIC_APOPT | GOBP_INTRINSIC_APOPTOTIC_SIGNALING_F | 37 | 0.49241 | 1.768466 | 0.006666667 | 0.10744077 | 0.998 | 1394 | tags=32%, list=16%, si | MM_2 |
| GOBP_PROTEIN_LOCALIZ | GOBP_PROTEIN_LOCALIZATION_TO_MITOCI | 81 | 0.4124 | 1.761584 | 0 | 0.11123962 | 0.998 | 1959 | tags=40%, list=22%, si | MM_2 |
| GOBP_RESPONSE_TO_TE | GOBP_RESPONSE_TO_TEMPERATURE_STIM | 92 | 0.409506 | 1.760968 | 0 | 0.10936748 | 0.998 | 819 | tags=20%, list=9%, sig | MM_2 |
| GOBP_NEGATIVE_REGUL | GOBP_NEGATIVE_REGULATION_OF_TYPE_II | 22 | 0.547908 | 1.754846 | 0.018087855 | 0.112621754 | 0.999 | 2083 | tags=41%, list=24%, si | MM_2 |
| PROTEIN_PROCESSING_IN | PROTEIN_PROCESSING_IN_ENDOPLASMIC_F | 145 | 0.371268 | 1.738529 | 0 | 0.12537235 | 0.999 | 1402 | tags=27%, list=16%, si | MM_2 |
| GOBP_SPERM_EGG_REC | GOBP_SPERM_EGG_RECOGNITION | 17 | 0.584679 | 1.731042 | 0.010869565 | 0.13048238 | 1 | 56 | tags=12%, list=1%, sig | MM_2 |
| GOBP_POSITIVE_REGULA | GOBP_POSITIVE_REGULATION_OF_CALCIUM | 54 | 0.438947 | 1.728856 | 0.006849315 | 0.13018952 | 1 | 1430 | tags=35%, list=16%, si | MM_2 |
| GOBP_CELLULAR_RESPON | GOBP_CELLULAR_RESPONSE_TO_HEAT | 37 | 0.494848 | 1.728622 | 0.006289308 | 0.12783982 | 1 | 819 | tags=24%, list=9%, sig | MM_2 |
| HUNTINGTON_DISEASE | HUNTINGTON_DISEASE | 139 | 0.367489 | 1.725242 | 0 | 0.1287262 | 1 | 2555 | tags=52%, list=29%, si | MM_2 |
| GOBP_REGULATION_OF_ | GOBP_REGULATION_OF_INTERLEUKIN_4_PF | 15 | 0.599412 | 1.721449 | 0.021052632 | 0.1297006 | 1 | 2341 | tags=73%, list=27%, si | MM_2 |
| GOBP_AEROBIC_ELECTRO | GOBP_AEROBIC_ELECTRON_TRANSPORT_CH | 56 | 0.43451 | 1.718872 | 0 | 0.13015035 | 1 | 2679 | tags=61%, list=31%, si | MM_2 |
| GOBP_ACTIVATION_OF_C | GOBP_ACTIVATION_OF_CYSTEINE_TYPE_EN | 55 | 0.435665 | 1.718433 | 0.006666667 | 0.12826303 | 1 | 1268 | tags=31%, list=14%, si | MM_2 |
| GOBP_COLLAGEN_FIBRIL | GOBP_COLLAGEN_FIBRIL_ORGANIZATION | 21 | 0.552538 | 1.717569 | 0.014409222 | 0.1267706 | 1 | 528 | tags=33%, list=6%, sig | MM_2 |
| GOBP_PROTEIN_TARGETI | GOBP_PROTEIN_TARGETING_TO_MITOCHO | 61 | 0.431762 | 1.713453 | 0.00295858 | 0.12853141 | 1 | 1959 | tags=43%, list=22%, si | MM_2 |
| GOBP_ANTIGEN_PROCES | GOBP_ANTIGEN_PROCESSING_AND_PRESEN | 27 | 0.527256 | 1.711617 | 0.014925373 | 0.12798104 | 1 | 1819 | tags=52%, list=21%, si | MM_2 |
| GOBP_SYMBIONT_ENTRY | GOBP_SYMBIONT_ENTRY_INTO_HOST | 60 | 0.419252 | 1.687588 | 0.009174312 | 0.15171164 | 1 | 1555 | tags=28%, list=18%, si | MM_2 |
| GOBP_ATP_SYNTHESIS_C | GOBP_ATP_SYNTHESIS_COUPLED_ELECTRO | 66 | 0.414716 | 1.668861 | 0 | 0.1707787 | 1 | 2679 | tags=58%, list=31%, si | MM_2 |
| CELL_CYCLE | CELL_CYCLE | 88 | 0.386103 | 1.66669 | 0.003344482 | 0.17088103 | 1 | 1045 | tags=22%, list=12%, si | MM_2 |
| DNA_REPLICATION | DNA_REPLICATION | 28 | 0.490251 | 1.653865 | 0.026706232 | 0.18440628 | 1 | 1892 | tags=50%, list=22%, si | MM_2 |
| GOBP_RESPONSE_TO_MU | GOBP_RESPONSE_TO_MURAMYL_DIPEPTID | 17 | 0.552816 | 1.653403 | 0.020618556 | 0.18212637 | 1 | 651 | tags=24%, list=7%, sig | MM_2 |
| GOBP_POSITIVE_REGULA | GOBP_POSITIVE_REGULATION_OF_CALCIUM | 16 | 0.581897 | 1.645423 | 0.025423728 | 0.18997228 | 1 | 777 | tags=25%, list=9%, sig | MM_2 |
| GOBP_VIRAL_LIFE_CYCLE | GOBP_VIRAL_LIFE_CYCLE | 167 | 0.340247 | 1.637747 | 0 | 0.19731505 | 1 | 1409 | tags=23%, list=16%, si | MM_2 |
| GOBP_NEGATIVE_REGUL | GOBP_NEGATIVE_REGULATION_OF_EPITHE | 35 | 0.450453 | 1.633651 | 0.017804155 | 0.19973809 | 1 | 1232 | tags=26%, list=14%, si | MM_2 |
| INFLAMMATORY_BOWEL | INFLAMMATORY_BOWEL_DISEASE_(IBD) | 31 | 0.465832 | 1.621057 | 0.048543688 | 0.21443078 | 1 | 1873 | tags=55%, list=21%, si | MM_2 |
| GOBP_RESPONSE_TO_INT | GOBP_RESPONSE_TO_INTERFERON_BETA | 43 | 0.43445 | 1.619932 | 0.015822785 | 0.21281241 | 1 | 792 | tags=30%, list=9%, sig | MM_2 |
| GOBP_POSITIVE_REGULA | GOBP_POSITIVE_REGULATION_OF_FIBROBL | 41 | 0.45115 | 1.618025 | 0.012345679 | 0.21243547 | 1 | 1437 | tags=34%, list=16%, si | MM_2 |
| GOBP_BINDING_OF_SPER | GOBP_BINDING_OF_SPERM_TO_ZONA_PEL | 15 | 0.564762 | 1.617116 | 0.050531916 | 0.21083419 | 1 | 2411 | tags=47%, list=28%, si | MM_2 |
| GOBP_VIRAL_GENOME_R | GOBP_VIRAL_GENOME_REPLICATION | 88 | 0.378687 | 1.614403 | 0.003717472 | 0.21175532 | 1 | 1368 | tags=27%, list=16%, si | MM_2 |
| GOBP_KILLING_OF_CELLS | GOBP_KILLING_OF_CELLS_OF_ANOTHER_OF | 29 | 0.47506 | 1.612969 | 0.020231213 | 0.21085979 | 1 | 778 | tags=28%, list=9%, sig | MM_2 |
| GOBP_ESTABLISHMENT_C | GOBP_ESTABLISHMENT_OF_PROTEIN_LOCA | 21 | 0.503802 | 1.593152 | 0.037037037 | 0.23669972 | 1 | 1941 | tags=52%, list=22%, si | MM_2 |
| GOBP_RESPONSE_TO_TY | GOBP_RESPONSE_TO_TYPE_I_INTERFERON | 48 | 0.412627 | 1.592368 | 0.021943573 | 0.23463897 | 1 | 1462 | tags=29%, list=17%, si | MM_2 |
| GOBP_REGULATION_OF_ | GOBP_REGULATION_OF_AEROBIC_RESPIRAT | 28 | 0.453595 | 1.583457 | 0.015060241 | 0.24537534 | 1 | 453 | tags=18%, list=5%, sig | MM_2 |
| NON-ALCOHOLIC_FATTY_ | NON-ALCOHOLIC_FATTY_LIVER_DISEASE_(N | 118 | 0.341671 | 1.582132 | 0 | 0.24417777 | 1 | 2448 | tags=47%, list=28%, si | MM_2 |
| GOBP_ANTIGEN_PROCES | GOBP_ANTIGEN_PROCESSING_AND_PRESEN | 45 | 0.419803 | 1.580841 | 0.015479876 | 0.2433764 | 1 | 1884 | tags=51%, list=22%, si | MM_2 |
| GOBP_INCLUSION_BODY_ | GOBP_INCLUSION_BODY_ASSEMBLY | 18 | 0.533088 | 1.578651 | 0.02549575 | 0.24408701 | 1 | 1410 | tags=44%, list=16%, si | MM_2 |
| GOBP_NEGATIVE_REGUL | GOBP_NEGATIVE_REGULATION_OF_RHO_PI | 15 | -0.79351 | -2.11211 | 0 | 0.07310724 | 0.067 | 1227 | tags=53%, list=14%, si | MM_2 |
| GOBP_POSITIVE_REGULA | GOBP_POSITIVE_REGULATION_OF_CELLULA | 18 | -0.663 | -2.14994 | 0 | 0.11354039 | 0.073 | 1052 | tags=50%, list=20%, si | MM_3 |
| GOBP_CELLULAR_RESPON | GOBP_CELLULAR_RESPONSE_TO_TYPE_II_IN | 39 | 0.644408 | 2.048492 | 0 | 0.018497823 | 0.026 | 323 | tags=31%, list=7%, sig | EN_4 |
| GOBP_RESPONSE_TO_TY | GOBP_RESPONSE_TO_TYPE_II_INTERFERON | 43 | 0.628574 | 2.010613 | 0 | 0.021966796 | 0.059 | 323 | tags=28%, list=7%, sig | EN_4 |
| CALCIUM_SIGNALING_PA | CALCIUM_SIGNALING_PATHWAY | 51 | -0.63102 | -2.25471 | 0 | 0.007414431 | 0.005 | 845 | tags=35%, list=18%, si | EN_4 |
| INSULIN_SECRETION | INSULIN_SECRETION | 28 | -0.6844 | -2.18087 | 0 | 0.008165275 | 0.011 | 1160 | tags=71%, list=25%, si | EN_4 |
| GOBP_REGULATION_OF_ | GOBP_REGULATION_OF_HEART_RATE | 30 | -0.63758 | -2.04849 | 0 | 0.0570929 | 0.111 | 624 | tags=37%, list=13%, si | EN_4 |
| GOBP_CARDIAC_CONDUC | GOBP_CARDIAC_CONDUCTION | 20 | -0.69096 | -2.01212 | 0 | 0.072882004 | 0.18 | 208 | tags=30%, list=4%, sig | EN_4 |
| GOBP_REGULATION_OF_ | GOBP_REGULATION_OF_ACTION_POTENTIA | 21 | -0.68455 | -1.99801 | 0 | 0.06993333 | 0.215 | 295 | tags=24%, list=6%, sig | EN_4 |
| GOBP_CARDIAC_MUSCLE | GOBP_CARDIAC_MUSCLE_CELL_CONTRACTI | 30 | -0.61618 | -1.97201 | 0.002364066 | 0.081730075 | 0.281 | 208 | tags=20%, list=4%, sig | EN_4 |
| GOBP_OUTFLOW_TRACT_ | GOBP_OUTFLOW_TRACT_MORPHOGENESIS | 35 | -0.59107 | -1.96037 | 0 | 0.08411596 | 0.327 | 591 | tags=29%, list=12%, si | EN_4 |
| GOBP_REGULATION_OF_ | GOBP_REGULATION_OF_STRIATED_MUSCLE | 30 | -0.59518 | -1.93127 | 0 | 0.107474424 | 0.433 | 1301 | tags=53%, list=27%, si | EN_4 |
| GOBP_HEART_MORPHOG | GOBP_HEART_MORPHOGENESIS | 99 | -0.48136 | -1.92426 | 0 | 0.10310952 | 0.459 | 511 | tags=23%, list=11%, si | EN_4 |
| GOBP_CARDIAC_SEPTUM | GOBP_CARDIAC_SEPTUM_MORPHOGENESIS | 33 | -0.60011 | -1.92272 | 0 | 0.09443584 | 0.461 | 591 | tags=30%, list=12%, si | EN_4 |
| GOBP_CARDIAC_VENTRIC | GOBP_CARDIAC_VENTRICLE_DEVELOPMENT | 53 | -0.52722 | -1.90741 | 0 | 0.10272265 | 0.524 | 563 | tags=25%, list=12%, si | EN_4 |
| CIRCADIAN_ENTRAINMEN | CIRCADIAN_ENTRAINMENT | 37 | -0.5689 | -1.8953 | 0 | 0.10761249 | 0.579 | 1270 | tags=49%, list=27%, si | EN_4 |
| GOBP_STRIATED_MUSCL | GOBP_STRIATED_MUSCLE_CONTRACTION | 54 | -0.52783 | -1.89405 | 0 | 0.10035534 | 0.583 | 1250 | tags=44%, list=26%, si | EN_4 |
| GOBP_CARDIAC_MUSCLE | GOBP_CARDIAC_MUSCLE_CONTRACTION | 46 | -0.53056 | -1.89256 | 0 | 0.094591536 | 0.586 | 282 | tags=17%, list=6%, sig | EN_4 |
| GOBP_ACTIN_FILAMENT_ | GOBP_ACTIN_FILAMENT_BASED_MOVEMEN | 51 | -0.52297 | -1.88718 | 0 | 0.093515806 | 0.606 | 658 | tags=25%, list=14%, si | EN_4 |
| RENIN_SECRETION | RENIN_SECRETION | 24 | -0.63856 | -1.88166 | 0.006896552 | 0.092773736 | 0.632 | 1270 | tags=63%, list=27%, si | EN_4 |
| GOBP_ACTIN_MEDIATED_ | GOBP_ACTIN_MEDIATED_CELL_CONTRACTI | 42 | -0.53883 | -1.8647 | 0 | 0.103417456 | 0.69 | 1305 | tags=45%, list=28%, si | EN_4 |
| GOBP_VISUAL_BEHAVIOR | GOBP_VISUAL_BEHAVIOR | 26 | -0.60009 | -1.84006 | 0 | 0.12641768 | 0.782 | 793 | tags=46%, list=17%, si | EN_4 |
| GOBP_ENDOCARDIAL_CU | GOBP_ENDOCARDIAL_CUSHION_MORPHOG | 15 | -0.70165 | -1.83996 | 0 | 0.119842 | 0.782 | 511 | tags=33%, list=11%, si | EN_4 |
| GOBP_REGULATION_OF_ | GOBP_REGULATION_OF_TISSUE_REMODELI | 25 | -0.59788 | -1.83785 | 0 | 0.11637954 | 0.789 | 580 | tags=36%, list=12%, si | EN_4 |
| GOBP_REGULATION_OF_ | GOBP_REGULATION_OF_MUSCLE_CONTRAC | 51 | -0.50462 | -1.82439 | 0 | 0.12670504 | 0.827 | 816 | tags=33%, list=17%, si | EN_4 |

| | | | | | | | | | | |
|---|---|---|---|---|---|---|---|---|---|---|
| GOBP_ENDOCARDIAL_CU | GOBP_ENDOCARDIAL_CUSHION_DEVELOPM | 20 | -0.62289 | -1.81852 | 0.006880734 | 0.12830058 | 0.845 | 511 | tags=30%, list=11%, si | EN_4 |
| GOBP_MUSCLE_CONTRA | GOBP_MUSCLE_CONTRACTION | 95 | -0.44478 | -1.81472 | 0 | 0.1273012 | 0.858 | 816 | tags=28%, list=17%, si | EN_4 |
| GOBP_CARDIAC_MUSCLE | GOBP_CARDIAC_MUSCLE_CELL_ACTION_PO | 22 | -0.60759 | -1.80949 | 0.004662005 | 0.12972927 | 0.872 | 1309 | tags=59%, list=28%, si | EN_4 |
| GOBP_REGULATION_OF_ | GOBP_REGULATION_OF_CARDIAC_MUSCLE | 27 | -0.58545 | -1.79954 | 0.009779952 | 0.13803531 | 0.889 | 1301 | tags=52%, list=27%, si | EN_4 |
| GOBP_REGULATION_OF_ | GOBP_REGULATION_OF_BLOOD_CIRCULATI | 83 | -0.46193 | -1.798 | 0 | 0.13518265 | 0.89 | 1253 | tags=46%, list=26%, si | EN_4 |
| GOBP_POSITIVE_REGULA | GOBP_POSITIVE_REGULATION_OF_NEURAL_ | 25 | -0.58574 | -1.79793 | 0.004962779 | 0.13023227 | 0.89 | 438 | tags=28%, list=9%, sig | EN_4 |
| GOBP_TRANSFORMING_( | GOBP_TRANSFORMING_GROWTH_FACTOR_ | 17 | -0.64851 | -1.79182 | 0.004494382 | 0.1337109 | 0.896 | 418 | tags=35%, list=9%, sig | EN_4 |
| GOBP_RESPONSE_TO_PU | GOBP_RESPONSE_TO_PURINE_CONTAINING | 34 | -0.54305 | -1.78764 | 0 | 0.13448054 | 0.906 | 1249 | tags=50%, list=26%, si | EN_4 |
| GOBP_CELL_FATE_SPECIF | GOBP_CELL_FATE_SPECIFICATION | 32 | -0.54105 | -1.77864 | 0.005089059 | 0.14171425 | 0.934 | 722 | tags=28%, list=15%, si | EN_4 |
| GOBP_AMINO_ACID_BIO | GOBP_AMINO_ACID_BIOSYNTHETIC_PROCE | 15 | -0.6839 | -1.77639 | 0.004454343 | 0.13953173 | 0.938 | 744 | tags=47%, list=16%, si | EN_4 |
| GOBP_REGULATION_OF_ | GOBP_REGULATION_OF_HEART_CONTRACT | 63 | -0.4777 | -1.77613 | 0.005050505 | 0.13585976 | 0.939 | 697 | tags=29%, list=15%, si | EN_4 |
| GOBP_SPINAL_CORD_DE | GOBP_SPINAL_CORD_DEVELOPMENT | 17 | -0.63476 | -1.77444 | 0.00896861 | 0.13421172 | 0.945 | 833 | tags=41%, list=18%, si | EN_4 |
| GOBP_POSITIVE_REGULA | GOBP_POSITIVE_REGULATION_OF_CALCIUN | 29 | -0.56103 | -1.77442 | 0.009389672 | 0.13030611 | 0.945 | 619 | tags=24%, list=13%, si | EN_4 |
| GOBP_SENSORY_PERCEP | GOBP_SENSORY_PERCEPTION_OF_MECHAN | 54 | -0.48939 | -1.7715 | 0 | 0.13053845 | 0.949 | 670 | tags=31%, list=14%, si | EN_4 |
| GOBP_REGULATION_OF_ | GOBP_REGULATION_OF_NEUROTRANSMITT | 21 | -0.6038 | -1.77009 | 0.004651163 | 0.12930547 | 0.953 | 580 | tags=43%, list=12%, si | EN_4 |
| GOBP_CARDIAC_CHAMBE | GOBP_CARDIAC_CHAMBER_MORPHOGENES | 50 | -0.49992 | -1.76925 | 0 | 0.1270052 | 0.954 | 591 | tags=24%, list=12%, si | EN_4 |
| GOBP_DEVELOPMENTAL_ | GOBP_DEVELOPMENTAL_PIGMENTATION | 17 | -0.63909 | -1.75782 | 0.004694836 | 0.13908061 | 0.964 | 933 | tags=47%, list=20%, si | EN_4 |
| GOBP_CELL_COMMUNIC | GOBP_CELL_COMMUNICATION_INVOLVED_ | 15 | -0.67281 | -1.75668 | 0.006666667 | 0.13737246 | 0.964 | 208 | tags=27%, list=4%, sig | EN_4 |
| GOBP_REGULATION_OF_ | GOBP_REGULATION_OF_SYSTEM_PROCESS | 183 | -0.40294 | -1.75626 | 0 | 0.13453285 | 0.964 | 833 | tags=28%, list=18%, si | EN_4 |
| ALDOSTERONE_SYNTHES | ALDOSTERONE_SYNTHESIS_AND_SECRETION | 37 | -0.53088 | -1.75201 | 0.002631579 | 0.13674255 | 0.968 | 1223 | tags=54%, list=26%, si | EN_4 |
| GLUTAMATERGIC_SYNAP | GLUTAMATERGIC_SYNAPSE | 33 | -0.54293 | -1.75159 | 0.002415459 | 0.13390495 | 0.968 | 1071 | tags=42%, list=23%, si | EN_4 |
| GOBP_REGULATION_OF_ | GOBP_REGULATION_OF_OSSIFICATION | 43 | -0.51262 | -1.75024 | 0 | 0.13265263 | 0.97 | 487 | tags=21%, list=10%, si | EN_4 |
| GOBP_HEAD_DEVELOPM | GOBP_HEAD_DEVELOPMENT | 249 | -0.37931 | -1.75001 | 0 | 0.13008274 | 0.97 | 865 | tags=29%, list=18%, si | EN_4 |
| PANCREATIC_SECRETION | PANCREATIC_SECRETION | 26 | -0.56153 | -1.74963 | 0.002288329 | 0.12758794 | 0.971 | 1090 | tags=50%, list=23%, si | EN_4 |
| GOBP_VENTRICULAR_SEI | GOBP_VENTRICULAR_SEPTUM_MORPHOGE | 22 | -0.58455 | -1.7453 | 0.011441648 | 0.12938893 | 0.971 | 487 | tags=27%, list=10%, si | EN_4 |
| GOBP_ESTABLISHMENT_ | GOBP_ESTABLISHMENT_OF_MITOTIC_SPINI | 18 | -0.62007 | -1.74197 | 0.004739337 | 0.13059832 | 0.974 | 504 | tags=44%, list=11%, si | EN_4 |
| GOBP_REGULATION_OF_ | GOBP_REGULATION_OF_CARDIAC_MUSCLE_ | 17 | -0.62673 | -1.74187 | 0.013793103 | 0.12797134 | 0.974 | 1301 | tags=59%, list=27%, si | EN_4 |
| SALIVARY_SECRETION | SALIVARY_SECRETION | 28 | -0.55587 | -1.73684 | 0.01330377 | 0.13163522 | 0.979 | 1090 | tags=57%, list=23%, si | EN_4 |
| GOBP_NEGATIVE_REGUL | GOBP_NEGATIVE_REGULATION_OF_SMALL_ | 24 | -0.57479 | -1.73666 | 0.004672897 | 0.12921081 | 0.979 | 1240 | tags=46%, list=26%, si | EN_4 |
| GOBP_PIGMENT_CELL_D | GOBP_PIGMENT_CELL_DIFFERENTIATION | 15 | -0.64428 | -1.73617 | 0.006864989 | 0.12722407 | 0.98 | 933 | tags=47%, list=20%, si | EN_4 |
| GOBP_REGULATION_OF_ | GOBP_REGULATION_OF_ACTIN_FILAMENT_ | 19 | -0.59761 | -1.73268 | 0.011185682 | 0.12905946 | 0.983 | 1301 | tags=53%, list=27%, si | EN_4 |
| GOBP_REGULATION_OF_ | GOBP_REGULATION_OF_STEM_CELL_DIFFEI | 45 | -0.49696 | -1.72937 | 0.005037783 | 0.13076155 | 0.986 | 779 | tags=36%, list=16%, si | EN_4 |
| GASTRIC_ACID_SECRETIO | GASTRIC_ACID_SECRETION | 25 | -0.5609 | -1.72802 | 0.00257732 | 0.13008447 | 0.987 | 1223 | tags=56%, list=26%, si | EN_4 |
| GOBP_REGULATION_OF_ | GOBP_REGULATION_OF_RELEASE_OF_SEQU | 26 | -0.55104 | -1.72573 | 0.002463054 | 0.13052344 | 0.989 | 1250 | tags=42%, list=26%, si | EN_4 |
| GOBP_PRIMARY_NEURAL | GOBP_PRIMARY_NEURAL_TUBE_FORMATIO | 42 | -0.4941 | -1.72428 | 0 | 0.12976436 | 0.99 | 728 | tags=29%, list=15%, si | EN_4 |
| GOBP_ESTABLISHMENT_ | GOBP_ESTABLISHMENT_OF_SPINDLE_ORIEN | 17 | -0.62282 | -1.72397 | 0.011415525 | 0.12793447 | 0.99 | 484 | tags=41%, list=10%, si | EN_4 |
| LEUKOCYTE_TRANSENDO | LEUKOCYTE_TRANSENDOTHELIAL_MIGRATI | 41 | -0.49853 | -1.72393 | 0.002457002 | 0.12580393 | 0.99 | 1157 | tags=39%, list=24%, si | EN_4 |
| GOBP_NEUROBLAST_PRO | GOBP_NEUROBLAST_PROLIFERATION | 36 | -0.52005 | -1.72207 | 0.004962779 | 0.12607786 | 0.99 | 660 | tags=28%, list=14%, si | EN_4 |
| GOBP_L_ALPHA_AMINO_ | GOBP_L_ALPHA_AMINO_ACID_TRANSMEM | 15 | -0.65403 | -1.7175 | 0.016055046 | 0.12933058 | 0.996 | 236 | tags=33%, list=5%, sig | EN_4 |
| GOBP_ASSOCIATIVE_LEA | GOBP_ASSOCIATIVE_LEARNING | 32 | -0.52491 | -1.71503 | 0.004842615 | 0.13007765 | 0.996 | 793 | tags=41%, list=17%, si | EN_4 |
| GOBP_VENTRICULAR_SEI | GOBP_VENTRICULAR_SEPTUM_DEVELOPME | 35 | -0.51971 | -1.71481 | 0.004889976 | 0.12812015 | 0.996 | 563 | tags=26%, list=12%, si | EN_4 |
| GOBP_POSITIVE_REGULA | GOBP_POSITIVE_REGULATION_OF_EPITHELI | 16 | -0.62866 | -1.71307 | 0.012315271 | 0.12825263 | 0.997 | 660 | tags=38%, list=14%, si | EN_4 |
| GOBP_NEGATIVE_REGUL | GOBP_NEGATIVE_REGULATION_OF_MAP_K | 16 | -0.61964 | -1.71202 | 0.006578947 | 0.12760918 | 0.998 | 580 | tags=38%, list=12%, si | EN_4 |
| GOBP_BODY_MORPHOG | GOBP_BODY_MORPHOGENESIS | 37 | -0.51685 | -1.71107 | 0.005167959 | 0.12652838 | 0.998 | 1460 | tags=54%, list=31%, si | EN_4 |
| ARRHYTHMOGENIC_RIGH | ARRHYTHMOGENIC_RIGHT_VENTRICULAR_ | 35 | -0.52243 | -1.71057 | 0.01010101 | 0.12523788 | 0.998 | 780 | tags=31%, list=16%, si | EN_4 |
| GOBP_LUNG_ALVEOLUS_ | GOBP_LUNG_ALVEOLUS_DEVELOPMENT | 26 | -0.54877 | -1.70945 | 0.006578947 | 0.12471861 | 0.998 | 820 | tags=46%, list=17%, si | EN_4 |
| GOBP_REGULATION_OF_ | GOBP_REGULATION_OF_TRANSMEMBRANE | 155 | -0.38795 | -1.70814 | 0 | 0.12439322 | 0.998 | 649 | tags=22%, list=14%, si | EN_4 |
| GOBP_EPITHELIAL_TUBE | GOBP_EPITHELIAL_TUBE_MORPHOGENESIS | 127 | -0.41183 | -1.70499 | 0 | 0.12606254 | 0.998 | 660 | tags=25%, list=14%, si | EN_4 |
| GOBP_REGULATION_OF_ | GOBP_REGULATION_OF_MULTICELLULAR_C | 33 | -0.52011 | -1.7019 | 0.010256411 | 0.12745921 | 0.998 | 800 | tags=33%, list=17%, si | EN_4 |
| GOBP_RESPONSE_TO_OR | GOBP_RESPONSE_TO_ORGANOPHOSPHORU | 28 | -0.54838 | -1.69743 | 0.011655011 | 0.13029854 | 0.999 | 1468 | tags=61%, list=31%, si | EN_4 |
| GOBP_EMBRYONIC_ORG | GOBP_EMBRYONIC_ORGAN_MORPHOGENE | 85 | -0.42576 | -1.69099 | 0 | 0.13563024 | 1 | 760 | tags=31%, list=16%, si | EN_4 |
| ADRENERGIC_SIGNALING | ADRENERGIC_SIGNALING_IN_CARDIOMYOC | 57 | -0.46122 | -1.68361 | 0.012254902 | 0.14253503 | 1 | 1160 | tags=40%, list=25%, si | EN_4 |
| GABAERGIC_SYNAPSE | GABAERGIC_SYNAPSE | 25 | -0.55672 | -1.68202 | 0.013513514 | 0.14277294 | 1 | 657 | tags=32%, list=14%, si | EN_4 |
| GOBP_REGULATION_OF_ | GOBP_REGULATION_OF_ORGANIC_ACID_TF | 22 | -0.57157 | -1.67337 | 0.018691588 | 0.15147048 | 1 | 758 | tags=36%, list=16%, si | EN_4 |
| GOBP_BIOLOGICAL_PRO | GOBP_BIOLOGICAL_PROCESS_INVOLVED_IN | 16 | -0.60827 | -1.67259 | 0.02247191 | 0.15049143 | 1 | 825 | tags=38%, list=17%, si | EN_4 |
| GOBP_BRANCHING_MOR | GOBP_BRANCHING_MORPHOGENESIS_OF_ | 61 | -0.44577 | -1.67117 | 0.0025 | 0.15017526 | 1 | 660 | tags=31%, list=14%, si | EN_4 |
| GOBP_MESENCHYME_M | GOBP_MESENCHYME_MORPHOGENESIS | 20 | -0.56868 | -1.66951 | 0.005037783 | 0.15013409 | 1 | 511 | tags=25%, list=11%, si | EN_4 |
| GOBP_REGULATION_OF_ | GOBP_REGULATION_OF_NEUROBLAST_PRO | 21 | -0.57733 | -1.66873 | 0.018181818 | 0.1491729 | 1 | 660 | tags=33%, list=14%, si | EN_4 |
| GOBP_SENSORY_ORGAN_ | GOBP_SENSORY_ORGAN_MORPHOGENESIS | 77 | -0.42556 | -1.66799 | 0 | 0.14821838 | 1 | 880 | tags=39%, list=19%, si | EN_4 |
| GOBP_SPINDLE_LOCALIZ | GOBP_SPINDLE_LOCALIZATION | 25 | -0.55039 | -1.66768 | 0.004950495 | 0.14669749 | 1 | 504 | tags=32%, list=11%, si | EN_4 |
| GOBP_SECRETION_BY_TI | GOBP_SECRETION_BY_TISSUE | 17 | -0.61145 | -1.66452 | 0.02027027 | 0.14891398 | 1 | 253 | tags=24%, list=5%, sig | EN_4 |
| CHOLINERGIC_SYNAPSE | CHOLINERGIC_SYNAPSE | 42 | -0.47541 | -1.65915 | 0.01038961 | 0.15425485 | 1 | 1160 | tags=48%, list=25%, si | EN_4 |
| GOBP_SENSORY_PERCEP | GOBP_SENSORY_PERCEPTION | 90 | -0.41228 | -1.65721 | 0 | 0.15478465 | 1 | 670 | tags=29%, list=14%, si | EN_4 |
| GOBP_RELAXATION_OF_ | GOBP_RELAXATION_OF_MUSCLE | 16 | -0.60903 | -1.65534 | 0.024752475 | 0.15517867 | 1 | 622 | tags=38%, list=13%, si | EN_4 |
| GOBP_HEART_DEVELOPM | GOBP_HEART_DEVELOPMENT | 241 | -0.35961 | -1.65343 | 0 | 0.1560793 | 1 | 857 | tags=24%, list=18%, si | EN_4 |
| GOBP_POTASSIUM_ION_ | GOBP_POTASSIUM_ION_TRANSPORT | 54 | -0.46227 | -1.65336 | 0.005128205 | 0.1543521 | 1 | 1249 | tags=52%, list=26%, si | EN_4 |
| GOBP_REGULATION_OF_ | GOBP_REGULATION_OF_MAP_KINASE_ACTI | 49 | -0.45979 | -1.65232 | 0.002415459 | 0.15391058 | 1 | 933 | tags=37%, list=20%, si | EN_4 |
| GOBP_AORTA_DEVELOPM | GOBP_AORTA_DEVELOPMENT | 28 | -0.5267 | -1.65141 | 0.009389672 | 0.1534857 | 1 | 323 | tags=21%, list=7%, sig | EN_4 |
| CGMP-PKG_SIGNALING_ | CGMP-PKG_SIGNALING_PATHWAY | 71 | -0.43262 | -1.64926 | 0.005509642 | 0.15419284 | 1 | 1326 | tags=46%, list=28%, si | EN_4 |
| GOBP_POSITIVE_REGULA | GOBP_POSITIVE_REGULATION_OF_MICROT | 17 | -0.59755 | -1.6481 | 0.013761468 | 0.15420972 | 1 | 950 | tags=47%, list=20%, si | EN_4 |
| GOBP_COLUMNAR_CUBO | GOBP_COLUMNAR_CUBOIDAL_EPITHELIAL_ | 19 | -0.56836 | -1.64741 | 0.02347418 | 0.15332362 | 1 | 638 | tags=32%, list=13%, si | EN_4 |
| GOBP_CALCIUM_ION_TR | GOBP_CALCIUM_ION_TRANSMEMBRANE_T | 91 | -0.41341 | -1.64694 | 0.002873563 | 0.15235288 | 1 | 969 | tags=31%, list=20%, si | EN_4 |
| GOBP_NEURAL_PRECURS | GOBP_NEURAL_PRECURSOR_CELL_PROLIFE | 79 | -0.41992 | -1.64586 | 0.002770083 | 0.15222813 | 1 | 660 | tags=23%, list=14%, si | EN_4 |
| GOBP_CALCIUM_ION_TR | GOBP_CALCIUM_ION_TRANSMEMBRANE_IN | 53 | -0.44978 | -1.64315 | 0.005449591 | 0.1538693 | 1 | 667 | tags=25%, list=14%, si | EN_4 |
| GOBP_LOCOMOTORY_BE | GOBP_LOCOMOTORY_BEHAVIOR | 89 | -0.41117 | -1.64274 | 0.005319149 | 0.15284428 | 1 | 833 | tags=30%, list=18%, si | EN_4 |
| GOBP_EMBRYONIC_HEAF | GOBP_EMBRYONIC_HEART_TUBE_MORPHO | 24 | -0.53657 | -1.64259 | 0.015625 | 0.15155824 | 1 | 640 | tags=25%, list=14%, si | EN_4 |
| GOBP_NEURAL_TUBE_FO | GOBP_NEURAL_TUBE_FORMATION | 45 | -0.46855 | -1.63717 | 0.002557545 | 0.1565153 | 1 | 728 | tags=27%, list=15%, si | EN_4 |
| GOBP_L_AMINO_ACID_T | GOBP_L_AMINO_ACID_TRANSPORT | 18 | -0.5757 | -1.63618 | 0.018223235 | 0.15608601 | 1 | 593 | tags=39%, list=13%, si | EN_4 |
| GOBP_TELENCEPHALON_ | GOBP_TELENCEPHALON_DEVELOPMENT | 84 | -0.41101 | -1.63429 | 0 | 0.15684848 | 1 | 898 | tags=31%, list=19%, si | EN_4 |
| GOBP_NEGATIVE_REGUL | GOBP_NEGATIVE_REGULATION_OF_STEM_ | 20 | -0.56728 | -1.63273 | 0.012315271 | 0.15707082 | 1 | 1328 | tags=60%, list=28%, si | EN_4 |
| DILATED_CARDIOMYOPA | DILATED_CARDIOMYOPATHY_(DCM) | 35 | -0.49323 | -1.63013 | 0.00511509 | 0.15880527 | 1 | 1332 | tags=46%, list=28%, si | EN_4 |
| GOBP_RESPONSE_TO_CA | GOBP_RESPONSE_TO_CAMP | 20 | -0.57246 | -1.62695 | 0.013986014 | 0.16118224 | 1 | 1468 | tags=65%, list=31%, si | EN_4 |

| | | | | | | | | | | |
|---|---|---|---|---|---|---|---|---|---|---|
| GOBP_CELL_FATE_COMM | GOBP_CELL_FATE_COMMITMENT | 80 | -0.41724 | -1.62654 | 0.002538071 | 0.16007465 | 1 | 831 | tags=29%, list=18%, si | EN_4 |
| GOBP_BODY_FLUID_SECR | GOBP_BODY_FLUID_SECRETION | 31 | -0.49904 | -1.62607 | 0.004796163 | 0.15908459 | 1 | 298 | tags=23%, list=6%, sig | EN_4 |
| GOBP_ANIMAL_ORGAN_ | GOBP_ANIMAL_ORGAN_MORPHOGENESIS | 323 | -0.34187 | -1.6186 | 0 | 0.16805945 | 1 | 834 | tags=27%, list=18%, si | EN_4 |
| GOBP_HEART_PROCESS | GOBP_HEART_PROCESS | 86 | -0.39824 | -1.61338 | 0.002873563 | 0.17317662 | 1 | 624 | tags=21%, list=13%, si | EN_4 |
| GOBP_REGULATION_OF_ | GOBP_REGULATION_OF_OLIGODENDROCYT | 17 | -0.58339 | -1.61246 | 0.02027027 | 0.17292808 | 1 | 681 | tags=29%, list=14%, si | EN_4 |
| GOBP_REGULATION_OF_ | GOBP_REGULATION_OF_POTASSIUM_ION_ | 33 | -0.49972 | -1.60914 | 0.009592326 | 0.17590155 | 1 | 1249 | tags=52%, list=26%, si | EN_4 |
| GOBP_METANEPHROS_D | GOBP_METANEPHROS_DEVELOPMENT | 24 | -0.51409 | -1.60586 | 0.017621145 | 0.17878446 | 1 | 677 | tags=25%, list=14%, si | EN_4 |
| GOBP_POSITIVE_REGULA | GOBP_POSITIVE_REGULATION_OF_CYTOSKE | 82 | -0.41838 | -1.60496 | 0.002717391 | 0.17838246 | 1 | 628 | tags=24%, list=13%, si | EN_4 |
| GOBP_TISSUE_MORPHOG | GOBP_TISSUE_MORPHOGENESIS | 220 | -0.35377 | -1.60476 | 0 | 0.1770795 | 1 | 679 | tags=24%, list=14%, si | EN_4 |
| GOBP_REGULATION_OF_ | GOBP_REGULATION_OF_GENERATION_OF_ | 56 | -0.43183 | -1.60413 | 0.00921659 | 0.17645444 | 1 | 837 | tags=30%, list=18%, si | EN_4 |
| GOBP_BONE_REMODELIN | GOBP_BONE_REMODELING | 29 | -0.50304 | -1.60229 | 0.006976744 | 0.17747584 | 1 | 580 | tags=31%, list=12%, si | EN_4 |
| GOBP_CEREBRAL_CORTEX | GOBP_CEREBRAL_CORTEX_DEVELOPMENT | 45 | -0.46207 | -1.60196 | 0.009569378 | 0.17636234 | 1 | 1103 | tags=40%, list=23%, si | EN_4 |
| GOBP_POSITIVE_REGULA | GOBP_POSITIVE_REGULATION_OF_INSULIN_ | 20 | -0.55523 | -1.59987 | 0.023255814 | 0.17792961 | 1 | 158 | tags=20%, list=3%, sig | EN_4 |
| GOBP_ARTERY_DEVELOPI | GOBP_ARTERY_DEVELOPMENT | 46 | -0.4541 | -1.59758 | 0.007281554 | 0.17993288 | 1 | 415 | tags=22%, list=9%, sig | EN_4 |
| GOBP_REGULATION_OF_ | GOBP_REGULATION_OF_CHROMATIN_ORG/ | 17 | -0.5781 | -1.597 | 0.025229357 | 0.17929678 | 1 | 1145 | tags=71%, list=24%, si | EN_4 |
| GOBP_NEGATIVE_REGUL | GOBP_NEGATIVE_REGULATION_OF_GROWT | 86 | -0.39787 | -1.58894 | 0 | 0.18877682 | 1 | 764 | tags=23%, list=16%, si | EN_4 |
| GOBP_LEARNING | GOBP_LEARNING | 51 | -0.44715 | -1.5881 | 0 | 0.18836777 | 1 | 883 | tags=39%, list=19%, si | EN_4 |
| GOBP_OVULATION_CYCL | GOBP_OVULATION_CYCLE | 18 | -0.56208 | -1.58665 | 0.029953917 | 0.1889429 | 1 | 776 | tags=39%, list=16%, si | EN_4 |
| GOBP_CENTRAL_NERVOU | GOBP_CENTRAL_NERVOUS_SYSTEM_DEVEL | 307 | -0.3427 | -1.58557 | 0 | 0.18885973 | 1 | 845 | tags=26%, list=18%, si | EN_4 |
| GOBP_CARBOHYDRATE_( | GOBP_CARBOHYDRATE_CATABOLIC_PROCES | 56 | -0.43568 | -1.58336 | 0.010810811 | 0.19051582 | 1 | 787 | tags=32%, list=17%, si | EN_4 |
| HYPERTROPHIC_CARDION | HYPERTROPHIC_CARDIOMYOPATHY_(HCM) | 35 | -0.47302 | -1.5833 | 0.005102041 | 0.189015 | 1 | 1421 | tags=49%, list=30%, si | EN_4 |
| GOBP_FOREBRAIN_DEVEI | GOBP_FOREBRAIN_DEVELOPMENT | 126 | -0.37523 | -1.57468 | 0.002849003 | 0.19975977 | 1 | 898 | tags=29%, list=19%, si | EN_4 |
| GOBP_POTASSIUM_ION_ | GOBP_POTASSIUM_ION_TRANSMEMBRANE | 45 | -0.4495 | -1.56978 | 0.017857144 | 0.2051237 | 1 | 1249 | tags=53%, list=26%, si | EN_4 |
| AMPHETAMINE_ADDICTI( | AMPHETAMINE_ADDICTION | 28 | -0.50352 | -1.56703 | 0.021686748 | 0.2078204 | 1 | 681 | tags=32%, list=14%, si | EN_4 |
| GOBP_BONE_RESORPTIO | GOBP_BONE_RESORPTION | 17 | -0.5602 | -1.56565 | 0.03529412 | 0.20809183 | 1 | 580 | tags=35%, list=12%, si | EN_4 |
| GOBP_MUSCLE_SYSTEM_ | GOBP_MUSCLE_SYSTEM_PROCESS | 148 | -0.36377 | -1.56522 | 0 | 0.20718901 | 1 | 826 | tags=25%, list=17%, si | EN_4 |
| GOBP_REGULATION_OF_ | GOBP_REGULATION_OF_FIBROBLAST_MIGR | 28 | -0.49423 | -1.55854 | 0.0369515 | 0.21617602 | 1 | 876 | tags=29%, list=19%, si | EN_4 |
| GOBP_NEGATIVE_REGUL | GOBP_NEGATIVE_REGULATION_OF_FAT_CE | 25 | -0.51571 | -1.55705 | 0.030516433 | 0.21673308 | 1 | 594 | tags=24%, list=13%, si | EN_4 |
| GOBP_HEAD_MORPHOGI | GOBP_HEAD_MORPHOGENESIS | 27 | -0.50521 | -1.55617 | 0.01754386 | 0.2164163 | 1 | 1636 | tags=67%, list=35%, si | EN_4 |
| GOBP_CARDIAC_CHAMBI | GOBP_CARDIAC_CHAMBER_DEVELOPMENT | 68 | -0.41049 | -1.55589 | 0.013157895 | 0.21529217 | 1 | 591 | tags=24%, list=12%, si | EN_4 |
| GOBP_AMINO_ACID_TRA | GOBP_AMINO_ACID_TRANSPORT | 31 | -0.48709 | -1.55342 | 0.019607844 | 0.21791512 | 1 | 593 | tags=35%, list=13%, si | EN_4 |
| GOBP_REGULATION_OF_ | GOBP_REGULATION_OF_STEM_CELL_PROLI | 35 | -0.46773 | -1.5529 | 0.030927835 | 0.21703663 | 1 | 638 | tags=31%, list=13%, si | EN_4 |
| GOBP_REGULATION_OF_ | GOBP_REGULATION_OF_BONE_MINERALIZ/ | 23 | -0.52894 | -1.55205 | 0.029885057 | 0.21683426 | 1 | 669 | tags=26%, list=14%, si | EN_4 |
| GOBP_CELLULAR_RESPOI | GOBP_CELLULAR_RESPONSE_TO_CAMP | 15 | -0.58228 | -1.54799 | 0.05755396 | 0.22191568 | 1 | 1354 | tags=67%, list=29%, si | EN_4 |
| GOBP_STEM_CELL_DIFFE | GOBP_STEM_CELL_DIFFERENTIATION | 103 | -0.38151 | -1.54729 | 0 | 0.22159865 | 1 | 1021 | tags=37%, list=22%, si | EN_4 |
| GOBP_REGULATION_OF_ | GOBP_REGULATION_OF_NEURON_MIGRATI | 19 | -0.53776 | -1.5452 | 0.044444446 | 0.22341207 | 1 | 970 | tags=58%, list=20%, si | EN_4 |
| GOBP_EMBRYONIC_PATT | GOBP_EMBRYONIC_PATTERN_SPECIFICATIO | 19 | -0.53933 | -1.54445 | 0.038116593 | 0.22298448 | 1 | 723 | tags=42%, list=15%, si | EN_4 |
| GOBP_BONE_GROWTH | GOBP_BONE_GROWTH | 15 | -0.57641 | -1.5442 | 0.040632054 | 0.22179289 | 1 | 638 | tags=33%, list=13%, si | EN_4 |
| GOBP_ACTION_POTENTI/ | GOBP_ACTION_POTENTIAL | 43 | -0.44832 | -1.54372 | 0.024213076 | 0.22096 | 1 | 1309 | tags=47%, list=28%, si | EN_4 |
| GOBP_EMBRYONIC_MOR | GOBP_EMBRYONIC_MORPHOGENESIS | 203 | -0.34525 | -1.54326 | 0 | 0.22009726 | 1 | 780 | tags=25%, list=16%, si | EN_4 |
| GOBP_OLIGODENDROCY1 | GOBP_OLIGODENDROCYTE_DIFFERENTIATIC | 39 | -0.45988 | -1.54016 | 0.018276762 | 0.22352915 | 1 | 328 | tags=18%, list=7%, sig | EN_4 |
| GOBP_REGULATION_OF_ | GOBP_REGULATION_OF_MONOATOMIC_IO | 107 | -0.37518 | -1.5391 | 0.005797102 | 0.22378777 | 1 | 649 | tags=21%, list=14%, si | EN_4 |
| GOBP_CELLULAR_RESPOI | GOBP_CELLULAR_RESPONSE_TO_NITROGEN | 221 | -0.33833 | -1.53809 | 0 | 0.22409263 | 1 | 845 | tags=30%, list=18%, si | EN_4 |
| GOBP_ENERGY_RESERVE_ | GOBP_ENERGY_RESERVE_METABOLIC_PRO( | 36 | -0.45931 | -1.53766 | 0.02970297 | 0.22335954 | 1 | 837 | tags=31%, list=18%, si | EN_4 |
| GOBP_VASODILATION | GOBP_VASODILATION | 22 | -0.50958 | -1.53759 | 0.040632054 | 0.22202115 | 1 | 880 | tags=36%, list=19%, si | EN_4 |
| GOBP_CARDIAC_VENTRI( | GOBP_CARDIAC_VENTRICLE_MORPHOGENE | 28 | -0.49545 | -1.53739 | 0.021686748 | 0.22072972 | 1 | 376 | tags=14%, list=8%, sig | EN_4 |
| GOBP_REGULATION_OF_ | GOBP_REGULATION_OF_NEURAL_PRECURS( | 46 | -0.43517 | -1.53151 | 0.03 | 0.22883819 | 1 | 660 | tags=24%, list=14%, si | EN_4 |
| GOBP_ADIPOSE_TISSUE_ | GOBP_ADIPOSE_TISSUE_DEVELOPMENT | 31 | -0.47377 | -1.5305 | 0.029411765 | 0.22896838 | 1 | 1283 | tags=45%, list=27%, si | EN_4 |
| GOBP_MESODERM_MOR | GOBP_MESODERM_MORPHOGENESIS | 20 | -0.53135 | -1.52342 | 0.040723983 | 0.23880635 | 1 | 535 | tags=30%, list=11%, si | EN_4 |
| GOBP_REGULATION_OF_ | GOBP_REGULATION_OF_CELL_CYCLE_G2_M | 36 | -0.45011 | -1.51943 | 0.027568921 | 0.24398758 | 1 | 413 | tags=25%, list=9%, sig | EN_4 |
| GOBP_METENCEPHALON_ | GOBP_METENCEPHALON_DEVELOPMENT | 39 | -0.44589 | -1.51917 | 0.02244389 | 0.2428255 | 1 | 619 | tags=38%, list=13%, si | EN_4 |
| GOBP_MORPHOGENESIS_ | GOBP_MORPHOGENESIS_OF_AN_EPITHELIU | 188 | -0.34508 | -1.51733 | 0 | 0.24422269 | 1 | 679 | tags=23%, list=14%, si | EN_4 |
| GOBP_INORGANIC_ION_ | GOBP_INORGANIC_ION_TRANSMEMBRANE | 179 | -0.34414 | -1.51661 | 0 | 0.243708 | 1 | 704 | tags=22%, list=15%, si | EN_4 |
| GOBP_EAR_MORPHOGEN | GOBP_EAR_MORPHOGENESIS | 29 | -0.48194 | -1.51642 | 0.034398034 | 0.2425735 | 1 | 760 | tags=38%, list=16%, si | EN_4 |
| GOBP_ADULT_LOCOMOT | GOBP_ADULT_LOCOMOTORY_BEHAVIOR | 40 | -0.44368 | -1.51598 | 0.020100502 | 0.24170288 | 1 | 833 | tags=33%, list=18%, si | EN_4 |
| GOBP_ADULT_BEHAVIOR | GOBP_ADULT_BEHAVIOR | 50 | -0.4247 | -1.51524 | 0.033333335 | 0.24131075 | 1 | 861 | tags=34%, list=18%, si | EN_4 |
| GOBP_DIGESTIVE_TRACT | GOBP_DIGESTIVE_TRACT_MORPHOGENESIS | 15 | -0.59129 | -1.51495 | 0.03255814 | 0.24029566 | 1 | 530 | tags=27%, list=11%, si | EN_4 |
| GOBP_FACULTATIVE_HET | GOBP_FACULTATIVE_HETEROCHROMATIN_ | 21 | -0.51575 | -1.51435 | 0.037220843 | 0.23999642 | 1 | 764 | tags=43%, list=16%, si | EN_4 |
| GOBP_ORGAN_GROWTH | GOBP_ORGAN_GROWTH | 83 | -0.38537 | -1.51162 | 0.010695187 | 0.2433981 | 1 | 826 | tags=25%, list=17%, si | EN_4 |
| GOBP_POSITIVE_REGULA | GOBP_POSITIVE_REGULATION_OF_PROTEIN | 41 | -0.43511 | -1.50983 | 0.013020833 | 0.24469528 | 1 | 950 | tags=39%, list=20%, si | EN_4 |
| GOBP_LYMPHOCYTE_HOI | GOBP_LYMPHOCYTE_HOMEOSTASIS | 31 | -0.46331 | -1.5081 | 0.03167421 | 0.24615748 | 1 | 849 | tags=26%, list=18%, si | EN_4 |
| GOBP_EMBRYONIC_HEAF | GOBP_EMBRYONIC_HEART_TUBE_DEVELOP | 29 | -0.47754 | -1.50643 | 0.048387095 | 0.24761033 | 1 | 640 | tags=21%, list=14%, si | EN_4 |
| GOBP_SENSORY_PERCEP1 | GOBP_SENSORY_PERCEPTION_OF_PAIN | 20 | -0.512 | -1.50448 | 0.061784897 | 0.24917112 | 1 | 670 | tags=40%, list=14%, si | EN_4 |
| DOPAMINERGIC_SYNAPS | DOPAMINERGIC_SYNAPSE | 57 | -0.40033 | -1.50097 | 0.01728395 | 0.2493844 | 1 | 1223 | tags=42%, list=26%, si | EN_4 |
| GOBP_MULTICELLULAR_( | GOBP_MULTICELLULAR_ORGANISM_GROW | 95 | -0.37262 | -1.5009 | 0.005714286 | 0.2479948 | 1 | 800 | tags=25%, list=17%, si | EN_4 |
| GOBP_PRESYNAPTIC_MO | GOBP_PRESYNAPTIC_MODULATION_OF_CH | 17 | -0.54291 | -1.50073 | 0.0389016 | 0.24679342 | 1 | 799 | tags=41%, list=17%, si | EN_4 |
| GOBP_SYNAPSE_ASSEMB | GOBP_SYNAPSE_ASSEMBLY | 81 | -0.37844 | -1.49948 | 0.013623978 | 0.247639 | 1 | 898 | tags=37%, list=19%, si | EN_4 |
| GOBP_HINDBRAIN_DEVEI | GOBP_HINDBRAIN_DEVELOPMENT | 59 | -0.40543 | -1.49731 | 0.014962593 | 0.24993126 | 1 | 619 | tags=31%, list=13%, si | EN_4 |
| GOBP_CAMERA_TYPE_EY | GOBP_CAMERA_TYPE_EYE_MORPHOGENES | 42 | -0.42974 | -1.49689 | 0.020997375 | 0.2493266 | 1 | 779 | tags=38%, list=16%, si | EN_4 |
| GOBP_MOLTING_CYCLE_ | GOBP_MOLTING_CYCLE_PROCESS | 36 | -0.45894 | -1.49645 | 0.028301887 | 0.2487176 | 1 | 681 | tags=28%, list=14%, si | EN_4 |
| GOBP_RESPONSE_TO_CH | GOBP_RESPONSE_TO_CHEMOKINE | 42 | 0.592906 | 2.024292 | 0 | 0.08881848 | 0.084 | 839 | tags=40%, list=10%, si | MM_4 |
| GOBP_T_CELL_MIGRATIO | GOBP_T_CELL_MIGRATION | 44 | 0.599602 | 2.018755 | 0 | 0.048586443 | 0.092 | 1296 | tags=34%, list=16%, si | MM_4 |
| GOBP_NEUTROPHIL_CHE | GOBP_NEUTROPHIL_CHEMOTAXIS | 49 | 0.584452 | 2.01545 | 0 | 0.03307668 | 0.094 | 869 | tags=31%, list=11%, si | MM_4 |
| GOBP_LYMPHOCYTE_CHE | GOBP_LYMPHOCYTE_CHEMOTAXIS | 23 | 0.6641 | 1.941036 | 0 | 0.090174735 | 0.294 | 1012 | tags=39%, list=13%, si | MM_4 |
| STAPHYLOCOCCUS_AURE | STAPHYLOCOCCUS_AUREUS_INFECTION | 20 | 0.669246 | 1.937581 | 0 | 0.07548899 | 0.305 | 886 | tags=45%, list=11%, si | MM_4 |
| GOBP_GRANULOCYTE_CH | GOBP_GRANULOCYTE_CHEMOTAXIS | 67 | 0.522404 | 1.916128 | 0 | 0.08846015 | 0.404 | 869 | tags=31%, list=11%, si | MM_4 |
| GOBP_REGULATION_OF_ | GOBP_REGULATION_OF_NEUTROPHIL_CHEI | 20 | 0.676783 | 1.895725 | 0 | 0.098460294 | 0.494 | 597 | tags=30%, list=7%, sig | MM_4 |
| GOBP_POSITIVE_REGULA | GOBP_POSITIVE_REGULATION_OF_LEUKOC | 48 | 0.538239 | 1.874842 | 0.001633987 | 0.114564605 | 0.594 | 1026 | tags=29%, list=13%, si | MM_4 |
| GOBP_NEGATIVE_REGUL | GOBP_NEGATIVE_REGULATION_OF_DNA_R | 26 | 0.627762 | 1.874279 | 0 | 0.102647714 | 0.599 | 1296 | tags=42%, list=16%, si | MM_4 |
| GOBP_HUMORAL_IMMU | GOBP_HUMORAL_IMMUNE_RESPONSE | 63 | 0.508494 | 1.860183 | 0 | 0.11356967 | 0.676 | 765 | tags=29%, list=10%, si | MM_4 |
| GOBP_T_CELL_CHEMOTA | GOBP_T_CELL_CHEMOTAXIS | 15 | 0.713983 | 1.857901 | 0.001760563 | 0.10676207 | 0.685 | 1012 | tags=47%, list=13%, si | MM_4 |
| GOBP_REGULATION_OF_ | GOBP_REGULATION_OF_T_CELL_MIGRATIO | 32 | 0.583576 | 1.849238 | 0.001779359 | 0.11161194 | 0.737 | 1296 | tags=38%, list=16%, si | MM_4 |

| | | | | | | | | | | |
|---|---|---|---|---|---|---|---|---|---|---|
| GOBP_RESPONSE_TO_INT | GOBP_RESPONSE_TO_INTERFERON_BETA | 36 | 0.567803 | 1.839601 | 0.001848429 | 0.11913211 | 0.794 | 874 | tags=33%, list=11%, si | MM_4 |
| GOBP_POSITIVE_REGULA | GOBP_POSITIVE_REGULATION_OF_STEROL_ | 19 | 0.652974 | 1.83822 | 0 | 0.11241774 | 0.802 | 1318 | tags=53%, list=16%, si | MM_4 |
| GOBP_POSITIVE_REGULA | GOBP_POSITIVE_REGULATION_OF_GRANUL | 15 | 0.698338 | 1.834351 | 0 | 0.10985507 | 0.82 | 597 | tags=33%, list=7%, sig | MM_4 |
| GOBP_VASCULAR_ENDOT | GOBP_VASCULAR_ENDOTHELIAL_GROWTH_ | 20 | 0.631138 | 1.825056 | 0.001736111 | 0.113816425 | 0.839 | 1480 | tags=55%, list=19%, si | MM_4 |
| GOBP_FATTY_ACID_DERI | GOBP_FATTY_ACID_DERIVATIVE_METABOLI | 19 | 0.657168 | 1.824498 | 0 | 0.10755532 | 0.84 | 1269 | tags=47%, list=16%, si | MM_4 |
| GOBP_POSITIVE_REGULA | GOBP_POSITIVE_REGULATION_OF_TUMOR_ | 76 | 0.486044 | 1.823854 | 0 | 0.10233573 | 0.841 | 2161 | tags=45%, list=27%, si | MM_4 |
| GOBP_MITOCHONDRIAL_ | GOBP_MITOCHONDRIAL_TRANSMEMBRANE | 65 | 0.496947 | 1.823759 | 0.00166113 | 0.09694964 | 0.841 | 1198 | tags=32%, list=15%, si | MM_4 |
| GOBP_NEGATIVE_REGUL | GOBP_NEGATIVE_REGULATION_OF_VIRAL_ | 24 | 0.614683 | 1.82003 | 0.001692047 | 0.09648786 | 0.852 | 889 | tags=33%, list=11%, si | MM_4 |
| GOBP_POSITIVE_REGULA | GOBP_POSITIVE_REGULATION_OF_MONON | 55 | 0.517298 | 1.819919 | 0 | 0.09214219 | 0.852 | 2152 | tags=47%, list=27%, si | MM_4 |
| GOBP_POSITIVE_REGULA | GOBP_POSITIVE_REGULATION_OF_LEUKOC | 15 | 0.678038 | 1.811739 | 0.009451796 | 0.09731134 | 0.879 | 1480 | tags=53%, list=19%, si | MM_4 |
| GOBP_REGULATION_OF_ | GOBP_REGULATION_OF_LEUKOCYTE_CHEM | 64 | 0.496247 | 1.810734 | 0.001694915 | 0.09457637 | 0.882 | 1026 | tags=27%, list=13%, si | MM_4 |
| GOBP_CELLULAR_EXTRAV | GOBP_CELLULAR_EXTRAVASATION | 37 | 0.552414 | 1.809959 | 0.001831502 | 0.09146429 | 0.885 | 1137 | tags=35%, list=14%, si | MM_4 |
| GOBP_NUCLEOBASE_ME | GOBP_NUCLEOBASE_METABOLIC_PROCESS | 19 | 0.64486 | 1.809391 | 0.005586592 | 0.08834099 | 0.886 | 1761 | tags=53%, list=22%, si | MM_4 |
| GOBP_CHOLESTEROL_EF | GOBP_CHOLESTEROL_EFFLUX | 28 | 0.589469 | 1.804352 | 0.001824818 | 0.090201415 | 0.902 | 1244 | tags=39%, list=16%, si | MM_4 |
| GOBP_REGULATION_OF_ | GOBP_REGULATION_OF_GRANULOCYTE_CH | 30 | 0.576613 | 1.802699 | 0.001748252 | 0.08874925 | 0.905 | 686 | tags=30%, list=9%, sig | MM_4 |
| GOBP_REGULATION_OF_ | GOBP_REGULATION_OF_VASCULATURE_DE | 137 | 0.428612 | 1.786591 | 0 | 0.106748 | 0.949 | 1480 | tags=28%, list=19%, si | MM_4 |
| GOBP_NUCLEOSIDE_BISP | GOBP_NUCLEOSIDE_BISPHOSPHATE_BIOSY | 22 | 0.616934 | 1.78652 | 0.001757469 | 0.103102714 | 0.949 | 1269 | tags=36%, list=16%, si | MM_4 |
| GOBP_POSITIVE_REGULA | GOBP_POSITIVE_REGULATION_OF_LYMPHO | 27 | 0.586765 | 1.782537 | 0.005555556 | 0.10425498 | 0.954 | 1296 | tags=33%, list=16%, si | MM_4 |
| GOBP_REGULATION_OF_ | GOBP_REGULATION_OF_CELLULAR_EXTRAV | 18 | 0.631085 | 1.768266 | 0.005454545 | 0.12088016 | 0.967 | 1137 | tags=50%, list=14%, si | MM_4 |
| GOBP_NEUTROPHIL_MIG | GOBP_NEUTROPHIL_MIGRATION | 62 | 0.49421 | 1.75383 | 0 | 0.13855252 | 0.983 | 869 | tags=24%, list=11%, si | MM_4 |
| GOBP_GMP_METABOLIC_ | GOBP_GMP_METABOLIC_PROCESS | 16 | 0.655466 | 1.751455 | 0.005328597 | 0.13780983 | 0.985 | 1492 | tags=50%, list=19%, si | MM_4 |
| GOBP_LEUKOCYTE_ADHE | GOBP_LEUKOCYTE_ADHESION_TO_VASCUL | 27 | 0.578725 | 1.750764 | 0.005319149 | 0.13492559 | 0.985 | 1037 | tags=37%, list=13%, si | MM_4 |
| GOBP_NEGATIVE_REGUL | GOBP_NEGATIVE_REGULATION_OF_EPITHE | 31 | 0.558722 | 1.746605 | 0.003533569 | 0.1375381 | 0.988 | 1909 | tags=45%, list=24%, si | MM_4 |
| GOBP_GRANULOCYTE_M | GOBP_GRANULOCYTE_MIGRATION | 82 | 0.460193 | 1.743268 | 0 | 0.13878495 | 0.989 | 869 | tags=26%, list=11%, si | MM_4 |
| GOBP_LEUKOCYTE_CHEM | GOBP_LEUKOCYTE_CHEMOTAXIS | 121 | 0.431672 | 1.740395 | 0.001610306 | 0.13906558 | 0.992 | 1026 | tags=25%, list=13%, si | MM_4 |
| GOBP_REGULATION_OF_ | GOBP_REGULATION_OF_MONOCYTE_CHEM | 18 | 0.616422 | 1.738594 | 0.005474452 | 0.137678 | 0.992 | 1788 | tags=56%, list=22%, si | MM_4 |
| GOBP_LYMPHOCYTE_MI | GOBP_LYMPHOCYTE_MIGRATION | 58 | 0.479699 | 1.737689 | 0 | 0.13554764 | 0.993 | 1296 | tags=28%, list=16%, si | MM_4 |
| GOBP_REGULATION_OF_ | GOBP_REGULATION_OF_LYMPHOCYTE_MIG | 43 | 0.519334 | 1.737 | 0.001703578 | 0.13364561 | 0.993 | 1296 | tags=30%, list=16%, si | MM_4 |
| CYTOSOLIC_DNA-SENSIN | CYTOSOLIC_DNA-SENSING_PATHWAY | 40 | 0.522574 | 1.736881 | 0 | 0.13064086 | 0.993 | 701 | tags=28%, list=9%, sig | MM_4 |
| GOBP_REGULATION_OF_ | GOBP_REGULATION_OF_MONONUCLEAR_C | 82 | 0.456516 | 1.727118 | 0.001712329 | 0.142067 | 0.997 | 1296 | tags=28%, list=16%, si | MM_4 |
| GOBP_NEGATIVE_REGUL | GOBP_NEGATIVE_REGULATION_OF_VASCUL | 44 | 0.515058 | 1.716208 | 0 | 0.15680788 | 1 | 1012 | tags=30%, list=13%, si | MM_4 |
| GOBP_REGULATION_OF_ | GOBP_REGULATION_OF_STEROL_TRANSPO | 28 | 0.548253 | 1.706589 | 0.005154639 | 0.16984913 | 1 | 1318 | tags=39%, list=16%, si | MM_4 |
| GOBP_ANTIMICROBIAL_H | GOBP_ANTIMICROBIAL_HUMORAL_IMMUN | 29 | 0.551155 | 1.705913 | 0.003508772 | 0.16732517 | 1 | 765 | tags=38%, list=10%, si | MM_4 |
| GOBP_POSITIVE_REGULA | GOBP_POSITIVE_REGULATION_OF_MACRO | 17 | 0.626775 | 1.701139 | 0.007181329 | 0.17189725 | 1 | 1244 | tags=53%, list=16%, si | MM_4 |
| GOBP_REGULATION_OF_ | GOBP_REGULATION_OF_PLASMA_LIPOPRO | 34 | 0.531383 | 1.69865 | 0.005145798 | 0.17206295 | 1 | 1867 | tags=47%, list=23%, si | MM_4 |
| GOBP_REGULATION_OF_ | GOBP_REGULATION_OF_MACROPHAGE_CH | 17 | 0.624585 | 1.69607 | 0.005357143 | 0.17330681 | 1 | 1026 | tags=47%, list=13%, si | MM_4 |
| PPAR_SIGNALING_PATH | PPAR_SIGNALING_PATHWAY | 27 | 0.54418 | 1.695161 | 0.008605852 | 0.17159893 | 1 | 1475 | tags=48%, list=18%, si | MM_4 |
| GOBP_REGULATION_OF_ | GOBP_REGULATION_OF_LEUKOCYTE_ADHE | 18 | 0.612194 | 1.691899 | 0.005385996 | 0.17405976 | 1 | 1037 | tags=44%, list=13%, si | MM_4 |
| GOBP_LEUKOCYTE_TETH | GOBP_LEUKOCYTE_TETHERING_OR_ROLLIN | 15 | 0.659578 | 1.690269 | 0.012280702 | 0.17359415 | 1 | 1037 | tags=40%, list=13%, si | MM_4 |
| GOBP_REGULATION_OF_ | GOBP_REGULATION_OF_MACROPHAGE_MI | 27 | 0.559111 | 1.690167 | 0.009140768 | 0.17039585 | 1 | 1244 | tags=44%, list=16%, si | MM_4 |
| SALIVARY_SECRETION | SALIVARY_SECRETION | 33 | 0.524157 | 1.683414 | 0.004983389 | 0.17855458 | 1 | 1252 | tags=27%, list=16%, si | MM_4 |
| CARBOHYDRATE_DIGESTI | CARBOHYDRATE_DIGESTION_AND_ABSORP | 19 | 0.587837 | 1.680143 | 0.014842301 | 0.18073562 | 1 | 1227 | tags=42%, list=15%, si | MM_4 |
| GOBP_IMPORT_INTO_TH | GOBP_IMPORT_INTO_THE_MITOCHONDRIO | 43 | 0.493711 | 1.670265 | 0.003502627 | 0.19593689 | 1 | 1466 | tags=35%, list=18%, si | MM_4 |
| GOBP_ANTIMICROBIAL_H | GOBP_ANTIMICROBIAL_HUMORAL_RESPON | 36 | 0.512377 | 1.669586 | 0.008403362 | 0.19372566 | 1 | 765 | tags=33%, list=10%, si | MM_4 |
| SYSTEMIC_LUPUS_ERYTH | SYSTEMIC_LUPUS_ERYTHEMATOSUS | 32 | 0.525245 | 1.665096 | 0.001821494 | 0.1994982 | 1 | 548 | tags=22%, list=7%, sig | MM_4 |
| GOBP_COLLAGEN_BIOSY | GOBP_COLLAGEN_BIOSYNTHETIC_PROCESS | 24 | 0.555674 | 1.658445 | 0.012345679 | 0.20872296 | 1 | 1482 | tags=42%, list=19%, si | MM_4 |
| GOBP_THIOESTER_BIOSY | GOBP_THIOESTER_BIOSYNTHETIC_PROCESS | 16 | 0.627587 | 1.657941 | 0.012237762 | 0.20623079 | 1 | 1201 | tags=31%, list=15%, si | MM_4 |
| GOBP_TELOMERE_MAIN | GOBP_TELOMERE_MAINTENANCE_IN_RESP | 25 | 0.55198 | 1.656917 | 0.01459854 | 0.20489421 | 1 | 1310 | tags=40%, list=16%, si | MM_4 |
| GOBP_CELL_CHEMOTAXI | GOBP_CELL_CHEMOTAXIS | 150 | 0.395556 | 1.645931 | 0.001579779 | 0.22477488 | 1 | 879 | tags=21%, list=11%, si | MM_4 |
| GOBP_DEFENSE_RESPON | GOBP_DEFENSE_RESPONSE_TO_GRAM_POS | 50 | 0.468985 | 1.643548 | 0.003389831 | 0.22610092 | 1 | 937 | tags=20%, list=12%, si | MM_4 |
| ALDOSTERONE-REGULAT | ALDOSTERONE-REGULATED_SODIUM_REAB | 18 | 0.599857 | 1.642677 | 0.01607143 | 0.2244993 | 1 | 1227 | tags=44%, list=15%, si | MM_4 |
| GOBP_REGULATION_OF_ | GOBP_REGULATION_OF_LYMPHOCYTE_CHE | 15 | 0.628762 | 1.64093 | 0.012681159 | 0.22490637 | 1 | 1012 | tags=40%, list=13%, si | MM_4 |
| GOBP_REGULATION_OF_ | GOBP_REGULATION_OF_NEUTROPHIL_MIG | 30 | 0.525306 | 1.636298 | 0.008787346 | 0.23153353 | 1 | 597 | tags=20%, list=7%, sig | MM_4 |
| GOBP_MACROPHAGE_CH | GOBP_MACROPHAGE_CHEMOTAXIS | 26 | 0.543421 | 1.634685 | 0.01831502 | 0.2318115 | 1 | 1271 | tags=42%, list=16%, si | MM_4 |
| PRION_DISEASES | PRION_DISEASES | 20 | 0.582669 | 1.634047 | 0.010869565 | 0.22963944 | 1 | 1223 | tags=40%, list=15%, si | MM_4 |
| PANCREATIC_SECRETION | PANCREATIC_SECRETION | 33 | 0.516802 | 1.628937 | 0.008802817 | 0.23786111 | 1 | 1482 | tags=33%, list=19%, si | MM_4 |
| GOBP_MONOCYTE_CHEM | GOBP_MONOCYTE_CHEMOTAXIS | 24 | 0.548665 | 1.627879 | 0.016245488 | 0.23676854 | 1 | 1788 | tags=50%, list=22%, si | MM_4 |
| GOBP_TRANSLATION_AT_ | GOBP_TRANSLATION_AT_SYNAPSE | 48 | -0.59912 | -2.12267 | 0 | 0.043442074 | 0.046 | 2953 | tags=85%, list=37%, si | MM_4 |
| **C1qaCKOvsiC1qaCKO** | | | | | | | | | | |
| GOBP_STEROL_BIOSYNTH | GOBP_STEROL_BIOSYNTHETIC_PROCESS | 58 | 0.543044 | 2.5594156 | 0.0 | 0 | 0 | 2864 | tags=40%, list=12%, si | EN_1 |
| GOBP_NADH_METABOLI | GOBP_NADH_METABOLIC_PROCESS | 39 | 0.607106 | 2.5455122 | 0.0 | 0 | 0 | 4470 | tags=67%, list=19%, si | EN_1 |
| FERROPTOSIS | FERROPTOSIS | 37 | 0.608282 | 2.4682345 | 0.0 | 0.002385722 | 0.002 | 5208 | tags=49%, list=22%, si | EN_1 |
| GOBP_NEGATIVE_REGUL | GOBP_NEGATIVE_REGULATION_OF_PROGR | 17 | 0.707899 | 2.4022624 | 0.0 | 0.007125821 | 0.008 | 5115 | tags=59%, list=22%, si | EN_1 |
| NON-SMALL_CELL_LUNG_ | NON-SMALL_CELL_LUNG_CANCER | 65 | 0.502252 | 2.3461833 | 0.0 | 0.012921074 | 0.018 | 6482 | tags=52%, list=27%, si | EN_1 |
| PEROXISOME | PEROXISOME | 81 | 0.477583 | 2.3418946 | 0.0 | 0.011350781 | 0.019 | 4149 | tags=40%, list=18%, si | EN_1 |
| GOBP_PHOSPHOLIPID_BI | GOBP_PHOSPHOLIPID_BIOSYNTHETIC_PRO | 214 | 0.418778 | 2.3386726 | 0.0 | 0.010219775 | 0.02 | 4619 | tags=38%, list=20%, si | EN_1 |
| GOBP_GLYCOLYTIC_PRO | GOBP_GLYCOLYTIC_PROCESS_THROUGH_G | 20 | 0.662509 | 2.3349013 | 0.0038910506 | 0.008942302 | 0.02 | 4212 | tags=75%, list=18%, si | EN_1 |
| GOBP_REGULATION_OF_ | GOBP_REGULATION_OF_PROGRAMMED_N | 27 | 0.608321 | 2.3298063 | 0.0 | 0.007948713 | 0.02 | 2888 | tags=33%, list=12%, si | EN_1 |
| CHRONIC_MYELOID_LEU | CHRONIC_MYELOID_LEUKEMIA | 75 | 0.483103 | 2.306371 | 0.0 | 0.008992722 | 0.025 | 6482 | tags=51%, list=27%, si | EN_1 |
| RIBOSOME | RIBOSOME | 128 | 0.432766 | 2.2866786 | 0.0 | 0.009143351 | 0.028 | 7882 | tags=51%, list=33%, si | EN_1 |
| GOBP_NADH_DEHYDROG | GOBP_NADH_DEHYDROGENASE_COMPLEX_ | 58 | 0.502905 | 2.2828043 | 0.0 | 0.009574836 | 0.032 | 6395 | tags=57%, list=27%, si | EN_1 |
| GOBP_GLYCOLYTIC_PRO | GOBP_GLYCOLYTIC_PROCESS_THROUGH_F | 25 | 0.591727 | 2.273676 | 0.0 | 0.009404174 | 0.034 | 4212 | tags=64%, list=18%, si | EN_1 |
| STEROID_BIOSYNTHESIS | STEROID_BIOSYNTHESIS | 17 | 0.677511 | 2.2500143 | 0.0 | 0.013857547 | 0.054 | 1747 | tags=53%, list=7%, sig | EN_1 |
| PARKINSON_DISEASE | PARKINSON_DISEASE | 122 | 0.424887 | 2.2447007 | 0.0 | 0.013181896 | 0.055 | 7583 | tags=71%, list=32%, si | EN_1 |
| GOBP_NAD_CATABOLIC_ | GOBP_NAD_CATABOLIC_PROCESS | 21 | 0.628454 | 2.243991 | 0.0 | 0.012358028 | 0.055 | 3874 | tags=67%, list=16%, si | EN_1 |
| GOBP_GLYCEROPHOSPH | GOBP_GLYCEROPHOSPHOLIPID_BIOSYNTHE | 160 | 0.400778 | 2.2065027 | 0.0 | 0.01685581 | 0.078 | 4610 | tags=38%, list=20%, si | EN_1 |
| GOBP_TRANSLATION_AT_ | GOBP_TRANSLATION_AT_SYNAPSE | 48 | 0.489678 | 2.206448 | 0.0 | 0.015919374 | 0.078 | 7882 | tags=58%, list=33%, si | EN_1 |
| OXIDATIVE_PHOSPHORYL | OXIDATIVE_PHOSPHORYLATION | 115 | 0.418496 | 2.1835837 | 0.0 | 0.017870117 | 0.093 | 6982 | tags=64%, list=30%, si | EN_1 |
| GOBP_PEROXISOME_ORG | GOBP_PEROXISOME_ORGANIZATION | 38 | 0.532287 | 2.1539152 | 0.0 | 0.022712212 | 0.12 | 5353 | tags=47%, list=23%, si | EN_1 |
| PPAR_SIGNALING_PATH | PPAR_SIGNALING_PATHWAY | 70 | 0.446484 | 2.153397 | 0.0 | 0.021630678 | 0.12 | 3043 | tags=40%, list=13%, si | EN_1 |
| GOBP_REGULATION_OF_ | GOBP_REGULATION_OF_RESPONSE_TO_EN | 80 | 0.433601 | 2.1510618 | 0.0 | 0.021776492 | 0.126 | 6262 | tags=45%, list=27%, si | EN_1 |
| GOBP_CELLULAR_RESPO | GOBP_CELLULAR_RESPONSE_TO_INTERLEU | 62 | 0.45382 | 2.1417568 | 0.0 | 0.023316247 | 0.139 | 3115 | tags=34%, list=13%, si | EN_1 |

| | | | | | | | | | | |
|---|---|---|---|---|---|---|---|---|---|---|
| GOBP_CELLULAR_RESPON | GOBP_CELLULAR_RESPONSE_TO_OXYGEN_I | 20 | 0.593726 | 2.1325178 | 0.0 | 0.02397164 | 0.15 | 5529 | tags=65%, list=23%, si | EN_1 |
| GOBP_REGULATION_OF_ | GOBP_REGULATION_OF_RESPONSE_TO_TYI | 15 | 0.641242 | 2.1169207 | 0.0 | 0.027298769 | 0.176 | 3013 | tags=47%, list=13%, si | EN_1 |
| GOBP_RESPONSE_TO_INT | GOBP_RESPONSE_TO_INTERLEUKIN_1 | 76 | 0.434292 | 2.1143699 | 0.0 | 0.026784906 | 0.178 | 3115 | tags=33%, list=13%, si | EN_1 |
| GOBP_TRNA_METHYLATI | GOBP_TRNA_METHYLATION | 39 | 0.501553 | 2.1094747 | 0.0 | 0.027253393 | 0.187 | 6556 | tags=62%, list=28%, si | EN_1 |
| GOBP_NUCLEOSIDE_BISP | GOBP_NUCLEOSIDE_BISPHOSPHATE_BIOSYN | 39 | 0.515576 | 2.0898283 | 0.0 | 0.031121947 | 0.218 | 3853 | tags=33%, list=16%, si | EN_1 |
| GOBP_LYSOSOMAL_LUM | GOBP_LYSOSOMAL_LUMEN_ACIDIFICATION | 15 | 0.631403 | 2.0895736 | 0.0 | 0.030302312 | 0.219 | 5364 | tags=53%, list=23%, si | EN_1 |
| GOBP_PHOSPHATIDYLCH | GOBP_PHOSPHATIDYLCHOLINE_METABOLIC | 59 | 0.460486 | 2.0852768 | 0.0 | 0.030938555 | 0.231 | 3189 | tags=39%, list=14%, si | EN_1 |
| GOBP_PIGMENT_GRANU | GOBP_PIGMENT_GRANULE_ORGANIZATION | 39 | 0.495456 | 2.080139 | 0.0 | 0.03165399 | 0.241 | 1282 | tags=18%, list=5%, sig | EN_1 |
| GOBP_COPII_COATED_VE | GOBP_COPII_COATED_VESICLE_CARGO_LO/ | 17 | 0.624427 | 2.0717392 | 0.0 | 0.03267998 | 0.257 | 6392 | tags=59%, list=27%, si | EN_1 |
| GOBP_REGULATION_OF_ | GOBP_REGULATION_OF_NECROPTOTIC_PR( | 22 | 0.581613 | 2.0714657 | 0.0 | 0.031795003 | 0.258 | 5115 | tags=45%, list=22%, si | EN_1 |
| GOBP_NUCLEOTIDE_BINI | GOBP_NUCLEOTIDE_BINDING_DOMAIN_LEU | 22 | 0.563463 | 2.0679455 | 0.0 | 0.031918414 | 0.266 | 4239 | tags=55%, list=18%, si | EN_1 |
| GOBP_RESPONSE_TO_OX | GOBP_RESPONSE_TO_OXYGEN_RADICAL | 24 | 0.559123 | 2.064358 | 0.0 | 0.03213595 | 0.274 | 5529 | tags=58%, list=23%, si | EN_1 |
| GOBP_GLYCEROLIPID_BIC | GOBP_GLYCEROLIPID_BIOSYNTHETIC_PROC | 210 | 0.35899 | 2.0569627 | 0.0 | 0.03272033 | 0.284 | 4610 | tags=34%, list=20%, si | EN_1 |
| GOBP_PROTEIN_MATUR/ | GOBP_PROTEIN_MATURATION_BY_IRON_SU | 16 | 0.62172 | 2.0532508 | 0.0 | 0.032908123 | 0.292 | 4945 | tags=56%, list=21%, si | EN_1 |
| GLYCOSAMINOGLYCAN_I | GLYCOSAMINOGLYCAN_DEGRADATION | 17 | 0.608995 | 2.0530455 | 0.0038610038 | 0.03222614 | 0.293 | 5299 | tags=59%, list=22%, si | EN_1 |
| GOBP_G0_TO_G1_TRANS | GOBP_G0_TO_G1_TRANSITION | 38 | 0.488484 | 2.0515523 | 0.0 | 0.032136124 | 0.299 | 5081 | tags=37%, list=22%, si | EN_1 |
| GLYCOSYLPHOSPHATIDYL | GLYCOSYLPHOSPHATIDYLINOSITOL_(GPI)-AN | 24 | 0.57779 | 2.0434058 | 0.004 | 0.03359387 | 0.315 | 4128 | tags=42%, list=17%, si | EN_1 |
| GLIOMA | GLIOMA | 71 | 0.435961 | 2.0403335 | 0.0 | 0.033995215 | 0.324 | 6482 | tags=48%, list=27%, si | EN_1 |
| GOBP_RESPONSE_TO_GR | GOBP_RESPONSE_TO_GROWTH_HORMONE | 20 | 0.585535 | 2.037978 | 0.0 | 0.034399964 | 0.333 | 5695 | tags=60%, list=24%, si | EN_1 |
| GOBP_NEGATIVE_REGUL | GOBP_NEGATIVE_REGULATION_OF_RESPON | 40 | 0.475792 | 2.0183384 | 0.0 | 0.040624354 | 0.391 | 5965 | tags=43%, list=25%, si | EN_1 |
| GOBP_COPII_COATED_VE | GOBP_COPII_COATED_VESICLE_BUDDING | 37 | 0.489736 | 2.017889 | 0.0 | 0.03970107 | 0.391 | 6747 | tags=57%, list=29%, si | EN_1 |
| GOBP_MITOCHONDRIAL_ | GOBP_MITOCHONDRIAL_RESPIRATORY_CHA | 91 | 0.410492 | 2.017458 | 0.0 | 0.038896587 | 0.391 | 6395 | tags=48%, list=27%, si | EN_1 |
| GOBP_PHOSPHATIDYLCH | GOBP_PHOSPHATIDYLCHOLINE_BIOSYNTHE | 21 | 0.547183 | 2.0082293 | 0.0 | 0.040912054 | 0.417 | 3189 | tags=52%, list=14%, si | EN_1 |
| GOBP_POSITIVE_REGULA | GOBP_POSITIVE_REGULATION_OF_COLLAGI | 39 | 0.469445 | 2.007134 | 0.005319149 | 0.040351633 | 0.419 | 3919 | tags=44%, list=17%, si | EN_1 |
| GOBP_FATTY_ACID_TRAN | GOBP_FATTY_ACID_TRANSMEMBRANE_TRA | 16 | 0.628151 | 1.9894361 | 0.0 | 0.04591854 | 0.461 | 4292 | tags=63%, list=18%, si | EN_1 |
| GOBP_ATP_BIOSYNTHETI | GOBP_ATP_BIOSYNTHETIC_PROCESS | 80 | 0.405069 | 1.9860312 | 0.0 | 0.04622225 | 0.471 | 6951 | tags=60%, list=29%, si | EN_1 |
| GOBP_REGULATION_OF_ | GOBP_REGULATION_OF_MONOCYTE_CHEM | 29 | 0.509888 | 1.9819723 | 0.0 | 0.046354868 | 0.478 | 3076 | tags=34%, list=13%, si | EN_1 |
| GOBP_TYPE_II_INTERFER | GOBP_TYPE_II_INTERFERON_MEDIATED_SIG | 23 | 0.548666 | 1.9793211 | 0.0 | 0.04622218 | 0.486 | 3576 | tags=43%, list=15%, si | EN_1 |
| GOBP_NEGATIVE_REGUL | GOBP_NEGATIVE_REGULATION_OF_EXTRIN | 26 | 0.528967 | 1.9790585 | 0.0 | 0.0455372 | 0.487 | 4561 | tags=50%, list=19%, si | EN_1 |
| GOBP_MAINTENANCE_OI | GOBP_MAINTENANCE_OF_PROTEIN_LOCAL | 43 | 0.450154 | 1.978582 | 0.0 | 0.044878203 | 0.487 | 6184 | tags=49%, list=26%, si | EN_1 |
| GOBP_NEGATIVE_REGUL | GOBP_NEGATIVE_REGULATION_OF_INSULIN | 17 | 0.573483 | 1.9688188 | 0.003717472 | 0.0480983 | 0.516 | 3413 | tags=47%, list=14%, si | EN_1 |
| GOBP_FATTY_ACYL_COA_ | GOBP_FATTY_ACYL_COA_METABOLIC_PROC | 18 | 0.56063 | 1.96643 | 0.0 | 0.047878396 | 0.519 | 4087 | tags=39%, list=17%, si | EN_1 |
| GOBP_CHAPERONE_MED | GOBP_CHAPERONE_MEDIATED_PROTEIN_C | 18 | 0.579908 | 1.9646926 | 0.0 | 0.047599174 | 0.522 | 1790 | tags=28%, list=8%, sig | EN_1 |
| ENDOMETRIAL_CANCER | ENDOMETRIAL_CANCER | 57 | 0.435356 | 1.9643146 | 0.0 | 0.046886798 | 0.522 | 6358 | tags=47%, list=27%, si | EN_1 |
| GOBP_PHOSPHOLIPID_M | GOBP_PHOSPHOLIPID_METABOLIC_PROCES | 352 | 0.328744 | 1.9624928 | 0.0 | 0.04676208 | 0.525 | 4723 | tags=34%, list=20%, si | EN_1 |
| GOBP_NAD_METABOLIC_ | GOBP_NAD_METABOLIC_PROCESS | 58 | 0.426245 | 1.9609044 | 0.0 | 0.04680764 | 0.53 | 3895 | tags=47%, list=16%, si | EN_1 |
| GOBP_NUCLEOTIDE_BINI | GOBP_NUCLEOTIDE_BINDING_OLIGOMERIZ | 18 | 0.566721 | 1.9551337 | 0.0038022813 | 0.048517756 | 0.546 | 3615 | tags=50%, list=15%, si | EN_1 |
| GOBP_NEGATIVE_REGUL | GOBP_NEGATIVE_REGULATION_OF_REACTI | 20 | 0.527532 | 1.9510477 | 0.0037313432 | 0.049075488 | 0.555 | 6126 | tags=45%, list=26%, si | EN_1 |
| GOBP_TRNA_MODIFICAT | GOBP_TRNA_MODIFICATION | 84 | 0.398892 | 1.9479914 | 0.0 | 0.049212497 | 0.562 | 6625 | tags=51%, list=28%, si | EN_1 |
| GOBP_NEGATIVE_REGUL | GOBP_NEGATIVE_REGULATION_OF_FIBROB | 44 | 0.449054 | 1.9434173 | 0.0 | 0.049842704 | 0.573 | 2813 | tags=27%, list=12%, si | EN_1 |
| GOBP_NEGATIVE_REGUL | GOBP_NEGATIVE_REGULATION_OF_PROTEI | 25 | 0.514916 | 1.941141 | 0.0 | 0.05016647 | 0.584 | 3866 | tags=44%, list=16%, si | EN_1 |
| GOBP_PROTEIN_STABILIZ | GOBP_PROTEIN_STABILIZATION | 190 | 0.34439 | 1.9235101 | 0.0 | 0.058325373 | 0.648 | 5422 | tags=34%, list=23%, si | EN_1 |
| GOBP_INFLAMMATORY_ | GOBP_INFLAMMATORY_RESPONSE_TO_WC | 30 | 0.484581 | 1.9229937 | 0.0 | 0.057608828 | 0.65 | 2667 | tags=33%, list=11%, si | EN_1 |
| GOBP_DNA_REPAIR_DEP | GOBP_DNA_REPAIR_DEPENDENT_CHROMA | 23 | 0.535659 | 1.9181894 | 0.0 | 0.059748553 | 0.666 | 2727 | tags=26%, list=12%, si | EN_1 |
| HEPATITIS_B | HEPATITIS_B | 147 | 0.353546 | 1.9168376 | 0.0 | 0.059436765 | 0.67 | 6358 | tags=41%, list=27%, si | EN_1 |
| GOBP_RESPONSE_TO_DE | GOBP_RESPONSE_TO_DEXAMETHASONE | 27 | 0.493192 | 1.9121597 | 0.0 | 0.060598608 | 0.686 | 4699 | tags=59%, list=20%, si | EN_1 |
| GOBP_PROTON_MOTIVE_ | GOBP_PROTON_MOTIVE_FORCE_DRIVEN_A | 52 | 0.430742 | 1.910728 | 0.0 | 0.06049373 | 0.691 | 7143 | tags=63%, list=30%, si | EN_1 |
| GOBP_MODULATION_BY_ | GOBP_MODULATION_BY_HOST_OF_VIRAL_ | 22 | 0.52754 | 1.9069082 | 0.0 | 0.06185258 | 0.704 | 1869 | tags=23%, list=8%, sig | EN_1 |
| FATTY_ACID_DEGRADATI | FATTY_ACID_DEGRADATION | 42 | 0.436931 | 1.9066955 | 0.0 | 0.06104405 | 0.705 | 4195 | tags=43%, list=18%, si | EN_1 |
| GOBP_REGULATION_OF_ | GOBP_REGULATION_OF_SUPEROXIDE_MET/ | 34 | 0.461199 | 1.8998008 | 0.0 | 0.06310398 | 0.719 | 1839 | tags=24%, list=8%, sig | EN_1 |
| ALZHEIMER_DISEASE | ALZHEIMER_DISEASE | 159 | 0.350644 | 1.8968078 | 0.0 | 0.06388619 | 0.725 | 6669 | tags=55%, list=28%, si | EN_1 |
| GOBP_COLLAGEN_FIBRIL | GOBP_COLLAGEN_FIBRIL_ORGANIZATION | 63 | 0.407675 | 1.8938112 | 0.0 | 0.06428225 | 0.734 | 5166 | tags=41%, list=22%, si | EN_1 |
| GOBP_GLUCOSE_CATABO | GOBP_GLUCOSE_CATABOLIC_PROCESS | 26 | 0.51095 | 1.8918979 | 0.004048583 | 0.064378105 | 0.738 | 3874 | tags=50%, list=16%, si | EN_1 |
| GOBP_ALCOHOL_BIOSYN | GOBP_ALCOHOL_BIOSYNTHETIC_PROCESS | 129 | 0.356519 | 1.891101 | 0.0 | 0.06386326 | 0.74 | 3296 | tags=29%, list=14%, si | EN_1 |
| GOBP_CELLULAR_RESPON | GOBP_CELLULAR_RESPONSE_TO_TOPOLOGI | 95 | 0.382408 | 1.8902148 | 0.0 | 0.06377786 | 0.746 | 6410 | tags=49%, list=27%, si | EN_1 |
| GOBP_REGULATION_OF_ | GOBP_REGULATION_OF_SULFUR_METABOL | 15 | 0.590461 | 1.8885416 | 0.014545455 | 0.063337214 | 0.746 | 4030 | tags=47%, list=17%, si | EN_1 |
| GOBP_TELOMERE_MAINT | GOBP_TELOMERE_MAINTENANCE_IN_RESP | 32 | 0.478841 | 1.8834084 | 0.0 | 0.06512235 | 0.765 | 3544 | tags=28%, list=15%, si | EN_1 |
| GOBP_HYALURONAN_CA | GOBP_HYALURONAN_CATABOLIC_PROCESS | 16 | 0.575037 | 1.881056 | 0.0 | 0.06540935 | 0.77 | 4473 | tags=56%, list=19%, si | EN_1 |
| GOBP_NEGATIVE_REGUL | GOBP_NEGATIVE_REGULATION_OF_FATTY_ | 17 | 0.575295 | 1.8806865 | 0.0034364262 | 0.064916335 | 0.771 | 3240 | tags=41%, list=14%, si | EN_1 |
| GOBP_CHONDROITIN_SU | GOBP_CHONDROITIN_SULFATE_METABOLIC | 22 | 0.524269 | 1.8764827 | 0.0 | 0.06677078 | 0.786 | 6417 | tags=59%, list=27%, si | EN_1 |
| GOBP_MAINTENANCE_OI | GOBP_MAINTENANCE_OF_PROTEIN_LOCAT | 97 | 0.368514 | 1.8762676 | 0.0 | 0.06610041 | 0.787 | 6184 | tags=45%, list=26%, si | EN_1 |
| GOBP_CELLULAR_RESPON | GOBP_CELLULAR_RESPONSE_TO_EXTERNAL | 36 | 0.452408 | 1.8753395 | 0.0049751243 | 0.06582693 | 0.789 | 5377 | tags=50%, list=23%, si | EN_1 |
| GOBP_RESPONSE_TO_INT | GOBP_RESPONSE_TO_INTERLEUKIN_4 | 35 | 0.464727 | 1.8747507 | 0.0 | 0.06543198 | 0.792 | 2195 | tags=34%, list=9%, sig | EN_1 |
| GOBP_REGULATION_OF_ | GOBP_REGULATION_OF_MACROPHAGE_DIF | 21 | 0.517653 | 1.8741298 | 0.01171875 | 0.06509327 | 0.793 | 4209 | tags=48%, list=18%, si | EN_1 |
| GOBP_MELANOSOME_AS | GOBP_MELANOSOME_ASSEMBLY | 19 | 0.552248 | 1.8739475 | 0.003787879 | 0.064478494 | 0.794 | 1282 | tags=21%, list=5%, sig | EN_1 |
| GOBP_REGULATION_OF_ | GOBP_REGULATION_OF_CELL_SUBSTRATE_ | 69 | 0.393592 | 1.8700631 | 0.0 | 0.06555159 | 0.802 | 6912 | tags=46%, list=29%, si | EN_1 |
| GOBP_ER_NUCLEUS_SIGN | GOBP_ER_NUCLEUS_SIGNALING_PATHWAY | 43 | 0.429741 | 1.8690335 | 0.0 | 0.06545955 | 0.805 | 6644 | tags=53%, list=28%, si | EN_1 |
| GOBP_REGULATION_OF_ | GOBP_REGULATION_OF_MITOCHONDRIAL_ | 23 | 0.506339 | 1.8672706 | 0.0 | 0.0656103 | 0.811 | 4695 | tags=39%, list=20%, si | EN_1 |
| GOBP_TAIL_ANCHORED_ | GOBP_TAIL_ANCHORED_MEMBRANE_PROT | 15 | 0.565972 | 1.8670604 | 0.0 | 0.064974606 | 0.811 | 6495 | tags=60%, list=28%, si | EN_1 |
| GOBP_TELOMERE_ORGAI | GOBP_TELOMERE_ORGANIZATION | 150 | 0.339908 | 1.865703 | 0.0 | 0.06473637 | 0.812 | 4276 | tags=27%, list=18%, si | EN_1 |
| GOBP_NEGATIVE_REGUL | GOBP_NEGATIVE_REGULATION_OF_MAP_K | 49 | 0.420265 | 1.8655424 | 0.0 | 0.06412245 | 0.812 | 3294 | tags=35%, list=14%, si | EN_1 |
| GOBP_PHOSPHATIDYLINC | GOBP_PHOSPHATIDYLINOSITOL_BIOSYNTHE | 95 | 0.366074 | 1.8654538 | 0.0 | 0.06348519 | 0.812 | 4546 | tags=35%, list=19%, si | EN_1 |
| THERMOGENESIS | THERMOGENESIS | 210 | 0.331996 | 1.8603715 | 0.0 | 0.06570305 | 0.825 | 5514 | tags=41%, list=23%, si | EN_1 |
| GOBP_MAINTENANCE_OI | GOBP_MAINTENANCE_OF_PROTEIN_LOCAT | 70 | 0.388281 | 1.8599788 | 0.0 | 0.065100454 | 0.825 | 5565 | tags=40%, list=24%, si | EN_1 |
| GOBP_MONOSACCHARID | GOBP_MONOSACCHARIDE_CATABOLIC_PRC | 40 | 0.431065 | 1.8588302 | 0.0 | 0.06520434 | 0.828 | 4468 | tags=45%, list=19%, si | EN_1 |
| SMALL_CELL_LUNG_CAN | SMALL_CELL_LUNG_CANCER | 91 | 0.364393 | 1.8513259 | 0.0 | 0.06882159 | 0.857 | 3275 | tags=25%, list=14%, si | EN_1 |
| GOBP_MONOCYTE_CHEM | GOBP_MONOCYTE_CHEMOTAXIS | 40 | 0.435889 | 1.8474852 | 0.0 | 0.06987405 | 0.865 | 3254 | tags=33%, list=14%, si | EN_1 |
| GOBP_POSITIVE_REGULA | GOBP_POSITIVE_REGULATION_OF_ERAD_P/ | 20 | 0.53019 | 1.847424 | 0.008658009 | 0.06925278 | 0.866 | 5741 | tags=55%, list=24%, si | EN_1 |
| GOBP_INTERFERON_MED | GOBP_INTERFERON_MEDIATED_SIGNALING | 76 | 0.385231 | 1.8471632 | 0.0 | 0.06874286 | 0.868 | 4393 | tags=41%, list=19%, si | EN_1 |
| GOBP_ACETYL_COA_MET | GOBP_ACETYL_COA_METABOLIC_PROCESS | 34 | 0.454353 | 1.8458811 | 0.0 | 0.06880043 | 0.869 | 3889 | tags=38%, list=16%, si | EN_1 |
| GOBP_VERY_LONG_CHAI | GOBP_VERY_LONG_CHAIN_FATTY_ACID_MI | 33 | 0.456309 | 1.8446736 | 0.0 | 0.06892285 | 0.873 | 3050 | tags=39%, list=13%, si | EN_1 |
| GOBP_METALLO_SULFUR | GOBP_METALLO_SULFUR_CLUSTER_ASSEMI | 25 | 0.478695 | 1.8422289 | 0.0 | 0.06951741 | 0.877 | 4635 | tags=40%, list=20%, si | EN_1 |

| | | | | | | | | | | |
|---|---|---|---|---|---|---|---|---|---|---|
| GOBP_PROTEIN_DESTABI | GOBP_PROTEIN_DESTABILIZATION | 56 | 0.405065 | 1.8419096 | 0.0 | 0.069029644 | 0.877 | 3417 | tags=23%, list=14%, si | EN_1 |
| GOBP_RESPONSE_TO_TY | GOBP_RESPONSE_TO_TYPE_II_INTERFERON | 122 | 0.346302 | 1.8413317 | 0.0 | 0.06868712 | 0.878 | 4172 | tags=36%, list=18%, si | EN_1 |
| GOBP_RETROGRADE_VES | GOBP_RETROGRADE_VESICLE_MEDIATED_T | 45 | 0.425616 | 1.8402712 | 0.0057803467 | 0.06874302 | 0.882 | 5849 | tags=38%, list=25%, si | EN_1 |
| BASE_EXCISION_REPAIR | BASE_EXCISION_REPAIR | 32 | 0.455407 | 1.8353047 | 0.0 | 0.07102873 | 0.889 | 4343 | tags=44%, list=18%, si | EN_1 |
| GOBP_POSITIVE_REGULA | GOBP_POSITIVE_REGULATION_OF_MONOC | 21 | 0.513862 | 1.8342314 | 0.004016064 | 0.071099035 | 0.89 | 628 | tags=19%, list=3%, sig | EN_1 |
| GOBP_POSITIVE_REGULA | GOBP_POSITIVE_REGULATION_OF_INFLAMI | 22 | 0.491877 | 1.8295226 | 0.004016064 | 0.07260039 | 0.901 | 2146 | tags=36%, list=9%, sig | EN_1 |
| TERPENOID_BACKBONE_ | TERPENOID_BACKBONE_BIOSYNTHESIS | 21 | 0.510082 | 1.8272886 | 0.008 | 0.07344909 | 0.904 | 3489 | tags=33%, list=15%, si | EN_1 |
| GOBP_MACROMOLECULE | GOBP_MACROMOLECULE_METHYLATION | 121 | 0.349107 | 1.8247591 | 0.0 | 0.07387706 | 0.908 | 6422 | tags=44%, list=27%, si | EN_1 |
| GOBP_REGULATION_OF_ | GOBP_REGULATION_OF_POST_TRANSLATIO | 242 | 0.32078 | 1.8246142 | 0.0 | 0.073290855 | 0.908 | 4741 | tags=29%, list=20%, si | EN_1 |
| GOBP_REGULATION_OF_ | GOBP_REGULATION_OF_CHOLESTEROL_BIO | 19 | 0.529807 | 1.8237967 | 0.003649635 | 0.07327732 | 0.909 | 2265 | tags=32%, list=10%, si | EN_1 |
| GOBP_REGULATION_OF_ | GOBP_REGULATION_OF_MYOBLAST_FUSIO | 22 | 0.506062 | 1.8233098 | 0.016528925 | 0.07292301 | 0.911 | 4118 | tags=50%, list=17%, si | EN_1 |
| ADIPOCYTOKINE_SIGNAL | ADIPOCYTOKINE_SIGNALING_PATHWAY | 68 | 0.387122 | 1.821717 | 0.0 | 0.07348954 | 0.914 | 5514 | tags=46%, list=23%, si | EN_1 |
| TNF_SIGNALING_PATHW | TNF_SIGNALING_PATHWAY | 108 | 0.355814 | 1.8168789 | 0.0 | 0.07522175 | 0.92 | 3254 | tags=28%, list=14%, si | EN_1 |
| GOBP_NADP_METABOLIC | GOBP_NADP_METABOLIC_PROCESS | 49 | 0.409906 | 1.8164685 | 0.0 | 0.07477094 | 0.92 | 3402 | tags=35%, list=14%, si | EN_1 |
| GOBP_GPI_ANCHOR_MET | GOBP_GPI_ANCHOR_METABOLIC_PROCESS | 33 | 0.451027 | 1.8158394 | 0.0 | 0.07447541 | 0.92 | 4317 | tags=36%, list=18%, si | EN_1 |
| GOBP_ORGANOPHOSPHA | GOBP_ORGANOPHOSPHATE_BIOSYNTHETIC | 500 | 0.300915 | 1.8134892 | 0.0 | 0.07507406 | 0.923 | 5519 | tags=39%, list=23%, si | EN_1 |
| GOBP_STEROL_METABOL | GOBP_STEROL_METABOLIC_PROCESS | 140 | 0.345098 | 1.8085176 | 0.0 | 0.07676541 | 0.925 | 2871 | tags=27%, list=12%, si | EN_1 |
| GOBP_ANTIGEN_PROCES | GOBP_ANTIGEN_PROCESSING_AND_PRESEN | 20 | 0.520007 | 1.8055449 | 0.008163265 | 0.07766055 | 0.929 | 1852 | tags=35%, list=8%, sig | EN_1 |
| GOBP_NEGATIVE_REGUL | GOBP_NEGATIVE_REGULATION_OF_REACTI | 52 | 0.401612 | 1.8053508 | 0.0 | 0.07709109 | 0.929 | 5841 | tags=38%, list=25%, si | EN_1 |
| GOBP_LIPOPROTEIN_BIO | GOBP_LIPOPROTEIN_BIOSYNTHETIC_PROCE | 104 | 0.356732 | 1.8045938 | 0.0 | 0.07695633 | 0.93 | 6052 | tags=44%, list=26%, si | EN_1 |
| GOBP_REGULATION_OF_ | GOBP_REGULATION_OF_ERAD_PATHWAY | 27 | 0.468959 | 1.8035233 | 0.004032258 | 0.0770844 | 0.931 | 5741 | tags=44%, list=24%, si | EN_1 |
| GOBP_NEGATIVE_REGUL | GOBP_NEGATIVE_REGULATION_OF_PEPTID | 52 | 0.404471 | 1.8028978 | 0.0 | 0.07692817 | 0.932 | 2420 | tags=37%, list=10%, si | EN_1 |
| EPSTEIN-BARR_VIRUS_IN | EPSTEIN-BARR_VIRUS_INFECTION | 190 | 0.317101 | 1.8023012 | 0.0 | 0.07674455 | 0.933 | 3973 | tags=26%, list=17%, si | EN_1 |
| GOBP_MUCOPOLYSACCH | GOBP_MUCOPOLYSACCHARIDE_METABOLIC | 73 | 0.387045 | 1.801496 | 0.0 | 0.07656906 | 0.934 | 4473 | tags=36%, list=19%, si | EN_1 |
| GOBP_NEGATIVE_REGUL | GOBP_NEGATIVE_REGULATION_OF_RECEPT | 27 | 0.460899 | 1.8001037 | 0.0045454544 | 0.07666054 | 0.936 | 2288 | tags=37%, list=10%, si | EN_1 |
| GOBP_THIOESTER_BIOSY | GOBP_THIOESTER_BIOSYNTHETIC_PROCESS | 27 | 0.469166 | 1.7998415 | 0.011952192 | 0.076211095 | 0.936 | 5917 | tags=48%, list=25%, si | EN_1 |
| GOBP_NEGATIVE_REGUL | GOBP_NEGATIVE_REGULATION_OF_TRANSF | 222 | 0.32213 | 1.7964396 | 0.0 | 0.07769313 | 0.944 | 3604 | tags=25%, list=15%, si | EN_1 |
| GOBP_POSITIVE_REGULA | GOBP_POSITIVE_REGULATION_OF_FATTY_A | 20 | 0.51388 | 1.7961416 | 0.0038610038 | 0.07727128 | 0.944 | 5514 | tags=70%, list=23%, si | EN_1 |
| THYROID_CANCER | THYROID_CANCER | 36 | 0.442633 | 1.7943507 | 0.0 | 0.077729486 | 0.945 | 4210 | tags=36%, list=18%, si | EN_1 |
| GOBP_REGULATION_OF_ | GOBP_REGULATION_OF_AUTOPHAGOSOME | 18 | 0.519104 | 1.7929232 | 0.007662835 | 0.07810883 | 0.948 | 6560 | tags=56%, list=28%, si | EN_1 |
| GOBP_COENZYME_A_ME | GOBP_COENZYME_A_METABOLIC_PROCESS | 19 | 0.519732 | 1.7910897 | 0.007518797 | 0.078509316 | 0.949 | 3853 | tags=42%, list=16%, si | EN_1 |
| GOBP_METHYLATION | GOBP_METHYLATION | 199 | 0.31402 | 1.7880777 | 0.0 | 0.07979616 | 0.954 | 6556 | tags=41%, list=28%, si | EN_1 |
| GOBP_GLYCEROPHOSPH | GOBP_GLYCEROPHOSPHOLIPID_METABOLIC | 267 | 0.315884 | 1.786949 | 0.0 | 0.0802553 | 0.955 | 4546 | tags=32%, list=19%, si | EN_1 |
| GOBP_MACROAUTOPHAG | GOBP_MACROAUTOPHAGY | 287 | 0.301292 | 1.7866182 | 0.0 | 0.07973061 | 0.955 | 6361 | tags=40%, list=27%, si | EN_1 |
| GOBP_NEGATIVE_REGUL | GOBP_NEGATIVE_REGULATION_OF_VASCUL | 33 | 0.43366 | 1.7833965 | 0.0 | 0.08104919 | 0.957 | 4935 | tags=48%, list=21%, si | EN_1 |
| GOBP_REGULATION_OF_ | GOBP_REGULATION_OF_EXTRINSIC_APOPT | 45 | 0.40794 | 1.7826917 | 0.019607844 | 0.081085734 | 0.959 | 4762 | tags=36%, list=20%, si | EN_1 |
| PANCREATIC_CANCER | PANCREATIC_CANCER | 74 | 0.362768 | 1.7812667 | 0.0 | 0.08133932 | 0.961 | 6524 | tags=47%, list=28%, si | EN_1 |
| GOBP_AEROBIC_ELECTRO | GOBP_AEROBIC_ELECTRON_TRANSPORT_CH | 63 | 0.37665 | 1.778629 | 0.0 | 0.082423784 | 0.963 | 5841 | tags=51%, list=25%, si | EN_1 |
| GOBP_COLLAGEN_BIOSYI | GOBP_COLLAGEN_BIOSYNTHETIC_PROCESS | 63 | 0.374711 | 1.776845 | 0.0 | 0.08279336 | 0.963 | 4930 | tags=43%, list=21%, si | EN_1 |
| GOBP_REGULATION_OF_ | GOBP_REGULATION_OF_MITOCHONDRIAL_ | 39 | 0.418968 | 1.7755721 | 0.0 | 0.08310916 | 0.965 | 4225 | tags=33%, list=18%, si | EN_1 |
| SPLICEOSOME | SPLICEOSOME | 127 | 0.343213 | 1.774476 | 0.0 | 0.08298238 | 0.966 | 14893 | tags=99%, list=63%, si | EN_1 |
| GOBP_TRICARBOXYLIC_A | GOBP_TRICARBOXYLIC_ACID_METABOLIC_P | 16 | 0.536365 | 1.7735646 | 0.011363637 | 0.083169706 | 0.967 | 3898 | tags=50%, list=17%, si | EN_1 |
| GOBP_CELLULAR_OXIDAN | GOBP_CELLULAR_OXIDANT_DETOXIFICATIO | 29 | 0.469417 | 1.7721629 | 0.0 | 0.08340607 | 0.968 | 3956 | tags=45%, list=17%, si | EN_1 |
| GOBP_FATTY_ACID_BIOS | GOBP_FATTY_ACID_BIOSYNTHETIC_PROCES | 145 | 0.328619 | 1.7694281 | 0.0 | 0.08469787 | 0.97 | 3240 | tags=30%, list=14%, si | EN_1 |
| GOBP_POSITIVE_REGULA | GOBP_POSITIVE_REGULATION_OF_PROTEIN | 24 | 0.460468 | 1.7685083 | 0.004464286 | 0.08492016 | 0.971 | 1473 | tags=21%, list=6%, sig | EN_1 |
| PROSTATE_CANCER | PROSTATE_CANCER | 93 | 0.354991 | 1.7670931 | 0.009803922 | 0.08511276 | 0.972 | 4546 | tags=31%, list=19%, si | EN_1 |
| GOBP_NEGATIVE_REGUL | GOBP_NEGATIVE_REGULATION_OF_KINASE | 188 | 0.319962 | 1.76639 | 0.0 | 0.08497405 | 0.972 | 3294 | tags=24%, list=14%, si | EN_1 |
| GOBP_ARGININE_METAB | GOBP_ARGININE_METABOLIC_PROCESS | 17 | 0.532878 | 1.7635999 | 0.011363637 | 0.08603479 | 0.975 | 3701 | tags=59%, list=16%, si | EN_1 |
| GOBP_NEGATIVE_REGUL | GOBP_NEGATIVE_REGULATION_OF_EPITHEI | 66 | 0.378428 | 1.7621168 | 0.0 | 0.08640621 | 0.976 | 2667 | tags=26%, list=11%, si | EN_1 |
| GOBP_CHONDROITIN_SU | GOBP_CHONDROITIN_SULFATE_PROTEOGLY | 29 | 0.453622 | 1.7619795 | 0.0 | 0.085917264 | 0.976 | 5299 | tags=52%, list=22%, si | EN_1 |
| NON-ALCOHOLIC_FATTY_ | NON-ALCOHOLIC_FATTY_LIVER_DISEASE_(N | 141 | 0.33125 | 1.7615287 | 0.0 | 0.085734695 | 0.976 | 6719 | tags=53%, list=28%, si | EN_1 |
| GOBP_POSITIVE_REGULA | GOBP_POSITIVE_REGULATION_OF_RESPON | 37 | 0.430479 | 1.7610993 | 0.005319149 | 0.08537266 | 0.976 | 5741 | tags=46%, list=24%, si | EN_1 |
| GOBP_TRNA_PROCESSIN | GOBP_TRNA_PROCESSING | 122 | 0.336642 | 1.7550514 | 0.0 | 0.08890697 | 0.978 | 6625 | tags=44%, list=28%, si | EN_1 |
| COLORECTAL_CANCER | COLORECTAL_CANCER | 86 | 0.35204 | 1.7539965 | 0.0 | 0.08920373 | 0.978 | 6358 | tags=42%, list=27%, si | EN_1 |
| GOBP_REGULATION_OF_ | GOBP_REGULATION_OF_PROTEIN_LOCALIZ | 44 | 0.408404 | 1.7531549 | 0.004901961 | 0.08918395 | 0.979 | 6224 | tags=43%, list=26%, si | EN_1 |
| GOBP_CELLULAR_RESPOI | GOBP_CELLULAR_RESPONSE_TO_DEXAMETI | 23 | 0.472531 | 1.7501334 | 0.0 | 0.09033684 | 0.98 | 4699 | tags=57%, list=20%, si | EN_1 |
| GOBP_CELLULAR_PIGMEI | GOBP_CELLULAR_PIGMENTATION | 57 | 0.386032 | 1.7500434 | 0.0 | 0.0898464 | 0.98 | 2193 | tags=18%, list=9%, sig | EN_1 |
| GOBP_ETHER_METABOLI | GOBP_ETHER_METABOLIC_PROCESS | 23 | 0.458867 | 1.7494475 | 0.004385965 | 0.08955674 | 0.98 | 2958 | tags=30%, list=13%, si | EN_1 |
| GOBP_REGULATION_OF_ | GOBP_REGULATION_OF_TELOMERE_MAINT | 98 | 0.337742 | 1.7489688 | 0.0 | 0.08929657 | 0.98 | 4276 | tags=24%, list=18%, si | EN_1 |
| GOBP_RNA_METHYLATIO | GOBP_RNA_METHYLATION | 65 | 0.382148 | 1.7463542 | 0.0 | 0.09047377 | 0.982 | 7377 | tags=65%, list=31%, si | EN_1 |
| GOBP_NEGATIVE_REGUL | GOBP_NEGATIVE_REGULATION_OF_AMIDE_ | 22 | 0.472554 | 1.7453543 | 0.004237288 | 0.090530686 | 0.982 | 5297 | tags=59%, list=22%, si | EN_1 |
| GOBP_REGULATION_OF_ | GOBP_REGULATION_OF_SUPEROXIDE_ANIC | 28 | 0.447724 | 1.738695 | 0.0045454544 | 0.094396286 | 0.985 | 1748 | tags=21%, list=7%, sig | EN_1 |
| GOBP_NUCLEOSIDE_TRIP | GOBP_NUCLEOSIDE_TRIPHOSPHATE_BIOSY | 100 | 0.343676 | 1.7385676 | 0.0 | 0.09387793 | 0.985 | 6395 | tags=50%, list=27%, si | EN_1 |
| GOBP_POLYAMINE_META | GOBP_POLYAMINE_METABOLIC_PROCESS | 19 | 0.502439 | 1.7370211 | 0.016949153 | 0.09417126 | 0.985 | 1862 | tags=26%, list=8%, sig | EN_1 |
| GOBP_GLYCERALDEHYDE | GOBP_GLYCERALDEHYDE_3_PHOSPHATE_M | 19 | 0.498487 | 1.7355877 | 0.012448133 | 0.094399236 | 0.986 | 5924 | tags=68%, list=25%, si | EN_1 |
| GOBP_RESPONSE_TO_TY | GOBP_RESPONSE_TO_TYPE_I_INTERFERON | 64 | 0.371291 | 1.7328187 | 0.0 | 0.09528188 | 0.987 | 4393 | tags=41%, list=19%, si | EN_1 |
| GOBP_POSITIVE_REGULA | GOBP_POSITIVE_REGULATION_OF_PROTEIN | 20 | 0.484197 | 1.731184 | 0.01724138 | 0.09606806 | 0.987 | 6484 | tags=50%, list=27%, si | EN_1 |
| GOBP_B_CELL_APOPTOTI | GOBP_B_CELL_APOPTOTIC_PROCESS | 29 | 0.441661 | 1.7288225 | 0.004608295 | 0.09728087 | 0.989 | 2627 | tags=34%, list=11%, si | EN_1 |
| GOBP_REGULATION_OF_ | GOBP_REGULATION_OF_TRANSLATIONAL_I | 17 | 0.521854 | 1.7274958 | 0.014084507 | 0.09775714 | 0.989 | 4150 | tags=35%, list=18%, si | EN_1 |
| HIF-1_SIGNALING_PATHV | HIF-1_SIGNALING_PATHWAY | 98 | 0.339377 | 1.7273141 | 0.0 | 0.09726141 | 0.989 | 4859 | tags=39%, list=21%, si | EN_1 |
| BLADDER_CANCER | BLADDER_CANCER | 37 | 0.413222 | 1.7257557 | 0.0056497175 | 0.09766317 | 0.99 | 6482 | tags=51%, list=27%, si | EN_1 |
| GOBP_RESPONSE_TO_AR | GOBP_RESPONSE_TO_ARSENIC_CONTAININ | 25 | 0.451434 | 1.7251748 | 0.004524887 | 0.09757884 | 0.991 | 6879 | tags=64%, list=29%, si | EN_1 |
| GOBP_QUINONE_METAB | GOBP_QUINONE_METABOLIC_PROCESS | 23 | 0.464626 | 1.7240818 | 0.009049774 | 0.097990215 | 0.993 | 5650 | tags=61%, list=24%, si | EN_1 |
| GOBP_NEGATIVE_REGUL | GOBP_NEGATIVE_REGULATION_OF_SMOOT | 36 | 0.414451 | 1.7234302 | 0.0 | 0.09814528 | 0.993 | 3516 | tags=31%, list=15%, si | EN_1 |
| GOBP_MONOCYTE_DIFFE | GOBP_MONOCYTE_DIFFERENTIATION | 28 | 0.446265 | 1.7220882 | 0.004464286 | 0.09847435 | 0.993 | 3169 | tags=36%, list=13%, si | EN_1 |
| GOBP_SULFUR_COMPOU | GOBP_SULFUR_COMPOUND_CATABOLIC_P | 35 | 0.419306 | 1.7191514 | 0.004901961 | 0.09988714 | 0.993 | 5299 | tags=51%, list=22%, si | EN_1 |
| GOBP_MITOTIC_G2_DNA | GOBP_MITOTIC_G2_DNA_DAMAGE_CHECK | 39 | 0.404342 | 1.7189109 | 0.0 | 0.09943724 | 0.993 | 3961 | tags=31%, list=17%, si | EN_1 |
| GOBP_CHAPERONE_COF | GOBP_CHAPERONE_COFACTOR_DEPENDEN | 31 | 0.42631 | 1.7178016 | 0.0050251256 | 0.09963653 | 0.994 | 4504 | tags=35%, list=19%, si | EN_1 |
| GOBP_PHOSPHATIDYLCH | GOBP_PHOSPHATIDYLCHOLINE_CATABOLIC_ | 15 | 0.54289 | 1.7169644 | 0.010416667 | 0.09986728 | 0.994 | 2608 | tags=40%, list=11%, si | EN_1 |
| GOBP_SUPEROXIDE_MET | GOBP_SUPEROXIDE_METABOLIC_PROCESS | 68 | 0.362862 | 1.7164105 | 0.0 | 0.099769734 | 0.994 | 3956 | tags=34%, list=17%, si | EN_1 |
| INFLUENZA_A | INFLUENZA_A | 139 | 0.316356 | 1.7150421 | 0.0 | 0.10031294 | 0.994 | 4779 | tags=34%, list=20%, si | EN_1 |
| GOBP_REGULATION_OF_ | GOBP_REGULATION_OF_FATTY_ACID_META | 99 | 0.329886 | 1.7149994 | 0.0 | 0.099795416 | 0.994 | 4131 | tags=31%, list=17%, si | EN_1 |

| | | | | | | | | | | |
|---|---|---|---|---|---|---|---|---|---|---|
| PROTEIN_PROCESSING_IN | PROTEIN_PROCESSING_IN_ENDOPLASMIC_R | 158 | 0.310004 | 1.7146071 | 0.0 | 0.09945324 | 0.994 | 7296 | tags=47%, list=31%, si | EN_1 |
| GOBP_BASE_EXCISION_R | GOBP_BASE_EXCISION_REPAIR | 41 | 0.403517 | 1.7070577 | 0.0 | 0.104218684 | 0.996 | 4272 | tags=34%, list=18%, si | EN_1 |
| GOBP_REGULATION_OF_ | GOBP_REGULATION_OF_B_CELL_APOPTOTI | 23 | 0.463759 | 1.7051347 | 0.013888889 | 0.105338685 | 0.997 | 2186 | tags=35%, list=9%, sig | EN_1 |
| FLUID_SHEAR_STRESS_AN | FLUID_SHEAR_STRESS_AND_ATHEROSCLERO | 135 | 0.317476 | 1.7028416 | 0.0 | 0.10642248 | 0.997 | 3647 | tags=27%, list=15%, si | EN_1 |
| GOBP_ANTIGEN_PROCES | GOBP_ANTIGEN_PROCESSING_AND_PRESEN | 28 | 0.420184 | 1.7025802 | 0.00456621 | 0.10605524 | 0.997 | 3154 | tags=32%, list=13%, si | EN_1 |
| GOBP_POSITIVE_REGULA | GOBP_POSITIVE_REGULATION_OF_TELOME | 15 | 0.536274 | 1.6994011 | 0.010135135 | 0.108084746 | 0.998 | 6452 | tags=47%, list=27%, si | EN_1 |
| GOBP_NEGATIVE_REGUL | GOBP_NEGATIVE_REGULATION_OF_AMYLO | 20 | 0.485041 | 1.6974423 | 0.00754717 | 0.10926107 | 0.999 | 3370 | tags=40%, list=14%, si | EN_1 |
| GOBP_NEGATIVE_REGUL | GOBP_NEGATIVE_REGULATION_OF_CANON | 57 | 0.372072 | 1.6956908 | 0.0 | 0.11015156 | 0.999 | 4637 | tags=37%, list=20%, si | EN_1 |
| GOBP_REGULATION_OF_ | GOBP_REGULATION_OF_LYSOSOMAL_LUME | 23 | 0.473877 | 1.6955453 | 0.012658228 | 0.109681904 | 0.999 | 5364 | tags=39%, list=23%, si | EN_1 |
| GOBP_ERAD_PATHWAY | GOBP_ERAD_PATHWAY | 108 | 0.326007 | 1.6953367 | 0.0 | 0.10934286 | 0.999 | 5741 | tags=34%, list=24%, si | EN_1 |
| GOBP_PROTEIN_LOCALIZ | GOBP_PROTEIN_LOCALIZATION_TO_GOLGI_ | 30 | 0.425449 | 1.6951708 | 0.0045454544 | 0.10897367 | 0.999 | 5277 | tags=33%, list=22%, si | EN_1 |
| GOBP_POSITIVE_REGULA | GOBP_POSITIVE_REGULATION_OF_TORC1_S | 52 | 0.381838 | 1.6944156 | 0.0 | 0.10885555 | 0.999 | 5377 | tags=35%, list=23%, si | EN_1 |
| GOBP_POSITIVE_REGULA | GOBP_POSITIVE_REGULATION_OF_PROTEAS | 102 | 0.333633 | 1.6933119 | 0.0 | 0.109149255 | 0.999 | 5929 | tags=41%, list=25%, si | EN_1 |
| GOBP_PURINE_CONTAIN | GOBP_PURINE_CONTAINING_COMPOUND_ | 150 | 0.315308 | 1.6932 | 0.0 | 0.108674295 | 0.999 | 3924 | tags=32%, list=17%, si | EN_1 |
| GOBP_PROGRAMMED_N | GOBP_PROGRAMMED_NECROTIC_CELL_DEA | 50 | 0.370089 | 1.6905708 | 0.0057803467 | 0.11055589 | 0.999 | 5726 | tags=44%, list=24%, si | EN_1 |
| GOBP_GLYCEROLIPID_ME | GOBP_GLYCEROLIPID_METABOLIC_PROCESS | 361 | 0.277204 | 1.6905696 | 0.0 | 0.11001127 | 0.999 | 3719 | tags=25%, list=16%, si | EN_1 |
| GOBP_REGULATION_OF_ | GOBP_REGULATION_OF_B_CELL_DIFFERENT | 37 | 0.404653 | 1.6899117 | 0.0 | 0.10998206 | 0.999 | 2625 | tags=32%, list=11%, si | EN_1 |
| GOBP_SMOOTH_MUSCLE | GOBP_SMOOTH_MUSCLE_CELL_PROLIFERAT | 191 | 0.29434 | 1.6897713 | 0.0 | 0.10955044 | 0.999 | 4307 | tags=34%, list=18%, si | EN_1 |
| GOBP_MITOCHONDRIAL_ | GOBP_MITOCHONDRIAL_OUTER_MEMBRAN | 34 | 0.406857 | 1.6897539 | 0.0049261083 | 0.10903642 | 0.999 | 6115 | tags=47%, list=26%, si | EN_1 |
| GOBP_REGULATION_OF_ | GOBP_REGULATION_OF_INFLAMMASOME_ | 40 | 0.395855 | 1.6895676 | 0.0 | 0.10859552 | 0.999 | 4138 | tags=38%, list=18%, si | EN_1 |
| MELANOMA | MELANOMA | 69 | 0.352967 | 1.6893119 | 0.0 | 0.108334325 | 0.999 | 4551 | tags=32%, list=19%, si | EN_1 |
| GOBP_POSITIVE_REGULA | GOBP_POSITIVE_REGULATION_OF_CELL_SU | 35 | 0.406021 | 1.6891481 | 0.009756098 | 0.10802008 | 0.999 | 7608 | tags=63%, list=32%, si | EN_1 |
| GOBP_RESPONSE_TO_EN | GOBP_RESPONSE_TO_ENDOPLASMIC_RETIC | 261 | 0.295912 | 1.6890409 | 0.0 | 0.10757502 | 0.999 | 5841 | tags=37%, list=25%, si | EN_1 |
| GOBP_POSITIVE_REGULA | GOBP_POSITIVE_REGULATION_OF_MYELOID | 21 | 0.477589 | 1.6881211 | 0.004115226 | 0.107912555 | 0.999 | 1643 | tags=29%, list=7%, sig | EN_1 |
| GOBP_REGULATION_OF_ | GOBP_REGULATION_OF_CYTOPLASMIC_PAT | 104 | 0.329044 | 1.6878996 | 0.0 | 0.10750635 | 0.999 | 4697 | tags=33%, list=20%, si | EN_1 |
| GOBP_SUBSTRATE_ADHE | GOBP_SUBSTRATE_ADHESION_DEPENDENT_ | 99 | 0.33804 | 1.6871893 | 0.0 | 0.107489966 | 0.999 | 3275 | tags=26%, list=14%, si | EN_1 |
| GOBP_FIBROBLAST_MIGR | GOBP_FIBROBLAST_MIGRATION | 68 | 0.351218 | 1.6866764 | 0.0 | 0.10734537 | 0.999 | 5095 | tags=34%, list=22%, si | EN_1 |
| GOBP_POSITIVE_REGULA | GOBP_POSITIVE_REGULATION_OF_RECEPTO | 17 | 0.500637 | 1.6864555 | 0.014545455 | 0.10696373 | 0.999 | 2824 | tags=35%, list=12%, si | EN_1 |
| CYSTEINE_AND_METHION | CYSTEINE_AND_METHIONINE_METABOLISM | 44 | 0.39851 | 1.6862398 | 0.011976048 | 0.106667824 | 0.999 | 4622 | tags=45%, list=20%, si | EN_1 |
| GOBP_RIBOSE_PHOSPHA | GOBP_RIBOSE_PHOSPHATE_BIOSYNTHETIC_ | 185 | 0.307114 | 1.6860772 | 0.0 | 0.106278025 | 0.999 | 5357 | tags=36%, list=23%, si | EN_1 |
| GOBP_UBIQUINONE_ME | GOBP_UBIQUINONE_METABOLIC_PROCESS | 19 | 0.491858 | 1.6856233 | 0.008196721 | 0.10613185 | 0.999 | 5650 | tags=58%, list=24%, si | EN_1 |
| GOBP_PURINE_NUCLEOS | GOBP_PURINE_NUCLEOSIDE_MONOPHOSPH | 22 | 0.464467 | 1.6849447 | 0.019762846 | 0.10612149 | 0.999 | 1569 | tags=23%, list=7%, sig | EN_1 |
| GOBP_REGULATION_OF_ | GOBP_REGULATION_OF_PROTEIN_UBIQUIT | 202 | 0.310075 | 1.6844226 | 0.0 | 0.10593109 | 0.999 | 4537 | tags=27%, list=19%, si | EN_1 |
| GOBP_VASCULAR_ASSOC | GOBP_VASCULAR_ASSOCIATED_SMOOTH_M | 80 | 0.342842 | 1.6840774 | 0.0 | 0.10577585 | 0.999 | 4951 | tags=40%, list=21%, si | EN_1 |
| GOBP_VESICLE_BUDDING | GOBP_VESICLE_BUDDING_FROM_MEMBRA | 79 | 0.329595 | 1.6838387 | 0.0 | 0.10550754 | 0.999 | 7130 | tags=51%, list=30%, si | EN_1 |
| GALACTOSE_METABOLISM | GALACTOSE_METABOLISM | 29 | 0.437289 | 1.68325 | 0.0049751243 | 0.10553125 | 0.999 | 4468 | tags=41%, list=19%, si | EN_1 |
| GOBP_PROTEIN_K63_LIN | GOBP_PROTEIN_K63_LINKED_DEUBIQUITIN | 29 | 0.437084 | 1.6816069 | 0.014018691 | 0.10628945 | 1 | 2129 | tags=17%, list=9%, sig | EN_1 |
| GOBP_PROTEIN_LOCALIZ | GOBP_PROTEIN_LOCALIZATION_TO_ENDOP | 76 | 0.344275 | 1.6801816 | 0.007874016 | 0.106835574 | 1 | 5734 | tags=42%, list=24%, si | EN_1 |
| GOBP_FOCAL_ADHESION | GOBP_FOCAL_ADHESION_ASSEMBLY | 87 | 0.332505 | 1.6790416 | 0.0 | 0.10717046 | 1 | 6912 | tags=47%, list=29%, si | EN_1 |
| GOBP_BONE_CELL_DEVE | GOBP_BONE_CELL_DEVELOPMENT | 17 | 0.492831 | 1.677987 | 0.033834588 | 0.107316 | 1 | 4254 | tags=47%, list=18%, si | EN_1 |
| GOBP_RESPONSE_TO_GA | GOBP_RESPONSE_TO_GAMMA_RADIATION | 48 | 0.378291 | 1.6768795 | 0.0 | 0.107599586 | 1 | 3379 | tags=29%, list=14%, si | EN_1 |
| GOBP_GOLGI_TO_PLASM | GOBP_GOLGI_TO_PLASMA_MEMBRANE_PR | 38 | 0.40195 | 1.6766042 | 0.010928961 | 0.1073483 | 1 | 4487 | tags=32%, list=19%, si | EN_1 |
| GOBP_CELLULAR_RESPON | GOBP_CELLULAR_RESPONSE_TO_TYPE_II_IN | 95 | 0.33199 | 1.6756644 | 0.0 | 0.107483536 | 1 | 4584 | tags=35%, list=19%, si | EN_1 |
| GOBP_LONG_CHAIN_FAT | GOBP_LONG_CHAIN_FATTY_ACID_BIOSYNT | 19 | 0.484386 | 1.673507 | 0.011857707 | 0.10876747 | 1 | 1690 | tags=32%, list=7%, sig | EN_1 |
| GOBP_RESPONSE_TO_MI | GOBP_RESPONSE_TO_MISFOLDED_PROTEIN | 18 | 0.488045 | 1.672943 | 0.0040650405 | 0.10879441 | 1 | 6356 | tags=56%, list=27%, si | EN_1 |
| GOBP_POSITIVE_REGULA | GOBP_POSITIVE_REGULATION_OF_MACROP | 15 | 0.53663 | 1.6706145 | 0.018656716 | 0.10985093 | 1 | 4209 | tags=47%, list=18%, si | EN_1 |
| ARGININE_BIOSYNTHESIS | ARGININE_BIOSYNTHESIS | 17 | 0.495454 | 1.669637 | 0.022304833 | 0.10998277 | 1 | 1751 | tags=35%, list=7%, sig | EN_1 |
| GOBP_MITOCHONDRIAL_ | GOBP_MITOCHONDRIAL_TRANSLATION | 128 | 0.31079 | 1.6677916 | 0.0 | 0.111136794 | 1 | 6187 | tags=38%, list=26%, si | EN_1 |
| GOBP_POSITIVE_REGULA | GOBP_POSITIVE_REGULATION_OF_TYPE_I_I | 19 | 0.474898 | 1.6670235 | 0.008032128 | 0.11125364 | 1 | 4393 | tags=58%, list=19%, si | EN_1 |
| GOBP_DE_NOVO_PROTEI | GOBP_DE_NOVO_PROTEIN_FOLDING | 38 | 0.405741 | 1.6656102 | 0.0053475937 | 0.111817524 | 1 | 5085 | tags=37%, list=22%, si | EN_1 |
| GOBP_NEGATIVE_REGUL | GOBP_NEGATIVE_REGULATION_OF_ENDOT | 30 | 0.422739 | 1.66385 | 0.005 | 0.112616204 | 1 | 2341 | tags=30%, list=10%, si | EN_1 |
| GOBP_POST_TRANSLATIO | GOBP_POST_TRANSLATIONAL_PROTEIN_TA | 18 | 0.487761 | 1.6633084 | 0.008097166 | 0.11247353 | 1 | 2816 | tags=33%, list=12%, si | EN_1 |
| GOBP_REGULATION_OF_ | GOBP_REGULATION_OF_ENDOPLASMIC_RE | 24 | 0.450275 | 1.6619167 | 0.00862069 | 0.113160715 | 1 | 4328 | tags=38%, list=18%, si | EN_1 |
| GOBP_POSTREPLICATION | GOBP_POSTREPLICATION_REPAIR | 29 | 0.436735 | 1.6610051 | 0.013043478 | 0.113269545 | 1 | 2546 | tags=28%, list=11%, si | EN_1 |
| GOBP_ELECTRON_TRANS | GOBP_ELECTRON_TRANSPORT_CHAIN | 107 | 0.323147 | 1.6591609 | 0.0 | 0.11437598 | 1 | 5841 | tags=47%, list=25%, si | EN_1 |
| GOBP_POSITIVE_REGULA | GOBP_POSITIVE_REGULATION_OF_GLYCOP | 26 | 0.433361 | 1.6551797 | 0.013100437 | 0.11733363 | 1 | 4742 | tags=38%, list=20%, si | EN_1 |
| GOBP_ENDOSOME_ORGA | GOBP_ENDOSOME_ORGANIZATION | 68 | 0.347088 | 1.6543167 | 0.0 | 0.11754398 | 1 | 3534 | tags=25%, list=15%, si | EN_1 |
| GOBP_ANTIGEN_PROCES | GOBP_ANTIGEN_PROCESSING_AND_PRESEN | 34 | 0.404247 | 1.6524886 | 0.014851485 | 0.11828194 | 1 | 3154 | tags=32%, list=13%, si | EN_1 |
| GOBP_GLYCOPROTEIN_C | GOBP_GLYCOPROTEIN_CATABOLIC_PROCES | 31 | 0.42014 | 1.6521833 | 0.0 | 0.118078545 | 1 | 2751 | tags=29%, list=12%, si | EN_1 |
| GOBP_REGULATION_OF_ | GOBP_REGULATION_OF_COLLAGEN_METAE | 61 | 0.35576 | 1.6510469 | 0.0 | 0.118557386 | 1 | 3919 | tags=38%, list=17%, si | EN_1 |
| GOBP_REGULATION_OF_ | GOBP_REGULATION_OF_CELL_MATRIX_ADH | 121 | 0.313969 | 1.6505371 | 0.0 | 0.11844836 | 1 | 5050 | tags=31%, list=21%, si | EN_1 |
| GOBP_POSITIVE_REGULA | GOBP_POSITIVE_REGULATION_OF_INTRINS | 69 | 0.344297 | 1.6502455 | 0.0 | 0.11826068 | 1 | 3523 | tags=29%, list=15%, si | EN_1 |
| GOBP_ATP_SYNTHESIS_C | GOBP_ATP_SYNTHESIS_COUPLED_ELECTRO | 76 | 0.339768 | 1.6489613 | 0.0 | 0.1187456 | 1 | 5841 | tags=49%, list=25%, si | EN_1 |
| GOBP_NUCLEOSIDE_BISP | GOBP_NUCLEOSIDE_BISPHOSPHATE_METAE | 102 | 0.321695 | 1.6483462 | 0.0 | 0.11869714 | 1 | 4149 | tags=32%, list=18%, si | EN_1 |
| GOBP_PEROXISOMAL_TR | GOBP_PEROXISOMAL_TRANSPORT | 21 | 0.458467 | 1.6472559 | 0.01754386 | 0.118935674 | 1 | 6779 | tags=57%, list=29%, si | EN_1 |
| GOBP_PYRIDINE_CONTAI | GOBP_PYRIDINE_CONTAINING_COMPOUND | 109 | 0.313937 | 1.6463345 | 0.0 | 0.119047195 | 1 | 3924 | tags=32%, list=17%, si | EN_1 |
| HEPATITIS_C | HEPATITIS_C | 138 | 0.312147 | 1.6462817 | 0.0 | 0.11859202 | 1 | 5180 | tags=36%, list=22%, si | EN_1 |
| GOBP_PROTEIN_LOCALIZ | GOBP_PROTEIN_LOCALIZATION_TO_NUCLEA | 15 | 0.51116 | 1.6457307 | 0.025362318 | 0.118553646 | 1 | 4802 | tags=47%, list=20%, si | EN_1 |
| GOBP_AMINOGLYCAN_B | GOBP_AMINOGLYCAN_BIOSYNTHETIC_PROC | 60 | 0.356714 | 1.6440479 | 0.0 | 0.11929241 | 1 | 5026 | tags=43%, list=21%, si | EN_1 |
| GOBP_RESPONSE_TO_MU | GOBP_RESPONSE_TO_MURAMYL_DIPEPTID | 20 | 0.462504 | 1.6422819 | 0.0125 | 0.12027532 | 1 | 3768 | tags=40%, list=16%, si | EN_1 |
| GOBP_ENDOPLASMIC_RE | GOBP_ENDOPLASMIC_RETICULUM_TO_GOL | 120 | 0.317526 | 1.6420183 | 0.0 | 0.119907096 | 1 | 6438 | tags=40%, list=27%, si | EN_1 |
| GOBP_CELLULAR_MODIFI | GOBP_CELLULAR_MODIFIED_AMINO_ACID_ | 40 | 0.374305 | 1.6414752 | 0.005263158 | 0.120007135 | 1 | 3790 | tags=40%, list=16%, si | EN_1 |
| GOBP_CANONICAL_NF_K | GOBP_CANONICAL_NF_KAPPAB_SIGNAL_TR | 247 | 0.276009 | 1.6406993 | 0.0 | 0.12016671 | 1 | 4209 | tags=30%, list=18%, si | EN_1 |
| GOBP_VESICLE_MEDIATE | GOBP_VESICLE_MEDIATED_TRANSPORT_BET | 46 | 0.378594 | 1.6386182 | 0.005235602 | 0.12142358 | 1 | 6583 | tags=46%, list=28%, si | EN_1 |
| GOBP_MITOCHONDRIAL_ | GOBP_MITOCHONDRIAL_GENE_EXPRESSION | 164 | 0.305021 | 1.6375816 | 0.0 | 0.12177238 | 1 | 6187 | tags=38%, list=26%, si | EN_1 |
| GOBP_ORGANOPHOSPHA | GOBP_ORGANOPHOSPHATE_CATABOLIC_PF | 227 | 0.280894 | 1.6369956 | 0.0 | 0.12164688 | 1 | 2669 | tags=24%, list=11%, si | EN_1 |
| GOBP_TORC1_SIGNALING | GOBP_TORC1_SIGNALING | 100 | 0.324728 | 1.6363211 | 0.0 | 0.1216771 | 1 | 15976 | tags=100%, list=68%, | EN_1 |
| GOBP_POSITIVE_REGULA | GOBP_POSITIVE_REGULATION_BY_HOST_OI | 22 | 0.458891 | 1.6361742 | 0.019762846 | 0.12129835 | 1 | 7456 | tags=64%, list=32%, si | EN_1 |
| GOBP_REGULATION_OF_ | GOBP_REGULATION_OF_FATTY_ACID_OXIDA | 41 | 0.400162 | 1.635951 | 0.010810811 | 0.121048614 | 1 | 4131 | tags=41%, list=17%, si | EN_1 |
| GOBP_NEGATIVE_REGUL | GOBP_NEGATIVE_REGULATION_OF_HEART_ | 15 | 0.491849 | 1.6334193 | 0.014184397 | 0.12257708 | 1 | 6067 | tags=53%, list=26%, si | EN_1 |
| GOBP_NEGATIVE_REGUL | GOBP_NEGATIVE_REGULATION_OF_INTRAC | 35 | 0.392391 | 1.6326607 | 0.0046948357 | 0.122803144 | 1 | 5841 | tags=37%, list=25%, si | EN_1 |
| GOBP_MITOTIC_G2_M_T | GOBP_MITOTIC_G2_M_TRANSITION_CHECK | 56 | 0.356792 | 1.6323562 | 0.0 | 0.12259861 | 1 | 5276 | tags=36%, list=22%, si | EN_1 |

| | | | | | | | | | | |
|---|---|---|---|---|---|---|---|---|---|---|
| GOBP_PEPTIDYL_LYSINE_ | GOBP_PEPTIDYL_LYSINE_MODIFICATION | 128 | 0.30476 | 1.6316228 | 0.0 | 0.12283934 | 1 | 4590 | tags=26%, list=19%, si | EN_1 |
| GOBP_SULFUR_COMPOU | GOBP_SULFUR_COMPOUND_METABOLIC_P | 292 | 0.284244 | 1.6314642 | 0.0 | 0.12247942 | 1 | 4284 | tags=31%, list=18%, si | EN_1 |
| GOBP_NEGATIVE_REGUL | GOBP_NEGATIVE_REGULATION_OF_VASCUL | 109 | 0.312797 | 1.6305984 | 0.010638298 | 0.122676 | 1 | 3441 | tags=28%, list=15%, si | EN_1 |
| GOBP_NEGATIVE_REGUL | GOBP_NEGATIVE_REGULATION_OF_SMALL_ | 61 | 0.347731 | 1.6281358 | 0.0074074073 | 0.12425476 | 1 | 3926 | tags=33%, list=17%, si | EN_1 |
| GOBP_MODULATION_BY_ | GOBP_MODULATION_BY_HOST_OF_VIRAL_ | 44 | 0.377761 | 1.6272541 | 0.01923077 | 0.124556296 | 1 | 3103 | tags=23%, list=13%, si | EN_1 |
| GOBP_POSITIVE_REGULA | GOBP_POSITIVE_REGULATION_OF_B_CELL_ | 18 | 0.474163 | 1.6267776 | 0.013636364 | 0.12454502 | 1 | 2043 | tags=33%, list=9%, sig | EN_1 |
| GOBP_SNRNA_METABOL | GOBP_SNRNA_METABOLIC_PROCESS | 50 | 0.361584 | 1.6247784 | 0.005952381 | 0.12562777 | 1 | 5092 | tags=34%, list=22%, si | EN_1 |
| GOBP_CGAS_STING_SIGN | GOBP_CGAS_STING_SIGNALING_PATHWAY | 16 | 0.483758 | 1.6243463 | 0.020905923 | 0.1255839 | 1 | 5377 | tags=38%, list=23%, si | EN_1 |
| GOBP_PHOSPHOLIPID_CA | GOBP_PHOSPHOLIPID_CATABOLIC_PROCESS | 52 | 0.35719 | 1.6237384 | 0.0 | 0.1256369 | 1 | 2608 | tags=29%, list=11%, si | EN_1 |
| OTHER_GLYCAN_DEGRAD | OTHER_GLYCAN_DEGRADATION | 16 | 0.487688 | 1.6216059 | 0.032520324 | 0.12683356 | 1 | 4761 | tags=44%, list=20%, si | EN_1 |
| FATTY_ACID_ELONGATIO | FATTY_ACID_ELONGATION | 27 | 0.422544 | 1.62031 | 0.011904762 | 0.12778744 | 1 | 3239 | tags=33%, list=14%, si | EN_1 |
| GOBP_REGULATION_OF_ | GOBP_REGULATION_OF_TRANSLATION_IN_ | 24 | 0.44207 | 1.6200341 | 0.021551725 | 0.12756383 | 1 | 5168 | tags=38%, list=22%, si | EN_1 |
| AUTOPHAGY | AUTOPHAGY | 126 | 0.31096 | 1.6199728 | 0.0 | 0.12712502 | 1 | 6802 | tags=43%, list=29%, si | EN_1 |
| GOBP_REGULATION_OF_ | GOBP_REGULATION_OF_PROTEIN_STABILIT | 305 | 0.265708 | 1.6199216 | 0.0 | 0.12675288 | 1 | 4715 | tags=27%, list=20%, si | EN_1 |
| GOBP_PLATELET_DERIVE | GOBP_PLATELET_DERIVED_GROWTH_FACTO | 63 | 0.351195 | 1.6191634 | 0.0 | 0.12693629 | 1 | 4891 | tags=32%, list=21%, si | EN_1 |
| GOBP_CARBOHYDRATE_D | GOBP_CARBOHYDRATE_DERIVATIVE_CATAB | 245 | 0.283141 | 1.6185769 | 0.0 | 0.12683022 | 1 | 3924 | tags=28%, list=17%, si | EN_1 |
| GOBP_RESPONSE_TO_TO | GOBP_RESPONSE_TO_TOPOLOGICALLY_INC | 127 | 0.314123 | 1.6180564 | 0.0 | 0.1266745 | 1 | 6513 | tags=46%, list=28%, si | EN_1 |
| GOBP_ANTIMICROBIAL_H | GOBP_ANTIMICROBIAL_HUMORAL_RESPON | 98 | 0.315613 | 1.6154782 | 0.0 | 0.12837036 | 1 | 1434 | tags=17%, list=6%, sig | EN_1 |
| GOBP_ESTABLISHMENT_( | GOBP_ESTABLISHMENT_OF_LYMPHOCYTE_I | 15 | 0.491726 | 1.6146353 | 0.016835017 | 0.12869872 | 1 | 3809 | tags=33%, list=16%, si | EN_1 |
| GOBP_REGULATION_OF_ | GOBP_REGULATION_OF_VACUOLE_ORGANI | 54 | 0.353595 | 1.614457 | 0.0 | 0.12837799 | 1 | 15281 | tags=100%, list=65%, | EN_1 |
| CYTOSOLIC_DNA-SENSING | CYTOSOLIC_DNA-SENSING_PATHWAY | 46 | 0.379912 | 1.6141967 | 0.011627907 | 0.12818095 | 1 | 4209 | tags=33%, list=18%, si | EN_1 |
| GOBP_CYTOPLASMIC_PA | GOBP_CYTOPLASMIC_PATTERN_RECOGNITI | 159 | 0.289162 | 1.6130518 | 0.0 | 0.12864688 | 1 | 4697 | tags=31%, list=20%, si | EN_1 |
| GOBP_REGULATION_OF_ | GOBP_REGULATION_OF_TELOMERE_MAINT | 20 | 0.470177 | 1.612382 | 0.027777778 | 0.12883459 | 1 | 3257 | tags=25%, list=14%, si | EN_1 |
| GOBP_REGULATION_OF_ | GOBP_REGULATION_OF_PATTERN_RECOGN | 157 | 0.291931 | 1.6110598 | 0.0 | 0.12957296 | 1 | 3079 | tags=23%, list=13%, si | EN_1 |
| GOBP_CELLULAR_RESPON | GOBP_CELLULAR_RESPONSE_TO_OXYGEN_L | 133 | 0.295021 | 1.6092514 | 0.0 | 0.13076603 | 1 | 3929 | tags=28%, list=17%, si | EN_1 |
| GOBP_VIRAL_TRANSLATI | GOBP_VIRAL_TRANSLATION | 18 | 0.468709 | 1.6088969 | 0.024 | 0.1306489 | 1 | 7786 | tags=67%, list=33%, si | EN_1 |
| GOBP_REGULATION_OF_ | GOBP_REGULATION_OF_TYPE_B_PANCREAT | 16 | 0.504424 | 1.6081876 | 0.023809524 | 0.13069877 | 1 | 1997 | tags=31%, list=8%, sig | EN_1 |
| GOBP_POSITIVE_REGULA | GOBP_POSITIVE_REGULATION_OF_SUBSTRA | 40 | 0.381113 | 1.6068832 | 0.01724138 | 0.13134581 | 1 | 3876 | tags=30%, list=16%, si | EN_1 |
| GOBP_PYRIDINE_CONTAI | GOBP_PYRIDINE_CONTAINING_COMPOUND | 182 | 0.30364 | 1.606284 | 0.0 | 0.13138324 | 1 | 3924 | tags=31%, list=17%, si | EN_1 |
| CITRATE_CYCLE_(TCA_CY | CITRATE_CYCLE_(TCA_CYCLE) | 28 | 0.408418 | 1.6062815 | 0.021929825 | 0.13094383 | 1 | 4822 | tags=50%, list=20%, si | EN_1 |
| GOBP_NEGATIVE_REGUL | GOBP_NEGATIVE_REGULATION_OF_FATTY_ | 33 | 0.389419 | 1.6045625 | 0.005181347 | 0.13175821 | 1 | 3898 | tags=30%, list=17%, si | EN_1 |
| BACTERIAL_INVASION_OI | BACTERIAL_INVASION_OF_EPITHELIAL_CELL | 72 | 0.327006 | 1.6042717 | 0.008403362 | 0.13152376 | 1 | 6323 | tags=40%, list=27%, si | EN_1 |
| GOBP_ANTIGEN_PROCES | GOBP_ANTIGEN_PROCESSING_AND_PRESEN | 24 | 0.436783 | 1.6034062 | 0.022123894 | 0.13190645 | 1 | 1852 | tags=29%, list=8%, sig | EN_1 |
| GOBP_REGULATION_OF_ | GOBP_REGULATION_OF_UBIQUITIN_PROTE | 39 | 0.380694 | 1.6030705 | 0.009661836 | 0.13171738 | 1 | 3661 | tags=28%, list=16%, si | EN_1 |
| GOBP_VESICLE_CARGO_L | GOBP_VESICLE_CARGO_LOADING | 27 | 0.417626 | 1.6027185 | 0.009478673 | 0.13162862 | 1 | 6392 | tags=41%, list=27%, si | EN_1 |
| PYRUVATE_METABOLISM | PYRUVATE_METABOLISM | 37 | 0.379565 | 1.6017485 | 0.0154639175 | 0.1321124 | 1 | 6179 | tags=57%, list=26%, si | EN_1 |
| GOBP_ENDOPLASMIC_RE | GOBP_ENDOPLASMIC_RETICULUM_UNFOLD | 69 | 0.338568 | 1.6002293 | 0.0 | 0.13296157 | 1 | 4459 | tags=33%, list=19%, si | EN_1 |
| GOBP_CELLULAR_RESPON | GOBP_CELLULAR_RESPONSE_TO_BIOTIC_ST | 224 | 0.284923 | 1.5990863 | 0.0 | 0.13351817 | 1 | 2264 | tags=20%, list=10%, si | EN_1 |
| GOBP_POSITIVE_REGULA | GOBP_POSITIVE_REGULATION_OF_MITOCH | 22 | 0.445052 | 1.5984122 | 0.013824885 | 0.13364111 | 1 | 5303 | tags=41%, list=22%, si | EN_1 |
| GOBP_POSITIVE_REGULA | GOBP_POSITIVE_REGULATION_OF_SIGNAL_ | 32 | 0.403044 | 1.5982442 | 0.015306123 | 0.13345413 | 1 | 4219 | tags=34%, list=18%, si | EN_1 |
| GOBP_GOLGI_TO_VACUO | GOBP_GOLGI_TO_VACUOLE_TRANSPORT | 21 | 0.444737 | 1.5979381 | 0.032786883 | 0.13324232 | 1 | 3818 | tags=29%, list=16%, si | EN_1 |
| BIOSYNTHESIS_OF_UNSA | BIOSYNTHESIS_OF_UNSATURATED_FATTY_A | 31 | 0.393925 | 1.5976658 | 0.017777778 | 0.13300848 | 1 | 3719 | tags=42%, list=16%, si | EN_1 |
| GOBP_NEGATIVE_REGUL | GOBP_NEGATIVE_REGULATION_OF_STRIATE | 39 | 0.380284 | 1.5972625 | 0.021978023 | 0.13287109 | 1 | 3898 | tags=31%, list=17%, si | EN_1 |
| HEPATOCELLULAR_CARCI | HEPATOCELLULAR_CARCINOMA | 164 | 0.291479 | 1.5971802 | 0.0 | 0.132458 | 1 | 4584 | tags=28%, list=19%, si | EN_1 |
| GOBP_RIBOSOME_ASSEM | GOBP_RIBOSOME_ASSEMBLY | 60 | 0.333135 | 1.5958968 | 0.0 | 0.13314985 | 1 | 15765 | tags=100%, list=67%, | EN_1 |
| GOBP_POSITIVE_REGULA | GOBP_POSITIVE_REGULATION_OF_PROTEIN | 192 | 0.276666 | 1.5953803 | 0.0 | 0.13318339 | 1 | 5987 | tags=37%, list=25%, si | EN_1 |
| GOBP_GLYCEROL_ETHER_ | GOBP_GLYCEROL_ETHER_METABOLIC_PROC | 19 | 0.469253 | 1.5925709 | 0.026415095 | 0.1353741 | 1 | 2958 | tags=32%, list=13%, si | EN_1 |
| GOBP_POSITIVE_REGULA | GOBP_POSITIVE_REGULATION_OF_DNA_RE | 135 | 0.307975 | 1.5923026 | 0.0 | 0.13516237 | 1 | 3896 | tags=21%, list=17%, si | EN_1 |
| GOBP_AEROBIC_RESPIRA | GOBP_AEROBIC_RESPIRATION | 170 | 0.294131 | 1.5910864 | 0.0 | 0.13589132 | 1 | 5841 | tags=45%, list=25%, si | EN_1 |
| GOBP_GOLGI_VESICLE_TI | GOBP_GOLGI_VESICLE_TRANSPORT | 276 | 0.28698 | 1.5898393 | 0.0 | 0.13676552 | 1 | 6438 | tags=37%, list=27%, si | EN_1 |
| GOBP_NEGATIVE_REGUL | GOBP_NEGATIVE_REGULATION_OF_NUCLEO | 20 | 0.461671 | 1.5885698 | 0.012875536 | 0.13748695 | 1 | 2392 | tags=35%, list=10%, si | EN_1 |
| GOBP_REGULATION_OF_ | GOBP_REGULATION_OF_FATTY_ACID_BIOSY | 47 | 0.36433 | 1.5861311 | 0.010204081 | 0.13928494 | 1 | 3240 | tags=28%, list=14%, si | EN_1 |
| GOBP_NEUTROPHIL_ACTI | GOBP_NEUTROPHIL_ACTIVATION_INVOLVEI | 18 | 0.453105 | 1.5858119 | 0.019305019 | 0.13907437 | 1 | 1981 | tags=28%, list=8%, sig | EN_1 |
| GOBP_POSITIVE_REGULA | GOBP_POSITIVE_REGULATION_OF_MEMBRA | 39 | 0.36798 | 1.5856652 | 0.005524862 | 0.13877437 | 1 | 6597 | tags=49%, list=28%, si | EN_1 |
| GOBP_REGULATION_OF_ | GOBP_REGULATION_OF_RRNA_PROCESSING | 16 | 0.500801 | 1.5854858 | 0.03731343 | 0.1385109 | 1 | 790 | tags=13%, list=3%, sig | EN_1 |
| GOBP_POSITIVE_REGULA | GOBP_POSITIVE_REGULATION_OF_CELLULA | 22 | 0.440594 | 1.5835444 | 0.033613447 | 0.13980162 | 1 | 2879 | tags=36%, list=12%, si | EN_1 |
| GOBP_NEGATIVE_REGUL | GOBP_NEGATIVE_REGULATION_OF_TYPE_I_ | 42 | 0.361484 | 1.5831301 | 0.016216217 | 0.13978612 | 1 | 5198 | tags=33%, list=22%, si | EN_1 |
| GOBP_CELLULAR_RESPON | GOBP_CELLULAR_RESPONSE_TO_GLUCOSE_ | 42 | 0.373017 | 1.5809442 | 0.023255814 | 0.14120431 | 1 | 14819 | tags=100%, list=63%, | EN_1 |
| BASAL_TRANSCRIPTION_ | BASAL_TRANSCRIPTION_FACTORS | 40 | 0.375396 | 1.5798187 | 0.011049724 | 0.14183164 | 1 | 4448 | tags=35%, list=19%, si | EN_1 |
| GOBP_NEGATIVE_REGUL | GOBP_NEGATIVE_REGULATION_OF_PROTEI | 93 | 0.309628 | 1.5789497 | 0.0 | 0.14226991 | 1 | 3294 | tags=25%, list=14%, si | EN_1 |
| GOBP_REGULATION_OF_ | GOBP_REGULATION_OF_TYPE_I_INTERFERO | 40 | 0.380442 | 1.5785824 | 0.011235955 | 0.14220126 | 1 | 4393 | tags=43%, list=19%, si | EN_1 |
| GOBP_MITOCHONDRIAL_ | GOBP_MITOCHONDRIAL_CYTOCHROME_C_ | 19 | 0.452481 | 1.5781174 | 0.027777778 | 0.14210609 | 1 | 3410 | tags=26%, list=14%, si | EN_1 |
| GOBP_NEGATIVE_REGUL | GOBP_NEGATIVE_REGULATION_OF_LEUKOC | 65 | 0.337856 | 1.5779636 | 0.0 | 0.14177485 | 1 | 5011 | tags=42%, list=21%, si | EN_1 |
| GOBP_REGULATION_OF_ | GOBP_REGULATION_OF_FIBROBLAST_PROL | 122 | 0.302658 | 1.577371 | 0.0 | 0.14199375 | 1 | 2576 | tags=18%, list=11%, si | EN_1 |
| GOBP_HEME_BIOSYNTHE | GOBP_HEME_BIOSYNTHETIC_PROCESS | 24 | 0.434499 | 1.5769906 | 0.03211009 | 0.14197308 | 1 | 7445 | tags=71%, list=32%, si | EN_1 |
| GOBP_LYTIC_VACUOLE_C | GOBP_LYTIC_VACUOLE_ORGANIZATION | 95 | 0.312501 | 1.5760475 | 0.0 | 0.14242254 | 1 | 4744 | tags=32%, list=20%, si | EN_1 |
| GOBP_NEGATIVE_REGUL | GOBP_NEGATIVE_REGULATION_OF_PROTEI | 17 | 0.464446 | 1.5758427 | 0.03041825 | 0.14211561 | 1 | 4509 | tags=47%, list=19%, si | EN_1 |
| GOBP_PROTEIN_TARGETI | GOBP_PROTEIN_TARGETING_TO_MITOCHO | 76 | 0.319975 | 1.5705442 | 0.0 | 0.14631782 | 1 | 6689 | tags=43%, list=28%, si | EN_1 |
| GOBP_MAINTENANCE_OI | GOBP_MAINTENANCE_OF_PROTEIN_LOCAT | 25 | 0.411841 | 1.5704641 | 0.016260162 | 0.1459675 | 1 | 6184 | tags=52%, list=26%, si | EN_1 |
| GOBP_REGULATION_OF_ | GOBP_REGULATION_OF_AUTOPHAGOSOME | 44 | 0.365453 | 1.5703948 | 0.0 | 0.1456124 | 1 | 14998 | tags=100%, list=64%, | EN_1 |
| GOBP_REGULATION_OF_ | GOBP_REGULATION_OF_DNA_STRAND_ELO | 18 | 0.458597 | 1.5686877 | 0.023529412 | 0.1467344 | 1 | 6836 | tags=44%, list=29%, si | EN_1 |
| GOBP_PROTEIN_K63_LIN | GOBP_PROTEIN_K63_LINKED_UBIQUITINAT | 68 | 0.330859 | 1.5679443 | 0.0 | 0.14694516 | 1 | 5316 | tags=38%, list=23%, si | EN_1 |
| GOBP_NEGATIVE_REGUL | GOBP_NEGATIVE_REGULATION_OF_VIRAL_ | 36 | 0.38863 | 1.5677465 | 0.029411765 | 0.1467341 | 1 | 4172 | tags=42%, list=18%, si | EN_1 |
| GOBP_POSITIVE_REGULA | GOBP_POSITIVE_REGULATION_OF_TELOME | 71 | 0.330154 | 1.5676104 | 0.0074074073 | 0.14638896 | 1 | 4214 | tags=23%, list=18%, si | EN_1 |
| GOBP_IMPORT_INTO_NU | GOBP_IMPORT_INTO_NUCLEUS | 180 | 0.286478 | 1.5671111 | 0.0 | 0.14648445 | 1 | 4509 | tags=29%, list=19%, si | EN_1 |
| GOBP_FATTY_ACID_ELON | GOBP_FATTY_ACID_ELONGATION | 15 | 0.489348 | 1.5670173 | 0.05514706 | 0.14617473 | 1 | 4812 | tags=53%, list=20%, si | EN_1 |
| GOBP_QUINONE_BIOSYN | GOBP_QUINONE_BIOSYNTHETIC_PROCESS | 16 | 0.479269 | 1.566761 | 0.04511278 | 0.14599732 | 1 | 5650 | tags=56%, list=24%, si | EN_1 |
| GOBP_POSITIVE_REGULA | GOBP_POSITIVE_REGULATION_OF_LIPID_BI | 104 | 0.305415 | 1.5662639 | 0.0 | 0.14621535 | 1 | 2002 | tags=17%, list=8%, sig | EN_1 |
| GOBP_CELLULAR_RESPON | GOBP_CELLULAR_RESPONSE_TO_HYPOXIA | 112 | 0.306033 | 1.5659498 | 0.0 | 0.14613412 | 1 | 3929 | tags=30%, list=17%, si | EN_1 |
| GOBP_SPHINGOID_META | GOBP_SPHINGOID_METABOLIC_PROCESS | 21 | 0.440415 | 1.5656426 | 0.02 | 0.14605263 | 1 | 1994 | tags=24%, list=8%, sig | EN_1 |
| GOBP_VACUOLE_ORGAN | GOBP_VACUOLE_ORGANIZATION | 223 | 0.281469 | 1.5651065 | 0.0 | 0.14624907 | 1 | 5198 | tags=32%, list=22%, si | EN_1 |
| GOBP_RNA_MODIFICATI( | GOBP_RNA_MODIFICATION | 141 | 0.285862 | 1.564133 | 0.0 | 0.14653404 | 1 | 6477 | tags=42%, list=27%, si | EN_1 |

| | | | | | | | | | | |
|---|---|---|---|---|---|---|---|---|---|---|
| GOBP_REGULATION_OF_ | GOBP_REGULATION_OF_DENDRITIC_SPINE_ | 60 | 0.347932 | 1.5636728 | 0.0 | 0.14654467 | 1 | 2087 | tags=17%, list=9%, sig | EN_1 |
| GOBP_REGULATION_OF_ | GOBP_REGULATION_OF_AMYLOID_BETA_F( | 37 | 0.377085 | 1.5633838 | 0.01058201 | 0.14644355 | 1 | 5983 | tags=49%, list=25%, si | EN_1 |
| GOBP_VACUOLAR_ACIDII | GOBP_VACUOLAR_ACIDIFICATION | 27 | 0.413781 | 1.5627311 | 0.018348623 | 0.14682554 | 1 | 5976 | tags=48%, list=25%, si | EN_1 |
| GOBP_DIOL_METABOLIC_ | GOBP_DIOL_METABOLIC_PROCESS | 29 | 0.397804 | 1.5623853 | 0.04090909 | 0.14668491 | 1 | 2473 | tags=24%, list=10%, si | EN_1 |
| GOBP_REGULATION_OF_ | GOBP_REGULATION_OF_BIOLOGICAL_PROC | 64 | 0.334712 | 1.5612881 | 0.006993007 | 0.14729021 | 1 | 3526 | tags=36%, list=15%, si | EN_1 |
| GOBP_POSITIVE_REGULA | GOBP_POSITIVE_REGULATION_OF_PROTEO | 120 | 0.304525 | 1.5602611 | 0.0 | 0.14772892 | 1 | 5929 | tags=39%, list=25%, si | EN_1 |
| PROPANOATE_METABOLI | PROPANOATE_METABOLISM | 29 | 0.400181 | 1.5597044 | 0.022727273 | 0.14779806 | 1 | 4822 | tags=41%, list=20%, si | EN_1 |
| GOBP_PSEUDOURIDINE_S | GOBP_PSEUDOURIDINE_SYNTHESIS | 18 | 0.463398 | 1.5590099 | 0.036885247 | 0.14807159 | 1 | 2729 | tags=28%, list=12%, si | EN_1 |
| GOBP_REGULATION_OF_ | GOBP_REGULATION_OF_MITOCHONDRIAL_ | 20 | 0.446146 | 1.5582637 | 0.041841004 | 0.14836472 | 1 | 2888 | tags=35%, list=12%, si | EN_1 |
| GOBP_RESPONSE_TO_STI | GOBP_RESPONSE_TO_STEROL_DEPLETION | 15 | 0.485404 | 1.5578331 | 0.027777778 | 0.1483297 | 1 | 6602 | tags=60%, list=28%, si | EN_1 |
| GOBP_NEGATIVE_REGUL | GOBP_NEGATIVE_REGULATION_OF_TYPE_I_ | 23 | 0.427303 | 1.5573999 | 0.026785715 | 0.1483651 | 1 | 2305 | tags=35%, list=10%, si | EN_1 |
| GOBP_AUTOPHAGOSOMI | GOBP_AUTOPHAGOSOME_MATURATION | 58 | 0.338162 | 1.5555155 | 0.0 | 0.14992777 | 1 | 4213 | tags=24%, list=18%, si | EN_1 |
| GOBP_AUTOPHAGOSOMI | GOBP_AUTOPHAGOSOME_ORGANIZATION | 128 | 0.28935 | 1.5545639 | 0.0 | 0.15020166 | 1 | 6361 | tags=42%, list=27%, si | EN_1 |
| GOBP_INTRINSIC_APOPT( | GOBP_INTRINSIC_APOPTOTIC_SIGNALING_I | 63 | 0.334003 | 1.5536052 | 0.0 | 0.15080287 | 1 | 6951 | tags=49%, list=29%, si | EN_1 |
| GOBP_TRANSLATIONAL_I | GOBP_TRANSLATIONAL_INITIATION | 112 | 0.298778 | 1.5535103 | 0.0 | 0.15045798 | 1 | 15763 | tags=99%, list=67%, si | EN_1 |
| MISMATCH_REPAIR | MISMATCH_REPAIR | 21 | 0.439846 | 1.5533249 | 0.046413504 | 0.15024374 | 1 | 3227 | tags=33%, list=14%, si | EN_1 |
| GOBP_NEGATIVE_REGUL | GOBP_NEGATIVE_REGULATION_OF_RESPON | 112 | 0.305585 | 1.5532922 | 0.01 | 0.14983548 | 1 | 3013 | tags=26%, list=13%, si | EN_1 |
| GOBP_NEGATIVE_REGUL | GOBP_NEGATIVE_REGULATION_OF_HEMOP | 121 | 0.302511 | 1.5523885 | 0.0 | 0.15016696 | 1 | 3417 | tags=28%, list=14%, si | EN_1 |
| GOBP_PHOSPHATIDYLIN( | GOBP_PHOSPHATIDYLINOSITOL_PHOSPHAT | 50 | 0.348666 | 1.551427 | 0.01724138 | 0.15059017 | 1 | 4834 | tags=34%, list=20%, si | EN_1 |
| GOBP_NEGATIVE_REGUL | GOBP_NEGATIVE_REGULATION_OF_CYTOPL | 50 | 0.339832 | 1.551116 | 0.006369427 | 0.15048184 | 1 | 4668 | tags=30%, list=20%, si | EN_1 |
| GOBP_REGULATION_OF_ | GOBP_REGULATION_OF_FIBROBLAST_MIGR | 47 | 0.345307 | 1.5506543 | 0.0 | 0.15038492 | 1 | 5377 | tags=38%, list=23%, si | EN_1 |
| GOBP_MONOCARBOXYLI( | GOBP_MONOCARBOXYLIC_ACID_CATABOLI( | 126 | 0.299779 | 1.5502576 | 0.0 | 0.1503563 | 1 | 4227 | tags=33%, list=18%, si | EN_1 |
| GOBP_REGULATION_OF_ | GOBP_REGULATION_OF_FATTY_ACID_BETA | 21 | 0.433589 | 1.5498476 | 0.026217228 | 0.15020368 | 1 | 5427 | tags=48%, list=23%, si | EN_1 |
| GOBP_PYRUVATE_METAE | GOBP_PYRUVATE_METABOLIC_PROCESS | 127 | 0.299505 | 1.5496751 | 0.0 | 0.14994611 | 1 | 3924 | tags=30%, list=17%, si | EN_1 |
| GOBP_MEMBRANE_BIOG | GOBP_MEMBRANE_BIOGENESIS | 63 | 0.323988 | 1.5492184 | 0.0074626864 | 0.14990757 | 1 | 2816 | tags=21%, list=12%, si | EN_1 |
| GOBP_AMIDE_METABOLI | GOBP_AMIDE_METABOLIC_PROCESS | 398 | 0.255848 | 1.5488575 | 0.0 | 0.14984904 | 1 | 3727 | tags=25%, list=16%, si | EN_1 |
| GOBP_REGULATION_OF_ | GOBP_REGULATION_OF_VASCULATURE_DE\ | 295 | 0.259939 | 1.5463941 | 0.0 | 0.15205844 | 1 | 5171 | tags=34%, list=22%, si | EN_1 |
| GOBP_PODOSOME_ASSE | GOBP_PODOSOME_ASSEMBLY | 25 | 0.408493 | 1.5463431 | 0.03587444 | 0.15174232 | 1 | 3306 | tags=32%, list=14%, si | EN_1 |
| GOBP_SIGNAL_PEPTIDE_ | GOBP_SIGNAL_PEPTIDE_PROCESSING | 15 | 0.473253 | 1.5461305 | 0.03533569 | 0.15151167 | 1 | 4632 | tags=60%, list=20%, si | EN_1 |
| GOBP_EXTRACELLULAR_V | GOBP_EXTRACELLULAR_VESICLE_BIOGENES | 24 | 0.416401 | 1.545414 | 0.04587156 | 0.1517336 | 1 | 7316 | tags=63%, list=31%, si | EN_1 |
| GOBP_REGULATION_OF_ | GOBP_REGULATION_OF_REACTIVE_OXYGEN | 160 | 0.280273 | 1.5449399 | 0.0 | 0.15167913 | 1 | 5937 | tags=39%, list=25%, si | EN_1 |
| GOBP_OXIDATIVE_PHOSF | GOBP_OXIDATIVE_PHOSPHORYLATION | 124 | 0.301456 | 1.5448421 | 0.0 | 0.15137745 | 1 | 6669 | tags=55%, list=28%, si | EN_1 |
| GOBP_NEGATIVE_REGUL | GOBP_NEGATIVE_REGULATION_OF_POST_T | 94 | 0.312671 | 1.5439848 | 0.0 | 0.15181026 | 1 | 5841 | tags=39%, list=25%, si | EN_1 |
| GOBP_CELLULAR_RESPOI | GOBP_CELLULAR_RESPONSE_TO_ARSENIC_( | 17 | 0.452221 | 1.543428 | 0.027586207 | 0.15185171 | 1 | 6021 | tags=53%, list=25%, si | EN_1 |
| GOBP_UBIQUITIN_DEPEN | GOBP_UBIQUITIN_DEPENDENT_PROTEIN_C/ | 29 | 0.391153 | 1.5434089 | 0.03196347 | 0.15147589 | 1 | 6701 | tags=41%, list=28%, si | EN_1 |
| HUNTINGTON_DISEASE | HUNTINGTON_DISEASE | 180 | 0.278611 | 1.5429853 | 0.0 | 0.15159261 | 1 | 6942 | tags=53%, list=29%, si | EN_1 |
| GOBP_XENOPHAGY | GOBP_XENOPHAGY | 17 | 0.449384 | 1.5424943 | 0.021052632 | 0.15174703 | 1 | 5316 | tags=53%, list=23%, si | EN_1 |
| DNA_REPLICATION | DNA_REPLICATION | 34 | 0.374251 | 1.5418589 | 0.025125628 | 0.15192619 | 1 | 4343 | tags=35%, list=18%, si | EN_1 |
| GOBP_VASCULAR_ASSOC | GOBP_VASCULAR_ASSOCIATED_SMOOTH_N | 16 | 0.473168 | 1.5407258 | 0.022304833 | 0.15262528 | 1 | 4245 | tags=50%, list=18%, si | EN_1 |
| GOBP_POSITIVE_REGULA | GOBP_POSITIVE_REGULATION_OF_CHEMOK | 15 | 0.469887 | 1.5404134 | 0.025089607 | 0.15263763 | 1 | 7086 | tags=60%, list=30%, si | EN_1 |
| GOBP_MITOCHONDRIAL_ | GOBP_MITOCHONDRIAL_OUTER_MEMBRAN | 28 | 0.401057 | 1.5403599 | 0.028301887 | 0.15230347 | 1 | 5793 | tags=46%, list=25%, si | EN_1 |
| GOBP_RESPONSE_TO_M( | GOBP_RESPONSE_TO_MOLECULE_OF_BACT | 296 | 0.261179 | 1.5402899 | 0.0 | 0.15196095 | 1 | 4567 | tags=32%, list=19%, si | EN_1 |
| GOBP_REGULATION_OF_ | GOBP_REGULATION_OF_LEUKOCYTE_APOPT | 103 | 0.307027 | 1.539572 | 0.011764706 | 0.15227443 | 1 | 4872 | tags=36%, list=21%, si | EN_1 |
| GOBP_NEGATIVE_REGUL | GOBP_NEGATIVE_REGULATION_OF_LYMPH( | 62 | 0.334207 | 1.539521 | 0.0 | 0.15193452 | 1 | 2625 | tags=27%, list=11%, si | EN_1 |
| GOBP_RESPONSE_TO_DE | GOBP_RESPONSE_TO_DECREASED_OXYGEN | 228 | 0.266292 | 1.5386256 | 0.0 | 0.15225701 | 1 | 3939 | tags=28%, list=17%, si | EN_1 |
| GOBP_PLATELET_DENSE_ | GOBP_PLATELET_DENSE_GRANULE_ORGAN | 21 | 0.433559 | 1.5374185 | 0.020833334 | 0.15320721 | 1 | 1282 | tags=19%, list=5%, sig | EN_1 |
| GOBP_PROTEIN_EXPORT_ | GOBP_PROTEIN_EXPORT_FROM_NUCLEUS | 64 | 0.332752 | 1.5359806 | 0.016 | 0.15436336 | 1 | 6709 | tags=42%, list=28%, si | EN_1 |
| GOBP_PROTEIN_REFOLDI | GOBP_PROTEIN_REFOLDING | 23 | 0.424229 | 1.5359231 | 0.008298756 | 0.15403871 | 1 | 6188 | tags=39%, list=26%, si | EN_1 |
| GOBP_PIGMENT_BIOSYN | GOBP_PIGMENT_BIOSYNTHETIC_PROCESS | 49 | 0.346027 | 1.5356925 | 0.004901961 | 0.15387562 | 1 | 5190 | tags=39%, list=22%, si | EN_1 |
| RIG-I-LIKE_RECEPTOR_SIG | RIG-I-LIKE_RECEPTOR_SIGNALING_PATHWA | 52 | 0.337205 | 1.5350509 | 0.014184397 | 0.15410182 | 1 | 2873 | tags=17%, list=12%, si | EN_1 |
| GOBP_VIRION_ASSEMBLY | GOBP_VIRION_ASSEMBLY | 26 | 0.412569 | 1.5347542 | 0.037209302 | 0.15389575 | 1 | 3953 | tags=31%, list=17%, si | EN_1 |
| GLUTATHIONE_METABOL | GLUTATHIONE_METABOLISM | 57 | 0.335935 | 1.5347005 | 0.0 | 0.15357538 | 1 | 4577 | tags=37%, list=19%, si | EN_1 |
| MITOPHAGY | MITOPHAGY | 62 | 0.337477 | 1.5340707 | 0.0 | 0.15380608 | 1 | 7050 | tags=52%, list=30%, si | EN_1 |
| GOBP_ESTABLISHMENT_( | GOBP_ESTABLISHMENT_OF_PROTEIN_LOCA | 435 | 0.251141 | 1.5312206 | 0.0 | 0.15612304 | 1 | 5398 | tags=30%, list=23%, si | EN_1 |
| GOBP_REGULATION_OF_ | GOBP_REGULATION_OF_ENDOPLASMIC_RE | 30 | 0.38656 | 1.5290035 | 0.009756098 | 0.15783937 | 1 | 6951 | tags=47%, list=29%, si | EN_1 |
| GOBP_TRNA_THREONYL( | GOBP_TRNA_THREONYLCARBAMOYLADEN( | 15 | 0.473978 | 1.5287971 | 0.023255814 | 0.15763575 | 1 | 3110 | tags=33%, list=13%, si | EN_1 |
| GOBP_CELL_SUBSTRATE_ | GOBP_CELL_SUBSTRATE_JUNCTION_ORGAN | 105 | 0.298958 | 1.5287163 | 0.0 | 0.15732025 | 1 | 7004 | tags=45%, list=30%, si | EN_1 |
| GOBP_NEGATIVE_REGUL | GOBP_NEGATIVE_REGULATION_OF_CELLUL | 51 | 0.343547 | 1.5285662 | 0.01875 | 0.15708503 | 1 | 4307 | tags=35%, list=18%, si | EN_1 |
| GOBP_LIPOPOLYSACCHAI | GOBP_LIPOPOLYSACCHARIDE_MEDIATED_SI | 57 | 0.330219 | 1.5276047 | 0.0124223605 | 0.1577746 | 1 | 3991 | tags=33%, list=17%, si | EN_1 |
| GOBP_PROTEIN_O_LINKE | GOBP_PROTEIN_O_LINKED_MANNOSYLATI( | 16 | 0.458462 | 1.5273777 | 0.041666668 | 0.15767579 | 1 | 6274 | tags=50%, list=27%, si | EN_1 |
| ARGININE_AND_PROLINE | ARGININE_AND_PROLINE_METABOLISM | 47 | 0.349151 | 1.5264349 | 0.0057803467 | 0.15822071 | 1 | 5913 | tags=51%, list=25%, si | EN_1 |
| GOBP_FIBROBLAST_APOI | GOBP_FIBROBLAST_APOPTOTIC_PROCESS | 31 | 0.377175 | 1.5261194 | 0.018604651 | 0.15811211 | 1 | 6358 | tags=48%, list=27%, si | EN_1 |
| GOBP_TRNA_METABOLIC | GOBP_TRNA_METABOLIC_PROCESS | 166 | 0.274158 | 1.5252643 | 0.0 | 0.15859663 | 1 | 6904 | tags=42%, list=29%, si | EN_1 |
| GOBP_EXTRINSIC_APOPT | GOBP_EXTRINSIC_APOPTOTIC_SIGNALING_I | 80 | 0.310296 | 1.5248455 | 0.0 | 0.15864146 | 1 | 4292 | tags=28%, list=18%, si | EN_1 |
| GOBP_REGULATION_OF_ | GOBP_REGULATION_OF_MACROAUTOPHA( | 129 | 0.288841 | 1.52439 | 0.0 | 0.15881166 | 1 | 6597 | tags=40%, list=28%, si | EN_1 |
| GOBP_AMYLOID_PRECUR | GOBP_AMYLOID_PRECURSOR_PROTEIN_ME | 75 | 0.316704 | 1.5238926 | 0.014492754 | 0.15900183 | 1 | 5018 | tags=33%, list=21%, si | EN_1 |
| GOBP_COPPER_ION_HON | GOBP_COPPER_ION_HOMEOSTASIS | 18 | 0.444751 | 1.5222464 | 0.040650405 | 0.16029605 | 1 | 5878 | tags=39%, list=25%, si | EN_1 |
| GOBP_ACYL_COA_METAE | GOBP_ACYL_COA_METABOLIC_PROCESS | 82 | 0.313738 | 1.5186512 | 0.0 | 0.16342543 | 1 | 4149 | tags=32%, list=18%, si | EN_1 |
| GOBP_REGULATION_OF_ | GOBP_REGULATION_OF_DOUBLE_STRAND_ | 28 | 0.402508 | 1.518621 | 0.016129032 | 0.16306269 | 1 | 3896 | tags=32%, list=17%, si | EN_1 |
| GOBP_INTRACELLULAR_I | GOBP_INTRACELLULAR_IRON_ION_HOMEO: | 64 | 0.320658 | 1.5172017 | 0.0 | 0.16413231 | 1 | 3620 | tags=31%, list=15%, si | EN_1 |
| GOBP_GLYCOSAMINOGL\ | GOBP_GLYCOSAMINOGLYCAN_CATABOLIC_ | 27 | 0.399668 | 1.5171509 | 0.03125 | 0.16379456 | 1 | 4692 | tags=41%, list=20%, si | EN_1 |
| GOBP_REGULATION_OF_ | GOBP_REGULATION_OF_AMIDE_METABOLI( | 69 | 0.331297 | 1.5169011 | 0.008196721 | 0.16363543 | 1 | 2824 | tags=25%, list=12%, si | EN_1 |
| GOBP_FIBROBLAST_ACTI\ | GOBP_FIBROBLAST_ACTIVATION | 22 | 0.418923 | 1.5164196 | 0.041666668 | 0.16375878 | 1 | 2406 | tags=32%, list=10%, si | EN_1 |
| GOBP_HYALURONAN_ME | GOBP_HYALURONAN_METABOLIC_PROCESS | 36 | 0.380081 | 1.5164038 | 0.028436018 | 0.16339815 | 1 | 4473 | tags=33%, list=19%, si | EN_1 |
| GOBP_REGULATION_OF_ | GOBP_REGULATION_OF_SYNCYTIUM_FORM | 31 | 0.38882 | 1.5155985 | 0.03153153 | 0.16388077 | 1 | 4118 | tags=45%, list=17%, si | EN_1 |
| RENAL_CELL_CARCINOM/ | RENAL_CELL_CARCINOMA | 67 | 0.320618 | 1.515498 | 0.012820513 | 0.163632 | 1 | 6293 | tags=40%, list=27%, si | EN_1 |
| GOBP_MODIFICATION_O | GOBP_MODIFICATION_OF_POSTSYNAPTIC_/ | 24 | 0.402342 | 1.5141361 | 0.042145595 | 0.16468547 | 1 | 6126 | tags=58%, list=26%, si | EN_1 |
| GOBP_CELLULAR_SENESC | GOBP_CELLULAR_SENESCENCE | 88 | 0.300239 | 1.5138309 | 0.018867925 | 0.16460936 | 1 | 3528 | tags=24%, list=15%, si | EN_1 |
| GOBP_NEGATIVE_REGUL | GOBP_NEGATIVE_REGULATION_OF_CELL_C | 69 | 0.314475 | 1.5137821 | 0.021276595 | 0.16427676 | 1 | 5276 | tags=33%, list=22%, si | EN_1 |
| GOBP_TUMOR_NECROSIS | GOBP_TUMOR_NECROSIS_FACTOR_SUPERF | 193 | 0.272017 | 1.5136545 | 0.0 | 0.16402777 | 1 | 4864 | tags=35%, list=21%, si | EN_1 |
| GOBP_HEPATIC_STELLATI | GOBP_HEPATIC_STELLATE_CELL_ACTIVATIO | 16 | 0.450454 | 1.5136242 | 0.044609666 | 0.1636559 | 1 | 2406 | tags=38%, list=10%, si | EN_1 |
| GOBP_MONOCARBOXYLI( | GOBP_MONOCARBOXYLIC_ACID_BIOSYNTH | 203 | 0.27946 | 1.5132208 | 0.0 | 0.16374978 | 1 | 3240 | tags=29%, list=14%, si | EN_1 |

| | | | | | | | | | | |
|---|---|---|---|---|---|---|---|---|---|---|
| GOBP_RESPIRATORY_CHA | GOBP_RESPIRATORY_CHAIN_COMPLEX_IV_ | 24 | 0.40665 | 1.5129501 | 0.021459227 | 0.16366738 | 1 | 3506 | tags=25%, list=15%, si | EN_1 |
| GOBP_PROTEIN_SUMOYL | GOBP_PROTEIN_SUMOYLATION | 42 | 0.364958 | 1.5126725 | 0.032608695 | 0.16361964 | 1 | 4560 | tags=26%, list=19%, si | EN_1 |
| GOBP_REGULATION_OF_ | GOBP_REGULATION_OF_PHOSPHOLIPID_BIO | 20 | 0.423871 | 1.5111785 | 0.027888447 | 0.16472049 | 1 | 2002 | tags=30%, list=8%, sig | EN_1 |
| GOBP_LIPOPROTEIN_MET | GOBP_LIPOPROTEIN_METABOLIC_PROCESS | 135 | 0.284859 | 1.5097027 | 0.0 | 0.16596915 | 1 | 4844 | tags=34%, list=21%, si | EN_1 |
| N-GLYCAN_BIOSYNTHESIS | N-GLYCAN_BIOSYNTHESIS | 48 | 0.334875 | 1.5087373 | 0.012578616 | 0.1667146 | 1 | 15720 | tags=100%, list=67%, | EN_1 |
| GOBP_CELLULAR_RESPON | GOBP_CELLULAR_RESPONSE_TO_TOXIC_SU | 63 | 0.3255 | 1.5086505 | 0.028985508 | 0.16644216 | 1 | 3956 | tags=37%, list=17%, si | EN_1 |
| GOBP_CELLULAR_RESPON | GOBP_CELLULAR_RESPONSE_TO_CADMIUM | 24 | 0.398764 | 1.5075049 | 0.03652968 | 0.16726445 | 1 | 5377 | tags=50%, list=23%, si | EN_1 |
| LYSOSOME | LYSOSOME | 118 | 0.295465 | 1.5072465 | 0.0 | 0.1671786 | 1 | 6296 | tags=42%, list=27%, si | EN_1 |
| GOBP_HEME_METABOLIC | GOBP_HEME_METABOLIC_PROCESS | 34 | 0.368979 | 1.5069524 | 0.044692736 | 0.16706899 | 1 | 7445 | tags=62%, list=32%, si | EN_1 |
| GOBP_LATE_ENDOSOME_ | GOBP_LATE_ENDOSOME_TO_LYSOSOME_TR | 19 | 0.438467 | 1.5058739 | 0.03265306 | 0.16773278 | 1 | 6701 | tags=53%, list=28%, si | EN_1 |
| GOBP_NUCLEOSIDE_DIPH | GOBP_NUCLEOSIDE_DIPHOSPHATE_CATABO | 111 | 0.297794 | 1.5056465 | 0.0 | 0.16758952 | 1 | 3924 | tags=32%, list=17%, si | EN_1 |
| GOBP_PURINE_CONTAIN | GOBP_PURINE_CONTAINING_COMPOUND_ | 19 | 0.43607 | 1.5049753 | 0.032128513 | 0.16795756 | 1 | 3751 | tags=32%, list=16%, si | EN_1 |
| GOBP_NEGATIVE_REGUL | GOBP_NEGATIVE_REGULATION_OF_EXTRIN | 95 | 0.297377 | 1.5048621 | 0.0 | 0.1677334 | 1 | 4225 | tags=28%, list=18%, si | EN_1 |
| GOBP_REGULATION_OF_ | GOBP_REGULATION_OF_PROTEIN_EXPORT_ | 37 | 0.352074 | 1.5038908 | 0.020725388 | 0.16836227 | 1 | 6709 | tags=46%, list=28%, si | EN_1 |
| GOBP_POSITIVE_REGULA | GOBP_POSITIVE_REGULATION_OF_CELL_M | 61 | 0.326489 | 1.5025532 | 0.0 | 0.16937642 | 1 | 6354 | tags=46%, list=27%, si | EN_1 |
| GOBP_AMYLOID_PRECUR | GOBP_AMYLOID_PRECURSOR_PROTEIN_CAT | 59 | 0.326875 | 1.5025319 | 0.013986014 | 0.16903089 | 1 | 5983 | tags=41%, list=25%, si | EN_1 |
| GOBP_GENERATION_OF_ | GOBP_GENERATION_OF_PRECURSOR_META | 439 | 0.237834 | 1.5023892 | 0.0 | 0.16881424 | 1 | 5779 | tags=38%, list=24%, si | EN_1 |
| GOBP_REGULATION_OF_ | GOBP_REGULATION_OF_CYCLIN_DEPENDEN | 67 | 0.317865 | 1.5016824 | 0.015873017 | 0.1693012 | 1 | 1876 | tags=21%, list=8%, sig | EN_1 |
| GOBP_REGULATION_OF_ | GOBP_REGULATION_OF_VASCULAR_ENDOT | 33 | 0.370852 | 1.5016812 | 0.033980582 | 0.16892663 | 1 | 4891 | tags=39%, list=21%, si | EN_1 |
| GOBP_RUFFLE_ORGANIZA | GOBP_RUFFLE_ORGANIZATION | 54 | 0.329499 | 1.5011799 | 0.01369863 | 0.1689693 | 1 | 4750 | tags=33%, list=20%, si | EN_1 |
| GOBP_UNSATURATED_FA | GOBP_UNSATURATED_FATTY_ACID_BIOSYN | 41 | 0.35319 | 1.5004092 | 0.01754386 | 0.1691852 | 1 | 1965 | tags=27%, list=8%, sig | EN_1 |
| GOBP_NUCLEOSIDE_TRIP | GOBP_NUCLEOSIDE_TRIPHOSPHATE_METAB | 250 | 0.256357 | 1.4998332 | 0.0 | 0.16946839 | 1 | 5514 | tags=37%, list=23%, si | EN_1 |
| GOBP_STEROID_BIOSYNT | GOBP_STEROID_BIOSYNTHETIC_PROCESS | 153 | 0.281129 | 1.4987838 | 0.0 | 0.17006092 | 1 | 2864 | tags=24%, list=12%, si | EN_1 |
| GOBP_GLYCEROPHOSPHO | GOBP_GLYCEROPHOSPHOLIPID_CATABOLIC_ | 31 | 0.37978 | 1.4977646 | 0.02631579 | 0.17076099 | 1 | 2608 | tags=32%, list=11%, si | EN_1 |
| SPHINGOLIPID_METABOL | SPHINGOLIPID_METABOLISM | 44 | 0.351591 | 1.4969255 | 0.020942409 | 0.1713251 | 1 | 2569 | tags=23%, list=11%, si | EN_1 |
| GOBP_PROTEIN_DEGLYCO | GOBP_PROTEIN_DEGLYCOSYLATION | 15 | 0.455015 | 1.4967816 | 0.05860806 | 0.17103675 | 1 | 12874 | tags=100%, list=55%, | EN_1 |
| GOBP_REGULATION_OF_ | GOBP_REGULATION_OF_MONOCYTE_DIFFE | 17 | 0.441823 | 1.4957165 | 0.05645161 | 0.17183384 | 1 | 4551 | tags=41%, list=19%, si | EN_1 |
| GOBP_SNRNA_TRANSCRI | GOBP_SNRNA_TRANSCRIPTION | 18 | 0.429788 | 1.4935502 | 0.057251908 | 0.17354144 | 1 | 5060 | tags=50%, list=21%, si | EN_1 |
| GOBP_INTEGRIN_MEDIAT | GOBP_INTEGRIN_MEDIATED_SIGNALING_P | 98 | 0.29267 | 1.4933482 | 0.0 | 0.17342757 | 1 | 4820 | tags=31%, list=20%, si | EN_1 |
| GOBP_RECEPTOR_RECYC | GOBP_RECEPTOR_RECYCLING | 54 | 0.332672 | 1.4932836 | 0.0 | 0.17316169 | 1 | 3534 | tags=31%, list=15%, si | EN_1 |
| GOBP_NEGATIVE_REGUL | GOBP_NEGATIVE_REGULATION_OF_MYELO | 89 | 0.308668 | 1.4919887 | 0.02020202 | 0.174178 | 1 | 4567 | tags=36%, list=19%, si | EN_1 |
| GOBP_ESTABLISHMENT_( | GOBP_ESTABLISHMENT_OF_PROTEIN_LOCA | 67 | 0.310229 | 1.4915137 | 0.01459854 | 0.17415726 | 1 | 4782 | tags=30%, list=20%, si | EN_1 |
| GOBP_REGULATION_OF_ | GOBP_REGULATION_OF_MACROPHAGE_DE | 25 | 0.39133 | 1.490518 | 0.0349345 | 0.17481638 | 1 | 2288 | tags=32%, list=10%, si | EN_1 |
| GOBP_POSITIVE_REGULA | GOBP_POSITIVE_REGULATION_OF_MEMBRA | 16 | 0.453182 | 1.4897361 | 0.03422053 | 0.17539094 | 1 | 5565 | tags=44%, list=24%, si | EN_1 |
| AMPK_SIGNALING_PATHW | AMPK_SIGNALING_PATHWAY | 120 | 0.292229 | 1.489637 | 0.024096385 | 0.17507015 | 1 | 5514 | tags=38%, list=23%, si | EN_1 |
| GOBP_VESICLE_TARGETIN | GOBP_VESICLE_TARGETING_TO_FROM_OR_ | 29 | 0.378295 | 1.4894017 | 0.051643193 | 0.17492431 | 1 | 7130 | tags=59%, list=30%, si | EN_1 |
| GOBP_TELOMERE_CAPPI | GOBP_TELOMERE_CAPPING | 38 | 0.356636 | 1.4885263 | 0.015 | 0.17554684 | 1 | 4697 | tags=34%, list=20%, si | EN_1 |
| GOBP_POSITIVE_REGULA | GOBP_POSITIVE_REGULATION_OF_TELOME | 31 | 0.378405 | 1.4881052 | 0.014778325 | 0.17567414 | 1 | 5052 | tags=35%, list=21%, si | EN_1 |
| GOBP_REGULATION_OF_ | GOBP_REGULATION_OF_CHOLESTEROL_ME | 41 | 0.360671 | 1.4869756 | 0.032608695 | 0.17664497 | 1 | 3112 | tags=32%, list=13%, si | EN_1 |
| GOBP_NEGATIVE_REGUL | GOBP_NEGATIVE_REGULATION_OF_SMOOT | 68 | 0.303879 | 1.4862498 | 0.0070422534 | 0.17711245 | 1 | 4286 | tags=34%, list=18%, si | EN_1 |
| GOBP_RIBOSOMAL_SMAI | GOBP_RIBOSOMAL_SMALL_SUBUNIT_BIOGE | 101 | 0.289505 | 1.4848458 | 0.0 | 0.17840973 | 1 | 15898 | tags=99%, list=67%, si | EN_1 |
| GOBP_TRANSLESION_SYN | GOBP_TRANSLESION_SYNTHESIS | 22 | 0.419156 | 1.4845694 | 0.07359307 | 0.17825219 | 1 | 2546 | tags=27%, list=11%, si | EN_1 |
| GOBP_POSITIVE_REGULA | GOBP_POSITIVE_REGULATION_OF_MYOBLA | 18 | 0.438528 | 1.4845654 | 0.055555556 | 0.1778777 | 1 | 4118 | tags=50%, list=17%, si | EN_1 |
| GOBP_NEGATIVE_REGUL | GOBP_NEGATIVE_REGULATION_OF_CELL_M | 36 | 0.359298 | 1.4820036 | 0.018604651 | 0.18008174 | 1 | 4292 | tags=33%, list=18%, si | EN_1 |
| GOBP_POSITIVE_REGULA | GOBP_POSITIVE_REGULATION_OF_PATTERN | 68 | 0.308156 | 1.4818497 | 0.01438849 | 0.17994414 | 1 | 2648 | tags=25%, list=11%, si | EN_1 |
| INSULIN_RESISTANCE | INSULIN_RESISTANCE | 105 | 0.290127 | 1.4809808 | 0.0125 | 0.18049698 | 1 | 5514 | tags=43%, list=23%, si | EN_1 |
| GOBP_DNA_TEMPLATED_ | GOBP_DNA_TEMPLATED_DNA_REPLICATION | 147 | 0.271602 | 1.4809319 | 0.0 | 0.18015759 | 1 | 4133 | tags=27%, list=18%, si | EN_1 |
| GOBP_HYALURONAN_BIO | GOBP_HYALURONAN_BIOSYNTHETIC_PROC | 15 | 0.451427 | 1.4798614 | 0.06153846 | 0.1811186 | 1 | 4297 | tags=33%, list=18%, si | EN_1 |
| GOBP_POSITIVE_REGULA | GOBP_POSITIVE_REGULATION_OF_POST_TR | 135 | 0.274203 | 1.4789346 | 0.0 | 0.18184821 | 1 | 4560 | tags=26%, list=19%, si | EN_1 |
| GOBP_LINOLEIC_ACID_M | GOBP_LINOLEIC_ACID_METABOLIC_PROCES | 18 | 0.422108 | 1.4787593 | 0.04494382 | 0.1817007 | 1 | 677 | tags=22%, list=3%, sig | EN_1 |
| GOBP_GLUTATHIONE_ME | GOBP_GLUTATHIONE_METABOLIC_PROCESS | 58 | 0.322341 | 1.4786434 | 0.025806451 | 0.18147355 | 1 | 4275 | tags=34%, list=18%, si | EN_1 |
| GOBP_NEGATIVE_REGUL | GOBP_NEGATIVE_REGULATION_OF_CELLUL | 106 | 0.284397 | 1.4773315 | 0.0 | 0.18264681 | 1 | 3964 | tags=25%, list=17%, si | EN_1 |
| GOBP_RIBOSOMAL_SMAI | GOBP_RIBOSOMAL_SMALL_SUBUNIT_ASSEM | 18 | 0.427736 | 1.4772627 | 0.060150377 | 0.18233697 | 1 | 13519 | tags=100%, list=57%, | EN_1 |
| GOBP_MYOBLAST_DIFFEI | GOBP_MYOBLAST_DIFFERENTIATION | 114 | 0.284718 | 1.4770663 | 0.0 | 0.18224175 | 1 | 4100 | tags=26%, list=17%, si | EN_1 |
| GOBP_REGULATION_OF_ | GOBP_REGULATION_OF_PROTEIN_TYROSIN | 60 | 0.318147 | 1.4737599 | 0.013513514 | 0.18553853 | 1 | 3902 | tags=30%, list=17%, si | EN_1 |
| GOBP_SMOOTH_MUSCLE | GOBP_SMOOTH_MUSCLE_CELL_MIGRATION | 115 | 0.282413 | 1.473679 | 0.0 | 0.18525366 | 1 | 4750 | tags=31%, list=20%, si | EN_1 |
| GOBP_PIGMENT_METABO | GOBP_PIGMENT_METABOLIC_PROCESS | 65 | 0.316453 | 1.4733832 | 0.015037594 | 0.18521309 | 1 | 5749 | tags=38%, list=24%, si | EN_1 |
| GOBP_ENDOPLASMIC_RE | GOBP_ENDOPLASMIC_RETICULUM_ORGANI | 90 | 0.291032 | 1.4731321 | 0.0 | 0.1851683 | 1 | 5774 | tags=39%, list=24%, si | EN_1 |
| NUCLEOTIDE_EXCISION_F | NUCLEOTIDE_EXCISION_REPAIR | 42 | 0.34424 | 1.4729023 | 0.02366864 | 0.18506035 | 1 | 4382 | tags=31%, list=19%, si | EN_1 |
| GOBP_MEMBRANE_LIPID | GOBP_MEMBRANE_LIPID_BIOSYNTHETIC_PI | 129 | 0.282974 | 1.4728917 | 0.0 | 0.18468498 | 1 | 3050 | tags=23%, list=13%, si | EN_1 |
| GOBP_NEGATIVE_REGUL | GOBP_NEGATIVE_REGULATION_OF_DNA_TI | 17 | 0.45074 | 1.4716688 | 0.06968641 | 0.18563329 | 1 | 7753 | tags=65%, list=33%, si | EN_1 |
| GOBP_PROTEIN_DNA_CO | GOBP_PROTEIN_DNA_COMPLEX_DISASSEM | 19 | 0.416079 | 1.4705011 | 0.036734693 | 0.18657358 | 1 | 5293 | tags=32%, list=22%, si | EN_1 |
| GOBP_REACTIVE_OXYGEI | GOBP_REACTIVE_OXYGEN_SPECIES_METAB | 239 | 0.258583 | 1.4697655 | 0.0 | 0.18697645 | 1 | 4297 | tags=27%, list=18%, si | EN_1 |
| GOBP_GLIAL_CELL_MIGR | GOBP_GLIAL_CELL_MIGRATION | 70 | 0.307749 | 1.469726 | 0.0 | 0.18662187 | 1 | 6305 | tags=41%, list=27%, si | EN_1 |
| GOBP_REGULATION_OF_ | GOBP_REGULATION_OF_ATP_BIOSYNTHETIC | 20 | 0.420808 | 1.4692876 | 0.052 | 0.18686679 | 1 | 6104 | tags=55%, list=26%, si | EN_1 |
| GOBP_POSITIVE_REGULA | GOBP_POSITIVE_REGULATION_OF_PROTEIN | 107 | 0.281676 | 1.4684405 | 0.0 | 0.18735906 | 1 | 3963 | tags=24%, list=17%, si | EN_1 |
| GOBP_MITOCHONDRIAL_ | GOBP_MITOCHONDRIAL_ELECTRON_TRANS | 29 | 0.375177 | 1.4679319 | 0.018779343 | 0.18746404 | 1 | 6533 | tags=55%, list=28%, si | EN_1 |
| GOBP_ENDOPLASMIC_RE | GOBP_ENDOPLASMIC_RETICULUM_TUBULA | 21 | 0.398938 | 1.4660623 | 0.060240965 | 0.18938522 | 1 | 7091 | tags=52%, list=30%, si | EN_1 |
| TYPE_II_DIABETES_MELLI | TYPE_II_DIABETES_MELLITUS | 45 | 0.327991 | 1.4656475 | 0.030303031 | 0.18952921 | 1 | 3155 | tags=29%, list=13%, si | EN_1 |
| GOBP_DNA_REPLICATION | GOBP_DNA_REPLICATION_INITIATION | 31 | 0.369406 | 1.4639195 | 0.051886793 | 0.19100173 | 1 | 3968 | tags=39%, list=17%, si | EN_1 |
| GOBP_POSITIVE_REGULA | GOBP_POSITIVE_REGULATION_OF_UBIQUIT | 18 | 0.42423 | 1.4638487 | 0.046332046 | 0.1907077 | 1 | 3661 | tags=39%, list=16%, si | EN_1 |
| GOBP_PROTEIN_LOCALIZ | GOBP_PROTEIN_LOCALIZATION_TO_VACUO | 78 | 0.30072 | 1.4631567 | 0.0 | 0.19117412 | 1 | 3620 | tags=19%, list=15%, si | EN_1 |
| GOBP_MATURATION_OF_ | GOBP_MATURATION_OF_SSU_RRNA | 48 | 0.333899 | 1.4629656 | 0.010989011 | 0.19106573 | 1 | 15743 | tags=100%, list=67%, | EN_1 |
| GOBP_FATTY_ACID_CATA | GOBP_FATTY_ACID_CATABOLIC_PROCESS | 100 | 0.29109 | 1.4628747 | 0.0091743115 | 0.19081506 | 1 | 4227 | tags=32%, list=18%, si | EN_1 |
| GOBP_INTERLEUKIN_1_M | GOBP_INTERLEUKIN_1_MEDIATED_SIGNALI | 28 | 0.386035 | 1.4618363 | 0.052863438 | 0.19163932 | 1 | 2780 | tags=32%, list=12%, si | EN_1 |
| GOBP_MODULATION_BY_ | GOBP_MODULATION_BY_SYMBIONT_OF_EN | 56 | 0.322196 | 1.4601649 | 0.021276595 | 0.193202 | 1 | 3526 | tags=34%, list=15%, si | EN_1 |
| GOBP_RESPONSE_TO_FLU | GOBP_RESPONSE_TO_FLUID_SHEAR_STRESS | 24 | 0.380845 | 1.459637 | 0.030837005 | 0.19335389 | 1 | 667 | tags=13%, list=3%, sig | EN_1 |
| GOBP_INTRACELLULAR_P | GOBP_INTRACELLULAR_PH_REDUCTION | 48 | 0.330263 | 1.4579636 | 0.040268455 | 0.19488066 | 1 | 5364 | tags=33%, list=23%, si | EN_1 |
| GOBP_REGULATION_OF_ | GOBP_REGULATION_OF_PROTEASOMAL_PR | 179 | 0.264374 | 1.4579424 | 0.0 | 0.19452035 | 1 | 5983 | tags=37%, list=25%, si | EN_1 |
| GOBP_REGULATION_OF_ | GOBP_REGULATION_OF_LIPOPOLYSACCHAR | 29 | 0.370409 | 1.4573938 | 0.046728972 | 0.19478403 | 1 | 1377 | tags=21%, list=6%, sig | EN_1 |
| GOBP_REGULATION_OF_ | GOBP_REGULATION_OF_PROTEIN_IMPORT_ | 76 | 0.306897 | 1.4573519 | 0.008474576 | 0.19444774 | 1 | 3725 | tags=29%, list=16%, si | EN_1 |
| TOLL-LIKE_RECEPTOR_SIG | TOLL-LIKE_RECEPTOR_SIGNALING_PATHWA | 84 | 0.298022 | 1.4564176 | 0.009090909 | 0.19514214 | 1 | 4209 | tags=29%, list=18%, si | EN_1 |

| | | | | | | | | | | |
|---|---|---|---|---|---|---|---|---|---|---|
| GOBP_CELL_AGGREGATI | GOBP_CELL_AGGREGATION | 19 | 0.427413 | 1.4562148 | 0.060240965 | 0.19501029 | 1 | 689 | tags=16%, list=3%, sig | EN_1 |
| GOBP_NEUROBLAST_DIV | GOBP_NEUROBLAST_DIVISION | 16 | 0.436752 | 1.4543141 | 0.05511811 | 0.19704649 | 1 | 3995 | tags=38%, list=17%, si | EN_1 |
| GOBP_ESTABLISHMENT_ | GOBP_ESTABLISHMENT_OF_PROTEIN_LOCA | 59 | 0.316063 | 1.4541235 | 0.012987013 | 0.19689177 | 1 | 3620 | tags=20%, list=15%, si | EN_1 |
| GOBP_POSITIVE_REGULA | GOBP_POSITIVE_REGULATION_OF_UBIQUIT | 93 | 0.29238 | 1.453023 | 0.010416667 | 0.19773957 | 1 | 5929 | tags=38%, list=25%, si | EN_1 |
| GOBP_LONG_CHAIN_FAT | GOBP_LONG_CHAIN_FATTY_ACID_IMPORT_ | 19 | 0.423482 | 1.4529495 | 0.061594203 | 0.19742808 | 1 | 4292 | tags=47%, list=18%, si | EN_1 |
| GOBP_ORGANIC_HYDRO | GOBP_ORGANIC_HYDROXY_COMPOUND_BI | 221 | 0.267962 | 1.4521612 | 0.0 | 0.19810872 | 1 | 2864 | tags=23%, list=12%, si | EN_1 |
| GOBP_LIPID_OXIDATION | GOBP_LIPID_OXIDATION | 118 | 0.280263 | 1.4518405 | 0.0 | 0.19812752 | 1 | 4227 | tags=31%, list=18%, si | EN_1 |
| GOBP_REGULATION_OF_ | GOBP_REGULATION_OF_SUBSTRATE_ADHES | 58 | 0.319727 | 1.4514452 | 0.01863354 | 0.19812231 | 1 | 4018 | tags=28%, list=17%, si | EN_1 |
| GOBP_TOLL_LIKE_RECEPT | GOBP_TOLL_LIKE_RECEPTOR_4_SIGNALING_ | 60 | 0.32105 | 1.4505005 | 0.021276595 | 0.19882716 | 1 | 3963 | tags=33%, list=17%, si | EN_1 |
| GOBP_MUSCLE_CELL_MI | GOBP_MUSCLE_CELL_MIGRATION | 129 | 0.275466 | 1.4501568 | 0.0 | 0.19891308 | 1 | 4935 | tags=34%, list=21%, si | EN_1 |
| GOBP_INNER_MITOCHON | GOBP_INNER_MITOCHONDRIAL_MEMBRAN | 36 | 0.347811 | 1.4499509 | 0.035714287 | 0.19874792 | 1 | 15411 | tags=100%, list=65%, | EN_1 |
| GOBP_PROTEIN_TARGETI | GOBP_PROTEIN_TARGETING_TO_VACUOLE | 46 | 0.320784 | 1.4490844 | 0.04191617 | 0.19942953 | 1 | 3620 | tags=20%, list=15%, si | EN_1 |
| GOBP_GERMINAL_CENTE | GOBP_GERMINAL_CENTER_FORMATION | 16 | 0.439533 | 1.4489796 | 0.06225681 | 0.1991727 | 1 | 3233 | tags=38%, list=14%, si | EN_1 |
| GOBP_EOSINOPHIL_CHEM | GOBP_EOSINOPHIL_CHEMOTAXIS | 22 | 0.385435 | 1.4484949 | 0.04347826 | 0.1994571 | 1 | 4151 | tags=50%, list=18%, si | EN_1 |
| GOBP_RIBOSOME_BIOGE | GOBP_RIBOSOME_BIOGENESIS | 306 | 0.247327 | 1.4466095 | 0.0 | 0.2013156 | 1 | 15910 | tags=97%, list=67%, si | EN_1 |
| GOBP_SNRNA_3_END_PR | GOBP_SNRNA_3_END_PROCESSING | 20 | 0.411827 | 1.4459381 | 0.048034936 | 0.20176412 | 1 | 2508 | tags=20%, list=11%, si | EN_1 |
| GOBP_REGULATION_OF_ | GOBP_REGULATION_OF_TELOMERASE_ACT | 42 | 0.341644 | 1.4451193 | 0.025 | 0.20254758 | 1 | 6253 | tags=45%, list=26%, si | EN_1 |
| GOBP_REGULATION_OF_ | GOBP_REGULATION_OF_LYMPHOCYTE_APO | 71 | 0.301532 | 1.4449393 | 0.0 | 0.20236808 | 1 | 5011 | tags=39%, list=21%, si | EN_1 |
| GOBP_LYMPHOCYTE_CHE | GOBP_LYMPHOCYTE_CHEMOTAXIS | 38 | 0.350422 | 1.4443308 | 0.009049774 | 0.20284912 | 1 | 1176 | tags=18%, list=5%, sig | EN_1 |
| GOBP_ESTABLISHMENT_ | GOBP_ESTABLISHMENT_OF_PROTEIN_LOCA | 262 | 0.251497 | 1.4438435 | 0.0 | 0.20310983 | 1 | 5734 | tags=34%, list=24%, si | EN_1 |
| GOBP_DNA_BIOSYNTHET | GOBP_DNA_BIOSYNTHETIC_PROCESS | 175 | 0.250926 | 1.4437588 | 0.0 | 0.2028384 | 1 | 5263 | tags=31%, list=22%, si | EN_1 |
| GOBP_PROTEIN_METHYL | GOBP_PROTEIN_METHYLATION | 57 | 0.313635 | 1.4432904 | 0.025477707 | 0.20303513 | 1 | 6326 | tags=37%, list=27%, si | EN_1 |
| GOBP_NEGATIVE_REGUL | GOBP_NEGATIVE_REGULATION_OF_INNATE | 86 | 0.2915 | 1.440457 | 0.036363635 | 0.20612824 | 1 | 2609 | tags=23%, list=11%, si | EN_1 |
| SNARE_INTERACTIONS_IN | SNARE_INTERACTIONS_IN_VESICULAR_TRAI | 32 | 0.364829 | 1.4403142 | 0.04901961 | 0.20590551 | 1 | 6851 | tags=44%, list=29%, si | EN_1 |
| GOBP_POSITIVE_REGULA | GOBP_POSITIVE_REGULATION_OF_PROTEIN | 35 | 0.362495 | 1.4400555 | 0.05 | 0.205814 | 1 | 4657 | tags=40%, list=20%, si | EN_1 |
| GOBP_NUCLEOSIDE_PHO | GOBP_NUCLEOSIDE_PHOSPHATE_CATABOLI | 170 | 0.265809 | 1.439775 | 0.0 | 0.20574512 | 1 | 3924 | tags=29%, list=17%, si | EN_1 |
| GOBP_REGULATION_OF_ | GOBP_REGULATION_OF_AUTOPHAGY | 295 | 0.240295 | 1.4395105 | 0.0 | 0.20571479 | 1 | 6379 | tags=40%, list=27%, si | EN_1 |
| LEISHMANIASIS | LEISHMANIASIS | 60 | 0.316441 | 1.4388797 | 0.023076924 | 0.20613909 | 1 | 2447 | tags=27%, list=10%, si | EN_1 |
| GOBP_REGULATION_OF_ | GOBP_REGULATION_OF_SPROUTING_ANGI | 42 | 0.339289 | 1.4388753 | 0.026595745 | 0.20577365 | 1 | 4544 | tags=43%, list=19%, si | EN_1 |
| GOBP_POSITIVE_REGULA | GOBP_POSITIVE_REGULATION_OF_CELL_SU | 134 | 0.275644 | 1.4378214 | 0.0 | 0.20671645 | 1 | 4820 | tags=31%, list=20%, si | EN_1 |
| GOBP_REGULATION_OF_ | GOBP_REGULATION_OF_PROTON_TRANSPO | 22 | 0.40243 | 1.4373091 | 0.059322033 | 0.20718278 | 1 | 941 | tags=18%, list=4%, sig | EN_1 |
| GOBP_REGULATION_OF_ | GOBP_REGULATION_OF_ACTIN_NUCLEATIO | 35 | 0.341891 | 1.4366424 | 0.033707865 | 0.20782627 | 1 | 2572 | tags=26%, list=11%, si | EN_1 |
| GOBP_RRNA_MODIFICAT | GOBP_RRNA_MODIFICATION | 29 | 0.365261 | 1.4366366 | 0.048245613 | 0.20744702 | 1 | 6472 | tags=52%, list=27%, si | EN_1 |
| GOBP_ATP_METABOLIC_ | GOBP_ATP_METABOLIC_PROCESS | 206 | 0.260846 | 1.4364167 | 0.0 | 0.20732495 | 1 | 5372 | tags=37%, list=23%, si | EN_1 |
| GOBP_MACROPHAGE_DI | GOBP_MACROPHAGE_DIFFERENTIATION | 61 | 0.316004 | 1.4358491 | 0.021582734 | 0.20760536 | 1 | 3725 | tags=31%, list=16%, si | EN_1 |
| GOBP_LATE_ENDOSOME_ | GOBP_LATE_ENDOSOME_TO_VACUOLE_TR | 25 | 0.391572 | 1.4356775 | 0.079831935 | 0.20742108 | 1 | 14375 | tags=100%, list=61%, | EN_1 |
| GLYCEROPHOSPHOLIPID_ | GLYCEROPHOSPHOLIPID_METABOLISM | 92 | 0.283766 | 1.4355006 | 0.0 | 0.20721948 | 1 | 3189 | tags=27%, list=14%, si | EN_1 |
| GOBP_CARDIOBLAST_DIF | GOBP_CARDIOBLAST_DIFFERENTIATION | 15 | 0.42843 | 1.4341285 | 0.05357143 | 0.2085319 | 1 | 2966 | tags=27%, list=13%, si | EN_1 |
| GOBP_RESPONSE_TO_MI | GOBP_RESPONSE_TO_MITOCHONDRIAL_DE | 21 | 0.392945 | 1.4334947 | 0.07594936 | 0.20872708 | 1 | 4534 | tags=29%, list=19%, si | EN_1 |
| GOBP_RESPONSE_TO_IN | GOBP_RESPONSE_TO_INTERLEUKIN_17 | 25 | 0.385751 | 1.4331086 | 0.058035713 | 0.208803 | 1 | 1434 | tags=20%, list=6%, sig | EN_1 |
| GOBP_NEGATIVE_REGUL | GOBP_NEGATIVE_REGULATION_OF_MYELO | 57 | 0.313151 | 1.4330448 | 0.014925373 | 0.20848669 | 1 | 3695 | tags=32%, list=16%, si | EN_1 |
| GOBP_REGULATION_OF_ | GOBP_REGULATION_OF_PROTEIN_AUTOPH | 41 | 0.3421 | 1.4330264 | 0.033707865 | 0.20812535 | 1 | 4546 | tags=29%, list=19%, si | EN_1 |
| GOBP_S_ADENOSYLMETH | GOBP_S_ADENOSYLMETHIONINE_METABOL | 17 | 0.43528 | 1.4328132 | 0.08058608 | 0.20802087 | 1 | 2066 | tags=24%, list=9%, sig | EN_1 |
| GOBP_CHONDROITIN_SU | GOBP_CHONDROITIN_SULFATE_PROTEOGLY | 21 | 0.411302 | 1.4326402 | 0.057377048 | 0.20781484 | 1 | 5026 | tags=52%, list=21%, si | EN_1 |
| GOBP_NEGATIVE_REGUL | GOBP_NEGATIVE_REGULATION_OF_CYCLIN | 25 | 0.376976 | 1.4321291 | 0.06467662 | 0.20811737 | 1 | 1433 | tags=20%, list=6%, sig | EN_1 |
| GOBP_KILLING_OF_CELLS | GOBP_KILLING_OF_CELLS_OF_ANOTHER_OI | 71 | 0.291883 | 1.4299592 | 0.008196721 | 0.21036322 | 1 | 2764 | tags=24%, list=12%, si | EN_1 |
| GOBP_NEGATIVE_REGUL | GOBP_NEGATIVE_REGULATION_OF_SPROUT | 15 | 0.440525 | 1.4292526 | 0.06137184 | 0.21087326 | 1 | 4506 | tags=47%, list=19%, si | EN_1 |
| GOBP_RETICULOPHAGY | GOBP_RETICULOPHAGY | 19 | 0.41111 | 1.4292519 | 0.08730159 | 0.21049869 | 1 | 5661 | tags=63%, list=24%, si | EN_1 |
| GOBP_PROTEIN_LOCALIZ | GOBP_PROTEIN_LOCALIZATION_TO_CHROM | 31 | 0.368984 | 1.4291115 | 0.035353534 | 0.21023956 | 1 | 14910 | tags=100%, list=63%, | EN_1 |
| GOBP_POSITIVE_REGULA | GOBP_POSITIVE_REGULATION_OF_SODIUM | 38 | 0.343343 | 1.4288785 | 0.046511628 | 0.21015392 | 1 | 1434 | tags=16%, list=6%, sig | EN_1 |
| GOBP_COLLAGEN_METAB | GOBP_COLLAGEN_METABOLIC_PROCESS | 119 | 0.278644 | 1.4282961 | 0.011904762 | 0.2104264 | 1 | 3768 | tags=28%, list=16%, si | EN_1 |
| GOBP_SYNAPTIC_VESICLE | GOBP_SYNAPTIC_VESICLE_TRANSPORT | 49 | 0.320702 | 1.4281317 | 0.031746034 | 0.21030799 | 1 | 5504 | tags=39%, list=23%, si | EN_1 |
| GOBP_FIBROBLAST_PROL | GOBP_FIBROBLAST_PROLIFERATION | 147 | 0.262629 | 1.4250304 | 0.0 | 0.21381557 | 1 | 3115 | tags=19%, list=13%, si | EN_1 |
| GOBP_MATURATION_OF_ | GOBP_MATURATION_OF_SSU_RRNA_FROM | 33 | 0.35099 | 1.4246298 | 0.026455026 | 0.2140061 | 1 | 15335 | tags=100%, list=65%, | EN_1 |
| GOBP_NEGATIVE_REGUL | GOBP_NEGATIVE_REGULATION_OF_RAS_PR | 26 | 0.380823 | 1.423854 | 0.053061225 | 0.21463342 | 1 | 3926 | tags=31%, list=17%, si | EN_1 |
| GOBP_NEGATIVE_REGUL | GOBP_NEGATIVE_REGULATION_OF_TELOM | 36 | 0.343018 | 1.4236801 | 0.028571429 | 0.21451502 | 1 | 15524 | tags=100%, list=66%, | EN_1 |
| GOBP_REGULATION_OF_ | GOBP_REGULATION_OF_MITOCHONDRIAL_ | 58 | 0.307356 | 1.422219 | 0.02173913 | 0.21592416 | 1 | 2888 | tags=24%, list=12%, si | EN_1 |
| GOBP_AUTOPHAGY_OF_ | GOBP_AUTOPHAGY_OF_MITOCHONDRION | 89 | 0.291257 | 1.4217663 | 0.0 | 0.21611434 | 1 | 5852 | tags=42%, list=25%, si | EN_1 |
| GOBP_NEGATIVE_REGUL | GOBP_NEGATIVE_REGULATION_OF_PHOSPH | 332 | 0.244716 | 1.4207805 | 0.0 | 0.21687849 | 1 | 4018 | tags=24%, list=17%, si | EN_1 |
| GOBP_RESPONSE_TO_TU | GOBP_RESPONSE_TO_TUMOR_NECROSIS_F | 167 | 0.257192 | 1.4204268 | 0.0 | 0.21699804 | 1 | 2503 | tags=18%, list=11%, si | EN_1 |
| GOBP_REGULATION_OF_ | GOBP_REGULATION_OF_CELL_SUBSTRATE_ | 219 | 0.256622 | 1.4201871 | 0.0 | 0.2169552 | 1 | 4820 | tags=29%, list=20%, si | EN_1 |
| GOBP_POSITIVE_REGULA | GOBP_POSITIVE_REGULATION_OF_SYNAPTI | 17 | 0.433084 | 1.4192388 | 0.064150944 | 0.21772589 | 1 | 1934 | tags=29%, list=8%, sig | EN_1 |
| GOBP_POSITIVE_REGULA | GOBP_POSITIVE_REGULATION_OF_PROTEAS | 72 | 0.300964 | 1.4187102 | 0.03508772 | 0.21800834 | 1 | 5929 | tags=38%, list=25%, si | EN_1 |
| MAPK_SIGNALING_PATH | MAPK_SIGNALING_PATHWAY | 287 | 0.24129 | 1.4175304 | 0.0 | 0.2191082 | 1 | 3725 | tags=25%, list=16%, si | EN_1 |
| GOBP_DEOXYRIBONUCLE | GOBP_DEOXYRIBONUCLEOTIDE_BIOSYNTHE | 17 | 0.43267 | 1.4164917 | 0.061594203 | 0.2199423 | 1 | 5509 | tags=47%, list=23%, si | EN_1 |
| GOBP_SNRNA_PROCESSI | GOBP_SNRNA_PROCESSING | 24 | 0.38314 | 1.4163141 | 0.05263158 | 0.21983564 | 1 | 7251 | tags=50%, list=31%, si | EN_1 |
| GOBP_MONOSACCHARID | GOBP_MONOSACCHARIDE_METABOLIC_PR | 255 | 0.240569 | 1.4156718 | 0.0 | 0.22037235 | 1 | 4131 | tags=27%, list=17%, si | EN_1 |
| GOBP_T_CELL_CHEMOTA | GOBP_T_CELL_CHEMOTAXIS | 23 | 0.374745 | 1.415264 | 0.053097345 | 0.22062477 | 1 | 1176 | tags=17%, list=5%, sig | EN_1 |
| GOBP_LIPID_MODIFICATI | GOBP_LIPID_MODIFICATION | 197 | 0.2453 | 1.4147736 | 0.0 | 0.22085944 | 1 | 4227 | tags=28%, list=18%, si | EN_1 |
| GOBP_REGULATION_OF_ | GOBP_REGULATION_OF_DNA_REPAIR | 219 | 0.247196 | 1.4130176 | 0.0 | 0.22292954 | 1 | 3544 | tags=18%, list=15%, si | EN_1 |
| GOBP_RESPONSE_TO_OX | GOBP_RESPONSE_TO_OXYGEN_LEVELS | 247 | 0.258211 | 1.4125795 | 0.0 | 0.22317092 | 1 | 3939 | tags=27%, list=17%, si | EN_1 |
| GOBP_REGULATION_OF_ | GOBP_REGULATION_OF_GLYCOPROTEIN_M | 47 | 0.318238 | 1.4119602 | 0.05 | 0.22355133 | 1 | 5490 | tags=34%, list=23%, si | EN_1 |
| AMINO_SUGAR_AND_NU | AMINO_SUGAR_AND_NUCLEOTIDE_SUGAR_ | 46 | 0.326296 | 1.4107742 | 0.03141361 | 0.22475316 | 1 | 4212 | tags=30%, list=18%, si | EN_1 |
| GOBP_NEGATIVE_REGUL | GOBP_NEGATIVE_REGULATION_OF_TOR_SI | 61 | 0.307588 | 1.4106668 | 0.03448276 | 0.22451155 | 1 | 16367 | tags=100%, list=69%, | EN_1 |
| GOBP_REGULATION_OF_ | GOBP_REGULATION_OF_CELLULAR_SENESC | 47 | 0.317029 | 1.410549 | 0.02793296 | 0.22427775 | 1 | 5617 | tags=38%, list=24%, si | EN_1 |
| GOBP_ALCOHOL_METAB | GOBP_ALCOHOL_METABOLIC_PROCESS | 308 | 0.250018 | 1.4105222 | 0.0 | 0.22394632 | 1 | 3499 | tags=24%, list=15%, si | EN_1 |
| GOBP_NEGATIVE_REGUL | GOBP_NEGATIVE_REGULATION_OF_APOPT | 241 | 0.251437 | 1.4070084 | 0.0 | 0.22823621 | 1 | 3253 | tags=19%, list=14%, si | EN_1 |
| GOBP_REGULATION_OF_ | GOBP_REGULATION_OF_INTRINSIC_APOPT | 185 | 0.249072 | 1.4060104 | 0.0 | 0.22926949 | 1 | 3523 | tags=20%, list=15%, si | EN_1 |
| GOBP_NUCLEAR_ENVELO | GOBP_NUCLEAR_ENVELOPE_ORGANIZATIO | 52 | 0.313221 | 1.4054214 | 0.023121387 | 0.22969976 | 1 | 6859 | tags=38%, list=29%, si | EN_1 |
| GOBP_ORGANIC_ACID_B | GOBP_ORGANIC_ACID_BIOSYNTHETIC_PRO | 287 | 0.236036 | 1.4047918 | 0.0 | 0.23015338 | 1 | 3719 | tags=28%, list=16%, si | EN_1 |
| GOBP_PROTEIN_LOCALIZ | GOBP_PROTEIN_LOCALIZATION_TO_MITOC | 101 | 0.275061 | 1.4043851 | 0.01 | 0.23033027 | 1 | 6689 | tags=40%, list=28%, si | EN_1 |
| GOBP_TETRAPYRROLE_BI | GOBP_TETRAPYRROLE_BIOSYNTHETIC_PRO | 29 | 0.360607 | 1.4043068 | 0.05479452 | 0.23007776 | 1 | 7445 | tags=62%, list=32%, si | EN_1 |

| | | | | | | | | | | |
|---|---|---|---|---|---|---|---|---|---|---|
| GOBP_CARBOHYDRATE_C | GOBP_CARBOHYDRATE_CATABOLIC_PROCES | 161 | 0.25513 | 1.404263 | 0.0 | 0.22974652 | 1 | 4859 | tags=33%, list=21%, si | EN_1 |
| GOBP_VASCULAR_ENDOT | GOBP_VASCULAR_ENDOTHELIAL_GROWTH_ | 50 | 0.310761 | 1.4036151 | 0.029940119 | 0.23018879 | 1 | 3476 | tags=30%, list=15%, si | EN_1 |
| GOBP_REGULATION_OF_ | GOBP_REGULATION_OF_HETEROTYPIC_CELI | 21 | 0.395357 | 1.4034586 | 0.06827309 | 0.22995509 | 1 | 4118 | tags=33%, list=17%, si | EN_1 |
| IL-17_SIGNALING_PATHW | IL-17_SIGNALING_PATHWAY | 85 | 0.283659 | 1.402844 | 0.0 | 0.23035747 | 1 | 4225 | tags=31%, list=18%, si | EN_1 |
| GOBP_NEGATIVE_REGUL | GOBP_NEGATIVE_REGULATION_OF_CELLUL | 18 | 0.409839 | 1.402455 | 0.0754717 | 0.23040265 | 1 | 4506 | tags=33%, list=19%, si | EN_1 |
| GOBP_MITOCHONDRIAL_ | GOBP_MITOCHONDRIAL_DNA_REPLICATION | 17 | 0.422755 | 1.4021469 | 0.10218978 | 0.23046316 | 1 | 4876 | tags=29%, list=21%, si | EN_1 |
| MICRORNAS_IN_CANCER | MICRORNAS_IN_CANCER | 139 | 0.261935 | 1.401671 | 0.0 | 0.23071164 | 1 | 5193 | tags=32%, list=22%, si | EN_1 |
| GOBP_PROTEIN_TRANSM | GOBP_PROTEIN_TRANSMEMBRANE_IMPOR | 36 | 0.335341 | 1.4014524 | 0.0591133 | 0.23063493 | 1 | 15705 | tags=100%, list=67%, | EN_1 |
| GOBP_POSITIVE_REGULA | GOBP_POSITIVE_REGULATION_OF_MYELOID | 38 | 0.340204 | 1.4003422 | 0.056338027 | 0.23178974 | 1 | 2630 | tags=26%, list=11%, si | EN_1 |
| GOBP_NECROPTOTIC_PR | GOBP_NECROPTOTIC_PROCESS | 37 | 0.341402 | 1.3987477 | 0.04494382 | 0.23357818 | 1 | 5726 | tags=43%, list=24%, si | EN_1 |
| GOBP_REGULATION_OF_ | GOBP_REGULATION_OF_PROTEIN_SUMOYL | 25 | 0.387607 | 1.3985281 | 0.072072074 | 0.23346515 | 1 | 4560 | tags=32%, list=19%, si | EN_1 |
| GOBP_MITOCHONDRIAL_ | GOBP_MITOCHONDRIAL_MEMBRANE_ORGA | 107 | 0.272956 | 1.3976597 | 0.0 | 0.23423329 | 1 | 6139 | tags=37%, list=26%, si | EN_1 |
| GOBP_REGULATION_OF_ | GOBP_REGULATION_OF_MITOCHONDRIAL_ | 32 | 0.350297 | 1.3974634 | 0.047826085 | 0.23408204 | 1 | 15351 | tags=100%, list=65%, | EN_1 |
| GOBP_POST_GOLGI_VESI | GOBP_POST_GOLGI_VESICLE_MEDIATED_TF | 101 | 0.282294 | 1.3957778 | 0.0 | 0.23622574 | 1 | 4487 | tags=25%, list=19%, si | EN_1 |
| GOBP_PHOSPHATIDYLINC | GOBP_PHOSPHATIDYLINOSITOL_METABOLI( | 148 | 0.259069 | 1.3957593 | 0.0 | 0.2358515 | 1 | 4546 | tags=28%, list=19%, si | EN_1 |
| GOBP_RESPONSE_TO_CH | GOBP_RESPONSE_TO_CHEMOKINE | 74 | 0.284076 | 1.3944882 | 0.008928572 | 0.23710525 | 1 | 2681 | tags=24%, list=11%, si | EN_1 |
| GOBP_NEUTRAL_LIPID_B | GOBP_NEUTRAL_LIPID_BIOSYNTHETIC_PRO( | 58 | 0.303571 | 1.3942415 | 0.03125 | 0.2370276 | 1 | 2989 | tags=21%, list=13%, si | EN_1 |
| GOBP_RESPONSE_TO_IOI | GOBP_RESPONSE_TO_IONIZING_RADIATION | 127 | 0.263637 | 1.3941492 | 0.0 | 0.23675749 | 1 | 5276 | tags=31%, list=22%, si | EN_1 |
| GOBP_SULFUR_COMPOU | GOBP_SULFUR_COMPOUND_BIOSYNTHETIC | 126 | 0.262391 | 1.3937716 | 0.025641026 | 0.23683025 | 1 | 3681 | tags=26%, list=16%, si | EN_1 |
| GOBP_REGULATION_OF_ | GOBP_REGULATION_OF_MYOBLAST_DIFFER | 81 | 0.28266 | 1.3932203 | 0.008403362 | 0.23723592 | 1 | 4100 | tags=27%, list=17%, si | EN_1 |
| GOBP_REGULATION_OF_ | GOBP_REGULATION_OF_INTRACELLULAR_P | 168 | 0.254685 | 1.3925402 | 0.0 | 0.23778336 | 1 | 6076 | tags=36%, list=26%, si | EN_1 |
| GOBP_REGULATION_OF_ | GOBP_REGULATION_OF_NUCLEOTIDE_BIOS | 32 | 0.342255 | 1.3921882 | 0.0456621 | 0.23787957 | 1 | 3870 | tags=38%, list=16%, si | EN_1 |
| GOBP_NUCLEOSIDE_DIPH | GOBP_NUCLEOSIDE_DIPHOSPHATE_METAB( | 127 | 0.261136 | 1.3918266 | 0.033333335 | 0.23796244 | 1 | 3924 | tags=29%, list=17%, si | EN_1 |
| GOBP_SCF_DEPENDENT_ | GOBP_SCF_DEPENDENT_PROTEASOMAL_UE | 47 | 0.320892 | 1.3911968 | 0.040462427 | 0.23847951 | 1 | 3911 | tags=30%, list=17%, si | EN_1 |
| GOBP_REGULATION_OF_ | GOBP_REGULATION_OF_PROTEIN_MATURA | 71 | 0.290494 | 1.3909364 | 0.032258064 | 0.23841383 | 1 | 4292 | tags=32%, list=18%, si | EN_1 |
| GOBP_VESICLE_TARGETIN | GOBP_VESICLE_TARGETING | 60 | 0.299851 | 1.3901048 | 0.0472973 | 0.23926471 | 1 | 7139 | tags=48%, list=30%, si | EN_1 |
| GOBP_NEGATIVE_REGUL | GOBP_NEGATIVE_REGULATION_OF_MUSCL | 69 | 0.287583 | 1.3899661 | 0.040650405 | 0.23908418 | 1 | 4245 | tags=33%, list=18%, si | EN_1 |
| GOBP_POSITIVE_REGULA | GOBP_POSITIVE_REGULATION_OF_NUCLEO | 74 | 0.283619 | 1.3894395 | 0.007751938 | 0.2394766 | 1 | 3725 | tags=24%, list=16%, si | EN_1 |
| GOBP_NEGATIVE_REGUL | GOBP_NEGATIVE_REGULATION_OF_TORC1_ | 41 | 0.323915 | 1.3884943 | 0.06779661 | 0.24047612 | 1 | 15976 | tags=100%, list=68%, | EN_1 |
| GOBP_RIBOSE_PHOSPHAT | GOBP_RIBOSE_PHOSPHATE_METABOLIC_PR | 419 | 0.216182 | 1.3882511 | 0.0 | 0.24044791 | 1 | 5372 | tags=33%, list=23%, si | EN_1 |
| GOBP_RRNA_PROCESSIN | GOBP_RRNA_PROCESSING | 211 | 0.248485 | 1.3881708 | 0.0 | 0.24015538 | 1 | 4952 | tags=21%, list=21%, si | EN_1 |
| GOBP_NEGATIVE_REGUL | GOBP_NEGATIVE_REGULATION_OF_DNA_B | 148 | 0.257589 | 1.3880204 | 0.0 | 0.23999453 | 1 | 6154 | tags=41%, list=26%, si | EN_1 |
| GOBP_PHOSPHOLIPID_DE | GOBP_PHOSPHOLIPID_DEPHOSPHORYLATIO | 43 | 0.321744 | 1.3878528 | 0.050847456 | 0.23994985 | 1 | 2010 | tags=21%, list=9%, sig | EN_1 |
| GOBP_POSITIVE_REGULA | GOBP_POSITIVE_REGULATION_OF_STEROID | 25 | 0.377396 | 1.3872024 | 0.07239819 | 0.24047568 | 1 | 2265 | tags=24%, list=10%, si | EN_1 |
| VALINE,_LEUCINE_AND_I | VALINE,_LEUCINE_AND_ISOLEUCINE_DEGRA | 49 | 0.307766 | 1.3865341 | 0.022988506 | 0.24103406 | 1 | 4195 | tags=33%, list=18%, si | EN_1 |
| GOBP_ORGANELLE_DISAS | GOBP_ORGANELLE_DISASSEMBLY | 155 | 0.256785 | 1.3859379 | 0.0 | 0.24148779 | 1 | 6597 | tags=43%, list=28%, si | EN_1 |
| GOBP_VACUOLAR_TRANS | GOBP_VACUOLAR_TRANSPORT | 161 | 0.254387 | 1.3856964 | 0.018181818 | 0.24143142 | 1 | 6296 | tags=34%, list=27%, si | EN_1 |
| GOBP_RESPONSE_TO_AN | GOBP_RESPONSE_TO_AMINO_ACID_STARV/ | 54 | 0.319487 | 1.3851017 | 0.07586207 | 0.2419141 | 1 | 5045 | tags=35%, list=21%, si | EN_1 |
| GOBP_DNA_REPLICATION | GOBP_DNA_REPLICATION | 263 | 0.231058 | 1.3846381 | 0.0 | 0.24213341 | 1 | 4133 | tags=22%, list=18%, si | EN_1 |
| GOBP_REGULATION_OF_ | GOBP_REGULATION_OF_CELLULAR_RESPON | 89 | 0.274136 | 1.3840624 | 0.0 | 0.24256966 | 1 | 4483 | tags=30%, list=19%, si | EN_1 |
| GOBP_MYOBLAST_FUSIO | GOBP_MYOBLAST_FUSION | 45 | 0.325124 | 1.3837246 | 0.051813472 | 0.24267228 | 1 | 4354 | tags=38%, list=18%, si | EN_1 |
| GOBP_POSITIVE_REGULA | GOBP_POSITIVE_REGULATION_OF_INTRACE | 108 | 0.272122 | 1.3836225 | 0.0 | 0.24243197 | 1 | 3737 | tags=24%, list=16%, si | EN_1 |
| GOBP_POSITIVE_REGULA | GOBP_POSITIVE_REGULATION_OF_LIPID_M | 147 | 0.256885 | 1.3836125 | 0.014492754 | 0.24205883 | 1 | 3112 | tags=22%, list=13%, si | EN_1 |
| GOBP_CELL_SURFACE_PA | GOBP_CELL_SURFACE_PATTERN_RECOGNIT | 70 | 0.287089 | 1.3829887 | 0.020689655 | 0.24253681 | 1 | 2112 | tags=21%, list=9%, sig | EN_1 |
| GLUCAGON_SIGNALING_ | GLUCAGON_SIGNALING_PATHWAY | 91 | 0.274291 | 1.3826424 | 0.019607844 | 0.24264944 | 1 | 5647 | tags=36%, list=24%, si | EN_1 |
| GOBP_RESPONSE_TO_AT | GOBP_RESPONSE_TO_ATP | 31 | 0.359171 | 1.3808811 | 0.06355932 | 0.2447816 | 1 | 3635 | tags=35%, list=15%, si | EN_1 |
| GOBP_REGULATION_OF_ | GOBP_REGULATION_OF_MAP_KINASE_ACTI | 159 | 0.254203 | 1.3800024 | 0.019607844 | 0.2456646 | 1 | 3317 | tags=24%, list=14%, si | EN_1 |
| PROTEIN_EXPORT | PROTEIN_EXPORT | 26 | 0.36493 | 1.3795016 | 0.0883721 | 0.24594782 | 1 | 5718 | tags=58%, list=24%, si | EN_1 |
| GOBP_NUCLEAR_MEMBR | GOBP_NUCLEAR_MEMBRANE_REASSEMBLY | 20 | 0.391489 | 1.3783543 | 0.08494209 | 0.24715206 | 1 | 14375 | tags=100%, list=61%, | EN_1 |
| GOBP_HEPATOCYTE_APO | GOBP_HEPATOCYTE_APOPTOTIC_PROCESS | 26 | 0.357792 | 1.3780032 | 0.09134615 | 0.24721345 | 1 | 6198 | tags=35%, list=26%, si | EN_1 |
| GOBP_REGULATION_OF_ | GOBP_REGULATION_OF_PROTEIN_CATABOI | 346 | 0.228627 | 1.3774269 | 0.0 | 0.24755268 | 1 | 5987 | tags=34%, list=25%, si | EN_1 |
| GOBP_NEGATIVE_REGUL | GOBP_NEGATIVE_REGULATION_OF_OXIDOF | 17 | 0.408684 | 1.3768568 | 0.09219858 | 0.24805897 | 1 | 4949 | tags=47%, list=21%, si | EN_1 |
| GOBP_DETOXIFICATION_( | GOBP_DETOXIFICATION_OF_INORGANIC_C( | 15 | 0.431449 | 1.3754519 | 0.113074206 | 0.24973354 | 1 | 4348 | tags=40%, list=18%, si | EN_1 |
| GOBP_NEGATIVE_REGUL | GOBP_NEGATIVE_REGULATION_OF_AUTOPI | 81 | 0.284889 | 1.3752311 | 0.025423728 | 0.24964772 | 1 | 4213 | tags=26%, list=18%, si | EN_1 |
| GOBP_MACROMOLECULE | GOBP_MACROMOLECULE_DEACYLATION | 59 | 0.296154 | 1.3751464 | 0.01369863 | 0.24938072 | 1 | 4744 | tags=27%, list=20%, si | EN_1 |
| GOBP_MEMBRANE_LIPID | GOBP_MEMBRANE_LIPID_METABOLIC_PRO | 186 | 0.245648 | 1.3750188 | 0.0 | 0.24917452 | 1 | 3050 | tags=22%, list=13%, si | EN_1 |
| GOBP_ISOPRENOID_BIOS | GOBP_ISOPRENOID_BIOSYNTHETIC_PROCES | 36 | 0.334015 | 1.3743194 | 0.061452515 | 0.24969928 | 1 | 2995 | tags=31%, list=13%, si | EN_1 |
| GOBP_RIBONUCLEOPROT | GOBP_RIBONUCLEOPROTEIN_COMPLEX_BI( | 417 | 0.221534 | 1.3740009 | 0.0 | 0.24974844 | 1 | 15910 | tags=96%, list=67%, si | EN_1 |
| GOBP_POSITIVE_REGULA | GOBP_POSITIVE_REGULATION_OF_MACROF | 29 | 0.359871 | 1.373923 | 0.04761905 | 0.24955851 | 1 | 2630 | tags=28%, list=11%, si | EN_1 |
| GOBP_SIGNALING_RECEP | GOBP_SIGNALING_RECEPTOR_LIGAND_PRE( | 29 | 0.353316 | 1.3738403 | 0.0787037 | 0.24934253 | 1 | 3851 | tags=28%, list=16%, si | EN_1 |
| GOBP_CONVERGENT_EXT | GOBP_CONVERGENT_EXTENSION | 15 | 0.429934 | 1.3734237 | 0.10309278 | 0.24945822 | 1 | 5481 | tags=53%, list=23%, si | EN_1 |
| GOBP_NEGATIVE_REGUL | GOBP_NEGATIVE_REGULATION_OF_PROTEI | 80 | 0.284245 | 1.3729695 | 0.031746034 | 0.24963863 | 1 | 3116 | tags=21%, list=13%, si | EN_1 |
| GOBP_NEGATIVE_REGUL | GOBP_NEGATIVE_REGULATION_OF_RELEAS | 22 | 0.373685 | 1.3728832 | 0.056 | 0.24939072 | 1 | 5517 | tags=41%, list=23%, si | EN_1 |
| PHENYLALANINE_METAB( | PHENYLALANINE_METABOLISM | 16 | 0.416929 | 1.372579 | 0.09293681 | 0.24949627 | 1 | 1756 | tags=25%, list=7%, sig | EN_1 |
| GOBP_REGULATION_OF_ | GOBP_REGULATION_OF_LIPID_CATABOLIC_ | 63 | 0.28279 | 1.3720425 | 0.0472973 | 0.24985968 | 1 | 2174 | tags=17%, list=9%, sig | EN_1 |
| GOBP_MITOTIC_DNA_INT | GOBP_MITOTIC_DNA_INTEGRITY_CHECKPO | 84 | 0.277664 | 1.3717057 | 0.0 | 0.24997853 | 1 | 3968 | tags=27%, list=17%, si | EN_1 |
| GOBP_REGULATION_OF_ | GOBP_REGULATION_OF_OXIDOREDUCTASE_ | 63 | 0.288644 | 1.3715239 | 0.04511278 | 0.24986911 | 1 | 2231 | tags=21%, list=9%, sig | EN_1 |
| GOBP_DISRUPTION_OF_A | GOBP_DISRUPTION_OF_ANATOMICAL_STRU | 73 | 0.288814 | 1.3708888 | 0.048387095 | 0.24998748 | 1 | 2764 | tags=23%, list=12%, si | EN_1 |
| OLFACTORY_TRANSDUCT | OLFACTORY_TRANSDUCTION | 186 | -0.5691 | -2.392394 | 0 | 0 | 0 | 7028 | tags=73%, list=30%, si | EN_1 |
| NEUROACTIVE_LIGAND-R | NEUROACTIVE_LIGAND-RECEPTOR_INTERA( | 274 | -0.44679 | -1.900511 | 0 | 0.039884962 | 0.279 | 7494 | tags=60%, list=32%, si | EN_1 |
| GOBP_RESPONSE_TO_PH | GOBP_RESPONSE_TO_PHEROMONE | 17 | -0.66351 | -1.884979 | 0.0013908206 | 0.043111842 | 0.355 | 4580 | tags=71%, list=19%, si | EN_1 |
| GOBP_ICOSANOID_TRAN | GOBP_ICOSANOID_TRANSPORT | 60 | -0.49347 | -1.812591 | 0 | 0.11327419 | 0.74 | 8168 | tags=60%, list=35%, si | EN_1 |
| OXIDATIVE_PHOSPHORYL | OXIDATIVE_PHOSPHORYLATION | 115 | 0.64164 | 3.396705 | 0 | 0 | 0 | 4105 | tags=70%, list=17%, si | MM_1 |
| GOBP_NADH_DEHYDROG | GOBP_NADH_DEHYDROGENASE_COMPLEX_ | 58 | 0.700112 | 3.3295238 | 0 | 0 | 0 | 4001 | tags=69%, list=17%, si | MM_1 |
| GOBP_MITOCHONDRIAL_ | GOBP_MITOCHONDRIAL_RESPIRATORY_CHA | 91 | 0.654221 | 3.3076682 | 0 | 0 | 0 | 6190 | tags=77%, list=26%, si | MM_1 |
| PARKINSON_DISEASE | PARKINSON_DISEASE | 122 | 0.607856 | 3.1695807 | 0 | 0 | 0 | 4088 | tags=64%, list=17%, si | MM_1 |
| GOBP_ATP_BIOSYNTHETI | GOBP_ATP_BIOSYNTHETIC_PROCESS | 80 | 0.648221 | 3.136742 | 0 | 0 | 0 | 3955 | tags=61%, list=17%, si | MM_1 |
| GOBP_AEROBIC_RESPIRA | GOBP_AEROBIC_RESPIRATION | 170 | 0.549264 | 3.130119 | 0 | 0 | 0 | 4873 | tags=56%, list=21%, si | MM_1 |
| GOBP_NUCLEOSIDE_TRIP | GOBP_NUCLEOSIDE_TRIPHOSPHATE_BIOSYI | 100 | 0.593444 | 3.0832527 | 0 | 0 | 0 | 3955 | tags=55%, list=17%, si | MM_1 |
| THERMOGENESIS | THERMOGENESIS | 210 | 0.526986 | 3.0478077 | 0 | 0 | 0 | 4974 | tags=53%, list=21%, si | MM_1 |
| RIBOSOME | RIBOSOME | 128 | 0.563912 | 3.0387163 | 0 | 0 | 0 | 8565 | tags=77%, list=36%, si | MM_1 |
| GOBP_MITOCHONDRIAL_ | GOBP_MITOCHONDRIAL_TRANSLATION | 128 | 0.565177 | 3.037871 | 0 | 0 | 0 | 6638 | tags=66%, list=28%, si | MM_1 |

| | | | | | | | | | | |
|---|---|---|---|---|---|---|---|---|---|---|
| GOBP_PROTON_MOTIVE_ | GOBP_PROTON_MOTIVE_FORCE_DRIVEN_A | 52 | 0.691683 | 3.032854 | 0 | 0 | 0 | 3955 | tags=73%, list=17%, si | MM_1 |
| ALZHEIMER_DISEASE | ALZHEIMER_DISEASE | 159 | 0.542757 | 2.966667 | 0 | 0 | 0 | 3654 | tags=52%, list=15%, si | MM_1 |
| HUNTINGTON_DISEASE | HUNTINGTON_DISEASE | 180 | 0.522001 | 2.9320657 | 0 | 0 | 0 | 3709 | tags=49%, list=16%, si | MM_1 |
| GOBP_CELLULAR_RESPIRA | GOBP_CELLULAR_RESPIRATION | 218 | 0.508713 | 2.926355 | 0 | 0 | 0 | 4873 | tags=53%, list=21%, si | MM_1 |
| GOBP_OXIDATIVE_PHOSP | GOBP_OXIDATIVE_PHOSPHORYLATION | 124 | 0.536942 | 2.8472924 | 0 | 0 | 0 | 3955 | tags=52%, list=17%, si | MM_1 |
| GOBP_MITOCHONDRIAL_ | GOBP_MITOCHONDRIAL_GENE_EXPRESSION | 164 | 0.510086 | 2.838428 | 0 | 0 | 0 | 6638 | tags=62%, list=28%, si | MM_1 |
| GOBP_ELECTRON_TRANS | GOBP_ELECTRON_TRANSPORT_CHAIN | 107 | 0.538287 | 2.828336 | 0 | 0 | 0 | 4873 | tags=58%, list=21%, si | MM_1 |
| SPLICEOSOME | SPLICEOSOME | 127 | 0.513984 | 2.7808032 | 0 | 0 | 0 | 7594 | tags=58%, list=32%, si | MM_1 |
| GOBP_RIBONUCLEOPROT | GOBP_RIBONUCLEOPROTEIN_COMPLEX_BIC | 417 | 0.436946 | 2.7719822 | 0 | 0 | 0 | 6641 | tags=45%, list=28%, si | MM_1 |
| GOBP_RIBOSOME_BIOGE | GOBP_RIBOSOME_BIOGENESIS | 306 | 0.446394 | 2.7311387 | 0 | 0 | 0 | 6641 | tags=46%, list=28%, si | MM_1 |
| GOBP_AEROBIC_ELECTRO | GOBP_AEROBIC_ELECTRON_TRANSPORT_CH | 63 | 0.577996 | 2.697869 | 0 | 0 | 0 | 3878 | tags=59%, list=16%, si | MM_1 |
| LYSOSOME | LYSOSOME | 118 | 0.50615 | 2.6845112 | 0 | 0 | 0 | 7059 | tags=71%, list=30%, si | MM_1 |
| GOBP_RIBOSE_PHOSPHAT | GOBP_RIBOSE_PHOSPHATE_BIOSYNTHETIC_ | 185 | 0.474324 | 2.6561728 | 0 | 0 | 0 | 6190 | tags=57%, list=26%, si | MM_1 |
| GOBP_PROTEIN_TARGETI | GOBP_PROTEIN_TARGETING_TO_MITOCHO | 76 | 0.542764 | 2.6491537 | 0 | 8.94524E-05 | 0.001 | 5153 | tags=51%, list=22%, si | MM_1 |
| GOBP_ATP_SYNTHESIS_C | GOBP_ATP_SYNTHESIS_COUPLED_ELECTRO | 76 | 0.5496 | 2.615396 | 0 | 8.58743E-05 | 0.001 | 3878 | tags=54%, list=16%, si | MM_1 |
| GOBP_AUTOPHAGOSOMI | GOBP_AUTOPHAGOSOME_MATURATION | 58 | 0.567104 | 2.592007 | 0 | 8.25715E-05 | 0.001 | 6569 | tags=66%, list=28%, si | MM_1 |
| CITRATE_CYCLE_(TCA_CY | CITRATE_CYCLE_(TCA_CYCLE) | 28 | 0.668661 | 2.5746841 | 0 | 7.95133E-05 | 0.001 | 4093 | tags=64%, list=17%, si | MM_1 |
| GOBP_PROTEIN_LOCALIZ | GOBP_PROTEIN_LOCALIZATION_TO_MITOC | 101 | 0.503665 | 2.5554037 | 0 | 0.000155803 | 0.002 | 5243 | tags=49%, list=22%, si | MM_1 |
| GOBP_RRNA_PROCESSIN | GOBP_RRNA_PROCESSING | 211 | 0.441965 | 2.5550861 | 0 | 0.00015043 | 0.002 | 6641 | tags=45%, list=28%, si | MM_1 |
| GOBP_HEME_METABOLIC | GOBP_HEME_METABOLIC_PROCESS | 34 | 0.627157 | 2.5535586 | 0 | 0.000145416 | 0.002 | 4566 | tags=62%, list=19%, si | MM_1 |
| GOBP_METALLO_SULFUR | GOBP_METALLO_SULFUR_CLUSTER_ASSEMI | 25 | 0.690999 | 2.5473151 | 0 | 0.000140725 | 0.002 | 4903 | tags=72%, list=21%, si | MM_1 |
| GOBP_TRICARBOXYLIC_A | GOBP_TRICARBOXYLIC_ACID_CYCLE | 33 | 0.622289 | 2.54462 | 0 | 0.000136328 | 0.002 | 4093 | tags=61%, list=17%, si | MM_1 |
| GOBP_INNER_MITOCHON | GOBP_INNER_MITOCHONDRIAL_MEMBRAN | 36 | 0.625844 | 2.5333087 | 0 | 0.00019805 | 0.003 | 5665 | tags=61%, list=24%, si | MM_1 |
| PROTEASOME | PROTEASOME | 45 | 0.593237 | 2.5332978 | 0 | 0.000192225 | 0.003 | 6686 | tags=84%, list=28%, si | MM_1 |
| GOBP_RIBOSOMAL_LARG | GOBP_RIBOSOMAL_LARGE_SUBUNIT_BIOGE | 64 | 0.530154 | 2.5288334 | 0 | 0.00031316 | 0.005 | 6798 | tags=56%, list=29%, si | MM_1 |
| GOBP_GENERATION_OF_ | GOBP_GENERATION_OF_PRECURSOR_META | 439 | 0.395325 | 2.506657 | 0 | 0.000304461 | 0.005 | 5284 | tags=44%, list=22%, si | MM_1 |
| GOBP_CYTOPLASMIC_TR | GOBP_CYTOPLASMIC_TRANSLATION | 143 | 0.463759 | 2.4976249 | 0 | 0.00041013 | 0.007 | 8834 | tags=66%, list=37%, si | MM_1 |
| GOBP_RRNA_METABOLIC | GOBP_RRNA_METABOLIC_PROCESS | 241 | 0.424829 | 2.4902775 | 0 | 0.000454661 | 0.008 | 6641 | tags=46%, list=28%, si | MM_1 |
| GOBP_TRNA_METABOLIC | GOBP_TRNA_METABOLIC_PROCESS | 166 | 0.438094 | 2.4874825 | 0 | 0.00055475 | 0.01 | 7162 | tags=55%, list=30%, si | MM_1 |
| GOBP_MITOCHONDRIAL_ | GOBP_MITOCHONDRIAL_ELECTRON_TRANS | 29 | 0.642753 | 2.474907 | 0 | 0.000646119 | 0.012 | 3611 | tags=62%, list=15%, si | MM_1 |
| GOBP_ENDOSOME_ORGA | GOBP_ENDOSOME_ORGANIZATION | 68 | 0.511959 | 2.4711373 | 0 | 0.000681636 | 0.013 | 6859 | tags=59%, list=29%, si | MM_1 |
| GOBP_CRISTAE_FORMAT | GOBP_CRISTAE_FORMATION | 17 | 0.736736 | 2.4681187 | 0 | 0.000665406 | 0.013 | 4404 | tags=65%, list=19%, si | MM_1 |
| GOBP_ENERGY_DERIVATI | GOBP_ENERGY_DERIVATION_BY_OXIDATIO | 321 | 0.412321 | 2.4676538 | 0 | 0.000649932 | 0.013 | 4873 | tags=45%, list=21%, si | MM_1 |
| GOBP_NEGATIVE_REGUL | GOBP_NEGATIVE_REGULATION_OF_TORC1_ | 41 | 0.573548 | 2.4613795 | 0 | 0.000685087 | 0.014 | 6606 | tags=59%, list=28%, si | MM_1 |
| GOBP_MACROAUTOPHAG | GOBP_MACROAUTOPHAGY | 287 | 0.415036 | 2.4610057 | 0 | 0.000669862 | 0.014 | 7443 | tags=55%, list=32%, si | MM_1 |
| NON-ALCOHOLIC_FATTY_ | NON-ALCOHOLIC_FATTY_LIVER_DISEASE_(N | 141 | 0.447815 | 2.4454348 | 0 | 0.000845248 | 0.018 | 3743 | tags=46%, list=16%, si | MM_1 |
| GOBP_TORC1_SIGNALING | GOBP_TORC1_SIGNALING | 100 | 0.485297 | 2.4431784 | 0 | 0.000827264 | 0.018 | 6606 | tags=52%, list=28%, si | MM_1 |
| GOBP_PROTEIN_INSERTI | GOBP_PROTEIN_INSERTION_INTO_ER_MEM | 26 | 0.641235 | 2.436561 | 0 | 0.000854353 | 0.019 | 5132 | tags=73%, list=22%, si | MM_1 |
| GOBP_MACROMOLECULE | GOBP_MACROMOLECULE_METHYLATION | 121 | 0.454519 | 2.4250329 | 0 | 0.000968296 | 0.022 | 5688 | tags=45%, list=24%, si | MM_1 |
| GOBP_RESPIRATORY_CHA | GOBP_RESPIRATORY_CHAIN_COMPLEX_IV_ | 24 | 0.676781 | 2.4242904 | 0 | 0.00094893 | 0.022 | 5940 | tags=75%, list=25%, si | MM_1 |
| GOBP_RNA_SPLICING | GOBP_RNA_SPLICING | 417 | 0.388844 | 2.414587 | 0 | 0.000930323 | 0.022 | 7609 | tags=48%, list=32%, si | MM_1 |
| RNA_POLYMERASE | RNA_POLYMERASE | 27 | 0.625478 | 2.408276 | 0 | 0.000995314 | 0.024 | 7253 | tags=74%, list=31%, si | MM_1 |
| GOBP_TRNA_PROCESSIN | GOBP_TRNA_PROCESSING | 122 | 0.451694 | 2.4051893 | 0 | 0.000976534 | 0.024 | 6752 | tags=52%, list=29%, si | MM_1 |
| GOBP_TRANSLATION_AT_ | GOBP_TRANSLATION_AT_SYNAPSE | 48 | 0.540818 | 2.4013603 | 0 | 0.00095845 | 0.024 | 9392 | tags=81%, list=40%, si | MM_1 |
| GOBP_NUCLEOSIDE_PHO | GOBP_NUCLEOSIDE_PHOSPHATE_BIOSYNTH | 234 | 0.404418 | 2.3866 | 0 | 0.001060656 | 0.027 | 6190 | tags=53%, list=26%, si | MM_1 |
| GOBP_NUCLEOSIDE_TRIP | GOBP_NUCLEOSIDE_TRIPHOSPHATE_METAE | 250 | 0.401451 | 2.385514 | 0 | 0.001041716 | 0.027 | 5198 | tags=42%, list=22%, si | MM_1 |
| GOBP_ORGANELLE_DISAS | GOBP_ORGANELLE_DISASSEMBLY | 155 | 0.435126 | 2.3852687 | 0 | 0.00102344 | 0.027 | 5223 | tags=41%, list=22%, si | MM_1 |
| GOBP_MITOCHONDRIAL_ | GOBP_MITOCHONDRIAL_CYTOCHROME_C_ | 19 | 0.69722 | 2.382862 | 0 | 0.001043292 | 0.028 | 5940 | tags=74%, list=25%, si | MM_1 |
| GOBP_ATP_METABOLIC_ | GOBP_ATP_METABOLIC_PROCESS | 206 | 0.422609 | 2.37177 | 0 | 0.00117387 | 0.032 | 5118 | tags=45%, list=22%, si | MM_1 |
| GOBP_VACUOLE_ORGAN | GOBP_VACUOLE_ORGANIZATION | 223 | 0.404587 | 2.3688672 | 0 | 0.001154306 | 0.032 | 7435 | tags=54%, list=31%, si | MM_1 |
| GOBP_TELOMERE_MAINT | GOBP_TELOMERE_MAINTENANCE_IN_RESP | 32 | 0.580483 | 2.3684375 | 0 | 0.001171173 | 0.033 | 7300 | tags=72%, list=31%, si | MM_1 |
| GOBP_MRNA_PROCESSIN | GOBP_MRNA_PROCESSING | 459 | 0.374101 | 2.3671885 | 0 | 0.001152284 | 0.033 | 7622 | tags=47%, list=32%, si | MM_1 |
| GOBP_METHYLATION | GOBP_METHYLATION | 199 | 0.414561 | 2.3666406 | 0 | 0.001167916 | 0.034 | 5783 | tags=42%, list=24%, si | MM_1 |
| GOBP_PROTEIN_RNA_CO | GOBP_PROTEIN_RNA_COMPLEX_ORGANIZA | 173 | 0.422627 | 2.3643868 | 0 | 0.00118365 | 0.034 | 7893 | tags=54%, list=33%, si | MM_1 |
| GOBP_CYTOCHROME_CO | GOBP_CYTOCHROME_COMPLEX_ASSEMBLY | 32 | 0.580852 | 2.3499253 | 0 | 0.001397408 | 0.041 | 5940 | tags=66%, list=25%, si | MM_1 |
| GOBP_TAIL_ANCHORED_ | GOBP_TAIL_ANCHORED_MEMBRANE_PROT | 15 | 0.741309 | 2.3498511 | 0 | 0.001376235 | 0.041 | 5132 | tags=87%, list=22%, si | MM_1 |
| GOBP_REGULATION_OF_ | GOBP_REGULATION_OF_TELOMERE_MAINT | 98 | 0.459733 | 2.3476765 | 0 | 0.001386659 | 0.042 | 6762 | tags=46%, list=29%, si | MM_1 |
| GOBP_POSITIVE_REGULA | GOBP_POSITIVE_REGULATION_OF_DNA_RE | 77 | 0.485713 | 2.3363795 | 0 | 0.001619797 | 0.049 | 5267 | tags=43%, list=22%, si | MM_1 |
| GOBP_POSITIVE_REGULA | GOBP_POSITIVE_REGULATION_OF_TELOME | 71 | 0.479479 | 2.3194184 | 0 | 0.001719159 | 0.053 | 6762 | tags=49%, list=29%, si | MM_1 |
| GOBP_PROTEIN_MATURA | GOBP_PROTEIN_MATURATION_BY_IRON_SU | 16 | 0.710526 | 2.3117301 | 0 | 0.001972425 | 0.06 | 6798 | tags=94%, list=29%, si | MM_1 |
| GOBP_GLIAL_CELL_MIGR | GOBP_GLIAL_CELL_MIGRATION | 70 | 0.480251 | 2.3107462 | 0 | 0.002005701 | 0.062 | 5073 | tags=44%, list=21%, si | MM_1 |
| MITOPHAGY | MITOPHAGY | 62 | 0.490647 | 2.302088 | 0 | 0.002217608 | 0.069 | 6792 | tags=52%, list=29%, si | MM_1 |
| GOBP_ORGANOPHOSPHA | GOBP_ORGANOPHOSPHATE_BIOSYNTHETIC | 500 | 0.364475 | 2.300549 | 0 | 0.002217322 | 0.07 | 6190 | tags=49%, list=26%, si | MM_1 |
| GOBP_GLYCERALDEHYDE | GOBP_GLYCERALDEHYDE_3_PHOSPHATE_M | 19 | 0.669392 | 2.295379 | 0 | 0.002391959 | 0.076 | 4709 | tags=79%, list=20%, si | MM_1 |
| GOBP_RNA_TEMPLATED_ | GOBP_RNA_TEMPLATED_DNA_BIOSYNTHET | 63 | 0.487782 | 2.2944212 | 0 | 0.002360066 | 0.076 | 7653 | tags=60%, list=32%, si | MM_1 |
| GOBP_REGULATION_OF_ | GOBP_REGULATION_OF_HEPATOCYTE_PRO | 25 | 0.61013 | 2.2941296 | 0 | 0.002329012 | 0.076 | 1258 | tags=28%, list=5%, sig | MM_1 |
| GOBP_ESTABLISHMENT_O | GOBP_ESTABLISHMENT_OF_PROTEIN_LOCA | 25 | 0.610718 | 2.2923207 | 0.0028735632 | 0.002383155 | 0.079 | 5580 | tags=60%, list=24%, si | MM_1 |
| GOBP_GOLGI_VESICLE_TI | GOBP_GOLGI_VESICLE_TRANSPORT | 276 | 0.392402 | 2.2912786 | 0 | 0.002380549 | 0.08 | 6378 | tags=44%, list=27%, si | MM_1 |
| GOBP_PORPHYRIN_CONT | GOBP_PORPHYRIN_CONTAINING_COMPOU | 40 | 0.547341 | 2.286651 | 0 | 0.002567073 | 0.088 | 4780 | tags=58%, list=20%, si | MM_1 |
| GOBP_RNA_SPLICING_VI | GOBP_RNA_SPLICING_VIA_TRANSESTERIFIC | 274 | 0.382447 | 2.2816238 | 0 | 0.002773146 | 0.097 | 7609 | tags=49%, list=32%, si | MM_1 |
| GOBP_PROTEIN_FOLDING | GOBP_PROTEIN_FOLDING | 162 | 0.412138 | 2.2778823 | 0 | 0.002818028 | 0.1 | 5078 | tags=41%, list=22%, si | MM_1 |
| GOBP_T_CELL_MIGRATIO | GOBP_T_CELL_MIGRATION | 64 | 0.492203 | 2.2758152 | 0 | 0.00283575 | 0.101 | 4139 | tags=39%, list=18%, si | MM_1 |
| GOBP_LYSOSOMAL_LUM | GOBP_LYSOSOMAL_LUMEN_ACIDIFICATION | 15 | 0.701445 | 2.2724462 | 0 | 0.002904699 | 0.105 | 7059 | tags=100%, list=30%, | MM_1 |
| GOBP_TRNA_MODIFICAT | GOBP_TRNA_MODIFICATION | 84 | 0.453954 | 2.26969 | 0 | 0.00297248 | 0.108 | 5515 | tags=44%, list=23%, si | MM_1 |
| GOBP_LYTIC_VACUOLE_O | GOBP_LYTIC_VACUOLE_ORGANIZATION | 95 | 0.439946 | 2.2647963 | 0 | 0.003162578 | 0.115 | 6544 | tags=57%, list=28%, si | MM_1 |
| GOBP_POSITIVE_REGULA | GOBP_POSITIVE_REGULATION_OF_TORC1_S | 52 | 0.510717 | 2.2606606 | 0 | 0.003175717 | 0.116 | 6321 | tags=50%, list=27%, si | MM_1 |
| GOBP_LATE_ENDOSOME_ | GOBP_LATE_ENDOSOME_TO_LYSOSOME_TI | 19 | 0.640026 | 2.2599084 | 0.0028089888 | 0.003213987 | 0.119 | 6569 | tags=79%, list=28%, si | MM_1 |
| GOBP_REGULATION_OF_ | GOBP_REGULATION_OF_MACROAUTOPHAG | 129 | 0.421477 | 2.2594323 | 0 | 0.003177464 | 0.119 | 7508 | tags=58%, list=32%, si | MM_1 |
| GOBP_REGULATION_OF_ | GOBP_REGULATION_OF_SUPEROXIDE_ANIC | 28 | 0.582894 | 2.2473488 | 0 | 0.003549509 | 0.134 | 4495 | tags=43%, list=19%, si | MM_1 |
| VALINE,_LEUCINE_AND_I | VALINE,_LEUCINE_AND_ISOLEUCINE_DEGRA | 49 | 0.500993 | 2.2467935 | 0 | 0.003582534 | 0.137 | 6152 | tags=65%, list=26%, si | MM_1 |
| GOBP_REGULATION_OF_ | GOBP_REGULATION_OF_SUPEROXIDE_META | 34 | 0.562045 | 2.2459228 | 0 | 0.003543166 | 0.137 | 4495 | tags=38%, list=19%, si | MM_1 |
| GOBP_TRANSCRIPTION_B | GOBP_TRANSCRIPTION_BY_RNA_POLYMERA | 60 | 0.481188 | 2.2349367 | 0 | 0.003922597 | 0.152 | 7594 | tags=60%, list=32%, si | MM_1 |

| | | | | | | | | | | |
|---|---|---|---|---|---|---|---|---|---|---|
| GOBP_TRANSLATIONAL_I | GOBP_TRANSLATIONAL_INITIATION | 112 | 0.421614 | 2.2140012 | 0 | 0.004847853 | 0.186 | 7961 | tags=50%, list=34%, si | MM_1 |
| GOBP_REGULATION_OF_ | GOBP_REGULATION_OF_T_CELL_MIGRATIO | 49 | 0.492729 | 2.2131963 | 0 | 0.004819667 | 0.187 | 4139 | tags=39%, list=18%, si | MM_1 |
| GOBP_NEGATIVE_REGUL | GOBP_NEGATIVE_REGULATION_OF_TRANSL | 20 | 0.631858 | 2.2063725 | 0 | 0.005152016 | 0.197 | 6474 | tags=55%, list=27%, si | MM_1 |
| GOBP_PROTEIN_K11_LIN | GOBP_PROTEIN_K11_LINKED_UBIQUITINAT | 29 | 0.573371 | 2.205803 | 0.003003003 | 0.005143995 | 0.199 | 7484 | tags=66%, list=32%, si | MM_1 |
| GOBP_TRNA_METHYLATI | GOBP_TRNA_METHYLATION | 39 | 0.527577 | 2.205111 | 0 | 0.005113368 | 0.2 | 5145 | tags=51%, list=22%, si | MM_1 |
| GOBP_HEME_BIOSYNTHE | GOBP_HEME_BIOSYNTHETIC_PROCESS | 24 | 0.607235 | 2.2049127 | 0 | 0.005061192 | 0.2 | 4281 | tags=58%, list=18%, si | MM_1 |
| GOBP_TRNA_THREONYLC | GOBP_TRNA_THREONYLCARBAMOYLADENC | 15 | 0.668541 | 2.1986477 | 0 | 0.005312556 | 0.21 | 6107 | tags=67%, list=26%, si | MM_1 |
| GOBP_POSITIVE_REGULA | GOBP_POSITIVE_REGULATION_OF_PROTEAS | 102 | 0.420193 | 2.193726 | 0 | 0.005583008 | 0.22 | 4955 | tags=37%, list=21%, si | MM_1 |
| GOBP_REGULATION_OF_ | GOBP_REGULATION_OF_RESPONSE_TO_EN | 80 | 0.445024 | 2.1918905 | 0 | 0.005570003 | 0.221 | 7328 | tags=56%, list=31%, si | MM_1 |
| GOBP_VIRAL_GENE_EXPR | GOBP_VIRAL_GENE_EXPRESSION | 75 | 0.450174 | 2.1904068 | 0 | 0.005556496 | 0.222 | 7511 | tags=53%, list=32%, si | MM_1 |
| GOBP_MATURATION_OF_ | GOBP_MATURATION_OF_LSU_RRNA | 25 | 0.577354 | 2.1845703 | 0 | 0.005876584 | 0.237 | 6758 | tags=60%, list=29%, si | MM_1 |
| GOBP_PROTEIN_HYDROX | GOBP_PROTEIN_HYDROXYLATION | 25 | 0.584271 | 2.183627 | 0 | 0.006003469 | 0.246 | 3595 | tags=44%, list=15%, si | MM_1 |
| GOBP_MEMBRANE_BIOG | GOBP_MEMBRANE_BIOGENESIS | 63 | 0.462716 | 2.17904 | 0 | 0.006193824 | 0.256 | 3452 | tags=32%, list=15%, si | MM_1 |
| GOBP_2_OXOGLUTARATE | GOBP_2_OXOGLUTARATE_METABOLIC_PRO | 18 | 0.633716 | 2.1758955 | 0.0026595744 | 0.006377013 | 0.264 | 2545 | tags=44%, list=11%, si | MM_1 |
| GOBP_POSITIVE_REGULA | GOBP_POSITIVE_REGULATION_OF_LYMPHO | 42 | 0.507646 | 2.1733692 | 0 | 0.006398505 | 0.268 | 4139 | tags=36%, list=18%, si | MM_1 |
| GOBP_AUTOPHAGY_OF_I | GOBP_AUTOPHAGY_OF_MITOCHONDRION | 89 | 0.43337 | 2.165009 | 0 | 0.006992688 | 0.293 | 5055 | tags=43%, list=21%, si | MM_1 |
| GOBP_RIBOSOMAL_SMAI | GOBP_RIBOSOMAL_SMALL_SUBUNIT_BIOGI | 101 | 0.418413 | 2.1647103 | 0 | 0.006947966 | 0.294 | 6560 | tags=44%, list=28%, si | MM_1 |
| AMINO_SUGAR_AND_NU | AMINO_SUGAR_AND_NUCLEOTIDE_SUGAR_ | 46 | 0.500512 | 2.160738 | 0 | 0.007080009 | 0.3 | 5342 | tags=52%, list=23%, si | MM_1 |
| GOBP_RESPONSE_TO_AM | GOBP_RESPONSE_TO_AMINO_ACID_STARV | 54 | 0.475049 | 2.1597145 | 0 | 0.007035278 | 0.3 | 5694 | tags=41%, list=24%, si | MM_1 |
| GOBP_PROTEIN_IMPORT_ | GOBP_PROTEIN_IMPORT_INTO_MITOCHON | 19 | 0.630838 | 2.1595762 | 0.003012048 | 0.006991659 | 0.301 | 5095 | tags=58%, list=22%, si | MM_1 |
| GOBP_TETRAPYRROLE_BI | GOBP_TETRAPYRROLE_BIOSYNTHETIC_PROC | 29 | 0.549574 | 2.1564186 | 0 | 0.007176394 | 0.31 | 4780 | tags=59%, list=20%, si | MM_1 |
| GOBP_BRANCHED_CHAIN | GOBP_BRANCHED_CHAIN_AMINO_ACID_CA | 21 | 0.615968 | 2.156238 | 0 | 0.007132303 | 0.31 | 6141 | tags=76%, list=26%, si | MM_1 |
| GOBP_TETRAPYRROLE_M | GOBP_TETRAPYRROLE_METABOLIC_PROCES | 49 | 0.491374 | 2.1555817 | 0 | 0.007106949 | 0.311 | 5453 | tags=55%, list=23%, si | MM_1 |
| GOBP_POSITIVE_REGULA | GOBP_POSITIVE_REGULATION_OF_MACROA | 81 | 0.426571 | 2.1552794 | 0 | 0.007045682 | 0.311 | 6051 | tags=43%, list=26%, si | MM_1 |
| GOBP_POSITIVE_REGULA | GOBP_POSITIVE_REGULATION_OF_TRANSLA | 125 | 0.404235 | 2.1536498 | 0 | 0.007075707 | 0.314 | 5291 | tags=34%, list=22%, si | MM_1 |
| GOBP_PROTEIN_QUALITY | GOBP_PROTEIN_QUALITY_CONTROL_FOR_M | 17 | 0.651714 | 2.1521335 | 0 | 0.007051939 | 0.314 | 2791 | tags=35%, list=12%, si | MM_1 |
| GOBP_PROTEIN_MATURA | GOBP_PROTEIN_MATURATION | 475 | 0.335847 | 2.1474113 | 0 | 0.007280648 | 0.326 | 5078 | tags=36%, list=22%, si | MM_1 |
| GOBP_TRANSLATIONAL_E | GOBP_TRANSLATIONAL_ELONGATION | 58 | 0.470502 | 2.136926 | 0 | 0.008248854 | 0.368 | 6840 | tags=50%, list=29%, si | MM_1 |
| GOBP_RETROGRADE_TRA | GOBP_RETROGRADE_TRANSPORT_ENDOSO | 87 | 0.422133 | 2.1369183 | 0 | 0.008180681 | 0.368 | 7735 | tags=61%, list=33%, si | MM_1 |
| GOBP_REGULATION_OF_ | GOBP_REGULATION_OF_TRANSLATION | 329 | 0.345652 | 2.1351767 | 0 | 0.008166345 | 0.371 | 5291 | tags=31%, list=22%, si | MM_1 |
| GOBP_RIBOSE_PHOSPHA | GOBP_RIBOSE_PHOSPHATE_METABOLIC_PR | 419 | 0.342281 | 2.1328826 | 0 | 0.008258493 | 0.376 | 5032 | tags=38%, list=21%, si | MM_1 |
| GOBP_VIRAL_TRANSLATI | GOBP_VIRAL_TRANSLATION | 18 | 0.631977 | 2.132732 | 0 | 0.008208334 | 0.377 | 7511 | tags=78%, list=32%, si | MM_1 |
| GOBP_RNA_METHYLATIO | GOBP_RNA_METHYLATION | 65 | 0.44255 | 2.1270814 | 0 | 0.008486087 | 0.393 | 6641 | tags=51%, list=28%, si | MM_1 |
| GOBP_REGULATION_OF_ | GOBP_REGULATION_OF_RESPONSE_TO_OX | 39 | 0.497935 | 2.126047 | 0 | 0.008452534 | 0.395 | 6194 | tags=44%, list=26%, si | MM_1 |
| GOBP_BRANCHED_CHAIN | GOBP_BRANCHED_CHAIN_AMINO_ACID_M | 24 | 0.577085 | 2.1231945 | 0 | 0.008553742 | 0.398 | 5573 | tags=71%, list=24%, si | MM_1 |
| GOBP_PROTEIN_INSERTI | GOBP_PROTEIN_INSERTION_INTO_MEMBRA | 39 | 0.512354 | 2.120191 | 0 | 0.008657888 | 0.404 | 5580 | tags=67%, list=24%, si | MM_1 |
| GOBP_POSITIVE_REGULA | GOBP_POSITIVE_REGULATION_OF_VIRAL_LI | 28 | 0.54462 | 2.1180556 | 0 | 0.008888962 | 0.416 | 2201 | tags=29%, list=9%, sig | MM_1 |
| GOBP_POSITIVE_REGULA | GOBP_POSITIVE_REGULATION_OF_TELOME | 38 | 0.503286 | 2.1177487 | 0 | 0.00885349 | 0.416 | 914 | tags=16%, list=4%, sig | MM_1 |
| GOBP_PROTEIN_TRANSM | GOBP_PROTEIN_TRANSMEMBRANE_IMPOR | 36 | 0.526794 | 2.1119204 | 0 | 0.009261726 | 0.435 | 5095 | tags=47%, list=22%, si | MM_1 |
| GOBP_NUCLEAR_ENVELO | GOBP_NUCLEAR_ENVELOPE_ORGANIZATIO | 52 | 0.46822 | 2.1112928 | 0 | 0.009256717 | 0.438 | 6958 | tags=46%, list=29%, si | MM_1 |
| GOBP_MITOCHONDRIAL_ | GOBP_MITOCHONDRIAL_MEMBRANE_ORGA | 107 | 0.407417 | 2.1085122 | 0 | 0.009492228 | 0.453 | 5741 | tags=50%, list=24%, si | MM_1 |
| GOBP_REGULATION_OF_ | GOBP_REGULATION_OF_INTEGRIN_MEDIAT | 19 | 0.595591 | 2.1039975 | 0 | 0.009739148 | 0.465 | 959 | tags=32%, list=4%, sig | MM_1 |
| GOBP_LATE_ENDOSOME_ | GOBP_LATE_ENDOSOME_TO_VACUOLE_TRA | 25 | 0.556491 | 2.1038566 | 0 | 0.009682464 | 0.466 | 6569 | tags=60%, list=28%, si | MM_1 |
| GOBP_REGULATION_OF_ | GOBP_REGULATION_OF_LYMPHOCYTE_MIG | 64 | 0.442608 | 2.0991924 | 0 | 0.009909304 | 0.476 | 4713 | tags=38%, list=20%, si | MM_1 |
| GOBP_REGULATION_OF_ | GOBP_REGULATION_OF_TELOMERE_CAPPIN | 26 | 0.55287 | 2.0988815 | 0.0028901733 | 0.009883588 | 0.478 | 4247 | tags=35%, list=18%, si | MM_1 |
| GOBP_NEGATIVE_REGUL | GOBP_NEGATIVE_REGULATION_OF_CYCLIN_ | 25 | 0.568277 | 2.0980818 | 0 | 0.009904693 | 0.481 | 3824 | tags=40%, list=16%, si | MM_1 |
| GOBP_NEGATIVE_REGUL | GOBP_NEGATIVE_REGULATION_OF_RESPON | 40 | 0.498262 | 2.0979757 | 0 | 0.009848893 | 0.482 | 6969 | tags=60%, list=30%, si | MM_1 |
| GOBP_SNRNA_PROCESSI | GOBP_SNRNA_PROCESSING | 24 | 0.55811 | 2.0942204 | 0 | 0.010085545 | 0.492 | 3738 | tags=33%, list=16%, si | MM_1 |
| GOBP_AUTOPHAGOSOMI | GOBP_AUTOPHAGOSOME_ORGANIZATION | 128 | 0.389064 | 2.089753 | 0 | 0.010333105 | 0.505 | 8037 | tags=57%, list=34%, si | MM_1 |
| GOBP_POSITIVE_REGULA | GOBP_POSITIVE_REGULATION_OF_TOR_SIG | 71 | 0.436646 | 2.0891612 | 0 | 0.010349674 | 0.509 | 6600 | tags=48%, list=28%, si | MM_1 |
| GOBP_PIGMENT_GRANU | GOBP_PIGMENT_GRANULE_ORGANIZATION | 39 | 0.491422 | 2.0839832 | 0 | 0.010813976 | 0.528 | 5913 | tags=46%, list=25%, si | MM_1 |
| GOBP_REGULATION_OF_ | GOBP_REGULATION_OF_VESICLE_SIZE | 21 | 0.568194 | 2.083467 | 0.0030395137 | 0.010813453 | 0.53 | 6105 | tags=62%, list=26%, si | MM_1 |
| GLYCOSYLPHOSPHATIDYL | GLYCOSYLPHOSPHATIDYLINOSITOL_(GPI)-AN | 24 | 0.56637 | 2.0832896 | 0.008928572 | 0.010783315 | 0.532 | 6120 | tags=63%, list=26%, si | MM_1 |
| PROTEIN_PROCESSING_IN | PROTEIN_PROCESSING_IN_ENDOPLASMIC_F | 158 | 0.377256 | 2.082391 | 0 | 0.010797073 | 0.535 | 6628 | tags=42%, list=28%, si | MM_1 |
| GOBP_GOLGI_TO_VACUC | GOBP_GOLGI_TO_VACUOLE_TRANSPORT | 21 | 0.585483 | 2.0798817 | 0 | 0.010882626 | 0.54 | 6378 | tags=57%, list=27%, si | MM_1 |
| GOBP_PROTEIN_METHYL | GOBP_PROTEIN_METHYLATION | 57 | 0.462113 | 2.0786233 | 0 | 0.010939832 | 0.545 | 5671 | tags=46%, list=24%, si | MM_1 |
| GOBP_GPI_ANCHOR_MET | GOBP_GPI_ANCHOR_METABOLIC_PROCESS | 33 | 0.507661 | 2.076871 | 0 | 0.010967489 | 0.548 | 4708 | tags=48%, list=20%, si | MM_1 |
| GOBP_RRNA_TRANSCRIPT | GOBP_RRNA_TRANSCRIPTION | 33 | 0.529101 | 2.0768344 | 0 | 0.010894373 | 0.548 | 7253 | tags=61%, list=31%, si | MM_1 |
| GOBP_REGULATION_OF_ | GOBP_REGULATION_OF_PROTEASOMAL_PR | 179 | 0.370819 | 2.0741084 | 0 | 0.011193385 | 0.56 | 5293 | tags=37%, list=22%, si | MM_1 |
| GOBP_RIBOSOME_ASSEM | GOBP_RIBOSOME_ASSEMBLY | 60 | 0.449473 | 2.0702467 | 0 | 0.011330356 | 0.565 | 8650 | tags=73%, list=37%, si | MM_1 |
| GOBP_REGULATION_OF_ | GOBP_REGULATION_OF_TELOMERE_MAINT | 61 | 0.451025 | 2.0700343 | 0 | 0.011256301 | 0.565 | 5917 | tags=39%, list=25%, si | MM_1 |
| GOBP_ACETYL_COA_BIOS | GOBP_ACETYL_COA_BIOSYNTHETIC_PROCES | 19 | 0.606754 | 2.06777 | 0.002739726 | 0.011364725 | 0.574 | 5476 | tags=63%, list=23%, si | MM_1 |
| GOBP_PEPTIDYL_LYSINE_ | GOBP_PEPTIDYL_LYSINE_MODIFICATION | 128 | 0.384391 | 2.067616 | 0 | 0.011318632 | 0.576 | 5358 | tags=37%, list=23%, si | MM_1 |
| GOBP_PYRUVATE_METAE | GOBP_PYRUVATE_METABOLIC_PROCESS | 127 | 0.384784 | 2.0654814 | 0 | 0.011518966 | 0.581 | 5198 | tags=40%, list=22%, si | MM_1 |
| GOBP_REGULATION_OF_ | GOBP_REGULATION_OF_MITOTIC_SPINDLE_ | 18 | 0.618838 | 2.064838 | 0.0027027028 | 0.011513397 | 0.583 | 3049 | tags=44%, list=13%, si | MM_1 |
| GOBP_GLYCOLYTIC_PROC | GOBP_GLYCOLYTIC_PROCESS_THROUGH_FR | 25 | 0.544561 | 2.0614216 | 0.0028735632 | 0.011709522 | 0.595 | 5076 | tags=56%, list=21%, si | MM_1 |
| GOBP_ERAD_PATHWAY | GOBP_ERAD_PATHWAY | 108 | 0.389645 | 2.0613356 | 0 | 0.011635876 | 0.595 | 6924 | tags=48%, list=29%, si | MM_1 |
| GOBP_PROTEASOME_ME | GOBP_PROTEASOME_MEDIATED_UBIQUITIN | 368 | 0.330904 | 2.061197 | 0 | 0.011563153 | 0.595 | 7487 | tags=49%, list=32%, si | MM_1 |
| GOBP_BASEMENT_MEME | GOBP_BASEMENT_MEMBRANE_ORGANIZAT | 40 | 0.486255 | 2.0603464 | 0 | 0.01159936 | 0.598 | 5262 | tags=50%, list=22%, si | MM_1 |
| GOBP_PROTEIN_TARGETI | GOBP_PROTEIN_TARGETING | 285 | 0.341614 | 2.05972 | 0 | 0.011594319 | 0.6 | 6547 | tags=47%, list=28%, si | MM_1 |
| GOBP_POSITIVE_REGULA | GOBP_POSITIVE_REGULATION_OF_DOUBLE_ | 43 | 0.478856 | 2.057243 | 0 | 0.011708746 | 0.606 | 4303 | tags=35%, list=18%, si | MM_1 |
| GOBP_TELOMERE_MAINT | GOBP_TELOMERE_MAINTENANCE_VIA_TEL( | 76 | 0.420255 | 2.0568252 | 0 | 0.01171401 | 0.609 | 7523 | tags=55%, list=32%, si | MM_1 |
| GOBP_POSITIVE_REGULA | GOBP_POSITIVE_REGULATION_OF_PROTEAS | 72 | 0.423994 | 2.054451 | 0 | 0.01190136 | 0.617 | 4849 | tags=38%, list=21%, si | MM_1 |
| GOBP_REGULATION_OF_ | GOBP_REGULATION_OF_T_CELL_APOPTOTI | 48 | 0.4696 | 2.054039 | 0 | 0.011921071 | 0.621 | 3053 | tags=33%, list=13%, si | MM_1 |
| GOBP_TRANSCRIPTION_P | GOBP_TRANSCRIPTION_PREINITIATION_CON | 63 | 0.439407 | 2.0505316 | 0 | 0.012246937 | 0.63 | 7167 | tags=46%, list=30%, si | MM_1 |
| GOBP_NUCLEOTIDE_SUG | GOBP_NUCLEOTIDE_SUGAR_METABOLIC_PI | 34 | 0.500584 | 2.0503278 | 0 | 0.012224596 | 0.632 | 6354 | tags=56%, list=27%, si | MM_1 |
| GOBP_ENDOPLASMIC_RE | GOBP_ENDOPLASMIC_RETICULUM_TO_GOL | 120 | 0.383946 | 2.0479352 | 0 | 0.012518456 | 0.643 | 6356 | tags=46%, list=27%, si | MM_1 |
| GOBP_VIRAL_PROCESS | GOBP_VIRAL_PROCESS | 303 | 0.339581 | 2.0466974 | 0 | 0.012595314 | 0.644 | 5972 | tags=39%, list=25%, si | MM_1 |
| GOBP_POSITIVE_REGULA | GOBP_POSITIVE_REGULATION_OF_CELL_MA | 61 | 0.448637 | 2.042174 | 0 | 0.013070807 | 0.656 | 3730 | tags=33%, list=16%, si | MM_1 |
| GOBP_MATURATION_OF_ | GOBP_MATURATION_OF_LSU_RRNA_FROM | 17 | 0.610681 | 2.0417414 | 0.003003003 | 0.013082692 | 0.658 | 5326 | tags=53%, list=23%, si | MM_1 |
| GOBP_NADH_METABOLI | GOBP_NADH_METABOLIC_PROCESS | 39 | 0.488837 | 2.0411606 | 0 | 0.013068767 | 0.661 | 5076 | tags=51%, list=21%, si | MM_1 |
| GOBP_REGULATION_OF_ | GOBP_REGULATION_OF_EARLY_ENDOSOME | 18 | 0.597781 | 2.039203 | 0 | 0.013154022 | 0.664 | 6070 | tags=44%, list=26%, si | MM_1 |

| | | | | | | | | | | |
|---|---|---|---|---|---|---|---|---|---|---|
| GOBP_ESTABLISHMENT_( | GOBP_ESTABLISHMENT_OF_LYMPHOCYTE_I | 15 | 0.638781 | 2.0366123 | 0.0026809652 | 0.013419457 | 0.672 | 1702 | tags=27%, list=7%, sig | MM_1 |
| GOBP_VIRAL_RELEASE_FI | GOBP_VIRAL_RELEASE_FROM_HOST_CELL | 24 | 0.544421 | 2.0350509 | 0 | 0.013560928 | 0.68 | 7471 | tags=71%, list=32%, si | MM_1 |
| GOBP_COENZYME_A_ME | GOBP_COENZYME_A_METABOLIC_PROCESS | 19 | 0.585242 | 2.0340261 | 0 | 0.013557329 | 0.681 | 2454 | tags=37%, list=10%, si | MM_1 |
| GOBP_LYMPHOCYTE_MIG | GOBP_LYMPHOCYTE_MIGRATION | 92 | 0.404176 | 2.0333636 | 0 | 0.013553328 | 0.683 | 4754 | tags=36%, list=20%, si | MM_1 |
| GOBP_REGULATION_OF_ | GOBP_REGULATION_OF_LYSOSOMAL_LUME | 23 | 0.558153 | 2.0320837 | 0 | 0.013680583 | 0.686 | 7059 | tags=83%, list=30%, si | MM_1 |
| GOBP_POSITIVE_REGULA | GOBP_POSITIVE_REGULATION_OF_PROTEO | 120 | 0.382869 | 2.0301893 | 0 | 0.013866349 | 0.692 | 4955 | tags=35%, list=21%, si | MM_1 |
| GOBP_TELOMERE_ORGAI | GOBP_TELOMERE_ORGANIZATION | 150 | 0.37349 | 2.0288112 | 0 | 0.013860563 | 0.694 | 6798 | tags=42%, list=29%, si | MM_1 |
| GOBP_RETROGRADE_VES | GOBP_RETROGRADE_VESICLE_MEDIATED_T | 45 | 0.475679 | 2.026056 | 0 | 0.01403059 | 0.699 | 8275 | tags=69%, list=35%, si | MM_1 |
| GOBP_REGULATION_OF_ | GOBP_REGULATION_OF_TRANSLATIONAL_I | 68 | 0.436753 | 2.025369 | 0 | 0.014036282 | 0.7 | 6474 | tags=35%, list=27%, si | MM_1 |
| GOBP_MEMBRANE_INVA | GOBP_MEMBRANE_INVAGINATION | 70 | 0.434857 | 2.025305 | 0 | 0.013959998 | 0.7 | 3673 | tags=31%, list=16%, si | MM_1 |
| FERROPTOSIS | FERROPTOSIS | 37 | 0.485986 | 2.0246944 | 0 | 0.013953264 | 0.702 | 5104 | tags=51%, list=22%, si | MM_1 |
| GOBP_UBIQUITIN_DEPEN | GOBP_UBIQUITIN_DEPENDENT_PROTEIN_C | 29 | 0.51227 | 2.0200295 | 0 | 0.014363398 | 0.713 | 6676 | tags=55%, list=28%, si | MM_1 |
| GOBP_PROTEIN_TRANSM | GOBP_PROTEIN_TRANSMEMBRANE_TRANS | 67 | 0.425882 | 2.0186994 | 0 | 0.0144236 | 0.718 | 5111 | tags=42%, list=22%, si | MM_1 |
| GOBP_REGULATION_OF_ | GOBP_REGULATION_OF_TRANSCRIPTION_B | 41 | 0.479388 | 2.0107808 | 0.003030303 | 0.015419823 | 0.746 | 7594 | tags=54%, list=32%, si | MM_1 |
| GOBP_REGULATION_OF_ | GOBP_REGULATION_OF_PROTEOLYSIS_INV | 215 | 0.350782 | 2.0077424 | 0 | 0.015632011 | 0.75 | 5293 | tags=34%, list=22%, si | MM_1 |
| GOBP_REGULATION_OF_ | GOBP_REGULATION_OF_TELOMERE_MAINT | 20 | 0.583067 | 2.0071692 | 0 | 0.01566077 | 0.752 | 6762 | tags=70%, list=29%, si | MM_1 |
| GOBP_CYTOPLASMIC_TR | GOBP_CYTOPLASMIC_TRANSLATIONAL_INIT | 33 | 0.504312 | 2.005578 | 0.0057636886 | 0.015790371 | 0.755 | 7895 | tags=61%, list=33%, si | MM_1 |
| GOBP_VASCULAR_ASSOC | GOBP_VASCULAR_ASSOCIATED_SMOOTH_M | 80 | 0.412738 | 2.0050263 | 0 | 0.01576381 | 0.756 | 5276 | tags=44%, list=22%, si | MM_1 |
| GOBP_IMPORT_INTO_TH | GOBP_IMPORT_INTO_THE_MITOCHONDRIO | 49 | 0.452436 | 2.0046635 | 0 | 0.015737833 | 0.757 | 5806 | tags=41%, list=25%, si | MM_1 |
| GOBP_REGULATION_OF_ | GOBP_REGULATION_OF_MACROPHAGE_MI | 47 | 0.453733 | 2.0032942 | 0 | 0.015901294 | 0.764 | 3779 | tags=34%, list=16%, si | MM_1 |
| GOBP_VESICLE_MEDIATE | GOBP_VESICLE_MEDIATED_TRANSPORT_BE | 46 | 0.460207 | 1.9996996 | 0 | 0.016320035 | 0.774 | 6070 | tags=33%, list=26%, si | MM_1 |
| GLYCOSAMINOGLYCAN_I | GLYCOSAMINOGLYCAN_DEGRADATION | 17 | 0.607107 | 1.9984572 | 0 | 0.016401518 | 0.779 | 6318 | tags=76%, list=27%, si | MM_1 |
| GOBP_RESPONSE_TO_MI | GOBP_RESPONSE_TO_MISFOLDED_PROTEIN | 18 | 0.587714 | 1.9969461 | 0.0051150895 | 0.0165025 | 0.78 | 5877 | tags=61%, list=25%, si | MM_1 |
| NUCLEOTIDE_EXCISION_ | NUCLEOTIDE_EXCISION_REPAIR | 42 | 0.475526 | 1.9966632 | 0 | 0.016473819 | 0.782 | 8320 | tags=69%, list=35%, si | MM_1 |
| GOBP_MITOCHONDRIAL_ | GOBP_MITOCHONDRIAL_TRANSPORT | 174 | 0.354293 | 1.9958384 | 0 | 0.016498344 | 0.786 | 5806 | tags=43%, list=25%, si | MM_1 |
| GOBP_REGULATION_OF_ | GOBP_REGULATION_OF_LYMPHOCYTE_APO | 71 | 0.414756 | 1.9928206 | 0 | 0.016865758 | 0.793 | 3053 | tags=30%, list=13%, si | MM_1 |
| GOBP_REGULATION_OF_ | GOBP_REGULATION_OF_MONONUCLEAR_C | 136 | 0.359798 | 1.9892166 | 0 | 0.017270869 | 0.8 | 4980 | tags=37%, list=21%, si | MM_1 |
| GOBP_NEGATIVE_REGUL | GOBP_NEGATIVE_REGULATION_OF_CILIUM | 21 | 0.56141 | 1.98864 | 0 | 0.01722706 | 0.801 | 5638 | tags=52%, list=24%, si | MM_1 |
| GOBP_TRNA_5_END_PRO | GOBP_TRNA_5_END_PROCESSING | 15 | 0.613999 | 1.9874188 | 0.0028089888 | 0.017321782 | 0.803 | 2443 | tags=40%, list=10%, si | MM_1 |
| RETROGRADE_ENDOCAN | RETROGRADE_ENDOCANNABINOID_SIGNAL | 133 | 0.369072 | 1.9871626 | 0 | 0.017247139 | 0.803 | 3624 | tags=35%, list=15%, si | MM_1 |
| GOBP_MITOCHONDRIAL_ | GOBP_MITOCHONDRIAL_DEPOLARIZATION | 29 | 0.51685 | 1.9871581 | 0.003012048 | 0.017163007 | 0.803 | 2034 | tags=21%, list=9%, sig | MM_1 |
| GOBP_ARP2_3_COMPLEX | GOBP_ARP2_3_COMPLEX_MEDIATED_ACTIN | 41 | 0.472526 | 1.986906 | 0.0030864198 | 0.017100902 | 0.803 | 8014 | tags=71%, list=34%, si | MM_1 |
| GOBP_VACUOLAR_TRANS | GOBP_VACUOLAR_TRANSPORT | 161 | 0.358626 | 1.9845262 | 0 | 0.017371982 | 0.808 | 7570 | tags=55%, list=32%, si | MM_1 |
| GOBP_POSITIVE_REGULA | GOBP_POSITIVE_REGULATION_OF_UBIQUIT | 93 | 0.388183 | 1.9793113 | 0 | 0.018018415 | 0.818 | 4849 | tags=35%, list=21%, si | MM_1 |
| GOBP_SYNAPTIC_VESICLE | GOBP_SYNAPTIC_VESICLE_MATURATION | 27 | 0.505163 | 1.9791304 | 0 | 0.017942552 | 0.818 | 3777 | tags=44%, list=16%, si | MM_1 |
| GOBP_EXTRACELLULAR_V | GOBP_EXTRACELLULAR_VESICLE_BIOGENES | 24 | 0.554493 | 1.978137 | 0 | 0.017960116 | 0.82 | 3485 | tags=29%, list=15%, si | MM_1 |
| GOBP_REGULATION_OF_ | GOBP_REGULATION_OF_POSITIVE_CHEMOT | 27 | 0.508656 | 1.9749898 | 0 | 0.018412951 | 0.829 | 3743 | tags=44%, list=16%, si | MM_1 |
| GOBP_ALDEHYDE_CATAB | GOBP_ALDEHYDE_CATABOLIC_PROCESS | 15 | 0.627247 | 1.9740402 | 0.007894737 | 0.018437047 | 0.831 | 5480 | tags=80%, list=23%, si | MM_1 |
| GOBP_MELANOSOME_AS | GOBP_MELANOSOME_ASSEMBLY | 19 | 0.569825 | 1.9730809 | 0.002770083 | 0.018471612 | 0.832 | 5148 | tags=53%, list=22%, si | MM_1 |
| GOBP_LYMPHOCYTE_APO | GOBP_LYMPHOCYTE_APOPTOTIC_PROCESS | 94 | 0.384517 | 1.9721371 | 0 | 0.018554851 | 0.835 | 3053 | tags=28%, list=13%, si | MM_1 |
| GOBP_SNRNA_3_END_PF | GOBP_SNRNA_3_END_PROCESSING | 20 | 0.565693 | 1.9698936 | 0 | 0.018797264 | 0.839 | 3738 | tags=35%, list=16%, si | MM_1 |
| PROTEIN_EXPORT | PROTEIN_EXPORT | 26 | 0.527323 | 1.9685938 | 0 | 0.018969055 | 0.842 | 6452 | tags=62%, list=27%, si | MM_1 |
| GOBP_T_CELL_APOPTOTI | GOBP_T_CELL_APOPTOTIC_PROCESS | 67 | 0.422701 | 1.9672389 | 0 | 0.019148618 | 0.845 | 4892 | tags=40%, list=21%, si | MM_1 |
| GOBP_PEROXISOME_ORG | GOBP_PEROXISOME_ORGANIZATION | 38 | 0.468974 | 1.9629668 | 0 | 0.019600637 | 0.852 | 6508 | tags=50%, list=28%, si | MM_1 |
| GOBP_ENDOSOMAL_TRA | GOBP_ENDOSOMAL_TRANSPORT | 272 | 0.326816 | 1.9627221 | 0 | 0.019560067 | 0.852 | 6909 | tags=44%, list=29%, si | MM_1 |
| GOBP_GOLGI_ORGANIZA | GOBP_GOLGI_ORGANIZATION | 135 | 0.366593 | 1.9624071 | 0 | 0.01949028 | 0.852 | 7735 | tags=51%, list=33%, si | MM_1 |
| GOBP_REGULATION_OF_ | GOBP_REGULATION_OF_LEUKOCYTE_APOP | 103 | 0.372787 | 1.9514297 | 0 | 0.020990383 | 0.875 | 4844 | tags=34%, list=21%, si | MM_1 |
| GOBP_RNA_MODIFICATI | GOBP_RNA_MODIFICATION | 141 | 0.351914 | 1.9512876 | 0 | 0.020915542 | 0.875 | 6107 | tags=40%, list=26%, si | MM_1 |
| GOBP_AMYLOID_FIBRIL_ | GOBP_AMYLOID_FIBRIL_FORMATION | 30 | 0.509743 | 1.949905 | 0.011904762 | 0.02098444 | 0.88 | 3673 | tags=43%, list=16%, si | MM_1 |
| GOBP_LAYER_FORMATIO | GOBP_LAYER_FORMATION_IN_CEREBRAL_C | 16 | 0.602027 | 1.949714 | 0 | 0.020909414 | 0.881 | 5073 | tags=69%, list=21%, si | MM_1 |
| GOBP_NEGATIVE_REGUL | GOBP_NEGATIVE_REGULATION_OF_PLASM | 40 | 0.469914 | 1.9484526 | 0 | 0.02107258 | 0.889 | 3282 | tags=33%, list=14%, si | MM_1 |
| GOBP_REGULATION_OF_ | GOBP_REGULATION_OF_VACUOLE_ORGANI | 54 | 0.42886 | 1.9469172 | 0 | 0.021329189 | 0.892 | 7937 | tags=61%, list=34%, si | MM_1 |
| GOBP_NUCLEOSIDE_DIPH | GOBP_NUCLEOSIDE_DIPHOSPHATE_METAB | 127 | 0.363932 | 1.9467506 | 0 | 0.021235228 | 0.892 | 5198 | tags=39%, list=22%, si | MM_1 |
| GOBP_ACETYL_COA_MET | GOBP_ACETYL_COA_METABOLIC_PROCESS | 34 | 0.480346 | 1.9447671 | 0 | 0.021544838 | 0.899 | 3877 | tags=44%, list=16%, si | MM_1 |
| GOBP_SULFUR_COMPOU | GOBP_SULFUR_COMPOUND_CATABOLIC_P | 35 | 0.479576 | 1.9419966 | 0.0030674846 | 0.021766473 | 0.903 | 5284 | tags=54%, list=22%, si | MM_1 |
| GOBP_POSITIVE_REGULA | GOBP_POSITIVE_REGULATION_OF_TELOME | 17 | 0.58859 | 1.9419365 | 0.0056657223 | 0.0216813 | 0.903 | 4247 | tags=41%, list=18%, si | MM_1 |
| GOBP_DNA_TEMPLATED_ | GOBP_DNA_TEMPLATED_TRANSCRIPTION_I | 112 | 0.3733 | 1.9396704 | 0 | 0.021976884 | 0.906 | 5811 | tags=37%, list=25%, si | MM_1 |
| GOBP_INTRA_GOLGI_VES | GOBP_INTRA_GOLGI_VESICLE_MEDIATED_T | 30 | 0.496086 | 1.9395659 | 0 | 0.02189141 | 0.906 | 6052 | tags=60%, list=26%, si | MM_1 |
| GOBP_RNA_3_END_PRO | GOBP_RNA_3_END_PROCESSING | 87 | 0.395687 | 1.9390756 | 0 | 0.021852635 | 0.906 | 7395 | tags=46%, list=31%, si | MM_1 |
| GOBP_T_CELL_CHEMOTA | GOBP_T_CELL_CHEMOTAXIS | 23 | 0.539628 | 1.9381073 | 0 | 0.021961212 | 0.908 | 3488 | tags=39%, list=15%, si | MM_1 |
| GOBP_TOR_SIGNALING | GOBP_TOR_SIGNALING | 165 | 0.345901 | 1.934132 | 0 | 0.022633081 | 0.916 | 6606 | tags=45%, list=28%, si | MM_1 |
| GOBP_GLYCOLYTIC_PRO | GOBP_GLYCOLYTIC_PROCESS_THROUGH_G | 20 | 0.555359 | 1.933466 | 0.0057142857 | 0.022628045 | 0.916 | 5076 | tags=55%, list=21%, si | MM_1 |
| GOBP_LEUKOCYTE_APOP | GOBP_LEUKOCYTE_APOPTOTIC_PROCESS | 131 | 0.356502 | 1.9331709 | 0 | 0.02258653 | 0.916 | 4908 | tags=34%, list=21%, si | MM_1 |
| GOBP_NEGATIVE_REGUL | GOBP_NEGATIVE_REGULATION_OF_RESPON | 15 | 0.599697 | 1.9323397 | 0.010610079 | 0.022572068 | 0.917 | 6127 | tags=53%, list=26%, si | MM_1 |
| GOBP_ALDEHYDE_BIOSYI | GOBP_ALDEHYDE_BIOSYNTHETIC_PROCESS | 26 | 0.520384 | 1.9319737 | 0 | 0.022557363 | 0.917 | 4617 | tags=50%, list=20%, si | MM_1 |
| GOBP_REGULATION_OF_ | GOBP_REGULATION_OF_LEUKOCYTE_MIGR | 230 | 0.323149 | 1.9314675 | 0 | 0.022544455 | 0.918 | 4035 | tags=29%, list=17%, si | MM_1 |
| GOBP_MEMBRANE_RAFT | GOBP_MEMBRANE_RAFT_ORGANIZATION | 24 | 0.518829 | 1.9304796 | 0 | 0.022609282 | 0.919 | 5902 | tags=54%, list=25%, si | MM_1 |
| GOBP_REGULATION_OF_ | GOBP_REGULATION_OF_TELOMERASE_ACT | 42 | 0.44372 | 1.9304215 | 0 | 0.022534411 | 0.919 | 5921 | tags=45%, list=25%, si | MM_1 |
| PHAGOSOME | PHAGOSOME | 146 | 0.347133 | 1.9287002 | 0 | 0.02273959 | 0.921 | 5000 | tags=32%, list=21%, si | MM_1 |
| GOBP_NEGATIVE_REGUL | GOBP_NEGATIVE_REGULATION_OF_TYPE_I_ | 42 | 0.456343 | 1.9285911 | 0 | 0.022672148 | 0.921 | 5668 | tags=48%, list=24%, si | MM_1 |
| RNA_TRANSPORT | RNA_TRANSPORT | 154 | 0.345272 | 1.9272897 | 0 | 0.022835158 | 0.925 | 5991 | tags=34%, list=25%, si | MM_1 |
| GOBP_MOTOR_BEHAVIO | GOBP_MOTOR_BEHAVIOR | 32 | 0.481901 | 1.9272721 | 0.0029325513 | 0.022742333 | 0.925 | 2248 | tags=31%, list=10%, si | MM_1 |
| GOBP_CELLULAR_ALDEH | GOBP_CELLULAR_ALDEHYDE_METABOLIC_P | 74 | 0.407202 | 1.926022 | 0 | 0.022884486 | 0.927 | 5598 | tags=54%, list=24%, si | MM_1 |
| GOBP_VIRION_ASSEMBLY | GOBP_VIRION_ASSEMBLY | 26 | 0.514187 | 1.9203607 | 0 | 0.023923017 | 0.936 | 7975 | tags=69%, list=34%, si | MM_1 |
| GOBP_PLATELET_MORPH | GOBP_PLATELET_MORPHOGENESIS | 25 | 0.509398 | 1.9160162 | 0 | 0.024491629 | 0.942 | 5963 | tags=60%, list=25%, si | MM_1 |
| GOBP_RESPONSE_TO_OX | GOBP_RESPONSE_TO_OXIDATIVE_STRESS | 364 | 0.31162 | 1.9152427 | 0 | 0.024522783 | 0.943 | 4910 | tags=30%, list=21%, si | MM_1 |
| GOBP_RIBOSOME_DISAS | GOBP_RIBOSOME_DISASSEMBLY | 31 | 0.479681 | 1.9151634 | 0.0056818184 | 0.024433667 | 0.943 | 6840 | tags=55%, list=29%, si | MM_1 |
| GOBP_REGULATION_OF_ | GOBP_REGULATION_OF_ATP_BIOSYNTHETI | 20 | 0.545744 | 1.9143338 | 0.019283747 | 0.024464898 | 0.943 | 3673 | tags=35%, list=16%, si | MM_1 |
| GOBP_REGULATION_OF_ | GOBP_REGULATION_OF_ERAD_PATHWAY | 27 | 0.502922 | 1.912375 | 0.0027472528 | 0.024748128 | 0.943 | 4955 | tags=48%, list=21%, si | MM_1 |
| GOBP_RETICULOPHAGY | GOBP_RETICULOPHAGY | 19 | 0.56158 | 1.9119916 | 0.002915452 | 0.02472627 | 0.944 | 4849 | tags=58%, list=21%, si | MM_1 |
| GOBP_NUCLEAR_MEMBR | GOBP_NUCLEAR_MEMBRANE_REASSEMBLY | 20 | 0.551687 | 1.9117723 | 0.0025906735 | 0.024679048 | 0.945 | 6958 | tags=55%, list=29%, si | MM_1 |
| GOBP_REGULATION_OF_ | GOBP_REGULATION_OF_GOLGI_ORGANIZAT | 17 | 0.559313 | 1.9105561 | 0.0026809652 | 0.024833169 | 0.95 | 8103 | tags=41%, list=34%, si | MM_1 |

| | | | | | | | | | | |
|---|---|---|---|---|---|---|---|---|---|---|
| GOBP_DE_NOVO_PROTEI | GOBP_DE_NOVO_PROTEIN_FOLDING | 38 | 0.463708 | 1.910062 | 0.0061919503 | 0.024828687 | 0.95 | 5051 | tags=42%, list=21%, si | MM_1 |
| GOBP_POSITIVE_REGULA | GOBP_POSITIVE_REGULATION_OF_ISOTYPE_ | 30 | 0.490885 | 1.9099683 | 0 | 0.024740413 | 0.95 | 4596 | tags=43%, list=19%, si | MM_1 |
| GOBP_MRNA_3_END_PR | GOBP_MRNA_3_END_PROCESSING | 41 | 0.453388 | 1.909279 | 0.00591716 | 0.024745125 | 0.95 | 1596 | tags=15%, list=7%, sig | MM_1 |
| GOBP_POST_TRANSCRIPT | GOBP_POST_TRANSCRIPTIONAL_REGULATIC | 480 | 0.296623 | 1.9064227 | 0 | 0.025211435 | 0.955 | 6129 | tags=33%, list=26%, si | MM_1 |
| GOBP_VIRAL_LIFE_CYCLE | GOBP_VIRAL_LIFE_CYCLE | 230 | 0.323397 | 1.9062957 | 0 | 0.025114838 | 0.955 | 5972 | tags=39%, list=25%, si | MM_1 |
| GOBP_POSITIVE_REGULA | GOBP_POSITIVE_REGULATION_OF_TELOME | 15 | 0.603819 | 1.9031662 | 0.010498688 | 0.025579019 | 0.957 | 6603 | tags=73%, list=28%, si | MM_1 |
| GOBP_MITOCHONDRION_ | GOBP_MITOCHONDRION_LOCALIZATION | 57 | 0.416002 | 1.9026169 | 0.003215434 | 0.025571529 | 0.957 | 3847 | tags=32%, list=16%, si | MM_1 |
| GOBP_VACUOLAR_ACIDI | GOBP_VACUOLAR_ACIDIFICATION | 27 | 0.499006 | 1.9004264 | 0.0056497175 | 0.025952114 | 0.959 | 4357 | tags=48%, list=18%, si | MM_1 |
| GOBP_PURINE_CONTAIN | GOBP_PURINE_CONTAINING_COMPOUND_ | 150 | 0.342932 | 1.8986895 | 0 | 0.0261945 | 0.961 | 5284 | tags=37%, list=22%, si | MM_1 |
| GOBP_TELOMERE_CAPPI | GOBP_TELOMERE_CAPPING | 38 | 0.448368 | 1.8974215 | 0 | 0.0263218 | 0.964 | 6762 | tags=47%, list=29%, si | MM_1 |
| GOBP_MATURATION_OF_ | GOBP_MATURATION_OF_5_8S_RRNA_FROM | 24 | 0.520859 | 1.8973454 | 0 | 0.026246725 | 0.964 | 5663 | tags=46%, list=24%, si | MM_1 |
| GOBP_REGULATION_OF_ | GOBP_REGULATION_OF_CYTOPLASMIC_PAT | 104 | 0.360057 | 1.8968447 | 0 | 0.026228728 | 0.965 | 4955 | tags=31%, list=21%, si | MM_1 |
| GOBP_ESTABLISHMENT_( | GOBP_ESTABLISHMENT_OF_PROTEIN_LOCA | 262 | 0.31964 | 1.891298 | 0 | 0.027159002 | 0.971 | 6325 | tags=45%, list=27%, si | MM_1 |
| GOBP_ESTABLISHMENT_( | GOBP_ESTABLISHMENT_OF_MITOCHONDRI( | 25 | 0.511223 | 1.8911116 | 0.0028985508 | 0.027090205 | 0.971 | 3847 | tags=44%, list=16%, si | MM_1 |
| GOBP_RNA_5_END_PRO( | GOBP_RNA_5_END_PROCESSING | 27 | 0.495713 | 1.8862252 | 0.002915452 | 0.02819162 | 0.975 | 5327 | tags=41%, list=23%, si | MM_1 |
| GOBP_NEGATIVE_REGUL | GOBP_NEGATIVE_REGULATION_OF_FIBROB | 44 | 0.436246 | 1.8859193 | 0.0032573289 | 0.028189981 | 0.976 | 3451 | tags=32%, list=15%, si | MM_1 |
| GOBP_NEGATIVE_REGUL | GOBP_NEGATIVE_REGULATION_OF_TELOM | 21 | 0.544222 | 1.88418 | 0.0028490028 | 0.028500713 | 0.977 | 5750 | tags=57%, list=24%, si | MM_1 |
| GOBP_DNA_DAMAGE_RE | GOBP_DNA_DAMAGE_RESPONSE_SIGNAL_T | 17 | 0.561262 | 1.8840556 | 0.01591512 | 0.02842058 | 0.977 | 4765 | tags=53%, list=20%, si | MM_1 |
| GOBP_REGULATION_OF_ | GOBP_REGULATION_OF_CELLULAR_RESPON | 26 | 0.499677 | 1.8810272 | 0 | 0.029066581 | 0.978 | 4732 | tags=46%, list=20%, si | MM_1 |
| GOBP_POSITIVE_REGULA | GOBP_POSITIVE_REGULATION_OF_MEMBRA | 16 | 0.568947 | 1.8807912 | 0.010471204 | 0.029030047 | 0.978 | 1245 | tags=25%, list=5%, sig | MM_1 |
| GOBP_POSITIVE_CHEMOT | GOBP_POSITIVE_CHEMOTAXIS | 45 | 0.431923 | 1.8799694 | 0 | 0.029126292 | 0.978 | 3743 | tags=33%, list=16%, si | MM_1 |
| GOBP_REGULATION_OF_ | GOBP_REGULATION_OF_CENTROSOME_DU | 46 | 0.42439 | 1.8797697 | 0.003012048 | 0.029036975 | 0.978 | 3289 | tags=30%, list=14%, si | MM_1 |
| GOBP_CARBOHYDRATE_D | GOBP_CARBOHYDRATE_DERIVATIVE_CATAB | 245 | 0.316203 | 1.8790324 | 0 | 0.029048052 | 0.98 | 5284 | tags=36%, list=22%, si | MM_1 |
| GOBP_MITOCHONDRIAL_ | GOBP_MITOCHONDRIAL_DNA_METABOLIC_ | 26 | 0.510906 | 1.8788316 | 0 | 0.028982228 | 0.98 | 2627 | tags=27%, list=11%, si | MM_1 |
| ARGININE_AND_PROLINE | ARGININE_AND_PROLINE_METABOLISM | 47 | 0.431176 | 1.8778374 | 0 | 0.02905384 | 0.98 | 4253 | tags=45%, list=18%, si | MM_1 |
| GOBP_NUCLEOSIDE_DIPH | GOBP_NUCLEOSIDE_DIPHOSPHATE_CATABO | 111 | 0.357779 | 1.8770132 | 0 | 0.029102542 | 0.98 | 5198 | tags=39%, list=22%, si | MM_1 |
| GOBP_GLYCOPROTEIN_C | GOBP_GLYCOPROTEIN_CATABOLIC_PROCES | 31 | 0.477056 | 1.875657 | 0 | 0.029225053 | 0.98 | 2864 | tags=35%, list=12%, si | MM_1 |
| GOBP_POSITIVE_REGULA | GOBP_POSITIVE_REGULATION_OF_CHROM( | 107 | 0.360266 | 1.8744726 | 0 | 0.029430892 | 0.982 | 6603 | tags=38%, list=28%, si | MM_1 |
| GOBP_REGULATION_OF_ | GOBP_REGULATION_OF_SPINDLE_ASSEMBL | 28 | 0.485693 | 1.8723197 | 0 | 0.02980953 | 0.982 | 3450 | tags=36%, list=15%, si | MM_1 |
| GOBP_SPLICEOSOMAL_SI | GOBP_SPLICEOSOMAL_SNRNP_ASSEMBLY | 35 | 0.47207 | 1.8722937 | 0.003012048 | 0.029720282 | 0.982 | 6776 | tags=57%, list=29%, si | MM_1 |
| GOBP_QUINONE_BIOSYN | GOBP_QUINONE_BIOSYNTHETIC_PROCESS | 16 | 0.55703 | 1.8702852 | 0.005263158 | 0.030105056 | 0.984 | 5824 | tags=63%, list=25%, si | MM_1 |
| GOBP_VIRAL_TRANSCRIP | GOBP_VIRAL_TRANSCRIPTION | 55 | 0.411019 | 1.8684454 | 0 | 0.030380165 | 0.985 | 6197 | tags=40%, list=26%, si | MM_1 |
| GOBP_TELENCEPHALON_ | GOBP_TELENCEPHALON_GLIAL_CELL_MIGR | 27 | 0.4949 | 1.8680758 | 0.0061919503 | 0.030356681 | 0.985 | 5868 | tags=48%, list=25%, si | MM_1 |
| GOBP_NEGATIVE_REGUL | GOBP_NEGATIVE_REGULATION_OF_RELEAS | 22 | 0.531539 | 1.8650827 | 0.0028653296 | 0.031028239 | 0.988 | 4350 | tags=50%, list=18%, si | MM_1 |
| GOBP_NEGATIVE_REGUL | GOBP_NEGATIVE_REGULATION_OF_CYTOPL | 50 | 0.419464 | 1.8649426 | 0 | 0.03093608 | 0.988 | 5300 | tags=38%, list=22%, si | MM_1 |
| GOBP_REGULATION_OF_ | GOBP_REGULATION_OF_VIRAL_PROCESS | 167 | 0.334907 | 1.8640813 | 0 | 0.03101382 | 0.988 | 5795 | tags=40%, list=25%, si | MM_1 |
| GOBP_HYPEROSMOTIC_R | GOBP_HYPEROSMOTIC_RESPONSE | 19 | 0.548369 | 1.8635787 | 0.005509642 | 0.031001823 | 0.988 | 4908 | tags=42%, list=21%, si | MM_1 |
| BLADDER_CANCER | BLADDER_CANCER | 37 | 0.456307 | 1.862552 | 0.005952381 | 0.031203354 | 0.988 | 2973 | tags=27%, list=13%, si | MM_1 |
| GOBP_TRNA_WOBBLE_U | GOBP_TRNA_WOBBLE_URIDINE_MODIFICAT | 16 | 0.573269 | 1.8621527 | 0.013089005 | 0.031199276 | 0.988 | 5264 | tags=63%, list=22%, si | MM_1 |
| PORPHYRIN_AND_CHLOR | PORPHYRIN_AND_CHLOROPHYLL_METABOL | 31 | 0.480324 | 1.8611683 | 0.003076923 | 0.03131842 | 0.988 | 5401 | tags=55%, list=23%, si | MM_1 |
| GOBP_ENDOSOME_TO_L | GOBP_ENDOSOME_TO_LYSOSOME_TRANSP | 16 | 0.57206 | 1.857574 | 0.010840109 | 0.03204916 | 0.988 | 6569 | tags=69%, list=28%, si | MM_1 |
| GOBP_MACROPHAGE_MI | GOBP_MACROPHAGE_MIGRATION | 65 | 0.398853 | 1.8574156 | 0 | 0.031984255 | 0.988 | 3779 | tags=32%, list=16%, si | MM_1 |
| GOBP_POSITIVE_REGULA | GOBP_POSITIVE_REGULATION_BY_HOST_OI | 22 | 0.527777 | 1.8559421 | 0.0054945056 | 0.032177962 | 0.989 | 7317 | tags=73%, list=31%, si | MM_1 |
| GOBP_CARBOHYDRATE_( | GOBP_CARBOHYDRATE_CATABOLIC_PROCES | 161 | 0.328013 | 1.8544976 | 0 | 0.032465175 | 0.99 | 5770 | tags=41%, list=24%, si | MM_1 |
| GOBP_NEGATIVE_REGUL | GOBP_NEGATIVE_REGULATION_OF_VASCUL | 33 | 0.465612 | 1.8537453 | 0.006289308 | 0.03255635 | 0.99 | 3261 | tags=39%, list=14%, si | MM_1 |
| GOBP_NUCLEOTIDE_SUG | GOBP_NUCLEOTIDE_SUGAR_BIOSYNTHETIC_ | 17 | 0.567688 | 1.8517694 | 0.0029239766 | 0.032882478 | 0.99 | 7037 | tags=59%, list=30%, si | MM_1 |
| GOBP_POSITIVE_REGULA | GOBP_POSITIVE_REGULATION_OF_TRANSCI | 32 | 0.460805 | 1.8504422 | 0.003076923 | 0.03318981 | 0.994 | 7594 | tags=56%, list=32%, si | MM_1 |
| GOBP_PHOSPHOLIPID_BI | GOBP_PHOSPHOLIPID_BIOSYNTHETIC_PROC | 214 | 0.318082 | 1.8501136 | 0 | 0.03317945 | 0.994 | 6120 | tags=44%, list=26%, si | MM_1 |
| GOBP_MITOCHONDRIAL_ | GOBP_MITOCHONDRIAL_TRANSMEMBRANE | 92 | 0.370683 | 1.8490039 | 0 | 0.033273615 | 0.994 | 5806 | tags=43%, list=25%, si | MM_1 |
| GOBP_NEGATIVE_REGUL | GOBP_NEGATIVE_REGULATION_OF_CHEMO | 39 | 0.441167 | 1.8479259 | 0.0031055901 | 0.033437226 | 0.994 | 3282 | tags=36%, list=14%, si | MM_1 |
| GOBP_RESPONSE_TO_EN | GOBP_RESPONSE_TO_ENDOPLASMIC_RETIC | 261 | 0.31496 | 1.8469558 | 0 | 0.033537135 | 0.994 | 6969 | tags=44%, list=30%, si | MM_1 |
| CARDIAC_MUSCLE_CONT | CARDIAC_MUSCLE_CONTRACTION | 72 | 0.384235 | 1.8462749 | 0 | 0.033601478 | 0.994 | 4906 | tags=47%, list=21%, si | MM_1 |
| GOBP_REGULATION_OF_ | GOBP_REGULATION_OF_NUCLEASE_ACTIVIT | 22 | 0.504652 | 1.8444895 | 0.003012048 | 0.033829696 | 0.994 | 6513 | tags=73%, list=28%, si | MM_1 |
| GOBP_T_CELL_HOMEOST | GOBP_T_CELL_HOMEOSTASIS | 52 | 0.419451 | 1.843333 | 0 | 0.033984035 | 0.994 | 4892 | tags=40%, list=21%, si | MM_1 |
| GOBP_RESPONSE_TO_TY | GOBP_RESPONSE_TO_TYPE_I_INTERFERON | 64 | 0.39759 | 1.8422718 | 0 | 0.034075856 | 0.994 | 4676 | tags=31%, list=20%, si | MM_1 |
| GOBP_REGULATION_OF_ | GOBP_REGULATION_OF_CELL_MATRIX_ADH | 121 | 0.34436 | 1.840721 | 0 | 0.03432233 | 0.994 | 4059 | tags=27%, list=17%, si | MM_1 |
| GOBP_MATURATION_OF_ | GOBP_MATURATION_OF_5_8S_RRNA | 35 | 0.460614 | 1.8398907 | 0 | 0.03444637 | 0.994 | 6046 | tags=43%, list=26%, si | MM_1 |
| RNA_DEGRADATION | RNA_DEGRADATION | 75 | 0.373924 | 1.8373787 | 0 | 0.034908466 | 0.996 | 7307 | tags=44%, list=31%, si | MM_1 |
| GOBP_REGULATION_OF_ | GOBP_REGULATION_OF_UBIQUITIN_DEPEN | 157 | 0.330054 | 1.8373533 | 0 | 0.034804516 | 0.996 | 5293 | tags=32%, list=22%, si | MM_1 |
| GOBP_REGULATION_OF_ | GOBP_REGULATION_OF_PROGRAMMED_NE | 27 | 0.491397 | 1.8365501 | 0.0032258064 | 0.034876544 | 0.996 | 3909 | tags=37%, list=17%, si | MM_1 |
| GOBP_VESICLE_TARGETI | GOBP_VESICLE_TARGETING | 60 | 0.395817 | 1.8363309 | 0 | 0.034834146 | 0.996 | 6853 | tags=47%, list=29%, si | MM_1 |
| GOBP_NUCLEOTIDE_EXCI | GOBP_NUCLEOTIDE_EXCISION_REPAIR | 76 | 0.371353 | 1.8346707 | 0.0066225166 | 0.035062276 | 0.996 | 5260 | tags=39%, list=22%, si | MM_1 |
| GOBP_LYSOSOMAL_TRAN | GOBP_LYSOSOMAL_TRANSPORT | 124 | 0.3471 | 1.8341067 | 0 | 0.03510049 | 0.996 | 6378 | tags=45%, list=27%, si | MM_1 |
| GOBP_RESPONSE_TO_HE | GOBP_RESPONSE_TO_HEAT | 86 | 0.367805 | 1.8339089 | 0 | 0.03500454 | 0.996 | 5256 | tags=36%, list=22%, si | MM_1 |
| GOBP_LIPOPROTEIN_BIO | GOBP_LIPOPROTEIN_BIOSYNTHETIC_PROCE | 104 | 0.354023 | 1.8309205 | 0 | 0.035655778 | 0.998 | 4439 | tags=34%, list=19%, si | MM_1 |
| GOBP_PROTEIN_STABILIZ | GOBP_PROTEIN_STABILIZATION | 190 | 0.32428 | 1.8308908 | 0 | 0.035545044 | 0.998 | 5381 | tags=32%, list=23%, si | MM_1 |
| GOBP_REGULATION_OF_ | GOBP_REGULATION_OF_VASCULOGENESIS | 16 | 0.567408 | 1.8305956 | 0.0026737968 | 0.035541445 | 0.998 | 1994 | tags=31%, list=8%, sig | MM_1 |
| GOBP_INTRACELLULAR_I | GOBP_INTRACELLULAR_IRON_ION_HOMEOS | 64 | 0.397212 | 1.8302411 | 0 | 0.03552435 | 0.998 | 4357 | tags=39%, list=18%, si | MM_1 |
| GOBP_POSITIVE_REGULA | GOBP_POSITIVE_REGULATION_OF_PROTEOI | 325 | 0.298262 | 1.8292054 | 0 | 0.03570568 | 0.998 | 4892 | tags=30%, list=21%, si | MM_1 |
| GOBP_NUCLEAR_MEMBR | GOBP_NUCLEAR_MEMBRANE_ORGANIZATI( | 35 | 0.442088 | 1.8288378 | 0.002994012 | 0.035701774 | 0.998 | 7656 | tags=54%, list=32%, si | MM_1 |
| GOBP_PROTEIN_REFOLDI | GOBP_PROTEIN_REFOLDING | 23 | 0.5054 | 1.8284551 | 0.011764706 | 0.03567701 | 0.998 | 4092 | tags=39%, list=17%, si | MM_1 |
| GOBP_POSITIVE_REGULA | GOBP_POSITIVE_REGULATION_OF_ATP_BIO | 15 | 0.573955 | 1.8277662 | 0.008219178 | 0.035752464 | 0.998 | 3673 | tags=40%, list=16%, si | MM_1 |
| GOBP_NUCLEOSIDE_BISP | GOBP_NUCLEOSIDE_BISPHOSPHATE_BIOSYN | 39 | 0.433705 | 1.8267308 | 0.0032894737 | 0.035917807 | 0.998 | 5660 | tags=49%, list=24%, si | MM_1 |
| GOBP_REGULATION_OF_ | GOBP_REGULATION_OF_AUTOPHAGY | 295 | 0.296707 | 1.8236673 | 0 | 0.036654156 | 0.998 | 7654 | tags=50%, list=32%, si | MM_1 |
| GOBP_NUCLEOLAR_LARG | GOBP_NUCLEOLAR_LARGE_RRNA_TRANSCR | 21 | 0.52554 | 1.8234884 | 0.010666667 | 0.03660821 | 0.998 | 7253 | tags=62%, list=31%, si | MM_1 |
| GOBP_VESICLE_TETHERIN | GOBP_VESICLE_TETHERING | 31 | 0.465631 | 1.8225988 | 0.0028571428 | 0.036755677 | 0.998 | 5762 | tags=39%, list=24%, si | MM_1 |
| GOBP_REGULATION_OF_ | GOBP_REGULATION_OF_ISOTYPE_SWITCHIN | 38 | 0.440788 | 1.8206583 | 0.002915452 | 0.037042983 | 0.998 | 4692 | tags=42%, list=20%, si | MM_1 |
| GOBP_PYRIDINE_CONTAI | GOBP_PYRIDINE_CONTAINING_COMPOUND | 182 | 0.323862 | 1.8193197 | 0 | 0.037194915 | 0.998 | 5198 | tags=38%, list=22%, si | MM_1 |
| GOBP_CELLULAR_SENESC | GOBP_CELLULAR_SENESCENCE | 88 | 0.357499 | 1.8190417 | 0 | 0.037185594 | 0.998 | 2930 | tags=19%, list=12%, si | MM_1 |
| GOBP_NEGATIVE_REGUL | GOBP_NEGATIVE_REGULATION_OF_TRANSL | 117 | 0.348407 | 1.8178132 | 0 | 0.03736386 | 0.998 | 5219 | tags=28%, list=22%, si | MM_1 |
| GOBP_POSITIVE_REGULA | GOBP_POSITIVE_REGULATION_OF_DNA_RE | 135 | 0.333141 | 1.8161936 | 0 | 0.037651952 | 0.998 | 6655 | tags=42%, list=28%, si | MM_1 |
| GOBP_POSITIVE_REGULA | GOBP_POSITIVE_REGULATION_OF_DOUBLE_ | 91 | 0.364411 | 1.814435 | 0.0034246575 | 0.038057804 | 0.998 | 4360 | tags=30%, list=18%, si | MM_1 |

| | | | | | | | | | | |
|---|---|---|---|---|---|---|---|---|---|---|
| GOBP_POSITIVE_REGULA | GOBP_POSITIVE_REGULATION_OF_LEUKOC | 159 | 0.322681 | 1.8138347 | 0 | 0.038057927 | 0.998 | 3779 | tags=27%, list=16%, si | MM_1 |
| GOBP_CELLULAR_COMPO | GOBP_CELLULAR_COMPONENT_DISASSEME | 472 | 0.283017 | 1.8115727 | 0 | 0.038578317 | 0.999 | 5286 | tags=33%, list=22%, si | MM_1 |
| GOBP_EXTRACELLULAR_N | GOBP_EXTRACELLULAR_MATRIX_ASSEMBLY | 48 | 0.411059 | 1.811176 | 0 | 0.038545907 | 0.999 | 3477 | tags=31%, list=15%, si | MM_1 |
| GOBP_RESPONSE_TO_ST | GOBP_RESPONSE_TO_STEROL_DEPLETION | 15 | 0.55451 | 1.8110065 | 0.018421052 | 0.038445678 | 0.999 | 7458 | tags=60%, list=32%, si | MM_1 |
| GOBP_MONOCARBOXYLI | GOBP_MONOCARBOXYLIC_ACID_CATABOLI | 126 | 0.335197 | 1.8108579 | 0 | 0.0383522 | 0.999 | 5379 | tags=42%, list=23%, si | MM_1 |
| MTOR_SIGNALING_PATH | MTOR_SIGNALING_PATHWAY | 146 | 0.328698 | 1.8096926 | 0 | 0.038495455 | 0.999 | 6167 | tags=38%, list=26%, si | MM_1 |
| GOBP_REGULATION_OF_ | GOBP_REGULATION_OF_AUTOPHAGOSOME | 44 | 0.424795 | 1.8078902 | 0.0062305294 | 0.038966 | 0.999 | 7937 | tags=59%, list=34%, si | MM_1 |
| GOBP_POSITIVE_REGULA | GOBP_POSITIVE_REGULATION_OF_MITOCH | 22 | 0.507158 | 1.8077666 | 0.014492754 | 0.038859636 | 0.999 | 6706 | tags=59%, list=28%, si | MM_1 |
| GOBP_CHAPERONE_MED | GOBP_CHAPERONE_MEDIATED_PROTEIN_C | 18 | 0.542046 | 1.8056738 | 0.00295858 | 0.039284684 | 0.999 | 3439 | tags=39%, list=15%, si | MM_1 |
| GOBP_POSITIVE_REGULA | GOBP_POSITIVE_REGULATION_OF_MONON | 94 | 0.360064 | 1.8038709 | 0 | 0.039607663 | 0.999 | 4980 | tags=35%, list=21%, si | MM_1 |
| GOBP_POSITIVE_REGULA | GOBP_POSITIVE_REGULATION_OF_DNA_TE | 72 | 0.380244 | 1.8034996 | 0 | 0.039542537 | 0.999 | 7428 | tags=43%, list=31%, si | MM_1 |
| GOBP_VASCULAR_ENDOT | GOBP_VASCULAR_ENDOTHELIAL_GROWTH_ | 40 | 0.432512 | 1.8030342 | 0.006060606 | 0.039559793 | 0.999 | 3840 | tags=33%, list=16%, si | MM_1 |
| GOBP_REGULATION_OF_ | GOBP_REGULATION_OF_TYPE_I_INTERFERO | 40 | 0.432552 | 1.8026898 | 0.002915452 | 0.03958769 | 0.999 | 4676 | tags=35%, list=20%, si | MM_1 |
| GOBP_PROTEIN_DESTABI | GOBP_PROTEIN_DESTABILIZATION | 56 | 0.405568 | 1.8017226 | 0 | 0.039796438 | 0.999 | 2280 | tags=20%, list=10%, si | MM_1 |
| GOBP_REGULATION_OF_ | GOBP_REGULATION_OF_ORGANELLE_ASSEN | 228 | 0.303594 | 1.8013288 | 0 | 0.03978704 | 0.999 | 6065 | tags=34%, list=26%, si | MM_1 |
| GOBP_REGULATION_OF_ | GOBP_REGULATION_OF_PROTEIN_TARGETI | 20 | 0.515008 | 1.800429 | 0.005063291 | 0.039957937 | 0.999 | 4259 | tags=45%, list=18%, si | MM_1 |
| GOBP_NUCLEOSIDE_PHO | GOBP_NUCLEOSIDE_PHOSPHATE_CATABOLI | 170 | 0.308708 | 1.799932 | 0 | 0.04000751 | 0.999 | 5284 | tags=35%, list=22%, si | MM_1 |
| GOBP_LYMPHOCYTE_COS | GOBP_LYMPHOCYTE_COSTIMULATION | 33 | 0.446668 | 1.7977265 | 0.0030487804 | 0.040465742 | 0.999 | 3053 | tags=33%, list=13%, si | MM_1 |
| GOBP_POSITIVE_REGULA | GOBP_POSITIVE_REGULATION_OF_MEMBRA | 17 | 0.538345 | 1.7972296 | 0.008595988 | 0.040484034 | 0.999 | 3673 | tags=47%, list=16%, si | MM_1 |
| GOBP_NEGATIVE_REGUL | GOBP_NEGATIVE_REGULATION_OF_VIRAL_ | 50 | 0.399915 | 1.7971245 | 0.0034129692 | 0.040394474 | 0.999 | 4676 | tags=38%, list=20%, si | MM_1 |
| GOBP_POSITIVE_REGULA | GOBP_POSITIVE_REGULATION_OF_PROTEIN | 192 | 0.313482 | 1.7962048 | 0 | 0.040531147 | 0.999 | 4955 | tags=31%, list=21%, si | MM_1 |
| GOBP_NEGATIVE_REGUL | GOBP_NEGATIVE_REGULATION_OF_LEUKO | 46 | 0.41074 | 1.7937909 | 0.0031545742 | 0.041044958 | 0.999 | 1607 | tags=26%, list=7%, sig | MM_1 |
| GOBP_POSITIVE_REGULA | GOBP_POSITIVE_REGULATION_OF_TYPE_I_I | 19 | 0.521929 | 1.7936534 | 0.014367816 | 0.040990233 | 0.999 | 2100 | tags=26%, list=9%, sig | MM_1 |
| GOBP_NEGATIVE_REGUL | GOBP_NEGATIVE_REGULATION_OF_B_CELL | 20 | 0.522363 | 1.7921156 | 0.013297873 | 0.041424014 | 0.999 | 1057 | tags=15%, list=4%, sig | MM_1 |
| GOBP_LIVER_MORPHOGE | GOBP_LIVER_MORPHOGENESIS | 36 | 0.430159 | 1.7908447 | 0.006134969 | 0.041662768 | 0.999 | 1499 | tags=22%, list=6%, sig | MM_1 |
| GOBP_REGULATION_OF_ | GOBP_REGULATION_OF_CHEMOTAXIS | 206 | 0.317055 | 1.790331 | 0 | 0.041720197 | 0.999 | 3786 | tags=29%, list=16%, si | MM_1 |
| GOBP_POSITIVE_REGULA | GOBP_POSITIVE_REGULATION_OF_FATTY_A | 20 | 0.509347 | 1.7854266 | 0.005602241 | 0.04297929 | 0.999 | 4804 | tags=50%, list=20%, si | MM_1 |
| GOBP_RUFFLE_ORGANIZ | GOBP_RUFFLE_ORGANIZATION | 54 | 0.392962 | 1.7850064 | 0.0031746032 | 0.042925403 | 0.999 | 5915 | tags=43%, list=25%, si | MM_1 |
| GOBP_REGULATION_OF_ | GOBP_REGULATION_OF_ARP2_3_COMPLEX | 27 | 0.468196 | 1.7846782 | 0 | 0.04287293 | 0.999 | 8548 | tags=81%, list=36%, si | MM_1 |
| GOBP_REGULATION_OF_ | GOBP_REGULATION_OF_TRANSLATION_IN_ | 24 | 0.482788 | 1.7821826 | 0.019830028 | 0.043385733 | 0.999 | 5219 | tags=33%, list=22%, si | MM_1 |
| GOBP_CELLULAR_RESPO | GOBP_CELLULAR_RESPONSE_TO_HEAT | 50 | 0.398432 | 1.7820143 | 0 | 0.043319725 | 0.999 | 5256 | tags=38%, list=22%, si | MM_1 |
| GOBP_REGULATION_OF_ | GOBP_REGULATION_OF_DNA_RECOMBINAT | 127 | 0.325237 | 1.7809169 | 0 | 0.043467607 | 0.999 | 4692 | tags=31%, list=20%, si | MM_1 |
| VASOPRESSIN-REGULATE | VASOPRESSIN-REGULATED_WATER_REABSO | 40 | 0.434385 | 1.7800431 | 0.009404388 | 0.043605447 | 0.999 | 5696 | tags=50%, list=24%, si | MM_1 |
| GOBP_POSITIVE_REGULA | GOBP_POSITIVE_REGULATION_OF_DNA_ME | 314 | 0.293615 | 1.7787267 | 0 | 0.043852314 | 0.999 | 4390 | tags=25%, list=19%, si | MM_1 |
| GOBP_MONOSACCHARID | GOBP_MONOSACCHARIDE_CATABOLIC_PRO | 40 | 0.427842 | 1.7782813 | 0.009868421 | 0.04386807 | 0.999 | 5343 | tags=52%, list=23%, si | MM_1 |
| GOBP_NEGATIVE_REGUL | GOBP_NEGATIVE_REGULATION_OF_SMOOT | 68 | 0.371187 | 1.7775567 | 0 | 0.04403731 | 0.999 | 3261 | tags=31%, list=14%, si | MM_1 |
| GOBP_SYMBIONT_ENTRY | GOBP_SYMBIONT_ENTRY_INTO_HOST | 97 | 0.347892 | 1.7771337 | 0 | 0.044052463 | 0.999 | 3192 | tags=26%, list=14%, si | MM_1 |
| GOBP_REGULATION_OF_ | GOBP_REGULATION_OF_PROTEIN_STABILIT | 305 | 0.294809 | 1.7769197 | 0 | 0.04402646 | 0.999 | 7376 | tags=48%, list=31%, si | MM_1 |
| GOBP_POSITIVE_REGULA | GOBP_POSITIVE_REGULATION_OF_B_CELL_ | 18 | 0.544694 | 1.7768961 | 0.008356546 | 0.043915443 | 0.999 | 5929 | tags=50%, list=25%, si | MM_1 |
| GOBP_SMALL_MOLECULE | GOBP_SMALL_MOLECULE_CATABOLIC_PRO | 318 | 0.289672 | 1.7766088 | 0 | 0.043896113 | 0.999 | 5573 | tags=42%, list=24%, si | MM_1 |
| GOBP_REGULATION_OF_ | GOBP_REGULATION_OF_ESTABLISHMENT_C | 25 | 0.478015 | 1.7754955 | 0.010695187 | 0.044103034 | 0.999 | 4259 | tags=40%, list=18%, si | MM_1 |
| GOBP_NEGATIVE_REGUL | GOBP_NEGATIVE_REGULATION_OF_VASCU | 21 | 0.501638 | 1.7742857 | 0.010989011 | 0.044410318 | 0.999 | 3730 | tags=48%, list=16%, si | MM_1 |
| GOBP_POSITIVE_REGULA | GOBP_POSITIVE_REGULATION_OF_VIRAL_P | 64 | 0.383895 | 1.7724032 | 0 | 0.044925854 | 0.999 | 5972 | tags=42%, list=25%, si | MM_1 |
| GOBP_NEGATIVE_REGUL | GOBP_NEGATIVE_REGULATION_OF_TOR_SI | 61 | 0.380641 | 1.772311 | 0.0032573289 | 0.04483083 | 0.999 | 6606 | tags=46%, list=28%, si | MM_1 |
| GOBP_MONOSACCHARID | GOBP_MONOSACCHARIDE_METABOLIC_PRO | 255 | 0.298543 | 1.7719316 | 0 | 0.04487547 | 0.999 | 5157 | tags=38%, list=22%, si | MM_1 |
| GOBP_MONONUCLEAR_C | GOBP_MONONUCLEAR_CELL_MIGRATION | 199 | 0.30545 | 1.7692171 | 0 | 0.04571292 | 0.999 | 3779 | tags=26%, list=16%, si | MM_1 |
| GOBP_REGULATION_OF_ | GOBP_REGULATION_OF_NUCLEOTIDE_BIOS | 32 | 0.442605 | 1.7691158 | 0.017441861 | 0.04561056 | 0.999 | 3673 | tags=34%, list=16%, si | MM_1 |
| GOBP_HISTONE_MRNA_N | GOBP_HISTONE_MRNA_METABOLIC_PROCE | 20 | 0.514049 | 1.7684201 | 0.01907357 | 0.045720946 | 0.999 | 7609 | tags=55%, list=32%, si | MM_1 |
| GOBP_IRON_ION_TRANS | GOBP_IRON_ION_TRANSPORT | 54 | 0.386887 | 1.7680879 | 0 | 0.04565225 | 0.999 | 4376 | tags=41%, list=19%, si | MM_1 |
| GOBP_GLYCOSYLATION | GOBP_GLYCOSYLATION | 218 | 0.303411 | 1.7675161 | 0 | 0.045710146 | 0.999 | 3130 | tags=24%, list=13%, si | MM_1 |
| GOBP_REGULATION_OF_ | GOBP_REGULATION_OF_PATTERN_RECOGN | 157 | 0.323986 | 1.7672635 | 0 | 0.045675613 | 0.999 | 6830 | tags=43%, list=29%, si | MM_1 |
| GOBP_SNO_S_RNA_MET | GOBP_SNO_S_RNA_METABOLIC_PROCESS | 17 | 0.526903 | 1.7664255 | 0.014367816 | 0.045817766 | 0.999 | 5900 | tags=47%, list=25%, si | MM_1 |
| GOBP_MATURATION_OF_ | GOBP_MATURATION_OF_SSU_RRNA | 48 | 0.393772 | 1.7649854 | 0.0100334445 | 0.04613247 | 0.999 | 6496 | tags=48%, list=28%, si | MM_1 |
| GOBP_PURINE_NUCLEOS | GOBP_PURINE_NUCLEOSIDE_MONOPHOSPH | 22 | 0.484044 | 1.7649459 | 0.005449591 | 0.046025794 | 0.999 | 8135 | tags=73%, list=34%, si | MM_1 |
| GOBP_REGULATION_OF_ | GOBP_REGULATION_OF_MEMBRANE_INVA | 21 | 0.50578 | 1.7649342 | 0.021978023 | 0.045908675 | 0.999 | 3673 | tags=43%, list=16%, si | MM_1 |
| GOBP_PHAGOCYTOSIS_E | GOBP_PHAGOCYTOSIS_ENGULFMENT | 53 | 0.396459 | 1.7640469 | 0.0032051282 | 0.04608564 | 0.999 | 3673 | tags=28%, list=16%, si | MM_1 |
| GOBP_REGULATION_OF_ | GOBP_REGULATION_OF_SEQUESTERING_OF | 15 | 0.553831 | 1.7636266 | 0.019283747 | 0.046066247 | 0.999 | 4110 | tags=40%, list=17%, si | MM_1 |
| GOBP_PROTEIN_NEDDYL | GOBP_PROTEIN_NEDDYLATION | 21 | 0.508744 | 1.762999 | 0.008264462 | 0.046155367 | 0.999 | 6323 | tags=52%, list=27%, si | MM_1 |
| GOBP_GLYCEROPHOSPHO | GOBP_GLYCEROPHOSPHOLIPID_BIOSYNTHE | 160 | 0.313803 | 1.7627472 | 0 | 0.046087414 | 0.999 | 6120 | tags=43%, list=26%, si | MM_1 |
| AUTOPHAGY | AUTOPHAGY | 126 | 0.329303 | 1.7612326 | 0 | 0.04645016 | 0.999 | 8036 | tags=50%, list=34%, si | MM_1 |
| GOBP_REGULATION_OF_ | GOBP_REGULATION_OF_UBIQUITIN_PROTE | 20 | 0.494073 | 1.7605333 | 0.016759777 | 0.046596456 | 0.999 | 318 | tags=10%, list=1%, sig | MM_1 |
| GOBP_INSULIN_LIKE_GR | GOBP_INSULIN_LIKE_GROWTH_FACTOR_RE | 49 | 0.399338 | 1.7554623 | 0.006060606 | 0.047956295 | 1 | 2041 | tags=20%, list=9%, sig | MM_1 |
| GOBP_NEGATIVE_REGUL | GOBP_NEGATIVE_REGULATION_OF_T_CELL | 29 | 0.453347 | 1.7542802 | 0.010238908 | 0.04812436 | 1 | 1469 | tags=21%, list=6%, sig | MM_1 |
| GOBP_CYTOKINETIC_PRO | GOBP_CYTOKINETIC_PROCESS | 45 | 0.404442 | 1.7529355 | 0.003215434 | 0.048399504 | 1 | 3261 | tags=33%, list=14%, si | MM_1 |
| GOBP_REGULATORY_T_C | GOBP_REGULATORY_T_CELL_DIFFERENTIAT | 41 | 0.407696 | 1.752303 | 0.005952381 | 0.04847205 | 1 | 3938 | tags=32%, list=17%, si | MM_1 |
| GOBP_POSITIVE_REGULA | GOBP_POSITIVE_REGULATION_OF_EXECUTI | 19 | 0.507617 | 1.7516768 | 0.013623978 | 0.048542157 | 1 | 2652 | tags=37%, list=11%, si | MM_1 |
| GOBP_PIECEMEAL_MICRO | GOBP_PIECEMEAL_MICROAUTOPHAGY_OF_ | 19 | 0.5083 | 1.7497994 | 0.0057306592 | 0.048912693 | 1 | 4799 | tags=47%, list=20%, si | MM_1 |
| GOBP_RESPONSE_TO_RE | GOBP_RESPONSE_TO_REACTIVE_OXYGEN_S | 181 | 0.30905 | 1.7477404 | 0 | 0.0494935 | 1 | 4697 | tags=28%, list=20%, si | MM_1 |
| GOBP_EMBRYONIC_BRAI | GOBP_EMBRYONIC_BRAIN_DEVELOPMENT | 24 | 0.480889 | 1.7465975 | 0.014245015 | 0.049708173 | 1 | 2510 | tags=33%, list=11%, si | MM_1 |
| GOBP_POSITIVE_REGULA | GOBP_POSITIVE_REGULATION_OF_VACUOL | 25 | 0.46814 | 1.7458959 | 0.017045455 | 0.049817428 | 1 | 6811 | tags=60%, list=29%, si | MM_1 |
| GOBP_NEGATIVE_REGUL | GOBP_NEGATIVE_REGULATION_OF_TELOM | 36 | 0.434711 | 1.745394 | 0.01529052 | 0.049925864 | 1 | 7523 | tags=56%, list=32%, si | MM_1 |
| GOBP_NEGATIVE_REGUL | GOBP_NEGATIVE_REGULATION_OF_TELOM | 27 | 0.45532 | 1.7448153 | 0.014662757 | 0.04997638 | 1 | 5750 | tags=48%, list=24%, si | MM_1 |
| GOBP_NUCLEUS_ORGAN | GOBP_NUCLEUS_ORGANIZATION | 136 | 0.323763 | 1.7429482 | 0 | 0.05050377 | 1 | 6076 | tags=36%, list=26%, si | MM_1 |
| GOBP_REGULATION_OF_ | GOBP_REGULATION_OF_MACROPHAGE_CH | 32 | 0.432281 | 1.7423012 | 0.015479876 | 0.050610736 | 1 | 1607 | tags=22%, list=7%, sig | MM_1 |
| GOBP_VESICLE_COATING | GOBP_VESICLE_COATING | 33 | 0.435362 | 1.7402937 | 0.009036144 | 0.05111064 | 1 | 7698 | tags=61%, list=33%, si | MM_1 |
| PYRUVATE_METABOLISM | PYRUVATE_METABOLISM | 37 | 0.412144 | 1.738991 | 0.0028735632 | 0.05154685 | 1 | 4133 | tags=49%, list=18%, si | MM_1 |
| GOBP_REGULATION_OF_ | GOBP_REGULATION_OF_MITOCHONDRIAL_ | 23 | 0.482594 | 1.7382895 | 0.0054644807 | 0.05170814 | 1 | 1245 | tags=17%, list=5%, sig | MM_1 |
| GOBP_POSITIVE_REGULA | GOBP_POSITIVE_REGULATION_OF_MACROF | 31 | 0.442223 | 1.7374312 | 0.0063492064 | 0.051846206 | 1 | 1194 | tags=19%, list=5%, sig | MM_1 |
| GOBP_SPHINGOMYELIN_ | GOBP_SPHINGOMYELIN_METABOLIC_PROC | 21 | 0.493699 | 1.7355478 | 0.011235955 | 0.052375346 | 1 | 7713 | tags=67%, list=33%, si | MM_1 |
| GOBP_AXONAL_TRANSPO | GOBP_AXONAL_TRANSPORT_OF_MITOCHO | 17 | 0.516003 | 1.7350675 | 0.0234375 | 0.052444857 | 1 | 3396 | tags=47%, list=14%, si | MM_1 |
| GOBP_PROTEIN_LOCALIZ | GOBP_PROTEIN_LOCALIZATION_TO_ENDOP | 76 | 0.359609 | 1.7349975 | 0 | 0.052370254 | 1 | 7396 | tags=58%, list=31%, si | MM_1 |
| GOBP_REGULATION_OF_ | GOBP_REGULATION_OF_AMYLOID_FIBRIL_F | 16 | 0.53695 | 1.7347418 | 0.019283747 | 0.05231179 | 1 | 5527 | tags=56%, list=23%, si | MM_1 |

| | | | | | | | | | | |
|---|---|---|---|---|---|---|---|---|---|---|
| GOBP_REGULATION_OF_ | GOBP_REGULATION_OF_DNA_STRAND_ELO | 18 | 0.518307 | 1.7330917 | 0.017441861 | 0.05273707 | 1 | 6603 | tags=67%, list=28%, si | MM_1 |
| GOBP_BENZENE_CONTAI | GOBP_BENZENE_CONTAINING_COMPOUND | 23 | 0.477248 | 1.7316984 | 0.017094018 | 0.053031806 | 1 | 2202 | tags=35%, list=9%, sig | MM_1 |
| GOBP_POSITIVE_REGULA | GOBP_POSITIVE_REGULATION_OF_CELL_SU | 134 | 0.318709 | 1.7304283 | 0 | 0.053372987 | 1 | 4129 | tags=31%, list=17%, si | MM_1 |
| GOBP_HEMATOPOIETIC_ | GOBP_HEMATOPOIETIC_STEM_CELL_PROLI | 37 | 0.422203 | 1.7298303 | 0.0056497175 | 0.05342427 | 1 | 4915 | tags=32%, list=21%, si | MM_1 |
| GOBP_ESTABLISHMENT_ | GOBP_ESTABLISHMENT_OF_PROTEIN_LOCA | 435 | 0.276561 | 1.7295008 | 0 | 0.05338927 | 1 | 6715 | tags=40%, list=28%, si | MM_1 |
| GOBP_VIRAL_GENOME_ | GOBP_VIRAL_GENOME_REPLICATION | 110 | 0.332589 | 1.7292657 | 0 | 0.05337352 | 1 | 5972 | tags=41%, list=25%, si | MM_1 |
| GOBP_MUSCLE_CELL_AP | GOBP_MUSCLE_CELL_APOPTOTIC_PROCESS | 124 | 0.318715 | 1.7279935 | 0 | 0.0536299 | 1 | 3553 | tags=23%, list=15%, si | MM_1 |
| GOBP_MEMBRANE_LIPID | GOBP_MEMBRANE_LIPID_BIOSYNTHETIC_P | 129 | 0.323678 | 1.7249655 | 0 | 0.05447 | 1 | 4708 | tags=34%, list=20%, si | MM_1 |
| GOBP_POSITIVE_REGULA | GOBP_POSITIVE_REGULATION_OF_B_CELL_ | 47 | 0.395788 | 1.7249569 | 0.003058104 | 0.05434303 | 1 | 4844 | tags=40%, list=21%, si | MM_1 |
| GOBP_REGULATION_OF_ | GOBP_REGULATION_OF_RELEASE_OF_CYTO | 49 | 0.389609 | 1.7248999 | 0.009708738 | 0.05423172 | 1 | 6116 | tags=49%, list=26%, si | MM_1 |
| GOBP_PROTEIN_CATABO | GOBP_PROTEIN_CATABOLIC_PROCESS_IN_T | 22 | 0.457038 | 1.7244686 | 0.008356546 | 0.054284062 | 1 | 7059 | tags=73%, list=30%, si | MM_1 |
| OTHER_GLYCAN_DEGRAD | OTHER_GLYCAN_DEGRADATION | 16 | 0.529464 | 1.7242978 | 0.017441861 | 0.05422275 | 1 | 6220 | tags=63%, list=26%, si | MM_1 |
| GOBP_MITOPHAGY | GOBP_MITOPHAGY | 58 | 0.371349 | 1.7240939 | 0.009146341 | 0.054162122 | 1 | 7418 | tags=57%, list=31%, si | MM_1 |
| GOBP_PEPTIDYL_LYSINE_ | GOBP_PEPTIDYL_LYSINE_METHYLATION | 24 | 0.477771 | 1.7238075 | 0.01907357 | 0.054165747 | 1 | 2553 | tags=25%, list=11%, si | MM_1 |
| GOBP_ESTABLISHMENT_ | GOBP_ESTABLISHMENT_OF_MITOCHONDRI | 34 | 0.425174 | 1.7236487 | 0.015384615 | 0.054104898 | 1 | 3847 | tags=35%, list=16%, si | MM_1 |
| GOBP_NAD_METABOLIC_ | GOBP_NAD_METABOLIC_PROCESS | 58 | 0.372414 | 1.7233876 | 0 | 0.05406926 | 1 | 5133 | tags=45%, list=22%, si | MM_1 |
| GOBP_RESPONSE_TO_ST | GOBP_RESPONSE_TO_STARVATION | 181 | 0.306841 | 1.7226332 | 0 | 0.054234978 | 1 | 6356 | tags=38%, list=27%, si | MM_1 |
| GOBP_ORGANIC_ACID_C | GOBP_ORGANIC_ACID_CATABOLIC_PROCES | 224 | 0.296625 | 1.7216194 | 0 | 0.05441822 | 1 | 5573 | tags=42%, list=24%, si | MM_1 |
| GOBP_NEGATIVE_REGUL | GOBP_NEGATIVE_REGULATION_OF_CELLUL | 18 | 0.494511 | 1.7214857 | 0.014492754 | 0.05433279 | 1 | 4313 | tags=44%, list=18%, si | MM_1 |
| GOBP_REGULATION_OF_ | GOBP_REGULATION_OF_MITOCHONDRIAL_ | 35 | 0.419398 | 1.7206286 | 0.008902078 | 0.05447614 | 1 | 6706 | tags=51%, list=28%, si | MM_1 |
| GOBP_COLLAGEN_FIBRIL | GOBP_COLLAGEN_FIBRIL_ORGANIZATION | 63 | 0.372252 | 1.719577 | 0.009646302 | 0.054798122 | 1 | 5128 | tags=43%, list=22%, si | MM_1 |
| GOBP_REGULATION_OF_ | GOBP_REGULATION_OF_PROTEASOMAL_UE | 118 | 0.322488 | 1.7185961 | 0 | 0.055001885 | 1 | 5293 | tags=34%, list=22%, si | MM_1 |
| GOBP_PEPTIDYL_LYSINE_ | GOBP_PEPTIDYL_LYSINE_ACETYLATION | 25 | 0.460201 | 1.7184471 | 0.0057306592 | 0.054926816 | 1 | 3888 | tags=36%, list=16%, si | MM_1 |
| SALIVARY_SECRETION | SALIVARY_SECRETION | 70 | 0.365022 | 1.7180374 | 0 | 0.05490974 | 1 | 2346 | tags=19%, list=10%, si | MM_1 |
| GOBP_NEGATIVE_REGUL | GOBP_NEGATIVE_REGULATION_OF_LYMPH | 44 | 0.4076 | 1.7178524 | 0.015337423 | 0.054843985 | 1 | 1607 | tags=20%, list=7%, sig | MM_1 |
| GOBP_DOUBLE_STRAND_ | GOBP_DOUBLE_STRAND_BREAK_REPAIR_VI | 67 | 0.365426 | 1.7175838 | 0.0033898305 | 0.054792836 | 1 | 4596 | tags=31%, list=19%, si | MM_1 |
| GOBP_TRNA_WOBBLE_B | GOBP_TRNA_WOBBLE_BASE_MODIFICATIO | 19 | 0.493564 | 1.716292 | 0.010869565 | 0.055130817 | 1 | 5264 | tags=53%, list=22%, si | MM_1 |
| GOBP_REGULATION_OF_ | GOBP_REGULATION_OF_PODOSOME_ASSE | 16 | 0.541705 | 1.7160774 | 0.0149700595 | 0.05509321 | 1 | 6875 | tags=63%, list=29%, si | MM_1 |
| GOBP_NEGATIVE_REGUL | GOBP_NEGATIVE_REGULATION_OF_INTRIN | 105 | 0.324767 | 1.7160031 | 0 | 0.054994464 | 1 | 5104 | tags=35%, list=22%, si | MM_1 |
| GOBP_CELLULAR_PIGME | GOBP_CELLULAR_PIGMENTATION | 57 | 0.377126 | 1.7156159 | 0.0034129692 | 0.05499634 | 1 | 5913 | tags=44%, list=25%, si | MM_1 |
| GOBP_NEGATIVE_REGUL | GOBP_NEGATIVE_REGULATION_OF_LYMPH | 62 | 0.362396 | 1.714986 | 0.0033670033 | 0.055101156 | 1 | 3665 | tags=31%, list=16%, si | MM_1 |
| GOBP_PROTEIN_LOCALIZ | GOBP_PROTEIN_LOCALIZATION_TO_CELL_S | 70 | 0.364674 | 1.7135391 | 0.0034602077 | 0.055477243 | 1 | 5589 | tags=41%, list=24%, si | MM_1 |
| GOBP_REGULATION_OF_ | GOBP_REGULATION_OF_ENDOTHELIAL_CEL | 21 | 0.487481 | 1.7128112 | 0.02259887 | 0.05565813 | 1 | 3786 | tags=38%, list=16%, si | MM_1 |
| GOBP_REGULATION_OF_ | GOBP_REGULATION_OF_NECROPTOTIC_PR | 22 | 0.485045 | 1.7126122 | 0.021621622 | 0.05559261 | 1 | 3909 | tags=36%, list=17%, si | MM_1 |
| GOBP_CYTOSOLIC_TRAN | GOBP_CYTOSOLIC_TRANSPORT | 124 | 0.322275 | 1.7119794 | 0 | 0.055654448 | 1 | 7735 | tags=53%, list=33%, si | MM_1 |
| GOBP_POSITIVE_REGULA | GOBP_POSITIVE_REGULATION_OF_ORGANE | 91 | 0.345976 | 1.71191 | 0.004048583 | 0.055555534 | 1 | 6811 | tags=40%, list=29%, si | MM_1 |
| GOBP_PIGMENT_CELL_D | GOBP_PIGMENT_CELL_DIFFERENTIATION | 37 | 0.414365 | 1.7116517 | 0.020114943 | 0.055527736 | 1 | 5111 | tags=41%, list=22%, si | MM_1 |
| GOBP_POSITIVE_REGULA | GOBP_POSITIVE_REGULATION_OF_MACRO | 25 | 0.452081 | 1.7109103 | 0.0060975607 | 0.05565572 | 1 | 1194 | tags=20%, list=5%, sig | MM_1 |
| GOBP_SMOOTH_MUSCLE | GOBP_SMOOTH_MUSCLE_CELL_APOPTOTIC | 39 | 0.416471 | 1.710056 | 0.002967359 | 0.05589765 | 1 | 3760 | tags=31%, list=16%, si | MM_1 |
| GOBP_POSITIVE_REGULA | GOBP_POSITIVE_REGULATION_OF_TRANSF | 18 | 0.503531 | 1.7089125 | 0.012953368 | 0.05620991 | 1 | 540 | tags=22%, list=2%, sig | MM_1 |
| GOBP_REGULATION_OF_ | GOBP_REGULATION_OF_PROTEIN_TARGETI | 59 | 0.371124 | 1.7084376 | 0.006802721 | 0.056245074 | 1 | 4682 | tags=41%, list=20%, si | MM_1 |
| GOBP_PHOSPHATIDIC_A | GOBP_PHOSPHATIDIC_ACID_BIOSYNTHETIC | 23 | 0.464082 | 1.7081871 | 0.010869565 | 0.05626273 | 1 | 6234 | tags=57%, list=26%, si | MM_1 |
| GOBP_NEGATIVE_REGUL | GOBP_NEGATIVE_REGULATION_OF_INSULI | 17 | 0.518612 | 1.7063979 | 0.022160664 | 0.05688118 | 1 | 35 | tags=12%, list=0%, sig | MM_1 |
| GOBP_REGULATION_OF_ | GOBP_REGULATION_OF_APOPTOTIC_SIGNA | 409 | 0.272179 | 1.7062683 | 0 | 0.05683276 | 1 | 5797 | tags=34%, list=25%, si | MM_1 |
| GOBP_NUCLEOPHAGY | GOBP_NUCLEOPHAGY | 24 | 0.471497 | 1.7050741 | 0.018018018 | 0.05717684 | 1 | 4799 | tags=42%, list=20%, si | MM_1 |
| FLUID_SHEAR_STRESS_A | FLUID_SHEAR_STRESS_AND_ATHEROSCLER | 135 | 0.314437 | 1.7041936 | 0 | 0.0573949 | 1 | 4665 | tags=35%, list=20%, si | MM_1 |
| GOBP_TYPE_I_INTERFER | GOBP_TYPE_I_INTERFERON_PRODUCTION | 127 | 0.320158 | 1.7034955 | 0 | 0.057501797 | 1 | 7011 | tags=47%, list=30%, si | MM_1 |
| GOBP_POSITIVE_REGULA | GOBP_POSITIVE_REGULATION_OF_CHEMOT | 150 | 0.310373 | 1.7031077 | 0 | 0.05746601 | 1 | 3786 | tags=28%, list=16%, si | MM_1 |
| GOBP_RESPONSE_TO_UV | GOBP_RESPONSE_TO_UV | 153 | 0.314631 | 1.7024943 | 0 | 0.057548046 | 1 | 4526 | tags=28%, list=19%, si | MM_1 |
| GOBP_LYMPHOCYTE_CHE | GOBP_LYMPHOCYTE_CHEMOTAXIS | 38 | 0.406042 | 1.7018276 | 0.0029325513 | 0.057678748 | 1 | 3736 | tags=29%, list=16%, si | MM_1 |
| GOBP_RESPONSE_TO_EP | GOBP_RESPONSE_TO_EPIDERMAL_GROWTH | 42 | 0.399428 | 1.7016889 | 0.012048192 | 0.057620082 | 1 | 7535 | tags=48%, list=32%, si | MM_1 |
| GOBP_POSITIVE_REGULA | GOBP_POSITIVE_REGULATION_OF_PROTEIN | 201 | 0.292256 | 1.7009649 | 0 | 0.057747256 | 1 | 6072 | tags=35%, list=26%, si | MM_1 |
| GOBP_DNA_REPAIR_DEP | GOBP_DNA_REPAIR_DEPENDENT_CHROMA | 23 | 0.465573 | 1.7006006 | 0.024657534 | 0.057774086 | 1 | 3350 | tags=30%, list=14%, si | MM_1 |
| GOBP_NEGATIVE_REGUL | GOBP_NEGATIVE_REGULATION_OF_LEUKO | 65 | 0.350569 | 1.699652 | 0.0034129692 | 0.058091596 | 1 | 4844 | tags=32%, list=21%, si | MM_1 |
| BASE_EXCISION_REPAIR | BASE_EXCISION_REPAIR | 32 | 0.428128 | 1.6988363 | 0.011627907 | 0.058312126 | 1 | 6340 | tags=41%, list=27%, si | MM_1 |
| GOBP_RESPONSE_TO_EL | GOBP_RESPONSE_TO_ELECTRICAL_STIMULU | 20 | 0.485923 | 1.6979337 | 0.008547009 | 0.05861278 | 1 | 4393 | tags=40%, list=19%, si | MM_1 |
| GOBP_PROTEIN_LOCALIZ | GOBP_PROTEIN_LOCALIZATION_TO_NUCLE | 17 | 0.502044 | 1.6966796 | 0.022857143 | 0.0589375 | 1 | 6593 | tags=53%, list=28%, si | MM_1 |
| GOBP_REGULATION_OF_ | GOBP_REGULATION_OF_CENTROSOME_CY | 55 | 0.375847 | 1.6956226 | 0.006369427 | 0.059217747 | 1 | 3289 | tags=25%, list=14%, si | MM_1 |
| GOBP_POSITIVE_REGULA | GOBP_POSITIVE_REGULATION_OF_PROTEIN | 87 | 0.336316 | 1.6951004 | 0 | 0.059313793 | 1 | 6473 | tags=41%, list=27%, si | MM_1 |
| GOBP_REGULATION_OF_ | GOBP_REGULATION_OF_NON_CANONICAL_ | 26 | 0.453585 | 1.6926585 | 0.012539185 | 0.060151763 | 1 | 3651 | tags=38%, list=15%, si | MM_1 |
| GOBP_ERYTHROCYTE_DE | GOBP_ERYTHROCYTE_DEVELOPMENT | 50 | 0.377451 | 1.692431 | 0.0031055901 | 0.060129497 | 1 | 1731 | tags=22%, list=7%, sig | MM_1 |
| GOBP_PIGMENT_METAB | GOBP_PIGMENT_METABOLIC_PROCESS | 65 | 0.360545 | 1.6921765 | 0.0069444445 | 0.0600938 | 1 | 2869 | tags=31%, list=12%, si | MM_1 |
| GOBP_POSITIVE_REGULA | GOBP_POSITIVE_REGULATION_OF_GLYCOP | 26 | 0.441615 | 1.690391 | 0.014084507 | 0.060729653 | 1 | 4390 | tags=35%, list=19%, si | MM_1 |
| GOBP_REGULATION_OF_ | GOBP_REGULATION_OF_VIRAL_LIFE_CYCLE | 139 | 0.313387 | 1.6902875 | 0 | 0.0606483 | 1 | 5657 | tags=37%, list=24%, si | MM_1 |
| GOBP_REGULATION_OF_ | GOBP_REGULATION_OF_MITOCHONDRION_ | 139 | 0.311632 | 1.6901463 | 0 | 0.060611628 | 1 | 5742 | tags=38%, list=24%, si | MM_1 |
| FRUCTOSE_AND_MANNO | FRUCTOSE_AND_MANNOSE_METABOLISM | 33 | 0.413834 | 1.6893892 | 0.00867052 | 0.060759846 | 1 | 3827 | tags=36%, list=16%, si | MM_1 |
| GOBP_CYTOSKELETON_D | GOBP_CYTOSKELETON_DEPENDENT_CYTOK | 97 | 0.323971 | 1.6881163 | 0 | 0.061162647 | 1 | 3428 | tags=27%, list=15%, si | MM_1 |
| GOBP_POSITIVE_REGULA | GOBP_POSITIVE_REGULATION_OF_INTERLE | 15 | 0.553272 | 1.6874328 | 0.013157895 | 0.061264083 | 1 | 2029 | tags=33%, list=9%, sig | MM_1 |
| GOBP_POSITIVE_REGULA | GOBP_POSITIVE_REGULATION_OF_ERAD_P | 20 | 0.486244 | 1.6862867 | 0.010810811 | 0.0616292 | 1 | 4955 | tags=50%, list=21%, si | MM_1 |
| GOBP_POSITIVE_REGULA | GOBP_POSITIVE_REGULATION_OF_PROTEIN | 35 | 0.412901 | 1.6860155 | 0.01179941 | 0.06159044 | 1 | 4868 | tags=40%, list=21%, si | MM_1 |
| GOBP_MAMMARY_GLAN | GOBP_MAMMARY_GLAND_MORPHOGENES | 58 | 0.361393 | 1.685525 | 0.0097402595 | 0.061677534 | 1 | 823 | tags=14%, list=3%, sig | MM_1 |
| OTHER_TYPES_OF_O-GLY | OTHER_TYPES_OF_O-GLYCAN_BIOSYNTHESI | 21 | 0.460082 | 1.6854669 | 0.03314917 | 0.061578255 | 1 | 5085 | tags=43%, list=22%, si | MM_1 |
| GOBP_MRNA_EXPORT_F | GOBP_MRNA_EXPORT_FROM_NUCLEUS | 60 | 0.35544 | 1.685381 | 0 | 0.06149261 | 1 | 4443 | tags=27%, list=19%, si | MM_1 |
| GOBP_POSITIVE_REGULA | GOBP_POSITIVE_REGULATION_OF_DNA_TE | 59 | 0.36299 | 1.6853688 | 0.0065789474 | 0.06136813 | 1 | 5727 | tags=32%, list=24%, si | MM_1 |
| GOBP_LIPOPROTEIN_ME | GOBP_LIPOPROTEIN_METABOLIC_PROCESS | 135 | 0.309782 | 1.6849337 | 0 | 0.06141334 | 1 | 4439 | tags=33%, list=19%, si | MM_1 |
| GOBP_NEGATIVE_REGUL | GOBP_NEGATIVE_REGULATION_OF_GLIAL_ | 23 | 0.46746 | 1.6842707 | 0.015384615 | 0.061528116 | 1 | 2627 | tags=30%, list=11%, si | MM_1 |
| GOBP_POSITIVE_REGULA | GOBP_POSITIVE_REGULATION_OF_ACTIN_F | 52 | 0.373841 | 1.6835161 | 0.019933555 | 0.061744362 | 1 | 6473 | tags=44%, list=27%, si | MM_1 |
| MANNOSE_TYPE_O-GLYC | MANNOSE_TYPE_O-GLYCAN_BIOSYNTHESIS | 21 | 0.472498 | 1.682962 | 0.007853403 | 0.06186458 | 1 | 3360 | tags=38%, list=14%, si | MM_1 |
| GOBP_CELLULAR_RESPO | GOBP_CELLULAR_RESPONSE_TO_UV | 90 | 0.333217 | 1.6827058 | 0.0037313432 | 0.06185599 | 1 | 4715 | tags=31%, list=20%, si | MM_1 |
| GOBP_PYRIDINE_CONTAI | GOBP_PYRIDINE_CONTAINING_COMPOUND | 109 | 0.323099 | 1.6807379 | 0.0038022813 | 0.062444028 | 1 | 5198 | tags=36%, list=22%, si | MM_1 |
| GOBP_CELLULAR_RESPO | GOBP_CELLULAR_RESPONSE_TO_VASCULAR | 60 | 0.363322 | 1.6805499 | 0.0032573289 | 0.06239705 | 1 | 3840 | tags=30%, list=16%, si | MM_1 |
| GOBP_THIOESTER_BIOSY | GOBP_THIOESTER_BIOSYNTHETIC_PROCESS | 27 | 0.444135 | 1.6794623 | 0.02346041 | 0.06285802 | 1 | 1885 | tags=30%, list=8%, sig | MM_1 |

| | | | | | | | | | | |
|---|---|---|---|---|---|---|---|---|---|---|
| GOBP_REGULATION_OF_ | GOBP_REGULATION_OF_CELLULAR_RESPON | 25 | 0.443077 | 1.6794608 | 0.02346041 | 0.06273736 | 1 | 3776 | tags=36%, list=16%, si | MM_1 |
| GOBP_PROTEIN_POLYME | GOBP_PROTEIN_POLYMERIZATION | 269 | 0.281745 | 1.6790401 | 0 | 0.062766 | 1 | 7694 | tags=49%, list=33%, si | MM_1 |
| GOBP_INTRACELLULAR_P | GOBP_INTRACELLULAR_PH_REDUCTION | 48 | 0.381934 | 1.6789919 | 0.0063492064 | 0.062650345 | 1 | 4357 | tags=35%, list=18%, si | MM_1 |
| GOBP_MITOCHONDRIAL_ | GOBP_MITOCHONDRIAL_ELECTRON_TRANS | 18 | 0.49546 | 1.6786053 | 0.028409092 | 0.06269551 | 1 | 3353 | tags=50%, list=14%, si | MM_1 |
| GOBP_NAD_CATABOLIC_ | GOBP_NAD_CATABOLIC_PROCESS | 21 | 0.463574 | 1.6780365 | 0.02616279 | 0.06278366 | 1 | 1075 | tags=24%, list=5%, sig | MM_1 |
| GOBP_PHOSPHATIDYLINC | GOBP_PHOSPHATIDYLINOSITOL_BIOSYNTHE | 95 | 0.326049 | 1.6776836 | 0.0036363637 | 0.06281978 | 1 | 6120 | tags=44%, list=26%, si | MM_1 |
| GOBP_THYMOCYTE_APOI | GOBP_THYMOCYTE_APOPTOTIC_PROCESS | 25 | 0.451075 | 1.676224 | 0.00867052 | 0.063301794 | 1 | 2878 | tags=36%, list=12%, si | MM_1 |
| GOBP_POSITIVE_REGULA | GOBP_POSITIVE_REGULATION_OF_LEUKOCY | 33 | 0.412543 | 1.6757715 | 0.0116959065 | 0.063366674 | 1 | 3053 | tags=27%, list=13%, si | MM_1 |
| GOBP_MITOTIC_INTRA_S | GOBP_MITOTIC_INTRA_S_DNA_DAMAGE_C | 16 | 0.51853 | 1.6739031 | 0.02617801 | 0.06396411 | 1 | 3896 | tags=44%, list=17%, si | MM_1 |
| GOBP_CELLULAR_RESPON | GOBP_CELLULAR_RESPONSE_TO_NUTRIENT | 213 | 0.293067 | 1.6735101 | 0 | 0.06402266 | 1 | 6116 | tags=36%, list=26%, si | MM_1 |
| GOBP_APOPTOTIC_MITO | GOBP_APOPTOTIC_MITOCHONDRIAL_CHAN | 118 | 0.315931 | 1.6734833 | 0 | 0.06390201 | 1 | 6204 | tags=44%, list=26%, si | MM_1 |
| GOBP_LONG_CHAIN_FAT | GOBP_LONG_CHAIN_FATTY_ACID_BIOSYNT | 19 | 0.487971 | 1.6729287 | 0.014367816 | 0.06396853 | 1 | 2636 | tags=26%, list=11%, si | MM_1 |
| GOBP_MACROPHAGE_CH | GOBP_MACROPHAGE_CHEMOTAXIS | 45 | 0.38445 | 1.6712532 | 0.017391304 | 0.064555116 | 1 | 3292 | tags=29%, list=14%, si | MM_1 |
| GOBP_CELL_MATRIX_ADI | GOBP_CELL_MATRIX_ADHESION | 225 | 0.285153 | 1.6709896 | 0 | 0.06453334 | 1 | 3945 | tags=25%, list=17%, si | MM_1 |
| GOBP_POSITIVE_REGULA | GOBP_POSITIVE_REGULATION_OF_IMMUN( | 55 | 0.370044 | 1.6708496 | 0.003508772 | 0.064486764 | 1 | 5547 | tags=44%, list=23%, si | MM_1 |
| ENDOCYTOSIS | ENDOCYTOSIS | 246 | 0.283238 | 1.6700282 | 0 | 0.064667694 | 1 | 7539 | tags=53%, list=32%, si | MM_1 |
| GOBP_MULTIVESICULAR_ | GOBP_MULTIVESICULAR_BODY_SORTING_P | 34 | 0.417566 | 1.6682191 | 0.0116959065 | 0.0653104 | 1 | 6569 | tags=50%, list=28%, si | MM_1 |
| GOBP_POSITIVE_REGULA | GOBP_POSITIVE_REGULATION_OF_AUTOPH | 19 | 0.490558 | 1.6681687 | 0.0359116 | 0.06522575 | 1 | 6811 | tags=58%, list=29%, si | MM_1 |
| GOBP_REGULATION_OF_ | GOBP_REGULATION_OF_AUTOPHAGOSOME | 18 | 0.497532 | 1.6669853 | 0.028985508 | 0.06562312 | 1 | 4300 | tags=50%, list=18%, si | MM_1 |
| GOBP_POSITIVE_REGULA | GOBP_POSITIVE_REGULATION_BY_HOST_OI | 17 | 0.497747 | 1.6664417 | 0.028571429 | 0.06572348 | 1 | 6197 | tags=53%, list=26%, si | MM_1 |
| HEPATOCELLULAR_CARCI | HEPATOCELLULAR_CARCINOMA | 164 | 0.294008 | 1.6662571 | 0 | 0.06569155 | 1 | 3743 | tags=23%, list=16%, si | MM_1 |
| GOBP_MITOCHONDRIAL_ | GOBP_MITOCHONDRIAL_GENOME_MAINTE | 35 | 0.406273 | 1.6652648 | 0.01734104 | 0.06595044 | 1 | 6207 | tags=40%, list=26%, si | MM_1 |
| TERPENOID_BACKBONE_ | TERPENOID_BACKBONE_BIOSYNTHESIS | 21 | 0.471352 | 1.6637833 | 0.013623978 | 0.06637461 | 1 | 2849 | tags=29%, list=12%, si | MM_1 |
| GOBP_VIRAL_BUDDING | GOBP_VIRAL_BUDDING | 18 | 0.496986 | 1.6632342 | 0.025139665 | 0.06646894 | 1 | 8493 | tags=83%, list=36%, si | MM_1 |
| GOBP_POSITIVE_REGULA | GOBP_POSITIVE_REGULATION_OF_MYELOID | 21 | 0.466914 | 1.6622523 | 0.017291067 | 0.0667497 | 1 | 7015 | tags=71%, list=30%, si | MM_1 |
| GOBP_REGULATION_OF_ | GOBP_REGULATION_OF_CELLULAR_EXTRAV | 43 | 0.391273 | 1.6611422 | 0.012232416 | 0.06707346 | 1 | 1194 | tags=19%, list=5%, sig | MM_1 |
| GOBP_MITOTIC_CYTOKIN | GOBP_MITOTIC_CYTOKINESIS | 83 | 0.334648 | 1.6607232 | 0.0068728523 | 0.06715679 | 1 | 3261 | tags=28%, list=14%, si | MM_1 |
| GLYOXYLATE_AND_DICAR | GLYOXYLATE_AND_DICARBOXYLATE_METAE | 25 | 0.435343 | 1.6600991 | 0.0057636886 | 0.06730964 | 1 | 7111 | tags=56%, list=30%, si | MM_1 |
| GOBP_REGULATION_OF_ | GOBP_REGULATION_OF_MEMBRANE_DEPO | 53 | 0.374227 | 1.659976 | 0.017006803 | 0.067215174 | 1 | 1245 | tags=17%, list=5%, sig | MM_1 |
| GOBP_S_ADENOSYLMETH | GOBP_S_ADENOSYLMETHIONINE_METABOL | 17 | 0.50005 | 1.6598326 | 0.02 | 0.06715248 | 1 | 5609 | tags=47%, list=24%, si | MM_1 |
| GOBP_POSITIVE_REGULA | GOBP_POSITIVE_REGULATION_OF_B_CELL_ | 90 | 0.329511 | 1.6591576 | 0 | 0.067328595 | 1 | 5569 | tags=37%, list=24%, si | MM_1 |
| GOBP_EXTRACELLULAR_S | GOBP_EXTRACELLULAR_STRUCTURE_ORGAN | 309 | 0.274144 | 1.6590134 | 0 | 0.06727891 | 1 | 3881 | tags=27%, list=16%, si | MM_1 |
| GOBP_REGULATION_OF_ | GOBP_REGULATION_OF_ATP_METABOLIC_F | 77 | 0.340749 | 1.6582204 | 0 | 0.06747755 | 1 | 5198 | tags=34%, list=22%, si | MM_1 |
| GOBP_NEGATIVE_REGUL | GOBP_NEGATIVE_REGULATION_OF_CELL_SU | 16 | 0.523943 | 1.6581893 | 0.023622047 | 0.06735575 | 1 | 6791 | tags=75%, list=29%, si | MM_1 |
| GOBP_CELLULAR_RESPON | GOBP_CELLULAR_RESPONSE_TO_CHEMICAL | 290 | 0.274578 | 1.6566797 | 0 | 0.06798752 | 1 | 4910 | tags=28%, list=21%, si | MM_1 |
| GOBP_INCLUSION_BODY_ | GOBP_INCLUSION_BODY_ASSEMBLY | 27 | 0.443954 | 1.6557517 | 0.031791907 | 0.068315916 | 1 | 7221 | tags=63%, list=31%, si | MM_1 |
| GOBP_MITOCHONDRIAL_ | GOBP_MITOCHONDRIAL_FISSION | 48 | 0.378661 | 1.6551639 | 0.012578616 | 0.068455584 | 1 | 6706 | tags=46%, list=28%, si | MM_1 |
| GOBP_MEMBRANE_FISSI | GOBP_MEMBRANE_FISSION | 31 | 0.427682 | 1.6537338 | 0.019444445 | 0.06891515 | 1 | 7840 | tags=61%, list=33%, si | MM_1 |
| GOBP_REGULATION_OF_ | GOBP_REGULATION_OF_GENERATION_OF_ | 148 | 0.30487 | 1.6536944 | 0.0044247787 | 0.0688037 | 1 | 5276 | tags=37%, list=22%, si | MM_1 |
| GOBP_VASCULOGENESIS | GOBP_VASCULOGENESIS | 92 | 0.327068 | 1.6535641 | 0.0036231885 | 0.06874769 | 1 | 1416 | tags=15%, list=6%, sig | MM_1 |
| GOBP_POST_GOLGI_VESI | GOBP_POST_GOLGI_VESICLE_MEDIATED_TF | 101 | 0.315824 | 1.6527367 | 0 | 0.0690001 | 1 | 6062 | tags=36%, list=26%, si | MM_1 |
| GOBP_ORGANOPHOSPHA | GOBP_ORGANOPHOSPHATE_CATABOLIC_PF | 227 | 0.282251 | 1.6525658 | 0 | 0.06897656 | 1 | 4804 | tags=31%, list=20%, si | MM_1 |
| GOBP_PLATELET_DENSE_ | GOBP_PLATELET_DENSE_GRANULE_ORGAN | 21 | 0.464866 | 1.6491629 | 0.023255814 | 0.070344634 | 1 | 5148 | tags=48%, list=22%, si | MM_1 |
| GOBP_NEGATIVE_REGUL | GOBP_NEGATIVE_REGULATION_OF_T_HELP | 22 | 0.459822 | 1.6489382 | 0.022222223 | 0.070279114 | 1 | 2145 | tags=27%, list=9%, sig | MM_1 |
| GOBP_REGULATION_OF_ | GOBP_REGULATION_OF_PLASMINOGEN_AC | 17 | 0.494341 | 1.6480424 | 0.012886598 | 0.07052795 | 1 | 1812 | tags=35%, list=8%, sig | MM_1 |
| GOBP_APOPTOTIC_CELL_ | GOBP_APOPTOTIC_CELL_CLEARANCE | 40 | 0.386312 | 1.6461232 | 0.014662757 | 0.07133417 | 1 | 6920 | tags=55%, list=29%, si | MM_1 |
| GOBP_NUCLEAR_EXPORT | GOBP_NUCLEAR_EXPORT | 160 | 0.298277 | 1.6459498 | 0 | 0.07125094 | 1 | 5991 | tags=34%, list=25%, si | MM_1 |
| TUBERCULOSIS | TUBERCULOSIS | 152 | 0.300508 | 1.6450894 | 0 | 0.07144845 | 1 | 4634 | tags=27%, list=20%, si | MM_1 |
| GOBP_MITOCHONDRIAL_ | GOBP_MITOCHONDRIAL_RNA_METABOLIC_ | 51 | 0.365411 | 1.6450485 | 0.013289036 | 0.07133052 | 1 | 6107 | tags=51%, list=26%, si | MM_1 |
| GOBP_REGULATION_OF_ | GOBP_REGULATION_OF_NEURON_MATURA | 16 | 0.503508 | 1.6450154 | 0.022038568 | 0.071213156 | 1 | 7138 | tags=69%, list=30%, si | MM_1 |
| GOBP_CELLULAR_RESPON | GOBP_CELLULAR_RESPONSE_TO_OXYGEN_L | 133 | 0.303014 | 1.64312 | 0 | 0.071981035 | 1 | 4382 | tags=30%, list=19%, si | MM_1 |
| GOBP_POSITIVE_REGULA | GOBP_POSITIVE_REGULATION_OF_DNA_DA | 15 | 0.52998 | 1.6430883 | 0.025069637 | 0.07187064 | 1 | 771 | tags=13%, list=3%, sig | MM_1 |
| GOBP_NEGATIVE_REGUL | GOBP_NEGATIVE_REGULATION_OF_AUTOPI | 81 | 0.328598 | 1.6418273 | 0.008064516 | 0.07236281 | 1 | 3781 | tags=26%, list=16%, si | MM_1 |
| GOBP_NEGATIVE_REGUL | GOBP_NEGATIVE_REGULATION_OF_UBIQUI | 18 | 0.49549 | 1.6416785 | 0.02617801 | 0.07231385 | 1 | 5276 | tags=39%, list=22%, si | MM_1 |
| GOBP_BIOLOGICAL_PROC | GOBP_BIOLOGICAL_PROCESS_INVOLVED_IN | 140 | 0.303538 | 1.6415198 | 0 | 0.072264604 | 1 | 3192 | tags=24%, list=14%, si | MM_1 |
| GOBP_REGULATION_OF_ | GOBP_REGULATION_OF_EXECUTION_PHASE | 26 | 0.435721 | 1.6403666 | 0.020114943 | 0.07258417 | 1 | 2878 | tags=31%, list=12%, si | MM_1 |
| GOBP_POSITIVE_REGULA | GOBP_POSITIVE_REGULATION_OF_PATTERN | 68 | 0.352307 | 1.6401477 | 0.006666667 | 0.072569735 | 1 | 4831 | tags=32%, list=20%, si | MM_1 |
| OOCYTE_MEIOSIS | OOCYTE_MEIOSIS | 108 | 0.315625 | 1.6399988 | 0 | 0.07250116 | 1 | 4628 | tags=29%, list=20%, si | MM_1 |
| GOBP_ACTIVATION_OF_( | GOBP_ACTIVATION_OF_GTPASE_ACTIVITY | 86 | 0.329813 | 1.6375989 | 0.0037313432 | 0.073482156 | 1 | 5490 | tags=42%, list=23%, si | MM_1 |
| GOBP_REGULATION_OF_ | GOBP_REGULATION_OF_ACTIN_NUCLEATIO | 35 | 0.400559 | 1.6375874 | 0.0091743115 | 0.07335516 | 1 | 8014 | tags=63%, list=34%, si | MM_1 |
| GOBP_POSITIVE_REGULA | GOBP_POSITIVE_REGULATION_OF_VASCUL/ | 44 | 0.372709 | 1.6348261 | 0.018404908 | 0.07449502 | 1 | 5114 | tags=39%, list=22%, si | MM_1 |
| GOBP_REGULATION_OF_ | GOBP_REGULATION_OF_INTRINSIC_APOPT( | 185 | 0.290238 | 1.633022 | 0 | 0.07519021 | 1 | 5374 | tags=34%, list=23%, si | MM_1 |
| GOBP_PROTEIN_PROCESS | GOBP_PROTEIN_PROCESSING | 237 | 0.28 | 1.6329265 | 0 | 0.07511782 | 1 | 4769 | tags=30%, list=20%, si | MM_1 |
| PROTEOGLYCANS_IN_CAI | PROTEOGLYCANS_IN_CANCER | 193 | 0.285334 | 1.6328878 | 0 | 0.0749926 | 1 | 4608 | tags=26%, list=20%, si | MM_1 |
| GOBP_POSITIVE_REGULA | GOBP_POSITIVE_REGULATION_OF_TELOME | 31 | 0.417525 | 1.6327361 | 0.023809524 | 0.07492085 | 1 | 914 | tags=16%, list=4%, sig | MM_1 |
| GOBP_CYTOPLASMIC_SE( | GOBP_CYTOPLASMIC_SEQUESTERING_OF_P | 17 | 0.490963 | 1.6322672 | 0.026706232 | 0.07499611 | 1 | 7613 | tags=59%, list=32%, si | MM_1 |
| GOBP_CELLULAR_RESPON | GOBP_CELLULAR_RESPONSE_TO_OXIDATIVE | 229 | 0.278273 | 1.6309569 | 0 | 0.07551764 | 1 | 4910 | tags=28%, list=21%, si | MM_1 |
| GOBP_POSITIVE_REGULA | GOBP_POSITIVE_REGULATION_OF_SMOOTH | 17 | 0.481252 | 1.6266347 | 0.023936171 | 0.077413455 | 1 | 3553 | tags=35%, list=15%, si | MM_1 |
| GOBP_COTRANSLATIONA | GOBP_COTRANSLATIONAL_PROTEIN_TARGE | 20 | 0.453047 | 1.6256379 | 0.030303031 | 0.07772323 | 1 | 6325 | tags=60%, list=27%, si | MM_1 |
| GOBP_NEGATIVE_REGUL | GOBP_NEGATIVE_REGULATION_OF_PROTEI | 82 | 0.333685 | 1.6249732 | 0 | 0.07793921 | 1 | 4350 | tags=35%, list=18%, si | MM_1 |
| GOBP_POSITIVE_REGULA | GOBP_POSITIVE_REGULATION_OF_PROTEIN | 16 | 0.49099 | 1.6240635 | 0.020253165 | 0.07821768 | 1 | 3 | tags=6%, list=0%, sign | MM_1 |
| GOBP_SNRNA_METABOL | GOBP_SNRNA_METABOLIC_PROCESS | 50 | 0.374329 | 1.6234999 | 0.015060241 | 0.07838013 | 1 | 3738 | tags=26%, list=16%, si | MM_1 |
| GOBP_AXONAL_TRANSP( | GOBP_AXONAL_TRANSPORT | 69 | 0.34364 | 1.6234348 | 0.0033783785 | 0.078277566 | 1 | 5332 | tags=45%, list=23%, si | MM_1 |
| GOBP_GTP_METABOLIC_ | GOBP_GTP_METABOLIC_PROCESS | 26 | 0.426804 | 1.6231142 | 0.025423728 | 0.07830102 | 1 | 6941 | tags=58%, list=29%, si | MM_1 |
| GOBP_AMINO_ACID_ACT | GOBP_AMINO_ACID_ACTIVATION | 26 | 0.429399 | 1.6229476 | 0.019886363 | 0.078250654 | 1 | 5261 | tags=50%, list=22%, si | MM_1 |
| GOBP_POSITIVE_REGULA | GOBP_POSITIVE_REGULATION_OF_LIPID_ST | 25 | 0.430843 | 1.6227812 | 0.025280898 | 0.07816737 | 1 | 2450 | tags=28%, list=10%, si | MM_1 |
| UBIQUITIN_MEDIATED_P | UBIQUITIN_MEDIATED_PROTEOLYSIS | 134 | 0.299114 | 1.6205003 | 0 | 0.07908651 | 1 | 7484 | tags=50%, list=32%, si | MM_1 |
| GOBP_RRNA_MODIFICAT | GOBP_RRNA_MODIFICATION | 29 | 0.414982 | 1.619037 | 0.026706232 | 0.07966901 | 1 | 6862 | tags=52%, list=29%, si | MM_1 |
| GOBP_POSITIVE_REGULA | GOBP_POSITIVE_REGULATION_OF_NUCLEO | 20 | 0.454081 | 1.6184547 | 0.032163743 | 0.07983355 | 1 | 3673 | tags=35%, list=16%, si | MM_1 |
| GOBP_REGULATION_OF_ | GOBP_REGULATION_OF_PROTEIN_CATABOI | 346 | 0.263535 | 1.6184087 | 0 | 0.07970397 | 1 | 5078 | tags=31%, list=22%, si | MM_1 |
| GOBP_REGULATION_OF_ | GOBP_REGULATION_OF_CELL_SUBSTRATE_ | 219 | 0.275508 | 1.6172106 | 0 | 0.08024726 | 1 | 4129 | tags=27%, list=17%, si | MM_1 |
| GOBP_WALKING_BEHAVI | GOBP_WALKING_BEHAVIOR | 58 | 0.36247 | 1.6166574 | 0.024024025 | 0.08040431 | 1 | 4935 | tags=40%, list=21%, si | MM_1 |

| | | | | | | | | | | |
|---|---|---|---|---|---|---|---|---|---|---|
| GOBP_POSITIVE_REGULA | GOBP_POSITIVE_REGULATION_OF_MITOCH | 17 | 0.482525 | 1.6163315 | 0.018766755 | 0.08042781 | 1 | 6277 | tags=53%, list=27%, si | MM_1 |
| GOBP_NEGATIVE_REGUL | GOBP_NEGATIVE_REGULATION_OF_ANOIKI | 15 | 0.492341 | 1.6152949 | 0.032663316 | 0.080901295 | 1 | 1541 | tags=27%, list=7%, sig | MM_1 |
| GOBP_NEGATIVE_REGUL | GOBP_NEGATIVE_REGULATION_OF_PROGR | 17 | 0.477207 | 1.6152648 | 0.03264095 | 0.08077078 | 1 | 3909 | tags=35%, list=17%, si | MM_1 |
| GOBP_LEUKOTRIENE_ME | GOBP_LEUKOTRIENE_METABOLIC_PROCESS | 20 | 0.455004 | 1.6149656 | 0.030812325 | 0.08075317 | 1 | 4066 | tags=40%, list=17%, si | MM_1 |
| GOBP_PHOSPHOLIPID_M | GOBP_PHOSPHOLIPID_METABOLIC_PROCES | 352 | 0.264634 | 1.6131126 | 0 | 0.08156869 | 1 | 6120 | tags=42%, list=26%, si | MM_1 |
| GOBP_POSITIVE_REGULA | GOBP_POSITIVE_REGULATION_OF_CELLULA | 22 | 0.455927 | 1.612632 | 0.027777778 | 0.08164197 | 1 | 1194 | tags=27%, list=5%, sig | MM_1 |
| GOBP_BASE_EXCISION_R | GOBP_BASE_EXCISION_REPAIR | 41 | 0.3795 | 1.612191 | 0.005988024 | 0.081714496 | 1 | 4715 | tags=27%, list=20%, si | MM_1 |
| GOBP_POSITIVE_REGULA | GOBP_POSITIVE_REGULATION_OF_CYTOSKE | 185 | 0.280115 | 1.6117208 | 0 | 0.081797384 | 1 | 6072 | tags=37%, list=26%, si | MM_1 |
| GOBP_NEGATIVE_REGUL | GOBP_NEGATIVE_REGULATION_OF_ALPHA_ | 30 | 0.414482 | 1.6114904 | 0.03380282 | 0.08182485 | 1 | 2145 | tags=23%, list=9%, sig | MM_1 |
| GOBP_TRNA_DECAY | GOBP_TRNA_DECAY | 16 | 0.49652 | 1.6113648 | 0.033240996 | 0.08175948 | 1 | 4111 | tags=31%, list=17%, si | MM_1 |
| GOBP_REGULATION_OF_ | GOBP_REGULATION_OF_INCLUSION_BODY_ | 19 | 0.465145 | 1.6113219 | 0.033426184 | 0.081632994 | 1 | 5527 | tags=47%, list=23%, si | MM_1 |
| GOBP_POSITIVE_REGULA | GOBP_POSITIVE_REGULATION_OF_AUTOPH | 157 | 0.292214 | 1.611277 | 0 | 0.08152844 | 1 | 7707 | tags=48%, list=33%, si | MM_1 |
| GOBP_ORGANELLE_TRAN | GOBP_ORGANELLE_TRANSPORT_ALONG_M | 86 | 0.322694 | 1.6111805 | 0.0036101083 | 0.08144566 | 1 | 5579 | tags=41%, list=24%, si | MM_1 |
| GOBP_NEGATIVE_REGUL | GOBP_NEGATIVE_REGULATION_OF_EPITHE | 159 | 0.293408 | 1.6111071 | 0 | 0.08133473 | 1 | 2976 | tags=21%, list=13%, si | MM_1 |
| GOBP_POSITIVE_REGULA | GOBP_POSITIVE_REGULATION_OF_PROTEIN | 16 | 0.490241 | 1.6106229 | 0.04735376 | 0.081462964 | 1 | 4611 | tags=56%, list=20%, si | MM_1 |
| GOBP_POSITIVE_REGULA | GOBP_POSITIVE_REGULATION_OF_PROTEIN | 28 | 0.419253 | 1.6097447 | 0.024390243 | 0.08183928 | 1 | 2158 | tags=29%, list=9%, sig | MM_1 |
| GOBP_PROTEIN_K48_LIN | GOBP_PROTEIN_K48_LINKED_DEUBIQUITIN | 29 | 0.422808 | 1.6087633 | 0.019337017 | 0.08214348 | 1 | 5276 | tags=38%, list=22%, si | MM_1 |
| GOBP_RESPONSE_TO_PR | GOBP_RESPONSE_TO_PROSTAGLANDIN_E | 19 | 0.469455 | 1.608404 | 0.024539877 | 0.08216023 | 1 | 1967 | tags=21%, list=8%, sig | MM_1 |
| GOBP_GLYCEROLIPID_BI | GOBP_GLYCEROLIPID_BIOSYNTHETIC_PROC | 210 | 0.274547 | 1.6068704 | 0 | 0.08280955 | 1 | 6120 | tags=40%, list=26%, si | MM_1 |
| GOBP_RESPONSE_TO_NU | GOBP_RESPONSE_TO_NUTRIENT | 75 | 0.328447 | 1.6050744 | 0.014760148 | 0.083544366 | 1 | 1963 | tags=21%, list=8%, sig | MM_1 |
| GOBP_SULFUR_COMPOU | GOBP_SULFUR_COMPOUND_METABOLIC_P | 292 | 0.265161 | 1.6042293 | 0 | 0.08387183 | 1 | 4093 | tags=29%, list=17%, si | MM_1 |
| GOBP_NUCLEAR_BODY_( | GOBP_NUCLEAR_BODY_ORGANIZATION | 19 | 0.468548 | 1.603994 | 0.026595745 | 0.08385728 | 1 | 2930 | tags=21%, list=12%, si | MM_1 |
| GOBP_SMOOTHENED_SIG | GOBP_SMOOTHENED_SIGNALING_PATHWA | 154 | 0.293628 | 1.6039455 | 0 | 0.08372981 | 1 | 2513 | tags=19%, list=11%, si | MM_1 |
| GOBP_PEROXISOMAL_TR | GOBP_PEROXISOMAL_TRANSPORT | 21 | 0.444846 | 1.602922 | 0.02710027 | 0.08405669 | 1 | 6508 | tags=48%, list=28%, si | MM_1 |
| PEROXISOME | PEROXISOME | 81 | 0.321342 | 1.6003418 | 0.0037453184 | 0.085350916 | 1 | 5150 | tags=36%, list=22%, si | MM_1 |
| GOBP_REGULATION_OF_ | GOBP_REGULATION_OF_LEUKOCYTE_TETHE | 19 | 0.454772 | 1.5992849 | 0.028947368 | 0.08581368 | 1 | 4015 | tags=32%, list=17%, si | MM_1 |
| GOBP_REGULATION_OF_ | GOBP_REGULATION_OF_CELLULAR_CATABO | 391 | 0.257939 | 1.5986251 | 0 | 0.08598787 | 1 | 6073 | tags=36%, list=26%, si | MM_1 |
| GOBP_POSITIVE_REGULA | GOBP_POSITIVE_REGULATION_OF_VASCUL/ | 172 | 0.279205 | 1.5977529 | 0 | 0.086292356 | 1 | 4133 | tags=26%, list=18%, si | MM_1 |
| GOBP_PROTEIN_MONOU | GOBP_PROTEIN_MONOUBIQUITINATION | 47 | 0.363573 | 1.5973284 | 0.0130293155 | 0.08635421 | 1 | 6334 | tags=45%, list=27%, si | MM_1 |
| GOBP_GOLGI_TO_PLASM | GOBP_GOLGI_TO_PLASMA_MEMBRANE_TR | 62 | 0.342809 | 1.5967817 | 0.010452962 | 0.086507216 | 1 | 6062 | tags=34%, list=26%, si | MM_1 |
| GOBP_DICARBOXYLIC_AC | GOBP_DICARBOXYLIC_ACID_METABOLIC_PF | 92 | 0.319904 | 1.5966606 | 0.0038461538 | 0.086428866 | 1 | 5428 | tags=45%, list=23%, si | MM_1 |
| GOBP_RESCUE_OF_STALL | GOBP_RESCUE_OF_STALLED_RIBOSOME | 25 | 0.436509 | 1.5958244 | 0.021276595 | 0.086772196 | 1 | 6840 | tags=52%, list=29%, si | MM_1 |
| GOBP_REGULATION_OF_ | GOBP_REGULATION_OF_MITOCHONDRIAL_ | 32 | 0.402466 | 1.5944138 | 0.0028735632 | 0.087370135 | 1 | 6277 | tags=38%, list=27%, si | MM_1 |
| GOBP_REGULATION_OF_ | GOBP_REGULATION_OF_GONAD_DEVELOPI | 16 | 0.50307 | 1.5943999 | 0.02688172 | 0.08723214 | 1 | 3609 | tags=31%, list=15%, si | MM_1 |
| GOBP_RESPONSE_TO_TO | GOBP_RESPONSE_TO_TOPOLOGICALLY_INC | 127 | 0.29942 | 1.5942982 | 0.003984064 | 0.08715743 | 1 | 6195 | tags=34%, list=26%, si | MM_1 |
| GOBP_SIGNAL_TRANSDU | GOBP_SIGNAL_TRANSDUCTION_BY_P53_CL | 151 | 0.287244 | 1.5942965 | 0 | 0.08701685 | 1 | 3960 | tags=25%, list=17%, si | MM_1 |
| GOBP_POSITIVE_REGULA | GOBP_POSITIVE_REGULATION_OF_ENDOTH | 107 | 0.309565 | 1.5934093 | 0.003787879 | 0.08736581 | 1 | 3786 | tags=25%, list=16%, si | MM_1 |
| GOBP_VESICLE_TARGETIN | GOBP_VESICLE_TARGETING_TO_FROM_OR_ | 29 | 0.418716 | 1.5917286 | 0.03405573 | 0.08816764 | 1 | 7182 | tags=52%, list=30%, si | MM_1 |
| GOBP_PROTEIN_LOCALIZ | GOBP_PROTEIN_LOCALIZATION_TO_PHAGO | 16 | 0.484501 | 1.5911257 | 0.040697675 | 0.08835293 | 1 | 7978 | tags=69%, list=34%, si | MM_1 |
| GOBP_DNA_TEMPLATED_ | GOBP_DNA_TEMPLATED_TRANSCRIPTION_I | 183 | 0.277813 | 1.5894557 | 0 | 0.089133166 | 1 | 7428 | tags=44%, list=31%, si | MM_1 |
| GOBP_POSITIVE_REGULA | GOBP_POSITIVE_REGULATION_OF_HYDROL | 427 | 0.254634 | 1.5894347 | 0 | 0.08901124 | 1 | 6221 | tags=36%, list=26%, si | MM_1 |
| GOBP_POSITIVE_REGULA | GOBP_POSITIVE_REGULATION_OF_CARTILA | 31 | 0.402354 | 1.5892085 | 0.023054754 | 0.08899159 | 1 | 1643 | tags=26%, list=7%, sig | MM_1 |
| GOBP_ANTEROGRADE_A) | GOBP_ANTEROGRADE_AXONAL_TRANSPOR | 53 | 0.355156 | 1.5883533 | 0.012307692 | 0.089289844 | 1 | 5332 | tags=47%, list=23%, si | MM_1 |
| GOBP_POSITIVE_REGULA | GOBP_POSITIVE_REGULATION_OF_RESPONS | 37 | 0.392023 | 1.587913 | 0.031073445 | 0.08938918 | 1 | 4955 | tags=35%, list=21%, si | MM_1 |
| GOBP_SMOOTH_MUSCLE | GOBP_SMOOTH_MUSCLE_CELL_PROLIFERAT | 191 | 0.277321 | 1.5878981 | 0 | 0.08925062 | 1 | 3945 | tags=26%, list=17%, si | MM_1 |
| GOBP_LYMPHOCYTE_HOI | GOBP_LYMPHOCYTE_HOMEOSTASIS | 92 | 0.313596 | 1.5877703 | 0.0039525693 | 0.08918336 | 1 | 5110 | tags=33%, list=22%, si | MM_1 |
| GOBP_REGULATION_OF_ | GOBP_REGULATION_OF_NUCLEOTIDE_MET/ | 96 | 0.313476 | 1.587656 | 0.01438849 | 0.0890824 | 1 | 5198 | tags=33%, list=22%, si | MM_1 |
| GOBP_EMBRYONIC_HEM | GOBP_EMBRYONIC_HEMOPOIESIS | 31 | 0.400112 | 1.5873073 | 0.026239067 | 0.08913129 | 1 | 3913 | tags=35%, list=17%, si | MM_1 |
| GOBP_POSITIVE_REGULA | GOBP_POSITIVE_REGULATION_OF_TUMOR_ | 113 | 0.300923 | 1.5872842 | 0.004310345 | 0.088997446 | 1 | 4676 | tags=28%, list=20%, si | MM_1 |
| GOBP_CEREBRAL_CORTE) | GOBP_CEREBRAL_CORTEX_RADIALLY_ORIEN | 38 | 0.380011 | 1.5862335 | 0.027272727 | 0.089462414 | 1 | 5982 | tags=39%, list=25%, si | MM_1 |
| GOBP_REGULATION_OF_ | GOBP_REGULATION_OF_MITOCHONDRIAL_ | 87 | 0.318361 | 1.5861621 | 0.010791367 | 0.08935534 | 1 | 4350 | tags=25%, list=18%, si | MM_1 |
| GOBP_PROTEIN_LOCALIZ | GOBP_PROTEIN_LOCALIZATION_TO_VACUO | 78 | 0.32251 | 1.5859122 | 0.0037453184 | 0.08933599 | 1 | 7174 | tags=49%, list=30%, si | MM_1 |
| GOBP_PLASMINOGEN_A( | GOBP_PLASMINOGEN_ACTIVATION | 23 | 0.434945 | 1.5858 | 0.023255814 | 0.08923646 | 1 | 1812 | tags=30%, list=8%, sig | MM_1 |
| GOBP_CELL_SUBSTRATE_ | GOBP_CELL_SUBSTRATE_ADHESION | 354 | 0.251612 | 1.5855334 | 0 | 0.08925093 | 1 | 3975 | tags=24%, list=17%, si | MM_1 |
| GOBP_POSITIVE_REGULA | GOBP_POSITIVE_REGULATION_OF_LYMPHO | 21 | 0.451257 | 1.5850898 | 0.038043477 | 0.089361876 | 1 | 3053 | tags=38%, list=13%, si | MM_1 |
| GOBP_REGULATION_OF_ | GOBP_REGULATION_OF_DNA_METABOLIC_ | 491 | 0.251817 | 1.5849447 | 0 | 0.08928545 | 1 | 4692 | tags=25%, list=20%, si | MM_1 |
| GOBP_TRANSCRIPTION_E | GOBP_TRANSCRIPTION_BY_RNA_POLYMER/ | 51 | 0.364297 | 1.5835906 | 0.012698413 | 0.08986975 | 1 | 7956 | tags=51%, list=34%, si | MM_1 |
| GOBP_MIRNA_MEDIATED | GOBP_MIRNA_MEDIATED_GENE_SILENCING | 17 | 0.481732 | 1.583529 | 0.051282052 | 0.08976316 | 1 | 5219 | tags=35%, list=22%, si | MM_1 |
| GOBP_NEGATIVE_REGUL | GOBP_NEGATIVE_REGULATION_OF_VIRAL_ | 18 | 0.452292 | 1.5818235 | 0.02849741 | 0.09054295 | 1 | 2825 | tags=28%, list=12%, si | MM_1 |
| GOBP_GLUTATHIONE_MI | GOBP_GLUTATHIONE_METABOLIC_PROCESS | 58 | 0.340601 | 1.5816745 | 0.010169491 | 0.090475686 | 1 | 4173 | tags=33%, list=18%, si | MM_1 |
| GLUTATHIONE_METABOL | GLUTATHIONE_METABOLISM | 57 | 0.344453 | 1.5808799 | 0 | 0.090726465 | 1 | 2636 | tags=26%, list=11%, si | MM_1 |
| GOBP_REGULATION_OF_ | GOBP_REGULATION_OF_CILIUM_ASSEMBLY | 73 | 0.332067 | 1.580735 | 0.013513514 | 0.0906956 | 1 | 4389 | tags=29%, list=19%, si | MM_1 |
| GOBP_PROTEIN_DEGLYC | GOBP_PROTEIN_DEGLYCOSYLATION | 15 | 0.484581 | 1.5801526 | 0.034759358 | 0.090954535 | 1 | 6545 | tags=47%, list=28%, si | MM_1 |
| GOBP_REGULATION_OF_ | GOBP_REGULATION_OF_SUPRAMOLECULAF | 378 | 0.254481 | 1.5792979 | 0 | 0.09126677 | 1 | 6174 | tags=34%, list=26%, si | MM_1 |
| GOBP_LEUKOCYTE_MIGR | GOBP_LEUKOCYTE_MIGRATION | 363 | 0.254349 | 1.5791261 | 0 | 0.09126235 | 1 | 3779 | tags=24%, list=16%, si | MM_1 |
| GOBP_PROTEIN_CONTAII | GOBP_PROTEIN_CONTAINING_COMPLEX_D | 247 | 0.264225 | 1.5779831 | 0 | 0.09179966 | 1 | 5360 | tags=31%, list=23%, si | MM_1 |
| GOBP_EXTRACELLULAR_I | GOBP_EXTRACELLULAR_MATRIX_CONSTITU | 16 | 0.489345 | 1.5773065 | 0.044619422 | 0.09200722 | 1 | 3302 | tags=25%, list=14%, si | MM_1 |
| GOBP_CELL_SUBSTRATE_ | GOBP_CELL_SUBSTRATE_JUNCTION_ORGAN | 105 | 0.303501 | 1.5767248 | 0.003717472 | 0.09215179 | 1 | 6791 | tags=44%, list=29%, si | MM_1 |
| GOBP_SKELETAL_MUSCLI | GOBP_SKELETAL_MUSCLE_ADAPTATION | 21 | 0.45028 | 1.5758814 | 0.037463978 | 0.09247429 | 1 | 2074 | tags=24%, list=9%, sig | MM_1 |
| GOBP_REGULATION_OF_ | GOBP_REGULATION_OF_T_CELL_CHEMOTA | 16 | 0.482468 | 1.5754962 | 0.03814714 | 0.0925761 | 1 | 2870 | tags=31%, list=12%, si | MM_1 |
| GOBP_REGULATION_OF_ | GOBP_REGULATION_OF_MEMBRANE_LIPID | 19 | 0.450094 | 1.5739795 | 0.041237112 | 0.09334671 | 1 | 7208 | tags=63%, list=31%, si | MM_1 |
| GOBP_CHAPERONE_MED | GOBP_CHAPERONE_MEDIATED_PROTEIN_F( | 62 | 0.339507 | 1.5737573 | 0.0063492064 | 0.09333261 | 1 | 4963 | tags=32%, list=21%, si | MM_1 |
| GOBP_POST_TRANSLATIO | GOBP_POST_TRANSLATIONAL_PROTEIN_TA | 18 | 0.468258 | 1.5735016 | 0.02849741 | 0.093336955 | 1 | 6278 | tags=61%, list=27%, si | MM_1 |
| GOBP_POSITIVE_REGULA | GOBP_POSITIVE_REGULATION_OF_PROTEIN | 120 | 0.296988 | 1.5728332 | 0.003921569 | 0.09360851 | 1 | 5016 | tags=37%, list=21%, si | MM_1 |
| GOBP_TOLL_LIKE_RECEPT | GOBP_TOLL_LIKE_RECEPTOR_9_SIGNALING_ | 16 | 0.477722 | 1.5727229 | 0.045584045 | 0.09352857 | 1 | 5138 | tags=44%, list=22%, si | MM_1 |
| GOBP_POSITIVE_REGULA | GOBP_POSITIVE_REGULATION_OF_APOPTO | 161 | 0.283991 | 1.5724746 | 0 | 0.09352611 | 1 | 6537 | tags=40%, list=28%, si | MM_1 |
| GOBP_GOLGI_TO_PLASM | GOBP_GOLGI_TO_PLASMA_MEMBRANE_PR | 38 | 0.385199 | 1.5718496 | 0.014245015 | 0.09374175 | 1 | 6062 | tags=39%, list=26%, si | MM_1 |
| GOBP_AGGREPHAGY | GOBP_AGGREPHAGY | 17 | 0.485572 | 1.5708214 | 0.031914894 | 0.09423946 | 1 | 4955 | tags=41%, list=21%, si | MM_1 |
| GOBP_VESICLE_DOCKING | GOBP_VESICLE_DOCKING_INVOLVED_IN_EX | 36 | 0.381173 | 1.5705411 | 0.019292604 | 0.09426212 | 1 | 2884 | tags=28%, list=12%, si | MM_1 |
| GLUCAGON_SIGNALING_ | GLUCAGON_SIGNALING_PATHWAY | 91 | 0.305073 | 1.570448 | 0.010526316 | 0.0941751 | 1 | 4350 | tags=30%, list=18%, si | MM_1 |
| GOBP_REGULATION_OF_ | GOBP_REGULATION_OF_PROTEIN_EXPORT_ | 37 | 0.382782 | 1.5692027 | 0.03448276 | 0.09476516 | 1 | 5465 | tags=49%, list=23%, si | MM_1 |
| N-GLYCAN_BIOSYNTHESIS | N-GLYCAN_BIOSYNTHESIS | 48 | 0.363013 | 1.5687118 | 0.01510574 | 0.09494417 | 1 | 4309 | tags=27%, list=18%, si | MM_1 |

| | | | | | | | | | | |
|---|---|---|---|---|---|---|---|---|---|---|
| GOBP_HEPATICOBILIARY_ | GOBP_HEPATICOBILIARY_SYSTEM_DEVELOP | 137 | 0.288494 | 1.568305 | 0 | 0.09497839 | 1 | 4910 | tags=29%, list=21%, si | MM_1 |
| GOBP_POSITIVE_REGULA | GOBP_POSITIVE_REGULATION_OF_CILIUM_ | 32 | 0.392344 | 1.5676354 | 0.021021022 | 0.0952336 | 1 | 3883 | tags=31%, list=16%, si | MM_1 |
| GOBP_REGULATION_OF_ | GOBP_REGULATION_OF_CHROMOSOME_OI | 242 | 0.267897 | 1.56741 | 0 | 0.09524145 | 1 | 5917 | tags=31%, list=25%, si | MM_1 |
| GOBP_DNA_DAMAGE_RE | GOBP_DNA_DAMAGE_RESPONSE_SIGNAL_T | 67 | 0.332162 | 1.5672243 | 0.013937282 | 0.09518256 | 1 | 3350 | tags=24%, list=14%, si | MM_1 |
| GOBP_PINOCYTOSIS | GOBP_PINOCYTOSIS | 22 | 0.444345 | 1.5672103 | 0.023188407 | 0.0950534 | 1 | 6845 | tags=55%, list=29%, si | MM_1 |
| GOBP_PROTEIN_K63_LIN | GOBP_PROTEIN_K63_LINKED_UBIQUITINAT | 68 | 0.330069 | 1.56669 | 0.013651877 | 0.09523715 | 1 | 6129 | tags=40%, list=26%, si | MM_1 |
| GOBP_TRANSCRIPTION_I | GOBP_TRANSCRIPTION_INITIATION_AT_RNA | 148 | 0.288149 | 1.5662339 | 0.0044247787 | 0.095337205 | 1 | 7167 | tags=40%, list=30%, si | MM_1 |
| GOBP_NEGATIVE_REGUL | GOBP_NEGATIVE_REGULATION_OF_CD4_PO | 26 | 0.420323 | 1.5657914 | 0.019498607 | 0.095469736 | 1 | 2145 | tags=23%, list=9%, sig | MM_1 |
| GOBP_COLLAGEN_ACTIVA | GOBP_COLLAGEN_ACTIVATED_SIGNALING_ | 17 | 0.468243 | 1.5651919 | 0.033505153 | 0.09573059 | 1 | 1809 | tags=29%, list=8%, sig | MM_1 |
| GOBP_GLYCOPROTEIN_M | GOBP_GLYCOPROTEIN_METABOLIC_PROCES | 342 | 0.25149 | 1.5649799 | 0 | 0.09569363 | 1 | 4478 | tags=28%, list=19%, si | MM_1 |
| GOBP_REGULATION_OF_ | GOBP_REGULATION_OF_SPINDLE_ORGANIZ | 42 | 0.366281 | 1.5638659 | 0.021875 | 0.0962454 | 1 | 7716 | tags=57%, list=33%, si | MM_1 |
| GOBP_REGULATION_OF_ | GOBP_REGULATION_OF_PHOSPHOLIPID_MI | 37 | 0.382365 | 1.5637774 | 0.017699115 | 0.096163124 | 1 | 2169 | tags=24%, list=9%, sig | MM_1 |
| GOBP_NEGATIVE_REGUL | GOBP_NEGATIVE_REGULATION_OF_RHO_PI | 25 | 0.416152 | 1.5613233 | 0.04037267 | 0.097471416 | 1 | 5118 | tags=44%, list=22%, si | MM_1 |
| GOBP_MUSCLE_CELL_CEL | GOBP_MUSCLE_CELL_CELLULAR_HOMEOST/ | 26 | 0.412326 | 1.56006 | 0.027108435 | 0.09814942 | 1 | 1486 | tags=27%, list=6%, sig | MM_1 |
| GOBP_REGULATION_OF_ | GOBP_REGULATION_OF_PROTEIN_LOCALIZ/ | 117 | 0.292418 | 1.5599711 | 0.004385965 | 0.098067984 | 1 | 5016 | tags=32%, list=21%, si | MM_1 |
| BASAL_TRANSCRIPTION_ | BASAL_TRANSCRIPTION_FACTORS | 40 | 0.356025 | 1.559636 | 0.022435898 | 0.098131254 | 1 | 7428 | tags=55%, list=31%, si | MM_1 |
| GOBP_MODULATION_BY_ | GOBP_MODULATION_BY_HOST_OF_VIRAL_ | 22 | 0.438846 | 1.5593586 | 0.035928145 | 0.09816066 | 1 | 5843 | tags=41%, list=25%, si | MM_1 |
| GOBP_NEGATIVE_REGUL | GOBP_NEGATIVE_REGULATION_OF_LEUKOC | 142 | 0.28653 | 1.5592669 | 0 | 0.09807001 | 1 | 4837 | tags=31%, list=20%, si | MM_1 |
| GLYCOLYSIS_/_GLUCONE | GLYCOLYSIS_/_GLUCONEOGENESIS | 60 | 0.341601 | 1.5588973 | 0.0067340066 | 0.09814129 | 1 | 4441 | tags=40%, list=19%, si | MM_1 |
| GOBP_AXO_DENDRITIC_T | GOBP_AXO_DENDRITIC_TRANSPORT | 92 | 0.308687 | 1.5583485 | 0.007246377 | 0.098416634 | 1 | 5332 | tags=39%, list=23%, si | MM_1 |
| GOBP_REGULATION_OF_ | GOBP_REGULATION_OF_AXON_EXTENSION | 15 | 0.490093 | 1.5580394 | 0.04812834 | 0.09847923 | 1 | 4326 | tags=47%, list=18%, si | MM_1 |
| GOBP_REGULATION_OF_ | GOBP_REGULATION_OF_NUCLEOTIDE_EXCIS | 29 | 0.402312 | 1.5575354 | 0.03561644 | 0.09860388 | 1 | 4360 | tags=34%, list=18%, si | MM_1 |
| GOBP_REGULATION_OF_ | GOBP_REGULATION_OF_PHOSPHOLIPID_BIO | 20 | 0.451966 | 1.5566065 | 0.036827195 | 0.0990484 | 1 | 2169 | tags=25%, list=9%, sig | MM_1 |
| GOBP_GLYCOLIPID_BIOSY | GOBP_GLYCOLIPID_BIOSYNTHETIC_PROCESS | 66 | 0.325769 | 1.5562924 | 0.02341137 | 0.09908827 | 1 | 4439 | tags=35%, list=19%, si | MM_1 |
| GOBP_IRON_ION_TRANS | GOBP_IRON_ION_TRANSMEMBRANE_TRAN | 24 | 0.419611 | 1.5560945 | 0.02253521 | 0.09908466 | 1 | 3104 | tags=33%, list=13%, si | MM_1 |
| ECM-RECEPTOR_INTERAC | ECM-RECEPTOR_INTERACTION | 79 | 0.321977 | 1.5555437 | 0.010791367 | 0.09928084 | 1 | 2192 | tags=25%, list=9%, sig | MM_1 |
| GOBP_REGULATION_OF_ | GOBP_REGULATION_OF_CENTRIOLE_REPLIC | 22 | 0.436295 | 1.5550466 | 0.044077136 | 0.09945211 | 1 | 3289 | tags=32%, list=14%, si | MM_1 |
| GOBP_RESPONSE_TO_X_ | GOBP_RESPONSE_TO_X_RAY | 24 | 0.42635 | 1.5548091 | 0.031161472 | 0.099460036 | 1 | 2797 | tags=33%, list=12%, si | MM_1 |
| GOBP_REGULATION_OF_ | GOBP_REGULATION_OF_CHOLESTEROL_BIO | 19 | 0.450326 | 1.5546876 | 0.036516853 | 0.09941278 | 1 | 5920 | tags=53%, list=25%, si | MM_1 |
| GOBP_PROTEIN_N_LINKE | GOBP_PROTEIN_N_LINKED_GLYCOSYLATION | 61 | 0.341428 | 1.5535656 | 0.009677419 | 0.099888355 | 1 | 6712 | tags=43%, list=28%, si | MM_1 |
| GOBP_RESPONSE_TO_TE | GOBP_RESPONSE_TO_TEMPERATURE_STIM | 159 | 0.280917 | 1.55351 | 0 | 0.099775895 | 1 | 5551 | tags=35%, list=24%, si | MM_1 |
| RIG-I-LIKE_RECEPTOR_SIG | RIG-I-LIKE_RECEPTOR_SIGNALING_PATHWA | 52 | 0.351351 | 1.5530185 | 0.016286645 | 0.099887125 | 1 | 5030 | tags=31%, list=21%, si | MM_1 |
| GOBP_DEFINITIVE_HEMO | GOBP_DEFINITIVE_HEMOPOIESIS | 25 | 0.415117 | 1.5518967 | 0.032608695 | 0.10045782 | 1 | 3605 | tags=32%, list=15%, si | MM_1 |
| GOBP_SPINDLE_ASSEMBL | GOBP_SPINDLE_ASSEMBLY | 122 | 0.295167 | 1.5514542 | 0 | 0.100595966 | 1 | 6064 | tags=38%, list=26%, si | MM_1 |
| GOBP_REGULATION_OF_ | GOBP_REGULATION_OF_ESTABLISHMENT_C | 25 | 0.415247 | 1.550869 | 0.032432433 | 0.10080998 | 1 | 6473 | tags=36%, list=27%, si | MM_1 |
| GOBP_ENDOTHELIAL_CEL | GOBP_ENDOTHELIAL_CELL_MIGRATION | 219 | 0.268469 | 1.5506645 | 0 | 0.100781746 | 1 | 3824 | tags=23%, list=16%, si | MM_1 |
| GOBP_LIPID_MODIFICATI | GOBP_LIPID_MODIFICATION | 197 | 0.268359 | 1.549173 | 0 | 0.1015218 | 1 | 5744 | tags=37%, list=24%, si | MM_1 |
| GOBP_REGULATION_OF_ | GOBP_REGULATION_OF_B_CELL_MEDIATED | 66 | 0.327191 | 1.5488102 | 0.013745705 | 0.10158128 | 1 | 4844 | tags=36%, list=21%, si | MM_1 |
| GOBP_NEGATIVE_REGUL | GOBP_NEGATIVE_REGULATION_OF_CATION | 32 | 0.386638 | 1.5486039 | 0.023880597 | 0.10155563 | 1 | 1336 | tags=19%, list=6%, sig | MM_1 |
| GOBP_CYTOPLASMIC_PA | GOBP_CYTOPLASMIC_PATTERN_RECOGNITI | 159 | 0.277181 | 1.5477927 | 0 | 0.101899624 | 1 | 5590 | tags=30%, list=24%, si | MM_1 |
| GOBP_REGULATION_OF_ | GOBP_REGULATION_OF_MITOCHONDRIAL_ | 26 | 0.404938 | 1.5475111 | 0.026392963 | 0.101964794 | 1 | 6277 | tags=38%, list=27%, si | MM_1 |
| GALACTOSE_METABOLISM | GALACTOSE_METABOLISM | 29 | 0.401377 | 1.5475059 | 0.032835823 | 0.101820774 | 1 | 6304 | tags=55%, list=27%, si | MM_1 |
| GOBP_CHEMOKINE_C_X_ | GOBP_CHEMOKINE_C_X_C_MOTIF_LIGAND | 25 | 0.415028 | 1.5466149 | 0.03076923 | 0.1022566 | 1 | 4676 | tags=36%, list=20%, si | MM_1 |
| GOBP_PROTON_TRANSPO | GOBP_PROTON_TRANSPORTING_TWO_SECT | 15 | 0.477634 | 1.5465121 | 0.037406486 | 0.102154694 | 1 | 5786 | tags=67%, list=25%, si | MM_1 |
| GOBP_CELLULAR_LIPID_C | GOBP_CELLULAR_LIPID_CATABOLIC_PROCES | 210 | 0.260781 | 1.546489 | 0 | 0.1020262 | 1 | 5455 | tags=38%, list=23%, si | MM_1 |
| GOBP_REGULATION_OF_ | GOBP_REGULATION_OF_GLYCOPROTEIN_M | 47 | 0.355139 | 1.5463172 | 0.015015015 | 0.1020008 | 1 | 4390 | tags=30%, list=19%, si | MM_1 |
| GOBP_NEGATIVE_REGUL | GOBP_NEGATIVE_REGULATION_OF_FIBROB | 19 | 0.450293 | 1.5462348 | 0.046272494 | 0.1019233 | 1 | 5598 | tags=42%, list=24%, si | MM_1 |
| GOBP_CELLULAR_RESPON | GOBP_CELLULAR_RESPONSE_TO_DSRNA | 19 | 0.453542 | 1.5461341 | 0.033994336 | 0.10183458 | 1 | 2100 | tags=26%, list=9%, sig | MM_1 |
| AMINOACYL-TRNA_BIOSY | AMINOACYL-TRNA_BIOSYNTHESIS | 43 | 0.360109 | 1.5433564 | 0.018348623 | 0.103283726 | 1 | 8117 | tags=63%, list=34%, si | MM_1 |
| GOBP_DEFENSE_RESPON | GOBP_DEFENSE_RESPONSE_TO_GRAM_NEG | 68 | 0.325063 | 1.5431849 | 0.01010101 | 0.1032416 | 1 | 1438 | tags=16%, list=6%, sig | MM_1 |
| GOBP_RESPONSE_TO_OX | GOBP_RESPONSE_TO_OXYGEN_LEVELS | 247 | 0.262032 | 1.5427551 | 0 | 0.10339961 | 1 | 5006 | tags=31%, list=21%, si | MM_1 |
| GOBP_REGULATION_OF_ | GOBP_REGULATION_OF_ACTIN_FILAMENT_ | 158 | 0.280016 | 1.5426561 | 0.0046948357 | 0.10331524 | 1 | 7480 | tags=44%, list=32%, si | MM_1 |
| GOBP_FOCAL_ADHESION | GOBP_FOCAL_ADHESION_ASSEMBLY | 87 | 0.315468 | 1.5418543 | 0.013745705 | 0.10362986 | 1 | 6791 | tags=46%, list=29%, si | MM_1 |
| GOBP_REGULATION_OF_ | GOBP_REGULATION_OF_VIRAL_GENOME_R | 78 | 0.31335 | 1.5414591 | 0.02749141 | 0.103693575 | 1 | 5657 | tags=38%, list=24%, si | MM_1 |
| GOBP_FATTY_ACID_TRAN | GOBP_FATTY_ACID_TRANSMEMBRANE_TRA | 16 | 0.466281 | 1.5413755 | 0.03058104 | 0.10360023 | 1 | 4804 | tags=50%, list=20%, si | MM_1 |
| GOBP_REGULATION_OF_ | GOBP_REGULATION_OF_ACTIN_FILAMENT_ | 266 | 0.257536 | 1.5400753 | 0 | 0.104254246 | 1 | 6982 | tags=41%, list=30%, si | MM_1 |
| GOBP_PROTEIN_TARGETI | GOBP_PROTEIN_TARGETING_TO_VACUOLE | 46 | 0.359456 | 1.5400404 | 0.024767801 | 0.10414567 | 1 | 7059 | tags=50%, list=30%, si | MM_1 |
| GOBP_G0_TO_G1_TRANS | GOBP_G0_TO_G1_TRANSITION | 38 | 0.370411 | 1.5397785 | 0.024539877 | 0.10415863 | 1 | 4360 | tags=34%, list=18%, si | MM_1 |
| GOBP_NON_CANONICAL_ | GOBP_NON_CANONICAL_WNT_SIGNALING_ | 59 | 0.329636 | 1.5393484 | 0.031347964 | 0.10429864 | 1 | 3731 | tags=31%, list=16%, si | MM_1 |
| GOBP_NEGATIVE_REGUL | GOBP_NEGATIVE_REGULATION_OF_NUCLEO | 29 | 0.390588 | 1.5391439 | 0.036697246 | 0.10430031 | 1 | 2627 | tags=24%, list=11%, si | MM_1 |
| GOBP_PROTEIN_K48_LIN | GOBP_PROTEIN_K48_LINKED_UBIQUITINAT | 78 | 0.319654 | 1.5387 | 0.007142857 | 0.10445251 | 1 | 6393 | tags=38%, list=27%, si | MM_1 |
| GOBP_INDUCTION_OF_PO | GOBP_INDUCTION_OF_POSITIVE_CHEMOTA | 15 | 0.467939 | 1.5384374 | 0.038251366 | 0.10449688 | 1 | 3730 | tags=40%, list=16%, si | MM_1 |
| GOBP_PROTEIN_O_LINKE | GOBP_PROTEIN_O_LINKED_GLYCOSYLATION | 79 | 0.309284 | 1.5380503 | 0.0035714286 | 0.104621604 | 1 | 3528 | tags=28%, list=15%, si | MM_1 |
| GOBP_REGULATION_OF_ | GOBP_REGULATION_OF_DNA_BIOSYNTHETI | 121 | 0.28916 | 1.5377258 | 0 | 0.1046646 | 1 | 5994 | tags=35%, list=25%, si | MM_1 |
| GOBP_RESPONSE_TO_INT | GOBP_RESPONSE_TO_INTERLEUKIN_4 | 35 | 0.373887 | 1.5367539 | 0.04191617 | 0.10518631 | 1 | 3556 | tags=29%, list=15%, si | MM_1 |
| GOBP_FIBROBLAST_PROL | GOBP_FIBROBLAST_PROLIFERATION | 147 | 0.279237 | 1.536741 | 0.007905139 | 0.1050543 | 1 | 4007 | tags=24%, list=17%, si | MM_1 |
| AMOEBIASIS | AMOEBIASIS | 89 | 0.31209 | 1.5364668 | 0.01532567 | 0.10511463 | 1 | 2648 | tags=24%, list=11%, si | MM_1 |
| GOBP_BIOLOGICAL_PROC | GOBP_BIOLOGICAL_PROCESS_INVOLVED_IN | 235 | 0.263906 | 1.5354195 | 0 | 0.10560829 | 1 | 5885 | tags=37%, list=25%, si | MM_1 |
| GOBP_REGULATION_OF_ | GOBP_REGULATION_OF_ENDOTHELIAL_CEL | 168 | 0.270907 | 1.5348262 | 0 | 0.10584867 | 1 | 3824 | tags=22%, list=16%, si | MM_1 |
| GNRH_SIGNALING_PATH | GNRH_SIGNALING_PATHWAY | 83 | 0.308734 | 1.534262 | 0.01724138 | 0.10606568 | 1 | 6197 | tags=35%, list=26%, si | MM_1 |
| GOBP_REGULATION_OF_ | GOBP_REGULATION_OF_VIRAL_TRANSCRIPT | 25 | 0.414245 | 1.533824 | 0.04532578 | 0.106185496 | 1 | 3321 | tags=28%, list=14%, si | MM_1 |
| GOBP_REGULATION_OF_ | GOBP_REGULATION_OF_CELLULAR_RESPIRA | 52 | 0.337213 | 1.5330178 | 0.027777778 | 0.10650886 | 1 | 5877 | tags=46%, list=25%, si | MM_1 |
| GOBP_ENDOLYSOSOMAL | GOBP_ENDOLYSOSOMAL_TOLL_LIKE_RECEP | 47 | 0.355029 | 1.5329882 | 0.026239067 | 0.10637055 | 1 | 5138 | tags=32%, list=22%, si | MM_1 |
| GOBP_INTERFERON_MED | GOBP_INTERFERON_MEDIATED_SIGNALING | 76 | 0.308763 | 1.5328743 | 0.017793594 | 0.10629607 | 1 | 4676 | tags=28%, list=20%, si | MM_1 |
| GOBP_REGULATION_OF_ | GOBP_REGULATION_OF_DNA_TEMPLATED_ | 84 | 0.312039 | 1.532842 | 0.022900764 | 0.10616702 | 1 | 7428 | tags=42%, list=31%, si | MM_1 |
| GOBP_MODULATION_BY_ | GOBP_MODULATION_BY_HOST_OF_SYMBIC | 79 | 0.311322 | 1.5320396 | 0.007490637 | 0.10663131 | 1 | 7591 | tags=53%, list=32%, si | MM_1 |
| GOBP_PROTEIN_LOCALIZ | GOBP_PROTEIN_LOCALIZATION_TO_CHROM | 31 | 0.397161 | 1.5311137 | 0.034883723 | 0.107050955 | 1 | 7653 | tags=45%, list=32%, si | MM_1 |
| GOBP_N_ACETYLGLUCOS | GOBP_N_ACETYLGLUCOSAMINE_METABOLI | 16 | 0.464383 | 1.5305438 | 0.045822103 | 0.107321516 | 1 | 4694 | tags=50%, list=20%, si | MM_1 |
| GOBP_ACTIN_POLYMERIZ | GOBP_ACTIN_POLYMERIZATION_OR_DEPOL | 187 | 0.269778 | 1.530195 | 0 | 0.107439585 | 1 | 6982 | tags=40%, list=30%, si | MM_1 |
| GOBP_RESPONSE_TO_INT | GOBP_RESPONSE_TO_INTERLEUKIN_7 | 17 | 0.464598 | 1.5300472 | 0.047887325 | 0.10737873 | 1 | 8333 | tags=76%, list=35%, si | MM_1 |
| GOBP_ASPARTATE_FAMI | GOBP_ASPARTATE_FAMILY_AMINO_ACID_M | 36 | 0.372022 | 1.5297798 | 0.036111113 | 0.10740469 | 1 | 5221 | tags=47%, list=22%, si | MM_1 |
| GOBP_NEGATIVE_REGUL | GOBP_NEGATIVE_REGULATION_OF_APOPTO | 241 | 0.258982 | 1.5296687 | 0 | 0.10733064 | 1 | 5157 | tags=30%, list=22%, si | MM_1 |

| | | | | | | | | | | |
|---|---|---|---|---|---|---|---|---|---|---|
| GOBP_RNA_EXPORT_FRO | GOBP_RNA_EXPORT_FROM_NUCLEUS | 78 | 0.318735 | 1.5293199 | 0.014234875 | 0.107444175 | 1 | 4443 | tags=23%, list=19%, si | MM_1 |
| GOBP_TRANSPORT_OF_V | GOBP_TRANSPORT_OF_VIRUS | 15 | 0.491536 | 1.5284785 | 0.055851065 | 0.107745305 | 1 | 4148 | tags=33%, list=18%, si | MM_1 |
| GOBP_MEMBRANE_LIPID | GOBP_MEMBRANE_LIPID_METABOLIC_PRO | 186 | 0.266732 | 1.5278826 | 0.005076142 | 0.107980944 | 1 | 4708 | tags=32%, list=20%, si | MM_1 |
| GOBP_MITOCHONDRIAL_ | GOBP_MITOCHONDRIAL_DNA_REPLICATION | 17 | 0.443298 | 1.5268717 | 0.040935673 | 0.10851899 | 1 | 2559 | tags=24%, list=11%, si | MM_1 |
| GOBP_DENDRITIC_SPINE | GOBP_DENDRITIC_SPINE_DEVELOPMENT | 133 | 0.285019 | 1.5267693 | 0.0040983604 | 0.10845977 | 1 | 3593 | tags=21%, list=15%, si | MM_1 |
| GOBP_TRANSFORMING_ | GOBP_TRANSFORMING_GROWTH_FACTOR_ | 35 | 0.376933 | 1.5260158 | 0.04761905 | 0.10886084 | 1 | 1371 | tags=20%, list=6%, sig | MM_1 |
| GOBP_REGULATION_OF_ | GOBP_REGULATION_OF_PROTEIN_BINDING | 188 | 0.272411 | 1.525392 | 0.005263158 | 0.10909742 | 1 | 4350 | tags=27%, list=18%, si | MM_1 |
| GOBP_FATTY_ACID_CATA | GOBP_FATTY_ACID_CATABOLIC_PROCESS | 100 | 0.30024 | 1.5244786 | 0.017921148 | 0.10958143 | 1 | 5379 | tags=40%, list=23%, si | MM_1 |
| GOBP_REGULATION_OF_ | GOBP_REGULATION_OF_PROTEIN_MATURA | 71 | 0.313566 | 1.5236678 | 0.0064102565 | 0.109978855 | 1 | 4305 | tags=32%, list=18%, si | MM_1 |
| GOBP_LIPOSACCHARIDE_ | GOBP_LIPOSACCHARIDE_METABOLIC_PROC | 93 | 0.300597 | 1.5236219 | 0.011627907 | 0.10986181 | 1 | 4439 | tags=33%, list=19%, si | MM_1 |
| GOBP_REGULATION_OF_ | GOBP_REGULATION_OF_STEM_CELL_DIFFE | 82 | 0.303372 | 1.5233535 | 0.010909091 | 0.10990683 | 1 | 4071 | tags=22%, list=17%, si | MM_1 |
| GOBP_NEGATIVE_REGUL | GOBP_NEGATIVE_REGULATION_OF_ALPHA_ | 47 | 0.353805 | 1.5231987 | 0.027777778 | 0.109877765 | 1 | 3667 | tags=30%, list=16%, si | MM_1 |
| GOBP_REGULATION_OF_ | GOBP_REGULATION_OF_TRANSLATIONAL_I | 17 | 0.446428 | 1.5230415 | 0.057534248 | 0.10986323 | 1 | 4734 | tags=29%, list=20%, si | MM_1 |
| SELENOCOMPOUND_ME | SELENOCOMPOUND_METABOLISM | 16 | 0.464188 | 1.5226588 | 0.05376344 | 0.109941766 | 1 | 5245 | tags=44%, list=22%, si | MM_1 |
| GOBP_ANGIOTENSIN_AC | GOBP_ANGIOTENSIN_ACTIVATED_SIGNALIN | 19 | 0.434036 | 1.5223168 | 0.039660055 | 0.110019535 | 1 | 1350 | tags=21%, list=6%, sig | MM_1 |
| GOBP_DNA_BIOSYNTHET | GOBP_DNA_BIOSYNTHETIC_PROCESS | 175 | 0.270967 | 1.5209371 | 0.0042735045 | 0.11076246 | 1 | 5994 | tags=35%, list=25%, si | MM_1 |
| GOBP_CELLULAR_RESPO | GOBP_CELLULAR_RESPONSE_TO_TOPOLOG | 95 | 0.303737 | 1.5203146 | 0 | 0.11108581 | 1 | 7328 | tags=43%, list=31%, si | MM_1 |
| GOBP_RIBOSOMAL_SMA | GOBP_RIBOSOMAL_SMALL_SUBUNIT_ASSE | 18 | 0.459077 | 1.520297 | 0.04896907 | 0.110946395 | 1 | 8508 | tags=78%, list=36%, si | MM_1 |
| GOBP_NEGATIVE_REGUL | GOBP_NEGATIVE_REGULATION_OF_SMALL_ | 61 | 0.325231 | 1.5198404 | 0.022727273 | 0.111143865 | 1 | 5723 | tags=38%, list=24%, si | MM_1 |
| GOBP_REGULATION_OF_ | GOBP_REGULATION_OF_ALCOHOL_BIOSYN | 48 | 0.341294 | 1.5187033 | 0.035820894 | 0.111797124 | 1 | 3272 | tags=27%, list=14%, si | MM_1 |
| GOBP_MORPHOGENESIS_ | GOBP_MORPHOGENESIS_OF_AN_ENDOTHE | 19 | 0.441249 | 1.5185158 | 0.049382716 | 0.111793496 | 1 | 6130 | tags=58%, list=26%, si | MM_1 |
| GOBP_RESPONSE_TO_MI | GOBP_RESPONSE_TO_MITOCHONDRIAL_DE | 21 | 0.409842 | 1.5185124 | 0.04644809 | 0.11165645 | 1 | 5631 | tags=43%, list=24%, si | MM_1 |
| GOBP_GLYCEROPHOSPH | GOBP_GLYCEROPHOSPHOLIPID_METABOLIC | 267 | 0.254727 | 1.5180959 | 0 | 0.11176992 | 1 | 6120 | tags=40%, list=26%, si | MM_1 |
| GOBP_REGULATION_OF_ | GOBP_REGULATION_OF_OXIDATIVE_STRESS | 39 | 0.359713 | 1.5177726 | 0.021943573 | 0.11185237 | 1 | 5276 | tags=41%, list=22%, si | MM_1 |
| GOBP_RESPONSE_TO_PR | GOBP_RESPONSE_TO_PROGESTERONE | 19 | 0.437608 | 1.5176475 | 0.042666666 | 0.11178013 | 1 | 4613 | tags=37%, list=20%, si | MM_1 |
| GOBP_ANTIGEN_PROCES | GOBP_ANTIGEN_PROCESSING_AND_PRESE | 105 | 0.29003 | 1.517463 | 0.016528925 | 0.111752145 | 1 | 5137 | tags=36%, list=22%, si | MM_1 |
| GOBP_VESICLE_ORGANIZ | GOBP_VESICLE_ORGANIZATION | 353 | 0.250067 | 1.5166936 | 0 | 0.1121256 | 1 | 5527 | tags=32%, list=23%, si | MM_1 |
| GOBP_NEGATIVE_REGUL | GOBP_NEGATIVE_REGULATION_OF_CHEMO | 29 | 0.399528 | 1.5166166 | 0.043010753 | 0.1120199 | 1 | 5016 | tags=41%, list=21%, si | MM_1 |
| GOBP_ACTIVATION_OF_ | GOBP_ACTIVATION_OF_PROTEIN_KINASE_A | 71 | 0.309534 | 1.5163455 | 0.020833334 | 0.11206539 | 1 | 3645 | tags=25%, list=15%, si | MM_1 |
| GOBP_MODULATION_BY_ | GOBP_MODULATION_BY_HOST_OF_VIRAL_ | 44 | 0.356222 | 1.5162685 | 0.021084337 | 0.111971 | 1 | 7591 | tags=55%, list=32%, si | MM_1 |
| GOBP_CELLULAR_RESPO | GOBP_CELLULAR_RESPONSE_TO_CADMIUM | 24 | 0.410288 | 1.5161604 | 0.04819277 | 0.11188493 | 1 | 4350 | tags=29%, list=18%, si | MM_1 |
| INSULIN_SECRETION | INSULIN_SECRETION | 81 | 0.310621 | 1.5157479 | 0.011450382 | 0.11202153 | 1 | 2569 | tags=23%, list=11%, si | MM_1 |
| GOBP_APOPTOTIC_PROC | GOBP_APOPTOTIC_PROCESS_INVOLVED_IN_ | 42 | 0.359223 | 1.515564 | 0.027863776 | 0.11201206 | 1 | 3907 | tags=36%, list=17%, si | MM_1 |
| GOBP_NUCLEAR_TRANSP | GOBP_NUCLEAR_TRANSPORT | 337 | 0.245714 | 1.5153741 | 0 | 0.11199571 | 1 | 5820 | tags=32%, list=25%, si | MM_1 |
| GOBP_MAINTENANCE_O | GOBP_MAINTENANCE_OF_PROTEIN_LOCAT | 97 | 0.30025 | 1.5149645 | 0 | 0.112133995 | 1 | 3360 | tags=25%, list=14%, si | MM_1 |
| GOBP_POSITIVE_REGULA | GOBP_POSITIVE_REGULATION_OF_T_CELL_ | 16 | 0.473673 | 1.5148987 | 0.04032258 | 0.11202894 | 1 | 4148 | tags=50%, list=18%, si | MM_1 |
| GOBP_POSITIVE_REGULA | GOBP_POSITIVE_REGULATION_OF_CHEMOK | 15 | 0.464882 | 1.5148174 | 0.04011461 | 0.111932755 | 1 | 1607 | tags=20%, list=7%, sig | MM_1 |
| CUSHING_SYNDROME | CUSHING_SYNDROME | 147 | 0.277866 | 1.5143781 | 0.008810572 | 0.11209691 | 1 | 2346 | tags=14%, list=10%, si | MM_1 |
| GOBP_STEROL_BIOSYNTH | GOBP_STEROL_BIOSYNTHETIC_PROCESS | 58 | 0.338602 | 1.5142957 | 0.029315962 | 0.11200364 | 1 | 2849 | tags=26%, list=12%, si | MM_1 |
| GOBP_NEGATIVE_REGUL | GOBP_NEGATIVE_REGULATION_OF_OSTEO | 37 | 0.365161 | 1.5130781 | 0.03416149 | 0.11266331 | 1 | 7797 | tags=57%, list=33%, si | MM_1 |
| GOBP_REGULATION_OF_ | GOBP_REGULATION_OF_GLIAL_CELL_MIGR | 26 | 0.401934 | 1.5128264 | 0.048991356 | 0.11265604 | 1 | 4990 | tags=35%, list=21%, si | MM_1 |
| GOBP_MELANOCYTE_DIF | GOBP_MELANOCYTE_DIFFERENTIATION | 26 | 0.405438 | 1.5125161 | 0.03846154 | 0.11271948 | 1 | 3856 | tags=35%, list=16%, si | MM_1 |
| GOBP_POSITIVE_REGULA | GOBP_POSITIVE_REGULATION_OF_CYSTEIN | 130 | 0.288228 | 1.5125136 | 0.0046296297 | 0.11257698 | 1 | 3186 | tags=21%, list=13%, si | MM_1 |
| GOBP_POSITIVE_REGULA | GOBP_POSITIVE_REGULATION_OF_MICROT | 34 | 0.372275 | 1.5103124 | 0.047169812 | 0.113894664 | 1 | 7406 | tags=56%, list=31%, si | MM_1 |
| GOBP_REGULATION_OF_ | GOBP_REGULATION_OF_DNA_DAMAGE_RE | 36 | 0.358943 | 1.5088377 | 0.040697675 | 0.11485979 | 1 | 2785 | tags=19%, list=12%, si | MM_1 |
| GOBP_LONG_CHAIN_FAT | GOBP_LONG_CHAIN_FATTY_ACID_IMPORT_ | 19 | 0.428398 | 1.5088097 | 0.056497175 | 0.114726074 | 1 | 4633 | tags=42%, list=20%, si | MM_1 |
| GOBP_NEGATIVE_REGUL | GOBP_NEGATIVE_REGULATION_OF_VIRAL_ | 95 | 0.299109 | 1.5083234 | 0.013377926 | 0.11492392 | 1 | 5788 | tags=38%, list=25%, si | MM_1 |
| GOBP_SPLICEOSOMAL_C | GOBP_SPLICEOSOMAL_COMPLEX_ASSEMBL | 49 | 0.336814 | 1.5072391 | 0.025236594 | 0.11546072 | 1 | 13104 | tags=96%, list=55%, si | MM_1 |
| GOBP_REGULATION_OF_ | GOBP_REGULATION_OF_INTRACELLULAR_E | 34 | 0.3671 | 1.5065384 | 0.035714287 | 0.115715474 | 1 | 5359 | tags=35%, list=23%, si | MM_1 |
| GOBP_REGULATION_OF_ | GOBP_REGULATION_OF_VASCULATURE_DE | 295 | 0.252202 | 1.506516 | 0 | 0.115607955 | 1 | 4176 | tags=24%, list=18%, si | MM_1 |
| GOBP_DNA_STRAND_ELO | GOBP_DNA_STRAND_ELONGATION | 36 | 0.368993 | 1.506342 | 0.055718474 | 0.11559 | 1 | 6603 | tags=53%, list=28%, si | MM_1 |
| GOBP_NEGATIVE_REGUL | GOBP_NEGATIVE_REGULATION_OF_ORGAN | 375 | 0.241758 | 1.5058221 | 0 | 0.11584915 | 1 | 4129 | tags=22%, list=17%, si | MM_1 |
| GOBP_REGULATION_OF_ | GOBP_REGULATION_OF_DNA_TEMPLATED_ | 79 | 0.311212 | 1.5053626 | 0.024475524 | 0.11601735 | 1 | 4390 | tags=23%, list=19%, si | MM_1 |
| GOBP_REGULATION_OF_ | GOBP_REGULATION_OF_UBIQUITIN_PROTE | 39 | 0.366572 | 1.505133 | 0.031446543 | 0.1160357 | 1 | 5276 | tags=28%, list=22%, si | MM_1 |
| GOBP_RNA_LOCALIZATIO | GOBP_RNA_LOCALIZATION | 164 | 0.270716 | 1.5048856 | 0.004830918 | 0.11609953 | 1 | 7651 | tags=43%, list=32%, si | MM_1 |
| GOBP_POSITIVE_REGULA | GOBP_POSITIVE_REGULATION_OF_DNA_BI | 76 | 0.306899 | 1.5043894 | 0.014492754 | 0.11634184 | 1 | 4390 | tags=24%, list=19%, si | MM_1 |
| GOBP_AMINO_ACID_ME | GOBP_AMINO_ACID_METABOLIC_PROCESS | 245 | 0.250912 | 1.504048 | 0 | 0.116438344 | 1 | 5609 | tags=38%, list=24%, si | MM_1 |
| GOBP_MICROAUTOPHAG | GOBP_MICROAUTOPHAGY | 24 | 0.417956 | 1.503851 | 0.06267806 | 0.11641626 | 1 | 4799 | tags=38%, list=20%, si | MM_1 |
| GLYCEROLIPID_METABOL | GLYCEROLIPID_METABOLISM | 57 | 0.328536 | 1.5033565 | 0.013986014 | 0.11661394 | 1 | 4189 | tags=37%, list=18%, si | MM_1 |
| GOBP_POSITIVE_REGULA | GOBP_POSITIVE_REGULATION_OF_TRANSL | 21 | 0.415208 | 1.5029316 | 0.053072624 | 0.116742425 | 1 | 6019 | tags=33%, list=25%, si | MM_1 |
| GOBP_CELLULAR_RESPO | GOBP_CELLULAR_RESPONSE_TO_GAMMA_ | 26 | 0.399926 | 1.5027868 | 0.049079753 | 0.11670907 | 1 | 5752 | tags=46%, list=24%, si | MM_1 |
| GOBP_NEGATIVE_REGUL | GOBP_NEGATIVE_REGULATION_OF_HEMOP | 121 | 0.287048 | 1.5023263 | 0.011904762 | 0.1168974 | 1 | 3665 | tags=23%, list=16%, si | MM_1 |
| GOBP_NEGATIVE_REGUL | GOBP_NEGATIVE_REGULATION_OF_CELL_S | 59 | 0.335461 | 1.5009484 | 0.024767801 | 0.11763982 | 1 | 3945 | tags=32%, list=17%, si | MM_1 |
| GOBP_PHOSPHATIDYLIN | GOBP_PHOSPHATIDYLINOSITOL_METABOLI | 148 | 0.278449 | 1.5000238 | 0.0042918455 | 0.11812682 | 1 | 6120 | tags=40%, list=26%, si | MM_1 |
| GOBP_APOPTOTIC_PROC | GOBP_APOPTOTIC_PROCESS_INVOLVED_IN_ | 30 | 0.384016 | 1.4997811 | 0.04761905 | 0.11815412 | 1 | 3907 | tags=37%, list=17%, si | MM_1 |
| GOBP_NEGATIVE_REGUL | GOBP_NEGATIVE_REGULATION_OF_SUPRA | 173 | 0.266865 | 1.4993339 | 0 | 0.11833834 | 1 | 4129 | tags=23%, list=17%, si | MM_1 |
| GOBP_ENDOPLASMIC_RE | GOBP_ENDOPLASMIC_RETICULUM_ORGANI | 90 | 0.299401 | 1.4993163 | 0.008 | 0.11820355 | 1 | 5497 | tags=36%, list=23%, si | MM_1 |
| GOBP_ADULT_WALKING_ | GOBP_ADULT_WALKING_BEHAVIOR | 48 | 0.338087 | 1.4991314 | 0.03416149 | 0.118208505 | 1 | 4749 | tags=38%, list=20%, si | MM_1 |
| GOBP_ENDOSOME_TO_L | GOBP_ENDOSOME_TO_LYSOSOME_TRANSP | 68 | 0.318124 | 1.4982924 | 0.029032258 | 0.11869499 | 1 | 6329 | tags=43%, list=27%, si | MM_1 |
| GOBP_MITOCHONDRIAL_ | GOBP_MITOCHONDRIAL_CALCIUM_ION_HO | 29 | 0.399327 | 1.4982772 | 0.048850574 | 0.11856575 | 1 | 4686 | tags=38%, list=20%, si | MM_1 |
| GOBP_REGULATION_OF_ | GOBP_REGULATION_OF_ENDOPLASMIC_RE | 24 | 0.413872 | 1.497499 | 0.059490085 | 0.11893638 | 1 | 7694 | tags=50%, list=33%, si | MM_1 |
| MEASLES | MEASLES | 128 | 0.276605 | 1.4974939 | 0.0038461538 | 0.11879134 | 1 | 4704 | tags=27%, list=20%, si | MM_1 |
| GOBP_VESICLE_BUDDING | GOBP_VESICLE_BUDDING_FROM_MEMBRA | 79 | 0.299092 | 1.497046 | 0.007874016 | 0.11891818 | 1 | 6329 | tags=39%, list=27%, si | MM_1 |
| GOBP_REGULATION_OF_ | GOBP_REGULATION_OF_PROTEIN_SUMOYL | 25 | 0.407676 | 1.495782 | 0.049723756 | 0.119647995 | 1 | 5836 | tags=36%, list=25%, si | MM_1 |
| GOBP_PROTEIN_EXPORT | GOBP_PROTEIN_EXPORT_FROM_NUCLEUS | 64 | 0.320859 | 1.4952934 | 0.021978023 | 0.119885355 | 1 | 5607 | tags=39%, list=24%, si | MM_1 |
| GOBP_POSITIVE_REGULA | GOBP_POSITIVE_REGULATION_OF_GENE_E | 54 | 0.328339 | 1.4952059 | 0.029801324 | 0.11978621 | 1 | 4150 | tags=28%, list=18%, si | MM_1 |
| GOBP_NEGATIVE_REGUL | GOBP_NEGATIVE_REGULATION_OF_GTPASE | 22 | 0.412973 | 1.4947712 | 0.04597701 | 0.12001555 | 1 | 7651 | tags=59%, list=32%, si | MM_1 |
| GOBP_REGULATION_OF_ | GOBP_REGULATION_OF_CYTOPLASMIC_TRA | 30 | 0.386848 | 1.4939317 | 0.035820894 | 0.1204017 | 1 | 7694 | tags=47%, list=33%, si | MM_1 |
| GOBP_REGULATION_OF_ | GOBP_REGULATION_OF_CELL_PROJECTION_ | 205 | 0.257078 | 1.4938949 | 0.004854369 | 0.12027684 | 1 | 3584 | tags=21%, list=15%, si | MM_1 |
| GOBP_MATURATION_OF_ | GOBP_MATURATION_OF_SSU_RRNA_FROM | 33 | 0.366144 | 1.4934855 | 0.044510387 | 0.120444514 | 1 | 6496 | tags=48%, list=28%, si | MM_1 |
| GOBP_REGULATION_OF_ | GOBP_REGULATION_OF_CELLULAR_RESPON | 27 | 0.401607 | 1.4929467 | 0.048780486 | 0.1207275 | 1 | 7660 | tags=52%, list=32%, si | MM_1 |
| GOBP_REGULATION_OF_ | GOBP_REGULATION_OF_TYPE_2_IMMUNE_ | 34 | 0.370341 | 1.4929336 | 0.0397351 | 0.120592356 | 1 | 1728 | tags=21%, list=7%, sig | MM_1 |

| | | | | | | | | | | |
|---|---|---|---|---|---|---|---|---|---|---|
| GOBP_REGULATION_OF_ | GOBP_REGULATION_OF_PROTEIN_POLYME | 194 | 0.261129 | 1.492692 | 0 | 0.12062182 | 1 | 7480 | tags=45%, list=32%, si | MM_1 |
| GOBP_CELL_CYCLE_G2_M | GOBP_CELL_CYCLE_G2_M_PHASE_TRANSITI | 152 | 0.269137 | 1.4918832 | 0.00913242 | 0.12103512 | 1 | 2973 | tags=18%, list=13%, si | MM_1 |
| GOBP_CELLULAR_RESPON | GOBP_CELLULAR_RESPONSE_TO_REACTIVE_ | 136 | 0.274447 | 1.4917883 | 0.013452915 | 0.12095403 | 1 | 4350 | tags=25%, list=18%, si | MM_1 |
| GOBP_GLUCOSE_CATABO | GOBP_GLUCOSE_CATABOLIC_PROCESS | 26 | 0.396636 | 1.4908379 | 0.045698926 | 0.12150346 | 1 | 5343 | tags=46%, list=23%, si | MM_1 |
| GOBP_POLARIZED_EPITHI | GOBP_POLARIZED_EPITHELIAL_CELL_DIFFER | 25 | 0.393689 | 1.489612 | 0.05029586 | 0.122232355 | 1 | 3261 | tags=24%, list=14%, si | MM_1 |
| GOBP_SIGNAL_PEPTIDE_ | GOBP_SIGNAL_PEPTIDE_PROCESSING | 15 | 0.454849 | 1.4882091 | 0.06906077 | 0.12315432 | 1 | 5576 | tags=53%, list=24%, si | MM_1 |
| GOBP_NEGATIVE_REGUL | GOBP_NEGATIVE_REGULATION_OF_MONO | 28 | 0.393175 | 1.4880173 | 0.05 | 0.123158425 | 1 | 4900 | tags=46%, list=21%, si | MM_1 |
| GOBP_MITOCHONDRIAL_ | GOBP_MITOCHONDRIAL_RNA_PROCESSING | 20 | 0.422482 | 1.4872212 | 0.05319149 | 0.123566665 | 1 | 7395 | tags=55%, list=31%, si | MM_1 |
| GOBP_POSITIVE_REGULA | GOBP_POSITIVE_REGULATION_OF_MUSCLE | 43 | 0.338832 | 1.4864719 | 0.034700315 | 0.12398323 | 1 | 3553 | tags=23%, list=15%, si | MM_1 |
| GOBP_DETERMINATION_ | GOBP_DETERMINATION_OF_ADULT_LIFESP | 48 | 0.338179 | 1.4856291 | 0.037617553 | 0.12448246 | 1 | 3552 | tags=27%, list=15%, si | MM_1 |
| GOBP_CYTOSKELETON_D | GOBP_CYTOSKELETON_DEPENDENT_INTRA | 196 | 0.261262 | 1.4854201 | 0 | 0.124471754 | 1 | 5999 | tags=39%, list=25%, si | MM_1 |
| FC_GAMMA_R-MEDIATE | FC_GAMMA_R-MEDIATED_PHAGOCYTOSIS | 84 | 0.304372 | 1.4846156 | 0.021352313 | 0.12491942 | 1 | 8036 | tags=55%, list=34%, si | MM_1 |
| GOBP_RESPONSE_TO_CA | GOBP_RESPONSE_TO_CADMIUM_ION | 35 | 0.365695 | 1.484345 | 0.03785489 | 0.12493667 | 1 | 7221 | tags=49%, list=31%, si | MM_1 |
| GOBP_GLUCOSE_METAB | GOBP_GLUCOSE_METABOLIC_PROCESS | 201 | 0.253619 | 1.4832735 | 0 | 0.12555079 | 1 | 5157 | tags=34%, list=22%, si | MM_1 |
| GOBP_T_HELPER_1_CELL | GOBP_T_HELPER_1_CELL_DIFFERENTIATION | 28 | 0.383145 | 1.4823035 | 0.029940119 | 0.12606564 | 1 | 392 | tags=11%, list=2%, sig | MM_1 |
| GOBP_REGULATION_OF_ | GOBP_REGULATION_OF_MITOCHONDRIAL_ | 58 | 0.324029 | 1.4815156 | 0.020066889 | 0.12643772 | 1 | 5741 | tags=43%, list=24%, si | MM_1 |
| GOBP_REGULATION_OF_ | GOBP_REGULATION_OF_BINDING | 331 | 0.243206 | 1.4805789 | 0 | 0.12697269 | 1 | 4350 | tags=25%, list=18%, si | MM_1 |
| GOBP_RIBOSOMAL_SUBU | GOBP_RIBOSOMAL_SUBUNIT_EXPORT_FRO | 18 | 0.433588 | 1.4804825 | 0.05927835 | 0.12690356 | 1 | 6440 | tags=56%, list=27%, si | MM_1 |
| GOBP_RNA_SURVEILLAN | GOBP_RNA_SURVEILLANCE | 17 | 0.449847 | 1.4801755 | 0.05899705 | 0.12696368 | 1 | 4111 | tags=24%, list=17%, si | MM_1 |
| GOBP_REGULATION_OF_ | GOBP_REGULATION_OF_THYMOCYTE_APO | 18 | 0.435992 | 1.4799987 | 0.08469945 | 0.12693514 | 1 | 2627 | tags=33%, list=11%, si | MM_1 |
| GOBP_ASPARTATE_FAMI | GOBP_ASPARTATE_FAMILY_AMINO_ACID_ | 21 | 0.422669 | 1.4799947 | 0.061971832 | 0.12678857 | 1 | 2227 | tags=33%, list=9%, sig | MM_1 |
| GOBP_POSITIVE_REGULA | GOBP_POSITIVE_REGULATION_OF_DENDRIT | 67 | 0.307422 | 1.4798573 | 0.023890786 | 0.12673283 | 1 | 5665 | tags=33%, list=24%, si | MM_1 |
| GOBP_MYELOID_CELL_D | GOBP_MYELOID_CELL_DEVELOPMENT | 98 | 0.289312 | 1.4796993 | 0.017482517 | 0.12672195 | 1 | 4869 | tags=30%, list=21%, si | MM_1 |
| GOBP_REGULATION_OF_ | GOBP_REGULATION_OF_PROTEIN_CONTAIN | 420 | 0.234808 | 1.4792976 | 0 | 0.12688355 | 1 | 7480 | tags=44%, list=32%, si | MM_1 |
| GOBP_CELLULAR_RESPON | GOBP_CELLULAR_RESPONSE_TO_LIPOPROT | 38 | 0.356901 | 1.4791522 | 0.055045873 | 0.12684272 | 1 | 4844 | tags=39%, list=21%, si | MM_1 |
| GOBP_AMIDE_METABOLI | GOBP_AMIDE_METABOLIC_PROCESS | 398 | 0.238171 | 1.4782759 | 0 | 0.12735505 | 1 | 4187 | tags=28%, list=18%, si | MM_1 |
| GOBP_ACYL_COA_METAE | GOBP_ACYL_COA_METABOLIC_PROCESS | 82 | 0.295894 | 1.4779377 | 0.022813689 | 0.12745391 | 1 | 4804 | tags=37%, list=20%, si | MM_1 |
| GOBP_PROTEIN_TARGETI | GOBP_PROTEIN_TARGETING_TO_MEMBRAN | 116 | 0.280472 | 1.4778916 | 0.0072202166 | 0.12733255 | 1 | 6325 | tags=46%, list=27%, si | MM_1 |
| RIBOSOME_BIOGENESIS_ | RIBOSOME_BIOGENESIS_IN_EUKARYOTES | 76 | 0.305046 | 1.476459 | 0.020905923 | 0.12821126 | 1 | 6496 | tags=49%, list=28%, si | MM_1 |
| BETA-ALANINE_METABOL | BETA-ALANINE_METABOLISM | 29 | 0.391686 | 1.4764501 | 0.05015674 | 0.12806946 | 1 | 3635 | tags=38%, list=15%, si | MM_1 |
| GOBP_NEGATIVE_REGUL | GOBP_NEGATIVE_REGULATION_OF_CELL_P | 208 | 0.256418 | 1.4763044 | 0 | 0.12802799 | 1 | 4918 | tags=30%, list=21%, si | MM_1 |
| GOBP_SYNAPTIC_VESICLE | GOBP_SYNAPTIC_VESICLE_LUMEN_ACIDIFIC | 19 | 0.427882 | 1.476207 | 0.038123168 | 0.12794413 | 1 | 4815 | tags=47%, list=20%, si | MM_1 |
| GOBP_POSITIVE_REGULA | GOBP_POSITIVE_REGULATION_OF_EXTRINS | 60 | 0.314256 | 1.4759358 | 0.01607717 | 0.12795696 | 1 | 4502 | tags=30%, list=19%, si | MM_1 |
| GOBP_REGULATION_OF_ | GOBP_REGULATION_OF_ACTIVATED_T_CEL | 41 | 0.342503 | 1.4736463 | 0.040625 | 0.12949762 | 1 | 4703 | tags=34%, list=20%, si | MM_1 |
| GOBP_MAGNESIUM_ION | GOBP_MAGNESIUM_ION_HOMEOSTASIS | 15 | 0.459066 | 1.4732921 | 0.07277628 | 0.12957305 | 1 | 1924 | tags=33%, list=8%, sig | MM_1 |
| GOBP_CHAPERONE_COF | GOBP_CHAPERONE_COFACTOR_DEPENDEN | 31 | 0.376938 | 1.4732307 | 0.04956268 | 0.12947525 | 1 | 4963 | tags=35%, list=21%, si | MM_1 |
| GOBP_REGULATION_OF_ | GOBP_REGULATION_OF_LEUKOCYTE_CHEM | 122 | 0.282108 | 1.4730942 | 0.016 | 0.12943551 | 1 | 3736 | tags=27%, list=16%, si | MM_1 |
| GOBP_POSITIVE_REGULA | GOBP_POSITIVE_REGULATION_OF_PEPTIDA | 161 | 0.259061 | 1.4725097 | 0.009345794 | 0.12972833 | 1 | 3016 | tags=20%, list=13%, si | MM_1 |
| GOBP_RESPONSE_TO_AN | GOBP_RESPONSE_TO_ANGIOTENSIN | 33 | 0.379459 | 1.472487 | 0.048710603 | 0.12960136 | 1 | 1350 | tags=15%, list=6%, sig | MM_1 |
| GOBP_PROTEIN_LOCALIZ | GOBP_PROTEIN_LOCALIZATION_TO_LYSOSO | 54 | 0.325395 | 1.4722278 | 0.023880597 | 0.12961455 | 1 | 7174 | tags=52%, list=30%, si | MM_1 |
| GOBP_GLIAL_CELL_APOP | GOBP_GLIAL_CELL_APOPTOTIC_PROCESS | 20 | 0.414085 | 1.4721408 | 0.054441262 | 0.12949783 | 1 | 4754 | tags=45%, list=20%, si | MM_1 |
| GOBP_IMMUNOLOGICAL_ | GOBP_IMMUNOLOGICAL_SYNAPSE_FORMA | 18 | 0.435701 | 1.4716328 | 0.07027027 | 0.12976696 | 1 | 5138 | tags=44%, list=22%, si | MM_1 |
| GOBP_PIGMENT_BIOSYN | GOBP_PIGMENT_BIOSYNTHETIC_PROCESS | 49 | 0.335052 | 1.471521 | 0.03521127 | 0.12968455 | 1 | 2869 | tags=31%, list=12%, si | MM_1 |
| GOBP_NUCLEOSIDE_BISP | GOBP_NUCLEOSIDE_BISPHOSPHATE_METAE | 102 | 0.284904 | 1.4712019 | 0.016194332 | 0.12979083 | 1 | 2454 | tags=23%, list=10%, si | MM_1 |
| GOBP_PODOSOME_ASSE | GOBP_PODOSOME_ASSEMBLY | 25 | 0.395149 | 1.4711752 | 0.0591716 | 0.12965229 | 1 | 6875 | tags=56%, list=29%, si | MM_1 |
| GOBP_POSITIVE_REGULA | GOBP_POSITIVE_REGULATION_OF_SMALL_ | 92 | 0.289671 | 1.4711014 | 0.010526316 | 0.12957753 | 1 | 1655 | tags=14%, list=7%, sig | MM_1 |
| GOBP_SUPEROXIDE_ANI | GOBP_SUPEROXIDE_ANION_GENERATION | 43 | 0.345979 | 1.4709836 | 0.041666668 | 0.12950847 | 1 | 4495 | tags=28%, list=19%, si | MM_1 |
| GOBP_REGULATION_OF_ | GOBP_REGULATION_OF_LYMPHOCYTE_CHE | 22 | 0.411909 | 1.4704827 | 0.060240965 | 0.12972526 | 1 | 3736 | tags=27%, list=16%, si | MM_1 |
| SNARE_INTERACTIONS_IN | SNARE_INTERACTIONS_IN_VESICULAR_TRA | 32 | 0.375968 | 1.470224 | 0.03880597 | 0.12977082 | 1 | 7570 | tags=59%, list=32%, si | MM_1 |
| GOBP_C_TERMINAL_PRO | GOBP_C_TERMINAL_PROTEIN_AMINO_ACID | 20 | 0.420511 | 1.4687049 | 0.053370785 | 0.13080887 | 1 | 5103 | tags=40%, list=22%, si | MM_1 |
| GOBP_GASTRULATION_W | GOBP_GASTRULATION_WITH_MOUTH_FOR | 34 | 0.364678 | 1.4685079 | 0.040935673 | 0.13080603 | 1 | 2510 | tags=24%, list=11%, si | MM_1 |
| GOBP_ACTIVATION_OF_P | GOBP_ACTIVATION_OF_PROTEIN_KINASE_B | 18 | 0.435599 | 1.4678528 | 0.0698324 | 0.13116512 | 1 | 3361 | tags=33%, list=14%, si | MM_1 |
| GOBP_PSEUDOURIDINE_S | GOBP_PSEUDOURIDINE_SYNTHESIS | 18 | 0.431593 | 1.4674009 | 0.06557377 | 0.13133965 | 1 | 13428 | tags=100%, list=57%, | MM_1 |
| GOBP_T_HELPER_2_CELL | GOBP_T_HELPER_2_CELL_DIFFERENTIATION | 21 | 0.415681 | 1.4665076 | 0.07082153 | 0.13186705 | 1 | 1728 | tags=19%, list=7%, sig | MM_1 |
| GOBP_LAMELLIPODIUM_ | GOBP_LAMELLIPODIUM_ASSEMBLY | 73 | 0.302883 | 1.4663534 | 0.018181818 | 0.1318686 | 1 | 6024 | tags=38%, list=26%, si | MM_1 |
| GOBP_ENDONUCLEOLYTI | GOBP_ENDONUCLEOLYTIC_CLEAVAGE_OF_ | 16 | 0.458243 | 1.466196 | 0.07837838 | 0.1317947 | 1 | 4575 | tags=38%, list=19%, si | MM_1 |
| GOBP_NEGATIVE_REGUL | GOBP_NEGATIVE_REGULATION_OF_LOCOM | 328 | 0.237052 | 1.4661605 | 0 | 0.13167048 | 1 | 4350 | tags=25%, list=18%, si | MM_1 |
| GOBP_POSITIVE_REGULA | GOBP_POSITIVE_REGULATION_OF_CELL_AD | 487 | 0.231335 | 1.4645835 | 0 | 0.13276695 | 1 | 3788 | tags=22%, list=16%, si | MM_1 |
| GOBP_RIBOSOMAL_LARG | GOBP_RIBOSOMAL_LARGE_SUBUNIT_ASSEM | 23 | 0.405087 | 1.4645177 | 0.0398773 | 0.13267085 | 1 | 6440 | tags=48%, list=27%, si | MM_1 |
| GOBP_PHAGOLYSOSOME | GOBP_PHAGOLYSOSOME_ASSEMBLY | 17 | 0.429946 | 1.4633228 | 0.05420054 | 0.13342838 | 1 | 6544 | tags=47%, list=28%, si | MM_1 |
| GOBP_NEGATIVE_REGUL | GOBP_NEGATIVE_REGULATION_OF_GLUCO | 20 | 0.43007 | 1.4631613 | 0.062322944 | 0.13337941 | 1 | 2420 | tags=25%, list=10%, si | MM_1 |
| GOBP_CGAS_STING_SIGN | GOBP_CGAS_STING_SIGNALING_PATHWAY | 16 | 0.448359 | 1.4621593 | 0.07417583 | 0.13394259 | 1 | 4350 | tags=38%, list=18%, si | MM_1 |
| GOBP_PHOSPHATIDYLIN | GOBP_PHOSPHATIDYLINOSITOL_DEPHOSPH | 29 | 0.374087 | 1.4608794 | 0.048913043 | 0.13478953 | 1 | 2348 | tags=21%, list=10%, si | MM_1 |
| GOBP_REGULATION_OF_ | GOBP_REGULATION_OF_AEROBIC_RESPIRAT | 33 | 0.357824 | 1.4605078 | 0.05345912 | 0.13491906 | 1 | 5877 | tags=45%, list=25%, si | MM_1 |
| GOBP_ALCOHOL_BIOSYN | GOBP_ALCOHOL_BIOSYNTHETIC_PROCESS | 129 | 0.270868 | 1.4602332 | 0.008474576 | 0.13498779 | 1 | 3272 | tags=22%, list=14%, si | MM_1 |
| GOBP_COPPER_ION_HON | GOBP_COPPER_ION_HOMEOSTASIS | 18 | 0.440918 | 1.4598794 | 0.0754717 | 0.13510914 | 1 | 5046 | tags=44%, list=21%, si | MM_1 |
| GOBP_REGULATION_OF_ | GOBP_REGULATION_OF_TOLL_LIKE_RECEPT | 28 | 0.380239 | 1.459879 | 0.0625 | 0.1349585 | 1 | 6948 | tags=57%, list=29%, si | MM_1 |
| GOBP_PORE_COMPLEX_A | GOBP_PORE_COMPLEX_ASSEMBLY | 20 | 0.415188 | 1.4596875 | 0.05691057 | 0.13495322 | 1 | 5553 | tags=35%, list=24%, si | MM_1 |
| GOBP_MODULATION_BY_ | GOBP_MODULATION_BY_SYMBIONT_OF_EN | 56 | 0.323269 | 1.4593413 | 0.03154574 | 0.13508898 | 1 | 3076 | tags=30%, list=13%, si | MM_1 |
| GOBP_NEURON_PROJECT | GOBP_NEURON_PROJECTION_EXTENSION_I | 19 | 0.417492 | 1.4593186 | 0.057377048 | 0.13495795 | 1 | 4776 | tags=42%, list=20%, si | MM_1 |
| GOBP_SIGNAL_TRANSDU | GOBP_SIGNAL_TRANSDUCTION_IN_RESPON | 176 | 0.257908 | 1.4592556 | 0 | 0.13488418 | 1 | 4335 | tags=24%, list=18%, si | MM_1 |
| GOBP_PARAXIAL_MESOD | GOBP_PARAXIAL_MESODERM_DEVELOPME | 21 | 0.40803 | 1.4582839 | 0.08571429 | 0.1354996 | 1 | 4164 | tags=38%, list=18%, si | MM_1 |
| GOBP_RUFFLE_ASSEMBLY | GOBP_RUFFLE_ASSEMBLY | 43 | 0.3375 | 1.4579644 | 0.043613706 | 0.13560264 | 1 | 5915 | tags=42%, list=25%, si | MM_1 |
| GOBP_MANNOSYLATION | GOBP_MANNOSYLATION | 22 | 0.403775 | 1.457779 | 0.05882353 | 0.1356142 | 1 | 6077 | tags=55%, list=26%, si | MM_1 |
| GOBP_MICROTUBULE_OF | GOBP_MICROTUBULE_ORGANIZING_CENTE | 34 | 0.357181 | 1.4577364 | 0.051829267 | 0.13549745 | 1 | 6174 | tags=35%, list=26%, si | MM_1 |
| GOBP_NEGATIVE_REGUL | GOBP_NEGATIVE_REGULATION_OF_MEIOTI | 21 | 0.419611 | 1.4577278 | 0.0694864 | 0.13535506 | 1 | 2456 | tags=29%, list=10%, si | MM_1 |
| GOBP_ENDOTHELIAL_CEL | GOBP_ENDOTHELIAL_CELL_PROLIFERATION | 164 | 0.260769 | 1.4575791 | 0.004608295 | 0.13533065 | 1 | 3905 | tags=23%, list=17%, si | MM_1 |
| GOBP_REGULATION_OF_ | GOBP_REGULATION_OF_GTPASE_ACTIVITY | 287 | 0.242844 | 1.4573495 | 0 | 0.13534659 | 1 | 5502 | tags=29%, list=23%, si | MM_1 |
| GOBP_REGULATION_OF_ | GOBP_REGULATION_OF_NUCLEAR_TRANSC | 15 | 0.457346 | 1.4568361 | 0.07821229 | 0.13558133 | 1 | 12819 | tags=100%, list=54%, | MM_1 |
| GOBP_REGULATION_OF_ | GOBP_REGULATION_OF_MEMBRANE_PERM | 70 | 0.304301 | 1.4564997 | 0.01724138 | 0.13569076 | 1 | 5741 | tags=40%, list=24%, si | MM_1 |
| GOBP_REGULATION_OF_ | GOBP_REGULATION_OF_ANOIKIS | 22 | 0.408024 | 1.4564301 | 0.05929919 | 0.13560088 | 1 | 2999 | tags=27%, list=13%, si | MM_1 |
| MRNA_SURVEILLANCE_P | MRNA_SURVEILLANCE_PATHWAY | 87 | 0.294812 | 1.4558709 | 0.010714286 | 0.13588983 | 1 | 4789 | tags=25%, list=20%, si | MM_1 |

| | | | | | | | | | | |
|---|---|---|---|---|---|---|---|---|---|---|
| GOBP_INFLAMMATORY_( | GOBP_INFLAMMATORY_CELL_APOPTOTIC_I | 28 | 0.383382 | 1.4553264 | 0.057803467 | 0.13615625 | 1 | 658 | tags=11%, list=3%, sig | MM_1 |
| GOBP_NUCLEOLUS_ORGA | GOBP_NUCLEOLUS_ORGANIZATION | 17 | 0.431496 | 1.4542831 | 0.068767905 | 0.1368249 | 1 | 5783 | tags=41%, list=24%, si | MM_1 |
| GOBP_TRICARBOXYLIC_A | GOBP_TRICARBOXYLIC_ACID_METABOLIC_P | 16 | 0.442292 | 1.4541976 | 0.06451613 | 0.13671751 | 1 | 6541 | tags=63%, list=28%, si | MM_1 |
| GOBP_POSITIVE_REGULA | GOBP_POSITIVE_REGULATION_OF_REACTIV | 81 | 0.29663 | 1.4537746 | 0.0114068445 | 0.13690978 | 1 | 4228 | tags=27%, list=18%, si | MM_1 |
| PANTOTHENATE_AND_CO | PANTOTHENATE_AND_COA_BIOSYNTHESIS | 17 | 0.442997 | 1.4532958 | 0.06371191 | 0.13713527 | 1 | 2454 | tags=35%, list=10%, si | MM_1 |
| GOBP_POSITIVE_REGULA | GOBP_POSITIVE_REGULATION_OF_CELLULA | 227 | 0.246393 | 1.4529605 | 0 | 0.1372567 | 1 | 7575 | tags=46%, list=32%, si | MM_1 |
| GOBP_REGULATION_OF_ | GOBP_REGULATION_OF_FIBROBLAST_PROL | 122 | 0.274541 | 1.4522433 | 0.030434782 | 0.13764693 | 1 | 3645 | tags=23%, list=15%, si | MM_1 |
| GOBP_RESPIRATORY_BUF | GOBP_RESPIRATORY_BURST | 27 | 0.393284 | 1.452032 | 0.056179777 | 0.13768172 | 1 | 3905 | tags=33%, list=17%, si | MM_1 |
| GOBP_AMINO_ACID_CAT | GOBP_AMINO_ACID_CATABOLIC_PROCESS | 85 | 0.292824 | 1.4513309 | 0.0234375 | 0.13803543 | 1 | 5573 | tags=45%, list=24%, si | MM_1 |
| GOBP_POSITIVE_REGULA | GOBP_POSITIVE_REGULATION_OF_DEPHOS | 53 | 0.314687 | 1.4512883 | 0.046666667 | 0.137911 | 1 | 3205 | tags=23%, list=14%, si | MM_1 |
| GOBP_PROTEIN_TARGETI | GOBP_PROTEIN_TARGETING_TO_LYSOSOMI | 29 | 0.378339 | 1.4492394 | 0.05263158 | 0.13932551 | 1 | 7059 | tags=48%, list=30%, si | MM_1 |
| GOBP_REGULATION_OF_ | GOBP_REGULATION_OF_NON_MOTILE_CILI | 15 | 0.442253 | 1.4474893 | 0.080729164 | 0.14050078 | 1 | 2405 | tags=33%, list=10%, si | MM_1 |
| GOBP_CELLULAR_RESPON | GOBP_CELLULAR_RESPONSE_TO_AMYLOID_ | 33 | 0.359026 | 1.4461465 | 0.060126584 | 0.14144129 | 1 | 4459 | tags=36%, list=19%, si | MM_1 |
| GOBP_NEGATIVE_REGUL | GOBP_NEGATIVE_REGULATION_OF_PROTEI | 93 | 0.288889 | 1.4451847 | 0.013793103 | 0.14210992 | 1 | 3115 | tags=18%, list=13%, si | MM_1 |
| GOBP_MYOBLAST_FUSIO | GOBP_MYOBLAST_FUSION | 45 | 0.338499 | 1.4437475 | 0.05944056 | 0.14310229 | 1 | 3710 | tags=29%, list=16%, si | MM_1 |
| GOBP_GLYCEROLIPID_ME | GOBP_GLYCEROLIPID_METABOLIC_PROCESS | 361 | 0.235257 | 1.4420035 | 0 | 0.1444438 | 1 | 6120 | tags=38%, list=26%, si | MM_1 |
| GOBP_NEGATIVE_REGUL | GOBP_NEGATIVE_REGULATION_OF_CELL_M | 36 | 0.357606 | 1.4417226 | 0.060509555 | 0.14452614 | 1 | 4325 | tags=36%, list=18%, si | MM_1 |
| GOBP_LATE_ENDOSOME_ | GOBP_LATE_ENDOSOME_TO_VACUOLE_TR | 16 | 0.446112 | 1.4401295 | 0.07506702 | 0.14562802 | 1 | 7282 | tags=50%, list=31%, si | MM_1 |
| GOBP_NEGATIVE_REGUL | GOBP_NEGATIVE_REGULATION_OF_ESTABL | 148 | 0.259479 | 1.4399596 | 0.0038610038 | 0.14560868 | 1 | 3461 | tags=22%, list=15%, si | MM_1 |
| GOBP_REGULATION_OF_ | GOBP_REGULATION_OF_CELLULAR_COMPO | 380 | 0.234496 | 1.4398497 | 0 | 0.1455443 | 1 | 5527 | tags=29%, list=23%, si | MM_1 |
| GOBP_LEUKOCYTE_CELL_ | GOBP_LEUKOCYTE_CELL_CELL_ADHESION | 390 | 0.22977 | 1.4396297 | 0 | 0.14556895 | 1 | 3788 | tags=21%, list=16%, si | MM_1 |
| GOBP_CELL_REDOX_HON | GOBP_CELL_REDOX_HOMEOSTASIS | 28 | 0.385723 | 1.4396261 | 0.05322129 | 0.14541775 | 1 | 3825 | tags=32%, list=16%, si | MM_1 |
| GOBP_DETOXIFICATION | GOBP_DETOXIFICATION | 70 | 0.303314 | 1.4386064 | 0.029508196 | 0.14613417 | 1 | 4661 | tags=34%, list=20%, si | MM_1 |
| GOBP_MACROPHAGE_AC | GOBP_MACROPHAGE_ACTIVATION | 107 | 0.273969 | 1.4382874 | 0.011764706 | 0.1462436 | 1 | 4831 | tags=31%, list=20%, si | MM_1 |
| GOBP_NEUROMUSCULAR | GOBP_NEUROMUSCULAR_PROCESS_CONTR | 69 | 0.303103 | 1.4379886 | 0.03514377 | 0.1463207 | 1 | 5046 | tags=33%, list=21%, si | MM_1 |
| GOBP_REGULATION_OF_ | GOBP_REGULATION_OF_LYMPHOCYTE_DIFF | 216 | 0.244482 | 1.4378258 | 0 | 0.1462964 | 1 | 3970 | tags=25%, list=17%, si | MM_1 |
| GOBP_INTRINSIC_APOPT | GOBP_INTRINSIC_APOPTOTIC_SIGNALING_I | 59 | 0.312494 | 1.4377309 | 0.049833886 | 0.14620502 | 1 | 6227 | tags=41%, list=26%, si | MM_1 |
| GOBP_NECROPTOTIC_PR | GOBP_NECROPTOTIC_PROCESS | 37 | 0.353401 | 1.4376985 | 0.043227665 | 0.14607242 | 1 | 3909 | tags=30%, list=17%, si | MM_1 |
| GOBP_MICROTUBULE_PO | GOBP_MICROTUBULE_POLYMERIZATION_O | 133 | 0.265564 | 1.4366976 | 0.004385965 | 0.14672337 | 1 | 3778 | tags=26%, list=16%, si | MM_1 |
| GOBP_POSITIVE_REGULA | GOBP_POSITIVE_REGULATION_OF_LEUKOC | 97 | 0.276898 | 1.4364327 | 0.02930403 | 0.14682639 | 1 | 3736 | tags=26%, list=16%, si | MM_1 |
| GOBP_ACTIN_FILAMENT_ | GOBP_ACTIN_FILAMENT_ORGANIZATION | 450 | 0.226833 | 1.4363965 | 0 | 0.14670028 | 1 | 6473 | tags=35%, list=27%, si | MM_1 |
| GOBP_POSITIVE_REGULA | GOBP_POSITIVE_REGULATION_OF_SUBSTR | 40 | 0.337401 | 1.4342862 | 0.044510387 | 0.14839694 | 1 | 6024 | tags=40%, list=26%, si | MM_1 |
| GOBP_INTERFERON_BETA | GOBP_INTERFERON_BETA_PRODUCTION | 66 | 0.304411 | 1.434227 | 0.031746034 | 0.14828719 | 1 | 6948 | tags=42%, list=29%, si | MM_1 |
| GOBP_CELLULAR_RESPON | GOBP_CELLULAR_RESPONSE_TO_LIGHT_STI | 106 | 0.276946 | 1.4335092 | 0.0038610038 | 0.14869802 | 1 | 4715 | tags=29%, list=20%, si | MM_1 |
| GOBP_PROTEIN_POLYUBI | GOBP_PROTEIN_POLYUBIQUITINATION | 250 | 0.24549 | 1.4333247 | 0.0057803467 | 0.14867657 | 1 | 7575 | tags=45%, list=32%, si | MM_1 |
| GOBP_MYELOID_LEUKOC | GOBP_MYELOID_LEUKOCYTE_MIGRATION | 222 | 0.246052 | 1.4323409 | 0 | 0.14930947 | 1 | 3779 | tags=23%, list=16%, si | MM_1 |
| GOBP_MEMBRANE_DOCI | GOBP_MEMBRANE_DOCKING | 78 | 0.294999 | 1.4322112 | 0.026143791 | 0.14927532 | 1 | 3121 | tags=22%, list=13%, si | MM_1 |
| GOBP_POSITIVE_REGULA | GOBP_POSITIVE_REGULATION_OF_CELLULA | 35 | 0.354921 | 1.4318676 | 0.051359516 | 0.14945321 | 1 | 2566 | tags=26%, list=11%, si | MM_1 |
| GOBP_NEGATIVE_REGUL | GOBP_NEGATIVE_REGULATION_OF_TYPE_I_ | 23 | 0.396836 | 1.4315896 | 0.07065217 | 0.14951383 | 1 | 4676 | tags=39%, list=20%, si | MM_1 |
| GOBP_NEGATIVE_REGUL | GOBP_NEGATIVE_REGULATION_OF_OXIDAT | 26 | 0.38132 | 1.4314367 | 0.07536232 | 0.14950238 | 1 | 5276 | tags=42%, list=22%, si | MM_1 |
| GOBP_POSITIVE_REGULA | GOBP_POSITIVE_REGULATION_OF_CELL_AD | 21 | 0.398096 | 1.430848 | 0.07506702 | 0.14987427 | 1 | 3645 | tags=29%, list=15%, si | MM_1 |
| GOBP_REGULATION_OF_ | GOBP_REGULATION_OF_RRNA_PROCESSIN | 16 | 0.434095 | 1.4306561 | 0.08055556 | 0.1498791 | 1 | 7089 | tags=56%, list=30%, si | MM_1 |
| GOBP_POSITIVE_REGULA | GOBP_POSITIVE_REGULATION_OF_PROTEIN | 100 | 0.277063 | 1.4306386 | 0.026717557 | 0.14972888 | 1 | 5638 | tags=36%, list=24%, si | MM_1 |
| GOBP_QUINONE_METAB | GOBP_QUINONE_METABOLIC_PROCESS | 23 | 0.388793 | 1.428495 | 0.06896552 | 0.15141033 | 1 | 5824 | tags=57%, list=25%, si | MM_1 |
| GOBP_POSITIVE_REGULA | GOBP_POSITIVE_REGULATION_OF_ENDOTH | 15 | 0.441586 | 1.4275203 | 0.08743169 | 0.15212959 | 1 | 6406 | tags=53%, list=27%, si | MM_1 |
| GOBP_POSITIVE_REGULA | GOBP_POSITIVE_REGULATION_OF_RNA_SPI | 50 | 0.318406 | 1.427145 | 0.055172414 | 0.1523332 | 1 | 8447 | tags=50%, list=36%, si | MM_1 |
| GOBP_REGULATION_OF_ | GOBP_REGULATION_OF_GLUCOSE_IMPORT | 66 | 0.299892 | 1.4268352 | 0.02919708 | 0.15245144 | 1 | 5114 | tags=33%, list=22%, si | MM_1 |
| GOBP_REGULATION_OF_ | GOBP_REGULATION_OF_RESPONSE_TO_NU | 34 | 0.355993 | 1.4265882 | 0.041791044 | 0.15250869 | 1 | 2535 | tags=26%, list=11%, si | MM_1 |
| GOBP_HEMATOPOIETIC_ | GOBP_HEMATOPOIETIC_STEM_CELL_DIFFER | 31 | 0.365124 | 1.426553 | 0.045714285 | 0.15237452 | 1 | 3840 | tags=26%, list=16%, si | MM_1 |
| GOBP_RESPONSE_TO_DE | GOBP_RESPONSE_TO_DECREASED_OXYGEN | 228 | 0.245488 | 1.426469 | 0 | 0.15227184 | 1 | 5006 | tags=29%, list=21%, si | MM_1 |
| GOBP_REGULATION_OF_ | GOBP_REGULATION_OF_NUCLEOCYTOPLAS | 130 | 0.26873 | 1.4261532 | 0.011583012 | 0.1523828 | 1 | 5276 | tags=34%, list=22%, si | MM_1 |
| GOBP_NEGATIVE_REGUL | GOBP_NEGATIVE_REGULATION_OF_CELLUL | 51 | 0.316332 | 1.4259437 | 0.037414964 | 0.15241595 | 1 | 1728 | tags=16%, list=7%, sig | MM_1 |
| MELANOGENESIS | MELANOGENESIS | 91 | 0.27938 | 1.4257687 | 0.017421603 | 0.15241586 | 1 | 2346 | tags=16%, list=10%, si | MM_1 |
| GOBP_NON_MEMBRANE | GOBP_NON_MEMBRANE_BOUNDED_ORGA | 405 | 0.224568 | 1.4256749 | 0 | 0.15236463 | 1 | 4485 | tags=23%, list=19%, si | MM_1 |
| GOBP_CYTOKINESIS | GOBP_CYTOKINESIS | 160 | 0.258936 | 1.4253343 | 0 | 0.15248242 | 1 | 3524 | tags=23%, list=15%, si | MM_1 |
| GOBP_POSITIVE_REGULA | GOBP_POSITIVE_REGULATION_OF_ACTIVAT | 27 | 0.376288 | 1.4247802 | 0.06798867 | 0.15281026 | 1 | 4703 | tags=37%, list=20%, si | MM_1 |
| GOBP_INTRACELLULAR_P | GOBP_INTRACELLULAR_PROTEIN_TRANSME | 32 | 0.361338 | 1.424107 | 0.07317073 | 0.15330589 | 1 | 4970 | tags=38%, list=21%, si | MM_1 |
| GOBP_REGULATION_OF_ | GOBP_REGULATION_OF_CYCLIN_DEPENDEN | 67 | 0.300271 | 1.4238378 | 0.015576324 | 0.15336384 | 1 | 4713 | tags=30%, list=20%, si | MM_1 |
| GOBP_EXECUTION_PHAS | GOBP_EXECUTION_PHASE_OF_APOPTOSIS | 78 | 0.291483 | 1.420634 | 0.034129694 | 0.15606166 | 1 | 3550 | tags=26%, list=15%, si | MM_1 |
| GOBP_PHOSPHOLIPID_DE | GOBP_PHOSPHOLIPID_DEPHOSPHORYLATIO | 43 | 0.333378 | 1.4198802 | 0.06382979 | 0.15657221 | 1 | 2411 | tags=23%, list=10%, si | MM_1 |
| GOBP_PLASMA_MEMBRA | GOBP_PLASMA_MEMBRANE_REPAIR | 30 | 0.367632 | 1.4197065 | 0.07232705 | 0.15656702 | 1 | 3049 | tags=30%, list=13%, si | MM_1 |
| GOBP_MYELOID_CELL_DI | GOBP_MYELOID_CELL_DIFFERENTIATION | 455 | 0.224083 | 1.4196154 | 0 | 0.1564992 | 1 | 5071 | tags=27%, list=21%, si | MM_1 |
| GOBP_PEPTIDYL_PROLINI | GOBP_PEPTIDYL_PROLINE_MODIFICATION | 39 | 0.339768 | 1.4196012 | 0.047477745 | 0.156352 | 1 | 5535 | tags=41%, list=23%, si | MM_1 |
| GOBP_POSITIVE_REGULA | GOBP_POSITIVE_REGULATION_OF_GTPASE_ | 203 | 0.249998 | 1.419454 | 0 | 0.1563346 | 1 | 5490 | tags=30%, list=23%, si | MM_1 |
| ENDOCRINE_AND_OTHER | ENDOCRINE_AND_OTHER_FACTOR-REGULAT | 42 | 0.333858 | 1.4194373 | 0.040133778 | 0.15619689 | 1 | 7093 | tags=52%, list=30%, si | MM_1 |
| GOBP_VESICLE_DOCKING | GOBP_VESICLE_DOCKING | 55 | 0.309449 | 1.4191628 | 0.03548387 | 0.1563063 | 1 | 3121 | tags=25%, list=13%, si | MM_1 |
| GOBP_IMMUNOGLOBULI | GOBP_IMMUNOGLOBULIN_PRODUCTION_II | 67 | 0.29323 | 1.4191253 | 0.024475524 | 0.15618172 | 1 | 4692 | tags=31%, list=20%, si | MM_1 |
| GOBP_REGULATION_OF_ | GOBP_REGULATION_OF_FATTY_ACID_OXID | 41 | 0.335477 | 1.4188579 | 0.050139274 | 0.15627114 | 1 | 5744 | tags=46%, list=24%, si | MM_1 |
| GOBP_TRIPARTITE_REGIO | GOBP_TRIPARTITE_REGIONAL_SUBDIVISION | 16 | 0.435444 | 1.4186599 | 0.09470752 | 0.15630156 | 1 | 4048 | tags=38%, list=17%, si | MM_1 |
| GOBP_FOAM_CELL_DIFFE | GOBP_FOAM_CELL_DIFFERENTIATION | 32 | 0.357637 | 1.4176489 | 0.066066064 | 0.1570502 | 1 | 1808 | tags=16%, list=8%, sig | MM_1 |
| GOBP_REGULATION_OF_ | GOBP_REGULATION_OF_MRNA_PROCESSIN | 131 | 0.262942 | 1.4169025 | 0.015873017 | 0.15755627 | 1 | 7719 | tags=39%, list=33%, si | MM_1 |
| GOBP_SEQUESTERING_O | GOBP_SEQUESTERING_OF_TRIGLYCERIDE | 18 | 0.407805 | 1.4162425 | 0.07084469 | 0.15799735 | 1 | 2253 | tags=28%, list=10%, si | MM_1 |
| HUMAN_PAPILLOMAVIRU | HUMAN_PAPILLOMAVIRUS_INFECTION | 315 | 0.231367 | 1.4156806 | 0 | 0.15836793 | 1 | 5188 | tags=28%, list=22%, si | MM_1 |
| GOBP_REGULATION_OF_ | GOBP_REGULATION_OF_CELL_CYCLE_G2_M | 117 | 0.270422 | 1.4151367 | 0.019157087 | 0.158716 | 1 | 2627 | tags=16%, list=11%, si | MM_1 |
| HERPES_SIMPLEX_VIRUS_ | HERPES_SIMPLEX_VIRUS_1_INFECTION | 373 | 0.227235 | 1.415097 | 0 | 0.15859425 | 1 | 5185 | tags=30%, list=22%, si | MM_1 |
| GOBP_RAC_PROTEIN_SIG | GOBP_RAC_PROTEIN_SIGNAL_TRANSDUCTI | 58 | 0.305185 | 1.4149938 | 0.05015674 | 0.15850975 | 1 | 5412 | tags=29%, list=23%, si | MM_1 |
| GOBP_ACTIVATION_OF_( | GOBP_ACTIVATION_OF_CYSTEINE_TYPE_EN | 81 | 0.286239 | 1.4141197 | 0.051194537 | 0.15912491 | 1 | 3186 | tags=22%, list=13%, si | MM_1 |
| GOBP_DEOXYRIBONUCLE | GOBP_DEOXYRIBONUCLEOTIDE_METABOLIC | 44 | 0.328924 | 1.4139 | 0.060606062 | 0.15917803 | 1 | 3155 | tags=25%, list=13%, si | MM_1 |
| GOBP_ALPHA_AMINO_AC | GOBP_ALPHA_AMINO_ACID_METABOLIC_P | 189 | 0.245796 | 1.4136289 | 0.004901961 | 0.15920076 | 1 | 5609 | tags=36%, list=24%, si | MM_1 |
| GOBP_POSITIVE_REGULA | GOBP_POSITIVE_REGULATION_OF_INTERFE | 45 | 0.326132 | 1.413499 | 0.057877813 | 0.15918766 | 1 | 4756 | tags=33%, list=20%, si | MM_1 |
| GOBP_REGULATION_OF_ | GOBP_REGULATION_OF_ANTIGEN_PROCESS | 21 | 0.391699 | 1.4128572 | 0.08695652 | 0.15957575 | 1 | 2720 | tags=24%, list=12%, si | MM_1 |
| GOBP_ENDOTHELIAL_CEL | GOBP_ENDOTHELIAL_CELL_APOPTOTIC_PRO | 58 | 0.309214 | 1.4125572 | 0.04262295 | 0.15967526 | 1 | 966 | tags=12%, list=4%, sig | MM_1 |

| | | | | | | | | | | |
|---|---|---|---|---|---|---|---|---|---|---|
| GOBP_REGULATION_OF_ | GOBP_REGULATION_OF_B_CELL_ACTIVATIO | 141 | 0.260012 | 1.4124823 | 0.02991453 | 0.15959658 | 1 | 5600 | tags=33%, list=24%, si | MM_1 |
| GOBP_SPHINGOID_META | GOBP_SPHINGOID_METABOLIC_PROCESS | 21 | 0.406858 | 1.4121943 | 0.07105263 | 0.15970087 | 1 | 2411 | tags=33%, list=10%, si | MM_1 |
| GOBP_ENDOTHELIAL_CEL | GOBP_ENDOTHELIAL_CELL_CHEMOTAXIS | 30 | 0.362074 | 1.4115423 | 0.056818184 | 0.16012745 | 1 | 4158 | tags=30%, list=18%, si | MM_1 |
| GOBP_TAXIS | GOBP_TAXIS | 411 | 0.22647 | 1.4113532 | 0 | 0.16013668 | 1 | 3786 | tags=23%, list=16%, si | MM_1 |
| GOBP_RESPONSE_TO_NU | GOBP_RESPONSE_TO_NUTRIENT_LEVELS | 371 | 0.228166 | 1.4108338 | 0 | 0.16043301 | 1 | 5976 | tags=33%, list=25%, si | MM_1 |
| GOBP_GLYCOPROTEIN_BI | GOBP_GLYCOPROTEIN_BIOSYNTHETIC_PRO( | 282 | 0.234333 | 1.4097188 | 0 | 0.16126513 | 1 | 3130 | tags=20%, list=13%, si | MM_1 |
| GOBP_ROOF_OF_MOUTH | GOBP_ROOF_OF_MOUTH_DEVELOPMENT | 95 | 0.272254 | 1.4092875 | 0.025547445 | 0.16156615 | 1 | 4620 | tags=25%, list=20%, si | MM_1 |
| GOBP_PROTEIN_N_LINKE | GOBP_PROTEIN_N_LINKED_GLYCOSYLATION | 18 | 0.41205 | 1.4085566 | 0.098314606 | 0.16207847 | 1 | 5996 | tags=39%, list=25%, si | MM_1 |
| GOBP_NEGATIVE_REGUL | GOBP_NEGATIVE_REGULATION_OF_NUCLE( | 20 | 0.406492 | 1.4083791 | 0.06267806 | 0.16209275 | 1 | 2521 | tags=25%, list=11%, si | MM_1 |
| GOBP_NEGATIVE_REGUL | GOBP_NEGATIVE_REGULATION_OF_CELL_M | 16 | 0.436354 | 1.4078557 | 0.09322034 | 0.16239934 | 1 | 3261 | tags=38%, list=14%, si | MM_1 |
| GOBP_PHOSPHATIDIC_AC | GOBP_PHOSPHATIDIC_ACID_METABOLIC_PI | 27 | 0.370372 | 1.4075491 | 0.04857143 | 0.16249399 | 1 | 4703 | tags=41%, list=20%, si | MM_1 |
| GOBP_REGULATION_OF_ | GOBP_REGULATION_OF_EXOSOMAL_SECRE | 17 | 0.413818 | 1.407053 | 0.08857143 | 0.16274446 | 1 | 8493 | tags=65%, list=36%, si | MM_1 |
| GOBP_REGULATION_OF_ | GOBP_REGULATION_OF_PROTEIN_LOCALIZ/ | 221 | 0.239049 | 1.4068748 | 0.005586592 | 0.16274658 | 1 | 5016 | tags=31%, list=21%, si | MM_1 |
| GOBP_REGULATION_OF_ | GOBP_REGULATION_OF_T_HELPER_CELL_D | 42 | 0.32622 | 1.4065846 | 0.04 | 0.16286755 | 1 | 2145 | tags=19%, list=9%, sig | MM_1 |
| GOBP_NUCLEAR_TRANSC | GOBP_NUCLEAR_TRANSCRIBED_MRNA_CAT | 118 | 0.267837 | 1.4063781 | 0.018691588 | 0.16290164 | 1 | 8084 | tags=47%, list=34%, si | MM_1 |
| GOBP_FATTY_ACID_BETA | GOBP_FATTY_ACID_BETA_OXIDATION | 75 | 0.28778 | 1.4050021 | 0.040145986 | 0.16398959 | 1 | 5741 | tags=43%, list=24%, si | MM_1 |
| GOBP_CELLULAR_RESPON | GOBP_CELLULAR_RESPONSE_TO_NUTRIENT | 38 | 0.346026 | 1.4043146 | 0.07058824 | 0.1643886 | 1 | 4754 | tags=29%, list=20%, si | MM_1 |
| GOBP_MEMBRANE_DEPO | GOBP_MEMBRANE_DEPOLARIZATION_DURI | 17 | 0.424325 | 1.4037385 | 0.08635098 | 0.16476238 | 1 | 5063 | tags=53%, list=21%, si | MM_1 |
| GOBP_MITOTIC_SISTER_( | GOBP_MITOTIC_SISTER_CHROMATID_SEGRI | 178 | 0.252213 | 1.4036566 | 0.023148147 | 0.1646858 | 1 | 3899 | tags=24%, list=17%, si | MM_1 |
| PENTOSE_AND_GLUCURO | PENTOSE_AND_GLUCURONATE_INTERCONV | 22 | 0.386645 | 1.4031807 | 0.070680626 | 0.16499361 | 1 | 3827 | tags=36%, list=16%, si | MM_1 |
| GOBP_NEGATIVE_REGUL | GOBP_NEGATIVE_REGULATION_OF_ORGAN | 47 | 0.32138 | 1.4030335 | 0.051446944 | 0.16494682 | 1 | 4452 | tags=30%, list=19%, si | MM_1 |
| HISTIDINE_METABOLISM | HISTIDINE_METABOLISM | 21 | 0.391053 | 1.4025089 | 0.10185185 | 0.16525914 | 1 | 3590 | tags=38%, list=15%, si | MM_1 |
| GOBP_ESTABLISHMENT_( | GOBP_ESTABLISHMENT_OF_PROTEIN_LOCA | 67 | 0.30327 | 1.4022559 | 0.05263158 | 0.16532202 | 1 | 6256 | tags=39%, list=26%, si | MM_1 |
| LONGEVITY_REGULATING | LONGEVITY_REGULATING_PATHWAY | 99 | 0.27382 | 1.4018688 | 0.023728814 | 0.16550495 | 1 | 4373 | tags=25%, list=19%, si | MM_1 |
| GOBP_CHROMATIN_REM | GOBP_CHROMATIN_REMODELING | 482 | 0.219149 | 1.400328 | 0 | 0.16682039 | 1 | 4491 | tags=22%, list=19%, si | MM_1 |
| GOBP_PROGRAMMED_N | GOBP_PROGRAMMED_NECROTIC_CELL_DE/ | 50 | 0.314215 | 1.4000918 | 0.053333335 | 0.16684428 | 1 | 3909 | tags=26%, list=17%, si | MM_1 |
| GOBP_GLAND_MORPHO( | GOBP_GLAND_MORPHOGENESIS | 148 | 0.258038 | 1.3991969 | 0.008810572 | 0.16746718 | 1 | 1688 | tags=14%, list=7%, sig | MM_1 |
| GOBP_MUSCLE_CELL_PR | GOBP_MUSCLE_CELL_PROLIFERATION | 262 | 0.236373 | 1.3988662 | 0.006535948 | 0.16758871 | 1 | 3945 | tags=23%, list=17%, si | MM_1 |
| GOBP_NEGATIVE_REGUL | GOBP_NEGATIVE_REGULATION_OF_BINDIN | 137 | 0.259435 | 1.3987756 | 0.008733625 | 0.16751038 | 1 | 4350 | tags=28%, list=18%, si | MM_1 |
| GOBP_REGULATION_OF_ | GOBP_REGULATION_OF_CELLULAR_RESPON | 89 | 0.277955 | 1.3987417 | 0.022222223 | 0.16736974 | 1 | 1890 | tags=16%, list=8%, sig | MM_1 |
| GOBP_REGULATION_OF_ | GOBP_REGULATION_OF_DNA_REPAIR | 219 | 0.240776 | 1.3984753 | 0.009433962 | 0.16749138 | 1 | 7400 | tags=44%, list=31%, si | MM_1 |
| GOBP_MRNA_TRANSPOR | GOBP_MRNA_TRANSPORT | 108 | 0.273754 | 1.3981644 | 0.015503876 | 0.16762689 | 1 | 5991 | tags=30%, list=25%, si | MM_1 |
| GOBP_SOMATIC_DIVERSI | GOBP_SOMATIC_DIVERSIFICATION_OF_IMN | 55 | 0.307405 | 1.3978751 | 0.038194444 | 0.16774382 | 1 | 4692 | tags=31%, list=20%, si | MM_1 |
| GOBP_POSITIVE_REGULA | GOBP_POSITIVE_REGULATION_OF_ALCOHO | 27 | 0.362374 | 1.3976012 | 0.06371191 | 0.16783732 | 1 | 3272 | tags=30%, list=14%, si | MM_1 |
| GOBP_ERYTHROCYTE_HO | GOBP_ERYTHROCYTE_HOMEOSTASIS | 163 | 0.247736 | 1.3970093 | 0.009756098 | 0.16829154 | 1 | 6390 | tags=37%, list=27%, si | MM_1 |
| GOBP_FATTY_ACID_ELON | GOBP_FATTY_ACID_ELONGATION | 15 | 0.436598 | 1.3966287 | 0.08743169 | 0.1684839 | 1 | 6273 | tags=60%, list=27%, si | MM_1 |
| GOBP_INSULIN_METABO | GOBP_INSULIN_METABOLIC_PROCESS | 18 | 0.412859 | 1.3955449 | 0.09366391 | 0.16932036 | 1 | 1542 | tags=22%, list=7%, sig | MM_1 |
| GOBP_MULTICELLULAR_( | GOBP_MULTICELLULAR_ORGANISM_GROW | 218 | 0.240995 | 1.3952578 | 0 | 0.16942677 | 1 | 2845 | tags=16%, list=12%, si | MM_1 |
| GOBP_INTEGRIN_MEDIAT | GOBP_INTEGRIN_MEDIATED_SIGNALING_P/ | 98 | 0.275735 | 1.3948456 | 0.030303031 | 0.16963795 | 1 | 3660 | tags=26%, list=16%, si | MM_1 |
| GOBP_CARBOHYDRATE_I | GOBP_CARBOHYDRATE_DERIVATIVE_TRANS | 76 | 0.290601 | 1.3941084 | 0.04609929 | 0.17018005 | 1 | 6079 | tags=42%, list=26%, si | MM_1 |
| GOBP_CERAMIDE_METAE | GOBP_CERAMIDE_METABOLIC_PROCESS | 94 | 0.283475 | 1.3938957 | 0.033210333 | 0.17024067 | 1 | 3727 | tags=30%, list=16%, si | MM_1 |
| GOBP_MAMMARY_GLAN | GOBP_MAMMARY_GLAND_DUCT_MORPHO | 40 | 0.332961 | 1.3935359 | 0.06460674 | 0.17045212 | 1 | 6223 | tags=45%, list=26%, si | MM_1 |
| GOBP_NEGATIVE_REGUL | GOBP_NEGATIVE_REGULATION_OF_CHROM | 96 | 0.273984 | 1.3933156 | 0.03984064 | 0.17046314 | 1 | 5750 | tags=34%, list=24%, si | MM_1 |
| GOBP_CELLULAR_RESPON | GOBP_CELLULAR_RESPONSE_TO_INSULIN_S | 207 | 0.23699 | 1.3928657 | 0.0054644807 | 0.17074803 | 1 | 2034 | tags=13%, list=9%, sig | MM_1 |
| GOBP_POSITIVE_REGULA | GOBP_POSITIVE_REGULATION_OF_NEURON | 15 | 0.437362 | 1.3928078 | 0.104 | 0.17064543 | 1 | 6043 | tags=40%, list=26%, si | MM_1 |
| GOBP_MYELOID_CELL_AF | GOBP_MYELOID_CELL_APOPTOTIC_PROCES | 45 | 0.313989 | 1.3919761 | 0.052941177 | 0.17129993 | 1 | 5386 | tags=33%, list=23%, si | MM_1 |
| GOBP_REGULATION_OF_ | GOBP_REGULATION_OF_FIBROBLAST_GROV | 33 | 0.341447 | 1.3914845 | 0.06940063 | 0.17162406 | 1 | 5598 | tags=36%, list=24%, si | MM_1 |
| GOBP_ALPHA_AMINO_AC | GOBP_ALPHA_AMINO_ACID_BIOSYNTHETIC | 54 | 0.30915 | 1.390693 | 0.053003535 | 0.17229721 | 1 | 5410 | tags=44%, list=23%, si | MM_1 |
| GOBP_REGULATION_OF_ | GOBP_REGULATION_OF_MACROPHAGE_AC | 57 | 0.294588 | 1.3902422 | 0.05862069 | 0.17254005 | 1 | 4831 | tags=37%, list=20%, si | MM_1 |
| GOBP_POSITIVE_REGULA | GOBP_POSITIVE_REGULATION_OF_ENDOTH | 99 | 0.272094 | 1.3901365 | 0.014869888 | 0.17249581 | 1 | 4035 | tags=25%, list=17%, si | MM_1 |
| GOBP_PROTEIN_LOCALIZ | GOBP_PROTEIN_LOCALIZATION_TO_CELL_C | 24 | 0.376928 | 1.3896832 | 0.07204611 | 0.17269954 | 1 | 3009 | tags=21%, list=13%, si | MM_1 |
| GOBP_VESICLE_TRANSPO | GOBP_VESICLE_TRANSPORT_ALONG_MICRO | 47 | 0.320013 | 1.3895657 | 0.0776699 | 0.17264669 | 1 | 5160 | tags=38%, list=22%, si | MM_1 |
| GOBP_EOSINOPHIL_CHEN | GOBP_EOSINOPHIL_CHEMOTAXIS | 22 | 0.387194 | 1.3892729 | 0.09116022 | 0.17273852 | 1 | 5850 | tags=50%, list=25%, si | MM_1 |
| GOBP_REGULATION_OF_ | GOBP_REGULATION_OF_EXTRINSIC_APOPT( | 161 | 0.246841 | 1.388705 | 0.008403362 | 0.17311281 | 1 | 3730 | tags=22%, list=16%, si | MM_1 |
| DNA_REPLICATION | DNA_REPLICATION | 34 | 0.34265 | 1.3886095 | 0.08383234 | 0.17305367 | 1 | 15532 | tags=100%, list=66%, | MM_1 |
| GOBP_SMALL_GTPASE_M | GOBP_SMALL_GTPASE_MEDIATED_SIGNAL_ | 434 | 0.21996 | 1.388397 | 0.008547009 | 0.17309919 | 1 | 4910 | tags=24%, list=21%, si | MM_1 |
| GOBP_ESTABLISHMENT_( | GOBP_ESTABLISHMENT_OF_PROTEIN_LOCA | 47 | 0.320529 | 1.3881025 | 0.061919503 | 0.1732326 | 1 | 6477 | tags=51%, list=27%, si | MM_1 |
| GOBP_VITAMIN_BIOSYNT | GOBP_VITAMIN_BIOSYNTHETIC_PROCESS | 25 | 0.369663 | 1.387813 | 0.08797654 | 0.17334624 | 1 | 2423 | tags=28%, list=10%, si | MM_1 |
| CYSTEINE_AND_METHION | CYSTEINE_AND_METHIONINE_METABOLISM | 44 | 0.323112 | 1.3872942 | 0.05732484 | 0.17371169 | 1 | 3803 | tags=27%, list=16%, si | MM_1 |
| GOBP_REGULATION_OF_ | GOBP_REGULATION_OF_CELL_GROWTH | 415 | 0.22182 | 1.3871101 | 0 | 0.17371085 | 1 | 4373 | tags=25%, list=19%, si | MM_1 |
| GOBP_REGULATION_OF_ | GOBP_REGULATION_OF_LEUKOCYTE_DIFFEI | 342 | 0.224219 | 1.3862959 | 0.006535948 | 0.1743388 | 1 | 3788 | tags=21%, list=16%, si | MM_1 |
| GOBP_LIPID_OXIDATION | GOBP_LIPID_OXIDATION | 118 | 0.266869 | 1.3862933 | 0.027667984 | 0.17417574 | 1 | 5744 | tags=38%, list=24%, si | MM_1 |
| GOBP_PROTEIN_LOCALIZ | GOBP_PROTEIN_LOCALIZATION_TO_NUCLEU | 326 | 0.225669 | 1.3859867 | 0.006756757 | 0.17430264 | 1 | 5175 | tags=28%, list=22%, si | MM_1 |
| PHOSPHATIDYLINOSITOL_ | PHOSPHATIDYLINOSITOL_SIGNALING_SYSTE | 94 | 0.275571 | 1.3857393 | 0.040892195 | 0.17437798 | 1 | 6234 | tags=35%, list=26%, si | MM_1 |
| GOBP_LEUKOCYTE_PROLI | GOBP_LEUKOCYTE_PROLIFERATION | 361 | 0.224972 | 1.3854915 | 0 | 0.17446423 | 1 | 3710 | tags=21%, list=16%, si | MM_1 |
| GOBP_LIPOPROTEIN_CAT | GOBP_LIPOPROTEIN_CATABOLIC_PROCESS | 17 | 0.42282 | 1.3844457 | 0.11716621 | 0.17533061 | 1 | 3927 | tags=35%, list=17%, si | MM_1 |
| GOBP_TISSUE_MIGRATIO | GOBP_TISSUE_MIGRATION | 314 | 0.225828 | 1.3839601 | 0.0068965517 | 0.17558244 | 1 | 3824 | tags=21%, list=16%, si | MM_1 |
| GOBP_CELLULAR_RESPON | GOBP_CELLULAR_RESPONSE_TO_RADIATIOI | 165 | 0.251938 | 1.3836995 | 0.018348623 | 0.17566739 | 1 | 4526 | tags=27%, list=19%, si | MM_1 |
| GOBP_CELLULAR_RESPON | GOBP_CELLULAR_RESPONSE_TO_HYDROGEI | 59 | 0.299968 | 1.3834121 | 0.053691275 | 0.17576641 | 1 | 2964 | tags=22%, list=13%, si | MM_1 |
| GOBP_REGULATION_OF_ | GOBP_REGULATION_OF_LEUKOCYTE_ADHES | 33 | 0.354894 | 1.3824241 | 0.077151336 | 0.17659776 | 1 | 4015 | tags=30%, list=17%, si | MM_1 |
| GOBP_REGULATION_OF_ | GOBP_REGULATION_OF_WNT_SIGNALING_ | 16 | 0.425602 | 1.3818733 | 0.101604275 | 0.17694318 | 1 | 3651 | tags=38%, list=15%, si | MM_1 |
| GOBP_RNA_CATABOLIC_ | GOBP_RNA_CATABOLIC_PROCESS | 278 | 0.231406 | 1.381294 | 0.006756757 | 0.17732762 | 1 | 5991 | tags=29%, list=25%, si | MM_1 |
| GOBP_INTEGRATED_STRE | GOBP_INTEGRATED_STRESS_RESPONSE_SIG | 23 | 0.381915 | 1.3805972 | 0.08888889 | 0.17780164 | 1 | 14602 | tags=100%, list=62%, | MM_1 |
| GOBP_CELLULAR_RESPON | GOBP_CELLULAR_RESPONSE_TO_ACID_CHE | 87 | 0.274263 | 1.3803377 | 0.036363635 | 0.17789984 | 1 | 4489 | tags=25%, list=19%, si | MM_1 |
| GOBP_REGULATION_OF_ | GOBP_REGULATION_OF_INTRACELLULAR_P | 168 | 0.249478 | 1.3800242 | 0.014354067 | 0.17798962 | 1 | 4756 | tags=30%, list=20%, si | MM_1 |
| GOBP_NEGATIVE_REGUL | GOBP_NEGATIVE_REGULATION_OF_PROTEI | 236 | 0.237274 | 1.3794435 | 0.0056497175 | 0.17838952 | 1 | 3461 | tags=19%, list=15%, si | MM_1 |
| GOBP_NEGATIVE_REGUL | GOBP_NEGATIVE_REGULATION_OF_INSULIN | 57 | 0.298576 | 1.3794003 | 0.06354515 | 0.17827299 | 1 | 1134 | tags=12%, list=5%, sig | MM_1 |
| GOBP_NEGATIVE_REGUL | GOBP_NEGATIVE_REGULATION_OF_CELL_C | 368 | 0.224932 | 1.37912 | 0 | 0.17835069 | 1 | 5569 | tags=31%, list=24%, si | MM_1 |
| GOBP_INNER_CELL_MAS! | GOBP_INNER_CELL_MASS_CELL_PROLIFERA | 16 | 0.419037 | 1.3789345 | 0.107344635 | 0.1783686 | 1 | 3522 | tags=38%, list=15%, si | MM_1 |
| GOBP_SISTER_CHROMAT | GOBP_SISTER_CHROMATID_SEGREGATION | 219 | 0.238646 | 1.3783181 | 0.005154639 | 0.17887662 | 1 | 3970 | tags=23%, list=17%, si | MM_1 |
| GOBP_REGULATION_OF_ | GOBP_REGULATION_OF_ADAPTIVE_IMMUN | 215 | 0.239046 | 1.3781072 | 0.012121212 | 0.17887227 | 1 | 4335 | tags=27%, list=18%, si | MM_1 |
| GOBP_POSITIVE_REGULA | GOBP_POSITIVE_REGULATION_OF_ALPHA_E | 27 | 0.360987 | 1.3776755 | 0.075418994 | 0.17914957 | 1 | 1310 | tags=19%, list=6%, sig | MM_1 |

| | | | | | | | | | | |
|---|---|---|---|---|---|---|---|---|---|---|
| TRYPTOPHAN_METABOLI | TRYPTOPHAN_METABOLISM | 43 | 0.316822 | 1.3776276 | 0.063253015 | 0.17902677 | 1 | 3280 | tags=30%, list=14%, si | MM_1 |
| GOBP_POSITIVE_REGULA | GOBP_POSITIVE_REGULATION_OF_LEUKOCY | 169 | 0.250202 | 1.3775353 | 0.022624435 | 0.17893249 | 1 | 3710 | tags=22%, list=16%, si | MM_1 |
| GOBP_MITOTIC_SPINDLE | GOBP_MITOTIC_SPINDLE_ASSEMBLY | 71 | 0.285773 | 1.37741 | 0.06690141 | 0.17890733 | 1 | 3883 | tags=25%, list=16%, si | MM_1 |
| GOBP_SMOOTH_MUSCLE | GOBP_SMOOTH_MUSCLE_CELL_MIGRATION | 115 | 0.258243 | 1.3770936 | 0.03305785 | 0.17906325 | 1 | 3261 | tags=22%, list=14%, si | MM_1 |
| GOBP_SPINDLE_ORGANIZ | GOBP_SPINDLE_ORGANIZATION | 190 | 0.238993 | 1.3757175 | 0.010471204 | 0.180273 | 1 | 6064 | tags=33%, list=26%, si | MM_1 |
| GOBP_ESTABLISHMENT_( | GOBP_ESTABLISHMENT_OR_MAINTENANCE | 223 | 0.23341 | 1.3756123 | 0.005154639 | 0.18018968 | 1 | 6174 | tags=33%, list=26%, si | MM_1 |
| GOBP_LAMELLIPODIUM_ | GOBP_LAMELLIPODIUM_ORGANIZATION | 92 | 0.269694 | 1.374869 | 0.040677965 | 0.1806654 | 1 | 5579 | tags=37%, list=24%, si | MM_1 |
| GOBP_CELLULAR_RESPON | GOBP_CELLULAR_RESPONSE_TO_HYPOXIA | 112 | 0.256885 | 1.3739724 | 0.046153847 | 0.18141781 | 1 | 5114 | tags=31%, list=22%, si | MM_1 |
| GOBP_WNT_SIGNALING_ | GOBP_WNT_SIGNALING_PATHWAY_PLANA | 38 | 0.330899 | 1.3737274 | 0.093247585 | 0.18147199 | 1 | 3731 | tags=29%, list=16%, si | MM_1 |
| GOBP_POSITIVE_REGULA | GOBP_POSITIVE_REGULATION_OF_MYOBLA | 45 | 0.310351 | 1.3735557 | 0.053412464 | 0.18145242 | 1 | 3970 | tags=29%, list=17%, si | MM_1 |
| GOBP_REGULATION_OF_ | GOBP_REGULATION_OF_ENDOCYTOSIS | 325 | 0.226137 | 1.3728911 | 0 | 0.18193725 | 1 | 4920 | tags=28%, list=21%, si | MM_1 |
| GOBP_ER_NUCLEUS_SIGN | GOBP_ER_NUCLEUS_SIGNALING_PATHWAY | 43 | 0.321229 | 1.372823 | 0.06603774 | 0.1818354 | 1 | 7694 | tags=44%, list=33%, si | MM_1 |
| GOBP_ENDOPLASMIC_RE | GOBP_ENDOPLASMIC_RETICULUM_TO_CYT | 27 | 0.360908 | 1.3727605 | 0.0922619 | 0.18174134 | 1 | 15099 | tags=100%, list=64%, | MM_1 |
| GOBP_POSITIVE_REGULA | GOBP_POSITIVE_REGULATION_OF_NUCLEO | 74 | 0.284179 | 1.3724474 | 0.044368602 | 0.1819101 | 1 | 4756 | tags=36%, list=20%, si | MM_1 |
| GOBP_PYRIMIDINE_NUCL | GOBP_PYRIMIDINE_NUCLEOSIDE_TRIPHOSP | 18 | 0.414136 | 1.3723247 | 0.109792285 | 0.18188854 | 1 | 3155 | tags=39%, list=13%, si | MM_1 |
| CHRONIC_MYELOID_LEUK | CHRONIC_MYELOID_LEUKEMIA | 75 | 0.278778 | 1.3719813 | 0.054237287 | 0.1820672 | 1 | 2974 | tags=19%, list=13%, si | MM_1 |
| CORTISOL_SYNTHESIS_AN | CORTISOL_SYNTHESIS_AND_SECRETION | 62 | 0.297517 | 1.3719662 | 0.04901961 | 0.18192211 | 1 | 2346 | tags=18%, list=10%, si | MM_1 |
| GOBP_PROTEIN_LOCALIZ | GOBP_PROTEIN_LOCALIZATION_TO_PLASM | 296 | 0.226833 | 1.3711516 | 0 | 0.18249853 | 1 | 5488 | tags=29%, list=23%, si | MM_1 |
| GOBP_CELLULAR_RESPON | GOBP_CELLULAR_RESPONSE_TO_GLUCOCO | 47 | 0.305779 | 1.3706896 | 0.055737704 | 0.18281573 | 1 | 1652 | tags=15%, list=7%, sig | MM_1 |
| GOBP_VESICLE_MEDIATE | GOBP_VESICLE_MEDIATED_TRANSPORT_TO | 158 | 0.246342 | 1.3701354 | 0.027027028 | 0.18325967 | 1 | 6955 | tags=41%, list=29%, si | MM_1 |
| GOBP_REGULATION_OF_ | GOBP_REGULATION_OF_DOUBLE_STRAND_ | 139 | 0.250043 | 1.370104 | 0.03773585 | 0.18312557 | 1 | 4360 | tags=24%, list=18%, si | MM_1 |
| GOBP_PLASMA_MEMBRA | GOBP_PLASMA_MEMBRANE_ORGANIZATIO | 149 | 0.250073 | 1.3700227 | 0.012987013 | 0.18301345 | 1 | 3185 | tags=21%, list=13%, si | MM_1 |
| PURINE_METABOLISM | PURINE_METABOLISM | 126 | 0.252882 | 1.3699719 | 0.028 | 0.18291329 | 1 | 6188 | tags=40%, list=26%, si | MM_1 |
| PROGESTERONE-MEDIATI | PROGESTERONE-MEDIATED_OOCYTE_MATU | 87 | 0.274975 | 1.3693029 | 0.043824703 | 0.18349431 | 1 | 4586 | tags=29%, list=19%, si | MM_1 |
| GOBP_POSITIVE_REGULA | GOBP_POSITIVE_REGULATION_OF_PHOSPH | 16 | 0.414924 | 1.3683611 | 0.11733333 | 0.1843216 | 1 | 5746 | tags=44%, list=24%, si | MM_1 |
| GOBP_CENTROSOME_DU | GOBP_CENTROSOME_DUPLICATION | 74 | 0.285557 | 1.3682578 | 0.035714287 | 0.18425362 | 1 | 5024 | tags=32%, list=21%, si | MM_1 |
| GOBP_POSITIVE_REGULA | GOBP_POSITIVE_REGULATION_OF_CELL_SU | 35 | 0.339105 | 1.3677437 | 0.0862069 | 0.18456192 | 1 | 6763 | tags=43%, list=29%, si | MM_1 |
| GOBP_GMP_BIOSYNTHET | GOBP_GMP_BIOSYNTHETIC_PROCESS | 15 | 0.42008 | 1.3670721 | 0.10743801 | 0.18503225 | 1 | 8135 | tags=67%, list=34%, si | MM_1 |
| GOBP_PROTEIN_LOCALIZ | GOBP_PROTEIN_LOCALIZATION_TO_NUCLE | 15 | 0.424263 | 1.365718 | 0.106770836 | 0.18616913 | 1 | 3238 | tags=27%, list=14%, si | MM_1 |
| GOBP_STRIATED_MUSCLE | GOBP_STRIATED_MUSCLE_CELL_APOPTOTIC | 73 | 0.274748 | 1.3645321 | 0.06122449 | 0.18724787 | 1 | 4733 | tags=29%, list=20%, si | MM_1 |
| GOBP_MICROTUBULE_PO | GOBP_MICROTUBULE_POLYMERIZATION | 89 | 0.272476 | 1.3644773 | 0.029411765 | 0.18714847 | 1 | 3778 | tags=25%, list=16%, si | MM_1 |
| GOBP_VESICLE_TARGETIN | GOBP_VESICLE_TARGETING_ROUGH_ER_TO | 20 | 0.402218 | 1.3643175 | 0.10597826 | 0.18716255 | 1 | 7182 | tags=50%, list=30%, si | MM_1 |
| GOBP_BLASTODERM_SEG | GOBP_BLASTODERM_SEGMENTATION | 26 | 0.365267 | 1.3642929 | 0.12011173 | 0.18702102 | 1 | 4048 | tags=31%, list=17%, si | MM_1 |
| GOBP_REGULATION_OF_ | GOBP_REGULATION_OF_PROTEIN_LOCALIZ | 158 | 0.245527 | 1.3642515 | 0.008230452 | 0.18690649 | 1 | 4468 | tags=26%, list=19%, si | MM_1 |
| GOBP_REGULATION_OF_ | GOBP_REGULATION_OF_EPITHELIAL_CELL_ | 238 | 0.230162 | 1.3639781 | 0.010416667 | 0.1870198 | 1 | 3824 | tags=21%, list=16%, si | MM_1 |
| GOBP_LEUKOCYTE_TETHE | GOBP_LEUKOCYTE_TETHERING_OR_ROLLIN | 35 | 0.327136 | 1.3635197 | 0.07886435 | 0.18736905 | 1 | 4015 | tags=26%, list=17%, si | MM_1 |
| GOBP_GLIOGENESIS | GOBP_GLIOGENESIS | 377 | 0.218815 | 1.3631865 | 0.008695652 | 0.18751192 | 1 | 3786 | tags=23%, list=16%, si | MM_1 |
| GOBP_TOLL_LIKE_RECEPT | GOBP_TOLL_LIKE_RECEPTOR_3_SIGNALING_ | 29 | 0.341541 | 1.3630221 | 0.09065156 | 0.1875668 | 1 | 5138 | tags=38%, list=22%, si | MM_1 |
| GOBP_T_CELL_DIFFERENT | GOBP_T_CELL_DIFFERENTIATION | 327 | 0.221354 | 1.3625891 | 0 | 0.1878693 | 1 | 3993 | tags=23%, list=17%, si | MM_1 |
| PROPANOATE_METABOLI | PROPANOATE_METABOLISM | 29 | 0.356387 | 1.3619272 | 0.08918919 | 0.18839315 | 1 | 6432 | tags=55%, list=27%, si | MM_1 |
| GOBP_POSITIVE_REGULA | GOBP_POSITIVE_REGULATION_OF_BINDING | 147 | 0.250902 | 1.3615822 | 0.016129032 | 0.18858528 | 1 | 5276 | tags=31%, list=22%, si | MM_1 |
| GOBP_BLASTOCYST_GRO | GOBP_BLASTOCYST_GROWTH | 29 | 0.349732 | 1.3603352 | 0.09439528 | 0.1896762 | 1 | 3713 | tags=38%, list=16%, si | MM_1 |
| GOBP_MICROTUBULE_BA | GOBP_MICROTUBULE_BASED_TRANSPORT | 217 | 0.230911 | 1.3589783 | 0.021052632 | 0.19090503 | 1 | 5967 | tags=38%, list=25%, si | MM_1 |
| GOBP_ESTABLISHMENT_( | GOBP_ESTABLISHMENT_OF_PROTEIN_LOCA | 20 | 0.402605 | 1.3577429 | 0.112021856 | 0.19210377 | 1 | 4614 | tags=25%, list=20%, si | MM_1 |
| PANCREATIC_CANCER | PANCREATIC_CANCER | 74 | 0.28278 | 1.3567923 | 0.059602648 | 0.19292246 | 1 | 5580 | tags=34%, list=24%, si | MM_1 |
| GOBP_PROTEIN_MODIFIC | GOBP_PROTEIN_MODIFICATION_BY_SMALL | 117 | 0.257778 | 1.3565613 | 0.014652015 | 0.19295932 | 1 | 5895 | tags=36%, list=25%, si | MM_1 |
| GOBP_ESTABLISHMENT_( | GOBP_ESTABLISHMENT_OF_RNA_LOCALIZA | 138 | 0.246507 | 1.3565375 | 0.04661017 | 0.1928067 | 1 | 5991 | tags=28%, list=25%, si | MM_1 |
| GOBP_NEGATIVE_REGUL | GOBP_NEGATIVE_REGULATION_OF_SMOOT | 36 | 0.333907 | 1.3563254 | 0.085365854 | 0.19287206 | 1 | 3261 | tags=31%, list=14%, si | MM_1 |
| GOBP_DEOXYRIBONUCLE | GOBP_DEOXYRIBONUCLEOTIDE_BIOSYNTHE | 17 | 0.40193 | 1.3561728 | 0.10344828 | 0.19289902 | 1 | 6745 | tags=59%, list=29%, si | MM_1 |
| GOBP_NUCLEOSOME_AS | GOBP_NUCLEOSOME_ASSEMBLY | 50 | 0.311983 | 1.3558455 | 0.06845238 | 0.19306679 | 1 | 3757 | tags=20%, list=16%, si | MM_1 |
| GOBP_REGULATION_OF_ | GOBP_REGULATION_OF_CELL_SUBSTRATE_ | 69 | 0.287101 | 1.355369 | 0.06666667 | 0.19338934 | 1 | 6791 | tags=45%, list=29%, si | MM_1 |
| GOBP_PROTEIN_O_LINKE | GOBP_PROTEIN_O_LINKED_MANNOSYLATIC | 16 | 0.421518 | 1.3535776 | 0.12912089 | 0.19518442 | 1 | 4309 | tags=44%, list=18%, si | MM_1 |
| GOBP_LEUKOCYTE_CHEM | GOBP_LEUKOCYTE_CHEMOTAXIS | 213 | 0.231026 | 1.3520682 | 0.014354067 | 0.19669822 | 1 | 3736 | tags=24%, list=16%, si | MM_1 |
| GOBP_NEGATIVE_REGUL | GOBP_NEGATIVE_REGULATION_OF_TRANSF | 222 | 0.231973 | 1.3516123 | 0 | 0.19700894 | 1 | 3525 | tags=18%, list=15%, si | MM_1 |
| GOBP_MITOTIC_METAPH | GOBP_MITOTIC_METAPHASE_CHROMOSOM | 58 | 0.289388 | 1.3510433 | 0.06031746 | 0.1974223 | 1 | 3350 | tags=24%, list=14%, si | MM_1 |
| GOBP_ALCOHOL_METAB( | GOBP_ALCOHOL_METABOLIC_PROCESS | 308 | 0.222298 | 1.3510095 | 0 | 0.19728374 | 1 | 3740 | tags=24%, list=16%, si | MM_1 |
| GOBP_AMINO_SUGAR_C/ | GOBP_AMINO_SUGAR_CATABOLIC_PROCES | 16 | 0.419388 | 1.3509358 | 0.09686609 | 0.19717228 | 1 | 5274 | tags=50%, list=22%, si | MM_1 |
| GOBP_REGULATION_OF_ | GOBP_REGULATION_OF_T_CELL_ACTIVATIO | 364 | 0.219527 | 1.3502749 | 0.0068965517 | 0.19770373 | 1 | 3667 | tags=21%, list=16%, si | MM_1 |
| GOBP_NEGATIVE_REGUL | GOBP_NEGATIVE_REGULATION_OF_SMOOT | 39 | 0.321739 | 1.3501455 | 0.089673914 | 0.19768058 | 1 | 4021 | tags=28%, list=17%, si | MM_1 |
| GOBP_DOUBLE_STRAND_ | GOBP_DOUBLE_STRAND_BREAK_REPAIR | 299 | 0.223618 | 1.3496646 | 0.012738854 | 0.19803663 | 1 | 4501 | tags=24%, list=19%, si | MM_1 |
| GOBP_RESPONSE_TO_IOI | GOBP_RESPONSE_TO_IONIZING_RADIATION | 127 | 0.250082 | 1.3495235 | 0.025531914 | 0.1980229 | 1 | 3350 | tags=23%, list=14%, si | MM_1 |
| GOBP_REGULATION_OF_ | GOBP_REGULATION_OF_B_CELL_DIFFERENT | 37 | 0.326887 | 1.3493515 | 0.0797546 | 0.1980184 | 1 | 6310 | tags=49%, list=27%, si | MM_1 |
| RELAXIN_SIGNALING_PAT | RELAXIN_SIGNALING_PATHWAY | 125 | 0.249884 | 1.3489311 | 0.027559055 | 0.19830339 | 1 | 4574 | tags=25%, list=19%, si | MM_1 |
| GOBP_REGULATION_OF_ | GOBP_REGULATION_OF_CALCIUM_ION_TR/ | 21 | 0.37802 | 1.3484305 | 0.11614731 | 0.19863743 | 1 | 4380 | tags=38%, list=19%, si | MM_1 |
| GOBP_REGULATION_OF_ | GOBP_REGULATION_OF_CYSTEINE_TYPE_EN | 197 | 0.233982 | 1.3475747 | 0.004761905 | 0.1993705 | 1 | 3730 | tags=21%, list=16%, si | MM_1 |
| GOBP_POSITIVE_REGULA | GOBP_POSITIVE_REGULATION_OF_EPITHELI | 158 | 0.244697 | 1.3475718 | 0.01793722 | 0.19919969 | 1 | 3194 | tags=20%, list=14%, si | MM_1 |
| GOBP_NEGATIVE_REGUL | GOBP_NEGATIVE_REGULATION_OF_CD4_PC | 37 | 0.32436 | 1.3463775 | 0.083076924 | 0.20026746 | 1 | 2145 | tags=19%, list=9%, sig | MM_1 |
| GOBP_MIDBODY_ABSCIS | GOBP_MIDBODY_ABSCISSION | 16 | 0.411733 | 1.3460276 | 0.11081081 | 0.20048103 | 1 | 3163 | tags=44%, list=13%, si | MM_1 |
| GOBP_COPII_COATED_VE | GOBP_COPII_COATED_VESICLE_BUDDING | 37 | 0.320854 | 1.345599 | 0.087878786 | 0.2008186 | 1 | 14397 | tags=97%, list=61%, si | MM_1 |
| GOBP_PROTEIN_KINASE_ | GOBP_PROTEIN_KINASE_C_ACTIVATING_G_ | 25 | 0.359201 | 1.345215 | 0.10755814 | 0.2010566 | 1 | 1190 | tags=20%, list=5%, sig | MM_1 |
| GOBP_REGULATION_OF_ | GOBP_REGULATION_OF_IMMUNOGLOBULI | 81 | 0.271399 | 1.345047 | 0.043165468 | 0.20108253 | 1 | 5192 | tags=36%, list=22%, si | MM_1 |
| GOBP_CALMODULIN_DEF | GOBP_CALMODULIN_DEPENDENT_KINASE_ | 16 | 0.407705 | 1.3447058 | 0.123324394 | 0.2012371 | 1 | 4170 | tags=38%, list=18%, si | MM_1 |
| GOBP_RESPONSE_TO_VIF | GOBP_RESPONSE_TO_VIRUS | 331 | 0.220905 | 1.3446932 | 0.007246377 | 0.20107493 | 1 | 4811 | tags=26%, list=20%, si | MM_1 |
| GOBP_NEGATIVE_REGUL | GOBP_NEGATIVE_REGULATION_OF_NEURO | 152 | 0.245566 | 1.344583 | 0.022222223 | 0.20101744 | 1 | 6113 | tags=38%, list=26%, si | MM_1 |
| METABOLISM_OF_XENOE | METABOLISM_OF_XENOBIOTICS_BY_CYTOC | 48 | 0.302881 | 1.3439641 | 0.073619634 | 0.20150147 | 1 | 3811 | tags=35%, list=16%, si | MM_1 |
| GOBP_NEGATIVE_REGUL | GOBP_NEGATIVE_REGULATION_OF_VASCUI | 109 | 0.256697 | 1.3436148 | 0.03787879 | 0.20168091 | 1 | 5052 | tags=29%, list=21%, si | MM_1 |
| GOBP_RESPONSE_TO_PL/ | GOBP_RESPONSE_TO_PLATELET_DERIVED_( | 22 | 0.372625 | 1.3435458 | 0.122507125 | 0.20157112 | 1 | 707 | tags=14%, list=3%, sig | MM_1 |
| GOBP_PROTEIN_LOCALIZ | GOBP_PROTEIN_LOCALIZATION_TO_CHROM | 112 | 0.254372 | 1.3434805 | 0.044354837 | 0.20147635 | 1 | 5945 | tags=33%, list=25%, si | MM_1 |
| GOBP_RECEPTOR_CLUSTI | GOBP_RECEPTOR_CLUSTERING | 79 | 0.279249 | 1.3425661 | 0.06418919 | 0.20229563 | 1 | 3134 | tags=24%, list=13%, si | MM_1 |
| GOBP_POSITIVE_REGULA | GOBP_POSITIVE_REGULATION_OF_TYPE_2_ | 18 | 0.408221 | 1.3421603 | 0.13586956 | 0.20257539 | 1 | 2487 | tags=28%, list=11%, si | MM_1 |
| SPHINGOLIPID_METABOL | SPHINGOLIPID_METABOLISM | 44 | 0.311818 | 1.3417344 | 0.09427609 | 0.20295767 | 1 | 3026 | tags=27%, list=13%, si | MM_1 |
| GOBP_FATTY_ACID_BIOS | GOBP_FATTY_ACID_BIOSYNTHETIC_PROCES | 145 | 0.242288 | 1.3412849 | 0.044334974 | 0.20330076 | 1 | 3708 | tags=24%, list=16%, si | MM_1 |

| | | | | | | | | | | |
|---|---|---|---|---|---|---|---|---|---|---|
| GOBP_ZYMOGEN_ACTIVA | GOBP_ZYMOGEN_ACTIVATION | 51 | 0.300268 | 1.3412212 | 0.07073955 | 0.20319746 | 1 | 1812 | tags=18%, list=8%, sig | MM_1 |
| GOBP_NEGATIVE_REGUL | GOBP_NEGATIVE_REGULATION_OF_TYPE_2 | 15 | 0.409133 | 1.3409082 | 0.13119534 | 0.2033397 | 1 | 572 | tags=20%, list=2%, sig | MM_1 |
| DRUG_METABOLISM | DRUG_METABOLISM | 93 | 0.262407 | 1.3402455 | 0.05836576 | 0.20395148 | 1 | 3685 | tags=30%, list=16%, si | MM_1 |
| GOBP_POSITIVE_REGULA | GOBP_POSITIVE_REGULATION_OF_T_CELL_ | 108 | 0.260551 | 1.3400444 | 0.030042918 | 0.20399185 | 1 | 3107 | tags=20%, list=13%, si | MM_1 |
| GOBP_RESPONSE_TO_GA | GOBP_RESPONSE_TO_GAMMA_RADIATION | 48 | 0.309056 | 1.3399317 | 0.090625 | 0.20395987 | 1 | 4034 | tags=27%, list=17%, si | MM_1 |
| GOBP_REGULATION_OF_ | GOBP_REGULATION_OF_LEUKOCYTE_DEGR/ | 48 | 0.30096 | 1.3397831 | 0.08383234 | 0.20395407 | 1 | 3660 | tags=25%, list=16%, si | MM_1 |
| GOBP_GLIAL_CELL_PROLI | GOBP_GLIAL_CELL_PROLIFERATION | 78 | 0.273684 | 1.3391492 | 0.07633588 | 0.2044494 | 1 | 4749 | tags=31%, list=20%, si | MM_1 |
| GOBP_POSITIVE_REGULA | GOBP_POSITIVE_REGULATION_OF_LYMPHO | 18 | 0.393647 | 1.3391117 | 0.14367816 | 0.20432179 | 1 | 3736 | tags=28%, list=16%, si | MM_1 |
| NON-SMALL_CELL_LUNG | NON-SMALL_CELL_LUNG_CANCER | 65 | 0.282145 | 1.3386315 | 0.055762082 | 0.2046754 | 1 | 3743 | tags=22%, list=16%, si | MM_1 |
| GOBP_PYRIMIDINE_DEOX | GOBP_PYRIMIDINE_DEOXYRIBONUCLEOTIDE | 20 | 0.378151 | 1.3384625 | 0.1278409 | 0.20468445 | 1 | 3155 | tags=35%, list=13%, si | MM_1 |
| GOBP_ACTIN_NUCLEATIO | GOBP_ACTIN_NUCLEATION | 58 | 0.286724 | 1.3382678 | 0.070063695 | 0.20470276 | 1 | 6691 | tags=45%, list=28%, si | MM_1 |
| GOBP_MITOTIC_CYTOKIN | GOBP_MITOTIC_CYTOKINETIC_PROCESS | 30 | 0.343229 | 1.3379176 | 0.097765364 | 0.2049182 | 1 | 3261 | tags=33%, list=14%, si | MM_1 |
| HIF-1_SIGNALING_PATHV | HIF-1_SIGNALING_PATHWAY | 98 | 0.262978 | 1.3370465 | 0.05357143 | 0.20568071 | 1 | 3743 | tags=23%, list=16%, si | MM_1 |
| GOBP_GRANULOCYTE_M | GOBP_GRANULOCYTE_MIGRATION | 146 | 0.24315 | 1.336633 | 0.020512821 | 0.20593144 | 1 | 4066 | tags=23%, list=17%, si | MM_1 |
| GOBP_ESTABLISHMENT_ | GOBP_ESTABLISHMENT_OR_MAINTENANCE | 51 | 0.298869 | 1.3365657 | 0.08615384 | 0.20581794 | 1 | 6125 | tags=39%, list=26%, si | MM_1 |
| GOBP_CELLULAR_COMPO | GOBP_CELLULAR_COMPONENT_DISASSEMB | 31 | 0.329415 | 1.336385 | 0.10619469 | 0.20583951 | 1 | 3550 | tags=32%, list=15%, si | MM_1 |
| GOBP_REGULATION_OF_ | GOBP_REGULATION_OF_MITOCHONDRIAL_ | 39 | 0.31792 | 1.3363303 | 0.075075075 | 0.20571879 | 1 | 5741 | tags=41%, list=24%, si | MM_1 |
| GOBP_NUCLEOSOME_OR | GOBP_NUCLEOSOME_ORGANIZATION | 67 | 0.28037 | 1.3342237 | 0.06440678 | 0.20791405 | 1 | 3970 | tags=18%, list=17%, si | MM_1 |
| GOBP_MITOTIC_NUCLEAI | GOBP_MITOTIC_NUCLEAR_DIVISION | 259 | 0.21932 | 1.3340415 | 0 | 0.20793338 | 1 | 3524 | tags=21%, list=15%, si | MM_1 |
| GOBP_DICARBOXYLIC_AC | GOBP_DICARBOXYLIC_ACID_CATABOLIC_PR | 17 | 0.401071 | 1.3329314 | 0.12983425 | 0.20900941 | 1 | 3579 | tags=35%, list=15%, si | MM_1 |
| GOBP_XENOBIOTIC_MET/ | GOBP_XENOBIOTIC_METABOLIC_PROCESS | 87 | 0.262921 | 1.3326837 | 0.07446808 | 0.20908867 | 1 | 2669 | tags=24%, list=11%, si | MM_1 |
| GOBP_PURINE_RIBONUC | GOBP_PURINE_RIBONUCLEOSIDE_METABOL | 24 | 0.357846 | 1.3325906 | 0.104986876 | 0.20901866 | 1 | 5913 | tags=42%, list=25%, si | MM_1 |
| GOBP_POSITIVE_REGULA | GOBP_POSITIVE_REGULATION_OF_INTERLE | 24 | 0.361409 | 1.3325512 | 0.12096774 | 0.20889471 | 1 | 5138 | tags=42%, list=22%, si | MM_1 |
| GOBP_REGULATION_OF_ | GOBP_REGULATION_OF_NEURONAL_SYNAF | 76 | 0.275059 | 1.3324915 | 0.056478404 | 0.2087906 | 1 | 2314 | tags=18%, list=10%, si | MM_1 |
| GOBP_INTRINSIC_APOPT | GOBP_INTRINSIC_APOPTOTIC_SIGNALING_F | 314 | 0.219551 | 1.3308047 | 0.011627907 | 0.2105987 | 1 | 5374 | tags=28%, list=23%, si | MM_1 |
| GOBP_B_CELL_MEDIATEI | GOBP_B_CELL_MEDIATED_IMMUNITY | 141 | 0.238361 | 1.3305198 | 0.034334764 | 0.21077521 | 1 | 4692 | tags=29%, list=20%, si | MM_1 |
| GOBP_NEGATIVE_REGUL | GOBP_NEGATIVE_REGULATION_OF_CELL_A | 301 | 0.219817 | 1.3299063 | 0 | 0.21130738 | 1 | 4613 | tags=26%, list=20%, si | MM_1 |
| GOBP_MEMBRANE_DEPO | GOBP_MEMBRANE_DEPOLARIZATION | 105 | 0.259602 | 1.329841 | 0.061068702 | 0.21119042 | 1 | 1245 | tags=12%, list=5%, sig | MM_1 |
| GOBP_INTERLEUKIN_13_ | GOBP_INTERLEUKIN_13_PRODUCTION | 21 | 0.386578 | 1.3291394 | 0.11363637 | 0.21169426 | 1 | 2029 | tags=24%, list=9%, sig | MM_1 |
| GOBP_RESPONSE_TO_AN | GOBP_RESPONSE_TO_AMYLOID_BETA | 39 | 0.327863 | 1.3289232 | 0.11111111 | 0.21178317 | 1 | 4459 | tags=33%, list=19%, si | MM_1 |
| GOBP_REGULATION_OF_ | GOBP_REGULATION_OF_INTRACELLULAR_T | 278 | 0.224574 | 1.3284053 | 0 | 0.21218023 | 1 | 5610 | tags=31%, list=24%, si | MM_1 |
| GOBP_POSITIVE_REGULA | GOBP_POSITIVE_REGULATION_OF_PLASMA | 115 | 0.252267 | 1.3279985 | 0.052434456 | 0.21245447 | 1 | 3883 | tags=21%, list=16%, si | MM_1 |
| GOBP_POSITIVE_REGULA | GOBP_POSITIVE_REGULATION_OF_INTRACE | 147 | 0.241962 | 1.3275559 | 0.028225806 | 0.21278216 | 1 | 4756 | tags=31%, list=20%, si | MM_1 |
| GOBP_REGULATION_OF_ | GOBP_REGULATION_OF_CELL_CYCLE_PHAS | 424 | 0.209054 | 1.3275205 | 0.019417476 | 0.21265386 | 1 | 5520 | tags=29%, list=23%, si | MM_1 |
| GOBP_MITOTIC_SISTER_ | GOBP_MITOTIC_SISTER_CHROMATID_SEPAF | 61 | 0.28865 | 1.3273401 | 0.074204944 | 0.21267332 | 1 | 3899 | tags=26%, list=17%, si | MM_1 |
| GOBP_RESPONSE_TO_AC | GOBP_RESPONSE_TO_ACID_CHEMICAL | 109 | 0.25399 | 1.3271273 | 0.052419353 | 0.21274996 | 1 | 4489 | tags=24%, list=19%, si | MM_1 |
| GOBP_REGULATION_OF_ | GOBP_REGULATION_OF_DOUBLE_STRAND_ | 80 | 0.273815 | 1.3262231 | 0.072413795 | 0.21365476 | 1 | 5267 | tags=31%, list=22%, si | MM_1 |
| GOBP_RECEPTOR_MEDIA | GOBP_RECEPTOR_MEDIATED_ENDOCYTOSIS | 269 | 0.224102 | 1.3257021 | 0.012578616 | 0.2140371 | 1 | 4871 | tags=28%, list=21%, si | MM_1 |
| GOBP_POSITIVE_REGULA | GOBP_POSITIVE_REGULATION_OF_PHOSPH | 26 | 0.363165 | 1.325186 | 0.121212125 | 0.21446039 | 1 | 5769 | tags=42%, list=24%, si | MM_1 |
| GOBP_REGULATION_OF_ | GOBP_REGULATION_OF_MITOTIC_CELL_CY | 331 | 0.216815 | 1.3250101 | 0.006289308 | 0.21450679 | 1 | 5758 | tags=31%, list=24%, si | MM_1 |
| GOBP_PROTEIN_SUMOYL | GOBP_PROTEIN_SUMOYLATION | 42 | 0.311883 | 1.3238897 | 0.10062893 | 0.21569861 | 1 | 5276 | tags=31%, list=22%, si | MM_1 |
| GOBP_REGULATION_OF_ | GOBP_REGULATION_OF_BIOLOGICAL_PROC | 64 | 0.287286 | 1.3237388 | 0.079617836 | 0.21567892 | 1 | 3076 | tags=28%, list=13%, si | MM_1 |
| GOBP_NUCLEOBASE_CON | GOBP_NUCLEOBASE_CONTAINING_COMPO | 195 | 0.231517 | 1.3226271 | 0.02347418 | 0.21675979 | 1 | 4443 | tags=22%, list=19%, si | MM_1 |
| GOBP_REGULATION_OF_ | GOBP_REGULATION_OF_INSULIN_LIKE_GRO | 22 | 0.373706 | 1.3225921 | 0.128 | 0.21660432 | 1 | 3313 | tags=32%, list=14%, si | MM_1 |
| GOBP_REGULATION_OF_ | GOBP_REGULATION_OF_LIPID_STORAGE | 55 | 0.289585 | 1.3224368 | 0.08588957 | 0.21658085 | 1 | 2450 | tags=20%, list=10%, si | MM_1 |
| GOBP_RESPONSE_TO_PR | GOBP_RESPONSE_TO_PROSTAGLANDIN | 25 | 0.349243 | 1.3219944 | 0.10481586 | 0.21694174 | 1 | 1967 | tags=16%, list=8%, sig | MM_1 |
| GOBP_REGULATION_OF_ | GOBP_REGULATION_OF_CELL_CYCLE_G1_S_ | 185 | 0.232441 | 1.32144 | 0.01843318 | 0.21743254 | 1 | 5744 | tags=32%, list=24%, si | MM_1 |
| GOBP_UBIQUINONE_MET | GOBP_UBIQUINONE_METABOLIC_PROCESS | 19 | 0.391893 | 1.3202873 | 0.14705883 | 0.21861829 | 1 | 5824 | tags=53%, list=25%, si | MM_1 |
| GOBP_MYOTUBE_DIFFER | GOBP_MYOTUBE_DIFFERENTIATION | 133 | 0.24566 | 1.3200536 | 0.022900764 | 0.21869615 | 1 | 3779 | tags=23%, list=16%, si | MM_1 |
| DOPAMINERGIC_SYNAPS | DOPAMINERGIC_SYNAPSE | 122 | 0.247567 | 1.3199255 | 0.04330709 | 0.21866553 | 1 | 4574 | tags=24%, list=19%, si | MM_1 |
| GOBP_REGULATION_OF_ | GOBP_REGULATION_OF_RESPONSE_TO_OSI | 18 | 0.396576 | 1.319082 | 0.1369863 | 0.2195506 | 1 | 5343 | tags=50%, list=23%, si | MM_1 |
| GOBP_ORGANELLE_INHE | GOBP_ORGANELLE_INHERITANCE | 15 | 0.424415 | 1.3179461 | 0.12755102 | 0.22072987 | 1 | 8103 | tags=53%, list=34%, si | MM_1 |
| GOBP_AXONAL_FASCICU | GOBP_AXONAL_FASCICULATION | 24 | 0.35138 | 1.3175402 | 0.118644066 | 0.22098503 | 1 | 5462 | tags=29%, list=23%, si | MM_1 |
| GOBP_L_AMINO_ACID_M | GOBP_L_AMINO_ACID_METABOLIC_PROCES | 154 | 0.235259 | 1.3175241 | 0.025125628 | 0.22082216 | 1 | 5276 | tags=33%, list=22%, si | MM_1 |
| GOBP_TYPE_2_IMMUNE_ | GOBP_TYPE_2_IMMUNE_RESPONSE | 46 | 0.302588 | 1.3174639 | 0.07185629 | 0.22070009 | 1 | 3570 | tags=26%, list=15%, si | MM_1 |
| GOBP_POSITIVE_REGULA | GOBP_POSITIVE_REGULATION_OF_CALCIUN | 17 | 0.396861 | 1.3170111 | 0.114369504 | 0.22104272 | 1 | 512 | tags=18%, list=2%, sig | MM_1 |
| GOBP_PERK_MEDIATED_ | GOBP_PERK_MEDIATED_UNFOLDED_PROTE | 16 | 0.400673 | 1.3169172 | 0.14095744 | 0.22096154 | 1 | 1620 | tags=13%, list=7%, sig | MM_1 |
| GOBP_REGULATION_OF_ | GOBP_REGULATION_OF_MITOTIC_CELL_CY | 490 | 0.205518 | 1.3167892 | 0.008928572 | 0.22093128 | 1 | 5114 | tags=27%, list=22%, si | MM_1 |
| GOBP_POSITIVE_REGULA | GOBP_POSITIVE_REGULATION_OF_ENDOCY | 182 | 0.230779 | 1.3167827 | 0.015544041 | 0.22075847 | 1 | 4920 | tags=28%, list=21%, si | MM_1 |
| GOBP_REGULATION_OF_ | GOBP_REGULATION_OF_DEPHOSPHORYLAT | 106 | 0.252943 | 1.3166287 | 0.03875969 | 0.22074088 | 1 | 3224 | tags=21%, list=14%, si | MM_1 |
| GOBP_RESPONSE_TO_XE | GOBP_RESPONSE_TO_XENOBIOTIC_STIMUL | 291 | 0.218926 | 1.3157337 | 0.0121951215 | 0.22156288 | 1 | 3360 | tags=22%, list=14%, si | MM_1 |
| GOBP_SUPEROXIDE_MET | GOBP_SUPEROXIDE_METABOLIC_PROCESS | 68 | 0.279009 | 1.3148817 | 0.089605734 | 0.22232032 | 1 | 4661 | tags=26%, list=20%, si | MM_1 |
| PENTOSE_PHOSPHATE_P/ | PENTOSE_PHOSPHATE_PATHWAY | 29 | 0.332476 | 1.3147407 | 0.11309524 | 0.22229126 | 1 | 5554 | tags=45%, list=24%, si | MM_1 |
| GOBP_NEGATIVE_REGUL | GOBP_NEGATIVE_REGULATION_OF_PROTEI | 87 | 0.263385 | 1.3143861 | 0.05533597 | 0.22251439 | 1 | 1313 | tags=13%, list=6%, sig | MM_1 |
| GOBP_FATTY_ACID_MET/ | GOBP_FATTY_ACID_METABOLIC_PROCESS | 399 | 0.211226 | 1.3137491 | 0.015503876 | 0.22308184 | 1 | 2874 | tags=20%, list=12%, si | MM_1 |
| GOBP_POSITIVE_REGULA | GOBP_POSITIVE_REGULATION_OF_PRODUC | 136 | 0.240608 | 1.3126628 | 0.04761905 | 0.22419237 | 1 | 5251 | tags=32%, list=22%, si | MM_1 |
| GOBP_STRESS_FIBER_AS | GOBP_STRESS_FIBER_ASSEMBLY | 117 | 0.24938 | 1.3123175 | 0.058577407 | 0.22438908 | 1 | 6982 | tags=44%, list=30%, si | MM_1 |
| GOBP_NEUTRAL_LIPID_M | GOBP_NEUTRAL_LIPID_METABOLIC_PROCES | 138 | 0.237656 | 1.3122846 | 0.03982301 | 0.22425787 | 1 | 3961 | tags=27%, list=17%, si | MM_1 |
| GOBP_ESTABLISHMENT_ | GOBP_ESTABLISHMENT_OR_MAINTENANCE | 25 | 0.353104 | 1.3118544 | 0.113259666 | 0.22459932 | 1 | 1926 | tags=16%, list=8%, sig | MM_1 |
| GOBP_PROTEIN_K63_LIN | GOBP_PROTEIN_K63_LINKED_DEUBIQUITIN | 29 | 0.342508 | 1.311803 | 0.09943182 | 0.22447503 | 1 | 6255 | tags=34%, list=26%, si | MM_1 |
| GOBP_REGULATION_OF_ | GOBP_REGULATION_OF_INTERLEUKIN_4_PF | 30 | 0.333677 | 1.3117787 | 0.106017195 | 0.22432223 | 1 | 5138 | tags=40%, list=22%, si | MM_1 |
| ERBB_SIGNALING_PATHV | ERBB_SIGNALING_PATHWAY | 83 | 0.264943 | 1.310137 | 0.06713781 | 0.22609903 | 1 | 2974 | tags=17%, list=13%, si | MM_1 |
| PPAR_SIGNALING_PATHV | PPAR_SIGNALING_PATHWAY | 70 | 0.275408 | 1.3098838 | 0.0777027 | 0.22618882 | 1 | 3961 | tags=29%, list=17%, si | MM_1 |
| GOBP_NEGATIVE_REGUL | GOBP_NEGATIVE_REGULATION_OF_MYELO | 57 | 0.283857 | 1.3095471 | 0.07954545 | 0.22639376 | 1 | 7797 | tags=53%, list=33%, si | MM_1 |
| GOBP_REGULATION_OF_ | GOBP_REGULATION_OF_LEUKOCYTE_PROLI | 273 | 0.216173 | 1.3084158 | 0 | 0.22763047 | 1 | 3710 | tags=21%, list=16%, si | MM_1 |
| GLYCEROPHOSPHOLIPID_ | GLYCEROPHOSPHOLIPID_METABOLISM | 92 | 0.258335 | 1.3058591 | 0.057471264 | 0.23053022 | 1 | 4703 | tags=35%, list=20%, si | MM_1 |
| GOBP_REGULATION_OF_ | GOBP_REGULATION_OF_ANIMAL_ORGAN_F | 29 | 0.33836 | 1.3053521 | 0.11904762 | 0.23095497 | 1 | 1643 | tags=17%, list=7%, sig | MM_1 |
| FOCAL_ADHESION | FOCAL_ADHESION | 194 | 0.225885 | 1.3053454 | 0.013333334 | 0.23078017 | 1 | 4514 | tags=26%, list=19%, si | MM_1 |
| GOBP_EXTRINSIC_APOPT | GOBP_EXTRINSIC_APOPTOTIC_SIGNALING_I | 233 | 0.221812 | 1.3043785 | 0.021621622 | 0.23183945 | 1 | 3730 | tags=20%, list=16%, si | MM_1 |
| GOBP_FIBROBLAST_MIGF | GOBP_FIBROBLAST_MIGRATION | 68 | 0.276227 | 1.303649 | 0.07284768 | 0.23254827 | 1 | 4897 | tags=28%, list=21%, si | MM_1 |
| GOBP_SPHINGOLIPID_ME | GOBP_SPHINGOLIPID_METABOLIC_PROCESS | 137 | 0.237344 | 1.3034676 | 0.050925925 | 0.23256865 | 1 | 3727 | tags=26%, list=16%, si | MM_1 |
| GOBP_CELLULAR_RESPOI | GOBP_CELLULAR_RESPONSE_TO_DEXAMETI | 23 | 0.359682 | 1.3026103 | 0.14409222 | 0.23344599 | 1 | 1652 | tags=17%, list=7%, sig | MM_1 |

| | | | | | | | | | | |
|---|---|---|---|---|---|---|---|---|---|---|
| GOBP_POSITIVE_REGULA | GOBP_POSITIVE_REGULATION_OF_DNA_BIN | 45 | 0.298304 | 1.3024035 | 0.06832298 | 0.23351218 | 1 | 5508 | tags=40%, list=23%, si | MM_1 |
| GOBP_NEURAL_TUBE_FO | GOBP_NEURAL_TUBE_FORMATION | 127 | 0.246178 | 1.3023008 | 0.050847456 | 0.23342557 | 1 | 2346 | tags=14%, list=10%, si | MM_1 |
| PYRIMIDINE_METABOLISI | PYRIMIDINE_METABOLISM | 53 | 0.282701 | 1.3022257 | 0.11111111 | 0.23334324 | 1 | 3627 | tags=30%, list=15%, si | MM_1 |
| GOBP_RIG_I_SIGNALING_ | GOBP_RIG_I_SIGNALING_PATHWAY | 24 | 0.354793 | 1.3019103 | 0.13150685 | 0.23361526 | 1 | 6004 | tags=42%, list=25%, si | MM_1 |
| GOBP_MORPHOGENESIS_ | GOBP_MORPHOGENESIS_OF_A_POLARIZED_ | 68 | 0.269616 | 1.3016299 | 0.08421053 | 0.2337559 | 1 | 3261 | tags=18%, list=14%, si | MM_1 |
| INOSITOL_PHOSPHATE_M | INOSITOL_PHOSPHATE_METABOLISM | 72 | 0.269492 | 1.3007431 | 0.10622711 | 0.23461294 | 1 | 6198 | tags=35%, list=26%, si | MM_1 |
| BACTERIAL_INVASION_OI | BACTERIAL_INVASION_OF_EPITHELIAL_CELL | 72 | 0.270543 | 1.300528 | 0.07482993 | 0.2346839 | 1 | 7388 | tags=53%, list=31%, si | MM_1 |
| ALDOSTERONE_SYNTHESI | ALDOSTERONE_SYNTHESIS_AND_SECRETION | 94 | 0.252125 | 1.2995473 | 0.05923345 | 0.23567861 | 1 | 2346 | tags=17%, list=10%, si | MM_1 |
| DILATED_CARDIOMYOPAT | DILATED_CARDIOMYOPATHY_(DCM) | 88 | 0.259622 | 1.2989653 | 0.09386282 | 0.23624131 | 1 | 4050 | tags=28%, list=17%, si | MM_1 |
| GOBP_CELL_CHEMOTAXIS | GOBP_CELL_CHEMOTAXIS | 295 | 0.216679 | 1.2987363 | 0.006451613 | 0.23632255 | 1 | 3786 | tags=22%, list=16%, si | MM_1 |
| RENIN_SECRETION | RENIN_SECRETION | 66 | 0.276756 | 1.2987313 | 0.07317073 | 0.23614383 | 1 | 4228 | tags=27%, list=18%, si | MM_1 |
| GOBP_REGULATION_OF_ | GOBP_REGULATION_OF_POSTSYNAPTIC_SP | 25 | 0.336351 | 1.2986367 | 0.13477089 | 0.23605227 | 1 | 2096 | tags=24%, list=9%, sig | MM_1 |
| GOBP_POSITIVE_REGULA | GOBP_POSITIVE_REGULATION_OF_PROTEIN | 63 | 0.276811 | 1.2982037 | 0.06666667 | 0.23641886 | 1 | 5016 | tags=35%, list=21%, si | MM_1 |
| GOBP_NEGATIVE_REGUL | GOBP_NEGATIVE_REGULATION_OF_T_CELL_ | 23 | 0.353491 | 1.297689 | 0.13390313 | 0.23693234 | 1 | 6649 | tags=52%, list=28%, si | MM_1 |
| GOBP_REGULATION_OF_ | GOBP_REGULATION_OF_FATTY_ACID_BETA | 21 | 0.371114 | 1.2976042 | 0.1402439 | 0.23683892 | 1 | 5666 | tags=52%, list=24%, si | MM_1 |
| GOBP_POSITIVE_REGULA | GOBP_POSITIVE_REGULATION_OF_INTERLE | 65 | 0.27522 | 1.2975171 | 0.06375839 | 0.23675732 | 1 | 2915 | tags=22%, list=12%, si | MM_1 |
| GOBP_DENDRITE_DEVELO | GOBP_DENDRITE_DEVELOPMENT | 319 | 0.210374 | 1.297242 | 0 | 0.2369072 | 1 | 2964 | tags=17%, list=13%, si | MM_1 |
| GOBP_MITOTIC_CELL_CY | GOBP_MITOTIC_CELL_CYCLE_PHASE_TRANS | 425 | 0.205953 | 1.2965024 | 0.01923077 | 0.23761939 | 1 | 6069 | tags=31%, list=26%, si | MM_1 |
| GOBP_DNA_CATABOLIC_ | GOBP_DNA_CATABOLIC_PROCESS | 29 | 0.335202 | 1.2961338 | 0.10086455 | 0.23788694 | 1 | 3550 | tags=38%, list=15%, si | MM_1 |
| GOBP_LIPID_DROPLET_O | GOBP_LIPID_DROPLET_ORGANIZATION | 36 | 0.323303 | 1.2949984 | 0.1319648 | 0.239074 | 1 | 4632 | tags=33%, list=20%, si | MM_1 |
| GOBP_PRE_MIRNA_PROC | GOBP_PRE_MIRNA_PROCESSING | 15 | 0.404987 | 1.2944509 | 0.14794521 | 0.23956564 | 1 | 6609 | tags=40%, list=28%, si | MM_1 |
| GOBP_REGULATION_OF_ | GOBP_REGULATION_OF_PROTEIN_CONTAIN | 131 | 0.240487 | 1.2940575 | 0.042735044 | 0.2398834 | 1 | 4868 | tags=28%, list=21%, si | MM_1 |
| GOBP_REGULATION_OF_ | GOBP_REGULATION_OF_EMBRYONIC_DEVE | 87 | 0.261242 | 1.2935866 | 0.074074075 | 0.24031045 | 1 | 3890 | tags=21%, list=16%, si | MM_1 |
| GOBP_CILIUM_ORGANIZA | GOBP_CILIUM_ORGANIZATION | 381 | 0.208579 | 1.293286 | 0.02238806 | 0.2404925 | 1 | 4548 | tags=25%, list=19%, si | MM_1 |
| GOBP_REGULATION_OF_ | GOBP_REGULATION_OF_VOLTAGE_GATED_ | 26 | 0.342923 | 1.293281 | 0.1260997 | 0.24030401 | 1 | 1457 | tags=15%, list=6%, sig | MM_1 |
| GOBP_LEUKOCYTE_MIGR | GOBP_LEUKOCYTE_MIGRATION_INVOLVED_ | 23 | 0.355289 | 1.2922928 | 0.13829787 | 0.2413768 | 1 | 3660 | tags=39%, list=16%, si | MM_1 |
| GOBP_REGULATION_OF_ | GOBP_REGULATION_OF_TOLL_LIKE_RECEPT | 32 | 0.327002 | 1.2918159 | 0.12084592 | 0.24180801 | 1 | 6255 | tags=47%, list=26%, si | MM_1 |
| GOBP_NEGATIVE_REGUL | GOBP_NEGATIVE_REGULATION_OF_DNA_R | 37 | 0.31043 | 1.2908363 | 0.11111111 | 0.2428987 | 1 | 8050 | tags=62%, list=34%, si | MM_1 |
| GOBP_CORONARY_VASCI | GOBP_CORONARY_VASCULATURE_MORPHO | 30 | 0.332654 | 1.2904737 | 0.12990937 | 0.24317206 | 1 | 4390 | tags=33%, list=19%, si | MM_1 |
| GOBP_FATTY_ACID_DERI | GOBP_FATTY_ACID_DERIVATIVE_METABOLI | 39 | 0.306237 | 1.2904286 | 0.1097561 | 0.24304828 | 1 | 5284 | tags=41%, list=22%, si | MM_1 |
| GOBP_POSITIVE_REGULA | GOBP_POSITIVE_REGULATION_OF_PROTEIN | 24 | 0.34721 | 1.2901117 | 0.14520548 | 0.24326938 | 1 | 5462 | tags=46%, list=23%, si | MM_1 |
| GOBP_POSTSYNAPTIC_SP | GOBP_POSTSYNAPTIC_SPECIALIZATION_ASS | 43 | 0.301771 | 1.2900075 | 0.106583074 | 0.2432124 | 1 | 1329 | tags=16%, list=6%, sig | MM_1 |
| GOBP_CELLULAR_EXTRAV | GOBP_CELLULAR_EXTRAVASATION | 74 | 0.271193 | 1.2898645 | 0.07063197 | 0.24316594 | 1 | 3691 | tags=23%, list=16%, si | MM_1 |
| GOBP_POSITIVE_REGULA | GOBP_POSITIVE_REGULATION_OF_CALCIUN | 30 | 0.328023 | 1.2895561 | 0.12154696 | 0.24337839 | 1 | 4877 | tags=27%, list=21%, si | MM_1 |
| GOBP_NEGATIVE_REGUL | GOBP_NEGATIVE_REGULATION_OF_MEMBF | 22 | 0.35933 | 1.2891811 | 0.122507125 | 0.24369307 | 1 | 3461 | tags=32%, list=15%, si | MM_1 |
| GOBP_XENOBIOTIC_CATA | GOBP_XENOBIOTIC_CATABOLIC_PROCESS | 21 | 0.366115 | 1.288914 | 0.15041783 | 0.24380113 | 1 | 2336 | tags=33%, list=10%, si | MM_1 |
| GOBP_PEPTIDYL_CYSTEIN | GOBP_PEPTIDYL_CYSTEINE_MODIFICATION | 44 | 0.298419 | 1.2887911 | 0.11490683 | 0.24377407 | 1 | 5419 | tags=39%, list=23%, si | MM_1 |
| GOBP_NEGATIVE_REGUL | GOBP_NEGATIVE_REGULATION_OF_ENDOC | 68 | 0.268881 | 1.2878464 | 0.09122807 | 0.2448115 | 1 | 3674 | tags=24%, list=16%, si | MM_1 |
| GOBP_REACTIVE_OXYGEI | GOBP_REACTIVE_OXYGEN_SPECIES_METAB | 239 | 0.219875 | 1.287064 | 0.051813472 | 0.24567144 | 1 | 3714 | tags=22%, list=16%, si | MM_1 |
| GOBP_EXOCYTIC_PROCES | GOBP_EXOCYTIC_PROCESS | 97 | 0.252756 | 1.2869852 | 0.062043797 | 0.24560003 | 1 | 2884 | tags=23%, list=12%, si | MM_1 |
| GOBP_POSITIVE_REGULA | GOBP_POSITIVE_REGULATION_OF_ESTABLIS | 345 | 0.208854 | 1.2869514 | 0.014705882 | 0.24544987 | 1 | 4756 | tags=28%, list=20%, si | MM_1 |
| GOBP_SPHINGOLIPID_BI | GOBP_SPHINGOLIPID_BIOSYNTHETIC_PROC | 94 | 0.253682 | 1.2861906 | 0.047445256 | 0.24631256 | 1 | 3582 | tags=24%, list=15%, si | MM_1 |
| GOBP_DEPHOSPHORYLAT | GOBP_DEPHOSPHORYLATION | 303 | 0.211086 | 1.2854911 | 0.005882353 | 0.24702042 | 1 | 5929 | tags=31%, list=25%, si | MM_1 |
| GOBP_ACYLGLYCEROL_H | GOBP_ACYLGLYCEROL_HOMEOSTASIS | 34 | 0.316868 | 1.2844175 | 0.111428574 | 0.24835387 | 1 | 2436 | tags=21%, list=10%, si | MM_1 |
| GOBP_NEGATIVE_REGUL | GOBP_NEGATIVE_REGULATION_OF_POST_T | 94 | 0.252735 | 1.2843868 | 0.0625 | 0.24819414 | 1 | 5276 | tags=31%, list=22%, si | MM_1 |
| GOBP_REGULATION_OF_ | GOBP_REGULATION_OF_AUTOPHAGY_OF_M | 32 | 0.31594 | 1.2840126 | 0.13611111 | 0.24851273 | 1 | 7508 | tags=59%, list=32%, si | MM_1 |
| GOBP_MUSCLE_CELL_DIF | GOBP_MUSCLE_CELL_DIFFERENTIATION | 444 | 0.20435 | 1.2837054 | 0.00862069 | 0.24874361 | 1 | 3806 | tags=21%, list=16%, si | MM_1 |
| GOBP_NEGATIVE_REGUL | GOBP_NEGATIVE_REGULATION_OF_CELL_C | 197 | 0.226814 | 1.283515 | 0.04347826 | 0.24880823 | 1 | 4918 | tags=27%, list=21%, si | MM_1 |
| GOBP_COPII_COATED_VE | GOBP_COPII_COATED_VESICLE_CARGO_LOA | 17 | 0.390507 | 1.2832376 | 0.14088398 | 0.24899861 | 1 | 14397 | tags=100%, list=61%, | MM_1 |
| GOBP_NEGATIVE_REGUL | GOBP_NEGATIVE_REGULATION_OF_DNA_M | 127 | 0.241829 | 1.2830328 | 0.0651341 | 0.24906962 | 1 | 4692 | tags=25%, list=20%, si | MM_1 |
| OLFACTORY_TRANSDUCT | OLFACTORY_TRANSDUCTION | 186 | -0.47714 | -2.355661 | 0 | 0.000505593 | 0.001 | 6032 | tags=62%, list=26%, si | MM_1 |
| GOBP_RESPONSE_TO_PH | GOBP_RESPONSE_TO_PHEROMONE | 17 | -0.6899 | -2.118717 | 0 | 0.009219206 | 0.035 | 4991 | tags=82%, list=21%, si | MM_1 |
| GOBP_ENTRAINMENT_OF | GOBP_ENTRAINMENT_OF_CIRCADIAN_CLO | 26 | -0.5779 | -2.022458 | 0 | 0.037716147 | 0.197 | 2539 | tags=35%, list=11%, si | MM_1 |
| GOBP_GANGLIOSIDE_BIO | GOBP_GANGLIOSIDE_BIOSYNTHETIC_PROCE | 17 | -0.60744 | -1.890778 | 0.001633987 | 0.17246613 | 0.732 | 2390 | tags=35%, list=10%, si | MM_1 |
| GOBP_CIRCADIAN_REGUI | GOBP_CIRCADIAN_REGULATION_OF_GENE_ | 69 | -0.44666 | -1.890348 | 0 | 0.13909045 | 0.737 | 1134 | tags=19%, list=5%, sig | MM_1 |
| GOBP_NEUTROPHIL_MED | GOBP_NEUTROPHIL_MEDIATED_CYTOTOXIC | 17 | -0.59808 | -1.838845 | 0 | 0.22457117 | 0.926 | 5887 | tags=65%, list=25%, si | MM_1 |
| GOBP_LIPID_EXPORT_FR | GOBP_LIPID_EXPORT_FROM_CELL | 56 | -0.45001 | -1.838366 | 0 | 0.19365542 | 0.927 | 766 | tags=18%, list=3%, sig | MM_1 |
| GOBP_NEUROPEPTIDE_SI | GOBP_NEUROPEPTIDE_SIGNALING_PATHW | 69 | -0.43128 | -1.834992 | 0 | 0.17687677 | 0.935 | 6255 | tags=52%, list=26%, si | MM_1 |
| GOBP_SPONTANEOUS_SY | GOBP_SPONTANEOUS_SYNAPTIC_TRANSMI | 18 | -0.59158 | -1.827699 | 0.001607717 | 0.17035125 | 0.949 | 7204 | tags=56%, list=31%, si | MM_1 |
| ASTHMA | ASTHMA | 16 | 0.746872 | 1.9437428 | 0 | 0.22449893 | 0.398 | 2105 | tags=50%, list=9%, sig | EN_2 |
| STAPHYLOCOCCUS_AURE | STAPHYLOCOCCUS_AUREUS_INFECTION | 44 | 0.59447 | 1.9400783 | 0 | 0.15970378 | 0.417 | 2105 | tags=32%, list=9%, sig | EN_2 |
| GOBP_REGULATION_OF_ | GOBP_REGULATION_OF_CELLULAR_EXTRAV | 43 | 0.595161 | 1.9328207 | 0 | 0.13552107 | 0.46 | 1873 | tags=30%, list=8%, sig | EN_2 |
| GOBP_POSITIVE_REGULA | GOBP_POSITIVE_REGULATION_OF_SYNCYTI | 25 | 0.660286 | 1.9272192 | 0 | 0.118261255 | 0.492 | 4821 | tags=64%, list=20%, si | EN_2 |
| GOBP_ANTIGEN_PROCES | GOBP_ANTIGEN_PROCESSING_AND_PRESEN | 24 | 0.657788 | 1.903114 | 0.0012626263 | 0.14722683 | 0.634 | 2105 | tags=38%, list=9%, sig | EN_2 |
| GOBP_NEGATIVE_REGUL | GOBP_NEGATIVE_REGULATION_OF_OSTEO | 37 | 0.605724 | 1.8913685 | 0 | 0.15670788 | 0.71 | 2261 | tags=27%, list=10%, si | EN_2 |
| GOBP_REGULATION_OF_ | GOBP_REGULATION_OF_SYNCYTIUM_FORM | 31 | 0.620767 | 1.8900027 | 0 | 0.14122525 | 0.726 | 4821 | tags=55%, list=20%, si | EN_2 |
| GOBP_REGULATION_OF_ | GOBP_REGULATION_OF_NUCLEOTIDE_BIOS | 32 | 0.605886 | 1.8706814 | 0 | 0.17015728 | 0.837 | 5196 | tags=56%, list=22%, si | EN_2 |
| GOBP_STEROL_BIOSYNTH | GOBP_STEROL_BIOSYNTHETIC_PROCESS | 58 | 0.541168 | 1.8670331 | 0 | 0.16255 | 0.856 | 5582 | tags=47%, list=24%, si | EN_2 |
| STEROID_BIOSYNTHESIS | STEROID_BIOSYNTHESIS | 17 | 0.695731 | 1.863692 | 0.0013089005 | 0.15615773 | 0.879 | 5114 | tags=71%, list=22%, si | EN_2 |
| PPAR_SIGNALING_PATHV | PPAR_SIGNALING_PATHWAY | 70 | 0.523427 | 1.842504 | 0 | 0.1943243 | 0.937 | 4913 | tags=40%, list=21%, si | EN_2 |
| GOBP_MACROPHAGE_CH | GOBP_MACROPHAGE_CHEMOTAXIS | 45 | 0.55042 | 1.8367745 | 0.0012077295 | 0.1946155 | 0.948 | 2800 | tags=31%, list=12%, si | EN_2 |
| GOBP_EOSINOPHIL_CHEN | GOBP_EOSINOPHIL_CHEMOTAXIS | 22 | 0.647033 | 1.8257699 | 0.0013003901 | 0.2133595 | 0.966 | 4667 | tags=55%, list=20%, si | EN_2 |
| GOBP_POSITIVE_REGULA | GOBP_POSITIVE_REGULATION_OF_NUCLEO | 20 | 0.653354 | 1.820709 | 0.0027247956 | 0.2115412 | 0.974 | 5196 | tags=60%, list=22%, si | EN_2 |
| GOBP_REGULATION_OF_ | GOBP_REGULATION_OF_MORPHOGENESIS_ | 71 | 0.518533 | 1.8172218 | 0 | 0.20750317 | 0.977 | 7114 | tags=61%, list=30%, si | EN_2 |
| GOBP_NEGATIVE_REGUL | GOBP_NEGATIVE_REGULATION_OF_MYELO | 57 | 0.54064 | 1.8079233 | 0 | 0.22122931 | 0.986 | 2261 | tags=25%, list=10%, si | EN_2 |
| GOBP_REGULATION_OF_ | GOBP_REGULATION_OF_MYOBLAST_FUSIOI | 22 | 0.630721 | 1.8040982 | 0.0026737968 | 0.22040057 | 0.986 | 4821 | tags=55%, list=20%, si | EN_2 |
| GOBP_POSITIVE_REGULA | GOBP_POSITIVE_REGULATION_OF_CELLULA | 22 | 0.630396 | 1.8011655 | 0.006451613 | 0.21807177 | 0.987 | 1873 | tags=36%, list=8%, sig | EN_2 |
| GOBP_POSITIVE_REGULA | GOBP_POSITIVE_REGULATION_OF_MYOBLA | 18 | 0.655022 | 1.7975097 | 0.0027322404 | 0.21870606 | 0.991 | 4821 | tags=61%, list=20%, si | EN_2 |
| GOBP_CELLULAR_MODIFI | GOBP_CELLULAR_MODIFIED_AMINO_ACID_ | 40 | 0.569905 | 1.7925256 | 0.0012062726 | 0.22371843 | 0.996 | 3548 | tags=33%, list=15%, si | EN_2 |
| GOBP_PROGRAMMED_N | GOBP_PROGRAMMED_NECROTIC_CELL_DEA | 50 | 0.537794 | 1.7862754 | 0.0011976048 | 0.23222016 | 0.997 | 3933 | tags=34%, list=17%, si | EN_2 |
| GOBP_SYNAPSE_PRUNIN | GOBP_SYNAPSE_PRUNING | 17 | 0.670542 | 1.7853023 | 0.0067476383 | 0.22474173 | 0.997 | 3010 | tags=47%, list=13%, si | EN_2 |
| GOBP_REGULATION_OF_ | GOBP_REGULATION_OF_MACROPHAGE_MI | 47 | 0.548982 | 1.7849036 | 0.0035971224 | 0.21644183 | 0.997 | 4692 | tags=43%, list=20%, si | EN_2 |

| | | | | | | | | | | |
|---|---|---|---|---|---|---|---|---|---|---|
| GOBP_LYMPHOCYTE_COS | GOBP_LYMPHOCYTE_COSTIMULATION | 33 | 0.572431 | 1.7790993 | 0 | 0.22384907 | 0.997 | 3834 | tags=39%, list=16%, si | EN_2 |
| GOBP_NUCLEAR_PORE_O | GOBP_NUCLEAR_PORE_ORGANIZATION | 15 | -0.74212 | -2.4260783 | 0 | 0.004178837 | 0.003 | 3314 | tags=67%, list=14%, si | EN_2 |
| GOBP_MITOTIC_CHROMO | GOBP_MITOTIC_CHROMOSOME_CONDENSA | 15 | -0.70844 | -2.2966354 | 0 | 0.025063826 | 0.034 | 3820 | tags=53%, list=16%, si | EN_2 |
| GOBP_BRANCHED_CHAIN | GOBP_BRANCHED_CHAIN_AMINO_ACID_CA | 21 | -0.63918 | -2.287693 | 0 | 0.018595254 | 0.038 | 3874 | tags=57%, list=16%, si | EN_2 |
| GOBP_CHROMOSOME_CO | GOBP_CHROMOSOME_CONDENSATION | 30 | -0.59541 | -2.2605639 | 0 | 0.018795496 | 0.05 | 3212 | tags=40%, list=14%, si | EN_2 |
| GOBP_NUCLEAR_TRANSC | GOBP_NUCLEAR_TRANSCRIBED_MRNA_CAT | 43 | -0.53183 | -2.2342207 | 0 | 0.02179708 | 0.071 | 5090 | tags=56%, list=22%, si | EN_2 |
| GOBP_BRANCHED_CHAIN | GOBP_BRANCHED_CHAIN_AMINO_ACID_M | 24 | -0.60929 | -2.211853 | 0 | 0.023071827 | 0.089 | 3149 | tags=50%, list=13%, si | EN_2 |
| GOBP_PROTEIN_LOCALIZ | GOBP_PROTEIN_LOCALIZATION_TO_MICRO | 39 | -0.50484 | -2.1455688 | 0 | 0.041135214 | 0.181 | 4381 | tags=46%, list=19%, si | EN_2 |
| GOBP_CENTRIOLE_ASSEM | GOBP_CENTRIOLE_ASSEMBLY | 46 | -0.48624 | -2.0946221 | 0 | 0.061268214 | 0.274 | 3433 | tags=46%, list=15%, si | EN_2 |
| GOBP_SECONDARY_PALA | GOBP_SECONDARY_PALATE_DEVELOPMENT | 25 | -0.54983 | -2.0522354 | 0 | 0.08445164 | 0.403 | 3950 | tags=44%, list=17%, si | EN_2 |
| GOBP_REGULATION_OF_ | GOBP_REGULATION_OF_CENTRIOLE_REPLIC | 22 | -0.57308 | -2.0270865 | 0 | 0.09674715 | 0.479 | 3433 | tags=55%, list=15%, si | EN_2 |
| GOBP_ENTRAINMENT_OF | GOBP_ENTRAINMENT_OF_CIRCADIAN_CLOC | 26 | -0.55034 | -2.0261335 | 0 | 0.088620275 | 0.484 | 4206 | tags=42%, list=18%, si | EN_2 |
| GOBP_MODIFICATION_O | GOBP_MODIFICATION_OF_POSTSYNAPTIC_S | 42 | -0.48197 | -2.00388 | 0 | 0.09882741 | 0.556 | 3341 | tags=29%, list=14%, si | EN_2 |
| PHOSPHATIDYLINOSITOL_ | PHOSPHATIDYLINOSITOL_SIGNALING_SYSTE | 94 | -0.39736 | -1.9946638 | 0 | 0.09937586 | 0.582 | 4525 | tags=45%, list=19%, si | EN_2 |
| GOBP_PORE_COMPLEX_A | GOBP_PORE_COMPLEX_ASSEMBLY | 20 | -0.58322 | -1.9789398 | 0 | 0.1066722 | 0.64 | 3314 | tags=55%, list=14%, si | EN_2 |
| GOBP_NUCLEAR_TRANSC | GOBP_NUCLEAR_TRANSCRIBED_MRNA_CAT | 118 | -0.39808 | -1.9773637 | 0 | 0.10092845 | 0.646 | 5090 | tags=47%, list=22%, si | EN_2 |
| GOBP_REGULATION_OF_ | GOBP_REGULATION_OF_NUCLEOTIDE_EXCIS | 29 | -0.49173 | -1.9743187 | 0.0045871558 | 0.0965483 | 0.656 | 4333 | tags=41%, list=18%, si | EN_2 |
| GOBP_PHOSPHATIDYLINO | GOBP_PHOSPHATIDYLINOSITOL_DEPHOSPH | 29 | -0.51662 | -1.9685704 | 0 | 0.093899816 | 0.665 | 4525 | tags=52%, list=19%, si | EN_2 |
| GOBP_NUCLEAR_TRANSC | GOBP_NUCLEAR_TRANSCRIBED_MRNA_CAT | 59 | -0.43255 | -1.9430636 | 0 | 0.10974929 | 0.744 | 4424 | tags=41%, list=19%, si | EN_2 |
| GOBP_CELLULAR_RESPON | GOBP_CELLULAR_RESPONSE_TO_GAMMA_ | 26 | -0.49858 | -1.9285148 | 0 | 0.116498336 | 0.783 | 9584 | tags=92%, list=41%, si | EN_2 |
| GOBP_NEGATIVE_REGULA | GOBP_NEGATIVE_REGULATION_OF_PROTEA | 51 | -0.43357 | -1.9088604 | 0 | 0.12868261 | 0.829 | 4126 | tags=29%, list=17%, si | EN_2 |
| GOBP_STRESS_GRANULE_ | GOBP_STRESS_GRANULE_ASSEMBLY | 35 | -0.47762 | -1.907884 | 0 | 0.123803794 | 0.835 | 5223 | tags=60%, list=22%, si | EN_2 |
| GOBP_NUCLEOLUS_ORGA | GOBP_NUCLEOLUS_ORGANIZATION | 17 | -0.57578 | -1.89792 | 0.008733625 | 0.12821773 | 0.858 | 3942 | tags=47%, list=17%, si | EN_2 |
| GOBP_MICROTUBULE_OR | GOBP_MICROTUBULE_ORGANIZING_CENTE | 149 | -0.35648 | -1.8923613 | 0 | 0.1266289 | 0.863 | 3538 | tags=33%, list=15%, si | EN_2 |
| GOBP_MRNA_METHYLGU | GOBP_MRNA_METHYLGUANOSINE_CAP_DE | 16 | -0.57481 | -1.8893609 | 0.0037453184 | 0.12404612 | 0.87 | 2518 | tags=25%, list=11%, si | EN_2 |
| GOBP_CEREBROSPINAL_F | GOBP_CEREBROSPINAL_FLUID_CIRCULATIO | 15 | -0.58171 | -1.8878042 | 0.0035842294 | 0.12073169 | 0.873 | 2565 | tags=47%, list=11%, si | EN_2 |
| GOBP_CENTROSOME_DU | GOBP_CENTROSOME_DUPLICATION | 74 | -0.40427 | -1.8854584 | 0 | 0.11818497 | 0.877 | 4401 | tags=47%, list=19%, si | EN_2 |
| GOBP_REGULATION_OF_ | GOBP_REGULATION_OF_DNA_DAMAGE_CH | 22 | -0.52218 | -1.872519 | 0.0042918455 | 0.12714237 | 0.903 | 3139 | tags=32%, list=13%, si | EN_2 |
| GOBP_MODIFICATION_O | GOBP_MODIFICATION_OF_SYNAPTIC_STRU | 51 | -0.42514 | -1.868631 | 0.0061728396 | 0.12686758 | 0.908 | 3341 | tags=25%, list=14%, si | EN_2 |
| GOBP_POSITIVE_REGULA | GOBP_POSITIVE_REGULATION_OF_CHROMO | 29 | -0.48383 | -1.8546475 | 0.005 | 0.13567786 | 0.928 | 3212 | tags=34%, list=14%, si | EN_2 |
| GOBP_POSITIVE_REGULA | GOBP_POSITIVE_REGULATION_OF_NOTCH_ | 42 | -0.4411 | -1.8501782 | 0 | 0.13588278 | 0.932 | 2466 | tags=29%, list=10%, si | EN_2 |
| GOBP_PROTEIN_DEGLYCO | GOBP_PROTEIN_DEGLYCOSYLATION | 15 | -0.58554 | -1.8454735 | 0 | 0.13608304 | 0.938 | 3728 | tags=47%, list=16%, si | EN_2 |
| GOBP_RESPONSE_TO_HE | GOBP_RESPONSE_TO_HEPATOCYTE_GROWT | 17 | -0.55079 | -1.8289672 | 0.0111524165 | 0.14884989 | 0.962 | 2498 | tags=29%, list=11%, si | EN_2 |
| GOBP_REGULATION_OF_ | GOBP_REGULATION_OF_CENTROSOME_CYC | 55 | -0.40596 | -1.8217674 | 0 | 0.15247189 | 0.972 | 3464 | tags=38%, list=15%, si | EN_2 |
| INOSITOL_PHOSPHATE_M | INOSITOL_PHOSPHATE_METABOLISM | 72 | -0.37555 | -1.8119165 | 0 | 0.15820289 | 0.977 | 4525 | tags=44%, list=19%, si | EN_2 |
| GOBP_NEGATIVE_REGULA | GOBP_NEGATIVE_REGULATION_OF_PROTEA | 36 | -0.4626 | -1.8040993 | 0 | 0.16278186 | 0.984 | 3553 | tags=25%, list=15%, si | EN_2 |
| VEGF_SIGNALING_PATHW | VEGF_SIGNALING_PATHWAY | 55 | -0.40435 | -1.7803904 | 0 | 0.18869154 | 0.995 | 2813 | tags=29%, list=12%, si | EN_2 |
| GOBP_MICROTUBULE_NU | GOBP_MICROTUBULE_NUCLEATION | 35 | -0.43611 | -1.7755947 | 0.004672897 | 0.1896201 | 0.995 | 4401 | tags=51%, list=19%, si | EN_2 |
| GOBP_MICROTUBULE_OR | GOBP_MICROTUBULE_ORGANIZING_CENTE | 34 | -0.4313 | -1.764972 | 0.005235602 | 0.19996004 | 0.997 | 4416 | tags=44%, list=19%, si | EN_2 |
| GOBP_GLANDULAR_EPITH | GOBP_GLANDULAR_EPITHELIAL_CELL_DEVE | 33 | -0.43969 | -1.7607602 | 0 | 0.20058896 | 0.997 | 3606 | tags=39%, list=15%, si | EN_2 |
| GOBP_POSITIVE_REGULA | GOBP_POSITIVE_REGULATION_OF_PROTEIN | 15 | -0.54568 | -1.7432266 | 0.014084507 | 0.22099526 | 0.999 | 4797 | tags=53%, list=20%, si | EN_2 |
| GOBP_REGULATION_OF_ | GOBP_REGULATION_OF_NEURON_MATURA | 16 | -0.5285 | -1.7409447 | 0.02919708 | 0.21884544 | 0.999 | 977 | tags=25%, list=4%, sig | EN_2 |
| GOBP_OLFACTORY_LOBE_ | GOBP_OLFACTORY_LOBE_DEVELOPMENT | 36 | -0.42036 | -1.7392831 | 0 | 0.21642584 | 0.999 | 1835 | tags=25%, list=8%, sig | EN_2 |
| GOBP_POSITIVE_REGULA | GOBP_POSITIVE_REGULATION_OF_MRNA_C | 95 | -0.35652 | -1.7341464 | 0 | 0.2192399 | 0.999 | 4651 | tags=40%, list=20%, si | EN_2 |
| GOBP_REGULATION_OF_ | GOBP_REGULATION_OF_CYTOKINESIS | 83 | -0.35175 | -1.7329991 | 0 | 0.21605186 | 0.999 | 5191 | tags=47%, list=22%, si | EN_2 |
| ADHERENS_JUNCTION | ADHERENS_JUNCTION | 70 | -0.36891 | -1.7321129 | 0 | 0.21248399 | 0.999 | 4099 | tags=37%, list=17%, si | EN_2 |
| GOBP_SISTER_CHROMAT | GOBP_SISTER_CHROMATID_SEGREGATION | 219 | -0.29903 | -1.7233804 | 0 | 0.2215666 | 1 | 4393 | tags=32%, list=19%, si | EN_2 |
| GOBP_VESICLE_TARGETIN | GOBP_VESICLE_TARGETING_TO_FROM_OR_ | 29 | -0.44709 | -1.7233136 | 0.00862069 | 0.21694762 | 1 | 1279 | tags=17%, list=5%, sig | EN_2 |
| GOBP_RESPONSE_TO_PLA | GOBP_RESPONSE_TO_PLATELET_DERIVED_C | 22 | -0.48929 | -1.7187537 | 0.004405286 | 0.21873195 | 1 | 4628 | tags=50%, list=20%, si | EN_2 |
| GOBP_PEPTIDYL_THREON | GOBP_PEPTIDYL_THREONINE_DEPHOSPHOR | 15 | -0.53395 | -1.7050575 | 0.014492754 | 0.23507695 | 1 | 3914 | tags=40%, list=17%, si | EN_2 |
| GOBP_DNA_INTEGRITY_C | GOBP_DNA_INTEGRITY_CHECKPOINT_SIGNA | 125 | -0.32483 | -1.6968405 | 0 | 0.24290013 | 1 | 3483 | tags=28%, list=15%, si | EN_2 |
| GOBP_MOTILE_CILIUM_A | GOBP_MOTILE_CILIUM_ASSEMBLY | 75 | -0.36271 | -1.693738 | 0 | 0.2427938 | 1 | 2577 | tags=28%, list=11%, si | EN_2 |
| GOBP_HIPPO_SIGNALING | GOBP_HIPPO_SIGNALING | 40 | -0.40321 | -1.692227 | 0.012738854 | 0.24016389 | 1 | 2714 | tags=33%, list=11%, si | EN_2 |
| GOBP_SPERM_FLAGELLU | GOBP_SPERM_FLAGELLUM_ASSEMBLY | 52 | -0.38377 | -1.6895427 | 0.01369863 | 0.23981258 | 1 | 2577 | tags=31%, list=11%, si | EN_2 |
| GOBP_TYPE_B_PANCREAT | GOBP_TYPE_B_PANCREATIC_CELL_DEVELOP | 19 | -0.49269 | -1.6877017 | 0.03731343 | 0.23802368 | 1 | 3336 | tags=42%, list=14%, si | EN_2 |
| GOBP_NEGATIVE_REGULA | GOBP_NEGATIVE_REGULATION_OF_CELL_C | 69 | -0.35431 | -1.6873242 | 0.0074626864 | 0.23441021 | 1 | 3465 | tags=30%, list=15%, si | EN_2 |
| GOBP_MEMBRANE_FISSIO | GOBP_MEMBRANE_FISSION | 31 | -0.4256 | -1.6869707 | 0.0091743115 | 0.23082697 | 1 | 5066 | tags=52%, list=21%, si | EN_2 |
| GOBP_ANOIKIS | GOBP_ANOIKIS | 31 | -0.42459 | -1.6816655 | 0.014925373 | 0.23532915 | 1 | 3582 | tags=39%, list=15%, si | EN_2 |
| GOBP_OSTEOBLAST_DEV | GOBP_OSTEOBLAST_DEVELOPMENT | 16 | -0.49435 | -1.6799301 | 0.014492754 | 0.23416784 | 1 | 1711 | tags=19%, list=7%, sig | EN_2 |
| GOBP_POSITIVE_REGULA | GOBP_POSITIVE_REGULATION_OF_VASCULA | 25 | -0.43872 | -1.677344 | 0.009259259 | 0.23411873 | 1 | 1393 | tags=24%, list=6%, sig | EN_2 |
| GOBP_FLUID_TRANSPORT | GOBP_FLUID_TRANSPORT | 28 | -0.44565 | -1.6711339 | 0.01746725 | 0.23970118 | 1 | 1722 | tags=32%, list=7%, sig | EN_2 |
| GOBP_MITOCHONDRIAL_ | GOBP_MITOCHONDRIAL_FISSION | 48 | -0.39881 | -1.6700543 | 0 | 0.23745203 | 1 | 4856 | tags=46%, list=21%, si | EN_2 |
| GOBP_MODIFICATION_O | GOBP_MODIFICATION_OF_POSTSYNAPTIC_A | 24 | -0.47018 | -1.6697663 | 0.009302326 | 0.2341586 | 1 | 2655 | tags=29%, list=11%, si | EN_2 |
| GOBP_INTERLEUKIN_6_M | GOBP_INTERLEUKIN_6_MEDIATED_SIGNALI | 19 | -0.49401 | -1.6687653 | 0.026717557 | 0.23202184 | 1 | 2785 | tags=32%, list=12%, si | EN_2 |
| GOBP_MICROAUTOPHAG | GOBP_MICROAUTOPHAGY | 24 | -0.45313 | -1.6595967 | 0.008 | 0.24174553 | 1 | 3361 | tags=38%, list=14%, si | EN_2 |
| GOBP_CELL_CYCLE_DNA_ | GOBP_CELL_CYCLE_DNA_REPLICATION | 40 | -0.40024 | -1.6564372 | 0.021978023 | 0.24346426 | 1 | 1935 | tags=20%, list=8%, sig | EN_2 |
| GOBP_NEGATIVE_REGULA | GOBP_NEGATIVE_REGULATION_OF_KERATI | 27 | -0.43289 | -1.652007 | 0.012048192 | 0.24630536 | 1 | 3804 | tags=44%, list=16%, si | EN_2 |
| GOBP_REGULATION_OF_ | GOBP_REGULATION_OF_GLUCONEOGENESI | 52 | -0.36445 | -1.6493309 | 0 | 0.24703442 | 1 | 2500 | tags=25%, list=11%, si | EN_2 |
| GOBP_TRANSLATION_AT_ | GOBP_TRANSLATION_AT_SYNAPSE | 48 | 0.638752 | 3.143856 | 0.0 | 0 | 0 | 6544 | tags=75%, list=28%, si | MM_2 |
| GOBP_RESPONSE_TO_INT | GOBP_RESPONSE_TO_INTERFERON_BETA | 64 | 0.588147 | 3.1171045 | 0.0 | 0 | 0 | 3297 | tags=45%, list=14%, si | MM_2 |
| CYTOSOLIC_DNA-SENSING | CYTOSOLIC_DNA-SENSING_PATHWAY | 46 | 0.653572 | 3.036163 | 0.0 | 0 | 0 | 3897 | tags=39%, list=17%, si | MM_2 |
| GOBP_NEGATIVE_REGULA | GOBP_NEGATIVE_REGULATION_OF_VIRAL_ | 50 | 0.611893 | 2.9368992 | 0.0 | 0 | 0 | 2379 | tags=44%, list=10%, si | MM_2 |
| GOBP_RESPONSE_TO_TYP | GOBP_RESPONSE_TO_TYPE_I_INTERFERON | 64 | 0.558849 | 2.8029294 | 0.0 | 0 | 0 | 3272 | tags=39%, list=14%, si | MM_2 |
| RIBOSOME | RIBOSOME | 128 | 0.489156 | 2.793752 | 0.0 | 0 | 0 | 6657 | tags=66%, list=28%, si | MM_2 |
| GOBP_DEFENSE_RESPON | GOBP_DEFENSE_RESPONSE_TO_VIRUS | 276 | 0.429581 | 2.7902582 | 0.0 | 0 | 0 | 2916 | tags=29%, list=12%, si | MM_2 |
| INFLUENZA_A | INFLUENZA_A | 139 | 0.455186 | 2.7542336 | 0.0 | 0 | 0 | 3435 | tags=35%, list=15%, si | MM_2 |
| MEASLES | MEASLES | 128 | 0.500404 | 2.7416468 | 0.0 | 0 | 0 | 3373 | tags=34%, list=14%, si | MM_2 |
| GOBP_RESPONSE_TO_INT | GOBP_RESPONSE_TO_INTERFERON_ALPHA | 28 | 0.638436 | 2.682579 | 0.0 | 0 | 0 | 2341 | tags=50%, list=10%, si | MM_2 |
| GOBP_NEGATIVE_REGULA | GOBP_NEGATIVE_REGULATION_OF_VIRAL_ | 95 | 0.504613 | 2.6794074 | 0.0 | 0 | 0 | 2379 | tags=35%, list=10%, si | MM_2 |
| GOBP_REGULATION_OF_ | GOBP_REGULATION_OF_VIRAL_GENOME_R | 78 | 0.508636 | 2.670124 | 0.0 | 0 | 0 | 2379 | tags=33%, list=10%, si | MM_2 |
| GOBP_CELLULAR_RESPON | GOBP_CELLULAR_RESPONSE_TO_INTERFERO | 16 | 0.772023 | 2.658967 | 0.0 | 0 | 0 | 2341 | tags=56%, list=10%, si | MM_2 |
| GOBP_REGULATION_OF_ | GOBP_REGULATION_OF_VIRAL_LIFE_CYCLE | 139 | 0.448984 | 2.658378 | 0.0 | 0 | 0 | 4357 | tags=47%, list=18%, si | MM_2 |

| | | | | | | | | | | |
|---|---|---|---|---|---|---|---|---|---|---|
| GOBP_RESPONSE_TO_VII | GOBP_RESPONSE_TO_VIRUS | 331 | 0.395353 | 2.6305034 | 0.0 | 0.000310256 | 0.001 | 2916 | tags=27%, list=12%, si | MM_2 |
| TOLL-LIKE_RECEPTOR_SIG | TOLL-LIKE_RECEPTOR_SIGNALING_PATHWA | 84 | 0.485483 | 2.612733 | 0.0 | 0.000290865 | 0.001 | 3189 | tags=30%, list=14%, si | MM_2 |
| GOBP_POSITIVE_REGULA | GOBP_POSITIVE_REGULATION_OF_INTERFE | 45 | 0.554634 | 2.5880475 | 0.0 | 0.001095577 | 0.004 | 2379 | tags=27%, list=10%, si | MM_2 |
| GOBP_ANTIGEN_PROCES | GOBP_ANTIGEN_PROCESSING_AND_PRESEN | 28 | 0.59424 | 2.5661228 | 0.0 | 0.001306802 | 0.005 | 4400 | tags=64%, list=19%, si | MM_2 |
| GOBP_REGULATION_OF_ | GOBP_REGULATION_OF_VIRAL_PROCESS | 167 | 0.442312 | 2.5276308 | 0.0 | 0.002448877 | 0.01 | 4357 | tags=45%, list=18%, si | MM_2 |
| GOBP_POSITIVE_REGULA | GOBP_POSITIVE_REGULATION_OF_LYMPHO | 21 | 0.642557 | 2.4870274 | 0.0 | 0.003938914 | 0.017 | 2279 | tags=38%, list=10%, si | MM_2 |
| GOBP_POSITIVE_REGULA | GOBP_POSITIVE_REGULATION_OF_CELL_AD | 21 | 0.64463 | 2.486093 | 0.0 | 0.003751347 | 0.017 | 729 | tags=24%, list=3%, sig | MM_2 |
| GOBP_REGULATION_OF_ | GOBP_REGULATION_OF_T_CELL_MIGRATIO | 49 | 0.510237 | 2.4653904 | 0.0 | 0.004851096 | 0.022 | 980 | tags=22%, list=4%, sig | MM_2 |
| TUBERCULOSIS | TUBERCULOSIS | 152 | 0.417737 | 2.4592702 | 0.0 | 0.005448029 | 0.025 | 3189 | tags=27%, list=14%, si | MM_2 |
| GOBP_T_CELL_CHEMOTA | GOBP_T_CELL_CHEMOTAXIS | 23 | 0.5952 | 2.4569812 | 0.0 | 0.005221028 | 0.025 | 432 | tags=26%, list=2%, sig | MM_2 |
| GOBP_POSITIVE_REGULA | GOBP_POSITIVE_REGULATION_OF_TYPE_I_I | 77 | 0.468556 | 2.4568062 | 0.0 | 0.005012187 | 0.025 | 2567 | tags=29%, list=11%, si | MM_2 |
| GOBP_T_CELL_MIGRATIO | GOBP_T_CELL_MIGRATION | 64 | 0.48562 | 2.456722 | 0.0 | 0.004819411 | 0.025 | 1967 | tags=25%, list=8%, sig | MM_2 |
| GOBP_REGULATION_OF_ | GOBP_REGULATION_OF_TYPE_I_INTERFERO | 40 | 0.539658 | 2.4512014 | 0.0 | 0.005326971 | 0.029 | 4281 | tags=45%, list=18%, si | MM_2 |
| GOBP_LYMPH_NODE_DE | GOBP_LYMPH_NODE_DEVELOPMENT | 32 | 0.579332 | 2.4349124 | 0.0 | 0.005809217 | 0.033 | 3326 | tags=44%, list=14%, si | MM_2 |
| GOBP_INTERFERON_BETA | GOBP_INTERFERON_BETA_PRODUCTION | 66 | 0.473831 | 2.413819 | 0.0 | 0.006432138 | 0.038 | 2950 | tags=29%, list=12%, si | MM_2 |
| OXIDATIVE_PHOSPHORYL | OXIDATIVE_PHOSPHORYLATION | 115 | 0.444533 | 2.401231 | 0.0 | 0.00698361 | 0.043 | 5841 | tags=64%, list=25%, si | MM_2 |
| GOBP_REGULATION_OF_ | GOBP_REGULATION_OF_NUCLEASE_ACTIVIT | 22 | 0.62105 | 2.395898 | 0.0 | 0.006909827 | 0.044 | 3674 | tags=59%, list=16%, si | MM_2 |
| GOBP_POSITIVE_REGULA | GOBP_POSITIVE_REGULATION_OF_LEUKOC | 33 | 0.547375 | 2.38786 | 0.0 | 0.006990557 | 0.046 | 2279 | tags=27%, list=10%, si | MM_2 |
| GOBP_ANTIGEN_PROCES | GOBP_ANTIGEN_PROCESSING_AND_PRESEN | 34 | 0.551039 | 2.3805823 | 0.0 | 0.007063641 | 0.048 | 4400 | tags=59%, list=19%, si | MM_2 |
| GOBP_VIRAL_GENOME_F | GOBP_VIRAL_GENOME_REPLICATION | 110 | 0.438336 | 2.3777027 | 0.0 | 0.007251467 | 0.051 | 4357 | tags=39%, list=18%, si | MM_2 |
| GOBP_INTERFERON_MED | GOBP_INTERFERON_MEDIATED_SIGNALING | 76 | 0.457512 | 2.373271 | 0.0 | 0.007436271 | 0.054 | 3272 | tags=33%, list=14%, si | MM_2 |
| GOBP_REGULATION_OF_ | GOBP_REGULATION_OF_LYMPHOCYTE_CHE | 22 | 0.591307 | 2.36119 | 0.0 | 0.008011012 | 0.059 | 1635 | tags=32%, list=7%, sig | MM_2 |
| GOBP_ANTIVIRAL_INNAT | GOBP_ANTIVIRAL_INNATE_IMMUNE_RESPO | 47 | 0.500028 | 2.3560548 | 0.0 | 0.008046437 | 0.061 | 2835 | tags=38%, list=12%, si | MM_2 |
| RIG-I-LIKE_RECEPTOR_SIG | RIG-I-LIKE_RECEPTOR_SIGNALING_PATHWA | 52 | 0.492723 | 2.3507206 | 0.0 | 0.008068871 | 0.063 | 2411 | tags=27%, list=10%, si | MM_2 |
| GOBP_REGULATION_OF_ | GOBP_REGULATION_OF_T_CELL_CHEMOTA | 16 | 0.65434 | 2.3417563 | 0.0 | 0.008475182 | 0.068 | 432 | tags=31%, list=2%, sig | MM_2 |
| GOBP_ANTIGEN_PROCES | GOBP_ANTIGEN_PROCESSING_AND_PRESEN | 20 | 0.616898 | 2.330743 | 0.0 | 0.009326878 | 0.076 | 4364 | tags=70%, list=18%, si | MM_2 |
| GOBP_REGULATION_OF_ | GOBP_REGULATION_OF_TRANSLATIONAL_I | 68 | 0.455969 | 2.3233292 | 0.0 | 0.010119774 | 0.085 | 4965 | tags=40%, list=21%, si | MM_2 |
| GOBP_POSITIVE_REGULA | GOBP_POSITIVE_REGULATION_OF_VIRAL_LI | 28 | 0.542429 | 2.3232286 | 0.0 | 0.009878827 | 0.085 | 4680 | tags=68%, list=20%, si | MM_2 |
| GOBP_POSITIVE_REGULA | GOBP_POSITIVE_REGULATION_OF_LYMPHO | 42 | 0.500681 | 2.2994215 | 0.0 | 0.011897611 | 0.105 | 1635 | tags=24%, list=7%, sig | MM_2 |
| GOBP_PROTEIN_K63_LIN | GOBP_PROTEIN_K63_LINKED_UBIQUITINAT | 68 | 0.439394 | 2.2928677 | 0.0 | 0.012576145 | 0.114 | 4715 | tags=43%, list=20%, si | MM_2 |
| GOBP_POSITIVE_REGULA | GOBP_POSITIVE_REGULATION_OF_PATTERN | 68 | 0.450606 | 2.290358 | 0.0 | 0.012799053 | 0.118 | 4643 | tags=40%, list=20%, si | MM_2 |
| GOBP_CYTOPLASMIC_SE | GOBP_CYTOPLASMIC_SEQUESTERING_OF_P | 17 | 0.634276 | 2.28981 | 0.0 | 0.012837442 | 0.12 | 3593 | tags=53%, list=15%, si | MM_2 |
| GOBP_CHEMOKINE_C_X_ | GOBP_CHEMOKINE_C_X_C_MOTIF_LIGAND_ | 25 | 0.562694 | 2.2818341 | 0.0 | 0.013515064 | 0.129 | 2341 | tags=40%, list=10%, si | MM_2 |
| GOBP_RESPONSE_TO_CH | GOBP_RESPONSE_TO_CHEMOKINE | 74 | 0.419828 | 2.280064 | 0.0 | 0.013426981 | 0.13 | 614 | tags=16%, list=3%, sig | MM_2 |
| TNF_SIGNALING_PATHW | TNF_SIGNALING_PATHWAY | 108 | 0.430482 | 2.2724314 | 0.0 | 0.013814056 | 0.137 | 4144 | tags=35%, list=18%, si | MM_2 |
| GOBP_GLIAL_CELL_MIGR | GOBP_GLIAL_CELL_MIGRATION | 70 | 0.439215 | 2.2648065 | 0.0 | 0.014165248 | 0.144 | 4964 | tags=47%, list=21%, si | MM_2 |
| HEPATITIS_B | HEPATITIS_B | 147 | 0.38161 | 2.2639558 | 0.0 | 0.013887499 | 0.144 | 3735 | tags=28%, list=16%, si | MM_2 |
| GOBP_REGULATION_OF_ | GOBP_REGULATION_OF_LYMPHOCYTE_MIG | 64 | 0.451454 | 2.247234 | 0.0 | 0.016048238 | 0.168 | 3558 | tags=34%, list=15%, si | MM_2 |
| GOBP_CELL_CELL_ADHES | GOBP_CELL_CELL_ADHESION_MEDIATED_B | 18 | 0.595731 | 2.1922097 | 0.005235602 | 0.023983143 | 0.246 | 15 | tags=11%, list=0%, sig | MM_2 |
| GOBP_CELLULAR_RESPOI | GOBP_CELLULAR_RESPONSE_TO_DSRNA | 19 | 0.586762 | 2.1833463 | 0.0 | 0.025655786 | 0.261 | 2379 | tags=32%, list=10%, si | MM_2 |
| GOBP_POSITIVE_REGULA | GOBP_POSITIVE_REGULATION_OF_FATTY_A | 20 | 0.580767 | 2.1828613 | 0.005319149 | 0.025189316 | 0.261 | 3877 | tags=60%, list=16%, si | MM_2 |
| PARKINSON_DISEASE | PARKINSON_DISEASE | 122 | 0.387549 | 2.1764278 | 0.0 | 0.025329499 | 0.264 | 4701 | tags=53%, list=20%, si | MM_2 |
| GOBP_POSITIVE_REGULA | GOBP_POSITIVE_REGULATION_OF_VIRAL_P | 64 | 0.444967 | 2.170731 | 0.0 | 0.02554108 | 0.268 | 4680 | tags=48%, list=20%, si | MM_2 |
| GOBP_LYMPHOCYTE_CHE | GOBP_LYMPHOCYTE_CHEMOTAXIS | 38 | 0.480902 | 2.1653874 | 0.0 | 0.026142731 | 0.279 | 1967 | tags=24%, list=8%, sig | MM_2 |
| GOBP_REGULATION_OF_ | GOBP_REGULATION_OF_TRANSLATIONAL_I | 17 | 0.599322 | 2.1487072 | 0.0 | 0.028608896 | 0.305 | 2003 | tags=29%, list=8%, si | MM_2 |
| GOBP_VIRAL_LIFE_CYCLE | GOBP_VIRAL_LIFE_CYCLE | 230 | 0.379224 | 2.1439953 | 0.0 | 0.029286046 | 0.318 | 4702 | tags=42%, list=20%, si | MM_2 |
| GOBP_LYMPHOCYTE_MIG | GOBP_LYMPHOCYTE_MIGRATION | 92 | 0.384738 | 2.1400938 | 0.0 | 0.029337665 | 0.322 | 1635 | tags=20%, list=7%, sig | MM_2 |
| GOBP_MACROPHAGE_MI | GOBP_MACROPHAGE_MIGRATION | 65 | 0.419541 | 2.1390293 | 0.0 | 0.029176183 | 0.324 | 4068 | tags=38%, list=17%, si | MM_2 |
| GOBP_POSITIVE_REGULA | GOBP_POSITIVE_REGULATION_OF_EXTRINS | 60 | 0.413032 | 2.1390088 | 0.0 | 0.02871307 | 0.324 | 3654 | tags=40%, list=15%, si | MM_2 |
| EPSTEIN-BARR_VIRUS_INI | EPSTEIN-BARR_VIRUS_INFECTION | 190 | 0.371543 | 2.1384602 | 0.0 | 0.02841138 | 0.326 | 3414 | tags=30%, list=14%, si | MM_2 |
| PHAGOSOME | PHAGOSOME | 146 | 0.368614 | 2.1267838 | 0.0 | 0.030475315 | 0.351 | 3357 | tags=30%, list=14%, si | MM_2 |
| GOBP_POSITIVE_REGULA | GOBP_POSITIVE_REGULATION_OF_CHEMOK | 15 | 0.594525 | 2.1107955 | 0.0 | 0.03362338 | 0.386 | 2198 | tags=40%, list=9%, sig | MM_2 |
| GOBP_REGULATION_OF_ | GOBP_REGULATION_OF_MYOBLAST_FUSIOI | 22 | 0.558171 | 2.1086056 | 0.00625 | 0.033533927 | 0.39 | 530 | tags=23%, list=2%, sig | MM_2 |
| GOBP_RESPONSE_TO_DS | GOBP_RESPONSE_TO_DSRNA | 43 | 0.447637 | 2.099892 | 0.0 | 0.034963414 | 0.409 | 4016 | tags=35%, list=17%, si | MM_2 |
| GOBP_PROTEIN_K48_LIN | GOBP_PROTEIN_K48_LINKED_DEUBIQUITIN | 29 | 0.514564 | 2.0990314 | 0.0 | 0.03472468 | 0.41 | 4163 | tags=38%, list=18%, si | MM_2 |
| GOBP_DENDRITIC_CELL_ | GOBP_DENDRITIC_CELL_CYTOKINE_PRODU | 16 | 0.605366 | 2.096564 | 0.0 | 0.034485016 | 0.412 | 3120 | tags=44%, list=13%, si | MM_2 |
| GOBP_OXIDATIVE_PHOSP | GOBP_OXIDATIVE_PHOSPHORYLATION | 124 | 0.380513 | 2.0958066 | 0.0 | 0.034253135 | 0.413 | 5351 | tags=52%, list=23%, si | MM_2 |
| GOBP_POSITIVE_REGULA | GOBP_POSITIVE_REGULATION_OF_LYMPHO | 18 | 0.558734 | 2.0946743 | 0.0 | 0.03410361 | 0.416 | 1635 | tags=28%, list=7%, sig | MM_2 |
| GOBP_ANTIGEN_PROCES | GOBP_ANTIGEN_PROCESSING_AND_PRESEN | 59 | 0.404821 | 2.0912948 | 0.0 | 0.034651097 | 0.428 | 4400 | tags=53%, list=19%, si | MM_2 |
| GOBP_POSITIVE_REGULA | GOBP_POSITIVE_REGULATION_OF_T_CELL_ | 16 | 0.596399 | 2.0840786 | 0.0 | 0.035868324 | 0.441 | 2279 | tags=38%, list=10%, si | MM_2 |
| GOBP_POSITIVE_REGULA | GOBP_POSITIVE_REGULATION_OF_TYPE_I_I | 19 | 0.544091 | 2.0781112 | 0.0 | 0.036765266 | 0.454 | 4614 | tags=53%, list=20%, si | MM_2 |
| NF-KAPPA_B_SIGNALING | NF-KAPPA_B_SIGNALING_PATHWAY | 93 | 0.387643 | 2.0740376 | 0.0 | 0.03739177 | 0.465 | 2995 | tags=30%, list=13%, si | MM_2 |
| GOBP_CYTOKINE_MEDIA | GOBP_CYTOKINE_MEDIATED_SIGNALING_P | 421 | 0.28471 | 2.0703335 | 0.0 | 0.037815224 | 0.47 | 3272 | tags=24%, list=14%, si | MM_2 |
| GOBP_POSITIVE_REGULA | GOBP_POSITIVE_REGULATION_OF_MYOBLA | 18 | 0.556275 | 2.0651174 | 0.0 | 0.038806036 | 0.484 | 530 | tags=22%, list=2%, sig | MM_2 |
| HEPATITIS_C | HEPATITIS_C | 138 | 0.363058 | 2.0579302 | 0.0 | 0.04025959 | 0.504 | 3374 | tags=26%, list=14%, si | MM_2 |
| GOBP_TYPE_I_INTERFER | GOBP_TYPE_I_INTERFERON_PRODUCTION | 127 | 0.371727 | 2.0567496 | 0.0 | 0.040163577 | 0.509 | 4416 | tags=35%, list=19%, si | MM_2 |
| GOBP_VIRAL_PROCESS | GOBP_VIRAL_PROCESS | 303 | 0.351055 | 2.055376 | 0.0 | 0.039787035 | 0.511 | 4749 | tags=39%, list=20%, si | MM_2 |
| GOBP_ANTIMICROBIAL_H | GOBP_ANTIMICROBIAL_HUMORAL_RESPON | 98 | 0.374084 | 2.0524487 | 0.0 | 0.04013678 | 0.517 | 2229 | tags=28%, list=9%, sig | MM_2 |
| GOBP_NEGATIVE_REGUL | GOBP_NEGATIVE_REGULATION_OF_TRANSL | 20 | 0.550048 | 2.0502486 | 0.00591716 | 0.040321656 | 0.525 | 4346 | tags=45%, list=18%, si | MM_2 |
| GOBP_ACTIVATION_OF_I | GOBP_ACTIVATION_OF_INNATE_IMMUNE_ | 290 | 0.305742 | 2.0501192 | 0.0 | 0.039959334 | 0.526 | 2889 | tags=21%, list=12%, si | MM_2 |
| GOBP_COPPER_ION_HOM | GOBP_COPPER_ION_HOMEOSTASIS | 18 | 0.544655 | 2.048155 | 0.0 | 0.04034942 | 0.532 | 3042 | tags=44%, list=13%, si | MM_2 |
| C-TYPE_LECTIN_RECEPTO | C-TYPE_LECTIN_RECEPTOR_SIGNALING_PAT | 104 | 0.375044 | 2.0456638 | 0.0 | 0.040691257 | 0.54 | 3189 | tags=27%, list=14%, si | MM_2 |
| GOBP_REGULATION_OF_ | GOBP_REGULATION_OF_B_CELL_DIFFERENT | 37 | 0.447007 | 2.0438871 | 0.0 | 0.040864877 | 0.546 | 2814 | tags=24%, list=12%, si | MM_2 |
| GOBP_ANTIMICROBIAL_H | GOBP_ANTIMICROBIAL_HUMORAL_IMMUN | 79 | 0.387216 | 2.0425494 | 0.0 | 0.040719498 | 0.548 | 1887 | tags=25%, list=8%, sig | MM_2 |
| GOBP_INTRACELLULAR_M | GOBP_INTRACELLULAR_MRNA_LOCALIZATI | 15 | 0.581709 | 2.038733 | 0.01 | 0.04145481 | 0.557 | 4995 | tags=53%, list=21%, si | MM_2 |
| GOBP_LYMPHOCYTE_APC | GOBP_LYMPHOCYTE_APOPTOTIC_PROCESS | 94 | 0.362887 | 2.036624 | 0.0 | 0.041614167 | 0.562 | 3238 | tags=30%, list=14%, si | MM_2 |
| GOBP_REGULATION_OF_ | GOBP_REGULATION_OF_T_CELL_APOPTOTI | 48 | 0.407947 | 2.0264738 | 0.0 | 0.04417549 | 0.574 | 2618 | tags=27%, list=11%, si | MM_2 |
| GOBP_ANTIGEN_PROCES | GOBP_ANTIGEN_PROCESSING_AND_PRESEN | 105 | 0.357451 | 2.0238233 | 0.0 | 0.04424925 | 0.58 | 4400 | tags=45%, list=19%, si | MM_2 |
| GOBP_REGULATION_OF_ | GOBP_REGULATION_OF_MONONUCLEAR_C | 136 | 0.341261 | 2.0225248 | 0.0 | 0.044314094 | 0.585 | 3739 | tags=30%, list=16%, si | MM_2 |
| GOBP_REGULATION_OF_ | GOBP_REGULATION_OF_LYMPHOCYTE_APO | 71 | 0.384925 | 2.0185742 | 0.0 | 0.04501564 | 0.596 | 3120 | tags=30%, list=13%, si | MM_2 |
| GOBP_RESPONSE_TO_TY | GOBP_RESPONSE_TO_TYPE_II_INTERFERON | 122 | 0.347396 | 2.0123177 | 0.0 | 0.045814324 | 0.606 | 2937 | tags=29%, list=12%, si | MM_2 |
| GOBP_NEGATIVE_REGUL | GOBP_NEGATIVE_REGULATION_OF_TYPE_I_ | 23 | 0.511115 | 2.0097463 | 0.0 | 0.04591523 | 0.612 | 2779 | tags=39%, list=12%, si | MM_2 |

| | | | | | | | | | | |
|---|---|---|---|---|---|---|---|---|---|---|
| GOBP_POSITIVE_REGULA | GOBP_POSITIVE_REGULATION_OF_NEURON | 24 | 0.488357 | 2.0078669 | 0.0 | 0.04597408 | 0.614 | 1724 | tags=21%, list=7%, sig | MM_2 |
| GOBP_CYTOPLASMIC_PA | GOBP_CYTOPLASMIC_PATTERN_RECOGNITI | 159 | 0.333612 | 2.0026155 | 0.0 | 0.0470125 | 0.626 | 2889 | tags=22%, list=12%, si | MM_2 |
| ALZHEIMER_DISEASE | ALZHEIMER_DISEASE | 159 | 0.346744 | 1.9960196 | 0.0 | 0.049400713 | 0.644 | 5351 | tags=52%, list=23%, si | MM_2 |
| GOBP_POSITIVE_REGULA | GOBP_POSITIVE_REGULATION_OF_RESPONS | 62 | 0.373828 | 1.9897867 | 0.0 | 0.05061833 | 0.654 | 5001 | tags=45%, list=21%, si | MM_2 |
| NOD-LIKE_RECEPTOR_SIG | NOD-LIKE_RECEPTOR_SIGNALING_PATHWA | 151 | 0.351396 | 1.9829462 | 0.0 | 0.05273497 | 0.672 | 3272 | tags=26%, list=14%, si | MM_2 |
| HUMAN_CYTOMEGALOVI | HUMAN_CYTOMEGALOVIRUS_INFECTION | 216 | 0.321443 | 1.9807136 | 0.0 | 0.052992754 | 0.677 | 3754 | tags=29%, list=16%, si | MM_2 |
| GOBP_REGULATION_OF_ | GOBP_REGULATION_OF_OXIDATIVE_PHOSP | 23 | 0.505294 | 1.9802686 | 0.0061728396 | 0.05261178 | 0.678 | 3925 | tags=43%, list=17%, si | MM_2 |
| GOBP_CELLULAR_RESPON | GOBP_CELLULAR_RESPONSE_TO_BIOTIC_ST | 224 | 0.306882 | 1.9676191 | 0.0 | 0.056075845 | 0.707 | 3739 | tags=27%, list=16%, si | MM_2 |
| LEISHMANIASIS | LEISHMANIASIS | 60 | 0.38825 | 1.9671909 | 0.0 | 0.055541787 | 0.707 | 2633 | tags=25%, list=11%, si | MM_2 |
| GOBP_ANTIGEN_PROCES | GOBP_ANTIGEN_PROCESSING_AND_PRESEN | 24 | 0.501026 | 1.961433 | 0.00625 | 0.057109762 | 0.718 | 4364 | tags=58%, list=18%, si | MM_2 |
| SPLICEOSOME | SPLICEOSOME | 127 | 0.358068 | 1.9578935 | 0.0 | 0.05848352 | 0.728 | 14619 | tags=99%, list=62%, si | MM_2 |
| GOBP_AEROBIC_RESPIRA | GOBP_AEROBIC_RESPIRATION | 170 | 0.337832 | 1.9562591 | 0.0 | 0.05841623 | 0.733 | 5058 | tags=46%, list=21%, si | MM_2 |
| GOBP_POSITIVE_REGULA | GOBP_POSITIVE_REGULATION_OF_TUMOR_ | 113 | 0.355424 | 1.9518125 | 0.0 | 0.059432328 | 0.739 | 2468 | tags=24%, list=10%, si | MM_2 |
| GOBP_T_CELL_APOPTOTI | GOBP_T_CELL_APOPTOTIC_PROCESS | 67 | 0.386866 | 1.9516829 | 0.0 | 0.058892034 | 0.739 | 3238 | tags=31%, list=14%, si | MM_2 |
| GOBP_MITOCHONDRIAL_ | GOBP_MITOCHONDRIAL_TRANSLATION | 128 | 0.33442 | 1.9509776 | 0.0 | 0.05856954 | 0.741 | 5034 | tags=41%, list=21%, si | MM_2 |
| GOBP_POSITIVE_REGULA | GOBP_POSITIVE_REGULATION_OF_B_CELL_ | 18 | 0.543593 | 1.9458996 | 0.0 | 0.059483968 | 0.747 | 4919 | tags=44%, list=21%, si | MM_2 |
| GOBP_GOLGI_TO_VACUO | GOBP_GOLGI_TO_VACUOLE_TRANSPORT | 21 | 0.485355 | 1.9407599 | 0.0058139535 | 0.06036087 | 0.757 | 12161 | tags=100%, list=52%, | MM_2 |
| LYSOSOME | LYSOSOME | 118 | 0.345231 | 1.9335861 | 0.0 | 0.06286732 | 0.775 | 5282 | tags=52%, list=22%, si | MM_2 |
| MISMATCH_REPAIR | MISMATCH_REPAIR | 21 | 0.50354 | 1.9309062 | 0.0 | 0.06328565 | 0.781 | 4698 | tags=52%, list=20%, si | MM_2 |
| GOBP_CELLULAR_RESPIR | GOBP_CELLULAR_RESPIRATION | 218 | 0.323542 | 1.928333 | 0.0 | 0.06415943 | 0.79 | 4784 | tags=40%, list=20%, si | MM_2 |
| KAPOSI_SARCOMA-ASSO | KAPOSI_SARCOMA-ASSOCIATED_HERPESVIR | 176 | 0.328435 | 1.9233443 | 0.0 | 0.06544443 | 0.798 | 3373 | tags=25%, list=14%, si | MM_2 |
| OSTEOCLAST_DIFFERENTI | OSTEOCLAST_DIFFERENTIATION | 121 | 0.344766 | 1.9222807 | 0.0 | 0.065280065 | 0.799 | 3484 | tags=26%, list=15%, si | MM_2 |
| STAPHYLOCOCCUS_AURE | STAPHYLOCOCCUS_AUREUS_INFECTION | 44 | 0.406429 | 1.9208536 | 0.0 | 0.065472685 | 0.802 | 1455 | tags=27%, list=6%, sig | MM_2 |
| GOBP_CHAPERONE_COFA | GOBP_CHAPERONE_COFACTOR_DEPENDENT | 31 | 0.433589 | 1.9185877 | 0.007874016 | 0.065740824 | 0.806 | 3089 | tags=32%, list=13%, si | MM_2 |
| GOBP_MACROPHAGE_AC | GOBP_MACROPHAGE_ACTIVATION_INVOLV | 20 | 0.521426 | 1.9133431 | 0.0 | 0.06728348 | 0.818 | 2579 | tags=30%, list=11%, si | MM_2 |
| GOBP_NON_CANONICAL_ | GOBP_NON_CANONICAL_NF_KAPPAB_SIGN | 115 | 0.330161 | 1.8983324 | 0.0 | 0.07357743 | 0.841 | 3669 | tags=30%, list=16%, si | MM_2 |
| GOBP_TRANSLATIONAL_I | GOBP_TRANSLATIONAL_INITIATION | 112 | 0.348298 | 1.8972123 | 0.0 | 0.073613 | 0.844 | 2779 | tags=20%, list=12%, si | MM_2 |
| GOBP_REGULATION_OF_ | GOBP_REGULATION_OF_PROGRAMMED_NE | 27 | 0.465759 | 1.8892444 | 0.0 | 0.076923974 | 0.862 | 4754 | tags=52%, list=20%, si | MM_2 |
| GOBP_POSITIVE_REGULA | GOBP_POSITIVE_REGULATION_OF_TORC1_S | 52 | 0.392876 | 1.8882848 | 0.0 | 0.076710485 | 0.863 | 3793 | tags=31%, list=16%, si | MM_2 |
| GOBP_LATE_ENDOSOME_ | GOBP_LATE_ENDOSOME_TO_VACUOLE_TRA | 16 | 0.53051 | 1.8875183 | 0.0 | 0.076544195 | 0.865 | 5323 | tags=63%, list=23%, si | MM_2 |
| GOBP_MACROPHAGE_CH | GOBP_MACROPHAGE_CHEMOTAXIS | 45 | 0.397139 | 1.8855183 | 0.0 | 0.076608635 | 0.867 | 3975 | tags=38%, list=17%, si | MM_2 |
| GOBP_PIGMENT_CELL_D | GOBP_PIGMENT_CELL_DIFFERENTIATION | 37 | 0.419439 | 1.8802366 | 0.0 | 0.07846782 | 0.876 | 1513 | tags=27%, list=6%, sig | MM_2 |
| GOBP_B_CELL_APOPTOTI | GOBP_B_CELL_APOPTOTIC_PROCESS | 29 | 0.441222 | 1.8796635 | 0.0 | 0.078110136 | 0.877 | 3866 | tags=38%, list=16%, si | MM_2 |
| GOBP_KILLING_OF_CELLS | GOBP_KILLING_OF_CELLS_OF_ANOTHER_OF | 71 | 0.350285 | 1.8785444 | 0.0 | 0.07819433 | 0.878 | 1489 | tags=17%, list=6%, sig | MM_2 |
| GOBP_REGULATION_OF_ | GOBP_REGULATION_OF_SUPEROXIDE_MET | 34 | 0.430078 | 1.8760284 | 0.0 | 0.079003595 | 0.883 | 4410 | tags=47%, list=19%, si | MM_2 |
| GOBP_REGULATION_OF_ | GOBP_REGULATION_OF_MITOCHONDRIAL_ | 26 | 0.446873 | 1.8696461 | 0.0 | 0.080732614 | 0.892 | 5306 | tags=54%, list=22%, si | MM_2 |
| GOBP_LIPOPOLYSACCHAR | GOBP_LIPOPOLYSACCHARIDE_MEDIATED_SI | 57 | 0.380087 | 1.8681414 | 0.0 | 0.080783375 | 0.896 | 5039 | tags=46%, list=21%, si | MM_2 |
| GOBP_POSITIVE_REGULA | GOBP_POSITIVE_REGULATION_OF_DEFENSE | 31 | 0.439983 | 1.867694 | 0.0 | 0.080456235 | 0.896 | 2289 | tags=29%, list=10%, si | MM_2 |
| GOBP_POSITIVE_REGULA | GOBP_POSITIVE_REGULATION_OF_EXECUTI | 19 | 0.50687 | 1.8612659 | 0.023121387 | 0.082515284 | 0.904 | 2995 | tags=42%, list=13%, si | MM_2 |
| GOBP_REGULATION_OF_ | GOBP_REGULATION_OF_B_CELL_APOPTOTI | 23 | 0.465911 | 1.8597646 | 0.0 | 0.08254862 | 0.906 | 3866 | tags=39%, list=16%, si | MM_2 |
| GOBP_REGULATION_OF_ | GOBP_REGULATION_OF_TYPE_2_IMMUNE_ | 34 | 0.443127 | 1.8585303 | 0.0 | 0.08243789 | 0.908 | 884 | tags=18%, list=4%, sig | MM_2 |
| GOBP_POSITIVE_REGULA | GOBP_POSITIVE_REGULATION_OF_MEMBR | 39 | 0.40515 | 1.8561671 | 0.0 | 0.08289611 | 0.911 | 5169 | tags=56%, list=22%, si | MM_2 |
| GOBP_DE_NOVO_PROTE | GOBP_DE_NOVO_PROTEIN_FOLDING | 38 | 0.41746 | 1.8500426 | 0.008130081 | 0.08502525 | 0.918 | 3089 | tags=29%, list=13%, si | MM_2 |
| GOBP_MORPHOGENESIS_ | GOBP_MORPHOGENESIS_OF_AN_ENDOTHE | 19 | 0.517152 | 1.8496407 | 0.01604278 | 0.08458131 | 0.918 | 5404 | tags=58%, list=23%, si | MM_2 |
| GOBP_NEGATIVE_REGUL | GOBP_NEGATIVE_REGULATION_OF_MACRO | 33 | 0.435476 | 1.8463027 | 0.0 | 0.0858496 | 0.925 | 5153 | tags=48%, list=22%, si | MM_2 |
| GOBP_T_HELPER_2_CELL | GOBP_T_HELPER_2_CELL_CYTOKINE_PRODU | 17 | 0.503332 | 1.8399225 | 0.0051282053 | 0.088196844 | 0.93 | 434 | tags=18%, list=2%, sig | MM_2 |
| GOBP_PROTEIN_INSERTIO | GOBP_PROTEIN_INSERTION_INTO_ER_MEM | 26 | 0.456378 | 1.8394535 | 0.0 | 0.087837294 | 0.93 | 5214 | tags=50%, list=22%, si | MM_2 |
| CYTOKINE-CYTOKINE_RE | CYTOKINE-CYTOKINE_RECEPTOR_INTERACTI | 239 | 0.287425 | 1.8388872 | 0.0 | 0.087389976 | 0.93 | 2413 | tags=21%, list=10%, si | MM_2 |
| CHEMOKINE_SIGNALING_ | CHEMOKINE_SIGNALING_PATHWAY | 177 | 0.331404 | 1.834534 | 0.0 | 0.08863377 | 0.935 | 2493 | tags=20%, list=11%, si | MM_2 |
| GOBP_POSITIVE_REGULA | GOBP_POSITIVE_REGULATION_OF_CHEMOK | 79 | 0.366378 | 1.8338106 | 0.0 | 0.08850092 | 0.935 | 2341 | tags=28%, list=10%, si | MM_2 |
| GOBP_GTP_METABOLIC_ | GOBP_GTP_METABOLIC_PROCESS | 26 | 0.432997 | 1.8298967 | 0.012738854 | 0.09001489 | 0.943 | 1365 | tags=23%, list=6%, sig | MM_2 |
| GOBP_POSITIVE_REGULA | GOBP_POSITIVE_REGULATION_OF_CELLULA | 22 | 0.475004 | 1.829183 | 0.005988024 | 0.08956132 | 0.943 | 1402 | tags=27%, list=6%, sig | MM_2 |
| NON-ALCOHOLIC_FATTY_ | NON-ALCOHOLIC_FATTY_LIVER_DISEASE_(N | 141 | 0.30436 | 1.8236742 | 0.0 | 0.09181515 | 0.951 | 5351 | tags=45%, list=23%, si | MM_2 |
| GOBP_POSITIVE_REGULA | GOBP_POSITIVE_REGULATION_OF_INTERFE | 26 | 0.437593 | 1.8231537 | 0.0 | 0.09145579 | 0.952 | 2379 | tags=27%, list=10%, si | MM_2 |
| GOBP_REGULATION_OF_ | GOBP_REGULATION_OF_CYSTEINE_TYPE_EN | 16 | 0.517494 | 1.8219345 | 0.014563107 | 0.09124797 | 0.953 | 3242 | tags=50%, list=14%, si | MM_2 |
| GOBP_MONONUCLEAR_C | GOBP_MONONUCLEAR_CELL_MIGRATION | 199 | 0.311355 | 1.8185414 | 0.0 | 0.092130676 | 0.955 | 3739 | tags=29%, list=16%, si | MM_2 |
| GOBP_NOTOCHORD_DEV | GOBP_NOTOCHORD_DEVELOPMENT | 22 | 0.447894 | 1.8183947 | 0.013245033 | 0.09158861 | 0.955 | 2334 | tags=36%, list=10%, si | MM_2 |
| GOBP_AEROBIC_ELECTRO | GOBP_AEROBIC_ELECTRON_TRANSPORT_CH | 63 | 0.363042 | 1.8182955 | 0.0 | 0.09105114 | 0.955 | 4879 | tags=48%, list=21%, si | MM_2 |
| GOBP_NEGATIVE_REGUL | GOBP_NEGATIVE_REGULATION_OF_CELL_S | 16 | 0.517516 | 1.8178465 | 0.024752475 | 0.0907045 | 0.955 | 5683 | tags=69%, list=24%, si | MM_2 |
| GOBP_NEGATIVE_REGUL | GOBP_NEGATIVE_REGULATION_OF_LEUKO | 46 | 0.389242 | 1.8176916 | 0.0 | 0.09015242 | 0.955 | 2616 | tags=30%, list=11%, si | MM_2 |
| GOBP_UBIQUITIN_DEPEN | GOBP_UBIQUITIN_DEPENDENT_PROTEIN_C | 29 | 0.422296 | 1.8163966 | 0.0074074073 | 0.090439096 | 0.956 | 13652 | tags=100%, list=58%, | MM_2 |
| GOBP_PROTEIN_IMPORT_ | GOBP_PROTEIN_IMPORT_INTO_MITOCHON | 19 | 0.492893 | 1.8131452 | 0.0048076925 | 0.09133956 | 0.957 | 4219 | tags=47%, list=18%, si | MM_2 |
| GOBP_REGULATION_OF_ | GOBP_REGULATION_OF_CELLULAR_RESPIRA | 52 | 0.374495 | 1.8119924 | 0.0 | 0.091290414 | 0.959 | 3925 | tags=37%, list=17%, si | MM_2 |
| GOBP_REGULATION_OF_ | GOBP_REGULATION_OF_CELL_ADHESION_M | 47 | 0.386678 | 1.8045576 | 0.0 | 0.09431571 | 0.962 | 729 | tags=13%, list=3%, sig | MM_2 |
| GOBP_POSITIVE_REGULA | GOBP_POSITIVE_REGULATION_OF_APOPTO | 161 | 0.328123 | 1.8022623 | 0.0 | 0.09482371 | 0.962 | 4141 | tags=33%, list=18%, si | MM_2 |
| GOBP_DISRUPTION_OF_A | GOBP_DISRUPTION_OF_ANATOMICAL_STRU | 73 | 0.34796 | 1.802024 | 0.0 | 0.09429463 | 0.962 | 1489 | tags=16%, list=6%, sig | MM_2 |
| GOBP_LYMPHOCYTE_HO | GOBP_LYMPHOCYTE_HOMEOSTASIS | 92 | 0.328588 | 1.8016998 | 0.0 | 0.09403293 | 0.962 | 3383 | tags=28%, list=14%, si | MM_2 |
| GOBP_CELL_REDOX_HOM | GOBP_CELL_REDOX_HOMEOSTASIS | 28 | 0.450014 | 1.8006821 | 0.014492754 | 0.09387934 | 0.963 | 4993 | tags=50%, list=21%, si | MM_2 |
| GOBP_RELEASE_OF_CYTO | GOBP_RELEASE_OF_CYTOCHROME_C_FROM | 66 | 0.365346 | 1.8004444 | 0.0 | 0.093421206 | 0.963 | 3574 | tags=35%, list=15%, si | MM_2 |
| GOBP_TYPE_2_IMMUNE_ | GOBP_TYPE_2_IMMUNE_RESPONSE | 46 | 0.378718 | 1.8000543 | 0.0 | 0.09296831 | 0.963 | 3498 | tags=30%, list=15%, si | MM_2 |
| GOBP_CELLULAR_EXTRAV | GOBP_CELLULAR_EXTRAVASATION | 74 | 0.339982 | 1.7978684 | 0.0 | 0.09343874 | 0.964 | 3163 | tags=28%, list=13%, si | MM_2 |
| GOBP_REGULATION_OF_ | GOBP_REGULATION_OF_PATTERN_RECOGN | 157 | 0.316175 | 1.7973707 | 0.0 | 0.09299457 | 0.964 | 2616 | tags=18%, list=11%, si | MM_2 |
| GOBP_NEUTROPHIL_MIG | GOBP_NEUTROPHIL_MIGRATION | 104 | 0.326279 | 1.7967589 | 0.0 | 0.09283292 | 0.964 | 1027 | tags=14%, list=4%, sig | MM_2 |
| GOBP_NEGATIVE_REGUL | GOBP_NEGATIVE_REGULATION_OF_STRIATE | 39 | 0.400993 | 1.7949309 | 0.018867925 | 0.093274884 | 0.964 | 4117 | tags=36%, list=17%, si | MM_2 |
| GOBP_INTERFERON_ALPH | GOBP_INTERFERON_ALPHA_PRODUCTION | 33 | 0.406365 | 1.7933841 | 0.0 | 0.09359266 | 0.964 | 2579 | tags=24%, list=11%, si | MM_2 |
| GOBP_NEGATIVE_REGUL | GOBP_NEGATIVE_REGULATION_OF_VASCUL | 21 | 0.458937 | 1.792945 | 0.0 | 0.09318908 | 0.964 | 545 | tags=19%, list=2%, sig | MM_2 |
| GOBP_POSITIVE_REGULA | GOBP_POSITIVE_REGULATION_OF_NUCLEO | 74 | 0.332901 | 1.7922764 | 0.0 | 0.09289477 | 0.964 | 4416 | tags=31%, list=19%, si | MM_2 |
| GOBP_PROTON_MOTIVE_ | GOBP_PROTON_MOTIVE_FORCE_DRIVEN_A | 52 | 0.377257 | 1.7893529 | 0.0 | 0.093930475 | 0.966 | 5841 | tags=60%, list=25%, si | MM_2 |
| GOBP_NEGATIVE_REGUL | GOBP_NEGATIVE_REGULATION_OF_CELLUL | 51 | 0.366488 | 1.7864127 | 0.011111111 | 0.09488509 | 0.967 | 3025 | tags=29%, list=13%, si | MM_2 |
| GOBP_MULTIVESICULAR_ | GOBP_MULTIVESICULAR_BODY_SORTING_P | 34 | 0.411195 | 1.7857693 | 0.023255814 | 0.094764344 | 0.967 | 3881 | tags=41%, list=16%, si | MM_2 |
| GOBP_RIG_I_SIGNALING_ | GOBP_RIG_I_SIGNALING_PATHWAY | 24 | 0.440566 | 1.7855368 | 0.017441861 | 0.09428053 | 0.967 | 2379 | tags=29%, list=10%, si | MM_2 |
| GOBP_SEQUESTERING_O | GOBP_SEQUESTERING_OF_TRIGLYCERIDE | 18 | 0.485793 | 1.7852992 | 0.011235955 | 0.09382372 | 0.967 | 1357 | tags=28%, list=6%, sig | MM_2 |

| | | | | | | | | | | |
|---|---|---|---|---|---|---|---|---|---|---|
| HUMAN_T-CELL_LEUKEM | HUMAN_T-CELL_LEUKEMIA_VIRUS_1_INFEC | 215 | 0.294796 | 1.7847787 | 0.0 | 0.09366441 | 0.967 | 4150 | tags=33%, list=18%, si | MM_2 |
| GOBP_REGULATION_OF_ | GOBP_REGULATION_OF_GLIAL_CELL_MIGRA | 26 | 0.440018 | 1.7835621 | 0.012903226 | 0.09375999 | 0.968 | 4572 | tags=42%, list=19%, si | MM_2 |
| GOBP_DEVELOPMENTAL_ | GOBP_DEVELOPMENTAL_PIGMENTATION | 50 | 0.368625 | 1.7821445 | 0.010869565 | 0.09401353 | 0.969 | 2279 | tags=26%, list=10%, si | MM_2 |
| GOBP_POSITIVE_REGULA | GOBP_POSITIVE_REGULATION_OF_MONON | 94 | 0.31705 | 1.7797917 | 0.0 | 0.094778255 | 0.971 | 3307 | tags=24%, list=14%, si | MM_2 |
| GOBP_POSITIVE_REGULA | GOBP_POSITIVE_REGULATION_OF_CELL_MA | 61 | 0.365187 | 1.7781767 | 0.0 | 0.09511819 | 0.974 | 3484 | tags=31%, list=15%, si | MM_2 |
| GOBP_RETROGRADE_VES | GOBP_RETROGRADE_VESICLE_MEDIATED_T | 45 | 0.368684 | 1.7779448 | 0.009259259 | 0.094679825 | 0.974 | 11868 | tags=96%, list=50%, si | MM_2 |
| GOBP_REGULATION_OF_ | GOBP_REGULATION_OF_EXECUTION_PHASE | 26 | 0.415261 | 1.7765692 | 0.0 | 0.094800964 | 0.975 | 3162 | tags=38%, list=13%, si | MM_2 |
| GOBP_BRANCHED_CHAIN | GOBP_BRANCHED_CHAIN_AMINO_ACID_CA | 21 | 0.453557 | 1.7758888 | 0.011560693 | 0.09471158 | 0.975 | 4036 | tags=43%, list=17%, si | MM_2 |
| GOBP_RIBOSOME_ASSEM | GOBP_RIBOSOME_ASSEMBLY | 60 | 0.336215 | 1.7757714 | 0.0 | 0.09433009 | 0.975 | 6063 | tags=53%, list=26%, si | MM_2 |
| GOBP_SYMBIONT_ENTRY | GOBP_SYMBIONT_ENTRY_INTO_HOST | 97 | 0.336027 | 1.7743293 | 0.0 | 0.09442048 | 0.975 | 4680 | tags=44%, list=20%, si | MM_2 |
| GOBP_REGULATION_OF_ | GOBP_REGULATION_OF_ANTIGEN_PROCESS | 21 | 0.469487 | 1.7730742 | 0.02793296 | 0.09450906 | 0.975 | 2130 | tags=33%, list=9%, sig | MM_2 |
| GOBP_PIGMENT_GRANU | GOBP_PIGMENT_GRANULE_ORGANIZATION | 39 | 0.39992 | 1.7705072 | 0.018181818 | 0.09514118 | 0.977 | 2392 | tags=28%, list=10%, si | MM_2 |
| INFLAMMATORY_BOWEL | INFLAMMATORY_BOWEL_DISEASE_(IBD) | 52 | 0.370819 | 1.7688965 | 0.0 | 0.09544042 | 0.977 | 3619 | tags=37%, list=15%, si | MM_2 |
| GOBP_REGULATION_OF_ | GOBP_REGULATION_OF_LEUKOCYTE_MIGRA | 230 | 0.276247 | 1.7671854 | 0.0 | 0.09581151 | 0.979 | 3130 | tags=25%, list=13%, si | MM_2 |
| GOBP_REGULATION_OF_ | GOBP_REGULATION_OF_EARLY_ENDOSOME | 18 | 0.472959 | 1.7668188 | 0.0054347827 | 0.095533095 | 0.979 | 12452 | tags=100%, list=53%, | MM_2 |
| GOBP_REGULATION_OF_ | GOBP_REGULATION_OF_INCLUSION_BODY_ | 19 | 0.457284 | 1.7606606 | 0.011111111 | 0.09842947 | 0.98 | 3478 | tags=32%, list=15%, si | MM_2 |
| OTHER_GLYCAN_DEGRAD | OTHER_GLYCAN_DEGRADATION | 16 | 0.494478 | 1.7581449 | 0.028089888 | 0.09976946 | 0.981 | 5263 | tags=63%, list=22%, si | MM_2 |
| GOBP_LEUKOCYTE_APOP | GOBP_LEUKOCYTE_APOPTOTIC_PROCESS | 131 | 0.298865 | 1.7559736 | 0.0 | 0.100177966 | 0.982 | 4466 | tags=33%, list=19%, si | MM_2 |
| GOBP_TAIL_ANCHORED_ | GOBP_TAIL_ANCHORED_MEMBRANE_PROT | 15 | 0.48142 | 1.7540096 | 0.015228426 | 0.10092 | 0.982 | 5056 | tags=53%, list=21%, si | MM_2 |
| GOBP_REGULATION_OF_ | GOBP_REGULATION_OF_AEROBIC_RESPIRAT | 33 | 0.394785 | 1.7533417 | 0.0070422534 | 0.10090512 | 0.983 | 3925 | tags=39%, list=17%, si | MM_2 |
| GOBP_BIOLOGICAL_PRO | GOBP_BIOLOGICAL_PROCESS_INVOLVED_IN | 140 | 0.313673 | 1.7525818 | 0.0 | 0.10083836 | 0.984 | 4680 | tags=42%, list=20%, si | MM_2 |
| GOBP_REGULATION_OF_ | GOBP_REGULATION_OF_DEFENSE_RESPONS | 67 | 0.34117 | 1.7511373 | 0.0 | 0.101482406 | 0.986 | 2633 | tags=25%, list=11%, si | MM_2 |
| GOBP_POSITIVE_REGULA | GOBP_POSITIVE_REGULATION_OF_PROTEIN | 52 | 0.35091 | 1.7486961 | 0.0 | 0.102096386 | 0.986 | 4416 | tags=33%, list=19%, si | MM_2 |
| GOBP_ESTABLISHMENT_ | GOBP_ESTABLISHMENT_OF_PROTEIN_LOCA | 25 | 0.423798 | 1.7475203 | 0.005524862 | 0.10237318 | 0.986 | 5686 | tags=48%, list=24%, si | MM_2 |
| GOBP_MYELOID_LEUKOC | GOBP_MYELOID_LEUKOCYTE_MIGRATION | 222 | 0.27868 | 1.746531 | 0.0 | 0.102353424 | 0.986 | 3026 | tags=23%, list=13%, si | MM_2 |
| GOBP_REGULATION_OF_ | GOBP_REGULATION_OF_TRANSLATION_IN_ | 24 | 0.442669 | 1.7460802 | 0.006369427 | 0.10210183 | 0.986 | 2003 | tags=21%, list=8%, sig | MM_2 |
| GOBP_INCLUSION_BODY | GOBP_INCLUSION_BODY_ASSEMBLY | 27 | 0.412872 | 1.7449743 | 0.016260162 | 0.10217533 | 0.986 | 5154 | tags=41%, list=22%, si | MM_2 |
| APOPTOSIS | APOPTOSIS | 137 | 0.288419 | 1.7422205 | 0.0 | 0.102926135 | 0.986 | 4069 | tags=33%, list=17%, si | MM_2 |
| GOBP_NADH_DEHYDROG | GOBP_NADH_DEHYDROGENASE_COMPLEX_ | 58 | 0.358728 | 1.7402089 | 0.0 | 0.103318974 | 0.986 | 5618 | tags=52%, list=24%, si | MM_2 |
| GOBP_POSITIVE_REGULA | GOBP_POSITIVE_REGULATION_OF_RESPONS | 381 | 0.250467 | 1.7372453 | 0.0 | 0.10441104 | 0.987 | 3077 | tags=21%, list=13%, si | MM_2 |
| GOBP_EXTRACELLULAR_ | GOBP_EXTRACELLULAR_MATRIX_ASSEMBLY | 48 | 0.36494 | 1.7360897 | 0.0 | 0.10464283 | 0.987 | 1657 | tags=21%, list=7%, sig | MM_2 |
| GOBP_LATE_ENDOSOME_ | GOBP_LATE_ENDOSOME_TO_VACUOLE_TRA | 25 | 0.436728 | 1.7333874 | 0.0 | 0.10593138 | 0.987 | 3881 | tags=40%, list=16%, si | MM_2 |
| GOBP_POSITIVE_REGULA | GOBP_POSITIVE_REGULATION_OF_CANONI | 148 | 0.292498 | 1.7319915 | 0.0 | 0.1060922 | 0.988 | 3125 | tags=26%, list=13%, si | MM_2 |
| GOBP_VIRION_ASSEMBLY | GOBP_VIRION_ASSEMBLY | 26 | 0.4332 | 1.7306017 | 0.0070422534 | 0.106383696 | 0.988 | 5938 | tags=65%, list=25%, si | MM_2 |
| GOBP_RETROGRADE_TRA | GOBP_RETROGRADE_TRANSPORT_ENDOSO | 87 | 0.319649 | 1.729463 | 0.0 | 0.10649019 | 0.988 | 14066 | tags=97%, list=60%, si | MM_2 |
| GOBP_APOPTOTIC_MITO | GOBP_APOPTOTIC_MITOCHONDRIAL_CHAN | 118 | 0.322335 | 1.7290511 | 0.0 | 0.106166594 | 0.988 | 4722 | tags=36%, list=20%, si | MM_2 |
| GOBP_LYSOSOMAL_LUM | GOBP_LYSOSOMAL_LUMEN_ACIDIFICATION | 15 | 0.490147 | 1.7281555 | 0.005376344 | 0.105953686 | 0.988 | 5028 | tags=67%, list=21%, si | MM_2 |
| GOBP_POSITIVE_REGULA | GOBP_POSITIVE_REGULATION_OF_MITOCH | 17 | 0.47046 | 1.7268871 | 0.015706806 | 0.105975874 | 0.99 | 5034 | tags=59%, list=21%, si | MM_2 |
| GOBP_REGULATION_OF_ | GOBP_REGULATION_OF_RELEASE_OF_CYTO | 49 | 0.355783 | 1.723058 | 0.010204081 | 0.10770978 | 0.99 | 3574 | tags=35%, list=15%, si | MM_2 |
| GOBP_MITOCHONDRIAL_ | GOBP_MITOCHONDRIAL_CYTOCHROME_C_ | 19 | 0.456422 | 1.7217398 | 0.021052632 | 0.108109646 | 0.991 | 3022 | tags=32%, list=13%, si | MM_2 |
| GOBP_MRNA_TRANSCRIP | GOBP_MRNA_TRANSCRIPTION | 50 | 0.357108 | 1.721004 | 0.01010101 | 0.10798354 | 0.993 | 13845 | tags=98%, list=59%, si | MM_2 |
| GOBP_VESICLE_TETHERIN | GOBP_VESICLE_TETHERING | 31 | 0.399898 | 1.7204111 | 0.0074626864 | 0.107845284 | 0.993 | 14181 | tags=100%, list=60%, | MM_2 |
| GOBP_REGULATION_OF_ | GOBP_REGULATION_OF_LEUKOCYTE_APOPT | 103 | 0.320629 | 1.7173007 | 0.0 | 0.109323196 | 0.993 | 736 | tags=13%, list=3%, sig | MM_2 |
| GOBP_REGULATION_OF_ | GOBP_REGULATION_OF_MITOCHONDRIAL_ | 39 | 0.367544 | 1.7166972 | 0.0 | 0.109180816 | 0.993 | 5169 | tags=49%, list=22%, si | MM_2 |
| GOBP_REGULATION_OF_ | GOBP_REGULATION_OF_BIOLOGICAL_PROC | 64 | 0.340795 | 1.7162416 | 0.0 | 0.10892152 | 0.994 | 1752 | tags=27%, list=7%, sig | MM_2 |
| GOBP_ENDOLYSOSOMAL | GOBP_ENDOLYSOSOMAL_TOLL_LIKE_RECEP | 47 | 0.358377 | 1.7159556 | 0.0 | 0.108625636 | 0.994 | 4331 | tags=36%, list=18%, si | MM_2 |
| GOBP_REGULATION_OF_ | GOBP_REGULATION_OF_LIPOPOLYSACCHAR | 29 | 0.406845 | 1.7150149 | 0.014814815 | 0.1086388 | 0.995 | 1733 | tags=28%, list=7%, sig | MM_2 |
| GOBP_CD4_POSITIVE_AL | GOBP_CD4_POSITIVE_ALPHA_BETA_T_CELL | 26 | 0.415028 | 1.7148064 | 0.0124223605 | 0.10830321 | 0.995 | 434 | tags=15%, list=2%, sig | MM_2 |
| GOBP_REGULATION_OF_ | GOBP_REGULATION_OF_HEAT_GENERATION | 17 | 0.470785 | 1.714163 | 0.025125628 | 0.108195424 | 0.995 | 610 | tags=18%, list=3%, sig | MM_2 |
| GOBP_GLYCOLYTIC_PRO | GOBP_GLYCOLYTIC_PROCESS_THROUGH_GL | 20 | 0.448367 | 1.7115227 | 0.011111111 | 0.109163955 | 0.996 | 2308 | tags=40%, list=10%, si | MM_2 |
| GOBP_POSITIVE_REGULA | GOBP_POSITIVE_REGULATION_OF_TYPE_2_ | 18 | 0.470323 | 1.7110972 | 0.026041666 | 0.10892798 | 0.996 | 3435 | tags=44%, list=15%, si | MM_2 |
| GOBP_NEGATIVE_REGUL | GOBP_NEGATIVE_REGULATION_OF_EXTRIN | 26 | 0.421178 | 1.7100959 | 0.013333334 | 0.10913238 | 0.996 | 4393 | tags=50%, list=19%, si | MM_2 |
| GOBP_VIRAL_BUDDING | GOBP_VIRAL_BUDDING | 18 | 0.470679 | 1.7073306 | 0.021978023 | 0.11024661 | 0.996 | 4859 | tags=50%, list=21%, si | MM_2 |
| GOBP_ENDOSOME_ORGA | GOBP_ENDOSOME_ORGANIZATION | 68 | 0.331328 | 1.7068899 | 0.0 | 0.110131204 | 0.996 | 4354 | tags=40%, list=18%, si | MM_2 |
| GOBP_POSITIVE_REGULA | GOBP_POSITIVE_REGULATION_OF_MONOC | 21 | 0.432582 | 1.7039604 | 0.0055555557 | 0.111159936 | 0.996 | 3307 | tags=33%, list=14%, si | MM_2 |
| GOBP_PROTEIN_K11_LIN | GOBP_PROTEIN_K11_LINKED_UBIQUITINAT | 29 | 0.410593 | 1.7031915 | 0.013793103 | 0.111085124 | 0.997 | 13928 | tags=100%, list=59%, | MM_2 |
| GOBP_MIRNA_MEDIATED | GOBP_MIRNA_MEDIATED_GENE_SILENCING | 17 | 0.468363 | 1.7010913 | 0.018181818 | 0.11177578 | 0.997 | 3996 | tags=41%, list=17%, si | MM_2 |
| GOBP_SUPPRESSION_OF_ | GOBP_SUPPRESSION_OF_VIRAL_RELEASE_B | 22 | 0.430159 | 1.6981269 | 0.005952381 | 0.113098934 | 0.997 | 1559 | tags=27%, list=7%, sig | MM_2 |
| GOBP_GRANULOCYTE_M | GOBP_GRANULOCYTE_MIGRATION | 146 | 0.288187 | 1.6979853 | 0.0 | 0.11275703 | 0.997 | 1489 | tags=14%, list=6%, sig | MM_2 |
| GOBP_NEGATIVE_REGUL | GOBP_NEGATIVE_REGULATION_OF_UBIQUI | 18 | 0.457038 | 1.6966338 | 0.016216217 | 0.113058664 | 0.997 | 3913 | tags=39%, list=17%, si | MM_2 |
| GOBP_POSITIVE_REGULA | GOBP_POSITIVE_REGULATION_OF_ACTIN_F | 52 | 0.340943 | 1.695891 | 0.011111111 | 0.113087386 | 0.997 | 2777 | tags=23%, list=12%, si | MM_2 |
| RHEUMATOID_ARTHRITIS | RHEUMATOID_ARTHRITIS | 73 | 0.322911 | 1.6938722 | 0.0 | 0.11370062 | 0.998 | 3962 | tags=36%, list=17%, si | MM_2 |
| ANTIGEN_PROCESSING_A | ANTIGEN_PROCESSING_AND_PRESENTATIO | 67 | 0.326667 | 1.6927565 | 0.0 | 0.11386438 | 0.998 | 3414 | tags=37%, list=14%, si | MM_2 |
| GOBP_NEGATIVE_REGUL | GOBP_NEGATIVE_REGULATION_OF_GTPASE | 22 | 0.443872 | 1.6918423 | 0.010752688 | 0.113915116 | 0.998 | 4000 | tags=32%, list=17%, si | MM_2 |
| GOBP_EMBRYONIC_BRAI | GOBP_EMBRYONIC_BRAIN_DEVELOPMENT | 24 | 0.413758 | 1.690507 | 0.0067114094 | 0.114363186 | 0.998 | 1168 | tags=29%, list=5%, sig | MM_2 |
| GOBP_REGULATION_OF_ | GOBP_REGULATION_OF_DEFENSE_RESPONS | 42 | 0.367511 | 1.6846521 | 0.009345794 | 0.11733782 | 0.998 | 2633 | tags=29%, list=11%, si | MM_2 |
| GOBP_REGULATION_OF_ | GOBP_REGULATION_OF_VACUOLE_ORGANI | 54 | 0.334954 | 1.6845102 | 0.013513514 | 0.116937906 | 0.998 | 5650 | tags=50%, list=24%, si | MM_2 |
| GOBP_MAINTENANCE_OI | GOBP_MAINTENANCE_OF_PROTEIN_LOCAT | 97 | 0.310987 | 1.6833409 | 0.0 | 0.1171497 | 0.998 | 3668 | tags=35%, list=16%, si | MM_2 |
| GOBP_REGULATION_OF_ | GOBP_REGULATION_OF_IMMUNOGLOBULI | 81 | 0.319691 | 1.6827286 | 0.0 | 0.11710723 | 0.998 | 2910 | tags=27%, list=12%, si | MM_2 |
| GOBP_COLLAGEN_FIBRIL | GOBP_COLLAGEN_FIBRIL_ORGANIZATION | 63 | 0.341772 | 1.6826663 | 0.0 | 0.11665383 | 0.998 | 1627 | tags=19%, list=7%, sig | MM_2 |
| GOBP_RESPONSE_TO_GR | GOBP_RESPONSE_TO_GROWTH_HORMONE | 20 | 0.43937 | 1.682363 | 0.005882353 | 0.11637459 | 0.998 | 3498 | tags=35%, list=15%, si | MM_2 |
| GOBP_REGULATION_OF_ | GOBP_REGULATION_OF_CELLULAR_RESPON | 26 | 0.405844 | 1.6813018 | 0.013513514 | 0.11648642 | 0.998 | 4155 | tags=35%, list=18%, si | MM_2 |
| GOBP_PROGRAMMED_N | GOBP_PROGRAMMED_NECROTIC_CELL_DEA | 50 | 0.347216 | 1.6808215 | 0.0 | 0.11629887 | 0.998 | 4863 | tags=42%, list=21%, si | MM_2 |
| GOBP_EOSINOPHIL_CHEM | GOBP_EOSINOPHIL_CHEMOTAXIS | 22 | 0.4198 | 1.6803583 | 0.013513514 | 0.116072826 | 0.998 | 3695 | tags=41%, list=16%, si | MM_2 |
| GOBP_ATP_SYNTHESIS_C | GOBP_ATP_SYNTHESIS_COUPLED_ELECTRO | 76 | 0.320219 | 1.6764159 | 0.0 | 0.118123114 | 0.998 | 5341 | tags=49%, list=23%, si | MM_2 |
| GOBP_POSITIVE_REGULA | GOBP_POSITIVE_REGULATION_OF_VASCULA | 172 | 0.272748 | 1.675549 | 0.0 | 0.11824677 | 0.998 | 2633 | tags=23%, list=11%, si | MM_2 |
| GOBP_REGULATION_OF_ | GOBP_REGULATION_OF_MITOCHONDRIAL_ | 58 | 0.329483 | 1.6701875 | 0.04054054 | 0.12126949 | 0.998 | 5169 | tags=47%, list=22%, si | MM_2 |
| GOBP_NEUTROPHIL_CHE | GOBP_NEUTROPHIL_CHEMOTAXIS | 81 | 0.322901 | 1.6697068 | 0.018867925 | 0.12117671 | 0.998 | 2775 | tags=25%, list=12%, si | MM_2 |
| GOBP_NATURAL_KILLER_ | GOBP_NATURAL_KILLER_CELL_ACTIVATION_ | 21 | 0.433198 | 1.6649312 | 0.010869565 | 0.1237264 | 0.999 | 629 | tags=19%, list=3%, sig | MM_2 |
| GOBP_REGULATION_OF_ | GOBP_REGULATION_OF_NEUROINFLAMMA | 30 | 0.382005 | 1.664392 | 0.0 | 0.12358192 | 1 | 2282 | tags=23%, list=10%, si | MM_2 |
| GOBP_PROTEIN_LOCALIZ | GOBP_PROTEIN_LOCALIZATION_TO_MITOC | 101 | 0.307038 | 1.6639751 | 0.0 | 0.123354256 | 1 | 4882 | tags=40%, list=21%, si | MM_2 |
| GOBP_VIRAL_TRANSLATI | GOBP_VIRAL_TRANSLATION | 18 | 0.443672 | 1.660706 | 0.021390375 | 0.12531121 | 1 | 13143 | tags=100%, list=56%, | MM_2 |

| | | | | | | | | | | |
|---|---|---|---|---|---|---|---|---|---|---|
| GOBP_T_CELL_HOMEOST | GOBP_T_CELL_HOMEOSTASIS | 52 | 0.343971 | 1.659074 | 0.0 | 0.1260348 | 1 | 3374 | tags=29%, list=14%, si | MM_2 |
| GOBP_INTRINSIC_APOPT | GOBP_INTRINSIC_APOPTOTIC_SIGNALING_F | 44 | 0.359079 | 1.6574855 | 0.00990099 | 0.12668477 | 1 | 2634 | tags=27%, list=11%, si | MM_2 |
| GOBP_NUCLEOTIDE_BIND | GOBP_NUCLEOTIDE_BINDING_DOMAIN_LEU | 22 | 0.414417 | 1.656696 | 0.022346368 | 0.12668249 | 1 | 2497 | tags=32%, list=11%, si | MM_2 |
| GOBP_POSITIVE_REGULA | GOBP_POSITIVE_REGULATION_OF_DEFENSE | 480 | 0.242892 | 1.6566447 | 0.0 | 0.1262393 | 1 | 3251 | tags=22%, list=14%, si | MM_2 |
| PANCREATIC_CANCER | PANCREATIC_CANCER | 74 | 0.310007 | 1.6543272 | 0.0 | 0.12724797 | 1 | 3619 | tags=22%, list=15%, si | MM_2 |
| PRION_DISEASES | PRION_DISEASES | 31 | 0.388616 | 1.6464099 | 0.013986014 | 0.13222691 | 1 | 3274 | tags=35%, list=14%, si | MM_2 |
| GOBP_LEUKOCYTE_CHEM | GOBP_LEUKOCYTE_CHEMOTAXIS | 213 | 0.26164 | 1.6453661 | 0.0 | 0.1324147 | 1 | 2979 | tags=23%, list=13%, si | MM_2 |
| GOBP_GMP_BIOSYNTHET | GOBP_GMP_BIOSYNTHETIC_PROCESS | 15 | 0.488654 | 1.6447104 | 0.027173912 | 0.13247222 | 1 | 1217 | tags=27%, list=5%, sig | MM_2 |
| GOBP_POSITIVE_REGULA | GOBP_POSITIVE_REGULATION_OF_MONOC | 16 | 0.458255 | 1.6439395 | 0.034285713 | 0.13259141 | 1 | 2633 | tags=38%, list=11%, si | MM_2 |
| GOBP_REGULATION_OF_ | GOBP_REGULATION_OF_ENDOPLASMIC_RE | 24 | 0.406096 | 1.6409863 | 0.017964073 | 0.13416253 | 1 | 14032 | tags=100%, list=59%, | MM_2 |
| GOBP_REGULATION_OF_ | GOBP_REGULATION_OF_MEMBRANE_PERM | 70 | 0.311953 | 1.6387228 | 0.0 | 0.13542625 | 1 | 5169 | tags=46%, list=22%, si | MM_2 |
| GOBP_NEGATIVE_REGUL | GOBP_NEGATIVE_REGULATION_OF_TELOM | 21 | 0.418397 | 1.6385409 | 0.04278075 | 0.13506338 | 1 | 3892 | tags=38%, list=16%, si | MM_2 |
| GOBP_NEGATIVE_REGUL | GOBP_NEGATIVE_REGULATION_OF_TYPE_2 | 15 | 0.469507 | 1.6371983 | 0.020408163 | 0.13551746 | 1 | 691 | tags=13%, list=3%, sig | MM_2 |
| GOBP_PROTEIN_DEGLYC | GOBP_PROTEIN_DEGLYCOSYLATION | 15 | 0.46311 | 1.6355883 | 0.015625 | 0.13610588 | 1 | 12683 | tags=100%, list=54%, | MM_2 |
| GOBP_CELL_ADHESION_M | GOBP_CELL_ADHESION_MEDIATED_BY_INT | 86 | 0.297261 | 1.6340525 | 0.018867925 | 0.13687809 | 1 | 801 | tags=10%, list=3%, sig | MM_2 |
| GOBP_VASCULAR_ASSOC | GOBP_VASCULAR_ASSOCIATED_SMOOTH_M | 15 | 0.455884 | 1.6337916 | 0.032258064 | 0.13656601 | 1 | 1123 | tags=27%, list=5%, sig | MM_2 |
| GOBP_PURINE_NUCLEOS | GOBP_PURINE_NUCLEOSIDE_MONOPHOSPH | 22 | 0.411157 | 1.6328137 | 0.010752688 | 0.13679297 | 1 | 3911 | tags=36%, list=17%, si | MM_2 |
| GOBP_MODULATION_BY_ | GOBP_MODULATION_BY_SYMBIONT_OF_EN | 56 | 0.33457 | 1.6326524 | 0.0 | 0.13648503 | 1 | 1752 | tags=27%, list=7%, sig | MM_2 |
| GOBP_MITOTIC_DNA_RE | GOBP_MITOTIC_DNA_REPLICATION | 16 | 0.469771 | 1.6325852 | 0.020618556 | 0.13608369 | 1 | 2900 | tags=25%, list=12%, si | MM_2 |
| GOBP_POSITIVE_REGULA | GOBP_POSITIVE_REGULATION_OF_INTERLE | 106 | 0.291788 | 1.6311531 | 0.0 | 0.13653606 | 1 | 3307 | tags=27%, list=14%, si | MM_2 |
| GOBP_REGULATION_OF_ | GOBP_REGULATION_OF_MACROPHAGE_MI | 47 | 0.346464 | 1.6307214 | 0.009433962 | 0.13632548 | 1 | 4068 | tags=34%, list=17%, si | MM_2 |
| GOBP_REGULATION_OF_ | GOBP_REGULATION_OF_T_CELL_CYTOKINE_ | 36 | 0.357277 | 1.6298566 | 0.009259259 | 0.13662855 | 1 | 2013 | tags=22%, list=9%, sig | MM_2 |
| GOBP_NEGATIVE_REGUL | GOBP_NEGATIVE_REGULATION_OF_AUTOP | 81 | 0.323602 | 1.6297144 | 0.0 | 0.13619307 | 1 | 4593 | tags=41%, list=19%, si | MM_2 |
| GOBP_MYOBLAST_FUSIO | GOBP_MYOBLAST_FUSION | 45 | 0.343471 | 1.628176 | 0.0 | 0.13696033 | 1 | 1395 | tags=18%, list=6%, sig | MM_2 |
| GOBP_FATTY_ACID_CATA | GOBP_FATTY_ACID_CATABOLIC_PROCESS | 100 | 0.296715 | 1.6268721 | 0.0 | 0.1373122 | 1 | 3933 | tags=37%, list=17%, si | MM_2 |
| GOBP_GRANULOCYTE_CH | GOBP_GRANULOCYTE_CHEMOTAXIS | 119 | 0.288527 | 1.6251628 | 0.0 | 0.13799746 | 1 | 897 | tags=12%, list=4%, sig | MM_2 |
| GOBP_REGULATION_OF_ | GOBP_REGULATION_OF_INTEGRIN_MEDIAT | 19 | 0.428728 | 1.624816 | 0.021857923 | 0.137724 | 1 | 263 | tags=11%, list=1%, sig | MM_2 |
| GOBP_INFLAMMATORY_C | GOBP_INFLAMMATORY_CELL_APOPTOTIC_F | 28 | 0.393481 | 1.6243268 | 0.012269938 | 0.1377389 | 1 | 342 | tags=11%, list=1%, sig | MM_2 |
| GOBP_CHEMOKINE_PROI | GOBP_CHEMOKINE_PRODUCTION | 119 | 0.287007 | 1.6242547 | 0.0 | 0.13732769 | 1 | 3554 | tags=29%, list=15%, si | MM_2 |
| GOBP_POSITIVE_REGULA | GOBP_POSITIVE_REGULATION_OF_CELLULA | 20 | 0.427924 | 1.6237028 | 0.02259887 | 0.13717301 | 1 | 3472 | tags=35%, list=15%, si | MM_2 |
| HUMAN_PAPILLOMAVIRU | HUMAN_PAPILLOMAVIRUS_INFECTION | 315 | 0.24534 | 1.622937 | 0.0 | 0.13729717 | 1 | 3405 | tags=23%, list=14%, si | MM_2 |
| GOBP_REGULATION_OF_ | GOBP_REGULATION_OF_NON_CANONICAL_ | 92 | 0.300608 | 1.620935 | 0.0 | 0.13841736 | 1 | 4947 | tags=38%, list=21%, si | MM_2 |
| GOBP_REGULATION_OF_ | GOBP_REGULATION_OF_MITOCHONDRIAL_ | 32 | 0.367772 | 1.6197451 | 0.02 | 0.13883752 | 1 | 5034 | tags=47%, list=21%, si | MM_2 |
| GOBP_DNA_UNWINDING | GOBP_DNA_UNWINDING_INVOLVED_IN_DN | 20 | 0.420609 | 1.619655 | 0.029239766 | 0.1384462 | 1 | 13688 | tags=100%, list=58%, | MM_2 |
| GOBP_REGULATION_OF_ | GOBP_REGULATION_OF_CYTOPLASMIC_PAT | 104 | 0.292346 | 1.6138939 | 0.0 | 0.14197259 | 1 | 4068 | tags=28%, list=17%, si | MM_2 |
| GOBP_POSITIVE_REGULA | GOBP_POSITIVE_REGULATION_OF_MACROF | 31 | 0.365928 | 1.6137757 | 0.022900764 | 0.14163469 | 1 | 159 | tags=10%, list=1%, sig | MM_2 |
| GOBP_REGULATION_OF_ | GOBP_REGULATION_OF_PROTEIN_SUMOYL | 25 | 0.413278 | 1.6134123 | 0.006289308 | 0.14148554 | 1 | 13863 | tags=100%, list=59%, | MM_2 |
| GOBP_REGULATION_OF_ | GOBP_REGULATION_OF_PHOSPHOLIPID_BIO | 20 | 0.409324 | 1.6127095 | 0.02173913 | 0.14155771 | 1 | 4104 | tags=40%, list=17%, si | MM_2 |
| GOBP_EOSINOPHIL_MIGR | GOBP_EOSINOPHIL_MIGRATION | 26 | 0.395901 | 1.6124263 | 0.01863354 | 0.14125808 | 1 | 3695 | tags=38%, list=16%, si | MM_2 |
| GOBP_RESPIRATORY_CHA | GOBP_RESPIRATORY_CHAIN_COMPLEX_IV_ | 24 | 0.393623 | 1.6096058 | 0.018404908 | 0.14290352 | 1 | 3022 | tags=29%, list=13%, si | MM_2 |
| GOBP_MITOCHONDRIAL_ | GOBP_MITOCHONDRIAL_GENE_EXPRESSION | 164 | 0.278235 | 1.6084471 | 0.0 | 0.1433181 | 1 | 4583 | tags=34%, list=19%, si | MM_2 |
| GOBP_POSITIVE_REGULA | GOBP_POSITIVE_REGULATION_OF_MACROF | 25 | 0.399916 | 1.608315 | 0.02 | 0.14293727 | 1 | 159 | tags=12%, list=1%, sig | MM_2 |
| CHAGAS_DISEASE_(AMER | CHAGAS_DISEASE_(AMERICAN_TRYPANOSO | 101 | 0.295918 | 1.6069176 | 0.0 | 0.1435255 | 1 | 3724 | tags=30%, list=16%, si | MM_2 |
| GOBP_REGULATION_OF_ | GOBP_REGULATION_OF_VESICLE_SIZE | 21 | 0.412638 | 1.6053724 | 0.0116959065 | 0.14410196 | 1 | 4291 | tags=43%, list=18%, si | MM_2 |
| GOBP_SPLICEOSOMAL_C | GOBP_SPLICEOSOMAL_COMPLEX_ASSEMBL | 49 | 0.33629 | 1.6052833 | 0.039215688 | 0.14370571 | 1 | 15687 | tags=100%, list=66%, | MM_2 |
| GOBP_LATE_ENDOSOME_ | GOBP_LATE_ENDOSOME_TO_LYSOSOME_TI | 19 | 0.429365 | 1.6039716 | 0.028089888 | 0.14445825 | 1 | 13481 | tags=100%, list=57%, | MM_2 |
| GOBP_PROTEIN_RNA_CO | GOBP_PROTEIN_RNA_COMPLEX_ORGANIZA | 173 | 0.273293 | 1.6039046 | 0.0 | 0.14404911 | 1 | 14569 | tags=96%, list=62%, si | MM_2 |
| GOBP_REGULATION_OF_ | GOBP_REGULATION_OF_SUPEROXIDE_ANIC | 28 | 0.390949 | 1.6032076 | 0.028368793 | 0.1440801 | 1 | 4410 | tags=43%, list=19%, si | MM_2 |
| GOBP_MITOCHONDRIAL_ | GOBP_MITOCHONDRIAL_TRANSPORT | 174 | 0.272761 | 1.6019182 | 0.0 | 0.14463033 | 1 | 4541 | tags=32%, list=19%, si | MM_2 |
| GOBP_ACETYL_COA_BIOS | GOBP_ACETYL_COA_BIOSYNTHETIC_PROCES | 19 | 0.421269 | 1.601601 | 0.019512195 | 0.14453728 | 1 | 13672 | tags=100%, list=58%, | MM_2 |
| IL-17_SIGNALING_PATHW | IL-17_SIGNALING_PATHWAY | 85 | 0.295479 | 1.6015489 | 0.0 | 0.14411817 | 1 | 3189 | tags=27%, list=14%, si | MM_2 |
| GOBP_TYPE_II_INTERFER | GOBP_TYPE_II_INTERFERON_PRODUCTION | 132 | 0.280579 | 1.6014307 | 0.0 | 0.14373128 | 1 | 2914 | tags=26%, list=12%, si | MM_2 |
| GOBP_PURINE_NUCLEOS | GOBP_PURINE_NUCLEOSIDE_MONOPHOSPH | 36 | 0.359043 | 1.5980111 | 0.0 | 0.14546172 | 1 | 2512 | tags=28%, list=11%, si | MM_2 |
| GOBP_TOLL_LIKE_RECEPT | GOBP_TOLL_LIKE_RECEPTOR_4_SIGNALING_ | 60 | 0.322321 | 1.5967408 | 0.0 | 0.14607836 | 1 | 2579 | tags=20%, list=11%, si | MM_2 |
| GOBP_G0_TO_G1_TRANS | GOBP_G0_TO_G1_TRANSITION | 38 | 0.347732 | 1.5956804 | 0.018691588 | 0.14632083 | 1 | 13785 | tags=97%, list=58%, si | MM_2 |
| GOBP_B_CELL_HOMEOST | GOBP_B_CELL_HOMEOSTASIS | 42 | 0.348563 | 1.5956411 | 0.031578947 | 0.1458578 | 1 | 3875 | tags=33%, list=16%, si | MM_2 |
| GOBP_ENDOSOME_TO_L | GOBP_ENDOSOME_TO_LYSOSOME_TRANSP | 16 | 0.446389 | 1.5915155 | 0.02538071 | 0.14839043 | 1 | 13078 | tags=100%, list=55%, | MM_2 |
| GOBP_POSITIVE_REGULA | GOBP_POSITIVE_REGULATION_OF_PROTEIN | 100 | 0.285721 | 1.5908438 | 0.0 | 0.1483786 | 1 | 3351 | tags=23%, list=14%, si | MM_2 |
| GOBP_REGULATION_OF_ | GOBP_REGULATION_OF_CELL_SUBSTRATE_ | 69 | 0.304141 | 1.5897579 | 0.013333334 | 0.14872734 | 1 | 3669 | tags=30%, list=16%, si | MM_2 |
| GOBP_POSITIVE_REGULA | GOBP_POSITIVE_REGULATION_OF_HOMOTY | 15 | 0.452788 | 1.5894605 | 0.032432433 | 0.148567 | 1 | 3189 | tags=33%, list=14%, si | MM_2 |
| GOBP_MRNA_CIS_SPLICI | GOBP_MRNA_CIS_SPLICING_VIA_SPLICEOS | 26 | 0.38438 | 1.5880661 | 0.006369427 | 0.14953387 | 1 | 14545 | tags=100%, list=62%, | MM_2 |
| GOBP_INTRINSIC_APOPT | GOBP_INTRINSIC_APOPTOTIC_SIGNALING_F | 112 | 0.277947 | 1.5873109 | 0.0 | 0.14959201 | 1 | 2652 | tags=22%, list=11%, si | MM_2 |
| GOBP_REGULATION_OF_ | GOBP_REGULATION_OF_DNA_DAMAGE_RE | 36 | 0.341755 | 1.5872413 | 0.016393442 | 0.14921318 | 1 | 1894 | tags=14%, list=8%, sig | MM_2 |
| GOBP_PORE_COMPLEX_A | GOBP_PORE_COMPLEX_ASSEMBLY | 20 | 0.421075 | 1.5871941 | 0.03314917 | 0.14875266 | 1 | 13677 | tags=100%, list=58%, | MM_2 |
| GOBP_METALLO_SULFUR | GOBP_METALLO_SULFUR_CLUSTER_ASSEMI | 25 | 0.38924 | 1.5854917 | 0.013986014 | 0.14951868 | 1 | 14430 | tags=100%, list=61%, | MM_2 |
| GOBP_NEGATIVE_REGUL | GOBP_NEGATIVE_REGULATION_OF_PHAGO | 24 | 0.386259 | 1.5844426 | 0.026737968 | 0.14973193 | 1 | 5385 | tags=46%, list=23%, si | MM_2 |
| GOBP_POSITIVE_REGULA | GOBP_POSITIVE_REGULATION_OF_INFLAMI | 22 | 0.391946 | 1.5809076 | 0.012658228 | 0.15215227 | 1 | 14365 | tags=100%, list=61%, | MM_2 |
| AGE-RAGE_SIGNALING_P | AGE-RAGE_SIGNALING_PATHWAY_IN_DIABE | 100 | 0.29172 | 1.5801194 | 0.0 | 0.15242158 | 1 | 4061 | tags=33%, list=17%, si | MM_2 |
| GOBP_NEGATIVE_REGUL | GOBP_NEGATIVE_REGULATION_OF_PROGR | 17 | 0.432219 | 1.5794098 | 0.020725388 | 0.1524611 | 1 | 4590 | tags=47%, list=19%, si | MM_2 |
| GOBP_NEGATIVE_REGUL | GOBP_NEGATIVE_REGULATION_OF_FIBROB | 44 | 0.332383 | 1.5787767 | 0.018018018 | 0.15239678 | 1 | 4016 | tags=32%, list=17%, si | MM_2 |
| GOBP_TELOMERE_MAINT | GOBP_TELOMERE_MAINTENANCE_VIA_REC | 15 | 0.444548 | 1.5786167 | 0.057291668 | 0.15207577 | 1 | 13121 | tags=100%, list=56%, | MM_2 |
| GOBP_IMMUNOGLOBULI | GOBP_IMMUNOGLOBULIN_PRODUCTION | 132 | 0.279985 | 1.5756465 | 0.0 | 0.15420674 | 1 | 3234 | tags=25%, list=14%, si | MM_2 |
| GOBP_POSITIVE_REGULA | GOBP_POSITIVE_REGULATION_OF_NF_KAP | 127 | 0.265753 | 1.5746707 | 0.0 | 0.1546848 | 1 | 3226 | tags=26%, list=14%, si | MM_2 |
| GOBP_PROTEIN_REFOLDI | GOBP_PROTEIN_REFOLDING | 23 | 0.397304 | 1.5734334 | 0.0058139535 | 0.15524702 | 1 | 14239 | tags=100%, list=60%, | MM_2 |
| GOBP_INTEGRATED_STRE | GOBP_INTEGRATED_STRESS_RESPONSE_SIG | 23 | 0.39582 | 1.5728127 | 0.0067114094 | 0.15523782 | 1 | 14274 | tags=100%, list=60%, | MM_2 |
| GOBP_CELL_SURFACE_PA | GOBP_CELL_SURFACE_PATTERN_RECOGNIT | 70 | 0.305091 | 1.5714604 | 0.0 | 0.15577844 | 1 | 2579 | tags=20%, list=11%, si | MM_2 |
| GOBP_SYNAPTIC_VESICLE | GOBP_SYNAPTIC_VESICLE_CYTOSKELETAL_T | 21 | 0.40538 | 1.571134 | 0.033898305 | 0.15565109 | 1 | 2392 | tags=33%, list=10%, si | MM_2 |
| GOBP_LEUKOCYTE_HOMI | GOBP_LEUKOCYTE_HOMEOSTASIS | 137 | 0.273695 | 1.5704393 | 0.0 | 0.15577778 | 1 | 3739 | tags=26%, list=16%, si | MM_2 |
| GOBP_TRANSPORT_OF_V | GOBP_TRANSPORT_OF_VIRUS | 15 | 0.450735 | 1.5702165 | 0.04040404 | 0.15552364 | 1 | 4471 | tags=67%, list=19%, si | MM_2 |
| GOBP_NEGATIVE_REGUL | GOBP_NEGATIVE_REGULATION_OF_VIRAL_ | 36 | 0.360967 | 1.5701733 | 0.037037037 | 0.15510602 | 1 | 3147 | tags=36%, list=13%, si | MM_2 |
| GOBP_CELLULAR_RESPON | GOBP_CELLULAR_RESPONSE_TO_TYPE_II_IN | 95 | 0.287172 | 1.5686665 | 0.0 | 0.15583225 | 1 | 3111 | tags=23%, list=13%, si | MM_2 |
| GOBP_TUMOR_NECROSIS | GOBP_TUMOR_NECROSIS_FACTOR_SUPERF | 193 | 0.262171 | 1.5677102 | 0.0 | 0.1560768 | 1 | 3739 | tags=27%, list=16%, si | MM_2 |

| | | | | | | | | | | |
|---|---|---|---|---|---|---|---|---|---|---|
| HUMAN_IMMUNODEFICI | HUMAN_IMMUNODEFICIENCY_VIRUS_1_IN | 201 | 0.251526 | 1.566864 | 0.0 | 0.15611206 | 1 | 3563 | tags=24%, list=15%, si | MM_2 |
| HUNTINGTON_DISEASE | HUNTINGTON_DISEASE | 180 | 0.253034 | 1.566566 | 0.0 | 0.1559025 | 1 | 5369 | tags=47%, list=23%, si | MM_2 |
| GOBP_REGULATION_OF_ | GOBP_REGULATION_OF_CHROMATIN_BIND | 20 | 0.417002 | 1.5659853 | 0.053892214 | 0.15587631 | 1 | 2000 | tags=25%, list=8%, sig | MM_2 |
| GOBP_AUTOPHAGOSOMI | GOBP_AUTOPHAGOSOME_MATURATION | 58 | 0.319856 | 1.5657446 | 0.0 | 0.1555595 | 1 | 5154 | tags=43%, list=22%, si | MM_2 |
| GOBP_HEAT_GENERATIO | GOBP_HEAT_GENERATION | 23 | 0.392867 | 1.5645485 | 0.044025157 | 0.15608016 | 1 | 2813 | tags=30%, list=12%, si | MM_2 |
| FERROPTOSIS | FERROPTOSIS | 37 | 0.357061 | 1.5645318 | 0.008264462 | 0.15567029 | 1 | 4697 | tags=41%, list=20%, si | MM_2 |
| GOBP_REGULATION_OF_ | GOBP_REGULATION_OF_T_CELL_PROLIFERA | 188 | 0.238742 | 1.5634661 | 0.0 | 0.15602884 | 1 | 3439 | tags=26%, list=15%, si | MM_2 |
| GOBP_REGULATION_OF_ | GOBP_REGULATION_OF_RESPONSE_TO_EN | 80 | 0.303005 | 1.5621355 | 0.0 | 0.1567995 | 1 | 14168 | tags=96%, list=60%, si | MM_2 |
| GOBP_RESPONSE_TO_MI | GOBP_RESPONSE_TO_MITOCHONDRIAL_DE | 21 | 0.397694 | 1.5560381 | 0.02 | 0.16197392 | 1 | 14229 | tags=100%, list=60%, | MM_2 |
| GOBP_SUBSTRATE_ADHE | GOBP_SUBSTRATE_ADHESION_DEPENDENT_ | 99 | 0.277115 | 1.5557382 | 0.0 | 0.16168256 | 1 | 4616 | tags=33%, list=20%, si | MM_2 |
| GOBP_REGULATION_OF_ | GOBP_REGULATION_OF_AUTOPHAGOSOME | 44 | 0.330545 | 1.555675 | 0.0 | 0.16129127 | 1 | 5247 | tags=43%, list=22%, si | MM_2 |
| GOBP_DEOXYRIBONUCLE | GOBP_DEOXYRIBONUCLEOTIDE_BIOSYNTHE | 17 | 0.437068 | 1.555169 | 0.033816423 | 0.1612025 | 1 | 5466 | tags=53%, list=23%, si | MM_2 |
| GOBP_REGULATION_OF_ | GOBP_REGULATION_OF_CELLULAR_EXTRAV | 43 | 0.330065 | 1.5530022 | 0.021052632 | 0.16257231 | 1 | 1887 | tags=21%, list=8%, sig | MM_2 |
| GOBP_REGULATION_OF_ | GOBP_REGULATION_OF_SYNCYTIUM_FORM | 31 | 0.372159 | 1.5522679 | 0.020547945 | 0.16269101 | 1 | 530 | tags=16%, list=2%, sig | MM_2 |
| GOBP_AMP_METABOLIC_ | GOBP_AMP_METABOLIC_PROCESS | 24 | 0.394326 | 1.5521901 | 0.04225352 | 0.1623109 | 1 | 1556 | tags=29%, list=7%, sig | MM_2 |
| GOBP_PROTEIN_N_LINKE | GOBP_PROTEIN_N_LINKED_GLYCOSYLATION | 18 | 0.437315 | 1.5513691 | 0.052380953 | 0.16262138 | 1 | 13293 | tags=100%, list=56%, | MM_2 |
| GOBP_POSITIVE_REGULA | GOBP_POSITIVE_REGULATION_OF_SIGNAL_ | 32 | 0.364766 | 1.5510273 | 0.040983606 | 0.1624629 | 1 | 698 | tags=13%, list=3%, sig | MM_2 |
| GOBP_MITOCHONDRIAL_ | GOBP_MITOCHONDRIAL_OUTER_MEMBRAN | 34 | 0.341748 | 1.5506727 | 0.023809524 | 0.16231935 | 1 | 5169 | tags=53%, list=22%, si | MM_2 |
| GOBP_POSITIVE_REGULA | GOBP_POSITIVE_REGULATION_OF_LEUKOC | 159 | 0.250844 | 1.5505626 | 0.0 | 0.16199706 | 1 | 3307 | tags=25%, list=14%, si | MM_2 |
| GOBP_ACTIVATION_OF_C | GOBP_ACTIVATION_OF_CYSTEINE_TYPE_EN | 81 | 0.295546 | 1.5495712 | 0.0 | 0.16227381 | 1 | 3167 | tags=25%, list=13%, si | MM_2 |
| GOBP_POSITIVE_REGULA | GOBP_POSITIVE_REGULATION_OF_TRANSL | 21 | 0.404673 | 1.5490824 | 0.04 | 0.16207998 | 1 | 2779 | tags=19%, list=12%, si | MM_2 |
| GOBP_LEUKOCYTE_ADHE | GOBP_LEUKOCYTE_ADHESION_TO_VASCUL | 51 | 0.320264 | 1.5475067 | 0.044444446 | 0.16310456 | 1 | 3189 | tags=29%, list=14%, si | MM_2 |
| GOBP_REGULATION_OF_ | GOBP_REGULATION_OF_HOMOTYPIC_CELL_ | 36 | 0.335476 | 1.5471795 | 0.022058824 | 0.16303311 | 1 | 2505 | tags=19%, list=11%, si | MM_2 |
| GOBP_CELLULAR_SENESC | GOBP_CELLULAR_SENESCENCE | 88 | 0.290003 | 1.5468895 | 0.024390243 | 0.16283579 | 1 | 3129 | tags=23%, list=13%, si | MM_2 |
| GOBP_RESPONSE_TO_INT | GOBP_RESPONSE_TO_INTERLEUKIN_17 | 25 | 0.387109 | 1.5468861 | 0.020408163 | 0.16239211 | 1 | 3189 | tags=40%, list=14%, si | MM_2 |
| GOBP_ENERGY_DERIVATI | GOBP_ENERGY_DERIVATION_BY_OXIDATIO | 321 | 0.233556 | 1.5468214 | 0.0 | 0.16196373 | 1 | 3868 | tags=30%, list=16%, si | MM_2 |
| INTESTINAL_IMMUNE_NE | INTESTINAL_IMMUNE_NETWORK_FOR_IGA_ | 37 | 0.344178 | 1.5467018 | 0.022900764 | 0.16166174 | 1 | 3525 | tags=41%, list=15%, si | MM_2 |
| GOBP_REGULATION_OF_ | GOBP_REGULATION_OF_VASCULOGENESIS | 16 | 0.443656 | 1.5465944 | 0.047872342 | 0.16134992 | 1 | 2430 | tags=31%, list=10%, si | MM_2 |
| GOBP_REGULATION_OF_ | GOBP_REGULATION_OF_TOLL_LIKE_RECEPT | 28 | 0.385283 | 1.5435051 | 0.04347826 | 0.1636666 | 1 | 5025 | tags=43%, list=21%, si | MM_2 |
| GOBP_TOLL_LIKE_RECEPT | GOBP_TOLL_LIKE_RECEPTOR_3_SIGNALING_ | 29 | 0.357138 | 1.5426023 | 0.023076924 | 0.16382599 | 1 | 4590 | tags=45%, list=19%, si | MM_2 |
| GOBP_POSITIVE_REGULA | GOBP_POSITIVE_REGULATION_OF_SYNCYTI | 25 | 0.378832 | 1.5409192 | 0.03311258 | 0.16488107 | 1 | 530 | tags=16%, list=2%, sig | MM_2 |
| GOBP_EXTRINSIC_APOPT | GOBP_EXTRINSIC_APOPTOTIC_SIGNALING_ | 233 | 0.243919 | 1.5390606 | 0.0 | 0.166202 | 1 | 4016 | tags=32%, list=17%, si | MM_2 |
| GOBP_REGULATION_OF_ | GOBP_REGULATION_OF_RESPONSE_TO_CYT | 153 | 0.260729 | 1.5372092 | 0.0 | 0.16762292 | 1 | 4633 | tags=35%, list=20%, si | MM_2 |
| GOBP_TRNA_WOBBLE_U | GOBP_TRNA_WOBBLE_URIDINE_MODIFICAT | 16 | 0.429902 | 1.5370722 | 0.05820106 | 0.16732441 | 1 | 4418 | tags=44%, list=19%, si | MM_2 |
| GOBP_T_CELL_CYTOKINE | GOBP_T_CELL_CYTOKINE_PRODUCTION | 47 | 0.314593 | 1.5362864 | 0.010309278 | 0.16764353 | 1 | 3435 | tags=32%, list=15%, si | MM_2 |
| GOBP_CYTOPLASMIC_TR | GOBP_CYTOPLASMIC_TRANSLATION | 143 | 0.272785 | 1.5351881 | 0.0 | 0.16800672 | 1 | 6544 | tags=55%, list=28%, si | MM_2 |
| GOBP_P_BODY_ASSEMBL | GOBP_P_BODY_ASSEMBLY | 20 | 0.400771 | 1.5350281 | 0.030303031 | 0.1677603 | 1 | 14156 | tags=100%, list=60%, | MM_2 |
| GOBP_REGULATION_OF_ | GOBP_REGULATION_OF_TOLL_LIKE_RECEPT | 32 | 0.363484 | 1.5343257 | 0.040983606 | 0.16800751 | 1 | 1438 | tags=13%, list=6%, sig | MM_2 |
| GOBP_FATTY_ACID_ELON | GOBP_FATTY_ACID_ELONGATION | 15 | 0.46435 | 1.5332745 | 0.027472528 | 0.16837986 | 1 | 3860 | tags=53%, list=16%, si | MM_2 |
| GOBP_NEGATIVE_REGUL | GOBP_NEGATIVE_REGULATION_OF_INNATE | 86 | 0.291294 | 1.5317073 | 0.0 | 0.16929172 | 1 | 4334 | tags=40%, list=18%, si | MM_2 |
| GOBP_DNA_TEMPLATED_ | GOBP_DNA_TEMPLATED_DNA_REPLICATIOI | 147 | 0.27244 | 1.5308951 | 0.0 | 0.16961603 | 1 | 3799 | tags=27%, list=16%, si | MM_2 |
| GOBP_PROTEIN_MATUR | GOBP_PROTEIN_MATURATION_BY_IRON_S | 16 | 0.435761 | 1.5306337 | 0.05263158 | 0.16943651 | 1 | 3651 | tags=44%, list=15%, si | MM_2 |
| GOBP_BRANCHED_CHAIN | GOBP_BRANCHED_CHAIN_AMINO_ACID_M | 24 | 0.386216 | 1.5303268 | 0.034285713 | 0.1692032 | 1 | 4036 | tags=42%, list=17%, si | MM_2 |
| GOBP_NEGATIVE_REGUL | GOBP_NEGATIVE_REGULATION_OF_T_CELL_ | 29 | 0.362438 | 1.5296255 | 0.03164557 | 0.16943468 | 1 | 691 | tags=14%, list=3%, sig | MM_2 |
| GOBP_POSITIVE_REGULA | GOBP_POSITIVE_REGULATION_OF_PROTEIN | 16 | 0.428025 | 1.5276566 | 0.05027933 | 0.17076857 | 1 | 5033 | tags=44%, list=21%, si | MM_2 |
| GOBP_GASTRULATION_W | GOBP_GASTRULATION_WITH_MOUTH_FOR | 34 | 0.348206 | 1.5262685 | 0.025 | 0.17160949 | 1 | 2583 | tags=26%, list=11%, si | MM_2 |
| GOBP_MITOCHONDRIAL_ | GOBP_MITOCHONDRIAL_RESPIRATORY_CH | 91 | 0.286747 | 1.5260364 | 0.023255814 | 0.1713482 | 1 | 5628 | tags=42%, list=24%, si | MM_2 |
| GOBP_NUCLEAR_MEMBR | GOBP_NUCLEAR_MEMBRANE_ORGANIZATI | 35 | 0.344106 | 1.5258539 | 0.026785715 | 0.17106456 | 1 | 15498 | tags=100%, list=66%, | MM_2 |
| GOBP_NEGATIVE_REGUL | GOBP_NEGATIVE_REGULATION_OF_LYMPH | 44 | 0.322204 | 1.5226462 | 0.050847456 | 0.1735792 | 1 | 736 | tags=14%, list=3%, sig | MM_2 |
| GOBP_NECROPTOTIC_PR | GOBP_NECROPTOTIC_PROCESS | 37 | 0.339764 | 1.5219069 | 0.00990099 | 0.17391227 | 1 | 4016 | tags=35%, list=17%, si | MM_2 |
| GOBP_TRANSLATIONAL_ | GOBP_TRANSLATIONAL_ELONGATION | 58 | 0.317167 | 1.521812 | 0.011904762 | 0.17356642 | 1 | 13494 | tags=97%, list=57%, si | MM_2 |
| GOBP_HUMORAL_IMMU | GOBP_HUMORAL_IMMUNE_RESPONSE | 184 | 0.268384 | 1.5205851 | 0.0 | 0.17445761 | 1 | 2232 | tags=21%, list=9%, sig | MM_2 |
| STEROID_BIOSYNTHESIS | STEROID_BIOSYNTHESIS | 17 | 0.419547 | 1.5203508 | 0.029761905 | 0.17423809 | 1 | 3545 | tags=47%, list=15%, si | MM_2 |
| GOBP_POSITIVE_REGULA | GOBP_POSITIVE_REGULATION_OF_MYELOID | 21 | 0.397122 | 1.5185018 | 0.033519555 | 0.1754927 | 1 | 4976 | tags=52%, list=21%, si | MM_2 |
| GOBP_IMPORT_INTO_TH | GOBP_IMPORT_INTO_THE_MITOCHONDRIO | 49 | 0.321394 | 1.518029 | 0.036697246 | 0.1755684 | 1 | 16038 | tags=100%, list=68%, | MM_2 |
| CHRONIC_MYELOID_LEU | CHRONIC_MYELOID_LEUKEMIA | 75 | 0.303622 | 1.5156282 | 0.018181818 | 0.17760605 | 1 | 3735 | tags=23%, list=16%, si | MM_2 |
| GOBP_PROTEIN_TARGETI | GOBP_PROTEIN_TARGETING_TO_MITOCHO | 76 | 0.299211 | 1.5146898 | 0.016949153 | 0.17791702 | 1 | 4465 | tags=34%, list=19%, si | MM_2 |
| GOBP_PIGMENTATION | GOBP_PIGMENTATION | 112 | 0.261953 | 1.5129595 | 0.0 | 0.17908978 | 1 | 2392 | tags=21%, list=10%, si | MM_2 |
| GOBP_RESPONSE_TO_AN | GOBP_RESPONSE_TO_ANTIBIOTIC | 32 | 0.358 | 1.5120112 | 0.045454547 | 0.17947008 | 1 | 3135 | tags=25%, list=13%, si | MM_2 |
| CELL_CYCLE | CELL_CYCLE | 121 | 0.2682 | 1.5106881 | 0.0 | 0.18010461 | 1 | 4784 | tags=32%, list=20%, si | MM_2 |
| GOBP_MRNA_3_END_PR | GOBP_MRNA_3_END_PROCESSING | 41 | 0.318926 | 1.5103506 | 0.023076924 | 0.17999056 | 1 | 14616 | tags=98%, list=62%, si | MM_2 |
| PROTEASOME | PROTEASOME | 45 | 0.310009 | 1.5100319 | 0.019607844 | 0.17978543 | 1 | 6635 | tags=49%, list=28%, si | MM_2 |
| UBIQUITIN_MEDIATED_P | UBIQUITIN_MEDIATED_PROTEOLYSIS | 134 | 0.25767 | 1.5081975 | 0.0 | 0.18110651 | 1 | 15149 | tags=96%, list=64%, si | MM_2 |
| GOBP_GLYCERALDEHYDE | GOBP_GLYCERALDEHYDE_3_PHOSPHATE_M | 19 | 0.404479 | 1.507932 | 0.040697675 | 0.18093112 | 1 | 4824 | tags=63%, list=20%, si | MM_2 |
| GOBP_ER_NUCLEUS_SIGN | GOBP_ER_NUCLEUS_SIGNALING_PATHWAY | 43 | 0.321946 | 1.5057869 | 0.028301887 | 0.18240577 | 1 | 14556 | tags=98%, list=62%, si | MM_2 |
| GOBP_RESPONSE_TO_PR | GOBP_RESPONSE_TO_PROGESTERONE | 19 | 0.406365 | 1.5056765 | 0.06043956 | 0.18204968 | 1 | 4016 | tags=42%, list=17%, si | MM_2 |
| GOBP_REGULATION_OF_ | GOBP_REGULATION_OF_ADAPTIVE_IMMUN | 215 | 0.231917 | 1.5045706 | 0.0 | 0.18278112 | 1 | 3326 | tags=25%, list=14%, si | MM_2 |
| GOBP_T_CELL_ACTIVATIO | GOBP_T_CELL_ACTIVATION_INVOLVED_IN_ | 112 | 0.275633 | 1.5045239 | 0.0 | 0.18235782 | 1 | 3682 | tags=27%, list=16%, si | MM_2 |
| GOBP_PRE_MIRNA_PROC | GOBP_PRE_MIRNA_PROCESSING | 15 | 0.434408 | 1.5040126 | 0.06511628 | 0.18228613 | 1 | 5508 | tags=67%, list=23%, si | MM_2 |
| GOBP_NEGATIVE_REGUL | GOBP_NEGATIVE_REGULATION_OF_ESTABL | 148 | 0.253823 | 1.5037937 | 0.0 | 0.18207377 | 1 | 3242 | tags=26%, list=14%, si | MM_2 |
| GOBP_NUCLEAR_PORE_C | GOBP_NUCLEAR_PORE_ORGANIZATION | 15 | 0.420986 | 1.5014392 | 0.057416268 | 0.18361118 | 1 | 13677 | tags=100%, list=58%, | MM_2 |
| GOBP_NUCLEAR_BODY_C | GOBP_NUCLEAR_BODY_ORGANIZATION | 19 | 0.400835 | 1.5003738 | 0.06486487 | 0.18414022 | 1 | 1191 | tags=16%, list=5%, sig | MM_2 |
| GOBP_NUCLEOSIDE_MON | GOBP_NUCLEOSIDE_MONOPHOSPHATE_ME | 62 | 0.296758 | 1.5000987 | 0.0 | 0.18397443 | 1 | 1556 | tags=18%, list=7%, sig | MM_2 |
| GOBP_CANONICAL_NF_K | GOBP_CANONICAL_NF_KAPPAB_SIGNAL_TF | 247 | 0.227273 | 1.4995505 | 0.0 | 0.1840254 | 1 | 3166 | tags=21%, list=13%, si | MM_2 |
| GOBP_STEM_CELL_DIVISI | GOBP_STEM_CELL_DIVISION | 43 | 0.314783 | 1.4994907 | 0.019607844 | 0.18368344 | 1 | 1585 | tags=16%, list=7%, sig | MM_2 |
| GOBP_NUCLEAR_MEMBR | GOBP_NUCLEAR_MEMBRANE_REASSEMBLY | 20 | 0.393057 | 1.499318 | 0.038251366 | 0.18336493 | 1 | 14338 | tags=100%, list=61%, | MM_2 |
| GOBP_RIBOSOMAL_SMA | GOBP_RIBOSOMAL_SMALL_SUBUNIT_ASSEI | 18 | 0.405156 | 1.4982649 | 0.04812834 | 0.18389623 | 1 | 6415 | tags=67%, list=27%, si | MM_2 |
| GOBP_TORC1_SIGNALING | GOBP_TORC1_SIGNALING | 100 | 0.277564 | 1.4977548 | 0.025 | 0.18383378 | 1 | 14492 | tags=96%, list=61%, si | MM_2 |
| GOBP_REGULATION_OF_ | GOBP_REGULATION_OF_PODOSOME_ASSEI | 16 | 0.426175 | 1.4976178 | 0.06417112 | 0.1835389 | 1 | 1752 | tags=19%, list=7%, sig | MM_2 |
| GOBP_REGULATION_OF_ | GOBP_REGULATION_OF_LEUKOCYTE_ADHE | 33 | 0.343024 | 1.4969765 | 0.02189781 | 0.18366395 | 1 | 1907 | tags=21%, list=8%, sig | MM_2 |
| GOBP_MAINTENANCE_O | GOBP_MAINTENANCE_OF_PROTEIN_LOCAL | 43 | 0.306017 | 1.4961439 | 0.06481481 | 0.18416345 | 1 | 4478 | tags=44%, list=19%, si | MM_2 |
| GOBP_MITOCHONDRIAL_ | GOBP_MITOCHONDRIAL_DNA_REPLICATION | 17 | 0.415981 | 1.4949137 | 0.048543688 | 0.18481994 | 1 | 1055 | tags=18%, list=4%, sig | MM_2 |

| | | | | | | | | | | |
|---|---|---|---|---|---|---|---|---|---|---|
| GOBP_RNA_MODIFICATIO | GOBP_RNA_MODIFICATION | 141 | 0.254241 | 1.4948432 | 0.0 | 0.18445009 | 1 | 4508 | tags=34%, list=19%, si | MM_2 |
| GOBP_TOLL_LIKE_RECEPT | GOBP_TOLL_LIKE_RECEPTOR_9_SIGNALING_ | 16 | 0.425458 | 1.493969 | 0.065656565 | 0.1849196 | 1 | 4331 | tags=44%, list=18%, si | MM_2 |
| GOBP_NEGATIVE_REGUL | GOBP_NEGATIVE_REGULATION_OF_RELEAS | 22 | 0.369728 | 1.4932927 | 0.038674034 | 0.18510714 | 1 | 4950 | tags=45%, list=21%, si | MM_2 |
| GOBP_ELECTRON_TRANS | GOBP_ELECTRON_TRANSPORT_CHAIN | 107 | 0.269935 | 1.4919099 | 0.037037037 | 0.18605411 | 1 | 5519 | tags=44%, list=23%, si | MM_2 |
| GOBP_LEUKOCYTE_CELL_ | GOBP_LEUKOCYTE_CELL_CELL_ADHESION | 390 | 0.208503 | 1.4910231 | 0.0 | 0.18645199 | 1 | 3523 | tags=23%, list=15%, si | MM_2 |
| GOBP_NEGATIVE_REGUL | GOBP_NEGATIVE_REGULATION_OF_CYCLIN | 25 | 0.363062 | 1.4900827 | 0.061643835 | 0.18689705 | 1 | 2632 | tags=28%, list=11%, si | MM_2 |
| GOBP_NEGATIVE_REGUL | GOBP_NEGATIVE_REGULATION_OF_MONOI | 28 | 0.356401 | 1.4888393 | 0.028169014 | 0.18759148 | 1 | 3739 | tags=36%, list=16%, si | MM_2 |
| GOBP_NUCLEOSIDE_MON | GOBP_NUCLEOSIDE_MONOPHOSPHATE_BIC | 40 | 0.33496 | 1.4887753 | 0.00877193 | 0.18721102 | 1 | 2512 | tags=23%, list=11%, si | MM_2 |
| GOBP_POSITIVE_REGULA | GOBP_POSITIVE_REGULATION_OF_ERAD_PA | 20 | 0.400263 | 1.4869967 | 0.042168673 | 0.18875931 | 1 | 14168 | tags=100%, list=60%, | MM_2 |
| GOBP_POSITIVE_REGULA | GOBP_POSITIVE_REGULATION_OF_SMOOTH | 69 | 0.293439 | 1.4856706 | 0.025974026 | 0.18979119 | 1 | 729 | tags=10%, list=3%, sig | MM_2 |
| SALMONELLA_INFECTION | SALMONELLA_INFECTION | 76 | 0.277278 | 1.485442 | 0.0 | 0.1896116 | 1 | 3633 | tags=22%, list=15%, si | MM_2 |
| GOBP_NEGATIVE_REGUL | GOBP_NEGATIVE_REGULATION_OF_NON_C | 26 | 0.37216 | 1.4845902 | 0.061643835 | 0.18992916 | 1 | 3510 | tags=31%, list=15%, si | MM_2 |
| GOBP_REGULATION_OF_ | GOBP_REGULATION_OF_NUCLEOCYTOPLAS | 130 | 0.261632 | 1.4841245 | 0.0 | 0.18995742 | 1 | 3519 | tags=22%, list=15%, si | MM_2 |
| GOBP_TRNA_WOBBLE_BA | GOBP_TRNA_WOBBLE_BASE_MODIFICATIO | 19 | 0.390633 | 1.4830962 | 0.06358381 | 0.19040728 | 1 | 4945 | tags=42%, list=21%, si | MM_2 |
| GOBP_NEGATIVE_REGUL | GOBP_NEGATIVE_REGULATION_OF_CHEMO | 29 | 0.358832 | 1.4809412 | 0.069767445 | 0.1922125 | 1 | 3554 | tags=41%, list=15%, si | MM_2 |
| GOBP_REGULATION_OF_ | GOBP_REGULATION_OF_CELLULAR_RESPON | 89 | 0.265333 | 1.4800683 | 0.0 | 0.19259538 | 1 | 3105 | tags=22%, list=13%, si | MM_2 |
| GOBP_NEGATIVE_REGUL | GOBP_NEGATIVE_REGULATION_OF_RESPON | 112 | 0.27143 | 1.480036 | 0.022222223 | 0.19218002 | 1 | 4343 | tags=37%, list=18%, si | MM_2 |
| GOBP_POSITIVE_REGULA | GOBP_POSITIVE_REGULATION_OF_FATTY_A | 45 | 0.317499 | 1.4798082 | 0.035714287 | 0.19195263 | 1 | 2889 | tags=36%, list=12%, si | MM_2 |
| GOBP_REGULATION_OF_ | GOBP_REGULATION_OF_NECROPTOTIC_PRO | 22 | 0.386492 | 1.4787484 | 0.06626506 | 0.19237092 | 1 | 4016 | tags=36%, list=17%, si | MM_2 |
| GOBP_NEGATIVE_REGUL | GOBP_NEGATIVE_REGULATION_OF_DNA_TI | 17 | 0.420682 | 1.47819 | 0.06422018 | 0.19242829 | 1 | 13685 | tags=100%, list=58%, | MM_2 |
| GOBP_NEGATIVE_REGUL | GOBP_NEGATIVE_REGULATION_OF_IMMUN | 15 | 0.430637 | 1.4781263 | 0.048076924 | 0.19203782 | 1 | 2615 | tags=33%, list=11%, si | MM_2 |
| GOBP_RESPONSE_TO_TU | GOBP_RESPONSE_TO_TUMOR_NECROSIS_F | 167 | 0.266748 | 1.477723 | 0.0 | 0.19202282 | 1 | 3189 | tags=24%, list=14%, si | MM_2 |
| GOBP_VASCULAR_ASSOC | GOBP_VASCULAR_ASSOCIATED_SMOOTH_M | 80 | 0.283412 | 1.4777057 | 0.03846154 | 0.1916428 | 1 | 4016 | tags=33%, list=17%, si | MM_2 |
| GOBP_POSITIVE_REGULA | GOBP_POSITIVE_REGULATION_OF_TYPE_II_ | 80 | 0.276265 | 1.4764308 | 0.0 | 0.19248329 | 1 | 2891 | tags=29%, list=12%, si | MM_2 |
| TOXOPLASMOSIS | TOXOPLASMOSIS | 103 | 0.275594 | 1.4763993 | 0.025 | 0.19206469 | 1 | 3724 | tags=25%, list=16%, si | MM_2 |
| GOBP_REGULATION_OF_ | GOBP_REGULATION_OF_SEQUESTERING_OF | 15 | 0.417546 | 1.4752686 | 0.054455444 | 0.1927606 | 1 | 1227 | tags=20%, list=5%, sig | MM_2 |
| NECROPTOSIS | NECROPTOSIS | 138 | 0.264333 | 1.4751909 | 0.0 | 0.19243743 | 1 | 3835 | tags=30%, list=16%, si | MM_2 |
| GOBP_REGULATION_OF_ | GOBP_REGULATION_OF_MEMBRANE_LIPID_ | 19 | 0.405201 | 1.4750488 | 0.056179777 | 0.19211227 | 1 | 2198 | tags=21%, list=9%, sig | MM_2 |
| GOBP_SURFACTANT_HON | GOBP_SURFACTANT_HOMEOSTASIS | 22 | 0.372058 | 1.4738628 | 0.03726708 | 0.1928269 | 1 | 2341 | tags=36%, list=10%, si | MM_2 |
| GOBP_CELL_KILLING | GOBP_CELL_KILLING | 202 | 0.241551 | 1.4735148 | 0.0 | 0.19277954 | 1 | 1535 | tags=13%, list=7%, sig | MM_2 |
| GOBP_SPINAL_CORD_PA | GOBP_SPINAL_CORD_PATTERNING | 17 | 0.409415 | 1.4720293 | 0.04712042 | 0.1936846 | 1 | 2319 | tags=41%, list=10%, si | MM_2 |
| GOBP_RIBOSOMAL_SMAI | GOBP_RIBOSOMAL_SMALL_SUBUNIT_BIOGI | 101 | 0.260621 | 1.471461 | 0.0 | 0.19385852 | 1 | 14435 | tags=96%, list=61%, si | MM_2 |
| GOBP_RESPONSE_TO_PL | GOBP_RESPONSE_TO_PLATELET_DERIVED_ | 22 | 0.374945 | 1.4692266 | 0.04 | 0.19569771 | 1 | 1357 | tags=18%, list=6%, sig | MM_2 |
| GOBP_PRODUCTION_OF_ | GOBP_PRODUCTION_OF_MOLECULAR_MED | 270 | 0.211671 | 1.4690461 | 0.0 | 0.19551213 | 1 | 2273 | tags=18%, list=10%, si | MM_2 |
| GOBP_POSITIVE_REGULA | GOBP_POSITIVE_REGULATION_OF_INTERLE | 55 | 0.290964 | 1.4687142 | 0.0 | 0.19544946 | 1 | 3435 | tags=33%, list=15%, si | MM_2 |
| GOBP_TOR_SIGNALING | GOBP_TOR_SIGNALING | 165 | 0.242686 | 1.4682304 | 0.0 | 0.19545029 | 1 | 3213 | tags=22%, list=14%, si | MM_2 |
| GOBP_REGULATION_OF_ | GOBP_REGULATION_OF_LEUKOCYTE_PROLI | 273 | 0.241626 | 1.4681815 | 0.0 | 0.19508518 | 1 | 3741 | tags=29%, list=16%, si | MM_2 |
| THERMOGENESIS | THERMOGENESIS | 210 | 0.25752 | 1.4676685 | 0.0 | 0.19512112 | 1 | 4701 | tags=39%, list=20%, si | MM_2 |
| GOBP_NUCLEOSIDE_BISP | GOBP_NUCLEOSIDE_BISPHOSPHATE_BIOSYN | 39 | 0.323848 | 1.4660851 | 0.029411765 | 0.19626825 | 1 | 4810 | tags=38%, list=20%, si | MM_2 |
| GOBP_REGULATION_OF_ | GOBP_REGULATION_OF_ESTABLISHMENT_O | 25 | 0.374698 | 1.463438 | 0.07092199 | 0.19887355 | 1 | 14773 | tags=100%, list=63%, | MM_2 |
| GRAFT-VERSUS-HOST_DIS | GRAFT-VERSUS-HOST_DISEASE | 42 | 0.316537 | 1.4620082 | 0.028301887 | 0.19971085 | 1 | 2633 | tags=36%, list=11%, si | MM_2 |
| GOBP_REGULATION_OF_ | GOBP_REGULATION_OF_FATTY_ACID_BETA | 21 | 0.374854 | 1.4613793 | 0.03846154 | 0.2000252 | 1 | 3877 | tags=48%, list=16%, si | MM_2 |
| GOBP_POSITIVE_REGULA | GOBP_POSITIVE_REGULATION_OF_INTRACE | 108 | 0.272802 | 1.4609555 | 0.0 | 0.19998035 | 1 | 2792 | tags=18%, list=12%, si | MM_2 |
| GOBP_REGULATION_OF_ | GOBP_REGULATION_OF_EXTRACELLULAR_N | 66 | 0.286797 | 1.4597896 | 0.0 | 0.2007822 | 1 | 1977 | tags=17%, list=8%, sig | MM_2 |
| LEGIONELLOSIS | LEGIONELLOSIS | 55 | 0.304172 | 1.4590499 | 0.025316456 | 0.20100571 | 1 | 1246 | tags=15%, list=5%, sig | MM_2 |
| GOBP_SIGNAL_TRANSDU | GOBP_SIGNAL_TRANSDUCTION_IN_ABSENC | 78 | 0.274915 | 1.4587842 | 0.0 | 0.2008969 | 1 | 2787 | tags=26%, list=12%, si | MM_2 |
| GOBP_NEUTROPHIL_ACTI | GOBP_NEUTROPHIL_ACTIVATION | 36 | 0.32973 | 1.4577072 | 0.046296295 | 0.20152318 | 1 | 2302 | tags=19%, list=10%, si | MM_2 |
| GOBP_TOLL_LIKE_RECEPT | GOBP_TOLL_LIKE_RECEPTOR_SIGNALING_PA | 71 | 0.281648 | 1.4570595 | 0.018867925 | 0.20172316 | 1 | 1438 | tags=11%, list=6%, sig | MM_2 |
| GOBP_REGULATION_OF_ | GOBP_REGULATION_OF_RELEASE_OF_SEQU | 80 | 0.266879 | 1.4561198 | 0.016949153 | 0.20230095 | 1 | 1152 | tags=15%, list=5%, sig | MM_2 |
| GOBP_NEGATIVE_REGUL | GOBP_NEGATIVE_REGULATION_OF_FATTY_ | 17 | 0.400963 | 1.4558724 | 0.060606062 | 0.20213811 | 1 | 4688 | tags=47%, list=20%, si | MM_2 |
| GOBP_REGULATION_OF_ | GOBP_REGULATION_OF_MACROPHAGE_CH | 32 | 0.336605 | 1.4544259 | 0.048 | 0.20323767 | 1 | 159 | tags=9%, list=1%, sign | MM_2 |
| GOBP_REGULATION_OF_ | GOBP_REGULATION_OF_OXIDOREDUCTASE | 63 | 0.278118 | 1.4539226 | 0.015625 | 0.20340782 | 1 | 3459 | tags=32%, list=15%, si | MM_2 |
| GOBP_NEGATIVE_REGUL | GOBP_NEGATIVE_REGULATION_OF_LYMPH | 62 | 0.286848 | 1.4537789 | 0.0 | 0.20316708 | 1 | 915 | tags=11%, list=4%, sig | MM_2 |
| GOBP_REGULATION_OF_ | GOBP_REGULATION_OF_INTEGRIN_ACTIVAT | 17 | 0.408202 | 1.4529756 | 0.072164945 | 0.20353395 | 1 | 2296 | tags=24%, list=10%, si | MM_2 |
| GOBP_RIBOSOMAL_LARG | GOBP_RIBOSOMAL_LARGE_SUBUNIT_ASSEM | 23 | 0.358444 | 1.4513587 | 0.039215688 | 0.20489521 | 1 | 6024 | tags=61%, list=26%, si | MM_2 |
| GOBP_MONOCYTE_CHEM | GOBP_MONOCYTE_CHEMOTAXIS | 40 | 0.326426 | 1.4509382 | 0.072916664 | 0.2049123 | 1 | 3669 | tags=30%, list=16%, si | MM_2 |
| GOBP_REGULATION_OF_ | GOBP_REGULATION_OF_DNA_STRAND_ELO | 18 | 0.382767 | 1.4501566 | 0.05882353 | 0.20549339 | 1 | 14580 | tags=100%, list=62%, | MM_2 |
| GOBP_REGULATION_OF_ | GOBP_REGULATION_OF_AUTOPHAGOSOME | 18 | 0.384845 | 1.4501343 | 0.053475935 | 0.20507644 | 1 | 4161 | tags=39%, list=18%, si | MM_2 |
| GOBP_CELLULAR_RESPON | GOBP_CELLULAR_RESPONSE_TO_LIPOPROT | 38 | 0.326589 | 1.449726 | 0.018518519 | 0.20509608 | 1 | 2792 | tags=24%, list=12%, si | MM_2 |
| GOBP_EXTRINSIC_APOPT | GOBP_EXTRINSIC_APOPTOTIC_SIGNALING_ | 80 | 0.272517 | 1.4482692 | 0.0 | 0.20634814 | 1 | 3922 | tags=33%, list=17%, si | MM_2 |
| PROTEIN_PROCESSING_IN | PROTEIN_PROCESSING_IN_ENDOPLASMIC_ | 158 | 0.250586 | 1.4482393 | 0.0 | 0.2060085 | 1 | 4978 | tags=33%, list=21%, si | MM_2 |
| GOBP_REGULATION_OF_ | GOBP_REGULATION_OF_CYTOKINE_PRODU | 122 | 0.24981 | 1.4481368 | 0.0 | 0.20567138 | 1 | 2078 | tags=19%, list=9%, sig | MM_2 |
| GOBP_ESTABLISHMENT_ | GOBP_ESTABLISHMENT_OF_LYMPHOCYTE_ | 15 | 0.421562 | 1.4478527 | 0.049261082 | 0.20559995 | 1 | 1752 | tags=27%, list=7%, sig | MM_2 |
| GOBP_NUCLEOSIDE_TRIP | GOBP_NUCLEOSIDE_TRIPHOSPHATE_BIOSYN | 100 | 0.265387 | 1.4477996 | 0.048780486 | 0.20523717 | 1 | 5661 | tags=45%, list=24%, si | MM_2 |
| GOBP_POSITIVE_REGULA | GOBP_POSITIVE_REGULATION_OF_RESPONS | 37 | 0.323971 | 1.4471796 | 0.03305785 | 0.20551091 | 1 | 14168 | tags=97%, list=60%, si | MM_2 |
| GOBP_RESPONSE_TO_HE | GOBP_RESPONSE_TO_HEAT | 86 | 0.268208 | 1.4465784 | 0.03448276 | 0.20580497 | 1 | 4416 | tags=31%, list=19%, si | MM_2 |
| GOBP_COLLAGEN_CATAB | GOBP_COLLAGEN_CATABOLIC_PROCESS | 37 | 0.323293 | 1.4460354 | 0.03968254 | 0.20595433 | 1 | 2814 | tags=30%, list=12%, si | MM_2 |
| GOBP_REGULATION_OF_ | GOBP_REGULATION_OF_ENDOTHELIAL_CEL | 18 | 0.380606 | 1.4458076 | 0.04812834 | 0.205801 | 1 | 5545 | tags=56%, list=23%, si | MM_2 |
| GOBP_REGULATION_OF_ | GOBP_REGULATION_OF_PHOSPHOLIPID_MI | 37 | 0.317212 | 1.4441692 | 0.017391304 | 0.20712869 | 1 | 4446 | tags=41%, list=19%, si | MM_2 |
| GOBP_REGULATION_OF_ | GOBP_REGULATION_OF_CYSTEINE_TYPE_EN | 197 | 0.233108 | 1.4431454 | 0.0 | 0.20781809 | 1 | 2724 | tags=22%, list=12%, si | MM_2 |
| GOBP_REGULATION_OF_ | GOBP_REGULATION_OF_SUBSTRATE_ADHES | 58 | 0.294593 | 1.4430398 | 0.023529412 | 0.20752202 | 1 | 4616 | tags=36%, list=20%, si | MM_2 |
| GOBP_NEGATIVE_REGUL | GOBP_NEGATIVE_REGULATION_OF_LEUKOC | 65 | 0.289069 | 1.4428947 | 0.013157895 | 0.20732723 | 1 | 736 | tags=11%, list=3%, sig | MM_2 |
| GOBP_MONOCYTE_CHEM | GOBP_MONOCYTE_CHEMOTACTIC_PROTEIN | 27 | 0.341686 | 1.4414543 | 0.07189543 | 0.20834388 | 1 | 3554 | tags=37%, list=15%, si | MM_2 |
| GOBP_POSITIVE_REGULA | GOBP_POSITIVE_REGULATION_OF_LYMPHO | 318 | 0.236278 | 1.4407173 | 0.0 | 0.20878002 | 1 | 3238 | tags=23%, list=14%, si | MM_2 |
| GOBP_ANTIGEN_PROCES | GOBP_ANTIGEN_PROCESSING_AND_PRESEN | 38 | 0.302285 | 1.4405466 | 0.024390243 | 0.20849258 | 1 | 4400 | tags=47%, list=19%, si | MM_2 |
| GOBP_GRANULOCYTE_AC | GOBP_GRANULOCYTE_ACTIVATION | 41 | 0.302221 | 1.4403454 | 0.06 | 0.20824537 | 1 | 1703 | tags=15%, list=7%, sig | MM_2 |
| GOBP_RESPONSE_TO_INT | GOBP_RESPONSE_TO_INTERLEUKIN_7 | 17 | 0.38376 | 1.4394372 | 0.077319585 | 0.20883161 | 1 | 5238 | tags=59%, list=22%, si | MM_2 |
| GOBP_CELLULAR_RESPON | GOBP_CELLULAR_RESPONSE_TO_AMYLOID_ | 33 | 0.328514 | 1.4384364 | 0.054263566 | 0.20950131 | 1 | 2723 | tags=18%, list=12%, si | MM_2 |
| GOBP_PEPTIDYL_LYSINE_ | GOBP_PEPTIDYL_LYSINE_ACETYLATION | 25 | 0.35601 | 1.4382938 | 0.05263158 | 0.20923078 | 1 | 4638 | tags=40%, list=20%, si | MM_2 |
| GOBP_PROTEIN_TRANSM | GOBP_PROTEIN_TRANSMEMBRANE_IMPOR | 36 | 0.315112 | 1.4375658 | 0.06557377 | 0.209527 | 1 | 4219 | tags=33%, list=18%, si | MM_2 |
| GOBP_MITOCHONDRIAL_ | GOBP_MITOCHONDRIAL_MEMBRANE_ORGA | 107 | 0.26826 | 1.4372411 | 0.0 | 0.20948409 | 1 | 4465 | tags=32%, list=19%, si | MM_2 |
| GOBP_POSITIVE_REGULA | GOBP_POSITIVE_REGULATION_OF_TOR_SIG | 71 | 0.279557 | 1.4372394 | 0.013888889 | 0.20907007 | 1 | 3163 | tags=25%, list=13%, si | MM_2 |

| | | | | | | | | | | |
|---|---|---|---|---|---|---|---|---|---|---|
| GOBP_REGULATION_OF_ | GOBP_REGULATION_OF_MONOCYTE_CHEM | 29 | 0.345835 | 1.4370612 | 0.059602648 | 0.2088288 | 1 | 3669 | tags=31%, list=16%, si | MM_2 |
| GOBP_ENDOPLASMIC_RE | GOBP_ENDOPLASMIC_RETICULUM_ORGANI | 90 | 0.263651 | 1.4366652 | 0.0 | 0.20890498 | 1 | 2965 | tags=21%, list=13%, si | MM_2 |
| GOBP_NEGATIVE_REGUL | GOBP_NEGATIVE_REGULATION_OF_CALCIU | 28 | 0.338988 | 1.4366202 | 0.04819277 | 0.20850375 | 1 | 921 | tags=14%, list=4%, sig | MM_2 |
| GOBP_NEGATIVE_REGUL | GOBP_NEGATIVE_REGULATION_OF_TELOM | 36 | 0.321354 | 1.4354272 | 0.028037382 | 0.20943376 | 1 | 14374 | tags=97%, list=61%, si | MM_2 |
| GOBP_INNER_MITOCHON | GOBP_INNER_MITOCHONDRIAL_MEMBRAN | 36 | 0.315417 | 1.434794 | 0.05 | 0.20983656 | 1 | 14599 | tags=97%, list=62%, si | MM_2 |
| GOBP_POSITIVE_REGULA | GOBP_POSITIVE_REGULATION_OF_LIPID_CA | 26 | 0.349948 | 1.4344224 | 0.051612902 | 0.20977215 | 1 | 3373 | tags=42%, list=14%, si | MM_2 |
| GOBP_HISTONE_MRNA_I | GOBP_HISTONE_MRNA_METABOLIC_PROCE | 20 | 0.381273 | 1.4337165 | 0.061349694 | 0.21012485 | 1 | 14616 | tags=100%, list=62%, | MM_2 |
| GOBP_NEGATIVE_REGUL | GOBP_NEGATIVE_REGULATION_OF_PLATEL | 16 | 0.414298 | 1.4325713 | 0.08585858 | 0.21108274 | 1 | 2505 | tags=25%, list=11%, si | MM_2 |
| GOBP_ORGANELLE_INHE | GOBP_ORGANELLE_INHERITANCE | 15 | 0.418485 | 1.4324404 | 0.054187194 | 0.21084523 | 1 | 13736 | tags=100%, list=58%, | MM_2 |
| GOBP_CELLULAR_RESPON | GOBP_CELLULAR_RESPONSE_TO_INTERLEUI | 62 | 0.281768 | 1.4307528 | 0.023255814 | 0.21255477 | 1 | 4251 | tags=32%, list=18%, si | MM_2 |
| GOBP_REGULATION_OF_ | GOBP_REGULATION_OF_PROTEIN_EXPORT_ | 37 | 0.308237 | 1.4307305 | 0.055555556 | 0.21214364 | 1 | 3242 | tags=27%, list=14%, si | MM_2 |
| GOBP_REGULATION_OF_ | GOBP_REGULATION_OF_ERAD_PATHWAY | 27 | 0.344965 | 1.4302113 | 0.046511628 | 0.21234326 | 1 | 15475 | tags=100%, list=66%, | MM_2 |
| GOBP_GLYCOLYTIC_PROC | GOBP_GLYCOLYTIC_PROCESS_THROUGH_FR | 25 | 0.345224 | 1.4298809 | 0.074324325 | 0.21234578 | 1 | 2308 | tags=36%, list=10%, si | MM_2 |
| GOBP_POSITIVE_REGULA | GOBP_POSITIVE_REGULATION_OF_LEUKOCY | 169 | 0.252331 | 1.4290974 | 0.0 | 0.21287616 | 1 | 3741 | tags=28%, list=16%, si | MM_2 |
| GOBP_REGULATION_OF_ | GOBP_REGULATION_OF_ESTABLISHMENT_C | 20 | 0.374619 | 1.425667 | 0.072222225 | 0.21620812 | 1 | 14773 | tags=100%, list=63%, | MM_2 |
| GOBP_NEGATIVE_REGUL | GOBP_NEGATIVE_REGULATION_OF_TOR_SI | 61 | 0.289044 | 1.425411 | 0.02631579 | 0.21606077 | 1 | 14492 | tags=97%, list=61%, si | MM_2 |
| GOBP_PROTEIN_LOCALIZ | GOBP_PROTEIN_LOCALIZATION_TO_CHROM | 41 | 0.320558 | 1.4247396 | 0.04032258 | 0.21632197 | 1 | 1795 | tags=20%, list=8%, sig | MM_2 |
| GOBP_PROTEIN_POLYUBI | GOBP_PROTEIN_POLYUBIQUITINATION | 250 | 0.24336 | 1.4246131 | 0.0 | 0.21605575 | 1 | 4756 | tags=33%, list=20%, si | MM_2 |
| GOBP_TRANSFORMING_C | GOBP_TRANSFORMING_GROWTH_FACTOR_ | 35 | 0.309966 | 1.4234396 | 0.06363636 | 0.216886 | 1 | 936 | tags=14%, list=4%, sig | MM_2 |
| GOBP_PROTEIN_QUALITY | GOBP_PROTEIN_QUALITY_CONTROL_FOR_N | 17 | 0.395719 | 1.4227998 | 0.10638298 | 0.21715191 | 1 | 14274 | tags=100%, list=60%, | MM_2 |
| GOBP_POSITIVE_REGULA | GOBP_POSITIVE_REGULATION_OF_RELEASE | 27 | 0.353142 | 1.4227126 | 0.055944055 | 0.21682885 | 1 | 2413 | tags=30%, list=10%, si | MM_2 |
| GOBP_CELLULAR_PIGMEI | GOBP_CELLULAR_PIGMENTATION | 57 | 0.279625 | 1.4220606 | 0.03448276 | 0.2172275 | 1 | 2392 | tags=21%, list=10%, si | MM_2 |
| GOBP_LYMPHOCYTE_ACT | GOBP_LYMPHOCYTE_ACTIVATION_INVOLVE | 202 | 0.234474 | 1.4218065 | 0.0 | 0.21713492 | 1 | 3189 | tags=21%, list=14%, si | MM_2 |
| GOBP_NUCLEOLUS_ORGA | GOBP_NUCLEOLUS_ORGANIZATION | 17 | 0.395126 | 1.4210176 | 0.07653061 | 0.21760724 | 1 | 14288 | tags=100%, list=61%, | MM_2 |
| GOBP_ADAPTIVE_IMMUN | GOBP_ADAPTIVE_IMMUNE_RESPONSE_BAS | 322 | 0.215375 | 1.4204282 | 0.0 | 0.21783432 | 1 | 3383 | tags=25%, list=14%, si | MM_2 |
| ALLOGRAFT_REJECTION | ALLOGRAFT_REJECTION | 40 | 0.304508 | 1.4196974 | 0.008196721 | 0.2183145 | 1 | 2313 | tags=33%, list=10%, si | MM_2 |
| HEMATOPOIETIC_CELL_LI | HEMATOPOIETIC_CELL_LINEAGE | 83 | 0.258945 | 1.4189285 | 0.02 | 0.21873511 | 1 | 3238 | tags=29%, list=14%, si | MM_2 |
| GOBP_REGULATION_OF_ | GOBP_REGULATION_OF_NEURON_MATURA | 16 | 0.404723 | 1.4186084 | 0.085427135 | 0.21878606 | 1 | 1365 | tags=25%, list=6%, sig | MM_2 |
| GOBP_CERAMIDE_BIOSYI | GOBP_CERAMIDE_BIOSYNTHETIC_PROCESS | 65 | 0.2808 | 1.4182369 | 0.06349207 | 0.21878196 | 1 | 2198 | tags=23%, list=9%, sig | MM_2 |
| GOBP_RESPONSE_TO_GA | GOBP_RESPONSE_TO_GAMMA_RADIATION | 48 | 0.296812 | 1.4166865 | 0.03539823 | 0.22027758 | 1 | 14549 | tags=96%, list=62%, si | MM_2 |
| GOBP_PROTEIN_MODIFIC | GOBP_PROTEIN_MODIFICATION_BY_SMALL | 117 | 0.260034 | 1.4163768 | 0.021276595 | 0.22029692 | 1 | 4163 | tags=27%, list=18%, si | MM_2 |
| GOBP_DNA_REPLICATION | GOBP_DNA_REPLICATION | 263 | 0.218158 | 1.4163728 | 0.0 | 0.21988745 | 1 | 4003 | tags=25%, list=17%, si | MM_2 |
| GOBP_RESPONSE_TO_MU | GOBP_RESPONSE_TO_MURAMYL_DIPEPTID | 20 | 0.376865 | 1.4152106 | 0.070063695 | 0.22087012 | 1 | 14720 | tags=100%, list=62%, | MM_2 |
| GOBP_NEGATIVE_REGUL | GOBP_NEGATIVE_REGULATION_OF_VASCUI | 109 | 0.253232 | 1.415085 | 0.030303031 | 0.22060837 | 1 | 1433 | tags=15%, list=6%, sig | MM_2 |
| GOBP_POSITIVE_REGULA | GOBP_POSITIVE_REGULATION_OF_T_CELL_ | 26 | 0.340388 | 1.4150236 | 0.06849315 | 0.22028734 | 1 | 3435 | tags=38%, list=15%, si | MM_2 |
| GOBP_POSITIVE_REGULA | GOBP_POSITIVE_REGULATION_OF_CELL_CE | 309 | 0.21812 | 1.4137743 | 0.0 | 0.22114228 | 1 | 3238 | tags=21%, list=14%, si | MM_2 |
| GOBP_INSULIN_LIKE_GRC | GOBP_INSULIN_LIKE_GROWTH_FACTOR_RE | 49 | 0.290625 | 1.4129782 | 0.055555556 | 0.22160017 | 1 | 607 | tags=10%, list=3%, sig | MM_2 |
| GOBP_PROTEIN_LOCALIZ | GOBP_PROTEIN_LOCALIZATION_TO_NUCLE | 17 | 0.386311 | 1.4115669 | 0.07317073 | 0.2226312 | 1 | 1014 | tags=12%, list=4%, sig | MM_2 |
| GOBP_HEME_METABOLIC | GOBP_HEME_METABOLIC_PROCESS | 34 | 0.324409 | 1.4104754 | 0.06481481 | 0.22353427 | 1 | 4397 | tags=41%, list=19%, si | MM_2 |
| GOBP_CHAPERONE_MED | GOBP_CHAPERONE_MEDIATED_PROTEIN_F | 62 | 0.278482 | 1.4081254 | 0.04054054 | 0.22578646 | 1 | 4193 | tags=31%, list=18%, si | MM_2 |
| GOBP_POSITIVE_REGULA | GOBP_POSITIVE_REGULATION_OF_EXTRACE | 32 | 0.343879 | 1.4078441 | 0.08955224 | 0.22563115 | 1 | 1657 | tags=19%, list=7%, sig | MM_2 |
| GOBP_DNA_DAMAGE_RE | GOBP_DNA_DAMAGE_RESPONSE_SIGNAL_T | 17 | 0.38957 | 1.40779 | 0.101604275 | 0.2252523 | 1 | 2461 | tags=29%, list=10%, si | MM_2 |
| GOBP_NEGATIVE_REGUL | GOBP_NEGATIVE_REGULATION_OF_LEUKOC | 142 | 0.247715 | 1.4076674 | 0.05263158 | 0.22491737 | 1 | 3682 | tags=27%, list=16%, si | MM_2 |
| GOBP_MACROAUTOPHAG | GOBP_MACROAUTOPHAGY | 287 | 0.215953 | 1.4066157 | 0.0 | 0.22574922 | 1 | 5279 | tags=37%, list=22%, si | MM_2 |
| GOBP_NUCLEAR_ENVELO | GOBP_NUCLEAR_ENVELOPE_ORGANIZATION | 52 | 0.289161 | 1.406388 | 0.031578947 | 0.22557685 | 1 | 4859 | tags=38%, list=21%, si | MM_2 |
| GOBP_RUFFLE_ORGANIZA | GOBP_RUFFLE_ORGANIZATION | 54 | 0.29115 | 1.4061203 | 0.029411765 | 0.22546877 | 1 | 5114 | tags=43%, list=22%, si | MM_2 |
| GOBP_RESPONSE_TO_SA | GOBP_RESPONSE_TO_SALT_STRESS | 20 | 0.353874 | 1.405091 | 0.04812834 | 0.22632802 | 1 | 2853 | tags=30%, list=12%, si | MM_2 |
| GOBP_RNA_METHYLATIO | GOBP_RNA_METHYLATION | 65 | 0.283641 | 1.4045084 | 0.06666667 | 0.2265373 | 1 | 3261 | tags=29%, list=14%, si | MM_2 |
| GOBP_REGULATION_OF_ | GOBP_REGULATION_OF_INTRINSIC_APOPTC | 15 | 0.395876 | 1.4044565 | 0.094527364 | 0.22617134 | 1 | 808 | tags=20%, list=3%, sig | MM_2 |
| GOBP_TELOMERE_ORGAI | GOBP_TELOMERE_ORGANIZATION | 150 | 0.242678 | 1.4040813 | 0.0 | 0.22628875 | 1 | 4473 | tags=30%, list=19%, si | MM_2 |
| GOBP_TRNA_METABOLIC | GOBP_TRNA_METABOLIC_PROCESS | 166 | 0.240896 | 1.4036388 | 0.0 | 0.22633561 | 1 | 3529 | tags=22%, list=15%, si | MM_2 |
| GOBP_DNA_REPAIR_DEPI | GOBP_DNA_REPAIR_DEPENDENT_CHROMAT | 23 | 0.36401 | 1.4028986 | 0.065476194 | 0.22683753 | 1 | 1304 | tags=17%, list=6%, sig | MM_2 |
| GOBP_PROTEIN_INSERTIC | GOBP_PROTEIN_INSERTION_INTO_MEMBRA | 39 | 0.313438 | 1.4017047 | 0.077586204 | 0.227778 | 1 | 3725 | tags=33%, list=16%, si | MM_2 |
| GOBP_ACETYL_COA_MET | GOBP_ACETYL_COA_METABOLIC_PROCESS | 34 | 0.3161 | 1.4013016 | 0.05 | 0.22790265 | 1 | 16158 | tags=100%, list=68%, | MM_2 |
| GOBP_MHC_CLASS_II_BI | GOBP_MHC_CLASS_II_BIOSYNTHETIC_PROC | 16 | 0.37494 | 1.4005731 | 0.07537688 | 0.22840479 | 1 | 5405 | tags=63%, list=23%, si | MM_2 |
| GOBP_POSITIVE_REGULA | GOBP_POSITIVE_REGULATION_OF_CYSTEIN | 130 | 0.248987 | 1.4000833 | 0.035714287 | 0.22863576 | 1 | 3167 | tags=24%, list=13%, si | MM_2 |
| GOBP_REGULATION_OF_ | GOBP_REGULATION_OF_INNATE_IMMUNE_ | 427 | 0.239404 | 1.3994464 | 0.0 | 0.22901416 | 1 | 4416 | tags=29%, list=19%, si | MM_2 |
| GOBP_DNA_STRAND_ELC | GOBP_DNA_STRAND_ELONGATION | 36 | 0.316508 | 1.3993773 | 0.056603774 | 0.2286823 | 1 | 16149 | tags=100%, list=68%, | MM_2 |
| GOBP_REGULATION_OF_ | GOBP_REGULATION_OF_DNA_TEMPLATED_ | 79 | 0.273509 | 1.3986036 | 0.015384615 | 0.22924553 | 1 | 14392 | tags=96%, list=61%, si | MM_2 |
| GOBP_MITOCHONDRIAL_ | GOBP_MITOCHONDRIAL_ELECTRON_TRANS | 29 | 0.331818 | 1.3975122 | 0.02962963 | 0.23030263 | 1 | 5341 | tags=48%, list=23%, si | MM_2 |
| GOBP_MELANOCYTE_DIF | GOBP_MELANOCYTE_DIFFERENTIATION | 26 | 0.346836 | 1.3974072 | 0.058441557 | 0.2299935 | 1 | 2151 | tags=23%, list=9%, sig | MM_2 |
| GOBP_HEMATOPOIETIC_ | GOBP_HEMATOPOIETIC_STEM_CELL_HOME | 29 | 0.345157 | 1.3969904 | 0.10294118 | 0.23003283 | 1 | 1362 | tags=17%, list=6%, sig | MM_2 |
| GOBP_POSITIVE_REGULA | GOBP_POSITIVE_REGULATION_OF_CELL_SU | 35 | 0.306475 | 1.3960365 | 0.08461539 | 0.23089883 | 1 | 3196 | tags=29%, list=14%, si | MM_2 |
| GOBP_FIBROBLAST_PROL | GOBP_FIBROBLAST_PROLIFERATION | 147 | 0.24011 | 1.3957548 | 0.0 | 0.23082323 | 1 | 4027 | tags=29%, list=17%, si | MM_2 |
| GOBP_NEGATIVE_REGUL | GOBP_NEGATIVE_REGULATION_OF_PEPTID | 29 | 0.329135 | 1.3953539 | 0.08450704 | 0.23104316 | 1 | 15849 | tags=100%, list=67%, | MM_2 |
| GOBP_CELLULAR_RESPON | GOBP_CELLULAR_RESPONSE_TO_UV | 90 | 0.25924 | 1.3940827 | 0.018518519 | 0.23202717 | 1 | 4115 | tags=28%, list=17%, si | MM_2 |
| ASTHMA | ASTHMA | 16 | 0.391128 | 1.3934115 | 0.0726257 | 0.2324181 | 1 | 2868 | tags=44%, list=12%, si | MM_2 |
| GOBP_REGULATION_OF_ | GOBP_REGULATION_OF_INSULIN_LIKE_GRC | 22 | 0.368497 | 1.3917177 | 0.07772021 | 0.23418301 | 1 | 362 | tags=14%, list=2%, sig | MM_2 |
| GOBP_LEUKOCYTE_MIGR | GOBP_LEUKOCYTE_MIGRATION_INVOLVED_ | 23 | 0.359481 | 1.3909475 | 0.098684214 | 0.23465568 | 1 | 1402 | tags=22%, list=6%, sig | MM_2 |
| GOBP_REGULATION_OF_ | GOBP_REGULATION_OF_RECEPTOR_LOCALI | 38 | 0.297864 | 1.3905238 | 0.042016808 | 0.23485501 | 1 | 1146 | tags=13%, list=5%, sig | MM_2 |
| GOBP_POSITIVE_REGULA | GOBP_POSITIVE_REGULATION_OF_OXIDORE | 37 | 0.303653 | 1.3904126 | 0.05 | 0.23460823 | 1 | 3357 | tags=35%, list=14%, si | MM_2 |
| GOBP_PERK_MEDIATED_ | GOBP_PERK_MEDIATED_UNFOLDED_PROTE | 16 | 0.405959 | 1.3898461 | 0.109947644 | 0.23483983 | 1 | 14032 | tags=100%, list=59%, | MM_2 |
| GOBP_NUCLEOTIDE_BINI | GOBP_NUCLEOTIDE_BINDING_OLIGOMERIZ | 18 | 0.376833 | 1.386428 | 0.075675674 | 0.23860082 | 1 | 14720 | tags=100%, list=62%, | MM_2 |
| GOBP_REGULATION_OF_ | GOBP_REGULATION_OF_TELOMERE_MAINT | 98 | 0.247801 | 1.3846964 | 0.0 | 0.24037974 | 1 | 4473 | tags=30%, list=19%, si | MM_2 |
| GOBP_LOCOMOTORY_EX | GOBP_LOCOMOTORY_EXPLORATION_BEHAV | 19 | 0.375216 | 1.3833327 | 0.09289618 | 0.24150653 | 1 | 2597 | tags=32%, list=11%, si | MM_2 |
| GOBP_NEGATIVE_REGUL | GOBP_NEGATIVE_REGULATION_OF_ATP_DE | 20 | 0.363493 | 1.3803893 | 0.09550562 | 0.24517433 | 1 | 4470 | tags=40%, list=19%, si | MM_2 |
| GOBP_REGULATION_OF_ | GOBP_REGULATION_OF_B_CELL_PROLIFERA | 71 | 0.259154 | 1.3802736 | 0.028169014 | 0.24491213 | 1 | 1521 | tags=18%, list=6%, sig | MM_2 |
| GOBP_TRNA_MODIFICAT | GOBP_TRNA_MODIFICATION | 84 | 0.258217 | 1.3796955 | 0.014925373 | 0.24515037 | 1 | 4979 | tags=37%, list=21%, si | MM_2 |
| GOBP_PEPTIDYL_GLUTAN | GOBP_PEPTIDYL_GLUTAMIC_ACID_MODIFIC | 25 | 0.341913 | 1.3792114 | 0.062111802 | 0.24534906 | 1 | 2088 | tags=28%, list=9%, sig | MM_2 |
| GOBP_ANTIFUNGAL_INN | GOBP_ANTIFUNGAL_INNATE_IMMUNE_RES | 19 | 0.36844 | 1.3789366 | 0.0880829 | 0.2453184 | 1 | 2995 | tags=37%, list=13%, si | MM_2 |
| GOBP_PEPTIDYL_THREON | GOBP_PEPTIDYL_THREONINE_DEPHOSPHOR | 15 | 0.39341 | 1.3786076 | 0.09615385 | 0.24534781 | 1 | 3257 | tags=40%, list=14%, si | MM_2 |
| GOBP_INFLAMMATORY_I | GOBP_INFLAMMATORY_RESPONSE_TO_WC | 30 | 0.317684 | 1.3782551 | 0.07801419 | 0.24535666 | 1 | 1027 | tags=13%, list=4%, sig | MM_2 |

| | | | | | | | | | | |
|---|---|---|---|---|---|---|---|---|---|---|
| GOBP_LAYER_FORMATIO | GOBP_LAYER_FORMATION_IN_CEREBRAL_C | 16 | 0.381606 | 1.3777316 | 0.0855615 | 0.24558511 | 1 | 4884 | tags=50%, list=21%, si | MM_2 |
| GOBP_NEGATIVE_REGUL | GOBP_NEGATIVE_REGULATION_OF_CELL_JU | 35 | 0.312491 | 1.3773739 | 0.05645161 | 0.24562697 | 1 | 3446 | tags=31%, list=15%, si | MM_2 |
| GOBP_REGULATION_OF_ | GOBP_REGULATION_OF_ESTABLISHMENT_C | 25 | 0.329402 | 1.3768506 | 0.09375 | 0.24587075 | 1 | 2493 | tags=32%, list=11%, si | MM_2 |
| GOBP_ENDOPLASMIC_RE | GOBP_ENDOPLASMIC_RETICULUM_TUBULA | 21 | 0.351321 | 1.3758925 | 0.072289154 | 0.24674657 | 1 | 15323 | tags=100%, list=65%, | MM_2 |
| GOBP_NEGATIVE_REGUL | GOBP_NEGATIVE_REGULATION_OF_B_CELL | 16 | 0.388256 | 1.3757408 | 0.10471204 | 0.24658027 | 1 | 3866 | tags=44%, list=16%, si | MM_2 |
| GOBP_INTEGRIN_ACTIVA | GOBP_INTEGRIN_ACTIVATION | 26 | 0.338914 | 1.3725891 | 0.070512824 | 0.24985033 | 1 | 2296 | tags=19%, list=10%, si | MM_2 |
| GOBP_DNA_REPLICATION | GOBP_DNA_REPLICATION_INITIATION | 31 | 0.323018 | 1.3724145 | 0.09375 | 0.24960524 | 1 | 3224 | tags=26%, list=14%, si | MM_2 |
| GOBP_NEGATIVE_REGUL | GOBP_NEGATIVE_REGULATION_OF_TELOM | 27 | 0.336586 | 1.3722862 | 0.062893085 | 0.2493494 | 1 | 3892 | tags=30%, list=16%, si | MM_2 |
| GOBP_REGULATION_OF_ | GOBP_REGULATION_OF_TELOMERE_MAINT | 20 | 0.382799 | 1.371987 | 0.10982659 | 0.24934524 | 1 | 14580 | tags=100%, list=62%, | MM_2 |
| GOBP_NEGATIVE_REGUL | GOBP_NEGATIVE_REGULATION_OF_CELL_A | 301 | 0.188689 | 1.3716476 | 0.0 | 0.2494147 | 1 | 2618 | tags=20%, list=11%, si | MM_2 |
| SMALL_CELL_LUNG_CAN | SMALL_CELL_LUNG_CANCER | 91 | 0.255942 | 1.3715019 | 0.0 | 0.24915506 | 1 | 3035 | tags=22%, list=13%, si | MM_2 |
| OLFACTORY_TRANSDUCT | OLFACTORY_TRANSDUCTION | 186 | -0.53646 | -2.212798 | 0 | 0 | 0 | 6665 | tags=72%, list=28%, si | MM_2 |
| NICOTINE_ADDICTION | NICOTINE_ADDICTION | 35 | -0.56281 | -1.940577 | 0 | 0.001958219 | 0.143 | 8569 | tags=74%, list=36%, si | MM_2 |
| STEROID_HORMONE_BIO | STEROID_HORMONE_BIOSYNTHESIS | 45 | -0.5433 | -1.911585 | 0 | 0.00321675 | 0.237 | 8695 | tags=73%, list=37%, si | MM_2 |
| PHOTOTRANSDUCTION | PHOTOTRANSDUCTION | 21 | -0.59468 | -1.821329 | 0.0011933175 | 0.011841587 | 0.643 | 6164 | tags=57%, list=26%, si | MM_2 |
| GOBP_DNA_TEMPLATED_ | GOBP_DNA_TEMPLATED_TRANSCRIPTION_ | 25 | -0.57682 | -1.800258 | 0 | 0.016196676 | 0.762 | 516 | tags=16%, list=2%, sig | MM_2 |
| GOBP_INTERMEDIATE_FI | GOBP_INTERMEDIATE_FILAMENT_ORGANIZ | 46 | -0.5075 | -1.786346 | 0 | 0.01902167 | 0.818 | 4568 | tags=57%, list=19%, si | MM_2 |
| RETINOL_METABOLISM | RETINOL_METABOLISM | 52 | -0.49749 | -1.769699 | 0 | 0.02373202 | 0.887 | 9302 | tags=75%, list=39%, si | MM_2 |
| SYSTEMIC_LUPUS_ERYTH | SYSTEMIC_LUPUS_ERYTHEMATOSUS | 105 | -0.44986 | -1.752821 | 0 | 0.029695183 | 0.941 | 3784 | tags=35%, list=16%, si | MM_2 |
| GOBP_CELL_FATE_DETER | GOBP_CELL_FATE_DETERMINATION | 33 | -0.50849 | -1.699408 | 0 | 0.05985871 | 0.995 | 8873 | tags=64%, list=38%, si | MM_2 |
| GOBP_POSITIVE_REGULA | GOBP_POSITIVE_REGULATION_OF_CATECH | 18 | -0.58857 | -1.680531 | 0.004889976 | 0.0746497 | 0.998 | 888 | tags=28%, list=4%, sig | MM_2 |
| GOBP_NEUROPEPTIDE_SI | GOBP_NEUROPEPTIDE_SIGNALING_PATHW | 69 | -0.44892 | -1.664821 | 0 | 0.09048898 | 1 | 6982 | tags=54%, list=30%, si | MM_2 |
| GOBP_GROOMING_BEHA | GOBP_GROOMING_BEHAVIOR | 23 | -0.53481 | -1.661753 | 0.0072639226 | 0.0925416 | 1 | 7773 | tags=57%, list=33%, si | MM_2 |
| GOBP_HEART_FIELD_SPE | GOBP_HEART_FIELD_SPECIFICATION | 15 | -0.60358 | -1.656310 | 0.004920049 | 0.097536296 | 1 | 1340 | tags=33%, list=6%, sig | MM_2 |
| GOBP_POSITIVE_REGULA | GOBP_POSITIVE_REGULATION_OF_NEUROT | 24 | -0.52525 | -1.648104 | 0.0024038462 | 0.10639027 | 1 | 1609 | tags=25%, list=7%, sig | MM_2 |
| GOBP_RESPONSE_TO_TE | GOBP_RESPONSE_TO_TESTOSTERONE | 15 | -0.59644 | -1.647983 | 0.0037593986 | 0.10483083 | 1 | 1441 | tags=27%, list=6%, sig | MM_2 |
| GOBP_NOREPINEPHRINE_ | GOBP_NOREPINEPHRINE_TRANSPORT | 30 | -0.50299 | -1.642211 | 0.005773672 | 0.11045813 | 1 | 8296 | tags=57%, list=35%, si | MM_2 |
| GOBP_FEEDING_BEHAVI | GOBP_FEEDING_BEHAVIOR | 117 | -0.41645 | -1.638471 | 0.0010277493 | 0.11401399 | 1 | 8771 | tags=56%, list=37%, si | MM_2 |
| GOBP_POSITIVE_REGULA | GOBP_POSITIVE_REGULATION_OF_CALCIUM | 18 | -0.55832 | -1.630213 | 0.0048367595 | 0.12407121 | 1 | 8129 | tags=61%, list=34%, si | MM_2 |
| GOBP_REGULATION_OF_ | GOBP_REGULATION_OF_NEUROTRANSMITT | 20 | -0.53938 | -1.620795 | 0.007134364 | 0.13712925 | 1 | 1616 | tags=30%, list=7%, sig | MM_2 |
| GOBP_CORONARY_VASCU | GOBP_CORONARY_VASCULATURE_MORPHO | 30 | -0.50093 | -1.610138 | 0.0023337223 | 0.15428923 | 1 | 1264 | tags=23%, list=5%, sig | MM_2 |
| GOBP_RESPONSE_TO_PH | GOBP_RESPONSE_TO_PHEROMONE | 17 | -0.55524 | -1.608972 | 0.013664597 | 0.15397283 | 1 | 5525 | tags=71%, list=23%, si | MM_2 |
| GOBP_NEURONAL_ACTIO | GOBP_NEURONAL_ACTION_POTENTIAL | 56 | -0.44385 | -1.606337 | 0.002202643 | 0.15618946 | 1 | 8802 | tags=59%, list=37%, si | MM_2 |
| GOBP_SEX_DETERMINAT | GOBP_SEX_DETERMINATION | 23 | -0.52368 | -1.602591 | 0.013157895 | 0.16092892 | 1 | 7569 | tags=57%, list=32%, si | MM_2 |
| GOBP_ATRIAL_CARDIAC_ | GOBP_ATRIAL_CARDIAC_MUSCLE_TISSUE_D | 16 | -0.56353 | -1.591982 | 0.013513514 | 0.17936887 | 1 | 9537 | tags=75%, list=40%, si | MM_2 |
| NEUROACTIVE_LIGAND-R | NEUROACTIVE_LIGAND-RECEPTOR_INTERAC | 274 | -0.38254 | -1.591447 | 0 | 0.1778308 | 1 | 8702 | tags=59%, list=37%, si | MM_2 |
| GOBP_REGULATION_OF_ | GOBP_REGULATION_OF_CATECHOLAMINE_ | 55 | -0.4389 | -1.586721 | 0.0032432433 | 0.1855047 | 1 | 8474 | tags=51%, list=36%, si | MM_2 |
| GOBP_ORGAN_OR_TISSU | GOBP_ORGAN_OR_TISSUE_SPECIFIC_IMMU | 25 | -0.51248 | -1.582035 | 0.009456265 | 0.19321579 | 1 | 6686 | tags=56%, list=28%, si | MM_2 |
| CIRCADIAN_RHYTHM | CIRCADIAN_RHYTHM | 29 | -0.4957 | -1.580462 | 0.012702079 | 0.19421618 | 1 | 1355 | tags=21%, list=6%, sig | MM_2 |
| GOBP_EPOXYGENASE_P4 | GOBP_EPOXYGENASE_P450_PATHWAY | 20 | -0.53809 | -1.577459 | 0.013563502 | 0.1981912 | 1 | 7003 | tags=70%, list=30%, si | MM_2 |
| GOBP_RETINOIC_ACID_M | GOBP_RETINOIC_ACID_METABOLIC_PROCES | 28 | -0.49841 | -1.577448 | 0.0035128805 | 0.1955642 | 1 | 11349 | tags=93%, list=48%, si | MM_2 |
| GOBP_SYNAPTIC_VESICLE | GOBP_SYNAPTIC_VESICLE_CLUSTERING | 20 | -0.52827 | -1.574886 | 0.015700484 | 0.19880255 | 1 | 1609 | tags=30%, list=7%, sig | MM_2 |
| GOBP_POSITIVE_REGULA | GOBP_POSITIVE_REGULATION_OF_NEUROT | 30 | -0.48382 | -1.563374 | 0.0070505287 | 0.22375453 | 1 | 1616 | tags=23%, list=7%, sig | MM_2 |
| LINOLEIC_ACID_METABO | LINOLEIC_ACID_METABOLISM | 33 | -0.46611 | -1.555782 | 0.011614402 | 0.24068023 | 1 | 9096 | tags=73%, list=39%, si | MM_2 |
| GOBP_PROTEIN_AUTOPR | GOBP_PROTEIN_AUTOPROCESSING | 23 | -0.50535 | -1.554745 | 0.021201413 | 0.24065803 | 1 | 771 | tags=22%, list=3%, sig | MM_2 |
| GOBP_NEUROTRANSMIT | GOBP_NEUROTRANSMITTER_REUPTAKE | 29 | -0.47841 | -1.551379 | 0.013969732 | 0.24697375 | 1 | 3517 | tags=34%, list=15%, si | MM_2 |
| GOBP_MITOCHONDRIAL_ | GOBP_MITOCHONDRIAL_MEMBRANE_ORG | 107 | 0.555959 | 2.1606212 | 0 | 0.01138874 | 0.011 | 5978 | tags=51%, list=25%, si | MM_3 |
| GOBP_SUPPRESSION_OF_ | GOBP_SUPPRESSION_OF_VIRAL_RELEASE_B | 22 | 0.732216 | 2.1443903 | 0 | 0.006718132 | 0.013 | 3652 | tags=50%, list=15%, si | MM_3 |
| GOBP_MONOCYTE_DIFFE | GOBP_MONOCYTE_DIFFERENTIATION | 28 | 0.667906 | 2.1036434 | 0 | 0.010693863 | 0.031 | 5262 | tags=57%, list=22%, si | MM_3 |
| LYSOSOME | LYSOSOME | 118 | 0.540545 | 2.078298 | 0 | 0.0160425 | 0.058 | 7225 | tags=66%, list=31%, si | MM_3 |
| GOBP_TRANSLATION_AT_ | GOBP_TRANSLATION_AT_SYNAPSE | 48 | 0.60419 | 2.0708995 | 0 | 0.01572703 | 0.07 | 9132 | tags=90%, list=39%, si | MM_3 |
| GOBP_MACROPHAGE_MI | GOBP_MACROPHAGE_MIGRATION | 65 | 0.570252 | 2.0650735 | 0 | 0.014310877 | 0.077 | 6411 | tags=60%, list=27%, si | MM_3 |
| GOBP_POSITIVE_REGULA | GOBP_POSITIVE_REGULATION_OF_DNA_TE | 72 | 0.563687 | 2.063577 | 0 | 0.01241378 | 0.078 | 5637 | tags=44%, list=24%, si | MM_3 |
| GOBP_CYTOSKELETON_D | GOBP_CYTOSKELETON_DEPENDENT_CYTOK | 97 | 0.547848 | 2.062326 | 0 | 0.011382124 | 0.081 | 7255 | tags=63%, list=31%, si | MM_3 |
| GOBP_MITOCHONDRIAL_ | GOBP_MITOCHONDRIAL_FUSION | 36 | 0.624202 | 2.0592527 | 0 | 0.010805985 | 0.087 | 4974 | tags=44%, list=21%, si | MM_3 |
| GOBP_MITOCHONDRIAL_ | GOBP_MITOCHONDRIAL_TRANSLATION | 128 | 0.526738 | 2.0501928 | 0 | 0.010659294 | 0.095 | 5225 | tags=48%, list=22%, si | MM_3 |
| GOBP_REGULATION_OF_ | GOBP_REGULATION_OF_DNA_TEMPLATED_ | 84 | 0.54228 | 2.0469565 | 0 | 0.01015811 | 0.1 | 5637 | tags=42%, list=24%, si | MM_3 |
| GOBP_INNER_MITOCHON | GOBP_INNER_MITOCHONDRIAL_MEMBRAN | 36 | 0.628364 | 2.039175 | 0 | 0.011123221 | 0.115 | 6208 | tags=58%, list=26%, si | MM_3 |
| GOBP_GLYCOLYTIC_PRO | GOBP_GLYCOLYTIC_PROCESS_THROUGH_FR | 25 | 0.665294 | 2.035561 | 0.0012755102 | 0.010987824 | 0.122 | 3972 | tags=52%, list=17%, si | MM_3 |
| GOBP_CHAPERONE_COF | GOBP_CHAPERONE_COFACTOR_DEPENDEN | 31 | 0.634767 | 2.0354395 | 0 | 0.01020298 | 0.122 | 7216 | tags=77%, list=31%, si | MM_3 |
| GOBP_CYTOKINETIC_PRO | GOBP_CYTOKINETIC_PROCESS | 45 | 0.588583 | 2.0318048 | 0 | 0.010146241 | 0.13 | 6931 | tags=64%, list=29%, si | MM_3 |
| GOBP_NEGATIVE_REGUL | GOBP_NEGATIVE_REGULATION_OF_MYELO | 57 | 0.555261 | 2.0139537 | 0 | 0.012875119 | 0.176 | 5762 | tags=47%, list=24%, si | MM_3 |
| GOBP_NEGATIVE_REGUL | GOBP_NEGATIVE_REGULATION_OF_B_CELL | 36 | 0.603742 | 2.007241 | 0 | 0.014068165 | 0.205 | 4794 | tags=47%, list=20%, si | MM_3 |
| GOBP_REGULATION_OF_ | GOBP_REGULATION_OF_INTEGRIN_MEDIAT | 19 | 0.690956 | 2.0064352 | 0 | 0.013402235 | 0.207 | 3124 | tags=53%, list=13%, si | MM_3 |
| GOBP_SNO_S_RNA_MET | GOBP_SNO_S_RNA_METABOLIC_PROCESS | 17 | 0.708451 | 2.005496 | 0 | 0.012860732 | 0.209 | 1513 | tags=35%, list=6%, sig | MM_3 |
| PANTOTHENATE_AND_C | PANTOTHENATE_AND_COA_BIOSYNTHESIS | 17 | 0.705733 | 2.0053575 | 0 | 0.012268757 | 0.209 | 2426 | tags=47%, list=10%, si | MM_3 |
| GOBP_MEMBRANE_RAFT | GOBP_MEMBRANE_RAFT_ORGANIZATION | 24 | 0.66344 | 2.0046701 | 0 | 0.011881238 | 0.213 | 6124 | tags=71%, list=26%, si | MM_3 |
| GOBP_PROTEIN_K48_LIN | GOBP_PROTEIN_K48_LINKED_DEUBIQUITIN | 29 | 0.636282 | 1.9936776 | 0 | 0.013837328 | 0.253 | 2764 | tags=34%, list=12%, si | MM_3 |
| GOBP_TRANSCRIPTION_P | GOBP_TRANSCRIPTION_PREINITIATION_CO | 63 | 0.555892 | 1.9862254 | 0 | 0.014855007 | 0.282 | 7520 | tags=60%, list=32%, si | MM_3 |
| GOBP_SNRNA_METABOL | GOBP_SNRNA_METABOLIC_PROCESS | 50 | 0.576224 | 1.9787767 | 0 | 0.016044581 | 0.312 | 4330 | tags=44%, list=18%, si | MM_3 |
| GOBP_MITOCHONDRIAL_ | GOBP_MITOCHONDRIAL_TRANSPORT | 174 | 0.492163 | 1.9764028 | 0 | 0.016231244 | 0.327 | 5552 | tags=44%, list=24%, si | MM_3 |
| GOBP_MITOTIC_CYTOKIN | GOBP_MITOTIC_CYTOKINESIS | 83 | 0.536165 | 1.975278 | 0 | 0.015926752 | 0.334 | 7254 | tags=63%, list=31%, si | MM_3 |
| GOBP_MONOSACCHARID | GOBP_MONOSACCHARIDE_CATABOLIC_PRO | 40 | 0.589871 | 1.974149 | 0 | 0.015528033 | 0.338 | 3972 | tags=43%, list=17%, si | MM_3 |
| GOBP_REGULATION_OF_ | GOBP_REGULATION_OF_MACROPHAGE_MI | 47 | 0.577349 | 1.9720889 | 0 | 0.015564713 | 0.348 | 6411 | tags=62%, list=27%, si | MM_3 |
| PYRUVATE_METABOLISM | PYRUVATE_METABOLISM | 37 | 0.593273 | 1.9633889 | 0 | 0.01756277 | 0.393 | 4994 | tags=57%, list=21%, si | MM_3 |
| RIBOSOME | RIBOSOME | 128 | 0.506123 | 1.9609438 | 0 | 0.017666077 | 0.404 | 8011 | tags=66%, list=34%, si | MM_3 |
| GOBP_DNA_TEMPLATED_ | GOBP_DNA_TEMPLATED_TRANSCRIPTION_I | 183 | 0.491454 | 1.9564135 | 0 | 0.018532552 | 0.426 | 7590 | tags=56%, list=32%, si | MM_3 |
| CYTOSOLIC_DNA-SENSIN | CYTOSOLIC_DNA-SENSING_PATHWAY | 46 | 0.568683 | 1.9550552 | 0 | 0.018309893 | 0.433 | 3422 | tags=39%, list=14%, si | MM_3 |
| GOBP_REGULATION_OF_ | GOBP_REGULATION_OF_MEMBRANE_PERM | 70 | 0.533364 | 1.9537143 | 0 | 0.018351316 | 0.442 | 5930 | tags=49%, list=25%, si | MM_3 |
| GOBP_MITOCHONDRIAL_ | GOBP_MITOCHONDRIAL_FISSION | 48 | 0.564416 | 1.9513736 | 0 | 0.018752962 | 0.461 | 4679 | tags=50%, list=20%, si | MM_3 |
| GOBP_DNA_TEMPLATED_ | GOBP_DNA_TEMPLATED_TRANSCRIPTION_ | 112 | 0.508376 | 1.9512272 | 0 | 0.018246666 | 0.462 | 6950 | tags=50%, list=29%, si | MM_3 |
| MITOPHAGY | MITOPHAGY | 62 | 0.54158 | 1.9467459 | 0 | 0.018802991 | 0.48 | 5939 | tags=52%, list=25%, si | MM_3 |

| | | | | | | | | | | |
|---|---|---|---|---|---|---|---|---|---|---|
| GOBP_CHAPERONE_MED | GOBP_CHAPERONE_MEDIATED_PROTEIN_F | 62 | 0.542895 | 1.9451977 | 0 | 0.018713966 | 0.487 | 7216 | tags=68%, list=31%, si | MM_3 |
| GOBP_NUCLEOSIDE_DIPH | GOBP_NUCLEOSIDE_DIPHOSPHATE_METAB | 127 | 0.498115 | 1.9441189 | 0 | 0.018575044 | 0.492 | 4327 | tags=35%, list=18%, si | MM_3 |
| GOBP_EOSINOPHIL_MIGR | GOBP_EOSINOPHIL_MIGRATION | 26 | 0.644371 | 1.9438494 | 0.00125 | 0.018257579 | 0.494 | 3662 | tags=54%, list=16%, si | MM_3 |
| GOBP_GLYCOLYTIC_PROC | GOBP_GLYCOLYTIC_PROCESS_THROUGH_GL | 20 | 0.656839 | 1.9438379 | 0.0026773761 | 0.017801138 | 0.494 | 3972 | tags=50%, list=17%, si | MM_3 |
| GOBP_REGULATION_OF_ | GOBP_REGULATION_OF_MITOCHONDRIAL_ | 39 | 0.584228 | 1.9428184 | 0 | 0.017543774 | 0.499 | 5019 | tags=54%, list=21%, si | MM_3 |
| GOBP_NUCLEOSIDE_DIPH | GOBP_NUCLEOSIDE_DIPHOSPHATE_CATABO | 111 | 0.504822 | 1.9337955 | 0 | 0.019267851 | 0.537 | 4327 | tags=34%, list=18%, si | MM_3 |
| AMINO_SUGAR_AND_NU | AMINO_SUGAR_AND_NUCLEOTIDE_SUGAR_ | 46 | 0.573108 | 1.9326845 | 0 | 0.01922916 | 0.546 | 5886 | tags=48%, list=25%, si | MM_3 |
| GOBP_TELOMERE_MAINT | GOBP_TELOMERE_MAINTENANCE_IN_RESP | 32 | 0.594867 | 1.9188924 | 0 | 0.02300158 | 0.62 | 3755 | tags=41%, list=16%, si | MM_3 |
| GOBP_CELL_REDOX_HOM | GOBP_CELL_REDOX_HOMEOSTASIS | 28 | 0.613056 | 1.9129798 | 0 | 0.024903651 | 0.661 | 5033 | tags=50%, list=21%, si | MM_3 |
| GOBP_REGULATION_OF_ | GOBP_REGULATION_OF_MITOCHONDRIAL_ | 58 | 0.545811 | 1.9082732 | 0 | 0.026365433 | 0.689 | 5930 | tags=52%, list=25%, si | MM_3 |
| GOBP_CYTOPLASMIC_TRA | GOBP_CYTOPLASMIC_TRANSLATION | 143 | 0.484934 | 1.9079875 | 0 | 0.026002567 | 0.69 | 8944 | tags=76%, list=38%, si | MM_3 |
| GOBP_GLIAL_CELL_MIGR | GOBP_GLIAL_CELL_MIGRATION | 70 | 0.522721 | 1.9053018 | 0 | 0.026258413 | 0.697 | 5307 | tags=41%, list=22%, si | MM_3 |
| GOBP_MACROPHAGE_CH | GOBP_MACROPHAGE_CHEMOTAXIS | 45 | 0.548064 | 1.9025339 | 0 | 0.026651824 | 0.71 | 6134 | tags=56%, list=26%, si | MM_3 |
| GOBP_AEROBIC_RESPIRA | GOBP_AEROBIC_RESPIRATION | 170 | 0.477127 | 1.9002126 | 0 | 0.026969556 | 0.723 | 7012 | tags=52%, list=30%, si | MM_3 |
| GOBP_PYRIDINE_CONTAI | GOBP_PYRIDINE_CONTAINING_COMPOUND | 182 | 0.474303 | 1.899445 | 0 | 0.026725585 | 0.727 | 4572 | tags=35%, list=19%, si | MM_3 |
| GOBP_NADP_METABOLIC | GOBP_NADP_METABOLIC_PROCESS | 49 | 0.542387 | 1.8984826 | 0 | 0.026549736 | 0.728 | 4852 | tags=41%, list=21%, si | MM_3 |
| GOBP_PROTEIN_HYDROX | GOBP_PROTEIN_HYDROXYLATION | 25 | 0.620823 | 1.8970044 | 0 | 0.026733873 | 0.738 | 3429 | tags=40%, list=15%, si | MM_3 |
| GOBP_MITOCHONDRIAL_ | GOBP_MITOCHONDRIAL_GENE_EXPRESSION | 164 | 0.48079 | 1.8940667 | 0 | 0.027370222 | 0.752 | 5950 | tags=48%, list=25%, si | MM_3 |
| GOBP_INTRINSIC_APOPT | GOBP_INTRINSIC_APOPTOTIC_SIGNALING_F | 83 | 0.504611 | 1.8891045 | 0 | 0.028868042 | 0.772 | 7217 | tags=57%, list=31%, si | MM_3 |
| GOBP_ADIPOSE_TISSUE_ | GOBP_ADIPOSE_TISSUE_DEVELOPMENT | 64 | 0.522992 | 1.884701 | 0 | 0.029940767 | 0.791 | 4948 | tags=42%, list=21%, si | MM_3 |
| GOBP_MRNA_EXPORT_FI | GOBP_MRNA_EXPORT_FROM_NUCLEUS | 60 | 0.526796 | 1.8845547 | 0 | 0.029451784 | 0.791 | 6445 | tags=52%, list=27%, si | MM_3 |
| PARKINSON_DISEASE | PARKINSON_DISEASE | 122 | 0.483306 | 1.8837565 | 0 | 0.029194085 | 0.791 | 7484 | tags=61%, list=32%, si | MM_3 |
| GOBP_ANTIGEN_PROCES | GOBP_ANTIGEN_PROCESSING_AND_PRESEN | 105 | 0.492693 | 1.8817244 | 0 | 0.029365206 | 0.799 | 6797 | tags=54%, list=29%, si | MM_3 |
| GOBP_NEGATIVE_REGUL | GOBP_NEGATIVE_REGULATION_OF_NUCLEO | 20 | 0.648832 | 1.8795421 | 0.0013386881 | 0.02961637 | 0.806 | 4327 | tags=50%, list=18%, si | MM_3 |
| GOBP_RESPONSE_TO_CH | GOBP_RESPONSE_TO_CHEMOKINE | 74 | 0.513376 | 1.8792229 | 0 | 0.02923239 | 0.806 | 3726 | tags=36%, list=16%, si | MM_3 |
| GOBP_PYRUVATE_METAB | GOBP_PYRUVATE_METABOLIC_PROCESS | 127 | 0.487798 | 1.8790319 | 0 | 0.028877761 | 0.807 | 4327 | tags=35%, list=18%, si | MM_3 |
| GOBP_APOPTOTIC_MITO | GOBP_APOPTOTIC_MITOCHONDRIAL_CHAN | 118 | 0.4864 | 1.878781 | 0 | 0.028534245 | 0.808 | 5484 | tags=43%, list=23%, si | MM_3 |
| GOBP_POSITIVE_REGULA | GOBP_POSITIVE_REGULATION_OF_MEMBRA | 39 | 0.56066 | 1.8779271 | 0 | 0.028573437 | 0.812 | 5470 | tags=54%, list=23%, si | MM_3 |
| GOBP_PROTEIN_K63_LIN | GOBP_PROTEIN_K63_LINKED_UBIQUITINAT | 68 | 0.508466 | 1.8772268 | 0 | 0.02835678 | 0.813 | 7075 | tags=50%, list=30%, si | MM_3 |
| GOBP_NADH_DEHYDROG | GOBP_NADH_DEHYDROGENASE_COMPLEX_ | 58 | 0.529581 | 1.8729118 | 0 | 0.029964654 | 0.834 | 6996 | tags=57%, list=30%, si | MM_3 |
| GOBP_TRNA_WOBBLE_BA | GOBP_TRNA_WOBBLE_BASE_MODIFICATIO | 19 | 0.651398 | 1.8660043 | 0.0013495276 | 0.032484047 | 0.864 | 4594 | tags=63%, list=19%, si | MM_3 |
| GOBP_EOSINOPHIL_CHEM | GOBP_EOSINOPHIL_CHEMOTAXIS | 22 | 0.633467 | 1.8652055 | 0.0013089005 | 0.032432012 | 0.865 | 3662 | tags=55%, list=16%, si | MM_3 |
| GOBP_PURINE_CONTAIN | GOBP_PURINE_CONTAINING_COMPOUND_ | 150 | 0.474183 | 1.8598753 | 0 | 0.034617454 | 0.885 | 5010 | tags=35%, list=21%, si | MM_3 |
| GOBP_GRANULOCYTE_M | GOBP_GRANULOCYTE_MIGRATION | 146 | 0.469811 | 1.8596454 | 0 | 0.034211714 | 0.885 | 6172 | tags=43%, list=26%, si | MM_3 |
| GOBP_LYTIC_VACUOLE_C | GOBP_LYTIC_VACUOLE_ORGANIZATION | 95 | 0.493762 | 1.8593218 | 0 | 0.03394884 | 0.887 | 7583 | tags=60%, list=32%, si | MM_3 |
| GOBP_DE_NOVO_PROTEI | GOBP_DE_NOVO_PROTEIN_FOLDING | 38 | 0.559907 | 1.8580927 | 0.0012033695 | 0.033995517 | 0.888 | 6496 | tags=61%, list=28%, si | MM_3 |
| GOBP_TRANSCRIPTION_I | GOBP_TRANSCRIPTION_INITIATION_AT_RNA | 148 | 0.473733 | 1.8575704 | 0 | 0.03384198 | 0.892 | 7590 | tags=55%, list=32%, si | MM_3 |
| GOBP_HEMATOPOIETIC_S | GOBP_HEMATOPOIETIC_STEM_CELL_PROLIF | 37 | 0.567929 | 1.8575511 | 0.0012019231 | 0.033384655 | 0.892 | 6117 | tags=54%, list=26%, si | MM_3 |
| GOBP_NAD_CATABOLIC_ | GOBP_NAD_CATABOLIC_PROCESS | 21 | 0.634314 | 1.8569453 | 0.0013280213 | 0.03318851 | 0.895 | 3972 | tags=43%, list=17%, si | MM_3 |
| NOTCH_SIGNALING_PATH | NOTCH_SIGNALING_PATHWAY | 47 | 0.538236 | 1.8560882 | 0 | 0.03324192 | 0.899 | 4975 | tags=40%, list=21%, si | MM_3 |
| GOBP_CELLULAR_RESPON | GOBP_CELLULAR_RESPONSE_TO_OXYGEN_L | 133 | 0.473692 | 1.855477 | 0 | 0.033038978 | 0.899 | 6906 | tags=56%, list=29%, si | MM_3 |
| GOBP_MITOCHONDRIAL_ | GOBP_MITOCHONDRIAL_RESPIRATORY_CHA | 91 | 0.491607 | 1.8544843 | 0 | 0.033119254 | 0.906 | 6996 | tags=57%, list=30%, si | MM_3 |
| GOBP_NEGATIVE_REGUL | GOBP_NEGATIVE_REGULATION_OF_VIRAL_ | 18 | 0.657727 | 1.8539999 | 0.003916449 | 0.03293577 | 0.907 | 3827 | tags=44%, list=16%, si | MM_3 |
| GOBP_SNRNA_3_END_PF | GOBP_SNRNA_3_END_PROCESSING | 20 | 0.628808 | 1.8529892 | 0.0026490067 | 0.033002652 | 0.91 | 4084 | tags=50%, list=17%, si | MM_3 |
| GOBP_REGULATION_OF_ | GOBP_REGULATION_OF_MITOCHONDRIAL_ | 35 | 0.577374 | 1.8525796 | 0 | 0.032774277 | 0.911 | 4679 | tags=49%, list=20%, si | MM_3 |
| GOBP_IRON_ION_TRANS | GOBP_IRON_ION_TRANSMEMBRANE_TRAN | 24 | 0.622312 | 1.8522078 | 0.0026490067 | 0.03250089 | 0.912 | 4729 | tags=50%, list=20%, si | MM_3 |
| GOBP_OXIDATIVE_PHOSP | GOBP_OXIDATIVE_PHOSPHORYLATION | 124 | 0.476795 | 1.8506664 | 0 | 0.032869533 | 0.917 | 7393 | tags=56%, list=31%, si | MM_3 |
| GOBP_TRNA_WOBBLE_U | GOBP_TRNA_WOBBLE_URIDINE_MODIFICAT | 16 | 0.67239 | 1.8466406 | 0 | 0.03446258 | 0.928 | 5957 | tags=81%, list=25%, si | MM_3 |
| CHRONIC_MYELOID_LEUK | CHRONIC_MYELOID_LEUKEMIA | 75 | 0.500582 | 1.8451896 | 0 | 0.034642085 | 0.933 | 8145 | tags=68%, list=34%, si | MM_3 |
| GOBP_HEMATOPOIETIC_S | GOBP_HEMATOPOIETIC_STEM_CELL_HOME | 29 | 0.58517 | 1.8430005 | 0 | 0.03523864 | 0.936 | 4758 | tags=45%, list=20%, si | MM_3 |
| GOBP_RNA_EXPORT_FRO | GOBP_RNA_EXPORT_FROM_NUCLEUS | 78 | 0.494757 | 1.8403499 | 0.0010857764 | 0.036142536 | 0.94 | 7083 | tags=50%, list=30%, si | MM_3 |
| GOBP_CELLULAR_RESPON | GOBP_CELLULAR_RESPONSE_TO_OSMOTIC_ | 52 | 0.529616 | 1.8402805 | 0 | 0.035802357 | 0.94 | 6558 | tags=52%, list=28%, si | MM_3 |
| PROTEIN_PROCESSING_IN | PROTEIN_PROCESSING_IN_ENDOPLASMIC_F | 158 | 0.460525 | 1.8395805 | 0 | 0.03569053 | 0.941 | 7162 | tags=52%, list=30%, si | MM_3 |
| GOBP_ENDOSOME_ORGA | GOBP_ENDOSOME_ORGANIZATION | 68 | 0.500094 | 1.8389422 | 0 | 0.03575366 | 0.943 | 7951 | tags=65%, list=34%, si | MM_3 |
| GOBP_SNRNA_PROCESSI | GOBP_SNRNA_PROCESSING | 24 | 0.602085 | 1.8386228 | 0.0026845639 | 0.03566803 | 0.945 | 4084 | tags=46%, list=17%, si | MM_3 |
| GOBP_RESPONSE_TO_OX | GOBP_RESPONSE_TO_OXYGEN_LEVELS | 247 | 0.450175 | 1.838026 | 0 | 0.03557283 | 0.945 | 6906 | tags=52%, list=29%, si | MM_3 |
| GOBP_PROTEIN_DNA_CO | GOBP_PROTEIN_DNA_COMPLEX_ASSEMBLY | 154 | 0.461824 | 1.8379883 | 0 | 0.035212632 | 0.945 | 5104 | tags=38%, list=22%, si | MM_3 |
| GOBP_ANTIGEN_PROCES | GOBP_ANTIGEN_PROCESSING_AND_PRESEN | 59 | 0.512631 | 1.8375703 | 0.0011273957 | 0.035058025 | 0.946 | 6692 | tags=63%, list=28%, si | MM_3 |
| GOBP_NUCLEAR_TRANSC | GOBP_NUCLEAR_TRANSCRIBED_MRNA_CAT | 43 | 0.544378 | 1.8365086 | 0.0011750881 | 0.035211757 | 0.953 | 7909 | tags=65%, list=33%, si | MM_3 |
| GOBP_REGULATION_OF_ | GOBP_REGULATION_OF_MITOCHONDRION_ | 139 | 0.468654 | 1.8343647 | 0 | 0.035740014 | 0.958 | 5470 | tags=40%, list=23%, si | MM_3 |
| GOBP_PYRIDINE_CONTAI | GOBP_PYRIDINE_CONTAINING_COMPOUND | 109 | 0.472646 | 1.8336953 | 0 | 0.035734426 | 0.958 | 4327 | tags=31%, list=18%, si | MM_3 |
| GOBP_GRANULOCYTE_CH | GOBP_GRANULOCYTE_CHEMOTAXIS | 119 | 0.477765 | 1.8322164 | 0 | 0.036088564 | 0.963 | 5889 | tags=43%, list=25%, si | MM_3 |
| GOBP_POSITIVE_REGULA | GOBP_POSITIVE_REGULATION_OF_MACROF | 31 | 0.574262 | 1.8317734 | 0.0025094103 | 0.035954203 | 0.963 | 6411 | tags=61%, list=27%, si | MM_3 |
| GOBP_ESTABLISHMENT_( | GOBP_ESTABLISHMENT_OF_PROTEIN_LOCA | 25 | 0.611252 | 1.8313988 | 0.0026007802 | 0.03581188 | 0.963 | 6208 | tags=52%, list=26%, si | MM_3 |
| RNA_POLYMERASE | RNA_POLYMERASE | 27 | 0.600813 | 1.8311739 | 0 | 0.03560061 | 0.964 | 4258 | tags=52%, list=18%, si | MM_3 |
| GOBP_CELLULAR_RESPON | GOBP_CELLULAR_RESPONSE_TO_HYPOXIA | 112 | 0.475048 | 1.8309125 | 0 | 0.03541392 | 0.964 | 5491 | tags=46%, list=23%, si | MM_3 |
| GOBP_NEGATIVE_REGUL | GOBP_NEGATIVE_REGULATION_OF_TYPE_I_ | 42 | 0.53885 | 1.8305912 | 0 | 0.035300843 | 0.964 | 6055 | tags=48%, list=26%, si | MM_3 |
| GOBP_NEGATIVE_REGUL | GOBP_NEGATIVE_REGULATION_OF_RESPON | 112 | 0.471961 | 1.8275955 | 0 | 0.036483377 | 0.971 | 6246 | tags=45%, list=26%, si | MM_3 |
| GOBP_RIBOSE_PHOSPHA | GOBP_RIBOSE_PHOSPHATE_BIOSYNTHETIC_ | 185 | 0.454819 | 1.8273916 | 0 | 0.03624463 | 0.971 | 7066 | tags=52%, list=30%, si | MM_3 |
| GOBP_REGULATION_OF_ | GOBP_REGULATION_OF_MONOCYTE_DIFFE | 17 | 0.642236 | 1.8271087 | 0.0027586208 | 0.036009833 | 0.971 | 7051 | tags=71%, list=30%, si | MM_3 |
| GOBP_MAINTENANCE_OI | GOBP_MAINTENANCE_OF_PROTEIN_LOCAT | 97 | 0.477901 | 1.8252437 | 0 | 0.03658238 | 0.975 | 7627 | tags=54%, list=32%, si | MM_3 |
| GOBP_NEGATIVE_REGUL | GOBP_NEGATIVE_REGULATION_OF_HEMOP | 121 | 0.471978 | 1.8234199 | 0 | 0.037048057 | 0.977 | 5872 | tags=40%, list=25%, si | MM_3 |
| GOBP_NEGATIVE_REGUL | GOBP_NEGATIVE_REGULATION_OF_OSTEOC | 37 | 0.552322 | 1.8231618 | 0.0012239902 | 0.03686942 | 0.977 | 3847 | tags=32%, list=16%, si | MM_3 |
| GOBP_NEGATIVE_REGUL | GOBP_NEGATIVE_REGULATION_OF_MYELO | 89 | 0.484565 | 1.8204622 | 0 | 0.038021103 | 0.982 | 5762 | tags=43%, list=24%, si | MM_3 |
| GOBP_REGULATION_OF_ | GOBP_REGULATION_OF_T_CELL_CHEMOTA | 16 | 0.64921 | 1.8182424 | 0.002688172 | 0.038891386 | 0.986 | 3486 | tags=38%, list=15%, si | MM_3 |
| GOBP_VACUOLE_ORGAN | GOBP_VACUOLE_ORGANIZATION | 223 | 0.45045 | 1.8178043 | 0 | 0.038765986 | 0.987 | 7782 | tags=61%, list=33%, si | MM_3 |
| GOBP_REGULATION_OF_ | GOBP_REGULATION_OF_UBIQUITIN_DEPEN | 157 | 0.459688 | 1.8174146 | 0 | 0.038670454 | 0.987 | 7506 | tags=54%, list=32%, si | MM_3 |
| GOBP_CRANIOFACIAL_SU | GOBP_CRANIOFACIAL_SUTURE_MORPHOGE | 15 | 0.660282 | 1.8165411 | 0.004207574 | 0.038884316 | 0.988 | 883 | tags=33%, list=4%, sig | MM_3 |
| GOBP_CELLULAR_COMPO | GOBP_CELLULAR_COMPONENT_DISASSEMB | 31 | 0.57616 | 1.8163022 | 0.0025348542 | 0.03862709 | 0.988 | 2403 | tags=32%, list=10%, si | MM_3 |
| GOBP_PROTEIN_K6_LINK | GOBP_PROTEIN_K6_LINKED_UBIQUITINATIO | 15 | 0.685382 | 1.8136978 | 0.0013550136 | 0.039678197 | 0.991 | 6677 | tags=67%, list=28%, si | MM_3 |
| GOBP_PEPTIDYL_THREON | GOBP_PEPTIDYL_THREONINE_DEPHOSPHOR | 15 | 0.680975 | 1.811392 | 0.0041436465 | 0.040400997 | 0.995 | 6567 | tags=73%, list=28%, si | MM_3 |
| GOBP_REGULATION_OF_ | GOBP_REGULATION_OF_LYSOSOMAL_LUME | 23 | 0.609658 | 1.8113378 | 0.0012970169 | 0.04010264 | 0.995 | 6481 | tags=61%, list=27%, si | MM_3 |

| | | | | | | | | | |
|---|---|---|---|---|---|---|---|---|---|
| GOBP_CYTOKINESIS | GOBP_CYTOKINESIS | 160 | 0.45531 | 1.8065258 | 0 | 0.042218376 | 0.996 | 7255 | tags=54%, list=31%, si | MM_3 |
| GOBP_RIBOSOME_ASSEM | GOBP_RIBOSOME_ASSEMBLY | 60 | 0.501282 | 1.8055907 | 0.0022421526 | 0.04227228 | 0.997 | 7202 | tags=60%, list=31%, si | MM_3 |
| GOBP_ANTIGEN_PROCES | GOBP_ANTIGEN_PROCESSING_AND_PRESEN | 38 | 0.54965 | 1.8055027 | 0 | 0.041982874 | 0.997 | 6109 | tags=61%, list=26%, si | MM_3 |
| GLYCOLYSIS_/_GLUCONE | GLYCOLYSIS_/_GLUCONEOGENESIS | 60 | 0.508803 | 1.804913 | 0 | 0.041909795 | 0.997 | 4224 | tags=37%, list=18%, si | MM_3 |
| GOBP_CRISTAE_FORMAT | GOBP_CRISTAE_FORMATION | 17 | 0.647466 | 1.8043158 | 0.0067658997 | 0.041897584 | 0.997 | 4217 | tags=47%, list=18%, si | MM_3 |
| GOBP_POSITIVE_REGULA | GOBP_POSITIVE_REGULATION_OF_CIRCADI/ | 19 | 0.639592 | 1.8031961 | 0.0013140604 | 0.04209469 | 0.997 | 3561 | tags=53%, list=15%, si | MM_3 |
| ARGININE_AND_PROLINE | ARGININE_AND_PROLINE_METABOLISM | 47 | 0.514761 | 1.8019828 | 0.0011778563 | 0.042370625 | 0.997 | 5181 | tags=47%, list=22%, si | MM_3 |
| GOBP_POSITIVE_REGULA | GOBP_POSITIVE_REGULATION_OF_PATTERN | 68 | 0.49596 | 1.8019737 | 0.0011273957 | 0.04204261 | 0.997 | 6324 | tags=43%, list=27%, si | MM_3 |
| GOBP_REGULATION_OF_ | GOBP_REGULATION_OF_DNA_TEMPLATED_ | 79 | 0.481206 | 1.8011097 | 0 | 0.04217606 | 0.997 | 6832 | tags=47%, list=29%, si | MM_3 |
| GOBP_RESPONSE_TO_CA | GOBP_RESPONSE_TO_CADMIUM_ION | 35 | 0.549036 | 1.8006856 | 0.0012468828 | 0.042146076 | 0.997 | 6968 | tags=63%, list=30%, si | MM_3 |
| GOBP_NUCLEOSIDE_TRIP | GOBP_NUCLEOSIDE_TRIPHOSPHATE_METAE | 250 | 0.44282 | 1.7988445 | 0 | 0.042813998 | 0.999 | 7036 | tags=48%, list=30%, si | MM_3 |
| GOBP_RIBOSOME_BIOGE | GOBP_RIBOSOME_BIOGENESIS | 306 | 0.440891 | 1.797228 | 0 | 0.04332883 | 0.999 | 7507 | tags=55%, list=32%, si | MM_3 |
| GOBP_COENZYME_A_ME | GOBP_COENZYME_A_METABOLIC_PROCESS | 19 | 0.621069 | 1.7959944 | 0.008253095 | 0.043725397 | 0.999 | 4537 | tags=58%, list=19%, si | MM_3 |
| GOBP_RESPONSE_TO_DE | GOBP_RESPONSE_TO_DECREASED_OXYGEN | 228 | 0.444802 | 1.7954804 | 0 | 0.04363724 | 0.999 | 5491 | tags=42%, list=23%, si | MM_3 |
| GOBP_MITOTIC_CYTOKIN | GOBP_MITOTIC_CYTOKINETIC_PROCESS | 30 | 0.568116 | 1.7952752 | 0.0025188916 | 0.04342577 | 0.999 | 7254 | tags=67%, list=31%, si | MM_3 |
| GOBP_REGULATION_OF_ | GOBP_REGULATION_OF_UBIQUITIN_PROTE | 39 | 0.536726 | 1.7904634 | 0.0012315271 | 0.045937557 | 0.999 | 5943 | tags=49%, list=25%, si | MM_3 |
| GOBP_VASCULAR_ASSOC | GOBP_VASCULAR_ASSOCIATED_SMOOTH_N | 16 | 0.638691 | 1.7902869 | 0.004297994 | 0.045720287 | 0.999 | 4601 | tags=56%, list=19%, si | MM_3 |
| PANCREATIC_CANCER | PANCREATIC_CANCER | 74 | 0.490146 | 1.7902081 | 0 | 0.045429826 | 0.999 | 6953 | tags=57%, list=29%, si | MM_3 |
| GOBP_RESPONSE_TO_LEI | GOBP_RESPONSE_TO_LEPTIN | 19 | 0.62987 | 1.7900909 | 0.0013280213 | 0.04518131 | 0.999 | 2539 | tags=32%, list=11%, si | MM_3 |
| GOBP_SNRNA_TRANSCRI | GOBP_SNRNA_TRANSCRIPTION | 18 | 0.628599 | 1.7899493 | 0.0026007802 | 0.044929165 | 0.999 | 5269 | tags=56%, list=22%, si | MM_3 |
| GOBP_POSITIVE_REGULA | GOBP_POSITIVE_REGULATION_OF_FATTY_A | 20 | 0.608744 | 1.7895948 | 0.0013175231 | 0.04473985 | 0.999 | 4463 | tags=50%, list=19%, si | MM_3 |
| GOBP_POSITIVE_REGULA | GOBP_POSITIVE_REGULATION_OF_VIRAL_LI | 28 | 0.57362 | 1.7887381 | 0.0025575447 | 0.04488638 | 1 | 5358 | tags=50%, list=23%, si | MM_3 |
| GOBP_MEMBRANE_BIOG | GOBP_MEMBRANE_BIOGENESIS | 63 | 0.492181 | 1.7870787 | 0 | 0.045631584 | 1 | 6931 | tags=57%, list=29%, si | MM_3 |
| GOBP_POSITIVE_REGULA | GOBP_POSITIVE_REGULATION_OF_MACROF | 25 | 0.579556 | 1.7868274 | 0.005050505 | 0.045470547 | 1 | 5872 | tags=56%, list=25%, si | MM_3 |
| GOBP_ANTIGEN_PROCES | GOBP_ANTIGEN_PROCESSING_AND_PRESEN | 34 | 0.548327 | 1.7859911 | 0.004884005 | 0.04562989 | 1 | 6692 | tags=68%, list=28%, si | MM_3 |
| GOBP_POSITIVE_REGULA | GOBP_POSITIVE_REGULATION_OF_MITOTIC | 126 | 0.459792 | 1.7857585 | 0 | 0.045485504 | 1 | 6386 | tags=45%, list=27%, si | MM_3 |
| GOBP_POSITIVE_REGULA | GOBP_POSITIVE_REGULATION_OF_MEMBRA | 22 | 0.594884 | 1.7853851 | 0.003916449 | 0.045392767 | 1 | 5262 | tags=45%, list=22%, si | MM_3 |
| GOBP_RESPONSE_TO_OS | GOBP_RESPONSE_TO_OSMOTIC_STRESS | 82 | 0.477712 | 1.7848498 | 0 | 0.04536579 | 1 | 6558 | tags=49%, list=28%, si | MM_3 |
| GOBP_REGULATION_OF_ | GOBP_REGULATION_OF_B_CELL_ACTIVATIC | 141 | 0.461787 | 1.7841325 | 0 | 0.04552888 | 1 | 4794 | tags=34%, list=20%, si | MM_3 |
| GOBP_PROTEIN_DEPHOS | GOBP_PROTEIN_DEPHOSPHORYLATION | 220 | 0.439119 | 1.7838712 | 0 | 0.04538206 | 1 | 6438 | tags=42%, list=27%, si | MM_3 |
| GOBP_PROTEASOME_ME | GOBP_PROTEASOME_MEDIATED_UBIQUITIN | 368 | 0.433642 | 1.7837137 | 0 | 0.045119118 | 1 | 6462 | tags=42%, list=27%, si | MM_3 |
| GOBP_NEGATIVE_REGUL | GOBP_NEGATIVE_REGULATION_OF_DNA_B | 148 | 0.450234 | 1.7832242 | 0 | 0.045059733 | 1 | 5501 | tags=39%, list=23%, si | MM_3 |
| GOBP_LYMPHOCYTE_CHE | GOBP_LYMPHOCYTE_CHEMOTAXIS | 38 | 0.532844 | 1.7830684 | 0.003533569 | 0.04485711 | 1 | 3662 | tags=37%, list=16%, si | MM_3 |
| GOBP_REGULATION_OF_ | GOBP_REGULATION_OF_TELOMERE_CAPPIN | 26 | 0.581009 | 1.7827554 | 0.0051612905 | 0.04474576 | 1 | 6250 | tags=58%, list=26%, si | MM_3 |
| GOBP_DNA_CATABOLIC_ | GOBP_DNA_CATABOLIC_PROCESS | 29 | 0.573272 | 1.7787184 | 0.0061500617 | 0.046841826 | 1 | 2403 | tags=34%, list=10%, si | MM_3 |
| GOBP_NEGATIVE_REGUL | GOBP_NEGATIVE_REGULATION_OF_INNATE | 86 | 0.468108 | 1.7783674 | 0 | 0.046753313 | 1 | 5518 | tags=41%, list=23%, si | MM_3 |
| GOBP_ENDOPLASMIC_RE | GOBP_ENDOPLASMIC_RETICULUM_TO_GOL | 120 | 0.460371 | 1.7779553 | 0 | 0.046752375 | 1 | 6583 | tags=51%, list=28%, si | MM_3 |
| GOBP_RESPIRATORY_BUF | GOBP_RESPIRATORY_BURST | 27 | 0.567173 | 1.7756916 | 0.0038709678 | 0.047753666 | 1 | 6192 | tags=59%, list=26%, si | MM_3 |
| GOBP_NEGATIVE_REGUL | GOBP_NEGATIVE_REGULATION_OF_VIRAL_ | 95 | 0.468192 | 1.7753234 | 0 | 0.047700077 | 1 | 5685 | tags=43%, list=24%, si | MM_3 |
| GOBP_T_CELL_CHEMOTA | GOBP_T_CELL_CHEMOTAXIS | 23 | 0.591458 | 1.7741203 | 0.0038809832 | 0.048086468 | 1 | 3486 | tags=35%, list=15%, si | MM_3 |
| GOBP_GLOMERULAR_EPI | GOBP_GLOMERULAR_EPITHELIUM_DEVELO | 21 | 0.59412 | 1.7732862 | 0.0065789474 | 0.048259176 | 1 | 3338 | tags=38%, list=14%, si | MM_3 |
| GOBP_GLUCOSE_CATABO | GOBP_GLUCOSE_CATABOLIC_PROCESS | 26 | 0.581681 | 1.7731909 | 0.00635324 | 0.048028894 | 1 | 5886 | tags=54%, list=25%, si | MM_3 |
| GOBP_MITOCHONDRIAL_ | GOBP_MITOCHONDRIAL_OUTER_MEMBRAN | 34 | 0.553935 | 1.7727393 | 0.0012674271 | 0.04796861 | 1 | 5019 | tags=50%, list=21%, si | MM_3 |
| GOBP_RNA_SURVEILLAN | GOBP_RNA_SURVEILLANCE | 17 | 0.626329 | 1.7727263 | 0.0054869684 | 0.047678865 | 1 | 3398 | tags=41%, list=14%, si | MM_3 |
| GOBP_CYTOPLASMIC_TR | GOBP_CYTOPLASMIC_TRANSLATIONAL_INIT | 33 | 0.545349 | 1.7724379 | 0.004813478 | 0.04758326 | 1 | 7222 | tags=61%, list=31%, si | MM_3 |
| GOBP_NEGATIVE_REGUL | GOBP_NEGATIVE_REGULATION_OF_MIRNA_ | 30 | 0.565909 | 1.7723483 | 0.0025157232 | 0.047324687 | 1 | 4520 | tags=40%, list=19%, si | MM_3 |
| GOBP_REGULATION_OF_ | GOBP_REGULATION_OF_PROTEASOMAL_UE | 118 | 0.457212 | 1.7719488 | 0 | 0.04729467 | 1 | 7490 | tags=55%, list=32%, si | MM_3 |
| GOBP_PROTEIN_FOLDING | GOBP_PROTEIN_FOLDING | 162 | 0.443269 | 1.7708803 | 0 | 0.047677144 | 1 | 7238 | tags=54%, list=31%, si | MM_3 |
| GOBP_REGULATION_OF_ | GOBP_REGULATION_OF_VIRAL_TRANSCRIPT | 25 | 0.586742 | 1.7706246 | 0.0012531328 | 0.047540635 | 1 | 4088 | tags=36%, list=17%, si | MM_3 |
| GOBP_CELLULAR_RESPIR | GOBP_CELLULAR_RESPIRATION | 218 | 0.436691 | 1.7703323 | 0 | 0.04748585 | 1 | 7012 | tags=48%, list=30%, si | MM_3 |
| GOBP_ATP_METABOLIC_ | GOBP_ATP_METABOLIC_PROCESS | 206 | 0.438186 | 1.7692807 | 0 | 0.047823627 | 1 | 7036 | tags=45%, list=30%, si | MM_3 |
| GOBP_NEUTROPHIL_MIG | GOBP_NEUTROPHIL_MIGRATION | 104 | 0.460317 | 1.7683984 | 0 | 0.04812048 | 1 | 6424 | tags=43%, list=27%, si | MM_3 |
| GOBP_LYSOSOMAL_LUM | GOBP_LYSOSOMAL_LUMEN_ACIDIFICATION | 15 | 0.646956 | 1.7677426 | 0.0055632824 | 0.04822014 | 1 | 6481 | tags=67%, list=27%, si | MM_3 |
| GOBP_NEGATIVE_REGUL | GOBP_NEGATIVE_REGULATION_OF_TRANSF | 222 | 0.431695 | 1.7660692 | 0 | 0.04893933 | 1 | 5501 | tags=35%, list=23%, si | MM_3 |
| GOBP_REGULATION_OF_ | GOBP_REGULATION_OF_GLIAL_CELL_MIGRA | 26 | 0.579909 | 1.7655638 | 0.0025873221 | 0.04894955 | 1 | 6781 | tags=58%, list=29%, si | MM_3 |
| GOBP_NEGATIVE_REGUL | GOBP_NEGATIVE_REGULATION_OF_PROTEI | 27 | 0.570703 | 1.764526 | 0.005082592 | 0.049310375 | 1 | 7521 | tags=67%, list=32%, si | MM_3 |
| GOBP_NADPH_REGENERA | GOBP_NADPH_REGENERATION | 23 | 0.588192 | 1.7640923 | 0.011904762 | 0.049247175 | 1 | 5470 | tags=52%, list=23%, si | MM_3 |
| GOBP_NEGATIVE_REGUL | GOBP_NEGATIVE_REGULATION_OF_REACTI | 52 | 0.500497 | 1.7633088 | 0.0011695906 | 0.049367018 | 1 | 5939 | tags=42%, list=25%, si | MM_3 |
| GOBP_MATURATION_OF_ | GOBP_MATURATION_OF_5_8S_RRNA | 35 | 0.541804 | 1.7624204 | 0.0012195122 | 0.04959097 | 1 | 6935 | tags=57%, list=29%, si | MM_3 |
| GOBP_METALLO_SULFUR | GOBP_METALLO_SULFUR_CLUSTER_ASSEMI | 25 | 0.580052 | 1.7622777 | 0.0038560412 | 0.049445815 | 1 | 4729 | tags=48%, list=20%, si | MM_3 |
| GOBP_MIRNA_MEDIATED | GOBP_MIRNA_MEDIATED_GENE_SILENCING | 17 | 0.627463 | 1.7606319 | 0.0070921984 | 0.050129276 | 1 | 3041 | tags=41%, list=13%, si | MM_3 |
| GOBP_MITOCHONDRIAL_ | GOBP_MITOCHONDRIAL_TRANSMEMBRANE | 92 | 0.467757 | 1.7596442 | 0 | 0.050408315 | 1 | 5552 | tags=41%, list=24%, si | MM_3 |
| GOBP_NEGATIVE_REGUL | GOBP_NEGATIVE_REGULATION_OF_DNA_R | 42 | 0.521156 | 1.7594991 | 0.0035046728 | 0.050198358 | 1 | 4668 | tags=38%, list=20%, si | MM_3 |
| COMPLEMENT_AND_COA | COMPLEMENT_AND_COAGULATION_CASCA | 72 | 0.471147 | 1.7581728 | 0.002219756 | 0.05068446 | 1 | 4577 | tags=38%, list=19%, si | MM_3 |
| GOBP_POSITIVE_REGULA | GOBP_POSITIVE_REGULATION_OF_AMYLOII | 20 | 0.601179 | 1.7575749 | 0.0066489363 | 0.050758246 | 1 | 3135 | tags=30%, list=13%, si | MM_3 |
| GOBP_INTRACILIARY_AN | GOBP_INTRACILIARY_ANTEROGRADE_TRAN | 15 | 0.635988 | 1.7568387 | 0.0013850415 | 0.050921444 | 1 | 2758 | tags=47%, list=12%, si | MM_3 |
| GOBP_NEUTROPHIL_CHE | GOBP_NEUTROPHIL_CHEMOTAXIS | 81 | 0.471196 | 1.7567668 | 0.0010857764 | 0.050685335 | 1 | 4387 | tags=33%, list=19%, si | MM_3 |
| GOBP_POSITIVE_REGULA | GOBP_POSITIVE_REGULATION_OF_LYMPHO | 18 | 0.611717 | 1.7567537 | 0.007978723 | 0.050423976 | 1 | 1766 | tags=28%, list=7%, sig | MM_3 |
| GOBP_CARBOHYDRATE_C | GOBP_CARBOHYDRATE_CATABOLIC_PROCES | 161 | 0.44909 | 1.7563698 | 0.0010384216 | 0.050387062 | 1 | 7036 | tags=50%, list=30%, si | MM_3 |
| GOBP_TRANSCRIPTION_B | GOBP_TRANSCRIPTION_BY_RNA_POLYMER/ | 51 | 0.506971 | 1.7561045 | 0 | 0.050251495 | 1 | 7218 | tags=55%, list=31%, si | MM_3 |
| GOBP_HYDROGEN_PERO | GOBP_HYDROGEN_PEROXIDE_CATABOLIC_F | 22 | 0.588852 | 1.7558595 | 0.009126467 | 0.05013905 | 1 | 4382 | tags=45%, list=19%, si | MM_3 |
| GOBP_INTRACELLULAR_I | GOBP_INTRACELLULAR_IRON_ION_HOMEO | 64 | 0.482448 | 1.7554381 | 0.0045924224 | 0.05016384 | 1 | 6675 | tags=47%, list=28%, si | MM_3 |
| GOBP_RESPONSE_TO_FO | GOBP_RESPONSE_TO_FOOD | 35 | 0.538859 | 1.7538759 | 0.0050955415 | 0.050745837 | 1 | 4558 | tags=40%, list=19%, si | MM_3 |
| GOBP_MITOCHONDRIAL_ | GOBP_MITOCHONDRIAL_OUTER_MEMBRAN | 28 | 0.559438 | 1.7537894 | 0.0025477707 | 0.050562218 | 1 | 5019 | tags=50%, list=21%, si | MM_3 |
| GOBP_ANTIMICROBIAL_H | GOBP_ANTIMICROBIAL_HUMORAL_RESPON | 98 | 0.458617 | 1.7522 | 0 | 0.0511742 | 1 | 5172 | tags=41%, list=22%, si | MM_3 |
| GOBP_ELECTRON_TRANS | GOBP_ELECTRON_TRANSPORT_CHAIN | 107 | 0.463766 | 1.7518146 | 0 | 0.05110809 | 1 | 7437 | tags=51%, list=31%, si | MM_3 |
| GOBP_PROTEIN_LOCALIZ | GOBP_PROTEIN_LOCALIZATION_TO_NUCLE/ | 15 | 0.648885 | 1.7510281 | 0.007173601 | 0.051307935 | 1 | 2945 | tags=47%, list=12%, si | MM_3 |
| GOBP_MACROAUTOPHAG | GOBP_MACROAUTOPHAGY | 287 | 0.427312 | 1.7507404 | 0 | 0.05123097 | 1 | 7782 | tags=58%, list=33%, si | MM_3 |
| GOBP_REGULATION_OF_ | GOBP_REGULATION_OF_MAMMARY_GLANI | 19 | 0.605756 | 1.7493787 | 0.0053619305 | 0.05171719 | 1 | 5067 | tags=53%, list=21%, si | MM_3 |
| GOBP_MODULATION_BY_ | GOBP_MODULATION_BY_SYMBIONT_OF_EN | 56 | 0.492488 | 1.7491548 | 0 | 0.051617954 | 1 | 6393 | tags=52%, list=27%, si | MM_3 |
| GOBP_TRNA_METABOLIC | GOBP_TRNA_METABOLIC_PROCESS | 166 | 0.438759 | 1.7463183 | 0 | 0.05316913 | 1 | 6906 | tags=52%, list=29%, si | MM_3 |
| GOBP_CEREBRAL_CORTE | GOBP_CEREBRAL_CORTEX_RADIALLY_ORIEN | 38 | 0.526715 | 1.7461964 | 0.0012239902 | 0.052980773 | 1 | 5307 | tags=47%, list=22%, si | MM_3 |

| | | | | | | | | | | |
|---|---|---|---|---|---|---|---|---|---|---|
| GOBP_NEUTROPHIL_MED | GOBP_NEUTROPHIL_MEDIATED_CYTOTOXIC | 17 | 0.62591 | 1.7447699 | 0.010796221 | 0.05378333 | 1 | 941 | tags=24%, list=4%, sig | MM_3 |
| GOBP_REGULATION_OF_ | GOBP_REGULATION_OF_CD8_POSITIVE_ALP | 18 | 0.607899 | 1.7445384 | 0.005284016 | 0.05365544 | 1 | 4364 | tags=50%, list=18%, si | MM_3 |
| GOBP_ATP_BIOSYNTHETI | GOBP_ATP_BIOSYNTHETIC_PROCESS | 80 | 0.469102 | 1.7431808 | 0.0010952903 | 0.054324634 | 1 | 7012 | tags=51%, list=30%, si | MM_3 |
| GOBP_AMYLOID_PRECUR | GOBP_AMYLOID_PRECURSOR_PROTEIN_CAT | 59 | 0.493508 | 1.7430003 | 0 | 0.054200284 | 1 | 7235 | tags=49%, list=31%, si | MM_3 |
| GOBP_POSITIVE_REGULA | GOBP_POSITIVE_REGULATION_OF_MITOTIC | 93 | 0.458572 | 1.7411768 | 0 | 0.054976314 | 1 | 6906 | tags=49%, list=29%, si | MM_3 |
| GOBP_REGULATION_OF_ | GOBP_REGULATION_OF_NUCLEOBASE_CON | 16 | 0.632773 | 1.7404242 | 0.0055710305 | 0.055141665 | 1 | 3581 | tags=38%, list=15%, si | MM_3 |
| GOBP_REGULATION_OF_ | GOBP_REGULATION_OF_MACROPHAGE_CH | 32 | 0.532589 | 1.7397615 | 0.007434944 | 0.055205517 | 1 | 6134 | tags=53%, list=26%, si | MM_3 |
| GOBP_POSITIVE_REGULA | GOBP_POSITIVE_REGULATION_OF_DNA_TE | 59 | 0.490666 | 1.7395093 | 0 | 0.055139028 | 1 | 5725 | tags=39%, list=24%, si | MM_3 |
| GOBP_REGULATION_OF_ | GOBP_REGULATION_OF_ADENYLATE_CYCLA | 24 | 0.569503 | 1.738574 | 0.008 | 0.05542974 | 1 | 5032 | tags=42%, list=21%, si | MM_3 |
| GOBP_REGULATION_OF_ | GOBP_REGULATION_OF_B_CELL_DIFFERENT | 37 | 0.525087 | 1.7385219 | 0.002538071 | 0.05521504 | 1 | 6594 | tags=51%, list=28%, si | MM_3 |
| GOBP_ANTIMICROBIAL_H | GOBP_ANTIMICROBIAL_HUMORAL_IMMUN | 79 | 0.474708 | 1.7380154 | 0.0011037528 | 0.055301663 | 1 | 3784 | tags=35%, list=16%, si | MM_3 |
| GOBP_NUCLEAR_TRANSC | GOBP_NUCLEAR_TRANSCRIBED_MRNA_CAT | 118 | 0.447576 | 1.7370155 | 0 | 0.055582743 | 1 | 7411 | tags=50%, list=31%, si | MM_3 |
| ALZHEIMER_DISEASE | ALZHEIMER_DISEASE | 159 | 0.435354 | 1.7368102 | 0 | 0.055452585 | 1 | 7393 | tags=57%, list=31%, si | MM_3 |
| GOBP_REGULATION_OF_ | GOBP_REGULATION_OF_PATTERN_RECOGN | 157 | 0.43356 | 1.7352078 | 0 | 0.056209933 | 1 | 7075 | tags=46%, list=30%, si | MM_3 |
| GOBP_REGULATION_OF_ | GOBP_REGULATION_OF_BIOLOGICAL_PROC | 64 | 0.482397 | 1.7351727 | 0.0011494253 | 0.0559629 | 1 | 5903 | tags=47%, list=25%, si | MM_3 |
| GOBP_POSITIVE_REGULA | GOBP_POSITIVE_REGULATION_OF_SMOOTH | 17 | 0.615394 | 1.7343347 | 0.009536785 | 0.056163576 | 1 | 3173 | tags=41%, list=13%, si | MM_3 |
| GOBP_NEGATIVE_REGUL | GOBP_NEGATIVE_REGULATION_OF_UBIQUI | 18 | 0.607476 | 1.7334247 | 0.006811989 | 0.05643865 | 1 | 3755 | tags=44%, list=16%, si | MM_3 |
| GOBP_RESPIRATORY_CHA | GOBP_RESPIRATORY_CHAIN_COMPLEX_IV_ | 24 | 0.578413 | 1.733301 | 0.002538071 | 0.056246094 | 1 | 5820 | tags=63%, list=25%, si | MM_3 |
| GOBP_PROTEIN_REFOLDI | GOBP_PROTEIN_REFOLDING | 23 | 0.569713 | 1.7331538 | 0.0067114094 | 0.056050703 | 1 | 6463 | tags=65%, list=27%, si | MM_3 |
| GOBP_REGULATION_OF_ | GOBP_REGULATION_OF_MITOCHONDRIAL_ | 87 | 0.462654 | 1.7327993 | 0 | 0.056078155 | 1 | 7322 | tags=54%, list=31%, si | MM_3 |
| GOBP_NUCLEOSIDE_PHO | GOBP_NUCLEOSIDE_PHOSPHATE_BIOSYNTH | 234 | 0.425429 | 1.7326503 | 0 | 0.05590862 | 1 | 6211 | tags=44%, list=26%, si | MM_3 |
| GOBP_PROTON_MOTIVE_ | GOBP_PROTON_MOTIVE_FORCE_DRIVEN_A | 52 | 0.495847 | 1.7323787 | 0.0011376564 | 0.055815313 | 1 | 7393 | tags=58%, list=31%, si | MM_3 |
| OXIDATIVE_PHOSPHORYL | OXIDATIVE_PHOSPHORYLATION | 115 | 0.453218 | 1.7317926 | 0.001059322 | 0.05597359 | 1 | 7393 | tags=62%, list=31%, si | MM_3 |
| GOBP_B_CELL_MEDIATED | GOBP_B_CELL_MEDIATED_IMMUNITY | 141 | 0.443363 | 1.7312534 | 0 | 0.05606565 | 1 | 5466 | tags=36%, list=23%, si | MM_3 |
| GOBP_REGULATION_OF_ | GOBP_REGULATION_OF_NEUTROPHIL_CHEM | 36 | 0.524681 | 1.7299274 | 0.0062421975 | 0.056621797 | 1 | 4387 | tags=36%, list=19%, si | MM_3 |
| GOBP_REGULATION_OF_ | GOBP_REGULATION_OF_INCLUSION_BODY_ | 19 | 0.602283 | 1.72885 | 0.0067385444 | 0.05715377 | 1 | 4191 | tags=47%, list=18%, si | MM_3 |
| GOBP_ATP_SYNTHESIS_C | GOBP_ATP_SYNTHESIS_COUPLED_ELECTRO | 76 | 0.47781 | 1.7288457 | 0.0011223345 | 0.056906562 | 1 | 7393 | tags=55%, list=31%, si | MM_3 |
| GOBP_RIBONUCLEOPROT | GOBP_RIBONUCLEOPROTEIN_COMPLEX_BIO | 417 | 0.418882 | 1.7283672 | 0 | 0.05703844 | 1 | 7454 | tags=52%, list=32%, si | MM_3 |
| GOBP_CYTOPLASMIC_SE | GOBP_CYTOPLASMIC_SEQUESTERING_OF_P | 17 | 0.620305 | 1.7282006 | 0.0055710305 | 0.056893386 | 1 | 5414 | tags=53%, list=23%, si | MM_3 |
| GOBP_REGULATION_OF_ | GOBP_REGULATION_OF_TRANSLATION | 329 | 0.420034 | 1.7280627 | 0 | 0.05672699 | 1 | 7357 | tags=51%, list=31%, si | MM_3 |
| GOBP_ANTIGEN_PROCES | GOBP_ANTIGEN_PROCESSING_AND_PRESEN | 27 | 0.552438 | 1.7268951 | 0.0074257427 | 0.057283696 | 1 | 5938 | tags=56%, list=25%, si | MM_3 |
| GOBP_REGULATION_OF_ | GOBP_REGULATION_OF_MEMBRANE_INVA | 21 | 0.583634 | 1.7259712 | 0.008805032 | 0.05770164 | 1 | 6192 | tags=52%, list=26%, si | MM_3 |
| GOBP_NUCLEOSIDE_PHO | GOBP_NUCLEOSIDE_PHOSPHATE_CATABOLI | 170 | 0.431088 | 1.7247005 | 0 | 0.058334142 | 1 | 7101 | tags=46%, list=30%, si | MM_3 |
| GOBP_REGULATION_OF_ | GOBP_REGULATION_OF_RELEASE_OF_CYTO | 49 | 0.501286 | 1.7243897 | 0.0046838406 | 0.058306113 | 1 | 6558 | tags=51%, list=28%, si | MM_3 |
| GOBP_REGULATION_OF_ | GOBP_REGULATION_OF_PHAGOCYTOSIS | 106 | 0.448805 | 1.7242631 | 0 | 0.058132906 | 1 | 7430 | tags=51%, list=31%, si | MM_3 |
| GOBP_FIBROBLAST_MIGR | GOBP_FIBROBLAST_MIGRATION | 68 | 0.479051 | 1.7235991 | 0.001109878 | 0.058382392 | 1 | 7955 | tags=63%, list=34%, si | MM_3 |
| GOBP_NUCLEOSIDE_BISP | GOBP_NUCLEOSIDE_BISPHOSPHATE_BIOSYN | 39 | 0.524395 | 1.7234344 | 0.0023894862 | 0.05825833 | 1 | 5444 | tags=46%, list=23%, si | MM_3 |
| KAPOSI_SARCOMA-ASSO | KAPOSI_SARCOMA-ASSOCIATED_HERPESVIR | 176 | 0.431748 | 1.7220055 | 0 | 0.058931552 | 1 | 5939 | tags=40%, list=25%, si | MM_3 |
| GOBP_REGULATION_OF_ | GOBP_REGULATION_OF_INFLAMMASOME_ | 40 | 0.508061 | 1.7217553 | 0.0084951455 | 0.058844972 | 1 | 4253 | tags=33%, list=18%, si | MM_3 |
| GOBP_REGULATION_OF_ | GOBP_REGULATION_OF_FIBROBLAST_MIGR | 47 | 0.497736 | 1.7211059 | 0.0035046728 | 0.059018057 | 1 | 6906 | tags=57%, list=29%, si | MM_3 |
| GOBP_COTRANSLATIONA | GOBP_COTRANSLATIONAL_PROTEIN_TARGE | 20 | 0.591125 | 1.7197258 | 0.01193634 | 0.059742957 | 1 | 6379 | tags=60%, list=27%, si | MM_3 |
| GOBP_HUMORAL_IMMU | GOBP_HUMORAL_IMMUNE_RESPONSE | 184 | 0.430109 | 1.7196605 | 0 | 0.059547476 | 1 | 4227 | tags=29%, list=18%, si | MM_3 |
| GOBP_POSITIVE_REGULA | GOBP_POSITIVE_REGULATION_OF_MUSCLE_ | 76 | 0.465249 | 1.7195799 | 0 | 0.059340782 | 1 | 4259 | tags=34%, list=18%, si | MM_3 |
| GOBP_POSITIVE_REGULA | GOBP_POSITIVE_REGULATION_OF_ENDOCY | 182 | 0.428661 | 1.717305 | 0 | 0.06064233 | 1 | 6781 | tags=46%, list=29%, si | MM_3 |
| GOBP_POSITIVE_REGULA | GOBP_POSITIVE_REGULATION_OF_MACROP | 29 | 0.547753 | 1.7161536 | 0.010322581 | 0.061130557 | 1 | 4917 | tags=41%, list=21%, si | MM_3 |
| GOBP_POSITIVE_REGULA | GOBP_POSITIVE_REGULATION_OF_MUSCLE_ | 43 | 0.508976 | 1.7155435 | 0.004784689 | 0.061361626 | 1 | 5470 | tags=42%, list=23%, si | MM_3 |
| GOBP_TRICARBOXYLIC_A | GOBP_TRICARBOXYLIC_ACID_CYCLE | 33 | 0.530655 | 1.7154253 | 0.0050188205 | 0.061222244 | 1 | 6124 | tags=52%, list=26%, si | MM_3 |
| GOBP_CANONICAL_NF_K | GOBP_CANONICAL_NF_KAPPAB_SIGNAL_TR | 247 | 0.418063 | 1.714503 | 0 | 0.061630882 | 1 | 7295 | tags=48%, list=31%, si | MM_3 |
| GOBP_POSITIVE_REGULA | GOBP_POSITIVE_REGULATION_OF_TRANSLA | 125 | 0.443753 | 1.71445 | 0.002120891 | 0.06142499 | 1 | 7253 | tags=54%, list=31%, si | MM_3 |
| GOBP_POSITIVE_REGULA | GOBP_POSITIVE_REGULATION_OF_DENDRIT | 31 | 0.533385 | 1.7142427 | 0.006142506 | 0.061340846 | 1 | 4281 | tags=39%, list=18%, si | MM_3 |
| GOBP_POSITIVE_REGULA | GOBP_POSITIVE_REGULATION_OF_PHAGOC | 74 | 0.463814 | 1.7142285 | 0 | 0.061104745 | 1 | 7359 | tags=54%, list=31%, si | MM_3 |
| GOBP_NEGATIVE_REGUL | GOBP_NEGATIVE_REGULATION_OF_NF_KAP | 78 | 0.460733 | 1.714162 | 0.003267974 | 0.06094438 | 1 | 5501 | tags=38%, list=23%, si | MM_3 |
| GOBP_MYELOID_LEUKOC | GOBP_MYELOID_LEUKOCYTE_MIGRATION | 222 | 0.424021 | 1.7141594 | 0 | 0.06070349 | 1 | 6424 | tags=42%, list=27%, si | MM_3 |
| GOBP_POSITIVE_REGULA | GOBP_POSITIVE_REGULATION_OF_MYELOID | 21 | 0.585406 | 1.7138101 | 0.006501951 | 0.0606843 | 1 | 7218 | tags=67%, list=31%, si | MM_3 |
| TNF_SIGNALING_PATHW | TNF_SIGNALING_PATHWAY | 108 | 0.45019 | 1.7134597 | 0 | 0.060693786 | 1 | 5833 | tags=44%, list=25%, si | MM_3 |
| GOBP_POSITIVE_REGULA | GOBP_POSITIVE_REGULATION_OF_UBIQUIT | 93 | 0.453719 | 1.7134573 | 0 | 0.06046069 | 1 | 6369 | tags=46%, list=27%, si | MM_3 |
| GOBP_REGULATION_OF_ | GOBP_REGULATION_OF_HEMATOPOIETIC_S | 19 | 0.593981 | 1.712522 | 0.009345794 | 0.060938023 | 1 | 4491 | tags=42%, list=19%, si | MM_3 |
| GOBP_MITOCHONDRIAL_ | GOBP_MITOCHONDRIAL_CYTOCHROME_C_ | 19 | 0.594889 | 1.7124709 | 0.005284016 | 0.06075384 | 1 | 5820 | tags=63%, list=25%, si | MM_3 |
| GOBP_PLASMA_MEMBRA | GOBP_PLASMA_MEMBRANE_TUBULATION | 19 | 0.596234 | 1.7120376 | 0.010666667 | 0.06087897 | 1 | 6678 | tags=63%, list=28%, si | MM_3 |
| GOBP_ENDOLYSOSOMAL_ | GOBP_ENDOLYSOSOMAL_TOLL_LIKE_RECEP | 47 | 0.49537 | 1.7109785 | 0.0035971224 | 0.061334036 | 1 | 5903 | tags=47%, list=25%, si | MM_3 |
| GOBP_INTRINSIC_APOPT | GOBP_INTRINSIC_APOPTOTIC_SIGNALING_P | 44 | 0.50584 | 1.7106916 | 0.0072202166 | 0.06127362 | 1 | 8226 | tags=68%, list=35%, si | MM_3 |
| GOBP_AMYLOID_FIBRIL_ | GOBP_AMYLOID_FIBRIL_FORMATION | 30 | 0.533495 | 1.7102516 | 0.007352941 | 0.06133195 | 1 | 6546 | tags=53%, list=28%, si | MM_3 |
| GOBP_LYSOSOMAL_TRAN | GOBP_LYSOSOMAL_TRANSPORT | 124 | 0.433305 | 1.7094613 | 0 | 0.06168113 | 1 | 7245 | tags=50%, list=31%, si | MM_3 |
| GOBP_CEREBRAL_CORTEX | GOBP_CEREBRAL_CORTEX_CELL_MIGRATIO | 50 | 0.498901 | 1.7079931 | 0.0034722222 | 0.062458903 | 1 | 5855 | tags=48%, list=25%, si | MM_3 |
| GOBP_REGULATION_OF_ | GOBP_REGULATION_OF_PROTEIN_LOCALIZA | 44 | 0.503681 | 1.7061887 | 0.0011655012 | 0.063418165 | 1 | 6906 | tags=57%, list=29%, si | MM_3 |
| GOBP_REGULATION_OF_ | GOBP_REGULATION_OF_NEUTROPHIL_MIGR | 52 | 0.487418 | 1.7055964 | 0.0034364262 | 0.063588336 | 1 | 6424 | tags=44%, list=27%, si | MM_3 |
| GOBP_AUTOPHAGOSOMI | GOBP_AUTOPHAGOSOME_MATURATION | 58 | 0.476531 | 1.7053368 | 0.0022346368 | 0.06348602 | 1 | 7657 | tags=64%, list=32%, si | MM_3 |
| GOBP_NEGATIVE_REGUL | GOBP_NEGATIVE_REGULATION_OF_VIRAL_ | 36 | 0.516268 | 1.7053096 | 0.0024125453 | 0.06327227 | 1 | 6706 | tags=64%, list=28%, si | MM_3 |
| GOBP_REGULATION_OF_ | GOBP_REGULATION_OF_GRANULOCYTE_CH | 59 | 0.474062 | 1.7049521 | 0.004459309 | 0.06323323 | 1 | 5889 | tags=41%, list=25%, si | MM_3 |
| GOBP_POSITIVE_REGULA | GOBP_POSITIVE_REGULATION_OF_INFLAMM | 22 | 0.572009 | 1.7032453 | 0.0026845639 | 0.06425621 | 1 | 4253 | tags=32%, list=18%, si | MM_3 |
| GOBP_REGULATION_OF_ | GOBP_REGULATION_OF_PROTEOLYSIS_INVO | 215 | 0.420785 | 1.7016394 | 0 | 0.06521838 | 1 | 7506 | tags=52%, list=32%, si | MM_3 |
| GOBP_MITOTIC_DNA_RE | GOBP_MITOTIC_DNA_REPLICATION | 16 | 0.614763 | 1.6996819 | 0.011034483 | 0.06641309 | 1 | 5929 | tags=69%, list=25%, si | MM_3 |
| GOBP_POSITIVE_REGULA | GOBP_POSITIVE_REGULATION_OF_MONON | 94 | 0.452472 | 1.6978006 | 0 | 0.067446716 | 1 | 6543 | tags=48%, list=28%, si | MM_3 |
| GOBP_REGULATION_OF_ | GOBP_REGULATION_OF_SUPEROXIDE_META | 34 | 0.528939 | 1.6973575 | 0.003773585 | 0.067503035 | 1 | 5033 | tags=41%, list=21%, si | MM_3 |
| GOBP_NEGATIVE_REGUL | GOBP_NEGATIVE_REGULATION_OF_RELEAS | 22 | 0.578709 | 1.6971354 | 0.014230272 | 0.0674646 | 1 | 5985 | tags=55%, list=25%, si | MM_3 |
| GOBP_SOMATIC_DIVERSI | GOBP_SOMATIC_DIVERSIFICATION_OF_IMN | 17 | 0.606827 | 1.6969668 | 0.0055632824 | 0.067328714 | 1 | 1709 | tags=35%, list=7%, sig | MM_3 |
| GOBP_CELLULAR_RESPON | GOBP_CELLULAR_RESPONSE_TO_LEPTIN_ST | 16 | 0.60868 | 1.6966549 | 0.011004127 | 0.067287326 | 1 | 2539 | tags=31%, list=11%, si | MM_3 |
| GOBP_NUCLEAR_TRANSC | GOBP_NUCLEAR_TRANSCRIBED_MRNA_CAT | 59 | 0.46524 | 1.6966385 | 0.005733945 | 0.06706778 | 1 | 7354 | tags=49%, list=31%, si | MM_3 |
| GOBP_MITOCHONDRIAL_ | GOBP_MITOCHONDRIAL_CALCIUM_ION_HO | 29 | 0.54059 | 1.6962796 | 0.0075 | 0.06712408 | 1 | 5230 | tags=45%, list=22%, si | MM_3 |
| GOBP_REGULATION_OF_ | GOBP_REGULATION_OF_NUCLEOTIDE_CATA | 56 | 0.471335 | 1.6954601 | 0.0023041475 | 0.06741653 | 1 | 6761 | tags=43%, list=29%, si | MM_3 |
| GOBP_PROTEIN_LOCALIZ | GOBP_PROTEIN_LOCALIZATION_TO_MITOC | 101 | 0.44188 | 1.6952778 | 0 | 0.067294426 | 1 | 7279 | tags=49%, list=31%, si | MM_3 |
| GOBP_NEGATIVE_REGUL | GOBP_NEGATIVE_REGULATION_OF_NUCLEO | 29 | 0.529363 | 1.6941117 | 0.008962868 | 0.06785546 | 1 | 6163 | tags=45%, list=26%, si | MM_3 |

| | | | | | | | | | | |
|---|---|---|---|---|---|---|---|---|---|---|
| GOBP_MACROPHAGE_DI | GOBP_MACROPHAGE_DIFFERENTIATION | 61 | 0.470284 | 1.6936613 | 0.0011312218 | 0.06793804 | 1 | 5993 | tags=39%, list=25%, si | MM_3 |
| GOBP_NAD_METABOLIC_ | GOBP_NAD_METABOLIC_PROCESS | 58 | 0.481353 | 1.6925353 | 0.007900678 | 0.06853699 | 1 | 4566 | tags=38%, list=19%, si | MM_3 |
| GOBP_PROTEIN_LOCALIZ | GOBP_PROTEIN_LOCALIZATION_TO_ENDOP | 76 | 0.454806 | 1.6919819 | 0.0022172949 | 0.06863445 | 1 | 8500 | tags=67%, list=36%, si | MM_3 |
| HUMAN_T-CELL_LEUKEM | HUMAN_T-CELL_LEUKEMIA_VIRUS_1_INFEC | 215 | 0.412783 | 1.6916819 | 0 | 0.06860093 | 1 | 7206 | tags=51%, list=31%, si | MM_3 |
| GOBP_POSITIVE_REGULA | GOBP_POSITIVE_REGULATION_OF_ISOTYPE_ | 30 | 0.535351 | 1.691531 | 0.012437811 | 0.068473555 | 1 | 4592 | tags=37%, list=19%, si | MM_3 |
| GOBP_SMALL_MOLECULE | GOBP_SMALL_MOLECULE_CATABOLIC_PRO | 318 | 0.414062 | 1.6915158 | 0 | 0.06825739 | 1 | 5181 | tags=35%, list=22%, si | MM_3 |
| GOBP_REGULATION_OF_ | GOBP_REGULATION_OF_PROTEIN_CATABOL | 346 | 0.410291 | 1.691379 | 0 | 0.068146504 | 1 | 7521 | tags=51%, list=32%, si | MM_3 |
| GOBP_INFLAMMATORY_ | GOBP_INFLAMMATORY_RESPONSE_TO_WO | 30 | 0.537321 | 1.691155 | 0.004938272 | 0.06805407 | 1 | 4403 | tags=40%, list=19%, si | MM_3 |
| GOBP_REGULATION_OF_ | GOBP_REGULATION_OF_CYTOPLASMIC_PAT | 104 | 0.441872 | 1.6908792 | 0.0010660981 | 0.06806577 | 1 | 6192 | tags=38%, list=26%, si | MM_3 |
| CHEMOKINE_SIGNALING_ | CHEMOKINE_SIGNALING_PATHWAY | 177 | 0.424304 | 1.6899986 | 0 | 0.06848845 | 1 | 5905 | tags=38%, list=25%, si | MM_3 |
| HUNTINGTON_DISEASE | HUNTINGTON_DISEASE | 180 | 0.423403 | 1.6895218 | 0 | 0.068561874 | 1 | 7149 | tags=51%, list=30%, si | MM_3 |
| GOBP_LYMPHOID_PROGE | GOBP_LYMPHOID_PROGENITOR_CELL_DIFF | 24 | 0.556856 | 1.6894358 | 0.0026525198 | 0.06839196 | 1 | 4913 | tags=50%, list=21%, si | MM_3 |
| GOBP_BASEMENT_MEMB | GOBP_BASEMENT_MEMBRANE_ORGANIZAT | 40 | 0.513023 | 1.6888083 | 0.0035460992 | 0.06860196 | 1 | 2453 | tags=30%, list=10%, si | MM_3 |
| GOBP_POSITIVE_REGULA | GOBP_POSITIVE_REGULATION_OF_CELL_CY | 31 | 0.535397 | 1.6876938 | 0.012121212 | 0.069164194 | 1 | 5711 | tags=52%, list=24%, si | MM_3 |
| GOBP_PROTEIN_MATURA | GOBP_PROTEIN_MATURATION | 475 | 0.405824 | 1.6871215 | 0 | 0.069335975 | 1 | 5562 | tags=37%, list=24%, si | MM_3 |
| GOBP_REGULATION_OF_ | GOBP_REGULATION_OF_TOLL_LIKE_RECEPT | 32 | 0.519686 | 1.6863356 | 0.0075662043 | 0.06973929 | 1 | 4539 | tags=41%, list=19%, si | MM_3 |
| GOBP_PROTEIN_KINASE_ | GOBP_PROTEIN_KINASE_C_SIGNALING | 36 | 0.518243 | 1.6845287 | 0.009852217 | 0.070887506 | 1 | 3330 | tags=33%, list=14%, si | MM_3 |
| GOBP_REGULATION_OF_ | GOBP_REGULATION_OF_DNA_BINDING_TR | 362 | 0.407132 | 1.6838366 | 0 | 0.07119993 | 1 | 5501 | tags=36%, list=23%, si | MM_3 |
| GOBP_RENAL_FILTRATIO | GOBP_RENAL_FILTRATION | 29 | 0.526389 | 1.6830664 | 0.0113350125 | 0.071417525 | 1 | 5778 | tags=48%, list=24%, si | MM_3 |
| GOBP_ENDOSOME_TO_L | GOBP_ENDOSOME_TO_LYSOSOME_TRANSP | 68 | 0.460601 | 1.6826248 | 0 | 0.07160248 | 1 | 8049 | tags=62%, list=34%, si | MM_3 |
| GOBP_REGULATION_OF_ | GOBP_REGULATION_OF_ACTIN_NUCLEATIO | 35 | 0.50555 | 1.6823118 | 0.004920049 | 0.07158474 | 1 | 2941 | tags=29%, list=12%, si | MM_3 |
| GOBP_REGULATION_OF_ | GOBP_REGULATION_OF_DEFENSE_RESPONS | 20 | 0.576463 | 1.6818414 | 0.009247027 | 0.071689986 | 1 | 6246 | tags=60%, list=26%, si | MM_3 |
| GOBP_REGULATION_OF_ | GOBP_REGULATION_OF_ANTIGEN_PROCESS | 21 | 0.577563 | 1.6818241 | 0.012379643 | 0.07146516 | 1 | 6192 | tags=62%, list=26%, si | MM_3 |
| GOBP_RRNA_PROCESSIN | GOBP_RRNA_PROCESSING | 211 | 0.414844 | 1.6816022 | 0 | 0.07138392 | 1 | 7507 | tags=53%, list=32%, si | MM_3 |
| GOBP_TRNA_MODIFICAT | GOBP_TRNA_MODIFICATION | 84 | 0.448319 | 1.6806527 | 0.002244669 | 0.07192993 | 1 | 4049 | tags=36%, list=17%, si | MM_3 |
| GOBP_TOLL_LIKE_RECEP | GOBP_TOLL_LIKE_RECEPTOR_SIGNALING_P | 71 | 0.457913 | 1.6805948 | 0.0011160715 | 0.071736895 | 1 | 4539 | tags=32%, list=19%, si | MM_3 |
| GOBP_NEGATIVE_REGUL | GOBP_NEGATIVE_REGULATION_OF_SIGNAL | 29 | 0.533349 | 1.6803969 | 0.009840098 | 0.071662165 | 1 | 4235 | tags=38%, list=18%, si | MM_3 |
| THERMOGENESIS | THERMOGENESIS | 210 | 0.42176 | 1.6803831 | 0 | 0.071454376 | 1 | 7407 | tags=52%, list=31%, si | MM_3 |
| GOBP_SIGNAL_TRANSDU | GOBP_SIGNAL_TRANSDUCTION_BY_P53_CL | 151 | 0.424632 | 1.6795971 | 0 | 0.07185338 | 1 | 7217 | tags=54%, list=31%, si | MM_3 |
| GOBP_REGULATION_OF_ | GOBP_REGULATION_OF_AMYLOID_FIBRIL_F | 16 | 0.611773 | 1.6795894 | 0.015827337 | 0.07162975 | 1 | 6546 | tags=69%, list=28%, si | MM_3 |
| GOBP_POSITIVE_REGULA | GOBP_POSITIVE_REGULATION_OF_CHROM | 29 | 0.536907 | 1.6795805 | 0.0062578223 | 0.07141086 | 1 | 7052 | tags=59%, list=30%, si | MM_3 |
| GOBP_MITOCHONDRION_ | GOBP_MITOCHONDRION_LOCALIZATION | 57 | 0.466198 | 1.6792926 | 0.0034129692 | 0.071430795 | 1 | 6285 | tags=46%, list=27%, si | MM_3 |
| GOBP_POSITIVE_REGULA | GOBP_POSITIVE_REGULATION_OF_TUMOR_ | 113 | 0.438027 | 1.6787703 | 0 | 0.071607724 | 1 | 5794 | tags=37%, list=25%, si | MM_3 |
| GOBP_FOREBRAIN_CELL_ | GOBP_FOREBRAIN_CELL_MIGRATION | 66 | 0.468268 | 1.6782556 | 0.0055309734 | 0.07174475 | 1 | 5855 | tags=44%, list=25%, si | MM_3 |
| GOBP_POSITIVE_REGULA | GOBP_POSITIVE_REGULATION_OF_RECEPTO | 61 | 0.470397 | 1.6779197 | 0.0057142857 | 0.07178267 | 1 | 6159 | tags=46%, list=26%, si | MM_3 |
| GOBP_PEPTIDYL_PROLIN | GOBP_PEPTIDYL_PROLINE_MODIFICATION | 39 | 0.499512 | 1.6772729 | 0.0036101083 | 0.0721008 | 1 | 4206 | tags=33%, list=18%, si | MM_3 |
| GOBP_POSITIVE_REGULA | GOBP_POSITIVE_REGULATION_OF_GLUCON | 26 | 0.543952 | 1.6771551 | 0.0088495575 | 0.07195598 | 1 | 5983 | tags=50%, list=25%, si | MM_3 |
| GOBP_NEGATIVE_REGUL | GOBP_NEGATIVE_REGULATION_OF_REACTI | 20 | 0.582844 | 1.6769857 | 0.007894737 | 0.07187352 | 1 | 1632 | tags=20%, list=7%, sig | MM_3 |
| GOBP_RESPONSE_TO_HE | GOBP_RESPONSE_TO_HEAT | 86 | 0.445057 | 1.6768081 | 0 | 0.07178499 | 1 | 7012 | tags=55%, list=30%, si | MM_3 |
| GOBP_POSITIVE_REGULA | GOBP_POSITIVE_REGULATION_OF_CELLULA | 227 | 0.415598 | 1.6762012 | 0 | 0.07201215 | 1 | 7383 | tags=49%, list=31%, si | MM_3 |
| GOBP_CELLULAR_RESPON | GOBP_CELLULAR_RESPONSE_TO_HYDROGE | 59 | 0.464744 | 1.6760051 | 0.005605381 | 0.07194001 | 1 | 3164 | tags=29%, list=13%, si | MM_3 |
| GOBP_PEPTIDYL_LYSINE_ | GOBP_PEPTIDYL_LYSINE_MODIFICATION | 128 | 0.430567 | 1.6748159 | 0 | 0.07262208 | 1 | 7313 | tags=48%, list=31%, si | MM_3 |
| GOBP_LACTATE_METABO | GOBP_LACTATE_METABOLIC_PROCESS | 22 | 0.563041 | 1.674076 | 0.013245033 | 0.07293715 | 1 | 3949 | tags=50%, list=17%, si | MM_3 |
| GOBP_ANTIGEN_PROCES | GOBP_ANTIGEN_PROCESSING_AND_PRESEN | 28 | 0.530181 | 1.6739857 | 0.0077021825 | 0.07280232 | 1 | 6692 | tags=71%, list=28%, si | MM_3 |
| GOBP_REGULATION_OF_ | GOBP_REGULATION_OF_REACTIVE_OXYGEN | 160 | 0.422191 | 1.6739722 | 0 | 0.07258602 | 1 | 7557 | tags=48%, list=32%, si | MM_3 |
| GOBP_POST_TRANSCRIPT | GOBP_POST_TRANSCRIPTIONAL_REGULATIO | 480 | 0.403295 | 1.6732358 | 0 | 0.072866075 | 1 | 7262 | tags=48%, list=31%, si | MM_3 |
| GOBP_REGULATION_OF_ | GOBP_REGULATION_OF_LYMPHOCYTE_CHE | 22 | 0.563426 | 1.6729518 | 0.017127799 | 0.072871245 | 1 | 3486 | tags=36%, list=15%, si | MM_3 |
| GOBP_THIOESTER_BIOSY | GOBP_THIOESTER_BIOSYNTHETIC_PROCESS | 27 | 0.539692 | 1.6729181 | 0.009020618 | 0.07267234 | 1 | 6841 | tags=59%, list=29%, si | MM_3 |
| GOBP_NEGATIVE_REGUL | GOBP_NEGATIVE_REGULATION_OF_DEPHO | 38 | 0.49962 | 1.672081 | 0.0072376356 | 0.073028356 | 1 | 5310 | tags=42%, list=22%, si | MM_3 |
| GOBP_VESICLE_TETHERIN | GOBP_VESICLE_TETHERING | 31 | 0.518684 | 1.6715045 | 0.0126103405 | 0.07323552 | 1 | 7360 | tags=71%, list=31%, si | MM_3 |
| GOBP_NUCLEOSIDE_TRIP | GOBP_NUCLEOSIDE_TRIPHOSPHATE_BIOSYN | 100 | 0.438003 | 1.6713506 | 0.0021413276 | 0.07313381 | 1 | 7012 | tags=53%, list=30%, si | MM_3 |
| GOBP_HEME_METABOLIC | GOBP_HEME_METABOLIC_PROCESS | 34 | 0.51848 | 1.6706094 | 0.007308161 | 0.07344744 | 1 | 6442 | tags=53%, list=27%, si | MM_3 |
| GOBP_RIBOSE_PHOSPHA | GOBP_RIBOSE_PHOSPHATE_METABOLIC_PR | 419 | 0.406033 | 1.670236 | 0 | 0.07348788 | 1 | 7066 | tags=45%, list=30%, si | MM_3 |
| GOBP_POSITIVE_REGULA | GOBP_POSITIVE_REGULATION_OF_TELOME | 71 | 0.451835 | 1.6692362 | 0.0010940919 | 0.07410132 | 1 | 6932 | tags=49%, list=29%, si | MM_3 |
| GOBP_POSITIVE_REGULA | GOBP_POSITIVE_REGULATION_OF_PROTEAS | 72 | 0.456167 | 1.6687578 | 0.0011235955 | 0.07421651 | 1 | 7490 | tags=57%, list=32%, si | MM_3 |
| GOBP_REGULATION_OF_ | GOBP_REGULATION_OF_PROTEIN_UBIQUIT | 202 | 0.410579 | 1.6686548 | 0 | 0.074094854 | 1 | 7228 | tags=50%, list=31%, si | MM_3 |
| GOBP_TYPE_I_INTERFER | GOBP_TYPE_I_INTERFERON_PRODUCTION | 127 | 0.434621 | 1.667275 | 0 | 0.07497869 | 1 | 5732 | tags=35%, list=24%, si | MM_3 |
| GOBP_REGULATION_OF_ | GOBP_REGULATION_OF_PHOSPHOPROTEIN | 42 | 0.486638 | 1.666916 | 0.007134364 | 0.07504446 | 1 | 8340 | tags=62%, list=35%, si | MM_3 |
| GOBP_PROTEIN_LOCALIZ | GOBP_PROTEIN_LOCALIZATION_TO_CELL_S | 70 | 0.447715 | 1.6663312 | 0.0044101435 | 0.0752799 | 1 | 6992 | tags=53%, list=30%, si | MM_3 |
| GOBP_MEMBRANE_PROT | GOBP_MEMBRANE_PROTEIN_ECTODOMAIN | 47 | 0.480607 | 1.6653678 | 0.005882353 | 0.0758047 | 1 | 6634 | tags=49%, list=28%, si | MM_3 |
| GOBP_IMPORT_INTO_TH | GOBP_IMPORT_INTO_THE_MITOCHONDRIO | 49 | 0.47683 | 1.665189 | 0.009501188 | 0.075746715 | 1 | 6993 | tags=53%, list=30%, si | MM_3 |
| GOBP_REGULATION_OF_ | GOBP_REGULATION_OF_PROTEIN_DEPHOSI | 76 | 0.451833 | 1.6642555 | 0.0032644179 | 0.07623969 | 1 | 7299 | tags=50%, list=31%, si | MM_3 |
| GOBP_FATTY_ACID_BETA | GOBP_FATTY_ACID_BETA_OXIDATION | 75 | 0.459896 | 1.6631178 | 0.0022123894 | 0.07691275 | 1 | 4743 | tags=32%, list=20%, si | MM_3 |
| GOBP_REGULATION_OF_ | GOBP_REGULATION_OF_RESPIRATORY_SYST | 15 | 0.612852 | 1.6627195 | 0.010796221 | 0.076956585 | 1 | 4363 | tags=47%, list=18%, si | MM_3 |
| EPSTEIN-BARR_VIRUS_IN | EPSTEIN-BARR_VIRUS_INFECTION | 190 | 0.419469 | 1.662429 | 0 | 0.07695821 | 1 | 7522 | tags=51%, list=32%, si | MM_3 |
| GOBP_CELLULAR_RESPON | GOBP_CELLULAR_RESPONSE_TO_REACTIVE_ | 136 | 0.428258 | 1.6623578 | 0.0010526315 | 0.076790534 | 1 | 5583 | tags=39%, list=24%, si | MM_3 |
| GOBP_VESICLE_BUDDING | GOBP_VESICLE_BUDDING_FROM_MEMBRA | 79 | 0.44816 | 1.6617239 | 0 | 0.077089444 | 1 | 6720 | tags=58%, list=28%, si | MM_3 |
| GOBP_NUCLEOTIDE_SUG | GOBP_NUCLEOTIDE_SUGAR_BIOSYNTHETIC | 17 | 0.595153 | 1.661655 | 0.014905149 | 0.076922335 | 1 | 6483 | tags=59%, list=27%, si | MM_3 |
| GOBP_REGULATION_OF_ | GOBP_REGULATION_OF_PROTEASOMAL_PR | 179 | 0.417343 | 1.6613032 | 0 | 0.07698644 | 1 | 7506 | tags=53%, list=32%, si | MM_3 |
| GOBP_RESPONSE_TO_MI | GOBP_RESPONSE_TO_MISFOLDED_PROTEIN | 18 | 0.588476 | 1.6605666 | 0.021857923 | 0.0773563 | 1 | 3271 | tags=39%, list=14%, si | MM_3 |
| GOBP_PERICARDIUM_DE | GOBP_PERICARDIUM_DEVELOPMENT | 27 | 0.533509 | 1.6598899 | 0.014906832 | 0.07768253 | 1 | 3410 | tags=30%, list=14%, si | MM_3 |
| GOBP_POSITIVE_REGULA | GOBP_POSITIVE_REGULATION_OF_NF_KAPP | 127 | 0.428454 | 1.6592507 | 0.0031413613 | 0.07803321 | 1 | 6329 | tags=43%, list=27%, si | MM_3 |
| GOBP_REGULATION_OF_ | GOBP_REGULATION_OF_NUCLEOTIDE_MET | 96 | 0.435224 | 1.6592314 | 0.0043149944 | 0.07783088 | 1 | 7012 | tags=44%, list=30%, si | MM_3 |
| GLUTATHIONE_METABOL | GLUTATHIONE_METABOLISM | 57 | 0.46506 | 1.6588143 | 0 | 0.077885166 | 1 | 5248 | tags=40%, list=22%, si | MM_3 |
| GOBP_NEGATIVE_REGUL | GOBP_NEGATIVE_REGULATION_OF_CELL_A | 207 | 0.415104 | 1.6584378 | 0 | 0.07797456 | 1 | 5055 | tags=32%, list=21%, si | MM_3 |
| GOBP_POSITIVE_REGULA | GOBP_POSITIVE_REGULATION_OF_MITOCH | 22 | 0.554437 | 1.6581572 | 0.01055409 | 0.07795327 | 1 | 4351 | tags=50%, list=18%, si | MM_3 |
| GOBP_PROTEIN_K48_LIN | GOBP_PROTEIN_K48_LINKED_UBIQUITINAT | 78 | 0.44162 | 1.6581366 | 0.0010917031 | 0.07774193 | 1 | 7311 | tags=51%, list=31%, si | MM_3 |
| GOBP_PEPTIDYL_GLUTAM | GOBP_PEPTIDYL_GLUTAMIC_ACID_MODIFIC | 25 | 0.541985 | 1.6577877 | 0.008997429 | 0.077798985 | 1 | 4616 | tags=44%, list=20%, si | MM_3 |
| GOBP_T_CELL_DIFFEREN | GOBP_T_CELL_DIFFERENTIATION_IN_THYM | 103 | 0.435131 | 1.6575797 | 0.0021141649 | 0.0777068 | 1 | 4283 | tags=31%, list=18%, si | MM_3 |
| GOBP_MYELOID_LEUKOC | GOBP_MYELOID_LEUKOCYTE_DIFFERENTIAT | 258 | 0.405145 | 1.6571623 | 0 | 0.077786624 | 1 | 6223 | tags=38%, list=26%, si | MM_3 |
| GOBP_MACROPHAGE_CY | GOBP_MACROPHAGE_CYTOKINE_PRODUCT | 51 | 0.476523 | 1.6571567 | 0.0057670125 | 0.0775752 | 1 | 5794 | tags=41%, list=25%, si | MM_3 |
| GOBP_NEGATIVE_REGUL | GOBP_NEGATIVE_REGULATION_OF_PEPTID | 52 | 0.471948 | 1.6569223 | 0.007017544 | 0.077552564 | 1 | 7299 | tags=54%, list=31%, si | MM_3 |

| | | | | | | | | | | |
|---|---|---|---|---|---|---|---|---|---|---|
| GOBP_GLUCOSE_6_PHOS | GOBP_GLUCOSE_6_PHOSPHATE_METABOLI | 28 | 0.537179 | 1.6568748 | 0.010204081 | 0.07737963 | 1 | 6330 | tags=46%, list=27%, si | MM_3 |
| GOBP_GENERATION_OF_ | GOBP_GENERATION_OF_PRECURSOR_META | 439 | 0.398426 | 1.6560749 | 0 | 0.07774228 | 1 | 6220 | tags=40%, list=26%, si | MM_3 |
| GOBP_REGULATION_OF_ | GOBP_REGULATION_OF_VACUOLE_ORGANI | 54 | 0.467945 | 1.6555984 | 0.005854801 | 0.07788271 | 1 | 7782 | tags=65%, list=33%, si | MM_3 |
| GOBP_MEMBRANE_FISSI | GOBP_MEMBRANE_FISSION | 31 | 0.519517 | 1.6554027 | 0.010088272 | 0.07779484 | 1 | 7068 | tags=65%, list=30%, si | MM_3 |
| GOBP_DEPHOSPHORYLAT | GOBP_DEPHOSPHORYLATION | 303 | 0.407172 | 1.6547587 | 0 | 0.07808663 | 1 | 6438 | tags=40%, list=27%, si | MM_3 |
| VIRAL_CARCINOGENESIS | VIRAL_CARCINOGENESIS | 201 | 0.412419 | 1.6546147 | 0 | 0.077979095 | 1 | 7175 | tags=48%, list=30%, si | MM_3 |
| GOBP_COPPER_ION_HON | GOBP_COPPER_ION_HOMEOSTASIS | 18 | 0.586573 | 1.6544987 | 0.010738255 | 0.077885866 | 1 | 5077 | tags=39%, list=22%, si | MM_3 |
| GOBP_MITOCHONDRION | GOBP_MITOCHONDRION_DISTRIBUTION | 27 | 0.529081 | 1.653942 | 0.014267186 | 0.07808834 | 1 | 3699 | tags=30%, list=16%, si | MM_3 |
| GOBP_IMMUNOGLOBULI | GOBP_IMMUNOGLOBULIN_PRODUCTION_II | 67 | 0.450875 | 1.6535933 | 0.0055991043 | 0.07815079 | 1 | 5024 | tags=37%, list=21%, si | MM_3 |
| GOBP_NEGATIVE_REGUL | GOBP_NEGATIVE_REGULATION_OF_PHOSPH | 398 | 0.401287 | 1.6534691 | 0 | 0.078027464 | 1 | 7313 | tags=44%, list=31%, si | MM_3 |
| GOBP_REGULATION_OF_ | GOBP_REGULATION_OF_B_CELL_RECEPTOR | 18 | 0.575676 | 1.653141 | 0.018445322 | 0.078043126 | 1 | 3518 | tags=33%, list=15%, si | MM_3 |
| GOBP_REGULATION_OF_ | GOBP_REGULATION_OF_TELOMERE_MAINT | 98 | 0.435331 | 1.652376 | 0.0010787486 | 0.0784962 | 1 | 7177 | tags=50%, list=30%, si | MM_3 |
| GOBP_PROTEIN_TRANSM | GOBP_PROTEIN_TRANSMEMBRANE_TRANS | 67 | 0.457982 | 1.6512607 | 0.0044792835 | 0.079156056 | 1 | 6470 | tags=46%, list=27%, si | MM_3 |
| GOBP_FEMALE_MEIOTIC_ | GOBP_FEMALE_MEIOTIC_NUCLEAR_DIVISIO | 41 | 0.495246 | 1.6511086 | 0.007290401 | 0.079072535 | 1 | 4612 | tags=37%, list=20%, si | MM_3 |
| GOBP_CELLULAR_RESPON | GOBP_CELLULAR_RESPONSE_TO_CADMIUM | 24 | 0.540063 | 1.6510992 | 0.012919896 | 0.0788775 | 1 | 6968 | tags=67%, list=30%, si | MM_3 |
| GOBP_NEGATIVE_REGUL | GOBP_NEGATIVE_REGULATION_OF_PROTEI | 431 | 0.397253 | 1.650862 | 0 | 0.07889611 | 1 | 7322 | tags=45%, list=31%, si | MM_3 |
| GOBP_HYDROGEN_PERO | GOBP_HYDROGEN_PEROXIDE_METABOLIC_ | 48 | 0.48561 | 1.6506709 | 0.00820633 | 0.07883302 | 1 | 6159 | tags=42%, list=26%, si | MM_3 |
| GOBP_AUTOPHAGOSOMI | GOBP_AUTOPHAGOSOME_ORGANIZATION | 128 | 0.427937 | 1.6498352 | 0 | 0.07925004 | 1 | 7782 | tags=62%, list=33%, si | MM_3 |
| MRNA_SURVEILLANCE_P/ | MRNA_SURVEILLANCE_PATHWAY | 87 | 0.434417 | 1.6495268 | 0.0033076075 | 0.07931366 | 1 | 7738 | tags=54%, list=33%, si | MM_3 |
| GOBP_ENDOSOMAL_TRA | GOBP_ENDOSOMAL_TRANSPORT | 272 | 0.403689 | 1.6492904 | 0 | 0.07929298 | 1 | 7012 | tags=45%, list=30%, si | MM_3 |
| GOBP_REGULATION_OF_ | GOBP_REGULATION_OF_PROTEIN_SUMOYL | 25 | 0.530627 | 1.6487273 | 0.0025641026 | 0.07949835 | 1 | 7413 | tags=64%, list=31%, si | MM_3 |
| GOBP_REGULATION_OF_ | GOBP_REGULATION_OF_DNA_CATABOLIC_I | 15 | 0.610728 | 1.6474268 | 0.013227513 | 0.08035411 | 1 | 2345 | tags=40%, list=10%, si | MM_3 |
| GOBP_POSITIVE_REGULA | GOBP_POSITIVE_REGULATION_OF_AUTOPH | 157 | 0.416565 | 1.6470189 | 0 | 0.08051826 | 1 | 7383 | tags=51%, list=31%, si | MM_3 |
| GOBP_POSITIVE_REGULA | GOBP_POSITIVE_REGULATION_OF_DNA_BIN | 217 | 0.40805 | 1.6468096 | 0 | 0.080486864 | 1 | 5395 | tags=34%, list=23%, si | MM_3 |
| GOBP_CD40_SIGNALING_ | GOBP_CD40_SIGNALING_PATHWAY | 16 | 0.610651 | 1.6454786 | 0.012587412 | 0.08130181 | 1 | 7374 | tags=75%, list=31%, si | MM_3 |
| GOBP_ORGANOPHOSPHA | GOBP_ORGANOPHOSPHATE_BIOSYNTHETIC | 500 | 0.394021 | 1.6446438 | 0 | 0.08174614 | 1 | 6239 | tags=40%, list=26%, si | MM_3 |
| GOBP_RETROGRADE_VES | GOBP_RETROGRADE_VESICLE_MEDIATED_T | 45 | 0.480285 | 1.6429265 | 0.008254717 | 0.08288976 | 1 | 8033 | tags=71%, list=34%, si | MM_3 |
| GOBP_ORGANIC_ACID_C | GOBP_ORGANIC_ACID_CATABOLIC_PROCES | 224 | 0.405513 | 1.6412703 | 0.0010121458 | 0.08408357 | 1 | 5181 | tags=35%, list=22%, si | MM_3 |
| GOBP_POSITIVE_REGULA | GOBP_POSITIVE_REGULATION_OF_LEUKOCY | 159 | 0.418566 | 1.6412299 | 0 | 0.0839143 | 1 | 6009 | tags=40%, list=25%, si | MM_3 |
| GOBP_GOLGI_VESICLE_TI | GOBP_GOLGI_VESICLE_TRANSPORT | 276 | 0.405763 | 1.6410162 | 0 | 0.08389562 | 1 | 7571 | tags=54%, list=32%, si | MM_3 |
| GOBP_REGULATION_OF_ | GOBP_REGULATION_OF_POST_TRANSLATIO | 242 | 0.405144 | 1.6406624 | 0 | 0.08399522 | 1 | 7506 | tags=52%, list=32%, si | MM_3 |
| GOBP_NEGATIVE_REGUL | GOBP_NEGATIVE_REGULATION_OF_T_CELL | 73 | 0.446038 | 1.6401627 | 0.00440044 | 0.08419604 | 1 | 4813 | tags=36%, list=20%, si | MM_3 |
| GOBP_NUCLEAR_ENVELO | GOBP_NUCLEAR_ENVELOPE_ORGANIZATIOI | 52 | 0.474531 | 1.6400039 | 0.006993007 | 0.08411956 | 1 | 6931 | tags=60%, list=29%, si | MM_3 |
| GOBP_POSITIVE_REGULA | GOBP_POSITIVE_REGULATION_OF_LYMPHO | 42 | 0.485753 | 1.6395634 | 0.010650887 | 0.084227905 | 1 | 3849 | tags=31%, list=16%, si | MM_3 |
| GOBP_ADAPTIVE_IMMUN | GOBP_ADAPTIVE_IMMUNE_RESPONSE_BAS | 322 | 0.40078 | 1.638108 | 0 | 0.08527759 | 1 | 6594 | tags=41%, list=28%, si | MM_3 |
| GOBP_POSITIVE_REGULA | GOBP_POSITIVE_REGULATION_OF_PROTEIN | 16 | 0.593584 | 1.6379728 | 0.017615177 | 0.08517041 | 1 | 6383 | tags=63%, list=27%, si | MM_3 |
| GOBP_RESPONSE_TO_RE | GOBP_RESPONSE_TO_REACTIVE_OXYGEN_S | 181 | 0.408495 | 1.6379076 | 0.0020597323 | 0.08501744 | 1 | 7204 | tags=51%, list=31%, si | MM_3 |
| GOBP_T_CELL_MIGRATIO | GOBP_T_CELL_MIGRATION | 64 | 0.446608 | 1.6362008 | 0.011160715 | 0.08619931 | 1 | 6933 | tags=48%, list=29%, si | MM_3 |
| GOBP_NEGATIVE_REGUL | GOBP_NEGATIVE_REGULATION_OF_FIBROB | 44 | 0.479296 | 1.6359023 | 0.010477299 | 0.08626277 | 1 | 5974 | tags=43%, list=25%, si | MM_3 |
| GOBP_POSITIVE_REGULA | GOBP_POSITIVE_REGULATION_OF_PHOSPH | 16 | 0.583876 | 1.6356546 | 0.026098901 | 0.086221 | 1 | 8195 | tags=75%, list=35%, si | MM_3 |
| GOBP_REGULATION_OF_ | GOBP_REGULATION_OF_SUPEROXIDE_ANIC | 28 | 0.526498 | 1.6354289 | 0.011392405 | 0.08621 | 1 | 5778 | tags=43%, list=24%, si | MM_3 |
| GOBP_ORGANELLE_DISAS | GOBP_ORGANELLE_DISASSEMBLY | 155 | 0.413366 | 1.6346116 | 0 | 0.086617276 | 1 | 7389 | tags=53%, list=31%, si | MM_3 |
| GOBP_PHOSPHATIDYLINC | GOBP_PHOSPHATIDYLINOSITOL_BIOSYNTHE | 95 | 0.429507 | 1.6345658 | 0.004338395 | 0.08642738 | 1 | 4421 | tags=34%, list=19%, si | MM_3 |
| GOBP_BROWN_FAT_CELL | GOBP_BROWN_FAT_CELL_DIFFERENTIATION | 63 | 0.456345 | 1.6345475 | 0.0045402953 | 0.08623839 | 1 | 5327 | tags=33%, list=23%, si | MM_3 |
| GOBP_TELENCEPHALON_ | GOBP_TELENCEPHALON_GLIAL_CELL_MIGR/ | 27 | 0.539299 | 1.6342739 | 0.015364917 | 0.08627042 | 1 | 5855 | tags=52%, list=25%, si | MM_3 |
| GOBP_LEUKOCYTE_ACTIV | GOBP_LEUKOCYTE_ACTIVATION_INVOLVED | 53 | 0.460353 | 1.6338975 | 0.009291521 | 0.08637314 | 1 | 5180 | tags=30%, list=22%, si | MM_3 |
| ENDOCYTOSIS | ENDOCYTOSIS | 246 | 0.399869 | 1.6338897 | 0 | 0.08616797 | 1 | 7601 | tags=52%, list=32%, si | MM_3 |
| GOBP_NEGATIVE_REGUL | GOBP_NEGATIVE_REGULATION_OF_MULTIC | 17 | 0.587596 | 1.6338406 | 0.014204546 | 0.086009115 | 1 | 2886 | tags=41%, list=12%, si | MM_3 |
| GOBP_POSITIVE_REGULA | GOBP_POSITIVE_REGULATION_OF_NEUTRO | 41 | 0.489182 | 1.6336662 | 0.0071174377 | 0.08592871 | 1 | 4917 | tags=37%, list=21%, si | MM_3 |
| GOBP_CARBOHYDRATE_F | GOBP_CARBOHYDRATE_PHOSPHORYLATION | 19 | 0.562775 | 1.6335964 | 0.02008032 | 0.08576871 | 1 | 7046 | tags=53%, list=30%, si | MM_3 |
| SPLICEOSOME | SPLICEOSOME | 127 | 0.421843 | 1.6321284 | 0 | 0.086898975 | 1 | 7323 | tags=44%, list=31%, si | MM_3 |
| GOBP_POSITIVE_REGULA | GOBP_POSITIVE_REGULATION_OF_CHROMO | 28 | 0.522564 | 1.6320485 | 0.0114068445 | 0.08676216 | 1 | 7052 | tags=57%, list=30%, si | MM_3 |
| GOBP_SMOOTH_MUSCLE | GOBP_SMOOTH_MUSCLE_CELL_MIGRATION | 115 | 0.420995 | 1.6320099 | 0 | 0.08659384 | 1 | 5906 | tags=35%, list=25%, si | MM_3 |
| GOBP_TUMOR_NECROSIS | GOBP_TUMOR_NECROSIS_FACTOR_SUPERF | 193 | 0.404208 | 1.631463 | 0 | 0.08680999 | 1 | 6518 | tags=40%, list=28%, si | MM_3 |
| GOBP_NEGATIVE_REGUL | GOBP_NEGATIVE_REGULATION_OF_LYMPH | 164 | 0.411829 | 1.6307927 | 0 | 0.08718312 | 1 | 5026 | tags=32%, list=21%, si | MM_3 |
| GOBP_PEPTIDYL_TYROSIN | GOBP_PEPTIDYL_TYROSINE_DEPHOSPHORYI | 33 | 0.503377 | 1.6307442 | 0.018248174 | 0.087010056 | 1 | 6567 | tags=45%, list=28%, si | MM_3 |
| GOBP_AEROBIC_ELECTRO | GOBP_AEROBIC_ELECTRON_TRANSPORT_CH | 63 | 0.457099 | 1.6304371 | 0.007856341 | 0.08709103 | 1 | 7393 | tags=54%, list=31%, si | MM_3 |
| GOBP_STEM_CELL_DIVISI | GOBP_STEM_CELL_DIVISION | 43 | 0.479961 | 1.629786 | 0.014101057 | 0.08742674 | 1 | 4567 | tags=28%, list=19%, si | MM_3 |
| GOBP_SMOOTH_MUSCLE | GOBP_SMOOTH_MUSCLE_CELL_APOPTOTIC | 39 | 0.496273 | 1.6293299 | 0.013317191 | 0.08758947 | 1 | 5977 | tags=49%, list=25%, si | MM_3 |
| GOBP_POSITIVE_REGULA | GOBP_POSITIVE_REGULATION_OF_GRANUL | 31 | 0.513832 | 1.6291238 | 0.016149068 | 0.08757591 | 1 | 4387 | tags=35%, list=19%, si | MM_3 |
| GOBP_HEMATOPOIETIC_S | GOBP_HEMATOPOIETIC_STEM_CELL_DIFFEF | 31 | 0.514629 | 1.6285433 | 0.017073171 | 0.08791527 | 1 | 6263 | tags=58%, list=27%, si | MM_3 |
| PROTEASOME | PROTEASOME | 45 | 0.477408 | 1.6260432 | 0.0058892816 | 0.08976719 | 1 | 8809 | tags=69%, list=37%, si | MM_3 |
| GOBP_ENERGY_DERIVATI | GOBP_ENERGY_DERIVATION_BY_OXIDATIOI | 321 | 0.395736 | 1.6252974 | 0 | 0.090192206 | 1 | 7393 | tags=49%, list=31%, si | MM_3 |
| GOBP_TELOMERE_ORGAI | GOBP_TELOMERE_ORGANIZATION | 150 | 0.413571 | 1.6251674 | 0 | 0.090088 | 1 | 7400 | tags=49%, list=31%, si | MM_3 |
| GOBP_NEGATIVE_REGUL | GOBP_NEGATIVE_REGULATION_OF_PROTEI | 93 | 0.426776 | 1.6250312 | 0.0021645022 | 0.089996375 | 1 | 7020 | tags=45%, list=30%, si | MM_3 |
| GOBP_HEMATOPOIETIC_I | GOBP_HEMATOPOIETIC_PROGENITOR_CELL | 158 | 0.408938 | 1.6238533 | 0 | 0.09076681 | 1 | 5323 | tags=39%, list=23%, si | MM_3 |
| GOBP_REGULATION_OF_ | GOBP_REGULATION_OF_T_CELL_MIGRATIO | 49 | 0.470233 | 1.623611 | 0.009237875 | 0.09077004 | 1 | 6543 | tags=47%, list=28%, si | MM_3 |
| LEGIONELLOSIS | LEGIONELLOSIS | 55 | 0.460774 | 1.6235889 | 0.013574661 | 0.090593465 | 1 | 6137 | tags=44%, list=26%, si | MM_3 |
| GOBP_EXECUTION_PHAS | GOBP_EXECUTION_PHASE_OF_APOPTOSIS | 78 | 0.443318 | 1.6232299 | 0.0044444446 | 0.09075477 | 1 | 3864 | tags=28%, list=16%, si | MM_3 |
| GOBP_MEMBRANE_INVA | GOBP_MEMBRANE_INVAGINATION | 70 | 0.446378 | 1.6224053 | 0.0022296545 | 0.09122291 | 1 | 6709 | tags=47%, list=28%, si | MM_3 |
| GOBP_POSITIVE_REGULA | GOBP_POSITIVE_REGULATION_OF_POST_TF | 135 | 0.414272 | 1.6223156 | 0.00524109 | 0.09110849 | 1 | 7506 | tags=55%, list=32%, si | MM_3 |
| GOBP_LIPOPOLYSACCHAF | GOBP_LIPOPOLYSACCHARIDE_MEDIATED_SI | 57 | 0.460362 | 1.6214216 | 0.0068728523 | 0.09165217 | 1 | 6329 | tags=46%, list=27%, si | MM_3 |
| GOBP_TELOMERE_CAPPI | GOBP_TELOMERE_CAPPING | 38 | 0.492782 | 1.6210654 | 0.013365735 | 0.091707475 | 1 | 6250 | tags=50%, list=26%, si | MM_3 |
| GOBP_POSITIVE_REGULA | GOBP_POSITIVE_REGULATION_OF_LIPID_CA | 26 | 0.521852 | 1.6207248 | 0.012484395 | 0.09180305 | 1 | 6035 | tags=46%, list=26%, si | MM_3 |
| GOBP_GLIAL_CELL_ACTIV | GOBP_GLIAL_CELL_ACTIVATION | 59 | 0.450522 | 1.6206564 | 0.006833713 | 0.09163627 | 1 | 5180 | tags=31%, list=22%, si | MM_3 |
| GLIOMA | GLIOMA | 71 | 0.436414 | 1.6204354 | 0.006644518 | 0.09159729 | 1 | 5583 | tags=44%, list=24%, si | MM_3 |
| GOBP_POSITIVE_REGULA | GOBP_POSITIVE_REGULATION_OF_PHOSPH | 182 | 0.402156 | 1.6199224 | 0 | 0.09189447 | 1 | 5994 | tags=40%, list=25%, si | MM_3 |
| GOBP_MAINTENANCE_OI | GOBP_MAINTENANCE_OF_PROTEIN_LOCAT | 70 | 0.441748 | 1.6197165 | 0.006615215 | 0.09191831 | 1 | 7627 | tags=51%, list=32%, si | MM_3 |
| GOBP_POSITIVE_REGULA | GOBP_POSITIVE_REGULATION_OF_GLIOGEN | 87 | 0.436352 | 1.6191345 | 0.0010952903 | 0.092271455 | 1 | 7329 | tags=51%, list=31%, si | MM_3 |
| GOBP_NEGATIVE_REGUL | GOBP_NEGATIVE_REGULATION_OF_UBIQUI | 55 | 0.455027 | 1.6182802 | 0.011350738 | 0.092835665 | 1 | 8099 | tags=60%, list=34%, si | MM_3 |
| GOBP_NEGATIVE_REGUL | GOBP_NEGATIVE_REGULATION_OF_BMP_SI | 53 | 0.463479 | 1.6181664 | 0.012585812 | 0.09269442 | 1 | 1968 | tags=21%, list=8%, sig | MM_3 |
| GOBP_POSITIVE_REGULA | GOBP_POSITIVE_REGULATION_OF_MYELOID | 38 | 0.490327 | 1.617793 | 0.014616322 | 0.09285344 | 1 | 4917 | tags=34%, list=21%, si | MM_3 |

| | | | | | | | | | | |
|---|---|---|---|---|---|---|---|---|---|---|
| GOBP_POSITIVE_REGULA | GOBP_POSITIVE_REGULATION_OF_CANONI | 148 | 0.414104 | 1.6174799 | 0.001044932 | 0.092930496 | 1 | 6413 | tags=41%, list=27%, si | MM_3 |
| GOBP_REGULATION_OF_ | GOBP_REGULATION_OF_HEMATOPOIETIC_ | 39 | 0.487321 | 1.6172279 | 0.020656137 | 0.09292182 | 1 | 6580 | tags=59%, list=28%, si | MM_3 |
| GOBP_POSITIVE_REGULA | GOBP_POSITIVE_REGULATION_OF_VACUOL | 25 | 0.524033 | 1.6166664 | 0.015810277 | 0.093213044 | 1 | 7130 | tags=68%, list=30%, si | MM_3 |
| GOBP_REGULATION_OF_ | GOBP_REGULATION_OF_RESPONSE_TO_FO | 21 | 0.549125 | 1.6161627 | 0.013140605 | 0.09347497 | 1 | 4558 | tags=43%, list=19%, si | MM_3 |
| GOBP_RNA_3_END_PRO | GOBP_RNA_3_END_PROCESSING | 87 | 0.432729 | 1.616053 | 0.0065645515 | 0.09337106 | 1 | 6500 | tags=41%, list=28%, si | MM_3 |
| GOBP_MEIOTIC_SPINDLE | GOBP_MEIOTIC_SPINDLE_ORGANIZATION | 21 | 0.548563 | 1.615005 | 0.029372497 | 0.09412385 | 1 | 3332 | tags=33%, list=14%, si | MM_3 |
| GOBP_POSITIVE_REGULA | GOBP_POSITIVE_REGULATION_OF_B_CELL_ | 47 | 0.469753 | 1.6149644 | 0.0083234245 | 0.093961835 | 1 | 4592 | tags=32%, list=19%, si | MM_3 |
| GOBP_REGULATION_OF_ | GOBP_REGULATION_OF_FIBROBLAST_PROL | 122 | 0.412424 | 1.6142411 | 0.002120891 | 0.094354756 | 1 | 5974 | tags=41%, list=25%, si | MM_3 |
| GOBP_SIGNAL_TRANSDU | GOBP_SIGNAL_TRANSDUCTION_IN_RESPON | 176 | 0.403923 | 1.6126982 | 0 | 0.09550574 | 1 | 5711 | tags=41%, list=24%, si | MM_3 |
| GOBP_INTRACILIARY_TRA | GOBP_INTRACILIARY_TRANSPORT | 43 | 0.473869 | 1.6121638 | 0.013431014 | 0.0957617 | 1 | 3828 | tags=40%, list=16%, si | MM_3 |
| GOBP_INTERFERON_BETA | GOBP_INTERFERON_BETA_PRODUCTION | 66 | 0.435259 | 1.6120998 | 0.0067873304 | 0.09562015 | 1 | 7383 | tags=44%, list=31%, si | MM_3 |
| HUMAN_CYTOMEGALOVI | HUMAN_CYTOMEGALOVIRUS_INFECTION | 216 | 0.395749 | 1.6118003 | 0 | 0.095653504 | 1 | 5938 | tags=38%, list=25%, si | MM_3 |
| GOBP_PROTEIN_PROCESS | GOBP_PROTEIN_PROCESSING | 237 | 0.401009 | 1.611743 | 0 | 0.095488034 | 1 | 5788 | tags=36%, list=25%, si | MM_3 |
| GOBP_DETECTION_OF_M | GOBP_DETECTION_OF_MECHANICAL_STIMU | 25 | 0.533617 | 1.6104295 | 0.015852047 | 0.09648404 | 1 | 5500 | tags=44%, list=23%, si | MM_3 |
| GOBP_KILLING_OF_CELLS | GOBP_KILLING_OF_CELLS_OF_ANOTHER_OF | 71 | 0.440134 | 1.6101786 | 0.007751938 | 0.096501656 | 1 | 3733 | tags=28%, list=16%, si | MM_3 |
| GOBP_ENDOPLASMIC_RE | GOBP_ENDOPLASMIC_RETICULUM_CALCIUM | 28 | 0.509663 | 1.6095637 | 0.017902814 | 0.096823886 | 1 | 6806 | tags=61%, list=29%, si | MM_3 |
| GOBP_CYTOPLASMIC_PA | GOBP_CYTOPLASMIC_PATTERN_RECOGNITI | 159 | 0.404885 | 1.6090705 | 0 | 0.09706406 | 1 | 7581 | tags=49%, list=32%, si | MM_3 |
| NON-ALCOHOLIC_FATTY_ | NON-ALCOHOLIC_FATTY_LIVER_DISEASE_(N | 141 | 0.412435 | 1.6090133 | 0.003121748 | 0.09689281 | 1 | 7521 | tags=55%, list=32%, si | MM_3 |
| GOBP_NEGATIVE_REGUL | GOBP_NEGATIVE_REGULATION_OF_PHOSPH | 17 | 0.560501 | 1.6087351 | 0.02173913 | 0.096940294 | 1 | 8340 | tags=82%, list=35%, si | MM_3 |
| GOBP_VACUOLAR_TRANS | GOBP_VACUOLAR_TRANSPORT | 161 | 0.405104 | 1.6087154 | 0.0020639836 | 0.09675453 | 1 | 7162 | tags=45%, list=30%, si | MM_3 |
| GOBP_NEGATIVE_REGUL | GOBP_NEGATIVE_REGULATION_OF_KINASE | 188 | 0.403651 | 1.6083978 | 0 | 0.09683762 | 1 | 6708 | tags=41%, list=28%, si | MM_3 |
| GOBP_REGULATION_OF_ | GOBP_REGULATION_OF_MAST_CELL_ACTIV | 24 | 0.531416 | 1.6081288 | 0.024804177 | 0.09685604 | 1 | 5027 | tags=33%, list=21%, si | MM_3 |
| GOBP_TRANSLATIONAL_E | GOBP_TRANSLATIONAL_ELONGATION | 58 | 0.44865 | 1.6073245 | 0.009237875 | 0.097325094 | 1 | 4931 | tags=38%, list=21%, si | MM_3 |
| GOBP_NUCLEOTIDE_SUG | GOBP_NUCLEOTIDE_SUGAR_METABOLIC_PI | 34 | 0.498448 | 1.6070317 | 0.02125 | 0.09737135 | 1 | 6787 | tags=53%, list=29%, si | MM_3 |
| GOBP_UNSATURATED_FA | GOBP_UNSATURATED_FATTY_ACID_BIOSYN | 41 | 0.476938 | 1.6068034 | 0.008423586 | 0.09731856 | 1 | 4203 | tags=29%, list=18%, si | MM_3 |
| PEROXISOME | PEROXISOME | 81 | 0.42961 | 1.6065848 | 0.004385965 | 0.09732293 | 1 | 5313 | tags=38%, list=23%, si | MM_3 |
| GOBP_MRNA_CATABOLIC | GOBP_MRNA_CATABOLIC_PROCESS | 233 | 0.395573 | 1.6064621 | 0.002042901 | 0.097228795 | 1 | 7545 | tags=47%, list=32%, si | MM_3 |
| GOBP_NEGATIVE_REGUL | GOBP_NEGATIVE_REGULATION_OF_TRANSL | 117 | 0.419092 | 1.6061426 | 0.0010548523 | 0.09727679 | 1 | 7354 | tags=51%, list=31%, si | MM_3 |
| GOBP_MYELOID_CELL_DI | GOBP_MYELOID_CELL_DIFFERENTIATION | 455 | 0.386339 | 1.6050452 | 0 | 0.09805012 | 1 | 6956 | tags=42%, list=29%, si | MM_3 |
| GOBP_RESPONSE_TO_MI | GOBP_RESPONSE_TO_MITOCHONDRIAL_DE | 21 | 0.53634 | 1.6048576 | 0.025606468 | 0.098015964 | 1 | 1678 | tags=24%, list=7%, sig | MM_3 |
| GOBP_REGULATION_OF_ | GOBP_REGULATION_OF_PROTEIN_DEACETY | 33 | 0.491532 | 1.6041408 | 0.020286396 | 0.09848512 | 1 | 5619 | tags=45%, list=24%, si | MM_3 |
| GOBP_PROTEIN_POLYUB | GOBP_PROTEIN_POLYUBIQUITINATION | 250 | 0.395798 | 1.6039933 | 0 | 0.098398216 | 1 | 6254 | tags=39%, list=26%, si | MM_3 |
| GOBP_VIRAL_GENE_EXPR | GOBP_VIRAL_GENE_EXPRESSION | 75 | 0.436877 | 1.6039544 | 0.011049724 | 0.09821879 | 1 | 7405 | tags=48%, list=31%, si | MM_3 |
| GOBP_ADRENAL_GLAND_ | GOBP_ADRENAL_GLAND_DEVELOPMENT | 22 | 0.538958 | 1.6039087 | 0.031887755 | 0.09805937 | 1 | 919 | tags=18%, list=4%, sig | MM_3 |
| GOBP_REGULATION_OF_ | GOBP_REGULATION_OF_DEPHOSPHORYLAT | 106 | 0.424468 | 1.6038665 | 0.0021299254 | 0.09790498 | 1 | 7299 | tags=50%, list=31%, si | MM_3 |
| GOBP_POSITIVE_REGULA | GOBP_POSITIVE_REGULATION_OF_CHROM | 107 | 0.423989 | 1.6036242 | 0.0021390375 | 0.09795296 | 1 | 6932 | tags=46%, list=29%, si | MM_3 |
| GOBP_PROGRAMMED_N | GOBP_PROGRAMMED_NECROTIC_CELL_DEA | 50 | 0.460097 | 1.6032088 | 0.011560693 | 0.09811433 | 1 | 5974 | tags=42%, list=25%, si | MM_3 |
| GOBP_MEMBRANE_PROT | GOBP_MEMBRANE_PROTEIN_PROTEOLYSIS | 60 | 0.447258 | 1.6029713 | 0.010344828 | 0.09811096 | 1 | 6634 | tags=48%, list=28%, si | MM_3 |
| GOBP_POSITIVE_REGULA | GOBP_POSITIVE_REGULATION_OF_TELOME | 15 | 0.596134 | 1.6026825 | 0.019525802 | 0.09814128 | 1 | 580 | tags=27%, list=2%, sig | MM_3 |
| GOBP_NADH_METABOLIC | GOBP_NADH_METABOLIC_PROCESS | 39 | 0.483001 | 1.6016028 | 0.015494636 | 0.098938465 | 1 | 4566 | tags=38%, list=19%, si | MM_3 |
| GOBP_REGULATION_OF_ | GOBP_REGULATION_OF_TELOMERE_MAINT | 20 | 0.562737 | 1.6003426 | 0.022193212 | 0.09986573 | 1 | 580 | tags=25%, list=2%, sig | MM_3 |
| GOBP_POSITIVE_REGULA | GOBP_POSITIVE_REGULATION_OF_AMYLOID | 26 | 0.524242 | 1.599611 | 0.024783147 | 0.100346245 | 1 | 3135 | tags=23%, list=13%, si | MM_3 |
| GOBP_RNA_MODIFICATIO | GOBP_RNA_MODIFICATION | 141 | 0.404714 | 1.5990958 | 0.0020811656 | 0.10065117 | 1 | 6935 | tags=50%, list=29%, si | MM_3 |
| GOBP_NUCLEAR_EXPORT | GOBP_NUCLEAR_EXPORT | 160 | 0.402799 | 1.5986392 | 0.0020682523 | 0.1008787 | 1 | 6445 | tags=40%, list=27%, si | MM_3 |
| GOBP_REGULATION_OF_ | GOBP_REGULATION_OF_MONONUCLEAR_C | 136 | 0.407826 | 1.5984923 | 0.002105263 | 0.10083763 | 1 | 7056 | tags=49%, list=30%, si | MM_3 |
| GOBP_BRANCHED_CHAIN | GOBP_BRANCHED_CHAIN_AMINO_ACID_CA | 21 | 0.540732 | 1.597991 | 0.02141901 | 0.10108912 | 1 | 4600 | tags=52%, list=19%, si | MM_3 |
| GOBP_DEFINITIVE_HEMO | GOBP_DEFINITIVE_HEMOPOIESIS | 25 | 0.527809 | 1.5974566 | 0.031725887 | 0.10142763 | 1 | 4020 | tags=44%, list=17%, si | MM_3 |
| CITRATE_CYCLE_(TCA_CY | CITRATE_CYCLE_(TCA_CYCLE) | 28 | 0.512239 | 1.5973592 | 0.026548672 | 0.101335414 | 1 | 6006 | tags=57%, list=25%, si | MM_3 |
| GOBP_RNA_CATABOLIC_ | GOBP_RNA_CATABOLIC_PROCESS | 278 | 0.393802 | 1.5970588 | 0 | 0.10143173 | 1 | 7545 | tags=49%, list=32%, si | MM_3 |
| GOBP_TRANSLATIONAL_I | GOBP_TRANSLATIONAL_INITIATION | 112 | 0.412674 | 1.5968724 | 0.0010582011 | 0.101396225 | 1 | 7536 | tags=52%, list=32%, si | MM_3 |
| RNA_TRANSPORT | RNA_TRANSPORT | 154 | 0.400357 | 1.5966353 | 0 | 0.10142124 | 1 | 7733 | tags=54%, list=33%, si | MM_3 |
| GOBP_NEGATIVE_REGUL | GOBP_NEGATIVE_REGULATION_OF_NEURO | 21 | 0.547527 | 1.5943389 | 0.023529412 | 0.10333721 | 1 | 5716 | tags=43%, list=24%, si | MM_3 |
| GOBP_ONE_CARBON_CO | GOBP_ONE_CARBON_COMPOUND_TRANSP | 16 | 0.56768 | 1.5933038 | 0.028649386 | 0.10419487 | 1 | 2146 | tags=25%, list=9%, sig | MM_3 |
| GOBP_PORPHYRIN_CONT | GOBP_PORPHYRIN_CONTAINING_COMPOU | 40 | 0.480057 | 1.5928749 | 0.020262217 | 0.104423285 | 1 | 6442 | tags=50%, list=27%, si | MM_3 |
| GOBP_RENAL_ABSORPTIO | GOBP_RENAL_ABSORPTION | 28 | 0.5058 | 1.592825 | 0.020942409 | 0.104270734 | 1 | 6009 | tags=46%, list=25%, si | MM_3 |
| GOBP_GLUCOSE_METAB | GOBP_GLUCOSE_METABOLIC_PROCESS | 201 | 0.396132 | 1.5922129 | 0.0010224949 | 0.10465727 | 1 | 5701 | tags=38%, list=24%, si | MM_3 |
| GOBP_MITOPHAGY | GOBP_MITOPHAGY | 58 | 0.446919 | 1.5918568 | 0.006849315 | 0.10480101 | 1 | 7533 | tags=53%, list=32%, si | MM_3 |
| RNA_DEGRADATION | RNA_DEGRADATION | 75 | 0.429801 | 1.591599 | 0.014145811 | 0.10481277 | 1 | 4408 | tags=28%, list=19%, si | MM_3 |
| GOBP_RESPONSE_TO_VI | GOBP_RESPONSE_TO_VIRUS | 331 | 0.387519 | 1.5901463 | 0 | 0.106011115 | 1 | 4317 | tags=27%, list=18%, si | MM_3 |
| GOBP_POSITIVE_REGULA | GOBP_POSITIVE_REGULATION_OF_AXON_E | 54 | 0.455965 | 1.5897032 | 0.011494253 | 0.10627621 | 1 | 7570 | tags=57%, list=32%, si | MM_3 |
| GOBP_KILLING_BY_HOST | GOBP_KILLING_BY_HOST_OF_SYMBIONT_C | 20 | 0.536169 | 1.5893921 | 0.025537634 | 0.10638929 | 1 | 941 | tags=20%, list=4%, sig | MM_3 |
| GOBP_HOMEOSTASIS_OF | GOBP_HOMEOSTASIS_OF_NUMBER_OF_CEI | 388 | 0.382877 | 1.5891933 | 0 | 0.10637751 | 1 | 7020 | tags=43%, list=30%, si | MM_3 |
| GOBP_RESPONSE_TO_LIT | GOBP_RESPONSE_TO_LITHIUM_ION | 16 | 0.583075 | 1.588946 | 0.027322404 | 0.10642273 | 1 | 1167 | tags=31%, list=5%, sig | MM_3 |
| GOBP_POSITIVE_REGULA | GOBP_POSITIVE_REGULATION_OF_TYPE_I_I | 77 | 0.429937 | 1.5888201 | 0.006615215 | 0.10633362 | 1 | 4303 | tags=27%, list=18%, si | MM_3 |
| GOBP_MEMBRANE_DOCI | GOBP_MEMBRANE_DOCKING | 78 | 0.430046 | 1.5887208 | 0.004474273 | 0.10622498 | 1 | 7118 | tags=55%, list=30%, si | MM_3 |
| GOBP_AMYLOID_BETA_M | GOBP_AMYLOID_BETA_METABOLIC_PROCES | 55 | 0.444919 | 1.5886284 | 0.012429379 | 0.10609834 | 1 | 4108 | tags=29%, list=17%, si | MM_3 |
| GOBP_SMOOTH_MUSCLE | GOBP_SMOOTH_MUSCLE_CELL_PROLIFERAT | 191 | 0.398725 | 1.5880854 | 0.0010152285 | 0.10638404 | 1 | 5730 | tags=36%, list=24%, si | MM_3 |
| GOBP_TRIPARTITE_REGIO | GOBP_TRIPARTITE_REGIONAL_SUBDIVISION | 16 | 0.566852 | 1.5879805 | 0.039492242 | 0.106247574 | 1 | 682 | tags=25%, list=3%, sig | MM_3 |
| GOBP_RRNA_METABOLIC | GOBP_RRNA_METABOLIC_PROCESS | 241 | 0.390784 | 1.5874772 | 0 | 0.10650979 | 1 | 7577 | tags=51%, list=32%, si | MM_3 |
| GOBP_REGULATION_OF_ | GOBP_REGULATION_OF_MACROAUTOPHAG | 129 | 0.409655 | 1.5867107 | 0.0030959751 | 0.107007295 | 1 | 7506 | tags=57%, list=32%, si | MM_3 |
| GOBP_REGULATION_OF_ | GOBP_REGULATION_OF_DNA_STRAND_ELO | 18 | 0.561811 | 1.5865456 | 0.017931035 | 0.10695049 | 1 | 580 | tags=22%, list=2%, sig | MM_3 |
| GOBP_POSITIVE_REGULA | GOBP_POSITIVE_REGULATION_OF_MACRO | 81 | 0.429344 | 1.5865092 | 0.0054112556 | 0.10677397 | 1 | 7377 | tags=57%, list=31%, si | MM_3 |
| GOBP_POST_TRANSCRIPT | GOBP_POST_TRANSCRIPTIONAL_GENE_SILE | 64 | 0.441371 | 1.5861372 | 0.0077951 | 0.10692331 | 1 | 2349 | tags=22%, list=10%, si | MM_3 |
| GOBP_CARBOHYDRATE_D | GOBP_CARBOHYDRATE_DERIVATIVE_CATAB | 245 | 0.3882 | 1.5859011 | 0 | 0.10693426 | 1 | 6163 | tags=38%, list=26%, si | MM_3 |
| GOBP_MRNA_3_END_PR | GOBP_MRNA_3_END_PROCESSING | 41 | 0.473318 | 1.5856893 | 0.016627079 | 0.10689792 | 1 | 6500 | tags=44%, list=28%, si | MM_3 |
| GOBP_NEGATIVE_REGUL | GOBP_NEGATIVE_REGULATION_OF_PROTEI | 82 | 0.429533 | 1.5852482 | 0.005446623 | 0.107119195 | 1 | 5764 | tags=41%, list=24%, si | MM_3 |
| GOBP_NUCLEUS_ORGAN | GOBP_NUCLEUS_ORGANIZATION | 136 | 0.402026 | 1.5851268 | 0.0010298662 | 0.10703309 | 1 | 7379 | tags=49%, list=31%, si | MM_3 |
| GOBP_CELL_CHEMOTAXIS | GOBP_CELL_CHEMOTAXIS | 295 | 0.387755 | 1.5850918 | 0 | 0.1068565 | 1 | 6056 | tags=39%, list=26%, si | MM_3 |
| GOBP_B_CELL_ACTIVATIO | GOBP_B_CELL_ACTIVATION | 276 | 0.386468 | 1.5850408 | 0 | 0.1066865 | 1 | 5120 | tags=30%, list=22%, si | MM_3 |
| GOBP_CELL_CYCLE_G2_N | GOBP_CELL_CYCLE_G2_M_PHASE_TRANSITI | 152 | 0.401587 | 1.5849755 | 0.0020661156 | 0.10653485 | 1 | 5809 | tags=41%, list=25%, si | MM_3 |
| GOBP_POSITIVE_REGULA | GOBP_POSITIVE_REGULATION_OF_CYTOKIN | 40 | 0.476583 | 1.5849338 | 0.019300362 | 0.10638958 | 1 | 6730 | tags=48%, list=29%, si | MM_3 |
| GOBP_NEURAL_TUBE_PA | GOBP_NEURAL_TUBE_PATTERNING | 38 | 0.475772 | 1.5826151 | 0.011933174 | 0.10842513 | 1 | 2758 | tags=32%, list=12%, si | MM_3 |

| | | | | | | | | | | |
|---|---|---|---|---|---|---|---|---|---|---|
| GOBP_SMALL_GTPASE_M | GOBP_SMALL_GTPASE_MEDIATED_SIGNAL_ | 434 | 0.381581 | 1.5825264 | 0 | 0.108304426 | 1 | 6730 | tags=41%, list=29%, si | MM_3 |
| GOBP_TRNA_PROCESSIN | GOBP_TRNA_PROCESSING | 122 | 0.408471 | 1.5825111 | 0 | 0.10811964 | 1 | 6906 | tags=50%, list=29%, si | MM_3 |
| GOBP_GLIAL_CELL_APOP | GOBP_GLIAL_CELL_APOPTOTIC_PROCESS | 20 | 0.533107 | 1.5824537 | 0.017264277 | 0.10797248 | 1 | 8009 | tags=75%, list=34%, si | MM_3 |
| GOBP_THYMOCYTE_APOI | GOBP_THYMOCYTE_APOPTOTIC_PROCESS | 25 | 0.524 | 1.582295 | 0.020698577 | 0.107909605 | 1 | 5470 | tags=44%, list=23%, si | MM_3 |
| GOBP_CELLULAR_RESPON | GOBP_CELLULAR_RESPONSE_TO_GLUCOSE_ | 42 | 0.464141 | 1.5820688 | 0.016241299 | 0.10790711 | 1 | 5939 | tags=48%, list=25%, si | MM_3 |
| GOBP_CEREBRAL_CORTE | GOBP_CEREBRAL_CORTEX_DEVELOPMENT | 113 | 0.408139 | 1.5820591 | 0.004223865 | 0.10771085 | 1 | 5855 | tags=40%, list=25%, si | MM_3 |
| GOBP_HEME_BIOSYNTHE | GOBP_HEME_BIOSYNTHETIC_PROCESS | 24 | 0.521688 | 1.5812086 | 0.027166883 | 0.108324096 | 1 | 4144 | tags=42%, list=18%, si | MM_3 |
| MICRORNAS_IN_CANCER | MICRORNAS_IN_CANCER | 139 | 0.403951 | 1.5798734 | 0.0010384216 | 0.109462336 | 1 | 7169 | tags=49%, list=30%, si | MM_3 |
| GOBP_CELLULAR_COMPO | GOBP_CELLULAR_COMPONENT_DISASSEMB | 472 | 0.378336 | 1.579607 | 0 | 0.10953806 | 1 | 7533 | tags=50%, list=32%, si | MM_3 |
| GOBP_RELEASE_OF_CYTO | GOBP_RELEASE_OF_CYTOCHROME_C_FRON | 66 | 0.439818 | 1.5795946 | 0.011025358 | 0.10933828 | 1 | 5985 | tags=44%, list=25%, si | MM_3 |
| GOBP_SHORT_TERM_ME | GOBP_SHORT_TERM_MEMORY | 18 | 0.5561 | 1.5794231 | 0.043301757 | 0.10929099 | 1 | 2901 | tags=33%, list=12%, si | MM_3 |
| GOBP_HYPOTHALAMUS_ | GOBP_HYPOTHALAMUS_DEVELOPMENT | 17 | 0.563487 | 1.5789492 | 0.038770054 | 0.109562054 | 1 | 5906 | tags=59%, list=25%, si | MM_3 |
| GOBP_POSITIVE_REGULA | GOBP_POSITIVE_REGULATION_OF_EXECUTI | 19 | 0.549931 | 1.5786349 | 0.04432855 | 0.10968488 | 1 | 2871 | tags=32%, list=12%, si | MM_3 |
| GOBP_MISMATCH_REPAI | GOBP_MISMATCH_REPAIR | 25 | 0.523577 | 1.5778282 | 0.027568921 | 0.11030757 | 1 | 3741 | tags=40%, list=16%, si | MM_3 |
| GOBP_GLUTATHIONE_ME | GOBP_GLUTATHIONE_METABOLIC_PROCESS | 58 | 0.447279 | 1.5766989 | 0.009153318 | 0.11119302 | 1 | 5301 | tags=40%, list=22%, si | MM_3 |
| GOBP_MYELOID_LEUKOC | GOBP_MYELOID_LEUKOCYTE_CYTOKINE_PR | 64 | 0.435617 | 1.5765977 | 0.012318029 | 0.111112304 | 1 | 7295 | tags=47%, list=31%, si | MM_3 |
| TOLL-LIKE_RECEPTOR_SIG | TOLL-LIKE_RECEPTOR_SIGNALING_PATHWA | 84 | 0.416138 | 1.5761027 | 0.007616975 | 0.111412294 | 1 | 5692 | tags=37%, list=24%, si | MM_3 |
| GOBP_CARBOHYDRATE_D | GOBP_CARBOHYDRATE_DERIVATIVE_TRANS | 76 | 0.430369 | 1.5760227 | 0.0066964286 | 0.111282274 | 1 | 5311 | tags=37%, list=22%, si | MM_3 |
| GOBP_QUINONE_BIOSYN | GOBP_QUINONE_BIOSYNTHETIC_PROCESS | 16 | 0.582153 | 1.5757415 | 0.025641026 | 0.111359105 | 1 | 4215 | tags=44%, list=18%, si | MM_3 |
| GOBP_NUCLEOSOME_OR | GOBP_NUCLEOSOME_ORGANIZATION | 67 | 0.428793 | 1.5749207 | 0.014412417 | 0.11202098 | 1 | 5104 | tags=34%, list=22%, si | MM_3 |
| GOBP_REGULATION_OF_ | GOBP_REGULATION_OF_MITOCHONDRIAL_ | 32 | 0.487528 | 1.5746386 | 0.02962963 | 0.112129785 | 1 | 6628 | tags=50%, list=28%, si | MM_3 |
| GOBP_REGULATION_OF_ | GOBP_REGULATION_OF_LEUKOCYTE_MIGR/ | 230 | 0.39114 | 1.5746151 | 0 | 0.111954145 | 1 | 6424 | tags=41%, list=27%, si | MM_3 |
| GOBP_POSITIVE_REGULA | GOBP_POSITIVE_REGULATION_OF_PROTEIN | 192 | 0.39926 | 1.5743605 | 0.001019368 | 0.11201015 | 1 | 7571 | tags=52%, list=32%, si | MM_3 |
| GOBP_CELLULAR_RESPON | GOBP_CELLULAR_RESPONSE_TO_CHEMICAL | 290 | 0.385253 | 1.5741074 | 0 | 0.1121091 | 1 | 7210 | tags=48%, list=31%, si | MM_3 |
| GOBP_POSITIVE_REGULA | GOBP_POSITIVE_REGULATION_OF_LEUKOCY | 97 | 0.420949 | 1.5740299 | 0.0064171124 | 0.111981265 | 1 | 6009 | tags=38%, list=25%, si | MM_3 |
| GOBP_NEGATIVE_REGUL | GOBP_NEGATIVE_REGULATION_OF_ANTIGE | 29 | 0.499162 | 1.5739784 | 0.03325123 | 0.111812726 | 1 | 6640 | tags=45%, list=28%, si | MM_3 |
| GOBP_CELLULAR_RESPON | GOBP_CELLULAR_RESPONSE_TO_TYPE_II_IN | 95 | 0.414731 | 1.5738204 | 0.0043431055 | 0.11179595 | 1 | 8133 | tags=59%, list=34%, si | MM_3 |
| GOBP_CALCIUM_ION_IM | GOBP_CALCIUM_ION_IMPORT | 75 | 0.425763 | 1.5737076 | 0.0066225166 | 0.1117141 | 1 | 5903 | tags=44%, list=25%, si | MM_3 |
| GOBP_NEGATIVE_REGUL | GOBP_NEGATIVE_REGULATION_OF_TRANSM | 142 | 0.401131 | 1.57364 | 0.0020768433 | 0.11159164 | 1 | 3707 | tags=25%, list=16%, si | MM_3 |
| GOBP_POSITIVE_REGULA | GOBP_POSITIVE_REGULATION_OF_PROTEIN | 24 | 0.523925 | 1.5732927 | 0.020860495 | 0.111743875 | 1 | 2817 | tags=33%, list=12%, si | MM_3 |
| GOBP_REGULATION_OF_ | GOBP_REGULATION_OF_BROWN_FAT_CELL | 27 | 0.503978 | 1.573244 | 0.024752475 | 0.11157547 | 1 | 5656 | tags=41%, list=24%, si | MM_3 |
| GOBP_POSITIVE_REGULA | GOBP_POSITIVE_REGULATION_OF_CHEMOT | 150 | 0.398271 | 1.5732057 | 0.0020533882 | 0.11140764 | 1 | 6192 | tags=38%, list=26%, si | MM_3 |
| GOBP_NEGATIVE_REGUL | GOBP_NEGATIVE_REGULATION_OF_CATALY | 487 | 0.376311 | 1.5728347 | 0 | 0.11160239 | 1 | 5643 | tags=33%, list=24%, si | MM_3 |
| GOBP_NEGATIVE_REGUL | GOBP_NEGATIVE_REGULATION_OF_INTERF | 16 | 0.577483 | 1.5723003 | 0.032520324 | 0.11191408 | 1 | 9009 | tags=69%, list=38%, si | MM_3 |
| GOBP_CELLULAR_RESPON | GOBP_CELLULAR_RESPONSE_TO_ABIOTIC_S | 276 | 0.385278 | 1.5722026 | 0.001019368 | 0.1117998 | 1 | 6761 | tags=45%, list=29%, si | MM_3 |
| GOBP_RNA_DECAPPING | GOBP_RNA_DECAPPING | 19 | 0.540683 | 1.5721447 | 0.040897097 | 0.11165482 | 1 | 2242 | tags=26%, list=9%, sig | MM_3 |
| GOBP_NEGATIVE_REGUL | GOBP_NEGATIVE_REGULATION_OF_PROTEI | 25 | 0.502336 | 1.5718099 | 0.03080873 | 0.111832716 | 1 | 6632 | tags=56%, list=28%, si | MM_3 |
| GOBP_ESTABLISHMENT_( | GOBP_ESTABLISHMENT_OF_PROTEIN_LOCA | 25 | 0.512585 | 1.5715699 | 0.028534371 | 0.11190032 | 1 | 3758 | tags=32%, list=16%, si | MM_3 |
| GOBP_LYMPHOCYTE_ME | GOBP_LYMPHOCYTE_MEDIATED_IMMUNITY | 300 | 0.388603 | 1.5713198 | 0.001004016 | 0.11193661 | 1 | 6059 | tags=36%, list=26%, si | MM_3 |
| GOBP_LEUKOCYTE_CHEM | GOBP_LEUKOCYTE_CHEMOTAXIS | 213 | 0.390006 | 1.5712521 | 0 | 0.111803465 | 1 | 5889 | tags=37%, list=25%, si | MM_3 |
| GOBP_REGULATION_OF_ | GOBP_REGULATION_OF_CELL_GROWTH | 415 | 0.378241 | 1.5710988 | 0 | 0.1117561 | 1 | 6907 | tags=40%, list=29%, si | MM_3 |
| GOBP_COPPER_ION_TRA | GOBP_COPPER_ION_TRANSPORT | 16 | 0.566126 | 1.571036 | 0.043417368 | 0.11161458 | 1 | 3528 | tags=38%, list=15%, si | MM_3 |
| GOBP_MONOSACCHARID | GOBP_MONOSACCHARIDE_METABOLIC_PR | 255 | 0.389503 | 1.5709995 | 0 | 0.11144637 | 1 | 5983 | tags=40%, list=25%, si | MM_3 |
| CELL_CYCLE | CELL_CYCLE | 121 | 0.408644 | 1.5708745 | 0.005285412 | 0.11135999 | 1 | 5999 | tags=39%, list=25%, si | MM_3 |
| BASAL_TRANSCRIPTION_ | BASAL_TRANSCRIPTION_FACTORS | 40 | 0.462501 | 1.5706304 | 0.01937046 | 0.11140224 | 1 | 3893 | tags=30%, list=16%, si | MM_3 |
| GOBP_ESTABLISHMENT_( | GOBP_ESTABLISHMENT_OF_RNA_LOCALIZA | 138 | 0.395196 | 1.5703504 | 0 | 0.11148421 | 1 | 7083 | tags=46%, list=30%, si | MM_3 |
| GOBP_REGULATION_OF_ | GOBP_REGULATION_OF_RESPONSE_TO_OS | 18 | 0.55652 | 1.5701064 | 0.025165563 | 0.11150828 | 1 | 6558 | tags=67%, list=28%, si | MM_3 |
| GOBP_AUTOPHAGY_OF_ | GOBP_AUTOPHAGY_OF_MITOCHONDRION | 89 | 0.420258 | 1.569147 | 0.008686211 | 0.11220583 | 1 | 7533 | tags=56%, list=32%, si | MM_3 |
| GOBP_NEGATIVE_REGUL | GOBP_NEGATIVE_REGULATION_OF_PHOSPH | 332 | 0.379103 | 1.5688069 | 0 | 0.11234328 | 1 | 5501 | tags=31%, list=23%, si | MM_3 |
| GOBP_REGULATION_OF_ | GOBP_REGULATION_OF_PHOSPHATASE_AC | 61 | 0.436458 | 1.568667 | 0.014755959 | 0.11228668 | 1 | 7299 | tags=49%, list=31%, si | MM_3 |
| GOBP_ESTABLISHMENT_( | GOBP_ESTABLISHMENT_OF_PROTEIN_LOCA | 435 | 0.376357 | 1.568232 | 0 | 0.11249848 | 1 | 7221 | tags=43%, list=31%, si | MM_3 |
| GOBP_DISRUPTION_OF_A | GOBP_DISRUPTION_OF_ANATOMICAL_STRU | 73 | 0.428872 | 1.5680406 | 0.0077348067 | 0.11249889 | 1 | 3733 | tags=27%, list=16%, si | MM_3 |
| GOBP_REACTIVE_OXYGEN | GOBP_REACTIVE_OXYGEN_SPECIES_METAB | 239 | 0.386848 | 1.5676717 | 0 | 0.112652786 | 1 | 5500 | tags=32%, list=23%, si | MM_3 |
| GOBP_REGULATION_OF_ | GOBP_REGULATION_OF_CELLULAR_CATABO | 391 | 0.380847 | 1.5675379 | 0 | 0.11259961 | 1 | 7383 | tags=47%, list=31%, si | MM_3 |
| GOBP_PROTEIN_TARGETI | GOBP_PROTEIN_TARGETING_TO_LYSOSOMI | 29 | 0.501541 | 1.5665109 | 0.024813896 | 0.113454275 | 1 | 7162 | tags=52%, list=30%, si | MM_3 |
| GOBP_MRNA_TRANSPOR | GOBP_MRNA_TRANSPORT | 108 | 0.409312 | 1.5664256 | 0.006322445 | 0.113361046 | 1 | 6628 | tags=44%, list=28%, si | MM_3 |
| GOBP_MRNA_TRANSCRIP | GOBP_MRNA_TRANSCRIPTION | 50 | 0.447124 | 1.5655698 | 0.019675925 | 0.11405411 | 1 | 5487 | tags=42%, list=23%, si | MM_3 |
| GOBP_COLLAGEN_CATAB | GOBP_COLLAGEN_CATABOLIC_PROCESS | 37 | 0.476236 | 1.5649348 | 0.023199024 | 0.1144496 | 1 | 6647 | tags=57%, list=28%, si | MM_3 |
| GOBP_RESPONSE_TO_HY | GOBP_RESPONSE_TO_HYDROGEN_PEROXID | 85 | 0.42334 | 1.5643549 | 0.009879254 | 0.114822395 | 1 | 5518 | tags=40%, list=23%, si | MM_3 |
| GOBP_ORGANOPHOSPHA | GOBP_ORGANOPHOSPHATE_CATABOLIC_PR | 227 | 0.383957 | 1.5641669 | 0 | 0.11485068 | 1 | 6624 | tags=38%, list=28%, si | MM_3 |
| GOBP_PROTEIN_SUMOYL | GOBP_PROTEIN_SUMOYLATION | 42 | 0.462462 | 1.5637354 | 0.022592152 | 0.11508807 | 1 | 7612 | tags=62%, list=32%, si | MM_3 |
| GOBP_POSITIVE_REGULA | GOBP_POSITIVE_REGULATION_OF_INTERLE | 65 | 0.430094 | 1.5631802 | 0.016949153 | 0.11544245 | 1 | 4655 | tags=26%, list=20%, si | MM_3 |
| GOBP_POSITIVE_REGULA | GOBP_POSITIVE_REGULATION_OF_PHOSPH | 26 | 0.512494 | 1.5620342 | 0.029299363 | 0.11645391 | 1 | 7299 | tags=65%, list=31%, si | MM_3 |
| GOBP_POSITIVE_REGULA | GOBP_POSITIVE_REGULATION_OF_SMOOTH | 114 | 0.407885 | 1.5618405 | 0.00524659 | 0.116427064 | 1 | 7761 | tags=54%, list=33%, si | MM_3 |
| GOBP_LEUKOCYTE_MIGR | GOBP_LEUKOCYTE_MIGRATION | 363 | 0.381027 | 1.5618151 | 0 | 0.1162534 | 1 | 5891 | tags=36%, list=25%, si | MM_3 |
| GOBP_REGULATION_OF_ | GOBP_REGULATION_OF_DNA_REPAIR | 219 | 0.385206 | 1.5613173 | 0.0010131713 | 0.11651951 | 1 | 4668 | tags=27%, list=20%, si | MM_3 |
| GOBP_NEGATIVE_REGUL | GOBP_NEGATIVE_REGULATION_OF_B_CELL | 20 | 0.541076 | 1.5605922 | 0.025956284 | 0.11701275 | 1 | 2571 | tags=30%, list=11%, si | MM_3 |
| GOBP_NEUROINFLAMMA | GOBP_NEUROINFLAMMATORY_RESPONSE | 84 | 0.421872 | 1.5605867 | 0.0076086954 | 0.11682682 | 1 | 5180 | tags=31%, list=22%, si | MM_3 |
| GOBP_ESTABLISHMENT_( | GOBP_ESTABLISHMENT_OF_EPITHELIAL_CE | 36 | 0.47343 | 1.5605181 | 0.032490976 | 0.116724774 | 1 | 7559 | tags=56%, list=32%, si | MM_3 |
| GOBP_GLYCERALDEHYDE | GOBP_GLYCERALDEHYDE_3_PHOSPHATE_M | 19 | 0.553022 | 1.5604348 | 0.046384722 | 0.116638675 | 1 | 4566 | tags=42%, list=19%, si | MM_3 |
| GOBP_REGULATION_OF_ | GOBP_REGULATION_OF_CELL_GROWTH_IN | 33 | 0.481888 | 1.5601765 | 0.022673031 | 0.11672212 | 1 | 3853 | tags=33%, list=16%, si | MM_3 |
| GOBP_NEGATIVE_REGUL | GOBP_NEGATIVE_REGULATION_OF_PROTE( | 69 | 0.42742 | 1.5595949 | 0.013348165 | 0.1171435 | 1 | 7217 | tags=45%, list=31%, si | MM_3 |
| GOBP_MATURATION_OF_ | GOBP_MATURATION_OF_5_8S_RRNA_FRON | 24 | 0.516312 | 1.5591823 | 0.02849741 | 0.117387906 | 1 | 7276 | tags=63%, list=31%, si | MM_3 |
| GOBP_REGULATION_OF_ | GOBP_REGULATION_OF_B_CELL_PROLIFERA | 71 | 0.43017 | 1.5586013 | 0.014492754 | 0.1177879 | 1 | 5024 | tags=31%, list=21%, si | MM_3 |
| GOBP_REGULATION_OF_ | GOBP_REGULATION_OF_MITOCHONDRIAL_ | 20 | 0.527464 | 1.5577608 | 0.047872342 | 0.11846303 | 1 | 5019 | tags=45%, list=21%, si | MM_3 |
| GOBP_RESPONSE_TO_IOI | GOBP_RESPONSE_TO_IONIZING_RADIATION | 127 | 0.402633 | 1.5575645 | 0.003125 | 0.11846397 | 1 | 8297 | tags=63%, list=35%, si | MM_3 |
| GOBP_PROTEIN_TARGETI | GOBP_PROTEIN_TARGETING | 285 | 0.378628 | 1.5573915 | 0 | 0.11846323 | 1 | 6609 | tags=39%, list=28%, si | MM_3 |
| GOBP_APOPTOTIC_DNA_ | GOBP_APOPTOTIC_DNA_FRAGMENTATION | 18 | 0.55344 | 1.5566897 | 0.03410641 | 0.11899228 | 1 | 2403 | tags=33%, list=10%, si | MM_3 |
| GOBP_DEFENSE_RESPON | GOBP_DEFENSE_RESPONSE_TO_VIRUS | 276 | 0.382108 | 1.5566506 | 0 | 0.1188337 | 1 | 4317 | tags=27%, list=18%, si | MM_3 |
| GOBP_REGULATION_OF_ | GOBP_REGULATION_OF_GLIOGENESIS | 133 | 0.398153 | 1.5559125 | 0.0021030493 | 0.119376905 | 1 | 5859 | tags=36%, list=25%, si | MM_3 |
| GOBP_ESTABLISHMENT_( | GOBP_ESTABLISHMENT_OF_EPITHELIAL_CEI | 18 | 0.546764 | 1.5557252 | 0.047683924 | 0.11938659 | 1 | 7297 | tags=61%, list=31%, si | MM_3 |
| GOBP_MYELOID_LEUKOC | GOBP_MYELOID_LEUKOCYTE_ACTIVATION | 247 | 0.379252 | 1.5556357 | 0 | 0.11929269 | 1 | 5087 | tags=28%, list=22%, si | MM_3 |

| | | | | | | | | | | |
|---|---|---|---|---|---|---|---|---|---|---|
| GOBP_MITOCHONDRIAL_ | GOBP_MITOCHONDRIAL_CALCIUM_ION_TR | 19 | 0.543021 | 1.5555826 | 0.043882977 | 0.11915845 | 1 | 6993 | tags=68%, list=30%, si | MM_3 |
| GOBP_REGULATION_OF_ | GOBP_REGULATION_OF_LIPOPOLYSACCHAR | 29 | 0.488527 | 1.5555608 | 0.0275 | 0.11898743 | 1 | 5501 | tags=45%, list=23%, si | MM_3 |
| GOBP_FIBROBLAST_PROL | GOBP_FIBROBLAST_PROLIFERATION | 147 | 0.393344 | 1.5548947 | 0.0030832477 | 0.1194673 | 1 | 5988 | tags=41%, list=25%, si | MM_3 |
| GOBP_NEGATIVE_REGUL | GOBP_NEGATIVE_REGULATION_OF_NEURO | 152 | 0.393573 | 1.5546409 | 0.0010319918 | 0.119518705 | 1 | 6781 | tags=45%, list=29%, si | MM_3 |
| GOBP_GLOMERULUS_DE | GOBP_GLOMERULUS_DEVELOPMENT | 70 | 0.427606 | 1.554255 | 0.010227272 | 0.11974677 | 1 | 3957 | tags=33%, list=17%, si | MM_3 |
| PHAGOSOME | PHAGOSOME | 146 | 0.397748 | 1.5539666 | 0.0020618557 | 0.1198549 | 1 | 7319 | tags=55%, list=31%, si | MM_3 |
| GOBP_POSITIVE_REGULA | GOBP_POSITIVE_REGULATION_OF_GLIAL_C | 31 | 0.490572 | 1.553822 | 0.036942676 | 0.11981334 | 1 | 5402 | tags=45%, list=23%, si | MM_3 |
| GOBP_B_CELL_HOMEOST | GOBP_B_CELL_HOMEOSTASIS | 42 | 0.458902 | 1.553779 | 0.019138755 | 0.11965136 | 1 | 4974 | tags=33%, list=21%, si | MM_3 |
| GOBP_SYNAPTIC_VESICLE | GOBP_SYNAPTIC_VESICLE_RECYCLING | 101 | 0.407912 | 1.5530488 | 0.0074231178 | 0.12018053 | 1 | 6972 | tags=47%, list=30%, si | MM_3 |
| GOBP_SIGNALING_RECEP | GOBP_SIGNALING_RECEPTOR_LIGAND_PRE | 29 | 0.489007 | 1.5529615 | 0.029776676 | 0.12007522 | 1 | 2190 | tags=28%, list=9%, sig | MM_3 |
| GOBP_LEUKOCYTE_MEDI | GOBP_LEUKOCYTE_MEDIATED_IMMUNITY | 400 | 0.375154 | 1.5529549 | 0 | 0.11988686 | 1 | 6518 | tags=38%, list=28%, si | MM_3 |
| GOBP_POSITIVE_REGULA | GOBP_POSITIVE_REGULATION_OF_CELL_GR | 185 | 0.388767 | 1.5520637 | 0.0020491802 | 0.120538004 | 1 | 6781 | tags=42%, list=29%, si | MM_3 |
| GOBP_DNA_DAMAGE_RE | GOBP_DNA_DAMAGE_RESPONSE_SIGNAL_T | 67 | 0.427336 | 1.5519937 | 0.012290503 | 0.12042911 | 1 | 5526 | tags=45%, list=23%, si | MM_3 |
| GOBP_RESPONSE_TO_AM | GOBP_RESPONSE_TO_AMINO_ACID_STARV | 54 | 0.43212 | 1.5518707 | 0.018539976 | 0.12035394 | 1 | 7203 | tags=48%, list=31%, si | MM_3 |
| GOBP_ACTIVATION_OF_I | GOBP_ACTIVATION_OF_INNATE_IMMUNE_ | 290 | 0.382228 | 1.5516768 | 0.001004016 | 0.12038155 | 1 | 7581 | tags=46%, list=32%, si | MM_3 |
| GOBP_MONOCARBOXYLI | GOBP_MONOCARBOXYLIC_ACID_CATABOLI | 126 | 0.403962 | 1.5516738 | 0.0010515247 | 0.1201909 | 1 | 5142 | tags=33%, list=22%, si | MM_3 |
| GOBP_REGULATION_OF_ | GOBP_REGULATION_OF_ENDOPLASMIC_RE | 24 | 0.513822 | 1.5516304 | 0.032051284 | 0.120032236 | 1 | 6457 | tags=63%, list=27%, si | MM_3 |
| GOBP_THYMIC_T_CELL_S | GOBP_THYMIC_T_CELL_SELECTION | 26 | 0.493559 | 1.5507923 | 0.031725887 | 0.120677084 | 1 | 1364 | tags=19%, list=6%, sig | MM_3 |
| GOBP_REGULATION_OF_ | GOBP_REGULATION_OF_UBIQUITIN_PROTE | 20 | 0.535818 | 1.5507876 | 0.029372497 | 0.120488524 | 1 | 7202 | tags=55%, list=31%, si | MM_3 |
| SNARE_INTERACTIONS_IN | SNARE_INTERACTIONS_IN_VESICULAR_TRA | 32 | 0.483713 | 1.550346 | 0.02774427 | 0.12078844 | 1 | 7089 | tags=66%, list=30%, si | MM_3 |
| GOBP_SMOOTH_MUSCLE | GOBP_SMOOTH_MUSCLE_CELL_DIFFERENT | 78 | 0.412524 | 1.5497782 | 0.015283843 | 0.12115448 | 1 | 4761 | tags=31%, list=20%, si | MM_3 |
| GOBP_REGULATION_OF_ | GOBP_REGULATION_OF_ATP_METABOLIC_ | 77 | 0.427815 | 1.5494889 | 0.011123471 | 0.12127896 | 1 | 6488 | tags=38%, list=27%, si | MM_3 |
| GOBP_PEPTIDYL_LYSINE_ | GOBP_PEPTIDYL_LYSINE_METHYLATION | 24 | 0.508335 | 1.5494858 | 0.043250326 | 0.12109063 | 1 | 5633 | tags=38%, list=24%, si | MM_3 |
| GOBP_CALCIUM_ION_IM | GOBP_CALCIUM_ION_IMPORT_ACROSS_PL | 34 | 0.478208 | 1.5494202 | 0.03638645 | 0.12097312 | 1 | 5903 | tags=53%, list=25%, si | MM_3 |
| GOBP_NEUROMUSCULAR | GOBP_NEUROMUSCULAR_PROCESS_CONTR | 69 | 0.428761 | 1.5493532 | 0.013422819 | 0.12084455 | 1 | 4042 | tags=30%, list=17%, si | MM_3 |
| FRUCTOSE_AND_MANNO | FRUCTOSE_AND_MANNOSE_METABOLISM | 33 | 0.488369 | 1.5492136 | 0.019680196 | 0.12082521 | 1 | 3972 | tags=27%, list=17%, si | MM_3 |
| GOBP_POSITIVE_REGULA | GOBP_POSITIVE_REGULATION_OF_PROTEIN | 201 | 0.383268 | 1.5489444 | 0.0030456854 | 0.12095403 | 1 | 7521 | tags=47%, list=32%, si | MM_3 |
| GOBP_NEGATIVE_REGUL | GOBP_NEGATIVE_REGULATION_OF_IMMUN | 186 | 0.392451 | 1.5487932 | 0.0030549897 | 0.12090733 | 1 | 6246 | tags=39%, list=26%, si | MM_3 |
| GOBP_REGULATION_OF_ | GOBP_REGULATION_OF_AUTOPHAGY | 295 | 0.379938 | 1.5479931 | 0 | 0.12150195 | 1 | 7506 | tags=51%, list=32%, si | MM_3 |
| GOBP_REGULATION_OF_ | GOBP_REGULATION_OF_VIRAL_PROCESS | 167 | 0.389453 | 1.5474972 | 0.002072539 | 0.12177942 | 1 | 7549 | tags=49%, list=32%, si | MM_3 |
| GOBP_ALDEHYDE_BIOSY | GOBP_ALDEHYDE_BIOSYNTHETIC_PROCESS | 26 | 0.496787 | 1.5471604 | 0.025094103 | 0.121930085 | 1 | 4997 | tags=46%, list=21%, si | MM_3 |
| GOBP_RESPONSE_TO_HY | GOBP_RESPONSE_TO_HYDROPEROXIDE | 18 | 0.54318 | 1.5470084 | 0.044959128 | 0.12186422 | 1 | 6356 | tags=50%, list=27%, si | MM_3 |
| GOBP_NEGATIVE_REGUL | GOBP_NEGATIVE_REGULATION_OF_LEUKO | 65 | 0.427388 | 1.5469778 | 0.013667426 | 0.12170526 | 1 | 4409 | tags=29%, list=19%, si | MM_3 |
| GOBP_HEPATOCYTE_APO | GOBP_HEPATOCYTE_APOPTOTIC_PROCESS | 26 | 0.50418 | 1.5468711 | 0.044757035 | 0.12164002 | 1 | 936 | tags=19%, list=4%, sig | MM_3 |
| HUMAN_IMMUNODEFICI | HUMAN_IMMUNODEFICIENCY_VIRUS_1_IN | 201 | 0.383869 | 1.5466627 | 0 | 0.12166806 | 1 | 5983 | tags=38%, list=25%, si | MM_3 |
| GOBP_REGULATION_OF_ | GOBP_REGULATION_OF_T_CELL_PROLIFERA | 188 | 0.387464 | 1.5464313 | 0 | 0.12170098 | 1 | 4871 | tags=30%, list=21%, si | MM_3 |
| PENTOSE_PHOSPHATE_P | PENTOSE_PHOSPHATE_PATHWAY | 29 | 0.501466 | 1.5460598 | 0.03241895 | 0.121906355 | 1 | 4019 | tags=31%, list=17%, si | MM_3 |
| GOBP_NEGATIVE_REGUL | GOBP_NEGATIVE_REGULATION_OF_ALPHA_ | 47 | 0.45003 | 1.5454466 | 0.03033839 | 0.12236198 | 1 | 6428 | tags=40%, list=27%, si | MM_3 |
| GOBP_NUCLEOBASE_CON | GOBP_NUCLEOBASE_CONTAINING_COMPO | 195 | 0.384297 | 1.5448053 | 0 | 0.12289459 | 1 | 6544 | tags=40%, list=28%, si | MM_3 |
| GOBP_PROTEIN_MONOU | GOBP_PROTEIN_MONOUBIQUITINATION | 47 | 0.44824 | 1.5444733 | 0.024390243 | 0.123056084 | 1 | 6843 | tags=51%, list=29%, si | MM_3 |
| GOBP_REGULATION_OF_ | GOBP_REGULATION_OF_CELL_CYCLE_CHEC | 48 | 0.449391 | 1.5443339 | 0.021052632 | 0.12301499 | 1 | 5401 | tags=38%, list=23%, si | MM_3 |
| GOBP_NEGATIVE_REGUL | GOBP_NEGATIVE_REGULATION_OF_DEFENS | 19 | 0.537071 | 1.5440402 | 0.036144577 | 0.12312972 | 1 | 5933 | tags=42%, list=25%, si | MM_3 |
| GOBP_REGULATION_OF_ | GOBP_REGULATION_OF_LYMPHOCYTE_MIG | 64 | 0.424911 | 1.5434313 | 0.020524515 | 0.123618044 | 1 | 3849 | tags=28%, list=16%, si | MM_3 |
| GOBP_REGULATION_OF_ | GOBP_REGULATION_OF_RNA_SPLICING | 178 | 0.388545 | 1.542985 | 0.0031023785 | 0.12391185 | 1 | 8563 | tags=58%, list=36%, si | MM_3 |
| OOCYTE_MEIOSIS | OOCYTE_MEIOSIS | 108 | 0.40509 | 1.5428239 | 0.004287245 | 0.123861656 | 1 | 5892 | tags=37%, list=25%, si | MM_3 |
| GOBP_PODOSOME_ASSE | GOBP_PODOSOME_ASSEMBLY | 25 | 0.501976 | 1.5427344 | 0.044699874 | 0.12376749 | 1 | 7600 | tags=64%, list=32%, si | MM_3 |
| BASE_EXCISION_REPAIR | BASE_EXCISION_REPAIR | 32 | 0.483809 | 1.5427296 | 0.040342297 | 0.12358383 | 1 | 4874 | tags=41%, list=21%, si | MM_3 |
| GOBP_REGULATION_OF_ | GOBP_REGULATION_OF_FIBROBLAST_APOP | 24 | 0.5099 | 1.542636 | 0.03698811 | 0.12350144 | 1 | 5974 | tags=46%, list=25%, si | MM_3 |
| GOBP_MAINTENANCE_OI | GOBP_MAINTENANCE_OF_PROTEIN_LOCAL | 43 | 0.457108 | 1.54256 | 0.027154664 | 0.12340358 | 1 | 7610 | tags=53%, list=32%, si | MM_3 |
| GOBP_REGULATION_OF_ | GOBP_REGULATION_OF_ENDOCYTOSIS | 325 | 0.372481 | 1.5422742 | 0 | 0.12352559 | 1 | 7490 | tags=46%, list=32%, si | MM_3 |
| GOBP_REGULATION_OF_ | GOBP_REGULATION_OF_MICROGLIAL_CELL | 17 | 0.561503 | 1.5422258 | 0.044414535 | 0.12338577 | 1 | 3518 | tags=35%, list=15%, si | MM_3 |
| GOBP_PROTEIN_TARGETI | GOBP_PROTEIN_TARGETING_TO_MITOCHO | 76 | 0.419893 | 1.5419122 | 0.0122358175 | 0.123552635 | 1 | 6208 | tags=39%, list=26%, si | MM_3 |
| GOBP_RESPONSE_TO_NI | GOBP_RESPONSE_TO_NITRIC_OXIDE | 17 | 0.566195 | 1.5418043 | 0.03888889 | 0.123495795 | 1 | 5977 | tags=47%, list=25%, si | MM_3 |
| GOBP_CELLULAR_EXTRAV | GOBP_CELLULAR_EXTRAVASATION | 74 | 0.41853 | 1.5415382 | 0.018619934 | 0.123588905 | 1 | 7118 | tags=47%, list=30%, si | MM_3 |
| GOBP_ENDOPLASMIC_RE | GOBP_ENDOPLASMIC_RETICULUM_ORGAN | 90 | 0.40721 | 1.5414083 | 0.015217391 | 0.123569585 | 1 | 6507 | tags=46%, list=28%, si | MM_3 |
| GOBP_POSITIVE_REGULA | GOBP_POSITIVE_REGULATION_OF_IMMUNE | 266 | 0.378054 | 1.541322 | 0 | 0.12347407 | 1 | 5242 | tags=29%, list=22%, si | MM_3 |
| GOBP_NEGATIVE_REGUL | GOBP_NEGATIVE_REGULATION_OF_INTERL | 24 | 0.508522 | 1.5412298 | 0.034793813 | 0.12336341 | 1 | 5501 | tags=50%, list=23%, si | MM_3 |
| GOBP_RNA_LOCALIZATIO | GOBP_RNA_LOCALIZATION | 164 | 0.386254 | 1.5406548 | 0.0031088083 | 0.12374111 | 1 | 7086 | tags=45%, list=30%, si | MM_3 |
| GOBP_ANTIGEN_PROCES | GOBP_ANTIGEN_PROCESSING_AND_PRESEN | 29 | 0.489446 | 1.5405768 | 0.029962547 | 0.123613395 | 1 | 6109 | tags=52%, list=26%, si | MM_3 |
| GOBP_REGULATION_OF_ | GOBP_REGULATION_OF_CELL_CYCLE_G2_M | 117 | 0.39746 | 1.5405688 | 0.011506276 | 0.12343198 | 1 | 5809 | tags=40%, list=25%, si | MM_3 |
| GOBP_POSITIVE_REGULA | GOBP_POSITIVE_REGULATION_OF_CELL_CY | 112 | 0.397672 | 1.5402768 | 0.005285412 | 0.12357832 | 1 | 6906 | tags=46%, list=29%, si | MM_3 |
| GOBP_NEGATIVE_REGUL | GOBP_NEGATIVE_REGULATION_OF_TRANSF | 94 | 0.408348 | 1.5399357 | 0.009635975 | 0.1237286 | 1 | 4943 | tags=33%, list=21%, si | MM_3 |
| GOBP_RESPONSE_TO_UV | GOBP_RESPONSE_TO_UV | 153 | 0.39173 | 1.539875 | 0.0010319918 | 0.123630874 | 1 | 5470 | tags=37%, list=23%, si | MM_3 |
| GOBP_FATTY_ACID_CATA | GOBP_FATTY_ACID_CATABOLIC_PROCESS | 100 | 0.405558 | 1.5395919 | 0.0096982755 | 0.12374988 | 1 | 5142 | tags=33%, list=22%, si | MM_3 |
| GOBP_REGULATION_OF_ | GOBP_REGULATION_OF_ISOTYPE_SWITCHIN | 38 | 0.471414 | 1.5391788 | 0.020334927 | 0.12398363 | 1 | 4592 | tags=39%, list=19%, si | MM_3 |
| BLADDER_CANCER | BLADDER_CANCER | 37 | 0.464543 | 1.5389725 | 0.031668697 | 0.124016136 | 1 | 6864 | tags=54%, list=29%, si | MM_3 |
| GOBP_NEGATIVE_REGUL | GOBP_NEGATIVE_REGULATION_OF_TYPE_I_ | 23 | 0.512625 | 1.5386798 | 0.038258575 | 0.12410523 | 1 | 1490 | tags=22%, list=6%, sig | MM_3 |
| GLYCINE,_SERINE_AND_T | GLYCINE,_SERINE_AND_THREONINE_META | 38 | 0.458261 | 1.53847 | 0.034400947 | 0.12416947 | 1 | 4166 | tags=32%, list=18%, si | MM_3 |
| GOBP_POSITIVE_REGULA | GOBP_POSITIVE_REGULATION_OF_VASCUL | 20 | 0.524814 | 1.5380721 | 0.043363996 | 0.12442855 | 1 | 4256 | tags=45%, list=18%, si | MM_3 |
| GOBP_INFLAMMASOME_ | GOBP_INFLAMMASOME_MEDIATED_SIGNA | 46 | 0.44432 | 1.5379331 | 0.030624263 | 0.12438724 | 1 | 4655 | tags=30%, list=20%, si | MM_3 |
| GOBP_REGULATION_OF_ | GOBP_REGULATION_OF_MITOCHONDRIAL_ | 26 | 0.496664 | 1.5376776 | 0.038119443 | 0.12445985 | 1 | 6628 | tags=50%, list=28%, si | MM_3 |
| GOBP_NEGATIVE_REGUL | GOBP_NEGATIVE_REGULATION_OF_LEUKO | 142 | 0.392698 | 1.5375445 | 0.006342495 | 0.12441108 | 1 | 5188 | tags=32%, list=22%, si | MM_3 |
| GOBP_POSITIVE_REGULA | GOBP_POSITIVE_REGULATION_OF_FATTY_A | 45 | 0.449927 | 1.535706 | 0.028673835 | 0.12609927 | 1 | 4463 | tags=36%, list=19%, si | MM_3 |
| GOBP_T_CELL_PROLIFER | GOBP_T_CELL_PROLIFERATION | 233 | 0.382408 | 1.5355351 | 0.0010070493 | 0.12608163 | 1 | 5500 | tags=31%, list=23%, si | MM_3 |
| GOBP_REGULATION_OF_ | GOBP_REGULATION_OF_EXTRACELLULAR_M | 66 | 0.424588 | 1.5352377 | 0.02 | 0.12618333 | 1 | 4581 | tags=36%, list=19%, si | MM_3 |
| GOBP_CYTOCHROME_CO | GOBP_CYTOCHROME_COMPLEX_ASSEMBLY | 32 | 0.475841 | 1.5351762 | 0.025454545 | 0.1260733 | 1 | 6365 | tags=59%, list=27%, si | MM_3 |
| GOBP_RESPONSE_TO_ES | GOBP_RESPONSE_TO_ESTRADIOL | 38 | 0.459875 | 1.5347271 | 0.027811367 | 0.12635851 | 1 | 5397 | tags=45%, list=23%, si | MM_3 |
| GOBP_LATE_ENDOSOME_ | GOBP_LATE_ENDOSOME_TO_LYSOSOME_T | 19 | 0.537889 | 1.5346835 | 0.031123139 | 0.12622298 | 1 | 7915 | tags=68%, list=34%, si | MM_3 |
| GOBP_INCLUSION_BODY_ | GOBP_INCLUSION_BODY_ASSEMBLY | 27 | 0.495541 | 1.5346344 | 0.039042823 | 0.12608935 | 1 | 7317 | tags=67%, list=31%, si | MM_3 |
| GOBP_SOMATIC_DIVERSI | GOBP_SOMATIC_DIVERSIFICATION_OF_IMM | 55 | 0.434285 | 1.5341002 | 0.017142856 | 0.12650093 | 1 | 4592 | tags=35%, list=19%, si | MM_3 |
| GOBP_REGULATION_OF_ | GOBP_REGULATION_OF_MUSCLE_CELL_DIF | 154 | 0.387857 | 1.5338755 | 0.004192872 | 0.1265936 | 1 | 3164 | tags=23%, list=13%, si | MM_3 |
| GOBP_POSITIVE_REGULA | GOBP_POSITIVE_REGULATION_OF_GLYCOP | 26 | 0.490699 | 1.5338552 | 0.033707865 | 0.12642562 | 1 | 6159 | tags=46%, list=26%, si | MM_3 |

| | | | | | | | | | | |
|---|---|---|---|---|---|---|---|---|---|---|
| GOBP_RESPONSE_TO_SA | GOBP_RESPONSE_TO_SALT_STRESS | 20 | 0.530558 | 1.53383 | 0.03504043 | 0.12626113 | 1 | 6500 | tags=60%, list=28%, si | MM_3 |
| GOBP_POSITIVE_REGULA | GOBP_POSITIVE_REGULATION_OF_PROTEIN | 107 | 0.403792 | 1.5337144 | 0.0053418805 | 0.12620014 | 1 | 7506 | tags=54%, list=32%, si | MM_3 |
| AMINOACYL-TRNA_BIOSY | AMINOACYL-TRNA_BIOSYNTHESIS | 43 | 0.45268 | 1.5331185 | 0.02361275 | 0.12661391 | 1 | 7423 | tags=63%, list=31%, si | MM_3 |
| GOBP_PRESYNAPTIC_END | GOBP_PRESYNAPTIC_ENDOCYTOSIS | 95 | 0.407189 | 1.5330652 | 0.0054406966 | 0.1264841 | 1 | 6972 | tags=44%, list=30%, si | MM_3 |
| GOBP_NEGATIVE_REGUL | GOBP_NEGATIVE_REGULATION_OF_DNA_R | 37 | 0.463824 | 1.5318689 | 0.036144577 | 0.12754123 | 1 | 4668 | tags=38%, list=20%, si | MM_3 |
| GOBP_RESPONSE_TO_MO | GOBP_RESPONSE_TO_MOLECULE_OF_BACT | 296 | 0.37618 | 1.5318323 | 0 | 0.127403 | 1 | 6682 | tags=40%, list=28%, si | MM_3 |
| GOBP_REGULATION_OF_ | GOBP_REGULATION_OF_THYMOCYTE_APOP | 18 | 0.5474 | 1.5313482 | 0.036262203 | 0.1277346 | 1 | 5470 | tags=44%, list=23%, si | MM_3 |
| GOBP_NEGATIVE_REGUL | GOBP_NEGATIVE_REGULATION_OF_TISSUE_ | 27 | 0.502522 | 1.5312208 | 0.030264817 | 0.12768812 | 1 | 1201 | tags=22%, list=5%, sig | MM_3 |
| GOBP_RENAL_SYSTEM_V | GOBP_RENAL_SYSTEM_VASCULATURE_DEV | 29 | 0.486585 | 1.530733 | 0.038265307 | 0.12800792 | 1 | 3680 | tags=34%, list=16%, si | MM_3 |
| VASOPRESSIN-REGULATE | VASOPRESSIN-REGULATED_WATER_REABSO | 40 | 0.461214 | 1.5305728 | 0.033333335 | 0.1279847 | 1 | 7243 | tags=52%, list=31%, si | MM_3 |
| GOBP_NEGATIVE_REGUL | GOBP_NEGATIVE_REGULATION_OF_DEFENS | 242 | 0.379051 | 1.5305668 | 0 | 0.1278085 | 1 | 5518 | tags=33%, list=23%, si | MM_3 |
| GOBP_NEGATIVE_REGUL | GOBP_NEGATIVE_REGULATION_OF_AMIDE_ | 22 | 0.51997 | 1.5305663 | 0.027954256 | 0.12762696 | 1 | 7235 | tags=59%, list=31%, si | MM_3 |
| GOBP_FIBROBLAST_APOP | GOBP_FIBROBLAST_APOPTOTIC_PROCESS | 31 | 0.484768 | 1.5302829 | 0.029666254 | 0.127753 | 1 | 6329 | tags=45%, list=27%, si | MM_3 |
| GOBP_SULFUR_COMPOU | GOBP_SULFUR_COMPOUND_CATABOLIC_PI | 35 | 0.46746 | 1.530218 | 0.042579077 | 0.12768058 | 1 | 5217 | tags=43%, list=22%, si | MM_3 |
| GOBP_PHOSPHATIDYLINC | GOBP_PHOSPHATIDYLINOSITOL_METABOLIC | 148 | 0.386293 | 1.5301583 | 0 | 0.127566 | 1 | 3381 | tags=25%, list=14%, si | MM_3 |
| GOBP_REGULATION_OF_ | GOBP_REGULATION_OF_DENDRITIC_SPINE_ | 60 | 0.424255 | 1.5293069 | 0.01793722 | 0.12825543 | 1 | 4545 | tags=32%, list=19%, si | MM_3 |
| GOBP_REGULATION_OF_ | GOBP_REGULATION_OF_INNATE_IMMUNE_ | 427 | 0.370329 | 1.5292734 | 0 | 0.12810513 | 1 | 6329 | tags=36%, list=27%, si | MM_3 |
| GOBP_POSITIVE_REGULA | GOBP_POSITIVE_REGULATION_OF_INTERLE | 15 | 0.564106 | 1.5283439 | 0.034387894 | 0.12891603 | 1 | 4655 | tags=47%, list=20%, si | MM_3 |
| GOBP_POSITIVE_REGULA | GOBP_POSITIVE_REGULATION_OF_GTPASE_ | 203 | 0.37871 | 1.5282739 | 0.0020470829 | 0.12881912 | 1 | 6037 | tags=36%, list=26%, si | MM_3 |
| GOBP_VASCULAR_ASSOC | GOBP_VASCULAR_ASSOCIATED_SMOOTH_M | 80 | 0.411908 | 1.5278374 | 0.0096982755 | 0.12915689 | 1 | 7741 | tags=50%, list=33%, si | MM_3 |
| GOBP_SUPEROXIDE_ANIC | GOBP_SUPEROXIDE_ANION_GENERATION | 43 | 0.449021 | 1.5278234 | 0.03199052 | 0.1289932 | 1 | 5778 | tags=35%, list=24%, si | MM_3 |
| GOBP_MYOBLAST_FUSIO | GOBP_MYOBLAST_FUSION | 45 | 0.443493 | 1.5277009 | 0.027122641 | 0.1289371 | 1 | 5715 | tags=38%, list=24%, si | MM_3 |
| GOBP_DETOXIFICATION | GOBP_DETOXIFICATION | 70 | 0.41978 | 1.5275766 | 0.016666668 | 0.12888996 | 1 | 4429 | tags=34%, list=19%, si | MM_3 |
| GOBP_POSITIVE_REGULA | GOBP_POSITIVE_REGULATION_OF_CELL_GR | 17 | 0.5539 | 1.5275291 | 0.04103967 | 0.12876925 | 1 | 5075 | tags=47%, list=21%, si | MM_3 |
| GOBP_REGULATION_OF_ | GOBP_REGULATION_OF_FILOPODIUM_ASSE | 54 | 0.428575 | 1.527428 | 0.033878505 | 0.12869217 | 1 | 7890 | tags=59%, list=33%, si | MM_3 |
| GOBP_REGULATION_OF_ | GOBP_REGULATION_OF_CHROMOSOME_OI | 242 | 0.375305 | 1.5267892 | 0 | 0.12921195 | 1 | 6963 | tags=41%, list=29%, si | MM_3 |
| GOBP_POSITIVE_REGULA | GOBP_POSITIVE_REGULATION_OF_MRNA_M | 131 | 0.391329 | 1.526493 | 0.0041237115 | 0.12933022 | 1 | 7545 | tags=51%, list=32%, si | MM_3 |
| GOBP_INNATE_IMMUNE_ | GOBP_INNATE_IMMUNE_RESPONSE_ACTIV | 254 | 0.37473 | 1.5259683 | 0 | 0.1297197 | 1 | 7581 | tags=46%, list=32%, si | MM_3 |
| GOBP_KETONE_BIOSYNTI | GOBP_KETONE_BIOSYNTHETIC_PROCESS | 45 | 0.436407 | 1.525792 | 0.018115941 | 0.1297249 | 1 | 4997 | tags=40%, list=21%, si | MM_3 |
| GOBP_LOCOMOTOR_RHY | GOBP_LOCOMOTOR_RHYTHM | 20 | 0.518455 | 1.5247594 | 0.04359313 | 0.13064738 | 1 | 1078 | tags=20%, list=5%, sig | MM_3 |
| GOBP_MITOTIC_RECOMB | GOBP_MITOTIC_RECOMBINATION | 24 | 0.502001 | 1.5246689 | 0.044795785 | 0.13057266 | 1 | 6905 | tags=63%, list=29%, si | MM_3 |
| GOBP_REGULATION_OF_ | GOBP_REGULATION_OF_DNA_METABOLIC_ | 491 | 0.367156 | 1.52443 | 0 | 0.13066971 | 1 | 7542 | tags=46%, list=32%, si | MM_3 |
| NF-KAPPA_B_SIGNALING_ | NF-KAPPA_B_SIGNALING_PATHWAY | 93 | 0.401948 | 1.5244136 | 0.00856531 | 0.13049662 | 1 | 6426 | tags=44%, list=27%, si | MM_3 |
| GOBP_RIBOSOMAL_LARG | GOBP_RIBOSOMAL_LARGE_SUBUNIT_BIOGE | 64 | 0.424712 | 1.5243952 | 0.018994413 | 0.13034253 | 1 | 7202 | tags=58%, list=31%, si | MM_3 |
| PROPANOATE_METABOLI | PROPANOATE_METABOLISM | 29 | 0.486215 | 1.5241739 | 0.034739453 | 0.13038115 | 1 | 4994 | tags=45%, list=21%, si | MM_3 |
| GLYOXYLATE_AND_DICAR | GLYOXYLATE_AND_DICARBOXYLATE_METAE | 25 | 0.501903 | 1.5239624 | 0.036363635 | 0.13045363 | 1 | 4994 | tags=52%, list=21%, si | MM_3 |
| GOBP_CHEMOKINE_PROI | GOBP_CHEMOKINE_PRODUCTION | 119 | 0.392677 | 1.5232208 | 0.009394572 | 0.13112654 | 1 | 4393 | tags=27%, list=19%, si | MM_3 |
| GOBP_REGULATION_OF_ | GOBP_REGULATION_OF_EXECUTION_PHASE | 26 | 0.480217 | 1.523115 | 0.037419356 | 0.13105462 | 1 | 2871 | tags=27%, list=12%, si | MM_3 |
| GOBP_POSITIVE_REGULA | GOBP_POSITIVE_REGULATION_OF_PROTEO | 325 | 0.370933 | 1.5230325 | 0 | 0.13096012 | 1 | 7534 | tags=50%, list=32%, si | MM_3 |
| GOBP_DEFENSE_RESPON | GOBP_DEFENSE_RESPONSE_TO_GRAM_NEG | 68 | 0.415818 | 1.5226468 | 0.018223235 | 0.1311389 | 1 | 3407 | tags=22%, list=14%, si | MM_3 |
| GOBP_RETROGRADE_TRA | GOBP_RETROGRADE_TRANSPORT_ENDOSO | 87 | 0.40847 | 1.5222679 | 0.00989011 | 0.13139078 | 1 | 7571 | tags=53%, list=32%, si | MM_3 |
| GOBP_REGULATION_OF_ | GOBP_REGULATION_OF_PROTEIN_BINDING | 188 | 0.382869 | 1.5220472 | 0.003051882 | 0.13142906 | 1 | 5764 | tags=35%, list=24%, si | MM_3 |
| GOBP_PEPTIDE_HORMON | GOBP_PEPTIDE_HORMONE_PROCESSING | 24 | 0.499642 | 1.5219848 | 0.037593983 | 0.13132358 | 1 | 4588 | tags=38%, list=19%, si | MM_3 |
| GOBP_POSITIVE_REGULA | GOBP_POSITIVE_REGULATION_OF_PROTEO | 120 | 0.393907 | 1.5217838 | 0.0062695923 | 0.13138835 | 1 | 6369 | tags=45%, list=27%, si | MM_3 |
| GOBP_REGULATION_OF_ | GOBP_REGULATION_OF_CELLULAR_RESPON | 26 | 0.491839 | 1.520698 | 0.027061855 | 0.1324196 | 1 | 5485 | tags=50%, list=23%, si | MM_3 |
| GOBP_RECEPTOR_MEDIA | GOBP_RECEPTOR_MEDIATED_ENDOCYTOSIS | 269 | 0.374476 | 1.5201424 | 0 | 0.132835 | 1 | 7149 | tags=43%, list=30%, si | MM_3 |
| GOBP_RAS_PROTEIN_SIG | GOBP_RAS_PROTEIN_SIGNAL_TRANSDUCTI | 112 | 0.390662 | 1.5187238 | 0.011727079 | 0.13419367 | 1 | 5840 | tags=36%, list=25%, si | MM_3 |
| GOBP_NEGATIVE_REGUL | GOBP_NEGATIVE_REGULATION_OF_CELL_G | 184 | 0.377343 | 1.5186733 | 0.0010266941 | 0.13405572 | 1 | 7289 | tags=43%, list=31%, si | MM_3 |
| GOBP_NEGATIVE_REGUL | GOBP_NEGATIVE_REGULATION_OF_APOPT | 241 | 0.373341 | 1.5184778 | 0.0010050251 | 0.13410664 | 1 | 6055 | tags=38%, list=26%, si | MM_3 |
| GOBP_POSITIVE_REGULA | GOBP_POSITIVE_REGULATION_OF_MEMBRA | 17 | 0.544269 | 1.5182414 | 0.046831954 | 0.13420062 | 1 | 6192 | tags=53%, list=26%, si | MM_3 |
| GOBP_ERYTHROCYTE_DE | GOBP_ERYTHROCYTE_DEVELOPMENT | 50 | 0.442008 | 1.5180683 | 0.02366864 | 0.13423178 | 1 | 4375 | tags=34%, list=19%, si | MM_3 |
| CELLULAR_SENESCENCE | CELLULAR_SENESCENCE | 159 | 0.38343 | 1.5177952 | 0.002074689 | 0.1343531 | 1 | 6953 | tags=48%, list=29%, si | MM_3 |
| GOBP_PROTON_TRANSP | GOBP_PROTON_TRANSPORTING_TWO_SEC | 15 | 0.557951 | 1.517549 | 0.05293276 | 0.13441719 | 1 | 2286 | tags=27%, list=10%, si | MM_3 |
| GOBP_REGULATION_OF_ | GOBP_REGULATION_OF_VASCULOGENESIS | 16 | 0.546827 | 1.5171938 | 0.056241427 | 0.13462402 | 1 | 5151 | tags=50%, list=22%, si | MM_3 |
| GOBP_RRNA_METHYLATI | GOBP_RRNA_METHYLATION | 20 | 0.511713 | 1.5169767 | 0.04605263 | 0.13469876 | 1 | 6935 | tags=60%, list=29%, si | MM_3 |
| GOBP_VIRAL_TRANSCRIP | GOBP_VIRAL_TRANSCRIPTION | 55 | 0.431745 | 1.516842 | 0.021372328 | 0.13465576 | 1 | 6991 | tags=44%, list=30%, si | MM_3 |
| GOBP_POSITIVE_REGULA | GOBP_POSITIVE_REGULATION_OF_DENDRIT | 67 | 0.421668 | 1.5166919 | 0.01793722 | 0.13462816 | 1 | 7337 | tags=45%, list=31%, si | MM_3 |
| GOBP_REGULATION_OF_ | GOBP_REGULATION_OF_ADAPTIVE_IMMUN | 215 | 0.37492 | 1.516161 | 0.0030800821 | 0.13506566 | 1 | 6059 | tags=38%, list=26%, si | MM_3 |
| GOBP_ASTROCYTE_ACTIV | GOBP_ASTROCYTE_ACTIVATION | 21 | 0.510367 | 1.5161234 | 0.046666667 | 0.13492435 | 1 | 6781 | tags=43%, list=29%, si | MM_3 |
| GOBP_CELLULAR_RESPON | GOBP_CELLULAR_RESPONSE_TO_TOPOLOG | 95 | 0.399124 | 1.5160154 | 0.010857764 | 0.13487704 | 1 | 6670 | tags=47%, list=28%, si | MM_3 |
| GOBP_LOCOMOTION_IN | GOBP_LOCOMOTION_INVOLVED_IN_LOCON | 16 | 0.544725 | 1.5152854 | 0.0585034 | 0.13548175 | 1 | 5892 | tags=50%, list=25%, si | MM_3 |
| GOBP_REGULATION_OF_ | GOBP_REGULATION_OF_MYELOID_LEUKOC | 140 | 0.387161 | 1.5150259 | 0.010351967 | 0.13560426 | 1 | 6330 | tags=39%, list=27%, si | MM_3 |
| GOBP_RETICULOPHAGY | GOBP_RETICULOPHAGY | 19 | 0.528027 | 1.514639 | 0.040789474 | 0.13585787 | 1 | 7912 | tags=68%, list=34%, si | MM_3 |
| GOBP_IMMUNE_RESPON | GOBP_IMMUNE_RESPONSE_TO_TUMOR_CE | 29 | 0.47248 | 1.5144705 | 0.04255319 | 0.13586694 | 1 | 4364 | tags=34%, list=18%, si | MM_3 |
| GOBP_SYMBIONT_ENTRY | GOBP_SYMBIONT_ENTRY_INTO_HOST | 97 | 0.402865 | 1.5144433 | 0.00855615 | 0.13572158 | 1 | 6526 | tags=46%, list=28%, si | MM_3 |
| GOBP_PROTEIN_AUTOPR | GOBP_PROTEIN_AUTOPROCESSING | 23 | 0.507816 | 1.5142481 | 0.05470738 | 0.13579234 | 1 | 2913 | tags=30%, list=12%, si | MM_3 |
| GOBP_NEGATIVE_REGUL | GOBP_NEGATIVE_REGULATION_OF_AXON_ | 16 | 0.551239 | 1.5141164 | 0.06469761 | 0.13575263 | 1 | 5716 | tags=44%, list=24%, si | MM_3 |
| GOBP_REGULATION_OF_ | GOBP_REGULATION_OF_ARP2_3_COMPLEX | 27 | 0.491683 | 1.5137284 | 0.0396927 | 0.13602343 | 1 | 4667 | tags=37%, list=20%, si | MM_3 |
| GOBP_RESPONSE_TO_TY | GOBP_RESPONSE_TO_TYPE_II_INTERFERON | 122 | 0.389132 | 1.5134878 | 0.0063626724 | 0.13611275 | 1 | 7649 | tags=53%, list=32%, si | MM_3 |
| GOBP_REGULATION_OF_ | GOBP_REGULATION_OF_CHROMOSOME_SE | 126 | 0.387279 | 1.5128145 | 0.0052687037 | 0.13676699 | 1 | 6708 | tags=39%, list=28%, si | MM_3 |
| GOBP_DNA_REPLICATION | GOBP_DNA_REPLICATION | 263 | 0.366093 | 1.5127326 | 0 | 0.1366814 | 1 | 6446 | tags=43%, list=27%, si | MM_3 |
| GOBP_ACTIN_NUCLEATIO | GOBP_ACTIN_NUCLEATION | 58 | 0.423263 | 1.5125166 | 0.019406393 | 0.13674766 | 1 | 7945 | tags=60%, list=34%, si | MM_3 |
| GOBP_REGULATION_OF_ | GOBP_REGULATION_OF_MUSCLE_HYPERTR | 80 | 0.407172 | 1.512434 | 0.013245033 | 0.13666904 | 1 | 7322 | tags=48%, list=31%, si | MM_3 |
| GOBP_REGULATION_OF_ | GOBP_REGULATION_OF_RECEPTOR_MEDIAT | 132 | 0.385686 | 1.5116653 | 0.008316008 | 0.13737443 | 1 | 6781 | tags=41%, list=29%, si | MM_3 |
| GOBP_LEUKOCYTE_HOMI | GOBP_LEUKOCYTE_HOMEOSTASIS | 137 | 0.38837 | 1.5116585 | 0.0031347962 | 0.1372047 | 1 | 4974 | tags=34%, list=21%, si | MM_3 |
| GOBP_ESTABLISHMENT_C | GOBP_ESTABLISHMENT_OF_PROTEIN_LOCA | 262 | 0.371171 | 1.5115969 | 0.001019368 | 0.13708675 | 1 | 6561 | tags=42%, list=28%, si | MM_3 |
| GOBP_RNA_SPLICING | GOBP_RNA_SPLICING | 417 | 0.367187 | 1.5112027 | 0 | 0.13736346 | 1 | 7551 | tags=46%, list=32%, si | MM_3 |
| GOBP_RESPONSE_TO_GR | GOBP_RESPONSE_TO_GROWTH_HORMONE | 20 | 0.518778 | 1.5107088 | 0.05 | 0.13770543 | 1 | 6158 | tags=40%, list=26%, si | MM_3 |
| GOBP_ERYTHROCYTE_HO | GOBP_ERYTHROCYTE_HOMEOSTASIS | 163 | 0.377377 | 1.5106387 | 0.0020597323 | 0.13760741 | 1 | 7574 | tags=47%, list=32%, si | MM_3 |
| GOBP_POSITIVE_REGULA | GOBP_POSITIVE_REGULATION_OF_DEFENSE | 480 | 0.361333 | 1.5103319 | 0 | 0.13776152 | 1 | 6329 | tags=36%, list=27%, si | MM_3 |
| GOBP_NEGATIVE_REGUL | GOBP_NEGATIVE_REGULATION_OF_POST_T | 94 | 0.405364 | 1.5097622 | 0.010683761 | 0.13825679 | 1 | 7206 | tags=50%, list=31%, si | MM_3 |
| GOBP_PHOSPHOLIPID_BI | GOBP_PHOSPHOLIPID_BIOSYNTHETIC_PROC | 214 | 0.373744 | 1.509516 | 0.002038736 | 0.1383498 | 1 | 4804 | tags=31%, list=20%, si | MM_3 |

| | | | | | | | | | | |
|---|---|---|---|---|---|---|---|---|---|---|
| GOBP_POSITIVE_REGULA | GOBP_POSITIVE_REGULATION_OF_RECEPTO | 19 | 0.522744 | 1.5092002 | 0.061842106 | 0.13851058 | 1 | 5518 | tags=53%, list=23%, si | MM_3 |
| GOBP_NON_CANONICAL_ | GOBP_NON_CANONICAL_NF_KAPPAB_SIGN | 115 | 0.396518 | 1.5091687 | 0.012711864 | 0.13837887 | 1 | 7953 | tags=54%, list=34%, si | MM_3 |
| GOBP_SULFUR_COMPOU | GOBP_SULFUR_COMPOUND_TRANSPORT | 49 | 0.440889 | 1.5090295 | 0.021101993 | 0.13837782 | 1 | 4687 | tags=31%, list=20%, si | MM_3 |
| APOPTOSIS | APOPTOSIS | 137 | 0.386735 | 1.5088121 | 0.00524109 | 0.13845818 | 1 | 5583 | tags=36%, list=24%, si | MM_3 |
| GOBP_NEURON_PROJECT | GOBP_NEURON_PROJECTION_REGENERATI | 68 | 0.409945 | 1.5080317 | 0.020594966 | 0.13913615 | 1 | 6810 | tags=43%, list=29%, si | MM_3 |
| GOBP_REGULATION_OF_ | GOBP_REGULATION_OF_MRNA_METABOLIC | 286 | 0.370399 | 1.5077399 | 0 | 0.13928969 | 1 | 7780 | tags=49%, list=33%, si | MM_3 |
| GOBP_REGULATION_OF_ | GOBP_REGULATION_OF_SYNAPTIC_VESICLE | 19 | 0.527314 | 1.5072662 | 0.04435995 | 0.13962561 | 1 | 4975 | tags=47%, list=21%, si | MM_3 |
| GOBP_NEGATIVE_REGUL | GOBP_NEGATIVE_REGULATION_OF_PROTEI | 236 | 0.371207 | 1.5070275 | 0.0010162601 | 0.1397129 | 1 | 7564 | tags=49%, list=32%, si | MM_3 |
| GOBP_CYTOKINE_PRODU | GOBP_CYTOKINE_PRODUCTION_INVOLVED_ | 63 | 0.422841 | 1.506757 | 0.019362187 | 0.139829 | 1 | 3253 | tags=21%, list=14%, si | MM_3 |
| GOBP_VACUOLAR_ACIDI | GOBP_VACUOLAR_ACIDIFICATION | 27 | 0.482896 | 1.5065403 | 0.044987146 | 0.13988969 | 1 | 6717 | tags=59%, list=28%, si | MM_3 |
| GOBP_NEGATIVE_REGUL | GOBP_NEGATIVE_REGULATION_OF_CATABO | 339 | 0.366297 | 1.5061042 | 0 | 0.14023642 | 1 | 7530 | tags=49%, list=32%, si | MM_3 |
| GOBP_TELOMERE_MAINT | GOBP_TELOMERE_MAINTENANCE_VIA_REC | 15 | 0.543506 | 1.5058964 | 0.051175658 | 0.14030555 | 1 | 5012 | tags=60%, list=21%, si | MM_3 |
| GOBP_RIBOSOMAL_SMAI | GOBP_RIBOSOMAL_SMALL_SUBUNIT_BIOGI | 101 | 0.400003 | 1.5054575 | 0.009625669 | 0.14056179 | 1 | 7577 | tags=53%, list=32%, si | MM_3 |
| GOBP_POSITIVE_REGULA | GOBP_POSITIVE_REGULATION_OF_HYDROL | 427 | 0.363906 | 1.5053872 | 0 | 0.14045821 | 1 | 7415 | tags=48%, list=31%, si | MM_3 |
| GOBP_POSITIVE_REGULA | GOBP_POSITIVE_REGULATION_OF_CYTOKIN | 27 | 0.489486 | 1.5048949 | 0.05429293 | 0.14083138 | 1 | 3253 | tags=26%, list=14%, si | MM_3 |
| GOBP_CRANIAL_SKELETA | GOBP_CRANIAL_SKELETAL_SYSTEM_DEVELO | 76 | 0.403307 | 1.5046221 | 0.025871767 | 0.14094797 | 1 | 2312 | tags=20%, list=10%, si | MM_3 |
| GOBP_INTRINSIC_APOPT | GOBP_INTRINSIC_APOPTOTIC_SIGNALING_I | 314 | 0.361639 | 1.5042105 | 0 | 0.14124116 | 1 | 7148 | tags=46%, list=30%, si | MM_3 |
| GOBP_MYELOID_DENDRI | GOBP_MYELOID_DENDRITIC_CELL_ACTIVAT | 37 | 0.454219 | 1.5040377 | 0.035151515 | 0.14126147 | 1 | 4975 | tags=38%, list=21%, si | MM_3 |
| GOBP_NUCLEOSOME_AS | GOBP_NUCLEOSOME_ASSEMBLY | 50 | 0.43567 | 1.5037512 | 0.038372092 | 0.1414006 | 1 | 5104 | tags=38%, list=22%, si | MM_3 |
| GOBP_AMYLOID_PRECUR | GOBP_AMYLOID_PRECURSOR_PROTEIN_ME | 75 | 0.408742 | 1.5032592 | 0.020765027 | 0.14181969 | 1 | 7235 | tags=45%, list=31%, si | MM_3 |
| GOBP_POSITIVE_REGULA | GOBP_POSITIVE_REGULATION_OF_ADAPTIV | 140 | 0.387153 | 1.5029416 | 0.0103412615 | 0.14200711 | 1 | 7028 | tags=45%, list=30%, si | MM_3 |
| GOBP_REGULATION_OF_ | GOBP_REGULATION_OF_LEUKOCYTE_PROLI | 273 | 0.363873 | 1.502598 | 0 | 0.14223313 | 1 | 4929 | tags=30%, list=21%, si | MM_3 |
| GOBP_ENDOPLASMIC_RE | GOBP_ENDOPLASMIC_RETICULUM_UNFOLD | 69 | 0.411636 | 1.502326 | 0.015555556 | 0.14238206 | 1 | 6670 | tags=48%, list=28%, si | MM_3 |
| GOBP_GLIAL_CELL_PROLI | GOBP_GLIAL_CELL_PROLIFERATION | 78 | 0.403513 | 1.502065 | 0.018518519 | 0.1425253 | 1 | 5751 | tags=41%, list=24%, si | MM_3 |
| GOBP_INTRACELLULAR_P | GOBP_INTRACELLULAR_PROTEIN_TRANSME | 32 | 0.469141 | 1.5019326 | 0.04045512 | 0.14247905 | 1 | 6379 | tags=47%, list=27%, si | MM_3 |
| FERROPTOSIS | FERROPTOSIS | 37 | 0.454283 | 1.5016303 | 0.039806996 | 0.14262038 | 1 | 7329 | tags=59%, list=31%, si | MM_3 |
| GOBP_POSITIVE_REGULA | GOBP_POSITIVE_REGULATION_OF_RESPONS | 381 | 0.361869 | 1.5015706 | 0 | 0.14250815 | 1 | 6329 | tags=34%, list=27%, si | MM_3 |
| GOBP_TAXIS | GOBP_TAXIS | 411 | 0.364668 | 1.5012801 | 0 | 0.14266497 | 1 | 6009 | tags=36%, list=25%, si | MM_3 |
| GOBP_STRESS_GRANULE_ | GOBP_STRESS_GRANULE_ASSEMBLY | 35 | 0.469536 | 1.4999642 | 0.038694076 | 0.14390405 | 1 | 8301 | tags=63%, list=35%, si | MM_3 |
| GOBP_ZINC_ION_TRANSF | GOBP_ZINC_ION_TRANSPORT | 28 | 0.479298 | 1.4988965 | 0.05108557 | 0.14494286 | 1 | 5884 | tags=43%, list=25%, si | MM_3 |
| B_CELL_RECEPTOR_SIGN/ | B_CELL_RECEPTOR_SIGNALING_PATHWAY | 68 | 0.403957 | 1.4988391 | 0.027964206 | 0.14482972 | 1 | 8684 | tags=65%, list=37%, si | MM_3 |
| GOBP_RESPONSE_TO_OX | GOBP_RESPONSE_TO_OXIDATIVE_STRESS | 364 | 0.363435 | 1.498794 | 0 | 0.14469245 | 1 | 6764 | tags=43%, list=29%, si | MM_3 |
| GOBP_ADULT_WALKING_ | GOBP_ADULT_WALKING_BEHAVIOR | 48 | 0.429076 | 1.4984555 | 0.03172738 | 0.14487343 | 1 | 5526 | tags=35%, list=23%, si | MM_3 |
| CHOLESTEROL_METABOL | CHOLESTEROL_METABOLISM | 44 | 0.43441 | 1.4980978 | 0.040767387 | 0.14510533 | 1 | 7206 | tags=50%, list=31%, si | MM_3 |
| GOBP_POSITIVE_REGULA | GOBP_POSITIVE_REGULATION_OF_MITOTIC | 18 | 0.530874 | 1.4976645 | 0.06754967 | 0.14541222 | 1 | 6905 | tags=61%, list=29%, si | MM_3 |
| GOBP_POSITIVE_REGULA | GOBP_POSITIVE_REGULATION_OF_INTERFE | 45 | 0.43357 | 1.4975786 | 0.04117647 | 0.14535163 | 1 | 5340 | tags=31%, list=23%, si | MM_3 |
| GOBP_AMINO_ACID_ACT | GOBP_AMINO_ACID_ACTIVATION | 26 | 0.487436 | 1.4972984 | 0.045918368 | 0.14547737 | 1 | 4095 | tags=38%, list=17%, si | MM_3 |
| GOBP_MIDBODY_ABSCIS | GOBP_MIDBODY_ABSCISSION | 16 | 0.540353 | 1.4969592 | 0.058583107 | 0.14564753 | 1 | 3916 | tags=38%, list=17%, si | MM_3 |
| GOBP_CELLULAR_ALDEH | GOBP_CELLULAR_ALDEHYDE_METABOLIC_P | 74 | 0.406079 | 1.496843 | 0.03181818 | 0.14561091 | 1 | 4166 | tags=32%, list=18%, si | MM_3 |
| GOBP_POSITIVE_REGULA | GOBP_POSITIVE_REGULATION_OF_LEUKOCY | 152 | 0.37643 | 1.495036 | 0.006243496 | 0.14751384 | 1 | 6192 | tags=36%, list=26%, si | MM_3 |
| GOBP_REGULATION_OF_ | GOBP_REGULATION_OF_APOPTOTIC_SIGNA | 409 | 0.363017 | 1.4949042 | 0 | 0.14748257 | 1 | 7534 | tags=49%, list=32%, si | MM_3 |
| GOBP_REGULATION_OF_ | GOBP_REGULATION_OF_AMYLOID_BETA_F | 37 | 0.448208 | 1.4947683 | 0.04572804 | 0.14745162 | 1 | 7983 | tags=51%, list=34%, si | MM_3 |
| GOBP_NEGATIVE_REGUL | GOBP_NEGATIVE_REGULATION_OF_DNA_M | 127 | 0.385312 | 1.494578 | 0.015957447 | 0.14749928 | 1 | 5012 | tags=31%, list=21%, si | MM_3 |
| GOBP_REGULATION_OF_ | GOBP_REGULATION_OF_IMMUNE_EFFECTO | 406 | 0.359972 | 1.4944626 | 0 | 0.14744036 | 1 | 6140 | tags=35%, list=26%, si | MM_3 |
| GOBP_PROTEIN_AUTOUB | GOBP_PROTEIN_AUTOUBIQUITINATION | 68 | 0.417442 | 1.4944624 | 0.029445074 | 0.14726286 | 1 | 6590 | tags=43%, list=28%, si | MM_3 |
| GOBP_REGULATION_OF_ | GOBP_REGULATION_OF_MRNA_CATABOLIC | 175 | 0.375921 | 1.4944526 | 0.0020618557 | 0.14709337 | 1 | 7545 | tags=46%, list=32%, si | MM_3 |
| GOBP_POSITIVE_REGULA | GOBP_POSITIVE_REGULATION_OF_DEPHOS | 53 | 0.425643 | 1.4937917 | 0.03490991 | 0.14760767 | 1 | 7299 | tags=55%, list=31%, si | MM_3 |
| GOBP_REGULATION_OF_ | GOBP_REGULATION_OF_CALCINEURIN_MEI | 40 | 0.441637 | 1.4936949 | 0.037214886 | 0.14753014 | 1 | 5892 | tags=43%, list=25%, si | MM_3 |
| GOBP_POSITIVE_REGULA | GOBP_POSITIVE_REGULATION_OF_FILOPOD | 35 | 0.456471 | 1.4935828 | 0.032019705 | 0.14748044 | 1 | 6853 | tags=49%, list=29%, si | MM_3 |
| HIF-1_SIGNALING_PATHV | HIF-1_SIGNALING_PATHWAY | 98 | 0.394421 | 1.4931338 | 0.010787486 | 0.14777884 | 1 | 5583 | tags=38%, list=24%, si | MM_3 |
| GOBP_MONONUCLEAR_C | GOBP_MONONUCLEAR_CELL_MIGRATION | 199 | 0.368823 | 1.4928864 | 0.0020325202 | 0.14787808 | 1 | 7056 | tags=45%, list=30%, si | MM_3 |
| GOBP_POSITIVE_REGULA | GOBP_POSITIVE_REGULATION_OF_DNA_ME | 314 | 0.3647 | 1.4923103 | 0 | 0.14832345 | 1 | 7435 | tags=46%, list=31%, si | MM_3 |
| GOBP_REGULATION_OF_ | GOBP_REGULATION_OF_OLIGODENDROCYT | 51 | 0.423592 | 1.4922436 | 0.028103044 | 0.14822042 | 1 | 1864 | tags=18%, list=8%, sig | MM_3 |
| GOBP_VESICLE_MEDIATE | GOBP_VESICLE_MEDIATED_TRANSPORT_TO | 158 | 0.373094 | 1.4922224 | 0.005165289 | 0.14807017 | 1 | 7111 | tags=46%, list=30%, si | MM_3 |
| GOBP_NEGATIVE_REGUL | GOBP_NEGATIVE_REGULATION_OF_TELOM | 36 | 0.45705 | 1.4920167 | 0.047503047 | 0.14814995 | 1 | 6708 | tags=47%, list=28%, si | MM_3 |
| GOBP_REGULATION_OF_ | GOBP_REGULATION_OF_DNA_DAMAGE_RE | 36 | 0.45707 | 1.4918576 | 0.058753 | 0.14814733 | 1 | 5526 | tags=56%, list=23%, si | MM_3 |
| GOBP_FC_RECEPTOR_ME | GOBP_FC_RECEPTOR_MEDIATED_STIMULAT | 17 | 0.531965 | 1.4915361 | 0.050561797 | 0.14834772 | 1 | 7473 | tags=59%, list=32%, si | MM_3 |
| GOBP_VIRAL_PROCESS | GOBP_VIRAL_PROCESS | 303 | 0.3645 | 1.4913924 | 0.001003009 | 0.1483212 | 1 | 7549 | tags=49%, list=32%, si | MM_3 |
| GOBP_INTERLEUKIN_1_P | GOBP_INTERLEUKIN_1_PRODUCTION | 114 | 0.387622 | 1.4910707 | 0.011640212 | 0.14849472 | 1 | 8291 | tags=53%, list=35%, si | MM_3 |
| GOBP_REGULATION_OF_ | GOBP_REGULATION_OF_CELLULAR_RESPON | 141 | 0.380946 | 1.490357 | 0.0062370063 | 0.14915591 | 1 | 6781 | tags=41%, list=29%, si | MM_3 |
| GOBP_CELLULAR_RESPO | GOBP_CELLULAR_RESPONSE_TO_BIOTIC_ST | 224 | 0.366912 | 1.4903127 | 0.0010090817 | 0.14902197 | 1 | 7534 | tags=47%, list=32%, si | MM_3 |
| GOBP_NEGATIVE_REGUL | GOBP_NEGATIVE_REGULATION_OF_PHOSPH | 25 | 0.484493 | 1.4894732 | 0.055837564 | 0.14988743 | 1 | 2065 | tags=24%, list=9%, sig | MM_3 |
| GOBP_NEGATIVE_REGUL | GOBP_NEGATIVE_REGULATION_OF_MAP_K | 49 | 0.420657 | 1.4891343 | 0.047281325 | 0.15009059 | 1 | 6567 | tags=47%, list=28%, si | MM_3 |
| GOBP_REGULATION_OF_ | GOBP_REGULATION_OF_CELL_CYCLE_PHAS | 424 | 0.357531 | 1.4890261 | 0 | 0.1500424 | 1 | 7175 | tags=44%, list=30%, si | MM_3 |
| GOBP_NEGATIVE_REGUL | GOBP_NEGATIVE_REGULATION_OF_PROTEI | 80 | 0.403449 | 1.4885793 | 0.018418202 | 0.1503709 | 1 | 6069 | tags=44%, list=26%, si | MM_3 |
| GOBP_AMINO_ACID_ME | GOBP_AMINO_ACID_METABOLIC_PROCESS | 245 | 0.367899 | 1.4877476 | 0 | 0.15124074 | 1 | 5181 | tags=35%, list=22%, si | MM_3 |
| GOBP_REGULATION_OF_ | GOBP_REGULATION_OF_NEURON_PROJECT | 43 | 0.436977 | 1.4877189 | 0.0405562 | 0.15109047 | 1 | 8710 | tags=63%, list=37%, si | MM_3 |
| GOBP_NEGATIVE_REGUL | GOBP_NEGATIVE_REGULATION_OF_CELL_D | 293 | 0.360456 | 1.4876792 | 0 | 0.15095776 | 1 | 5906 | tags=32%, list=25%, si | MM_3 |
| GOBP_RESPONSE_TO_TU | GOBP_RESPONSE_TO_TUMOR_NECROSIS_F | 167 | 0.376362 | 1.4874154 | 0.0010235414 | 0.15112491 | 1 | 6640 | tags=43%, list=28%, si | MM_3 |
| GOBP_REGULATION_OF_ | GOBP_REGULATION_OF_VASCULATURE_DE | 295 | 0.362822 | 1.4872695 | 0 | 0.15110894 | 1 | 5788 | tags=37%, list=25%, si | MM_3 |
| GOBP_REGULATION_OF_ | GOBP_REGULATION_OF_FATTY_ACID_OXID | 41 | 0.447651 | 1.4863547 | 0.051373955 | 0.15199743 | 1 | 3188 | tags=24%, list=14%, si | MM_3 |
| GOBP_POSITIVE_REGULA | GOBP_POSITIVE_REGULATION_OF_B_CELL_ | 90 | 0.397537 | 1.4853489 | 0.016304348 | 0.15300955 | 1 | 5024 | tags=29%, list=21%, si | MM_3 |
| GOBP_PRODUCTION_OF_ | GOBP_PRODUCTION_OF_MOLECULAR_MED | 270 | 0.364761 | 1.4852496 | 0 | 0.1529425 | 1 | 5024 | tags=28%, list=21%, si | MM_3 |
| GOBP_STRIATED_MUSCL | GOBP_STRIATED_MUSCLE_CELL_APOPTOTI | 73 | 0.405072 | 1.4852182 | 0.02724177 | 0.15279385 | 1 | 5470 | tags=33%, list=23%, si | MM_3 |
| GOBP_POSITIVE_REGULA | GOBP_POSITIVE_REGULATION_OF_VIRAL_P | 64 | 0.418341 | 1.4850343 | 0.020134227 | 0.1528491 | 1 | 7383 | tags=52%, list=31%, si | MM_3 |
| GOBP_INHIBITORY_POST | GOBP_INHIBITORY_POSTSYNAPTIC_POTENT | 26 | 0.490245 | 1.48444 | 0.061277706 | 0.15332823 | 1 | 3677 | tags=35%, list=16%, si | MM_3 |
| GOBP_POSITIVE_REGULA | GOBP_POSITIVE_REGULATION_OF_ENDOTH | 99 | 0.391793 | 1.4842993 | 0.01945946 | 0.15331332 | 1 | 4842 | tags=33%, list=21%, si | MM_3 |
| GOBP_ACETYL_COA_BIOS | GOBP_ACETYL_COA_BIOSYNTHETIC_PROCES | 19 | 0.529196 | 1.4837372 | 0.06084656 | 0.15379316 | 1 | 6841 | tags=58%, list=29%, si | MM_3 |
| GOBP_REGULATION_OF_ | GOBP_REGULATION_OF_CELL_SHAPE | 137 | 0.378944 | 1.4836808 | 0.0072916667 | 0.15368457 | 1 | 6406 | tags=41%, list=27%, si | MM_3 |
| GOBP_MITOCHONDRIAL_ | GOBP_MITOCHONDRIAL_ELECTRON_TRANS | 29 | 0.475715 | 1.4835472 | 0.05283505 | 0.15365969 | 1 | 8099 | tags=55%, list=34%, si | MM_3 |
| GOBP_INTEGRIN_ACTIVA | GOBP_INTEGRIN_ACTIVATION | 26 | 0.486988 | 1.4835376 | 0.05148005 | 0.15349086 | 1 | 3007 | tags=27%, list=13%, si | MM_3 |
| GOBP_LYMPHOCYTE_MIG | GOBP_LYMPHOCYTE_MIGRATION | 92 | 0.401084 | 1.4831582 | 0.014008621 | 0.1537709 | 1 | 5751 | tags=36%, list=24%, si | MM_3 |

| | | | | | | | | | | |
|---|---|---|---|---|---|---|---|---|---|---|
| UBIQUITIN_MEDIATED_P | UBIQUITIN_MEDIATED_PROTEOLYSIS | 134 | 0.382172 | 1.4828511 | 0.008333334 | 0.15392838 | 1 | 8310 | tags=54%, list=35%, si | MM_3 |
| GOBP_TRNA_DECAY | GOBP_TRNA_DECAY | 16 | 0.54005 | 1.4827746 | 0.048846677 | 0.15385863 | 1 | 3784 | tags=44%, list=16%, si | MM_3 |
| GOBP_TETRAPYRROLE_M | GOBP_TETRAPYRROLE_METABOLIC_PROCES | 49 | 0.430798 | 1.4825798 | 0.024475524 | 0.15391944 | 1 | 6684 | tags=47%, list=28%, si | MM_3 |
| GOBP_ISOTYPE_SWITCHI | GOBP_ISOTYPE_SWITCHING_TO_IGG_ISOTY | 18 | 0.515617 | 1.4823066 | 0.066390045 | 0.1540643 | 1 | 2916 | tags=33%, list=12%, si | MM_3 |
| GOBP_REGULATION_OF_ | GOBP_REGULATION_OF_LEUKOCYTE_CHEM | 122 | 0.383045 | 1.4821872 | 0.008403362 | 0.15403682 | 1 | 7056 | tags=44%, list=30%, si | MM_3 |
| GOBP_VENOUS_BLOOD_ | GOBP_VENOUS_BLOOD_VESSEL_DEVELOPM | 18 | 0.526414 | 1.4820805 | 0.0630749 | 0.15399627 | 1 | 5781 | tags=44%, list=24%, si | MM_3 |
| GALACTOSE_METABOLISN | GALACTOSE_METABOLISM | 29 | 0.475502 | 1.482033 | 0.060529634 | 0.15387782 | 1 | 3953 | tags=31%, list=17%, si | MM_3 |
| GOBP_MONOSACCHARID | GOBP_MONOSACCHARIDE_BIOSYNTHETIC_ | 93 | 0.390985 | 1.481653 | 0.027085591 | 0.1541357 | 1 | 5216 | tags=35%, list=22%, si | MM_3 |
| GOBP_REGULATION_OF_ | GOBP_REGULATION_OF_SMOOTH_MUSCLE | 37 | 0.451661 | 1.4814845 | 0.042475726 | 0.15415153 | 1 | 4601 | tags=30%, list=19%, si | MM_3 |
| PURINE_METABOLISM | PURINE_METABOLISM | 126 | 0.385338 | 1.4814466 | 0.010638298 | 0.1540261 | 1 | 4288 | tags=28%, list=18%, si | MM_3 |
| GOBP_ORGANELLE_INHE | GOBP_ORGANELLE_INHERITANCE | 15 | 0.550782 | 1.4813834 | 0.05263158 | 0.1539211 | 1 | 6479 | tags=60%, list=27%, si | MM_3 |
| GOBP_DNA_DAMAGE_RE | GOBP_DNA_DAMAGE_RESPONSE_SIGNAL_ | 17 | 0.533641 | 1.4811788 | 0.06775068 | 0.15397128 | 1 | 7135 | tags=71%, list=30%, si | MM_3 |
| GOBP_REGULATION_OF_ | GOBP_REGULATION_OF_CELLULAR_RESPIRA | 52 | 0.424144 | 1.4811608 | 0.03202847 | 0.15381911 | 1 | 6112 | tags=42%, list=26%, si | MM_3 |
| GOBP_RESPONSE_TO_TO | GOBP_RESPONSE_TO_TOPOLOGICALLY_INC | 127 | 0.379883 | 1.4807823 | 0.008412197 | 0.15409712 | 1 | 6027 | tags=39%, list=26%, si | MM_3 |
| GOBP_REGULATORY_NCR | GOBP_REGULATORY_NCRNA_MEDIATED_GI | 116 | 0.379698 | 1.4805373 | 0.013626864 | 0.15423624 | 1 | 6341 | tags=37%, list=27%, si | MM_3 |
| GOBP_HISTONE_MRNA_ | GOBP_HISTONE_MRNA_METABOLIC_PROCE | 20 | 0.515896 | 1.4804512 | 0.06447368 | 0.1541734 | 1 | 3091 | tags=30%, list=13%, si | MM_3 |
| GOBP_TOLL_LIKE_RECEPT | GOBP_TOLL_LIKE_RECEPTOR_3_SIGNALING_ | 29 | 0.471837 | 1.4802573 | 0.052434456 | 0.15421264 | 1 | 5903 | tags=48%, list=25%, si | MM_3 |
| GOBP_REGULATION_OF_ | GOBP_REGULATION_OF_GLIAL_CELL_PROLI | 56 | 0.415853 | 1.4795694 | 0.040677965 | 0.15486912 | 1 | 5751 | tags=39%, list=24%, si | MM_3 |
| GOBP_RESOLUTION_OF_ | GOBP_RESOLUTION_OF_MEIOTIC_RECOMBI | 17 | 0.520177 | 1.4794573 | 0.06097561 | 0.15482934 | 1 | 3759 | tags=35%, list=16%, si | MM_3 |
| GOBP_CELL_KILLING | GOBP_CELL_KILLING | 202 | 0.367291 | 1.4793929 | 0.0040983604 | 0.1547364 | 1 | 5255 | tags=32%, list=22%, si | MM_3 |
| GOBP_PEPTIDYL_LYSINE_ | GOBP_PEPTIDYL_LYSINE_ACETYLATION | 25 | 0.485749 | 1.479388 | 0.05321782 | 0.1545635 | 1 | 6761 | tags=44%, list=29%, si | MM_3 |
| GOBP_REGULATION_OF_ | GOBP_REGULATION_OF_PROTEIN_SERINE_ | 280 | 0.3604 | 1.4793812 | 0.002016129 | 0.15439336 | 1 | 7408 | tags=44%, list=31%, si | MM_3 |
| GOBP_GLOMERULAR_ME | GOBP_GLOMERULAR_MESANGIUM_DEVELC | 16 | 0.542215 | 1.479363 | 0.07027027 | 0.15423892 | 1 | 3680 | tags=38%, list=16%, si | MM_3 |
| GOBP_TRANSITION_MET | GOBP_TRANSITION_METAL_ION_TRANSPOR | 95 | 0.386365 | 1.4790894 | 0.016163792 | 0.15440318 | 1 | 6393 | tags=42%, list=27%, si | MM_3 |
| AFRICAN_TRYPANOSOMI | AFRICAN_TRYPANOSOMIASIS | 31 | 0.462178 | 1.4784827 | 0.048387095 | 0.15498306 | 1 | 215 | tags=13%, list=1%, sig | MM_3 |
| GLYCOSAMINOGLYCAN_I | GLYCOSAMINOGLYCAN_DEGRADATION | 17 | 0.523607 | 1.4780399 | 0.06374502 | 0.1553061 | 1 | 5217 | tags=53%, list=22%, si | MM_3 |
| GOBP_POLY_A_PLUS_MR | GOBP_POLY_A_PLUS_MRNA_EXPORT_FRON | 21 | 0.510743 | 1.4775724 | 0.06442953 | 0.1557306 | 1 | 2360 | tags=33%, list=10%, si | MM_3 |
| GOBP_CYTOSOLIC_TRANS | GOBP_CYTOSOLIC_TRANSPORT | 124 | 0.376174 | 1.4775587 | 0.0063829785 | 0.15557201 | 1 | 7571 | tags=51%, list=32%, si | MM_3 |
| GOBP_RETINA_VASCULAT | GOBP_RETINA_VASCULATURE_DEVELOPME | 22 | 0.488744 | 1.4772921 | 0.07539683 | 0.15571211 | 1 | 5018 | tags=41%, list=21%, si | MM_3 |
| GOBP_NEGATIVE_REGUL | GOBP_NEGATIVE_REGULATION_OF_GLUCO | 20 | 0.510645 | 1.4764383 | 0.07490145 | 0.15656006 | 1 | 6166 | tags=40%, list=26%, si | MM_3 |
| GOBP_NEURON_MIGRAT | GOBP_NEURON_MIGRATION | 190 | 0.368734 | 1.4762173 | 0 | 0.15664262 | 1 | 5577 | tags=35%, list=24%, si | MM_3 |
| GOBP_PHAGOCYTOSIS | GOBP_PHAGOCYTOSIS | 213 | 0.366991 | 1.4761556 | 0.0030487804 | 0.15652567 | 1 | 7508 | tags=46%, list=32%, si | MM_3 |
| GOBP_MYELOID_CELL_H | GOBP_MYELOID_CELL_HOMEOSTASIS | 209 | 0.366154 | 1.4751055 | 0.0060790274 | 0.15761538 | 1 | 7574 | tags=47%, list=32%, si | MM_3 |
| GOBP_POSITIVE_REGULA | GOBP_POSITIVE_REGULATION_OF_PROTEAS | 102 | 0.386474 | 1.4750032 | 0.019210245 | 0.15757197 | 1 | 7506 | tags=55%, list=32%, si | MM_3 |
| GOBP_CLATHRIN_DEPENI | GOBP_CLATHRIN_DEPENDENT_ENDOCYTOS | 48 | 0.421386 | 1.4749964 | 0.04311377 | 0.15739746 | 1 | 7244 | tags=50%, list=31%, si | MM_3 |
| GOBP_HOMEOSTASIS_OF | GOBP_HOMEOSTASIS_OF_NUMBER_OF_CEI | 48 | 0.421225 | 1.4748772 | 0.043632075 | 0.15737756 | 1 | 6992 | tags=46%, list=30%, si | MM_3 |
| GOBP_CELLULAR_RESPON | GOBP_CELLULAR_RESPONSE_TO_HEAT | 50 | 0.427447 | 1.47487 | 0.03211009 | 0.1572083 | 1 | 7177 | tags=52%, list=30%, si | MM_3 |
| GOBP_ADAPTIVE_IMMUN | GOBP_ADAPTIVE_IMMUNE_RESPONSE | 433 | 0.354257 | 1.47482 | 0.0010050251 | 0.15710196 | 1 | 5024 | tags=30%, list=21%, si | MM_3 |
| GOBP_CYTOKINE_MEDIAT | GOBP_CYTOKINE_MEDIATED_SIGNALING_P | 421 | 0.357677 | 1.4746828 | 0 | 0.1570895 | 1 | 6192 | tags=35%, list=26%, si | MM_3 |
| GOBP_POSITIVE_REGULA | GOBP_POSITIVE_REGULATION_OF_AUTOPH | 19 | 0.509517 | 1.4743905 | 0.06451613 | 0.15724705 | 1 | 1978 | tags=32%, list=8%, sig | MM_3 |
| GOBP_NON_MEMBRANE | GOBP_NON_MEMBRANE_BOUNDED_ORGA | 405 | 0.356806 | 1.473924 | 0 | 0.1576867 | 1 | 7088 | tags=42%, list=30%, si | MM_3 |
| GOBP_DNA_STRAND_ELC | GOBP_DNA_STRAND_ELONGATION | 36 | 0.440548 | 1.4734534 | 0.043956045 | 0.15806769 | 1 | 6689 | tags=53%, list=28%, si | MM_3 |
| GOBP_NUCLEAR_TRANSP | GOBP_NUCLEAR_TRANSPORT | 337 | 0.357661 | 1.4733535 | 0 | 0.15800013 | 1 | 7142 | tags=43%, list=30%, si | MM_3 |
| GOBP_NEGATIVE_REGUL | GOBP_NEGATIVE_REGULATION_OF_ALPHA_ | 16 | 0.534522 | 1.4730006 | 0.053521127 | 0.15829493 | 1 | 4515 | tags=44%, list=19%, si | MM_3 |
| GOBP_INTRINSIC_APOPT | GOBP_INTRINSIC_APOPTOTIC_SIGNALING_I | 112 | 0.381052 | 1.4727987 | 0.0148936175 | 0.15837109 | 1 | 7135 | tags=50%, list=30%, si | MM_3 |
| GOBP_EXTRACELLULAR_S | GOBP_EXTRACELLULAR_STRUCTURE_ORGAI | 309 | 0.361005 | 1.4725752 | 0 | 0.15844916 | 1 | 4635 | tags=31%, list=20%, si | MM_3 |
| IL-17_SIGNALING_PATHW | IL-17_SIGNALING_PATHWAY | 85 | 0.390183 | 1.472072 | 0.025945947 | 0.15885277 | 1 | 5583 | tags=35%, list=24%, si | MM_3 |
| GOBP_LONG_TERM_SYN | GOBP_LONG_TERM_SYNAPTIC_POTENTIATI | 116 | 0.376563 | 1.4717782 | 0.010615711 | 0.15902291 | 1 | 6548 | tags=38%, list=28%, si | MM_3 |
| GOBP_REGULATION_OF_ | GOBP_REGULATION_OF_INTRINSIC_APOPTC | 34 | 0.450884 | 1.4717628 | 0.06909091 | 0.15886202 | 1 | 7115 | tags=53%, list=30%, si | MM_3 |
| GOBP_REGULATION_OF_ | GOBP_REGULATION_OF_COAGULATION | 64 | 0.40864 | 1.4709117 | 0.041340783 | 0.15975133 | 1 | 5903 | tags=45%, list=25%, si | MM_3 |
| GOBP_PROTEIN_KINASE_ | GOBP_PROTEIN_KINASE_C_ACTIVATING_G_ | 25 | 0.487464 | 1.4705489 | 0.06878307 | 0.16000797 | 1 | 3164 | tags=36%, list=13%, si | MM_3 |
| GOBP_REGULATION_OF_ | GOBP_REGULATION_OF_SISTER_CHROMATI | 104 | 0.387473 | 1.4700372 | 0.020408163 | 0.16048834 | 1 | 6963 | tags=40%, list=29%, si | MM_3 |
| GOBP_DENDRITIC_CELL_ | GOBP_DENDRITIC_CELL_CHEMOTAXIS | 19 | 0.510658 | 1.4699311 | 0.060321715 | 0.16044168 | 1 | 6134 | tags=53%, list=26%, si | MM_3 |
| GOBP_PHOSPHATIDYLIN | GOBP_PHOSPHATIDYLINOSITOL_3_KINASE_ | 307 | 0.358498 | 1.4696603 | 0.001002004 | 0.16061457 | 1 | 4554 | tags=29%, list=19%, si | MM_3 |
| GOBP_REGULATION_OF_ | GOBP_REGULATION_OF_PROTEIN_CONTAIN | 420 | 0.356597 | 1.4695848 | 0.001003009 | 0.16054046 | 1 | 7521 | tags=48%, list=32%, si | MM_3 |
| GOBP_REGULATION_OF_ | GOBP_REGULATION_OF_PROTEIN_LOCALIZ | 158 | 0.373002 | 1.4693255 | 0.012371134 | 0.16065295 | 1 | 7322 | tags=42%, list=31%, si | MM_3 |
| GLYCOSYLPHOSPHATIDYL | GLYCOSYLPHOSPHATIDYLINOSITOL_(GPI)-AN | 24 | 0.479927 | 1.4693204 | 0.05699482 | 0.16048832 | 1 | 4421 | tags=50%, list=19%, si | MM_3 |
| GOBP_PROTEIN_DEUBIQU | GOBP_PROTEIN_DEUBIQUITINATION | 98 | 0.384135 | 1.4692802 | 0.024625268 | 0.1603603 | 1 | 2974 | tags=21%, list=13%, si | MM_3 |
| GOBP_POSITIVE_REGULA | GOBP_POSITIVE_REGULATION_OF_NEURON | 99 | 0.381414 | 1.4692453 | 0.014069264 | 0.16023377 | 1 | 7521 | tags=45%, list=32%, si | MM_3 |
| GOBP_REGULATION_OF_ | GOBP_REGULATION_OF_VIRAL_LIFE_CYCLE | 139 | 0.376085 | 1.46904 | 0.015560166 | 0.16030402 | 1 | 5669 | tags=35%, list=24%, si | MM_3 |
| GOBP_NEGATIVE_REGUL | GOBP_NEGATIVE_REGULATION_OF_PROTEI | 29 | 0.467229 | 1.4685376 | 0.06218274 | 0.1607635 | 1 | 3151 | tags=28%, list=13%, si | MM_3 |
| GOBP_REGULATION_OF_ | GOBP_REGULATION_OF_AUTOPHAGY_OF_N | 32 | 0.461601 | 1.4680173 | 0.05860806 | 0.16118827 | 1 | 7506 | tags=59%, list=32%, si | MM_3 |
| FANCONI_ANEMIA_PATH | FANCONI_ANEMIA_PATHWAY | 50 | 0.420559 | 1.4678996 | 0.045184303 | 0.16115233 | 1 | 5150 | tags=40%, list=22%, si | MM_3 |
| GOBP_MITOTIC_CHROM | GOBP_MITOTIC_CHROMOSOME_CONDENSA | 15 | 0.53883 | 1.4678867 | 0.06989247 | 0.1609929 | 1 | 3754 | tags=33%, list=16%, si | MM_3 |
| GOBP_ENERGY_HOMEOS | GOBP_ENERGY_HOMEOSTASIS | 92 | 0.392229 | 1.4677925 | 0.023630505 | 0.16094725 | 1 | 6141 | tags=37%, list=26%, si | MM_3 |
| GOBP_NEGATIVE_REGUL | GOBP_NEGATIVE_REGULATION_OF_INSULIN | 17 | 0.532853 | 1.4676194 | 0.06346968 | 0.16096593 | 1 | 7652 | tags=65%, list=32%, si | MM_3 |
| GOBP_POSITIVE_REGULA | GOBP_POSITIVE_REGULATION_OF_MRNA_C | 95 | 0.386552 | 1.4670883 | 0.021574972 | 0.16142659 | 1 | 7545 | tags=47%, list=32%, si | MM_3 |
| GOBP_ORGANELLE_FUSI | GOBP_ORGANELLE_FUSION | 167 | 0.3677 | 1.4669653 | 0.004149378 | 0.16139866 | 1 | 7664 | tags=44%, list=32%, si | MM_3 |
| GOBP_MUSCLE_CELL_AP | GOBP_MUSCLE_CELL_APOPTOTIC_PROCESS | 124 | 0.377823 | 1.4666146 | 0.018848168 | 0.16164044 | 1 | 5977 | tags=37%, list=25%, si | MM_3 |
| GOBP_RESPONSE_TO_TE | GOBP_RESPONSE_TO_TESTOSTERONE | 15 | 0.538928 | 1.4655352 | 0.08344924 | 0.16273822 | 1 | 1632 | tags=27%, list=7%, sig | MM_3 |
| GOBP_POSITIVE_REGULA | GOBP_POSITIVE_REGULATION_OF_DNA_RE | 135 | 0.375657 | 1.4647223 | 0.014814815 | 0.16353291 | 1 | 6787 | tags=39%, list=29%, si | MM_3 |
| GOBP_POSITIVE_REGULA | GOBP_POSITIVE_REGULATION_OF_PROTEIN | 35 | 0.445581 | 1.4644927 | 0.04342432 | 0.16364099 | 1 | 7521 | tags=51%, list=32%, si | MM_3 |
| GOBP_MRNA_CIS_SPLICI | GOBP_MRNA_CIS_SPLICING_VIA_SPLICEOSC | 26 | 0.467314 | 1.464168 | 0.065656565 | 0.16386254 | 1 | 7536 | tags=50%, list=32%, si | MM_3 |
| GOBP_REGULATION_OF_ | GOBP_REGULATION_OF_AUTOPHAGOSOME | 44 | 0.430547 | 1.4641166 | 0.04107981 | 0.16375868 | 1 | 7782 | tags=64%, list=33%, si | MM_3 |
| GOBP_BRANCHED_CHAIN | GOBP_BRANCHED_CHAIN_AMINO_ACID_M | 24 | 0.489869 | 1.4637746 | 0.061579652 | 0.1640029 | 1 | 4600 | tags=46%, list=19%, si | MM_3 |
| GOBP_CELLULAR_RESPON | GOBP_CELLULAR_RESPONSE_TO_TOXIC_SU | 63 | 0.408014 | 1.4630545 | 0.043624163 | 0.16467093 | 1 | 5033 | tags=35%, list=21%, si | MM_3 |
| GOBP_NEGATIVE_REGUL | GOBP_NEGATIVE_REGULATION_OF_LEUKOC | 66 | 0.402303 | 1.4629613 | 0.038636364 | 0.16461536 | 1 | 7331 | tags=52%, list=31%, si | MM_3 |
| GOBP_TORC1_SIGNALING | GOBP_TORC1_SIGNALING | 100 | 0.384146 | 1.4624641 | 0.020540541 | 0.165062 | 1 | 7468 | tags=48%, list=32%, si | MM_3 |
| GOBP_MIRNA_TRANSCRI | GOBP_MIRNA_TRANSCRIPTION | 78 | 0.39382 | 1.4624162 | 0.034103412 | 0.1649397 | 1 | 5641 | tags=32%, list=24%, si | MM_3 |
| GOBP_POSITIVE_REGULA | GOBP_POSITIVE_REGULATION_OF_PRODUC | 136 | 0.373974 | 1.4622931 | 0.009473684 | 0.16492602 | 1 | 4684 | tags=26%, list=20%, si | MM_3 |
| GOBP_RRNA_MODIFICAT | GOBP_RRNA_MODIFICATION | 29 | 0.461641 | 1.4622208 | 0.06377551 | 0.1648504 | 1 | 6935 | tags=55%, list=29%, si | MM_3 |
| GOBP_SUPEROXIDE_MET | GOBP_SUPEROXIDE_METABOLIC_PROCESS | 68 | 0.405855 | 1.4619619 | 0.03131991 | 0.16500886 | 1 | 7190 | tags=41%, list=30%, si | MM_3 |

| | | | | | | | | | | |
|---|---|---|---|---|---|---|---|---|---|---|
| GOBP_REGULATION_OF_ | GOBP_REGULATION_OF_TOLL_LIKE_RECEPT | 28 | 0.470562 | 1.4618949 | 0.058676653 | 0.16490817 | 1 | 7383 | tags=68%, list=31%, si | MM_3 |
| GOBP_CHAPERONE_MED | GOBP_CHAPERONE_MEDIATED_PROTEIN_C | 18 | 0.518384 | 1.4615084 | 0.06793478 | 0.1652362 | 1 | 4299 | tags=44%, list=18%, si | MM_3 |
| GOBP_NEGATIVE_REGUL | GOBP_NEGATIVE_REGULATION_OF_LEUKOC | 96 | 0.390523 | 1.4615062 | 0.01754386 | 0.16506389 | 1 | 4813 | tags=31%, list=20%, si | MM_3 |
| GOBP_POSITIVE_REGULA | GOBP_POSITIVE_REGULATION_OF_RNA_SPL | 50 | 0.420143 | 1.4614183 | 0.04181601 | 0.16499633 | 1 | 7536 | tags=56%, list=32%, si | MM_3 |
| GOBP_AMIDE_METABOLI | GOBP_AMIDE_METABOLIC_PROCESS | 398 | 0.355019 | 1.4611098 | 0.002006018 | 0.16520363 | 1 | 5556 | tags=32%, list=24%, si | MM_3 |
| GOBP_POSITIVE_REGULA | GOBP_POSITIVE_REGULATION_OF_MUSCLE | 43 | 0.430719 | 1.460906 | 0.042475726 | 0.16529049 | 1 | 7651 | tags=58%, list=32%, si | MM_3 |
| GOBP_PROTEIN_TETRAM | GOBP_PROTEIN_TETRAMERIZATION | 82 | 0.385322 | 1.4608374 | 0.023255814 | 0.16520648 | 1 | 6189 | tags=34%, list=26%, si | MM_3 |
| GOBP_POSITIVE_REGULA | GOBP_POSITIVE_REGULATION_OF_VASCULA | 172 | 0.368472 | 1.4606903 | 0.0081883315 | 0.16520038 | 1 | 5491 | tags=35%, list=23%, si | MM_3 |
| GOBP_REGULATION_OF_ | GOBP_REGULATION_OF_NEUROINFLAMMA | 30 | 0.464923 | 1.4601525 | 0.061173532 | 0.16564997 | 1 | 4391 | tags=37%, list=19%, si | MM_3 |
| GOBP_POSITIVE_REGULA | GOBP_POSITIVE_REGULATION_OF_PROTEIN | 100 | 0.386896 | 1.4601272 | 0.022508038 | 0.16551231 | 1 | 5993 | tags=35%, list=25%, si | MM_3 |
| GOBP_POSITIVE_REGULA | GOBP_POSITIVE_REGULATION_OF_CELL_CY | 330 | 0.354564 | 1.4598473 | 0.0020120724 | 0.16569553 | 1 | 6906 | tags=40%, list=29%, si | MM_3 |
| GOBP_POSITIVE_REGULA | GOBP_POSITIVE_REGULATION_OF_CHEMOK | 79 | 0.392674 | 1.459472 | 0.032222223 | 0.16595252 | 1 | 4649 | tags=28%, list=20%, si | MM_3 |
| GOBP_PROTEIN_MATURA | GOBP_PROTEIN_MATURATION_BY_IRON_SU | 16 | 0.528382 | 1.4594598 | 0.076071925 | 0.16579762 | 1 | 7420 | tags=63%, list=31%, si | MM_3 |
| GOBP_REGULATION_OF_ | GOBP_REGULATION_OF_PH | 104 | 0.384157 | 1.4593589 | 0.022245763 | 0.16577679 | 1 | 6867 | tags=45%, list=29%, si | MM_3 |
| GOBP_CELL_CYCLE_PHAS | GOBP_CELL_CYCLE_PHASE | 21 | 0.482285 | 1.4591521 | 0.05781866 | 0.1658461 | 1 | 5801 | tags=43%, list=25%, si | MM_3 |
| GOBP_SULFUR_COMPOU | GOBP_SULFUR_COMPOUND_METABOLIC_P | 292 | 0.354965 | 1.4591426 | 0.002016129 | 0.16568859 | 1 | 5307 | tags=32%, list=22%, si | MM_3 |
| GOBP_CIRCADIAN_REGUI | GOBP_CIRCADIAN_REGULATION_OF_GENE_ | 69 | 0.399628 | 1.459129 | 0.03405221 | 0.16553134 | 1 | 7389 | tags=58%, list=31%, si | MM_3 |
| GOBP_REGULATION_OF_ | GOBP_REGULATION_OF_MITOTIC_CELL_CY( | 490 | 0.351889 | 1.4591287 | 0 | 0.16535819 | 1 | 6906 | tags=41%, list=29%, si | MM_3 |
| GOBP_VESICLE_DOCKING | GOBP_VESICLE_DOCKING | 55 | 0.418721 | 1.458789 | 0.03337169 | 0.16563739 | 1 | 7071 | tags=55%, list=30%, si | MM_3 |
| GOBP_VENTRICULAR_SYS | GOBP_VENTRICULAR_SYSTEM_DEVELOPME | 36 | 0.444753 | 1.4587784 | 0.05582524 | 0.16547418 | 1 | 2539 | tags=28%, list=11%, si | MM_3 |
| GOBP_POSITIVE_REGULA | GOBP_POSITIVE_REGULATION_OF_T_CELL_ | 66 | 0.400553 | 1.4586848 | 0.036954086 | 0.16540293 | 1 | 7372 | tags=48%, list=31%, si | MM_3 |
| NECROPTOSIS | NECROPTOSIS | 138 | 0.375863 | 1.4586815 | 0.0062305294 | 0.1652339 | 1 | 6645 | tags=43%, list=28%, si | MM_3 |
| GOBP_NEGATIVE_REGUL | GOBP_NEGATIVE_REGULATION_OF_TUMOR | 67 | 0.398165 | 1.4586772 | 0.033860046 | 0.16506845 | 1 | 6246 | tags=42%, list=26%, si | MM_3 |
| GOBP_CELLULAR_KETONI | GOBP_CELLULAR_KETONE_METABOLIC_PRO | 222 | 0.363023 | 1.4585661 | 0.0030364373 | 0.16502923 | 1 | 4311 | tags=26%, list=18%, si | MM_3 |
| GOBP_EMBRYONIC_DIGE | GOBP_EMBRYONIC_DIGESTIVE_TRACT_DEV | 32 | 0.456418 | 1.4581227 | 0.06875754 | 0.16541658 | 1 | 1616 | tags=22%, list=7%, sig | MM_3 |
| GOBP_PRODUCTION_OF_ | GOBP_PRODUCTION_OF_MOLECULAR_MED | 97 | 0.378304 | 1.4578956 | 0.019210245 | 0.1655199 | 1 | 3329 | tags=20%, list=14%, si | MM_3 |
| GOBP_REGULATION_OF_ | GOBP_REGULATION_OF_HUMORAL_IMMUN | 47 | 0.424463 | 1.4576033 | 0.06272189 | 0.16568606 | 1 | 3957 | tags=26%, list=17%, si | MM_3 |
| GOBP_REGULATION_OF_ | GOBP_REGULATION_OF_CALCIUM_ION_IMI | 45 | 0.421156 | 1.4573452 | 0.058402862 | 0.165838 | 1 | 5903 | tags=40%, list=25%, si | MM_3 |
| GOBP_REGULATION_OF_ | GOBP_REGULATION_OF_NITRIC_OXIDE_SYN | 20 | 0.497056 | 1.4566581 | 0.061579652 | 0.16656552 | 1 | 5903 | tags=45%, list=25%, si | MM_3 |
| GOBP_RESPONSE_TO_TU | GOBP_RESPONSE_TO_TUMOR_CELL | 42 | 0.431023 | 1.4564818 | 0.038004752 | 0.16659771 | 1 | 4712 | tags=33%, list=20%, si | MM_3 |
| GOBP_REGULATION_OF_ | GOBP_REGULATION_OF_NUCLEOTIDE_EXCIS | 29 | 0.455231 | 1.4562198 | 0.056847546 | 0.16675882 | 1 | 8678 | tags=69%, list=37%, si | MM_3 |
| GOBP_POSITIVE_REGULA | GOBP_POSITIVE_REGULATION_OF_BEHAVIO | 33 | 0.452689 | 1.4561626 | 0.051660515 | 0.16665512 | 1 | 4599 | tags=30%, list=19%, si | MM_3 |
| GOBP_MIRNA_METABOL | GOBP_MIRNA_METABOLIC_PROCESS | 103 | 0.380932 | 1.4551854 | 0.02255639 | 0.16767997 | 1 | 6762 | tags=39%, list=29%, si | MM_3 |
| GOBP_POSITIVE_REGULA | GOBP_POSITIVE_REGULATION_OF_RESPONS | 19 | 0.504077 | 1.4550272 | 0.06473829 | 0.16771932 | 1 | 4364 | tags=37%, list=18%, si | MM_3 |
| GOBP_INTERLEUKIN_6_P | GOBP_INTERLEUKIN_6_PRODUCTION | 173 | 0.364291 | 1.4544933 | 0.0071868584 | 0.16821496 | 1 | 6192 | tags=36%, list=26%, si | MM_3 |
| GOBP_GTP_METABOLIC_ | GOBP_GTP_METABOLIC_PROCESS | 26 | 0.475856 | 1.4541634 | 0.067183465 | 0.16846004 | 1 | 4324 | tags=38%, list=18%, si | MM_3 |
| GOBP_DNA_INTEGRITY_C | GOBP_DNA_INTEGRITY_CHECKPOINT_SIGNA | 125 | 0.378466 | 1.4541632 | 0.01049318 | 0.16828726 | 1 | 7164 | tags=49%, list=30%, si | MM_3 |
| GOBP_RESPONSE_TO_CO | GOBP_RESPONSE_TO_COPPER_ION | 23 | 0.484474 | 1.4535685 | 0.087126136 | 0.168838 | 1 | 6026 | tags=43%, list=26%, si | MM_3 |
| GOBP_VIRAL_LIFE_CYCLE | GOBP_VIRAL_LIFE_CYCLE | 230 | 0.358979 | 1.4534745 | 0.004052685 | 0.1687743 | 1 | 7549 | tags=50%, list=32%, si | MM_3 |
| GOBP_NECROPTOTIC_PR | GOBP_NECROPTOTIC_PROCESS | 37 | 0.443213 | 1.4530811 | 0.06993865 | 0.16913633 | 1 | 5903 | tags=38%, list=25%, si | MM_3 |
| GOBP_MUSCLE_CELL_MI | GOBP_MUSCLE_CELL_MIGRATION | 129 | 0.373731 | 1.4530464 | 0.023231257 | 0.16901216 | 1 | 5906 | tags=33%, list=25%, si | MM_3 |
| GOBP_CELL_CYCLE_G1_S | GOBP_CELL_CYCLE_G1_S_PHASE_TRANSITIC | 236 | 0.359253 | 1.4523566 | 0.004040404 | 0.1696837 | 1 | 7568 | tags=47%, list=32%, si | MM_3 |
| GOBP_RECEPTOR_CLUST | GOBP_RECEPTOR_CLUSTERING | 79 | 0.394208 | 1.4523301 | 0.020787746 | 0.16953716 | 1 | 4062 | tags=32%, list=17%, si | MM_3 |
| GOBP_REGULATION_OF_ | GOBP_REGULATION_OF_ADIPOSE_TISSUE_[ | 16 | 0.523591 | 1.4518819 | 0.076398365 | 0.16993155 | 1 | 5397 | tags=44%, list=23%, si | MM_3 |
| GOBP_POSITIVE_REGULA | GOBP_POSITIVE_REGULATION_OF_REACTIV | 81 | 0.39351 | 1.4518037 | 0.036144577 | 0.16986705 | 1 | 5470 | tags=32%, list=23%, si | MM_3 |
| GOBP_REGULATION_OF_ | GOBP_REGULATION_OF_MRNA_SPLICING_V | 108 | 0.379604 | 1.4517353 | 0.020234292 | 0.1697703 | 1 | 8264 | tags=55%, list=35%, si | MM_3 |
| GOBP_NUCLEOTIDE_BINI | GOBP_NUCLEOTIDE_BINDING_DOMAIN_LEU | 22 | 0.481847 | 1.4516793 | 0.078457445 | 0.169662 | 1 | 6867 | tags=59%, list=29%, si | MM_3 |
| GOBP_RIBOSOMAL_LARG | GOBP_RIBOSOMAL_LARGE_SUBUNIT_ASSEN | 23 | 0.481393 | 1.4515312 | 0.079634465 | 0.16967475 | 1 | 819 | tags=22%, list=3%, sig | MM_3 |
| GOBP_SOMATIC_DIVERSI | GOBP_SOMATIC_DIVERSIFICATION_OF_IMN | 71 | 0.400256 | 1.451493 | 0.036544852 | 0.16955528 | 1 | 4592 | tags=32%, list=19%, si | MM_3 |
| SALMONELLA_INFECTION | SALMONELLA_INFECTION | 76 | 0.388335 | 1.4514173 | 0.046875 | 0.16947477 | 1 | 8279 | tags=66%, list=35%, si | MM_3 |
| GOBP_NEGATIVE_REGUL | GOBP_NEGATIVE_REGULATION_OF_DOUBL | 24 | 0.477275 | 1.4511706 | 0.072072074 | 0.16963738 | 1 | 4668 | tags=33%, list=20%, si | MM_3 |
| GOBP_NEGATIVE_REGUL | GOBP_NEGATIVE_REGULATION_OF_CALCIN | 16 | 0.523116 | 1.4503933 | 0.07713126 | 0.17040505 | 1 | 1887 | tags=25%, list=8%, sig | MM_3 |
| GOBP_POSITIVE_REGULA | GOBP_POSITIVE_REGULATION_OF_CELL_SU | 134 | 0.369726 | 1.4501265 | 0.014690451 | 0.17055793 | 1 | 4229 | tags=31%, list=18%, si | MM_3 |
| GOBP_TOLL_LIKE_RECEPT | GOBP_TOLL_LIKE_RECEPTOR_9_SIGNALING_ | 16 | 0.516777 | 1.4495219 | 0.07723577 | 0.17113855 | 1 | 5807 | tags=50%, list=25%, si | MM_3 |
| GOBP_REGULATION_OF_ | GOBP_REGULATION_OF_CARBOHYDRATE_C | 72 | 0.394797 | 1.4493474 | 0.041758243 | 0.17120379 | 1 | 6488 | tags=42%, list=27%, si | MM_3 |
| GOBP_REGULATION_OF_ | GOBP_REGULATION_OF_TRANSLATION_IN_ | 24 | 0.477767 | 1.4493208 | 0.058047492 | 0.17106691 | 1 | 7262 | tags=58%, list=31%, si | MM_3 |
| GOBP_STORE_OPERATED | GOBP_STORE_OPERATED_CALCIUM_ENTRY | 25 | 0.479477 | 1.4490594 | 0.07822686 | 0.17123489 | 1 | 3818 | tags=32%, list=16%, si | MM_3 |
| GOBP_REGULATION_OF_ | GOBP_REGULATION_OF_VESICLE_SIZE | 21 | 0.492542 | 1.448749 | 0.06818182 | 0.17143196 | 1 | 6579 | tags=52%, list=28%, si | MM_3 |
| FLUID_SHEAR_STRESS_AI | FLUID_SHEAR_STRESS_AND_ATHEROSCLERO | 135 | 0.371799 | 1.4482716 | 0.010471204 | 0.17189424 | 1 | 6572 | tags=45%, list=28%, si | MM_3 |
| GOBP_GLIOGENESIS | GOBP_GLIOGENESIS | 377 | 0.350763 | 1.4480147 | 0 | 0.17204866 | 1 | 5930 | tags=34%, list=25%, si | MM_3 |
| GOBP_POSITIVE_REGULA | GOBP_POSITIVE_REGULATION_OF_CYCLIN_ | 20 | 0.494904 | 1.4478185 | 0.062169313 | 0.17209512 | 1 | 2693 | tags=25%, list=11%, si | MM_3 |
| GOBP_REGULATION_OF_ | GOBP_REGULATION_OF_G_PROTEIN_COUP | 142 | 0.367829 | 1.4477731 | 0.013626834 | 0.17198604 | 1 | 5275 | tags=35%, list=22%, si | MM_3 |
| GOBP_REGULATION_OF_ | GOBP_REGULATION_OF_LEUKOCYTE_DIFFEI | 342 | 0.354537 | 1.4466269 | 0 | 0.1732329 | 1 | 6594 | tags=38%, list=28%, si | MM_3 |
| GOBP_ER_NUCLEUS_SIGI | GOBP_ER_NUCLEUS_SIGNALING_PATHWAY | 43 | 0.420945 | 1.4465846 | 0.05344418 | 0.17311473 | 1 | 6194 | tags=40%, list=26%, si | MM_3 |
| GOBP_PHYSIOLOGICAL_C | GOBP_PHYSIOLOGICAL_CARDIAC_MUSCLE_ | 46 | 0.420248 | 1.4459467 | 0.056338027 | 0.17375225 | 1 | 3164 | tags=24%, list=13%, si | MM_3 |
| GOBP_POSITIVE_REGULA | GOBP_POSITIVE_REGULATION_OF_UBIQUIT | 18 | 0.515148 | 1.4458228 | 0.080368906 | 0.17371128 | 1 | 7506 | tags=61%, list=32%, si | MM_3 |
| GOBP_REGULATION_OF_ | GOBP_REGULATION_OF_DENDRITIC_SPINE_ | 87 | 0.385803 | 1.4457537 | 0.021064302 | 0.17362793 | 1 | 7337 | tags=43%, list=31%, si | MM_3 |
| GOBP_SOMATIC_DIVERSI | GOBP_SOMATIC_DIVERSIFICATION_OF_IMN | 81 | 0.386185 | 1.4454798 | 0.027027028 | 0.17383914 | 1 | 3367 | tags=26%, list=14%, si | MM_3 |
| GOBP_P_BODY_ASSEMBL | GOBP_P_BODY_ASSEMBLY | 20 | 0.499794 | 1.4449446 | 0.071614586 | 0.1742913 | 1 | 8313 | tags=80%, list=35%, si | MM_3 |
| GOBP_NEGATIVE_REGUL | GOBP_NEGATIVE_REGULATION_OF_TOLL_L | 22 | 0.48911 | 1.4447439 | 0.077025235 | 0.17434534 | 1 | 5255 | tags=45%, list=22%, si | MM_3 |
| GOBP_POSITIVE_REGULA | GOBP_POSITIVE_REGULATION_OF_CELL_SU | 35 | 0.429782 | 1.4445214 | 0.0746988 | 0.17447305 | 1 | 7254 | tags=51%, list=31%, si | MM_3 |
| GOBP_REGULATION_OF_ | GOBP_REGULATION_OF_MYELOID_CELL_DI | 217 | 0.356649 | 1.4444916 | 0.0051124743 | 0.17433716 | 1 | 6330 | tags=37%, list=27%, si | MM_3 |
| GOBP_MICROGLIA_DIFFE | GOBP_MICROGLIA_DIFFERENTIATION | 16 | 0.533086 | 1.4442947 | 0.07630522 | 0.17443813 | 1 | 4391 | tags=31%, list=19%, si | MM_3 |
| GOBP_LYMPHOCYTE_HOI | GOBP_LYMPHOCYTE_HOMEOSTASIS | 92 | 0.383816 | 1.4441434 | 0.025806451 | 0.17446311 | 1 | 7020 | tags=45%, list=30%, si | MM_3 |
| GOBP_L_AMINO_ACID_M | GOBP_L_AMINO_ACID_METABOLIC_PROCES | 154 | 0.358383 | 1.4440534 | 0.016632017 | 0.17441246 | 1 | 5181 | tags=33%, list=22%, si | MM_3 |
| GOBP_REGULATION_OF_ | GOBP_REGULATION_OF_LIPID_CATABOLIC_ | 63 | 0.401948 | 1.4437904 | 0.04942529 | 0.17460784 | 1 | 6181 | tags=35%, list=26%, si | MM_3 |
| GOBP_CALCINEURIN_ME | GOBP_CALCINEURIN_MEDIATED_SIGNALING | 50 | 0.415738 | 1.4437561 | 0.04079254 | 0.17447966 | 1 | 5892 | tags=38%, list=25%, si | MM_3 |
| GOBP_REGULATION_OF_ | GOBP_REGULATION_OF_LYMPHOCYTE_ACT | 495 | 0.345225 | 1.4425461 | 0 | 0.17589855 | 1 | 5751 | tags=33%, list=24%, si | MM_3 |
| GOBP_POSITIVE_REGULA | GOBP_POSITIVE_REGULATION_OF_TORC1_S | 52 | 0.418008 | 1.4423481 | 0.052691866 | 0.17598718 | 1 | 7468 | tags=48%, list=32%, si | MM_3 |
| GOBP_REGULATION_OF_ | GOBP_REGULATION_OF_INFLAMMATORY_F | 33 | 0.448234 | 1.4423323 | 0.05507956 | 0.17583355 | 1 | 6692 | tags=45%, list=28%, si | MM_3 |
| GOBP_REGULATION_OF_ | GOBP_REGULATION_OF_FATTY_ACID_BETA | 21 | 0.489048 | 1.4421597 | 0.079427086 | 0.17586716 | 1 | 1101 | tags=19%, list=5%, sig | MM_3 |
| GOBP_NEGATIVE_REGUL | GOBP_NEGATIVE_REGULATION_OF_CELL_C | 197 | 0.359375 | 1.4418716 | 0.007135576 | 0.17608312 | 1 | 6040 | tags=34%, list=26%, si | MM_3 |

| | | | | | | | | | | |
|---|---|---|---|---|---|---|---|---|---|---|
| GOBP_POSITIVE_REGULA | GOBP_POSITIVE_REGULATION_OF_OLIGODE | 28 | 0.464426 | 1.4417745 | 0.06947891 | 0.17603123 | 1 | 3801 | tags=25%, list=16%, si | MM_3 |
| GOBP_POSITIVE_REGULA | GOBP_POSITIVE_REGULATION_OF_FIBROBL | 76 | 0.389112 | 1.4412676 | 0.03463687 | 0.17650612 | 1 | 5262 | tags=38%, list=22%, si | MM_3 |
| TRYPTOPHAN_METABOLI | TRYPTOPHAN_METABOLISM | 43 | 0.421745 | 1.4408296 | 0.06490385 | 0.1768797 | 1 | 2961 | tags=28%, list=13%, si | MM_3 |
| GOBP_POSITIVE_REGULA | GOBP_POSITIVE_REGULATION_OF_STRIATE | 50 | 0.410326 | 1.4407626 | 0.05704307 | 0.17680296 | 1 | 5274 | tags=38%, list=22%, si | MM_3 |
| GOBP_VESICLE_DOCKING | GOBP_VESICLE_DOCKING_INVOLVED_IN_EX | 36 | 0.436374 | 1.4406599 | 0.056580566 | 0.17674263 | 1 | 7071 | tags=53%, list=30%, si | MM_3 |
| GOBP_MITOTIC_CELL_CY | GOBP_MITOTIC_CELL_CYCLE_PHASE_TRANS | 425 | 0.349711 | 1.4406354 | 0 | 0.17660466 | 1 | 6931 | tags=42%, list=29%, si | MM_3 |
| GOBP_REGULATION_OF_ | GOBP_REGULATION_OF_DNA_RECOMBINAT | 127 | 0.372723 | 1.4404899 | 0.013655462 | 0.17661427 | 1 | 4668 | tags=28%, list=20%, si | MM_3 |
| GOBP_PROTEIN_INSERTI | GOBP_PROTEIN_INSERTION_INTO_MEMBRA | 39 | 0.427212 | 1.4403753 | 0.065398335 | 0.17658655 | 1 | 6806 | tags=46%, list=29%, si | MM_3 |
| GOBP_POSITIVE_REGULA | GOBP_POSITIVE_REGULATION_OF_SIGNAL_ | 32 | 0.446683 | 1.4400643 | 0.06273526 | 0.1768233 | 1 | 7202 | tags=63%, list=31%, si | MM_3 |
| GOBP_REGULATION_OF_ | GOBP_REGULATION_OF_SIGNAL_TRANSDU | 83 | 0.391417 | 1.43995 | 0.031624865 | 0.17679936 | 1 | 7202 | tags=54%, list=31%, si | MM_3 |
| GOBP_NEGATIVE_REGUL | GOBP_NEGATIVE_REGULATION_OF_CYCLIN | 25 | 0.469714 | 1.4394945 | 0.062015504 | 0.17719698 | 1 | 5501 | tags=36%, list=23%, si | MM_3 |
| GOBP_VIRAL_RELEASE_F | GOBP_VIRAL_RELEASE_FROM_HOST_CELL | 24 | 0.471378 | 1.439065 | 0.07721519 | 0.17759027 | 1 | 7508 | tags=67%, list=32%, si | MM_3 |
| GOBP_REGULATION_OF_ | GOBP_REGULATION_OF_LYMPHOCYTE_APO | 71 | 0.393306 | 1.4387497 | 0.04185022 | 0.17782709 | 1 | 5470 | tags=34%, list=23%, si | MM_3 |
| GOBP_POSITIVE_REGULA | GOBP_POSITIVE_REGULATION_OF_INFLAM | 134 | 0.369198 | 1.4385674 | 0.009473684 | 0.1778973 | 1 | 6912 | tags=43%, list=29%, si | MM_3 |
| GOBP_CELL_DIFFERENTIA | GOBP_CELL_DIFFERENTIATION_INVOLVED_ | 50 | 0.418282 | 1.4384407 | 0.0521327 | 0.17788646 | 1 | 3599 | tags=30%, list=15%, si | MM_3 |
| GOBP_BIOLOGICAL_PRO | GOBP_BIOLOGICAL_PROCESS_INVOLVED_IN | 235 | 0.354598 | 1.4384277 | 0.005045409 | 0.17772771 | 1 | 7525 | tags=49%, list=32%, si | MM_3 |
| GOBP_DNA_TEMPLATED_ | GOBP_DNA_TEMPLATED_DNA_REPLICATION | 54 | 0.409317 | 1.4380847 | 0.05740528 | 0.17796871 | 1 | 4488 | tags=31%, list=19%, si | MM_3 |
| GOBP_PERK_MEDIATED_ | GOBP_PERK_MEDIATED_UNFOLDED_PROTE | 16 | 0.523211 | 1.4376647 | 0.09090909 | 0.17835875 | 1 | 6194 | tags=44%, list=26%, si | MM_3 |
| GOBP_CELL_CYCLE_CHEC | GOBP_CELL_CYCLE_CHECKPOINT_SIGNALIN | 191 | 0.359767 | 1.4374797 | 0.008196721 | 0.17843395 | 1 | 7164 | tags=45%, list=30%, si | MM_3 |
| GOBP_REGULATION_OF_ | GOBP_REGULATION_OF_MYOBLAST_FUSIO | 22 | 0.482599 | 1.4369905 | 0.074122235 | 0.17888546 | 1 | 5715 | tags=41%, list=24%, si | MM_3 |
| GOBP_RNA_TEMPLATED_ | GOBP_RNA_TEMPLATED_DNA_BIOSYNTHET | 63 | 0.404608 | 1.4369483 | 0.03502825 | 0.17878023 | 1 | 6932 | tags=44%, list=29%, si | MM_3 |
| LYSINE_DEGRADATION | LYSINE_DEGRADATION | 57 | 0.403427 | 1.436554 | 0.057603687 | 0.17909162 | 1 | 7645 | tags=53%, list=32%, si | MM_3 |
| GOBP_LEUKOCYTE_MEDI | GOBP_LEUKOCYTE_MEDIATED_CYTOTOXICI | 135 | 0.36598 | 1.4363182 | 0.016649324 | 0.17923659 | 1 | 6013 | tags=37%, list=25%, si | MM_3 |
| GOBP_BASIC_AMINO_AC | GOBP_BASIC_AMINO_ACID_TRANSPORT | 16 | 0.525091 | 1.4361637 | 0.09090909 | 0.17927392 | 1 | 3160 | tags=38%, list=13%, si | MM_3 |
| GOBP_REGULATION_OF_ | GOBP_REGULATION_OF_PROTEIN_KINASE_ | 22 | 0.482219 | 1.4360079 | 0.068331145 | 0.17931847 | 1 | 5031 | tags=36%, list=21%, si | MM_3 |
| GOBP_REGULATION_OF_ | GOBP_REGULATION_OF_HIPPO_SIGNALING | 22 | 0.486227 | 1.4357907 | 0.08322497 | 0.1794668 | 1 | 3520 | tags=32%, list=15%, si | MM_3 |
| GOBP_REGULATION_OF_ | GOBP_REGULATION_OF_AEROBIC_RESPIRAT | 33 | 0.448496 | 1.434895 | 0.0682095 | 0.18045199 | 1 | 7521 | tags=58%, list=32%, si | MM_3 |
| GOBP_MACROMOLECULE | GOBP_MACROMOLECULE_METHYLATION | 121 | 0.370392 | 1.4348934 | 0.01898734 | 0.18028176 | 1 | 6935 | tags=45%, list=29%, si | MM_3 |
| GOBP_TETRAPYRROLE_BI | GOBP_TETRAPYRROLE_BIOSYNTHETIC_PRO | 29 | 0.445989 | 1.4348049 | 0.083333336 | 0.18023132 | 1 | 4144 | tags=34%, list=18%, si | MM_3 |
| GOBP_ALANINE_TRANSP | GOBP_ALANINE_TRANSPORT | 17 | 0.514714 | 1.434711 | 0.07317073 | 0.18018782 | 1 | 2884 | tags=29%, list=12%, si | MM_3 |
| GOBP_CELL_PROLIFERATI | GOBP_CELL_PROLIFERATION_INVOLVED_IN | 20 | 0.485988 | 1.434619 | 0.07591623 | 0.18013167 | 1 | 3680 | tags=40%, list=16%, si | MM_3 |
| GOBP_SYNCYTIUM_FORM | GOBP_SYNCYTIUM_FORMATION | 63 | 0.40003 | 1.4345855 | 0.053409092 | 0.17998996 | 1 | 6853 | tags=43%, list=29%, si | MM_3 |
| GOBP_REGULATION_OF_ | GOBP_REGULATION_OF_CELLULAR_PH | 88 | 0.380606 | 1.4344983 | 0.027956989 | 0.17991835 | 1 | 6867 | tags=45%, list=29%, si | MM_3 |
| GOBP_TRABECULA_FORM | GOBP_TRABECULA_FORMATION | 27 | 0.460789 | 1.4340862 | 0.07060334 | 0.18028875 | 1 | 3547 | tags=30%, list=15%, si | MM_3 |
| GOBP_REGULATION_OF_ | GOBP_REGULATION_OF_ATTACHMENT_OF_ | 20 | 0.494025 | 1.4340663 | 0.07850708 | 0.18013848 | 1 | 6709 | tags=50%, list=28%, si | MM_3 |
| GOBP_POSITIVE_REGULA | GOBP_POSITIVE_REGULATION_OF_INTRINS | 69 | 0.387584 | 1.4340228 | 0.039106146 | 0.18001986 | 1 | 7518 | tags=54%, list=32%, si | MM_3 |
| GOBP_PROTEIN_MODIFI | GOBP_PROTEIN_MODIFICATION_BY_SMALL | 117 | 0.376905 | 1.4338571 | 0.018828452 | 0.18006401 | 1 | 2974 | tags=21%, list=13%, si | MM_3 |
| GOBP_PURINE_DEOXYRIB | GOBP_PURINE_DEOXYRIBONUCLEOTIDE_M | 20 | 0.480758 | 1.4338318 | 0.08940853 | 0.17992416 | 1 | 6982 | tags=55%, list=30%, si | MM_3 |
| VIRAL_MYOCARDITIS | VIRAL_MYOCARDITIS | 63 | 0.39554 | 1.4337684 | 0.042841036 | 0.17983918 | 1 | 6533 | tags=44%, list=28%, si | MM_3 |
| GOBP_NEGATIVE_REGUL | GOBP_NEGATIVE_REGULATION_OF_IMMUN | 132 | 0.370384 | 1.433588 | 0.017689906 | 0.17988618 | 1 | 6140 | tags=39%, list=26%, si | MM_3 |
| CYSTEINE_AND_METHION | CYSTEINE_AND_METHIONINE_METABOLISM | 44 | 0.42028 | 1.4332818 | 0.051825676 | 0.18013245 | 1 | 4601 | tags=39%, list=19%, si | MM_3 |
| GOBP_DNA_BIOSYNTHET | GOBP_DNA_BIOSYNTHETIC_PROCESS | 175 | 0.36215 | 1.4329257 | 0.0030895984 | 0.18040133 | 1 | 4818 | tags=30%, list=20%, si | MM_3 |
| GOBP_COMPLEMENT_AC | GOBP_COMPLEMENT_ACTIVATION | 50 | 0.416875 | 1.4328822 | 0.050458714 | 0.18028422 | 1 | 4577 | tags=30%, list=19%, si | MM_3 |
| GOBP_NEGATIVE_REGUL | GOBP_NEGATIVE_REGULATION_OF_NEURO | 15 | 0.520012 | 1.4328381 | 0.07832168 | 0.1801712 | 1 | 1394 | tags=20%, list=6%, sig | MM_3 |
| GOBP_CELLULAR_RESPO | GOBP_CELLULAR_RESPONSE_TO_OXIDATIVE | 229 | 0.357681 | 1.432779 | 0.0060913707 | 0.18008465 | 1 | 7204 | tags=47%, list=31%, si | MM_3 |
| GOBP_POSITIVE_REGULA | GOBP_POSITIVE_REGULATION_OF_VASCUL | 44 | 0.42042 | 1.4323577 | 0.054413542 | 0.18048069 | 1 | 7731 | tags=52%, list=33%, si | MM_3 |
| GOBP_RESPONSE_TO_RA | GOBP_RESPONSE_TO_RADIATION | 411 | 0.347491 | 1.4318866 | 0.001002004 | 0.18090436 | 1 | 6761 | tags=43%, list=29%, si | MM_3 |
| GOBP_POSITIVE_REGULA | GOBP_POSITIVE_REGULATION_OF_CELLULA | 22 | 0.485457 | 1.4317893 | 0.08979592 | 0.18086198 | 1 | 7330 | tags=59%, list=31%, si | MM_3 |
| GOBP_MATURATION_OF_ | GOBP_MATURATION_OF_LSU_RRNA | 25 | 0.471183 | 1.4316204 | 0.077120826 | 0.18092926 | 1 | 7086 | tags=60%, list=30%, si | MM_3 |
| GOBP_REGULATION_OF_ | GOBP_REGULATION_OF_NON_CANONICAL_ | 92 | 0.380045 | 1.4315516 | 0.017278617 | 0.18086658 | 1 | 7953 | tags=53%, list=34%, si | MM_3 |
| STEROID_BIOSYNTHESIS | STEROID_BIOSYNTHESIS | 17 | 0.517779 | 1.4312904 | 0.09271523 | 0.18103206 | 1 | 7631 | tags=65%, list=32%, si | MM_3 |
| GOBP_MAMMARY_GLAN | GOBP_MAMMARY_GLAND_EPITHELIAL_CEL | 16 | 0.514299 | 1.4309978 | 0.0996016 | 0.1812736 | 1 | 5067 | tags=38%, list=21%, si | MM_3 |
| GOBP_REGULATION_OF_ | GOBP_REGULATION_OF_DNA_DAMAGE_CH | 22 | 0.478261 | 1.4308639 | 0.104336046 | 0.18127374 | 1 | 6252 | tags=45%, list=26%, si | MM_3 |
| GOBP_BIOLOGICAL_PRO | GOBP_BIOLOGICAL_PROCESS_INVOLVED_IN | 140 | 0.364947 | 1.4306015 | 0.01975052 | 0.18147787 | 1 | 7525 | tags=53%, list=32%, si | MM_3 |
| GOBP_NUCLEAR_MEMBR | GOBP_NUCLEAR_MEMBRANE_ORGANIZATI | 35 | 0.438381 | 1.4305049 | 0.08062575 | 0.18144487 | 1 | 6931 | tags=54%, list=29%, si | MM_3 |
| GOBP_REGULATION_OF_ | GOBP_REGULATION_OF_AXON_REGENERAT | 36 | 0.436194 | 1.430321 | 0.05882353 | 0.18149646 | 1 | 8586 | tags=61%, list=36%, si | MM_3 |
| GOBP_NUCLEOSIDE_BISP | GOBP_NUCLEOSIDE_BISPHOSPHATE_METAB | 102 | 0.371101 | 1.4300905 | 0.024520256 | 0.18163346 | 1 | 4994 | tags=32%, list=21%, si | MM_3 |
| GOBP_PROTEIN_LOCALIZ | GOBP_PROTEIN_LOCALIZATION_TO_CHROM | 112 | 0.370317 | 1.43004 | 0.015856236 | 0.18152258 | 1 | 6154 | tags=38%, list=26%, si | MM_3 |
| GOBP_REGULATION_OF_ | GOBP_REGULATION_OF_CALCIUM_MEDIAT | 93 | 0.376569 | 1.4296317 | 0.033369213 | 0.18192986 | 1 | 5509 | tags=33%, list=23%, si | MM_3 |
| GOBP_NEGATIVE_REGUL | GOBP_NEGATIVE_REGULATION_OF_VASOC | 17 | 0.508024 | 1.4296093 | 0.0880829 | 0.18179116 | 1 | 7570 | tags=53%, list=32%, si | MM_3 |
| GOBP_CORTICAL_CYTOSK | GOBP_CORTICAL_CYTOSKELETON_ORGANIZ | 53 | 0.407857 | 1.4294171 | 0.04367816 | 0.18190941 | 1 | 7254 | tags=51%, list=31%, si | MM_3 |
| GOBP_PEROXISOMAL_TR | GOBP_PEROXISOMAL_TRANSPORT | 21 | 0.492748 | 1.4293981 | 0.086065575 | 0.18176897 | 1 | 5313 | tags=43%, list=23%, si | MM_3 |
| STARCH_AND_SUCROSE_ | STARCH_AND_SUCROSE_METABOLISM | 26 | 0.465887 | 1.428629 | 0.06949807 | 0.18253487 | 1 | 6220 | tags=42%, list=26%, si | MM_3 |
| GOBP_GRANULOCYTE_AC | GOBP_GRANULOCYTE_ACTIVATION | 41 | 0.430125 | 1.4286108 | 0.06793802 | 0.18238558 | 1 | 7568 | tags=54%, list=32%, si | MM_3 |
| GOBP_NEGATIVE_REGUL | GOBP_NEGATIVE_REGULATION_OF_ESTABL | 148 | 0.361717 | 1.4285756 | 0.014403292 | 0.1822614 | 1 | 6779 | tags=43%, list=29%, si | MM_3 |
| GOBP_INFLAMMATORY_ | GOBP_INFLAMMATORY_RESPONSE_TO_ANT | 62 | 0.398085 | 1.4283131 | 0.05824176 | 0.18240105 | 1 | 4236 | tags=27%, list=18%, si | MM_3 |
| GOBP_POSITIVE_REGULA | GOBP_POSITIVE_REGULATION_OF_PROTEIN | 156 | 0.36057 | 1.4279568 | 0.010288066 | 0.18270728 | 1 | 7081 | tags=40%, list=30%, si | MM_3 |
| GOBP_TRANSFORMING_ | GOBP_TRANSFORMING_GROWTH_FACTOR_ | 202 | 0.352639 | 1.4277889 | 0.007142857 | 0.18276784 | 1 | 6781 | tags=41%, list=29%, si | MM_3 |
| GOBP_VASCULAR_ASSOC | GOBP_VASCULAR_ASSOCIATED_SMOOTH_M | 39 | 0.425137 | 1.427761 | 0.06769596 | 0.18263792 | 1 | 5307 | tags=38%, list=22%, si | MM_3 |
| GOBP_IRON_ION_TRANS | GOBP_IRON_ION_TRANSPORT | 54 | 0.403547 | 1.4277135 | 0.055813953 | 0.18253694 | 1 | 6393 | tags=44%, list=27%, si | MM_3 |
| GOBP_POSITIVE_REGULA | GOBP_POSITIVE_REGULATION_OF_PROTEIN | 78 | 0.384894 | 1.4276716 | 0.031868134 | 0.18243235 | 1 | 7322 | tags=45%, list=31%, si | MM_3 |
| GOBP_NEGATIVE_REGUL | GOBP_NEGATIVE_REGULATION_OF_STRIATE | 39 | 0.421081 | 1.4268558 | 0.06872038 | 0.1832549 | 1 | 1750 | tags=18%, list=7%, sig | MM_3 |
| GOBP_NEGATIVE_REGUL | GOBP_NEGATIVE_REGULATION_OF_SMOOT | 36 | 0.441187 | 1.4267827 | 0.07393939 | 0.18317533 | 1 | 5903 | tags=36%, list=25%, si | MM_3 |
| GOBP_PROTEIN_CATABO | GOBP_PROTEIN_CATABOLIC_PROCESS_IN_T | 22 | 0.476095 | 1.4265442 | 0.06298201 | 0.1833259 | 1 | 7147 | tags=68%, list=30%, si | MM_3 |
| GOBP_RECEPTOR_INTERN | GOBP_RECEPTOR_INTERNALIZATION | 154 | 0.35556 | 1.4262173 | 0.014537903 | 0.18360645 | 1 | 6781 | tags=38%, list=29%, si | MM_3 |
| GOBP_RESPONSE_TO_DI | GOBP_RESPONSE_TO_DIETARY_EXCESS | 44 | 0.416428 | 1.4260044 | 0.067058824 | 0.18369952 | 1 | 6124 | tags=41%, list=26%, si | MM_3 |
| GOBP_MITOCHONDRIAL_ | GOBP_MITOCHONDRIAL_GENOME_MAINTE | 35 | 0.435532 | 1.4257139 | 0.077970296 | 0.18391125 | 1 | 7230 | tags=51%, list=31%, si | MM_3 |
| GOBP_REGULATION_OF_ | GOBP_REGULATION_OF_MUSCLE_ADAPTAT | 102 | 0.37676 | 1.4256798 | 0.033405174 | 0.1837956 | 1 | 7339 | tags=46%, list=31%, si | MM_3 |
| GOBP_ESTABLISHMENT_ | GOBP_ESTABLISHMENT_OF_BLOOD_BRAIN_ | 17 | 0.502706 | 1.4255841 | 0.10854817 | 0.18376875 | 1 | 4152 | tags=41%, list=18%, si | MM_3 |
| GOBP_SPHINGOMYELIN_ | GOBP_SPHINGOMYELIN_METABOLIC_PROC | 21 | 0.4822 | 1.425576 | 0.09466667 | 0.18360923 | 1 | 7190 | tags=62%, list=30%, si | MM_3 |
| GOBP_QUINONE_METAB | GOBP_QUINONE_METABOLIC_PROCESS | 23 | 0.46836 | 1.4254158 | 0.090314135 | 0.18362667 | 1 | 4215 | tags=35%, list=18%, si | MM_3 |
| PORPHYRIN_AND_CHLOR | PORPHYRIN_AND_CHLOROPHYLL_METABOL | 31 | 0.447315 | 1.425266 | 0.07503075 | 0.18367049 | 1 | 5217 | tags=35%, list=22%, si | MM_3 |

| | | | | | | | | | | |
|---|---|---|---|---|---|---|---|---|---|---|
| GOBP_NEGATIVE_REGUL | GOBP_NEGATIVE_REGULATION_OF_EXOCYT | 37 | 0.428906 | 1.4252411 | 0.073405534 | 0.18353206 | 1 | 7377 | tags=54%, list=31%, si | MM_3 |
| GOBP_TRICARBOXYLIC_A | GOBP_TRICARBOXYLIC_ACID_METABOLIC_P | 16 | 0.512037 | 1.4251101 | 0.08855586 | 0.18352237 | 1 | 3872 | tags=38%, list=16%, si | MM_3 |
| GOBP_SERINE_FAMILY_A | GOBP_SERINE_FAMILY_AMINO_ACID_META | 30 | 0.449162 | 1.4249831 | 0.07481297 | 0.1835278 | 1 | 4000 | tags=33%, list=17%, si | MM_3 |
| GOBP_TELOMERE_MAINT | GOBP_TELOMERE_MAINTENANCE_VIA_TEL | 76 | 0.388391 | 1.4248022 | 0.0436205 | 0.18358468 | 1 | 7216 | tags=45%, list=31%, si | MM_3 |
| GOBP_REGULATION_OF_ | GOBP_REGULATION_OF_SIGNALING_RECEP | 109 | 0.3691 | 1.4245687 | 0.027484143 | 0.18373497 | 1 | 3571 | tags=23%, list=15%, si | MM_3 |
| GOBP_DOUBLE_STRAND_ | GOBP_DOUBLE_STRAND_BREAK_REPAIR_VI | 67 | 0.392443 | 1.4244306 | 0.04290429 | 0.18374947 | 1 | 5012 | tags=37%, list=21%, si | MM_3 |
| GOBP_CGAS_STING_SIGN | GOBP_CGAS_STING_SIGNALING_PATHWAY | 16 | 0.515207 | 1.4243804 | 0.09641873 | 0.18363413 | 1 | 4409 | tags=38%, list=19%, si | MM_3 |
| GOBP_UBIQUINONE_MET | GOBP_UBIQUINONE_METABOLIC_PROCESS | 19 | 0.49055 | 1.4238154 | 0.08695652 | 0.18421818 | 1 | 4215 | tags=37%, list=18%, si | MM_3 |
| GOBP_NEGATIVE_REGUL | GOBP_NEGATIVE_REGULATION_OF_CANON | 57 | 0.405678 | 1.4237872 | 0.05956472 | 0.18409525 | 1 | 6635 | tags=42%, list=28%, si | MM_3 |
| GOBP_IMMUNOGLOBULI | GOBP_IMMUNOGLOBULIN_PRODUCTION | 132 | 0.36635 | 1.4236497 | 0.017989418 | 0.18413618 | 1 | 5024 | tags=30%, list=21%, si | MM_3 |
| GOBP_NEGATIVE_REGUL | GOBP_NEGATIVE_REGULATION_OF_LYMPH | 44 | 0.420756 | 1.4232844 | 0.06720742 | 0.18446259 | 1 | 4409 | tags=30%, list=19%, si | MM_3 |
| GOBP_NEGATIVE_REGUL | GOBP_NEGATIVE_REGULATION_OF_CELLUL | 51 | 0.413439 | 1.4221971 | 0.0575374 | 0.18569256 | 1 | 6079 | tags=37%, list=26%, si | MM_3 |
| GOBP_POSITIVE_REGULA | GOBP_POSITIVE_REGULATION_OF_DNA_BI | 76 | 0.385554 | 1.4214327 | 0.04605263 | 0.18653479 | 1 | 7698 | tags=51%, list=33%, si | MM_3 |
| GOBP_PHOSPHATIDYLINO | GOBP_PHOSPHATIDYLINOSITOL_PHOSPHAT | 50 | 0.408309 | 1.421383 | 0.052878965 | 0.18643256 | 1 | 4302 | tags=30%, list=18%, si | MM_3 |
| GOBP_ESTABLISHMENT_ | GOBP_ESTABLISHMENT_OR_MAINTENANCE | 25 | 0.475508 | 1.4213485 | 0.093709886 | 0.18630004 | 1 | 7600 | tags=56%, list=32%, si | MM_3 |
| GOBP_NEGATIVE_REGUL | GOBP_NEGATIVE_REGULATION_OF_CALCIU | 22 | 0.486161 | 1.420834 | 0.096985586 | 0.18681012 | 1 | 4272 | tags=32%, list=18%, si | MM_3 |
| GOBP_PROTEIN_LOCALIZ | GOBP_PROTEIN_LOCALIZATION_TO_NUCLE | 326 | 0.346178 | 1.4207028 | 0.0020120724 | 0.18682458 | 1 | 7544 | tags=45%, list=32%, si | MM_3 |
| GOBP_SEQUESTERING_O | GOBP_SEQUESTERING_OF_TRIGLYCERIDE | 18 | 0.510022 | 1.4205667 | 0.07856192 | 0.18684344 | 1 | 5113 | tags=39%, list=22%, si | MM_3 |
| GOBP_NEGATIVE_REGUL | GOBP_NEGATIVE_REGULATION_OF_GROWT | 239 | 0.353427 | 1.4204761 | 0.005055612 | 0.18681175 | 1 | 4256 | tags=24%, list=18%, si | MM_3 |
| GOBP_PHOSPHOLIPID_M | GOBP_PHOSPHOLIPID_METABOLIC_PROCES | 352 | 0.34333 | 1.4199537 | 0 | 0.18733609 | 1 | 4804 | tags=28%, list=20%, si | MM_3 |
| GOBP_POSITIVE_REGULA | GOBP_POSITIVE_REGULATION_OF_PEPTIDA | 161 | 0.355453 | 1.4193276 | 0.021649485 | 0.18803056 | 1 | 7534 | tags=49%, list=32%, si | MM_3 |
| GOBP_NEGATIVE_REGUL | GOBP_NEGATIVE_REGULATION_OF_PROTEA | 51 | 0.404148 | 1.4192957 | 0.05854801 | 0.18791482 | 1 | 1369 | tags=14%, list=6%, sig | MM_3 |
| GOBP_LAYER_FORMATIO | GOBP_LAYER_FORMATION_IN_CEREBRAL_C | 16 | 0.521259 | 1.419141 | 0.093622796 | 0.18797979 | 1 | 5855 | tags=50%, list=25%, si | MM_3 |
| GOBP_PROTEIN_TARGETI | GOBP_PROTEIN_TARGETING_TO_VACUOLE | 46 | 0.41005 | 1.419071 | 0.06884481 | 0.18791488 | 1 | 7162 | tags=43%, list=30%, si | MM_3 |
| GOBP_POSITIVE_REGULA | GOBP_POSITIVE_REGULATION_OF_MYELINA | 25 | 0.462212 | 1.4189627 | 0.0786802 | 0.18787861 | 1 | 5985 | tags=44%, list=25%, si | MM_3 |
| GOBP_DOUBLE_STRAND_ | GOBP_DOUBLE_STRAND_BREAK_REPAIR | 299 | 0.346436 | 1.4188191 | 0.0020120724 | 0.18793601 | 1 | 4820 | tags=28%, list=20%, si | MM_3 |
| GOBP_GLYCEROPHOSPH | GOBP_GLYCEROPHOSPHOLIPID_BIOSYNTHE | 160 | 0.359628 | 1.4185992 | 0.011363637 | 0.1880574 | 1 | 4804 | tags=30%, list=20%, si | MM_3 |
| GOBP_NEURON_REMODE | GOBP_NEURON_REMODELING | 19 | 0.489724 | 1.4184508 | 0.112137206 | 0.18809168 | 1 | 3849 | tags=42%, list=16%, si | MM_3 |
| GOBP_SPLEEN_DEVELOPI | GOBP_SPLEEN_DEVELOPMENT | 46 | 0.409844 | 1.4184227 | 0.06928407 | 0.1879702 | 1 | 4794 | tags=35%, list=20%, si | MM_3 |
| GOBP_POSITIVE_REGULA | GOBP_POSITIVE_REGULATION_OF_SEQUEST | 16 | 0.51028 | 1.4182827 | 0.0941337 | 0.18797688 | 1 | 4256 | tags=38%, list=18%, si | MM_3 |
| GOBP_INTRACELLULAR_R | GOBP_INTRACELLULAR_RECEPTOR_SIGNALI | 325 | 0.34448 | 1.4181417 | 0.001001001 | 0.18801655 | 1 | 7082 | tags=40%, list=30%, si | MM_3 |
| GOBP_MICROTUBULE_NU | GOBP_MICROTUBULE_NUCLEATION | 35 | 0.434522 | 1.4176412 | 0.07655502 | 0.1884942 | 1 | 6981 | tags=51%, list=30%, si | MM_3 |
| GOBP_REGULATION_OF_ | GOBP_REGULATION_OF_VASCULAR_PERME | 48 | 0.404862 | 1.4176253 | 0.056287427 | 0.18834452 | 1 | 4256 | tags=33%, list=18%, si | MM_3 |
| GOBP_STEROL_TRANSPO | GOBP_STEROL_TRANSPORT | 109 | 0.373761 | 1.4173427 | 0.03315508 | 0.18860805 | 1 | 7182 | tags=48%, list=30%, si | MM_3 |
| GOBP_PURINE_NUCLEOS | GOBP_PURINE_NUCLEOSIDE_MONOPHOSP | 36 | 0.43912 | 1.4172298 | 0.07 | 0.18859884 | 1 | 4801 | tags=36%, list=20%, si | MM_3 |
| GOBP_NEGATIVE_REGUL | GOBP_NEGATIVE_REGULATION_OF_NON_C | 26 | 0.460834 | 1.4169825 | 0.09343434 | 0.1887631 | 1 | 5614 | tags=38%, list=24%, si | MM_3 |
| GOBP_MODULATION_BY_ | GOBP_MODULATION_BY_HOST_OF_VIRAL_ | 22 | 0.46995 | 1.4168835 | 0.074270554 | 0.18873008 | 1 | 7909 | tags=73%, list=33%, si | MM_3 |
| GOBP_CHROMOSOME_SE | GOBP_CHROMOSOME_SEGREGATION | 403 | 0.340592 | 1.4167062 | 0 | 0.18879588 | 1 | 7081 | tags=41%, list=30%, si | MM_3 |
| GOBP_ENDOSOME_TO_P | GOBP_ENDOSOME_TO_PLASMA_MEMBRAN | 21 | 0.482305 | 1.4165593 | 0.087012984 | 0.18882014 | 1 | 5729 | tags=52%, list=24%, si | MM_3 |
| GOBP_NEGATIVE_REGUL | GOBP_NEGATIVE_REGULATION_OF_CELLUL | 25 | 0.463345 | 1.416526 | 0.070876285 | 0.18870819 | 1 | 6636 | tags=44%, list=28%, si | MM_3 |
| GOBP_PROTEIN_LOCALIZ | GOBP_PROTEIN_LOCALIZATION_TO_LYSOSO | 54 | 0.404498 | 1.4164265 | 0.05499439 | 0.18866545 | 1 | 7214 | tags=54%, list=31%, si | MM_3 |
| GOBP_LIPOPROTEIN_BIOS | GOBP_LIPOPROTEIN_BIOSYNTHETIC_PROCE | 104 | 0.367947 | 1.4163846 | 0.035181236 | 0.18855481 | 1 | 4421 | tags=32%, list=19%, si | MM_3 |
| GOBP_MELANOSOME_AS | GOBP_MELANOSOME_ASSEMBLY | 19 | 0.498476 | 1.415761 | 0.0845839 | 0.18925999 | 1 | 4131 | tags=42%, list=17%, si | MM_3 |
| GOBP_REGULATION_OF_ | GOBP_REGULATION_OF_LYMPHOCYTE_ME | 190 | 0.35356 | 1.4156454 | 0.009137056 | 0.1892476 | 1 | 6059 | tags=36%, list=26%, si | MM_3 |
| GOBP_POSITIVE_REGULA | GOBP_POSITIVE_REGULATION_OF_MAPK_C | 496 | 0.339237 | 1.4155123 | 0 | 0.18926121 | 1 | 6030 | tags=35%, list=26%, si | MM_3 |
| GOBP_ALPHA_AMINO_A | GOBP_ALPHA_AMINO_ACID_METABOLIC_P | 189 | 0.355173 | 1.4153663 | 0.009211873 | 0.18930286 | 1 | 5181 | tags=33%, list=22%, si | MM_3 |
| GOBP_POSITIVE_REGULA | GOBP_POSITIVE_REGULATION_OF_AXONO | 98 | 0.371034 | 1.4153442 | 0.042826552 | 0.18917234 | 1 | 6781 | tags=42%, list=29%, si | MM_3 |
| CYTOKINE-CYTOKINE_RE | CYTOKINE-CYTOKINE_RECEPTOR_INTERACTI | 239 | 0.350661 | 1.4151281 | 0.004048583 | 0.18925461 | 1 | 3801 | tags=24%, list=16%, si | MM_3 |
| LONGEVITY_REGULATING | LONGEVITY_REGULATING_PATHWAY | 99 | 0.372284 | 1.4146876 | 0.031182796 | 0.18966213 | 1 | 7204 | tags=48%, list=31%, si | MM_3 |
| GOBP_POSITIVE_REGULA | GOBP_POSITIVE_REGULATION_OF_PROTEIN | 41 | 0.418473 | 1.4142512 | 0.07311321 | 0.19013377 | 1 | 7299 | tags=51%, list=31%, si | MM_3 |
| GOBP_ENSHEATHMENT_ | GOBP_ENSHEATHMENT_OF_NEURONS | 169 | 0.35495 | 1.4139891 | 0.016477859 | 0.19035397 | 1 | 6124 | tags=38%, list=26%, si | MM_3 |
| GOBP_PLASMINOGEN_A | GOBP_PLASMINOGEN_ACTIVATION | 23 | 0.469261 | 1.4137312 | 0.086563304 | 0.1905056 | 1 | 4183 | tags=39%, list=18%, si | MM_3 |
| HEPATITIS_B | HEPATITIS_B | 147 | 0.356408 | 1.4135312 | 0.017689906 | 0.1906024 | 1 | 8145 | tags=52%, list=34%, si | MM_3 |
| GOBP_ENDOCYTIC_RECY | GOBP_ENDOCYTIC_RECYCLING | 88 | 0.377352 | 1.4134705 | 0.03504929 | 0.19051911 | 1 | 7011 | tags=44%, list=30%, si | MM_3 |
| GOBP_MYELIN_MAINTEN | GOBP_MYELIN_MAINTENANCE | 16 | 0.514187 | 1.4127822 | 0.08309859 | 0.19131686 | 1 | 5115 | tags=38%, list=22%, si | MM_3 |
| GOBP_POSITIVE_REGULA | GOBP_POSITIVE_REGULATION_OF_BROWN_ | 17 | 0.508718 | 1.4126753 | 0.09226594 | 0.1912767 | 1 | 4190 | tags=35%, list=18%, si | MM_3 |
| ENDOCRINE_AND_OTHER | ENDOCRINE_AND_OTHER_FACTOR-REGULAT | 42 | 0.416618 | 1.4123083 | 0.07482185 | 0.19160709 | 1 | 6213 | tags=43%, list=26%, si | MM_3 |
| GOBP_RESPONSE_TO_X_ | GOBP_RESPONSE_TO_X_RAY | 24 | 0.464827 | 1.4120148 | 0.073834196 | 0.19182031 | 1 | 2345 | tags=25%, list=10%, si | MM_3 |
| GOBP_PLATELET_DERIVE | GOBP_PLATELET_DERIVED_GROWTH_FACT | 63 | 0.388573 | 1.4118788 | 0.057757646 | 0.19183342 | 1 | 6914 | tags=40%, list=29%, si | MM_3 |
| GOBP_POSITIVE_REGULA | GOBP_POSITIVE_REGULATION_OF_NEURON | 15 | 0.527511 | 1.4110798 | 0.10887097 | 0.19276136 | 1 | 8550 | tags=73%, list=36%, si | MM_3 |
| GOBP_PROTEIN_INSERTI | GOBP_PROTEIN_INSERTION_INTO_ER_MEM | 26 | 0.456465 | 1.4110109 | 0.08928572 | 0.19267935 | 1 | 6806 | tags=54%, list=29%, si | MM_3 |
| GOBP_MAST_CELL_MIGR | GOBP_MAST_CELL_MIGRATION | 17 | 0.513678 | 1.4106839 | 0.08769448 | 0.19298168 | 1 | 4170 | tags=29%, list=18%, si | MM_3 |
| GOBP_NEGATIVE_REGUL | GOBP_NEGATIVE_REGULATION_OF_INFLAM | 153 | 0.359348 | 1.4106778 | 0.011530398 | 0.19282222 | 1 | 5515 | tags=33%, list=23%, si | MM_3 |
| GOBP_REGENERATION | GOBP_REGENERATION | 161 | 0.35415 | 1.4105772 | 0.016528925 | 0.19279537 | 1 | 5054 | tags=30%, list=21%, si | MM_3 |
| GOBP_MUSCLE_HYPERTR | GOBP_MUSCLE_HYPERTROPHY | 115 | 0.366362 | 1.4104241 | 0.028723404 | 0.19283931 | 1 | 7322 | tags=42%, list=31%, si | MM_3 |
| GOBP_MYELOID_LEUKOC | GOBP_MYELOID_LEUKOCYTE_MEDIATED_IN | 108 | 0.371057 | 1.4100958 | 0.033862434 | 0.19313611 | 1 | 7218 | tags=42%, list=31%, si | MM_3 |
| GOBP_OSTEOCLAST_DIFF | GOBP_OSTEOCLAST_DIFFERENTIATION | 126 | 0.363598 | 1.4096904 | 0.023084994 | 0.19347313 | 1 | 6330 | tags=34%, list=27%, si | MM_3 |
| GOBP_REGULATION_OF_ | GOBP_REGULATION_OF_T_CELL_ACTIVATIO | 364 | 0.342765 | 1.4095689 | 0.002006018 | 0.19346869 | 1 | 5751 | tags=32%, list=24%, si | MM_3 |
| GOBP_VESICLE_ORGANIZ | GOBP_VESICLE_ORGANIZATION | 353 | 0.340538 | 1.4094164 | 0.0010050251 | 0.1935183 | 1 | 7089 | tags=44%, list=30%, si | MM_3 |
| GOBP_POSITIVE_REGULA | GOBP_POSITIVE_REGULATION_OF_CELL_DI | 79 | 0.383126 | 1.4090508 | 0.036026202 | 0.19383793 | 1 | 7731 | tags=52%, list=33%, si | MM_3 |
| GOBP_HIPPO_SIGNALING | GOBP_HIPPO_SIGNALING | 40 | 0.421252 | 1.4087305 | 0.08452381 | 0.1941096 | 1 | 3782 | tags=30%, list=16%, si | MM_3 |
| GOBP_POSITIVE_REGULA | GOBP_POSITIVE_REGULATION_OF_MITOCH | 17 | 0.504772 | 1.407933 | 0.1195795 | 0.19505684 | 1 | 6164 | tags=41%, list=26%, si | MM_3 |
| GOBP_REGULATION_OF_ | GOBP_REGULATION_OF_GTPASE_ACTIVITY | 287 | 0.344954 | 1.4077817 | 0.0030395137 | 0.19508989 | 1 | 5778 | tags=32%, list=24%, si | MM_3 |
| GOBP_REGULATION_OF_ | GOBP_REGULATION_OF_LEUKOCYTE_APOP | 103 | 0.372711 | 1.4077437 | 0.041304346 | 0.19497679 | 1 | 4409 | tags=27%, list=19%, si | MM_3 |
| GOBP_POSITIVE_REGULA | GOBP_POSITIVE_REGULATION_OF_SMOOTH | 69 | 0.394352 | 1.4075519 | 0.045146726 | 0.19504498 | 1 | 7542 | tags=48%, list=32%, si | MM_3 |
| GOBP_ARP2_3_COMPLEX | GOBP_ARP2_3_COMPLEX_MEDIATED_ACTI | 41 | 0.416841 | 1.4064308 | 0.07125604 | 0.19640766 | 1 | 7483 | tags=54%, list=32%, si | MM_3 |
| GOBP_CHROMOSOME_LO | GOBP_CHROMOSOME_LOCALIZATION | 111 | 0.369296 | 1.4061497 | 0.036208734 | 0.19662651 | 1 | 7083 | tags=43%, list=30%, si | MM_3 |
| GOBP_CELLULAR_RESPON | GOBP_CELLULAR_RESPONSE_TO_RADIATIO | 165 | 0.356994 | 1.4059656 | 0.014256619 | 0.19670023 | 1 | 5759 | tags=38%, list=24%, si | MM_3 |
| GOBP_METAPHASE_CHR | GOBP_METAPHASE_CHROMOSOME_ALIGN | 93 | 0.369629 | 1.4056062 | 0.04883227 | 0.19701046 | 1 | 6709 | tags=41%, list=28%, si | MM_3 |
| GOBP_NEGATIVE_REGUL | GOBP_NEGATIVE_REGULATION_OF_CYTOKI | 292 | 0.347637 | 1.405243 | 0.004016064 | 0.1973103 | 1 | 6442 | tags=38%, list=27%, si | MM_3 |
| GOBP_LEUKOCYTE_PROL | GOBP_LEUKOCYTE_PROLIFERATION | 361 | 0.341337 | 1.4047893 | 0.001004016 | 0.19775267 | 1 | 5711 | tags=32%, list=24%, si | MM_3 |
| AGE-RAGE_SIGNALING_P | AGE-RAGE_SIGNALING_PATHWAY_IN_DIAB | 100 | 0.368457 | 1.4046968 | 0.038751345 | 0.19770725 | 1 | 5977 | tags=36%, list=25%, si | MM_3 |

| | | | | | | | | | | |
|---|---|---|---|---|---|---|---|---|---|---|
| GOBP_NEGATIVE_REGUL | GOBP_NEGATIVE_REGULATION_OF_RESPON | 362 | 0.341411 | 1.4046425 | 0.002002002 | 0.19761117 | 1 | 5518 | tags=31%, list=23%, si | MM_3 |
| GOBP_NEGATIVE_REGUL | GOBP_NEGATIVE_REGULATION_OF_LYMPH | 62 | 0.390859 | 1.403767 | 0.06300115 | 0.19861327 | 1 | 6040 | tags=34%, list=26%, si | MM_3 |
| GOBP_REGULATION_OF_ | GOBP_REGULATION_OF_MAP_KINASE_ACTI | 159 | 0.354909 | 1.4035693 | 0.016427105 | 0.19869617 | 1 | 6634 | tags=40%, list=28%, si | MM_3 |
| GOBP_ORGANELLE_FISSI | GOBP_ORGANELLE_FISSION | 477 | 0.338284 | 1.4032892 | 0 | 0.19890794 | 1 | 6936 | tags=40%, list=29%, si | MM_3 |
| GOBP_NEGATIVE_REGUL | GOBP_NEGATIVE_REGULATION_OF_STEM_ | 25 | 0.459125 | 1.4030248 | 0.09395109 | 0.199089 | 1 | 2888 | tags=24%, list=12%, si | MM_3 |
| GOBP_POSITIVE_REGULA | GOBP_POSITIVE_REGULATION_OF_INTERLE | 106 | 0.368535 | 1.4028828 | 0.0393617 | 0.19910705 | 1 | 5807 | tags=32%, list=25%, si | MM_3 |
| GOBP_NEGATIVE_REGUL | GOBP_NEGATIVE_REGULATION_OF_CELL_C | 368 | 0.34315 | 1.4024918 | 0.003003003 | 0.19948561 | 1 | 6697 | tags=39%, list=28%, si | MM_3 |
| GOBP_PEROXISOME_ORG | GOBP_PEROXISOME_ORGANIZATION | 38 | 0.422946 | 1.4021974 | 0.08591885 | 0.19970219 | 1 | 5313 | tags=34%, list=23%, si | MM_3 |
| GOBP_REGULATION_OF_ | GOBP_REGULATION_OF_HEMOPOIESIS | 425 | 0.338525 | 1.4021425 | 0.001002004 | 0.199605 | 1 | 6594 | tags=36%, list=28%, si | MM_3 |
| GOBP_NEGATIVE_REGUL | GOBP_NEGATIVE_REGULATION_OF_PHAGO | 24 | 0.465567 | 1.4020166 | 0.08454426 | 0.19960782 | 1 | 7430 | tags=54%, list=31%, si | MM_3 |
| GOBP_INTERLEUKIN_8_P | GOBP_INTERLEUKIN_8_PRODUCTION | 73 | 0.385533 | 1.4019016 | 0.050055616 | 0.19959588 | 1 | 7521 | tags=47%, list=32%, si | MM_3 |
| GOBP_REGULATION_OF_ | GOBP_REGULATION_OF_SODIUM_ION_TRA | 86 | 0.37732 | 1.4016517 | 0.03969957 | 0.19979581 | 1 | 6906 | tags=45%, list=29%, si | MM_3 |
| GOBP_OLIGODENDROCYT | GOBP_OLIGODENDROCYTE_DIFFERENTIATIO | 112 | 0.361576 | 1.4015986 | 0.030494217 | 0.19970585 | 1 | 5930 | tags=31%, list=25%, si | MM_3 |
| GOBP_IMMUNOLOGICAL_ | GOBP_IMMUNOLOGICAL_SYNAPSE_FORMA | 18 | 0.494835 | 1.4014215 | 0.096945554 | 0.19979306 | 1 | 3486 | tags=39%, list=15%, si | MM_3 |
| GOBP_NEGATIVE_REGUL | GOBP_NEGATIVE_REGULATION_OF_MRNA_ | 92 | 0.366262 | 1.4012666 | 0.04120267 | 0.1998457 | 1 | 5323 | tags=33%, list=23%, si | MM_3 |
| GOBP_POSITIVE_REGULA | GOBP_POSITIVE_REGULATION_OF_APOPTO | 161 | 0.352645 | 1.4007716 | 0.018499486 | 0.20038456 | 1 | 7534 | tags=50%, list=32%, si | MM_3 |
| GOBP_RESPONSE_TO_GA | GOBP_RESPONSE_TO_GAMMA_RADIATION | 48 | 0.397426 | 1.400668 | 0.064327486 | 0.20036002 | 1 | 6864 | tags=52%, list=29%, si | MM_3 |
| GOBP_PLATELET_MORPH | GOBP_PLATELET_MORPHOGENESIS | 25 | 0.461486 | 1.4006345 | 0.11182519 | 0.20024075 | 1 | 6945 | tags=56%, list=29%, si | MM_3 |
| GAP_JUNCTION | GAP_JUNCTION | 83 | 0.373891 | 1.4005824 | 0.04600219 | 0.20014562 | 1 | 6131 | tags=35%, list=26%, si | MM_3 |
| GOBP_POSITIVE_REGULA | GOBP_POSITIVE_REGULATION_OF_LEUKOC | 169 | 0.350055 | 1.399929 | 0.021604938 | 0.20089756 | 1 | 5711 | tags=33%, list=24%, si | MM_3 |
| GOBP_LIPID_OXIDATION | GOBP_LIPID_OXIDATION | 118 | 0.364124 | 1.3996732 | 0.026123302 | 0.2010848 | 1 | 5029 | tags=29%, list=21%, si | MM_3 |
| GOBP_VASCULAR_ENDOT | GOBP_VASCULAR_ENDOTHELIAL_GROWTH_ | 50 | 0.399817 | 1.3994858 | 0.06666667 | 0.20116957 | 1 | 4264 | tags=32%, list=18%, si | MM_3 |
| GOBP_NEGATIVE_REGUL | GOBP_NEGATIVE_REGULATION_OF_CELL_A | 301 | 0.341137 | 1.3994603 | 0.004032258 | 0.2010346 | 1 | 5781 | tags=32%, list=24%, si | MM_3 |
| GOBP_POSITIVE_REGULA | GOBP_POSITIVE_REGULATION_OF_METAPH | 15 | 0.503843 | 1.3992152 | 0.10822511 | 0.2012075 | 1 | 6905 | tags=60%, list=29%, si | MM_3 |
| GOBP_POSITIVE_REGULA | GOBP_POSITIVE_REGULATION_OF_SMALL_ | 174 | 0.354535 | 1.3991942 | 0.015353122 | 0.20106934 | 1 | 5262 | tags=30%, list=22%, si | MM_3 |
| GOBP_MITOTIC_NUCLEA | GOBP_MITOTIC_NUCLEAR_DIVISION | 259 | 0.343417 | 1.3984917 | 0.003012048 | 0.20189779 | 1 | 7054 | tags=43%, list=30%, si | MM_3 |
| GOBP_POSITIVE_REGULA | GOBP_POSITIVE_REGULATION_OF_STRIATE | 26 | 0.453267 | 1.3983912 | 0.07780612 | 0.20187806 | 1 | 5470 | tags=38%, list=23%, si | MM_3 |
| NOD-LIKE_RECEPTOR_SIG | NOD-LIKE_RECEPTOR_SIGNALING_PATHWA | 151 | 0.355191 | 1.3982879 | 0.02376033 | 0.2018464 | 1 | 6125 | tags=37%, list=26%, si | MM_3 |
| GOBP_PHAGOCYTOSIS_E | GOBP_PHAGOCYTOSIS_ENGULFMENT | 53 | 0.394877 | 1.3977789 | 0.0804077 | 0.20235944 | 1 | 6919 | tags=45%, list=29%, si | MM_3 |
| GOBP_INTERLEUKIN_1_B | GOBP_INTERLEUKIN_1_BETA_PRODUCTION | 95 | 0.365748 | 1.3976804 | 0.043525573 | 0.20232227 | 1 | 7534 | tags=43%, list=32%, si | MM_3 |
| GOBP_BASE_EXCISION_R | GOBP_BASE_EXCISION_REPAIR | 41 | 0.414466 | 1.3976312 | 0.077844314 | 0.20222306 | 1 | 4874 | tags=32%, list=21%, si | MM_3 |
| GOBP_CELL_VOLUME_HO | GOBP_CELL_VOLUME_HOMEOSTASIS | 38 | 0.41535 | 1.3967612 | 0.0776583 | 0.20325355 | 1 | 6488 | tags=37%, list=27%, si | MM_3 |
| GOBP_POSITIVE_REGULA | GOBP_POSITIVE_REGULATION_OF_DNA_DA | 15 | 0.515901 | 1.3966187 | 0.11173184 | 0.20328912 | 1 | 4532 | tags=53%, list=19%, si | MM_3 |
| AMOEBIASIS | AMOEBIASIS | 89 | 0.370819 | 1.3966166 | 0.050108932 | 0.20312479 | 1 | 3208 | tags=24%, list=14%, si | MM_3 |
| GOBP_DNA_RECOMBINA | GOBP_DNA_RECOMBINATION | 313 | 0.340004 | 1.396232 | 0.0070422534 | 0.20353562 | 1 | 4746 | tags=28%, list=20%, si | MM_3 |
| GOBP_PROTEIN_CONTAI | GOBP_PROTEIN_CONTAINING_COMPLEX_L | 259 | 0.343363 | 1.395854 | 0.00907258 | 0.20389974 | 1 | 6809 | tags=40%, list=29%, si | MM_3 |
| TRANSCRIPTIONAL_MISR | TRANSCRIPTIONAL_MISREGULATION_IN_CA | 170 | 0.352815 | 1.3956283 | 0.014344262 | 0.20404312 | 1 | 5977 | tags=34%, list=25%, si | MM_3 |
| GOBP_MUSCLE_ADAPTAT | GOBP_MUSCLE_ADAPTATION | 126 | 0.360278 | 1.3950732 | 0.02620545 | 0.20464508 | 1 | 7339 | tags=44%, list=31%, si | MM_3 |
| GOBP_GOLGI_ORGANIZA | GOBP_GOLGI_ORGANIZATION | 135 | 0.356896 | 1.3948617 | 0.01875 | 0.20476052 | 1 | 6523 | tags=40%, list=28%, si | MM_3 |
| GOBP_REGULATION_OF_ | GOBP_REGULATION_OF_STORE_OPERATED_ | 18 | 0.497453 | 1.3946337 | 0.09838275 | 0.20490353 | 1 | 1811 | tags=28%, list=8%, sig | MM_3 |
| GOBP_ERAD_PATHWAY | GOBP_ERAD_PATHWAY | 108 | 0.363537 | 1.3945367 | 0.036247335 | 0.20484707 | 1 | 7070 | tags=44%, list=30%, si | MM_3 |
| GOBP_POSITIVE_REGULA | GOBP_POSITIVE_REGULATION_OF_IMMUN | 55 | 0.397122 | 1.3941486 | 0.079768784 | 0.20526177 | 1 | 4592 | tags=27%, list=19%, si | MM_3 |
| GOBP_MAINTENANCE_O | GOBP_MAINTENANCE_OF_CELL_NUMBER | 205 | 0.346354 | 1.3939469 | 0.0091743115 | 0.20537956 | 1 | 6263 | tags=35%, list=27%, si | MM_3 |
| RENAL_CELL_CARCINOM | RENAL_CELL_CARCINOMA | 67 | 0.38116 | 1.3938098 | 0.06087937 | 0.20538254 | 1 | 7064 | tags=55%, list=30%, si | MM_3 |
| GOBP_POSITIVE_REGULA | GOBP_POSITIVE_REGULATION_OF_CELLULA | 20 | 0.488649 | 1.3936509 | 0.09004093 | 0.205431 | 1 | 1273 | tags=20%, list=5%, sig | MM_3 |
| GOBP_PROTEIN_LOCALIZ | GOBP_PROTEIN_LOCALIZATION_TO_VACUO | 78 | 0.371551 | 1.3936256 | 0.05771365 | 0.205296 | 1 | 7214 | tags=47%, list=31%, si | MM_3 |
| GOBP_MITOCHONDRIAL_ | GOBP_MITOCHONDRIAL_DNA_METABOLIC_ | 26 | 0.44905 | 1.3935645 | 0.103846155 | 0.20521997 | 1 | 7230 | tags=58%, list=31%, si | MM_3 |
| GOBP_AXON_EXTENSION | GOBP_AXON_EXTENSION | 125 | 0.357004 | 1.3931509 | 0.036055144 | 0.20565501 | 1 | 7337 | tags=42%, list=31%, si | MM_3 |
| PENTOSE_AND_GLUCURO | PENTOSE_AND_GLUCURONATE_INTERCONV | 22 | 0.464183 | 1.3923796 | 0.112299465 | 0.20662436 | 1 | 3362 | tags=27%, list=14%, si | MM_3 |
| GOBP_ACTIN_POLYMERIZ | GOBP_ACTIN_POLYMERIZATION_OR_DEPOL | 187 | 0.346652 | 1.392202 | 0.014285714 | 0.20669441 | 1 | 8106 | tags=51%, list=34%, si | MM_3 |
| GOBP_CORTICAL_ACTIN_ | GOBP_CORTICAL_ACTIN_CYTOSKELETON_O | 42 | 0.407543 | 1.3915681 | 0.07783019 | 0.20744728 | 1 | 8273 | tags=62%, list=35%, si | MM_3 |
| GOBP_REGULATION_OF_ | GOBP_REGULATION_OF_CHEMOTAXIS | 206 | 0.3461 | 1.3915555 | 0.014155713 | 0.20730038 | 1 | 4790 | tags=27%, list=20%, si | MM_3 |
| HUMAN_PAPILLOMAVIRU | HUMAN_PAPILLOMAVIRUS_INFECTION | 315 | 0.340841 | 1.3913811 | 0.004016064 | 0.20736705 | 1 | 6131 | tags=36%, list=26%, si | MM_3 |
| GOBP_GOLGI_TO_PLASM | GOBP_GOLGI_TO_PLASMA_MEMBRANE_TR | 62 | 0.389191 | 1.3913035 | 0.06966292 | 0.20730741 | 1 | 7649 | tags=55%, list=32%, si | MM_3 |
| GOBP_REGULATION_OF_ | GOBP_REGULATION_OF_INTRINSIC_APOPT | 185 | 0.350548 | 1.3911961 | 0.017435897 | 0.20727046 | 1 | 7148 | tags=46%, list=30%, si | MM_3 |
| GOBP_REGULATION_OF_ | GOBP_REGULATION_OF_CIRCADIAN_RHYTH | 112 | 0.361591 | 1.3911601 | 0.037593983 | 0.20714429 | 1 | 7506 | tags=48%, list=32%, si | MM_3 |
| GOBP_RESPONSE_TO_EN | GOBP_RESPONSE_TO_ENDOPLASMIC_RETIC | 261 | 0.342029 | 1.3907543 | 0.0030549897 | 0.20752041 | 1 | 7363 | tags=47%, list=31%, si | MM_3 |
| INFLUENZA_A | INFLUENZA_A | 139 | 0.356998 | 1.3905265 | 0.041322313 | 0.20767751 | 1 | 7534 | tags=45%, list=32%, si | MM_3 |
| GOBP_DETECTION_OF_M | GOBP_DETECTION_OF_MECHANICAL_STIMU | 53 | 0.393725 | 1.3903948 | 0.06868452 | 0.20770265 | 1 | 5500 | tags=38%, list=23%, si | MM_3 |
| GOBP_POSITIVE_REGULA | GOBP_POSITIVE_REGULATION_OF_SMOOTH | 17 | 0.499316 | 1.3903095 | 0.10714286 | 0.20766711 | 1 | 5893 | tags=47%, list=25%, si | MM_3 |
| STAPHYLOCOCCUS_AURE | STAPHYLOCOCCUS_AUREUS_INFECTION | 44 | 0.41163 | 1.3900748 | 0.084295616 | 0.20784718 | 1 | 6533 | tags=48%, list=28%, si | MM_3 |
| GOBP_CHROMATIN_REM | GOBP_CHROMATIN_REMODELING | 482 | 0.335206 | 1.3898697 | 0 | 0.20800272 | 1 | 7645 | tags=46%, list=32%, si | MM_3 |
| GOBP_REGULATION_OF_ | GOBP_REGULATION_OF_URINE_VOLUME | 20 | 0.471988 | 1.3898658 | 0.10484928 | 0.20783676 | 1 | 3164 | tags=35%, list=13%, si | MM_3 |
| GOBP_EMBRYONIC_EYE_ | GOBP_EMBRYONIC_EYE_MORPHOGENESIS | 33 | 0.426223 | 1.3894439 | 0.07965686 | 0.2082616 | 1 | 2953 | tags=21%, list=13%, si | MM_3 |
| PROTEIN_EXPORT | PROTEIN_EXPORT | 26 | 0.456362 | 1.3894304 | 0.09875 | 0.20811054 | 1 | 8493 | tags=62%, list=36%, si | MM_3 |
| VALINE,_LEUCINE_AND_I | VALINE,_LEUCINE_AND_ISOLEUCINE_DEGRA | 49 | 0.400802 | 1.3894283 | 0.081850536 | 0.20794736 | 1 | 4743 | tags=33%, list=20%, si | MM_3 |
| GOBP_REGULATION_OF_ | GOBP_REGULATION_OF_GLUCONEOGENESI | 52 | 0.396095 | 1.3893586 | 0.06659012 | 0.20787294 | 1 | 5216 | tags=35%, list=22%, si | MM_3 |
| GOBP_REGULATION_OF_ | GOBP_REGULATION_OF_AMIDE_METABOLI | 69 | 0.384329 | 1.3887303 | 0.06050605 | 0.20862043 | 1 | 5470 | tags=32%, list=23%, si | MM_3 |
| GOBP_POSITIVE_REGULA | GOBP_POSITIVE_REGULATION_OF_DNA_BIN | 45 | 0.404212 | 1.3884773 | 0.08244994 | 0.20885123 | 1 | 5235 | tags=36%, list=22%, si | MM_3 |
| GOBP_TELENCEPHALON_ | GOBP_TELENCEPHALON_DEVELOPMENT | 245 | 0.341038 | 1.3884479 | 0.004060914 | 0.20872024 | 1 | 5391 | tags=32%, list=23%, si | MM_3 |
| GOBP_POSITIVE_REGULA | GOBP_POSITIVE_REGULATION_OF_NON_CA | 64 | 0.384161 | 1.3881812 | 0.066059224 | 0.20892529 | 1 | 7663 | tags=52%, list=32%, si | MM_3 |
| GOBP_REGULATION_OF_ | GOBP_REGULATION_OF_OSTEOCLAST_DIFF | 84 | 0.36918 | 1.3878629 | 0.04714912 | 0.2091974 | 1 | 6330 | tags=36%, list=27%, si | MM_3 |
| GOBP_POSITIVE_REGULA | GOBP_POSITIVE_REGULATION_OF_GLIAL_C | 51 | 0.40061 | 1.3862739 | 0.090694934 | 0.21127081 | 1 | 5822 | tags=33%, list=25%, si | MM_3 |
| GOBP_REGULATION_OF_ | GOBP_REGULATION_OF_AUTOPHAGOSOME | 18 | 0.490953 | 1.3858857 | 0.12068965 | 0.21165025 | 1 | 7599 | tags=56%, list=32%, si | MM_3 |
| GOBP_REGULATION_OF_ | GOBP_REGULATION_OF_RNA_STABILITY | 174 | 0.344886 | 1.3856581 | 0.01934827 | 0.21183154 | 1 | 7545 | tags=45%, list=32%, si | MM_3 |
| GOBP_REGULATION_OF_ | GOBP_REGULATION_OF_IMMATURE_T_CEL | 16 | 0.508334 | 1.3852571 | 0.10320781 | 0.21224108 | 1 | 1155 | tags=19%, list=5%, sig | MM_3 |
| GOBP_NEUTROPHIL_ACT | GOBP_NEUTROPHIL_ACTIVATION | 36 | 0.421793 | 1.3851787 | 0.08837772 | 0.21219021 | 1 | 5778 | tags=36%, list=24%, si | MM_3 |
| GOBP_B_CELL_PROLIFER | GOBP_B_CELL_PROLIFERATION | 97 | 0.36987 | 1.3847517 | 0.042964555 | 0.21261977 | 1 | 5024 | tags=27%, list=21%, si | MM_3 |
| GOBP_NEGATIVE_REGUL | GOBP_NEGATIVE_REGULATION_OF_REGULA | 25 | 0.451162 | 1.3846561 | 0.09317585 | 0.21258664 | 1 | 1581 | tags=16%, list=7%, sig | MM_3 |
| GOBP_C_TERMINAL_PRO | GOBP_C_TERMINAL_PROTEIN_AMINO_ACID | 20 | 0.474879 | 1.3845304 | 0.09517241 | 0.21258225 | 1 | 4616 | tags=45%, list=20%, si | MM_3 |
| METABOLISM_OF_XENOE | METABOLISM_OF_XENOBIOTICS_BY_CYTOC | 48 | 0.39602 | 1.3845128 | 0.09239766 | 0.2124355 | 1 | 5248 | tags=35%, list=22%, si | MM_3 |
| GOBP_REGULATION_OF_ | GOBP_REGULATION_OF_ESTABLISHMENT_O | 25 | 0.447143 | 1.3844912 | 0.10152284 | 0.21229872 | 1 | 4949 | tags=36%, list=21%, si | MM_3 |

| | | | | | | | | | | |
|---|---|---|---|---|---|---|---|---|---|---|
| GOBP_RHYTHMIC_BEHAV | GOBP_RHYTHMIC_BEHAVIOR | 61 | 0.381273 | 1.3842431 | 0.07386363 | 0.21246937 | 1 | 2368 | tags=20%, list=10%, si | MM_3 |
| GOBP_METHYLATION | GOBP_METHYLATION | 199 | 0.344583 | 1.3834772 | 0.024489796 | 0.21337497 | 1 | 7590 | tags=48%, list=32%, si | MM_3 |
| GOBP_GRANULOCYTE_DI | GOBP_GRANULOCYTE_DIFFERENTIATION | 50 | 0.401424 | 1.3831669 | 0.07582939 | 0.21367322 | 1 | 6172 | tags=40%, list=26%, si | MM_3 |
| GOBP_REGULATION_OF_ | GOBP_REGULATION_OF_ANTIGEN_RECEPTO | 64 | 0.386067 | 1.3829606 | 0.053511705 | 0.21379276 | 1 | 5026 | tags=30%, list=21%, si | MM_3 |
| GOBP_AMINO_ACID_CAT | GOBP_AMINO_ACID_CATABOLIC_PROCESS | 85 | 0.366855 | 1.3824648 | 0.059912853 | 0.21432191 | 1 | 5181 | tags=36%, list=22%, si | MM_3 |
| GOBP_ETHER_METABOLI | GOBP_ETHER_METABOLIC_PROCESS | 23 | 0.457711 | 1.382387 | 0.10671937 | 0.21428679 | 1 | 3888 | tags=30%, list=16%, si | MM_3 |
| GOBP_NEGATIVE_REGUL | GOBP_NEGATIVE_REGULATION_OF_RAS_PR | 26 | 0.447159 | 1.38232 | 0.098997496 | 0.21422005 | 1 | 1842 | tags=19%, list=8%, sig | MM_3 |
| GOBP_REGULATION_OF_ | GOBP_REGULATION_OF_GENERATION_OF_ | 148 | 0.352471 | 1.3821486 | 0.028183715 | 0.21429811 | 1 | 6488 | tags=37%, list=27%, si | MM_3 |
| GOBP_MUSCLE_CELL_DIF | GOBP_MUSCLE_CELL_DIFFERENTIATION | 444 | 0.330309 | 1.3819393 | 0.002 | 0.21443528 | 1 | 5526 | tags=30%, list=23%, si | MM_3 |
| GOBP_ESTABLISHMENT_( | GOBP_ESTABLISHMENT_OF_PROTEIN_LOCA | 67 | 0.380071 | 1.3815347 | 0.061179087 | 0.21485546 | 1 | 5863 | tags=43%, list=25%, si | MM_3 |
| GOBP_SISTER_CHROMAT | GOBP_SISTER_CHROMATID_SEGREGATION | 219 | 0.343798 | 1.3815335 | 0.012244898 | 0.21468803 | 1 | 7054 | tags=40%, list=30%, si | MM_3 |
| GOBP_REGULATION_OF_ | GOBP_REGULATION_OF_STEM_CELL_DIFFEI | 82 | 0.3729 | 1.3814445 | 0.04939627 | 0.214656 | 1 | 7322 | tags=44%, list=31%, si | MM_3 |
| GOBP_NEGATIVE_REGUL | GOBP_NEGATIVE_REGULATION_OF_SMOOT | 68 | 0.381313 | 1.3807354 | 0.07150838 | 0.21549648 | 1 | 5903 | tags=37%, list=25%, si | MM_3 |
| GOBP_LYMPHOCYTE_COS | GOBP_LYMPHOCYTE_COSTIMULATION | 33 | 0.440339 | 1.3802253 | 0.092364535 | 0.21603726 | 1 | 5903 | tags=52%, list=25%, si | MM_3 |
| GOBP_REGULATION_OF_ | GOBP_REGULATION_OF_B_CELL_MEDIATED | 66 | 0.380061 | 1.3800762 | 0.06873614 | 0.21608122 | 1 | 4592 | tags=30%, list=19%, si | MM_3 |
| GOBP_SMOOTHENED_SIG | GOBP_SMOOTHENED_SIGNALING_PATHWA | 154 | 0.348812 | 1.3799683 | 0.022657055 | 0.21605985 | 1 | 3327 | tags=24%, list=14%, si | MM_3 |
| GOBP_ESTABLISHMENT_( | GOBP_ESTABLISHMENT_OF_PROTEIN_LOCA | 59 | 0.386571 | 1.3791604 | 0.06395349 | 0.2170687 | 1 | 7162 | tags=41%, list=30%, si | MM_3 |
| GOBP_MITOTIC_SISTER_( | GOBP_MITOTIC_SISTER_CHROMATID_SEGRI | 178 | 0.344842 | 1.3790709 | 0.022703819 | 0.21703832 | 1 | 7054 | tags=42%, list=30%, si | MM_3 |
| GOBP_REGULATION_OF_ | GOBP_REGULATION_OF_AMYLOID_PRECUR | 44 | 0.412243 | 1.3789481 | 0.086448595 | 0.21703848 | 1 | 7983 | tags=48%, list=34%, si | MM_3 |
| GOBP_INTRACELLULAR_L | GOBP_INTRACELLULAR_LIPID_TRANSPORT | 42 | 0.407022 | 1.3787493 | 0.096736595 | 0.2171515 | 1 | 6992 | tags=43%, list=30%, si | MM_3 |
| GOBP_CENTROMERE_CO | GOBP_CENTROMERE_COMPLEX_ASSEMBLY | 27 | 0.445541 | 1.3783804 | 0.10566038 | 0.21751186 | 1 | 4422 | tags=33%, list=19%, si | MM_3 |
| GOBP_NEGATIVE_REGUL | GOBP_NEGATIVE_REGULATION_OF_PROTEI | 87 | 0.360762 | 1.3782289 | 0.051819183 | 0.21756564 | 1 | 6561 | tags=45%, list=28%, si | MM_3 |
| GOBP_REGULATION_OF_ | GOBP_REGULATION_OF_OXIDATIVE_PHOSP | 23 | 0.454735 | 1.3781774 | 0.117117114 | 0.21747397 | 1 | 7521 | tags=57%, list=32%, si | MM_3 |
| DNA_REPLICATION | DNA_REPLICATION | 34 | 0.419631 | 1.3781035 | 0.102402024 | 0.21740656 | 1 | 4035 | tags=35%, list=17%, si | MM_3 |
| GOBP_INTEGRATED_STRE | GOBP_INTEGRATED_STRESS_RESPONSE_SIG | 23 | 0.459378 | 1.3780241 | 0.10761155 | 0.21735124 | 1 | 6194 | tags=39%, list=26%, si | MM_3 |
| GOBP_CELLULAR_SENESC | GOBP_CELLULAR_SENESCENCE | 88 | 0.367382 | 1.3779029 | 0.050111357 | 0.21736816 | 1 | 7190 | tags=48%, list=30%, si | MM_3 |
| GOBP_POSITIVE_REGULA | GOBP_POSITIVE_REGULATION_OF_INTRACE | 147 | 0.348145 | 1.3770176 | 0.038381744 | 0.21847999 | 1 | 6965 | tags=44%, list=29%, si | MM_3 |
| GOBP_REGULATION_OF_ | GOBP_REGULATION_OF_ALPHA_BETA_T_CE | 119 | 0.354914 | 1.3765991 | 0.03043022 | 0.21896242 | 1 | 5001 | tags=28%, list=21%, si | MM_3 |
| GOBP_LYMPHOCYTE_APO | GOBP_LYMPHOCYTE_APOPTOTIC_PROCESS | 94 | 0.362157 | 1.3763773 | 0.052401748 | 0.21912336 | 1 | 4409 | tags=28%, list=19%, si | MM_3 |
| GOBP_NEGATIVE_REGUL | GOBP_NEGATIVE_REGULATION_OF_VIRAL_ | 50 | 0.396056 | 1.3762459 | 0.08963911 | 0.219158 | 1 | 5253 | tags=34%, list=22%, si | MM_3 |
| GOBP_NEGATIVE_REGUL | GOBP_NEGATIVE_REGULATION_OF_MUSCL | 35 | 0.415486 | 1.3761734 | 0.0995086 | 0.2190857 | 1 | 3571 | tags=23%, list=15%, si | MM_3 |
| GOBP_CORPUS_CALLOSU | GOBP_CORPUS_CALLOSUM_DEVELOPMENT | 27 | 0.446903 | 1.3759869 | 0.093709886 | 0.21921596 | 1 | 5213 | tags=33%, list=22%, si | MM_3 |
| GOBP_RESPONSE_TO_OR | GOBP_RESPONSE_TO_ORGANOPHOSPHORU | 93 | 0.363543 | 1.3753271 | 0.049092848 | 0.22005057 | 1 | 5426 | tags=29%, list=23%, si | MM_3 |
| GOBP_BLASTODERM_SEG | GOBP_BLASTODERM_SEGMENTATION | 26 | 0.43742 | 1.3751152 | 0.10673443 | 0.22019675 | 1 | 1472 | tags=23%, list=6%, sig | MM_3 |
| GOBP_RESPONSE_TO_TR | GOBP_RESPONSE_TO_TRANSFORMING_GRO | 246 | 0.339399 | 1.374956 | 0.01417004 | 0.2202527 | 1 | 6812 | tags=40%, list=29%, si | MM_3 |
| GOBP_POSITIVE_REGULA | GOBP_POSITIVE_REGULATION_OF_CALCIUN | 26 | 0.450654 | 1.3748533 | 0.112658225 | 0.22025773 | 1 | 3206 | tags=27%, list=14%, si | MM_3 |
| GOBP_ESTABLISHMENT_( | GOBP_ESTABLISHMENT_OF_PROTEIN_LOCA | 47 | 0.401412 | 1.374538 | 0.10161663 | 0.22054207 | 1 | 8493 | tags=62%, list=36%, si | MM_3 |
| GOBP_MACROPHAGE_AC | GOBP_MACROPHAGE_ACTIVATION | 107 | 0.361737 | 1.3743844 | 0.045502644 | 0.22058812 | 1 | 4391 | tags=23%, list=19%, si | MM_3 |
| GOBP_ANTERIOR_POSTE | GOBP_ANTERIOR_POSTERIOR_AXIS_SPECIFI | 46 | 0.403185 | 1.3741281 | 0.077192985 | 0.22077902 | 1 | 5307 | tags=33%, list=22%, si | MM_3 |
| GOBP_NEUTRAL_AMINO_ | GOBP_NEUTRAL_AMINO_ACID_TRANSPORT | 51 | 0.395511 | 1.3738976 | 0.07462686 | 0.22099026 | 1 | 2884 | tags=24%, list=12%, si | MM_3 |
| N-GLYCAN_BIOSYNTHESIS | N-GLYCAN_BIOSYNTHESIS | 48 | 0.400188 | 1.3735424 | 0.09501738 | 0.22131619 | 1 | 4705 | tags=38%, list=20%, si | MM_3 |
| GOBP_PROTEIN_CONTAI | GOBP_PROTEIN_CONTAINING_COMPLEX_D | 247 | 0.339082 | 1.3730148 | 0.012158055 | 0.2218785 | 1 | 7761 | tags=50%, list=33%, si | MM_3 |
| GOBP_NEGATIVE_REGUL | GOBP_NEGATIVE_REGULATION_OF_GLUCO | 24 | 0.453813 | 1.372948 | 0.11011524 | 0.22180377 | 1 | 6166 | tags=33%, list=26%, si | MM_3 |
| GOBP_REGULATION_OF_ | GOBP_REGULATION_OF_GLIAL_CELL_DIFFEI | 89 | 0.364637 | 1.3728274 | 0.0613563 | 0.22181465 | 1 | 5859 | tags=31%, list=25%, si | MM_3 |
| GOBP_SARCOMERE_ORG | GOBP_SARCOMERE_ORGANIZATION | 53 | 0.390596 | 1.3726412 | 0.07764706 | 0.22191058 | 1 | 3612 | tags=28%, list=15%, si | MM_3 |
| GOBP_PROTEIN_POLYME | GOBP_PROTEIN_POLYMERIZATION | 269 | 0.335981 | 1.3725661 | 0.0040201005 | 0.22183926 | 1 | 7602 | tags=46%, list=32%, si | MM_3 |
| GOBP_SPINDLE_ASSEMBL | GOBP_SPINDLE_ASSEMBLY | 122 | 0.353964 | 1.3724208 | 0.04197272 | 0.22189564 | 1 | 7169 | tags=44%, list=30%, si | MM_3 |
| GOBP_CARDIAC_ATRIUM | GOBP_CARDIAC_ATRIUM_DEVELOPMENT | 36 | 0.412141 | 1.372397 | 0.09296482 | 0.22175756 | 1 | 5428 | tags=31%, list=23%, si | MM_3 |
| GOBP_RIBONUCLEOSIDE_ | GOBP_RIBONUCLEOSIDE_MONOPHOSPHATI | 48 | 0.402404 | 1.3720341 | 0.06849315 | 0.22213538 | 1 | 4987 | tags=38%, list=21%, si | MM_3 |
| GOBP_REGULATION_OF_ | GOBP_REGULATION_OF_MRNA_PROCESSIN | 131 | 0.350583 | 1.3719648 | 0.03215768 | 0.22206502 | 1 | 8191 | tags=55%, list=35%, si | MM_3 |
| GOBP_VIRAL_TRANSLATI | GOBP_VIRAL_TRANSLATION | 18 | 0.480937 | 1.3717084 | 0.116504855 | 0.22226118 | 1 | 7222 | tags=50%, list=31%, si | MM_3 |
| GOBP_POSITIVE_REGULA | GOBP_POSITIVE_REGULATION_OF_CELL_AD | 487 | 0.330825 | 1.3713007 | 0 | 0.22271703 | 1 | 5725 | tags=32%, list=24%, si | MM_3 |
| GOBP_RNA_METHYLATIO | GOBP_RNA_METHYLATION | 65 | 0.376087 | 1.3712343 | 0.07777778 | 0.2226281 | 1 | 6935 | tags=49%, list=29%, si | MM_3 |
| TUBERCULOSIS | TUBERCULOSIS | 152 | 0.345897 | 1.3712145 | 0.027083334 | 0.22248377 | 1 | 8045 | tags=55%, list=34%, si | MM_3 |
| GOBP_RHO_PROTEIN_SIG | GOBP_RHO_PROTEIN_SIGNAL_TRANSDUCTI | 142 | 0.347437 | 1.3710973 | 0.038381744 | 0.22249232 | 1 | 7662 | tags=49%, list=32%, si | MM_3 |
| GOBP_ATTACHMENT_OF_ | GOBP_ATTACHMENT_OF_MITOTIC_SPINDLE | 24 | 0.455283 | 1.3710959 | 0.10755442 | 0.22232555 | 1 | 6708 | tags=54%, list=28%, si | MM_3 |
| GOBP_NEGATIVE_REGUL | GOBP_NEGATIVE_REGULATION_OF_HYDRO | 238 | 0.336818 | 1.3709599 | 0.010121457 | 0.2223899 | 1 | 4880 | tags=29%, list=21%, si | MM_3 |
| GOBP_POSITIVE_REGULA | GOBP_POSITIVE_REGULATION_OF_MYOBLA | 18 | 0.490068 | 1.3707103 | 0.10372341 | 0.22263025 | 1 | 5715 | tags=44%, list=24%, si | MM_3 |
| GOBP_REGULATION_OF_ | GOBP_REGULATION_OF_INFLAMMATORY_R | 348 | 0.329938 | 1.3701295 | 0.0060362173 | 0.22327094 | 1 | 5542 | tags=30%, list=23%, si | MM_3 |
| GOBP_POSITIVE_REGULA | GOBP_POSITIVE_REGULATION_OF_RELEASE | 27 | 0.446707 | 1.3699123 | 0.09355246 | 0.22341168 | 1 | 6558 | tags=48%, list=28%, si | MM_3 |
| GOBP_POSITIVE_REGULA | GOBP_POSITIVE_REGULATION_OF_CHEMOK | 15 | 0.503536 | 1.3698399 | 0.13483146 | 0.2233463 | 1 | 8133 | tags=87%, list=34%, si | MM_3 |
| GOBP_POSITIVE_REGULA | GOBP_POSITIVE_REGULATION_OF_SMAD_P | 38 | 0.413353 | 1.3698124 | 0.09169765 | 0.2232155 | 1 | 4997 | tags=34%, list=21%, si | MM_3 |
| GOBP_NEGATIVE_REGUL | GOBP_NEGATIVE_REGULATION_OF_OXIDAT | 26 | 0.445503 | 1.3695682 | 0.0996119 | 0.22342452 | 1 | 5643 | tags=50%, list=24%, si | MM_3 |
| GOBP_REGULATION_OF_ | GOBP_REGULATION_OF_LEUKOCYTE_MEDIA | 253 | 0.334306 | 1.3694834 | 0.014127145 | 0.22337887 | 1 | 7028 | tags=40%, list=30%, si | MM_3 |
| GOBP_MRNA_PROCESSIN | GOBP_MRNA_PROCESSING | 459 | 0.332511 | 1.3693596 | 0 | 0.22337961 | 1 | 7551 | tags=44%, list=32%, si | MM_3 |
| GOBP_POSITIVE_REGULA | GOBP_POSITIVE_REGULATION_OF_OXIDORE | 37 | 0.419029 | 1.36907 | 0.09529703 | 0.2236316 | 1 | 4473 | tags=32%, list=19%, si | MM_3 |
| GOBP_PROTEIN_RNA_CO | GOBP_PROTEIN_RNA_COMPLEX_ORGANIZA | 173 | 0.344036 | 1.3690434 | 0.032258064 | 0.22351117 | 1 | 7222 | tags=46%, list=31%, si | MM_3 |
| GOBP_POSITIVE_REGULA | GOBP_POSITIVE_REGULATION_OF_CELLULA | 29 | 0.440941 | 1.3688856 | 0.1011378 | 0.22359888 | 1 | 7175 | tags=45%, list=30%, si | MM_3 |
| GOBP_POSITIVE_REGULA | GOBP_POSITIVE_REGULATION_OF_LYMPHO | 318 | 0.334237 | 1.3686118 | 0.0060301507 | 0.22383246 | 1 | 5724 | tags=32%, list=24%, si | MM_3 |
| GOBP_SCF_DEPENDENT_ | GOBP_SCF_DEPENDENT_PROTEASOMAL_UE | 47 | 0.394559 | 1.368607 | 0.08382527 | 0.22367425 | 1 | 3183 | tags=26%, list=13%, si | MM_3 |
| GOBP_REGULATION_OF_ | GOBP_REGULATION_OF_CELL_SUBSTRATE_ | 219 | 0.340091 | 1.3685217 | 0.021341464 | 0.22361836 | 1 | 5822 | tags=35%, list=25%, si | MM_3 |
| GOBP_MONOCYTE_CHEM | GOBP_MONOCYTE_CHEMOTACTIC_PROTEIN | 27 | 0.44115 | 1.3677297 | 0.11235955 | 0.2246325 | 1 | 5872 | tags=37%, list=25%, si | MM_3 |
| GOBP_REGULATION_OF_ | GOBP_REGULATION_OF_SYNCYTIUM_FORM | 31 | 0.429727 | 1.3677098 | 0.11630847 | 0.22449253 | 1 | 6853 | tags=45%, list=29%, si | MM_3 |
| GOBP_PALLIUM_DEVELO | GOBP_PALLIUM_DEVELOPMENT | 165 | 0.345232 | 1.3672947 | 0.030991735 | 0.22490127 | 1 | 5733 | tags=35%, list=24%, si | MM_3 |
| GOBP_NEGATIVE_REGUL | GOBP_NEGATIVE_REGULATION_OF_NEURO | 83 | 0.368733 | 1.3668144 | 0.059912853 | 0.22542296 | 1 | 2690 | tags=18%, list=11%, si | MM_3 |
| GOBP_RIBONUCLEOSIDE_ | GOBP_RIBONUCLEOSIDE_MONOPHOSPHATI | 30 | 0.426103 | 1.3666302 | 0.112840466 | 0.22552352 | 1 | 4288 | tags=40%, list=18%, si | MM_3 |
| GOBP_REGULATION_OF_ | GOBP_REGULATION_OF_ACUTE_INFLAMMA | 48 | 0.397516 | 1.3664085 | 0.08215962 | 0.22570013 | 1 | 4236 | tags=33%, list=18%, si | MM_3 |
| GOBP_REGULATION_OF_ | GOBP_REGULATION_OF_LYMPHOCYTE_DIFF | 216 | 0.34038 | 1.3663137 | 0.016260162 | 0.2256616 | 1 | 6641 | tags=38%, list=28%, si | MM_3 |
| GOBP_POSITIVE_REGULA | GOBP_POSITIVE_REGULATION_OF_CELL_AC | 373 | 0.330178 | 1.3661544 | 0.006012024 | 0.22573312 | 1 | 5724 | tags=31%, list=24%, si | MM_3 |
| GOBP_NEUROMUSCULAR | GOBP_NEUROMUSCULAR_PROCESS | 204 | 0.343146 | 1.365439 | 0.015243903 | 0.22660512 | 1 | 4042 | tags=24%, list=17%, si | MM_3 |
| GOBP_REGULATION_OF_ | GOBP_REGULATION_OF_NUCLEASE_ACTIVIT | 22 | 0.462647 | 1.3653771 | 0.109947644 | 0.22652993 | 1 | 7684 | tags=68%, list=33%, si | MM_3 |
| GOBP_REGULATION_OF_ | GOBP_REGULATION_OF_CLATHRIN_DEPENI | 18 | 0.478711 | 1.3651404 | 0.14324693 | 0.22672293 | 1 | 7244 | tags=56%, list=31%, si | MM_3 |

| | | | | | | | | | | |
|---|---|---|---|---|---|---|---|---|---|---|
| GOBP_IMMUNE_RESPON | GOBP_IMMUNE_RESPONSE_REGULATING_S | 457 | 0.330303 | 1.3647928 | 0 | 0.22707854 | 1 | 6329 | tags=33%, list=27%, si | MM_3 |
| GOBP_POSTREPLICATION | GOBP_POSTREPLICATION_REPAIR | 29 | 0.429705 | 1.3647773 | 0.12136536 | 0.22692868 | 1 | 3755 | tags=31%, list=16%, si | MM_3 |
| GOBP_ICOSANOID_BIOSY | GOBP_ICOSANOID_BIOSYNTHETIC_PROCESS | 43 | 0.39784 | 1.3645451 | 0.1017964 | 0.22710803 | 1 | 4203 | tags=26%, list=18%, si | MM_3 |
| GOBP_REGULATION_OF_ | GOBP_REGULATION_OF_STEM_CELL_POPU | 74 | 0.37449 | 1.3643205 | 0.060908083 | 0.22729054 | 1 | 6655 | tags=38%, list=28%, si | MM_3 |
| GOBP_NEUTROPHIL_MED | GOBP_NEUTROPHIL_MEDIATED_IMMUNITY | 37 | 0.409734 | 1.36428 | 0.10866911 | 0.2271845 | 1 | 1322 | tags=16%, list=6%, sig | MM_3 |
| GOBP_NEGATIVE_REGUL | GOBP_NEGATIVE_REGULATION_OF_CYTOKI | 22 | 0.462499 | 1.3642532 | 0.13753213 | 0.22705811 | 1 | 3202 | tags=23%, list=14%, si | MM_3 |
| GOBP_MITOTIC_SISTER_C | GOBP_MITOTIC_SISTER_CHROMATID_SEPAR | 61 | 0.380918 | 1.3640432 | 0.0703125 | 0.22719546 | 1 | 6521 | tags=38%, list=28%, si | MM_3 |
| PROSTATE_CANCER | PROSTATE_CANCER | 93 | 0.3621 | 1.3639395 | 0.060671724 | 0.22717597 | 1 | 8093 | tags=54%, list=34%, si | MM_3 |
| GOBP_CELLULAR_RESPON | GOBP_CELLULAR_RESPONSE_TO_ESTRADIO | 19 | 0.478775 | 1.3638535 | 0.10569106 | 0.22713517 | 1 | 5353 | tags=37%, list=23%, si | MM_3 |
| GOBP_LIPID_HOMEOSTA | GOBP_LIPID_HOMEOSTASIS | 167 | 0.343181 | 1.3637633 | 0.039054472 | 0.22711122 | 1 | 7206 | tags=40%, list=31%, si | MM_3 |
| GOBP_PRIMARY_NEURAL | GOBP_PRIMARY_NEURAL_TUBE_FORMATIO | 115 | 0.354516 | 1.3636605 | 0.0516333 | 0.22708508 | 1 | 3973 | tags=20%, list=17%, si | MM_3 |
| GOBP_NITRIC_OXIDE_ME | GOBP_NITRIC_OXIDE_MEDIATED_SIGNAL_T | 26 | 0.443803 | 1.3630327 | 0.1185567 | 0.22785653 | 1 | 4629 | tags=38%, list=20%, si | MM_3 |
| GOBP_POSITIVE_REGULA | GOBP_POSITIVE_REGULATION_OF_GROWTH | 289 | 0.331912 | 1.3629564 | 0.006012024 | 0.2278122 | 1 | 6781 | tags=36%, list=29%, si | MM_3 |
| GOBP_ATTACHMENT_OF_ | GOBP_ATTACHMENT_OF_SPINDLE_MICROT | 48 | 0.394605 | 1.3629421 | 0.0893067 | 0.22766706 | 1 | 6709 | tags=46%, list=28%, si | MM_3 |
| GOBP_REGULATION_OF_ | GOBP_REGULATION_OF_ACTIN_FILAMENT_ | 158 | 0.345766 | 1.3627647 | 0.031860225 | 0.22778188 | 1 | 7483 | tags=45%, list=32%, si | MM_3 |
| GOBP_INTRACELLULAR_P | GOBP_INTRACELLULAR_PH_REDUCTION | 48 | 0.39799 | 1.3625501 | 0.083908044 | 0.22795506 | 1 | 6731 | tags=52%, list=29%, si | MM_3 |
| GOBP_POSITIVE_REGULA | GOBP_POSITIVE_REGULATION_OF_INTERLE | 42 | 0.402697 | 1.3625472 | 0.105199516 | 0.22779338 | 1 | 7383 | tags=48%, list=31%, si | MM_3 |
| GOBP_INTRACELLULAR_Z | GOBP_INTRACELLULAR_ZINC_ION_HOMEOS | 24 | 0.450391 | 1.3623631 | 0.118489586 | 0.22789249 | 1 | 7553 | tags=54%, list=32%, si | MM_3 |
| GOBP_GMP_METABOLIC_ | GOBP_GMP_METABOLIC_PROCESS | 21 | 0.465798 | 1.3620449 | 0.12696335 | 0.22820218 | 1 | 4288 | tags=38%, list=18%, si | MM_3 |
| GOBP_REGULATION_OF_ | GOBP_REGULATION_OF_INTRACELLULAR_T | 278 | 0.335339 | 1.3609514 | 0.012133468 | 0.22963993 | 1 | 6779 | tags=40%, list=29%, si | MM_3 |
| GOBP_MITOCHONDRIAL_ | GOBP_MITOCHONDRIAL_DNA_REPLICATION | 17 | 0.489769 | 1.3607504 | 0.12696335 | 0.22980615 | 1 | 5468 | tags=47%, list=23%, si | MM_3 |
| GOBP_ORGANELLE_TRAN | GOBP_ORGANELLE_TRANSPORT_ALONG_M | 86 | 0.362003 | 1.360728 | 0.0595883 | 0.22966719 | 1 | 6709 | tags=41%, list=28%, si | MM_3 |
| GOBP_POSITIVE_REGULA | GOBP_POSITIVE_REGULATION_OF_SYNAPTI | 42 | 0.398138 | 1.3605642 | 0.10653753 | 0.22975416 | 1 | 5946 | tags=38%, list=25%, si | MM_3 |
| GOBP_ACYLGLYCEROL_H | GOBP_ACYLGLYCEROL_HOMEOSTASIS | 34 | 0.417626 | 1.3600881 | 0.10617284 | 0.2303047 | 1 | 2115 | tags=24%, list=9%, sig | MM_3 |
| GOBP_NEGATIVE_REGUL | GOBP_NEGATIVE_REGULATION_OF_CARDIA | 30 | 0.431083 | 1.359851 | 0.1037037 | 0.23048793 | 1 | 3474 | tags=30%, list=15%, si | MM_3 |
| GOBP_POLYOL_METABOL | GOBP_POLYOL_METABOLIC_PROCESS | 99 | 0.360998 | 1.3595862 | 0.06903991 | 0.23068927 | 1 | 6171 | tags=35%, list=26%, si | MM_3 |
| GOBP_INTERLEUKIN_17_ | GOBP_INTERLEUKIN_17_PRODUCTION | 42 | 0.400502 | 1.3592324 | 0.10769231 | 0.23107119 | 1 | 4256 | tags=26%, list=18%, si | MM_3 |
| GOBP_REGULATION_OF_ | GOBP_REGULATION_OF_HEART_GROWTH | 85 | 0.365693 | 1.3591897 | 0.06555556 | 0.23097087 | 1 | 3474 | tags=24%, list=15%, si | MM_3 |
| GOBP_INTRACELLULAR_S | GOBP_INTRACELLULAR_STEROL_TRANSPOR | 34 | 0.418096 | 1.3591019 | 0.10918114 | 0.23093924 | 1 | 6992 | tags=47%, list=30%, si | MM_3 |
| GOBP_RESPONSE_TO_TE | GOBP_RESPONSE_TO_TEMPERATURE_STIM | 159 | 0.343665 | 1.3590202 | 0.03205791 | 0.2309144 | 1 | 5502 | tags=37%, list=23%, si | MM_3 |
| GOBP_CELLULAR_OXIDAN | GOBP_CELLULAR_OXIDANT_DETOXIFICATIO | 29 | 0.433284 | 1.3587898 | 0.10125 | 0.23109832 | 1 | 6764 | tags=45%, list=29%, si | MM_3 |
| GOBP_CGMP_MEDIATED | GOBP_CGMP_MEDIATED_SIGNALING | 32 | 0.420881 | 1.3584983 | 0.11742892 | 0.23139137 | 1 | 2727 | tags=22%, list=12%, si | MM_3 |
| GOBP_REGULATION_OF_ | GOBP_REGULATION_OF_DNA_REPLICATION | 118 | 0.353527 | 1.3584579 | 0.059134107 | 0.23127858 | 1 | 6709 | tags=43%, list=28%, si | MM_3 |
| MAPK_SIGNALING_PATH | MAPK_SIGNALING_PATHWAY | 287 | 0.328397 | 1.3582956 | 0.005050505 | 0.2313524 | 1 | 6953 | tags=41%, list=29%, si | MM_3 |
| GOBP_POST_GOLGI_VESI | GOBP_POST_GOLGI_VESICLE_MEDIATED_TF | 101 | 0.356481 | 1.3581926 | 0.061068702 | 0.23132063 | 1 | 5514 | tags=36%, list=23%, si | MM_3 |
| GOBP_REGULATION_OF_ | GOBP_REGULATION_OF_WNT_SIGNALING_ | 16 | 0.480147 | 1.3581339 | 0.12990528 | 0.2312305 | 1 | 4997 | tags=44%, list=21%, si | MM_3 |
| GOBP_REGULATION_OF_ | GOBP_REGULATION_OF_RESPONSE_TO_WC | 161 | 0.339472 | 1.3574353 | 0.035379812 | 0.23217122 | 1 | 7383 | tags=45%, list=31%, si | MM_3 |
| GOBP_REGULATION_OF_ | GOBP_REGULATION_OF_MULTICELLULAR_C | 83 | 0.365033 | 1.3574312 | 0.06263737 | 0.23200892 | 1 | 2886 | tags=19%, list=12%, si | MM_3 |
| GOBP_MUSCLE_CELL_PR | GOBP_MUSCLE_CELL_PROLIFERATION | 262 | 0.334462 | 1.3573512 | 0.0130916415 | 0.23196551 | 1 | 5730 | tags=32%, list=24%, si | MM_3 |
| DRUG_METABOLISM | DRUG_METABOLISM | 93 | 0.360908 | 1.3573407 | 0.073993474 | 0.2318163 | 1 | 5248 | tags=30%, list=22%, si | MM_3 |
| GOBP_PROTEIN_O_LINKE | GOBP_PROTEIN_O_LINKED_GLYCOSYLATION | 79 | 0.365789 | 1.3573127 | 0.059275523 | 0.23169938 | 1 | 3592 | tags=25%, list=15%, si | MM_3 |
| GOBP_NEGATIVE_REGUL | GOBP_NEGATIVE_REGULATION_OF_RECEPT | 27 | 0.43557 | 1.3570682 | 0.12716049 | 0.23188424 | 1 | 3381 | tags=33%, list=14%, si | MM_3 |
| GOBP_POSITIVE_REGULA | GOBP_POSITIVE_REGULATION_OF_CELL_CY | 252 | 0.334491 | 1.356962 | 0.011178861 | 0.23187093 | 1 | 6906 | tags=39%, list=29%, si | MM_3 |
| GOBP_POSITIVE_REGULA | GOBP_POSITIVE_REGULATION_OF_MACROF | 15 | 0.502484 | 1.3565524 | 0.12133891 | 0.23231845 | 1 | 4649 | tags=40%, list=20%, si | MM_3 |
| GOBP_REGULATION_OF_ | GOBP_REGULATION_OF_PROTEIN_TARGETI | 59 | 0.379203 | 1.3564433 | 0.090492554 | 0.23231654 | 1 | 4201 | tags=29%, list=18%, si | MM_3 |
| GOBP_DICARBOXYLIC_AC | GOBP_DICARBOXYLIC_ACID_CATABOLIC_PR | 17 | 0.482804 | 1.3563892 | 0.12947658 | 0.23224871 | 1 | 5700 | tags=47%, list=24%, si | MM_3 |
| GOBP_POSITIVE_REGULA | GOBP_POSITIVE_REGULATION_OF_SYNCYTI | 25 | 0.444191 | 1.3562627 | 0.107830554 | 0.23227811 | 1 | 6853 | tags=48%, list=29%, si | MM_3 |
| TGF-BETA_SIGNALING_PA | TGF-BETA_SIGNALING_PATHWAY | 85 | 0.361632 | 1.3560761 | 0.06352683 | 0.23240462 | 1 | 7175 | tags=46%, list=30%, si | MM_3 |
| GOBP_SEROTONIN_SECR | GOBP_SEROTONIN_SECRETION | 15 | 0.497228 | 1.3560404 | 0.14345405 | 0.232305 | 1 | 3206 | tags=27%, list=14%, si | MM_3 |
| GOBP_CELLULAR_RESPON | GOBP_CELLULAR_RESPONSE_TO_IONIZING_ | 69 | 0.372529 | 1.3560132 | 0.084269665 | 0.23218328 | 1 | 6864 | tags=45%, list=29%, si | MM_3 |
| MISMATCH_REPAIR | MISMATCH_REPAIR | 21 | 0.464136 | 1.3558868 | 0.10353866 | 0.23220375 | 1 | 2407 | tags=33%, list=10%, si | MM_3 |
| GOBP_POSITIVE_REGULA | GOBP_POSITIVE_REGULATION_OF_RESPONS | 73 | 0.366823 | 1.3555111 | 0.069922306 | 0.23262054 | 1 | 5297 | tags=34%, list=22%, si | MM_3 |
| GOBP_CHEMOKINE_C_X_ | GOBP_CHEMOKINE_C_X_C_MOTIF_LIGAND | 25 | 0.44274 | 1.3554331 | 0.118863046 | 0.23258094 | 1 | 7521 | tags=64%, list=32%, si | MM_3 |
| GOBP_B_CELL_DIFFEREN | GOBP_B_CELL_DIFFERENTIATION | 151 | 0.344785 | 1.3550484 | 0.036082473 | 0.23296 | 1 | 7299 | tags=41%, list=31%, si | MM_3 |
| GOBP_LYMPH_NODE_DE | GOBP_LYMPH_NODE_DEVELOPMENT | 32 | 0.43007 | 1.3548245 | 0.12469136 | 0.23313628 | 1 | 5942 | tags=53%, list=25%, si | MM_3 |
| GOBP_MULTICELLULAR_C | GOBP_MULTICELLULAR_ORGANISMAL_LEVE | 69 | 0.370871 | 1.3548108 | 0.07399103 | 0.23299007 | 1 | 5774 | tags=36%, list=24%, si | MM_3 |
| C-TYPE_LECTIN_RECEPTO | C-TYPE_LECTIN_RECEPTOR_SIGNALING_PAT | 104 | 0.354275 | 1.3545264 | 0.057359308 | 0.23328461 | 1 | 6906 | tags=42%, list=29%, si | MM_3 |
| GOBP_COPII_COATED_VE | GOBP_COPII_COATED_VESICLE_BUDDING | 37 | 0.416207 | 1.354515 | 0.08782936 | 0.2331393 | 1 | 6720 | tags=54%, list=28%, si | MM_3 |
| GOBP_REGULATION_OF_ | GOBP_REGULATION_OF_GLIAL_CELL_APOPT | 15 | 0.505714 | 1.3534693 | 0.1232493 | 0.23454618 | 1 | 8009 | tags=73%, list=34%, si | MM_3 |
| GOBP_RESPONSE_TO_DE | GOBP_RESPONSE_TO_DEXAMETHASONE | 27 | 0.432597 | 1.3534026 | 0.12403101 | 0.2344884 | 1 | 2220 | tags=22%, list=9%, sig | MM_3 |
| GOBP_CELLULAR_RESPON | GOBP_CELLULAR_RESPONSE_TO_NUTRIENT | 213 | 0.335396 | 1.3531852 | 0.022403259 | 0.23467389 | 1 | 7203 | tags=43%, list=31%, si | MM_3 |
| GOBP_SPINDLE_ORGANIZ | GOBP_SPINDLE_ORGANIZATION | 190 | 0.336652 | 1.3526565 | 0.042008195 | 0.23530228 | 1 | 7081 | tags=41%, list=30%, si | MM_3 |
| COLORECTAL_CANCER | COLORECTAL_CANCER | 86 | 0.361115 | 1.3523767 | 0.08478261 | 0.2355758 | 1 | 6975 | tags=50%, list=30%, si | MM_3 |
| GOBP_NEGATIVE_REGUL | GOBP_NEGATIVE_REGULATION_OF_GLIAL_( | 34 | 0.414732 | 1.3522823 | 0.11524164 | 0.23553947 | 1 | 6192 | tags=38%, list=26%, si | MM_3 |
| GOBP_POSITIVE_REGULA | GOBP_POSITIVE_REGULATION_OF_CYSTEIN | 130 | 0.346178 | 1.3522528 | 0.052742615 | 0.23540932 | 1 | 7135 | tags=45%, list=30%, si | MM_3 |
| GOBP_FILOPODIUM_ASS | GOBP_FILOPODIUM_ASSEMBLY | 74 | 0.372775 | 1.3521038 | 0.07079646 | 0.23546745 | 1 | 7600 | tags=51%, list=32%, si | MM_3 |
| GOBP_ANIMAL_ORGAN_ | GOBP_ANIMAL_ORGAN_REGENERATION | 28 | 0.426366 | 1.3519808 | 0.11801242 | 0.23546934 | 1 | 6442 | tags=50%, list=27%, si | MM_3 |
| GOBP_NUCLEAR_TRANSC | GOBP_NUCLEAR_TRANSCRIBED_MRNA_POL | 26 | 0.436426 | 1.3515061 | 0.116071425 | 0.23601939 | 1 | 5917 | tags=38%, list=25%, si | MM_3 |
| GOBP_REGULATION_OF_ | GOBP_REGULATION_OF_EXTRINSIC_APOPT( | 45 | 0.394931 | 1.3514706 | 0.11004785 | 0.23589508 | 1 | 7564 | tags=60%, list=32%, si | MM_3 |
| GOBP_GASTRO_INTESTIN | GOBP_GASTRO_INTESTINAL_SYSTEM_SMOC | 15 | 0.492231 | 1.350998 | 0.13387978 | 0.2364637 | 1 | 4713 | tags=40%, list=20%, si | MM_3 |
| GOBP_AMYLOID_BETA_C | GOBP_AMYLOID_BETA_CLEARANCE | 37 | 0.41385 | 1.3507618 | 0.10513739 | 0.2366684 | 1 | 7118 | tags=46%, list=30%, si | MM_3 |
| GOBP_PROTEIN_LOCALIZ | GOBP_PROTEIN_LOCALIZATION_TO_CHROM | 41 | 0.393298 | 1.35062 | 0.10550997 | 0.23672774 | 1 | 2731 | tags=22%, list=12%, si | MM_3 |
| GOBP_POSITIVE_REGULA | GOBP_POSITIVE_REGULATION_OF_CARBOH | 94 | 0.35869 | 1.3502964 | 0.058315333 | 0.23705786 | 1 | 6488 | tags=36%, list=27%, si | MM_3 |
| GOBP_PROTEIN_LOCALIZ | GOBP_PROTEIN_LOCALIZATION_TO_CHROM | 31 | 0.423669 | 1.3500743 | 0.10828026 | 0.23723698 | 1 | 6932 | tags=52%, list=29%, si | MM_3 |
| GOBP_NEUROTROPHIN_T | GOBP_NEUROTROPHIN_TRK_RECEPTOR_SIG | 33 | 0.410766 | 1.3498257 | 0.12332112 | 0.2374885 | 1 | 2430 | tags=27%, list=10%, si | MM_3 |
| GOBP_REGULATION_OF_ | GOBP_REGULATION_OF_CELLULAR_EXTRAV | 43 | 0.386487 | 1.3497933 | 0.11575179 | 0.23736897 | 1 | 4713 | tags=35%, list=20%, si | MM_3 |
| GOBP_RESPONSE_TO_INT | GOBP_RESPONSE_TO_INTERLEUKIN_7 | 17 | 0.479155 | 1.3487993 | 0.14056225 | 0.23871124 | 1 | 8680 | tags=82%, list=37%, si | MM_3 |
| GOBP_NEGATIVE_REGUL | GOBP_NEGATIVE_REGULATION_OF_RNA_CA | 82 | 0.363335 | 1.3487612 | 0.0852459 | 0.23860627 | 1 | 7186 | tags=46%, list=30%, si | MM_3 |
| GOBP_PROTEIN_DNA_CO | GOBP_PROTEIN_DNA_COMPLEX_DISASSEM | 19 | 0.461668 | 1.3486872 | 0.15289256 | 0.23855944 | 1 | 8907 | tags=74%, list=38%, si | MM_3 |
| GOBP_ACTIVATION_OF_C | GOBP_ACTIVATION_OF_CYSTEINE_TYPE_EN | 81 | 0.364092 | 1.3485672 | 0.079434164 | 0.23858741 | 1 | 7135 | tags=48%, list=30%, si | MM_3 |
| GOBP_REGULATION_OF_ | GOBP_REGULATION_OF_OXIDATIVE_STRESS | 39 | 0.407649 | 1.3483591 | 0.10722891 | 0.23874407 | 1 | 5643 | tags=44%, list=24%, si | MM_3 |
| GOBP_GLYCOPROTEIN_C | GOBP_GLYCOPROTEIN_CATABOLIC_PROCES | 31 | 0.424825 | 1.3480121 | 0.104477614 | 0.23911466 | 1 | 5587 | tags=35%, list=24%, si | MM_3 |

| | | | | | | | | | | |
|---|---|---|---|---|---|---|---|---|---|---|
| GOBP_REGULATION_OF_ | GOBP_REGULATION_OF_EARLY_ENDOSOME | 18 | 0.478793 | 1.3477426 | 0.125855 | 0.23935047 | 1 | 6933 | tags=50%, list=29%, si | MM_3 |
| AMPK_SIGNALING_PATH\ | AMPK_SIGNALING_PATHWAY | 120 | 0.350801 | 1.3476713 | 0.05122732 | 0.2392838 | 1 | 6468 | tags=37%, list=27%, si | MM_3 |
| GOBP_INTRINSIC_APOPT( | GOBP_INTRINSIC_APOPTOTIC_SIGNALING_F | 59 | 0.383335 | 1.3474909 | 0.08135593 | 0.23937677 | 1 | 7534 | tags=53%, list=32%, si | MM_3 |
| GOBP_REGULATION_OF_ | GOBP_REGULATION_OF_MACROPHAGE_DIF | 21 | 0.461899 | 1.34727 | 0.11686587 | 0.23955855 | 1 | 5993 | tags=43%, list=25%, si | MM_3 |
| GOBP_SPINAL_CORD_MC | GOBP_SPINAL_CORD_MOTOR_NEURON_DII | 28 | 0.4329 | 1.3471918 | 0.122347064 | 0.23950219 | 1 | 2530 | tags=21%, list=11%, si | MM_3 |
| GOBP_POSITIVE_REGULA | GOBP_POSITIVE_REGULATION_OF_MONOC | 21 | 0.45499 | 1.3470016 | 0.12873863 | 0.23962443 | 1 | 3888 | tags=33%, list=16%, si | MM_3 |
| GOBP_DENDRITIC_CELL_I | GOBP_DENDRITIC_CELL_MIGRATION | 26 | 0.440236 | 1.3465475 | 0.12005277 | 0.24016926 | 1 | 7028 | tags=50%, list=30%, si | MM_3 |
| GOBP_REGULATION_OF_ | GOBP_REGULATION_OF_PLASMINOGEN_AC | 17 | 0.483268 | 1.3462827 | 0.13580246 | 0.24044627 | 1 | 3578 | tags=41%, list=15%, si | MM_3 |
| GOBP_POSITIVE_REGULA | GOBP_POSITIVE_REGULATION_OF_NUCLEO | 49 | 0.385578 | 1.3462367 | 0.09815242 | 0.24035116 | 1 | 6488 | tags=37%, list=27%, si | MM_3 |
| GOBP_NEGATIVE_REGUL | GOBP_NEGATIVE_REGULATION_OF_PROTEI | 110 | 0.348922 | 1.3461499 | 0.0508658 | 0.24032082 | 1 | 7235 | tags=45%, list=31%, si | MM_3 |
| GOBP_ISOPRENOID_BIOS | GOBP_ISOPRENOID_BIOSYNTHETIC_PROCES | 36 | 0.416487 | 1.3459398 | 0.120987654 | 0.24046198 | 1 | 4551 | tags=39%, list=19%, si | MM_3 |
| GOBP_POSITIVE_REGULA | GOBP_POSITIVE_REGULATION_OF_COAGUL | 28 | 0.427839 | 1.3458974 | 0.110275686 | 0.24036701 | 1 | 5788 | tags=57%, list=25%, si | MM_3 |
| GOBP_REGULATION_OF_ | GOBP_REGULATION_OF_SMALL_MOLECULE | 356 | 0.327893 | 1.3457915 | 0.009018037 | 0.24036676 | 1 | 6192 | tags=33%, list=26%, si | MM_3 |
| GOBP_ENDONUCLEOLYTI | GOBP_ENDONUCLEOLYTIC_CLEAVAGE_OF_ | 16 | 0.488133 | 1.3451809 | 0.15641712 | 0.24114798 | 1 | 7391 | tags=69%, list=31%, si | MM_3 |
| GOBP_EMBRYONIC_CAM | GOBP_EMBRYONIC_CAMERA_TYPE_EYE_MC | 28 | 0.428266 | 1.3451725 | 0.13823164 | 0.24099472 | 1 | 2953 | tags=21%, list=13%, si | MM_3 |
| GOBP_REGULATION_OF_ | GOBP_REGULATION_OF_DEFENSE_RESPONS | 67 | 0.375269 | 1.3448477 | 0.08906426 | 0.24133414 | 1 | 5933 | tags=30%, list=25%, si | MM_3 |
| GOBP_RIBOSOMAL_SUBU | GOBP_RIBOSOMAL_SUBUNIT_EXPORT_FROI | 18 | 0.485852 | 1.3448082 | 0.15208614 | 0.24122386 | 1 | 5966 | tags=50%, list=25%, si | MM_3 |
| GOBP_NEGATIVE_REGUL | GOBP_NEGATIVE_REGULATION_OF_PLATEL | 16 | 0.476297 | 1.3445146 | 0.14084508 | 0.24149196 | 1 | 7190 | tags=56%, list=30%, si | MM_3 |
| GOBP_NEGATIVE_REGUL | GOBP_NEGATIVE_REGULATION_OF_PROTE( | 266 | 0.331221 | 1.3443955 | 0.011099899 | 0.2415141 | 1 | 5577 | tags=29%, list=24%, si | MM_3 |
| GOBP_REGULATION_OF_ | GOBP_REGULATION_OF_INTRACELLULAR_S | 72 | 0.368065 | 1.3442864 | 0.08676307 | 0.2415265 | 1 | 7071 | tags=40%, list=30%, si | MM_3 |
| GOBP_NUCLEOTIDE_EXCI | GOBP_NUCLEOTIDE_EXCISION_REPAIR | 76 | 0.366231 | 1.3442572 | 0.08093126 | 0.24141848 | 1 | 6641 | tags=41%, list=28%, si | MM_3 |
| GOBP_MAMMARY_GLAN | GOBP_MAMMARY_GLAND_EPITHELIAL_CEL | 32 | 0.415113 | 1.3441366 | 0.13167702 | 0.2414176 | 1 | 5067 | tags=44%, list=21%, si | MM_3 |
| GOBP_GLYCEROPHOSPH( | GOBP_GLYCEROPHOSPHOLIPID_METABOLIC | 267 | 0.327884 | 1.3441207 | 0.011066399 | 0.2412828 | 1 | 4804 | tags=27%, list=20%, si | MM_3 |
| GOBP_B_CELL_APOPTOTI | GOBP_B_CELL_APOPTOTIC_PROCESS | 29 | 0.42856 | 1.344018 | 0.11547912 | 0.24127467 | 1 | 7468 | tags=59%, list=32%, si | MM_3 |
| GOBP_POSITIVE_REGULA | GOBP_POSITIVE_REGULATION_OF_FIBROBL | 24 | 0.437723 | 1.3440154 | 0.1118421 | 0.24111383 | 1 | 6906 | tags=54%, list=29%, si | MM_3 |
| GOBP_CARDIAC_CELL_DE | GOBP_CARDIAC_CELL_DEVELOPMENT | 114 | 0.350853 | 1.3439293 | 0.04726101 | 0.24107984 | 1 | 3330 | tags=22%, list=14%, si | MM_3 |
| GOBP_RESPONSE_TO_ST/ | GOBP_RESPONSE_TO_STARVATION | 181 | 0.332008 | 1.3439102 | 0.036960986 | 0.2409378 | 1 | 5977 | tags=36%, list=25%, si | MM_3 |
| GOBP_GPI_ANCHOR_ME | GOBP_GPI_ANCHOR_METABOLIC_PROCESS | 33 | 0.418019 | 1.3438078 | 0.111817025 | 0.24095424 | 1 | 4421 | tags=36%, list=19%, si | MM_3 |
| GOBP_RESPONSE_TO_PE | GOBP_RESPONSE_TO_PEPTIDE_HORMONE | 361 | 0.324256 | 1.3432978 | 0.01003009 | 0.24159274 | 1 | 6159 | tags=33%, list=26%, si | MM_3 |
| GOBP_ALPHA_BETA_T_C | GOBP_ALPHA_BETA_T_CELL_PROLIFERATIO | 54 | 0.382455 | 1.3431095 | 0.10689655 | 0.24170437 | 1 | 4515 | tags=33%, list=19%, si | MM_3 |
| GOBP_REGULATION_OF_ | GOBP_REGULATION_OF_NUCLEOCYTOPLAS | 130 | 0.344702 | 1.3430961 | 0.058885384 | 0.24156874 | 1 | 6779 | tags=40%, list=29%, si | MM_3 |
| GOBP_REGULATION_OF_ | GOBP_REGULATION_OF_CELLULAR_SENESC | 47 | 0.38897 | 1.3430098 | 0.109283194 | 0.24153803 | 1 | 6636 | tags=47%, list=28%, si | MM_3 |
| GOBP_VESICLE_LOCALIZA | GOBP_VESICLE_LOCALIZATION | 218 | 0.330573 | 1.3427718 | 0.024539877 | 0.24175425 | 1 | 6720 | tags=43%, list=28%, si | MM_3 |
| GOBP_VESICLE_MEDIATE | GOBP_VESICLE_MEDIATED_TRANSPORT_BE | 46 | 0.389536 | 1.3426586 | 0.117508814 | 0.24175042 | 1 | 8945 | tags=67%, list=38%, si | MM_3 |
| GOBP_POSITIVE_REGULA | GOBP_POSITIVE_REGULATION_OF_LYMPHO | 21 | 0.456237 | 1.3424604 | 0.14095744 | 0.2418934 | 1 | 5470 | tags=43%, list=23%, si | MM_3 |
| GOBP_N_TERMINAL_PRO | GOBP_N_TERMINAL_PROTEIN_AMINO_ACII | 25 | 0.432976 | 1.3424437 | 0.12041885 | 0.24176227 | 1 | 5614 | tags=44%, list=24%, si | MM_3 |
| GOBP_FOREBRAIN_DEVE | GOBP_FOREBRAIN_DEVELOPMENT | 374 | 0.32605 | 1.3423117 | 0.008024072 | 0.24180971 | 1 | 5402 | tags=30%, list=23%, si | MM_3 |
| GOBP_POSITIVE_REGULA | GOBP_POSITIVE_REGULATION_OF_T_CELL_ | 108 | 0.350549 | 1.342056 | 0.061755147 | 0.24204424 | 1 | 4871 | tags=26%, list=21%, si | MM_3 |
| GOBP_REGULATION_OF_ | GOBP_REGULATION_OF_CARBOHYDRATE_N | 196 | 0.333019 | 1.3419253 | 0.03169734 | 0.24206986 | 1 | 6569 | tags=37%, list=28%, si | MM_3 |
| GOBP_PIGMENT_METAB( | GOBP_PIGMENT_METABOLIC_PROCESS | 65 | 0.371634 | 1.3409787 | 0.093573846 | 0.24339116 | 1 | 6660 | tags=46%, list=28%, si | MM_3 |
| GOBP_POSITIVE_REGULA | GOBP_POSITIVE_REGULATION_OF_DNA_RE | 77 | 0.360043 | 1.3408467 | 0.08314856 | 0.24341775 | 1 | 6076 | tags=31%, list=26%, si | MM_3 |
| GOBP_NEGATIVE_REGUL | GOBP_NEGATIVE_REGULATION_OF_NERVO | 22 | 0.44987 | 1.3406286 | 0.123697914 | 0.24359481 | 1 | 1190 | tags=18%, list=5%, sig | MM_3 |
| GOBP_REGULATION_OF_ | GOBP_REGULATION_OF_PEROXISOME_PRO | 17 | 0.479567 | 1.3406266 | 0.12867133 | 0.24343154 | 1 | 4881 | tags=41%, list=21%, si | MM_3 |
| GOBP_POSITIVE_REGULA | GOBP_POSITIVE_REGULATION_OF_CYTOKIN | 488 | 0.325814 | 1.3405931 | 0 | 0.2433191 | 1 | 4986 | tags=25%, list=21%, si | MM_3 |
| GOBP_REGULATION_OF_ | GOBP_REGULATION_OF_OXIDOREDUCTASE | 63 | 0.370164 | 1.3405396 | 0.08117249 | 0.24323833 | 1 | 4679 | tags=30%, list=20%, si | MM_3 |
| GOBP_POSITIVE_REGULA | GOBP_POSITIVE_REGULATION_OF_ALPHA_E | 75 | 0.362975 | 1.3403848 | 0.07 | 0.24330556 | 1 | 4655 | tags=24%, list=20%, si | MM_3 |
| GOBP_VESICLE_COATING | GOBP_VESICLE_COATING | 33 | 0.409521 | 1.3402836 | 0.114285715 | 0.24329095 | 1 | 6720 | tags=55%, list=28%, si | MM_3 |
| P53_SIGNALING_PATHW/ | P53_SIGNALING_PATHWAY | 70 | 0.361784 | 1.3400611 | 0.09492274 | 0.2434891 | 1 | 5926 | tags=37%, list=25%, si | MM_3 |
| GOBP_CELLULAR_MODIF | GOBP_CELLULAR_MODIFIED_AMINO_ACID_ | 181 | 0.335688 | 1.3399693 | 0.044102564 | 0.24345553 | 1 | 5301 | tags=32%, list=22%, si | MM_3 |
| GOBP_DETOXIFICATION_( | GOBP_DETOXIFICATION_OF_INORGANIC_C( | 15 | 0.495638 | 1.3399009 | 0.14420485 | 0.24338754 | 1 | 2449 | tags=40%, list=10%, si | MM_3 |
| GOBP_ASYMMETRIC_CEL | GOBP_ASYMMETRIC_CELL_DIVISION | 21 | 0.460568 | 1.339873 | 0.124675326 | 0.2432631 | 1 | 882 | tags=19%, list=4%, sig | MM_3 |
| GOBP_MACROPHAGE_AC | GOBP_MACROPHAGE_ACTIVATION_INVOLV | 20 | 0.459939 | 1.3398368 | 0.129771 | 0.24315622 | 1 | 2564 | tags=20%, list=11%, si | MM_3 |
| GOBP_BONE_TRABECULA | GOBP_BONE_TRABECULA_MORPHOGENESIS | 17 | 0.480245 | 1.339576 | 0.13903743 | 0.24342771 | 1 | 5724 | tags=47%, list=24%, si | MM_3 |
| GOBP_RESPONSE_TO_IN( | GOBP_RESPONSE_TO_INORGANIC_SUBSTAN | 432 | 0.325682 | 1.3394736 | 0.003003003 | 0.24342757 | 1 | 4738 | tags=25%, list=20%, si | MM_3 |
| GOBP_POSITIVE_REGULA | GOBP_POSITIVE_REGULATION_OF_GLUCOS | 54 | 0.382449 | 1.3393642 | 0.104118995 | 0.24342962 | 1 | 6159 | tags=39%, list=26%, si | MM_3 |
| GOBP_INSULIN_RECEPTO | GOBP_INSULIN_RECEPTOR_SIGNALING_PAT | 133 | 0.346163 | 1.3392999 | 0.04831933 | 0.24336468 | 1 | 6159 | tags=34%, list=26%, si | MM_3 |
| GOBP_PEROXISOME_PRC | GOBP_PEROXISOME_PROLIFERATOR_ACTIV/ | 27 | 0.425012 | 1.3389411 | 0.1376721 | 0.24376544 | 1 | 5397 | tags=33%, list=23%, si | MM_3 |
| FATTY_ACID_DEGRADATI | FATTY_ACID_DEGRADATION | 42 | 0.398798 | 1.338899 | 0.11216056 | 0.24366358 | 1 | 4743 | tags=31%, list=20%, si | MM_3 |
| GOBP_REGULATION_OF_ | GOBP_REGULATION_OF_MITOTIC_CELL_CY( | 331 | 0.323579 | 1.3388054 | 0.012024048 | 0.2436432 | 1 | 6906 | tags=38%, list=29%, si | MM_3 |
| GOBP_REGULATION_OF_ | GOBP_REGULATION_OF_ODONTOGENESIS | 19 | 0.462254 | 1.3387233 | 0.13769124 | 0.24359497 | 1 | 5057 | tags=42%, list=21%, si | MM_3 |
| GOBP_REGULATION_OF_ | GOBP_REGULATION_OF_GLUCOSE_METAB( | 109 | 0.347522 | 1.3387175 | 0.057815846 | 0.24344 | 1 | 5983 | tags=36%, list=25%, si | MM_3 |
| GOBP_NUCLEAR_CHROM | GOBP_NUCLEAR_CHROMOSOME_SEGREGAT | 305 | 0.325632 | 1.3386519 | 0.0090452265 | 0.24337868 | 1 | 7067 | tags=40%, list=30%, si | MM_3 |
| GOBP_REGULATION_OF_ | GOBP_REGULATION_OF_NON_CANONICAL_ | 26 | 0.43423 | 1.3377801 | 0.1259542 | 0.2445708 | 1 | 4997 | tags=31%, list=21%, si | MM_3 |
| GOBP_CELLULAR_RESPON | GOBP_CELLULAR_RESPONSE_TO_VASCULAF | 60 | 0.374306 | 1.3376024 | 0.09859155 | 0.24469668 | 1 | 3694 | tags=25%, list=16%, si | MM_3 |
| GOBP_RESPONSE_TO_FU | GOBP_RESPONSE_TO_FUNGUS | 59 | 0.38229 | 1.3375975 | 0.09885058 | 0.24453996 | 1 | 4256 | tags=24%, list=18%, si | MM_3 |
| GOBP_REGULATION_OF_ | GOBP_REGULATION_OF_RESPONSE_TO_NU | 34 | 0.409486 | 1.3375936 | 0.11590629 | 0.24438131 | 1 | 5192 | tags=35%, list=22%, si | MM_3 |
| GOBP_NUCLEOTIDE_BINI | GOBP_NUCLEOTIDE_BINDING_OLIGOMERIZ | 18 | 0.469054 | 1.3372036 | 0.15552524 | 0.2448246 | 1 | 6867 | tags=61%, list=29%, si | MM_3 |
| GOBP_CELLULAR_RESPON | GOBP_CELLULAR_RESPONSE_TO_CAMP | 38 | 0.401498 | 1.3367801 | 0.11311673 | 0.24533407 | 1 | 5317 | tags=32%, list=23%, si | MM_3 |
| GOBP_NEGATIVE_REGUL | GOBP_NEGATIVE_REGULATION_OF_MRNA_ | 69 | 0.3682 | 1.3365798 | 0.08305648 | 0.24550767 | 1 | 5323 | tags=32%, list=23%, si | MM_3 |
| GOBP_CELLULAR_RESPON | GOBP_CELLULAR_RESPONSE_TO_PEPTIDE_H | 276 | 0.326449 | 1.3362651 | 0.016129032 | 0.24581471 | 1 | 6159 | tags=33%, list=26%, si | MM_3 |
| GOBP_POSITIVE_REGULA | GOBP_POSITIVE_REGULATION_OF_KINASE_ | 329 | 0.321796 | 1.3358626 | 0.006024096 | 0.24628204 | 1 | 5031 | tags=26%, list=21%, si | MM_3 |
| GOBP_MITOCHONDRIAL_ | GOBP_MITOCHONDRIAL_TRANSCRIPTION | 21 | 0.452957 | 1.3355571 | 0.1296061 | 0.2465666 | 1 | 7204 | tags=57%, list=31%, si | MM_3 |
| GOBP_ARF_PROTEIN_SIG | GOBP_ARF_PROTEIN_SIGNAL_TRANSDUCTI( | 23 | 0.442171 | 1.3355197 | 0.13289474 | 0.24646723 | 1 | 5346 | tags=35%, list=23%, si | MM_3 |
| GOBP_MYELOID_CELL_DE | GOBP_MYELOID_CELL_DEVELOPMENT | 98 | 0.353541 | 1.3352715 | 0.06878307 | 0.24668981 | 1 | 7051 | tags=45%, list=30%, si | MM_3 |
| GOBP_CELL_MIGRATION_ | GOBP_CELL_MIGRATION_INVOLVED_IN_HE | 19 | 0.457586 | 1.3351719 | 0.17001338 | 0.2466959 | 1 | 4761 | tags=37%, list=20%, si | MM_3 |
| GOBP_GLYCEROLIPID_BI( | GOBP_GLYCEROLIPID_BIOSYNTHETIC_PROC | 210 | 0.329674 | 1.3349527 | 0.030612245 | 0.24689592 | 1 | 5055 | tags=31%, list=21%, si | MM_3 |
| GOBP_RESPONSE_TO_TO | GOBP_RESPONSE_TO_TOXIC_SUBSTANCE | 136 | 0.340088 | 1.3343215 | 0.049525816 | 0.24771686 | 1 | 4679 | tags=30%, list=20%, si | MM_3 |
| GOBP_REGULATION_OF_ | GOBP_REGULATION_OF_ACUTE_INFLAMMA | 17 | 0.472272 | 1.3341819 | 0.16487935 | 0.2477785 | 1 | 6692 | tags=47%, list=28%, si | MM_3 |
| GOBP_INTRINSIC_APOPT( | GOBP_INTRINSIC_APOPTOTIC_SIGNALING_F | 63 | 0.372638 | 1.3341225 | 0.09662921 | 0.24770771 | 1 | 5485 | tags=38%, list=23%, si | MM_3 |
| GOBP_NEGATIVE_REGUL | GOBP_NEGATIVE_REGULATION_OF_AUTOPI | 81 | 0.360331 | 1.3341119 | 0.08039647 | 0.24755879 | 1 | 7599 | tags=53%, list=32%, si | MM_3 |
| GOBP_POSITIVE_REGULA | GOBP_POSITIVE_REGULATION_OF_TELOME | 17 | 0.47254 | 1.3340712 | 0.14247312 | 0.24744838 | 1 | 6250 | tags=41%, list=26%, si | MM_3 |
| GOBP_REGULATION_OF_ | GOBP_REGULATION_OF_HYDROGEN_PERO) | 17 | 0.475738 | 1.3340192 | 0.14986376 | 0.2473621 | 1 | 1273 | tags=18%, list=5%, sig | MM_3 |

| | | | | | | | | | | |
|---|---|---|---|---|---|---|---|---|---|---|
| GOBP_REGULATION_OF_ | GOBP_REGULATION_OF_MEMBRANE_PROT | 25 | 0.438157 | 1.3335752 | 0.123655915 | 0.24785326 | 1 | 7521 | tags=48%, list=32%, si | MM_3 |
| GOBP_RNA_SPLICING_VI | GOBP_RNA_SPLICING_VIA_TRANSESTERIFIC | 274 | 0.324701 | 1.3331307 | 0.011055277 | 0.248361 | 1 | 8269 | tags=51%, list=35%, si | MM_3 |
| GOBP_CALCIUM_ION_TR | GOBP_CALCIUM_ION_TRANSPORT_INTO_CY | 30 | 0.420195 | 1.3328013 | 0.11153846 | 0.24874514 | 1 | 6697 | tags=50%, list=28%, si | MM_3 |
| GOBP_NEGATIVE_REGUL | GOBP_NEGATIVE_REGULATION_OF_T_CELL | 23 | 0.447903 | 1.3327478 | 0.1484375 | 0.24866363 | 1 | 6489 | tags=43%, list=27%, si | MM_3 |
| GOBP_REGULATION_OF_ | GOBP_REGULATION_OF_CELL_MORPHOGEN | 267 | 0.327363 | 1.332747 | 0.019172553 | 0.24850109 | 1 | 5559 | tags=32%, list=24%, si | MM_3 |
| GOBP_REGULATION_OF_ | GOBP_REGULATION_OF_BINDING | 331 | 0.323401 | 1.3323905 | 0.010070493 | 0.24888521 | 1 | 6906 | tags=41%, list=29%, si | MM_3 |
| GOBP_RESPONSE_TO_PR | GOBP_RESPONSE_TO_PROGESTERONE | 19 | 0.465425 | 1.3320957 | 0.15680881 | 0.24920648 | 1 | 5903 | tags=47%, list=25%, si | MM_3 |
| GOBP_EXTRINSIC_APOPT | GOBP_EXTRINSIC_APOPTOTIC_SIGNALING_I | 80 | 0.357008 | 1.3320789 | 0.07717751 | 0.24906568 | 1 | 7564 | tags=54%, list=32%, si | MM_3 |
| GOBP_IMMATURE_T_CEL | GOBP_IMMATURE_T_CELL_PROLIFERATION | 18 | 0.476365 | 1.3319721 | 0.14498645 | 0.24906331 | 1 | 1155 | tags=17%, list=5%, sig | MM_3 |
| GOBP_PROTEIN_HOMOT | GOBP_PROTEIN_HOMOTETRAMERIZATION | 49 | 0.389628 | 1.3319153 | 0.11355736 | 0.24899344 | 1 | 5318 | tags=33%, list=23%, si | MM_3 |
| GOBP_REGULATION_OF_ | GOBP_REGULATION_OF_DOUBLE_STRAND_ | 139 | 0.337477 | 1.3318005 | 0.056962024 | 0.24900796 | 1 | 4668 | tags=24%, list=20%, si | MM_3 |
| GOBP_VIRION_ASSEMBLY | GOBP_VIRION_ASSEMBLY | 26 | 0.435308 | 1.3316816 | 0.14231259 | 0.24904366 | 1 | 7229 | tags=62%, list=31%, si | MM_3 |
| GOBP_REGULATION_OF_ | GOBP_REGULATION_OF_CELLULAR_COMPO | 380 | 0.32178 | 1.3312776 | 0.006024096 | 0.2495055 | 1 | 7570 | tags=42%, list=32%, si | MM_3 |
| GOBP_NEGATIVE_REGUL | GOBP_NEGATIVE_REGULATION_OF_ADENY | 17 | 0.471041 | 1.3312742 | 0.14945652 | 0.24934538 | 1 | 1690 | tags=18%, list=7%, sig | MM_3 |
| GOBP_STEM_CELL_PROLI | GOBP_STEM_CELL_PROLIFERATION | 128 | 0.343521 | 1.3308271 | 0.05549738 | 0.24990788 | 1 | 5507 | tags=30%, list=23%, si | MM_3 |
| GOBP_REGULATION_OF_ | GOBP_REGULATION_OF_MYELINATION | 58 | 0.379773 | 1.3308219 | 0.08924485 | 0.24975452 | 1 | 7521 | tags=50%, list=32%, si | MM_3 |
| GOBP_MULTICELLULAR_( | GOBP_MULTICELLULAR_ORGANISMAL_LEVE | 20 | 0.454495 | 1.3307931 | 0.14496644 | 0.24963649 | 1 | 880 | tags=15%, list=4%, sig | MM_3 |
| GOBP_PSEUDOURIDINE_S | GOBP_PSEUDOURIDINE_SYNTHESIS | 18 | 0.468153 | 1.3306937 | 0.14114514 | 0.24962546 | 1 | 6341 | tags=50%, list=27%, si | MM_3 |
| GOBP_REGULATION_OF_ | GOBP_REGULATION_OF_CELL_PROLIFERATI | 15 | 0.485514 | 1.3306559 | 0.1438451 | 0.24951303 | 1 | 3680 | tags=33%, list=16%, si | MM_3 |
| GOBP_CELL_ACTIVATION | GOBP_CELL_ACTIVATION_INVOLVED_IN_IM | 291 | 0.324569 | 1.3306164 | 0.014084507 | 0.24941163 | 1 | 5012 | tags=26%, list=21%, si | MM_3 |
| GOBP_NEGATIVE_REGUL | GOBP_NEGATIVE_REGULATION_OF_T_CELL | 29 | 0.418156 | 1.3301699 | 0.14338236 | 0.24996822 | 1 | 2935 | tags=24%, list=12%, si | MM_3 |
| GOBP_CONNECTIVE_TISS | GOBP_CONNECTIVE_TISSUE_DEVELOPMENT | 292 | 0.32425 | 1.3301048 | 0.022199798 | 0.24990304 | 1 | 5583 | tags=31%, list=24%, si | MM_3 |
| GOBP_REGULATION_OF_ | GOBP_REGULATION_OF_CELL_KILLING | 116 | 0.34565 | 1.3301008 | 0.06503198 | 0.24974564 | 1 | 5938 | tags=34%, list=25%, si | MM_3 |
| GOBP_INTEGRIN_MEDIAT | GOBP_INTEGRIN_MEDIATED_SIGNALING_P | 98 | 0.354282 | 1.3300854 | 0.08145767 | 0.24960256 | 1 | 7219 | tags=46%, list=31%, si | MM_3 |
| GOBP_PROTEIN_LOCALIZ | GOBP_PROTEIN_LOCALIZATION_TO_PLASM | 296 | 0.324593 | 1.3299166 | 0.012158055 | 0.24970677 | 1 | 7649 | tags=50%, list=32%, si | MM_3 |
| GOBP_ESTABLISHMENT_( | GOBP_ESTABLISHMENT_OF_ORGANELLE_LO | 472 | 0.320206 | 1.3291962 | 0.003003003 | 0.24997081 | 1 | 6931 | tags=39%, list=29%, si | MM_3 |
| OLFACTORY_TRANSDUCT | OLFACTORY_TRANSDUCTION | 186 | -0.38624 | -2.220427 | 0 | 0.07281068 | 0.089 | 7745 | tags=74%, list=33%, si | MM_3 |
| GOBP_RENAL_VESICLE_D | GOBP_RENAL_VESICLE_DEVELOPMENT | 18 | -0.60787 | -2.068948 | 0 | 0.18671498 | 0.371 | 934 | tags=28%, list=4%, sig | MM_3 |
| GOBP_REGULATION_OF_ | GOBP_REGULATION_OF_SKELETAL_MUSCLE | 21 | -0.52849 | -2.000707 | 0.0046296297 | 0.2346745 | 0.583 | 1621 | tags=33%, list=7%, sig | MM_3 |
| RIBOSOME | RIBOSOME | 128 | 0.64121 | 3.114226 | 0 | 0 | 0 | 5223 | tags=67%, list=22%, si | EN_3 |
| GOBP_CYTOPLASMIC_TR | GOBP_CYTOPLASMIC_TRANSLATION | 143 | 0.590112 | 2.9275281 | 0 | 0 | 0 | 6141 | tags=64%, list=26%, si | EN_3 |
| GOBP_TRANSLATION_AT_ | GOBP_TRANSLATION_AT_SYNAPSE | 48 | 0.696777 | 2.8099957 | 0 | 0 | 0 | 5958 | tags=83%, list=25%, si | EN_3 |
| OXIDATIVE_PHOSPHORYL | OXIDATIVE_PHOSPHORYLATION | 115 | 0.576614 | 2.7561688 | 0 | 0 | 0 | 4894 | tags=53%, list=21%, si | EN_3 |
| PARKINSON_DISEASE | PARKINSON_DISEASE | 122 | 0.543979 | 2.6161716 | 0 | 0 | 0 | 4894 | tags=50%, list=21%, si | EN_3 |
| GOBP_OXIDATIVE_PHOSP | GOBP_OXIDATIVE_PHOSPHORYLATION | 124 | 0.527476 | 2.5284312 | 0 | 0 | 0 | 5859 | tags=54%, list=25%, si | EN_3 |
| GOBP_AEROBIC_RESPIRA | GOBP_AEROBIC_RESPIRATION | 170 | 0.491153 | 2.485345 | 0 | 0 | 0 | 5897 | tags=49%, list=25%, si | EN_3 |
| GOBP_ATP_BIOSYNTHETI | GOBP_ATP_BIOSYNTHETIC_PROCESS | 80 | 0.54261 | 2.467636 | 0 | 0 | 0 | 5859 | tags=54%, list=25%, si | EN_3 |
| HUNTINGTON_DISEASE | HUNTINGTON_DISEASE | 180 | 0.480809 | 2.4451027 | 0 | 0 | 0 | 4894 | tags=42%, list=21%, si | EN_3 |
| GOBP_NUCLEOSIDE_TRIP | GOBP_NUCLEOSIDE_TRIPHOSPHATE_BIOSYN | 100 | 0.525134 | 2.4369886 | 0 | 0 | 0 | 4894 | tags=43%, list=21%, si | EN_3 |
| NON-ALCOHOLIC_FATTY_ | NON-ALCOHOLIC_FATTY_LIVER_DISEASE_(N | 141 | 0.492144 | 2.4321294 | 0 | 0 | 0 | 5000 | tags=45%, list=21%, si | EN_3 |
| ALZHEIMER_DISEASE | ALZHEIMER_DISEASE | 159 | 0.474286 | 2.384569 | 0 | 0 | 0 | 5000 | tags=43%, list=21%, si | EN_3 |
| GOBP_MITOCHONDRIAL_ | GOBP_MITOCHONDRIAL_RESPIRATORY_CHA | 91 | 0.52125 | 2.3803012 | 0 | 9.02381E-05 | 0.001 | 5971 | tags=51%, list=25%, si | EN_3 |
| THERMOGENESIS | THERMOGENESIS | 210 | 0.454608 | 2.3429704 | 0 | 0.000337107 | 0.004 | 7074 | tags=54%, list=30%, si | EN_3 |
| GOBP_CELLULAR_RESPIR | GOBP_CELLULAR_RESPIRATION | 218 | 0.451443 | 2.3345902 | 0 | 0.000553256 | 0.007 | 5897 | tags=46%, list=25%, si | EN_3 |
| GOBP_AEROBIC_ELECTRO | GOBP_AEROBIC_ELECTRON_TRANSPORT_CH | 63 | 0.539966 | 2.3204644 | 0 | 0.00074212 | 0.01 | 5837 | tags=59%, list=25%, si | EN_3 |
| GOBP_PROTON_MOTIVE_ | GOBP_PROTON_MOTIVE_FORCE_DRIVEN_A | 52 | 0.558248 | 2.2832375 | 0 | 0.001463825 | 0.021 | 4894 | tags=48%, list=21%, si | EN_3 |
| GOBP_RIBOSE_PHOSPHA | GOBP_RIBOSE_PHOSPHATE_BIOSYNTHETIC_ | 185 | 0.441547 | 2.2642756 | 0 | 0.001647874 | 0.025 | 5859 | tags=43%, list=25%, si | EN_3 |
| GOBP_NUCLEOSIDE_TRIP | GOBP_NUCLEOSIDE_TRIPHOSPHATE_METAB | 250 | 0.42774 | 2.2638764 | 0 | 0.001561144 | 0.025 | 5097 | tags=37%, list=22%, si | EN_3 |
| GOBP_GLYCOLYTIC_PROC | GOBP_GLYCOLYTIC_PROCESS_THROUGH_FR | 25 | 0.634432 | 2.2063682 | 0 | 0.004985701 | 0.083 | 5019 | tags=48%, list=21%, si | EN_3 |
| GOBP_NADH_DEHYDROG | GOBP_NADH_DEHYDROGENASE_COMPLEX_ | 58 | 0.516368 | 2.1825082 | 0 | 0.006781936 | 0.117 | 5886 | tags=48%, list=25%, si | EN_3 |
| GOBP_GENERATION_OF_ | GOBP_GENERATION_OF_PRECURSOR_META | 439 | 0.393355 | 2.1779094 | 0 | 0.007065312 | 0.128 | 5423 | tags=37%, list=23%, si | EN_3 |
| GOBP_INNER_MITOCHON | GOBP_INNER_MITOCHONDRIAL_MEMBRAN | 36 | 0.567794 | 2.155882 | 0 | 0.00974638 | 0.178 | 4033 | tags=36%, list=17%, si | EN_3 |
| GOBP_ATP_SYNTHESIS_C | GOBP_ATP_SYNTHESIS_COUPLED_ELECTRO | 76 | 0.492411 | 2.154306 | 0 | 0.009588433 | 0.182 | 4914 | tags=46%, list=21%, si | EN_3 |
| GOBP_PROTEIN_STABILIZ | GOBP_PROTEIN_STABILIZATION | 190 | 0.422259 | 2.1500933 | 0 | 0.009630967 | 0.191 | 5484 | tags=35%, list=23%, si | EN_3 |
| GOBP_ELECTRON_TRANS | GOBP_ELECTRON_TRANSPORT_CHAIN | 107 | 0.456533 | 2.147845 | 0 | 0.00953436 | 0.196 | 4914 | tags=43%, list=21%, si | EN_3 |
| GOBP_MAINTENANCE_OI | GOBP_MAINTENANCE_OF_PROTEIN_LOCAL | 43 | 0.548729 | 2.1433742 | 0 | 0.009663356 | 0.207 | 4934 | tags=53%, list=21%, si | EN_3 |
| BASE_EXCISION_REPAIR | BASE_EXCISION_REPAIR | 32 | 0.583413 | 2.1398718 | 0 | 0.009657259 | 0.213 | 4836 | tags=50%, list=20%, si | EN_3 |
| GOBP_CARBOHYDRATE_F | GOBP_CARBOHYDRATE_PHOSPHORYLATION | 19 | 0.655429 | 2.1254117 | 0 | 0.011289915 | 0.25 | 3092 | tags=47%, list=13%, si | EN_3 |
| GOBP_RIBONUCLEOPROT | GOBP_RIBONUCLEOPROTEIN_COMPLEX_BIC | 417 | 0.387549 | 2.1227503 | 0 | 0.011389213 | 0.259 | 6100 | tags=42%, list=26%, si | EN_3 |
| GOBP_NUCLEOSIDE_PHO | GOBP_NUCLEOSIDE_PHOSPHATE_BIOSYNTH | 234 | 0.409212 | 2.1180952 | 0 | 0.011866669 | 0.274 | 4894 | tags=35%, list=21%, si | EN_3 |
| GOBP_PYRIMIDINE_NUCL | GOBP_PYRIMIDINE_NUCLEOTIDE_BIOSYNTH | 24 | 0.621172 | 2.1142979 | 0 | 0.011978628 | 0.284 | 4815 | tags=50%, list=20%, si | EN_3 |
| GOBP_RIBOSOME_BIOGE | GOBP_RIBOSOME_BIOGENESIS | 306 | 0.391976 | 2.1071177 | 0 | 0.012550608 | 0.304 | 5227 | tags=39%, list=22%, si | EN_3 |
| GOBP_CELLULAR_RESPOI | GOBP_CELLULAR_RESPONSE_TO_ACID_CHE | 87 | 0.463461 | 2.102876 | 0 | 0.013019554 | 0.316 | 5721 | tags=43%, list=24%, si | EN_3 |
| GOBP_ENERGY_DERIVATI | GOBP_ENERGY_DERIVATION_BY_OXIDATIO | 321 | 0.389246 | 2.100413 | 0 | 0.012952531 | 0.321 | 5916 | tags=40%, list=25%, si | EN_3 |
| GOBP_PYRIMIDINE_NUCL | GOBP_PYRIMIDINE_NUCLEOSIDE_TRIPHOSP | 18 | 0.6583 | 2.0988932 | 0 | 0.012757955 | 0.326 | 1287 | tags=28%, list=5%, sig | EN_3 |
| GOBP_MONOSACCHARID | GOBP_MONOSACCHARIDE_METABOLIC_PR | 255 | 0.395572 | 2.0974112 | 0 | 0.012638491 | 0.327 | 4833 | tags=32%, list=20%, si | EN_3 |
| GOBP_MITOCHONDRIAL_ | GOBP_MITOCHONDRIAL_TRANSLATION | 128 | 0.430146 | 2.0967834 | 0 | 0.012367944 | 0.329 | 3789 | tags=33%, list=16%, si | EN_3 |
| GOBP_ATP_METABOLIC_ | GOBP_ATP_METABOLIC_PROCESS | 206 | 0.412195 | 2.0956542 | 0 | 0.012172762 | 0.332 | 5097 | tags=37%, list=22%, si | EN_3 |
| GOBP_NEGATIVE_REGUL | GOBP_NEGATIVE_REGULATION_OF_RHO_P | 25 | 0.607366 | 2.0899544 | 0.0016051364 | 0.012759283 | 0.347 | 5740 | tags=48%, list=24%, si | EN_3 |
| GOBP_RIBOSE_PHOSPHA | GOBP_RIBOSE_PHOSPHATE_METABOLIC_PR | 419 | 0.376541 | 2.0876899 | 0 | 0.012940903 | 0.354 | 5160 | tags=33%, list=22%, si | EN_3 |
| GOBP_MITOTIC_DNA_IN | GOBP_MITOTIC_DNA_INTEGRITY_CHECKPO | 84 | 0.456949 | 2.0850942 | 0 | 0.012885348 | 0.361 | 5653 | tags=40%, list=24%, si | EN_3 |
| GOBP_FIBROBLAST_MIGR | GOBP_FIBROBLAST_MIGRATION | 68 | 0.478006 | 2.0839846 | 0 | 0.012778136 | 0.368 | 4995 | tags=41%, list=21%, si | EN_3 |
| GOBP_PROTEIN_LOCALIZ | GOBP_PROTEIN_LOCALIZATION_TO_MITOC | 101 | 0.442138 | 2.0834966 | 0 | 0.012514683 | 0.369 | 7767 | tags=58%, list=33%, si | EN_3 |
| GOBP_RIBOSOMAL_SMAI | GOBP_RIBOSOMAL_SMALL_SUBUNIT_BIOGE | 101 | 0.444213 | 2.0788984 | 0 | 0.01297837 | 0.386 | 6275 | tags=50%, list=27%, si | EN_3 |
| GOBP_REGULATION_OF_ | GOBP_REGULATION_OF_ERAD_PATHWAY | 27 | 0.5941 | 2.0745285 | 0 | 0.013134801 | 0.398 | 3656 | tags=37%, list=15%, si | EN_3 |
| GOBP_GLUCOSE_METABO | GOBP_GLUCOSE_METABOLIC_PROCESS | 201 | 0.397917 | 2.0637724 | 0 | 0.014522497 | 0.437 | 4386 | tags=31%, list=19%, si | EN_3 |
| GOBP_PYRIMIDINE_NUCL | GOBP_PYRIMIDINE_NUCLEOTIDE_METABOL | 38 | 0.539346 | 2.0631216 | 0 | 0.014294058 | 0.44 | 4815 | tags=39%, list=20%, si | EN_3 |
| GOBP_GLYCOLYTIC_PROC | GOBP_GLYCOLYTIC_PROCESS_THROUGH_GL | 20 | 0.641927 | 2.0606108 | 0.0016778524 | 0.014414161 | 0.451 | 5019 | tags=50%, list=21%, si | EN_3 |
| GOBP_STEROL_BIOSYNTH | GOBP_STEROL_BIOSYNTHETIC_PROCESS | 58 | 0.482508 | 2.0592763 | 0 | 0.014411054 | 0.457 | 2896 | tags=34%, list=12%, si | EN_3 |
| GOBP_POSITIVE_REGULA | GOBP_POSITIVE_REGULATION_OF_TORC1_S | 52 | 0.503385 | 2.0588262 | 0 | 0.014221875 | 0.457 | 4852 | tags=44%, list=21%, si | EN_3 |
| GOBP_MAINTENANCE_OI | GOBP_MAINTENANCE_OF_PROTEIN_LOCAT | 97 | 0.445621 | 2.0564055 | 0 | 0.014358089 | 0.469 | 4934 | tags=42%, list=21%, si | EN_3 |
| GOBP_ENDOPLASMIC_RE | GOBP_ENDOPLASMIC_RETICULUM_TO_GOL | 120 | 0.427965 | 2.0547862 | 0 | 0.014468137 | 0.476 | 5542 | tags=36%, list=23%, si | EN_3 |

| | | | | | | | | | | |
|---|---|---|---|---|---|---|---|---|---|---|
| GOBP_RETINOIC_ACID_R | GOBP_RETINOIC_ACID_RECEPTOR_SIGNALIN | 34 | 0.552379 | 2.0490353 | 0 | 0.015320265 | 0.506 | 4660 | tags=41%, list=20%, si | EN_3 |
| GOBP_COLLAGEN_FIBRIL | GOBP_COLLAGEN_FIBRIL_ORGANIZATION | 63 | 0.476134 | 2.045664 | 0 | 0.01579746 | 0.525 | 5692 | tags=41%, list=24%, si | EN_3 |
| GOBP_RIBOSOME_ASSEM | GOBP_RIBOSOME_ASSEMBLY | 60 | 0.477445 | 2.0401742 | 0 | 0.01663928 | 0.549 | 4834 | tags=47%, list=20%, si | EN_3 |
| GLYCOLYSIS_/_GLUCONE | GLYCOLYSIS_/_GLUCONEOGENESIS | 60 | 0.489126 | 2.0388544 | 0 | 0.016638035 | 0.553 | 5103 | tags=45%, list=22%, si | EN_3 |
| GOBP_NEGATIVE_REGUL | GOBP_NEGATIVE_REGULATION_OF_CYTOPL | 50 | 0.500693 | 2.0368345 | 0 | 0.016618075 | 0.556 | 3321 | tags=34%, list=14%, si | EN_3 |
| GOBP_PROTEIN_RNA_CO | GOBP_PROTEIN_RNA_COMPLEX_ORGANIZA | 173 | 0.40007 | 2.0169575 | 0 | 0.021387538 | 0.654 | 5290 | tags=37%, list=22%, si | EN_3 |
| GOBP_DEFENSE_RESPON | GOBP_DEFENSE_RESPONSE_TO_VIRUS | 276 | 0.377425 | 2.0084846 | 0 | 0.023145447 | 0.693 | 2997 | tags=24%, list=13%, si | EN_3 |
| GOBP_RESPONSE_TO_AC | GOBP_RESPONSE_TO_ACID_CHEMICAL | 109 | 0.426407 | 2.006622 | 0 | 0.023233641 | 0.701 | 5721 | tags=39%, list=24%, si | EN_3 |
| GOBP_NEGATIVE_REGUL | GOBP_NEGATIVE_REGULATION_OF_VIRAL_ | 95 | 0.434938 | 2.0052152 | 0 | 0.023413902 | 0.709 | 2487 | tags=26%, list=11%, si | EN_3 |
| GOBP_REGULATION_OF_ | GOBP_REGULATION_OF_PROGRAMMED_NE | 27 | 0.561287 | 2.0041354 | 0 | 0.023325566 | 0.714 | 5583 | tags=48%, list=24%, si | EN_3 |
| GOBP_ISOPRENOID_MET | GOBP_ISOPRENOID_METABOLIC_PROCESS | 87 | 0.442302 | 1.9987609 | 0 | 0.024462998 | 0.734 | 4023 | tags=33%, list=17%, si | EN_3 |
| GOBP_PROTEIN_TARGETI | GOBP_PROTEIN_TARGETING_TO_MITOCHO | 76 | 0.453369 | 1.9966879 | 0 | 0.02467009 | 0.74 | 7304 | tags=54%, list=31%, si | EN_3 |
| GOBP_REGULATION_OF_ | GOBP_REGULATION_OF_RESPONSE_TO_EN | 80 | 0.439082 | 1.9959296 | 0 | 0.024583498 | 0.743 | 4076 | tags=28%, list=17%, si | EN_3 |
| GOBP_MITOCHONDRIAL_ | GOBP_MITOCHONDRIAL_GENE_EXPRESSION | 164 | 0.399316 | 1.9928973 | 0 | 0.025031254 | 0.758 | 3789 | tags=31%, list=16%, si | EN_3 |
| GOBP_DNA_BIOSYNTHET | GOBP_DNA_BIOSYNTHETIC_PROCESS | 175 | 0.386673 | 1.9914521 | 0 | 0.024942327 | 0.762 | 5201 | tags=33%, list=22%, si | EN_3 |
| GOBP_NEGATIVE_REGUL | GOBP_NEGATIVE_REGULATION_OF_RELEAS | 22 | 0.600523 | 1.987354 | 0 | 0.025700195 | 0.776 | 3418 | tags=32%, list=14%, si | EN_3 |
| GOBP_MACROMOLECULE | GOBP_MACROMOLECULE_DEACYLATION | 59 | 0.474767 | 1.986188 | 0 | 0.025586965 | 0.778 | 5253 | tags=39%, list=22%, si | EN_3 |
| GOBP_REGULATION_OF_ | GOBP_REGULATION_OF_FIBROBLAST_MIGR | 47 | 0.492476 | 1.9860013 | 0 | 0.025377238 | 0.78 | 4672 | tags=38%, list=20%, si | EN_3 |
| GOBP_PROTEIN_QUALITY | GOBP_PROTEIN_QUALITY_CONTROL_FOR_M | 17 | 0.64201 | 1.985195 | 0.0016778524 | 0.025304945 | 0.784 | 8070 | tags=82%, list=34%, si | EN_3 |
| GOBP_MACROAUTOPHAG | GOBP_MACROAUTOPHAGY | 287 | 0.37312 | 1.9842293 | 0 | 0.025315864 | 0.789 | 6301 | tags=35%, list=27%, si | EN_3 |
| GOBP_POSITIVE_REGULA | GOBP_POSITIVE_REGULATION_OF_SUBSTRA | 40 | 0.503909 | 1.9800094 | 0 | 0.026285665 | 0.802 | 4672 | tags=38%, list=20%, si | EN_3 |
| GOBP_MAINTENANCE_OF | GOBP_MAINTENANCE_OF_PROTEIN_LOCAT | 70 | 0.45075 | 1.9762353 | 0 | 0.0270246 | 0.811 | 4934 | tags=47%, list=21%, si | EN_3 |
| GOBP_ISOPRENOID_BIOS | GOBP_ISOPRENOID_BIOSYNTHETIC_PROCES | 36 | 0.517835 | 1.9731576 | 0 | 0.027669128 | 0.82 | 3983 | tags=39%, list=17%, si | EN_3 |
| GOBP_HEME_METABOLIC | GOBP_HEME_METABOLIC_PROCESS | 34 | 0.523016 | 1.9724756 | 0 | 0.027556099 | 0.824 | 4522 | tags=41%, list=19%, si | EN_3 |
| GOBP_PYRIMIDINE_CONT | GOBP_PYRIMIDINE_CONTAINING_COMPOU | 60 | 0.462354 | 1.970146 | 0.0013280213 | 0.027978819 | 0.834 | 3814 | tags=27%, list=16%, si | EN_3 |
| GOBP_RRNA_METABOLIC | GOBP_RRNA_METABOLIC_PROCESS | 241 | 0.376945 | 1.9701097 | 0 | 0.027654782 | 0.835 | 4783 | tags=35%, list=20%, si | EN_3 |
| GOBP_POSITIVE_REGULA | GOBP_POSITIVE_REGULATION_OF_ACTIN_F | 52 | 0.476839 | 1.9688858 | 0.0014144272 | 0.027591974 | 0.838 | 2908 | tags=33%, list=12%, si | EN_3 |
| DNA_REPLICATION | DNA_REPLICATION | 34 | 0.522484 | 1.9681674 | 0.001529052 | 0.027485272 | 0.839 | 3249 | tags=35%, list=14%, si | EN_3 |
| GOBP_HYDROGEN_PERO | GOBP_HYDROGEN_PEROXIDE_METABOLIC_ | 48 | 0.49016 | 1.9679292 | 0 | 0.027193254 | 0.839 | 3634 | tags=33%, list=15%, si | EN_3 |
| GOBP_ORGANOPHOSPHA | GOBP_ORGANOPHOSPHATE_BIOSYNTHETIC | 500 | 0.357606 | 1.9646609 | 0 | 0.02798004 | 0.852 | 5888 | tags=38%, list=25%, si | EN_3 |
| GOBP_MITOCHONDRIAL_ | GOBP_MITOCHONDRIAL_MEMBRANE_ORGA | 107 | 0.414756 | 1.9641473 | 0 | 0.02787389 | 0.853 | 3635 | tags=30%, list=15%, si | EN_3 |
| GOBP_GLUCOSE_IMPORT | GOBP_GLUCOSE_IMPORT | 84 | 0.435804 | 1.9629476 | 0 | 0.027938873 | 0.86 | 4986 | tags=35%, list=21%, si | EN_3 |
| GOBP_RRNA_PROCESSIN | GOBP_RRNA_PROCESSING | 211 | 0.382432 | 1.9607118 | 0 | 0.028415835 | 0.869 | 4772 | tags=36%, list=20%, si | EN_3 |
| GOBP_TORC1_SIGNALING | GOBP_TORC1_SIGNALING | 100 | 0.42731 | 1.9589666 | 0 | 0.028512824 | 0.87 | 4537 | tags=36%, list=19%, si | EN_3 |
| GOBP_G0_TO_G1_TRANS | GOBP_G0_TO_G1_TRANSITION | 38 | 0.511497 | 1.9588679 | 0.0015698588 | 0.028202308 | 0.87 | 3339 | tags=32%, list=14%, si | EN_3 |
| GOBP_MYELOID_LEUKOC | GOBP_MYELOID_LEUKOCYTE_MEDIATED_IM | 108 | 0.41099 | 1.952101 | 0 | 0.030270027 | 0.89 | 3448 | tags=28%, list=15%, si | EN_3 |
| GOBP_NUCLEOTIDE_EXCI | GOBP_NUCLEOTIDE_EXCISION_REPAIR | 76 | 0.441422 | 1.951898 | 0 | 0.030013343 | 0.89 | 3625 | tags=30%, list=15%, si | EN_3 |
| GOBP_TELOMERE_ORGAI | GOBP_TELOMERE_ORGANIZATION | 150 | 0.392603 | 1.9514124 | 0 | 0.02986629 | 0.89 | 5418 | tags=33%, list=23%, si | EN_3 |
| GOBP_REGULATION_OF_ | GOBP_REGULATION_OF_MITOCHONDRION_ | 139 | 0.396496 | 1.9492184 | 0 | 0.030211879 | 0.895 | 4024 | tags=28%, list=17%, si | EN_3 |
| N-GLYCAN_BIOSYNTHESIS | N-GLYCAN_BIOSYNTHESIS | 48 | 0.479273 | 1.9469488 | 0.0014858841 | 0.030551689 | 0.898 | 4715 | tags=31%, list=20%, si | EN_3 |
| GOBP_MITOCHONDRIAL_ | GOBP_MITOCHONDRIAL_ELECTRON_TRANS | 18 | 0.619699 | 1.945893 | 0.011627907 | 0.030554712 | 0.9 | 4758 | tags=61%, list=20%, si | EN_3 |
| GOBP_NEGATIVE_REGUL | GOBP_NEGATIVE_REGULATION_OF_VIRAL_ | 36 | 0.513233 | 1.9434192 | 0.0030816642 | 0.030969203 | 0.904 | 2208 | tags=36%, list=9%, sig | EN_3 |
| GOBP_POSITIVE_REGULA | GOBP_POSITIVE_REGULATION_OF_RESPONS | 37 | 0.508561 | 1.9419179 | 0.0015174507 | 0.031288203 | 0.908 | 3635 | tags=32%, list=15%, si | EN_3 |
| GOBP_FIBROBLAST_APOP | GOBP_FIBROBLAST_APOPTOTIC_PROCESS | 31 | 0.54272 | 1.9416374 | 0 | 0.03103896 | 0.909 | 4135 | tags=42%, list=18%, si | EN_3 |
| GOBP_CYTOPLASMIC_TR | GOBP_CYTOPLASMIC_TRANSLATIONAL_INIT | 33 | 0.516714 | 1.9404505 | 0.0015898251 | 0.031133676 | 0.911 | 6141 | tags=52%, list=26%, si | EN_3 |
| GOBP_NUCLEOSIDE_DIPH | GOBP_NUCLEOSIDE_DIPHOSPHATE_METAB | 127 | 0.398108 | 1.9377583 | 0 | 0.031827003 | 0.918 | 5371 | tags=37%, list=23%, si | EN_3 |
| GOBP_CHOLESTEROL_ST | GOBP_CHOLESTEROL_STORAGE | 27 | 0.550997 | 1.9364471 | 0.006153846 | 0.031876124 | 0.918 | 1508 | tags=22%, list=6%, sig | EN_3 |
| GOBP_EXTRINSIC_APOPT | GOBP_EXTRINSIC_APOPTOTIC_SIGNALING_ | 80 | 0.430844 | 1.9362068 | 0 | 0.031619538 | 0.918 | 4514 | tags=35%, list=19%, si | EN_3 |
| GOBP_REGULATION_OF_ | GOBP_REGULATION_OF_FIBROBLAST_PROL | 122 | 0.402359 | 1.9344463 | 0 | 0.031855766 | 0.92 | 5489 | tags=37%, list=23%, si | EN_3 |
| SPLICEOSOME | SPLICEOSOME | 127 | 0.400947 | 1.9339451 | 0 | 0.03180087 | 0.921 | 7081 | tags=43%, list=30%, si | EN_3 |
| AMPK_SIGNALING_PATH | AMPK_SIGNALING_PATHWAY | 120 | 0.406904 | 1.933939 | 0 | 0.031495094 | 0.921 | 3876 | tags=25%, list=16%, si | EN_3 |
| GOBP_LIPOPROTEIN_BIO | GOBP_LIPOPROTEIN_BIOSYNTHETIC_PROCE | 104 | 0.411904 | 1.9319932 | 0 | 0.031952456 | 0.926 | 5882 | tags=39%, list=25%, si | EN_3 |
| GOBP_REGULATION_OF_ | GOBP_REGULATION_OF_MITOCHONDRIAL_ | 35 | 0.515848 | 1.9294672 | 0.001510574 | 0.03261469 | 0.93 | 2847 | tags=31%, list=12%, si | EN_3 |
| GOBP_INDUCTION_OF_P | GOBP_INDUCTION_OF_POSITIVE_CHEMOTA | 15 | 0.635608 | 1.927039 | 0.0033167496 | 0.033230443 | 0.931 | 3345 | tags=47%, list=14%, si | EN_3 |
| GOBP_VASCULAR_ASSOC | GOBP_VASCULAR_ASSOCIATED_SMOOTH_M | 80 | 0.431579 | 1.925675 | 0 | 0.033517234 | 0.934 | 4129 | tags=29%, list=17%, si | EN_3 |
| GOBP_POSITIVE_REGULA | GOBP_POSITIVE_REGULATION_OF_MYOBLA | 18 | 0.607658 | 1.92387 | 0.006329114 | 0.033917014 | 0.937 | 3525 | tags=44%, list=15%, si | EN_3 |
| GOBP_POSITIVE_REGULA | GOBP_POSITIVE_REGULATION_OF_PROTEIN | 192 | 0.374667 | 1.9237161 | 0 | 0.03367363 | 0.938 | 5369 | tags=32%, list=23%, si | EN_3 |
| GOBP_SPLICEOSOMAL_SI | GOBP_SPLICEOSOMAL_SNRNP_ASSEMBLY | 35 | 0.506155 | 1.9230763 | 0.0030395137 | 0.033700746 | 0.94 | 6100 | tags=51%, list=26%, si | EN_3 |
| GOBP_AUTOPHAGOSOMI | GOBP_AUTOPHAGOSOME_MATURATION | 58 | 0.457326 | 1.9229434 | 0 | 0.03343182 | 0.94 | 5015 | tags=34%, list=21%, si | EN_3 |
| GOBP_SIGNAL_TRANSDU | GOBP_SIGNAL_TRANSDUCTION_BY_P53_CL | 151 | 0.384014 | 1.9225379 | 0 | 0.033282887 | 0.941 | 6046 | tags=40%, list=26%, si | EN_3 |
| GOBP_NEGATIVE_REGUL | GOBP_NEGATIVE_REGULATION_OF_SMOOT | 36 | 0.503543 | 1.9212354 | 0 | 0.033542305 | 0.945 | 3154 | tags=31%, list=13%, si | EN_3 |
| GOBP_STEROL_METABOL | GOBP_STEROL_METABOLIC_PROCESS | 140 | 0.393321 | 1.9211252 | 0 | 0.033281762 | 0.945 | 4085 | tags=32%, list=17%, si | EN_3 |
| GOBP_REGULATION_OF_ | GOBP_REGULATION_OF_CHOLESTEROL_STC | 21 | 0.579779 | 1.9204503 | 0.0032573289 | 0.03323022 | 0.948 | 1508 | tags=24%, list=6%, sig | EN_3 |
| GOBP_REGULATION_OF_ | GOBP_REGULATION_OF_AMYLOID_FIBRIL_F | 16 | 0.63686 | 1.9202383 | 0.0033840947 | 0.032976512 | 0.948 | 5632 | tags=50%, list=24%, si | EN_3 |
| GOBP_POSITIVE_REGULA | GOBP_POSITIVE_REGULATION_OF_FIBROBL | 24 | 0.561096 | 1.9182233 | 0.0032733225 | 0.03337151 | 0.951 | 4498 | tags=42%, list=19%, si | EN_3 |
| GOBP_PYRIMIDINE_CONT | GOBP_PYRIMIDINE_CONTAINING_COMPOU | 32 | 0.518225 | 1.9178106 | 0.00155521 | 0.03323175 | 0.952 | 4815 | tags=41%, list=20%, si | EN_3 |
| GOBP_RESPONSE_TO_TY | GOBP_RESPONSE_TO_TYPE_II_INTERFERON | 122 | 0.395063 | 1.9175892 | 0 | 0.032994628 | 0.952 | 5806 | tags=40%, list=25%, si | EN_3 |
| GOBP_BASE_EXCISION_R | GOBP_BASE_EXCISION_REPAIR | 41 | 0.494059 | 1.9135022 | 0 | 0.03411645 | 0.959 | 5281 | tags=51%, list=22%, si | EN_3 |
| PENTOSE_PHOSPHATE_P | PENTOSE_PHOSPHATE_PATHWAY | 29 | 0.526252 | 1.909815 | 0 | 0.035345666 | 0.962 | 4233 | tags=41%, list=18%, si | EN_3 |
| GOBP_POSITIVE_REGULA | GOBP_POSITIVE_REGULATION_OF_GLUCOS | 42 | 0.492843 | 1.9089072 | 0 | 0.035405885 | 0.964 | 4986 | tags=43%, list=21%, si | EN_3 |
| GOBP_POSITIVE_REGULA | GOBP_POSITIVE_REGULATION_OF_DNA_BIO | 76 | 0.429542 | 1.9080518 | 0 | 0.03544566 | 0.967 | 3784 | tags=28%, list=16%, si | EN_3 |
| GOBP_RESPONSE_TO_RE | GOBP_RESPONSE_TO_REACTIVE_OXYGEN_S | 181 | 0.370103 | 1.9066592 | 0 | 0.035608 | 0.969 | 6409 | tags=40%, list=27%, si | EN_3 |
| GOBP_VACUOLAR_LOCAL | GOBP_VACUOLAR_LOCALIZATION | 85 | 0.418322 | 1.9062923 | 0.0012970169 | 0.03550448 | 0.969 | 4428 | tags=29%, list=19%, si | EN_3 |
| GOBP_HEPATIC_STELLATI | GOBP_HEPATIC_STELLATE_CELL_ACTIVATIO | 16 | 0.619013 | 1.904019 | 0.003539823 | 0.03605676 | 0.97 | 2001 | tags=38%, list=8%, sig | EN_3 |
| GOBP_MITOCHONDRION | GOBP_MITOCHONDRION_LOCALIZATION | 57 | 0.449919 | 1.9000988 | 0 | 0.037518866 | 0.976 | 6341 | tags=47%, list=27%, si | EN_3 |
| GOBP_CELLULAR_RESPON | GOBP_CELLULAR_RESPONSE_TO_TYPE_II_IN | 95 | 0.410989 | 1.9000672 | 0.0012836971 | 0.03722802 | 0.976 | 5806 | tags=40%, list=25%, si | EN_3 |
| GOBP_REGULATION_OF_ | GOBP_REGULATION_OF_DNA_BIOSYNTHETI | 121 | 0.402747 | 1.8983386 | 0 | 0.03766211 | 0.979 | 5470 | tags=35%, list=23%, si | EN_3 |
| GOBP_COPII_COATED_VE | GOBP_COPII_COATED_VESICLE_BUDDING | 37 | 0.497363 | 1.8961359 | 0.0015037594 | 0.03820863 | 0.98 | 5409 | tags=46%, list=23%, si | EN_3 |
| GOBP_REGULATION_OF_ | GOBP_REGULATION_OF_GLUCOSE_IMPORT | 66 | 0.435521 | 1.895745 | 0.0013831259 | 0.038072206 | 0.981 | 4986 | tags=36%, list=21%, si | EN_3 |
| GOBP_RECOMBINATIONA | GOBP_RECOMBINATIONAL_REPAIR | 177 | 0.369131 | 1.8952582 | 0 | 0.037972864 | 0.981 | 5408 | tags=36%, list=23%, si | EN_3 |
| GOBP_VESICLE_TARGETIN | GOBP_VESICLE_TARGETING | 60 | 0.448239 | 1.8941635 | 0.0013888889 | 0.03808827 | 0.981 | 6019 | tags=40%, list=25%, si | EN_3 |
| GOBP_PYRIMIDINE_DEOX | GOBP_PYRIMIDINE_DEOXYRIBONUCLEOTIDE | 20 | 0.566842 | 1.8921269 | 0.0032840723 | 0.038757104 | 0.981 | 4815 | tags=45%, list=20%, si | EN_3 |

| | | | | | | | | | | |
|---|---|---|---|---|---|---|---|---|---|---|
| FANCONI_ANEMIA_PATH | FANCONI_ANEMIA_PATHWAY | 50 | 0.468313 | 1.8910758 | 0 | 0.038838938 | 0.981 | 6120 | tags=42%, list=26%, si | EN_3 |
| GOBP_RESPONSE_TO_OX | GOBP_RESPONSE_TO_OXIDATIVE_STRESS | 364 | 0.346271 | 1.8893827 | 0 | 0.03905813 | 0.981 | 6409 | tags=38%, list=27%, si | EN_3 |
| GOBP_AUTOPHAGY_OF_ | GOBP_AUTOPHAGY_OF_MITOCHONDRION | 89 | 0.416899 | 1.8870995 | 0 | 0.039626896 | 0.981 | 6685 | tags=42%, list=28%, si | EN_3 |
| GOBP_CELLULAR_RESPON | GOBP_CELLULAR_RESPONSE_TO_DSRNA | 19 | 0.591366 | 1.8862728 | 0 | 0.03980225 | 0.982 | 6185 | tags=68%, list=26%, si | EN_3 |
| GOBP_MITOCHONDRIAL_ | GOBP_MITOCHONDRIAL_FISSION | 48 | 0.466246 | 1.8846002 | 0.0014164306 | 0.040255707 | 0.985 | 6801 | tags=52%, list=29%, si | EN_3 |
| GOBP_CELLULAR_RESPON | GOBP_CELLULAR_RESPONSE_TO_OXIDATIVE | 229 | 0.358451 | 1.881721 | 0 | 0.04111634 | 0.989 | 5269 | tags=29%, list=22%, si | EN_3 |
| GOBP_RESPONSE_TO_INT | GOBP_RESPONSE_TO_INTERLEUKIN_4 | 35 | 0.493913 | 1.8807777 | 0 | 0.041212056 | 0.99 | 2963 | tags=34%, list=13%, si | EN_3 |
| GOBP_VACUOLE_ORGAN | GOBP_VACUOLE_ORGANIZATION | 223 | 0.361579 | 1.8800977 | 0 | 0.041231878 | 0.99 | 6720 | tags=40%, list=28%, si | EN_3 |
| GOBP_REGULATION_OF_ | GOBP_REGULATION_OF_RELEASE_OF_CYTO | 49 | 0.45749 | 1.8798238 | 0.002915452 | 0.04104441 | 0.99 | 4024 | tags=37%, list=17%, si | EN_3 |
| GOBP_REGULATION_OF_ | GOBP_REGULATION_OF_CELLULAR_RESPIRA | 52 | 0.460171 | 1.8772194 | 0.001459854 | 0.04179387 | 0.99 | 5859 | tags=50%, list=25%, si | EN_3 |
| GOBP_PROTEIN_FOLDING | GOBP_PROTEIN_FOLDING | 162 | 0.373753 | 1.876851 | 0 | 0.041726086 | 0.99 | 5530 | tags=38%, list=23%, si | EN_3 |
| GOBP_ALCOHOL_BIOSYN | GOBP_ALCOHOL_BIOSYNTHETIC_PROCESS | 129 | 0.385655 | 1.8733277 | 0 | 0.043050207 | 0.992 | 2983 | tags=25%, list=13%, si | EN_3 |
| GOBP_LIPOPROTEIN_ME | GOBP_LIPOPROTEIN_METABOLIC_PROCESS | 135 | 0.381987 | 1.8722805 | 0 | 0.04320043 | 0.992 | 3816 | tags=27%, list=16%, si | EN_3 |
| GOBP_VIRAL_TRANSLATI | GOBP_VIRAL_TRANSLATION | 18 | 0.591744 | 1.872259 | 0.0016920473 | 0.042926393 | 0.992 | 7985 | tags=78%, list=34%, si | EN_3 |
| GOBP_PYRIDINE_CONTAI | GOBP_PYRIDINE_CONTAINING_COMPOUND | 182 | 0.367981 | 1.8721324 | 0 | 0.042711858 | 0.992 | 5418 | tags=37%, list=23%, si | EN_3 |
| GOBP_REGULATION_OF_ | GOBP_REGULATION_OF_INFLAMMASOME_ | 40 | 0.485501 | 1.8716073 | 0 | 0.04269568 | 0.992 | 3292 | tags=35%, list=14%, si | EN_3 |
| GOBP_HEME_BIOSYNTHE | GOBP_HEME_BIOSYNTHETIC_PROCESS | 24 | 0.540877 | 1.8713968 | 0.008064516 | 0.042485055 | 0.992 | 5579 | tags=54%, list=24%, si | EN_3 |
| GOBP_POSITIVE_REGULA | GOBP_POSITIVE_REGULATION_OF_TOR_SIG | 71 | 0.428602 | 1.870116 | 0.0013513514 | 0.042781737 | 0.992 | 4934 | tags=39%, list=21%, si | EN_3 |
| GOBP_DENDRITIC_TRANS | GOBP_DENDRITIC_TRANSPORT | 19 | 0.573487 | 1.8685105 | 0.005 | 0.04308974 | 0.992 | 5100 | tags=53%, list=22%, si | EN_3 |
| GOBP_GRANULOCYTE_AC | GOBP_GRANULOCYTE_ACTIVATION | 41 | 0.478088 | 1.8660715 | 0 | 0.04389192 | 0.993 | 3053 | tags=34%, list=13%, si | EN_3 |
| GOBP_ALCOHOL_METAB | GOBP_ALCOHOL_METABOLIC_PROCESS | 308 | 0.348805 | 1.8644211 | 0 | 0.044372506 | 0.993 | 4125 | tags=27%, list=17%, si | EN_3 |
| GOBP_TUBULIN_DEACET | GOBP_TUBULIN_DEACETYLATION | 24 | 0.537007 | 1.8634866 | 0.0015873016 | 0.044543542 | 0.993 | 4253 | tags=38%, list=18%, si | EN_3 |
| PORPHYRIN_AND_CHLOR | PORPHYRIN_AND_CHLOROPHYLL_METABOL | 31 | 0.51605 | 1.8632069 | 0.0015151515 | 0.044419546 | 0.993 | 4522 | tags=42%, list=19%, si | EN_3 |
| GOBP_REGULATION_OF_ | GOBP_REGULATION_OF_PROTEIN_STABILIT | 305 | 0.34617 | 1.8630016 | 0 | 0.044245157 | 0.993 | 5817 | tags=33%, list=25%, si | EN_3 |
| GOBP_POSITIVE_REGULA | GOBP_POSITIVE_REGULATION_OF_CELL_SU | 134 | 0.384948 | 1.860841 | 0 | 0.04495588 | 0.993 | 6236 | tags=37%, list=26%, si | EN_3 |
| GOBP_POSITIVE_REGULA | GOBP_POSITIVE_REGULATION_OF_FIBROBL | 76 | 0.428126 | 1.8585066 | 0 | 0.04560534 | 0.993 | 5419 | tags=37%, list=23%, si | EN_3 |
| GOBP_ALDITOL_METABO | GOBP_ALDITOL_METABOLIC_PROCESS | 22 | 0.554248 | 1.8582512 | 0.0031847134 | 0.045433793 | 0.993 | 3983 | tags=45%, list=17%, si | EN_3 |
| GOBP_PROTEIN_DEACET | GOBP_PROTEIN_DEACETYLATION | 46 | 0.471468 | 1.8572214 | 0.0029542097 | 0.0455843 | 0.993 | 5253 | tags=37%, list=22%, si | EN_3 |
| GOBP_AUTOPHAGOSOM | GOBP_AUTOPHAGOSOME_ORGANIZATION | 128 | 0.381341 | 1.8566054 | 0 | 0.04559608 | 0.994 | 5484 | tags=34%, list=23%, si | EN_3 |
| GOBP_ACETYL_COA_MET | GOBP_ACETYL_COA_METABOLIC_PROCESS | 34 | 0.498657 | 1.8560972 | 0.0015337423 | 0.04553522 | 0.994 | 6561 | tags=50%, list=28%, si | EN_3 |
| GOBP_RESPONSE_TO_LIT | GOBP_RESPONSE_TO_LITHIUM_ION | 16 | 0.606033 | 1.8558314 | 0.0034305318 | 0.045411382 | 0.994 | 1330 | tags=31%, list=6%, sig | EN_3 |
| GOBP_RESPONSE_TO_HY | GOBP_RESPONSE_TO_HYDROGEN_PEROXID | 85 | 0.412073 | 1.8545343 | 0.0013227513 | 0.04562949 | 0.994 | 2265 | tags=24%, list=10%, si | EN_3 |
| GOBP_VASCULAR_ENDOT | GOBP_VASCULAR_ENDOTHELIAL_GROWTH_ | 40 | 0.487042 | 1.854446 | 0.0015243902 | 0.045393426 | 0.994 | 6236 | tags=48%, list=26%, si | EN_3 |
| VALINE,_LEUCINE_AND_I | VALINE,_LEUCINE_AND_ISOLEUCINE_DEGRA | 49 | 0.453084 | 1.8540038 | 0.0028328612 | 0.045363687 | 0.995 | 3959 | tags=31%, list=17%, si | EN_3 |
| GOBP_PYRIMIDINE_NUCL | GOBP_PYRIMIDINE_NUCLEOSIDE_MONOPH | 21 | 0.567051 | 1.8528069 | 0 | 0.045648832 | 0.997 | 4383 | tags=43%, list=19%, si | EN_3 |
| GOBP_DNA_DAMAGE_RE | GOBP_DNA_DAMAGE_RESPONSE_SIGNAL_T | 67 | 0.433011 | 1.8516352 | 0.0013831259 | 0.04593775 | 0.997 | 3050 | tags=25%, list=13%, si | EN_3 |
| GOBP_POSITIVE_REGULA | GOBP_POSITIVE_REGULATION_OF_ERAD_P | 20 | 0.56244 | 1.8515475 | 0.008237232 | 0.045732826 | 0.997 | 6029 | tags=50%, list=26%, si | EN_3 |
| BASAL_TRANSCRIPTION_ | BASAL_TRANSCRIPTION_FACTORS | 40 | 0.475549 | 1.8513229 | 0.0014619883 | 0.045509692 | 0.997 | 7193 | tags=50%, list=30%, si | EN_3 |
| GOBP_TRIGLYCERIDE_ME | GOBP_TRIGLYCERIDE_METABOLIC_PROCESS | 105 | 0.395353 | 1.8508786 | 0.001270648 | 0.04537758 | 0.997 | 4386 | tags=30%, list=19%, si | EN_3 |
| GOBP_FIBROBLAST_ACTI | GOBP_FIBROBLAST_ACTIVATION | 22 | 0.563314 | 1.8507894 | 0.008090615 | 0.045159113 | 0.997 | 2001 | tags=32%, list=8%, sig | EN_3 |
| GOBP_POSITIVE_REGULA | GOBP_POSITIVE_REGULATION_OF_CELLULA | 20 | 0.561289 | 1.8501742 | 0.0046875 | 0.045266252 | 0.997 | 3177 | tags=50%, list=13%, si | EN_3 |
| GOBP_PYRIDINE_CONTAI | GOBP_PYRIDINE_CONTAINING_COMPOUND | 109 | 0.395546 | 1.8490462 | 0 | 0.045413207 | 0.997 | 5371 | tags=37%, list=23%, si | EN_3 |
| GOBP_DEOXYRIBONUCLE | GOBP_DEOXYRIBONUCLEOTIDE_BIOSYNTHE | 17 | 0.609535 | 1.8489457 | 0.0033557047 | 0.04522468 | 0.997 | 4815 | tags=47%, list=20%, si | EN_3 |
| GOBP_REGULATION_OF_ | GOBP_REGULATION_OF_NECROPTOTIC_PR | 22 | 0.548787 | 1.8483832 | 0.0094191525 | 0.045204576 | 0.997 | 4485 | tags=41%, list=19%, si | EN_3 |
| GOBP_NEGATIVE_REGUL | GOBP_NEGATIVE_REGULATION_OF_CELL_C | 69 | 0.428207 | 1.8482327 | 0.0027063598 | 0.045039028 | 0.997 | 5457 | tags=42%, list=23%, si | EN_3 |
| GOBP_VESICLE_TARGETI | GOBP_VESICLE_TARGETING_TO_FROM_OR_ | 29 | 0.511502 | 1.847756 | 0.0049261083 | 0.04499976 | 0.997 | 5409 | tags=52%, list=23%, si | EN_3 |
| GOBP_REGULATION_OF_ | GOBP_REGULATION_OF_OXIDATIVE_PHOSP | 23 | 0.548474 | 1.8474317 | 0.008474576 | 0.044928756 | 0.997 | 5859 | tags=65%, list=25%, si | EN_3 |
| GOBP_RESPONSE_TO_RE | GOBP_RESPONSE_TO_RETINOIC_ACID | 83 | 0.412425 | 1.8470907 | 0.002642008 | 0.044845294 | 0.997 | 3004 | tags=24%, list=13%, si | EN_3 |
| GOBP_DEOXYRIBONUCLE | GOBP_DEOXYRIBONUCLEOTIDE_METABOLIC | 44 | 0.47206 | 1.8468767 | 0.004398827 | 0.04467888 | 0.997 | 4815 | tags=36%, list=20%, si | EN_3 |
| GOBP_LIGAND_GATED_I | GOBP_LIGAND_GATED_ION_CHANNEL_SIGN | 33 | 0.500172 | 1.8453882 | 0.0015197569 | 0.04500863 | 0.997 | 2696 | tags=24%, list=11%, si | EN_3 |
| GOBP_DNA_REPLICATION | GOBP_DNA_REPLICATION | 263 | 0.34796 | 1.8450471 | 0 | 0.04487554 | 0.997 | 5450 | tags=32%, list=23%, si | EN_3 |
| GOBP_POSITIVE_REGULA | GOBP_POSITIVE_REGULATION_OF_TELOME | 71 | 0.422781 | 1.8448983 | 0.002688172 | 0.04470559 | 0.997 | 4200 | tags=27%, list=18%, si | EN_3 |
| PROTEASOME | PROTEASOME | 45 | 0.466264 | 1.8442808 | 0 | 0.044802062 | 0.997 | 7880 | tags=64%, list=33%, si | EN_3 |
| GOBP_REGULATION_OF_ | GOBP_REGULATION_OF_CELL_CYCLE_G2_M | 117 | 0.391356 | 1.8440272 | 0 | 0.044702925 | 0.997 | 5457 | tags=36%, list=23%, si | EN_3 |
| GOBP_DNA_TEMPLATED_ | GOBP_DNA_TEMPLATED_DNA_REPLICATION | 147 | 0.37463 | 1.8431506 | 0 | 0.044924296 | 0.997 | 5450 | tags=33%, list=23%, si | EN_3 |
| GOBP_TRANSPORT_OF_V | GOBP_TRANSPORT_OF_VIRUS | 15 | 0.617106 | 1.8422475 | 0.0034602077 | 0.045036297 | 0.997 | 3947 | tags=40%, list=17%, si | EN_3 |
| GOBP_SIGNAL_TRANSDU | GOBP_SIGNAL_TRANSDUCTION_IN_RESPON | 176 | 0.364101 | 1.8379972 | 0 | 0.04671131 | 0.997 | 6141 | tags=35%, list=26%, si | EN_3 |
| GOBP_REGULATION_OF_ | GOBP_REGULATION_OF_VACUOLE_ORGANI | 54 | 0.436944 | 1.8379524 | 0 | 0.046481453 | 0.997 | 5484 | tags=41%, list=23%, si | EN_3 |
| GOBP_RRNA_MODIFICAT | GOBP_RRNA_MODIFICATION | 29 | 0.50919 | 1.8354139 | 0.003003003 | 0.047386106 | 0.999 | 3731 | tags=38%, list=16%, si | EN_3 |
| GOBP_TELOMERE_MAINT | GOBP_TELOMERE_MAINTENANCE_VIA_REC | 15 | 0.604441 | 1.8334272 | 0.008695652 | 0.048189275 | 0.999 | 5201 | tags=47%, list=22%, si | EN_3 |
| GOBP_PYRIMIDINE_RIBO | GOBP_PYRIMIDINE_RIBONUCLEOTIDE_BIOS | 17 | 0.58982 | 1.8328521 | 0.011744967 | 0.048240744 | 0.999 | 4815 | tags=47%, list=20%, si | EN_3 |
| GOBP_REGULATION_OF_ | GOBP_REGULATION_OF_PROTEIN_LOCALIZ | 117 | 0.381272 | 1.8312788 | 0 | 0.048676345 | 0.999 | 4041 | tags=27%, list=17%, si | EN_3 |
| GOBP_NUCLEOSIDE_PHO | GOBP_NUCLEOSIDE_PHOSPHATE_CATABOL | 170 | 0.363432 | 1.8308598 | 0 | 0.04864031 | 0.999 | 5371 | tags=34%, list=23%, si | EN_3 |
| GOBP_NEGATIVE_REGUL | GOBP_NEGATIVE_REGULATION_OF_VIRAL_ | 50 | 0.44581 | 1.8285637 | 0.004219409 | 0.049470566 | 0.999 | 2487 | tags=24%, list=11%, si | EN_3 |
| GOBP_CELLULAR_RESPON | GOBP_CELLULAR_RESPONSE_TO_REACTIVE_ | 136 | 0.378289 | 1.8284578 | 0 | 0.049294583 | 0.999 | 2935 | tags=23%, list=12%, si | EN_3 |
| GOBP_ORGANELLE_DISAS | GOBP_ORGANELLE_DISASSEMBLY | 155 | 0.370133 | 1.8281672 | 0 | 0.049208365 | 0.999 | 5418 | tags=31%, list=23%, si | EN_3 |
| GOBP_IONOTROPIC_GLU | GOBP_IONOTROPIC_GLUTAMATE_RECEPTO | 20 | 0.558245 | 1.8261744 | 0.0094191525 | 0.0499658 | 0.999 | 2696 | tags=30%, list=11%, si | EN_3 |
| GOBP_REGULATION_OF_ | GOBP_REGULATION_OF_FIBROBLAST_APOP | 24 | 0.536834 | 1.825723 | 0.0064935065 | 0.04994833 | 0.999 | 4029 | tags=42%, list=17%, si | EN_3 |
| GOBP_REGULATION_OF_ | GOBP_REGULATION_OF_DEFENSE_RESPONS | 67 | 0.419071 | 1.8229052 | 0.0013661202 | 0.051000267 | 0.999 | 2997 | tags=28%, list=13%, si | EN_3 |
| GOBP_REGULATION_OF_ | GOBP_REGULATION_OF_TELOMERE_MAINT | 98 | 0.39756 | 1.821892 | 0 | 0.051296107 | 1 | 5418 | tags=30%, list=23%, si | EN_3 |
| GOBP_NEGATIVE_REGUL | GOBP_NEGATIVE_REGULATION_OF_NUCLE | 27 | 0.513276 | 1.8207397 | 0.0063593006 | 0.051674113 | 1 | 5231 | tags=52%, list=22%, si | EN_3 |
| GOBP_GOLGI_VESICLE_T | GOBP_GOLGI_VESICLE_TRANSPORT | 276 | 0.339934 | 1.8198571 | 0 | 0.051815547 | 1 | 8097 | tags=47%, list=34%, si | EN_3 |
| GOBP_REGULATION_OF_ | GOBP_REGULATION_OF_CELL_CYCLE_G1_S_ | 185 | 0.355897 | 1.8193924 | 0 | 0.051765855 | 1 | 4998 | tags=31%, list=21%, si | EN_3 |
| GOBP_FIBROBLAST_PROL | GOBP_FIBROBLAST_PROLIFERATION | 147 | 0.370293 | 1.8184692 | 0 | 0.05196707 | 1 | 4668 | tags=32%, list=20%, si | EN_3 |
| GOBP_NEUTROPHIL_ACT | GOBP_NEUTROPHIL_ACTIVATION | 36 | 0.474953 | 1.8179308 | 0.0077160494 | 0.051973756 | 1 | 3053 | tags=33%, list=13%, si | EN_3 |
| GOBP_REGULATION_OF_ | GOBP_REGULATION_OF_AEROBIC_RESPIRAT | 33 | 0.491555 | 1.817679 | 0.0029411765 | 0.05182348 | 1 | 5859 | tags=58%, list=25%, si | EN_3 |
| GOBP_DNA_REPLICATION | GOBP_DNA_REPLICATION_INITIATION | 31 | 0.50158 | 1.8175019 | 0.0112 | 0.05165747 | 1 | 3614 | tags=35%, list=15%, si | EN_3 |
| GOBP_NEGATIVE_REGUL | GOBP_NEGATIVE_REGULATION_OF_DEFENS | 19 | 0.570625 | 1.8164412 | 0.0051369863 | 0.05203988 | 1 | 2260 | tags=37%, list=10%, si | EN_3 |
| GOBP_CELLULAR_RESPON | GOBP_CELLULAR_RESPONSE_TO_VASCULAR | 60 | 0.42898 | 1.8162448 | 0.0014204546 | 0.051874198 | 1 | 4477 | tags=35%, list=19%, si | EN_3 |
| LONGEVITY_REGULATING | LONGEVITY_REGULATING_PATHWAY | 99 | 0.390774 | 1.815387 | 0.0012690355 | 0.051992066 | 1 | 7102 | tags=43%, list=30%, si | EN_3 |
| GOBP_PINOCYTOSIS | GOBP_PINOCYTOSIS | 22 | 0.545465 | 1.8143965 | 0.005059022 | 0.052350424 | 1 | 4502 | tags=50%, list=19%, si | EN_3 |
| GOBP_REGULATION_OF_ | GOBP_REGULATION_OF_CYTOPLASMIC_PAT | 104 | 0.387365 | 1.8128178 | 0 | 0.052841544 | 1 | 3321 | tags=25%, list=14%, si | EN_3 |

| | | | | | | | | | | |
|---|---|---|---|---|---|---|---|---|---|---|
| GOBP_ACTIVATION_OF_C | GOBP_ACTIVATION_OF_CYSTEINE_TYPE_EN | 81 | 0.401077 | 1.8117127 | 0 | 0.053111184 | 1 | 3177 | tags=25%, list=13%, si | EN_3 |
| GOBP_SMOOTH_MUSCLE | GOBP_SMOOTH_MUSCLE_CELL_MIGRATION | 115 | 0.382232 | 1.8117024 | 0 | 0.05287944 | 1 | 3811 | tags=24%, list=16%, si | EN_3 |
| GOBP_CARBOHYDRATE_C | GOBP_CARBOHYDRATE_CATABOLIC_PROCES | 161 | 0.361609 | 1.8111138 | 0 | 0.052909005 | 1 | 5418 | tags=36%, list=23%, si | EN_3 |
| TERPENOID_BACKBONE_I | TERPENOID_BACKBONE_BIOSYNTHESIS | 21 | 0.548759 | 1.8110844 | 0.014128729 | 0.052675016 | 1 | 3800 | tags=38%, list=16%, si | EN_3 |
| GOBP_NEGATIVE_REGUL | GOBP_NEGATIVE_REGULATION_OF_INTRINS | 21 | 0.553527 | 1.8107059 | 0.008156607 | 0.05266274 | 1 | 7214 | tags=62%, list=31%, si | EN_3 |
| GOBP_CELLULAR_RESPON | GOBP_CELLULAR_RESPONSE_TO_HYDROGE | 59 | 0.435779 | 1.810038 | 0.0026990552 | 0.052751213 | 1 | 2265 | tags=25%, list=10%, si | EN_3 |
| GOBP_REGULATION_OF_ | GOBP_REGULATION_OF_MYOBLAST_FUSION | 22 | 0.541766 | 1.8088499 | 0.012461059 | 0.053103857 | 1 | 3525 | tags=36%, list=15%, si | EN_3 |
| GOBP_MONOSACCHARID | GOBP_MONOSACCHARIDE_BIOSYNTHETIC_I | 93 | 0.394161 | 1.808522 | 0 | 0.05303165 | 1 | 4833 | tags=34%, list=20%, si | EN_3 |
| GOBP_RESPONSE_TO_UV | GOBP_RESPONSE_TO_UV | 153 | 0.359937 | 1.8082997 | 0.0011682243 | 0.05293884 | 1 | 6141 | tags=39%, list=26%, si | EN_3 |
| GOBP_REGULATION_OF_ | GOBP_REGULATION_OF_REGULATED_SECRE | 139 | 0.365198 | 1.8078864 | 0 | 0.05294106 | 1 | 3780 | tags=25%, list=16%, si | EN_3 |
| GOBP_MUSCLE_CELL_MI | GOBP_MUSCLE_CELL_MIGRATION | 129 | 0.372571 | 1.8075786 | 0.0012004802 | 0.052818533 | 1 | 3811 | tags=23%, list=16%, si | EN_3 |
| GOBP_CELLULAR_RESPON | GOBP_CELLULAR_RESPONSE_TO_LIPOPROT | 38 | 0.463748 | 1.8064967 | 0.004470939 | 0.053080887 | 1 | 3321 | tags=29%, list=14%, si | EN_3 |
| GOBP_NUCLEOSIDE_MON | GOBP_NUCLEOSIDE_MONOPHOSPHATE_BIC | 40 | 0.465028 | 1.8053849 | 0.008695652 | 0.053412676 | 1 | 4383 | tags=38%, list=19%, si | EN_3 |
| GOBP_POSITIVE_REGULA | GOBP_POSITIVE_REGULATION_OF_PROTEOI | 120 | 0.381628 | 1.8047961 | 0 | 0.05344369 | 1 | 4076 | tags=25%, list=17%, si | EN_3 |
| RNA_POLYMERASE | RNA_POLYMERASE | 27 | 0.518239 | 1.8046014 | 0.0063091484 | 0.05332063 | 1 | 2788 | tags=37%, list=12%, si | EN_3 |
| GOBP_MATURATION_OF_ | GOBP_MATURATION_OF_5_8S_RRNA | 35 | 0.477948 | 1.802771 | 0.0045871558 | 0.053987294 | 1 | 5205 | tags=46%, list=22%, si | EN_3 |
| GOBP_RIBONUCLEOSIDE_ | GOBP_RIBONUCLEOSIDE_MONOPHOSPHATI | 30 | 0.500045 | 1.8023796 | 0.009360374 | 0.053944048 | 1 | 4140 | tags=40%, list=18%, si | EN_3 |
| GOBP_RIBONUCLEOSIDE_ | GOBP_RIBONUCLEOSIDE_MONOPHOSPHATI | 48 | 0.448637 | 1.8010137 | 0.0014749262 | 0.054396447 | 1 | 4140 | tags=33%, list=18%, si | EN_3 |
| GOBP_PROTEIN_REFOLDI | GOBP_PROTEIN_REFOLDING | 23 | 0.538723 | 1.8007803 | 0 | 0.054252263 | 1 | 6559 | tags=57%, list=28%, si | EN_3 |
| GOBP_ESTABLISHMENT_C | GOBP_ESTABLISHMENT_OF_PROTEIN_LOCA | 435 | 0.326468 | 1.8007603 | 0 | 0.054028403 | 1 | 7212 | tags=41%, list=31%, si | EN_3 |
| GOBP_ENDOSOME_TO_L | GOBP_ENDOSOME_TO_LYSOSOME_TRANSP | 16 | 0.588133 | 1.7976979 | 0.0102214655 | 0.05544757 | 1 | 4771 | tags=63%, list=20%, si | EN_3 |
| GOBP_PURINE_CONTAIN | GOBP_PURINE_CONTAINING_COMPOUND_ | 150 | 0.365722 | 1.7972387 | 0.0012285012 | 0.055439245 | 1 | 6317 | tags=41%, list=27%, si | EN_3 |
| GOBP_TETRAPYRROLE_BI | GOBP_TETRAPYRROLE_BIOSYNTHETIC_PROC | 29 | 0.50147 | 1.7965893 | 0.006134969 | 0.05549566 | 1 | 5579 | tags=52%, list=24%, si | EN_3 |
| HOMOLOGOUS_RECOMB | HOMOLOGOUS_RECOMBINATION | 40 | 0.45784 | 1.7959532 | 0.0060975607 | 0.05564999 | 1 | 4514 | tags=38%, list=19%, si | EN_3 |
| GOBP_REGULATION_OF_ | GOBP_REGULATION_OF_ENDOTHELIAL_CEL | 18 | 0.561223 | 1.7943319 | 0.007886435 | 0.05628693 | 1 | 5040 | tags=33%, list=21%, si | EN_3 |
| GOBP_ORGANELLE_TRAN | GOBP_ORGANELLE_TRANSPORT_ALONG_M | 86 | 0.389639 | 1.793415 | 0.0012642225 | 0.056558862 | 1 | 6290 | tags=40%, list=27%, si | EN_3 |
| GOBP_REGULATION_OF_ | GOBP_REGULATION_OF_VIRAL_TRANSCRIPT | 25 | 0.521468 | 1.792177 | 0.0064102565 | 0.05696006 | 1 | 3411 | tags=40%, list=14%, si | EN_3 |
| GOBP_LEUKOCYTE_DEGR | GOBP_LEUKOCYTE_DEGRANULATION | 77 | 0.400588 | 1.7915812 | 0.003875969 | 0.0570327 | 1 | 3780 | tags=29%, list=16%, si | EN_3 |
| GOBP_MONOSACCHARID | GOBP_MONOSACCHARIDE_CATABOLIC_PRC | 40 | 0.454976 | 1.7892973 | 0.0014858841 | 0.057958692 | 1 | 5418 | tags=38%, list=23%, si | EN_3 |
| GOBP_RIBOSOMAL_LARG | GOBP_RIBOSOMAL_LARGE_SUBUNIT_BIOGE | 64 | 0.414511 | 1.7891097 | 0 | 0.057829473 | 1 | 5205 | tags=47%, list=22%, si | EN_3 |
| GOBP_SMOOTH_MUSCLE | GOBP_SMOOTH_MUSCLE_CELL_PROLIFERAT | 191 | 0.349102 | 1.7888513 | 0 | 0.057711273 | 1 | 4149 | tags=25%, list=18%, si | EN_3 |
| GOBP_REGULATION_OF_ | GOBP_REGULATION_OF_TELOMERE_MAINT | 61 | 0.422131 | 1.7872107 | 0.0014204546 | 0.05837503 | 1 | 1956 | tags=16%, list=8%, sig | EN_3 |
| GOBP_REGULATION_OF_ | GOBP_REGULATION_OF_BILE_ACID_METAB | 17 | 0.579289 | 1.7869071 | 0.0016806723 | 0.05829363 | 1 | 3253 | tags=41%, list=14%, si | EN_3 |
| GOBP_RESPONSE_TO_VIR | GOBP_RESPONSE_TO_VIRUS | 331 | 0.331335 | 1.7859728 | 0 | 0.05853916 | 1 | 5452 | tags=32%, list=23%, si | EN_3 |
| GOBP_REGULATION_OF_ | GOBP_REGULATION_OF_MICROGLIAL_CELL_ | 17 | 0.583787 | 1.7838614 | 0.0082236845 | 0.059460618 | 1 | 3592 | tags=47%, list=15%, si | EN_3 |
| GOBP_CELLULAR_RESPON | GOBP_CELLULAR_RESPONSE_TO_EXTERNAL | 36 | 0.475379 | 1.7834996 | 0.0059790732 | 0.0594181 | 1 | 4672 | tags=39%, list=20%, si | EN_3 |
| GOBP_RNA_SPLICING | GOBP_RNA_SPLICING | 417 | 0.325508 | 1.7834377 | 0 | 0.05920278 | 1 | 7575 | tags=43%, list=32%, si | EN_3 |
| GOBP_PEPTIDYL_LYSINE_ | GOBP_PEPTIDYL_LYSINE_ACETYLATION | 25 | 0.515237 | 1.7829125 | 0.006557377 | 0.05926865 | 1 | 5089 | tags=36%, list=22%, si | EN_3 |
| GOBP_BARBED_END_ACT | GOBP_BARBED_END_ACTIN_FILAMENT_CAI | 25 | 0.51045 | 1.782771 | 0.009646302 | 0.05910674 | 1 | 6240 | tags=48%, list=26%, si | EN_3 |
| GOBP_LYSOSOMAL_LUM | GOBP_LYSOSOMAL_LUMEN_ACIDIFICATION | 15 | 0.583869 | 1.7815874 | 0.008833922 | 0.05949088 | 1 | 4463 | tags=40%, list=19%, si | EN_3 |
| GOBP_REGULATION_OF_ | GOBP_REGULATION_OF_AUTOPHAGOSOME | 18 | 0.563301 | 1.7802755 | 0.009966778 | 0.060015257 | 1 | 3020 | tags=33%, list=13%, si | EN_3 |
| GOBP_NADH_METABOLIC | GOBP_NADH_METABOLIC_PROCESS | 39 | 0.465639 | 1.7790211 | 0.004464286 | 0.060398154 | 1 | 6125 | tags=54%, list=26%, si | EN_3 |
| GOBP_GLYCEROLIPID_ME | GOBP_GLYCEROLIPID_METABOLIC_PROCESS | 361 | 0.328171 | 1.7780616 | 0 | 0.060704883 | 1 | 4386 | tags=25%, list=19%, si | EN_3 |
| GOBP_CYTOCHROME_CO | GOBP_CYTOCHROME_COMPLEX_ASSEMBLY | 32 | 0.484861 | 1.7771554 | 0.01529052 | 0.061004277 | 1 | 3729 | tags=34%, list=16%, si | EN_3 |
| GOBP_TERPENOID_META | GOBP_TERPENOID_METABOLIC_PROCESS | 60 | 0.422384 | 1.7766889 | 0.0013477089 | 0.060975138 | 1 | 4022 | tags=35%, list=17%, si | EN_3 |
| GOBP_TRNA_PROCESSIN | GOBP_TRNA_PROCESSING | 122 | 0.371832 | 1.7762313 | 0.0012391574 | 0.061005224 | 1 | 3826 | tags=29%, list=16%, si | EN_3 |
| GOBP_REGULATION_OF_ | GOBP_REGULATION_OF_MYELOID_LEUKOC | 61 | 0.417413 | 1.7757252 | 0.0013422819 | 0.06112918 | 1 | 3780 | tags=30%, list=16%, si | EN_3 |
| GOBP_REGULATION_OF_ | GOBP_REGULATION_OF_NON_CANONICAL_ | 92 | 0.389448 | 1.7756927 | 0.0026595744 | 0.060921017 | 1 | 5561 | tags=40%, list=24%, si | EN_3 |
| GOBP_PROTEIN_LOCALIZ | GOBP_PROTEIN_LOCALIZATION_TO_CELL_S | 70 | 0.41415 | 1.7727771 | 0 | 0.062428236 | 1 | 5759 | tags=43%, list=24%, si | EN_3 |
| GOBP_RNA_MODIFICATIO | GOBP_RNA_MODIFICATION | 141 | 0.364727 | 1.7721081 | 0 | 0.06256774 | 1 | 4312 | tags=31%, list=18%, si | EN_3 |
| TYPE_II_DIABETES_MELLI | TYPE_II_DIABETES_MELLITUS | 45 | 0.445866 | 1.7719178 | 0.0056577087 | 0.062436443 | 1 | 4360 | tags=36%, list=18%, si | EN_3 |
| GOBP_ER_NUCLEUS_SIGN | GOBP_ER_NUCLEUS_SIGNALING_PATHWAY | 43 | 0.446849 | 1.7718172 | 0.0014619883 | 0.062261686 | 1 | 2963 | tags=26%, list=13%, si | EN_3 |
| GOBP_POSITIVE_REGULA | GOBP_POSITIVE_REGULATION_OF_CALCIUN | 26 | 0.512377 | 1.771588 | 0.014827018 | 0.062149715 | 1 | 3182 | tags=42%, list=13%, si | EN_3 |
| GOBP_MYELOID_CELL_AC | GOBP_MYELOID_CELL_ACTIVATION_INVOLV | 98 | 0.378172 | 1.7696301 | 0.0012787724 | 0.063084096 | 1 | 3363 | tags=28%, list=14%, si | EN_3 |
| GOBP_ESTABLISHMENT_C | GOBP_ESTABLISHMENT_OF_MITOCHONDRI | 34 | 0.480305 | 1.7695954 | 0.00466563 | 0.06285656 | 1 | 6290 | tags=47%, list=27%, si | EN_3 |
| GOBP_STEROID_BIOSYNT | GOBP_STEROID_BIOSYNTHETIC_PROCESS | 153 | 0.354803 | 1.7689697 | 0 | 0.06299216 | 1 | 4386 | tags=31%, list=19%, si | EN_3 |
| GOBP_POSITIVE_REGULA | GOBP_POSITIVE_REGULATION_OF_REGULAT | 55 | 0.423418 | 1.7689549 | 0.0028490028 | 0.062766545 | 1 | 2841 | tags=29%, list=12%, si | EN_3 |
| GOBP_REGULATION_OF_ | GOBP_REGULATION_OF_GLIAL_CELL_MIGRA | 26 | 0.508683 | 1.7655743 | 0.0152207 | 0.06457377 | 1 | 4660 | tags=42%, list=20%, si | EN_3 |
| GOBP_NECROPTOTIC_PR | GOBP_NECROPTOTIC_PROCESS | 37 | 0.461632 | 1.7655386 | 0.009259259 | 0.06438277 | 1 | 4485 | tags=32%, list=19%, si | EN_3 |
| GOBP_REGULATION_OF_ | GOBP_REGULATION_OF_EXTRINSIC_APOPT( | 45 | 0.435708 | 1.765206 | 0.0074850298 | 0.064343534 | 1 | 4514 | tags=33%, list=19%, si | EN_3 |
| GOBP_REGULATION_OF_ | GOBP_REGULATION_OF_PROTEOLYSIS_INV( | 215 | 0.343287 | 1.7648953 | 0 | 0.06435946 | 1 | 4076 | tags=24%, list=17%, si | EN_3 |
| GOBP_ESTABLISHMENT_C | GOBP_ESTABLISHMENT_OF_PROTEIN_LOCA | 25 | 0.526671 | 1.7634883 | 0.0078125 | 0.06489855 | 1 | 4033 | tags=44%, list=17%, si | EN_3 |
| GOBP_AMYLOID_FIBRIL_ | GOBP_AMYLOID_FIBRIL_FORMATION | 30 | 0.486022 | 1.7627654 | 0.013761468 | 0.065111585 | 1 | 5632 | tags=37%, list=24%, si | EN_3 |
| GOBP_NEGATIVE_REGUL | GOBP_NEGATIVE_REGULATION_OF_SMOOT | 68 | 0.40581 | 1.7621664 | 0 | 0.0651587 | 1 | 3569 | tags=28%, list=15%, si | EN_3 |
| GOBP_NUCLEOSIDE_DIPH | GOBP_NUCLEOSIDE_DIPHOSPHATE_CATABC | 111 | 0.378873 | 1.7617533 | 0.0024600246 | 0.06513771 | 1 | 5371 | tags=35%, list=23%, si | EN_3 |
| GALACTOSE_METABOLISN | GALACTOSE_METABOLISM | 29 | 0.492885 | 1.7616996 | 0.0063492064 | 0.064941026 | 1 | 4233 | tags=38%, list=18%, si | EN_3 |
| GOBP_APOPTOTIC_MITO | GOBP_APOPTOTIC_MITOCHONDRIAL_CHAN | 118 | 0.367503 | 1.7607805 | 0 | 0.065310396 | 1 | 3635 | tags=26%, list=15%, si | EN_3 |
| GOBP_MONOCYTE_CHEM | GOBP_MONOCYTE_CHEMOTAXIS | 40 | 0.448883 | 1.7606792 | 0.0076569677 | 0.06511881 | 1 | 3182 | tags=30%, list=13%, si | EN_3 |
| GOBP_POSITIVE_REGULA | GOBP_POSITIVE_REGULATION_OF_PROTEIN | 171 | 0.348558 | 1.760514 | 0 | 0.06502393 | 1 | 3345 | tags=21%, list=14%, si | EN_3 |
| GOBP_NEGATIVE_REGUL | GOBP_NEGATIVE_REGULATION_OF_VASCUI | 33 | 0.46913 | 1.759019 | 0.004651163 | 0.06563262 | 1 | 3569 | tags=27%, list=15%, si | EN_3 |
| P53_SIGNALING_PATHW/ | P53_SIGNALING_PATHWAY | 70 | 0.399482 | 1.7588755 | 0.00979021 | 0.06547076 | 1 | 3635 | tags=26%, list=15%, si | EN_3 |
| GOBP_RNA_SPLICING_VI | GOBP_RNA_SPLICING_VIA_TRANSESTERIFIC | 274 | 0.331426 | 1.7571173 | 0 | 0.06620908 | 1 | 7575 | tags=44%, list=32%, si | EN_3 |
| GOBP_PROTEIN_K11_LIN | GOBP_PROTEIN_K11_LINKED_UBIQUITINAT | 29 | 0.494806 | 1.7570137 | 0.00952381 | 0.06606678 | 1 | 6853 | tags=55%, list=29%, si | EN_3 |
| GOBP_RESPONSE_TO_MI | GOBP_RESPONSE_TO_MITOCHONDRIAL_DE | 21 | 0.538271 | 1.755781 | 0.010118044 | 0.066520736 | 1 | 2725 | tags=29%, list=12%, si | EN_3 |
| GOBP_REGULATION_OF_ | GOBP_REGULATION_OF_INTRINSIC_APOPT( | 34 | 0.469026 | 1.7553779 | 0.010736196 | 0.06652071 | 1 | 7214 | tags=59%, list=31%, si | EN_3 |
| GOBP_MITOTIC_G1_S_TR | GOBP_MITOTIC_G1_S_TRANSITION_CHECKI | 24 | 0.506534 | 1.7550985 | 0.003322259 | 0.06645575 | 1 | 5418 | tags=42%, list=23%, si | EN_3 |
| RETROGRADE_ENDOCANI | RETROGRADE_ENDOCANNABINOID_SIGNAL | 133 | 0.359236 | 1.7547578 | 0 | 0.06645613 | 1 | 5839 | tags=37%, list=25%, si | EN_3 |
| GOBP_POSITIVE_REGULA | GOBP_POSITIVE_REGULATION_OF_LEUKOC | 29 | 0.489798 | 1.7529147 | 0.0124223605 | 0.067433946 | 1 | 2841 | tags=34%, list=12%, si | EN_3 |
| GOBP_CELLULAR_RESPON | GOBP_CELLULAR_RESPONSE_TO_OXYGEN_I | 20 | 0.539052 | 1.7523528 | 0.006779661 | 0.06747457 | 1 | 2571 | tags=35%, list=11%, si | EN_3 |
| GOBP_PYRUVATE_METAE | GOBP_PYRUVATE_METABOLIC_PROCESS | 127 | 0.365273 | 1.7520331 | 0.0012515645 | 0.06741111 | 1 | 5371 | tags=34%, list=23%, si | EN_3 |
| GOBP_POSITIVE_REGULA | GOBP_POSITIVE_REGULATION_OF_PROTEIN | 201 | 0.33927 | 1.7515779 | 0 | 0.067448065 | 1 | 4081 | tags=25%, list=17%, si | EN_3 |
| GOBP_DNA_INTEGRITY_C | GOBP_DNA_INTEGRITY_CHECKPOINT_SIGN/ | 125 | 0.365262 | 1.7492448 | 0 | 0.06848425 | 1 | 4966 | tags=29%, list=21%, si | EN_3 |

| | | | | | | | | | | |
|---|---|---|---|---|---|---|---|---|---|---|
| GOBP_NEGATIVE_REGUL | GOBP_NEGATIVE_REGULATION_OF_UBIQUI | 18 | 0.548552 | 1.7487243 | 0.014950166 | 0.06857288 | 1 | 4343 | tags=44%, list=18%, si | EN_3 |
| GOBP_CORTICAL_CYTOSK | GOBP_CORTICAL_CYTOSKELETON_ORGANIZ | 53 | 0.429323 | 1.7482905 | 0.0042918455 | 0.068589486 | 1 | 4774 | tags=32%, list=20%, si | EN_3 |
| GOBP_REGULATION_OF_ | GOBP_REGULATION_OF_SIGNAL_TRANSDU | 83 | 0.388382 | 1.7482027 | 0.0039421814 | 0.068421386 | 1 | 5859 | tags=43%, list=25%, si | EN_3 |
| GOBP_ENDOSOMAL_TRA | GOBP_ENDOSOMAL_TRANSPORT | 272 | 0.331543 | 1.746316 | 0 | 0.069406524 | 1 | 4662 | tags=26%, list=20%, si | EN_3 |
| GOBP_POSITIVE_REGULA | GOBP_POSITIVE_REGULATION_OF_INFLAMI | 22 | 0.526005 | 1.7460325 | 0.0033840947 | 0.06938495 | 1 | 3060 | tags=36%, list=13%, si | EN_3 |
| GOBP_POSITIVE_REGULA | GOBP_POSITIVE_REGULATION_OF_PEPTIDY | 28 | 0.481017 | 1.7459306 | 0.0096 | 0.069216006 | 1 | 1080 | tags=18%, list=5%, sig | EN_3 |
| GOBP_REGULATION_OF_ | GOBP_REGULATION_OF_DEFENSE_RESPONS | 42 | 0.447806 | 1.7445085 | 0.010432191 | 0.069815576 | 1 | 2841 | tags=26%, list=12%, si | EN_3 |
| GOBP_REGULATION_OF_ | GOBP_REGULATION_OF_MACROAUTOPHAG | 129 | 0.360663 | 1.7444094 | 0.0012210013 | 0.069658004 | 1 | 5681 | tags=32%, list=24%, si | EN_3 |
| GOBP_REGULATION_OF_ | GOBP_REGULATION_OF_MONOCYTE_CHEM | 29 | 0.488338 | 1.7429489 | 0.010852713 | 0.07021798 | 1 | 3182 | tags=31%, list=13%, si | EN_3 |
| GOBP_REGULATION_OF_ | GOBP_REGULATION_OF_NEUROINFLAMMA | 30 | 0.477864 | 1.7404317 | 0.006557377 | 0.07173033 | 1 | 3592 | tags=37%, list=15%, si | EN_3 |
| GOBP_EOSINOPHIL_CHEM | GOBP_EOSINOPHIL_CHEMOTAXIS | 22 | 0.509919 | 1.740163 | 0.010016695 | 0.07169012 | 1 | 3118 | tags=41%, list=13%, si | EN_3 |
| GOBP_VESICLE_TARGETIN | GOBP_VESICLE_TARGETING_ROUGH_ER_TO | 20 | 0.529099 | 1.7397321 | 0.011532125 | 0.071696326 | 1 | 6019 | tags=65%, list=25%, si | EN_3 |
| GOBP_LIPID_STORAGE | GOBP_LIPID_STORAGE | 97 | 0.37753 | 1.7388332 | 0.0012594458 | 0.07200724 | 1 | 3764 | tags=24%, list=16%, si | EN_3 |
| GOBP_POSITIVE_REGULA | GOBP_POSITIVE_REGULATION_OF_DNA_ME | 314 | 0.324148 | 1.7380619 | 0 | 0.072239734 | 1 | 5457 | tags=29%, list=23%, si | EN_3 |
| GOBP_PROTEIN_ACETYLA | GOBP_PROTEIN_ACETYLATION | 56 | 0.41676 | 1.737961 | 0.006868132 | 0.07208132 | 1 | 5492 | tags=30%, list=23%, si | EN_3 |
| GOBP_REGULATION_OF_ | GOBP_REGULATION_OF_CALCIUM_ION_IMI | 45 | 0.439969 | 1.7378759 | 0.00877193 | 0.0718828 | 1 | 3182 | tags=31%, list=13%, si | EN_3 |
| GOBP_MITOTIC_G2_M_T | GOBP_MITOTIC_G2_M_TRANSITION_CHECK | 56 | 0.414473 | 1.7375227 | 0.007072136 | 0.07186937 | 1 | 5457 | tags=39%, list=23%, si | EN_3 |
| GOBP_NEGATIVE_REGUL | GOBP_NEGATIVE_REGULATION_OF_PROGR | 17 | 0.560765 | 1.7358755 | 0.011627907 | 0.072724834 | 1 | 4485 | tags=41%, list=19%, si | EN_3 |
| GOBP_REGULATION_OF_ | GOBP_REGULATION_OF_ENDOPLASMIC_RE | 30 | 0.472381 | 1.7350237 | 0.00764526 | 0.073077805 | 1 | 7975 | tags=57%, list=34%, si | EN_3 |
| GOBP_GLYCEROLIPID_BIC | GOBP_GLYCEROLIPID_BIOSYNTHETIC_PROC | 210 | 0.334701 | 1.7348489 | 0 | 0.07294552 | 1 | 4311 | tags=27%, list=18%, si | EN_3 |
| GOBP_PORPHYRIN_CONT | GOBP_PORPHYRIN_CONTAINING_COMPOU | 40 | 0.449357 | 1.7346784 | 0.0044182623 | 0.07283998 | 1 | 4522 | tags=38%, list=19%, si | EN_3 |
| GOBP_DISRUPTION_OF_A | GOBP_DISRUPTION_OF_ANATOMICAL_STRU | 73 | 0.394695 | 1.7340059 | 0.00794702 | 0.07299728 | 1 | 3182 | tags=27%, list=13%, si | EN_3 |
| PPAR_SIGNALING_PATHV | PPAR_SIGNALING_PATHWAY | 70 | 0.394543 | 1.7331958 | 0.0026007802 | 0.073311925 | 1 | 3314 | tags=24%, list=14%, si | EN_3 |
| GOBP_REGULATION_OF_ | GOBP_REGULATION_OF_PROTEIN_CATABOI | 346 | 0.318462 | 1.7320355 | 0 | 0.07378755 | 1 | 5087 | tags=28%, list=22%, si | EN_3 |
| GOBP_MITOCHONDRIAL_ | GOBP_MITOCHONDRIAL_GENOME_MAINTE | 35 | 0.469707 | 1.7308718 | 0.0045941807 | 0.074394815 | 1 | 5418 | tags=40%, list=23%, si | EN_3 |
| GOBP_REGULATION_OF_ | GOBP_REGULATION_OF_CYCLIN_DEPENDEN | 67 | 0.399596 | 1.7306217 | 0.004115226 | 0.074286535 | 1 | 4187 | tags=34%, list=18%, si | EN_3 |
| GOBP_POSITIVE_REGULA | GOBP_POSITIVE_REGULATION_OF_TRANSLA | 125 | 0.35516 | 1.7301928 | 0.0012468828 | 0.07435721 | 1 | 4859 | tags=30%, list=21%, si | EN_3 |
| GOBP_PHOSPHOLIPID_BI | GOBP_PHOSPHOLIPID_BIOSYNTHETIC_PROC | 214 | 0.336776 | 1.7301248 | 0 | 0.074162446 | 1 | 4378 | tags=27%, list=19%, si | EN_3 |
| GOBP_NEGATIVE_REGUL | GOBP_NEGATIVE_REGULATION_OF_VIRAL_ | 18 | 0.543623 | 1.7270627 | 0.016806724 | 0.07590591 | 1 | 2176 | tags=39%, list=9%, sig | EN_3 |
| GOBP_REGULATION_OF_ | GOBP_REGULATION_OF_MITOCHONDRIAL_ | 39 | 0.447629 | 1.726402 | 0.0091047045 | 0.07610516 | 1 | 3635 | tags=33%, list=15%, si | EN_3 |
| GOBP_DNA_TEMPLATED_ | GOBP_DNA_TEMPLATED_TRANSCRIPTION_ | 112 | 0.360629 | 1.7260523 | 0.00124533 | 0.07605803 | 1 | 3709 | tags=22%, list=16%, si | EN_3 |
| GOBP_REGULATION_OF_ | GOBP_REGULATION_OF_SUPEROXIDE_ANIC | 28 | 0.484325 | 1.7243837 | 0.00931677 | 0.076908484 | 1 | 3321 | tags=32%, list=14%, si | EN_3 |
| GOBP_LATE_ENDOSOME_ | GOBP_LATE_ENDOSOME_TO_LYSOSOME_T | 19 | 0.540367 | 1.723756 | 0.010169491 | 0.07713083 | 1 | 5091 | tags=53%, list=22%, si | EN_3 |
| GOBP_VENTRICULAR_SYS | GOBP_VENTRICULAR_SYSTEM_DEVELOPME | 36 | 0.464119 | 1.7233703 | 0.010204081 | 0.07719489 | 1 | 1689 | tags=19%, list=7%, sig | EN_3 |
| GOBP_REGULATION_OF_ | GOBP_REGULATION_OF_TRIGLYCERIDE_BIO | 25 | 0.503301 | 1.7216549 | 0.0127795525 | 0.07806917 | 1 | 4092 | tags=36%, list=17%, si | EN_3 |
| GOBP_REGULATION_OF_ | GOBP_REGULATION_OF_SUPEROXIDE_MET | 34 | 0.462526 | 1.7215261 | 0.0106870225 | 0.07792492 | 1 | 3879 | tags=32%, list=16%, si | EN_3 |
| GOBP_REGULATION_OF_ | GOBP_REGULATION_OF_SMALL_MOLECULE | 356 | 0.31847 | 1.7206254 | 0 | 0.07825508 | 1 | 4399 | tags=26%, list=19%, si | EN_3 |
| GOBP_CELLULAR_RESPON | GOBP_CELLULAR_RESPONSE_TO_RETINOIC_ | 44 | 0.442847 | 1.7197587 | 0.0060698027 | 0.07852962 | 1 | 3004 | tags=25%, list=13%, si | EN_3 |
| GOBP_REGULATION_OF_ | GOBP_REGULATION_OF_RECEPTOR_RECYCL | 30 | 0.472158 | 1.7190522 | 0.0128410915 | 0.0787087 | 1 | 5191 | tags=40%, list=22%, si | EN_3 |
| GOBP_POSITIVE_REGULA | GOBP_POSITIVE_REGULATION_OF_PROTEO | 325 | 0.318433 | 1.7185202 | 0 | 0.07881272 | 1 | 3411 | tags=21%, list=14%, si | EN_3 |
| GOBP_ASPARTATE_FAMII | GOBP_ASPARTATE_FAMILY_AMINO_ACID_N | 36 | 0.453273 | 1.7183628 | 0.013157895 | 0.078678615 | 1 | 3532 | tags=36%, list=15%, si | EN_3 |
| SNARE_INTERACTIONS_IN | SNARE_INTERACTIONS_IN_VESICULAR_TRAI | 32 | 0.471415 | 1.718198 | 0.017991005 | 0.07855205 | 1 | 3617 | tags=31%, list=15%, si | EN_3 |
| GOBP_REGULATION_OF_ | GOBP_REGULATION_OF_PROTEIN_CONTAIN | 420 | 0.312655 | 1.7180408 | 0 | 0.07844047 | 1 | 5484 | tags=30%, list=23%, si | EN_3 |
| GOBP_CGAS_STING_SIGN | GOBP_CGAS_STING_SIGNALING_PATHWAY | 16 | 0.568182 | 1.7180146 | 0.011532125 | 0.07823597 | 1 | 3321 | tags=38%, list=14%, si | EN_3 |
| GOBP_MITOCHONDRIAL_ | GOBP_MITOCHONDRIAL_TRANSPORT | 174 | 0.341359 | 1.7180058 | 0 | 0.078015454 | 1 | 6376 | tags=41%, list=27%, si | EN_3 |
| GOBP_MRNA_TRANSCRIP | GOBP_MRNA_TRANSCRIPTION | 50 | 0.424842 | 1.7171729 | 0.008547009 | 0.078367405 | 1 | 8463 | tags=58%, list=36%, si | EN_3 |
| GOBP_INTRINSIC_APOPT | GOBP_INTRINSIC_APOPTOTIC_SIGNALING_ | 83 | 0.382852 | 1.7169304 | 0.0052980133 | 0.07828841 | 1 | 5977 | tags=40%, list=25%, si | EN_3 |
| GOBP_NEURON_APOPTO | GOBP_NEURON_APOPTOTIC_PROCESS | 369 | 0.315078 | 1.7167139 | 0 | 0.07818922 | 1 | 6290 | tags=35%, list=27%, si | EN_3 |
| STEROID_BIOSYNTHESIS | STEROID_BIOSYNTHESIS | 17 | 0.540615 | 1.7165828 | 0.017482517 | 0.078077205 | 1 | 5219 | tags=53%, list=22%, si | EN_3 |
| GOBP_REGULATION_OF_ | GOBP_REGULATION_OF_MONONUCLEAR_C | 136 | 0.348787 | 1.715954 | 0 | 0.07821758 | 1 | 3851 | tags=26%, list=16%, si | EN_3 |
| GOBP_VESICLE_COATING | GOBP_VESICLE_COATING | 33 | 0.457328 | 1.7136368 | 0.01857585 | 0.07944528 | 1 | 6954 | tags=55%, list=29%, si | EN_3 |
| GOBP_KILLING_OF_CELLS | GOBP_KILLING_OF_CELLS_OF_ANOTHER_OI | 71 | 0.391974 | 1.712373 | 0.004048583 | 0.08002435 | 1 | 3182 | tags=27%, list=13%, si | EN_3 |
| GOBP_CELLULAR_RESPON | GOBP_CELLULAR_RESPONSE_TO_BIOTIC_ST | 224 | 0.32796 | 1.7108415 | 0 | 0.08089349 | 1 | 4015 | tags=24%, list=17%, si | EN_3 |
| GOBP_VIRAL_PROCESS | GOBP_VIRAL_PROCESS | 303 | 0.320802 | 1.7095805 | 0 | 0.08152525 | 1 | 4771 | tags=28%, list=20%, si | EN_3 |
| GOBP_NEGATIVE_REGUL | GOBP_NEGATIVE_REGULATION_OF_PROTEI | 151 | 0.342419 | 1.7092835 | 0 | 0.08145933 | 1 | 6320 | tags=38%, list=27%, si | EN_3 |
| GOBP_TRNA_METABOLIC | GOBP_TRNA_METABOLIC_PROCESS | 166 | 0.343512 | 1.7091454 | 0.0011454754 | 0.081340276 | 1 | 3007 | tags=25%, list=13%, si | EN_3 |
| GOBP_VESICLE_BUDDING | GOBP_VESICLE_BUDDING_FROM_MEMBRA | 79 | 0.380757 | 1.7087173 | 0 | 0.08136843 | 1 | 5579 | tags=39%, list=24%, si | EN_3 |
| GLUTATHIONE_METABOL | GLUTATHIONE_METABOLISM | 57 | 0.404632 | 1.7073243 | 0.007102273 | 0.082107715 | 1 | 3307 | tags=32%, list=14%, si | EN_3 |
| PEROXISOME | PEROXISOME | 81 | 0.380032 | 1.7061965 | 0.006729475 | 0.082630605 | 1 | 5510 | tags=36%, list=23%, si | EN_3 |
| GOBP_REGULATION_OF_ | GOBP_REGULATION_OF_LIPID_BIOSYNTHET | 197 | 0.331291 | 1.7061298 | 0.0011494253 | 0.08245346 | 1 | 4300 | tags=27%, list=18%, si | EN_3 |
| GOBP_NUCLEOSIDE_MON | GOBP_NUCLEOSIDE_MONOPHOSPHATE_ME | 62 | 0.403587 | 1.7061137 | 0.0027510317 | 0.08223433 | 1 | 4383 | tags=31%, list=19%, si | EN_3 |
| GOBP_REGULATION_OF_ | GOBP_REGULATION_OF_EXOCYTOSIS | 194 | 0.331891 | 1.7046617 | 0 | 0.08313026 | 1 | 4158 | tags=24%, list=18%, si | EN_3 |
| GOBP_ORGANIC_HYDRO) | GOBP_ORGANIC_HYDROXY_COMPOUND_BI | 221 | 0.327011 | 1.7044102 | 0.0034129692 | 0.0830417 | 1 | 2983 | tags=22%, list=13%, si | EN_3 |
| GOBP_CELLULAR_OXIDAN | GOBP_CELLULAR_OXIDANT_DETOXIFICATIO | 29 | 0.475914 | 1.7042902 | 0.015576324 | 0.082914524 | 1 | 2571 | tags=31%, list=11%, si | EN_3 |
| GOBP_MAINTENANCE_OI | GOBP_MAINTENANCE_OF_LOCATION | 341 | 0.314852 | 1.704122 | 0 | 0.082817316 | 1 | 4608 | tags=27%, list=20%, si | EN_3 |
| GOBP_NEGATIVE_REGUL | GOBP_NEGATIVE_REGULATION_OF_CYCLIN | 25 | 0.486968 | 1.7037362 | 0.011290323 | 0.08285072 | 1 | 6035 | tags=52%, list=26%, si | EN_3 |
| GOBP_CELLULAR_RESPON | GOBP_CELLULAR_RESPONSE_TO_CHEMICAL | 290 | 0.318305 | 1.7037237 | 0.0010881393 | 0.082630746 | 1 | 4168 | tags=22%, list=18%, si | EN_3 |
| GOBP_REGULATION_OF_ | GOBP_REGULATION_OF_GLUCONEOGENESI | 52 | 0.416931 | 1.70264 | 0.0070921984 | 0.083210334 | 1 | 6290 | tags=40%, list=27%, si | EN_3 |
| GOBP_NUCLEAR_BODY_C | GOBP_NUCLEAR_BODY_ORGANIZATION | 19 | 0.52562 | 1.7007167 | 0.01986755 | 0.084322825 | 1 | 3195 | tags=32%, list=14%, si | EN_3 |
| GOBP_POSITIVE_REGULA | GOBP_POSITIVE_REGULATION_OF_PROTEAS | 102 | 0.364573 | 1.6991848 | 0 | 0.0851674 | 1 | 6029 | tags=33%, list=26%, si | EN_3 |
| GOBP_TETRAPYRROLE_M | GOBP_TETRAPYRROLE_METABOLIC_PROCES | 49 | 0.417374 | 1.6987733 | 0.013196481 | 0.08520679 | 1 | 5243 | tags=41%, list=22%, si | EN_3 |
| GOBP_ACTIN_POLYMERIZ | GOBP_ACTIN_POLYMERIZATION_OR_DEPOL | 187 | 0.335883 | 1.6980778 | 0.0011494253 | 0.08546995 | 1 | 4798 | tags=28%, list=20%, si | EN_3 |
| GOBP_REGULATION_OF_ | GOBP_REGULATION_OF_LEUKOCYTE_DEGR | 48 | 0.423097 | 1.6980281 | 0.013719512 | 0.08528799 | 1 | 3780 | tags=31%, list=16%, si | EN_3 |
| GOBP_PEPTIDYL_LYSINE_ | GOBP_PEPTIDYL_LYSINE_MODIFICATION | 128 | 0.355285 | 1.6977174 | 0.0024783148 | 0.08528852 | 1 | 7159 | tags=43%, list=30%, si | EN_3 |
| GOBP_NEGATIVE_REGUL | GOBP_NEGATIVE_REGULATION_OF_LIPID_S | 27 | 0.478608 | 1.6971446 | 0.0125 | 0.08551123 | 1 | 3616 | tags=22%, list=15%, si | EN_3 |
| GOBP_NEGATIVE_REGUL | GOBP_NEGATIVE_REGULATION_OF_CELL_M | 36 | 0.452694 | 1.6955214 | 0.013740458 | 0.08645126 | 1 | 4995 | tags=33%, list=21%, si | EN_3 |
| GOBP_POSITIVE_REGULA | GOBP_POSITIVE_REGULATION_OF_EXOCYT | 90 | 0.370088 | 1.6953185 | 0.0025974025 | 0.08641068 | 1 | 2841 | tags=22%, list=12%, si | EN_3 |
| GOBP_PROTEIN_ACYLATI | GOBP_PROTEIN_ACYLATION | 94 | 0.370927 | 1.6952379 | 0 | 0.08626007 | 1 | 5089 | tags=29%, list=22%, si | EN_3 |
| GOBP_NEGATIVE_REGUL | GOBP_NEGATIVE_REGULATION_OF_NON_C | 26 | 0.483085 | 1.6951504 | 0.022950819 | 0.08608233 | 1 | 5561 | tags=46%, list=24%, si | EN_3 |
| GLYCOSYLPHOSPHATIDYL | GLYCOSYLPHOSPHATIDYLINOSITOL_(GPI)-AN | 24 | 0.510783 | 1.6950831 | 0.01453958 | 0.08589908 | 1 | 1514 | tags=29%, list=6%, sig | EN_3 |
| GOBP_REGULATION_OF_ | GOBP_REGULATION_OF_STEROID_METABO | 107 | 0.359192 | 1.6939571 | 0.0012755102 | 0.086469606 | 1 | 4085 | tags=31%, list=17%, si | EN_3 |
| PHAGOSOME | PHAGOSOME | 146 | 0.344218 | 1.6938652 | 0 | 0.08628315 | 1 | 5341 | tags=34%, list=23%, si | EN_3 |

| | | | | | | | | | | |
|---|---|---|---|---|---|---|---|---|---|---|
| GOBP_POSITIVE_REGULA | GOBP_POSITIVE_REGULATION_OF_DNA_BIN | 217 | 0.329733 | 1.693683 | 0 | 0.08616264 | 1 | 4168 | tags=27%, list=18%, si | EN_3 |
| GOBP_REGULATION_OF_ | GOBP_REGULATION_OF_PROTEIN_UBIQUIT | 202 | 0.331468 | 1.693634 | 0.0011389522 | 0.08597799 | 1 | 5318 | tags=32%, list=23%, si | EN_3 |
| GOBP_EXTRACELLULAR_S | GOBP_EXTRACELLULAR_STRUCTURE_ORGAN | 309 | 0.312984 | 1.6936208 | 0 | 0.085769504 | 1 | 5496 | tags=31%, list=23%, si | EN_3 |
| GOBP_MUSCLE_CELL_CEI | GOBP_MUSCLE_CELL_CELLULAR_HOMEOST/ | 26 | 0.481771 | 1.6934557 | 0.012861736 | 0.08567326 | 1 | 3719 | tags=31%, list=16%, si | EN_3 |
| GOBP_ORGANOPHOSPHA | GOBP_ORGANOPHOSPHATE_CATABOLIC_PF | 227 | 0.324897 | 1.6933241 | 0 | 0.08554078 | 1 | 4579 | tags=28%, list=19%, si | EN_3 |
| GOBP_NEGATIVE_REGUL | GOBP_NEGATIVE_REGULATION_OF_PEPTID | 52 | 0.406103 | 1.6922718 | 0.010884354 | 0.08600688 | 1 | 4923 | tags=37%, list=21%, si | EN_3 |
| GOBP_VASCULAR_ENDOT | GOBP_VASCULAR_ENDOTHELIAL_CELL_PRO | 29 | 0.477859 | 1.6917745 | 0.0048701297 | 0.08616721 | 1 | 577 | tags=17%, list=2%, sig | EN_3 |
| GOBP_NEUTRAL_LIPID_M | GOBP_NEUTRAL_LIPID_METABOLIC_PROCES | 138 | 0.348194 | 1.6902151 | 0.0012077295 | 0.08705937 | 1 | 4386 | tags=26%, list=19%, si | EN_3 |
| GOBP_REGULATION_OF_ | GOBP_REGULATION_OF_AMIDE_METABOLI | 69 | 0.388757 | 1.6902096 | 0.008275862 | 0.086836144 | 1 | 6470 | tags=39%, list=27%, si | EN_3 |
| GOBP_GPI_ANCHOR_MET | GOBP_GPI_ANCHOR_METABOLIC_PROCESS | 33 | 0.463125 | 1.6892111 | 0.01529052 | 0.087266706 | 1 | 3864 | tags=36%, list=16%, si | EN_3 |
| LEGIONELLOSIS | LEGIONELLOSIS | 55 | 0.406697 | 1.6881412 | 0.008534851 | 0.087858476 | 1 | 7060 | tags=55%, list=30%, si | EN_3 |
| GOBP_CELLULAR_RESPON | GOBP_CELLULAR_RESPONSE_TO_UV | 90 | 0.366978 | 1.6878324 | 0.002541296 | 0.08785568 | 1 | 6141 | tags=38%, list=26%, si | EN_3 |
| GOBP_PROTEIN_K63_LIN | GOBP_PROTEIN_K63_LINKED_DEUBIQUITIN | 29 | 0.472344 | 1.6859051 | 0.018404908 | 0.088974245 | 1 | 4785 | tags=41%, list=20%, si | EN_3 |
| GOBP_RESPONSE_TO_DE | GOBP_RESPONSE_TO_DEXAMETHASONE | 27 | 0.478499 | 1.685473 | 0.015479876 | 0.08902834 | 1 | 2829 | tags=26%, list=12%, si | EN_3 |
| GOBP_REGULATION_OF_ | GOBP_REGULATION_OF_CELL_MATRIX_ADH | 121 | 0.3531 | 1.6853606 | 0.0024420025 | 0.088887595 | 1 | 7590 | tags=45%, list=32%, si | EN_3 |
| GOBP_REGULATION_OF_ | GOBP_REGULATION_OF_INTRINSIC_APOPT( | 185 | 0.327635 | 1.6834099 | 0.0011467889 | 0.090079166 | 1 | 6057 | tags=35%, list=26%, si | EN_3 |
| GOBP_RELEASE_OF_CYTO | GOBP_RELEASE_OF_CYTOCHROME_C_FRON | 66 | 0.391236 | 1.682938 | 0.006887052 | 0.09016322 | 1 | 5792 | tags=41%, list=25%, si | EN_3 |
| GOBP_REGULATION_OF_ | GOBP_REGULATION_OF_PATTERN_RECOGN | 157 | 0.335353 | 1.6825054 | 0.0023310024 | 0.09026756 | 1 | 3467 | tags=23%, list=15%, si | EN_3 |
| GOBP_REGULATION_OF_ | GOBP_REGULATION_OF_CELL_SUBSTRATE_ | 219 | 0.327031 | 1.6824722 | 0 | 0.09005373 | 1 | 6263 | tags=34%, list=27%, si | EN_3 |
| GOBP_POSITIVE_REGULA | GOBP_POSITIVE_REGULATION_OF_VASCUL/ | 172 | 0.334253 | 1.6821474 | 0.0011587485 | 0.090065874 | 1 | 3851 | tags=23%, list=16%, si | EN_3 |
| GOBP_REGULATION_OF_ | GOBP_REGULATION_OF_PROTEASOMAL_PR | 179 | 0.329478 | 1.6812575 | 0.0011261262 | 0.09059561 | 1 | 4898 | tags=26%, list=21%, si | EN_3 |
| GOBP_NEGATIVE_REGUL | GOBP_NEGATIVE_REGULATION_OF_RNA_SF | 29 | 0.47453 | 1.6811816 | 0.010819165 | 0.090438336 | 1 | 7216 | tags=62%, list=31%, si | EN_3 |
| GOBP_MUSCLE_CELL_PR( | GOBP_MUSCLE_CELL_PROLIFERATION | 262 | 0.317272 | 1.6811092 | 0 | 0.09027039 | 1 | 4149 | tags=23%, list=18%, si | EN_3 |
| GOBP_NEGATIVE_REGUL | GOBP_NEGATIVE_REGULATION_OF_APOPT( | 241 | 0.322603 | 1.6806191 | 0 | 0.09041118 | 1 | 4195 | tags=25%, list=18%, si | EN_3 |
| GOBP_CELLULAR_RESPON | GOBP_CELLULAR_RESPONSE_TO_LIPID | 498 | 0.303949 | 1.6805298 | 0 | 0.090270266 | 1 | 3402 | tags=19%, list=14%, si | EN_3 |
| GOBP_REGULATION_OF_ | GOBP_REGULATION_OF_PROTEIN_LOCALIZ/ | 44 | 0.433434 | 1.6805038 | 0.014409222 | 0.09006604 | 1 | 3659 | tags=30%, list=15%, si | EN_3 |
| GOBP_RETROGRADE_VES | GOBP_RETROGRADE_VESICLE_MEDIATED_T | 45 | 0.419848 | 1.6789769 | 0.008450705 | 0.09087811 | 1 | 9253 | tags=76%, list=39%, si | EN_3 |
| GOBP_MAST_CELL_ACTIV | GOBP_MAST_CELL_ACTIVATION_INVOLVED_ | 56 | 0.402501 | 1.6789677 | 0.006887052 | 0.090661734 | 1 | 3363 | tags=27%, list=14%, si | EN_3 |
| GOBP_RESPONSE_TO_AM | GOBP_RESPONSE_TO_AMYLOID_BETA | 39 | 0.434601 | 1.6788392 | 0.010401188 | 0.09051865 | 1 | 3074 | tags=28%, list=13%, si | EN_3 |
| ENDOCYTOSIS | ENDOCYTOSIS | 246 | 0.315205 | 1.6759361 | 0 | 0.092396274 | 1 | 6637 | tags=39%, list=28%, si | EN_3 |
| GOBP_REGULATION_OF_ | GOBP_REGULATION_OF_MITOCHONDRIAL_ | 58 | 0.396152 | 1.6757355 | 0.0028735632 | 0.09229033 | 1 | 3635 | tags=31%, list=15%, si | EN_3 |
| GOBP_PROTEIN_MATURA | GOBP_PROTEIN_MATURATION | 475 | 0.30443 | 1.6755028 | 0 | 0.09223926 | 1 | 5799 | tags=33%, list=25%, si | EN_3 |
| GOBP_INTRINSIC_APOPT( | GOBP_INTRINSIC_APOPTOTIC_SIGNALING_F | 59 | 0.401573 | 1.6748465 | 0.007032349 | 0.0925292 | 1 | 3814 | tags=25%, list=16%, si | EN_3 |
| GOBP_NEGATIVE_REGUL | GOBP_NEGATIVE_REGULATION_OF_PROTEI | 80 | 0.370832 | 1.6745068 | 0.010695187 | 0.092517935 | 1 | 5762 | tags=34%, list=24%, si | EN_3 |
| GOBP_NEGATIVE_REGUL | GOBP_NEGATIVE_REGULATION_OF_CELL_C | 67 | 0.389747 | 1.6742812 | 0.009575923 | 0.09245256 | 1 | 4966 | tags=30%, list=21%, si | EN_3 |
| GOBP_RESPONSE_TO_NE | GOBP_RESPONSE_TO_NERVE_GROWTH_FA( | 42 | 0.425966 | 1.6740565 | 0.01161103 | 0.092416026 | 1 | 3321 | tags=26%, list=14%, si | EN_3 |
| GOBP_VESICLE_ORGANIZ | GOBP_VESICLE_ORGANIZATION | 353 | 0.307155 | 1.6719244 | 0 | 0.09384142 | 1 | 5806 | tags=32%, list=25%, si | EN_3 |
| GOBP_MODULATION_BY_ | GOBP_MODULATION_BY_HOST_OF_VIRAL_ | 22 | 0.503227 | 1.671583 | 0.013605442 | 0.09386368 | 1 | 6559 | tags=41%, list=28%, si | EN_3 |
| GOBP_POSITIVE_REGULA | GOBP_POSITIVE_REGULATION_OF_CYTOSKE | 185 | 0.326589 | 1.6710523 | 0 | 0.09400712 | 1 | 2941 | tags=19%, list=12%, si | EN_3 |
| GOBP_NEGATIVE_REGUL | GOBP_NEGATIVE_REGULATION_OF_INTRAC | 35 | 0.449002 | 1.6704336 | 0.010574019 | 0.094223544 | 1 | 5231 | tags=46%, list=22%, si | EN_3 |
| GOBP_POSITIVE_REGULA | GOBP_POSITIVE_REGULATION_OF_ENDOTH | 107 | 0.353423 | 1.6701404 | 0.006385696 | 0.09421137 | 1 | 4085 | tags=24%, list=17%, si | EN_3 |
| GOBP_NEUTROPHIL_ACTI | GOBP_NEUTROPHIL_ACTIVATION_INVOLVEI | 18 | 0.524076 | 1.6700637 | 0.014705882 | 0.094033495 | 1 | 2453 | tags=33%, list=10%, si | EN_3 |
| GOBP_REGULATION_OF_ | GOBP_REGULATION_OF_POST_TRANSLATIO | 242 | 0.315962 | 1.669957 | 0 | 0.09386218 | 1 | 5446 | tags=31%, list=23%, si | EN_3 |
| ASCORBATE_AND_ALDAR | ASCORBATE_AND_ALDARATE_METABOLISM | 16 | 0.543409 | 1.6699435 | 0.011608624 | 0.09365798 | 1 | 3393 | tags=50%, list=14%, si | EN_3 |
| GOBP_REGULATION_OF_ | GOBP_REGULATION_OF_SUBSTRATE_ADHES | 58 | 0.403835 | 1.6697726 | 0.012517385 | 0.09355507 | 1 | 4672 | tags=33%, list=20%, si | EN_3 |
| GOBP_POSITIVE_REGULA | GOBP_POSITIVE_REGULATION_OF_CELL_GR | 17 | 0.530082 | 1.669104 | 0.024752475 | 0.09380853 | 1 | 3586 | tags=47%, list=15%, si | EN_3 |
| GOBP_POSITIVE_REGULA | GOBP_POSITIVE_REGULATION_OF_PROTEIN | 107 | 0.356116 | 1.6689013 | 0.0024660912 | 0.09373109 | 1 | 4934 | tags=31%, list=21%, si | EN_3 |
| GOBP_IMPORT_INTO_TH | GOBP_IMPORT_INTO_THE_MITOCHONDRIO | 49 | 0.419178 | 1.6685262 | 0.010071943 | 0.093792014 | 1 | 7511 | tags=53%, list=32%, si | EN_3 |
| GOBP_PHOSPHATIDYLCH( | GOBP_PHOSPHATIDYLCHOLINE_METABOLIC | 59 | 0.396072 | 1.667601 | 0.009433962 | 0.0942885 | 1 | 1532 | tags=19%, list=6%, sig | EN_3 |
| FC_GAMMA_R-MEDIATEI | FC_GAMMA_R-MEDIATED_PHAGOCYTOSIS | 84 | 0.372207 | 1.6675997 | 0.009296149 | 0.09406973 | 1 | 4672 | tags=30%, list=20%, si | EN_3 |
| GOBP_POSITIVE_REGULA | GOBP_POSITIVE_REGULATION_OF_GLUCOS | 50 | 0.414436 | 1.6665727 | 0.011267605 | 0.09467398 | 1 | 4986 | tags=36%, list=21%, si | EN_3 |
| GOBP_POSITIVE_REGULA | GOBP_POSITIVE_REGULATION_OF_MUSCLE | 76 | 0.374428 | 1.6663395 | 0.0038659794 | 0.09461962 | 1 | 3586 | tags=25%, list=15%, si | EN_3 |
| GOBP_REGULATION_OF_ | GOBP_REGULATION_OF_PRODUCTION_OF_ | 195 | 0.319649 | 1.6662757 | 0.0011428571 | 0.094467275 | 1 | 3080 | tags=22%, list=13%, si | EN_3 |
| GOBP_PRIMARY_AMINO_ | GOBP_PRIMARY_AMINO_COMPOUND_MET | 17 | 0.557219 | 1.6649072 | 0.023688663 | 0.09518604 | 1 | 6563 | tags=65%, list=28%, si | EN_3 |
| GOBP_COTRANSLATIONA | GOBP_COTRANSLATIONAL_PROTEIN_TARGE | 20 | 0.513859 | 1.6632501 | 0.025931928 | 0.096342966 | 1 | 6949 | tags=60%, list=29%, si | EN_3 |
| GOBP_SNRNA_PROCESSII | GOBP_SNRNA_PROCESSING | 24 | 0.484676 | 1.6626797 | 0.02488336 | 0.09655482 | 1 | 5628 | tags=50%, list=24%, si | EN_3 |
| GOBP_REGULATION_OF_ | GOBP_REGULATION_OF_GLUCOSE_METABO | 109 | 0.350344 | 1.6623816 | 0.002528445 | 0.096526906 | 1 | 6457 | tags=38%, list=27%, si | EN_3 |
| PROTEIN_PROCESSING_IN | PROTEIN_PROCESSING_IN_ENDOPLASMIC_I | 158 | 0.33244 | 1.6620747 | 0.0011750881 | 0.09656654 | 1 | 6689 | tags=37%, list=28%, si | EN_3 |
| GOBP_REGULATION_OF_ | GOBP_REGULATION_OF_GENERATION_OF_ | 148 | 0.335326 | 1.6617883 | 0.0011947431 | 0.09653836 | 1 | 5859 | tags=36%, list=25%, si | EN_3 |
| GOBP_MYELOID_LEUKOC | GOBP_MYELOID_LEUKOCYTE_MIGRATION | 222 | 0.318567 | 1.6615409 | 0 | 0.096491754 | 1 | 3851 | tags=23%, list=16%, si | EN_3 |
| GOBP_COMPLEMENT_AC | GOBP_COMPLEMENT_ACTIVATION_CLASSIC | 29 | 0.460689 | 1.6614999 | 0.020280812 | 0.09630822 | 1 | 3500 | tags=34%, list=15%, si | EN_3 |
| GOBP_PROGRAMMED_N | GOBP_PROGRAMMED_NECROTIC_CELL_DE/ | 50 | 0.411008 | 1.6608111 | 0.00896861 | 0.09659239 | 1 | 6174 | tags=40%, list=26%, si | EN_3 |
| GOBP_RESPONSE_TO_AN | GOBP_RESPONSE_TO_AMINO_ACID_STARV/ | 54 | 0.403099 | 1.6605898 | 0.01746725 | 0.0965567 | 1 | 6373 | tags=44%, list=27%, si | EN_3 |
| GOBP_TRIGLYCERIDE_BIO | GOBP_TRIGLYCERIDE_BIOSYNTHETIC_PROCE | 46 | 0.416258 | 1.6604911 | 0.012912482 | 0.09641717 | 1 | 4311 | tags=30%, list=18%, si | EN_3 |
| GOBP_CHAPERONE_MED | GOBP_CHAPERONE_MEDIATED_PROTEIN_C | 18 | 0.532392 | 1.660443 | 0.016722407 | 0.09624357 | 1 | 5467 | tags=61%, list=23%, si | EN_3 |
| GOBP_MITOTIC_DNA_RE | GOBP_MITOTIC_DNA_REPLICATION | 16 | 0.544926 | 1.6603175 | 0.028475711 | 0.09613712 | 1 | 5093 | tags=56%, list=22%, si | EN_3 |
| GOBP_NEGATIVE_REGUL | GOBP_NEGATIVE_REGULATION_OF_AMIDE_ | 22 | 0.490292 | 1.6602749 | 0.014354067 | 0.09595169 | 1 | 6770 | tags=59%, list=29%, si | EN_3 |
| GOBP_CELLULAR_RESPON | GOBP_CELLULAR_RESPONSE_TO_TOXIC_SU | 63 | 0.386745 | 1.6595958 | 0.012482663 | 0.09634308 | 1 | 2725 | tags=29%, list=12%, si | EN_3 |
| GOBP_REGULATION_OF_ | GOBP_REGULATION_OF_CELL_CYCLE_PHASI | 424 | 0.302616 | 1.6591707 | 0 | 0.096490964 | 1 | 4998 | tags=28%, list=21%, si | EN_3 |
| GOBP_VESICLE_TETHERIN | GOBP_VESICLE_TETHERING | 31 | 0.454334 | 1.6589848 | 0.018404908 | 0.0964084 | 1 | 6019 | tags=35%, list=25%, si | EN_3 |
| GOBP_RESPONSE_TO_ISC | GOBP_RESPONSE_TO_ISCHEMIA | 48 | 0.408917 | 1.6589164 | 0.00877193 | 0.09625795 | 1 | 3115 | tags=27%, list=13%, si | EN_3 |
| GOBP_REGULATION_OF_ | GOBP_REGULATION_OF_CHOLESTEROL_ME | 41 | 0.43009 | 1.6588479 | 0.018489985 | 0.09611112 | 1 | 4085 | tags=34%, list=17%, si | EN_3 |
| GOBP_DNA_UNWINDING | GOBP_DNA_UNWINDING_INVOLVED_IN_DI | 20 | 0.501228 | 1.6583499 | 0.026229508 | 0.09623969 | 1 | 5324 | tags=60%, list=23%, si | EN_3 |
| GOBP_PROTEIN_LOCALIZ | GOBP_PROTEIN_LOCALIZATION_TO_NUCLE( | 17 | 0.539854 | 1.6582699 | 0.026711185 | 0.09607255 | 1 | 2296 | tags=35%, list=10%, si | EN_3 |
| GOBP_REGULATION_OF_ | GOBP_REGULATION_OF_LIPOPROTEIN_MET | 18 | 0.528009 | 1.6571584 | 0.04139073 | 0.09666135 | 1 | 3743 | tags=39%, list=16%, si | EN_3 |
| GOBP_REACTIVE_OXYGEI | GOBP_REACTIVE_OXYGEN_SPECIES_METAB | 239 | 0.315407 | 1.6559361 | 0 | 0.09740025 | 1 | 5751 | tags=33%, list=24%, si | EN_3 |
| GOBP_AXO_DENDRITIC_T | GOBP_AXO_DENDRITIC_TRANSPORT | 92 | 0.36247 | 1.6555287 | 0.0025445293 | 0.09752452 | 1 | 5416 | tags=32%, list=23%, si | EN_3 |
| GOBP_EPITHELIAL_TUBE_ | GOBP_EPITHELIAL_TUBE_BRANCHING_INVO | 37 | 0.435842 | 1.6550702 | 0.012084592 | 0.09763293 | 1 | 2842 | tags=27%, list=12%, si | EN_3 |
| GOBP_RESPONSE_TO_CH | GOBP_RESPONSE_TO_CHEMOKINE | 74 | 0.371986 | 1.6544585 | 0.0026773761 | 0.097836345 | 1 | 3182 | tags=30%, list=13%, si | EN_3 |
| GOBP_RIBOSOMAL_SMAI | GOBP_RIBOSOMAL_SMALL_SUBUNIT_ASSEI | 18 | 0.524965 | 1.653749 | 0.028346457 | 0.09817784 | 1 | 6557 | tags=61%, list=28%, si | EN_3 |
| GOBP_INTRINSIC_APOPT( | GOBP_INTRINSIC_APOPTOTIC_SIGNALING_F | 314 | 0.307333 | 1.6537126 | 0 | 0.097996145 | 1 | 6480 | tags=36%, list=27%, si | EN_3 |
| GOBP_REGULATION_OF_ | GOBP_REGULATION_OF_RESPONSE_TO_TYF | 15 | 0.546439 | 1.6536136 | 0.023255814 | 0.09785625 | 1 | 1080 | tags=27%, list=5%, sig | EN_3 |

| | | | | | | | | | | |
|---|---|---|---|---|---|---|---|---|---|---|
| GOBP_SYNAPTIC_VESICLE | GOBP_SYNAPTIC_VESICLE_MATURATION | 27 | 0.467674 | 1.6535847 | 0.031397175 | 0.097663194 | 1 | 7781 | tags=52%, list=33%, si | EN_3 |
| GOBP_HUMORAL_IMMUI | GOBP_HUMORAL_IMMUNE_RESPONSE | 184 | 0.323326 | 1.652989 | 0.0022779044 | 0.09787687 | 1 | 3182 | tags=23%, list=13%, si | EN_3 |
| GOBP_SNRNA_METABOL | GOBP_SNRNA_METABOLIC_PROCESS | 50 | 0.408274 | 1.6527299 | 0.0088626295 | 0.09781206 | 1 | 5769 | tags=44%, list=24%, si | EN_3 |
| GOBP_CARBOHYDRATE_T | GOBP_CARBOHYDRATE_TRANSMEMBRANE_ | 122 | 0.342792 | 1.6526582 | 0.0036809817 | 0.097663715 | 1 | 4986 | tags=30%, list=21%, si | EN_3 |
| GOBP_MRNA_PROCESSIN | GOBP_MRNA_PROCESSING | 459 | 0.298521 | 1.6525304 | 0 | 0.09754121 | 1 | 6661 | tags=34%, list=28%, si | EN_3 |
| GOBP_PROXIMAL_DISTAL | GOBP_PROXIMAL_DISTAL_PATTERN_FORM/ | 33 | 0.450641 | 1.6510468 | 0.013657056 | 0.09845158 | 1 | 1421 | tags=21%, list=6%, sig | EN_3 |
| GOBP_NEGATIVE_REGUL/ | GOBP_NEGATIVE_REGULATION_OF_OSTEO( | 37 | 0.438747 | 1.65094 | 0.016393442 | 0.09833311 | 1 | 4923 | tags=41%, list=21%, si | EN_3 |
| GOBP_TRICARBOXYLIC_A | GOBP_TRICARBOXYLIC_ACID_CYCLE | 33 | 0.439498 | 1.6502222 | 0.022012578 | 0.09864054 | 1 | 6125 | tags=48%, list=26%, si | EN_3 |
| GOBP_REGULATION_OF_ | GOBP_REGULATION_OF_DNA_METABOLIC_ | 491 | 0.297064 | 1.6495124 | 0 | 0.09894471 | 1 | 5470 | tags=29%, list=23%, si | EN_3 |
| GOBP_NEGATIVE_REGUL/ | GOBP_NEGATIVE_REGULATION_OF_TELOM | 36 | 0.433072 | 1.6491215 | 0.013293944 | 0.09903926 | 1 | 5418 | tags=33%, list=23%, si | EN_3 |
| GOBP_REGULATION_OF_ | GOBP_REGULATION_OF_ANIMAL_ORGAN_I | 29 | 0.45406 | 1.6487136 | 0.014240506 | 0.09912334 | 1 | 5266 | tags=41%, list=22%, si | EN_3 |
| GOBP_NEGATIVE_REGUL/ | GOBP_NEGATIVE_REGULATION_OF_TYPE_I_ | 42 | 0.418511 | 1.648512 | 0.017699115 | 0.09906476 | 1 | 7711 | tags=60%, list=33%, si | EN_3 |
| GOBP_POSITIVE_REGULA | GOBP_POSITIVE_REGULATION_OF_TRIGLYC | 18 | 0.516762 | 1.6481032 | 0.017571885 | 0.0991627 | 1 | 3864 | tags=33%, list=16%, si | EN_3 |
| RENAL_CELL_CARCINOM/ | RENAL_CELL_CARCINOMA | 67 | 0.386806 | 1.6479352 | 0.010680908 | 0.09907432 | 1 | 7961 | tags=61%, list=34%, si | EN_3 |
| GOBP_NEGATIVE_REGUL/ | GOBP_NEGATIVE_REGULATION_OF_REACTI\ | 52 | 0.401986 | 1.6471705 | 0.014204546 | 0.09946016 | 1 | 6290 | tags=46%, list=27%, si | EN_3 |
| GOBP_POSITIVE_REGULA | GOBP_POSITIVE_REGULATION_OF_CYTOKIN | 82 | 0.367394 | 1.6465731 | 0.0077021825 | 0.09971077 | 1 | 2936 | tags=24%, list=12%, si | EN_3 |
| GOBP_ENDOPLASMIC_RE | GOBP_ENDOPLASMIC_RETICULUM_CALCIUI | 28 | 0.456999 | 1.6463975 | 0.029457364 | 0.09962909 | 1 | 4044 | tags=29%, list=17%, si | EN_3 |
| GOBP_NEGATIVE_REGUL/ | GOBP_NEGATIVE_REGULATION_OF_INTRAC | 36 | 0.434357 | 1.6460856 | 0.013372957 | 0.09966876 | 1 | 5165 | tags=39%, list=22%, si | EN_3 |
| GOBP_BLASTOCYST_GRO | GOBP_BLASTOCYST_GROWTH | 29 | 0.465472 | 1.6446213 | 0.013473053 | 0.10065423 | 1 | 2288 | tags=24%, list=10%, si | EN_3 |
| GOBP_PROTEIN_POLYUBI | GOBP_PROTEIN_POLYUBIQUITINATION | 250 | 0.3104 | 1.6444281 | 0.0011025359 | 0.10060784 | 1 | 5269 | tags=30%, list=22%, si | EN_3 |
| MRNA_SURVEILLANCE_P/ | MRNA_SURVEILLANCE_PATHWAY | 87 | 0.362984 | 1.6443677 | 0.0026595744 | 0.10043169 | 1 | 7216 | tags=46%, list=31%, si | EN_3 |
| GOBP_NEGATIVE_REGUL/ | GOBP_NEGATIVE_REGULATION_OF_PROTEI | 80 | 0.370664 | 1.6424451 | 0.013531799 | 0.10184248 | 1 | 5223 | tags=34%, list=22%, si | EN_3 |
| GOBP_REGULATION_OF_ | GOBP_REGULATION_OF_ACTIN_FILAMENT_ | 266 | 0.308374 | 1.6422851 | 0 | 0.10176966 | 1 | 5306 | tags=28%, list=22%, si | EN_3 |
| GOBP_PROTEIN_LOCALIZ | GOBP_PROTEIN_LOCALIZATION_TO_ENDOP | 76 | 0.370446 | 1.6420572 | 0.00795756 | 0.10171917 | 1 | 4909 | tags=34%, list=21%, si | EN_3 |
| GOBP_REGULATION_OF_ | GOBP_REGULATION_OF_APOPTOTIC_SIGNA | 409 | 0.30045 | 1.6394348 | 0 | 0.1035586 | 1 | 4424 | tags=26%, list=19%, si | EN_3 |
| APOPTOSIS | APOPTOSIS | 137 | 0.336785 | 1.6390702 | 0.0011904762 | 0.103621155 | 1 | 4417 | tags=26%, list=19%, si | EN_3 |
| GOBP_LUNG_CELL_DIFFE | GOBP_LUNG_CELL_DIFFERENTIATION | 38 | 0.428428 | 1.6387316 | 0.010432191 | 0.10366888 | 1 | 5371 | tags=32%, list=23%, si | EN_3 |
| GOBP_PHOSPHOLIPID_M | GOBP_PHOSPHOLIPID_METABOLIC_PROCES | 352 | 0.304578 | 1.6386318 | 0 | 0.10354221 | 1 | 4228 | tags=24%, list=18%, si | EN_3 |
| GOBP_RESPONSE_TO_FA | GOBP_RESPONSE_TO_FATTY_ACID | 46 | 0.412563 | 1.637598 | 0.022889843 | 0.10421224 | 1 | 3743 | tags=35%, list=16%, si | EN_3 |
| GOBP_GLYCEROL_METAB | GOBP_GLYCEROL_METABOLIC_PROCESS | 19 | 0.513203 | 1.6368324 | 0.029268293 | 0.104595445 | 1 | 3983 | tags=42%, list=17%, si | EN_3 |
| GOBP_AMYLOID_BETA_C | GOBP_AMYLOID_BETA_CLEARANCE | 37 | 0.424713 | 1.6364743 | 0.015873017 | 0.10463602 | 1 | 4076 | tags=32%, list=17%, si | EN_3 |
| GOBP_POSITIVE_REGULA | GOBP_POSITIVE_REGULATION_OF_MYOBLA | 45 | 0.416344 | 1.6363134 | 0.011577424 | 0.10459792 | 1 | 3647 | tags=24%, list=15%, si | EN_3 |
| GOBP_METANEPHROS_D | GOBP_METANEPHROS_DEVELOPMENT | 82 | 0.361373 | 1.635058 | 0.009485095 | 0.10535344 | 1 | 3285 | tags=24%, list=14%, si | EN_3 |
| GOBP_TOR_SIGNALING | GOBP_TOR_SIGNALING | 165 | 0.323772 | 1.6325649 | 0.0023781213 | 0.107234396 | 1 | 4537 | tags=28%, list=19%, si | EN_3 |
| GOBP_REGULATION_OF_ | GOBP_REGULATION_OF_CELL_SUBSTRATE_ | 69 | 0.378353 | 1.6325256 | 0.011283498 | 0.1070619 | 1 | 6942 | tags=46%, list=29%, si | EN_3 |
| GOBP_NEUTRAL_LIPID_C/ | GOBP_NEUTRAL_LIPID_CATABOLIC_PROCES | 34 | 0.433054 | 1.6323162 | 0.02560241 | 0.1069992 | 1 | 964 | tags=15%, list=4%, sig | EN_3 |
| NEUROTROPHIN_SIGNALI | NEUROTROPHIN_SIGNALING_PATHWAY | 116 | 0.344683 | 1.6321682 | 0.0012135922 | 0.1069065 | 1 | 7876 | tags=50%, list=33%, si | EN_3 |
| GOBP_NEGATIVE_REGUL/ | GOBP_NEGATIVE_REGULATION_OF_MYELO | 57 | 0.386382 | 1.6321206 | 0.015895953 | 0.106738195 | 1 | 4975 | tags=37%, list=21%, si | EN_3 |
| GOBP_MITOCHONDRIAL_ | GOBP_MITOCHONDRIAL_OUTER_MEMBRAI | 34 | 0.433939 | 1.6316907 | 0.0116959065 | 0.10688968 | 1 | 3635 | tags=35%, list=15%, si | EN_3 |
| GOBP_POSITIVE_REGULA | GOBP_POSITIVE_REGULATION_OF_CHEMOT | 150 | 0.331383 | 1.6314052 | 0.0024213076 | 0.10692027 | 1 | 3851 | tags=25%, list=16%, si | EN_3 |
| GLIOMA | GLIOMA | 71 | 0.372032 | 1.630533 | 0.0069156294 | 0.107461534 | 1 | 7044 | tags=41%, list=30%, si | EN_3 |
| GOBP_ENDOPLASMIC_RE | GOBP_ENDOPLASMIC_RETICULUM_TUBULA | 21 | 0.487466 | 1.6303034 | 0.025641026 | 0.10743215 | 1 | 7949 | tags=57%, list=34%, si | EN_3 |
| GOBP_POSITIVE_REGULA | GOBP_POSITIVE_REGULATION_OF_ESTABLIS | 345 | 0.298609 | 1.6299337 | 0 | 0.10754793 | 1 | 5134 | tags=27%, list=22%, si | EN_3 |
| GOBP_POSITIVE_REGULA | GOBP_POSITIVE_REGULATION_OF_CELL_GR | 185 | 0.322412 | 1.6298131 | 0.0034324944 | 0.10742484 | 1 | 5919 | tags=34%, list=25%, si | EN_3 |
| GOBP_POSITIVE_REGULA | GOBP_POSITIVE_REGULATION_OF_CELL_M/ | 61 | 0.382733 | 1.6298013 | 0.012379643 | 0.107220374 | 1 | 6236 | tags=38%, list=26%, si | EN_3 |
| GOBP_INTERFERON_MED | GOBP_INTERFERON_MEDIATED_SIGNALING | 76 | 0.369675 | 1.62914 | 0.009320905 | 0.10744816 | 1 | 5423 | tags=36%, list=23%, si | EN_3 |
| GOBP_REGULATION_OF_ | GOBP_REGULATION_OF_ACTIN_FILAMENT_ | 158 | 0.326765 | 1.627917 | 0.0011862396 | 0.10829423 | 1 | 5011 | tags=28%, list=21%, si | EN_3 |
| GOBP_NEGATIVE_REGUL/ | GOBP_NEGATIVE_REGULATION_OF_ENDOP | 18 | 0.507982 | 1.6271237 | 0.025764896 | 0.10873632 | 1 | 1822 | tags=17%, list=8%, sig | EN_3 |
| GOBP_VIRAL_GENOME_R | GOBP_VIRAL_GENOME_REPLICATION | 110 | 0.348001 | 1.6268451 | 0.0077021825 | 0.10878167 | 1 | 4129 | tags=24%, list=17%, si | EN_3 |
| GOBP_NEGATIVE_REGUL/ | GOBP_NEGATIVE_REGULATION_OF_POST_T | 94 | 0.351468 | 1.6256045 | 0.003821656 | 0.109596185 | 1 | 4983 | tags=32%, list=21%, si | EN_3 |
| GOBP_REGULATION_OF_ | GOBP_REGULATION_OF_MITOTIC_CELL_CY( | 331 | 0.301505 | 1.6254678 | 0 | 0.10948914 | 1 | 5457 | tags=31%, list=23%, si | EN_3 |
| GOBP_REGULATION_OF_ | GOBP_REGULATION_OF_ACTIN_NUCLEATIO | 35 | 0.429358 | 1.6250755 | 0.013931888 | 0.109603815 | 1 | 5091 | tags=43%, list=22%, si | EN_3 |
| GOBP_TRNA_MODIFICAT | GOBP_TRNA_MODIFICATION | 84 | 0.364897 | 1.624086 | 0.009090909 | 0.11019 | 1 | 3826 | tags=29%, list=16%, si | EN_3 |
| GOBP_SPLEEN_DEVELOPI | GOBP_SPLEEN_DEVELOPMENT | 46 | 0.403636 | 1.6239712 | 0.013043478 | 0.110073015 | 1 | 4743 | tags=28%, list=20%, si | EN_3 |
| GOBP_POSITIVE_REGULA | GOBP_POSITIVE_REGULATION_OF_BINDING | 147 | 0.330713 | 1.6235754 | 0 | 0.110238835 | 1 | 6068 | tags=33%, list=26%, si | EN_3 |
| GOBP_MITOCHONDRIAL_ | GOBP_MITOCHONDRIAL_FUSION | 36 | 0.423834 | 1.6230543 | 0.020537125 | 0.11045194 | 1 | 3635 | tags=28%, list=15%, si | EN_3 |
| GOBP_PHOSPHATIDYLETH | GOBP_PHOSPHATIDYLETHANOLAMINE_MET | 25 | 0.476464 | 1.6218668 | 0.016025642 | 0.111214094 | 1 | 2822 | tags=36%, list=12%, si | EN_3 |
| GOBP_GLUTAMATE_RECE | GOBP_GLUTAMATE_RECEPTOR_SIGNALING_ | 49 | 0.403788 | 1.6216543 | 0.017366135 | 0.11118277 | 1 | 2696 | tags=20%, list=11%, si | EN_3 |
| GOBP_REGULATED_EXOC | GOBP_REGULATED_EXOCYTOSIS | 243 | 0.308264 | 1.6213579 | 0.002224694 | 0.11118322 | 1 | 4182 | tags=24%, list=18%, si | EN_3 |
| GOBP_MEMBRANE_RAFT | GOBP_MEMBRANE_RAFT_ORGANIZATION | 24 | 0.47559 | 1.6205491 | 0.019417476 | 0.11165591 | 1 | 3068 | tags=29%, list=13%, si | EN_3 |
| GOBP_POSITIVE_REGULA | GOBP_POSITIVE_REGULATION_OF_NON_CA | 64 | 0.380396 | 1.6184673 | 0.015027323 | 0.11321627 | 1 | 5223 | tags=39%, list=22%, si | EN_3 |
| GOBP_SNRNA_3_END_PF | GOBP_SNRNA_3_END_PROCESSING | 20 | 0.489691 | 1.6162109 | 0.033167496 | 0.11498242 | 1 | 2267 | tags=30%, list=10%, si | EN_3 |
| GOBP_REGULATION_OF_ | GOBP_REGULATION_OF_VASCULATURE_DE\ | 295 | 0.304264 | 1.6156847 | 0 | 0.115140736 | 1 | 4168 | tags=23%, list=18%, si | EN_3 |
| GOBP_REGULATION_OF_ | GOBP_REGULATION_OF_CELL_GROWTH | 415 | 0.2948 | 1.6146482 | 0 | 0.11579434 | 1 | 5849 | tags=32%, list=25%, si | EN_3 |
| GOBP_MITOCHONDRIAL_ | GOBP_MITOCHONDRIAL_DNA_METABOLIC_ | 26 | 0.469828 | 1.6142157 | 0.02283105 | 0.11596633 | 1 | 4693 | tags=38%, list=20%, si | EN_3 |
| GOBP_INTRACILIARY_ANT | GOBP_INTRACILIARY_ANTEROGRADE_TRAN | 15 | 0.550678 | 1.6141812 | 0.04310345 | 0.115785256 | 1 | 9346 | tags=80%, list=40%, si | EN_3 |
| GOBP_PROTEIN_COMPLE | GOBP_PROTEIN_COMPLEX_OLIGOMERIZATI | 249 | 0.305701 | 1.6139884 | 0.0022075055 | 0.115761556 | 1 | 5418 | tags=33%, list=23%, si | EN_3 |
| GOBP_TERPENOID_BIOSY | GOBP_TERPENOID_BIOSYNTHETIC_PROCESS | 20 | 0.493336 | 1.6138527 | 0.033444814 | 0.11568224 | 1 | 3856 | tags=40%, list=16%, si | EN_3 |
| GOBP_POSITIVE_REGULA | GOBP_POSITIVE_REGULATION_OF_MAST_C | 21 | 0.485899 | 1.6136249 | 0.02631579 | 0.115670174 | 1 | 2841 | tags=33%, list=12%, si | EN_3 |
| GOBP_CEREBELLAR_CORT | GOBP_CEREBELLAR_CORTEX_FORMATION | 28 | 0.452568 | 1.6134604 | 0.033639144 | 0.11558681 | 1 | 7896 | tags=57%, list=33%, si | EN_3 |
| GOBP_POSITIVE_REGULA | GOBP_POSITIVE_REGULATION_OF_PROTEIN | 288 | 0.304049 | 1.6126873 | 0 | 0.116052456 | 1 | 5134 | tags=27%, list=22%, si | EN_3 |
| GOBP_TISSUE_REGENERA | GOBP_TISSUE_REGENERATION | 68 | 0.369356 | 1.6126307 | 0.014666666 | 0.11591583 | 1 | 5371 | tags=34%, list=23%, si | EN_3 |
| GOBP_NEGATIVE_REGUL/ | GOBP_NEGATIVE_REGULATION_OF_OXIDOF | 17 | 0.523393 | 1.611503 | 0.025559105 | 0.11664916 | 1 | 1536 | tags=24%, list=7%, sig | EN_3 |
| GOBP_VIRAL_GENE_EXPF | GOBP_VIRAL_GENE_EXPRESSION | 75 | 0.36532 | 1.6110667 | 0.010989011 | 0.11682074 | 1 | 4177 | tags=27%, list=18%, si | EN_3 |
| GOBP_ANIMAL_ORGAN_ | GOBP_ANIMAL_ORGAN_MATURATION | 33 | 0.436348 | 1.6110486 | 0.023659306 | 0.11662339 | 1 | 1506 | tags=18%, list=6%, sig | EN_3 |
| GOBP_PYRIMIDINE_RIBO | GOBP_PYRIMIDINE_RIBONUCLEOTIDE_MET/ | 26 | 0.456089 | 1.6109593 | 0.020866774 | 0.11650415 | 1 | 4815 | tags=35%, list=20%, si | EN_3 |
| HISTIDINE_METABOLISM | HISTIDINE_METABOLISM | 21 | 0.490083 | 1.6100415 | 0.039123632 | 0.117079765 | 1 | 2280 | tags=33%, list=10%, si | EN_3 |
| GOBP_POSITIVE_REGULA | GOBP_POSITIVE_REGULATION_OF_MACRO/ | 81 | 0.354073 | 1.6100025 | 0.010610079 | 0.1168897 | 1 | 8015 | tags=54%, list=34%, si | EN_3 |
| GOBP_POSITIVE_REGULA | GOBP_POSITIVE_REGULATION_OF_TYPE_2_ | 18 | 0.513059 | 1.6096935 | 0.0397351 | 0.11694148 | 1 | 4572 | tags=50%, list=19%, si | EN_3 |
| SMALL_CELL_LUNG_CAN( | SMALL_CELL_LUNG_CANCER | 91 | 0.345244 | 1.608799 | 0.009370817 | 0.11755489 | 1 | 6024 | tags=34%, list=26%, si | EN_3 |
| GOBP_ARP2_3_COMPLEX | GOBP_ARP2_3_COMPLEX_MEDIATED_ACTII | 41 | 0.413712 | 1.6086426 | 0.019145804 | 0.11746741 | 1 | 5091 | tags=37%, list=22%, si | EN_3 |
| GOBP_ESTABLISHMENT_( | GOBP_ESTABLISHMENT_OF_MITOCHONDRI( | 25 | 0.475943 | 1.6085004 | 0.030015798 | 0.11739599 | 1 | 6290 | tags=48%, list=27%, si | EN_3 |

| | | | | | | | | | | |
|---|---|---|---|---|---|---|---|---|---|---|
| GOBP_REGULATION_OF_ | GOBP_REGULATION_OF_PROTEIN_DEACETY | 33 | 0.433956 | 1.6083262 | 0.031296574 | 0.11734849 | 1 | 5253 | tags=33%, list=22%, si | EN_3 |
| GOBP_REGULATION_OF_ | GOBP_REGULATION_OF_MEMBRANE_PERM | 70 | 0.37361 | 1.607739 | 0.011095701 | 0.1176721 | 1 | 3635 | tags=29%, list=15%, si | EN_3 |
| PYRIMIDINE_METABOLISI | PYRIMIDINE_METABOLISM | 53 | 0.390497 | 1.6070281 | 0.015625 | 0.11815442 | 1 | 3666 | tags=28%, list=16%, si | EN_3 |
| GOBP_INFLAMMASOME_ | GOBP_INFLAMMASOME_MEDIATED_SIGNA | 46 | 0.405555 | 1.6070126 | 0.023977432 | 0.11795434 | 1 | 3292 | tags=33%, list=14%, si | EN_3 |
| GOBP_VESICLE_CARGO_L | GOBP_VESICLE_CARGO_LOADING | 27 | 0.453929 | 1.6068578 | 0.028301887 | 0.1178563 | 1 | 6861 | tags=44%, list=29%, si | EN_3 |
| GOBP_REGULATION_OF_ | GOBP_REGULATION_OF_MEMBRANE_LIPID_ | 19 | 0.503933 | 1.6062313 | 0.033167496 | 0.118163794 | 1 | 6470 | tags=47%, list=27%, si | EN_3 |
| GOBP_MACROPHAGE_MI | GOBP_MACROPHAGE_MIGRATION | 65 | 0.376084 | 1.6061374 | 0.017333332 | 0.11803804 | 1 | 2230 | tags=22%, list=9%, sig | EN_3 |
| GOBP_ACTIN_NUCLEATIC | GOBP_ACTIN_NUCLEATION | 58 | 0.385592 | 1.6052675 | 0.014784946 | 0.118568785 | 1 | 5091 | tags=38%, list=22%, si | EN_3 |
| GOBP_GLYCOLIPID_BIOSY | GOBP_GLYCOLIPID_BIOSYNTHETIC_PROCESS | 66 | 0.367388 | 1.604375 | 0.012605042 | 0.11912346 | 1 | 3913 | tags=29%, list=17%, si | EN_3 |
| GOBP_REGULATION_OF_ | GOBP_REGULATION_OF_MYOBLAST_DIFFER | 81 | 0.354435 | 1.6039196 | 0.014608234 | 0.11928943 | 1 | 3647 | tags=23%, list=15%, si | EN_3 |
| GOBP_NEGATIVE_REGUL | GOBP_NEGATIVE_REGULATION_OF_INTRAC | 54 | 0.381121 | 1.6033914 | 0.012838801 | 0.119527854 | 1 | 5843 | tags=39%, list=25%, si | EN_3 |
| GOBP_POSITIVE_REGULA | GOBP_POSITIVE_REGULATION_OF_VASCUL | 44 | 0.407126 | 1.6032618 | 0.020679468 | 0.119426414 | 1 | 4129 | tags=32%, list=17%, si | EN_3 |
| GOBP_RESPONSE_TO_TO | GOBP_RESPONSE_TO_TOPOLOGICALLY_INC | 127 | 0.33165 | 1.6025071 | 0.0037974683 | 0.119863495 | 1 | 7785 | tags=50%, list=33%, si | EN_3 |
| GOBP_POSITIVE_REGULA | GOBP_POSITIVE_REGULATION_OF_OLIGODE | 28 | 0.452194 | 1.6017098 | 0.026153846 | 0.12036302 | 1 | 4541 | tags=39%, list=19%, si | EN_3 |
| GOBP_CELLULAR_RESPON | GOBP_CELLULAR_RESPONSE_TO_TOPOLOGI | 95 | 0.348492 | 1.6009319 | 0.007915568 | 0.120860055 | 1 | 7785 | tags=52%, list=33%, si | EN_3 |
| GOBP_ESTABLISHMENT_( | GOBP_ESTABLISHMENT_OF_PROTEIN_LOCA | 262 | 0.302177 | 1.6008141 | 0.002247191 | 0.12075053 | 1 | 5021 | tags=27%, list=21%, si | EN_3 |
| GOBP_NEGATIVE_REGUL | GOBP_NEGATIVE_REGULATION_OF_RESPON | 40 | 0.41873 | 1.600776 | 0.010852713 | 0.12058016 | 1 | 7975 | tags=50%, list=34%, si | EN_3 |
| GOBP_NEGATIVE_REGUL | GOBP_NEGATIVE_REGULATION_OF_INTERLI | 18 | 0.505877 | 1.6005305 | 0.037037037 | 0.1205871 | 1 | 5389 | tags=56%, list=23%, si | EN_3 |
| GOBP_STEROL_HOMEOST | GOBP_STEROL_HOMEOSTASIS | 94 | 0.345926 | 1.6002779 | 0.011734028 | 0.12060495 | 1 | 4078 | tags=27%, list=17%, si | EN_3 |
| GOBP_RUFFLE_ORGANIZ | GOBP_RUFFLE_ORGANIZATION | 54 | 0.382532 | 1.5993346 | 0.014556041 | 0.12119357 | 1 | 3470 | tags=30%, list=15%, si | EN_3 |
| GOBP_DIOL_BIOSYNTHET | GOBP_DIOL_BIOSYNTHETIC_PROCESS | 21 | 0.496603 | 1.5992727 | 0.030351438 | 0.12104692 | 1 | 1786 | tags=24%, list=8%, sig | EN_3 |
| GOBP_CORTICAL_ACTIN_ | GOBP_CORTICAL_ACTIN_CYTOSKELETON_O | 42 | 0.41152 | 1.5991092 | 0.01754386 | 0.120960966 | 1 | 4774 | tags=31%, list=20%, si | EN_3 |
| GOBP_RESPONSE_TO_CO | GOBP_RESPONSE_TO_COPPER_ION | 23 | 0.466942 | 1.5987825 | 0.022617124 | 0.12105326 | 1 | 6168 | tags=57%, list=26%, si | EN_3 |
| GOBP_REGULATION_OF_ | GOBP_REGULATION_OF_PROTEIN_LOCALIZ | 158 | 0.318484 | 1.5979576 | 0.0058343057 | 0.12157443 | 1 | 6046 | tags=32%, list=26%, si | EN_3 |
| GOBP_RECEPTOR_RECYCI | GOBP_RECEPTOR_RECYCLING | 54 | 0.387839 | 1.5975584 | 0.018156424 | 0.12174443 | 1 | 6105 | tags=41%, list=26%, si | EN_3 |
| GOBP_GLYCEROPHOSPHO | GOBP_GLYCEROPHOSPHOLIPID_METABOLIC | 267 | 0.301426 | 1.597447 | 0 | 0.121620394 | 1 | 3022 | tags=19%, list=13%, si | EN_3 |
| GOBP_MEMBRANE_LIPID | GOBP_MEMBRANE_LIPID_BIOSYNTHETIC_PI | 129 | 0.330749 | 1.596835 | 0.0049566296 | 0.12191864 | 1 | 6672 | tags=41%, list=28%, si | EN_3 |
| GOBP_PROTEIN_TARGETI | GOBP_PROTEIN_TARGETING | 285 | 0.300015 | 1.5967518 | 0.0032292788 | 0.12179513 | 1 | 7210 | tags=40%, list=31%, si | EN_3 |
| GOBP_RESPONSE_TO_OX | GOBP_RESPONSE_TO_OXYGEN_RADICAL | 24 | 0.467291 | 1.5959014 | 0.03267974 | 0.12229844 | 1 | 2571 | tags=29%, list=11%, si | EN_3 |
| GOBP_POSITIVE_REGULA | GOBP_POSITIVE_REGULATION_OF_MYELOID | 38 | 0.427999 | 1.5955232 | 0.031866465 | 0.122426435 | 1 | 2936 | tags=24%, list=12%, si | EN_3 |
| GOBP_NEGATIVE_REGUL | GOBP_NEGATIVE_REGULATION_OF_CELL_SI | 59 | 0.377031 | 1.5949882 | 0.016304348 | 0.12268604 | 1 | 5011 | tags=36%, list=21%, si | EN_3 |
| GOBP_COPII_COATED_VE | GOBP_COPII_COATED_VESICLE_CARGO_LOA | 17 | 0.507358 | 1.5947684 | 0.024590164 | 0.122664765 | 1 | 4795 | tags=35%, list=20%, si | EN_3 |
| GOBP_CELL_CYCLE_G1_S | GOBP_CELL_CYCLE_G1_S_PHASE_TRANSITIC | 236 | 0.305381 | 1.5947629 | 0.0010940919 | 0.122452535 | 1 | 4998 | tags=28%, list=21%, si | EN_3 |
| GOBP_REGULATION_OF_ | GOBP_REGULATION_OF_VIRAL_PROCESS | 167 | 0.316948 | 1.594657 | 0.007025761 | 0.12233325 | 1 | 2250 | tags=18%, list=10%, si | EN_3 |
| GOBP_STEROID_METABO | GOBP_STEROID_METABOLIC_PROCESS | 287 | 0.299997 | 1.5946001 | 0 | 0.12218364 | 1 | 4197 | tags=27%, list=18%, si | EN_3 |
| GOBP_POSITIVE_REGULA | GOBP_POSITIVE_REGULATION_OF_MEMBRA | 16 | 0.515995 | 1.594089 | 0.032367975 | 0.12243295 | 1 | 6236 | tags=75%, list=26%, si | EN_3 |
| GOBP_RESPONSE_TO_ETI | GOBP_RESPONSE_TO_ETHANOL | 60 | 0.37511 | 1.593715 | 0.015006822 | 0.12253477 | 1 | 2936 | tags=22%, list=12%, si | EN_3 |
| GOBP_VESICLE_DOCKING | GOBP_VESICLE_DOCKING | 55 | 0.383754 | 1.5937082 | 0.013947002 | 0.12233068 | 1 | 3617 | tags=25%, list=15%, si | EN_3 |
| GOBP_REGULATION_OF_ | GOBP_REGULATION_OF_NEURON_APOPTOT | 313 | 0.294451 | 1.5936568 | 0.0010729614 | 0.12218026 | 1 | 6290 | tags=33%, list=27%, si | EN_3 |
| GOBP_CELLULAR_KETONI | GOBP_CELLULAR_KETONE_METABOLIC_PRC | 222 | 0.307863 | 1.593526 | 0.0022624435 | 0.122075066 | 1 | 4734 | tags=29%, list=20%, si | EN_3 |
| GOBP_PROTEIN_LOCALIZ | GOBP_PROTEIN_LOCALIZATION_TO_PLASM | 296 | 0.296378 | 1.5920384 | 0 | 0.12318311 | 1 | 5012 | tags=27%, list=21%, si | EN_3 |
| GOBP_NEGATIVE_REGUL | GOBP_NEGATIVE_REGULATION_OF_TELOM | 27 | 0.461113 | 1.5919883 | 0.024038462 | 0.12302366 | 1 | 1956 | tags=19%, list=8%, sig | EN_3 |
| ADIPOCYTOKINE_SIGNALI | ADIPOCYTOKINE_SIGNALING_PATHWAY | 68 | 0.361818 | 1.5910355 | 0.024657534 | 0.12361605 | 1 | 5000 | tags=29%, list=21%, si | EN_3 |
| GOBP_ANTIMICROBIAL_H | GOBP_ANTIMICROBIAL_HUMORAL_RESPON | 98 | 0.337227 | 1.5906261 | 0.0115681235 | 0.123767205 | 1 | 3182 | tags=26%, list=13%, si | EN_3 |
| GOBP_POSITIVE_REGULA | GOBP_POSITIVE_REGULATION_OF_MEMBRA | 39 | 0.409546 | 1.5902331 | 0.01627219 | 0.12389738 | 1 | 3635 | tags=33%, list=15%, si | EN_3 |
| GOBP_NEGATIVE_REGUL | GOBP_NEGATIVE_REGULATION_OF_ACTIN_ | 64 | 0.371713 | 1.5900383 | 0.016483517 | 0.12386226 | 1 | 5762 | tags=36%, list=24%, si | EN_3 |
| GOBP_SYNAPTIC_VESICLE | GOBP_SYNAPTIC_VESICLE_LUMEN_ACIDIFIC | 19 | 0.488549 | 1.5895872 | 0.041254126 | 0.12409866 | 1 | 5923 | tags=42%, list=25%, si | EN_3 |
| GOBP_REGULATION_OF_ | GOBP_REGULATION_OF_AUTOPHAGY | 295 | 0.298297 | 1.5895466 | 0 | 0.12391745 | 1 | 5484 | tags=28%, list=23%, si | EN_3 |
| GOBP_HYDROGEN_PERO | GOBP_HYDROGEN_PEROXIDE_CATABOLIC_I | 22 | 0.478124 | 1.5869694 | 0.034201954 | 0.12607728 | 1 | 3475 | tags=32%, list=15%, si | EN_3 |
| GOBP_POSITIVE_REGULA | GOBP_POSITIVE_REGULATION_OF_CELL_CY | 61 | 0.370308 | 1.5860107 | 0.012311902 | 0.1267807 | 1 | 3321 | tags=26%, list=14%, si | EN_3 |
| GOBP_LIVER_MORPHOGI | GOBP_LIVER_MORPHOGENESIS | 36 | 0.421524 | 1.5855178 | 0.02587519 | 0.12702014 | 1 | 4386 | tags=33%, list=19%, si | EN_3 |
| GOBP_MELANOSOME_AS | GOBP_MELANOSOME_ASSEMBLY | 19 | 0.500891 | 1.5845965 | 0.043269232 | 0.12763053 | 1 | 2600 | tags=26%, list=11%, si | EN_3 |
| GOBP_VIRAL_LIFE_CYCLE | GOBP_VIRAL_LIFE_CYCLE | 230 | 0.304412 | 1.5840981 | 0 | 0.12796275 | 1 | 4771 | tags=27%, list=20%, si | EN_3 |
| GOBP_REGULATION_OF_ | GOBP_REGULATION_OF_STORE_OPERATED_ | 18 | 0.493122 | 1.5838125 | 0.02617801 | 0.12802032 | 1 | 7796 | tags=67%, list=33%, si | EN_3 |
| GOBP_GLYCEROPHOSPHO | GOBP_GLYCEROPHOSPHOLIPID_BIOSYNTHE | 160 | 0.32034 | 1.5827013 | 0.0011614402 | 0.12882677 | 1 | 3624 | tags=23%, list=15%, si | EN_3 |
| GOBP_PIGMENT_METABO | GOBP_PIGMENT_METABOLIC_PROCESS | 65 | 0.370058 | 1.5825585 | 0.023160763 | 0.12875642 | 1 | 2261 | tags=22%, list=10%, si | EN_3 |
| GOBP_POSITIVE_REGULA | GOBP_POSITIVE_REGULATION_OF_TELOME | 31 | 0.440077 | 1.5823057 | 0.02021773 | 0.12875175 | 1 | 6373 | tags=45%, list=27%, si | EN_3 |
| GOBP_SUPEROXIDE_MET | GOBP_SUPEROXIDE_METABOLIC_PROCESS | 68 | 0.360424 | 1.581856 | 0.012379643 | 0.12894802 | 1 | 3879 | tags=28%, list=16%, si | EN_3 |
| GOBP_LYTIC_VACUOLE_C | GOBP_LYTIC_VACUOLE_ORGANIZATION | 95 | 0.343163 | 1.5814365 | 0.010322581 | 0.12919822 | 1 | 6711 | tags=42%, list=28%, si | EN_3 |
| GOBP_POSITIVE_REGULA | GOBP_POSITIVE_REGULATION_OF_MEMBRA | 17 | 0.516879 | 1.580967 | 0.034188036 | 0.12946199 | 1 | 3926 | tags=41%, list=17%, si | EN_3 |
| GOBP_NEGATIVE_REGUL | GOBP_NEGATIVE_REGULATION_OF_MONOI | 28 | 0.438592 | 1.5808594 | 0.032786883 | 0.12932484 | 1 | 3321 | tags=29%, list=14%, si | EN_3 |
| GOBP_REGULATION_OF_ | GOBP_REGULATION_OF_MUSCLE_CELL_DIF | 154 | 0.312495 | 1.5807217 | 0.0011792453 | 0.12925442 | 1 | 3586 | tags=22%, list=15%, si | EN_3 |
| GOBP_REGULATION_OF_ | GOBP_REGULATION_OF_NUCLEOCYTOPLAS | 130 | 0.331311 | 1.580504 | 0.010909091 | 0.12928604 | 1 | 5363 | tags=32%, list=23%, si | EN_3 |
| GOBP_EMBRYONIC_CAM | GOBP_EMBRYONIC_CAMERA_TYPE_EYE_DE | 40 | 0.407884 | 1.5801786 | 0.022955524 | 0.12937757 | 1 | 3856 | tags=33%, list=16%, si | EN_3 |
| GOBP_MAST_CELL_ACTIV | GOBP_MAST_CELL_ACTIVATION | 77 | 0.351707 | 1.5795437 | 0.018396847 | 0.12972589 | 1 | 3402 | tags=26%, list=14%, si | EN_3 |
| GOBP_LIPID_HOMEOSTA | GOBP_LIPID_HOMEOSTASIS | 167 | 0.310034 | 1.5789222 | 0.0035211267 | 0.13005987 | 1 | 4537 | tags=26%, list=19%, si | EN_3 |
| GOBP_CELLULAR_ALDEHY | GOBP_CELLULAR_ALDEHYDE_METABOLIC_P | 74 | 0.359871 | 1.5772971 | 0.020297699 | 0.13130012 | 1 | 4300 | tags=32%, list=18%, si | EN_3 |
| GOBP_PROTEIN_TRANSM | GOBP_PROTEIN_TRANSMEMBRANE_TRANS | 67 | 0.361638 | 1.5770831 | 0.019178081 | 0.13132046 | 1 | 6804 | tags=42%, list=29%, si | EN_3 |
| GOBP_NEGATIVE_REGUL | GOBP_NEGATIVE_REGULATION_OF_TRANSI | 222 | 0.302541 | 1.5766547 | 0.0022779044 | 0.1314954 | 1 | 6110 | tags=36%, list=26%, si | EN_3 |
| GOBP_REGULATION_OF_ | GOBP_REGULATION_OF_GLUCOSE_TRANSM | 84 | 0.349461 | 1.5764222 | 0.017994858 | 0.13151097 | 1 | 4986 | tags=31%, list=21%, si | EN_3 |
| GOBP_REGULATION_OF_ | GOBP_REGULATION_OF_NUCLEOTIDE_MET | 96 | 0.342034 | 1.5762779 | 0.010126582 | 0.13143253 | 1 | 6348 | tags=38%, list=27%, si | EN_3 |
| GOBP_INDOLE_CONTAIN | GOBP_INDOLE_CONTAINING_COMPOUND_ | 25 | 0.450373 | 1.5762182 | 0.035603717 | 0.13126372 | 1 | 6563 | tags=56%, list=28%, si | EN_3 |
| GOBP_POSITIVE_REGULA | GOBP_POSITIVE_REGULATION_OF_POST_TF | 135 | 0.320938 | 1.5746449 | 0.0062578223 | 0.13252828 | 1 | 4934 | tags=27%, list=21%, si | EN_3 |
| GOBP_GLIAL_CELL_APOP | GOBP_GLIAL_CELL_APOPTOTIC_PROCESS | 20 | 0.482606 | 1.5746051 | 0.034539472 | 0.1323525 | 1 | 5684 | tags=50%, list=24%, si | EN_3 |
| PYRUVATE_METABOLISM | PYRUVATE_METABOLISM | 37 | 0.407744 | 1.573969 | 0.027859237 | 0.13274977 | 1 | 2381 | tags=24%, list=10%, si | EN_3 |
| BUTANOATE_METABOLIS | BUTANOATE_METABOLISM | 23 | 0.469937 | 1.5732429 | 0.04265403 | 0.13325562 | 1 | 6895 | tags=57%, list=29%, si | EN_3 |
| GOBP_NEGATIVE_REGUL | GOBP_NEGATIVE_REGULATION_OF_CELLUL | 51 | 0.383565 | 1.5732154 | 0.024926687 | 0.13306056 | 1 | 4228 | tags=29%, list=18%, si | EN_3 |
| GOBP_POSITIVE_REGULA | GOBP_POSITIVE_REGULATION_OF_MITOCH | 22 | 0.461387 | 1.5730363 | 0.044728436 | 0.1330279 | 1 | 7737 | tags=64%, list=33%, si | EN_3 |
| GOBP_REGULATION_OF_ | GOBP_REGULATION_OF_CELLULAR_KETONE | 154 | 0.311752 | 1.5728757 | 0.005966587 | 0.13297226 | 1 | 4300 | tags=27%, list=18%, si | EN_3 |
| GOBP_DENDRITIC_CELL_( | GOBP_DENDRITIC_CELL_CYTOKINE_PRODU( | 16 | 0.516683 | 1.5727235 | 0.045 | 0.1329116 | 1 | 2936 | tags=38%, list=12%, si | EN_3 |
| GOBP_MULTIVESICULAR_ | GOBP_MULTIVESICULAR_BODY_SORTING_P | 34 | 0.426929 | 1.5722753 | 0.02472952 | 0.13310492 | 1 | 4771 | tags=41%, list=20%, si | EN_3 |
| GOBP_REGULATION_OF_ | GOBP_REGULATION_OF_CYTOKINE_PRODU( | 122 | 0.326474 | 1.5705566 | 0.0037453184 | 0.13462324 | 1 | 2936 | tags=22%, list=12%, si | EN_3 |

| | | | | | | | | | | |
|---|---|---|---|---|---|---|---|---|---|---|
| GOBP_POSITIVE_REGULA | GOBP_POSITIVE_REGULATION_OF_PRODUC | 136 | 0.320956 | 1.5704151 | 0.005868545 | 0.13454147 | 1 | 3080 | tags=21%, list=13%, si | EN_3 |
| GOBP_REGULATION_OF_ | GOBP_REGULATION_OF_CELLULAR_CATABO | 391 | 0.288567 | 1.570194 | 0.0021008404 | 0.13455024 | 1 | 5484 | tags=28%, list=23%, si | EN_3 |
| GOBP_REGULATION_OF_ | GOBP_REGULATION_OF_PLASMINOGEN_AC | 17 | 0.503365 | 1.5701172 | 0.034201954 | 0.13441195 | 1 | 3017 | tags=29%, list=13%, si | EN_3 |
| GOBP_INTRACELLULAR_P | GOBP_INTRACELLULAR_PROTEIN_TRANSME | 32 | 0.427789 | 1.5698521 | 0.04718417 | 0.13450375 | 1 | 5269 | tags=34%, list=22%, si | EN_3 |
| GOBP_REGULATION_OF_ | GOBP_REGULATION_OF_TOLL_LIKE_RECEPT | 28 | 0.443042 | 1.5696436 | 0.041411042 | 0.13448405 | 1 | 4750 | tags=36%, list=20%, si | EN_3 |
| GOBP_REGULATION_OF_ | GOBP_REGULATION_OF_REACTIVE_OXYGEN | 160 | 0.312938 | 1.5692574 | 0.008343265 | 0.1346444 | 1 | 5751 | tags=32%, list=24%, si | EN_3 |
| GOBP_MAMMARY_GLAN | GOBP_MAMMARY_GLAND_ALVEOLUS_DEV | 21 | 0.47446 | 1.5691582 | 0.035353534 | 0.13454646 | 1 | 2715 | tags=33%, list=11%, si | EN_3 |
| GOBP_RESPONSE_TO_ST | GOBP_RESPONSE_TO_STEROL_DEPLETION | 15 | 0.51976 | 1.5687898 | 0.04467354 | 0.1346788 | 1 | 2840 | tags=33%, list=12%, si | EN_3 |
| GOBP_NEGATIVE_REGUL | GOBP_NEGATIVE_REGULATION_OF_MRNA_ | 24 | 0.469459 | 1.5672883 | 0.0375 | 0.13595714 | 1 | 7216 | tags=58%, list=31%, si | EN_3 |
| GOBP_REGULATION_OF_ | GOBP_REGULATION_OF_PROTEIN_LOCALIZA | 160 | 0.312504 | 1.5669858 | 0.0036319613 | 0.13601792 | 1 | 4774 | tags=26%, list=20%, si | EN_3 |
| GOBP_REGULATION_OF_ | GOBP_REGULATION_OF_STEROID_BIOSYNT | 81 | 0.350743 | 1.5666918 | 0.018229166 | 0.13612665 | 1 | 4300 | tags=32%, list=18%, si | EN_3 |
| GOBP_PROTEIN_POLYME | GOBP_PROTEIN_POLYMERIZATION | 269 | 0.296547 | 1.566573 | 0 | 0.136014 | 1 | 5823 | tags=31%, list=25%, si | EN_3 |
| GOBP_PROTEIN_DNA_CO | GOBP_PROTEIN_DNA_COMPLEX_ASSEMBLY | 154 | 0.314881 | 1.5664679 | 0.011918951 | 0.13590886 | 1 | 6098 | tags=35%, list=26%, si | EN_3 |
| GOBP_REGULATION_OF_ | GOBP_REGULATION_OF_LEUKOCYTE_MIGRA | 230 | 0.297231 | 1.5663387 | 0.0022421526 | 0.1358134 | 1 | 3851 | tags=23%, list=16%, si | EN_3 |
| PURINE_METABOLISM | PURINE_METABOLISM | 126 | 0.320215 | 1.5656503 | 0.012531328 | 0.1362527 | 1 | 4415 | tags=27%, list=19%, si | EN_3 |
| GOBP_MEMBRANE_DOCI | GOBP_MEMBRANE_DOCKING | 78 | 0.3558 | 1.5652705 | 0.014745308 | 0.13641387 | 1 | 4431 | tags=24%, list=19%, si | EN_3 |
| GOBP_NEGATIVE_REGUL | GOBP_NEGATIVE_REGULATION_OF_SIGNAL | 31 | 0.432333 | 1.564469 | 0.039936103 | 0.13705002 | 1 | 5790 | tags=48%, list=25%, si | EN_3 |
| GOBP_REGULATION_OF_ | GOBP_REGULATION_OF_POSITIVE_CHEMOT | 27 | 0.453032 | 1.5641822 | 0.031007752 | 0.1371464 | 1 | 3345 | tags=30%, list=14%, si | EN_3 |
| GOBP_MITOTIC_G2_DNA | GOBP_MITOTIC_G2_DNA_DAMAGE_CHECKI | 39 | 0.403539 | 1.5641721 | 0.032738097 | 0.13693774 | 1 | 5457 | tags=38%, list=23%, si | EN_3 |
| GOBP_MATURATION_OF_ | GOBP_MATURATION_OF_LSU_RRNA | 25 | 0.454531 | 1.5637671 | 0.037398376 | 0.13716838 | 1 | 1659 | tags=28%, list=7%, sig | EN_3 |
| GOBP_COMPLEMENT_AC | GOBP_COMPLEMENT_ACTIVATION | 50 | 0.385791 | 1.5635706 | 0.021802325 | 0.13712695 | 1 | 3641 | tags=28%, list=15%, si | EN_3 |
| GOBP_MATURATION_OF_ | GOBP_MATURATION_OF_5_8S_RRNA_FROM | 24 | 0.465194 | 1.5635697 | 0.031446543 | 0.13691565 | 1 | 4772 | tags=46%, list=20%, si | EN_3 |
| GOBP_POSITIVE_REGULA | GOBP_POSITIVE_REGULATION_OF_PATTERN | 68 | 0.355044 | 1.5634781 | 0.017173052 | 0.136769 | 1 | 3467 | tags=24%, list=15%, si | EN_3 |
| GOBP_ACYL_COA_META | GOBP_ACYL_COA_METABOLIC_PROCESS | 82 | 0.350829 | 1.563163 | 0.010610079 | 0.13685034 | 1 | 5590 | tags=34%, list=24%, si | EN_3 |
| GOBP_REGULATION_OF_ | GOBP_REGULATION_OF_AUTOPHAGOSOME | 44 | 0.3941 | 1.5630174 | 0.029154519 | 0.136786 | 1 | 5484 | tags=41%, list=23%, si | EN_3 |
| GOBP_MITOTIC_INTRA_S | GOBP_MITOTIC_INTRA_S_DNA_DAMAGE_C | 16 | 0.505104 | 1.562526 | 0.05756579 | 0.13705316 | 1 | 3976 | tags=38%, list=17%, si | EN_3 |
| GOBP_PROTEOGLYCAN_M | GOBP_PROTEOGLYCAN_METABOLIC_PROCE | 84 | 0.347895 | 1.5624956 | 0.012903226 | 0.1368708 | 1 | 5365 | tags=31%, list=23%, si | EN_3 |
| GOBP_POSITIVE_REGULA | GOBP_POSITIVE_REGULATION_OF_ENDOTH | 99 | 0.338506 | 1.5622181 | 0.0127551025 | 0.13695881 | 1 | 3345 | tags=22%, list=14%, si | EN_3 |
| GOBP_DEOXYRIBONUCLE | GOBP_DEOXYRIBONUCLEOTIDE_CATABOLIC | 24 | 0.457989 | 1.5621048 | 0.044481054 | 0.13685142 | 1 | 3914 | tags=33%, list=17%, si | EN_3 |
| GOBP_POSITIVE_REGULA | GOBP_POSITIVE_REGULATION_OF_PROTEIN | 87 | 0.341421 | 1.5619843 | 0.018158237 | 0.13676569 | 1 | 2941 | tags=22%, list=12%, si | EN_3 |
| GOBP_NEGATIVE_REGUL | GOBP_NEGATIVE_REGULATION_OF_PEPTID | 29 | 0.438484 | 1.5618535 | 0.032407407 | 0.13668595 | 1 | 4129 | tags=31%, list=17%, si | EN_3 |
| GOBP_REGULATION_OF_ | GOBP_REGULATION_OF_RUFFLE_ASSEMBLY | 29 | 0.43071 | 1.5616528 | 0.04225352 | 0.13668182 | 1 | 3167 | tags=31%, list=13%, si | EN_3 |
| GOBP_DOUBLE_STRAND_ | GOBP_DOUBLE_STRAND_BREAK_REPAIR | 299 | 0.291836 | 1.5616524 | 0 | 0.13647473 | 1 | 5498 | tags=30%, list=23%, si | EN_3 |
| GOBP_REGULATION_OF_ | GOBP_REGULATION_OF_INTRACELLULAR_T | 278 | 0.291614 | 1.5616388 | 0.0010764262 | 0.13628443 | 1 | 7899 | tags=47%, list=33%, si | EN_3 |
| GOBP_REGULATION_OF_ | GOBP_REGULATION_OF_DNA_REPLICATION | 118 | 0.330882 | 1.5615342 | 0.0061576352 | 0.13620603 | 1 | 5450 | tags=32%, list=23%, si | EN_3 |
| GOBP_REGULATION_OF_ | GOBP_REGULATION_OF_CYTOPLASMIC_TRA | 30 | 0.437461 | 1.5615027 | 0.038880248 | 0.13602735 | 1 | 7722 | tags=63%, list=33%, si | EN_3 |
| GOBP_REGULATION_OF_ | GOBP_REGULATION_OF_TYPE_B_PANCREAT | 16 | 0.513775 | 1.5600991 | 0.05308219 | 0.13714854 | 1 | 2879 | tags=31%, list=12%, si | EN_3 |
| GOBP_SKELETAL_MUSCLI | GOBP_SKELETAL_MUSCLE_TISSUE_REGENER | 43 | 0.395101 | 1.559701 | 0.034139402 | 0.13734767 | 1 | 5785 | tags=33%, list=25%, si | EN_3 |
| GOBP_CHAPERONE_MED | GOBP_CHAPERONE_MEDIATED_PROTEIN_F | 62 | 0.363561 | 1.558688 | 0.01591512 | 0.13813232 | 1 | 4381 | tags=32%, list=19%, si | EN_3 |
| GOBP_SNRNA_TRANSCRI | GOBP_SNRNA_TRANSCRIPTION | 18 | 0.48883 | 1.5584406 | 0.036006548 | 0.1381849 | 1 | 7443 | tags=67%, list=32%, si | EN_3 |
| GOBP_ORGANIC_HYDRO | GOBP_ORGANIC_HYDROXY_COMPOUND_M | 492 | 0.284082 | 1.5581825 | 0 | 0.13822155 | 1 | 4156 | tags=24%, list=18%, si | EN_3 |
| GOBP_DETOXIFICATION | GOBP_DETOXIFICATION | 70 | 0.356466 | 1.5579247 | 0.016371077 | 0.13824382 | 1 | 2682 | tags=26%, list=11%, si | EN_3 |
| GOBP_REGULATION_OF_ | GOBP_REGULATION_OF_VESICLE_SIZE | 21 | 0.473798 | 1.5576876 | 0.046774194 | 0.13826796 | 1 | 3288 | tags=38%, list=14%, si | EN_3 |
| GOBP_CARBOHYDRATE_T | GOBP_CARBOHYDRATE_TRANSPORT | 148 | 0.31545 | 1.5569109 | 0.0058275056 | 0.13880482 | 1 | 4363 | tags=26%, list=18%, si | EN_3 |
| CENTRAL_CARBON_META | CENTRAL_CARBON_METABOLISM_IN_CANC | 61 | 0.365162 | 1.5567168 | 0.018544937 | 0.1387608 | 1 | 2433 | tags=16%, list=10%, si | EN_3 |
| GOBP_POSITIVE_REGULA | GOBP_POSITIVE_REGULATION_OF_PEPTIDA | 161 | 0.3121 | 1.5566283 | 0.009512485 | 0.1386428 | 1 | 3284 | tags=21%, list=14%, si | EN_3 |
| LYSOSOME | LYSOSOME | 118 | 0.327886 | 1.5562264 | 0.008706467 | 0.13882485 | 1 | 7210 | tags=41%, list=31%, si | EN_3 |
| MANNOSE_TYPE_O-GLYC | MANNOSE_TYPE_O-GLYCAN_BIOSYNTHESIS | 21 | 0.47574 | 1.5561147 | 0.03974563 | 0.13872483 | 1 | 2982 | tags=33%, list=13%, si | EN_3 |
| GOBP_NEGATIVE_REGUL | GOBP_NEGATIVE_REGULATION_OF_PROTEI | 431 | 0.28164 | 1.5556691 | 0 | 0.13891993 | 1 | 6110 | tags=32%, list=26%, si | EN_3 |
| GOBP_EOSINOPHIL_MIGR | GOBP_EOSINOPHIL_MIGRATION | 26 | 0.450585 | 1.554308 | 0.052050475 | 0.1400057 | 1 | 3118 | tags=35%, list=13%, si | EN_3 |
| GOBP_POSITIVE_REGULA | GOBP_POSITIVE_REGULATION_OF_CELL_CY | 330 | 0.291073 | 1.554126 | 0.0010706638 | 0.1399725 | 1 | 5011 | tags=28%, list=21%, si | EN_3 |
| GOBP_NEUTRAL_LIPID_B | GOBP_NEUTRAL_LIPID_BIOSYNTHETIC_PRO | 58 | 0.365476 | 1.5531496 | 0.03208556 | 0.14075054 | 1 | 4311 | tags=28%, list=18%, si | EN_3 |
| GOBP_INTRINSIC_APOPT | GOBP_INTRINSIC_APOPTOTIC_SIGNALING_F | 112 | 0.325389 | 1.552944 | 0.012953368 | 0.1407651 | 1 | 5859 | tags=37%, list=25%, si | EN_3 |
| GOBP_ORGANELLE_FUSI | GOBP_ORGANELLE_FUSION | 167 | 0.310051 | 1.5522258 | 0.009345794 | 0.14126828 | 1 | 3797 | tags=23%, list=16%, si | EN_3 |
| GOBP_REGULATION_OF_ | GOBP_REGULATION_OF_VIRAL_GENOME_R | 78 | 0.351865 | 1.5515643 | 0.017639078 | 0.1416795 | 1 | 2793 | tags=19%, list=12%, si | EN_3 |
| GOBP_POSITIVE_REGULA | GOBP_POSITIVE_REGULATION_OF_SIGNAL_ | 32 | 0.424451 | 1.5505607 | 0.036050156 | 0.14246704 | 1 | 7903 | tags=63%, list=33%, si | EN_3 |
| GOBP_REGULATION_OF_ | GOBP_REGULATION_OF_PROTEIN_POLYMEI | 194 | 0.302496 | 1.5503554 | 0.0045714285 | 0.14244305 | 1 | 5467 | tags=28%, list=23%, si | EN_3 |
| GOBP_RESPONSE_TO_M | GOBP_RESPONSE_TO_MOLECULE_OF_BACT | 296 | 0.287318 | 1.5503416 | 0 | 0.14225075 | 1 | 5490 | tags=30%, list=23%, si | EN_3 |
| GOBP_REGULATION_OF_ | GOBP_REGULATION_OF_TYPE_2_IMMUNE_ | 34 | 0.411356 | 1.5501806 | 0.049382716 | 0.14218384 | 1 | 2870 | tags=29%, list=12%, si | EN_3 |
| GOBP_FC_RECEPTOR_ME | GOBP_FC_RECEPTOR_MEDIATED_STIMULAT | 17 | 0.492017 | 1.5499405 | 0.03898305 | 0.14219502 | 1 | 4961 | tags=41%, list=21%, si | EN_3 |
| GOBP_REGULATION_OF_ | GOBP_REGULATION_OF_SYNAPTIC_VESICLE | 19 | 0.49163 | 1.5498179 | 0.049099836 | 0.14210214 | 1 | 5684 | tags=58%, list=24%, si | EN_3 |
| GOBP_AMEBOIDAL_TYPE | GOBP_AMEBOIDAL_TYPE_CELL_MIGRATION | 443 | 0.28292 | 1.5494317 | 0 | 0.14228174 | 1 | 3776 | tags=21%, list=16%, si | EN_3 |
| GOBP_FATTY_ACYL_COA | GOBP_FATTY_ACYL_COA_METABOLIC_PRO | 18 | 0.481656 | 1.5491394 | 0.048309177 | 0.14237317 | 1 | 4311 | tags=50%, list=18%, si | EN_3 |
| GOBP_NEGATIVE_REGUL | GOBP_NEGATIVE_REGULATION_OF_GLUCO | 16 | 0.498041 | 1.549001 | 0.055737704 | 0.1423251 | 1 | 2153 | tags=31%, list=9%, sig | EN_3 |
| NECROPTOSIS | NECROPTOSIS | 138 | 0.317639 | 1.5486449 | 0.013157895 | 0.14243153 | 1 | 5021 | tags=33%, list=21%, si | EN_3 |
| GOBP_PROTEIN_TRANSM | GOBP_PROTEIN_TRANSMEMBRANE_IMPOR | 36 | 0.411409 | 1.5486243 | 0.025525525 | 0.1422585 | 1 | 7450 | tags=53%, list=32%, si | EN_3 |
| GOBP_NEGATIVE_REGUL | GOBP_NEGATIVE_REGULATION_OF_KINASE | 188 | 0.303132 | 1.5483236 | 0.0068965517 | 0.14234796 | 1 | 6110 | tags=36%, list=26%, si | EN_3 |
| GOBP_REGULATION_OF_ | GOBP_REGULATION_OF_LIPID_METABOLIC_ | 343 | 0.287188 | 1.5481641 | 0.0010683761 | 0.14229842 | 1 | 4300 | tags=23%, list=18%, si | EN_3 |
| GOBP_RENAL_SYSTEM_V | GOBP_RENAL_SYSTEM_VASCULATURE_DEV | 29 | 0.427895 | 1.5481515 | 0.04107425 | 0.1421042 | 1 | 2316 | tags=28%, list=10%, si | EN_3 |
| GOBP_NEGATIVE_REGUL | GOBP_NEGATIVE_REGULATION_OF_MYELO | 89 | 0.3404 | 1.5481056 | 0.021333333 | 0.1419411 | 1 | 4975 | tags=30%, list=21%, si | EN_3 |
| GOBP_REGULATION_OF_ | GOBP_REGULATION_OF_SMOOTH_MUSCLE | 67 | 0.359068 | 1.5477598 | 0.021008404 | 0.14209305 | 1 | 4214 | tags=28%, list=18%, si | EN_3 |
| GOBP_MICROTUBULE_BA | GOBP_MICROTUBULE_BASED_TRANSPORT | 217 | 0.296498 | 1.5472848 | 0.0011223345 | 0.1423691 | 1 | 6418 | tags=33%, list=27%, si | EN_3 |
| GOBP_PROTEIN_LOCALIZ | GOBP_PROTEIN_LOCALIZATION_TO_NUCLEU | 326 | 0.284258 | 1.5471457 | 0 | 0.14229448 | 1 | 6121 | tags=31%, list=26%, si | EN_3 |
| GOBP_SNO_S_RNA_MET | GOBP_SNO_S_RNA_METABOLIC_PROCESS | 17 | 0.499556 | 1.546069 | 0.03752039 | 0.14308381 | 1 | 4917 | tags=47%, list=21%, si | EN_3 |
| GOBP_POSITIVE_REGULA | GOBP_POSITIVE_REGULATION_OF_EPITHELI | 158 | 0.310599 | 1.5454369 | 0.0035714286 | 0.14346041 | 1 | 3864 | tags=21%, list=16%, si | EN_3 |
| HUMAN_CYTOMEGALOVI | HUMAN_CYTOMEGALOVIRUS_INFECTION | 216 | 0.296318 | 1.5453454 | 0.007856341 | 0.1433476 | 1 | 7044 | tags=39%, list=30%, si | EN_3 |
| GOBP_NEGATIVE_REGUL | GOBP_NEGATIVE_REGULATION_OF_MITOTI | 230 | 0.292671 | 1.5452626 | 0.005605381 | 0.14322847 | 1 | 5457 | tags=30%, list=23%, si | EN_3 |
| GOBP_CELL_CYCLE_G2_N | GOBP_CELL_CYCLE_G2_M_PHASE_TRANSITI | 152 | 0.313977 | 1.5450518 | 0.0058962265 | 0.14327788 | 1 | 5457 | tags=31%, list=23%, si | EN_3 |
| GOBP_REGULATION_OF_ | GOBP_REGULATION_OF_MAST_CELL_DEGRA | 33 | 0.415049 | 1.5449252 | 0.05279503 | 0.14320281 | 1 | 3363 | tags=27%, list=14%, si | EN_3 |
| GOBP_POSITIVE_REGULA | GOBP_POSITIVE_REGULATION_OF_SYNCYTI | 25 | 0.445223 | 1.5448732 | 0.04784689 | 0.14304721 | 1 | 3525 | tags=36%, list=15%, si | EN_3 |
| GOBP_CRISTAE_FORMAT | GOBP_CRISTAE_FORMATION | 17 | 0.496238 | 1.5434799 | 0.043333333 | 0.14430705 | 1 | 2859 | tags=24%, list=12%, si | EN_3 |
| GOBP_REGULATION_OF_ | GOBP_REGULATION_OF_MITOCHONDRIAL_ | 87 | 0.335311 | 1.5425869 | 0.023968043 | 0.14501615 | 1 | 6341 | tags=41%, list=27%, si | EN_3 |

| | | | | | | | | | | |
|---|---|---|---|---|---|---|---|---|---|---|
| GOBP_NUCLEAR_TRANSC | GOBP_NUCLEAR_TRANSCRIBED_MRNA_CAT | 43 | 0.392711 | 1.5424167 | 0.034795765 | 0.145001 | 1 | 7291 | tags=47%, list=31%, si | EN_3 |
| GOBP_ADULT_LOCOMOT | GOBP_ADULT_LOCOMOTORY_BEHAVIOR | 109 | 0.327059 | 1.541998 | 0.0163522 | 0.1451741 | 1 | 4232 | tags=27%, list=18%, si | EN_3 |
| GOBP_FOCAL_ADHESION | GOBP_FOCAL_ADHESION_ASSEMBLY | 87 | 0.338249 | 1.5415109 | 0.018181818 | 0.14545378 | 1 | 7534 | tags=49%, list=32%, si | EN_3 |
| GOBP_REGULATION_OF_ | GOBP_REGULATION_OF_MITOTIC_CELL_CY | 490 | 0.279012 | 1.541396 | 0 | 0.14536293 | 1 | 5470 | tags=29%, list=23%, si | EN_3 |
| GOBP_POSITIVE_REGULA | GOBP_POSITIVE_REGULATION_OF_VESICLE_ | 19 | 0.484327 | 1.5413125 | 0.03305785 | 0.14523579 | 1 | 1838 | tags=21%, list=8%, sig | EN_3 |
| GOBP_VESICLE_TRANSPO | GOBP_VESICLE_TRANSPORT_ALONG_MICR | 47 | 0.381564 | 1.5404598 | 0.031111112 | 0.14593595 | 1 | 3531 | tags=23%, list=15%, si | EN_3 |
| FOCAL_ADHESION | FOCAL_ADHESION | 194 | 0.301476 | 1.5400803 | 0.00921659 | 0.14615919 | 1 | 6875 | tags=39%, list=29%, si | EN_3 |
| GOBP_POSITIVE_REGULA | GOBP_POSITIVE_REGULATION_OF_SMOOTH | 34 | 0.417683 | 1.5397938 | 0.039274924 | 0.14622194 | 1 | 945 | tags=18%, list=4%, sig | EN_3 |
| GOBP_PROTEIN_TRIMERI | GOBP_PROTEIN_TRIMERIZATION | 16 | 0.499305 | 1.5395304 | 0.04769737 | 0.14629094 | 1 | 6269 | tags=50%, list=27%, si | EN_3 |
| GOBP_MYELOID_LEUKOC | GOBP_MYELOID_LEUKOCYTE_ACTIVATION | 247 | 0.292123 | 1.539472 | 0.006651885 | 0.14615028 | 1 | 3402 | tags=22%, list=14%, si | EN_3 |
| GOBP_MITOTIC_CELL_CY | GOBP_MITOTIC_CELL_CYCLE_CHECKPOINT_ | 147 | 0.312581 | 1.5391529 | 0.008353222 | 0.14631832 | 1 | 5457 | tags=34%, list=23%, si | EN_3 |
| GOBP_POSITIVE_REGULA | GOBP_POSITIVE_REGULATION_OF_CELLULA | 22 | 0.457852 | 1.5389934 | 0.055910543 | 0.14627494 | 1 | 3714 | tags=36%, list=16%, si | EN_3 |
| GOBP_REGULATION_OF_ | GOBP_REGULATION_OF_TRANSLATION | 329 | 0.285387 | 1.538857 | 0.001059322 | 0.14621383 | 1 | 7554 | tags=45%, list=32%, si | EN_3 |
| GOBP_NEGATIVE_REGUL | GOBP_NEGATIVE_REGULATION_OF_MACRO | 33 | 0.423072 | 1.53879 | 0.04379562 | 0.14605922 | 1 | 5418 | tags=39%, list=23%, si | EN_3 |
| GOBP_MUSCLE_CELL_DIF | GOBP_MUSCLE_CELL_DIFFERENTIATION | 444 | 0.277289 | 1.5385762 | 0 | 0.14608535 | 1 | 4986 | tags=26%, list=21%, si | EN_3 |
| GOBP_POSITIVE_REGULA | GOBP_POSITIVE_REGULATION_OF_TELOME | 38 | 0.401098 | 1.5383425 | 0.029239766 | 0.1461507 | 1 | 4200 | tags=21%, list=18%, si | EN_3 |
| GOBP_NEGATIVE_REGUL | GOBP_NEGATIVE_REGULATION_OF_NEURO | 211 | 0.296605 | 1.5366966 | 0.007838746 | 0.14769356 | 1 | 3208 | tags=18%, list=14%, si | EN_3 |
| GOBP_POSITIVE_REGULA | GOBP_POSITIVE_REGULATION_OF_MUSCLE | 43 | 0.388549 | 1.5362046 | 0.033430234 | 0.14801152 | 1 | 7578 | tags=56%, list=32%, si | EN_3 |
| GOBP_NEGATIVE_REGUL | GOBP_NEGATIVE_REGULATION_OF_INTRINS | 105 | 0.331406 | 1.5361893 | 0.021573603 | 0.14782286 | 1 | 7214 | tags=43%, list=31%, si | EN_3 |
| GOBP_POSITIVE_REGULA | GOBP_POSITIVE_REGULATION_OF_MITOTIC | 126 | 0.318387 | 1.5360146 | 0.015970515 | 0.14779915 | 1 | 4944 | tags=26%, list=21%, si | EN_3 |
| GOBP_MEMBRANE_PROT | GOBP_MEMBRANE_PROTEIN_PROTEOLYSIS | 60 | 0.358071 | 1.5359092 | 0.034965035 | 0.14769427 | 1 | 7208 | tags=47%, list=31%, si | EN_3 |
| GOBP_REGULATION_OF_ | GOBP_REGULATION_OF_BICELLULAR_TIGHT | 26 | 0.439109 | 1.5355524 | 0.03733766 | 0.14788324 | 1 | 4112 | tags=31%, list=17%, si | EN_3 |
| GOBP_NEGATIVE_REGUL | GOBP_NEGATIVE_REGULATION_OF_TORC1_ | 41 | 0.392384 | 1.5352578 | 0.03776435 | 0.14798598 | 1 | 6373 | tags=41%, list=27%, si | EN_3 |
| GOBP_POSITIVE_REGULA | GOBP_POSITIVE_REGULATION_OF_GLUCON | 26 | 0.43756 | 1.5352135 | 0.032206118 | 0.14782463 | 1 | 6290 | tags=38%, list=27%, si | EN_3 |
| GOBP_REGULATION_OF_ | GOBP_REGULATION_OF_MEMBRANE_INVA | 21 | 0.4629 | 1.5348088 | 0.03858521 | 0.14800815 | 1 | 3926 | tags=38%, list=17%, si | EN_3 |
| GOBP_NEGATIVE_REGUL | GOBP_NEGATIVE_REGULATION_OF_BMP_S | 53 | 0.368193 | 1.5347075 | 0.021489972 | 0.1479083 | 1 | 4029 | tags=28%, list=17%, si | EN_3 |
| GOBP_RIBOSOME_DISASS | GOBP_RIBOSOME_DISASSEMBLY | 31 | 0.415016 | 1.5344118 | 0.043010753 | 0.14799605 | 1 | 5170 | tags=32%, list=22%, si | EN_3 |
| GOBP_RESPONSE_TO_GA | GOBP_RESPONSE_TO_GAMMA_RADIATION | 48 | 0.379958 | 1.534289 | 0.037037037 | 0.1479129 | 1 | 7066 | tags=44%, list=30%, si | EN_3 |
| GOBP_TYPE_II_INTERFER | GOBP_TYPE_II_INTERFERON_MEDIATED_SIG | 23 | 0.449748 | 1.5339234 | 0.052202284 | 0.14808086 | 1 | 1080 | tags=17%, list=5%, sig | EN_3 |
| GOBP_POSITIVE_REGULA | GOBP_POSITIVE_REGULATION_OF_PLASMA | 115 | 0.323288 | 1.5336889 | 0.008641975 | 0.14813249 | 1 | 2908 | tags=17%, list=12%, si | EN_3 |
| GOBP_REGULATION_OF_ | GOBP_REGULATION_OF_CHOLESTEROL_EFF | 41 | 0.385913 | 1.5335364 | 0.03357664 | 0.14808947 | 1 | 1432 | tags=15%, list=6%, sig | EN_3 |
| GOBP_MYELOID_LEUKOC | GOBP_MYELOID_LEUKOCYTE_CYTOKINE_PR | 64 | 0.35845 | 1.5331837 | 0.019914651 | 0.14817017 | 1 | 3699 | tags=25%, list=16%, si | EN_3 |
| GOBP_REGULATION_OF_ | GOBP_REGULATION_OF_UBIQUITIN_PROTE | 39 | 0.398027 | 1.5326769 | 0.041297935 | 0.14848731 | 1 | 4343 | tags=36%, list=18%, si | EN_3 |
| GOBP_PEPTIDYL_CYSTEIN | GOBP_PEPTIDYL_CYSTEINE_MODIFICATION | 44 | 0.382937 | 1.532439 | 0.032208588 | 0.148535 | 1 | 4152 | tags=30%, list=18%, si | EN_3 |
| GOBP_GAMMA_AMINOB | GOBP_GAMMA_AMINOBUTYRIC_ACID_SIGN | 30 | 0.421477 | 1.5324012 | 0.04477612 | 0.14837848 | 1 | 786 | tags=17%, list=3%, sig | EN_3 |
| GOBP_HEART_GROWTH | GOBP_HEART_GROWTH | 116 | 0.321812 | 1.532361 | 0.013732834 | 0.14821756 | 1 | 3649 | tags=21%, list=15%, si | EN_3 |
| GOBP_INTERLEUKIN_8_P | GOBP_INTERLEUKIN_8_PRODUCTION | 73 | 0.346987 | 1.532349 | 0.02406417 | 0.14802524 | 1 | 3017 | tags=25%, list=13%, si | EN_3 |
| GOBP_POSITIVE_REGULA | GOBP_POSITIVE_REGULATION_OF_SMOOTH | 114 | 0.326748 | 1.5323476 | 0.0125 | 0.1478287 | 1 | 4129 | tags=24%, list=17%, si | EN_3 |
| GOBP_CD4_POSITIVE_AL | GOBP_CD4_POSITIVE_ALPHA_BETA_T_CELL | 28 | 0.426472 | 1.5322514 | 0.045454547 | 0.14772461 | 1 | 845 | tags=14%, list=4%, sig | EN_3 |
| GOBP_POSITIVE_REGULA | GOBP_POSITIVE_REGULATION_OF_SMOOTH | 69 | 0.353122 | 1.5321716 | 0.02853261 | 0.14761138 | 1 | 4097 | tags=25%, list=17%, si | EN_3 |
| GOBP_TYPE_I_INTERFER | GOBP_TYPE_I_INTERFERON_PRODUCTION | 127 | 0.318542 | 1.5318019 | 0.0121951215 | 0.1478053 | 1 | 6185 | tags=38%, list=26%, si | EN_3 |
| GOBP_MITOCHONDRIAL_ | GOBP_MITOCHONDRIAL_OUTER_MEMBRAN | 28 | 0.431975 | 1.5317736 | 0.053291537 | 0.14763066 | 1 | 3586 | tags=36%, list=15%, si | EN_3 |
| GOBP_POSITIVE_REGULA | GOBP_POSITIVE_REGULATION_OF_MONON | 94 | 0.337258 | 1.5316426 | 0.016971279 | 0.14757179 | 1 | 3851 | tags=27%, list=16%, si | EN_3 |
| GOBP_NUCLEOSIDE_BISP | GOBP_NUCLEOSIDE_BISPHOSPHATE_META | 102 | 0.325208 | 1.531539 | 0.0163522 | 0.14749712 | 1 | 5590 | tags=32%, list=24%, si | EN_3 |
| GOBP_NEGATIVE_REGUL | GOBP_NEGATIVE_REGULATION_OF_SMALL_ | 61 | 0.353888 | 1.5313816 | 0.032212887 | 0.1474715 | 1 | 6224 | tags=39%, list=26%, si | EN_3 |
| FLUID_SHEAR_STRESS_AI | FLUID_SHEAR_STRESS_AND_ATHEROSCLER | 135 | 0.313589 | 1.5312864 | 0.01438849 | 0.1473753 | 1 | 7007 | tags=41%, list=30%, si | EN_3 |
| GOBP_REGULATION_OF_ | GOBP_REGULATION_OF_ENDOTHELIAL_CEL | 168 | 0.309149 | 1.530684 | 0.009467456 | 0.1477816 | 1 | 6395 | tags=34%, list=27%, si | EN_3 |
| GOBP_CELL_CYCLE_DNA_ | GOBP_CELL_CYCLE_DNA_REPLICATION | 40 | 0.400593 | 1.5305363 | 0.02427921 | 0.14775753 | 1 | 5450 | tags=40%, list=23%, si | EN_3 |
| GOBP_POSITIVE_REGULA | GOBP_POSITIVE_REGULATION_OF_NF_KAP | 127 | 0.318442 | 1.5300275 | 0.013496933 | 0.14804639 | 1 | 4326 | tags=26%, list=18%, si | EN_3 |
| GOBP_CELLULAR_RESPON | GOBP_CELLULAR_RESPONSE_TO_AMYLOID_ | 33 | 0.420776 | 1.5297587 | 0.034795765 | 0.14812824 | 1 | 3074 | tags=27%, list=13%, si | EN_3 |
| GOBP_REGULATION_OF_ | GOBP_REGULATION_OF_CELLULAR_RESPON | 25 | 0.445761 | 1.5293595 | 0.051805336 | 0.1483502 | 1 | 6110 | tags=44%, list=26%, si | EN_3 |
| CITRATE_CYCLE_(TCA_CY | CITRATE_CYCLE_(TCA_CYCLE) | 28 | 0.430637 | 1.5292268 | 0.039184954 | 0.14827232 | 1 | 6125 | tags=46%, list=26%, si | EN_3 |
| MALARIA | MALARIA | 41 | 0.389445 | 1.5291972 | 0.03539823 | 0.14811519 | 1 | 3058 | tags=32%, list=13%, si | EN_3 |
| GOBP_ERAD_PATHWAY | GOBP_ERAD_PATHWAY | 108 | 0.328805 | 1.5291636 | 0.011612903 | 0.1479491 | 1 | 6053 | tags=32%, list=26%, si | EN_3 |
| GOBP_PROTEIN_LOCALIZ | GOBP_PROTEIN_LOCALIZATION_TO_CHROM | 31 | 0.424603 | 1.5291047 | 0.04769737 | 0.14782986 | 1 | 4200 | tags=29%, list=18%, si | EN_3 |
| GOBP_SMOOTH_MUSCLE | GOBP_SMOOTH_MUSCLE_CONTRACTION | 115 | 0.318353 | 1.5284657 | 0.014760148 | 0.14827305 | 1 | 4761 | tags=28%, list=20%, si | EN_3 |
| GOBP_CARDIOCYTE_DIFF | GOBP_CARDIOCYTE_DIFFERENTIATION | 184 | 0.300011 | 1.5275149 | 0.008092485 | 0.14899826 | 1 | 3795 | tags=22%, list=16%, si | EN_3 |
| GOBP_CELLULAR_RESPON | GOBP_CELLULAR_RESPONSE_TO_IONIZING_ | 69 | 0.348568 | 1.5266984 | 0.033103447 | 0.14965568 | 1 | 5457 | tags=32%, list=23%, si | EN_3 |
| GOBP_ANTIGEN_PROCES | GOBP_ANTIGEN_PROCESSING_AND_PRESEN | 20 | 0.469645 | 1.5263759 | 0.050080776 | 0.14980836 | 1 | 3066 | tags=35%, list=13%, si | EN_3 |
| GOBP_POSITIVE_REGULA | GOBP_POSITIVE_REGULATION_OF_DNA_BIN | 45 | 0.383449 | 1.5259312 | 0.044992745 | 0.15003844 | 1 | 2591 | tags=20%, list=11%, si | EN_3 |
| GOBP_NEGATIVE_REGUL | GOBP_NEGATIVE_REGULATION_OF_CELL_A | 207 | 0.299683 | 1.5249381 | 0.0011286682 | 0.15089858 | 1 | 3363 | tags=23%, list=14%, si | EN_3 |
| GOBP_POLYOL_METABOL | GOBP_POLYOL_METABOLIC_PROCESS | 99 | 0.326802 | 1.5248207 | 0.019035533 | 0.15083237 | 1 | 4125 | tags=24%, list=17%, si | EN_3 |
| GOBP_REGULATION_OF_ | GOBP_REGULATION_OF_NUCLEOTIDE_EXCI | 29 | 0.421216 | 1.5246508 | 0.051036682 | 0.15082009 | 1 | 3339 | tags=24%, list=14%, si | EN_3 |
| GOBP_ANTIGEN_PROCES | GOBP_ANTIGEN_PROCESSING_AND_PRESEN | 24 | 0.445241 | 1.5245885 | 0.056198347 | 0.15069574 | 1 | 3074 | tags=33%, list=13%, si | EN_3 |
| GOBP_PLATELET_AGGRE | GOBP_PLATELET_AGGREGATION | 57 | 0.360805 | 1.5241005 | 0.03159341 | 0.15099004 | 1 | 3429 | tags=25%, list=15%, si | EN_3 |
| GOBP_PIGMENT_BIOSYN | GOBP_PIGMENT_BIOSYNTHETIC_PROCESS | 49 | 0.383582 | 1.5228561 | 0.029126214 | 0.15206634 | 1 | 1994 | tags=22%, list=8%, sig | EN_3 |
| GOBP_HEMATOPOIETIC_ | GOBP_HEMATOPOIETIC_OR_LYMPHOID_OR | 118 | 0.317383 | 1.5225754 | 0.017369727 | 0.1521353 | 1 | 6385 | tags=36%, list=27%, si | EN_3 |
| GOBP_ENDOSOME_ORGA | GOBP_ENDOSOME_ORGANIZATION | 68 | 0.345506 | 1.5219793 | 0.036061026 | 0.15255223 | 1 | 5797 | tags=34%, list=25%, si | EN_3 |
| HERPES_SIMPLEX_VIRUS_ | HERPES_SIMPLEX_VIRUS_1_INFECTION | 373 | 0.28065 | 1.5213883 | 0.0020920502 | 0.1530013 | 1 | 3989 | tags=23%, list=17%, si | EN_3 |
| GOBP_NON_CANONICAL_ | GOBP_NON_CANONICAL_NF_KAPPAB_SIGN | 115 | 0.316854 | 1.5213225 | 0.019631902 | 0.15287207 | 1 | 4844 | tags=30%, list=21%, si | EN_3 |
| CELL_CYCLE | CELL_CYCLE | 121 | 0.31708 | 1.5208541 | 0.021924483 | 0.15314673 | 1 | 6619 | tags=38%, list=28%, si | EN_3 |
| GOBP_SULFUR_COMPOU | GOBP_SULFUR_COMPOUND_METABOLIC_P | 292 | 0.284343 | 1.5207402 | 0.0021621622 | 0.1530631 | 1 | 5590 | tags=30%, list=24%, si | EN_3 |
| GOBP_PHAGOSOME_MA | GOBP_PHAGOSOME_MATURATION | 29 | 0.422853 | 1.5200291 | 0.047169812 | 0.15359034 | 1 | 7606 | tags=55%, list=32%, si | EN_3 |
| GOBP_AMYLOID_PRECUR | GOBP_AMYLOID_PRECURSOR_PROTEIN_CA | 59 | 0.359752 | 1.5192173 | 0.04378531 | 0.15421571 | 1 | 6170 | tags=34%, list=26%, si | EN_3 |
| GOBP_TELOMERE_MAIN | GOBP_TELOMERE_MAINTENANCE_VIA_TEL | 76 | 0.345964 | 1.5191976 | 0.023746701 | 0.15403117 | 1 | 4359 | tags=24%, list=18%, si | EN_3 |
| GOBP_INTERLEUKIN_17_ | GOBP_INTERLEUKIN_17_PRODUCTION | 42 | 0.396622 | 1.51865 | 0.043541364 | 0.15439898 | 1 | 2936 | tags=26%, list=12%, si | EN_3 |
| GOBP_POSITIVE_REGULA | GOBP_POSITIVE_REGULATION_OF_G1_S_TR | 48 | 0.37639 | 1.517789 | 0.03768116 | 0.15512224 | 1 | 4258 | tags=31%, list=18%, si | EN_3 |
| GOBP_NEGATIVE_REGUL | GOBP_NEGATIVE_REGULATION_OF_PROTEI | 93 | 0.332805 | 1.5174977 | 0.017038008 | 0.1552584 | 1 | 6110 | tags=40%, list=26%, si | EN_3 |
| NUCLEOTIDE_EXCISION_ | NUCLEOTIDE_EXCISION_REPAIR | 42 | 0.38317 | 1.5168306 | 0.04511278 | 0.15574956 | 1 | 7521 | tags=48%, list=32%, si | EN_3 |
| GOBP_CARDIAC_MUSCLE | GOBP_CARDIAC_MUSCLE_CELL_DIFFERENTI | 147 | 0.310899 | 1.5167603 | 0.015550239 | 0.1556305 | 1 | 3740 | tags=22%, list=16%, si | EN_3 |
| GOBP_REGULATION_OF_ | GOBP_REGULATION_OF_PLASMA_LIPOPRO | 66 | 0.352339 | 1.5167071 | 0.029530201 | 0.15548769 | 1 | 1678 | tags=17%, list=7%, sig | EN_3 |
| GOBP_TRNA_THREONYL | GOBP_TRNA_THREONYLCARBAMOYLADEN | 15 | 0.499267 | 1.5163045 | 0.07575758 | 0.15572512 | 1 | 3619 | tags=40%, list=15%, si | EN_3 |

| | | | | | | | | | | |
|---|---|---|---|---|---|---|---|---|---|---|
| GOBP_REGULATION_OF_ | GOBP_REGULATION_OF_BMP_SIGNALING_ | 96 | 0.322552 | 1.5157926 | 0.03137255 | 0.1561084 | 1 | 4029 | tags=24%, list=17%, si | EN_3 |
| GOBP_REGULATION_OF_ | GOBP_REGULATION_OF_MACROPHAGE_AC | 57 | 0.360704 | 1.5154303 | 0.030985916 | 0.15633371 | 1 | 3592 | tags=26%, list=15%, si | EN_3 |
| GOBP_NEGATIVE_REGUL | GOBP_NEGATIVE_REGULATION_OF_DNA_M | 127 | 0.312364 | 1.5152911 | 0.012150669 | 0.15629388 | 1 | 5418 | tags=31%, list=23%, si | EN_3 |
| GOBP_REGULATION_OF_ | GOBP_REGULATION_OF_EPITHELIAL_CELL_ | 118 | 0.318005 | 1.5150218 | 0.013563502 | 0.15640494 | 1 | 4934 | tags=26%, list=21%, si | EN_3 |
| GOBP_REGULATION_OF_ | GOBP_REGULATION_OF_RESPONSE_TO_CY | 153 | 0.305187 | 1.5145985 | 0.01911589 | 0.15666334 | 1 | 3441 | tags=22%, list=15%, si | EN_3 |
| GOBP_NEGATIVE_REGUL | GOBP_NEGATIVE_REGULATION_OF_PROTEI | 17 | 0.479914 | 1.5144576 | 0.061769616 | 0.15665591 | 1 | 7508 | tags=71%, list=32%, si | EN_3 |
| GOBP_NEGATIVE_REGUL | GOBP_NEGATIVE_REGULATION_OF_PHOSPI | 332 | 0.280389 | 1.5135016 | 0.0031847134 | 0.15748163 | 1 | 5843 | tags=31%, list=25%, si | EN_3 |
| GOBP_MITOCHONDRION | GOBP_MITOCHONDRION_DISTRIBUTION | 27 | 0.42918 | 1.5129448 | 0.05351682 | 0.15783563 | 1 | 3432 | tags=30%, list=15%, si | EN_3 |
| GOBP_EXTRINSIC_APOPT | GOBP_EXTRINSIC_APOPTOTIC_SIGNALING_ | 233 | 0.291033 | 1.5129113 | 0.0088495575 | 0.15766354 | 1 | 4541 | tags=26%, list=19%, si | EN_3 |
| GOBP_POSITIVE_REGULA | GOBP_POSITIVE_REGULATION_OF_INTRINS | 69 | 0.351747 | 1.5128173 | 0.025469169 | 0.1575556 | 1 | 5418 | tags=38%, list=23%, si | EN_3 |
| GOBP_TRANSLATIONAL_I | GOBP_TRANSLATIONAL_INITIATION | 112 | 0.316872 | 1.5125841 | 0.016149068 | 0.15759152 | 1 | 7308 | tags=46%, list=31%, si | EN_3 |
| GOBP_REGULATION_OF_ | GOBP_REGULATION_OF_PROTEIN_EXPORT_ | 37 | 0.393776 | 1.5125675 | 0.041420117 | 0.15741597 | 1 | 4934 | tags=38%, list=21%, si | EN_3 |
| GOBP_NEGATIVE_REGUL | GOBP_NEGATIVE_REGULATION_OF_CD4_P | 37 | 0.394009 | 1.5124394 | 0.04239766 | 0.1573397 | 1 | 2841 | tags=24%, list=12%, si | EN_3 |
| GOBP_NEGATIVE_REGUL | GOBP_NEGATIVE_REGULATION_OF_CELL_C | 368 | 0.277219 | 1.5123552 | 0.0010471204 | 0.157246 | 1 | 5473 | tags=29%, list=23%, si | EN_3 |
| GOBP_MYELOID_CELL_H | GOBP_MYELOID_CELL_HOMEOSTASIS | 209 | 0.292523 | 1.5119338 | 0.004530011 | 0.15751728 | 1 | 5684 | tags=30%, list=24%, si | EN_3 |
| GOBP_VIRAL_TRANSCRIP | GOBP_VIRAL_TRANSCRIPTION | 55 | 0.365341 | 1.5116804 | 0.032440055 | 0.15760458 | 1 | 3411 | tags=24%, list=14%, si | EN_3 |
| GOBP_CHRONIC_INFLAM | GOBP_CHRONIC_INFLAMMATORY_RESPONS | 18 | 0.484152 | 1.5111 | 0.05882353 | 0.15804948 | 1 | 2841 | tags=39%, list=12%, si | EN_3 |
| GOBP_LIPOPROTEIN_LOC | GOBP_LIPOPROTEIN_LOCALIZATION | 20 | 0.469717 | 1.5108602 | 0.051409617 | 0.15809456 | 1 | 5783 | tags=45%, list=24%, si | EN_3 |
| GOBP_EXOCYTOSIS | GOBP_EXOCYTOSIS | 367 | 0.279023 | 1.5105101 | 0.004237288 | 0.15826891 | 1 | 4322 | tags=22%, list=18%, si | EN_3 |
| GLUTAMATERGIC_SYNAP | GLUTAMATERGIC_SYNAPSE | 108 | 0.320488 | 1.5104053 | 0.027166883 | 0.15818344 | 1 | 4274 | tags=25%, list=18%, si | EN_3 |
| GOBP_REGULATION_OF_ | GOBP_REGULATION_OF_CARTILAGE_DEVEL | 71 | 0.342048 | 1.5098988 | 0.021680217 | 0.1585441 | 1 | 4321 | tags=28%, list=18%, si | EN_3 |
| GOBP_REGULATION_OF_ | GOBP_REGULATION_OF_INTRACELLULAR_P | 168 | 0.300558 | 1.5094568 | 0.012820513 | 0.1588443 | 1 | 7368 | tags=45%, list=31%, si | EN_3 |
| INFLUENZA_A | INFLUENZA_A | 139 | 0.3058 | 1.5091251 | 0.015795868 | 0.15899485 | 1 | 5812 | tags=32%, list=25%, si | EN_3 |
| FOXO_SIGNALING_PATHV | FOXO_SIGNALING_PATHWAY | 127 | 0.316147 | 1.5076613 | 0.013398294 | 0.1604417 | 1 | 3321 | tags=19%, list=14%, si | EN_3 |
| GOBP_REGULATION_OF_ | GOBP_REGULATION_OF_LIPID_STORAGE | 55 | 0.362361 | 1.5074621 | 0.035663337 | 0.1604737 | 1 | 4092 | tags=20%, list=17%, si | EN_3 |
| GOBP_INTERSTRAND_CR | GOBP_INTERSTRAND_CROSS_LINK_REPAIR | 31 | 0.418128 | 1.5074311 | 0.047244094 | 0.16030623 | 1 | 5653 | tags=52%, list=24%, si | EN_3 |
| GOBP_POSITIVE_REGULA | GOBP_POSITIVE_REGULATION_OF_SECRETI | 385 | 0.279244 | 1.5073025 | 0.003177966 | 0.16025242 | 1 | 3345 | tags=19%, list=14%, si | EN_3 |
| GOBP_POSITIVE_REGULA | GOBP_POSITIVE_REGULATION_OF_TYPE_I_I | 77 | 0.335679 | 1.5072526 | 0.031209363 | 0.16011024 | 1 | 6185 | tags=38%, list=26%, si | EN_3 |
| GOBP_I_KAPPAB_PHOSPI | GOBP_I_KAPPAB_PHOSPHORYLATION | 17 | 0.484989 | 1.5068234 | 0.06335616 | 0.16035919 | 1 | 3321 | tags=35%, list=14%, si | EN_3 |
| GOBP_NEGATIVE_REGUL | GOBP_NEGATIVE_REGULATION_OF_CELLUL | 106 | 0.318943 | 1.5065792 | 0.018726591 | 0.16043563 | 1 | 3864 | tags=25%, list=16%, si | EN_3 |
| GOBP_NUCLEOTIDE_BINI | GOBP_NUCLEOTIDE_BINDING_OLIGOMERIZ | 18 | 0.480743 | 1.5064349 | 0.058430716 | 0.1604096 | 1 | 4168 | tags=39%, list=18%, si | EN_3 |
| GOBP_CHONDROCYTE_D | GOBP_CHONDROCYTE_DIFFERENTIATION | 115 | 0.315453 | 1.5056288 | 0.022304833 | 0.1610515 | 1 | 4467 | tags=24%, list=19%, si | EN_3 |
| GOBP_REGULATION_OF_ | GOBP_REGULATION_OF_OXIDOREDUCTASE_ | 63 | 0.34733 | 1.504784 | 0.025920874 | 0.16183518 | 1 | 3672 | tags=25%, list=16%, si | EN_3 |
| GOBP_ANTEROGRADE_A | GOBP_ANTEROGRADE_AXONAL_TRANSPOR | 53 | 0.366206 | 1.5045631 | 0.041608877 | 0.16189656 | 1 | 7781 | tags=49%, list=33%, si | EN_3 |
| ANTIGEN_PROCESSING_A | ANTIGEN_PROCESSING_AND_PRESENTATIO | 67 | 0.347713 | 1.5042189 | 0.023743017 | 0.1620673 | 1 | 5341 | tags=42%, list=23%, si | EN_3 |
| GOBP_NEGATIVE_REGUL | GOBP_NEGATIVE_REGULATION_OF_LOCOM | 328 | 0.27686 | 1.5041217 | 0.0021367522 | 0.16197205 | 1 | 3649 | tags=20%, list=15%, si | EN_3 |
| GOBP_THYMUS_DEVELOI | GOBP_THYMUS_DEVELOPMENT | 53 | 0.363004 | 1.5041065 | 0.040616248 | 0.16178791 | 1 | 7884 | tags=51%, list=33%, si | EN_3 |
| GOBP_ESTABLISHMENT_ | GOBP_ESTABLISHMENT_OF_PROTEIN_LOCA | 19 | 0.470895 | 1.5036136 | 0.058267716 | 0.16215554 | 1 | 4200 | tags=32%, list=18%, si | EN_3 |
| GOBP_FRUCTOSE_6_PHO | GOBP_FRUCTOSE_6_PHOSPHATE_METABOL | 16 | 0.503234 | 1.5035709 | 0.065743946 | 0.16201597 | 1 | 2433 | tags=25%, list=10%, si | EN_3 |
| GOBP_SKELETAL_MUSCLI | GOBP_SKELETAL_MUSCLE_ACETYLCHOLINE_ | 21 | 0.460584 | 1.5034001 | 0.055464927 | 0.16201115 | 1 | 6824 | tags=57%, list=29%, si | EN_3 |
| GOBP_DNA_TEMPLATED_ | GOBP_DNA_TEMPLATED_TRANSCRIPTION_I | 183 | 0.297523 | 1.5027876 | 0.010309278 | 0.1624596 | 1 | 5869 | tags=32%, list=25%, si | EN_3 |
| GOBP_SHORT_TERM_ME | GOBP_SHORT_TERM_MEMORY | 18 | 0.476885 | 1.5018307 | 0.062711865 | 0.16328725 | 1 | 459 | tags=17%, list=2%, sig | EN_3 |
| GOBP_RESPONSE_TO_IN | GOBP_RESPONSE_TO_INORGANIC_SUBSTAI | 432 | 0.27046 | 1.5015761 | 0.0010362695 | 0.1633803 | 1 | 5792 | tags=29%, list=25%, si | EN_3 |
| GOBP_ACETYL_COA_BIOS | GOBP_ACETYL_COA_BIOSYNTHETIC_PROCES | 19 | 0.476618 | 1.5007985 | 0.0608 | 0.16407457 | 1 | 6561 | tags=47%, list=28%, si | EN_3 |
| GOBP_POSITIVE_REGULA | GOBP_POSITIVE_REGULATION_OF_PROTEIN | 41 | 0.388566 | 1.5002738 | 0.046062406 | 0.16441411 | 1 | 3266 | tags=24%, list=14%, si | EN_3 |
| GOBP_STRIATED_MUSCLI | GOBP_STRIATED_MUSCLE_TISSUE_DEVELOI | 281 | 0.283758 | 1.5000707 | 0.0077262693 | 0.16444737 | 1 | 3963 | tags=21%, list=17%, si | EN_3 |
| GOBP_LOW_DENSITY_LIP | GOBP_LOW_DENSITY_LIPOPROTEIN_PARTIC | 23 | 0.450928 | 1.4999028 | 0.044374008 | 0.16442244 | 1 | 1508 | tags=22%, list=6%, sig | EN_3 |
| GOBP_POSITIVE_REGULA | GOBP_POSITIVE_REGULATION_OF_HYDROL | 427 | 0.275739 | 1.4998165 | 0 | 0.16431998 | 1 | 3321 | tags=18%, list=14%, si | EN_3 |
| GOBP_REGULATION_OF_ | GOBP_REGULATION_OF_DNA_TEMPLATED_ | 79 | 0.338251 | 1.49924 | 0.036 | 0.16477187 | 1 | 5846 | tags=30%, list=25%, si | EN_3 |
| AGE-RAGE_SIGNALING_P | AGE-RAGE_SIGNALING_PATHWAY_IN_DIABI | 100 | 0.324621 | 1.4991554 | 0.019851116 | 0.16466765 | 1 | 3837 | tags=23%, list=16%, si | EN_3 |
| GOBP_NEGATIVE_REGUL | GOBP_NEGATIVE_REGULATION_OF_MEMBI | 22 | 0.450636 | 1.4988872 | 0.051612902 | 0.16475417 | 1 | 3586 | tags=32%, list=15%, si | EN_3 |
| GOBP_REGULATION_OF_ | GOBP_REGULATION_OF_PROTEIN_SECRETI | 300 | 0.281684 | 1.4987595 | 0.0043243244 | 0.16468965 | 1 | 3729 | tags=20%, list=16%, si | EN_3 |
| GOBP_DNA_REPLICATION | GOBP_DNA_REPLICATION_CHECKPOINT_SIG | 18 | 0.483055 | 1.4986228 | 0.06472492 | 0.16463648 | 1 | 3625 | tags=39%, list=15%, si | EN_3 |
| GOBP_NAD_METABOLIC_ | GOBP_NAD_METABOLIC_PROCESS | 58 | 0.356092 | 1.4984572 | 0.033379693 | 0.16463138 | 1 | 6125 | tags=45%, list=26%, si | EN_3 |
| GOBP_NEGATIVE_REGUL | GOBP_NEGATIVE_REGULATION_OF_STRIATI | 39 | 0.389097 | 1.4984403 | 0.0456621 | 0.16446486 | 1 | 4017 | tags=33%, list=17%, si | EN_3 |
| VASOPRESSIN-REGULATEI | VASOPRESSIN-REGULATED_WATER_REABSO | 40 | 0.386681 | 1.4982458 | 0.041990668 | 0.16446544 | 1 | 6374 | tags=43%, list=27%, si | EN_3 |
| GOBP_LATE_ENDOSOME_ | GOBP_LATE_ENDOSOME_TO_VACUOLE_TR | 25 | 0.438769 | 1.4977885 | 0.06549521 | 0.16475429 | 1 | 4771 | tags=40%, list=20%, si | EN_3 |
| METABOLISM_OF_XENOB | METABOLISM_OF_XENOBIOTICS_BY_CYTOC | 48 | 0.374867 | 1.4974097 | 0.03954802 | 0.1649365 | 1 | 4796 | tags=40%, list=20%, si | EN_3 |
| GOBP_MAINTENANCE_O | GOBP_MAINTENANCE_OF_LOCATION_IN_C | 224 | 0.29241 | 1.4972705 | 0.0054884744 | 0.16486101 | 1 | 4934 | tags=29%, list=21%, si | EN_3 |
| GOBP_BONE_DEVELOPM | GOBP_BONE_DEVELOPMENT | 222 | 0.28783 | 1.497138 | 0.0122631 | 0.1648066 | 1 | 4849 | tags=27%, list=21%, si | EN_3 |
| GOBP_MUSCLE_ADAPTAT | GOBP_MUSCLE_ADAPTATION | 126 | 0.314496 | 1.4971275 | 0.017094018 | 0.16462721 | 1 | 5000 | tags=26%, list=21%, si | EN_3 |
| GOBP_PROTEIN_TARGETI | GOBP_PROTEIN_TARGETING_TO_MEMBRAI | 116 | 0.312851 | 1.496977 | 0.023370234 | 0.1646163 | 1 | 4950 | tags=28%, list=21%, si | EN_3 |
| GOBP_AXONAL_TRANSPO | GOBP_AXONAL_TRANSPORT | 69 | 0.349563 | 1.4967899 | 0.024456521 | 0.16465023 | 1 | 7873 | tags=49%, list=33%, si | EN_3 |
| GOBP_POSITIVE_REGULA | GOBP_POSITIVE_REGULATION_OF_RELEASE | 27 | 0.426949 | 1.4964802 | 0.06260032 | 0.16479634 | 1 | 4024 | tags=44%, list=17%, si | EN_3 |
| GOBP_GLYCEROLIPID_CA | GOBP_GLYCEROLIPID_CATABOLIC_PROCESS | 58 | 0.356513 | 1.4963573 | 0.036414567 | 0.16474105 | 1 | 1563 | tags=16%, list=7%, sig | EN_3 |
| GOBP_ESTABLISHMENT_ | GOBP_ESTABLISHMENT_OF_EPITHELIAL_CEI | 36 | 0.39095 | 1.4962856 | 0.04427481 | 0.16460708 | 1 | 5011 | tags=31%, list=21%, si | EN_3 |
| GOBP_MUSCLE_HYPERTR | GOBP_MUSCLE_HYPERTROPHY | 115 | 0.313316 | 1.4960567 | 0.017135862 | 0.16466697 | 1 | 5000 | tags=25%, list=21%, si | EN_3 |
| GOBP_SPECIFICATION_OI | GOBP_SPECIFICATION_OF_SYMMETRY | 123 | 0.310811 | 1.4960066 | 0.019607844 | 0.16451952 | 1 | 5217 | tags=24%, list=22%, si | EN_3 |
| GOBP_RUFFLE_ASSEMBLY | GOBP_RUFFLE_ASSEMBLY | 43 | 0.377274 | 1.4956359 | 0.042735044 | 0.16473526 | 1 | 3470 | tags=30%, list=15%, si | EN_3 |
| GOBP_IMPORT_INTO_NU | GOBP_IMPORT_INTO_NUCLEUS | 180 | 0.293149 | 1.4952805 | 0.011587486 | 0.16490355 | 1 | 7637 | tags=44%, list=32%, si | EN_3 |
| GOBP_VASODILATION | GOBP_VASODILATION | 64 | 0.347604 | 1.4952204 | 0.03314917 | 0.16477425 | 1 | 4129 | tags=30%, list=17%, si | EN_3 |
| GOBP_NEGATIVE_REGUL | GOBP_NEGATIVE_REGULATION_OF_ACTIN_ | 45 | 0.373207 | 1.4948957 | 0.046511628 | 0.16493823 | 1 | 7278 | tags=44%, list=31%, si | EN_3 |
| GOBP_EXTRACELLULAR_I | GOBP_EXTRACELLULAR_MATRIX_CONSTITU | 16 | 0.484928 | 1.4945962 | 0.06623586 | 0.16504538 | 1 | 1506 | tags=19%, list=6%, sig | EN_3 |
| GOBP_POSITIVE_REGULA | GOBP_POSITIVE_REGULATION_OF_STRIATE | 50 | 0.367645 | 1.4945374 | 0.044604316 | 0.16492064 | 1 | 3586 | tags=28%, list=15%, si | EN_3 |
| GOBP_TRANSCRIPTION_P | GOBP_TRANSCRIPTION_PREINITIATION_CO | 63 | 0.344296 | 1.4944377 | 0.033602152 | 0.1648441 | 1 | 6098 | tags=38%, list=26%, si | EN_3 |
| GOBP_REGULATION_OF_ | GOBP_REGULATION_OF_MACROPHAGE_MI | 47 | 0.373478 | 1.4939026 | 0.04040404 | 0.16526048 | 1 | 3851 | tags=28%, list=16%, si | EN_3 |
| GOBP_POSITIVE_REGULA | GOBP_POSITIVE_REGULATION_OF_DEFENSE | 31 | 0.409332 | 1.4938112 | 0.056923077 | 0.16515906 | 1 | 2841 | tags=26%, list=12%, si | EN_3 |
| GOBP_REGULATION_OF_ | GOBP_REGULATION_OF_DNA_DAMAGE_RE | 36 | 0.38905 | 1.4934944 | 0.039647575 | 0.1653551 | 1 | 6046 | tags=42%, list=26%, si | EN_3 |
| GOBP_POSITIVE_REGULA | GOBP_POSITIVE_REGULATION_OF_MACROI | 31 | 0.411622 | 1.4932435 | 0.044585988 | 0.16543414 | 1 | 3851 | tags=32%, list=16%, si | EN_3 |
| GOBP_ENDOCYTIC_RECYC | GOBP_ENDOCYTIC_RECYCLING | 88 | 0.326789 | 1.4927019 | 0.027704485 | 0.16584112 | 1 | 4139 | tags=25%, list=18%, si | EN_3 |
| GOBP_POSITIVE_REGULA | GOBP_POSITIVE_REGULATION_OF_VACUOL | 25 | 0.428357 | 1.4923133 | 0.048361935 | 0.16609746 | 1 | 5484 | tags=48%, list=23%, si | EN_3 |
| GOBP_NEGATIVE_REGUL | GOBP_NEGATIVE_REGULATION_OF_INFLAM | 153 | 0.301981 | 1.4918813 | 0.015294118 | 0.16639431 | 1 | 3184 | tags=21%, list=13%, si | EN_3 |

| | | | | | | | | | | |
|---|---|---|---|---|---|---|---|---|---|---|
| GOBP_DIOL_METABOLIC_ | GOBP_DIOL_METABOLIC_PROCESS | 29 | 0.417513 | 1.4916844 | 0.053680982 | 0.1664106 | 1 | 1786 | tags=17%, list=8%, sig | EN_3 |
| GOBP_REGULATION_OF_ | GOBP_REGULATION_OF_MUSCLE_ADAPTAT | 102 | 0.316484 | 1.4916501 | 0.03375 | 0.16624753 | 1 | 5000 | tags=27%, list=21%, si | EN_3 |
| GOBP_REGULATION_OF_ | GOBP_REGULATION_OF_CELL_SHAPE | 137 | 0.302269 | 1.4914864 | 0.024390243 | 0.16625297 | 1 | 3243 | tags=20%, list=14%, si | EN_3 |
| GOBP_UBIQUITIN_DEPEN | GOBP_UBIQUITIN_DEPENDENT_PROTEIN_C | 29 | 0.414589 | 1.4914503 | 0.048818897 | 0.16611211 | 1 | 4771 | tags=38%, list=20%, si | EN_3 |
| GOBP_REGULATION_OF_ | GOBP_REGULATION_OF_PHAGOCYTOSIS | 106 | 0.316083 | 1.491056 | 0.021464646 | 0.16635184 | 1 | 3926 | tags=24%, list=17%, si | EN_3 |
| GOBP_REGULATION_OF_ | GOBP_REGULATION_OF_INTEGRIN_MEDIAT | 19 | 0.462615 | 1.4907471 | 0.06532663 | 0.16649596 | 1 | 4486 | tags=32%, list=19%, si | EN_3 |
| GOBP_PHYSIOLOGICAL_C | GOBP_PHYSIOLOGICAL_CARDIAC_MUSCLE_ | 46 | 0.37632 | 1.490556 | 0.039647575 | 0.16650799 | 1 | 3586 | tags=26%, list=15%, si | EN_3 |
| GOBP_CELLULAR_COMPO | GOBP_CELLULAR_COMPONENT_DISASSEMB | 472 | 0.267921 | 1.4890826 | 0.0020639836 | 0.16794331 | 1 | 5418 | tags=27%, list=23%, si | EN_3 |
| GOBP_MITOPHAGY | GOBP_MITOPHAGY | 58 | 0.35503 | 1.488962 | 0.033898305 | 0.16787797 | 1 | 4372 | tags=22%, list=19%, si | EN_3 |
| GOBP_PHOSPHATIDYLCH | GOBP_PHOSPHATIDYLCHOLINE_BIOSYNTHE | 21 | 0.445597 | 1.4889144 | 0.067993365 | 0.16774438 | 1 | 1036 | tags=19%, list=4%, sig | EN_3 |
| GOBP_POSITIVE_REGULA | GOBP_POSITIVE_REGULATION_OF_ERYTHRO | 28 | 0.418782 | 1.4888511 | 0.046616543 | 0.16763788 | 1 | 1994 | tags=14%, list=8%, sig | EN_3 |
| GOBP_CYTOKINE_PRODU | GOBP_CYTOKINE_PRODUCTION_INVOLVED_ | 145 | 0.297248 | 1.488419 | 0.018181818 | 0.16792874 | 1 | 2936 | tags=21%, list=12%, si | EN_3 |
| GOBP_COENZYME_A_ME | GOBP_COENZYME_A_METABOLIC_PROCESS | 19 | 0.463165 | 1.4883807 | 0.056692913 | 0.16777661 | 1 | 4552 | tags=37%, list=19%, si | EN_3 |
| GOBP_NEGATIVE_REGUL | GOBP_NEGATIVE_REGULATION_OF_MITOTI | 174 | 0.296046 | 1.4880961 | 0.018539976 | 0.16790886 | 1 | 5457 | tags=32%, list=23%, si | EN_3 |
| GOBP_POSITIVE_REGULA | GOBP_POSITIVE_REGULATION_OF_CYSTEIN | 130 | 0.307898 | 1.4880418 | 0.02389937 | 0.16776529 | 1 | 3184 | tags=20%, list=13%, si | EN_3 |
| GOBP_RESPONSE_TO_TY | GOBP_RESPONSE_TO_TYPE_I_INTERFERON | 64 | 0.351006 | 1.4875927 | 0.03009576 | 0.16808389 | 1 | 2972 | tags=22%, list=13%, si | EN_3 |
| GOBP_ORGAN_INDUCTIO | GOBP_ORGAN_INDUCTION | 24 | 0.443486 | 1.4874918 | 0.07219662 | 0.16799101 | 1 | 5266 | tags=42%, list=22%, si | EN_3 |
| GOBP_RRNA_METHYLATI | GOBP_RRNA_METHYLATION | 20 | 0.459944 | 1.4871042 | 0.07754443 | 0.16822773 | 1 | 3731 | tags=35%, list=16%, si | EN_3 |
| GOBP_CGMP_MEDIATED_ | GOBP_CGMP_MEDIATED_SIGNALING | 32 | 0.396716 | 1.4868561 | 0.044822257 | 0.1683108 | 1 | 2849 | tags=28%, list=12%, si | EN_3 |
| GOBP_REGULATION_OF_ | GOBP_REGULATION_OF_DNA_BINDING_TRA | 362 | 0.274998 | 1.4865801 | 0.0010582011 | 0.16843608 | 1 | 4367 | tags=23%, list=18%, si | EN_3 |
| GOBP_NEGATIVE_REGUL | GOBP_NEGATIVE_REGULATION_OF_EPITHEL | 66 | 0.343984 | 1.486427 | 0.03888889 | 0.16843483 | 1 | 6475 | tags=38%, list=27%, si | EN_3 |
| AUTOPHAGY | AUTOPHAGY | 126 | 0.310157 | 1.4855148 | 0.019512195 | 0.16934745 | 1 | 7005 | tags=37%, list=30%, si | EN_3 |
| GOBP_TUMOR_NECROSIS | GOBP_TUMOR_NECROSIS_FACTOR_SUPERF | 193 | 0.290061 | 1.4854188 | 0.010215664 | 0.16926043 | 1 | 5000 | tags=28%, list=21%, si | EN_3 |
| GOBP_GLIAL_CELL_MIGR | GOBP_GLIAL_CELL_MIGRATION | 70 | 0.337532 | 1.4845458 | 0.032831736 | 0.17005064 | 1 | 5182 | tags=30%, list=22%, si | EN_3 |
| GOBP_REGULATION_OF_ | GOBP_REGULATION_OF_MUSCLE_HYPERTR | 80 | 0.336283 | 1.4844735 | 0.028683182 | 0.16994713 | 1 | 5462 | tags=31%, list=23%, si | EN_3 |
| GOBP_NEGATIVE_REGUL | GOBP_NEGATIVE_REGULATION_OF_RESPON | 79 | 0.33916 | 1.4828088 | 0.029972753 | 0.1717112 | 1 | 3441 | tags=24%, list=15%, si | EN_3 |
| GOBP_POSITIVE_REGULA | GOBP_POSITIVE_REGULATION_OF_MYELOID | 21 | 0.45129 | 1.482691 | 0.06414474 | 0.1716355 | 1 | 3780 | tags=33%, list=16%, si | EN_3 |
| GOBP_PHOSPHATIDYLCH | GOBP_PHOSPHATIDYLCHOLINE_CATABOLIC_ | 15 | 0.501027 | 1.4822774 | 0.08443709 | 0.17194948 | 1 | 1532 | tags=27%, list=6%, sig | EN_3 |
| GOBP_ANTIGEN_PROCES | GOBP_ANTIGEN_PROCESSING_AND_PRESEN | 34 | 0.396378 | 1.4817451 | 0.06849315 | 0.17239055 | 1 | 5263 | tags=38%, list=22%, si | EN_3 |
| GOBP_MULTICELLULAR_C | GOBP_MULTICELLULAR_ORGANISM_GROW | 218 | 0.285134 | 1.4816854 | 0.012600229 | 0.17227066 | 1 | 4232 | tags=22%, list=18%, si | EN_3 |
| GOBP_ESTABLISHMENT_C | GOBP_ESTABLISHMENT_OF_PROTEIN_LOCA | 47 | 0.371894 | 1.4815971 | 0.038406827 | 0.17219564 | 1 | 4909 | tags=36%, list=21%, si | EN_3 |
| GOBP_ANTIGEN_PROCES | GOBP_ANTIGEN_PROCESSING_AND_PRESEN | 105 | 0.320701 | 1.4812605 | 0.024050632 | 0.17242368 | 1 | 5341 | tags=32%, list=23%, si | EN_3 |
| GOBP_REGULATION_OF_ | GOBP_REGULATION_OF_CELLULAR_COMPO | 380 | 0.272536 | 1.4812117 | 0.0031512605 | 0.17227525 | 1 | 5801 | tags=30%, list=25%, si | EN_3 |
| GOBP_POSITIVE_REGULA | GOBP_POSITIVE_REGULATION_OF_MONOC | 21 | 0.446827 | 1.4809102 | 0.06138934 | 0.17242812 | 1 | 3182 | tags=29%, list=13%, si | EN_3 |
| GOBP_POSITIVE_THYMIC | GOBP_POSITIVE_THYMIC_T_CELL_SELECTIO | 15 | 0.489664 | 1.4808035 | 0.08169935 | 0.17234917 | 1 | 5128 | tags=53%, list=22%, si | EN_3 |
| GOBP_REGULATION_OF_ | GOBP_REGULATION_OF_EPITHELIAL_CELL_ | 238 | 0.28312 | 1.4798515 | 0.013468013 | 0.17322606 | 1 | 3864 | tags=19%, list=16%, si | EN_3 |
| GOBP_NEGATIVE_REGUL | GOBP_NEGATIVE_REGULATION_OF_SMOOT | 18 | 0.472799 | 1.4798285 | 0.06478405 | 0.17305264 | 1 | 3811 | tags=39%, list=16%, si | EN_3 |
| GOBP_PROTEIN_CONTAI | GOBP_PROTEIN_CONTAINING_COMPLEX_D | 247 | 0.280123 | 1.4796607 | 0.00874317 | 0.17302942 | 1 | 3177 | tags=19%, list=13%, si | EN_3 |
| GOBP_NUCLEOLUS_ORGA | GOBP_NUCLEOLUS_ORGANIZATION | 17 | 0.478493 | 1.4795539 | 0.068292685 | 0.17296074 | 1 | 1925 | tags=24%, list=8%, sig | EN_3 |
| GOBP_POSITIVE_REGULA | GOBP_POSITIVE_REGULATION_OF_TRIGLYC | 24 | 0.430881 | 1.4793037 | 0.074484944 | 0.17305093 | 1 | 3864 | tags=25%, list=16%, si | EN_3 |
| HUMAN_T-CELL_LEUKEM | HUMAN_T-CELL_LEUKEMIA_VIRUS_1_INFEC | 215 | 0.286873 | 1.4787761 | 0.014348786 | 0.17346203 | 1 | 7026 | tags=41%, list=30%, si | EN_3 |
| GOBP_GLUCOSE_6_PHOS | GOBP_GLUCOSE_6_PHOSPHATE_METABOLI | 28 | 0.411734 | 1.4785248 | 0.072013095 | 0.17354631 | 1 | 3307 | tags=25%, list=14%, si | EN_3 |
| GOBP_REGULATION_OF_ | GOBP_REGULATION_OF_GASTRULATION | 16 | 0.498766 | 1.4784542 | 0.07717042 | 0.17343836 | 1 | 1489 | tags=19%, list=6%, sig | EN_3 |
| GOBP_POSITIVE_REGULA | GOBP_POSITIVE_REGULATION_OF_LEUKOC | 159 | 0.298056 | 1.4782765 | 0.011862396 | 0.17342937 | 1 | 3851 | tags=25%, list=16%, si | EN_3 |
| GOBP_AMINOGLYCAN_B | GOBP_AMINOGLYCAN_BIOSYNTHETIC_PRO | 60 | 0.351852 | 1.4781849 | 0.035856575 | 0.17333496 | 1 | 4348 | tags=28%, list=18%, si | EN_3 |
| GOBP_MAINTENANCE_O | GOBP_MAINTENANCE_OF_PROTEIN_LOCAT | 25 | 0.433562 | 1.4779208 | 0.056634303 | 0.17344108 | 1 | 4934 | tags=44%, list=21%, si | EN_3 |
| GOBP_STRIATED_MUSCL | GOBP_STRIATED_MUSCLE_CELL_APOPTOTI | 73 | 0.335424 | 1.4777696 | 0.035763413 | 0.17342654 | 1 | 4017 | tags=25%, list=17%, si | EN_3 |
| GOBP_ACTIN_FILAMENT_ | GOBP_ACTIN_FILAMENT_DEPOLYMERIZATI | 57 | 0.353945 | 1.477749 | 0.056497175 | 0.17327552 | 1 | 7578 | tags=47%, list=32%, si | EN_3 |
| GOBP_CEREBELLAR_PUR | GOBP_CEREBELLAR_PURKINJE_CELL_LAYER_ | 16 | 0.471207 | 1.4775444 | 0.06324787 | 0.17331544 | 1 | 249 | tags=13%, list=1%, sig | EN_3 |
| PANCREATIC_CANCER | PANCREATIC_CANCER | 74 | 0.336756 | 1.4774554 | 0.030872483 | 0.17323443 | 1 | 5419 | tags=28%, list=23%, si | EN_3 |
| GOBP_GOLGI_TO_PLASM | GOBP_GOLGI_TO_PLASMA_MEMBRANE_PR | 38 | 0.383776 | 1.4771807 | 0.06576981 | 0.17336956 | 1 | 3331 | tags=21%, list=14%, si | EN_3 |
| GOBP_REGULATION_OF_ | GOBP_REGULATION_OF_LIPASE_ACTIVITY | 50 | 0.36279 | 1.476671 | 0.04674221 | 0.17374912 | 1 | 2990 | tags=22%, list=13%, si | EN_3 |
| GOBP_ENDOTHELIAL_CE | GOBP_ENDOTHELIAL_CELL_MIGRATION | 219 | 0.285317 | 1.4765642 | 0.0055555557 | 0.17366387 | 1 | 7106 | tags=39%, list=30%, si | EN_3 |
| GOBP_POSITIVE_REGULA | GOBP_POSITIVE_REGULATION_OF_ACTIN_F | 67 | 0.344861 | 1.4756321 | 0.031914894 | 0.17457007 | 1 | 5302 | tags=31%, list=22%, si | EN_3 |
| GOBP_NEGATIVE_REGUL | GOBP_NEGATIVE_REGULATION_OF_AUTOP | 81 | 0.329204 | 1.4753898 | 0.034528553 | 0.1746326 | 1 | 3791 | tags=25%, list=16%, si | EN_3 |
| GOBP_LAMELLIPODIUM_ | GOBP_LAMELLIPODIUM_ASSEMBLY | 73 | 0.330315 | 1.475319 | 0.037760418 | 0.17454037 | 1 | 5189 | tags=29%, list=22%, si | EN_3 |
| GOBP_PROTEIN_K48_LIN | GOBP_PROTEIN_K48_LINKED_DEUBIQUITIN | 29 | 0.412999 | 1.4751022 | 0.062992126 | 0.17459129 | 1 | 6029 | tags=48%, list=26%, si | EN_3 |
| GOBP_POSITIVE_REGULA | GOBP_POSITIVE_REGULATION_OF_PROTEIN | 75 | 0.332069 | 1.4748929 | 0.03230148 | 0.17464848 | 1 | 3670 | tags=23%, list=16%, si | EN_3 |
| GOBP_NON_MEMBRANE | GOBP_NON_MEMBRANE_BOUNDED_ORGA | 405 | 0.267223 | 1.4747186 | 0.0010559662 | 0.17464444 | 1 | 5302 | tags=29%, list=22%, si | EN_3 |
| GOBP_LONG_TERM_MEM | GOBP_LONG_TERM_MEMORY | 44 | 0.376618 | 1.47388 | 0.04950495 | 0.17540509 | 1 | 6432 | tags=45%, list=27%, si | EN_3 |
| GOBP_ANGIOGENESIS_IN | GOBP_ANGIOGENESIS_INVOLVED_IN_WOU | 25 | 0.429424 | 1.473599 | 0.06635802 | 0.17552388 | 1 | 3017 | tags=24%, list=13%, si | EN_3 |
| GOBP_VIRAL_BUDDING | GOBP_VIRAL_BUDDING | 18 | 0.460644 | 1.473498 | 0.086319216 | 0.17545034 | 1 | 4771 | tags=44%, list=20%, si | EN_3 |
| GOBP_TRNA_WOBBLE_B | GOBP_TRNA_WOBBLE_BASE_MODIFICATIO | 19 | 0.454632 | 1.4733077 | 0.07560138 | 0.17544928 | 1 | 2890 | tags=32%, list=12%, si | EN_3 |
| GLYCEROPHOSPHOLIPID_ | GLYCEROPHOSPHOLIPID_METABOLISM | 92 | 0.32137 | 1.4729887 | 0.03267974 | 0.17563874 | 1 | 4378 | tags=27%, list=19%, si | EN_3 |
| GOBP_REGULATION_OF_ | GOBP_REGULATION_OF_T_HELPER_17_TYP | 33 | 0.403475 | 1.4727819 | 0.060559005 | 0.17566672 | 1 | 2870 | tags=24%, list=12%, si | EN_3 |
| GOBP_REGULATION_OF_ | GOBP_REGULATION_OF_SUPRAMOLECULAR | 378 | 0.270189 | 1.4724752 | 0.0010570824 | 0.1758212 | 1 | 5467 | tags=26%, list=23%, si | EN_3 |
| GOBP_REGULATION_OF_ | GOBP_REGULATION_OF_NUCLEOTIDE_BIOS | 32 | 0.402247 | 1.4719484 | 0.054263566 | 0.17622408 | 1 | 6348 | tags=41%, list=27%, si | EN_3 |
| GOBP_POSITIVE_REGULA | GOBP_POSITIVE_REGULATION_OF_PROTEIN | 24 | 0.4274 | 1.4719334 | 0.05970149 | 0.17605193 | 1 | 3321 | tags=29%, list=14%, si | EN_3 |
| GOBP_HUMORAL_IMMU | GOBP_HUMORAL_IMMUNE_RESPONSE_ME | 44 | 0.373524 | 1.4717516 | 0.052857142 | 0.1760843 | 1 | 3500 | tags=30%, list=15%, si | EN_3 |
| GOBP_MAST_CELL_MIGR | GOBP_MAST_CELL_MIGRATION | 17 | 0.473416 | 1.471737 | 0.06785137 | 0.175919 | 1 | 1097 | tags=18%, list=5%, sig | EN_3 |
| GOBP_NUCLEAR_TRANSC | GOBP_NUCLEAR_TRANSCRIBED_MRNA_CAT | 118 | 0.308365 | 1.4708112 | 0.025188917 | 0.1768568 | 1 | 7539 | tags=43%, list=32%, si | EN_3 |
| GOBP_PROTEIN_DESTABI | GOBP_PROTEIN_DESTABILIZATION | 56 | 0.347401 | 1.4706205 | 0.04494382 | 0.17691313 | 1 | 7467 | tags=46%, list=32%, si | EN_3 |
| RNA_TRANSPORT | RNA_TRANSPORT | 154 | 0.293904 | 1.4701707 | 0.016110472 | 0.17725544 | 1 | 7771 | tags=49%, list=33%, si | EN_3 |
| GOBP_MUSCLE_TISSUE_D | GOBP_MUSCLE_TISSUE_DEVELOPMENT | 464 | 0.265283 | 1.4701265 | 0.0041067763 | 0.17711969 | 1 | 3963 | tags=19%, list=17%, si | EN_3 |
| GOBP_C_TERMINAL_PRO | GOBP_C_TERMINAL_PROTEIN_AMINO_ACID | 20 | 0.457758 | 1.4699464 | 0.07072368 | 0.17714591 | 1 | 1080 | tags=20%, list=5%, sig | EN_3 |
| MORPHINE_ADDICTION | MORPHINE_ADDICTION | 85 | 0.323793 | 1.4687233 | 0.029411765 | 0.1784512 | 1 | 6251 | tags=39%, list=26%, si | EN_3 |
| GOBP_POSITIVE_REGULA | GOBP_POSITIVE_REGULATION_OF_DNA_TE | 59 | 0.348717 | 1.4683892 | 0.045519203 | 0.17861967 | 1 | 4040 | tags=24%, list=17%, si | EN_3 |
| GOBP_CARBOHYDRATE_D | GOBP_CARBOHYDRATE_DERIVATIVE_CATAB | 245 | 0.280639 | 1.4678713 | 0.008830022 | 0.17906608 | 1 | 5375 | tags=30%, list=23%, si | EN_3 |
| GOBP_NUCLEOPHAGY | GOBP_NUCLEOPHAGY | 24 | 0.434187 | 1.4675821 | 0.0686901 | 0.17921107 | 1 | 6659 | tags=46%, list=28%, si | EN_3 |
| GOBP_NEGATIVE_REGUL | GOBP_NEGATIVE_REGULATION_OF_DNA_B | 37 | 0.383027 | 1.4673593 | 0.059970014 | 0.1792946 | 1 | 4743 | tags=32%, list=20%, si | EN_3 |
| GOBP_NEGATIVE_REGUL | GOBP_NEGATIVE_REGULATION_OF_SMALL_ | 108 | 0.312432 | 1.4673191 | 0.031094527 | 0.17916015 | 1 | 4514 | tags=31%, list=19%, si | EN_3 |
| GOBP_INTRINSIC_APOPT | GOBP_INTRINSIC_APOPTOTIC_SIGNALING_ | 63 | 0.341101 | 1.4667101 | 0.047954865 | 0.17965308 | 1 | 7975 | tags=48%, list=34%, si | EN_3 |

| | | | | | | | | | | |
|---|---|---|---|---|---|---|---|---|---|---|
| GOBP_DNA_DUPLEX_UN\ | GOBP_DNA_DUPLEX_UNWINDING | 43 | 0.373273 | 1.466202 | 0.05652174 | 0.18002142 | 1 | 5387 | tags=44%, list=23%, si | EN_3 |
| GOBP_LEUKOCYTE_ACTIV | GOBP_LEUKOCYTE_ACTIVATION_INVOLVED_ | 53 | 0.356474 | 1.4658542 | 0.04964539 | 0.18027104 | 1 | 4076 | tags=32%, list=17%, si | EN_3 |
| GOBP_REGULATION_OF_ | GOBP_REGULATION_OF_OXIDATIVE_STRESS | 39 | 0.378467 | 1.4653897 | 0.05909091 | 0.18061414 | 1 | 3814 | tags=26%, list=16%, si | EN_3 |
| GOBP_CYTOSKELETON_D | GOBP_CYTOSKELETON_DEPENDENT_INTRA( | 196 | 0.286065 | 1.4652919 | 0.015204678 | 0.18053332 | 1 | 8313 | tags=49%, list=35%, si | EN_3 |
| GOBP_OSSIFICATION | GOBP_OSSIFICATION | 400 | 0.267199 | 1.4649605 | 0.0021030493 | 0.18073353 | 1 | 4329 | tags=23%, list=18%, si | EN_3 |
| INSULIN_SIGNALING_PAT | INSULIN_SIGNALING_PATHWAY | 131 | 0.29819 | 1.4647149 | 0.024390243 | 0.18081594 | 1 | 7597 | tags=41%, list=32%, si | EN_3 |
| INSULIN_RESISTANCE | INSULIN_RESISTANCE | 105 | 0.316463 | 1.464709 | 0.025510205 | 0.18063658 | 1 | 5460 | tags=26%, list=23%, si | EN_3 |
| GOBP_POSITIVE_REGULA | GOBP_POSITIVE_REGULATION_OF_LIPID_BI | 104 | 0.308307 | 1.4636766 | 0.027918782 | 0.18161966 | 1 | 4257 | tags=26%, list=18%, si | EN_3 |
| GOBP_CARBOHYDRATE_B | GOBP_CARBOHYDRATE_BIOSYNTHETIC_PRC | 197 | 0.284369 | 1.4628513 | 0.018223235 | 0.18238817 | 1 | 5160 | tags=26%, list=22%, si | EN_3 |
| GOBP_REGULATION_OF_ | GOBP_REGULATION_OF_ADAPTIVE_IMMUN | 215 | 0.280007 | 1.4623312 | 0.012571429 | 0.18283378 | 1 | 5393 | tags=31%, list=23%, si | EN_3 |
| GOBP_RIBOSOMAL_LARG | GOBP_RIBOSOMAL_LARGE_SUBUNIT_ASSEN | 23 | 0.42739 | 1.4620411 | 0.07073955 | 0.18299955 | 1 | 4509 | tags=48%, list=19%, si | EN_3 |
| GOBP_POSITIVE_REGULA | GOBP_POSITIVE_REGULATION_OF_PROTEIN | 63 | 0.344386 | 1.4608448 | 0.047954865 | 0.18425643 | 1 | 3670 | tags=25%, list=16%, si | EN_3 |
| GOBP_POSITIVE_REGULA | GOBP_POSITIVE_REGULATION_OF_DNA_RE | 135 | 0.298899 | 1.4608194 | 0.03151515 | 0.18409678 | 1 | 3339 | tags=19%, list=14%, si | EN_3 |
| GOBP_PLASMINOGEN_A( | GOBP_PLASMINOGEN_ACTIVATION | 23 | 0.431854 | 1.4606756 | 0.08386076 | 0.18410414 | 1 | 3017 | tags=26%, list=13%, si | EN_3 |
| GOBP_BILE_ACID_METAB | GOBP_BILE_ACID_METABOLIC_PROCESS | 41 | 0.379238 | 1.4602219 | 0.049707603 | 0.18446282 | 1 | 3253 | tags=24%, list=14%, si | EN_3 |
| GOBP_REGULATION_OF_ | GOBP_REGULATION_OF_ACTIN_FILAMENT_ | 387 | 0.268585 | 1.4602027 | 0.0041884817 | 0.18429734 | 1 | 5011 | tags=25%, list=21%, si | EN_3 |
| GOBP_ANOIKIS | GOBP_ANOIKIS | 31 | 0.409921 | 1.4601291 | 0.06554878 | 0.18420056 | 1 | 3615 | tags=29%, list=15%, si | EN_3 |
| GOBP_REGULATION_OF_ | GOBP_REGULATION_OF_PEPTIDYL_THREON | 45 | 0.369335 | 1.4600765 | 0.057142857 | 0.18407716 | 1 | 3266 | tags=20%, list=14%, si | EN_3 |
| GOBP_INTRACELLULAR_R | GOBP_INTRACELLULAR_RECEPTOR_SIGNALI | 325 | 0.269879 | 1.4599985 | 0.0074074073 | 0.18397972 | 1 | 3321 | tags=18%, list=14%, si | EN_3 |
| GOBP_REGULATION_OF_ | GOBP_REGULATION_OF_SEQUESTERING_OF | 15 | 0.480219 | 1.4599581 | 0.0794702 | 0.18383963 | 1 | 4092 | tags=27%, list=17%, si | EN_3 |
| HUMAN_PAPILLOMAVIRU | HUMAN_PAPILLOMAVIRUS_INFECTION | 315 | 0.27334 | 1.4599165 | 0.008519702 | 0.18371195 | 1 | 5496 | tags=27%, list=23%, si | EN_3 |
| GOBP_LEUKOCYTE_MEDI | GOBP_LEUKOCYTE_MEDIATED_IMMUNITY | 400 | 0.26595 | 1.459766 | 0.0020833334 | 0.18367437 | 1 | 3547 | tags=21%, list=15%, si | EN_3 |
| GOBP_NEGATIVE_REGUL | GOBP_NEGATIVE_REGULATION_OF_LEUKO( | 46 | 0.36167 | 1.4595509 | 0.067349926 | 0.1837483 | 1 | 4612 | tags=30%, list=20%, si | EN_3 |
| CYTOSOLIC_DNA-SENSIN( | CYTOSOLIC_DNA-SENSING_PATHWAY | 46 | 0.365187 | 1.4585793 | 0.057909604 | 0.18473938 | 1 | 2736 | tags=20%, list=12%, si | EN_3 |
| GOBP_POSITIVE_REGULA | GOBP_POSITIVE_REGULATION_OF_MACROF | 29 | 0.406889 | 1.4584141 | 0.06726457 | 0.18475284 | 1 | 2936 | tags=24%, list=12%, si | EN_3 |
| GOBP_POSITIVE_REGULA | GOBP_POSITIVE_REGULATION_OF_INTERLE | 55 | 0.352082 | 1.4583722 | 0.04895105 | 0.1846179 | 1 | 3017 | tags=25%, list=13%, si | EN_3 |
| GOBP_BILE_ACID_BIOSYN | GOBP_BILE_ACID_BIOSYNTHETIC_PROCESS | 27 | 0.41533 | 1.4582888 | 0.06373293 | 0.18455102 | 1 | 1334 | tags=19%, list=6%, sig | EN_3 |
| GOBP_REGULATION_OF_ | GOBP_REGULATION_OF_PROTEIN_CONTAIN | 131 | 0.298605 | 1.4580653 | 0.026410565 | 0.1846313 | 1 | 3167 | tags=20%, list=13%, si | EN_3 |
| GOBP_MUSCLE_CELL_AP( | GOBP_MUSCLE_CELL_APOPTOTIC_PROCESS | 124 | 0.296497 | 1.4577588 | 0.024844721 | 0.18478025 | 1 | 4129 | tags=22%, list=17%, si | EN_3 |
| GOBP_NEGATIVE_REGUL | GOBP_NEGATIVE_REGULATION_OF_MUSCL | 69 | 0.332514 | 1.4576938 | 0.04489796 | 0.18466152 | 1 | 4017 | tags=28%, list=17%, si | EN_3 |
| GOBP_PROTEIN_N_LINKE | GOBP_PROTEIN_N_LINKED_GLYCOSYLATION | 61 | 0.344448 | 1.4575956 | 0.04931507 | 0.18459454 | 1 | 5646 | tags=33%, list=24%, si | EN_3 |
| GOBP_CELLULAR_RESPON | GOBP_CELLULAR_RESPONSE_TO_RADIATION | 165 | 0.290532 | 1.457461 | 0.010309278 | 0.18457012 | 1 | 5457 | tags=28%, list=23%, si | EN_3 |
| GOBP_INNER_CELL_MASS | GOBP_INNER_CELL_MASS_CELL_PROLIFERA | 16 | 0.472894 | 1.456864 | 0.07952622 | 0.1850817 | 1 | 1256 | tags=25%, list=5%, sig | EN_3 |
| GOBP_RESPONSE_TO_MI | GOBP_RESPONSE_TO_MISFOLDED_PROTEIN | 18 | 0.456732 | 1.4564333 | 0.086319216 | 0.18543936 | 1 | 4135 | tags=39%, list=18%, si | EN_3 |
| GOBP_ANTIGEN_PROCES | GOBP_ANTIGEN_PROCESSING_AND_PRESEN | 28 | 0.407382 | 1.4561311 | 0.05203252 | 0.18561497 | 1 | 3448 | tags=29%, list=15%, si | EN_3 |
| GOBP_ORGANELLE_MEM | GOBP_ORGANELLE_MEMBRANE_FUSION | 123 | 0.301441 | 1.455957 | 0.03404792 | 0.18567444 | 1 | 5806 | tags=33%, list=25%, si | EN_3 |
| GOBP_RETINAL_METABO | GOBP_RETINAL_METABOLIC_PROCESS | 16 | 0.482585 | 1.4559377 | 0.08117444 | 0.18550403 | 1 | 3856 | tags=38%, list=16%, si | EN_3 |
| GOBP_CYTOKINE_MEDIAT | GOBP_CYTOKINE_MEDIATED_SIGNALING_P | 421 | 0.263694 | 1.4557508 | 0.0041753654 | 0.1855322 | 1 | 3441 | tags=20%, list=15%, si | EN_3 |
| GOBP_POSITIVE_REGULA | GOBP_POSITIVE_REGULATION_OF_STEROID | 33 | 0.389684 | 1.4549439 | 0.057142857 | 0.18633188 | 1 | 2847 | tags=30%, list=12%, si | EN_3 |
| GOBP_SEQUESTERING_O | GOBP_SEQUESTERING_OF_TRIGLYCERIDE | 18 | 0.461532 | 1.4545616 | 0.066450566 | 0.18659571 | 1 | 4092 | tags=28%, list=17%, si | EN_3 |
| GOBP_POSITIVE_REGULA | GOBP_POSITIVE_REGULATION_OF_CYCLIN_ | 20 | 0.442256 | 1.4536206 | 0.087458745 | 0.1875768 | 1 | 3321 | tags=30%, list=14%, si | EN_3 |
| GOBP_GLYCOPROTEIN_M | GOBP_GLYCOPROTEIN_METABOLIC_PROCES | 342 | 0.26816 | 1.4531163 | 0.0031678986 | 0.18803032 | 1 | 5937 | tags=31%, list=25%, si | EN_3 |
| GOBP_PLASMA_MEMBRA | GOBP_PLASMA_MEMBRANE_TUBULATION | 19 | 0.453916 | 1.4529513 | 0.0732899 | 0.18802136 | 1 | 4594 | tags=42%, list=19%, si | EN_3 |
| GOBP_POSITIVE_REGULA | GOBP_POSITIVE_REGULATION_OF_INTRACE | 147 | 0.294393 | 1.4529448 | 0.020481927 | 0.18784148 | 1 | 7899 | tags=48%, list=33%, si | EN_3 |
| GOBP_STEROL_TRANSPO | GOBP_STEROL_TRANSPORT | 109 | 0.308012 | 1.4527013 | 0.03 | 0.18793851 | 1 | 1784 | tags=14%, list=8%, sig | EN_3 |
| GOBP_MYOBLAST_DIFFEI | GOBP_MYOBLAST_DIFFERENTIATION | 114 | 0.30628 | 1.4525858 | 0.031407036 | 0.18789332 | 1 | 3647 | tags=19%, list=15%, si | EN_3 |
| GOBP_REGULATION_OF_ | GOBP_REGULATION_OF_NEUROTROPHIN_T | 17 | 0.475302 | 1.4520774 | 0.068292685 | 0.18831973 | 1 | 6110 | tags=53%, list=26%, si | EN_3 |
| GOBP_MACROPHAGE_CY | GOBP_MACROPHAGE_CYTOKINE_PRODUCT | 51 | 0.354789 | 1.451727 | 0.057522126 | 0.18859604 | 1 | 3699 | tags=27%, list=16%, si | EN_3 |
| GOBP_POSITIVE_REGULA | GOBP_POSITIVE_REGULATION_OF_ERK1_AN | 201 | 0.28258 | 1.4506751 | 0.019384265 | 0.18965417 | 1 | 5136 | tags=30%, list=22%, si | EN_3 |
| GOBP_CYTOPLASMIC_PA | GOBP_CYTOPLASMIC_PATTERN_RECOGNITI | 159 | 0.287442 | 1.4502395 | 0.021226415 | 0.1899839 | 1 | 3321 | tags=21%, list=14%, si | EN_3 |
| GOBP_REGULATION_OF_ | GOBP_REGULATION_OF_INFLAMMATORY_F | 348 | 0.267076 | 1.4501498 | 0.0031712474 | 0.18991065 | 1 | 3185 | tags=19%, list=13%, si | EN_3 |
| GOBP_FORELIMB_MORPI | GOBP_FORELIMB_MORPHOGENESIS | 40 | 0.367288 | 1.4497472 | 0.06525038 | 0.1902183 | 1 | 3761 | tags=30%, list=16%, si | EN_3 |
| GOBP_REGULATION_OF_ | GOBP_REGULATION_OF_FAT_CELL_DIFFERE | 143 | 0.292956 | 1.4495598 | 0.021924483 | 0.19026543 | 1 | 2250 | tags=13%, list=10%, si | EN_3 |
| GOBP_NEUTROPHIL_MEC | GOBP_NEUTROPHIL_MEDIATED_IMMUNITY | 37 | 0.381676 | 1.4494848 | 0.064615384 | 0.19017325 | 1 | 2453 | tags=24%, list=10%, si | EN_3 |
| GOBP_REGULATION_OF_ | GOBP_REGULATION_OF_STRIATED_MUSCLE | 97 | 0.314892 | 1.449438 | 0.027603513 | 0.19004017 | 1 | 3586 | tags=25%, list=15%, si | EN_3 |
| GOBP_RESPONSE_TO_SA | GOBP_RESPONSE_TO_SALT_STRESS | 20 | 0.450081 | 1.4492573 | 0.067708336 | 0.1900997 | 1 | 1810 | tags=25%, list=8%, sig | EN_3 |
| PROTEIN_EXPORT | PROTEIN_EXPORT | 26 | 0.41211 | 1.4490215 | 0.07436709 | 0.19021223 | 1 | 4909 | tags=42%, list=21%, si | EN_3 |
| GOBP_METHYLATION | GOBP_METHYLATION | 199 | 0.285036 | 1.4488275 | 0.009237875 | 0.1902777 | 1 | 5887 | tags=33%, list=25%, si | EN_3 |
| BIOSYNTHESIS_OF_UNSA | BIOSYNTHESIS_OF_UNSATURATED_FATTY_A | 31 | 0.399244 | 1.4481764 | 0.074132495 | 0.1909009 | 1 | 4777 | tags=42%, list=20%, si | EN_3 |
| GLYCEROLIPID_METABOL | GLYCEROLIPID_METABOLISM | 57 | 0.344734 | 1.4475253 | 0.053900708 | 0.19153194 | 1 | 4125 | tags=28%, list=17%, si | EN_3 |
| OTHER_TYPES_OF_O-GLY | OTHER_TYPES_OF_O-GLYCAN_BIOSYNTHESI | 21 | 0.439672 | 1.4471675 | 0.08056872 | 0.19172534 | 1 | 4710 | tags=38%, list=20%, si | EN_3 |
| GOBP_ANTIMICROBIAL_H | GOBP_ANTIMICROBIAL_HUMORAL_IMMUN | 79 | 0.325199 | 1.4470512 | 0.051282052 | 0.1917072 | 1 | 3182 | tags=25%, list=13%, si | EN_3 |
| GOBP_NEGATIVE_REGUL | GOBP_NEGATIVE_REGULATION_OF_PHOSPI | 398 | 0.264012 | 1.4462194 | 0.0041710115 | 0.19262168 | 1 | 6110 | tags=31%, list=26%, si | EN_3 |
| GOBP_ACTIN_FILAMENT_ | GOBP_ACTIN_FILAMENT_ORGANIZATION | 450 | 0.263585 | 1.4459937 | 0.003051882 | 0.19274837 | 1 | 5042 | tags=26%, list=21%, si | EN_3 |
| GOBP_LIPOPROTEIN_CAT | GOBP_LIPOPROTEIN_CATABOLIC_PROCESS | 17 | 0.465481 | 1.4458973 | 0.08471761 | 0.19267581 | 1 | 4386 | tags=35%, list=19%, si | EN_3 |
| GOBP_REGULATION_OF_ | GOBP_REGULATION_OF_MITOTIC_SPINDLE_ | 18 | 0.45395 | 1.445731 | 0.07109005 | 0.19270591 | 1 | 4771 | tags=39%, list=20%, si | EN_3 |
| GOBP_POSITIVE_REGULA | GOBP_POSITIVE_REGULATION_OF_T_HELPE | 15 | 0.493079 | 1.445627 | 0.07907743 | 0.19264609 | 1 | 1625 | tags=20%, list=7%, sig | EN_3 |
| GOBP_RNA_METHYLATIO | GOBP_RNA_METHYLATION | 65 | 0.339264 | 1.4455378 | 0.060522698 | 0.1925682 | 1 | 3878 | tags=31%, list=16%, si | EN_3 |
| GOBP_SUPEROXIDE_ANI( | GOBP_SUPEROXIDE_ANION_GENERATION | 43 | 0.369337 | 1.4451138 | 0.07204611 | 0.1928568 | 1 | 3634 | tags=28%, list=15%, si | EN_3 |
| GOBP_SUPPRESSION_OF_ | GOBP_SUPPRESSION_OF_VIRAL_RELEASE_B | 22 | 0.432778 | 1.4444351 | 0.093699515 | 0.19347562 | 1 | 4117 | tags=36%, list=17%, si | EN_3 |
| GOBP_PROTEIN_O_LINKE | GOBP_PROTEIN_O_LINKED_MANNOSYLATIO | 16 | 0.463417 | 1.4439018 | 0.098615915 | 0.19401595 | 1 | 2982 | tags=38%, list=13%, si | EN_3 |
| GOBP_DE_NOVO_PROTEI | GOBP_DE_NOVO_PROTEIN_FOLDING | 38 | 0.373727 | 1.4438142 | 0.08256881 | 0.19394033 | 1 | 5314 | tags=37%, list=23%, si | EN_3 |
| GOBP_REGULATION_OF_ | GOBP_REGULATION_OF_CALCIUM_ION_DEI | 41 | 0.370891 | 1.4437354 | 0.06518518 | 0.19384636 | 1 | 1049 | tags=17%, list=4%, sig | EN_3 |
| GOBP_CARDIAC_MYOFIB | GOBP_CARDIAC_MYOFIBRIL_ASSEMBLY | 17 | 0.464139 | 1.4428629 | 0.10183639 | 0.19475298 | 1 | 3129 | tags=35%, list=13%, si | EN_3 |
| GOBP_REGULATION_OF_ | GOBP_REGULATION_OF_ANTIGEN_PROCESS | 21 | 0.441696 | 1.4424871 | 0.08072488 | 0.19501436 | 1 | 3074 | tags=29%, list=13%, si | EN_3 |
| GOBP_SYNAPTIC_VESICLE | GOBP_SYNAPTIC_VESICLE_LOCALIZATION | 64 | 0.338668 | 1.4423095 | 0.047945205 | 0.19503674 | 1 | 5764 | tags=33%, list=24%, si | EN_3 |
| GOBP_VESICLE_MEDIATE | GOBP_VESICLE_MEDIATED_TRANSPORT_TO | 158 | 0.2863 | 1.4420747 | 0.027972028 | 0.1951062 | 1 | 4139 | tags=21%, list=18%, si | EN_3 |
| GOBP_TRIGLYCERIDE_CA | GOBP_TRIGLYCERIDE_CATABOLIC_PROCESS | 23 | 0.430694 | 1.4415284 | 0.0776699 | 0.19557786 | 1 | 964 | tags=17%, list=4%, sig | EN_3 |
| GOBP_REGULATION_OF_ | GOBP_REGULATION_OF_SODIUM_ION_TRA | 45 | 0.361143 | 1.441183 | 0.069503546 | 0.19584253 | 1 | 5521 | tags=40%, list=23%, si | EN_3 |
| GOBP_NEGATIVE_REGUL | GOBP_NEGATIVE_REGULATION_OF_SMAD_ | 23 | 0.436563 | 1.4410932 | 0.08589952 | 0.19579665 | 1 | 2467 | tags=22%, list=10%, si | EN_3 |
| GOBP_BONE_MATURATI( | GOBP_BONE_MATURATION | 29 | 0.410554 | 1.4403784 | 0.088188976 | 0.19645575 | 1 | 1506 | tags=17%, list=6%, sig | EN_3 |
| GOBP_NEGATIVE_REGUL | GOBP_NEGATIVE_REGULATION_OF_CATAB( | 339 | 0.26391 | 1.4403762 | 0.007415254 | 0.19626747 | 1 | 4413 | tags=22%, list=19%, si | EN_3 |

| | | | | | | | | | | |
|---|---|---|---|---|---|---|---|---|---|---|
| GOBP_TRANSLESION_SYN | GOBP_TRANSLESION_SYNTHESIS | 22 | 0.429393 | 1.4403268 | 0.100164205 | 0.19612294 | 1 | 6930 | tags=59%, list=29%, si | EN_3 |
| GOBP_REGULATION_OF_ | GOBP_REGULATION_OF_FATTY_ACID_META | 99 | 0.310598 | 1.440292 | 0.033942558 | 0.19597529 | 1 | 3926 | tags=24%, list=17%, si | EN_3 |
| GOBP_POLYOL_BIOSYNTH | GOBP_POLYOL_BIOSYNTHETIC_PROCESS | 58 | 0.338053 | 1.4402707 | 0.046767537 | 0.19581197 | 1 | 2983 | tags=21%, list=13%, si | EN_3 |
| GOBP_NEGATIVE_REGUL | GOBP_NEGATIVE_REGULATION_OF_CYTOKI | 292 | 0.269671 | 1.44023 | 0.0064655175 | 0.19565804 | 1 | 3791 | tags=22%, list=16%, si | EN_3 |
| GOBP_NEGATIVE_REGUL | GOBP_NEGATIVE_REGULATION_OF_CELLUL | 100 | 0.313198 | 1.4392091 | 0.03860523 | 0.19667889 | 1 | 4159 | tags=26%, list=18%, si | EN_3 |
| GOBP_STRIATED_MUSCLI | GOBP_STRIATED_MUSCLE_CELL_DIFFERENT | 330 | 0.265943 | 1.4391088 | 0.0075512407 | 0.19663131 | 1 | 3657 | tags=20%, list=15%, si | EN_3 |
| GOBP_POSITIVE_REGULA | GOBP_POSITIVE_REGULATION_OF_EPITHELI | 216 | 0.276194 | 1.4391057 | 0.012643678 | 0.19644755 | 1 | 5266 | tags=27%, list=22%, si | EN_3 |
| GOBP_DNA_DAMAGE_RE | GOBP_DNA_DAMAGE_RESPONSE_SIGNAL_T | 17 | 0.462904 | 1.4385989 | 0.087314665 | 0.19690415 | 1 | 6035 | tags=53%, list=26%, si | EN_3 |
| GOBP_MORPHOGENESIS_ | GOBP_MORPHOGENESIS_OF_AN_EPITHELIA | 17 | 0.467484 | 1.4374319 | 0.093587525 | 0.19824141 | 1 | 5266 | tags=41%, list=22%, si | EN_3 |
| GOBP_LAMELLIPODIUM_ | GOBP_LAMELLIPODIUM_ORGANIZATION | 92 | 0.310925 | 1.4373785 | 0.038809832 | 0.19812015 | 1 | 5189 | tags=26%, list=22%, si | EN_3 |
| GOBP_ENDOTHELIAL_CEL | GOBP_ENDOTHELIAL_CELL_PROLIFERATION | 164 | 0.284561 | 1.4373709 | 0.026442308 | 0.19793794 | 1 | 3567 | tags=19%, list=15%, si | EN_3 |
| GOBP_CELL_SUBSTRATE_ | GOBP_CELL_SUBSTRATE_ADHESION | 354 | 0.265001 | 1.4370558 | 0.0063829785 | 0.1981398 | 1 | 6263 | tags=32%, list=27%, si | EN_3 |
| GOBP_RESPONSE_TO_EN | GOBP_RESPONSE_TO_ENDOPLASMIC_RETIC | 261 | 0.273311 | 1.4370265 | 0.012061403 | 0.19798832 | 1 | 7785 | tags=43%, list=33%, si | EN_3 |
| GOBP_NEGATIVE_REGUL | GOBP_NEGATIVE_REGULATION_OF_ESTABL | 148 | 0.294074 | 1.4367833 | 0.019559903 | 0.1981103 | 1 | 5231 | tags=30%, list=22%, si | EN_3 |
| GOBP_POSITIVE_REGULA | GOBP_POSITIVE_REGULATION_OF_RNA_SPL | 50 | 0.355851 | 1.4367694 | 0.064896755 | 0.19794112 | 1 | 7056 | tags=42%, list=30%, si | EN_3 |
| GOBP_APPENDAGE_MOR | GOBP_APPENDAGE_MORPHOGENESIS | 164 | 0.287163 | 1.4366455 | 0.026589595 | 0.19791217 | 1 | 3761 | tags=20%, list=16%, si | EN_3 |
| GOBP_PROTEIN_PROCESS | GOBP_PROTEIN_PROCESSING | 237 | 0.272668 | 1.4363412 | 0.016519824 | 0.19805993 | 1 | 5877 | tags=32%, list=25%, si | EN_3 |
| GOBP_REGULATION_OF_ | GOBP_REGULATION_OF_COLLAGEN_METAB | 61 | 0.340102 | 1.4357698 | 0.039402176 | 0.19861123 | 1 | 3017 | tags=21%, list=13%, si | EN_3 |
| GOBP_LIPOPOLYSACCHAR | GOBP_LIPOPOLYSACCHARIDE_MEDIATED_SI | 57 | 0.34055 | 1.4357691 | 0.051460363 | 0.19842315 | 1 | 3949 | tags=25%, list=17%, si | EN_3 |
| GOBP_NEUROINFLAMMA | GOBP_NEUROINFLAMMATORY_RESPONSE | 84 | 0.322268 | 1.4355136 | 0.055033557 | 0.19856267 | 1 | 4076 | tags=27%, list=17%, si | EN_3 |
| GOBP_TISSUE_MIGRATIO | GOBP_TISSUE_MIGRATION | 314 | 0.267892 | 1.435475 | 0.004278075 | 0.19843335 | 1 | 3745 | tags=19%, list=16%, si | EN_3 |
| GOBP_NEGATIVE_REGUL | GOBP_NEGATIVE_REGULATION_OF_PROTEI | 236 | 0.276367 | 1.435056 | 0.014380531 | 0.1987461 | 1 | 5231 | tags=27%, list=22%, si | EN_3 |
| GOBP_MEMBRANE_PROT | GOBP_MEMBRANE_PROTEIN_ECTODOMAIN | 47 | 0.360333 | 1.4349058 | 0.058309037 | 0.1987367 | 1 | 6127 | tags=40%, list=26%, si | EN_3 |
| GOBP_REGULATION_OF_ | GOBP_REGULATION_OF_LYSOSOMAL_LUME | 23 | 0.422933 | 1.4346026 | 0.097288676 | 0.1989322 | 1 | 7200 | tags=48%, list=30%, si | EN_3 |
| GOBP_SYNAPTIC_VESICLE | GOBP_SYNAPTIC_VESICLE_CYCLE | 242 | 0.275734 | 1.4345775 | 0.0174482 | 0.198775 | 1 | 4256 | tags=22%, list=18%, si | EN_3 |
| GOBP_REGULATION_OF_ | GOBP_REGULATION_OF_GLIAL_CELL_APOPT | 15 | 0.487116 | 1.4343195 | 0.09201389 | 0.19892293 | 1 | 4774 | tags=47%, list=20%, si | EN_3 |
| GOBP_GLYCOSYLATION | GOBP_GLYCOSYLATION | 218 | 0.277417 | 1.4342059 | 0.011198209 | 0.1988824 | 1 | 4822 | tags=26%, list=20%, si | EN_3 |
| GOBP_REGULATION_OF_ | GOBP_REGULATION_OF_CALCIUM_ION_TRA | 265 | 0.271514 | 1.4341927 | 0.01655629 | 0.19871573 | 1 | 3363 | tags=20%, list=14%, si | EN_3 |
| GOBP_METANEPHRIC_NE | GOBP_METANEPHRIC_NEPHRON_DEVELOPI | 35 | 0.371044 | 1.433669 | 0.06896552 | 0.19913258 | 1 | 3208 | tags=26%, list=14%, si | EN_3 |
| GOBP_PLATELET_ACTIVAT | GOBP_PLATELET_ACTIVATION | 94 | 0.308215 | 1.4331859 | 0.03889586 | 0.19944441 | 1 | 3429 | tags=21%, list=15%, si | EN_3 |
| GOBP_RESPIRATORY_SYS | GOBP_RESPIRATORY_SYSTEM_DEVELOPMEN | 256 | 0.270116 | 1.4328591 | 0.011049724 | 0.1996666 | 1 | 5425 | tags=27%, list=23%, si | EN_3 |
| GOBP_DEVELOPMENTAL_ | GOBP_DEVELOPMENTAL_INDUCTION | 37 | 0.381984 | 1.4327735 | 0.06240487 | 0.19957092 | 1 | 5266 | tags=38%, list=22%, si | EN_3 |
| GOBP_NEGATIVE_REGUL | GOBP_NEGATIVE_REGULATION_OF_DEFENS | 242 | 0.271436 | 1.4325308 | 0.017660044 | 0.19968815 | 1 | 3184 | tags=19%, list=13%, si | EN_3 |
| GOBP_FATTY_ACID_ELON | GOBP_FATTY_ACID_ELONGATION | 15 | 0.484502 | 1.4323785 | 0.09043478 | 0.19971651 | 1 | 4777 | tags=53%, list=20%, si | EN_3 |
| GOBP_REGULATION_OF_ | GOBP_REGULATION_OF_SULFUR_METABOL | 15 | 0.474927 | 1.4321908 | 0.10963455 | 0.1997916 | 1 | 249 | tags=13%, list=1%, sig | EN_3 |
| GOBP_ESTABLISHMENT_( | GOBP_ESTABLISHMENT_OR_MAINTENANCE | 25 | 0.410891 | 1.4315959 | 0.07915994 | 0.20035563 | 1 | 6301 | tags=44%, list=27%, si | EN_3 |
| GOBP_REGULATION_OF_ | GOBP_REGULATION_OF_BINDING | 331 | 0.264153 | 1.4312894 | 0.00857449 | 0.2005207 | 1 | 4056 | tags=21%, list=17%, si | EN_3 |
| GOBP_POSITIVE_REGULA | GOBP_POSITIVE_REGULATION_OF_PHAGOC | 74 | 0.327599 | 1.4307779 | 0.03960396 | 0.20096672 | 1 | 3926 | tags=26%, list=17%, si | EN_3 |
| GOBP_POST_TRANSCRIPT | GOBP_POST_TRANSCRIPTIONAL_REGULATIO | 480 | 0.259808 | 1.4305233 | 0.001025641 | 0.20112297 | 1 | 7554 | tags=42%, list=32%, si | EN_3 |
| GOBP_MEMBRANE_LIPID | GOBP_MEMBRANE_LIPID_METABOLIC_PRO | 186 | 0.280572 | 1.4304042 | 0.017301038 | 0.20109496 | 1 | 5720 | tags=32%, list=24%, si | EN_3 |
| GOBP_LEUKOTRIENE_ME | GOBP_LEUKOTRIENE_METABOLIC_PROCESS | 20 | 0.446971 | 1.4303505 | 0.09178744 | 0.20099004 | 1 | 3755 | tags=35%, list=16%, si | EN_3 |
| GOBP_LEUKOCYTE_MIGR | GOBP_LEUKOCYTE_MIGRATION | 363 | 0.266955 | 1.430119 | 0.0020986358 | 0.2010942 | 1 | 3851 | tags=21%, list=16%, si | EN_3 |
| GOBP_PROTEIN_CATABO | GOBP_PROTEIN_CATABOLIC_PROCESS_IN_T | 22 | 0.428489 | 1.4294585 | 0.079617836 | 0.20170243 | 1 | 7391 | tags=41%, list=31%, si | EN_3 |
| GOBP_POSITIVE_REGULA | GOBP_POSITIVE_REGULATION_OF_APOPTO | 161 | 0.289741 | 1.4289808 | 0.033254158 | 0.20210887 | 1 | 4975 | tags=32%, list=21%, si | EN_3 |
| GOBP_ENDOPLASMIC_RE | GOBP_ENDOPLASMIC_RETICULUM_ORGANI | 90 | 0.311214 | 1.4287001 | 0.047244094 | 0.20226412 | 1 | 7393 | tags=38%, list=31%, si | EN_3 |
| GOBP_NEGATIVE_REGUL | GOBP_NEGATIVE_REGULATION_OF_ALPHA_ | 16 | 0.469727 | 1.4285785 | 0.112026356 | 0.20224954 | 1 | 845 | tags=19%, list=4%, sig | EN_3 |
| GOBP_REGULATION_OF_ | GOBP_REGULATION_OF_CELLULAR_RESPON | 308 | 0.264084 | 1.4283339 | 0.010775862 | 0.20237269 | 1 | 5418 | tags=26%, list=23%, si | EN_3 |
| GOBP_INFLAMMATORY_ | GOBP_INFLAMMATORY_RESPONSE_TO_WO | 30 | 0.393746 | 1.4281595 | 0.075235106 | 0.20240846 | 1 | 3795 | tags=27%, list=16%, si | EN_3 |
| GOBP_PROTEIN_LOCALIZ | GOBP_PROTEIN_LOCALIZATION_TO_PHAGO | 16 | 0.455163 | 1.4279834 | 0.109298535 | 0.20244846 | 1 | 6659 | tags=56%, list=28%, si | EN_3 |
| GOBP_POSITIVE_REGULA | GOBP_POSITIVE_REGULATION_OF_PROTEIN | 78 | 0.32458 | 1.4274548 | 0.05148005 | 0.20296744 | 1 | 6068 | tags=32%, list=26%, si | EN_3 |
| GOBP_GLYCOPROTEIN_B | GOBP_GLYCOPROTEIN_BIOSYNTHETIC_PRO | 282 | 0.268146 | 1.4272658 | 0.014084507 | 0.203009 | 1 | 5713 | tags=30%, list=24%, si | EN_3 |
| GOBP_BONE_REMODELIN | GOBP_BONE_REMODELING | 102 | 0.305209 | 1.4266789 | 0.04918033 | 0.20358953 | 1 | 2708 | tags=18%, list=11%, si | EN_3 |
| GOBP_REGULATION_OF_ | GOBP_REGULATION_OF_RNA_SPLICING | 178 | 0.281639 | 1.4259068 | 0.03128621 | 0.20443466 | 1 | 7216 | tags=40%, list=31%, si | EN_3 |
| GOBP_CELL_COMMUNIC | GOBP_CELL_COMMUNICATION_BY_ELECTRI | 19 | 0.43638 | 1.4258581 | 0.08846761 | 0.20429733 | 1 | 2384 | tags=26%, list=10%, si | EN_3 |
| GOBP_POSITIVE_REGULA | GOBP_POSITIVE_REGULATION_OF_RECEPTO | 19 | 0.448346 | 1.425823 | 0.08399366 | 0.20417117 | 1 | 6824 | tags=53%, list=29%, si | EN_3 |
| GOBP_NEGATIVE_REGUL | GOBP_NEGATIVE_REGULATION_OF_TRANSF | 65 | 0.32776 | 1.4257655 | 0.043723553 | 0.20405166 | 1 | 1878 | tags=15%, list=8%, sig | EN_3 |
| GOBP_GLIAL_CELL_ACTIV | GOBP_GLIAL_CELL_ACTIVATION | 59 | 0.337371 | 1.4257284 | 0.059065934 | 0.20392267 | 1 | 4076 | tags=31%, list=17%, si | EN_3 |
| GOBP_PRODUCTION_OF_ | GOBP_PRODUCTION_OF_MOLECULAR_MED | 270 | 0.26774 | 1.4251627 | 0.00990099 | 0.20448628 | 1 | 3080 | tags=19%, list=13%, si | EN_3 |
| GOBP_VESICLE_LOCALIZA | GOBP_VESICLE_LOCALIZATION | 218 | 0.2757 | 1.425089 | 0.019079685 | 0.20438768 | 1 | 6019 | tags=31%, list=25%, si | EN_3 |
| GOBP_VASCULAR_PROCE | GOBP_VASCULAR_PROCESS_IN_CIRCULATO | 238 | 0.27065 | 1.4249516 | 0.013392857 | 0.20436457 | 1 | 4835 | tags=26%, list=20%, si | EN_3 |
| GOBP_RESCUE_OF_STALL | GOBP_RESCUE_OF_STALLED_RIBOSOME | 25 | 0.416937 | 1.4244919 | 0.0829421 | 0.20475356 | 1 | 7518 | tags=44%, list=32%, si | EN_3 |
| GOBP_VASCULAR_WOUN | GOBP_VASCULAR_WOUND_HEALING | 18 | 0.448148 | 1.4244252 | 0.08922559 | 0.20465575 | 1 | 3017 | tags=28%, list=13%, si | EN_3 |
| GOBP_ERYTHROCYTE_DE | GOBP_ERYTHROCYTE_DEVELOPMENT | 50 | 0.347757 | 1.4235083 | 0.08213256 | 0.20566313 | 1 | 2408 | tags=22%, list=10%, si | EN_3 |
| GOBP_CELL_CHEMOTAXIS | GOBP_CELL_CHEMOTAXIS | 295 | 0.264442 | 1.4233251 | 0.010787486 | 0.20567681 | 1 | 5220 | tags=28%, list=22%, si | EN_3 |
| GOBP_EMBRYONIC_APPE | GOBP_EMBRYONIC_APPENDAGE_MORPHO( | 134 | 0.290786 | 1.422725 | 0.025240384 | 0.2062387 | 1 | 3761 | tags=20%, list=16%, si | EN_3 |
| GOBP_REGULATION_OF_ | GOBP_REGULATION_OF_PROTEIN_SERINE_ | 280 | 0.270257 | 1.422723 | 0.009814613 | 0.206056 | 1 | 7237 | tags=43%, list=31%, si | EN_3 |
| GOBP_RESPONSE_TO_ME | GOBP_RESPONSE_TO_MECHANICAL_STIMU | 160 | 0.281894 | 1.4214858 | 0.026963657 | 0.20743407 | 1 | 5467 | tags=30%, list=23%, si | EN_3 |
| GOBP_PHOSPHATIDYLINC | GOBP_PHOSPHATIDYLINOSITOL_BIOSYNTHE | 95 | 0.305615 | 1.4214218 | 0.054637864 | 0.20733334 | 1 | 3794 | tags=24%, list=16%, si | EN_3 |
| GOBP_NUCLEAR_TRANSP | GOBP_NUCLEAR_TRANSPORT | 337 | 0.264834 | 1.4212539 | 0.013993542 | 0.20734875 | 1 | 5843 | tags=30%, list=25%, si | EN_3 |
| GOBP_MYELOID_CELL_DE | GOBP_MYELOID_CELL_DEVELOPMENT | 98 | 0.301774 | 1.4206789 | 0.044720497 | 0.20789494 | 1 | 2489 | tags=19%, list=11%, si | EN_3 |
| GOBP_CALMODULIN_DEF | GOBP_CALMODULIN_DEPENDENT_KINASE_ | 16 | 0.467413 | 1.4205052 | 0.113793105 | 0.20794345 | 1 | 3115 | tags=38%, list=13%, si | EN_3 |
| GOBP_POSITIVE_REGULA | GOBP_POSITIVE_REGULATION_OF_COLLAGI | 39 | 0.370311 | 1.4204824 | 0.06518283 | 0.20778269 | 1 | 2058 | tags=21%, list=9%, sig | EN_3 |
| GOBP_NEGATIVE_REGUL | GOBP_NEGATIVE_REGULATION_OF_ORGAN | 47 | 0.351264 | 1.4203439 | 0.05693431 | 0.20775923 | 1 | 4701 | tags=32%, list=20%, si | EN_3 |
| GOBP_REGULATION_OF_ | GOBP_REGULATION_OF_MITOCHONDRIAL_ | 20 | 0.436032 | 1.4202774 | 0.11130137 | 0.20765674 | 1 | 3586 | tags=30%, list=15%, si | EN_3 |
| GOBP_POSITIVE_REGULA | GOBP_POSITIVE_REGULATION_OF_PHOSPH | 182 | 0.278437 | 1.4200237 | 0.04090909 | 0.20778167 | 1 | 4833 | tags=26%, list=20%, si | EN_3 |
| GOBP_MORPHOGENESIS_ | GOBP_MORPHOGENESIS_OF_AN_ENDOTHE | 19 | 0.448168 | 1.4196322 | 0.0968254 | 0.20810899 | 1 | 6236 | tags=53%, list=26%, si | EN_3 |
| GOBP_REGULATION_OF_ | GOBP_REGULATION_OF_AMYLOID_BETA_CI | 18 | 0.449143 | 1.4193668 | 0.093650796 | 0.2082811 | 1 | 4076 | tags=39%, list=17%, si | EN_3 |
| GOBP_CELLULAR_RESPON | GOBP_CELLULAR_RESPONSE_TO_FATTY_AC | 30 | 0.390381 | 1.4193062 | 0.07817109 | 0.20816146 | 1 | 3743 | tags=33%, list=16%, si | EN_3 |
| GOBP_AMYLOID_PRECUR | GOBP_AMYLOID_PRECURSOR_PROTEIN_ME | 75 | 0.322618 | 1.4192871 | 0.062251657 | 0.20800047 | 1 | 5870 | tags=32%, list=25%, si | EN_3 |
| GOBP_POSITIVE_REGULA | GOBP_POSITIVE_REGULATION_OF_IMMUNE | 266 | 0.266424 | 1.4192051 | 0.015317286 | 0.20791845 | 1 | 3080 | tags=18%, list=13%, si | EN_3 |
| GOBP_B_CELL_MEDIATED | GOBP_B_CELL_MEDIATED_IMMUNITY | 141 | 0.292825 | 1.4191903 | 0.032926828 | 0.2077643 | 1 | 3500 | tags=21%, list=15%, si | EN_3 |
| GOBP_CELLULAR_RESPON | GOBP_CELLULAR_RESPONSE_TO_DEXAMETI | 23 | 0.419662 | 1.4188684 | 0.08728943 | 0.20800708 | 1 | 2829 | tags=22%, list=12%, si | EN_3 |

| | | | | | | | | | | |
|---|---|---|---|---|---|---|---|---|---|---|
| GOBP_PYRIMIDINE_CONT | GOBP_PYRIMIDINE_CONTAINING_COMPOU | 26 | 0.411853 | 1.4177715 | 0.09878419 | 0.20918019 | 1 | 3666 | tags=23%, list=16%, si | EN_3 |
| GOBP_CARDIAC_CELL_DE | GOBP_CARDIAC_CELL_DEVELOPMENT | 114 | 0.297673 | 1.4176563 | 0.05 | 0.20913446 | 1 | 3588 | tags=21%, list=15%, si | EN_3 |
| GOBP_REGULATION_OF_ | GOBP_REGULATION_OF_ENDOPLASMIC_RE | 24 | 0.41069 | 1.4175985 | 0.09259259 | 0.20902427 | 1 | 4034 | tags=33%, list=17%, si | EN_3 |
| GOBP_RESPONSE_TO_ST | GOBP_RESPONSE_TO_STARVATION | 181 | 0.27439 | 1.4167153 | 0.02640643 | 0.20999983 | 1 | 6373 | tags=36%, list=27%, si | EN_3 |
| GOBP_REGULATION_OF_ | GOBP_REGULATION_OF_ATP_BIOSYNTHETI | 20 | 0.440031 | 1.4166505 | 0.09328969 | 0.20989646 | 1 | 5859 | tags=40%, list=25%, si | EN_3 |
| GOBP_PHAGOCYTOSIS_R | GOBP_PHAGOCYTOSIS_RECOGNITION | 24 | 0.417941 | 1.4165859 | 0.08227848 | 0.20978154 | 1 | 4314 | tags=33%, list=18%, si | EN_3 |
| GOBP_PIGMENT_GRANU | GOBP_PIGMENT_GRANULE_ORGANIZATION | 39 | 0.369088 | 1.4158748 | 0.07692308 | 0.21051776 | 1 | 3061 | tags=23%, list=13%, si | EN_3 |
| FRUCTOSE_AND_MANNO | FRUCTOSE_AND_MANNOSE_METABOLISM | 33 | 0.382822 | 1.4158312 | 0.07936508 | 0.21039419 | 1 | 5323 | tags=36%, list=23%, si | EN_3 |
| GOBP_REGULATION_OF_ | GOBP_REGULATION_OF_PHOSPHATASE_AC | 61 | 0.329578 | 1.4157777 | 0.06364922 | 0.21026966 | 1 | 2326 | tags=16%, list=10%, si | EN_3 |
| TUBERCULOSIS | TUBERCULOSIS | 152 | 0.289321 | 1.4155822 | 0.036384977 | 0.21032524 | 1 | 5115 | tags=28%, list=22%, si | EN_3 |
| GOBP_REGULATION_OF_ | GOBP_REGULATION_OF_ATP_METABOLIC_ | 77 | 0.317941 | 1.415556 | 0.060565274 | 0.21018209 | 1 | 6685 | tags=39%, list=28%, si | EN_3 |
| GOBP_INNATE_IMMUNE_ | GOBP_INNATE_IMMUNE_RESPONSE_ACTIV | 254 | 0.268098 | 1.4155085 | 0.010055866 | 0.21004865 | 1 | 3467 | tags=20%, list=15%, si | EN_3 |
| GOBP_REGULATION_OF_ | GOBP_REGULATION_OF_CARBOHYDRATE_ | 196 | 0.276521 | 1.4154803 | 0.020524515 | 0.2098935 | 1 | 5418 | tags=27%, list=23%, si | EN_3 |
| GOBP_RIG_I_SIGNALING_ | GOBP_RIG_I_SIGNALING_PATHWAY | 24 | 0.418114 | 1.4154665 | 0.07371795 | 0.20972505 | 1 | 6335 | tags=46%, list=27%, si | EN_3 |
| GOBP_PROTEIN_LOCALIZ | GOBP_PROTEIN_LOCALIZATION_TO_CELL_C | 24 | 0.414056 | 1.4150388 | 0.0732899 | 0.21010034 | 1 | 7373 | tags=50%, list=31%, si | EN_3 |
| GOBP_VASCULAR_ASSOC | GOBP_VASCULAR_ASSOCIATED_SMOOTH_M | 39 | 0.368806 | 1.4146031 | 0.097201765 | 0.21049806 | 1 | 1864 | tags=18%, list=8%, sig | EN_3 |
| GOBP_CANONICAL_NF_K | GOBP_CANONICAL_NF_KAPPAB_SIGNAL_TF | 247 | 0.266708 | 1.4145854 | 0.013259669 | 0.21033059 | 1 | 5011 | tags=28%, list=21%, si | EN_3 |
| GOBP_REACTIVE_NITROG | GOBP_REACTIVE_NITROGEN_SPECIES_MET | 92 | 0.310688 | 1.4141849 | 0.05026455 | 0.21064766 | 1 | 3791 | tags=21%, list=16%, si | EN_3 |
| GOBP_ADAPTIVE_THERM | GOBP_ADAPTIVE_THERMOGENESIS | 169 | 0.284657 | 1.4140145 | 0.034924332 | 0.21067134 | 1 | 3151 | tags=20%, list=13%, si | EN_3 |
| GOBP_PROTEIN_K63_LIN | GOBP_PROTEIN_K63_LINKED_UBIQUITINAT | 68 | 0.326201 | 1.4138417 | 0.061842106 | 0.21071365 | 1 | 4417 | tags=26%, list=19%, si | EN_3 |
| GOBP_WHITE_FAT_CELL_ | GOBP_WHITE_FAT_CELL_DIFFERENTIATION | 22 | 0.425113 | 1.4136028 | 0.1095008 | 0.21081202 | 1 | 2842 | tags=23%, list=12%, si | EN_3 |
| GOBP_CELL_CYCLE_CHEC | GOBP_CELL_CYCLE_CHECKPOINT_SIGNALIN | 191 | 0.280702 | 1.413198 | 0.026558891 | 0.21114045 | 1 | 5457 | tags=30%, list=23%, si | EN_3 |
| HIF-1_SIGNALING_PATHV | HIF-1_SIGNALING_PATHWAY | 98 | 0.306134 | 1.4131256 | 0.047919292 | 0.2110492 | 1 | 5530 | tags=30%, list=23%, si | EN_3 |
| FATTY_ACID_ELONGATIO | FATTY_ACID_ELONGATION | 27 | 0.407474 | 1.4125854 | 0.0755287 | 0.21155632 | 1 | 6022 | tags=41%, list=26%, si | EN_3 |
| GOBP_NEGATIVE_REGUL | GOBP_NEGATIVE_REGULATION_OF_AMYLO | 20 | 0.438219 | 1.4124528 | 0.098726116 | 0.21154042 | 1 | 6170 | tags=50%, list=26%, si | EN_3 |
| GOBP_LUNG_MORPHOGI | GOBP_LUNG_MORPHOGENESIS | 70 | 0.324853 | 1.4112614 | 0.070866145 | 0.21291824 | 1 | 6572 | tags=41%, list=28%, si | EN_3 |
| GOBP_TRANSCRIPTION_I | GOBP_TRANSCRIPTION_INITIATION_AT_RN | 148 | 0.287089 | 1.411246 | 0.048019208 | 0.21275106 | 1 | 6927 | tags=39%, list=29%, si | EN_3 |
| HUMAN_IMMUNODEFICI | HUMAN_IMMUNODEFICIENCY_VIRUS_1_IN | 201 | 0.275659 | 1.4111303 | 0.037931036 | 0.21270023 | 1 | 5802 | tags=31%, list=25%, si | EN_3 |
| GOBP_POSITIVE_REGULA | GOBP_POSITIVE_REGULATION_OF_VASCUL | 19 | 0.433932 | 1.4107965 | 0.10823909 | 0.21294597 | 1 | 577 | tags=16%, list=2%, sig | EN_3 |
| GOBP_POSITIVE_REGULA | GOBP_POSITIVE_REGULATION_OF_REACTIV | 81 | 0.31204 | 1.4107777 | 0.061923582 | 0.21279463 | 1 | 4209 | tags=25%, list=18%, si | EN_3 |
| NOTCH_SIGNALING_PATH | NOTCH_SIGNALING_PATHWAY | 47 | 0.352618 | 1.4095095 | 0.08797654 | 0.21427871 | 1 | 5668 | tags=36%, list=24%, si | EN_3 |
| GOBP_NEGATIVE_REGUL | GOBP_NEGATIVE_REGULATION_OF_FATTY_ | 16 | 0.469065 | 1.4090556 | 0.09967846 | 0.21472584 | 1 | 3926 | tags=44%, list=17%, si | EN_3 |
| GOBP_CELL_SUBSTRATE_ | GOBP_CELL_SUBSTRATE_JUNCTION_ORGAN | 105 | 0.299828 | 1.4090455 | 0.04109589 | 0.21455286 | 1 | 6942 | tags=40%, list=29%, si | EN_3 |
| GOBP_POSITIVE_REGULA | GOBP_POSITIVE_REGULATION_OF_MRNA_ | 37 | 0.376168 | 1.4088782 | 0.08748115 | 0.21459769 | 1 | 3881 | tags=24%, list=16%, si | EN_3 |
| GOBP_MAMMARY_GLAN | GOBP_MAMMARY_GLAND_EPITHELIAL_CEL | 32 | 0.386222 | 1.4084206 | 0.09259259 | 0.21502556 | 1 | 1468 | tags=16%, list=6%, sig | EN_3 |
| GOBP_REGULATION_OF_ | GOBP_REGULATION_OF_CELL_GROWTH_IN | 33 | 0.378602 | 1.4081368 | 0.08296296 | 0.21517146 | 1 | 3586 | tags=30%, list=15%, si | EN_3 |
| GOBP_REGULATION_OF_ | GOBP_REGULATION_OF_MITOCHONDRIAL_ | 23 | 0.416074 | 1.4080178 | 0.095846646 | 0.21514341 | 1 | 4799 | tags=39%, list=20%, si | EN_3 |
| GOBP_CELL_SURFACE_PA | GOBP_CELL_SURFACE_PATTERN_RECOGNIT | 70 | 0.32894 | 1.407889 | 0.053547524 | 0.21512783 | 1 | 4168 | tags=26%, list=18%, si | EN_3 |
| GOBP_MEMBRANE_BIOG | GOBP_MEMBRANE_BIOGENESIS | 63 | 0.327078 | 1.4075459 | 0.05298913 | 0.21540107 | 1 | 5371 | tags=33%, list=23%, si | EN_3 |
| GOBP_REGULATION_OF_ | GOBP_REGULATION_OF_IMMUNE_EFFECTO | 406 | 0.255532 | 1.4074368 | 0.007322176 | 0.21534419 | 1 | 3780 | tags=21%, list=16%, si | EN_3 |
| GOBP_INTERLEUKIN_6_P | GOBP_INTERLEUKIN_6_PRODUCTION | 173 | 0.274871 | 1.4071689 | 0.030267753 | 0.21549955 | 1 | 4271 | tags=25%, list=18%, si | EN_3 |
| GOBP_FC_RECEPTOR_SIG | GOBP_FC_RECEPTOR_SIGNALING_PATHWA | 16 | 0.465655 | 1.4068447 | 0.108474575 | 0.21570659 | 1 | 4961 | tags=38%, list=21%, si | EN_3 |
| GOBP_VASCULAR_TRANS | GOBP_VASCULAR_TRANSPORT | 22 | 0.419395 | 1.4067248 | 0.108843535 | 0.21569556 | 1 | 2917 | tags=27%, list=12%, si | EN_3 |
| GOBP_REGULATION_OF_ | GOBP_REGULATION_OF_CHEMOTAXIS | 206 | 0.26919 | 1.4064404 | 0.025056947 | 0.21589655 | 1 | 3851 | tags=20%, list=16%, si | EN_3 |
| GOBP_PROTEIN_LOCALIZ | GOBP_PROTEIN_LOCALIZATION_TO_CELL_P | 391 | 0.258161 | 1.4061474 | 0.010537407 | 0.21609457 | 1 | 5012 | tags=25%, list=21%, si | EN_3 |
| GOBP_REGULATION_OF_ | GOBP_REGULATION_OF_AMYLOID_BETA_F | 37 | 0.368874 | 1.4061356 | 0.092675634 | 0.21592236 | 1 | 6170 | tags=38%, list=26%, si | EN_3 |
| GOBP_MUCUS_SECRETIO | GOBP_MUCUS_SECRETION | 22 | 0.425023 | 1.406045 | 0.11093248 | 0.21585506 | 1 | 3336 | tags=27%, list=14%, si | EN_3 |
| NON-SMALL_CELL_LUNG | NON-SMALL_CELL_LUNG_CANCER | 65 | 0.325703 | 1.4058474 | 0.06207827 | 0.21593562 | 1 | 3635 | tags=18%, list=15%, si | EN_3 |
| GOBP_SYMBIONT_ENTRY | GOBP_SYMBIONT_ENTRY_INTO_HOST | 97 | 0.299826 | 1.405703 | 0.056962024 | 0.21595822 | 1 | 4129 | tags=29%, list=17%, si | EN_3 |
| GOBP_DNA_TEMPLATED_ | GOBP_DNA_TEMPLATED_DNA_REPLICATION | 54 | 0.342696 | 1.4047 | 0.06926407 | 0.2170585 | 1 | 2029 | tags=17%, list=9%, sig | EN_3 |
| GOBP_REGULATION_OF_ | GOBP_REGULATION_OF_PHOSPHOLIPASE_A | 36 | 0.368614 | 1.404594 | 0.09050445 | 0.2170301 | 1 | 2990 | tags=22%, list=13%, si | EN_3 |
| GOBP_RESOLUTION_OF_ | GOBP_RESOLUTION_OF_MEIOTIC_RECOMB | 17 | 0.454435 | 1.4042811 | 0.10787671 | 0.21724036 | 1 | 289 | tags=18%, list=1%, sig | EN_3 |
| GOBP_NEGATIVE_REGUL | GOBP_NEGATIVE_REGULATION_OF_OSTEO | 58 | 0.337625 | 1.4040768 | 0.06993007 | 0.21731341 | 1 | 5761 | tags=38%, list=24%, si | EN_3 |
| GOBP_NEGATIVE_REGUL | GOBP_NEGATIVE_REGULATION_OF_EXOCYT | 37 | 0.370812 | 1.4031975 | 0.076576576 | 0.21820658 | 1 | 3980 | tags=24%, list=17%, si | EN_3 |
| GOBP_TISSUE_REMODELI | GOBP_TISSUE_REMODELING | 202 | 0.274073 | 1.4021289 | 0.029816514 | 0.21938202 | 1 | 2842 | tags=15%, list=12%, si | EN_3 |
| GLYCOSAMINOGLYCAN_I | GLYCOSAMINOGLYCAN_DEGRADATION | 17 | 0.446377 | 1.4018084 | 0.10016695 | 0.21967423 | 1 | 5365 | tags=41%, list=23%, si | EN_3 |
| GOBP_PSEUDOURIDINE_ | GOBP_PSEUDOURIDINE_SYNTHESIS | 18 | 0.438175 | 1.4017617 | 0.11423841 | 0.21955606 | 1 | 3442 | tags=33%, list=15%, si | EN_3 |
| CHOLESTEROL_METABOL | CHOLESTEROL_METABOLISM | 44 | 0.350256 | 1.4008698 | 0.08421053 | 0.2206039 | 1 | 4906 | tags=34%, list=21%, si | EN_3 |
| RNA_DEGRADATION | RNA_DEGRADATION | 75 | 0.322525 | 1.4006447 | 0.07506702 | 0.22070469 | 1 | 5072 | tags=24%, list=21%, si | EN_3 |
| GOBP_POSITIVE_REGULA | GOBP_POSITIVE_REGULATION_OF_RESPON | 73 | 0.317817 | 1.4004229 | 0.066755675 | 0.22083338 | 1 | 6290 | tags=36%, list=27%, si | EN_3 |
| GOBP_NEGATIVE_REGUL | GOBP_NEGATIVE_REGULATION_OF_FIBROB | 44 | 0.356495 | 1.4001802 | 0.08588064 | 0.22097707 | 1 | 4029 | tags=30%, list=17%, si | EN_3 |
| GOBP_BIOLOGICAL_PRO | GOBP_BIOLOGICAL_PROCESS_INVOLVED_IN | 140 | 0.290532 | 1.3995528 | 0.045614034 | 0.22160438 | 1 | 4771 | tags=31%, list=20%, si | EN_3 |
| GOBP_NEGATIVE_REGUL | GOBP_NEGATIVE_REGULATION_OF_PROTEI | 29 | 0.387169 | 1.3995394 | 0.11217949 | 0.221439 | 1 | 3672 | tags=21%, list=16%, si | EN_3 |
| GOBP_ERYTHROCYTE_HO | GOBP_ERYTHROCYTE_HOMEOSTASIS | 163 | 0.278159 | 1.399214 | 0.03868032 | 0.22169706 | 1 | 5684 | tags=28%, list=24%, si | EN_3 |
| PENTOSE_AND_GLUCURO | PENTOSE_AND_GLUCURONATE_INTERCONV | 22 | 0.422352 | 1.3990452 | 0.0968254 | 0.22173241 | 1 | 4820 | tags=45%, list=20%, si | EN_3 |
| GOBP_CELL_ACTIVATION | GOBP_CELL_ACTIVATION_INVOLVED_IN_IM | 291 | 0.26036 | 1.3986993 | 0.018220793 | 0.22203907 | 1 | 3365 | tags=20%, list=14%, si | EN_3 |
| GOBP_CELLULAR_RESPON | GOBP_CELLULAR_RESPONSE_TO_GLUCOCO | 47 | 0.34454 | 1.3984134 | 0.08479532 | 0.22224733 | 1 | 2914 | tags=21%, list=12%, si | EN_3 |
| GOBP_CELLULAR_RESPON | GOBP_CELLULAR_RESPONSE_TO_ABIOTIC_S | 276 | 0.264739 | 1.3982326 | 0.017410228 | 0.22229417 | 1 | 5457 | tags=28%, list=23%, si | EN_3 |
| GOBP_POSITIVE_REGULA | GOBP_POSITIVE_REGULATION_OF_PROTEIN | 120 | 0.289617 | 1.3982089 | 0.042079207 | 0.22213794 | 1 | 5721 | tags=33%, list=24%, si | EN_3 |
| GOBP_RESPONSE_TO_INT | GOBP_RESPONSE_TO_INTERFERON_BETA | 64 | 0.320368 | 1.3980085 | 0.06911142 | 0.22222869 | 1 | 2487 | tags=20%, list=11%, si | EN_3 |
| GOBP_MYOTUBE_DIFFER | GOBP_MYOTUBE_DIFFERENTIATION | 133 | 0.287821 | 1.3967011 | 0.046398047 | 0.22381234 | 1 | 3199 | tags=18%, list=14%, si | EN_3 |
| GOBP_POSITIVE_REGULA | GOBP_POSITIVE_REGULATION_OF_VASCUL | 25 | 0.408445 | 1.3961831 | 0.1107595 | 0.22428791 | 1 | 4097 | tags=24%, list=17%, si | EN_3 |
| GOBP_NON_PROTEINOGI | GOBP_NON_PROTEINOGENIC_AMINO_ACID | 15 | 0.466686 | 1.3960631 | 0.11352254 | 0.22427413 | 1 | 1917 | tags=27%, list=8%, sig | EN_3 |
| GOBP_ESTABLISHMENT_ | GOBP_ESTABLISHMENT_OR_MAINTENANCE | 223 | 0.264736 | 1.3958038 | 0.018952062 | 0.22443622 | 1 | 7371 | tags=39%, list=31%, si | EN_3 |
| TRYPTOPHAN_METABOLI | TRYPTOPHAN_METABOLISM | 43 | 0.351221 | 1.3955815 | 0.072289154 | 0.22454514 | 1 | 3747 | tags=28%, list=16%, si | EN_3 |
| GOBP_NEGATIVE_REGUL | GOBP_NEGATIVE_REGULATION_OF_PROTEI | 90 | 0.30792 | 1.3953588 | 0.06451613 | 0.22463043 | 1 | 3167 | tags=20%, list=13%, si | EN_3 |
| GOBP_REGULATION_OF_ | GOBP_REGULATION_OF_SPINDLE_ASSEMBL | 28 | 0.39857 | 1.3951949 | 0.0968254 | 0.22466987 | 1 | 4771 | tags=32%, list=20%, si | EN_3 |
| GOBP_MONONUCLEAR_ | GOBP_MONONUCLEAR_CELL_MIGRATION | 199 | 0.274624 | 1.3948718 | 0.028058361 | 0.22490455 | 1 | 3851 | tags=23%, list=16%, si | EN_3 |
| GOBP_POSITIVE_REGULA | GOBP_POSITIVE_REGULATION_OF_TRANSL | 21 | 0.422236 | 1.394537 | 0.1028481 | 0.22519737 | 1 | 2874 | tags=29%, list=12%, si | EN_3 |
| GOBP_GOLGI_TO_PLASM | GOBP_GOLGI_TO_PLASMA_MEMBRANE_TR | 62 | 0.330928 | 1.3945208 | 0.0877193 | 0.22502834 | 1 | 3541 | tags=18%, list=15%, si | EN_3 |
| GOBP_POSITIVE_REGULA | GOBP_POSITIVE_REGULATION_OF_VASCUL | 20 | 0.433666 | 1.3942723 | 0.115 | 0.22521007 | 1 | 2058 | tags=25%, list=9%, sig | EN_3 |
| GOBP_RNA_TEMPLATED_ | GOBP_RNA_TEMPLATED_DNA_BIOSYNTHET | 63 | 0.325636 | 1.3941683 | 0.06824513 | 0.22517185 | 1 | 5418 | tags=25%, list=23%, si | EN_3 |

| | | | | | | | | | | |
|---|---|---|---|---|---|---|---|---|---|---|
| BACTERIAL_INVASION_OI | BACTERIAL_INVASION_OF_EPITHELIAL_CELL | 72 | 0.317619 | 1.3940954 | 0.05548038 | 0.22509213 | 1 | 6865 | tags=39%, list=29%, si | EN_3 |
| GOBP_SYNAPTIC_VESICLE | GOBP_SYNAPTIC_VESICLE_CYTOSKELETAL_T | 21 | 0.425159 | 1.3940129 | 0.12096774 | 0.2250215 | 1 | 3155 | tags=29%, list=13%, si | EN_3 |
| GOBP_CONNECTIVE_TISS | GOBP_CONNECTIVE_TISSUE_DEVELOPMENT | 292 | 0.258146 | 1.3939743 | 0.013071896 | 0.22489005 | 1 | 4467 | tags=22%, list=19%, si | EN_3 |
| GOBP_HEMOSTASIS | GOBP_HEMOSTASIS | 168 | 0.278554 | 1.3939213 | 0.04117647 | 0.22477236 | 1 | 3441 | tags=21%, list=15%, si | EN_3 |
| ASTHMA | ASTHMA | 16 | 0.450115 | 1.3937321 | 0.11666667 | 0.2248717 | 1 | 3066 | tags=38%, list=13%, si | EN_3 |
| GOBP_POSITIVE_REGULA | GOBP_POSITIVE_REGULATION_OF_INFLAMI | 134 | 0.286632 | 1.3937082 | 0.05269461 | 0.2247119 | 1 | 3074 | tags=21%, list=13%, si | EN_3 |
| GOBP_POSITIVE_REGULA | GOBP_POSITIVE_REGULATION_OF_STEROID | 25 | 0.400662 | 1.3937078 | 0.10737179 | 0.22452685 | 1 | 2847 | tags=36%, list=12%, si | EN_3 |
| GOBP_PROTEIN_LOCALIZ | GOBP_PROTEIN_LOCALIZATION_TO_EXTRAC | 411 | 0.255352 | 1.3936739 | 0.012565445 | 0.22437437 | 1 | 3729 | tags=19%, list=16%, si | EN_3 |
| GOBP_TYROSINE_PHOSPI | GOBP_TYROSINE_PHOSPHORYLATION_OF_S | 69 | 0.318067 | 1.3931468 | 0.08264463 | 0.22489995 | 1 | 3263 | tags=22%, list=14%, si | EN_3 |
| GOBP_LEUKOCYTE_APOP | GOBP_LEUKOCYTE_APOPTOTIC_PROCESS | 131 | 0.28578 | 1.3931127 | 0.047272727 | 0.22476223 | 1 | 5418 | tags=31%, list=23%, si | EN_3 |
| GOBP_AXONEME_ASSEM | GOBP_AXONEME_ASSEMBLY | 83 | 0.310822 | 1.3930832 | 0.04853129 | 0.22461691 | 1 | 3962 | tags=25%, list=17%, si | EN_3 |
| GOBP_MONOCARBOXYLI | GOBP_MONOCARBOXYLIC_ACID_BIOSYNTH | 203 | 0.269895 | 1.3928121 | 0.038106237 | 0.22478732 | 1 | 4796 | tags=26%, list=20%, si | EN_3 |
| GOBP_DNA_CONFORMAT | GOBP_DNA_CONFORMATION_CHANGE | 58 | 0.330807 | 1.392611 | 0.08891929 | 0.2248482 | 1 | 5387 | tags=38%, list=23%, si | EN_3 |
| GOBP_POSITIVE_REGULA | GOBP_POSITIVE_REGULATION_OF_LIPASE_ | 35 | 0.37225 | 1.3913182 | 0.090771556 | 0.2264628 | 1 | 2990 | tags=23%, list=13%, si | EN_3 |
| GOBP_REGULATION_OF_ | GOBP_REGULATION_OF_PROTEIN_LOCALIZ | 221 | 0.270032 | 1.3910737 | 0.026845638 | 0.22659373 | 1 | 5253 | tags=28%, list=22%, si | EN_3 |
| GOBP_NEGATIVE_REGUL | GOBP_NEGATIVE_REGULATION_OF_MULTIC | 17 | 0.447238 | 1.3906316 | 0.11111111 | 0.22700346 | 1 | 3139 | tags=29%, list=13%, si | EN_3 |
| GOBP_RECEPTOR_METAE | GOBP_RECEPTOR_METABOLIC_PROCESS | 80 | 0.306719 | 1.390492 | 0.07480315 | 0.22702485 | 1 | 6105 | tags=35%, list=26%, si | EN_3 |
| GOBP_ORGAN_GROWTH | GOBP_ORGAN_GROWTH | 201 | 0.272067 | 1.3902158 | 0.022274327 | 0.22716391 | 1 | 3740 | tags=19%, list=16%, si | EN_3 |
| GOBP_MAINTENANCE_OI | GOBP_MAINTENANCE_OF_BLOOD_BRAIN_E | 22 | 0.413913 | 1.3894339 | 0.08658744 | 0.22809336 | 1 | 885 | tags=18%, list=4%, sig | EN_3 |
| THYROID_CANCER | THYROID_CANCER | 36 | 0.371544 | 1.3893802 | 0.09559939 | 0.2279688 | 1 | 7005 | tags=44%, list=30%, si | EN_3 |
| GOBP_RESPONSE_TO_OX | GOBP_RESPONSE_TO_OXYGEN_LEVELS | 247 | 0.264601 | 1.3889025 | 0.029126214 | 0.22844271 | 1 | 4484 | tags=25%, list=19%, si | EN_3 |
| GOBP_POSITIVE_REGULA | GOBP_POSITIVE_REGULATION_OF_INTERFE | 45 | 0.352943 | 1.3887577 | 0.08392603 | 0.22844045 | 1 | 6185 | tags=42%, list=26%, si | EN_3 |
| GOBP_REGULATION_OF_ | GOBP_REGULATION_OF_OLIGODENDROCYT | 51 | 0.337945 | 1.388547 | 0.07380608 | 0.2285205 | 1 | 4541 | tags=25%, list=19%, si | EN_3 |
| GOBP_ANTIVIRAL_INNAT | GOBP_ANTIVIRAL_INNATE_IMMUNE_RESPO | 47 | 0.348013 | 1.388392 | 0.083215795 | 0.22854003 | 1 | 5178 | tags=32%, list=22%, si | EN_3 |
| CHRONIC_MYELOID_LEUK | CHRONIC_MYELOID_LEUKEMIA | 75 | 0.317585 | 1.3882595 | 0.070680626 | 0.22851801 | 1 | 7905 | tags=48%, list=33%, si | EN_3 |
| GOBP_MITOTIC_CELL_CY | GOBP_MITOTIC_CELL_CYCLE_PHASE_TRANS | 425 | 0.254076 | 1.3880683 | 0.008333334 | 0.22860666 | 1 | 5457 | tags=28%, list=23%, si | EN_3 |
| GOBP_PROTEOGLYCAN_E | GOBP_PROTEOGLYCAN_BIOSYNTHETIC_PRO | 61 | 0.327656 | 1.387496 | 0.08695652 | 0.22925127 | 1 | 5155 | tags=30%, list=22%, si | EN_3 |
| GOBP_POSITIVE_REGULA | GOBP_POSITIVE_REGULATION_OF_RUFFLE_ | 15 | 0.467054 | 1.3872925 | 0.122349106 | 0.22932947 | 1 | 2908 | tags=33%, list=12%, si | EN_3 |
| GOBP_NEGATIVE_REGUL | GOBP_NEGATIVE_REGULATION_OF_TOR_SI | 61 | 0.329366 | 1.3869771 | 0.07094134 | 0.22962284 | 1 | 6373 | tags=36%, list=27%, si | EN_3 |
| GOBP_REGULATION_OF_ | GOBP_REGULATION_OF_PROTEIN_IMPORT_ | 76 | 0.313396 | 1.386285 | 0.08918919 | 0.23037086 | 1 | 5279 | tags=30%, list=22%, si | EN_3 |
| GOBP_CALCIUM_ION_IM | GOBP_CALCIUM_ION_IMPORT | 75 | 0.318933 | 1.3862593 | 0.068 | 0.23022048 | 1 | 3182 | tags=24%, list=13%, si | EN_3 |
| GOBP_POSITIVE_REGULA | GOBP_POSITIVE_REGULATION_OF_RECEPTO | 17 | 0.453694 | 1.3862181 | 0.11627907 | 0.23009929 | 1 | 5191 | tags=47%, list=22%, si | EN_3 |
| GOBP_POSITIVE_REGULA | GOBP_POSITIVE_REGULATION_OF_LEUKOCY | 97 | 0.302608 | 1.3860737 | 0.0634715 | 0.23012842 | 1 | 3851 | tags=25%, list=16%, si | EN_3 |
| GOBP_PROTEIN_DEPOLYI | GOBP_PROTEIN_DEPOLYMERIZATION | 125 | 0.283618 | 1.3858986 | 0.039615847 | 0.23016886 | 1 | 3167 | tags=20%, list=13%, si | EN_3 |
| GOBP_POSITIVE_REGULA | GOBP_POSITIVE_REGULATION_OF_LIPID_M | 147 | 0.282871 | 1.3856386 | 0.048076924 | 0.23037556 | 1 | 2847 | tags=18%, list=12%, si | EN_3 |
| GOBP_POSITIVE_REGULA | GOBP_POSITIVE_REGULATION_OF_HEART_( | 49 | 0.343589 | 1.385604 | 0.07102273 | 0.23023345 | 1 | 3586 | tags=24%, list=15%, si | EN_3 |
| GOBP_NAD_CATABOLIC_ | GOBP_NAD_CATABOLIC_PROCESS | 21 | 0.41864 | 1.3851216 | 0.11308562 | 0.23071818 | 1 | 5019 | tags=38%, list=21%, si | EN_3 |
| GOBP_POSITIVE_REGULA | GOBP_POSITIVE_REGULATION_OF_CHROM( | 107 | 0.293041 | 1.3848324 | 0.052238807 | 0.23089808 | 1 | 4251 | tags=19%, list=18%, si | EN_3 |
| GOBP_POSITIVE_REGULA | GOBP_POSITIVE_REGULATION_OF_TYPE_II_ | 80 | 0.311445 | 1.3846306 | 0.057692308 | 0.23100503 | 1 | 2936 | tags=23%, list=12%, si | EN_3 |
| FERROPTOSIS | FERROPTOSIS | 37 | 0.368003 | 1.3846061 | 0.0962406 | 0.23085707 | 1 | 5418 | tags=41%, list=23%, si | EN_3 |
| GOBP_REGULATION_OF_ | GOBP_REGULATION_OF_MAMMARY_GLANI | 19 | 0.42965 | 1.3842964 | 0.11755233 | 0.23110852 | 1 | 3635 | tags=26%, list=15%, si | EN_3 |
| GOBP_REGULATION_OF_ | GOBP_REGULATION_OF_LEUKOCYTE_CHEM | 122 | 0.287402 | 1.3835021 | 0.04213135 | 0.23197506 | 1 | 4750 | tags=27%, list=20%, si | EN_3 |
| GOBP_REGULATION_OF_ | GOBP_REGULATION_OF_LEUKOCYTE_MEDI/ | 253 | 0.260839 | 1.383181 | 0.018418202 | 0.23223367 | 1 | 5341 | tags=30%, list=23%, si | EN_3 |
| GOBP_PROTEASOME_ME | GOBP_PROTEASOME_MEDIATED_UBIQUITIN | 368 | 0.254337 | 1.3829756 | 0.009404388 | 0.23233594 | 1 | 5546 | tags=26%, list=23%, si | EN_3 |
| GOBP_VASCULAR_ASSOC | GOBP_VASCULAR_ASSOCIATED_SMOOTH_N | 16 | 0.4529 | 1.3829068 | 0.11945392 | 0.23224668 | 1 | 2829 | tags=25%, list=12%, si | EN_3 |
| GOBP_PROTEIN_KINASE_ | GOBP_PROTEIN_KINASE_C_SIGNALING | 36 | 0.363492 | 1.3825202 | 0.09687034 | 0.23259823 | 1 | 3351 | tags=25%, list=14%, si | EN_3 |
| GOBP_PROTEIN_DNA_CO | GOBP_PROTEIN_DNA_COMPLEX_DISASSEM | 19 | 0.426186 | 1.3821223 | 0.113175675 | 0.23296756 | 1 | 13556 | tags=100%, list=57%, | EN_3 |
| GOBP_REGULATION_OF_ | GOBP_REGULATION_OF_SMOOTH_MUSCLE | 37 | 0.361662 | 1.3818733 | 0.08320493 | 0.2331222 | 1 | 1924 | tags=16%, list=8%, sig | EN_3 |
| GOBP_ZYMOGEN_ACTIVA | GOBP_ZYMOGEN_ACTIVATION | 51 | 0.333833 | 1.3816242 | 0.07692308 | 0.23326512 | 1 | 3017 | tags=24%, list=13%, si | EN_3 |
| GOBP_POSITIVE_REGULA | GOBP_POSITIVE_REGULATION_OF_TYROSIN | 51 | 0.339152 | 1.3814301 | 0.08695652 | 0.23336151 | 1 | 2922 | tags=22%, list=12%, si | EN_3 |
| GOBP_REGULATION_OF_ | GOBP_REGULATION_OF_DNA_BINDING | 99 | 0.295929 | 1.3812109 | 0.061946902 | 0.23347773 | 1 | 2868 | tags=16%, list=12%, si | EN_3 |
| GOBP_REGULATION_OF_ | GOBP_REGULATION_OF_VIRAL_LIFE_CYCLE | 139 | 0.282251 | 1.381197 | 0.052884616 | 0.23331408 | 1 | 2487 | tags=17%, list=11%, si | EN_3 |
| GOBP_VACUOLAR_ACIDII | GOBP_VACUOLAR_ACIDIFICATION | 27 | 0.400071 | 1.3807951 | 0.10708402 | 0.23368378 | 1 | 5923 | tags=33%, list=25%, si | EN_3 |
| GOBP_REGENERATION | GOBP_REGENERATION | 161 | 0.277153 | 1.3805125 | 0.042725172 | 0.23390648 | 1 | 3937 | tags=20%, list=17%, si | EN_3 |
| GOBP_AUDITORY_RECEPT | GOBP_AUDITORY_RECEPTOR_CELL_MORPH | 26 | 0.396418 | 1.3799659 | 0.10509031 | 0.2344883 | 1 | 6563 | tags=54%, list=28%, si | EN_3 |
| GOBP_PROTEIN_LOCALIZ | GOBP_PROTEIN_LOCALIZATION_TO_EARLY_ | 15 | 0.450619 | 1.379871 | 0.12231405 | 0.23444253 | 1 | 5817 | tags=47%, list=25%, si | EN_3 |
| GOBP_RETINA_VASCULAT | GOBP_RETINA_VASCULATURE_DEVELOPME | 22 | 0.422236 | 1.3797214 | 0.12026359 | 0.23445244 | 1 | 6904 | tags=50%, list=29%, si | EN_3 |
| GOBP_POST_TRANSLATIC | GOBP_POST_TRANSLATIONAL_PROTEIN_TA | 18 | 0.434149 | 1.3797114 | 0.12357724 | 0.23427756 | 1 | 4909 | tags=39%, list=21%, si | EN_3 |
| GOBP_NUCLEOTIDE_SUG | GOBP_NUCLEOTIDE_SUGAR_METABOLIC_PI | 34 | 0.369335 | 1.3796325 | 0.10990712 | 0.23419994 | 1 | 7064 | tags=44%, list=30%, si | EN_3 |
| GOBP_MITOTIC_RECOMB | GOBP_MITOTIC_RECOMBINATION | 24 | 0.405942 | 1.3795588 | 0.114779875 | 0.23411869 | 1 | 4320 | tags=33%, list=18%, si | EN_3 |
| GOBP_POSITIVE_REGULA | GOBP_POSITIVE_REGULATION_OF_MITOTIC | 93 | 0.298477 | 1.3791881 | 0.058746736 | 0.23444434 | 1 | 5318 | tags=30%, list=23%, si | EN_3 |
| GOBP_POSITIVE_REGULA | GOBP_POSITIVE_REGULATION_OF_TRANSM | 97 | 0.296905 | 1.3791637 | 0.06480305 | 0.23429722 | 1 | 4738 | tags=25%, list=20%, si | EN_3 |
| GOBP_POSITIVE_REGULA | GOBP_POSITIVE_REGULATION_OF_ENDOCY | 182 | 0.269709 | 1.3785158 | 0.04157044 | 0.23496106 | 1 | 4076 | tags=21%, list=17%, si | EN_3 |
| GOBP_MAINTENANCE_OI | GOBP_MAINTENANCE_OF_CELL_POLARITY | 17 | 0.444192 | 1.3785132 | 0.12186978 | 0.23478 | 1 | 7124 | tags=53%, list=30%, si | EN_3 |
| GOBP_REGULATION_OF_ | GOBP_REGULATION_OF_NITRIC_OXIDE_SYN | 20 | 0.431716 | 1.3784072 | 0.1097973 | 0.23475014 | 1 | 1432 | tags=15%, list=6%, sig | EN_3 |
| GOBP_RESPONSE_TO_HE | GOBP_RESPONSE_TO_HEAT | 86 | 0.303898 | 1.3782932 | 0.068601586 | 0.23472433 | 1 | 5435 | tags=30%, list=23%, si | EN_3 |
| GOBP_SUBSTRATE_ADHE | GOBP_SUBSTRATE_ADHESION_DEPENDENT_ | 99 | 0.296196 | 1.3779608 | 0.07317073 | 0.23498948 | 1 | 4672 | tags=27%, list=20%, si | EN_3 |
| GOBP_REGULATION_OF_ | GOBP_REGULATION_OF_HETEROTYPIC_CELI | 21 | 0.421548 | 1.3779225 | 0.11221122 | 0.23485772 | 1 | 5077 | tags=38%, list=22%, si | EN_3 |
| GOBP_CELLULAR_RESPOI | GOBP_CELLULAR_RESPONSE_TO_INSULIN_S | 207 | 0.266488 | 1.3773829 | 0.029612755 | 0.2354089 | 1 | 4401 | tags=22%, list=19%, si | EN_3 |
| GOBP_CELLULAR_SENESC | GOBP_CELLULAR_SENESCENCE | 88 | 0.298693 | 1.3773005 | 0.07850708 | 0.23534139 | 1 | 3780 | tags=22%, list=16%, si | EN_3 |
| SALMONELLA_INFECTION | SALMONELLA_INFECTION | 76 | 0.311204 | 1.3772304 | 0.07692308 | 0.23526461 | 1 | 4672 | tags=28%, list=20%, si | EN_3 |
| GOBP_VIRION_ASSEMBLY | GOBP_VIRION_ASSEMBLY | 26 | 0.396763 | 1.3770196 | 0.11735537 | 0.23536317 | 1 | 6720 | tags=46%, list=28%, si | EN_3 |
| GOBP_NOREPINEPHRINE_ | GOBP_NOREPINEPHRINE_METABOLIC_PROC | 17 | 0.444563 | 1.3767985 | 0.13057852 | 0.23548968 | 1 | 1585 | tags=29%, list=7%, sig | EN_3 |
| GLYCOSAMINOGLYCAN_E | GLYCOSAMINOGLYCAN_BIOSYNTHESIS | 51 | 0.338308 | 1.3765891 | 0.07737226 | 0.2355919 | 1 | 5155 | tags=31%, list=22%, si | EN_3 |
| GOBP_TYPE_B_PANCREAT | GOBP_TYPE_B_PANCREATIC_CELL_APOPTO | 15 | 0.456007 | 1.3764241 | 0.12295082 | 0.23564455 | 1 | 4934 | tags=40%, list=21%, si | EN_3 |
| GOBP_NEGATIVE_REGUL | GOBP_NEGATIVE_REGULATION_OF_CELL_C | 290 | 0.255637 | 1.3763199 | 0.02056277 | 0.23561157 | 1 | 5457 | tags=28%, list=23%, si | EN_3 |
| GOBP_LOCOMOTORY_EX | GOBP_LOCOMOTORY_EXPLORATION_BEHA | 19 | 0.434938 | 1.3763152 | 0.10508475 | 0.23543374 | 1 | 5632 | tags=47%, list=24%, si | EN_3 |
| GOBP_INTRINSIC_APOPT | GOBP_INTRINSIC_APOPTOTIC_SIGNALING_I | 44 | 0.349881 | 1.3762566 | 0.076358296 | 0.23533943 | 1 | 3080 | tags=20%, list=13%, si | EN_3 |
| GOBP_TETRAHYDROFOLA | GOBP_TETRAHYDROFOLATE_METABOLIC_PI | 20 | 0.416376 | 1.3760647 | 0.1248 | 0.23540694 | 1 | 6687 | tags=50%, list=28%, si | EN_3 |
| GOBP_POSITIVE_REGULA | GOBP_POSITIVE_REGULATION_OF_ADAPTIV | 140 | 0.280701 | 1.3759731 | 0.056140352 | 0.23535115 | 1 | 3632 | tags=21%, list=15%, si | EN_3 |
| GOBP_POSITIVE_REGULA | GOBP_POSITIVE_REGULATION_OF_NEURON | 99 | 0.300855 | 1.3759503 | 0.06683805 | 0.23519985 | 1 | 5914 | tags=33%, list=25%, si | EN_3 |
| GOBP_ENDONUCLEOLYTI | GOBP_ENDONUCLEOLYTIC_CLEAVAGE_OF_ | 16 | 0.4449 | 1.3755302 | 0.13504274 | 0.23560265 | 1 | 4087 | tags=44%, list=17%, si | EN_3 |

| | | | | | | | | | | |
|---|---|---|---|---|---|---|---|---|---|---|
| CARDIAC_MUSCLE_CONT | CARDIAC_MUSCLE_CONTRACTION | 72 | 0.315242 | 1.3754275 | 0.0731383 | 0.23555873 | 1 | 5042 | tags=40%, list=21%, si | EN_3 |
| GOBP_POSITIVE_REGULA | GOBP_POSITIVE_REGULATION_OF_SMOOTH | 17 | 0.437763 | 1.3751662 | 0.12745099 | 0.23573956 | 1 | 3337 | tags=24%, list=14%, si | EN_3 |
| GOBP_NITRIC_OXIDE_ME | GOBP_NITRIC_OXIDE_MEDIATED_SIGNAL_T | 26 | 0.397071 | 1.3750622 | 0.10130719 | 0.2356966 | 1 | 4579 | tags=35%, list=19%, si | EN_3 |
| EPSTEIN-BARR_VIRUS_IN | EPSTEIN-BARR_VIRUS_INFECTION | 190 | 0.267038 | 1.3742942 | 0.03686636 | 0.23657322 | 1 | 5892 | tags=32%, list=25%, si | EN_3 |
| GOBP_NEGATIVE_REGUL | GOBP_NEGATIVE_REGULATION_OF_HEMOP | 121 | 0.288846 | 1.3742564 | 0.058080807 | 0.23643284 | 1 | 4975 | tags=29%, list=21%, si | EN_3 |
| GOBP_TRANSLATIONAL_E | GOBP_TRANSLATIONAL_ELONGATION | 58 | 0.324943 | 1.3740392 | 0.09312321 | 0.23656707 | 1 | 7782 | tags=57%, list=33%, si | EN_3 |
| GOBP_MRNA_CATABOLIC | GOBP_MRNA_CATABOLIC_PROCESS | 233 | 0.262204 | 1.3739358 | 0.033632286 | 0.23652546 | 1 | 7387 | tags=38%, list=31%, si | EN_3 |
| GOBP_ANTIBACTERIAL_H | GOBP_ANTIBACTERIAL_HUMORAL_RESPONS | 36 | 0.364377 | 1.3734295 | 0.09800919 | 0.23706445 | 1 | 3920 | tags=33%, list=17%, si | EN_3 |
| GOBP_T_CELL_APOPTOTI | GOBP_T_CELL_APOPTOTIC_PROCESS | 67 | 0.31882 | 1.3723496 | 0.075177304 | 0.23838739 | 1 | 4386 | tags=28%, list=19%, si | EN_3 |
| GOBP_REGULATION_OF_ | GOBP_REGULATION_OF_CELLULAR_EXTRAV | 43 | 0.34049 | 1.3721459 | 0.108308606 | 0.23850042 | 1 | 3714 | tags=26%, list=16%, si | EN_3 |
| GOBP_REGULATION_OF_ | GOBP_REGULATION_OF_SYNAPTIC_VESICLE | 55 | 0.338244 | 1.3720828 | 0.09248555 | 0.23841615 | 1 | 3545 | tags=20%, list=15%, si | EN_3 |
| GOBP_NEGATIVE_REGUL | GOBP_NEGATIVE_REGULATION_OF_ANDRO | 15 | 0.459331 | 1.3720807 | 0.12881356 | 0.23823476 | 1 | 4792 | tags=33%, list=20%, si | EN_3 |
| GOBP_CELL_PROLIFERATI | GOBP_CELL_PROLIFERATION_INVOLVED_IN | 20 | 0.423961 | 1.3719807 | 0.11611785 | 0.23818196 | 1 | 2817 | tags=25%, list=12%, si | EN_3 |
| GOBP_CHONDROCYTE_PI | GOBP_CHONDROCYTE_PROLIFERATION | 28 | 0.383411 | 1.3716127 | 0.107361965 | 0.23850583 | 1 | 2605 | tags=25%, list=11%, si | EN_3 |
| GOBP_NEGATIVE_REGUL | GOBP_NEGATIVE_REGULATION_OF_INTRINS | 25 | 0.399722 | 1.3706964 | 0.11490683 | 0.2395826 | 1 | 5790 | tags=48%, list=25%, si | EN_3 |
| GOBP_POSITIVE_REGULA | GOBP_POSITIVE_REGULATION_OF_COLD_IN | 100 | 0.294576 | 1.3704745 | 0.06779661 | 0.23971336 | 1 | 2859 | tags=22%, list=12%, si | EN_3 |
| DOPAMINERGIC_SYNAPS | DOPAMINERGIC_SYNAPSE | 122 | 0.287564 | 1.3703753 | 0.06356968 | 0.23965245 | 1 | 7410 | tags=38%, list=31%, si | EN_3 |
| GOBP_POSITIVE_REGULA | GOBP_POSITIVE_REGULATION_OF_CYTOKIN | 488 | 0.244284 | 1.3697596 | 0.009155646 | 0.2403288 | 1 | 5263 | tags=26%, list=22%, si | EN_3 |
| GOBP_SYNCYTIUM_FORM | GOBP_SYNCYTIUM_FORMATION | 63 | 0.316321 | 1.3696308 | 0.08240224 | 0.24032216 | 1 | 3525 | tags=25%, list=15%, si | EN_3 |
| GOBP_MITOCHONDRIAL_ | GOBP_MITOCHONDRIAL_CYTOCHROME_C_ | 19 | 0.422064 | 1.3692851 | 0.13300492 | 0.24060875 | 1 | 6268 | tags=58%, list=27%, si | EN_3 |
| GOBP_POSITIVE_REGULA | GOBP_POSITIVE_REGULATION_OF_OXIDORE | 37 | 0.360639 | 1.3691801 | 0.111600585 | 0.2405779 | 1 | 3655 | tags=24%, list=15%, si | EN_3 |
| GOBP_POSITIVE_REGULA | GOBP_POSITIVE_REGULATION_OF_CELL_CY | 112 | 0.289559 | 1.369088 | 0.062901154 | 0.24051438 | 1 | 4944 | tags=28%, list=21%, si | EN_3 |
| GOBP_PYRIMIDINE_DEOX | GOBP_PYRIMIDINE_DEOXYRIBONUCLEOSIDE | 15 | 0.45837 | 1.3690294 | 0.10016978 | 0.24041837 | 1 | 6961 | tags=53%, list=29%, si | EN_3 |
| GOBP_RESPONSE_TO_ME | GOBP_RESPONSE_TO_METAL_ION | 278 | 0.259445 | 1.3688866 | 0.024678111 | 0.24047324 | 1 | 5583 | tags=27%, list=24%, si | EN_3 |
| GOBP_REGULATION_OF_ | GOBP_REGULATION_OF_SMAD_PROTEIN_S | 61 | 0.322889 | 1.3686602 | 0.08707483 | 0.24062689 | 1 | 2467 | tags=16%, list=10%, si | EN_3 |
| GOBP_REGULATION_OF_ | GOBP_REGULATION_OF_SYNCYTIUM_FORM | 31 | 0.379406 | 1.3686391 | 0.11480363 | 0.24047203 | 1 | 3525 | tags=29%, list=15%, si | EN_3 |
| PROTEOGLYCANS_IN_CA | PROTEOGLYCANS_IN_CANCER | 193 | 0.26951 | 1.3686228 | 0.040137615 | 0.24031207 | 1 | 7099 | tags=38%, list=30%, si | EN_3 |
| GOBP_RESPONSE_TO_W | GOBP_RESPONSE_TO_WOUNDING | 492 | 0.24882 | 1.3677024 | 0.009307136 | 0.24142285 | 1 | 5216 | tags=26%, list=22%, si | EN_3 |
| GOBP_POSITIVE_REGULA | GOBP_POSITIVE_REGULATION_OF_CELL_SU | 35 | 0.368409 | 1.367692 | 0.09516381 | 0.24125063 | 1 | 7475 | tags=49%, list=32%, si | EN_3 |
| GOBP_REGULATION_OF_ | GOBP_REGULATION_OF_TELOMERASE_ACT | 42 | 0.351805 | 1.3673296 | 0.108938545 | 0.24156965 | 1 | 4200 | tags=26%, list=18%, si | EN_3 |
| GOBP_CHAPERONE_COF | GOBP_CHAPERONE_COFACTOR_DEPENDEN | 31 | 0.367381 | 1.3671597 | 0.105511814 | 0.24164143 | 1 | 5314 | tags=35%, list=23%, si | EN_3 |
| GOBP_CELL_MATRIX_AD | GOBP_CELL_MATRIX_ADHESION | 225 | 0.262404 | 1.3666847 | 0.040044494 | 0.24212863 | 1 | 7590 | tags=41%, list=32%, si | EN_3 |
| GOBP_REGULATION_OF_ | GOBP_REGULATION_OF_NITRIC_OXIDE_ME | 76 | 0.311114 | 1.3662542 | 0.07989348 | 0.24250786 | 1 | 4076 | tags=22%, list=17%, si | EN_3 |
| GOBP_ESTABLISHMENT_ | GOBP_ESTABLISHMENT_OF_PROTEIN_LOCA | 20 | 0.42756 | 1.3659986 | 0.122719735 | 0.2427168 | 1 | 13524 | tags=100%, list=57%, | EN_3 |
| GOBP_CELLULAR_RESPO | GOBP_CELLULAR_RESPONSE_TO_ALKALOID | 30 | 0.385887 | 1.3658593 | 0.10461538 | 0.2427398 | 1 | 5684 | tags=43%, list=24%, si | EN_3 |
| GOBP_NEGATIVE_REGUL | GOBP_NEGATIVE_REGULATION_OF_ORGAN | 375 | 0.249813 | 1.3655244 | 0.015739769 | 0.24309829 | 1 | 5467 | tags=27%, list=23%, si | EN_3 |
| GOBP_INTERLEUKIN_1_P | GOBP_INTERLEUKIN_1_PRODUCTION | 114 | 0.285872 | 1.3645778 | 0.06426735 | 0.24428532 | 1 | 4934 | tags=29%, list=21%, si | EN_3 |
| GOBP_GLIOGENESIS | GOBP_GLIOGENESIS | 377 | 0.251654 | 1.3643945 | 0.019128587 | 0.24436173 | 1 | 5698 | tags=27%, list=24%, si | EN_3 |
| GOBP_POSITIVE_REGULA | GOBP_POSITIVE_REGULATION_OF_NEURON | 24 | 0.397544 | 1.3637419 | 0.11598746 | 0.24513426 | 1 | 7371 | tags=46%, list=31%, si | EN_3 |
| GOBP_PROSTATE_GLAND | GOBP_PROSTATE_GLAND_MORPHOGENESIS | 30 | 0.371066 | 1.3634223 | 0.1276252 | 0.24540472 | 1 | 2842 | tags=23%, list=12%, si | EN_3 |
| GOBP_REGULATION_OF_ | GOBP_REGULATION_OF_REPRODUCTIVE_PF | 186 | 0.266575 | 1.3626698 | 0.039414413 | 0.24634181 | 1 | 4214 | tags=23%, list=18%, si | EN_3 |
| GOBP_REGULATION_OF_ | GOBP_REGULATION_OF_DNA_REPAIR | 219 | 0.263493 | 1.3621132 | 0.030542986 | 0.24698323 | 1 | 5709 | tags=28%, list=24%, si | EN_3 |
| AMYOTROPHIC_LATERAL | AMYOTROPHIC_LATERAL_SCLEROSIS_(ALS) | 50 | 0.33667 | 1.3620821 | 0.10197869 | 0.24685071 | 1 | 6044 | tags=38%, list=26%, si | EN_3 |
| GOBP_DNA_RECOMBINA | GOBP_DNA_RECOMBINATION | 313 | 0.250405 | 1.3618257 | 0.02826087 | 0.24702895 | 1 | 5408 | tags=29%, list=23%, si | EN_3 |
| GOBP_POSITIVE_REGULA | GOBP_POSITIVE_REGULATION_OF_SIGNALI | 28 | 0.38245 | 1.361467 | 0.11111111 | 0.2473585 | 1 | 3084 | tags=25%, list=13%, si | EN_3 |
| GOBP_ANTIGEN_PROCES | GOBP_ANTIGEN_PROCESSING_AND_PRESEN | 59 | 0.327212 | 1.3614182 | 0.08 | 0.24724913 | 1 | 5341 | tags=36%, list=23%, si | EN_3 |
| GOBP_GLIAL_CELL_DEVE | GOBP_GLIAL_CELL_DEVELOPMENT | 134 | 0.275008 | 1.3613225 | 0.061728396 | 0.247192 | 1 | 4086 | tags=22%, list=17%, si | EN_3 |
| GOBP_GLAND_MORPHO | GOBP_GLAND_MORPHOGENESIS | 148 | 0.275931 | 1.3612949 | 0.05443787 | 0.24705175 | 1 | 2369 | tags=15%, list=10%, si | EN_3 |
| GOBP_LYMPHOCYTE_APO | GOBP_LYMPHOCYTE_APOPTOTIC_PROCESS | 94 | 0.299328 | 1.3611256 | 0.070680626 | 0.24712178 | 1 | 5418 | tags=33%, list=23%, si | EN_3 |
| GOBP_NEGATIVE_REGUL | GOBP_NEGATIVE_REGULATION_OF_INFLAM | 17 | 0.446283 | 1.3610753 | 0.15772358 | 0.2469979 | 1 | 3292 | tags=35%, list=14%, si | EN_3 |
| GOBP_COLLAGEN_META | GOBP_COLLAGEN_METABOLIC_PROCESS | 119 | 0.285833 | 1.3606377 | 0.06634499 | 0.24743123 | 1 | 5649 | tags=29%, list=24%, si | EN_3 |
| GOBP_MITOCHONDRIAL_ | GOBP_MITOCHONDRIAL_DNA_REPLICATION | 17 | 0.447025 | 1.3605983 | 0.13188648 | 0.24730368 | 1 | 3276 | tags=29%, list=14%, si | EN_3 |
| LONG-TERM_DEPRESSION | LONG-TERM_DEPRESSION | 55 | 0.325692 | 1.3604034 | 0.089108914 | 0.2473912 | 1 | 633 | tags=9%, list=3%, sign | EN_3 |
| GOBP_REGULATION_OF_ | GOBP_REGULATION_OF_RAC_PROTEIN_SIG | 35 | 0.365577 | 1.360237 | 0.1012461 | 0.24744862 | 1 | 6770 | tags=51%, list=29%, si | EN_3 |
| GOBP_RESPONSE_TO_IO | GOBP_RESPONSE_TO_IONIZING_RADIATION | 127 | 0.281515 | 1.360065 | 0.06297229 | 0.24749805 | 1 | 7098 | tags=39%, list=30%, si | EN_3 |
| GOBP_OSTEOBLAST_PRO | GOBP_OSTEOBLAST_PROLIFERATION | 38 | 0.351394 | 1.3600643 | 0.0953125 | 0.247315 | 1 | 6139 | tags=42%, list=26%, si | EN_3 |
| OOCYTE_MEIOSIS | OOCYTE_MEIOSIS | 108 | 0.28679 | 1.3600364 | 0.061173532 | 0.24717642 | 1 | 6853 | tags=38%, list=29%, si | EN_3 |
| GOBP_REGULATION_OF_ | GOBP_REGULATION_OF_PEPTIDYL_TYROSIN | 238 | 0.257364 | 1.3598431 | 0.042410713 | 0.2472796 | 1 | 3306 | tags=19%, list=14%, si | EN_3 |
| GOBP_RRNA_TRANSCRIP | GOBP_RRNA_TRANSCRIPTION | 33 | 0.377171 | 1.3596635 | 0.1102719 | 0.24735607 | 1 | 2990 | tags=21%, list=13%, si | EN_3 |
| PROSTATE_CANCER | PROSTATE_CANCER | 93 | 0.296466 | 1.3593688 | 0.06815203 | 0.24759124 | 1 | 7986 | tags=44%, list=34%, si | EN_3 |
| GOBP_VACUOLAR_TRANS | GOBP_VACUOLAR_TRANSPORT | 161 | 0.2773 | 1.3590152 | 0.04481132 | 0.24791053 | 1 | 4555 | tags=23%, list=19%, si | EN_3 |
| GOBP_LIPOSACCHARIDE_ | GOBP_LIPOSACCHARIDE_METABOLIC_PROC | 93 | 0.294447 | 1.3582786 | 0.06363636 | 0.24879988 | 1 | 3913 | tags=26%, list=17%, si | EN_3 |
| GOBP_POSITIVE_REGULA | GOBP_POSITIVE_REGULATION_OF_CELL_PR | 453 | 0.245335 | 1.3579806 | 0.013457557 | 0.24904227 | 1 | 4209 | tags=18%, list=18%, si | EN_3 |
| GOBP_REGULATION_OF_ | GOBP_REGULATION_OF_ADIPOSE_TISSUE_ | 16 | 0.448255 | 1.3573895 | 0.11675127 | 0.24971639 | 1 | 3780 | tags=25%, list=16%, si | EN_3 |
| GOBP_CELLULAR_RESPO | GOBP_CELLULAR_RESPONSE_TO_PEPTIDE_ | 276 | 0.254332 | 1.3571745 | 0.02860286 | 0.24982268 | 1 | 4878 | tags=23%, list=21%, si | EN_3 |
| GOBP_WOUND_HEALING | GOBP_WOUND_HEALING | 350 | 0.252359 | 1.356976 | 0.01577287 | 0.2498998 | 1 | 3582 | tags=19%, list=15%, si | EN_3 |
| GOBP_NEGATIVE_REGUL | GOBP_NEGATIVE_REGULATION_OF_FATTY_ | 33 | 0.362338 | 1.3563119 | 0.1095679 | 0.24999255 | 1 | 3926 | tags=36%, list=17%, si | EN_3 |
| OLFACTORY_TRANSDUCT | OLFACTORY_TRANSDUCTION | 186 | -0.43677 | -2.745003 | 0 | 0 | 0 | 7135 | tags=71%, list=30%, si | EN_3 |
| GOBP_REGULATION_OF_ | GOBP_REGULATION_OF_CLATHRIN_DEPEND | 18 | -0.62663 | -2.169060 | 0 | 0.045415524 | 0.156 | 1566 | tags=28%, list=7%, sig | EN_3 |
| GOBP_CLATHRIN_DEPEN | GOBP_CLATHRIN_DEPENDENT_ENDOCYTOS | 48 | -0.4112 | -1.964256 | 0 | 0.24390821 | 0.753 | 1566 | tags=17%, list=7%, sig | EN_3 |
| GOBP_GLYCOSPHINGOLI | GOBP_GLYCOSPHINGOLIPID_METABOLIC_PI | 51 | 0.65604 | 1.874934 | 0 | 0.111440696 | 0.097 | 3400 | tags=37%, list=14%, si | EN_4 |
| GOBP_OLIGOSACCHARID | GOBP_OLIGOSACCHARIDE_METABOLIC_PRO | 48 | 0.666807 | 1.8598349 | 0 | 0.080312125 | 0.139 | 3084 | tags=35%, list=13%, si | EN_4 |
| GOBP_GANGLIOSIDE_ME | GOBP_GANGLIOSIDE_METABOLIC_PROCESS | 27 | 0.715575 | 1.8234243 | 0.0016474464 | 0.13071223 | 0.295 | 3400 | tags=44%, list=14%, si | EN_4 |
| GOBP_REGULATION_OF_ | GOBP_REGULATION_OF_BILE_ACID_METAB | 17 | 0.783407 | 1.8000997 | 0 | 0.16432242 | 0.444 | 1507 | tags=29%, list=6%, sig | EN_4 |
| GOBP_DETECTION_OF_M | GOBP_DETECTION_OF_MECHANICAL_STIMU | 25 | 0.714851 | 1.79555 | 0.0016025641 | 0.14585382 | 0.475 | 2420 | tags=36%, list=10%, si | EN_4 |
| GOBP_SEQUESTERING_O | GOBP_SEQUESTERING_OF_TRIGLYCERIDE | 18 | 0.753988 | 1.7826968 | 0 | 0.1568857 | 0.576 | 3382 | tags=50%, list=14%, si | EN_4 |
| GOBP_MODIFICATION_O | GOBP_MODIFICATION_OF_SYNAPTIC_STRU | 51 | 0.62099 | 1.7732813 | 0.0013793104 | 0.16410153 | 0.643 | 4925 | tags=55%, list=21%, si | EN_4 |
| GOBP_MODIFICATION_O | GOBP_MODIFICATION_OF_POSTSYNAPTIC_ | 24 | 0.723436 | 1.7723715 | 0.0015873016 | 0.14590767 | 0.644 | 4925 | tags=67%, list=21%, si | EN_4 |
| FOCAL_ADHESION | FOCAL_ADHESION | 194 | 0.523114 | 1.7554557 | 0 | 0.18500818 | 0.788 | 5575 | tags=52%, list=24%, si | EN_4 |
| GOBP_SENSORY_PERCEP | GOBP_SENSORY_PERCEPTION_OF_MECHAN | 195 | 0.527285 | 1.7523683 | 0 | 0.17599317 | 0.805 | 4093 | tags=29%, list=17%, si | EN_4 |
| STAPHYLOCOCCUS_AURE | STAPHYLOCOCCUS_AUREUS_INFECTION | 44 | 0.630363 | 1.7428124 | 0 | 0.18989629 | 0.863 | 2131 | tags=23%, list=9%, sig | EN_4 |
| GOBP_POSITIVE_REGULA | GOBP_POSITIVE_REGULATION_OF_AXON_E | 54 | 0.602154 | 1.7321199 | 0 | 0.2099208 | 0.9 | 5575 | tags=57%, list=24%, si | EN_4 |

| | | | | | | | | | | |
|---|---|---|---|---|---|---|---|---|---|---|
| GOBP_NEURON_PROJECT | GOBP_NEURON_PROJECTION_EXTENSION_I | 19 | 0.724693 | 1.7266546 | 0 | 0.21359617 | 0.924 | 3835 | tags=63%, list=16%, si | EN_4 |
| GOBP_REGULATION_OF_ | GOBP_REGULATION_OF_AXON_EXTENSION | 15 | 0.75994 | 1.7263236 | 0 | 0.19989717 | 0.925 | 3835 | tags=60%, list=16%, si | EN_4 |
| GOBP_REGULATION_OF_ | GOBP_REGULATION_OF_NITRIC_OXIDE_SYN | 20 | 0.717908 | 1.7250756 | 0.0032626428 | 0.19108509 | 0.927 | 2934 | tags=35%, list=12%, si | EN_4 |
| GOBP_REGULATION_OF_ | GOBP_REGULATION_OF_MAST_CELL_ACTIV | 24 | 0.675581 | 1.7139488 | 0.003236246 | 0.2173695 | 0.959 | 3849 | tags=46%, list=16%, si | EN_4 |
| GOBP_NEGATIVE_REGUL | GOBP_NEGATIVE_REGULATION_OF_AMINO | 19 | 0.71808 | 1.7103593 | 0.0016393443 | 0.21685456 | 0.962 | 1708 | tags=26%, list=7%, sig | EN_4 |
| GOBP_GLYCOSYLATION | GOBP_GLYCOSYLATION | 218 | 0.502721 | 1.7094831 | 0 | 0.20765719 | 0.963 | 3455 | tags=30%, list=15%, si | EN_4 |
| GOBP_REGULATION_OF_ | GOBP_REGULATION_OF_HETEROTYPIC_CELI | 21 | 0.711319 | 1.7087705 | 0.0031397175 | 0.19845682 | 0.965 | 5121 | tags=62%, list=22%, si | EN_4 |
| GOBP_AMYLOID_PRECUR | GOBP_AMYLOID_PRECURSOR_PROTEIN_CAT | 59 | 0.589394 | 1.7074993 | 0 | 0.19247103 | 0.969 | 5044 | tags=46%, list=21%, si | EN_4 |
| GOBP_MAINTENANCE_OI | GOBP_MAINTENANCE_OF_BLOOD_BRAIN_E | 22 | 0.699617 | 1.7061034 | 0 | 0.18720895 | 0.972 | 3104 | tags=36%, list=13%, si | EN_4 |
| GOBP_MODIFICATION_O | GOBP_MODIFICATION_OF_POSTSYNAPTIC_S | 42 | 0.627644 | 1.7049394 | 0 | 0.18227324 | 0.975 | 4925 | tags=55%, list=21%, si | EN_4 |
| GOBP_POSITIVE_REGULA | GOBP_POSITIVE_REGULATION_OF_MACROF | 29 | 0.65142 | 1.7003182 | 0.0015384615 | 0.18763585 | 0.982 | 4269 | tags=55%, list=18%, si | EN_4 |
| GOBP_REGULATION_OF_ | GOBP_REGULATION_OF_AMYLOID_BETA_FO | 37 | 0.623372 | 1.6965414 | 0 | 0.19045916 | 0.986 | 2994 | tags=35%, list=13%, si | EN_4 |
| GOBP_POSITIVE_REGULA | GOBP_POSITIVE_REGULATION_OF_INTERLEU | 55 | 0.585784 | 1.6923579 | 0 | 0.1951027 | 0.99 | 3770 | tags=36%, list=16%, si | EN_4 |
| ECM-RECEPTOR_INTERAC | ECM-RECEPTOR_INTERACTION | 79 | 0.559419 | 1.6907561 | 0.0014409221 | 0.1924228 | 0.99 | 5443 | tags=51%, list=23%, si | EN_4 |
| GOBP_POSITIVE_REGULA | GOBP_POSITIVE_REGULATION_OF_MONOO | 22 | 0.681044 | 1.6880263 | 0.004651163 | 0.19386916 | 0.991 | 1812 | tags=32%, list=8%, sig | EN_4 |
| GOBP_PINOCYTOSIS | GOBP_PINOCYTOSIS | 22 | 0.680918 | 1.681086 | 0.0032258064 | 0.20913057 | 0.998 | 3931 | tags=55%, list=17%, si | EN_4 |
| SALMONELLA_INFECTION | SALMONELLA_INFECTION | 76 | 0.550673 | 1.6723021 | 0 | 0.22964257 | 0.999 | 4594 | tags=38%, list=19%, si | EN_4 |
| GOBP_TYPE_B_PANCREAT | GOBP_TYPE_B_PANCREATIC_CELL_PROLIFEF | 35 | 0.629745 | 1.6709187 | 0.0015503876 | 0.22661044 | 1 | 3633 | tags=31%, list=15%, si | EN_4 |
| GOBP_EXTRACELLULAR_N | GOBP_EXTRACELLULAR_MATRIX_ASSEMBLY | 48 | 0.591399 | 1.6681834 | 0 | 0.22834946 | 1 | 4470 | tags=44%, list=19%, si | EN_4 |
| GOBP_CELLULAR_RESPON | GOBP_CELLULAR_RESPONSE_TO_VITAMIN | 16 | 0.715127 | 1.6636908 | 0.006644518 | 0.2369486 | 1 | 3012 | tags=50%, list=13%, si | EN_4 |
| GOBP_VASCULAR_ASSOC | GOBP_VASCULAR_ASSOCIATED_SMOOTH_M | 16 | 0.731523 | 1.659172 | 0.0048 | 0.24687746 | 1 | 1865 | tags=31%, list=8%, sig | EN_4 |
| GOBP_POSITIVE_REGULA | GOBP_POSITIVE_REGULATION_OF_ACTIN_F | 52 | 0.572926 | 1.6248198 | 0.008645534 | 0.24942668 | 1 | 2865 | tags=25%, list=12%, si | EN_4 |
| GOBP_POSITIVE_REGULA | GOBP_POSITIVE_REGULATION_OF_MUSCLE | 52 | 0.56688 | 1.624298 | 0.001497006 | 0.24634545 | 1 | 3561 | tags=33%, list=15%, si | EN_4 |
| GOBP_GLYCOLYTIC_PROC | GOBP_GLYCOLYTIC_PROCESS_THROUGH_FR | 25 | 0.656662 | 1.623544 | 0.0015015015 | 0.24420883 | 1 | 5116 | tags=56%, list=22%, si | EN_4 |
| GOBP_POSITIVE_REGULA | GOBP_POSITIVE_REGULATION_OF_GLIAL_C | 31 | 0.636203 | 1.6182292 | 0.0077279755 | 0.24926262 | 1 | 4311 | tags=45%, list=18%, si | EN_4 |
| GOBP_POSITIVE_REGULA | GOBP_POSITIVE_REGULATION_OF_INTERLEU | 65 | 0.540753 | 1.617896 | 0.007215007 | 0.2458713 | 1 | 4290 | tags=35%, list=18%, si | EN_4 |
| GOBP_POSITIVE_REGULA | GOBP_POSITIVE_REGULATION_OF_NEUROG | 312 | 0.469146 | 1.617352 | 0 | 0.24359493 | 1 | 4341 | tags=34%, list=18%, si | EN_4 |
| TOXOPLASMOSIS | TOXOPLASMOSIS | 103 | 0.522423 | 1.6167684 | 0 | 0.2411591 | 1 | 5390 | tags=44%, list=23%, si | EN_4 |
| GOBP_LEUKOCYTE_AGGR | GOBP_LEUKOCYTE_AGGREGATION | 16 | 0.699435 | 1.6134804 | 0.011744967 | 0.2482187 | 1 | 5604 | tags=63%, list=24%, si | EN_4 |
| GOBP_GLYCOSYLCERAMI | GOBP_GLYCOSYLCERAMIDE_METABOLIC_PF | 15 | 0.723293 | 1.6133355 | 0.0067114094 | 0.24477969 | 1 | 3084 | tags=47%, list=13%, si | EN_4 |
| GLYCOSPHINGOLIPID_BIC | GLYCOSPHINGOLIPID_BIOSYNTHESIS | 44 | 0.579042 | 1.6130916 | 0.0030165913 | 0.24150115 | 1 | 3400 | tags=34%, list=14%, si | EN_4 |
| GOBP_REGULATION_OF_ | GOBP_REGULATION_OF_JNK_CASCADE | 149 | 0.497441 | 1.6103102 | 0 | 0.2467344 | 1 | 4306 | tags=36%, list=18%, si | EN_4 |
| GOBP_CELL_SURFACE_PA | GOBP_CELL_SURFACE_PATTERN_RECOGNIT | 70 | 0.539652 | 1.6093575 | 0.005873715 | 0.24599138 | 1 | 4776 | tags=39%, list=20%, si | EN_4 |
| GOBP_LIVER_MORPHOGE | GOBP_LIVER_MORPHOGENESIS | 36 | 0.595978 | 1.6082743 | 0.0057803467 | 0.24572457 | 1 | 5719 | tags=50%, list=24%, si | EN_4 |
| PERTUSSIS | PERTUSSIS | 69 | 0.542388 | 1.6048677 | 0.0014104373 | 0.24943775 | 1 | 3444 | tags=29%, list=15%, si | EN_4 |
| GOBP_CAMERA_TYPE_EY | GOBP_CAMERA_TYPE_EYE_PHOTORECEPTO | 41 | 0.587871 | 1.6030607 | 0.0089552235 | 0.24825548 | 1 | 3039 | tags=29%, list=13%, si | EN_4 |
| GOBP_EPIBOLY | GOBP_EPIBOLY | 32 | 0.61424 | 1.6015387 | 0.009756098 | 0.24944703 | 1 | 4612 | tags=50%, list=20%, si | EN_4 |
| GOBP_NEUROMUSCULAR | GOBP_NEUROMUSCULAR_JUNCTION_DEVE | 66 | 0.548676 | 1.6013223 | 0.0028530671 | 0.2468385 | 1 | 4158 | tags=41%, list=18%, si | EN_4 |
| GOBP_POSITIVE_REGULA | GOBP_POSITIVE_REGULATION_OF_VASCULA | 25 | 0.645518 | 1.5995984 | 0.0190779 | 0.2487437 | 1 | 4206 | tags=52%, list=18%, si | EN_4 |
| GOBP_GAMMA_AMINOB | GOBP_GAMMA_AMINOBUTYRIC_ACID_SIGN | 30 | 0.615239 | 1.599506 | 0.009202454 | 0.24568826 | 1 | 2348 | tags=20%, list=10%, si | EN_4 |
| GOBP_RESPONSE_TO_VIT | GOBP_RESPONSE_TO_VITAMIN_D | 15 | 0.711777 | 1.5983905 | 0.015410959 | 0.24604836 | 1 | 3012 | tags=47%, list=13%, si | EN_4 |
| GOBP_REGULATION_OF_ | GOBP_REGULATION_OF_NEURON_PROJECT | 43 | 0.570252 | 1.5972788 | 0.007824726 | 0.24618992 | 1 | 4520 | tags=51%, list=19%, si | EN_4 |
| GOBP_INNATE_IMMUNE_ | GOBP_INNATE_IMMUNE_RESPONSE_ACTIV | 82 | 0.524199 | 1.5967928 | 0.0013947001 | 0.24445964 | 1 | 4776 | tags=34%, list=20%, si | EN_4 |
| GOBP_PREPULSE_INHIBIT | GOBP_PREPULSE_INHIBITION | 18 | 0.664845 | 1.5951408 | 0.012638231 | 0.24650723 | 1 | 4520 | tags=44%, list=19%, si | EN_4 |
| GOBP_GLYCERALDEHYDE | GOBP_GLYCERALDEHYDE_3_PHOSPHATE_M | 19 | 0.671103 | 1.5942614 | 0.0176 | 0.2461451 | 1 | 3861 | tags=42%, list=16%, si | EN_4 |
| GOBP_POSITIVE_REGULA | GOBP_POSITIVE_REGULATION_OF_TYPE_I_I | 19 | 0.649274 | 1.5941793 | 0.015254238 | 0.24333508 | 1 | 3472 | tags=42%, list=15%, si | EN_4 |
| GOBP_CARBOHYDRATE_D | GOBP_CARBOHYDRATE_DERIVATIVE_TRANS | 76 | 0.53034 | 1.5941521 | 0 | 0.24050057 | 1 | 4739 | tags=39%, list=20%, si | EN_4 |
| GOBP_DETECTION_OF_M | GOBP_DETECTION_OF_MECHANICAL_STIMU | 53 | 0.554079 | 1.5911362 | 0.0014164306 | 0.24742971 | 1 | 2420 | tags=21%, list=10%, si | EN_4 |
| GOBP_GANGLIOSIDE_BIO | GOBP_GANGLIOSIDE_BIOSYNTHETIC_PROCE | 17 | 0.681477 | 1.5901204 | 0.021172639 | 0.24803218 | 1 | 3084 | tags=35%, list=13%, si | EN_4 |
| GOBP_POSITIVE_REGULA | GOBP_POSITIVE_REGULATION_OF_NERVOU | 374 | 0.456062 | 1.5890261 | 0 | 0.2482587 | 1 | 4341 | tags=33%, list=18%, si | EN_4 |
| GOBP_MICROVILLUS_ASS | GOBP_MICROVILLUS_ASSEMBLY | 20 | 0.658502 | 1.5887774 | 0.01309329 | 0.24611928 | 1 | 3291 | tags=40%, list=14%, si | EN_4 |
| TNF_SIGNALING_PATHW. | TNF_SIGNALING_PATHWAY | 108 | 0.50623 | 1.5884013 | 0 | 0.24462202 | 1 | 3683 | tags=34%, list=16%, si | EN_4 |
| GOBP_DETECTION_OF_BI | GOBP_DETECTION_OF_BIOTIC_STIMULUS | 37 | 0.591829 | 1.5881373 | 0.007575758 | 0.24257424 | 1 | 3451 | tags=27%, list=15%, si | EN_4 |
| GOBP_REGULATION_OF_ | GOBP_REGULATION_OF_TYPE_B_PANCREAT | 16 | 0.676106 | 1.577341 | 0.02173913 | 0.24941511 | 1 | 3633 | tags=38%, list=15%, si | EN_4 |
| GOBP_POSITIVE_REGULA | GOBP_POSITIVE_REGULATION_OF_INTERLE | 15 | 0.690419 | 1.5770688 | 0.013490725 | 0.24766158 | 1 | 2796 | tags=40%, list=12%, si | EN_4 |
| GOBP_JNK_CASCADE | GOBP_JNK_CASCADE | 179 | 0.473744 | 1.5765047 | 0 | 0.24689253 | 1 | 4311 | tags=35%, list=18%, si | EN_4 |
| GOBP_MEGAKARYOCYTE_ | GOBP_MEGAKARYOCYTE_DEVELOPMENT | 25 | 0.629174 | 1.576294 | 0.014446228 | 0.24500197 | 1 | 4294 | tags=40%, list=18%, si | EN_4 |
| GOBP_POSITIVE_REGULA | GOBP_POSITIVE_REGULATION_OF_JNK_CAS | 101 | 0.509356 | 1.5758158 | 0 | 0.24384117 | 1 | 4306 | tags=38%, list=18%, si | EN_4 |
| GOBP_WATER_TRANSPOI | GOBP_WATER_TRANSPORT | 17 | 0.690209 | 1.5728571 | 0.008264462 | 0.2494617 | 1 | 3174 | tags=35%, list=13%, si | EN_4 |
| RIBOSOME | RIBOSOME | 128 | -0.53785 | -1.973488 | 0 | 0.10236795 | 0.122 | 4590 | tags=53%, list=19%, si | EN_4 |
| BASE_EXCISION_REPAIR | BASE_EXCISION_REPAIR | 32 | -0.66387 | -1.971789 | 0.0028248588 | 0.053999383 | 0.127 | 2376 | tags=41%, list=10%, si | EN_4 |
| GOBP_GLUCOCORTICOID | GOBP_GLUCOCORTICOID_METABOLIC_PRO | 21 | -0.7053 | -1.893585 | 0 | 0.14012752 | 0.409 | 1573 | tags=24%, list=7%, sig | EN_4 |
| GOBP_BASE_EXCISION_R | GOBP_BASE_EXCISION_REPAIR | 41 | -0.63483 | -1.882000 | 0 | 0.12518652 | 0.465 | 2058 | tags=29%, list=9%, sig | EN_4 |
| GOBP_LATE_ENDOSOME_ | GOBP_LATE_ENDOSOME_TO_VACUOLE_TR | 25 | -0.66955 | -1.870688 | 0 | 0.118247256 | 0.519 | 3255 | tags=48%, list=14%, si | EN_4 |
| GOBP_SPLICEOSOMAL_SI | GOBP_SPLICEOSOMAL_SNRNP_ASSEMBLY | 35 | -0.62599 | -1.852773 | 0 | 0.12596656 | 0.6 | 4205 | tags=60%, list=18%, si | EN_4 |
| GOBP_ESTABLISHMENT_( | GOBP_ESTABLISHMENT_OF_PROTEIN_LOCA | 25 | -0.6766 | -1.845873 | 0.0027100272 | 0.11718072 | 0.627 | 4443 | tags=60%, list=19%, si | EN_4 |
| GOBP_GAMMA_DELTA_T | GOBP_GAMMA_DELTA_T_CELL_DIFFERENTI | 17 | -0.72818 | -1.83504 | 0.0052083335 | 0.11855301 | 0.679 | 1819 | tags=41%, list=8%, sig | EN_4 |
| GOBP_REGULATION_OF_ | GOBP_REGULATION_OF_MITOTIC_SPINDLE_ | 18 | -0.72721 | -1.834834 | 0.002538071 | 0.10546725 | 0.68 | 2534 | tags=44%, list=11%, si | EN_4 |
| GOBP_ATTACHMENT_OF_ | GOBP_ATTACHMENT_OF_MITOTIC_SPINDLE | 24 | -0.65842 | -1.799116 | 0 | 0.14927673 | 0.84 | 2761 | tags=38%, list=12%, si | EN_4 |
| GOBP_TRANSLATION_AT_ | GOBP_TRANSLATION_AT_SYNAPSE | 48 | 0.722612 | 3.8591926 | 0 | 0 | 0 | 5691 | tags=83%, list=24%, si | MM_4 |
| RIBOSOME | RIBOSOME | 128 | 0.536381 | 3.5031 | 0 | 0 | 0 | 5691 | tags=70%, list=24%, si | MM_4 |
| PROTEASOME | PROTEASOME | 45 | 0.640639 | 3.2674563 | 0 | 0 | 0 | 4874 | tags=69%, list=21%, si | MM_4 |
| GOBP_ANTIGEN_PROCES | GOBP_ANTIGEN_PROCESSING_AND_PRESEN | 34 | 0.663611 | 3.09633 | 0 | 0 | 0 | 4073 | tags=68%, list=17%, si | MM_4 |
| GOBP_ANTIGEN_PROCES | GOBP_ANTIGEN_PROCESSING_AND_PRESEN | 105 | 0.491035 | 3.0735114 | 0 | 0 | 0 | 2979 | tags=34%, list=13%, si | MM_4 |
| GOBP_ANTIGEN_PROCES | GOBP_ANTIGEN_PROCESSING_AND_PRESEN | 28 | 0.657206 | 3.0479264 | 0 | 0 | 0 | 4073 | tags=68%, list=17%, si | MM_4 |
| OXIDATIVE_PHOSPHORYL | OXIDATIVE_PHOSPHORYLATION | 115 | 0.462157 | 2.9916317 | 0 | 0 | 0 | 4772 | tags=57%, list=20%, si | MM_4 |
| ALZHEIMER_DISEASE | ALZHEIMER_DISEASE | 159 | 0.43438 | 2.9501038 | 0 | 0 | 0 | 5204 | tags=55%, list=22%, si | MM_4 |
| LYSOSOME | LYSOSOME | 118 | 0.444772 | 2.8261306 | 0 | 0 | 0 | 4120 | tags=45%, list=17%, si | MM_4 |
| PARKINSON_DISEASE | PARKINSON_DISEASE | 122 | 0.434078 | 2.7993479 | 0 | 0.000235262 | 0.001 | 5478 | tags=59%, list=23%, si | MM_4 |
| GOBP_ANTIGEN_PROCES | GOBP_ANTIGEN_PROCESSING_AND_PRESEN | 24 | 0.656917 | 2.7441583 | 0 | 0.000438799 | 0.002 | 4073 | tags=67%, list=17%, si | MM_4 |
| GOBP_RIBOSOME_ASSEM | GOBP_RIBOSOME_ASSEMBLY | 60 | 0.463285 | 2.7064514 | 0 | 0.000598284 | 0.003 | 5506 | tags=60%, list=23%, si | MM_4 |
| GOBP_REGULATION_OF_ | GOBP_REGULATION_OF_TELOMERE_CAPPIN | 26 | 0.604411 | 2.6990037 | 0 | 0.000552262 | 0.003 | 4454 | tags=62%, list=19%, si | MM_4 |

| | | | | | | | | | | |
|---|---|---|---|---|---|---|---|---|---|---|
| GOBP_AEROBIC_RESPIRA | GOBP_AEROBIC_RESPIRATION | 170 | 0.394142 | 2.6879854 | 0 | 0.001011736 | 0.006 | 4824 | tags=46%, list=20%, si | MM_4 |
| GOBP_REGULATION_OF_ | GOBP_REGULATION_OF_MACROPHAGE_MI | 47 | 0.525146 | 2.6461477 | 0 | 0.00208492 | 0.013 | 3882 | tags=45%, list=16%, si | MM_4 |
| GOBP_REGULATION_OF_ | GOBP_REGULATION_OF_ANTIGEN_PROCESS | 21 | 0.66641 | 2.6450253 | 0 | 0.001954613 | 0.013 | 2499 | tags=48%, list=11%, si | MM_4 |
| GOBP_TAIL_ANCHORED_ | GOBP_TAIL_ANCHORED_MEMBRANE_PROT | 15 | 0.698984 | 2.6310534 | 0 | 0.002264796 | 0.015 | 2720 | tags=47%, list=12%, si | MM_4 |
| GOBP_ANTIGEN_PROCES | GOBP_ANTIGEN_PROCESSING_AND_PRESEN | 59 | 0.472897 | 2.624465 | 0 | 0.002270249 | 0.016 | 4073 | tags=47%, list=17%, si | MM_4 |
| GOBP_POSITIVE_REGULA | GOBP_POSITIVE_REGULATION_OF_MACROF | 31 | 0.560752 | 2.6071858 | 0 | 0.002644428 | 0.02 | 3681 | tags=48%, list=16%, si | MM_4 |
| GOBP_ANTIGEN_PROCES | GOBP_ANTIGEN_PROCESSING_AND_PRESEN | 20 | 0.662107 | 2.6070633 | 0 | 0.002512207 | 0.02 | 4468 | tags=80%, list=19%, si | MM_4 |
| GOBP_RIBOSOMAL_LARG | GOBP_RIBOSOMAL_LARGE_SUBUNIT_ASSEN | 23 | 0.604074 | 2.6058943 | 0 | 0.002392578 | 0.02 | 5702 | tags=78%, list=24%, si | MM_4 |
| GOBP_CYTOPLASMIC_TR/ | GOBP_CYTOPLASMIC_TRANSLATION | 143 | 0.378875 | 2.591646 | 0 | 0.002396805 | 0.021 | 5691 | tags=59%, list=24%, si | MM_4 |
| GOBP_PHAGOCYTOSIS_EI | GOBP_PHAGOCYTOSIS_ENGULFMENT | 53 | 0.486368 | 2.5720081 | 0 | 0.002716308 | 0.025 | 2094 | tags=28%, list=9%, sig | MM_4 |
| GOBP_MEMBRANE_INVA | GOBP_MEMBRANE_INVAGINATION | 70 | 0.452357 | 2.5714107 | 0 | 0.002603129 | 0.025 | 2094 | tags=24%, list=9%, sig | MM_4 |
| GOBP_CHAPERONE_COF/ | GOBP_CHAPERONE_COFACTOR_DEPENDENT | 31 | 0.537127 | 2.5114787 | 0 | 0.004148768 | 0.041 | 10945 | tags=100%, list=46%, | MM_4 |
| GOBP_CELL_KILLING | GOBP_CELL_KILLING | 202 | 0.351063 | 2.5081234 | 0 | 0.0039892 | 0.041 | 2880 | tags=24%, list=12%, si | MM_4 |
| GOBP_TELOMERE_MAINT | GOBP_TELOMERE_MAINTENANCE_IN_RESP | 32 | 0.546908 | 2.504819 | 0 | 0.004019029 | 0.043 | 2865 | tags=44%, list=12%, si | MM_4 |
| GOBP_SNRNA_PROCESSII | GOBP_SNRNA_PROCESSING | 24 | 0.593191 | 2.4898193 | 0 | 0.004903406 | 0.055 | 3686 | tags=50%, list=16%, si | MM_4 |
| GOBP_CELLULAR_RESPIR/ | GOBP_CELLULAR_RESPIRATION | 218 | 0.353243 | 2.488844 | 0 | 0.004734323 | 0.055 | 4824 | tags=43%, list=20%, si | MM_4 |
| GOBP_SNRNA_3_END_PF | GOBP_SNRNA_3_END_PROCESSING | 20 | 0.616267 | 2.474492 | 0.004048583 | 0.005382385 | 0.064 | 3686 | tags=55%, list=16%, si | MM_4 |
| GOBP_REGULATION_OF_ | GOBP_REGULATION_OF_MACROPHAGE_CH | 32 | 0.530224 | 2.4569626 | 0 | 0.00590495 | 0.073 | 3681 | tags=44%, list=16%, si | MM_4 |
| GOBP_REGULATION_OF_ | GOBP_REGULATION_OF_MONONUCLEAR_C | 136 | 0.364416 | 2.4568722 | 0 | 0.00572042 | 0.073 | 3681 | tags=32%, list=16%, si | MM_4 |
| GOBP_POSITIVE_REGULA | GOBP_POSITIVE_REGULATION_OF_TRANSLA | 125 | 0.386049 | 2.4561877 | 0 | 0.005547074 | 0.073 | 4540 | tags=40%, list=19%, si | MM_4 |
| GOBP_POSITIVE_REGULA | GOBP_POSITIVE_REGULATION_OF_TELOME | 17 | 0.662622 | 2.4543302 | 0 | 0.005599534 | 0.076 | 3745 | tags=59%, list=16%, si | MM_4 |
| GOBP_TELOMERE_CAPPII | GOBP_TELOMERE_CAPPING | 38 | 0.507914 | 2.4436533 | 0 | 0.005988468 | 0.081 | 4454 | tags=53%, list=19%, si | MM_4 |
| SPLICEOSOME | SPLICEOSOME | 127 | 0.383931 | 2.4342017 | 0 | 0.006284636 | 0.087 | 11317 | tags=94%, list=48%, si | MM_4 |
| GOBP_T_CELL_MIGRATIO | GOBP_T_CELL_MIGRATION | 64 | 0.441243 | 2.433892 | 0 | 0.006114781 | 0.087 | 3572 | tags=36%, list=15%, si | MM_4 |
| GOBP_T_CELL_CHEMOTA | GOBP_T_CELL_CHEMOTAXIS | 23 | 0.548825 | 2.42449 | 0.003937008 | 0.006263062 | 0.092 | 1090 | tags=26%, list=5%, sig | MM_4 |
| GOBP_CELLULAR_EXTRAV | GOBP_CELLULAR_EXTRAVASATION | 74 | 0.412075 | 2.422924 | 0 | 0.006102471 | 0.092 | 3307 | tags=34%, list=14%, si | MM_4 |
| GOBP_KILLING_OF_CELLS | GOBP_KILLING_OF_CELLS_OF_ANOTHER_OI | 71 | 0.415734 | 2.4221003 | 0 | 0.006009838 | 0.093 | 348 | tags=11%, list=1%, sig | MM_4 |
| GOBP_PROTON_MOTIVE_ | GOBP_PROTON_MOTIVE_FORCE_DRIVEN_A | 52 | 0.455296 | 2.4215643 | 0 | 0.005922379 | 0.094 | 11228 | tags=98%, list=48%, si | MM_4 |
| GOBP_LYMPHOCYTE_MIG | GOBP_LYMPHOCYTE_MIGRATION | 92 | 0.39291 | 2.4197624 | 0 | 0.005892792 | 0.095 | 3614 | tags=34%, list=15%, si | MM_4 |
| ANTIGEN_PROCESSING_A | ANTIGEN_PROCESSING_AND_PRESENTATIO | 67 | 0.42417 | 2.414785 | 0 | 0.006036223 | 0.1 | 2958 | tags=37%, list=13%, si | MM_4 |
| GOBP_AEROBIC_ELECTRO | GOBP_AEROBIC_ELECTRON_TRANSPORT_CH | 63 | 0.430416 | 2.3997746 | 0 | 0.006609893 | 0.112 | 11986 | tags=97%, list=51%, si | MM_4 |
| GOBP_FIBROBLAST_MIGF | GOBP_FIBROBLAST_MIGRATION | 68 | 0.413709 | 2.391775 | 0 | 0.006896525 | 0.12 | 3562 | tags=34%, list=15%, si | MM_4 |
| GOBP_POSITIVE_REGULA | GOBP_POSITIVE_REGULATION_OF_CELL_AD | 21 | 0.583174 | 2.3747656 | 0 | 0.008256409 | 0.142 | 1982 | tags=43%, list=8%, sig | MM_4 |
| GOBP_RESCUE_OF_STALL | GOBP_RESCUE_OF_STALLED_RIBOSOME | 25 | 0.560029 | 2.3723645 | 0 | 0.008337317 | 0.147 | 5300 | tags=60%, list=22%, si | MM_4 |
| N-GLYCAN_BIOSYNTHESIS | N-GLYCAN_BIOSYNTHESIS | 48 | 0.438732 | 2.3678343 | 0 | 0.008668196 | 0.155 | 3355 | tags=40%, list=14%, si | MM_4 |
| GOBP_DISRUPTION_OF_A | GOBP_DISRUPTION_OF_ANATOMICAL_STRU | 73 | 0.411442 | 2.3651147 | 0 | 0.008638826 | 0.156 | 348 | tags=11%, list=1%, sig | MM_4 |
| GOBP_POSITIVE_REGULA | GOBP_POSITIVE_REGULATION_OF_MACROF | 25 | 0.549472 | 2.359149 | 0 | 0.008845067 | 0.162 | 3681 | tags=48%, list=16%, si | MM_4 |
| GOBP_TRANSLATIONAL_E | GOBP_TRANSLATIONAL_ELONGATION | 58 | 0.441861 | 2.3564274 | 0 | 0.008814253 | 0.165 | 12127 | tags=98%, list=51%, si | MM_4 |
| GOBP_POSITIVE_REGULA | GOBP_POSITIVE_REGULATION_OF_TELOME | 71 | 0.411658 | 2.3519943 | 0 | 0.008878805 | 0.17 | 4454 | tags=41%, list=19%, si | MM_4 |
| GOBP_REGULATION_OF_ | GOBP_REGULATION_OF_TELOMERE_MAINT | 98 | 0.372917 | 2.3467042 | 0 | 0.009162277 | 0.179 | 4454 | tags=39%, list=19%, si | MM_4 |
| GOBP_PROTEIN_DEGLYC | GOBP_PROTEIN_DEGLYCOSYLATION | 15 | 0.639403 | 2.3458874 | 0 | 0.009127495 | 0.182 | 8523 | tags=100%, list=36%, | MM_4 |
| GOBP_REGULATION_OF_ | GOBP_REGULATION_OF_TRANSLATIONAL_I | 68 | 0.411542 | 2.3442152 | 0 | 0.0090908 | 0.185 | 11125 | tags=94%, list=47%, si | MM_4 |
| CHEMOKINE_SIGNALING_ | CHEMOKINE_SIGNALING_PATHWAY | 177 | 0.348207 | 2.3398066 | 0 | 0.009187506 | 0.19 | 2915 | tags=23%, list=12%, si | MM_4 |
| GOBP_TELOMERE_ORGAI | GOBP_TELOMERE_ORGANIZATION | 150 | 0.338822 | 2.3339741 | 0 | 0.009620257 | 0.202 | 4454 | tags=38%, list=19%, si | MM_4 |
| GOBP_GLYCOPROTEIN_C | GOBP_GLYCOPROTEIN_CATABOLIC_PROCES | 31 | 0.515884 | 2.333953 | 0.004716981 | 0.00945439 | 0.202 | 3385 | tags=45%, list=14%, si | MM_4 |
| GOBP_RIBOSOME_DISAS | GOBP_RIBOSOME_DISASSEMBLY | 31 | 0.514115 | 2.331642 | 0 | 0.009376556 | 0.204 | 4638 | tags=52%, list=20%, si | MM_4 |
| GOBP_RIBOSOMAL_LARG | GOBP_RIBOSOMAL_LARGE_SUBUNIT_BIOGE | 64 | 0.399528 | 2.3305814 | 0 | 0.009298545 | 0.206 | 5702 | tags=59%, list=24%, si | MM_4 |
| GOBP_IMMUNOLOGICAL_ | GOBP_IMMUNOLOGICAL_SYNAPSE_FORMA | 18 | 0.610921 | 2.3297842 | 0 | 0.00914611 | 0.206 | 1561 | tags=28%, list=7%, sig | MM_4 |
| GOBP_MEMBRANE_RAFT | GOBP_MEMBRANE_RAFT_ORGANIZATION | 24 | 0.537211 | 2.3290603 | 0 | 0.008998592 | 0.206 | 4446 | tags=54%, list=19%, si | MM_4 |
| GOBP_PROTEIN_IMPORT | GOBP_PROTEIN_IMPORT_INTO_MITOCHON | 19 | 0.602417 | 2.3227477 | 0 | 0.009308878 | 0.216 | 5376 | tags=68%, list=23%, si | MM_4 |
| GOBP_OXIDATIVE_PHOSF | GOBP_OXIDATIVE_PHOSPHORYLATION | 124 | 0.365132 | 2.3219657 | 0 | 0.009240572 | 0.218 | 5456 | tags=52%, list=23%, si | MM_4 |
| HUNTINGTON_DISEASE | HUNTINGTON_DISEASE | 180 | 0.331486 | 2.3217185 | 0 | 0.00909841 | 0.218 | 4772 | tags=48%, list=20%, si | MM_4 |
| GOBP_POSITIVE_REGULA | GOBP_POSITIVE_REGULATION_OF_TELOME | 31 | 0.503662 | 2.3190851 | 0 | 0.009215194 | 0.225 | 5057 | tags=55%, list=21%, si | MM_4 |
| GOBP_REGULATION_OF_ | GOBP_REGULATION_OF_DEFENSE_RESPONS | 67 | 0.401552 | 2.2966294 | 0 | 0.010834698 | 0.261 | 2538 | tags=30%, list=11%, si | MM_4 |
| GOBP_GLYCOLYTIC_PROC | GOBP_GLYCOLYTIC_PROCESS_THROUGH_GI | 20 | 0.56522 | 2.296499 | 0.0035211267 | 0.010712043 | 0.262 | 3613 | tags=55%, list=15%, si | MM_4 |
| GOBP_NEGATIVE_REGUL | GOBP_NEGATIVE_REGULATION_OF_GLUCO | 24 | 0.539494 | 2.287475 | 0 | 0.011494608 | 0.281 | 2815 | tags=33%, list=12%, si | MM_4 |
| GOBP_NEGATIVE_REGUL | GOBP_NEGATIVE_REGULATION_OF_PHAGO | 24 | 0.533378 | 2.2814105 | 0 | 0.011878721 | 0.293 | 4679 | tags=54%, list=20%, si | MM_4 |
| GOBP_ATP_SYNTHESIS_C | GOBP_ATP_SYNTHESIS_COUPLED_ELECTROI | 76 | 0.394955 | 2.2804756 | 0 | 0.011846933 | 0.297 | 4772 | tags=49%, list=20%, si | MM_4 |
| GOBP_REGULATION_OF_ | GOBP_REGULATION_OF_DEFENSE_RESPONS | 42 | 0.463294 | 2.2664518 | 0 | 0.012910424 | 0.322 | 2538 | tags=33%, list=11%, si | MM_4 |
| MISMATCH_REPAIR | MISMATCH_REPAIR | 21 | 0.560025 | 2.2661312 | 0 | 0.012799869 | 0.324 | 4128 | tags=52%, list=17%, si | MM_4 |
| GOBP_LYMPHOCYTE_CHE | GOBP_LYMPHOCYTE_CHEMOTAXIS | 38 | 0.464943 | 2.2634552 | 0 | 0.012888044 | 0.33 | 1495 | tags=21%, list=6%, sig | MM_4 |
| GOBP_RESPONSE_TO_CH | GOBP_RESPONSE_TO_CHEMOKINE | 74 | 0.383793 | 2.2538774 | 0 | 0.013775566 | 0.355 | 917 | tags=14%, list=4%, sig | MM_4 |
| CITRATE_CYCLE_(TCA_CY | CITRATE_CYCLE_(TCA_CYCLE) | 28 | 0.507724 | 2.2515996 | 0 | 0.013783666 | 0.358 | 4729 | tags=54%, list=20%, si | MM_4 |
| GOBP_LYSOSOMAL_LUM | GOBP_LYSOSOMAL_LUMEN_ACIDIFICATION | 15 | 0.604866 | 2.251235 | 0 | 0.013604658 | 0.358 | 4249 | tags=73%, list=18%, si | MM_4 |
| GOBP_NUCLEAR_TRANSC | GOBP_NUCLEAR_TRANSCRIBED_MRNA_POL | 26 | 0.508174 | 2.2508488 | 0.004 | 0.013430239 | 0.358 | 4133 | tags=46%, list=18%, si | MM_4 |
| GOBP_VIRION_ASSEMBLY | GOBP_VIRION_ASSEMBLY | 26 | 0.522429 | 2.2465625 | 0 | 0.013836443 | 0.37 | 11289 | tags=100%, list=48%, | MM_4 |
| GOBP_CYTOPLASMIC_SE | GOBP_CYTOPLASMIC_SEQUESTERING_OF_P | 17 | 0.589312 | 2.246284 | 0 | 0.013663488 | 0.37 | 5023 | tags=65%, list=21%, si | MM_4 |
| GOBP_MITOCHONDRIAL_ | GOBP_MITOCHONDRIAL_ELECTRON_TRANS | 29 | 0.49294 | 2.239646 | 0 | 0.013883167 | 0.379 | 11986 | tags=100%, list=51%, | MM_4 |
| GOBP_LEUKOCYTE_CELL_ | GOBP_LEUKOCYTE_CELL_CELL_ADHESION | 390 | 0.292621 | 2.2370222 | 0 | 0.013915152 | 0.385 | 3042 | tags=25%, list=13%, si | MM_4 |
| GOBP_RESPONSE_TO_VII | GOBP_RESPONSE_TO_VIRUS | 331 | 0.296331 | 2.2313018 | 0 | 0.014240206 | 0.396 | 2656 | tags=24%, list=11%, si | MM_4 |
| GOBP_POST_TRANSCRIPT | GOBP_POST_TRANSCRIPTIONAL_REGULATIO | 480 | 0.280748 | 2.2111366 | 0 | 0.016816664 | 0.448 | 4133 | tags=31%, list=18%, si | MM_4 |
| GOBP_CHAPERONE_MED | GOBP_CHAPERONE_MEDIATED_PROTEIN_F | 62 | 0.399006 | 2.2049284 | 0 | 0.01754757 | 0.46 | 3267 | tags=31%, list=14%, si | MM_4 |
| GOBP_GRANULOCYTE_CH | GOBP_GRANULOCYTE_CHEMOTAXIS | 119 | 0.342512 | 2.2043438 | 0 | 0.017371248 | 0.461 | 2096 | tags=19%, list=9%, sig | MM_4 |
| CYTOSOLIC_DNA-SENSING | CYTOSOLIC_DNA-SENSING_PATHWAY | 46 | 0.434144 | 2.2026932 | 0 | 0.017441785 | 0.468 | 2880 | tags=30%, list=12%, si | MM_4 |
| GOBP_NEGATIVE_REGUL | GOBP_NEGATIVE_REGULATION_OF_LEUKO | 46 | 0.442193 | 2.2008538 | 0 | 0.017545583 | 0.475 | 3653 | tags=35%, list=15%, si | MM_4 |
| GOBP_POSITIVE_REGULA | GOBP_POSITIVE_REGULATION_OF_DEFENSE | 31 | 0.486882 | 2.1984787 | 0 | 0.017728593 | 0.479 | 2471 | tags=35%, list=10%, si | MM_4 |
| GOBP_REGULATION_OF_ | GOBP_REGULATION_OF_FIBROBLAST_MIGR | 47 | 0.421947 | 2.1878045 | 0 | 0.018790117 | 0.501 | 2544 | tags=26%, list=11%, si | MM_4 |
| NUCLEOTIDE_EXCISION_F | NUCLEOTIDE_EXCISION_REPAIR | 42 | 0.438241 | 2.187451 | 0 | 0.018661518 | 0.503 | 2851 | tags=33%, list=12%, si | MM_4 |
| GOBP_RIBONUCLEOPROT | GOBP_RIBONUCLEOPROTEIN_COMPLEX_BIC | 417 | 0.272896 | 2.1774135 | 0 | 0.01989558 | 0.528 | 3877 | tags=30%, list=16%, si | MM_4 |
| GOBP_REGULATION_OF_ | GOBP_REGULATION_OF_DNA_TEMPLATED_ | 79 | 0.373816 | 2.1761723 | 0 | 0.01973241 | 0.53 | 4439 | tags=46%, list=19%, si | MM_4 |
| GOBP_RNA_METHYLATIO | GOBP_RNA_METHYLATION | 65 | 0.406796 | 2.1761377 | 0 | 0.01952249 | 0.53 | 4316 | tags=49%, list=18%, si | MM_4 |
| GOBP_I_KAPPAB_PHOSPI | GOBP_I_KAPPAB_PHOSPHORYLATION | 17 | 0.575903 | 2.1744912 | 0.0038022813 | 0.019468125 | 0.532 | 4082 | tags=59%, list=17%, si | MM_4 |

| | | | | | | | | | | |
|---|---|---|---|---|---|---|---|---|---|---|
| RNA_DEGRADATION | RNA_DEGRADATION | 75 | 0.382558 | 2.1729012 | 0 | 0.019739145 | 0.543 | 3854 | tags=35%, list=16%, si | MM_4 |
| GOBP_DE_NOVO_PROTEI | GOBP_DE_NOVO_PROTEIN_FOLDING | 38 | 0.451368 | 2.1710265 | 0 | 0.0197063 | 0.545 | 2719 | tags=32%, list=12%, si | MM_4 |
| GOBP_TRNA_METABOLIC | GOBP_TRNA_METABOLIC_PROCESS | 166 | 0.31493 | 2.169216 | 0 | 0.019678451 | 0.55 | 4103 | tags=38%, list=17%, si | MM_4 |
| GOBP_T_CELL_PROLIFERA | GOBP_T_CELL_PROLIFERATION | 233 | 0.299046 | 2.1676567 | 0 | 0.019794768 | 0.559 | 3042 | tags=27%, list=13%, si | MM_4 |
| GOBP_POSITIVE_REGULA | GOBP_POSITIVE_REGULATION_OF_TELOME | 38 | 0.447556 | 2.1621494 | 0 | 0.020367917 | 0.572 | 4454 | tags=45%, list=19%, si | MM_4 |
| GOBP_REGULATION_OF_ | GOBP_REGULATION_OF_GRANULOCYTE_CH | 59 | 0.386126 | 2.1619406 | 0 | 0.020166254 | 0.572 | 3681 | tags=34%, list=16%, si | MM_4 |
| GOBP_INNER_MITOCHON | GOBP_INNER_MITOCHONDRIAL_MEMBRAN | 36 | 0.444444 | 2.1611395 | 0 | 0.02004033 | 0.572 | 11581 | tags=97%, list=49%, si | MM_4 |
| GOBP_NEGATIVE_REGUL | GOBP_NEGATIVE_REGULATION_OF_T_CELL_ | 73 | 0.372952 | 2.1602695 | 0 | 0.020032318 | 0.572 | 1528 | tags=23%, list=6%, sig | MM_4 |
| GOBP_RNA_3_END_PROC | GOBP_RNA_3_END_PROCESSING | 87 | 0.360766 | 2.159644 | 0 | 0.019952716 | 0.573 | 4470 | tags=44%, list=19%, si | MM_4 |
| GOBP_LEUKOCYTE_MIGR | GOBP_LEUKOCYTE_MIGRATION | 363 | 0.274176 | 2.1586778 | 0 | 0.019944403 | 0.576 | 3681 | tags=27%, list=16%, si | MM_4 |
| GOBP_TRANSLATIONAL_I | GOBP_TRANSLATIONAL_INITIATION | 112 | 0.337298 | 2.1580505 | 0 | 0.01977837 | 0.577 | 5121 | tags=40%, list=22%, si | MM_4 |
| EPSTEIN-BARR_VIRUS_IN | EPSTEIN-BARR_VIRUS_INFECTION | 190 | 0.303413 | 2.1569862 | 0 | 0.019771554 | 0.581 | 4931 | tags=40%, list=21%, si | MM_4 |
| GOBP_NEGATIVE_REGUL | GOBP_NEGATIVE_REGULATION_OF_OSTEO | 37 | 0.446444 | 2.1565711 | 0 | 0.019633174 | 0.582 | 3364 | tags=41%, list=14%, si | MM_4 |
| GOBP_PROTEIN_TRANSM | GOBP_PROTEIN_TRANSMEMBRANE_IMPOR | 36 | 0.43654 | 2.1545935 | 0 | 0.01969588 | 0.585 | 3196 | tags=39%, list=14%, si | MM_4 |
| GOBP_ORGANELLE_DISAS | GOBP_ORGANELLE_DISASSEMBLY | 155 | 0.320745 | 2.1522913 | 0 | 0.019909833 | 0.592 | 3412 | tags=31%, list=14%, si | MM_4 |
| GOBP_REGULATION_OF_ | GOBP_REGULATION_OF_NUCLEAR_TRANSC | 15 | 0.577192 | 2.1465483 | 0 | 0.02052978 | 0.606 | 4133 | tags=53%, list=18%, si | MM_4 |
| CHRONIC_MYELOID_LEUK | CHRONIC_MYELOID_LEUKEMIA | 75 | 0.372805 | 2.1419039 | 0 | 0.021092622 | 0.617 | 5064 | tags=47%, list=21%, si | MM_4 |
| GOBP_DEFENSE_RESPON | GOBP_DEFENSE_RESPONSE_TO_VIRUS | 276 | 0.294954 | 2.1416333 | 0 | 0.020971037 | 0.618 | 2656 | tags=24%, list=11%, si | MM_4 |
| GOBP_POSITIVE_REGULA | GOBP_POSITIVE_REGULATION_OF_LYMPHO | 18 | 0.542023 | 2.137404 | 0 | 0.021590097 | 0.629 | 53 | tags=17%, list=0%, sig | MM_4 |
| GOBP_BROWN_FAT_CELL | GOBP_BROWN_FAT_CELL_DIFFERENTIATION | 63 | 0.38098 | 2.1356964 | 0 | 0.021712068 | 0.638 | 975 | tags=14%, list=4%, sig | MM_4 |
| GOBP_REGULATION_OF_ | GOBP_REGULATION_OF_MYELOID_LEUKOC | 61 | 0.379507 | 2.134909 | 0 | 0.021668768 | 0.64 | 3842 | tags=41%, list=16%, si | MM_4 |
| GOBP_ANTIGEN_PROCES | GOBP_ANTIGEN_PROCESSING_AND_PRESEN | 38 | 0.422544 | 2.1339097 | 0 | 0.02162759 | 0.643 | 2817 | tags=34%, list=12%, si | MM_4 |
| GOBP_MYELOID_LEUKOC | GOBP_MYELOID_LEUKOCYTE_MEDIATED_IM | 108 | 0.34988 | 2.1334946 | 0 | 0.021504968 | 0.644 | 3842 | tags=35%, list=16%, si | MM_4 |
| GOBP_RNA_CATABOLIC_ | GOBP_RNA_CATABOLIC_PROCESS | 278 | 0.285623 | 2.1325982 | 0 | 0.021426506 | 0.647 | 4402 | tags=34%, list=19%, si | MM_4 |
| GOBP_REGULATION_OF_ | GOBP_REGULATION_OF_TRANSLATION | 329 | 0.281117 | 2.1323977 | 0 | 0.02126714 | 0.647 | 4550 | tags=35%, list=19%, si | MM_4 |
| GOBP_GRANULOCYTE_M | GOBP_GRANULOCYTE_MIGRATION | 146 | 0.323869 | 2.1321225 | 0 | 0.021091377 | 0.647 | 3157 | tags=26%, list=13%, si | MM_4 |
| GOBP_POSITIVE_REGULA | GOBP_POSITIVE_REGULATION_OF_DNA_TE | 59 | 0.3859 | 2.1316738 | 0 | 0.021035265 | 0.649 | 4439 | tags=47%, list=19%, si | MM_4 |
| GOBP_PHAGOCYTOSIS | GOBP_PHAGOCYTOSIS | 213 | 0.291992 | 2.1309395 | 0 | 0.020944772 | 0.649 | 3842 | tags=29%, list=16%, si | MM_4 |
| GOBP_POSITIVE_REGULA | GOBP_POSITIVE_REGULATION_OF_MONON | 94 | 0.361534 | 2.1301389 | 0 | 0.020832788 | 0.65 | 3681 | tags=33%, list=16%, si | MM_4 |
| GOBP_MAINTENANCE_OI | GOBP_MAINTENANCE_OF_PROTEIN_LOCAT | 97 | 0.34369 | 2.1272323 | 0 | 0.02106738 | 0.659 | 4000 | tags=35%, list=17%, si | MM_4 |
| GOBP_MRNA_CATABOLIC | GOBP_MRNA_CATABOLIC_PROCESS | 233 | 0.291279 | 2.1259289 | 0 | 0.021014092 | 0.662 | 4184 | tags=31%, list=18%, si | MM_4 |
| GOBP_NEGATIVE_REGUL | GOBP_NEGATIVE_REGULATION_OF_GLUCO | 20 | 0.537798 | 2.1255288 | 0 | 0.020848626 | 0.662 | 2321 | tags=30%, list=10%, si | MM_4 |
| GOBP_CD4_POSITIVE_AL | GOBP_CD4_POSITIVE_ALPHA_BETA_T_CELL_ | 28 | 0.463218 | 2.1200671 | 0.012096774 | 0.021601997 | 0.674 | 2217 | tags=32%, list=9%, sig | MM_4 |
| GOBP_MATERNAL_PROC | GOBP_MATERNAL_PROCESS_INVOLVED_IN_ | 48 | 0.403332 | 2.1170788 | 0 | 0.021847092 | 0.684 | 3518 | tags=40%, list=15%, si | MM_4 |
| GOBP_REGULATION_OF_ | GOBP_REGULATION_OF_LYMPHOCYTE_MIG | 64 | 0.378409 | 2.1081986 | 0 | 0.022904662 | 0.703 | 3614 | tags=33%, list=15%, si | MM_4 |
| GOBP_PROTEIN_INSERTI | GOBP_PROTEIN_INSERTION_INTO_ER_MEM | 26 | 0.494651 | 2.105135 | 0 | 0.02314905 | 0.709 | 4525 | tags=50%, list=19%, si | MM_4 |
| LEISHMANIASIS | LEISHMANIASIS | 60 | 0.383576 | 2.104849 | 0 | 0.02301068 | 0.709 | 2815 | tags=35%, list=12%, si | MM_4 |
| GOBP_NEGATIVE_REGUL | GOBP_NEGATIVE_REGULATION_OF_INNATE | 86 | 0.356798 | 2.1008391 | 0 | 0.023307156 | 0.717 | 3850 | tags=38%, list=16%, si | MM_4 |
| GOBP_RIBOSOME_BIOGE | GOBP_RIBOSOME_BIOGENESIS | 306 | 0.279078 | 2.0950317 | 0 | 0.024157675 | 0.732 | 3877 | tags=31%, list=16%, si | MM_4 |
| GOBP_CELLULAR_RESPON | GOBP_CELLULAR_RESPONSE_TO_INTERFER | 16 | 0.582803 | 2.0922267 | 0.006779661 | 0.02453318 | 0.739 | 1488 | tags=31%, list=6%, sig | MM_4 |
| GOBP_REGULATION_OF_ | GOBP_REGULATION_OF_NEUROINFLAMMA | 30 | 0.446684 | 2.0918732 | 0 | 0.024369717 | 0.74 | 2987 | tags=40%, list=13%, si | MM_4 |
| GOBP_POLYPRENOL_MET | GOBP_POLYPRENOL_METABOLIC_PROCESS | 17 | 0.556337 | 2.0917408 | 0.0039525693 | 0.024191836 | 0.74 | 3109 | tags=59%, list=13%, si | MM_4 |
| GOBP_NEGATIVE_REGUL | GOBP_NEGATIVE_REGULATION_OF_CELL_K | 28 | 0.490406 | 2.0912912 | 0 | 0.02406881 | 0.741 | 1982 | tags=36%, list=8%, sig | MM_4 |
| GOBP_NEGATIVE_REGUL | GOBP_NEGATIVE_REGULATION_OF_INTERL | 51 | 0.401944 | 2.0910037 | 0 | 0.023913475 | 0.741 | 2140 | tags=24%, list=9%, sig | MM_4 |
| GOBP_DNA_TEMPLATED_ | GOBP_DNA_TEMPLATED_TRANSCRIPTION_ | 112 | 0.319521 | 2.0898623 | 0 | 0.023844784 | 0.742 | 4786 | tags=46%, list=20%, si | MM_4 |
| GOBP_PROTEIN_TARGETI | GOBP_PROTEIN_TARGETING_TO_MITOCHO | 76 | 0.359874 | 2.085106 | 0 | 0.024216292 | 0.752 | 3196 | tags=29%, list=14%, si | MM_4 |
| GOBP_REGULATION_OF_ | GOBP_REGULATION_OF_T_CELL_PROLIFERA | 188 | 0.30058 | 2.0848749 | 0 | 0.024080051 | 0.753 | 3024 | tags=27%, list=13%, si | MM_4 |
| GOBP_REGULATION_OF_ | GOBP_REGULATION_OF_LYMPHOCYTE_CHE | 22 | 0.488866 | 2.0842512 | 0.0037313432 | 0.024077931 | 0.758 | 1090 | tags=18%, list=5%, sig | MM_4 |
| GOBP_SNO_S_RNA_MET | GOBP_SNO_S_RNA_METABOLIC_PROCESS | 17 | 0.549078 | 2.0840452 | 0 | 0.023910724 | 0.758 | 4041 | tags=59%, list=17%, si | MM_4 |
| GOBP_MELANOSOME_AS | GOBP_MELANOSOME_ASSEMBLY | 19 | 0.544596 | 2.0822968 | 0.0035971224 | 0.024092767 | 0.763 | 974 | tags=32%, list=4%, sig | MM_4 |
| GOBP_POSITIVE_REGULA | GOBP_POSITIVE_REGULATION_OF_MYELOID | 21 | 0.513922 | 2.0813005 | 0 | 0.024125488 | 0.765 | 3842 | tags=62%, list=16%, si | MM_4 |
| NON-ALCOHOLIC_FATTY_ | NON-ALCOHOLIC_FATTY_LIVER_DISEASE_(N | 141 | 0.311376 | 2.0724766 | 0 | 0.025340784 | 0.786 | 4772 | tags=45%, list=20%, si | MM_4 |
| GOBP_NUCLEOSIDE_TRIP | GOBP_NUCLEOSIDE_TRIPHOSPHATE_BIOSYN | 100 | 0.335679 | 2.0709264 | 0 | 0.025591912 | 0.794 | 11575 | tags=92%, list=49%, si | MM_4 |
| GOBP_POSITIVE_REGULA | GOBP_POSITIVE_REGULATION_OF_PROTEIN | 16 | 0.542957 | 2.069066 | 0.007168459 | 0.025644235 | 0.797 | 1322 | tags=31%, list=6%, sig | MM_4 |
| GOBP_REGULATION_OF_ | GOBP_REGULATION_OF_B_CELL_DIFFERENT | 37 | 0.421529 | 2.068661 | 0 | 0.02548915 | 0.797 | 3328 | tags=43%, list=14%, si | MM_4 |
| GOBP_POSITIVE_REGULA | GOBP_POSITIVE_REGULATION_BY_HOST_OI | 22 | 0.498552 | 2.0676396 | 0 | 0.025495913 | 0.799 | 2296 | tags=36%, list=10%, si | MM_4 |
| GOBP_NUCLEAR_TRANSC | GOBP_NUCLEAR_TRANSCRIBED_MRNA_CAT | 118 | 0.319256 | 2.067407 | 0 | 0.025344662 | 0.799 | 4184 | tags=32%, list=18%, si | MM_4 |
| GOBP_REGULATION_OF_ | GOBP_REGULATION_OF_LEUKOCYTE_MEDI | 253 | 0.277237 | 2.0642648 | 0 | 0.025805693 | 0.805 | 3636 | tags=30%, list=15%, si | MM_4 |
| ECM-RECEPTOR_INTERAC | ECM-RECEPTOR_INTERACTION | 79 | 0.353248 | 2.0636203 | 0 | 0.025715796 | 0.805 | 1170 | tags=19%, list=5%, sig | MM_4 |
| GOBP_MONONUCLEAR_C | GOBP_MONONUCLEAR_CELL_MIGRATION | 199 | 0.297633 | 2.063261 | 0 | 0.02556585 | 0.805 | 3681 | tags=29%, list=16%, si | MM_4 |
| GOBP_CHEMOKINE_PROI | GOBP_CHEMOKINE_PRODUCTION | 119 | 0.317329 | 2.0627975 | 0 | 0.025510162 | 0.805 | 2657 | tags=23%, list=11%, si | MM_4 |
| GOBP_MRNA_3_END_PR | GOBP_MRNA_3_END_PROCESSING | 41 | 0.405475 | 2.0601852 | 0.0039525693 | 0.025854021 | 0.812 | 4819 | tags=46%, list=20%, si | MM_4 |
| GOBP_PROTEIN_DNA_CO | GOBP_PROTEIN_DNA_COMPLEX_DISASSEM | 19 | 0.51049 | 2.0583029 | 0 | 0.025842538 | 0.816 | 11567 | tags=100%, list=49%, | MM_4 |
| GOBP_PROTON_TRANSP | GOBP_PROTON_TRANSPORTING_TWO_SEC | 15 | 0.562744 | 2.0573728 | 0.0073260074 | 0.025939206 | 0.818 | 3776 | tags=73%, list=16%, si | MM_4 |
| GOBP_REGULATION_OF_ | GOBP_REGULATION_OF_T_CELL_MIGRATIO | 49 | 0.391166 | 2.0572581 | 0 | 0.025836986 | 0.819 | 3572 | tags=33%, list=15%, si | MM_4 |
| GOBP_TUMOR_NECROSIS | GOBP_TUMOR_NECROSIS_FACTOR_SUPERF | 193 | 0.289584 | 2.0571902 | 0 | 0.025676508 | 0.819 | 3157 | tags=23%, list=13%, si | MM_4 |
| GOBP_VIRAL_BUDDING | GOBP_VIRAL_BUDDING | 18 | 0.540137 | 2.0566473 | 0.010989011 | 0.025622312 | 0.821 | 10867 | tags=100%, list=46%, | MM_4 |
| GOBP_MRNA_PROCESSIN | GOBP_MRNA_PROCESSING | 459 | 0.260493 | 2.0535567 | 0 | 0.025865244 | 0.828 | 4743 | tags=33%, list=20%, si | MM_4 |
| GOBP_PROTEIN_INSERTI | GOBP_PROTEIN_INSERTION_INTO_MEMBRA | 39 | 0.423698 | 2.052821 | 0 | 0.025809534 | 0.829 | 3572 | tags=33%, list=15%, si | MM_4 |
| HUMAN_T-CELL_LEUKEM | HUMAN_T-CELL_LEUKEMIA_VIRUS_1_INFEC | 215 | 0.291608 | 2.052119 | 0 | 0.025812482 | 0.83 | 2885 | tags=26%, list=12%, si | MM_4 |
| GOBP_LATE_ENDOSOME_ | GOBP_LATE_ENDOSOME_TO_LYSOSOME_T | 19 | 0.526766 | 2.0503237 | 0 | 0.025933104 | 0.833 | 11183 | tags=100%, list=47%, | MM_4 |
| GOBP_NEGATIVE_REGUL | GOBP_NEGATIVE_REGULATION_OF_ENDOT | 54 | 0.391302 | 2.0502627 | 0 | 0.025777815 | 0.833 | 2526 | tags=24%, list=11%, si | MM_4 |
| GOBP_LEUKOCYTE_ADHE | GOBP_LEUKOCYTE_ADHESION_TO_VASCUL | 51 | 0.380187 | 2.046157 | 0.0045871558 | 0.02639252 | 0.842 | 4232 | tags=43%, list=18%, si | MM_4 |
| GOBP_REGULATION_OF_ | GOBP_REGULATION_OF_TELOMERASE_ACT | 42 | 0.418462 | 2.0455241 | 0 | 0.026334563 | 0.843 | 3765 | tags=40%, list=16%, si | MM_4 |
| GOBP_POSITIVE_REGULA | GOBP_POSITIVE_REGULATION_OF_LEUKOC | 261 | 0.279222 | 2.0447497 | 0 | 0.026307136 | 0.846 | 2811 | tags=23%, list=12%, si | MM_4 |
| GOBP_RNA_SPLICING | GOBP_RNA_SPLICING | 417 | 0.269106 | 2.043273 | 0 | 0.026431378 | 0.849 | 3451 | tags=25%, list=15%, si | MM_4 |
| GOBP_TRNA_WOBBLE_B | GOBP_TRNA_WOBBLE_BASE_MODIFICATIO | 19 | 0.515594 | 2.041016 | 0.003921569 | 0.026667114 | 0.852 | 3058 | tags=42%, list=13%, si | MM_4 |
| GOBP_NEGATIVE_REGUL | GOBP_NEGATIVE_REGULATION_OF_LEUKO | 96 | 0.332872 | 2.04044 | 0 | 0.026652252 | 0.854 | 3371 | tags=34%, list=14%, si | MM_4 |
| GOBP_REGULATION_OF_ | GOBP_REGULATION_OF_MEMBRANE_INVA | 21 | 0.4877 | 2.0397823 | 0.003787879 | 0.026554557 | 0.855 | 3636 | tags=48%, list=15%, si | MM_4 |
| GOBP_REGULATION_OF_ | GOBP_REGULATION_OF_TRANSLATION_IN_ | 24 | 0.483146 | 2.0372446 | 0 | 0.026884092 | 0.858 | 3553 | tags=46%, list=15%, si | MM_4 |
| GOBP_TRANSCRIPTION_P | GOBP_TRANSCRIPTION_PREINITIATION_CO | 63 | 0.365913 | 2.037145 | 0.0056179776 | 0.026745614 | 0.859 | 3987 | tags=43%, list=17%, si | MM_4 |
| GOBP_CELLULAR_RESPON | GOBP_CELLULAR_RESPONSE_TO_DSRNA | 19 | 0.506071 | 2.032803 | 0.0036630037 | 0.027161006 | 0.862 | 3144 | tags=42%, list=13%, si | MM_4 |

| | | | | | | | | | | |
|---|---|---|---|---|---|---|---|---|---|---|
| GOBP_PROTEIN_QUALITY | GOBP_PROTEIN_QUALITY_CONTROL_FOR_N | 17 | 0.52829 | 2.0326042 | 0.01048951 | 0.027021267 | 0.862 | 11146 | tags=100%, list=47%, | MM_4 |
| GOBP_REGULATION_OF_ | GOBP_REGULATION_OF_T_CELL_ACTIVATIO | 364 | 0.269633 | 2.0325396 | 0 | 0.026897619 | 0.862 | 3042 | tags=23%, list=13%, si | MM_4 |
| TERPENOID_BACKBONE_I | TERPENOID_BACKBONE_BIOSYNTHESIS | 21 | 0.498282 | 2.0323153 | 0.003787879 | 0.026761819 | 0.862 | 2785 | tags=43%, list=12%, si | MM_4 |
| GOBP_POSITIVE_REGULA | GOBP_POSITIVE_REGULATION_OF_MRNA_C | 95 | 0.328584 | 2.0317283 | 0 | 0.026627539 | 0.862 | 4133 | tags=32%, list=18%, si | MM_4 |
| GOBP_NEGATIVE_REGUL | GOBP_NEGATIVE_REGULATION_OF_ALPHA_ | 16 | 0.564572 | 2.029898 | 0 | 0.026746493 | 0.866 | 2569 | tags=38%, list=11%, si | MM_4 |
| GOBP_CELL_ADHESION_N | GOBP_CELL_ADHESION_MEDIATED_BY_INTI | 86 | 0.34183 | 2.0280147 | 0 | 0.027008235 | 0.874 | 2423 | tags=26%, list=10%, si | MM_4 |
| GOBP_ENDOSOME_TO_L | GOBP_ENDOSOME_TO_LYSOSOME_TRANSP | 16 | 0.540092 | 2.026405 | 0.003508772 | 0.027110709 | 0.877 | 10867 | tags=100%, list=46%, | MM_4 |
| GOBP_MITOCHONDRIAL_ | GOBP_MITOCHONDRIAL_GENE_EXPRESSION | 164 | 0.30501 | 2.0226345 | 0 | 0.027584694 | 0.886 | 4322 | tags=35%, list=18%, si | MM_4 |
| GOBP_REGULATION_OF_ | GOBP_REGULATION_OF_T_CELL_CHEMOTA | 16 | 0.550473 | 2.0217397 | 0.0068728523 | 0.02763113 | 0.889 | 1090 | tags=25%, list=5%, sig | MM_4 |
| GOBP_REGULATION_OF_ | GOBP_REGULATION_OF_LYMPHOCYTE_MEI | 190 | 0.281082 | 2.01963 | 0 | 0.027754745 | 0.893 | 3042 | tags=26%, list=13%, si | MM_4 |
| GOBP_LEUKOCYTE_PROLI | GOBP_LEUKOCYTE_PROLIFERATION | 361 | 0.260759 | 2.0174966 | 0 | 0.027925111 | 0.897 | 3042 | tags=25%, list=13%, si | MM_4 |
| GOBP_POSITIVE_REGULA | GOBP_POSITIVE_REGULATION_OF_TUMOR_ | 113 | 0.306066 | 2.016724 | 0 | 0.027954817 | 0.9 | 3527 | tags=24%, list=15%, si | MM_4 |
| GOBP_RNA_SPLICING_VI | GOBP_RNA_SPLICING_VIA_TRANSESTERIFIC | 274 | 0.26934 | 2.016583 | 0 | 0.027857479 | 0.9 | 4820 | tags=33%, list=20%, si | MM_4 |
| GOBP_REGULATION_OF_ | GOBP_REGULATION_OF_LYMPHOCYTE_ACT | 495 | 0.257696 | 2.012825 | 0 | 0.028533123 | 0.909 | 3066 | tags=23%, list=13%, si | MM_4 |
| GOBP_CYTOSOLIC_TRANS | GOBP_CYTOSOLIC_TRANSPORT | 124 | 0.316819 | 2.01196 | 0 | 0.028460024 | 0.909 | 4694 | tags=37%, list=20%, si | MM_4 |
| GOBP_VIRAL_TRANSLATI | GOBP_VIRAL_TRANSLATION | 18 | 0.510876 | 2.0107708 | 0.0036101083 | 0.028650865 | 0.91 | 2151 | tags=33%, list=9%, sig | MM_4 |
| GOBP_POSITIVE_REGULA | GOBP_POSITIVE_REGULATION_OF_GTPASE_ | 203 | 0.282786 | 2.0105138 | 0 | 0.028564734 | 0.91 | 3277 | tags=23%, list=14%, si | MM_4 |
| GOBP_NEUTROPHIL_MIG | GOBP_NEUTROPHIL_MIGRATION | 104 | 0.32202 | 2.0102637 | 0 | 0.028480733 | 0.911 | 2899 | tags=24%, list=12%, si | MM_4 |
| GOBP_ENERGY_DERIVATI | GOBP_ENERGY_DERIVATION_BY_OXIDATIOI | 321 | 0.267571 | 2.0089989 | 0 | 0.028553952 | 0.915 | 3835 | tags=31%, list=16%, si | MM_4 |
| GOBP_RIBOSOMAL_SMA | GOBP_RIBOSOMAL_SMALL_SUBUNIT_ASSEI | 18 | 0.50623 | 2.0079665 | 0.0034246575 | 0.028531862 | 0.915 | 11667 | tags=100%, list=49%, | MM_4 |
| GOBP_MACROPHAGE_MI | GOBP_MACROPHAGE_MIGRATION | 65 | 0.358248 | 2.0051544 | 0.004878049 | 0.029032478 | 0.917 | 3681 | tags=35%, list=16%, si | MM_4 |
| HUMAN_CYTOMEGALOVI | HUMAN_CYTOMEGALOVIRUS_INFECTION | 216 | 0.277058 | 2.0048819 | 0 | 0.028959423 | 0.917 | 3744 | tags=27%, list=16%, si | MM_4 |
| GOBP_INTRA_GOLGI_VES | GOBP_INTRA_GOLGI_VESICLE_MEDIATED_T | 30 | 0.455847 | 2.0031507 | 0 | 0.029159471 | 0.919 | 4210 | tags=40%, list=18%, si | MM_4 |
| GOBP_NEGATIVE_REGUL | GOBP_NEGATIVE_REGULATION_OF_MONOI | 28 | 0.446603 | 2.003062 | 0 | 0.029037848 | 0.919 | 2350 | tags=25%, list=10%, si | MM_4 |
| GOBP_CELL_CELL_ADHES | GOBP_CELL_CELL_ADHESION_MEDIATED_B\ | 18 | 0.515052 | 2.0024755 | 0.006802721 | 0.028964043 | 0.919 | 3727 | tags=39%, list=16%, si | MM_4 |
| GOBP_TOLL_LIKE_RECEPT | GOBP_TOLL_LIKE_RECEPTOR_9_SIGNALING_ | 16 | 0.540426 | 2.002104 | 0.0034965035 | 0.028939145 | 0.921 | 3953 | tags=50%, list=17%, si | MM_4 |
| GOBP_PROTEASOME_ME | GOBP_PROTEASOME_MEDIATED_UBIQUITIN | 368 | 0.270308 | 1.9994612 | 0 | 0.02934742 | 0.923 | 4340 | tags=34%, list=18%, si | MM_4 |
| GOBP_REGULATION_OF_ | GOBP_REGULATION_OF_INNATE_IMMUNE_ | 427 | 0.251279 | 1.9992756 | 0 | 0.029215876 | 0.923 | 3465 | tags=26%, list=15%, si | MM_4 |
| GLYCOSAMINOGLYCAN_I | GLYCOSAMINOGLYCAN_DEGRADATION | 17 | 0.522319 | 1.9979856 | 0.0035842294 | 0.029365905 | 0.925 | 3385 | tags=59%, list=14%, si | MM_4 |
| GOBP_REGULATION_OF_ | GOBP_REGULATION_OF_PATTERN_RECOGN | 157 | 0.291454 | 1.9942138 | 0 | 0.030059593 | 0.93 | 4070 | tags=31%, list=17%, si | MM_4 |
| GOBP_PROTEIN_FOLDING | GOBP_PROTEIN_FOLDING | 162 | 0.304181 | 1.9927517 | 0 | 0.030089045 | 0.93 | 3592 | tags=28%, list=15%, si | MM_4 |
| GOBP_REGULATION_OF_ | GOBP_REGULATION_OF_LEUKOCYTE_MIGR/ | 230 | 0.269216 | 1.9914501 | 0 | 0.03021065 | 0.932 | 3727 | tags=29%, list=16%, si | MM_4 |
| GOBP_POSITIVE_REGULA | GOBP_POSITIVE_REGULATION_OF_CELLULA | 29 | 0.435883 | 1.9908146 | 0.003937008 | 0.03020351 | 0.932 | 1075 | tags=14%, list=5%, sig | MM_4 |
| GOBP_LEUKOCYTE_MEDI | GOBP_LEUKOCYTE_MEDIATED_IMMUNITY | 400 | 0.254502 | 1.9905759 | 0 | 0.030118175 | 0.932 | 3636 | tags=28%, list=15%, si | MM_4 |
| GOBP_ELECTRON_TRANS | GOBP_ELECTRON_TRANSPORT_CHAIN | 107 | 0.314987 | 1.9889414 | 0 | 0.030177902 | 0.932 | 4871 | tags=44%, list=21%, si | MM_4 |
| GOBP_MITOCHONDRIAL_ | GOBP_MITOCHONDRIAL_RNA_PROCESSING | 20 | 0.487957 | 1.988705 | 0.007246377 | 0.030081863 | 0.933 | 4322 | tags=60%, list=18%, si | MM_4 |
| GOBP_RNA_SURVEILLAN | GOBP_RNA_SURVEILLANCE | 17 | 0.52161 | 1.9886572 | 0.0032786885 | 0.029952582 | 0.933 | 4601 | tags=59%, list=19%, si | MM_4 |
| GOBP_LIPOPROTEIN_LOC | GOBP_LIPOPROTEIN_LOCALIZATION | 20 | 0.485607 | 1.9886172 | 0.010909091 | 0.029813267 | 0.933 | 1689 | tags=25%, list=7%, sig | MM_4 |
| GOBP_MYELOID_LEUKOC | GOBP_MYELOID_LEUKOCYTE_MIGRATION | 222 | 0.274215 | 1.9882337 | 0 | 0.029754605 | 0.934 | 2140 | tags=18%, list=9%, sig | MM_4 |
| GOBP_POSITIVE_REGULA | GOBP_POSITIVE_REGULATION_OF_CELL_CE | 309 | 0.267959 | 1.9873121 | 0 | 0.029852757 | 0.935 | 2811 | tags=22%, list=12%, si | MM_4 |
| BACTERIAL_INVASION_OI | BACTERIAL_INVASION_OF_EPITHELIAL_CELL | 72 | 0.338506 | 1.9870685 | 0 | 0.02979288 | 0.935 | 2799 | tags=25%, list=12%, si | MM_4 |
| GOBP_PROTEIN_LOCALIZ | GOBP_PROTEIN_LOCALIZATION_TO_MITOCI | 101 | 0.322416 | 1.9868693 | 0 | 0.029689796 | 0.936 | 3196 | tags=27%, list=14%, si | MM_4 |
| GOBP_RUFFLE_ORGANIZA | GOBP_RUFFLE_ORGANIZATION | 54 | 0.361171 | 1.9862804 | 0 | 0.02963177 | 0.936 | 2638 | tags=26%, list=11%, si | MM_4 |
| GOBP_CARBOHYDRATE_I | GOBP_CARBOHYDRATE_DERIVATIVE_CATAB | 245 | 0.272835 | 1.9815155 | 0 | 0.030388523 | 0.94 | 3189 | tags=29%, list=14%, si | MM_4 |
| GOBP_CRISTAE_FORMAT | GOBP_CRISTAE_FORMATION | 17 | 0.509854 | 1.9803909 | 0 | 0.030478166 | 0.941 | 11581 | tags=100%, list=49%, | MM_4 |
| GOBP_REGULATION_OF_ | GOBP_REGULATION_OF_TELOMERE_MAINT | 20 | 0.51703 | 1.9799626 | 0.011538462 | 0.030449104 | 0.941 | 2696 | tags=40%, list=11%, si | MM_4 |
| AMINOACYL-TRNA_BIOSY | AMINOACYL-TRNA_BIOSYNTHESIS | 43 | 0.40268 | 1.9795785 | 0.004608295 | 0.03038641 | 0.942 | 3909 | tags=47%, list=17%, si | MM_4 |
| GOBP_ALDEHYDE_CATAB | GOBP_ALDEHYDE_CATABOLIC_PROCESS | 15 | 0.544239 | 1.9759057 | 0.0070921984 | 0.030807322 | 0.945 | 2888 | tags=47%, list=12%, si | MM_4 |
| GOBP_REGULATION_OF_ | GOBP_REGULATION_OF_MRNA_METABOLIC | 286 | 0.259177 | 1.9756193 | 0 | 0.030736038 | 0.945 | 3320 | tags=23%, list=14%, si | MM_4 |
| GOBP_RIBOSE_PHOSPHA | GOBP_RIBOSE_PHOSPHATE_BIOSYNTHETIC_ | 185 | 0.280512 | 1.9711947 | 0 | 0.031385884 | 0.949 | 4733 | tags=42%, list=20%, si | MM_4 |
| GOBP_RESPONSE_TO_TU | GOBP_RESPONSE_TO_TUMOR_NECROSIS_F | 167 | 0.287342 | 1.9709044 | 0 | 0.031299062 | 0.949 | 3846 | tags=29%, list=16%, si | MM_4 |
| GOBP_SNRNA_METABOL | GOBP_SNRNA_METABOLIC_PROCESS | 50 | 0.380849 | 1.9702926 | 0 | 0.031268675 | 0.949 | 4721 | tags=42%, list=20%, si | MM_4 |
| GOBP_POSITIVE_REGULA | GOBP_POSITIVE_REGULATION_OF_FIBROBL | 24 | 0.458985 | 1.9693187 | 0.004 | 0.031268895 | 0.949 | 872 | tags=17%, list=4%, sig | MM_4 |
| GOBP_MATERNAL_PLACE | GOBP_MATERNAL_PLACENTA_DEVELOPMEI | 34 | 0.422289 | 1.9686882 | 0 | 0.031289633 | 0.949 | 3518 | tags=38%, list=15%, si | MM_4 |
| GOBP_INTEGRATED_STRE | GOBP_INTEGRATED_STRESS_RESPONSE_SIG | 23 | 0.463686 | 1.9684886 | 0.0038910506 | 0.031185465 | 0.95 | 2459 | tags=30%, list=10%, si | MM_4 |
| GOBP_GOLGI_TO_VACUC | GOBP_GOLGI_TO_VACUOLE_TRANSPORT | 21 | 0.466932 | 1.9673524 | 0.0035211267 | 0.03138323 | 0.951 | 5230 | tags=57%, list=22%, si | MM_4 |
| GOBP_REGULATION_OF_ | GOBP_REGULATION_OF_EXECUTION_PHASE | 26 | 0.448514 | 1.9646721 | 0 | 0.031730074 | 0.954 | 11912 | tags=96%, list=50%, si | MM_4 |
| GOBP_REGULATION_OF_ | GOBP_REGULATION_OF_MRNA_CATABOLIC | 175 | 0.281699 | 1.9646075 | 0 | 0.031615324 | 0.954 | 4133 | tags=31%, list=18%, si | MM_4 |
| GOBP_CYTOCHROME_CO | GOBP_CYTOCHROME_COMPLEX_ASSEMBLY | 32 | 0.416404 | 1.963852 | 0.008438818 | 0.031645995 | 0.956 | 3249 | tags=38%, list=14%, si | MM_4 |
| GOBP_TRICARBOXYLIC_A | GOBP_TRICARBOXYLIC_ACID_CYCLE | 33 | 0.413359 | 1.9623001 | 0.009803922 | 0.031795144 | 0.959 | 4729 | tags=52%, list=20%, si | MM_4 |
| GOBP_OSTEOCLAST_DIFF | GOBP_OSTEOCLAST_DIFFERENTIATION | 126 | 0.291016 | 1.9614782 | 0 | 0.031742632 | 0.959 | 3882 | tags=33%, list=16%, si | MM_4 |
| GOBP_NEUTROPHIL_CHE | GOBP_NEUTROPHIL_CHEMOTAXIS | 81 | 0.340759 | 1.9607235 | 0 | 0.03172065 | 0.96 | 3287 | tags=26%, list=14%, si | MM_4 |
| GOBP_ATP_BIOSYNTHETI | GOBP_ATP_BIOSYNTHETIC_PROCESS | 80 | 0.331098 | 1.9593067 | 0 | 0.031869568 | 0.961 | 11575 | tags=93%, list=49%, si | MM_4 |
| GOBP_PIGMENT_GRANU | GOBP_PIGMENT_GRANULE_ORGANIZATION | 39 | 0.403075 | 1.9550258 | 0.008810572 | 0.032623608 | 0.966 | 1194 | tags=23%, list=5%, sig | MM_4 |
| GOBP_POSITIVE_REGULA | GOBP_POSITIVE_REGULATION_OF_LYMPHO | 42 | 0.38977 | 1.9548339 | 0 | 0.032508735 | 0.966 | 1742 | tags=21%, list=7%, sig | MM_4 |
| GOBP_POSITIVE_REGULA | GOBP_POSITIVE_REGULATION_OF_EXECUTI | 19 | 0.495867 | 1.9543895 | 0.014705882 | 0.032464076 | 0.967 | 11912 | tags=100%, list=50%, | MM_4 |
| GOBP_B_CELL_RECEPTOR | GOBP_B_CELL_RECEPTOR_SIGNALING_PATH | 55 | 0.355285 | 1.9540235 | 0.004672897 | 0.032389056 | 0.967 | 3882 | tags=33%, list=16%, si | MM_4 |
| GOBP_VESICLE_COATING | GOBP_VESICLE_COATING | 33 | 0.408705 | 1.9519409 | 0 | 0.03271542 | 0.969 | 2079 | tags=27%, list=9%, sig | MM_4 |
| GOBP_POSITIVE_REGULA | GOBP_POSITIVE_REGULATION_OF_PATTERN | 68 | 0.347962 | 1.9514525 | 0 | 0.032660633 | 0.969 | 3612 | tags=28%, list=15%, si | MM_4 |
| GOBP_VIRAL_PROCESS | GOBP_VIRAL_PROCESS | 303 | 0.260531 | 1.9502699 | 0 | 0.032808386 | 0.971 | 4575 | tags=35%, list=19%, si | MM_4 |
| GOBP_VACUOLAR_LOCAL | GOBP_VACUOLAR_LOCALIZATION | 85 | 0.315609 | 1.9498347 | 0 | 0.03281259 | 0.971 | 3842 | tags=32%, list=16%, si | MM_4 |
| GOBP_NEGATIVE_REGUL | GOBP_NEGATIVE_REGULATION_OF_DEFENS | 242 | 0.265614 | 1.9478912 | 0 | 0.033076137 | 0.972 | 2302 | tags=21%, list=10%, si | MM_4 |
| GOBP_POSITIVE_REGULA | GOBP_POSITIVE_REGULATION_OF_CELL_AD | 487 | 0.24108 | 1.9476664 | 0 | 0.03298233 | 0.972 | 2755 | tags=20%, list=12%, si | MM_4 |
| GOBP_INTERFERON_ALPH | GOBP_INTERFERON_ALPHA_PRODUCTION | 33 | 0.40628 | 1.9469702 | 0 | 0.03300301 | 0.972 | 2891 | tags=33%, list=12%, si | MM_4 |
| GOBP_NEGATIVE_REGUL | GOBP_NEGATIVE_REGULATION_OF_CHEMO | 39 | 0.396566 | 1.9458526 | 0.0041841003 | 0.033042993 | 0.972 | 3653 | tags=33%, list=15%, si | MM_4 |
| GOBP_TRANSCRIPTION_I | GOBP_TRANSCRIPTION_INITIATION_AT_RN/ | 148 | 0.29153 | 1.9433978 | 0 | 0.033331722 | 0.973 | 4257 | tags=36%, list=18%, si | MM_4 |
| GOBP_TRNA_MODIFICAT | GOBP_TRNA_MODIFICATION | 84 | 0.334332 | 1.9417725 | 0 | 0.033455625 | 0.974 | 4103 | tags=40%, list=17%, si | MM_4 |
| GOBP_REGULATION_OF_ | GOBP_REGULATION_OF_TELOMERE_MAINT | 61 | 0.357729 | 1.9367803 | 0 | 0.034224253 | 0.975 | 3939 | tags=38%, list=17%, si | MM_4 |
| THERMOGENESIS | THERMOGENESIS | 210 | 0.281516 | 1.9361515 | 0 | 0.03420417 | 0.975 | 5279 | tags=48%, list=22%, si | MM_4 |
| GOBP_REGULATION_OF_ | GOBP_REGULATION_OF_OSTEOCLAST_DIFF | 84 | 0.338317 | 1.9361395 | 0 | 0.03407108 | 0.975 | 3528 | tags=33%, list=15%, si | MM_4 |
| GOBP_CARBOHYDRATE_C | GOBP_CARBOHYDRATE_CATABOLIC_PROCE | 161 | 0.293512 | 1.9329437 | 0 | 0.034593005 | 0.975 | 3412 | tags=30%, list=14%, si | MM_4 |
| GOBP_RNA_MODIFICATI | GOBP_RNA_MODIFICATION | 141 | 0.288855 | 1.9316878 | 0 | 0.0347022 | 0.975 | 4397 | tags=38%, list=19%, si | MM_4 |

| | | | | | | | | | | |
|---|---|---|---|---|---|---|---|---|---|---|
| GOBP_REGULATION_OF_ | GOBP_REGULATION_OF_TRANSLATIONAL_I | 17 | 0.510972 | 1.9314399 | 0.01 | 0.034632154 | 0.975 | 2081 | tags=35%, list=9%, sig | MM_4 |
| GOBP_NEGATIVE_REGUL | GOBP_NEGATIVE_REGULATION_OF_MULTIC | 17 | 0.508991 | 1.9307898 | 0.0033333334 | 0.034628324 | 0.975 | 10 | tags=12%, list=0%, sig | MM_4 |
| GOBP_GENERATION_OF_ | GOBP_GENERATION_OF_PRECURSOR_META | 439 | 0.245112 | 1.9306673 | 0 | 0.034523804 | 0.976 | 3835 | tags=30%, list=16%, si | MM_4 |
| GOBP_GERMINAL_CENTE | GOBP_GERMINAL_CENTER_FORMATION | 16 | 0.512515 | 1.9298608 | 0.0035842294 | 0.03456455 | 0.977 | 2719 | tags=38%, list=12%, si | MM_4 |
| GOBP_TRNA_METHYLATI | GOBP_TRNA_METHYLATION | 39 | 0.400939 | 1.9288841 | 0 | 0.034570985 | 0.977 | 4314 | tags=51%, list=18%, si | MM_4 |
| GOBP_ALPHA_BETA_T_C | GOBP_ALPHA_BETA_T_CELL_PROLIFERATIO | 54 | 0.353542 | 1.9253349 | 0.0047393367 | 0.03512064 | 0.981 | 2569 | tags=22%, list=11%, si | MM_4 |
| GOBP_REGULATION_OF_ | GOBP_REGULATION_OF_LEUKOCYTE_CHEM | 122 | 0.302428 | 1.9251113 | 0 | 0.035050426 | 0.981 | 3727 | tags=30%, list=16%, si | MM_4 |
| GOBP_REGULATION_OF_ | GOBP_REGULATION_OF_B_CELL_RECEPTOR | 18 | 0.497318 | 1.925103 | 0.018656716 | 0.03491915 | 0.981 | 3882 | tags=50%, list=16%, si | MM_4 |
| GOBP_RESPIRATORY_CH | GOBP_RESPIRATORY_CHAIN_COMPLEX_IV_ | 24 | 0.445895 | 1.9221436 | 0 | 0.035417825 | 0.983 | 3249 | tags=42%, list=14%, si | MM_4 |
| GOBP_POSITIVE_REGULA | GOBP_POSITIVE_REGULATION_OF_LYMPHO | 318 | 0.253945 | 1.92163 | 0 | 0.03541137 | 0.983 | 3042 | tags=23%, list=13%, si | MM_4 |
| SALIVARY_SECRETION | SALIVARY_SECRETION | 70 | 0.341484 | 1.9213551 | 0 | 0.03530614 | 0.983 | 1013 | tags=10%, list=4%, sig | MM_4 |
| GOBP_NEGATIVE_REGUL | GOBP_NEGATIVE_REGULATION_OF_LEUKO | 142 | 0.291556 | 1.9201932 | 0 | 0.035433304 | 0.983 | 2986 | tags=25%, list=13%, si | MM_4 |
| GOBP_POLYSACCHARIDE_ | GOBP_POLYSACCHARIDE_CATABOLIC_PROC | 35 | 0.41357 | 1.9194959 | 0.008474576 | 0.035407744 | 0.983 | 2695 | tags=34%, list=11%, si | MM_4 |
| GOBP_REGULATION_OF_ | GOBP_REGULATION_OF_GTPASE_ACTIVITY | 287 | 0.259034 | 1.9166033 | 0 | 0.035895333 | 0.984 | 2685 | tags=20%, list=11%, si | MM_4 |
| GOBP_TELOMERE_MAINT | GOBP_TELOMERE_MAINTENANCE_VIA_REC | 15 | 0.515488 | 1.9109592 | 0.02166065 | 0.037126854 | 0.989 | 2640 | tags=47%, list=11%, si | MM_4 |
| GOBP_REGULATION_OF_ | GOBP_REGULATION_OF_UBIQUITIN_DEPEN | 157 | 0.281733 | 1.9100114 | 0 | 0.03715897 | 0.989 | 3628 | tags=29%, list=15%, si | MM_4 |
| GOBP_REGULATION_OF_ | GOBP_REGULATION_OF_PHAGOCYTOSIS | 106 | 0.301536 | 1.9077555 | 0 | 0.03748392 | 0.99 | 3905 | tags=32%, list=17%, si | MM_4 |
| GOBP_MACROPHAGE_CH | GOBP_MACROPHAGE_CHEMOTAXIS | 45 | 0.371045 | 1.9069668 | 0 | 0.037555683 | 0.99 | 3681 | tags=36%, list=16%, si | MM_4 |
| OSTEOCLAST_DIFFERENTI | OSTEOCLAST_DIFFERENTIATION | 121 | 0.297311 | 1.9047157 | 0 | 0.03783479 | 0.99 | 3658 | tags=26%, list=15%, si | MM_4 |
| GOBP_POSITIVE_REGULA | GOBP_POSITIVE_REGULATION_OF_DNA_RE | 135 | 0.290384 | 1.9039068 | 0 | 0.03779292 | 0.991 | 2913 | tags=23%, list=12%, si | MM_4 |
| GOBP_MEMBRANE_BIOG | GOBP_MEMBRANE_BIOGENESIS | 63 | 0.338647 | 1.9029534 | 0.0053475937 | 0.037830237 | 0.991 | 4213 | tags=35%, list=18%, si | MM_4 |
| GOBP_SPLICEOSOMAL_S | GOBP_SPLICEOSOMAL_SNRNP_ASSEMBLY | 35 | 0.394161 | 1.8999391 | 0.009009009 | 0.038345892 | 0.991 | 3673 | tags=43%, list=16%, si | MM_4 |
| GOBP_MITOCHONDRIAL_ | GOBP_MITOCHONDRIAL_TRANSMEMBRANE | 92 | 0.310199 | 1.8992122 | 0 | 0.038356315 | 0.991 | 4266 | tags=38%, list=18%, si | MM_4 |
| GOBP_RESPONSE_TO_TR | GOBP_RESPONSE_TO_TRANSFORMING_GR | 246 | 0.263071 | 1.8974574 | 0 | 0.03852747 | 0.992 | 3223 | tags=24%, list=14%, si | MM_4 |
| GOBP_POSITIVE_REGULA | GOBP_POSITIVE_REGULATION_OF_HYDROL | 427 | 0.238378 | 1.8970853 | 0 | 0.038433664 | 0.992 | 3377 | tags=24%, list=14%, si | MM_4 |
| GOBP_LYMPHOCYTE_ME | GOBP_LYMPHOCYTE_MEDIATED_IMMUNITY | 300 | 0.251559 | 1.8968452 | 0 | 0.03833323 | 0.992 | 3437 | tags=26%, list=15%, si | MM_4 |
| GOBP_CELL_CHEMOTAXIS | GOBP_CELL_CHEMOTAXIS | 295 | 0.254624 | 1.8942612 | 0 | 0.038709782 | 0.992 | 2096 | tags=16%, list=9%, sig | MM_4 |
| GOBP_REGULATION_OF_ | GOBP_REGULATION_OF_STRESS_ACTIVATED | 41 | 0.371027 | 1.8898371 | 0.0088495575 | 0.039621606 | 0.992 | 4467 | tags=49%, list=19%, si | MM_4 |
| GOBP_POSITIVE_REGULA | GOBP_POSITIVE_REGULATION_OF_STEM_C | 50 | 0.365151 | 1.8888725 | 0.004761905 | 0.039756566 | 0.992 | 5167 | tags=54%, list=22%, si | MM_4 |
| GOBP_POSITIVE_REGULA | GOBP_POSITIVE_REGULATION_OF_CELL_AC | 373 | 0.243082 | 1.8874589 | 0 | 0.03994105 | 0.993 | 3051 | tags=22%, list=13%, si | MM_4 |
| GOBP_GOLGI_VESICLE_T | GOBP_GOLGI_VESICLE_TRANSPORT | 276 | 0.258224 | 1.886754 | 0 | 0.039952237 | 0.993 | 4101 | tags=31%, list=17%, si | MM_4 |
| GOBP_NUCLEOTIDE_EXCI | GOBP_NUCLEOTIDE_EXCISION_REPAIR | 76 | 0.321639 | 1.8860005 | 0 | 0.040006448 | 0.993 | 3800 | tags=37%, list=16%, si | MM_4 |
| GOBP_MITOCHONDRIAL_ | GOBP_MITOCHONDRIAL_TRANSLATION | 128 | 0.286995 | 1.8829991 | 0 | 0.040554125 | 0.994 | 4103 | tags=31%, list=17%, si | MM_4 |
| GOBP_PROTEIN_N_LINKE | GOBP_PROTEIN_N_LINKED_GLYCOSYLATION | 61 | 0.34083 | 1.882766 | 0.004950495 | 0.04046523 | 0.994 | 3109 | tags=31%, list=13%, si | MM_4 |
| GOBP_RNA_TEMPLATED_ | GOBP_RNA_TEMPLATED_DNA_BIOSYNTHET | 63 | 0.342115 | 1.8827412 | 0 | 0.04032759 | 0.994 | 4454 | tags=40%, list=19%, si | MM_4 |
| GOBP_VACUOLE_ORGAN | GOBP_VACUOLE_ORGANIZATION | 223 | 0.26109 | 1.8821534 | 0 | 0.040329885 | 0.994 | 4249 | tags=35%, list=18%, si | MM_4 |
| GOBP_ESTABLISHMENT_ | GOBP_ESTABLISHMENT_OF_PROTEIN_LOCA | 25 | 0.426823 | 1.8813297 | 0.0033333334 | 0.040448282 | 0.994 | 5614 | tags=52%, list=24%, si | MM_4 |
| GOBP_REGULATION_OF_ | GOBP_REGULATION_OF_LEUKOCYTE_PROLI | 273 | 0.253054 | 1.8803233 | 0 | 0.040554997 | 0.994 | 3025 | tags=25%, list=13%, si | MM_4 |
| GOBP_NEGATIVE_REGUL | GOBP_NEGATIVE_REGULATION_OF_DNA_T | 17 | 0.510172 | 1.8765222 | 0.01986755 | 0.041385822 | 0.995 | 5311 | tags=65%, list=22%, si | MM_4 |
| GOBP_REGULATION_OF_ | GOBP_REGULATION_OF_DNA_STRAND_ELO | 18 | 0.471326 | 1.8762805 | 0.0073260074 | 0.041312285 | 0.995 | 2696 | tags=33%, list=11%, si | MM_4 |
| GOBP_TRNA_PROCESSIN | GOBP_TRNA_PROCESSING | 122 | 0.290484 | 1.876154 | 0 | 0.041182443 | 0.995 | 4397 | tags=40%, list=19%, si | MM_4 |
| GOBP_NEGATIVE_REGUL | GOBP_NEGATIVE_REGULATION_OF_IMMUN | 186 | 0.273248 | 1.8758165 | 0 | 0.04115854 | 0.995 | 2302 | tags=23%, list=10%, si | MM_4 |
| GOBP_POSITIVE_REGULA | GOBP_POSITIVE_REGULATION_OF_LEUKOC | 97 | 0.308399 | 1.8741925 | 0 | 0.041491017 | 0.995 | 3727 | tags=31%, list=16%, si | MM_4 |
| GOBP_REGULATION_OF_ | GOBP_REGULATION_OF_PROTEOLYSIS_INV | 215 | 0.260125 | 1.873508 | 0 | 0.04150413 | 0.995 | 3628 | tags=28%, list=15%, si | MM_4 |
| GOBP_RIBOSOMAL_SMA | GOBP_RIBOSOMAL_SMALL_SUBUNIT_BIOG | 101 | 0.299374 | 1.8721051 | 0 | 0.041667722 | 0.995 | 11667 | tags=91%, list=49%, si | MM_4 |
| RNA_POLYMERASE | RNA_POLYMERASE | 27 | 0.41884 | 1.8696345 | 0.003968254 | 0.04224691 | 0.996 | 3444 | tags=41%, list=15%, si | MM_4 |
| GOBP_NEGATIVE_REGUL | GOBP_NEGATIVE_REGULATION_OF_RELEAS | 22 | 0.439365 | 1.8690609 | 0.014184397 | 0.042281643 | 0.996 | 11575 | tags=95%, list=49%, si | MM_4 |
| GOBP_REGULATION_OF_ | GOBP_REGULATION_OF_NUCLEOBASE_CON | 16 | 0.503094 | 1.8680675 | 0.01056338 | 0.042353984 | 0.996 | 11740 | tags=100%, list=50%, | MM_4 |
| PROTEIN_PROCESSING_I | PROTEIN_PROCESSING_IN_ENDOPLASMIC_ | 158 | 0.273327 | 1.8680003 | 0 | 0.042231858 | 0.996 | 11267 | tags=89%, list=48%, si | MM_4 |
| GOBP_NAD_CATABOLIC_ | GOBP_NAD_CATABOLIC_PROCESS | 21 | 0.467631 | 1.8668747 | 0.02811245 | 0.042444073 | 0.996 | 3613 | tags=52%, list=15%, si | MM_4 |
| GOBP_IMPORT_INTO_TH | GOBP_IMPORT_INTO_THE_MITOCHONDRIO | 49 | 0.366965 | 1.8653299 | 0.004608295 | 0.04270283 | 0.996 | 4266 | tags=41%, list=18%, si | MM_4 |
| VIRAL_MYOCARDITIS | VIRAL_MYOCARDITIS | 63 | 0.334375 | 1.864734 | 0 | 0.042672664 | 0.996 | 3523 | tags=33%, list=15%, si | MM_4 |
| GOBP_DEFENSE_RESPON | GOBP_DEFENSE_RESPONSE_TO_GRAM_POS | 95 | 0.296598 | 1.863721 | 0.006849315 | 0.042756744 | 0.997 | 2271 | tags=19%, list=10%, si | MM_4 |
| GOBP_LYTIC_VACUOLE_ | GOBP_LYTIC_VACUOLE_ORGANIZATION | 95 | 0.3046 | 1.8613328 | 0 | 0.043304373 | 0.997 | 4249 | tags=38%, list=18%, si | MM_4 |
| GOBP_TELOMERE_MAINT | GOBP_TELOMERE_MAINTENANCE_VIA_TEL | 76 | 0.325233 | 1.8583194 | 0 | 0.04386137 | 0.997 | 4454 | tags=38%, list=19%, si | MM_4 |
| GOBP_ANTIMICROBIAL_H | GOBP_ANTIMICROBIAL_HUMORAL_RESPON | 98 | 0.298226 | 1.8559506 | 0 | 0.04439844 | 0.997 | 2271 | tags=20%, list=10%, si | MM_4 |
| GOBP_LEUKOCYTE_CHEM | GOBP_LEUKOCYTE_CHEMOTAXIS | 213 | 0.266174 | 1.8558887 | 0 | 0.044287626 | 0.997 | 2096 | tags=16%, list=9%, sig | MM_4 |
| GOBP_NEGATIVE_REGUL | GOBP_NEGATIVE_REGULATION_OF_LEUKO | 66 | 0.335378 | 1.8545727 | 0 | 0.044400383 | 0.998 | 2986 | tags=33%, list=13%, si | MM_4 |
| GOBP_NEGATIVE_REGUL | GOBP_NEGATIVE_REGULATION_OF_VASCU | 109 | 0.289199 | 1.8528167 | 0.00625 | 0.04470775 | 0.998 | 3078 | tags=26%, list=13%, si | MM_4 |
| GOBP_NATURAL_KILLER_ | GOBP_NATURAL_KILLER_CELL_MEDIATED_I | 73 | 0.32726 | 1.8526871 | 0.004854369 | 0.04460464 | 0.998 | 1562 | tags=22%, list=7%, sig | MM_4 |
| GOBP_TRNA_THREONYL | GOBP_TRNA_THREONYLCARBAMOYLADEN | 15 | 0.506332 | 1.8518203 | 0.014869888 | 0.04464479 | 0.998 | 4397 | tags=60%, list=19%, si | MM_4 |
| GOBP_REGULATION_OF_ | GOBP_REGULATION_OF_DNA_REPAIR | 219 | 0.255284 | 1.8509686 | 0 | 0.044678293 | 0.998 | 2963 | tags=22%, list=13%, si | MM_4 |
| GOBP_POSITIVE_REGULA | GOBP_POSITIVE_REGULATION_OF_DNA_BI | 76 | 0.317708 | 1.8501666 | 0 | 0.04474179 | 0.998 | 3765 | tags=33%, list=16%, si | MM_4 |
| GOBP_RESPONSE_TO_IN | GOBP_RESPONSE_TO_INTERFERON_ALPHA | 28 | 0.412908 | 1.8498381 | 0.003968254 | 0.044698905 | 0.998 | 1488 | tags=25%, list=6%, sig | MM_4 |
| GOBP_REGULATION_OF_ | GOBP_REGULATION_OF_EXTRINSIC_APOPT | 45 | 0.35001 | 1.8496028 | 0.0050251256 | 0.044613052 | 0.998 | 2556 | tags=24%, list=11%, si | MM_4 |
| HUMAN_PAPILLOMAVIRU | HUMAN_PAPILLOMAVIRUS_INFECTION | 315 | 0.248614 | 1.8493192 | 0 | 0.04454216 | 0.998 | 3082 | tags=23%, list=13%, si | MM_4 |
| GOBP_TRANSFORMING_ | GOBP_TRANSFORMING_GROWTH_FACTOR_ | 202 | 0.264992 | 1.8485473 | 0 | 0.044610742 | 0.998 | 3223 | tags=24%, list=14%, si | MM_4 |
| GOBP_NEGATIVE_REGUL | GOBP_NEGATIVE_REGULATION_OF_B_CELL | 36 | 0.386472 | 1.846114 | 0.008097166 | 0.0452245 | 0.999 | 3371 | tags=39%, list=14%, si | MM_4 |
| GOBP_LYMPH_NODE_DE | GOBP_LYMPH_NODE_DEVELOPMENT | 32 | 0.4007 | 1.8452982 | 0.016393442 | 0.045319807 | 0.999 | 18 | tags=9%, list=0%, sign | MM_4 |
| GOBP_MACROMOLECULE | GOBP_MACROMOLECULE_METHYLATION | 121 | 0.287396 | 1.8449749 | 0 | 0.045254845 | 0.999 | 4316 | tags=37%, list=18%, si | MM_4 |
| GOBP_DNA_STRAND_EL | GOBP_DNA_STRAND_ELONGATION | 36 | 0.375264 | 1.8448993 | 0.00456621 | 0.045124877 | 0.999 | 2701 | tags=28%, list=11%, si | MM_4 |
| FOCAL_ADHESION | FOCAL_ADHESION | 194 | 0.267077 | 1.8438587 | 0 | 0.045252025 | 0.999 | 3334 | tags=24%, list=14%, si | MM_4 |
| GOBP_PHOSPHATIDIC_A | GOBP_PHOSPHATIDIC_ACID_BIOSYNTHETIC_ | 23 | 0.441846 | 1.843316 | 0.015503876 | 0.04521649 | 0.999 | 2032 | tags=30%, list=9%, sig | MM_4 |
| GOBP_NEGATIVE_REGUL | GOBP_NEGATIVE_REGULATION_OF_TRANS | 127 | 0.285789 | 1.841291 | 0 | 0.04559494 | 0.999 | 2815 | tags=24%, list=12%, si | MM_4 |
| GOBP_LEUKOCYTE_MEDI | GOBP_LEUKOCYTE_MEDIATED_CYTOTOXICI | 135 | 0.287773 | 1.840718 | 0 | 0.04567439 | 0.999 | 3435 | tags=29%, list=15%, si | MM_4 |
| GOBP_NEGATIVE_REGUL | GOBP_NEGATIVE_REGULATION_OF_NATUR | 19 | 0.467137 | 1.838388 | 0.018115941 | 0.046207484 | 0.999 | 1562 | tags=37%, list=7%, sig | MM_4 |
| GOBP_GLIAL_CELL_MIGR | GOBP_GLIAL_CELL_MIGRATION | 70 | 0.32069 | 1.8371686 | 0.0057142857 | 0.04637073 | 0.999 | 5281 | tags=50%, list=22%, si | MM_4 |
| GOBP_REGULATION_OF_ | GOBP_REGULATION_OF_LEUKOCYTE_DIFFE | 342 | 0.239819 | 1.8367999 | 0 | 0.046305634 | 0.999 | 3364 | tags=24%, list=14%, si | MM_4 |
| GOBP_RESPONSE_TO_EP | GOBP_RESPONSE_TO_EPIDERMAL_GROWTH | 42 | 0.367365 | 1.8359263 | 0.0045454544 | 0.04638117 | 0.999 | 2882 | tags=33%, list=12%, si | MM_4 |
| HEMATOPOIETIC_CELL_L | HEMATOPOIETIC_CELL_LINEAGE | 83 | 0.300628 | 1.8351622 | 0.0057471264 | 0.046457883 | 0.999 | 786 | tags=13%, list=3%, sig | MM_4 |
| GOBP_REGULATION_OF_ | GOBP_REGULATION_OF_IMMUNE_EFFECTO | 406 | 0.235591 | 1.8333538 | 0 | 0.04684514 | 0.999 | 3641 | tags=27%, list=15%, si | MM_4 |
| GOBP_REGULATION_OF_ | GOBP_REGULATION_OF_VASCULAR_ENDOT | 33 | 0.386038 | 1.8326672 | 0 | 0.04687006 | 0.999 | 892 | tags=18%, list=4%, sig | MM_4 |

| | | | | | | | | | | |
|---|---|---|---|---|---|---|---|---|---|---|
| GOBP_NEGATIVE_REGUL | GOBP_NEGATIVE_REGULATION_OF_RESPON | 112 | 0.297611 | 1.8326164 | 0 | 0.046761114 | 0.999 | 3944 | tags=34%, list=17%, si | MM_4 |
| GOBP_APOPTOTIC_CELL_ | GOBP_APOPTOTIC_CELL_CLEARANCE | 40 | 0.374962 | 1.8325607 | 0 | 0.04663879 | 0.999 | 2387 | tags=28%, list=10%, si | MM_4 |
| STAPHYLOCOCCUS_AURE | STAPHYLOCOCCUS_AUREUS_INFECTION | 44 | 0.360886 | 1.8316951 | 0 | 0.046719976 | 0.999 | 4035 | tags=43%, list=17%, si | MM_4 |
| NOD-LIKE_RECEPTOR_SIG | NOD-LIKE_RECEPTOR_SIGNALING_PATHWA | 151 | 0.267079 | 1.8314731 | 0 | 0.04663989 | 0.999 | 3732 | tags=27%, list=16%, si | MM_4 |
| GOBP_MEMBRANE_FISSI | GOBP_MEMBRANE_FISSION | 31 | 0.400362 | 1.8243164 | 0.011857707 | 0.048265196 | 0.999 | 12431 | tags=97%, list=53%, si | MM_4 |
| GOBP_VIRAL_GENE_EXPR | GOBP_VIRAL_GENE_EXPRESSION | 75 | 0.319421 | 1.8226786 | 0 | 0.04850265 | 0.999 | 4279 | tags=36%, list=18%, si | MM_4 |
| GOBP_PROTEIN_TRANSM | GOBP_PROTEIN_TRANSMEMBRANE_TRANS | 67 | 0.319106 | 1.821056 | 0.005586592 | 0.048777726 | 0.999 | 3289 | tags=34%, list=14%, si | MM_4 |
| GOBP_POSITIVE_REGULA | GOBP_POSITIVE_REGULATION_OF_MRNA_M | 131 | 0.278595 | 1.8203803 | 0 | 0.04882985 | 0.999 | 4133 | tags=29%, list=18%, si | MM_4 |
| GOBP_REGULATION_OF_ | GOBP_REGULATION_OF_CELL_CELL_ADHESI | 475 | 0.23417 | 1.8197708 | 0 | 0.048881512 | 0.999 | 3042 | tags=22%, list=13%, si | MM_4 |
| GOBP_NEGATIVE_REGUL | GOBP_NEGATIVE_REGULATION_OF_TYPE_I_ | 23 | 0.432404 | 1.8191098 | 0 | 0.048927337 | 0.999 | 4931 | tags=57%, list=21%, si | MM_4 |
| GOBP_NUCLEOSIDE_PHO | GOBP_NUCLEOSIDE_PHOSPHATE_BIOSYNTH | 234 | 0.250376 | 1.8169676 | 0 | 0.049502656 | 0.999 | 4841 | tags=39%, list=21%, si | MM_4 |
| GOBP_TRNA_WOBBLE_U | GOBP_TRNA_WOBBLE_URIDINE_MODIFICAT | 16 | 0.483987 | 1.8168751 | 0.02166065 | 0.04938294 | 0.999 | 3058 | tags=44%, list=13%, si | MM_4 |
| GOBP_NEGATIVE_REGUL | GOBP_NEGATIVE_REGULATION_OF_LYMPH | 53 | 0.337002 | 1.8165447 | 0 | 0.04933807 | 0.999 | 2217 | tags=28%, list=9%, sig | MM_4 |
| GOBP_RESPONSE_TO_CO | GOBP_RESPONSE_TO_COLD | 44 | 0.350151 | 1.8153031 | 0.0091743115 | 0.04945615 | 0.999 | 2561 | tags=30%, list=11%, si | MM_4 |
| GOBP_POSITIVE_REGULA | GOBP_POSITIVE_REGULATION_OF_DNA_ME | 314 | 0.237811 | 1.8146188 | 0 | 0.049499616 | 0.999 | 2913 | tags=20%, list=12%, si | MM_4 |
| GOBP_LEUKOCYTE_HOM | GOBP_LEUKOCYTE_HOMEOSTASIS | 137 | 0.267441 | 1.8139819 | 0 | 0.04956209 | 0.999 | 3596 | tags=28%, list=15%, si | MM_4 |
| GOBP_AMYLOID_FIBRIL_ | GOBP_AMYLOID_FIBRIL_FORMATION | 30 | 0.390622 | 1.8132734 | 0.008064516 | 0.049585402 | 0.999 | 3233 | tags=33%, list=14%, si | MM_4 |
| GOBP_MITOCHONDRIAL_ | GOBP_MITOCHONDRIAL_ELECTRON_TRANS | 18 | 0.481331 | 1.8127605 | 0.017985612 | 0.049628273 | 0.999 | 4755 | tags=61%, list=20%, si | MM_4 |
| GOBP_NEGATIVE_REGUL | GOBP_NEGATIVE_REGULATION_OF_LYMPH | 164 | 0.269871 | 1.8100586 | 0 | 0.050184976 | 0.999 | 1765 | tags=18%, list=7%, sig | MM_4 |
| GOBP_POSITIVE_REGULA | GOBP_POSITIVE_REGULATION_OF_LIPID_ST | 25 | 0.421124 | 1.8097978 | 0.007246377 | 0.050112918 | 0.999 | 2346 | tags=28%, list=10%, si | MM_4 |
| GOBP_DNA_TEMPLATED_ | GOBP_DNA_TEMPLATED_TRANSCRIPTION_I | 183 | 0.260589 | 1.8078923 | 0 | 0.050505318 | 0.999 | 4185 | tags=34%, list=18%, si | MM_4 |
| GOBP_POSITIVE_REGULA | GOBP_POSITIVE_REGULATION_OF_T_CELL_ | 108 | 0.282923 | 1.8054161 | 0.006369427 | 0.051165376 | 0.999 | 1982 | tags=21%, list=8%, sig | MM_4 |
| GOBP_MACROAUTOPHA | GOBP_MACROAUTOPHAGY | 287 | 0.243742 | 1.8041091 | 0 | 0.05130862 | 0.999 | 3425 | tags=24%, list=15%, si | MM_4 |
| GOBP_NEGATIVE_REGUL | GOBP_NEGATIVE_REGULATION_OF_CELL_M | 16 | 0.473236 | 1.8036081 | 0.014925373 | 0.05133902 | 0.999 | 2526 | tags=31%, list=11%, si | MM_4 |
| GOBP_POSITIVE_REGULA | GOBP_POSITIVE_REGULATION_OF_CHEMOK | 15 | 0.489492 | 1.8027952 | 0.014134276 | 0.051473696 | 0.999 | 2599 | tags=47%, list=11%, si | MM_4 |
| GOBP_REGULATION_OF_ | GOBP_REGULATION_OF_CYTOPLASMIC_PAT | 104 | 0.280818 | 1.8013165 | 0 | 0.051707305 | 0.999 | 3287 | tags=25%, list=14%, si | MM_4 |
| GOBP_REGULATION_OF_ | GOBP_REGULATION_OF_CELL_CYCLE_G1_S_ | 185 | 0.264625 | 1.801083 | 0 | 0.05163917 | 0.999 | 3259 | tags=27%, list=14%, si | MM_4 |
| GOBP_MITOCHONDRIAL_ | GOBP_MITOCHONDRIAL_CYTOCHROME_C_ | 19 | 0.450888 | 1.800806 | 0.014440433 | 0.051596694 | 0.999 | 3249 | tags=42%, list=14%, si | MM_4 |
| GOBP_ORGANELLE_INHE | GOBP_ORGANELLE_INHERITANCE | 15 | 0.505343 | 1.8007754 | 0.017985612 | 0.051470462 | 0.999 | 4421 | tags=47%, list=19%, si | MM_4 |
| GOBP_GLIAL_CELL_APOP | GOBP_GLIAL_CELL_APOPTOTIC_PROCESS | 20 | 0.437508 | 1.800218 | 0.013986014 | 0.051543947 | 0.999 | 11145 | tags=95%, list=47%, si | MM_4 |
| GOBP_GPI_ANCHOR_ME | GOBP_GPI_ANCHOR_METABOLIC_PROCESS | 33 | 0.383318 | 1.7989511 | 0.012096774 | 0.051733647 | 0.999 | 1643 | tags=27%, list=7%, sig | MM_4 |
| GOBP_AMYLOID_BETA_C | GOBP_AMYLOID_BETA_CLEARANCE | 37 | 0.370426 | 1.796526 | 0.013513514 | 0.052396778 | 0.999 | 2723 | tags=24%, list=12%, si | MM_4 |
| GOBP_NEGATIVE_REGUL | GOBP_NEGATIVE_REGULATION_OF_CHEMO | 29 | 0.39647 | 1.7960567 | 0.013574661 | 0.0523989 | 0.999 | 3216 | tags=34%, list=14%, si | MM_4 |
| GOBP_TRANSPORT_OF_V | GOBP_TRANSPORT_OF_VIRUS | 15 | 0.487441 | 1.7933549 | 0.006968641 | 0.053058952 | 0.999 | 3098 | tags=47%, list=13%, si | MM_4 |
| GOBP_POSITIVE_REGULA | GOBP_POSITIVE_REGULATION_OF_CHROM | 107 | 0.284186 | 1.7916064 | 0 | 0.053358316 | 0.999 | 4465 | tags=36%, list=19%, si | MM_4 |
| GOBP_NEGATIVE_REGUL | GOBP_NEGATIVE_REGULATION_OF_TRANSL | 20 | 0.450513 | 1.7902982 | 0.01923077 | 0.053590726 | 0.999 | 10977 | tags=95%, list=46%, si | MM_4 |
| GOBP_MITOCHONDRIAL_ | GOBP_MITOCHONDRIAL_TRANSPORT | 174 | 0.25795 | 1.7890625 | 0 | 0.05385437 | 0.999 | 4012 | tags=32%, list=17%, si | MM_4 |
| GOBP_REGULATION_OF_ | GOBP_REGULATION_OF_MYELOID_LEUKOC | 140 | 0.27416 | 1.7877449 | 0 | 0.054079145 | 0.999 | 3364 | tags=27%, list=14%, si | MM_4 |
| GOBP_B_CELL_HOMEOST | GOBP_B_CELL_HOMEOSTASIS | 42 | 0.350423 | 1.7871755 | 0.018348623 | 0.05408831 | 0.999 | 11435 | tags=90%, list=48%, si | MM_4 |
| GOBP_RESPONSE_TO_TY | GOBP_RESPONSE_TO_TYPE_II_INTERFERON | 122 | 0.272854 | 1.78407 | 0 | 0.054777563 | 0.999 | 4035 | tags=36%, list=17%, si | MM_4 |
| GOBP_CELLULAR_RESPO | GOBP_CELLULAR_RESPONSE_TO_VITAMIN | 16 | 0.468188 | 1.7815034 | 0.016025642 | 0.055401802 | 0.999 | 1811 | tags=31%, list=8%, sig | MM_4 |
| GOBP_RESPONSE_TO_MI | GOBP_RESPONSE_TO_MISFOLDED_PROTEIN | 18 | 0.462796 | 1.7809613 | 0.003875969 | 0.05543273 | 0.999 | 11153 | tags=94%, list=47%, si | MM_4 |
| GOBP_NADH_METABOLI | GOBP_NADH_METABOLIC_PROCESS | 39 | 0.363222 | 1.780843 | 0.004149378 | 0.05533167 | 0.999 | 4292 | tags=46%, list=18%, si | MM_4 |
| GOBP_PROTEIN_RNA_CO | GOBP_PROTEIN_RNA_COMPLEX_ORGANIZA | 173 | 0.262852 | 1.779131 | 0 | 0.05562383 | 0.999 | 5026 | tags=40%, list=21%, si | MM_4 |
| GOBP_LEUKOCYTE_TETH | GOBP_LEUKOCYTE_TETHERING_OR_ROLLIN | 35 | 0.375452 | 1.7773945 | 0.008064516 | 0.056042492 | 0.999 | 3573 | tags=40%, list=15%, si | MM_4 |
| GOBP_ACTIN_POLYMERIZ | GOBP_ACTIN_POLYMERIZATION_OR_DEPOL | 187 | 0.257934 | 1.7770104 | 0 | 0.056069747 | 0.999 | 3905 | tags=29%, list=17%, si | MM_4 |
| GOBP_POSITIVE_REGULA | GOBP_POSITIVE_REGULATION_OF_DEFENSE | 480 | 0.230107 | 1.7768589 | 0 | 0.05595611 | 0.999 | 3465 | tags=25%, list=15%, si | MM_4 |
| GOBP_PROTEIN_CATABO | GOBP_PROTEIN_CATABOLIC_PROCESS_IN_T | 22 | 0.427162 | 1.7768394 | 0.014440433 | 0.055818267 | 0.999 | 4249 | tags=59%, list=18%, si | MM_4 |
| GOBP_REGULATION_OF_ | GOBP_REGULATION_OF_JUN_KINASE_ACTIV | 62 | 0.321646 | 1.7753218 | 0 | 0.056067318 | 0.999 | 3953 | tags=31%, list=17%, si | MM_4 |
| GOBP_POSITIVE_REGULA | GOBP_POSITIVE_REGULATION_OF_AUTOPH | 19 | 0.447724 | 1.7738677 | 0.018518519 | 0.05631856 | 0.999 | 3144 | tags=32%, list=13%, si | MM_4 |
| GOBP_ACTIVATION_OF_I | GOBP_ACTIVATION_OF_INNATE_IMMUNE_ | 290 | 0.240772 | 1.7735176 | 0 | 0.05624111 | 0.999 | 3465 | tags=25%, list=15%, si | MM_4 |
| TUBERCULOSIS | TUBERCULOSIS | 152 | 0.268491 | 1.7735121 | 0 | 0.056098003 | 0.999 | 2693 | tags=22%, list=11%, si | MM_4 |
| GOBP_CELL_CYCLE_G1_S | GOBP_CELL_CYCLE_G1_S_PHASE_TRANSITI | 236 | 0.244744 | 1.7733849 | 0 | 0.055967744 | 0.999 | 3377 | tags=26%, list=14%, si | MM_4 |
| GOBP_NEGATIVE_REGUL | GOBP_NEGATIVE_REGULATION_OF_CYTOKI | 292 | 0.231832 | 1.7720002 | 0 | 0.056206565 | 0.999 | 3372 | tags=25%, list=14%, si | MM_4 |
| GOBP_MITOTIC_G2_M_T | GOBP_MITOTIC_G2_M_TRANSITION_CHECK | 56 | 0.324788 | 1.7707766 | 0 | 0.056480955 | 0.999 | 2866 | tags=30%, list=12%, si | MM_4 |
| GOBP_PYRIMIDINE_DEOX | GOBP_PYRIMIDINE_DEOXYRIBONUCLEOTIDE | 20 | 0.431812 | 1.7704172 | 0.012145749 | 0.056424867 | 0.999 | 3106 | tags=45%, list=13%, si | MM_4 |
| GOBP_POSITIVE_REGULA | GOBP_POSITIVE_REGULATION_OF_LEUKOC | 26 | 0.398274 | 1.7681137 | 0.0042735045 | 0.05708593 | 0.999 | 3307 | tags=35%, list=14%, si | MM_4 |
| GOBP_REGULATION_OF_ | GOBP_REGULATION_OF_T_CELL_MEDIATED | 92 | 0.296307 | 1.7670125 | 0 | 0.057302024 | 0.999 | 3561 | tags=30%, list=15%, si | MM_4 |
| GOBP_DENSE_CORE_GRA | GOBP_DENSE_CORE_GRANULE_LOCALIZATI | 28 | 0.397171 | 1.766443 | 0.01826484 | 0.057342898 | 0.999 | 1531 | tags=21%, list=6%, sig | MM_4 |
| GOBP_ORGANOPHOSPHA | GOBP_ORGANOPHOSPHATE_BIOSYNTHETIC | 500 | 0.215482 | 1.765768 | 0 | 0.05735491 | 0.999 | 3370 | tags=27%, list=14%, si | MM_4 |
| GOBP_VESICLE_TARGETI | GOBP_VESICLE_TARGETING | 60 | 0.319104 | 1.765648 | 0.010256411 | 0.057241865 | 1 | 3982 | tags=35%, list=17%, si | MM_4 |
| GOBP_GMP_BIOSYNTHET | GOBP_GMP_BIOSYNTHETIC_PROCESS | 15 | 0.486333 | 1.763088 | 0.024734983 | 0.057948306 | 1 | 2803 | tags=47%, list=12%, si | MM_4 |
| GOBP_CELLULAR_RESPO | GOBP_CELLULAR_RESPONSE_TO_VIRUS | 47 | 0.342882 | 1.7630509 | 0.014778325 | 0.05780487 | 1 | 2538 | tags=28%, list=11%, si | MM_4 |
| GOBP_PROTEIN_AUTOUB | GOBP_PROTEIN_AUTOUBIQUITINATION | 68 | 0.318587 | 1.7616235 | 0.0052083335 | 0.058035906 | 1 | 2128 | tags=21%, list=9%, sig | MM_4 |
| GOBP_QUINONE_BIOSYN | GOBP_QUINONE_BIOSYNTHETIC_PROCESS | 16 | 0.46584 | 1.7604796 | 0.033444814 | 0.058271077 | 1 | 3486 | tags=44%, list=15%, si | MM_4 |
| GOBP_CELLULAR_COMPO | GOBP_CELLULAR_COMPONENT_DISASSEMB | 472 | 0.220197 | 1.7601379 | 0 | 0.058197863 | 1 | 3727 | tags=25%, list=16%, si | MM_4 |
| GOBP_RESPONSE_TO_AT | GOBP_RESPONSE_TO_ATP | 31 | 0.396421 | 1.7592789 | 0.027237354 | 0.058320824 | 1 | 2093 | tags=29%, list=9%, sig | MM_4 |
| GOBP_GMP_METABOLIC_ | GOBP_GMP_METABOLIC_PROCESS | 21 | 0.436258 | 1.7590626 | 0.016736401 | 0.058248613 | 1 | 3757 | tags=48%, list=16%, si | MM_4 |
| GOBP_RESPONSE_TO_LIT | GOBP_RESPONSE_TO_LITHIUM_ION | 16 | 0.462482 | 1.758118 | 0.014184397 | 0.058398377 | 1 | 3051 | tags=38%, list=13%, si | MM_4 |
| GOBP_NEGATIVE_REGUL | GOBP_NEGATIVE_REGULATION_OF_CELL_S | 59 | 0.323508 | 1.7581074 | 0.0099502485 | 0.0582682 | 1 | 2672 | tags=27%, list=11%, si | MM_4 |
| GOBP_NEGATIVE_REGUL | GOBP_NEGATIVE_REGULATION_OF_SPROUT | 15 | 0.485326 | 1.7562004 | 0.010791367 | 0.058736768 | 1 | 3268 | tags=53%, list=14%, si | MM_4 |
| GOBP_RESPONSE_TO_ST | GOBP_RESPONSE_TO_STEROL_DEPLETION | 15 | 0.48582 | 1.7561586 | 0.01831502 | 0.058606073 | 1 | 5266 | tags=60%, list=22%, si | MM_4 |
| GOBP_REGULATION_OF_ | GOBP_REGULATION_OF_NATURAL_KILLER_ | 55 | 0.327297 | 1.7561176 | 0.0052083335 | 0.058487974 | 1 | 1562 | tags=24%, list=7%, sig | MM_4 |
| GOBP_EOSINOPHIL_CHEM | GOBP_EOSINOPHIL_CHEMOTAXIS | 22 | 0.415102 | 1.7559093 | 0.01992032 | 0.058422673 | 1 | 154 | tags=14%, list=1%, sig | MM_4 |
| GOBP_MIRNA_MEDIATED | GOBP_MIRNA_MEDIATED_GENE_SILENCING | 17 | 0.46865 | 1.7555875 | 0.022364218 | 0.058391087 | 1 | 4133 | tags=53%, list=18%, si | MM_4 |
| GOBP_COENZYME_A_ME | GOBP_COENZYME_A_METABOLIC_PROCESS | 19 | 0.443097 | 1.7551206 | 0.017421603 | 0.058400456 | 1 | 998 | tags=26%, list=4%, sig | MM_4 |
| GOBP_REGULATION_OF_ | GOBP_REGULATION_OF_ACTIN_FILAMENT_ | 158 | 0.261772 | 1.7550908 | 0 | 0.05826632 | 1 | 3905 | tags=30%, list=17%, si | MM_4 |
| GOBP_REGULATION_OF_ | GOBP_REGULATION_OF_TYPE_I_INTERFERO | 40 | 0.360171 | 1.7548069 | 0.018181818 | 0.05822564 | 1 | 4282 | tags=40%, list=18%, si | MM_4 |
| GOBP_NEGATIVE_REGUL | GOBP_NEGATIVE_REGULATION_OF_VIRAL_ | 50 | 0.340387 | 1.7537566 | 0.00952381 | 0.058350626 | 1 | 4506 | tags=40%, list=19%, si | MM_4 |
| GOBP_MITOCHONDRIAL_ | GOBP_MITOCHONDRIAL_RNA_METABOLIC_ | 51 | 0.339429 | 1.7531639 | 0 | 0.058382414 | 1 | 4322 | tags=43%, list=18%, si | MM_4 |
| GOBP_AMINO_SUGAR_C | GOBP_AMINO_SUGAR_CATABOLIC_PROCES | 16 | 0.477031 | 1.7529233 | 0.010638298 | 0.058323193 | 1 | 800 | tags=19%, list=3%, sig | MM_4 |
| GOBP_POSITIVE_REGULA | GOBP_POSITIVE_REGULATION_OF_RESPON | 381 | 0.236974 | 1.7526066 | 0 | 0.05831512 | 1 | 3465 | tags=25%, list=15%, si | MM_4 |

| | | | | | | | | | | |
|---|---|---|---|---|---|---|---|---|---|---|
| GOBP_NEGATIVE_REGUL | GOBP_NEGATIVE_REGULATION_OF_EPITHEI | 71 | 0.306126 | 1.7524058 | 0.011111111 | 0.05821219 | 1 | 2526 | tags=18%, list=11%, si | MM_4 |
| GOBP_REGULATION_OF_ | GOBP_REGULATION_OF_VASCULATURE_DE | 295 | 0.23373 | 1.7523835 | 0 | 0.05808641 | 1 | 3095 | tags=22%, list=13%, si | MM_4 |
| GOBP_REGULATION_OF_ | GOBP_REGULATION_OF_PROTEIN_SUMOYL | 25 | 0.423638 | 1.7513386 | 0.008196721 | 0.05831148 | 1 | 2977 | tags=28%, list=13%, si | MM_4 |
| GOBP_LYSOSOMAL_TRAN | GOBP_LYSOSOMAL_TRANSPORT | 124 | 0.269879 | 1.7499634 | 0 | 0.058568675 | 1 | 4315 | tags=35%, list=18%, si | MM_4 |
| GOBP_POSITIVE_REGULA | GOBP_POSITIVE_REGULATION_OF_PHAGOC | 74 | 0.301026 | 1.7486488 | 0 | 0.05880398 | 1 | 2014 | tags=20%, list=9%, sig | MM_4 |
| GOBP_REGULATION_OF_ | GOBP_REGULATION_OF_INFLAMMATORY_F | 348 | 0.229547 | 1.7475502 | 0 | 0.05899091 | 1 | 3073 | tags=23%, list=13%, si | MM_4 |
| GOBP_ORGANELLE_TRAN | GOBP_ORGANELLE_TRANSPORT_ALONG_M | 86 | 0.291249 | 1.7472988 | 0.0061728396 | 0.058941867 | 1 | 4101 | tags=34%, list=17%, si | MM_4 |
| GOBP_NEGATIVE_REGUL | GOBP_NEGATIVE_REGULATION_OF_INFLAM | 153 | 0.264666 | 1.7466575 | 0 | 0.05897847 | 1 | 2140 | tags=20%, list=9%, sig | MM_4 |
| GOBP_VIRAL_GENOME_R | GOBP_VIRAL_GENOME_REPLICATION | 110 | 0.283143 | 1.7454385 | 0.0064102565 | 0.059225246 | 1 | 3780 | tags=31%, list=16%, si | MM_4 |
| GOBP_AMYLOID_BETA_M | GOBP_AMYLOID_BETA_METABOLIC_PROCES | 55 | 0.328339 | 1.7452959 | 0.005263158 | 0.059121557 | 1 | 2833 | tags=27%, list=12%, si | MM_4 |
| GOBP_GLYCOLYTIC_PROC | GOBP_GLYCOLYTIC_PROCESS_THROUGH_FR | 25 | 0.398941 | 1.7448028 | 0.028688524 | 0.059151366 | 1 | 3613 | tags=48%, list=15%, si | MM_4 |
| GOBP_ENDOLYSOSOMAL | GOBP_ENDOLYSOSOMAL_TOLL_LIKE_RECEP | 47 | 0.336967 | 1.744663 | 0.0045454544 | 0.05907591 | 1 | 2498 | tags=26%, list=11%, si | MM_4 |
| GOBP_AMIDE_METABOLI | GOBP_AMIDE_METABOLIC_PROCESS | 398 | 0.218813 | 1.7438726 | 0 | 0.059189454 | 1 | 3638 | tags=28%, list=15%, si | MM_4 |
| GOBP_ANTIMICROBIAL_H | GOBP_ANTIMICROBIAL_HUMORAL_IMMUN | 79 | 0.304484 | 1.7436879 | 0 | 0.059108626 | 1 | 2450 | tags=20%, list=10%, si | MM_4 |
| GOBP_UBIQUINONE_MET | GOBP_UBIQUINONE_METABOLIC_PROCESS | 19 | 0.440433 | 1.7435703 | 0.0111524165 | 0.05901206 | 1 | 3486 | tags=42%, list=15%, si | MM_4 |
| GOBP_REGULATION_OF_ | GOBP_REGULATION_OF_LEUKOCYTE_DEGR | 48 | 0.350243 | 1.7427576 | 0.010256411 | 0.059140485 | 1 | 3842 | tags=35%, list=16%, si | MM_4 |
| GOBP_PROTEIN_TARGETI | GOBP_PROTEIN_TARGETING | 285 | 0.236781 | 1.7414322 | 0 | 0.059410162 | 1 | 4386 | tags=30%, list=19%, si | MM_4 |
| TOLL-LIKE_RECEPTOR_SIG | TOLL-LIKE_RECEPTOR_SIGNALING_PATHWA | 84 | 0.294911 | 1.7408817 | 0 | 0.05940146 | 1 | 4150 | tags=32%, list=18%, si | MM_4 |
| GOBP_NUCLEOBASE_MET | GOBP_NUCLEOBASE_METABOLIC_PROCESS | 34 | 0.364989 | 1.7406462 | 0.021551725 | 0.059327345 | 1 | 3774 | tags=44%, list=16%, si | MM_4 |
| GOBP_TOR_SIGNALING | GOBP_TOR_SIGNALING | 165 | 0.254552 | 1.7403253 | 0 | 0.059317466 | 1 | 3776 | tags=30%, list=16%, si | MM_4 |
| GOBP_AMINO_ACID_ACT | GOBP_AMINO_ACID_ACTIVATION | 26 | 0.398452 | 1.7383386 | 0.020325202 | 0.059808727 | 1 | 3909 | tags=46%, list=17%, si | MM_4 |
| GOBP_POSITIVE_REGULA | GOBP_POSITIVE_REGULATION_OF_NEUTRO | 41 | 0.344797 | 1.736863 | 0.0045454544 | 0.0601461 | 1 | 4146 | tags=37%, list=18%, si | MM_4 |
| GOBP_MITOCHONDRIAL_ | GOBP_MITOCHONDRIAL_RESPIRATORY_CHA | 91 | 0.287768 | 1.7347759 | 0.0125 | 0.060695704 | 1 | 5332 | tags=46%, list=23%, si | MM_4 |
| GOBP_ESTABLISHMENT_ | GOBP_ESTABLISHMENT_OF_PROTEIN_LOCA | 435 | 0.221568 | 1.7340391 | 0 | 0.060790576 | 1 | 3196 | tags=22%, list=14%, si | MM_4 |
| GOBP_REGULATION_OF_ | GOBP_REGULATION_OF_ADAPTIVE_IMMUN | 215 | 0.238512 | 1.7334474 | 0 | 0.060789436 | 1 | 2986 | tags=25%, list=13%, si | MM_4 |
| GOBP_PROTEIN_LOCALIZ | GOBP_PROTEIN_LOCALIZATION_TO_CHROM | 31 | 0.381841 | 1.7315484 | 0.017021276 | 0.06134954 | 1 | 4127 | tags=32%, list=17%, si | MM_4 |
| GOBP_GLUCOSAMINE_C | GOBP_GLUCOSAMINE_CONTAINING_COMP | 24 | 0.413781 | 1.7307025 | 0.011904762 | 0.061480336 | 1 | 800 | tags=13%, list=3%, sig | MM_4 |
| GOBP_NEGATIVE_REGUL | GOBP_NEGATIVE_REGULATION_OF_MYELO | 57 | 0.313735 | 1.7306718 | 0 | 0.061344016 | 1 | 3364 | tags=30%, list=14%, si | MM_4 |
| GOBP_MODULATION_BY_ | GOBP_MODULATION_BY_HOST_OF_SYMBIC | 79 | 0.302677 | 1.7292825 | 0 | 0.061626483 | 1 | 2519 | tags=25%, list=11%, si | MM_4 |
| GOBP_ANTIGEN_PROCES | GOBP_ANTIGEN_PROCESSING_AND_PRESEN | 29 | 0.391586 | 1.7291921 | 0.036 | 0.06151794 | 1 | 2817 | tags=34%, list=12%, si | MM_4 |
| GOBP_REGULATION_OF_ | GOBP_REGULATION_OF_INTEGRIN_ACTIVAT | 17 | 0.461781 | 1.7284384 | 0.024647888 | 0.06164048 | 1 | 1281 | tags=24%, list=5%, sig | MM_4 |
| GOBP_MAINTENANCE_O | GOBP_MAINTENANCE_OF_LOCATION | 341 | 0.227793 | 1.7279229 | 0 | 0.061748553 | 1 | 3037 | tags=22%, list=13%, si | MM_4 |
| GOBP_REGULATION_OF_ | GOBP_REGULATION_OF_NATURAL_KILLER_ | 17 | 0.453511 | 1.727678 | 0.0034722222 | 0.061681017 | 1 | 3452 | tags=47%, list=15%, si | MM_4 |
| GOBP_NUCLEAR_TRANSP | GOBP_NUCLEAR_TRANSPORT | 337 | 0.227515 | 1.7276355 | 0 | 0.061561666 | 1 | 3986 | tags=26%, list=17%, si | MM_4 |
| GOBP_INNATE_IMMUNE_ | GOBP_INNATE_IMMUNE_RESPONSE_ACTIV | 254 | 0.228802 | 1.7269926 | 0 | 0.061658323 | 1 | 3406 | tags=23%, list=14%, si | MM_4 |
| GOBP_POSITIVE_REGULA | GOBP_POSITIVE_REGULATION_OF_JUN_KIN | 47 | 0.337727 | 1.7267839 | 0.013392857 | 0.06162928 | 1 | 3783 | tags=30%, list=16%, si | MM_4 |
| GOBP_IMP_METABOLIC_ | GOBP_IMP_METABOLIC_PROCESS | 19 | 0.430258 | 1.7263935 | 0.021276595 | 0.061614912 | 1 | 3189 | tags=42%, list=14%, si | MM_4 |
| ACUTE_MYELOID_LEUKEM | ACUTE_MYELOID_LEUKEMIA | 66 | 0.302355 | 1.7248033 | 0.0055555557 | 0.062013544 | 1 | 3163 | tags=26%, list=13%, si | MM_4 |
| GOBP_CHEMOKINE_C_X_ | GOBP_CHEMOKINE_C_X_C_MOTIF_LIGAND | 25 | 0.404511 | 1.7238721 | 0.02892562 | 0.062154874 | 1 | 2599 | tags=32%, list=11%, si | MM_4 |
| DNA_REPLICATION | DNA_REPLICATION | 34 | 0.352534 | 1.7237407 | 0.020920502 | 0.062093567 | 1 | 2851 | tags=29%, list=12%, si | MM_4 |
| GOBP_REGULATION_OF_ | GOBP_REGULATION_OF_B_CELL_ACTIVATIO | 141 | 0.260256 | 1.7235311 | 0 | 0.062021524 | 1 | 3025 | tags=26%, list=13%, si | MM_4 |
| GOBP_ERAD_PATHWAY | GOBP_ERAD_PATHWAY | 108 | 0.273256 | 1.7208263 | 0 | 0.06274262 | 1 | 4172 | tags=32%, list=18%, si | MM_4 |
| GOBP_POSITIVE_REGULA | GOBP_POSITIVE_REGULATION_OF_CYTOKIN | 488 | 0.221077 | 1.7207453 | 0 | 0.06263307 | 1 | 2633 | tags=19%, list=11%, si | MM_4 |
| PROTEIN_DIGESTION_AN | PROTEIN_DIGESTION_AND_ABSORPTION | 78 | 0.30038 | 1.7185868 | 0.010638298 | 0.06336615 | 1 | 1509 | tags=21%, list=6%, sig | MM_4 |
| GOBP_MYELOID_LEUKOC | GOBP_MYELOID_LEUKOCYTE_DIFFERENTIAT | 258 | 0.231072 | 1.7181228 | 0 | 0.06336824 | 1 | 3385 | tags=26%, list=14%, si | MM_4 |
| GOBP_LEUKOCYTE_DEGR | GOBP_LEUKOCYTE_DEGRANULATION | 77 | 0.299271 | 1.7175694 | 0 | 0.063428536 | 1 | 3842 | tags=32%, list=16%, si | MM_4 |
| GOBP_POSITIVE_REGULA | GOBP_POSITIVE_REGULATION_OF_MEMBRA | 17 | 0.449849 | 1.7175307 | 0.010600707 | 0.06329358 | 1 | 3636 | tags=41%, list=15%, si | MM_4 |
| GOBP_VIRAL_LIFE_CYCLE | GOBP_VIRAL_LIFE_CYCLE | 230 | 0.236602 | 1.7170402 | 0 | 0.06333789 | 1 | 4575 | tags=33%, list=19%, si | MM_4 |
| NOTCH_SIGNALING_PATH | NOTCH_SIGNALING_PATHWAY | 47 | 0.346818 | 1.7156012 | 0.010050251 | 0.06379254 | 1 | 3082 | tags=32%, list=13%, si | MM_4 |
| GOBP_EXTRINSIC_APOPT | GOBP_EXTRINSIC_APOPTOTIC_SIGNALING_ | 80 | 0.289832 | 1.7155957 | 0 | 0.06365767 | 1 | 2556 | tags=21%, list=11%, si | MM_4 |
| ENDOCYTOSIS | ENDOCYTOSIS | 246 | 0.236035 | 1.7155863 | 0 | 0.063523374 | 1 | 3078 | tags=21%, list=13%, si | MM_4 |
| GOBP_NEGATIVE_REGUL | GOBP_NEGATIVE_REGULATION_OF_LEUKOC | 16 | 0.45533 | 1.7146889 | 0.02238806 | 0.0637495 | 1 | 4000 | tags=44%, list=17%, si | MM_4 |
| GOBP_MYELOID_LEUKOC | GOBP_MYELOID_LEUKOCYTE_ACTIVATION | 247 | 0.241021 | 1.714013 | 0 | 0.06392859 | 1 | 3051 | tags=23%, list=13%, si | MM_4 |
| GOBP_NEGATIVE_REGUL | GOBP_NEGATIVE_REGULATION_OF_CELL_M | 36 | 0.354452 | 1.7132244 | 0 | 0.06405075 | 1 | 2672 | tags=28%, list=11%, si | MM_4 |
| GOBP_NEGATIVE_REGUL | GOBP_NEGATIVE_REGULATION_OF_LOCOM | 328 | 0.219214 | 1.7131011 | 0 | 0.0639372 | 1 | 3231 | tags=22%, list=14%, si | MM_4 |
| GOBP_TYPE_I_INTERFER | GOBP_TYPE_I_INTERFERON_PRODUCTION | 127 | 0.27009 | 1.7122779 | 0.013245033 | 0.06408772 | 1 | 2891 | tags=24%, list=12%, si | MM_4 |
| GOBP_ENDOPLASMIC_RE | GOBP_ENDOPLASMIC_RETICULUM_ORGANI | 90 | 0.277929 | 1.7118338 | 0.0061728396 | 0.06407354 | 1 | 3087 | tags=26%, list=13%, si | MM_4 |
| GOBP_NEUTROPHIL_MED | GOBP_NEUTROPHIL_MEDIATED_IMMUNITY | 37 | 0.36163 | 1.7116249 | 0.013274336 | 0.06404892 | 1 | 3218 | tags=35%, list=14%, si | MM_4 |
| GOBP_NUCLEOSIDE_MO | GOBP_NUCLEOSIDE_MONOPHOSPHATE_ME | 62 | 0.30739 | 1.7112457 | 0 | 0.06404552 | 1 | 3189 | tags=34%, list=14%, si | MM_4 |
| GOBP_BASEMENT_MEMB | GOBP_BASEMENT_MEMBRANE_ORGANIZAT | 40 | 0.349563 | 1.7106411 | 0.02739726 | 0.064210966 | 1 | 3027 | tags=33%, list=13%, si | MM_4 |
| GOBP_NEGATIVE_REGUL | GOBP_NEGATIVE_REGULATION_OF_TYPE_2 | 15 | 0.467869 | 1.7105519 | 0.024647888 | 0.06412224 | 1 | 347 | tags=20%, list=1%, sig | MM_4 |
| CELLULAR_SENESCENCE | CELLULAR_SENESCENCE | 159 | 0.250789 | 1.7102109 | 0 | 0.064104624 | 1 | 4237 | tags=31%, list=18%, si | MM_4 |
| GOBP_BIOLOGICAL_PRO | GOBP_BIOLOGICAL_PROCESS_INVOLVED_IN | 235 | 0.236624 | 1.709904 | 0 | 0.06407636 | 1 | 2549 | tags=21%, list=11%, si | MM_4 |
| GOBP_TRICARBOXYLIC_A | GOBP_TRICARBOXYLIC_ACID_METABOLIC_P | 16 | 0.458389 | 1.7093163 | 0.028985508 | 0.06413612 | 1 | 1172 | tags=31%, list=5%, sig | MM_4 |
| GOBP_REGULATION_OF_ | GOBP_REGULATION_OF_CELL_ADHESION_N | 47 | 0.337875 | 1.7089126 | 0.009615385 | 0.06412825 | 1 | 1281 | tags=19%, list=5%, sig | MM_4 |
| GOBP_POSITIVE_REGULA | GOBP_POSITIVE_REGULATION_OF_DNA_TE | 72 | 0.30938 | 1.7084273 | 0 | 0.06417355 | 1 | 3987 | tags=40%, list=17%, si | MM_4 |
| GOBP_GLYCOPROTEIN_M | GOBP_GLYCOPROTEIN_METABOLIC_PROCES | 342 | 0.218309 | 1.7078556 | 0 | 0.06424723 | 1 | 3550 | tags=27%, list=15%, si | MM_4 |
| GOBP_TRANSFORMING_ | GOBP_TRANSFORMING_GROWTH_FACTOR_ | 35 | 0.347647 | 1.7075984 | 0.012765957 | 0.06420393 | 1 | 879 | tags=17%, list=4%, sig | MM_4 |
| GOBP_NUCLEOSIDE_TRIP | GOBP_NUCLEOSIDE_TRIPHOSPHATE_METAE | 250 | 0.227124 | 1.7062727 | 0 | 0.06454252 | 1 | 4901 | tags=38%, list=21%, si | MM_4 |
| GOBP_RETROGRADE_VES | GOBP_RETROGRADE_VESICLE_MEDIATED_T | 45 | 0.340604 | 1.7059523 | 0.009852217 | 0.06454855 | 1 | 4068 | tags=36%, list=17%, si | MM_4 |
| GOBP_AMP_METABOLIC_ | GOBP_AMP_METABOLIC_PROCESS | 24 | 0.398255 | 1.7055031 | 0.020325202 | 0.064574055 | 1 | 3189 | tags=42%, list=14%, si | MM_4 |
| GOBP_DEFENSE_RESPON | GOBP_DEFENSE_RESPONSE_TO_BACTERIUM | 252 | 0.234105 | 1.7054662 | 0 | 0.06446287 | 1 | 2271 | tags=17%, list=10%, si | MM_4 |
| GOBP_REGULATION_OF_ | GOBP_REGULATION_OF_CELLULAR_RESPON | 141 | 0.258951 | 1.7048203 | 0 | 0.06457317 | 1 | 3223 | tags=23%, list=14%, si | MM_4 |
| PHAGOSOME | PHAGOSOME | 146 | 0.264436 | 1.7023298 | 0 | 0.065260924 | 1 | 4249 | tags=32%, list=18%, si | MM_4 |
| GOBP_REGULATION_OF_ | GOBP_REGULATION_OF_CELLULAR_EXTRAV | 43 | 0.331351 | 1.7019255 | 0.014492754 | 0.06527866 | 1 | 3307 | tags=28%, list=14%, si | MM_4 |
| GOBP_POSITIVE_REGULA | GOBP_POSITIVE_REGULATION_OF_UBIQUIT | 93 | 0.279415 | 1.7013701 | 0.005988024 | 0.065334454 | 1 | 3583 | tags=30%, list=15%, si | MM_4 |
| GOBP_POSITIVE_REGULA | GOBP_POSITIVE_REGULATION_OF_CHEMOT | 150 | 0.256369 | 1.6991743 | 0 | 0.06602982 | 1 | 3731 | tags=26%, list=16%, si | MM_4 |
| GOBP_COLLAGEN_ACTIV | GOBP_COLLAGEN_ACTIVATED_SIGNALING_ | 17 | 0.456804 | 1.6991088 | 0.026119404 | 0.065907635 | 1 | 3236 | tags=41%, list=14%, si | MM_4 |
| GOBP_POSITIVE_REGULA | GOBP_POSITIVE_REGULATION_OF_INFLAMI | 134 | 0.264584 | 1.6978866 | 0 | 0.06620456 | 1 | 1741 | tags=18%, list=7%, sig | MM_4 |
| GOBP_PYRIMIDINE_NUCL | GOBP_PYRIMIDINE_NUCLEOSIDE_TRIPHOSP | 18 | 0.431322 | 1.6977905 | 0.021276595 | 0.06608734 | 1 | 11014 | tags=94%, list=47%, si | MM_4 |
| GOBP_NEGATIVE_REGUL | GOBP_NEGATIVE_REGULATION_OF_PROTEC | 69 | 0.291118 | 1.6962194 | 0 | 0.06645643 | 1 | 2807 | tags=22%, list=12%, si | MM_4 |
| GOBP_T_CELL_APOPTOTI | GOBP_T_CELL_APOPTOTIC_PROCESS | 67 | 0.300657 | 1.6949685 | 0 | 0.066833116 | 1 | 2755 | tags=22%, list=12%, si | MM_4 |

| | | | | | | | | | | |
|---|---|---|---|---|---|---|---|---|---|---|
| GOBP_REGULATION_OF_ | GOBP_REGULATION_OF_MULTICELLULAR_C | 83 | 0.29527 | 1.6948384 | 0.0056818184 | 0.06673855 | 1 | 2338 | tags=17%, list=10%, si | MM_4 |
| GOBP_PROTEIN_LOCALIZ | GOBP_PROTEIN_LOCALIZATION_TO_LYSOSC | 54 | 0.305042 | 1.6940544 | 0.004830918 | 0.0668951 | 1 | 4315 | tags=35%, list=18%, si | MM_4 |
| GOBP_NEGATIVE_REGUL | GOBP_NEGATIVE_REGULATION_OF_DNA_R | 42 | 0.333052 | 1.6938607 | 0 | 0.06683838 | 1 | 3586 | tags=36%, list=15%, si | MM_4 |
| GOBP_RESPONSE_TO_NI | GOBP_RESPONSE_TO_NITRIC_OXIDE | 17 | 0.45552 | 1.6928163 | 0.025925925 | 0.06703703 | 1 | 783 | tags=12%, list=3%, sig | MM_4 |
| GOBP_NEGATIVE_REGUL | GOBP_NEGATIVE_REGULATION_OF_CELL_C | 69 | 0.298467 | 1.6919413 | 0.009756098 | 0.0672724 | 1 | 2866 | tags=26%, list=12%, si | MM_4 |
| GOBP_LYMPHOID_PROGI | GOBP_LYMPHOID_PROGENITOR_CELL_DIFF | 24 | 0.392133 | 1.6915975 | 0.02892562 | 0.06728629 | 1 | 2972 | tags=29%, list=13%, si | MM_4 |
| GOBP_CLATHRIN_COAT_ | GOBP_CLATHRIN_COAT_ASSEMBLY | 21 | 0.40863 | 1.6912367 | 0.014981274 | 0.067344144 | 1 | 13971 | tags=100%, list=59%, | MM_4 |
| GOBP_NEGATIVE_REGUL | GOBP_NEGATIVE_REGULATION_OF_T_CELL_ | 22 | 0.402181 | 1.6911764 | 0.015209125 | 0.06723594 | 1 | 3512 | tags=36%, list=15%, si | MM_4 |
| GOBP_AUTOPHAGY_OF_I | GOBP_AUTOPHAGY_OF_PEROXISOME | 15 | 0.456191 | 1.6909758 | 0.016835017 | 0.0671563 | 1 | 3178 | tags=40%, list=13%, si | MM_4 |
| GOBP_CELL_MATRIX_ADI | GOBP_CELL_MATRIX_ADHESION | 225 | 0.239046 | 1.6908472 | 0 | 0.06705388 | 1 | 3059 | tags=22%, list=13%, si | MM_4 |
| GOBP_REGULATION_OF_ | GOBP_REGULATION_OF_LYSOSOMAL_LUMI | 23 | 0.403305 | 1.6903306 | 0.034351144 | 0.067109756 | 1 | 4754 | tags=57%, list=20%, si | MM_4 |
| GOBP_MYELOID_CELL_AI | GOBP_MYELOID_CELL_APOPTOTIC_PROCES | 45 | 0.337033 | 1.6898049 | 0.017021276 | 0.067179345 | 1 | 2662 | tags=24%, list=11%, si | MM_4 |
| GOBP_PHOSPHATIDIC_AC | GOBP_PHOSPHATIDIC_ACID_METABOLIC_PI | 27 | 0.376747 | 1.687195 | 0.032258064 | 0.06794466 | 1 | 2032 | tags=26%, list=9%, sig | MM_4 |
| GOBP_REGULATION_OF_ | GOBP_REGULATION_OF_LYMPHOCYTE_DIFF | 216 | 0.240026 | 1.6867808 | 0 | 0.06795725 | 1 | 3042 | tags=22%, list=13%, si | MM_4 |
| GOBP_PROTEIN_K63_LIN | GOBP_PROTEIN_K63_LINKED_UBIQUITINAT | 68 | 0.291415 | 1.6866101 | 0.004784689 | 0.06787293 | 1 | 2776 | tags=24%, list=12%, si | MM_4 |
| GOBP_MYELOID_CELL_DI | GOBP_MYELOID_CELL_DIFFERENTIATION | 455 | 0.210758 | 1.6864746 | 0 | 0.06779398 | 1 | 3385 | tags=23%, list=14%, si | MM_4 |
| GOBP_B_CELL_DIFFEREN | GOBP_B_CELL_DIFFERENTIATION | 151 | 0.253697 | 1.6859344 | 0 | 0.067841925 | 1 | 2972 | tags=25%, list=13%, si | MM_4 |
| GOBP_POSITIVE_REGULA | GOBP_POSITIVE_REGULATION_OF_LEUKOC | 159 | 0.243645 | 1.6854346 | 0 | 0.06792261 | 1 | 1742 | tags=16%, list=7%, sig | MM_4 |
| GOBP_PLATELET_DENSE_ | GOBP_PLATELET_DENSE_GRANULE_ORGAN | 21 | 0.421625 | 1.6849841 | 0.012048192 | 0.06790379 | 1 | 974 | tags=29%, list=4%, sig | MM_4 |
| GOBP_POSITIVE_REGULA | GOBP_POSITIVE_REGULATION_OF_FATTY_A | 45 | 0.333354 | 1.6836793 | 0.01746725 | 0.06825061 | 1 | 3638 | tags=31%, list=15%, si | MM_4 |
| GOBP_NUCLEAR_EXPORT | GOBP_NUCLEAR_EXPORT | 160 | 0.247407 | 1.6828539 | 0 | 0.068502635 | 1 | 3986 | tags=26%, list=17%, si | MM_4 |
| GOBP_REGULATION_OF_ | GOBP_REGULATION_OF_LEUKOCYTE_ADHE | 33 | 0.361299 | 1.6825829 | 0.012396694 | 0.06851386 | 1 | 3307 | tags=30%, list=14%, si | MM_4 |
| GOBP_BIOLOGICAL_PROC | GOBP_BIOLOGICAL_PROCESS_INVOLVED_IN | 104 | 0.271327 | 1.6824061 | 0.0057471264 | 0.068493076 | 1 | 2519 | tags=24%, list=11%, si | MM_4 |
| GOBP_EXECUTION_PHASI | GOBP_EXECUTION_PHASE_OF_APOPTOSIS | 78 | 0.286968 | 1.6821725 | 0.006666667 | 0.068494126 | 1 | 2432 | tags=19%, list=10%, si | MM_4 |
| GOBP_REGULATION_OF_ | GOBP_REGULATION_OF_SUPRAMOLECULAR | 378 | 0.21477 | 1.6816025 | 0 | 0.0685143 | 1 | 3740 | tags=23%, list=16%, si | MM_4 |
| GOBP_REGULATION_OF_ | GOBP_REGULATION_OF_ENDOTHELIAL_CEL | 21 | 0.415162 | 1.6801413 | 0.030612245 | 0.06888564 | 1 | 892 | tags=14%, list=4%, sig | MM_4 |
| GOBP_EOSINOPHIL_MIGI | GOBP_EOSINOPHIL_MIGRATION | 26 | 0.385645 | 1.6754872 | 0.029850746 | 0.07052664 | 1 | 154 | tags=12%, list=1%, sig | MM_4 |
| GOBP_VESICLE_BUDDING | GOBP_VESICLE_BUDDING_FROM_MEMBRA | 79 | 0.275006 | 1.6750869 | 0.0058139535 | 0.07056525 | 1 | 3825 | tags=29%, list=16%, si | MM_4 |
| GOBP_REGULATION_OF_ | GOBP_REGULATION_OF_AMYLOID_FIBRIL_F | 16 | 0.443576 | 1.6750125 | 0.053691275 | 0.07045599 | 1 | 3233 | tags=38%, list=14%, si | MM_4 |
| GOBP_MYELOID_CELL_AC | GOBP_MYELOID_CELL_ACTIVATION_INVOLV | 98 | 0.270743 | 1.6743125 | 0.007194245 | 0.07059386 | 1 | 3042 | tags=23%, list=13%, si | MM_4 |
| GOBP_REGULATION_OF_ | GOBP_REGULATION_OF_DNA_BIOSYNTHETI | 121 | 0.262343 | 1.6733626 | 0 | 0.070834 | 1 | 4028 | tags=31%, list=17%, si | MM_4 |
| GOBP_PROTEIN_LOCALIZ | GOBP_PROTEIN_LOCALIZATION_TO_PHAGO | 16 | 0.439834 | 1.6731378 | 0.017985612 | 0.070777245 | 1 | 2022 | tags=31%, list=9%, sig | MM_4 |
| GOBP_NUCLEAR_TRANSC | GOBP_NUCLEAR_TRANSCRIBED_MRNA_CAT | 59 | 0.309523 | 1.6719049 | 0.0054644807 | 0.071083404 | 1 | 4133 | tags=32%, list=18%, si | MM_4 |
| GOBP_RRNA_PROCESSIN | GOBP_RRNA_PROCESSING | 211 | 0.239089 | 1.6716992 | 0 | 0.071013846 | 1 | 3854 | tags=30%, list=16%, si | MM_4 |
| GOBP_POSITIVE_REGULA | GOBP_POSITIVE_REGULATION_OF_PROTEIN | 192 | 0.231894 | 1.6715851 | 0 | 0.070931494 | 1 | 2927 | tags=20%, list=12%, si | MM_4 |
| GOBP_CYTOKINE_PRODU | GOBP_CYTOKINE_PRODUCTION_INVOLVED_ | 145 | 0.244151 | 1.6711588 | 0 | 0.07099598 | 1 | 2498 | tags=22%, list=11%, si | MM_4 |
| GOBP_ACTIN_FILAMENT_ | GOBP_ACTIN_FILAMENT_SEVERING | 16 | 0.442123 | 1.6704512 | 0.035587188 | 0.071100265 | 1 | 4503 | tags=56%, list=19%, si | MM_4 |
| GOBP_PROTEIN_EXPORT_ | GOBP_PROTEIN_EXPORT_FROM_NUCLEUS | 64 | 0.29485 | 1.667721 | 0.005291005 | 0.07199972 | 1 | 11382 | tags=86%, list=48%, si | MM_4 |
| GOBP_PURINE_NUCLEOB | GOBP_PURINE_NUCLEOBASE_METABOLIC_F | 18 | 0.440543 | 1.6662029 | 0.041198503 | 0.07244464 | 1 | 3189 | tags=39%, list=14%, si | MM_4 |
| GOBP_POSITIVE_REGULA | GOBP_POSITIVE_REGULATION_OF_CHEMOK | 79 | 0.287135 | 1.6661094 | 0.027027028 | 0.07235086 | 1 | 2599 | tags=20%, list=11%, si | MM_4 |
| GOBP_REGULATION_OF_ | GOBP_REGULATION_OF_RNA_STABILITY | 174 | 0.241281 | 1.6650072 | 0 | 0.0725888 | 1 | 3259 | tags=24%, list=14%, si | MM_4 |
| GOBP_POSITIVE_REGULA | GOBP_POSITIVE_REGULATION_OF_DOUBLE_ | 15 | 0.446259 | 1.6620562 | 0.021818181 | 0.07355823 | 1 | 3916 | tags=47%, list=17%, si | MM_4 |
| GOBP_PROTEIN_SUMOYL | GOBP_PROTEIN_SUMOYLATION | 42 | 0.343 | 1.6610289 | 0.022727273 | 0.073817864 | 1 | 2977 | tags=26%, list=13%, si | MM_4 |
| GOBP_NEGATIVE_REGUL | GOBP_NEGATIVE_REGULATION_OF_MYELO | 23 | 0.385567 | 1.6593902 | 0.02846975 | 0.07438686 | 1 | 1176 | tags=22%, list=5%, sig | MM_4 |
| GOBP_PYRUVATE_METAE | GOBP_PYRUVATE_METABOLIC_PROCESS | 127 | 0.2591 | 1.6589651 | 0.0067114094 | 0.074468836 | 1 | 3746 | tags=31%, list=16%, si | MM_4 |
| THYROID_CANCER | THYROID_CANCER | 36 | 0.352968 | 1.6584979 | 0.008583691 | 0.07448925 | 1 | 5515 | tags=58%, list=23%, si | MM_4 |
| GOBP_REGULATION_OF_ | GOBP_REGULATION_OF_TOLL_LIKE_RECEPT | 28 | 0.375085 | 1.6583635 | 0.044354837 | 0.07441514 | 1 | 11452 | tags=93%, list=49%, si | MM_4 |
| GOBP_INTERLEUKIN_1_P | GOBP_INTERLEUKIN_1_PRODUCTION | 114 | 0.260117 | 1.6573229 | 0.014814815 | 0.07472262 | 1 | 2031 | tags=18%, list=9%, sig | MM_4 |
| GOBP_NEGATIVE_REGUL | GOBP_NEGATIVE_REGULATION_OF_CELL_A | 207 | 0.232956 | 1.6564428 | 0 | 0.07490925 | 1 | 3066 | tags=23%, list=13%, si | MM_4 |
| GOBP_NEGATIVE_REGUL | GOBP_NEGATIVE_REGULATION_OF_TRANSF | 222 | 0.229993 | 1.6563803 | 0 | 0.07479138 | 1 | 2544 | tags=22%, list=11%, si | MM_4 |
| GOBP_NEGATIVE_REGUL | GOBP_NEGATIVE_REGULATION_OF_CELLUL | 51 | 0.314253 | 1.6562184 | 0.019607844 | 0.07471704 | 1 | 3013 | tags=27%, list=13%, si | MM_4 |
| GOBP_NEGATIVE_REGUL | GOBP_NEGATIVE_REGULATION_OF_UBIQUI | 55 | 0.312708 | 1.6537185 | 0.014218009 | 0.075580895 | 1 | 2709 | tags=22%, list=11%, si | MM_4 |
| GOBP_POSITIVE_REGULA | GOBP_POSITIVE_REGULATION_OF_CYTOSKE | 185 | 0.23768 | 1.6518291 | 0 | 0.07628048 | 1 | 2799 | tags=19%, list=12%, si | MM_4 |
| GOBP_RIBONUCLEOSIDE_ | GOBP_RIBONUCLEOSIDE_MONOPHOSPHATI | 48 | 0.323162 | 1.6505227 | 0.004878049 | 0.07676345 | 1 | 2803 | tags=33%, list=12%, si | MM_4 |
| GOBP_NEGATIVE_REGUL | GOBP_NEGATIVE_REGULATION_OF_LEUKOC | 65 | 0.290216 | 1.6498319 | 0.009433962 | 0.07690826 | 1 | 1611 | tags=17%, list=7%, sig | MM_4 |
| GOBP_CGAS_STING_SIGN | GOBP_CGAS_STING_SIGNALING_PATHWAY | 16 | 0.43964 | 1.6496326 | 0.019933555 | 0.07686521 | 1 | 3148 | tags=50%, list=13%, si | MM_4 |
| GOBP_SECRETORY_GRAN | GOBP_SECRETORY_GRANULE_LOCALIZATIOI | 34 | 0.35503 | 1.6495552 | 0.024096385 | 0.076749355 | 1 | 1531 | tags=18%, list=6%, sig | MM_4 |
| GOBP_NAD_METABOLIC_ | GOBP_NAD_METABOLIC_PROCESS | 58 | 0.3043 | 1.649293 | 0.0054347827 | 0.076723404 | 1 | 3660 | tags=34%, list=16%, si | MM_4 |
| GOBP_REGULATION_OF_ | GOBP_REGULATION_OF_RESPONSE_TO_EN | 80 | 0.2775 | 1.6492742 | 0.0065789474 | 0.07659571 | 1 | 2525 | tags=23%, list=11%, si | MM_4 |
| GOBP_ENDOPLASMIC_RE | GOBP_ENDOPLASMIC_RETICULUM_TO_GOL | 120 | 0.25797 | 1.6489782 | 0.0069444445 | 0.07658723 | 1 | 4072 | tags=31%, list=17%, si | MM_4 |
| RNA_TRANSPORT | RNA_TRANSPORT | 154 | 0.238975 | 1.6481932 | 0 | 0.076770805 | 1 | 4337 | tags=31%, list=18%, si | MM_4 |
| GOBP_REGULATION_OF_ | GOBP_REGULATION_OF_DNA_TEMPLATED_ | 84 | 0.275965 | 1.6473598 | 0.0057803467 | 0.076999046 | 1 | 4257 | tags=39%, list=18%, si | MM_4 |
| INFLUENZA_A | INFLUENZA_A | 139 | 0.251571 | 1.6463054 | 0 | 0.07733925 | 1 | 3722 | tags=27%, list=16%, si | MM_4 |
| GOBP_REGULATION_OF_ | GOBP_REGULATION_OF_CELLULAR_RESPIRA | 52 | 0.313175 | 1.6426085 | 0.015151516 | 0.07875991 | 1 | 4282 | tags=33%, list=18%, si | MM_4 |
| GOBP_NEUTRAL_LIPID_M | GOBP_NEUTRAL_LIPID_METABOLIC_PROCES | 138 | 0.248744 | 1.6425823 | 0.006535948 | 0.07862174 | 1 | 3083 | tags=27%, list=13%, si | MM_4 |
| KAPOSI_SARCOMA-ASSO | KAPOSI_SARCOMA-ASSOCIATED_HERPESVIR | 176 | 0.23982 | 1.6424898 | 0 | 0.07851327 | 1 | 3744 | tags=23%, list=16%, si | MM_4 |
| GOBP_POSITIVE_REGULA | GOBP_POSITIVE_REGULATION_OF_PROTEOI | 325 | 0.21788 | 1.6414801 | 0 | 0.07878332 | 1 | 3409 | tags=25%, list=14%, si | MM_4 |
| GOBP_REGULATION_OF_ | GOBP_REGULATION_OF_DNA_METABOLIC_ | 491 | 0.21515 | 1.6400855 | 0 | 0.079162024 | 1 | 2963 | tags=20%, list=13%, si | MM_4 |
| GOBP_ADAPTIVE_IMMUN | GOBP_ADAPTIVE_IMMUNE_RESPONSE | 433 | 0.204246 | 1.6396804 | 0 | 0.07920065 | 1 | 2930 | tags=20%, list=12%, si | MM_4 |
| GOBP_B_CELL_ACTIVATIC | GOBP_B_CELL_ACTIVATION | 276 | 0.221515 | 1.6369381 | 0 | 0.080230154 | 1 | 3042 | tags=23%, list=13%, si | MM_4 |
| GOBP_CELLULAR_PIGMEI | GOBP_CELLULAR_PIGMENTATION | 57 | 0.302952 | 1.6364292 | 0.016304348 | 0.080387615 | 1 | 1692 | tags=19%, list=7%, sig | MM_4 |
| GOBP_NUCLEAR_ENVELC | GOBP_NUCLEAR_ENVELOPE_ORGANIZATIOI | 52 | 0.308849 | 1.6362809 | 0.018518519 | 0.080319345 | 1 | 4650 | tags=46%, list=20%, si | MM_4 |
| GOBP_REGULATION_OF_ | GOBP_REGULATION_OF_ALTERNATIVE_MRI | 56 | 0.311528 | 1.6359973 | 0.004716981 | 0.08032274 | 1 | 15339 | tags=98%, list=65%, si | MM_4 |
| GOBP_REGULATION_OF_ | GOBP_REGULATION_OF_PROTEASOMAL_PR | 179 | 0.239923 | 1.635748 | 0 | 0.080351025 | 1 | 3551 | tags=27%, list=15%, si | MM_4 |
| GOBP_INTEGRIN_ACTIVA | GOBP_INTEGRIN_ACTIVATION | 26 | 0.377542 | 1.6355147 | 0.023255814 | 0.08032906 | 1 | 2423 | tags=23%, list=10%, si | MM_4 |
| GOBP_PINOCYTOSIS | GOBP_PINOCYTOSIS | 22 | 0.395816 | 1.6348945 | 0.03065134 | 0.0804356 | 1 | 4679 | tags=50%, list=20%, si | MM_4 |
| PANCREATIC_CANCER | PANCREATIC_CANCER | 74 | 0.27369 | 1.6348287 | 0 | 0.0803263 | 1 | 3523 | tags=34%, list=15%, si | MM_4 |
| GOBP_RESPONSE_TO_GR | GOBP_RESPONSE_TO_GROWTH_HORMONE | 20 | 0.406185 | 1.6342711 | 0.033834588 | 0.08045925 | 1 | 756 | tags=15%, list=3%, sig | MM_4 |
| GOBP_NEGATIVE_REGUL | GOBP_NEGATIVE_REGULATION_OF_PROTEI | 80 | 0.275619 | 1.6340119 | 0.010869565 | 0.08044846 | 1 | 3163 | tags=31%, list=13%, si | MM_4 |
| GOBP_PROTEIN_POLYUBI | GOBP_PROTEIN_POLYUBIQUITINATION | 250 | 0.226768 | 1.6335102 | 0 | 0.080548234 | 1 | 2776 | tags=20%, list=12%, si | MM_4 |
| GOBP_COPPER_ION_HOM | GOBP_COPPER_ION_HOMEOSTASIS | 18 | 0.412123 | 1.632905 | 0.022058824 | 0.080686994 | 1 | 5219 | tags=61%, list=22%, si | MM_4 |
| GOBP_RESPONSE_TO_IN | GOBP_RESPONSE_TO_INTERLEUKIN_7 | 17 | 0.421629 | 1.632858 | 0.039426524 | 0.0805821 | 1 | 11116 | tags=94%, list=47%, si | MM_4 |

| | | | | | | | | | | |
|---|---|---|---|---|---|---|---|---|---|---|
| HUMAN_IMMUNODEFICI | HUMAN_IMMUNODEFICIENCY_VIRUS_1_IN | 201 | 0.235429 | 1.6320673 | 0 | 0.080785714 | 1 | 4123 | tags=29%, list=17%, si | MM_4 |
| GOBP_POSITIVE_REGULA | GOBP_POSITIVE_REGULATION_OF_ACTIN_F | 52 | 0.308658 | 1.631468 | 0.009852217 | 0.080951095 | 1 | 2698 | tags=23%, list=11%, si | MM_4 |
| AUTOPHAGY | AUTOPHAGY | 126 | 0.252901 | 1.6287656 | 0 | 0.08185175 | 1 | 4249 | tags=33%, list=18%, si | MM_4 |
| FERROPTOSIS | FERROPTOSIS | 37 | 0.344421 | 1.6286634 | 0.008810572 | 0.08174639 | 1 | 3847 | tags=41%, list=16%, si | MM_4 |
| GOBP_NUCLEOSIDE_BISP | GOBP_NUCLEOSIDE_BISPHOSPHATE_BIOSYN | 39 | 0.338885 | 1.6285092 | 0.020661157 | 0.08165682 | 1 | 3497 | tags=41%, list=15%, si | MM_4 |
| GOBP_LEUKOCYTE_APOP | GOBP_LEUKOCYTE_APOPTOTIC_PROCESS | 131 | 0.249241 | 1.6281301 | 0 | 0.08168474 | 1 | 2755 | tags=20%, list=12%, si | MM_4 |
| GOBP_NUCLEAR_BODY_C | GOBP_NUCLEAR_BODY_ORGANIZATION | 19 | 0.414386 | 1.6267406 | 0.04868914 | 0.08213569 | 1 | 3728 | tags=42%, list=16%, si | MM_4 |
| GOBP_RETROGRADE_TRA | GOBP_RETROGRADE_TRANSPORT_ENDOSO | 87 | 0.261024 | 1.6262519 | 0.0064935065 | 0.082246274 | 1 | 4076 | tags=30%, list=17%, si | MM_4 |
| GOBP_NEGATIVE_REGUL | GOBP_NEGATIVE_REGULATION_OF_RESPON | 362 | 0.213274 | 1.6243079 | 0 | 0.08299085 | 1 | 2140 | tags=16%, list=9%, sig | MM_4 |
| GOBP_MODULATION_BY_ | GOBP_MODULATION_BY_HOST_OF_VIRAL_ | 44 | 0.313695 | 1.6234037 | 0.036199097 | 0.08321305 | 1 | 2384 | tags=25%, list=10%, si | MM_4 |
| GOBP_MYELOID_LEUKOC | GOBP_MYELOID_LEUKOCYTE_CYTOKINE_PR | 64 | 0.285787 | 1.6229017 | 0.015706806 | 0.083318464 | 1 | 3905 | tags=34%, list=17%, si | MM_4 |
| GOBP_PURINE_NUCLEOS | GOBP_PURINE_NUCLEOSIDE_MONOPHOSPI | 22 | 0.393112 | 1.622317 | 0.01119403 | 0.083439365 | 1 | 2803 | tags=36%, list=12%, si | MM_4 |
| GOBP_POSITIVE_REGULA | GOBP_POSITIVE_REGULATION_OF_LEUKOCY | 152 | 0.244939 | 1.6221881 | 0 | 0.08334787 | 1 | 3636 | tags=28%, list=15%, si | MM_4 |
| GOBP_CYTOPLASMIC_PA | GOBP_CYTOPLASMIC_PATTERN_RECOGNITI | 159 | 0.233598 | 1.618833 | 0 | 0.08476941 | 1 | 2538 | tags=19%, list=11%, si | MM_4 |
| GOBP_ENDOSOME_TO_L | GOBP_ENDOSOME_TO_LYSOSOME_TRANSP | 68 | 0.282684 | 1.6182443 | 0.0044247787 | 0.08486706 | 1 | 4315 | tags=34%, list=18%, si | MM_4 |
| GOBP_AUTOPHAGOSOMI | GOBP_AUTOPHAGOSOME_ORGANIZATION | 128 | 0.248395 | 1.617744 | 0 | 0.08495732 | 1 | 3258 | tags=27%, list=14%, si | MM_4 |
| GOBP_REGULATION_OF_ | GOBP_REGULATION_OF_MONOCYTE_DIFFE | 17 | 0.429371 | 1.6166817 | 0.033707865 | 0.085332595 | 1 | 13480 | tags=100%, list=57%, | MM_4 |
| GOBP_POSITIVE_REGULA | GOBP_POSITIVE_REGULATION_OF_PROTEOI | 120 | 0.259896 | 1.6153915 | 0.0064102565 | 0.08581389 | 1 | 3583 | tags=28%, list=15%, si | MM_4 |
| GOBP_SULFUR_COMPOU | GOBP_SULFUR_COMPOUND_CATABOLIC_PI | 35 | 0.343046 | 1.6143776 | 0.017391304 | 0.08610648 | 1 | 2821 | tags=34%, list=12%, si | MM_4 |
| GOBP_HUMORAL_IMMU | GOBP_HUMORAL_IMMUNE_RESPONSE | 184 | 0.233619 | 1.6129335 | 0.009615385 | 0.08665648 | 1 | 2271 | tags=17%, list=10%, si | MM_4 |
| GOBP_RESPONSE_TO_VI | GOBP_RESPONSE_TO_VITAMIN | 33 | 0.348714 | 1.611905 | 0.035856575 | 0.086923376 | 1 | 1811 | tags=24%, list=8%, sig | MM_4 |
| OTHER_GLYCAN_DEGRAD | OTHER_GLYCAN_DEGRADATION | 16 | 0.431129 | 1.6118017 | 0.03787879 | 0.0868401 | 1 | 3263 | tags=50%, list=14%, si | MM_4 |
| GOBP_NEGATIVE_REGUL | GOBP_NEGATIVE_REGULATION_OF_IMMUN | 132 | 0.256367 | 1.6103989 | 0.007874016 | 0.08735166 | 1 | 2217 | tags=23%, list=9%, sig | MM_4 |
| GOBP_RESPONSE_TO_EN | GOBP_RESPONSE_TO_ENDOPLASMIC_RETIC | 261 | 0.216348 | 1.610369 | 0 | 0.08722427 | 1 | 4193 | tags=28%, list=18%, si | MM_4 |
| GOBP_NEGATIVE_REGUL | GOBP_NEGATIVE_REGULATION_OF_POST_T | 94 | 0.267696 | 1.6099393 | 0.006289308 | 0.087353714 | 1 | 3163 | tags=31%, list=13%, si | MM_4 |
| GOBP_GLUCOSE_CATABO | GOBP_GLUCOSE_CATABOLIC_PROCESS | 26 | 0.371833 | 1.6080726 | 0.04797048 | 0.08809113 | 1 | 3613 | tags=46%, list=15%, si | MM_4 |
| GOBP_NEGATIVE_REGUL | GOBP_NEGATIVE_REGULATION_OF_DNA_B | 148 | 0.239118 | 1.6077596 | 0 | 0.088089235 | 1 | 4274 | tags=33%, list=18%, si | MM_4 |
| GOBP_RRNA_METHYLATI | GOBP_RRNA_METHYLATION | 20 | 0.409499 | 1.6050909 | 0.045454547 | 0.08917877 | 1 | 2272 | tags=30%, list=10%, si | MM_4 |
| GOBP_MEMBRANE_LIPID | GOBP_MEMBRANE_LIPID_METABOLIC_PRO | 186 | 0.235672 | 1.603506 | 0 | 0.08980409 | 1 | 3483 | tags=27%, list=15%, si | MM_4 |
| GOBP_NEUROTRANSMIT | GOBP_NEUROTRANSMITTER_RECEPTOR_TR | 27 | 0.352427 | 1.6026567 | 0.01746725 | 0.08999798 | 1 | 15299 | tags=100%, list=65%, | MM_4 |
| GOBP_POSITIVE_REGULA | GOBP_POSITIVE_REGULATION_OF_ACTIVAT | 27 | 0.362871 | 1.6015004 | 0.05323194 | 0.09034801 | 1 | 1427 | tags=26%, list=6%, sig | MM_4 |
| GOBP_CELLULAR_RESPOI | GOBP_CELLULAR_RESPONSE_TO_BIOTIC_ST | 224 | 0.226381 | 1.6006389 | 0 | 0.090575695 | 1 | 3963 | tags=29%, list=17%, si | MM_4 |
| GOBP_DEPHOSPHORYLAT | GOBP_DEPHOSPHORYLATION | 303 | 0.217786 | 1.5999058 | 0 | 0.09073087 | 1 | 4164 | tags=30%, list=18%, si | MM_4 |
| GOBP_TAXIS | GOBP_TAXIS | 411 | 0.204384 | 1.5994287 | 0 | 0.09077897 | 1 | 2693 | tags=16%, list=11%, si | MM_4 |
| GOBP_RIBOSE_PHOSPHA | GOBP_RIBOSE_PHOSPHATE_METABOLIC_PR | 419 | 0.201828 | 1.5993844 | 0 | 0.090655625 | 1 | 3835 | tags=29%, list=16%, si | MM_4 |
| GOBP_INTERLEUKIN_1_B | GOBP_INTERLEUKIN_1_BETA_PRODUCTION | 95 | 0.255003 | 1.5979216 | 0.012658228 | 0.09116701 | 1 | 2031 | tags=18%, list=9%, sig | MM_4 |
| GOBP_REGULATION_OF_ | GOBP_REGULATION_OF_FATTY_ACID_META | 99 | 0.255381 | 1.5975411 | 0.0058479533 | 0.09121776 | 1 | 3483 | tags=26%, list=15%, si | MM_4 |
| GOBP_REGULATION_OF_ | GOBP_REGULATION_OF_NITRIC_OXIDE_ME | 76 | 0.277471 | 1.5974898 | 0.015151516 | 0.09110647 | 1 | 3583 | tags=30%, list=15%, si | MM_4 |
| GOBP_INTRACELLULAR_E | GOBP_INTRACELLULAR_ESTROGEN_RECEPT | 47 | 0.308527 | 1.5971085 | 0.009615385 | 0.0911642 | 1 | 12805 | tags=91%, list=54%, si | MM_4 |
| GOBP_IMMUNE_RESPON | GOBP_IMMUNE_RESPONSE_REGULATING_S | 457 | 0.203027 | 1.5962118 | 0 | 0.09149927 | 1 | 3722 | tags=23%, list=16%, si | MM_4 |
| GOBP_STRESS_ACTIVATE | GOBP_STRESS_ACTIVATED_PROTEIN_KINAS | 59 | 0.286729 | 1.5956753 | 0.014285714 | 0.09162085 | 1 | 3783 | tags=36%, list=16%, si | MM_4 |
| GOBP_NEGATIVE_REGUL | GOBP_NEGATIVE_REGULATION_OF_ENDOC | 68 | 0.280863 | 1.5950313 | 0.015228426 | 0.09174361 | 1 | 13491 | tags=90%, list=57%, si | MM_4 |
| GOBP_DEFENSE_RESPON | GOBP_DEFENSE_RESPONSE_TO_GRAM_NE | 68 | 0.28704 | 1.5948188 | 0.016216217 | 0.0916978 | 1 | 2037 | tags=18%, list=9%, sig | MM_4 |
| GOBP_POSITIVE_REGULA | GOBP_POSITIVE_REGULATION_OF_PHOSPH | 182 | 0.231767 | 1.5938977 | 0 | 0.09199389 | 1 | 2646 | tags=21%, list=11%, si | MM_4 |
| GOBP_REGULATION_OF_ | GOBP_REGULATION_OF_HEMOPOIESIS | 425 | 0.208025 | 1.5938438 | 0 | 0.09187492 | 1 | 3364 | tags=22%, list=14%, si | MM_4 |
| GOBP_REGULATION_OF_ | GOBP_REGULATION_OF_CELL_KILLING | 116 | 0.262141 | 1.593709 | 0 | 0.09180986 | 1 | 3435 | tags=28%, list=15%, si | MM_4 |
| GOBP_REGULATION_OF_ | GOBP_REGULATION_OF_AEROBIC_RESPIRAT | 33 | 0.338398 | 1.5936077 | 0.015625 | 0.091703296 | 1 | 3638 | tags=27%, list=15%, si | MM_4 |
| GOBP_NUCLEOPHAGY | GOBP_NUCLEOPHAGY | 24 | 0.372331 | 1.5935687 | 0.05109489 | 0.09156263 | 1 | 2311 | tags=33%, list=10%, si | MM_4 |
| GOBP_CYTOKINE_MEDIAT | GOBP_CYTOKINE_MEDIATED_SIGNALING_P | 421 | 0.209544 | 1.5921944 | 0 | 0.09213028 | 1 | 3636 | tags=25%, list=15%, si | MM_4 |
| GOBP_LIPOSACCHARIDE_ | GOBP_LIPOSACCHARIDE_METABOLIC_PROC | 93 | 0.262701 | 1.5918595 | 0.006535948 | 0.0921437 | 1 | 3439 | tags=31%, list=15%, si | MM_4 |
| GOBP_DNA_INTEGRITY_C | GOBP_DNA_INTEGRITY_CHECKPOINT_SIGN | 125 | 0.250287 | 1.5917958 | 0 | 0.092021324 | 1 | 2926 | tags=26%, list=12%, si | MM_4 |
| GOBP_ENDOTHELIAL_CEL | GOBP_ENDOTHELIAL_CELL_CHEMOTAXIS | 30 | 0.355171 | 1.59129 | 0.0373444 | 0.092143826 | 1 | 892 | tags=10%, list=4%, sig | MM_4 |
| GOBP_REGULATION_OF_ | GOBP_REGULATION_OF_ACTIN_FILAMENT_ | 266 | 0.219678 | 1.5909948 | 0 | 0.092179835 | 1 | 2799 | tags=19%, list=12%, si | MM_4 |
| GOBP_NEGATIVE_REGUL | GOBP_NEGATIVE_REGULATION_OF_CELL_A | 301 | 0.2084 | 1.5901436 | 0 | 0.0924834 | 1 | 2387 | tags=18%, list=10%, si | MM_4 |
| GOBP_CEREBELLAR_PURI | GOBP_CEREBELLAR_PURKINJE_CELL_LAYER_ | 16 | 0.417496 | 1.5897571 | 0.041984733 | 0.09251613 | 1 | 2563 | tags=31%, list=11%, si | MM_4 |
| GOBP_PROTEIN_LOCALIZ | GOBP_PROTEIN_LOCALIZATION_TO_NUCLE | 326 | 0.205645 | 1.5885937 | 0 | 0.09295802 | 1 | 3169 | tags=21%, list=13%, si | MM_4 |
| GOBP_PROTEIN_MATURA | GOBP_PROTEIN_MATURATION | 475 | 0.201121 | 1.5884098 | 0 | 0.09289114 | 1 | 3422 | tags=22%, list=14%, si | MM_4 |
| GOBP_HEPATICOBILIARY_ | GOBP_HEPATICOBILIARY_SYSTEM_DEVELOP | 137 | 0.237422 | 1.587648 | 0 | 0.09316214 | 1 | 3689 | tags=30%, list=16%, si | MM_4 |
| ENDOCRINE_AND_OTHER | ENDOCRINE_AND_OTHER_FACTOR-REGULA | 42 | 0.319585 | 1.5876065 | 0.008695652 | 0.09303274 | 1 | 2815 | tags=31%, list=12%, si | MM_4 |
| GOBP_REGULATION_OF_ | GOBP_REGULATION_OF_MRNA_PROCESSIN | 131 | 0.241582 | 1.5873814 | 0.0074074073 | 0.09303742 | 1 | 3559 | tags=23%, list=15%, si | MM_4 |
| GOBP_PHOSPHATIDYLINC | GOBP_PHOSPHATIDYLINOSITOL_3_KINASE_ | 307 | 0.211644 | 1.5871896 | 0 | 0.09295325 | 1 | 2662 | tags=19%, list=11%, si | MM_4 |
| NON-SMALL_CELL_LUNG | NON-SMALL_CELL_LUNG_CANCER | 65 | 0.287714 | 1.586105 | 0.015151516 | 0.093326844 | 1 | 2867 | tags=29%, list=12%, si | MM_4 |
| GOBP_GRANULOCYTE_AC | GOBP_GRANULOCYTE_ACTIVATION | 41 | 0.309153 | 1.5819783 | 0.033333335 | 0.09521301 | 1 | 3218 | tags=29%, list=14%, si | MM_4 |
| GOBP_ALTERNATIVE_MR | GOBP_ALTERNATIVE_MRNA_SPLICING_VIA_ | 75 | 0.271009 | 1.5813502 | 0.016666668 | 0.09539865 | 1 | 15339 | tags=96%, list=65%, si | MM_4 |
| GOBP_POSITIVE_REGULA | GOBP_POSITIVE_REGULATION_OF_DOUBLE_ | 91 | 0.26177 | 1.5810297 | 0.011764706 | 0.09541314 | 1 | 2580 | tags=21%, list=11%, si | MM_4 |
| GOBP_GLYCEROLIPID_BIC | GOBP_GLYCEROLIPID_BIOSYNTHETIC_PROC | 210 | 0.222752 | 1.5790917 | 0 | 0.096250154 | 1 | 3370 | tags=25%, list=14%, si | MM_4 |
| GOBP_NUCLEAR_MEMBR | GOBP_NUCLEAR_MEMBRANE_REASSEMBLY | 20 | 0.395659 | 1.577422 | 0.025089607 | 0.09704395 | 1 | 10867 | tags=95%, list=46%, si | MM_4 |
| GOBP_POSITIVE_REGULA | GOBP_POSITIVE_REGULATION_OF_INTERFE | 26 | 0.360896 | 1.5766537 | 0.041198503 | 0.09726663 | 1 | 2538 | tags=27%, list=11%, si | MM_4 |
| GOBP_CELLULAR_RESPOI | GOBP_CELLULAR_RESPONSE_TO_OXIDATIVE | 229 | 0.215811 | 1.5760952 | 0 | 0.09740466 | 1 | 3477 | tags=27%, list=15%, si | MM_4 |
| GOBP_REGULATION_OF_ | GOBP_REGULATION_OF_TOLL_LIKE_RECEPT | 32 | 0.345247 | 1.5755012 | 0.04366812 | 0.097577915 | 1 | 4003 | tags=44%, list=17%, si | MM_4 |
| GOBP_MAINTENANCE_OI | GOBP_MAINTENANCE_OF_PROTEIN_LOCAT | 70 | 0.273626 | 1.5750116 | 0 | 0.097717926 | 1 | 3156 | tags=29%, list=13%, si | MM_4 |
| GOBP_VACUOLAR_ACIDII | GOBP_VACUOLAR_ACIDIFICATION | 27 | 0.36197 | 1.5740249 | 0.02 | 0.098099716 | 1 | 4343 | tags=56%, list=18%, si | MM_4 |
| GOBP_NEGATIVE_REGUL | GOBP_NEGATIVE_REGULATION_OF_CELLUL | 100 | 0.264413 | 1.573975 | 0.0057803467 | 0.09797677 | 1 | 1899 | tags=18%, list=8%, sig | MM_4 |
| GOBP_NEGATIVE_REGUL | GOBP_NEGATIVE_REGULATION_OF_RECEPT | 16 | 0.428886 | 1.57365 | 0.028880866 | 0.097992584 | 1 | 13491 | tags=100%, list=57%, | MM_4 |
| GOBP_POSITIVE_REGULA | GOBP_POSITIVE_REGULATION_OF_NATURA | 26 | 0.366487 | 1.5735083 | 0.04477612 | 0.09791738 | 1 | 3452 | tags=42%, list=15%, si | MM_4 |
| GOBP_POSITIVE_REGULA | GOBP_POSITIVE_REGULATION_OF_ACUTE_I | 27 | 0.350722 | 1.5714723 | 0.040441178 | 0.09887406 | 1 | 1713 | tags=22%, list=7%, sig | MM_4 |
| GOBP_NEGATIVE_REGUL | GOBP_NEGATIVE_REGULATION_OF_PROTEI | 431 | 0.199321 | 1.5710676 | 0 | 0.09890934 | 1 | 3364 | tags=25%, list=14%, si | MM_4 |
| GOBP_RESPONSE_TO_OX | GOBP_RESPONSE_TO_OXIDATIVE_STRESS | 364 | 0.204501 | 1.5710378 | 0 | 0.09877505 | 1 | 3698 | tags=27%, list=16%, si | MM_4 |
| GOBP_REGULATION_OF_ | GOBP_REGULATION_OF_MYELOID_CELL_AP | 37 | 0.330441 | 1.5705056 | 0.025210084 | 0.09893119 | 1 | 1176 | tags=16%, list=5%, sig | MM_4 |
| GOBP_NEUTROPHIL_ACTI | GOBP_NEUTROPHIL_ACTIVATION | 36 | 0.324845 | 1.5694891 | 0.053097345 | 0.09933176 | 1 | 3218 | tags=31%, list=14%, si | MM_4 |
| GOBP_REGULATION_OF_ | GOBP_REGULATION_OF_GLIAL_CELL_APOPT | 15 | 0.423949 | 1.5692095 | 0.039087947 | 0.09930092 | 1 | 11145 | tags=93%, list=47%, si | MM_4 |
| GOBP_POST_GOLGI_VESI | GOBP_POST_GOLGI_VESICLE_MEDIATED_TF | 101 | 0.259788 | 1.5687096 | 0.006134969 | 0.09941435 | 1 | 4101 | tags=33%, list=17%, si | MM_4 |

| | | | | | | | | | | |
|---|---|---|---|---|---|---|---|---|---|---|
| GOBP_AMINOGLYCAN_C | GOBP_AMINOGLYCAN_CATABOLIC_PROCES | 33 | 0.339214 | 1.5685563 | 0.04366812 | 0.09933769 | 1 | 2821 | tags=30%, list=12%, si | MM_4 |
| GOBP_NEGATIVE_REGUL | GOBP_NEGATIVE_REGULATION_OF_B_CELL | 20 | 0.384742 | 1.5682542 | 0.03649635 | 0.09934612 | 1 | 3328 | tags=50%, list=14%, si | MM_4 |
| GOBP_MITOCHONDRIAL_ | GOBP_MITOCHONDRIAL_MEMBRANE_ORG/ | 107 | 0.246716 | 1.5682462 | 0.014184397 | 0.09919829 | 1 | 13126 | tags=90%, list=56%, si | MM_4 |
| GOBP_LEUKOCYTE_AGGR | GOBP_LEUKOCYTE_AGGREGATION | 16 | 0.418047 | 1.5679371 | 0.04347826 | 0.09926845 | 1 | 3523 | tags=38%, list=15%, si | MM_4 |
| GOBP_FC_RECEPTOR_ME | GOBP_FC_RECEPTOR_MEDIATED_STIMULAT | 17 | 0.396914 | 1.5674492 | 0.039568346 | 0.099376164 | 1 | 11227 | tags=94%, list=48%, si | MM_4 |
| GOBP_POSITIVE_REGULA | GOBP_POSITIVE_REGULATION_OF_GLYCOPI | 26 | 0.357951 | 1.5674255 | 0.031496063 | 0.09923617 | 1 | 3550 | tags=35%, list=15%, si | MM_4 |
| GOBP_POSITIVE_REGULA | GOBP_POSITIVE_REGULATION_OF_ACTIN_F | 67 | 0.280351 | 1.5667534 | 0.024271844 | 0.09949099 | 1 | 1744 | tags=16%, list=7%, sig | MM_4 |
| GOBP_POSITIVE_REGULA | GOBP_POSITIVE_REGULATION_OF_RECEPTC | 17 | 0.416232 | 1.5663483 | 0.05263158 | 0.09956791 | 1 | 13790 | tags=100%, list=58%, | MM_4 |
| GOBP_FATTY_ACID_TRAN | GOBP_FATTY_ACID_TRANSMEMBRANE_TRA | 16 | 0.426977 | 1.5659233 | 0.048442908 | 0.099675775 | 1 | 2941 | tags=38%, list=12%, si | MM_4 |
| GOBP_AUTOPHAGY_OF_I | GOBP_AUTOPHAGY_OF_MITOCHONDRION | 89 | 0.263897 | 1.565848 | 0.0054945056 | 0.09956403 | 1 | 3412 | tags=27%, list=14%, si | MM_4 |
| GOBP_RESPONSE_TO_IN | GOBP_RESPONSE_TO_INORGANIC_SUBSTAI | 432 | 0.199156 | 1.5657473 | 0 | 0.09946359 | 1 | 3737 | tags=25%, list=16%, si | MM_4 |
| GOBP_RESPONSE_TO_MI | GOBP_RESPONSE_TO_METAL_ION | 278 | 0.210878 | 1.5657134 | 0 | 0.09933184 | 1 | 2350 | tags=18%, list=10%, si | MM_4 |
| GOBP_POSITIVE_REGULA | GOBP_POSITIVE_REGULATION_OF_TELOME | 15 | 0.438686 | 1.5650582 | 0.039215688 | 0.0994912 | 1 | 2696 | tags=33%, list=11%, si | MM_4 |
| GOBP_POSITIVE_REGULA | GOBP_POSITIVE_REGULATION_OF_ADAPTIV | 140 | 0.240868 | 1.5643026 | 0 | 0.09973196 | 1 | 2817 | tags=22%, list=12%, si | MM_4 |
| GOBP_DIACYLGLYCEROL_ | GOBP_DIACYLGLYCEROL_METABOLIC_PROC | 28 | 0.352482 | 1.5640829 | 0.038314175 | 0.09967418 | 1 | 3037 | tags=39%, list=13%, si | MM_4 |
| GOBP_POSITIVE_REGULA | GOBP_POSITIVE_REGULATION_OF_VACUOL | 25 | 0.365582 | 1.5638684 | 0.032128513 | 0.099609226 | 1 | 1207 | tags=20%, list=5%, sig | MM_4 |
| GOBP_POSITIVE_REGULA | GOBP_POSITIVE_REGULATION_OF_CELL_SU | 134 | 0.23943 | 1.5621243 | 0 | 0.10037321 | 1 | 2377 | tags=15%, list=10%, si | MM_4 |
| GOBP_NUCLEAR_TRANSC | GOBP_NUCLEAR_TRANSCRIBED_MRNA_CAT | 43 | 0.312671 | 1.5617968 | 0.030973451 | 0.100431785 | 1 | 3929 | tags=28%, list=17%, si | MM_4 |
| GOBP_VACUOLAR_TRANS | GOBP_VACUOLAR_TRANSPORT | 161 | 0.226919 | 1.5617354 | 0.007142857 | 0.10032009 | 1 | 4694 | tags=35%, list=20%, si | MM_4 |
| GOBP_CELL_SUBSTRATE_ | GOBP_CELL_SUBSTRATE_ADHESION | 354 | 0.20505 | 1.5616107 | 0 | 0.10023735 | 1 | 3528 | tags=23%, list=15%, si | MM_4 |
| GOBP_LYMPHOCYTE_APO | GOBP_LYMPHOCYTE_APOPTOTIC_PROCESS | 94 | 0.251903 | 1.5608977 | 0.0056818184 | 0.100482196 | 1 | 2755 | tags=20%, list=12%, si | MM_4 |
| GOBP_REGULATION_OF_ | GOBP_REGULATION_OF_LEUKOCYTE_TETHE | 19 | 0.401273 | 1.5607978 | 0.039215688 | 0.10038588 | 1 | 3307 | tags=37%, list=14%, si | MM_4 |
| GOBP_NEGATIVE_REGUL | GOBP_NEGATIVE_REGULATION_OF_UBIQUI | 18 | 0.399551 | 1.5603251 | 0.033210333 | 0.100490384 | 1 | 830 | tags=22%, list=4%, sig | MM_4 |
| GOBP_POSITIVE_REGULA | GOBP_POSITIVE_REGULATION_OF_TRIGLYC | 24 | 0.358576 | 1.5598931 | 0.04743083 | 0.100563034 | 1 | 3037 | tags=38%, list=13%, si | MM_4 |
| GOBP_ANTIGEN_PROCES | GOBP_ANTIGEN_PROCESSING_AND_PRESEN | 27 | 0.351215 | 1.5589129 | 0.045627378 | 0.1009235 | 1 | 2817 | tags=30%, list=12%, si | MM_4 |
| GOBP_FATTY_ACID_MET/ | GOBP_FATTY_ACID_METABOLIC_PROCESS | 399 | 0.207074 | 1.5585687 | 0 | 0.100916214 | 1 | 2350 | tags=18%, list=10%, si | MM_4 |
| GOBP_CELLULAR_COMPO | GOBP_CELLULAR_COMPONENT_DISASSEME | 31 | 0.341547 | 1.5573565 | 0.049586777 | 0.10140231 | 1 | 630 | tags=16%, list=3%, sig | MM_4 |
| GOBP_REGULATION_OF_ | GOBP_REGULATION_OF_CHROMATIN_BIND | 20 | 0.388776 | 1.5572413 | 0.064150944 | 0.10130115 | 1 | 1403 | tags=25%, list=6%, sig | MM_4 |
| PARATHYROID_HORMON | PARATHYROID_HORMONE_SYNTHESIS,_SEC | 97 | 0.253362 | 1.5559685 | 0.031446543 | 0.101800054 | 1 | 4158 | tags=36%, list=18%, si | MM_4 |
| GOBP_PYRIMIDINE_RIBO | GOBP_PYRIMIDINE_RIBONUCLEOTIDE_MET/ | 26 | 0.352767 | 1.5555974 | 0.05982906 | 0.10185136 | 1 | 4140 | tags=46%, list=18%, si | MM_4 |
| GOBP_VESICLE_TETHERIN | GOBP_VESICLE_TETHERING | 31 | 0.341758 | 1.5553943 | 0.016528925 | 0.101822495 | 1 | 3570 | tags=35%, list=15%, si | MM_4 |
| GOBP_ATP_METABOLIC_ | GOBP_ATP_METABOLIC_PROCESS | 206 | 0.216981 | 1.5553677 | 0 | 0.10169437 | 1 | 4901 | tags=37%, list=21%, si | MM_4 |
| GOBP_POSITIVE_REGULA | GOBP_POSITIVE_REGULATION_OF_FATTY_A | 20 | 0.393029 | 1.5549673 | 0.048780486 | 0.101760745 | 1 | 3638 | tags=35%, list=15%, si | MM_4 |
| GOBP_RESPONSE_TO_UV | GOBP_RESPONSE_TO_UV | 153 | 0.234461 | 1.5543158 | 0 | 0.10195354 | 1 | 2544 | tags=24%, list=11%, si | MM_4 |
| GOBP_POSITIVE_REGULA | GOBP_POSITIVE_REGULATION_OF_TOR_SIG | 71 | 0.277819 | 1.553376 | 0.0051282053 | 0.10226891 | 1 | 2553 | tags=24%, list=11%, si | MM_4 |
| GOBP_RESPONSE_TO_MC | GOBP_RESPONSE_TO_MOLECULE_OF_BACT | 296 | 0.205485 | 1.5526713 | 0 | 0.102477476 | 1 | 2880 | tags=20%, list=12%, si | MM_4 |
| GOBP_NEGATIVE_REGUL | GOBP_NEGATIVE_REGULATION_OF_MYELO | 89 | 0.2693 | 1.5520859 | 0.026143791 | 0.10259438 | 1 | 3583 | tags=28%, list=15%, si | MM_4 |
| GOBP_PROTEIN_MONOU | GOBP_PROTEIN_MONOUBIQUITINATION | 47 | 0.286012 | 1.5515054 | 0.03196347 | 0.102795824 | 1 | 3178 | tags=30%, list=13%, si | MM_4 |
| GOBP_POSITIVE_REGULA | GOBP_POSITIVE_REGULATION_OF_FILOPOD | 35 | 0.331008 | 1.5514798 | 0.060085837 | 0.10265739 | 1 | 12302 | tags=91%, list=52%, si | MM_4 |
| GOBP_POSITIVE_REGULA | GOBP_POSITIVE_REGULATION_OF_HOMOTY | 15 | 0.417866 | 1.5506178 | 0.050541516 | 0.10296118 | 1 | 1760 | tags=27%, list=7%, sig | MM_4 |
| GOBP_REGULATION_OF_ | GOBP_REGULATION_OF_CELLULAR_RESPON | 27 | 0.341056 | 1.5499158 | 0.038022812 | 0.10318903 | 1 | 715 | tags=15%, list=3%, sig | MM_4 |
| GOBP_POSITIVE_REGULA | GOBP_POSITIVE_REGULATION_OF_PROTEIN | 201 | 0.21479 | 1.5498912 | 0 | 0.103057645 | 1 | 3780 | tags=25%, list=16%, si | MM_4 |
| GOBP_ACTIVATION_OF_C | GOBP_ACTIVATION_OF_GTPASE_ACTIVITY | 86 | 0.261151 | 1.5489712 | 0.0056179776 | 0.10338242 | 1 | 3982 | tags=29%, list=17%, si | MM_4 |
| GOBP_NEGATIVE_REGUL | GOBP_NEGATIVE_REGULATION_OF_TELOM | 36 | 0.318469 | 1.5485336 | 0.022624435 | 0.10345993 | 1 | 4237 | tags=39%, list=18%, si | MM_4 |
| GOBP_REGULATION_OF_ | GOBP_REGULATION_OF_PROTEIN_EXPORT_ | 37 | 0.308509 | 1.5476438 | 0.039473683 | 0.1038163 | 1 | 15318 | tags=97%, list=65%, si | MM_4 |
| GOBP_POSITIVE_REGULA | GOBP_POSITIVE_REGULATION_OF_TYPE_I_I | 77 | 0.269837 | 1.547467 | 0.0060975607 | 0.10375145 | 1 | 2538 | tags=22%, list=11%, si | MM_4 |
| GOBP_REGULATION_OF_ | GOBP_REGULATION_OF_PROTEASOMAL_UI | 118 | 0.247465 | 1.5468565 | 0.025974026 | 0.103939675 | 1 | 3551 | tags=27%, list=15%, si | MM_4 |
| COMPLEMENT_AND_COA | COMPLEMENT_AND_COAGULATION_CASCA | 72 | 0.27287 | 1.5465986 | 0.010989011 | 0.103939295 | 1 | 464 | tags=13%, list=2%, sig | MM_4 |
| GOBP_REGULATION_OF_ | GOBP_REGULATION_OF_CELL_MATRIX_ADH | 121 | 0.243819 | 1.5458083 | 0.006289308 | 0.10419374 | 1 | 2672 | tags=19%, list=11%, si | MM_4 |
| GOBP_N_ACETYLGLUCOS | GOBP_N_ACETYLGLUCOSAMINE_METABOLI | 16 | 0.415791 | 1.545556 | 0.04659498 | 0.104178816 | 1 | 13800 | tags=100%, list=58%, | MM_4 |
| GOBP_REGULATION_OF_ | GOBP_REGULATION_OF_PROTEIN_POLYMEI | 194 | 0.218583 | 1.5452071 | 0 | 0.10418111 | 1 | 3905 | tags=27%, list=17%, si | MM_4 |
| GOBP_RESPONSE_TO_IN | GOBP_RESPONSE_TO_INTERFERON_BETA | 64 | 0.267522 | 1.544939 | 0.0056818184 | 0.10417602 | 1 | 3245 | tags=30%, list=14%, si | MM_4 |
| GOBP_NADH_DEHYDROG | GOBP_NADH_DEHYDROGENASE_COMPLEX_ | 58 | 0.287611 | 1.5435755 | 0.026595745 | 0.10470691 | 1 | 5651 | tags=53%, list=24%, si | MM_4 |
| GOBP_PROTEIN_DEPHOS | GOBP_PROTEIN_DEPHOSPHORYLATION | 220 | 0.217399 | 1.5428704 | 0 | 0.104924634 | 1 | 4164 | tags=29%, list=18%, si | MM_4 |
| GOBP_REGULATION_OF_ | GOBP_REGULATION_OF_PROTEIN_MATURA | 71 | 0.267004 | 1.5424391 | 0.027472528 | 0.10499445 | 1 | 1469 | tags=14%, list=6%, sig | MM_4 |
| GOBP_RESPONSE_TO_VI | GOBP_RESPONSE_TO_VITAMIN_D | 15 | 0.429657 | 1.542154 | 0.053956833 | 0.105018325 | 1 | 1811 | tags=27%, list=8%, sig | MM_4 |
| GOBP_NEGATIVE_REGUL | GOBP_NEGATIVE_REGULATION_OF_LIPID_T | 30 | 0.339434 | 1.5407878 | 0.041984733 | 0.10555091 | 1 | 3526 | tags=33%, list=15%, si | MM_4 |
| GOBP_REGULATION_OF_ | GOBP_REGULATION_OF_PROGRAMMED_NE | 27 | 0.346093 | 1.539844 | 0.048387095 | 0.10590229 | 1 | 11555 | tags=93%, list=49%, si | MM_4 |
| GOBP_IMPORT_INTO_NU | GOBP_IMPORT_INTO_NUCLEUS | 180 | 0.219899 | 1.5390037 | 0.009615385 | 0.10619456 | 1 | 3098 | tags=23%, list=13%, si | MM_4 |
| GOBP_MATURATION_OF_ | GOBP_MATURATION_OF_SSU_RRNA | 48 | 0.299328 | 1.5387797 | 0.028037382 | 0.10614768 | 1 | 4331 | tags=40%, list=18%, si | MM_4 |
| GOBP_REGULATION_OF_ | GOBP_REGULATION_OF_ENDOTHELIAL_CEL | 168 | 0.228992 | 1.5364884 | 0.007936508 | 0.107259646 | 1 | 2662 | tags=18%, list=11%, si | MM_4 |
| GOBP_GLYCOSYL_COMPO | GOBP_GLYCOSYL_COMPOUND_BIOSYNTHET | 18 | 0.394915 | 1.5360975 | 0.052447554 | 0.10732862 | 1 | 2758 | tags=39%, list=12%, si | MM_4 |
| GOBP_REGULATION_OF_ | GOBP_REGULATION_OF_ACUTE_INFLAMMA | 17 | 0.398383 | 1.5350538 | 0.052805282 | 0.10773599 | 1 | 981 | tags=24%, list=4%, sig | MM_4 |
| GOBP_CELL_ACTIVATION | GOBP_CELL_ACTIVATION_INVOLVED_IN_IM | 291 | 0.209193 | 1.5348022 | 0 | 0.10777213 | 1 | 2945 | tags=21%, list=12%, si | MM_4 |
| GOBP_DENDRITIC_CELL_ | GOBP_DENDRITIC_CELL_CHEMOTAXIS | 19 | 0.404346 | 1.5331595 | 0.040650405 | 0.108638346 | 1 | 1522 | tags=21%, list=6%, sig | MM_4 |
| GOBP_POSITIVE_REGULA | GOBP_POSITIVE_REGULATION_OF_SYNAPTI | 17 | 0.409536 | 1.532398 | 0.040677965 | 0.10896294 | 1 | 13948 | tags=100%, list=59%, | MM_4 |
| GOBP_REGULATION_OF_ | GOBP_REGULATION_OF_SYNAPTIC_VESICLE | 42 | 0.309431 | 1.5321337 | 0.02173913 | 0.10894853 | 1 | 15281 | tags=98%, list=65%, si | MM_4 |
| GOBP_REGULATION_OF_ | GOBP_REGULATION_OF_CHEMOTAXIS | 206 | 0.216709 | 1.5319835 | 0 | 0.108891375 | 1 | 3731 | tags=24%, list=16%, si | MM_4 |
| GOBP_REGULATION_OF_ | GOBP_REGULATION_OF_VIRAL_PROCESS | 167 | 0.226819 | 1.5315179 | 0 | 0.108974695 | 1 | 3592 | tags=26%, list=15%, si | MM_4 |
| GOBP_RUFFLE_ASSEMBL | GOBP_RUFFLE_ASSEMBLY | 43 | 0.293954 | 1.5312485 | 0.061032865 | 0.10895724 | 1 | 2449 | tags=23%, list=10%, si | MM_4 |
| GOBP_PROTEIN_K48_LIN | GOBP_PROTEIN_K48_LINKED_DEUBIQUITIN | 29 | 0.35021 | 1.5299346 | 0.035856575 | 0.10954682 | 1 | 15352 | tags=100%, list=65%, | MM_4 |
| GOBP_ADAPTIVE_IMMUN | GOBP_ADAPTIVE_IMMUNE_RESPONSE_BAS | 322 | 0.200922 | 1.529147 | 0 | 0.10990836 | 1 | 3019 | tags=21%, list=13%, si | MM_4 |
| GOBP_RESPONSE_TO_TY | GOBP_RESPONSE_TO_TYPE_I_INTERFERON | 64 | 0.273549 | 1.52813 | 0.00952381 | 0.110317975 | 1 | 4373 | tags=38%, list=19%, si | MM_4 |
| AGE-RAGE_SIGNALING_P | AGE-RAGE_SIGNALING_PATHWAY_IN_DIABI | 100 | 0.247737 | 1.5278947 | 0.01986755 | 0.11033159 | 1 | 3317 | tags=26%, list=14%, si | MM_4 |
| GOBP_PROTEIN_N_LINKE | GOBP_PROTEIN_N_LINKED_GLYCOSYLATION | 18 | 0.413499 | 1.5277721 | 0.05685619 | 0.11023846 | 1 | 11588 | tags=94%, list=49%, si | MM_4 |
| GOBP_NEGATIVE_REGUL | GOBP_NEGATIVE_REGULATION_OF_GROWT | 239 | 0.211029 | 1.5277464 | 0 | 0.110097036 | 1 | 3175 | tags=22%, list=13%, si | MM_4 |
| ADHERENS_JUNCTION | ADHERENS_JUNCTION | 70 | 0.260307 | 1.527528 | 0.01980198 | 0.11006901 | 1 | 14865 | tags=96%, list=63%, si | MM_4 |
| GOBP_REGULATION_OF_ | GOBP_REGULATION_OF_LEUKOCYTE_APOP1 | 103 | 0.256172 | 1.526538 | 0.014814815 | 0.110529505 | 1 | 1611 | tags=15%, list=7%, sig | MM_4 |
| MALARIA | MALARIA | 41 | 0.302466 | 1.5251386 | 0.023148147 | 0.11127531 | 1 | 3024 | tags=27%, list=13%, si | MM_4 |
| GOBP_DEOXYRIBONUCLE | GOBP_DEOXYRIBONUCLEOTIDE_BIOSYNTHE | 17 | 0.412642 | 1.5249562 | 0.045454547 | 0.11127875 | 1 | 4220 | tags=47%, list=18%, si | MM_4 |
| GOBP_PRODUCTION_OF_ | GOBP_PRODUCTION_OF_MOLECULAR_MED | 270 | 0.20382 | 1.5235609 | 0 | 0.11192972 | 1 | 2672 | tags=20%, list=11%, si | MM_4 |
| GOBP_REGULATION_OF_ | GOBP_REGULATION_OF_NEUTROPHIL_CHEI | 36 | 0.322252 | 1.5229708 | 0.034188036 | 0.1120824 | 1 | 3681 | tags=31%, list=16%, si | MM_4 |

| | | | | | | | | | | |
|---|---|---|---|---|---|---|---|---|---|---|
| GOBP_POSITIVE_REGULA | GOBP_POSITIVE_REGULATION_OF_EXTRINS | 60 | 0.269651 | 1.5224704 | 0.030612245 | 0.1122436 | 1 | 2555 | tags=20%, list=11%, si | MM_4 |
| GOBP_APOPTOTIC_DNA_ | GOBP_APOPTOTIC_DNA_FRAGMENTATION | 18 | 0.382962 | 1.5223404 | 0.06792453 | 0.1121747 | 1 | 595 | tags=17%, list=3%, sig | MM_4 |
| RENAL_CELL_CARCINOMA | RENAL_CELL_CARCINOMA | 67 | 0.270516 | 1.52225 | 0.035714287 | 0.112082876 | 1 | 2693 | tags=22%, list=11%, si | MM_4 |
| GOBP_REGULATION_OF_ | GOBP_REGULATION_OF_DEPHOSPHORYLAT | 106 | 0.241102 | 1.52223 | 0.006756757 | 0.11195357 | 1 | 4063 | tags=34%, list=17%, si | MM_4 |
| GOBP_REGULATION_OF_ | GOBP_REGULATION_OF_CELL_SUBSTRATE_ | 219 | 0.217822 | 1.5208274 | 0 | 0.11263107 | 1 | 2418 | tags=16%, list=10%, si | MM_4 |
| GOBP_DEOXYRIBONUCLE | GOBP_DEOXYRIBONUCLEOTIDE_METABOLIC | 44 | 0.294625 | 1.5196885 | 0.020746889 | 0.11313558 | 1 | 3189 | tags=34%, list=14%, si | MM_4 |
| GOBP_LIPOPROTEIN_CAT | GOBP_LIPOPROTEIN_CATABOLIC_PROCESS | 17 | 0.394331 | 1.5193462 | 0.061594203 | 0.11320216 | 1 | 2425 | tags=41%, list=10%, si | MM_4 |
| GOBP_ORGANOPHOSPHA | GOBP_ORGANOPHOSPHATE_CATABOLIC_PR | 227 | 0.210082 | 1.5191296 | 0 | 0.11322066 | 1 | 3189 | tags=24%, list=14%, si | MM_4 |
| GOBP_SYMBIONT_ENTRY | GOBP_SYMBIONT_ENTRY_INTO_HOST | 97 | 0.246933 | 1.5190828 | 0.025477707 | 0.113093875 | 1 | 4575 | tags=38%, list=19%, si | MM_4 |
| GOBP_POSITIVE_REGULA | GOBP_POSITIVE_REGULATION_OF_VASCUL | 172 | 0.216787 | 1.5179111 | 0.008333334 | 0.1134929 | 1 | 2414 | tags=17%, list=10%, si | MM_4 |
| GOBP_CERAMIDE_METAB | GOBP_CERAMIDE_METABOLIC_PROCESS | 94 | 0.246112 | 1.5170962 | 0.00625 | 0.11385191 | 1 | 3439 | tags=29%, list=15%, si | MM_4 |
| GOBP_CHOLESTEROL_STO | GOBP_CHOLESTEROL_STORAGE | 27 | 0.367893 | 1.5164784 | 0.0375 | 0.11406761 | 1 | 2987 | tags=33%, list=13%, si | MM_4 |
| NATURAL_KILLER_CELL_M | NATURAL_KILLER_CELL_MEDIATED_CYTOTO | 97 | 0.24945 | 1.5160925 | 0.006024096 | 0.11415787 | 1 | 2817 | tags=24%, list=12%, si | MM_4 |
| GOBP_POSITIVE_REGULA | GOBP_POSITIVE_REGULATION_OF_FATTY_A | 25 | 0.353777 | 1.5156535 | 0.05691057 | 0.11426637 | 1 | 3638 | tags=36%, list=15%, si | MM_4 |
| GOBP_INDOLE_CONTAIN | GOBP_INDOLE_CONTAINING_COMPOUND_ | 25 | 0.354788 | 1.5151569 | 0.06557377 | 0.11438394 | 1 | 3097 | tags=28%, list=13%, si | MM_4 |
| GOBP_POSITIVE_REGULA | GOBP_POSITIVE_REGULATION_OF_TRANSLA | 21 | 0.36977 | 1.5148643 | 0.073800735 | 0.11439159 | 1 | 4124 | tags=33%, list=17%, si | MM_4 |
| GOBP_HEMATOPOIETIC_S | GOBP_HEMATOPOIETIC_STEM_CELL_PROLIF | 37 | 0.307537 | 1.5148 | 0.04368932 | 0.11426416 | 1 | 1414 | tags=14%, list=6%, sig | MM_4 |
| GOBP_RIBONUCLEOSIDE_ | GOBP_RIBONUCLEOSIDE_MONOPHOSPHATE | 30 | 0.339818 | 1.5145439 | 0.057377048 | 0.11425016 | 1 | 3999 | tags=47%, list=17%, si | MM_4 |
| GOBP_TRIGLYCERIDE_ME | GOBP_TRIGLYCERIDE_METABOLIC_PROCESS | 105 | 0.246759 | 1.5132723 | 0.020833334 | 0.114875846 | 1 | 3037 | tags=27%, list=13%, si | MM_4 |
| GOBP_REGULATION_OF_ | GOBP_REGULATION_OF_T_CELL_APOPTOTI | 48 | 0.279713 | 1.5128164 | 0.018099548 | 0.11501075 | 1 | 1611 | tags=15%, list=7%, sig | MM_4 |
| GOBP_NEGATIVE_REGUL | GOBP_NEGATIVE_REGULATION_OF_DNA_M | 127 | 0.236197 | 1.5121279 | 0.023076924 | 0.11528572 | 1 | 3704 | tags=28%, list=16%, si | MM_4 |
| GOBP_REGULATION_OF_ | GOBP_REGULATION_OF_RNA_SPLICING | 178 | 0.21874 | 1.5107034 | 0.008333334 | 0.11603019 | 1 | 3451 | tags=22%, list=15%, si | MM_4 |
| GOBP_ESTABLISHMENT_( | GOBP_ESTABLISHMENT_OF_LYMPHOCYTE_ | 15 | 0.41707 | 1.5099672 | 0.053156145 | 0.1163552 | 1 | 1138 | tags=13%, list=5%, sig | MM_4 |
| GOBP_REGULATION_OF_ | GOBP_REGULATION_OF_DOUBLE_STRAND_ | 139 | 0.228349 | 1.50963 | 0.0074074073 | 0.11642102 | 1 | 4491 | tags=32%, list=19%, si | MM_4 |
| GOBP_NEGATIVE_REGUL | GOBP_NEGATIVE_REGULATION_OF_VIRAL_ | 95 | 0.247781 | 1.5094799 | 0.01910828 | 0.11635181 | 1 | 4575 | tags=37%, list=19%, si | MM_4 |
| GOBP_MAINTENANCE_OI | GOBP_MAINTENANCE_OF_LOCATION_IN_C | 224 | 0.211341 | 1.5090175 | 0 | 0.116440274 | 1 | 3583 | tags=25%, list=15%, si | MM_4 |
| GOBP_POSITIVE_REGULA | GOBP_POSITIVE_REGULATION_OF_TRANSF | 18 | 0.399186 | 1.5085757 | 0.09057971 | 0.11656283 | 1 | 263 | tags=17%, list=1%, sig | MM_4 |
| GOBP_VASCULAR_ENDOT | GOBP_VASCULAR_ENDOTHELIAL_GROWTH_ | 39 | 0.305183 | 1.5083094 | 0.056179777 | 0.11655304 | 1 | 918 | tags=13%, list=4%, sig | MM_4 |
| GOBP_INTERLEUKIN_6_P | GOBP_INTERLEUKIN_6_PRODUCTION | 173 | 0.222847 | 1.5081913 | 0.011363637 | 0.116487086 | 1 | 3527 | tags=23%, list=15%, si | MM_4 |
| GOBP_3_UTR_MEDIATED | GOBP_3_UTR_MEDIATED_MRNA_STABILIZA | 21 | 0.377564 | 1.5078322 | 0.05185185 | 0.116508976 | 1 | 4613 | tags=52%, list=20%, si | MM_4 |
| LONG-TERM_DEPRESSION | LONG-TERM_DEPRESSION | 55 | 0.274365 | 1.507547 | 0.030456852 | 0.11651072 | 1 | 4123 | tags=35%, list=17%, si | MM_4 |
| TGF-BETA_SIGNALING_PA | TGF-BETA_SIGNALING_PATHWAY | 85 | 0.250075 | 1.5072864 | 0.010526316 | 0.11647835 | 1 | 3040 | tags=21%, list=13%, si | MM_4 |
| GOBP_GLYCOSIDE_META | GOBP_GLYCOSIDE_METABOLIC_PROCESS | 18 | 0.385382 | 1.5072218 | 0.06719368 | 0.11635139 | 1 | 3108 | tags=44%, list=13%, si | MM_4 |
| GOBP_REGULATION_OF_ | GOBP_REGULATION_OF_ACTIN_FILAMENT_ | 387 | 0.194672 | 1.506396 | 0 | 0.11675472 | 1 | 2799 | tags=18%, list=12%, si | MM_4 |
| GOBP_CELLULAR_RESPON | GOBP_CELLULAR_RESPONSE_TO_NUTRIENT | 213 | 0.212632 | 1.5062107 | 0 | 0.11671 | 1 | 3740 | tags=26%, list=16%, si | MM_4 |
| GOBP_NUCLEOSIDE_MON | GOBP_NUCLEOSIDE_MONOPHOSPHATE_BIC | 40 | 0.310031 | 1.5036345 | 0.042016808 | 0.1182041 | 1 | 4155 | tags=43%, list=18%, si | MM_4 |
| GOBP_MISMATCH_REPAI | GOBP_MISMATCH_REPAIR | 25 | 0.338354 | 1.5031272 | 0.049808428 | 0.118349925 | 1 | 1914 | tags=24%, list=8%, sig | MM_4 |
| GOBP_RIBOSOMAL_SUBU | GOBP_RIBOSOMAL_SUBUNIT_EXPORT_FRO | 18 | 0.382399 | 1.5030018 | 0.069078945 | 0.11825515 | 1 | 3868 | tags=39%, list=16%, si | MM_4 |
| GOBP_VESICLE_CARGO_L | GOBP_VESICLE_CARGO_LOADING | 27 | 0.342022 | 1.5021693 | 0.034188036 | 0.118552975 | 1 | 3946 | tags=37%, list=17%, si | MM_4 |
| GOBP_CELLULAR_SENESC | GOBP_CELLULAR_SENESCENCE | 88 | 0.247654 | 1.5003304 | 0.024096385 | 0.119446784 | 1 | 2867 | tags=24%, list=12%, si | MM_4 |
| GOBP_RIG_I_SIGNALING_ | GOBP_RIG_I_SIGNALING_PATHWAY | 24 | 0.355037 | 1.5002588 | 0.062962964 | 0.11933858 | 1 | 5058 | tags=50%, list=21%, si | MM_4 |
| GOBP_REGULATION_OF_ | GOBP_REGULATION_OF_NUCLEOCYTOPLAS | 130 | 0.235015 | 1.5002402 | 0.0066225166 | 0.11919698 | 1 | 3028 | tags=23%, list=13%, si | MM_4 |
| APELIN_SIGNALING_PATH | APELIN_SIGNALING_PATHWAY | 131 | 0.229054 | 1.4997227 | 0 | 0.119337335 | 1 | 4389 | tags=34%, list=19%, si | MM_4 |
| VASOPRESSIN-REGULATE | VASOPRESSIN-REGULATED_WATER_REABSO | 40 | 0.304631 | 1.4997187 | 0.04784689 | 0.119187236 | 1 | 4531 | tags=48%, list=19%, si | MM_4 |
| GOBP_NEGATIVE_REGUL | GOBP_NEGATIVE_REGULATION_OF_INTERF | 16 | 0.410449 | 1.4996791 | 0.08208955 | 0.1190617 | 1 | 1383 | tags=19%, list=6%, sig | MM_4 |
| GOBP_POSITIVE_REGULA | GOBP_POSITIVE_REGULATION_OF_PROTEAS | 102 | 0.23896 | 1.4988844 | 0.005882353 | 0.11945464 | 1 | 3551 | tags=26%, list=15%, si | MM_4 |
| GOBP_GLYCOLIPID_BIOSY | GOBP_GLYCOLIPID_BIOSYNTHETIC_PROCESS | 66 | 0.266449 | 1.4985054 | 0.02173913 | 0.119454786 | 1 | 1778 | tags=21%, list=8%, sig | MM_4 |
| GOBP_NEUTRAL_LIPID_B | GOBP_NEUTRAL_LIPID_BIOSYNTHETIC_PRO | 58 | 0.275408 | 1.4979903 | 0.035532996 | 0.11961077 | 1 | 2743 | tags=28%, list=12%, si | MM_4 |
| GOBP_NEGATIVE_REGUL | GOBP_NEGATIVE_REGULATION_OF_SMALL_ | 61 | 0.269529 | 1.4975226 | 0.043209877 | 0.11976185 | 1 | 2672 | tags=23%, list=11%, si | MM_4 |
| AMINO_SUGAR_AND_NU | AMINO_SUGAR_AND_NUCLEOTIDE_SUGAR_ | 46 | 0.293803 | 1.49714 | 0.029045643 | 0.11989147 | 1 | 2967 | tags=28%, list=13%, si | MM_4 |
| HYPERTROPHIC_CARDIOM | HYPERTROPHIC_CARDIOMYOPATHY_(HCM) | 84 | 0.24833 | 1.4953871 | 0.021276595 | 0.12085807 | 1 | 1127 | tags=15%, list=5%, sig | MM_4 |
| GOBP_PERK_MEDIATED_ | GOBP_PERK_MEDIATED_UNFOLDED_PROTE | 16 | 0.413195 | 1.4952384 | 0.07885305 | 0.12078818 | 1 | 1561 | tags=19%, list=7%, sig | MM_4 |
| GOBP_REGULATION_OF_ | GOBP_REGULATION_OF_DEFENSE_RESPONS | 20 | 0.364395 | 1.4942955 | 0.047138046 | 0.12125168 | 1 | 1760 | tags=30%, list=7%, sig | MM_4 |
| BASAL_TRANSCRIPTION_ | BASAL_TRANSCRIPTION_FACTORS | 40 | 0.301177 | 1.4939128 | 0.036036037 | 0.12135466 | 1 | 3987 | tags=35%, list=17%, si | MM_4 |
| PLATELET_ACTIVATION | PLATELET_ACTIVATION | 116 | 0.234843 | 1.4903312 | 0.0064102565 | 0.12343451 | 1 | 4235 | tags=34%, list=18%, si | MM_4 |
| GOBP_CD40_SIGNALING_ | GOBP_CD40_SIGNALING_PATHWAY | 16 | 0.39189 | 1.4889235 | 0.06514658 | 0.124191344 | 1 | 11801 | tags=94%, list=50%, si | MM_4 |
| GOBP_RESPONSE_TO_AM | GOBP_RESPONSE_TO_AMINO_ACID_STARV | 54 | 0.286622 | 1.4888318 | 0.05 | 0.12409088 | 1 | 3306 | tags=31%, list=14%, si | MM_4 |
| GOBP_REGULATION_OF_ | GOBP_REGULATION_OF_NEUTROPHIL_MIG | 52 | 0.289246 | 1.4885547 | 0.04639175 | 0.1240648 | 1 | 3905 | tags=31%, list=17%, si | MM_4 |
| GOBP_REGULATION_OF_ | GOBP_REGULATION_OF_AMYLOID_BETA_C | 18 | 0.38271 | 1.4881035 | 0.049822062 | 0.12417312 | 1 | 2387 | tags=28%, list=10%, si | MM_4 |
| GOBP_NEGATIVE_REGUL | GOBP_NEGATIVE_REGULATION_OF_RESPON | 40 | 0.300375 | 1.4879527 | 0.03524229 | 0.12412307 | 1 | 2525 | tags=25%, list=11%, si | MM_4 |
| GOBP_PEPTIDE_CROSS_L | GOBP_PEPTIDE_CROSS_LINKING | 22 | 0.348998 | 1.4870654 | 0.06866953 | 0.124486215 | 1 | 879 | tags=14%, list=4%, sig | MM_4 |
| GOBP_POSITIVE_REGULA | GOBP_POSITIVE_REGULATION_OF_LEUKOC | 169 | 0.221817 | 1.4863406 | 0 | 0.12475123 | 1 | 2755 | tags=21%, list=12%, si | MM_4 |
| GOBP_REGULATION_OF_ | GOBP_REGULATION_OF_PROTEIN_CONTAIN | 420 | 0.190255 | 1.4851295 | 0 | 0.12541284 | 1 | 3609 | tags=23%, list=15%, si | MM_4 |
| GOBP_REACTIVE_NITROG | GOBP_REACTIVE_NITROGEN_SPECIES_MET | 92 | 0.242097 | 1.4841994 | 0.023952097 | 0.12592055 | 1 | 3698 | tags=28%, list=16%, si | MM_4 |
| GOBP_VESICLE_LOCALIZA | GOBP_VESICLE_LOCALIZATION | 218 | 0.202838 | 1.4824144 | 0.00990099 | 0.12693042 | 1 | 3042 | tags=20%, list=13%, si | MM_4 |
| GOBP_GLYCEROPHOSPHO | GOBP_GLYCEROPHOSPHOLIPID_BIOSYNTHET | 160 | 0.214513 | 1.4821373 | 0 | 0.12694095 | 1 | 3741 | tags=26%, list=16%, si | MM_4 |
| GOBP_STEROL_TRANSPO | GOBP_STEROL_TRANSPORT | 109 | 0.240373 | 1.48175 | 0.012658228 | 0.12704368 | 1 | 2506 | tags=23%, list=11%, si | MM_4 |
| GOBP_PEPTIDYL_CYSTEIN | GOBP_PEPTIDYL_CYSTEINE_MODIFICATION | 44 | 0.282003 | 1.4808275 | 0.037383176 | 0.12756361 | 1 | 2789 | tags=25%, list=12%, si | MM_4 |
| NECROPTOSIS | NECROPTOSIS | 138 | 0.223765 | 1.4808083 | 0.013245033 | 0.1274228 | 1 | 4186 | tags=32%, list=18%, si | MM_4 |
| TOXOPLASMOSIS | TOXOPLASMOSIS | 103 | 0.243617 | 1.4807143 | 0.02027027 | 0.12732291 | 1 | 2693 | tags=22%, list=11%, si | MM_4 |
| GOBP_NUCLEOTIDE_SUG | GOBP_NUCLEOTIDE_SUGAR_BIOSYNTHETIC_ | 17 | 0.389438 | 1.4806893 | 0.0703125 | 0.12718248 | 1 | 3618 | tags=41%, list=15%, si | MM_4 |
| GOBP_REGULATION_OF_ | GOBP_REGULATION_OF_ACTIN_FILAMENT_ | 113 | 0.229876 | 1.480457 | 0.02739726 | 0.12716247 | 1 | 2799 | tags=19%, list=12%, si | MM_4 |
| GOBP_REGULATION_OF_ | GOBP_REGULATION_OF_MRNA_SPLICING_V | 108 | 0.235638 | 1.4802344 | 0.013513514 | 0.1271881 | 1 | 3320 | tags=21%, list=14%, si | MM_4 |
| GOBP_NEGATIVE_REGUL | GOBP_NEGATIVE_REGULATION_OF_TOLL_L | 22 | 0.362818 | 1.4801819 | 0.05106383 | 0.12706575 | 1 | 4003 | tags=45%, list=17%, si | MM_4 |
| GOBP_AMYLOID_PRECUR | GOBP_AMYLOID_PRECURSOR_PROTEIN_CAT | 59 | 0.271365 | 1.4800885 | 0.04232804 | 0.12696147 | 1 | 2833 | tags=24%, list=12%, si | MM_4 |
| GOBP_INTRACELLULAR_R | GOBP_INTRACELLULAR_RECEPTOR_SIGNALI | 325 | 0.198558 | 1.4784263 | 0 | 0.12789173 | 1 | 3973 | tags=26%, list=17%, si | MM_4 |
| GOBP_RRNA_MODIFICAT | GOBP_RRNA_MODIFICATION | 29 | 0.328165 | 1.4783945 | 0.04743083 | 0.12775427 | 1 | 4613 | tags=41%, list=20%, si | MM_4 |
| GOBP_T_CELL_DIFFERENT | GOBP_T_CELL_DIFFERENTIATION_IN_THYM | 103 | 0.230175 | 1.4779186 | 0.01910828 | 0.12792432 | 1 | 3209 | tags=25%, list=14%, si | MM_4 |
| GOBP_PYRIDINE_CONTAI | GOBP_PYRIDINE_CONTAINING_COMPOUND | 109 | 0.233392 | 1.4769149 | 0.01875 | 0.1283931 | 1 | 3708 | tags=29%, list=16%, si | MM_4 |
| GOBP_NEGATIVE_REGUL | GOBP_NEGATIVE_REGULATION_OF_RNA_SI | 29 | 0.320082 | 1.475802 | 0.05283019 | 0.12891616 | 1 | 11317 | tags=90%, list=48%, si | MM_4 |
| GOBP_REGULATION_OF_ | GOBP_REGULATION_OF_ANOIKIS | 22 | 0.361516 | 1.4757271 | 0.057142857 | 0.12880686 | 1 | 3662 | tags=45%, list=16%, si | MM_4 |
| GOBP_DENDRITIC_CELL_ | GOBP_DENDRITIC_CELL_MIGRATION | 26 | 0.332863 | 1.4743934 | 0.07083333 | 0.12950045 | 1 | 1522 | tags=15%, list=6%, sig | MM_4 |

| | | | | | | | | | | |
|---|---|---|---|---|---|---|---|---|---|---|
| GOBP_SYNAPTIC_VESICLE | GOBP_SYNAPTIC_VESICLE_MATURATION | 27 | 0.34279 | 1.4741225 | 0.0513834 | 0.12953305 | 1 | 4343 | tags=41%, list=18%, si | MM_4 |
| GOBP_CANONICAL_NF_K | GOBP_CANONICAL_NF_KAPPAB_SIGNAL_TR | 247 | 0.198411 | 1.4740881 | 0 | 0.12938923 | 1 | 2538 | tags=19%, list=11%, si | MM_4 |
| GOBP_COLLAGEN_FIBRIL | GOBP_COLLAGEN_FIBRIL_ORGANIZATION | 63 | 0.270393 | 1.4732878 | 0.015228426 | 0.12982294 | 1 | 1464 | tags=19%, list=6%, sig | MM_4 |
| GOBP_NUCLEOSIDE_PHO | GOBP_NUCLEOSIDE_PHOSPHATE_CATABOLI | 170 | 0.213349 | 1.4730635 | 0 | 0.12982602 | 1 | 3189 | tags=26%, list=14%, si | MM_4 |
| GOBP_LYMPHOCYTE_HOI | GOBP_LYMPHOCYTE_HOMEOSTASIS | 92 | 0.241333 | 1.4716715 | 0.012658228 | 0.13059369 | 1 | 3755 | tags=29%, list=16%, si | MM_4 |
| GOBP_T_CELL_DIFFERENT | GOBP_T_CELL_DIFFERENTIATION | 327 | 0.19284 | 1.4716007 | 0 | 0.13048318 | 1 | 2930 | tags=19%, list=12%, si | MM_4 |
| GOBP_NEGATIVE_REGUL | GOBP_NEGATIVE_REGULATION_OF_TRANSF | 94 | 0.241309 | 1.4715817 | 0.0064935065 | 0.13034487 | 1 | 3160 | tags=28%, list=13%, si | MM_4 |
| HEPATOCELLULAR_CARCI | HEPATOCELLULAR_CARCINOMA | 164 | 0.222012 | 1.4714794 | 0.009090909 | 0.13025522 | 1 | 2867 | tags=21%, list=12%, si | MM_4 |
| GOBP_PHOSPHOLIPID_BI | GOBP_PHOSPHOLIPID_BIOSYNTHETIC_PROC | 214 | 0.202403 | 1.4713664 | 0.01904762 | 0.13015746 | 1 | 3370 | tags=25%, list=14%, si | MM_4 |
| GOBP_CELLULAR_RESPON | GOBP_CELLULAR_RESPONSE_TO_NUTRIENT | 38 | 0.303576 | 1.4711461 | 0.069869 | 0.13017261 | 1 | 4008 | tags=29%, list=17%, si | MM_4 |
| GOBP_PURINE_NUCLEOS | GOBP_PURINE_NUCLEOSIDE_MONOPHOSPH | 36 | 0.311532 | 1.4708409 | 0.03984064 | 0.13021664 | 1 | 3189 | tags=31%, list=14%, si | MM_4 |
| GOBP_POSITIVE_REGULA | GOBP_POSITIVE_REGULATION_OF_PEPTIDA | 161 | 0.220938 | 1.4693911 | 0 | 0.1309598 | 1 | 3371 | tags=25%, list=14%, si | MM_4 |
| GOBP_RNA_EXPORT_FRO | GOBP_RNA_EXPORT_FROM_NUCLEUS | 78 | 0.245687 | 1.4688121 | 0.021621622 | 0.13113348 | 1 | 5909 | tags=49%, list=25%, si | MM_4 |
| GOBP_NEGATIVE_REGUL | GOBP_NEGATIVE_REGULATION_OF_TUMOR | 67 | 0.259099 | 1.4673979 | 0.03448276 | 0.13187423 | 1 | 2917 | tags=24%, list=12%, si | MM_4 |
| GOBP_ZINC_ION_TRANSF | GOBP_ZINC_ION_TRANSPORT | 28 | 0.338802 | 1.4667028 | 0.06849315 | 0.1321562 | 1 | 2227 | tags=29%, list=9%, sig | MM_4 |
| GOBP_DECIDUALIZATION | GOBP_DECIDUALIZATION | 24 | 0.345191 | 1.4664446 | 0.08064516 | 0.13217618 | 1 | 3518 | tags=33%, list=15%, si | MM_4 |
| GOBP_NEGATIVE_REGUL | GOBP_NEGATIVE_REGULATION_OF_HEMOP | 121 | 0.230206 | 1.4663849 | 0 | 0.13205193 | 1 | 3364 | tags=22%, list=14%, si | MM_4 |
| GOBP_RESPONSE_TO_TO | GOBP_RESPONSE_TO_TOPOLOGICALLY_INC | 127 | 0.228061 | 1.4662161 | 0.02238806 | 0.13201791 | 1 | 4631 | tags=34%, list=20%, si | MM_4 |
| GOBP_REGULATION_OF_ | GOBP_REGULATION_OF_MYELOID_CELL_DI | 217 | 0.213158 | 1.4654552 | 0.011111111 | 0.13243283 | 1 | 3780 | tags=26%, list=16%, si | MM_4 |
| GOBP_CELL_REDOX_HON | GOBP_CELL_REDOX_HOMEOSTASIS | 28 | 0.335469 | 1.4647974 | 0.07377049 | 0.13273633 | 1 | 3944 | tags=36%, list=17%, si | MM_4 |
| GOBP_DNA_REPLICATION | GOBP_DNA_REPLICATION_CHECKPOINT_SIG | 18 | 0.382587 | 1.4639113 | 0.09966777 | 0.13316655 | 1 | 2552 | tags=39%, list=11%, si | MM_4 |
| GOBP_TORC1_SIGNALING | GOBP_TORC1_SIGNALING | 100 | 0.238024 | 1.4631544 | 0.01764706 | 0.13350514 | 1 | 3776 | tags=30%, list=16%, si | MM_4 |
| GOBP_REGULATION_OF_ | GOBP_REGULATION_OF_INTRACELLULAR_E | 34 | 0.320984 | 1.4629579 | 0.063025214 | 0.13352567 | 1 | 14523 | tags=97%, list=62%, si | MM_4 |
| GOBP_REGULATION_OF_ | GOBP_REGULATION_OF_ALPHA_BETA_T_CE | 119 | 0.227284 | 1.4626498 | 0.0067114094 | 0.13353036 | 1 | 2850 | tags=19%, list=12%, si | MM_4 |
| DILATED_CARDIOMYOPAT | DILATED_CARDIOMYOPATHY_(DCM) | 88 | 0.237964 | 1.4617832 | 0.036585364 | 0.13399495 | 1 | 3141 | tags=27%, list=13%, si | MM_4 |
| GOBP_NEUROINFLAMMA | GOBP_NEUROINFLAMMATORY_RESPONSE | 84 | 0.257039 | 1.4615015 | 0.046242774 | 0.13400339 | 1 | 3681 | tags=31%, list=16%, si | MM_4 |
| GOBP_REGULATION_OF_ | GOBP_REGULATION_OF_AUTOPHAGY | 295 | 0.200571 | 1.4609917 | 0.015625 | 0.13421011 | 1 | 3258 | tags=23%, list=14%, si | MM_4 |
| GOBP_POSITIVE_REGULA | GOBP_POSITIVE_REGULATION_OF_PROTEIN | 20 | 0.359571 | 1.4597808 | 0.070866145 | 0.13483194 | 1 | 15128 | tags=100%, list=64%, | MM_4 |
| GOBP_CEREBELLAR_PURK | GOBP_CEREBELLAR_PURKINJE_CELL_LAYER_ | 21 | 0.357746 | 1.4596609 | 0.08108108 | 0.13475402 | 1 | 4489 | tags=48%, list=19%, si | MM_4 |
| GOBP_RESPONSE_TO_CA | GOBP_RESPONSE_TO_CALCIUM_ION | 130 | 0.227866 | 1.4594858 | 0.030303031 | 0.13473137 | 1 | 2283 | tags=18%, list=10%, si | MM_4 |
| GOBP_REGULATION_OF_ | GOBP_REGULATION_OF_ENDOCYTOSIS | 325 | 0.194267 | 1.4587191 | 0 | 0.13510592 | 1 | 3982 | tags=25%, list=17%, si | MM_4 |
| GOBP_ORGANIC_CATION | GOBP_ORGANIC_CATION_TRANSPORT | 122 | 0.220133 | 1.4582742 | 0.006666667 | 0.13528456 | 1 | 3336 | tags=26%, list=14%, si | MM_4 |
| GOBP_RESPONSE_TO_NE | GOBP_RESPONSE_TO_NERVE_GROWTH_FA | 42 | 0.29396 | 1.4580888 | 0.052173913 | 0.13521986 | 1 | 14459 | tags=95%, list=61%, si | MM_4 |
| GOBP_DNA_DAMAGE_RE | GOBP_DNA_DAMAGE_RESPONSE_SIGNAL_ | 17 | 0.376057 | 1.4580342 | 0.094736844 | 0.13509205 | 1 | 4846 | tags=47%, list=21%, si | MM_4 |
| GOBP_T_CELL_HOMEOST | GOBP_T_CELL_HOMEOSTASIS | 52 | 0.272563 | 1.4578538 | 0.04901961 | 0.13506393 | 1 | 3512 | tags=31%, list=15%, si | MM_4 |
| GOBP_PYRIMIDINE_DEOX | GOBP_PYRIMIDINE_DEOXYRIBONUCLEOSIDE | 15 | 0.395578 | 1.4575208 | 0.06312292 | 0.13514036 | 1 | 3106 | tags=40%, list=13%, si | MM_4 |
| GOBP_ORGANIC_ACID_TI | GOBP_ORGANIC_ACID_TRANSMEMBRANE_ | 135 | 0.220462 | 1.4574581 | 0.014084507 | 0.13501811 | 1 | 2986 | tags=23%, list=13%, si | MM_4 |
| GOBP_SEQUESTERING_O | GOBP_SEQUESTERING_OF_CALCIUM_ION | 129 | 0.220698 | 1.4574007 | 0.013888889 | 0.13488539 | 1 | 2539 | tags=17%, list=11%, si | MM_4 |
| GOBP_REGULATION_OF_ | GOBP_REGULATION_OF_CHROMATIN_ORG | 36 | 0.309069 | 1.4569076 | 0.05485232 | 0.13506702 | 1 | 2777 | tags=25%, list=12%, si | MM_4 |
| GOBP_NEGATIVE_REGUL | GOBP_NEGATIVE_REGULATION_OF_RHO_P | 25 | 0.350054 | 1.4567206 | 0.04819277 | 0.13505195 | 1 | 2672 | tags=28%, list=11%, si | MM_4 |
| GOBP_DENSE_CORE_GRA | GOBP_DENSE_CORE_GRANULE_EXOCYTOSIS | 15 | 0.403995 | 1.4565706 | 0.08214286 | 0.13497236 | 1 | 1304 | tags=20%, list=6%, sig | MM_4 |
| GOBP_ESTABLISHMENT_ | GOBP_ESTABLISHMENT_OF_RNA_LOCALIZA | 138 | 0.224241 | 1.456246 | 0.014705882 | 0.1350478 | 1 | 3572 | tags=23%, list=15%, si | MM_4 |
| GOBP_NEGATIVE_REGUL | GOBP_NEGATIVE_REGULATION_OF_VASCU | 33 | 0.306869 | 1.4561977 | 0.047169812 | 0.13490719 | 1 | 3543 | tags=39%, list=15%, si | MM_4 |
| GOBP_AGGREPHAGY | GOBP_AGGREPHAGY | 17 | 0.382988 | 1.4557604 | 0.062730625 | 0.13501354 | 1 | 2086 | tags=24%, list=9%, sig | MM_4 |
| CELL_ADHESION_MOLECU | CELL_ADHESION_MOLECULES_(CAMS) | 139 | 0.220986 | 1.4555777 | 0.016806724 | 0.13496834 | 1 | 1666 | tags=14%, list=7%, sig | MM_4 |
| GOBP_COPII_COATED_VE | GOBP_COPII_COATED_VESICLE_BUDDING | 37 | 0.304298 | 1.455481 | 0.05263158 | 0.1348718 | 1 | 3503 | tags=32%, list=15%, si | MM_4 |
| GOBP_QUINONE_METAB | GOBP_QUINONE_METABOLIC_PROCESS | 23 | 0.340941 | 1.4551798 | 0.06530612 | 0.1349217 | 1 | 3486 | tags=35%, list=15%, si | MM_4 |
| GOBP_CHOLESTEROL_EFF | GOBP_CHOLESTEROL_EFFLUX | 59 | 0.269027 | 1.4543428 | 0.056818184 | 0.13534185 | 1 | 2387 | tags=27%, list=10%, si | MM_4 |
| GOBP_NUCLEAR_MEMBR | GOBP_NUCLEAR_MEMBRANE_ORGANIZATI | 35 | 0.299708 | 1.4540007 | 0.06363636 | 0.13543253 | 1 | 4486 | tags=43%, list=19%, si | MM_4 |
| GOBP_PYRIMIDINE_NUCL | GOBP_PYRIMIDINE_NUCLEOTIDE_METABOL | 38 | 0.305399 | 1.4536724 | 0.05676856 | 0.13547753 | 1 | 4220 | tags=47%, list=18%, si | MM_4 |
| OTHER_TYPES_OF_O-GLY | OTHER_TYPES_OF_O-GLYCAN_BIOSYNTHESI | 21 | 0.364042 | 1.4530398 | 0.074074075 | 0.13572736 | 1 | 2153 | tags=29%, list=9%, sig | MM_4 |
| GOBP_LONG_CHAIN_FAT | GOBP_LONG_CHAIN_FATTY_ACID_IMPORT_ | 19 | 0.373755 | 1.4516617 | 0.083333336 | 0.13647969 | 1 | 2743 | tags=26%, list=12%, si | MM_4 |
| GOBP_NATURAL_KILLER_ | GOBP_NATURAL_KILLER_CELL_ACTIVATION_ | 21 | 0.355291 | 1.4507711 | 0.06944445 | 0.13694264 | 1 | 1090 | tags=19%, list=5%, sig | MM_4 |
| GOBP_LUNG_ALVEOLUS_ | GOBP_LUNG_ALVEOLUS_DEVELOPMENT | 67 | 0.259587 | 1.45042 | 0.04368932 | 0.13703685 | 1 | 2270 | tags=25%, list=10%, si | MM_4 |
| GOBP_MATURATION_OF_ | GOBP_MATURATION_OF_5_8S_RRNA | 35 | 0.301126 | 1.450371 | 0.060344826 | 0.13690203 | 1 | 3686 | tags=37%, list=16%, si | MM_4 |
| PRIMARY_IMMUNODEFIC | PRIMARY_IMMUNODEFICIENCY | 33 | 0.301374 | 1.4498595 | 0.07359307 | 0.13709103 | 1 | 1931 | tags=24%, list=8%, sig | MM_4 |
| GOBP_POSITIVE_REGULA | GOBP_POSITIVE_REGULATION_OF_TYPE_I_I | 19 | 0.35708 | 1.4498469 | 0.07835821 | 0.13694765 | 1 | 3722 | tags=37%, list=16%, si | MM_4 |
| PROGESTERONE-MEDIATI | PROGESTERONE-MEDIATED_OOCYTE_MATU | 87 | 0.238995 | 1.4482557 | 0.011976048 | 0.13783088 | 1 | 2718 | tags=28%, list=12%, si | MM_4 |
| GOBP_P_BODY_ASSEMBL | GOBP_P_BODY_ASSEMBLY | 20 | 0.354097 | 1.4465386 | 0.062962964 | 0.13873813 | 1 | 10883 | tags=95%, list=46%, si | MM_4 |
| CHOLESTEROL_METABOL | CHOLESTEROL_METABOLISM | 44 | 0.288666 | 1.4462882 | 0.07920792 | 0.13872865 | 1 | 3207 | tags=30%, list=14%, si | MM_4 |
| GOBP_REGULATION_OF_ | GOBP_REGULATION_OF_GLYCOPROTEIN_M | 47 | 0.277526 | 1.4460044 | 0.051282052 | 0.13876186 | 1 | 3255 | tags=30%, list=14%, si | MM_4 |
| GOBP_VIRAL_TRANSCRIP | GOBP_VIRAL_TRANSCRIPTION | 55 | 0.276717 | 1.4451938 | 0.03211009 | 0.13916403 | 1 | 4279 | tags=35%, list=18%, si | MM_4 |
| RHEUMATOID_ARTHRITIS | RHEUMATOID_ARTHRITIS | 73 | 0.250121 | 1.4451338 | 0.04 | 0.13904348 | 1 | 4137 | tags=36%, list=18%, si | MM_4 |
| GOBP_NEGATIVE_REGUL | GOBP_NEGATIVE_REGULATION_OF_PHOSPH | 398 | 0.18496 | 1.4447436 | 0 | 0.13914281 | 1 | 3364 | tags=23%, list=14%, si | MM_4 |
| GOBP_PHOSPHOLIPID_M | GOBP_PHOSPHOLIPID_METABOLIC_PROCES | 352 | 0.19137 | 1.4442258 | 0 | 0.13936563 | 1 | 3377 | tags=23%, list=14%, si | MM_4 |
| GOBP_LIPOPROTEIN_BIO | GOBP_LIPOPROTEIN_BIOSYNTHETIC_PROCE | 104 | 0.227961 | 1.4435421 | 0.023255814 | 0.13972154 | 1 | 2115 | tags=20%, list=9%, sig | MM_4 |
| GOBP_BIOLOGICAL_PROC | GOBP_BIOLOGICAL_PROCESS_INVOLVED_IN | 140 | 0.222692 | 1.4426584 | 0.006289308 | 0.14020419 | 1 | 3570 | tags=26%, list=15%, si | MM_4 |
| GOBP_NEGATIVE_REGUL | GOBP_NEGATIVE_REGULATION_OF_T_CELL_ | 29 | 0.325397 | 1.4414921 | 0.07569721 | 0.14077246 | 1 | 1611 | tags=17%, list=7%, sig | MM_4 |
| MRNA_SURVEILLANCE_P | MRNA_SURVEILLANCE_PATHWAY | 87 | 0.239781 | 1.4412265 | 0.018292682 | 0.14080119 | 1 | 4470 | tags=29%, list=19%, si | MM_4 |
| GOBP_RESPIRATORY_BUR | GOBP_RESPIRATORY_BURST | 27 | 0.316611 | 1.4410025 | 0.052208837 | 0.14083992 | 1 | 3944 | tags=33%, list=17%, si | MM_4 |
| GOBP_ENDOTHELIAL_CEL | GOBP_ENDOTHELIAL_CELL_MIGRATION | 219 | 0.200077 | 1.4406815 | 0 | 0.14088893 | 1 | 2350 | tags=15%, list=10%, si | MM_4 |
| GOBP_LATE_ENDOSOME_ | GOBP_LATE_ENDOSOME_TO_VACUOLE_TR | 25 | 0.338763 | 1.4404695 | 0.093023255 | 0.14086291 | 1 | 11250 | tags=92%, list=48%, si | MM_4 |
| GOBP_GLYCOPROTEIN_B | GOBP_GLYCOPROTEIN_BIOSYNTHETIC_PRO | 282 | 0.194291 | 1.4386088 | 0 | 0.1419729 | 1 | 2818 | tags=21%, list=12%, si | MM_4 |
| GOBP_MAINTENANCE_OI | GOBP_MAINTENANCE_OF_PROTEIN_LOCAT | 25 | 0.333707 | 1.4379209 | 0.08921933 | 0.14226972 | 1 | 1774 | tags=20%, list=8%, sig | MM_4 |
| GOBP_POSITIVE_REGULA | GOBP_POSITIVE_REGULATION_OF_IMMUNE | 266 | 0.195419 | 1.4377412 | 0 | 0.14224039 | 1 | 3641 | tags=25%, list=15%, si | MM_4 |
| GOBP_POSITIVE_REGULA | GOBP_POSITIVE_REGULATION_OF_GRANUL | 31 | 0.312931 | 1.4372989 | 0.054347824 | 0.1424015 | 1 | 3681 | tags=29%, list=16%, si | MM_4 |
| GOBP_NEUROTRANSMITT | GOBP_NEUROTRANSMITTER_RECEPTOR_TR | 40 | 0.288615 | 1.4327155 | 0.071428575 | 0.14546297 | 1 | 15299 | tags=95%, list=65%, si | MM_4 |
| GOBP_GLYCOSYLATION | GOBP_GLYCOSYLATION | 218 | 0.198681 | 1.4321984 | 0 | 0.14566231 | 1 | 3609 | tags=28%, list=15%, si | MM_4 |
| GOBP_PURINE_RIBONUC | GOBP_PURINE_RIBONUCLEOSIDE_METABOL | 24 | 0.344753 | 1.4316727 | 0.093023255 | 0.1458886 | 1 | 3378 | tags=42%, list=14%, si | MM_4 |
| FATTY_ACID_ELONGATIO | FATTY_ACID_ELONGATION | 27 | 0.324142 | 1.4316089 | 0.049822062 | 0.14577122 | 1 | 2425 | tags=30%, list=10%, si | MM_4 |
| GOBP_REGULATION_OF_ | GOBP_REGULATION_OF_EARLY_ENDOSOME | 18 | 0.375867 | 1.431147 | 0.0625 | 0.14595586 | 1 | 12075 | tags=94%, list=51%, si | MM_4 |
| GOBP_LYMPHOCYTE_COS | GOBP_LYMPHOCYTE_COSTIMULATION | 33 | 0.306767 | 1.431118 | 0.07048458 | 0.14581986 | 1 | 1931 | tags=24%, list=8%, sig | MM_4 |

| | | | | | | | | | | |
|---|---|---|---|---|---|---|---|---|---|---|
| GOBP_SPHINGOLIPID_ME | GOBP_SPHINGOLIPID_METABOLIC_PROCESS | 137 | 0.219537 | 1.4310781 | 0.007936508 | 0.14567877 | 1 | 3483 | tags=26%, list=15%, si | MM_4 |
| GOBP_CELLULAR_RESPON | GOBP_CELLULAR_RESPONSE_TO_INORGANI | 177 | 0.208911 | 1.4305284 | 0.009259259 | 0.14591575 | 1 | 3052 | tags=22%, list=13%, si | MM_4 |
| GOBP_DNA_BIOSYNTHET | GOBP_DNA_BIOSYNTHETIC_PROCESS | 175 | 0.20125 | 1.4300596 | 0.010526316 | 0.14610235 | 1 | 4454 | tags=31%, list=19%, si | MM_4 |
| GOBP_POSITIVE_REGULA | GOBP_POSITIVE_REGULATION_OF_SYSTEMI | 23 | 0.338656 | 1.4296247 | 0.06367041 | 0.14629869 | 1 | 331 | tags=13%, list=1%, sig | MM_4 |
| GOBP_PYRIMIDINE_NUCL | GOBP_PYRIMIDINE_NUCLEOSIDE_MONOPH | 21 | 0.347503 | 1.4295341 | 0.094736844 | 0.14620546 | 1 | 3999 | tags=48%, list=17%, si | MM_4 |
| C-TYPE_LECTIN_RECEPTO | C-TYPE_LECTIN_RECEPTOR_SIGNALING_PAT | 104 | 0.230372 | 1.4291072 | 0.018867925 | 0.14634117 | 1 | 1511 | tags=10%, list=6%, sig | MM_4 |
| GOBP_POSITIVE_REGULA | GOBP_POSITIVE_REGULATION_OF_VASCULA | 28 | 0.322262 | 1.4287652 | 0.094017096 | 0.14643578 | 1 | 311 | tags=11%, list=1%, sig | MM_4 |
| CENTRAL_CARBON_META | CENTRAL_CARBON_METABOLISM_IN_CANC | 61 | 0.250359 | 1.4282454 | 0.05102041 | 0.1466669 | 1 | 3301 | tags=25%, list=14%, si | MM_4 |
| GOBP_POSITIVE_REGULA | GOBP_POSITIVE_REGULATION_OF_HEMOPO | 202 | 0.202494 | 1.4281232 | 0.011235955 | 0.1466228 | 1 | 3769 | tags=25%, list=16%, si | MM_4 |
| GOBP_NEGATIVE_REGUL | GOBP_NEGATIVE_REGULATION_OF_MEMBR | 22 | 0.343974 | 1.4272933 | 0.09774436 | 0.14706463 | 1 | 3985 | tags=41%, list=17%, si | MM_4 |
| GOBP_NEGATIVE_REGUL | GOBP_NEGATIVE_REGULATION_OF_BLOOD | 35 | 0.297359 | 1.4272164 | 0.0921659 | 0.14697586 | 1 | 2987 | tags=23%, list=13%, si | MM_4 |
| GOBP_INTERFERON_BETA | GOBP_INTERFERON_BETA_PRODUCTION | 66 | 0.251748 | 1.4271419 | 0.03448276 | 0.1468535 | 1 | 3953 | tags=30%, list=17%, si | MM_4 |
| GOBP_PYRIDINE_CONTAI | GOBP_PYRIDINE_CONTAINING_COMPOUND | 182 | 0.2099 | 1.4255188 | 0.030612245 | 0.14791901 | 1 | 3815 | tags=27%, list=16%, si | MM_4 |
| GOBP_CELLULAR_RESPON | GOBP_CELLULAR_RESPONSE_TO_CHEMICAL | 290 | 0.194298 | 1.4253901 | 0.014925373 | 0.14784516 | 1 | 3705 | tags=26%, list=16%, si | MM_4 |
| GOBP_POSITIVE_REGULA | GOBP_POSITIVE_REGULATION_OF_SIGNAL_ | 32 | 0.309667 | 1.42482 | 0.09417041 | 0.1481225 | 1 | 5534 | tags=59%, list=23%, si | MM_4 |
| GOBP_CELLULAR_LIPID_C | GOBP_CELLULAR_LIPID_CATABOLIC_PROCES | 210 | 0.20289 | 1.4245836 | 0 | 0.14812529 | 1 | 3178 | tags=24%, list=13%, si | MM_4 |
| GOBP_REGULATION_OF_ | GOBP_REGULATION_OF_DNA_REPLICATION | 118 | 0.222024 | 1.4244639 | 0.02247191 | 0.14808728 | 1 | 3099 | tags=20%, list=13%, si | MM_4 |
| GOBP_NEGATIVE_REGUL | GOBP_NEGATIVE_REGULATION_OF_CYTOPL | 50 | 0.26948 | 1.4242903 | 0.04712042 | 0.14802022 | 1 | 4352 | tags=36%, list=18%, si | MM_4 |
| GOBP_NEGATIVE_REGUL | GOBP_NEGATIVE_REGULATION_OF_CELL_C | 197 | 0.203539 | 1.4232856 | 0.00862069 | 0.14855957 | 1 | 1528 | tags=14%, list=6%, sig | MM_4 |
| GOBP_T_CELL_MEDIATED | GOBP_T_CELL_MEDIATED_IMMUNITY | 131 | 0.228552 | 1.4228851 | 0.008403362 | 0.14874499 | 1 | 3561 | tags=27%, list=15%, si | MM_4 |
| GOBP_MORPHOGENESIS_ | GOBP_MORPHOGENESIS_OF_AN_EPITHELIA | 17 | 0.382284 | 1.4228219 | 0.094339624 | 0.14862435 | 1 | 14591 | tags=100%, list=62%, | MM_4 |
| GOBP_HEME_BIOSYNTHE | GOBP_HEME_BIOSYNTHETIC_PROCESS | 24 | 0.337683 | 1.422522 | 0.08368201 | 0.14869301 | 1 | 2181 | tags=29%, list=9%, sig | MM_4 |
| GOBP_NEGATIVE_REGUL | GOBP_NEGATIVE_REGULATION_OF_MICRO | 15 | 0.410137 | 1.4216803 | 0.110738255 | 0.14921023 | 1 | 13933 | tags=100%, list=59%, | MM_4 |
| GOBP_DEOXYRIBONUCLE | GOBP_DEOXYRIBONUCLEOSIDE_MONOPHO | 28 | 0.318566 | 1.41969 | 0.075187966 | 0.15045162 | 1 | 3189 | tags=32%, list=14%, si | MM_4 |
| GOBP_NEGATIVE_REGUL | GOBP_NEGATIVE_REGULATION_OF_LYMPH | 44 | 0.279914 | 1.4186732 | 0.060344826 | 0.15106645 | 1 | 1611 | tags=16%, list=7%, sig | MM_4 |
| GOBP_POSITIVE_REGULA | GOBP_POSITIVE_REGULATION_OF_ENDOCY | 182 | 0.204262 | 1.4177077 | 0.01010101 | 0.1516181 | 1 | 2037 | tags=15%, list=9%, sig | MM_4 |
| GOBP_PEROXISOME_PRO | GOBP_PEROXISOME_PROLIFERATOR_ACTIV | 27 | 0.319861 | 1.4165165 | 0.08300395 | 0.1523947 | 1 | 3461 | tags=37%, list=15%, si | MM_4 |
| GOBP_MACROPHAGE_CY | GOBP_MACROPHAGE_CYTOKINE_PRODUCT | 51 | 0.268321 | 1.4160709 | 0.03723404 | 0.15252434 | 1 | 2498 | tags=24%, list=11%, si | MM_4 |
| GOBP_POSITIVE_REGULA | GOBP_POSITIVE_REGULATION_OF_NITRIC_ | 51 | 0.262113 | 1.4142574 | 0.069565214 | 0.15369825 | 1 | 4350 | tags=37%, list=18%, si | MM_4 |
| GOBP_REGULATION_OF_ | GOBP_REGULATION_OF_VESICLE_SIZE | 21 | 0.347596 | 1.4139351 | 0.111524165 | 0.15379149 | 1 | 2814 | tags=24%, list=12%, si | MM_4 |
| GOBP_NEGATIVE_REGUL | GOBP_NEGATIVE_REGULATION_OF_AUTOP | 81 | 0.244443 | 1.4138142 | 0.06741573 | 0.15371494 | 1 | 4846 | tags=40%, list=21%, si | MM_4 |
| GOBP_NEGATIVE_REGUL | GOBP_NEGATIVE_REGULATION_OF_AMYLO | 20 | 0.357337 | 1.4135839 | 0.097122304 | 0.15377176 | 1 | 2790 | tags=30%, list=12%, si | MM_4 |
| GOBP_MACROPHAGE_AC | GOBP_MACROPHAGE_ACTIVATION | 107 | 0.227199 | 1.4131465 | 0.036585364 | 0.15391424 | 1 | 3681 | tags=28%, list=16%, si | MM_4 |
| GOBP_FOAM_CELL_DIFFE | GOBP_FOAM_CELL_DIFFERENTIATION | 32 | 0.297981 | 1.4114754 | 0.061904762 | 0.15500525 | 1 | 3528 | tags=38%, list=15%, si | MM_4 |
| GOBP_REGULATION_OF_ | GOBP_REGULATION_OF_T_CELL_CYTOKINE_ | 36 | 0.286633 | 1.4113486 | 0.06511628 | 0.15493566 | 1 | 3882 | tags=31%, list=16%, si | MM_4 |
| GOBP_RESPONSE_TO_CA | GOBP_RESPONSE_TO_CADMIUM_ION | 35 | 0.299367 | 1.4111803 | 0.08695652 | 0.15489677 | 1 | 3368 | tags=40%, list=14%, si | MM_4 |
| GOBP_REGULATION_OF_ | GOBP_REGULATION_OF_CYTOKINE_PRODU | 122 | 0.221124 | 1.4111363 | 0.0064935065 | 0.15475206 | 1 | 2498 | tags=20%, list=11%, si | MM_4 |
| GOBP_POSITIVE_REGULA | GOBP_POSITIVE_REGULATION_OF_PROTEAS | 72 | 0.243619 | 1.4107919 | 0.04494382 | 0.15487939 | 1 | 3551 | tags=28%, list=15%, si | MM_4 |
| GOBP_NEGATIVE_REGUL | GOBP_NEGATIVE_REGULATION_OF_TYPE_I_ | 42 | 0.275452 | 1.4107854 | 0.07359307 | 0.1547195 | 1 | 3157 | tags=29%, list=13%, si | MM_4 |
| GOBP_MITOCHONDRIAL_ | GOBP_MITOCHONDRIAL_DNA_METABOLIC_ | 26 | 0.330333 | 1.4100865 | 0.091240875 | 0.1551177 | 1 | 3556 | tags=38%, list=15%, si | MM_4 |
| GOBP_ALPHA_BETA_T_CI | GOBP_ALPHA_BETA_T_CELL_ACTIVATION | 191 | 0.196039 | 1.4098165 | 0 | 0.15518345 | 1 | 2850 | tags=18%, list=12%, si | MM_4 |
| GOBP_REGULATION_OF_ | GOBP_REGULATION_OF_PRODUCTION_OF_ | 195 | 0.202629 | 1.4073344 | 0.012345679 | 0.15691847 | 1 | 3953 | tags=28%, list=17%, si | MM_4 |
| GOBP_LYMPHOCYTE_ACT | GOBP_LYMPHOCYTE_ACTIVATION_INVOLVE | 202 | 0.200715 | 1.4073157 | 0 | 0.15677205 | 1 | 2685 | tags=18%, list=11%, si | MM_4 |
| GOBP_REGULATION_OF_ | GOBP_REGULATION_OF_PROTEIN_CATABOL | 346 | 0.18003 | 1.4071752 | 0 | 0.1567211 | 1 | 2718 | tags=18%, list=12%, si | MM_4 |
| GOBP_MAST_CELL_ACTIV | GOBP_MAST_CELL_ACTIVATION_INVOLVED | 56 | 0.265032 | 1.4068733 | 0.04950495 | 0.1567953 | 1 | 3842 | tags=29%, list=16%, si | MM_4 |
| TH17_CELL_DIFFERENTIA | TH17_CELL_DIFFERENTIATION | 94 | 0.235951 | 1.4064454 | 0.056497175 | 0.15693837 | 1 | 2693 | tags=19%, list=11%, si | MM_4 |
| GOBP_TOLL_LIKE_RECEPT | GOBP_TOLL_LIKE_RECEPTOR_2_SIGNALING | 17 | 0.364365 | 1.4059565 | 0.12244898 | 0.1571012 | 1 | 4919 | tags=53%, list=21%, si | MM_4 |
| GOBP_ESTABLISHMENT_( | GOBP_ESTABLISHMENT_OF_BLOOD_BRAIN_ | 17 | 0.370019 | 1.4057162 | 0.088435374 | 0.15710436 | 1 | 1134 | tags=18%, list=5%, sig | MM_4 |
| GOBP_MITOCHONDRIAL_ | GOBP_MITOCHONDRIAL_GENOME_MAINTE | 35 | 0.301929 | 1.4056671 | 0.088 | 0.15697876 | 1 | 4097 | tags=37%, list=17%, si | MM_4 |
| GOBP_ESTABLISHMENT_( | GOBP_ESTABLISHMENT_OF_PROTEIN_LOCA | 262 | 0.190664 | 1.404938 | 0 | 0.15741411 | 1 | 3210 | tags=21%, list=14%, si | MM_4 |
| GOBP_REGULATION_OF_ | GOBP_REGULATION_OF_LYMPHOCYTE_APO | 71 | 0.243579 | 1.4040653 | 0.044554457 | 0.15793967 | 1 | 1611 | tags=14%, list=7%, sig | MM_4 |
| GOBP_PROSTATE_GLAND | GOBP_PROSTATE_GLAND_MORPHOGENESIS | 30 | 0.308289 | 1.4040118 | 0.07826087 | 0.15783826 | 1 | 14591 | tags=93%, list=62%, si | MM_4 |
| GOBP_POSITIVE_REGULA | GOBP_POSITIVE_REGULATION_OF_APOPTO | 161 | 0.210648 | 1.4029689 | 0.032786883 | 0.15847966 | 1 | 2656 | tags=17%, list=11%, si | MM_4 |
| GOBP_LIPID_MODIFICATI | GOBP_LIPID_MODIFICATION | 197 | 0.196504 | 1.4026401 | 0 | 0.15853715 | 1 | 3328 | tags=25%, list=14%, si | MM_4 |
| GOBP_INTRINSIC_APOPT( | GOBP_INTRINSIC_APOPTOTIC_SIGNALING_F | 112 | 0.220023 | 1.4025413 | 0.026666667 | 0.15844038 | 1 | 11575 | tags=84%, list=49%, si | MM_4 |
| GOBP_EPITHELIAL_TO_M | GOBP_EPITHELIAL_TO_MESENCHYMAL_TRA | 16 | 0.370656 | 1.4015952 | 0.12237762 | 0.15898518 | 1 | 14865 | tags=100%, list=63%, | MM_4 |
| GOBP_POSITIVE_REGULA | GOBP_POSITIVE_REGULATION_OF_MYELOID | 71 | 0.244884 | 1.40095 | 0.06878307 | 0.15926462 | 1 | 3769 | tags=28%, list=16%, si | MM_4 |
| GOBP_REGULATION_OF_ | GOBP_REGULATION_OF_CELLULAR_CATABO | 391 | 0.181585 | 1.4008589 | 0 | 0.15917726 | 1 | 3258 | tags=22%, list=14%, si | MM_4 |
| GOBP_GLYCOSYL_COMPO | GOBP_GLYCOSYL_COMPOUND_CATABOLIC_ | 26 | 0.325385 | 1.4005417 | 0.09448819 | 0.15923862 | 1 | 3189 | tags=38%, list=14%, si | MM_4 |
| GOBP_RESPONSE_TO_GA | GOBP_RESPONSE_TO_GAMMA_RADIATION | 48 | 0.273805 | 1.4004503 | 0.066079296 | 0.15912184 | 1 | 4867 | tags=46%, list=21%, si | MM_4 |
| GOBP_REGULATION_OF_ | GOBP_REGULATION_OF_SYNCYTIUM_FORM | 31 | 0.308366 | 1.4000673 | 0.115079366 | 0.15926477 | 1 | 3769 | tags=32%, list=16%, si | MM_4 |
| PI3K-AKT_SIGNALING_PA | PI3K-AKT_SIGNALING_PATHWAY | 324 | 0.186379 | 1.3994226 | 0 | 0.15960135 | 1 | 1518 | tags=11%, list=6%, sig | MM_4 |
| GOBP_NEUTROPHIL_ACTI | GOBP_NEUTROPHIL_ACTIVATION_INVOLVE | 18 | 0.360873 | 1.3993825 | 0.09665427 | 0.15946516 | 1 | 2791 | tags=33%, list=12%, si | MM_4 |
| GOBP_RESPONSE_TO_IOI | GOBP_RESPONSE_TO_IONIZING_RADIATION | 127 | 0.211517 | 1.3993589 | 0.021428572 | 0.1593144 | 1 | 2424 | tags=20%, list=10%, si | MM_4 |
| GOBP_REGULATION_OF_ | GOBP_REGULATION_OF_MACROAUTOPHAG | 129 | 0.217269 | 1.3986257 | 0.04225352 | 0.15966198 | 1 | 3258 | tags=22%, list=14%, si | MM_4 |
| GOBP_ANOIKIS | GOBP_ANOIKIS | 31 | 0.304654 | 1.3979839 | 0.08778626 | 0.15995927 | 1 | 3662 | tags=39%, list=16%, si | MM_4 |
| GOBP_REGULATION_OF_ | GOBP_REGULATION_OF_NATURAL_KILLER_( | 46 | 0.282435 | 1.395562 | 0.07692308 | 0.1617573 | 1 | 3616 | tags=35%, list=15%, si | MM_4 |
| GOBP_BASE_EXCISION_R | GOBP_BASE_EXCISION_REPAIR | 41 | 0.287433 | 1.3955299 | 0.09090909 | 0.16160984 | 1 | 3800 | tags=32%, list=16%, si | MM_4 |
| GOBP_CELL_SURFACE_RE | GOBP_CELL_SURFACE_RECEPTOR_PROTEIN_ | 348 | 0.186822 | 1.3955041 | 0.023255814 | 0.16146974 | 1 | 3223 | tags=20%, list=14%, si | MM_4 |
| GOBP_PROTEIN_K11_LIN | GOBP_PROTEIN_K11_LINKED_UBIQUITINAT | 29 | 0.315313 | 1.3952273 | 0.06910569 | 0.16152762 | 1 | 4647 | tags=45%, list=20%, si | MM_4 |
| GOBP_INFLAMMATORY_I | GOBP_INFLAMMATORY_RESPONSE_TO_ANT | 62 | 0.247188 | 1.394604 | 0.07106599 | 0.16184768 | 1 | 3042 | tags=27%, list=13%, si | MM_4 |
| GOBP_NEGATIVE_REGUL | GOBP_NEGATIVE_REGULATION_OF_PHOSPI | 332 | 0.186286 | 1.3945042 | 0.015873017 | 0.16175158 | 1 | 3364 | tags=23%, list=14%, si | MM_4 |
| GOBP_REGULATION_OF_ | GOBP_REGULATION_OF_GOLGI_ORGANIZA | 17 | 0.361221 | 1.3934219 | 0.10996564 | 0.16249378 | 1 | 15088 | tags=100%, list=64%, | MM_4 |
| GOBP_REGULATION_OF_ | GOBP_REGULATION_OF_AUTOPHAGOSOME | 44 | 0.278906 | 1.3928511 | 0.043902438 | 0.16276215 | 1 | 11265 | tags=89%, list=48%, si | MM_4 |
| GOBP_GLYCEROLIPID_ME | GOBP_GLYCEROLIPID_METABOLIC_PROCESS | 361 | 0.180834 | 1.3922127 | 0 | 0.16313232 | 1 | 3377 | tags=22%, list=14%, si | MM_4 |
| GOBP_NEGATIVE_REGUL | GOBP_NEGATIVE_REGULATION_OF_CALCIU | 51 | 0.258324 | 1.3919494 | 0.050632913 | 0.16319071 | 1 | 3839 | tags=31%, list=16%, si | MM_4 |
| GOBP_REGULATION_OF_ | GOBP_REGULATION_OF_CELLULAR_RESPON | 308 | 0.188947 | 1.3908291 | 0 | 0.1640072 | 1 | 3223 | tags=20%, list=14%, si | MM_4 |
| GOBP_CHONDROITIN_SU | GOBP_CHONDROITIN_SULFATE_METABOLIC | 22 | 0.340297 | 1.3893801 | 0.09090909 | 0.16497208 | 1 | 2019 | tags=27%, list=9%, sig | MM_4 |
| GOBP_AMYLOID_PRECUR | GOBP_AMYLOID_PRECURSOR_PROTEIN_ME | 75 | 0.241616 | 1.3890362 | 0.036842104 | 0.1650909 | 1 | 2833 | tags=23%, list=12%, si | MM_4 |
| GOBP_NEGATIVE_REGUL | GOBP_NEGATIVE_REGULATION_OF_DOUBL | 24 | 0.327004 | 1.3883474 | 0.0875 | 0.16549444 | 1 | 3586 | tags=38%, list=15%, si | MM_4 |
| GOBP_DENDRITIC_CELL_I | GOBP_DENDRITIC_CELL_DIFFERENTIATION | 42 | 0.271163 | 1.3876133 | 0.08888889 | 0.16585392 | 1 | 1522 | tags=14%, list=6%, sig | MM_4 |
| GOBP_NUCLEOSIDE_DIPH | GOBP_NUCLEOSIDE_DIPHOSPHATE_METAB( | 127 | 0.209826 | 1.3875793 | 0.03125 | 0.16570646 | 1 | 3774 | tags=28%, list=16%, si | MM_4 |

| | | | | | | | | | | |
|---|---|---|---|---|---|---|---|---|---|---|
| GOBP_REGULATION_OF_ | GOBP_REGULATION_OF_HYDROGEN_PERO | 17 | 0.365713 | 1.387519 | 0.118971065 | 0.16557649 | 1 | 14982 | tags=100%, list=63%, | MM_4 |
| GOBP_RESPONSE_TO_DS | GOBP_RESPONSE_TO_DSRNA | 43 | 0.263426 | 1.386552 | 0.074418604 | 0.16622448 | 1 | 2693 | tags=26%, list=11%, si | MM_4 |
| GOBP_NEGATIVE_REGUL | GOBP_NEGATIVE_REGULATION_OF_TELOM | 27 | 0.319691 | 1.3835943 | 0.094339624 | 0.16847192 | 1 | 3939 | tags=41%, list=17%, si | MM_4 |
| GOBP_REGULATION_OF_ | GOBP_REGULATION_OF_EXTRINSIC_APOPT | 161 | 0.199348 | 1.382298 | 0.015037594 | 0.16937532 | 1 | 2556 | tags=17%, list=11%, si | MM_4 |
| GOBP_REGULATION_OF_ | GOBP_REGULATION_OF_STEM_CELL_POPU | 74 | 0.239693 | 1.3822963 | 0.05172414 | 0.16920628 | 1 | 5167 | tags=41%, list=22%, si | MM_4 |
| GOBP_ACUTE_INFLAMM | GOBP_ACUTE_INFLAMMATORY_RESPONSE | 97 | 0.222096 | 1.3809624 | 0.027624309 | 0.16996263 | 1 | 1747 | tags=16%, list=7%, sig | MM_4 |
| PROTEOGLYCANS_IN_CA | PROTEOGLYCANS_IN_CANCER | 193 | 0.198493 | 1.3809543 | 0.029126214 | 0.16979335 | 1 | 2693 | tags=18%, list=11%, si | MM_4 |
| GOBP_REGULATION_OF_ | GOBP_REGULATION_OF_VIRAL_GENOME_R | 78 | 0.242466 | 1.3799518 | 0.037837837 | 0.1705481 | 1 | 3592 | tags=27%, list=15%, si | MM_4 |
| GOBP_REGULATION_OF_ | GOBP_REGULATION_OF_CELLULAR_RESPON | 26 | 0.313449 | 1.378897 | 0.115226336 | 0.17125109 | 1 | 5313 | tags=50%, list=23%, si | MM_4 |
| GOBP_NEGATIVE_REGUL | GOBP_NEGATIVE_REGULATION_OF_TRANSL | 117 | 0.219056 | 1.3786119 | 0.03448276 | 0.17131442 | 1 | 3982 | tags=30%, list=17%, si | MM_4 |
| GOBP_REGULATION_OF_ | GOBP_REGULATION_OF_PROTEIN_TARGETI | 20 | 0.348096 | 1.3784999 | 0.12 | 0.17122109 | 1 | 1965 | tags=25%, list=8%, sig | MM_4 |
| GOBP_VESICLE_TRANSPO | GOBP_VESICLE_TRANSPORT_ALONG_MICR | 47 | 0.275374 | 1.3767685 | 0.08465608 | 0.17250545 | 1 | 2398 | tags=23%, list=10%, si | MM_4 |
| GOBP_ER_NUCLEUS_SIG | GOBP_ER_NUCLEUS_SIGNALING_PATHWAY | 43 | 0.271089 | 1.3764415 | 0.070754714 | 0.17257722 | 1 | 4384 | tags=35%, list=19%, si | MM_4 |
| GOBP_REGULATION_OF_ | GOBP_REGULATION_OF_GLIAL_CELL_MIGR | 26 | 0.317016 | 1.376437 | 0.09615385 | 0.17241594 | 1 | 3955 | tags=31%, list=17%, si | MM_4 |
| GOBP_NEGATIVE_REGUL | GOBP_NEGATIVE_REGULATION_OF_CELL_S | 16 | 0.37294 | 1.3758012 | 0.12631579 | 0.17286295 | 1 | 2387 | tags=31%, list=10%, si | MM_4 |
| GOBP_ACTIN_FILAMENT_ | GOBP_ACTIN_FILAMENT_ORGANIZATION | 450 | 0.172884 | 1.3745756 | 0 | 0.17362553 | 1 | 2698 | tags=17%, list=11%, si | MM_4 |
| GOBP_INTRACELLULAR_P | GOBP_INTRACELLULAR_PH_REDUCTION | 48 | 0.26367 | 1.3744913 | 0.092307694 | 0.17352943 | 1 | 4343 | tags=42%, list=18%, si | MM_4 |
| GOBP_REGULATION_OF_ | GOBP_REGULATION_OF_VIRAL_LIFE_CYCLE | 139 | 0.208019 | 1.3741823 | 0.0234375 | 0.17362301 | 1 | 3592 | tags=24%, list=15%, si | MM_4 |
| GOBP_POSITIVE_REGULA | GOBP_POSITIVE_REGULATION_OF_MACRO | 15 | 0.388354 | 1.373953 | 0.10526316 | 0.17367598 | 1 | 14447 | tags=100%, list=61%, | MM_4 |
| GOBP_ANTEROGRADE_A | GOBP_ANTEROGRADE_AXONAL_TRANSPOR | 53 | 0.253367 | 1.3736312 | 0.08520179 | 0.17373434 | 1 | 2686 | tags=25%, list=11%, si | MM_4 |
| GOBP_RESPONSE_TO_IN | GOBP_RESPONSE_TO_INTERLEUKIN_1 | 76 | 0.236419 | 1.3734725 | 0.03846154 | 0.17368159 | 1 | 3270 | tags=26%, list=14%, si | MM_4 |
| GOBP_PROTEIN_PROCES | GOBP_PROTEIN_PROCESSING | 237 | 0.191994 | 1.3730978 | 0.012345679 | 0.17385592 | 1 | 2989 | tags=19%, list=13%, si | MM_4 |
| GOBP_NEGATIVE_REGUL | GOBP_NEGATIVE_REGULATION_OF_CALCIU | 69 | 0.24366 | 1.3724134 | 0.06382979 | 0.1742893 | 1 | 2756 | tags=23%, list=12%, si | MM_4 |
| GOBP_PROTEIN_MATUR | GOBP_PROTEIN_MATURATION_BY_IRON_S | 16 | 0.37139 | 1.3720299 | 0.13357401 | 0.174481 | 1 | 2754 | tags=38%, list=12%, si | MM_4 |
| GOBP_NEGATIVE_REGUL | GOBP_NEGATIVE_REGULATION_OF_ANOIKI | 15 | 0.37064 | 1.3717308 | 0.115384616 | 0.17451812 | 1 | 14865 | tags=100%, list=63%, | MM_4 |
| GOBP_RESPONSE_TO_NU | GOBP_RESPONSE_TO_NUTRIENT | 75 | 0.241828 | 1.3715082 | 0.10059172 | 0.1745374 | 1 | 3163 | tags=21%, list=13%, si | MM_4 |
| GOBP_INSULIN_LIKE_GR | GOBP_INSULIN_LIKE_GROWTH_FACTOR_RE | 49 | 0.259767 | 1.3706337 | 0.0625 | 0.17512618 | 1 | 2693 | tags=24%, list=11%, si | MM_4 |
| GOBP_PIGMENTATION | GOBP_PIGMENTATION | 112 | 0.21157 | 1.3701924 | 0.042424243 | 0.17527647 | 1 | 1692 | tags=13%, list=7%, sig | MM_4 |
| GOBP_POSITIVE_REGULA | GOBP_POSITIVE_REGULATION_OF_STRESS_ | 24 | 0.325737 | 1.3701662 | 0.115523465 | 0.17513621 | 1 | 2220 | tags=25%, list=9%, sig | MM_4 |
| GOBP_NEGATIVE_REGUL | GOBP_NEGATIVE_REGULATION_OF_MACRO | 33 | 0.3042 | 1.3701384 | 0.082987554 | 0.17498907 | 1 | 3163 | tags=27%, list=13%, si | MM_4 |
| GOBP_NEGATIVE_REGUL | GOBP_NEGATIVE_REGULATION_OF_CELL_J | 35 | 0.287127 | 1.3701348 | 0.09170306 | 0.17481886 | 1 | 2387 | tags=17%, list=10%, si | MM_4 |
| GOBP_VESICLE_TARGETI | GOBP_VESICLE_TARGETING_TO_FROM_OR_ | 29 | 0.297842 | 1.370058 | 0.10373444 | 0.1747072 | 1 | 4480 | tags=38%, list=19%, si | MM_4 |
| GOBP_VESICLE_CYTOSKE | GOBP_VESICLE_CYTOSKELETAL_TRAFFICKIN | 61 | 0.252929 | 1.3690209 | 0.058252428 | 0.17538087 | 1 | 2398 | tags=21%, list=10%, si | MM_4 |
| GOBP_POSITIVE_REGULA | GOBP_POSITIVE_REGULATION_OF_CELL_M | 61 | 0.249125 | 1.3679001 | 0.08743169 | 0.17614157 | 1 | 3583 | tags=23%, list=15%, si | MM_4 |
| GOBP_ENDOSOME_ORG | GOBP_ENDOSOME_ORGANIZATION | 68 | 0.24606 | 1.3676591 | 0.06358381 | 0.17615429 | 1 | 11525 | tags=87%, list=49%, si | MM_4 |
| GOBP_POSITIVE_REGULA | GOBP_POSITIVE_REGULATION_OF_MEMBR | 39 | 0.273547 | 1.3661464 | 0.08796296 | 0.1772623 | 1 | 12552 | tags=90%, list=53%, si | MM_4 |
| GOBP_RETROGRADE_AX | GOBP_RETROGRADE_AXONAL_TRANSPORT | 24 | 0.315072 | 1.3651851 | 0.08646616 | 0.17788441 | 1 | 2445 | tags=21%, list=10%, si | MM_4 |
| GOBP_NEGATIVE_REGUL | GOBP_NEGATIVE_REGULATION_OF_ANDRO | 15 | 0.379056 | 1.364951 | 0.09558824 | 0.17792518 | 1 | 1602 | tags=33%, list=7%, sig | MM_4 |
| GOBP_REGULATION_OF_ | GOBP_REGULATION_OF_APOPTOTIC_SIGNA | 409 | 0.17278 | 1.3649423 | 0 | 0.17776027 | 1 | 2690 | tags=17%, list=11%, si | MM_4 |
| GOBP_NEGATIVE_REGUL | GOBP_NEGATIVE_REGULATION_OF_EPITHE | 159 | 0.205159 | 1.3642399 | 0.028301887 | 0.17821738 | 1 | 3387 | tags=23%, list=14%, si | MM_4 |
| GOBP_POSITIVE_REGULA | GOBP_POSITIVE_REGULATION_OF_PROTEIN | 87 | 0.228885 | 1.3636866 | 0.042682927 | 0.17854102 | 1 | 3740 | tags=25%, list=16%, si | MM_4 |
| GOBP_LIPID_HOMEOSTA | GOBP_LIPID_HOMEOSTASIS | 167 | 0.19459 | 1.3630593 | 0.033333335 | 0.17897417 | 1 | 4029 | tags=30%, list=17%, si | MM_4 |
| GOBP_CHONDROITIN_SU | GOBP_CHONDROITIN_SULFATE_PROTEOGL | 29 | 0.304793 | 1.3629483 | 0.079831935 | 0.17888999 | 1 | 2019 | tags=24%, list=9%, sig | MM_4 |
| GOBP_CELLULAR_RESPO | GOBP_CELLULAR_RESPONSE_TO_INTERLEU | 62 | 0.249675 | 1.362939 | 0.06842105 | 0.17872512 | 1 | 3270 | tags=27%, list=14%, si | MM_4 |
| GOBP_GRANULOCYTE_DI | GOBP_GRANULOCYTE_DIFFERENTIATION | 50 | 0.254857 | 1.3623693 | 0.082417585 | 0.17899704 | 1 | 3364 | tags=30%, list=14%, si | MM_4 |
| GOBP_PRODUCTION_OF_ | GOBP_PRODUCTION_OF_MOLECULAR_MED | 97 | 0.224085 | 1.3620586 | 0.052910052 | 0.1791053 | 1 | 3981 | tags=29%, list=17%, si | MM_4 |
| GOBP_MONOCARBOXYLI | GOBP_MONOCARBOXYLIC_ACID_CATABOLI | 126 | 0.211241 | 1.3615372 | 0.03076923 | 0.17937781 | 1 | 3178 | tags=25%, list=13%, si | MM_4 |
| GOBP_POSITIVE_REGULA | GOBP_POSITIVE_REGULATION_OF_BLOOD_ | 58 | 0.250432 | 1.361222 | 0.06382979 | 0.17950775 | 1 | 958 | tags=14%, list=4%, sig | MM_4 |
| GOBP_POSITIVE_REGULA | GOBP_POSITIVE_REGULATION_OF_MONOC | 21 | 0.332161 | 1.3606029 | 0.11363637 | 0.17994502 | 1 | 3134 | tags=29%, list=13%, si | MM_4 |
| CHAGAS_DISEASE_(AME | CHAGAS_DISEASE_(AMERICAN_TRYPANOSO | 101 | 0.221548 | 1.3605062 | 0.048611112 | 0.17984454 | 1 | 2693 | tags=19%, list=11%, si | MM_4 |
| GOBP_MAST_CELL_ACTI | GOBP_MAST_CELL_ACTIVATION | 77 | 0.236563 | 1.3602085 | 0.05263158 | 0.17998932 | 1 | 3842 | tags=26%, list=16%, si | MM_4 |
| HEPATITIS_C | HEPATITIS_C | 138 | 0.211013 | 1.3595454 | 0.03787879 | 0.18046407 | 1 | 5064 | tags=33%, list=21%, si | MM_4 |
| GOBP_GLYCOSYL_COMP | GOBP_GLYCOSYL_COMPOUND_METABOLIC | 68 | 0.240369 | 1.3583906 | 0.070754714 | 0.18131532 | 1 | 3378 | tags=34%, list=14%, si | MM_4 |
| GOBP_GLYCOSYLCERAMI | GOBP_GLYCOSYLCERAMIDE_METABOLIC_P | 15 | 0.370787 | 1.3575972 | 0.125 | 0.18184376 | 1 | 2704 | tags=40%, list=11%, si | MM_4 |
| GOBP_ARGININE_METAB | GOBP_ARGININE_METABOLIC_PROCESS | 17 | 0.362703 | 1.3575227 | 0.13780919 | 0.18175298 | 1 | 685 | tags=18%, list=3%, sig | MM_4 |
| GOBP_POSITIVE_REGULA | GOBP_POSITIVE_REGULATION_OF_CYSTEIN | 130 | 0.206982 | 1.357497 | 0.04195804 | 0.18160327 | 1 | 3802 | tags=26%, list=16%, si | MM_4 |
| GOBP_NEGATIVE_REGUL | GOBP_NEGATIVE_REGULATION_OF_RECEPT | 28 | 0.298661 | 1.3564444 | 0.08810572 | 0.18231668 | 1 | 14147 | tags=93%, list=60%, si | MM_4 |
| GOBP_REGULATION_OF_ | GOBP_REGULATION_OF_SYNAPTIC_VESICLE | 33 | 0.295598 | 1.3554615 | 0.09795918 | 0.18303123 | 1 | 15281 | tags=97%, list=65%, si | MM_4 |
| GOBP_CELLULAR_KETON | GOBP_CELLULAR_KETONE_METABOLIC_PR | 222 | 0.191242 | 1.3554088 | 0.0121951215 | 0.18292199 | 1 | 1935 | tags=15%, list=8%, sig | MM_4 |
| GOBP_NEGATIVE_REGUL | GOBP_NEGATIVE_REGULATION_OF_MRNA_ | 92 | 0.223445 | 1.3552163 | 0.03649635 | 0.18293425 | 1 | 3287 | tags=23%, list=14%, si | MM_4 |
| GOBP_AMIDE_BIOSYNTH | GOBP_AMIDE_BIOSYNTHETIC_PROCESS | 128 | 0.212545 | 1.3550206 | 0.044444446 | 0.18291394 | 1 | 3497 | tags=28%, list=15%, si | MM_4 |
| GOBP_NATURAL_KILLER_ | GOBP_NATURAL_KILLER_CELL_ACTIVATION | 86 | 0.233453 | 1.3549377 | 0.05263158 | 0.18284419 | 1 | 3452 | tags=28%, list=15%, si | MM_4 |
| UBIQUITIN_MEDIATED_P | UBIQUITIN_MEDIATED_PROTEOLYSIS | 134 | 0.208917 | 1.3547455 | 0.020408163 | 0.18283014 | 1 | 4686 | tags=33%, list=20%, si | MM_4 |
| GOBP_REGULATION_OF_ | GOBP_REGULATION_OF_OXIDATIVE_PHOSP | 23 | 0.33163 | 1.354395 | 0.12554112 | 0.18301374 | 1 | 3638 | tags=30%, list=15%, si | MM_4 |
| INTESTINAL_IMMUNE_N | INTESTINAL_IMMUNE_NETWORK_FOR_IGA_ | 37 | 0.277761 | 1.3542782 | 0.08267716 | 0.18295902 | 1 | 1931 | tags=24%, list=8%, sig | MM_4 |
| GOBP_MONOSACCHARID | GOBP_MONOSACCHARIDE_CATABOLIC_PR | 40 | 0.271118 | 1.3535448 | 0.074418604 | 0.18339884 | 1 | 3708 | tags=38%, list=16%, si | MM_4 |
| GOBP_ESTABLISHMENT_ | GOBP_ESTABLISHMENT_OF_PROTEIN_LOCA | 19 | 0.342482 | 1.3532513 | 0.1024735 | 0.18348011 | 1 | 3577 | tags=32%, list=15%, si | MM_4 |
| GOBP_LIPID_STORAGE | GOBP_LIPID_STORAGE | 97 | 0.217589 | 1.3532472 | 0.06666667 | 0.18331008 | 1 | 3037 | tags=24%, list=13%, si | MM_4 |
| GOBP_PHOSPHATIDYLIN | GOBP_PHOSPHATIDYLINOSITOL_BIOSYNTHE | 95 | 0.223436 | 1.3513278 | 0.061349694 | 0.18491556 | 1 | 3370 | tags=27%, list=14%, si | MM_4 |
| GOBP_NEGATIVE_REGUL | GOBP_NEGATIVE_REGULATION_OF_LYMPH | 62 | 0.240689 | 1.3509398 | 0.085 | 0.1850937 | 1 | 347 | tags=8%, list=1%, sign | MM_4 |
| GOBP_REGULATION_OF_ | GOBP_REGULATION_OF_RRNA_PROCESSIN | 16 | 0.374615 | 1.3507577 | 0.13148789 | 0.1850738 | 1 | 11784 | tags=94%, list=50%, si | MM_4 |
| GOBP_POSITIVE_REGULA | GOBP_POSITIVE_REGULATION_OF_PROTEIN | 100 | 0.216847 | 1.3499576 | 0.0729927 | 0.18557832 | 1 | 2693 | tags=19%, list=11%, si | MM_4 |
| GOBP_POSITIVE_REGULA | GOBP_POSITIVE_REGULATION_OF_SUBSTR | 40 | 0.273558 | 1.3498557 | 0.10152284 | 0.18548505 | 1 | 14726 | tags=95%, list=62%, si | MM_4 |
| GOBP_ACETYL_COA_MET | GOBP_ACETYL_COA_METABOLIC_PROCESS | 34 | 0.287277 | 1.3493687 | 0.09053498 | 0.18573308 | 1 | 3601 | tags=41%, list=15%, si | MM_4 |
| GOBP_TELENCEPHALON_ | GOBP_TELENCEPHALON_GLIAL_CELL_MIGR | 27 | 0.306407 | 1.3485808 | 0.11885246 | 0.18634729 | 1 | 2076 | tags=26%, list=9%, sig | MM_4 |
| AMOEBIASIS | AMOEBIASIS | 89 | 0.22209 | 1.3480387 | 0.06936416 | 0.18665719 | 1 | 3031 | tags=25%, list=13%, si | MM_4 |
| GOBP_GLUTATHIONE_M | GOBP_GLUTATHIONE_METABOLIC_PROCES | 58 | 0.248036 | 1.3478963 | 0.08629441 | 0.18661834 | 1 | 3386 | tags=33%, list=14%, si | MM_4 |
| GOBP_BEHAVIORAL_RES | GOBP_BEHAVIORAL_RESPONSE_TO_PAIN | 29 | 0.30401 | 1.347552 | 0.13944224 | 0.18678924 | 1 | 607 | tags=10%, list=3%, sig | MM_4 |
| GOBP_POSITIVE_REGULA | GOBP_POSITIVE_REGULATION_OF_NUCLEO | 74 | 0.230738 | 1.3473629 | 0.06010929 | 0.18678275 | 1 | 4462 | tags=32%, list=19%, si | MM_4 |
| GOBP_CHAPERONE_MED | GOBP_CHAPERONE_MEDIATED_PROTEIN_C | 18 | 0.346741 | 1.3469254 | 0.12142857 | 0.1869813 | 1 | 15430 | tags=100%, list=65%, | MM_4 |
| GOBP_NEGATIVE_REGUL | GOBP_NEGATIVE_REGULATION_OF_AMIDE_ | 22 | 0.320869 | 1.3461092 | 0.101886794 | 0.18755966 | 1 | 1906 | tags=23%, list=8%, sig | MM_4 |
| GOBP_REGULATION_OF_ | GOBP_REGULATION_OF_RESPONSE_TO_NU | 34 | 0.286674 | 1.3444675 | 0.11792453 | 0.18897612 | 1 | 1891 | tags=18%, list=8%, sig | MM_4 |

| | | | | | | | | | | |
|---|---|---|---|---|---|---|---|---|---|---|
| GOBP_TRIGLYCERIDE_BIC | GOBP_TRIGLYCERIDE_BIOSYNTHETIC_PROCE | 46 | 0.266996 | 1.3443636 | 0.1 | 0.18889731 | 1 | 3037 | tags=28%, list=13%, si | MM_4 |
| GOBP_MAINTENANCE_O | GOBP_MAINTENANCE_OF_PROTEIN_LOCAL | 43 | 0.261546 | 1.3443311 | 0.071428575 | 0.18874021 | 1 | 2885 | tags=23%, list=12%, si | MM_4 |
| GOBP_RESPONSE_TO_ST/ | GOBP_RESPONSE_TO_STARVATION | 181 | 0.190624 | 1.3437188 | 0.021505376 | 0.18907191 | 1 | 3652 | tags=26%, list=15%, si | MM_4 |
| GOBP_REGULATION_OF_ | GOBP_REGULATION_OF_GLUCOSE_TRANSM | 84 | 0.225988 | 1.3432226 | 0.042424243 | 0.18937594 | 1 | 2815 | tags=20%, list=12%, si | MM_4 |
| PANTOTHENATE_AND_C( | PANTOTHENATE_AND_COA_BIOSYNTHESIS | 17 | 0.352333 | 1.3423905 | 0.1220339 | 0.18998438 | 1 | 1220 | tags=24%, list=5%, sig | MM_4 |
| GOBP_RECEPTOR_CATAB | GOBP_RECEPTOR_CATABOLIC_PROCESS | 29 | 0.30455 | 1.3413892 | 0.12307692 | 0.19070561 | 1 | 12463 | tags=90%, list=53%, si | MM_4 |
| GOBP_POSITIVE_REGULA | GOBP_POSITIVE_REGULATION_OF_DNA_RE | 77 | 0.227386 | 1.341143 | 0.07027027 | 0.1907133 | 1 | 2566 | tags=18%, list=11%, si | MM_4 |
| GOBP_POSITIVE_REGULA | GOBP_POSITIVE_REGULATION_OF_B_CELL_ | 18 | 0.346915 | 1.340742 | 0.14693877 | 0.19092709 | 1 | 3900 | tags=50%, list=17%, si | MM_4 |
| GOBP_MOTOR_BEHAVIO | GOBP_MOTOR_BEHAVIOR | 32 | 0.288449 | 1.3403991 | 0.112 | 0.19109121 | 1 | 4242 | tags=41%, list=18%, si | MM_4 |
| GOBP_AXONAL_TRANSP( | GOBP_AXONAL_TRANSPORT | 69 | 0.239238 | 1.3396059 | 0.06896552 | 0.19164397 | 1 | 2686 | tags=22%, list=11%, si | MM_4 |
| GOBP_LIPID_OXIDATION | GOBP_LIPID_OXIDATION | 118 | 0.210355 | 1.3393685 | 0.044871796 | 0.19170325 | 1 | 3178 | tags=26%, list=13%, si | MM_4 |
| GOBP_LOCOMOTORY_EX | GOBP_LOCOMOTORY_EXPLORATION_BEHA\ | 19 | 0.345489 | 1.3390436 | 0.16853933 | 0.191847 | 1 | 3730 | tags=42%, list=16%, si | MM_4 |
| GOBP_POSITIVE_REGULA | GOBP_POSITIVE_REGULATION_OF_ALPHA_E | 27 | 0.306851 | 1.3379239 | 0.15510204 | 0.19261976 | 1 | 1641 | tags=19%, list=7%, sig | MM_4 |
| GOBP_POST_TRANSLATIC | GOBP_POST_TRANSLATIONAL_PROTEIN_TA | 18 | 0.344273 | 1.3375099 | 0.101123594 | 0.19286394 | 1 | 10945 | tags=89%, list=46%, si | MM_4 |
| GOBP_PROTEIN_POLYME | GOBP_PROTEIN_POLYMERIZATION | 269 | 0.180161 | 1.3374301 | 0.01010101 | 0.19275084 | 1 | 3905 | tags=24%, list=17%, si | MM_4 |
| GOBP_POSITIVE_REGULA | GOBP_POSITIVE_REGULATION_OF_DNA_DA | 15 | 0.367178 | 1.3366045 | 0.13993174 | 0.19340949 | 1 | 4519 | tags=47%, list=19%, si | MM_4 |
| GOBP_ENUCLEATE_ERYTH | GOBP_ENUCLEATE_ERYTHROCYTE_DIFFEREI | 15 | 0.374091 | 1.3363227 | 0.13240418 | 0.19344561 | 1 | 5017 | tags=53%, list=21%, si | MM_4 |
| GOBP_ACTIVATED_T_CEL | GOBP_ACTIVATED_T_CELL_PROLIFERATION | 50 | 0.245007 | 1.3359995 | 0.08080808 | 0.19360486 | 1 | 2986 | tags=24%, list=13%, si | MM_4 |
| GOBP_PROTEIN_PEPTIDY | GOBP_PROTEIN_PEPTIDYL_PROLYL_ISOMER | 25 | 0.310596 | 1.3359454 | 0.105932206 | 0.19347206 | 1 | 4272 | tags=32%, list=18%, si | MM_4 |
| PHOSPHATIDYLINOSITOL_ | PHOSPHATIDYLINOSITOL_SIGNALING_SYSTE | 94 | 0.222616 | 1.3348471 | 0.080213904 | 0.1943214 | 1 | 3776 | tags=30%, list=16%, si | MM_4 |
| GOBP_NEGATIVE_REGUL | GOBP_NEGATIVE_REGULATION_OF_CANON | 57 | 0.251126 | 1.3346145 | 0.098445594 | 0.19436598 | 1 | 2516 | tags=23%, list=11%, si | MM_4 |
| THYROID_HORMONE_SIG | THYROID_HORMONE_SIGNALING_PATHWA\ | 113 | 0.213987 | 1.3336487 | 0.08552632 | 0.19499305 | 1 | 3490 | tags=27%, list=15%, si | MM_4 |
| GOBP_ESTABLISHMENT_( | GOBP_ESTABLISHMENT_OF_PROTEIN_LOCA | 20 | 0.331897 | 1.3336283 | 0.12765957 | 0.19483577 | 1 | 1987 | tags=25%, list=8%, sig | MM_4 |
| MEASLES | MEASLES | 128 | 0.206992 | 1.333251 | 0.055555556 | 0.19502169 | 1 | 2662 | tags=20%, list=11%, si | MM_4 |
| GOBP_MYELOID_CELL_DE | GOBP_MYELOID_CELL_DEVELOPMENT | 98 | 0.218686 | 1.3330051 | 0.0647482 | 0.19509023 | 1 | 4619 | tags=33%, list=20%, si | MM_4 |
| GOBP_G0_TO_G1_TRANS | GOBP_G0_TO_G1_TRANSITION | 38 | 0.268264 | 1.332807 | 0.11374407 | 0.19511625 | 1 | 5256 | tags=45%, list=22%, si | MM_4 |
| GOBP_CELL_DIFFERENTIA | GOBP_CELL_DIFFERENTIATION_INVOLVED_I | 30 | 0.305912 | 1.3323882 | 0.11885246 | 0.1953104 | 1 | 2555 | tags=27%, list=11%, si | MM_4 |
| GOBP_REGULATION_OF_ | GOBP_REGULATION_OF_MITOCHONDRIAL_ | 87 | 0.227429 | 1.331928 | 0.093333334 | 0.1955565 | 1 | 3798 | tags=28%, list=16%, si | MM_4 |
| GOBP_PROTEIN_LOCALIZ | GOBP_PROTEIN_LOCALIZATION_TO_ENDOP | 76 | 0.229774 | 1.33191 | 0.08 | 0.19540584 | 1 | 4158 | tags=30%, list=18%, si | MM_4 |
| GOBP_LIPID_IMPORT_IN | GOBP_LIPID_IMPORT_INTO_CELL | 24 | 0.317495 | 1.3283418 | 0.09338521 | 0.1986887 | 1 | 2743 | tags=25%, list=12%, si | MM_4 |
| GOBP_MITOTIC_DNA_IN | GOBP_MITOTIC_DNA_INTEGRITY_CHECKPO | 84 | 0.220094 | 1.3280907 | 0.04819277 | 0.19875638 | 1 | 2926 | tags=24%, list=12%, si | MM_4 |
| GOBP_CELLULAR_RESPOI | GOBP_CELLULAR_RESPONSE_TO_OXYGEN_I | 20 | 0.32742 | 1.3275983 | 0.11673152 | 0.19905703 | 1 | 4376 | tags=45%, list=19%, si | MM_4 |
| GOBP_REGULATION_OF_ | GOBP_REGULATION_OF_PHOSPHOLIPID_TR | 15 | 0.363845 | 1.3275938 | 0.13915858 | 0.1988846 | 1 | 1977 | tags=33%, list=8%, sig | MM_4 |
| GOBP_POSITIVE_REGULA | GOBP_POSITIVE_REGULATION_OF_LYMPHO | 21 | 0.326865 | 1.3275051 | 0.107407406 | 0.19880694 | 1 | 613 | tags=14%, list=3%, sig | MM_4 |
| GOBP_T_HELPER_2_CELL | GOBP_T_HELPER_2_CELL_CYTOKINE_PRODU | 17 | 0.345183 | 1.3267946 | 0.13978495 | 0.19932015 | 1 | 3882 | tags=41%, list=16%, si | MM_4 |
| GOBP_POSITIVE_REGULA | GOBP_POSITIVE_REGULATION_OF_T_CELL_ | 66 | 0.236851 | 1.3265568 | 0.08465608 | 0.19937961 | 1 | 3561 | tags=26%, list=15%, si | MM_4 |
| GOBP_NEGATIVE_REGUL | GOBP_NEGATIVE_REGULATION_OF_CYTOKI | 34 | 0.288468 | 1.3265043 | 0.10236221 | 0.1992465 | 1 | 1290 | tags=24%, list=5%, sig | MM_4 |
| GLYCEROLIPID_METABOL | GLYCEROLIPID_METABOLISM | 57 | 0.244006 | 1.3263049 | 0.11891892 | 0.1992788 | 1 | 2206 | tags=21%, list=9%, sig | MM_4 |
| GOBP_BRANCHED_CHAIN | GOBP_BRANCHED_CHAIN_AMINO_ACID_CA | 21 | 0.319514 | 1.3260306 | 0.13108614 | 0.19937125 | 1 | 13650 | tags=95%, list=58%, si | MM_4 |
| GOBP_POSITIVE_REGULA | GOBP_POSITIVE_REGULATION_OF_LEUKOC\ | 33 | 0.285289 | 1.3250084 | 0.12556054 | 0.20024018 | 1 | 613 | tags=12%, list=3%, sig | MM_4 |
| GOBP_REGULATION_OF_ | GOBP_REGULATION_OF_TRIGLYCERIDE_ME | 47 | 0.258356 | 1.3241644 | 0.08490566 | 0.20078923 | 1 | 2714 | tags=30%, list=11%, si | MM_4 |
| GOBP_NEGATIVE_REGUL | GOBP_NEGATIVE_REGULATION_OF_ACTIN_ | 39 | 0.257354 | 1.3241187 | 0.10313901 | 0.20065099 | 1 | 15373 | tags=95%, list=65%, si | MM_4 |
| GOBP_ISOPRENOID_MET. | GOBP_ISOPRENOID_METABOLIC_PROCESS | 87 | 0.216593 | 1.3241055 | 0.06818182 | 0.20049152 | 1 | 3113 | tags=25%, list=13%, si | MM_4 |
| GOBP_REGULATION_OF_ | GOBP_REGULATION_OF_PROTEIN_DEPHOSI | 76 | 0.222263 | 1.3231146 | 0.039325844 | 0.20124882 | 1 | 4063 | tags=33%, list=17%, si | MM_4 |
| GOBP_DNA_TEMPLATED_ | GOBP_DNA_TEMPLATED_TRANSCRIPTION_ | 25 | 0.307086 | 1.3218757 | 0.118320614 | 0.2022121 | 1 | 3572 | tags=36%, list=15%, si | MM_4 |
| GOBP_VASCULAR_ASSOC | GOBP_VASCULAR_ASSOCIATED_SMOOTH_N | 16 | 0.358179 | 1.3216373 | 0.12727273 | 0.20226519 | 1 | 3386 | tags=44%, list=14%, si | MM_4 |
| GOBP_MEMBRANE_LIPID | GOBP_MEMBRANE_LIPID_CATABOLIC_PRO( | 33 | 0.284071 | 1.3209592 | 0.12711865 | 0.2028098 | 1 | 2676 | tags=27%, list=11%, si | MM_4 |
| GOBP_REGULATION_OF_ | GOBP_REGULATION_OF_CELL_GROWTH_IN | 33 | 0.277382 | 1.3207047 | 0.12295082 | 0.2028769 | 1 | 3163 | tags=30%, list=13%, si | MM_4 |
| GOBP_REGULATION_OF_ | GOBP_REGULATION_OF_BLOOD_VESSEL_EN | 94 | 0.217752 | 1.320673 | 0.045161292 | 0.20272917 | 1 | 2526 | tags=19%, list=11%, si | MM_4 |
| GOBP_OLIGOSACCHARIDI | GOBP_OLIGOSACCHARIDE_METABOLIC_PRO | 48 | 0.264291 | 1.3206398 | 0.10729614 | 0.2025858 | 1 | 3439 | tags=33%, list=15%, si | MM_4 |
| GOBP_DEOXYRIBONUCLE | GOBP_DEOXYRIBONUCLEOSIDE_MONOPHO | 17 | 0.342518 | 1.3201169 | 0.14509805 | 0.2029348 | 1 | 3189 | tags=35%, list=14%, si | MM_4 |
| GOBP_PHOSPHOLIPID_CA | GOBP_PHOSPHOLIPID_CATABOLIC_PROCESS | 52 | 0.247634 | 1.3185486 | 0.08252427 | 0.20429006 | 1 | 2618 | tags=19%, list=11%, si | MM_4 |
| GOBP_NEGATIVE_REGUL | GOBP_NEGATIVE_REGULATION_OF_CATAB( | 339 | 0.172993 | 1.3175061 | 0.016666668 | 0.20508246 | 1 | 2890 | tags=19%, list=12%, si | MM_4 |
| GOBP_NEGATIVE_REGUL | GOBP_NEGATIVE_REGULATION_OF_MUSCL | 29 | 0.28923 | 1.316695 | 0.13360325 | 0.20570149 | 1 | 3232 | tags=24%, list=14%, si | MM_4 |
| GOBP_ENDOPLASMIC_RE | GOBP_ENDOPLASMIC_RETICULUM_TO_CYT( | 27 | 0.296634 | 1.3165367 | 0.13358779 | 0.20568913 | 1 | 11153 | tags=89%, list=47%, si | MM_4 |
| GOBP_MATURATION_OF_ | GOBP_MATURATION_OF_5_8S_RRNA_FRON | 24 | 0.309874 | 1.3159876 | 0.13653137 | 0.20602094 | 1 | 3686 | tags=42%, list=16%, si | MM_4 |
| GOBP_TYPE_2_IMMUNE_ | GOBP_TYPE_2_IMMUNE_RESPONSE | 46 | 0.253162 | 1.315649 | 0.0969163 | 0.2061842 | 1 | 1375 | tags=15%, list=6%, sig | MM_4 |
| GOBP_COPII_COATED_VE | GOBP_COPII_COATED_VESICLE_CARGO_LO/ | 17 | 0.344308 | 1.3153765 | 0.1368421 | 0.20626585 | 1 | 3503 | tags=41%, list=15%, si | MM_4 |
| GOBP_REGULATION_OF_ | GOBP_REGULATION_OF_CHOLESTEROL_STC | 21 | 0.323668 | 1.3153373 | 0.119521916 | 0.20613724 | 1 | 2987 | tags=29%, list=13%, si | MM_4 |
| GOBP_SEQUESTERING_O | GOBP_SEQUESTERING_OF_TRIGLYCERIDE | 18 | 0.336728 | 1.3151156 | 0.13620071 | 0.20616758 | 1 | 2987 | tags=28%, list=13%, si | MM_4 |
| GOBP_REGULATION_OF_ | GOBP_REGULATION_OF_FIBROBLAST_GROV | 33 | 0.285297 | 1.3150347 | 0.128 | 0.20604765 | 1 | 2320 | tags=21%, list=10%, si | MM_4 |
| GOBP_NUCLEOBASE_COI | GOBP_NUCLEOBASE_CONTAINING_COMPO | 195 | 0.186521 | 1.3148625 | 0.046296295 | 0.20608038 | 1 | 4117 | tags=26%, list=17%, si | MM_4 |
| GOBP_REGULATION_OF_ | GOBP_REGULATION_OF_INFLAMMATORY_F | 33 | 0.28414 | 1.3144294 | 0.12552302 | 0.20632409 | 1 | 1817 | tags=24%, list=8%, sig | MM_4 |
| GOBP_REGULATION_OF_ | GOBP_REGULATION_OF_DNA_RECOMBINAT | 127 | 0.198226 | 1.314209 | 0.031007752 | 0.20634283 | 1 | 2929 | tags=21%, list=12%, si | MM_4 |
| GOBP_PROTEIN_TARGETI | GOBP_PROTEIN_TARGETING_TO_LYSOSOMI | 29 | 0.291629 | 1.3132551 | 0.13178295 | 0.20714855 | 1 | 4694 | tags=38%, list=20%, si | MM_4 |
| GOBP_SULFUR_COMPOU | GOBP_SULFUR_COMPOUND_METABOLIC_P | 292 | 0.177544 | 1.3121529 | 0 | 0.20803249 | 1 | 3638 | tags=27%, list=15%, si | MM_4 |
| GOBP_NEGATIVE_REGUL | GOBP_NEGATIVE_REGULATION_OF_TRANSF | 463 | 0.17172 | 1.311799 | 0 | 0.20819093 | 1 | 3839 | tags=24%, list=16%, si | MM_4 |
| GOBP_MACROPHAGE_AC | GOBP_MACROPHAGE_ACTIVATION_INVOLV | 20 | 0.342444 | 1.3114426 | 0.12878788 | 0.20837899 | 1 | 1418 | tags=20%, list=6%, sig | MM_4 |
| GOBP_TRANSITION_MET/ | GOBP_TRANSITION_METAL_ION_TRANSPOF | 95 | 0.217437 | 1.3111387 | 0.087628864 | 0.20846717 | 1 | 2470 | tags=22%, list=10%, si | MM_4 |
| GOBP_POSITIVE_REGULA | GOBP_POSITIVE_REGULATION_OF_T_CELL_/ | 16 | 0.353246 | 1.3106533 | 0.16551724 | 0.20880014 | 1 | 561 | tags=13%, list=2%, sig | MM_4 |
| GOBP_INTRACELLULAR_I | GOBP_INTRACELLULAR_IRON_ION_HOMEO: | 64 | 0.235328 | 1.3100802 | 0.07177033 | 0.20920183 | 1 | 3575 | tags=27%, list=15%, si | MM_4 |
| GOBP_NEGATIVE_REGUL | GOBP_NEGATIVE_REGULATION_OF_DEPHO: | 38 | 0.270183 | 1.3099521 | 0.12068965 | 0.20915665 | 1 | 3839 | tags=34%, list=16%, si | MM_4 |
| GOBP_NEGATIVE_REGUL | GOBP_NEGATIVE_REGULATION_OF_PEPTID\ | 18 | 0.328595 | 1.3090981 | 0.15209125 | 0.2098072 | 1 | 3553 | tags=33%, list=15%, si | MM_4 |
| CUSHING_SYNDROME | CUSHING_SYNDROME | 147 | 0.197344 | 1.3089532 | 0.046979867 | 0.20978986 | 1 | 2932 | tags=21%, list=12%, si | MM_4 |
| GOBP_RESPONSE_TO_NU | GOBP_RESPONSE_TO_NUTRIENT_LEVELS | 371 | 0.170805 | 1.3087965 | 0.023255814 | 0.20975527 | 1 | 3474 | tags=22%, list=15%, si | MM_4 |
| GLYCOSPHINGOLIPID_BIC | GLYCOSPHINGOLIPID_BIOSYNTHESIS | 44 | 0.258775 | 1.3087837 | 0.0862069 | 0.20958622 | 1 | 2811 | tags=30%, list=12%, si | MM_4 |
| GOBP_REGULATION_OF_ | GOBP_REGULATION_OF_MAST_CELL_DEGR/ | 33 | 0.287259 | 1.3087268 | 0.12 | 0.20945027 | 1 | 1262 | tags=15%, list=5%, sig | MM_4 |
| GOBP_REGULATION_OF_ | GOBP_REGULATION_OF_SUBSTRATE_ADHE: | 58 | 0.243456 | 1.3082811 | 0.11111111 | 0.2097073 | 1 | 3721 | tags=26%, list=16%, si | MM_4 |
| GOBP_REGULATION_OF_ | GOBP_REGULATION_OF_TYPE_2_IMMUNE_ | 34 | 0.287091 | 1.3082671 | 0.13004485 | 0.20954038 | 1 | 1375 | tags=18%, list=6%, sig | MM_4 |
| GOBP_NEGATIVE_REGUL | GOBP_NEGATIVE_REGULATION_OF_PEPTID\ | 29 | 0.286967 | 1.3082008 | 0.11764706 | 0.20942137 | 1 | 3805 | tags=31%, list=16%, si | MM_4 |
| GOBP_POSITIVE_REGULA | GOBP_POSITIVE_REGULATION_OF_AMIDE_I | 33 | 0.278465 | 1.3078759 | 0.11155379 | 0.2095666 | 1 | 2228 | tags=21%, list=9%, sig | MM_4 |
| GOBP_MITOCHONDRIAL_ | GOBP_MITOCHONDRIAL_OUTER_MEMBRAN | 28 | 0.290458 | 1.3077226 | 0.12970711 | 0.20956296 | 1 | 12094 | tags=89%, list=51%, si | MM_4 |

| | | | | | | | | | | |
|---|---|---|---|---|---|---|---|---|---|---|
| GOBP_ANTIGEN_RECEPTO | GOBP_ANTIGEN_RECEPTOR_MEDIATED_SIG | 171 | 0.187517 | 1.3072233 | 0.025210084 | 0.20989059 | 1 | 3636 | tags=21%, list=15%, si | MM_4 |
| GOBP_HEME_METABOLIC | GOBP_HEME_METABOLIC_PROCESS | 34 | 0.276711 | 1.3054398 | 0.097345136 | 0.21154742 | 1 | 3552 | tags=32%, list=15%, si | MM_4 |
| GOBP_INTEGRIN_MEDIAT | GOBP_INTEGRIN_MEDIATED_SIGNALING_PA | 98 | 0.214849 | 1.3053929 | 0.090277776 | 0.21139687 | 1 | 2797 | tags=19%, list=12%, si | MM_4 |
| GOBP_POSITIVE_REGULA | GOBP_POSITIVE_REGULATION_OF_INTERLEU | 65 | 0.233585 | 1.3053092 | 0.082051285 | 0.2113016 | 1 | 4180 | tags=31%, list=18%, si | MM_4 |
| GOBP_REGULATION_OF_ | GOBP_REGULATION_OF_ESTABLISHMENT_C | 25 | 0.296676 | 1.3043176 | 0.14741036 | 0.21211095 | 1 | 1965 | tags=20%, list=8%, sig | MM_4 |
| GOBP_POSITIVE_REGULA | GOBP_POSITIVE_REGULATION_OF_EXCITATO | 40 | 0.257199 | 1.3037708 | 0.10619469 | 0.21249181 | 1 | 15408 | tags=95%, list=65%, si | MM_4 |
| GOBP_TOLL_LIKE_RECEPT | GOBP_TOLL_LIKE_RECEPTOR_SIGNALING_PA | 71 | 0.222218 | 1.3035268 | 0.1017964 | 0.21256219 | 1 | 4003 | tags=32%, list=17%, si | MM_4 |
| GOBP_MEMBRANE_LIPID | GOBP_MEMBRANE_LIPID_BIOSYNTHETIC_PI | 129 | 0.198582 | 1.3032973 | 0.033333335 | 0.21263267 | 1 | 1876 | tags=16%, list=8%, sig | MM_4 |
| GOBP_IMMUNE_RESPON | GOBP_IMMUNE_RESPONSE_REGULATING_C | 278 | 0.177393 | 1.3023763 | 0.055555556 | 0.21328968 | 1 | 3681 | tags=22%, list=16%, si | MM_4 |
| GOBP_NEGATIVE_REGUL | GOBP_NEGATIVE_REGULATION_OF_STEM_C | 25 | 0.296349 | 1.3022426 | 0.1392405 | 0.21324274 | 1 | 5003 | tags=52%, list=21%, si | MM_4 |
| GOBP_CELLULAR_RESPON | GOBP_CELLULAR_RESPONSE_TO_OXYGEN_L | 133 | 0.197532 | 1.301706 | 0.047945205 | 0.21358617 | 1 | 3590 | tags=26%, list=15%, si | MM_4 |
| GOBP_VASCULAR_ENDOT | GOBP_VASCULAR_ENDOTHELIAL_GROWTH_ | 50 | 0.243956 | 1.3014921 | 0.11442786 | 0.2136026 | 1 | 892 | tags=12%, list=4%, sig | MM_4 |
| GOBP_DEFINITIVE_HEMO | GOBP_DEFINITIVE_HEMOPOIESIS | 25 | 0.314616 | 1.301033 | 0.13229571 | 0.21386433 | 1 | 2882 | tags=28%, list=12%, si | MM_4 |
| GOBP_NATURAL_KILLER_ | GOBP_NATURAL_KILLER_CELL_DIFFERENTIA | 34 | 0.276301 | 1.3010005 | 0.13043478 | 0.21370898 | 1 | 3452 | tags=35%, list=15%, si | MM_4 |
| GOBP_ACTIN_FILAMENT_ | GOBP_ACTIN_FILAMENT_BUNDLE_ORGANIZ | 180 | 0.191407 | 1.3001012 | 0.069565214 | 0.2144671 | 1 | 2799 | tags=18%, list=12%, si | MM_4 |
| GOBP_NEGATIVE_REGUL | GOBP_NEGATIVE_REGULATION_OF_PROGRA | 17 | 0.341182 | 1.299543 | 0.15384616 | 0.21486033 | 1 | 11555 | tags=94%, list=49%, si | MM_4 |
| GOBP_REGULATION_OF_ | GOBP_REGULATION_OF_SULFUR_METABOL | 15 | 0.361642 | 1.2986997 | 0.13986014 | 0.21550314 | 1 | 4340 | tags=47%, list=18%, si | MM_4 |
| GOBP_ESTABLISHMENT_C | GOBP_ESTABLISHMENT_OF_MITOCHONDRIC | 25 | 0.306335 | 1.298199 | 0.14079422 | 0.21582098 | 1 | 14720 | tags=96%, list=62%, si | MM_4 |
| GOBP_NEGATIVE_REGUL | GOBP_NEGATIVE_REGULATION_OF_REGULA | 25 | 0.311861 | 1.2981011 | 0.115879826 | 0.21573417 | 1 | 3641 | tags=36%, list=15%, si | MM_4 |
| GOBP_FILOPODIUM_ASSI | GOBP_FILOPODIUM_ASSEMBLY | 74 | 0.228483 | 1.2979285 | 0.08092485 | 0.21575741 | 1 | 4655 | tags=32%, list=20%, si | MM_4 |
| GOBP_REGULATION_OF_ | GOBP_REGULATION_OF_PLASMA_LIPOPROT | 66 | 0.230234 | 1.2976134 | 0.10810811 | 0.21592648 | 1 | 2346 | tags=20%, list=10%, si | MM_4 |
| GOBP_EXTRINSIC_APOPT | GOBP_EXTRINSIC_APOPTOTIC_SIGNALING_I | 233 | 0.182857 | 1.2971306 | 0.023809524 | 0.21628504 | 1 | 2556 | tags=16%, list=11%, si | MM_4 |
| GOBP_WOUND_HEALING | GOBP_WOUND_HEALING_SPREADING_OF_I | 17 | 0.344792 | 1.2962395 | 0.14046822 | 0.21699463 | 1 | 3163 | tags=29%, list=13%, si | MM_4 |
| GOBP_CHONDROITIN_SU | GOBP_CHONDROITIN_SULFATE_PROTEOGLY | 21 | 0.320643 | 1.2958592 | 0.15412186 | 0.21722502 | 1 | 950 | tags=19%, list=4%, sig | MM_4 |
| GOBP_CELLULAR_RESPON | GOBP_CELLULAR_RESPONSE_TO_COLD | 16 | 0.345392 | 1.2955127 | 0.16334662 | 0.21736367 | 1 | 3383 | tags=44%, list=14%, si | MM_4 |
| GOBP_NEGATIVE_REGUL | GOBP_NEGATIVE_REGULATION_OF_INTERLI | 30 | 0.279655 | 1.2951728 | 0.11764706 | 0.21753915 | 1 | 2031 | tags=23%, list=9%, sig | MM_4 |
| GOBP_NEGATIVE_REGUL | GOBP_NEGATIVE_REGULATION_OF_SUPRAI | 173 | 0.183754 | 1.2948735 | 0.06451613 | 0.21763597 | 1 | 4619 | tags=32%, list=20%, si | MM_4 |
| GOBP_POSITIVE_REGULA | GOBP_POSITIVE_REGULATION_OF_INTRACE | 147 | 0.192327 | 1.2945698 | 0.08527132 | 0.21774161 | 1 | 4484 | tags=33%, list=19%, si | MM_4 |
| GOBP_NEGATIVE_REGUL | GOBP_NEGATIVE_REGULATION_OF_ENDOT | 43 | 0.258643 | 1.2941247 | 0.14741036 | 0.21801376 | 1 | 2471 | tags=19%, list=10%, si | MM_4 |
| GOBP_POSITIVE_REGULA | GOBP_POSITIVE_REGULATION_OF_MACROI | 15 | 0.35393 | 1.294089 | 0.1641791 | 0.21788579 | 1 | 136 | tags=13%, list=1%, sig | MM_4 |
| GOBP_POSITIVE_REGULA | GOBP_POSITIVE_REGULATION_OF_RESPONS | 73 | 0.220966 | 1.293168 | 0.10714286 | 0.21862161 | 1 | 2448 | tags=18%, list=10%, si | MM_4 |
| GOBP_REGULATION_OF_ | GOBP_REGULATION_OF_SEQUESTERING_OI | 15 | 0.351579 | 1.2925413 | 0.16187051 | 0.21908997 | 1 | 2987 | tags=33%, list=13%, si | MM_4 |
| GOBP_PEPTIDE_CATABOL | GOBP_PEPTIDE_CATABOLIC_PROCESS | 20 | 0.332425 | 1.291856 | 0.14642857 | 0.21960586 | 1 | 2485 | tags=25%, list=11%, si | MM_4 |
| B_CELL_RECEPTOR_SIGN/ | B_CELL_RECEPTOR_SIGNALING_PATHWAY | 68 | 0.233211 | 1.291853 | 0.1122449 | 0.2194221 | 1 | 4029 | tags=28%, list=17%, si | MM_4 |
| GOBP_HYDROGEN_PERO) | GOBP_HYDROGEN_PEROXIDE_CATABOLIC_I | 22 | 0.310899 | 1.2913053 | 0.14 | 0.21981426 | 1 | 1817 | tags=18%, list=8%, sig | MM_4 |
| GOBP_FATTY_ACID_BETA | GOBP_FATTY_ACID_BETA_OXIDATION | 75 | 0.218032 | 1.2908136 | 0.09836066 | 0.22012703 | 1 | 3178 | tags=28%, list=13%, si | MM_4 |
| GOBP_REGULATION_OF_ | GOBP_REGULATION_OF_RESPONSE_TO_CYT | 153 | 0.195303 | 1.2904315 | 0.04385965 | 0.22038715 | 1 | 4282 | tags=32%, list=18%, si | MM_4 |
| GOBP_REGULATION_OF_ | GOBP_REGULATION_OF_NUCLEOTIDE_EXCIS | 29 | 0.283022 | 1.289384 | 0.15789473 | 0.2213173 | 1 | 5256 | tags=41%, list=22%, si | MM_4 |
| GOBP_COPPER_ION_TRA | GOBP_COPPER_ION_TRANSPORT | 16 | 0.361984 | 1.2879399 | 0.16479401 | 0.2226406 | 1 | 3067 | tags=44%, list=13%, si | MM_4 |
| GOBP_REGULATION_OF_ | GOBP_REGULATION_OF_NECROPTOTIC_PRC | 22 | 0.317036 | 1.287609 | 0.15625 | 0.22278532 | 1 | 11555 | tags=91%, list=49%, si | MM_4 |
| GOBP_POSITIVE_REGULA | GOBP_POSITIVE_REGULATION_OF_RAC_PRC | 20 | 0.330059 | 1.2871023 | 0.1594203 | 0.22313018 | 1 | 3741 | tags=35%, list=16%, si | MM_4 |
| SALMONELLA_INFECTION | SALMONELLA_INFECTION | 76 | 0.220947 | 1.2868888 | 0.081871346 | 0.2231246 | 1 | 2799 | tags=21%, list=12%, si | MM_4 |
| GOBP_POSITIVE_REGULA | GOBP_POSITIVE_REGULATION_OF_LYMPHO | 134 | 0.199664 | 1.286435 | 0.07092199 | 0.2234084 | 1 | 3534 | tags=24%, list=15%, si | MM_4 |
| GOBP_HOMEOSTASIS_OF | GOBP_HOMEOSTASIS_OF_NUMBER_OF_CEI | 388 | 0.164285 | 1.2858789 | 0 | 0.22385342 | 1 | 3780 | tags=23%, list=16%, si | MM_4 |
| GOBP_MITOTIC_G1_S_TR | GOBP_MITOTIC_G1_S_TRANSITION_CHECKF | 24 | 0.292248 | 1.2858568 | 0.12669684 | 0.2236818 | 1 | 4389 | tags=38%, list=19%, si | MM_4 |
| GOBP_FATTY_ACID_CATA | GOBP_FATTY_ACID_CATABOLIC_PROCESS | 100 | 0.208742 | 1.2857239 | 0.07065217 | 0.2236456 | 1 | 3178 | tags=26%, list=13%, si | MM_4 |
| GOBP_INTERFERON_MED | GOBP_INTERFERON_MEDIATED_SIGNALING | 76 | 0.219372 | 1.2856861 | 0.10638298 | 0.22349821 | 1 | 4373 | tags=36%, list=19%, si | MM_4 |
| GOBP_CALCIUM_ION_TR | GOBP_CALCIUM_ION_TRANSMEMBRANE_IN | 180 | 0.182156 | 1.2852422 | 0.018867925 | 0.223787 | 1 | 2258 | tags=14%, list=10%, si | MM_4 |
| GOBP_SUBSTRATE_ADHE | GOBP_SUBSTRATE_ADHESION_DEPENDENT_ | 99 | 0.209139 | 1.285205 | 0.06962025 | 0.22365141 | 1 | 2423 | tags=18%, list=10%, si | MM_4 |
| GOBP_CELLULAR_RESPON | GOBP_CELLULAR_RESPONSE_TO_METAL_IO | 155 | 0.192956 | 1.2851799 | 0.07438017 | 0.22348501 | 1 | 3052 | tags=23%, list=13%, si | MM_4 |
| REGULATION_OF_ACTIN_ | REGULATION_OF_ACTIN_CYTOSKELETON | 208 | 0.18549 | 1.2842896 | 0.03809524 | 0.22433133 | 1 | 3543 | tags=23%, list=15%, si | MM_4 |
| GOBP_POSITIVE_REGULA | GOBP_POSITIVE_REGULATION_OF_PRODUC | 136 | 0.194823 | 1.2836862 | 0.045801528 | 0.224761 | 1 | 3953 | tags=29%, list=17%, si | MM_4 |
| GOBP_REGULATION_OF_ | GOBP_REGULATION_OF_CYSTEINE_TYPE_EN | 197 | 0.182931 | 1.2815385 | 0.055555556 | 0.22679637 | 1 | 2483 | tags=17%, list=11%, si | MM_4 |
| GOBP_MITOCHONDRIAL_ | GOBP_MITOCHONDRIAL_OUTER_MEMBRAN | 34 | 0.260105 | 1.2814168 | 0.13671875 | 0.22672805 | 1 | 3985 | tags=35%, list=17%, si | MM_4 |
| GOBP_NEGATIVE_REGUL | GOBP_NEGATIVE_REGULATION_OF_DEVELC | 110 | 0.200251 | 1.2813263 | 0.083333336 | 0.22664763 | 1 | 2338 | tags=15%, list=10%, si | MM_4 |
| GOBP_MICROGLIA_DIFFE | GOBP_MICROGLIA_DIFFERENTIATION | 16 | 0.340718 | 1.2812732 | 0.14705883 | 0.22652435 | 1 | 3943 | tags=44%, list=17%, si | MM_4 |
| GOBP_BONE_RESORPTIO | GOBP_BONE_RESORPTION | 76 | 0.218825 | 1.2809724 | 0.12755102 | 0.22663762 | 1 | 3769 | tags=34%, list=16%, si | MM_4 |
| GOBP_GLYCEROLIPID_CA | GOBP_GLYCEROLIPID_CATABOLIC_PROCESS | 58 | 0.233068 | 1.280793 | 0.116161615 | 0.22663882 | 1 | 3134 | tags=24%, list=13%, si | MM_4 |
| GLYCEROPHOSPHOLIPID_ | GLYCEROPHOSPHOLIPID_METABOLISM | 92 | 0.208753 | 1.2807585 | 0.07692308 | 0.22649609 | 1 | 1851 | tags=16%, list=8%, sig | MM_4 |
| GOBP_REGULATION_OF_ | GOBP_REGULATION_OF_AMYLOID_BETA_FC | 37 | 0.263857 | 1.2803757 | 0.15246637 | 0.2267071 | 1 | 2228 | tags=22%, list=9%, sig | MM_4 |
| GOBP_MITOTIC_RECOMB | GOBP_MITOTIC_RECOMBINATION | 24 | 0.297718 | 1.2794058 | 0.16226415 | 0.22753602 | 1 | 2640 | tags=33%, list=11%, si | MM_4 |
| GOBP_PYRIMIDINE_CONT | GOBP_PYRIMIDINE_CONTAINING_COMPOU | 32 | 0.275556 | 1.2793498 | 0.16386555 | 0.22741258 | 1 | 4220 | tags=47%, list=18%, si | MM_4 |
| GOBP_REGULATION_OF_ | GOBP_REGULATION_OF_CELLULAR_SENESC | 47 | 0.249544 | 1.2783016 | 0.13551402 | 0.2284406 | 1 | 3477 | tags=30%, list=15%, si | MM_4 |
| GOBP_POSITIVE_REGULA | GOBP_POSITIVE_REGULATION_OF_MITOCH | 17 | 0.330054 | 1.2782747 | 0.15646258 | 0.22828573 | 1 | 4505 | tags=53%, list=19%, si | MM_4 |
| GOBP_RESPONSE_TO_KE | GOBP_RESPONSE_TO_KETONE | 134 | 0.197195 | 1.2774712 | 0.065789476 | 0.22894366 | 1 | 2599 | tags=19%, list=11%, si | MM_4 |
| RELAXIN_SIGNALING_PAT | RELAXIN_SIGNALING_PATHWAY | 125 | 0.197581 | 1.2771906 | 0.06382979 | 0.22903764 | 1 | 4099 | tags=30%, list=17%, si | MM_4 |
| GOBP_RECEPTOR_CLUST | GOBP_RECEPTOR_CLUSTERING | 79 | 0.216361 | 1.2771802 | 0.08064516 | 0.22886269 | 1 | 2445 | tags=18%, list=10%, si | MM_4 |
| GOBP_REGULATION_OF_ | GOBP_REGULATION_OF_EXTRACELLULAR_M | 66 | 0.228859 | 1.2769188 | 0.11578947 | 0.22895588 | 1 | 1004 | tags=14%, list=4%, sig | MM_4 |
| GOBP_POST_TRANSCRIPT | GOBP_POST_TRANSCRIPTIONAL_GENE_SILE | 64 | 0.219556 | 1.2765955 | 0.10989011 | 0.22915623 | 1 | 4133 | tags=28%, list=18%, si | MM_4 |
| PYRIMIDINE_METABOLISI | PYRIMIDINE_METABOLISM | 53 | 0.235972 | 1.2760994 | 0.08585858 | 0.2294635 | 1 | 4220 | tags=38%, list=18%, si | MM_4 |
| GOBP_ESTABLISHMENT_C | GOBP_ESTABLISHMENT_OR_MAINTENANCE | 51 | 0.241702 | 1.2758 | 0.125 | 0.22964402 | 1 | 15856 | tags=96%, list=67%, si | MM_4 |
| GOBP_CELL_CELL_RECOG | GOBP_CELL_CELL_RECOGNITION | 69 | 0.226066 | 1.2741412 | 0.13407822 | 0.23111926 | 1 | 1599 | tags=13%, list=7%, sig | MM_4 |
| GOBP_CGMP_MEDIATED | GOBP_CGMP_MEDIATED_SIGNALING | 32 | 0.268836 | 1.2738483 | 0.14096916 | 0.23124342 | 1 | 1470 | tags=13%, list=6%, sig | MM_4 |
| GOBP_POSITIVE_REGULA | GOBP_POSITIVE_REGULATION_OF_CANONI | 148 | 0.193654 | 1.2738105 | 0.05645161 | 0.23108533 | 1 | 2538 | tags=18%, list=11%, si | MM_4 |
| GOBP_RESPONSE_TO_PR | GOBP_RESPONSE_TO_PROSTAGLANDIN_E | 19 | 0.323135 | 1.2730563 | 0.18382353 | 0.23178844 | 1 | 2350 | tags=21%, list=10%, si | MM_4 |
| GOBP_PURINE_CONTAIN | GOBP_PURINE_CONTAINING_COMPOUND_ | 150 | 0.189871 | 1.2721733 | 0.08510638 | 0.23268722 | 1 | 3708 | tags=27%, list=16%, si | MM_4 |
| GOBP_NEGATIVE_REGUL | GOBP_NEGATIVE_REGULATION_OF_CYTOSC | 23 | 0.295168 | 1.2721578 | 0.15648855 | 0.23251083 | 1 | 2809 | tags=30%, list=12%, si | MM_4 |
| GOBP_REGULATION_OF_ | GOBP_REGULATION_OF_MITOCHONDRIAL_ | 58 | 0.229559 | 1.2713498 | 0.0959596 | 0.23323445 | 1 | 3985 | tags=31%, list=17%, si | MM_4 |
| GOBP_LONG_CHAIN_FAT | GOBP_LONG_CHAIN_FATTY_ACID_TRANSPC | 63 | 0.232631 | 1.2707213 | 0.12921348 | 0.233674 | 1 | 2987 | tags=21%, list=13%, si | MM_4 |
| GOBP_INTRINSIC_APOPT | GOBP_INTRINSIC_APOPTOTIC_SIGNALING_F | 44 | 0.250237 | 1.2684921 | 0.13675214 | 0.2360172 | 1 | 11362 | tags=86%, list=48%, si | MM_4 |
| GOBP_CELLULAR_RESPON | GOBP_CELLULAR_RESPONSE_TO_HYPOXIA | 112 | 0.201645 | 1.2676927 | 0.089655176 | 0.23670454 | 1 | 3590 | tags=26%, list=15%, si | MM_4 |
| GOBP_NUCLEOSIDE_DIPH | GOBP_NUCLEOSIDE_DIPHOSPHATE_CATABC | 111 | 0.200373 | 1.267316 | 0.1183432 | 0.23688433 | 1 | 3708 | tags=27%, list=16%, si | MM_4 |

| | | | | | | | | | | |
|---|---|---|---|---|---|---|---|---|---|---|
| GOBP_POSITIVE_REGULA | GOBP_POSITIVE_REGULATION_OF_TORC1_S | 52 | 0.240744 | 1.26607 | 0.12616822 | 0.23806639 | 1 | 2454 | tags=21%, list=10%, si | MM_4 |
| GOBP_DNA_CATABOLIC_ | GOBP_DNA_CATABOLIC_PROCESS | 29 | 0.269574 | 1.265638 | 0.16205534 | 0.23834257 | 1 | 1147 | tags=14%, list=5%, sig | MM_4 |
| GOBP_PROTEOGLYCAN_M | GOBP_PROTEOGLYCAN_METABOLIC_PROCE | 84 | 0.212799 | 1.2649896 | 0.086021505 | 0.23888448 | 1 | 2019 | tags=19%, list=9%, sig | MM_4 |
| GOBP_IMMUNOGLOBULI | GOBP_IMMUNOGLOBULIN_PRODUCTION_II | 67 | 0.222217 | 1.2639225 | 0.09424084 | 0.23988858 | 1 | 2928 | tags=22%, list=12%, si | MM_4 |
| GOBP_CELLULAR_RESPON | GOBP_CELLULAR_RESPONSE_TO_TOPOLOGI | 95 | 0.209408 | 1.2632154 | 0.07317073 | 0.24049032 | 1 | 5007 | tags=42%, list=21%, si | MM_4 |
| GOBP_PROTEIN_STABILIZ | GOBP_PROTEIN_STABILIZATION | 190 | 0.179114 | 1.2628641 | 0.042735044 | 0.24071647 | 1 | 3667 | tags=21%, list=16%, si | MM_4 |
| GOBP_HYDROGEN_PERO) | GOBP_HYDROGEN_PEROXIDE_BIOSYNTHETI | 18 | 0.323091 | 1.262625 | 0.15517241 | 0.2408044 | 1 | 15988 | tags=100%, list=68%, | MM_4 |
| GOBP_NEGATIVE_REGUL | GOBP_NEGATIVE_REGULATION_OF_PROTEI | 29 | 0.288191 | 1.2623079 | 0.18584071 | 0.24088052 | 1 | 373 | tags=10%, list=2%, sig | MM_4 |
| GOBP_REGULATION_OF_ | GOBP_REGULATION_OF_CD4_POSITIVE_ALF | 82 | 0.2117 | 1.2613895 | 0.09677419 | 0.24173236 | 1 | 2377 | tags=17%, list=10%, si | MM_4 |
| GOBP_REGULATION_OF_ | GOBP_REGULATION_OF_NUCLEOTIDE_CATA | 56 | 0.236311 | 1.2612698 | 0.12903225 | 0.24166754 | 1 | 3163 | tags=21%, list=13%, si | MM_4 |
| GOBP_CLATHRIN_DEPENI | GOBP_CLATHRIN_DEPENDENT_ENDOCYTOS | 48 | 0.241317 | 1.261082 | 0.12135922 | 0.2416795 | 1 | 2917 | tags=23%, list=12%, si | MM_4 |
| PROTEIN_EXPORT | PROTEIN_EXPORT | 26 | 0.282357 | 1.2608873 | 0.20242915 | 0.24173194 | 1 | 4362 | tags=31%, list=18%, si | MM_4 |
| GOBP_NEGATIVE_REGUL | GOBP_NEGATIVE_REGULATION_OF_BINDIN | 137 | 0.187889 | 1.2606778 | 0.06944445 | 0.24174394 | 1 | 3160 | tags=21%, list=13%, si | MM_4 |
| ALDOSTERONE_SYNTHESI | ALDOSTERONE_SYNTHESIS_AND_SECRETION | 94 | 0.207732 | 1.2592348 | 0.1017964 | 0.2431474 | 1 | 2815 | tags=22%, list=12%, si | MM_4 |
| GOBP_POSITIVE_REGULA | GOBP_POSITIVE_REGULATION_OF_PROTEIN | 28 | 0.287402 | 1.2589962 | 0.15 | 0.24320982 | 1 | 2946 | tags=25%, list=12%, si | MM_4 |
| GOBP_REGULATION_OF_ | GOBP_REGULATION_OF_GENERATION_OF_ | 148 | 0.185744 | 1.2589284 | 0.07936508 | 0.24310778 | 1 | 3638 | tags=22%, list=15%, si | MM_4 |
| GOBP_STRESS_FIBER_ASS | GOBP_STRESS_FIBER_ASSEMBLY | 117 | 0.199354 | 1.2581449 | 0.10526316 | 0.24383321 | 1 | 1762 | tags=14%, list=7%, sig | MM_4 |
| GOBP_NEGATIVE_REGUL | GOBP_NEGATIVE_REGULATION_OF_PROTEA | 51 | 0.234549 | 1.2576655 | 0.14155251 | 0.2442059 | 1 | 2807 | tags=20%, list=12%, si | MM_4 |
| GOBP_REGULATION_OF_ | GOBP_REGULATION_OF_NON_CANONICAL_ | 26 | 0.287133 | 1.2574562 | 0.1627907 | 0.24423483 | 1 | 15321 | tags=96%, list=65%, si | MM_4 |
| GOBP_PROTEIN_LOCALIZ | GOBP_PROTEIN_LOCALIZATION_TO_VACUO | 78 | 0.216437 | 1.2573771 | 0.0867052 | 0.24411786 | 1 | 4393 | tags=31%, list=19%, si | MM_4 |
| GOBP_PHAGOSOME_MA | GOBP_PHAGOSOME_MATURATION | 29 | 0.283517 | 1.2568635 | 0.15040651 | 0.24450791 | 1 | 15360 | tags=97%, list=65%, si | MM_4 |
| GOBP_FEMALE_PREGNAI | GOBP_FEMALE_PREGNANCY | 140 | 0.19281 | 1.2567251 | 0.07438017 | 0.24446267 | 1 | 3287 | tags=27%, list=14%, si | MM_4 |
| GOBP_MITOCHONDRIAL_ | GOBP_MITOCHONDRIAL_TRANSCRIPTION | 21 | 0.308654 | 1.2553225 | 0.1904762 | 0.24592045 | 1 | 3564 | tags=38%, list=15%, si | MM_4 |
| GOBP_CD4_POSITIVE_AL | GOBP_CD4_POSITIVE_ALPHA_BETA_T_CELL | 132 | 0.189441 | 1.2549428 | 0.096296296 | 0.24612948 | 1 | 2811 | tags=17%, list=12%, si | MM_4 |
| GOBP_REGULATION_OF_ | GOBP_REGULATION_OF_EXTRACELLULAR_M | 16 | 0.343084 | 1.2537951 | 0.15679443 | 0.2472004 | 1 | 2899 | tags=31%, list=12%, si | MM_4 |
| MELANOGENESIS | MELANOGENESIS | 91 | 0.219569 | 1.2536335 | 0.160221 | 0.24716339 | 1 | 1744 | tags=15%, list=7%, sig | MM_4 |
| GOBP_PIGMENT_BIOSYN | GOBP_PIGMENT_BIOSYNTHETIC_PROCESS | 49 | 0.238448 | 1.2534329 | 0.1352459 | 0.24719445 | 1 | 2181 | tags=20%, list=9%, sig | MM_4 |
| GOBP_REGULATION_OF_ | GOBP_REGULATION_OF_CELLULAR_KETONE | 154 | 0.184122 | 1.2529529 | 0.056074765 | 0.24750729 | 1 | 1847 | tags=13%, list=8%, sig | MM_4 |
| GOBP_NATURAL_KILLER_ | GOBP_NATURAL_KILLER_CELL_PROLIFERATI | 16 | 0.334411 | 1.2524718 | 0.17132868 | 0.24789919 | 1 | 912 | tags=19%, list=4%, sig | MM_4 |
| GLUTATHIONE_METABOL | GLUTATHIONE_METABOLISM | 57 | 0.227369 | 1.2510443 | 0.114583336 | 0.2494017 | 1 | 2356 | tags=25%, list=10%, si | MM_4 |
| GOBP_POSITIVE_REGULA | GOBP_POSITIVE_REGULATION_OF_SMALL_I | 174 | 0.176785 | 1.2508037 | 0.05645161 | 0.24950892 | 1 | 3336 | tags=21%, list=14%, si | MM_4 |
| GOBP_SPINDLE_CHECKPC | GOBP_SPINDLE_CHECKPOINT_SIGNALING | 53 | -0.68404 | -2.9953718 | 0 | 0 | 0 | 1087 | tags=40%, list=5%, sig | MM_4 |
| GOBP_REGULATION_OF_ | GOBP_REGULATION_OF_MITOTIC_SISTER_C | 56 | -0.65423 | -2.9780362 | 0 | 0 | 0 | 1170 | tags=39%, list=5%, sig | MM_4 |
| GOBP_MITOTIC_SISTER_C | GOBP_MITOTIC_SISTER_CHROMATID_SEPAF | 61 | -0.61594 | -2.831533 | 0 | 0 | 0 | 1144 | tags=34%, list=5%, sig | MM_4 |
| GOBP_MITOTIC_SISTER_C | GOBP_MITOTIC_SISTER_CHROMATID_SEGRI | 178 | -0.48027 | -2.6919255 | 0 | 0 | 0 | 1841 | tags=28%, list=8%, sig | MM_4 |
| GOBP_REGULATION_OF_ | GOBP_REGULATION_OF_MITOTIC_NUCLEAF | 114 | -0.50768 | -2.6335022 | 0 | 0 | 0 | 2123 | tags=32%, list=9%, sig | MM_4 |
| GOBP_NEGATIVE_REGUL | GOBP_NEGATIVE_REGULATION_OF_NUCLEA | 66 | -0.56468 | -2.6301161 | 0 | 0 | 0 | 1794 | tags=35%, list=8%, sig | MM_4 |
| GOBP_CHROMOSOME_SE | GOBP_CHROMOSOME_SEPARATION | 80 | -0.5339 | -2.5914328 | 0 | 0 | 0 | 1807 | tags=33%, list=8%, sig | MM_4 |
| OLFACTORY_TRANSDUCT | OLFACTORY_TRANSDUCTION | 186 | -0.45557 | -2.5621395 | 0 | 0 | 0 | 7871 | tags=72%, list=33%, si | MM_4 |
| GOBP_NEGATIVE_REGUL | GOBP_NEGATIVE_REGULATION_OF_CHROM | 96 | -0.50575 | -2.532422 | 0 | 0 | 0 | 1334 | tags=25%, list=6%, sig | MM_4 |
| GOBP_MITOTIC_NUCLEAI | GOBP_MITOTIC_NUCLEAR_DIVISION | 259 | -0.43366 | -2.5277138 | 0 | 0 | 0 | 1841 | tags=24%, list=8%, sig | MM_4 |
| GOBP_SISTER_CHROMAT | GOBP_SISTER_CHROMATID_SEGREGATION | 219 | -0.41845 | -2.4204035 | 0 | 5.80658E-05 | 0.001 | 1841 | tags=23%, list=8%, sig | MM_4 |
| GOBP_REGULATION_OF_ | GOBP_REGULATION_OF_CHROMOSOME_SE | 126 | -0.46021 | -2.4162931 | 0 | 5.3227E-05 | 0.001 | 1807 | tags=25%, list=8%, sig | MM_4 |
| GOBP_FEMALE_MEIOTIC_ | GOBP_FEMALE_MEIOTIC_NUCLEAR_DIVISIO | 41 | -0.58374 | -2.4093454 | 0 | 9.90114E-05 | 0.002 | 1934 | tags=39%, list=8%, sig | MM_4 |
| GOBP_CENTROSOME_DU | GOBP_CENTROSOME_DUPLICATION | 74 | -0.48791 | -2.3356473 | 0 | 0.000229397 | 0.005 | 2364 | tags=28%, list=10%, si | MM_4 |
| GOBP_METAPHASE_CHRO | GOBP_METAPHASE_CHROMOSOME_ALIGNI | 93 | -0.4651 | -2.3216321 | 0 | 0.000257184 | 0.006 | 1794 | tags=26%, list=8%, sig | MM_4 |
| GOBP_PROTEIN_LOCALIZ | GOBP_PROTEIN_LOCALIZATION_TO_CHROM | 24 | -0.64682 | -2.307102 | 0 | 0.000320962 | 0.008 | 1891 | tags=42%, list=8%, sig | MM_4 |
| GOBP_REGULATION_OF_ | GOBP_REGULATION_OF_NUCLEAR_DIVISIOI | 144 | -0.42265 | -2.3051305 | 0 | 0.000302082 | 0.008 | 2123 | tags=26%, list=9%, sig | MM_4 |
| GOBP_REGULATION_OF_ | GOBP_REGULATION_OF_SISTER_CHROMATI | 104 | -0.44143 | -2.2593079 | 0 | 0.000856047 | 0.024 | 1170 | tags=22%, list=5%, sig | MM_4 |
| GOBP_NUCLEAR_CHROM | GOBP_NUCLEAR_CHROMOSOME_SEGREGAT | 305 | -0.37163 | -2.2218838 | 0 | 0.001421289 | 0.042 | 1857 | tags=21%, list=8%, sig | MM_4 |
| GOBP_CHROMOSOME_LO | GOBP_CHROMOSOME_LOCALIZATION | 111 | -0.43127 | -2.2103593 | 0 | 0.001543026 | 0.047 | 1794 | tags=24%, list=8%, sig | MM_4 |
| GOBP_PROTEIN_LOCALIZ | GOBP_PROTEIN_LOCALIZATION_TO_CONDE | 16 | -0.68222 | -2.1976783 | 0 | 0.001714428 | 0.055 | 995 | tags=44%, list=4%, sig | MM_4 |
| GOBP_MICROTUBULE_CY | GOBP_MICROTUBULE_CYTOSKELETON_ORG | 158 | -0.40045 | -2.184286 | 0 | 0.002015526 | 0.067 | 2302 | tags=25%, list=10%, si | MM_4 |
| GOBP_CHROMOSOME_SE | GOBP_CHROMOSOME_SEGREGATION | 403 | -0.35652 | -2.1771364 | 0 | 0.002123775 | 0.074 | 1891 | tags=19%, list=8%, sig | MM_4 |
| GOBP_SPINDLE_ASSEMBL | GOBP_SPINDLE_ASSEMBLY | 122 | -0.42283 | -2.1751891 | 0 | 0.002088936 | 0.076 | 2278 | tags=23%, list=10%, si | MM_4 |
| GOBP_REGULATION_OF_ | GOBP_REGULATION_OF_KERATINOCYTE_MI | 15 | -0.70062 | -2.1710122 | 0 | 0.00215953 | 0.081 | 431 | tags=27%, list=2%, sig | MM_4 |
| GOBP_MICROTUBULE_OF | GOBP_MICROTUBULE_ORGANIZING_CENTE | 149 | -0.39496 | -2.1377943 | 0 | 0.00316292 | 0.122 | 1782 | tags=19%, list=8%, sig | MM_4 |
| GOBP_MITOTIC_METAPH | GOBP_MITOTIC_METAPHASE_CHROMOSOM | 58 | -0.46695 | -2.1189556 | 0 | 0.003926682 | 0.151 | 1619 | tags=26%, list=7%, sig | MM_4 |
| GOBP_MITOTIC_SPINDLE | GOBP_MITOTIC_SPINDLE_ORGANIZATION | 125 | -0.39867 | -2.1047955 | 0 | 0.004452987 | 0.175 | 2302 | tags=25%, list=10%, si | MM_4 |
| GOBP_SPINDLE_ORGANIZ | GOBP_SPINDLE_ORGANIZATION | 190 | -0.3708 | -2.0814686 | 0 | 0.006094351 | 0.239 | 2302 | tags=21%, list=10%, si | MM_4 |
| GOBP_ORGANELLE_FISSI | GOBP_ORGANELLE_FISSION | 477 | -0.33494 | -2.078218 | 0 | 0.006212106 | 0.249 | 2278 | tags=20%, list=10%, si | MM_4 |
| GOBP_ERBB2_SIGNALING | GOBP_ERBB2_SIGNALING_PATHWAY | 23 | -0.58191 | -2.0723811 | 0 | 0.006301502 | 0.263 | 709 | tags=26%, list=3%, sig | MM_4 |
| GOBP_POSITIVE_REGULA | GOBP_POSITIVE_REGULATION_OF_CELL_DIV | 79 | -0.43044 | -2.0641298 | 0 | 0.006808001 | 0.29 | 2278 | tags=27%, list=10%, si | MM_4 |
| NEUROACTIVE_LIGAND-R | NEUROACTIVE_LIGAND-RECEPTOR_INTERAC | 274 | -0.35064 | -2.0546768 | 0 | 0.007399354 | 0.319 | 11657 | tags=72%, list=49%, si | MM_4 |
| GOBP_ATTACHMENT_OF | GOBP_ATTACHMENT_OF_SPINDLE_MICROT | 48 | -0.47309 | -2.0543451 | 0 | 0.007200618 | 0.32 | 1170 | tags=27%, list=5%, sig | MM_4 |
| GOBP_MITOTIC_CYTOKIN | GOBP_MITOTIC_CYTOKINESIS | 83 | -0.41208 | -2.0346431 | 0 | 0.008666475 | 0.378 | 2278 | tags=25%, list=10%, si | MM_4 |
| GOBP_POSITIVE_REGULA | GOBP_POSITIVE_REGULATION_OF_NITRIC_( | 20 | -0.59194 | -2.0331566 | 0 | 0.00865784 | 0.386 | 1262 | tags=25%, list=5%, sig | MM_4 |
| GOBP_REGULATION_OF_ | GOBP_REGULATION_OF_LIPOPROTEIN_MET | 18 | -0.6093 | -2.0206738 | 0 | 0.010022668 | 0.435 | 897 | tags=33%, list=4%, sig | MM_4 |
| GOBP_MITOTIC_SPINDLE | GOBP_MITOTIC_SPINDLE_ASSEMBLY | 71 | -0.42925 | -2.0074781 | 0 | 0.01143413 | 0.493 | 2278 | tags=25%, list=10%, si | MM_4 |
| GOBP_NITRIC_OXIDE_SYI | GOBP_NITRIC_OXIDE_SYNTHASE_BIOSYNTH | 27 | -0.53001 | -1.9967003 | 0 | 0.01290286 | 0.546 | 1718 | tags=26%, list=7%, sig | MM_4 |
| GOBP_REGULATION_OF_ | GOBP_REGULATION_OF_SPINDLE_CHECKPC | 24 | -0.551 | -1.992025 | 0.0013679891 | 0.013207244 | 0.564 | 640 | tags=25%, list=3%, sig | MM_4 |
| GOBP_MITOTIC_CELL_CY | GOBP_MITOTIC_CELL_CYCLE_CHECKPOINT_ | 147 | -0.36331 | -1.9812888 | 0 | 0.014607852 | 0.604 | 2239 | tags=21%, list=9%, sig | MM_4 |
| GOBP_CELL_CYCLE_CHEC | GOBP_CELL_CYCLE_CHECKPOINT_SIGNALIN | 191 | -0.35149 | -1.9641786 | 0 | 0.017764188 | 0.685 | 2322 | tags=21%, list=10%, si | MM_4 |
| GOBP_POSITIVE_REGULA | GOBP_POSITIVE_REGULATION_OF_MITOTIC | 44 | -0.45422 | -1.9402742 | 0 | 0.022821603 | 0.781 | 2818 | tags=34%, list=12%, si | MM_4 |
| GOBP_NEGATIVE_REGUL | GOBP_NEGATIVE_REGULATION_OF_MITOTI | 174 | -0.3488 | -1.9363389 | 0 | 0.023411004 | 0.801 | 2356 | tags=22%, list=10%, si | MM_4 |
| GOBP_KINETOCHORE_OR | GOBP_KINETOCHORE_ORGANIZATION | 20 | -0.56399 | -1.9288954 | 0 | 0.024819115 | 0.822 | 802 | tags=20%, list=3%, sig | MM_4 |
| GOBP_NEGATIVE_REGUL | GOBP_NEGATIVE_REGULATION_OF_CELL_C | 290 | -0.32648 | -1.9267249 | 0 | 0.025007064 | 0.829 | 2364 | tags=22%, list=10%, si | MM_4 |
| GOBP_CENTRIOLE_ASSEN | GOBP_CENTRIOLE_ASSEMBLY | 46 | -0.44218 | -1.9259082 | 0 | 0.024735963 | 0.833 | 2364 | tags=28%, list=10%, si | MM_4 |
| GOBP_CILIUM_MOVEMEI | GOBP_CILIUM_MOVEMENT | 202 | -0.33955 | -1.9204842 | 0 | 0.025745992 | 0.853 | 12268 | tags=69%, list=52%, si | MM_4 |
| GOBP_REGULATION_OF_ | GOBP_REGULATION_OF_CELL_DIVISION | 168 | -0.34556 | -1.9204159 | 0 | 0.025259666 | 0.854 | 2364 | tags=23%, list=10%, si | MM_4 |
| GOBP_POSITIVE_REGULA | GOBP_POSITIVE_REGULATION_OF_PROTEIN | 24 | -0.52917 | -1.917203 | 0 | 0.02573168 | 0.864 | 2312 | tags=29%, list=10%, si | MM_4 |
| GOBP_POSITIVE_REGULA | GOBP_POSITIVE_REGULATION_OF_CELL_CY | 17 | -0.59012 | -1.9168428 | 0.0013192613 | 0.02529007 | 0.864 | 559 | tags=29%, list=2%, sig | MM_4 |

| | | | | | | | | | | |
|---|---|---|---|---|---|---|---|---|---|---|
| GOBP_MEIOTIC_SPINDLE | GOBP_MEIOTIC_SPINDLE_ORGANIZATION | 21 | -0.55191 | -1.9140531 | 0 | 0.025557445 | 0.871 | 1857 | tags=33%, list=8%, sig | MM_4 |
| GOBP_REGULATION_OF_ | GOBP_REGULATION_OF_CILIUM_MOVEMEN | 33 | -0.47744 | -1.907863 | 0 | 0.027283072 | 0.892 | 11457 | tags=76%, list=49%, si | MM_4 |
| GOBP_CYTOSKELETON_D | GOBP_CYTOSKELETON_DEPENDENT_CYTOK | 97 | -0.37665 | -1.8966852 | 0 | 0.030371072 | 0.918 | 2278 | tags=23%, list=10%, si | MM_4 |
| GOBP_POSITIVE_REGULA | GOBP_POSITIVE_REGULATION_OF_CELLULA | 35 | -0.47091 | -1.8818077 | 0.001285347 | 0.035066415 | 0.955 | 2813 | tags=31%, list=12%, si | MM_4 |
| GOBP_MICROTUBULE_OF | GOBP_MICROTUBULE_ORGANIZING_CENTE | 34 | -0.46519 | -1.8789672 | 0 | 0.035484098 | 0.96 | 2169 | tags=24%, list=9%, sig | MM_4 |
| GOBP_CYTOKINESIS | GOBP_CYTOKINESIS | 160 | -0.34555 | -1.8771234 | 0 | 0.035504334 | 0.961 | 2278 | tags=22%, list=10%, si | MM_4 |
| GOBP_KERATINOCYTE_M | GOBP_KERATINOCYTE_MIGRATION | 19 | -0.54913 | -1.8756416 | 0.004076087 | 0.035524756 | 0.964 | 431 | tags=21%, list=2%, sig | MM_4 |
| GOBP_REGULATION_OF_ | GOBP_REGULATION_OF_CENTROSOME_DU | 46 | -0.43683 | -1.8718307 | 0 | 0.03634996 | 0.967 | 2364 | tags=24%, list=10%, si | MM_4 |
| GOBP_NEUROPEPTIDE_SI | GOBP_NEUROPEPTIDE_SIGNALING_PATHWA | 69 | -0.39946 | -1.8712292 | 0 | 0.035947252 | 0.968 | 9613 | tags=68%, list=41%, si | MM_4 |
| GOBP_POSITIVE_REGULA | GOBP_POSITIVE_REGULATION_OF_PEPTIDY | 28 | -0.49237 | -1.8681407 | 0.0012953368 | 0.03642189 | 0.969 | 1606 | tags=29%, list=7%, sig | MM_4 |
| NICOTINE_ADDICTION | NICOTINE_ADDICTION | 35 | -0.45359 | -1.8649956 | 0.0013089005 | 0.036901414 | 0.973 | 11419 | tags=80%, list=48%, si | MM_4 |
| GOBP_REGULATION_OF_ | GOBP_REGULATION_OF_CYTOKINESIS | 83 | -0.38303 | -1.8589965 | 0 | 0.038497366 | 0.976 | 2278 | tags=27%, list=10%, si | MM_4 |
| GOBP_INTERMEDIATE_FI | GOBP_INTERMEDIATE_FILAMENT_ORGANIZ | 46 | -0.42896 | -1.8566866 | 0 | 0.038699374 | 0.979 | 5141 | tags=50%, list=22%, si | MM_4 |
| GOBP_REGULATION_OF_ | GOBP_REGULATION_OF_ENDOTHELIAL_CEL | 18 | -0.55366 | -1.8525864 | 0.0013831259 | 0.039795455 | 0.981 | 2039 | tags=28%, list=9%, sig | MM_4 |
| GOBP_CILIUM_OR_FLAGE | GOBP_CILIUM_OR_FLAGELLUM_DEPENDEN | 173 | -0.33337 | -1.8491944 | 0 | 0.040538397 | 0.984 | 12268 | tags=69%, list=52%, si | MM_4 |
| GOBP_POSITIVE_REGULA | GOBP_POSITIVE_REGULATION_OF_UBIQUIT | 18 | -0.55106 | -1.8450171 | 0.0027434842 | 0.04159311 | 0.986 | 990 | tags=28%, list=4%, sig | MM_4 |
| GOBP_REGULATION_OF_ | GOBP_REGULATION_OF_CILIUM_DEPENDEN | 23 | -0.511 | -1.8394253 | 0 | 0.043429792 | 0.991 | 11007 | tags=74%, list=47%, si | MM_4 |
| GOBP_MAMMARY_GLAN | GOBP_MAMMARY_GLAND_ALVEOLUS_DEV | 21 | -0.53687 | -1.8378657 | 0.0027247956 | 0.04354638 | 0.992 | 1261 | tags=29%, list=5%, sig | MM_4 |
| GOBP_MHC_CLASS_II_BI | GOBP_MHC_CLASS_II_BIOSYNTHETIC_PROC | 16 | -0.57119 | -1.8262897 | 0.0041724616 | 0.048028808 | 0.995 | 1262 | tags=31%, list=5%, sig | MM_4 |
| GOBP_OVULATION | GOBP_OVULATION | 22 | -0.52122 | -1.8249125 | 0.0026560426 | 0.04809579 | 0.995 | 1170 | tags=27%, list=5%, sig | MM_4 |
| GOBP_RESPONSE_TO_AM | GOBP_RESPONSE_TO_AMYLOID_BETA | 39 | -0.44175 | -1.8172139 | 0 | 0.05127508 | 0.997 | 1673 | tags=21%, list=7%, sig | MM_4 |
| GOBP_POSITIVE_REGULA | GOBP_POSITIVE_REGULATION_OF_CYTOKIN | 40 | -0.44072 | -1.8147041 | 0.0012836971 | 0.051815696 | 0.998 | 2278 | tags=30%, list=10%, si | MM_4 |
| MICRORNAS_IN_CANCER | MICRORNAS_IN_CANCER | 139 | -0.33814 | -1.8123295 | 0 | 0.052253287 | 0.999 | 2053 | tags=17%, list=9%, sig | MM_4 |
| GOBP_SYNAPTIC_TRANSM | GOBP_SYNAPTIC_TRANSMISSION_CHOLINEI | 33 | -0.45559 | -1.811441 | 0.001310616 | 0.052044947 | 0.999 | 12246 | tags=85%, list=52%, si | MM_4 |
| GOBP_NEGATIVE_REGUL | GOBP_NEGATIVE_REGULATION_OF_CELL_C | 368 | -0.29829 | -1.8105158 | 0 | 0.05191853 | 0.999 | 2364 | tags=20%, list=10%, si | MM_4 |
| GOBP_NEGATIVE_REGUL | GOBP_NEGATIVE_REGULATION_OF_MITOTI | 230 | -0.31498 | -1.8041431 | 0 | 0.05464166 | 1 | 2356 | tags=20%, list=10%, si | MM_4 |
| FANCONI_ANEMIA_PATH | FANCONI_ANEMIA_PATHWAY | 50 | -0.41945 | -1.7985474 | 0 | 0.05716992 | 1 | 2043 | tags=26%, list=9%, sig | MM_4 |
| GOBP_NEURONAL_ACTIO | GOBP_NEURONAL_ACTION_POTENTIAL | 56 | -0.40215 | -1.7961401 | 0.0012722646 | 0.057950214 | 1 | 11695 | tags=79%, list=50%, si | MM_4 |
| PROSTATE_CANCER | PROSTATE_CANCER | 93 | -0.35989 | -1.7942855 | 0 | 0.05835111 | 1 | 1153 | tags=12%, list=5%, sig | MM_4 |
| GOBP_NEGATIVE_REGUL | GOBP_NEGATIVE_REGULATION_OF_INTERLI | 19 | -0.51984 | -1.7873179 | 0.007002801 | 0.061797824 | 1 | 918 | tags=32%, list=4%, sig | MM_4 |
| GOBP_ATTACHMENT_OF_ | GOBP_ATTACHMENT_OF_MITOTIC_SPINDLE | 24 | -0.49231 | -1.778371 | 0.0027210885 | 0.06661211 | 1 | 1170 | tags=29%, list=5%, sig | MM_4 |
| GOBP_CELLULAR_RESPON | GOBP_CELLULAR_RESPONSE_TO_ACID_CHE | 87 | -0.35442 | -1.7767155 | 0.0011961722 | 0.06687848 | 1 | 1679 | tags=18%, list=7%, sig | MM_4 |
| GOBP_POSITIVE_REGULA | GOBP_POSITIVE_REGULATION_OF_CELL_CY | 252 | -0.30605 | -1.775225 | 0 | 0.06696238 | 1 | 2415 | tags=19%, list=10%, si | MM_4 |
| GOBP_REGULATION_OF_ | GOBP_REGULATION_OF_CYSTEINE_TYPE_EN | 16 | -0.54784 | -1.7692994 | 0.0042253523 | 0.069919005 | 1 | 2032 | tags=19%, list=9%, sig | MM_4 |
| GOBP_NUCLEOLAR_LARG | GOBP_NUCLEOLAR_LARGE_RRNA_TRANSCR | 21 | -0.51412 | -1.7646275 | 0.0053547523 | 0.07237141 | 1 | 2032 | tags=29%, list=9%, sig | MM_4 |
| GOBP_REGULATION_OF_ | GOBP_REGULATION_OF_CELL_CYCLE_CHECI | 48 | -0.40383 | -1.7625921 | 0.0078023407 | 0.07310625 | 1 | 1772 | tags=21%, list=8%, sig | MM_4 |
| GOBP_PROTEIN_LOCALIZ | GOBP_PROTEIN_LOCALIZATION_TO_CHROM | 112 | -0.33651 | -1.7606263 | 0 | 0.07356881 | 1 | 2322 | tags=18%, list=10%, si | MM_4 |
| GOBP_RESPONSE_TO_MI | GOBP_RESPONSE_TO_MITOCHONDRIAL_DE | 21 | -0.50975 | -1.7598804 | 0.006756757 | 0.0731757 | 1 | 477 | tags=14%, list=2%, sig | MM_4 |
| GOBP_REGULATION_OF_ | GOBP_REGULATION_OF_UBIQUITIN_PROTE | 20 | -0.51889 | -1.7576272 | 0.0027510317 | 0.07406216 | 1 | 1144 | tags=30%, list=5%, sig | MM_4 |
| GOBP_POSITIVE_REGULA | GOBP_POSITIVE_REGULATION_OF_INTERLEU | 15 | -0.56361 | -1.7558329 | 0.011111111 | 0.074484326 | 1 | 2225 | tags=47%, list=9%, sig | MM_4 |
| GOBP_POSITIVE_REGULA | GOBP_POSITIVE_REGULATION_OF_SMOOTH | 34 | -0.43278 | -1.748219 | 0.008985879 | 0.079166785 | 1 | 11590 | tags=71%, list=49%, si | MM_4 |
| GOBP_REGULATION_OF_ | GOBP_REGULATION_OF_CHROMOSOME_OI | 242 | -0.30048 | -1.7453704 | 0 | 0.08041758 | 1 | 2307 | tags=17%, list=10%, si | MM_4 |
| GOBP_REGULATION_OF_ | GOBP_REGULATION_OF_MICROTUBULE_BA | 48 | -0.40446 | -1.7451677 | 0.006105006 | 0.07971283 | 1 | 12463 | tags=69%, list=53%, si | MM_4 |
| GOBP_S_ADENOSYLMETH | GOBP_S_ADENOSYLMETHIONINE_METABOL | 17 | -0.53822 | -1.7438493 | 0.011331445 | 0.0800103 | 1 | 1679 | tags=29%, list=7%, sig | MM_4 |
| GOBP_KINETOCHORE_AS | GOBP_KINETOCHORE_ASSEMBLY | 16 | -0.53999 | -1.7381433 | 0.006821282 | 0.08385824 | 1 | 802 | tags=19%, list=3%, sig | MM_4 |
| GOBP_CILIUM_ORGANIZA | GOBP_CILIUM_ORGANIZATION | 381 | -0.28467 | -1.7380741 | 0 | 0.08303995 | 1 | 2232 | tags=18%, list=9%, sig | MM_4 |
| GOBP_CENTROMERE_CO | GOBP_CENTROMERE_COMPLEX_ASSEMBLY | 27 | -0.47045 | -1.7372863 | 0.004 | 0.08272399 | 1 | 1986 | tags=26%, list=8%, sig | MM_4 |
| GOBP_REGULATION_OF_ | GOBP_REGULATION_OF_CENTROSOME_CYC | 55 | -0.38947 | -1.736618 | 0 | 0.0824074 | 1 | 2364 | tags=20%, list=10%, si | MM_4 |
| GOBP_SPERM_MOTILITY | GOBP_SPERM_MOTILITY | 151 | -0.31676 | -1.7357926 | 0 | 0.082245335 | 1 | 12268 | tags=69%, list=52%, si | MM_4 |
| GOBP_MANNOSYLATION | GOBP_MANNOSYLATION | 22 | -0.49171 | -1.7227091 | 0.015256588 | 0.09282908 | 1 | 2767 | tags=36%, list=12%, si | MM_4 |
| GOBP_MITOPHAGY | GOBP_MITOPHAGY | 58 | -0.37343 | -1.7153798 | 0.002469136 | 0.098252244 | 1 | 1606 | tags=16%, list=7%, sig | MM_4 |
| GOBP_PROTEIN_LOCALIZ | GOBP_PROTEIN_LOCALIZATION_TO_EARLY_ | 15 | -0.53661 | -1.7118958 | 0.011204482 | 0.10079873 | 1 | 709 | tags=20%, list=3%, sig | MM_4 |
| GOBP_POSITIVE_REGULA | GOBP_POSITIVE_REGULATION_OF_NUCLEAI | 62 | -0.36638 | -1.7077744 | 0.008684863 | 0.10374287 | 1 | 3838 | tags=31%, list=16%, si | MM_4 |
| GOBP_REGULATION_OF_ | GOBP_REGULATION_OF_MICROTUBULE_BA | 248 | -0.29165 | -1.7077069 | 0 | 0.102809764 | 1 | 2427 | tags=18%, list=10%, si | MM_4 |
| GOBP_REGULATION_OF_ | GOBP_REGULATION_OF_PROTEIN_AUTOPH | 41 | -0.41172 | -1.6988688 | 0.011363637 | 0.11074467 | 1 | 1633 | tags=20%, list=7%, sig | MM_4 |
| GOBP_AXONAL_FASCICU | GOBP_AXONAL_FASCICULATION | 24 | -0.46928 | -1.6897231 | 0.0066489363 | 0.11961606 | 1 | 1589 | tags=25%, list=7%, sig | MM_4 |
| GOBP_PROTEIN_O_LINKE | GOBP_PROTEIN_O_LINKED_MANNOSYLATIO | 16 | -0.53115 | -1.6873955 | 0.00405954 | 0.12108785 | 1 | 2233 | tags=38%, list=9%, sig | MM_4 |
| GOBP_ACTION_POTENTIA | GOBP_ACTION_POTENTIAL | 171 | -0.30638 | -1.6857688 | 0 | 0.12161046 | 1 | 11695 | tags=60%, list=50%, si | MM_4 |
| GOBP_NEGATIVE_REGUL | GOBP_NEGATIVE_REGULATION_OF_MEMBI | 18 | -0.49989 | -1.6834726 | 0.0043290043 | 0.1231506 | 1 | 2328 | tags=28%, list=10%, si | MM_4 |
| GOBP_FIBROBLAST_PROL | GOBP_FIBROBLAST_PROLIFERATION | 147 | -0.30949 | -1.6821243 | 0.003382187 | 0.12349385 | 1 | 2364 | tags=20%, list=10%, si | MM_4 |
| GOBP_REGULATION_OF_ | GOBP_REGULATION_OF_RECEPTOR_BINDIN | 20 | -0.4826 | -1.6770103 | 0.012178619 | 0.12846504 | 1 | 2103 | tags=25%, list=9%, sig | MM_4 |
| GLYCINE,_SERINE_AND_T | GLYCINE,_SERINE_AND_THREONINE_METAE | 38 | -0.40642 | -1.676741 | 0.006451613 | 0.12770376 | 1 | 5139 | tags=47%, list=22%, si | MM_4 |
| GOBP_PEPTIDYL_THREON | GOBP_PEPTIDYL_THREONINE_MODIFICATIO | 110 | -0.32495 | -1.67284 | 0 | 0.13122949 | 1 | 2026 | tags=17%, list=9%, sig | MM_4 |
| GOBP_STRIATUM_DEVEL | GOBP_STRIATUM_DEVELOPMENT | 17 | -0.51063 | -1.6707093 | 0.017191976 | 0.13269788 | 1 | 1684 | tags=29%, list=7%, sig | MM_4 |
| GOBP_REGULATION_OF_ | GOBP_REGULATION_OF_ADENYLATE_CYCLA | 36 | -0.41635 | -1.6706264 | 0.0038910506 | 0.13166995 | 1 | 10256 | tags=75%, list=43%, si | MM_4 |
| GOBP_POSITIVE_REGULA | GOBP_POSITIVE_REGULATION_OF_CHROMO | 28 | -0.44719 | -1.669522 | 0.010854817 | 0.13190687 | 1 | 1807 | tags=29%, list=8%, sig | MM_4 |
| GOBP_TERTIARY_ALCOHO | GOBP_TERTIARY_ALCOHOL_METABOLIC_PR | 20 | -0.4858 | -1.6687298 | 0.015492958 | 0.13175733 | 1 | 4661 | tags=50%, list=20%, si | MM_4 |
| GOBP_PROTEIN_NEDDYL | GOBP_PROTEIN_NEDDYLATION | 21 | -0.47452 | -1.6681908 | 0.01650619 | 0.1313963 | 1 | 1849 | tags=29%, list=8%, sig | MM_4 |
| GOBP_EXIT_FROM_MITO | GOBP_EXIT_FROM_MITOSIS | 26 | -0.45521 | -1.6681378 | 0.010854817 | 0.13038181 | 1 | 957 | tags=15%, list=4%, sig | MM_4 |
| GOBP_NEGATIVE_REGUL | GOBP_NEGATIVE_REGULATION_OF_ORGAN | 375 | -0.27445 | -1.6664515 | 0 | 0.13119006 | 1 | 2364 | tags=17%, list=10%, si | MM_4 |
| GOBP_REGULATION_OF_ | GOBP_REGULATION_OF_PEPTIDYL_THREON | 45 | -0.39737 | -1.6656563 | 0.011292347 | 0.13109447 | 1 | 1606 | tags=20%, list=7%, sig | MM_4 |
| GOBP_POSITIVE_REGULA | GOBP_POSITIVE_REGULATION_OF_MITOTIC | 18 | -0.49588 | -1.6655505 | 0.02010724 | 0.13013332 | 1 | 559 | tags=28%, list=2%, sig | MM_4 |
| GOBP_PEPTIDYL_GLUTAN | GOBP_PEPTIDYL_GLUTAMIC_ACID_MODIFIC | 25 | -0.46126 | -1.665033 | 0.013477089 | 0.12968536 | 1 | 1365 | tags=24%, list=6%, sig | MM_4 |
| GOBP_ESTABLISHMENT_ | GOBP_ESTABLISHMENT_OF_MITOTIC_SPIND | 41 | -0.39638 | -1.6609919 | 0.011378002 | 0.13347659 | 1 | 2202 | tags=24%, list=9%, sig | MM_4 |
| GOBP_SPINDLE_LOCALIZ | GOBP_SPINDLE_LOCALIZATION | 61 | -0.36263 | -1.6524322 | 0.0051020407 | 0.14266531 | 1 | 2202 | tags=21%, list=9%, sig | MM_4 |
| GOBP_BILE_ACID_BIOSYN | GOBP_BILE_ACID_BIOSYNTHETIC_PROCESS | 27 | -0.43584 | -1.6515901 | 0.010781671 | 0.14253472 | 1 | 9163 | tags=63%, list=39%, si | MM_4 |
| GOBP_MESENCHYMAL_C | GOBP_MESENCHYMAL_CELL_PROLIFERATIO | 55 | -0.36723 | -1.6463445 | 0.0075662043 | 0.14780027 | 1 | 760 | tags=16%, list=3%, sig | MM_4 |
| GOBP_REGULATION_OF_ | GOBP_REGULATION_OF_CYCLIN_DEPENDEN | 67 | -0.34963 | -1.643199 | 0.00973236 | 0.15069309 | 1 | 1794 | tags=19%, list=8%, sig | MM_4 |
| GOBP_GLUTAMATERGIC_ | GOBP_GLUTAMATERGIC_NEURON_DIFFERE | 25 | -0.45336 | -1.6412562 | 0.014550265 | 0.15219499 | 1 | 7658 | tags=60%, list=32%, si | MM_4 |
| GOBP_REGULATION_OF_ | GOBP_REGULATION_OF_MITOTIC_CYTOKIN | 15 | -0.50797 | -1.6370597 | 0.01724138 | 0.1566169 | 1 | 2278 | tags=40%, list=10%, si | MM_4 |
| GOBP_AXONEME_ASSEM | GOBP_AXONEME_ASSEMBLY | 83 | -0.32991 | -1.6335721 | 0.0012135922 | 0.16028039 | 1 | 12202 | tags=70%, list=52%, si | MM_4 |
| GOBP_POSITIVE_REGULA | GOBP_POSITIVE_REGULATION_OF_CELL_CY | 330 | -0.27069 | -1.6335138 | 0 | 0.15912353 | 1 | 2415 | tags=17%, list=10%, si | MM_4 |

| | | | | | | | | | | |
|---|---|---|---|---|---|---|---|---|---|---|
| GOBP_NEGATIVE_REGUL | GOBP_NEGATIVE_REGULATION_OF_ORGAN | 47 | -0.37837 | -1.631656 | 0.012578616 | 0.16051587 | 1 | 1782 | tags=19%, list=8%, sig | MM_4 |
| GOBP_REGULATION_OF_ | GOBP_REGULATION_OF_ATTACHMENT_OF_ | 20 | -0.46107 | -1.630758 | 0.021156559 | 0.1604842 | 1 | 1794 | tags=30%, list=8%, sig | MM_4 |
| GOBP_POSITIVE_REGULA | GOBP_POSITIVE_REGULATION_OF_P38MAP | 30 | -0.42011 | -1.629980 | 0.021978023 | 0.16033505 | 1 | 1730 | tags=20%, list=7%, sig | MM_4 |
| GOBP_MITOTIC_CELL_CY | GOBP_MITOTIC_CELL_CYCLE_PHASE_TRANS | 425 | -0.26364 | -1.628453 | 0 | 0.16130961 | 1 | 2415 | tags=17%, list=10%, si | MM_4 |
| GOBP_REGULATION_OF_ | GOBP_REGULATION_OF_FIBROBLAST_PROL | 122 | -0.31303 | -1.627816 | 0.0011627907 | 0.16107732 | 1 | 2356 | tags=20%, list=10%, si | MM_4 |
| GOBP_SENSORY_PERCEPT | GOBP_SENSORY_PERCEPTION_OF_TASTE | 33 | -0.41205 | -1.626890 | 0.006418485 | 0.16109313 | 1 | 11295 | tags=73%, list=48%, si | MM_4 |
| GOBP_EYE_PHOTORECEP | GOBP_EYE_PHOTORECEPTOR_CELL_DIFFERE | 58 | -0.35292 | -1.625487 | 0.014814815 | 0.16193873 | 1 | 12871 | tags=74%, list=55%, si | MM_4 |
| GOBP_SISTER_CHROMAT | GOBP_SISTER_CHROMATID_COHESION | 54 | -0.36217 | -1.623082 | 0.01511335 | 0.16413207 | 1 | 2415 | tags=20%, list=10%, si | MM_4 |
| GOBP_REGULATION_OF_ | GOBP_REGULATION_OF_MITOTIC_CELL_CY | 490 | -0.25988 | -1.622704 | 0 | 0.16354214 | 1 | 2356 | tags=17%, list=10%, si | MM_4 |
| GOBP_MEIOTIC_CHROMO | GOBP_MEIOTIC_CHROMOSOME_SEGREGAT | 94 | -0.32833 | -1.618414 | 0.007380074 | 0.1686861 | 1 | 1857 | tags=19%, list=8%, sig | MM_4 |
| GOBP_MEIOTIC_CELL_CY | GOBP_MEIOTIC_CELL_CYCLE_PROCESS | 220 | -0.28159 | -1.616833 | 0 | 0.16971904 | 1 | 1993 | tags=17%, list=8%, sig | MM_4 |
| LINOLEIC_ACID_METABO | LINOLEIC_ACID_METABOLISM | 33 | -0.40632 | -1.616473 | 0.0131752305 | 0.16908425 | 1 | 12133 | tags=82%, list=51%, si | MM_4 |
| GOBP_CAMERA_TYPE_EY | GOBP_CAMERA_TYPE_EYE_PHOTORECEPTO | 41 | -0.38076 | -1.614587 | 0.01058201 | 0.17081922 | 1 | 10954 | tags=68%, list=46%, si | MM_4 |
| GOBP_REGULATION_OF_ | GOBP_REGULATION_OF_CILIUM_BEAT_FRE | 15 | -0.50353 | -1.612987 | 0.031724136 | 0.17199163 | 1 | 9536 | tags=73%, list=40%, si | MM_4 |
| GOBP_MICROTUBULE_BA | GOBP_MICROTUBULE_BASED_PROTEIN_TRA | 16 | -0.50542 | -1.606117 | 0.017142856 | 0.18123008 | 1 | 762 | tags=25%, list=3%, sig | MM_4 |
| GOBP_SEGMENT_SPECIFI | GOBP_SEGMENT_SPECIFICATION | 17 | -0.49968 | -1.602133 | 0.021947874 | 0.18597974 | 1 | 10717 | tags=82%, list=45%, si | MM_4 |
| GOBP_NEPHRON_TUBULI | GOBP_NEPHRON_TUBULE_FORMATION | 18 | -0.48079 | -1.601472 | 0.017857144 | 0.1857765 | 1 | 10178 | tags=83%, list=43%, si | MM_4 |
| GOBP_XENOPHAGY | GOBP_XENOPHAGY | 17 | -0.49347 | -1.599037 | 0.021276595 | 0.18844105 | 1 | 1606 | tags=24%, list=7%, sig | MM_4 |
| GOBP_REGULATION_OF_ | GOBP_REGULATION_OF_MITOTIC_CELL_CY | 331 | -0.26546 | -1.596179 | 0 | 0.19182354 | 1 | 2415 | tags=17%, list=10%, si | MM_4 |
| GOBP_INTRACELLULAR_N | GOBP_INTRACELLULAR_MRNA_LOCALIZATI | 15 | -0.50953 | -1.596115 | 0.023255814 | 0.19070846 | 1 | 3307 | tags=27%, list=14%, si | MM_4 |
| GOBP_ASTROCYTE_DEVE | GOBP_ASTROCYTE_DEVELOPMENT | 36 | -0.39572 | -1.593656 | 0.014102564 | 0.19318616 | 1 | 1262 | tags=17%, list=5%, sig | MM_4 |
| GOBP_NON_MEMBRANE | GOBP_NON_MEMBRANE_BOUNDED_ORGA | 405 | -0.26167 | -1.589249 | 0 | 0.19920084 | 1 | 2364 | tags=15%, list=10%, si | MM_4 |
| GOBP_INNER_EAR_MORF | GOBP_INNER_EAR_MORPHOGENESIS | 108 | -0.31157 | -1.586783 | 0.004716981 | 0.20182262 | 1 | 12298 | tags=67%, list=52%, si | MM_4 |
| GOBP_POLYSACCHARIDE | GOBP_POLYSACCHARIDE_BIOSYNTHETIC_PR | 65 | -0.34217 | -1.585373 | 0.01875 | 0.2028305 | 1 | 1458 | tags=15%, list=6%, sig | MM_4 |
| GOBP_SENSORY_PERCEPT | GOBP_SENSORY_PERCEPTION_OF_CHEMICA | 69 | -0.34102 | -1.584664 | 0.012345679 | 0.202751 | 1 | 11429 | tags=70%, list=48%, si | MM_4 |
| GOBP_POSITIVE_REGULA | GOBP_POSITIVE_REGULATION_OF_CELL_CY | 31 | -0.40211 | -1.583872 | 0.017402945 | 0.20294312 | 1 | 1993 | tags=23%, list=8%, sig | MM_4 |
| GOBP_PHOSPHATE_ION_ | GOBP_PHOSPHATE_ION_TRANSPORT | 25 | -0.43073 | -1.580077 | 0.02734375 | 0.20804633 | 1 | 9643 | tags=56%, list=41%, si | MM_4 |
| GOBP_REGULATION_OF_ | GOBP_REGULATION_OF_PRESYNAPSE_ORG | 43 | -0.3711 | -1.578899 | 0.024675325 | 0.2087023 | 1 | 10699 | tags=70%, list=45%, si | MM_4 |
| GOBP_PROTEIN_LOCALIZ | GOBP_PROTEIN_LOCALIZATION_TO_NUCLE | 15 | -0.50158 | -1.577581 | 0.02367688 | 0.20942159 | 1 | 718 | tags=20%, list=3%, sig | MM_4 |
| GOBP_REGULATION_OF_ | GOBP_REGULATION_OF_FLAGELLATED_SPE | 15 | -0.50482 | -1.577095 | 0.031636864 | 0.20903642 | 1 | 11007 | tags=73%, list=47%, si | MM_4 |
| GOBP_NON_MOTILE_CILI | GOBP_NON_MOTILE_CILIUM_ASSEMBLY | 73 | -0.32991 | -1.576781 | 0.013142174 | 0.20843585 | 1 | 2187 | tags=16%, list=9%, sig | MM_4 |
| GOBP_SULFATION | GOBP_SULFATION | 20 | -0.45115 | -1.576057 | 0.02112676 | 0.20833367 | 1 | 10926 | tags=65%, list=46%, si | MM_4 |
| GOBP_NEGATIVE_REGUL | GOBP_NEGATIVE_REGULATION_OF_CILIUM | 21 | -0.44728 | -1.566159 | 0.026737968 | 0.22451851 | 1 | 917 | tags=24%, list=4%, sig | MM_4 |
| GOBP_MRNA_TRANSCRIP | GOBP_MRNA_TRANSCRIPTION | 50 | -0.35574 | -1.565792 | 0.014322917 | 0.22389735 | 1 | 2319 | tags=18%, list=10%, si | MM_4 |
| GOBP_T_HELPER_1_TYPE | GOBP_T_HELPER_1_TYPE_IMMUNE_RESPOI | 53 | -0.35094 | -1.563489 | 0.016434893 | 0.22676016 | 1 | 2676 | tags=25%, list=11%, si | MM_4 |
| GOBP_POSITIVE_REGULA | GOBP_POSITIVE_REGULATION_OF_HEART_ | 29 | -0.4043 | -1.563029 | 0.024707412 | 0.22635327 | 1 | 11821 | tags=76%, list=50%, si | MM_4 |
| GOBP_CELL_CELL_SIGNA | GOBP_CELL_CELL_SIGNALING_INVOLVED_IN | 25 | -0.42611 | -1.562704 | 0.017663043 | 0.22560787 | 1 | 11363 | tags=76%, list=48%, si | MM_4 |
| GOBP_REGULATION_OF_ | GOBP_REGULATION_OF_GLUCONEOGENESI | 52 | -0.35106 | -1.558307 | 0.018726591 | 0.23237927 | 1 | 1611 | tags=17%, list=7%, sig | MM_4 |
| GOBP_GLUTAMATE_MET | GOBP_GLUTAMATE_METABOLIC_PROCESS | 30 | -0.40548 | -1.558018 | 0.025232404 | 0.23150998 | 1 | 1794 | tags=23%, list=8%, sig | MM_4 |
| GOBP_POSITIVE_REGULA | GOBP_POSITIVE_REGULATION_OF_RELEASE | 27 | -0.41544 | -1.557362 | 0.03137255 | 0.23152049 | 1 | 2328 | tags=19%, list=10%, si | MM_4 |
| GOBP_REGULATION_OF_ | GOBP_REGULATION_OF_PROTEIN_LOCALIZ | 15 | -0.50061 | -1.555323 | 0.03301238 | 0.233907 | 1 | 2196 | tags=27%, list=9%, sig | MM_4 |
| GOBP_REGULATION_OF_ | GOBP_REGULATION_OF_CELL_CYCLE_PHAS | 424 | -0.25214 | -1.553755 | 0 | 0.23551616 | 1 | 2415 | tags=16%, list=10%, si | MM_4 |
| GOBP_POSITIVE_REGULA | GOBP_POSITIVE_REGULATION_OF_FIBROBL | 76 | -0.32092 | -1.552104 | 0.011083744 | 0.23745531 | 1 | 1962 | tags=18%, list=8%, sig | MM_4 |
| GOBP_PIRNA_PROCESSIN | GOBP_PIRNA_PROCESSING | 18 | -0.46885 | -1.551829 | 0.027816411 | 0.2365746 | 1 | 10250 | tags=78%, list=43%, si | MM_4 |
| GOBP_REGULATION_OF_ | GOBP_REGULATION_OF_CILIUM_ASSEMBLY | 73 | -0.32539 | -1.551623 | 0.014510278 | 0.23557058 | 1 | 1541 | tags=15%, list=7%, sig | MM_4 |
| GOBP_EPOXYGENASE_P4 | GOBP_EPOXYGENASE_P450_PATHWAY | 20 | -0.45592 | -1.549645 | 0.027285129 | 0.23785536 | 1 | 8281 | tags=70%, list=35%, si | MM_4 |
| GOBP_REGULATION_OF_ | GOBP_REGULATION_OF_PEPTIDYL_TYROSIN | 238 | -0.26935 | -1.548735 | 0.0021715527 | 0.23829795 | 1 | 2495 | tags=18%, list=11%, si | MM_4 |
| GOBP_G_PROTEIN_COUP | GOBP_G_PROTEIN_COUPLED_RECEPTOR_IN | 19 | -0.46176 | -1.548349 | 0.038081806 | 0.23773842 | 1 | 3838 | tags=42%, list=16%, si | MM_4 |
| GOBP_VENTRICULAR_SYS | GOBP_VENTRICULAR_SYSTEM_DEVELOPME | 36 | -0.38161 | -1.547693 | 0.025469169 | 0.23764424 | 1 | 1783 | tags=19%, list=8%, sig | MM_4 |
| GOBP_NEGATIVE_REGUL | GOBP_NEGATIVE_REGULATION_OF_COAGU | 43 | -0.36812 | -1.545460 | 0.022020726 | 0.24053495 | 1 | 5021 | tags=40%, list=21%, si | MM_4 |
| GOBP_NUCLEOLUS_ORGA | GOBP_NUCLEOLUS_ORGANIZATION | 17 | -0.45897 | -1.543755 | 0.032033425 | 0.2426153 | 1 | 2248 | tags=41%, list=10%, si | MM_4 |
| GOBP_MEIOTIC_CELL_CY | GOBP_MEIOTIC_CELL_CYCLE | 282 | -0.25974 | -1.540903 | 0.0010787486 | 0.24684198 | 1 | 3369 | tags=21%, list=14%, si | MM_4 |
| GOBP_SENSORY_PERCEPT | GOBP_SENSORY_PERCEPTION | 446 | -0.25223 | -1.540318 | 0 | 0.24662665 | 1 | 11917 | tags=59%, list=50%, si | MM_4 |
| GOBP_NEGATIVE_REGUL | GOBP_NEGATIVE_REGULATION_OF_CHOND | 25 | -0.42127 | -1.537700 | 0.036339164 | 0.24932165 | 1 | 12109 | tags=72%, list=51%, si | MM_4 |
| **iC1qaCKOvsC1qaFL** | | | | | | | | | | |
| GOBP_REGULATION_OF_ | GOBP_REGULATION_OF_MITOCHONDRIAL_ | 39 | -0.59555 | -2.56313 | 0.0 | 0.004022648 | 0.001 | 5440 | tags=54%, list=23%, si | EN_1 |
| GOBP_RESPONSE_TO_GR | GOBP_RESPONSE_TO_GROWTH_HORMONE | 20 | -0.70768 | -2.47578 | 0.0 | 0.007534674 | 0.004 | 5414 | tags=75%, list=23%, si | EN_1 |
| GOBP_CHONDROITIN_SU | GOBP_CHONDROITIN_SULFATE_METABOLIC | 22 | -0.62671 | -2.369 | 0.0 | 0.014857627 | 0.012 | 4651 | tags=59%, list=20%, si | EN_1 |
| NON-SMALL_CELL_LUNG | NON-SMALL_CELL_LUNG_CANCER | 65 | -0.49357 | -2.33686 | 0.0 | 0.016714094 | 0.016 | 5442 | tags=43%, list=23%, si | EN_1 |
| GOBP_MITOCHONDRIAL_ | GOBP_MITOCHONDRIAL_OUTER_MEMBRAN | 34 | -0.55245 | -2.25684 | 0.0 | 0.037268076 | 0.048 | 5440 | tags=56%, list=23%, si | EN_1 |
| STEROID_BIOSYNTHESIS | STEROID_BIOSYNTHESIS | 17 | -0.67434 | -2.25534 | 0.0 | 0.031056728 | 0.048 | 2242 | tags=53%, list=9%, sig | EN_1 |
| GOBP_NEGATIVE_REGUL | GOBP_NEGATIVE_REGULATION_OF_PROGR | 17 | -0.65302 | -2.25473 | 0.0 | 0.026620053 | 0.048 | 3717 | tags=53%, list=16%, si | EN_1 |
| GOBP_MITOCHONDRIAL_ | GOBP_MITOCHONDRIAL_OUTER_MEMBRAN | 28 | -0.58848 | -2.25327 | 0.0 | 0.023292545 | 0.048 | 5440 | tags=54%, list=23%, si | EN_1 |
| GOBP_PHOSPHATIDYLCH | GOBP_PHOSPHATIDYLCHOLINE_METABOLIC | 59 | -0.47831 | -2.24598 | 0.0 | 0.021979451 | 0.051 | 2684 | tags=36%, list=11%, si | EN_1 |
| GOBP_PHOSPHOLIPID_BI | GOBP_PHOSPHOLIPID_BIOSYNTHETIC_PROC | 214 | -0.38883 | -2.21188 | 0.0 | 0.030175034 | 0.076 | 4514 | tags=36%, list=19%, si | EN_1 |
| GOBP_TAIL_ANCHORED_ | GOBP_TAIL_ANCHORED_MEMBRANE_PROT | 15 | -0.67588 | -2.18366 | 0.0 | 0.03317768 | 0.093 | 6466 | tags=73%, list=27%, si | EN_1 |
| GOBP_MAINTENANCE_O | GOBP_MAINTENANCE_OF_PROTEIN_LOCAL | 43 | -0.49485 | -2.14107 | 0.0 | 0.041883223 | 0.122 | 5229 | tags=44%, list=22%, si | EN_1 |
| CHRONIC_MYELOID_LEUK | CHRONIC_MYELOID_LEUKEMIA | 75 | -0.44268 | -2.13886 | 0.0 | 0.0392224 | 0.124 | 5442 | tags=40%, list=23%, si | EN_1 |
| GOBP_NEGATIVE_REGUL | GOBP_NEGATIVE_REGULATION_OF_NUCLE | 20 | -0.58998 | -2.13625 | 0.0 | 0.036959793 | 0.126 | 3051 | tags=45%, list=13%, si | EN_1 |
| GOBP_POSITIVE_REGULA | GOBP_POSITIVE_REGULATION_OF_MEMBR | 39 | -0.5067 | -2.12373 | 0.0 | 0.038964763 | 0.142 | 5440 | tags=51%, list=23%, si | EN_1 |
| GOBP_NEGATIVE_REGUL | GOBP_NEGATIVE_REGULATION_OF_REACTI | 20 | -0.58832 | -2.12335 | 0.0 | 0.036529463 | 0.142 | 6008 | tags=55%, list=25%, si | EN_1 |
| GOBP_MODULATION_BY_ | GOBP_MODULATION_BY_HOST_OF_VIRAL_ | 22 | -0.57418 | -2.10714 | 0.0 | 0.039410494 | 0.163 | 4803 | tags=36%, list=20%, si | EN_1 |
| GOBP_ENDOPLASMIC_RE | GOBP_ENDOPLASMIC_RETICULUM_CALCIU | 28 | -0.53262 | -2.10073 | 0.0 | 0.041176915 | 0.18 | 2766 | tags=32%, list=12%, si | EN_1 |
| GOBP_PEROXISOME_ORG | GOBP_PEROXISOME_ORGANIZATION | 38 | -0.49858 | -2.09731 | 0.0 | 0.03981242 | 0.184 | 4355 | tags=39%, list=18%, si | EN_1 |
| GOBP_REGULATION_OF_ | GOBP_REGULATION_OF_PROGRAMMED_NE | 27 | -0.54405 | -2.09681 | 0.0 | 0.038014214 | 0.185 | 3717 | tags=37%, list=16%, si | EN_1 |
| GOBP_CHONDROITIN_SU | GOBP_CHONDROITIN_SULFATE_PROTEOGLY | 29 | -0.52714 | -2.09345 | 0.0 | 0.037811 | 0.193 | 4651 | tags=59%, list=20%, si | EN_1 |
| PEROXISOME | PEROXISOME | 81 | -0.42532 | -2.08277 | 0.0 | 0.040172234 | 0.214 | 4939 | tags=40%, list=21%, si | EN_1 |
| GOBP_DEOXYRIBONUCLE | GOBP_DEOXYRIBONUCLEOTIDE_BIOSYNTHE | 17 | -0.63909 | -2.0827 | 0.0 | 0.038425613 | 0.214 | 5325 | tags=65%, list=23%, si | EN_1 |
| GOBP_MAINTENANCE_O | GOBP_MAINTENANCE_OF_PROTEIN_LOCAT | 70 | -0.42128 | -2.076 | 0.0 | 0.039023776 | 0.222 | 5280 | tags=40%, list=22%, si | EN_1 |
| GOBP_GLYCEROPHOSPH | GOBP_GLYCEROPHOSPHOLIPID_BIOSYNTHE | 160 | -0.37005 | -2.06564 | 0.0 | 0.04196608 | 0.246 | 4492 | tags=35%, list=19%, si | EN_1 |
| AUTOPHAGY | AUTOPHAGY | 126 | -0.38291 | -2.06024 | 0.0 | 0.041926455 | 0.254 | 5499 | tags=40%, list=23%, si | EN_1 |
| GOBP_STEROL_BIOSYNTH | GOBP_STEROL_BIOSYNTHETIC_PROCESS | 58 | -0.44616 | -2.05892 | 0.0 | 0.040641088 | 0.255 | 2704 | tags=38%, list=11%, si | EN_1 |

| | | | | | | | | | | |
|---|---|---|---|---|---|---|---|---|---|---|
| GOBP_MAINTENANCE_OI | GOBP_MAINTENANCE_OF_PROTEIN_LOCAT | 97 | -0.39724 | -2.01723 | 0.0 | 0.05652814 | 0.345 | 5280 | tags=40%, list=22%, si | EN_1 |
| GOBP_FATTY_ACID_TRAN | GOBP_FATTY_ACID_TRANSMEMBRANE_TRA | 16 | -0.62035 | -2.01351 | 0.0 | 0.056012277 | 0.353 | 5880 | tags=75%, list=25%, si | EN_1 |
| ENDOMETRIAL_CANCER | ENDOMETRIAL_CANCER | 57 | -0.44042 | -1.99902 | 0.0 | 0.061832897 | 0.386 | 6300 | tags=47%, list=27%, si | EN_1 |
| GOBP_ACETYL_COA_MET | GOBP_ACETYL_COA_METABOLIC_PROCESS | 34 | -0.48104 | -1.99888 | 0.0 | 0.059953947 | 0.387 | 3952 | tags=38%, list=17%, si | EN_1 |
| GOBP_PINOCYTOSIS | GOBP_PINOCYTOSIS | 22 | -0.55319 | -1.99163 | 0.0 | 0.061803184 | 0.407 | 3944 | tags=41%, list=17%, si | EN_1 |
| GOBP_MIRNA_PROCESSII | GOBP_MIRNA_PROCESSING | 42 | -0.46792 | -1.98082 | 0.0 | 0.06682387 | 0.438 | 4982 | tags=33%, list=21%, si | EN_1 |
| GOBP_MUCOPOLYSACCH | GOBP_MUCOPOLYSACCHARIDE_METABOLIC | 73 | -0.40863 | -1.98018 | 0.0 | 0.06541137 | 0.44 | 4278 | tags=40%, list=18%, si | EN_1 |
| GOBP_LIPOPROTEIN_BIO: | GOBP_LIPOPROTEIN_BIOSYNTHETIC_PROCE | 104 | -0.373 | -1.97707 | 0.0 | 0.06546992 | 0.455 | 4157 | tags=37%, list=18%, si | EN_1 |
| GOBP_REGULATION_OF_ | GOBP_REGULATION_OF_MITOCHONDRIAL_ | 58 | -0.42797 | -1.9733 | 0.0 | 0.06531563 | 0.462 | 5440 | tags=47%, list=23%, si | EN_1 |
| GOBP_REGULATION_OF_ | GOBP_REGULATION_OF_MITOCHONDRIAL_ | 20 | -0.55643 | -1.97252 | 0.0 | 0.06373989 | 0.463 | 5440 | tags=55%, list=23%, si | EN_1 |
| GOBP_POSITIVE_REGULA | GOBP_POSITIVE_REGULATION_OF_DNA_RE | 135 | -0.35742 | -1.96302 | 0.0 | 0.067111984 | 0.489 | 5858 | tags=36%, list=25%, si | EN_1 |
| GOBP_REGULATION_OF_ | GOBP_REGULATION_OF_CYTOPLASMIC_PAT | 104 | -0.38129 | -1.96237 | 0.0 | 0.06577829 | 0.491 | 4803 | tags=40%, list=20%, si | EN_1 |
| GOBP_REGULATION_OF_ | GOBP_REGULATION_OF_TRANSLATION_IN_ | 24 | -0.52254 | -1.94035 | 0.0 | 0.07660977 | 0.555 | 3906 | tags=42%, list=17%, si | EN_1 |
| GOBP_REGULATION_OF_ | GOBP_REGULATION_OF_CELLULAR_RESPON | 26 | -0.50641 | -1.93992 | 0.00990099 | 0.07474123 | 0.555 | 1705 | tags=31%, list=7%, sig | EN_1 |
| GOBP_REGULATION_OF_ | GOBP_REGULATION_OF_SULFUR_METABOL | 15 | -0.59189 | -1.93288 | 0.004132231 | 0.07664181 | 0.572 | 4511 | tags=47%, list=19%, si | EN_1 |
| GOBP_REGULATION_OF_ | GOBP_REGULATION_OF_RESPONSE_TO_EN | 80 | -0.38529 | -1.92467 | 0.0 | 0.080138706 | 0.604 | 4775 | tags=34%, list=20%, si | EN_1 |
| GLIOMA | GLIOMA | 71 | -0.39927 | -1.92035 | 0.0 | 0.08053105 | 0.614 | 6381 | tags=48%, list=27%, si | EN_1 |
| GOBP_POSTREPLICATION | GOBP_POSTREPLICATION_REPAIR | 29 | -0.48473 | -1.91336 | 0.0 | 0.08385194 | 0.639 | 2568 | tags=21%, list=11%, si | EN_1 |
| GOBP_TRANSLATION_AT_ | GOBP_TRANSLATION_AT_SYNAPSE | 48 | -0.43737 | -1.90242 | 0.0 | 0.08897294 | 0.669 | 12188 | tags=98%, list=52%, si | EN_1 |
| GOBP_NEGATIVE_REGUL | GOBP_NEGATIVE_REGULATION_OF_SMOOT | 36 | -0.47162 | -1.88994 | 0.0 | 0.096778534 | 0.706 | 4471 | tags=44%, list=19%, si | EN_1 |
| GOBP_NUCLEOTIDE_BINI | GOBP_NUCLEOTIDE_BINDING_OLIGOMERIZ | 18 | -0.53935 | -1.88798 | 0.008032128 | 0.09614687 | 0.71 | 2056 | tags=44%, list=9%, sig | EN_1 |
| GOBP_MACROAUTOPHAG | GOBP_MACROAUTOPHAGY | 287 | -0.31867 | -1.88669 | 0.0 | 0.09479408 | 0.713 | 5607 | tags=35%, list=24%, si | EN_1 |
| GOBP_GLYCEROPHOSPH | GOBP_GLYCEROPHOSPHOLIPID_METABOLIC | 267 | -0.32436 | -1.88493 | 0.0 | 0.09431005 | 0.72 | 3678 | tags=28%, list=16%, si | EN_1 |
| GOBP_PHOSPHATIDYLCH | GOBP_PHOSPHATIDYLCHOLINE_BIOSYNTHET | 21 | -0.51614 | -1.88214 | 0.003984064 | 0.0944789 | 0.726 | 1856 | tags=38%, list=8%, sig | EN_1 |
| GLYCOSYLPHOSPHATIDYL | GLYCOSYLPHOSPHATIDYLINOSITOL_(GPI)-AN | 24 | -0.49568 | -1.87942 | 0.0 | 0.09451805 | 0.731 | 5277 | tags=46%, list=22%, si | EN_1 |
| ONE_CARBON_POOL_BY_ | ONE_CARBON_POOL_BY_FOLATE | 18 | -0.54176 | -1.8747 | 0.004310345 | 0.09731702 | 0.745 | 5293 | tags=56%, list=22%, si | EN_1 |
| GOBP_REGULATION_OF_ | GOBP_REGULATION_OF_MACROAUTOPHAG | 129 | -0.35056 | -1.85988 | 0.0 | 0.10687335 | 0.786 | 6374 | tags=43%, list=27%, si | EN_1 |
| GOBP_MAINTENANCE_OI | GOBP_MAINTENANCE_OF_PROTEIN_LOCAT | 25 | -0.47444 | -1.85959 | 0.0 | 0.10519492 | 0.786 | 5595 | tags=60%, list=24%, si | EN_1 |
| PHOSPHATIDYLINOSITOL_ | PHOSPHATIDYLINOSITOL_SIGNALING_SYSTE | 94 | -0.36776 | -1.85945 | 0.0 | 0.10338315 | 0.787 | 4542 | tags=35%, list=19%, si | EN_1 |
| GOBP_REGULATION_OF_ | GOBP_REGULATION_OF_MYOBLAST_FUSIOI | 22 | -0.50896 | -1.85797 | 0.004761905 | 0.10255567 | 0.791 | 4650 | tags=55%, list=20%, si | EN_1 |
| GOBP_HYALURONAN_BI | GOBP_HYALURONAN_BIOSYNTHETIC_PROC | 15 | -0.57171 | -1.85795 | 0.004048583 | 0.10085035 | 0.791 | 4278 | tags=53%, list=18%, si | EN_1 |
| GOBP_TRANSLESION_SYI | GOBP_TRANSLESION_SYNTHESIS | 22 | -0.50417 | -1.85551 | 0.0049751243 | 0.10058999 | 0.795 | 6031 | tags=50%, list=26%, si | EN_1 |
| GOBP_NEGATIVE_REGUL | GOBP_NEGATIVE_REGULATION_OF_CELLUL | 18 | -0.54681 | -1.84851 | 0.016460905 | 0.10445788 | 0.812 | 1705 | tags=39%, list=7%, sig | EN_1 |
| GOBP_NEGATIVE_REGUL | GOBP_NEGATIVE_REGULATION_OF_INSULIN | 17 | -0.55779 | -1.84531 | 0.008695652 | 0.10606178 | 0.823 | 4956 | tags=53%, list=21%, si | EN_1 |
| GOBP_PHOSPHATIDYLIN | GOBP_PHOSPHATIDYLINOSITOL_BIOSYNTHE | 95 | -0.37681 | -1.84452 | 0.0 | 0.10488622 | 0.824 | 4492 | tags=35%, list=19%, si | EN_1 |
| GOBP_REGULATION_OF_ | GOBP_REGULATION_OF_TRANSLATIONAL_I | 17 | -0.53744 | -1.84452 | 0.0 | 0.10322136 | 0.824 | 3906 | tags=35%, list=17%, si | EN_1 |
| BIOSYNTHESIS_OF_UNSA | BIOSYNTHESIS_OF_UNSATURATED_FATTY_A | 31 | -0.45149 | -1.84266 | 0.006329114 | 0.102951825 | 0.826 | 4579 | tags=42%, list=19%, si | EN_1 |
| GOBP_ATP_BIOSYNTHETI | GOBP_ATP_BIOSYNTHETIC_PROCESS | 80 | -0.37619 | -1.84034 | 0.0 | 0.103103824 | 0.831 | 7507 | tags=55%, list=32%, si | EN_1 |
| GOBP_NUCLEOTIDE_BINI | GOBP_NUCLEOTIDE_BINDING_DOMAIN_LEU | 22 | -0.50466 | -1.84006 | 0.0 | 0.10165294 | 0.832 | 2964 | tags=45%, list=13%, si | EN_1 |
| GOBP_AMINOGLYCAN_BI | GOBP_AMINOGLYCAN_BIOSYNTHETIC_PRO | 60 | -0.39145 | -1.83064 | 0.0 | 0.106990516 | 0.853 | 4651 | tags=43%, list=20%, si | EN_1 |
| GOBP_REGULATION_OF_ | GOBP_REGULATION_OF_ATP_BIOSYNTHETI | 20 | -0.52902 | -1.82797 | 0.0076923077 | 0.10865793 | 0.866 | 3051 | tags=40%, list=13%, si | EN_1 |
| GOBP_POSITIVE_REGULA | GOBP_POSITIVE_REGULATION_OF_UBIQUIT | 18 | -0.53782 | -1.8247 | 0.00952381 | 0.11010049 | 0.871 | 6306 | tags=61%, list=27%, si | EN_1 |
| GOBP_REGULATION_OF_ | GOBP_REGULATION_OF_AUTOPHAGOSOME | 44 | -0.41864 | -1.82271 | 0.0 | 0.11010368 | 0.873 | 5571 | tags=41%, list=24%, si | EN_1 |
| GOBP_HYDROGEN_PERO | GOBP_HYDROGEN_PEROXIDE_CATABOLIC_I | 22 | -0.48309 | -1.82004 | 0.0 | 0.1104268 | 0.875 | 1826 | tags=23%, list=8%, sig | EN_1 |
| GOBP_MITOTIC_G2_DNA | GOBP_MITOTIC_G2_DNA_DAMAGE_CHECKI | 39 | -0.43072 | -1.81884 | 0.005882353 | 0.10993042 | 0.876 | 4711 | tags=38%, list=20%, si | EN_1 |
| GOBP_NEGATIVE_REGUL | GOBP_NEGATIVE_REGULATION_OF_SMOOT | 68 | -0.37421 | -1.81652 | 0.0 | 0.109865524 | 0.88 | 4332 | tags=41%, list=18%, si | EN_1 |
| GOBP_GLYCEROPHOSPH | GOBP_GLYCEROPHOSPHOLIPID_CATABOLIC | 31 | -0.45591 | -1.81567 | 0.0 | 0.10927687 | 0.88 | 3005 | tags=32%, list=13%, si | EN_1 |
| GOBP_REGULATION_OF_ | GOBP_REGULATION_OF_CHOLESTEROL_BIO | 19 | -0.49251 | -1.81525 | 0.0045454544 | 0.10822176 | 0.881 | 4649 | tags=47%, list=20%, si | EN_1 |
| GOBP_CHONDROITIN_SU | GOBP_CHONDROITIN_SULFATE_PROTEOGLY | 21 | -0.50642 | -1.81161 | 0.004385965 | 0.11006322 | 0.889 | 3024 | tags=52%, list=13%, si | EN_1 |
| GOBP_NEGATIVE_REGUL | GOBP_NEGATIVE_REGULATION_OF_VASCU | 33 | -0.44172 | -1.811 | 0.0 | 0.109118745 | 0.892 | 5107 | tags=52%, list=22%, si | EN_1 |
| GOBP_CHAPERONE_MED | GOBP_CHAPERONE_MEDIATED_PROTEIN_C | 18 | -0.52665 | -1.80824 | 0.007968128 | 0.10965816 | 0.895 | 2698 | tags=39%, list=11%, si | EN_1 |
| GOBP_TETRAHYDROFOLA | GOBP_TETRAHYDROFOLATE_METABOLIC_PI | 20 | -0.5015 | -1.80627 | 0.0 | 0.110074 | 0.901 | 5293 | tags=60%, list=22%, si | EN_1 |
| GLYCOSAMINOGLYCAN_I | GLYCOSAMINOGLYCAN_DEGRADATION | 17 | -0.53107 | -1.80602 | 0.0074626864 | 0.108790025 | 0.901 | 2503 | tags=41%, list=11%, si | EN_1 |
| GOBP_POSITIVE_REGULA | GOBP_POSITIVE_REGULATION_OF_PROTEO | 120 | -0.34528 | -1.803 | 0.012345679 | 0.11029694 | 0.908 | 5526 | tags=38%, list=23%, si | EN_1 |
| GOBP_NADH_DEHYDROG | GOBP_NADH_DEHYDROGENASE_COMPLEX_ | 58 | -0.38243 | -1.79924 | 0.0 | 0.112426184 | 0.914 | 6734 | tags=47%, list=29%, si | EN_1 |
| GOBP_REGULATION_OF_ | GOBP_REGULATION_OF_TELOMERE_MAINT | 20 | -0.49903 | -1.7982 | 0.0 | 0.11178758 | 0.915 | 5858 | tags=45%, list=25%, si | EN_1 |
| GOBP_NEGATIVE_REGUL | GOBP_NEGATIVE_REGULATION_OF_CELL_SI | 16 | -0.52794 | -1.79422 | 0.0038910506 | 0.1138176 | 0.919 | 3616 | tags=50%, list=15%, si | EN_1 |
| GOBP_NADH_METABOLI | GOBP_NADH_METABOLIC_PROCESS | 39 | -0.42679 | -1.79402 | 0.0 | 0.1128307 | 0.92 | 5003 | tags=49%, list=21%, si | EN_1 |
| GOBP_DIOL_BIOSYNTHET | GOBP_DIOL_BIOSYNTHETIC_PROCESS | 21 | -0.50214 | -1.79329 | 0.008658009 | 0.11203794 | 0.921 | 4909 | tags=48%, list=21%, si | EN_1 |
| PPAR_SIGNALING_PATHV | PPAR_SIGNALING_PATHWAY | 70 | -0.39012 | -1.79124 | 0.0 | 0.11211496 | 0.926 | 3451 | tags=39%, list=15%, si | EN_1 |
| GOBP_PROTEIN_DESTABI | GOBP_PROTEIN_DESTABILIZATION | 56 | -0.40057 | -1.79047 | 0.0 | 0.11122302 | 0.926 | 4814 | tags=30%, list=20%, si | EN_1 |
| GOBP_REGULATION_OF_ | GOBP_REGULATION_OF_FIBROBLAST_MIGR | 47 | -0.39462 | -1.78819 | 0.0 | 0.112320416 | 0.927 | 3426 | tags=32%, list=15%, si | EN_1 |
| GOBP_RESPONSE_TO_OX | GOBP_RESPONSE_TO_OXYGEN_RADICAL | 24 | -0.47526 | -1.78787 | 0.004784689 | 0.11132296 | 0.927 | 3939 | tags=38%, list=17%, si | EN_1 |
| GOBP_PHOSPHOLIPID_M | GOBP_PHOSPHOLIPID_METABOLIC_PROCES | 352 | -0.32168 | -1.78775 | 0.0 | 0.11009963 | 0.927 | 4542 | tags=34%, list=19%, si | EN_1 |
| GOBP_GLYCOLYTIC_PROC | GOBP_GLYCOLYTIC_PROCESS_THROUGH_GI | 20 | -0.50718 | -1.78427 | 0.004484305 | 0.11181734 | 0.934 | 4679 | tags=60%, list=20%, si | EN_1 |
| GOBP_NUCLEOSIDE_BISP | GOBP_NUCLEOSIDE_BISPHOSPHATE_BIOSYN | 39 | -0.42134 | -1.7826 | 0.0 | 0.112055175 | 0.935 | 4396 | tags=33%, list=19%, si | EN_1 |
| GOBP_POSITIVE_REGULA | GOBP_POSITIVE_REGULATION_OF_RESPONS | 37 | -0.43451 | -1.7807 | 0.006289308 | 0.1128084 | 0.937 | 4775 | tags=43%, list=20%, si | EN_1 |
| GOBP_PROGRAMMED_N | GOBP_PROGRAMMED_NECROTIC_CELL_DE/ | 50 | -0.40217 | -1.78043 | 0.0 | 0.11193973 | 0.938 | 4171 | tags=34%, list=18%, si | EN_1 |
| GOBP_REGULATION_OF_ | GOBP_REGULATION_OF_NECROPTOTIC_PRO | 22 | -0.48005 | -1.77889 | 0.009661836 | 0.1118978 | 0.939 | 3717 | tags=36%, list=16%, si | EN_1 |
| GOBP_PEPTIDYL_TYROSIN | GOBP_PEPTIDYL_TYROSINE_DEPHOSPHORYI | 33 | -0.44591 | -1.77717 | 0.0 | 0.112168536 | 0.94 | 3983 | tags=45%, list=17%, si | EN_1 |
| GOBP_NEGATIVE_REGUL | GOBP_NEGATIVE_REGULATION_OF_NUCLEC | 29 | -0.47282 | -1.77672 | 0.0 | 0.11144153 | 0.941 | 3051 | tags=38%, list=13%, si | EN_1 |
| GOBP_NEGATIVE_REGUL | GOBP_NEGATIVE_REGULATION_OF_CELL_M | 36 | -0.41498 | -1.77667 | 0.0 | 0.110354036 | 0.941 | 4332 | tags=42%, list=18%, si | EN_1 |
| GOBP_POSITIVE_REGULA | GOBP_POSITIVE_REGULATION_OF_PROTEAS | 102 | -0.35277 | -1.77583 | 0.0 | 0.11018672 | 0.942 | 5506 | tags=36%, list=23%, si | EN_1 |
| GOBP_REGULATION_OF_ | GOBP_REGULATION_OF_PHOSPHOLIPID_BI | 20 | -0.49562 | -1.77448 | 0.014423077 | 0.11012851 | 0.944 | 1364 | tags=30%, list=6%, sig | EN_1 |
| GOBP_AUTOPHAGOSOMI | GOBP_AUTOPHAGOSOME_ORGANIZATION | 128 | -0.33338 | -1.77336 | 0.0 | 0.1098556 | 0.944 | 5797 | tags=41%, list=25%, si | EN_1 |
| GOBP_NEGATIVE_REGUL | GOBP_NEGATIVE_REGULATION_OF_REACTI | 52 | -0.39017 | -1.76868 | 0.0 | 0.112750046 | 0.948 | 3563 | tags=25%, list=15%, si | EN_1 |
| GOBP_POSITIVE_REGULA | GOBP_POSITIVE_REGULATION_OF_TYPE_I_I | 19 | -0.50196 | -1.76423 | 0.004484305 | 0.11537099 | 0.953 | 4483 | tags=58%, list=19%, si | EN_1 |
| GOBP_PHOSPHATIDYLIN | GOBP_PHOSPHATIDYLINOSITOL_PHOSPHAT | 50 | -0.40505 | -1.76297 | 0.0 | 0.11496465 | 0.954 | 4332 | tags=36%, list=18%, si | EN_1 |
| SNARE_INTERACTIONS_IN | SNARE_INTERACTIONS_IN_VESICULAR_TRAI | 32 | -0.44297 | -1.76292 | 0.0 | 0.11394675 | 0.954 | 3783 | tags=28%, list=16%, si | EN_1 |
| GOBP_POSITIVE_REGULA | GOBP_POSITIVE_REGULATION_BY_HOST_OI | 22 | -0.48853 | -1.76009 | 0.009661836 | 0.11519072 | 0.96 | 3736 | tags=36%, list=16%, si | EN_1 |
| GOBP_PROTEIN_LOCALIZ | GOBP_PROTEIN_LOCALIZATION_TO_PHAGO | 16 | -0.53402 | -1.75735 | 0.011235955 | 0.11653222 | 0.964 | 6086 | tags=75%, list=26%, si | EN_1 |
| GOBP_RESPONSE_TO_MI | GOBP_RESPONSE_TO_MISFOLDED_PROTEIN | 18 | -0.51104 | -1.75234 | 0.0 | 0.11960913 | 0.968 | 3166 | tags=33%, list=13%, si | EN_1 |

| | | | | | | | | | | |
|---|---|---|---|---|---|---|---|---|---|---|
| GOBP_POSITIVE_REGULA | GOBP_POSITIVE_REGULATION_OF_UBIQUIT | 93 | -0.33979 | -1.75161 | 0.0 | 0.11916392 | 0.969 | 3550 | tags=25%, list=15%, si | EN_1 |
| GOBP_CELLULAR_RESPON | GOBP_CELLULAR_RESPONSE_TO_AMYLOID_ | 33 | -0.42926 | -1.74914 | 0.0 | 0.1200524 | 0.974 | 3686 | tags=33%, list=16%, si | EN_1 |
| GOBP_TRNA_MODIFICAT | GOBP_TRNA_MODIFICATION | 84 | -0.34841 | -1.74343 | 0.0 | 0.1245835 | 0.976 | 5618 | tags=39%, list=24%, si | EN_1 |
| GOBP_PROTEIN_LOCALIZ | GOBP_PROTEIN_LOCALIZATION_TO_CELL_C | 24 | -0.46314 | -1.74152 | 0.0049751243 | 0.1249999 | 0.977 | 7372 | tags=75%, list=31%, si | EN_1 |
| GOBP_CYTOPLASMIC_PA | GOBP_CYTOPLASMIC_PATTERN_RECOGNITI | 159 | -0.31649 | -1.74039 | 0.0 | 0.124900624 | 0.978 | 4309 | tags=34%, list=18%, si | EN_1 |
| GOBP_PORE_COMPLEX_A | GOBP_PORE_COMPLEX_ASSEMBLY | 20 | -0.48278 | -1.7389 | 0.0093896715 | 0.124789424 | 0.979 | 5156 | tags=50%, list=22%, si | EN_1 |
| GOBP_CELLULAR_RESPON | GOBP_CELLULAR_RESPONSE_TO_OXYGEN_L | 133 | -0.31842 | -1.73811 | 0.0 | 0.12428618 | 0.979 | 4611 | tags=33%, list=20%, si | EN_1 |
| GOBP_POSITIVE_REGULA | GOBP_POSITIVE_REGULATION_OF_ERAD_PA | 20 | -0.48861 | -1.73626 | 0.02202643 | 0.12482285 | 0.98 | 4775 | tags=55%, list=20%, si | EN_1 |
| BACTERIAL_INVASION_OI | BACTERIAL_INVASION_OF_EPITHELIAL_CELL | 72 | -0.36527 | -1.7317 | 0.0 | 0.12780538 | 0.982 | 5682 | tags=39%, list=24%, si | EN_1 |
| GOBP_NEGATIVE_REGUL | GOBP_NEGATIVE_REGULATION_OF_CELL_C | 69 | -0.35862 | -1.73103 | 0.0 | 0.12710807 | 0.982 | 4370 | tags=30%, list=19%, si | EN_1 |
| INSULIN_RESISTANCE | INSULIN_RESISTANCE | 105 | -0.33543 | -1.73033 | 0.0 | 0.12676598 | 0.982 | 4332 | tags=35%, list=18%, si | EN_1 |
| GOBP_RETROGRADE_VES | GOBP_RETROGRADE_VESICLE_MEDIATED_T | 45 | -0.39965 | -1.72972 | 0.0 | 0.12621069 | 0.982 | 5603 | tags=36%, list=24%, si | EN_1 |
| GOBP_CELLULAR_PIGMEI | GOBP_CELLULAR_PIGMENTATION | 57 | -0.37646 | -1.72777 | 0.008 | 0.12666279 | 0.982 | 4379 | tags=33%, list=19%, si | EN_1 |
| GOBP_ER_NUCLEUS_SIGN | GOBP_ER_NUCLEUS_SIGNALING_PATHWAY | 43 | -0.40128 | -1.72675 | 0.0066225166 | 0.12669408 | 0.982 | 5609 | tags=42%, list=24%, si | EN_1 |
| GOBP_REGULATION_OF_ | GOBP_REGULATION_OF_VACUOLE_ORGANI | 54 | -0.37386 | -1.72436 | 0.0 | 0.12797706 | 0.984 | 5571 | tags=37%, list=24%, si | EN_1 |
| GOBP_PHOSPHOLIPID_DE | GOBP_PHOSPHOLIPID_DEPHOSPHORYLATIO | 43 | -0.39434 | -1.72395 | 0.007352941 | 0.12739766 | 0.984 | 4140 | tags=40%, list=18%, si | EN_1 |
| GOBP_MEMBRANE_BIOG | GOBP_MEMBRANE_BIOGENESIS | 63 | -0.36275 | -1.72275 | 0.0 | 0.1272679 | 0.984 | 4547 | tags=37%, list=19%, si | EN_1 |
| GOBP_DNA_REPAIR_DEPI | GOBP_DNA_REPAIR_DEPENDENT_CHROMA | 23 | -0.46961 | -1.72145 | 0.023364486 | 0.12737465 | 0.984 | 5595 | tags=39%, list=24%, si | EN_1 |
| GOBP_POSITIVE_REGULA | GOBP_POSITIVE_REGULATION_OF_INFLAMI | 22 | -0.47415 | -1.71634 | 0.013761468 | 0.13102952 | 0.989 | 2553 | tags=32%, list=11%, si | EN_1 |
| SPLICEOSOME | SPLICEOSOME | 127 | -0.32976 | -1.71557 | 0.0 | 0.1306706 | 0.99 | 15152 | tags=99%, list=64%, si | EN_1 |
| GOBP_MYELIN_MAINTEN | GOBP_MYELIN_MAINTENANCE | 16 | -0.5076 | -1.71435 | 0.021367522 | 0.13061525 | 0.99 | 360 | tags=13%, list=2%, sig | EN_1 |
| GOBP_STEROL_METABOL | GOBP_STEROL_METABOLIC_PROCESS | 140 | -0.31262 | -1.71141 | 0.0 | 0.13237122 | 0.992 | 2813 | tags=29%, list=12%, si | EN_1 |
| GOBP_PROTEIN_HETERO | GOBP_PROTEIN_HETEROOLIGOMERIZATION | 27 | -0.4387 | -1.70674 | 0.010695187 | 0.13558127 | 0.993 | 472 | tags=15%, list=2%, sig | EN_1 |
| GOBP_PIGMENT_GRANU | GOBP_PIGMENT_GRANULE_ORGANIZATION | 39 | -0.39192 | -1.70447 | 0.0 | 0.13643615 | 0.993 | 4101 | tags=28%, list=17%, si | EN_1 |
| GOBP_PROTEIN_INSERTI | GOBP_PROTEIN_INSERTION_INTO_ER_MEM | 26 | -0.44157 | -1.7028 | 0.0 | 0.13715355 | 0.994 | 6466 | tags=54%, list=27%, si | EN_1 |
| GOBP_RNA_SURVEILLAN | GOBP_RNA_SURVEILLANCE | 17 | -0.5068 | -1.70118 | 0.015625 | 0.13778844 | 0.994 | 11653 | tags=100%, list=49%, | EN_1 |
| GOBP_MODULATION_BY_ | GOBP_MODULATION_BY_HOST_OF_VIRAL_ | 44 | -0.40103 | -1.69984 | 0.0 | 0.1380578 | 0.996 | 3906 | tags=27%, list=17%, si | EN_1 |
| GOBP_ANATOMICAL_STR | GOBP_ANATOMICAL_STRUCTURE_ARRANGI | 22 | -0.46366 | -1.69916 | 0.024038462 | 0.13770095 | 0.996 | 1716 | tags=27%, list=7%, sig | EN_1 |
| GOBP_SPHINGOID_META | GOBP_SPHINGOID_METABOLIC_PROCESS | 21 | -0.48385 | -1.69897 | 0.00921659 | 0.13683572 | 0.996 | 1591 | tags=29%, list=7%, sig | EN_1 |
| GOBP_MORPHOGENESIS_ | GOBP_MORPHOGENESIS_OF_AN_ENDOTHE | 19 | -0.48591 | -1.69349 | 0.023809524 | 0.14137898 | 0.996 | 3988 | tags=42%, list=17%, si | EN_1 |
| GOBP_MITOCHONDRIAL_ | GOBP_MITOCHONDRIAL_TRANSPORT | 174 | -0.30192 | -1.69271 | 0.0 | 0.1411919 | 0.996 | 3670 | tags=23%, list=16%, si | EN_1 |
| GOBP_MEMBRANE_RAFT | GOBP_MEMBRANE_RAFT_ORGANIZATION | 24 | -0.4609 | -1.69185 | 0.014285714 | 0.14111397 | 0.996 | 6539 | tags=54%, list=28%, si | EN_1 |
| GOBP_CELL_SUBSTRATE_ | GOBP_CELL_SUBSTRATE_JUNCTION_ORGAN | 105 | -0.33709 | -1.69064 | 0.0 | 0.14143582 | 0.996 | 5224 | tags=41%, list=22%, si | EN_1 |
| GOBP_ERYTHROCYTE_DE | GOBP_ERYTHROCYTE_DEVELOPMENT | 50 | -0.38225 | -1.68976 | 0.0069444445 | 0.14151157 | 0.997 | 2330 | tags=24%, list=10%, si | EN_1 |
| GOBP_CELLULAR_RESPON | GOBP_CELLULAR_RESPONSE_TO_OXYGEN_ | 20 | -0.47057 | -1.68965 | 0.013824885 | 0.14068425 | 0.997 | 3939 | tags=35%, list=17%, si | EN_1 |
| GOBP_RESPONSE_TO_IN | GOBP_RESPONSE_TO_INTERLEUKIN_1 | 76 | -0.34135 | -1.68693 | 0.0 | 0.14207965 | 0.998 | 3812 | tags=33%, list=16%, si | EN_1 |
| GOBP_TRNA_PROCESSIN | GOBP_TRNA_PROCESSING | 122 | -0.31629 | -1.68657 | 0.0 | 0.14154996 | 0.998 | 5618 | tags=37%, list=24%, si | EN_1 |
| GOBP_REGULATION_OF_ | GOBP_REGULATION_OF_MEMBRANE_PERM | 70 | -0.36655 | -1.68635 | 0.0 | 0.14074048 | 0.998 | 5440 | tags=41%, list=23%, si | EN_1 |
| RIBOSOME | RIBOSOME | 128 | -0.3166 | -1.68361 | 0.0 | 0.14207587 | 0.998 | 14528 | tags=98%, list=62%, si | EN_1 |
| GOBP_POSITIVE_REGULA | GOBP_POSITIVE_REGULATION_OF_FATTY_A | 20 | -0.46879 | -1.68327 | 0.0045871558 | 0.14154588 | 0.998 | 3943 | tags=50%, list=17%, si | EN_1 |
| RENAL_CELL_CARCINOM | RENAL_CELL_CARCINOMA | 67 | -0.3603 | -1.68047 | 0.0 | 0.14329027 | 0.998 | 5666 | tags=37%, list=24%, si | EN_1 |
| GOBP_REGULATION_OF_ | GOBP_REGULATION_OF_UBIQUITIN_PROTE | 39 | -0.39177 | -1.67974 | 0.0 | 0.1432791 | 0.999 | 6473 | tags=54%, list=27%, si | EN_1 |
| GOBP_COPII_COATED_VE | GOBP_COPII_COATED_VESICLE_CARGO_LO | 17 | -0.49331 | -1.67819 | 0.01746725 | 0.14373615 | 0.999 | 2855 | tags=24%, list=12%, si | EN_1 |
| GOBP_CELLULAR_RESPON | GOBP_CELLULAR_RESPONSE_TO_HYPOXIA | 112 | -0.32807 | -1.67759 | 0.0 | 0.14338027 | 0.999 | 4325 | tags=32%, list=18%, si | EN_1 |
| GOBP_DNA_DUPLEX_UN | GOBP_DNA_DUPLEX_UNWINDING | 43 | -0.38694 | -1.67744 | 0.0 | 0.14261785 | 0.999 | 1767 | tags=23%, list=7%, sig | EN_1 |
| GOBP_MITOCHONDRIAL_ | GOBP_MITOCHONDRIAL_MEMBRANE_ORG | 107 | -0.32737 | -1.67615 | 0.0 | 0.14316696 | 0.999 | 5520 | tags=34%, list=23%, si | EN_1 |
| PANCREATIC_CANCER | PANCREATIC_CANCER | 74 | -0.34587 | -1.67192 | 0.0 | 0.14710459 | 0.999 | 5442 | tags=38%, list=23%, si | EN_1 |
| GOBP_HEME_BIOSYNTHE | GOBP_HEME_BIOSYNTHETIC_PROCESS | 24 | -0.43662 | -1.67085 | 0.0049751243 | 0.14729808 | 0.999 | 2594 | tags=29%, list=11%, si | EN_1 |
| GOBP_NUCLEAR_ENVELO | GOBP_NUCLEAR_ENVELOPE_ORGANIZATIO | 52 | -0.36767 | -1.67076 | 0.007246377 | 0.14648256 | 0.999 | 5109 | tags=38%, list=22%, si | EN_1 |
| GOBP_GLYCEROLIPID_BIO | GOBP_GLYCEROLIPID_BIOSYNTHETIC_PROC | 210 | -0.29824 | -1.67048 | 0.0 | 0.14589041 | 0.999 | 3597 | tags=27%, list=15%, si | EN_1 |
| GOBP_POSITIVE_REGULA | GOBP_POSITIVE_REGULATION_OF_STEROID | 25 | -0.44594 | -1.67043 | 0.004651163 | 0.14502482 | 0.999 | 4372 | tags=44%, list=19%, si | EN_1 |
| GOBP_NEGATIVE_REGUL | GOBP_NEGATIVE_REGULATION_OF_MACRO | 33 | -0.41796 | -1.66788 | 0.011428571 | 0.14672531 | 0.999 | 6272 | tags=48%, list=27%, si | EN_1 |
| GOBP_RESPONSE_TO_IN | GOBP_RESPONSE_TO_INTERLEUKIN_17 | 25 | -0.44904 | -1.66739 | 0.020942409 | 0.1462079 | 0.999 | 3051 | tags=28%, list=13%, si | EN_1 |
| GOBP_FIBROBLAST_MIGR | GOBP_FIBROBLAST_MIGRATION | 68 | -0.34615 | -1.66474 | 0.0 | 0.14812492 | 0.999 | 3882 | tags=31%, list=16%, si | EN_1 |
| GOBP_FOCAL_ADHESION | GOBP_FOCAL_ADHESION_ASSEMBLY | 87 | -0.33158 | -1.66401 | 0.0 | 0.14797139 | 0.999 | 5505 | tags=45%, list=23%, si | EN_1 |
| GOBP_POSTSYNAPTIC_M | GOBP_POSTSYNAPTIC_MEMBRANE_ORGAN | 53 | -0.37106 | -1.66204 | 0.0 | 0.14923385 | 0.999 | 3805 | tags=32%, list=16%, si | EN_1 |
| GOBP_POSITIVE_REGULA | GOBP_POSITIVE_REGULATION_OF_TELOME | 15 | -0.51764 | -1.65928 | 0.016194332 | 0.15090328 | 0.999 | 5858 | tags=47%, list=25%, si | EN_1 |
| GOBP_MAMMARY_GLAN | GOBP_MAMMARY_GLAND_EPITHELIAL_CEL | 16 | -0.49328 | -1.65909 | 0.021186441 | 0.15006863 | 0.999 | 1364 | tags=25%, list=6%, sig | EN_1 |
| GOBP_NUCLEAR_PORE_C | GOBP_NUCLEAR_PORE_ORGANIZATION | 15 | -0.51609 | -1.65907 | 0.015686275 | 0.14921905 | 0.999 | 4571 | tags=40%, list=19%, si | EN_1 |
| GOBP_CELLULAR_MODIFI | GOBP_CELLULAR_MODIFIED_AMINO_ACID_ | 40 | -0.38358 | -1.65801 | 0.0 | 0.14975049 | 0.999 | 3275 | tags=35%, list=14%, si | EN_1 |
| GOBP_NEGATIVE_REGUL | GOBP_NEGATIVE_REGULATION_OF_CYTOPL | 50 | -0.36228 | -1.65447 | 0.0064102565 | 0.15267822 | 1 | 4803 | tags=38%, list=20%, si | EN_1 |
| GOBP_NEGATIVE_REGUL | GOBP_NEGATIVE_REGULATION_OF_PEPTID | 29 | -0.42231 | -1.654 | 0.010050251 | 0.15222186 | 1 | 2351 | tags=28%, list=10%, si | EN_1 |
| GOBP_LIPID_MODIFICATI | GOBP_LIPID_MODIFICATION | 197 | -0.2941 | -1.65041 | 0.0 | 0.15533842 | 1 | 3988 | tags=30%, list=17%, si | EN_1 |
| GOBP_REGULATION_OF_ | GOBP_REGULATION_OF_SUBSTRATE_ADHE | 58 | -0.35465 | -1.65033 | 0.0 | 0.15452835 | 1 | 5321 | tags=43%, list=23%, si | EN_1 |
| GOBP_CELLULAR_RESPON | GOBP_CELLULAR_RESPONSE_TO_INTERLEU | 62 | -0.35245 | -1.65009 | 0.0 | 0.15383257 | 1 | 3812 | tags=34%, list=16%, si | EN_1 |
| GOBP_DIOL_METABOLIC_ | GOBP_DIOL_METABOLIC_PROCESS | 29 | -0.41704 | -1.64956 | 0.010638298 | 0.15357246 | 1 | 2915 | tags=31%, list=12%, si | EN_1 |
| GOBP_REGULATION_OF_ | GOBP_REGULATION_OF_DNA_STRAND_ELO | 18 | -0.47161 | -1.64765 | 0.016949153 | 0.15470502 | 1 | 5958 | tags=44%, list=25%, si | EN_1 |
| GOBP_UBIQUITIN_DEPEN | GOBP_UBIQUITIN_DEPENDENT_PROTEIN_C | 29 | -0.40948 | -1.64404 | 0.010416667 | 0.15789756 | 1 | 3996 | tags=28%, list=17%, si | EN_1 |
| GOBP_GLYCOLYTIC_PROC | GOBP_GLYCOLYTIC_PROCESS_THROUGH_FR | 25 | -0.42982 | -1.64007 | 0.005 | 0.16147906 | 1 | 4810 | tags=52%, list=20%, si | EN_1 |
| ADIPOCYTOKINE_SIGNALI | ADIPOCYTOKINE_SIGNALING_PATHWAY | 68 | -0.3486 | -1.63922 | 0.009009009 | 0.16149485 | 1 | 3983 | tags=34%, list=17%, si | EN_1 |
| GOBP_POSITIVE_REGULA | GOBP_POSITIVE_REGULATION_OF_ERYTHR | 28 | -0.40627 | -1.63554 | 0.010416667 | 0.1645012 | 1 | 5135 | tags=43%, list=22%, si | EN_1 |
| GOBP_MACROMOLECULE | GOBP_MACROMOLECULE_DEACYLATION | 59 | -0.35819 | -1.63364 | 0.0070422534 | 0.16588612 | 1 | 6511 | tags=49%, list=28%, si | EN_1 |
| GOBP_MONOSACCHARID | GOBP_MONOSACCHARIDE_CATABOLIC_PRO | 40 | -0.38795 | -1.62849 | 0.012738854 | 0.17091012 | 1 | 6247 | tags=58%, list=26%, si | EN_1 |
| GOBP_PEPTIDYL_LYSINE_ | GOBP_PEPTIDYL_LYSINE_ACETYLATION | 25 | -0.43375 | -1.62715 | 0.008928572 | 0.17150982 | 1 | 2015 | tags=20%, list=9%, sig | EN_1 |
| GOBP_PROTON_MOTIVE_ | GOBP_PROTON_MOTIVE_FORCE_DRIVEN_A | 52 | -0.35989 | -1.62302 | 0.0070422534 | 0.17543054 | 1 | 15132 | tags=100%, list=64%, | EN_1 |
| GOBP_NEGATIVE_REGUL | GOBP_NEGATIVE_REGULATION_OF_FATTY_ | 17 | -0.46608 | -1.62292 | 0.016129032 | 0.17464645 | 1 | 7606 | tags=76%, list=32%, si | EN_1 |
| GOBP_RESPONSE_TO_STI | GOBP_RESPONSE_TO_STEROL_DEPLETION | 15 | -0.49462 | -1.62157 | 0.011363637 | 0.17501329 | 1 | 5609 | tags=53%, list=24%, si | EN_1 |
| GOBP_SIGNAL_TRANSDU | GOBP_SIGNAL_TRANSDUCTION_IN_RESPON | 176 | -0.2891 | -1.62152 | 0.0 | 0.17416082 | 1 | 5305 | tags=34%, list=22%, si | EN_1 |
| GOBP_POSITIVE_REGULA | GOBP_POSITIVE_REGULATION_OF_MACRO | 81 | -0.32686 | -1.62084 | 0.0 | 0.17384815 | 1 | 5571 | tags=36%, list=24%, si | EN_1 |
| GOBP_PHOSPHATIDYLIN | GOBP_PHOSPHATIDYLINOSITOL_METABOLI | 148 | -0.29489 | -1.61894 | 0.0 | 0.17531317 | 1 | 4542 | tags=32%, list=19%, si | EN_1 |
| GOBP_DNA_TEMPLATED_ | GOBP_DNA_TEMPLATED_DNA_REPLICATIO | 54 | -0.35159 | -1.61844 | 0.0 | 0.17498411 | 1 | 1808 | tags=20%, list=8%, sig | EN_1 |
| GOBP_REGULATION_OF_ | GOBP_REGULATION_OF_NUCLEOTIDE_BIOS | 32 | -0.4107 | -1.61688 | 0.0 | 0.17609082 | 1 | 3293 | tags=38%, list=14%, si | EN_1 |

| | | | | | | | | | | |
|---|---|---|---|---|---|---|---|---|---|---|
| GOBP_REGULATION_OF_ | GOBP_REGULATION_OF_PATTERN_RECOGN | 157 | -0.29429 | -1.61604 | 0.0 | 0.17594364 | 1 | 4332 | tags=33%, list=18%, si | EN_1 |
| GOBP_REGULATION_OF_ | GOBP_REGULATION_OF_AUTOPHAGY | 295 | -0.26748 | -1.61442 | 0.0 | 0.17694217 | 1 | 5406 | tags=35%, list=23%, si | EN_1 |
| GOBP_INTRINSIC_APOPT | GOBP_INTRINSIC_APOPTOTIC_SIGNALING_F | 63 | -0.33837 | -1.61417 | 0.009090909 | 0.17621739 | 1 | 5477 | tags=33%, list=23%, si | EN_1 |
| GOBP_NUCLEOSIDE_TRIP | GOBP_NUCLEOSIDE_TRIPHOSPHATE_BIOSYN | 100 | -0.31608 | -1.61125 | 0.0 | 0.17883395 | 1 | 6734 | tags=45%, list=29%, si | EN_1 |
| GOBP_NEGATIVE_REGUL | GOBP_NEGATIVE_REGULATION_OF_RECEPT | 27 | -0.42438 | -1.6057 | 0.029411765 | 0.18418297 | 1 | 3644 | tags=41%, list=15%, si | EN_1 |
| GOBP_PIGMENT_BIOSYN | GOBP_PIGMENT_BIOSYNTHETIC_PROCESS | 49 | -0.36741 | -1.60468 | 0.0 | 0.18469964 | 1 | 3804 | tags=33%, list=16%, si | EN_1 |
| GOBP_ORGANOPHOSPHA | GOBP_ORGANOPHOSPHATE_BIOSYNTHETIC | 500 | -0.25472 | -1.60456 | 0.0 | 0.18393295 | 1 | 4835 | tags=30%, list=20%, si | EN_1 |
| GOBP_MITOCHONDRIAL_ | GOBP_MITOCHONDRIAL_RNA_PROCESSING | 20 | -0.45499 | -1.60452 | 0.03301887 | 0.1830648 | 1 | 6026 | tags=65%, list=26%, si | EN_1 |
| GOBP_PEROXISOMAL_TR | GOBP_PEROXISOMAL_TRANSPORT | 21 | -0.44788 | -1.60393 | 0.008733625 | 0.1830037 | 1 | 4842 | tags=33%, list=21%, si | EN_1 |
| GOBP_PYRUVATE_METAB | GOBP_PYRUVATE_METABOLIC_PROCESS | 127 | -0.30913 | -1.60319 | 0.0 | 0.18313383 | 1 | 4138 | tags=30%, list=18%, si | EN_1 |
| GOBP_RESPONSE_TO_AM | GOBP_RESPONSE_TO_AMYLOID_BETA | 39 | -0.36731 | -1.60092 | 0.011299435 | 0.18515746 | 1 | 3686 | tags=31%, list=16%, si | EN_1 |
| GOBP_HYALURONAN_CA | GOBP_HYALURONAN_CATABOLIC_PROCESS | 16 | -0.4924 | -1.5985 | 0.037815128 | 0.18732631 | 1 | 2838 | tags=38%, list=12%, si | EN_1 |
| GOBP_REGULATION_OF_ | GOBP_REGULATION_OF_NITRIC_OXIDE_SYN | 20 | -0.45175 | -1.59811 | 0.007843138 | 0.18684293 | 1 | 3146 | tags=35%, list=13%, si | EN_1 |
| GOBP_CELL_REDOX_HOM | GOBP_CELL_REDOX_HOMEOSTASIS | 28 | -0.4152 | -1.5979 | 0.02347418 | 0.18620452 | 1 | 13819 | tags=100%, list=59%, | EN_1 |
| GOBP_METHYLATION | GOBP_METHYLATION | 199 | -0.28653 | -1.59764 | 0.0 | 0.18561032 | 1 | 5547 | tags=34%, list=23%, si | EN_1 |
| GOBP_MYOBLAST_FUSIO | GOBP_MYOBLAST_FUSION | 45 | -0.369 | -1.59445 | 0.0 | 0.18853857 | 1 | 4708 | tags=40%, list=20%, si | EN_1 |
| GOBP_GLUCOSE_CATABO | GOBP_GLUCOSE_CATABOLIC_PROCESS | 26 | -0.40767 | -1.59347 | 0.024154589 | 0.18872719 | 1 | 4679 | tags=46%, list=20%, si | EN_1 |
| GOBP_MITOTIC_G2_M_T | GOBP_MITOTIC_G2_M_TRANSITION_CHECK | 56 | -0.35811 | -1.59292 | 0.008 | 0.18845423 | 1 | 4370 | tags=29%, list=19%, si | EN_1 |
| VALINE,_LEUCINE_AND_I | VALINE,_LEUCINE_AND_ISOLEUCINE_DEGRA | 49 | -0.35385 | -1.59194 | 0.007352941 | 0.1886554 | 1 | 3451 | tags=27%, list=15%, si | EN_1 |
| GOBP_NEGATIVE_REGUL | GOBP_NEGATIVE_REGULATION_OF_EXTRIN | 26 | -0.42843 | -1.59089 | 0.014354067 | 0.18917212 | 1 | 5440 | tags=58%, list=23%, si | EN_1 |
| GOBP_POSITIVE_REGULA | GOBP_POSITIVE_REGULATION_OF_EXCITAT | 40 | -0.37257 | -1.59008 | 0.006369427 | 0.18893659 | 1 | 3911 | tags=33%, list=17%, si | EN_1 |
| GOBP_MITOCHONDRIAL_ | GOBP_MITOCHONDRIAL_TRANSLATION | 128 | -0.30792 | -1.58942 | 0.0 | 0.18882123 | 1 | 15530 | tags=99%, list=66%, si | EN_1 |
| TERPENOID_BACKBONE_ | TERPENOID_BACKBONE_BIOSYNTHESIS | 21 | -0.45525 | -1.58798 | 0.035353534 | 0.18949524 | 1 | 5114 | tags=48%, list=22%, si | EN_1 |
| GOBP_AUTOPHAGY_OF_ | GOBP_AUTOPHAGY_OF_PEROXISOME | 15 | -0.50353 | -1.58604 | 0.03292181 | 0.19111703 | 1 | 5499 | tags=60%, list=23%, si | EN_1 |
| GOBP_PHOSPHATIDYLCH | GOBP_PHOSPHATIDYLCHOLINE_CATABOLIC_ | 15 | -0.49761 | -1.58584 | 0.034883723 | 0.1905454 | 1 | 5053 | tags=60%, list=21%, si | EN_1 |
| GOBP_FIBROBLAST_APOP | GOBP_FIBROBLAST_APOPTOTIC_PROCESS | 31 | -0.38762 | -1.58557 | 0.03 | 0.18995793 | 1 | 4563 | tags=29%, list=19%, si | EN_1 |
| GOBP_GLYCERALDEHYDE | GOBP_GLYCERALDEHYDE_3_PHOSPHATE_M | 19 | -0.45588 | -1.58371 | 0.0349345 | 0.19149664 | 1 | 5497 | tags=53%, list=23%, si | EN_1 |
| GOBP_REGULATION_OF_ | GOBP_REGULATION_OF_RESPIRATORY_SYST | 15 | -0.47658 | -1.58055 | 0.030075189 | 0.19456969 | 1 | 5821 | tags=60%, list=25%, si | EN_1 |
| GOBP_REGULATION_OF_ | GOBP_REGULATION_OF_ORGANELLE_ASSEN | 228 | -0.274 | -1.58033 | 0.0 | 0.19400601 | 1 | 4627 | tags=31%, list=20%, si | EN_1 |
| GOBP_VACUOLE_ORGAN | GOBP_VACUOLE_ORGANIZATION | 223 | -0.27689 | -1.57714 | 0.0 | 0.19689238 | 1 | 4440 | tags=29%, list=19%, si | EN_1 |
| GOBP_CARBOHYDRATE_C | GOBP_CARBOHYDRATE_CATABOLIC_PROCES | 161 | -0.27964 | -1.57668 | 0.0 | 0.19680633 | 1 | 5000 | tags=35%, list=21%, si | EN_1 |
| GOBP_POSITIVE_REGULA | GOBP_POSITIVE_REGULATION_OF_RECEPTC | 17 | -0.4685 | -1.57505 | 0.03875969 | 0.19811235 | 1 | 5218 | tags=53%, list=22%, si | EN_1 |
| GOBP_NEGATIVE_REGUL | GOBP_NEGATIVE_REGULATION_OF_FIBROB | 44 | -0.35633 | -1.57433 | 0.022222223 | 0.19796178 | 1 | 3190 | tags=25%, list=14%, si | EN_1 |
| GOBP_FATTY_ACID_BIOS | GOBP_FATTY_ACID_BIOSYNTHETIC_PROCES | 145 | -0.29224 | -1.57424 | 0.0 | 0.19717877 | 1 | 2901 | tags=25%, list=12%, si | EN_1 |
| GOBP_REGULATION_OF_ | GOBP_REGULATION_OF_TYPE_I_INTERFERC | 40 | -0.3706 | -1.5736 | 0.013245033 | 0.19707754 | 1 | 4483 | tags=45%, list=19%, si | EN_1 |
| GOBP_REGULATION_OF_ | GOBP_REGULATION_OF_PROTEOLYSIS_INV( | 215 | -0.27288 | -1.5732 | 0.0 | 0.19661497 | 1 | 5526 | tags=33%, list=23%, si | EN_1 |
| GOBP_REGULATION_OF_ | GOBP_REGULATION_OF_MONOCYTE_CHEM | 29 | -0.40049 | -1.57203 | 0.015384615 | 0.19715525 | 1 | 913 | tags=17%, list=4%, sig | EN_1 |
| GOBP_SULFUR_COMPOU | GOBP_SULFUR_COMPOUND_METABOLIC_P | 292 | -0.27655 | -1.5718 | 0.0 | 0.19656418 | 1 | 3299 | tags=25%, list=14%, si | EN_1 |
| GOBP_HYALURONAN_ME | GOBP_HYALURONAN_METABOLIC_PROCESS | 36 | -0.37779 | -1.57167 | 0.016759777 | 0.19598958 | 1 | 4278 | tags=39%, list=18%, si | EN_1 |
| GOBP_TRANSLATIONAL_ | GOBP_TRANSLATIONAL_ELONGATION | 58 | -0.3447 | -1.57138 | 0.00862069 | 0.19544668 | 1 | 15492 | tags=100%, list=66%, | EN_1 |
| GOBP_PROTEIN_O_LINKE | GOBP_PROTEIN_O_LINKED_MANNOSYLATIC | 16 | -0.47108 | -1.57061 | 0.04 | 0.19558243 | 1 | 3194 | tags=44%, list=14%, si | EN_1 |
| GOBP_RESPONSE_TO_AM | GOBP_RESPONSE_TO_AMINO_ACID_STARV. | 54 | -0.34362 | -1.57001 | 0.022222223 | 0.19555679 | 1 | 5469 | tags=39%, list=23%, si | EN_1 |
| FATTY_ACID_DEGRADATI | FATTY_ACID_DEGRADATION | 42 | -0.36384 | -1.56992 | 0.006451613 | 0.19481653 | 1 | 3858 | tags=33%, list=16%, si | EN_1 |
| GOBP_NEGATIVE_REGUL | GOBP_NEGATIVE_REGULATION_OF_CATECH | 17 | -0.46684 | -1.56928 | 0.031111112 | 0.19485293 | 1 | 1734 | tags=29%, list=7%, sig | EN_1 |
| GOBP_VESICLE_TETHERIN | GOBP_VESICLE_TETHERING | 31 | -0.38556 | -1.56739 | 0.017142856 | 0.19628382 | 1 | 6120 | tags=45%, list=26%, si | EN_1 |
| GOBP_REGULATION_OF_ | GOBP_REGULATION_OF_ASTROCYTE_DIFFEI | 34 | -0.38477 | -1.56192 | 0.021052632 | 0.20246397 | 1 | 991 | tags=15%, list=4%, sig | EN_1 |
| GOBP_CYTOPLASMIC_SE | GOBP_CYTOPLASMIC_SEQUESTERING_OF_P | 17 | -0.44832 | -1.56049 | 0.033898305 | 0.20336862 | 1 | 3717 | tags=41%, list=16%, si | EN_1 |
| GOBP_LIPOPROTEIN_ME | GOBP_LIPOPROTEIN_METABOLIC_PROCESS | 135 | -0.28287 | -1.55882 | 0.0 | 0.20478868 | 1 | 4157 | tags=31%, list=18%, si | EN_1 |
| GOBP_LONG_CHAIN_FAT | GOBP_LONG_CHAIN_FATTY_ACID_BIOSYNT | 19 | -0.43774 | -1.55851 | 0.040650405 | 0.20430923 | 1 | 1700 | tags=32%, list=7%, sig | EN_1 |
| HEPATITIS_B | HEPATITIS_B | 147 | -0.29378 | -1.55782 | 0.0 | 0.20438094 | 1 | 5476 | tags=37%, list=23%, si | EN_1 |
| GOBP_NEGATIVE_REGUL | GOBP_NEGATIVE_REGULATION_OF_EXTRIN | 95 | -0.30432 | -1.55656 | 0.0 | 0.20501587 | 1 | 3663 | tags=31%, list=16%, si | EN_1 |
| GOBP_G0_TO_G1_TRANS | GOBP_G0_TO_G1_TRANSITION | 38 | -0.36739 | -1.55518 | 0.0125 | 0.20595388 | 1 | 2367 | tags=18%, list=10%, si | EN_1 |
| GOBP_NAD_CATABOLIC_ | GOBP_NAD_CATABOLIC_PROCESS | 21 | -0.43976 | -1.55391 | 0.012931035 | 0.20680241 | 1 | 4679 | tags=57%, list=20%, si | EN_1 |
| GOBP_AUTOPHAGY_OF_ | GOBP_AUTOPHAGY_OF_MITOCHONDRION | 89 | -0.3128 | -1.5538 | 0.0 | 0.20604752 | 1 | 6374 | tags=47%, list=27%, si | EN_1 |
| GOBP_ENDOSOME_ORGA | GOBP_ENDOSOME_ORGANIZATION | 68 | -0.33172 | -1.55289 | 0.009803922 | 0.20625857 | 1 | 4308 | tags=35%, list=18%, si | EN_1 |
| GOBP_ALCOHOL_BIOSYN | GOBP_ALCOHOL_BIOSYNTHETIC_PROCESS | 129 | -0.29693 | -1.55107 | 0.0 | 0.20759758 | 1 | 2915 | tags=26%, list=12%, si | EN_1 |
| GOBP_SULFUR_COMPOU | GOBP_SULFUR_COMPOUND_BIOSYNTHETIC | 126 | -0.30175 | -1.55017 | 0.0 | 0.20791961 | 1 | 3299 | tags=25%, list=14%, si | EN_1 |
| GOBP_REGULATION_OF_ | GOBP_REGULATION_OF_CELL_SUBSTRATE_ | 69 | -0.33197 | -1.54967 | 0.01010101 | 0.20773079 | 1 | 5388 | tags=39%, list=23%, si | EN_1 |
| GOBP_MONOCYTE_CHEM | GOBP_MONOCYTE_CHEMOTAXIS | 40 | -0.36311 | -1.54779 | 0.007751938 | 0.2093163 | 1 | 913 | tags=15%, list=4%, sig | EN_1 |
| GOBP_REGULATION_OF_ | GOBP_REGULATION_OF_REGULATORY_NCR | 15 | -0.48014 | -1.54759 | 0.046511628 | 0.20874886 | 1 | 4982 | tags=40%, list=21%, si | EN_1 |
| GOBP_TETRAPYRROLE_BI | GOBP_TETRAPYRROLE_BIOSYNTHETIC_PROC | 29 | -0.39388 | -1.54699 | 0.009569378 | 0.20869145 | 1 | 2594 | tags=28%, list=11%, si | EN_1 |
| GOBP_POSITIVE_REGULA | GOBP_POSITIVE_REGULATION_OF_DOUBLE_ | 91 | -0.30566 | -1.54381 | 0.0 | 0.21174328 | 1 | 4251 | tags=24%, list=18%, si | EN_1 |
| GOBP_REGULATION_OF_ | GOBP_REGULATION_OF_ACTIN_NUCLEATIO | 35 | -0.37764 | -1.54289 | 0.018292682 | 0.21233948 | 1 | 513 | tags=14%, list=2%, sig | EN_1 |
| GOBP_NEUROBLAST_DIV | GOBP_NEUROBLAST_DIVISION | 16 | -0.45396 | -1.5426 | 0.05179283 | 0.21180001 | 1 | 2487 | tags=31%, list=11%, si | EN_1 |
| GOBP_REGULATION_OF_ | GOBP_REGULATION_OF_ARP2_3_COMPLEX | 27 | -0.39541 | -1.54116 | 0.033816423 | 0.21287492 | 1 | 513 | tags=15%, list=2%, sig | EN_1 |
| GOBP_TRNA_METHYLATI | GOBP_TRNA_METHYLATION | 39 | -0.37112 | -1.53979 | 0.012738854 | 0.21378164 | 1 | 3586 | tags=28%, list=15%, si | EN_1 |
| GOBP_POSITIVE_REGULA | GOBP_POSITIVE_REGULATION_OF_PROTEAS | 72 | -0.32016 | -1.53925 | 0.008474576 | 0.21355806 | 1 | 3550 | tags=22%, list=15%, si | EN_1 |
| GOBP_VERY_LONG_CHAI | GOBP_VERY_LONG_CHAIN_FATTY_ACID_MI | 33 | -0.37901 | -1.53905 | 0.010256411 | 0.21303672 | 1 | 3772 | tags=33%, list=16%, si | EN_1 |
| GOBP_REGULATION_OF_ | GOBP_REGULATION_OF_SYNCYTIUM_FORM | 31 | -0.38568 | -1.53838 | 0.010989011 | 0.21316403 | 1 | 4926 | tags=52%, list=21%, si | EN_1 |
| GOBP_ORGANELLE_DISAS | GOBP_ORGANELLE_DISASSEMBLY | 155 | -0.28295 | -1.53681 | 0.0 | 0.21450639 | 1 | 6374 | tags=39%, list=27%, si | EN_1 |
| GOBP_MODULATION_OF | GOBP_MODULATION_OF_EXCITATORY_POS | 59 | -0.32022 | -1.53658 | 0.0 | 0.21398893 | 1 | 3911 | tags=29%, list=17%, si | EN_1 |
| GOBP_REGULATION_OF_ | GOBP_REGULATION_OF_DNA_REPAIR | 219 | -0.27869 | -1.53385 | 0.0 | 0.21687146 | 1 | 4370 | tags=24%, list=19%, si | EN_1 |
| GOBP_POSITIVE_REGULA | GOBP_POSITIVE_REGULATION_OF_DOUBLE_ | 15 | -0.48293 | -1.53371 | 0.043824703 | 0.21627556 | 1 | 5530 | tags=60%, list=23%, si | EN_1 |
| GOBP_RIBOSOME_DISASS | GOBP_RIBOSOME_DISASSEMBLY | 31 | -0.37361 | -1.53367 | 0.021621622 | 0.21550354 | 1 | 14801 | tags=100%, list=63%, | EN_1 |
| GOBP_S_ADENOSYLMETH | GOBP_S_ADENOSYLMETHIONINE_METABOL | 17 | -0.44451 | -1.53272 | 0.0456621 | 0.215742 | 1 | 3667 | tags=35%, list=16%, si | EN_1 |
| GOBP_NEGATIVE_REGUL | GOBP_NEGATIVE_REGULATION_OF_TRANSL | 117 | -0.28937 | -1.53256 | 0.0 | 0.21510325 | 1 | 4364 | tags=27%, list=18%, si | EN_1 |
| GOBP_N_ACETYLGLUCOS | GOBP_N_ACETYLGLUCOSAMINE_METABOLI | 16 | -0.45884 | -1.5305 | 0.06437768 | 0.21703885 | 1 | 2006 | tags=31%, list=8%, sig | EN_1 |
| GOBP_REGULATION_OF_ | GOBP_REGULATION_OF_REACTIVE_OXYGEN | 50 | -0.35122 | -1.53045 | 0.0 | 0.21628614 | 1 | 5135 | tags=40%, list=22%, si | EN_1 |
| GOBP_POSITIVE_REGULA | GOBP_POSITIVE_REGULATION_OF_AUTOPH | 157 | -0.27799 | -1.52939 | 0.0 | 0.21697278 | 1 | 3515 | tags=25%, list=15%, si | EN_1 |
| GOBP_REGULATION_OF_ | GOBP_REGULATION_OF_PROTEASOMAL_PR | 179 | -0.27213 | -1.52924 | 0.0 | 0.21633562 | 1 | 5506 | tags=33%, list=23%, si | EN_1 |
| GOBP_REACTIVE_OXYGEI | GOBP_REACTIVE_OXYGEN_SPECIES_METAB | 239 | -0.24885 | -1.52404 | 0.0 | 0.22309767 | 1 | 4054 | tags=24%, list=17%, si | EN_1 |
| GOBP_POSITIVE_REGULA | GOBP_POSITIVE_REGULATION_OF_MYOBLA | 18 | -0.44408 | -1.52399 | 0.037453182 | 0.22234961 | 1 | 4650 | tags=56%, list=20%, si | EN_1 |

| | | | | | | | | | | |
|---|---|---|---|---|---|---|---|---|---|---|
| GOBP_NUCLEOSIDE_DIPH | GOBP_NUCLEOSIDE_DIPHOSPHATE_CATABO | 111 | -0.29349 | -1.52228 | 0.015625 | 0.22378945 | 1 | 4881 | tags=37%, list=21%, si | EN_1 |
| GOBP_POSITIVE_REGULA | GOBP_POSITIVE_REGULATION_OF_ORGANE | 91 | -0.29514 | -1.52084 | 0.0 | 0.22488692 | 1 | 4570 | tags=32%, list=19%, si | EN_1 |
| GOBP_MITOTIC_DNA_INT | GOBP_MITOTIC_DNA_INTEGRITY_CHECKPO | 84 | -0.31647 | -1.51969 | 0.011494253 | 0.22590086 | 1 | 4711 | tags=33%, list=20%, si | EN_1 |
| GOBP_PROTEIN_ACYLATI | GOBP_PROTEIN_ACYLATION | 94 | -0.30672 | -1.51943 | 0.015384615 | 0.22543317 | 1 | 4210 | tags=29%, list=18%, si | EN_1 |
| GOBP_POSITIVE_REGULA | GOBP_POSITIVE_REGULATION_OF_B_CELL_ | 18 | -0.44138 | -1.51932 | 0.04888889 | 0.22472703 | 1 | 2611 | tags=33%, list=11%, si | EN_1 |
| GOBP_REGULATION_OF_ | GOBP_REGULATION_OF_STEM_CELL_DIFFEI | 82 | -0.30453 | -1.51917 | 0.011363637 | 0.22425197 | 1 | 4846 | tags=38%, list=21%, si | EN_1 |
| GOBP_NUCLEOSIDE_DIPH | GOBP_NUCLEOSIDE_DIPHOSPHATE_METABO | 127 | -0.29223 | -1.51828 | 0.0 | 0.22453542 | 1 | 4881 | tags=37%, list=21%, si | EN_1 |
| GOBP_NUCLEOPHAGY | GOBP_NUCLEOPHAGY | 24 | -0.4086 | -1.51815 | 0.033980582 | 0.22394907 | 1 | 4288 | tags=54%, list=18%, si | EN_1 |
| GOBP_REGULATION_OF_ | GOBP_REGULATION_OF_TRANSCRIPTION_B | 41 | -0.34576 | -1.51624 | 0.027586207 | 0.2257725 | 1 | 15461 | tags=100%, list=65%, | EN_1 |
| GOBP_MODIFICATION_O | GOBP_MODIFICATION_OF_POSTSYNAPTIC_ | 24 | -0.40752 | -1.5153 | 0.03314917 | 0.2261981 | 1 | 3701 | tags=42%, list=16%, si | EN_1 |
| GOBP_PURINE_CONTAIN | GOBP_PURINE_CONTAINING_COMPOUND_ | 150 | -0.27748 | -1.51442 | 0.0 | 0.22676647 | 1 | 4881 | tags=37%, list=21%, si | EN_1 |
| GOBP_REGULATION_OF_ | GOBP_REGULATION_OF_PROTEIN_EXPORT_ | 37 | -0.36222 | -1.51393 | 0.030864198 | 0.22664848 | 1 | 6579 | tags=54%, list=28%, si | EN_1 |
| GOBP_REGULATION_OF_ | GOBP_REGULATION_OF_DOUBLE_STRAND_ | 28 | -0.39617 | -1.51366 | 0.04712042 | 0.22628422 | 1 | 4707 | tags=36%, list=20%, si | EN_1 |
| GOBP_PIGMENT_CELL_DI | GOBP_PIGMENT_CELL_DIFFERENTIATION | 37 | -0.37054 | -1.5133 | 0.0 | 0.22594751 | 1 | 4138 | tags=32%, list=18%, si | EN_1 |
| GOBP_NUCLEAR_TRANSC | GOBP_NUCLEAR_TRANSCRIBED_MRNA_POL | 26 | -0.39981 | -1.51295 | 0.057142857 | 0.2255798 | 1 | 14181 | tags=100%, list=60%, | EN_1 |
| GOBP_REGULATION_OF_ | GOBP_REGULATION_OF_NUCLEOTIDE_MET | 96 | -0.29134 | -1.51211 | 0.0 | 0.22607227 | 1 | 3577 | tags=28%, list=15%, si | EN_1 |
| GOBP_PHOSPHATIDYLETH | GOBP_PHOSPHATIDYLETHANOLAMINE_MET | 25 | -0.39825 | -1.51028 | 0.03809524 | 0.22790731 | 1 | 5053 | tags=52%, list=21%, si | EN_1 |
| GOBP_REGULATION_OF_ | GOBP_REGULATION_OF_CELL_MATRIX_ADH | 121 | -0.28114 | -1.50686 | 0.0 | 0.232031 | 1 | 4875 | tags=37%, list=21%, si | EN_1 |
| GOBP_POSITIVE_REGULA | GOBP_POSITIVE_REGULATION_OF_LIPID_BI | 104 | -0.29042 | -1.50632 | 0.0 | 0.23187076 | 1 | 3846 | tags=30%, list=16%, si | EN_1 |
| GOBP_NADP_METABOLIC | GOBP_NADP_METABOLIC_PROCESS | 49 | -0.34616 | -1.50569 | 0.02097902 | 0.23212437 | 1 | 3596 | tags=24%, list=15%, si | EN_1 |
| GOBP_GOLGI_TO_VACUC | GOBP_GOLGI_TO_VACUOLE_TRANSPORT | 21 | -0.433 | -1.50508 | 0.049327355 | 0.23245496 | 1 | 4617 | tags=29%, list=20%, si | EN_1 |
| GOBP_PROTEIN_LOCALIZ | GOBP_PROTEIN_LOCALIZATION_TO_ENDOP | 76 | -0.30326 | -1.50394 | 0.0 | 0.2333908 | 1 | 5390 | tags=32%, list=23%, si | EN_1 |
| GOBP_VESICLE_TARGETII | GOBP_VESICLE_TARGETING | 60 | -0.3176 | -1.50362 | 0.0 | 0.23303168 | 1 | 14924 | tags=98%, list=63%, si | EN_1 |
| GOBP_PROTEIN_LOCALIZ | GOBP_PROTEIN_LOCALIZATION_TO_VACUO | 78 | -0.30168 | -1.50359 | 0.0 | 0.2322728 | 1 | 3195 | tags=18%, list=14%, si | EN_1 |
| GOBP_REGULATION_OF_ | GOBP_REGULATION_OF_DOUBLE_STRAND_ | 80 | -0.30529 | -1.50313 | 0.027777778 | 0.23219785 | 1 | 1901 | tags=15%, list=8%, sig | EN_1 |
| GOBP_REGULATION_OF_ | GOBP_REGULATION_OF_MOTOR_NEURON_ | 19 | -0.44026 | -1.50273 | 0.06410257 | 0.23216261 | 1 | 13224 | tags=100%, list=56%, | EN_1 |
| GOBP_RESPONSE_TO_SA | GOBP_RESPONSE_TO_SALT_STRESS | 20 | -0.40753 | -1.5026 | 0.030973451 | 0.23150067 | 1 | 6249 | tags=45%, list=26%, si | EN_1 |
| GOBP_RIG_I_SIGNALING_ | GOBP_RIG_I_SIGNALING_PATHWAY | 24 | -0.4106 | -1.50216 | 0.03448276 | 0.23145993 | 1 | 2507 | tags=25%, list=11%, si | EN_1 |
| PROLACTIN_SIGNALING_ | PROLACTIN_SIGNALING_PATHWAY | 64 | -0.31782 | -1.50205 | 0.007575758 | 0.23081674 | 1 | 5442 | tags=44%, list=23%, si | EN_1 |
| GOBP_LIPID_DROPLET_O | GOBP_LIPID_DROPLET_ORGANIZATION | 36 | -0.36174 | -1.50078 | 0.030612245 | 0.23182447 | 1 | 2242 | tags=22%, list=9%, sig | EN_1 |
| GOBP_MANNOSYLATION | GOBP_MANNOSYLATION | 22 | -0.41895 | -1.50054 | 0.043269232 | 0.2314794 | 1 | 3194 | tags=36%, list=14%, si | EN_1 |
| GOBP_VIRAL_TRANSLATI | GOBP_VIRAL_TRANSLATION | 18 | -0.45003 | -1.49965 | 0.05327869 | 0.23177734 | 1 | 12993 | tags=100%, list=55%, | EN_1 |
| GOBP_INNER_MITOCHON | GOBP_INNER_MITOCHONDRIAL_MEMBRAN | 36 | -0.36189 | -1.49821 | 0.040935673 | 0.23318037 | 1 | 15079 | tags=100%, list=64%, | EN_1 |
| GOBP_POSITIVE_REGULA | GOBP_POSITIVE_REGULATION_OF_AUTOPH | 19 | -0.42589 | -1.49806 | 0.047008548 | 0.23263787 | 1 | 5571 | tags=53%, list=24%, si | EN_1 |
| GOBP_RESPONSE_TO_MI | GOBP_RESPONSE_TO_MITOCHONDRIAL_DE | 21 | -0.42223 | -1.49755 | 0.04910714 | 0.23262255 | 1 | 3482 | tags=24%, list=15%, si | EN_1 |
| GOBP_PYRIDINE_CONTAI | GOBP_PYRIDINE_CONTAINING_COMPOUND | 109 | -0.29017 | -1.49669 | 0.013157895 | 0.2332566 | 1 | 4881 | tags=37%, list=21%, si | EN_1 |
| GOBP_POSITIVE_REGULA | GOBP_POSITIVE_REGULATION_OF_TELOME | 71 | -0.30831 | -1.49566 | 0.0 | 0.23396696 | 1 | 6638 | tags=41%, list=28%, si | EN_1 |
| BASAL_TRANSCRIPTION_ | BASAL_TRANSCRIPTION_FACTORS | 40 | -0.34827 | -1.49509 | 0.017441861 | 0.23403904 | 1 | 2385 | tags=20%, list=10%, si | EN_1 |
| GOBP_GLYCOLIPID_BIOSY | GOBP_GLYCOLIPID_BIOSYNTHETIC_PROCESS | 66 | -0.31264 | -1.49507 | 0.010309278 | 0.23333667 | 1 | 3213 | tags=27%, list=14%, si | EN_1 |
| GOBP_MEMBRANE_LIPID | GOBP_MEMBRANE_LIPID_BIOSYNTHETIC_PI | 129 | -0.28036 | -1.49418 | 0.0 | 0.23391537 | 1 | 3213 | tags=26%, list=14%, si | EN_1 |
| GOBP_FATTY_ACID_ELON | GOBP_FATTY_ACID_ELONGATION | 15 | -0.45682 | -1.49266 | 0.033707865 | 0.2355116 | 1 | 2118 | tags=33%, list=9%, sig | EN_1 |
| GOBP_REGULATION_OF_ | GOBP_REGULATION_OF_POST_TRANSLATIO | 242 | -0.25453 | -1.49245 | 0.0 | 0.23506995 | 1 | 3395 | tags=21%, list=14%, si | EN_1 |
| GOBP_RUFFLE_ORGANIZ | GOBP_RUFFLE_ORGANIZATION | 54 | -0.33509 | -1.49179 | 0.017094018 | 0.23523219 | 1 | 5219 | tags=37%, list=22%, si | EN_1 |
| GOBP_NUCLEUS_ORGAN | GOBP_NUCLEUS_ORGANIZATION | 136 | -0.28576 | -1.49127 | 0.0 | 0.23519024 | 1 | 5329 | tags=34%, list=23%, si | EN_1 |
| GOBP_REGULATION_OF_ | GOBP_REGULATION_OF_AUTOPHAGOSOME | 18 | -0.42939 | -1.49113 | 0.05485232 | 0.23460358 | 1 | 3264 | tags=28%, list=14%, si | EN_1 |
| GOBP_POSITIVE_REGULA | GOBP_POSITIVE_REGULATION_OF_MEMBRA | 16 | -0.44963 | -1.4909 | 0.05529954 | 0.23416965 | 1 | 4881 | tags=44%, list=21%, si | EN_1 |
| HEPATITIS_C | HEPATITIS_C | 138 | -0.27475 | -1.49084 | 0.0 | 0.23351923 | 1 | 4759 | tags=32%, list=20%, si | EN_1 |
| TYPE_II_DIABETES_MELLI | TYPE_II_DIABETES_MELLITUS | 45 | -0.3413 | -1.49067 | 0.01923077 | 0.2329765 | 1 | 2690 | tags=22%, list=11%, si | EN_1 |
| MITOPHAGY | MITOPHAGY | 62 | -0.32208 | -1.49002 | 0.0091743115 | 0.23322631 | 1 | 7236 | tags=48%, list=31%, si | EN_1 |
| GOBP_REGULATION_OF_ | GOBP_REGULATION_OF_VASCULAR_ENDOT | 33 | -0.35557 | -1.4895 | 0.03468208 | 0.23313774 | 1 | 2160 | tags=24%, list=9%, sig | EN_1 |
| GOBP_REGULATION_OF_ | GOBP_REGULATION_OF_EXTRINSIC_APOPT | 45 | -0.34796 | -1.48898 | 0.012345679 | 0.23319921 | 1 | 3699 | tags=36%, list=16%, si | EN_1 |
| GOBP_REGULATION_OF_ | GOBP_REGULATION_OF_DENDRITIC_SPINE_ | 60 | -0.31577 | -1.48736 | 0.009259259 | 0.23506077 | 1 | 3703 | tags=27%, list=16%, si | EN_1 |
| GOBP_POSITIVE_REGULA | GOBP_POSITIVE_REGULATION_OF_ACTIN_F | 52 | -0.32902 | -1.48666 | 0.014705882 | 0.23529133 | 1 | 1124 | tags=15%, list=5%, sig | EN_1 |
| GOBP_CYTOPLASMIC_TR | GOBP_CYTOPLASMIC_TRANSLATIONAL_INIT | 33 | -0.37003 | -1.48618 | 0.02259887 | 0.23524503 | 1 | 14886 | tags=100%, list=63%, | EN_1 |
| GOBP_PROTEIN_TRANSM | GOBP_PROTEIN_TRANSMEMBRANE_IMPOR | 36 | -0.36529 | -1.48569 | 0.03125 | 0.23508716 | 1 | 14999 | tags=100%, list=64%, | EN_1 |
| CYTOSOLIC_DNA-SENSING | CYTOSOLIC_DNA-SENSING_PATHWAY | 46 | -0.33181 | -1.48152 | 0.025316456 | 0.24039991 | 1 | 2293 | tags=22%, list=10%, si | EN_1 |
| GOBP_SUBSTRATE_ADHE | GOBP_SUBSTRATE_ADHESION_DEPENDENT_ | 99 | -0.28961 | -1.48083 | 0.014084507 | 0.24075998 | 1 | 4120 | tags=29%, list=17%, si | EN_1 |
| GOBP_PEPTIDYL_LYSINE_ | GOBP_PEPTIDYL_LYSINE_MODIFICATION | 128 | -0.28067 | -1.47917 | 0.0 | 0.242453 | 1 | 4210 | tags=23%, list=18%, si | EN_1 |
| GOBP_REGULATION_OF_ | GOBP_REGULATION_OF_EXTRINSIC_APOPT | 161 | -0.2721 | -1.47738 | 0.0 | 0.24435094 | 1 | 3717 | tags=29%, list=16%, si | EN_1 |
| GOBP_INSULIN_RECEPTO | GOBP_INSULIN_RECEPTOR_SIGNALING_PAT | 133 | -0.27457 | -1.47721 | 0.0 | 0.24387252 | 1 | 4430 | tags=33%, list=19%, si | EN_1 |
| GOBP_ENUCLEATE_ERYTI | GOBP_ENUCLEATE_ERYTHROCYTE_DIFFEREI | 15 | -0.46663 | -1.47658 | 0.07662835 | 0.24413872 | 1 | 1050 | tags=20%, list=4%, sig | EN_1 |
| GOBP_PROTEIN_SUMOYL | GOBP_PROTEIN_SUMOYLATION | 42 | -0.34247 | -1.47463 | 0.02259887 | 0.24619398 | 1 | 2769 | tags=21%, list=12%, si | EN_1 |
| GOBP_NECROPTOTIC_PR | GOBP_NECROPTOTIC_PROCESS | 37 | -0.3477 | -1.47403 | 0.05732484 | 0.24637362 | 1 | 3717 | tags=30%, list=16%, si | EN_1 |
| GOBP_NEGATIVE_REGUL | GOBP_NEGATIVE_REGULATION_OF_ANOIKI | 15 | -0.45439 | -1.47376 | 0.05755396 | 0.24603914 | 1 | 3426 | tags=33%, list=15%, si | EN_1 |
| GOBP_MEMBRANE_FISSI | GOBP_MEMBRANE_FISSION | 31 | -0.35931 | -1.47169 | 0.02116402 | 0.2482199 | 1 | 15138 | tags=100%, list=64%, | EN_1 |
| OLFACTORY_TRANSDUCT | OLFACTORY_TRANSDUCTION | 186 | 0.539906 | 2.167154 | 0 | 0 | 0 | 5788 | tags=67%, list=25%, si | EN_1 |
| GOBP_INTERMEDIATE_FI | GOBP_INTERMEDIATE_FILAMENT_ORGANIZ | 46 | 0.53751 | 1.861755 | 0 | 0.03142632 | 0.391 | 3720 | tags=52%, list=16%, si | EN_1 |
| GOBP_RESPONSE_TO_PH | GOBP_RESPONSE_TO_PHEROMONE | 17 | 0.656257 | 1.826318 | 0 | 0.048588496 | 0.569 | 4676 | tags=82%, list=20%, si | EN_1 |
| GOBP_CHLORIDE_TRANS | GOBP_CHLORIDE_TRANSMEMBRANE_TRAN | 56 | 0.492475 | 1.745372 | 0 | 0.14634748 | 0.943 | 7666 | tags=55%, list=32%, si | EN_1 |
| GOBP_SENSORY_PERCEP | GOBP_SENSORY_PERCEPTION_OF_TASTE | 33 | 0.533708 | 1.744034 | 0.0012269939 | 0.13723956 | 0.948 | 7762 | tags=67%, list=33%, si | EN_1 |
| GOBP_POSITIVE_REGULA | GOBP_POSITIVE_REGULATION_OF_NATURA | 35 | 0.509927 | 1.717556 | 0 | 0.1871687 | 0.985 | 8451 | tags=71%, list=36%, si | EN_1 |
| SYSTEMIC_LUPUS_ERYTH | SYSTEMIC_LUPUS_ERYTHEMATOSUS | 105 | 0.44202 | 1.711407 | 0 | 0.18905126 | 0.991 | 8870 | tags=58%, list=38%, si | EN_1 |
| GOBP_REGULATION_OF_ | GOBP_REGULATION_OF_NATURAL_KILLER_ | 55 | 0.476689 | 1.699083 | 0.0011261262 | 0.20909561 | 0.997 | 7975 | tags=60%, list=34%, si | EN_1 |
| NEUROACTIVE_LIGAND-R | NEUROACTIVE_LIGAND-RECEPTOR_INTERAC | 274 | 0.413732 | 1.694739 | 0 | 0.20833828 | 0.998 | 7914 | tags=57%, list=34%, si | EN_1 |
| GOBP_INTERMEDIATE_FI | GOBP_INTERMEDIATE_FILAMENT_BASED_P | 67 | 0.462482 | 1.687919 | 0 | 0.2137262 | 1 | 3720 | tags=39%, list=16%, si | EN_1 |
| GOBP_NEUROPEPTIDE_SI | GOBP_NEUROPEPTIDE_SIGNALING_PATHW | 69 | 0.45569 | 1.675383 | 0 | 0.23788984 | 1 | 7699 | tags=57%, list=33%, si | EN_1 |
| GOBP_RETINOIC_ACID_M | GOBP_RETINOIC_ACID_METABOLIC_PROCES | 28 | 0.535015 | 1.667866 | 0.003726708 | 0.24956216 | 1 | 8343 | tags=64%, list=35%, si | EN_1 |
| GOBP_REGULATION_OF_ | GOBP_REGULATION_OF_MACROPHAGE_MI | 47 | -0.62428 | -1.84627 | 0.0010989011 | 0.18583845 | 0.431 | 2898 | tags=32%, list=12%, si | MM_1 |
| GOBP_NEGATIVE_REGUL | GOBP_NEGATIVE_REGULATION_OF_COLLAG | 18 | 0.574863 | 1.90398 | 0 | 0.21109883 | 0.679 | 261 | tags=22%, list=1%, sig | MM_1 |
| GOBP_RESPONSE_TO_VIT | GOBP_RESPONSE_TO_VITAMIN_D | 15 | 0.588716 | 1.900057 | 0.0045045046 | 0.17991036 | 0.686 | 528 | tags=20%, list=2%, sig | MM_1 |
| GOBP_VENTRICULAR_CA | GOBP_VENTRICULAR_CARDIAC_MUSCLE_CE | 15 | 0.64054 | 1.89961 | 0 | 0.15489826 | 0.689 | 1115 | tags=33%, list=5%, sig | MM_1 |
| GOBP_POSITIVE_REGULA | GOBP_POSITIVE_REGULATION_OF_PROTEIN | 24 | 0.519481 | 1.895967 | 0.00591716 | 0.14020061 | 0.699 | 3089 | tags=33%, list=13%, si | MM_1 |

| | | | | | | | | | | |
|---|---|---|---|---|---|---|---|---|---|---|
| GOBP_ENTRAINMENT_OF | GOBP_ENTRAINMENT_OF_CIRCADIAN_CLO | 26 | 0.51098 | 1.80062 | 0.0061728396 | 0.24804842 | 0.901 | 1607 | tags=27%, list=7%, sig | MM_1 |
| GOBP_NEGATIVE_REGUL | GOBP_NEGATIVE_REGULATION_OF_VIRAL_ | 50 | -0.57411 | -2.55135 | 0 | 0.001460335 | 0.001 | 2864 | tags=40%, list=12%, si | MM_2 |
| GOBP_CHEMOKINE_C_X_ | GOBP_CHEMOKINE_C_X_C_MOTIF_LIGAND | 25 | -0.67361 | -2.51168 | 0 | 0.002920532 | 0.004 | 1704 | tags=32%, list=7%, sig | MM_2 |
| GOBP_ANTIGEN_PROCES | GOBP_ANTIGEN_PROCESSING_AND_PRESEN | 28 | -0.66146 | -2.49173 | 0.0024390244 | 0.004888768 | 0.01 | 5246 | tags=57%, list=22%, si | MM_2 |
| GOBP_RESPONSE_TO_INT | GOBP_RESPONSE_TO_INTERFERON_BETA | 64 | -0.53902 | -2.46132 | 0 | 0.004745569 | 0.013 | 4622 | tags=45%, list=20%, si | MM_2 |
| GOBP_REGULATION_OF_ | GOBP_REGULATION_OF_NUCLEASE_ACTIVIT | 22 | -0.69071 | -2.45724 | 0 | 0.004090426 | 0.014 | 3865 | tags=55%, list=16%, si | MM_2 |
| GOBP_ANTIGEN_PROCES | GOBP_ANTIGEN_PROCESSING_AND_PRESEN | 20 | -0.69686 | -2.41964 | 0 | 0.006109877 | 0.024 | 5246 | tags=65%, list=22%, si | MM_2 |
| GOBP_RESPONSE_TO_TY | GOBP_RESPONSE_TO_TYPE_I_INTERFERON | 64 | -0.52484 | -2.39723 | 0 | 0.007106361 | 0.033 | 5819 | tags=55%, list=25%, si | MM_2 |
| GOBP_MITOTIC_SISTER_ | GOBP_MITOTIC_SISTER_CHROMATID_SEPAR | 61 | -0.51128 | -2.38096 | 0 | 0.007848762 | 0.041 | 4413 | tags=41%, list=19%, si | MM_2 |
| CYTOSOLIC_DNA-SENSIN | CYTOSOLIC_DNA-SENSING_PATHWAY | 46 | -0.55297 | -2.37899 | 0 | 0.006976678 | 0.041 | 4207 | tags=37%, list=18%, si | MM_2 |
| GOBP_ANTIGEN_PROCES | GOBP_ANTIGEN_PROCESSING_AND_PRESEN | 34 | -0.6095 | -2.35874 | 0 | 0.008909876 | 0.056 | 5465 | tags=56%, list=23%, si | MM_2 |
| GOBP_REGULATION_OF_ | GOBP_REGULATION_OF_TRANSLATIONAL_I | 68 | -0.50007 | -2.35588 | 0 | 0.008500111 | 0.059 | 5819 | tags=44%, list=25%, si | MM_2 |
| C-TYPE_LECTIN_RECEPTO | C-TYPE_LECTIN_RECEPTOR_SIGNALING_PAT | 104 | -0.45566 | -2.35364 | 0 | 0.007911334 | 0.06 | 3775 | tags=30%, list=16%, si | MM_2 |
| GOBP_LYMPH_NODE_DE | GOBP_LYMPH_NODE_DEVELOPMENT | 32 | -0.60907 | -2.35116 | 0 | 0.007413548 | 0.061 | 4239 | tags=44%, list=18%, si | MM_2 |
| GOBP_CHROMOSOME_SE | GOBP_CHROMOSOME_SEPARATION | 80 | -0.48479 | -2.3493 | 0 | 0.007089807 | 0.063 | 4413 | tags=38%, list=19%, si | MM_2 |
| GOBP_POSITIVE_REGULA | GOBP_POSITIVE_REGULATION_OF_PATTERN | 68 | -0.49974 | -2.33253 | 0 | 0.008182682 | 0.078 | 4446 | tags=40%, list=19%, si | MM_2 |
| HEPATITIS_B | HEPATITIS_B | 147 | -0.43373 | -2.3174 | 0 | 0.009046272 | 0.093 | 5002 | tags=41%, list=21%, si | MM_2 |
| MEASLES | MEASLES | 128 | -0.43878 | -2.31482 | 0 | 0.008950076 | 0.098 | 3921 | tags=27%, list=17%, si | MM_2 |
| GOBP_CHROMOSOME_C | GOBP_CHROMOSOME_CONDENSATION | 30 | -0.61367 | -2.30669 | 0 | 0.009101098 | 0.105 | 2053 | tags=37%, list=9%, sig | MM_2 |
| GOBP_CELLULAR_RESPON | GOBP_CELLULAR_RESPONSE_TO_INTERFER | 16 | -0.72819 | -2.27574 | 0 | 0.01217644 | 0.147 | 1873 | tags=38%, list=8%, sig | MM_2 |
| TOLL-LIKE_RECEPTOR_SIG | TOLL-LIKE_RECEPTOR_SIGNALING_PATHWA | 84 | -0.45651 | -2.27301 | 0 | 0.011859455 | 0.148 | 5628 | tags=43%, list=24%, si | MM_2 |
| GOBP_MITOTIC_SISTER_ | GOBP_MITOTIC_SISTER_CHROMATID_SEGR | 178 | -0.41188 | -2.26908 | 0 | 0.011852236 | 0.156 | 4270 | tags=33%, list=18%, si | MM_2 |
| GOBP_POST_TRANSCRIPT | GOBP_POST_TRANSCRIPTIONAL_GENE_SILE | 64 | -0.48962 | -2.25995 | 0 | 0.012654389 | 0.174 | 4370 | tags=44%, list=19%, si | MM_2 |
| LYSOSOME | LYSOSOME | 118 | -0.4285 | -2.24183 | 0 | 0.015351931 | 0.214 | 4935 | tags=42%, list=21%, si | MM_2 |
| TUBERCULOSIS | TUBERCULOSIS | 152 | -0.40774 | -2.22611 | 0 | 0.017042704 | 0.246 | 5095 | tags=33%, list=22%, si | MM_2 |
| GOBP_REGULATION_OF_ | GOBP_REGULATION_OF_VIRAL_GENOME_R | 78 | -0.45741 | -2.2226 | 0 | 0.01707442 | 0.258 | 3123 | tags=29%, list=13%, si | MM_2 |
| INFLUENZA_A | INFLUENZA_A | 139 | -0.41515 | -2.21297 | 0 | 0.018832764 | 0.288 | 5615 | tags=43%, list=24%, si | MM_2 |
| TNF_SIGNALING_PATHW | TNF_SIGNALING_PATHWAY | 108 | -0.4293 | -2.21221 | 0 | 0.018189313 | 0.289 | 5042 | tags=40%, list=21%, si | MM_2 |
| GOBP_MIRNA_MEDIATED | GOBP_MIRNA_MEDIATED_GENE_SILENCING | 17 | -0.6952 | -2.21121 | 0 | 0.017904835 | 0.295 | 4205 | tags=59%, list=18%, si | MM_2 |
| GOBP_NEGATIVE_REGUL | GOBP_NEGATIVE_REGULATION_OF_RHO_P | 25 | -0.59693 | -2.20355 | 0 | 0.018811187 | 0.317 | 1825 | tags=36%, list=8%, sig | MM_2 |
| GOBP_REGULATION_OF_ | GOBP_REGULATION_OF_MITOTIC_SISTER_C | 56 | -0.48647 | -2.1994 | 0 | 0.019160844 | 0.328 | 2002 | tags=25%, list=8%, sig | MM_2 |
| GOBP_POSITIVE_REGULA | GOBP_POSITIVE_REGULATION_OF_FATTY_A | 20 | -0.64082 | -2.19926 | 0.0023923444 | 0.018542752 | 0.328 | 4077 | tags=60%, list=17%, si | MM_2 |
| GOBP_PROTEIN_LOCALIZ | GOBP_PROTEIN_LOCALIZATION_TO_CHROM | 112 | -0.42016 | -2.1882 | 0 | 0.019743156 | 0.357 | 5176 | tags=33%, list=22%, si | MM_2 |
| GOBP_REGULATION_OF_ | GOBP_REGULATION_OF_VACUOLE_ORGANI | 54 | -0.49622 | -2.18692 | 0 | 0.019545805 | 0.363 | 5856 | tags=52%, list=25%, si | MM_2 |
| GOBP_PHOSPHATIDYLINO | GOBP_PHOSPHATIDYLINOSITOL_BIOSYNTHE | 95 | -0.43984 | -2.17357 | 0 | 0.021765836 | 0.404 | 4203 | tags=35%, list=18%, si | MM_2 |
| GOBP_PROTEIN_LOCALIZ | GOBP_PROTEIN_LOCALIZATION_TO_CHROM | 41 | -0.51516 | -2.17131 | 0 | 0.02177268 | 0.413 | 3874 | tags=32%, list=16%, si | MM_2 |
| GOBP_REGULATION_OF_ | GOBP_REGULATION_OF_TYPE_I_INTERFERO | 40 | -0.53004 | -2.16918 | 0.002444988 | 0.021494163 | 0.419 | 2174 | tags=30%, list=9%, sig | MM_2 |
| GOBP_POSITIVE_REGULA | GOBP_POSITIVE_REGULATION_OF_TYPE_I_I | 77 | -0.45649 | -2.16205 | 0 | 0.02242411 | 0.441 | 4010 | tags=38%, list=17%, si | MM_2 |
| GOBP_SPINDLE_CHECKPO | GOBP_SPINDLE_CHECKPOINT_SIGNALING | 53 | -0.49069 | -2.14657 | 0 | 0.025672931 | 0.497 | 4413 | tags=38%, list=19%, si | MM_2 |
| GOBP_REGULATION_OF_ | GOBP_REGULATION_OF_AUTOPHAGOSOME | 44 | -0.50637 | -2.12985 | 0 | 0.029748013 | 0.548 | 5856 | tags=50%, list=25%, si | MM_2 |
| GOBP_REGULATION_OF_ | GOBP_REGULATION_OF_LYMPHOCYTE_MIG | 64 | -0.4571 | -2.12658 | 0 | 0.029740622 | 0.556 | 4175 | tags=33%, list=18%, si | MM_2 |
| GOBP_NUCLEAR_CHROM | GOBP_NUCLEAR_CHROMOSOME_SEGREGAT | 305 | -0.34467 | -2.12218 | 0 | 0.03041175 | 0.577 | 4270 | tags=30%, list=18%, si | MM_2 |
| GOBP_POSITIVE_REGULA | GOBP_POSITIVE_REGULATION_OF_CHROMO | 28 | -0.57614 | -2.12015 | 0 | 0.030104838 | 0.581 | 2727 | tags=36%, list=12%, si | MM_2 |
| GOBP_REGULATION_OF_ | GOBP_REGULATION_OF_LYMPHOCYTE_CHE | 22 | -0.60555 | -2.11836 | 0.0023640662 | 0.029881893 | 0.589 | 3602 | tags=41%, list=15%, si | MM_2 |
| GOBP_AUTOPHAGOSOMI | GOBP_AUTOPHAGOSOME_ORGANIZATION | 128 | -0.39803 | -2.11485 | 0 | 0.030305352 | 0.601 | 6278 | tags=45%, list=27%, si | MM_2 |
| GOBP_POSITIVE_REGULA | GOBP_POSITIVE_REGULATION_OF_CHEMOK | 15 | -0.66846 | -2.11324 | 0.004842615 | 0.029890483 | 0.603 | 3755 | tags=40%, list=16%, si | MM_2 |
| GOBP_ATTACHMENT_OF_ | GOBP_ATTACHMENT_OF_SPINDLE_MICROT | 48 | -0.4954 | -2.10803 | 0 | 0.030447789 | 0.613 | 4413 | tags=46%, list=19%, si | MM_2 |
| GOBP_REGULATION_OF_ | GOBP_REGULATION_OF_T_CELL_MIGRATIO | 49 | -0.48626 | -2.1064 | 0 | 0.03011191 | 0.617 | 4175 | tags=35%, list=18%, si | MM_2 |
| FC_GAMMA_R-MEDIATED | FC_GAMMA_R-MEDIATED_PHAGOCYTOSIS | 84 | -0.43073 | -2.10399 | 0 | 0.030275667 | 0.625 | 6225 | tags=44%, list=26%, si | MM_2 |
| GOBP_POSITIVE_REGULA | GOBP_POSITIVE_REGULATION_OF_LYMPHO | 42 | -0.50207 | -2.0971 | 0.002352941 | 0.03150989 | 0.654 | 3868 | tags=31%, list=16%, si | MM_2 |
| GOBP_NEGATIVE_REGUL | GOBP_NEGATIVE_REGULATION_OF_TRANSL | 20 | -0.6155 | -2.09476 | 0 | 0.031816848 | 0.667 | 3339 | tags=35%, list=14%, si | MM_2 |
| GOBP_TRANSLATIONAL_I | GOBP_TRANSLATIONAL_INITIATION | 112 | -0.41199 | -2.09433 | 0 | 0.031250115 | 0.668 | 3793 | tags=26%, list=16%, si | MM_2 |
| EPSTEIN-BARR_VIRUS_IN | EPSTEIN-BARR_VIRUS_INFECTION | 190 | -0.36707 | -2.08507 | 0 | 0.033269607 | 0.697 | 3921 | tags=29%, list=17%, si | MM_2 |
| GOBP_POSITIVE_REGULA | GOBP_POSITIVE_REGULATION_OF_INTERFE | 45 | -0.50121 | -2.08316 | 0.0024875621 | 0.033084527 | 0.702 | 3959 | tags=38%, list=17%, si | MM_2 |
| GOBP_T_CELL_MIGRATIO | GOBP_T_CELL_MIGRATION | 64 | -0.46213 | -2.0791 | 0 | 0.033639375 | 0.719 | 4858 | tags=36%, list=21%, si | MM_2 |
| GOBP_VIRAL_GENOME_F | GOBP_VIRAL_GENOME_REPLICATION | 110 | -0.40068 | -2.07854 | 0 | 0.033162158 | 0.72 | 2864 | tags=23%, list=12%, si | MM_2 |
| GOBP_CHROMOSOME_SE | GOBP_CHROMOSOME_SEGREGATION | 403 | -0.3283 | -2.07732 | 0 | 0.033042453 | 0.721 | 4361 | tags=29%, list=18%, si | MM_2 |
| GOBP_REGULATION_OF_ | GOBP_REGULATION_OF_DNA_TEMPLATED_ | 40 | -0.50528 | -2.07385 | 0.0022675737 | 0.03354613 | 0.733 | 3943 | tags=43%, list=17%, si | MM_2 |
| GOBP_CYTOSKELETON_D | GOBP_CYTOSKELETON_DEPENDENT_CYTOK | 97 | -0.41944 | -2.07105 | 0 | 0.033497404 | 0.738 | 3795 | tags=27%, list=16%, si | MM_2 |
| RIG-I-LIKE_RECEPTOR_SIG | RIG-I-LIKE_RECEPTOR_SIGNALING_PATHWA | 52 | -0.47743 | -2.06952 | 0.0026595744 | 0.033201605 | 0.741 | 4288 | tags=35%, list=18%, si | MM_2 |
| GOBP_POSITIVE_REGULA | GOBP_POSITIVE_REGULATION_OF_ACTIN_F | 52 | -0.47641 | -2.06012 | 0.0022172949 | 0.035552934 | 0.77 | 3875 | tags=31%, list=16%, si | MM_2 |
| GOBP_POSITIVE_REGULA | GOBP_POSITIVE_REGULATION_OF_B_CELL_ | 18 | -0.64027 | -2.05916 | 0 | 0.035329226 | 0.772 | 3025 | tags=28%, list=13%, si | MM_2 |
| OSTEOCLAST_DIFFERENTI | OSTEOCLAST_DIFFERENTIATION | 121 | -0.39058 | -2.05774 | 0 | 0.035209864 | 0.775 | 5357 | tags=37%, list=23%, si | MM_2 |
| GOBP_NEGATIVE_REGUL | GOBP_NEGATIVE_REGULATION_OF_CHROM | 96 | -0.41349 | -2.05541 | 0 | 0.03532724 | 0.782 | 2696 | tags=24%, list=11%, si | MM_2 |
| GOBP_FATTY_ACID_CATA | GOBP_FATTY_ACID_CATABOLIC_PROCESS | 100 | -0.40031 | -2.05289 | 0 | 0.035486918 | 0.788 | 4470 | tags=37%, list=19%, si | MM_2 |
| GOBP_MITOTIC_SPINDLE | GOBP_MITOTIC_SPINDLE_ORGANIZATION | 125 | -0.39032 | -2.0483 | 0 | 0.036341354 | 0.803 | 4230 | tags=30%, list=18%, si | MM_2 |
| GOBP_SEQUESTERING_O | GOBP_SEQUESTERING_OF_TRIGLYCERIDE | 18 | -0.62048 | -2.04737 | 0 | 0.03599049 | 0.804 | 4943 | tags=61%, list=21%, si | MM_2 |
| GOBP_PROTEIN_LOCALIZ | GOBP_PROTEIN_LOCALIZATION_TO_NUCLE | 15 | -0.65805 | -2.04725 | 0.004484305 | 0.03545332 | 0.804 | 1039 | tags=27%, list=4%, sig | MM_2 |
| GOBP_TYPE_I_INTERFERO | GOBP_TYPE_I_INTERFERON_PRODUCTION | 127 | -0.39129 | -2.04676 | 0 | 0.035082906 | 0.806 | 4338 | tags=34%, list=18%, si | MM_2 |
| GOBP_MITOTIC_CYTOKIN | GOBP_MITOTIC_CYTOKINESIS | 83 | -0.41969 | -2.04639 | 0 | 0.034680292 | 0.809 | 4788 | tags=33%, list=20%, si | MM_2 |
| GOBP_SPINDLE_ORGANIZ | GOBP_SPINDLE_ORGANIZATION | 190 | -0.36083 | -2.04547 | 0 | 0.034477107 | 0.812 | 4863 | tags=34%, list=21%, si | MM_2 |
| GOBP_TRANSLATION_AT_ | GOBP_TRANSLATION_AT_SYNAPSE | 48 | -0.48264 | -2.04246 | 0 | 0.034830093 | 0.822 | 12238 | tags=100%, list=52%, | MM_2 |
| HUMAN_T-CELL_LEUKEM | HUMAN_T-CELL_LEUKEMIA_VIRUS_1_INFEC | 215 | -0.35601 | -2.0424 | 0 | 0.03438748 | 0.822 | 5211 | tags=40%, list=22%, si | MM_2 |
| GOBP_CYTOKINESIS | GOBP_CYTOKINESIS | 160 | -0.37146 | -2.04181 | 0 | 0.034035083 | 0.823 | 4788 | tags=30%, list=20%, si | MM_2 |
| GOBP_RUFFLE_ORGANIZA | GOBP_RUFFLE_ORGANIZATION | 54 | -0.46217 | -2.04079 | 0.0023255814 | 0.033872925 | 0.827 | 4318 | tags=41%, list=18%, si | MM_2 |
| GOBP_DEFENSE_RESPON | GOBP_DEFENSE_RESPONSE_TO_VIRUS | 276 | -0.33951 | -2.03991 | 0 | 0.03361612 | 0.828 | 4023 | tags=30%, list=17%, si | MM_2 |
| GOBP_REGULATION_OF_ | GOBP_REGULATION_OF_PATTERN_RECOGN | 157 | -0.36835 | -2.03972 | 0 | 0.033211768 | 0.829 | 4261 | tags=32%, list=18%, si | MM_2 |
| GOBP_REGULATION_OF_ | GOBP_REGULATION_OF_T_CELL_CHEMOTA | 16 | -0.64332 | -2.03925 | 0.004716981 | 0.0329149 | 0.831 | 3602 | tags=44%, list=15%, si | MM_2 |
| STAPHYLOCOCCUS_AURE | STAPHYLOCOCCUS_AUREUS_INFECTION | 44 | -0.48396 | -2.03456 | 0 | 0.033654526 | 0.845 | 3276 | tags=34%, list=14%, si | MM_2 |
| GOBP_INTERFERON_MED | GOBP_INTERFERON_MEDIATED_SIGNALING | 76 | -0.41971 | -2.02465 | 0 | 0.036245324 | 0.861 | 5399 | tags=47%, list=23%, si | MM_2 |
| GOBP_INTERFERON_BETA | GOBP_INTERFERON_BETA_PRODUCTION | 66 | -0.43834 | -2.02452 | 0 | 0.035829097 | 0.862 | 4272 | tags=36%, list=18%, si | MM_2 |
| GOBP_ANTIGEN_PROCES | GOBP_ANTIGEN_PROCESSING_AND_PRESEN | 24 | -0.57393 | -2.02193 | 0.00243309 | 0.03616092 | 0.867 | 5246 | tags=58%, list=22%, si | MM_2 |

| | | | | | | | | | | |
|---|---|---|---|---|---|---|---|---|---|---|
| GOBP_SISTER_CHROMAT | GOBP_SISTER_CHROMATID_SEGREGATION | 219 | -0.35049 | -2.02131 | 0 | 0.03593244 | 0.87 | 4270 | tags=29%, list=18%, si | MM_2 |
| GOBP_POSITIVE_REGULA | GOBP_POSITIVE_REGULATION_OF_CHEMOK | 79 | -0.42151 | -2.0212 | 0 | 0.035517197 | 0.87 | 2772 | tags=25%, list=12%, si | MM_2 |
| GOBP_T_CELL_CHEMOTA | GOBP_T_CELL_CHEMOTAXIS | 23 | -0.57356 | -2.0185 | 0.008830022 | 0.035792205 | 0.874 | 3602 | tags=35%, list=15%, si | MM_2 |
| GOBP_POST_TRANSCRIPT | GOBP_POST_TRANSCRIPTIONAL_REGULATIO | 480 | -0.31532 | -2.01822 | 0 | 0.03540532 | 0.874 | 5234 | tags=33%, list=22%, si | MM_2 |
| GOBP_NEGATIVE_REGUL | GOBP_NEGATIVE_REGULATION_OF_NUCLEA | 66 | -0.44357 | -2.01745 | 0 | 0.035163984 | 0.876 | 4413 | tags=39%, list=19%, si | MM_2 |
| GOBP_RESPONSE_TO_INT | GOBP_RESPONSE_TO_INTERFERON_ALPHA | 28 | -0.54244 | -2.01589 | 0.0024570024 | 0.035216082 | 0.881 | 4252 | tags=39%, list=18%, si | MM_2 |
| GOBP_CELLULAR_RESPON | GOBP_CELLULAR_RESPONSE_TO_LIPOPROT | 38 | -0.50161 | -2.01446 | 0 | 0.035264578 | 0.885 | 4355 | tags=32%, list=18%, si | MM_2 |
| GOBP_CERAMIDE_BIOSYN | GOBP_CERAMIDE_BIOSYNTHETIC_PROCESS | 65 | -0.43265 | -2.01353 | 0.0025062656 | 0.035098046 | 0.887 | 3755 | tags=34%, list=16%, si | MM_2 |
| GOBP_RESPONSE_TO_VIR | GOBP_RESPONSE_TO_VIRUS | 331 | -0.32738 | -2.00987 | 0 | 0.0358466 | 0.893 | 4290 | tags=31%, list=18%, si | MM_2 |
| GOBP_NEGATIVE_REGUL | GOBP_NEGATIVE_REGULATION_OF_DOUBL | 24 | -0.56341 | -2.00738 | 0 | 0.03619253 | 0.9 | 4308 | tags=42%, list=18%, si | MM_2 |
| GOBP_MEIOTIC_CHROMO | GOBP_MEIOTIC_CHROMOSOME_SEGREGAT | 94 | -0.39732 | -1.99751 | 0 | 0.038794313 | 0.916 | 3014 | tags=32%, list=13%, si | MM_2 |
| CHEMOKINE_SIGNALING_ | CHEMOKINE_SIGNALING_PATHWAY | 177 | -0.36112 | -1.99485 | 0 | 0.039289616 | 0.922 | 5254 | tags=39%, list=22%, si | MM_2 |
| GOBP_PORE_COMPLEX_A | GOBP_PORE_COMPLEX_ASSEMBLY | 20 | -0.58631 | -1.99157 | 0.004901961 | 0.039949838 | 0.925 | 4467 | tags=50%, list=19%, si | MM_2 |
| GOBP_REGULATION_OF_ | GOBP_REGULATION_OF_CELL_SUBSTRATE_ | 69 | -0.42645 | -1.99062 | 0 | 0.039869808 | 0.925 | 3783 | tags=35%, list=16%, si | MM_2 |
| GOBP_NEGATIVE_REGUL | GOBP_NEGATIVE_REGULATION_OF_DNA_R | 37 | -0.50653 | -1.98533 | 0 | 0.041359805 | 0.934 | 5101 | tags=38%, list=22%, si | MM_2 |
| GOBP_PROTEIN_LOCALIZ | GOBP_PROTEIN_LOCALIZATION_TO_CHROM | 24 | -0.5548 | -1.98168 | 0.004796163 | 0.042245686 | 0.935 | 5158 | tags=50%, list=22%, si | MM_2 |
| GOBP_POSITIVE_REGULA | GOBP_POSITIVE_REGULATION_OF_MITOTIC | 18 | -0.60202 | -1.97878 | 0.005181347 | 0.04282936 | 0.941 | 965 | tags=28%, list=4%, sig | MM_2 |
| GOBP_MEIOSIS_I_CELL_C | GOBP_MEIOSIS_I_CELL_CYCLE_PROCESS | 128 | -0.36785 | -1.97876 | 0 | 0.042396743 | 0.941 | 3014 | tags=29%, list=13%, si | MM_2 |
| GOBP_REGULATION_OF_ | GOBP_REGULATION_OF_ATTACHMENT_OF_ | 20 | -0.58538 | -1.97667 | 0.004705882 | 0.042883772 | 0.944 | 4413 | tags=55%, list=19%, si | MM_2 |
| GOBP_NEGATIVE_REGUL | GOBP_NEGATIVE_REGULATION_OF_CELL_SI | 16 | -0.61922 | -1.97187 | 0.0025125628 | 0.04414972 | 0.947 | 3783 | tags=50%, list=16%, si | MM_2 |
| GOBP_CORPUS_CALLOSU | GOBP_CORPUS_CALLOSUM_DEVELOPMENT | 27 | -0.53418 | -1.9664 | 0.0047393367 | 0.045471586 | 0.95 | 1815 | tags=26%, list=8%, sig | MM_2 |
| GOBP_TOLL_LIKE_RECEPT | GOBP_TOLL_LIKE_RECEPTOR_3_SIGNALING_ | 29 | -0.51364 | -1.96203 | 0 | 0.046721876 | 0.959 | 3308 | tags=38%, list=14%, si | MM_2 |
| GOBP_POSITIVE_REGULA | GOBP_POSITIVE_REGULATION_OF_PHOSPH | 26 | -0.53598 | -1.96108 | 0.006976744 | 0.046581358 | 0.959 | 4918 | tags=46%, list=21%, si | MM_2 |
| HEPATITIS_C | HEPATITIS_C | 138 | -0.36361 | -1.96003 | 0 | 0.046500646 | 0.959 | 5002 | tags=34%, list=21%, si | MM_2 |
| GOBP_ENDOSOME_ORGA | GOBP_ENDOSOME_ORGANIZATION | 68 | -0.42044 | -1.95978 | 0 | 0.046145055 | 0.959 | 5645 | tags=41%, list=24%, si | MM_2 |
| GOBP_POSITIVE_REGULA | GOBP_POSITIVE_REGULATION_OF_PROTEIN | 52 | -0.452 | -1.95939 | 0 | 0.04576817 | 0.959 | 2318 | tags=25%, list=10%, si | MM_2 |
| GOBP_REGULATION_OF_ | GOBP_REGULATION_OF_VIRAL_LIFE_CYCLE | 139 | -0.36484 | -1.95368 | 0 | 0.047489334 | 0.963 | 3123 | tags=27%, list=13%, si | MM_2 |
| PHAGOSOME | PHAGOSOME | 146 | -0.35781 | -1.94952 | 0 | 0.04875629 | 0.964 | 3721 | tags=23%, list=16%, si | MM_2 |
| GOBP_REGULATION_OF_ | GOBP_REGULATION_OF_CHROMATIN_BIND | 20 | -0.57125 | -1.94412 | 0 | 0.05047345 | 0.968 | 7232 | tags=80%, list=31%, si | MM_2 |
| GOBP_VIRAL_PROCESS | GOBP_VIRAL_PROCESS | 303 | -0.31782 | -1.94093 | 0 | 0.051386554 | 0.972 | 5727 | tags=39%, list=24%, si | MM_2 |
| GOBP_MEIOTIC_SPINDLE | GOBP_MEIOTIC_SPINDLE_ORGANIZATION | 21 | -0.55877 | -1.93389 | 0 | 0.05396341 | 0.977 | 3646 | tags=43%, list=15%, si | MM_2 |
| GOBP_SPINDLE_ASSEMBL | GOBP_SPINDLE_ASSEMBLY | 122 | -0.36805 | -1.93116 | 0 | 0.05458489 | 0.978 | 4361 | tags=32%, list=18%, si | MM_2 |
| GOBP_LYMPHOCYTE_MIG | GOBP_LYMPHOCYTE_MIGRATION | 92 | -0.39324 | -1.92935 | 0 | 0.054772492 | 0.978 | 3602 | tags=26%, list=15%, si | MM_2 |
| GOBP_REGULATION_OF_ | GOBP_REGULATION_OF_VIRAL_PROCESS | 167 | -0.34594 | -1.92904 | 0 | 0.054462314 | 0.978 | 4590 | tags=34%, list=19%, si | MM_2 |
| GOBP_POSITIVE_REGULA | GOBP_POSITIVE_REGULATION_OF_CELL_MA | 61 | -0.42392 | -1.9285 | 0 | 0.054258473 | 0.978 | 3649 | tags=30%, list=15%, si | MM_2 |
| GOBP_METAPHASE_CHRO | GOBP_METAPHASE_CHROMOSOME_ALIGNI | 93 | -0.38906 | -1.92787 | 0 | 0.054120917 | 0.98 | 4413 | tags=34%, list=19%, si | MM_2 |
| GOBP_REGULATION_OF_ | GOBP_REGULATION_OF_CYTOKINESIS | 83 | -0.39341 | -1.92504 | 0 | 0.054754622 | 0.984 | 4918 | tags=36%, list=21%, si | MM_2 |
| GOBP_DENDRITIC_CELL_ | GOBP_DENDRITIC_CELL_CYTOKINE_PRODUC | 16 | -0.61359 | -1.92418 | 0.0050125313 | 0.05465172 | 0.985 | 5674 | tags=56%, list=24%, si | MM_2 |
| GOBP_NATURAL_KILLER_ | GOBP_NATURAL_KILLER_CELL_DIFFERENTIA | 34 | -0.48891 | -1.92328 | 0.002352941 | 0.05449933 | 0.986 | 5109 | tags=47%, list=22%, si | MM_2 |
| GOBP_MITOTIC_CHROMO | GOBP_MITOTIC_CHROMOSOME_CONDENSA | 15 | -0.61737 | -1.9223 | 0.005063291 | 0.054450624 | 0.987 | 2053 | tags=40%, list=9%, sig | MM_2 |
| GOBP_POSITIVE_REGULA | GOBP_POSITIVE_REGULATION_OF_SMOOTH | 69 | -0.40758 | -1.91782 | 0 | 0.055745434 | 0.989 | 4572 | tags=30%, list=19%, si | MM_2 |
| GOBP_REGULATION_OF_ | GOBP_REGULATION_OF_TRANSLATIONAL_I | 17 | -0.5913 | -1.91668 | 0.007194245 | 0.0556857 | 0.989 | 1929 | tags=24%, list=8%, sig | MM_2 |
| GOBP_REGULATION_OF_ | GOBP_REGULATION_OF_CHROMOSOME_SE | 126 | -0.36267 | -1.91425 | 0 | 0.05627854 | 0.99 | 4413 | tags=31%, list=19%, si | MM_2 |
| GOBP_MITOTIC_NUCLEA | GOBP_MITOTIC_NUCLEAR_DIVISION | 259 | -0.32458 | -1.90372 | 0 | 0.06012264 | 0.993 | 4271 | tags=31%, list=18%, si | MM_2 |
| GOBP_PRE_MIRNA_PROC | GOBP_PRE_MIRNA_PROCESSING | 15 | -0.60078 | -1.89969 | 0.0069444445 | 0.06141151 | 0.993 | 4917 | tags=60%, list=21%, si | MM_2 |
| GOBP_SMOOTH_MUSCLE | GOBP_SMOOTH_MUSCLE_CELL_MIGRATION | 115 | -0.37159 | -1.89156 | 0.0024390244 | 0.06463055 | 0.995 | 4887 | tags=34%, list=21%, si | MM_2 |
| UBIQUITIN_MEDIATED_P | UBIQUITIN_MEDIATED_PROTEOLYSIS | 134 | -0.35651 | -1.88966 | 0 | 0.06501928 | 0.995 | 5873 | tags=37%, list=25%, si | MM_2 |
| GOBP_NEGATIVE_REGUL | GOBP_NEGATIVE_REGULATION_OF_CELLUL | 51 | -0.43956 | -1.88876 | 0.0023041475 | 0.06499115 | 0.995 | 3521 | tags=35%, list=15%, si | MM_2 |
| KAPOSI_SARCOMA-ASSO | KAPOSI_SARCOMA-ASSOCIATED_HERPESVIR | 176 | -0.34314 | -1.88853 | 0 | 0.06460454 | 0.995 | 5252 | tags=38%, list=22%, si | MM_2 |
| GOBP_POSITIVE_REGULA | GOBP_POSITIVE_REGULATION_OF_CHROMO | 29 | -0.49977 | -1.88838 | 0.009478673 | 0.06414527 | 0.995 | 4649 | tags=48%, list=20%, si | MM_2 |
| GOBP_CELL_VOLUME_HO | GOBP_CELL_VOLUME_HOMEOSTASIS | 38 | -0.47499 | -1.88552 | 0.0024271845 | 0.06521411 | 0.995 | 1769 | tags=29%, list=7%, sig | MM_2 |
| GOBP_CELLULAR_RESPON | GOBP_CELLULAR_RESPONSE_TO_DSRNA | 19 | -0.55744 | -1.88409 | 0 | 0.065419815 | 0.995 | 1902 | tags=21%, list=8%, sig | MM_2 |
| GOBP_RESPONSE_TO_MA | GOBP_RESPONSE_TO_MAGNESIUM_ION | 15 | -0.60351 | -1.8805 | 0.0121654505 | 0.066470824 | 0.995 | 882 | tags=33%, list=4%, sig | MM_2 |
| GOBP_VESICLE_DOCKING | GOBP_VESICLE_DOCKING | 55 | -0.42468 | -1.88041 | 0 | 0.066054255 | 0.995 | 2105 | tags=22%, list=9%, sig | MM_2 |
| GOBP_REGULATION_OF_ | GOBP_REGULATION_OF_PROTEIN_STABILIT | 305 | -0.30989 | -1.87952 | 0 | 0.065945305 | 0.995 | 4338 | tags=30%, list=18%, si | MM_2 |
| GOBP_REGULATION_OF_ | GOBP_REGULATION_OF_CELL_DIVISION | 168 | -0.33245 | -1.87892 | 0 | 0.06572101 | 0.995 | 4929 | tags=36%, list=21%, si | MM_2 |
| GOBP_MITOTIC_CELL_CY | GOBP_MITOTIC_CELL_CYCLE_PHASE_TRANS | 425 | -0.29624 | -1.87617 | 0 | 0.06652152 | 0.995 | 4986 | tags=32%, list=21%, si | MM_2 |
| FERROPTOSIS | FERROPTOSIS | 37 | -0.47181 | -1.87426 | 0.004842615 | 0.066874236 | 0.996 | 6982 | tags=65%, list=30%, si | MM_2 |
| GOBP_GLYCEROPHOSPHO | GOBP_GLYCEROPHOSPHOLIPID_BIOSYNTHE | 160 | -0.34125 | -1.87389 | 0 | 0.0666563 | 0.996 | 4203 | tags=31%, list=18%, si | MM_2 |
| GOBP_POSITIVE_REGULA | GOBP_POSITIVE_REGULATION_OF_NUCLEO | 74 | -0.40368 | -1.87344 | 0.0025445293 | 0.06639254 | 0.996 | 4220 | tags=30%, list=18%, si | MM_2 |
| GOBP_POSITIVE_REGULA | GOBP_POSITIVE_REGULATION_OF_TYPE_I_I | 19 | -0.54338 | -1.87172 | 0.016627079 | 0.0669378 | 0.996 | 2174 | tags=32%, list=9%, sig | MM_2 |
| NF-KAPPA_B_SIGNALING | NF-KAPPA_B_SIGNALING_PATHWAY | 93 | -0.37385 | -1.86966 | 0.0023584906 | 0.06741913 | 0.996 | 5104 | tags=38%, list=22%, si | MM_2 |
| GOBP_POSITIVE_REGULA | GOBP_POSITIVE_REGULATION_OF_TUMOR_ | 113 | -0.36888 | -1.86946 | 0 | 0.06712332 | 0.996 | 4220 | tags=28%, list=18%, si | MM_2 |
| GOBP_ESTABLISHMENT_ | GOBP_ESTABLISHMENT_OF_PROTEIN_LOCA | 20 | -0.54104 | -1.86873 | 0.0025252525 | 0.06704445 | 0.997 | 3901 | tags=45%, list=17%, si | MM_2 |
| GOBP_ESTABLISHMENT_ | GOBP_ESTABLISHMENT_OR_MAINTENANCE | 25 | -0.51185 | -1.86763 | 0.0051150895 | 0.0670858 | 0.997 | 2390 | tags=32%, list=10%, si | MM_2 |
| GOBP_PHOSPHOLIPID_BI | GOBP_PHOSPHOLIPID_BIOSYNTHETIC_PROC | 214 | -0.32046 | -1.86746 | 0 | 0.06665955 | 0.997 | 4259 | tags=29%, list=18%, si | MM_2 |
| GOBP_NEGATIVE_REGUL | GOBP_NEGATIVE_REGULATION_OF_DNA_R | 42 | -0.45507 | -1.86681 | 0.002386635 | 0.066426665 | 0.997 | 4463 | tags=36%, list=19%, si | MM_2 |
| GOBP_POSITIVE_REGULA | GOBP_POSITIVE_REGULATION_OF_DEPHOS | 53 | -0.43064 | -1.86507 | 0.0023364485 | 0.06678164 | 0.998 | 4918 | tags=38%, list=21%, si | MM_2 |
| GOBP_FEMALE_MEIOTIC_ | GOBP_FEMALE_MEIOTIC_NUCLEAR_DIVISIO | 41 | -0.45063 | -1.86221 | 0.007125891 | 0.067566745 | 0.999 | 3646 | tags=44%, list=15%, si | MM_2 |
| GOBP_ENDOLYSOSOMAL_ | GOBP_ENDOLYSOSOMAL_TOLL_LIKE_RECEP | 47 | -0.44119 | -1.85969 | 0.004576659 | 0.068376645 | 1 | 3308 | tags=30%, list=14%, si | MM_2 |
| GOBP_VACUOLE_ORGAN | GOBP_VACUOLE_ORGANIZATION | 223 | -0.32222 | -1.85896 | 0 | 0.068293065 | 1 | 6092 | tags=42%, list=26%, si | MM_2 |
| GOBP_VESICLE_DOCKING | GOBP_VESICLE_DOCKING_INVOLVED_IN_EX | 36 | -0.46597 | -1.85884 | 0 | 0.06791223 | 1 | 1675 | tags=22%, list=7%, sig | MM_2 |
| GOBP_POSITIVE_REGULA | GOBP_POSITIVE_REGULATION_OF_DEFENSE | 31 | -0.47692 | -1.85683 | 0 | 0.06849583 | 1 | 3720 | tags=42%, list=16%, si | MM_2 |
| GOBP_VIRAL_LIFE_CYCLE | GOBP_VIRAL_LIFE_CYCLE | 230 | -0.32078 | -1.85467 | 0 | 0.06917019 | 1 | 5727 | tags=38%, list=24%, si | MM_2 |
| GOBP_CYTOKINE_MEDIAT | GOBP_CYTOKINE_MEDIATED_SIGNALING_P | 421 | -0.29336 | -1.85424 | 0 | 0.068962805 | 1 | 4618 | tags=30%, list=20%, si | MM_2 |
| GOBP_NUCLEAR_ENVELO | GOBP_NUCLEAR_ENVELOPE_ORGANIZATION | 52 | -0.42407 | -1.85276 | 0 | 0.06926106 | 1 | 3960 | tags=25%, list=17%, si | MM_2 |
| GOBP_REGULATION_OF_ | GOBP_REGULATION_OF_DEFENSE_RESPONS | 67 | -0.39826 | -1.8516 | 0 | 0.06943841 | 1 | 3720 | tags=36%, list=16%, si | MM_2 |
| GOBP_POSITIVE_REGULA | GOBP_POSITIVE_REGULATION_OF_MEMBRA | 39 | -0.45197 | -1.84882 | 0.0073710075 | 0.07026379 | 1 | 4842 | tags=41%, list=21%, si | MM_2 |
| SPHINGOLIPID_METABOL | SPHINGOLIPID_METABOLISM | 44 | -0.45078 | -1.84871 | 0.002538071 | 0.06989779 | 1 | 3317 | tags=32%, list=14%, si | MM_2 |
| CELL_CYCLE | CELL_CYCLE | 121 | -0.35162 | -1.84854 | 0 | 0.06949984 | 1 | 4954 | tags=34%, list=21%, si | MM_2 |
| GOBP_HIPPO_SIGNALING | GOBP_HIPPO_SIGNALING | 40 | -0.44803 | -1.84712 | 0.0046620048 | 0.06965872 | 1 | 4999 | tags=48%, list=21%, si | MM_2 |
| GOBP_ANTIGEN_PROCES | GOBP_ANTIGEN_PROCESSING_AND_PRESEN | 105 | -0.36226 | -1.84379 | 0 | 0.070900194 | 1 | 5465 | tags=39%, list=23%, si | MM_2 |

| | | | | | | | | | | |
|---|---|---|---|---|---|---|---|---|---|---|
| GOBP_VASCULAR_ASSOC | GOBP_VASCULAR_ASSOCIATED_SMOOTH_N | 80 | -0.39033 | -1.84359 | 0 | 0.07056645 | 1 | 4833 | tags=38%, list=20%, si | MM_2 |
| GOBP_TOLL_LIKE_RECEPT | GOBP_TOLL_LIKE_RECEPTOR_4_SIGNALING_ | 60 | -0.4086 | -1.84051 | 0 | 0.07170161 | 1 | 4244 | tags=33%, list=18%, si | MM_2 |
| GOBP_RESPONSE_TO_PL | GOBP_RESPONSE_TO_PLATELET_DERIVED_C | 22 | -0.52891 | -1.83921 | 0.004784689 | 0.072106145 | 1 | 2631 | tags=27%, list=11%, si | MM_2 |
| GOBP_REGULATION_OF_ | GOBP_REGULATION_OF_NATURAL_KILLER_C | 17 | -0.57142 | -1.83688 | 0.0046189376 | 0.07279045 | 1 | 5109 | tags=53%, list=22%, si | MM_2 |
| GOBP_COPPER_ION_HON | GOBP_COPPER_ION_HOMEOSTASIS | 18 | -0.57561 | -1.83604 | 0.012953368 | 0.07274974 | 1 | 3039 | tags=39%, list=13%, si | MM_2 |
| GOBP_MONONUCLEAR_C | GOBP_MONONUCLEAR_CELL_MIGRATION | 199 | -0.3192 | -1.83374 | 0 | 0.07345294 | 1 | 3926 | tags=26%, list=17%, si | MM_2 |
| GOBP_SMOOTH_MUSCLE | GOBP_SMOOTH_MUSCLE_CELL_PROLIFERAT | 191 | -0.3225 | -1.83354 | 0 | 0.07307247 | 1 | 4887 | tags=34%, list=21%, si | MM_2 |
| GOBP_MICROTUBULE_CY | GOBP_MICROTUBULE_CYTOSKELETON_ORG | 158 | -0.337 | -1.83171 | 0 | 0.073654994 | 1 | 4230 | tags=28%, list=18%, si | MM_2 |
| GOBP_REGULATION_OF_ | GOBP_REGULATION_OF_TRANSLATION | 329 | -0.30014 | -1.8317 | 0 | 0.073235124 | 1 | 5181 | tags=31%, list=22%, si | MM_2 |
| GOBP_REGULATION_OF_ | GOBP_REGULATION_OF_B_CELL_DIFFERENT | 37 | -0.46311 | -1.83013 | 0.0025316456 | 0.07358284 | 1 | 5647 | tags=41%, list=24%, si | MM_2 |
| GOBP_MACROPHAGE_MI | GOBP_MACROPHAGE_MIGRATION | 65 | -0.40298 | -1.82796 | 0.0070588235 | 0.07431356 | 1 | 3883 | tags=34%, list=16%, si | MM_2 |
| GOBP_NEGATIVE_REGUL | GOBP_NEGATIVE_REGULATION_OF_TRANSL | 117 | -0.35243 | -1.82591 | 0 | 0.07483434 | 1 | 4923 | tags=33%, list=21%, si | MM_2 |
| GOBP_MICROTUBULE_OF | GOBP_MICROTUBULE_ORGANIZING_CENTE | 34 | -0.47012 | -1.82574 | 0.0073349634 | 0.07445938 | 1 | 1737 | tags=21%, list=7%, sig | MM_2 |
| GOBP_LYMPHOCYTE_CHE | GOBP_LYMPHOCYTE_CHEMOTAXIS | 38 | -0.46574 | -1.82556 | 0.004555809 | 0.074162245 | 1 | 3602 | tags=29%, list=15%, si | MM_2 |
| GOBP_MITOTIC_SISTER_C | GOBP_MITOTIC_SISTER_CHROMATID_COHE | 26 | -0.49366 | -1.82554 | 0.006772009 | 0.07375381 | 1 | 3874 | tags=31%, list=16%, si | MM_2 |
| GOBP_REGULATION_OF_ | GOBP_REGULATION_OF_LEUKOCYTE_APOPT | 103 | -0.35445 | -1.8249 | 0 | 0.073766135 | 1 | 5104 | tags=31%, list=22%, si | MM_2 |
| GOBP_NEGATIVE_REGUL | GOBP_NEGATIVE_REGULATION_OF_VIRAL_ | 95 | -0.36908 | -1.82408 | 0.0024096386 | 0.073698334 | 1 | 4023 | tags=37%, list=17%, si | MM_2 |
| GOBP_REGULATION_OF_ | GOBP_REGULATION_OF_TRANSLATION_IN_ | 24 | -0.5058 | -1.82201 | 0.009345794 | 0.074440576 | 1 | 2288 | tags=21%, list=10%, si | MM_2 |
| GOBP_POSITIVE_REGULA | GOBP_POSITIVE_REGULATION_OF_CELL_AD | 487 | -0.28445 | -1.82019 | 0 | 0.07498853 | 1 | 3931 | tags=25%, list=17%, si | MM_2 |
| GOBP_NUCLEAR_PORE_C | GOBP_NUCLEAR_PORE_ORGANIZATION | 15 | -0.58456 | -1.81956 | 0.013513514 | 0.0749241 | 1 | 4467 | tags=60%, list=19%, si | MM_2 |
| GOBP_CERAMIDE_METAE | GOBP_CERAMIDE_METABOLIC_PROCESS | 94 | -0.37384 | -1.81711 | 0 | 0.07584635 | 1 | 4373 | tags=35%, list=19%, si | MM_2 |
| GOBP_MUSCLE_CELL_MI | GOBP_MUSCLE_CELL_MIGRATION | 129 | -0.34055 | -1.81692 | 0 | 0.075586446 | 1 | 4887 | tags=33%, list=21%, si | MM_2 |
| GOBP_PLATELET_DENSE_ | GOBP_PLATELET_DENSE_GRANULE_ORGAN | 21 | -0.52151 | -1.81679 | 0.011520738 | 0.07520395 | 1 | 6967 | tags=71%, list=30%, si | MM_2 |
| GOBP_GLIAL_CELL_APOP | GOBP_GLIAL_CELL_APOPTOTIC_PROCESS | 20 | -0.52181 | -1.81605 | 0.00954654 | 0.07520169 | 1 | 4831 | tags=45%, list=20%, si | MM_2 |
| GOBP_POSITIVE_REGULA | GOBP_POSITIVE_REGULATION_OF_CELL_CY | 330 | -0.29007 | -1.81394 | 0 | 0.07593291 | 1 | 4345 | tags=30%, list=18%, si | MM_2 |
| GOBP_ANTIGEN_PROCES | GOBP_ANTIGEN_PROCESSING_AND_PRESEN | 59 | -0.40136 | -1.81383 | 0.006960557 | 0.07557017 | 1 | 5246 | tags=41%, list=22%, si | MM_2 |
| GOBP_POSITIVE_REGULA | GOBP_POSITIVE_REGULATION_OF_PROTEIN | 100 | -0.35559 | -1.81291 | 0 | 0.07561029 | 1 | 4383 | tags=33%, list=19%, si | MM_2 |
| GOBP_DNA_TEMPLATED_ | GOBP_DNA_TEMPLATED_DNA_REPLICATION | 147 | -0.33251 | -1.81282 | 0 | 0.07526061 | 1 | 5048 | tags=37%, list=21%, si | MM_2 |
| GOBP_FOCAL_ADHESION | GOBP_FOCAL_ADHESION_ASSEMBLY | 87 | -0.36346 | -1.81022 | 0 | 0.07626279 | 1 | 3824 | tags=33%, list=16%, si | MM_2 |
| MISMATCH_REPAIR | MISMATCH_REPAIR | 21 | -0.52673 | -1.80646 | 0.013544018 | 0.07788389 | 1 | 5275 | tags=48%, list=22%, si | MM_2 |
| GOBP_NEURON_PROJECT | GOBP_NEURON_PROJECTION_REGENERATIO | 68 | -0.38564 | -1.80437 | 0.0049261083 | 0.07860052 | 1 | 3602 | tags=34%, list=15%, si | MM_2 |
| GOBP_REGULATION_OF_ | GOBP_REGULATION_OF_SEQUESTERING_OF | 15 | -0.58104 | -1.80281 | 0.019950125 | 0.079164706 | 1 | 4943 | tags=53%, list=21%, si | MM_2 |
| GOBP_REGULATION_OF_ | GOBP_REGULATION_OF_EXECUTION_PHASE | 26 | -0.49404 | -1.80201 | 0.01686747 | 0.07922457 | 1 | 4523 | tags=46%, list=19%, si | MM_2 |
| GOBP_PIGMENT_GRANU | GOBP_PIGMENT_GRANULE_ORGANIZATION | 39 | -0.44187 | -1.8012 | 0.0024213076 | 0.07932167 | 1 | 4034 | tags=38%, list=17%, si | MM_2 |
| GOBP_CELLULAR_LIPID_C | GOBP_CELLULAR_LIPID_CATABOLIC_PROCES | 210 | -0.30736 | -1.79792 | 0 | 0.08098364 | 1 | 4470 | tags=31%, list=19%, si | MM_2 |
| GOBP_LIPID_STORAGE | GOBP_LIPID_STORAGE | 97 | -0.35712 | -1.79545 | 0.0049875313 | 0.08190253 | 1 | 4972 | tags=41%, list=21%, si | MM_2 |
| GOBP_NEGATIVE_REGUL | GOBP_NEGATIVE_REGULATION_OF_SMOOT | 36 | -0.44969 | -1.79301 | 0.006880734 | 0.0828233 | 1 | 4833 | tags=36%, list=20%, si | MM_2 |
| GOBP_PROTEIN_K48_LIN | GOBP_PROTEIN_K48_LINKED_UBIQUITINAT | 78 | -0.37291 | -1.79019 | 0.0022123894 | 0.08402685 | 1 | 7447 | tags=58%, list=32%, si | MM_2 |
| GOBP_CYTOPLASMIC_PA | GOBP_CYTOPLASMIC_PATTERN_RECOGNITI | 159 | -0.32286 | -1.78985 | 0 | 0.08380626 | 1 | 3509 | tags=25%, list=15%, si | MM_2 |
| GOBP_ESTABLISHMENT_C | GOBP_ESTABLISHMENT_OF_LYMPHOCYTE_ | 15 | -0.56664 | -1.78741 | 0.015665796 | 0.0848791 | 1 | 2519 | tags=33%, list=11%, si | MM_2 |
| GOBP_NEGATIVE_REGUL | GOBP_NEGATIVE_REGULATION_OF_DNA_M | 127 | -0.34328 | -1.78684 | 0.002386635 | 0.0848447 | 1 | 5176 | tags=32%, list=22%, si | MM_2 |
| APOPTOSIS | APOPTOSIS | 137 | -0.33348 | -1.78593 | 0 | 0.08518185 | 1 | 5086 | tags=34%, list=22%, si | MM_2 |
| GOBP_MACROAUTOPHAG | GOBP_MACROAUTOPHAGY | 287 | -0.29974 | -1.78473 | 0 | 0.085393764 | 1 | 6278 | tags=41%, list=27%, si | MM_2 |
| GOBP_REGULATION_OF_ | GOBP_REGULATION_OF_LEUKOCYTE_PROLI | 273 | -0.29851 | -1.78297 | 0 | 0.08591277 | 1 | 3351 | tags=22%, list=14%, si | MM_2 |
| GOBP_REGULATION_OF_ | GOBP_REGULATION_OF_PODOSOME_ASSEI | 16 | -0.5582 | -1.78176 | 0.011876484 | 0.08613337 | 1 | 6062 | tags=63%, list=26%, si | MM_2 |
| GOBP_CHROMOSOME_LC | GOBP_CHROMOSOME_LOCALIZATION | 111 | -0.34656 | -1.77905 | 0 | 0.087387465 | 1 | 5752 | tags=40%, list=24%, si | MM_2 |
| GOBP_MYELOID_CELL_DE | GOBP_MYELOID_CELL_DEVELOPMENT | 98 | -0.35565 | -1.77748 | 0 | 0.087866254 | 1 | 2917 | tags=22%, list=12%, si | MM_2 |
| GOBP_COLLAGEN_METAE | GOBP_COLLAGEN_METABOLIC_PROCESS | 119 | -0.33744 | -1.77566 | 0 | 0.08847461 | 1 | 1507 | tags=16%, list=6%, sig | MM_2 |
| GOBP_NEGATIVE_REGUL | GOBP_NEGATIVE_REGULATION_OF_STEM_C | 28 | -0.46335 | -1.7739 | 0.00954654 | 0.08911436 | 1 | 4604 | tags=46%, list=19%, si | MM_2 |
| GOBP_NECROPTOTIC_PR | GOBP_NECROPTOTIC_PROCESS | 37 | -0.44171 | -1.77183 | 0.012468828 | 0.08997396 | 1 | 3638 | tags=32%, list=15%, si | MM_2 |
| GOBP_NEGATIVE_REGUL | GOBP_NEGATIVE_REGULATION_OF_MACRO | 33 | -0.44956 | -1.77163 | 0.004819277 | 0.08969765 | 1 | 3826 | tags=36%, list=16%, si | MM_2 |
| GOBP_CELL_SUBSTRATE_ | GOBP_CELL_SUBSTRATE_JUNCTION_ORGAN | 105 | -0.34656 | -1.77008 | 0 | 0.090259545 | 1 | 3931 | tags=30%, list=17%, si | MM_2 |
| GOBP_POSITIVE_REGULA | GOBP_POSITIVE_REGULATION_OF_PHAGOC | 74 | -0.37255 | -1.7696 | 0.0024813896 | 0.090133466 | 1 | 3025 | tags=20%, list=13%, si | MM_2 |
| GOBP_REGULATION_OF_ | GOBP_REGULATION_OF_UBIQUITIN_PROTE | 20 | -0.52156 | -1.76949 | 0.012077294 | 0.089772165 | 1 | 2472 | tags=30%, list=10%, si | MM_2 |
| GOBP_TRNA_METHYLATI | GOBP_TRNA_METHYLATION | 39 | -0.43414 | -1.76886 | 0.013513514 | 0.08965563 | 1 | 4169 | tags=38%, list=18%, si | MM_2 |
| GOBP_NEGATIVE_REGUL | GOBP_NEGATIVE_REGULATION_OF_TYPE_I_ | 23 | -0.50986 | -1.76779 | 0.013986014 | 0.08983506 | 1 | 4543 | tags=52%, list=19%, si | MM_2 |
| GOBP_LIPOPROTEIN_BIO | GOBP_LIPOPROTEIN_BIOSYNTHETIC_PROCE | 104 | -0.35058 | -1.76771 | 0.0022675737 | 0.089500174 | 1 | 4356 | tags=32%, list=18%, si | MM_2 |
| HERPES_SIMPLEX_VIRUS_ | HERPES_SIMPLEX_VIRUS_1_INFECTION | 373 | -0.28444 | -1.76759 | 0 | 0.08916116 | 1 | 3244 | tags=23%, list=14%, si | MM_2 |
| GOBP_PLATELET_MORPH | GOBP_PLATELET_MORPHOGENESIS | 25 | -0.49521 | -1.76749 | 0.016431924 | 0.08883169 | 1 | 4495 | tags=48%, list=19%, si | MM_2 |
| GOBP_DEPHOSPHORYLAT | GOBP_DEPHOSPHORYLATION | 303 | -0.29384 | -1.7666 | 0 | 0.08893387 | 1 | 5111 | tags=33%, list=22%, si | MM_2 |
| GOBP_POSITIVE_REGULA | GOBP_POSITIVE_REGULATION_OF_MONOC | 16 | -0.55862 | -1.76251 | 0.015486726 | 0.090907045 | 1 | 1704 | tags=31%, list=7%, sig | MM_2 |
| GOBP_DOUBLE_STRAND_ | GOBP_DOUBLE_STRAND_BREAK_REPAIR_VI | 67 | -0.37616 | -1.76202 | 0.0024271845 | 0.09075051 | 1 | 3206 | tags=22%, list=14%, si | MM_2 |
| GOBP_POSITIVE_REGULA | GOBP_POSITIVE_REGULATION_OF_CYTOKIN | 40 | -0.43181 | -1.76171 | 0.011904762 | 0.09045881 | 1 | 1624 | tags=23%, list=7%, sig | MM_2 |
| GOBP_GLYCEROLIPID_BIC | GOBP_GLYCEROLIPID_BIOSYNTHETIC_PROC | 210 | -0.30732 | -1.76136 | 0 | 0.090247646 | 1 | 4203 | tags=28%, list=18%, si | MM_2 |
| GOBP_PHOSPHOLIPID_M | GOBP_PHOSPHOLIPID_METABOLIC_PROCES | 352 | -0.28038 | -1.76058 | 0 | 0.09034521 | 1 | 4265 | tags=27%, list=18%, si | MM_2 |
| GOBP_NON_MEMBRANE | GOBP_NON_MEMBRANE_BOUNDED_ORGA | 405 | -0.27833 | -1.76022 | 0 | 0.090156354 | 1 | 4242 | tags=27%, list=18%, si | MM_2 |
| GOBP_REGULATION_OF_ | GOBP_REGULATION_OF_CYSTEINE_TYPE_EN | 16 | -0.54141 | -1.76004 | 0.016431924 | 0.089898564 | 1 | 3904 | tags=44%, list=17%, si | MM_2 |
| GOBP_EMBRYONIC_BRAI | GOBP_EMBRYONIC_BRAIN_DEVELOPMENT | 24 | -0.48424 | -1.75958 | 0.022421524 | 0.08970625 | 1 | 3808 | tags=38%, list=16%, si | MM_2 |
| HUMAN_CYTOMEGALOVI | HUMAN_CYTOMEGALOVIRUS_INFECTION | 216 | -0.30647 | -1.75875 | 0 | 0.089740835 | 1 | 5279 | tags=36%, list=22%, si | MM_2 |
| GOBP_LYMPHOCYTE_APC | GOBP_LYMPHOCYTE_APOPTOTIC_PROCESS | 94 | -0.35541 | -1.75874 | 0 | 0.08936175 | 1 | 4172 | tags=29%, list=18%, si | MM_2 |
| GOBP_POSITIVE_REGULA | GOBP_POSITIVE_REGULATION_OF_MITOTIC | 126 | -0.33473 | -1.75806 | 0 | 0.089417316 | 1 | 5515 | tags=40%, list=23%, si | MM_2 |
| GOBP_REGULATION_OF_ | GOBP_REGULATION_OF_CHROMOSOME_OI | 242 | -0.29737 | -1.75804 | 0 | 0.08904305 | 1 | 4563 | tags=26%, list=19%, si | MM_2 |
| GOBP_LEUKOCYTE_APOP | GOBP_LEUKOCYTE_APOPTOTIC_PROCESS | 131 | -0.32787 | -1.75462 | 0 | 0.09079234 | 1 | 5104 | tags=32%, list=22%, si | MM_2 |
| GOBP_ACETYL_COA_MET | GOBP_ACETYL_COA_METABOLIC_PROCESS | 34 | -0.44946 | -1.75352 | 0.009876544 | 0.09101451 | 1 | 5054 | tags=47%, list=21%, si | MM_2 |
| GOBP_CELLULAR_PIGMEI | GOBP_CELLULAR_PIGMENTATION | 57 | -0.39476 | -1.75162 | 0.010075566 | 0.091691196 | 1 | 6412 | tags=51%, list=27%, si | MM_2 |
| GOBP_REGULATION_OF_ | GOBP_REGULATION_OF_MONONUCLEAR_C | 136 | -0.33474 | -1.75116 | 0 | 0.091643974 | 1 | 3926 | tags=27%, list=17%, si | MM_2 |
| GOBP_REPLACEMENT_OS | GOBP_REPLACEMENT_OSSIFICATION | 34 | -0.45394 | -1.74962 | 0.011574074 | 0.09231481 | 1 | 3374 | tags=35%, list=14%, si | MM_2 |
| ACUTE_MYELOID_LEUKEI | ACUTE_MYELOID_LEUKEMIA | 66 | -0.38351 | -1.74853 | 0 | 0.092588164 | 1 | 4982 | tags=44%, list=21%, si | MM_2 |
| GOBP_MONOCARBOXYLI | GOBP_MONOCARBOXYLIC_ACID_CATABOLI | 126 | -0.33429 | -1.74811 | 0.0025316456 | 0.092466526 | 1 | 4514 | tags=36%, list=19%, si | MM_2 |
| GOBP_REGULATION_OF_ | GOBP_REGULATION_OF_RESPONSE_TO_CYT | 153 | -0.32172 | -1.74512 | 0 | 0.09415863 | 1 | 4543 | tags=32%, list=19%, si | MM_2 |
| GOBP_REGULATION_OF_ | GOBP_REGULATION_OF_SISTER_CHROMATI | 104 | -0.34208 | -1.74455 | 0.0025316456 | 0.09414487 | 1 | 1318 | tags=14%, list=6%, sig | MM_2 |
| GOBP_POSITIVE_REGULA | GOBP_POSITIVE_REGULATION_OF_INTRACE | 108 | -0.34063 | -1.74375 | 0 | 0.09420838 | 1 | 5012 | tags=30%, list=21%, si | MM_2 |

| | | | | | | | | | | |
|---|---|---|---|---|---|---|---|---|---|---|
| GOBP_POSITIVE_REGULA | GOBP_POSITIVE_REGULATION_OF_MITOTIC | 93 | -0.35153 | -1.74343 | 0.0045871558 | 0.09402274 | 1 | 5515 | tags=43%, list=23%, si | MM_2 |
| GOBP_GLYCEROLIPID_ME | GOBP_GLYCEROLIPID_METABOLIC_PROCESS | 361 | -0.28131 | -1.74298 | 0 | 0.09399232 | 1 | 4203 | tags=26%, list=18%, si | MM_2 |
| GOBP_RNA_MODIFICATIO | GOBP_RNA_MODIFICATION | 141 | -0.3245 | -1.74243 | 0 | 0.09401945 | 1 | 4377 | tags=31%, list=19%, si | MM_2 |
| GOBP_PHOSPHATIDYLINO | GOBP_PHOSPHATIDYLINOSITOL_METABOLIC | 148 | -0.32601 | -1.74176 | 0 | 0.0941489 | 1 | 5747 | tags=39%, list=24%, si | MM_2 |
| GOBP_REGULATION_OF_ | GOBP_REGULATION_OF_LYMPHOCYTE_APO | 71 | -0.37356 | -1.74167 | 0.0049751243 | 0.09381892 | 1 | 3384 | tags=23%, list=14%, si | MM_2 |
| GOBP_REGULATION_OF_ | GOBP_REGULATION_OF_CYTOPLASMIC_PAT | 104 | -0.34668 | -1.74153 | 0 | 0.09352644 | 1 | 3138 | tags=26%, list=13%, si | MM_2 |
| GOBP_REGULATION_OF_ | GOBP_REGULATION_OF_ANTIGEN_PROCESS | 21 | -0.4997 | -1.73947 | 0.02238806 | 0.09427487 | 1 | 3131 | tags=38%, list=13%, si | MM_2 |
| GOBP_POSITIVE_REGULA | GOBP_POSITIVE_REGULATION_OF_SMOOTH | 114 | -0.33195 | -1.73932 | 0.002375297 | 0.094012484 | 1 | 4724 | tags=32%, list=20%, si | MM_2 |
| GOBP_POSITIVE_REGULA | GOBP_POSITIVE_REGULATION_OF_INTRACE | 147 | -0.32413 | -1.73877 | 0.0023094688 | 0.09399421 | 1 | 5127 | tags=30%, list=22%, si | MM_2 |
| GOBP_REGULATION_OF_ | GOBP_REGULATION_OF_NUCLEOCYTOPLAS | 130 | -0.33051 | -1.73791 | 0 | 0.094180964 | 1 | 4893 | tags=32%, list=21%, si | MM_2 |
| GOBP_RIBOSOME_ASSEM | GOBP_RIBOSOME_ASSEMBLY | 60 | -0.3835 | -1.73708 | 0.0024937657 | 0.09433947 | 1 | 4315 | tags=32%, list=18%, si | MM_2 |
| GOBP_MAINTENANCE_OI | GOBP_MAINTENANCE_OF_PROTEIN_LOCAT | 97 | -0.35006 | -1.7364 | 0 | 0.094285026 | 1 | 4920 | tags=33%, list=21%, si | MM_2 |
| GOBP_REGULATION_OF_ | GOBP_REGULATION_OF_LYMPHOCYTE_ACT | 495 | -0.26942 | -1.73512 | 0 | 0.094776124 | 1 | 3675 | tags=22%, list=16%, si | MM_2 |
| GOBP_RESPONSE_TO_TU | GOBP_RESPONSE_TO_TUMOR_NECROSIS_F | 167 | -0.31123 | -1.73478 | 0 | 0.0946411 | 1 | 4023 | tags=28%, list=17%, si | MM_2 |
| RIBOSOME | RIBOSOME | 128 | -0.32414 | -1.73292 | 0 | 0.09543731 | 1 | 13156 | tags=96%, list=56%, si | MM_2 |
| GOBP_POSITIVE_REGULA | GOBP_POSITIVE_REGULATION_OF_CELL_DIV | 79 | -0.35919 | -1.7326 | 0.0024509805 | 0.095252156 | 1 | 4561 | tags=34%, list=19%, si | MM_2 |
| GOBP_PEROXISOME_ORG | GOBP_PEROXISOME_ORGANIZATION | 38 | -0.42872 | -1.73217 | 0.012106538 | 0.09515204 | 1 | 3969 | tags=37%, list=17%, si | MM_2 |
| GOBP_POSITIVE_REGULA | GOBP_POSITIVE_REGULATION_OF_CELL_SU | 134 | -0.32735 | -1.73128 | 0 | 0.09529596 | 1 | 3931 | tags=28%, list=17%, si | MM_2 |
| GOBP_ATTACHMENT_OF_ | GOBP_ATTACHMENT_OF_MITOTIC_SPINDLE | 24 | -0.47987 | -1.73123 | 0.018604651 | 0.09496827 | 1 | 4230 | tags=42%, list=18%, si | MM_2 |
| GLYCOSYLPHOSPHATIDYL | GLYCOSYLPHOSPHATIDYLINOSITOL_(GPI)-AN | 24 | -0.48053 | -1.73112 | 0.0077319588 | 0.09466481 | 1 | 3732 | tags=33%, list=16%, si | MM_2 |
| GOBP_POSITIVE_REGULA | GOBP_POSITIVE_REGULATION_OF_ERYTHRO | 28 | -0.4585 | -1.73039 | 0.016241299 | 0.09481405 | 1 | 6728 | tags=54%, list=28%, si | MM_2 |
| GOBP_RIBOSOME_DISASS | GOBP_RIBOSOME_DISASSEMBLY | 31 | -0.44341 | -1.72825 | 0.009615385 | 0.09564169 | 1 | 13155 | tags=100%, list=56%, | MM_2 |
| GOBP_POSITIVE_REGULA | GOBP_POSITIVE_REGULATION_OF_LYMPHO | 18 | -0.52392 | -1.7267 | 0.016990291 | 0.09628465 | 1 | 3602 | tags=28%, list=15%, si | MM_2 |
| GOBP_CELL_CYCLE_CHEC | GOBP_CELL_CYCLE_CHECKPOINT_SIGNALIN | 191 | -0.30979 | -1.7261 | 0 | 0.096274614 | 1 | 4524 | tags=30%, list=19%, si | MM_2 |
| GOBP_LEUKOCYTE_PROLI | GOBP_LEUKOCYTE_PROLIFERATION | 361 | -0.27809 | -1.72512 | 0 | 0.096644215 | 1 | 3351 | tags=20%, list=14%, si | MM_2 |
| GOBP_POSITIVE_REGULA | GOBP_POSITIVE_REGULATION_OF_CYTOKIN | 488 | -0.26543 | -1.72487 | 0 | 0.09647649 | 1 | 4023 | tags=26%, list=17%, si | MM_2 |
| GOBP_MONOCYTE_CHEM | GOBP_MONOCYTE_CHEMOTACTIC_PROTEIN | 27 | -0.47 | -1.72465 | 0.011709602 | 0.0962997 | 1 | 1704 | tags=22%, list=7%, sig | MM_2 |
| GOBP_LIPID_MODIFICATI | GOBP_LIPID_MODIFICATION | 197 | -0.3007 | -1.72405 | 0 | 0.096359886 | 1 | 4508 | tags=34%, list=19%, si | MM_2 |
| GOBP_DNA_REPAIR_DEP | GOBP_DNA_REPAIR_DEPENDENT_CHROMA | 23 | -0.48539 | -1.72344 | 0.016129032 | 0.096382596 | 1 | 2944 | tags=26%, list=12%, si | MM_2 |
| GOBP_T_CELL_DIFFERENT | GOBP_T_CELL_DIFFERENTIATION | 327 | -0.27852 | -1.72338 | 0 | 0.0960638 | 1 | 3776 | tags=26%, list=16%, si | MM_2 |
| GOBP_REGULATION_OF_ | GOBP_REGULATION_OF_T_CELL_ACTIVATIO | 364 | -0.27925 | -1.72328 | 0 | 0.09580669 | 1 | 3670 | tags=24%, list=16%, si | MM_2 |
| GOBP_POSITIVE_REGULA | GOBP_POSITIVE_REGULATION_OF_LYMPHO | 318 | -0.28688 | -1.72298 | 0 | 0.0955936 | 1 | 4276 | tags=26%, list=18%, si | MM_2 |
| GOBP_MATURATION_OF_ | GOBP_MATURATION_OF_SSU_RRNA_FROM | 33 | -0.43673 | -1.72186 | 0.014634146 | 0.09597584 | 1 | 4451 | tags=27%, list=19%, si | MM_2 |
| GOBP_POSITIVE_REGULA | GOBP_POSITIVE_REGULATION_OF_EXTRINS | 60 | -0.38043 | -1.72144 | 0.009661836 | 0.095940046 | 1 | 4260 | tags=30%, list=18%, si | MM_2 |
| GOBP_TELOMERE_CAPPI | GOBP_TELOMERE_CAPPING | 38 | -0.4268 | -1.72136 | 0.012437811 | 0.09564962 | 1 | 5368 | tags=42%, list=23%, si | MM_2 |
| GOBP_MIRNA_METABOL | GOBP_MIRNA_METABOLIC_PROCESS | 103 | -0.34013 | -1.71971 | 0.0023364485 | 0.09619403 | 1 | 6839 | tags=52%, list=29%, si | MM_2 |
| GOBP_REGULATION_OF_ | GOBP_REGULATION_OF_TOLL_LIKE_RECEPT | 28 | -0.4518 | -1.71905 | 0.026004728 | 0.09628964 | 1 | 4023 | tags=36%, list=17%, si | MM_2 |
| GOBP_ORGANELLE_FISSI | GOBP_ORGANELLE_FISSION | 477 | -0.26782 | -1.71784 | 0 | 0.096723184 | 1 | 3970 | tags=26%, list=17%, si | MM_2 |
| GOBP_REGULATION_OF_ | GOBP_REGULATION_OF_MITOTIC_CELL_CY | 331 | -0.27883 | -1.71766 | 0 | 0.09651133 | 1 | 4986 | tags=32%, list=21%, si | MM_2 |
| GOBP_POSITIVE_REGULA | GOBP_POSITIVE_REGULATION_OF_NUCLEA | 15 | -0.54526 | -1.71539 | 0.027027028 | 0.09776985 | 1 | 5797 | tags=67%, list=25%, si | MM_2 |
| GOBP_TUMOR_NECROSIS | GOBP_TUMOR_NECROSIS_FACTOR_SUPERF | 193 | -0.30366 | -1.71485 | 0.004842615 | 0.09777643 | 1 | 4272 | tags=27%, list=18%, si | MM_2 |
| GOBP_PROTEASOME_ME | GOBP_PROTEASOME_MEDIATED_UBIQUITIN | 368 | -0.27388 | -1.71457 | 0 | 0.09760394 | 1 | 5812 | tags=32%, list=25%, si | MM_2 |
| GOBP_CRANIOFACIAL_SU | GOBP_CRANIOFACIAL_SUTURE_MORPHOGE | 15 | -0.53801 | -1.71453 | 0.033936653 | 0.09729052 | 1 | 1988 | tags=33%, list=8%, sig | MM_2 |
| GOBP_REGULATION_OF_ | GOBP_REGULATION_OF_CELL_CYCLE_PHAS | 424 | -0.26999 | -1.71432 | 0 | 0.09710068 | 1 | 5228 | tags=33%, list=22%, si | MM_2 |
| GOBP_INTRACELLULAR_N | GOBP_INTRACELLULAR_MRNA_LOCALIZATIO | 15 | -0.54551 | -1.71379 | 0.017114915 | 0.09701318 | 1 | 5174 | tags=67%, list=22%, si | MM_2 |
| GOBP_PROTEIN_POLYUB | GOBP_PROTEIN_POLYUBIQUITINATION | 250 | -0.29272 | -1.71295 | 0 | 0.09720248 | 1 | 5755 | tags=37%, list=24%, si | MM_2 |
| GOBP_EXOCYTIC_PROCES | GOBP_EXOCYTIC_PROCESS | 97 | -0.34218 | -1.71229 | 0 | 0.09725513 | 1 | 1741 | tags=20%, list=7%, sig | MM_2 |
| GOBP_CELLULAR_COMPO | GOBP_CELLULAR_COMPONENT_DISASSEMB | 472 | -0.26609 | -1.71163 | 0 | 0.09736668 | 1 | 5965 | tags=37%, list=25%, si | MM_2 |
| GOBP_REGULATION_OF_ | GOBP_REGULATION_OF_MITOTIC_NUCLEAR | 114 | -0.32877 | -1.71117 | 0 | 0.09733518 | 1 | 4413 | tags=34%, list=19%, si | MM_2 |
| GOBP_RETROGRADE_TRA | GOBP_RETROGRADE_TRANSPORT_ENDOSO | 87 | -0.36018 | -1.71041 | 0.0024937657 | 0.09750133 | 1 | 6686 | tags=45%, list=28%, si | MM_2 |
| GOBP_CELL_SURFACE_PA | GOBP_CELL_SURFACE_PATTERN_RECOGNIT | 70 | -0.36843 | -1.71035 | 0.0050125313 | 0.097196646 | 1 | 4244 | tags=31%, list=18%, si | MM_2 |
| GOBP_PIGMENTATION | GOBP_PIGMENTATION | 112 | -0.3359 | -1.70927 | 0.0023980816 | 0.097461544 | 1 | 2267 | tags=20%, list=10%, si | MM_2 |
| GOBP_NEGATIVE_REGUL | GOBP_NEGATIVE_REGULATION_OF_SMALL_ | 61 | -0.38085 | -1.70545 | 0.0050125313 | 0.099527 | 1 | 2663 | tags=25%, list=11%, si | MM_2 |
| GOBP_RESPONSE_TO_INT | GOBP_RESPONSE_TO_INTERLEUKIN_17 | 25 | -0.47205 | -1.70485 | 0.016990291 | 0.099581584 | 1 | 4023 | tags=44%, list=17%, si | MM_2 |
| GOBP_NEGATIVE_REGUL | GOBP_NEGATIVE_REGULATION_OF_PLATEL | 16 | -0.54063 | -1.70472 | 0.01909308 | 0.09929374 | 1 | 4011 | tags=38%, list=17%, si | MM_2 |
| PANCREATIC_CANCER | PANCREATIC_CANCER | 74 | -0.35531 | -1.70174 | 0.0077120825 | 0.10102256 | 1 | 4982 | tags=38%, list=21%, si | MM_2 |
| GOBP_POSITIVE_REGULA | GOBP_POSITIVE_REGULATION_OF_VASCULA | 44 | -0.40089 | -1.70144 | 0.0073710075 | 0.100852974 | 1 | 4572 | tags=39%, list=19%, si | MM_2 |
| GOBP_MELANOSOME_AS | GOBP_MELANOSOME_ASSEMBLY | 19 | -0.5266 | -1.70057 | 0.027459955 | 0.101167925 | 1 | 6967 | tags=74%, list=30%, si | MM_2 |
| GOBP_MATURATION_OF_ | GOBP_MATURATION_OF_SSU_RRNA | 48 | -0.39557 | -1.69997 | 0.014705882 | 0.101167426 | 1 | 6266 | tags=40%, list=27%, si | MM_2 |
| GOBP_REGULATION_OF_ | GOBP_REGULATION_OF_MITOTIC_CYTOKIN | 15 | -0.54786 | -1.69966 | 0.028037382 | 0.10102362 | 1 | 3296 | tags=47%, list=14%, si | MM_2 |
| GOBP_VASCULAR_ASSOC | GOBP_VASCULAR_ASSOCIATED_SMOOTH_M | 15 | -0.54973 | -1.6993 | 0.021791767 | 0.10096693 | 1 | 4268 | tags=60%, list=18%, si | MM_2 |
| GOBP_VIRION_ASSEMBLY | GOBP_VIRION_ASSEMBLY | 26 | -0.4576 | -1.69867 | 0.021582734 | 0.10108719 | 1 | 5900 | tags=50%, list=25%, si | MM_2 |
| GOBP_POSITIVE_REGULA | GOBP_POSITIVE_REGULATION_OF_EXECUTI | 19 | -0.51365 | -1.69739 | 0.024630541 | 0.10160351 | 1 | 4243 | tags=42%, list=18%, si | MM_2 |
| GOBP_EXTRACELLULAR_N | GOBP_EXTRACELLULAR_MATRIX_ASSEMBLY | 48 | -0.38635 | -1.69478 | 0.012077294 | 0.102990374 | 1 | 2646 | tags=25%, list=11%, si | MM_2 |
| GOBP_MITOTIC_CELL_CY | GOBP_MITOTIC_CELL_CYCLE_CHECKPOINT_ | 147 | -0.31311 | -1.6946 | 0 | 0.10282354 | 1 | 4524 | tags=30%, list=19%, si | MM_2 |
| GOBP_MEMBRANE_DOCI | GOBP_MEMBRANE_DOCKING | 78 | -0.35667 | -1.69244 | 0.009975063 | 0.103857405 | 1 | 2105 | tags=18%, list=9%, sig | MM_2 |
| PLATELET_ACTIVATION | PLATELET_ACTIVATION | 116 | -0.32512 | -1.6912 | 0.002375297 | 0.10440279 | 1 | 5279 | tags=36%, list=22%, si | MM_2 |
| GOBP_ACTIVATION_OF_I | GOBP_ACTIVATION_OF_INNATE_IMMUNE_ | 290 | -0.28253 | -1.6902 | 0 | 0.104757726 | 1 | 4261 | tags=29%, list=18%, si | MM_2 |
| GOBP_REGULATION_OF_ | GOBP_REGULATION_OF_LIPID_STORAGE | 55 | -0.38139 | -1.68951 | 0.0072815535 | 0.10487698 | 1 | 4943 | tags=38%, list=21%, si | MM_2 |
| GOBP_MUSCLE_CELL_PR | GOBP_MUSCLE_CELL_PROLIFERATION | 262 | -0.28427 | -1.68883 | 0 | 0.10504062 | 1 | 4658 | tags=30%, list=20%, si | MM_2 |
| GOBP_CELL_CYCLE_G2_N | GOBP_CELL_CYCLE_G2_M_PHASE_TRANSITI | 152 | -0.30859 | -1.68825 | 0.0023364485 | 0.1051481 | 1 | 3105 | tags=24%, list=13%, si | MM_2 |
| GOBP_NEGATIVE_REGUL | GOBP_NEGATIVE_REGULATION_OF_CELL_C | 368 | -0.27284 | -1.68571 | 0 | 0.106692776 | 1 | 4469 | tags=28%, list=19%, si | MM_2 |
| GOBP_POSITIVE_REGULA | GOBP_POSITIVE_REGULATION_OF_CELL_SU | 35 | -0.42403 | -1.68417 | 0.018823529 | 0.10753688 | 1 | 3649 | tags=34%, list=15%, si | MM_2 |
| GOBP_POSITIVE_REGULA | GOBP_POSITIVE_REGULATION_OF_FAT_CEL | 71 | -0.36152 | -1.68402 | 0.004901961 | 0.10733795 | 1 | 4262 | tags=30%, list=18%, si | MM_2 |
| GOBP_PHOSPHATIDYLGLY | GOBP_PHOSPHATIDYLGLYCEROL_METABOLI | 25 | -0.46332 | -1.68358 | 0.011792453 | 0.10728638 | 1 | 3387 | tags=36%, list=14%, si | MM_2 |
| GOBP_REGULATION_OF_ | GOBP_REGULATION_OF_HOMOTYPIC_CELL_ | 36 | -0.4152 | -1.68337 | 0.013953488 | 0.10710295 | 1 | 5104 | tags=33%, list=22%, si | MM_2 |
| CHRONIC_MYELOID_LEUK | CHRONIC_MYELOID_LEUKEMIA | 75 | -0.35806 | -1.68328 | 0.0046620048 | 0.10683831 | 1 | 5999 | tags=48%, list=25%, si | MM_2 |
| GOBP_ENDOSOMAL_TRA | GOBP_ENDOSOMAL_TRANSPORT | 272 | -0.27969 | -1.68094 | 0 | 0.108115464 | 1 | 6724 | tags=42%, list=28%, si | MM_2 |
| GOBP_CHEMOKINE_PROI | GOBP_CHEMOKINE_PRODUCTION | 119 | -0.32251 | -1.68091 | 0.004784689 | 0.10780449 | 1 | 4023 | tags=26%, list=17%, si | MM_2 |
| GOBP_RNA_METHYLATIO | GOBP_RNA_METHYLATION | 65 | -0.36616 | -1.67922 | 0.007633588 | 0.10870341 | 1 | 4169 | tags=34%, list=18%, si | MM_2 |
| GOBP_GLIAL_CELL_MIGR | GOBP_GLIAL_CELL_MIGRATION | 70 | -0.36153 | -1.67753 | 0.0121951215 | 0.1095653 | 1 | 4003 | tags=37%, list=17%, si | MM_2 |
| GOBP_NEGATIVE_REGUL | GOBP_NEGATIVE_REGULATION_OF_CYCLIN | 25 | -0.46136 | -1.67747 | 0.013729977 | 0.109248176 | 1 | 4413 | tags=52%, list=19%, si | MM_2 |

| | | | | | | | | | | |
|---|---|---|---|---|---|---|---|---|---|---|
| GOBP_POSITIVE_REGULA | GOBP_POSITIVE_REGULATION_OF_INFLAMI | 22 | -0.47046 | -1.67562 | 0.03233831 | 0.1102724 | 1 | 5241 | tags=41%, list=22%, si | MM_2 |
| GOBP_RESPONSE_TO_PR | GOBP_RESPONSE_TO_PROGESTERONE | 19 | -0.49778 | -1.67484 | 0.02910053 | 0.110471256 | 1 | 3228 | tags=37%, list=14%, si | MM_2 |
| GOBP_CD4_POSITIVE_AL | GOBP_CD4_POSITIVE_ALPHA_BETA_T_CELL_ | 100 | -0.34017 | -1.67468 | 0.009638554 | 0.110220894 | 1 | 3548 | tags=25%, list=15%, si | MM_2 |
| GOBP_MIRNA_TRANSCRI | GOBP_MIRNA_TRANSCRIPTION | 78 | -0.35153 | -1.67462 | 0.004819277 | 0.109931685 | 1 | 3762 | tags=32%, list=16%, si | MM_2 |
| GOBP_FATTY_ACID_BETA | GOBP_FATTY_ACID_BETA_OXIDATION | 75 | -0.35841 | -1.67449 | 0.0024096386 | 0.10971537 | 1 | 4470 | tags=36%, list=19%, si | MM_2 |
| GOBP_ERYTHROCYTE_DE | GOBP_ERYTHROCYTE_DEVELOPMENT | 50 | -0.38409 | -1.67208 | 0.004796163 | 0.111000404 | 1 | 2917 | tags=26%, list=12%, si | MM_2 |
| COLORECTAL_CANCER | COLORECTAL_CANCER | 86 | -0.33992 | -1.67206 | 0.0074074073 | 0.11069861 | 1 | 4982 | tags=35%, list=21%, si | MM_2 |
| GOBP_ORGANOPHOSPHA | GOBP_ORGANOPHOSPHATE_BIOSYNTHETIC_ | 500 | -0.26054 | -1.67181 | 0 | 0.1105297 | 1 | 4376 | tags=26%, list=19%, si | MM_2 |
| GOBP_PIGMENT_CELL_DI | GOBP_PIGMENT_CELL_DIFFERENTIATION | 37 | -0.41152 | -1.6713 | 0.027272727 | 0.110637255 | 1 | 6259 | tags=49%, list=27%, si | MM_2 |
| GOBP_T_CELL_PROLIFERA | GOBP_T_CELL_PROLIFERATION | 233 | -0.28536 | -1.67039 | 0 | 0.110922106 | 1 | 3244 | tags=21%, list=14%, si | MM_2 |
| GOBP_REGULATION_OF_ | GOBP_REGULATION_OF_PHAGOCYTOSIS | 106 | -0.32636 | -1.66997 | 0.009090909 | 0.110922575 | 1 | 3863 | tags=22%, list=16%, si | MM_2 |
| GOBP_REGULATION_OF_ | GOBP_REGULATION_OF_PROTEIN_IMPORT_ | 76 | -0.35681 | -1.66978 | 0.007633588 | 0.11070818 | 1 | 2318 | tags=21%, list=10%, si | MM_2 |
| GOBP_ANTIVIRAL_INNAT | GOBP_ANTIVIRAL_INNATE_IMMUNE_RESPO | 47 | -0.39431 | -1.6697 | 0.014527845 | 0.1104044 | 1 | 2736 | tags=32%, list=12%, si | MM_2 |
| GOBP_REGULATION_OF_ | GOBP_REGULATION_OF_ACTIN_FILAMENT_ | 158 | -0.30655 | -1.66869 | 0 | 0.11084972 | 1 | 4706 | tags=27%, list=20%, si | MM_2 |
| GOBP_REGULATION_OF_ | GOBP_REGULATION_OF_HIPPO_SIGNALING | 22 | -0.48752 | -1.66761 | 0.027777778 | 0.111256406 | 1 | 4762 | tags=50%, list=20%, si | MM_2 |
| GOBP_SNRNA_3_END_PF | GOBP_SNRNA_3_END_PROCESSING | 20 | -0.4974 | -1.66759 | 0.020050125 | 0.110953145 | 1 | 2765 | tags=30%, list=12%, si | MM_2 |
| GOBP_ANTIMICROBIAL_H | GOBP_ANTIMICROBIAL_HUMORAL_IMMUN | 79 | -0.34651 | -1.66749 | 0.014251782 | 0.11072017 | 1 | 3996 | tags=30%, list=17%, si | MM_2 |
| GOBP_REGULATION_OF_ | GOBP_REGULATION_OF_REGULATORY_NCR | 15 | -0.52114 | -1.66695 | 0.025943397 | 0.11079939 | 1 | 3655 | tags=47%, list=15%, si | MM_2 |
| GOBP_RUFFLE_ASSEMBLY | GOBP_RUFFLE_ASSEMBLY | 43 | -0.39949 | -1.66678 | 0.004796163 | 0.11057646 | 1 | 4318 | tags=40%, list=18%, si | MM_2 |
| GOBP_POSITIVE_REGULA | GOBP_POSITIVE_REGULATION_OF_DEFENSE | 480 | -0.25865 | -1.66668 | 0 | 0.1103882 | 1 | 3509 | tags=23%, list=15%, si | MM_2 |
| GOBP_REGULATION_OF_ | GOBP_REGULATION_OF_DEFENSE_RESPONS | 42 | -0.39927 | -1.66623 | 0.015037594 | 0.110387266 | 1 | 3720 | tags=36%, list=16%, si | MM_2 |
| GOBP_POSITIVE_REGULA | GOBP_POSITIVE_REGULATION_OF_CELL_AC | 373 | -0.26815 | -1.66582 | 0 | 0.110338815 | 1 | 3675 | tags=23%, list=16%, si | MM_2 |
| GOBP_CYTOPLASMIC_SE | GOBP_CYTOPLASMIC_SEQUESTERING_OF_P | 17 | -0.51706 | -1.66549 | 0.030444965 | 0.11020416 | 1 | 4920 | tags=41%, list=21%, si | MM_2 |
| GOBP_INNATE_IMMUNE_ | GOBP_INNATE_IMMUNE_RESPONSE_ACTIV | 254 | -0.28167 | -1.66385 | 0 | 0.110993475 | 1 | 4261 | tags=27%, list=18%, si | MM_2 |
| GOBP_TELOMERE_ORGAI | GOBP_TELOMERE_ORGANIZATION | 150 | -0.30245 | -1.66202 | 0.004716981 | 0.11198069 | 1 | 4626 | tags=27%, list=20%, si | MM_2 |
| GOBP_DNA_REPLICATION | GOBP_DNA_REPLICATION | 263 | -0.27581 | -1.66183 | 0 | 0.11177546 | 1 | 4974 | tags=31%, list=21%, si | MM_2 |
| GOBP_POSITIVE_REGULA | GOBP_POSITIVE_REGULATION_OF_LEUKOC | 261 | -0.27676 | -1.66174 | 0 | 0.11152594 | 1 | 4276 | tags=28%, list=18%, si | MM_2 |
| IL-17_SIGNALING_PATHW | IL-17_SIGNALING_PATHWAY | 85 | -0.33983 | -1.65848 | 0.00913242 | 0.11349116 | 1 | 2771 | tags=22%, list=12%, si | MM_2 |
| GOBP_POSITIVE_REGULA | GOBP_POSITIVE_REGULATION_OF_MONON | 94 | -0.33368 | -1.65789 | 0.0048076925 | 0.11361904 | 1 | 3926 | tags=26%, list=17%, si | MM_2 |
| OOCYTE_MEIOSIS | OOCYTE_MEIOSIS | 108 | -0.32278 | -1.65785 | 0.0025974025 | 0.11332537 | 1 | 5211 | tags=33%, list=22%, si | MM_2 |
| GOBP_REGULATION_OF_ | GOBP_REGULATION_OF_SUPEROXIDE_ANIC | 28 | -0.45336 | -1.65725 | 0.013921114 | 0.113470055 | 1 | 919 | tags=14%, list=4%, sig | MM_2 |
| GOBP_TRNA_METABOLIC | GOBP_TRNA_METABOLIC_PROCESS | 166 | -0.30217 | -1.65614 | 0.0022883294 | 0.11402226 | 1 | 4449 | tags=31%, list=19%, si | MM_2 |
| GOBP_CGAS_STING_SIGN | GOBP_CGAS_STING_SIGNALING_PATHWAY | 16 | -0.52118 | -1.65611 | 0.026634382 | 0.11372991 | 1 | 3458 | tags=38%, list=15%, si | MM_2 |
| GOBP_NEGATIVE_REGUL | GOBP_NEGATIVE_REGULATION_OF_FIBROB | 44 | -0.3985 | -1.65557 | 0.02690583 | 0.113819495 | 1 | 6629 | tags=52%, list=28%, si | MM_2 |
| GOBP_POSITIVE_REGULA | GOBP_POSITIVE_REGULATION_OF_PROTEIN | 28 | -0.44225 | -1.65479 | 0.016587678 | 0.114058524 | 1 | 2598 | tags=36%, list=11%, si | MM_2 |
| GOBP_POSITIVE_REGULA | GOBP_POSITIVE_REGULATION_OF_PHOSPH | 16 | -0.51968 | -1.65469 | 0.03782506 | 0.11382074 | 1 | 2325 | tags=31%, list=10%, si | MM_2 |
| GOBP_CELL_SUBSTRATE_ | GOBP_CELL_SUBSTRATE_ADHESION | 354 | -0.26801 | -1.65335 | 0 | 0.11456783 | 1 | 3931 | tags=25%, list=17%, si | MM_2 |
| GOBP_PROTEIN_POLYME | GOBP_PROTEIN_POLYMERIZATION | 269 | -0.27833 | -1.65328 | 0 | 0.11430203 | 1 | 4744 | tags=28%, list=20%, si | MM_2 |
| GOBP_REGULATION_OF_ | GOBP_REGULATION_OF_NEUROINFLAMMA | 30 | -0.43319 | -1.65318 | 0.01891253 | 0.11405372 | 1 | 2784 | tags=33%, list=12%, si | MM_2 |
| HUMAN_PAPILLOMAVIRU | HUMAN_PAPILLOMAVIRUS_INFECTION | 315 | -0.27228 | -1.65211 | 0 | 0.11446675 | 1 | 3660 | tags=23%, list=16%, si | MM_2 |
| GOBP_REGULATION_OF_ | GOBP_REGULATION_OF_PROTEIN_MATURA | 71 | -0.35553 | -1.6521 | 0.0070422534 | 0.11415971 | 1 | 3087 | tags=30%, list=13%, si | MM_2 |
| GOBP_DNA_BIOSYNTHET | GOBP_DNA_BIOSYNTHETIC_PROCESS | 175 | -0.29363 | -1.65174 | 0 | 0.11409196 | 1 | 4369 | tags=26%, list=19%, si | MM_2 |
| GOBP_AORTA_DEVELOPN | GOBP_AORTA_DEVELOPMENT | 70 | -0.34738 | -1.65152 | 0.0050251256 | 0.113922015 | 1 | 3429 | tags=29%, list=15%, si | MM_2 |
| GOBP_CELL_CYCLE_G1_S | GOBP_CELL_CYCLE_G1_S_PHASE_TRANSITIC | 236 | -0.28659 | -1.65144 | 0 | 0.11366216 | 1 | 4954 | tags=34%, list=21%, si | MM_2 |
| GOBP_REGULATION_OF_ | GOBP_REGULATION_OF_CELL_SUBSTRATE_ | 219 | -0.28411 | -1.65085 | 0 | 0.11381695 | 1 | 3931 | tags=26%, list=17%, si | MM_2 |
| GOBP_NEGATIVE_REGUL | GOBP_NEGATIVE_REGULATION_OF_RESPON | 79 | -0.34566 | -1.65054 | 0.004830918 | 0.113735214 | 1 | 4543 | tags=34%, list=19%, si | MM_2 |
| GOBP_REGULATION_OF_ | GOBP_REGULATION_OF_DNA_METABOLIC_ | 491 | -0.25349 | -1.65027 | 0 | 0.113626845 | 1 | 5009 | tags=28%, list=21%, si | MM_2 |
| GOBP_REGULATION_OF_ | GOBP_REGULATION_OF_T_CELL_PROLIFERA | 188 | -0.2945 | -1.65022 | 0.00243309 | 0.1133554 | 1 | 3244 | tags=22%, list=14%, si | MM_2 |
| GOBP_REGULATION_OF_ | GOBP_REGULATION_OF_CELL_MATRIX_ADH | 121 | -0.31452 | -1.64902 | 0.00243309 | 0.11390674 | 1 | 3783 | tags=27%, list=16%, si | MM_2 |
| GOBP_POSITIVE_REGULA | GOBP_POSITIVE_REGULATION_OF_VACUOL | 25 | -0.45724 | -1.64897 | 0.034883723 | 0.113635726 | 1 | 5691 | tags=60%, list=24%, si | MM_2 |
| GOBP_GLYCEROPHOSPHO | GOBP_GLYCEROPHOSPHOLIPID_METABOLIC | 267 | -0.27804 | -1.64836 | 0.002538071 | 0.11376152 | 1 | 4203 | tags=26%, list=18%, si | MM_2 |
| GOBP_REGULATION_OF_ | GOBP_REGULATION_OF_CELLULAR_RESPON | 89 | -0.33398 | -1.64744 | 0.0073710075 | 0.11411544 | 1 | 3521 | tags=28%, list=15%, si | MM_2 |
| GOBP_NEGATIVE_REGUL | GOBP_NEGATIVE_REGULATION_OF_ORGAN | 375 | -0.26172 | -1.64665 | 0 | 0.11435249 | 1 | 4585 | tags=27%, list=19%, si | MM_2 |
| GOBP_ALPHA_BETA_T_CI | GOBP_ALPHA_BETA_T_CELL_DIFFERENTIATI | 137 | -0.30906 | -1.64596 | 0.0046838406 | 0.11454405 | 1 | 5682 | tags=39%, list=24%, si | MM_2 |
| GOBP_REGULATION_OF_ | GOBP_REGULATION_OF_OSTEOCLAST_DIFF | 84 | -0.33269 | -1.64533 | 0.0024570024 | 0.114659816 | 1 | 4125 | tags=25%, list=17%, si | MM_2 |
| GOBP_POST_GOLGI_VESI | GOBP_POST_GOLGI_VESICLE_MEDIATED_TF | 101 | -0.32911 | -1.64293 | 0.0025188916 | 0.11620016 | 1 | 5132 | tags=31%, list=22%, si | MM_2 |
| GOBP_COLLAGEN_CATAB | GOBP_COLLAGEN_CATABOLIC_PROCESS | 37 | -0.40715 | -1.64064 | 0.019950125 | 0.11762272 | 1 | 1507 | tags=16%, list=6%, sig | MM_2 |
| GOBP_KINETOCHORE_AS | GOBP_KINETOCHORE_ASSEMBLY | 16 | -0.50189 | -1.63838 | 0.03626943 | 0.119005494 | 1 | 1916 | tags=25%, list=8%, sig | MM_2 |
| GOBP_POSITIVE_REGULA | GOBP_POSITIVE_REGULATION_OF_CELL_AD | 21 | -0.47301 | -1.63818 | 0.03782506 | 0.11885649 | 1 | 1355 | tags=19%, list=6%, sig | MM_2 |
| GOBP_NEGATIVE_REGUL | GOBP_NEGATIVE_REGULATION_OF_LEUKOC | 65 | -0.35809 | -1.63696 | 0.014888338 | 0.11951487 | 1 | 4746 | tags=29%, list=20%, si | MM_2 |
| GOBP_POSITIVE_REGULA | GOBP_POSITIVE_REGULATION_OF_CELL_CY | 112 | -0.31353 | -1.63555 | 0.002444988 | 0.12015874 | 1 | 4744 | tags=35%, list=20%, si | MM_2 |
| GOBP_NEUTROPHIL_ACTI | GOBP_NEUTROPHIL_ACTIVATION | 36 | -0.41026 | -1.6342 | 0.023584906 | 0.12075153 | 1 | 3276 | tags=28%, list=14%, si | MM_2 |
| GOBP_CYTOPLASMIC_TR | GOBP_CYTOPLASMIC_TRANSLATION | 143 | -0.30332 | -1.63352 | 0.0047393367 | 0.120940916 | 1 | 12611 | tags=94%, list=53%, si | MM_2 |
| GOBP_RESPONSE_TO_AX | GOBP_RESPONSE_TO_AXON_INJURY | 75 | -0.3421 | -1.63155 | 0.014814815 | 0.12206767 | 1 | 3754 | tags=31%, list=16%, si | MM_2 |
| GOBP_POSITIVE_REGULA | GOBP_POSITIVE_REGULATION_OF_RECEPTC | 17 | -0.49187 | -1.63114 | 0.039906103 | 0.122023925 | 1 | 2552 | tags=29%, list=11%, si | MM_2 |
| GOBP_POSITIVE_REGULA | GOBP_POSITIVE_REGULATION_OF_INTERLE | 106 | -0.32406 | -1.62971 | 0.0025575447 | 0.12275968 | 1 | 4618 | tags=31%, list=20%, si | MM_2 |
| GOBP_HOMEOSTASIS_OF | GOBP_HOMEOSTASIS_OF_NUMBER_OF_CEI | 388 | -0.26389 | -1.62878 | 0 | 0.123114176 | 1 | 5969 | tags=37%, list=25%, si | MM_2 |
| GOBP_NATURAL_KILLER_ | GOBP_NATURAL_KILLER_CELL_ACTIVATION | 86 | -0.32737 | -1.62853 | 0.00968523 | 0.12304733 | 1 | 4065 | tags=29%, list=17%, si | MM_2 |
| GOBP_NEGATIVE_REGUL | GOBP_NEGATIVE_REGULATION_OF_CELL_C | 290 | -0.27243 | -1.62818 | 0 | 0.122987665 | 1 | 4524 | tags=28%, list=19%, si | MM_2 |
| GOBP_POSITIVE_REGULA | GOBP_POSITIVE_REGULATION_OF_PROTEIN | 16 | -0.51156 | -1.62756 | 0.030952381 | 0.12318278 | 1 | 5956 | tags=56%, list=25%, si | MM_2 |
| GOBP_REGULATION_OF_ | GOBP_REGULATION_OF_ERYTHROCYTE_DIF | 48 | -0.3795 | -1.6262 | 0.00982801 | 0.1239401 | 1 | 5160 | tags=44%, list=22%, si | MM_2 |
| GOBP_SUBSTRATE_ADHE | GOBP_SUBSTRATE_ADHESION_DEPENDENT_ | 99 | -0.32512 | -1.6261 | 0.0046948357 | 0.123717375 | 1 | 5745 | tags=40%, list=24%, si | MM_2 |
| GOBP_PROTEIN_LOCALIZ | GOBP_PROTEIN_LOCALIZATION_TO_NUCLEI | 326 | -0.26639 | -1.62481 | 0 | 0.12447045 | 1 | 4610 | tags=27%, list=20%, si | MM_2 |
| GOBP_POSITIVE_REGULA | GOBP_POSITIVE_REGULATION_OF_LEUKOC | 169 | -0.29449 | -1.62444 | 0 | 0.1243941 | 1 | 3351 | tags=22%, list=14%, si | MM_2 |
| GOBP_VESICLE_MEDIATE | GOBP_VESICLE_MEDIATED_TRANSPORT_BE | 46 | -0.37338 | -1.62308 | 0.014018691 | 0.12519448 | 1 | 6508 | tags=46%, list=28%, si | MM_2 |
| GOBP_REGULATION_OF_ | GOBP_REGULATION_OF_FAT_CELL_DIFFERE | 143 | -0.30097 | -1.62233 | 0.0025062656 | 0.12543356 | 1 | 4383 | tags=27%, list=19%, si | MM_2 |
| GOBP_PROTEIN_DEPHOS | GOBP_PROTEIN_DEPHOSPHORYLATION | 220 | -0.27797 | -1.62229 | 0 | 0.12514108 | 1 | 5085 | tags=32%, list=22%, si | MM_2 |
| GOBP_NEGATIVE_REGUL | GOBP_NEGATIVE_REGULATION_OF_NUCLE( | 20 | -0.47209 | -1.62171 | 0.041463416 | 0.12529759 | 1 | 4795 | tags=45%, list=20%, si | MM_2 |
| GOBP_RESCUE_OF_STALL | GOBP_RESCUE_OF_STALLED_RIBOSOME | 25 | -0.4433 | -1.62128 | 0.047169812 | 0.12533867 | 1 | 13155 | tags=100%, list=56%, | MM_2 |
| GOBP_CD40_SIGNALING_ | GOBP_CD40_SIGNALING_PATHWAY | 16 | -0.50397 | -1.62067 | 0.037861917 | 0.12549238 | 1 | 1698 | tags=25%, list=7%, sig | MM_2 |
| GOBP_INNER_EAR_RECEI | GOBP_INNER_EAR_RECEPTOR_CELL_STERE( | 44 | -0.38096 | -1.62063 | 0.023529412 | 0.12520675 | 1 | 3347 | tags=25%, list=14%, si | MM_2 |
| GOBP_REGULATION_OF_ | GOBP_REGULATION_OF_DNA_REPLICATION | 118 | -0.31138 | -1.6202 | 0.009367681 | 0.1252146 | 1 | 4874 | tags=33%, list=21%, si | MM_2 |

| | | | | | | | | | | |
|---|---|---|---|---|---|---|---|---|---|---|
| GOBP_VIRAL_TRANSLATI | GOBP_VIRAL_TRANSLATION | 18 | -0.48534 | -1.62018 | 0.035460994 | 0.124952056 | 1 | 12160 | tags=100%, list=51%, | MM_2 |
| GOBP_REGULATION_OF_ | GOBP_REGULATION_OF_NUCLEAR_DIVISIO | 144 | -0.29934 | -1.61963 | 0.0022935779 | 0.12511118 | 1 | 2088 | tags=20%, list=9%, sig | MM_2 |
| GOBP_POSITIVE_REGULA | GOBP_POSITIVE_REGULATION_OF_HOMOTY | 15 | -0.52463 | -1.61884 | 0.036876354 | 0.12550665 | 1 | 1149 | tags=20%, list=5%, sig | MM_2 |
| GOBP_INTEGRIN_MEDIAT | GOBP_INTEGRIN_MEDIATED_SIGNALING_P | 98 | -0.32227 | -1.61831 | 0.0074074073 | 0.1255759 | 1 | 6684 | tags=43%, list=28%, si | MM_2 |
| NON-SMALL_CELL_LUNG_ | NON-SMALL_CELL_LUNG_CANCER | 65 | -0.35506 | -1.618 | 0.0070422534 | 0.12549561 | 1 | 5968 | tags=48%, list=25%, si | MM_2 |
| GOBP_CHONDROITIN_SU | GOBP_CHONDROITIN_SULFATE_METABOLIC | 22 | -0.466 | -1.61768 | 0.03233831 | 0.12543668 | 1 | 5178 | tags=36%, list=22%, si | MM_2 |
| GOBP_PEPTIDYL_GLUTAN | GOBP_PEPTIDYL_GLUTAMIC_ACID_MODIFIC | 25 | -0.4289 | -1.61641 | 0.02247191 | 0.1262094 | 1 | 2639 | tags=32%, list=11%, si | MM_2 |
| GOBP_RESPONSE_TO_TY | GOBP_RESPONSE_TO_TYPE_II_INTERFERON | 122 | -0.30815 | -1.61491 | 0.0091743115 | 0.12707587 | 1 | 4773 | tags=37%, list=20%, si | MM_2 |
| GOBP_MITOTIC_SPINDLE | GOBP_MITOTIC_SPINDLE_ASSEMBLY | 71 | -0.34136 | -1.61457 | 0.0046838406 | 0.12698144 | 1 | 3879 | tags=27%, list=16%, si | MM_2 |
| GOBP_REGULATION_OF_ | GOBP_REGULATION_OF_TOLL_LIKE_RECEPT | 32 | -0.41038 | -1.61369 | 0.0247191 | 0.1273066 | 1 | 2301 | tags=22%, list=10%, si | MM_2 |
| GOBP_POSITIVE_REGULA | GOBP_POSITIVE_REGULATION_OF_CELL_CY | 252 | -0.27267 | -1.61235 | 0 | 0.12808056 | 1 | 4744 | tags=32%, list=20%, si | MM_2 |
| GOBP_GASTRULATION_W | GOBP_GASTRULATION_WITH_MOUTH_FOR | 34 | -0.40124 | -1.61104 | 0.025125628 | 0.12878586 | 1 | 4190 | tags=29%, list=18%, si | MM_2 |
| GOBP_NEGATIVE_REGUL | GOBP_NEGATIVE_REGULATION_OF_LYMPH | 44 | -0.37965 | -1.60905 | 0.0154639175 | 0.13004382 | 1 | 3384 | tags=25%, list=14%, si | MM_2 |
| GOBP_MONOATOMIC_AN | GOBP_MONOATOMIC_ANION_HOMEOSTAS | 25 | -0.44639 | -1.60898 | 0.023980815 | 0.12978116 | 1 | 1499 | tags=24%, list=6%, sig | MM_2 |
| GOBP_POSITIVE_REGULA | GOBP_POSITIVE_REGULATION_OF_RESPON | 62 | -0.35416 | -1.60776 | 0.025839793 | 0.1305264 | 1 | 5649 | tags=47%, list=24%, si | MM_2 |
| GOBP_TRNA_PROCESSIN | GOBP_TRNA_PROCESSING | 122 | -0.31273 | -1.60744 | 0.014634146 | 0.13049778 | 1 | 5688 | tags=41%, list=24%, si | MM_2 |
| GOBP_DEMETHYLATION | GOBP_DEMETHYLATION | 34 | -0.40331 | -1.60728 | 0.026190476 | 0.13032877 | 1 | 5210 | tags=41%, list=22%, si | MM_2 |
| GOBP_UBIQUITIN_DEPEN | GOBP_UBIQUITIN_DEPENDENT_PROTEIN_C | 29 | -0.42124 | -1.60667 | 0.032745592 | 0.13054101 | 1 | 4919 | tags=38%, list=21%, si | MM_2 |
| GOBP_COLLAGEN_FIBRIL | GOBP_COLLAGEN_FIBRIL_ORGANIZATION | 63 | -0.34507 | -1.60624 | 0.017456358 | 0.13057129 | 1 | 2175 | tags=19%, list=9%, sig | MM_2 |
| GOBP_VESICLE_CARGO_L | GOBP_VESICLE_CARGO_LOADING | 27 | -0.42728 | -1.6053 | 0.026378896 | 0.13095185 | 1 | 7143 | tags=59%, list=30%, si | MM_2 |
| GOBP_REGULATION_OF_ | GOBP_REGULATION_OF_EARLY_ENDOSOME | 18 | -0.49525 | -1.60394 | 0.021226415 | 0.1318464 | 1 | 11926 | tags=100%, list=51%, | MM_2 |
| GOBP_B_CELL_APOPTOTI | GOBP_B_CELL_APOPTOTIC_PROCESS | 29 | -0.428 | -1.60381 | 0.035377357 | 0.1316594 | 1 | 4172 | tags=31%, list=18%, si | MM_2 |
| GOBP_MULTIVESICULAR_ | GOBP_MULTIVESICULAR_BODY_SORTING_P | 34 | -0.40625 | -1.60341 | 0.028508771 | 0.13168018 | 1 | 4995 | tags=32%, list=21%, si | MM_2 |
| GOBP_CARDIAC_LEFT_VE | GOBP_CARDIAC_LEFT_VENTRICLE_MORPHO | 20 | -0.46746 | -1.60235 | 0.040572792 | 0.13223903 | 1 | 3413 | tags=30%, list=14%, si | MM_2 |
| GOBP_TRNA_MODIFICAT | GOBP_TRNA_MODIFICATION | 84 | -0.3357 | -1.60221 | 0.009925558 | 0.13204889 | 1 | 4169 | tags=32%, list=18%, si | MM_2 |
| GOBP_RIG_I_SIGNALING_ | GOBP_RIG_I_SIGNALING_PATHWAY | 24 | -0.44594 | -1.60082 | 0.03820225 | 0.13278577 | 1 | 2954 | tags=29%, list=13%, si | MM_2 |
| GOBP_SPHINGOMYELIN_ | GOBP_SPHINGOMYELIN_METABOLIC_PROC | 21 | -0.45595 | -1.59998 | 0.03406326 | 0.13319106 | 1 | 2490 | tags=33%, list=11%, si | MM_2 |
| GOBP_POSITIVE_REGULA | GOBP_POSITIVE_REGULATION_OF_AUTOPH | 19 | -0.47772 | -1.59886 | 0.04589372 | 0.13384931 | 1 | 5625 | tags=58%, list=24%, si | MM_2 |
| GOBP_REGULATION_OF_ | GOBP_REGULATION_OF_UBIQUITIN_DEPEN | 157 | -0.29089 | -1.59871 | 0.004796163 | 0.133704 | 1 | 4904 | tags=28%, list=21%, si | MM_2 |
| GOBP_ESTABLISHMENT_ | GOBP_ESTABLISHMENT_OF_CELL_POLARITY | 156 | -0.29056 | -1.59849 | 0.004938272 | 0.13359249 | 1 | 4674 | tags=28%, list=20%, si | MM_2 |
| GOBP_REGULATION_OF_ | GOBP_REGULATION_OF_UBIQUITIN_PROTE | 39 | -0.39428 | -1.59783 | 0.0069124424 | 0.13381432 | 1 | 4271 | tags=31%, list=18%, si | MM_2 |
| GOBP_NEGATIVE_REGUL | GOBP_NEGATIVE_REGULATION_OF_AUTOP | 81 | -0.32877 | -1.59675 | 0.014084507 | 0.1343878 | 1 | 5998 | tags=46%, list=25%, si | MM_2 |
| GOBP_LEUKOCYTE_ADHE | GOBP_LEUKOCYTE_ADHESION_TO_VASCUL | 51 | -0.36273 | -1.59635 | 0.030379746 | 0.13439794 | 1 | 4061 | tags=33%, list=17%, si | MM_2 |
| GOBP_AXON_REGENERAT | GOBP_AXON_REGENERATION | 56 | -0.35272 | -1.59604 | 0.019559903 | 0.1343227 | 1 | 3602 | tags=32%, list=15%, si | MM_2 |
| GOBP_POSITIVE_REGULA | GOBP_POSITIVE_REGULATION_OF_TORC1_ | 52 | -0.36634 | -1.59595 | 0.020100502 | 0.13408627 | 1 | 4761 | tags=38%, list=20%, si | MM_2 |
| GOBP_MRNA_TRANSCRIP | GOBP_MRNA_TRANSCRIPTION | 50 | -0.36512 | -1.59361 | 0.028571429 | 0.13559124 | 1 | 5244 | tags=38%, list=22%, si | MM_2 |
| GOBP_REGULATION_OF_ | GOBP_REGULATION_OF_T_CELL_APOPTOTI | 48 | -0.37769 | -1.59327 | 0.017429193 | 0.13552018 | 1 | 3105 | tags=21%, list=13%, si | MM_2 |
| GOBP_NEGATIVE_REGUL | GOBP_NEGATIVE_REGULATION_OF_IMMUN | 15 | -0.51439 | -1.59284 | 0.025641026 | 0.1355617 | 1 | 840 | tags=20%, list=4%, sig | MM_2 |
| GOBP_REGULATION_OF_ | GOBP_REGULATION_OF_B_CELL_APOPTOTI | 23 | -0.44271 | -1.59246 | 0.03341289 | 0.13556929 | 1 | 5593 | tags=39%, list=24%, si | MM_2 |
| GOBP_MISMATCH_REPAI | GOBP_MISMATCH_REPAIR | 25 | -0.44012 | -1.59199 | 0.037470724 | 0.13560912 | 1 | 3159 | tags=24%, list=13%, si | MM_2 |
| GOBP_AMIDE_BIOSYNTH | GOBP_AMIDE_BIOSYNTHETIC_PROCESS | 128 | -0.29774 | -1.59162 | 0.012376238 | 0.13557447 | 1 | 3819 | tags=27%, list=16%, si | MM_2 |
| GOBP_MIRNA_PROCESSI | GOBP_MIRNA_PROCESSING | 42 | -0.38253 | -1.59149 | 0.016548464 | 0.13538103 | 1 | 4917 | tags=40%, list=21%, si | MM_2 |
| GOBP_NUCLEOSIDE_DIPH | GOBP_NUCLEOSIDE_DIPHOSPHATE_METAB | 127 | -0.30634 | -1.59121 | 0.011876484 | 0.13532087 | 1 | 4935 | tags=36%, list=21%, si | MM_2 |
| GOBP_RESPONSE_TO_AN | GOBP_RESPONSE_TO_AMINO_ACID_STARV | 54 | -0.35738 | -1.59038 | 0.019607844 | 0.13569967 | 1 | 4221 | tags=35%, list=18%, si | MM_2 |
| GOBP_POSITIVE_REGULA | GOBP_POSITIVE_REGULATION_OF_FATTY_A | 45 | -0.38162 | -1.59036 | 0.01965602 | 0.13542336 | 1 | 5037 | tags=47%, list=21%, si | MM_2 |
| GOBP_LEUKOCYTE_MIGR | GOBP_LEUKOCYTE_MIGRATION_INVOLVED_ | 23 | -0.44279 | -1.58724 | 0.02117647 | 0.13775593 | 1 | 201 | tags=13%, list=1%, sig | MM_2 |
| GOBP_ENDOPLASMIC_RE | GOBP_ENDOPLASMIC_RETICULUM_TUBULA | 21 | -0.45549 | -1.58549 | 0.04400978 | 0.13880971 | 1 | 4998 | tags=33%, list=21%, si | MM_2 |
| GOBP_LYSOSOMAL_LUM | GOBP_LYSOSOMAL_LUMEN_ACIDIFICATION | 15 | -0.49308 | -1.58545 | 0.036764707 | 0.13856065 | 1 | 6243 | tags=60%, list=26%, si | MM_2 |
| GOBP_REGULATION_OF_ | GOBP_REGULATION_OF_TELOMERASE_ACT | 42 | -0.38702 | -1.5847 | 0.01946472 | 0.13887747 | 1 | 4474 | tags=29%, list=19%, si | MM_2 |
| GOBP_NAD_METABOLIC_ | GOBP_NAD_METABOLIC_PROCESS | 58 | -0.35334 | -1.5846 | 0.020134227 | 0.13867077 | 1 | 4983 | tags=40%, list=21%, si | MM_2 |
| GOBP_REGULATION_OF_ | GOBP_REGULATION_OF_SUPEROXIDE_MET | 34 | -0.4016 | -1.5845 | 0.04408353 | 0.13845886 | 1 | 1757 | tags=15%, list=7%, sig | MM_2 |
| GOBP_REGULATION_OF_ | GOBP_REGULATION_OF_MITOTIC_CELL_CY | 490 | -0.24747 | -1.58449 | 0 | 0.13816804 | 1 | 4993 | tags=30%, list=21%, si | MM_2 |
| GOBP_LEUKOCYTE_CELL_ | GOBP_LEUKOCYTE_CELL_CELL_ADHESION | 390 | -0.25171 | -1.58447 | 0 | 0.13789116 | 1 | 3670 | tags=22%, list=16%, si | MM_2 |
| GOBP_POSITIVE_REGULA | GOBP_POSITIVE_REGULATION_OF_VIRAL_L | 28 | -0.41984 | -1.58427 | 0.050772626 | 0.13776143 | 1 | 5538 | tags=43%, list=23%, si | MM_2 |
| GOBP_POSITIVE_REGULA | GOBP_POSITIVE_REGULATION_OF_RESPON | 381 | -0.25248 | -1.58384 | 0 | 0.13784373 | 1 | 3509 | tags=23%, list=15%, si | MM_2 |
| GOBP_NON_CANONICAL_ | GOBP_NON_CANONICAL_NF_KAPPAB_SIGN | 115 | -0.30083 | -1.5832 | 0.009661836 | 0.13810699 | 1 | 5022 | tags=30%, list=21%, si | MM_2 |
| GOBP_NUCLEOSIDE_MON | GOBP_NUCLEOSIDE_MONOPHOSPHATE_BIO | 40 | -0.38162 | -1.58219 | 0.024691358 | 0.13860384 | 1 | 3576 | tags=25%, list=15%, si | MM_2 |
| GOBP_ACTIN_POLYMERIZ | GOBP_ACTIN_POLYMERIZATION_OR_DEPOL | 187 | -0.28294 | -1.58145 | 0.0024509805 | 0.13887669 | 1 | 4706 | tags=25%, list=20%, si | MM_2 |
| GOBP_REGULATION_OF_ | GOBP_REGULATION_OF_MITOCHONDRIAL_ | 32 | -0.41121 | -1.58057 | 0.027638191 | 0.1393078 | 1 | 3219 | tags=28%, list=14%, si | MM_2 |
| GOBP_NEGATIVE_REGUL | GOBP_NEGATIVE_REGULATION_OF_VASCU | 21 | -0.45561 | -1.58001 | 0.03827751 | 0.13949215 | 1 | 4383 | tags=48%, list=19%, si | MM_2 |
| GOBP_POSITIVE_REGULA | GOBP_POSITIVE_REGULATION_OF_TRANSL | 125 | -0.30244 | -1.57996 | 0.0072639226 | 0.13923566 | 1 | 5004 | tags=30%, list=21%, si | MM_2 |
| GOBP_PHOSPHOLIPID_DE | GOBP_PHOSPHOLIPID_DEPHOSPHORYLATIO | 43 | -0.38126 | -1.57917 | 0.033573143 | 0.13965221 | 1 | 5692 | tags=47%, list=24%, si | MM_2 |
| GOBP_POLARIZED_EPITH | GOBP_POLARIZED_EPITHELIAL_CELL_DIFFER | 25 | -0.43997 | -1.57895 | 0.028301887 | 0.13952984 | 1 | 2934 | tags=28%, list=12%, si | MM_2 |
| GOBP_REGULATION_OF_ | GOBP_REGULATION_OF_RNA_STABILITY | 174 | -0.28641 | -1.57866 | 0.002415459 | 0.13949506 | 1 | 5234 | tags=32%, list=22%, si | MM_2 |
| GOBP_SNRNA_PROCESSI | GOBP_SNRNA_PROCESSING | 24 | -0.43577 | -1.57839 | 0.03892944 | 0.13947308 | 1 | 2765 | tags=25%, list=12%, si | MM_2 |
| GOBP_POSITIVE_REGULA | GOBP_POSITIVE_REGULATION_OF_ORGANE | 91 | -0.31698 | -1.57806 | 0.0071090045 | 0.13943212 | 1 | 5625 | tags=37%, list=24%, si | MM_2 |
| GOBP_LEUKOCYTE_MIGR | GOBP_LEUKOCYTE_MIGRATION | 363 | -0.25247 | -1.57706 | 0 | 0.13989791 | 1 | 3967 | tags=23%, list=17%, si | MM_2 |
| PHOSPHATIDYLINOSITOL_ | PHOSPHATIDYLINOSITOL_SIGNALING_SYSTE | 94 | -0.31551 | -1.57626 | 0.019950125 | 0.14026858 | 1 | 5968 | tags=43%, list=25%, si | MM_2 |
| GOBP_MYELOID_CELL_DI | GOBP_MYELOID_CELL_DIFFERENTIATION | 455 | -0.24701 | -1.57547 | 0 | 0.14064266 | 1 | 5402 | tags=31%, list=23%, si | MM_2 |
| GOBP_CELLULAR_RESPO | GOBP_CELLULAR_RESPONSE_TO_DEXAMET | 23 | -0.43694 | -1.57506 | 0.03818616 | 0.14070429 | 1 | 3521 | tags=30%, list=15%, si | MM_2 |
| GOBP_NEGATIVE_REGUL | GOBP_NEGATIVE_REGULATION_OF_MITOTI | 174 | -0.28053 | -1.57358 | 0.0051282053 | 0.14171943 | 1 | 4524 | tags=28%, list=19%, si | MM_2 |
| GOBP_TOR_SIGNALING | GOBP_TOR_SIGNALING | 165 | -0.28601 | -1.57314 | 0.0025188916 | 0.14186274 | 1 | 4935 | tags=32%, list=21%, si | MM_2 |
| GOBP_CYTOKINETIC_PRO | GOBP_CYTOKINETIC_PROCESS | 45 | -0.37411 | -1.57238 | 0.02606635 | 0.14225419 | 1 | 2018 | tags=16%, list=9%, sig | MM_2 |
| GOBP_GRANULOCYTE_AC | GOBP_GRANULOCYTE_ACTIVATION | 41 | -0.3798 | -1.57204 | 0.03311258 | 0.14218038 | 1 | 3276 | tags=27%, list=14%, si | MM_2 |
| GOBP_ENDONUCLEOLYTI | GOBP_ENDONUCLEOLYTIC_CLEAVAGE_OF_ | 16 | -0.499 | -1.57133 | 0.04694836 | 0.14250112 | 1 | 6171 | tags=63%, list=26%, si | MM_2 |
| GOBP_REGULATION_OF_ | GOBP_REGULATION_OF_PROTEIN_CONTAIN | 420 | -0.24882 | -1.57089 | 0 | 0.14253958 | 1 | 3901 | tags=22%, list=17%, si | MM_2 |
| GOBP_LYMPHOCYTE_HO | GOBP_LYMPHOCYTE_HOMEOSTASIS | 92 | -0.31399 | -1.57068 | 0.0093896715 | 0.1424181 | 1 | 4172 | tags=28%, list=18%, si | MM_2 |
| GOBP_POSITIVE_REGULA | GOBP_POSITIVE_REGULATION_OF_APOPTO | 161 | -0.28618 | -1.57067 | 0.0024630541 | 0.14212564 | 1 | 4276 | tags=24%, list=18%, si | MM_2 |
| GOBP_PODOSOME_ASSE | GOBP_PODOSOME_ASSEMBLY | 25 | -0.4307 | -1.57057 | 0.021276595 | 0.1419038 | 1 | 6217 | tags=52%, list=26%, si | MM_2 |
| GOBP_POSITIVE_REGULA | GOBP_POSITIVE_REGULATION_OF_PROTEIN | 41 | -0.37835 | -1.57056 | 0.033254158 | 0.14162256 | 1 | 4918 | tags=34%, list=21%, si | MM_2 |
| GOBP_POSITIVE_REGULA | GOBP_POSITIVE_REGULATION_OF_ACUTE_I | 27 | -0.42045 | -1.57017 | 0.043778803 | 0.14165036 | 1 | 3315 | tags=26%, list=14%, si | MM_2 |
| GOBP_MEIOTIC_CELL_CY | GOBP_MEIOTIC_CELL_CYCLE_PROCESS | 220 | -0.27188 | -1.57013 | 0.0049875313 | 0.14138281 | 1 | 3014 | tags=24%, list=13%, si | MM_2 |

| | | | | | | | | | | |
|---|---|---|---|---|---|---|---|---|---|---|
| GOBP_REGULATION_OF_ | GOBP_REGULATION_OF_GENE_SILENCING_ | 29 | -0.41453 | -1.56943 | 0.04524887 | 0.14173797 | 1 | 4370 | tags=45%, list=19%, si | MM_2 |
| GOBP_NICOTINAMIDE_N | GOBP_NICOTINAMIDE_NUCLEOTIDE_BIOSYI | 16 | -0.47669 | -1.56872 | 0.05399568 | 0.14206493 | 1 | 4983 | tags=50%, list=21%, si | MM_2 |
| GOBP_DETECTION_OF_BI | GOBP_DETECTION_OF_BIOTIC_STIMULUS | 37 | -0.38568 | -1.56862 | 0.026004728 | 0.14187233 | 1 | 3627 | tags=27%, list=15%, si | MM_2 |
| GOBP_POSITIVE_REGULA | GOBP_POSITIVE_REGULATION_OF_DNA_BI( | 76 | -0.3247 | -1.56831 | 0.020512821 | 0.1418686 | 1 | 1510 | tags=14%, list=6%, sig | MM_2 |
| GOBP_INTRACELLULAR_R | GOBP_INTRACELLULAR_RECEPTOR_SIGNALI | 325 | -0.25836 | -1.56779 | 0.0024752475 | 0.14207003 | 1 | 3513 | tags=22%, list=15%, si | MM_2 |
| GOBP_REGULATION_OF_ | GOBP_REGULATION_OF_INTRACELLULAR_P | 168 | -0.28228 | -1.56704 | 0.004819277 | 0.14241363 | 1 | 4536 | tags=26%, list=19%, si | MM_2 |
| AMINOACYL-TRNA_BIOSY | AMINOACYL-TRNA_BIOSYNTHESIS | 43 | -0.37752 | -1.56676 | 0.03640777 | 0.14230022 | 1 | 3483 | tags=33%, list=15%, si | MM_2 |
| GOBP_REGULATION_OF_ | GOBP_REGULATION_OF_PHOSPHATASE_AC | 61 | -0.3393 | -1.5658 | 0.009345794 | 0.14273645 | 1 | 5218 | tags=36%, list=22%, si | MM_2 |
| GOBP_REGULATION_OF_ | GOBP_REGULATION_OF_INTEGRIN_MEDIAT | 19 | -0.45676 | -1.56513 | 0.029953917 | 0.14297861 | 1 | 3462 | tags=32%, list=15%, si | MM_2 |
| GOBP_RESPONSE_TO_CH | GOBP_RESPONSE_TO_CHEMOKINE | 74 | -0.33105 | -1.56492 | 0.026506023 | 0.14285395 | 1 | 3571 | tags=27%, list=15%, si | MM_2 |
| NECROPTOSIS | NECROPTOSIS | 138 | -0.29729 | -1.56466 | 0.007125891 | 0.14281924 | 1 | 5086 | tags=36%, list=22%, si | MM_2 |
| GOBP_REGULATION_OF_ | GOBP_REGULATION_OF_LEUKOCYTE_DIFFEI | 342 | -0.25163 | -1.56365 | 0 | 0.14339294 | 1 | 4317 | tags=24%, list=18%, si | MM_2 |
| PEROXISOME | PEROXISOME | 81 | -0.32693 | -1.56341 | 0.024213076 | 0.14332965 | 1 | 4470 | tags=33%, list=19%, si | MM_2 |
| GOBP_SEROTONIN_SECRI | GOBP_SEROTONIN_SECRETION | 15 | -0.50019 | -1.56338 | 0.06097561 | 0.14309199 | 1 | 1382 | tags=27%, list=6%, sig | MM_2 |
| GOBP_KINETOCHORE_OR | GOBP_KINETOCHORE_ORGANIZATION | 20 | -0.46715 | -1.56199 | 0.053398058 | 0.14400509 | 1 | 1916 | tags=25%, list=8%, sig | MM_2 |
| GLYCEROLIPID_METABOL | GLYCEROLIPID_METABOLISM | 57 | -0.33927 | -1.56133 | 0.012135922 | 0.14431413 | 1 | 2557 | tags=19%, list=11%, si | MM_2 |
| GOBP_NATURAL_KILLER_ | GOBP_NATURAL_KILLER_CELL_ACTIVATION_ | 21 | -0.45041 | -1.56122 | 0.04589372 | 0.1441365 | 1 | 1427 | tags=24%, list=6%, sig | MM_2 |
| GOBP_REGULATION_OF_ | GOBP_REGULATION_OF_ACTIN_FILAMENT_ | 266 | -0.26338 | -1.56002 | 0 | 0.1448698 | 1 | 4825 | tags=27%, list=20%, si | MM_2 |
| GOBP_NEGATIVE_REGUL | GOBP_NEGATIVE_REGULATION_OF_CATABO | 339 | -0.25286 | -1.55913 | 0 | 0.14535277 | 1 | 4886 | tags=29%, list=21%, si | MM_2 |
| GOBP_REGULATION_OF_ | GOBP_REGULATION_OF_DEPHOSPHORYLAT | 106 | -0.30797 | -1.55892 | 0.011627907 | 0.14520913 | 1 | 5218 | tags=32%, list=22%, si | MM_2 |
| GOBP_MITOTIC_METAPH | GOBP_MITOTIC_METAPHASE_CHROMOSOM | 58 | -0.34529 | -1.55836 | 0.02494331 | 0.14535718 | 1 | 4230 | tags=33%, list=18%, si | MM_2 |
| GOBP_POSITIVE_REGULA | GOBP_POSITIVE_REGULATION_OF_BLOOD_ | 42 | -0.37541 | -1.55741 | 0.023364486 | 0.1458787 | 1 | 2683 | tags=31%, list=11%, si | MM_2 |
| GOBP_LATE_ENDOSOME_ | GOBP_LATE_ENDOSOME_TO_VACUOLE_TR/ | 25 | -0.42413 | -1.55679 | 0.029545454 | 0.14615859 | 1 | 13607 | tags=100%, list=58%, | MM_2 |
| GOBP_NEGATIVE_REGUL | GOBP_NEGATIVE_REGULATION_OF_NOTCH | 39 | -0.38256 | -1.55626 | 0.018390805 | 0.1463266 | 1 | 2764 | tags=28%, list=12%, si | MM_2 |
| GOBP_POSITIVE_REGULA | GOBP_POSITIVE_REGULATION_OF_TELOME | 71 | -0.33298 | -1.55609 | 0.026506023 | 0.14617658 | 1 | 5368 | tags=31%, list=23%, si | MM_2 |
| GOBP_REGULATION_OF_ | GOBP_REGULATION_OF_MYELOID_LEUKOC | 61 | -0.35216 | -1.55591 | 0.02538071 | 0.14605074 | 1 | 3276 | tags=30%, list=14%, si | MM_2 |
| GOBP_T_CELL_ACTIVATIO | GOBP_T_CELL_ACTIVATION_INVOLVED_IN_ | 112 | -0.30302 | -1.55578 | 0.0121951215 | 0.14589895 | 1 | 3872 | tags=24%, list=16%, si | MM_2 |
| GOBP_LEUKOCYTE_CHEM | GOBP_LEUKOCYTE_CHEMOTAXIS | 213 | -0.26906 | -1.55421 | 0 | 0.1469478 | 1 | 3967 | tags=23%, list=17%, si | MM_2 |
| GOBP_ATRIOVENTRICULA | GOBP_ATRIOVENTRICULAR_VALVE_DEVELO | 30 | -0.40508 | -1.55397 | 0.040669855 | 0.14689282 | 1 | 3755 | tags=37%, list=16%, si | MM_2 |
| GOBP_STRESS_FIBER_ASS | GOBP_STRESS_FIBER_ASSEMBLY | 117 | -0.29836 | -1.55393 | 0.012224939 | 0.14663908 | 1 | 3783 | tags=27%, list=16%, si | MM_2 |
| GOBP_EXTRACELLULAR_S | GOBP_EXTRACELLULAR_STRUCTURE_ORGAI | 309 | -0.25335 | -1.55382 | 0 | 0.14648397 | 1 | 4009 | tags=25%, list=17%, si | MM_2 |
| GOBP_REGULATION_OF_ | GOBP_REGULATION_OF_RHO_PROTEIN_SIG | 91 | -0.30961 | -1.55235 | 0.016949153 | 0.1475461 | 1 | 3807 | tags=24%, list=16%, si | MM_2 |
| GOBP_NEGATIVE_REGUL | GOBP_NEGATIVE_REGULATION_OF_LEUKO( | 142 | -0.28744 | -1.55234 | 0.009411764 | 0.14726453 | 1 | 3571 | tags=23%, list=15%, si | MM_2 |
| GOBP_REGULATION_OF_ | GOBP_REGULATION_OF_GASTRULATION | 16 | -0.49084 | -1.55225 | 0.054245282 | 0.14706235 | 1 | 3783 | tags=44%, list=16%, si | MM_2 |
| GOBP_REGULATION_OF_ | GOBP_REGULATION_OF_LEUKOCYTE_MIGR/ | 230 | -0.26724 | -1.55204 | 0.005037783 | 0.14691329 | 1 | 3754 | tags=23%, list=16%, si | MM_2 |
| GOBP_NEGATIVE_REGUL | GOBP_NEGATIVE_REGULATION_OF_TRANSF | 222 | -0.26761 | -1.55135 | 0 | 0.14721292 | 1 | 4513 | tags=29%, list=19%, si | MM_2 |
| GOBP_LEUKOCYTE_HOMI | GOBP_LEUKOCYTE_HOMEOSTASIS | 137 | -0.28891 | -1.55128 | 0.002352941 | 0.14698407 | 1 | 5884 | tags=38%, list=25%, si | MM_2 |
| GOBP_MICROGLIA_DIFFE | GOBP_MICROGLIA_DIFFERENTIATION | 16 | -0.50576 | -1.55115 | 0.05172414 | 0.14680582 | 1 | 3942 | tags=31%, list=17%, si | MM_2 |
| N-GLYCAN_BIOSYNTHESIS | N-GLYCAN_BIOSYNTHESIS | 48 | -0.36517 | -1.55105 | 0.026252983 | 0.14658968 | 1 | 4945 | tags=29%, list=21%, si | MM_2 |
| GOBP_ORGANELLE_DISAS | GOBP_ORGANELLE_DISASSEMBLY | 155 | -0.28343 | -1.55095 | 0.012437811 | 0.1463857 | 1 | 7118 | tags=48%, list=30%, si | MM_2 |
| GOBP_CELL_MATRIX_ADI | GOBP_CELL_MATRIX_ADHESION | 225 | -0.26503 | -1.55051 | 0.0023696683 | 0.14652053 | 1 | 3824 | tags=24%, list=16%, si | MM_2 |
| GOBP_MICROTUBULE_DE | GOBP_MICROTUBULE_DEPOLYMERIZATION | 58 | -0.34842 | -1.54932 | 0.03125 | 0.14721757 | 1 | 4489 | tags=41%, list=19%, si | MM_2 |
| GOBP_NUCLEOSIDE_DIPH | GOBP_NUCLEOSIDE_DIPHOSPHATE_CATABO | 111 | -0.29923 | -1.54923 | 0.009756098 | 0.14703487 | 1 | 4935 | tags=36%, list=21%, si | MM_2 |
| GOBP_NEGATIVE_REGUL | GOBP_NEGATIVE_REGULATION_OF_TOR_SI | 61 | -0.34457 | -1.54872 | 0.02948403 | 0.1472275 | 1 | 3598 | tags=30%, list=15%, si | MM_2 |
| GOBP_POSITIVE_REGULA | GOBP_POSITIVE_REGULATION_OF_MACROF | 15 | -0.49212 | -1.5486 | 0.048076924 | 0.14704317 | 1 | 1047 | tags=20%, list=4%, sig | MM_2 |
| GOBP_NUCLEOSOME_AS | GOBP_NUCLEOSOME_ASSEMBLY | 50 | -0.35677 | -1.54843 | 0.03301887 | 0.14690292 | 1 | 6598 | tags=46%, list=28%, si | MM_2 |
| GOBP_MITOCHONDRIAL_ | GOBP_MITOCHONDRIAL_TRANSPORT | 174 | -0.27735 | -1.54825 | 0.00990099 | 0.14677961 | 1 | 6000 | tags=34%, list=25%, si | MM_2 |
| GOBP_DEFENSE_RESPON | GOBP_DEFENSE_RESPONSE_TO_GRAM_POS | 95 | -0.30788 | -1.54761 | 0.023584906 | 0.14707962 | 1 | 2877 | tags=20%, list=12%, si | MM_2 |
| HUMAN_IMMUNODEFICI | HUMAN_IMMUNODEFICIENCY_VIRUS_1_IN | 201 | -0.26877 | -1.54747 | 0 | 0.14695077 | 1 | 5254 | tags=32%, list=22%, si | MM_2 |
| GOBP_NUCLEOPHAGY | GOBP_NUCLEOPHAGY | 24 | -0.42994 | -1.54661 | 0.032482598 | 0.14744507 | 1 | 6848 | tags=63%, list=29%, si | MM_2 |
| GOBP_POSITIVE_REGULA | GOBP_POSITIVE_REGULATION_OF_CELL_CE | 309 | -0.2558 | -1.54639 | 0 | 0.14732714 | 1 | 4276 | tags=26%, list=18%, si | MM_2 |
| GOBP_LIPID_OXIDATION | GOBP_LIPID_OXIDATION | 118 | -0.29964 | -1.54543 | 0.014962593 | 0.14796054 | 1 | 3901 | tags=30%, list=17%, si | MM_2 |
| GOBP_NEGATIVE_REGUL | GOBP_NEGATIVE_REGULATION_OF_MIRNA | 30 | -0.40247 | -1.54542 | 0.03117506 | 0.14769405 | 1 | 3762 | tags=40%, list=16%, si | MM_2 |
| GOBP_REGULATION_OF_ | GOBP_REGULATION_OF_MEMBRANE_PERM | 70 | -0.32815 | -1.54426 | 0.030092593 | 0.14841604 | 1 | 4842 | tags=34%, list=21%, si | MM_2 |
| GOBP_ESTABLISHMENT_( | GOBP_ESTABLISHMENT_OF_PROTEIN_LOCA | 435 | -0.24352 | -1.54237 | 0 | 0.14967258 | 1 | 4538 | tags=24%, list=19%, si | MM_2 |
| GOBP_DEVELOPMENTAL_ | GOBP_DEVELOPMENTAL_PIGMENTATION | 50 | -0.36902 | -1.54195 | 0.016746411 | 0.1497366 | 1 | 6259 | tags=48%, list=27%, si | MM_2 |
| GOBP_STEM_CELL_DIVISI | GOBP_STEM_CELL_DIVISION | 43 | -0.36858 | -1.5415 | 0.034642033 | 0.14979976 | 1 | 6665 | tags=53%, list=28%, si | MM_2 |
| GOBP_SULFUR_COMPOU | GOBP_SULFUR_COMPOUND_CATABOLIC_PI | 35 | -0.39307 | -1.54044 | 0.028871391 | 0.15046015 | 1 | 5131 | tags=49%, list=22%, si | MM_2 |
| REGULATION_OF_LIPOLY | REGULATION_OF_LIPOLYSIS_IN_ADIPOCYTE | 53 | -0.34973 | -1.53908 | 0.015981736 | 0.15129164 | 1 | 3458 | tags=28%, list=15%, si | MM_2 |
| GOBP_MEMBRANE_LIPID | GOBP_MEMBRANE_LIPID_CATABOLIC_PRO( | 33 | -0.38147 | -1.53727 | 0.038004752 | 0.15262651 | 1 | 4297 | tags=36%, list=18%, si | MM_2 |
| GOBP_PHOSPHATIDYLINO | GOBP_PHOSPHATIDYLINOSITOL_PHOSPHAT | 50 | -0.35913 | -1.53662 | 0.03422983 | 0.15291564 | 1 | 5868 | tags=42%, list=25%, si | MM_2 |
| GOBP_CD4_POSITIVE_AL | GOBP_CD4_POSITIVE_ALPHA_BETA_T_CELL | 132 | -0.29481 | -1.53561 | 0.009259259 | 0.15356568 | 1 | 3548 | tags=23%, list=15%, si | MM_2 |
| GOBP_REGULATION_OF_ | GOBP_REGULATION_OF_MRNA_METABOLI( | 286 | -0.25559 | -1.53486 | 0 | 0.15403903 | 1 | 5963 | tags=35%, list=25%, si | MM_2 |
| GOBP_CELLULAR_RESPON | GOBP_CELLULAR_RESPONSE_TO_NUTRIENT | 38 | -0.3787 | -1.5345 | 0.03087886 | 0.15406278 | 1 | 4184 | tags=32%, list=18%, si | MM_2 |
| GOBP_REGULATION_OF_ | GOBP_REGULATION_OF_TELOMERE_MAINT | 98 | -0.31268 | -1.53442 | 0.009803922 | 0.15384635 | 1 | 5368 | tags=32%, list=23%, si | MM_2 |
| GOBP_GOLGI_VESICLE_TI | GOBP_GOLGI_VESICLE_TRANSPORT | 276 | -0.25349 | -1.53393 | 0.0024875621 | 0.15400656 | 1 | 5900 | tags=32%, list=25%, si | MM_2 |
| GOBP_MYELOID_CELL_HO | GOBP_MYELOID_CELL_HOMEOSTASIS | 209 | -0.26612 | -1.53328 | 0.00477327 | 0.1543187 | 1 | 5898 | tags=37%, list=25%, si | MM_2 |
| GOBP_COPII_COATED_VE | GOBP_COPII_COATED_VESICLE_CARGO_LO/ | 17 | -0.47951 | -1.53324 | 0.06787331 | 0.1540743 | 1 | 12297 | tags=100%, list=52%, | MM_2 |
| GOBP_REGULATION_OF_ | GOBP_REGULATION_OF_HEMOPOIESIS | 425 | -0.24056 | -1.53273 | 0 | 0.15424323 | 1 | 4203 | tags=24%, list=18%, si | MM_2 |
| GOBP_ACTIN_FILAMENT_ | GOBP_ACTIN_FILAMENT_ORGANIZATION | 450 | -0.2394 | -1.53173 | 0 | 0.15490024 | 1 | 4637 | tags=26%, list=20%, si | MM_2 |
| GOBP_PROTEIN_MODIFIC | GOBP_PROTEIN_MODIFICATION_BY_SMALL | 117 | -0.29388 | -1.53168 | 0.014705882 | 0.15466878 | 1 | 6576 | tags=39%, list=28%, si | MM_2 |
| GOBP_ERYTHROCYTE_HO | GOBP_ERYTHROCYTE_HOMEOSTASIS | 163 | -0.28098 | -1.53121 | 0.013824885 | 0.15480766 | 1 | 5303 | tags=34%, list=22%, si | MM_2 |
| GOBP_PROTEIN_TRIMERI | GOBP_PROTEIN_TRIMERIZATION | 16 | -0.48241 | -1.53039 | 0.07692308 | 0.15526268 | 1 | 3438 | tags=44%, list=15%, | MM_2 |
| GOBP_CELLULAR_RESPON | GOBP_CELLULAR_RESPONSE_TO_METAL_IO | 155 | -0.28244 | -1.53017 | 0.00968523 | 0.15521799 | 1 | 4240 | tags=30%, list=18%, si | MM_2 |
| ETHER_LIPID_METABOLIS | ETHER_LIPID_METABOLISM | 42 | -0.36952 | -1.52958 | 0.04157044 | 0.15545578 | 1 | 2673 | tags=24%, list=11%, si | MM_2 |
| GOBP_PROTEIN_LOCALIZ | GOBP_PROTEIN_LOCALIZATION_TO_CONDE | 16 | -0.48407 | -1.52927 | 0.06733167 | 0.15547103 | 1 | 5158 | tags=50%, list=22%, si | MM_2 |
| GOBP_REGULATION_OF_ | GOBP_REGULATION_OF_ADIPOSE_TISSUE_I | 16 | -0.4678 | -1.52893 | 0.06526806 | 0.1554968 | 1 | 2631 | tags=19%, list=11%, si | MM_2 |
| GOBP_CELLULAR_RESPON | GOBP_CELLULAR_RESPONSE_TO_INORGANI | 177 | -0.27491 | -1.52889 | 0.0076530613 | 0.15526459 | 1 | 3931 | tags=27%, list=17%, si | MM_2 |
| GOBP_NEGATIVE_REGUL | GOBP_NEGATIVE_REGULATION_OF_INNATE | 86 | -0.31887 | -1.5285 | 0.009615385 | 0.15535896 | 1 | 3000 | tags=24%, list=13%, si | MM_2 |
| GOBP_REGULATION_OF_ | GOBP_REGULATION_OF_GTPASE_ACTIVITY | 287 | -0.25582 | -1.52847 | 0.0045871558 | 0.15509704 | 1 | 4125 | tags=24%, list=17%, si | MM_2 |
| GOBP_REGULATION_OF_ | GOBP_REGULATION_OF_INNATE_IMMUNE_ | 427 | -0.24249 | -1.52829 | 0 | 0.15498914 | 1 | 3138 | tags=21%, list=13%, si | MM_2 |
| GOBP_POSITIVE_REGULA | GOBP_POSITIVE_REGULATION_OF_DNA_ME | 314 | -0.25219 | -1.52784 | 0 | 0.15513559 | 1 | 4616 | tags=25%, list=20%, si | MM_2 |

| | | | | | | | | | | |
|---|---|---|---|---|---|---|---|---|---|---|
| GOBP_CHROMATIN_REM | GOBP_CHROMATIN_REMODELING | 482 | -0.23987 | -1.52686 | 0.002469136 | 0.15578201 | 1 | 5243 | tags=29%, list=22%, si | MM_2 |
| GOBP_ALPHA_BETA_T_C | GOBP_ALPHA_BETA_T_CELL_ACTIVATION | 191 | -0.2749 | -1.52667 | 0.0022222223 | 0.15566643 | 1 | 3660 | tags=25%, list=16%, si | MM_2 |
| GOBP_PROTEIN_K48_LIN | GOBP_PROTEIN_K48_LINKED_DEUBIQUITIN | 29 | -0.40056 | -1.52637 | 0.053164557 | 0.15566467 | 1 | 5833 | tags=45%, list=25%, si | MM_2 |
| GOBP_POSITIVE_REGULA | GOBP_POSITIVE_REGULATION_OF_DENDRIT | 67 | -0.33175 | -1.52599 | 0.034739453 | 0.15576264 | 1 | 4150 | tags=25%, list=18%, si | MM_2 |
| GOBP_HEMATOPOIETIC_( | GOBP_HEMATOPOIETIC_OR_LYMPHOID_OR | 118 | -0.29672 | -1.52521 | 0.013986014 | 0.15624063 | 1 | 4986 | tags=31%, list=21%, si | MM_2 |
| GOBP_IMPORT_INTO_NU | GOBP_IMPORT_INTO_NUCLEUS | 180 | -0.26934 | -1.52474 | 0.0071090045 | 0.15642336 | 1 | 5332 | tags=33%, list=23%, si | MM_2 |
| GOBP_CELLULAR_RESPON | GOBP_CELLULAR_RESPONSE_TO_KETONE | 88 | -0.31131 | -1.52407 | 0.011961723 | 0.1567986 | 1 | 3521 | tags=25%, list=15%, si | MM_2 |
| GOBP_ADAPTIVE_IMMUN | GOBP_ADAPTIVE_IMMUNE_RESPONSE_BAS | 322 | -0.24847 | -1.5239 | 0 | 0.15667012 | 1 | 3776 | tags=21%, list=16%, si | MM_2 |
| NOD-LIKE_RECEPTOR_SIG | NOD-LIKE_RECEPTOR_SIGNALING_PATHWA | 151 | -0.27914 | -1.52348 | 0.010309278 | 0.15679358 | 1 | 3720 | tags=24%, list=16%, si | MM_2 |
| GOBP_POSITIVE_REGULA | GOBP_POSITIVE_REGULATION_OF_SYNAPTI | 17 | -0.45781 | -1.52303 | 0.06367925 | 0.15691176 | 1 | 1971 | tags=29%, list=8%, sig | MM_2 |
| GOBP_REGULATION_OF_ | GOBP_REGULATION_OF_MACROPHAGE_MI | 47 | -0.35717 | -1.52294 | 0.03341289 | 0.15671547 | 1 | 3351 | tags=30%, list=14%, si | MM_2 |
| GOBP_REGULATION_OF_ | GOBP_REGULATION_OF_DNA_REPAIR | 219 | -0.25917 | -1.5229 | 0.002375297 | 0.15649763 | 1 | 5176 | tags=30%, list=22%, si | MM_2 |
| GOBP_T_CELL_HOMEOST | GOBP_T_CELL_HOMEOSTASIS | 52 | -0.34488 | -1.52205 | 0.031746034 | 0.1569715 | 1 | 3989 | tags=31%, list=17%, si | MM_2 |
| GOBP_ORGANELLE_INHE | GOBP_ORGANELLE_INHERITANCE | 15 | -0.48201 | -1.52131 | 0.06122449 | 0.15739395 | 1 | 12237 | tags=100%, list=52%, | MM_2 |
| GOBP_POSITIVE_REGULA | GOBP_POSITIVE_REGULATION_OF_CALCINE | 19 | -0.45757 | -1.52117 | 0.04245283 | 0.15727048 | 1 | 4501 | tags=53%, list=19%, si | MM_2 |
| GOBP_REGULATION_OF_ | GOBP_REGULATION_OF_PROTEIN_LOCALIZ/ | 158 | -0.27776 | -1.52116 | 0.009367681 | 0.15700346 | 1 | 4383 | tags=28%, list=19%, si | MM_2 |
| GOBP_NEGATIVE_REGUL | GOBP_NEGATIVE_REGULATION_OF_CELL_JU | 35 | -0.38225 | -1.52093 | 0.038990825 | 0.15698758 | 1 | 3783 | tags=31%, list=16%, si | MM_2 |
| GOBP_SNRNA_METABOL | GOBP_SNRNA_METABOLIC_PROCESS | 50 | -0.3469 | -1.52054 | 0.025125628 | 0.1570784 | 1 | 3954 | tags=24%, list=17%, si | MM_2 |
| GOBP_NEGATIVE_REGUL | GOBP_NEGATIVE_REGULATION_OF_CELL_G | 184 | -0.27376 | -1.52031 | 0.0069444445 | 0.15708853 | 1 | 4003 | tags=30%, list=17%, si | MM_2 |
| GOBP_ACETYL_COA_BIOS | GOBP_ACETYL_COA_BIOSYNTHETIC_PROCES | 19 | -0.45499 | -1.51913 | 0.07363421 | 0.15791203 | 1 | 5613 | tags=53%, list=24%, si | MM_2 |
| GOBP_NEGATIVE_REGUL | GOBP_NEGATIVE_REGULATION_OF_B_CELL | 16 | -0.48612 | -1.51862 | 0.065727696 | 0.15809102 | 1 | 4172 | tags=38%, list=18%, si | MM_2 |
| GOBP_APOPTOTIC_MITO | GOBP_APOPTOTIC_MITOCHONDRIAL_CHAN | 118 | -0.29106 | -1.51831 | 0.01511335 | 0.15813076 | 1 | 4893 | tags=29%, list=21%, si | MM_2 |
| GLYCOSAMINOGLYCAN_I | GLYCOSAMINOGLYCAN_DEGRADATION | 17 | -0.47301 | -1.518 | 0.07476635 | 0.15813571 | 1 | 1951 | tags=35%, list=8%, sig | MM_2 |
| GOBP_NEGATIVE_REGUL | GOBP_NEGATIVE_REGULATION_OF_CELL_A | 207 | -0.26322 | -1.51705 | 0 | 0.15872493 | 1 | 3342 | tags=21%, list=14%, si | MM_2 |
| B_CELL_RECEPTOR_SIGN/ | B_CELL_RECEPTOR_SIGNALING_PATHWAY | 68 | -0.32433 | -1.51644 | 0.024038462 | 0.15900128 | 1 | 5254 | tags=34%, list=22%, si | MM_2 |
| GOBP_NEGATIVE_REGUL | GOBP_NEGATIVE_REGULATION_OF_SMOOT | 17 | -0.46373 | -1.51568 | 0.068292685 | 0.15945074 | 1 | 4557 | tags=35%, list=19%, si | MM_2 |
| GOBP_REGULATION_OF_ | GOBP_REGULATION_OF_CELL_CYCLE_G1_S_ | 185 | -0.26753 | -1.51525 | 0.0072289156 | 0.15962236 | 1 | 4873 | tags=32%, list=21%, si | MM_2 |
| GOBP_REGULATION_OF_ | GOBP_REGULATION_OF_MONOCYTE_CHEM | 29 | -0.40397 | -1.51524 | 0.052896727 | 0.15936394 | 1 | 5635 | tags=52%, list=24%, si | MM_2 |
| GOBP_ESTABLISHMENT_( | GOBP_ESTABLISHMENT_OR_MAINTENANCE | 51 | -0.34891 | -1.51458 | 0.03301887 | 0.15968807 | 1 | 4765 | tags=31%, list=20%, si | MM_2 |
| GOBP_RESPONSE_TO_HY | GOBP_RESPONSE_TO_HYDROGEN_PEROXID | 85 | -0.30552 | -1.51445 | 0.021791767 | 0.15957807 | 1 | 6784 | tags=49%, list=29%, si | MM_2 |
| GOBP_SOMITOGENESIS | GOBP_SOMITOGENESIS | 63 | -0.32906 | -1.51422 | 0.024038462 | 0.15953442 | 1 | 4018 | tags=30%, list=17%, si | MM_2 |
| AGE-RAGE_SIGNALING_P | AGE-RAGE_SIGNALING_PATHWAY_IN_DIABI | 100 | -0.29928 | -1.51355 | 0.011574074 | 0.159935 | 1 | 3659 | tags=28%, list=15%, si | MM_2 |
| GOBP_TRANSLATIONAL_I | GOBP_TRANSLATIONAL_ELONGATION | 58 | -0.3342 | -1.5123 | 0.017721519 | 0.16087198 | 1 | 5703 | tags=36%, list=24%, si | MM_2 |
| GOBP_PROTEIN_COMPLE | GOBP_PROTEIN_COMPLEX_OLIGOMERIZATI | 249 | -0.25436 | -1.51219 | 0.004938272 | 0.16071521 | 1 | 4843 | tags=33%, list=21%, si | MM_2 |
| GOBP_RESOLUTION_OF_ | GOBP_RESOLUTION_OF_MEIOTIC_RECOMBI | 17 | -0.46523 | -1.51171 | 0.07126437 | 0.16090511 | 1 | 4836 | tags=53%, list=20%, si | MM_2 |
| GOBP_HEMATOPOIETIC_! | GOBP_HEMATOPOIETIC_STEM_CELL_HOME | 29 | -0.40549 | -1.51164 | 0.050239235 | 0.1607155 | 1 | 6668 | tags=62%, list=28%, si | MM_2 |
| GOBP_T_CELL_APOPTOTI | GOBP_T_CELL_APOPTOTIC_PROCESS | 67 | -0.32945 | -1.51083 | 0.03640777 | 0.16119422 | 1 | 4172 | tags=28%, list=18%, si | MM_2 |
| GOBP_REGULATION_OF_ | GOBP_REGULATION_OF_CYCLIN_DEPENDEN | 67 | -0.32653 | -1.51023 | 0.026506023 | 0.16147853 | 1 | 4954 | tags=43%, list=21%, si | MM_2 |
| GOBP_REGULATION_OF_ | GOBP_REGULATION_OF_NECROPTOTIC_PR( | 22 | -0.42474 | -1.50977 | 0.07160494 | 0.16163759 | 1 | 1698 | tags=18%, list=7%, sig | MM_2 |
| GOBP_VESICLE_ORGANIZ | GOBP_VESICLE_ORGANIZATION | 353 | -0.24668 | -1.50899 | 0 | 0.16201432 | 1 | 5132 | tags=30%, list=22%, si | MM_2 |
| GOBP_PROTEIN_HETERO( | GOBP_PROTEIN_HETEROOLIGOMERIZATION | 27 | -0.40438 | -1.50872 | 0.059101656 | 0.16197883 | 1 | 3086 | tags=30%, list=13%, si | MM_2 |
| GOBP_EXTRACELLULAR_I | GOBP_EXTRACELLULAR_MATRIX_DISASSEM | 32 | -0.39329 | -1.50844 | 0.050251257 | 0.16199 | 1 | 1829 | tags=22%, list=8%, sig | MM_2 |
| GOBP_POSITIVE_REGULA | GOBP_POSITIVE_REGULATION_OF_CALCIUN | 52 | -0.34296 | -1.5083 | 0.032786883 | 0.16186999 | 1 | 4501 | tags=35%, list=19%, si | MM_2 |
| GOBP_NEGATIVE_REGUL | GOBP_NEGATIVE_REGULATION_OF_MUSCL | 67 | -0.32412 | -1.50797 | 0.0275 | 0.16191134 | 1 | 4557 | tags=34%, list=19%, si | MM_2 |
| GOBP_REGULATION_OF_ | GOBP_REGULATION_OF_MRNA_CATABOLIC | 175 | -0.27505 | -1.50783 | 0.0049261083 | 0.16178225 | 1 | 5232 | tags=31%, list=22%, si | MM_2 |
| GOBP_ESTABLISHMENT_( | GOBP_ESTABLISHMENT_OF_ORGANELLE_LC | 472 | -0.23592 | -1.50769 | 0 | 0.16164212 | 1 | 4290 | tags=24%, list=18%, si | MM_2 |
| GOBP_LIPID_CATABOLIC_ | GOBP_LIPID_CATABOLIC_PROCESS | 304 | -0.24943 | -1.50751 | 0.002375297 | 0.16153298 | 1 | 4156 | tags=26%, list=18%, si | MM_2 |
| GOBP_DEFENSE_RESPON | GOBP_DEFENSE_RESPONSE_TO_GRAM_NE( | 68 | -0.32002 | -1.50658 | 0.03373494 | 0.16211693 | 1 | 2877 | tags=19%, list=12%, si | MM_2 |
| GOBP_REGULATION_OF_ | GOBP_REGULATION_OF_CELL_CELL_ADHESI | 475 | -0.23702 | -1.50524 | 0.0023696683 | 0.16310745 | 1 | 3670 | tags=21%, list=16%, si | MM_2 |
| GOBP_VACUOLAR_LOCAL | GOBP_VACUOLAR_LOCALIZATION | 85 | -0.30859 | -1.50506 | 0.021686748 | 0.16302986 | 1 | 5254 | tags=33%, list=22%, si | MM_2 |
| GOBP_CELL_ADHESION_I | GOBP_CELL_ADHESION_MEDIATED_BY_INTI | 86 | -0.31487 | -1.50475 | 0.030373832 | 0.16305687 | 1 | 3602 | tags=23%, list=15%, si | MM_2 |
| GOBP_BASEMENT_MEMI | GOBP_BASEMENT_MEMBRANE_ORGANIZAT | 40 | -0.36903 | -1.50466 | 0.04157044 | 0.1628637 | 1 | 3979 | tags=33%, list=17%, si | MM_2 |
| GOBP_NUCLEAR_TRANSP | GOBP_NUCLEAR_TRANSPORT | 337 | -0.24853 | -1.50374 | 0.002415459 | 0.16341752 | 1 | 5351 | tags=31%, list=23%, si | MM_2 |
| GOBP_POSITIVE_REGULA | GOBP_POSITIVE_REGULATION_OF_METAPH | 15 | -0.48276 | -1.50373 | 0.06407323 | 0.16317084 | 1 | 3916 | tags=33%, list=17%, si | MM_2 |
| GOBP_SMALL_GTPASE_M | GOBP_SMALL_GTPASE_MEDIATED_SIGNAL_ | 434 | -0.2367 | -1.50327 | 0 | 0.1633396 | 1 | 5372 | tags=31%, list=23%, si | MM_2 |
| GOBP_REGULATORY_NCF | GOBP_REGULATORY_NCRNA_MEDIATED_GI | 116 | -0.29363 | -1.50321 | 0.024886878 | 0.16313803 | 1 | 5232 | tags=34%, list=22%, si | MM_2 |
| GOBP_REGULATION_OF_ | GOBP_REGULATION_OF_PROTEIN_POLYMEI | 194 | -0.26411 | -1.50279 | 0.009070295 | 0.16323718 | 1 | 4706 | tags=27%, list=20%, si | MM_2 |
| GOBP_REGULATION_OF_ | GOBP_REGULATION_OF_SMALL_GTPASE_M | 242 | -0.25574 | -1.50261 | 0.009478673 | 0.16314065 | 1 | 5310 | tags=33%, list=22%, si | MM_2 |
| GOBP_REGULATION_OF_ | GOBP_REGULATION_OF_RUFFLE_ASSEMBLY | 29 | -0.39542 | -1.50121 | 0.07 | 0.16414572 | 1 | 3484 | tags=31%, list=15%, si | MM_2 |
| GOBP_REGULATION_OF_ | GOBP_REGULATION_OF_GLIAL_CELL_APOPT | 15 | -0.48158 | -1.50072 | 0.059241705 | 0.16431516 | 1 | 4831 | tags=47%, list=20%, si | MM_2 |
| GOBP_NEGATIVE_REGUL | GOBP_NEGATIVE_REGULATION_OF_UBIQUI | 18 | -0.44889 | -1.50004 | 0.07865169 | 0.16468827 | 1 | 2472 | tags=28%, list=10%, si | MM_2 |
| GOBP_CELLULAR_RESPON | GOBP_CELLULAR_RESPONSE_TO_TYPE_II_IN | 95 | -0.29968 | -1.49986 | 0.011792453 | 0.16462274 | 1 | 4773 | tags=36%, list=20%, si | MM_2 |
| GOBP_POSITIVE_REGULA | GOBP_POSITIVE_REGULATION_OF_PROTEIN | 201 | -0.26534 | -1.49958 | 0.0024271845 | 0.16464247 | 1 | 3875 | tags=23%, list=16%, si | MM_2 |
| GOBP_HOMOTYPIC_CELL | GOBP_HOMOTYPIC_CELL_CELL_ADHESION | 85 | -0.3054 | -1.4994 | 0.011904762 | 0.16452032 | 1 | 3017 | tags=19%, list=13%, si | MM_2 |
| GOBP_RRNA_MODIFICAT | GOBP_RRNA_MODIFICATION | 29 | -0.40413 | -1.49908 | 0.042986427 | 0.16458853 | 1 | 1678 | tags=21%, list=7%, sig | MM_2 |
| GOBP_REGULATION_OF_ | GOBP_REGULATION_OF_FATTY_ACID_OXID/ | 41 | -0.36304 | -1.49863 | 0.041463416 | 0.16475353 | 1 | 4077 | tags=44%, list=17%, si | MM_2 |
| GOBP_NEGATIVE_REGUL | GOBP_NEGATIVE_REGULATION_OF_PEPTID | 18 | -0.45329 | -1.49802 | 0.07760532 | 0.16511609 | 1 | 2821 | tags=28%, list=12%, si | MM_2 |
| GOBP_MEMBRANE_LIPID | GOBP_MEMBRANE_LIPID_METABOLIC_PRO | 186 | -0.2659 | -1.49788 | 0.009569378 | 0.165009 | 1 | 4373 | tags=28%, list=19%, si | MM_2 |
| GOBP_NEGATIVE_REGUL | GOBP_NEGATIVE_REGULATION_OF_NON_C | 26 | -0.40993 | -1.49755 | 0.06904762 | 0.16506036 | 1 | 6702 | tags=58%, list=28%, si | MM_2 |
| GOBP_TOLL_LIKE_RECEPT | GOBP_TOLL_LIKE_RECEPTOR_SIGNALING_P/ | 71 | -0.32043 | -1.49755 | 0.035714287 | 0.16480815 | 1 | 2448 | tags=17%, list=10%, si | MM_2 |
| GOBP_POTASSIUM_ION_ | GOBP_POTASSIUM_ION_HOMEOSTASIS | 30 | -0.40046 | -1.49717 | 0.042713568 | 0.16491804 | 1 | 2166 | tags=30%, list=9%, sig | MM_2 |
| GOBP_NEGATIVE_REGUL | GOBP_NEGATIVE_REGULATION_OF_MUSCL | 35 | -0.37915 | -1.49532 | 0.050480768 | 0.16653278 | 1 | 4655 | tags=46%, list=20%, si | MM_2 |
| GOBP_ESTABLISHMENT_( | GOBP_ESTABLISHMENT_OF_EPITHELIAL_CEI | 36 | -0.37717 | -1.49444 | 0.046296295 | 0.16709551 | 1 | 4674 | tags=33%, list=20%, si | MM_2 |
| GOBP_GMP_BIOSYNTHET | GOBP_GMP_BIOSYNTHETIC_PROCESS | 15 | -0.47417 | -1.49371 | 0.06407323 | 0.16749848 | 1 | 3359 | tags=27%, list=14%, si | MM_2 |
| GOBP_POSITIVE_REGULA | GOBP_POSITIVE_REGULATION_OF_VIRAL_P | 64 | -0.32419 | -1.49335 | 0.033653848 | 0.16757822 | 1 | 5900 | tags=38%, list=25%, si | MM_2 |
| GOBP_POSITIVE_REGULA | GOBP_POSITIVE_REGULATION_OF_NATURA | 26 | -0.40649 | -1.49193 | 0.049763035 | 0.168708 | 1 | 4065 | tags=38%, list=17%, si | MM_2 |
| GOBP_INSULIN_RECEPTO | GOBP_INSULIN_RECEPTOR_SIGNALING_PAT | 133 | -0.2852 | -1.49158 | 0.015267176 | 0.16878621 | 1 | 4413 | tags=29%, list=19%, si | MM_2 |
| GOBP_KILLING_OF_CELLS | GOBP_KILLING_OF_CELLS_OF_ANOTHER_OI | 71 | -0.31891 | -1.49158 | 0.02189781 | 0.16853145 | 1 | 1415 | tags=11%, list=6%, sig | MM_2 |
| GOBP_NUCLEUS_ORGAN | GOBP_NUCLEUS_ORGANIZATION | 136 | -0.27719 | -1.49147 | 0.021686748 | 0.16837446 | 1 | 3970 | tags=24%, list=17%, si | MM_2 |
| GOBP_INTERSTRAND_CR( | GOBP_INTERSTRAND_CROSS_LINK_REPAIR | 31 | -0.39808 | -1.49098 | 0.056790125 | 0.16859268 | 1 | 2480 | tags=26%, list=11%, si | MM_2 |
| GOBP_PEPTIDYL_SERINE_ | GOBP_PEPTIDYL_SERINE_MODIFICATION | 286 | -0.25051 | -1.49085 | 0.004819277 | 0.16847356 | 1 | 3013 | tags=17%, list=13%, si | MM_2 |
| GOBP_CELLULAR_RESPON | GOBP_CELLULAR_RESPONSE_TO_NUTRIENT | 213 | -0.25808 | -1.49064 | 0.008810572 | 0.16841589 | 1 | 4337 | tags=28%, list=18%, si | MM_2 |

| | | | | | | | | | | |
|---|---|---|---|---|---|---|---|---|---|---|
| GOBP_POSITIVE_REGULA | GOBP_POSITIVE_REGULATION_OF_LIPID_ST | 25 | -0.4206 | -1.49046 | 0.059382424 | 0.16834015 | 1 | 4943 | tags=40%, list=21%, si | MM_2 |
| GOBP_REGULATION_OF_ | GOBP_REGULATION_OF_ACTIN_FILAMENT_ | 387 | -0.2392 | -1.49036 | 0 | 0.16819249 | 1 | 2977 | tags=18%, list=13%, si | MM_2 |
| GOBP_ADIPOSE_TISSUE_ | GOBP_ADIPOSE_TISSUE_DEVELOPMENT | 64 | -0.32623 | -1.48992 | 0.025761124 | 0.16839513 | 1 | 5201 | tags=33%, list=22%, si | MM_2 |
| GOBP_REGULATION_OF_ | GOBP_REGULATION_OF_LYMPHOCYTE_DIFF | 216 | -0.25786 | -1.48991 | 0 | 0.16815074 | 1 | 3670 | tags=23%, list=16%, si | MM_2 |
| GOBP_MELANOCYTE_DIF | GOBP_MELANOCYTE_DIFFERENTIATION | 26 | -0.40625 | -1.48944 | 0.053398058 | 0.16836132 | 1 | 1984 | tags=23%, list=8%, sig | MM_2 |
| GOBP_REGULATION_OF_ | GOBP_REGULATION_OF_PROTEASOMAL_UE | 118 | -0.28532 | -1.489 | 0.021077283 | 0.16854134 | 1 | 4904 | tags=27%, list=21%, si | MM_2 |
| GOBP_MICROTUBULE_PO | GOBP_MICROTUBULE_POLYMERIZATION_O | 133 | -0.28051 | -1.48882 | 0.009852217 | 0.16849284 | 1 | 4489 | tags=34%, list=19%, si | MM_2 |
| GOBP_NEGATIVE_REGUL | GOBP_NEGATIVE_REGULATION_OF_CELL_C | 197 | -0.26013 | -1.48759 | 0.0073710075 | 0.16941488 | 1 | 3571 | tags=22%, list=15%, si | MM_2 |
| GOBP_PROTEIN_CATABO | GOBP_PROTEIN_CATABOLIC_PROCESS_IN_T | 22 | -0.43484 | -1.48709 | 0.06082725 | 0.16969258 | 1 | 6076 | tags=45%, list=26%, si | MM_2 |
| PROTEASOME | PROTEASOME | 45 | -0.35696 | -1.48706 | 0.05882353 | 0.16945936 | 1 | 12780 | tags=96%, list=54%, si | MM_2 |
| GOBP_INTEGRIN_ACTIVA | GOBP_INTEGRIN_ACTIVATION | 26 | -0.40488 | -1.48681 | 0.054117646 | 0.16945973 | 1 | 4383 | tags=31%, list=19%, si | MM_2 |
| GOBP_PROTEIN_STABILIZ | GOBP_PROTEIN_STABILIZATION | 190 | -0.26243 | -1.48666 | 0.009615385 | 0.16934797 | 1 | 3611 | tags=25%, list=15%, si | MM_2 |
| GOBP_NEGATIVE_REGUL | GOBP_NEGATIVE_REGULATION_OF_ERBB_S | 32 | -0.39003 | -1.48657 | 0.039387308 | 0.16915767 | 1 | 4513 | tags=41%, list=19%, si | MM_2 |
| GOBP_MACROPHAGE_AC | GOBP_MACROPHAGE_ACTIVATION_INVOLV | 20 | -0.43302 | -1.48641 | 0.060096152 | 0.1690847 | 1 | 4244 | tags=30%, list=18%, si | MM_2 |
| GOBP_MITOTIC_CYTOKIN | GOBP_MITOTIC_CYTOKINETIC_PROCESS | 30 | -0.38309 | -1.48545 | 0.042154565 | 0.16980486 | 1 | 3795 | tags=23%, list=16%, si | MM_2 |
| GOBP_PROTEIN_QUALITY | GOBP_PROTEIN_QUALITY_CONTROL_FOR_M | 17 | -0.45624 | -1.48511 | 0.08674699 | 0.1698723 | 1 | 12846 | tags=100%, list=54%, | MM_2 |
| GOBP_POSITIVE_REGULA | GOBP_POSITIVE_REGULATION_OF_OSTEOCI | 36 | -0.36388 | -1.48458 | 0.04589372 | 0.17017987 | 1 | 4017 | tags=25%, list=17%, si | MM_2 |
| GOBP_RESPONSE_TO_PR | GOBP_RESPONSE_TO_PROTOZOAN | 44 | -0.34578 | -1.4836 | 0.04918033 | 0.17090145 | 1 | 1959 | tags=27%, list=8%, sig | MM_2 |
| GOBP_REGULATION_OF_ | GOBP_REGULATION_OF_EPIDERMAL_GROW | 20 | -0.43343 | -1.48356 | 0.06382979 | 0.17068477 | 1 | 4432 | tags=40%, list=19%, si | MM_2 |
| GOBP_REGULATION_OF_ | GOBP_REGULATION_OF_POSTSYNAPSE_ASS | 41 | -0.3639 | -1.48286 | 0.058536585 | 0.17119129 | 1 | 3127 | tags=24%, list=13%, si | MM_2 |
| GOBP_OSSIFICATION | GOBP_OSSIFICATION | 400 | -0.23849 | -1.48154 | 0 | 0.17216739 | 1 | 3755 | tags=23%, list=16%, si | MM_2 |
| GOBP_CELLULAR_RESPON | GOBP_CELLULAR_RESPONSE_TO_FATTY_AC | 30 | -0.38375 | -1.48112 | 0.066176474 | 0.17232409 | 1 | 380 | tags=13%, list=2%, sig | MM_2 |
| PROGESTERONE-MEDIATI | PROGESTERONE-MEDIATED_OOCYTE_MATU | 87 | -0.30313 | -1.48073 | 0.018779343 | 0.17244542 | 1 | 5211 | tags=32%, list=22%, si | MM_2 |
| GOBP_LYMPHOCYTE_ACT | GOBP_LYMPHOCYTE_ACTIVATION_INVOLVE | 202 | -0.25966 | -1.48047 | 0.0024570024 | 0.17240055 | 1 | 3232 | tags=18%, list=14%, si | MM_2 |
| GOBP_MYELOID_LEUKOC | GOBP_MYELOID_LEUKOCYTE_MIGRATION | 222 | -0.25403 | -1.47909 | 0.004750594 | 0.17344712 | 1 | 3967 | tags=24%, list=17%, si | MM_2 |
| GOBP_CELLULAR_EXTRAV | GOBP_CELLULAR_EXTRAVASATION | 74 | -0.31527 | -1.47909 | 0.055023924 | 0.1731945 | 1 | 3276 | tags=28%, list=14%, si | MM_2 |
| GOBP_SMOOTHENED_SIG | GOBP_SMOOTHENED_SIGNALING_PATHWA | 154 | -0.27124 | -1.47883 | 0.009638554 | 0.17319827 | 1 | 3316 | tags=25%, list=14%, si | MM_2 |
| GOBP_NUCLEAR_MIGRAT | GOBP_NUCLEAR_MIGRATION | 32 | -0.37815 | -1.47839 | 0.058139537 | 0.17330633 | 1 | 455 | tags=9%, list=2%, sign | MM_2 |
| GOBP_ACTOMYOSIN_STR | GOBP_ACTOMYOSIN_STRUCTURE_ORGANIZ | 219 | -0.25559 | -1.47787 | 0.0069444445 | 0.17354673 | 1 | 3783 | tags=25%, list=16%, si | MM_2 |
| GOBP_PEROXISOMAL_TR | GOBP_PEROXISOMAL_TRANSPORT | 21 | -0.43045 | -1.47693 | 0.0883721 | 0.17417909 | 1 | 3901 | tags=38%, list=17%, si | MM_2 |
| GOBP_CHROMOSOME_O | GOBP_CHROMOSOME_ORGANIZATION_INV | 65 | -0.32542 | -1.47683 | 0.025252525 | 0.17401932 | 1 | 3014 | tags=26%, list=13%, si | MM_2 |
| GOBP_MACROPHAGE_CH | GOBP_MACROPHAGE_CHEMOTAXIS | 45 | -0.34628 | -1.47616 | 0.037914693 | 0.17443119 | 1 | 3351 | tags=24%, list=14%, si | MM_2 |
| GOBP_POSITIVE_REGULA | GOBP_POSITIVE_REGULATION_OF_NEURON | 15 | -0.46629 | -1.47548 | 0.09245742 | 0.17488295 | 1 | 12608 | tags=100%, list=53%, | MM_2 |
| JAK-STAT_SIGNALING_PA | JAK-STAT_SIGNALING_PATHWAY | 138 | -0.27278 | -1.47514 | 0.0275 | 0.17498326 | 1 | 3514 | tags=25%, list=15%, si | MM_2 |
| GOBP_REGULATION_OF_ | GOBP_REGULATION_OF_SUPRAMOLECULAR | 378 | -0.23909 | -1.4747 | 0.0024937657 | 0.17520936 | 1 | 4637 | tags=25%, list=20%, si | MM_2 |
| GOBP_REGULATION_OF_ | GOBP_REGULATION_OF_SYNAPTIC_VESICLE | 55 | -0.33277 | -1.47417 | 0.0275 | 0.1754783 | 1 | 2556 | tags=25%, list=11%, si | MM_2 |
| GOBP_DISRUPTION_OF_A | GOBP_DISRUPTION_OF_ANATOMICAL_STRU | 73 | -0.31483 | -1.4739 | 0.037128713 | 0.17552558 | 1 | 1415 | tags=11%, list=6%, sig | MM_2 |
| GOBP_REGULATION_OF_ | GOBP_REGULATION_OF_MEMBRANE_LIPID_ | 19 | -0.43225 | -1.47329 | 0.079822615 | 0.17594992 | 1 | 4220 | tags=37%, list=18%, si | MM_2 |
| GOBP_REGULATION_OF_ | GOBP_REGULATION_OF_IMMUNE_EFFECTO | 406 | -0.23262 | -1.47328 | 0.0022883294 | 0.1757197 | 1 | 3276 | tags=19%, list=14%, si | MM_2 |
| GOBP_TYPE_II_INTERFER | GOBP_TYPE_II_INTERFERON_PRODUCTION | 132 | -0.27576 | -1.47311 | 0.015037594 | 0.17561886 | 1 | 3534 | tags=23%, list=15%, si | MM_2 |
| GOBP_DETECTION_OF_EX | GOBP_DETECTION_OF_EXTERNAL_BIOTIC_S | 27 | -0.40128 | -1.47305 | 0.050724637 | 0.17543381 | 1 | 3627 | tags=26%, list=15%, si | MM_2 |
| GOBP_REGULATION_OF_ | GOBP_REGULATION_OF_NOTCH_SIGNALING | 92 | -0.29167 | -1.47298 | 0.03521127 | 0.17526168 | 1 | 3452 | tags=27%, list=15%, si | MM_2 |
| GOBP_CELLULAR_SENESC | GOBP_CELLULAR_SENESCENCE | 88 | -0.2999 | -1.47282 | 0.02606635 | 0.1751702 | 1 | 3743 | tags=26%, list=16%, si | MM_2 |
| GOBP_REGULATORY_T_C | GOBP_REGULATORY_T_CELL_DIFFERENTIAT | 41 | -0.35642 | -1.47195 | 0.046189375 | 0.17572653 | 1 | 4917 | tags=44%, list=21%, si | MM_2 |
| GOBP_NEGATIVE_REGUL | GOBP_NEGATIVE_REGULATION_OF_RESPON | 112 | -0.29031 | -1.47174 | 0.027989822 | 0.17570819 | 1 | 3123 | tags=24%, list=13%, si | MM_2 |
| GOBP_REGULATION_OF_ | GOBP_REGULATION_OF_DNA_BIOSYNTHETI | 121 | -0.27951 | -1.4715 | 0.009638554 | 0.17571022 | 1 | 4572 | tags=26%, list=19%, si | MM_2 |
| GOBP_EXECUTION_PHAS | GOBP_EXECUTION_PHASE_OF_APOPTOSIS | 78 | -0.31048 | -1.46933 | 0.026442308 | 0.17756698 | 1 | 4243 | tags=27%, list=18%, si | MM_2 |
| GOBP_LEUKOCYTE_MEDI | GOBP_LEUKOCYTE_MEDIATED_IMMUNITY | 400 | -0.23197 | -1.46925 | 0.0023148148 | 0.17740983 | 1 | 3308 | tags=18%, list=14%, si | MM_2 |
| GOBP_POSITIVE_REGULA | GOBP_POSITIVE_REGULATION_OF_PROTEIN | 87 | -0.30241 | -1.46878 | 0.04157044 | 0.17762719 | 1 | 3875 | tags=25%, list=16%, si | MM_2 |
| PYRIMIDINE_METABOLISI | PYRIMIDINE_METABOLISM | 53 | -0.33202 | -1.46806 | 0.0311804 | 0.1781033 | 1 | 4531 | tags=36%, list=19%, si | MM_2 |
| GOBP_RHO_PROTEIN_SIG | GOBP_RHO_PROTEIN_SIGNAL_TRANSDUCTI | 142 | -0.27154 | -1.46791 | 0.024570024 | 0.17799993 | 1 | 3807 | tags=23%, list=16%, si | MM_2 |
| GOBP_MEMBRANE_LIPID | GOBP_MEMBRANE_LIPID_BIOSYNTHETIC_P | 129 | -0.27724 | -1.46714 | 0.00921659 | 0.17844674 | 1 | 3755 | tags=26%, list=16%, si | MM_2 |
| GOBP_FIBROBLAST_PROL | GOBP_FIBROBLAST_PROLIFERATION | 147 | -0.26894 | -1.4666 | 0.013793103 | 0.1786596 | 1 | 4109 | tags=26%, list=17%, si | MM_2 |
| GOBP_T_CELL_DIFFERENT | GOBP_T_CELL_DIFFERENTIATION_IN_THYM | 103 | -0.29164 | -1.46579 | 0.030444965 | 0.17927195 | 1 | 4887 | tags=37%, list=21%, si | MM_2 |
| GOBP_GASTRULATION | GOBP_GASTRULATION | 172 | -0.26399 | -1.46538 | 0.009411764 | 0.17947942 | 1 | 5965 | tags=38%, list=25%, si | MM_2 |
| GOBP_POSITIVE_REGULA | GOBP_POSITIVE_REGULATION_OF_MIRNA_ | 64 | -0.31645 | -1.46464 | 0.037037037 | 0.17993523 | 1 | 3726 | tags=27%, list=16%, si | MM_2 |
| GOBP_PLATELET_ACTIVAT | GOBP_PLATELET_ACTIVATION | 94 | -0.29787 | -1.46458 | 0.037470724 | 0.17972738 | 1 | 5104 | tags=32%, list=22%, si | MM_2 |
| GOBP_NEGATIVE_REGUL | GOBP_NEGATIVE_REGULATION_OF_HOMOT | 16 | -0.46566 | -1.4638 | 0.06511628 | 0.1802545 | 1 | 4982 | tags=44%, list=21%, si | MM_2 |
| GOBP_RESPONSE_TO_DE | GOBP_RESPONSE_TO_DEXAMETHASONE | 27 | -0.38939 | -1.46319 | 0.062211983 | 0.18058652 | 1 | 3521 | tags=26%, list=15%, si | MM_2 |
| CELLULAR_SENESCENCE | CELLULAR_SENESCENCE | 159 | -0.26782 | -1.46305 | 0.022222223 | 0.1804815 | 1 | 5054 | tags=33%, list=21%, si | MM_2 |
| GOBP_REGULATION_OF_ | GOBP_REGULATION_OF_SPINDLE_ORGANIZ | 42 | -0.36065 | -1.46297 | 0.046403714 | 0.18028435 | 1 | 3795 | tags=26%, list=16%, si | MM_2 |
| GOBP_T_HELPER_2_CELL | GOBP_T_HELPER_2_CELL_CYTOKINE_PRODU | 17 | -0.44837 | -1.46265 | 0.06954437 | 0.1803563 | 1 | 811 | tags=18%, list=3%, sig | MM_2 |
| GLYCEROPHOSPHOLIPID_ | GLYCEROPHOSPHOLIPID_METABOLISM | 92 | -0.29604 | -1.46254 | 0.025316456 | 0.18022323 | 1 | 4265 | tags=27%, list=18%, si | MM_2 |
| GOBP_POSITIVE_REGULA | GOBP_POSITIVE_REGULATION_OF_MYELOID | 21 | -0.43294 | -1.46154 | 0.07263923 | 0.18107662 | 1 | 3276 | tags=33%, list=14%, si | MM_2 |
| GOBP_POSITIVE_REGULA | GOBP_POSITIVE_REGULATION_OF_BINDING | 147 | -0.26838 | -1.46121 | 0.00729927 | 0.18115143 | 1 | 5092 | tags=31%, list=22%, si | MM_2 |
| GOBP_HOMOLOGOUS_CI | GOBP_HOMOLOGOUS_CHROMOSOME_SEG | 66 | -0.31606 | -1.4603 | 0.04513064 | 0.18185496 | 1 | 3014 | tags=26%, list=13%, si | MM_2 |
| GOBP_PEPTIDYL_THREON | GOBP_PEPTIDYL_THREONINE_MODIFICATIO | 110 | -0.27939 | -1.4598 | 0.01909308 | 0.18216302 | 1 | 3166 | tags=23%, list=13%, si | MM_2 |
| RELAXIN_SIGNALING_PAT | RELAXIN_SIGNALING_PATHWAY | 125 | -0.28203 | -1.45979 | 0.031325303 | 0.1819232 | 1 | 3555 | tags=26%, list=15%, si | MM_2 |
| GOBP_REGULATION_OF_ | GOBP_REGULATION_OF_NON_CANONICAL_ | 92 | -0.29288 | -1.45939 | 0.021276595 | 0.18203564 | 1 | 6591 | tags=41%, list=28%, si | MM_2 |
| GOBP_NEGATIVE_REGUL | GOBP_NEGATIVE_REGULATION_OF_TYPE_I_ | 42 | -0.34719 | -1.45886 | 0.05882353 | 0.18236555 | 1 | 4272 | tags=33%, list=18%, si | MM_2 |
| GOBP_RESPONSE_TO_AN | GOBP_RESPONSE_TO_ANGIOTENSIN | 33 | -0.36705 | -1.45862 | 0.062650606 | 0.18236415 | 1 | 3509 | tags=27%, list=15%, si | MM_2 |
| GOBP_NEGATIVE_REGUL | GOBP_NEGATIVE_REGULATION_OF_VASCUL | 109 | -0.28322 | -1.45855 | 0.029545454 | 0.18219034 | 1 | 3560 | tags=27%, list=15%, si | MM_2 |
| PARKINSON_DISEASE | PARKINSON_DISEASE | 122 | -0.27815 | -1.45818 | 0.009433962 | 0.18227598 | 1 | 7142 | tags=45%, list=30%, si | MM_2 |
| GOBP_POSITIVE_REGULA | GOBP_POSITIVE_REGULATION_OF_PROTEIN | 24 | -0.41733 | -1.45812 | 0.055690072 | 0.1820681 | 1 | 4816 | tags=38%, list=20%, si | MM_2 |
| GOBP_DEFENSE_RESPON | GOBP_DEFENSE_RESPONSE_TO_BACTERIUM | 252 | -0.24835 | -1.45799 | 0.004750594 | 0.18197073 | 1 | 2772 | tags=18%, list=12%, si | MM_2 |
| GOBP_RESPONSE_TO_EP | GOBP_RESPONSE_TO_EPIDERMAL_GROWTH | 42 | -0.35021 | -1.45796 | 0.05299539 | 0.18174002 | 1 | 6259 | tags=48%, list=27%, si | MM_2 |
| GOBP_REGULATION_OF_ | GOBP_REGULATION_OF_CELL_CYCLE_G2_M | 117 | -0.27766 | -1.45793 | 0.018867925 | 0.18152964 | 1 | 3105 | tags=22%, list=13%, si | MM_2 |
| GOBP_POSITIVE_REGULA | GOBP_POSITIVE_REGULATION_OF_LYMPHO | 21 | -0.42212 | -1.45676 | 0.07191011 | 0.1824766 | 1 | 4677 | tags=29%, list=20%, si | MM_2 |
| CYTOKINE-CYTOKINE_REC | CYTOKINE-CYTOKINE_RECEPTOR_INTERACTI | 239 | -0.24843 | -1.45624 | 0.00913242 | 0.18278316 | 1 | 3938 | tags=24%, list=17%, si | MM_2 |
| GOBP_SPHINGOLIPID_ME | GOBP_SPHINGOLIPID_METABOLIC_PROCESS | 137 | -0.26252 | -1.45584 | 0.009975063 | 0.18297131 | 1 | 3755 | tags=25%, list=16%, si | MM_2 |
| GOBP_CELL_ACTIVATION | GOBP_CELL_ACTIVATION_INVOLVED_IN_IM | 291 | -0.24076 | -1.45541 | 0 | 0.18318217 | 1 | 3776 | tags=20%, list=16%, si | MM_2 |
| GOBP_REGULATION_OF_ | GOBP_REGULATION_OF_MACROPHAGE_DE | 25 | -0.39709 | -1.45502 | 0.060324825 | 0.1833511 | 1 | 480 | tags=12%, list=2%, sig | MM_2 |

| | | | | | | | | | | |
|---|---|---|---|---|---|---|---|---|---|---|
| GLYCOSAMINOGLYCAN_E | GLYCOSAMINOGLYCAN_BIOSYNTHESIS | 51 | -0.33966 | -1.45476 | 0.046004843 | 0.1833642 | 1 | 3741 | tags=27%, list=16%, si | MM_2 |
| GOBP_REGULATION_OF_ | GOBP_REGULATION_OF_VESICLE_SIZE | 21 | -0.42535 | -1.45382 | 0.06904232 | 0.1841403 | 1 | 6399 | tags=48%, list=27%, si | MM_2 |
| BACTERIAL_INVASION_OI | BACTERIAL_INVASION_OF_EPITHELIAL_CELL | 72 | -0.31163 | -1.4529 | 0.02676399 | 0.18485548 | 1 | 6847 | tags=50%, list=29%, si | MM_2 |
| GOBP_POSITIVE_REGULA | GOBP_POSITIVE_REGULATION_OF_MACROF | 31 | -0.38254 | -1.45158 | 0.07619048 | 0.18607289 | 1 | 3754 | tags=32%, list=16%, si | MM_2 |
| GOBP_POSITIVE_REGULA | GOBP_POSITIVE_REGULATION_OF_HEMOP( | 202 | -0.25301 | -1.45154 | 0.0072289156 | 0.18586376 | 1 | 4276 | tags=24%, list=18%, si | MM_2 |
| GOBP_REGULATION_OF_ | GOBP_REGULATION_OF_LEUKOCYTE_MEDIA | 253 | -0.24386 | -1.45131 | 0.00486618 | 0.18585916 | 1 | 3276 | tags=20%, list=14%, si | MM_2 |
| GOBP_ACYL_COA_METAE | GOBP_ACYL_COA_METABOLIC_PROCESS | 82 | -0.2965 | -1.45104 | 0.031476997 | 0.1858853 | 1 | 5054 | tags=35%, list=21%, si | MM_2 |
| GOBP_POTASSIUM_ION_ | GOBP_POTASSIUM_ION_IMPORT_ACROSS_ | 44 | -0.34144 | -1.45042 | 0.056603774 | 0.18628827 | 1 | 2166 | tags=25%, list=9%, sig | MM_2 |
| GOBP_NEUROINFLAMMA | GOBP_NEUROINFLAMMATORY_RESPONSE | 84 | -0.29458 | -1.44997 | 0.033816423 | 0.186449 | 1 | 4262 | tags=31%, list=18%, si | MM_2 |
| GOBP_PYRIMIDINE_CONT | GOBP_PYRIMIDINE_CONTAINING_COMPOU | 32 | -0.37063 | -1.44989 | 0.054245282 | 0.18629162 | 1 | 2501 | tags=25%, list=11%, si | MM_2 |
| GOBP_ANTIMICROBIAL_H | GOBP_ANTIMICROBIAL_HUMORAL_RESPON | 98 | -0.28495 | -1.44931 | 0.03341289 | 0.18660092 | 1 | 2877 | tags=21%, list=12%, si | MM_2 |
| GOBP_REGULATION_OF_ | GOBP_REGULATION_OF_MITOCHONDRIAL_ | 26 | -0.39187 | -1.44926 | 0.089330025 | 0.18640052 | 1 | 3219 | tags=27%, list=14%, si | MM_2 |
| GOBP_POSITIVE_REGULA | GOBP_POSITIVE_REGULATION_OF_DNA_TE | 72 | -0.31228 | -1.4489 | 0.055684455 | 0.18650153 | 1 | 5479 | tags=38%, list=23%, si | MM_2 |
| GOBP_PROTON_TRANSP( | GOBP_PROTON_TRANSPORTING_TWO_SECT | 15 | -0.45679 | -1.44846 | 0.07692308 | 0.18672717 | 1 | 4890 | tags=53%, list=21%, si | MM_2 |
| GOBP_BONE_DEVELOPM | GOBP_BONE_DEVELOPMENT | 222 | -0.24455 | -1.44805 | 0.0074074073 | 0.18689527 | 1 | 1988 | tags=14%, list=8%, sig | MM_2 |
| GOBP_REGULATION_OF_ | GOBP_REGULATION_OF_ACTOMYOSIN_STR | 111 | -0.28192 | -1.44803 | 0.021226415 | 0.18666269 | 1 | 2145 | tags=19%, list=9%, sig | MM_2 |
| GOBP_PROTEIN_TETRAM | GOBP_PROTEIN_TETRAMERIZATION | 82 | -0.29673 | -1.4479 | 0.027027028 | 0.18655728 | 1 | 4263 | tags=34%, list=18%, si | MM_2 |
| GOBP_ENDOPLASMIC_RE | GOBP_ENDOPLASMIC_RETICULUM_ORGANI | 90 | -0.29635 | -1.44759 | 0.023584906 | 0.18666741 | 1 | 4195 | tags=26%, list=18%, si | MM_2 |
| GOBP_NEGATIVE_REGUL | GOBP_NEGATIVE_REGULATION_OF_MITOTI | 230 | -0.24959 | -1.44615 | 0.0069444445 | 0.1878549 | 1 | 4524 | tags=27%, list=19%, si | MM_2 |
| GOBP_PROTEIN_LOCALIZ | GOBP_PROTEIN_LOCALIZATION_TO_VACUO | 78 | -0.3035 | -1.4452 | 0.043583535 | 0.18864465 | 1 | 6076 | tags=37%, list=26%, si | MM_2 |
| SMALL_CELL_LUNG_CANC | SMALL_CELL_LUNG_CANCER | 91 | -0.29516 | -1.44504 | 0.020356234 | 0.18859524 | 1 | 5220 | tags=34%, list=22%, si | MM_2 |
| GOBP_ORGANIC_ACID_C | GOBP_ORGANIC_ACID_CATABOLIC_PROCES | 224 | -0.24862 | -1.44483 | 0.0048076925 | 0.18855731 | 1 | 4514 | tags=30%, list=19%, si | MM_2 |
| GOBP_POSITIVE_REGULA | GOBP_POSITIVE_REGULATION_OF_NF_KAPI | 127 | -0.27341 | -1.44381 | 0.024663677 | 0.1894219 | 1 | 5176 | tags=38%, list=22%, si | MM_2 |
| GOBP_POSITIVE_REGULA | GOBP_POSITIVE_REGULATION_OF_T_CELL_ | 108 | -0.28378 | -1.44355 | 0.032745592 | 0.18946388 | 1 | 3218 | tags=22%, list=14%, si | MM_2 |
| GOBP_REGULATION_OF_ | GOBP_REGULATION_OF_PLATELET_AGGREC | 25 | -0.40669 | -1.44324 | 0.06235566 | 0.1895807 | 1 | 4011 | tags=32%, list=17%, si | MM_2 |
| GOBP_MITOCHONDRIAL_ | GOBP_MITOCHONDRIAL_RNA_PROCESSING | 20 | -0.42322 | -1.44228 | 0.08232445 | 0.1903047 | 1 | 5686 | tags=50%, list=24%, si | MM_2 |
| GOBP_POSITIVE_REGULA | GOBP_POSITIVE_REGULATION_OF_MACROF | 15 | -0.46377 | -1.44196 | 0.08454106 | 0.19038957 | 1 | 480 | tags=13%, list=2%, sig | MM_2 |
| GOBP_PROTEIN_LOCALIZ | GOBP_PROTEIN_LOCALIZATION_TO_ENDOS | 31 | -0.36934 | -1.4414 | 0.06930693 | 0.19080693 | 1 | 6116 | tags=48%, list=26%, si | MM_2 |
| GOBP_BIOMINERAL_TISS | GOBP_BIOMINERAL_TISSUE_DEVELOPMENT | 172 | -0.26006 | -1.44024 | 0.012077294 | 0.19182195 | 1 | 3755 | tags=24%, list=16%, si | MM_2 |
| GOBP_EXTRINSIC_APOPT | GOBP_EXTRINSIC_APOPTOTIC_SIGNALING_I | 233 | -0.24565 | -1.43984 | 0.0071599046 | 0.1920053 | 1 | 5002 | tags=31%, list=21%, si | MM_2 |
| GOBP_NEGATIVE_REGUL | GOBP_NEGATIVE_REGULATION_OF_CELL_A | 301 | -0.23456 | -1.43856 | 0 | 0.19302848 | 1 | 3039 | tags=20%, list=13%, si | MM_2 |
| GOBP_MONOSACCHARID | GOBP_MONOSACCHARIDE_METABOLIC_PR( | 255 | -0.24303 | -1.4374 | 0.012562814 | 0.19399022 | 1 | 4262 | tags=30%, list=18%, si | MM_2 |
| GOBP_CELLULAR_RESPON | GOBP_CELLULAR_RESPONSE_TO_REACTIVE_ | 136 | -0.26659 | -1.43721 | 0.018518519 | 0.19392854 | 1 | 4272 | tags=25%, list=18%, si | MM_2 |
| GOBP_RELEASE_OF_CYTO | GOBP_RELEASE_OF_CYTOCHROME_C_FROM | 66 | -0.3185 | -1.43697 | 0.05352798 | 0.19395569 | 1 | 4893 | tags=33%, list=21%, si | MM_2 |
| GOBP_PRODUCTION_OF_ | GOBP_PRODUCTION_OF_MOLECULAR_MED | 270 | -0.23797 | -1.43695 | 0.012077294 | 0.1937293 | 1 | 3232 | tags=19%, list=14%, si | MM_2 |
| GOBP_NUCLEAR_MEMBR | GOBP_NUCLEAR_MEMBRANE_ORGANIZATI( | 35 | -0.3661 | -1.43693 | 0.07380953 | 0.19349949 | 1 | 3960 | tags=23%, list=17%, si | MM_2 |
| GOBP_REGULATION_OF_ | GOBP_REGULATION_OF_PRODUCTION_OF_ | 195 | -0.25109 | -1.43648 | 0.004555809 | 0.1937116 | 1 | 3232 | tags=19%, list=14%, si | MM_2 |
| GOBP_VENTRICULAR_SYS | GOBP_VENTRICULAR_SYSTEM_DEVELOPME | 36 | -0.35993 | -1.43638 | 0.06896552 | 0.1935728 | 1 | 2166 | tags=19%, list=9%, sig | MM_2 |
| GOBP_REGULATION_OF_ | GOBP_REGULATION_OF_ERBB_SIGNALING_ | 71 | -0.3045 | -1.43588 | 0.043037973 | 0.19379325 | 1 | 4513 | tags=34%, list=19%, si | MM_2 |
| GOBP_MAINTENANCE_OI | GOBP_MAINTENANCE_OF_LOCATION | 341 | -0.23549 | -1.43577 | 0.002570694 | 0.1936422 | 1 | 4972 | tags=31%, list=21%, si | MM_2 |
| GOBP_SISTER_CHROMAT | GOBP_SISTER_CHROMATID_COHESION | 54 | -0.3305 | -1.43563 | 0.03373494 | 0.19351009 | 1 | 2826 | tags=20%, list=12%, si | MM_2 |
| GOBP_CELLULAR_RESPON | GOBP_CELLULAR_RESPONSE_TO_PEPTIDE_I | 276 | -0.24082 | -1.43555 | 0.004672897 | 0.19333501 | 1 | 4337 | tags=25%, list=18%, si | MM_2 |
| GOBP_REGULATION_OF_ | GOBP_REGULATION_OF_PROTEIN_SUMOYL | 25 | -0.39181 | -1.43554 | 0.08728179 | 0.19310221 | 1 | 6918 | tags=52%, list=29%, si | MM_2 |
| GOBP_POSITIVE_REGULA | GOBP_POSITIVE_REGULATION_OF_GTPASE_ | 203 | -0.2491 | -1.4352 | 0.013513514 | 0.19325249 | 1 | 5946 | tags=33%, list=25%, si | MM_2 |
| GOBP_INTERFERON_ALPI | GOBP_INTERFERON_ALPHA_PRODUCTION | 33 | -0.37535 | -1.43458 | 0.07606264 | 0.19368537 | 1 | 1902 | tags=18%, list=8%, sig | MM_2 |
| GOBP_REGULATION_OF_ | GOBP_REGULATION_OF_MITOCHONDRIAL_ | 58 | -0.32197 | -1.43458 | 0.06812652 | 0.19344738 | 1 | 4842 | tags=33%, list=21%, si | MM_2 |
| GOBP_REGULATION_OF_ | GOBP_REGULATION_OF_CHOLESTEROL_STO | 21 | -0.41246 | -1.43442 | 0.098280095 | 0.19336353 | 1 | 3563 | tags=29%, list=15%, si | MM_2 |
| GOBP_REGULATION_OF_ | GOBP_REGULATION_OF_CELL_GROWTH | 415 | -0.2277 | -1.4344 | 0.009367681 | 0.1931339 | 1 | 4035 | tags=26%, list=17%, si | MM_2 |
| GOBP_REGULATION_OF_ | GOBP_REGULATION_OF_CYTOPLASMIC_TRA | 30 | -0.36477 | -1.43426 | 0.07341772 | 0.19299999 | 1 | 15009 | tags=100%, list=64%, | MM_2 |
| GOBP_BONE_TRABECULA | GOBP_BONE_TRABECULA_MORPHOGENESIS | 17 | -0.45117 | -1.43334 | 0.08695652 | 0.19376694 | 1 | 1391 | tags=29%, list=6%, sig | MM_2 |
| GOBP_N_TERMINAL_PRO | GOBP_N_TERMINAL_PROTEIN_AMINO_ACII | 25 | -0.39545 | -1.43308 | 0.0817757 | 0.19376932 | 1 | 6186 | tags=48%, list=26%, si | MM_2 |
| GOBP_DETECTION_OF_O | GOBP_DETECTION_OF_OTHER_ORGANISM | 18 | -0.43799 | -1.43303 | 0.09192825 | 0.19358107 | 1 | 3627 | tags=33%, list=15%, si | MM_2 |
| GOBP_PROTEIN_DNA_CO | GOBP_PROTEIN_DNA_COMPLEX_ASSEMBLY | 154 | -0.2646 | -1.43303 | 0.01703163 | 0.1933489 | 1 | 5507 | tags=33%, list=23%, si | MM_2 |
| GOBP_CEREBRAL_CORTE> | GOBP_CEREBRAL_CORTEX_DEVELOPMENT | 113 | -0.27562 | -1.43166 | 0.026570048 | 0.19455576 | 1 | 3304 | tags=21%, list=14%, si | MM_2 |
| GOBP_REGULATION_OF_ | GOBP_REGULATION_OF_DOUBLE_STRAND_ | 80 | -0.29123 | -1.43108 | 0.062801935 | 0.19496338 | 1 | 6389 | tags=43%, list=27%, si | MM_2 |
| GOBP_MEMBRANE_FISSI | GOBP_MEMBRANE_FISSION | 31 | -0.37318 | -1.43106 | 0.082914576 | 0.1947323 | 1 | 1320 | tags=13%, list=6%, sig | MM_2 |
| GOBP_CELLULAR_RESPON | GOBP_CELLULAR_RESPONSE_TO_ORGANIC_ | 443 | -0.22225 | -1.43103 | 0 | 0.19452737 | 1 | 4017 | tags=23%, list=17%, si | MM_2 |
| GOBP_VASCULAR_ASSOC | GOBP_VASCULAR_ASSOCIATED_SMOOTH_N | 39 | -0.35428 | -1.42983 | 0.08980583 | 0.1956251 | 1 | 4557 | tags=36%, list=19%, si | MM_2 |
| GOBP_REGULATION_OF_ | GOBP_REGULATION_OF_MYELOID_LEUKOC | 140 | -0.26873 | -1.42957 | 0.03241895 | 0.19567005 | 1 | 4413 | tags=24%, list=19%, si | MM_2 |
| GOBP_PROTEIN_DEGLYC | GOBP_PROTEIN_DEGLYCOSYLATION | 15 | -0.45582 | -1.42952 | 0.110311754 | 0.19547927 | 1 | 12855 | tags=100%, list=54%, | MM_2 |
| GOBP_NEGATIVE_REGUL | GOBP_NEGATIVE_REGULATION_OF_POST_T | 94 | -0.28823 | -1.42908 | 0.036585364 | 0.19571596 | 1 | 4778 | tags=32%, list=20%, si | MM_2 |
| GOBP_ESTABLISHMENT_( | GOBP_ESTABLISHMENT_OR_MAINTENANCE | 223 | -0.2448 | -1.42832 | 0.007075472 | 0.19636771 | 1 | 4674 | tags=26%, list=20%, si | MM_2 |
| GOBP_REGULATION_OF_ | GOBP_REGULATION_OF_ACTIN_FILAMENT_ | 113 | -0.28031 | -1.42766 | 0.02244389 | 0.19685206 | 1 | 2145 | tags=19%, list=9%, sig | MM_2 |
| GOBP_LIPOPROTEIN_MET | GOBP_LIPOPROTEIN_METABOLIC_PROCESS | 135 | -0.26311 | -1.42749 | 0.027160494 | 0.19679126 | 1 | 4356 | tags=29%, list=18%, si | MM_2 |
| GOBP_NEGATIVE_REGUL | GOBP_NEGATIVE_REGULATION_OF_OSTEO( | 37 | -0.34908 | -1.42741 | 0.05035971 | 0.19664086 | 1 | 1095 | tags=14%, list=5%, sig | MM_2 |
| GOBP_REGULATION_OF_ | GOBP_REGULATION_OF_MITOCHONDRION_ | 139 | -0.26589 | -1.42692 | 0.018691588 | 0.19694802 | 1 | 4842 | tags=29%, list=21%, si | MM_2 |
| GOBP_POSITIVE_REGULA | GOBP_POSITIVE_REGULATION_OF_NON_CA | 64 | -0.30551 | -1.42685 | 0.046620045 | 0.19678843 | 1 | 5176 | tags=31%, list=22%, si | MM_2 |
| GOBP_REGULATION_OF_ | GOBP_REGULATION_OF_MYELOID_CELL_DI | 217 | -0.24538 | -1.42603 | 0.011792453 | 0.19741388 | 1 | 4203 | tags=23%, list=18%, si | MM_2 |
| GOBP_PROTEIN_LOCALIZ | GOBP_PROTEIN_LOCALIZATION_TO_EARLY_ | 15 | -0.43614 | -1.42474 | 0.10869565 | 0.19858047 | 1 | 2145 | tags=27%, list=9%, sig | MM_2 |
| GOBP_PHOSPHATIDYLIN( | GOBP_PHOSPHATIDYLINOSITOL_3_KINASE_ | 307 | -0.23438 | -1.42439 | 0.0051150895 | 0.19871715 | 1 | 4870 | tags=31%, list=21%, si | MM_2 |
| GOBP_REGULATION_OF_ | GOBP_REGULATION_OF_CD8_POSITIVE_ALI | 18 | -0.42222 | -1.42405 | 0.10268949 | 0.19884288 | 1 | 6406 | tags=56%, list=27%, si | MM_2 |
| PROPANOATE_METABOLI | PROPANOATE_METABOLISM | 29 | -0.37895 | -1.42356 | 0.061320756 | 0.19918044 | 1 | 3643 | tags=28%, list=15%, si | MM_2 |
| GOBP_NEGATIVE_REGUL | GOBP_NEGATIVE_REGULATION_OF_ESTABL | 148 | -0.2645 | -1.4222 | 0.032407407 | 0.2004382 | 1 | 4471 | tags=28%, list=19%, si | MM_2 |
| GOBP_MEIOTIC_CELL_CY | GOBP_MEIOTIC_CELL_CYCLE | 282 | -0.23883 | -1.42202 | 0.011961723 | 0.20042935 | 1 | 3826 | tags=26%, list=16%, si | MM_2 |
| GOBP_TELOMERE_MAINT | GOBP_TELOMERE_MAINTENANCE_IN_RESP | 32 | -0.36514 | -1.42194 | 0.05952381 | 0.20026177 | 1 | 5541 | tags=41%, list=23%, si | MM_2 |
| GOBP_NOTOCHORD_DEV | GOBP_NOTOCHORD_DEVELOPMENT | 22 | -0.40856 | -1.42169 | 0.1 | 0.20032921 | 1 | 611 | tags=18%, list=3%, sig | MM_2 |
| GOBP_RESPONSE_TO_M | GOBP_RESPONSE_TO_METAL_ION | 278 | -0.23486 | -1.42084 | 0.014527845 | 0.20102905 | 1 | 4602 | tags=27%, list=19%, si | MM_2 |
| GOBP_PROTEIN_CONTAI | GOBP_PROTEIN_CONTAINING_COMPLEX_D | 247 | -0.24196 | -1.41974 | 0.004750594 | 0.20198856 | 1 | 4495 | tags=28%, list=19%, si | MM_2 |
| GOBP_CELLULAR_RESPON | GOBP_CELLULAR_RESPONSE_TO_GLUCOCO | 47 | -0.3326 | -1.41935 | 0.07537688 | 0.20218045 | 1 | 3851 | tags=28%, list=16%, si | MM_2 |
| GOBP_TUMOR_NECROSIS | GOBP_TUMOR_NECROSIS_FACTOR_MEDIAT | 91 | -0.28306 | -1.41896 | 0.035885166 | 0.20236112 | 1 | 4904 | tags=31%, list=21%, si | MM_2 |
| GOBP_REGULATION_OF_ | GOBP_REGULATION_OF_ALPHA_BETA_T_CE | 75 | -0.30094 | -1.41845 | 0.03217822 | 0.2026702 | 1 | 5682 | tags=40%, list=24%, si | MM_2 |
| GOBP_DENDRITIC_CELL_( | GOBP_DENDRITIC_CELL_CHEMOTAXIS | 19 | -0.42481 | -1.41777 | 0.09006929 | 0.2031864 | 1 | 3542 | tags=26%, list=15%, si | MM_2 |

| | | | | | | | | | | |
|---|---|---|---|---|---|---|---|---|---|---|
| GOBP_RIBOSOME_BIOGE | GOBP_RIBOSOME_BIOGENESIS | 306 | -0.23419 | -1.41763 | 0.009615385 | 0.20309429 | 1 | 6203 | tags=31%, list=26%, si | MM_2 |
| GOBP_FATTY_ACID_ELON | GOBP_FATTY_ACID_ELONGATION | 15 | -0.45022 | -1.41688 | 0.087677725 | 0.20359068 | 1 | 3029 | tags=40%, list=13%, si | MM_2 |
| GOBP_DNA_MODIFICATI | GOBP_DNA_MODIFICATION | 26 | -0.39099 | -1.41664 | 0.09090909 | 0.20364408 | 1 | 2864 | tags=23%, list=12%, si | MM_2 |
| GOBP_VIRAL_BUDDING | GOBP_VIRAL_BUDDING | 18 | -0.43 | -1.41655 | 0.086124405 | 0.20349431 | 1 | 5645 | tags=44%, list=24%, si | MM_2 |
| GOBP_B_CELL_DIFFEREN | GOBP_B_CELL_DIFFERENTIATION | 151 | -0.26109 | -1.41647 | 0.028985508 | 0.20334996 | 1 | 3768 | tags=21%, list=16%, si | MM_2 |
| GOBP_LATE_ENDOSOME_ | GOBP_LATE_ENDOSOME_TO_VACUOLE_TR | 16 | -0.45206 | -1.41629 | 0.10199005 | 0.20333593 | 1 | 4995 | tags=38%, list=21%, si | MM_2 |
| GOBP_CELLULAR_COMPO | GOBP_CELLULAR_COMPONENT_DISASSEMB | 31 | -0.37468 | -1.41587 | 0.08851675 | 0.20356493 | 1 | 3865 | tags=29%, list=16%, si | MM_2 |
| GOBP_RIBOSOMAL_SMA | GOBP_RIBOSOMAL_SMALL_SUBUNIT_BIOG | 101 | -0.28235 | -1.41575 | 0.032786883 | 0.20345339 | 1 | 13219 | tags=93%, list=56%, si | MM_2 |
| GOBP_NEGATIVE_REGUL | GOBP_NEGATIVE_REGULATION_OF_PHOSP | 398 | -0.22741 | -1.41538 | 0.0073891627 | 0.20364974 | 1 | 4459 | tags=25%, list=19%, si | MM_2 |
| GOBP_REGULATION_OF_ | GOBP_REGULATION_OF_NUCLEAR_TRANSC | 26 | -0.39098 | -1.41535 | 0.08533333 | 0.20343412 | 1 | 5797 | tags=54%, list=25%, si | MM_2 |
| GOBP_REGULATION_OF_ | GOBP_REGULATION_OF_VASCULOGENESIS | 16 | -0.44406 | -1.41494 | 0.10864745 | 0.20361367 | 1 | 4011 | tags=44%, list=17%, si | MM_2 |
| GOBP_BODY_MORPHOGE | GOBP_BODY_MORPHOGENESIS | 63 | -0.30635 | -1.41487 | 0.06356968 | 0.20343195 | 1 | 4752 | tags=35%, list=20%, si | MM_2 |
| GOBP_NEUTRAL_LIPID_C | GOBP_NEUTRAL_LIPID_CATABOLIC_PROCES | 34 | -0.36694 | -1.41484 | 0.075 | 0.2032207 | 1 | 3085 | tags=26%, list=13%, si | MM_2 |
| GOBP_EXOCYTOSIS | GOBP_EXOCYTOSIS | 367 | -0.22651 | -1.41399 | 0.0070093456 | 0.20395763 | 1 | 3276 | tags=19%, list=14%, si | MM_2 |
| GOBP_PROTEIN_CONTAI | GOBP_PROTEIN_CONTAINING_COMPLEX_L | 259 | -0.23599 | -1.41376 | 0.016826924 | 0.20398006 | 1 | 2824 | tags=18%, list=12%, si | MM_2 |
| GOBP_CLATHRIN_COAT_ | GOBP_CLATHRIN_COAT_ASSEMBLY | 21 | -0.40169 | -1.4137 | 0.087759815 | 0.20380777 | 1 | 2845 | tags=24%, list=12%, si | MM_2 |
| GOBP_POSITIVE_REGULA | GOBP_POSITIVE_REGULATION_OF_TOR_SIG | 71 | -0.2991 | -1.41305 | 0.050480768 | 0.2043019 | 1 | 4761 | tags=34%, list=20%, si | MM_2 |
| GOBP_POSITIVE_REGULA | GOBP_POSITIVE_REGULATION_OF_CHROM | 107 | -0.27615 | -1.41258 | 0.024122806 | 0.20460999 | 1 | 4649 | tags=24%, list=20%, si | MM_2 |
| GOBP_NEGATIVE_REGUL | GOBP_NEGATIVE_REGULATION_OF_LYMPH | 164 | -0.25766 | -1.41244 | 0.030805686 | 0.20452099 | 1 | 3465 | tags=21%, list=15%, si | MM_2 |
| GOBP_RAC_PROTEIN_SIG | GOBP_RAC_PROTEIN_SIGNAL_TRANSDUCTI | 58 | -0.31729 | -1.41174 | 0.059382424 | 0.20509852 | 1 | 4830 | tags=40%, list=20%, si | MM_2 |
| GOBP_POSITIVE_REGULA | GOBP_POSITIVE_REGULATION_OF_NITRIC_ | 20 | -0.41182 | -1.41086 | 0.09468822 | 0.20579785 | 1 | 3272 | tags=30%, list=14%, si | MM_2 |
| GOBP_CELLULAR_RESPO | GOBP_CELLULAR_RESPONSE_TO_CORTICOS | 55 | -0.31902 | -1.41028 | 0.04901961 | 0.20617825 | 1 | 3851 | tags=27%, list=16%, si | MM_2 |
| GOBP_NEGATIVE_REGUL | GOBP_NEGATIVE_REGULATION_OF_HEMOP | 121 | -0.26739 | -1.40982 | 0.040816326 | 0.20647478 | 1 | 2583 | tags=18%, list=11%, si | MM_2 |
| GOBP_REGULATION_OF_ | GOBP_REGULATION_OF_DNA_TEMPLATED_ | 84 | -0.29099 | -1.40958 | 0.050480768 | 0.20652768 | 1 | 5479 | tags=33%, list=23%, si | MM_2 |
| GOBP_RNA_EXPORT_FRO | GOBP_RNA_EXPORT_FROM_NUCLEUS | 78 | -0.28885 | -1.40878 | 0.054320987 | 0.2072025 | 1 | 4667 | tags=32%, list=20%, si | MM_2 |
| GOBP_NEGATIVE_REGUL | GOBP_NEGATIVE_REGULATION_OF_PHOSP | 62 | -0.30512 | -1.40834 | 0.03782506 | 0.20744027 | 1 | 4829 | tags=29%, list=20%, si | MM_2 |
| GOBP_REGULATION_OF_ | GOBP_REGULATION_OF_INSULIN_RECEPTO | 81 | -0.28296 | -1.40819 | 0.05491991 | 0.20734578 | 1 | 3521 | tags=26%, list=15%, si | MM_2 |
| GOBP_HEMATOPOIETIC_ | GOBP_HEMATOPOIETIC_STEM_CELL_PROLI | 37 | -0.3519 | -1.40807 | 0.07746479 | 0.2072479 | 1 | 4543 | tags=32%, list=19%, si | MM_2 |
| GOBP_REGULATION_OF_ | GOBP_REGULATION_OF_NUCLEAR_TRANSC | 15 | -0.46557 | -1.40787 | 0.12686567 | 0.20722839 | 1 | 5797 | tags=53%, list=25%, si | MM_2 |
| GOBP_TRACHEA_DEVELO | GOBP_TRACHEA_DEVELOPMENT | 21 | -0.40588 | -1.40683 | 0.08988764 | 0.20819011 | 1 | 1344 | tags=19%, list=6%, sig | MM_2 |
| GOBP_POSITIVE_REGULA | GOBP_POSITIVE_REGULATION_OF_UBIQUIT | 93 | -0.28001 | -1.40675 | 0.048899755 | 0.20804344 | 1 | 6639 | tags=42%, list=28%, si | MM_2 |
| GOBP_REGULATION_OF_ | GOBP_REGULATION_OF_PROTEIN_DEPHOS | 76 | -0.29207 | -1.4066 | 0.051044084 | 0.20796965 | 1 | 3723 | tags=22%, list=16%, si | MM_2 |
| MICRORNAS_IN_CANCER | MICRORNAS_IN_CANCER | 139 | -0.26402 | -1.40622 | 0.03309693 | 0.20815758 | 1 | 3659 | tags=20%, list=15%, si | MM_2 |
| GOBP_GLIAL_CELL_DEVE | GOBP_GLIAL_CELL_DEVELOPMENT | 134 | -0.26439 | -1.40586 | 0.025462963 | 0.20832485 | 1 | 3539 | tags=24%, list=15%, si | MM_2 |
| GOBP_NEGATIVE_REGUL | GOBP_NEGATIVE_REGULATION_OF_PHAGO | 24 | -0.39903 | -1.4053 | 0.08530806 | 0.20872083 | 1 | 7848 | tags=63%, list=33%, si | MM_2 |
| GOBP_POSITIVE_REGULA | GOBP_POSITIVE_REGULATION_OF_UBIQUIT | 18 | -0.42887 | -1.40456 | 0.10909091 | 0.20935693 | 1 | 714 | tags=17%, list=3%, sig | MM_2 |
| GOBP_REGULATION_OF_ | GOBP_REGULATION_OF_SUBSTRATE_ADHE | 58 | -0.3075 | -1.40451 | 0.04691358 | 0.20918895 | 1 | 4825 | tags=34%, list=20%, si | MM_2 |
| GOBP_IMMUNOGLOBULI | GOBP_IMMUNOGLOBULIN_PRODUCTION | 132 | -0.26706 | -1.4044 | 0.03440367 | 0.20904921 | 1 | 3232 | tags=18%, list=14%, si | MM_2 |
| GOBP_POSITIVE_REGULA | GOBP_POSITIVE_REGULATION_OF_CANONI | 148 | -0.25829 | -1.40381 | 0.024038462 | 0.20946681 | 1 | 5674 | tags=41%, list=24%, si | MM_2 |
| GOBP_REGULATION_OF_ | GOBP_REGULATION_OF_FATTY_ACID_BETA | 21 | -0.40816 | -1.40375 | 0.10526316 | 0.20930806 | 1 | 4077 | tags=43%, list=17%, si | MM_2 |
| GOBP_REGULATION_OF_ | GOBP_REGULATION_OF_SPINDLE_CHECKPO | 24 | -0.39496 | -1.4032 | 0.0879397 | 0.20971958 | 1 | 409 | tags=13%, list=2%, sig | MM_2 |
| GOBP_DOUBLE_STRAND_ | GOBP_DOUBLE_STRAND_BREAK_REPAIR | 299 | -0.23337 | -1.40301 | 0.011876484 | 0.2097088 | 1 | 5228 | tags=30%, list=22%, si | MM_2 |
| GOBP_TOOTH_MINERALI | GOBP_TOOTH_MINERALIZATION | 30 | -0.36758 | -1.40284 | 0.09090909 | 0.20966415 | 1 | 2010 | tags=20%, list=9%, sig | MM_2 |
| GOBP_REGULATION_OF_ | GOBP_REGULATION_OF_ANDROGEN_RECE | 25 | -0.38366 | -1.40249 | 0.085784316 | 0.20984034 | 1 | 6558 | tags=56%, list=28%, si | MM_2 |
| GOBP_SULFUR_COMPOU | GOBP_SULFUR_COMPOUND_METABOLIC_P | 292 | -0.23024 | -1.40146 | 0.0121951215 | 0.2107971 | 1 | 5223 | tags=35%, list=22%, si | MM_2 |
| GOBP_REGENERATION | GOBP_REGENERATION | 161 | -0.2504 | -1.40099 | 0.021479713 | 0.2110757 | 1 | 3718 | tags=25%, list=16%, si | MM_2 |
| GOBP_AUTOPHAGY_OF_ | GOBP_AUTOPHAGY_OF_MITOCHONDRION | 89 | -0.28336 | -1.4007 | 0.027431421 | 0.21117164 | 1 | 7092 | tags=48%, list=30%, si | MM_2 |
| GOBP_MITOCHONDRIAL_ | GOBP_MITOCHONDRIAL_OUTER_MEMBRAN | 34 | -0.36011 | -1.4006 | 0.0783848 | 0.21104456 | 1 | 7073 | tags=53%, list=30%, si | MM_2 |
| GOBP_INTERLEUKIN_6_P | GOBP_INTERLEUKIN_6_PRODUCTION | 173 | -0.24959 | -1.40033 | 0.023980815 | 0.2110533 | 1 | 4618 | tags=27%, list=20%, si | MM_2 |
| GOBP_PROTEIN_DEUBIQ | GOBP_PROTEIN_DEUBIQUITINATION | 98 | -0.27733 | -1.39997 | 0.044496488 | 0.21121304 | 1 | 4925 | tags=27%, list=21%, si | MM_2 |
| GOBP_HYDROGEN_PERO | GOBP_HYDROGEN_PEROXIDE_CATABOLIC_ | 22 | -0.3951 | -1.39986 | 0.101895735 | 0.21111342 | 1 | 2006 | tags=18%, list=8%, sig | MM_2 |
| GOBP_RESPONSE_TO_IO | GOBP_RESPONSE_TO_IONIZING_RADIATION | 127 | -0.26517 | -1.39975 | 0.027160494 | 0.21101886 | 1 | 3879 | tags=24%, list=16%, si | MM_2 |
| GOBP_GLYCEROLIPID_CA | GOBP_GLYCEROLIPID_CATABOLIC_PROCESS | 58 | -0.31264 | -1.39974 | 0.07178218 | 0.21079363 | 1 | 3161 | tags=26%, list=13%, si | MM_2 |
| GOBP_VACUOLAR_ACIDI | GOBP_VACUOLAR_ACIDIFICATION | 27 | -0.37732 | -1.39971 | 0.09443099 | 0.21057396 | 1 | 6243 | tags=48%, list=26%, si | MM_2 |
| GOBP_LATE_ENDOSOME_ | GOBP_LATE_ENDOSOME_TO_LYSOSOME_T | 19 | -0.42402 | -1.39955 | 0.11600928 | 0.21052894 | 1 | 13607 | tags=100%, list=58%, | MM_2 |
| AUTOPHAGY | AUTOPHAGY | 126 | -0.26338 | -1.399 | 0.0450237 | 0.21095736 | 1 | 6848 | tags=42%, list=29%, si | MM_2 |
| CHAGAS_DISEASE_(AMER | CHAGAS_DISEASE_(AMERICAN_TRYPANOSO | 101 | -0.28079 | -1.39804 | 0.050808314 | 0.21184634 | 1 | 3921 | tags=24%, list=17%, si | MM_2 |
| GOBP_PURINE_NUCLEOS | GOBP_PURINE_NUCLEOSIDE_MONOPHOSP | 22 | -0.39412 | -1.39798 | 0.07656612 | 0.21167473 | 1 | 3576 | tags=23%, list=15%, si | MM_2 |
| GOBP_CELL_KILLING | GOBP_CELL_KILLING | 202 | -0.23964 | -1.39782 | 0.012285012 | 0.2116086 | 1 | 3070 | tags=17%, list=13%, si | MM_2 |
| GOBP_TELENCEPHALON_ | GOBP_TELENCEPHALON_DEVELOPMENT | 245 | -0.23675 | -1.3977 | 0.012787724 | 0.21149607 | 1 | 2570 | tags=16%, list=11%, si | MM_2 |
| GOBP_POSITIVE_REGULA | GOBP_POSITIVE_REGULATION_OF_REACTIV | 81 | -0.29375 | -1.39752 | 0.04608295 | 0.21147662 | 1 | 2696 | tags=17%, list=11%, si | MM_2 |
| GOBP_PROTEIN_LOCALIZ | GOBP_PROTEIN_LOCALIZATION_TO_PLASM | 296 | -0.22923 | -1.39684 | 0.0091743115 | 0.21208088 | 1 | 5058 | tags=28%, list=21%, si | MM_2 |
| GOBP_RESPONSE_TO_ST | GOBP_RESPONSE_TO_STARVATION | 181 | -0.25106 | -1.39677 | 0.03836317 | 0.2119066 | 1 | 4221 | tags=27%, list=18%, si | MM_2 |
| GOBP_RETROGRADE_VES | GOBP_RETROGRADE_VESICLE_MEDIATED_T | 45 | -0.33145 | -1.39671 | 0.057142857 | 0.21172962 | 1 | 12980 | tags=96%, list=55%, si | MM_2 |
| GOBP_BONE_RESORPTIO | GOBP_BONE_RESORPTION | 76 | -0.29671 | -1.39661 | 0.061320756 | 0.21158578 | 1 | 4334 | tags=33%, list=18%, si | MM_2 |
| GOBP_NEGATIVE_REGUL | GOBP_NEGATIVE_REGULATION_OF_LEUKO | 16 | -0.44908 | -1.39502 | 0.10585585 | 0.21323878 | 1 | 5635 | tags=50%, list=24%, si | MM_2 |
| GOBP_REGULATION_OF_ | GOBP_REGULATION_OF_ADAPTIVE_IMMUN | 215 | -0.24222 | -1.3949 | 0.018779343 | 0.21315543 | 1 | 3232 | tags=18%, list=14%, si | MM_2 |
| GOBP_REGULATION_OF_ | GOBP_REGULATION_OF_LEUKOCYTE_ADHE | 33 | -0.36058 | -1.39486 | 0.09158416 | 0.21293996 | 1 | 4858 | tags=42%, list=21%, si | MM_2 |
| GOBP_STRIATED_MUSCL | GOBP_STRIATED_MUSCLE_ADAPTATION | 49 | -0.31861 | -1.39473 | 0.07451923 | 0.21287794 | 1 | 3551 | tags=27%, list=15%, si | MM_2 |
| GOBP_METHYLATION | GOBP_METHYLATION | 199 | -0.24276 | -1.39389 | 0.018518519 | 0.213581 | 1 | 4976 | tags=32%, list=21%, si | MM_2 |
| GOBP_REGULATION_OF_ | GOBP_REGULATION_OF_FIBROBLAST_PROL | 122 | -0.26863 | -1.39371 | 0.027295286 | 0.2135354 | 1 | 4789 | tags=29%, list=20%, si | MM_2 |
| GOBP_FIBROBLAST_MIGR | GOBP_FIBROBLAST_MIGRATION | 68 | -0.3013 | -1.39357 | 0.055421688 | 0.21344739 | 1 | 5074 | tags=35%, list=21%, si | MM_2 |
| GOBP_NEGATIVE_REGUL | GOBP_NEGATIVE_REGULATION_OF_PEPTID | 52 | -0.32055 | -1.39283 | 0.070776254 | 0.21410249 | 1 | 4513 | tags=31%, list=19%, si | MM_2 |
| GOBP_DNA_RECOMBINA | GOBP_DNA_RECOMBINATION | 313 | -0.23068 | -1.39243 | 0.00954654 | 0.2143411 | 1 | 4708 | tags=25%, list=20%, si | MM_2 |
| GOBP_INTERLEUKIN_8_P | GOBP_INTERLEUKIN_8_PRODUCTION | 73 | -0.29161 | -1.39235 | 0.07038835 | 0.21419328 | 1 | 4172 | tags=32%, list=18%, si | MM_2 |
| GOBP_REGULATION_OF_ | GOBP_REGULATION_OF_ACUTE_INFLAMMA | 48 | -0.32408 | -1.39153 | 0.06531531 | 0.2148828 | 1 | 2736 | tags=21%, list=12%, si | MM_2 |
| GOBP_REGULATION_OF_ | GOBP_REGULATION_OF_PROTEASOMAL_PR | 179 | -0.24775 | -1.39149 | 0.02173913 | 0.21469346 | 1 | 4932 | tags=26%, list=21%, si | MM_2 |
| GOBP_LEUKOCYTE_TETH | GOBP_LEUKOCYTE_TETHERING_OR_ROLLIN | 35 | -0.35654 | -1.39139 | 0.09951457 | 0.21454233 | 1 | 4858 | tags=40%, list=21%, si | MM_2 |
| GOBP_CELLULAR_RESPO | GOBP_CELLULAR_RESPONSE_TO_HYDROGE | 59 | -0.30486 | -1.3911 | 0.07025761 | 0.2146417 | 1 | 5579 | tags=37%, list=24%, si | MM_2 |
| GOBP_CELLULAR_RESPO | GOBP_CELLULAR_RESPONSE_TO_INSULIN_S | 207 | -0.24468 | -1.39077 | 0.022959184 | 0.21482518 | 1 | 4413 | tags=25%, list=19%, si | MM_2 |
| GOBP_SKELETAL_MUSCL | GOBP_SKELETAL_MUSCLE_ADAPTATION | 21 | -0.41099 | -1.39049 | 0.106235564 | 0.21493033 | 1 | 2149 | tags=19%, list=9%, sig | MM_2 |
| GOBP_AMIDE_METABOLI | GOBP_AMIDE_METABOLIC_PROCESS | 398 | -0.22016 | -1.38996 | 0.009592326 | 0.21527454 | 1 | 4386 | tags=27%, list=19%, si | MM_2 |

| | | | | | | | | | | |
|---|---|---|---|---|---|---|---|---|---|---|
| GOBP_RESPONSE_TO_VIT | GOBP_RESPONSE_TO_VITAMIN_D | 15 | -0.44062 | -1.3898 | 0.10576923 | 0.21519184 | 1 | 3431 | tags=33%, list=15%, si | MM_2 |
| GOBP_MRNA_3_END_PR | GOBP_MRNA_3_END_PROCESSING | 41 | -0.33741 | -1.38933 | 0.08461539 | 0.21550882 | 1 | 5665 | tags=37%, list=24%, si | MM_2 |
| GOBP_NEGATIVE_ADAPT | GOBP_NEGATIVE_ADAPTATION_OF_SIGNAL | 29 | -0.36494 | -1.38931 | 0.10273973 | 0.21530932 | 1 | 471 | tags=14%, list=2%, sig | MM_2 |
| GOBP_POSITIVE_REGULA | GOBP_POSITIVE_REGULATION_OF_DNA_RE | 47 | -0.32095 | -1.38908 | 0.07638889 | 0.21534134 | 1 | 2618 | tags=21%, list=11%, si | MM_2 |
| GOBP_REGULATION_OF_ | GOBP_REGULATION_OF_ORGANELLE_ASSEM | 228 | -0.23737 | -1.38896 | 0.02097902 | 0.2152426 | 1 | 5645 | tags=35%, list=24%, si | MM_2 |
| MITOPHAGY | MITOPHAGY | 62 | -0.30255 | -1.38859 | 0.0746988 | 0.21542378 | 1 | 7092 | tags=47%, list=30%, si | MM_2 |
| GOBP_ENUCLEATE_ERYTH | GOBP_ENUCLEATE_ERYTHROCYTE_DIFFEREN | 15 | -0.44474 | -1.3881 | 0.12740384 | 0.21577753 | 1 | 2996 | tags=27%, list=13%, si | MM_2 |
| GOBP_NEGATIVE_REGUL | GOBP_NEGATIVE_REGULATION_OF_LEUKOC | 96 | -0.27597 | -1.38796 | 0.048661802 | 0.21571735 | 1 | 3342 | tags=21%, list=14%, si | MM_2 |
| GOBP_POSITIVE_REGULA | GOBP_POSITIVE_REGULATION_OF_INTERFE | 26 | -0.38275 | -1.38764 | 0.12790698 | 0.21586077 | 1 | 1902 | tags=19%, list=8%, sig | MM_2 |
| GOBP_POSITIVE_REGULA | GOBP_POSITIVE_REGULATION_OF_PROTEIN | 288 | -0.2322 | -1.3873 | 0.014634146 | 0.21601772 | 1 | 4919 | tags=27%, list=21%, si | MM_2 |
| GOBP_COPII_COATED_VE | GOBP_COPII_COATED_VESICLE_BUDDING | 37 | -0.34729 | -1.38649 | 0.08549223 | 0.21675768 | 1 | 13205 | tags=95%, list=56%, si | MM_2 |
| GOBP_ACTIN_FILAMENT_ | GOBP_ACTIN_FILAMENT_BUNDLE_ORGANIZ | 180 | -0.24745 | -1.38589 | 0.03883495 | 0.21725002 | 1 | 3868 | tags=23%, list=16%, si | MM_2 |
| GOBP_RESPONSE_TO_DIE | GOBP_RESPONSE_TO_DIETARY_EXCESS | 44 | -0.32671 | -1.38563 | 0.07881773 | 0.21730709 | 1 | 3447 | tags=23%, list=15%, si | MM_2 |
| INOSITOL_PHOSPHATE_M | INOSITOL_PHOSPHATE_METABOLISM | 72 | -0.29043 | -1.38562 | 0.07017544 | 0.2170857 | 1 | 5692 | tags=38%, list=24%, si | MM_2 |
| GOBP_NEGATIVE_REGUL | GOBP_NEGATIVE_REGULATION_OF_T_CELL | 73 | -0.29279 | -1.3853 | 0.061611373 | 0.21722746 | 1 | 3244 | tags=22%, list=14%, si | MM_2 |
| GOBP_INTRACELLULAR_M | GOBP_INTRACELLULAR_MONOATOMIC_ANI | 15 | -0.43699 | -1.38509 | 0.11963882 | 0.2172095 | 1 | 915 | tags=20%, list=4%, sig | MM_2 |
| GOBP_REGULATION_OF_ | GOBP_REGULATION_OF_LEUKOCYTE_DEGR | 48 | -0.32431 | -1.3843 | 0.079404466 | 0.21787934 | 1 | 5254 | tags=35%, list=22%, si | MM_2 |
| CHOLINE_METABOLISM_I | CHOLINE_METABOLISM_IN_CANCER | 97 | -0.27535 | -1.38428 | 0.062068965 | 0.21766764 | 1 | 6188 | tags=39%, list=26%, si | MM_2 |
| GOBP_RESPONSE_TO_AN | GOBP_RESPONSE_TO_ANTIBIOTIC | 32 | -0.35776 | -1.38315 | 0.085995086 | 0.21883942 | 1 | 3943 | tags=28%, list=17%, si | MM_2 |
| GOBP_REGULATION_OF_ | GOBP_REGULATION_OF_RAC_PROTEIN_SIG | 35 | -0.35486 | -1.3831 | 0.10705596 | 0.21866181 | 1 | 4830 | tags=46%, list=20%, si | MM_2 |
| GOBP_NUCLEOSIDE_BISP | GOBP_NUCLEOSIDE_BISPHOSPHATE_METAB | 102 | -0.2775 | -1.38232 | 0.04556355 | 0.21931818 | 1 | 4305 | tags=29%, list=18%, si | MM_2 |
| GOBP_REGULATION_OF_ | GOBP_REGULATION_OF_IMMUNOGLOBULI | 81 | -0.28072 | -1.38227 | 0.05172414 | 0.2191502 | 1 | 3232 | tags=19%, list=14%, si | MM_2 |
| GOBP_REGULATION_OF_ | GOBP_REGULATION_OF_INTRACELLULAR_T | 278 | -0.2303 | -1.38214 | 0.00982801 | 0.2190586 | 1 | 5012 | tags=27%, list=21%, si | MM_2 |
| RENAL_CELL_CARCINOMA | RENAL_CELL_CARCINOMA | 67 | -0.29489 | -1.38196 | 0.06326034 | 0.21902733 | 1 | 6309 | tags=45%, list=27%, si | MM_2 |
| GOBP_EPIGENETIC_REGU | GOBP_EPIGENETIC_REGULATION_OF_GENE | 242 | -0.23667 | -1.3818 | 0.027568921 | 0.21897222 | 1 | 4264 | tags=23%, list=18%, si | MM_2 |
| NOTCH_SIGNALING_PATH | NOTCH_SIGNALING_PATHWAY | 47 | -0.32304 | -1.3815 | 0.0741688 | 0.21910149 | 1 | 6819 | tags=55%, list=29%, si | MM_2 |
| GOBP_POSITIVE_REGULA | GOBP_POSITIVE_REGULATION_OF_LEUKOCY | 33 | -0.35539 | -1.38125 | 0.09779951 | 0.2191291 | 1 | 5104 | tags=30%, list=22%, si | MM_2 |
| GOBP_RESPONSE_TO_UV | GOBP_RESPONSE_TO_UV | 153 | -0.25314 | -1.3808 | 0.041162226 | 0.21938743 | 1 | 3813 | tags=20%, list=16%, si | MM_2 |
| GOBP_REGULATION_OF_ | GOBP_REGULATION_OF_NITRIC_OXIDE_SYN | 20 | -0.40634 | -1.38079 | 0.10837439 | 0.21915463 | 1 | 6075 | tags=50%, list=26%, si | MM_2 |
| GOBP_NEGATIVE_REGUL | GOBP_NEGATIVE_REGULATION_OF_MYELO | 23 | -0.39116 | -1.38024 | 0.106024094 | 0.21958175 | 1 | 5039 | tags=30%, list=21%, si | MM_2 |
| GOBP_POSITIVE_REGULA | GOBP_POSITIVE_REGULATION_OF_CYTOKIN | 27 | -0.36763 | -1.3795 | 0.0920398 | 0.22020863 | 1 | 3452 | tags=30%, list=15%, si | MM_2 |
| GOBP_RESPONSE_TO_RE | GOBP_RESPONSE_TO_REACTIVE_OXYGEN_S | 181 | -0.24708 | -1.37904 | 0.024509804 | 0.2204899 | 1 | 4276 | tags=24%, list=18%, si | MM_2 |
| GOBP_NEGATIVE_REGUL | GOBP_NEGATIVE_REGULATION_OF_ANDRO | 15 | -0.42882 | -1.37863 | 0.1191067 | 0.22075649 | 1 | 6556 | tags=67%, list=28%, si | MM_2 |
| GOBP_LEUKOCYTE_DEGR | GOBP_LEUKOCYTE_DEGRANULATION | 77 | -0.28707 | -1.3782 | 0.060185187 | 0.22094013 | 1 | 3276 | tags=19%, list=14%, si | MM_2 |
| GOBP_ENDOSOME_TO_L | GOBP_ENDOSOME_TO_LYSOSOME_TRANSP | 16 | -0.42397 | -1.37802 | 0.11192214 | 0.2209184 | 1 | 13607 | tags=100%, list=58%, | MM_2 |
| GOBP_PHOSPHATIDIC_AC | GOBP_PHOSPHATIDIC_ACID_BIOSYNTHETIC | 23 | -0.38879 | -1.37781 | 0.12679426 | 0.22092532 | 1 | 5933 | tags=43%, list=25%, si | MM_2 |
| GOBP_MITOCHONDRIAL_ | GOBP_MITOCHONDRIAL_OUTER_MEMBRAN | 28 | -0.37316 | -1.37765 | 0.098173514 | 0.22086154 | 1 | 4842 | tags=32%, list=21%, si | MM_2 |
| GOBP_SYMBIONT_ENTRY | GOBP_SYMBIONT_ENTRY_INTO_HOST | 97 | -0.27192 | -1.3768 | 0.0425 | 0.22166994 | 1 | 4407 | tags=29%, list=19%, si | MM_2 |
| GOBP_NEGATIVE_REGUL | GOBP_NEGATIVE_REGULATION_OF_PROTEI | 431 | -0.21862 | -1.37652 | 0.009779952 | 0.22176935 | 1 | 4459 | tags=26%, list=19%, si | MM_2 |
| GOBP_POSITIVE_REGULA | GOBP_POSITIVE_REGULATION_OF_PROTEAS | 72 | -0.29322 | -1.37649 | 0.06153846 | 0.22156179 | 1 | 6639 | tags=42%, list=28%, si | MM_2 |
| GOBP_GLUCOSE_METABO | GOBP_GLUCOSE_METABOLIC_PROCESS | 201 | -0.23972 | -1.37643 | 0.02112676 | 0.22143058 | 1 | 4338 | tags=31%, list=18%, si | MM_2 |
| GOBP_RESPONSE_TO_OX | GOBP_RESPONSE_TO_OXIDATIVE_STRESS | 364 | -0.22234 | -1.37632 | 0.005 | 0.2213119 | 1 | 4381 | tags=23%, list=19%, si | MM_2 |
| GOBP_TRNA_WOBBLE_BA | GOBP_TRNA_WOBBLE_BASE_MODIFICATIO | 19 | -0.40777 | -1.37594 | 0.13286713 | 0.22156309 | 1 | 3499 | tags=26%, list=15%, si | MM_2 |
| GOBP_VESICLE_TRANSPO | GOBP_VESICLE_TRANSPORT_ALONG_MICRO | 47 | -0.32828 | -1.37584 | 0.096412554 | 0.22144768 | 1 | 2267 | tags=21%, list=10%, si | MM_2 |
| GOBP_NEGATIVE_REGUL | GOBP_NEGATIVE_REGULATION_OF_LEUKOC | 46 | -0.3304 | -1.37582 | 0.098130845 | 0.22123991 | 1 | 3571 | tags=28%, list=15%, si | MM_2 |
| GOBP_RESPONSE_TO_PE | GOBP_RESPONSE_TO_PEPTIDE_HORMONE | 361 | -0.22259 | -1.37494 | 0.013574661 | 0.22208579 | 1 | 3539 | tags=20%, list=15%, si | MM_2 |
| GOBP_NEGATIVE_REGUL | GOBP_NEGATIVE_REGULATION_OF_PROTEA | 51 | -0.32195 | -1.37466 | 0.08148148 | 0.22222383 | 1 | 4398 | tags=25%, list=19%, si | MM_2 |
| GOBP_POSITIVE_REGULA | GOBP_POSITIVE_REGULATION_OF_T_CELL_ | 16 | -0.42798 | -1.37367 | 0.13475177 | 0.22314748 | 1 | 4677 | tags=31%, list=20%, si | MM_2 |
| GOBP_CD8_POSITIVE_AL | GOBP_CD8_POSITIVE_ALPHA_BETA_T_CELL | 21 | -0.40003 | -1.37336 | 0.12206573 | 0.22328073 | 1 | 5682 | tags=52%, list=24%, si | MM_2 |
| GOBP_SMAD_PROTEIN_S | GOBP_SMAD_PROTEIN_SIGNAL_TRANSDUC | 83 | -0.28511 | -1.37289 | 0.062015504 | 0.22359757 | 1 | 1355 | tags=14%, list=6%, sig | MM_2 |
| GOBP_REGULATION_OF_ | GOBP_REGULATION_OF_INTEGRIN_ACTIVAT | 17 | -0.41426 | -1.37271 | 0.13636364 | 0.22360924 | 1 | 1500 | tags=18%, list=6%, sig | MM_2 |
| PROTEIN_DIGESTION_AN | PROTEIN_DIGESTION_AND_ABSORPTION | 78 | -0.28379 | -1.37251 | 0.07110092 | 0.22362098 | 1 | 3555 | tags=26%, list=15%, si | MM_2 |
| HIPPO_SIGNALING_PATH | HIPPO_SIGNALING_PATHWAY | 153 | -0.25212 | -1.37248 | 0.022058824 | 0.22341725 | 1 | 4238 | tags=27%, list=18%, si | MM_2 |
| GOBP_SNRNA_TRANSCRI | GOBP_SNRNA_TRANSCRIPTION | 18 | -0.41965 | -1.37221 | 0.12315271 | 0.22355632 | 1 | 4726 | tags=39%, list=20%, si | MM_2 |
| GOBP_REGULATION_OF_ | GOBP_REGULATION_OF_PEPTIDYL_THREON | 45 | -0.32546 | -1.37214 | 0.080684595 | 0.22341308 | 1 | 2821 | tags=24%, list=12%, si | MM_2 |
| GOBP_VESICLE_MEDIATE | GOBP_VESICLE_MEDIATED_TRANSPORT_TO | 158 | -0.2477 | -1.372 | 0.023923445 | 0.22334054 | 1 | 3889 | tags=22%, list=16%, si | MM_2 |
| GOBP_RRNA_PROCESSIN | GOBP_RRNA_PROCESSING | 211 | -0.23748 | -1.37114 | 0.01923077 | 0.22419128 | 1 | 3040 | tags=16%, list=13%, si | MM_2 |
| GOBP_IMMUNE_RESPON | GOBP_IMMUNE_RESPONSE_REGULATING_S | 457 | -0.21512 | -1.37088 | 0.0046296297 | 0.22426194 | 1 | 3793 | tags=20%, list=16%, si | MM_2 |
| GOBP_MEMBRANE_INVA | GOBP_MEMBRANE_INVAGINATION | 70 | -0.29118 | -1.37077 | 0.055690072 | 0.2241455 | 1 | 4584 | tags=26%, list=19%, si | MM_2 |
| GOBP_PYRUVATE_METAB | GOBP_PYRUVATE_METABOLIC_PROCESS | 127 | -0.25689 | -1.37064 | 0.061124694 | 0.22404042 | 1 | 5054 | tags=36%, list=21%, si | MM_2 |
| GOBP_POSITIVE_REGULA | GOBP_POSITIVE_REGULATION_OF_LYMPHO | 134 | -0.25914 | -1.37058 | 0.030534351 | 0.22388531 | 1 | 6021 | tags=38%, list=25%, si | MM_2 |
| GOBP_RESPONSE_TO_NU | GOBP_RESPONSE_TO_NUTRIENT_LEVELS | 371 | -0.22139 | -1.37053 | 0.016018307 | 0.22370303 | 1 | 4248 | tags=24%, list=18%, si | MM_2 |
| GOBP_REGULATION_OF_ | GOBP_REGULATION_OF_PROTEOLYSIS_INV | 215 | -0.2411 | -1.37034 | 0.02905569 | 0.22368638 | 1 | 4932 | tags=27%, list=21%, si | MM_2 |
| GOBP_NEGATIVE_REGUL | GOBP_NEGATIVE_REGULATION_OF_VASOC | 17 | -0.42583 | -1.37009 | 0.13194445 | 0.22374727 | 1 | 979 | tags=18%, list=4%, sig | MM_2 |
| GOBP_DEOXYRIBONUCLE | GOBP_DEOXYRIBONUCLEOTIDE_BIOSYNTHE | 17 | -0.42277 | -1.36972 | 0.09832134 | 0.22397113 | 1 | 2569 | tags=29%, list=11%, si | MM_2 |
| GOBP_IMMUNOLOGICAL_ | GOBP_IMMUNOLOGICAL_SYNAPSE_FORMA | 18 | -0.40996 | -1.36908 | 0.110311754 | 0.22445294 | 1 | 1095 | tags=11%, list=5%, sig | MM_2 |
| GOBP_EMBRYONIC_BODY | GOBP_EMBRYONIC_BODY_MORPHOGENESI | 15 | -0.44545 | -1.36863 | 0.095794395 | 0.22472693 | 1 | 1865 | tags=27%, list=8%, sig | MM_2 |
| GOBP_CENTROMERE_CO | GOBP_CENTROMERE_COMPLEX_ASSEMBLY | 27 | -0.3666 | -1.36825 | 0.10538641 | 0.22494696 | 1 | 1916 | tags=19%, list=8%, sig | MM_2 |
| GOBP_MONOCYTE_CHEM | GOBP_MONOCYTE_CHEMOTAXIS | 40 | -0.32859 | -1.36781 | 0.07674944 | 0.22529703 | 1 | 5635 | tags=40%, list=24%, si | MM_2 |
| GOBP_PROGRAMMED_N | GOBP_PROGRAMMED_NECROTIC_CELL_DEA | 50 | -0.30964 | -1.36713 | 0.08726415 | 0.22591494 | 1 | 3638 | tags=28%, list=15%, si | MM_2 |
| GOBP_POSITIVE_REGULA | GOBP_POSITIVE_REGULATION_OF_HYDROL | 427 | -0.21628 | -1.36685 | 0.014563107 | 0.22601089 | 1 | 4873 | tags=28%, list=21%, si | MM_2 |
| GOBP_VIRAL_GENE_EXPR | GOBP_VIRAL_GENE_EXPRESSION | 75 | -0.29252 | -1.36599 | 0.050251257 | 0.22679508 | 1 | 7063 | tags=53%, list=30%, si | MM_2 |
| GOBP_BRAIN_MORPHOG | GOBP_BRAIN_MORPHOGENESIS | 39 | -0.332 | -1.36572 | 0.1007371 | 0.22687006 | 1 | 3304 | tags=21%, list=14%, si | MM_2 |
| GOBP_ECTODERMAL_PLA | GOBP_ECTODERMAL_PLACODE_DEVELOPM | 16 | -0.42602 | -1.3657 | 0.117506 | 0.22666888 | 1 | 2697 | tags=19%, list=11%, si | MM_2 |
| RNA_TRANSPORT | RNA_TRANSPORT | 154 | -0.25248 | -1.36532 | 0.047029704 | 0.22687091 | 1 | 5820 | tags=37%, list=25%, si | MM_2 |
| GOBP_NUCLEOSIDE_BISP | GOBP_NUCLEOSIDE_BISPHOSPHATE_BIOSYN | 39 | -0.33833 | -1.36508 | 0.09876543 | 0.2269456 | 1 | 4184 | tags=36%, list=18%, si | MM_2 |
| GOBP_NEGATIVE_REGUL | GOBP_NEGATIVE_REGULATION_OF_T_CELL | 29 | -0.36377 | -1.36503 | 0.103044495 | 0.22677039 | 1 | 2071 | tags=17%, list=9%, sig | MM_2 |
| GOBP_REGULATION_OF_ | GOBP_REGULATION_OF_VASCULATURE_DE | 295 | -0.22113 | -1.3647 | 0.01703163 | 0.22699818 | 1 | 3560 | tags=22%, list=15%, si | MM_2 |
| GOBP_GOLGI_TO_PLASM | GOBP_GOLGI_TO_PLASMA_MEMBRANE_TR | 62 | -0.30156 | -1.36453 | 0.0691358 | 0.22698042 | 1 | 4788 | tags=26%, list=20%, si | MM_2 |
| GOBP_NEGATIVE_REGUL | GOBP_NEGATIVE_REGULATION_OF_PROTEI | 110 | -0.26814 | -1.36439 | 0.03686636 | 0.22694732 | 1 | 4845 | tags=28%, list=21%, si | MM_2 |
| GOBP_PROTEIN_K11_LIN | GOBP_PROTEIN_K11_LINKED_UBIQUITINAT | 29 | -0.36065 | -1.36391 | 0.09411765 | 0.22733204 | 1 | 5753 | tags=41%, list=24%, si | MM_2 |
| GOBP_TORC1_SIGNALING | GOBP_TORC1_SIGNALING | 100 | -0.27156 | -1.36322 | 0.037914693 | 0.22797029 | 1 | 4488 | tags=31%, list=19%, si | MM_2 |

| | | | | | | | | | | |
|---|---|---|---|---|---|---|---|---|---|---|
| GOBP_REGULATION_OF_ | GOBP_REGULATION_OF_TELOMERE_MAINT | 61 | -0.29255 | -1.36315 | 0.07424594 | 0.22782949 | 1 | 2696 | tags=16%, list=11%, si | MM_2 |
| GOBP_CELLULAR_RESPON | GOBP_CELLULAR_RESPONSE_TO_UV | 90 | -0.27345 | -1.36314 | 0.05472637 | 0.2276101 | 1 | 3813 | tags=22%, list=16%, si | MM_2 |
| GOBP_BRANCHED_CHAIN | GOBP_BRANCHED_CHAIN_AMINO_ACID_CA | 21 | -0.39499 | -1.36195 | 0.118329465 | 0.22887948 | 1 | 4894 | tags=38%, list=21%, si | MM_2 |
| GOBP_REGULATION_OF_ | GOBP_REGULATION_OF_T_CELL_CYTOKINE_ | 36 | -0.33883 | -1.36178 | 0.106024094 | 0.22881484 | 1 | 1147 | tags=14%, list=5%, sig | MM_2 |
| GOBP_EMBRYONIC_HEM | GOBP_EMBRYONIC_HEMOPOIESIS | 31 | -0.35702 | -1.36168 | 0.083538085 | 0.22870809 | 1 | 6728 | tags=55%, list=28%, si | MM_2 |
| GOBP_PRIMARY_NEURAL | GOBP_PRIMARY_NEURAL_TUBE_FORMATIO | 115 | -0.26103 | -1.3607 | 0.03448276 | 0.22960725 | 1 | 2946 | tags=21%, list=12%, si | MM_2 |
| GOBP_REGULATION_OF_ | GOBP_REGULATION_OF_MITOCHONDRIAL_ | 39 | -0.32921 | -1.35954 | 0.11057692 | 0.23079696 | 1 | 4842 | tags=31%, list=21%, si | MM_2 |
| GOBP_FATTY_ACYL_COA | GOBP_FATTY_ACYL_COA_METABOLIC_PRO | 18 | -0.41504 | -1.35947 | 0.12626262 | 0.23065397 | 1 | 4730 | tags=39%, list=20%, si | MM_2 |
| GOBP_POSITIVE_REGULA | GOBP_POSITIVE_REGULATION_OF_LEUKOC | 159 | -0.24646 | -1.35919 | 0.038369305 | 0.23079789 | 1 | 3926 | tags=22%, list=17%, si | MM_2 |
| GOBP_CELLULAR_RESPON | GOBP_CELLULAR_RESPONSE_TO_LIPID | 498 | -0.21206 | -1.35875 | 0.0024390244 | 0.23106492 | 1 | 3521 | tags=19%, list=15%, si | MM_2 |
| GOBP_ERYTHROCYTE_MA | GOBP_ERYTHROCYTE_MATURATION | 23 | -0.38226 | -1.35826 | 0.10144927 | 0.2315005 | 1 | 2917 | tags=30%, list=12%, si | MM_2 |
| GOBP_RESPONSE_TO_STI | GOBP_RESPONSE_TO_STEROID_HORMONE | 226 | -0.23348 | -1.35797 | 0.008988764 | 0.23160015 | 1 | 5218 | tags=30%, list=22%, si | MM_2 |
| GOBP_REGULATION_OF_ | GOBP_REGULATION_OF_BINDING | 331 | -0.22196 | -1.35748 | 0.014251782 | 0.23189631 | 1 | 5253 | tags=33%, list=22%, si | MM_2 |
| GOBP_GPI_ANCHOR_ME | GOBP_GPI_ANCHOR_METABOLIC_PROCESS | 33 | -0.35207 | -1.35623 | 0.08863636 | 0.2331529 | 1 | 4039 | tags=30%, list=17%, si | MM_2 |
| GOBP_GRANULOCYTE_CH | GOBP_GRANULOCYTE_CHEMOTAXIS | 119 | -0.25932 | -1.35615 | 0.03836317 | 0.23300335 | 1 | 3967 | tags=24%, list=17%, si | MM_2 |
| GOBP_NEGATIVE_REGUL | GOBP_NEGATIVE_REGULATION_OF_MICRO | 49 | -0.32015 | -1.35614 | 0.09151786 | 0.23277736 | 1 | 4413 | tags=39%, list=19%, si | MM_2 |
| GOBP_REGULATION_OF_ | GOBP_REGULATION_OF_FIBROBLAST_MIGR | 47 | -0.31697 | -1.35595 | 0.09738717 | 0.23282704 | 1 | 4833 | tags=34%, list=20%, si | MM_2 |
| GOBP_REGULATION_OF_ | GOBP_REGULATION_OF_ARP2_3_COMPLEX | 27 | -0.3719 | -1.35545 | 0.12718205 | 0.23318578 | 1 | 1928 | tags=19%, list=8%, sig | MM_2 |
| GOBP_PEPTIDYL_THREON | GOBP_PEPTIDYL_THREONINE_DEPHOSPHOR | 15 | -0.43429 | -1.35518 | 0.13776723 | 0.2332314 | 1 | 3567 | tags=33%, list=15%, si | MM_2 |
| GOBP_NEGATIVE_REGUL | GOBP_NEGATIVE_REGULATION_OF_TELOM | 21 | -0.39179 | -1.35512 | 0.12652068 | 0.23306899 | 1 | 4474 | tags=38%, list=19%, si | MM_2 |
| GOBP_PYRIDINE_NUCLEO | GOBP_PYRIDINE_NUCLEOTIDE_BIOSYNTHET | 19 | -0.39484 | -1.35498 | 0.13452914 | 0.2330326 | 1 | 4983 | tags=42%, list=21%, si | MM_2 |
| GOBP_POSITIVE_REGULA | GOBP_POSITIVE_REGULATION_OF_DNA_BIN | 217 | -0.23468 | -1.35472 | 0.013729977 | 0.2331041 | 1 | 5176 | tags=35%, list=22%, si | MM_2 |
| GOBP_INTRACELLULAR_P | GOBP_INTRACELLULAR_PH_REDUCTION | 48 | -0.31324 | -1.35461 | 0.103365384 | 0.23298162 | 1 | 6306 | tags=44%, list=27%, si | MM_2 |
| GOBP_NEGATIVE_REGUL | GOBP_NEGATIVE_REGULATION_OF_OSTEOB | 58 | -0.30188 | -1.35445 | 0.078282826 | 0.23298967 | 1 | 3424 | tags=28%, list=15%, si | MM_2 |
| GOBP_REGULATION_OF_ | GOBP_REGULATION_OF_CELLULAR_COMPO | 380 | -0.21734 | -1.35442 | 0.018648019 | 0.23279944 | 1 | 4655 | tags=24%, list=20%, si | MM_2 |
| GOBP_REGULATION_OF_ | GOBP_REGULATION_OF_CENTRIOLE_REPLIC | 22 | -0.37863 | -1.35436 | 0.124390244 | 0.23264267 | 1 | 5324 | tags=36%, list=23%, si | MM_2 |
| GOBP_BASE_EXCISION_R | GOBP_BASE_EXCISION_REPAIR | 41 | -0.32738 | -1.35417 | 0.08584687 | 0.23268236 | 1 | 4245 | tags=27%, list=18%, si | MM_2 |
| GOBP_POSITIVE_REGULA | GOBP_POSITIVE_REGULATION_OF_MONOC | 21 | -0.38837 | -1.35371 | 0.12085308 | 0.23301348 | 1 | 3926 | tags=38%, list=17%, si | MM_2 |
| GOBP_PROTEIN_TARGETI | GOBP_PROTEIN_TARGETING | 285 | -0.22781 | -1.35355 | 0.02347418 | 0.23298432 | 1 | 4337 | tags=24%, list=18%, si | MM_2 |
| GOBP_MRNA_PROCESSIN | GOBP_MRNA_PROCESSING | 459 | -0.21121 | -1.35343 | 0.004854369 | 0.23288599 | 1 | 6318 | tags=35%, list=27%, si | MM_2 |
| GOBP_RIBOSOMAL_SMA | GOBP_RIBOSOMAL_SMALL_SUBUNIT_ASSEM | 18 | -0.40952 | -1.3532 | 0.13145539 | 0.23296204 | 1 | 4315 | tags=28%, list=18%, si | MM_2 |
| GOBP_MRNA_CATABOLIC | GOBP_MRNA_CATABOLIC_PROCESS | 233 | -0.22959 | -1.35319 | 0.017766498 | 0.23274557 | 1 | 5232 | tags=29%, list=22%, si | MM_2 |
| GOBP_REGULATION_OF_ | GOBP_REGULATION_OF_REGULATED_SECRE | 139 | -0.25554 | -1.35298 | 0.047191013 | 0.232784 | 1 | 3276 | tags=22%, list=14%, si | MM_2 |
| GOBP_DNA_INTEGRITY_C | GOBP_DNA_INTEGRITY_CHECKPOINT_SIGNA | 125 | -0.25445 | -1.35281 | 0.038990825 | 0.23275216 | 1 | 4469 | tags=29%, list=19%, si | MM_2 |
| GOBP_RECEPTOR_RECYCL | GOBP_RECEPTOR_RECYCLING | 54 | -0.30597 | -1.35277 | 0.086419754 | 0.23256813 | 1 | 4208 | tags=26%, list=18%, si | MM_2 |
| GOBP_REGULATION_OF_ | GOBP_REGULATION_OF_DNA_BINDING_TRA | 362 | -0.21859 | -1.35244 | 0.014492754 | 0.23273066 | 1 | 5244 | tags=31%, list=22%, si | MM_2 |
| GOBP_BIOLOGICAL_PROC | GOBP_BIOLOGICAL_PROCESS_INVOLVED_IN | 235 | -0.23084 | -1.35201 | 0.040302265 | 0.23303917 | 1 | 5425 | tags=32%, list=23%, si | MM_2 |
| GOBP_POSITIVE_REGULA | GOBP_POSITIVE_REGULATION_OF_RELEASE | 27 | -0.36073 | -1.35201 | 0.11809045 | 0.23281524 | 1 | 3842 | tags=30%, list=16%, si | MM_2 |
| GOBP_ALDITOL_METABO | GOBP_ALDITOL_METABOLIC_PROCESS | 22 | -0.37882 | -1.35168 | 0.11320755 | 0.23301116 | 1 | 4889 | tags=41%, list=21%, si | MM_2 |
| GOBP_CANONICAL_NF_K | GOBP_CANONICAL_NF_KAPPAB_SIGNAL_TR | 247 | -0.22843 | -1.35137 | 0.03448276 | 0.23321088 | 1 | 5714 | tags=36%, list=24%, si | MM_2 |
| GOBP_NOTCH_SIGNALIN | GOBP_NOTCH_SIGNALING_PATHWAY | 179 | -0.23869 | -1.35133 | 0.040201005 | 0.23303746 | 1 | 3954 | tags=26%, list=17%, si | MM_2 |
| GOBP_RECEPTOR_CLUST | GOBP_RECEPTOR_CLUSTERING | 79 | -0.28084 | -1.3509 | 0.07434053 | 0.2333357 | 1 | 3348 | tags=23%, list=14%, si | MM_2 |
| GOBP_REGULATION_OF_ | GOBP_REGULATION_OF_MEMBRANE_INVA | 21 | -0.38966 | -1.35086 | 0.12235294 | 0.23313968 | 1 | 3447 | tags=33%, list=15%, si | MM_2 |
| GOBP_NEGATIVE_REGUL | GOBP_NEGATIVE_REGULATION_OF_UBIQUI | 55 | -0.30385 | -1.35084 | 0.08980583 | 0.23294544 | 1 | 4398 | tags=24%, list=19%, si | MM_2 |
| OTHER_TYPES_OF_O-GLY | OTHER_TYPES_OF_O-GLYCAN_BIOSYNTHESI | 21 | -0.38542 | -1.34975 | 0.106575966 | 0.23407514 | 1 | 3909 | tags=29%, list=17%, si | MM_2 |
| GOBP_POSITIVE_REGULA | GOBP_POSITIVE_REGULATION_BY_HOST_O | 22 | -0.37853 | -1.34925 | 0.14030612 | 0.234503 | 1 | 12814 | tags=95%, list=54%, si | MM_2 |
| GOBP_RESPONSE_TO_KE | GOBP_RESPONSE_TO_KETONE | 134 | -0.24982 | -1.34877 | 0.031026253 | 0.23486792 | 1 | 3521 | tags=22%, list=15%, si | MM_2 |
| GOBP_MICROTUBULE_OR | GOBP_MICROTUBULE_ORGANIZING_CENTE | 149 | -0.24763 | -1.34751 | 0.05936073 | 0.23617537 | 1 | 4225 | tags=23%, list=18%, si | MM_2 |
| GOBP_ARTERY_DEVELOP | GOBP_ARTERY_DEVELOPMENT | 117 | -0.25848 | -1.34748 | 0.06067961 | 0.23598227 | 1 | 3726 | tags=25%, list=16%, si | MM_2 |
| GOBP_INNATE_IMMUNE_ | GOBP_INNATE_IMMUNE_RESPONSE_ACTIV | 82 | -0.28145 | -1.34669 | 0.086448595 | 0.23675333 | 1 | 4244 | tags=27%, list=18%, si | MM_2 |
| GOBP_POSITIVE_REGULA | GOBP_POSITIVE_REGULATION_OF_ALPHA_B | 52 | -0.31041 | -1.34656 | 0.0928382 | 0.23667206 | 1 | 5682 | tags=42%, list=24%, si | MM_2 |
| GOBP_NEGATIVE_REGUL | GOBP_NEGATIVE_REGULATION_OF_TELOM | 36 | -0.33416 | -1.34647 | 0.105882354 | 0.23657489 | 1 | 5368 | tags=39%, list=23%, si | MM_2 |
| GOBP_AXONAL_TRANSPO | GOBP_AXONAL_TRANSPORT | 69 | -0.28466 | -1.34643 | 0.06699752 | 0.23638733 | 1 | 6967 | tags=59%, list=30%, si | MM_2 |
| GOBP_PURINE_CONTAIN | GOBP_PURINE_CONTAINING_COMPOUND_ | 150 | -0.24755 | -1.34574 | 0.04157044 | 0.23706257 | 1 | 4935 | tags=33%, list=21%, si | MM_2 |
| GOBP_TOLL_LIKE_RECEPT | GOBP_TOLL_LIKE_RECEPTOR_9_SIGNALING | 16 | -0.42338 | -1.34547 | 0.13711584 | 0.23717429 | 1 | 4181 | tags=38%, list=18%, si | MM_2 |
| GOBP_CELLULAR_RESPON | GOBP_CELLULAR_RESPONSE_TO_OXIDATIVE | 229 | -0.22934 | -1.34538 | 0.03529412 | 0.2370643 | 1 | 4272 | tags=24%, list=18%, si | MM_2 |
| APELIN_SIGNALING_PATH | APELIN_SIGNALING_PATHWAY | 131 | -0.25513 | -1.34472 | 0.04379562 | 0.23767862 | 1 | 3534 | tags=24%, list=15%, si | MM_2 |
| GOBP_THIOESTER_BIOSY | GOBP_THIOESTER_BIOSYNTHETIC_PROCESS | 27 | -0.35956 | -1.34472 | 0.112941176 | 0.23745304 | 1 | 5054 | tags=44%, list=21%, si | MM_2 |
| GOBP_MAGNESIUM_ION | GOBP_MAGNESIUM_ION_TRANSPORT | 17 | -0.41428 | -1.34446 | 0.15403423 | 0.23755108 | 1 | 5619 | tags=47%, list=24%, si | MM_2 |
| GOBP_SEGMENTATION | GOBP_SEGMENTATION | 97 | -0.26755 | -1.34403 | 0.060150377 | 0.23780717 | 1 | 4190 | tags=28%, list=18%, si | MM_2 |
| GOBP_LAYER_FORMATIO | GOBP_LAYER_FORMATION_IN_CEREBRAL_C | 16 | -0.42013 | -1.34398 | 0.12589073 | 0.23764625 | 1 | 2543 | tags=38%, list=11%, si | MM_2 |
| GOBP_REGULATION_OF_ | GOBP_REGULATION_OF_OSSIFICATION | 135 | -0.2527 | -1.34377 | 0.060606062 | 0.23765829 | 1 | 3755 | tags=23%, list=16%, si | MM_2 |
| GOBP_REGULATION_OF_ | GOBP_REGULATION_OF_POST_TRANSLATIO | 242 | -0.22948 | -1.34365 | 0.029850746 | 0.2375752 | 1 | 4778 | tags=24%, list=20%, si | MM_2 |
| STEROID_BIOSYNTHESIS | STEROID_BIOSYNTHESIS | 17 | -0.41651 | -1.34353 | 0.14153132 | 0.2375218 | 1 | 3253 | tags=35%, list=14%, si | MM_2 |
| GOBP_NEGATIVE_REGUL | GOBP_NEGATIVE_REGULATION_OF_PHOSPH | 332 | -0.22017 | -1.34332 | 0.012626262 | 0.23754369 | 1 | 4513 | tags=26%, list=19%, si | MM_2 |
| GOBP_SPINAL_CORD_PA | GOBP_SPINAL_CORD_PATTERNING | 17 | -0.40462 | -1.34149 | 0.13995485 | 0.23950708 | 1 | 1842 | tags=24%, list=8%, sig | MM_2 |
| GOBP_LYMPHOCYTE_ME | GOBP_LYMPHOCYTE_MEDIATED_IMMUNITY | 300 | -0.2227 | -1.34138 | 0.014814815 | 0.23941879 | 1 | 3232 | tags=17%, list=14%, si | MM_2 |
| GOBP_ADAPTIVE_IMMUN | GOBP_ADAPTIVE_IMMUNE_RESPONSE | 433 | -0.21053 | -1.34133 | 0.004784689 | 0.23924425 | 1 | 3813 | tags=19%, list=16%, si | MM_2 |
| GOBP_NEGATIVE_REGUL | GOBP_NEGATIVE_REGULATION_OF_LYMPH | 62 | -0.2966 | -1.34119 | 0.103797466 | 0.23917273 | 1 | 2905 | tags=24%, list=12%, si | MM_2 |
| GOBP_RESPONSE_TO_AR | GOBP_RESPONSE_TO_ARSENIC_CONTAININ | 25 | -0.36848 | -1.34112 | 0.118644066 | 0.23903167 | 1 | 2559 | tags=20%, list=11%, si | MM_2 |
| GOBP_REGULATION_OF_ | GOBP_REGULATION_OF_MYELOID_CELL_AP | 37 | -0.32569 | -1.34032 | 0.10401891 | 0.23978767 | 1 | 5039 | tags=30%, list=21%, si | MM_2 |
| GOBP_REGULATION_OF_ | GOBP_REGULATION_OF_PEPTIDYL_TYROSIN | 238 | -0.23059 | -1.3401 | 0.035799522 | 0.23983331 | 1 | 4432 | tags=25%, list=19%, si | MM_2 |
| GOBP_POSTREPLICATION | GOBP_POSTREPLICATION_REPAIR | 29 | -0.35906 | -1.33873 | 0.112709835 | 0.24137002 | 1 | 4463 | tags=41%, list=19%, si | MM_2 |
| GOBP_ODONTOGENESIS_ | GOBP_ODONTOGENESIS_OF_DENTIN_CONT | 82 | -0.28011 | -1.33839 | 0.0625 | 0.24157919 | 1 | 2409 | tags=17%, list=10%, si | MM_2 |
| GOBP_EPITHELIAL_TO_M | GOBP_EPITHELIAL_TO_MESENCHYMAL_TRA | 16 | -0.41953 | -1.33552 | 0.155902 | 0.24503249 | 1 | 7660 | tags=69%, list=32%, si | MM_2 |
| GOBP_LIPID_HOMEOSTA | GOBP_LIPID_HOMEOSTASIS | 167 | -0.24153 | -1.33517 | 0.041262135 | 0.24527276 | 1 | 3967 | tags=25%, list=17%, si | MM_2 |
| GOBP_PHAGOSOME_MA | GOBP_PHAGOSOME_MATURATION | 29 | -0.3535 | -1.33473 | 0.12672812 | 0.24559578 | 1 | 6526 | tags=59%, list=28%, si | MM_2 |
| GOBP_PIECEMEAL_MICR | GOBP_PIECEMEAL_MICROAUTOPHAGY_OF_ | 19 | -0.39084 | -1.33397 | 0.113425925 | 0.24634825 | 1 | 6848 | tags=63%, list=29%, si | MM_2 |
| GOBP_ORGANOPHOSPHA | GOBP_ORGANOPHOSPHATE_CATABOLIC_PR | 227 | -0.22702 | -1.33397 | 0.022779044 | 0.24611628 | 1 | 4951 | tags=30%, list=21%, si | MM_2 |
| GOBP_CELLULAR_RESPON | GOBP_CELLULAR_RESPONSE_TO_VITAMIN | 16 | -0.42355 | -1.33339 | 0.15116279 | 0.24665943 | 1 | 3431 | tags=38%, list=15%, si | MM_2 |
| GOBP_RECOMBINATIONA | GOBP_RECOMBINATIONAL_REPAIR | 177 | -0.23752 | -1.33338 | 0.050397877 | 0.24643171 | 1 | 5228 | tags=30%, list=22%, si | MM_2 |
| GOBP_PROTEIN_AUTOPH | GOBP_PROTEIN_AUTOPHOSPHORYLATION | 204 | -0.23149 | -1.33251 | 0.02849741 | 0.24737905 | 1 | 2946 | tags=18%, list=12%, si | MM_2 |

| | | | | | | | | | | |
|---|---|---|---|---|---|---|---|---|---|---|
| GOBP_MITOCHONDRIAL_ | GOBP_MITOCHONDRIAL_TRANSMEMBRANE | 92 | -0.26656 | -1.33233 | 0.070953436 | 0.24736837 | 1 | 6237 | tags=38%, list=26%, si | MM_2 |
| GOBP_VIRAL_TRANSCRIP | GOBP_VIRAL_TRANSCRIPTION | 55 | -0.29877 | -1.33195 | 0.08830549 | 0.2476173 | 1 | 5456 | tags=42%, list=23%, si | MM_2 |
| GOBP_RIBOSOMAL_LARG | GOBP_RIBOSOMAL_LARGE_SUBUNIT_ASSEN | 23 | -0.37003 | -1.33156 | 0.11764706 | 0.2478899 | 1 | 2196 | tags=22%, list=9%, sig | MM_2 |
| GOBP_PRESYNAPTIC_END | GOBP_PRESYNAPTIC_ENDOCYTOSIS | 95 | -0.26878 | -1.33135 | 0.059850372 | 0.24796206 | 1 | 2549 | tags=18%, list=11%, si | MM_2 |
| GOBP_MACROMOLECULE | GOBP_MACROMOLECULE_METHYLATION | 121 | -0.25271 | -1.3307 | 0.07289294 | 0.2485218 | 1 | 4231 | tags=27%, list=18%, si | MM_2 |
| OLFACTORY_TRANSDUCT | OLFACTORY_TRANSDUCTION | 186 | 0.440376 | 2.406622 | 0 | 0.001999845 | 0.003 | 7718 | tags=72%, list=33%, si | MM_2 |
| GOBP_HEART_FIELD_SPE | GOBP_HEART_FIELD_SPECIFICATION | 15 | 0.704634 | 2.113167 | 0.0017452007 | 0.07001468 | 0.185 | 1242 | tags=40%, list=5%, sig | MM_2 |
| GOBP_DNA_TEMPLATED_ | GOBP_DNA_TEMPLATED_TRANSCRIPTION_T | 25 | 0.597613 | 2.06335 | 0 | 0.09183906 | 0.345 | 2075 | tags=24%, list=9%, sig | MM_2 |
| CIRCADIAN_RHYTHM | CIRCADIAN_RHYTHM | 29 | 0.557877 | 2.016838 | 0.0017301039 | 0.11300759 | 0.497 | 2512 | tags=28%, list=11%, si | MM_2 |
| GOBP_LIGAND_GATED_IO | GOBP_LIGAND_GATED_ION_CHANNEL_SIGN | 33 | 0.52095 | 1.990335 | 0.0017361111 | 0.1258352 | 0.618 | 1635 | tags=21%, list=7%, sig | MM_2 |
| GOBP_REGULATION_OF_ | GOBP_REGULATION_OF_NON_CANONICAL_ | 26 | 0.564725 | 1.986132 | 0.0017421603 | 0.108958855 | 0.631 | 2067 | tags=31%, list=9%, sig | MM_2 |
| GOBP_G_PROTEIN_COUP | GOBP_G_PROTEIN_COUPLED_ACETYLCHOLI | 20 | 0.603102 | 1.971504 | 0 | 0.10948112 | 0.689 | 3676 | tags=40%, list=16%, si | MM_2 |
| SYSTEMIC_LUPUS_ERYTH | SYSTEMIC_LUPUS_ERYTHEMATOSUS | 105 | 0.393659 | 1.935262 | 0 | 0.14510423 | 0.826 | 1473 | tags=20%, list=6%, sig | MM_2 |
| GOBP_REGULATION_OF_ | GOBP_REGULATION_OF_ENDOTHELIAL_CEL | 21 | 0.575896 | 1.929601 | 0 | 0.13660254 | 0.845 | 2709 | tags=43%, list=11%, si | MM_2 |
| GOBP_ESTABLISHMENT_C | GOBP_ESTABLISHMENT_OF_PLANAR_POLAR | 18 | 0.600488 | 1.921631 | 0.0035149385 | 0.13446705 | 0.875 | 712 | tags=22%, list=3%, sig | MM_2 |
| GOBP_ESTABLISHMENT_C | GOBP_ESTABLISHMENT_OF_PLANAR_POLAR | 16 | 0.617786 | 1.914974 | 0.0034843206 | 0.13234735 | 0.891 | 712 | tags=25%, list=3%, sig | MM_2 |
| GOBP_POSITIVE_REGULA | GOBP_POSITIVE_REGULATION_OF_ANIMAL_ | 37 | 0.475272 | 1.896128 | 0.0017513135 | 0.15148094 | 0.938 | 1637 | tags=27%, list=7%, sig | MM_2 |
| GOBP_PROTEIN_FOLDING | GOBP_PROTEIN_FOLDING | 162 | 0.349057 | 1.889771 | 0 | 0.15059865 | 0.95 | 2573 | tags=22%, list=11%, si | MM_2 |
| GOBP_NEGATIVE_REGUL | GOBP_NEGATIVE_REGULATION_OF_LIPID_T | 30 | 0.511699 | 1.888119 | 0.0033500837 | 0.14265098 | 0.951 | 2016 | tags=33%, list=9%, sig | MM_2 |
| GOBP_ENDOTHELIAL_CEL | GOBP_ENDOTHELIAL_CELL_CHEMOTAXIS | 30 | 0.514644 | 1.880708 | 0.0016556291 | 0.14365974 | 0.957 | 1184 | tags=23%, list=5%, sig | MM_2 |
| GOBP_PROTEIN_REFOLDI | GOBP_PROTEIN_REFOLDING | 23 | 0.544799 | 1.875855 | 0.0051020407 | 0.14180796 | 0.96 | 2491 | tags=39%, list=11%, si | MM_2 |
| GOBP_REGULATION_OF_ | GOBP_REGULATION_OF_ANIMAL_ORGAN_F | 29 | 0.51614 | 1.873194 | 0.0017211704 | 0.13758704 | 0.962 | 1173 | tags=28%, list=5%, sig | MM_2 |
| GOBP_MYELIN_MAINTEN | GOBP_MYELIN_MAINTENANCE | 16 | 0.612072 | 1.871207 | 0 | 0.13212483 | 0.967 | 1179 | tags=31%, list=5%, sig | MM_2 |
| GOBP_ENTRAINMENT_OF | GOBP_ENTRAINMENT_OF_CIRCADIAN_CLOC | 26 | 0.519941 | 1.868366 | 0.0017513135 | 0.1290956 | 0.972 | 1838 | tags=27%, list=8%, sig | MM_2 |
| GOBP_RENAL_FILTRATIOI | GOBP_RENAL_FILTRATION | 29 | 0.518489 | 1.86684 | 0.0017182131 | 0.12434279 | 0.978 | 2026 | tags=28%, list=9%, sig | MM_2 |
| GOBP_POSITIVE_REGULA | GOBP_POSITIVE_REGULATION_OF_ENDOTH | 15 | 0.618396 | 1.852889 | 0.0069444445 | 0.13497531 | 0.991 | 2109 | tags=47%, list=9%, sig | MM_2 |
| RETINOL_METABOLISM | RETINOL_METABOLISM | 52 | 0.428728 | 1.852166 | 0 | 0.12977763 | 0.991 | 10322 | tags=75%, list=44%, si | MM_2 |
| GOBP_MODIFICATION_O | GOBP_MODIFICATION_OF_POSTSYNAPTIC_S | 42 | 0.461458 | 1.85156 | 0.0017699115 | 0.1251734 | 0.991 | 1201 | tags=19%, list=5%, sig | MM_2 |
| GOBP_MODIFICATION_O | GOBP_MODIFICATION_OF_SYNAPTIC_STRU | 51 | 0.4423 | 1.849197 | 0 | 0.12260232 | 0.992 | 1201 | tags=18%, list=5%, sig | MM_2 |
| GOBP_INTERMEDIATE_FI | GOBP_INTERMEDIATE_FILAMENT_ORGANIZ | 46 | 0.448567 | 1.848483 | 0 | 0.11830938 | 0.992 | 5522 | tags=61%, list=23%, si | MM_2 |
| GOBP_CIRCADIAN_REGUI | GOBP_CIRCADIAN_REGULATION_OF_GENE_ | 69 | 0.405233 | 1.819973 | 0 | 0.1504223 | 0.998 | 2348 | tags=20%, list=10%, si | MM_2 |
| ALCOHOLISM | ALCOHOLISM | 161 | 0.339567 | 1.812508 | 0 | 0.1553717 | 0.999 | 991 | tags=14%, list=4%, sig | MM_2 |
| GOBP_CRANIAL_NERVE_ | GOBP_CRANIAL_NERVE_DEVELOPMENT | 53 | 0.420759 | 1.802089 | 0 | 0.1661356 | 0.999 | 3437 | tags=36%, list=15%, si | MM_2 |
| GOBP_FAT_CELL_PROLIFE | GOBP_FAT_CELL_PROLIFERATION | 16 | 0.585235 | 1.794134 | 0.006968641 | 0.17278 | 0.999 | 2536 | tags=31%, list=11%, si | MM_2 |
| GOBP_CIRCADIAN_SLEEP | GOBP_CIRCADIAN_SLEEP_WAKE_CYCLE_PRO | 28 | 0.486917 | 1.779946 | 0.001669449 | 0.1915898 | 0.999 | 6898 | tags=54%, list=29%, si | MM_2 |
| GOBP_NON_CANONICAL_ | GOBP_NON_CANONICAL_WNT_SIGNALING_ | 59 | 0.415225 | 1.772457 | 0.0032733225 | 0.19946155 | 0.999 | 2067 | tags=24%, list=9%, sig | MM_2 |
| GOBP_REGULATION_OF_ | GOBP_REGULATION_OF_MODIFICATION_OF | 29 | 0.481749 | 1.771336 | 0.0069204154 | 0.19516331 | 0.999 | 1023 | tags=17%, list=4%, sig | MM_2 |
| PHOTOTRANSDUCTION | PHOTOTRANSDUCTION | 21 | 0.533919 | 1.769932 | 0.005226481 | 0.19155163 | 0.999 | 7207 | tags=57%, list=31%, si | MM_2 |
| GOBP_REGULATION_OF_ | GOBP_REGULATION_OF_NEUROTRANSMITT | 20 | 0.536057 | 1.766911 | 0.010544815 | 0.19063576 | 0.999 | 947 | tags=25%, list=4%, sig | MM_2 |
| GOBP_REGULATION_OF_ | GOBP_REGULATION_OF_PROTEIN_DEACETY | 33 | 0.468223 | 1.765226 | 0.0072072074 | 0.18792887 | 0.999 | 629 | tags=15%, list=3%, sig | MM_2 |
| GOBP_RESPONSE_TO_TE | GOBP_RESPONSE_TO_TESTOSTERONE | 15 | 0.586595 | 1.763209 | 0.011382114 | 0.1863192 | 0.999 | 1137 | tags=20%, list=5%, sig | MM_2 |
| GOBP_POSITIVE_REGULA | GOBP_POSITIVE_REGULATION_OF_SMOOTH | 17 | 0.565017 | 1.761519 | 0.010398613 | 0.18414675 | 0.999 | 439 | tags=18%, list=2%, sig | MM_2 |
| GOBP_REGULATION_OF_ | GOBP_REGULATION_OF_ESTABLISHMENT_C | 19 | 0.544133 | 1.757519 | 0.0049261083 | 0.18641877 | 1 | 712 | tags=21%, list=3%, sig | MM_2 |
| GOBP_PYROPTOSIS | GOBP_PYROPTOSIS | 35 | 0.463391 | 1.755056 | 0.0054644807 | 0.18572412 | 1 | 3048 | tags=23%, list=13%, si | MM_2 |
| GOBP_WNT_SIGNALING_ | GOBP_WNT_SIGNALING_PATHWAY_PLANA | 38 | 0.445058 | 1.749955 | 0.001669449 | 0.18996398 | 1 | 2067 | tags=24%, list=9%, sig | MM_2 |
| GOBP_PROTEIN_AUTOPR | GOBP_PROTEIN_AUTOPROCESSING | 23 | 0.511642 | 1.744718 | 0.008576329 | 0.19333889 | 1 | 1187 | tags=30%, list=5%, sig | MM_2 |
| STEROID_HORMONE_BIO | STEROID_HORMONE_BIOSYNTHESIS | 45 | 0.422033 | 1.737074 | 0.005119454 | 0.20195763 | 1 | 10393 | tags=80%, list=44%, si | MM_2 |
| ARGININE_BIOSYNTHESIS | ARGININE_BIOSYNTHESIS | 17 | 0.553579 | 1.736773 | 0.010676157 | 0.19764945 | 1 | 867 | tags=24%, list=4%, sig | MM_2 |
| GOBP_INFLAMMASOME_ | GOBP_INFLAMMASOME_MEDIATED_SIGNA | 46 | 0.424936 | 1.736518 | 0.003539823 | 0.19356608 | 1 | 1395 | tags=17%, list=6%, sig | MM_2 |
| GOBP_REGULATION_OF_ | GOBP_REGULATION_OF_INCLUSION_BODY_ | 19 | 0.543962 | 1.720909 | 0.008741259 | 0.21727435 | 1 | 2074 | tags=32%, list=9%, sig | MM_2 |
| GOBP_PARASYMPATHETI | GOBP_PARASYMPATHETIC_NERVOUS_SYSTE | 21 | 0.533359 | 1.714219 | 0.0072072074 | 0.2256592 | 1 | 2903 | tags=43%, list=12%, si | MM_2 |
| MATURITY_ONSET_DIABE | MATURITY_ONSET_DIABETES_OF_THE_YOU | 17 | 0.559177 | 1.714126 | 0.005319149 | 0.22099963 | 1 | 10251 | tags=94%, list=43%, si | MM_2 |
| GOBP_SPECIFICATION_OI | GOBP_SPECIFICATION_OF_ANIMAL_ORGAN | 37 | 0.442611 | 1.712707 | 0.006779661 | 0.21893312 | 1 | 1242 | tags=24%, list=5%, sig | MM_2 |
| GOBP_REGULATION_OF_ | GOBP_REGULATION_OF_AXONOGENESIS | 170 | 0.313005 | 1.702215 | 0 | 0.23420651 | 1 | 1730 | tags=15%, list=7%, sig | MM_2 |
| GOBP_POSITIVE_REGULA | GOBP_POSITIVE_REGULATION_OF_AXONOG | 98 | 0.351849 | 1.698276 | 0.0052724076 | 0.23803721 | 1 | 1730 | tags=16%, list=7%, sig | MM_2 |
| GOBP_ANIMAL_ORGAN_ | GOBP_ANIMAL_ORGAN_FORMATION | 70 | 0.377584 | 1.692571 | 0.0053667263 | 0.24506181 | 1 | 1246 | tags=20%, list=5%, sig | MM_2 |
| GOBP_TRANSLATION_AT_ | GOBP_TRANSLATION_AT_SYNAPSE | 48 | -0.65351 | -2.00065 | 0 | 0.014750369 | 0.014 | 7947 | tags=92%, list=34%, si | MM_3 |
| GOBP_AMYLOID_PRECUR | GOBP_AMYLOID_PRECURSOR_PROTEIN_CA | 59 | -0.60764 | -1.90006 | 0 | 0.10619598 | 0.19 | 7091 | tags=68%, list=30%, si | MM_3 |
| GOBP_POSITIVE_REGULA | GOBP_POSITIVE_REGULATION_OF_ISOTYPE | 30 | -0.66693 | -1.88517 | 0 | 0.09512084 | 0.244 | 3651 | tags=43%, list=15%, si | MM_3 |
| GOBP_NEGATIVE_REGUL | GOBP_NEGATIVE_REGULATION_OF_MIRNA | 30 | -0.67262 | -1.88004 | 0 | 0.07741121 | 0.257 | 4798 | tags=57%, list=20%, si | MM_3 |
| GOBP_MACROPHAGE_MI | GOBP_MACROPHAGE_MIGRATION | 65 | -0.58695 | -1.8672 | 0 | 0.085391484 | 0.338 | 5962 | tags=58%, list=25%, si | MM_3 |
| GOBP_NEGATIVE_REGUL | GOBP_NEGATIVE_REGULATION_OF_DOUBL | 24 | -0.68862 | -1.86603 | 0.0013513514 | 0.07308982 | 0.346 | 2356 | tags=33%, list=10%, si | MM_3 |
| GOBP_SNRNA_METABOL | GOBP_SNRNA_METABOLIC_PROCESS | 50 | -0.59927 | -1.81642 | 0 | 0.15688547 | 0.67 | 5504 | tags=56%, list=23%, si | MM_3 |
| GOBP_POSITIVE_REGULA | GOBP_POSITIVE_REGULATION_OF_CIRCADI | 19 | -0.70422 | -1.81261 | 0 | 0.14995843 | 0.702 | 2938 | tags=47%, list=12%, si | MM_3 |
| GOBP_AEROBIC_RESPIRA | GOBP_AEROBIC_RESPIRATION | 170 | -0.51797 | -1.81219 | 0 | 0.13400029 | 0.704 | 7071 | tags=57%, list=30%, si | MM_3 |
| GOBP_NEGATIVE_REGUL | GOBP_NEGATIVE_REGULATION_OF_AMIDE_ | 22 | -0.68382 | -1.80773 | 0 | 0.13105962 | 0.74 | 4285 | tags=59%, list=18%, si | MM_3 |
| GOBP_NEUROINFLAMMA | GOBP_NEUROINFLAMMATORY_RESPONSE | 84 | -0.54638 | -1.80487 | 0 | 0.12509584 | 0.755 | 6719 | tags=55%, list=28%, si | MM_3 |
| GOBP_AMYLOID_BETA_M | GOBP_AMYLOID_BETA_METABOLIC_PROCES | 55 | -0.57783 | -1.80369 | 0.0012254902 | 0.11722639 | 0.762 | 4458 | tags=42%, list=19%, si | MM_3 |
| GOBP_PROTEIN_AUTOUB | GOBP_PROTEIN_AUTOUBIQUITINATION | 68 | -0.57075 | -1.80334 | 0 | 0.10861515 | 0.764 | 6186 | tags=51%, list=26%, si | MM_3 |
| GOBP_REGULATION_OF_ | GOBP_REGULATION_OF_MACROPHAGE_MI | 47 | -0.59463 | -1.79743 | 0 | 0.11415345 | 0.809 | 5962 | tags=62%, list=25%, si | MM_3 |
| PANTOTHENATE_AND_CO | PANTOTHENATE_AND_COA_BIOSYNTHESIS | 17 | -0.6971 | -1.79171 | 0.0013245033 | 0.1180991 | 0.842 | 2446 | tags=35%, list=10%, si | MM_3 |
| GOBP_OXIDATIVE_PHOSP | GOBP_OXIDATIVE_PHOSPHORYLATION | 124 | -0.52043 | -1.7859 | 0 | 0.12168775 | 0.868 | 7071 | tags=58%, list=30%, si | MM_3 |
| GOBP_ONE_CARBON_CO | GOBP_ONE_CARBON_COMPOUND_TRANSP | 16 | -0.71045 | -1.78133 | 0.0014347202 | 0.12422471 | 0.881 | 400 | tags=25%, list=2%, sig | MM_3 |
| GOBP_SUPPRESSION_OF_ | GOBP_SUPPRESSION_OF_VIRAL_RELEASE_B | 22 | -0.6685 | -1.77988 | 0 | 0.120429136 | 0.886 | 892 | tags=32%, list=4%, sig | MM_3 |
| GOBP_SNRNA_PROCESSI | GOBP_SNRNA_PROCESSING | 24 | -0.6523 | -1.77813 | 0.00397878 | 0.11737297 | 0.891 | 3283 | tags=46%, list=14%, si | MM_3 |
| GOBP_IMPORT_INTO_TH | GOBP_IMPORT_INTO_THE_MITOCHONDRIO | 49 | -0.58534 | -1.77592 | 0 | 0.11567688 | 0.899 | 5448 | tags=49%, list=23%, si | MM_3 |
| GOBP_POSITIVE_REGULA | GOBP_POSITIVE_REGULATION_OF_INTERFE | 45 | -0.58287 | -1.77382 | 0 | 0.11434509 | 0.902 | 3515 | tags=33%, list=15%, si | MM_3 |
| GOBP_GLIAL_CELL_ACTIV | GOBP_GLIAL_CELL_ACTIVATION | 59 | -0.56172 | -1.77381 | 0 | 0.109195605 | 0.902 | 6268 | tags=56%, list=27%, si | MM_3 |
| GOBP_POSITIVE_REGULA | GOBP_POSITIVE_REGULATION_OF_BROWN_ | 17 | -0.70587 | -1.76872 | 0 | 0.1140437 | 0.923 | 2439 | tags=35%, list=10%, si | MM_3 |
| GOBP_ANTIGEN_PROCES | GOBP_ANTIGEN_PROCESSING_AND_PRESEN | 28 | -0.63378 | -1.76823 | 0 | 0.11012713 | 0.925 | 6253 | tags=79%, list=26%, si | MM_3 |
| GOBP_NUCLEOTIDE_SUG | GOBP_NUCLEOTIDE_SUGAR_METABOLIC_PI | 34 | -0.60411 | -1.7638 | 0 | 0.11367129 | 0.941 | 4595 | tags=47%, list=19%, si | MM_3 |
| GOBP_CYTOPLASMIC_TR | GOBP_CYTOPLASMIC_TRANSLATION | 143 | -0.50592 | -1.76139 | 0 | 0.11340917 | 0.949 | 8015 | tags=76%, list=34%, si | MM_3 |

| | | | | | | | | | | |
|---|---|---|---|---|---|---|---|---|---|---|
| GOBP_NEGATIVE_REGUL | GOBP_NEGATIVE_REGULATION_OF_B_CELL | 36 | -0.59745 | -1.76109 | 0.002484472 | 0.10975537 | 0.949 | 2920 | tags=36%, list=12%, si | MM_3 |
| GOBP_SNRNA_3_END_PF | GOBP_SNRNA_3_END_PROCESSING | 20 | -0.67772 | -1.76081 | 0 | 0.10632502 | 0.951 | 3283 | tags=50%, list=14%, si | MM_3 |
| GOBP_ANTIGEN_PROCES | GOBP_ANTIGEN_PROCESSING_AND_PRESEN | 34 | -0.60459 | -1.75818 | 0.0013020834 | 0.107252955 | 0.959 | 6830 | tags=76%, list=29%, si | MM_3 |
| GOBP_REGULATION_OF_ | GOBP_REGULATION_OF_OLIGODENDROCYT | 51 | -0.56102 | -1.7575 | 0 | 0.10501751 | 0.962 | 3606 | tags=27%, list=15%, si | MM_3 |
| GOBP_SOMATIC_DIVERSI | GOBP_SOMATIC_DIVERSIFICATION_OF_IMN | 17 | -0.69164 | -1.74946 | 0.0013755158 | 0.115612246 | 0.977 | 2772 | tags=41%, list=12%, si | MM_3 |
| GOBP_NEGATIVE_REGUL | GOBP_NEGATIVE_REGULATION_OF_TYPE_I_ | 23 | -0.65602 | -1.74867 | 0.0040983604 | 0.11341893 | 0.979 | 1655 | tags=30%, list=7%, sig | MM_3 |
| GOBP_TRICARBOXYLIC_A | GOBP_TRICARBOXYLIC_ACID_METABOLIC_P | 16 | -0.71523 | -1.74797 | 0 | 0.111262195 | 0.98 | 2791 | tags=50%, list=12%, si | MM_3 |
| CHRONIC_MYELOID_LEUK | CHRONIC_MYELOID_LEUKEMIA | 75 | -0.53704 | -1.74696 | 0 | 0.109511845 | 0.98 | 6338 | tags=53%, list=27%, si | MM_3 |
| GOBP_PROTON_MOTIVE_ | GOBP_PROTON_MOTIVE_FORCE_DRIVEN_A | 52 | -0.56197 | -1.73997 | 0.0011820331 | 0.11777291 | 0.986 | 7071 | tags=63%, list=30%, si | MM_3 |
| GOBP_REGULATION_OF_ | GOBP_REGULATION_OF_AMYLOID_BETA_FO | 37 | -0.59172 | -1.73705 | 0.0012919897 | 0.11958338 | 0.991 | 4458 | tags=43%, list=19%, si | MM_3 |
| PYRUVATE_METABOLISM | PYRUVATE_METABOLISM | 37 | -0.59376 | -1.73115 | 0.0012658228 | 0.12705919 | 0.995 | 5415 | tags=59%, list=23%, si | MM_3 |
| GOBP_PEPTIDYL_LYSINE_ | GOBP_PEPTIDYL_LYSINE_ACETYLATION | 25 | -0.62545 | -1.72736 | 0.004 | 0.1306135 | 0.997 | 3309 | tags=32%, list=14%, si | MM_3 |
| GOBP_HEMATOPOIETIC_S | GOBP_HEMATOPOIETIC_STEM_CELL_PROLIF | 37 | -0.5883 | -1.72532 | 0.0012578616 | 0.13130379 | 0.997 | 4549 | tags=41%, list=19%, si | MM_3 |
| GOBP_ATP_BIOSYNTHETI | GOBP_ATP_BIOSYNTHETIC_PROCESS | 80 | -0.52419 | -1.72464 | 0 | 0.12907769 | 0.997 | 7071 | tags=59%, list=30%, si | MM_3 |
| GOBP_NEGATIVE_REGUL | GOBP_NEGATIVE_REGULATION_OF_DNA_R | 37 | -0.58208 | -1.72321 | 0.0038659794 | 0.12889084 | 0.997 | 2470 | tags=30%, list=10%, si | MM_3 |
| GOBP_CELLULAR_RESPON | GOBP_CELLULAR_RESPONSE_TO_HYPOXIA | 112 | -0.50966 | -1.72274 | 0 | 0.12677996 | 0.997 | 4991 | tags=38%, list=21%, si | MM_3 |
| GOBP_RESPONSE_TO_FO | GOBP_RESPONSE_TO_FOOD | 35 | -0.59087 | -1.71977 | 0.0037926675 | 0.1297796 | 0.998 | 5334 | tags=46%, list=23%, si | MM_3 |
| GOBP_NEGATIVE_REGUL | GOBP_NEGATIVE_REGULATION_OF_DNA_R | 42 | -0.57294 | -1.71956 | 0.0038412292 | 0.12728731 | 0.998 | 3312 | tags=31%, list=14%, si | MM_3 |
| GOBP_MITOTIC_DNA_RE | GOBP_MITOTIC_DNA_REPLICATION | 16 | -0.68579 | -1.71891 | 0 | 0.12582228 | 0.998 | 2941 | tags=44%, list=12%, si | MM_3 |
| GOBP_AMYLOID_PRECUR | GOBP_AMYLOID_PRECURSOR_PROTEIN_ME | 75 | -0.52397 | -1.7182 | 0 | 0.12430589 | 0.998 | 6044 | tags=52%, list=26%, si | MM_3 |
| GOBP_EOSINOPHIL_MIGR | GOBP_EOSINOPHIL_MIGRATION | 26 | -0.62568 | -1.71693 | 0 | 0.123908006 | 0.998 | 5452 | tags=58%, list=23%, si | MM_3 |
| GOBP_PROTEIN_K6_LINK | GOBP_PROTEIN_K6_LINKED_UBIQUITINATIO | 15 | -0.6976 | -1.71643 | 0.0028129395 | 0.12242924 | 0.998 | 4323 | tags=60%, list=18%, si | MM_3 |
| GOBP_POSITIVE_REGULA | GOBP_POSITIVE_REGULATION_OF_CELL_CY | 31 | -0.60614 | -1.71614 | 0.0038412292 | 0.12038456 | 0.998 | 4987 | tags=55%, list=21%, si | MM_3 |
| GOBP_COENZYME_A_ME | GOBP_COENZYME_A_METABOLIC_PROCESS | 19 | -0.66701 | -1.71603 | 0.004237288 | 0.11810343 | 0.998 | 2446 | tags=37%, list=10%, si | MM_3 |
| GOBP_INFLAMMATORY_ | GOBP_INFLAMMATORY_RESPONSE_TO_WO | 30 | -0.60562 | -1.71508 | 0.0013495276 | 0.116970345 | 0.998 | 2075 | tags=30%, list=9%, sig | MM_3 |
| GOBP_PORPHYRIN_CONT | GOBP_PORPHYRIN_CONTAINING_COMPOU | 40 | -0.57862 | -1.71444 | 0.0012547051 | 0.11583913 | 0.998 | 4980 | tags=45%, list=21%, si | MM_3 |
| NOTCH_SIGNALING_PATH | NOTCH_SIGNALING_PATHWAY | 47 | -0.5555 | -1.71413 | 0.0012195122 | 0.11427144 | 0.998 | 5922 | tags=47%, list=25%, si | MM_3 |
| GOBP_NEGATIVE_REGUL | GOBP_NEGATIVE_REGULATION_OF_AMYLO | 20 | -0.65356 | -1.71364 | 0.0055632824 | 0.11319083 | 0.998 | 7091 | tags=80%, list=30%, si | MM_3 |
| GOBP_REGULATION_OF_ | GOBP_REGULATION_OF_BROWN_FAT_CELL | 27 | -0.61441 | -1.71081 | 0.0013736264 | 0.11580109 | 0.998 | 2439 | tags=30%, list=10%, si | MM_3 |
| CITRATE_CYCLE_(TCA_CY | CITRATE_CYCLE_(TCA_CYCLE) | 28 | -0.61178 | -1.70977 | 0.007782101 | 0.11580946 | 0.998 | 6449 | tags=68%, list=27%, si | MM_3 |
| GOBP_GOLGI_TO_PLASM | GOBP_GOLGI_TO_PLASMA_MEMBRANE_TR | 62 | -0.53132 | -1.70928 | 0.0011890606 | 0.11453661 | 0.998 | 7588 | tags=55%, list=32%, si | MM_3 |
| GOBP_REGULATION_OF_ | GOBP_REGULATION_OF_GLIAL_CELL_DIFFE | 89 | -0.51391 | -1.70686 | 0 | 0.116661504 | 0.999 | 3606 | tags=28%, list=15%, si | MM_3 |
| LYSOSOME | LYSOSOME | 118 | -0.49885 | -1.7041 | 0 | 0.119001925 | 0.999 | 6102 | tags=58%, list=26%, si | MM_3 |
| GOBP_REGULATION_OF_ | GOBP_REGULATION_OF_AMYLOID_PRECUR | 44 | -0.56466 | -1.70224 | 0.0024301338 | 0.11980298 | 0.999 | 7091 | tags=61%, list=30%, si | MM_3 |
| GOBP_UNSATURATED_FA | GOBP_UNSATURATED_FATTY_ACID_BIOSYN | 41 | -0.56591 | -1.70108 | 0.0012330456 | 0.12012543 | 0.999 | 5935 | tags=46%, list=25%, si | MM_3 |
| GOBP_GRANULOCYTE_CH | GOBP_GRANULOCYTE_CHEMOTAXIS | 119 | -0.49623 | -1.70056 | 0 | 0.119109355 | 0.999 | 5954 | tags=50%, list=25%, si | MM_3 |
| GOBP_TOLL_LIKE_RECEPT | GOBP_TOLL_LIKE_RECEPTOR_SIGNALING_P | 71 | -0.52903 | -1.6965 | 0 | 0.12413022 | 1 | 6746 | tags=56%, list=29%, si | MM_3 |
| GOBP_CELLULAR_RESPON | GOBP_CELLULAR_RESPONSE_TO_TOXIC_SU | 63 | -0.53222 | -1.69547 | 0.0011682243 | 0.12379188 | 1 | 6127 | tags=49%, list=26%, si | MM_3 |
| GOBP_MYELOID_LEUKOC | GOBP_MYELOID_LEUKOCYTE_MIGRATION | 222 | -0.4677 | -1.6944 | 0 | 0.12366031 | 1 | 6268 | tags=49%, list=27%, si | MM_3 |
| GOBP_POSITIVE_REGULA | GOBP_POSITIVE_REGULATION_OF_VASCULA | 20 | -0.64332 | -1.69256 | 0.006830601 | 0.12527886 | 1 | 5988 | tags=65%, list=25%, si | MM_3 |
| GOBP_GRANULOCYTE_M | GOBP_GRANULOCYTE_MIGRATION | 146 | -0.49061 | -1.6914 | 0 | 0.12565053 | 1 | 5954 | tags=48%, list=25%, si | MM_3 |
| SNARE_INTERACTIONS_IN | SNARE_INTERACTIONS_IN_VESICULAR_TRAN | 32 | -0.58418 | -1.68977 | 0.0037831021 | 0.12661672 | 1 | 6482 | tags=69%, list=27%, si | MM_3 |
| GOBP_HEME_METABOLIC | GOBP_HEME_METABOLIC_PROCESS | 34 | -0.59505 | -1.68962 | 0.0049937577 | 0.1250112 | 1 | 4425 | tags=44%, list=19%, si | MM_3 |
| GOBP_ERYTHROCYTE_DE | GOBP_ERYTHROCYTE_DEVELOPMENT | 50 | -0.54721 | -1.68855 | 0 | 0.12500542 | 1 | 6027 | tags=50%, list=26%, si | MM_3 |
| GOBP_CELLULAR_RESPON | GOBP_CELLULAR_RESPONSE_TO_OXYGEN_L | 133 | -0.49455 | -1.68747 | 0 | 0.1252255 | 1 | 5347 | tags=38%, list=23%, si | MM_3 |
| GOBP_ENDOPLASMIC_RE | GOBP_ENDOPLASMIC_RETICULUM_CALCIUM | 28 | -0.60572 | -1.6863 | 0.004048583 | 0.12560032 | 1 | 6718 | tags=61%, list=28%, si | MM_3 |
| DNA_REPLICATION | DNA_REPLICATION | 34 | -0.58623 | -1.68551 | 0.0012755102 | 0.12521106 | 1 | 2672 | tags=44%, list=11%, si | MM_3 |
| GOBP_IRON_ION_TRANS | GOBP_IRON_ION_TRANSMEMBRANE_TRAN | 24 | -0.61239 | -1.68515 | 0.0054200543 | 0.12407645 | 1 | 4999 | tags=42%, list=21%, si | MM_3 |
| GOBP_NEGATIVE_REGUL | GOBP_NEGATIVE_REGULATION_OF_DNA_TE | 17 | -0.65707 | -1.68473 | 0.0040816325 | 0.123351224 | 1 | 4283 | tags=35%, list=18%, si | MM_3 |
| GOBP_TOLL_LIKE_RECEPT | GOBP_TOLL_LIKE_RECEPTOR_3_SIGNALING_ | 29 | -0.5959 | -1.68464 | 0.0066313 | 0.12186688 | 1 | 3216 | tags=34%, list=14%, si | MM_3 |
| GOBP_MEMBRANE_RAFT | GOBP_MEMBRANE_RAFT_ORGANIZATION | 24 | -0.61291 | -1.68446 | 0.0053475937 | 0.12061345 | 1 | 7091 | tags=67%, list=30%, si | MM_3 |
| GOBP_ANTIGEN_PROCES | GOBP_ANTIGEN_PROCESSING_AND_PRESEN | 59 | -0.53955 | -1.68262 | 0.0012048193 | 0.12203402 | 1 | 6253 | tags=59%, list=26%, si | MM_3 |
| GOBP_RESPONSE_TO_LIT | GOBP_RESPONSE_TO_LITHIUM_ION | 16 | -0.66635 | -1.68252 | 0.0028490028 | 0.12066331 | 1 | 2822 | tags=44%, list=12%, si | MM_3 |
| GOBP_REGULATION_OF_ | GOBP_REGULATION_OF_TELOMERE_MAINT | 20 | -0.6389 | -1.68212 | 0.004103967 | 0.11973628 | 1 | 5074 | tags=55%, list=21%, si | MM_3 |
| GOBP_MITOCHONDRIAL_ | GOBP_MITOCHONDRIAL_CALCIUM_ION_TR | 19 | -0.64484 | -1.6798 | 0.0055172415 | 0.12236727 | 1 | 5413 | tags=58%, list=23%, si | MM_3 |
| GOBP_DNA_TEMPLATED_ | GOBP_DNA_TEMPLATED_TRANSCRIPTION_E | 112 | -0.49562 | -1.67582 | 0 | 0.12762986 | 1 | 6091 | tags=48%, list=26%, si | MM_3 |
| GOBP_CELLULAR_RESPIRA | GOBP_CELLULAR_RESPIRATION | 218 | -0.47107 | -1.67492 | 0 | 0.12768364 | 1 | 7071 | tags=55%, list=30%, si | MM_3 |
| GOBP_LEUKOCYTE_ACTIV | GOBP_LEUKOCYTE_ACTIVATION_INVOLVED | 53 | -0.53969 | -1.67373 | 0.001234568 | 0.12821287 | 1 | 4960 | tags=42%, list=21%, si | MM_3 |
| GOBP_POSITIVE_REGULA | GOBP_POSITIVE_REGULATION_OF_MUSCLE | 43 | -0.56253 | -1.67279 | 0.0037313432 | 0.12839438 | 1 | 5440 | tags=51%, list=23%, si | MM_3 |
| GOBP_NEUTROPHIL_MED | GOBP_NEUTROPHIL_MEDIATED_CYTOTOXIC | 17 | -0.66739 | -1.67084 | 0.0056818184 | 0.13016969 | 1 | 780 | tags=24%, list=3%, sig | MM_3 |
| GOBP_POSITIVE_REGULA | GOBP_POSITIVE_REGULATION_OF_VASCULA | 172 | -0.47683 | -1.6698 | 0 | 0.13076067 | 1 | 4715 | tags=35%, list=20%, si | MM_3 |
| GOBP_TRICARBOXYLIC_A | GOBP_TRICARBOXYLIC_ACID_CYCLE | 33 | -0.57159 | -1.66803 | 0.0025773195 | 0.13224246 | 1 | 6449 | tags=58%, list=27%, si | MM_3 |
| GOBP_NEGATIVE_REGUL | GOBP_NEGATIVE_REGULATION_OF_VIRAL_ | 95 | -0.50129 | -1.66678 | 0.0022522523 | 0.13296546 | 1 | 5582 | tags=42%, list=24%, si | MM_3 |
| GOBP_TYPE_I_INTERFER | GOBP_TYPE_I_INTERFERON_PRODUCTION | 127 | -0.48279 | -1.66511 | 0 | 0.13433988 | 1 | 4982 | tags=39%, list=21%, si | MM_3 |
| GOBP_REGULATION_OF_ | GOBP_REGULATION_OF_PEROXISOME_PRO | 17 | -0.658 | -1.6651 | 0.006821282 | 0.13289863 | 1 | 3183 | tags=35%, list=13%, si | MM_3 |
| GOBP_RESPONSE_TO_OX | GOBP_RESPONSE_TO_OXYGEN_LEVELS | 247 | -0.46283 | -1.66364 | 0 | 0.13411058 | 1 | 5372 | tags=38%, list=23%, si | MM_3 |
| GOBP_HEMATOPOIETIC_S | GOBP_HEMATOPOIETIC_STEM_CELL_HOME | 29 | -0.5901 | -1.66326 | 0.0052980133 | 0.13346404 | 1 | 5006 | tags=48%, list=21%, si | MM_3 |
| GOBP_NEGATIVE_REGUL | GOBP_NEGATIVE_REGULATION_OF_OLIGOD | 17 | -0.64607 | -1.66171 | 0.011049724 | 0.1347534 | 1 | 3606 | tags=35%, list=15%, si | MM_3 |
| GOBP_REGULATION_OF_ | GOBP_REGULATION_OF_CELL_GROWTH_IN | 33 | -0.57267 | -1.66139 | 0.0065616798 | 0.13382451 | 1 | 4567 | tags=48%, list=19%, si | MM_3 |
| GOBP_ORGANELLE_FUSIO | GOBP_ORGANELLE_FUSION | 167 | -0.46958 | -1.66135 | 0 | 0.13245247 | 1 | 5861 | tags=41%, list=25%, si | MM_3 |
| GOBP_MITOCHONDRIAL_ | GOBP_MITOCHONDRIAL_TRANSMEMBRANE | 92 | -0.50364 | -1.66082 | 0.0022547913 | 0.13199148 | 1 | 5448 | tags=37%, list=23%, si | MM_3 |
| GOBP_CHONDROITIN_SU | GOBP_CHONDROITIN_SULFATE_PROTEOGLY | 29 | -0.59172 | -1.66029 | 0.0051746443 | 0.13158236 | 1 | 5403 | tags=62%, list=23%, si | MM_3 |
| GOBP_CHAPERONE_COFA | GOBP_CHAPERONE_COFACTOR_DEPENDEN | 31 | -0.58207 | -1.65841 | 0.0038610038 | 0.13322163 | 1 | 4371 | tags=42%, list=19%, si | MM_3 |
| GOBP_MITOCHONDRIAL_ | GOBP_MITOCHONDRIAL_TRANSPORT | 174 | -0.47567 | -1.65791 | 0 | 0.13284013 | 1 | 4991 | tags=36%, list=21%, si | MM_3 |
| GOBP_POSITIVE_REGULA | GOBP_POSITIVE_REGULATION_OF_GLUCON | 26 | -0.60324 | -1.65728 | 0.0064683054 | 0.13246611 | 1 | 2026 | tags=27%, list=9%, sig | MM_3 |
| GOBP_RESPONSE_TO_DE | GOBP_RESPONSE_TO_DECREASED_OXYGEN | 228 | -0.46388 | -1.65521 | 0 | 0.13469072 | 1 | 5372 | tags=39%, list=23%, si | MM_3 |
| GOBP_MACROPHAGE_CH | GOBP_MACROPHAGE_CHEMOTAXIS | 45 | -0.54261 | -1.65492 | 0.0024479805 | 0.13411129 | 1 | 5954 | tags=53%, list=25%, si | MM_3 |
| GOBP_CELLULAR_RESPON | GOBP_CELLULAR_RESPONSE_TO_OSMOTIC_ | 52 | -0.53577 | -1.65418 | 0.0025608195 | 0.13396062 | 1 | 3970 | tags=31%, list=17%, si | MM_3 |
| GOBP_NEUTROPHIL_MIG | GOBP_NEUTROPHIL_MIGRATION | 104 | -0.48616 | -1.65346 | 0 | 0.1339235 | 1 | 5907 | tags=49%, list=25%, si | MM_3 |
| GOBP_REGULATION_OF_ | GOBP_REGULATION_OF_CD8_POSITIVE_ALF | 18 | -0.64074 | -1.65142 | 0.0054200543 | 0.13593021 | 1 | 5882 | tags=56%, list=25%, si | MM_3 |
| GOBP_NEUTROPHIL_CHE | GOBP_NEUTROPHIL_CHEMOTAXIS | 81 | -0.50185 | -1.64986 | 0.0011441647 | 0.13738713 | 1 | 5907 | tags=52%, list=25%, si | MM_3 |
| GOBP_MATURATION_OF_ | GOBP_MATURATION_OF_5_8S_RRNA_FRON | 24 | -0.60691 | -1.64954 | 0.0013850415 | 0.1365072 | 1 | 1782 | tags=38%, list=8%, sig | MM_3 |

| | | | | | | | | | | |
|---|---|---|---|---|---|---|---|---|---|---|
| GOBP_REGULATION_OF_ | GOBP_REGULATION_OF_HEMATOPOIETIC_S | 19 | -0.6343 | -1.6495 | 0.008344923 | 0.13525483 | 1 | 2439 | tags=32%, list=10%, si | MM_3 |
| GOBP_NEGATIVE_REGUL | GOBP_NEGATIVE_REGULATION_OF_RESPON | 112 | -0.48796 | -1.64879 | 0 | 0.13544795 | 1 | 3703 | tags=31%, list=16%, si | MM_3 |
| GOBP_REGULATION_OF_ | GOBP_REGULATION_OF_PROTEIN_SUMOYL | 25 | -0.59973 | -1.64644 | 0.007792208 | 0.13826634 | 1 | 6497 | tags=56%, list=28%, si | MM_3 |
| GOBP_VESICLE_DOCKING | GOBP_VESICLE_DOCKING_INVOLVED_IN_EX | 36 | -0.56777 | -1.64627 | 0.004968944 | 0.13726753 | 1 | 5612 | tags=53%, list=24%, si | MM_3 |
| GOBP_POSITIVE_REGULA | GOBP_POSITIVE_REGULATION_OF_MACROF | 31 | -0.57747 | -1.64553 | 0.001285347 | 0.13724095 | 1 | 5962 | tags=58%, list=25%, si | MM_3 |
| GOBP_METALLO_SULFUR | GOBP_METALLO_SULFUR_CLUSTER_ASSEMI | 25 | -0.59339 | -1.64552 | 0.0068587107 | 0.13603708 | 1 | 4979 | tags=52%, list=21%, si | MM_3 |
| PANCREATIC_CANCER | PANCREATIC_CANCER | 74 | -0.50531 | -1.64517 | 0.0022753128 | 0.13546035 | 1 | 6853 | tags=55%, list=29%, si | MM_3 |
| GOBP_SNRNA_TRANSCRI | GOBP_SNRNA_TRANSCRIPTION | 18 | -0.64492 | -1.64207 | 0.00660502 | 0.13904789 | 1 | 4577 | tags=61%, list=19%, si | MM_3 |
| GOBP_PROTEIN_SUMOYL | GOBP_PROTEIN_SUMOYLATION | 42 | -0.546 | -1.64129 | 0.006329114 | 0.13927661 | 1 | 6601 | tags=55%, list=28%, si | MM_3 |
| GOBP_MITOCHONDRIAL_ | GOBP_MITOCHONDRIAL_CALCIUM_ION_HC | 29 | -0.58143 | -1.64105 | 0.015768725 | 0.13856171 | 1 | 5227 | tags=45%, list=22%, si | MM_3 |
| GOBP_ARGININE_METAB | GOBP_ARGININE_METABOLIC_PROCESS | 17 | -0.65559 | -1.64053 | 0.008633094 | 0.13839315 | 1 | 3167 | tags=35%, list=13%, si | MM_3 |
| GOBP_INTERFERON_BETA | GOBP_INTERFERON_BETA_PRODUCTION | 66 | -0.50688 | -1.64007 | 0.0023501762 | 0.13811234 | 1 | 6346 | tags=45%, list=27%, si | MM_3 |
| GOBP_PROTEIN_K48_LIN | GOBP_PROTEIN_K48_LINKED_DEUBIQUITIN | 29 | -0.58345 | -1.63909 | 0.0052562417 | 0.13881364 | 1 | 1739 | tags=28%, list=7%, sig | MM_3 |
| GOBP_MEMBRANE_DOCI | GOBP_MEMBRANE_DOCKING | 78 | -0.50447 | -1.63792 | 0.0022883294 | 0.13961895 | 1 | 6237 | tags=54%, list=26%, si | MM_3 |
| GOBP_MANNOSYLATION | GOBP_MANNOSYLATION | 22 | -0.61669 | -1.6377 | 0.0054719565 | 0.13886195 | 1 | 6206 | tags=55%, list=26%, si | MM_3 |
| GOBP_ENDOLYSOSOMAL | GOBP_ENDOLYSOSOMAL_TOLL_LIKE_RECEP | 47 | -0.53138 | -1.63685 | 0.006234414 | 0.13908932 | 1 | 5034 | tags=40%, list=21%, si | MM_3 |
| GOBP_NEGATIVE_REGUL | GOBP_NEGATIVE_REGULATION_OF_BMP_SI | 53 | -0.53032 | -1.63589 | 0.0023980816 | 0.13967547 | 1 | 2241 | tags=23%, list=9%, sig | MM_3 |
| GOBP_NEGATIVE_REGUL | GOBP_NEGATIVE_REGULATION_OF_NUCLE( | 20 | -0.62841 | -1.63576 | 0.008207934 | 0.13876832 | 1 | 4842 | tags=50%, list=21%, si | MM_3 |
| GOBP_VESICLE_DOCKING | GOBP_VESICLE_DOCKING | 55 | -0.52448 | -1.63403 | 0.0046948357 | 0.14084075 | 1 | 5837 | tags=51%, list=25%, si | MM_3 |
| GOBP_MATURATION_OF_ | GOBP_MATURATION_OF_5_8S_RRNA | 35 | -0.55952 | -1.63247 | 0.006393862 | 0.14248931 | 1 | 2467 | tags=37%, list=10%, si | MM_3 |
| GOBP_NEGATIVE_REGUL | GOBP_NEGATIVE_REGULATION_OF_VIRAL_ | 18 | -0.63108 | -1.63164 | 0.01462766 | 0.14295717 | 1 | 3678 | tags=39%, list=16%, si | MM_3 |
| GOBP_POSITIVE_REGULA | GOBP_POSITIVE_REGULATION_OF_MYELOID | 21 | -0.62003 | -1.6316 | 0.0041958042 | 0.14195494 | 1 | 6965 | tags=71%, list=29%, si | MM_3 |
| TRANSCRIPTIONAL_MISRI | TRANSCRIPTIONAL_MISREGULATION_IN_CA | 170 | -0.46873 | -1.63045 | 0 | 0.14285566 | 1 | 6323 | tags=43%, list=27%, si | MM_3 |
| GOBP_NEGATIVE_REGUL | GOBP_NEGATIVE_REGULATION_OF_NUCLE( | 29 | -0.57985 | -1.63007 | 0.00397351 | 0.1424295 | 1 | 4842 | tags=41%, list=21%, si | MM_3 |
| GOBP_NUCLEAR_TRANSC | GOBP_NUCLEAR_TRANSCRIBED_MRNA_CAT | 118 | -0.47162 | -1.62863 | 0 | 0.14401281 | 1 | 6483 | tags=45%, list=27%, si | MM_3 |
| GOBP_NEGATIVE_REGUL | GOBP_NEGATIVE_REGULATION_OF_DEFENS | 242 | -0.45288 | -1.62768 | 0 | 0.14453965 | 1 | 4736 | tags=34%, list=20%, si | MM_3 |
| GOBP_NEGATIVE_REGUL | GOBP_NEGATIVE_REGULATION_OF_VIRAL_( | 50 | -0.52745 | -1.62716 | 0.006045949 | 0.14433749 | 1 | 5582 | tags=42%, list=24%, si | MM_3 |
| GOBP_RESPONSE_TO_IN | GOBP_RESPONSE_TO_INTERFERON_BETA | 64 | -0.5061 | -1.6264 | 0.002366864 | 0.14455065 | 1 | 4206 | tags=38%, list=18%, si | MM_3 |
| GOBP_GLOMERULAR_EPI | GOBP_GLOMERULAR_EPITHELIUM_DEVELO | 21 | -0.60826 | -1.62583 | 0.008152174 | 0.1445379 | 1 | 5457 | tags=52%, list=23%, si | MM_3 |
| GOBP_RESPIRATORY_CHA | GOBP_RESPIRATORY_CHAIN_COMPLEX_IV_ | 24 | -0.59558 | -1.6248 | 0.010723861 | 0.14535119 | 1 | 6597 | tags=67%, list=28%, si | MM_3 |
| GOBP_NEGATIVE_REGUL | GOBP_NEGATIVE_REGULATION_OF_RHO_PI | 25 | -0.58571 | -1.62458 | 0.0068399454 | 0.14458683 | 1 | 5028 | tags=52%, list=21%, si | MM_3 |
| GOBP_POSITIVE_REGULA | GOBP_POSITIVE_REGULATION_OF_STRIATE | 26 | -0.58989 | -1.62354 | 0.012096774 | 0.14545679 | 1 | 5440 | tags=50%, list=23%, si | MM_3 |
| GOBP_POSITIVE_REGULA | GOBP_POSITIVE_REGULATION_OF_INTERLE | 15 | -0.66045 | -1.62215 | 0.009817672 | 0.14677857 | 1 | 6363 | tags=73%, list=27%, si | MM_3 |
| GOBP_INTRACELLULAR_Z | GOBP_INTRACELLULAR_ZINC_ION_HOMEOS | 24 | -0.59869 | -1.62207 | 0.009536785 | 0.14587933 | 1 | 3702 | tags=38%, list=16%, si | MM_3 |
| GOBP_NEGATIVE_REGUL | GOBP_NEGATIVE_REGULATION_OF_INTRINS | 21 | -0.60981 | -1.62099 | 0.012430939 | 0.1469073 | 1 | 6165 | tags=57%, list=26%, si | MM_3 |
| GOBP_SNO_S_RNA_META | GOBP_SNO_S_RNA_METABOLIC_PROCESS | 17 | -0.64698 | -1.61981 | 0.012448133 | 0.14791979 | 1 | 2175 | tags=29%, list=9%, sig | MM_3 |
| GOBP_PYRIDINE_CONTAI | GOBP_PYRIDINE_CONTAINING_COMPOUND | 182 | -0.46097 | -1.61958 | 0 | 0.14724974 | 1 | 5936 | tags=43%, list=25%, si | MM_3 |
| GOBP_FATTY_ACID_BIOS | GOBP_FATTY_ACID_BIOSYNTHETIC_PROCES | 145 | -0.46423 | -1.61956 | 0 | 0.14631338 | 1 | 5702 | tags=38%, list=24%, si | MM_3 |
| GOBP_RESPONSE_TO_LEI | GOBP_RESPONSE_TO_LEPTIN | 19 | -0.6275 | -1.61803 | 0.016830295 | 0.1479867 | 1 | 4844 | tags=47%, list=21%, si | MM_3 |
| GOBP_NEGATIVE_REGUL | GOBP_NEGATIVE_REGULATION_OF_CELL_A | 207 | -0.45929 | -1.61785 | 0 | 0.14733005 | 1 | 4408 | tags=32%, list=19%, si | MM_3 |
| GOBP_ENERGY_HOMEOS | GOBP_ENERGY_HOMEOSTASIS | 92 | -0.49013 | -1.61749 | 0.0011273957 | 0.14701578 | 1 | 5328 | tags=35%, list=23%, si | MM_3 |
| OXIDATIVE_PHOSPHORYL | OXIDATIVE_PHOSPHORYLATION | 115 | -0.47705 | -1.61707 | 0 | 0.1467547 | 1 | 7071 | tags=65%, list=30%, si | MM_3 |
| N-GLYCAN_BIOSYNTHESIS | N-GLYCAN_BIOSYNTHESIS | 48 | -0.53279 | -1.61545 | 0.00243309 | 0.14870127 | 1 | 6969 | tags=63%, list=30%, si | MM_3 |
| GOBP_REGULATION_OF_ | GOBP_REGULATION_OF_PROTEIN_ACETYLA | 22 | -0.61308 | -1.61519 | 0.012552301 | 0.1482031 | 1 | 6626 | tags=59%, list=28%, si | MM_3 |
| GOBP_POSITIVE_REGULA | GOBP_POSITIVE_REGULATION_OF_GRANUL | 31 | -0.57209 | -1.61475 | 0.013003902 | 0.1478971 | 1 | 5896 | tags=61%, list=25%, si | MM_3 |
| GOBP_CHONDROITIN_SU | GOBP_CHONDROITIN_SULFATE_METABOLIC | 22 | -0.59782 | -1.61447 | 0.017264277 | 0.14728686 | 1 | 5403 | tags=64%, list=23%, si | MM_3 |
| GOBP_REGULATION_OF_ | GOBP_REGULATION_OF_RELEASE_OF_CYTO | 49 | -0.52716 | -1.61415 | 0.0061500617 | 0.14690278 | 1 | 4991 | tags=43%, list=21%, si | MM_3 |
| GOBP_RESPONSE_TO_HY | GOBP_RESPONSE_TO_HYDROPEROXIDE | 18 | -0.63634 | -1.61378 | 0.017615177 | 0.14648235 | 1 | 5946 | tags=56%, list=25%, si | MM_3 |
| GOBP_POSITIVE_REGULA | GOBP_POSITIVE_REGULATION_OF_MITOTIC | 126 | -0.46816 | -1.61254 | 0.0010940919 | 0.14769019 | 1 | 6346 | tags=48%, list=27%, si | MM_3 |
| RIBOSOME | RIBOSOME | 128 | -0.47402 | -1.61202 | 0 | 0.14776555 | 1 | 7947 | tags=79%, list=34%, si | MM_3 |
| GOBP_DNA_STRAND_ELC | GOBP_DNA_STRAND_ELONGATION | 36 | -0.54691 | -1.61198 | 0.0050251256 | 0.14688952 | 1 | 2967 | tags=33%, list=13%, si | MM_3 |
| GOBP_POSITIVE_REGULA | GOBP_POSITIVE_REGULATION_OF_MITOTIC | 93 | -0.48978 | -1.61084 | 0 | 0.14805295 | 1 | 6162 | tags=48%, list=26%, si | MM_3 |
| GOBP_NEGATIVE_REGUL | GOBP_NEGATIVE_REGULATION_OF_INNATE | 86 | -0.48973 | -1.61046 | 0 | 0.14777724 | 1 | 4466 | tags=35%, list=19%, si | MM_3 |
| GOBP_REGULATION_OF_ | GOBP_REGULATION_OF_MITOCHONDRIAL_ | 35 | -0.55714 | -1.60926 | 0.006393862 | 0.14891294 | 1 | 2575 | tags=37%, list=11%, si | MM_3 |
| GOBP_POSITIVE_REGULA | GOBP_POSITIVE_REGULATION_OF_MEMBRA | 22 | -0.59826 | -1.60902 | 0.007894737 | 0.14846021 | 1 | 4991 | tags=50%, list=21%, si | MM_3 |
| GOBP_CELLULAR_RESPOI | GOBP_CELLULAR_RESPONSE_TO_LEPTIN_ST | 16 | -0.63868 | -1.60896 | 0.018030513 | 0.14769626 | 1 | 4844 | tags=50%, list=21%, si | MM_3 |
| GOBP_TETRAPYRROLE_BI | GOBP_TETRAPYRROLE_BIOSYNTHETIC_PROC | 29 | -0.56681 | -1.60867 | 0.01344086 | 0.14732635 | 1 | 4980 | tags=45%, list=21%, si | MM_3 |
| GOBP_ATP_SYNTHESIS_C | GOBP_ATP_SYNTHESIS_COUPLED_ELECTRO | 76 | -0.49115 | -1.60665 | 0.0023255814 | 0.14995448 | 1 | 7269 | tags=59%, list=31%, si | MM_3 |
| GOBP_POSITIVE_REGULA | GOBP_POSITIVE_REGULATION_OF_TUMOR_ | 113 | -0.4706 | -1.60653 | 0 | 0.14921477 | 1 | 6556 | tags=55%, list=28%, si | MM_3 |
| GOBP_TELOMERE_MAINT | GOBP_TELOMERE_MAINTENANCE_IN_RESP | 32 | -0.55824 | -1.60605 | 0.010376135 | 0.14916341 | 1 | 4326 | tags=44%, list=18%, si | MM_3 |
| PARKINSON_DISEASE | PARKINSON_DISEASE | 122 | -0.47235 | -1.60472 | 0.0010787486 | 0.15075155 | 1 | 7071 | tags=61%, list=30%, si | MM_3 |
| GOBP_S_ADENOSYLMETH | GOBP_S_ADENOSYLMETHIONINE_METABOL | 17 | -0.63617 | -1.60422 | 0.017687075 | 0.15077898 | 1 | 6775 | tags=76%, list=29%, si | MM_3 |
| GOBP_MITOCHONDRIAL_ | GOBP_MITOCHONDRIAL_FISSION | 48 | -0.52409 | -1.60418 | 0.007246377 | 0.14993435 | 1 | 2575 | tags=33%, list=11%, si | MM_3 |
| GOBP_WHITE_FAT_CELL_ | GOBP_WHITE_FAT_CELL_DIFFERENTIATION | 22 | -0.60247 | -1.60258 | 0.012396694 | 0.15201262 | 1 | 2376 | tags=32%, list=10%, si | MM_3 |
| GOBP_PEPTIDYL_LYSINE_ | GOBP_PEPTIDYL_LYSINE_MODIFICATION | 128 | -0.46747 | -1.60194 | 0.00110011 | 0.1521358 | 1 | 6125 | tags=42%, list=26%, si | MM_3 |
| GOBP_NEGATIVE_REGUL | GOBP_NEGATIVE_REGULATION_OF_GLIAL_( | 34 | -0.54006 | -1.60088 | 0.01303781 | 0.15302372 | 1 | 3606 | tags=32%, list=15%, si | MM_3 |
| GOBP_TETRAPYRROLE_M | GOBP_TETRAPYRROLE_METABOLIC_PROCES | 49 | -0.53101 | -1.60043 | 0.0011990408 | 0.15291104 | 1 | 4980 | tags=41%, list=21%, si | MM_3 |
| GOBP_ENERGY_DERIVATI | GOBP_ENERGY_DERIVATION_BY_OXIDATIOI | 321 | -0.44019 | -1.60039 | 0 | 0.15209623 | 1 | 7071 | tags=51%, list=30%, si | MM_3 |
| GOBP_MEMBRANE_PROT | GOBP_MEMBRANE_PROTEIN_ECTODOMAIN | 47 | -0.51842 | -1.60021 | 0.00373599 | 0.15152954 | 1 | 6718 | tags=55%, list=28%, si | MM_3 |
| GOBP_POSITIVE_REGULA | GOBP_POSITIVE_REGULATION_OF_B_CELL_ | 47 | -0.52356 | -1.59985 | 0.0012376237 | 0.15130179 | 1 | 3809 | tags=38%, list=16%, si | MM_3 |
| GOBP_POSITIVE_REGULA | GOBP_POSITIVE_REGULATION_OF_PATTERN | 68 | -0.49976 | -1.59956 | 0.0023419203 | 0.15100004 | 1 | 6602 | tags=51%, list=28%, si | MM_3 |
| GOBP_REGULATION_OF_ | GOBP_REGULATION_OF_INTRINSIC_APOPT( | 34 | -0.55777 | -1.59936 | 0.005121639 | 0.15049629 | 1 | 7515 | tags=62%, list=32%, si | MM_3 |
| GOBP_ADRENAL_GLAND_ | GOBP_ADRENAL_GLAND_DEVELOPMENT | 22 | -0.59947 | -1.59772 | 0.01589404 | 0.15263806 | 1 | 1617 | tags=23%, list=7%, sig | MM_3 |
| GOBP_POSITIVE_REGULA | GOBP_POSITIVE_REGULATION_OF_TYPE_I_I | 77 | -0.49253 | -1.59667 | 0.003508772 | 0.15350655 | 1 | 3585 | tags=30%, list=15%, si | MM_3 |
| GOBP_PERICARDIUM_DE | GOBP_PERICARDIUM_DEVELOPMENT | 27 | -0.5722 | -1.5964 | 0.015706806 | 0.15311281 | 1 | 2846 | tags=30%, list=12%, si | MM_3 |
| GOBP_ASTROCYTE_ACTIV | GOBP_ASTROCYTE_ACTIVATION | 21 | -0.60536 | -1.5933 | 0.017105263 | 0.15774168 | 1 | 6142 | tags=67%, list=26%, si | MM_3 |
| GOBP_CRANIOFACIAL_SU | GOBP_CRANIOFACIAL_SUTURE_MORPHOGE | 15 | -0.64432 | -1.59324 | 0.024390243 | 0.15695775 | 1 | 2814 | tags=40%, list=12%, si | MM_3 |
| GOBP_PROTEIN_O_LINKE | GOBP_PROTEIN_O_LINKED_MANNOSYLATIC | 16 | -0.6438 | -1.5927 | 0.016783217 | 0.15715371 | 1 | 5942 | tags=56%, list=25%, si | MM_3 |
| GOBP_CELLULAR_OXIDAN | GOBP_CELLULAR_OXIDANT_DETOXIFICATIO | 29 | -0.56441 | -1.59145 | 0.0132626 | 0.15853463 | 1 | 5116 | tags=48%, list=22%, si | MM_3 |
| GOBP_NAD_METABOLIC_ | GOBP_NAD_METABOLIC_PROCESS | 58 | -0.50873 | -1.59132 | 0 | 0.15794405 | 1 | 5278 | tags=41%, list=22%, si | MM_3 |
| GOBP_CELLULAR_RESPOI | GOBP_CELLULAR_RESPONSE_TO_DSRNA | 19 | -0.61534 | -1.58871 | 0.01830986 | 0.16159973 | 1 | 1159 | tags=26%, list=5%, sig | MM_3 |
| GOBP_NEGATIVE_REGUL | GOBP_NEGATIVE_REGULATION_OF_CELLUL | 18 | -0.61475 | -1.58776 | 0.015384615 | 0.1625864 | 1 | 4739 | tags=50%, list=20%, si | MM_3 |

| | | | | | | | | | | |
|---|---|---|---|---|---|---|---|---|---|---|
| GOBP_MODULATION_OF_ | GOBP_MODULATION_OF_PROCESS_OF_AN | 17 | -0.63234 | -1.58677 | 0.017906336 | 0.16355594 | 1 | 1710 | tags=24%, list=7%, sig | MM_3 |
| GOBP_PROSTANOID_BIO | GOBP_PROSTANOID_BIOSYNTHETIC_PROCE | 30 | -0.55728 | -1.58655 | 0.009043927 | 0.16308922 | 1 | 5935 | tags=47%, list=25%, si | MM_3 |
| GOBP_APOPTOTIC_MITO | GOBP_APOPTOTIC_MITOCHONDRIAL_CHAN | 118 | -0.46503 | -1.5848 | 0.0022346368 | 0.16529347 | 1 | 4991 | tags=38%, list=21%, si | MM_3 |
| GOBP_SOMATIC_DIVERSI | GOBP_SOMATIC_DIVERSIFICATION_OF_IMN | 55 | -0.515 | -1.5842 | 0.0024271845 | 0.1655426 | 1 | 3651 | tags=33%, list=15%, si | MM_3 |
| GOBP_POSITIVE_REGULA | GOBP_POSITIVE_REGULATION_OF_GLIAL_C | 51 | -0.51461 | -1.58393 | 0.0060975607 | 0.1651004 | 1 | 3348 | tags=25%, list=14%, si | MM_3 |
| GOBP_MAMMARY_GLAN | GOBP_MAMMARY_GLAND_EPITHELIAL_CEL | 16 | -0.6392 | -1.58312 | 0.013908206 | 0.16573112 | 1 | 4951 | tags=56%, list=21%, si | MM_3 |
| GOBP_MITOCHONDRIAL_ | GOBP_MITOCHONDRIAL_FUSION | 36 | -0.54114 | -1.58245 | 0.010335918 | 0.16619805 | 1 | 3576 | tags=31%, list=15%, si | MM_3 |
| GOBP_REGULATION_OF_ | GOBP_REGULATION_OF_DNA_DAMAGE_RE | 36 | -0.54756 | -1.58159 | 0.010152284 | 0.1669008 | 1 | 5538 | tags=47%, list=23%, si | MM_3 |
| GOBP_REGULATION_OF_ | GOBP_REGULATION_OF_NEUROINFLAMMA | 30 | -0.55401 | -1.57939 | 0.023255814 | 0.17032045 | 1 | 3489 | tags=37%, list=15%, si | MM_3 |
| GOBP_REGULATION_OF_ | GOBP_REGULATION_OF_SUPEROXIDE_MET | 34 | -0.54131 | -1.57925 | 0.015132409 | 0.16972248 | 1 | 4991 | tags=50%, list=21%, si | MM_3 |
| GOBP_ADIPOSE_TISSUE_ | GOBP_ADIPOSE_TISSUE_DEVELOPMENT | 64 | -0.49933 | -1.57924 | 0.004716981 | 0.16889381 | 1 | 6133 | tags=44%, list=26%, si | MM_3 |
| GOBP_CELL_REDOX_HOM | GOBP_CELL_REDOX_HOMEOSTASIS | 28 | -0.57027 | -1.57915 | 0.021390375 | 0.1682249 | 1 | 5116 | tags=54%, list=22%, si | MM_3 |
| GOBP_ANTIGEN_PROCES | GOBP_ANTIGEN_PROCESSING_AND_PRESEN | 105 | -0.46822 | -1.57896 | 0.0011198209 | 0.16771875 | 1 | 6966 | tags=57%, list=30%, si | MM_3 |
| GOBP_MITOCHONDRIAL_ | GOBP_MITOCHONDRIAL_MEMBRANE_ORG | 107 | -0.46941 | -1.57792 | 0.0011111111 | 0.16867806 | 1 | 5448 | tags=41%, list=23%, si | MM_3 |
| GOBP_CARDIAC_MUSCLE | GOBP_CARDIAC_MUSCLE_ADAPTATION | 30 | -0.54864 | -1.5779 | 0.013157895 | 0.16788612 | 1 | 2473 | tags=20%, list=10%, si | MM_3 |
| GOBP_POSITIVE_REGULA | GOBP_POSITIVE_REGULATION_OF_ENDOTH | 99 | -0.47254 | -1.57767 | 0.0011111111 | 0.16747682 | 1 | 6209 | tags=47%, list=26%, si | MM_3 |
| GOBP_NADH_DEHYDROG | GOBP_NADH_DEHYDROGENASE_COMPLEX_ | 58 | -0.50511 | -1.57691 | 0.010869565 | 0.16805063 | 1 | 7153 | tags=64%, list=30%, si | MM_3 |
| GOBP_REGULATION_OF_ | GOBP_REGULATION_OF_SYNAPTIC_VESICLE | 19 | -0.60258 | -1.57649 | 0.018741634 | 0.16788757 | 1 | 5922 | tags=68%, list=25%, si | MM_3 |
| GOBP_NUCLEOSIDE_PHO | GOBP_NUCLEOSIDE_PHOSPHATE_BIOSYNTH | 234 | -0.43529 | -1.57629 | 0 | 0.1675093 | 1 | 5983 | tags=41%, list=25%, si | MM_3 |
| GOBP_CYTOPLASMIC_TR | GOBP_CYTOPLASMIC_TRANSLATIONAL_INIT | 33 | -0.55612 | -1.57536 | 0.016905071 | 0.16828671 | 1 | 5775 | tags=39%, list=24%, si | MM_3 |
| GOBP_COPPER_ION_HON | GOBP_COPPER_ION_HOMEOSTASIS | 18 | -0.61816 | -1.57431 | 0.021008404 | 0.16939744 | 1 | 4131 | tags=39%, list=17%, si | MM_3 |
| GOBP_RIBOSE_PHOSPHA | GOBP_RIBOSE_PHOSPHATE_BIOSYNTHETIC_ | 185 | -0.44726 | -1.574 | 0 | 0.16913716 | 1 | 5671 | tags=41%, list=24%, si | MM_3 |
| CYTOSOLIC_DNA-SENSING | CYTOSOLIC_DNA-SENSING_PATHWAY | 46 | -0.51444 | -1.57298 | 0.0073710075 | 0.17011844 | 1 | 6111 | tags=50%, list=26%, si | MM_3 |
| GOBP_LYMPHOCYTE_CHE | GOBP_LYMPHOCYTE_CHEMOTAXIS | 38 | -0.53761 | -1.57251 | 0.012578616 | 0.1702322 | 1 | 2464 | tags=26%, list=10%, si | MM_3 |
| GOBP_MITOCHONDRIAL_ | GOBP_MITOCHONDRIAL_CYTOCHROME_C_ | 19 | -0.59865 | -1.57211 | 0.019283747 | 0.17005576 | 1 | 6597 | tags=74%, list=28%, si | MM_3 |
| GOBP_POSITIVE_REGULA | GOBP_POSITIVE_REGULATION_OF_INFLAMI | 22 | -0.58779 | -1.57169 | 0.01897019 | 0.17006086 | 1 | 8111 | tags=73%, list=34%, si | MM_3 |
| GOBP_GENERATION_OF_ | GOBP_GENERATION_OF_PRECURSOR_MET | 439 | -0.42996 | -1.57162 | 0 | 0.16935982 | 1 | 7071 | tags=51%, list=30%, si | MM_3 |
| GOBP_REGULATION_OF_ | GOBP_REGULATION_OF_ISOTYPE_SWITCHIN | 38 | -0.53791 | -1.57142 | 0.010025063 | 0.16895162 | 1 | 3651 | tags=37%, list=15%, si | MM_3 |
| GOBP_REGULATION_OF_ | GOBP_REGULATION_OF_MUSCLE_HYPERTR | 80 | -0.47578 | -1.57115 | 0.0022701477 | 0.16866736 | 1 | 5491 | tags=40%, list=23%, si | MM_3 |
| GOBP_DEOXYRIBONUCLE | GOBP_DEOXYRIBONUCLEOTIDE_BIOSYNTHE | 17 | -0.61548 | -1.56932 | 0.02202643 | 0.17105177 | 1 | 5405 | tags=53%, list=23%, si | MM_3 |
| GOBP_NUCLEOSIDE_DIPH | GOBP_NUCLEOSIDE_DIPHOSPHATE_METAB | 127 | -0.45986 | -1.56926 | 0.0010976949 | 0.17038274 | 1 | 5278 | tags=39%, list=22%, si | MM_3 |
| GOBP_POSITIVE_REGULA | GOBP_POSITIVE_REGULATION_OF_FATTY_A | 45 | -0.52293 | -1.56911 | 0.0037783375 | 0.16989154 | 1 | 2458 | tags=24%, list=10%, si | MM_3 |
| GOBP_POSITIVE_REGULA | GOBP_POSITIVE_REGULATION_OF_CELLULA | 20 | -0.5952 | -1.56886 | 0.01650619 | 0.16960812 | 1 | 5686 | tags=50%, list=24%, si | MM_3 |
| GOBP_ANTIGEN_PROCES | GOBP_ANTIGEN_PROCESSING_AND_PRESEN | 20 | -0.59414 | -1.56856 | 0.022911051 | 0.16951154 | 1 | 6240 | tags=75%, list=26%, si | MM_3 |
| GOBP_MIRNA_METABOL | GOBP_MIRNA_METABOLIC_PROCESS | 103 | -0.46417 | -1.56756 | 0.0011210763 | 0.17059462 | 1 | 6718 | tags=48%, list=28%, si | MM_3 |
| GOBP_RESPONSE_TO_CH | GOBP_RESPONSE_TO_CHEMOKINE | 74 | -0.483 | -1.56669 | 0.0023228803 | 0.17147176 | 1 | 4529 | tags=31%, list=19%, si | MM_3 |
| GOBP_REGULATION_OF_ | GOBP_REGULATION_OF_TELOMERE_CAPPIN | 26 | -0.57892 | -1.56659 | 0.014765101 | 0.17087045 | 1 | 5125 | tags=54%, list=22%, si | MM_3 |
| GOBP_REGULATION_OF_ | GOBP_REGULATION_OF_RESPONSE_TO_OS | 18 | -0.60981 | -1.56619 | 0.020519836 | 0.17089088 | 1 | 4707 | tags=44%, list=20%, si | MM_3 |
| GOBP_ORGANELLE_MEM | GOBP_ORGANELLE_MEMBRANE_FUSION | 123 | -0.45876 | -1.56614 | 0 | 0.17023237 | 1 | 5861 | tags=42%, list=25%, si | MM_3 |
| MITOPHAGY | MITOPHAGY | 62 | -0.4944 | -1.56485 | 0.009603841 | 0.17189968 | 1 | 6338 | tags=45%, list=27%, si | MM_3 |
| GOBP_REGULATION_OF_ | GOBP_REGULATION_OF_B_CELL_ACTIVATIC | 141 | -0.45092 | -1.56456 | 0.0010706638 | 0.1716361 | 1 | 3764 | tags=32%, list=16%, si | MM_3 |
| GOBP_NEGATIVE_REGUL | GOBP_NEGATIVE_REGULATION_OF_RESPON | 79 | -0.48323 | -1.56444 | 0.0046189376 | 0.17109671 | 1 | 4568 | tags=34%, list=19%, si | MM_3 |
| GOBP_REGULATION_OF_ | GOBP_REGULATION_OF_SIGNAL_TRANSDU | 83 | -0.47994 | -1.5642 | 0 | 0.17088892 | 1 | 5886 | tags=41%, list=25%, si | MM_3 |
| GOBP_POSITIVE_REGULA | GOBP_POSITIVE_REGULATION_OF_PROTEIN | 16 | -0.63049 | -1.56281 | 0.029577464 | 0.17271852 | 1 | 6120 | tags=50%, list=26%, si | MM_3 |
| GOBP_ESTABLISHMENT_ | GOBP_ESTABLISHMENT_OF_PROTEIN_LOCA | 67 | -0.47868 | -1.56259 | 0.0023148148 | 0.1724737 | 1 | 5760 | tags=43%, list=24%, si | MM_3 |
| GOBP_POSITIVE_REGULA | GOBP_POSITIVE_REGULATION_OF_FAT_CEL | 71 | -0.48466 | -1.56244 | 0.0023201855 | 0.17200251 | 1 | 3518 | tags=28%, list=15%, si | MM_3 |
| GOBP_NEGATIVE_REGUL | GOBP_NEGATIVE_REGULATION_OF_REACTI | 52 | -0.49914 | -1.56219 | 0.012106538 | 0.17177671 | 1 | 5992 | tags=46%, list=25%, si | MM_3 |
| GOBP_REGULATION_OF_ | GOBP_REGULATION_OF_MACROPHAGE_CH | 32 | -0.54248 | -1.56168 | 0.010416667 | 0.17201853 | 1 | 5954 | tags=53%, list=25%, si | MM_3 |
| GOBP_REGULATION_OF_ | GOBP_REGULATION_OF_MACROPHAGE_DE | 25 | -0.57532 | -1.56162 | 0.019736841 | 0.17143594 | 1 | 3426 | tags=32%, list=15%, si | MM_3 |
| GOBP_FEMALE_MEIOTIC_ | GOBP_FEMALE_MEIOTIC_NUCLEAR_DIVISIO | 41 | -0.52349 | -1.56139 | 0.014962593 | 0.17108782 | 1 | 3762 | tags=41%, list=16%, si | MM_3 |
| GOBP_EXECUTION_PHAS | GOBP_EXECUTION_PHASE_OF_APOPTOSIS | 78 | -0.47958 | -1.56116 | 0.0034364262 | 0.17091313 | 1 | 4046 | tags=37%, list=17%, si | MM_3 |
| GOBP_PROTEIN_LOCALIZ | GOBP_PROTEIN_LOCALIZATION_TO_ENDOP | 76 | -0.48224 | -1.56102 | 0.0035211267 | 0.17049928 | 1 | 6391 | tags=50%, list=27%, si | MM_3 |
| GOBP_HEME_BIOSYNTHE | GOBP_HEME_BIOSYNTHETIC_PROCESS | 24 | -0.57447 | -1.56092 | 0.017333332 | 0.1699762 | 1 | 4980 | tags=46%, list=21%, si | MM_3 |
| GOBP_LEUKOCYTE_AGGR | GOBP_LEUKOCYTE_AGGREGATION | 16 | -0.62271 | -1.56068 | 0.024390243 | 0.16967364 | 1 | 5896 | tags=63%, list=25%, si | MM_3 |
| GOBP_ICOSANOID_BIOSY | GOBP_ICOSANOID_BIOSYNTHETIC_PROCESS | 43 | -0.51998 | -1.56061 | 0.019255456 | 0.16908456 | 1 | 5935 | tags=40%, list=25%, si | MM_3 |
| GOBP_CGAS_STING_SIGN | GOBP_CGAS_STING_SIGNALING_PATHWAY | 16 | -0.62436 | -1.56011 | 0.02086231 | 0.16925183 | 1 | 5663 | tags=63%, list=24%, si | MM_3 |
| GOBP_PEPTIDYL_THREON | GOBP_PEPTIDYL_THREONINE_DEPHOSPHOR | 15 | -0.63691 | -1.55903 | 0.020718232 | 0.17058639 | 1 | 4736 | tags=53%, list=20%, si | MM_3 |
| GOBP_REGULATION_OF_ | GOBP_REGULATION_OF_VASCULATURE_DE | 295 | -0.43209 | -1.5589 | 0 | 0.17012423 | 1 | 4783 | tags=33%, list=20%, si | MM_3 |
| GOBP_DNA_REPLICATION | GOBP_DNA_REPLICATION | 263 | -0.43414 | -1.55801 | 0 | 0.17106235 | 1 | 5651 | tags=40%, list=24%, si | MM_3 |
| GOBP_NEGATIVE_REGUL | GOBP_NEGATIVE_REGULATION_OF_HEMOP | 121 | -0.45493 | -1.55771 | 0.0021715527 | 0.17102116 | 1 | 4942 | tags=32%, list=21%, si | MM_3 |
| GOBP_SIGNALING_RECEP | GOBP_SIGNALING_RECEPTOR_LIGAND_PRE | 29 | -0.55235 | -1.55738 | 0.013404826 | 0.1709505 | 1 | 2246 | tags=31%, list=10%, si | MM_3 |
| GOBP_MUSCLE_HYPERTR | GOBP_MUSCLE_HYPERTROPHY | 115 | -0.45576 | -1.55638 | 0.0021551724 | 0.17223594 | 1 | 6172 | tags=43%, list=26%, si | MM_3 |
| GOBP_REGULATION_OF_ | GOBP_REGULATION_OF_GRANULOCYTE_CH | 59 | -0.49204 | -1.55566 | 0.007194245 | 0.17289169 | 1 | 5954 | tags=56%, list=25%, si | MM_3 |
| GOBP_NUCLEOSIDE_DIPH | GOBP_NUCLEOSIDE_DIPHOSPHATE_CATABO | 111 | -0.45926 | -1.55525 | 0.0033076075 | 0.17292649 | 1 | 5278 | tags=38%, list=22%, si | MM_3 |
| GOBP_DEFINITIVE_HEMO | GOBP_DEFINITIVE_HEMOPOIESIS | 25 | -0.5715 | -1.55517 | 0.021650879 | 0.17239326 | 1 | 4458 | tags=44%, list=19%, si | MM_3 |
| GOBP_RESPONSE_TO_DI | GOBP_RESPONSE_TO_DIETARY_EXCESS | 44 | -0.51011 | -1.55448 | 0.017521903 | 0.17300369 | 1 | 2198 | tags=20%, list=9%, sig | MM_3 |
| GOBP_REGULATION_OF_ | GOBP_REGULATION_OF_DNA_TEMPLATED_ | 79 | -0.47582 | -1.55381 | 0.0022497189 | 0.17359701 | 1 | 4738 | tags=37%, list=20%, si | MM_3 |
| GOBP_HYPEROSMOTIC_R | GOBP_HYPEROSMOTIC_RESPONSE | 19 | -0.60332 | -1.55206 | 0.031161472 | 0.17617305 | 1 | 6341 | tags=47%, list=27%, si | MM_3 |
| GOBP_INNER_MITOCHON | GOBP_INNER_MITOCHONDRIAL_MEMBRAN | 36 | -0.52435 | -1.55204 | 0.022900764 | 0.1755092 | 1 | 7043 | tags=58%, list=30%, si | MM_3 |
| INFLUENZA_A | INFLUENZA_A | 139 | -0.45003 | -1.55156 | 0.0010834236 | 0.17577371 | 1 | 6321 | tags=40%, list=27%, si | MM_3 |
| GOBP_EOSINOPHIL_CHEM | GOBP_EOSINOPHIL_CHEMOTAXIS | 22 | -0.58109 | -1.55144 | 0.022849463 | 0.17529859 | 1 | 5452 | tags=55%, list=23%, si | MM_3 |
| GOBP_NEGATIVE_REGUL | GOBP_NEGATIVE_REGULATION_OF_PHAGO | 24 | -0.58953 | -1.55118 | 0.021709634 | 0.17509744 | 1 | 7238 | tags=63%, list=31%, si | MM_3 |
| GOBP_CELLULAR_COMPO | GOBP_CELLULAR_COMPONENT_DISASSEME | 31 | -0.54625 | -1.54968 | 0.019710906 | 0.17721266 | 1 | 3950 | tags=48%, list=17%, si | MM_3 |
| GOBP_REGULATION_OF_ | GOBP_REGULATION_OF_VIRAL_TRANSCRIPT | 25 | -0.57348 | -1.54929 | 0.028263796 | 0.17720178 | 1 | 5309 | tags=40%, list=22%, si | MM_3 |
| GOBP_REGULATION_OF_ | GOBP_REGULATION_OF_MITOCHONDRIAL_ | 39 | -0.52991 | -1.54882 | 0.0127551025 | 0.1774658 | 1 | 4732 | tags=44%, list=20%, si | MM_3 |
| LEISHMANIASIS | LEISHMANIASIS | 60 | -0.49366 | -1.54843 | 0.0060313633 | 0.17751794 | 1 | 6540 | tags=48%, list=28%, si | MM_3 |
| GOBP_OLIGODENDROCYT | GOBP_OLIGODENDROCYTE_DIFFERENTIATIC | 112 | -0.46335 | -1.54838 | 0.0032537961 | 0.1769285 | 1 | 3606 | tags=25%, list=15%, si | MM_3 |
| GOBP_MITOCHONDRIAL_ | GOBP_MITOCHONDRIAL_RESPIRATORY_CHA | 91 | -0.47051 | -1.54836 | 0.0033333334 | 0.17632365 | 1 | 7153 | tags=63%, list=30%, si | MM_3 |
| GOBP_POSITIVE_REGULA | GOBP_POSITIVE_REGULATION_OF_FATTY_A | 20 | -0.60545 | -1.54754 | 0.018741634 | 0.17706682 | 1 | 2458 | tags=30%, list=10%, si | MM_3 |
| GOBP_NEUTROPHIL_ACTI | GOBP_NEUTROPHIL_ACTIVATION | 36 | -0.52995 | -1.54743 | 0.019157087 | 0.1765989 | 1 | 5455 | tags=50%, list=23%, si | MM_3 |
| GOBP_NEGATIVE_REGUL | GOBP_NEGATIVE_REGULATION_OF_MYELO | 57 | -0.49305 | -1.54719 | 0.009638554 | 0.17637631 | 1 | 6112 | tags=46%, list=26%, si | MM_3 |
| GOBP_REGULATION_OF_ | GOBP_REGULATION_OF_NUCLEASE_ACTIVIT | 22 | -0.58206 | -1.54712 | 0.030303031 | 0.17587961 | 1 | 4991 | tags=55%, list=21%, si | MM_3 |

| | | | | | | | | | | |
|---|---|---|---|---|---|---|---|---|---|---|
| GOBP_CHEMOKINE_C_X_ | GOBP_CHEMOKINE_C_X_C_MOTIF_LIGAND_ | 25 | -0.56926 | -1.54652 | 0.03006536 | 0.17635791 | 1 | 3620 | tags=40%, list=15%, si | MM_3 |
| GOBP_ERYTHROCYTE_HO | GOBP_ERYTHROCYTE_HOMEOSTASIS | 163 | -0.44317 | -1.54513 | 0.0010559662 | 0.1780909 | 1 | 6225 | tags=45%, list=26%, si | MM_3 |
| SPLICEOSOME | SPLICEOSOME | 127 | -0.44917 | -1.5446 | 0.0021482278 | 0.17849974 | 1 | 5848 | tags=37%, list=25%, si | MM_3 |
| GOBP_LEUKOCYTE_CHEM | GOBP_LEUKOCYTE_CHEMOTAXIS | 213 | -0.43318 | -1.54436 | 0 | 0.17827028 | 1 | 6048 | tags=44%, list=26%, si | MM_3 |
| GOBP_NEGATIVE_REGUL | GOBP_NEGATIVE_REGULATION_OF_DNA_B | 148 | -0.44418 | -1.5442 | 0.0010570824 | 0.17795068 | 1 | 6194 | tags=42%, list=26%, si | MM_3 |
| GOBP_ANTIGEN_PROCES | GOBP_ANTIGEN_PROCESSING_AND_PRESEN | 24 | -0.57194 | -1.54312 | 0.010943913 | 0.17942855 | 1 | 6240 | tags=71%, list=26%, si | MM_3 |
| GOBP_AEROBIC_ELECTRO | GOBP_AEROBIC_ELECTRON_TRANSPORT_CH | 63 | -0.48024 | -1.54305 | 0.0012033695 | 0.17893681 | 1 | 7269 | tags=60%, list=31%, si | MM_3 |
| GOBP_POSITIVE_REGULA | GOBP_POSITIVE_REGULATION_OF_NEUTRO | 41 | -0.52185 | -1.54291 | 0.015834348 | 0.17856577 | 1 | 6268 | tags=61%, list=27%, si | MM_3 |
| GOBP_RESPIRATORY_BUR | GOBP_RESPIRATORY_BURST | 27 | -0.55344 | -1.54251 | 0.028493894 | 0.17866038 | 1 | 6618 | tags=67%, list=28%, si | MM_3 |
| GOBP_REGULATION_OF_ | GOBP_REGULATION_OF_CYTOPLASMIC_PAT | 104 | -0.45691 | -1.54194 | 0.0011148272 | 0.17900436 | 1 | 4991 | tags=37%, list=21%, si | MM_3 |
| GOBP_ORGANELLE_INHE | GOBP_ORGANELLE_INHERITANCE | 15 | -0.62615 | -1.5419 | 0.028694404 | 0.17845014 | 1 | 2725 | tags=40%, list=12%, si | MM_3 |
| GOBP_REGULATION_OF_ | GOBP_REGULATION_OF_STEM_CELL_POPUL | 74 | -0.47833 | -1.54135 | 0.006984866 | 0.17897522 | 1 | 6461 | tags=50%, list=27%, si | MM_3 |
| GOBP_STEM_CELL_DIVISI | GOBP_STEM_CELL_DIVISION | 43 | -0.50907 | -1.54072 | 0.018337408 | 0.17957832 | 1 | 4611 | tags=35%, list=20%, si | MM_3 |
| GOBP_PYRIDINE_CONTAI | GOBP_PYRIDINE_CONTAINING_COMPOUND | 109 | -0.45713 | -1.54068 | 0 | 0.17902797 | 1 | 6718 | tags=47%, list=28%, si | MM_3 |
| GOBP_REGULATION_OF_ | GOBP_REGULATION_OF_VASCULAR_PERME | 48 | -0.50947 | -1.53898 | 0.009708738 | 0.18145369 | 1 | 5028 | tags=44%, list=21%, si | MM_3 |
| GOBP_SIGNAL_TRANSDU | GOBP_SIGNAL_TRANSDUCTION_BY_P53_CL | 151 | -0.44783 | -1.53895 | 0 | 0.18087505 | 1 | 5733 | tags=38%, list=24%, si | MM_3 |
| GOBP_CYTOPLASMIC_PA | GOBP_CYTOPLASMIC_PATTERN_RECOGNITI | 159 | -0.44015 | -1.53852 | 0.0042643924 | 0.18106449 | 1 | 5034 | tags=36%, list=21%, si | MM_3 |
| GOBP_DENDRITIC_CELL_ | GOBP_DENDRITIC_CELL_MIGRATION | 26 | -0.55431 | -1.53839 | 0.034528553 | 0.18069549 | 1 | 3586 | tags=38%, list=15%, si | MM_3 |
| GOBP_MYELOID_CELL_DI | GOBP_MYELOID_CELL_DIFFERENTIATION | 455 | -0.41712 | -1.53824 | 0 | 0.18033998 | 1 | 6323 | tags=42%, list=27%, si | MM_3 |
| GOBP_B_CELL_MEDIATED | GOBP_B_CELL_MEDIATED_IMMUNITY | 141 | -0.44796 | -1.5377 | 0.0031982942 | 0.18075737 | 1 | 5284 | tags=36%, list=22%, si | MM_3 |
| GOBP_MIRNA_TRANSCRI | GOBP_MIRNA_TRANSCRIPTION | 78 | -0.47603 | -1.53658 | 0.010440836 | 0.18231279 | 1 | 4942 | tags=37%, list=21%, si | MM_3 |
| GOBP_NEUTRAL_AMINO_ | GOBP_NEUTRAL_AMINO_ACID_TRANSPORT | 51 | -0.49361 | -1.53628 | 0.010895884 | 0.18223561 | 1 | 3744 | tags=27%, list=16%, si | MM_3 |
| GOBP_RELEASE_OF_CYTO | GOBP_RELEASE_OF_CYTOCHROME_C_FROM | 66 | -0.48495 | -1.53599 | 0.0034443168 | 0.1821555 | 1 | 4991 | tags=39%, list=21%, si | MM_3 |
| GOBP_NEGATIVE_REGUL | GOBP_NEGATIVE_REGULATION_OF_EXOCYT | 37 | -0.52214 | -1.53572 | 0.016169155 | 0.18205412 | 1 | 4617 | tags=41%, list=20%, si | MM_3 |
| GOBP_PROTEIN_ACYLATI | GOBP_PROTEIN_ACYLATION | 94 | -0.46219 | -1.535 | 0.0068104425 | 0.1828512 | 1 | 6436 | tags=44%, list=27%, si | MM_3 |
| GOBP_PROTEASOME_ME | GOBP_PROTEASOME_MEDIATED_UBIQUITIN | 368 | -0.419 | -1.534 | 0 | 0.18415597 | 1 | 6185 | tags=41%, list=26%, si | MM_3 |
| GOBP_COCHLEA_DEVELO | GOBP_COCHLEA_DEVELOPMENT | 51 | -0.49592 | -1.53378 | 0.013381995 | 0.18393953 | 1 | 4799 | tags=39%, list=20%, si | MM_3 |
| GOBP_CARDIAC_LEFT_VE | GOBP_CARDIAC_LEFT_VENTRICLE_MORPHO | 20 | -0.58214 | -1.53258 | 0.030812325 | 0.1855426 | 1 | 1199 | tags=25%, list=5%, sig | MM_3 |
| GOBP_REGULATION_OF_ | GOBP_REGULATION_OF_REACTIVE_OXYGEN | 160 | -0.44098 | -1.53251 | 0 | 0.18504918 | 1 | 6305 | tags=43%, list=27%, si | MM_3 |
| GOBP_RNA_3_END_PROC | GOBP_RNA_3_END_PROCESSING | 87 | -0.46525 | -1.53227 | 0.0056497175 | 0.18490878 | 1 | 4813 | tags=36%, list=20%, si | MM_3 |
| GOBP_NUCLEOSIDE_TRIP | GOBP_NUCLEOSIDE_TRIPHOSPHATE_BIOSYN | 100 | -0.45919 | -1.53182 | 0.0067873304 | 0.18528913 | 1 | 7071 | tags=58%, list=30%, si | MM_3 |
| GOBP_MYELOID_CELL_HO | GOBP_MYELOID_CELL_HOMEOSTASIS | 209 | -0.42618 | -1.53172 | 0 | 0.1848327 | 1 | 6254 | tags=44%, list=26%, si | MM_3 |
| GOBP_REGULATION_OF_ | GOBP_REGULATION_OF_MITOCHONDRIAL_ | 58 | -0.4901 | -1.53137 | 0.01911589 | 0.18485394 | 1 | 4754 | tags=40%, list=20%, si | MM_3 |
| GOBP_REGULATION_OF_ | GOBP_REGULATION_OF_MAMMARY_GLAN | 19 | -0.6002 | -1.53073 | 0.0332871 | 0.18551788 | 1 | 7036 | tags=74%, list=30%, si | MM_3 |
| GOBP_VACUOLE_ORGAN | GOBP_VACUOLE_ORGANIZATION | 223 | -0.43086 | -1.53023 | 0.0020920502 | 0.18586352 | 1 | 6114 | tags=47%, list=26%, si | MM_3 |
| GOBP_POSITIVE_REGULA | GOBP_POSITIVE_REGULATION_OF_LEUKOCY | 97 | -0.45566 | -1.52972 | 0.0022675737 | 0.18613572 | 1 | 6738 | tags=49%, list=29%, si | MM_3 |
| GOBP_NEGATIVE_REGUL | GOBP_NEGATIVE_REGULATION_OF_B_CELL | 20 | -0.56974 | -1.52889 | 0.033512063 | 0.18715124 | 1 | 2920 | tags=35%, list=12%, si | MM_3 |
| NON-ALCOHOLIC_FATTY_ | NON-ALCOHOLIC_FATTY_LIVER_DISEASE_(N | 141 | -0.4447 | -1.52765 | 0.0010764262 | 0.1888077 | 1 | 7108 | tags=58%, list=30%, si | MM_3 |
| GOBP_PEPTIDE_HORMON | GOBP_PEPTIDE_HORMONE_PROCESSING | 24 | -0.55613 | -1.52711 | 0.02710027 | 0.18912527 | 1 | 4016 | tags=38%, list=17%, si | MM_3 |
| GOBP_EMBRYONIC_DIGE | GOBP_EMBRYONIC_DIGESTIVE_TRACT_MOR | 19 | -0.57114 | -1.52676 | 0.03064067 | 0.18918996 | 1 | 1834 | tags=21%, list=8%, sig | MM_3 |
| GOBP_NEGATIVE_REGUL | GOBP_NEGATIVE_REGULATION_OF_MULTIC | 17 | -0.61303 | -1.52591 | 0.027434843 | 0.19030777 | 1 | 4844 | tags=47%, list=21%, si | MM_3 |
| GOBP_RESPONSE_TO_OS | GOBP_RESPONSE_TO_OSMOTIC_STRESS | 82 | -0.46561 | -1.52568 | 0.0056433408 | 0.19014283 | 1 | 6738 | tags=43%, list=29%, si | MM_3 |
| GOBP_GRANULOCYTE_AC | GOBP_GRANULOCYTE_ACTIVATION | 41 | -0.51576 | -1.5253 | 0.017699115 | 0.19020353 | 1 | 5455 | tags=49%, list=23%, si | MM_3 |
| GOBP_NEGATIVE_REGUL | GOBP_NEGATIVE_REGULATION_OF_LYMPH | 164 | -0.43465 | -1.52516 | 0.0010638298 | 0.18985584 | 1 | 5742 | tags=38%, list=24%, si | MM_3 |
| GOBP_NEGATIVE_REGUL | GOBP_NEGATIVE_REGULATION_OF_TYPE_I_ | 42 | -0.50596 | -1.52476 | 0.01891551 | 0.19002064 | 1 | 4960 | tags=50%, list=21%, si | MM_3 |
| GOBP_MACROPHAGE_AC | GOBP_MACROPHAGE_ACTIVATION | 107 | -0.45257 | -1.52435 | 0.0033594626 | 0.19015655 | 1 | 7418 | tags=54%, list=31%, si | MM_3 |
| GOBP_REGULATION_OF_ | GOBP_REGULATION_OF_TOLL_LIKE_RECEPT | 32 | -0.53317 | -1.52433 | 0.024453025 | 0.1896002 | 1 | 5287 | tags=44%, list=22%, si | MM_3 |
| GOBP_REGULATION_OF_ | GOBP_REGULATION_OF_NEUTROPHIL_MIGI | 52 | -0.49437 | -1.52315 | 0.012019231 | 0.19131938 | 1 | 3586 | tags=42%, list=15%, si | MM_3 |
| GOBP_BONE_TRABECULA | GOBP_BONE_TRABECULA_MORPHOGENESIS | 17 | -0.61141 | -1.52286 | 0.025 | 0.19125363 | 1 | 2487 | tags=29%, list=11%, si | MM_3 |
| GOBP_RNA_SURVEILLAN | GOBP_RNA_SURVEILLANCE | 17 | -0.61616 | -1.52238 | 0.027548209 | 0.19151726 | 1 | 3708 | tags=41%, list=16%, si | MM_3 |
| GOBP_RENAL_SYSTEM_V | GOBP_RENAL_SYSTEM_VASCULATURE_DEV | 29 | -0.53606 | -1.52213 | 0.024173029 | 0.19137302 | 1 | 5018 | tags=38%, list=21%, si | MM_3 |
| GOBP_VASCULAR_ASSOC | GOBP_VASCULAR_ASSOCIATED_SMOOTH_M | 80 | -0.46429 | -1.52191 | 0.0091743115 | 0.19117053 | 1 | 4987 | tags=36%, list=21%, si | MM_3 |
| GOBP_ANTIGEN_PROCES | GOBP_ANTIGEN_PROCESSING_AND_PRESEN | 38 | -0.51462 | -1.52163 | 0.018359853 | 0.19113536 | 1 | 5762 | tags=50%, list=24%, si | MM_3 |
| MALARIA | MALARIA | 41 | -0.5123 | -1.52091 | 0.020731708 | 0.19191386 | 1 | 6342 | tags=56%, list=27%, si | MM_3 |
| GOBP_NEGATIVE_REGUL | GOBP_NEGATIVE_REGULATION_OF_INFLAM | 153 | -0.44084 | -1.5205 | 0.004278075 | 0.19215378 | 1 | 4946 | tags=35%, list=21%, si | MM_3 |
| GOBP_GLYCOLYTIC_PROC | GOBP_GLYCOLYTIC_PROCESS_THROUGH_FR | 25 | -0.55655 | -1.52046 | 0.03576159 | 0.19162586 | 1 | 6396 | tags=56%, list=27%, si | MM_3 |
| GOBP_HYDROGEN_PERO | GOBP_HYDROGEN_PEROXIDE_CATABOLIC_F | 22 | -0.56618 | -1.52021 | 0.030955585 | 0.19148672 | 1 | 4984 | tags=55%, list=21%, si | MM_3 |
| GOBP_POSITIVE_REGULA | GOBP_POSITIVE_REGULATION_OF_MACROF | 25 | -0.54746 | -1.5201 | 0.021447722 | 0.19105676 | 1 | 6408 | tags=56%, list=27%, si | MM_3 |
| CELL_CYCLE | CELL_CYCLE | 121 | -0.4484 | -1.52009 | 0.0010834236 | 0.1905244 | 1 | 5407 | tags=42%, list=23%, si | MM_3 |
| GOBP_ASTROCYTE_DEVEI | GOBP_ASTROCYTE_DEVELOPMENT | 36 | -0.52408 | -1.52005 | 0.014175258 | 0.18998246 | 1 | 3470 | tags=36%, list=15%, si | MM_3 |
| GOBP_REGULATION_OF_ | GOBP_REGULATION_OF_CELLULAR_RESPON | 26 | -0.55203 | -1.51974 | 0.038512617 | 0.18996681 | 1 | 4739 | tags=38%, list=20%, si | MM_3 |
| GOBP_DETOXIFICATION | GOBP_DETOXIFICATION | 70 | -0.47341 | -1.51903 | 0.008254717 | 0.19075869 | 1 | 5678 | tags=43%, list=24%, si | MM_3 |
| KAPOSI_SARCOMA-ASSO | KAPOSI_SARCOMA-ASSOCIATED_HERPESVIR | 176 | -0.42992 | -1.51775 | 0.0021344717 | 0.19249314 | 1 | 6615 | tags=43%, list=28%, si | MM_3 |
| GOBP_NEGATIVE_REGUL | GOBP_NEGATIVE_REGULATION_OF_OSTEO | 37 | -0.51516 | -1.51773 | 0.0175 | 0.19197388 | 1 | 4879 | tags=38%, list=21%, si | MM_3 |
| GOBP_REGULATION_OF_ | GOBP_REGULATION_OF_AMIDE_METABOLI | 69 | -0.47984 | -1.51747 | 0.0081206495 | 0.19190893 | 1 | 6476 | tags=49%, list=27%, si | MM_3 |
| ALZHEIMER_DISEASE | ALZHEIMER_DISEASE | 159 | -0.4356 | -1.51571 | 0.0021459227 | 0.19469453 | 1 | 7071 | tags=60%, list=30%, si | MM_3 |
| MICRORNAS_IN_CANCER | MICRORNAS_IN_CANCER | 139 | -0.43963 | -1.51494 | 0.0010905125 | 0.19554505 | 1 | 6556 | tags=42%, list=28%, si | MM_3 |
| GOBP_DNA_TEMPLATED_ | GOBP_DNA_TEMPLATED_DNA_REPLICATION | 147 | -0.43262 | -1.51487 | 0 | 0.19509408 | 1 | 5651 | tags=40%, list=24%, si | MM_3 |
| GOBP_CHROMOSOME_C | GOBP_CHROMOSOME_CONDENSATION | 30 | -0.53182 | -1.51445 | 0.022049287 | 0.19543968 | 1 | 3981 | tags=40%, list=17%, si | MM_3 |
| GOBP_CELLULAR_RESPO | GOBP_CELLULAR_RESPONSE_TO_REACTIVE_ | 136 | -0.43588 | -1.51406 | 0.0054704593 | 0.19561341 | 1 | 5964 | tags=44%, list=25%, si | MM_3 |
| GOBP_POSITIVE_REGULA | GOBP_POSITIVE_REGULATION_OF_VIRAL_LI | 28 | -0.53847 | -1.51369 | 0.044058744 | 0.19572929 | 1 | 3491 | tags=32%, list=15%, si | MM_3 |
| GOBP_SPERMATID_NUCL | GOBP_SPERMATID_NUCLEUS_DIFFERENTIAT | 24 | -0.55897 | -1.51294 | 0.033738192 | 0.19662923 | 1 | 3809 | tags=38%, list=16%, si | MM_3 |
| GOBP_NADPH_REGENER | GOBP_NADPH_REGENERATION | 23 | -0.56503 | -1.51293 | 0.01907357 | 0.1960838 | 1 | 5846 | tags=61%, list=25%, si | MM_3 |
| GOBP_KILLING_BY_HOST | GOBP_KILLING_BY_HOST_OF_SYMBIONT_C | 20 | -0.58783 | -1.51266 | 0.036111113 | 0.1960303 | 1 | 780 | tags=20%, list=3%, sig | MM_3 |
| GOBP_REGULATION_OF_ | GOBP_REGULATION_OF_PHAGOCYTOSIS | 106 | -0.44979 | -1.51223 | 0.0033333334 | 0.19627841 | 1 | 7113 | tags=53%, list=30%, si | MM_3 |
| GOBP_COTRANSLATIONA | GOBP_COTRANSLATIONAL_PROTEIN_TARGE | 20 | -0.57735 | -1.51185 | 0.03950954 | 0.19644372 | 1 | 6588 | tags=55%, list=28%, si | MM_3 |
| GOBP_TRANSCRIPTION_E | GOBP_TRANSCRIPTION_BY_RNA_POLYMERA | 51 | -0.48985 | -1.51171 | 0.012150669 | 0.19615641 | 1 | 4986 | tags=47%, list=21%, si | MM_3 |
| GOBP_REGULATION_OF_ | GOBP_REGULATION_OF_PROTEIN_DEACETY | 33 | -0.52845 | -1.51136 | 0.02774275 | 0.19618237 | 1 | 3309 | tags=27%, list=14%, si | MM_3 |
| AMINOACYL-TRNA_BIOSY | AMINOACYL-TRNA_BIOSYNTHESIS | 43 | -0.50712 | -1.51131 | 0.02189781 | 0.19571699 | 1 | 2812 | tags=37%, list=12%, si | MM_3 |
| GOBP_CELLULAR_RESPO | GOBP_CELLULAR_RESPONSE_TO_CHEMICAL | 290 | -0.4188 | -1.51069 | 0 | 0.19632284 | 1 | 5964 | tags=39%, list=25%, si | MM_3 |
| GOBP_MONOCYTE_DIFFE | GOBP_MONOCYTE_DIFFERENTIATION | 28 | -0.53413 | -1.51028 | 0.027237354 | 0.19665872 | 1 | 4942 | tags=43%, list=21%, si | MM_3 |
| GOBP_REGULATION_OF_ | GOBP_REGULATION_OF_PATTERN_RECOGN | 157 | -0.4322 | -1.51003 | 0.0010626992 | 0.19661376 | 1 | 6126 | tags=42%, list=26%, si | MM_3 |

| | | | | | | | | | | |
|---|---|---|---|---|---|---|---|---|---|---|
| GOBP_NADH_METABOLIC | GOBP_NADH_METABOLIC_PROCESS | 39 | -0.51596 | -1.50987 | 0.017902814 | 0.19635475 | 1 | 5278 | tags=41%, list=22%, si | MM_3 |
| GOBP_INTRINSIC_APOPT | GOBP_INTRINSIC_APOPTOTIC_SIGNALING_F | 83 | -0.46345 | -1.50944 | 0.0078037903 | 0.19662203 | 1 | 7586 | tags=57%, list=32%, si | MM_3 |
| GOBP_REGULATION_OF_ | GOBP_REGULATION_OF_BIOLOGICAL_PROC | 64 | -0.47295 | -1.50924 | 0.017730497 | 0.19642635 | 1 | 3809 | tags=31%, list=16%, si | MM_3 |
| GOBP_REGULATION_OF_ | GOBP_REGULATION_OF_ENDOPLASMIC_RE | 24 | -0.55093 | -1.50905 | 0.030428769 | 0.19627333 | 1 | 7091 | tags=71%, list=30%, si | MM_3 |
| GOBP_PHAGOSOME_MAT | GOBP_PHAGOSOME_MATURATION | 29 | -0.54578 | -1.50824 | 0.026007803 | 0.19740772 | 1 | 6943 | tags=69%, list=29%, si | MM_3 |
| GOBP_MYELOID_LEUKOC | GOBP_MYELOID_LEUKOCYTE_ACTIVATION | 247 | -0.422 | -1.50782 | 0 | 0.19761123 | 1 | 6363 | tags=43%, list=27%, si | MM_3 |
| GOBP_DNA_SYNTHESIS_I | GOBP_DNA_SYNTHESIS_INVOLVED_IN_DNA | 35 | -0.5266 | -1.50759 | 0.016993465 | 0.19751157 | 1 | 4374 | tags=46%, list=19%, si | MM_3 |
| GOBP_FOAM_CELL_DIFFE | GOBP_FOAM_CELL_DIFFERENTIATION | 32 | -0.52982 | -1.50635 | 0.033591732 | 0.19919607 | 1 | 4055 | tags=31%, list=17%, si | MM_3 |
| GOBP_PROTEIN_DEPHOS | GOBP_PROTEIN_DEPHOSPHORYLATION | 220 | -0.4207 | -1.50618 | 0.0010351967 | 0.19897662 | 1 | 6606 | tags=45%, list=28%, si | MM_3 |
| GOBP_REGULATION_OF_ | GOBP_REGULATION_OF_T_CELL_CHEMOTA | 16 | -0.59979 | -1.50508 | 0.028288543 | 0.20040214 | 1 | 2003 | tags=25%, list=8%, sig | MM_3 |
| GOBP_CYTOSKELETON_D | GOBP_CYTOSKELETON_DEPENDENT_CYTOK | 97 | -0.45004 | -1.50482 | 0.0056179776 | 0.20036867 | 1 | 6406 | tags=48%, list=27%, si | MM_3 |
| RHEUMATOID_ARTHRITIS | RHEUMATOID_ARTHRITIS | 73 | -0.46821 | -1.50481 | 0.016431924 | 0.19985585 | 1 | 6579 | tags=59%, list=28%, si | MM_3 |
| GOBP_CELLULAR_RESPON | GOBP_CELLULAR_RESPONSE_TO_OXYGEN_F | 20 | -0.57391 | -1.50467 | 0.037921347 | 0.1995534 | 1 | 4984 | tags=50%, list=21%, si | MM_3 |
| GOBP_SUPEROXIDE_MET | GOBP_SUPEROXIDE_METABOLIC_PROCESS | 68 | -0.47098 | -1.50343 | 0.019473081 | 0.20140372 | 1 | 5916 | tags=46%, list=25%, si | MM_3 |
| GOBP_INTERFERON_MED | GOBP_INTERFERON_MEDIATED_SIGNALING | 76 | -0.46197 | -1.50297 | 0.008027523 | 0.2016802 | 1 | 4798 | tags=33%, list=20%, si | MM_3 |
| GOBP_MODULATION_BY_ | GOBP_MODULATION_BY_HOST_OF_VIRAL_ | 44 | -0.49868 | -1.50279 | 0.024783147 | 0.2014692 | 1 | 6357 | tags=48%, list=27%, si | MM_3 |
| GOBP_POSITIVE_REGULA | GOBP_POSITIVE_REGULATION_OF_LEUKOCY | 33 | -0.52579 | -1.5027 | 0.025477707 | 0.20108859 | 1 | 6929 | tags=64%, list=29%, si | MM_3 |
| GOBP_TUMOR_NECROSIS | GOBP_TUMOR_NECROSIS_FACTOR_SUPERF | 193 | -0.42592 | -1.5022 | 0 | 0.2014919 | 1 | 6718 | tags=50%, list=28%, si | MM_3 |
| GOBP_POSITIVE_REGULA | GOBP_POSITIVE_REGULATION_OF_LEUKOCY | 159 | -0.42899 | -1.50206 | 0.0021299254 | 0.20122062 | 1 | 6408 | tags=49%, list=27%, si | MM_3 |
| GLIOMA | GLIOMA | 71 | -0.46191 | -1.50183 | 0.011534025 | 0.20113216 | 1 | 5254 | tags=39%, list=22%, si | MM_3 |
| GOBP_N_TERMINAL_PRO | GOBP_N_TERMINAL_PROTEIN_AMINO_ACID | 25 | -0.54934 | -1.50139 | 0.04149933 | 0.20148036 | 1 | 5752 | tags=48%, list=24%, si | MM_3 |
| GOBP_REGULATION_OF_ | GOBP_REGULATION_OF_B_CELL_PROLIFERA | 71 | -0.46908 | -1.50135 | 0.009535161 | 0.20103161 | 1 | 6305 | tags=48%, list=27%, si | MM_3 |
| GOBP_NEGATIVE_REGUL | GOBP_NEGATIVE_REGULATION_OF_T_HELP | 22 | -0.55128 | -1.50088 | 0.033377837 | 0.20138745 | 1 | 3690 | tags=27%, list=16%, si | MM_3 |
| GOBP_LEUKOCYTE_MIGR | GOBP_LEUKOCYTE_MIGRATION | 363 | -0.41264 | -1.50015 | 0 | 0.20225556 | 1 | 6048 | tags=42%, list=26%, si | MM_3 |
| GOBP_RESPONSE_TO_UV | GOBP_RESPONSE_TO_UV | 153 | -0.43296 | -1.49979 | 0.0021276595 | 0.20245034 | 1 | 5239 | tags=39%, list=22%, si | MM_3 |
| PROPANOATE_METABOLI | PROPANOATE_METABOLISM | 29 | -0.53466 | -1.49898 | 0.025503356 | 0.20336662 | 1 | 6233 | tags=66%, list=26%, si | MM_3 |
| GOBP_ORGANOPHOSPHA | GOBP_ORGANOPHOSPHATE_BIOSYNTHETIC | 500 | -0.40643 | -1.49772 | 0 | 0.205175 | 1 | 5681 | tags=37%, list=24%, si | MM_3 |
| GOBP_SEQUESTERING_O | GOBP_SEQUESTERING_OF_TRIGLYCERIDE | 18 | -0.58246 | -1.49706 | 0.050847456 | 0.20591974 | 1 | 5927 | tags=50%, list=25%, si | MM_3 |
| GOBP_NEGATIVE_REGUL | GOBP_NEGATIVE_REGULATION_OF_NF_KAF | 78 | -0.45746 | -1.49694 | 0.014639639 | 0.2055947 | 1 | 6194 | tags=44%, list=26%, si | MM_3 |
| GOBP_SYNAPTIC_TRANSM | GOBP_SYNAPTIC_TRANSMISSION_DOPAMIN | 27 | -0.53193 | -1.49677 | 0.04296875 | 0.20538138 | 1 | 3763 | tags=33%, list=16%, si | MM_3 |
| GOBP_NEGATIVE_REGUL | GOBP_NEGATIVE_REGULATION_OF_RNA_SF | 29 | -0.53339 | -1.49648 | 0.037564766 | 0.20542188 | 1 | 7853 | tags=62%, list=33%, si | MM_3 |
| GOBP_ATP_METABOLIC_ | GOBP_ATP_METABOLIC_PROCESS | 206 | -0.42411 | -1.49642 | 0.0010438414 | 0.20502114 | 1 | 7071 | tags=49%, list=30%, si | MM_3 |
| GOBP_RESPONSE_TO_FU | GOBP_RESPONSE_TO_FUNGUS | 59 | -0.47213 | -1.49612 | 0.015568863 | 0.20503172 | 1 | 4046 | tags=29%, list=17%, si | MM_3 |
| GOBP_POSITIVE_REGULA | GOBP_POSITIVE_REGULATION_OF_MACROF | 15 | -0.61179 | -1.49586 | 0.03724138 | 0.2050567 | 1 | 4819 | tags=47%, list=20%, si | MM_3 |
| GOBP_REACTIVE_OXYGEI | GOBP_REACTIVE_OXYGEN_SPECIES_METAB | 239 | -0.41686 | -1.49539 | 0 | 0.20538537 | 1 | 4991 | tags=33%, list=21%, si | MM_3 |
| GOBP_POSITIVE_REGULA | GOBP_POSITIVE_REGULATION_BY_HOST_OI | 22 | -0.55026 | -1.49488 | 0.035460994 | 0.20581791 | 1 | 6357 | tags=55%, list=27%, si | MM_3 |
| GOBP_PHAGOLYSOSOME | GOBP_PHAGOLYSOSOME_ASSEMBLY | 17 | -0.59401 | -1.49478 | 0.03763441 | 0.20545122 | 1 | 6943 | tags=71%, list=29%, si | MM_3 |
| GOBP_POSITIVE_REGULA | GOBP_POSITIVE_REGULATION_OF_CHEMOT | 150 | -0.43042 | -1.49469 | 0.0032327587 | 0.20511316 | 1 | 6408 | tags=45%, list=27%, si | MM_3 |
| GOBP_REGULATION_OF_ | GOBP_REGULATION_OF_DNA_RECOMBINAT | 127 | -0.43953 | -1.49426 | 0.0042964555 | 0.20542476 | 1 | 6278 | tags=39%, list=27%, si | MM_3 |
| STAPHYLOCOCCUS_AURE | STAPHYLOCOCCUS_AUREUS_INFECTION | 44 | -0.49202 | -1.49341 | 0.029739777 | 0.20663056 | 1 | 6717 | tags=55%, list=28%, si | MM_3 |
| GOBP_GOLGI_TO_PLASM | GOBP_GOLGI_TO_PLASMA_MEMBRANE_PR | 38 | -0.50788 | -1.49326 | 0.01965602 | 0.20638825 | 1 | 7588 | tags=58%, list=32%, si | MM_3 |
| GOBP_MIRNA_PROCESSII | GOBP_MIRNA_PROCESSING | 42 | -0.49916 | -1.49203 | 0.019305019 | 0.20823489 | 1 | 6710 | tags=50%, list=28%, si | MM_3 |
| GOBP_CELL_CYCLE_G2_M | GOBP_CELL_CYCLE_G2_M_PHASE_TRANSITI | 152 | -0.43001 | -1.49201 | 0.00857449 | 0.20772901 | 1 | 4759 | tags=38%, list=20%, si | MM_3 |
| GOBP_PYRUVATE_METAB | GOBP_PYRUVATE_METABOLIC_PROCESS | 127 | -0.4383 | -1.49124 | 0.0065146578 | 0.20870322 | 1 | 5278 | tags=39%, list=22%, si | MM_3 |
| TOLL-LIKE_RECEPTOR_SIG | TOLL-LIKE_RECEPTOR_SIGNALING_PATHWA | 84 | -0.45577 | -1.49118 | 0.010285715 | 0.20826222 | 1 | 5511 | tags=39%, list=23%, si | MM_3 |
| GOBP_CARBOHYDRATE_C | GOBP_CARBOHYDRATE_CATABOLIC_PROCES | 161 | -0.42623 | -1.49088 | 0 | 0.20836796 | 1 | 5403 | tags=37%, list=23%, si | MM_3 |
| GOBP_REGULATION_OF_ | GOBP_REGULATION_OF_RESPONSE_TO_FO | 21 | -0.55686 | -1.49077 | 0.055330634 | 0.20805135 | 1 | 3523 | tags=33%, list=15%, si | MM_3 |
| HUNTINGTON_DISEASE | HUNTINGTON_DISEASE | 180 | -0.42461 | -1.49015 | 0.0010559662 | 0.20878287 | 1 | 7071 | tags=54%, list=30%, si | MM_3 |
| GOBP_ASTROCYTE_DIFFE | GOBP_ASTROCYTE_DIFFERENTIATION | 88 | -0.44996 | -1.48995 | 0.0056433408 | 0.20861374 | 1 | 4473 | tags=31%, list=19%, si | MM_3 |
| GOBP_PROTEIN_ACETYLA | GOBP_PROTEIN_ACETYLATION | 56 | -0.48003 | -1.48993 | 0.023980815 | 0.208134 | 1 | 6749 | tags=48%, list=29%, si | MM_3 |
| GOBP_LYTIC_VACUOLE_C | GOBP_LYTIC_VACUOLE_ORGANIZATION | 95 | -0.44914 | -1.48966 | 0.005592841 | 0.20813195 | 1 | 6073 | tags=54%, list=26%, si | MM_3 |
| GOBP_GLIAL_CELL_APOP | GOBP_GLIAL_CELL_APOPTOTIC_PROCESS | 20 | -0.57299 | -1.48913 | 0.047419805 | 0.20867158 | 1 | 6596 | tags=55%, list=28%, si | MM_3 |
| GOBP_PHAGOCYTOSIS | GOBP_PHAGOCYTOSIS | 213 | -0.41711 | -1.4883 | 0.0031282585 | 0.2097298 | 1 | 7420 | tags=54%, list=31%, si | MM_3 |
| GOBP_RESPONSE_TO_VII | GOBP_RESPONSE_TO_VIRUS | 331 | -0.40731 | -1.48789 | 0 | 0.20998245 | 1 | 4765 | tags=30%, list=20%, si | MM_3 |
| GOBP_POSITIVE_REGULA | GOBP_POSITIVE_REGULATION_OF_DNA_RE | 77 | -0.45449 | -1.48725 | 0.011709602 | 0.2107689 | 1 | 5422 | tags=36%, list=23%, si | MM_3 |
| MISMATCH_REPAIR | MISMATCH_REPAIR | 21 | -0.55243 | -1.48718 | 0.037138928 | 0.21040678 | 1 | 5694 | tags=48%, list=24%, si | MM_3 |
| COMPLEMENT_AND_COA | COMPLEMENT_AND_COAGULATION_CASCA | 72 | -0.46155 | -1.48714 | 0.010714286 | 0.20998512 | 1 | 5132 | tags=32%, list=22%, si | MM_3 |
| GOBP_MATURATION_OF_ | GOBP_MATURATION_OF_LSU_RRNA | 25 | -0.54079 | -1.48695 | 0.045393858 | 0.20984489 | 1 | 4920 | tags=48%, list=21%, si | MM_3 |
| GOBP_REGULATION_OF_ | GOBP_REGULATION_OF_DNA_METABOLIC_ | 491 | -0.4061 | -1.48681 | 0 | 0.2095619 | 1 | 6617 | tags=42%, list=28%, si | MM_3 |
| GOBP_EXIT_FROM_MITO | GOBP_EXIT_FROM_MITOSIS | 26 | -0.54232 | -1.48678 | 0.04149933 | 0.20912462 | 1 | 5367 | tags=46%, list=23%, si | MM_3 |
| GOBP_RESPONSE_TO_OX | GOBP_RESPONSE_TO_OXYGEN_RADICAL | 24 | -0.54143 | -1.48618 | 0.043715846 | 0.20977224 | 1 | 4984 | tags=46%, list=21%, si | MM_3 |
| GOBP_MUCOPOLYSACCH | GOBP_MUCOPOLYSACCHARIDE_METABOLIC | 73 | -0.46904 | -1.48529 | 0.010356732 | 0.21108827 | 1 | 6049 | tags=55%, list=26%, si | MM_3 |
| GOBP_CARDIAC_RIGHT_V | GOBP_CARDIAC_RIGHT_VENTRICLE_MORPH | 19 | -0.56713 | -1.48525 | 0.045152724 | 0.21065292 | 1 | 3243 | tags=37%, list=14%, si | MM_3 |
| GOBP_NUCLEAR_TRANSC | GOBP_NUCLEAR_TRANSCRIBED_MRNA_CAT | 59 | -0.47612 | -1.48513 | 0.021428572 | 0.2103963 | 1 | 8027 | tags=64%, list=34%, si | MM_3 |
| GOBP_REGULATION_OF_ | GOBP_REGULATION_OF_INFLAMMASOME_ | 40 | -0.49724 | -1.48472 | 0.033007335 | 0.2106785 | 1 | 1650 | tags=23%, list=7%, sig | MM_3 |
| GOBP_HISTONE_MRNA_M | GOBP_HISTONE_MRNA_METABOLIC_PROCE | 20 | -0.57178 | -1.48408 | 0.040616248 | 0.21140994 | 1 | 2920 | tags=35%, list=12%, si | MM_3 |
| GOBP_DE_NOVO_PROTEI | GOBP_DE_NOVO_PROTEIN_FOLDING | 38 | -0.50986 | -1.48374 | 0.0390625 | 0.2115309 | 1 | 4388 | tags=37%, list=19%, si | MM_3 |
| GOBP_NEGATIVE_REGUL | GOBP_NEGATIVE_REGULATION_OF_LEUKOC | 142 | -0.42843 | -1.4833 | 0.0032292788 | 0.2119314 | 1 | 5082 | tags=34%, list=22%, si | MM_3 |
| GOBP_THIOESTER_BIOSYI | GOBP_THIOESTER_BIOSYNTHETIC_PROCESS | 27 | -0.53458 | -1.48285 | 0.041884817 | 0.21232219 | 1 | 5223 | tags=48%, list=22%, si | MM_3 |
| GOBP_REGULATION_OF_ | GOBP_REGULATION_OF_MUSCLE_CELL_DIF | 154 | -0.42682 | -1.48283 | 0.005285412 | 0.2118514 | 1 | 3489 | tags=27%, list=15%, si | MM_3 |
| GOBP_POSITIVE_REGULA | GOBP_POSITIVE_REGULATION_OF_OLIGODE | 28 | -0.53078 | -1.482 | 0.050938338 | 0.21298687 | 1 | 2658 | tags=21%, list=11%, si | MM_3 |
| GOBP_DENDRITIC_CELL_C | GOBP_DENDRITIC_CELL_CHEMOTAXIS | 19 | -0.57325 | -1.48199 | 0.04258242 | 0.21249184 | 1 | 3586 | tags=42%, list=15%, si | MM_3 |
| GOBP_NEGATIVE_REGUL | GOBP_NEGATIVE_REGULATION_OF_TISSUE_ | 27 | -0.5416 | -1.4817 | 0.038007863 | 0.21252373 | 1 | 4766 | tags=44%, list=20%, si | MM_3 |
| GOBP_PHOSPHATIDYLINC | GOBP_PHOSPHATIDYLINOSITOL_3_KINASE_ | 307 | -0.40724 | -1.48127 | 0 | 0.21292304 | 1 | 6439 | tags=43%, list=27%, si | MM_3 |
| RENAL_CELL_CARCINOMA | RENAL_CELL_CARCINOMA | 67 | -0.46288 | -1.48115 | 0.01629802 | 0.21264106 | 1 | 5254 | tags=40%, list=22%, si | MM_3 |
| GOBP_REGULATION_OF_ | GOBP_REGULATION_OF_RESPONSE_TO_CYT | 153 | -0.42296 | -1.481 | 0.0010752688 | 0.21243416 | 1 | 4568 | tags=29%, list=19%, si | MM_3 |
| GOBP_HEMATOPOIETIC_ | GOBP_HEMATOPOIETIC_PROGENITOR_CELL | 158 | -0.4224 | -1.47954 | 0.0010493179 | 0.21481249 | 1 | 6023 | tags=46%, list=26%, si | MM_3 |
| GOBP_TRACHEA_DEVELO | GOBP_TRACHEA_DEVELOPMENT | 21 | -0.56311 | -1.47929 | 0.05114401 | 0.21484844 | 1 | 2725 | tags=33%, list=12%, si | MM_3 |
| GOBP_PHYSIOLOGICAL_C | GOBP_PHYSIOLOGICAL_CARDIAC_MUSCLE_ | 46 | -0.48923 | -1.47904 | 0.02345679 | 0.21483555 | 1 | 4567 | tags=41%, list=19%, si | MM_3 |
| GOBP_MONOSACCHARID | GOBP_MONOSACCHARIDE_CATABOLIC_PRO | 40 | -0.49773 | -1.47895 | 0.03860523 | 0.21451084 | 1 | 5278 | tags=38%, list=22%, si | MM_3 |
| GOBP_NEGATIVE_REGUL | GOBP_NEGATIVE_REGULATION_OF_DEFENS | 19 | -0.57724 | -1.47883 | 0.04264099 | 0.21425343 | 1 | 5673 | tags=47%, list=24%, si | MM_3 |
| GOBP_VIRAL_GENOME_R | GOBP_VIRAL_GENOME_REPLICATION | 110 | -0.43221 | -1.47848 | 0.007683864 | 0.21447754 | 1 | 6357 | tags=39%, list=27%, si | MM_3 |

| | | | | | | | | | | |
|---|---|---|---|---|---|---|---|---|---|---|
| GOBP_POSITIVE_REGULA | GOBP_POSITIVE_REGULATION_OF_B_CELL_ | 47 | -0.48299 | -1.47791 | 0.023199024 | 0.21523754 | 1 | 6305 | tags=47%, list=27%, si | MM_3 |
| NUCLEOTIDE_EXCISION_F | NUCLEOTIDE_EXCISION_REPAIR | 42 | -0.48829 | -1.47711 | 0.029003782 | 0.21635836 | 1 | 5520 | tags=48%, list=23%, si | MM_3 |
| GOBP_SULFUR_COMPOU | GOBP_SULFUR_COMPOUND_TRANSPORT | 49 | -0.48459 | -1.47705 | 0.026666667 | 0.21603632 | 1 | 4580 | tags=35%, list=19%, si | MM_3 |
| GOBP_MEMBRANE_FUSI( | GOBP_MEMBRANE_FUSION | 156 | -0.42498 | -1.47671 | 0.0031813362 | 0.21619673 | 1 | 4136 | tags=31%, list=18%, si | MM_3 |
| GOBP_SERTOLI_CELL_DE\ | GOBP_SERTOLI_CELL_DEVELOPMENT | 22 | -0.5507 | -1.47667 | 0.03994674 | 0.2157833 | 1 | 1500 | tags=23%, list=6%, sig | MM_3 |
| GOBP_POSITIVE_REGULA | GOBP_POSITIVE_REGULATION_OF_MITOCH | 22 | -0.55536 | -1.47663 | 0.043715846 | 0.21538135 | 1 | 2512 | tags=36%, list=11%, si | MM_3 |
| GOBP_REGULATION_OF_ | GOBP_REGULATION_OF_MITOCHONDRION_ | 139 | -0.42823 | -1.47525 | 0.0032608695 | 0.21760067 | 1 | 4991 | tags=36%, list=21%, si | MM_3 |
| GOBP_PROTEIN_CATABO | GOBP_PROTEIN_CATABOLIC_PROCESS_IN_T | 22 | -0.55348 | -1.47498 | 0.043835618 | 0.21760587 | 1 | 6473 | tags=64%, list=27%, si | MM_3 |
| GOBP_CORTICAL_CYTOSK | GOBP_CORTICAL_CYTOSKELETON_ORGANIZ | 53 | -0.47745 | -1.47474 | 0.024509804 | 0.21762764 | 1 | 7271 | tags=51%, list=31%, si | MM_3 |
| GOBP_REGULATION_OF_ | GOBP_REGULATION_OF_MAST_CELL_ACTIV | 24 | -0.54345 | -1.47441 | 0.044354837 | 0.2178908 | 1 | 6014 | tags=50%, list=25%, si | MM_3 |
| GOBP_DETECTION_OF_M | GOBP_DETECTION_OF_MECHANICAL_STIMU | 25 | -0.54025 | -1.47377 | 0.05263158 | 0.21857904 | 1 | 4172 | tags=36%, list=18%, si | MM_3 |
| GOBP_CELLULAR_RESPON | GOBP_CELLULAR_RESPONSE_TO_INTERLEU | 62 | -0.45922 | -1.47368 | 0.025059666 | 0.21829867 | 1 | 4326 | tags=32%, list=18%, si | MM_3 |
| GOBP_RESPONSE_TO_CO | GOBP_RESPONSE_TO_COPPER_ION | 23 | -0.54498 | -1.47366 | 0.056786705 | 0.2178454 | 1 | 4131 | tags=35%, list=17%, si | MM_3 |
| GOBP_POSITIVE_REGULA | GOBP_POSITIVE_REGULATION_OF_MORPHO | 39 | -0.50152 | -1.47333 | 0.029908972 | 0.21800259 | 1 | 4017 | tags=28%, list=17%, si | MM_3 |
| GOBP_POSITIVE_REGULA | GOBP_POSITIVE_REGULATION_OF_VASCUL/ | 44 | -0.48924 | -1.47271 | 0.028930817 | 0.21881028 | 1 | 4987 | tags=41%, list=21%, si | MM_3 |
| GOBP_DNA_CATABOLIC_ | GOBP_DNA_CATABOLIC_PROCESS | 29 | -0.52574 | -1.47265 | 0.028871391 | 0.21842453 | 1 | 3950 | tags=41%, list=17%, si | MM_3 |
| GOBP_DETECTION_OF_M | GOBP_DETECTION_OF_MECHANICAL_STIMU | 53 | -0.47478 | -1.47228 | 0.024330901 | 0.21863589 | 1 | 6453 | tags=42%, list=27%, si | MM_3 |
| GOBP_T_CELL_PROLIFERA | GOBP_T_CELL_PROLIFERATION | 233 | -0.41228 | -1.47215 | 0 | 0.2183802 | 1 | 6656 | tags=45%, list=28%, si | MM_3 |
| GOBP_VACUOLAR_ACIDII | GOBP_VACUOLAR_ACIDIFICATION | 27 | -0.52683 | -1.47208 | 0.038107753 | 0.21804658 | 1 | 6522 | tags=74%, list=28%, si | MM_3 |
| UBIQUITIN_MEDIATED_P | UBIQUITIN_MEDIATED_PROTEOLYSIS | 134 | -0.42934 | -1.47194 | 0.0065075923 | 0.2178605 | 1 | 7018 | tags=46%, list=30%, si | MM_3 |
| GOBP_PROTEIN_LOCALIZ | GOBP_PROTEIN_LOCALIZATION_TO_CELL_S | 70 | -0.45888 | -1.47178 | 0.01627907 | 0.21768676 | 1 | 5251 | tags=44%, list=22%, si | MM_3 |
| GOBP_SYNAPTIC_VESICLE | GOBP_SYNAPTIC_VESICLE_MEMBRANE_ORG | 35 | -0.5026 | -1.47141 | 0.04227213 | 0.21800771 | 1 | 2817 | tags=29%, list=12%, si | MM_3 |
| GOBP_POSITIVE_REGULA | GOBP_POSITIVE_REGULATION_OF_REACTIV | 81 | -0.46065 | -1.46889 | 0.013651877 | 0.2225605 | 1 | 4960 | tags=37%, list=21%, si | MM_3 |
| CELLULAR_SENESCENCE | CELLULAR_SENESCENCE | 159 | -0.42085 | -1.46876 | 0.0021482278 | 0.22235455 | 1 | 6439 | tags=46%, list=27%, si | MM_3 |
| GOBP_NEGATIVE_REGUL | GOBP_NEGATIVE_REGULATION_OF_LYMPH( | 62 | -0.46663 | -1.46831 | 0.014218009 | 0.222787 | 1 | 3690 | tags=27%, list=16%, si | MM_3 |
| GOBP_NEGATIVE_REGUL | GOBP_NEGATIVE_REGULATION_OF_RELEAS | 22 | -0.5499 | -1.46798 | 0.052770447 | 0.22300185 | 1 | 4991 | tags=55%, list=21%, si | MM_3 |
| GOBP_POSITIVE_REGULA | GOBP_POSITIVE_REGULATION_OF_NEURON | 99 | -0.43803 | -1.46753 | 0.010022271 | 0.22338988 | 1 | 7004 | tags=41%, list=30%, si | MM_3 |
| PHAGOSOME | PHAGOSOME | 146 | -0.42465 | -1.46732 | 0.0063829785 | 0.2233525 | 1 | 6540 | tags=55%, list=28%, si | MM_3 |
| GOBP_GLIAL_CELL_MIGR | GOBP_GLIAL_CELL_MIGRATION | 70 | -0.45555 | -1.46635 | 0.012716763 | 0.2248076 | 1 | 6011 | tags=46%, list=25%, si | MM_3 |
| GOBP_HOMEOSTASIS_OF | GOBP_HOMEOSTASIS_OF_NUMBER_OF_CE | 388 | -0.40284 | -1.46628 | 0 | 0.22444567 | 1 | 5921 | tags=40%, list=25%, si | MM_3 |
| GOBP_NEUTROPHIL_MED | GOBP_NEUTROPHIL_MEDIATED_IMMUNITY | 37 | -0.4947 | -1.4659 | 0.038314175 | 0.22470516 | 1 | 1875 | tags=22%, list=8%, sig | MM_3 |
| GOBP_ACETYL_COA_BIOS | GOBP_ACETYL_COA_BIOSYNTHETIC_PROCES | 19 | -0.55901 | -1.46584 | 0.05344585 | 0.22433569 | 1 | 5137 | tags=53%, list=22%, si | MM_3 |
| GOBP_ELECTRON_TRANS | GOBP_ELECTRON_TRANSPORT_CHAIN | 107 | -0.43918 | -1.4653 | 0.010989011 | 0.22494128 | 1 | 7048 | tags=52%, list=30%, si | MM_3 |
| GOBP_CHAPERONE_MED | GOBP_CHAPERONE_MEDIATED_PROTEIN_F( | 62 | -0.46218 | -1.46521 | 0.021608643 | 0.22463147 | 1 | 4715 | tags=37%, list=20%, si | MM_3 |
| GOBP_PEROXISOME_PRO | GOBP_PEROXISOME_PROLIFERATOR_ACTIV/ | 27 | -0.53703 | -1.46488 | 0.05019815 | 0.22482777 | 1 | 3183 | tags=26%, list=13%, si | MM_3 |
| GOBP_TRANSLATIONAL_E | GOBP_TRANSLATIONAL_ELONGATION | 58 | -0.46611 | -1.46465 | 0.02184466 | 0.2247949 | 1 | 6359 | tags=48%, list=27%, si | MM_3 |
| GOBP_MIRNA_MEDIATED | GOBP_MIRNA_MEDIATED_GENE_SILENCING | 17 | -0.58772 | -1.46436 | 0.04901961 | 0.22492059 | 1 | 408 | tags=24%, list=2%, sig | MM_3 |
| GOBP_REGULATION_OF_ | GOBP_REGULATION_OF_ACUTE_INFLAMMA | 17 | -0.5724 | -1.46401 | 0.053596616 | 0.22517745 | 1 | 5678 | tags=47%, list=24%, si | MM_3 |
| GOBP_RESPONSE_TO_TY | GOBP_RESPONSE_TO_TYPE_I_INTERFERON | 64 | -0.46583 | -1.4638 | 0.023201857 | 0.22518149 | 1 | 4798 | tags=31%, list=20%, si | MM_3 |
| GOBP_DEFENSE_RESPON | GOBP_DEFENSE_RESPONSE_TO_VIRUS | 276 | -0.40258 | -1.46343 | 0 | 0.2255616 | 1 | 4765 | tags=30%, list=20%, si | MM_3 |
| GOBP_POSITIVE_REGULA | GOBP_POSITIVE_REGULATION_OF_MUSCLE | 43 | -0.48586 | -1.4629 | 0.023370234 | 0.22626966 | 1 | 5352 | tags=44%, list=23%, si | MM_3 |
| GOBP_PROTEIN_POLYUB | GOBP_PROTEIN_POLYUBIQUITINATION | 250 | -0.41212 | -1.4626 | 0 | 0.22639567 | 1 | 6231 | tags=42%, list=26%, si | MM_3 |
| GOBP_FIBROBLAST_APOF | GOBP_FIBROBLAST_APOPTOTIC_PROCESS | 31 | -0.51499 | -1.4625 | 0.040575918 | 0.22611298 | 1 | 7113 | tags=58%, list=30%, si | MM_3 |
| GOBP_SMOOTH_MUSCLE | GOBP_SMOOTH_MUSCLE_CELL_PROLIFERAT | 191 | -0.41223 | -1.46233 | 0.001048218 | 0.22603835 | 1 | 5688 | tags=38%, list=24%, si | MM_3 |
| GOBP_TYROSINE_PHOSPI | GOBP_TYROSINE_PHOSPHORYLATION_OF_S | 69 | -0.45453 | -1.46195 | 0.019976499 | 0.22630675 | 1 | 3621 | tags=29%, list=15%, si | MM_3 |
| GOBP_ORGANIC_ACID_TI | GOBP_ORGANIC_ACID_TRANSMEMBRANE_ | 135 | -0.42689 | -1.46111 | 0.0043149944 | 0.22750767 | 1 | 5032 | tags=33%, list=21%, si | MM_3 |
| GOBP_NEUROTRANSMIT | GOBP_NEUROTRANSMITTER_RECEPTOR_TR | 27 | -0.52293 | -1.46103 | 0.051655628 | 0.2271921 | 1 | 2630 | tags=37%, list=11%, si | MM_3 |
| GOBP_INTRACELLULAR_I | GOBP_INTRACELLULAR_IRON_ION_HOMEO: | 64 | -0.45946 | -1.4608 | 0.027611045 | 0.22718987 | 1 | 7232 | tags=55%, list=31%, si | MM_3 |
| GOBP_REGULATION_OF_ | GOBP_REGULATION_OF_CIRCADIAN_RHYTH | 112 | -0.43341 | -1.46064 | 0.011025358 | 0.22701587 | 1 | 4166 | tags=29%, list=18%, si | MM_3 |
| THERMOGENESIS | THERMOGENESIS | 210 | -0.41561 | -1.46049 | 0 | 0.22684485 | 1 | 7153 | tags=54%, list=30%, si | MM_3 |
| TNF_SIGNALING_PATHW | TNF_SIGNALING_PATHWAY | 108 | -0.4348 | -1.46044 | 0.01430143 | 0.22647682 | 1 | 5717 | tags=40%, list=24%, si | MM_3 |
| GOBP_REGULATION_OF_ | GOBP_REGULATION_OF_HEART_GROWTH | 85 | -0.44569 | -1.46042 | 0.014823261 | 0.22608206 | 1 | 3489 | tags=32%, list=15%, si | MM_3 |
| GOBP_REGULATION_OF_ | GOBP_REGULATION_OF_DNA_BINDING_TR/ | 362 | -0.40198 | -1.4601 | 0 | 0.22622283 | 1 | 6287 | tags=40%, list=27%, si | MM_3 |
| GOBP_REGULATION_OF_ | GOBP_REGULATION_OF_FATTY_ACID_BIOSY | 47 | -0.48258 | -1.45996 | 0.01965602 | 0.22609757 | 1 | 1792 | tags=23%, list=8%, sig | MM_3 |
| GOBP_REGULATION_OF_ | GOBP_REGULATION_OF_GLIOGENESIS | 133 | -0.42736 | -1.45972 | 0.0065146578 | 0.2261405 | 1 | 6126 | tags=37%, list=26%, si | MM_3 |
| GOBP_POSITIVE_REGULA | GOBP_POSITIVE_REGULATION_OF_VASCUL/ | 19 | -0.56704 | -1.45964 | 0.057342656 | 0.22584192 | 1 | 7108 | tags=53%, list=30%, si | MM_3 |
| GOBP_NEGATIVE_REGUL | GOBP_NEGATIVE_REGULATION_OF_RESPON | 362 | -0.39914 | -1.45955 | 0 | 0.22555523 | 1 | 4502 | tags=30%, list=19%, si | MM_3 |
| GOBP_MODULATION_BY_ | GOBP_MODULATION_BY_HOST_OF_VIRAL_ | 22 | -0.5527 | -1.45912 | 0.048648648 | 0.2259459 | 1 | 6357 | tags=55%, list=27%, si | MM_3 |
| GOBP_MONONUCLEAR_( | GOBP_MONONUCLEAR_CELL_MIGRATION | 199 | -0.41323 | -1.45893 | 0.0021074815 | 0.225893 | 1 | 6408 | tags=45%, list=27%, si | MM_3 |
| GOBP_REGULATION_OF_ | GOBP_REGULATION_OF_ATP_BIOSYNTHETI( | 20 | -0.55953 | -1.45822 | 0.048846677 | 0.22690366 | 1 | 4942 | tags=40%, list=21%, si | MM_3 |
| GOBP_REGULATION_OF_ | GOBP_REGULATION_OF_VIRAL_LIFE_CYCLE | 139 | -0.42161 | -1.45795 | 0.004310345 | 0.22694455 | 1 | 5872 | tags=37%, list=25%, si | MM_3 |
| GOBP_POSITIVE_REGULA | GOBP_POSITIVE_REGULATION_OF_INTERLE | 42 | -0.49203 | -1.45789 | 0.028571429 | 0.22658886 | 1 | 5115 | tags=45%, list=22%, si | MM_3 |
| GOBP_NEGATIVE_REGUL | GOBP_NEGATIVE_REGULATION_OF_IMMUN | 186 | -0.41546 | -1.45734 | 0.002120891 | 0.22728728 | 1 | 4466 | tags=28%, list=19%, si | MM_3 |
| GOBP_GLUCOSE_METAB( | GOBP_GLUCOSE_METABOLIC_PROCESS | 201 | -0.41206 | -1.45726 | 0.0010384216 | 0.22701032 | 1 | 5153 | tags=36%, list=22%, si | MM_3 |
| GOBP_REGULATION_OF_ | GOBP_REGULATION_OF_VIRAL_PROCESS | 167 | -0.41452 | -1.45714 | 0.004250797 | 0.22679353 | 1 | 5872 | tags=36%, list=25%, si | MM_3 |
| GOBP_GLYCOLYTIC_PRO( | GOBP_GLYCOLYTIC_PROCESS_THROUGH_GI | 20 | -0.56218 | -1.45695 | 0.03797468 | 0.22672689 | 1 | 6396 | tags=60%, list=27%, si | MM_3 |
| GOBP_MITOTIC_CYTOKIN | GOBP_MITOTIC_CYTOKINESIS | 83 | -0.44183 | -1.45677 | 0.022909507 | 0.22667956 | 1 | 6406 | tags=51%, list=27%, si | MM_3 |
| GOBP_TRANSCRIPTION_F | GOBP_TRANSCRIPTION_PREINITIATION_CO | 63 | -0.45746 | -1.45672 | 0.025974026 | 0.226341 | 1 | 5860 | tags=46%, list=25%, si | MM_3 |
| GOBP_POSITIVE_REGULA | GOBP_POSITIVE_REGULATION_OF_DNA_ME | 314 | -0.40199 | -1.45651 | 0 | 0.22633617 | 1 | 5125 | tags=32%, list=22%, si | MM_3 |
| GOBP_NEGATIVE_REGUL | GOBP_NEGATIVE_REGULATION_OF_MUSCL | 67 | -0.44941 | -1.45617 | 0.022016222 | 0.22657356 | 1 | 3423 | tags=28%, list=14%, si | MM_3 |
| GOBP_REGULATION_OF_ | GOBP_REGULATION_OF_FATTY_ACID_OXID/ | 41 | -0.48868 | -1.45592 | 0.024360536 | 0.22659163 | 1 | 2458 | tags=22%, list=10%, si | MM_3 |
| GOBP_REGULATION_OF_ | GOBP_REGULATION_OF_LEUKOCYTE_MIGR/ | 230 | -0.41075 | -1.45589 | 0.0010384216 | 0.2262206 | 1 | 6408 | tags=47%, list=27%, si | MM_3 |
| GOBP_FATTY_ACID_ELON | GOBP_FATTY_ACID_ELONGATION | 15 | -0.59302 | -1.45584 | 0.05698006 | 0.22586767 | 1 | 4591 | tags=47%, list=19%, si | MM_3 |
| GOBP_RESPONSE_TO_CA | GOBP_RESPONSE_TO_CADMIUM_ION | 35 | -0.50328 | -1.45567 | 0.050128534 | 0.22574192 | 1 | 5321 | tags=43%, list=23%, si | MM_3 |
| GOBP_CELLULAR_RESPON | GOBP_CELLULAR_RESPONSE_TO_CADMIUM | 24 | -0.53215 | -1.45554 | 0.053262316 | 0.22556302 | 1 | 5321 | tags=46%, list=23%, si | MM_3 |
| GOBP_NEGATIVE_REGUL | GOBP_NEGATIVE_REGULATION_OF_MYELO | 89 | -0.44352 | -1.45535 | 0.014573991 | 0.22551046 | 1 | 6738 | tags=46%, list=29%, si | MM_3 |
| GOBP_NEGATIVE_REGUL | GOBP_NEGATIVE_REGULATION_OF_VASCU | 109 | -0.42755 | -1.45533 | 0.008733625 | 0.22512443 | 1 | 4502 | tags=31%, list=19%, si | MM_3 |
| GOBP_NUCLEOSIDE_BISP | GOBP_NUCLEOSIDE_BISPHOSPHATE_BIOSYN | 39 | -0.49956 | -1.45475 | 0.034653466 | 0.22583008 | 1 | 5223 | tags=44%, list=22%, si | MM_3 |
| GOBP_REGULATION_OF_ | GOBP_REGULATION_OF_MYELOID_LEUKOC | 61 | -0.4554 | -1.45427 | 0.028503563 | 0.22626995 | 1 | 7051 | tags=56%, list=30%, si | MM_3 |
| GOBP_REGULATION_OF_ | GOBP_REGULATION_OF_GLIAL_CELL_APOPT | 15 | -0.58932 | -1.45418 | 0.049861494 | 0.22603656 | 1 | 7113 | tags=60%, list=30%, si | MM_3 |
| GOBP_DEPHOSPHORYLAT | GOBP_DEPHOSPHORYLATION | 303 | -0.40558 | -1.45376 | 0 | 0.22634579 | 1 | 6606 | tags=41%, list=28%, si | MM_3 |
| GOBP_REGULATION_OF_ | GOBP_REGULATION_OF_PROTEIN_UBIQUIT | 202 | -0.40836 | -1.45369 | 0.0010416667 | 0.22604771 | 1 | 6948 | tags=50%, list=29%, si | MM_3 |

| | | | | | | | | | | |
|---|---|---|---|---|---|---|---|---|---|---|
| GOBP_REGULATION_OF_ | GOBP_REGULATION_OF_DNA_REPAIR | 219 | -0.40689 | -1.45355 | 0.0031185031 | 0.22590359 | 1 | 6258 | tags=39%, list=27%, si | MM_3 |
| GOBP_REGULATION_OF_ | GOBP_REGULATION_OF_ASTROCYTE_DIFFE | 34 | -0.50695 | -1.45299 | 0.031887755 | 0.22659318 | 1 | 3324 | tags=29%, list=14%, si | MM_3 |
| GOBP_SOMATIC_DIVERSI | GOBP_SOMATIC_DIVERSIFICATION_OF_IMN | 81 | -0.43934 | -1.4529 | 0.014823261 | 0.22638364 | 1 | 3651 | tags=30%, list=15%, si | MM_3 |
| PERTUSSIS | PERTUSSIS | 69 | -0.44848 | -1.45155 | 0.03804994 | 0.22845086 | 1 | 6254 | tags=36%, list=26%, si | MM_3 |
| HIF-1_SIGNALING_PATHV | HIF-1_SIGNALING_PATHWAY | 98 | -0.4377 | -1.45141 | 0.0067114094 | 0.22830716 | 1 | 6316 | tags=47%, list=27%, si | MM_3 |
| GOBP_NEUROMUSCULAR | GOBP_NEUROMUSCULAR_PROCESS_CONTR | 69 | -0.44813 | -1.4513 | 0.028935185 | 0.22809142 | 1 | 5403 | tags=41%, list=23%, si | MM_3 |
| GOBP_ENDOSOME_TO_P | GOBP_ENDOSOME_TO_PLASMA_MEMBRAN | 21 | -0.54693 | -1.45126 | 0.05235602 | 0.22773552 | 1 | 2682 | tags=43%, list=11%, si | MM_3 |
| GOBP_REGULATION_OF_ | GOBP_REGULATION_OF_TYPE_B_PANCREAT | 16 | -0.58048 | -1.4512 | 0.058487874 | 0.22743535 | 1 | 6083 | tags=50%, list=26%, si | MM_3 |
| GOBP_RIBOSOMAL_LARG | GOBP_RIBOSOMAL_LARGE_SUBUNIT_BIOGE | 64 | -0.46208 | -1.45103 | 0.02 | 0.22729258 | 1 | 5888 | tags=48%, list=25%, si | MM_3 |
| GOBP_ACTIN_CROSSLINK | GOBP_ACTIN_CROSSLINK_FORMATION | 15 | -0.58416 | -1.45102 | 0.07173601 | 0.22689608 | 1 | 5900 | tags=47%, list=25%, si | MM_3 |
| GOBP_DEFENSE_RESPON | GOBP_DEFENSE_RESPONSE_TO_GRAM_NEC | 68 | -0.44975 | -1.45078 | 0.019721577 | 0.22689675 | 1 | 4654 | tags=32%, list=20%, si | MM_3 |
| GOBP_REGULATION_OF_ | GOBP_REGULATION_OF_LEUKOCYTE_CHEM | 122 | -0.42604 | -1.45002 | 0.0076923077 | 0.22793235 | 1 | 6738 | tags=51%, list=29%, si | MM_3 |
| GOBP_GLIAL_CELL_DEVE | GOBP_GLIAL_CELL_DEVELOPMENT | 134 | -0.41627 | -1.44993 | 0.0075268815 | 0.22771004 | 1 | 3470 | tags=25%, list=15%, si | MM_3 |
| GOBP_T_CELL_CHEMOTA | GOBP_T_CELL_CHEMOTAXIS | 23 | -0.53739 | -1.44963 | 0.055181697 | 0.22789979 | 1 | 2003 | tags=22%, list=8%, sig | MM_3 |
| GOBP_INNATE_IMMUNE_ | GOBP_INNATE_IMMUNE_RESPONSE_ACTIV | 254 | -0.40243 | -1.44959 | 0 | 0.22757936 | 1 | 5034 | tags=32%, list=21%, si | MM_3 |
| GOBP_POSITIVE_REGULA | GOBP_POSITIVE_REGULATION_OF_IMMUNE | 266 | -0.40357 | -1.44947 | 0.0010204081 | 0.22737668 | 1 | 6988 | tags=48%, list=30%, si | MM_3 |
| BIOSYNTHESIS_OF_UNSA | BIOSYNTHESIS_OF_UNSATURATED_FATTY_A | 31 | -0.50881 | -1.44942 | 0.04232804 | 0.22707205 | 1 | 4591 | tags=39%, list=19%, si | MM_3 |
| GOBP_SKELETAL_MUSCLE | GOBP_SKELETAL_MUSCLE_TISSUE_REGENEF | 43 | -0.48971 | -1.44876 | 0.03670886 | 0.22791222 | 1 | 6181 | tags=40%, list=26%, si | MM_3 |
| GOBP_POSITIVE_REGULA | GOBP_POSITIVE_REGULATION_OF_IMMUNC | 55 | -0.46682 | -1.44872 | 0.025672372 | 0.22755079 | 1 | 6278 | tags=47%, list=27%, si | MM_3 |
| GOBP_MYELOID_LEUKOC | GOBP_MYELOID_LEUKOCYTE_CYTOKINE_PR | 64 | -0.44729 | -1.44854 | 0.024418605 | 0.22745791 | 1 | 7113 | tags=56%, list=30%, si | MM_3 |
| GOBP_PROTEIN_DNA_CO | GOBP_PROTEIN_DNA_COMPLEX_ASSEMBLY | 154 | -0.41645 | -1.4479 | 0.0063897762 | 0.22823977 | 1 | 5934 | tags=42%, list=25%, si | MM_3 |
| GOBP_CELLULAR_RESPON | GOBP_CELLULAR_RESPONSE_TO_OXIDATIVE | 229 | -0.40414 | -1.44786 | 0.0010298662 | 0.22787768 | 1 | 5964 | tags=40%, list=25%, si | MM_3 |
| GOBP_CHEMOKINE_PROI | GOBP_CHEMOKINE_PRODUCTION | 119 | -0.42794 | -1.44775 | 0.012127894 | 0.22769287 | 1 | 3651 | tags=26%, list=15%, si | MM_3 |
| GOBP_REGULATION_OF_ | GOBP_REGULATION_OF_CYSTEINE_TYPE_EN | 16 | -0.5741 | -1.44745 | 0.05284553 | 0.227854 | 1 | 5678 | tags=44%, list=24%, si | MM_3 |
| GOBP_REGULATION_OF_ | GOBP_REGULATION_OF_UBIQUITIN_DEPEN | 157 | -0.41597 | -1.44736 | 0.0042462847 | 0.2276329 | 1 | 6802 | tags=46%, list=29%, si | MM_3 |
| GOBP_REGULATION_OF_ | GOBP_REGULATION_OF_PROTEASOMAL_UE | 118 | -0.42803 | -1.447 | 0.010857764 | 0.22797456 | 1 | 6317 | tags=44%, list=27%, si | MM_3 |
| GOBP_REGULATION_OF_ | GOBP_REGULATION_OF_MORPHOGENESIS_ | 71 | -0.44936 | -1.44689 | 0.019813519 | 0.22776961 | 1 | 3426 | tags=25%, list=15%, si | MM_3 |
| GOBP_POSITIVE_REGULA | GOBP_POSITIVE_REGULATION_OF_VACUOL | 25 | -0.52618 | -1.44641 | 0.050938338 | 0.2282916 | 1 | 5724 | tags=56%, list=24%, si | MM_3 |
| GOBP_REGULATORY_T_C | GOBP_REGULATORY_T_CELL_DIFFERENTIAT | 41 | -0.48502 | -1.44575 | 0.037943695 | 0.22920175 | 1 | 4924 | tags=41%, list=21%, si | MM_3 |
| GOBP_NUCLEOTIDE_SUG | GOBP_NUCLEOTIDE_SUGAR_BIOSYNTHETIC_ | 17 | -0.57264 | -1.44552 | 0.049861494 | 0.2292732 | 1 | 6179 | tags=53%, list=26%, si | MM_3 |
| GOBP_REGULATION_OF_ | GOBP_REGULATION_OF_PH | 104 | -0.42502 | -1.44524 | 0.010169491 | 0.22941764 | 1 | 6523 | tags=47%, list=28%, si | MM_3 |
| GOBP_POSITIVE_REGULA | GOBP_POSITIVE_REGULATION_OF_GLIOGEN | 87 | -0.43832 | -1.44515 | 0.016835017 | 0.22917923 | 1 | 6268 | tags=40%, list=27%, si | MM_3 |
| AGE-RAGE_SIGNALING_P | AGE-RAGE_SIGNALING_PATHWAY_IN_DIABI | 100 | -0.42969 | -1.44498 | 0.012222222 | 0.22909725 | 1 | 6615 | tags=45%, list=28%, si | MM_3 |
| GOBP_MICROTUBULE_NU | GOBP_MICROTUBULE_NUCLEATION | 35 | -0.49987 | -1.44489 | 0.05019815 | 0.22886199 | 1 | 5253 | tags=46%, list=22%, si | MM_3 |
| GOBP_REGULATION_OF_ | GOBP_REGULATION_OF_DNA_REPLICATION | 118 | -0.42695 | -1.44479 | 0.010834237 | 0.22863528 | 1 | 5462 | tags=36%, list=23%, si | MM_3 |
| SPHINGOLIPID_SIGNALIN | SPHINGOLIPID_SIGNALING_PATHWAY | 118 | -0.42877 | -1.44471 | 0.01 | 0.2283849 | 1 | 6876 | tags=47%, list=29%, si | MM_3 |
| GOBP_SOMATIC_DIVERSI | GOBP_SOMATIC_DIVERSIFICATION_OF_IMN | 71 | -0.44849 | -1.44423 | 0.027058823 | 0.22892365 | 1 | 3651 | tags=31%, list=15%, si | MM_3 |
| GOBP_MONOSACCHARID | GOBP_MONOSACCHARIDE_METABOLIC_PRC | 255 | -0.40047 | -1.44412 | 0.00101833 | 0.2287401 | 1 | 5346 | tags=36%, list=23%, si | MM_3 |
| GOBP_POSITIVE_REGULA | GOBP_POSITIVE_REGULATION_OF_PROTEIN | 15 | -0.59424 | -1.44396 | 0.071532845 | 0.22867689 | 1 | 6626 | tags=53%, list=28%, si | MM_3 |
| GOBP_RESPONSE_TO_SA | GOBP_RESPONSE_TO_SALT_STRESS | 20 | -0.54832 | -1.44373 | 0.049928673 | 0.22863628 | 1 | 6738 | tags=55%, list=29%, si | MM_3 |
| GOBP_POSITIVE_REGULA | GOBP_POSITIVE_REGULATION_OF_B_CELL_ | 90 | -0.43213 | -1.44367 | 0.015819209 | 0.22834285 | 1 | 3764 | tags=30%, list=16%, si | MM_3 |
| GOBP_CELL_CHEMOTAXIS | GOBP_CELL_CHEMOTAXIS | 295 | -0.39905 | -1.44343 | 0.0020222447 | 0.22841817 | 1 | 6254 | tags=43%, list=26%, si | MM_3 |
| GOBP_REGULATION_OF_ | GOBP_REGULATION_OF_LIPID_CATABOLIC_ | 63 | -0.45654 | -1.44301 | 0.025821596 | 0.228761 | 1 | 4155 | tags=30%, list=18%, si | MM_3 |
| GOBP_COMPLEMENT_AC | GOBP_COMPLEMENT_ACTIVATION | 50 | -0.46432 | -1.44228 | 0.03508772 | 0.22976148 | 1 | 5919 | tags=36%, list=25%, si | MM_3 |
| GOBP_POSITIVE_REGULA | GOBP_POSITIVE_REGULATION_OF_CELL_CY | 330 | -0.39619 | -1.44157 | 0 | 0.23074216 | 1 | 6618 | tags=42%, list=28%, si | MM_3 |
| GOBP_POSITIVE_REGULA | GOBP_POSITIVE_REGULATION_OF_CHEMOK | 79 | -0.43885 | -1.44081 | 0.030127462 | 0.2318732 | 1 | 3975 | tags=29%, list=17%, si | MM_3 |
| GOBP_FIBROBLAST_PROL | GOBP_FIBROBLAST_PROLIFERATION | 147 | -0.41388 | -1.44036 | 0.008733625 | 0.23229712 | 1 | 6458 | tags=44%, list=27%, si | MM_3 |
| GOBP_REGULATION_OF_ | GOBP_REGULATION_OF_INTEGRIN_MEDIAT | 19 | -0.55065 | -1.44022 | 0.066390045 | 0.23218007 | 1 | 5058 | tags=47%, list=21%, si | MM_3 |
| GOBP_NEGATIVE_REGUL | GOBP_NEGATIVE_REGULATION_OF_LIPID_N | 104 | -0.42233 | -1.44009 | 0.017877094 | 0.2320316 | 1 | 3990 | tags=28%, list=17%, si | MM_3 |
| GOBP_OLIGODENDROCY | GOBP_OLIGODENDROCYTE_DEVELOPMENT | 47 | -0.47244 | -1.44008 | 0.033962265 | 0.23164177 | 1 | 2157 | tags=23%, list=9%, sig | MM_3 |
| GOBP_NEGATIVE_REGUL | GOBP_NEGATIVE_REGULATION_OF_REACTI | 20 | -0.5517 | -1.43987 | 0.061497327 | 0.2316803 | 1 | 1728 | tags=25%, list=7%, sig | MM_3 |
| GOBP_SERTOLI_CELL_DIF | GOBP_SERTOLI_CELL_DIFFERENTIATION | 28 | -0.51008 | -1.43972 | 0.048114434 | 0.23151688 | 1 | 1834 | tags=21%, list=8%, sig | MM_3 |
| GOBP_POSITIVE_REGULA | GOBP_POSITIVE_REGULATION_OF_PHOSPH | 182 | -0.41019 | -1.43972 | 0.0031746032 | 0.23112437 | 1 | 6195 | tags=43%, list=26%, si | MM_3 |
| GOBP_NADP_METABOLIC | GOBP_NADP_METABOLIC_PROCESS | 49 | -0.47043 | -1.43965 | 0.03606965 | 0.23084947 | 1 | 5867 | tags=49%, list=25%, si | MM_3 |
| GOBP_REGULATION_OF_ | GOBP_REGULATION_OF_INFLAMMATORY_F | 348 | -0.39774 | -1.4392 | 0 | 0.2313654 | 1 | 6137 | tags=38%, list=26%, si | MM_3 |
| GOBP_POSITIVE_REGULA | GOBP_POSITIVE_REGULATION_OF_STEM_CI | 50 | -0.46984 | -1.43878 | 0.044957474 | 0.23176341 | 1 | 6461 | tags=50%, list=27%, si | MM_3 |
| GOBP_POSITIVE_REGULA | GOBP_POSITIVE_REGULATION_OF_AMYLOII | 20 | -0.55199 | -1.43865 | 0.06777317 | 0.23166938 | 1 | 4458 | tags=35%, list=19%, si | MM_3 |
| GOBP_REGULATION_OF_ | GOBP_REGULATION_OF_ADENYLATE_CYCLA | 24 | -0.53618 | -1.43831 | 0.06925208 | 0.23194063 | 1 | 2743 | tags=25%, list=12%, si | MM_3 |
| GOBP_POSITIVE_REGULA | GOBP_POSITIVE_REGULATION_OF_TYROSIN | 51 | -0.46833 | -1.43783 | 0.025578562 | 0.2324469 | 1 | 3489 | tags=31%, list=15%, si | MM_3 |
| GOBP_POSITIVE_REGULA | GOBP_POSITIVE_REGULATION_OF_MYELOID | 38 | -0.48359 | -1.43778 | 0.04005168 | 0.23214892 | 1 | 6988 | tags=63%, list=30%, si | MM_3 |
| GOBP_REGULATION_OF_ | GOBP_REGULATION_OF_EXOSOMAL_SECRE | 17 | -0.5667 | -1.43712 | 0.06901409 | 0.23307855 | 1 | 6405 | tags=65%, list=27%, si | MM_3 |
| GOBP_POSITIVE_REGULA | GOBP_POSITIVE_REGULATION_OF_PROTEIN | 107 | -0.42626 | -1.43701 | 0.013274336 | 0.23287927 | 1 | 6948 | tags=53%, list=29%, si | MM_3 |
| GOBP_ESTABLISHMENT_ | GOBP_ESTABLISHMENT_OF_PROTEIN_LOCA | 47 | -0.47413 | -1.43693 | 0.045743328 | 0.23261303 | 1 | 6588 | tags=47%, list=28%, si | MM_3 |
| GOBP_REGULATION_OF_ | GOBP_REGULATION_OF_FIBROBLAST_PROL | 122 | -0.41861 | -1.43625 | 0.009782609 | 0.23356965 | 1 | 5239 | tags=34%, list=22%, si | MM_3 |
| GOBP_GLUCOSE_6_PHOS | GOBP_GLUCOSE_6_PHOSPHATE_METABOLI | 28 | -0.51734 | -1.4361 | 0.05242464 | 0.23351207 | 1 | 5846 | tags=54%, list=25%, si | MM_3 |
| GOBP_WATER_TRANSPOI | GOBP_WATER_TRANSPORT | 17 | -0.56092 | -1.43576 | 0.07260274 | 0.23377246 | 1 | 4790 | tags=35%, list=20%, si | MM_3 |
| GOBP_MITOCHONDRIAL_ | GOBP_MITOCHONDRIAL_ELECTRON_TRANS | 29 | -0.51267 | -1.43552 | 0.058517557 | 0.23385921 | 1 | 7045 | tags=55%, list=30%, si | MM_3 |
| GOBP_REGULATION_OF_ | GOBP_REGULATION_OF_NEUTROPHIL_CHEI | 36 | -0.50229 | -1.43541 | 0.042690814 | 0.23369281 | 1 | 3491 | tags=44%, list=15%, si | MM_3 |
| GOBP_ACUTE_INFLAMM | GOBP_ACUTE_INFLAMMATORY_RESPONSE_ | 28 | -0.52314 | -1.43531 | 0.063719116 | 0.23354508 | 1 | 5717 | tags=39%, list=24%, si | MM_3 |
| GOBP_REGULATORY_NCF | GOBP_REGULATORY_NCRNA_MEDIATED_GI | 116 | -0.424 | -1.43518 | 0.010964912 | 0.23342673 | 1 | 6710 | tags=42%, list=28%, si | MM_3 |
| GOBP_GLYCERALDEHYDE | GOBP_GLYCERALDEHYDE_3_PHOSPHATE_M | 19 | -0.56211 | -1.43463 | 0.061797753 | 0.23422143 | 1 | 4068 | tags=42%, list=17%, si | MM_3 |
| GOBP_REGULATION_OF_ | GOBP_REGULATION_OF_OXIDOREDUCTASE | 63 | -0.45665 | -1.43411 | 0.02997602 | 0.23486345 | 1 | 6206 | tags=48%, list=26%, si | MM_3 |
| GOBP_NUCLEOSIDE_PHO | GOBP_NUCLEOSIDE_PHOSPHATE_CATABOLI | 170 | -0.40246 | -1.43369 | 0.008324662 | 0.23533325 | 1 | 6720 | tags=42%, list=28%, si | MM_3 |
| GOBP_POSITIVE_REGULA | GOBP_POSITIVE_REGULATION_OF_DNA_TEI | 59 | -0.449 | -1.43348 | 0.0382387 | 0.23536777 | 1 | 6091 | tags=46%, list=26%, si | MM_3 |
| GOBP_REGULATION_OF_ | GOBP_REGULATION_OF_STEROL_TRANSPOI | 59 | -0.46263 | -1.43312 | 0.039806996 | 0.23567677 | 1 | 6321 | tags=41%, list=27%, si | MM_3 |
| GOBP_IMMUNOGLOBULI | GOBP_IMMUNOGLOBULIN_PRODUCTION_II | 67 | -0.44747 | -1.4331 | 0.028571429 | 0.23531742 | 1 | 3651 | tags=30%, list=15%, si | MM_3 |
| GOBP_PROSTANOID_ME | GOBP_PROSTANOID_METABOLIC_PROCESS | 53 | -0.46327 | -1.43286 | 0.027577939 | 0.23541379 | 1 | 6035 | tags=38%, list=26%, si | MM_3 |
| GOBP_NUCLEOSIDE_TRIP | GOBP_NUCLEOSIDE_TRIPHOSPHATE_METAE | 250 | -0.40061 | -1.43225 | 0.004085802 | 0.23631532 | 1 | 6720 | tags=46%, list=28%, si | MM_3 |
| GOBP_STEROL_TRANSPO | GOBP_STEROL_TRANSPORT | 109 | -0.42263 | -1.43203 | 0.01419214 | 0.23633203 | 1 | 6534 | tags=43%, list=28%, si | MM_3 |
| GOBP_NEGATIVE_REGUL | GOBP_NEGATIVE_REGULATION_OF_TRANSI | 117 | -0.41587 | -1.43181 | 0.019911505 | 0.2364017 | 1 | 6820 | tags=46%, list=29%, si | MM_3 |
| GOBP_LIPOPROTEIN_CAT | GOBP_LIPOPROTEIN_CATABOLIC_PROCESS | 17 | -0.56768 | -1.43161 | 0.075418994 | 0.23639143 | 1 | 5659 | tags=65%, list=24%, si | MM_3 |
| GOBP_PROTEIN_HOMOTI | GOBP_PROTEIN_HOMOTETRAMERIZATION | 49 | -0.46526 | -1.43102 | 0.044205494 | 0.23713315 | 1 | 5462 | tags=39%, list=23%, si | MM_3 |

| | | | | | | | | | | |
|---|---|---|---|---|---|---|---|---|---|---|
| GOBP_PEPTIDYL_CYSTEIN | GOBP_PEPTIDYL_CYSTEINE_MODIFICATION | 44 | -0.47729 | -1.43097 | 0.03960396 | 0.2368546 | 1 | 5868 | tags=41%, list=25%, si | MM_3 |
| GOBP_NEGATIVE_REGUL | GOBP_NEGATIVE_REGULATION_OF_CELL_D | 293 | -0.39714 | -1.43094 | 0.0010121458 | 0.23652087 | 1 | 5474 | tags=33%, list=23%, si | MM_3 |
| GOBP_REGULATION_OF_ | GOBP_REGULATION_OF_TRANSLATION | 329 | -0.39216 | -1.43086 | 0 | 0.23628972 | 1 | 6472 | tags=43%, list=27%, si | MM_3 |
| GOBP_MITOCHONDRIAL_ | GOBP_MITOCHONDRIAL_DNA_REPLICATION | 17 | -0.56685 | -1.43078 | 0.06039326 | 0.23605806 | 1 | 5462 | tags=59%, list=23%, si | MM_3 |
| GOBP_NEGATIVE_REGUL | GOBP_NEGATIVE_REGULATION_OF_DNA_M | 127 | -0.41872 | -1.43055 | 0.013071896 | 0.23610109 | 1 | 5432 | tags=34%, list=23%, si | MM_3 |
| GOBP_VESICLE_MEDIATE | GOBP_VESICLE_MEDIATED_TRANSPORT_TO | 158 | -0.40915 | -1.42992 | 0.0075675678 | 0.2369673 | 1 | 6054 | tags=39%, list=26%, si | MM_3 |
| GOBP_TRANSFORMING_G | GOBP_TRANSFORMING_GROWTH_FACTOR_ | 202 | -0.39849 | -1.42989 | 0.001039501 | 0.23663783 | 1 | 6326 | tags=43%, list=27%, si | MM_3 |
| GOBP_NEGATIVE_REGUL | GOBP_NEGATIVE_REGULATION_OF_CYTOKI | 292 | -0.39344 | -1.42963 | 0.0010235414 | 0.23681015 | 1 | 5018 | tags=34%, list=21%, si | MM_3 |
| GOBP_PEPTIDE_CATABOL | GOBP_PEPTIDE_CATABOLIC_PROCESS | 20 | -0.54602 | -1.42951 | 0.06327373 | 0.2366382 | 1 | 5399 | tags=40%, list=23%, si | MM_3 |
| GOBP_REGULATION_OF_ | GOBP_REGULATION_OF_DOUBLE_STRAND_ | 80 | -0.43448 | -1.42923 | 0.036488026 | 0.23684639 | 1 | 6258 | tags=36%, list=27%, si | MM_3 |
| GOBP_REGULATION_OF_ | GOBP_REGULATION_OF_STORE_OPERATED | 18 | -0.56239 | -1.42899 | 0.06509695 | 0.23689665 | 1 | 1806 | tags=33%, list=8%, sig | MM_3 |
| GOBP_ADULT_WALKING_ | GOBP_ADULT_WALKING_BEHAVIOR | 48 | -0.47051 | -1.42844 | 0.049200494 | 0.23758945 | 1 | 4222 | tags=27%, list=18%, si | MM_3 |
| GOBP_NEGATIVE_REGUL | GOBP_NEGATIVE_REGULATION_OF_CARBO | 58 | -0.45146 | -1.42794 | 0.041567694 | 0.23823142 | 1 | 4950 | tags=36%, list=21%, si | MM_3 |
| GOBP_RIBONUCLEOPROT | GOBP_RIBONUCLEOPROTEIN_COMPLEX_BIO | 417 | -0.38828 | -1.42793 | 0 | 0.23785898 | 1 | 6878 | tags=47%, list=29%, si | MM_3 |
| GOBP_SMOOTH_MUSCLE | GOBP_SMOOTH_MUSCLE_CELL_MIGRATION | 115 | -0.41876 | -1.42768 | 0.01659292 | 0.23798785 | 1 | 7297 | tags=54%, list=31%, si | MM_3 |
| GOBP_REGULATION_OF_ | GOBP_REGULATION_OF_VIRAL_GENOME_R | 78 | -0.43645 | -1.42764 | 0.03364269 | 0.23768392 | 1 | 5582 | tags=36%, list=24%, si | MM_3 |
| GOBP_POLYOL_METABOL | GOBP_POLYOL_METABOLIC_PROCESS | 99 | -0.42681 | -1.42737 | 0.014396456 | 0.23785228 | 1 | 4950 | tags=28%, list=21%, si | MM_3 |
| GOBP_NEGATIVE_REGUL | GOBP_NEGATIVE_REGULATION_OF_LEUKOC | 96 | -0.42958 | -1.42692 | 0.016816143 | 0.23836733 | 1 | 5034 | tags=35%, list=21%, si | MM_3 |
| PROGESTERONE-MEDIATI | PROGESTERONE-MEDIATED_OOCYTE_MATU | 87 | -0.43259 | -1.42632 | 0.01803833 | 0.23920938 | 1 | 5254 | tags=36%, list=22%, si | MM_3 |
| GOBP_REGULATION_OF_ | GOBP_REGULATION_OF_LEUKOCYTE_PROLI | 273 | -0.39476 | -1.42601 | 0.0010266941 | 0.23944429 | 1 | 4408 | tags=31%, list=19%, si | MM_3 |
| GOBP_MUSCLE_ADAPTAT | GOBP_MUSCLE_ADAPTATION | 126 | -0.41892 | -1.426 | 0.008648649 | 0.23908752 | 1 | 6172 | tags=41%, list=26%, si | MM_3 |
| GOBP_ORGANIC_ACID_C | GOBP_ORGANIC_ACID_CATABOLIC_PROCES | 224 | -0.39786 | -1.42564 | 0.0020470829 | 0.23940188 | 1 | 5482 | tags=34%, list=23%, si | MM_3 |
| GOBP_REGULATION_OF_ | GOBP_REGULATION_OF_SUPEROXIDE_ANIC | 28 | -0.51151 | -1.42547 | 0.051181104 | 0.23931779 | 1 | 4991 | tags=46%, list=21%, si | MM_3 |
| GOBP_CARBOHYDRATE_P | GOBP_CARBOHYDRATE_PHOSPHORYLATION | 19 | -0.55637 | -1.42536 | 0.072829135 | 0.23913786 | 1 | 3220 | tags=37%, list=14%, si | MM_3 |
| GOBP_REGULATION_OF_ | GOBP_REGULATION_OF_LYMPHOCYTE_CHE | 22 | -0.53749 | -1.42489 | 0.06901218 | 0.23971061 | 1 | 2003 | tags=23%, list=8%, sig | MM_3 |
| GOBP_REGULATION_OF_ | GOBP_REGULATION_OF_EARLY_ENDOSOME | 18 | -0.56529 | -1.42474 | 0.068306014 | 0.23966229 | 1 | 4549 | tags=39%, list=19%, si | MM_3 |
| GOBP_NEGATIVE_REGUL | GOBP_NEGATIVE_REGULATION_OF_TRANSM | 142 | -0.41681 | -1.42449 | 0.011866235 | 0.23980626 | 1 | 4768 | tags=30%, list=20%, si | MM_3 |
| GOBP_REGULATION_OF_ | GOBP_REGULATION_OF_UBIQUITIN_PROTE | 39 | -0.48055 | -1.42423 | 0.064020485 | 0.23999839 | 1 | 5922 | tags=49%, list=25%, si | MM_3 |
| GOBP_POSITIVE_REGULA | GOBP_POSITIVE_REGULATION_OF_EXECUTI | 19 | -0.55149 | -1.42419 | 0.056878306 | 0.23971263 | 1 | 4732 | tags=47%, list=20%, si | MM_3 |
| GOBP_NEGATIVE_REGUL | GOBP_NEGATIVE_REGULATION_OF_MAPK_ | 163 | -0.40752 | -1.424 | 0.0031982942 | 0.23966637 | 1 | 6625 | tags=43%, list=28%, si | MM_3 |
| GOBP_REGULATION_OF_ | GOBP_REGULATION_OF_POST_TRANSLATIO | 242 | -0.40089 | -1.4239 | 0.0030895984 | 0.23947573 | 1 | 6948 | tags=49%, list=29%, si | MM_3 |
| GOBP_POSITIVE_REGULA | GOBP_POSITIVE_REGULATION_OF_MONON | 94 | -0.4259 | -1.42337 | 0.017857144 | 0.24013211 | 1 | 6738 | tags=51%, list=29%, si | MM_3 |
| SALMONELLA_INFECTION | SALMONELLA_INFECTION | 76 | -0.43485 | -1.42261 | 0.031322505 | 0.24119234 | 1 | 6097 | tags=42%, list=26%, si | MM_3 |
| GOBP_MONOCARBOXYLIC | GOBP_MONOCARBOXYLIC_ACID_BIOSYNTH | 203 | -0.40117 | -1.42247 | 0.0052301255 | 0.24108689 | 1 | 5746 | tags=35%, list=24%, si | MM_3 |
| PENTOSE_AND_GLUCURO | PENTOSE_AND_GLUCURONATE_INTERCONV | 22 | -0.52968 | -1.4221 | 0.07537012 | 0.2414566 | 1 | 5817 | tags=45%, list=25%, si | MM_3 |
| GOBP_PROTEIN_DEGLYCO | GOBP_PROTEIN_DEGLYCOSYLATION | 15 | -0.57379 | -1.42201 | 0.08567416 | 0.24125317 | 1 | 2832 | tags=27%, list=12%, si | MM_3 |
| GOBP_REGULATION_OF_ | GOBP_REGULATION_OF_OSTEOCLAST_DIFFE | 84 | -0.42948 | -1.42199 | 0.020179372 | 0.24091256 | 1 | 4960 | tags=35%, list=21%, si | MM_3 |
| GOBP_POSITIVE_REGULA | GOBP_POSITIVE_REGULATION_OF_SMOOTH | 114 | -0.42182 | -1.42196 | 0.0174482 | 0.24059786 | 1 | 4991 | tags=36%, list=21%, si | MM_3 |
| GOBP_B_CELL_HOMEOST | GOBP_B_CELL_HOMEOSTASIS | 42 | -0.46681 | -1.42171 | 0.06140351 | 0.24070929 | 1 | 6014 | tags=45%, list=25%, si | MM_3 |
| GOBP_SUPEROXIDE_ANIO | GOBP_SUPEROXIDE_ANION_GENERATION | 43 | -0.47147 | -1.42138 | 0.051833123 | 0.24097368 | 1 | 7093 | tags=56%, list=30%, si | MM_3 |
| GOBP_POSITIVE_REGULA | GOBP_POSITIVE_REGULATION_OF_HEART_C | 49 | -0.46287 | -1.42127 | 0.04187192 | 0.24082848 | 1 | 2439 | tags=31%, list=10%, si | MM_3 |
| GOBP_NUCLEAR_ENVELO | GOBP_NUCLEAR_ENVELOPE_ORGANIZATION | 52 | -0.46648 | -1.42092 | 0.031476997 | 0.24114819 | 1 | 5075 | tags=48%, list=21%, si | MM_3 |
| GOBP_PROTEIN_REFOLDI | GOBP_PROTEIN_REFOLDING | 23 | -0.53443 | -1.42054 | 0.069798656 | 0.24153595 | 1 | 1538 | tags=22%, list=7%, sig | MM_3 |
| GOBP_REGULATION_OF_ | GOBP_REGULATION_OF_EXTRINSIC_APOPTO | 45 | -0.46971 | -1.42046 | 0.04375 | 0.2413576 | 1 | 6619 | tags=60%, list=28%, si | MM_3 |
| GOBP_POSITIVE_REGULA | GOBP_POSITIVE_REGULATION_OF_NEURON | 15 | -0.56877 | -1.42031 | 0.07982583 | 0.24131292 | 1 | 6001 | tags=53%, list=25%, si | MM_3 |
| GOBP_REGULATION_OF_ | GOBP_REGULATION_OF_CELLULAR_RESPON | 141 | -0.40807 | -1.42029 | 0.014130435 | 0.24098672 | 1 | 6326 | tags=44%, list=27%, si | MM_3 |
| GOBP_NEUROTRANSMITT | GOBP_NEUROTRANSMITTER_GATED_ION_C | 21 | -0.53946 | -1.42022 | 0.06232687 | 0.2407499 | 1 | 3987 | tags=38%, list=17%, si | MM_3 |
| GOBP_REGULATION_OF_ | GOBP_REGULATION_OF_DEFENSE_RESPONS | 20 | -0.53539 | -1.42009 | 0.07934336 | 0.24066561 | 1 | 3753 | tags=45%, list=16%, si | MM_3 |
| GOBP_REGULATION_OF_ | GOBP_REGULATION_OF_FATTY_ACID_META | 99 | -0.42571 | -1.4195 | 0.01803833 | 0.2414851 | 1 | 2458 | tags=19%, list=10%, si | MM_3 |
| GOBP_CELLULAR_RESPON | GOBP_CELLULAR_RESPONSE_TO_NUTRIENT | 38 | -0.47672 | -1.41916 | 0.04936709 | 0.24173637 | 1 | 5214 | tags=42%, list=22%, si | MM_3 |
| GOBP_ESTABLISHMENT_O | GOBP_ESTABLISHMENT_OF_PROTEIN_LOCA | 25 | -0.51783 | -1.41911 | 0.073076926 | 0.24147207 | 1 | 6691 | tags=56%, list=28%, si | MM_3 |
| GOBP_REGULATION_OF_ | GOBP_REGULATION_OF_MITOCHONDRIAL_ | 87 | -0.4334 | -1.41887 | 0.029378532 | 0.24159291 | 1 | 4991 | tags=40%, list=21%, si | MM_3 |
| GOBP_POSITIVE_REGULA | GOBP_POSITIVE_REGULATION_OF_FIBROBL | 76 | -0.43512 | -1.41885 | 0.032482598 | 0.241252 | 1 | 5130 | tags=36%, list=22%, si | MM_3 |
| GOBP_SMAD_PROTEIN_S | GOBP_SMAD_PROTEIN_SIGNAL_TRANSDUC | 83 | -0.44049 | -1.41841 | 0.021914648 | 0.24181929 | 1 | 4926 | tags=30%, list=21%, si | MM_3 |
| GOBP_NEGATIVE_REGUL | GOBP_NEGATIVE_REGULATION_OF_JNK_CA | 37 | -0.48002 | -1.4183 | 0.052564103 | 0.24167122 | 1 | 6602 | tags=41%, list=28%, si | MM_3 |
| GOBP_MATERNAL_PLACE | GOBP_MATERNAL_PLACENTA_DEVELOPME | 34 | -0.4908 | -1.41767 | 0.054361567 | 0.24250181 | 1 | 5568 | tags=41%, list=24%, si | MM_3 |
| GOBP_REGULATION_OF_ | GOBP_REGULATION_OF_ADIPOSE_TISSUE_D | 16 | -0.56851 | -1.41739 | 0.08108108 | 0.2426778 | 1 | 6133 | tags=63%, list=26%, si | MM_3 |
| GOBP_NUCLEOTIDE_EXCI | GOBP_NUCLEOTIDE_EXCISION_REPAIR | 76 | -0.43527 | -1.41718 | 0.026589595 | 0.24274933 | 1 | 6002 | tags=50%, list=25%, si | MM_3 |
| GOBP_PIGMENT_METABO | GOBP_PIGMENT_METABOLIC_PROCESS | 65 | -0.44783 | -1.41684 | 0.03846154 | 0.24302109 | 1 | 4980 | tags=34%, list=21%, si | MM_3 |
| GOBP_REGULATION_OF_ | GOBP_REGULATION_OF_CELL_CYCLE_G2_M | 117 | -0.41677 | -1.41653 | 0.01754386 | 0.24329011 | 1 | 4759 | tags=36%, list=20%, si | MM_3 |
| GOBP_POSITIVE_REGULA | GOBP_POSITIVE_REGULATION_OF_MEMBRA | 39 | -0.48326 | -1.4163 | 0.0479798 | 0.24338278 | 1 | 4754 | tags=41%, list=20%, si | MM_3 |
| GOBP_RESPONSE_TO_TU | GOBP_RESPONSE_TO_TUMOR_NECROSIS_F | 167 | -0.39915 | -1.41623 | 0.011615628 | 0.24314134 | 1 | 7317 | tags=51%, list=31%, si | MM_3 |
| GOBP_REGULATION_OF_ | GOBP_REGULATION_OF_RECEPTOR_BINDIN | 20 | -0.53932 | -1.41609 | 0.08891929 | 0.2430522 | 1 | 6721 | tags=50%, list=28%, si | MM_3 |
| GOBP_REGULATION_OF_ | GOBP_REGULATION_OF_MONOCYTE_CHEM | 29 | -0.50936 | -1.41601 | 0.06435006 | 0.24283576 | 1 | 7212 | tags=66%, list=31%, si | MM_3 |
| GOBP_NEGATIVE_REGUL | GOBP_NEGATIVE_REGULATION_OF_T_CELL_ | 73 | -0.43975 | -1.41499 | 0.032786883 | 0.2444017 | 1 | 4362 | tags=36%, list=18%, si | MM_3 |
| MANNOSE_TYPE_O-GLYC | MANNOSE_TYPE_O-GLYCAN_BIOSYNTHESIS | 21 | -0.52436 | -1.41479 | 0.07181572 | 0.24444416 | 1 | 3045 | tags=29%, list=13%, si | MM_3 |
| GOBP_DNA_DAMAGE_RE | GOBP_DNA_DAMAGE_RESPONSE_SIGNAL_T | 67 | -0.44251 | -1.41469 | 0.045296166 | 0.24428368 | 1 | 5637 | tags=39%, list=24%, si | MM_3 |
| CYTOKINE-CYTOKINE_REC | CYTOKINE-CYTOKINE_RECEPTOR_INTERACTI | 239 | -0.39995 | -1.41462 | 0.0010298662 | 0.2440653 | 1 | 4133 | tags=29%, list=18%, si | MM_3 |
| GOBP_REGULATION_OF_ | GOBP_REGULATION_OF_CELL_FATE_COMM | 41 | -0.47351 | -1.41449 | 0.045454547 | 0.24395163 | 1 | 4488 | tags=27%, list=19%, si | MM_3 |
| GOBP_QUINONE_METAB | GOBP_QUINONE_METABOLIC_PROCESS | 23 | -0.52283 | -1.41445 | 0.06334232 | 0.24366005 | 1 | 3254 | tags=26%, list=14%, si | MM_3 |
| GOBP_REGULATION_OF_ | GOBP_REGULATION_OF_T_CELL_ACTIVATIO | 364 | -0.38777 | -1.41422 | 0 | 0.24374284 | 1 | 4963 | tags=32%, list=21%, si | MM_3 |
| GOBP_TRABECULA_FORM | GOBP_TRABECULA_FORMATION | 27 | -0.50705 | -1.41415 | 0.07049608 | 0.24352483 | 1 | 3423 | tags=33%, list=14%, si | MM_3 |
| GOBP_POST_TRANSCRIPT | GOBP_POST_TRANSCRIPTIONAL_GENE_SILE | 64 | -0.44723 | -1.41334 | 0.038416762 | 0.24467549 | 1 | 6710 | tags=45%, list=28%, si | MM_3 |
| GOBP_GLOMERULAR_ME | GOBP_GLOMERULAR_MESANGIUM_DEVELO | 16 | -0.56989 | -1.41303 | 0.07616361 | 0.24493805 | 1 | 1783 | tags=25%, list=8%, sig | MM_3 |
| GOBP_POSITIVE_REGULA | GOBP_POSITIVE_REGULATION_OF_CARTILA | 31 | -0.49914 | -1.41268 | 0.064020485 | 0.24529332 | 1 | 2473 | tags=26%, list=10%, si | MM_3 |
| GOBP_TELOMERE_CAPPI | GOBP_TELOMERE_CAPPING | 38 | -0.47985 | -1.41244 | 0.05897436 | 0.24535304 | 1 | 5125 | tags=45%, list=22%, si | MM_3 |
| FLUID_SHEAR_STRESS_AN | FLUID_SHEAR_STRESS_AND_ATHEROSCLERO | 135 | -0.40894 | -1.4122 | 0.010881393 | 0.24547297 | 1 | 6571 | tags=44%, list=28%, si | MM_3 |
| GOBP_STORE_OPERATED | GOBP_STORE_OPERATED_CALCIUM_ENTRY | 25 | -0.51816 | -1.41193 | 0.07208388 | 0.24567561 | 1 | 3162 | tags=36%, list=13%, si | MM_3 |
| GOBP_INTRACELLULAR_R | GOBP_INTRACELLULAR_RECEPTOR_SIGNALI | 325 | -0.38915 | -1.41193 | 0 | 0.24532163 | 1 | 5866 | tags=36%, list=25%, si | MM_3 |
| GOBP_REGULATION_OF_ | GOBP_REGULATION_OF_T_CELL_PROLIFERA | 188 | -0.40306 | -1.41186 | 0.0062695923 | 0.24511626 | 1 | 4362 | tags=31%, list=18%, si | MM_3 |
| GOBP_LEUKOCYTE_MIGR | GOBP_LEUKOCYTE_MIGRATION_INVOLVED_ | 23 | -0.52206 | -1.41157 | 0.08672799 | 0.24535729 | 1 | 4339 | tags=43%, list=18%, si | MM_3 |
| GOBP_NEGATIVE_REGUL | GOBP_NEGATIVE_REGULATION_OF_PHOSPH | 62 | -0.44378 | -1.41153 | 0.045507584 | 0.24506721 | 1 | 6625 | tags=45%, list=28%, si | MM_3 |

| | | | | | | | | | | |
|---|---|---|---|---|---|---|---|---|---|---|
| GOBP_POSITIVE_REGULA | GOBP_POSITIVE_REGULATION_OF_RELEASE | 27 | -0.50808 | -1.41138 | 0.055128206 | 0.24501176 | 1 | 2720 | tags=26%, list=12%, si | MM_3 |
| GOBP_NEGATIVE_REGUL | GOBP_NEGATIVE_REGULATION_OF_TRANSF | 94 | -0.42713 | -1.41133 | 0.018888889 | 0.24476178 | 1 | 4768 | tags=34%, list=20%, si | MM_3 |
| GOBP_REGULATION_OF_ | GOBP_REGULATION_OF_MONONUCLEAR_C | 136 | -0.4073 | -1.41127 | 0.019438446 | 0.24450628 | 1 | 7212 | tags=55%, list=31%, si | MM_3 |
| GOBP_MYELOID_LEUKOC | GOBP_MYELOID_LEUKOCYTE_DIFFERENTIAT | 258 | -0.39168 | -1.41106 | 0.0030643514 | 0.24454567 | 1 | 6363 | tags=41%, list=27%, si | MM_3 |
| GOBP_NUCLEAR_TRANSC | GOBP_NUCLEAR_TRANSCRIBED_MRNA_POL | 26 | -0.51111 | -1.41101 | 0.072289154 | 0.24430056 | 1 | 8027 | tags=69%, list=34%, si | MM_3 |
| GOBP_POSITIVE_REGULA | GOBP_POSITIVE_REGULATION_OF_ORGAN_ | 60 | -0.44567 | -1.41094 | 0.035151515 | 0.2440623 | 1 | 3489 | tags=32%, list=15%, si | MM_3 |
| GOBP_DOPAMINE_RECEP | GOBP_DOPAMINE_RECEPTOR_SIGNALING_ | 35 | -0.48345 | -1.4107 | 0.07152145 | 0.24422675 | 1 | 3321 | tags=29%, list=14%, si | MM_3 |
| GOBP_MYELOID_CELL_DE | GOBP_MYELOID_CELL_DEVELOPMENT | 98 | -0.41823 | -1.4104 | 0.017837236 | 0.24444729 | 1 | 6781 | tags=48%, list=29%, si | MM_3 |
| GOBP_MYELOID_LEUKOC | GOBP_MYELOID_LEUKOCYTE_MEDIATED_IM | 108 | -0.42026 | -1.41035 | 0.0122631 | 0.24418068 | 1 | 6540 | tags=44%, list=28%, si | MM_3 |
| GOBP_POSITIVE_REGULA | GOBP_POSITIVE_REGULATION_OF_SIGNAL_ | 32 | -0.49314 | -1.41033 | 0.06435006 | 0.24385993 | 1 | 7515 | tags=59%, list=32%, si | MM_3 |
| GOBP_POSITIVE_REGULA | GOBP_POSITIVE_REGULATION_OF_FATTY_A | 25 | -0.50879 | -1.41026 | 0.078947365 | 0.24362394 | 1 | 1792 | tags=24%, list=8%, sig | MM_3 |
| GOBP_ENDOSOMAL_TRA | GOBP_ENDOSOMAL_TRANSPORT | 272 | -0.39149 | -1.41018 | 0.0010245901 | 0.24344051 | 1 | 6548 | tags=44%, list=28%, si | MM_3 |
| GOBP_DNA_DUPLEX_UN | GOBP_DNA_DUPLEX_UNWINDING | 43 | -0.46981 | -1.41013 | 0.054545455 | 0.24320129 | 1 | 3852 | tags=33%, list=16%, si | MM_3 |
| GOBP_ACTIVATION_OF_I | GOBP_ACTIVATION_OF_INNATE_IMMUNE_ | 290 | -0.39249 | -1.41012 | 0.0040983604 | 0.24288067 | 1 | 5034 | tags=32%, list=21%, si | MM_3 |
| GOBP_POSITIVE_REGULA | GOBP_POSITIVE_REGULATION_OF_INTERLE | 106 | -0.41708 | -1.40922 | 0.01323043 | 0.24430314 | 1 | 6746 | tags=47%, list=29%, si | MM_3 |
| GOBP_MAINTENANCE_O | GOBP_MAINTENANCE_OF_CELL_NUMBER | 205 | -0.39637 | -1.40919 | 0.006185567 | 0.24402577 | 1 | 6195 | tags=39%, list=26%, si | MM_3 |
| GOBP_PROSTATE_GLAND | GOBP_PROSTATE_GLAND_DEVELOPMENT | 47 | -0.46324 | -1.40918 | 0.060270604 | 0.24370244 | 1 | 4017 | tags=28%, list=17%, si | MM_3 |
| GOBP_GOLGI_VESICLE_T | GOBP_GOLGI_VESICLE_TRANSPORT | 276 | -0.38939 | -1.40847 | 0 | 0.24472503 | 1 | 7055 | tags=50%, list=30%, si | MM_3 |
| GOBP_RESPONSE_TO_PR | GOBP_RESPONSE_TO_PROGESTERONE | 19 | -0.53879 | -1.40845 | 0.082530946 | 0.24439855 | 1 | 1036 | tags=26%, list=4%, sig | MM_3 |
| GOBP_REGULATION_OF_ | GOBP_REGULATION_OF_MEMBRANE_PERM | 70 | -0.4377 | -1.40824 | 0.022432113 | 0.2444525 | 1 | 4754 | tags=36%, list=20%, si | MM_3 |
| GOBP_REGULATION_OF_ | GOBP_REGULATION_OF_GENERATION_OF_ | 148 | -0.40099 | -1.40791 | 0.011664899 | 0.24475126 | 1 | 5967 | tags=35%, list=25%, si | MM_3 |
| GOBP_PROTEIN_MATURA | GOBP_PROTEIN_MATURATION | 475 | -0.38444 | -1.40713 | 0 | 0.24590397 | 1 | 6055 | tags=37%, list=26%, si | MM_3 |
| FOXO_SIGNALING_PATHV | FOXO_SIGNALING_PATHWAY | 127 | -0.41013 | -1.40631 | 0.014130435 | 0.24719137 | 1 | 6429 | tags=39%, list=27%, si | MM_3 |
| GOBP_CYTOKINE_MEDIA | GOBP_CYTOKINE_MEDIATED_SIGNALING_P | 421 | -0.38493 | -1.406 | 0 | 0.24748017 | 1 | 5018 | tags=30%, list=21%, si | MM_3 |
| GOBP_PROTEIN_K11_LIN | GOBP_PROTEIN_K11_LINKED_UBIQUITINAT | 29 | -0.50138 | -1.40576 | 0.06382979 | 0.24759206 | 1 | 4323 | tags=34%, list=18%, si | MM_3 |
| GOBP_STRIATED_MUSCL | GOBP_STRIATED_MUSCLE_CELL_APOPTOTI | 73 | -0.43345 | -1.40574 | 0.037166085 | 0.24727824 | 1 | 4987 | tags=34%, list=21%, si | MM_3 |
| GOBP_VIRAL_LIFE_CYCLE | GOBP_VIRAL_LIFE_CYCLE | 230 | -0.39263 | -1.40557 | 0.0030737706 | 0.24726643 | 1 | 6484 | tags=42%, list=27%, si | MM_3 |
| GOBP_NEGATIVE_REGUL | GOBP_NEGATIVE_REGULATION_OF_ALPHA_ | 47 | -0.46455 | -1.40511 | 0.05036855 | 0.24780992 | 1 | 3972 | tags=28%, list=17%, si | MM_3 |
| GOBP_LEUKOCYTE_PROL | GOBP_LEUKOCYTE_PROLIFERATION | 361 | -0.38208 | -1.40501 | 0.0010090817 | 0.2476205 | 1 | 4408 | tags=29%, list=19%, si | MM_3 |
| GOBP_CARBOHYDRATE_D | GOBP_CARBOHYDRATE_DERIVATIVE_TRANS | 76 | -0.43766 | -1.40462 | 0.036342323 | 0.24803126 | 1 | 5411 | tags=41%, list=23%, si | MM_3 |
| APOPTOSIS | APOPTOSIS | 137 | -0.41184 | -1.40426 | 0.011789925 | 0.2483922 | 1 | 6917 | tags=47%, list=29%, si | MM_3 |
| GOBP_RNA_CATABOLIC_ | GOBP_RNA_CATABOLIC_PROCESS | 278 | -0.39116 | -1.40425 | 0.0010266941 | 0.24807839 | 1 | 6736 | tags=43%, list=29%, si | MM_3 |
| GOBP_REGULATION_OF_ | GOBP_REGULATION_OF_MEMBRANE_PROT | 25 | -0.50949 | -1.40421 | 0.090322584 | 0.24779063 | 1 | 6718 | tags=48%, list=28%, si | MM_3 |
| GOBP_POSITIVE_REGULA | GOBP_POSITIVE_REGULATION_OF_CELL_GR | 17 | -0.55464 | -1.4042 | 0.081575245 | 0.24745782 | 1 | 4567 | tags=53%, list=19%, si | MM_3 |
| GOBP_POSITIVE_REGULA | GOBP_POSITIVE_REGULATION_OF_OSTEOBI | 74 | -0.4298 | -1.40388 | 0.03076923 | 0.24771121 | 1 | 5568 | tags=35%, list=24%, si | MM_3 |
| GOBP_CELLULAR_RESPON | GOBP_CELLULAR_RESPONSE_TO_VASCULAR | 60 | -0.44537 | -1.40367 | 0.04211793 | 0.24780914 | 1 | 6378 | tags=40%, list=27%, si | MM_3 |
| GOBP_ORGANOPHOSPHA | GOBP_ORGANOPHOSPHATE_CATABOLIC_PR | 227 | -0.39251 | -1.4036 | 0.0020618557 | 0.24761751 | 1 | 6720 | tags=41%, list=28%, si | MM_3 |
| GOBP_HYDROGEN_PERO | GOBP_HYDROGEN_PEROXIDE_METABOLIC_ | 48 | -0.45908 | -1.40302 | 0.050251257 | 0.24840942 | 1 | 4984 | tags=40%, list=21%, si | MM_3 |
| GOBP_NAD_CATABOLIC_ | GOBP_NAD_CATABOLIC_PROCESS | 21 | -0.53204 | -1.4027 | 0.07744565 | 0.24868503 | 1 | 5278 | tags=43%, list=22%, si | MM_3 |
| GOBP_RENAL_ABSORPTI | GOBP_RENAL_ABSORPTION | 28 | -0.49702 | -1.40268 | 0.06586021 | 0.24838297 | 1 | 5457 | tags=36%, list=23%, si | MM_3 |
| GOBP_NEGATIVE_REGUL | GOBP_NEGATIVE_REGULATION_OF_CATABO | 339 | -0.3862 | -1.40253 | 0.003033367 | 0.24831142 | 1 | 6387 | tags=39%, list=27%, si | MM_3 |
| RETROGRADE_ENDOCAN | RETROGRADE_ENDOCANNABINOID_SIGNAL | 133 | -0.40951 | -1.40248 | 0.009825327 | 0.24805595 | 1 | 6124 | tags=36%, list=26%, si | MM_3 |
| GOBP_REGULATION_OF_ | GOBP_REGULATION_OF_LEUKOCYTE_DIFFE | 342 | -0.38532 | -1.40192 | 0 | 0.2487549 | 1 | 4960 | tags=32%, list=21%, si | MM_3 |
| GOBP_POSITIVE_REGULA | GOBP_POSITIVE_REGULATION_OF_TELOME | 15 | -0.55788 | -1.40132 | 0.09361702 | 0.24952297 | 1 | 6544 | tags=53%, list=28%, si | MM_3 |
| GOBP_RIBOSE_PHOSPHA | GOBP_RIBOSE_PHOSPHATE_METABOLIC_PR | 419 | -0.38239 | -1.40118 | 0 | 0.249459 | 1 | 5922 | tags=36%, list=25%, si | MM_3 |
| GOBP_HEART_GROWTH | GOBP_HEART_GROWTH | 116 | -0.4146 | -1.40107 | 0.022271715 | 0.24931684 | 1 | 4657 | tags=35%, list=20%, si | MM_3 |
| GOBP_REGULATION_OF_ | GOBP_REGULATION_OF_SKELETAL_MUSCLE | 21 | 0.634075 | 2.06915 | 0 | 0.15568073 | 0.184 | 815 | tags=29%, list=3%, sig | MM_3 |
| GOBP_GLYCOSYL_COMP | GOBP_GLYCOSYL_COMPOUND_CATABOLIC_ | 26 | 0.579115 | 1.993397 | 0.004385965 | 0.17467268 | 0.374 | 1041 | tags=19%, list=4%, sig | MM_3 |
| GOBP_RETINAL_METABO | GOBP_RETINAL_METABOLIC_PROCESS | 16 | 0.665906 | 1.991669 | 0 | 0.11892934 | 0.379 | 1747 | tags=44%, list=7%, sig | MM_3 |
| GOBP_PARAXIAL_MESOD | GOBP_PARAXIAL_MESODERM_DEVELOPME | 21 | 0.62641 | 1.986268 | 0 | 0.09596585 | 0.399 | 1809 | tags=38%, list=8%, sig | MM_3 |
| GOBP_POSITIVE_REGULA | GOBP_POSITIVE_REGULATION_OF_SYNAPTI | 17 | 0.633786 | 1.935717 | 0.010989011 | 0.13232717 | 0.573 | 2128 | tags=29%, list=9%, sig | MM_3 |
| GOBP_PARASYMPATHETI | GOBP_PARASYMPATHETIC_NERVOUS_SYSTE | 21 | 0.565076 | 1.801769 | 0 | 0.24114978 | 0.948 | 1490 | tags=29%, list=6%, sig | MM_3 |
| RIBOSOME | RIBOSOME | 128 | -0.65521 | -2.77974 | 0 | 0 | 0 | 5349 | tags=71%, list=23%, si | EN_3 |
| GOBP_CYTOPLASMIC_TR | GOBP_CYTOPLASMIC_TRANSLATION | 143 | -0.60884 | -2.63556 | 0 | 0 | 0 | 6593 | tags=70%, list=28%, si | EN_3 |
| GOBP_TRANSLATION_AT_ | GOBP_TRANSLATION_AT_SYNAPSE | 48 | -0.704 | -2.55833 | 0 | 0 | 0 | 6530 | tags=94%, list=28%, si | EN_3 |
| OXIDATIVE_PHOSPHORYL | OXIDATIVE_PHOSPHORYLATION | 115 | -0.56704 | -2.44653 | 0 | 0.000259421 | 0.001 | 6040 | tags=57%, list=26%, si | EN_3 |
| PARKINSON_DISEASE | PARKINSON_DISEASE | 122 | -0.54867 | -2.35176 | 0 | 0.000207537 | 0.001 | 4914 | tags=48%, list=21%, si | EN_3 |
| GOBP_OXIDATIVE_PHOSP | GOBP_OXIDATIVE_PHOSPHORYLATION | 124 | -0.54384 | -2.32394 | 0 | 0.000172947 | 0.001 | 5997 | tags=52%, list=25%, si | EN_3 |
| GOBP_AEROBIC_ELECTRO | GOBP_AEROBIC_ELECTRON_TRANSPORT_CH | 63 | -0.59235 | -2.26241 | 0 | 0.000735667 | 0.005 | 4898 | tags=44%, list=21%, si | EN_3 |
| ALZHEIMER_DISEASE | ALZHEIMER_DISEASE | 159 | -0.50735 | -2.22561 | 0 | 0.001156194 | 0.009 | 6464 | tags=53%, list=27%, si | EN_3 |
| THERMOGENESIS | THERMOGENESIS | 210 | -0.48335 | -2.20791 | 0 | 0.001368664 | 0.012 | 6433 | tags=50%, list=27%, si | EN_3 |
| GOBP_MITOCHONDRIAL_ | GOBP_MITOCHONDRIAL_RESPIRATORY_CHA | 91 | -0.54046 | -2.1961 | 0 | 0.001644046 | 0.016 | 6223 | tags=55%, list=26%, si | EN_3 |
| HUNTINGTON_DISEASE | HUNTINGTON_DISEASE | 180 | -0.49479 | -2.17304 | 0 | 0.002430617 | 0.026 | 5997 | tags=50%, list=25%, si | EN_3 |
| GOBP_PROTON_MOTIVE_ | GOBP_PROTON_MOTIVE_FORCE_DRIVEN_A | 52 | -0.58845 | -2.16301 | 0 | 0.002823539 | 0.033 | 5997 | tags=56%, list=25%, si | EN_3 |
| GOBP_AEROBIC_RESPIRA | GOBP_AEROBIC_RESPIRATION | 170 | -0.4805 | -2.13747 | 0 | 0.004267408 | 0.053 | 6128 | tags=51%, list=26%, si | EN_3 |
| GOBP_NADH_DEHYDROG | GOBP_NADH_DEHYDROGENASE_COMPLEX_ | 58 | -0.56401 | -2.13686 | 0 | 0.003962593 | 0.053 | 6223 | tags=52%, list=26%, si | EN_3 |
| GOBP_LIPOPROTEIN_LOC | GOBP_LIPOPROTEIN_LOCALIZATION | 20 | -0.70928 | -2.12858 | 0 | 0.004864834 | 0.069 | 3198 | tags=50%, list=14%, si | EN_3 |
| GOBP_MATURATION_OF_ | GOBP_MATURATION_OF_5_8S_RRNA | 35 | -0.61004 | -2.08782 | 0 | 0.009378733 | 0.139 | 2603 | tags=43%, list=11%, si | EN_3 |
| GOBP_CELLULAR_RESPIR | GOBP_CELLULAR_RESPIRATION | 218 | -0.4582 | -2.0869 | 0 | 0.009128886 | 0.142 | 5905 | tags=45%, list=25%, si | EN_3 |
| GOBP_ATP_SYNTHESIS_C | GOBP_ATP_SYNTHESIS_COUPLED_ELECTRO | 76 | -0.51385 | -2.06278 | 0 | 0.012559755 | 0.202 | 4905 | tags=39%, list=21%, si | EN_3 |
| GOBP_DNA_UNWINDING | GOBP_DNA_UNWINDING_INVOLVED_IN_DI | 20 | -0.67882 | -2.05233 | 0 | 0.014437866 | 0.235 | 3462 | tags=45%, list=15%, si | EN_3 |
| GOBP_PROTEIN_LOCALIZ | GOBP_PROTEIN_LOCALIZATION_TO_MITOC | 101 | -0.48696 | -2.03322 | 0 | 0.018494088 | 0.301 | 3458 | tags=28%, list=15%, si | EN_3 |
| GOBP_ATP_BIOSYNTHETI | GOBP_ATP_BIOSYNTHETIC_PROCESS | 80 | -0.50488 | -2.03295 | 0 | 0.017759517 | 0.302 | 6151 | tags=52%, list=26%, si | EN_3 |
| GOBP_ELECTRON_TRANS | GOBP_ELECTRON_TRANSPORT_CHAIN | 107 | -0.48293 | -2.02663 | 0 | 0.018680047 | 0.32 | 5084 | tags=39%, list=22%, si | EN_3 |
| GOBP_MITOTIC_DNA_RE | GOBP_MITOTIC_DNA_REPLICATION | 16 | -0.68757 | -2.00563 | 0 | 0.02608608 | 0.433 | 3045 | tags=44%, list=13%, si | EN_3 |
| GOBP_RIBOSOMAL_SMA | GOBP_RIBOSOMAL_SMALL_SUBUNIT_BIOGI | 101 | -0.47518 | -1.9952 | 0 | 0.028421821 | 0.475 | 4815 | tags=42%, list=20%, si | EN_3 |
| GOBP_NUCLEOSIDE_TRIP | GOBP_NUCLEOSIDE_TRIPHOSPHATE_BIOSY | 100 | -0.48165 | -1.98122 | 0 | 0.033036128 | 0.543 | 5997 | tags=48%, list=25%, si | EN_3 |
| GOBP_NEGATIVE_REGUL | GOBP_NEGATIVE_REGULATION_OF_NUCLEO | 27 | -0.60723 | -1.98011 | 0 | 0.032438 | 0.551 | 4152 | tags=37%, list=18%, si | EN_3 |
| GOBP_STEROL_BIOSYNTH | GOBP_STEROL_BIOSYNTHETIC_PROCESS | 58 | -0.51972 | -1.97235 | 0.0013297872 | 0.03455334 | 0.594 | 4246 | tags=45%, list=18%, si | EN_3 |
| GOBP_REGULATION_OF_ | GOBP_REGULATION_OF_NECROPTOTIC_PRO | 22 | -0.63124 | -1.97061 | 0.0015151515 | 0.034052756 | 0.604 | 5727 | tags=59%, list=24%, si | EN_3 |
| GOBP_DISRUPTION_OF_A | GOBP_DISRUPTION_OF_ANATOMICAL_STRU | 73 | -0.49595 | -1.96715 | 0 | 0.03419161 | 0.619 | 2577 | tags=29%, list=11%, si | EN_3 |
| GOBP_PROTEIN_TARGETI | GOBP_PROTEIN_TARGETING_TO_MITOCHO | 76 | -0.48323 | -1.95278 | 0 | 0.040479567 | 0.701 | 3458 | tags=28%, list=15%, si | EN_3 |

| | | | | | | | | | | |
|---|---|---|---|---|---|---|---|---|---|---|
| GOBP_HYDROGEN_PERO) | GOBP_HYDROGEN_PEROXIDE_METABOLIC_ | 48 | -0.53196 | -1.95213 | 0 | 0.03960386 | 0.704 | 4307 | tags=40%, list=18%, si | EN_3 |
| GOBP_REGULATION_OF_ | GOBP_REGULATION_OF_VIRAL_TRANSCRIPT | 25 | -0.62447 | -1.95198 | 0 | 0.03839852 | 0.704 | 4955 | tags=60%, list=21%, si | EN_3 |
| NON-ALCOHOLIC_FATTY_ | NON-ALCOHOLIC_FATTY_LIVER_DISEASE_(N | 141 | -0.45421 | -1.95027 | 0 | 0.03841897 | 0.718 | 5065 | tags=41%, list=21%, si | EN_3 |
| GOBP_RIBOSOME_BIOGE | GOBP_RIBOSOME_BIOGENESIS | 306 | -0.41669 | -1.94168 | 0 | 0.0416739 | 0.759 | 5135 | tags=39%, list=22%, si | EN_3 |
| GOBP_CELLULAR_RESPON | GOBP_CELLULAR_RESPONSE_TO_HEAT | 50 | -0.52617 | -1.93786 | 0.0026773761 | 0.04268518 | 0.774 | 4283 | tags=38%, list=18%, si | EN_3 |
| GOBP_RESPONSE_TO_HE | GOBP_RESPONSE_TO_HEAT | 86 | -0.47616 | -1.92569 | 0 | 0.049465414 | 0.82 | 4283 | tags=33%, list=18%, si | EN_3 |
| GOBP_BLASTOCYST_GRO | GOBP_BLASTOCYST_GROWTH | 29 | -0.59369 | -1.92314 | 0 | 0.04987892 | 0.831 | 5067 | tags=48%, list=21%, si | EN_3 |
| GLYCINE,_SERINE_AND_T | GLYCINE,_SERINE_AND_THREONINE_METAE | 38 | -0.5456 | -1.91952 | 0 | 0.05084036 | 0.846 | 2700 | tags=32%, list=11%, si | EN_3 |
| GOBP_CELL_REDOX_HON | GOBP_CELL_REDOX_HOMEOSTASIS | 28 | -0.58331 | -1.91765 | 0.0014662757 | 0.05061892 | 0.852 | 8157 | tags=89%, list=35%, si | EN_3 |
| GOBP_KILLING_OF_CELLS | GOBP_KILLING_OF_CELLS_OF_ANOTHER_OI | 71 | -0.49007 | -1.91647 | 0 | 0.05015082 | 0.857 | 2577 | tags=28%, list=11%, si | EN_3 |
| GOBP_CARBOHYDRATE_F | GOBP_CARBOHYDRATE_PHOSPHORYLATION | 19 | -0.64623 | -1.91584 | 0 | 0.049178004 | 0.857 | 1123 | tags=32%, list=5%, sig | EN_3 |
| GOBP_ENDOPLASMIC_RE | GOBP_ENDOPLASMIC_RETICULUM_CALCIUI | 28 | -0.57861 | -1.9104 | 0.0014430014 | 0.05118598 | 0.871 | 5301 | tags=39%, list=22%, si | EN_3 |
| GOBP_POSITIVE_REGULA | GOBP_POSITIVE_REGULATION_OF_CELLULA | 20 | -0.6311 | -1.90125 | 0 | 0.05606766 | 0.901 | 3708 | tags=50%, list=16%, si | EN_3 |
| GOBP_MAINTENANCE_OI | GOBP_MAINTENANCE_OF_PROTEIN_LOCAT | 97 | -0.45877 | -1.90083 | 0 | 0.055145685 | 0.901 | 6099 | tags=46%, list=26%, si | EN_3 |
| GOBP_NEGATIVE_REGUL | GOBP_NEGATIVE_REGULATION_OF_PROTEI | 17 | -0.65175 | -1.89654 | 0 | 0.056731265 | 0.917 | 4152 | tags=41%, list=18%, si | EN_3 |
| GOBP_REGULATION_OF_ | GOBP_REGULATION_OF_RESPONSE_TO_EN | 80 | -0.47113 | -1.8963 | 0 | 0.05556511 | 0.917 | 6940 | tags=48%, list=29%, si | EN_3 |
| GOBP_REGULATION_OF_ | GOBP_REGULATION_OF_PROGRAMMED_NE | 27 | -0.57858 | -1.89445 | 0.0043290043 | 0.055870004 | 0.92 | 5727 | tags=56%, list=24%, si | EN_3 |
| FANCONI_ANEMIA_PATH | FANCONI_ANEMIA_PATHWAY | 50 | -0.51039 | -1.89105 | 0 | 0.056889806 | 0.925 | 4794 | tags=46%, list=20%, si | EN_3 |
| GOBP_INNER_MITOCHON | GOBP_INNER_MITOCHONDRIAL_MEMBRAN | 36 | -0.54816 | -1.88738 | 0.0029282577 | 0.058812484 | 0.931 | 5823 | tags=42%, list=25%, si | EN_3 |
| GOBP_SPLICEOSOMAL_SI | GOBP_SPLICEOSOMAL_SNRNP_ASSEMBLY | 35 | -0.5517 | -1.88604 | 0 | 0.058849018 | 0.936 | 5280 | tags=49%, list=22%, si | EN_3 |
| GOBP_ACETYL_COA_MET | GOBP_ACETYL_COA_METABOLIC_PROCESS | 34 | -0.56019 | -1.88444 | 0 | 0.05872151 | 0.941 | 4572 | tags=44%, list=19%, si | EN_3 |
| GOBP_REGULATION_OF_ | GOBP_REGULATION_OF_CELLULAR_RESPIRA | 52 | -0.50257 | -1.88431 | 0.0013262599 | 0.057710618 | 0.941 | 3919 | tags=38%, list=17%, si | EN_3 |
| GOBP_NEGATIVE_REGUL | GOBP_NEGATIVE_REGULATION_OF_INTRAC | 54 | -0.49831 | -1.8837 | 0 | 0.057105716 | 0.944 | 7651 | tags=54%, list=32%, si | EN_3 |
| GOBP_AMMONIUM_ION_ | GOBP_AMMONIUM_ION_METABOLIC_PRO( | 24 | -0.59376 | -1.88125 | 0.0014684288 | 0.057744242 | 0.945 | 1725 | tags=29%, list=7%, sig | EN_3 |
| GOBP_EXTRINSIC_APOPT | GOBP_EXTRINSIC_APOPTOTIC_SIGNALING_ | 80 | -0.46985 | -1.87978 | 0 | 0.057776332 | 0.946 | 5656 | tags=44%, list=24%, si | EN_3 |
| GOBP_REGULATION_OF_ | GOBP_REGULATION_OF_SYNAPTIC_VESICLE | 19 | -0.62764 | -1.87941 | 0.0030674846 | 0.05700228 | 0.948 | 3680 | tags=53%, list=16%, si | EN_3 |
| GOBP_PROTEIN_QUALITY | GOBP_PROTEIN_QUALITY_CONTROL_FOR_N | 17 | -0.64808 | -1.87824 | 0 | 0.056919854 | 0.951 | 6996 | tags=65%, list=30%, si | EN_3 |
| GOBP_RIBOSOMAL_LARG | GOBP_RIBOSOMAL_LARGE_SUBUNIT_BIOGE | 64 | -0.4873 | -1.87779 | 0 | 0.05632806 | 0.951 | 5066 | tags=48%, list=21%, si | EN_3 |
| GOBP_NEGATIVE_REGUL | GOBP_NEGATIVE_REGULATION_OF_INTRAC | 35 | -0.5483 | -1.87473 | 0.004115226 | 0.05718123 | 0.953 | 4152 | tags=31%, list=18%, si | EN_3 |
| GOBP_INNER_CELL_MAS | GOBP_INNER_CELL_MASS_CELL_PROLIFERA | 16 | -0.66864 | -1.87357 | 0.0030674846 | 0.056861047 | 0.955 | 3517 | tags=50%, list=15%, si | EN_3 |
| GOBP_REGULATION_OF_ | GOBP_REGULATION_OF_FIBROBLAST_MIGR | 47 | -0.51088 | -1.87029 | 0.0026595744 | 0.058117047 | 0.959 | 5915 | tags=45%, list=25%, si | EN_3 |
| GOBP_MITOCHONDRIAL_ | GOBP_MITOCHONDRIAL_ELECTRON_TRANS | 18 | -0.63267 | -1.87019 | 0.00770416 | 0.057279453 | 0.959 | 4878 | tags=50%, list=21%, si | EN_3 |
| GOBP_FIBROBLAST_MIGF | GOBP_FIBROBLAST_MIGRATION | 68 | -0.47825 | -1.8671 | 0 | 0.058507092 | 0.962 | 6105 | tags=46%, list=26%, si | EN_3 |
| GOBP_POSITIVE_REGULA | GOBP_POSITIVE_REGULATION_OF_FIBROBL | 24 | -0.58706 | -1.86645 | 0 | 0.058234897 | 0.965 | 4381 | tags=46%, list=19%, si | EN_3 |
| GOBP_RIBOSOMAL_SMAI | GOBP_RIBOSOMAL_SMALL_SUBUNIT_ASSEI | 18 | -0.62677 | -1.86373 | 0.0029717681 | 0.059361175 | 0.97 | 5313 | tags=67%, list=23%, si | EN_3 |
| GOBP_REGULATION_OF_ | GOBP_REGULATION_OF_CYCLIN_DEPENDEN | 67 | -0.47546 | -1.85946 | 0.00124533 | 0.061512645 | 0.975 | 3722 | tags=34%, list=16%, si | EN_3 |
| GOBP_MATURATION_OF_ | GOBP_MATURATION_OF_5_8S_RRNA_FROM | 24 | -0.58638 | -1.85791 | 0.0015432099 | 0.06180674 | 0.976 | 2603 | tags=42%, list=11%, si | EN_3 |
| GOBP_RIBONUCLEOPROT | GOBP_RIBONUCLEOPROTEIN_COMPLEX_BI( | 417 | -0.39132 | -1.85404 | 0 | 0.06354276 | 0.982 | 5135 | tags=35%, list=22%, si | EN_3 |
| GOBP_DIOL_METABOLIC_ | GOBP_DIOL_METABOLIC_PROCESS | 29 | -0.54416 | -1.84871 | 0 | 0.06699815 | 0.986 | 4505 | tags=41%, list=19%, si | EN_3 |
| GOBP_DEOXYRIBONUCLE | GOBP_DEOXYRIBONUCLEOTIDE_BIOSYNTHE | 17 | -0.63201 | -1.84854 | 0.0065146578 | 0.06607072 | 0.986 | 3078 | tags=41%, list=13%, si | EN_3 |
| GOBP_VASCULAR_ENDOT | GOBP_VASCULAR_ENDOTHELIAL_GROWTH_ | 40 | -0.51752 | -1.8455 | 0.004166667 | 0.06735493 | 0.986 | 5612 | tags=48%, list=24%, si | EN_3 |
| GOBP_RESPONSE_TO_UV | GOBP_RESPONSE_TO_UV | 153 | -0.42144 | -1.84429 | 0 | 0.067475036 | 0.987 | 5403 | tags=35%, list=23%, si | EN_3 |
| GOBP_REGULATION_OF_ | GOBP_REGULATION_OF_BILE_ACID_METAB | 17 | -0.64043 | -1.84335 | 0.0047095763 | 0.06729724 | 0.987 | 931 | tags=29%, list=4%, sig | EN_3 |
| GOBP_RIBOSOME_ASSEM | GOBP_RIBOSOME_ASSEMBLY | 60 | -0.47603 | -1.84208 | 0 | 0.067207694 | 0.989 | 5751 | tags=55%, list=24%, si | EN_3 |
| GOBP_BASEMENT_MEME | GOBP_BASEMENT_MEMBRANE_ORGANIZAT | 40 | -0.52225 | -1.84061 | 0.004166667 | 0.06746228 | 0.989 | 6669 | tags=55%, list=28%, si | EN_3 |
| GOBP_REGULATION_OF_ | GOBP_REGULATION_OF_OXIDATIVE_PHOSP | 23 | -0.58628 | -1.83978 | 0 | 0.06719647 | 0.99 | 3442 | tags=43%, list=15%, si | EN_3 |
| GOBP_MAINTENANCE_OI | GOBP_MAINTENANCE_OF_PROTEIN_LOCAL | 43 | -0.50775 | -1.83846 | 0 | 0.067591876 | 0.991 | 6534 | tags=53%, list=28%, si | EN_3 |
| GOBP_REGULATION_OF_ | GOBP_REGULATION_OF_ERAD_PATHWAY | 27 | -0.56643 | -1.83532 | 0.0014245014 | 0.06893795 | 0.993 | 4330 | tags=41%, list=18%, si | EN_3 |
| GOBP_IMPORT_INTO_TH | GOBP_IMPORT_INTO_THE_MITOCHONDRIO | 49 | -0.49917 | -1.8345 | 0.004054054 | 0.06880612 | 0.993 | 4266 | tags=33%, list=18%, si | EN_3 |
| GOBP_CELLULAR_RESPON | GOBP_CELLULAR_RESPONSE_TO_VASCULAR | 60 | -0.47865 | -1.83264 | 0 | 0.06924457 | 0.994 | 5612 | tags=43%, list=24%, si | EN_3 |
| GOBP_REGULATION_OF_ | GOBP_REGULATION_OF_VACUOLE_ORGANI | 54 | -0.49332 | -1.82885 | 0.0013368984 | 0.071535036 | 0.997 | 2469 | tags=28%, list=10%, si | EN_3 |
| GOBP_RRNA_METABOLIC | GOBP_RRNA_METABOLIC_PROCESS | 241 | -0.40134 | -1.8288 | 0 | 0.07073831 | 0.997 | 5135 | tags=35%, list=22%, si | EN_3 |
| GOBP_PRIMARY_AMINO_ | GOBP_PRIMARY_AMINO_COMPOUND_MET | 17 | -0.61903 | -1.82785 | 0.00591716 | 0.07066726 | 0.997 | 1725 | tags=35%, list=7%, sig | EN_3 |
| GOBP_RETINOIC_ACID_R | GOBP_RETINOIC_ACID_RECEPTOR_SIGNALII | 34 | -0.53151 | -1.82655 | 0 | 0.07079248 | 0.997 | 4259 | tags=38%, list=18%, si | EN_3 |
| GOBP_REGULATION_OF_ | GOBP_REGULATION_OF_MONOCYTE_CHEM | 29 | -0.56306 | -1.82606 | 0.0042796005 | 0.07040643 | 0.997 | 2175 | tags=28%, list=9%, sig | EN_3 |
| GOBP_POSITIVE_REGULA | GOBP_POSITIVE_REGULATION_OF_INFLAMI | 22 | -0.57623 | -1.82306 | 0.0045731706 | 0.0722404 | 0.998 | 3138 | tags=32%, list=13%, si | EN_3 |
| GOBP_MITOCHONDRIAL_ | GOBP_MITOCHONDRIAL_TRANSPORT | 174 | -0.41654 | -1.82191 | 0 | 0.072259754 | 0.998 | 5136 | tags=36%, list=22%, si | EN_3 |
| GOBP_NEGATIVE_REGUL | GOBP_NEGATIVE_REGULATION_OF_VIRAL_ | 18 | -0.62331 | -1.81929 | 0.00147929 | 0.07364706 | 0.998 | 4955 | tags=61%, list=21%, si | EN_3 |
| GOBP_ESTABLISHMENT_( | GOBP_ESTABLISHMENT_OF_MITOCHONDRI( | 34 | -0.5307 | -1.81684 | 0.004491018 | 0.07503348 | 0.998 | 6391 | tags=53%, list=27%, si | EN_3 |
| GOBP_CELLULAR_RESPON | GOBP_CELLULAR_RESPONSE_TO_DSRNA | 19 | -0.61361 | -1.81239 | 0.001545595 | 0.07831194 | 0.999 | 5586 | tags=58%, list=24%, si | EN_3 |
| GOBP_RRNA_PROCESSIN( | GOBP_RRNA_PROCESSING | 211 | -0.40475 | -1.80591 | 0 | 0.083130255 | 0.999 | 5135 | tags=37%, list=22%, si | EN_3 |
| GOBP_INDOLE_CONTAIN | GOBP_INDOLE_CONTAINING_COMPOUND_ | 25 | -0.57248 | -1.80555 | 0.0060331826 | 0.08256057 | 0.999 | 5077 | tags=52%, list=22%, si | EN_3 |
| GOBP_POSITIVE_REGULA | GOBP_POSITIVE_REGULATION_OF_MONOC | 21 | -0.58808 | -1.8051 | 0.004538578 | 0.08197138 | 0.999 | 2175 | tags=29%, list=9%, sig | EN_3 |
| PROTEASOME | PROTEASOME | 45 | -0.49307 | -1.80203 | 0.005457026 | 0.08390973 | 0.999 | 7430 | tags=64%, list=31%, si | EN_3 |
| SMALL_CELL_LUNG_CAN( | SMALL_CELL_LUNG_CANCER | 91 | -0.443 | -1.80161 | 0 | 0.083394505 | 0.999 | 4154 | tags=31%, list=18%, si | EN_3 |
| GOBP_NEGATIVE_REGUL | GOBP_NEGATIVE_REGULATION_OF_RESPON | 40 | -0.50825 | -1.80068 | 0.0041436465 | 0.08354159 | 0.999 | 5090 | tags=33%, list=22%, si | EN_3 |
| GOBP_RESPONSE_TO_TY | GOBP_RESPONSE_TO_TYPE_II_INTERFERON | 122 | -0.42433 | -1.79924 | 0 | 0.084088735 | 0.999 | 4831 | tags=34%, list=20%, si | EN_3 |
| GOBP_CGAS_STING_SIGN | GOBP_CGAS_STING_SIGNALING_PATHWAY | 16 | -0.62501 | -1.79777 | 0.007974481 | 0.08448895 | 0.999 | 2573 | tags=38%, list=11%, si | EN_3 |
| GOBP_RIBOSE_PHOSPHA | GOBP_RIBOSE_PHOSPHATE_BIOSYNTHETIC_ | 185 | -0.40536 | -1.7976 | 0 | 0.0838848 | 0.999 | 5338 | tags=37%, list=23%, si | EN_3 |
| GOBP_GENERATION_OF_ | GOBP_GENERATION_OF_PRECURSOR_META | 439 | -0.3784 | -1.79605 | 0 | 0.08440186 | 0.999 | 5905 | tags=38%, list=25%, si | EN_3 |
| GOBP_TERPENOID_META | GOBP_TERPENOID_METABOLIC_PROCESS | 60 | -0.46046 | -1.79524 | 0 | 0.084187835 | 1 | 4205 | tags=37%, list=18%, si | EN_3 |
| GOBP_ENERGY_DERIVATI | GOBP_ENERGY_DERIVATION_BY_OXIDATIOI | 321 | -0.38283 | -1.79461 | 0 | 0.0839757 | 1 | 6337 | tags=42%, list=27%, si | EN_3 |
| GOBP_REGULATION_OF_ | GOBP_REGULATION_OF_UBIQUITIN_PROTE | 39 | -0.51363 | -1.79449 | 0.0014064697 | 0.08324024 | 1 | 5105 | tags=46%, list=22%, si | EN_3 |
| GOBP_POSITIVE_REGULA | GOBP_POSITIVE_REGULATION_OF_MYOBLA | 18 | -0.61295 | -1.79217 | 0.0091883615 | 0.084812805 | 1 | 3515 | tags=44%, list=15%, si | EN_3 |
| GOBP_PHOSPHATIDYLCH( | GOBP_PHOSPHATIDYLCHOLINE_BIOSYNTHE | 21 | -0.56656 | -1.79073 | 0.00927357 | 0.08527869 | 1 | 1598 | tags=29%, list=7%, sig | EN_3 |
| GOBP_HEME_METABOLI( | GOBP_HEME_METABOLIC_PROCESS | 34 | -0.52129 | -1.7895 | 0.005641749 | 0.0856176 | 1 | 4070 | tags=44%, list=17%, si | EN_3 |
| FERROPTOSIS | FERROPTOSIS | 37 | -0.50848 | -1.7863 | 0.004231312 | 0.08787956 | 1 | 5502 | tags=54%, list=23%, si | EN_3 |
| GOBP_POSITIVE_REGULA | GOBP_POSITIVE_REGULATION_OF_RESPONS | 37 | -0.50308 | -1.78206 | 0.005738881 | 0.09073838 | 1 | 6940 | tags=54%, list=29%, si | EN_3 |
| GOBP_DEFENSE_RESPON | GOBP_DEFENSE_RESPONSE_TO_VIRUS | 276 | -0.38232 | -1.77905 | 0 | 0.09277026 | 1 | 4325 | tags=28%, list=18%, si | EN_3 |
| GOBP_POSITIVE_REGULA | GOBP_POSITIVE_REGULATION_OF_TYPE_I_I | 19 | -0.5997 | -1.77524 | 0.0062015504 | 0.095843524 | 1 | 1015 | tags=21%, list=4%, sig | EN_3 |
| GOBP_PROTEIN_RNA_CO | GOBP_PROTEIN_RNA_COMPLEX_ORGANIZA | 173 | -0.39913 | -1.77437 | 0 | 0.09583189 | 1 | 5342 | tags=35%, list=23%, si | EN_3 |
| GOBP_PHOSPHATIDYLCH( | GOBP_PHOSPHATIDYLCHOLINE_METABOLIC | 59 | -0.45884 | -1.7705 | 0 | 0.098908864 | 1 | 1598 | tags=20%, list=7%, sig | EN_3 |

| | | | | | | | | | | |
|---|---|---|---|---|---|---|---|---|---|---|
| GOBP_PYRIMIDINE_NUCL | GOBP_PYRIMIDINE_NUCLEOTIDE_METABOL | 38 | -0.50901 | -1.76839 | 0.004178273 | 0.100244954 | 1 | 1811 | tags=24%, list=8%, sig | EN_3 |
| GOBP_INTRINSIC_APOPT | GOBP_INTRINSIC_APOPTOTIC_SIGNALING_F | 59 | -0.46092 | -1.76673 | 0.003968254 | 0.10088702 | 1 | 7654 | tags=58%, list=32%, si | EN_3 |
| GALACTOSE_METABOLISM | GALACTOSE_METABOLISM | 29 | -0.53028 | -1.76559 | 0.010159652 | 0.10114496 | 1 | 2202 | tags=28%, list=9%, sig | EN_3 |
| GOBP_SERINE_FAMILY_A | GOBP_SERINE_FAMILY_AMINO_ACID_META | 30 | -0.53682 | -1.7651 | 0.0014144272 | 0.10069829 | 1 | 1215 | tags=23%, list=5%, sig | EN_3 |
| GOBP_CELLULAR_RESPON | GOBP_CELLULAR_RESPONSE_TO_TYPE_II_IN | 95 | -0.42834 | -1.76457 | 0.0023980816 | 0.10040776 | 1 | 5003 | tags=36%, list=21%, si | EN_3 |
| GOBP_NEGATIVE_REGUL | GOBP_NEGATIVE_REGULATION_OF_VIRAL_ | 95 | -0.4337 | -1.7612 | 0.0012254902 | 0.1031887 | 1 | 4955 | tags=36%, list=21%, si | EN_3 |
| GOBP_PEPTIDYL_PROLIN | GOBP_PEPTIDYL_PROLINE_MODIFICATION | 39 | -0.50601 | -1.76055 | 0.0027855153 | 0.1030724 | 1 | 5327 | tags=46%, list=23%, si | EN_3 |
| GOBP_LATE_ENDOSOME_ | GOBP_LATE_ENDOSOME_TO_LYSOSOME_TI | 19 | -0.58971 | -1.75807 | 0.0078125 | 0.10476514 | 1 | 3843 | tags=53%, list=16%, si | EN_3 |
| GOBP_POSITIVE_REGULA | GOBP_POSITIVE_REGULATION_OF_SYSTEMI | 23 | -0.55558 | -1.7562 | 0.015527951 | 0.10604865 | 1 | 2307 | tags=30%, list=10%, si | EN_3 |
| GOBP_PYRIMIDINE_DEOX | GOBP_PYRIMIDINE_DEOXYRIBONUCLEOTIDE | 20 | -0.58559 | -1.75306 | 0.0015384615 | 0.10894344 | 1 | 2950 | tags=30%, list=12%, si | EN_3 |
| GOBP_HYDROGEN_PERO | GOBP_HYDROGEN_PEROXIDE_CATABOLIC_F | 22 | -0.56689 | -1.75217 | 0.010401188 | 0.10910952 | 1 | 4810 | tags=50%, list=20%, si | EN_3 |
| GOBP_NUCLEOSIDE_PHO | GOBP_NUCLEOSIDE_PHOSPHATE_BIOSYNTH | 234 | -0.38648 | -1.75181 | 0 | 0.108570166 | 1 | 5338 | tags=36%, list=23%, si | EN_3 |
| GOBP_NEGATIVE_REGUL | GOBP_NEGATIVE_REGULATION_OF_ENDOP | 18 | -0.59238 | -1.74983 | 0.0047095763 | 0.10985538 | 1 | 4975 | tags=39%, list=21%, si | EN_3 |
| GOBP_TROPHOBLAST_CE | GOBP_TROPHOBLAST_CELL_MIGRATION | 17 | -0.59814 | -1.74983 | 0.006042296 | 0.10898352 | 1 | 1515 | tags=24%, list=6%, sig | EN_3 |
| GOBP_POSITIVE_REGULA | GOBP_POSITIVE_REGULATION_OF_SUBSTRA | 40 | -0.49499 | -1.74844 | 0 | 0.1098001 | 1 | 5567 | tags=43%, list=24%, si | EN_3 |
| GOBP_ISOPRENOID_MET | GOBP_ISOPRENOID_METABOLIC_PROCESS | 87 | -0.42919 | -1.74805 | 0.0036674817 | 0.10932815 | 1 | 4205 | tags=34%, list=18%, si | EN_3 |
| GOBP_RESPONSE_TO_MI | GOBP_RESPONSE_TO_MITOCHONDRIAL_DE | 21 | -0.5715 | -1.74781 | 0.0059435363 | 0.10879105 | 1 | 2100 | tags=24%, list=9%, sig | EN_3 |
| GOBP_REGULATION_OF_ | GOBP_REGULATION_OF_AMYLOID_FIBRIL_F | 16 | -0.60654 | -1.74777 | 0.007849294 | 0.10801775 | 1 | 4764 | tags=44%, list=20%, si | EN_3 |
| GOBP_NEGATIVE_REGUL | GOBP_NEGATIVE_REGULATION_OF_SMOOT | 36 | -0.50074 | -1.74574 | 0.006954103 | 0.10942248 | 1 | 6815 | tags=53%, list=29%, si | EN_3 |
| HOMOLOGOUS_RECOMB | HOMOLOGOUS_RECOMBINATION | 40 | -0.48418 | -1.74288 | 0.004103967 | 0.11163033 | 1 | 4011 | tags=35%, list=17%, si | EN_3 |
| GOBP_NECROPTOTIC_PR | GOBP_NECROPTOTIC_PROCESS | 37 | -0.50106 | -1.74189 | 0.008097166 | 0.11194901 | 1 | 5727 | tags=43%, list=24%, si | EN_3 |
| GOBP_LYSOSOMAL_LUM | GOBP_LYSOSOMAL_LUMEN_ACIDIFICATION | 15 | -0.61276 | -1.74188 | 0.018489985 | 0.11111356 | 1 | 3460 | tags=47%, list=15%, si | EN_3 |
| GOBP_MONOCYTE_CHEM | GOBP_MONOCYTE_CHEMOTAXIS | 40 | -0.49048 | -1.74134 | 0.0026631157 | 0.11092249 | 1 | 2175 | tags=23%, list=9%, sig | EN_3 |
| GOBP_CELLULAR_RESPON | GOBP_CELLULAR_RESPONSE_TO_UV | 90 | -0.42563 | -1.74069 | 0 | 0.11083288 | 1 | 6141 | tags=38%, list=26%, si | EN_3 |
| GOBP_RNA_SURVEILLAN | GOBP_RNA_SURVEILLANCE | 17 | -0.58903 | -1.73868 | 0.0061633284 | 0.11217581 | 1 | 3724 | tags=47%, list=16%, si | EN_3 |
| GOBP_PROTEIN_STABILIZ | GOBP_PROTEIN_STABILIZATION | 190 | -0.38964 | -1.73509 | 0 | 0.11543588 | 1 | 6329 | tags=37%, list=27%, si | EN_3 |
| BASE_EXCISION_REPAIR | BASE_EXCISION_REPAIR | 32 | -0.5062 | -1.73443 | 0.0029112082 | 0.115263134 | 1 | 3335 | tags=41%, list=14%, si | EN_3 |
| GOBP_G0_TO_G1_TRANS | GOBP_G0_TO_G1_TRANSITION | 38 | -0.4962 | -1.73397 | 0.007002801 | 0.11488046 | 1 | 4698 | tags=37%, list=20%, si | EN_3 |
| GOBP_ISOPRENOID_BIOS | GOBP_ISOPRENOID_BIOSYNTHETIC_PROCES | 36 | -0.49568 | -1.73384 | 0.0057553956 | 0.114189506 | 1 | 4091 | tags=42%, list=17%, si | EN_3 |
| GOBP_NEGATIVE_REGUL | GOBP_NEGATIVE_REGULATION_OF_HEART_ | 15 | -0.60589 | -1.7327 | 0.011254019 | 0.11486005 | 1 | 2307 | tags=33%, list=10%, si | EN_3 |
| GOBP_ASPARTATE_FAMI | GOBP_ASPARTATE_FAMILY_AMINO_ACID_N | 36 | -0.50899 | -1.73175 | 0.0042253523 | 0.11521367 | 1 | 1704 | tags=25%, list=7%, sig | EN_3 |
| GOBP_REGULATION_OF_ | GOBP_REGULATION_OF_ENDOPLASMIC_RE | 30 | -0.51839 | -1.73074 | 0.008720931 | 0.115569934 | 1 | 6745 | tags=47%, list=29%, si | EN_3 |
| GOBP_REGULATION_OF_ | GOBP_REGULATION_OF_PROTEIN_KINASE_ | 22 | -0.56584 | -1.73041 | 0.011494253 | 0.11513456 | 1 | 4635 | tags=45%, list=20%, si | EN_3 |
| GOBP_POSITIVE_REGULA | GOBP_POSITIVE_REGULATION_OF_PATTERN | 68 | -0.44377 | -1.729 | 0.0012531328 | 0.11598665 | 1 | 4251 | tags=29%, list=18%, si | EN_3 |
| SPLICEOSOME | SPLICEOSOME | 127 | -0.4005 | -1.72542 | 0 | 0.11905843 | 1 | 6874 | tags=43%, list=29%, si | EN_3 |
| GOBP_INTERFERON_MED | GOBP_INTERFERON_MEDIATED_SIGNALING | 76 | -0.42989 | -1.72379 | 0 | 0.12054574 | 1 | 4325 | tags=32%, list=18%, si | EN_3 |
| GOBP_REGULATION_OF_ | GOBP_REGULATION_OF_MEMBRANE_LIPID | 19 | -0.575 | -1.72337 | 0.011976048 | 0.12008849 | 1 | 4995 | tags=47%, list=21%, si | EN_3 |
| GOBP_REGULATION_OF_ | GOBP_REGULATION_OF_PROTEOLYSIS_INV | 215 | -0.38171 | -1.72327 | 0 | 0.11941116 | 1 | 4859 | tags=27%, list=21%, si | EN_3 |
| GOBP_REGULATION_OF_ | GOBP_REGULATION_OF_CHOLESTEROL_ME | 41 | -0.47753 | -1.72255 | 0.004037685 | 0.11946458 | 1 | 3706 | tags=32%, list=16%, si | EN_3 |
| GOBP_PROTEIN_TRANSM | GOBP_PROTEIN_TRANSMEMBRANE_IMPOR | 36 | -0.5004 | -1.72126 | 0.008275862 | 0.12026808 | 1 | 3442 | tags=28%, list=15%, si | EN_3 |
| GOBP_MITOCHONDRIAL_ | GOBP_MITOCHONDRIAL_TRANSMEMBRANE | 92 | -0.41946 | -1.71859 | 0.0036407767 | 0.122746944 | 1 | 4817 | tags=36%, list=20%, si | EN_3 |
| GOBP_MYELOID_LEUKOC | GOBP_MYELOID_LEUKOCYTE_MEDIATED_IN | 108 | -0.40646 | -1.71335 | 0 | 0.12808667 | 1 | 2451 | tags=21%, list=10%, si | EN_3 |
| GOBP_REGULATION_OF_ | GOBP_REGULATION_OF_MYOBLAST_FUSIO | 22 | -0.55945 | -1.71079 | 0.004451039 | 0.13047458 | 1 | 3515 | tags=36%, list=15%, si | EN_3 |
| GOBP_NEGATIVE_REGUL | GOBP_NEGATIVE_REGULATION_OF_ESTABL | 148 | -0.39175 | -1.70878 | 0 | 0.1319749 | 1 | 4997 | tags=31%, list=21%, si | EN_3 |
| GOBP_COPPER_ION_HON | GOBP_COPPER_ION_HOMEOSTASIS | 18 | -0.57019 | -1.70746 | 0.018489985 | 0.13260703 | 1 | 4908 | tags=50%, list=21%, si | EN_3 |
| GOBP_MATURATION_OF_ | GOBP_MATURATION_OF_LSU_RRNA | 25 | -0.53175 | -1.70732 | 0.013293944 | 0.13193668 | 1 | 2494 | tags=36%, list=11%, si | EN_3 |
| GOBP_DNA_REPLICATION | GOBP_DNA_REPLICATION_INITIATION | 31 | -0.50962 | -1.70647 | 0.007062147 | 0.13202544 | 1 | 6595 | tags=52%, list=28%, si | EN_3 |
| CELL_CYCLE | CELL_CYCLE | 121 | -0.39906 | -1.70572 | 0 | 0.13206792 | 1 | 4859 | tags=34%, list=21%, si | EN_3 |
| GOBP_LIPOPROTEIN_CAT | GOBP_LIPOPROTEIN_CATABOLIC_PROCESS | 17 | -0.59192 | -1.7046 | 0.01076923 | 0.13245453 | 1 | 4457 | tags=41%, list=19%, si | EN_3 |
| GOBP_REGULATION_OF_ | GOBP_REGULATION_OF_GLIAL_CELL_MIGR | 26 | -0.5284 | -1.70388 | 0.0071123755 | 0.13255784 | 1 | 5800 | tags=50%, list=25%, si | EN_3 |
| GOBP_NEGATIVE_REGUL | GOBP_NEGATIVE_REGULATION_OF_CELLUL | 106 | -0.40599 | -1.70233 | 0.0012048193 | 0.13377477 | 1 | 2441 | tags=22%, list=10%, si | EN_3 |
| GOBP_RIG_I_SIGNALING_ | GOBP_RIG_I_SIGNALING_PATHWAY | 24 | -0.54087 | -1.70148 | 0.009202454 | 0.13410538 | 1 | 468 | tags=17%, list=2%, sig | EN_3 |
| GOBP_PROTEIN_FOLDING | GOBP_PROTEIN_FOLDING | 162 | -0.38774 | -1.70121 | 0 | 0.13369787 | 1 | 5801 | tags=33%, list=25%, si | EN_3 |
| GOBP_CELLULAR_RESPON | GOBP_CELLULAR_RESPONSE_TO_ALKALOID | 30 | -0.51186 | -1.69966 | 0.014306151 | 0.13485454 | 1 | 6275 | tags=50%, list=27%, si | EN_3 |
| GOBP_AUTOPHAGOSOMI | GOBP_AUTOPHAGOSOME_MATURATION | 58 | -0.44532 | -1.69956 | 0.0026560426 | 0.13415812 | 1 | 4910 | tags=41%, list=21%, si | EN_3 |
| GOBP_VACUOLAR_LOCAL | GOBP_VACUOLAR_LOCALIZATION | 85 | -0.41937 | -1.69825 | 0.0012360939 | 0.13521406 | 1 | 2451 | tags=21%, list=10%, si | EN_3 |
| GOBP_PINOCYTOSIS | GOBP_PINOCYTOSIS | 22 | -0.55274 | -1.69695 | 0.011093502 | 0.13593976 | 1 | 2967 | tags=36%, list=13%, si | EN_3 |
| GOBP_MACROAUTOPHAG | GOBP_MACROAUTOPHAGY | 287 | -0.36689 | -1.69592 | 0 | 0.13638598 | 1 | 4506 | tags=26%, list=19%, si | EN_3 |
| GOBP_POSITIVE_REGULA | GOBP_POSITIVE_REGULATION_OF_PEPTIDY | 28 | -0.5177 | -1.69503 | 0.008683068 | 0.13679664 | 1 | 2724 | tags=25%, list=12%, si | EN_3 |
| N-GLYCAN_BIOSYNTHESIS | N-GLYCAN_BIOSYNTHESIS | 48 | -0.45879 | -1.69368 | 0.008163265 | 0.13770989 | 1 | 4195 | tags=29%, list=18%, si | EN_3 |
| GOBP_VIRAL_TRANSCRIP | GOBP_VIRAL_TRANSCRIPTION | 55 | -0.44378 | -1.69347 | 0.009032258 | 0.13722274 | 1 | 4955 | tags=36%, list=21%, si | EN_3 |
| GOBP_PYRIMIDINE_CONT | GOBP_PYRIMIDINE_CONTAINING_COMPOU | 60 | -0.44502 | -1.6931 | 0.0025575447 | 0.13695985 | 1 | 2950 | tags=25%, list=12%, si | EN_3 |
| GOBP_PROTEIN_K11_LIN | GOBP_PROTEIN_K11_LINKED_UBIQUITINAT | 29 | -0.51031 | -1.69226 | 0.02124646 | 0.1372224 | 1 | 6674 | tags=55%, list=28%, si | EN_3 |
| GOBP_MEMBRANE_RAFT | GOBP_MEMBRANE_RAFT_ORGANIZATION | 24 | -0.5339 | -1.68997 | 0.012802276 | 0.13945523 | 1 | 3588 | tags=42%, list=15%, si | EN_3 |
| GOBP_PROTEIN_PEPTIDY | GOBP_PROTEIN_PEPTIDYL_PROLYL_ISOMER | 25 | -0.53165 | -1.68983 | 0.013719512 | 0.13887657 | 1 | 2259 | tags=32%, list=10%, si | EN_3 |
| GOBP_VESICLE_COATING | GOBP_VESICLE_COATING | 33 | -0.50274 | -1.68897 | 0.008547009 | 0.13922377 | 1 | 4204 | tags=39%, list=18%, si | EN_3 |
| GOBP_RESPONSE_TO_MI | GOBP_RESPONSE_TO_MISFOLDED_PROTEIN | 18 | -0.58294 | -1.68618 | 0.01925926 | 0.14216092 | 1 | 3075 | tags=28%, list=13%, si | EN_3 |
| GOBP_MAINTENANCE_OI | GOBP_MAINTENANCE_OF_PROTEIN_LOCAT | 70 | -0.43257 | -1.68212 | 0.0037926675 | 0.14681768 | 1 | 6099 | tags=47%, list=26%, si | EN_3 |
| GOBP_TETRAPYRROLE_M | GOBP_TETRAPYRROLE_METABOLIC_PROCES | 49 | -0.45374 | -1.68096 | 0.01058201 | 0.14750022 | 1 | 4070 | tags=37%, list=17%, si | EN_3 |
| GOBP_VESICLE_TARGETII | GOBP_VESICLE_TARGETING_ROUGH_ER_TO | 20 | -0.55054 | -1.68017 | 0.010324484 | 0.14784174 | 1 | 4204 | tags=50%, list=18%, si | EN_3 |
| GOBP_NEGATIVE_REGUL | GOBP_NEGATIVE_REGULATION_OF_UBIQUI | 18 | -0.56325 | -1.68014 | 0.016344724 | 0.14708433 | 1 | 4764 | tags=56%, list=20%, si | EN_3 |
| GOBP_REGULATION_OF_ | GOBP_REGULATION_OF_AUTOPHAGOSOME | 44 | -0.46452 | -1.68011 | 0.009395973 | 0.14631271 | 1 | 2442 | tags=27%, list=10%, si | EN_3 |
| GOBP_AMYLOID_PRECUR | GOBP_AMYLOID_PRECURSOR_PROTEIN_CAT | 59 | -0.43841 | -1.67939 | 0.0077720205 | 0.14645578 | 1 | 6287 | tags=49%, list=27%, si | EN_3 |
| GOBP_ACTIVATION_OF_( | GOBP_ACTIVATION_OF_CYSTEINE_TYPE_EN | 81 | -0.41295 | -1.67892 | 0.0037974683 | 0.14627607 | 1 | 5267 | tags=33%, list=22%, si | EN_3 |
| GOBP_PROTEIN_K63_LIN | GOBP_PROTEIN_K63_LINKED_UBIQUITINAT | 68 | -0.4245 | -1.67747 | 0.0064683054 | 0.14754313 | 1 | 2647 | tags=22%, list=11%, si | EN_3 |
| GOBP_VIRAL_GENE_EXPR | GOBP_VIRAL_GENE_EXPRESSION | 75 | -0.42428 | -1.67702 | 0.005076142 | 0.14732754 | 1 | 6527 | tags=45%, list=28%, si | EN_3 |
| GOBP_PROTEIN_HYDROX | GOBP_PROTEIN_HYDROXYLATION | 25 | -0.52506 | -1.6765 | 0.009146341 | 0.14710343 | 1 | 5327 | tags=44%, list=23%, si | EN_3 |
| GOBP_STEROL_METABOL | GOBP_STEROL_METABOLIC_PROCESS | 140 | -0.39346 | -1.67555 | 0 | 0.1474436 | 1 | 4868 | tags=36%, list=21%, si | EN_3 |
| GOBP_NEGATIVE_REGUL | GOBP_NEGATIVE_REGULATION_OF_MACRO | 33 | -0.49028 | -1.67519 | 0.012622721 | 0.14710167 | 1 | 5245 | tags=39%, list=22%, si | EN_3 |
| GOBP_PEPTIDYL_LYSINE_ | GOBP_PEPTIDYL_LYSINE_MODIFICATION | 128 | -0.3887 | -1.67232 | 0 | 0.15009513 | 1 | 5818 | tags=33%, list=25%, si | EN_3 |
| GOBP_NEGATIVE_REGUL | GOBP_NEGATIVE_REGULATION_OF_SMOOT | 68 | -0.41923 | -1.67216 | 0.0062421975 | 0.14953569 | 1 | 4036 | tags=31%, list=17%, si | EN_3 |
| GOBP_POSITIVE_REGULA | GOBP_POSITIVE_REGULATION_OF_TELOME | 71 | -0.41685 | -1.66893 | 0.0026075619 | 0.15315776 | 1 | 7334 | tags=54%, list=31%, si | EN_3 |

| | | | | | | | | | | |
|---|---|---|---|---|---|---|---|---|---|---|
| GOBP_NUCLEOLUS_ORGA | GOBP_NUCLEOLUS_ORGANIZATION | 17 | -0.57399 | -1.66823 | 0.024242423 | 0.15329392 | 1 | 7308 | tags=59%, list=31%, si | EN_3 |
| GOBP_MITOCHONDRION | GOBP_MITOCHONDRION_LOCALIZATION | 57 | -0.43346 | -1.66566 | 0.005291005 | 0.15600371 | 1 | 6391 | tags=42%, list=27%, si | EN_3 |
| GOBP_RESPONSE_TO_VII | GOBP_RESPONSE_TO_VIRUS | 331 | -0.35589 | -1.66351 | 0 | 0.15811147 | 1 | 4325 | tags=26%, list=18%, si | EN_3 |
| GOBP_REGULATION_OF_ | GOBP_REGULATION_OF_CELL_CYCLE_G2_M | 117 | -0.38462 | -1.66119 | 0.0023228803 | 0.16040711 | 1 | 6081 | tags=41%, list=26%, si | EN_3 |
| GOBP_REGULATION_OF_ | GOBP_REGULATION_OF_EXTRINSIC_APOPT( | 45 | -0.45265 | -1.66118 | 0.013642564 | 0.15960625 | 1 | 4282 | tags=36%, list=18%, si | EN_3 |
| GOBP_DIOL_BIOSYNTHET | GOBP_DIOL_BIOSYNTHETIC_PROCESS | 21 | -0.55347 | -1.66042 | 0.022288261 | 0.15989798 | 1 | 4505 | tags=43%, list=19%, si | EN_3 |
| GOBP_DENTATE_GYRUS_ | GOBP_DENTATE_GYRUS_DEVELOPMENT | 26 | -0.50862 | -1.66009 | 0.010494753 | 0.1595674 | 1 | 4776 | tags=42%, list=20%, si | EN_3 |
| GOBP_PEPTIDYL_CYSTEIN | GOBP_PEPTIDYL_CYSTEINE_MODIFICATION | 44 | -0.45579 | -1.65937 | 0.014245015 | 0.1598404 | 1 | 4893 | tags=34%, list=21%, si | EN_3 |
| GOBP_LEUKOCYTE_ACTIV | GOBP_LEUKOCYTE_ACTIVATION_INVOLVED | 53 | -0.44803 | -1.65904 | 0.013605442 | 0.15951438 | 1 | 3138 | tags=28%, list=13%, si | EN_3 |
| GOBP_PORPHYRIN_CONT | GOBP_PORPHYRIN_CONTAINING_COMPOU | 40 | -0.46733 | -1.6581 | 0.008450705 | 0.15996543 | 1 | 4070 | tags=40%, list=17%, si | EN_3 |
| GOBP_POSITIVE_REGULA | GOBP_POSITIVE_REGULATION_OF_PROTEO | 120 | -0.39033 | -1.65771 | 0.0023809525 | 0.15965652 | 1 | 4859 | tags=28%, list=21%, si | EN_3 |
| GOBP_NUCLEOSIDE_TRIP | GOBP_NUCLEOSIDE_TRIPHOSPHATE_METAE | 250 | -0.36057 | -1.65736 | 0 | 0.15942052 | 1 | 5997 | tags=38%, list=25%, si | EN_3 |
| PURINE_METABOLISM | PURINE_METABOLISM | 126 | -0.38867 | -1.6569 | 0.002361275 | 0.15943012 | 1 | 4423 | tags=30%, list=19%, si | EN_3 |
| GOBP_DENDRITIC_CELL_( | GOBP_DENDRITIC_CELL_CYTOKINE_PRODU( | 16 | -0.5701 | -1.65668 | 0.026143791 | 0.15898426 | 1 | 1127 | tags=25%, list=5%, sig | EN_3 |
| GOBP_PROTEIN_LOCALIZ | GOBP_PROTEIN_LOCALIZATION_TO_ENDOP | 76 | -0.41439 | -1.65631 | 0.002503129 | 0.15872402 | 1 | 4674 | tags=36%, list=20%, si | EN_3 |
| GOBP_PURINE_RIBONUC | GOBP_PURINE_RIBONUCLEOSIDE_METABOL | 24 | -0.5193 | -1.65465 | 0.02672956 | 0.16012594 | 1 | 5256 | tags=46%, list=22%, si | EN_3 |
| GOBP_PIGMENT_METAB( | GOBP_PIGMENT_METABOLIC_PROCESS | 65 | -0.42564 | -1.65323 | 0.005148005 | 0.1614658 | 1 | 3661 | tags=31%, list=16%, si | EN_3 |
| GOBP_ORGANELLE_TRAN | GOBP_ORGANELLE_TRANSPORT_ALONG_M | 86 | -0.40924 | -1.65222 | 0.0048250905 | 0.16211529 | 1 | 7109 | tags=49%, list=30%, si | EN_3 |
| GOBP_PROTEIN_IMPORT | GOBP_PROTEIN_IMPORT_INTO_MITOCHON | 19 | -0.54907 | -1.65179 | 0.0149700595 | 0.16195698 | 1 | 5714 | tags=37%, list=24%, si | EN_3 |
| GOBP_INTRINSIC_APOPT( | GOBP_INTRINSIC_APOPTOTIC_SIGNALING_F | 63 | -0.42965 | -1.65132 | 0.007874016 | 0.16192487 | 1 | 4315 | tags=27%, list=18%, si | EN_3 |
| GOBP_CYTOCHROME_CO | GOBP_CYTOCHROME_COMPLEX_ASSEMBLY | 32 | -0.48694 | -1.65066 | 0.016081871 | 0.1620228 | 1 | 3361 | tags=38%, list=14%, si | EN_3 |
| GOBP_BARBED_END_ACT | GOBP_BARBED_END_ACTIN_FILAMENT_CAF | 25 | -0.51197 | -1.65056 | 0.018544937 | 0.1614108 | 1 | 5941 | tags=52%, list=25%, si | EN_3 |
| HUMAN_CYTOMEGALOVI | HUMAN_CYTOMEGALOVIRUS_INFECTION | 216 | -0.36313 | -1.64974 | 0 | 0.16190779 | 1 | 6464 | tags=38%, list=27%, si | EN_3 |
| PORPHYRIN_AND_CHLOR | PORPHYRIN_AND_CHLOROPHYLL_METABOL | 31 | -0.48864 | -1.64967 | 0.012311902 | 0.16127792 | 1 | 4187 | tags=42%, list=18%, si | EN_3 |
| GOBP_ESTABLISHMENT_( | GOBP_ESTABLISHMENT_OF_PROTEIN_LOCA | 25 | -0.51019 | -1.64827 | 0.026946107 | 0.16256775 | 1 | 6697 | tags=52%, list=28%, si | EN_3 |
| GOBP_POSITIVE_REGULA | GOBP_POSITIVE_REGULATION_OF_PROTEO | 325 | -0.35535 | -1.64708 | 0 | 0.16342627 | 1 | 4859 | tags=27%, list=21%, si | EN_3 |
| GOBP_POSITIVE_REGULA | GOBP_POSITIVE_REGULATION_OF_AMYLOII | 26 | -0.5047 | -1.6456 | 0.021097047 | 0.16479684 | 1 | 6287 | tags=50%, list=27%, si | EN_3 |
| GOBP_POSITIVE_REGULA | GOBP_POSITIVE_REGULATION_OF_AMIDE_I | 33 | -0.48886 | -1.64544 | 0.010174419 | 0.16424881 | 1 | 6287 | tags=45%, list=27%, si | EN_3 |
| GOBP_REGULATION_OF_ | GOBP_REGULATION_OF_MACROAUTOPHAG | 129 | -0.38098 | -1.64499 | 0 | 0.16414319 | 1 | 6376 | tags=38%, list=27%, si | EN_3 |
| GOBP_TELOMERE_MAINT | GOBP_TELOMERE_MAINTENANCE_VIA_REC | 15 | -0.57994 | -1.64483 | 0.025356578 | 0.16363543 | 1 | 4011 | tags=40%, list=17%, si | EN_3 |
| GOBP_COPII_COATED_VE | GOBP_COPII_COATED_VESICLE_BUDDING | 37 | -0.47116 | -1.64347 | 0.018361581 | 0.16486727 | 1 | 4204 | tags=38%, list=18%, si | EN_3 |
| GOBP_REGULATION_OF_ | GOBP_REGULATION_OF_AEROBIC_RESPIRAT | 33 | -0.47764 | -1.6428 | 0.021156559 | 0.1651515 | 1 | 3919 | tags=42%, list=17%, si | EN_3 |
| GOBP_ENDOPLASMIC_RE | GOBP_ENDOPLASMIC_RETICULUM_TO_GOL | 120 | -0.38664 | -1.64206 | 0.0035545023 | 0.16543344 | 1 | 4947 | tags=33%, list=21%, si | EN_3 |
| GOBP_REGULATION_OF_ | GOBP_REGULATION_OF_FIBROBLAST_APOP | 24 | -0.51275 | -1.642 | 0.02467344 | 0.16476645 | 1 | 2541 | tags=25%, list=11%, si | EN_3 |
| GOBP_VESICLE_TETHERIN | GOBP_VESICLE_TETHERING | 31 | -0.48858 | -1.64194 | 0.01551481 | 0.1641458 | 1 | 4012 | tags=29%, list=17%, si | EN_3 |
| GOBP_RESPONSE_TO_DE | GOBP_RESPONSE_TO_DEXAMETHASONE | 27 | -0.49553 | -1.64101 | 0.017366135 | 0.16458043 | 1 | 3375 | tags=30%, list=14%, si | EN_3 |
| GOBP_REGULATION_OF_ | GOBP_REGULATION_OF_AMIDE_METABOLI( | 69 | -0.41931 | -1.63879 | 0.005319149 | 0.16711596 | 1 | 6287 | tags=43%, list=27%, si | EN_3 |
| GOBP_MITOCHONDRIAL_ | GOBP_MITOCHONDRIAL_MEMBRANE_ORG/ | 107 | -0.38862 | -1.63583 | 0.007398274 | 0.17063037 | 1 | 5092 | tags=33%, list=22%, si | EN_3 |
| STEROID_BIOSYNTHESIS | STEROID_BIOSYNTHESIS | 17 | -0.55701 | -1.63557 | 0.027231468 | 0.17032573 | 1 | 3556 | tags=53%, list=15%, si | EN_3 |
| GOBP_REGULATION_OF_ | GOBP_REGULATION_OF_INTEGRIN_MEDIAT | 19 | -0.54558 | -1.63384 | 0.02143951 | 0.17223734 | 1 | 3269 | tags=26%, list=14%, si | EN_3 |
| GOBP_REGULATION_OF_ | GOBP_REGULATION_OF_MYELOID_LEUKOC | 61 | -0.4348 | -1.63288 | 0.0065616798 | 0.1730396 | 1 | 2451 | tags=25%, list=10%, si | EN_3 |
| GOBP_AMYLOID_FIBRIL_I | GOBP_AMYLOID_FIBRIL_FORMATION | 30 | -0.49097 | -1.63284 | 0.025280898 | 0.17235413 | 1 | 7193 | tags=57%, list=30%, si | EN_3 |
| GOBP_RIBOSE_PHOSPHA | GOBP_RIBOSE_PHOSPHATE_METABOLIC_PR | 419 | -0.34505 | -1.63115 | 0 | 0.17392054 | 1 | 5902 | tags=35%, list=25%, si | EN_3 |
| GOBP_FIBROBLAST_APOF | GOBP_FIBROBLAST_APOPTOTIC_PROCESS | 31 | -0.4889 | -1.63103 | 0.011644833 | 0.17340086 | 1 | 4677 | tags=35%, list=20%, si | EN_3 |
| CARDIAC_MUSCLE_CONT | CARDIAC_MUSCLE_CONTRACTION | 72 | -0.40828 | -1.62846 | 0.007444169 | 0.17661504 | 1 | 5169 | tags=42%, list=22%, si | EN_3 |
| PENTOSE_PHOSPHATE_P | PENTOSE_PHOSPHATE_PATHWAY | 29 | -0.47915 | -1.62845 | 0.014577259 | 0.17587915 | 1 | 4545 | tags=38%, list=19%, si | EN_3 |
| FRUCTOSE_AND_MANNO | FRUCTOSE_AND_MANNOSE_METABOLISM | 33 | -0.48268 | -1.62825 | 0.011527377 | 0.17543063 | 1 | 4545 | tags=33%, list=19%, si | EN_3 |
| GOBP_REGULATION_OF_ | GOBP_REGULATION_OF_CYTOPLASMIC_PAT | 104 | -0.38882 | -1.62821 | 0.0012135922 | 0.17478608 | 1 | 5017 | tags=30%, list=21%, si | EN_3 |
| GOBP_PROSTANOID_BIO | GOBP_PROSTANOID_BIOSYNTHETIC_PROCE | 30 | -0.48792 | -1.62767 | 0.022796353 | 0.1747301 | 1 | 487 | tags=17%, list=2%, sig | EN_3 |
| GOBP_POSITIVE_REGULA | GOBP_POSITIVE_REGULATION_OF_GLUCOS | 42 | -0.45703 | -1.62766 | 0.017333332 | 0.17403089 | 1 | 7230 | tags=50%, list=31%, si | EN_3 |
| GOBP_PIGMENT_BIOSYN | GOBP_PIGMENT_BIOSYNTHETIC_PROCESS | 49 | -0.44005 | -1.62746 | 0.015748031 | 0.17357208 | 1 | 5346 | tags=43%, list=23%, si | EN_3 |
| GOBP_POSITIVE_REGULA | GOBP_POSITIVE_REGULATION_OF_TORC1_S | 52 | -0.43241 | -1.62703 | 0.016260162 | 0.17350148 | 1 | 4859 | tags=33%, list=21%, si | EN_3 |
| GOBP_INTRINSIC_APOPT( | GOBP_INTRINSIC_APOPTOTIC_SIGNALING_F | 314 | -0.34657 | -1.62625 | 0 | 0.17386435 | 1 | 6339 | tags=37%, list=27%, si | EN_3 |
| GOBP_SPHINGOMYELIN_ | GOBP_SPHINGOMYELIN_METABOLIC_PROC | 21 | -0.53339 | -1.62593 | 0.023564065 | 0.17369364 | 1 | 6623 | tags=52%, list=28%, si | EN_3 |
| GOBP_POSITIVE_REGULA | GOBP_POSITIVE_REGULATION_OF_CALCIUN | 26 | -0.49954 | -1.62491 | 0.025104603 | 0.17439927 | 1 | 3448 | tags=31%, list=15%, si | EN_3 |
| GOBP_ORGANOPHOSPHA | GOBP_ORGANOPHOSPHATE_BIOSYNTHETIC | 500 | -0.33866 | -1.62418 | 0 | 0.17474565 | 1 | 5187 | tags=32%, list=22%, si | EN_3 |
| GOBP_TERPENOID_BIOSY | GOBP_TERPENOID_BIOSYNTHETIC_PROCESS | 20 | -0.53493 | -1.62268 | 0.025757575 | 0.1761739 | 1 | 4091 | tags=45%, list=17%, si | EN_3 |
| GOBP_RESPONSE_TO_OX | GOBP_RESPONSE_TO_OXIDATIVE_STRESS | 364 | -0.34635 | -1.6221 | 0 | 0.17629814 | 1 | 6357 | tags=39%, list=27%, si | EN_3 |
| GOBP_VESICLE_TARGETIN | GOBP_VESICLE_TARGETING | 60 | -0.42452 | -1.61981 | 0.011749348 | 0.17906837 | 1 | 4941 | tags=32%, list=21%, si | EN_3 |
| GOBP_NEGATIVE_REGUL | GOBP_NEGATIVE_REGULATION_OF_CANON | 57 | -0.42387 | -1.61959 | 0.017994858 | 0.17869955 | 1 | 2346 | tags=23%, list=10%, si | EN_3 |
| GOBP_MITOTIC_DNA_INT | GOBP_MITOTIC_DNA_INTEGRITY_CHECKPO | 84 | -0.39819 | -1.61903 | 0.015037594 | 0.17884073 | 1 | 5267 | tags=37%, list=22%, si | EN_3 |
| GOBP_NEGATIVE_REGUL | GOBP_NEGATIVE_REGULATION_OF_BLOOD_ | 27 | -0.49706 | -1.61863 | 0.017518248 | 0.17886885 | 1 | 2307 | tags=22%, list=10%, si | EN_3 |
| GOBP_VESICLE_TARGETIN | GOBP_VESICLE_TARGETING_TO_FROM_OR_ | 29 | -0.49629 | -1.61844 | 0.017416546 | 0.1784367 | 1 | 4941 | tags=48%, list=21%, si | EN_3 |
| GOBP_SNRNA_METABOL | GOBP_SNRNA_METABOLIC_PROCESS | 50 | -0.43605 | -1.61806 | 0.011889036 | 0.17827955 | 1 | 4839 | tags=38%, list=20%, si | EN_3 |
| GOBP_POSITIVE_REGULA | GOBP_POSITIVE_REGULATION_OF_ERAD_P/ | 20 | -0.53524 | -1.61767 | 0.030627871 | 0.17816655 | 1 | 4315 | tags=40%, list=18%, si | EN_3 |
| GOBP_DE_NOVO_PROTEI | GOBP_DE_NOVO_PROTEIN_FOLDING | 38 | -0.45532 | -1.61743 | 0.014905149 | 0.17788547 | 1 | 5912 | tags=45%, list=25%, si | EN_3 |
| GOBP_CYTOPLASMIC_TR/ | GOBP_CYTOPLASMIC_TRANSLATIONAL_INIT | 33 | -0.47856 | -1.61615 | 0.012658228 | 0.17902328 | 1 | 6593 | tags=45%, list=28%, si | EN_3 |
| GOBP_ENDOSOME_TO_L | GOBP_ENDOSOME_TO_LYSOSOME_TRANSP | 16 | -0.568 | -1.61614 | 0.024806201 | 0.17835973 | 1 | 3075 | tags=44%, list=13%, si | EN_3 |
| GOBP_NEGATIVE_REGUL | GOBP_NEGATIVE_REGULATION_OF_TELOM | 36 | -0.47203 | -1.61572 | 0.021156559 | 0.17831838 | 1 | 7299 | tags=56%, list=31%, si | EN_3 |
| GOBP_POSITIVE_REGULA | GOBP_POSITIVE_REGULATION_OF_MYELOII | 38 | -0.46156 | -1.61271 | 0.009722223 | 0.18211652 | 1 | 1713 | tags=21%, list=7%, sig | EN_3 |
| GOBP_REGULATION_OF_ | GOBP_REGULATION_OF_MICROGLIAL_CELL_ | 17 | -0.55319 | -1.61225 | 0.026356589 | 0.18210459 | 1 | 3729 | tags=41%, list=16%, si | EN_3 |
| APOPTOSIS | APOPTOSIS | 137 | -0.37507 | -1.6112 | 0.0034762456 | 0.18289217 | 1 | 6339 | tags=38%, list=27%, si | EN_3 |
| GOBP_ACYL_COA_METAE | GOBP_ACYL_COA_METABOLIC_PROCESS | 82 | -0.39605 | -1.60961 | 0.010075566 | 0.18452793 | 1 | 5857 | tags=37%, list=25%, si | EN_3 |
| GOBP_MODULATION_BY_ | GOBP_MODULATION_BY_HOST_OF_VIRAL_ | 22 | -0.52196 | -1.60958 | 0.03397341 | 0.18387008 | 1 | 6319 | tags=45%, list=27%, si | EN_3 |
| GOBP_POSITIVE_REGULA | GOBP_POSITIVE_REGULATION_OF_PROTEAS | 102 | -0.38588 | -1.60917 | 0.0072202166 | 0.18381225 | 1 | 7160 | tags=44%, list=30%, si | EN_3 |
| GOBP_NEUROINFLAMMA | GOBP_NEUROINFLAMMATORY_RESPONSE | 84 | -0.40251 | -1.60898 | 0.0037831021 | 0.18348141 | 1 | 2775 | tags=21%, list=12%, si | EN_3 |
| GOBP_MEMBRANE_DOCI | GOBP_MEMBRANE_DOCKING | 78 | -0.39959 | -1.60886 | 0.012254902 | 0.18295634 | 1 | 4293 | tags=28%, list=18%, si | EN_3 |
| GOBP_T_CELL_APOPTOTI | GOBP_T_CELL_APOPTOTIC_PROCESS | 67 | -0.41072 | -1.60869 | 0.013853904 | 0.18257086 | 1 | 3462 | tags=30%, list=15%, si | EN_3 |
| RNA_DEGRADATION | RNA_DEGRADATION | 75 | -0.40679 | -1.60674 | 0.0050251256 | 0.1848031 | 1 | 6699 | tags=45%, list=28%, si | EN_3 |
| GOBP_AXO_DENDRITIC_T | GOBP_AXO_DENDRITIC_TRANSPORT | 92 | -0.38947 | -1.60653 | 0.008816121 | 0.18443576 | 1 | 5238 | tags=34%, list=22%, si | EN_3 |
| GOBP_RESPONSE_TO_TE | GOBP_RESPONSE_TO_TEMPERATURE_STIM | 159 | -0.36349 | -1.60614 | 0.0011198209 | 0.1844047 | 1 | 4283 | tags=27%, list=18%, si | EN_3 |
| GOBP_REGULATION_OF_ | GOBP_REGULATION_OF_CELLULAR_RESPON | 25 | -0.49791 | -1.60391 | 0.020989506 | 0.18708827 | 1 | 1945 | tags=24%, list=8%, sig | EN_3 |

| | | | | | | | | | | |
|---|---|---|---|---|---|---|---|---|---|---|
| GOBP_LATE_ENDOSOME_ | GOBP_LATE_ENDOSOME_TO_VACUOLE_TRA | 25 | -0.50004 | -1.60309 | 0.026988637 | 0.18762287 | 1 | 4088 | tags=40%, list=17%, si | EN_3 |
| NECROPTOSIS | NECROPTOSIS | 138 | -0.36352 | -1.60306 | 0.004555809 | 0.18700342 | 1 | 5354 | tags=34%, list=23%, si | EN_3 |
| GOBP_NEGATIVE_REGUL | GOBP_NEGATIVE_REGULATION_OF_LEUKOC | 46 | -0.44003 | -1.60292 | 0.01591512 | 0.18657987 | 1 | 2612 | tags=28%, list=11%, si | EN_3 |
| GOBP_LIPOPROTEIN_ME | GOBP_LIPOPROTEIN_METABOLIC_PROCESS | 135 | -0.37356 | -1.60262 | 0.0011723329 | 0.18632038 | 1 | 4893 | tags=33%, list=21%, si | EN_3 |
| GOBP_REGULATION_OF_ | GOBP_REGULATION_OF_NEUROINFLAMMA | 30 | -0.48368 | -1.60253 | 0.02691218 | 0.1857817 | 1 | 2775 | tags=27%, list=12%, si | EN_3 |
| GOBP_FRUCTOSE_6_PHO | GOBP_FRUCTOSE_6_PHOSPHATE_METABOL | 16 | -0.57092 | -1.60187 | 0.034865294 | 0.18615764 | 1 | 1436 | tags=25%, list=6%, sig | EN_3 |
| GOBP_RESPONSE_TO_CO | GOBP_RESPONSE_TO_COPPER_ION | 23 | -0.51205 | -1.60068 | 0.026548672 | 0.18720293 | 1 | 3272 | tags=35%, list=14%, si | EN_3 |
| GOBP_REGULATION_OF_ | GOBP_REGULATION_OF_CYTOPLASMIC_TRA | 30 | -0.46914 | -1.60017 | 0.026462395 | 0.18732858 | 1 | 6983 | tags=60%, list=30%, si | EN_3 |
| GLUTATHIONE_METABOL | GLUTATHIONE_METABOLISM | 57 | -0.41644 | -1.59929 | 0.007884363 | 0.18808176 | 1 | 6366 | tags=53%, list=27%, si | EN_3 |
| GOBP_POSITIVE_REGULA | GOBP_POSITIVE_REGULATION_OF_PEPTIDA | 161 | -0.36401 | -1.59911 | 0.0022701477 | 0.18769386 | 1 | 5362 | tags=32%, list=23%, si | EN_3 |
| GOBP_NEUTROPHIL_ACTI | GOBP_NEUTROPHIL_ACTIVATION_INVOLVE | 18 | -0.53681 | -1.59879 | 0.036036037 | 0.18756348 | 1 | 2243 | tags=33%, list=9%, sig | EN_3 |
| GOBP_REGULATION_OF_ | GOBP_REGULATION_OF_TELOMERE_MAINT | 98 | -0.38771 | -1.5985 | 0.0084951455 | 0.18731882 | 1 | 7334 | tags=51%, list=31%, si | EN_3 |
| GOBP_PEPTIDYL_THREON | GOBP_PEPTIDYL_THREONINE_DEPHOSPHOR | 15 | -0.56253 | -1.59772 | 0.03726708 | 0.1877625 | 1 | 7298 | tags=73%, list=31%, si | EN_3 |
| GOBP_CELLULAR_RESPO | GOBP_CELLULAR_RESPONSE_TO_INTERLEU | 62 | -0.41467 | -1.59618 | 0.01662404 | 0.18952833 | 1 | 5846 | tags=40%, list=25%, si | EN_3 |
| GOBP_CAMP_MEDIATED_ | GOBP_CAMP_MEDIATED_SIGNALING | 67 | -0.40493 | -1.59542 | 0.015037594 | 0.19007823 | 1 | 2074 | tags=19%, list=9%, sig | EN_3 |
| GOBP_PROTEIN_CATABO | GOBP_PROTEIN_CATABOLIC_PROCESS_IN_T | 22 | -0.50579 | -1.59517 | 0.026153846 | 0.18980734 | 1 | 3497 | tags=27%, list=15%, si | EN_3 |
| GOBP_REGULATION_OF_ | GOBP_REGULATION_OF_FIBROBLAST_PROL | 122 | -0.37516 | -1.59464 | 0.0058139535 | 0.18986057 | 1 | 4746 | tags=32%, list=20%, si | EN_3 |
| GOBP_ERAD_PATHWAY | GOBP_ERAD_PATHWAY | 108 | -0.37814 | -1.59287 | 0.0047562425 | 0.19172427 | 1 | 5793 | tags=35%, list=25%, si | EN_3 |
| GOBP_PROGRAMMED_N | GOBP_PROGRAMMED_NECROTIC_CELL_DE | 50 | -0.43509 | -1.59277 | 0.015364917 | 0.19118239 | 1 | 3024 | tags=24%, list=13%, si | EN_3 |
| GOBP_MYELOID_LEUKOC | GOBP_MYELOID_LEUKOCYTE_CYTOKINE_PR | 64 | -0.41321 | -1.5922 | 0.015228426 | 0.19142462 | 1 | 2115 | tags=20%, list=9%, sig | EN_3 |
| GOBP_SNO_S_RNA_MET | GOBP_SNO_S_RNA_METABOLIC_PROCESS | 17 | -0.54417 | -1.59219 | 0.029411765 | 0.19081119 | 1 | 3119 | tags=35%, list=13%, si | EN_3 |
| GOBP_RESPONSE_TO_CA | GOBP_RESPONSE_TO_CADMIUM_ION | 35 | -0.4616 | -1.59198 | 0.02827381 | 0.19047071 | 1 | 4908 | tags=37%, list=21%, si | EN_3 |
| GOBP_ESTABLISHMENT_ | GOBP_ESTABLISHMENT_OF_MITOCHONDRI | 25 | -0.50421 | -1.59195 | 0.01594203 | 0.18988493 | 1 | 6239 | tags=52%, list=26%, si | EN_3 |
| GOBP_NEGATIVE_REGUL | GOBP_NEGATIVE_REGULATION_OF_OSTEO | 37 | -0.44641 | -1.59103 | 0.026773762 | 0.19065984 | 1 | 2282 | tags=24%, list=10%, si | EN_3 |
| GOBP_MULTIVESICULAR_ | GOBP_MULTIVESICULAR_BODY_SORTING_P | 34 | -0.46113 | -1.59011 | 0.022284122 | 0.19142328 | 1 | 4088 | tags=41%, list=17%, si | EN_3 |
| PHENYLALANINE_METAB | PHENYLALANINE_METABOLISM | 16 | -0.5504 | -1.58931 | 0.042414356 | 0.19217446 | 1 | 2996 | tags=44%, list=13%, si | EN_3 |
| GOBP_REGULATION_OF_ | GOBP_REGULATION_OF_DEFENSE_RESPONS | 42 | -0.44996 | -1.58916 | 0.023383768 | 0.19180101 | 1 | 4283 | tags=31%, list=18%, si | EN_3 |
| TERPENOID_BACKBONE_ | TERPENOID_BACKBONE_BIOSYNTHESIS | 21 | -0.51561 | -1.58794 | 0.028213166 | 0.19305587 | 1 | 948 | tags=24%, list=4%, sig | EN_3 |
| GOBP_AUTOPHAGOSOMI | GOBP_AUTOPHAGOSOME_ORGANIZATION | 128 | -0.3696 | -1.58714 | 0.0034722222 | 0.19371015 | 1 | 4506 | tags=27%, list=19%, si | EN_3 |
| GOBP_REGULATION_OF_ | GOBP_REGULATION_OF_MITOCHONDRIAL_ | 35 | -0.4635 | -1.5868 | 0.028248588 | 0.19361398 | 1 | 4454 | tags=34%, list=19%, si | EN_3 |
| GOBP_PROTEIN_TARGETI | GOBP_PROTEIN_TARGETING | 285 | -0.34646 | -1.58661 | 0 | 0.19326101 | 1 | 5136 | tags=28%, list=22%, si | EN_3 |
| GOBP_ATP_METABOLIC_ | GOBP_ATP_METABOLIC_PROCESS | 206 | -0.35129 | -1.58531 | 0 | 0.19468947 | 1 | 6351 | tags=41%, list=27%, si | EN_3 |
| GOBP_NEGATIVE_REGUL | GOBP_NEGATIVE_REGULATION_OF_TELOM | 27 | -0.48484 | -1.58463 | 0.025373135 | 0.1950966 | 1 | 6895 | tags=52%, list=29%, si | EN_3 |
| GOBP_SIGNAL_TRANSDU | GOBP_SIGNAL_TRANSDUCTION_BY_P53_CL | 151 | -0.36683 | -1.58444 | 0 | 0.1947988 | 1 | 6081 | tags=40%, list=26%, si | EN_3 |
| GOBP_REGULATION_OF_ | GOBP_REGULATION_OF_PATTERN_RECOGN | 157 | -0.35998 | -1.58295 | 0.0011587485 | 0.19640851 | 1 | 4251 | tags=26%, list=18%, si | EN_3 |
| GOBP_NEGATIVE_REGUL | GOBP_NEGATIVE_REGULATION_OF_INTRINS | 105 | -0.37842 | -1.58257 | 0.011876484 | 0.19642466 | 1 | 6764 | tags=44%, list=29%, si | EN_3 |
| GOBP_NEGATIVE_REGUL | GOBP_NEGATIVE_REGULATION_OF_AUTOP | 81 | -0.39544 | -1.58227 | 0.008663367 | 0.19624338 | 1 | 2231 | tags=21%, list=9%, sig | EN_3 |
| GOBP_ERYTHROCYTE_MA | GOBP_ERYTHROCYTE_MATURATION | 23 | -0.4995 | -1.58129 | 0.031626508 | 0.19718002 | 1 | 2161 | tags=22%, list=9%, sig | EN_3 |
| GOBP_LIPOPROTEIN_BIO | GOBP_LIPOPROTEIN_BIOSYNTHETIC_PROCE | 104 | -0.37849 | -1.58084 | 0.007290401 | 0.19725819 | 1 | 6647 | tags=47%, list=28%, si | EN_3 |
| GOBP_NEGATIVE_REGUL | GOBP_NEGATIVE_REGULATION_OF_PROTEI | 80 | -0.39375 | -1.58025 | 0.009840098 | 0.19762906 | 1 | 6099 | tags=39%, list=26%, si | EN_3 |
| GOBP_PROTEIN_MONOU | GOBP_PROTEIN_MONOUBIQUITINATION | 47 | -0.43578 | -1.58021 | 0.023968043 | 0.19707046 | 1 | 7406 | tags=51%, list=31%, si | EN_3 |
| GOBP_REGULATION_OF_ | GOBP_REGULATION_OF_NEUROTROPHIN_T | 17 | -0.55305 | -1.57993 | 0.04017857 | 0.19692592 | 1 | 5692 | tags=53%, list=24%, si | EN_3 |
| HISTIDINE_METABOLISM | HISTIDINE_METABOLISM | 21 | -0.52267 | -1.57822 | 0.032884903 | 0.19909477 | 1 | 2996 | tags=33%, list=13%, si | EN_3 |
| AMPK_SIGNALING_PATH | AMPK_SIGNALING_PATHWAY | 120 | -0.36841 | -1.57798 | 0.009489917 | 0.19882871 | 1 | 4819 | tags=28%, list=20%, si | EN_3 |
| VIRAL_CARCINOGENESIS | VIRAL_CARCINOGENESIS | 201 | -0.35418 | -1.57583 | 0.0022271716 | 0.20153809 | 1 | 3741 | tags=23%, list=16%, si | EN_3 |
| GOBP_NEGATIVE_REGUL | GOBP_NEGATIVE_REGULATION_OF_CELL_C | 69 | -0.40592 | -1.57496 | 0.015564202 | 0.20235106 | 1 | 6334 | tags=46%, list=27%, si | EN_3 |
| GOBP_SMOOTH_MUSCLE | GOBP_SMOOTH_MUSCLE_CELL_MIGRATION | 115 | -0.36961 | -1.57374 | 0.0070422534 | 0.20358606 | 1 | 6980 | tags=43%, list=30%, si | EN_3 |
| GOBP_NEGATIVE_REGUL | GOBP_NEGATIVE_REGULATION_OF_ACTIN_ | 64 | -0.40743 | -1.5737 | 0.020075282 | 0.20300826 | 1 | 6099 | tags=41%, list=26%, si | EN_3 |
| GOBP_POSITIVE_REGULA | GOBP_POSITIVE_REGULATION_OF_MACROF | 29 | -0.46777 | -1.57359 | 0.033850495 | 0.20257318 | 1 | 3815 | tags=31%, list=16%, si | EN_3 |
| GOBP_MITOCHONDRIAL_ | GOBP_MITOCHONDRIAL_TRANSLATION | 128 | -0.36818 | -1.57276 | 0.0035545023 | 0.20328362 | 1 | 4523 | tags=32%, list=19%, si | EN_3 |
| GOBP_RESPONSE_TO_ST | GOBP_RESPONSE_TO_STEROL_DEPLETION | 15 | -0.55384 | -1.57232 | 0.044728436 | 0.20341928 | 1 | 6399 | tags=60%, list=27%, si | EN_3 |
| GOBP_REGULATION_OF_ | GOBP_REGULATION_OF_THE_FORCE_OF_H | 30 | -0.47072 | -1.57177 | 0.03372434 | 0.20370162 | 1 | 6562 | tags=43%, list=28%, si | EN_3 |
| GOBP_ETHER_METABOLI | GOBP_ETHER_METABOLIC_PROCESS | 23 | -0.5132 | -1.57096 | 0.034534536 | 0.20447303 | 1 | 943 | tags=17%, list=4%, sig | EN_3 |
| GOBP_PROTEIN_K6_LINK | GOBP_PROTEIN_K6_LINKED_UBIQUITINATIO | 15 | -0.54859 | -1.57045 | 0.050847456 | 0.20469399 | 1 | 1672 | tags=27%, list=7%, sig | EN_3 |
| GLYCOLYSIS_/_GLUCONE | GLYCOLYSIS_/_GLUCONEOGENESIS | 60 | -0.41388 | -1.57045 | 0.02 | 0.2040849 | 1 | 5724 | tags=43%, list=24%, si | EN_3 |
| GOBP_STEROID_BIOSYNT | GOBP_STEROID_BIOSYNTHETIC_PROCESS | 153 | -0.36075 | -1.56901 | 0.0033975085 | 0.20577788 | 1 | 4246 | tags=28%, list=18%, si | EN_3 |
| GOBP_RESPONSE_TO_TY | GOBP_RESPONSE_TO_TYPE_I_INTERFERON | 64 | -0.40298 | -1.56871 | 0.0144356955 | 0.20565027 | 1 | 4325 | tags=28%, list=18%, si | EN_3 |
| GOBP_RETINOL_METABO | GOBP_RETINOL_METABOLIC_PROCESS | 21 | -0.51409 | -1.56798 | 0.041604754 | 0.2061014 | 1 | 4051 | tags=38%, list=17%, si | EN_3 |
| GOBP_L_AMINO_ACID_M | GOBP_L_AMINO_ACID_METABOLIC_PROCE | 154 | -0.35717 | -1.56639 | 0.0033707866 | 0.20811819 | 1 | 4153 | tags=25%, list=18%, si | EN_3 |
| GOBP_TORC1_SIGNALIN | GOBP_TORC1_SIGNALING | 100 | -0.37859 | -1.56573 | 0.004878049 | 0.20856011 | 1 | 5309 | tags=33%, list=22%, si | EN_3 |
| GOBP_NEGATIVE_REGUL | GOBP_NEGATIVE_REGULATION_OF_BMP_S | 53 | -0.41315 | -1.56391 | 0.021680217 | 0.21087095 | 1 | 4651 | tags=32%, list=20%, si | EN_3 |
| GOBP_REGULATION_OF_ | GOBP_REGULATION_OF_PEPTIDYL_THREON | 45 | -0.43184 | -1.563 | 0.02035278 | 0.21172188 | 1 | 2724 | tags=20%, list=12%, si | EN_3 |
| GOBP_RRNA_TRANSCRIP | GOBP_RRNA_TRANSCRIPTION | 33 | -0.4635 | -1.56294 | 0.027104137 | 0.21118534 | 1 | 1600 | tags=15%, list=7%, sig | EN_3 |
| GOBP_DENDRITIC_TRANS | GOBP_DENDRITIC_TRANSPORT | 19 | -0.50965 | -1.56097 | 0.044977512 | 0.21385801 | 1 | 3310 | tags=37%, list=14%, si | EN_3 |
| EPSTEIN-BARR_VIRUS_IN | EPSTEIN-BARR_VIRUS_INFECTION | 190 | -0.35143 | -1.56074 | 0.0033482143 | 0.21366443 | 1 | 3914 | tags=23%, list=17%, si | EN_3 |
| GOBP_DEOXYRIBONUCLE | GOBP_DEOXYRIBONUCLEOTIDE_METABOLIC | 44 | -0.42987 | -1.55981 | 0.021361817 | 0.2144301 | 1 | 5256 | tags=36%, list=22%, si | EN_3 |
| GOBP_MONOSACCHARID | GOBP_MONOSACCHARIDE_METABOLIC_PR | 255 | -0.34093 | -1.55836 | 0 | 0.21619253 | 1 | 4147 | tags=24%, list=18%, si | EN_3 |
| GLYCEROPHOSPHOLIPID_ | GLYCEROPHOSPHOLIPID_METABOLISM | 92 | -0.38145 | -1.55803 | 0.006142506 | 0.21610443 | 1 | 4610 | tags=32%, list=20%, si | EN_3 |
| GOBP_NEGATIVE_REGUL | GOBP_NEGATIVE_REGULATION_OF_POST_T | 94 | -0.38083 | -1.55772 | 0.006024096 | 0.21600251 | 1 | 5287 | tags=36%, list=22%, si | EN_3 |
| GOBP_POSITIVE_REGULA | GOBP_POSITIVE_REGULATION_OF_UBIQUIT | 18 | -0.52188 | -1.55741 | 0.03894081 | 0.2159065 | 1 | 1911 | tags=28%, list=8%, sig | EN_3 |
| GOBP_NEGATIVE_REGUL | GOBP_NEGATIVE_REGULATION_OF_PROGR | 17 | -0.53864 | -1.55688 | 0.040971167 | 0.2161009 | 1 | 5727 | tags=53%, list=24%, si | EN_3 |
| GOBP_TELOMERE_ORGA | GOBP_TELOMERE_ORGANIZATION | 150 | -0.35397 | -1.55618 | 0.0034207525 | 0.2166462 | 1 | 6308 | tags=39%, list=27%, si | EN_3 |
| GOBP_DNA_TEMPLATED_ | GOBP_DNA_TEMPLATED_DNA_REPLICATION | 147 | -0.35439 | -1.55474 | 0.004514673 | 0.21854354 | 1 | 3492 | tags=25%, list=15%, si | EN_3 |
| GOBP_MUSCLE_CELL_MI | GOBP_MUSCLE_CELL_MIGRATION | 129 | -0.36648 | -1.55436 | 0.003440367 | 0.21850942 | 1 | 6980 | tags=43%, list=30%, si | EN_3 |
| GOBP_GLUCOSE_METAB | GOBP_GLUCOSE_METABOLIC_PROCESS | 201 | -0.34684 | -1.55316 | 0.0044345898 | 0.21997671 | 1 | 4147 | tags=24%, list=18%, si | EN_3 |
| GOBP_MRNA_TRANSCRIP | GOBP_MRNA_TRANSCRIPTION | 50 | -0.42474 | -1.55262 | 0.01864181 | 0.2202423 | 1 | 6942 | tags=50%, list=29%, si | EN_3 |
| GOBP_POSITIVE_REGULA | GOBP_POSITIVE_REGULATION_OF_PROTEIN | 41 | -0.44005 | -1.5525 | 0.030042918 | 0.21989247 | 1 | 3698 | tags=27%, list=16%, si | EN_3 |
| GOBP_POSITIVE_REGULA | GOBP_POSITIVE_REGULATION_OF_PROTEIN | 192 | -0.34548 | -1.55128 | 0.0011013215 | 0.22133058 | 1 | 6982 | tags=40%, list=30%, si | EN_3 |
| GOBP_IMMATURE_B_CEL | GOBP_IMMATURE_B_CELL_DIFFERENTIATIO | 17 | -0.52467 | -1.55063 | 0.03974563 | 0.22192352 | 1 | 499 | tags=18%, list=2%, sig | EN_3 |
| GOBP_HEME_BIOSYNTHE | GOBP_HEME_BIOSYNTHETIC_PROCESS | 24 | -0.49876 | -1.54993 | 0.038748138 | 0.22248383 | 1 | 5346 | tags=54%, list=23%, si | EN_3 |
| GOBP_SULFUR_COMPOU | GOBP_SULFUR_COMPOUND_METABOLIC_P | 292 | -0.33392 | -1.54974 | 0.0010537407 | 0.22218004 | 1 | 4624 | tags=29%, list=20%, si | EN_3 |
| GOBP_REGULATION_OF_ | GOBP_REGULATION_OF_INFLAMMASOME_ | 40 | -0.43676 | -1.54941 | 0.026836159 | 0.22205283 | 1 | 5017 | tags=33%, list=21%, si | EN_3 |

| | | | | | | | | | | |
|---|---|---|---|---|---|---|---|---|---|---|
| GOBP_REGULATION_OF_ | GOBP_REGULATION_OF_STRESS_ACTIVATED | 41 | -0.43619 | -1.54869 | 0.02351314 | 0.22269455 | 1 | 6624 | tags=44%, list=28%, si | EN_3 |
| GOBP_REGULATION_OF_ | GOBP_REGULATION_OF_PROTEIN_STABILIT | 305 | -0.33089 | -1.54863 | 0 | 0.22215578 | 1 | 6136 | tags=35%, list=26%, si | EN_3 |
| GOBP_NEGATIVE_REGUL | GOBP_NEGATIVE_REGULATION_OF_RECEPT | 27 | -0.47276 | -1.54825 | 0.04270987 | 0.22216953 | 1 | 4783 | tags=37%, list=20%, si | EN_3 |
| GOBP_PROTEIN_REFOLDI | GOBP_PROTEIN_REFOLDING | 23 | -0.4928 | -1.54818 | 0.03851852 | 0.22164378 | 1 | 8045 | tags=70%, list=34%, si | EN_3 |
| GOBP_POSITIVE_REGULA | GOBP_POSITIVE_REGULATION_OF_EPITHELI | 158 | -0.35517 | -1.54795 | 0.006880734 | 0.22142926 | 1 | 6694 | tags=39%, list=28%, si | EN_3 |
| GOBP_PURINE_NUCLEOS | GOBP_PURINE_NUCLEOSIDE_METABOLIC_P | 30 | -0.46254 | -1.54652 | 0.0335277 | 0.22328004 | 1 | 2150 | tags=23%, list=9%, sig | EN_3 |
| GOBP_POSITIVE_REGULA | GOBP_POSITIVE_REGULATION_OF_MACRO/ | 81 | -0.38145 | -1.54633 | 0.023929471 | 0.22297801 | 1 | 6349 | tags=40%, list=27%, si | EN_3 |
| GOBP_POSITIVE_REGULA | GOBP_POSITIVE_REGULATION_OF_GLUCOS | 50 | -0.41248 | -1.54605 | 0.02110818 | 0.22285742 | 1 | 7230 | tags=46%, list=31%, si | EN_3 |
| GOBP_UBIQUITIN_DEPEN | GOBP_UBIQUITIN_DEPENDENT_PROTEIN_C | 29 | -0.46174 | -1.54595 | 0.038737446 | 0.22241549 | 1 | 3278 | tags=34%, list=14%, si | EN_3 |
| GOBP_NEGATIVE_REGUL | GOBP_NEGATIVE_REGULATION_OF_AMIDE_ | 22 | -0.49373 | -1.54575 | 0.03697997 | 0.22213474 | 1 | 5837 | tags=55%, list=25%, si | EN_3 |
| GOBP_REGULATION_OF_ | GOBP_REGULATION_OF_VASCULATURE_DE' | 295 | -0.33304 | -1.54554 | 0 | 0.22187774 | 1 | 5292 | tags=32%, list=22%, si | EN_3 |
| GOBP_SNRNA_TRANSCRI | GOBP_SNRNA_TRANSCRIPTION | 18 | -0.5203 | -1.54511 | 0.057057057 | 0.22198394 | 1 | 6312 | tags=50%, list=27%, si | EN_3 |
| GOBP_ESTABLISHMENT_( | GOBP_ESTABLISHMENT_OF_PROTEIN_LOCA | 435 | -0.32722 | -1.54449 | 0 | 0.22251293 | 1 | 7473 | tags=43%, list=32%, si | EN_3 |
| GOBP_DNA_TEMPLATED_ | GOBP_DNA_TEMPLATED_TRANSCRIPTION_I | 112 | -0.3685 | -1.54449 | 0.010638298 | 0.22191478 | 1 | 7999 | tags=54%, list=34%, si | EN_3 |
| GOBP_POSITIVE_REGULA | GOBP_POSITIVE_REGULATION_OF_MEMBR/ | 39 | -0.44307 | -1.54417 | 0.026685392 | 0.22183257 | 1 | 4942 | tags=44%, list=21%, si | EN_3 |
| GOBP_INTRINSIC_APOPT( | GOBP_INTRINSIC_APOPTOTIC_SIGNALING_I | 44 | -0.43094 | -1.54257 | 0.02631579 | 0.22393064 | 1 | 2632 | tags=25%, list=11%, si | EN_3 |
| GOBP_RESPONSE_TO_LIT | GOBP_RESPONSE_TO_LITHIUM_ION | 16 | -0.53774 | -1.5411 | 0.034755133 | 0.22578256 | 1 | 595 | tags=25%, list=3%, sig | EN_3 |
| GOBP_RESPONSE_TO_INT | GOBP_RESPONSE_TO_INTERFERON_BETA | 64 | -0.40319 | -1.54025 | 0.023376623 | 0.22659297 | 1 | 4696 | tags=34%, list=20%, si | EN_3 |
| GOBP_MYELOID_CELL_A( | GOBP_MYELOID_CELL_ACTIVATION_INVOLV | 98 | -0.37521 | -1.53984 | 0.010676157 | 0.22667618 | 1 | 2510 | tags=22%, list=11%, si | EN_3 |
| GOBP_RESPONSE_TO_FA | GOBP_RESPONSE_TO_FATTY_ACID | 46 | -0.42137 | -1.53976 | 0.031746034 | 0.22621517 | 1 | 3787 | tags=33%, list=16%, si | EN_3 |
| GOBP_REGULATION_OF_ | GOBP_REGULATION_OF_PROTEASOMAL_PR | 179 | -0.34796 | -1.53973 | 0.0011135858 | 0.22565907 | 1 | 6982 | tags=38%, list=30%, si | EN_3 |
| GOBP_REGULATION_OF_ | GOBP_REGULATION_OF_MITOTIC_SPINDLE_ | 18 | -0.51359 | -1.53952 | 0.04658385 | 0.22541912 | 1 | 3294 | tags=33%, list=14%, si | EN_3 |
| GOBP_RIBONUCLEOSIDE_ | GOBP_RIBONUCLEOSIDE_METABOLIC_PRO( | 30 | -0.45886 | -1.53913 | 0.044992745 | 0.22552295 | 1 | 2150 | tags=23%, list=9%, sig | EN_3 |
| GOBP_MITOCHONDRIAL_ | GOBP_MITOCHONDRIAL_ELECTRON_TRANS | 29 | -0.47221 | -1.53891 | 0.040697675 | 0.2253546 | 1 | 5902 | tags=38%, list=25%, si | EN_3 |
| BUTANOATE_METABOLIS | BUTANOATE_METABOLISM | 23 | -0.49164 | -1.5384 | 0.05007364 | 0.22562146 | 1 | 981 | tags=22%, list=4%, sig | EN_3 |
| GOBP_POSITIVE_REGULA | GOBP_POSITIVE_REGULATION_OF_VACUOL | 25 | -0.48191 | -1.53839 | 0.046478875 | 0.22504456 | 1 | 2469 | tags=32%, list=10%, si | EN_3 |
| AMINO_SUGAR_AND_NU | AMINO_SUGAR_AND_NUCLEOTIDE_SUGAR_ | 46 | -0.41706 | -1.53799 | 0.03448276 | 0.22513558 | 1 | 3699 | tags=26%, list=16%, si | EN_3 |
| GOBP_POSITIVE_REGULA | GOBP_POSITIVE_REGULATION_OF_ENDOTH | 107 | -0.36509 | -1.53776 | 0.00973236 | 0.22495183 | 1 | 4528 | tags=26%, list=19%, si | EN_3 |
| GOBP_GLUCOSE_6_PHOS | GOBP_GLUCOSE_6_PHOSPHATE_METABOLI | 28 | -0.46593 | -1.53766 | 0.04101839 | 0.22458039 | 1 | 1436 | tags=21%, list=6%, sig | EN_3 |
| GOBP_NEGATIVE_REGUL | GOBP_NEGATIVE_REGULATION_OF_VIRAL_( | 50 | -0.42287 | -1.5376 | 0.030625831 | 0.22410771 | 1 | 2572 | tags=20%, list=11%, si | EN_3 |
| GOBP_TETRAPYRROLE_BI | GOBP_TETRAPYRROLE_BIOSYNTHETIC_PRO( | 29 | -0.46121 | -1.53719 | 0.042016808 | 0.22417648 | 1 | 5346 | tags=52%, list=23%, si | EN_3 |
| GOBP_VACUOLE_ORGAN | GOBP_VACUOLE_ORGANIZATION | 223 | -0.34038 | -1.53689 | 0.0022075055 | 0.22416316 | 1 | 4556 | tags=28%, list=19%, si | EN_3 |
| FOLATE_BIOSYNTHESIS | FOLATE_BIOSYNTHESIS | 20 | -0.51361 | -1.53633 | 0.04418262 | 0.22456262 | 1 | 3417 | tags=40%, list=14%, si | EN_3 |
| GOBP_CELLULAR_RESPON | GOBP_CELLULAR_RESPONSE_TO_LIGHT_STI | 106 | -0.36872 | -1.53536 | 0.01459854 | 0.22557032 | 1 | 5403 | tags=31%, list=23%, si | EN_3 |
| GOBP_GLYCOLYTIC_PRO( | GOBP_GLYCOLYTIC_PROCESS_THROUGH_FR | 25 | -0.48357 | -1.53526 | 0.05202312 | 0.22517164 | 1 | 5724 | tags=44%, list=24%, si | EN_3 |
| GOBP_INFLAMMATORY_I | GOBP_INFLAMMATORY_RESPONSE_TO_WO | 30 | -0.46109 | -1.53437 | 0.032967035 | 0.2261421 | 1 | 1127 | tags=23%, list=5%, sig | EN_3 |
| GOBP_CELLULAR_RESPON | GOBP_CELLULAR_RESPONSE_TO_TOPOLOG | 95 | -0.374 | -1.53401 | 0.018564356 | 0.22613148 | 1 | 7432 | tags=44%, list=31%, si | EN_3 |
| GOBP_CELLULAR_RESPON | GOBP_CELLULAR_RESPONSE_TO_TOXIC_SU | 63 | -0.39892 | -1.533 | 0.015768725 | 0.22737703 | 1 | 5243 | tags=41%, list=22%, si | EN_3 |
| GOBP_CHOLESTEROL_ST( | GOBP_CHOLESTEROL_STORAGE | 27 | -0.46083 | -1.53297 | 0.050847456 | 0.22685349 | 1 | 3286 | tags=30%, list=14%, si | EN_3 |
| GOBP_AUTOPHAGY_OF_I | GOBP_AUTOPHAGY_OF_MITOCHONDRION | 89 | -0.38139 | -1.53297 | 0.01743462 | 0.22628877 | 1 | 7405 | tags=51%, list=31%, si | EN_3 |
| GOBP_AMYLOID_PRECUR | GOBP_AMYLOID_PRECURSOR_PROTEIN_ME | 75 | -0.38202 | -1.53218 | 0.018018018 | 0.22709689 | 1 | 6287 | tags=45%, list=27%, si | EN_3 |
| GOBP_VIRAL_GENOME_F | GOBP_VIRAL_GENOME_REPLICATION | 110 | -0.36175 | -1.53212 | 0.011778563 | 0.22661142 | 1 | 5586 | tags=34%, list=24%, si | EN_3 |
| GOBP_INTRINSIC_APOPT( | GOBP_INTRINSIC_APOPTOTIC_SIGNALING_I | 83 | -0.37757 | -1.53211 | 0.01743462 | 0.22605914 | 1 | 4512 | tags=29%, list=19%, si | EN_3 |
| GOBP_CHAPERONE_MED | GOBP_CHAPERONE_MEDIATED_PROTEIN_C | 18 | -0.513 | -1.53092 | 0.048231512 | 0.22742832 | 1 | 3398 | tags=39%, list=14%, si | EN_3 |
| GOBP_INTRINSIC_APOPT( | GOBP_INTRINSIC_APOPTOTIC_SIGNALING_I | 112 | -0.36305 | -1.52847 | 0.019347038 | 0.23112781 | 1 | 6388 | tags=41%, list=27%, si | EN_3 |
| GOBP_NEGATIVE_REGUL | GOBP_NEGATIVE_REGULATION_OF_CELL_C | 67 | -0.3936 | -1.52816 | 0.02948718 | 0.23106469 | 1 | 3059 | tags=24%, list=13%, si | EN_3 |
| GOBP_TRANSLESION_SYN | GOBP_TRANSLESION_SYNTHESIS | 22 | -0.49714 | -1.52783 | 0.050974514 | 0.23097877 | 1 | 5402 | tags=55%, list=23%, si | EN_3 |
| GOBP_REGULATION_OF_ | GOBP_REGULATION_OF_STEROID_METABO | 107 | -0.36013 | -1.52749 | 0.015348288 | 0.23103975 | 1 | 3718 | tags=24%, list=16%, si | EN_3 |
| GOBP_REGULATION_OF_ | GOBP_REGULATION_OF_LEUKOCYTE_DEGR/ | 48 | -0.41607 | -1.52619 | 0.041208792 | 0.23275673 | 1 | 2451 | tags=23%, list=10%, si | EN_3 |
| GOBP_ENDOSOMAL_TRA | GOBP_ENDOSOMAL_TRANSPORT | 272 | -0.33182 | -1.52565 | 0.0031813362 | 0.23321642 | 1 | 5866 | tags=34%, list=25%, si | EN_3 |
| GOBP_REGULATION_OF_ | GOBP_REGULATION_OF_LONG_TERM_SYN/ | 19 | -0.50372 | -1.52447 | 0.054135337 | 0.23467936 | 1 | 4439 | tags=47%, list=19%, si | EN_3 |
| GOBP_POSITIVE_REGULA | GOBP_POSITIVE_REGULATION_OF_AMYLOI[ | 20 | -0.51186 | -1.5234 | 0.056296296 | 0.23602717 | 1 | 3706 | tags=35%, list=16%, si | EN_3 |
| GOBP_REGULATION_OF_ | GOBP_REGULATION_OF_DEFENSE_RESPON! | 67 | -0.38682 | -1.52252 | 0.025940338 | 0.23698257 | 1 | 4283 | tags=30%, list=18%, si | EN_3 |
| GOBP_REGULATION_OF_ | GOBP_REGULATION_OF_TELOMERE_MAINT | 61 | -0.39477 | -1.52183 | 0.031088082 | 0.23753473 | 1 | 7334 | tags=46%, list=31%, si | EN_3 |
| GOBP_ORGANELLE_DISA! | GOBP_ORGANELLE_DISASSEMBLY | 155 | -0.35249 | -1.52162 | 0.0068259384 | 0.2373232 | 1 | 4631 | tags=26%, list=20%, si | EN_3 |
| GOBP_POSITIVE_REGULA | GOBP_POSITIVE_REGULATION_OF_TRANSL/ | 125 | -0.35698 | -1.52134 | 0.013002364 | 0.23727114 | 1 | 6448 | tags=42%, list=27%, si | EN_3 |
| GOBP_EMBRYONIC_BODY | GOBP_EMBRYONIC_BODY_MORPHOGENESI | 15 | -0.53445 | -1.5207 | 0.04368175 | 0.23772994 | 1 | 6795 | tags=53%, list=29%, si | EN_3 |
| GOBP_NEGATIVE_REGUL | GOBP_NEGATIVE_REGULATION_OF_PROTE( | 69 | -0.38955 | -1.52045 | 0.024600247 | 0.23757328 | 1 | 4764 | tags=28%, list=20%, si | EN_3 |
| GOBP_VIRAL_RELEASE_FI | GOBP_VIRAL_RELEASE_FROM_HOST_CELL | 24 | -0.47774 | -1.51953 | 0.04261796 | 0.23868157 | 1 | 4910 | tags=46%, list=21%, si | EN_3 |
| GOBP_POSITIVE_REGULA | GOBP_POSITIVE_REGULATION_OF_T_CELL_ | 19 | -0.51316 | -1.51946 | 0.05648855 | 0.2382188 | 1 | 6364 | tags=63%, list=27%, si | EN_3 |
| GOBP_PYRIMIDINE_CONT | GOBP_PYRIMIDINE_CONTAINING_COMPOU | 32 | -0.45315 | -1.518 | 0.039244186 | 0.24036004 | 1 | 2950 | tags=28%, list=12%, si | EN_3 |
| GOBP_GOLGI_VESICLE_TI | GOBP_GOLGI_VESICLE_TRANSPORT | 276 | -0.33022 | -1.51761 | 0.0010416667 | 0.2404285 | 1 | 6901 | tags=41%, list=29%, si | EN_3 |
| GOBP_NEGATIVE_REGUL | GOBP_NEGATIVE_REGULATION_OF_CATAB( | 339 | -0.32523 | -1.5175 | 0.0020942409 | 0.24003296 | 1 | 4849 | tags=27%, list=21%, si | EN_3 |
| GOBP_PROTEIN_ACYLATI | GOBP_PROTEIN_ACYLATION | 94 | -0.36827 | -1.51725 | 0.0094451 | 0.23993413 | 1 | 5011 | tags=30%, list=21%, si | EN_3 |
| GOBP_PROTEIN_POLYUBI | GOBP_PROTEIN_POLYUBIQUITINATION | 250 | -0.3356 | -1.51663 | 0.0021574972 | 0.2404998 | 1 | 4872 | tags=26%, list=21%, si | EN_3 |
| GOBP_NEGATIVE_REGUL | GOBP_NEGATIVE_REGULATION_OF_VIRAL_I | 36 | -0.43215 | -1.51633 | 0.041608877 | 0.2404609 | 1 | 4955 | tags=44%, list=21%, si | EN_3 |
| GOBP_ALCOHOL_BIOSYN | GOBP_ALCOHOL_BIOSYNTHETIC_PROCESS | 129 | -0.35738 | -1.51606 | 0.007075472 | 0.24033017 | 1 | 4246 | tags=31%, list=18%, si | EN_3 |
| GOBP_AMINOGLYCAN_BI | GOBP_AMINOGLYCAN_BIOSYNTHETIC_PRO( | 60 | -0.4013 | -1.51595 | 0.02486911 | 0.2399784 | 1 | 4203 | tags=33%, list=18%, si | EN_3 |
| GOBP_REGULATION_OF_ | GOBP_REGULATION_OF_SENSORY_PERCEPT | 18 | -0.5231 | -1.5155 | 0.05808477 | 0.24021074 | 1 | 2613 | tags=33%, list=11%, si | EN_3 |
| GOBP_GLIAL_CELL_ACTIV | GOBP_GLIAL_CELL_ACTIVATION | 59 | -0.39823 | -1.51343 | 0.028534371 | 0.24335133 | 1 | 3138 | tags=24%, list=13%, si | EN_3 |
| GOBP_VIRAL_PROCESS | GOBP_VIRAL_PROCESS | 303 | -0.32431 | -1.51312 | 0 | 0.24335207 | 1 | 5666 | tags=33%, list=24%, si | EN_3 |
| GOBP_LENS_MORPHOGE | GOBP_LENS_MORPHOGENESIS_IN_CAMERA | 26 | -0.46087 | -1.51265 | 0.045123726 | 0.24357457 | 1 | 3375 | tags=27%, list=14%, si | EN_3 |
| GOBP_NEGATIVE_REGUL | GOBP_NEGATIVE_REGULATION_OF_VASCUI | 33 | -0.44514 | -1.51258 | 0.043296088 | 0.2431287 | 1 | 4036 | tags=27%, list=17%, si | EN_3 |
| GOBP_POSITIVE_REGULA | GOBP_POSITIVE_REGULATION_OF_DEFENSE | 31 | -0.45637 | -1.51196 | 0.038356163 | 0.24366309 | 1 | 4048 | tags=32%, list=17%, si | EN_3 |
| HERPES_SIMPLEX_VIRUS_ | HERPES_SIMPLEX_VIRUS_1_INFECTION | 373 | -0.31868 | -1.51148 | 0 | 0.24395718 | 1 | 4494 | tags=28%, list=19%, si | EN_3 |
| GOBP_ALCOHOL_METAB( | GOBP_ALCOHOL_METABOLIC_PROCESS | 308 | -0.32754 | -1.5099 | 0.0010537407 | 0.24636187 | 1 | 4974 | tags=31%, list=21%, si | EN_3 |
| GOBP_PHOSPHOLIPID_DI | GOBP_PHOSPHOLIPID_DEPHOSPHORYLATIO | 43 | -0.42309 | -1.50979 | 0.04295302 | 0.2459868 | 1 | 5309 | tags=28%, list=22%, si | EN_3 |
| GOBP_ICOSANOID_BIOSY | GOBP_ICOSANOID_BIOSYNTHETIC_PROCESS | 43 | -0.41863 | -1.5096 | 0.04109589 | 0.24582034 | 1 | 1595 | tags=19%, list=7%, sig | EN_3 |
| GOBP_NEGATIVE_REGUL | GOBP_NEGATIVE_REGULATION_OF_STEM_( | 28 | -0.45692 | -1.50935 | 0.037845705 | 0.24572833 | 1 | 7701 | tags=54%, list=33%, si | EN_3 |
| GOBP_REGULATION_OF_ | GOBP_REGULATION_OF_INTRINSIC_APOPT( | 34 | -0.43374 | -1.50915 | 0.059914406 | 0.24550503 | 1 | 5511 | tags=41%, list=23%, si | EN_3 |
| GOBP_REGULATION_OF_ | GOBP_REGULATION_OF_BMP_SIGNALING_I | 96 | -0.36435 | -1.50861 | 0.01726264 | 0.24594098 | 1 | 4394 | tags=29%, list=19%, si | EN_3 |
| GOBP_POSITIVE_REGULA | GOBP_POSITIVE_REGULATION_OF_UBIQUIT | 93 | -0.36723 | -1.50851 | 0.017073171 | 0.24554794 | 1 | 7160 | tags=42%, list=30%, si | EN_3 |

| | | | | | | | | | | |
|---|---|---|---|---|---|---|---|---|---|---|
| GOBP_PROTEIN_LOCALIZ | GOBP_PROTEIN_LOCALIZATION_TO_CELL_C | 24 | -0.4716 | -1.50727 | 0.04411765 | 0.24712995 | 1 | 4507 | tags=38%, list=19%, si | EN_3 |
| GOBP_SYNAPTIC_VESICLE | GOBP_SYNAPTIC_VESICLE_LUMEN_ACIDIFIC | 19 | -0.50245 | -1.50696 | 0.060029283 | 0.24716334 | 1 | 6050 | tags=53%, list=26%, si | EN_3 |
| GOBP_NEGATIVE_REGUL | GOBP_NEGATIVE_REGULATION_OF_CYCLIN | 25 | -0.46219 | -1.50693 | 0.057991512 | 0.24665879 | 1 | 3073 | tags=32%, list=13%, si | EN_3 |
| GOBP_REGULATION_OF_ | GOBP_REGULATION_OF_ACTIN_NUCLEATIO | 35 | -0.44216 | -1.50686 | 0.044295304 | 0.2462283 | 1 | 4869 | tags=46%, list=21%, si | EN_3 |
| GOBP_SNRNA_PROCESSI | GOBP_SNRNA_PROCESSING | 24 | -0.47626 | -1.50664 | 0.0637037 | 0.2460326 | 1 | 3629 | tags=42%, list=15%, si | EN_3 |
| GOBP_POSITIVE_REGULA | GOBP_POSITIVE_REGULATION_OF_OSTEOBI | 74 | -0.38028 | -1.5062 | 0.026717557 | 0.24629228 | 1 | 6656 | tags=50%, list=28%, si | EN_3 |
| GOBP_ANTIVIRAL_INNAT | GOBP_ANTIVIRAL_INNATE_IMMUNE_RESPO | 47 | -0.41422 | -1.50575 | 0.03133515 | 0.24651633 | 1 | 3708 | tags=19%, list=16%, si | EN_3 |
| GOBP_GAMMA_AMINOB | GOBP_GAMMA_AMINOBUTYRIC_ACID_SIGN | 30 | -0.44841 | -1.50523 | 0.04454023 | 0.24693927 | 1 | 2360 | tags=23%, list=10%, si | EN_3 |
| GOBP_NEGATIVE_REGUL | GOBP_NEGATIVE_REGULATION_OF_PROTEI | 87 | -0.3765 | -1.50427 | 0.02836005 | 0.24811059 | 1 | 4997 | tags=32%, list=21%, si | EN_3 |
| GOBP_RESPONSE_TO_SA | GOBP_RESPONSE_TO_SALT_STRESS | 20 | -0.49816 | -1.50375 | 0.06505295 | 0.24854565 | 1 | 7050 | tags=55%, list=30%, si | EN_3 |
| GOBP_ERYTHROCYTE_DE | GOBP_ERYTHROCYTE_DEVELOPMENT | 50 | -0.40143 | -1.50373 | 0.04032258 | 0.24803324 | 1 | 5475 | tags=42%, list=23%, si | EN_3 |
| GOBP_AXONAL_TRANSPO | GOBP_AXONAL_TRANSPORT | 69 | -0.38813 | -1.50333 | 0.026515152 | 0.24817573 | 1 | 7058 | tags=49%, list=30%, si | EN_3 |
| GOBP_POSITIVE_REGULA | GOBP_POSITIVE_REGULATION_OF_PROTEIN | 107 | -0.35554 | -1.50271 | 0.022249691 | 0.24877566 | 1 | 6007 | tags=35%, list=25%, si | EN_3 |
| HEPATITIS_C | HEPATITIS_C | 138 | -0.34402 | -1.50251 | 0.014789534 | 0.24861735 | 1 | 4325 | tags=23%, list=18%, si | EN_3 |
| GOBP_RNA_SPLICING_VI | GOBP_RNA_SPLICING_VIA_TRANSESTERIFIC | 274 | -0.3258 | -1.50237 | 0.0043243244 | 0.24829258 | 1 | 6896 | tags=37%, list=29%, si | EN_3 |
| GOBP_NEGATIVE_REGUL | GOBP_NEGATIVE_REGULATION_OF_PROTEI | 80 | -0.37057 | -1.50221 | 0.033333335 | 0.24810436 | 1 | 4764 | tags=34%, list=20%, si | EN_3 |
| GOBP_RECOMBINATIONA | GOBP_RECOMBINATIONAL_REPAIR | 177 | -0.33672 | -1.50194 | 0.0033370412 | 0.24800023 | 1 | 4486 | tags=28%, list=19%, si | EN_3 |
| GOBP_REGULATION_OF_ | GOBP_REGULATION_OF_POST_TRANSLATIO | 242 | -0.32949 | -1.50129 | 0.0032119914 | 0.24858256 | 1 | 5325 | tags=31%, list=23%, si | EN_3 |
| GOBP_CELLULAR_RESPON | GOBP_CELLULAR_RESPONSE_TO_CADMIUM | 24 | -0.46899 | -1.50093 | 0.05882353 | 0.24874145 | 1 | 2690 | tags=29%, list=11%, si | EN_3 |
| GOBP_RESPONSE_TO_INT | GOBP_RESPONSE_TO_INTERLEUKIN_4 | 35 | -0.43486 | -1.50081 | 0.04434907 | 0.2484239 | 1 | 2114 | tags=23%, list=9%, sig | EN_3 |
| GOBP_GLYCEROL_ETHER_ | GOBP_GLYCEROL_ETHER_METABOLIC_PRO | 19 | -0.49247 | -1.50012 | 0.055205047 | 0.24904665 | 1 | 943 | tags=16%, list=4%, sig | EN_3 |
| GOBP_NEGATIVE_REGUL | GOBP_NEGATIVE_REGULATION_OF_INSULIN | 17 | -0.51358 | -1.49876 | 0.05919003 | 0.24973628 | 1 | 4819 | tags=41%, list=20%, si | EN_3 |
| GOBP_MITOCHONDRIAL_ | GOBP_MITOCHONDRIAL_CYTOCHROME_C_ | 19 | -0.49689 | -1.49867 | 0.06481481 | 0.2493629 | 1 | 3361 | tags=37%, list=14%, si | EN_3 |
| GOBP_CENTRAL_NERVOU | GOBP_CENTRAL_NERVOUS_SYSTEM_NEURO | 41 | 0.51739 | 2.227364 | 0 | 0.07723793 | 0.078 | 2571 | tags=41%, list=11%, si | EN_3 |
| GOBP_TRNA_METHYLATI | GOBP_TRNA_METHYLATION | 39 | 0.508841 | 2.169182 | 0 | 0.08749803 | 0.172 | 2728 | tags=44%, list=12%, si | EN_3 |
| GOBP_REGULATION_OF_ | GOBP_REGULATION_OF_POSTSYNAPTIC_NE | 17 | 0.64755 | 2.149957 | 0 | 0.07264409 | 0.205 | 1247 | tags=35%, list=5%, sig | EN_3 |
| GOBP_CENTRAL_NERVOU | GOBP_CENTRAL_NERVOUS_SYSTEM_PROJE | 30 | 0.513449 | 2.006266 | 0 | 0.21929404 | 0.623 | 2571 | tags=47%, list=11%, si | EN_3 |
| GOBP_POSITIVE_REGULA | GOBP_POSITIVE_REGULATION_OF_CATION_ | 43 | 0.460651 | 1.936475 | 0.0036363637 | 0.23774731 | 0.847 | 2072 | tags=30%, list=9%, sig | EN_3 |
| GOBP_POSITIVE_REGULA | GOBP_POSITIVE_REGULATION_OF_TRANSP | 81 | 0.389874 | 1.928059 | 0 | 0.22465922 | 0.865 | 2072 | tags=23%, list=9%, sig | EN_3 |
| GOBP_REGULATION_OF_ | GOBP_REGULATION_OF_CILIUM_BEAT_FRE | 15 | 0.617373 | 1.920276 | 0.005449591 | 0.21426791 | 0.88 | 2848 | tags=47%, list=12%, si | EN_3 |
| GOBP_FOREBRAIN_CELL_ | GOBP_FOREBRAIN_CELL_MIGRATION | 66 | 0.401423 | 1.91238 | 0.004524887 | 0.20619544 | 0.895 | 2771 | tags=29%, list=12%, si | EN_3 |
| OLFACTORY_TRANSDUCT | OLFACTORY_TRANSDUCTION | 186 | 0.342764 | 1.911174 | 0 | 0.18926364 | 0.897 | 8224 | tags=74%, list=35%, si | EN_3 |
| GOBP_COCHLEA_MORPH | GOBP_COCHLEA_MORPHOGENESIS | 27 | -0.78057 | -1.80222 | 0 | 0.14664592 | 0.119 | 1829 | tags=26%, list=8%, sig | EN_4 |
| GOBP_COCHLEA_DEVELO | GOBP_COCHLEA_DEVELOPMENT | 51 | -0.69359 | -1.79716 | 0 | 0.08951801 | 0.146 | 1829 | tags=20%, list=8%, sig | EN_4 |
| GOBP_TRANSLATION_AT_ | GOBP_TRANSLATION_AT_SYNAPSE | 48 | -0.80905 | -4.41609 | 0.0 | 0 | 0 | 4101 | tags=94%, list=17%, si | MM_4 |
| RIBOSOME | RIBOSOME | 128 | -0.55806 | -3.59932 | 0.0 | 0 | 0 | 4101 | tags=69%, list=17%, si | MM_4 |
| GOBP_CYTOPLASMIC_TR | GOBP_CYTOPLASMIC_TRANSLATION | 143 | -0.5248 | -3.58726 | 0.0 | 0 | 0 | 4193 | tags=68%, list=18%, si | MM_4 |
| GOBP_ANTIGEN_PROCES | GOBP_ANTIGEN_PROCESSING_AND_PRESEN | 28 | -0.62739 | -3.00443 | 0.0 | 0 | 0 | 3833 | tags=68%, list=16%, si | MM_4 |
| GOBP_ANTIGEN_PROCES | GOBP_ANTIGEN_PROCESSING_AND_PRESEN | 34 | -0.59666 | -2.99658 | 0.0 | 0 | 0 | 3833 | tags=65%, list=16%, si | MM_4 |
| SPLICEOSOME | SPLICEOSOME | 127 | -0.4428 | -2.97206 | 0.0 | 0 | 0 | 11552 | tags=98%, list=49%, si | MM_4 |
| GOBP_ANTIGEN_PROCES | GOBP_ANTIGEN_PROCESSING_AND_PRESEN | 105 | -0.41599 | -2.71922 | 0.0 | 0.002033818 | 0.004 | 3970 | tags=46%, list=17%, si | MM_4 |
| GOBP_ANTIGEN_PROCES | GOBP_ANTIGEN_PROCESSING_AND_PRESEN | 24 | -0.60021 | -2.68655 | 0.0 | 0.001779591 | 0.004 | 3833 | tags=63%, list=16%, si | MM_4 |
| GOBP_ANTIGEN_PROCES | GOBP_ANTIGEN_PROCESSING_AND_PRESEN | 20 | -0.64198 | -2.67349 | 0.0 | 0.002007784 | 0.005 | 3833 | tags=70%, list=16%, si | MM_4 |
| GOBP_REGULATION_OF_ | GOBP_REGULATION_OF_ANTIGEN_PROCESS | 21 | -0.61488 | -2.66952 | 0.0 | 0.001807006 | 0.005 | 2693 | tags=52%, list=11%, si | MM_4 |
| GOBP_REGULATION_OF_ | GOBP_REGULATION_OF_B_CELL_DIFFERENT | 37 | -0.49213 | -2.60117 | 0.0 | 0.003254861 | 0.01 | 3782 | tags=51%, list=16%, si | MM_4 |
| GOBP_NUCLEAR_TRANSC | GOBP_NUCLEAR_TRANSCRIBED_MRNA_POL | 26 | -0.55058 | -2.58433 | 0.0 | 0.00386349 | 0.013 | 10625 | tags=100%, list=45%, | MM_4 |
| LYSOSOME | LYSOSOME | 118 | -0.37461 | -2.5382 | 0.0 | 0.005520863 | 0.02 | 3833 | tags=38%, list=16%, si | MM_4 |
| GOBP_LYMPHOID_PROGE | GOBP_LYMPHOID_PROGENITOR_CELL_DIFF | 24 | -0.55815 | -2.48133 | 0.0 | 0.009396956 | 0.035 | 1343 | tags=25%, list=6%, sig | MM_4 |
| GOBP_NEGATIVE_REGUL | GOBP_NEGATIVE_REGULATION_OF_LEUKO | 46 | -0.45215 | -2.46622 | 0.0 | 0.009720919 | 0.039 | 2579 | tags=39%, list=11%, si | MM_4 |
| GOBP_NEGATIVE_REGUL | GOBP_NEGATIVE_REGULATION_OF_MONO | 28 | -0.51465 | -2.454 | 0.0 | 0.010445421 | 0.045 | 2579 | tags=43%, list=11%, si | MM_4 |
| GOBP_MEMBRANE_INVA | GOBP_MEMBRANE_INVAGINATION | 70 | -0.39609 | -2.44517 | 0.0 | 0.010449828 | 0.048 | 885 | tags=16%, list=4%, sig | MM_4 |
| GOBP_RIBOSOME_ASSEM | GOBP_RIBOSOME_ASSEMBLY | 60 | -0.41286 | -2.41435 | 0.0 | 0.012625973 | 0.062 | 4758 | tags=62%, list=20%, si | MM_4 |
| GOBP_PHAGOCYTOSIS_E | GOBP_PHAGOCYTOSIS_ENGULFMENT | 53 | -0.41965 | -2.39103 | 0.0 | 0.013449339 | 0.07 | 868 | tags=17%, list=4%, sig | MM_4 |
| GOBP_TRANSLATIONAL_ | GOBP_TRANSLATIONAL_ELONGATION | 58 | -0.42315 | -2.38419 | 0.0 | 0.012776872 | 0.07 | 4301 | tags=48%, list=18%, si | MM_4 |
| GOBP_PROTON_MOTIVE_ | GOBP_PROTON_MOTIVE_FORCE_DRIVEN_A | 52 | -0.42702 | -2.38167 | 0.0 | 0.012493334 | 0.072 | 11098 | tags=98%, list=47%, si | MM_4 |
| GLYCOSYLPHOSPHATIDYL | GLYCOSYLPHOSPHATIDYLINOSITOL_(GPI)-AN | 24 | -0.51669 | -2.3419 | 0.0 | 0.017846338 | 0.104 | 2934 | tags=46%, list=12%, si | MM_4 |
| GOBP_RESCUE_OF_STALL | GOBP_RESCUE_OF_STALLED_RIBOSOME | 25 | -0.51405 | -2.32094 | 0.0 | 0.02002533 | 0.121 | 11486 | tags=100%, list=49%, | MM_4 |
| GOBP_RETROGRADE_VES | GOBP_RETROGRADE_VESICLE_MEDIATED_T | 45 | -0.41915 | -2.30615 | 0.0 | 0.021102587 | 0.133 | 11606 | tags=96%, list=49%, si | MM_4 |
| GOBP_ENDOPLASMIC_RE | GOBP_ENDOPLASMIC_RETICULUM_TO_CYT | 27 | -0.49956 | -2.28898 | 0.0 | 0.023228144 | 0.151 | 9519 | tags=96%, list=40%, si | MM_4 |
| GOBP_MODULATION_BY_ | GOBP_MODULATION_BY_HOST_OF_VIRAL_ | 22 | -0.51378 | -2.27263 | 0.0058479533 | 0.024659103 | 0.166 | 11491 | tags=100%, list=49%, | MM_4 |
| GOBP_GLYCOLYTIC_PRO | GOBP_GLYCOLYTIC_PROCESS_THROUGH_GI | 20 | -0.53285 | -2.2636 | 0.0 | 0.025886081 | 0.181 | 11040 | tags=100%, list=47%, | MM_4 |
| GOBP_REGULATION_OF_ | GOBP_REGULATION_OF_TRANSLATIONAL_I | 68 | -0.38311 | -2.24399 | 0.0 | 0.02875329 | 0.209 | 4502 | tags=49%, list=19%, si | MM_4 |
| GOBP_NEGATIVE_REGUL | GOBP_NEGATIVE_REGULATION_OF_TYPE_2 | 15 | -0.60102 | -2.23676 | 0.0 | 0.028880926 | 0.216 | 1031 | tags=33%, list=4%, sig | MM_4 |
| GOBP_REGULATION_OF_ | GOBP_REGULATION_OF_MACROPHAGE_MI | 47 | -0.41664 | -2.2182 | 0.0 | 0.031323485 | 0.237 | 2721 | tags=32%, list=12%, si | MM_4 |
| GOBP_REGULATION_OF_ | GOBP_REGULATION_OF_NUCLEAR_TRANSC | 15 | -0.56617 | -2.2158 | 0.0 | 0.031005735 | 0.242 | 10251 | tags=100%, list=43%, | MM_4 |
| GOBP_ANTIGEN_PROCES | GOBP_ANTIGEN_PROCESSING_AND_PRESEN | 59 | -0.37221 | -2.21539 | 0.0 | 0.030140739 | 0.243 | 3970 | tags=47%, list=17%, si | MM_4 |
| GOBP_RETICULOPHAGY | GOBP_RETICULOPHAGY | 19 | -0.51193 | -2.20153 | 0.0 | 0.03191246 | 0.257 | 11533 | tags=100%, list=49%, | MM_4 |
| GOBP_TRANSLATIONAL_I | GOBP_TRANSLATIONAL_INITIATION | 112 | -0.35074 | -2.19742 | 0.0 | 0.032034736 | 0.263 | 4502 | tags=47%, list=19%, si | MM_4 |
| GOBP_CGAS_STING_SIGN | GOBP_CGAS_STING_SIGNALING_PATHWAY | 16 | -0.58675 | -2.19649 | 0.0052083335 | 0.031630713 | 0.266 | 9766 | tags=100%, list=41%, | MM_4 |
| GOBP_RESPONSE_TO_NI | GOBP_RESPONSE_TO_NITRIC_OXIDE | 17 | -0.53505 | -2.19349 | 0.005 | 0.031234166 | 0.27 | 924 | tags=24%, list=4%, sig | MM_4 |
| GOBP_LATE_ENDOSOME_ | GOBP_LATE_ENDOSOME_TO_LYSOSOME_T | 19 | -0.51227 | -2.18449 | 0.0 | 0.03221878 | 0.284 | 11525 | tags=100%, list=49%, | MM_4 |
| GOBP_PROTEIN_DNA_CO | GOBP_PROTEIN_DNA_COMPLEX_DISASSEM | 19 | -0.51049 | -2.18188 | 0.0 | 0.032029454 | 0.29 | 11567 | tags=100%, list=49%, | MM_4 |
| GOBP_NATURAL_KILLER_ | GOBP_NATURAL_KILLER_CELL_PROLIFERATI | 16 | -0.54626 | -2.17523 | 0.0 | 0.032293804 | 0.299 | 458 | tags=25%, list=2%, sig | MM_4 |
| GOBP_NUCLEAR_MEMBR | GOBP_NUCLEAR_MEMBRANE_REASSEMBLY | 20 | -0.51229 | -2.17411 | 0.0 | 0.031663693 | 0.3 | 11525 | tags=100%, list=49%, | MM_4 |
| GOBP_RESPONSE_TO_ST | GOBP_RESPONSE_TO_STEROL_DEPLETION | 15 | -0.57115 | -2.17184 | 0.0 | 0.031585347 | 0.305 | 4917 | tags=73%, list=21%, si | MM_4 |
| GOBP_VIRAL_BUDDING | GOBP_VIRAL_BUDDING | 18 | -0.51225 | -2.16595 | 0.0057471264 | 0.032230325 | 0.316 | 11525 | tags=100%, list=49%, | MM_4 |
| GOBP_POSITIVE_REGULA | GOBP_POSITIVE_REGULATION_OF_FIBROBL | 24 | -0.48039 | -2.16314 | 0.0 | 0.032129895 | 0.32 | 435 | tags=21%, list=2%, sig | MM_4 |
| GOBP_ERAD_PATHWAY | GOBP_ERAD_PATHWAY | 108 | -0.32044 | -2.16233 | 0.0 | 0.03172191 | 0.324 | 13448 | tags=94%, list=57%, si | MM_4 |
| GOBP_POSITIVE_REGULA | GOBP_POSITIVE_REGULATION_OF_MACRO | 25 | -0.46272 | -2.162 | 0.0 | 0.031096557 | 0.324 | 272 | tags=20%, list=1%, sig | MM_4 |
| GOBP_GPI_ANCHOR_ME | GOBP_GPI_ANCHOR_METABOLIC_PROCESS | 33 | -0.42326 | -2.15375 | 0.0 | 0.0315797 | 0.335 | 2934 | tags=42%, list=12%, si | MM_4 |
| GOBP_TAIL_ANCHORED_ | GOBP_TAIL_ANCHORED_MEMBRANE_PROT | 15 | -0.55516 | -2.15353 | 0.005181347 | 0.03090779 | 0.335 | 10511 | tags=100%, list=45%, | MM_4 |
| GOBP_ACTIN_FILAMENT_ | GOBP_ACTIN_FILAMENT_SEVERING | 16 | -0.5242 | -2.14809 | 0.0050251256 | 0.030922988 | 0.342 | 11242 | tags=100%, list=48%, | MM_4 |

| | | | | | | | | | | |
|---|---|---|---|---|---|---|---|---|---|---|
| GOBP_MELANOSOME_AS | GOBP_MELANOSOME_ASSEMBLY | 19 | -0.50444 | -2.14775 | 0.0051020407 | 0.030291906 | 0.342 | 1789 | tags=37%, list=8%, sig | MM_4 |
| GOBP_POSITIVE_REGULA | GOBP_POSITIVE_REGULATION_OF_DEFENSE | 31 | -0.43479 | -2.14752 | 0.0 | 0.029686067 | 0.342 | 1866 | tags=26%, list=8%, sig | MM_4 |
| GOBP_REGULATION_OF_ | GOBP_REGULATION_OF_FIBROBLAST_MIGR | 47 | -0.40503 | -2.14673 | 0.0 | 0.029378781 | 0.345 | 2119 | tags=28%, list=9%, sig | MM_4 |
| GOBP_REGULATION_OF_ | GOBP_REGULATION_OF_MACROPHAGE_CH | 32 | -0.42249 | -2.14383 | 0.0 | 0.029653436 | 0.352 | 272 | tags=16%, list=1%, sig | MM_4 |
| GOBP_REGULATION_OF_ | GOBP_REGULATION_OF_TRANSLATION_IN_ | 24 | -0.48647 | -2.14318 | 0.0 | 0.029232092 | 0.354 | 4142 | tags=50%, list=18%, si | MM_4 |
| GOBP_RIBOSOMAL_LARG | GOBP_RIBOSOMAL_LARGE_SUBUNIT_BIOGE | 64 | -0.36158 | -2.13756 | 0.0 | 0.02967121 | 0.364 | 11616 | tags=94%, list=49%, si | MM_4 |
| PROTEASOME | PROTEASOME | 45 | -0.3878 | -2.13636 | 0.0 | 0.029318659 | 0.365 | 10843 | tags=93%, list=46%, si | MM_4 |
| GOBP_POSITIVE_REGULA | GOBP_POSITIVE_REGULATION_OF_TRANSLA | 125 | -0.32361 | -2.1328 | 0.0 | 0.029685738 | 0.376 | 4366 | tags=47%, list=18%, si | MM_4 |
| GOBP_MRNA_PROCESSIN | GOBP_MRNA_PROCESSING | 459 | -0.23582 | -2.1075 | 0.0 | 0.03391563 | 0.422 | 11661 | tags=88%, list=49%, si | MM_4 |
| GOBP_P_BODY_ASSEMBL | GOBP_P_BODY_ASSEMBLY | 20 | -0.51339 | -2.09747 | 0.006451613 | 0.035469245 | 0.442 | 11499 | tags=100%, list=49%, | MM_4 |
| GOBP_NEGATIVE_REGUL | GOBP_NEGATIVE_REGULATION_OF_RHO_P | 25 | -0.45138 | -2.09436 | 0.0058479533 | 0.03593461 | 0.452 | 2434 | tags=32%, list=10%, si | MM_4 |
| GOBP_PROTEIN_LOCALIZ | GOBP_PROTEIN_LOCALIZATION_TO_PHAGO | 16 | -0.51362 | -2.08433 | 0.0 | 0.037866935 | 0.475 | 2475 | tags=38%, list=10%, si | MM_4 |
| GOBP_REGULATION_OF_ | GOBP_REGULATION_OF_RRNA_PROCESSING | 16 | -0.53369 | -2.08306 | 0.009708738 | 0.03764582 | 0.481 | 11018 | tags=100%, list=47%, | MM_4 |
| GOBP_RNA_SPLICING | GOBP_RNA_SPLICING | 417 | -0.25365 | -2.07505 | 0.0 | 0.038794857 | 0.499 | 11661 | tags=88%, list=49%, si | MM_4 |
| GOBP_PEROXISOMAL_TR | GOBP_PEROXISOMAL_TRANSPORT | 21 | -0.48374 | -2.06759 | 0.0 | 0.039967302 | 0.514 | 12199 | tags=100%, list=52%, | MM_4 |
| GOBP_REGULATION_OF_ | GOBP_REGULATION_OF_NATURAL_KILLER_ | 55 | -0.37029 | -2.06495 | 0.0 | 0.03997096 | 0.52 | 637 | tags=18%, list=3%, sig | MM_4 |
| GOBP_RESPONSE_TO_TY | GOBP_RESPONSE_TO_TYPE_II_INTERFERON | 122 | -0.2989 | -2.05826 | 0.0 | 0.041633703 | 0.537 | 3004 | tags=26%, list=13%, si | MM_4 |
| GOBP_RNA_SPLICING_VI | GOBP_RNA_SPLICING_VIA_TRANSESTERIFIC | 274 | -0.26393 | -2.05646 | 0.0 | 0.041491844 | 0.541 | 11661 | tags=89%, list=49%, si | MM_4 |
| GOBP_ESTABLISHMENT_ | GOBP_ESTABLISHMENT_OF_PROTEIN_LOCA | 20 | -0.48372 | -2.0523 | 0.0 | 0.04219672 | 0.551 | 12199 | tags=100%, list=52%, | MM_4 |
| GOBP_NEGATIVE_REGUL | GOBP_NEGATIVE_REGULATION_OF_TISSUE_ | 27 | -0.44377 | -2.04608 | 0.006756757 | 0.04375086 | 0.569 | 2760 | tags=48%, list=12%, si | MM_4 |
| GOBP_POSITIVE_REGULA | GOBP_POSITIVE_REGULATION_OF_NATURA | 26 | -0.44361 | -2.04506 | 0.012345679 | 0.04327199 | 0.569 | 3672 | tags=54%, list=16%, si | MM_4 |
| GOBP_RIBOSOMAL_SMAI | GOBP_RIBOSOMAL_SMALL_SUBUNIT_BIOGI | 101 | -0.30405 | -2.04167 | 0.0 | 0.043610323 | 0.574 | 4106 | tags=45%, list=17%, si | MM_4 |
| RNA_TRANSPORT | RNA_TRANSPORT | 154 | -0.28428 | -2.04055 | 0.0 | 0.043448035 | 0.575 | 11602 | tags=90%, list=49%, si | MM_4 |
| GOBP_ENDOSOME_TO_L | GOBP_ENDOSOME_TO_LYSOSOME_TRANSP | 16 | -0.51221 | -2.04005 | 0.0 | 0.043149106 | 0.578 | 11525 | tags=100%, list=49%, | MM_4 |
| GOBP_NUCLEOTIDE_SUG | GOBP_NUCLEOTIDE_SUGAR_BIOSYNTHETIC_ | 17 | -0.51346 | -2.03886 | 0.005586592 | 0.04284487 | 0.58 | 11496 | tags=100%, list=49%, | MM_4 |
| GOBP_KILLING_OF_CELLS | GOBP_KILLING_OF_CELLS_OF_ANOTHER_OI | 71 | -0.3366 | -2.02635 | 0.0 | 0.046268765 | 0.618 | 708 | tags=11%, list=3%, sig | MM_4 |
| GOBP_IMMUNOLOGICAL_ | GOBP_IMMUNOLOGICAL_SYNAPSE_FORMA | 18 | -0.49138 | -2.01849 | 0.011494253 | 0.04857337 | 0.637 | 2591 | tags=39%, list=11%, si | MM_4 |
| GOBP_DISRUPTION_OF_A | GOBP_DISRUPTION_OF_ANATOMICAL_STRU | 73 | -0.33311 | -2.01656 | 0.0 | 0.048733864 | 0.64 | 708 | tags=11%, list=3%, sig | MM_4 |
| GOBP_NEGATIVE_REGUL | GOBP_NEGATIVE_REGULATION_OF_LIPID_T | 30 | -0.41131 | -2.00806 | 0.0 | 0.050149217 | 0.654 | 2553 | tags=30%, list=11%, si | MM_4 |
| GOBP_POSITIVE_REGULA | GOBP_POSITIVE_REGULATION_OF_MACROI | 31 | -0.41205 | -2.00655 | 0.0 | 0.04998133 | 0.657 | 272 | tags=16%, list=1%, sig | MM_4 |
| GOBP_RIBONUCLEOPROT | GOBP_RIBONUCLEOPROTEIN_COMPLEX_BIC | 417 | -0.24643 | -1.99177 | 0.0 | 0.054410398 | 0.698 | 11616 | tags=88%, list=49%, si | MM_4 |
| GOBP_NEGATIVE_REGUL | GOBP_NEGATIVE_REGULATION_OF_CELL_M | 16 | -0.48918 | -1.99091 | 0.01058201 | 0.053773608 | 0.699 | 945 | tags=13%, list=4%, sig | MM_4 |
| GOBP_POSITIVE_REGULA | GOBP_POSITIVE_REGULATION_OF_MRNA_ | 95 | -0.31182 | -1.98594 | 0.0 | 0.0550771 | 0.71 | 4723 | tags=47%, list=20%, si | MM_4 |
| GOBP_MEMBRANE_FISSI | GOBP_MEMBRANE_FISSION | 31 | -0.39142 | -1.98485 | 0.0 | 0.05500733 | 0.716 | 12477 | tags=97%, list=53%, si | MM_4 |
| CHEMOKINE_SIGNALING_ | CHEMOKINE_SIGNALING_PATHWAY | 177 | -0.26908 | -1.98443 | 0.0 | 0.054430384 | 0.718 | 1270 | tags=13%, list=5%, sig | MM_4 |
| GOBP_RIBOSOME_DISAS | GOBP_RIBOSOME_DISASSEMBLY | 31 | -0.40533 | -1.9814 | 0.0 | 0.054403614 | 0.723 | 11486 | tags=97%, list=49%, si | MM_4 |
| GOBP_RIBOSOME_BIOGE | GOBP_RIBOSOME_BIOGENESIS | 306 | -0.24949 | -1.967 | 0.0 | 0.059008878 | 0.76 | 11721 | tags=89%, list=50%, si | MM_4 |
| RNA_POLYMERASE | RNA_POLYMERASE | 27 | -0.41612 | -1.96412 | 0.012903226 | 0.05945717 | 0.768 | 2459 | tags=30%, list=10%, si | MM_4 |
| GOBP_PROTEIN_DEGLYC | GOBP_PROTEIN_DEGLYCOSYLATION | 15 | -0.51892 | -1.95942 | 0.0 | 0.06023445 | 0.775 | 11366 | tags=100%, list=48%, | MM_4 |
| GOBP_NEGATIVE_REGUL | GOBP_NEGATIVE_REGULATION_OF_ALPHA_ | 16 | -0.48897 | -1.95518 | 0.004830918 | 0.061142575 | 0.789 | 554 | tags=19%, list=2%, sig | MM_4 |
| GOBP_PLASMA_MEMBRA | GOBP_PLASMA_MEMBRANE_TUBULATION | 19 | -0.47192 | -1.95098 | 0.011627907 | 0.061961107 | 0.794 | 12477 | tags=100%, list=53%, | MM_4 |
| GOBP_NEGATIVE_REGUL | GOBP_NEGATIVE_REGULATION_OF_MULTIC | 17 | -0.4662 | -1.94814 | 0.0 | 0.062449977 | 0.805 | 897 | tags=18%, list=4%, sig | MM_4 |
| GOBP_REGULATION_OF_ | GOBP_REGULATION_OF_DEFENSE_RESPONS | 42 | -0.36866 | -1.94593 | 0.0 | 0.062579095 | 0.807 | 2039 | tags=21%, list=9%, sig | MM_4 |
| GOBP_POSITIVE_REGULA | GOBP_POSITIVE_REGULATION_OF_CHEMOK | 15 | -0.50576 | -1.93961 | 0.0048076925 | 0.064015836 | 0.815 | 918 | tags=40%, list=4%, sig | MM_4 |
| GOBP_AMINO_ACID_ACT | GOBP_AMINO_ACID_ACTIVATION | 26 | -0.40686 | -1.93491 | 0.0065789474 | 0.0651648 | 0.832 | 3136 | tags=38%, list=13%, si | MM_4 |
| GOBP_BASE_EXCISION_R | GOBP_BASE_EXCISION_REPAIR | 41 | -0.36353 | -1.93158 | 0.0 | 0.06564321 | 0.84 | 3601 | tags=39%, list=15%, si | MM_4 |
| GOBP_REGULATION_OF_ | GOBP_REGULATION_OF_INNATE_IMMUNE_ | 427 | -0.20016 | -1.93021 | 0.0 | 0.06562517 | 0.843 | 1876 | tags=16%, list=8%, sig | MM_4 |
| GOBP_VESICLE_TETHERIN | GOBP_VESICLE_TETHERING | 31 | -0.40063 | -1.92968 | 0.0 | 0.065125875 | 0.843 | 11150 | tags=94%, list=47%, si | MM_4 |
| GOBP_MRNA_SPLICE_SIT | GOBP_MRNA_SPLICE_SITE_RECOGNITION | 27 | -0.41145 | -1.9271 | 0.0 | 0.06575928 | 0.848 | 13907 | tags=100%, list=59%, | MM_4 |
| GOBP_RIBOSOMAL_SMAI | GOBP_RIBOSOMAL_SMALL_SUBUNIT_ASSEI | 18 | -0.47346 | -1.92632 | 0.016304348 | 0.06552922 | 0.849 | 3679 | tags=56%, list=16%, si | MM_4 |
| ALZHEIMER_DISEASE | ALZHEIMER_DISEASE | 159 | -0.26561 | -1.92529 | 0.0 | 0.06544407 | 0.852 | 3801 | tags=30%, list=16%, si | MM_4 |
| GOBP_POSITIVE_REGULA | GOBP_POSITIVE_REGULATION_OF_T_CELL_ | 16 | -0.50203 | -1.92451 | 0.0 | 0.06517145 | 0.855 | 11765 | tags=100%, list=50%, | MM_4 |
| GOBP_NATURAL_KILLER_ | GOBP_NATURAL_KILLER_CELL_MEDIATED_I | 73 | -0.31913 | -1.91975 | 0.0 | 0.066513024 | 0.863 | 1719 | tags=23%, list=7%, sig | MM_4 |
| GOBP_VIRION_ASSEMBLY | GOBP_VIRION_ASSEMBLY | 26 | -0.41781 | -1.9195 | 0.006802721 | 0.06596322 | 0.863 | 11525 | tags=96%, list=49%, si | MM_4 |
| GOBP_REGULATION_OF_ | GOBP_REGULATION_OF_TRANSLATION | 329 | -0.24275 | -1.91685 | 0.0 | 0.06662085 | 0.865 | 3957 | tags=35%, list=17%, si | MM_4 |
| GOBP_RNA_EXPORT_FRO | GOBP_RNA_EXPORT_FROM_NUCLEUS | 78 | -0.31879 | -1.91397 | 0.0 | 0.06720161 | 0.87 | 11581 | tags=90%, list=49%, si | MM_4 |
| CHRONIC_MYELOID_LEUI | CHRONIC_MYELOID_LEUKEMIA | 75 | -0.31006 | -1.91261 | 0.0 | 0.06711221 | 0.873 | 3222 | tags=28%, list=14%, si | MM_4 |
| GOBP_CELLULAR_RESPON | GOBP_CELLULAR_RESPONSE_TO_INTERFER | 16 | -0.48307 | -1.91255 | 0.005154639 | 0.066583276 | 0.873 | 3653 | tags=50%, list=15%, si | MM_4 |
| EPSTEIN-BARR_VIRUS_IN | EPSTEIN-BARR_VIRUS_INFECTION | 190 | -0.25791 | -1.90624 | 0.0 | 0.06863488 | 0.884 | 3972 | tags=33%, list=17%, si | MM_4 |
| GOBP_FATTY_ACID_TRAN | GOBP_FATTY_ACID_TRANSMEMBRANE_TRA | 16 | -0.48426 | -1.90436 | 0.010752688 | 0.06874534 | 0.887 | 2476 | tags=25%, list=10%, si | MM_4 |
| GOBP_NEGATIVE_REGUL | GOBP_NEGATIVE_REGULATION_OF_OSTEO | 37 | -0.36591 | -1.90234 | 0.0 | 0.06872902 | 0.889 | 2788 | tags=35%, list=12%, si | MM_4 |
| GOBP_LYMPHOCYTE_MIG | GOBP_LYMPHOCYTE_MIGRATION | 92 | -0.29795 | -1.90207 | 0.0 | 0.06820168 | 0.889 | 1198 | tags=18%, list=5%, sig | MM_4 |
| GOBP_NEGATIVE_REGUL | GOBP_NEGATIVE_REGULATION_OF_SMALL_ | 61 | -0.33767 | -1.90172 | 0.010869565 | 0.06784464 | 0.891 | 2577 | tags=28%, list=11%, si | MM_4 |
| GOBP_REGULATION_OF_ | GOBP_REGULATION_OF_CHOLESTEROL_BIO | 19 | -0.45822 | -1.8945 | 0.0056818184 | 0.06963832 | 0.901 | 2064 | tags=32%, list=9%, sig | MM_4 |
| GOBP_GOLGI_TO_VACUO | GOBP_GOLGI_TO_VACUOLE_TRANSPORT | 21 | -0.43706 | -1.87661 | 0.005952381 | 0.07692811 | 0.931 | 10969 | tags=95%, list=46%, si | MM_4 |
| GOBP_RESPONSE_TO_ZIN | GOBP_RESPONSE_TO_ZINC_ION | 34 | -0.37875 | -1.87639 | 0.0 | 0.07634403 | 0.931 | 2240 | tags=26%, list=9%, sig | MM_4 |
| GOBP_MONOCYTE_DIFFE | GOBP_MONOCYTE_DIFFERENTIATION | 28 | -0.39868 | -1.8752 | 0.007194245 | 0.07602022 | 0.931 | 3792 | tags=43%, list=16%, si | MM_4 |
| GOBP_NEGATIVE_REGUL | GOBP_NEGATIVE_REGULATION_OF_NATUR | 19 | -0.44531 | -1.86446 | 0.010256411 | 0.08088127 | 0.942 | 637 | tags=21%, list=3%, sig | MM_4 |
| GOBP_VESICLE_COATING | GOBP_VESICLE_COATING | 33 | -0.36933 | -1.85831 | 0.0 | 0.08328171 | 0.946 | 2987 | tags=39%, list=13%, si | MM_4 |
| GOBP_RRNA_PROCESSIN | GOBP_RRNA_PROCESSING | 211 | -0.23664 | -1.85727 | 0.0 | 0.08293868 | 0.946 | 11616 | tags=89%, list=49%, si | MM_4 |
| GOBP_CHEMOKINE_C_X_ | GOBP_CHEMOKINE_C_X_C_MOTIF_LIGAND_ | 25 | -0.40855 | -1.85534 | 0.0067114094 | 0.08310656 | 0.947 | 1118 | tags=28%, list=5%, sig | MM_4 |
| GOBP_B_CELL_DIFFEREN | GOBP_B_CELL_DIFFERENTIATION | 151 | -0.276 | -1.85392 | 0.0 | 0.08323271 | 0.949 | 3618 | tags=34%, list=15%, si | MM_4 |
| GOBP_CELL_KILLING | GOBP_CELL_KILLING | 202 | -0.26757 | -1.85261 | 0.0 | 0.08325138 | 0.95 | 1714 | tags=15%, list=7%, sig | MM_4 |
| GOBP_REGULATION_OF_ | GOBP_REGULATION_OF_DEFENSE_RESPONS | 67 | -0.3029 | -1.84758 | 0.0 | 0.084995165 | 0.954 | 2122 | tags=16%, list=9%, sig | MM_4 |
| GOBP_POSITIVE_REGULA | GOBP_POSITIVE_REGULATION_OF_DNA_TE | 59 | -0.33255 | -1.84548 | 0.012658228 | 0.085406914 | 0.958 | 3472 | tags=34%, list=15%, si | MM_4 |
| GOBP_REGULATION_OF_ | GOBP_REGULATION_OF_B_CELL_RECEPTOR | 18 | -0.43182 | -1.8432 | 0.016129032 | 0.08591572 | 0.959 | 3748 | tags=39%, list=16%, si | MM_4 |
| GOBP_PIGMENT_GRANU | GOBP_PIGMENT_GRANULE_ORGANIZATION | 39 | -0.34897 | -1.84018 | 0.0 | 0.086873226 | 0.96 | 1789 | tags=23%, list=8%, sig | MM_4 |
| GOBP_RIBOSOMAL_LARG | GOBP_RIBOSOMAL_LARGE_SUBUNIT_ASSEI | 23 | -0.41249 | -1.82997 | 0.006849315 | 0.09185749 | 0.968 | 4758 | tags=65%, list=20%, si | MM_4 |
| GOBP_REGULATION_OF_ | GOBP_REGULATION_OF_B_CELL_ACTIVATIC | 141 | -0.25972 | -1.82593 | 0.0 | 0.093213856 | 0.969 | 3782 | tags=33%, list=16%, si | MM_4 |
| GOBP_REGULATION_OF_ | GOBP_REGULATION_OF_PATTERN_RECOGN | 157 | -0.25201 | -1.82417 | 0.0 | 0.0934494 | 0.969 | 4010 | tags=33%, list=17%, si | MM_4 |
| GOBP_TRANSFORMING_ | GOBP_TRANSFORMING_GROWTH_FACTOR_ | 35 | -0.36043 | -1.82098 | 0.0 | 0.094625354 | 0.973 | 1368 | tags=26%, list=6%, sig | MM_4 |
| GOBP_TRICARBOXYLIC_A | GOBP_TRICARBOXYLIC_ACID_METABOLIC_P | 16 | -0.45888 | -1.81882 | 0.005076142 | 0.09492225 | 0.974 | 2649 | tags=44%, list=11%, si | MM_4 |

| | | | | | | | | | | |
|---|---|---|---|---|---|---|---|---|---|---|
| GOBP_CELLULAR_RESPON | GOBP_CELLULAR_RESPONSE_TO_TYPE_II_IN | 95 | -0.28987 | -1.81658 | 0.0 | 0.09538221 | 0.975 | 3004 | tags=26%, list=13%, si | MM_4 |
| GOBP_POSITIVE_REGULA | GOBP_POSITIVE_REGULATION_OF_GTPASE_ | 203 | -0.23576 | -1.81459 | 0.0 | 0.0958634 | 0.976 | 2300 | tags=18%, list=10%, si | MM_4 |
| GOBP_NUCLEAR_EXPORT | GOBP_NUCLEAR_EXPORT | 160 | -0.25298 | -1.81231 | 0.0 | 0.09641497 | 0.98 | 11581 | tags=86%, list=49%, si | MM_4 |
| TOLL-LIKE_RECEPTOR_SIG | TOLL-LIKE_RECEPTOR_SIGNALING_PATHWA | 84 | -0.28705 | -1.81082 | 0.0 | 0.09640266 | 0.98 | 2037 | tags=20%, list=9%, sig | MM_4 |
| PROTEIN_PROCESSING_IN | PROTEIN_PROCESSING_IN_ENDOPLASMIC_F | 158 | -0.24967 | -1.8097 | 0.0 | 0.09633947 | 0.981 | 11538 | tags=89%, list=49%, si | MM_4 |
| GOBP_POSITIVE_REGULA | GOBP_POSITIVE_REGULATION_OF_STEM_CI | 50 | -0.32114 | -1.80845 | 0.0 | 0.09611755 | 0.981 | 3567 | tags=38%, list=15%, si | MM_4 |
| GOBP_DEFENSE_RESPON | GOBP_DEFENSE_RESPONSE_TO_VIRUS | 276 | -0.20181 | -1.80219 | 0.0 | 0.099041246 | 0.983 | 2136 | tags=17%, list=9%, sig | MM_4 |
| GOBP_REGULATION_OF_ | GOBP_REGULATION_OF_CELL_KILLING | 116 | -0.25409 | -1.80169 | 0.0 | 0.09880213 | 0.983 | 1963 | tags=17%, list=8%, sig | MM_4 |
| GOBP_RETROGRADE_TRA | GOBP_RETROGRADE_TRANSPORT_ENDOSO | 87 | -0.27956 | -1.80015 | 0.0 | 0.09923081 | 0.985 | 11492 | tags=90%, list=49%, si | MM_4 |
| GOBP_POSITIVE_REGULA | GOBP_POSITIVE_REGULATION_OF_MRNA_N | 131 | -0.27017 | -1.79915 | 0.0 | 0.09922163 | 0.987 | 4723 | tags=42%, list=20%, si | MM_4 |
| GOBP_POST_TRANSCRIPT | GOBP_POST_TRANSCRIPTIONAL_REGULATIC | 480 | -0.22423 | -1.79691 | 0.0 | 0.09990416 | 0.987 | 4242 | tags=36%, list=18%, si | MM_4 |
| GOBP_POSITIVE_REGULA | GOBP_POSITIVE_REGULATION_OF_INTERLEU | 44 | -0.33467 | -1.79424 | 0.0 | 0.10059206 | 0.987 | 1163 | tags=16%, list=5%, sig | MM_4 |
| GOBP_TYPE_I_INTERFERO | GOBP_TYPE_I_INTERFERON_PRODUCTION | 127 | -0.27453 | -1.79402 | 0.0 | 0.10000863 | 0.987 | 2136 | tags=18%, list=9%, sig | MM_4 |
| GOBP_NEGATIVE_REGUL | GOBP_NEGATIVE_REGULATION_OF_CELL_K | 28 | -0.3769 | -1.79345 | 0.005586592 | 0.099609464 | 0.987 | 637 | tags=18%, list=3%, sig | MM_4 |
| GOBP_MRNA_3_END_PR | GOBP_MRNA_3_END_PROCESSING | 41 | -0.33881 | -1.78813 | 0.017094018 | 0.10191479 | 0.987 | 13182 | tags=95%, list=56%, si | MM_4 |
| GOBP_APOPTOTIC_CELL_ | GOBP_APOPTOTIC_CELL_CLEARANCE | 40 | -0.35184 | -1.7819 | 0.0088495575 | 0.104731366 | 0.988 | 1491 | tags=18%, list=6%, sig | MM_4 |
| GOBP_NEGATIVE_REGUL | GOBP_NEGATIVE_REGULATION_OF_BONE_I | 18 | -0.42534 | -1.78071 | 0.013824885 | 0.10459466 | 0.988 | 2760 | tags=50%, list=12%, si | MM_4 |
| GOBP_NAD_CATABOLIC_ | GOBP_NAD_CATABOLIC_PROCESS | 21 | -0.40005 | -1.77956 | 0.021505376 | 0.104488164 | 0.989 | 12287 | tags=95%, list=52%, si | MM_4 |
| ANTIGEN_PROCESSING_A | ANTIGEN_PROCESSING_AND_PRESENTATIO | 67 | -0.30381 | -1.77842 | 0.0 | 0.104501426 | 0.99 | 4080 | tags=45%, list=17%, si | MM_4 |
| GOBP_SPLICEOSOMAL_C | GOBP_SPLICEOSOMAL_COMPLEX_ASSEMBL | 49 | -0.31873 | -1.7736 | 0.0 | 0.10651694 | 0.991 | 13907 | tags=96%, list=59%, si | MM_4 |
| GOBP_REGULATION_OF_ | GOBP_REGULATION_OF_AEROBIC_RESPIRAT | 33 | -0.35754 | -1.77343 | 0.007142857 | 0.1058581 | 0.991 | 11456 | tags=91%, list=49%, si | MM_4 |
| GOBP_PROTEIN_INSERTIO | GOBP_PROTEIN_INSERTION_INTO_ER_MEM | 26 | -0.38651 | -1.77232 | 0.0 | 0.10575899 | 0.991 | 10511 | tags=92%, list=45%, si | MM_4 |
| GOBP_CYTOPLASMIC_SE | GOBP_CYTOPLASMIC_SEQUESTERING_OF_P | 17 | -0.44177 | -1.77214 | 0.0051282053 | 0.10518371 | 0.991 | 4270 | tags=47%, list=18%, si | MM_4 |
| GOBP_MITOCHONDRIAL_ | GOBP_MITOCHONDRIAL_CYTOCHROME_C_ | 19 | -0.41423 | -1.76901 | 0.0052083335 | 0.106216155 | 0.992 | 1706 | tags=32%, list=7%, sig | MM_4 |
| GOBP_POSITIVE_REGULA | GOBP_POSITIVE_REGULATION_OF_CELL_AD | 21 | -0.41314 | -1.76382 | 0.0062111802 | 0.109140076 | 0.993 | 2144 | tags=33%, list=9%, sig | MM_4 |
| GOBP_ARGININE_METAB | GOBP_ARGININE_METABOLIC_PROCESS | 17 | -0.42265 | -1.76356 | 0.020304568 | 0.10866841 | 0.993 | 2119 | tags=35%, list=9%, sig | MM_4 |
| NOTCH_SIGNALING_PATH | NOTCH_SIGNALING_PATHWAY | 47 | -0.31045 | -1.75892 | 0.0 | 0.11089239 | 0.994 | 4051 | tags=38%, list=17%, si | MM_4 |
| GOBP_PROTEIN_TARGETI | GOBP_PROTEIN_TARGETING_TO_VACUOLE | 46 | -0.3208 | -1.75774 | 0.010416667 | 0.11106591 | 0.994 | 3542 | tags=35%, list=15%, si | MM_4 |
| MISMATCH_REPAIR | MISMATCH_REPAIR | 21 | -0.42153 | -1.75439 | 0.01734104 | 0.11221761 | 0.994 | 3601 | tags=43%, list=15%, si | MM_4 |
| GOBP_RUFFLE_ORGANIZA | GOBP_RUFFLE_ORGANIZATION | 54 | -0.31518 | -1.75384 | 0.0 | 0.11182278 | 0.994 | 1895 | tags=22%, list=8%, sig | MM_4 |
| GOBP_ALDEHYDE_CATAB | GOBP_ALDEHYDE_CATABOLIC_PROCESS | 15 | -0.43227 | -1.7537 | 0.0 | 0.11125973 | 0.994 | 2829 | tags=40%, list=12%, si | MM_4 |
| GOBP_FIBROBLAST_MIGR | GOBP_FIBROBLAST_MIGRATION | 68 | -0.2875 | -1.75033 | 0.0 | 0.11282912 | 0.994 | 2794 | tags=24%, list=12%, si | MM_4 |
| GOBP_STEROL_BIOSYNTH | GOBP_STEROL_BIOSYNTHETIC_PROCESS | 58 | -0.3056 | -1.75015 | 0.0 | 0.11222023 | 0.994 | 2592 | tags=26%, list=11%, si | MM_4 |
| GOBP_POSITIVE_REGULA | GOBP_POSITIVE_REGULATION_OF_LYMPHO | 318 | -0.20903 | -1.74905 | 0.0 | 0.11206902 | 0.994 | 3564 | tags=29%, list=15%, si | MM_4 |
| GOBP_POSITIVE_REGULA | GOBP_POSITIVE_REGULATION_OF_EXTRINS | 60 | -0.29383 | -1.74714 | 0.0 | 0.112752326 | 0.994 | 2666 | tags=23%, list=11%, si | MM_4 |
| AUTOPHAGY | AUTOPHAGY | 126 | -0.25832 | -1.74637 | 0.0 | 0.112516426 | 0.994 | 11737 | tags=90%, list=50%, si | MM_4 |
| GOBP_REGULATION_OF_ | GOBP_REGULATION_OF_DNA_TEMPLATED_ | 79 | -0.29374 | -1.74192 | 0.0 | 0.11417397 | 0.996 | 11552 | tags=89%, list=49%, si | MM_4 |
| GOBP_REGULATION_OF_ | GOBP_REGULATION_OF_ERAD_PATHWAY | 27 | -0.37461 | -1.73452 | 0.012820513 | 0.11795075 | 0.997 | 13256 | tags=96%, list=56%, si | MM_4 |
| GOBP_BROWN_FAT_CELL | GOBP_BROWN_FAT_CELL_DIFFERENTIATION | 63 | -0.28843 | -1.73353 | 0.013888889 | 0.11788065 | 0.997 | 2301 | tags=21%, list=10%, si | MM_4 |
| GOBP_SKELETAL_MUSCLE | GOBP_SKELETAL_MUSCLE_ADAPTATION | 21 | -0.40707 | -1.73318 | 0.01910828 | 0.11733279 | 0.997 | 2433 | tags=38%, list=10%, si | MM_4 |
| OXIDATIVE_PHOSPHORYL | OXIDATIVE_PHOSPHORYLATION | 115 | -0.25022 | -1.72595 | 0.0 | 0.1211933 | 0.998 | 3632 | tags=26%, list=15%, si | MM_4 |
| GOBP_MACROPHAGE_MI | GOBP_MACROPHAGE_MIGRATION | 65 | -0.28378 | -1.72551 | 0.010869565 | 0.12078225 | 0.998 | 2500 | tags=25%, list=11%, si | MM_4 |
| GOBP_ORGANELLE_DISAS | GOBP_ORGANELLE_DISASSEMBLY | 155 | -0.2423 | -1.72377 | 0.0 | 0.1213123 | 0.999 | 2597 | tags=21%, list=11%, si | MM_4 |
| GOBP_SNRNA_3_END_PF | GOBP_SNRNA_3_END_PROCESSING | 20 | -0.39291 | -1.72315 | 0.036585364 | 0.121059306 | 0.999 | 3487 | tags=45%, list=15%, si | MM_4 |
| GOBP_PHAGOCYTOSIS | GOBP_PHAGOCYTOSIS | 213 | -0.23418 | -1.72273 | 0.0 | 0.12063116 | 0.999 | 2421 | tags=19%, list=10%, si | MM_4 |
| GOBP_LEUKOCYTE_AGGR | GOBP_LEUKOCYTE_AGGREGATION | 16 | -0.42992 | -1.72267 | 0.032258064 | 0.12004947 | 0.999 | 2108 | tags=31%, list=9%, sig | MM_4 |
| GOBP_CYTOSOLIC_TRANS | GOBP_CYTOSOLIC_TRANSPORT | 124 | -0.25475 | -1.7217 | 0.0 | 0.12001497 | 0.999 | 11716 | tags=89%, list=50%, si | MM_4 |
| GOBP_VESICLE_TARGETIN | GOBP_VESICLE_TARGETING_ROUGH_ER_TO | 20 | -0.39734 | -1.72122 | 0.0058139535 | 0.11955918 | 0.999 | 2887 | tags=45%, list=12%, si | MM_4 |
| GOBP_BEHAVIORAL_RESF | GOBP_BEHAVIORAL_RESPONSE_TO_PAIN | 29 | -0.34534 | -1.72058 | 0.013245033 | 0.11924813 | 0.999 | 2037 | tags=24%, list=9%, sig | MM_4 |
| GOBP_INTRA_GOLGI_VES | GOBP_INTRA_GOLGI_VESICLE_MEDIATED_T | 30 | -0.36546 | -1.72008 | 0.02097902 | 0.118898936 | 0.999 | 11474 | tags=93%, list=49%, si | MM_4 |
| GOBP_NEGATIVE_REGUL | GOBP_NEGATIVE_REGULATION_OF_MUSCL | 29 | -0.35672 | -1.71954 | 0.0 | 0.118632264 | 0.999 | 2178 | tags=31%, list=9%, sig | MM_4 |
| GOBP_POSITIVE_REGULA | GOBP_POSITIVE_REGULATION_OF_ACTIN_F | 52 | -0.30656 | -1.71943 | 0.011235955 | 0.11803747 | 0.999 | 3473 | tags=33%, list=15%, si | MM_4 |
| GOBP_B_CELL_ACTIVATIO | GOBP_B_CELL_ACTIVATION | 276 | -0.21368 | -1.71764 | 0.0 | 0.118870355 | 0.999 | 3848 | tags=31%, list=16%, si | MM_4 |
| GOBP_SNRNA_PROCESSI | GOBP_SNRNA_PROCESSING | 24 | -0.38979 | -1.71643 | 0.021428572 | 0.1189145 | 0.999 | 3487 | tags=46%, list=15%, si | MM_4 |
| GOBP_RETROGRADE_AXO | GOBP_RETROGRADE_AXONAL_TRANSPORT | 24 | -0.37769 | -1.71605 | 0.022988506 | 0.11854236 | 0.999 | 14702 | tags=100%, list=62%, | MM_4 |
| GOBP_INTERLEUKIN_12_ | GOBP_INTERLEUKIN_12_PRODUCTION | 65 | -0.28229 | -1.71377 | 0.0 | 0.11962405 | 0.999 | 732 | tags=12%, list=3%, sig | MM_4 |
| GOBP_MRNA_CATABOLIC | GOBP_MRNA_CATABOLIC_PROCESS | 233 | -0.23185 | -1.71355 | 0.0 | 0.11907953 | 0.999 | 11602 | tags=86%, list=49%, si | MM_4 |
| GOBP_MITOCHONDRIAL_ | GOBP_MITOCHONDRIAL_RNA_PROCESSING | 20 | -0.41756 | -1.71133 | 0.018404908 | 0.11955431 | 0.999 | 4573 | tags=65%, list=19%, si | MM_4 |
| TOXOPLASMOSIS | TOXOPLASMOSIS | 103 | -0.2594 | -1.71001 | 0.0 | 0.11976433 | 0.999 | 2801 | tags=22%, list=12%, si | MM_4 |
| GOBP_PROTEIN_TARGETI | GOBP_PROTEIN_TARGETING_TO_LYSOSOMI | 29 | -0.33372 | -1.70993 | 0.008333334 | 0.11919049 | 0.999 | 3542 | tags=38%, list=15%, si | MM_4 |
| HUMAN_T-CELL_LEUKEM | HUMAN_T-CELL_LEUKEMIA_VIRUS_1_INFEC | 215 | -0.22744 | -1.70926 | 0.0 | 0.11910342 | 0.999 | 3886 | tags=34%, list=16%, si | MM_4 |
| GOBP_POSITIVE_REGULA | GOBP_POSITIVE_REGULATION_OF_B_CELL_ | 18 | -0.4176 | -1.70384 | 0.020942409 | 0.12232611 | 0.999 | 2279 | tags=39%, list=10%, si | MM_4 |
| GOBP_LEUKOCYTE_CELL_ | GOBP_LEUKOCYTE_CELL_CELL_ADHESION | 390 | -0.20553 | -1.70381 | 0.0 | 0.12171055 | 0.999 | 3574 | tags=27%, list=15%, si | MM_4 |
| GOBP_OSTEOCLAST_DIFF | GOBP_OSTEOCLAST_DIFFERENTIATION | 126 | -0.23206 | -1.70367 | 0.0 | 0.12117446 | 0.999 | 2867 | tags=25%, list=12%, si | MM_4 |
| GOBP_TORC2_SIGNALING | GOBP_TORC2_SIGNALING | 15 | -0.42334 | -1.7023 | 0.027777778 | 0.12146854 | 0.999 | 1986 | tags=40%, list=8%, sig | MM_4 |
| GOBP_EXTRINSIC_APOPT | GOBP_EXTRINSIC_APOPTOTIC_SIGNALING_I | 80 | -0.26661 | -1.70031 | 0.0 | 0.12192471 | 0.999 | 2871 | tags=23%, list=12%, si | MM_4 |
| GOBP_POSITIVE_REGULA | GOBP_POSITIVE_REGULATION_OF_VACUOL | 25 | -0.39636 | -1.70008 | 0.02 | 0.12148273 | 0.999 | 10914 | tags=96%, list=46%, si | MM_4 |
| BACTERIAL_INVASION_OI | BACTERIAL_INVASION_OF_EPITHELIAL_CELL | 72 | -0.28627 | -1.69807 | 0.0 | 0.122138165 | 0.999 | 2788 | tags=28%, list=12%, si | MM_4 |
| GOBP_REGULATION_OF_ | GOBP_REGULATION_OF_MRNA_CATABOLIC | 175 | -0.2509 | -1.6979 | 0.0 | 0.12164873 | 0.999 | 4234 | tags=38%, list=18%, si | MM_4 |
| CENTRAL_CARBON_META | CENTRAL_CARBON_METABOLISM_IN_CANC | 61 | -0.28522 | -1.69715 | 0.010752688 | 0.12154642 | 0.999 | 3667 | tags=33%, list=16%, si | MM_4 |
| GOBP_POSITIVE_REGULA | GOBP_POSITIVE_REGULATION_OF_CELLULA | 29 | -0.34935 | -1.69664 | 0.0068965517 | 0.12129279 | 0.999 | 12358 | tags=93%, list=52%, si | MM_4 |
| GOBP_NATURAL_KILLER_ | GOBP_NATURAL_KILLER_CELL_ACTIVATION_ | 21 | -0.40109 | -1.6944 | 0.017441861 | 0.12231478 | 0.999 | 1423 | tags=24%, list=6%, sig | MM_4 |
| GOBP_POSITIVE_REGULA | GOBP_POSITIVE_REGULATION_OF_CARDIOC | 15 | -0.44924 | -1.69122 | 0.04411765 | 0.123906486 | 0.999 | 197 | tags=13%, list=1%, sig | MM_4 |
| GOBP_CELLULAR_PIGME | GOBP_CELLULAR_PIGMENTATION | 57 | -0.29104 | -1.69056 | 0.0 | 0.12360625 | 0.999 | 1789 | tags=21%, list=8%, sig | MM_4 |
| GOBP_ALTERNATIVE_MR | GOBP_ALTERNATIVE_MRNA_SPLICING_VIA_ | 75 | -0.26741 | -1.68901 | 0.0 | 0.12404261 | 0.999 | 15855 | tags=97%, list=67%, si | MM_4 |
| GOBP_REGULATION_OF_ | GOBP_REGULATION_OF_RNA_STABILITY | 174 | -0.22033 | -1.68771 | 0.0 | 0.12439993 | 0.999 | 4234 | tags=36%, list=18%, si | MM_4 |
| GOBP_LIPOPROTEIN_CAT | GOBP_LIPOPROTEIN_CATABOLIC_PROCESS | 17 | -0.42604 | -1.68614 | 0.015957447 | 0.12466829 | 0.999 | 2453 | tags=35%, list=10%, si | MM_4 |
| GOBP_PREPULSE_INHIBIT | GOBP_PREPULSE_INHIBITION | 18 | -0.41374 | -1.68098 | 0.027472528 | 0.12802532 | 0.999 | 1718 | tags=33%, list=7%, sig | MM_4 |
| GOBP_CYTOPLASMIC_TRA | GOBP_CYTOPLASMIC_TRANSLATIONAL_INIT | 33 | -0.33148 | -1.68073 | 0.0125 | 0.1276343 | 0.999 | 11552 | tags=94%, list=49%, si | MM_4 |
| GOBP_COPPER_ION_HON | GOBP_COPPER_ION_HOMEOSTASIS | 18 | -0.40295 | -1.68033 | 0.031746034 | 0.12734051 | 0.999 | 1733 | tags=28%, list=7%, sig | MM_4 |
| GOBP_INNER_MITOCHON | GOBP_INNER_MITOCHONDRIAL_MEMBRAN | 36 | -0.32201 | -1.67823 | 0.007194245 | 0.12812717 | 0.999 | 1724 | tags=19%, list=7%, sig | MM_4 |
| GOBP_MICROAUTOPHAG | GOBP_MICROAUTOPHAGY | 24 | -0.36534 | -1.67404 | 0.018518519 | 0.13044058 | 0.999 | 13100 | tags=96%, list=55%, si | MM_4 |

| | | | | | | | | | | |
|---|---|---|---|---|---|---|---|---|---|---|
| GOBP_REGULATION_OF_ | GOBP_REGULATION_OF_BLOOD_VESSEL_EN | 17 | -0.40855 | -1.67222 | 0.011363637 | 0.13129695 | 0.999 | 11 | tags=12%, list=0%, sig | MM_4 |
| GOBP_NEGATIVE_REGUL | GOBP_NEGATIVE_REGULATION_OF_CELLUL | 18 | -0.4098 | -1.67203 | 0.02631579 | 0.13078447 | 0.999 | 1953 | tags=33%, list=8%, sig | MM_4 |
| GOBP_REGULATION_OF_ | GOBP_REGULATION_OF_LEUKOCYTE_PROLI | 273 | -0.21273 | -1.67182 | 0.0 | 0.13029072 | 0.999 | 3574 | tags=30%, list=15%, si | MM_4 |
| GOBP_REGULATION_OF_ | GOBP_REGULATION_OF_CHROMATIN_BIND | 20 | -0.39604 | -1.67083 | 0.028735632 | 0.13045368 | 0.999 | 3134 | tags=45%, list=13%, si | MM_4 |
| GOBP_EXTRINSIC_APOPT | GOBP_EXTRINSIC_APOPTOTIC_SIGNALING_ | 233 | -0.21449 | -1.67067 | 0.0 | 0.12998477 | 0.999 | 2704 | tags=21%, list=11%, si | MM_4 |
| GOBP_LEUKOCYTE_PROLI | GOBP_LEUKOCYTE_PROLIFERATION | 361 | -0.20629 | -1.66987 | 0.0 | 0.12993312 | 0.999 | 2801 | tags=22%, list=12%, si | MM_4 |
| GOBP_MATURATION_OF_ | GOBP_MATURATION_OF_SSU_RRNA | 48 | -0.31291 | -1.66685 | 0.0 | 0.13158439 | 0.999 | 11485 | tags=92%, list=49%, si | MM_4 |
| GOBP_INTERFERON_ALPH | GOBP_INTERFERON_ALPHA_PRODUCTION | 33 | -0.34587 | -1.6661 | 0.01183432 | 0.13152039 | 0.999 | 2675 | tags=30%, list=11%, si | MM_4 |
| GOBP_REGULATION_OF_ | GOBP_REGULATION_OF_ACTIVATED_T_CELI | 41 | -0.31773 | -1.66211 | 0.008695652 | 0.1338436 | 0.999 | 2955 | tags=32%, list=13%, si | MM_4 |
| GOBP_RESPONSE_TO_TU | GOBP_RESPONSE_TO_TUMOR_NECROSIS_F | 167 | -0.23579 | -1.65975 | 0.0 | 0.13475035 | 1 | 2955 | tags=24%, list=13%, si | MM_4 |
| GOBP_NEGATIVE_REGUL | GOBP_NEGATIVE_REGULATION_OF_B_CELL_ | 36 | -0.32698 | -1.65942 | 0.01724138 | 0.13441145 | 1 | 3720 | tags=42%, list=16%, si | MM_4 |
| GOBP_MAST_CELL_MIGR | GOBP_MAST_CELL_MIGRATION | 17 | -0.41777 | -1.65847 | 0.040816326 | 0.13445511 | 1 | 2432 | tags=35%, list=10%, si | MM_4 |
| GOBP_REGULATION_OF_ | GOBP_REGULATION_OF_T_CELL_PROLIFERA | 188 | -0.21817 | -1.65632 | 0.0 | 0.1352284 | 1 | 3574 | tags=30%, list=15%, si | MM_4 |
| GOBP_ACTIVATION_OF_ | GOBP_ACTIVATION_OF_GTPASE_ACTIVITY | 86 | -0.26428 | -1.65413 | 0.015873017 | 0.135931 | 1 | 2687 | tags=26%, list=11%, si | MM_4 |
| GOBP_REGULATION_OF_ | GOBP_REGULATION_OF_ENDOTHELIAL_CEL | 21 | -0.36603 | -1.65303 | 0.036363635 | 0.1361245 | 1 | 343 | tags=14%, list=1%, sig | MM_4 |
| GOBP_DNA_TEMPLATED_ | GOBP_DNA_TEMPLATED_TRANSCRIPTION_ | 112 | -0.25069 | -1.65237 | 0.0 | 0.13586397 | 1 | 3023 | tags=26%, list=13%, si | MM_4 |
| GOBP_REGULATION_OF_ | GOBP_REGULATION_OF_OSTEOCLAST_DIFF | 84 | -0.255 | -1.65195 | 0.0 | 0.13544402 | 1 | 2788 | tags=25%, list=12%, si | MM_4 |
| GOBP_LYSOSOMAL_LUM | GOBP_LYSOSOMAL_LUMEN_ACIDIFICATION | 15 | -0.42677 | -1.65037 | 0.05263158 | 0.13617095 | 1 | 3573 | tags=47%, list=15%, si | MM_4 |
| GOBP_PIECEMEAL_MICR | GOBP_PIECEMEAL_MICROAUTOPHAGY_OF_ | 19 | -0.41344 | -1.65004 | 0.025252525 | 0.13587071 | 1 | 11533 | tags=95%, list=49%, si | MM_4 |
| GOBP_POSITIVE_REGULA | GOBP_POSITIVE_REGULATION_OF_NATURA | 35 | -0.33425 | -1.64785 | 0.024793388 | 0.13702214 | 1 | 1719 | tags=23%, list=7%, sig | MM_4 |
| GOBP_MITOCHONDRIAL_ | GOBP_MITOCHONDRIAL_DNA_METABOLIC_ | 26 | -0.35541 | -1.64785 | 0.02919708 | 0.13643406 | 1 | 3445 | tags=46%, list=15%, si | MM_4 |
| GOBP_NEGATIVE_REGUL | GOBP_NEGATIVE_REGULATION_OF_TRANSI | 127 | -0.24354 | -1.64744 | 0.0 | 0.13609421 | 1 | 2780 | tags=24%, list=12%, si | MM_4 |
| GOBP_POSITIVE_REGULA | GOBP_POSITIVE_REGULATION_OF_TYPE_I_I | 19 | -0.3982 | -1.64481 | 0.032786883 | 0.13723463 | 1 | 1388 | tags=26%, list=6%, sig | MM_4 |
| GOBP_POSITIVE_REGULA | GOBP_POSITIVE_REGULATION_OF_SYNAPTI | 17 | -0.40712 | -1.64402 | 0.02538071 | 0.13717787 | 1 | 14005 | tags=100%, list=59%, | MM_4 |
| GOBP_MODULATION_BY_ | GOBP_MODULATION_BY_HOST_OF_VIRAL_ | 44 | -0.30452 | -1.64319 | 0.017857144 | 0.13729933 | 1 | 13062 | tags=93%, list=55%, si | MM_4 |
| GOBP_RNA_SURVEILLAN | GOBP_RNA_SURVEILLANCE | 17 | -0.40996 | -1.64184 | 0.027322404 | 0.13777101 | 1 | 11009 | tags=94%, list=47%, si | MM_4 |
| GOBP_PHOSPHATIDIC_AC | GOBP_PHOSPHATIDIC_ACID_BIOSYNTHETIC_ | 23 | -0.36117 | -1.6414 | 0.022857143 | 0.13747707 | 1 | 2594 | tags=43%, list=11%, si | MM_4 |
| GOBP_RNA_METHYLATIO | GOBP_RNA_METHYLATION | 65 | -0.27437 | -1.64013 | 0.012987013 | 0.13773765 | 1 | 3936 | tags=40%, list=17%, si | MM_4 |
| GOBP_MRNA_CIS_SPLICI | GOBP_MRNA_CIS_SPLICING_VIA_SPLICEOS | 26 | -0.36461 | -1.64001 | 0.014084507 | 0.13725285 | 1 | 12421 | tags=96%, list=53%, si | MM_4 |
| GOBP_VACUOLE_ORGAN | GOBP_VACUOLE_ORGANIZATION | 223 | -0.20919 | -1.6394 | 0.0 | 0.13700819 | 1 | 3654 | tags=28%, list=15%, si | MM_4 |
| GOBP_REGULATION_OF_ | GOBP_REGULATION_OF_NATURAL_KILLER_ | 46 | -0.29146 | -1.6382 | 0.00952381 | 0.13735041 | 1 | 470 | tags=13%, list=2%, sig | MM_4 |
| GOBP_MEMBRANE_LIPID | GOBP_MEMBRANE_LIPID_METABOLIC_PRO | 186 | -0.21429 | -1.63764 | 0.0 | 0.13724847 | 1 | 2085 | tags=22%, list=9%, sig | MM_4 |
| GOBP_NEGATIVE_REGUL | GOBP_NEGATIVE_REGULATION_OF_INTERL | 51 | -0.28948 | -1.63722 | 0.010309278 | 0.13700421 | 1 | 1031 | tags=14%, list=4%, sig | MM_4 |
| GOBP_RNA_CATABOLIC_ | GOBP_RNA_CATABOLIC_PROCESS | 278 | -0.23529 | -1.63457 | 0.0 | 0.13822651 | 1 | 4234 | tags=35%, list=18%, si | MM_4 |
| GOBP_REGULATION_OF_ | GOBP_REGULATION_OF_MONONUCLEAR_C | 136 | -0.23888 | -1.63395 | 0.0 | 0.13796882 | 1 | 3723 | tags=32%, list=16%, si | MM_4 |
| GOBP_POSITIVE_REGULA | GOBP_POSITIVE_REGULATION_OF_TYPE_I_I | 77 | -0.26929 | -1.63365 | 0.0 | 0.13768473 | 1 | 2675 | tags=23%, list=11%, si | MM_4 |
| GOBP_NEUROTRANSMITT | GOBP_NEUROTRANSMITTER_RECEPTOR_TR | 27 | -0.34848 | -1.63184 | 0.034722224 | 0.13832128 | 1 | 15392 | tags=100%, list=65%, | MM_4 |
| GOBP_ESTABLISHMENT_ | GOBP_ESTABLISHMENT_OF_PROTEIN_LOCA | 25 | -0.34213 | -1.6295 | 0.024539877 | 0.13931993 | 1 | 10733 | tags=92%, list=45%, si | MM_4 |
| GOBP_MAINTENANCE_O | GOBP_MAINTENANCE_OF_PROTEIN_LOCAT | 97 | -0.26733 | -1.62885 | 0.021276595 | 0.13918915 | 1 | 3292 | tags=29%, list=14%, si | MM_4 |
| GOBP_GERMINAL_CENTE | GOBP_GERMINAL_CENTER_FORMATION | 16 | -0.40041 | -1.62728 | 0.023923445 | 0.13970122 | 1 | 11239 | tags=94%, list=48%, si | MM_4 |
| GOBP_REGULATION_OF_ | GOBP_REGULATION_OF_CYTOPLASMIC_PAT | 104 | -0.24208 | -1.62687 | 0.0 | 0.13945632 | 1 | 4092 | tags=32%, list=17%, si | MM_4 |
| GOBP_NATURAL_KILLER_ | GOBP_NATURAL_KILLER_CELL_ACTIVATION | 86 | -0.25384 | -1.62686 | 0.0 | 0.13890728 | 1 | 3672 | tags=36%, list=16%, si | MM_4 |
| GOBP_POSITIVE_REGULA | GOBP_POSITIVE_REGULATION_OF_EXECUTI | 19 | -0.39661 | -1.62683 | 0.032786883 | 0.13837574 | 1 | 11976 | tags=95%, list=51%, si | MM_4 |
| GOBP_MITOCHONDRIAL_ | GOBP_MITOCHONDRIAL_GENOME_MAINTE | 35 | -0.31604 | -1.62517 | 0.020408163 | 0.13914149 | 1 | 3445 | tags=40%, list=15%, si | MM_4 |
| GOBP_MYELOID_CELL_DI | GOBP_MYELOID_CELL_DIFFERENTIATION | 455 | -0.20124 | -1.62492 | 0.0 | 0.13865475 | 1 | 2867 | tags=22%, list=12%, si | MM_4 |
| GOBP_VACUOLAR_LOCAL | GOBP_VACUOLAR_LOCALIZATION | 85 | -0.26017 | -1.62482 | 0.0 | 0.13814521 | 1 | 1826 | tags=16%, list=8%, sig | MM_4 |
| GOBP_CARDIOBLAST_DIF | GOBP_CARDIOBLAST_DIFFERENTIATION | 15 | -0.41989 | -1.62317 | 0.036458332 | 0.13894093 | 1 | 1370 | tags=27%, list=6%, sig | MM_4 |
| GOBP_MRNA_EXPORT_F | GOBP_MRNA_EXPORT_FROM_NUCLEUS | 60 | -0.27987 | -1.62263 | 0.0 | 0.13880333 | 1 | 11465 | tags=88%, list=49%, si | MM_4 |
| GOBP_NEGATIVE_REGUL | GOBP_NEGATIVE_REGULATION_OF_LEUKO | 96 | -0.24729 | -1.62073 | 0.016666668 | 0.13966163 | 1 | 3782 | tags=35%, list=16%, si | MM_4 |
| AMINOACYL-TRNA_BIOSY | AMINOACYL-TRNA_BIOSYNTHESIS | 43 | -0.29366 | -1.61877 | 0.017699115 | 0.14035457 | 1 | 4081 | tags=37%, list=17%, si | MM_4 |
| GOBP_NEGATIVE_REGUL | GOBP_NEGATIVE_REGULATION_OF_T_CELL_ | 73 | -0.26105 | -1.61867 | 0.0 | 0.13988745 | 1 | 3574 | tags=32%, list=15%, si | MM_4 |
| GOBP_DIACYLGLYCEROL_ | GOBP_DIACYLGLYCEROL_METABOLIC_PROC | 28 | -0.33793 | -1.61861 | 0.01734104 | 0.13941167 | 1 | 2594 | tags=46%, list=11%, si | MM_4 |
| GOBP_COLLAGEN_ACTIV | GOBP_COLLAGEN_ACTIVATED_SIGNALING_ | 17 | -0.41444 | -1.6177 | 0.043243244 | 0.13953233 | 1 | 3221 | tags=41%, list=14%, si | MM_4 |
| GOBP_ERYTHROCYTE_HO | GOBP_ERYTHROCYTE_HOMEOSTASIS | 163 | -0.23621 | -1.61621 | 0.0 | 0.14013104 | 1 | 3679 | tags=31%, list=16%, si | MM_4 |
| GOBP_POSITIVE_REGULA | GOBP_POSITIVE_REGULATION_OF_INTERFE | 26 | -0.35754 | -1.61527 | 0.039215688 | 0.14017513 | 1 | 2675 | tags=31%, list=11%, si | MM_4 |
| GOBP_PROTEIN_N_LINKE | GOBP_PROTEIN_N_LINKED_GLYCOSYLATION | 18 | -0.3965 | -1.61443 | 0.04232804 | 0.14027406 | 1 | 11523 | tags=94%, list=49%, si | MM_4 |
| GOBP_NEGATIVE_REGUL | GOBP_NEGATIVE_REGULATION_OF_NUCLE | 27 | -0.33683 | -1.61406 | 0.028368793 | 0.14008275 | 1 | 3881 | tags=37%, list=16%, si | MM_4 |
| GOBP_LEUKOCYTE_MIGR | GOBP_LEUKOCYTE_MIGRATION | 363 | -0.20992 | -1.61224 | 0.0 | 0.14090449 | 1 | 1198 | tags=13%, list=5%, sig | MM_4 |
| GOBP_NEGATIVE_REGUL | GOBP_NEGATIVE_REGULATION_OF_INNATE | 86 | -0.26266 | -1.61212 | 0.0 | 0.14047314 | 1 | 1118 | tags=13%, list=5%, sig | MM_4 |
| C-TYPE_LECTIN_RECEPTO | C-TYPE_LECTIN_RECEPTOR_SIGNALING_PAT | 104 | -0.24968 | -1.61168 | 0.0 | 0.14026773 | 1 | 2037 | tags=14%, list=9%, sig | MM_4 |
| GOBP_CD4_POSITIVE_AL | GOBP_CD4_POSITIVE_ALPHA_BETA_T_CELL_ | 28 | -0.33096 | -1.61124 | 0.020408163 | 0.14000928 | 1 | 11834 | tags=89%, list=50%, si | MM_4 |
| GOBP_PROTEIN_LOCALIZ | GOBP_PROTEIN_LOCALIZATION_TO_LYSOS | 54 | -0.2774 | -1.60834 | 0.0 | 0.14148737 | 1 | 3545 | tags=31%, list=15%, si | MM_4 |
| GOBP_POSITIVE_REGULA | GOBP_POSITIVE_REGULATION_BY_HOST_O | 22 | -0.36115 | -1.60759 | 0.035928145 | 0.14158006 | 1 | 10254 | tags=91%, list=43%, si | MM_4 |
| GOBP_REGULATION_OF_ | GOBP_REGULATION_OF_EXTRINSIC_APOPT | 161 | -0.22364 | -1.60623 | 0.0 | 0.14224657 | 1 | 2704 | tags=20%, list=11%, si | MM_4 |
| GOBP_REGULATION_OF_ | GOBP_REGULATION_OF_GTPASE_ACTIVITY | 287 | -0.21028 | -1.60553 | 0.0 | 0.14232083 | 1 | 2757 | tags=20%, list=12%, si | MM_4 |
| GOBP_T_CELL_CHEMOTA | GOBP_T_CELL_CHEMOTAXIS | 23 | -0.36887 | -1.60347 | 0.031578947 | 0.14327757 | 1 | 272 | tags=22%, list=1%, sig | MM_4 |
| AMINO_SUGAR_AND_NU | AMINO_SUGAR_AND_NUCLEOTIDE_SUGAR_ | 46 | -0.28968 | -1.60222 | 0.009803922 | 0.14381541 | 1 | 2518 | tags=35%, list=11%, si | MM_4 |
| GOBP_ERYTHROCYTE_DE | GOBP_ERYTHROCYTE_DEVELOPMENT | 50 | -0.2939 | -1.60207 | 0.01904762 | 0.14347735 | 1 | 3472 | tags=34%, list=15%, si | MM_4 |
| GOBP_NUCLEAR_TRANSC | GOBP_NUCLEAR_TRANSCRIBED_MRNA_CAT | 118 | -0.24787 | -1.60176 | 0.0 | 0.14319259 | 1 | 11602 | tags=89%, list=49%, si | MM_4 |
| GOBP_T_CELL_MIGRATIO | GOBP_T_CELL_MIGRATION | 64 | -0.26466 | -1.60145 | 0.013333334 | 0.14293782 | 1 | 1198 | tags=17%, list=5%, sig | MM_4 |
| GOBP_GLYCOLYTIC_PROC | GOBP_GLYCOLYTIC_PROCESS_THROUGH_FR | 25 | -0.35956 | -1.59992 | 0.026845638 | 0.14352153 | 1 | 11213 | tags=92%, list=47%, si | MM_4 |
| GOBP_POSITIVE_REGULA | GOBP_POSITIVE_REGULATION_OF_ACTIVAT | 27 | -0.33868 | -1.5992 | 0.01754386 | 0.14362912 | 1 | 3295 | tags=41%, list=14%, si | MM_4 |
| GOBP_POSTSYNAPSE_TO | GOBP_POSTSYNAPSE_TO_NUCLEUS_SIGNAL | 16 | -0.40861 | -1.59904 | 0.022522522 | 0.1432123 | 1 | 2153 | tags=25%, list=9%, sig | MM_4 |
| GOBP_NEGATIVE_REGUL | GOBP_NEGATIVE_REGULATION_OF_IMMUN | 186 | -0.22173 | -1.59829 | 0.0 | 0.14320521 | 1 | 2291 | tags=18%, list=10%, si | MM_4 |
| GOBP_PROTEIN_TRANSM | GOBP_PROTEIN_TRANSMEMBRANE_IMPOR | 36 | -0.31875 | -1.59826 | 0.028301887 | 0.14273047 | 1 | 12199 | tags=92%, list=52%, si | MM_4 |
| GOBP_GLYCOPROTEIN_C | GOBP_GLYCOPROTEIN_CATABOLIC_PROCES | 31 | -0.33569 | -1.59779 | 0.035460994 | 0.14254276 | 1 | 15695 | tags=100%, list=66%, | MM_4 |
| GOBP_CELLULAR_RESPO | GOBP_CELLULAR_RESPONSE_TO_ARSENIC_ | 17 | -0.42177 | -1.59621 | 0.026737968 | 0.14327843 | 1 | 2273 | tags=29%, list=10%, si | MM_4 |
| GOBP_REGULATION_OF_ | GOBP_REGULATION_OF_TOLL_LIKE_RECEPT | 28 | -0.33002 | -1.59562 | 0.021276595 | 0.14328632 | 1 | 2421 | tags=25%, list=10%, si | MM_4 |
| GOBP_CHONDROITIN_SU | GOBP_CHONDROITIN_SULFATE_METABOLIC | 22 | -0.36431 | -1.59486 | 0.061349694 | 0.14343834 | 1 | 4329 | tags=64%, list=18%, si | MM_4 |
| GOBP_POSITIVE_REGULA | GOBP_POSITIVE_REGULATION_OF_LIPID_ST | 25 | -0.34695 | -1.59413 | 0.03773585 | 0.14346783 | 1 | 596 | tags=20%, list=3%, sig | MM_4 |
| GOBP_NEGATIVE_REGUL | GOBP_NEGATIVE_REGULATION_OF_LEUKO | 65 | -0.26611 | -1.59255 | 0.0 | 0.14427985 | 1 | 1553 | tags=18%, list=7%, sig | MM_4 |
| GOBP_PROTEIN_RNA_CO | GOBP_PROTEIN_RNA_COMPLEX_ORGANIZA | 173 | -0.23981 | -1.59246 | 0.0 | 0.14383525 | 1 | 11721 | tags=88%, list=50%, si | MM_4 |

| | | | | | | | | | | |
|---|---|---|---|---|---|---|---|---|---|---|
| GOBP_GLYCOSIDE_META | GOBP_GLYCOSIDE_METABOLIC_PROCESS | 18 | -0.38903 | -1.59 | 0.067307696 | 0.14543097 | 1 | 1615 | tags=33%, list=7%, sig | MM_4 |
| PROTEIN_EXPORT | PROTEIN_EXPORT | 26 | -0.34739 | -1.58848 | 0.03797468 | 0.14619131 | 1 | 11305 | tags=92%, list=48%, si | MM_4 |
| GOBP_NEGATIVE_REGUL | GOBP_NEGATIVE_REGULATION_OF_DEPHO | 38 | -0.30434 | -1.58617 | 0.02586207 | 0.14766736 | 1 | 3781 | tags=42%, list=16%, si | MM_4 |
| INFLUENZA_A | INFLUENZA_A | 139 | -0.23533 | -1.58567 | 0.0 | 0.14748555 | 1 | 3872 | tags=33%, list=16%, si | MM_4 |
| GOBP_CELL_ADHESION_M | GOBP_CELL_ADHESION_MEDIATED_BY_INT | 86 | -0.25742 | -1.58484 | 0.029411765 | 0.14751013 | 1 | 2357 | tags=23%, list=10%, si | MM_4 |
| GOBP_POSITIVE_REGULA | GOBP_POSITIVE_REGULATION_OF_PATTERN | 68 | -0.25687 | -1.58482 | 0.026666667 | 0.14705499 | 1 | 4092 | tags=35%, list=17%, si | MM_4 |
| GOBP_INTRACELLULAR_P | GOBP_INTRACELLULAR_PROTEIN_TRANSME | 32 | -0.33446 | -1.58457 | 0.04255319 | 0.14672028 | 1 | 13117 | tags=94%, list=56%, si | MM_4 |
| GOBP_PLATELET_DENSE_ | GOBP_PLATELET_DENSE_GRANULE_ORGAN | 21 | -0.36962 | -1.58327 | 0.02116402 | 0.1473335 | 1 | 1789 | tags=33%, list=8%, sig | MM_4 |
| GOBP_REGULATION_OF_ | GOBP_REGULATION_OF_PROTEIN_EXPORT_ | 37 | -0.29827 | -1.58268 | 0.044247787 | 0.14740442 | 1 | 15230 | tags=97%, list=65%, si | MM_4 |
| GOBP_SECRETORY_GRAN | GOBP_SECRETORY_GRANULE_LOCALIZATIO | 34 | -0.3153 | -1.58266 | 0.0234375 | 0.14693049 | 1 | 15227 | tags=97%, list=64%, si | MM_4 |
| GOBP_RETROGRADE_TRA | GOBP_RETROGRADE_TRANS_SYNAPTIC_SIG | 15 | -0.41521 | -1.58193 | 0.03902439 | 0.14703973 | 1 | 1632 | tags=33%, list=7%, sig | MM_4 |
| INFLAMMATORY_BOWEL | INFLAMMATORY_BOWEL_DISEASE_(IBD) | 52 | -0.28269 | -1.58165 | 0.0 | 0.14675713 | 1 | 643 | tags=15%, list=3%, sig | MM_4 |
| GOBP_POSITIVE_REGULA | GOBP_POSITIVE_REGULATION_OF_ERAD_P | 20 | -0.37462 | -1.58058 | 0.048648648 | 0.14711742 | 1 | 12785 | tags=95%, list=54%, si | MM_4 |
| GOBP_REGULATION_OF_ | GOBP_REGULATION_OF_LYMPHOCYTE_MIG | 64 | -0.26449 | -1.58052 | 0.0 | 0.14668502 | 1 | 1031 | tags=16%, list=4%, sig | MM_4 |
| GOBP_ACTIN_POLYMERIZ | GOBP_ACTIN_POLYMERIZATION_OR_DEPOL | 187 | -0.21376 | -1.5802 | 0.0 | 0.1464298 | 1 | 3477 | tags=27%, list=15%, si | MM_4 |
| RNA_DEGRADATION | RNA_DEGRADATION | 75 | -0.25611 | -1.57926 | 0.02631579 | 0.14662658 | 1 | 11499 | tags=89%, list=49%, si | MM_4 |
| GOBP_REGULATION_OF_ | GOBP_REGULATION_OF_IMMUNE_EFFECTO | 406 | -0.1865 | -1.57862 | 0.0 | 0.1466161 | 1 | 2291 | tags=17%, list=10%, si | MM_4 |
| GOBP_B_CELL_RECEPTOR | GOBP_B_CELL_RECEPTOR_SIGNALING_PATH | 55 | -0.27426 | -1.57728 | 0.010638298 | 0.14706363 | 1 | 3748 | tags=31%, list=16%, si | MM_4 |
| GOBP_TRNA_METHYLATI | GOBP_TRNA_METHYLATION | 39 | -0.3088 | -1.57261 | 0.016528925 | 0.15004855 | 1 | 2516 | tags=31%, list=11%, si | MM_4 |
| GOBP_RUFFLE_ASSEMBLY | GOBP_RUFFLE_ASSEMBLY | 43 | -0.28776 | -1.5699 | 0.036036037 | 0.15173274 | 1 | 1826 | tags=21%, list=8%, sig | MM_4 |
| GOBP_RNA_3_END_PROC | GOBP_RNA_3_END_PROCESSING | 87 | -0.25229 | -1.5688 | 0.0 | 0.15233186 | 1 | 11577 | tags=89%, list=49%, si | MM_4 |
| GOBP_RESPONSE_TO_NE | GOBP_RESPONSE_TO_NERVE_GROWTH_FA | 42 | -0.29852 | -1.56808 | 0.032520324 | 0.15241157 | 1 | 14520 | tags=95%, list=61%, si | MM_4 |
| GOBP_NEGATIVE_REGUL | GOBP_NEGATIVE_REGULATION_OF_TYPE_I_ | 42 | -0.30095 | -1.56732 | 0.016129032 | 0.15236582 | 1 | 470 | tags=10%, list=2%, sig | MM_4 |
| GOBP_NEGATIVE_REGUL | GOBP_NEGATIVE_REGULATION_OF_DEFENS | 242 | -0.20512 | -1.56672 | 0.0 | 0.1524022 | 1 | 2291 | tags=17%, list=10%, si | MM_4 |
| GOBP_MATURATION_OF_ | GOBP_MATURATION_OF_SSU_RRNA_FROM | 33 | -0.30261 | -1.56578 | 0.015384615 | 0.15265127 | 1 | 11485 | tags=91%, list=49%, si | MM_4 |
| GOBP_NEGATIVE_REGUL | GOBP_NEGATIVE_REGULATION_OF_CYTOKI | 292 | -0.2088 | -1.56489 | 0.0 | 0.15316074 | 1 | 1118 | tags=11%, list=5%, sig | MM_4 |
| GOBP_AUTOPHAGOSOMI | GOBP_AUTOPHAGOSOME_ORGANIZATION | 128 | -0.22669 | -1.56362 | 0.0 | 0.15369184 | 1 | 3635 | tags=27%, list=15%, si | MM_4 |
| GOBP_NEGATIVE_REGUL | GOBP_NEGATIVE_REGULATION_OF_CHEMO | 39 | -0.29523 | -1.5634 | 0.01923077 | 0.15340237 | 1 | 2432 | tags=21%, list=10%, si | MM_4 |
| GOBP_POSITIVE_REGULA | GOBP_POSITIVE_REGULATION_OF_RAC_PR | 20 | -0.37067 | -1.56266 | 0.031446543 | 0.15357232 | 1 | 662 | tags=15%, list=3%, sig | MM_4 |
| GOBP_ALPHA_BETA_T_CI | GOBP_ALPHA_BETA_T_CELL_PROLIFERATIO | 54 | -0.28287 | -1.56218 | 0.020408163 | 0.15349866 | 1 | 665 | tags=11%, list=3%, sig | MM_4 |
| GOBP_MONONUCLEAR_C | GOBP_MONONUCLEAR_CELL_MIGRATION | 199 | -0.2199 | -1.56107 | 0.0 | 0.15395896 | 1 | 2846 | tags=24%, list=12%, si | MM_4 |
| GOBP_RESPIRATORY_CHA | GOBP_RESPIRATORY_CHAIN_COMPLEX_IV_ | 24 | -0.34397 | -1.56038 | 0.049723756 | 0.15396687 | 1 | 1706 | tags=25%, list=7%, sig | MM_4 |
| GOBP_PROTEIN_LOCALIZ | GOBP_PROTEIN_LOCALIZATION_TO_VACUO | 78 | -0.25028 | -1.55982 | 0.0 | 0.15399593 | 1 | 3545 | tags=28%, list=15%, si | MM_4 |
| GOBP_REGULATION_OF_ | GOBP_REGULATION_OF_MRNA_METABOLI | 286 | -0.21288 | -1.55864 | 0.0 | 0.15437712 | 1 | 4234 | tags=33%, list=18%, si | MM_4 |
| GOBP_HEMATOPOIETIC_ | GOBP_HEMATOPOIETIC_STEM_CELL_DIFFER | 31 | -0.31997 | -1.55604 | 0.034722224 | 0.15583919 | 1 | 3472 | tags=39%, list=15%, si | MM_4 |
| GOBP_3_UTR_MEDIATED | GOBP_3_UTR_MEDIATED_MRNA_DESTABIL | 17 | -0.38841 | -1.55551 | 0.042682927 | 0.15584134 | 1 | 4681 | tags=53%, list=20%, si | MM_4 |
| GOBP_ENDOSOME_ORGA | GOBP_ENDOSOME_ORGANIZATION | 68 | -0.25319 | -1.55422 | 0.03797468 | 0.1562743 | 1 | 11950 | tags=88%, list=51%, si | MM_4 |
| GOBP_RESPONSE_TO_VI | GOBP_RESPONSE_TO_VIRUS | 331 | -0.19866 | -1.55323 | 0.0 | 0.15672952 | 1 | 1423 | tags=13%, list=6%, sig | MM_4 |
| GOBP_REGULATION_OF_ | GOBP_REGULATION_OF_LEUKOCYTE_APOP | 103 | -0.2317 | -1.55309 | 0.0 | 0.1563862 | 1 | 1553 | tags=16%, list=7%, sig | MM_4 |
| GOBP_POSITIVE_REGULA | GOBP_POSITIVE_REGULATION_OF_T_CELL_ | 108 | -0.22704 | -1.55249 | 0.018181818 | 0.15651442 | 1 | 3564 | tags=32%, list=15%, si | MM_4 |
| GOBP_NEGATIVE_REGUL | GOBP_NEGATIVE_REGULATION_OF_LEUKO | 16 | -0.39968 | -1.55224 | 0.03827751 | 0.15621707 | 1 | 9292 | tags=88%, list=39%, si | MM_4 |
| GOBP_REGULATION_OF_ | GOBP_REGULATION_OF_TRANSLATIONAL_I | 17 | -0.39357 | -1.55074 | 0.036458332 | 0.15695795 | 1 | 10677 | tags=94%, list=45%, si | MM_4 |
| GOBP_NEGATIVE_REGUL | GOBP_NEGATIVE_REGULATION_OF_RESPON | 112 | -0.23237 | -1.54769 | 0.0 | 0.15891738 | 1 | 4010 | tags=29%, list=17%, si | MM_4 |
| GOBP_REGULATION_OF_ | GOBP_REGULATION_OF_SYNCYTIUM_FORM | 31 | -0.31262 | -1.5468 | 0.024390243 | 0.1592758 | 1 | 411 | tags=16%, list=2%, sig | MM_4 |
| GOBP_MYELOID_CELL_HO | GOBP_MYELOID_CELL_HOMEOSTASIS | 209 | -0.20175 | -1.54678 | 0.0 | 0.15881589 | 1 | 2846 | tags=23%, list=12%, si | MM_4 |
| GOBP_MACROPHAGE_CH | GOBP_MACROPHAGE_CHEMOTAXIS | 45 | -0.29453 | -1.54618 | 0.0 | 0.1589172 | 1 | 272 | tags=11%, list=1%, sig | MM_4 |
| GOBP_REGULATION_OF_ | GOBP_REGULATION_OF_RAC_PROTEIN_SIG | 35 | -0.29272 | -1.5455 | 0.03846154 | 0.15919094 | 1 | 662 | tags=11%, list=3%, sig | MM_4 |
| GOBP_POLYPRENOL_MET | GOBP_POLYPRENOL_METABOLIC_PROCESS | 17 | -0.38165 | -1.54464 | 0.071428575 | 0.15953338 | 1 | 3265 | tags=53%, list=14%, si | MM_4 |
| GOBP_GLUCOSE_CATABO | GOBP_GLUCOSE_CATABOLIC_PROCESS | 26 | -0.32327 | -1.54302 | 0.030487806 | 0.16022453 | 1 | 3630 | tags=42%, list=15%, si | MM_4 |
| GOBP_PEROXISOME_PRO | GOBP_PEROXISOME_PROLIFERATOR_ACTIV | 27 | -0.32729 | -1.54284 | 0.028368793 | 0.15991376 | 1 | 1141 | tags=22%, list=5%, sig | MM_4 |
| GOBP_RAC_PROTEIN_SIG | GOBP_RAC_PROTEIN_SIGNAL_TRANSDUCTI | 58 | -0.26864 | -1.54277 | 0.0 | 0.15947121 | 1 | 2161 | tags=17%, list=9%, sig | MM_4 |
| GOBP_TRANSCRIPTION_P | GOBP_TRANSCRIPTION_PREINITIATION_CO | 63 | -0.26635 | -1.5425 | 0.013513514 | 0.15922557 | 1 | 11340 | tags=87%, list=48%, si | MM_4 |
| GOBP_NEGATIVE_REGUL | GOBP_NEGATIVE_REGULATION_OF_CYTOKI | 34 | -0.31322 | -1.54202 | 0.027586207 | 0.15921623 | 1 | 619 | tags=21%, list=3%, sig | MM_4 |
| GOBP_REGULATION_OF_ | GOBP_REGULATION_OF_ACTIN_FILAMENT_ | 158 | -0.22319 | -1.54192 | 0.0 | 0.1588499 | 1 | 3553 | tags=28%, list=15%, si | MM_4 |
| GOBP_POSITIVE_REGULA | GOBP_POSITIVE_REGULATION_OF_T_HELPE | 19 | -0.37212 | -1.54169 | 0.041025642 | 0.15857573 | 1 | 1763 | tags=21%, list=7%, sig | MM_4 |
| GOBP_REGULATION_OF_ | GOBP_REGULATION_OF_LYMPHOCYTE_APO | 71 | -0.24898 | -1.54159 | 0.0 | 0.15820414 | 1 | 918 | tags=13%, list=4%, sig | MM_4 |
| GOBP_NEGATIVE_REGUL | GOBP_NEGATIVE_REGULATION_OF_TELOM | 27 | -0.33078 | -1.54133 | 0.045751635 | 0.15801547 | 1 | 4693 | tags=63%, list=20%, si | MM_4 |
| TH17_CELL_DIFFERENTIA | TH17_CELL_DIFFERENTIATION | 94 | -0.24695 | -1.54118 | 0.016129032 | 0.15769917 | 1 | 2801 | tags=27%, list=12%, si | MM_4 |
| GOBP_REGULATION_OF_ | GOBP_REGULATION_OF_PROTEOLYSIS_INV | 215 | -0.20289 | -1.54058 | 0.0 | 0.15769164 | 1 | 3599 | tags=30%, list=15%, si | MM_4 |
| GOBP_PLATELET_MORPH | GOBP_PLATELET_MORPHOGENESIS | 25 | -0.33266 | -1.53936 | 0.026845638 | 0.158083 | 1 | 2377 | tags=32%, list=10%, si | MM_4 |
| GOBP_LYMPH_NODE_DE | GOBP_LYMPH_NODE_DEVELOPMENT | 32 | -0.31899 | -1.53872 | 0.041666668 | 0.1581841 | 1 | 158 | tags=9%, list=1%, sign | MM_4 |
| GOBP_REGULATION_OF_ | GOBP_REGULATION_OF_LEUKOCYTE_MIGR | 230 | -0.20032 | -1.53621 | 0.0 | 0.15989292 | 1 | 2077 | tags=17%, list=9%, sig | MM_4 |
| LEISHMANIASIS | LEISHMANIASIS | 60 | -0.27289 | -1.53565 | 0.011764706 | 0.1599613 | 1 | 2755 | tags=23%, list=12%, si | MM_4 |
| GOBP_BLOOD_VESSEL_E | GOBP_BLOOD_VESSEL_ENDOTHELIAL_CELL_ | 23 | -0.35098 | -1.53563 | 0.022727273 | 0.15954393 | 1 | 11 | tags=9%, list=0%, sign | MM_4 |
| GOBP_NEGATIVE_REGUL | GOBP_NEGATIVE_REGULATION_OF_TRANS | 463 | -0.17071 | -1.53434 | 0.0 | 0.16032307 | 1 | 2698 | tags=18%, list=11%, si | MM_4 |
| GOBP_CELLULAR_RESPO | GOBP_CELLULAR_RESPONSE_TO_DSRNA | 19 | -0.36612 | -1.53394 | 0.05154639 | 0.16021019 | 1 | 4494 | tags=47%, list=19%, si | MM_4 |
| GOBP_NEGATIVE_REGUL | GOBP_NEGATIVE_REGULATION_OF_LYMPH | 53 | -0.26965 | -1.53359 | 0.010869565 | 0.16000059 | 1 | 2270 | tags=23%, list=10%, si | MM_4 |
| ACUTE_MYELOID_LEUKEM | ACUTE_MYELOID_LEUKEMIA | 66 | -0.24593 | -1.53158 | 0.039473683 | 0.16129121 | 1 | 2284 | tags=18%, list=10%, si | MM_4 |
| GOBP_ESTABLISHMENT_ | GOBP_ESTABLISHMENT_OF_PROTEIN_LOCA | 24 | -0.33298 | -1.53018 | 0.029411765 | 0.16188346 | 1 | 13967 | tags=96%, list=59%, si | MM_4 |
| GOBP_DNA_STRAND_ELO | GOBP_DNA_STRAND_ELONGATION | 36 | -0.30526 | -1.52926 | 0.03649635 | 0.16215427 | 1 | 11197 | tags=89%, list=47%, si | MM_4 |
| GOBP_AEROBIC_RESPIRA | GOBP_AEROBIC_RESPIRATION | 170 | -0.21285 | -1.52818 | 0.0 | 0.16265318 | 1 | 12107 | tags=89%, list=51%, si | MM_4 |
| GOBP_REGULATION_OF_ | GOBP_REGULATION_OF_T_CELL_APOPTOTI | 48 | -0.28137 | -1.52762 | 0.010989011 | 0.16276844 | 1 | 343 | tags=10%, list=1%, sig | MM_4 |
| GOBP_TOLL_LIKE_RECEPT | GOBP_TOLL_LIKE_RECEPTOR_9_SIGNALING_ | 16 | -0.3898 | -1.52698 | 0.038647342 | 0.16289805 | 1 | 3770 | tags=50%, list=16%, si | MM_4 |
| GOBP_CYTOCHROME_CO | GOBP_CYTOCHROME_COMPLEX_ASSEMBLY | 32 | -0.31641 | -1.52635 | 0.04477612 | 0.16306247 | 1 | 1706 | tags=22%, list=7%, sig | MM_4 |
| GOBP_REGULATION_OF_ | GOBP_REGULATION_OF_EXTRINSIC_APOPT | 45 | -0.28458 | -1.52615 | 0.00862069 | 0.16275592 | 1 | 2666 | tags=22%, list=11%, si | MM_4 |
| GOBP_PROTEIN_QUALITY | GOBP_PROTEIN_QUALITY_CONTROL_FOR_M | 17 | -0.37266 | -1.52581 | 0.056410257 | 0.16261339 | 1 | 4166 | tags=47%, list=18%, si | MM_4 |
| GOBP_REGULATION_OF_ | GOBP_REGULATION_OF_NUCLEOBASE_CON | 16 | -0.38657 | -1.52406 | 0.039772727 | 0.16356085 | 1 | 2627 | tags=38%, list=11%, si | MM_4 |
| GOBP_VESICLE_TARGETIN | GOBP_VESICLE_TARGETING | 60 | -0.27218 | -1.52389 | 0.0 | 0.16332208 | 1 | 2987 | tags=35%, list=13%, si | MM_4 |
| GOBP_ESTABLISHMENT_ | GOBP_ESTABLISHMENT_OF_RNA_LOCALIZA | 138 | -0.21601 | -1.52087 | 0.0 | 0.16549423 | 1 | 11581 | tags=86%, list=49%, si | MM_4 |
| GOBP_ENDOTHELIAL_CEL | GOBP_ENDOTHELIAL_CELL_MORPHOGENES | 15 | -0.40694 | -1.52061 | 0.075 | 0.16517457 | 1 | 4925 | tags=60%, list=21%, si | MM_4 |
| GOBP_INTERFERON_BET | GOBP_INTERFERON_BETA_PRODUCTION | 66 | -0.25289 | -1.51961 | 0.022222223 | 0.16559723 | 1 | 2675 | tags=24%, list=11%, si | MM_4 |
| GOBP_RESPONSE_TO_AT | GOBP_RESPONSE_TO_ATP | 31 | -0.30415 | -1.51899 | 0.033557046 | 0.16557328 | 1 | 2758 | tags=29%, list=12%, si | MM_4 |

| | | | | | | | | | | |
|---|---|---|---|---|---|---|---|---|---|---|
| TGF-BETA_SIGNALING_PA | TGF-BETA_SIGNALING_PATHWAY | 85 | -0.23832 | -1.51894 | 0.0 | 0.16521746 | 1 | 1067 | tags=12%, list=5%, sig | MM_4 |
| GOBP_REGULATION_OF_ | GOBP_REGULATION_OF_TELOMERE_MAINT | 98 | -0.22863 | -1.51853 | 0.0 | 0.16512719 | 1 | 4745 | tags=45%, list=20%, si | MM_4 |
| TH1_AND_TH2_CELL_DIF | TH1_AND_TH2_CELL_DIFFERENTIATION | 81 | -0.23601 | -1.51821 | 0.016393442 | 0.1649454 | 1 | 2801 | tags=26%, list=12%, si | MM_4 |
| GOBP_LEUKOCYTE_DEGR | GOBP_LEUKOCYTE_DEGRANULATION | 77 | -0.24209 | -1.51805 | 0.027777778 | 0.16461366 | 1 | 1775 | tags=17%, list=8%, sig | MM_4 |
| GOBP_MITOCHONDRIAL_ | GOBP_MITOCHONDRIAL_RNA_METABOLIC_ | 51 | -0.27581 | -1.51743 | 0.021505376 | 0.1647622 | 1 | 4311 | tags=43%, list=18%, si | MM_4 |
| GOBP_PROTEIN_INSERTIO | GOBP_PROTEIN_INSERTION_INTO_MEMBRA | 39 | -0.29337 | -1.51528 | 0.028301887 | 0.16630635 | 1 | 4426 | tags=41%, list=19%, si | MM_4 |
| GOBP_GLYCEROLIPID_BIO | GOBP_GLYCEROLIPID_BIOSYNTHETIC_PROC | 210 | -0.20571 | -1.5139 | 0.0 | 0.16705209 | 1 | 3350 | tags=29%, list=14%, si | MM_4 |
| GOBP_ENDOPLASMIC_RE | GOBP_ENDOPLASMIC_RETICULUM_ORGANI | 90 | -0.23376 | -1.5133 | 0.0 | 0.16715413 | 1 | 4317 | tags=36%, list=18%, si | MM_4 |
| GOBP_T_CELL_APOPTOTI | GOBP_T_CELL_APOPTOTIC_PROCESS | 67 | -0.26102 | -1.51293 | 0.01369863 | 0.16708088 | 1 | 3703 | tags=30%, list=16%, si | MM_4 |
| GOBP_SEQUESTERING_O | GOBP_SEQUESTERING_OF_CALCIUM_ION | 129 | -0.2187 | -1.51249 | 0.030303031 | 0.16703458 | 1 | 1163 | tags=12%, list=5%, sig | MM_4 |
| GOBP_RESPONSE_TO_INT | GOBP_RESPONSE_TO_INTERFERON_BETA | 64 | -0.25627 | -1.51234 | 0.032967035 | 0.16670349 | 1 | 1135 | tags=17%, list=5%, sig | MM_4 |
| GOBP_REGULATION_OF_ | GOBP_REGULATION_OF_UBIQUITIN_DEPEN | 157 | -0.21864 | -1.51113 | 0.0 | 0.1672265 | 1 | 3599 | tags=30%, list=15%, si | MM_4 |
| GOBP_POSITIVE_REGULA | GOBP_POSITIVE_REGULATION_OF_TRANSLA | 21 | -0.34196 | -1.51041 | 0.06395349 | 0.16738012 | 1 | 10638 | tags=90%, list=45%, si | MM_4 |
| GOBP_REGULATION_OF_ | GOBP_REGULATION_OF_T_HELPER_1_TYPE_ | 33 | -0.30041 | -1.51028 | 0.024096385 | 0.1670415 | 1 | 1031 | tags=15%, list=4%, sig | MM_4 |
| GOBP_BONE_RESORPTIO | GOBP_BONE_RESORPTION | 76 | -0.25284 | -1.50909 | 0.0 | 0.16782397 | 1 | 3833 | tags=38%, list=16%, si | MM_4 |
| GOBP_NEUROTRANSMITT | GOBP_NEUROTRANSMITTER_RECEPTOR_TR | 40 | -0.27585 | -1.50909 | 0.036697246 | 0.16739585 | 1 | 15392 | tags=95%, list=65%, si | MM_4 |
| GOBP_SNRNA_METABOL | GOBP_SNRNA_METABOLIC_PROCESS | 50 | -0.26584 | -1.50821 | 0.035714287 | 0.16776137 | 1 | 11425 | tags=90%, list=48%, si | MM_4 |
| GOBP_NEGATIVE_REGUL | GOBP_NEGATIVE_REGULATION_OF_RESPON | 362 | -0.17932 | -1.50792 | 0.0 | 0.16761929 | 1 | 2432 | tags=17%, list=10%, si | MM_4 |
| GOBP_POSITIVE_REGULA | GOBP_POSITIVE_REGULATION_OF_CELL_KIL | 79 | -0.23578 | -1.50792 | 0.0 | 0.16720437 | 1 | 1963 | tags=16%, list=8%, sig | MM_4 |
| GOBP_NEGATIVE_REGUL | GOBP_NEGATIVE_REGULATION_OF_MYELO | 89 | -0.23447 | -1.5057 | 0.033898305 | 0.1687588 | 1 | 2788 | tags=24%, list=12%, si | MM_4 |
| GOBP_RESPONSE_TO_AR | GOBP_RESPONSE_TO_ARSENIC_CONTAININ | 25 | -0.32356 | -1.50526 | 0.041666668 | 0.16867997 | 1 | 2731 | tags=28%, list=12%, si | MM_4 |
| GOBP_SYNAPTIC_VESICLE | GOBP_SYNAPTIC_VESICLE_MATURATION | 27 | -0.31983 | -1.50517 | 0.046666667 | 0.16831891 | 1 | 3825 | tags=30%, list=16%, si | MM_4 |
| GOBP_LIPOSACCHARIDE_ | GOBP_LIPOSACCHARIDE_METABOLIC_PROC | 93 | -0.23377 | -1.50459 | 0.02173913 | 0.16832079 | 1 | 2934 | tags=27%, list=12%, si | MM_4 |
| GOBP_POSITIVE_REGULA | GOBP_POSITIVE_REGULATION_OF_CELL_AC | 373 | -0.18219 | -1.50427 | 0.0 | 0.16814624 | 1 | 3790 | tags=28%, list=16%, si | MM_4 |
| GLYCOSPHINGOLIPID_BIO | GLYCOSPHINGOLIPID_BIOSYNTHESIS | 44 | -0.27766 | -1.50387 | 0.03773585 | 0.16804706 | 1 | 1798 | tags=20%, list=8%, sig | MM_4 |
| GOBP_ENDOPLASMIC_RE | GOBP_ENDOPLASMIC_RETICULUM_TUBULA | 21 | -0.35501 | -1.50217 | 0.074285716 | 0.16906472 | 1 | 15236 | tags=100%, list=65%, | MM_4 |
| GOBP_MEMBRANE_RAFT | GOBP_MEMBRANE_RAFT_ORGANIZATION | 24 | -0.33079 | -1.5017 | 0.04117647 | 0.16901326 | 1 | 2607 | tags=33%, list=11%, si | MM_4 |
| GOBP_ENDONUCLEOLYTI | GOBP_ENDONUCLEOLYTIC_CLEAVAGE_OF_ | 16 | -0.37748 | -1.50111 | 0.05263158 | 0.16907576 | 1 | 11485 | tags=94%, list=49%, si | MM_4 |
| GOBP_RIG_I_SIGNALING_ | GOBP_RIG_I_SIGNALING_PATHWAY | 24 | -0.33843 | -1.50049 | 0.025974026 | 0.16914709 | 1 | 3817 | tags=33%, list=16%, si | MM_4 |
| GOBP_POSITIVE_REGULA | GOBP_POSITIVE_REGULATION_OF_PROTEIN | 201 | -0.20332 | -1.50032 | 0.0 | 0.1688088 | 1 | 3289 | tags=25%, list=14%, si | MM_4 |
| GOBP_POSITIVE_REGULA | GOBP_POSITIVE_REGULATION_OF_PROTEIN | 24 | -0.33733 | -1.49899 | 0.05421687 | 0.16944633 | 1 | 15654 | tags=100%, list=66%, | MM_4 |
| GOBP_REGULATION_OF_ | GOBP_REGULATION_OF_LYMPHOCYTE_MED | 190 | -0.20237 | -1.49786 | 0.0 | 0.1701584 | 1 | 2270 | tags=18%, list=10%, si | MM_4 |
| GOBP_POSITIVE_REGULA | GOBP_POSITIVE_REGULATION_OF_TELOME | 71 | -0.2454 | -1.49755 | 0.0 | 0.17006166 | 1 | 4013 | tags=38%, list=17%, si | MM_4 |
| GOBP_CD4_POSITIVE_AL | GOBP_CD4_POSITIVE_ALPHA_BETA_T_CELL | 26 | -0.32172 | -1.49567 | 0.030487806 | 0.17114075 | 1 | 1372 | tags=19%, list=6%, sig | MM_4 |
| GOBP_POSITIVE_REGULA | GOBP_POSITIVE_REGULATION_OF_LEUKOCY | 169 | -0.21842 | -1.4952 | 0.0 | 0.17116271 | 1 | 3564 | tags=31%, list=15%, si | MM_4 |
| GOBP_POSITIVE_REGULA | GOBP_POSITIVE_REGULATION_OF_LYMPHO | 21 | -0.3434 | -1.49446 | 0.06043956 | 0.171412 | 1 | 665 | tags=14%, list=3%, sig | MM_4 |
| RENAL_CELL_CARCINOMA | RENAL_CELL_CARCINOMA | 67 | -0.26485 | -1.49343 | 0.032608695 | 0.17181718 | 1 | 2037 | tags=19%, list=9%, sig | MM_4 |
| GOBP_RESPONSE_TO_KE | GOBP_RESPONSE_TO_KETONE | 134 | -0.21391 | -1.4922 | 0.0 | 0.1724987 | 1 | 2501 | tags=22%, list=11%, si | MM_4 |
| GOBP_REGULATION_OF_ | GOBP_REGULATION_OF_LEUKOCYTE_MEDIA | 253 | -0.20495 | -1.49151 | 0.0 | 0.1726326 | 1 | 2291 | tags=18%, list=10%, si | MM_4 |
| HYPERTROPHIC_CARDIOM | HYPERTROPHIC_CARDIOMYOPATHY_(HCM) | 84 | -0.23934 | -1.49016 | 0.030303031 | 0.1735216 | 1 | 1577 | tags=20%, list=7%, sig | MM_4 |
| GOBP_PROTEIN_K48_LIN | GOBP_PROTEIN_K48_LINKED_DEUBIQUITIN | 29 | -0.29968 | -1.48957 | 0.040268455 | 0.17369223 | 1 | 14397 | tags=97%, list=61%, si | MM_4 |
| GOBP_POSITIVE_REGULA | GOBP_POSITIVE_REGULATION_OF_FILOPOD | 35 | -0.29888 | -1.4889 | 0.05042017 | 0.1738505 | 1 | 2704 | tags=29%, list=11%, si | MM_4 |
| GOBP_PODOSOME_ASSE | GOBP_PODOSOME_ASSEMBLY | 25 | -0.33733 | -1.48849 | 0.07913669 | 0.17382881 | 1 | 1995 | tags=24%, list=8%, sig | MM_4 |
| GOBP_POSITIVE_REGULA | GOBP_POSITIVE_REGULATION_OF_PROTEIN | 87 | -0.23321 | -1.4879 | 0.0 | 0.17383741 | 1 | 3624 | tags=33%, list=15%, si | MM_4 |
| GOBP_NEGATIVE_REGUL | GOBP_NEGATIVE_REGULATION_OF_INTERF | 16 | -0.3679 | -1.48693 | 0.054347824 | 0.17427261 | 1 | 14930 | tags=100%, list=63%, | MM_4 |
| GOBP_ENDOCYTIC_RECYC | GOBP_ENDOCYTIC_RECYCLING | 88 | -0.24356 | -1.48557 | 0.018867925 | 0.17531298 | 1 | 15180 | tags=95%, list=64%, si | MM_4 |
| GOBP_REGULATION_OF_ | GOBP_REGULATION_OF_MULTICELLULAR_O | 83 | -0.25215 | -1.48414 | 0.03508772 | 0.17622262 | 1 | 1817 | tags=13%, list=8%, sig | MM_4 |
| GOBP_ENDOPLASMIC_RE | GOBP_ENDOPLASMIC_RETICULUM_TO_GOL | 120 | -0.22524 | -1.48373 | 0.0 | 0.17617503 | 1 | 2987 | tags=24%, list=13%, si | MM_4 |
| GOBP_POSITIVE_REGULA | GOBP_POSITIVE_REGULATION_OF_AUTOPH | 19 | -0.35858 | -1.48341 | 0.08121827 | 0.17599471 | 1 | 10914 | tags=95%, list=46%, si | MM_4 |
| GOBP_NUCLEOPHAGY | GOBP_NUCLEOPHAGY | 24 | -0.32029 | -1.48123 | 0.048484847 | 0.17758448 | 1 | 11533 | tags=92%, list=49%, si | MM_4 |
| GOBP_REGULATION_OF_ | GOBP_REGULATION_OF_TISSUE_REMODELI | 84 | -0.23556 | -1.48117 | 0.0 | 0.17725101 | 1 | 2760 | tags=26%, list=12%, si | MM_4 |
| GOBP_POSITIVE_REGULA | GOBP_POSITIVE_REGULATION_OF_DNA_TE | 72 | -0.23457 | -1.48079 | 0.0 | 0.17719556 | 1 | 11426 | tags=86%, list=48%, si | MM_4 |
| GOBP_T_HELPER_2_CELL | GOBP_T_HELPER_2_CELL_CYTOKINE_PRODU | 17 | -0.36397 | -1.48046 | 0.057291668 | 0.17709936 | 1 | 2469 | tags=35%, list=10%, si | MM_4 |
| GOBP_ENDOLYSOSOMAL_ | GOBP_ENDOLYSOSOMAL_TOLL_LIKE_RECEP | 47 | -0.26914 | -1.48042 | 0.067226894 | 0.17673545 | 1 | 3910 | tags=43%, list=17%, si | MM_4 |
| GOBP_NEGATIVE_REGUL | GOBP_NEGATIVE_REGULATION_OF_T_CELL_ | 22 | -0.33445 | -1.48036 | 0.05882353 | 0.17637442 | 1 | 1963 | tags=27%, list=8%, sig | MM_4 |
| GOBP_REGULATION_OF_ | GOBP_REGULATION_OF_GRANULOCYTE_CH | 59 | -0.25306 | -1.47992 | 0.0 | 0.17630471 | 1 | 918 | tags=14%, list=4%, sig | MM_4 |
| GOBP_MEMBRANE_BIOG | GOBP_MEMBRANE_BIOGENESIS | 63 | -0.24562 | -1.4772 | 0.013157895 | 0.17834012 | 1 | 13148 | tags=89%, list=56%, si | MM_4 |
| GOBP_RESPONSE_TO_TR | GOBP_RESPONSE_TO_TRANSFORMING_GRO | 246 | -0.19254 | -1.47593 | 0.0 | 0.17908196 | 1 | 1953 | tags=16%, list=8%, sig | MM_4 |
| GOBP_LYMPHOCYTE_APO | GOBP_LYMPHOCYTE_APOPTOTIC_PROCESS | 94 | -0.22142 | -1.47499 | 0.0 | 0.17956999 | 1 | 3720 | tags=28%, list=16%, si | MM_4 |
| GOBP_NEGATIVE_REGUL | GOBP_NEGATIVE_REGULATION_OF_LEUKOC | 66 | -0.24863 | -1.47424 | 0.033707865 | 0.17985925 | 1 | 2291 | tags=21%, list=10%, si | MM_4 |
| GOBP_CELLULAR_RESPON | GOBP_CELLULAR_RESPONSE_TO_VIRUS | 47 | -0.26859 | -1.47359 | 0.021505376 | 0.18012878 | 1 | 1039 | tags=13%, list=4%, sig | MM_4 |
| GOBP_REGULATION_OF_ | GOBP_REGULATION_OF_DNA_STRAND_ELO | 18 | -0.36792 | -1.47255 | 0.07476635 | 0.18074083 | 1 | 11169 | tags=94%, list=47%, si | MM_4 |
| GOBP_MAINTENANCE_OF | GOBP_MAINTENANCE_OF_LOCATION | 341 | -0.18897 | -1.47252 | 0.0 | 0.18037757 | 1 | 2564 | tags=18%, list=11%, si | MM_4 |
| GOBP_RADIAL_GLIAL_CEL | GOBP_RADIAL_GLIAL_CELL_DIFFERENTIATIO | 18 | -0.35535 | -1.471 | 0.07734807 | 0.1814591 | 1 | 512 | tags=17%, list=2%, sig | MM_4 |
| GOBP_NEGATIVE_REGUL | GOBP_NEGATIVE_REGULATION_OF_MYELO | 57 | -0.24394 | -1.47078 | 0.026666667 | 0.1812419 | 1 | 2975 | tags=28%, list=13%, si | MM_4 |
| ARGININE_BIOSYNTHESIS | ARGININE_BIOSYNTHESIS | 17 | -0.34772 | -1.46965 | 0.06532663 | 0.18176606 | 1 | 433 | tags=18%, list=2%, sig | MM_4 |
| GOBP_POSITIVE_REGULA | GOBP_POSITIVE_REGULATION_OF_RESPONS | 73 | -0.24034 | -1.46959 | 0.014925373 | 0.18144141 | 1 | 1294 | tags=14%, list=5%, sig | MM_4 |
| GOBP_CEREBELLAR_PURK | GOBP_CEREBELLAR_PURKINJE_CELL_LAYER_ | 16 | -0.36917 | -1.46823 | 0.07486631 | 0.18235022 | 1 | 14900 | tags=100%, list=63%, | MM_4 |
| GOBP_I_KAPPAB_PHOSPH | GOBP_I_KAPPAB_PHOSPHORYLATION | 17 | -0.36297 | -1.46715 | 0.06726457 | 0.18312316 | 1 | 4353 | tags=41%, list=18%, si | MM_4 |
| GOBP_POSITIVE_REGULA | GOBP_POSITIVE_REGULATION_OF_APOPTO | 161 | -0.2073 | -1.46556 | 0.0 | 0.18423611 | 1 | 2704 | tags=19%, list=11%, si | MM_4 |
| GOBP_REGULATION_OF_ | GOBP_REGULATION_OF_ALTERNATIVE_MRN | 56 | -0.25111 | -1.46456 | 0.05 | 0.18482567 | 1 | 15855 | tags=96%, list=67%, si | MM_4 |
| GOBP_ORGANIC_CATION | GOBP_ORGANIC_CATION_TRANSPORT | 122 | -0.21454 | -1.46443 | 0.0 | 0.18448459 | 1 | 1513 | tags=16%, list=6%, sig | MM_4 |
| PPAR_SIGNALING_PATHW | PPAR_SIGNALING_PATHWAY | 70 | -0.24066 | -1.46433 | 0.0 | 0.1841966 | 1 | 1076 | tags=14%, list=5%, sig | MM_4 |
| NUCLEOTIDE_EXCISION_R | NUCLEOTIDE_EXCISION_REPAIR | 42 | -0.28054 | -1.46427 | 0.019607844 | 0.18383385 | 1 | 11559 | tags=90%, list=49%, si | MM_4 |
| GOBP_TELOMERE_MAINT | GOBP_TELOMERE_MAINTENANCE_VIA_REC | 15 | -0.38918 | -1.46315 | 0.087628864 | 0.18438162 | 1 | 4477 | tags=47%, list=19%, si | MM_4 |
| GOBP_MYELOID_CELL_DE | GOBP_MYELOID_CELL_DEVELOPMENT | 98 | -0.23226 | -1.4608 | 0.019607844 | 0.18627025 | 1 | 3472 | tags=28%, list=15%, si | MM_4 |
| GOBP_RNA_LOCALIZATIO | GOBP_RNA_LOCALIZATION | 164 | -0.1992 | -1.46047 | 0.0 | 0.18616274 | 1 | 11581 | tags=85%, list=49%, si | MM_4 |
| GOBP_DNA_REPLICATION | GOBP_DNA_REPLICATION_CHECKPOINT_SIG | 18 | -0.35952 | -1.4591 | 0.06593407 | 0.18693523 | 1 | 3376 | tags=39%, list=14%, si | MM_4 |
| GOBP_SKELETAL_MUSCLE | GOBP_SKELETAL_MUSCLE_FIBER_DIFFERENT | 15 | -0.36666 | -1.45823 | 0.05612245 | 0.18733148 | 1 | 14959 | tags=100%, list=63%, | MM_4 |
| GOBP_POSITIVE_REGULA | GOBP_POSITIVE_REGULATION_OF_MYELOID | 21 | -0.34352 | -1.45787 | 0.051282052 | 0.18736054 | 1 | 11407 | tags=90%, list=48%, si | MM_4 |
| GOBP_NEGATIVE_REGUL | GOBP_NEGATIVE_REGULATION_OF_CELL_SI | 16 | -0.36477 | -1.45721 | 0.073446326 | 0.18761523 | 1 | 15004 | tags=100%, list=64%, | MM_4 |
| GOBP_NEGATIVE_REGUL | GOBP_NEGATIVE_REGULATION_OF_CYTOPL | 50 | -0.25777 | -1.45715 | 0.034782607 | 0.18727541 | 1 | 2421 | tags=18%, list=10%, si | MM_4 |

| | | | | | | | | | | |
|---|---|---|---|---|---|---|---|---|---|---|
| GOBP_NEGATIVE_REGUL | GOBP_NEGATIVE_REGULATION_OF_CELL_C | 67 | -0.2364 | -1.4565 | 0.019607844 | 0.18746142 | 1 | 2977 | tags=22%, list=13%, si | MM_4 |
| GOBP_STRESS_ACTIVATEI | GOBP_STRESS_ACTIVATED_PROTEIN_KINAS | 59 | -0.2592 | -1.4559 | 0.021052632 | 0.18763453 | 1 | 3667 | tags=34%, list=16%, si | MM_4 |
| DNA_REPLICATION | DNA_REPLICATION | 34 | -0.29095 | -1.4558 | 0.042857144 | 0.18730971 | 1 | 1440 | tags=21%, list=6%, sig | MM_4 |
| GOBP_REGULATION_OF_ | GOBP_REGULATION_OF_TELOMERE_CAPPIN | 26 | -0.31418 | -1.45567 | 0.056962024 | 0.18704107 | 1 | 4683 | tags=58%, list=20%, si | MM_4 |
| GOBP_NEGATIVE_REGUL | GOBP_NEGATIVE_REGULATION_OF_SPROUT | 15 | -0.38051 | -1.45546 | 0.08372093 | 0.18686461 | 1 | 662 | tags=20%, list=3%, sig | MM_4 |
| GOBP_RESPONSE_TO_IN( | GOBP_RESPONSE_TO_INORGANIC_SUBSTAN | 432 | -0.16742 | -1.45503 | 0.0 | 0.18684632 | 1 | 2548 | tags=18%, list=11%, si | MM_4 |
| GOBP_ACTIVATED_T_CEL | GOBP_ACTIVATED_T_CELL_PROLIFERATION | 50 | -0.26945 | -1.45422 | 0.03883495 | 0.1871649 | 1 | 2955 | tags=30%, list=13%, si | MM_4 |
| GOBP_CHOLESTEROL_STO | GOBP_CHOLESTEROL_STORAGE | 27 | -0.32229 | -1.45393 | 0.05925926 | 0.18708128 | 1 | 1682 | tags=26%, list=7%, sig | MM_4 |
| GOBP_POSITIVE_REGULA | GOBP_POSITIVE_REGULATION_OF_LEUKOCY | 261 | -0.19378 | -1.45337 | 0.0 | 0.18725286 | 1 | 3564 | tags=27%, list=15%, si | MM_4 |
| GOBP_GRANULOCYTE_M | GOBP_GRANULOCYTE_MIGRATION | 146 | -0.2005 | -1.45293 | 0.03846154 | 0.18727514 | 1 | 1178 | tags=12%, list=5%, sig | MM_4 |
| GOBP_AEROBIC_ELECTRO | GOBP_AEROBIC_ELECTRON_TRANSPORT_CH | 63 | -0.24078 | -1.45116 | 0.021276595 | 0.18853743 | 1 | 12107 | tags=90%, list=51%, si | MM_4 |
| GOBP_COTRANSLATIONA | GOBP_COTRANSLATIONAL_PROTEIN_TARGE | 20 | -0.34026 | -1.44918 | 0.0867052 | 0.19005258 | 1 | 12352 | tags=95%, list=52%, si | MM_4 |
| GOBP_REGULATION_OF_ | GOBP_REGULATION_OF_STRESS_ACTIVATED | 41 | -0.28733 | -1.44901 | 0.04347826 | 0.18974784 | 1 | 3667 | tags=34%, list=16%, si | MM_4 |
| PHAGOSOME | PHAGOSOME | 146 | -0.20423 | -1.44584 | 0.0 | 0.19219324 | 1 | 3604 | tags=25%, list=15%, si | MM_4 |
| GOBP_POSITIVE_REGULA | GOBP_POSITIVE_REGULATION_OF_CYTOKIN | 27 | -0.31037 | -1.44571 | 0.075 | 0.1918999 | 1 | 1876 | tags=26%, list=8%, sig | MM_4 |
| GOBP_LYMPHOCYTE_CHE | GOBP_LYMPHOCYTE_CHEMOTAXIS | 38 | -0.28425 | -1.44546 | 0.06557377 | 0.19171728 | 1 | 1158 | tags=21%, list=5%, sig | MM_4 |
| GOBP_REGULATION_OF_ | GOBP_REGULATION_OF_CELLULAR_RESPON | 26 | -0.3141 | -1.44412 | 0.061728396 | 0.19267417 | 1 | 1953 | tags=23%, list=8%, sig | MM_4 |
| GOBP_LEUKOCYTE_MEDI | GOBP_LEUKOCYTE_MEDIATED_CYTOTOXICIT | 135 | -0.20648 | -1.44341 | 0.032258064 | 0.193015 | 1 | 1963 | tags=16%, list=8%, sig | MM_4 |
| HEMATOPOIETIC_CELL_LI | HEMATOPOIETIC_CELL_LINEAGE | 83 | -0.23121 | -1.44332 | 0.013513514 | 0.19269136 | 1 | 1890 | tags=18%, list=8%, sig | MM_4 |
| GOBP_MACROAUTOPHAG | GOBP_MACROAUTOPHAGY | 287 | -0.18999 | -1.44217 | 0.0 | 0.19334088 | 1 | 3817 | tags=26%, list=16%, si | MM_4 |
| GOBP_NEGATIVE_REGUL | GOBP_NEGATIVE_REGULATION_OF_DNA_B | 148 | -0.21622 | -1.44209 | 0.0 | 0.19298153 | 1 | 1886 | tags=16%, list=8%, sig | MM_4 |
| GOBP_FILOPODIUM_ASSI | GOBP_FILOPODIUM_ASSEMBLY | 74 | -0.23533 | -1.44206 | 0.0 | 0.1926243 | 1 | 2704 | tags=24%, list=11%, si | MM_4 |
| GOBP_MITOCHONDRIAL_ | GOBP_MITOCHONDRIAL_DNA_REPLICATION | 17 | -0.35585 | -1.43994 | 0.07329843 | 0.19423312 | 1 | 11389 | tags=94%, list=48%, si | MM_4 |
| GOBP_DENSE_CORE_GRA | GOBP_DENSE_CORE_GRANULE_LOCALIZATI | 28 | -0.31072 | -1.43803 | 0.07586207 | 0.19570763 | 1 | 15227 | tags=96%, list=64%, si | MM_4 |
| GOBP_POSITIVE_REGULA | GOBP_POSITIVE_REGULATION_OF_TRANSF | 18 | -0.33751 | -1.43739 | 0.073033705 | 0.19594528 | 1 | 619 | tags=17%, list=3%, sig | MM_4 |
| GOBP_PROTEIN_PEPTIDY | GOBP_PROTEIN_PEPTIDYL_PROLYL_ISOMER | 25 | -0.32587 | -1.43734 | 0.09589041 | 0.19562116 | 1 | 11546 | tags=92%, list=49%, si | MM_4 |
| GOBP_REGULATION_OF_ | GOBP_REGULATION_OF_T_CELL_ACTIVATIO | 364 | -0.18578 | -1.43401 | 0.0 | 0.1984181 | 1 | 3574 | tags=26%, list=15%, si | MM_4 |
| GOBP_NEUTROPHIL_MED | GOBP_NEUTROPHIL_MEDIATED_IMMUNITY | 37 | -0.27693 | -1.43342 | 0.07964602 | 0.19860145 | 1 | 1178 | tags=19%, list=5%, sig | MM_4 |
| GOBP_REGULATION_OF_ | GOBP_REGULATION_OF_EXECUTION_PHASE | 26 | -0.31144 | -1.43314 | 0.08391608 | 0.19850078 | 1 | 14714 | tags=96%, list=62%, si | MM_4 |
| GOBP_POSITIVE_REGULA | GOBP_POSITIVE_REGULATION_OF_SMOOTH | 17 | -0.35767 | -1.43297 | 0.07272727 | 0.19825403 | 1 | 1273 | tags=18%, list=5%, sig | MM_4 |
| GOBP_INTRINSIC_APOPT( | GOBP_INTRINSIC_APOPTOTIC_SIGNALING_F | 112 | -0.2091 | -1.43281 | 0.0 | 0.1980164 | 1 | 3025 | tags=25%, list=13%, si | MM_4 |
| GOBP_REGULATION_OF_ | GOBP_REGULATION_OF_TYPE_2_IMMUNE_ | 34 | -0.27828 | -1.43109 | 0.05882353 | 0.19924803 | 1 | 2209 | tags=26%, list=9%, sig | MM_4 |
| GOBP_LYTIC_VACUOLE_C | GOBP_LYTIC_VACUOLE_ORGANIZATION | 95 | -0.21959 | -1.43075 | 0.023809524 | 0.19922405 | 1 | 3654 | tags=31%, list=15%, si | MM_4 |
| GOBP_REGULATION_OF_ | GOBP_REGULATION_OF_VESICLE_SIZE | 21 | -0.3373 | -1.43049 | 0.06395349 | 0.19907723 | 1 | 12654 | tags=95%, list=54%, si | MM_4 |
| GOBP_REGULATION_OF_ | GOBP_REGULATION_OF_JUN_KINASE_ACTIV | 62 | -0.23842 | -1.42833 | 0.025 | 0.20071867 | 1 | 3081 | tags=31%, list=13%, si | MM_4 |
| GOBP_PHOSPHATIDIC_AC | GOBP_PHOSPHATIDIC_ACID_METABOLIC_PI | 27 | -0.30431 | -1.42801 | 0.06622516 | 0.20068686 | 1 | 2594 | tags=37%, list=11%, si | MM_4 |
| GOBP_REGULATION_OF_ | GOBP_REGULATION_OF_GOLGI_ORGANIZAT | 17 | -0.35732 | -1.42777 | 0.060606062 | 0.20049736 | 1 | 15180 | tags=100%, list=64%, | MM_4 |
| GOBP_VESICLE_TARGETIN | GOBP_VESICLE_TARGETING_TO_FROM_OR_ | 29 | -0.2849 | -1.42768 | 0.062937066 | 0.20018682 | 1 | 2887 | tags=38%, list=12%, si | MM_4 |
| GOBP_NEGATIVE_REGUL | GOBP_NEGATIVE_REGULATION_OF_CALCIU | 69 | -0.24587 | -1.42669 | 0.0 | 0.2007863 | 1 | 2780 | tags=23%, list=12%, si | MM_4 |
| GOBP_MATURATION_OF_ | GOBP_MATURATION_OF_LSU_RRNA_FROM | 17 | -0.35291 | -1.42588 | 0.082125604 | 0.20129874 | 1 | 11590 | tags=94%, list=49%, si | MM_4 |
| RIG-I-LIKE_RECEPTOR_SIG | RIG-I-LIKE_RECEPTOR_SIGNALING_PATHWA | 52 | -0.25042 | -1.42572 | 0.06122449 | 0.20108755 | 1 | 1495 | tags=13%, list=6%, sig | MM_4 |
| GOBP_NEGATIVE_REGUL | GOBP_NEGATIVE_REGULATION_OF_PEPTID | 18 | -0.34603 | -1.42563 | 0.12290503 | 0.20076911 | 1 | 4785 | tags=61%, list=20%, si | MM_4 |
| GOBP_AXONAL_TRANSPO | GOBP_AXONAL_TRANSPORT_OF_MITOCHO | 17 | -0.37377 | -1.42494 | 0.07650273 | 0.20098957 | 1 | 14792 | tags=100%, list=63%, | MM_4 |
| GOBP_PROTEIN_TARGETI | GOBP_PROTEIN_TARGETING | 285 | -0.17955 | -1.42477 | 0.0 | 0.20075803 | 1 | 3099 | tags=21%, list=13%, si | MM_4 |
| GOBP_NEGATIVE_REGUL | GOBP_NEGATIVE_REGULATION_OF_LYMPH | 44 | -0.26199 | -1.42263 | 0.042735044 | 0.20259634 | 1 | 1904 | tags=18%, list=8%, sig | MM_4 |
| GOBP_RRNA_METABOLIC | GOBP_RRNA_METABOLIC_PROCESS | 241 | -0.20067 | -1.4224 | 0.0 | 0.20241071 | 1 | 11616 | tags=88%, list=49%, si | MM_4 |
| GOBP_DNA_REPAIR_DEP | GOBP_DNA_REPAIR_DEPENDENT_CHROMAT | 23 | -0.31667 | -1.42153 | 0.06369427 | 0.20296438 | 1 | 11402 | tags=91%, list=48%, si | MM_4 |
| GOBP_NEGATIVE_REGUL | GOBP_NEGATIVE_REGULATION_OF_INTERL | 18 | -0.34484 | -1.42146 | 0.08064516 | 0.20263335 | 1 | 1634 | tags=22%, list=7%, sig | MM_4 |
| GOBP_NEGATIVE_REGUL | GOBP_NEGATIVE_REGULATION_OF_UBIQUI | 55 | -0.23642 | -1.41713 | 0.036144577 | 0.20679997 | 1 | 14572 | tags=93%, list=62%, si | MM_4 |
| GOBP_AUTOPHAGY_OF_I | GOBP_AUTOPHAGY_OF_MITOCHONDRION | 89 | -0.22466 | -1.41579 | 0.020408163 | 0.20779215 | 1 | 2750 | tags=20%, list=12%, si | MM_4 |
| GOBP_VESICLE_MEDIATE | GOBP_VESICLE_MEDIATED_TRANSPORT_TO | 158 | -0.19799 | -1.4141 | 0.0 | 0.2089579 | 1 | 15626 | tags=95%, list=66%, si | MM_4 |
| GOBP_POSITIVE_REGULA | GOBP_POSITIVE_REGULATION_OF_LEUKOCY | 152 | -0.19448 | -1.41403 | 0.0 | 0.20861867 | 1 | 2209 | tags=17%, list=9%, sig | MM_4 |
| GOBP_NAD_METABOLIC_ | GOBP_NAD_METABOLIC_PROCESS | 58 | -0.24414 | -1.41351 | 0.019417476 | 0.20876554 | 1 | 3907 | tags=36%, list=17%, si | MM_4 |
| GOBP_ENDOSOMAL_TRA | GOBP_ENDOSOMAL_TRANSPORT | 272 | -0.17042 | -1.41335 | 0.0 | 0.20853148 | 1 | 12088 | tags=86%, list=51%, si | MM_4 |
| GOBP_LONG_CHAIN_FAT | GOBP_LONG_CHAIN_FATTY_ACID_IMPORT_ | 19 | -0.36156 | -1.4121 | 0.09090909 | 0.20946471 | 1 | 2476 | tags=26%, list=10%, si | MM_4 |
| GOBP_CELLULAR_RESPON | GOBP_CELLULAR_RESPONSE_TO_OXYGEN_F | 20 | -0.33587 | -1.41185 | 0.078947365 | 0.20939393 | 1 | 3307 | tags=40%, list=14%, si | MM_4 |
| GOBP_POSITIVE_REGULA | GOBP_POSITIVE_REGULATION_OF_MITOCH | 17 | -0.34905 | -1.41114 | 0.06111111 | 0.2097147 | 1 | 4240 | tags=53%, list=18%, si | MM_4 |
| GOBP_RESPONSE_TO_OX | GOBP_RESPONSE_TO_OXYGEN_RADICAL | 24 | -0.31042 | -1.40999 | 0.059210528 | 0.210595 | 1 | 2015 | tags=29%, list=9%, sig | MM_4 |
| GOBP_REGULATION_OF_ | GOBP_REGULATION_OF_PROGRAMMED_NE | 27 | -0.29816 | -1.40914 | 0.08 | 0.21123686 | 1 | 12532 | tags=93%, list=53%, si | MM_4 |
| GOBP_RESPONSE_TO_NU | GOBP_RESPONSE_TO_NUTRIENT | 75 | -0.22883 | -1.40895 | 0.027027028 | 0.21106131 | 1 | 4263 | tags=40%, list=18%, si | MM_4 |
| GOBP_POSITIVE_REGULA | GOBP_POSITIVE_REGULATION_OF_TOR_SIG | 71 | -0.23575 | -1.40696 | 0.016129032 | 0.2127471 | 1 | 4633 | tags=41%, list=20%, si | MM_4 |
| GOBP_REGULATION_OF_ | GOBP_REGULATION_OF_PHAGOCYTOSIS | 106 | -0.214 | -1.40633 | 0.04761905 | 0.21310768 | 1 | 1905 | tags=16%, list=8%, sig | MM_4 |
| GOBP_NEGATIVE_REGUL | GOBP_NEGATIVE_REGULATION_OF_INFLAM | 153 | -0.2061 | -1.40614 | 0.04761905 | 0.21293776 | 1 | 2291 | tags=19%, list=10%, si | MM_4 |
| GOBP_LEUKOCYTE_APOP | GOBP_LEUKOCYTE_APOPTOTIC_PROCESS | 131 | -0.20327 | -1.40582 | 0.032258064 | 0.21280745 | 1 | 2105 | tags=17%, list=9%, sig | MM_4 |
| GOBP_PORE_COMPLEX_A | GOBP_PORE_COMPLEX_ASSEMBLY | 20 | -0.33907 | -1.40573 | 0.09352518 | 0.21251893 | 1 | 3545 | tags=40%, list=15%, si | MM_4 |
| GOBP_REGULATION_OF_ | GOBP_REGULATION_OF_NUCLEASE_ACTIVIT | 22 | -0.29826 | -1.40521 | 0.09615385 | 0.21270686 | 1 | 4013 | tags=50%, list=17%, si | MM_4 |
| GOBP_POSITIVE_REGULA | GOBP_POSITIVE_REGULATION_OF_DNA_DA | 15 | -0.34788 | -1.40165 | 0.07368421 | 0.21609432 | 1 | 15402 | tags=100%, list=65%, | MM_4 |
| GOBP_ORGANELLE_TRAN | GOBP_ORGANELLE_TRANSPORT_ALONG_M | 86 | -0.22448 | -1.40113 | 0.023809524 | 0.21626623 | 1 | 2894 | tags=22%, list=12%, si | MM_4 |
| GOBP_TYPE_B_PANCREA | GOBP_TYPE_B_PANCREATIC_CELL_APOPTO | 15 | -0.35983 | -1.40063 | 0.093596056 | 0.2163779 | 1 | 15120 | tags=100%, list=64%, | MM_4 |
| GOBP_TELOMERE_CAPPI | GOBP_TELOMERE_CAPPING | 38 | -0.25789 | -1.40039 | 0.08130081 | 0.21624294 | 1 | 4213 | tags=47%, list=18%, si | MM_4 |
| GOBP_RESPONSE_TO_CA | GOBP_RESPONSE_TO_CADMIUM_ION | 35 | -0.28334 | -1.40036 | 0.075187966 | 0.21583416 | 1 | 2240 | tags=26%, list=9%, sig | MM_4 |
| GOBP_REGULATION_OF_ | GOBP_REGULATION_OF_PROTEIN_POLYME | 194 | -0.19693 | -1.40027 | 0.0 | 0.21552092 | 1 | 3626 | tags=29%, list=15%, si | MM_4 |
| GOBP_MYELOID_LEUKOC | GOBP_MYELOID_LEUKOCYTE_MEDIATED_IM | 108 | -0.20797 | -1.40024 | 0.02631579 | 0.21516137 | 1 | 2563 | tags=19%, list=11%, si | MM_4 |
| GOBP_SPHINGOLIPID_ME | GOBP_SPHINGOLIPID_METABOLIC_PROCESS | 137 | -0.19589 | -1.39886 | 0.0 | 0.2161878 | 1 | 2085 | tags=20%, list=9%, sig | MM_4 |
| GOBP_VASCULAR_ASSOC | GOBP_VASCULAR_ASSOCIATED_SMOOTH_N | 41 | -0.26562 | -1.39797 | 0.10344828 | 0.21684971 | 1 | 1767 | tags=20%, list=7%, sig | MM_4 |
| GOBP_INDOLE_CONTAIN | GOBP_INDOLE_CONTAINING_COMPOUND_ | 25 | -0.30972 | -1.3973 | 0.094339624 | 0.21721256 | 1 | 338 | tags=12%, list=1%, sig | MM_4 |
| GOBP_REGULATION_OF_ | GOBP_REGULATION_OF_OXIDATIVE_PHOSP | 23 | -0.31554 | -1.39665 | 0.062111802 | 0.21747483 | 1 | 2273 | tags=26%, list=10%, si | MM_4 |
| GOBP_COPPER_ION_TRA | GOBP_COPPER_ION_TRANSPORT | 16 | -0.35476 | -1.39623 | 0.108571425 | 0.21755087 | 1 | 1733 | tags=31%, list=7%, sig | MM_4 |
| FOCAL_ADHESION | FOCAL_ADHESION | 194 | -0.19334 | -1.3961 | 0.0 | 0.21725748 | 1 | 1628 | tags=14%, list=7%, sig | MM_4 |
| GOBP_NEGATIVE_REGUL | GOBP_NEGATIVE_REGULATION_OF_TUMOR | 67 | -0.24296 | -1.39605 | 0.03076923 | 0.21689363 | 1 | 1031 | tags=13%, list=4%, sig | MM_4 |
| GOBP_NEGATIVE_REGUL | GOBP_NEGATIVE_REGULATION_OF_RNA_SF | 29 | -0.28773 | -1.39538 | 0.05970149 | 0.21722868 | 1 | 11582 | tags=90%, list=49%, si | MM_4 |
| GOBP_TOLL_LIKE_RECEPT | GOBP_TOLL_LIKE_RECEPTOR_2_SIGNALING_ | 17 | -0.35152 | -1.39436 | 0.08510638 | 0.21787944 | 1 | 2133 | tags=29%, list=9%, sig | MM_4 |

| | | | | | | | | | | |
|---|---|---|---|---|---|---|---|---|---|---|
| GOBP_ENUCLEATE_ERYTH | GOBP_ENUCLEATE_ERYTHROCYTE_DIFFEREN | 15 | -0.36337 | -1.39377 | 0.11290322 | 0.21802905 | 1 | 3618 | tags=47%, list=15%, si | MM_4 |
| GOBP_ANTIGEN_PROCES | GOBP_ANTIGEN_PROCESSING_AND_PRESEN | 38 | -0.27161 | -1.39329 | 0.058333334 | 0.21817839 | 1 | 3970 | tags=42%, list=17%, si | MM_4 |
| GOBP_REGULATION_OF_ | GOBP_REGULATION_OF_POSTSYNAPTIC_NE | 34 | -0.28881 | -1.39283 | 0.08695652 | 0.21835195 | 1 | 2757 | tags=26%, list=12%, si | MM_4 |
| GOBP_POSITIVE_REGULA | GOBP_POSITIVE_REGULATION_OF_CELL_GR | 185 | -0.18359 | -1.39278 | 0.0 | 0.21801035 | 1 | 2846 | tags=22%, list=12%, si | MM_4 |
| GOBP_FOAM_CELL_DIFFE | GOBP_FOAM_CELL_DIFFERENTIATION | 32 | -0.27728 | -1.39265 | 0.0661157 | 0.21775453 | 1 | 1682 | tags=22%, list=7%, sig | MM_4 |
| GOBP_AUTOPHAGOSOMI | GOBP_AUTOPHAGOSOME_MATURATION | 58 | -0.25113 | -1.39162 | 0.03960396 | 0.21863584 | 1 | 11525 | tags=88%, list=49%, si | MM_4 |
| GOBP_NEGATIVE_REGUL | GOBP_NEGATIVE_REGULATION_OF_CELL_M | 36 | -0.26882 | -1.39161 | 0.05970149 | 0.21824911 | 1 | 1972 | tags=17%, list=8%, sig | MM_4 |
| GOBP_POSITIVE_REGULA | GOBP_POSITIVE_REGULATION_OF_VASCULA | 25 | -0.31233 | -1.39086 | 0.077922076 | 0.21857059 | 1 | 1729 | tags=20%, list=7%, sig | MM_4 |
| GOBP_HAIR_CYCLE_PHAS | GOBP_HAIR_CYCLE_PHASE | 17 | -0.34469 | -1.39084 | 0.1010101 | 0.21820395 | 1 | 4028 | tags=41%, list=17%, si | MM_4 |
| GOBP_T_CELL_PROLIFERA | GOBP_T_CELL_PROLIFERATION | 233 | -0.19867 | -1.38828 | 0.0 | 0.22085863 | 1 | 3574 | tags=27%, list=15%, si | MM_4 |
| GOBP_RESPONSE_TO_HY | GOBP_RESPONSE_TO_HYDROPEROXIDE | 18 | -0.34244 | -1.38801 | 0.086021505 | 0.22070651 | 1 | 3563 | tags=44%, list=15%, si | MM_4 |
| GOBP_CELL_CELL_ADHES | GOBP_CELL_CELL_ADHESION_MEDIATED_BY | 18 | -0.34315 | -1.38721 | 0.09859155 | 0.22123641 | 1 | 13640 | tags=94%, list=58%, si | MM_4 |
| GOBP_PROTEIN_EXIT_FR | GOBP_PROTEIN_EXIT_FROM_ENDOPLASMIC | 42 | -0.25549 | -1.38692 | 0.07258064 | 0.22107345 | 1 | 9519 | tags=86%, list=40%, si | MM_4 |
| GOBP_REGULATION_OF_ | GOBP_REGULATION_OF_RIG_I_SIGNALING_ | 18 | -0.34239 | -1.38601 | 0.10152284 | 0.2217746 | 1 | 10792 | tags=89%, list=46%, si | MM_4 |
| GOBP_POSITIVE_REGULA | GOBP_POSITIVE_REGULATION_OF_IMMUNE | 266 | -0.17289 | -1.38334 | 0.0 | 0.22431763 | 1 | 2209 | tags=16%, list=9%, sig | MM_4 |
| GOBP_BONE_REMODELIN | GOBP_BONE_REMODELING | 102 | -0.21245 | -1.38268 | 0.0 | 0.22468194 | 1 | 2820 | tags=26%, list=12%, si | MM_4 |
| GOBP_REGULATION_OF_ | GOBP_REGULATION_OF_TELOMERE_MAINT | 61 | -0.23911 | -1.38199 | 0.030927835 | 0.22498508 | 1 | 4318 | tags=46%, list=18%, si | MM_4 |
| CYTOKINE-CYTOKINE_REC | CYTOKINE-CYTOKINE_RECEPTOR_INTERACTI | 239 | -0.18328 | -1.38176 | 0.0 | 0.22477631 | 1 | 1046 | tags=10%, list=4%, sig | MM_4 |
| GOBP_PROTEIN_EXPORT_ | GOBP_PROTEIN_EXPORT_FROM_NUCLEUS | 64 | -0.22429 | -1.38151 | 0.025641026 | 0.22467987 | 1 | 15230 | tags=94%, list=65%, si | MM_4 |
| GOBP_POSITIVE_REGULA | GOBP_POSITIVE_REGULATION_OF_AXON_E | 54 | -0.2544 | -1.38112 | 0.030303031 | 0.22476235 | 1 | 2083 | tags=20%, list=9%, sig | MM_4 |
| GOBP_ENDOTHELIAL_CEL | GOBP_ENDOTHELIAL_CELL_CHEMOTAXIS | 30 | -0.28664 | -1.38087 | 0.07633588 | 0.22474736 | 1 | 343 | tags=10%, list=1%, sig | MM_4 |
| GOBP_REGULATION_OF_ | GOBP_REGULATION_OF_LYMPHOCYTE_CHE | 22 | -0.31083 | -1.37961 | 0.08522727 | 0.22562648 | 1 | 1016 | tags=23%, list=4%, sig | MM_4 |
| GOBP_REGULATION_OF_ | GOBP_REGULATION_OF_T_CELL_CHEMOTA | 16 | -0.36306 | -1.37801 | 0.09137056 | 0.22695695 | 1 | 272 | tags=19%, list=1%, sig | MM_4 |
| ECM-RECEPTOR_INTERAC | ECM-RECEPTOR_INTERACTION | 79 | -0.22662 | -1.37577 | 0.03846154 | 0.22907934 | 1 | 1444 | tags=16%, list=6%, sig | MM_4 |
| GOBP_RESPONSE_TO_ME | GOBP_RESPONSE_TO_METAL_ION | 278 | -0.18617 | -1.37556 | 0.0 | 0.22893219 | 1 | 2357 | tags=17%, list=10%, si | MM_4 |
| GOBP_MEGAKARYOCYTE_ | GOBP_MEGAKARYOCYTE_DEVELOPMENT | 25 | -0.30215 | -1.37502 | 0.0952381 | 0.22903766 | 1 | 2411 | tags=28%, list=10%, si | MM_4 |
| GOBP_REGULATION_OF_ | GOBP_REGULATION_OF_MYELOID_LEUKOCY | 61 | -0.2453 | -1.37418 | 0.037037037 | 0.22949024 | 1 | 2446 | tags=20%, list=10%, si | MM_4 |
| GOBP_NEGATIVE_REGUL | GOBP_NEGATIVE_REGULATION_OF_B_CELL | 16 | -0.35921 | -1.37389 | 0.10909091 | 0.22944742 | 1 | 11765 | tags=94%, list=50%, si | MM_4 |
| HUNTINGTON_DISEASE | HUNTINGTON_DISEASE | 180 | -0.18235 | -1.37255 | 0.05882353 | 0.23057902 | 1 | 3906 | tags=28%, list=17%, si | MM_4 |
| GOBP_NEUTRAL_LIPID_B | GOBP_NEUTRAL_LIPID_BIOSYNTHETIC_PRO | 58 | -0.23532 | -1.37198 | 0.05263158 | 0.23076339 | 1 | 2476 | tags=28%, list=10%, si | MM_4 |
| GOBP_NEGATIVE_REGUL | GOBP_NEGATIVE_REGULATION_OF_ENDOT | 43 | -0.25835 | -1.37187 | 0.08256881 | 0.23047607 | 1 | 411 | tags=9%, list=2%, sign | MM_4 |
| GOBP_NEGATIVE_REGUL | GOBP_NEGATIVE_REGULATION_OF_PROTEI | 17 | -0.34403 | -1.37174 | 0.1036036 | 0.23025899 | 1 | 1212 | tags=24%, list=5%, sig | MM_4 |
| GOBP_REGULATION_OF_ | GOBP_REGULATION_OF_CELL_CYCLE_G1_S_ | 185 | -0.19359 | -1.37157 | 0.0 | 0.23011316 | 1 | 3066 | tags=24%, list=13%, si | MM_4 |
| GOBP_NEGATIVE_REGUL | GOBP_NEGATIVE_REGULATION_OF_MICRO | 15 | -0.35805 | -1.37067 | 0.122994654 | 0.2307461 | 1 | 15162 | tags=100%, list=64%, | MM_4 |
| GOBP_NEGATIVE_REGUL | GOBP_NEGATIVE_REGULATION_OF_RNA_CA | 82 | -0.21722 | -1.37056 | 0.04477612 | 0.23047933 | 1 | 3497 | tags=29%, list=15%, si | MM_4 |
| GOBP_B_CELL_HOMEOST | GOBP_B_CELL_HOMEOSTASIS | 42 | -0.25586 | -1.36917 | 0.07368421 | 0.23171313 | 1 | 11219 | tags=88%, list=48%, si | MM_4 |
| GOBP_VIRAL_TRANSLATI | GOBP_VIRAL_TRANSLATION | 18 | -0.33158 | -1.36846 | 0.1097561 | 0.23210494 | 1 | 4662 | tags=44%, list=20%, si | MM_4 |
| GOBP_ORGANELLE_INHE | GOBP_ORGANELLE_INHERITANCE | 15 | -0.37524 | -1.36786 | 0.105820104 | 0.23245436 | 1 | 3667 | tags=53%, list=16%, si | MM_4 |
| GOBP_POSITIVE_REGULA | GOBP_POSITIVE_REGULATION_OF_LYMPHO | 18 | -0.33777 | -1.36721 | 0.10526316 | 0.23285763 | 1 | 593 | tags=22%, list=3%, sig | MM_4 |
| GOBP_NEGATIVE_REGUL | GOBP_NEGATIVE_REGULATION_OF_DNA_T | 17 | -0.34359 | -1.36698 | 0.11794872 | 0.23270498 | 1 | 11394 | tags=94%, list=48%, si | MM_4 |
| N-GLYCAN_BIOSYNTHESIS | N-GLYCAN_BIOSYNTHESIS | 48 | -0.24426 | -1.3659 | 0.07777778 | 0.2336302 | 1 | 2866 | tags=33%, list=12%, si | MM_4 |
| MITOPHAGY | MITOPHAGY | 62 | -0.22928 | -1.36581 | 0.047058824 | 0.23335707 | 1 | 3784 | tags=34%, list=16%, si | MM_4 |
| GOBP_MACROMOLECULE | GOBP_MACROMOLECULE_DEACYLATION | 59 | -0.23836 | -1.36579 | 0.071428575 | 0.23298697 | 1 | 2308 | tags=24%, list=10%, si | MM_4 |
| GOBP_MYELOID_LEUKOC | GOBP_MYELOID_LEUKOCYTE_MIGRATION | 222 | -0.18916 | -1.36492 | 0.0 | 0.23368111 | 1 | 2100 | tags=17%, list=9%, sig | MM_4 |
| GOBP_NEGATIVE_REGUL | GOBP_NEGATIVE_REGULATION_OF_CANON | 57 | -0.23673 | -1.36407 | 0.010989011 | 0.23433582 | 1 | 804 | tags=14%, list=3%, sig | MM_4 |
| GOBP_GLYCOLIPID_BIOSY | GOBP_GLYCOLIPID_BIOSYNTHETIC_PROCESS | 66 | -0.2238 | -1.36382 | 0.023529412 | 0.2342968 | 1 | 2934 | tags=26%, list=12%, si | MM_4 |
| GOBP_MATURATION_OF_ | GOBP_MATURATION_OF_LSU_RRNA | 25 | -0.29583 | -1.36149 | 0.069767445 | 0.23652299 | 1 | 11590 | tags=92%, list=49%, si | MM_4 |
| GOBP_RESPONSE_TO_UV | GOBP_RESPONSE_TO_UV_B | 21 | -0.31577 | -1.36129 | 0.08888889 | 0.23637341 | 1 | 499 | tags=14%, list=2%, sig | MM_4 |
| GOBP_NEGATIVE_REGUL | GOBP_NEGATIVE_REGULATION_OF_CHEMO | 29 | -0.2884 | -1.36124 | 0.087248325 | 0.2360182 | 1 | 2175 | tags=21%, list=9%, sig | MM_4 |
| GOBP_DEFENSE_RESPON | GOBP_DEFENSE_RESPONSE_TO_GRAM_NEG | 68 | -0.22986 | -1.361 | 0.07692308 | 0.23591644 | 1 | 1102 | tags=9%, list=5%, sign | MM_4 |
| GOBP_NICOTINAMIDE_N | GOBP_NICOTINAMIDE_NUCLEOTIDE_BIOSYN | 16 | -0.34988 | -1.36086 | 0.13679245 | 0.23564804 | 1 | 2458 | tags=31%, list=10%, si | MM_4 |
| GOBP_NEGATIVE_REGUL | GOBP_NEGATIVE_REGULATION_OF_RESPON | 40 | -0.26327 | -1.35982 | 0.10169491 | 0.23646845 | 1 | 3771 | tags=40%, list=16%, si | MM_4 |
| GOBP_POSITIVE_REGULA | GOBP_POSITIVE_REGULATION_OF_NEUTRO | 41 | -0.25337 | -1.35955 | 0.06 | 0.23638071 | 1 | 1634 | tags=22%, list=7%, sig | MM_4 |
| GOBP_REGULATION_OF_ | GOBP_REGULATION_OF_INFLAMMATORY_R | 348 | -0.16565 | -1.35936 | 0.0 | 0.23623982 | 1 | 2291 | tags=17%, list=10%, si | MM_4 |
| GOBP_TRANSPORT_OF_V | GOBP_TRANSPORT_OF_VIRUS | 15 | -0.36009 | -1.35911 | 0.12182741 | 0.23616473 | 1 | 1021 | tags=20%, list=4%, sig | MM_4 |
| GOBP_REGULATION_OF_ | GOBP_REGULATION_OF_LEUKOCYTE_DEGRA | 48 | -0.25181 | -1.35859 | 0.07476635 | 0.23645833 | 1 | 2446 | tags=21%, list=10%, si | MM_4 |
| GOBP_RESPONSE_TO_EP | GOBP_RESPONSE_TO_EPIDERMAL_GROWTH | 42 | -0.25944 | -1.35794 | 0.075 | 0.23679022 | 1 | 2489 | tags=26%, list=11%, si | MM_4 |
| GOBP_CELLULAR_RESPON | GOBP_CELLULAR_RESPONSE_TO_KETONE | 88 | -0.22045 | -1.35767 | 0.06349207 | 0.23666546 | 1 | 3325 | tags=31%, list=14%, si | MM_4 |
| DILATED_CARDIOMYOPA | DILATED_CARDIOMYOPATHY_(DCM) | 88 | -0.21607 | -1.35741 | 0.030303031 | 0.2366734 | 1 | 1817 | tags=19%, list=8%, sig | MM_4 |
| GOBP_REGULATION_OF_ | GOBP_REGULATION_OF_EXTRINSIC_APOPT | 44 | -0.25588 | -1.35663 | 0.097087376 | 0.2372983 | 1 | 363 | tags=9%, list=2%, sign | MM_4 |
| GOBP_REGULATION_OF_ | GOBP_REGULATION_OF_CELLULAR_RESPIRA | 52 | -0.23689 | -1.35527 | 0.01923077 | 0.23852888 | 1 | 3920 | tags=33%, list=17%, si | MM_4 |
| GOBP_REGULATION_OF_ | GOBP_REGULATION_OF_DNA_TEMPLATED_ | 84 | -0.21258 | -1.35469 | 0.0754717 | 0.23882754 | 1 | 2304 | tags=19%, list=10%, si | MM_4 |
| GOBP_NUCLEAR_TRANSC | GOBP_NUCLEAR_TRANSCRIBED_MRNA_CAT | 59 | -0.23969 | -1.354 | 0.047058824 | 0.23908037 | 1 | 10781 | tags=88%, list=46%, si | MM_4 |
| GOBP_MITOCHONDRIAL_ | GOBP_MITOCHONDRIAL_MEMBRANE_ORGA | 107 | -0.20254 | -1.35384 | 0.03773585 | 0.23889144 | 1 | 13165 | tags=88%, list=56%, si | MM_4 |
| GOBP_POSITIVE_REGULA | GOBP_POSITIVE_REGULATION_OF_CHOLEST | 19 | -0.33865 | -1.35268 | 0.11340206 | 0.2399012 | 1 | 2064 | tags=26%, list=9%, sig | MM_4 |
| GOBP_COPII_COATED_VE | GOBP_COPII_COATED_VESICLE_BUDDING | 37 | -0.2734 | -1.35262 | 0.07194245 | 0.2395817 | 1 | 2887 | tags=30%, list=12%, si | MM_4 |
| GOBP_NEGATIVE_REGUL | GOBP_NEGATIVE_REGULATION_OF_LEUKO | 142 | -0.19292 | -1.35252 | 0.0 | 0.23933072 | 1 | 2946 | tags=21%, list=12%, si | MM_4 |
| GOBP_IMMUNE_SYSTEM | GOBP_IMMUNE_SYSTEM_DEVELOPMENT | 216 | -0.17275 | -1.35223 | 0.0 | 0.2393394 | 1 | 2846 | tags=19%, list=12%, si | MM_4 |
| GOBP_REGULATION_OF_ | GOBP_REGULATION_OF_CELL_ADHESION_M | 47 | -0.25153 | -1.3512 | 0.07619048 | 0.2402317 | 1 | 2357 | tags=26%, list=10%, si | MM_4 |
| GOBP_ESTABLISHMENT_ | GOBP_ESTABLISHMENT_OF_MITOCHONDRI | 25 | -0.29212 | -1.35098 | 0.07096774 | 0.24001922 | 1 | 14792 | tags=96%, list=63%, si | MM_4 |
| GOBP_POSITIVE_REGULA | GOBP_POSITIVE_REGULATION_OF_GLYCOP | 26 | -0.29139 | -1.35081 | 0.1097561 | 0.2398103 | 1 | 3790 | tags=42%, list=16%, si | MM_4 |
| GOBP_POSITIVE_REGULA | GOBP_POSITIVE_REGULATION_OF_RECEPTO | 17 | -0.34301 | -1.35051 | 0.121212125 | 0.23987944 | 1 | 4028 | tags=41%, list=17%, si | MM_4 |
| GOBP_NUCLEAR_TRANSC | GOBP_NUCLEAR_TRANSCRIBED_MRNA_CAT | 43 | -0.24846 | -1.34999 | 0.091836736 | 0.24000667 | 1 | 11602 | tags=88%, list=49%, si | MM_4 |
| GOBP_REGULATION_OF_ | GOBP_REGULATION_OF_PROTEIN_SUMOYL | 25 | -0.30358 | -1.34903 | 0.10119048 | 0.24077214 | 1 | 14540 | tags=96%, list=62%, si | MM_4 |
| GOBP_REGULATION_OF_ | GOBP_REGULATION_OF_BONE_RESORPTIO | 51 | -0.23804 | -1.34821 | 0.054945055 | 0.24127848 | 1 | 2760 | tags=29%, list=12%, si | MM_4 |
| GOBP_STRIATED_MUSCLI | GOBP_STRIATED_MUSCLE_CONTRACTION | 167 | -0.1832 | -1.34722 | 0.0 | 0.2419328 | 1 | 2593 | tags=21%, list=11%, si | MM_4 |
| GOBP_GRANULOCYTE_CH | GOBP_GRANULOCYTE_CHEMOTAXIS | 119 | -0.20505 | -1.34711 | 0.057142857 | 0.24170174 | 1 | 918 | tags=10%, list=4%, sig | MM_4 |
| GOBP_REGULATION_OF_ | GOBP_REGULATION_OF_FILOPODIUM_ASSE | 54 | -0.24051 | -1.34658 | 0.04 | 0.24198617 | 1 | 2704 | tags=24%, list=11%, si | MM_4 |
| GOBP_LYMPHOCYTE_ME | GOBP_LYMPHOCYTE_MEDIATED_IMMUNITY | 300 | -0.17102 | -1.34654 | 0.0 | 0.24162886 | 1 | 2279 | tags=17%, list=10%, si | MM_4 |
| GOBP_REGULATION_OF_ | GOBP_REGULATION_OF_ACTIN_FILAMENT_ | 48 | -0.24893 | -1.34562 | 0.06930693 | 0.24234049 | 1 | 908 | tags=17%, list=4%, sig | MM_4 |
| GOBP_VACUOLAR_ACIDI | GOBP_VACUOLAR_ACIDIFICATION | 27 | -0.28458 | -1.3455 | 0.10135135 | 0.24209905 | 1 | 3573 | tags=37%, list=15%, si | MM_4 |
| GOBP_ESTABLISHMENT_ | GOBP_ESTABLISHMENT_OF_LYMPHOCYTE_ | 15 | -0.33823 | -1.34222 | 0.088541664 | 0.24557996 | 1 | 2256 | tags=33%, list=10%, si | MM_4 |

| | | | | | | | | | | |
|---|---|---|---|---|---|---|---|---|---|---|
| GOBP_VIRAL_GENOME_R | GOBP_VIRAL_GENOME_REPLICATION | 110 | -0.20395 | -1.34201 | 0.044444446 | 0.24545905 | 1 | 11887 | tags=86%, list=50%, si | MM_4 |
| GOBP_MITOCHONDRIAL_ | GOBP_MITOCHONDRIAL_FUSION | 36 | -0.26353 | -1.34153 | 0.10526316 | 0.24568042 | 1 | 11650 | tags=89%, list=49%, si | MM_4 |
| GOBP_REGULATION_OF_ | GOBP_REGULATION_OF_MONOCYTE_DIFFE | 17 | -0.34437 | -1.34119 | 0.12903225 | 0.24570154 | 1 | 3605 | tags=35%, list=15%, si | MM_4 |
| TUBERCULOSIS | TUBERCULOSIS | 152 | -0.1934 | -1.3395 | 0.0882353 | 0.2473208 | 1 | 2755 | tags=19%, list=12%, si | MM_4 |
| GOBP_EMBRYONIC_HEM | GOBP_EMBRYONIC_HEMOPOIESIS | 31 | -0.27375 | -1.33949 | 0.09016393 | 0.24694836 | 1 | 14305 | tags=94%, list=61%, si | MM_4 |
| GOBP_POSITIVE_REGULA | GOBP_POSITIVE_REGULATION_OF_TELOME | 31 | -0.27925 | -1.33834 | 0.10526316 | 0.24784921 | 1 | 4213 | tags=52%, list=18%, si | MM_4 |
| GOBP_T_CELL_MEDIATED | GOBP_T_CELL_MEDIATED_CYTOTOXICITY | 53 | -0.23927 | -1.33814 | 0.04255319 | 0.24767986 | 1 | 1963 | tags=17%, list=8%, sig | MM_4 |
| GOBP_MITOCHONDRIAL_ | GOBP_MITOCHONDRIAL_RESPIRATORY_CHA | 91 | -0.21689 | -1.3381 | 0.06666667 | 0.24733225 | 1 | 11456 | tags=90%, list=49%, si | MM_4 |
| GOBP_REGULATION_OF_ | GOBP_REGULATION_OF_CHOLESTEROL_STO | 21 | -0.3084 | -1.33768 | 0.13953489 | 0.24744324 | 1 | 1491 | tags=24%, list=6%, sig | MM_4 |
| GOBP_VACUOLAR_TRANS | GOBP_VACUOLAR_TRANSPORT | 161 | -0.18571 | -1.33738 | 0.0 | 0.24741293 | 1 | 4170 | tags=31%, list=18%, si | MM_4 |
| GOBP_HEMATOPOIETIC_S | GOBP_HEMATOPOIETIC_STEM_CELL_PROLIF | 37 | -0.26096 | -1.33726 | 0.07575758 | 0.24716519 | 1 | 3797 | tags=32%, list=16%, si | MM_4 |
| GOBP_RESPONSE_TO_CA | GOBP_RESPONSE_TO_CALCIUM_ION | 130 | -0.19184 | -1.33669 | 0.030303031 | 0.24745835 | 1 | 1644 | tags=13%, list=7%, sig | MM_4 |
| GOBP_POSITIVE_REGULA | GOBP_POSITIVE_REGULATION_OF_TORC1_S | 52 | -0.23915 | -1.33643 | 0.068627454 | 0.24740843 | 1 | 4633 | tags=42%, list=20%, si | MM_4 |
| GOBP_POSITIVE_REGULA | GOBP_POSITIVE_REGULATION_OF_TYROSIN | 51 | -0.23369 | -1.33496 | 0.089108914 | 0.2487283 | 1 | 2209 | tags=20%, list=9%, sig | MM_4 |
| GOBP_REGULATION_OF_ | GOBP_REGULATION_OF_CHROMATIN_ORGA | 36 | -0.26422 | -1.33396 | 0.09090909 | 0.24955425 | 1 | 12527 | tags=89%, list=53%, si | MM_4 |
| GOBP_LIPID_IMPORT_INT | GOBP_LIPID_IMPORT_INTO_CELL | 24 | -0.29025 | -1.33373 | 0.10135135 | 0.24944662 | 1 | 2476 | tags=25%, list=10%, si | MM_4 |
| GOBP_SPINDLE_CHECKPO | GOBP_SPINDLE_CHECKPOINT_SIGNALING | 53 | 0.698226 | 2.925449 | 0 | 0 | 0 | 2288 | tags=51%, list=10%, si | MM_4 |
| GOBP_REGULATION_OF_ | GOBP_REGULATION_OF_MITOTIC_SISTER_C | 56 | 0.665668 | 2.813424 | 0 | 0 | 0 | 2288 | tags=48%, list=10%, si | MM_4 |
| GOBP_MITOTIC_SISTER_C | GOBP_MITOTIC_SISTER_CHROMATID_SEPAR | 61 | 0.65266 | 2.782056 | 0 | 0 | 0 | 2288 | tags=43%, list=10%, si | MM_4 |
| GOBP_MITOTIC_SISTER_C | GOBP_MITOTIC_SISTER_CHROMATID_SEGRE | 178 | 0.562202 | 2.752264 | 0 | 0 | 0 | 2288 | tags=34%, list=10%, si | MM_4 |
| GOBP_NEGATIVE_REGUL | GOBP_NEGATIVE_REGULATION_OF_CHROM | 96 | 0.585641 | 2.724268 | 0 | 0 | 0 | 2288 | tags=34%, list=10%, si | MM_4 |
| GOBP_CHROMOSOME_SE | GOBP_CHROMOSOME_SEPARATION | 80 | 0.603394 | 2.694019 | 0 | 0 | 0 | 1756 | tags=38%, list=7%, sig | MM_4 |
| GOBP_ATTACHMENT_OF_ | GOBP_ATTACHMENT_OF_SPINDLE_MICROT | 48 | 0.647594 | 2.64561 | 0 | 0 | 0 | 1978 | tags=42%, list=8%, sig | MM_4 |
| GOBP_REGULATION_OF_ | GOBP_REGULATION_OF_MITOTIC_NUCLEAR | 114 | 0.552036 | 2.614773 | 0 | 0 | 0 | 2288 | tags=36%, list=10%, si | MM_4 |
| GOBP_MITOTIC_NUCLEAR | GOBP_MITOTIC_NUCLEAR_DIVISION | 259 | 0.503422 | 2.556676 | 0 | 0 | 0 | 2288 | tags=29%, list=10%, si | MM_4 |
| GOBP_NEGATIVE_REGUL | GOBP_NEGATIVE_REGULATION_OF_NUCLEA | 66 | 0.591188 | 2.554793 | 0 | 0 | 0 | 2288 | tags=39%, list=10%, si | MM_4 |
| GOBP_SISTER_CHROMAT | GOBP_SISTER_CHROMATID_SEGREGATION | 219 | 0.500129 | 2.520574 | 0 | 0 | 0 | 2288 | tags=28%, list=10%, si | MM_4 |
| GOBP_FEMALE_MEIOTIC_ | GOBP_FEMALE_MEIOTIC_NUCLEAR_DIVISIO | 41 | 0.631555 | 2.514751 | 0 | 0 | 0 | 1318 | tags=34%, list=6%, sig | MM_4 |
| GOBP_METAPHASE_CHRO | GOBP_METAPHASE_CHROMOSOME_ALIGNI | 93 | 0.546816 | 2.511123 | 0 | 0 | 0 | 2193 | tags=31%, list=9%, sig | MM_4 |
| GOBP_KINETOCHORE_OR | GOBP_KINETOCHORE_ORGANIZATION | 20 | 0.7264 | 2.450933 | 0 | 0 | 0 | 1736 | tags=45%, list=7%, sig | MM_4 |
| GOBP_MICROTUBULE_CY | GOBP_MICROTUBULE_CYTOSKELETON_ORG | 158 | 0.495508 | 2.447306 | 0 | 0 | 0 | 1940 | tags=26%, list=8%, sig | MM_4 |
| GOBP_MITOTIC_SPINDLE | GOBP_MITOTIC_SPINDLE_ORGANIZATION | 125 | 0.502019 | 2.411546 | 0 | 2.69863E-05 | 0.001 | 2273 | tags=27%, list=10%, si | MM_4 |
| GOBP_REGULATION_OF_ | GOBP_REGULATION_OF_SISTER_CHROMATI | 104 | 0.513854 | 2.396849 | 0 | 2.59868E-05 | 0.001 | 1756 | tags=27%, list=7%, sig | MM_4 |
| GOBP_CENTROSOME_DU | GOBP_CENTROSOME_DUPLICATION | 74 | 0.540987 | 2.388712 | 0 | 2.50587E-05 | 0.001 | 2193 | tags=31%, list=9%, sig | MM_4 |
| SYSTEMIC_LUPUS_ERYTH | SYSTEMIC_LUPUS_ERYTHEMATOSUS | 105 | 0.506553 | 2.37879 | 0 | 2.41946E-05 | 0.001 | 2510 | tags=33%, list=11%, si | MM_4 |
| GOBP_SPINDLE_ASSEMBL | GOBP_SPINDLE_ASSEMBLY | 122 | 0.499876 | 2.370058 | 0 | 2.33881E-05 | 0.001 | 2273 | tags=24%, list=10%, si | MM_4 |
| GOBP_REGULATION_OF_ | GOBP_REGULATION_OF_CHROMOSOME_SE | 126 | 0.498482 | 2.362695 | 0 | 2.26337E-05 | 0.001 | 1835 | tags=26%, list=8%, sig | MM_4 |
| GOBP_CENTROMERE_CO | GOBP_CENTROMERE_COMPLEX_ASSEMBLY | 27 | 0.643824 | 2.329949 | 0 | 2.19264E-05 | 0.001 | 1833 | tags=37%, list=8%, sig | MM_4 |
| GOBP_MITOTIC_SPINDLE | GOBP_MITOTIC_SPINDLE_ASSEMBLY | 71 | 0.523088 | 2.32548 | 0 | 2.12619E-05 | 0.001 | 2273 | tags=27%, list=10%, si | MM_4 |
| GOBP_REGULATION_OF_ | GOBP_REGULATION_OF_NUCLEAR_DIVISIO | 144 | 0.476178 | 2.307696 | 0 | 2.06366E-05 | 0.001 | 2288 | tags=28%, list=10%, si | MM_4 |
| GOBP_NUCLEAR_CHROM | GOBP_NUCLEAR_CHROMOSOME_SEGREGAT | 305 | 0.440469 | 2.299639 | 0 | 4.02009E-05 | 0.002 | 2288 | tags=25%, list=10%, si | MM_4 |
| GOBP_PROTEIN_LOCALIZ | GOBP_PROTEIN_LOCALIZATION_TO_CHROM | 24 | 0.653621 | 2.299128 | 0 | 5.87936E-05 | 0.003 | 1249 | tags=38%, list=5%, sig | MM_4 |
| GOBP_MICROTUBULE_OR | GOBP_MICROTUBULE_ORGANIZING_CENTE | 149 | 0.472408 | 2.289536 | 0 | 7.61727E-05 | 0.004 | 2271 | tags=27%, list=10%, si | MM_4 |
| GOBP_POSITIVE_REGULA | GOBP_POSITIVE_REGULATION_OF_MITOTIC | 18 | 0.696148 | 2.285256 | 0 | 0.000111135 | 0.006 | 265 | tags=33%, list=1%, sig | MM_4 |
| GOBP_ATTACHMENT_OF_ | GOBP_ATTACHMENT_OF_MITOTIC_SPINDLE | 24 | 0.646875 | 2.279181 | 0 | 0.000108286 | 0.006 | 1978 | tags=46%, list=8%, sig | MM_4 |
| GOBP_MITOTIC_METAPH | GOBP_MITOTIC_METAPHASE_CHROMOSOM | 58 | 0.527644 | 2.26896 | 0 | 0.000123124 | 0.007 | 2186 | tags=31%, list=9%, sig | MM_4 |
| GOBP_REGULATION_OF_ | GOBP_REGULATION_OF_SPINDLE_CHECKPO | 24 | 0.653463 | 2.267273 | 0 | 0.000137295 | 0.008 | 666 | tags=38%, list=3%, sig | MM_4 |
| OLFACTORY_TRANSDUCT | OLFACTORY_TRANSDUCTION | 186 | 0.458486 | 2.266463 | 0 | 0.000134026 | 0.008 | 7760 | tags=73%, list=33%, si | MM_4 |
| GOBP_REGULATION_OF_ | GOBP_REGULATION_OF_CELL_CYCLE_CHECI | 48 | 0.556794 | 2.264762 | 0 | 0.000130909 | 0.008 | 2193 | tags=33%, list=9%, sig | MM_4 |
| GOBP_CHROMOSOME_LO | GOBP_CHROMOSOME_LOCALIZATION | 111 | 0.480277 | 2.263347 | 0 | 0.000127934 | 0.008 | 2193 | tags=27%, list=9%, sig | MM_4 |
| GOBP_POSITIVE_REGULA | GOBP_POSITIVE_REGULATION_OF_CELL_CY | 17 | 0.708862 | 2.257556 | 0 | 0.000140703 | 0.009 | 666 | tags=41%, list=3%, sig | MM_4 |
| GOBP_SPINDLE_ORGANIZ | GOBP_SPINDLE_ORGANIZATION | 190 | 0.448727 | 2.24008 | 0 | 0.00015289 | 0.01 | 2273 | tags=23%, list=10%, si | MM_4 |
| GOBP_CHROMOSOME_SE | GOBP_CHROMOSOME_SEGREGATION | 403 | 0.4274 | 2.234004 | 0 | 0.00016446 | 0.011 | 2288 | tags=23%, list=10%, si | MM_4 |
| GOBP_KINETOCHORE_AS | GOBP_KINETOCHORE_ASSEMBLY | 16 | 0.715683 | 2.208237 | 0 | 0.000219355 | 0.015 | 1736 | tags=44%, list=7%, sig | MM_4 |
| GOBP_MITOTIC_CYTOKIN | GOBP_MITOTIC_CYTOKINESIS | 83 | 0.488959 | 2.199561 | 0 | 0.000229126 | 0.016 | 2273 | tags=28%, list=10%, si | MM_4 |
| GOBP_MITOTIC_CELL_CY | GOBP_MITOTIC_CELL_CYCLE_CHECKPOINT_ | 147 | 0.447963 | 2.196323 | 0 | 0.000238637 | 0.017 | 2068 | tags=24%, list=9%, sig | MM_4 |
| GOBP_MEIOTIC_SPINDLE | GOBP_MEIOTIC_SPINDLE_ORGANIZATION | 21 | 0.641024 | 2.180827 | 0 | 0.000330095 | 0.023 | 1410 | tags=33%, list=6%, sig | MM_4 |
| GOBP_POSITIVE_REGULA | GOBP_POSITIVE_REGULATION_OF_CHROMO | 28 | 0.595646 | 2.166643 | 0 | 0.000418464 | 0.03 | 1399 | tags=32%, list=6%, sig | MM_4 |
| GOBP_REGULATION_OF_ | GOBP_REGULATION_OF_ATTACHMENT_OF_ | 20 | 0.651949 | 2.15142 | 0 | 0.000516388 | 0.038 | 1855 | tags=45%, list=8%, sig | MM_4 |
| PRION_DISEASES | PRION_DISEASES | 31 | 0.575349 | 2.147417 | 0 | 0.00051975 | 0.039 | 2145 | tags=32%, list=9%, sig | MM_4 |
| GOBP_CELL_CYCLE_CHEC | GOBP_CELL_CYCLE_CHECKPOINT_SIGNALIN | 191 | 0.427177 | 2.111964 | 0 | 0.001008293 | 0.075 | 2288 | tags=24%, list=10%, si | MM_4 |
| GOBP_NEGATIVE_REGUL | GOBP_NEGATIVE_REGULATION_OF_MITOTI | 174 | 0.421162 | 2.11054 | 0 | 0.001027905 | 0.078 | 2139 | tags=23%, list=9%, sig | MM_4 |
| GOBP_POSITIVE_REGULA | GOBP_POSITIVE_REGULATION_OF_MITOTIC | 44 | 0.520656 | 2.088589 | 0 | 0.001551206 | 0.119 | 2717 | tags=36%, list=12%, si | MM_4 |
| GOBP_CYTOSKELETON_D | GOBP_CYTOSKELETON_DEPENDENT_CYTOK | 97 | 0.445379 | 2.054689 | 0 | 0.00225056 | 0.17 | 2273 | tags=25%, list=10%, si | MM_4 |
| GOBP_REGULATION_OF_ | GOBP_REGULATION_OF_CYTOKINESIS | 83 | 0.456127 | 2.054397 | 0 | 0.00222434 | 0.17 | 2273 | tags=28%, list=10%, si | MM_4 |
| GOBP_MICROTUBULE_OR | GOBP_MICROTUBULE_ORGANIZING_CENTE | 34 | 0.532016 | 2.043542 | 0 | 0.002514564 | 0.193 | 630 | tags=21%, list=3%, sig | MM_4 |
| GOBP_POSITIVE_REGULA | GOBP_POSITIVE_REGULATION_OF_CYTOKIN | 40 | 0.518017 | 2.033935 | 0 | 0.002898763 | 0.22 | 2273 | tags=33%, list=10%, si | MM_4 |
| GOBP_PROTEIN_LOCALIZ | GOBP_PROTEIN_LOCALIZATION_TO_CONDE | 16 | 0.643063 | 2.027318 | 0 | 0.003180473 | 0.242 | 1155 | tags=38%, list=5%, sig | MM_4 |
| GOBP_INTERMEDIATE_FI | GOBP_INTERMEDIATE_FILAMENT_ORGANIZ | 46 | 0.498373 | 2.009634 | 0 | 0.003830894 | 0.293 | 9427 | tags=74%, list=40%, si | MM_4 |
| GOBP_ORGANELLE_FISSI | GOBP_ORGANELLE_FISSION | 477 | 0.380502 | 1.99879 | 0 | 0.004396006 | 0.33 | 2291 | tags=22%, list=10%, si | MM_4 |
| GOBP_REGULATION_OF_ | GOBP_REGULATION_OF_MITOTIC_CYTOKIN | 15 | 0.651733 | 1.989931 | 0 | 0.004781138 | 0.355 | 797 | tags=40%, list=3%, sig | MM_4 |
| GOBP_REGULATION_OF_ | GOBP_REGULATION_OF_LIPOPOLYSACCHAR | 29 | 0.539876 | 1.985678 | 0 | 0.004953473 | 0.369 | 2234 | tags=31%, list=9%, sig | MM_4 |
| GOBP_CYTOKINETIC_PRO | GOBP_CYTOKINETIC_PROCESS | 45 | 0.488287 | 1.978663 | 0 | 0.005423904 | 0.403 | 1867 | tags=22%, list=8%, sig | MM_4 |
| GOBP_CYTOKINESIS | GOBP_CYTOKINESIS | 160 | 0.40475 | 1.971045 | 0 | 0.005880791 | 0.433 | 2273 | tags=23%, list=10%, si | MM_4 |
| GOBP_ESTABLISHMENT_C | GOBP_ESTABLISHMENT_OF_MITOTIC_SPIND | 41 | 0.495091 | 1.965124 | 0 | 0.006272949 | 0.456 | 1940 | tags=27%, list=8%, sig | MM_4 |
| GOBP_REGULATION_OF_ | GOBP_REGULATION_OF_CENTROSOME_CYC | 55 | 0.462249 | 1.96004 | 0.0010752688 | 0.006584985 | 0.478 | 2186 | tags=24%, list=9%, sig | MM_4 |
| GOBP_NEGATIVE_REGUL | GOBP_NEGATIVE_REGULATION_OF_CELL_C | 290 | 0.380399 | 1.955811 | 0 | 0.006877469 | 0.495 | 2193 | tags=21%, list=9%, sig | MM_4 |
| GOBP_VENTRICULAR_SYS | GOBP_VENTRICULAR_SYSTEM_DEVELOPME | 36 | 0.508079 | 1.951691 | 0 | 0.007122868 | 0.517 | 1177 | tags=22%, list=5%, sig | MM_4 |
| GOBP_CENTRIOLE_ASSEN | GOBP_CENTRIOLE_ASSEMBLY | 46 | 0.477455 | 1.95003 | 0 | 0.007149943 | 0.523 | 2153 | tags=26%, list=9%, sig | MM_4 |
| GOBP_NEGATIVE_REGUL | GOBP_NEGATIVE_REGULATION_OF_MITOTI | 230 | 0.381445 | 1.932998 | 0 | 0.008694066 | 0.6 | 2139 | tags=22%, list=9%, sig | MM_4 |
| GOBP_POSITIVE_REGULA | GOBP_POSITIVE_REGULATION_OF_CELL_CY | 252 | 0.377823 | 1.924589 | 0 | 0.009522876 | 0.641 | 1985 | tags=20%, list=8%, sig | MM_4 |
| GOBP_REGULATION_OF_ | GOBP_REGULATION_OF_CHROMOSOME_OI | 242 | 0.377171 | 1.917793 | 0 | 0.010284267 | 0.676 | 2111 | tags=19%, list=9%, sig | MM_4 |

| | | | | | | | | | | |
|---|---|---|---|---|---|---|---|---|---|---|
| GOBP_POSITIVE_REGULA | GOBP_POSITIVE_REGULATION_OF_CHROMO | 29 | 0.514748 | 1.916898 | 0 | 0.01032382 | 0.681 | 1399 | tags=28%, list=6%, sig | MM_4 |
| GOBP_CHROMOSOME_C | GOBP_CHROMOSOME_CONDENSATION | 30 | 0.516825 | 1.915589 | 0 | 0.010326451 | 0.685 | 2722 | tags=30%, list=12%, si | MM_4 |
| GOBP_REGULATION_OF_ | GOBP_REGULATION_OF_CENTROSOME_DU | 46 | 0.466819 | 1.911624 | 0 | 0.010772492 | 0.703 | 2186 | tags=24%, list=9%, sig | MM_4 |
| GOBP_POSITIVE_REGULA | GOBP_POSITIVE_REGULATION_OF_NUCLEA | 62 | 0.442754 | 1.899552 | 0 | 0.012171558 | 0.752 | 3729 | tags=32%, list=16%, si | MM_4 |
| GOBP_REGULATION_OF_ | GOBP_REGULATION_OF_EPIDERMAL_GROW | 20 | 0.564354 | 1.899114 | 0 | 0.012047351 | 0.753 | 1634 | tags=25%, list=7%, sig | MM_4 |
| GOBP_MITOCHONDRIAL_ | GOBP_MITOCHONDRIAL_CALCIUM_ION_HO | 29 | 0.52023 | 1.892158 | 0 | 0.01289303 | 0.779 | 2084 | tags=28%, list=9%, sig | MM_4 |
| RETINOL_METABOLISM | RETINOL_METABOLISM | 52 | 0.439126 | 1.860835 | 0 | 0.018484857 | 0.884 | 12012 | tags=87%, list=51%, si | MM_4 |
| GOBP_POSITIVE_REGULA | GOBP_POSITIVE_REGULATION_OF_CELL_DIV | 79 | 0.415458 | 1.853849 | 0 | 0.019851062 | 0.908 | 1915 | tags=23%, list=8%, sig | MM_4 |
| GOBP_REGULATION_OF_ | GOBP_REGULATION_OF_SPINDLE_ASSEMBL | 28 | 0.506907 | 1.849691 | 0.0048721074 | 0.020830812 | 0.917 | 1050 | tags=18%, list=4%, sig | MM_4 |
| GOBP_BRANCH_ELONGA | GOBP_BRANCH_ELONGATION_OF_AN_EPIT | 21 | 0.542761 | 1.838795 | 0.0036014405 | 0.023345893 | 0.942 | 2881 | tags=38%, list=12%, si | MM_4 |
| GOBP_MEIOTIC_CHROMO | GOBP_MEIOTIC_CHROMOSOME_SEGREGAT | 94 | 0.398177 | 1.833275 | 0 | 0.024600858 | 0.946 | 1736 | tags=22%, list=7%, sig | MM_4 |
| NEUROACTIVE_LIGAND-R | NEUROACTIVE_LIGAND-RECEPTOR_INTERAC | 274 | 0.357267 | 1.825046 | 0 | 0.0266646 | 0.963 | 11688 | tags=72%, list=50%, si | MM_4 |
| GOBP_RESPONSE_TO_PH | GOBP_RESPONSE_TO_PHEROMONE | 17 | 0.577934 | 1.821223 | 0.0025125628 | 0.027508039 | 0.967 | 6475 | tags=76%, list=27%, si | MM_4 |
| GOBP_EXIT_FROM_MITO | GOBP_EXIT_FROM_MITOSIS | 26 | 0.503744 | 1.812255 | 0.0035799523 | 0.030164856 | 0.976 | 2095 | tags=19%, list=9%, sig | MM_4 |
| GOBP_MODIFICATION_O | GOBP_MODIFICATION_OF_POSTSYNAPTIC_ | 24 | 0.521206 | 1.812026 | 0.0035714286 | 0.02991061 | 0.976 | 2149 | tags=33%, list=9%, sig | MM_4 |
| GOBP_MITOCHONDRIAL_ | GOBP_MITOCHONDRIAL_CALCIUM_ION_TR | 19 | 0.554287 | 1.80301 | 0.0036719707 | 0.03273448 | 0.985 | 2084 | tags=26%, list=9%, sig | MM_4 |
| GOBP_NEGATIVE_REGUL | GOBP_NEGATIVE_REGULATION_OF_VIRAL_ | 36 | 0.468032 | 1.789267 | 0 | 0.037504308 | 0.992 | 3788 | tags=36%, list=16%, si | MM_4 |
| GOBP_SPINDLE_LOCALIZ | GOBP_SPINDLE_LOCALIZATION | 61 | 0.416283 | 1.779386 | 0 | 0.041387487 | 0.995 | 1940 | tags=23%, list=8%, sig | MM_4 |
| GOBP_SUPPRESSION_OF_ | GOBP_SUPPRESSION_OF_VIRAL_RELEASE_B | 22 | 0.528346 | 1.77784 | 0.0012062726 | 0.04161647 | 0.995 | 1858 | tags=32%, list=8%, sig | MM_4 |
| GOBP_POSITIVE_REGULA | GOBP_POSITIVE_REGULATION_OF_METAPH | 15 | 0.573127 | 1.776551 | 0.0038461538 | 0.04168716 | 0.996 | 242 | tags=20%, list=1%, sig | MM_4 |
| GOBP_POSITIVE_REGULA | GOBP_POSITIVE_REGULATION_OF_UBIQUIT | 18 | 0.544516 | 1.771878 | 0.0038071065 | 0.043325856 | 0.996 | 601 | tags=22%, list=3%, sig | MM_4 |
| GOBP_ERBB2_SIGNALING | GOBP_ERBB2_SIGNALING_PATHWAY | 23 | 0.518457 | 1.770119 | 0.0036540804 | 0.043693062 | 0.997 | 264 | tags=17%, list=1%, sig | MM_4 |
| GOBP_NEGATIVE_REGUL | GOBP_NEGATIVE_REGULATION_OF_CELL_C | 368 | 0.339852 | 1.766642 | 0 | 0.04494417 | 0.998 | 2193 | tags=19%, list=9%, sig | MM_4 |
| GOBP_REGULATION_OF_ | GOBP_REGULATION_OF_DOUBLE_STRAND_ | 80 | 0.387241 | 1.750666 | 0.001059322 | 0.052575275 | 1 | 2291 | tags=23%, list=10%, si | MM_4 |
| GOBP_INTERLEUKIN_1_M | GOBP_INTERLEUKIN_1_MEDIATED_SIGNALI | 28 | 0.480674 | 1.749154 | 0.008196721 | 0.05286067 | 1 | 2234 | tags=32%, list=9%, sig | MM_4 |
| GOBP_POSITIVE_REGULA | GOBP_POSITIVE_REGULATION_OF_CELL_CY | 330 | 0.336301 | 1.742147 | 0 | 0.05607622 | 1 | 2273 | tags=20%, list=10%, si | MM_4 |
| GOBP_NEUROPEPTIDE_SI | GOBP_NEUROPEPTIDE_SIGNALING_PATHW | 69 | 0.39447 | 1.73355 | 0 | 0.061006933 | 1 | 9043 | tags=67%, list=38%, si | MM_4 |
| GOBP_KERATINIZATION | GOBP_KERATINIZATION | 31 | 0.470981 | 1.733115 | 0 | 0.060724057 | 1 | 11028 | tags=84%, list=47%, si | MM_4 |
| GOBP_REGULATION_OF_ | GOBP_REGULATION_OF_SPINDLE_ORGANIZ | 42 | 0.438506 | 1.732098 | 0 | 0.06076007 | 1 | 1806 | tags=19%, list=8%, sig | MM_4 |
| GOBP_REGULATION_OF_ | GOBP_REGULATION_OF_UBIQUITIN_PROTE | 20 | 0.516925 | 1.729146 | 0.009661836 | 0.0618804 | 1 | 1318 | tags=25%, list=6%, sig | MM_4 |
| GOBP_CELL_CYCLE_G2_M | GOBP_CELL_CYCLE_G2_M_PHASE_TRANSITI | 152 | 0.353441 | 1.727886 | 0 | 0.062173795 | 1 | 2111 | tags=17%, list=9%, sig | MM_4 |
| GOBP_MHC_CLASS_II_BI | GOBP_MHC_CLASS_II_BIOSYNTHETIC_PROC | 16 | 0.556223 | 1.720278 | 0.0114068445 | 0.06650916 | 1 | 1057 | tags=25%, list=4%, sig | MM_4 |
| GOBP_REGULATION_OF_ | GOBP_REGULATION_OF_INTERLEUKIN_4_P | 30 | 0.466166 | 1.715621 | 0.005760369 | 0.06902575 | 1 | 3236 | tags=43%, list=14%, si | MM_4 |
| GOBP_ESTABLISHMENT_ | GOBP_ESTABLISHMENT_OF_SPINDLE_ORIEN | 43 | 0.430878 | 1.707554 | 0.0011534025 | 0.074296914 | 1 | 1940 | tags=26%, list=8%, sig | MM_4 |
| GOBP_CELLULAR_RESPON | GOBP_CELLULAR_RESPONSE_TO_IONIZING_ | 69 | 0.3874 | 1.706999 | 0.0010683761 | 0.07404501 | 1 | 2717 | tags=23%, list=12%, si | MM_4 |
| GOBP_RESPONSE_TO_AM | GOBP_RESPONSE_TO_AMYLOID_BETA | 39 | 0.433165 | 1.705014 | 0.0034324944 | 0.07497444 | 1 | 2207 | tags=26%, list=9%, sig | MM_4 |
| GOBP_REGULATION_OF_ | GOBP_REGULATION_OF_MITOTIC_CELL_CY | 331 | 0.32773 | 1.699268 | 0 | 0.07874957 | 1 | 2139 | tags=16%, list=9%, sig | MM_4 |
| GOBP_REGULATION_OF_ | GOBP_REGULATION_OF_CELL_DIVISION | 168 | 0.345547 | 1.695621 | 0 | 0.0809756 | 1 | 2273 | tags=20%, list=10%, si | MM_4 |
| GOBP_PROTEIN_K6_LINK | GOBP_PROTEIN_K6_LINKED_UBIQUITINATI | 15 | 0.548199 | 1.695498 | 0.012150669 | 0.08036897 | 1 | 1947 | tags=20%, list=8%, sig | MM_4 |
| GOBP_INTERMEDIATE_FI | GOBP_INTERMEDIATE_FILAMENT_BASED_P | 67 | 0.387346 | 1.694791 | 0.0010810811 | 0.080395505 | 1 | 9427 | tags=61%, list=40%, si | MM_4 |
| GOBP_CILIUM_MOVEME | GOBP_CILIUM_MOVEMENT | 202 | 0.337857 | 1.692983 | 0 | 0.081201166 | 1 | 12049 | tags=67%, list=51%, si | MM_4 |
| GOBP_DETECTION_OF_V | GOBP_DETECTION_OF_VISIBLE_LIGHT | 25 | 0.481674 | 1.689855 | 0.004889976 | 0.08332443 | 1 | 10129 | tags=80%, list=43%, si | MM_4 |
| NICOTINE_ADDICTION | NICOTINE_ADDICTION | 35 | 0.44792 | 1.6897 | 0.0022753128 | 0.08275966 | 1 | 11688 | tags=86%, list=50%, si | MM_4 |
| GOBP_RECOMBINATIONA | GOBP_RECOMBINATIONAL_REPAIR | 177 | 0.336324 | 1.68655 | 0 | 0.08472337 | 1 | 2291 | tags=19%, list=10%, si | MM_4 |
| GOBP_LIPOPOLYSACCHA | GOBP_LIPOPOLYSACCHARIDE_MEDIATED_S | 57 | 0.397518 | 1.679173 | 0.0010952903 | 0.09023168 | 1 | 2234 | tags=23%, list=9%, sig | MM_4 |
| GOBP_NEGATIVE_REGUL | GOBP_NEGATIVE_REGULATION_OF_ORGAN | 47 | 0.415202 | 1.675312 | 0.003318584 | 0.093061864 | 1 | 2259 | tags=26%, list=10%, si | MM_4 |
| GOBP_REGULATION_OF_ | GOBP_REGULATION_OF_MITOTIC_CELL_CY | 490 | 0.317313 | 1.672943 | 0 | 0.09452495 | 1 | 2288 | tags=18%, list=10%, si | MM_4 |
| CHOLINE_METABOLISM_ | CHOLINE_METABOLISM_IN_CANCER | 97 | 0.365293 | 1.672561 | 0.0010548523 | 0.09404524 | 1 | 2206 | tags=21%, list=9%, sig | MM_4 |
| GOBP_NEGATIVE_REGUL | GOBP_NEGATIVE_REGULATION_OF_DNA_R | 42 | 0.416018 | 1.672548 | 0.004634994 | 0.09329849 | 1 | 2187 | tags=24%, list=9%, sig | MM_4 |
| GOBP_MITOTIC_CELL_CY | GOBP_MITOTIC_CELL_CYCLE_PHASE_TRANS | 425 | 0.320532 | 1.663367 | 0 | 0.101331495 | 1 | 2139 | tags=16%, list=9%, sig | MM_4 |
| GOBP_NEGATIVE_REGUL | GOBP_NEGATIVE_REGULATION_OF_PROTEI | 25 | 0.467783 | 1.660045 | 0.009638554 | 0.10388704 | 1 | 2191 | tags=32%, list=9%, sig | MM_4 |
| GOBP_REGULATION_OF_ | GOBP_REGULATION_OF_MYELINATION | 58 | 0.388185 | 1.659725 | 0.0010810811 | 0.10341552 | 1 | 2243 | tags=24%, list=9%, sig | MM_4 |
| GOBP_REGULATION_OF_ | GOBP_REGULATION_OF_ADENYLATE_CYCLA | 36 | 0.426379 | 1.655101 | 0.012629162 | 0.107305914 | 1 | 10211 | tags=72%, list=43%, si | MM_4 |
| GOBP_POSITIVE_REGULA | GOBP_POSITIVE_REGULATION_OF_FIBROBL | 76 | 0.375509 | 1.653839 | 0 | 0.107607774 | 1 | 2181 | tags=22%, list=9%, sig | MM_4 |
| ALCOHOLISM | ALCOHOLISM | 161 | 0.337135 | 1.653125 | 0 | 0.10752591 | 1 | 1810 | tags=18%, list=8%, sig | MM_4 |
| GOBP_REGULATION_OF_ | GOBP_REGULATION_OF_CENTRIOLE_REPLIC | 22 | 0.483832 | 1.650408 | 0.015568863 | 0.10952289 | 1 | 1546 | tags=23%, list=7%, sig | MM_4 |
| GOBP_REGULATION_OF_ | GOBP_REGULATION_OF_REGULATORY_NCR | 15 | 0.527134 | 1.640086 | 0.018891688 | 0.120502785 | 1 | 1910 | tags=20%, list=8%, sig | MM_4 |
| GOBP_REGULATION_OF_ | GOBP_REGULATION_OF_DNA_DAMAGE_CH | 22 | 0.483355 | 1.639092 | 0.0071856286 | 0.12069209 | 1 | 2193 | tags=27%, list=9%, sig | MM_4 |
| GOBP_NEGATIVE_REGUL | GOBP_NEGATIVE_REGULATION_OF_ORGAN | 375 | 0.314997 | 1.636522 | 0 | 0.12274381 | 1 | 2290 | tags=19%, list=10%, si | MM_4 |
| GOBP_NEGATIVE_REGUL | GOBP_NEGATIVE_REGULATION_OF_CILIUM | 21 | 0.485115 | 1.636389 | 0.01438849 | 0.122022115 | 1 | 2250 | tags=33%, list=10%, si | MM_4 |
| GOBP_INTERLEUKIN_4_P | GOBP_INTERLEUKIN_4_PRODUCTION | 38 | 0.420266 | 1.635151 | 0.007963595 | 0.12251892 | 1 | 3236 | tags=37%, list=14%, si | MM_4 |
| GOBP_DNA_DUPLEX_UN | GOBP_DNA_DUPLEX_UNWINDING | 43 | 0.408799 | 1.633082 | 0.004449388 | 0.12406466 | 1 | 2134 | tags=23%, list=9%, sig | MM_4 |
| GOBP_ODONTOBLAST_DI | GOBP_ODONTOBLAST_DIFFERENTIATION | 18 | 0.504748 | 1.632907 | 0.0111524165 | 0.12334861 | 1 | 2561 | tags=33%, list=11%, si | MM_4 |
| GOBP_LEUKOTRIENE_ME | GOBP_LEUKOTRIENE_METABOLIC_PROCESS | 20 | 0.485426 | 1.626785 | 0.011002445 | 0.1299536 | 1 | 3568 | tags=35%, list=15%, si | MM_4 |
| GOBP_REGULATION_OF_ | GOBP_REGULATION_OF_KERATINOCYTE_M | 15 | 0.518561 | 1.625688 | 0.021656051 | 0.13039541 | 1 | 1664 | tags=27%, list=7%, sig | MM_4 |
| GOBP_REGULATION_OF_ | GOBP_REGULATION_OF_NEUROTRANSMITT | 25 | 0.453016 | 1.623187 | 0.016509434 | 0.13263766 | 1 | 2871 | tags=28%, list=12%, si | MM_4 |
| GOBP_REGULATION_OF_ | GOBP_REGULATION_OF_LYASE_ACTIVITY | 48 | 0.389183 | 1.615563 | 0.005518764 | 0.14146544 | 1 | 9654 | tags=65%, list=41%, si | MM_4 |
| GOBP_FATTY_ACYL_COA_ | GOBP_FATTY_ACYL_COA_METABOLIC_PRO | 18 | 0.493664 | 1.614253 | 0.02354399 | 0.14216825 | 1 | 2276 | tags=28%, list=10%, si | MM_4 |
| GOBP_MITOTIC_CYTOKIN | GOBP_MITOTIC_CYTOKINETIC_PROCESS | 30 | 0.439497 | 1.613597 | 0.009090909 | 0.14200063 | 1 | 1828 | tags=17%, list=8%, sig | MM_4 |
| GOBP_PROTEIN_LOCALIZ | GOBP_PROTEIN_LOCALIZATION_TO_CHROM | 112 | 0.3412 | 1.610064 | 0.0010341262 | 0.14562245 | 1 | 1249 | tags=13%, list=5%, sig | MM_4 |
| GOBP_FATTY_ACID_HOM | GOBP_FATTY_ACID_HOMEOSTASIS | 15 | 0.527153 | 1.607929 | 0.017834395 | 0.14752497 | 1 | 2712 | tags=27%, list=11%, si | MM_4 |
| GOBP_REGULATION_OF_ | GOBP_REGULATION_OF_CYCLASE_ACTIVITY | 46 | 0.394105 | 1.60485 | 0.007829977 | 0.1509699 | 1 | 9654 | tags=65%, list=41%, si | MM_4 |
| GOBP_POSITIVE_REGULA | GOBP_POSITIVE_REGULATION_OF_MITOTIC | 93 | 0.349702 | 1.602931 | 0.0010582011 | 0.15276635 | 1 | 1581 | tags=15%, list=7%, sig | MM_4 |
| GOBP_CILIUM_OR_FLAG | GOBP_CILIUM_OR_FLAGELLUM_DEPENDEN | 173 | 0.323579 | 1.60053 | 0 | 0.15500796 | 1 | 13025 | tags=71%, list=55%, si | MM_4 |
| GOBP_REGULATION_OF_ | GOBP_REGULATION_OF_CELL_CYCLE_PHAS | 424 | 0.307004 | 1.600514 | 0 | 0.15401392 | 1 | 2288 | tags=16%, list=10%, si | MM_4 |
| GOBP_PRE_MIRNA_PRO | GOBP_PRE_MIRNA_PROCESSING | 15 | 0.521564 | 1.600065 | 0.022613065 | 0.15364654 | 1 | 1599 | tags=27%, list=7%, sig | MM_4 |
| GOBP_PERICARDIUM_DE | GOBP_PERICARDIUM_DEVELOPMENT | 27 | 0.437488 | 1.599533 | 0.015240328 | 0.15340742 | 1 | 1819 | tags=22%, list=8%, sig | MM_4 |
| GOBP_ASTROCYTE_ACTIV | GOBP_ASTROCYTE_ACTIVATION | 21 | 0.470872 | 1.595796 | 0.020807834 | 0.15773626 | 1 | 2207 | tags=29%, list=9%, sig | MM_4 |
| GOBP_DNA_CONFORMAT | GOBP_DNA_CONFORMATION_CHANGE | 58 | 0.379735 | 1.591159 | 0.0075431033 | 0.16339868 | 1 | 2134 | tags=22%, list=9%, sig | MM_4 |
| GOBP_SEMINIFEROUS_TU | GOBP_SEMINIFEROUS_TUBULE_DEVELOPM | 20 | 0.469576 | 1.590465 | 0.018094089 | 0.16339462 | 1 | 4171 | tags=40%, list=18%, si | MM_4 |
| GOBP_POSITIVE_REGULA | GOBP_POSITIVE_REGULATION_OF_NITRIC_ | 20 | 0.479403 | 1.588459 | 0.023337223 | 0.16543238 | 1 | 2179 | tags=30%, list=9%, sig | MM_4 |
| FANCONI_ANEMIA_PATH | FANCONI_ANEMIA_PATHWAY | 50 | 0.382971 | 1.588017 | 0.013318535 | 0.16511776 | 1 | 2193 | tags=26%, list=9%, sig | MM_4 |

| | | | | | | | | | | |
|---|---|---|---|---|---|---|---|---|---|---|
| GOBP_NUCLEOLAR_LARG | GOBP_NUCLEOLAR_LARGE_RRNA_TRANSCR | 21 | 0.464734 | 1.587187 | 0.021686748 | 0.165266 | 1 | 2072 | tags=19%, list=9%, sig | MM_4 |
| GOBP_POSITIVE_REGULA | GOBP_POSITIVE_REGULATION_OF_INTERLE | 15 | 0.51586 | 1.58637 | 0.022727273 | 0.16549486 | 1 | 2243 | tags=53%, list=9%, sig | MM_4 |
| GOBP_NEGATIVE_REGUL | GOBP_NEGATIVE_REGULATION_OF_REACTI | 20 | 0.466763 | 1.584737 | 0.015037594 | 0.16699854 | 1 | 2712 | tags=25%, list=11%, si | MM_4 |
| GOBP_REGULATION_OF_ | GOBP_REGULATION_OF_CYCLIN_DEPENDEN | 67 | 0.362328 | 1.578837 | 0.010869565 | 0.17537268 | 1 | 1806 | tags=18%, list=8%, sig | MM_4 |
| GOBP_MODIFICATION_O | GOBP_MODIFICATION_OF_POSTSYNAPTIC_ | 42 | 0.394789 | 1.57753 | 0.01033295 | 0.17637204 | 1 | 2149 | tags=24%, list=9%, sig | MM_4 |
| GOBP_AXIS_ELONGATION | GOBP_AXIS_ELONGATION | 36 | 0.409848 | 1.575526 | 0.020361992 | 0.17863353 | 1 | 2881 | tags=31%, list=12%, si | MM_4 |
| GOBP_INTERLEUKIN_6_M | GOBP_INTERLEUKIN_6_MEDIATED_SIGNALI | 19 | 0.48085 | 1.573495 | 0.020807834 | 0.18096758 | 1 | 2236 | tags=26%, list=9%, sig | MM_4 |
| GOBP_POSITIVE_REGULA | GOBP_POSITIVE_REGULATION_OF_PHOSPH | 26 | 0.440348 | 1.573424 | 0.016470589 | 0.17998317 | 1 | 1477 | tags=19%, list=6%, sig | MM_4 |
| GOBP_EYE_PHOTORECEP | GOBP_EYE_PHOTORECEPTOR_CELL_DIFFERE | 58 | 0.36851 | 1.571458 | 0.010845987 | 0.18205673 | 1 | 12463 | tags=72%, list=53%, si | MM_4 |
| GOBP_SISTER_CHROMAT | GOBP_SISTER_CHROMATID_COHESION | 54 | 0.377729 | 1.569015 | 0.013201321 | 0.18505694 | 1 | 1249 | tags=15%, list=5%, sig | MM_4 |
| GOBP_NEURON_INTRINS | GOBP_NEURON_INTRINSIC_APOPTOTIC_SIG | 21 | 0.459863 | 1.569013 | 0.02208589 | 0.18397026 | 1 | 2259 | tags=29%, list=10%, si | MM_4 |
| GOBP_REGULATION_OF_ | GOBP_REGULATION_OF_FIBROBLAST_APOP | 24 | 0.444315 | 1.568634 | 0.016847173 | 0.18360198 | 1 | 2158 | tags=25%, list=9%, sig | MM_4 |
| GOBP_POSITIVE_REGULA | GOBP_POSITIVE_REGULATION_OF_INTERLE | 24 | 0.451821 | 1.568141 | 0.01783591 | 0.18332446 | 1 | 3116 | tags=46%, list=13%, si | MM_4 |
| GOBP_REGULATION_OF_ | GOBP_REGULATION_OF_MITOCHONDRIAL_ | 32 | 0.419094 | 1.567057 | 0.014806378 | 0.18404922 | 1 | 2171 | tags=28%, list=9%, sig | MM_4 |
| GOBP_COMPLEMENT_AC | GOBP_COMPLEMENT_ACTIVATION_CLASSIC | 29 | 0.42429 | 1.566208 | 0.009280742 | 0.18441638 | 1 | 2207 | tags=24%, list=9%, sig | MM_4 |
| FOXO_SIGNALING_PATH | FOXO_SIGNALING_PATHWAY | 127 | 0.325951 | 1.565052 | 0 | 0.18527159 | 1 | 2712 | tags=20%, list=11%, si | MM_4 |
| GOBP_PROTEIN_LOCALIZ | GOBP_PROTEIN_LOCALIZATION_TO_MICRO | 39 | 0.397807 | 1.564664 | 0.009049774 | 0.18483798 | 1 | 1940 | tags=21%, list=8%, sig | MM_4 |
| GOBP_MITOCHONDRIAL_ | GOBP_MITOCHONDRIAL_TRANSLATION | 128 | 0.329011 | 1.564264 | 0.0031120332 | 0.18448572 | 1 | 2038 | tags=19%, list=9%, sig | MM_4 |
| GOBP_EXTRACELLULAR_T | GOBP_EXTRACELLULAR_TRANSPORT | 49 | 0.382453 | 1.564158 | 0.0121546965 | 0.18359779 | 1 | 11894 | tags=67%, list=50%, si | MM_4 |
| GOBP_REGULATION_OF_ | GOBP_REGULATION_OF_CELL_CYCLE_G2_M | 117 | 0.32844 | 1.561477 | 0.0010373445 | 0.18689324 | 1 | 748 | tags=12%, list=3%, sig | MM_4 |
| GOBP_REGULATION_OF_ | GOBP_REGULATION_OF_MODIFICATION_OF | 22 | 0.457664 | 1.561124 | 0.02256532 | 0.18649559 | 1 | 2149 | tags=27%, list=9%, sig | MM_4 |
| GOBP_NEURONAL_ACTIO | GOBP_NEURONAL_ACTION_POTENTIAL | 56 | 0.364948 | 1.557586 | 0.01416122 | 0.19165942 | 1 | 11634 | tags=70%, list=49%, si | MM_4 |
| GOBP_POSITIVE_REGULA | GOBP_POSITIVE_REGULATION_OF_NON_CA | 64 | 0.352832 | 1.557057 | 0.006521739 | 0.19152327 | 1 | 2207 | tags=22%, list=9%, sig | MM_4 |
| GOBP_REGULATION_OF_ | GOBP_REGULATION_OF_MICROTUBULE_BA | 248 | 0.304848 | 1.55653 | 0 | 0.19120297 | 1 | 2349 | tags=19%, list=10%, si | MM_4 |
| GOBP_C_TERMINAL_PRO | GOBP_C_TERMINAL_PROTEIN_AMINO_ACID | 20 | 0.46262 | 1.556365 | 0.020481927 | 0.19043419 | 1 | 2068 | tags=20%, list=9%, sig | MM_4 |
| GOBP_BRANCHED_CHAIN | GOBP_BRANCHED_CHAIN_AMINO_ACID_CA | 21 | 0.455952 | 1.552802 | 0.02 | 0.19583547 | 1 | 2262 | tags=33%, list=10%, si | MM_4 |
| GOBP_POSITIVE_REGULA | GOBP_POSITIVE_REGULATION_OF_CELL_CY | 112 | 0.329673 | 1.551611 | 0.004158004 | 0.19698095 | 1 | 1770 | tags=15%, list=7%, sig | MM_4 |
| GOBP_NEGATIVE_REGUL | GOBP_NEGATIVE_REGULATION_OF_MEMB | 18 | 0.4764 | 1.551175 | 0.02093596 | 0.19662353 | 1 | 1910 | tags=28%, list=8%, sig | MM_4 |
| GOBP_REGULATION_OF_ | GOBP_REGULATION_OF_MITOTIC_SPINDLE_ | 18 | 0.484038 | 1.550289 | 0.02631579 | 0.1971098 | 1 | 78 | tags=11%, list=0%, sig | MM_4 |
| CELL_CYCLE | CELL_CYCLE | 121 | 0.32567 | 1.548449 | 0.0010351967 | 0.19934033 | 1 | 2134 | tags=16%, list=9%, sig | MM_4 |
| GOBP_POSITIVE_REGULA | GOBP_POSITIVE_REGULATION_OF_CHOLES | 31 | 0.414112 | 1.548106 | 0.018779343 | 0.19884215 | 1 | 5713 | tags=45%, list=24%, si | MM_4 |
| GOBP_POSITIVE_REGULA | GOBP_POSITIVE_REGULATION_OF_PROTEIN | 15 | 0.509803 | 1.542495 | 0.029077118 | 0.20795919 | 1 | 2135 | tags=40%, list=9%, sig | MM_4 |
| GOBP_REGULATION_OF_ | GOBP_REGULATION_OF_PLATELET_ACTIVAT | 45 | 0.382391 | 1.537672 | 0.015503876 | 0.21601187 | 1 | 3775 | tags=31%, list=16%, si | MM_4 |
| GOBP_CAMERA_TYPE_EY | GOBP_CAMERA_TYPE_EYE_PHOTORECEPTO | 41 | 0.388175 | 1.536992 | 0.017857144 | 0.2160556 | 1 | 12463 | tags=78%, list=53%, si | MM_4 |
| GOBP_PROTEIN_K63_LIN | GOBP_PROTEIN_K63_LINKED_UBIQUITINAT | 68 | 0.352667 | 1.536024 | 0.009615385 | 0.2168623 | 1 | 2234 | tags=18%, list=9%, sig | MM_4 |
| GOBP_GROWTH_HORMO | GOBP_GROWTH_HORMONE_SECRETION | 20 | 0.458983 | 1.53553 | 0.033088237 | 0.2165762 | 1 | 2871 | tags=30%, list=12%, si | MM_4 |
| GOBP_RETINA_MORPHO | GOBP_RETINA_MORPHOGENESIS_IN_CAME | 77 | 0.342527 | 1.535173 | 0.004310345 | 0.21610226 | 1 | 12463 | tags=74%, list=53%, si | MM_4 |
| GOBP_SPERM_MOTILITY | GOBP_SPERM_MOTILITY | 151 | 0.315399 | 1.532622 | 0.002051282 | 0.21986035 | 1 | 11695 | tags=64%, list=50%, si | MM_4 |
| GOBP_REGULATION_OF_ | GOBP_REGULATION_OF_CILIUM_MOVEMEN | 33 | 0.410855 | 1.531669 | 0.022753129 | 0.2206316 | 1 | 11412 | tags=73%, list=48%, si | MM_4 |
| GOBP_PROTEIN_METHYL | GOBP_PROTEIN_METHYLATION | 57 | 0.363253 | 1.530006 | 0.009846827 | 0.2226991 | 1 | 2221 | tags=21%, list=9%, sig | MM_4 |
| GOBP_DETECTION_OF_LI | GOBP_DETECTION_OF_LIGHT_STIMULUS_IN | 19 | 0.462643 | 1.526253 | 0.03253012 | 0.2291031 | 1 | 10129 | tags=79%, list=43%, si | MM_4 |
| GOBP_CELLULAR_RESPON | GOBP_CELLULAR_RESPONSE_TO_CHOLESTE | 18 | 0.466325 | 1.523483 | 0.0404908 | 0.23356523 | 1 | 5713 | tags=56%, list=24%, si | MM_4 |
| GOBP_NUCLEOSIDE_MET | GOBP_NUCLEOSIDE_METABOLIC_PROCESS | 45 | 0.379671 | 1.52219 | 0.014444444 | 0.23497942 | 1 | 2228 | tags=24%, list=9%, sig | MM_4 |
| GOBP_POSITIVE_REGULA | GOBP_POSITIVE_REGULATION_OF_EXTRACE | 32 | 0.407474 | 1.521184 | 0.021965317 | 0.23598681 | 1 | 1910 | tags=25%, list=8%, sig | MM_4 |
| GOBP_MITOTIC_CHROM | GOBP_MITOTIC_CHROMOSOME_CONDENS | 15 | 0.485655 | 1.521173 | 0.033834588 | 0.23483473 | 1 | 1399 | tags=27%, list=6%, sig | MM_4 |
| GOBP_OLIGOPEPTIDE_TR | GOBP_OLIGOPEPTIDE_TRANSPORT | 15 | 0.497589 | 1.521044 | 0.03860523 | 0.2339172 | 1 | 4047 | tags=47%, list=17%, si | MM_4 |
| GOBP_FOREBRAIN_REGI | GOBP_FOREBRAIN_REGIONALIZATION | 18 | 0.464748 | 1.520314 | 0.024479805 | 0.23414151 | 1 | 9067 | tags=67%, list=38%, si | MM_4 |
| GOBP_CARDIAC_RIGHT_V | GOBP_CARDIAC_RIGHT_VENTRICLE_MORPH | 19 | 0.466895 | 1.518887 | 0.03151515 | 0.23588277 | 1 | 1469 | tags=32%, list=6%, sig | MM_4 |
| GOBP_NEGATIVE_REGUL | GOBP_NEGATIVE_REGULATION_OF_DOUBL | 24 | 0.43924 | 1.518407 | 0.031212484 | 0.23571582 | 1 | 2187 | tags=29%, list=9%, sig | MM_4 |
| GOBP_REGULATION_OF_ | GOBP_REGULATION_OF_PROTEIN_ACETYLA | 22 | 0.445478 | 1.518113 | 0.028436018 | 0.23515895 | 1 | 2135 | tags=32%, list=9%, sig | MM_4 |
| GOBP_MEIOTIC_CELL_CY | GOBP_MEIOTIC_CELL_CYCLE_PROCESS | 220 | 0.301497 | 1.515782 | 0 | 0.23878552 | 1 | 1736 | tags=15%, list=7%, sig | MM_4 |
| GOBP_RESPONSE_TO_ES | GOBP_RESPONSE_TO_ESTRADIOL | 38 | 0.389232 | 1.514782 | 0.020594966 | 0.23980273 | 1 | 638 | tags=13%, list=3%, sig | MM_4 |
| GOBP_REGULATION_OF_ | GOBP_REGULATION_OF_MODIFICATION_OF | 29 | 0.412541 | 1.514231 | 0.029988466 | 0.23987257 | 1 | 2149 | tags=24%, list=9%, sig | MM_4 |
| GOBP_DOPAMINE_TRAN | GOBP_DOPAMINE_TRANSPORT | 54 | 0.364515 | 1.512061 | 0.016611295 | 0.24318072 | 1 | 11352 | tags=63%, list=48%, si | MM_4 |
| GOBP_PROTEIN_ACETYLA | GOBP_PROTEIN_ACETYLATION | 56 | 0.356746 | 1.511373 | 0.02202643 | 0.2435474 | 1 | 2135 | tags=23%, list=9%, sig | MM_4 |
| GOBP_ORGAN_OR_TISSU | GOBP_ORGAN_OR_TISSUE_SPECIFIC_IMMU | 25 | 0.42942 | 1.510804 | 0.034117647 | 0.24361895 | 1 | 7785 | tags=64%, list=33%, si | MM_4 |
| GOBP_ERBB_SIGNALING_ | GOBP_ERBB_SIGNALING_PATHWAY | 125 | 0.31619 | 1.509779 | 0.0031023785 | 0.24463265 | 1 | 1985 | tags=17%, list=8%, sig | MM_4 |
| GOBP_ACTIVATION_OF_ | GOBP_ACTIVATION_OF_ADENYLATE_CYCLA | 19 | 0.459966 | 1.509391 | 0.03448276 | 0.24435383 | 1 | 10211 | tags=79%, list=43%, si | MM_4 |
