## Supplementary Table 5 for "C1qa□ muscularis macrophages maintain enteric synaptic homeostasis to regulate gastrointestinal motility"

| GO_term | FDR | Group | genes_present_in_term |
|---|---|---|---|
| anchored component of synaptic vesicle membrane | 2.59309E-09 | shared | RAB3A;RAB3B;RAB5A;RAB5C;RAB35;RAB11B;RAB10;RAB4A;RAB14;RAB7A;RAB2A;RAC1 |
| presynaptic ribosome | 4.10864E-43 | shared | RPL22;RPS6;RPL9;RPL10;RPL10A;RPL11;RPL12;RPL13;RPL13A;RPL14;RPL17;RPL15;RPL18;RPL23;RPL23A;RPL26;RPL24;RPL27;RPL27A;RPL28;RPL29;RPL32;RPL34;RPL35;RPL35A;RPL36;RPL36A;RPL37A;RPL38;RPL4;RPL5;RPL6;RPL7;RPL7A;RPL8;RPLP0;RPLP2;RPS10;RPS11;RPS12;RPS14;RPS15A;RPS16;RPS24;RPS26;RPS27A;RPS28;RPS5;RPS13;RPS27 |
| postsynaptic density membrane | 0.221919671 | shared | TRAPPC4;AKAP9;DGKB;NPTN;RTN4;VDAC1;STX1A;LRP1 |
| regulation of synapse organization | 2.09851E-05 | shared | NCKAP1;VCP;MECP2;CSNK2B;ARF4;SEPTIN11;TUBA1A;TUBB5;GPM6A |
| translation at presynapse | 7.67836E-42 | shared | RPL22;RPL9;RPL10;RPL10A;RPL11;RPL12;RPL13;RPL13A;RPL14;RPL17;RPL15;RPL18;RPL23;RPL23A;RPL26;RPL24;RPL27;RPL27A;RPL28;RPL29;RPL32;RPL34;RPL35;RPL35A;RPL36;RPL36A;RPL37A;RPL38;RPL4;RPL5;RPL6;RPL7;RPL7A;RPL8;RPLP0;RPLP2;RPS10;RPS11;RPS12;RPS14;RPS15A;RPS16;RPS24;RPS26;RPS27A;RPS28;RPS5;RPS13;RPS27 |
| translation at postsynapse | 9.06457E-42 | shared | EEF2;RPL9;RPL10;RPL10A;RPL11;RPL12;RPL13;RPL13A;RPL14;RPL17;RPL15;RPL18;RPL23;RPL22;RPL23A;RPL26;RPL24;RPL27;RPL27A;RPL28;RPL29;RPL32;RPL34;RPL35;RPL35A;RPL36;RPL36A;RPL37A;RPL38;RPL4;RPL5;RPL6;RPL7;RPL7A;RPL8;RPLP0;RPLP2;RPS10;RPS11;RPS12;RPS14;RPS15A;RPS16;RPS24;RPS26;RPS27A;RPS28;RPS5;RPS13;RPS27 |
| postsynaptic density membrane | 2.49755E-09 | Brain_specific | IQSEC2;ACTN2;PACSIN1;MLLT4;DLG2;DLG4;MPP2;NTM;IGSF21;CNKSR2;OLFM1;ATP2B2;ADCY1;ADAM22;GRIA1;GRIA2;GRIN1;GRIN2B;LRRC7;PTPRZ1;GRIN2A;GRIA3;KCNAB2;BAI3;GPR158;VANGL2 |
| regulation of postsynaptic membrane potential | 6.04129E-06 | Brain_specific | GABBR1;KCTD16;GRIN1;GRIN2A;GRIN2B;GRIA1;GRIA2;GABRA1;GABRB3;GABRA3;GABRG2;GRIA3;KCND2 |
| integral component of synaptic membrane | 0.33496598 | GI_specific | ITGB5;ITGB3;ITGB1;ANO6;NCAM2;EGFR |
| anchored component of synaptic vesicle membrane | 0.954630928 | GI_specific | RAB27B;DNAJC5;RAB5B |
| synaptic cleft | 0.189651401 | GI_specific | APOE;ACHE;LAMA5;LAMB2;LAMA2;LAMC1;LAMA4;SPARCL1 |
| regulation of postsynaptic membrane potential | 0.994432219 | GI_specific | NOS1;ANO6;DPP6 |
| regulation of synapse organization | 0.807325732 | GI_specific | ARF2;SPARC;APOE;DAG1;SPARCL1 |
| neurotransmitter receptor transport to postsynaptic membrane | 0.276498389 | GI_specific | RAB8A;SACM1L;MAPK10;MYLK |
