## Supplementary Table 6 for "C1qa□ muscularis macrophages maintain enteric synaptic homeostasis to regulate gastrointestinal motility"

| NAME | GS<br> follow link to MSigDB | SIZE | ES | NES | NOM p-val | FDR q-val | FWER p-val | RANK AT MAX | LEADING EDGE |
|---|---|---|---|---|---|---|---|---|---|
| COMPLEMENT AND COAGULATION CASCADES | COMPLEMENT AND COAGULATION CASCADES | 67 | 0.772645 | 1.810834 | 0 | 0.04705016 | 0.046 | 564 | tags=19%, list=3%, signal=20% |
| CELLULAR PROTEIN CATABOLIC PROCESS (GO:0044257) | CELLULAR PROTEIN CATABOLIC PROCESS (GO:0044257) | 55 | 0.79663 | 1.80734 | 0 | 0.025586067 | 0.049 | 1241 | tags=33%, list=6%, signal=35% |
| POSITIVE REGULATION OF LEUKOCYTE CHEMOTAXIS (GO:0002690) | POSITIVE REGULATION OF LEUKOCYTE CHEMOTAXIS (GO:0002690) | 50 | 0.803121 | 1.792562 | 0 | 0.031068569 | 0.09 | 1221 | tags=42%, list=6%, signal=44% |
| REGULATION OF PROTEIN ACTIVATION CASCADE (GO:2000257) | REGULATION OF PROTEIN ACTIVATION CASCADE (GO:2000257) | 35 | 0.825441 | 1.764428 | 0 | 0.051902983 | 0.189 | 564 | tags=34%, list=3%, signal=35% |
| REGULATION OF HUMORAL IMMUNE RESPONSE (GO:0002920) | REGULATION OF HUMORAL IMMUNE RESPONSE (GO:0002920) | 38 | 0.817931 | 1.751683 | 0 | 0.05866286 | 0.244 | 583 | tags=37%, list=3%, signal=38% |
| ARACHIDONIC ACID METABOLISM | ARACHIDONIC ACID METABOLISM | 57 | 0.769152 | 1.74549 | 0 | 0.0562224 | 0.275 | 1827 | tags=25%, list=8%, signal=27% |
| REGULATION OF COMPLEMENT ACTIVATION (GO:0030449) | REGULATION OF COMPLEMENT ACTIVATION (GO:0030449) | 36 | 0.820421 | 1.744964 | 0.001824818 | 0.048626494 | 0.277 | 564 | tags=33%, list=3%, signal=34% |
| SYNAPTIC TRANSMISSION, CHOLINERGIC (GO:0007271) | SYNAPTIC TRANSMISSION, CHOLINERGIC (GO:0007271) | 23 | 0.891883 | 1.743714 | 0 | 0.043956645 | 0.286 | 2 | tags=4%, list=0%, signal=4% |
| MINERAL ABSORPTION | MINERAL ABSORPTION | 39 | 0.800003 | 1.726809 | 0 | 0.063247316 | 0.412 | 1219 | tags=33%, list=6%, signal=35% |
| REGULATION OF ACUTE INFLAMMATORY RESPONSE (GO:0002673) | REGULATION OF ACUTE INFLAMMATORY RESPONSE (GO:0002673) | 46 | 0.773438 | 1.703456 | 0.001798561 | 0.09955903 | 0.597 | 944 | tags=30%, list=4%, signal=32% |
| REGULATION OF IMMUNE EFFECTOR PROCESS (GO:0002697) | REGULATION OF IMMUNE EFFECTOR PROCESS (GO:0002697) | 39 | 0.79937 | 1.69887 | 0 | 0.1018509 | 0.645 | 564 | tags=33%, list=3%, signal=34% |
| NEGATIVE REGULATION OF BLOOD VESSEL MORPHOGENESIS (GO:2000181) | NEGATIVE REGULATION OF BLOOD VESSEL MORPHOGENESIS (GO:2000181) | 59 | 0.735982 | 1.690873 | 0.00177305 | 0.11343288 | 0.724 | 1671 | tags=29%, list=8%, signal=31% |
| POSITIVE REGULATION OF OSSIFICATION (GO:0045778) | POSITIVE REGULATION OF OSSIFICATION (GO:0045778) | 46 | 0.757017 | 1.687347 | 0.00177305 | 0.11164735 | 0.741 | 2644 | tags=35%, list=12%, signal=39% |
| REGULATION OF LONG-TERM SYNAPTIC POTENTIATION (GO:1900271) | REGULATION OF LONG-TERM SYNAPTIC POTENTIATION (GO:1900271) | 16 | 0.912382 | 1.683443 | 0 | 0.11366469 | 0.777 | 2 | tags=6%, list=0%, signal=6% |
| LYSOSOME | LYSOSOME | 116 | 0.66966 | 1.681551 | 0 | 0.11038413 | 0.793 | 1688 | tags=48%, list=8%, signal=52% |
| CELLULAR RESPONSE TO AMYLOID-BETA (GO:1904646) | CELLULAR RESPONSE TO AMYLOID-BETA (GO:1904646) | 22 | 0.867703 | 1.673234 | 0.001776199 | 0.12225698 | 0.838 | 1527 | tags=55%, list=7%, signal=59% |
| SYNAPTIC VESICLE ENDOCYTOSIS (GO:0048488) | SYNAPTIC VESICLE ENDOCYTOSIS (GO:0048488) | 25 | 0.833362 | 1.663043 | 0.005395684 | 0.144077 | 0.904 | 1101 | tags=20%, list=5%, signal=21% |
| NEGATIVE REGULATION OF ANGIOGENESIS (GO:0016525) | NEGATIVE REGULATION OF ANGIOGENESIS (GO:0016525) | 63 | 0.719378 | 1.653218 | 0 | 0.16393217 | 0.939 | 1955 | tags=32%, list=9%, signal=35% |
| SENSORY PERCEPTION OF PAIN (GO:0019233) | SENSORY PERCEPTION OF PAIN (GO:0019233) | 19 | 0.872433 | 1.649446 | 0.00742115 | 0.16681187 | 0.952 | 541 | tags=11%, list=2%, signal=11% |
| REGULATION OF BONE MINERALIZATION (GO:0030500) | REGULATION OF BONE MINERALIZATION (GO:0030500) | 45 | 0.738178 | 1.63897 | 0.005190312 | 0.19206625 | 0.971 | 1510 | tags=33%, list=7%, signal=36% |
| CLATHRIN-DEPENDENT ENDOCYTOSIS (GO:0072583) | CLATHRIN-DEPENDENT ENDOCYTOSIS (GO:0072583) | 36 | 0.782439 | 1.638643 | 0.001782531 | 0.1841899 | 0.972 | 1038 | tags=25%, list=5%, signal=26% |
| NEGATIVE REGULATION OF CELLULAR METABOLIC PROCESS (GO:0031324) | NEGATIVE REGULATION OF CELLULAR METABOLIC PROCESS (GO:0031324) | 28 | 0.802992 | 1.637275 | 0.007490637 | 0.18055789 | 0.978 | 1892 | tags=39%, list=9%, signal=43% |
| POSITIVE REGULATION OF BONE MINERALIZATION (GO:0030501) | POSITIVE REGULATION OF BONE MINERALIZATION (GO:0030501) | 28 | 0.805981 | 1.636457 | 0.001858736 | 0.17493993 | 0.98 | 2644 | tags=36%, list=12%, signal=41% |
| SYNAPTIC VESICLE RECYCLING (GO:0036465) | SYNAPTIC VESICLE RECYCLING (GO:0036465) | 24 | 0.831947 | 1.629235 | 0.003610108 | 0.1916577 | 0.986 | 1354 | tags=21%, list=6%, signal=22% |
| REGULATION OF PROTEIN PROCESSING (GO:0070613) | REGULATION OF PROTEIN PROCESSING (GO:0070613) | 49 | 0.729503 | 1.624804 | 0 | 0.19797125 | 0.987 | 997 | tags=29%, list=5%, signal=30% |
| INORGANIC CATION TRANSMEMBRANE TRANSPORT (GO:0098662) | INORGANIC CATION TRANSMEMBRANE TRANSPORT (GO:0098662) | 122 | 0.64098 | 1.623967 | 0 | 0.19362977 | 0.989 | 2063 | tags=27%, list=9%, signal=30% |
| SECONDARY ALCOHOL METABOLIC PROCESS (GO:1902652) | SECONDARY ALCOHOL METABOLIC PROCESS (GO:1902652) | 23 | 0.824551 | 1.61965 | 0.00907441 | 0.20161892 | 0.993 | 1186 | tags=26%, list=5%, signal=28% |
| ANION HOMEOSTASIS (GO:0055081) | ANION HOMEOSTASIS (GO:0055081) | 15 | 0.89987 | 1.619145 | 0.005434783 | 0.1961723 | 0.994 | 26 | tags=13%, list=0%, signal=13% |
| CYTOKINE-CYTOKINE RECEPTOR INTERACTION | CYTOKINE-CYTOKINE RECEPTOR INTERACTION | 230 | 0.595993 | 1.612563 | 0 | 0.21071084 | 0.999 | 945 | tags=16%, list=4%, signal=17% |
| CHEMOKINE-MEDIATED SIGNALING PATHWAY (GO:0070098) | CHEMOKINE-MEDIATED SIGNALING PATHWAY (GO:0070098) | 33 | 0.776142 | 1.612084 | 0.003502627 | 0.20512067 | 0.999 | 686 | tags=39%, list=3%, signal=41% |
| POSITIVE REGULATION OF BIOMINERAL TISSUE DEVELOPMENT (GO:0070169) | POSITIVE REGULATION OF BIOMINERAL TISSUE DEVELOPMENT (GO:0070169) | 30 | 0.801635 | 1.611462 | 0.001858736 | 0.20041588 | 0.999 | 2644 | tags=33%, list=12%, signal=38% |
| POSITIVE REGULATION OF CALCIUM-MEDIATED SIGNALING (GO:0050850) | POSITIVE REGULATION OF CALCIUM-MEDIATED SIGNALING (GO:0050850) | 25 | 0.82586 | 1.609823 | 0.001727116 | 0.20044842 | 1 | 541 | tags=28%, list=2%, signal=29% |
| REGULATION OF OSSIFICATION (GO:0030278) | REGULATION OF OSSIFICATION (GO:0030278) | 35 | 0.772222 | 1.606119 | 0.007067138 | 0.2072269 | 1 | 2162 | tags=40%, list=10%, signal=44% |
| REGULATION OF NEURONAL SYNAPTIC PLASTICITY (GO:0048168) | REGULATION OF NEURONAL SYNAPTIC PLASTICITY (GO:0048168) | 22 | 0.830046 | 1.604313 | 0.003696858 | 0.20716636 | 1 | 307 | tags=14%, list=1%, signal=14% |
| POSITIVE REGULATION OF TRANSMEMBRANE RECEPTOR PROTEIN SERINE/THREONINE KINASE SIGNALING PATHWAY (GO:0090100) | POSITIVE REGULATION OF TRANSMEMBRANE RECEPTOR PROTEIN SERINE/THREONINE KINASE SIGNALING PATHWAY (GO:0090100) | 72 | 0.682689 | 1.598913 | 0.005347594 | 0.2203364 | 1 | 1355 | tags=22%, list=6%, signal=24% |
| POSITIVE REGULATION OF ION TRANSPORT (GO:0043270) | POSITIVE REGULATION OF ION TRANSPORT (GO:0043270) | 42 | 0.729246 | 1.595527 | 0.007326007 | 0.22575252 | 1 | 1340 | tags=31%, list=6%, signal=33% |
| REGULATION OF ERBB SIGNALING PATHWAY (GO:1901184) | REGULATION OF ERBB SIGNALING PATHWAY (GO:1901184) | 21 | 0.83574 | 1.58912 | 0.009225092 | 0.24129902 | 1 | 914 | tags=33%, list=4%, signal=35% |
| CHOLESTEROL TRANSPORT (GO:0030301) | CHOLESTEROL TRANSPORT (GO:0030301) | 37 | 0.739971 | 1.58897 | 0.007380074 | 0.235592 | 1 | 145 | tags=11%, list=1%, signal=11% |
| FERROPTOSIS | FERROPTOSIS | 37 | 0.749444 | 1.587925 | 0.003636364 | 0.2328035 | 1 | 1650 | tags=49%, list=7%, signal=52% |
| POSITIVE REGULATION OF DENDRITE DEVELOPMENT (GO:1900006) | POSITIVE REGULATION OF DENDRITE DEVELOPMENT (GO:1900006) | 30 | 0.784176 | 1.58659 | 0.005597015 | 0.23211631 | 1 | 127 | tags=7%, list=1%, signal=7% |
| PROTEOGLYCAN METABOLIC PROCESS (GO:0006029) | PROTEOGLYCAN METABOLIC PROCESS (GO:0006029) | 19 | 0.846191 | 1.584804 | 0.009225092 | 0.23252994 | 1 | 1539 | tags=21%, list=7%, signal=23% |
| LINOLEIC ACID METABOLISM | LINOLEIC ACID METABOLISM | 28 | 0.771579 | 1.584588 | 0.009242144 | 0.22784553 | 1 | 818 | tags=11%, list=4%, signal=11% |
| POSITIVE REGULATION OF ERK1 AND ERK2 CASCADE (GO:0070374) | POSITIVE REGULATION OF ERK1 AND ERK2 CASCADE (GO:0070374) | 173 | 0.603463 | 1.578182 | 0 | 0.24507926 | 1 | 818 | tags=16%, list=4%, signal=17% |
| STEROL METABOLIC PROCESS (GO:0016125) | STEROL METABOLIC PROCESS (GO:0016125) | 44 | 0.711721 | 1.577445 | 0.016014235 | 0.24187143 | 1 | 1186 | tags=16%, list=5%, signal=17% |
| NEGATIVE REGULATION OF PROTEIN LOCALIZATION TO PLASMA MEMBRANE (GO:1903077) | NEGATIVE REGULATION OF PROTEIN LOCALIZATION TO PLASMA MEMBRANE (GO:1903077) | 17 | 0.836375 | 1.576232 | 0.007352941 | 0.24097155 | 1 | 1690 | tags=47%, list=8%, signal=51% |
| STAPHYLOCOCCUS AUREUS INFECTION | STAPHYLOCOCCUS AUREUS INFECTION | 41 | 0.737954 | 1.575626 | 0.007407407 | 0.23794779 | 1 | 564 | tags=32%, list=3%, signal=32% |
| POSITIVE REGULATION OF PATHWAY-RESTRICTED SMAD PROTEIN PHOSPHORYLATION (GO:0010862) | POSITIVE REGULATION OF PATHWAY-RESTRICTED SMAD PROTEIN PHOSPHORYLATION (GO:0010862) | 40 | 0.726459 | 1.575065 | 0.016453382 | 0.23481841 | 1 | 1355 | tags=23%, list=6%, signal=24% |
| REGULATION OF BLOOD COAGULATION (GO:0030193) | REGULATION OF BLOOD COAGULATION (GO:0030193) | 25 | 0.790083 | 1.573699 | 0.003610108 | 0.23494686 | 1 | 1603 | tags=36%, list=7%, signal=39% |
| STRIATED MUSCLE CELL DEVELOPMENT (GO:0055002) | STRIATED MUSCLE CELL DEVELOPMENT (GO:0055002) | 18 | 0.841501 | 1.570549 | 0.003649635 | 0.24105509 | 1 | 754 | tags=22%, list=3%, signal=23% |
| LIPID TRANSPORT (GO:0006869) | LIPID TRANSPORT (GO:0006869) | 71 | 0.66776 | 1.569059 | 0.006956522 | 0.24173126 | 1 | 2305 | tags=28%, list=10%, signal=31% |
| PERTUSSIS | PERTUSSIS | 67 | 0.670574 | 1.568796 | 0.012195122 | 0.23783259 | 1 | 944 | tags=24%, list=4%, signal=25% |
| INTRACELLULAR PH REDUCTION (GO:0051452) | INTRACELLULAR PH REDUCTION (GO:0051452) | 24 | 0.793695 | 1.567978 | 0.014925373 | 0.23588704 | 1 | 2075 | tags=58%, list=9%, signal=64% |
| NEGATIVE REGULATION OF MYELOID LEUKOCYTE DIFFERENTIATION (GO:0002762) | NEGATIVE REGULATION OF MYELOID LEUKOCYTE DIFFERENTIATION (GO:0002762) | 21 | 0.823328 | 1.566948 | 0.01362862 | 0.23513527 | 1 | 385 | tags=24%, list=2%, signal=24% |
| PORPHYRIN AND CHLOROPHYLL METABOLISM | PORPHYRIN AND CHLOROPHYLL METABOLISM | 28 | 0.773205 | 1.562995 | 0.012345679 | 0.24411201 | 1 | 1939 | tags=29%, list=9%, signal=31% |
| NEGATIVE REGULATION OF BLOOD COAGULATION (GO:0030195) | NEGATIVE REGULATION OF BLOOD COAGULATION (GO:0030195) | 26 | 0.777573 | 1.562076 | 0.016917294 | 0.24307485 | 1 | 1603 | tags=35%, list=7%, signal=37% |
| ODONTOGENESIS (GO:0042476) | ODONTOGENESIS (GO:0042476) | 32 | 0.76047 | 1.560311 | 0.00907441 | 0.24467127 | 1 | 945 | tags=19%, list=4%, signal=20% |
| FATTY ACID CATABOLIC PROCESS (GO:0009062) | FATTY ACID CATABOLIC PROCESS (GO:0009062) | 58 | 0.685422 | 1.559085 | 0.008417509 | 0.24407329 | 1 | 3344 | tags=50%, list=15%, signal=59% |
| NEGATIVE REGULATION OF PROTEIN LOCALIZATION TO CELL PERIPHERY (GO:1904376) | NEGATIVE REGULATION OF PROTEIN LOCALIZATION TO CELL PERIPHERY (GO:1904376) | 16 | 0.823747 | 1.557955 | 0.005484461 | 0.24330132 | 1 | 1690 | tags=50%, list=8%, signal=54% |
| CALCIUM ION HOMEOSTASIS (GO:0055074) | CALCIUM ION HOMEOSTASIS (GO:0055074) | 74 | 0.656886 | 1.556752 | 0.005347594 | 0.24313968 | 1 | 1257 | tags=16%, list=6%, signal=17% |
| FERRIC IRON TRANSPORT (GO:0015682) | FERRIC IRON TRANSPORT (GO:0015682) | 30 | 0.758641 | 1.553628 | 0.012006861 | 0.24609067 | 1 | 2132 | tags=57%, list=10%, signal=63% |
| POSITIVE REGULATION OF INTERLEUKIN-1 BETA PRODUCTION (GO:0032731) | POSITIVE REGULATION OF INTERLEUKIN-1 BETA PRODUCTION (GO:0032731) | 22 | 0.79548 | 1.552717 | 0.012658228 | 0.2456671 | 1 | 1001 | tags=27%, list=5%, signal=29% |
| VIRAL GENE EXPRESSION (GO:0019080) | VIRAL GENE EXPRESSION (GO:0019080) | 105 | -0.90822 | -2.364 | 0 | 0 | 0 | 1048 | tags=70%, list=5%, signal=73% |
| VIRAL TRANSCRIPTION (GO:0019083) | VIRAL TRANSCRIPTION (GO:0019083) | 108 | -0.89497 | -2.3608 | 0 | 0 | 0 | 1048 | tags=67%, list=5%, signal=70% |
| NUCLEAR-TRANSCRIBED MRNA CATABOLIC PROCESS, NONSENSE-MEDIATED DECAY (GO:0000184) | NUCLEAR-TRANSCRIBED MRNA CATABOLIC PROCESS, NONSENSE-MEDIATED DECAY (GO:0000184) | 109 | -0.88333 | -2.33171 | 0 | 0 | 0 | 1530 | tags=71%, list=7%, signal=76% |
| COTRANSLATIONAL PROTEIN TARGETING TO MEMBRANE (GO:0006613) | COTRANSLATIONAL PROTEIN TARGETING TO MEMBRANE (GO:0006613) | 88 | -0.90289 | -2.32396 | 0 | 0 | 0 | 1048 | tags=81%, list=5%, signal=84% |
| SRP-DEPENDENT COTRANSLATIONAL PROTEIN TARGETING TO MEMBRANE (GO:0006614) | SRP-DEPENDENT COTRANSLATIONAL PROTEIN TARGETING TO MEMBRANE (GO:0006614) | 85 | -0.91793 | -2.31828 | 0 | 0 | 0 | 1048 | tags=85%, list=5%, signal=89% |
| RIBOSOME | RIBOSOME | 128 | -0.85397 | -2.30107 | 0 | 0 | 0 | 1048 | tags=60%, list=5%, signal=63% |
| PROTEIN TARGETING TO ER (GO:0045047) | PROTEIN TARGETING TO ER (GO:0045047) | 91 | -0.8882 | -2.26427 | 0 | 0 | 0 | 1116 | tags=78%, list=5%, signal=82% |
| RRNA PROCESSING (GO:0006364) | RRNA PROCESSING (GO:0006364) | 194 | -0.78911 | -2.23936 | 0 | 0 | 0 | 1991 | tags=46%, list=9%, signal=50% |
| PEPTIDE BIOSYNTHETIC PROCESS (GO:0043043) | PEPTIDE BIOSYNTHETIC PROCESS (GO:0043043) | 159 | -0.80119 | -2.22503 | 0 | 0 | 0 | 1048 | tags=46%, list=5%, signal=48% |
| RRNA METABOLIC PROCESS (GO:0016072) | RRNA METABOLIC PROCESS (GO:0016072) | 193 | -0.776 | -2.18916 | 0 | 0 | 0 | 1371 | tags=40%, list=6%, signal=43% |
| NUCLEAR-TRANSCRIBED MRNA CATABOLIC PROCESS (GO:0000956) | NUCLEAR-TRANSCRIBED MRNA CATABOLIC PROCESS (GO:0000956) | 165 | -0.79278 | -2.18001 | 0 | 0 | 0 | 1980 | tags=55%, list=9%, signal=59% |
| RIBOSOME BIOGENESIS (GO:0042254) | RIBOSOME BIOGENESIS (GO:0042254) | 216 | -0.76462 | -2.16952 | 0 | 0 | 0 | 1991 | tags=41%, list=9%, signal=44% |
| NCRNA PROCESSING (GO:0034470) | NCRNA PROCESSING (GO:0034470) | 219 | -0.74273 | -2.11769 | 0 | 0 | 0 | 1991 | tags=41%, list=9%, signal=44% |
| CYTOPLASMIC TRANSLATION (GO:0002181) | CYTOPLASMIC TRANSLATION (GO:0002181) | 49 | -0.89086 | -2.06273 | 0 | 0 | 0 | 1246 | tags=71%, list=6%, signal=76% |
| TRANSLATION (GO:0006412) | TRANSLATION (GO:0006412) | 220 | -0.72992 | -2.05725 | 0 | 0 | 0 | 1246 | tags=35%, list=6%, signal=37% |
| CELLULAR MACROMOLECULE BIOSYNTHETIC PROCESS (GO:0034645) | CELLULAR MACROMOLECULE BIOSYNTHETIC PROCESS (GO:0034645) | 348 | -0.62808 | -1.87827 | 0 | 0.00048594 | 0.008 | 2011 | tags=32%, list=9%, signal=35% |
| GENE EXPRESSION (GO:0010467) | GENE EXPRESSION (GO:0010467) | 391 | -0.61311 | -1.84132 | 0 | 0.001375494 | 0.024 | 2610 | tags=37%, list=12%, signal=42% |
| RIBOSOME ASSEMBLY (GO:0042255) | RIBOSOME ASSEMBLY (GO:0042255) | 50 | -0.79031 | -1.82227 | 0 | 0.002917721 | 0.054 | 1038 | tags=40%, list=5%, signal=42% |
| RIBOSOMAL SMALL SUBUNIT BIOGENESIS (GO:0042274) | RIBOSOMAL SMALL SUBUNIT BIOGENESIS (GO:0042274) | 39 | -0.80466 | -1.79133 | 0 | 0.007207729 | 0.133 | 1837 | tags=46%, list=8%, signal=50% |
| VIRAL PROCESS (GO:0016032) | VIRAL PROCESS (GO:0016032) | 208 | -0.62039 | -1.76617 | 0 | 0.012482396 | 0.223 | 1048 | tags=39%, list=5%, signal=41% |
| GRAFT-VERSUS-HOST DISEASE | GRAFT-VERSUS-HOST DISEASE | 36 | -0.78091 | -1.74003 | 0.002309469 | 0.021366095 | 0.37 | 903 | tags=31%, list=4%, signal=32% |
| TYPE I DIABETES MELLITUS | TYPE I DIABETES MELLITUS | 39 | -0.77062 | -1.73218 | 0 | 0.024417154 | 0.431 | 903 | tags=28%, list=4%, signal=29% |
| RIBOSOMAL SMALL SUBUNIT ASSEMBLY (GO:0000028) | RIBOSOMAL SMALL SUBUNIT ASSEMBLY (GO:0000028) | 16 | -0.88362 | -1.71865 | 0.002237137 | 0.03210418 | 0.533 | 764 | tags=56%, list=3%, signal=58% |
| RIBOSOMAL LARGE SUBUNIT BIOGENESIS (GO:0042273) | RIBOSOMAL LARGE SUBUNIT BIOGENESIS (GO:0042273) | 58 | -0.71791 | -1.70203 | 0.002304148 | 0.044450298 | 0.67 | 3225 | tags=53%, list=15%, signal=62% |
| ANTIGEN RECEPTOR-MEDIATED SIGNALING PATHWAY (GO:0050851) | ANTIGEN RECEPTOR-MEDIATED SIGNALING PATHWAY (GO:0050851) | 159 | -0.61993 | -1.7008 | 0 | 0.043764967 | 0.678 | 1971 | tags=32%, list=9%, signal=35% |
| T CELL RECEPTOR SIGNALING PATHWAY (GO:0050852) | T CELL RECEPTOR SIGNALING PATHWAY (GO:0050852) | 132 | -0.62056 | -1.6909 | 0.002450981 | 0.051389072 | 0.753 | 1971 | tags=29%, list=9%, signal=31% |
| NEGATIVE REGULATION OF LIPID METABOLIC PROCESS (GO:0045833) | NEGATIVE REGULATION OF LIPID METABOLIC PROCESS (GO:0045833) | 15 | -0.87146 | -1.68165 | 0.008948546 | 0.058895163 | 0.809 | 970 | tags=27%, list=4%, signal=28% |
| CHROMATIN ASSEMBLY (GO:0031497) | CHROMATIN ASSEMBLY (GO:0031497) | 46 | -0.71703 | -1.67413 | 0 | 0.06492101 | 0.847 | 2095 | tags=26%, list=9%, signal=29% |
| FC-GAMMA RECEPTOR SIGNALING PATHWAY INVOLVED IN PHAGOCYTOSIS (GO:0038096) | FC-GAMMA RECEPTOR SIGNALING PATHWAY INVOLVED IN PHAGOCYTOSIS (GO:0038096) | 70 | -0.67584 | -1.67188 | 0 | 0.06592484 | 0.86 | 1739 | tags=39%, list=8%, signal=42% |
| CELLULAR RESPONSE TO INTERLEUKIN-12 (GO:0071349) | CELLULAR RESPONSE TO INTERLEUKIN-12 (GO:0071349) | 44 | -0.7132 | -1.65805 | 0.007407407 | 0.08250905 | 0.919 | 1949 | tags=50%, list=9%, signal=55% |
| VIRAL MYOCARDITIS | VIRAL MYOCARDITIS | 59 | -0.69418 | -1.65019 | 0.004608295 | 0.09267421 | 0.948 | 1096 | tags=29%, list=5%, signal=30% |
| ALLOGRAFT REJECTION | ALLOGRAFT REJECTION | 35 | -0.75098 | -1.64636 | 0 | 0.09546033 | 0.957 | 903 | tags=29%, list=4%, signal=30% |
| NIK/NF-KAPPAB SIGNALING (GO:0038061) | NIK/NF-KAPPAB SIGNALING (GO:0038061) | 76 | -0.65552 | -1.64235 | 0.002469136 | 0.09881811 | 0.964 | 2637 | tags=39%, list=12%, signal=45% |
| NUCLEOSOME ASSEMBLY (GO:0006334) | NUCLEOSOME ASSEMBLY (GO:0006334) | 42 | -0.70687 | -1.63232 | 0.009501188 | 0.112918966 | 0.984 | 3111 | tags=40%, list=14%, signal=47% |
| FC RECEPTOR MEDIATED STIMULATORY SIGNALING PATHWAY (GO:0002431) | FC RECEPTOR MEDIATED STIMULATORY SIGNALING PATHWAY (GO:0002431) | 72 | -0.65312 | -1.61707 | 0 | 0.14025156 | 0.994 | 1739 | tags=39%, list=8%, signal=42% |
| INTERLEUKIN-12-MEDIATED SIGNALING PATHWAY (GO:0035722) | INTERLEUKIN-12-MEDIATED SIGNALING PATHWAY (GO:0035722) | 44 | -0.7132 | -1.61036 | 0.004524887 | 0.15218686 | 0.999 | 1949 | tags=50%, list=9%, signal=55% |
| NUCLEOSOME ORGANIZATION (GO:0034728) | NUCLEOSOME ORGANIZATION (GO:0034728) | 58 | -0.67791 | -1.59631 | 0.004694836 | 0.18458031 | 1 | 3133 | tags=41%, list=14%, signal=48% |
| CELL ADHESION MOLECULES (CAMS) | CELL ADHESION MOLECULES (CAMS) | 131 | -0.59349 | -1.58875 | 0.00248139 | 0.20053594 | 1 | 1041 | tags=21%, list=5%, signal=22% |
| FC-GAMMA RECEPTOR SIGNALING PATHWAY (GO:0038094) | FC-GAMMA RECEPTOR SIGNALING PATHWAY (GO:0038094) | 71 | -0.63945 | -1.58799 | 0.002450981 | 0.19770704 | 1 | 1739 | tags=38%, list=8%, signal=41% |
| INNATE IMMUNE RESPONSE ACTIVATING CELL SURFACE RECEPTOR SIGNALING PATHWAY (GO:0002220) | INNATE IMMUNE RESPONSE ACTIVATING CELL SURFACE RECEPTOR SIGNALING PATHWAY (GO:0002220) | 104 | -0.59699 | -1.58471 | 0.002252252 | 0.20168094 | 1 | 1945 | tags=30%, list=9%, signal=33% |
| REGULATION OF T CELL RECEPTOR SIGNALING PATHWAY (GO:0050856) | REGULATION OF T CELL RECEPTOR SIGNALING PATHWAY (GO:0050856) | 23 | -0.79414 | -1.57764 | 0.013215859 | 0.21849595 | 1 | 1528 | tags=39%, list=7%, signal=42% |
| REGULATION OF LIPOPROTEIN LIPASE ACTIVITY (GO:0051004) | REGULATION OF LIPOPROTEIN LIPASE ACTIVITY (GO:0051004) | 16 | -0.82862 | -1.57317 | 0.008830022 | 0.22700022 | 1 | 512 | tags=13%, list=2%, signal=13% |
